# ATF6 activation alters colonic lipid metabolism causing tumor-associated microbial adaptation

**DOI:** 10.1101/2023.11.03.565267

**Authors:** Olivia I. Coleman, Adam Sorbie, Alessandra Riva, Miriam von Stern, Stephanie Kuhls, Denise M. Selegato, Nikolai Köhler, Jakob Wirbel, Tim Kacprowski, Andreas Dunkel, Josh K. Pauling, Johannes Plagge, Diego Mediel-Cuadra, Sophia Wagner, Ines Chadly, Sandra Bierwith, Tingying Peng, Thomas Metzler, Clemens Schafmayer, Sebastian Hinz, Christian Röder, Christoph Röcken, Michael Zimmermann, Philip Rosenstiel, Katja Steiger, Moritz Jesinghaus, Gerhard Liebisch, Josef Ecker, Christina Schmidt, Georg Zeller, Klaus-Peter Janssen, Dirk Haller

## Abstract

Endoplasmic reticulum unfolded protein responses (UPR^ER^) contribute to cancer development and the activating transcription factor 6 (ATF6) is involved in microbiota-dependent tumorigenesis. Here, we substantiate the clinical relevance of ATF6 in early-onset and late colorectal cancer patient cohorts. Transcriptional analysis in intestinal epithelial cells (IEC) of ATF6 transgenic mice (nATF6^IEC^) identified bacteria-specific changes in cellular metabolism enriched for fatty acid biosynthesis. Untargeted metabolomics and isotype-labeling confirmed ATF6-related enrichment of long chain fatty acids in colonic tissue of patients, mice and organoid cultures. FASN inhibition and microbiota transfer in germ-free nATF6^IEC^ mice confirmed the causal involvement of ATF6-induced lipid alterations in tumorigenesis. The selective expansion of tumor-relevant microbial taxa was mechanistically linked to long chain fatty acid exposure, using bioorthogonal non-canonical amino acid tagging (BONCAT) and growth analysis of Desulfovibrio isolates. We postulate chronic ATF6 signaling in the epithelium to select for tumor-promoting microbiota by altering lipid metabolism.

Graphical Abstract:
Chronic ATF6 signaling in the colonic epithelium alters lipid metabolism to select a tumor-promoting microbiota

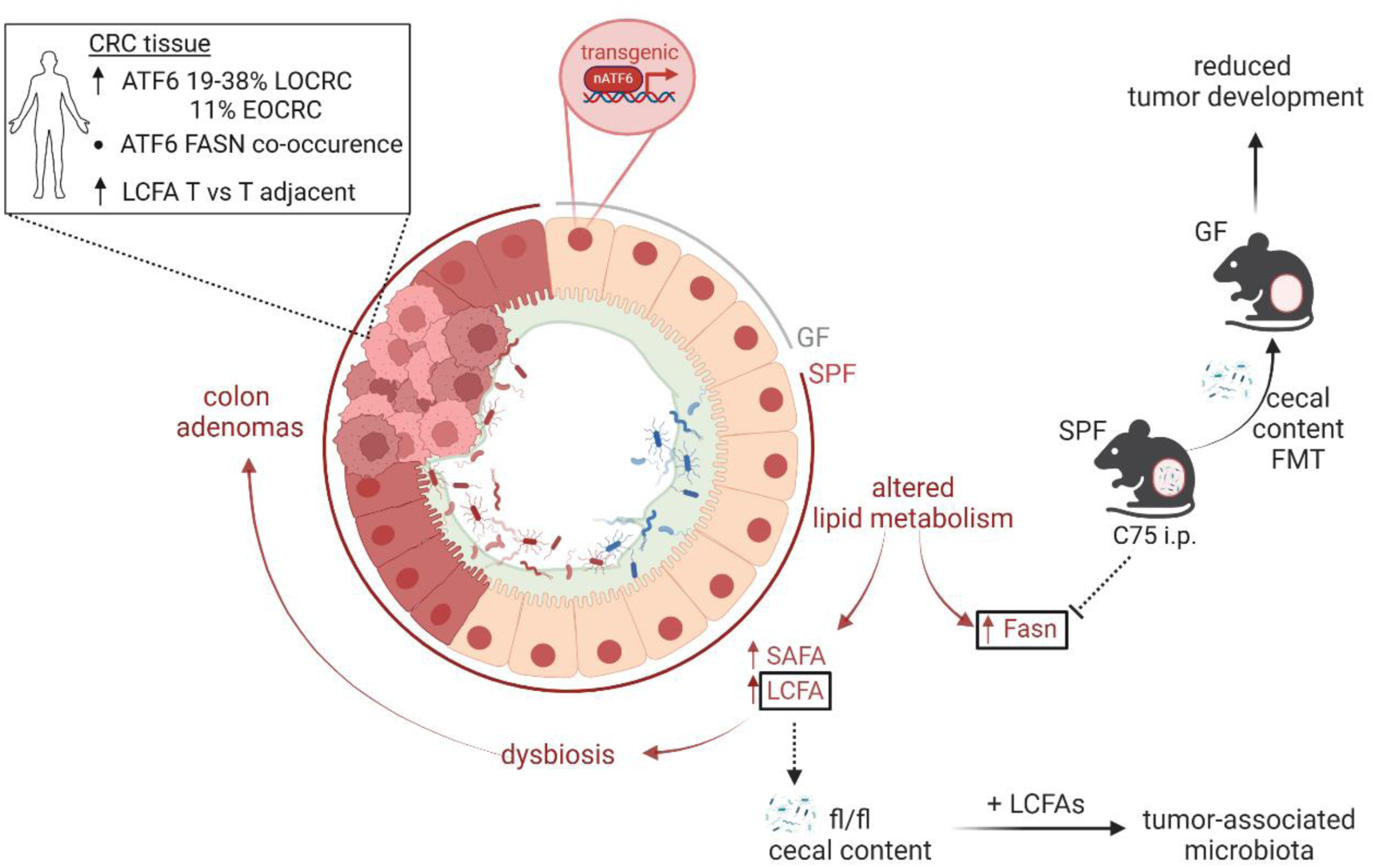

- Biallelic expression of activated ATF6 (p50 nuclear fragment) in intestinal epithelial cells (nATF6^IEC^) induces spontaneous colon tumors in SPF but not GF mice
- Mechanistically, biallelic SPF nATF6^IEC^ mice alter colonic lipid metabolism, including the upregulation of LCFAs and Fasn
- Inhibition of FASN prevents colon tumor formation in mice, and reduces the tumor-promoting potential of the intestinal microbiota (FMT)
- Exposure of fl/fl control mouse microbiota to LCFAs *ex vivo* translationally activates tumor-associated bacteria, including *Desulfovibrio fairfieldensis*
- Human CRC patients show ATF6 upregulation, FASN co-occurrence and increased LCFAs in T tissue
- ATF6 activity links with CRC-associated microbiota in patients, including *Desulfovibrio*

Created with BioRender.com

nATF6: activated activating transcription factor 6; LCFA: long-chain fatty acids; SAFA: saturated fatty acids; Fasn: fatty acid synthase; C75 i.p.: intraperitoneal injection of the Fasn inhibitor C75.

## Introduction

The endoplasmic reticulum (ER) in mammalian cells forms an extensive tubular-reticular network that acts as a gatekeeper controlling the synthesis and folding of proteins, and the synthesis of cellular lipids ^1,2^. The accumulation of un- and misfolded ER proteins activates the unfolded protein response (UPR^ER^), a highly conserved group of intricately regulated signaling pathways that aim to restore ER homeostasis. PKR-like ER kinase (PERK), inositol-requiring enzyme 1 (IRE1) and activating transcription factor 6 (ATF6) constitute the three main arms of the UPR^ER 3^. ER-stress results in the dissociation of membrane-bound ATF6 complexes from the ER chaperone glucose-regulating protein 78 (GRP78), subsequent cleavage by the site-1 and site-2 proteases at the Golgi apparatus, and translocation of the p50 active cytosolic N-terminal portion of the transcription factor (nATF6) into the nucleus to bind the ER-stress response element (ERSE) ^4,5^.

Numerous studies recognize the UPR^ER^ as a fundamental mediator in cellular physiology and the pathogenesis of inflammatory disorders, metabolic diseases and cancer, including colorectal cancer (CRC) ^6–12^. Enhanced expression of GRP78, a downstream target of nATF6, correlates with growth, invasion and metastasis of tumors ^13^. Furthermore, ATF6 mRNA expression and polymorphisms are associated with CRC metastasis and relapse ^14,15^, and hepatocellular carcinoma ^16^, respectively. We could show that chronic transgenic expression of nATF6 specifically in intestinal epithelial cells (IECs) induces spontaneous and microbiota-dependent colon adenoma formation in a murine model of early-onset CRC (nATF6^IEC^) ^11^. Accumulating evidence unarguably renders the intestinal microbiota as an important regulator of host health and a causal player in the development and progression of CRC. Our findings that dysbiosis precedes tumorigenesis in nATF6^IEC^ mice and that nATF6-expressing germ-free mice remain tumor-free ^11^, clearly indicate a mechanistic link between ATF6 signaling and the intestinal microbiota in the context of colon tumorigenesis. What remains to be elucidated, is how ATF6 signaling induces dysbiosis at the pre-tumor stage.

The multifaceted possibilities of physiological outcomes in response to the UPR^ER^ underlines the necessity to understand the mechanistic contribution of UPR^ER^ signaling to disease pathology. Here we firstly set out to identify the relevance of ATF6 in CRC patient populations, and secondly to understand the consequence of ATF6 signaling on the murine host. We characterize ATF6 expression in multiple independent German CRC cohorts and causally link chronic ATF6 activity to altered lipid metabolism and tumor-relevant microbiota changes.

## Results

### High ATF6 expression defines a subset of CRC patients

We previously identified increased *ATF6* expression to be associated with reduced disease-free survival in colorectal cancer (CRC) patients ^11^. To substantiate these findings, we quantified ATF6 protein expression via immunohistochemistry (IHC) staining in a German CRC patient cohort (cohort 1; *n=959*) using QuPath ^17,18^. ATF6 antibody specificity was confirmed through negative staining in human heart muscle, and the appropriate isotype control staining in CRC tissue (Extended Data Fig. 1a). ATF6 expression was compared between the tumor-center (ATF6-TC) and the tumor invasive front (ATF6-IF), showing similar expression levels between these two localizations (data not shown), and subsequently averaged to determine the ATF6 H-score (capturing both the intensity and the percentage of stained cells) (Fig. 1a). Findings show that ATF6 upregulation occurs in 38% of CRC patients that can be classed as ATF6-high in terms of their nuclear (NUC) and cytoplasmic (CYT) expression (Fig. 1a; Extended Data Fig. 1b,c). Stratification into all possible combinations of NUC and CYT ATF6 staining into low (L) and high (H) shows that all NUC H patients are also CYT H (38%), with 0% of NUC H patients being CYT L (Extended Data Fig. 1d). The other combinations of NUC L/CYTL and NUC L/CYT H make up 21% and 41% of the CRC patients, respectively (Extended Data Fig. 1d). In the same cohort 1, we assessed the global levels of endoplasmic reticulum (ER) stress through IHC staining of glucose regulating protein 78 (GRP78), the master chaperone of the ER. Interestingly, quantification of GRP78 using QuPath showed that, despite the wide range of GRP78 H-scores amongst the CRC cohort, there is no difference in GRP78 H-scores between ATF6-high and ATF6-low CRC patient groups (Extended Data Fig. 1e,f). To validate the above findings, we trained a deep learning model to predict ATF6 expression levels of CRC patient biopsy images. The model was trained on an independent CRC patient cohort (cohort 2; *n=50*) with classification and regression objectives using 5-fold cross validation. In line with our results, external testing results identified 39% (classification task) and 35.9% (regression task) of CRC patients in cohort 1 as ATF6-high (Fig. 1b-e, Extended Data Fig. 1g,h), confirming the robustness and applicability of our AI methodology. To further support our findings, we quantified ATF6 expression using QuPath in a third German CRC patient cohort (cohort 3; *n=256*). Here we identified a subpopulation of 19% of CRC patients to be ATF6-high (Fig. 1f). To investigate a potentially dissimilar effect of ATF6 expression in patients with early-onset CRC (age<50; EOCRC) versus late-onset CRC (age>50; LOCRC), we quantified ATF6 expression in additional patients from cohort 3 with EOCRC (cohort 3; *n=55* EOCRC). Findings show similar expression levels in EOCRC, with 11% of patients classified as ATF6-high (Fig. 1g). Together, these results clearly demonstrate that high ATF6 expression defines a subpopulation of CRC patients (19-38% LOCRC; 11% EOCRC).

**Fig. 1:**
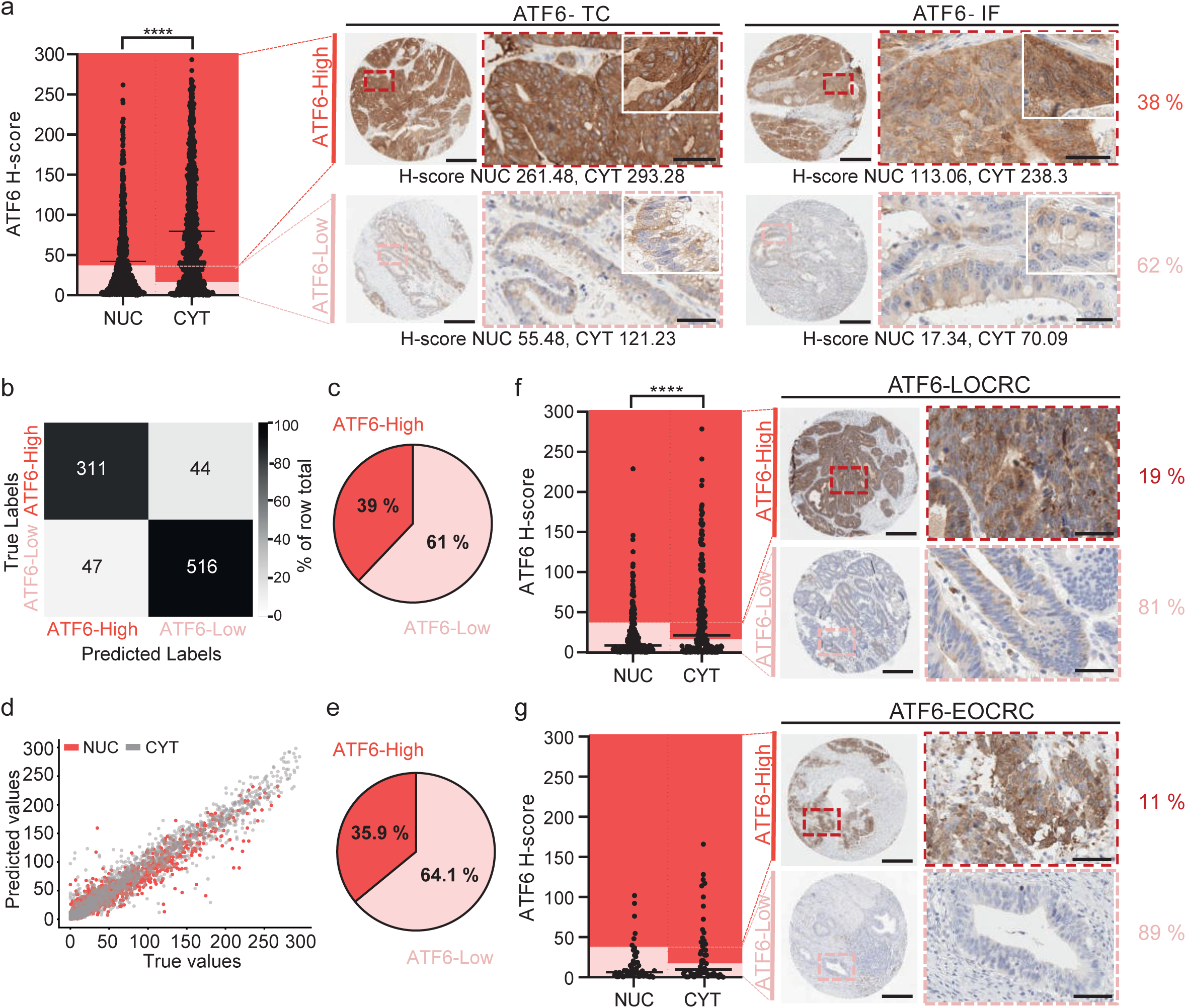
High ATF6 expression defines a subset of CRC patients. **a,** QuPath quantified ATF6 H-score of nuclear (NUC) and cytoplasmic (CYT) ATF6 expression in immunohistochemically stained CRC patient tissue samples (cohort 1, n=959). Significance was calculated using the Student’s two-tailed unpaired T-Test (*p*<0.0001), indicated is the mean. Representative images of ATF6-high and ATF6-low staining in the tumor centre (TC) and the invasive front (IF). Percentage shows subpopulation of CRC patients in each ATF6-stained category (ATF6-low, light red; ATF6-high, dark red). Scale bars: 300µm overview, 50µm zoom-in. **b,** Confusion matrix depicting the classification model of ATF6 H-scores quantified using QuPath (true labels) versus ATF6 H-scores predicted by the classification algorithm (predicted labels) for ATF6-high and ATF6-low CRC patients. **c,** Pie chart showing percentage of ATF6-high and ATF6-low CRC patients identified in CRC cohort 2 (n=50) using the classification model. **d,** Regression model of ATF6 H-scores quantified using QuPath (true values) versus ATF6 H-scores predicted by the regression algorithm (predicted values) for nuclear (NUC) and cytoplasmic (CYT) ATF6 expression. **e,** Pie chart showing percentage of ATF6-high and ATF6-low CRC patients identified in CRC cohort 2 (n=50) using the regression model. **f,** QuPath quantified ATF6 H-score of nuclear (NUC) and cytoplasmic (CYT) ATF6 expression in immunohistochemically stained CRC patient tissue samples (cohort 3, LOCRC, n=256). Significance was calculated using the Student’s two-tailed unpaired T-Test (*p*<0.0001), indicated is the mean. Representative images of ATF6-high and ATF6-low staining. Percentage shows subpopulation of CRC patients in each ATF6-stained category. Scale bars: 300µm overview, 50 µm zoom-in. **g,** QuPath quantified ATF6 H-score of nuclear (NUC) and cytoplasmic (CYT) ATF6 expression in immunohistochemically stained CRC patient tissue samples (cohort 3, EOCRC, n=55). Significance was calculated using the Student’s two-tailed unpaired T-Test, indicated is the mean. Representative images of ATF6-high and ATF6-low staining. Percentage shows subpopulation of CRC patients in each ATF6-stained category. Scale bars: 300µm overview, 50µm zoom-in. A significant *p*-value <0.05 is represented as an asterisk: \**p* < 0.05, \*\**p* < 0.01. \*\*\**p* < 0.001, \*\*\*\**p* < 0.0001.

### ATF6 alters colonic fatty acid metabolism in the presence of bacteria

In our ATF6 transgenic mouse model (nATF6^IEC^), we previously established a causal role of the microbiota in adenoma formation ^11^. In agreement with these findings, biallelic nATF6^IEC^ mice (tg/tg) show a reduced survival (Fig. 2a), and develop colonic tumors with an incidence of 100% under specific pathogen-free (SPF) conditions, while controls (fl/fl) and monoallelic (tg/wt) nATF6 expression, as well as germ-free (GF) tg/tg mice remain tumor-free (Fig. 2b). To characterize the transcriptional response induced by chronic ATF6 activity and to further understand the contribution of bacteria, we performed bulk mRNA sequencing of colonic epithelial cells in fl/fl and tg/tg mice at the pre-tumor time point (5wk, highlighted in orange) under SPF and GF conditions (Fig. 2c). Transgenic chronic activation of nATF6 in SPF tg/tg mice results in a total of 1941 differentially expressed genes (DEGs; threshold: -0.5 > log2FC > 0.5, p adj. < 0.05) compared to fl/fl controls, of which 1236 DEGs are upregulated and 705 DEGs are downregulated (Fig. 2d; Supplementary Table 1, sheet 1). Functional analysis using the Kyoto Encyclopedia of Genes and Genomes (KEGG) pathways showed that almost half (16/36; blue labels) of the top and bottom ranked pathways, according to the normalized enrichment score (NES), are related to metabolic pathways in SPF tg/tg mice (Fig. 2e, Supplementary Table 2, sheet 1). Under GF conditions, tg/tg mice showed a total of 706 DEGs, of which 491 DEGS are upregulated and 215 DEGs are downregulated (Extended Data Fig. 2a; Supplementary Table 1, sheet 2). Again, half of the top and bottom ranked KEGG pathways (14/28; blue labels) in GF tg/tg mice are linked to metabolic pathways (Extended Data Fig. 2b, Supplementary Table 2, sheet 2). Metabolic KEGG pathways associated with SPF tg/tg mice (tumor development) but not with GF tg/tg mice (no tumor development) are beta alanine metabolism, butanoate metabolism, fatty-acid (FA) metabolism, glycolysis gluconeogenesis, metabolism of xenobiotics by cytochrome P450 and nicotinate and nicotinamide metabolism (Fig. 2e). Of these, FA metabolism represents the highest ranked KEGG pathway upregulated in SPF tg/tg mice. To characterize the specificity of ATF6-related activation in the context of all UPR arms, we specifically looked at UPR-related genes, including the chronic SPF model, GF model, as well as our acute (tamoxifen-inducible transgenic model at 4 days post ATF6-activation; nATF6 Vil-Cre^ERT2Tg^) SPF model (Extended Data Fig. 2c). We see that eight out of the nine top regulated genes are also described to be ATF6 targets (marked in red), with considerable overlap in the other UPR^ER^ arms. The panel reveals that the expression level of target genes is amplified by chronicity of ATF6-activation and bacterial presence.

**Fig. 2:**
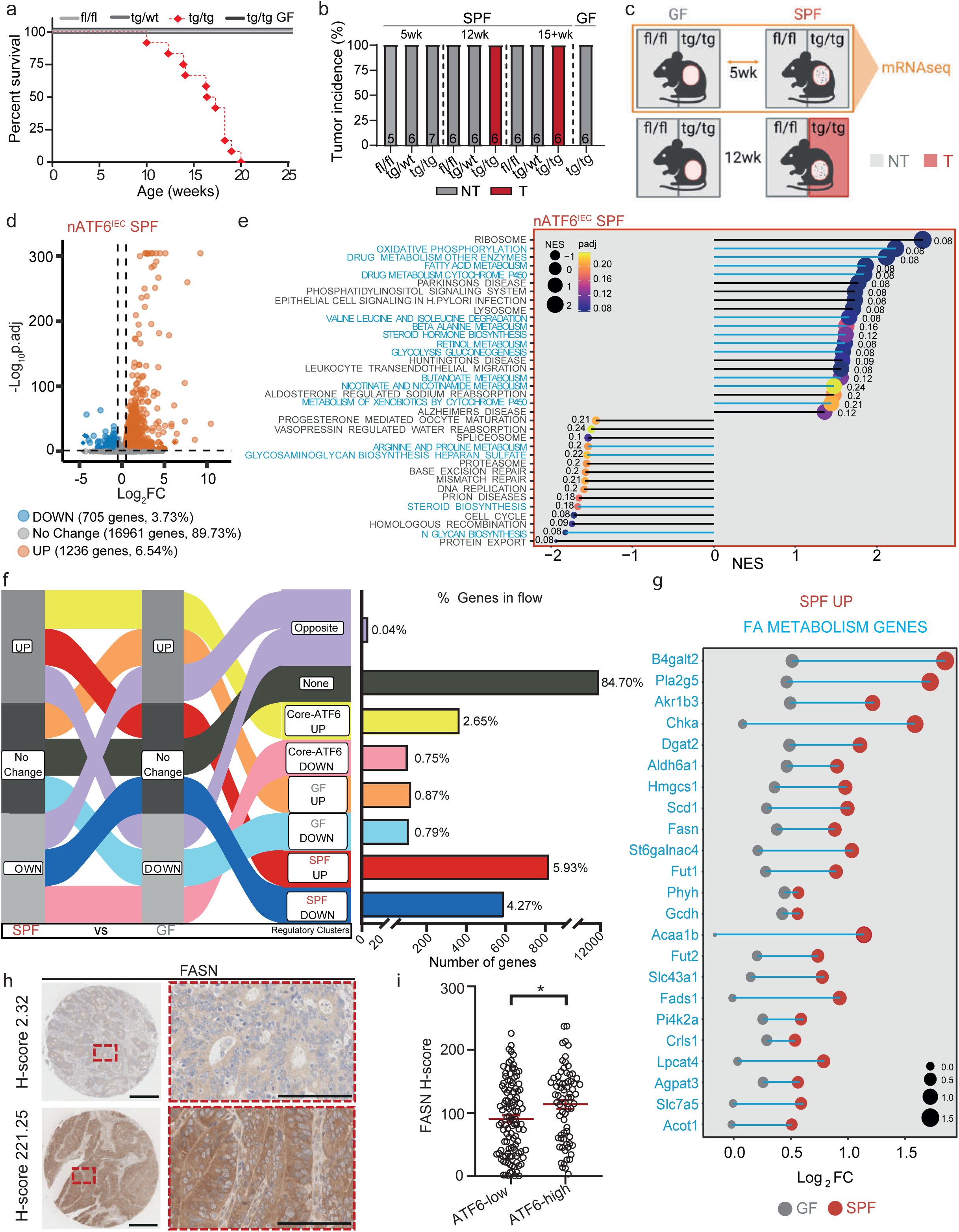
ATF6 alters colonic fatty acid metabolism in the presence of bacteria. **a,** Survival curve showing percent survival of fl/fl, tg/wt, tg/tg specific pathogen-free (SPF) and tg/tg germ-free (GF) mice between 0-25 weeks of age (n per genotype = 6-12). **b,** Tumor incidence (percentage) of SPF fl/fl, tg/wt and tg/tg mice and GF tg/tg mice at the pre-tumor (5wk), tumor (12wk) and late-tumor (15+wk) time points. Grey represents no tumor, red represents tumor. Number of mice is stated in each column. **c,** Schematic of non-tumor (NT) and tumor (T) phenotypes associated with GF and SPF fl/fl and tg/tg mice, highlighting the genotypes and colonization status used for the RNA sequencing analyses (orange box and arrow) (n=6). **d**, Volcano plot of detected genes in fl/fl versus tg/tg nATF6^IEC^ mice showing the number and percentage of genes that are unchanged (No Change), upregulated in tg/tg mice (UP) or downregulated in tg/tg mice (DOWN). Threshold: -0.5 > log2FC >0.5, *p* adj. <0.05. **e,** Lollipop graph of the top and bottom regulated KEGG pathways in tg/tg versus fl/fl nATF6^IEC^ SPF mice (*p* adj. <= 0.25, −1 =>NES>= 1, with a minimum of 50% of genes detected in the pathway). Metabolic pathways are highlighted in blue. NES, normalized enrichment score. **f,** Alluvial plot depicting the flows used to define SiRCle clusters in a comparison between SPF and GF mice. Each data type (SPF and GF) has been labelled as a column, with one of three states (UP, No Change, DOWN) defined for each column based on the results for differential analysis between fl/fl versus tg/tg in that data type. Shown are the flows between each data type and state, which define the regulatory clusters (third column). The number and percentage of genes in the flow are depicted next to each regulatory cluster. **g,** Lollipop graph of the FA metabolism related genes from the SPF UP cluster (blue) shown for GF (grey) and SPF (red) mice. **h,** Representative images of low H-score and high H-score FASN immunohistochemically stained in CRC patient tissue samples (cohort 1, n=181). Scale bars: 300µm overview, 50µm zoom-in. **i,** QuPath quantified FASN H-score with patients grouped into ATF6-low and ATF6-high (according to QuPath quantified ATF6 H-score). Significance was calculated using the Student’s two-tailed unpaired T-Test (*p*=0.0105), indicated is the mean. A significant *p*-value <0.05 is represented as an asterisk: \**p* < 0.05, \*\**p* < 0.01. \*\*\**p* < 0.001, \*\*\*\**p* < 0.0001.

To disentangle the transcriptional regulation between chronic tg/tg signaling under SPF and GF conditions, differentiating between the tumor-developing and non-tumor developing phenotype, we applied Signature Regulatory Clustering (SiRCle) ^19^. We defined three states of genes in tg/tg mice compared to fl/fl mice for each colonization status group, namely No Change (not regulated, -0.5 < log2FC < 0.5, p adj. > 0.05), UP (upregulated, log2FC > 0.5 and p. adj. <0.05) and DOWN (downregulated, log2FC <-0.5 and p adj. <0.05) (Fig. 2f, Supplementary Table 3, sheet 1). Following the flow of genes between SPF and GF conditions identified eight regulatory clusters named after the biological phenotype they include, for instance Core-ATF6 UP (Fig. 2f; Supplementary Table 3, sheet 2). Depicted next to the Alluvial plot are the number and percentage of genes found in each of the clusters. Additionally, we performed functional analysis using KEGG pathways (threshold: >5% of pathway genes detected, p adj. < 0.2, supplementary Table 4), and identified transcription factors that drive individual clusters (supplementary Table 5). After the regulatory cluster “none” (84.70%), the highest percentages of DEGs were found in the clusters “SPF-UP” (5.93 %), “SPF-DOWN” (4.27%) and “Core-ATF6 UP” (2.65%) (Fig. 2f). As anticipated, more than half (*P4hb*, *Atf6*, *Calr*, *Hspa5*, *Pdia4*, *Anxa6*, *Manf*, *Sdf2I1*, *Hsp90b1*, *CreId2*, *Nucb2*, *C1qtnf1*, *Herpud1* and *Dnajb11*) of the highly DEGs in the Core-ATF6 UP cluster (24 DEGs; black labels) are classically associated with ER protein folding and calcium homeostasis, and localized within the ER (Extended Data Fig. 2d). Transcription factors *Atf4* and *Atf6* were identified as the drivers of the CORE-ATF6 UP cluster (Extended Data Fig. 2d, Supplementary Table 5, sheet 1). Highly DEGs in the Core-ATF6 DOWN cluster (black labels) include *Cyba*, *Ces1g* and *Cyp4b1*, which are associated with intestinal ROS production and defense against bacteria at the colonic epithelium ^20^, hydrolysis of lipids and xenobiotics ^21^, and the hydroxylation of fatty acids and alcohols ^22^, respectively (Extended Data Fig. 2e). Functionally, the CORE-ATF6 DOWN cluster associates with five KEGG pathways, including the drug metabolism cytochrome p450 pathway, associated with cellular metabolism, homeostasis and the detoxification of drugs ^23^ (Extended Data Fig. 2e, Supplementary Table 4, sheet 1). *Arntl* is the identified transcription factor driving the CORE-ATF6 DOWN cluster, responsible for maintenance of the circadian rhythm and has been shown to be downregulated in cancers with an association to anti-cancer roles ^24^ (Extended Figure 2e, Supplementary Table 5, sheet 2). Highly regulated DEGs in the SPF DOWN cluster (black labels) include genes that have been associated with tumor suppression and CRC ^25–27^ (*Meg3*, *Cwh43, Naprt)*, lipid metabolism and detoxification ^28^ (*Ces1f*), and bactericidal/antimicrobial activity ^29^ (*Iapp*) (Extended Data Fig. 2f). The majority of KEGG pathways associated with the SPF DOWN cluster are metabolic pathways, including the drug metabolism cytochrome p450 pathway also identified in the CORE-ATF6 DOWN cluster (Extended Data Fig. 2f, Supplementary Table 4, sheet 2).

Most relevant to our murine model and the role of ATF6 signaling in colon adenoma development in tg/tg mice is the SPF UP cluster. The highest DEG in this cluster (black label) is ER localized *Serp2*, classically involved in the UPR and protein glycosylation (Extended Data Fig. 2g). Among the other highly DEGs (black labels) in this cluster, *Nup88*, *Dmxl1* and *Igf2bp3* are associated with dysplasia and cancer, including CRC ^30–32^. Importantly, and in line with our findings above (Fig. 2e), the SPF UP cluster contains 23 differentially expressed FA metabolism-related genes (blue labels) (Fig. 2g, Extended Data Fig. 2g). Of note, additional analysis of the pre-tumor time point using our inducible nATF6 Vil-Cre^ERT2Tg^ murine model that addresses the transcriptional response induced by acute (4d) activation of nATF6, demonstrates that the upregulation of the majority (13/23) of these FA metabolism-related genes (blue labels) occurs already early on after nATF6 activation (Extended Data Fig. 2h, Supplementary Table 1, sheet 3). In accordance with our tumor phenotype, one of the two identified KEGG pathways associated with the SPF UP cluster is cell cycle (Extended Data Fig. 2g, Supplementary Table 4, sheet 3). Three transcription factors were identified as drivers of the SPF UP cluster (Extended Data Fig. 2g, Supplementary Table 5, sheet 3), namely *E2f4* (control of cell cycle, increased expression in cancers^33^), *Foxm1* (oncogenic transcription factor, control of cell cycle, regulation of CRC metastasis and progression ^34^) and *NFyb* (promotes invasion, metastasis and differentiation, associated with multiple cancers, activator or repressor in a context dependent manner ^35^).

### ATF6 positively correlates with FASN in CRC patients

Alterations in the lipidomic profile have been observed in several cancer types ^36–38^, including CRC ^39^, which marks lipid metabolic rewiring as a phenotypic hallmark of cancer. Upregulation of fatty acid synthase (FASN), a key multi-enzyme complex of *de novo* lipid biosynthesis ^40^, is associated with aggressive disease and poor prognosis in CRC ^41,42^. We recently described a robust CRC-specific lipid signature based on quantitative lipidomics in a multi-cohort and cross-center study, identifying triacylglycerol species and FASN gene expression as differential markers of cancerous tissue and bad prognosis, respectively ^43^. *Fasn* was shown to be upregulated in response to both chronic and acute activation of nATF6 in our analyses (Fig 2.g, Extended Data Fig. 2g,h). Here, we stained a random subset (n=181) of tissue cores from our human CRC patient cohort 1 for FASN and correlated the QuPath quantified FASN H-score with our ATF6 classification (from Fig. 1a) in the same patients. Exemplary images show that patients clearly differed in their FASN expression levels (Fig. 2h). Strikingly, FASN H-scores were significantly higher in CRC patients classified as ATF6-high compared to those classified as ATF6-low (Fig. 2i), identifying a positive correlation between ATF6 and FASN expression in CRC patient tissue. In a further approach, the publicly available Pan-Cancer Atlas (*n=10.967* cases including approx. 600 CRC cases) and TCGA (*n=640* CRC cases) databases were screened for genomic amplifications/gains (amp/gain) in either *ATF6* alone (*ATF6*-only), *FASN* alone (*FASN*-only) or in both genes simultaneously (*ATF6*+*FASN*). The relationship between *ATF6*-high and *FASN*-high patients is evident through their association classified as co-occurrence, despite not reaching significance in the TCGA database (*p=0.297*), likely due to lower case numbers compared to the Pan-Cancer Atlas database (*p<0.001*) (Extended Data Fig. 3a). While the IHC-quantified ATF6 expression did not associate with CRC patient disease–free survival in our human cohorts (Extended Data Fig. 3b-d), Kaplan-Meier analysis in both databases showed a significantly reduced disease-free survival in cases that were classed as *ATF6*-only amp/gain (p<0.001 pan-cancer, p=0.0122 TCGA) and those classed as combined *ATF6* and *FASN* amp/gain (p<0.0001 pan-cancer, p<0.0001 TCGA), compared to those cases unaltered in either of the two genes (Extended Data Fig. 3e,f). Importantly, disease-free survival was significantly reduced in cases of combined *ATF6* and *FASN* amp/gain compared to those showing *ATF6*-only amp/gain in the Pan-Cancer Atlas (*p=0.0394*), with a trend observed in the TCGA database (*p=0.4046*) (Extended Data Fig. 3e,f). Amp/gain modifications were not associated with overall survival in the two databases (Extended Data Fig. 3g,h).

### ATF6 drives FA elongation and LCFAs accumulate in mice and CRC patients

ATF6 signaling clearly alters epithelial metabolism. To understand its effects on the metabolome, we performed untargeted metabolomics of nATF6^IEC^ cecal content at pre-tumor (5wk), tumor (12wk) and late-tumor (20wk) time points using LC-TOF-MS (left side schematic, Fig 3a). In parallel, we performed untargeted metabolomics of CRC patient tumor tissue (T) and patient-matched tumor adjacent (T adjacent) tissue in a cohort of 259 EOCRC and LOCRC patients (cohort 4; right side schematic, Fig. 3a). In our nATF6^IEC^ murine model, principal component analysis revealed a drastic difference in the metabolome between tumor-bearing mice (red) and non-tumor mice (grey), while metabolite profiles clustered in closer proximity for all three genotypes at the pre-tumor time point (Extended Data Fig. 4a). Differential enrichment analysis at the tumor time point demonstrated a depletion of peptide metabolites and sphingolipids, while long-chain fatty acids (LCFA) and other complex lipid species were enriched in T mice (Extended Data Fig. 4b). Principal component analysis showed that lipid profiles of non-tumor (NT) mice separate from tumor (T) mice already at the pre-tumor time point, with increasing diversity over time (Extended Data Fig. 4c). Differential metabolite analysis of the significantly regulated metabolites, shown between all genotypes and time points, demonstrated that tg/tg associated metabolites are mostly comprised of lysophospholipids and LCFAs (Extended Data Fig. 4d). In the CRC patient cohort 4, we identified 56 FAs that were significantly regulated in T tissue compared to T adjacent tissue (Table 1). We next examined the overlap between LCFAs (≥C20) which were significantly regulated in CRC patient T tissue and in T mice. Multiple fatty acids were significantly regulated in both, including Docosatetranoic acid (FA 22:4), Docosapentanoic acid (FA 22:5), Docosahexanoic acid (FA 22:6), and putative FAs: FA 20:3, C20 hydroxy FAs, FA 24:5 and FA 24:6 (Fig. 3b,c). Stratifying by timepoint, an unknown C20 hydroxy fatty acid was the only LCFA enriched in tg/tg mice at the pre-tumor time point, however the number of significant shared LCFAs increased at the tumor and late-tumor time points, indicating that tumor onset increases the LCFA pool (Fig 3b). When further stratifying significantly regulated FAs in CRC patient cohort 4 into EOCRC and LOCRC, we identified two LCFAs (C20:2 hydroxy FA and C20:3 FA) to also be significantly regulated in EOCRC T tissue compared to T adjacent tissue (Extended Data Fig. 4e). FAs 22:4, 22:5 and 22:6 have been validated by comparison with pure standards (Extended Data Fig. 4f).

**Fig. 3:**
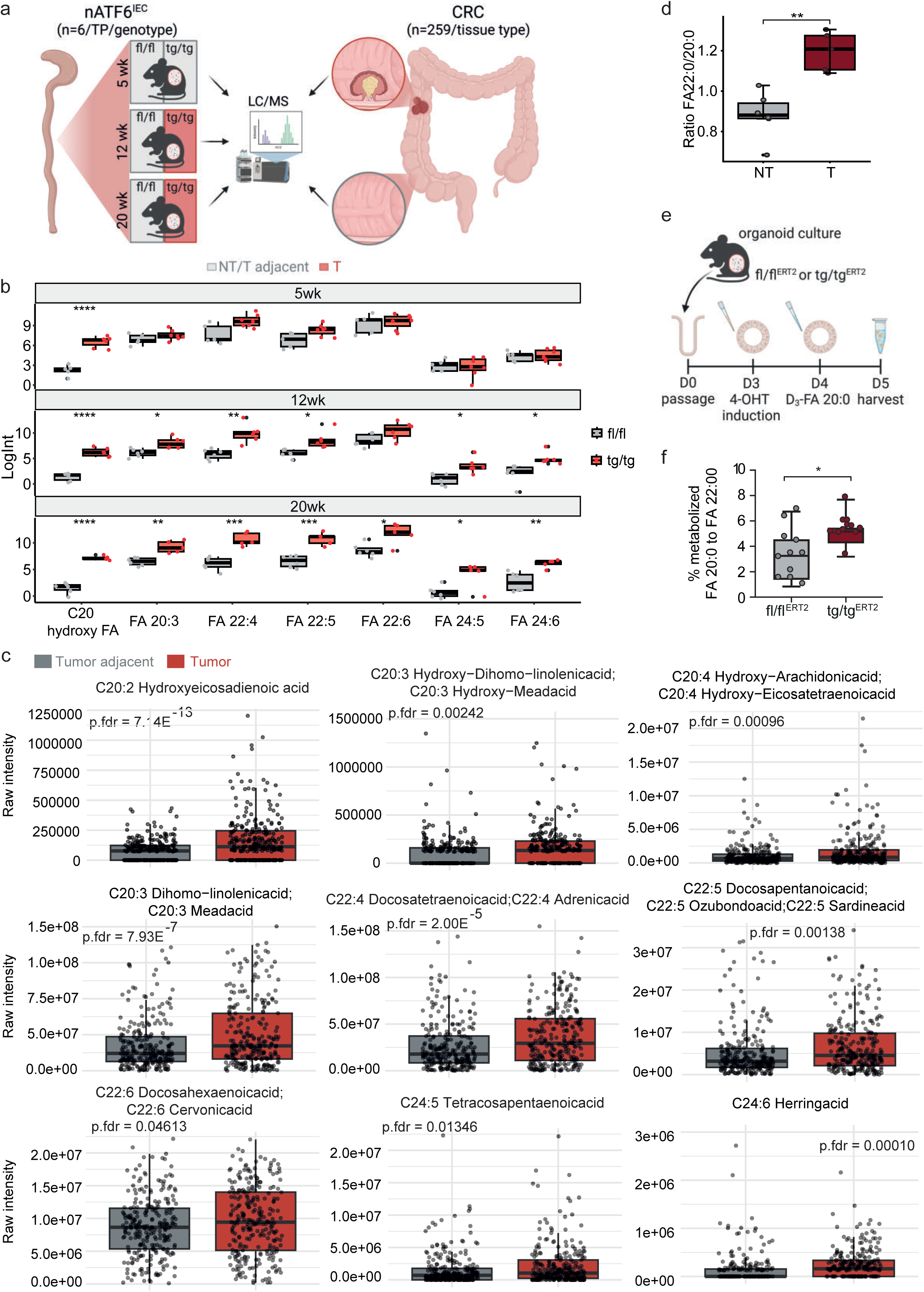
ATF6 drives FA elongation and LCFAs accumulate in mice and CRC patients. **a,** Schematic depicting metabolomic analysis strategy in nATF6^IEC^ mice and human CRC patients (cohort 4). Created with BioRender.com. **b,** Boxplots comparison of log-transformed fatty acid intensities in caecal content, comparing fl/fl to tg/tg mice, stratified by timepoint. Whiskers extend to the minimum and maximum values. Only fatty acid metabolites which were significant in mouse data and overlapped with CRC patient data are shown. Statistical significance was calculated using pairwise T-tests and adjusted for multiple comparisons using the Benjamini-Hochberg procedure. **c,** Boxplot comparison of fatty acid intensities in tumor tissue and tumor-adjacent tissue of samples from cohort 4. The plot displays the intensity differences for nine fatty acids that shown significant abundance in tumor samples (left to right): hydroxyeicosadenoic acid (FA 20:2), hydroxy-dihomo-linolenic acid (FA 20:3), hydroxy-arachidonic acid (FA 20:4), dihomo-linoleic acid (FA 20:3), docosatetranoic acid (FA 22:4), docosapentanoic acid (FA 22:5), docosahexanoic acid (FA 22:6), tetracosapentanoic acid (FA 24:5) and herring acid (FA 24:6). Whiskers extend to the minimum and maximum values, excluding outliers, which are shown as individual points. Paired T-Test was used to calculate *p*-values (and FDR corrected). Only fatty acids with p-values (FDR corrected) <0.05 are shown. **d,** Percentage of saturated fatty acids (SAFA) in NT and T tissue after quantification of total FA using GC-MS. Statistical analysis was perfomed using using an unpaired Wilcoxon test (*p*=0.0043). **e,** Scheme showing the treatment of intestinal organoids from nATF6 Vil-CreERT2 (fl/fl^ERT2^ and tg/tg^ERT2^) mice cultured *ex vivo* (n=11-12). Organoids were passaged at D0, ATF6 expression in all wells was induced at D3 with 500nM (Z)-4-hydroxytamoxifen (4-OHT) and treated with D3-FA 20:0 at D4, before harvesting at D5 (see also Supplementary Figure 1). Created with BioRender.com. **f,** Percentage of metabolized D_3_-FA20:0 to D_3_-FA:22:0 in organoids of tg/tg^ERT2^, compared to fl/fl^ERT2^ controls using GC-MS. Significance was calculated using the Student’s two-tailed unpaired T-Test (*p*=0.0184). A significant *p*-value <0.05 is represented as an asterisk: \**p* < 0.05, \*\**p* < 0.01. \*\*\**p* < 0.001, \*\*\*\**p* < 0.0001.

To further characterize the altered lipid environment in tg/tg mice, we quantified total FAs in colonic tissue using GC-MS. Analyses revealed an increase in saturated fatty acids (SAFAs) in T compared to NT mice, which also correlated with tumor number (Extended Data Fig. 4g,h). Furthermore, T mice show a clear elongation of SAFAs including FA 20:0 and FA 22:0 compared to NT mice (Fig. 3d). To determine whether elongation is a direct consequence of ATF6 signaling, we assessed ATF6-mediated elongation of FAs in *ex vivo* colon organoid cultures. We first phenotypically characterized colon organoids derived from fl/fl controls and tg/tg mice at the late-tumor time point (20wk). Both fl/fl and tg/tg crypts formed viable organoids in culture, including cyst formation (d1) and budding (d7) of cysts (Extended Data Fig. 4i). Strikingly, tg/tg organoids in culture did not show any significant increase in organoid size (Extended Data Fig. 4j) or the number of budding crypts per cyst (Extended Data Fig. 4k) compared to fl/fl controls, indicating that the hyper-proliferative phenotype observed *in vivo* in the presence of the microbiota is lost in this *ex vivo* culture system. In a separate experiment, intestinal organoids from nATF6 Vil-Cre^ERT2^ (fl/fl^ERT2^ and tg/tg^ERT2^) mice were cultured, and *ATF6* expression induced by adding 500nM (Z)-4-hydroxytamoxifen (4-OHT) to the culture medium for 24h. This was followed by a 24h exposure of organoids to deuterated eicoseinoic acid (D_3_-FA 20:0) (Fig. 3e, Supplementary Fig. 1). Importantly, tg/tg^ERT2^ intestinal organoids showed a significantly increased potential of FA elongation, quantified as the percentage of metabolized D_3_-FA20:0 to D_3_-FA:22:0, compared to fl/fl^ERT2^ controls (Fig. 3f). The above findings demonstrate that ATF6 activation is the driver of FA elongation in the absence of microbial and immune signaling.

### ATF6-related lipid profiles are causally linked to tumor-associated microbiota

We next sought to investigate whether chronic *Atf6* activation would alter the luminal and mucosa-associated microbiota. To isolate the impact of tumor formation, we first examined changes in microbiota composition at 5wks, before tumor onset. We observed augmented modulation of the mucosa-associated microbiota with increasing *Atf6* gene dose as shown by mean GUniFrac dissimilarity relative to control, between genotypes, an effect comparable to changes observed in the luminal environment (Fig. 4a). Global microbiota shifts were observed between all timepoints and genotypes, in both luminal and mucosal datasets, and appeared more distinct at later timepoints, particularly in tg/tg mice (Extended Data Fig. 5a,b; luminal PERMANOVA, p=0.001, mucosal PERMANOVA p=0.001). We applied machine learning models (Random forest (RF), Ridge regression (RR), LASSO (L), Elastic-net (E) and L1-penalized regression (LL)) to both sample types, demonstrating that the mucosal microbiota can better classify the tumorigenic phenotype compared to the luminal microbiota. Supporting increased utility of mucosal over luminal microbiota, four out of five models trained on mucosal microbiota data showed higher AUC values across all leave-one-out cross validation repetitions (Extended Data Fig. 5c). Moreover, we observed a significant correlation between tumor number and Shannon Effective diversity in mucosal data which was absent in luminal (Extended data Fig. 5d,e), further supporting this finding and suggesting a link between tumor burden and microbial diversity.

**Fig. 4:**
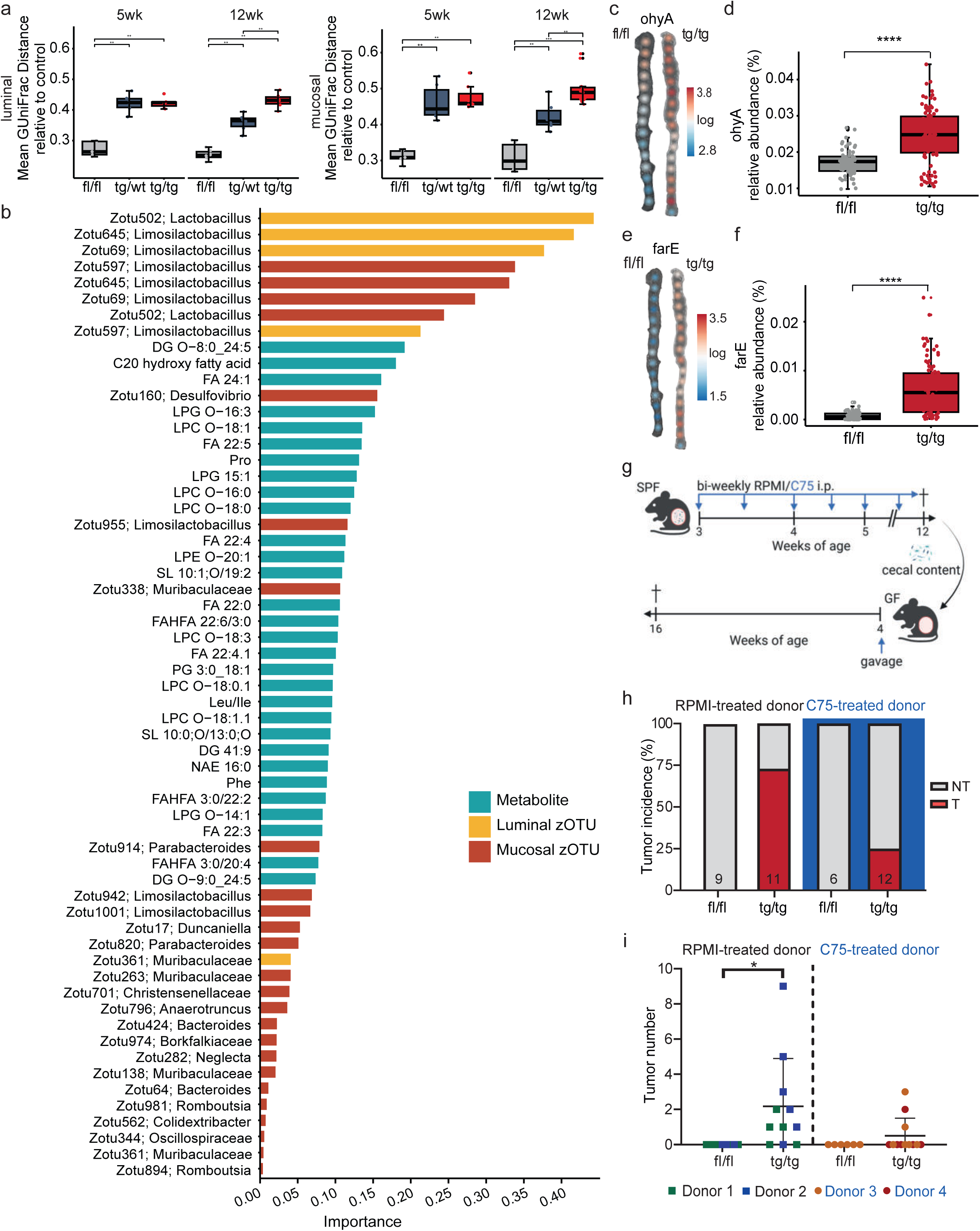
ATF6-related lipid profiles are causally linked to tumor-associated microbiota. **a,** Mean generalized UniFrac distance relative to control across fl/fl, tg/wt and tg/tg samples, at pre-tumor (5wk) and tumor-onset (12wk) in luminal (left) and mucosal communities. *P* values were calculated using a pairwise t-test and adjusted for multiple comparisons using the Benjamini-Hochberg procedure. Whiskers extend to the minimum and maximum values (n=6-7 mice per group). **b,** Loadings plots of omic features selected by the sPLS-DA model, discriminating tg/tg mice from other genotypes. Luminal zOTUs are shown in yellow, mucosal zOTUs in red and metabolites in blue. Features are sorted by importance (n = 17-18 per genotype). **c,** Spatial maps of Log10 transformed predicted relative abundance of oleate hydratase (ohyA) positive ASVs along the entire colon in fl/fl and tg/tg mice (n=6 mice per group). **i,** Predicted relative abundance of ohyA in fl/fl and tg/tg mice. **d,** Relative abundance of oleate hydratase (ohyA) in tg/tg and fl/fl mice *p*=4.7e-10). **e,** Spatial maps of Log10 transformed predicted relative abundance of FA efflux pump (farE) positive ASVs along the entire colon in fl/fl and tg/tg mice (n=6 mice per group). **f,** Predicted relative abundance of fatty acid efflux pump, farE in tg/tg and fl/fl mice (*p*=<2.26e-16). **g,** Schematic showing the experimental design for in vivo C75 FASN inhibitor intervention and transfer experiment. SPF mice were intraperitoneally (i.p.) injected either with the RPMI control or the C57 FASN inhibitor bi-weekly from the age of 3 weeks. After an intervention period of 9 weeks, mice were sacrificed at 12 weeks of age. Arrows indicate i.p. injections. **h,** Tumor incidence (percentage) of GF fl/fl and tg/tg mice after gavage with donor material from either RPMI-treated (controls) or C75 FASN inhibitor-treated mice. **i,** Tumor number of GF fl/fl and tg/tg mice after gavage with donor material from either RPMI-treated (controls) or C75 FASN inhibitor-treated mice. Points are colored by donor. Statistical significance was calculated using an ordinary one-way ANOVA with Tukey’s multiple comparison test (*p*=0.0205). A significant *p*-value <0.05 is represented as an asterisk: \**p* < 0.05, \*\**p* < 0.01. \*\*\**p* < 0.001, \*\*\*\**p* < 0.0001.

**Fig. 5:**
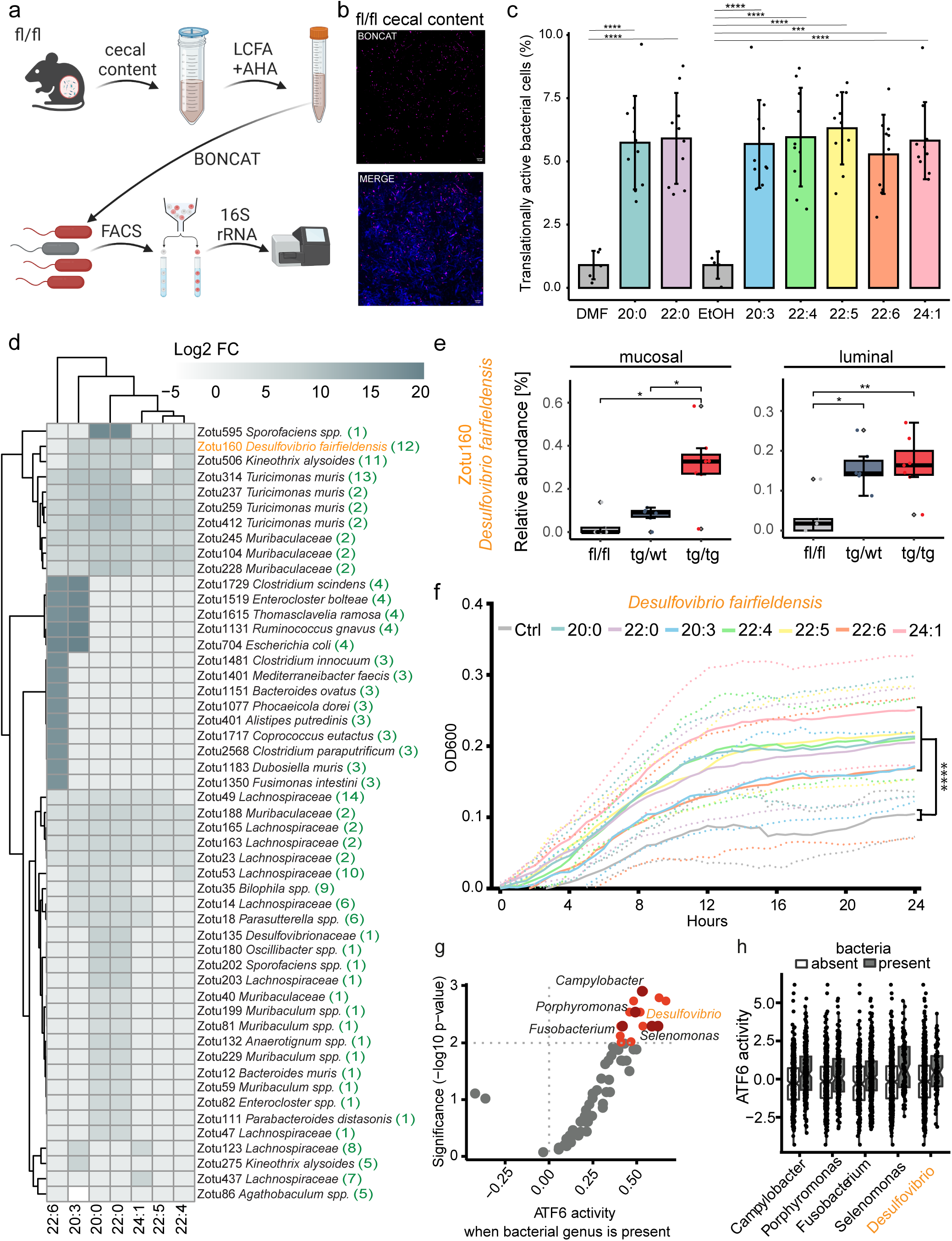
LCFAs selectively activate the growth of tumor-related bacteria. **a,** Schematic overview of bioorthogonal non-canonical amino acid tagging (BONCAT). Mouse cecal content was incubated in anaerobic conditions and stimulated with seven long-chain fatty acids (LCFAs), including arachidic acid (20:0), homo-γ-linolenic acid (20:3), docosanoic acid (22:0), docosatetraenoic acid (22:4), docosapentaenoic acid (22:5), docosahexaenoic acid (22:6), and nervonic acid (24:1) and along with the cellular activity marker L-azidohomoalanine (AHA). Translationally active bacterial cells were labeled by azide-alkyne click chemistry, sorted with FACS and sequenced by 16S rRNA gene amplicon sequencing. Created with BioRender.com. **b,** Representative confocal microscopic images of a mouse cecal content stimulated with nervonic acid (24:1). Pink: active cells (BONCAT-Cy5); blue: all cells (DAPI); merge image. Scale bar is 10 µm. **c,** Percentage of translationally active bacteria after LCFA amendment. LCFA are compared to their respective solvent control: 20:0 and 22:0 were compared to DMF (DMF), 20:3, 22:4, 22:5, 22:6 and 24:1 were compared to ethanol (Etoh). P values were calculated by ANOVA and Tukey’s test for multiple comparisons (mean ± sd: 5.8 ± 1.6% for LCFAs in total; 0.9 ± 0.51 for the controls; ANOVA, *p*<0.0001 for all comparisons, n=80). Error bars represent standard deviation of the mean. **d,** Heatmap displayed log2-fold changes of significantly enriched or depleted zOTUs in the translationally active fraction compared to the unsorted group for each LCFA as calculated with the Wald test (*p*<0.05, n=160). All the zOTUs (51) listed in the heatmap are significantly enriched or depleted. zOTUs were identified using the 16S-based ID tool of EzBioCloud. Numbers in parenthesis correspond to the barplots of Extended Data 7d where each barplot illustrate the shared and unique zOTUS between all LCFAs. **e,** Relative abundance of a *Desulfovibrio fairfieldensis* zOTU, in luminal (left) and mucosal (right) communities comparing fl/fl, tg/wt and tg/tg mice at the 5-week timepoint. Whiskers extend to the minimum and maximum values. Statistical significance was calculated using pairwise t-tests and adjusted for multiple comparisons using the Benjamini-Hochberg procedure . **f,** Growth curve of *Desulfovibrio fairfieldensis* in postgate medium, supplemented with different LCFAs. Each curve represents averaged values of n = 5 biological replicates in three technical replicates. Long-chain fatty acids (20:0, 22:0, 20:3, 22:4, 22:5, 22:6 and 24:1) were added to postgate medium to a concentration of 10µM each. The control (gray) contains only postgate medium with bacteria. All LCFA supplementations show significant increase of growth compared to the control (two-way ANOVA, Bonferroni test for multiple comparisons, *p* < 0,0001 for all comparisons, n=735). Error bars shown as dotted line represent standard deviation of the mean. **g,** Plot of ATF6 activity in CRC patients (TCGA Database) when bacterial genus is present (x-axis) and the significance (y-axis). Marked in orange/red are those genera significantly associated with ATF6 activity, with red dots representing classical CRC-associated genera. **h,** Plot of ATF6 activity in CRC patients (TCGA Database) when bacteria are absent (grey) or present (red) for the CRC-associated genera marked red in h (*p* values: Campylobacter = 0.00123, Desulfovibrio = 0.00507, Porphyromonas = 0.00287, Fusobacterium = 0.00507, Selenomonas = 0.00507). A significant *p*-value <0.05 is represented as an asterisk: \**p* < 0.05, \*\**p* < 0.01. \*\*\**p* < 0.001, \*\*\*\**p* < 0.0001.

To examine whether these changes in microbiota community structure were associated with shifts in metabolite profiles, we performed a multi-omic data integration using sPLS-DA analysis of luminal untargeted metabolomic profiles and luminal and mucosal 16S data ^44^. To delineate between genotype-driven and tumor-driven alterations, we performed two integrations that model genotype and phenotype separately. By sorting tg/tg discriminating features according to importance, we identified Lactobacillus/Limosilactobacillus and Desulfovibrio zOTUs as well as multiple LCFAs and lysophospholipids as microbial and metabolite contributions, respectively (Fig. 4b). Metabolite contributions to phenotype discrimination were similar and comprised mostly LCFAs and Lysophospholipids. However microbial contributions differed greatly, with mostly zOTUs classified as Parabacteroides and Romboutsia, indicating the presence of tumors differentially modulates microbiota, compared to *Atf6* activation alone (Extended data Fig. 6a). Next, to identify microbiota-correlated metabolites, we generated relevance networks using the network function of mixOmics DIABLO R package ^44^. Importantly, several fatty acids, including an unidentified C20 hydroxy fatty acid, nervonic acid (FA 24:1), Docosapentanoic acid (FA 22:5) and Docosatetranoic acid (FA22:4), correlated positively with tg/tg discriminative zOTUs, particularly those classified as Lactobacillus/Limosilactobacillus, and additionally with Desulfovibrio in the mucosal dataset (Extended data Fig. 6b,c, Supplementary Table 6). LCFA compounds are known to be toxic to bacteria, requiring adaptation mechanisms to allow growth in FA-rich milieus. Using PICRUSt2, we generated predictions of genomic content ^45,46^, following a spatially-resolved analysis of 16S rRNA amplicon sequencing-based bacterial profiles using individually defined 0.5 x 0.5 cm sites along the longitudinal colonic axis in fl/fl controls and tg/tg mice at the tumor (12wk) time point. We identified genes involved in bacterial responses to FAs in the proximal tumor-susceptible region of the colon, which are known to be involved in FA detoxification, catabolism and efflux (Extended Data Fig. 7a). Of this FA-response signature, oleate hydratase (*ohyA*) and the FA efflux pump (*farE*) were observed to be upregulated in tumor mice compared to controls (Extended Data Fig. 7a). *OhyA*-mediated fatty acid hydroxylation enables growth of certain bacteria in lipid-rich environments^47^. Spatial mapping of log_10_ transformed relative abundance showed clear enrichment of *ohyA* across all sites in tg/tg mice compared to fl/fl controls (Fig. 4c). The relative abundance of *ohyA* was increased in tumor-bearing tg/tg mice compared to fl/fl controls (Fig. 4d). Furthermore, the ratio of *ohyA*+ to *ohyA*-taxa was increased in all three mucosal phenotypes (non-tumor (NT), tumor adjacent (TA) and tumor (T)) in tg/tg mice, compared to fl/fl controls (Extended Data Fig. 7b). *FarE* constitutes a lipid efflux pump associated with antimicrobial FA resistance to limit the cellular accumulation of toxic FAs ^48^. Similar to *ohyA*, we observed an increased relative abundance across all tg/tg sites (Fig. 4e), and increased relative abundance of *farE* in tumor-bearing tg/tg mice (Fig. 4f). Furthermore, the ratio of *farE*+ to *farE*-taxa was increased in all tg/tg mucosal phenotypes, compared to fl/fl controls (Extended Data Fig. 7c). Taken together, our findings clearly indicate an early pre-tumor microbial adaptation to the ATF6-altered FA-rich intestinal environment, to shape a tumor-associated bacterial lipid-response signature.

To understand the causal role played by FASN in colonic tumorigenesis in our nATF6^IEC^ murine model, we exposed fl/fl and tg/tg SPF mice to bi-weekly intraperitoneal injections (i.p.) of the FASN inhibitor C75 from the age of 3 weeks (pre-tumor time point) until 12 weeks (tumor time point) (Fig. 4g). Interestingly, FASN inhibition reduced tumor incidence in tg/tg mice from 80% to 0% with no tumors observed in the tg/tg colon (Extended Data Fig. 7d,e). In line with this, FASN inhibition was able to reduce tumor number and tumor volume in the colon of tg/tg mice (Extended Data Fig. 7f,g). To validate the link between FA synthesis and microbiota-dependent tumorigenesis in nATF6^IEC^ mice, we performed microbiota transfer experiments from C75-treated mice and RPMI-treated controls into GF tg/tg mice and fl/fl controls (Fig. 4g). Compared to fl/fl recipient mice, all tg/tg recipient mice show reduced survival (Extended Data Fig. 7h). Strikingly, the tumor-promoting capacity of the C75-treatment-derived microbiota was reduced from a tumor incidence of 73% to 25% (Fig. 4h). In line with this, the microbiota from C75-treated mice reduced both tumor number and volume to levels comparable to tumor-free fl/fl controls (Fig. 4i, Extended Data Fig. 7i). Taken together, these findings clearly demonstrate that microbiota changes associated with ATF6-induced lipid alterations is causally involved in inducing a more aggressive tumorigenic phenotype in nATF6^IEC^ mice.

### LCFAs selectively activate the growth of tumor-related bacteria

To assess whether LCFAs directly impact bacteria, we exposed the complex microbiota of 5 week-old fl/fl mice to previously identified (Fig. 3b, Table 1) ATF6-associated LCFAs (24:1, 22:0, 20:0, 22:6, 22:5, 20:3 and 22:4). Our approach applied biorthogonal non-canonical amino acid tagging (BONCAT) combined with fluorescence-activated cell sorting (FACS) to identify translationally active bacteria in response to LCFA exposure (Fig. 5a). Confocal microscopy images displayed a significantly higher proportion of translationally active cells after LCFA supplementation compared to the controls (Fig. 5b, Supplementary Fig. 2). Microscopy images were confirmed with quantitative measure of translationally active bacterial cells exhibiting a higher percentage of translational activity compared to the controls (ANOVA, p<0.0001 for all comparisons) (Fig. 5c). A distinct separation between sorted and unsorted communities along with a significant decrease of zOTUs in the sorted fraction, demonstrates a specific microbial response to LCFA exposure (PERMANOVA, p<0.0001; Wilconox test: p<0.0001) (Extended Data Fig. 8 a,b). In total, we identified 50 zOTUs that were significantly enriched in response to one or multiple LCFAs, while only one zOTUs was depleted (Wald test, p<0.05) (Fig. 5d, Extended Data Fig. 8c). Strikingly, LCFA selectively activated tumor-associated bacteria including those of the Desulfovibrionaceae family (e.g., *Desulfovibrio fairfieldensis* and *Bilophila* spp.) (Fig 5 d), while the overall microbiota profiles were similar in response to the different LCFAs (Extended Data Fig. 8d; PERMANOVA, p=0.8045). Supporting the *in vivo* relevance of these findings, we examined the overlap between LCFA-enriched zOTUs and differentially abundant zOTUs at the pre-tumor timepoint, identifying *Desulfovibrio fairfieldensis* as one of three significantly enriched zOTUs in tg/tg mice at the pre-tumor timepoint in both the mucosal and luminal environments (Fig. 5e, Extended Data Fig. 8e). The higher translational activity in Desulfovibrionaceae was accompanied by a significant enhancement in bacterial growth when exposed to individual LCFAs, as indicated by the increased expansion of *Desulfovibrio fairfieldensis* (two-way ANOVA, p<0.0001 for all comparisons) (Fig. 5f, Extended Data Fig. 8f). A similar growth-promoting effect was observed for another Desulfovibrio isolate, *D. piger*, further confirming the ability of Desulfovibrio spp. to selectively grow in the presence of LCFAs (p<0.0001 for all comparisons; Extended Data 8g). Additional zOTUs responding to the LCFA-rich environment included Sutterella spp., Lachnospiraceae, Muribaculaceae, *Turicimonas muris*, *Ruminococcus gnavus*, Oscillibacter spp. (Fig. 5d). The above demonstrates that LCFAs translationally activate specific bacteria, including tumor-associated taxa, providing a direct link between tumor-associated fatty acid profiles and the selective expansion of bacterial taxa in mice.

Recent developments in identifying microbial signatures in the publicly available The Cancer Genome Atlas (TCGA) dataset ^49^, allowed us to compare computationally inferred transcription factor activity using previously validated transcription factor-target gene relationships (RNA-seq data) and microbial profiles (identified in whole-exome sequencing data from the same samples) in CRC patients. In a hypothesis-free approach, ATF6 was identified among the most significant transcription factors, as calculated using a linear mixed effects model, whose activities correlated positively with the presence of CRC-enriched genera, including Campylobacter, Porphyromonas, Fusobacterium, Selonomonas and Desulfovibrio (Fig. 5 g,h). Both *ATF6* activity and bacterial profiles are strongly associated with microsatellite instability (MSI) status of the cancer, which could be a potential confounding factor, but we observed a strong correlation of the CRC-enriched genera with ATF6 activity even after correcting our association models for MSI status (Extended Data Fig. 8h). Of note is the observed overlap in specific tumor- and ATF6-associated taxa between mouse and human microbiota (Fig. 5d,e,g,h and Extended Data Fig. 8h), namely *D. fairfieldensis*, Oscillibacter spp., Desulfovibrionaceae spp. Bilophila spp., supporting a causal link between ATF6-mediated LCFA synthesis and tumor-relevant changes in the microbiota.

## Discussion

As the second leading cause of deaths worldwide ^50^, early and accurate diagnosis as well as novel treatment strategies are imperative in the management of CRC. As a fundamental mediator in the pathogenesis of CRC, the UPR^ER^ represents a promising host cellular mechanism for preventive and therapeutic intervention. While our mechanistic understanding of the role of ATF6 signaling in CRC remains limited, its expression has been associated with poor prognosis in CRC patients ^14,15^. In the present study, we demonstrate the relevance of ATF6 expression in CRC patients across multiple independent German cohorts by identifying a clear sub-population (19-38% LOCRC; 11% EOCRC) of patients that present with high ATF6 expression, underlining the importance of ATF6 in CRC. Most importantly, we identified a mechanism through which ATF6-mediated changes in cellular lipid metabolism of the intestinal epithelium contribute to the development of tumor-promoting microbiota changes at an early disease stage.

We previously showed that chronic ATF6 activity specifically in murine intestinal epithelial cells (nATF6^IEC^) results in microbiota-dependent early-onset colonic adenomas ^11^. The necessity of microbial triggers in tumor induction was shown via GF mice, which remained tumor-free ^11^. Furthermore, the transfer of pre-tumor murine cecal content into GF mice revealed the dysbiotic microbiota from tumor-susceptible mice to have an increased capacity to induce tumors ^11^. Importantly, the development of dysbiosis in nATF6^IEC^ mice precedes tumor formation ^11^, suggesting that tumor-independent mechanisms at early stages of the ATF6-driven pathogenesis initiate microbial changes. To ascertain these mechanisms, we set out to characterize the transcriptional response induced by chronic ATF6 activation in the presence (SPF) and absence (GF) of bacteria at the pre-tumor stage. We identify the FA metabolism KEGG pathway as one of the top ranked metabolic pathways altered in tumor-susceptible SPF mice when compared to GF mice, and detect 23 upregulated FA metabolism-related genes. The majority are upregulated already in response to acute ATF6 activation, confirming FA alterations to be an early event occurring in direct response to ATF6 signaling in the presence of the microbiota. While the specific contribution of ATF6 to lipid metabolism is less well understood, the UPR^ER^ is known to play an important role in lipid metabolism ^51–54^. Lipid metabolic rewiring forms a phenotypic hallmark of cancer, with altered lipidome profiles identified in tumors of numerous organs ^37,55^. Ecker *et al.* recently identified a robust CRC tumor-specific triglyceride species signature that correlates with postoperative disease-free survival in a multicohort study spanning two clinical centres ^43^. Moreover, gene expression of *FASN*, which forms a multi-enzyme complex responsible for catalyzing FA synthesis, was prognostically detrimental ^43^. Strikingly, the *in vivo* inhibition of FASN in nATF6^IEC^ mice results in the prevention of colon adenoma development. We show that our tumor-associated cecal metabolite signature is mostly comprised of lysophospholipids and LCFAs, with an overlap of several significantly altered LCFAs between tg/tg mice and CRC patient tumor tissue, confirming the human relevance of the observations in our transgenic murine tumor model. Analyses of murine colonic tissue demonstrated increases in SAFAs and LCFAs. These findings are in line with a study profiling FAs in tissue and plasma of CRC patients and healthy controls, which identified a CRC-associated FA panel and found that LCFAs were the most increased FAs ^56^. Our experimental approach in *ex vivo* organoids demonstrates that FA elongation is a direct consequence of ATF6 signaling, occurring in the absence of microbial signals.

A handful of studies have identified pro-oncogenic bacteria enriched in adenoma samples, however, questions remain over whether these taxa initiate disease, since adenoma formation induces metabolic alterations that likely impact the local microenvironment and alter microbial composition ^57–59^. Our knowledge on which triggers may enable taxa to expand early on, and potentially become tumor-promoting, is sparse. We hypothesize that ATF6-driven lipid alterations impact the intestinal microbiota as an early event of tumorigenesis in nATF6^IEC^ mice. Our multi-omic data integration of 16S rRNA microbiota (luminal and mucosal) profiling and cecal metabolites identified several zOTUs, including those classified as Lactobacillus/Limosilactobacillus and Desulfovibrio, to be correlated with several LCFAs associated with our tumor-susceptible mice. Importantly, and in support of our hypothesis, PICRUSt2 predictions of genomic content confirm that the ATF6-driven FA-rich intestinal environment results in microbial adaptation in the form of a lipid-response signature that is associated with genes involved in FA detoxification, catabolism and efflux. Our data suggest a clear link between ATF6-induced lipid alterations and changes in the intestinal microbiota preceding tumor formation. Strikingly, FMT in GF mice using cecal content of FASN inhibitor-treated mice clearly demonstrated that the microbiota associated with ATF6-induced lipid alterations is causally involved in inducing a more aggressive tumorigenic phenotype in nATF6^IEC^ mice. To identify selective taxa of the intestinal microbiota responding to LCFAs, we here applied an approach to directly expose the microbiota of fl/fl control mice *ex vivo* to our identified LCFAs, showing that tumor-associated taxa are translationally activated in response to the LCFAs. Among these was *Desulfovibrio fairfieldensis*, which was significantly enriched in tg/tg mice at the pre-tumor time point in both the mucosal and luminal environments. In addition to CRC-associated lipid signatures, CRC-associated microbial signatures have also been identified ^60,61^. In a hypothesis-free approach analyzing transcription factor activity and microbial profiles in CRC patients, we show *ATF6* activity to positively associate with the presence of such a risk profile associated bacterial genera, identifying a clear link between ATF6 signaling and the microbiota in CRC pathogenesis. Interestingly, and in confirmation of our murine data, we identified Desulfovibrio among the ATF6-associated CRC genera.

In conclusion, we identify a clear mechanistic sequence of events in the initiation of colon tumorigenesis whereby ATF6 signaling drives a clinically relevant pathologic response by altering lipid metabolism to induce microbial adaptation, ultimately leading to a tumor-relevant bacterial dysbiosis. Although beyond the scope of this study, future work should focus on single-cell transcriptomics to characterize the ATF6 transcriptional response at the cellular level, as well as extending gnotobiotic mouse experiments with individual bacterial candidates, to identify causal tumor drivers and elucidate mechanisms of tumor initiation. Together, the above highlights novel mechanistic findings in an exciting new avenue of research bringing together cellular stress responses, lipid metabolism and the intestinal microbiota in the context of CRC. This evident triangular interplay presents new targetable vulnerabilities in CRC that warrant future research efforts to identify potential future prevention and treatment strategies.

## Author contributions

OC and DH wrote the manuscript. AS, AR, MS, DS and CS supported manuscript writing. OC, AS, AR, MS, SB, SK, IC, MS, DS and DH designed the experiments and performed data analysis. AS, MS and SK performed mouse experiments and respective mouse tissue and microbiota processing and analysis. KS performed IHC staining on human cohort TMAs. OC performed the quantification and data analysis of human IHC staining. TM supported quantification of human IHC staining. MJ provided CRC cohort 1 tissue and supported data analysis. KPJ provided CRC cohort 2 tissue and generated TCGA survival curves. CS, SH, CR (Christoph Röcken) and CR (Christian Röder) provided CRC cohort 3 tissue and supported data analysis. MZ provided untargeted metabolomics analyses of CRC cohort 4. DMC, SW and TP performed ATF6 quantification validation classification and regression models. JW and GZ performed microbial and transcription factor TCGA analyses. SB prepared mouse tissue for mRNA sequencing samples and supported mRNA sequencing analyses. PR performed mRNA sequencing. CS and TK performed mRNA sequencing data analyses. AD, DS and MZ performed untargeted metabolomics. JP (Josh Pauling) and NK performed metabolomics analysis and integration. JP (Johannes Plagge), JE and GL performed fatty acid analyses and data analysis.

## Data Availability

Raw and processed RNASeq data are available at GEO under accession GSE247122. Human CRC metabolomics data is available in the MetaboLights Database ^62^ (https://www.ebi.ac.uk/metabolights/) under accession number MTBLS7387. 16S rRNA gene amplicon sequence data have been deposited in the NCBI Short Read Archive under PRJNA1227744. Source data are provided with this paper.

## Acknowledgements

*Desulfovibrio piger* was kindly provided by Thomas Clavel. HiBC – Human intestinal bacterial collection: www.hibc.rwth-aachen.de.

## Online Methods

### Ethics statement regarding animal experiments

All animal experiments, as well as maintenance and breeding of mouse lines, were approved by the Committee on Animal Health Care and Use of the state of Upper Bavaria (Regierung von Oberbayern; AZ ROB-55.2-1-54-2532-217-2014, AZ ROB-55.2-2532.Vet_02-20-58, AZ TVA 55.2-2532.Vet_02-18-149, AZ TVA 55.2-2532.Vet_02-18-121, AZ TVA 55.2-2532.Vet_02-19-006) and performed in strict compliance with the EEC recommendations for the care and use of laboratory animals (Directive 2010/63/EU of the European Parliament and of the Council of 22 September 2010 on the protection of animals used for scientific purposes).

### Ethics statement regarding patient samples

Immunohistochemistry staining and quantification of ATF6 were performed in three independent human cohorts: a German large single-center cohort comprising 1004 colorectal cancer patients that underwent surgical resection spanning two decades, for which 959 patient tissue samples could be quantified (cohort 1) ^17^, a German cohort consisting of 104 CRC patients followed up over 20 years, for which 50 patient samples could be quantified (cohort 2), and a German single center cohort comprising more than 1,500 resected CRC patient samples spanning approximately 25 years, for which 55 EOCRC (age <50y) and 256 LOCRC (age >50y) were obtained and could be quantified (cohort 3). The use of surgically resected human tissue samples was approved by the local Ethics Committee of the Technical University of Munich (TUM) (ref.-no. 252/16 s, cohort 1), by the Ethics Committee of the Medical Faculty of TUM (ref.-no. 1926/7, 375/16S and 2022-169-S-KH, cohort 2), and by the local Ethics Committee of the University Hospital Schleswig-Holstein (ref.-no. A 110/99, cohort 3). CRC patient cohort 3 tissue specimens were supplied by the biobank TRIBanK (Translational Interdisciplinary Biobank Kiel) together with the Institute of Pathology, University Hospital Schleswig-Holstein, Kiel, Germany. LC-MS analyses were performed in a fourth German human cohort consisting of 259 patients diagnosed with EOCRC and LOCRC and included one tumor tissue (T) and one adjacent tissue (T adjacent) for each patient (cohort 4). The use of surgically resected human tissue samples was approved by the local Ethics Committee of the Medical School – The Christian-Albrechts-University of Kiel (ref.-no. A 156/03), the Ethics Committee of the Medical School of the University of Rostock (ref.-no. A 2019-0187) and the Ethics Committee of the EMBL (ref.- no. 2024/HE000062 - MiEOCRC). For all human cohorts in this study, both sexes were deliberately included, with slightly more male cases, corresponding to the incidence in the overall population. Sex-dependent differences were not observed in our data. Patient samples from all four cohorts were obtained after prior informed written consent, and information on sex was acquired by self-reporting.

### Immunohistochemical staining and quantification of ATF6

TMA cores were immunohistochemically stained with an anti-ATF6 antibody (Sigma-Aldrich, HPA-005935) diluted 1:100, following antigen retrieval in an EDTA buffer (30 min, pH 9), or an anti-FASN antibody (LS Bio, LS-B3636) or anti-GRP78 (Cell Signaling, 3177s) diluted 1:200 following antigen retrieval in a Citrate buffer (20 min, pH 6). Nuclei were counterstained with hematoxylin (Waldeck). Stainings were performed using the Leica Bond RXm (Leica) and scanned using the Aperio AT2 (Leica Biosystems). ATF6- and FASN-stained TMA cores were quantified using the QuPath Software ^18^. Cells were identified through positive cell detection of the hematoxylin-positive nucleus. ATF6 staining intensities for the nuclear and the cytoplasmic regions were classified according to defined thresholds of the DAB OD max and DAB OD mean, respectively. FASN staining was quantified based on cytoplasmic regions only. Furthermore, the object classifier was trained and applied to distinguish between epithelial cells and stromal cells. Exported epithelial histochemical-scores (H-scores, capturing both the intensity and the percentage of stained cells) for the nucleus and the cytoplasm were used for further patient cohort analyses

### ATF6 expression analysis using AI

Whole slide images were segmented using QuPath into individual cores and scaled down to 312×312 pixels. This pre-processing step yielded a dataset of 176 TMA cores from a cohort of 50 patients (cohort 2). For our deep learning model, a ResNet18 was trained (<u>arXiv:1512.03385v1</u> [cs.CV]), freezing all layers except the final convolutional and fully connected layers for the classification task and freezing the first layer for the regression task. This allowed leveraging pre-trained features while fine-tuning the model’s output layers for our specific task. To improve model generalization, data augmentation techniques were applied, including random horizontal and vertical flips and random rotations. In the final layer, dropout regularization was incorporated to prevent overfitting. A 5-fold cross-validation was conducted to evaluate our model’s performance. Each fold used data from 40 patients for training and 10 patients for validation. Throughout cross-validation, the best models were recorded based on their accuracy (classification task) or loss (regression task) in the validation dataset. Their weights were preserved for subsequent use. In our final prediction task of classifying samples as ATF6-high or ATF6-low, a weighted average ensemble strategy was employed by combining the predictions of the five best models. In a final evaluation step, our ensemble’s predictions were tested on an entirely independent cohort comprising 918 patients (cohort 1).

### Transgenic nATF6^IEC^ mice (SPF and GF)

nATF6^IEC^ mice were generated as previously described ^11^ and housed under SPF and GF conditions (12 h light/dark cycles and 24-26°C) at the Technical University of Munich (Weihenstephan, Germany). All mice were fed a standard chow diet (autoclaved V1124-300; Ssniff, Soest, Germany) *ad libitum*. Mice were cage-separated according to sex but not according to genotype. Both sexes were used for all experiments in this study. For the determination of the tumor phenotype, mice were sacrificed and the colon excised. The presence and number of tumors was recorded for each mouse as macroscopically observed.

### Isolation of murine primary colonic IECs

Colonic tissue was longitudinally opened, and transferred to 20 mL Dulbecco’s modified Eagle medium (Gibco) containing 10% fetal calf serum, 1% L-glutamine, and 0.8% antibiotics/antimycotics (IEC isolation medium) supplemented with 1 mmol/L dithiothreitol (Roth). The tissue was vigorously vortexed for 1 minute, incubated under continuous shaking (37°C, 15 minutes) and vortexed again for 1 minute. The cell suspension was centrifuged (7 minutes, 300xge) and the cell pellet was resuspended in 5 mL IEC isolation medium. After vortexing for 1 minute, the remaining tissue pieces were incubated in 20 mL PBS (10 minutes, 37°C) containing 1.5 mM EDTA under continuous shaking. The cell suspension was vortexed again followed by another centrifugation to pellet the cells. Both cell suspension fractions were combined and purified by centrifugation through a 20%/40% (in medium/PBS) discontinuous Percoll gradient (GE Healthcare Life Sciences) at 600xg for 30 minutes. The IEC cells at the interface between the two Percoll phases were collected and washed twice (with medium/PBS). The isolated IECs were lysed in RA1 RNA lysis buffer (Macherey-Nagel).

### RNA Isolation and library construction

RNA of colonic IECs was isolated according to the manufacturer’s instructions (NucleoSpin RNAII kit, Macherey-Nagel) and measured by NanoDrop ND-1000 spectrophotometer (Thermo Fisher Scientific). TruSeq stranded Total RNA kit (Illumina) was used for the construction of strand-specific sequencing libraries from the isolated colonic IEC RNA. Sequencing was performed on an Illumina HiSeq2500 (75-nucleotide (nt) paired-end reads for each run, 1/10 lanes per sample.

### RNASeq preprocessing

After the initial trimming of reads using trim_galore (version 0.6.4) to satisfy a Phred score of at least 30, RNASeq data was processed with the ‘nf-core/rnaseq‘ pipeline (version 1.4.2) and aligned to the GRCm38 genome using the ‘STAR‘ aligner (version 2.6.1d). Quantification of transcripts was performed using Salmon (version 0.14.1). All sequence properties were inspected in MultiQC and passed the quality thresholds.

### RNAseq functional analysis

Gene Set enrichment analysis (GSEA) was performed using the differential expression results ranked by t-value ^63^. KEGG gene set was downloaded from the Molecular Signatures Database ^63–65^. Gene names were translated into mouse gene names before the analysis using scibiomart (v.), a wrapper around the API from BioMart. GSEA was performed using the packages fgsea ^66^ (v1.24.0) and GSEABase ^67^ (v 1.60.0). Next we performed regulatory clustering based on regulatory rules (Supplementary Table 3, sheet 2) using <u>sircleR package</u> function sircleRCM_2Cond ^19^. In brief, each gene was assigned to “UP” (log2FC > 0.5, *p*.adj <0.05), “DOWN” (log2FC < -0.5, *p*.adj <0.05) or “No Change” (0.5 > log2FC > -0.5 and/or *p*.adj > 0.05) based on its expression change between fl/fl versus tg/tg. The background method chosen was “C1&C2”, which means a gene had to be detected in both conditions in order to be included in the clustering. Plots were generated using the EnhancedVolcano package ^68^ (v1.16.0) and ggplot2 ^69^ (v 3.4.2).

### Untargeted metabolomics of murine cecal content or tissue

Sample preparation was performed as previously described ^70^. Briefly, mouse cecal content (∼20mg) or tissue (∼25mg) was mixed with 1ml methanol-based extraction solvent in a 2ml bead beater tube (CKMix 2ml, Bertin Technologies) filled with ceramic beads (1.4 mm and 2.8 mm ceramic beads). Samples were then homogenized using a bead beater (Precellys Evolution, Bertin Technologies) supplied with a Cryolys cooling module (Bertin Technologies, cooled with liquid nitrogen) three times for a duration of 20 seconds, at a speed of 8000rpm. Subsequently, the resulting suspension was centrifuged for 10 minutes, at 6000rpm. Finally, 100μl of supernatant was mixed with 20μl internal standard solution (7 µmol/l) and injected into the Liquid chromatography time-of-flight mass spectrometer (LC-TOF-MS) system for untargeted analysis. Untargeted LC-TOF-MS analysis was carried out as described in Metwaly *et al.* ^70^. Briefly, untargeted analysis was performed on an ExionLC Ultra-high performance liquid chromatography (UHPLC) system (Sciex), connected to a 6600 TripleTOF instrument (Sciex) operating in positive and negative electrospray mode and calibrated using ESI positive and negative calibration solutions (Sciex). UHPLC phase separation was performed in reverse as well as hydrophilic interaction stationary phase (HILIC), on a Kinetex C18 column (Phenomenex) and ACQUITY BEH Amide column (Water) respectively. Mass spectrometry was performed using SWATH mode, with fragment spectra recorded in high-sensitivity mode. Metabolomics data processing was performed as described previously ^70^. Briefly, Raw data files were converted into Reifycs Abf (Analysis Base File) files and subsequent untargeted peak picking was performed using MS-DIAL software (version 3.5274) ^71^. Alignment was performed across all samples and the peak area of individual features were exported for further analysis using the R statistical software environment. Peak normalisation was based on QC samples, employing the method described in Wehrens *et al.* ^72^. All features were then combined into a single table and further normalised according to sample weight, before analysis.

### Fatty acid quantification

For FA quantification in mouse tissues and investigation of eicosanoic acid elongation in organoids, FA methyl esters (FAMEs) were generated by acetyl chloride and methanol treatment, and extracted with hexan ^73^. Total FA analysis was carried out using a Shimadzu 2010 GC-MS system (Duisburg, Germany). FAMEs were separated on a BPX70 column (10 m length, 0.10 mm diameter, 0.20 μm film thickness) from SGE using helium as the carrier gas. The initial oven temperature was 50°C and was programed to increase at 40°C/min to 155°C, 6°C/min to 210°C, and finally 15°C/min to 250°C. D3-eicosanoic acid and its elongation products in organoids were quantified by single-ion monitoring of specific fragment ions (D3-eicosanoic acid, 329 m/z; D3-docosanoic acid, 357 m/z). The internal standard was C21:0-iso.

### Untargeted metabolomics of CRC patient tissue sample preparation

Solid tissues were prepared for LC-MS analysis by organic solvent extraction. In brief, 200 μL of 0.1 mm zirconia/silica beads (BioSpec Products) and 500 µL of organic solvent (acetonitrile:methanol, 1:1) supplemented with an internal standard mixture were added to 70-100 mg of pre-weighed solid material. Internal standard mixture consisted of phenylalanine-*d5*, tryptophan-*d5*, ibuprofen-*d4*, tolfenamic acid-*d4*, estriol-*d3*, diclofenac-*d4*, warfarin-*d5*, oxfendazole-*d3*, chloramphenicol-*d5*, nafcillin-*d5* and caffeine-*d9*, each to a final concentration of 80 nM. The material was homogenized by mechanical disruption with a bead beater (BioSpec Products) set for 2 minutes and 30 seconds on the highest setting at room temperature. After incubation for at least 1 h at -20°C, samples were centrifuged (3220 rcf, 4°C) for 15 min. For C18 LC-MS acquisition, 15 µL of supernatant was diluted with 15 µL H2O.

### Untargeted metabolomics of CRC patient tissue LC-MS data acquisition

Chromatographic separation was performed by reversed-phase chromatography using an Agilent 1200 Infinity UHPLC system. The qTOF instrument (Agilent 6550) was operated in negative scanning mode (50–1,700 *m/z*). Online mass calibration was performed using a second ionization source and a constant flow of reference solution (*m/z* 112.9857 and 1033.9881). InfinityLab Poroshell HPH-C18 column (Agilent, 2.1 × 100 mm, 1.9 Micron particle size) was used for chromatography. LC solvent A was water with 0.1% formic acid (v/v) and solvent B was methanol with 0.1% formic acid (v/v). Column operated at 45 °C. 5 µL of sample were injected at the following gradients: 0 min 5% B; 0.1 min 5% B; 5.50 min 95% B; 6.49min 95% B; 6.5 min 5% B. The time between injections was 0.5 min. The qTOF was operated in negative scanning mode (50 - 1700 m/z) and the following source parameters: VCap: 3500 V, nozzle voltage: 2000 V, gas temp: 225 °C; drying gas 13 L/min; nebulizer: 20 psig; sheath gas temp 225 °C; sheath gas flow 12 L/min. Tandem mass spectrometry analysis (LC-MS/MS) was performed for fatty acids using the chromatographic separation and source parameters described above and the targeted-MS/MS mode of the instrument with a preferred inclusion list for parent ion with 20 ppm tolerance, Iso width set to ‘narrow width’ and collision energy to either 10, 20 or 40 eV.

### Semi-targeted metabolomics of CRC patient tissue LC-MS data processing

An *in-silico* library of 134 fatty acids was mined again C18-MS negative data for the extraction of area values of FA-putative features (Supplementary Table 7). The area under the curve from the targeted analysis was integrated using the MassHunter Quantitative Analysis Software (Agilent, version 7.0) based the accurate high-resolution mass of these reference analytes and the following parameters: signal threshold of 50,000; mass tolerances of 0.002 amu or 20 ppm; retention time tolerance of 0.2 min. The resulting table was further normalized in R 4.4.1 using RStudio/2023.06.1 according to established protocols ^74,75^.

### Validation of significant fatty acids of CRC patient tissue

To ensure the annotation of the significant fatty acids, we have performed level-1 metabolite annotation, which is the direct comparison of an authentic chemical standard analyzed under identical analytical conditions. For this, we have used standards of docotetranoic acid, docopentanoic acid and docohexanoic acid, as well as samples of CRC-patients that have shown high abundance of these metabolites. Correct annotation occurs when all three orthogonal properties of the metabolite of interest (retention time, high-resolution mass, and fragmentation mass spectrum) are similar between the analyte and the CRC-sample.

### Murine primary crypt isolation and intestinal organoid culture

Murine intestinal crypts were isolated following digestion with EDTA (30 min; 4°C) and mechanical dissociation. The supernatant was filtered through a 70μm cell strainer, pelleted by centrifugation, and resuspended in Matrigel (BD Biosciences). 25µl Matrigel^®^-organoid domes were plated in 48-well plates (Eppendorf) together with 250µl IntestiCult™ Organoid Growth Medium (STEMCELL Technologies) and maintained in a humidified 5% CO_2_ atmosphere at 37°C. Organoids were passaged every 7 days by mechanical disruption using a 1ml syringe with a 20G needle, pelleted by centrifugation, and embedded in fresh Matrigel.

### Induction of *ex vivo* recombination and fatty acid stimulation of intestinal organoids

Intestinal organoids from nATF6 Vil-Cre^ERT2^ (fl/fl^ERT2^ and tg/tg^ERT2^) mice ^11^ were passaged three days before the start of the experiment and cultivated in self-made crypt culture medium (CCM; advanced DMEM/F12 medium (Gibbco), 2mM GlutaMax (Gibbco), 10mM HEPES, penicillin, streptomycin and amphotericin (all Sigma-Aldrich) supplemented with N2, B27 (both Gibbco), 1mM N-acetylcysteine (Sigma-Aldrich), 50ng/ml EGF (ImmunoTools), 100ng/ml noggin and 0.5µg/ml R-spondin 1 (both PeproTec)). *ATF6* overexpression in intestinal organoids was induced by adding 500nM (Z)-4-hydroxytamoxifen (4-OHT; LKT) to the culture medium for 24h. After induction, organoids were exposed to Eicosanoic 20,20,20-D3 acid (30µM; Larodan) in CCM for 24h. Organoids were subsequently harvested in 0.2% SDS in ddH_2_0.

### Sampling of murine cecal content and colonic mucosa-associated microbiota

Cecal content was harvested immediately after euthanization and stored at -80°C before DNA extraction. The murine colon was excised and opened longitudinally before clearing of colonic content using a sterile needle and sterile PBS washes. Tissue sections were then excised using a sterile scalpel pre-treated with DNA Away™ (Fisher Scientific) to destroy contaminating DNA fragments. For spatial microbiota analyses, the entire colon was laid on a grid and dissected into individual 0.5 cm x 0.5 cm sections and subsequently frozen at -80°C before DNA extraction. All steps were carried out in a laminar flow cabinet.

### Microbial DNA extraction from murine cecal content and colonic tissue

DNA extraction from frozen cecal content was carried out using a modified version of the protocol described by Godon et al ^76^. Briefly, cecal samples were homogenized by vortexing and transferred to autoclaved screw-cap Eppendorf tubes containing 500mg silica beads (0.1mm Carl Roth) and kept on ice. To this, 600μl of DNA stabilizer (Macherey Nagel), 250μl 4M Guanidine thiocyanate and 500μl 5% N-laurolyl-sarcosine was added. Samples were incubated under moderate shaking (700rpm) for 1 hour at 70°C. Lysis was achieved via mechanical disruption with a FastPrep ®-24 bead beater (MP Biomedicals) using three 40-second cycles at a speed of 6.5 m/s. 15mg Polyvinylpolypyrrolidone (PVPP; Sigma-Aldrich) was added to homogenate to remove phenol contamination, before centrifugation at 15,000×g for 3 minutes at 4°C. The resulting supernatant was recovered, and centrifuged again under the same conditions, resulting in a clear supernatant containing lysed bacterial cells. To this mixture, 10mg/ml RNase was added to degrade bacterial RNA and incubated at 37°C for 30 minutes under constant shaking (700rpm). DNA extraction from colonic tissue was performed using enzymatic digestion. Where possible samples were extracted using the same kit batch, to limit differences in inherent kit contamination between different batches ^77^. Tissues were placed in 180μl sterile lysis buffer (20Mm Tris/HCL, 2Mm EDTA, 1% Triton-X100; pH 8 supplemented with 20mg/ml lysozyme) and incubated for 1 hour in a shaking incubator (950rpm) at 37°C. 10mg/ml Proteinase K (Macherey Nagel) was then added, and samples were incubated for 1-3 hours (until complete lysis of the tissue was obtained) at 56°C under moderate shaking (950rpm). The resulting genomic DNA was then purified using the NucleoSpin® gDNA clean-up kit (Macherey Nagel), according to the manufacturer’s instructions. The concentration and purity of extracted genomic DNA was determined using a Nanodrop ND-1000 Spectrophotometer. Samples were stored at -20°C before sequencing.

### 16S rRNA gene amplicon sequencing

High throughput amplicon sequencing of the 16S rRNA gene was carried out as previously described ^70^. In brief, the V3 and V4 hypervariable regions were amplified via a 2-step protocol, using the 341f and 785r primer pair (341f 5’-CCTACGGGNGGCWGCAG-3’, 785r 5’-GACTACHVGGGTATCTAATCC-3’). With 10×15 cycles for cecal content samples and 15×15 for mucosal samples. Samples were barcoded with a double index, according to Kozich *et al.* ^78^. The resulting amplicons were purified using AGENCOURT AMPure XP Beads (Beckman Coulter, Krefeld, Germany), pooled in equimolar ratios and then sequenced on an Illumina MiSeq system (Illumina), in paired-end mode (2 x 275bp). Raw reads were processed using IMNGS, which wraps the UPARSE/USEARCH software pipeline ^79,80^. Sequences were demultiplexed, trimmed to first base with a quality score <3 and merged. To remove spurious sequences, a length filter was applied based on the expected amplicon size of 444bp, removing those smaller than 300 and larger than 600bp. The resulting paired sequences were then dereplicated and denoised using UNOISE3, to generate zero-radius OTUs or zOTUs, reflecting true biological sequences ^81^. Taxonomy was assigned using the RDP classifier version 2.11 ^82^. Samples with less than 5000 total reads were excluded from further analysis. Since low biomass samples are susceptible to contamination which can lead to spurious conclusions, all mucosal samples were further processed with the R package decontam to remove putative contaminants ^83^.

### Supervised classification of 16S rRNA sequencing

Supervised classification of mucosal and luminal microbial profiles in non-tumor (fl/fl and tg/wt) and T (tg/tg) mice was performed using the SIAMCAT R package, utilising Random forest (RF), Ridge regression (RR), LASSO (L), Elastic-net (E) and L1-penalized regression (LL), which were chosen based on the ease of interpretation of model output ^84,85^. To account for differences in sample size between mucosal and luminal data, mucosal data was randomly subsampled to match luminal data. Prior to model training, low prevalent features - defined as those present in less than one-third of samples - were removed, and the data transformed using a centred log-ratio transform. All models were trained using five-fold CV, or in case of smaller sample size (<50 total samples), leave-one-out CV (LOOCV), each with five rounds of resampling.

### Multi-omics integration

Integration of metabolomic and microbiota data was performed on log-transformed, scaled, and centred log-ratio (clr) transformed metabolite and zOTU matrices, respectively. General associations between the microbiota (luminal and mucosal) and metabolites were identified using a multi-block sparse partial least squares discriminant analysis (sPLS-DA) model with seven components and 10-fold CV, as implemented in the DIABLO framework of the mixOmics R package ^44,86^..

### Functional potential prediction

To calculate predicted functional profiles based on 16S rRNA sequencing data, we utilised PICRUSt2 version 2.3 ^45^. FASTA files of representative sequences and minimum sum scaled zOTU tables were used as input for the command: *picrust2_pipeline.py*, which runs the full PICRUSt2 pipeline, aligning and placing the sequences into a reference tree, calculation of 16S copy number, Enzyme Commission (EC) and KEGG orthologs abundances, adjustment of these by 16S abundance and finally infers MetaCyc pathways by collapsing EC numbers according to their associated metabolic pathway. All PICRUSt2 data were generated in a high-performance computing environment, utilising the Linux-cluster system at the Leibniz Rechenzentrum, Garching.

### Intraperitoneal injections of mice with FASN inhibitor C75

The FASN inhibitor C75 (Biomol) was dissolved in RPMI 1640 (Sigma Aldrich) to a final concentration of 1.5 mg/mL, aliquoted and stored at -20°C until used. C75 or RPMI were injected intraperitoneally (i.p.; 10 ml/kg body weight) biweekly, starting at the age of three weeks until twelve weeks of age (tumor time point). When necessary, mice were given a subcutaneous depot (1:1 mixture (v/v) of 5% glucose and ringer’s lactate (B. Braun)) to alleviate potential side-effects of C75. Mice were euthanized with CO_2_ at the age of twelve weeks or when abortion criteria were met.

### Fecal microbiota transfer into GF mice

Cecal content from C75-treated mice and RPMI-treated controls was instantly suspended at 1:10 weight/volume in filter-sterilized phosphate-buffered saline (PBS)/40% glycerol and stored at –80°C. For gavage, cecal content solutions were centrifuged (3 minutes, 300*g*, 4°C) to pellet debris, followed by centrifugation (10 minutes, 8000*g*, 4°C) to collect microbes. This fraction was resuspended in an equal volume of sterile PBS. Each recipient mouse (tg/tg or fl/fl) was gavaged with 100 μL of the bacterial suspension at the age of 4 weeks. Recipient mice were housed in microbiota-specific isolators and euthanized after 12 weeks, or when abortion criteria (as defined in ethical proposals) were fulfilled.

### *Ex vivo* anaerobic incubations with LCFAs in cecal content of fl/fl mice

Cecal content was collected from 10 wild-type mice at 5 weeks of age and immediately transferred to an anaerobic chamber (10% H2, 90% N2). To ensure sufficient material, cecal contents of two mice of the same genotype and litter were combined. Cecal content (200-300 mg each) was homogenized in 1X PBS and filtered through a 40 μm filter (Corning, Germany) to remove larger particles. The samples were then transferred to sterile falcon tubes containing a final concentration of 50 µM of the cellular activity marker L-azidohomoalanine (AHA) (Baseclick GmbH, Germany) and between 400 pM and 50 nM of seven long-chain fatty acids (LCFAs), including FAs 20:0, 20:3, 22:0, 22:4, 22:5, 22:6, and 24:1, dissolved in either ethanol or DMF at a final concentration of 1%. Specifically, ethanol and DMF were included as solvent controls for FAs 20:3, 22:4, 22:5, 22:6, 24:1 and 20:0, 22:0, respectively, while 2 mg/mL glucose was the positive control for each experiment. Samples were incubated under anaerobic conditions for 6 hours. After incubation, a fraction of the samples was frozen for DNA extraction, and other aliquots were washed twice with PBS and fixed in 1:1 ethanol for further analysis by fluorescence-activated cell sorting (FACS) ^87^.

### Bioorthogonal non-canonical amino acid tagging (BONCAT) and bacterial cell quantification

To assess the ability of gut microbiota to utilize LCFAs, we applied BONCAT, a fluorescence-based single-cell labeling technique. BONCAT is based on the incorporation of the non-canonical amino acid AHA followed by fluorescent labeling of AHA-containing proteins via azide-alkyne click chemistry ^88^. The click chemistry reaction was carried out on microscopy slides, as previously described (Hatzenpichler et al. 2014, doi: 10.1111/1462-2920.12436). Samples were counterstained with 4′,6-diamidino-2-phenylindole (DAPI). Microscopy pictures were captured using a confocal microscope (Olympus, fluoview, FV10i, Germany) and processed with Fiji software ^89^ (**Supplementary** Figure 1). Translationally active cells were quantified with flow cytometry using absolute counting beads (CountBrightTM, Invitrogen, ThermoFisher Scientific, Germany) according to the manufacturer’s instructions ^87^ .

### BONCAT-FACS and DNA extraction

BONCAT combined with FACS was employed to identify bacteria capable of utilizing LCFAs. Samples were processed according to ^88^. Briefly, ethanol-fixed samples were first washed with 1X sterile PBS, resuspended in 96% ethanol, and centrifuged. The BONCAT dye solution was then added, and the samples were incubated for 30 minutes in the dark at room temperature ^88^. Following incubation, the samples were washed three times with 1X sterile PBS and filtered through a 35 mm nylon mesh using BD tubes (12 x 75 mm, Corning, Germany) prior to sorting. Cy5-positive bacterial cells were sorted using a FACS Melody instrument (BD, Germany), collected in sterile tubes, and stored at - 80°C until DNA extraction. The gating strategy is depicted in Extended Data Fig. 8 for fl/fl mice. DNA from both sorted and unsorted bacterial cells was extracted using the QIAamp DNA Mini Kit (Qiagen, Germany), following the manufacturer’s instructions and according to ^87^ .

### Desulfovibrio fairfieldensis and Desulfovibrio piger cultivation in LCFA supplemented media

*Desulfovibrio fairfieldensis* (CSUR P7329, from Collection de Souches de l’Unité des Rickettsies) and *Desulfovibrio piger* (CLA-AA-H201, from Human intestinal bacterial collection) were grown in postgate medium (PG; DSMZ 63). Strains were initially grown in PG medium broth at 37 °C in an anaerobic chamber (5 % CO2, 5 % H2, 90% N2). Once the cultures reached their maximum OD_600_, they were diluted to 0.05 OD_600_ into fresh PG medium broth supplemented with 10µM of each of the seven long-chain fatty acids (LCFAs), including arachidic acid (20:0), homo-γ-linolenic acid (20:3), docosanoic acid (22:0), docosatetraenoic acid (22:4), docosapentaenoic acid (22:5), docosahexaenoic acid (22:6), and nervonic acid (24:1). PG medium with *D. fairfieldensis* or *D. piger* was used as control. Bacteria suspensions were aliquoted into a sterile 96-well flat-bottom plate (BRANDplates, BRAND, Germany) and incubated for 24 h at 37 °C in a microplate reader (Cerillo, Germany) under anaerobic conditions. Growth data were measured every 30 minutes. Every experiment was conducted five times for *D. fairfieldensis* and tree times for *D. piger*, with three technical replicates. LCFA added to postgate medium in the same amount as to bacteria-inoculated tubes, served as blanks for each condition. The average OD-values of the blanks were subtracted from each technical replicate of the corresponding sample incubated with each LCFA. Of these, the average and standard deviation were calculated and plotted.

### TCGA Dataset

For both CRC-related TCGA projects (TCGA-COAD and TCGA-READ), gene expression profiles were downloaded from the firebrowse.org website (RSEM normalized gene expression). Using those tables, transcription factor activities were inferred using the DoRothEA R package (version 1.10.0) ^90^ with default parameters (using transcription factor-target interactions of medium- or higher confidence (A-C) and using the viper algorithm for statistical inference). Bacterial profiles derived from whole exome sequencing of the same samples were extracted from the supplemental material of Dohlman *et al.* ^49^. For all samples present in both data types (610 samples overlap: 451 COAD, 159 READ), the inferred activity of ATF6 was associated with bacterial presence, using the project information as random effect in a random effect model implemented in the lmerTest R package ^91^. Since we observed microsatellite instability (MSI) status to be very strongly associated with both bacterial profiles and ATF6 activity, we additionally included MSI-status as another random effect in a second association model. Bacterial genera were tested for CRC-enrichment with a random effects model for all patients with paired tumor and normal tissue samples and significantly CRC-enriched genera are highlighted in Figure 1 and Extended Data 1. All *p*-values were corrected for multiple hypothesis testing using the Benjamini-Hochberg procedure ^92^.

### Statistical analysis

For data analysis, unless otherwise stated below, statistical analyses were carried out using R (R software foundation, Vienna, Austria) or GraphPad Prism (version 9.00; GraphPad Software, San Diego, CA, USA). For flow cytometry analyses, data processing and analysis were performed with FlowJo™ v10.10.0 software (BD Life Sciences). (Reference: FlowJo™ Software (for Windows) Version v10.10.0. Ashland, OR: Becton, Dickinson and Company; 2023). For ATF6 quantification of IHC staining, the H-score cut-off was determined using the online *Cutoff Finder* Web Application for Biomarker cutoff optimization ^93^. For comparing the mean of two groups, unpaired student’s T-test or Wilcoxon test was used where appropriate. Differences between multiple groups were tested using ANOVA or Kruskal-Wallis tests. For testing differences in frequency, Fisher’s exact test was used. To test for differences in beta diversity, multivariate statistical testing was conducted using Permutational multivariate analysis of variance (PERMANOVA) ^94^.For hierarchical clustering of metabolite intensities, Ward’s method was used ^95^. Multiple comparisons were controlled using the method of Benjamini & Hochberg ^96^.

For semi-targeted metabolomics of CRC patient tissue, statistical analysis and plotting were performed in R 4.4.1 using RStudio/2023.06.1. The statistical significance of molecular feature in T versus T adjacent was assessed with paired t-test (t.test function in R), and p-values were FDR-corrected for multiple hypotheses testing using the Benjamini-Hochberg procedure (p.adjust function in R with ‘fdr’ parameter). Significant metabolites were selected when their intensity had a corrected p-value < 0.05.

For 16S rRNA gene amplicon sequencing, the decontam package utilises the relationship between pooled DNA concentration prior to sequencing and the prevalence and abundance of taxa in negative controls to identify putative contaminants. Identification of contaminants was performed with the isContaminant function using default parameters. Downstream analyses were performed using Rhea and phyloseq ^97,98^. Briefly, zOTU tables were normalized using minimum sum scaling or relative abundance. Alpha diversity was measured using Shannon Effective and diversity. To assess differences between groups, beta diversity was calculated based on generalized UniFrac distance (GUniFrac) ^99^. Mean GUniFrac dissimilarity for a given sample was calculated as the dissimilarity to the respective control mean. Differentially abundant taxa were identified using the Linear Discriminant analysis Effect size (LEfSe) algorithm, with a Linear discriminant analysis threshold (LDA) threshold of 3.0, to limit false positives ^100,101^.

For BONCAT-FACS experiments, statistical analysis was performed using R statistical software (https://www.r-project.org/). Raw sequencing data were pre-processed using the web platform IMNGS2 (https://www.imngs2.org/). Contaminants originating from PCR and kit reagents were detected and removed using the R package decontam ^102^. zOTU counts were subsampled to a number of reads smaller than the smallest library (1000 reads). Beta diversity was calculated using generalized UniFrac distances ^103^ and visualized using multidimensional scaling (MDS) ^97^. Permutational multivariate analysis of variance (perPERMANOVA) was performed to determine the significance of samples grouping. Alpha-diversity was evaluated based on taxa richness ^97^ and group comparison was calculated using Wilcoxon test. Significant enriched and depleted zOTUs between sorted and unsorted fractions were performed using DEseq2 ^104^.

Significant zOTUs sequences were identified using the 16S-based ID tool of EzBioCloud (https://www.microbiologyresearch.org/content/journal/ijsem/10.1099/ijsem.0.001755).

The variation in microbial communities across the seven LCFAs was further analyzed using the UpSetR package in R, which employs a matrix-based layout to represent set intersections and their sizes ^105^.

For quantifications of the percentage of translationally active cells, ANOVA and Tukey’s test for multiple comparisons were applied.

For growth curves, two-way ANOVA was used to compare *D. fairfieldensis* or *D. piger* stimulated with each LCFA with the control without LCFAs.Variables were expressed as mean ± standard deviation (SD), a probability value (p-value) less than 0.05 was considered statistically significant and Bonferroni method was used for multiple comparison. For single time point comparison one-way ANOVA was used and Dunnet’s test for multiple comparisons.

Unless otherwise specified all data are presented as the mean ± standard deviation, *p*-values < 0.05 are considered statistically significant (*p*<0.05=*, *p*<0.01=**, *p*<0.01=***, *p*<0.001=****).

