## Supplementary figures and images for "ATF6 activation alters colonic lipid metabolism causing tumor-associated microbial adaptation"

### Extended Data Fig 1

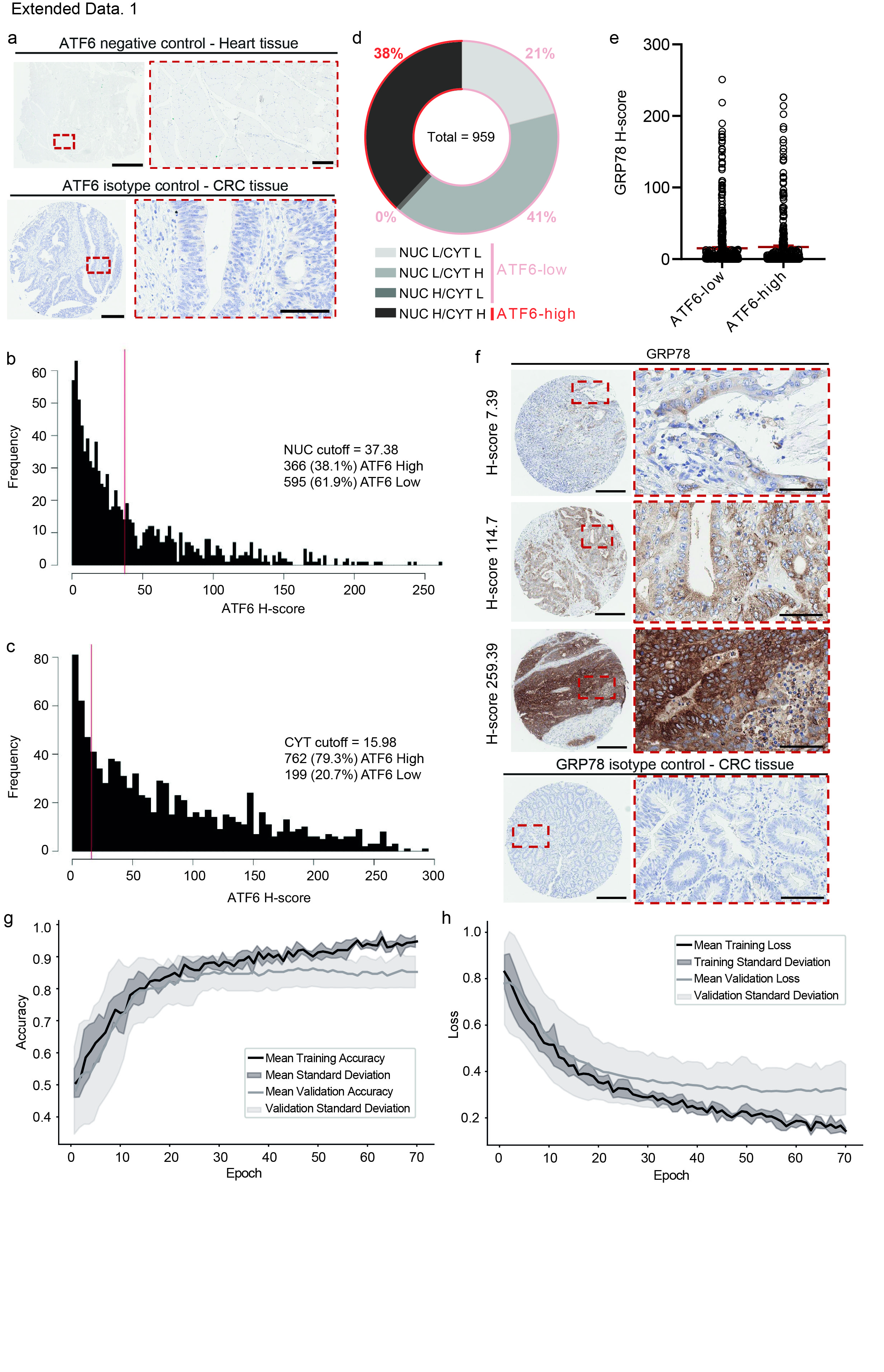

### Extended Data Fig 2

Extended Data 2.

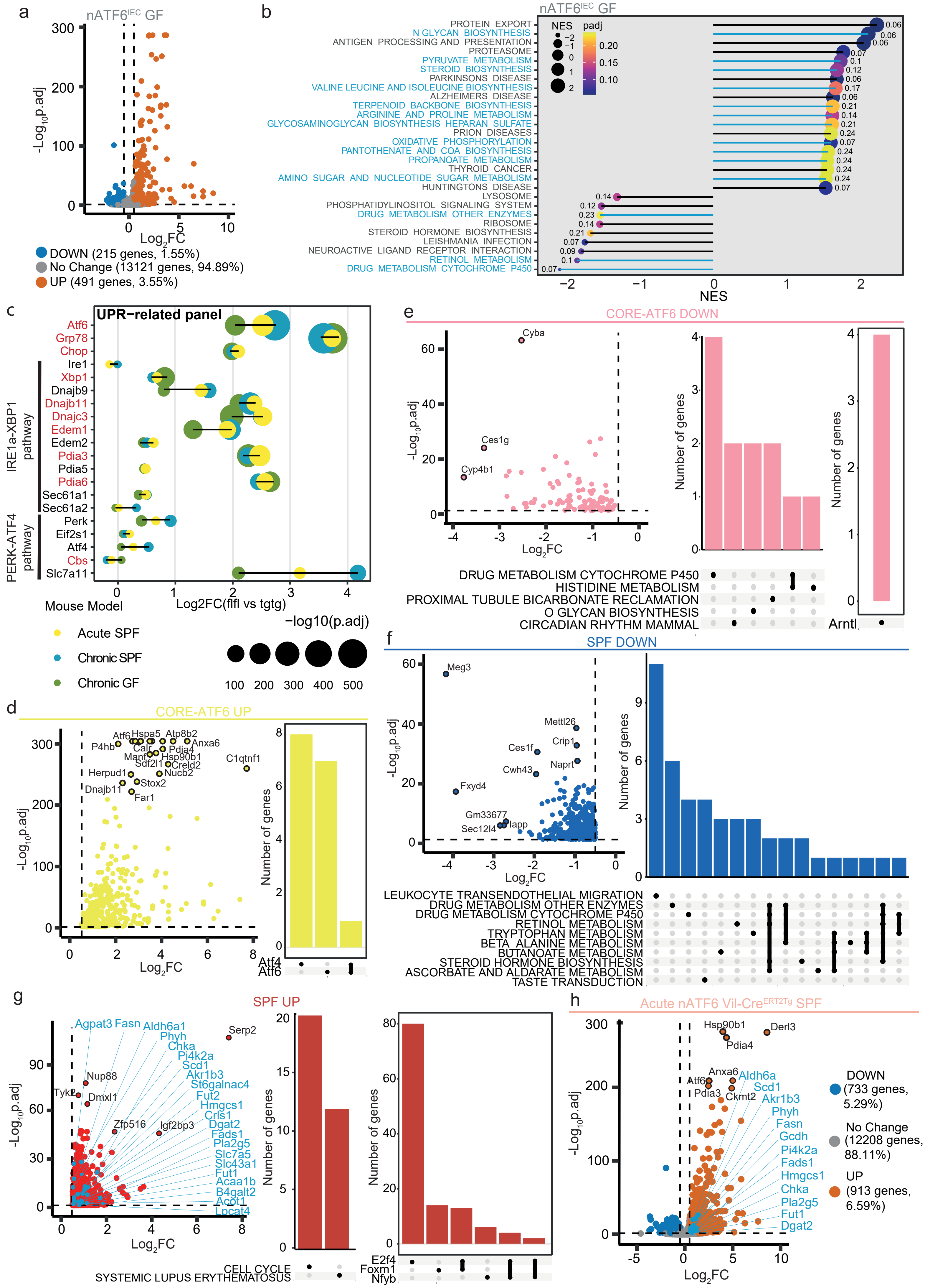

### Extended Data Fig 3

Extended Data 3.

a

| ATF6-FASN | Pan-cancer | CRC TCGA |
|---|---|---|
| Log2 OR | 1.459 | 0.640 |
| p-value | <0.001 | 0.297 |
| Association | Co-occurrence | |

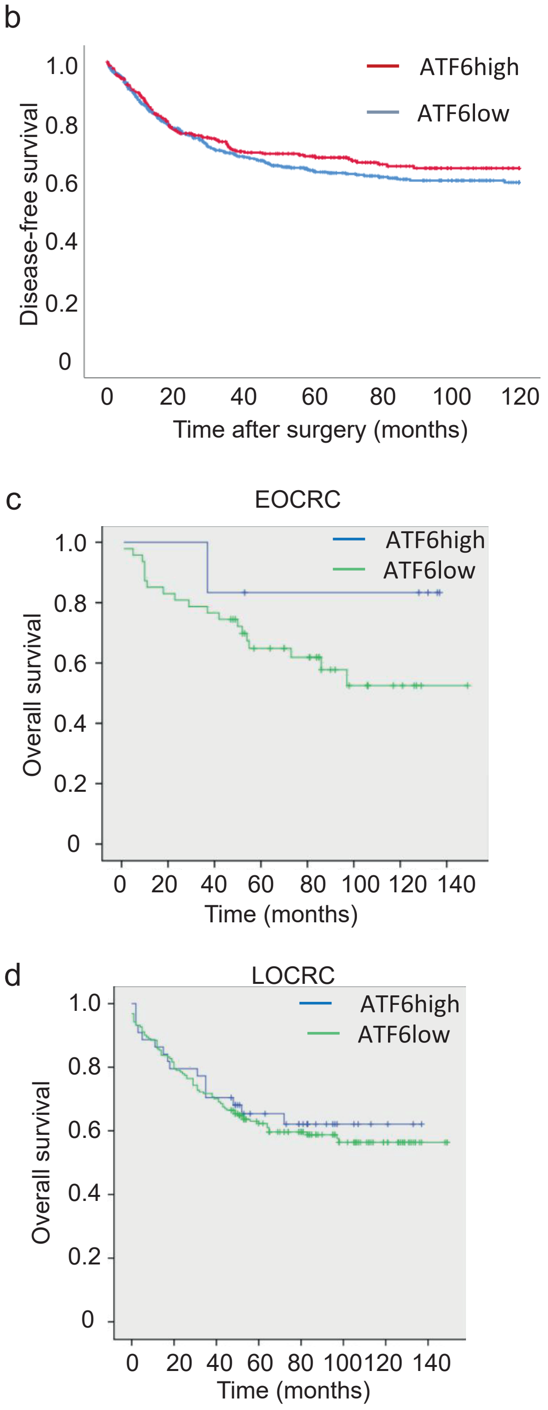


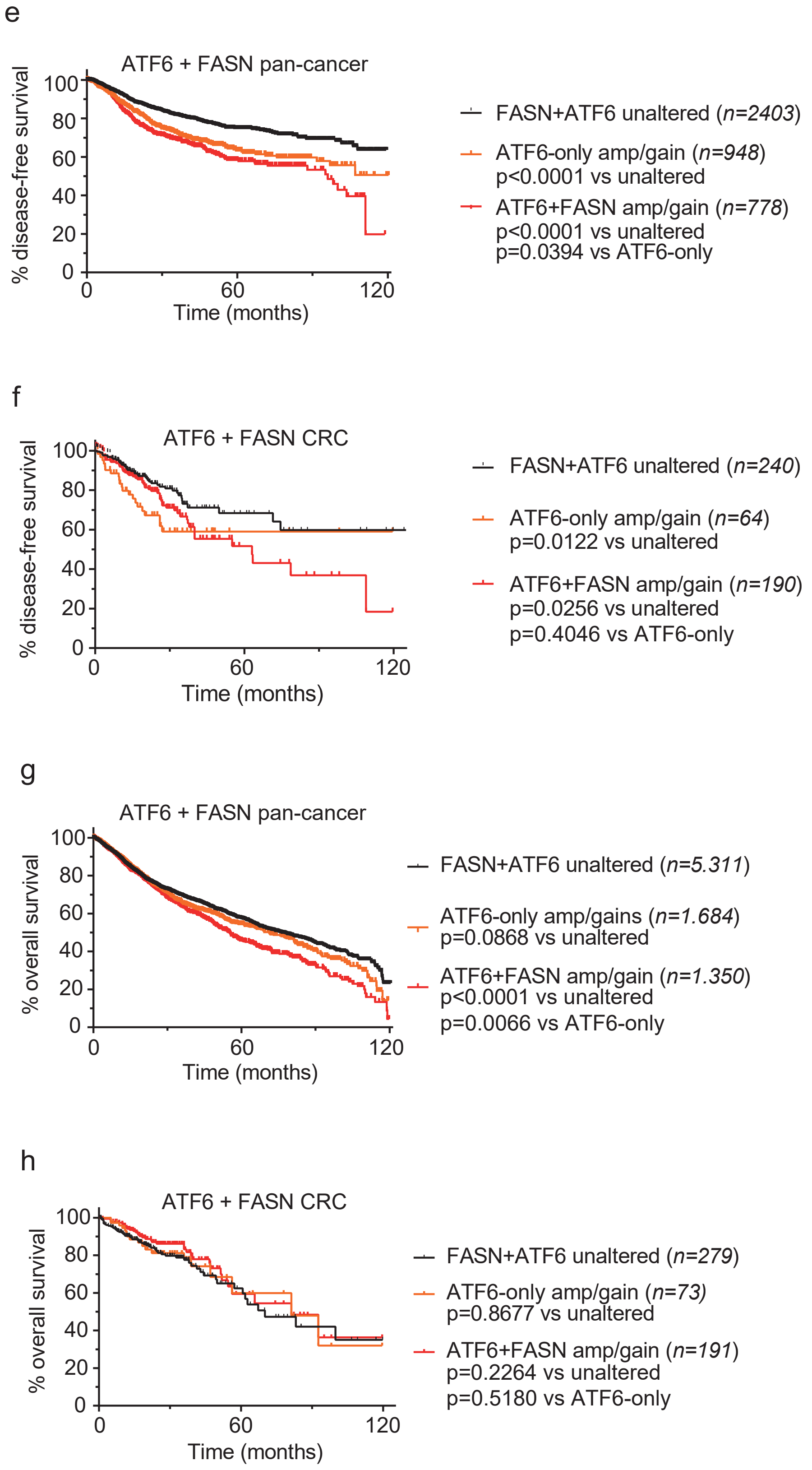

### Extended Data Fig 4

Extended Data 4.

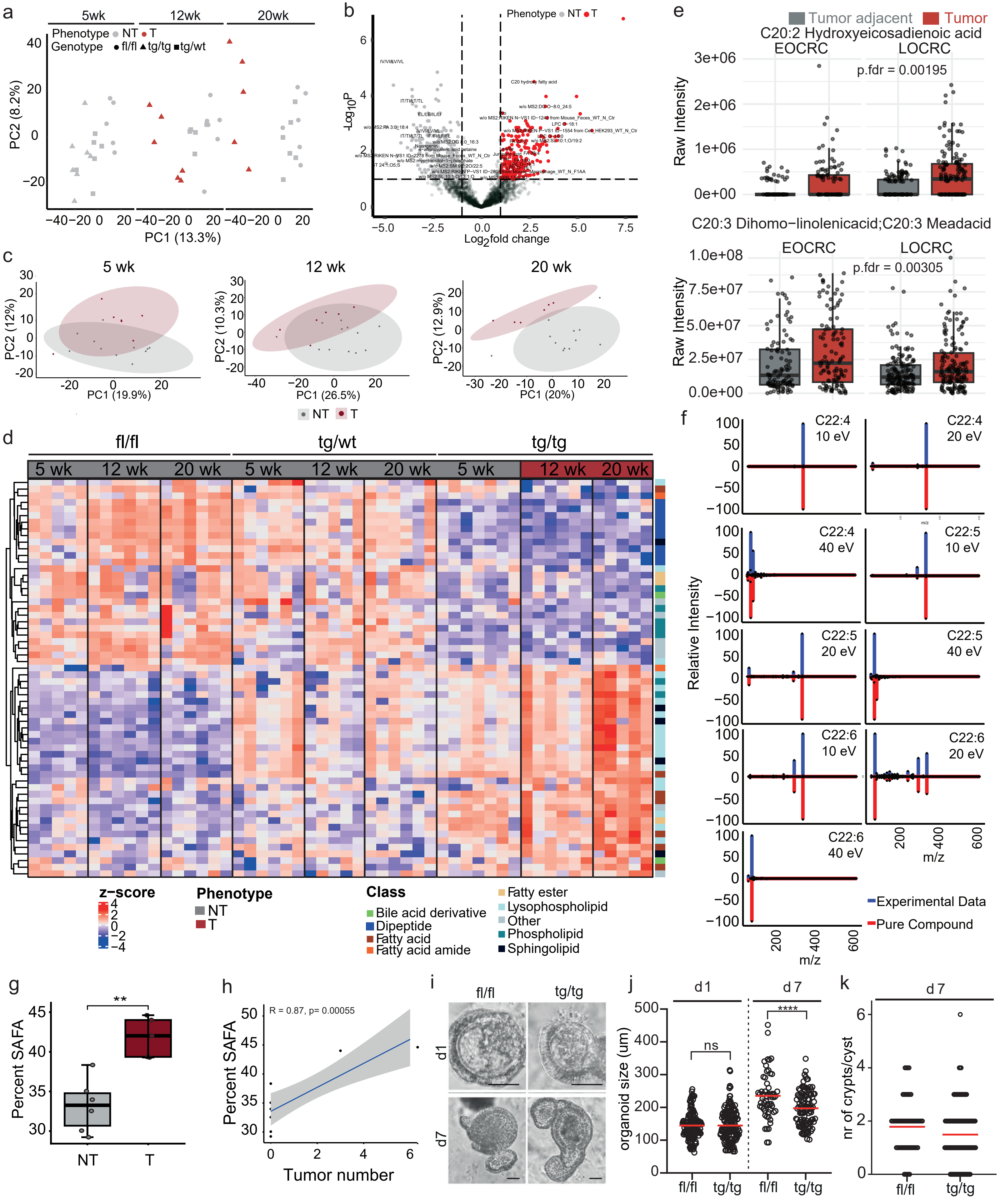

### Extended Data Fig 5

Extended Data 5.

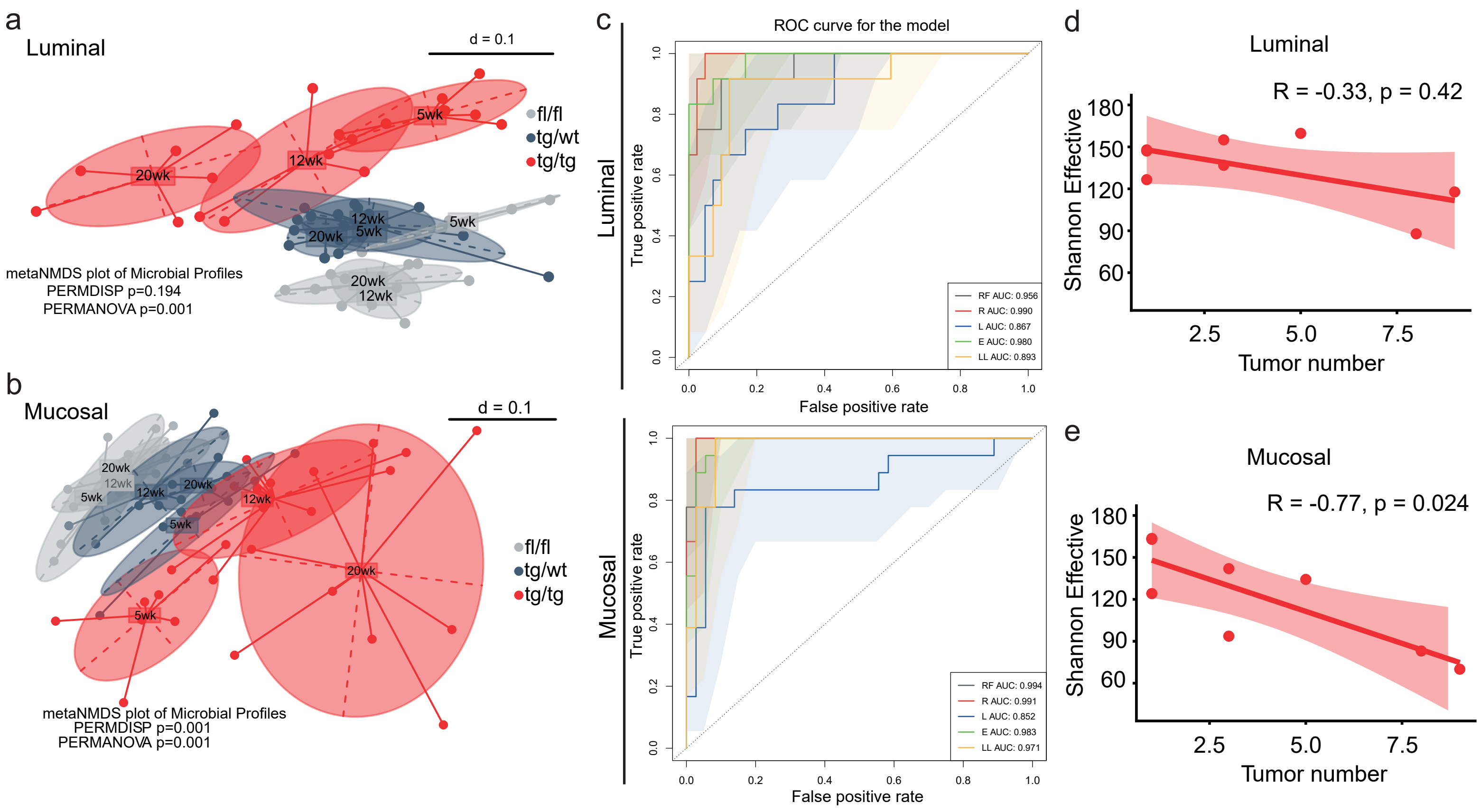

### Extended Data Fig 6

Extended Data 6.

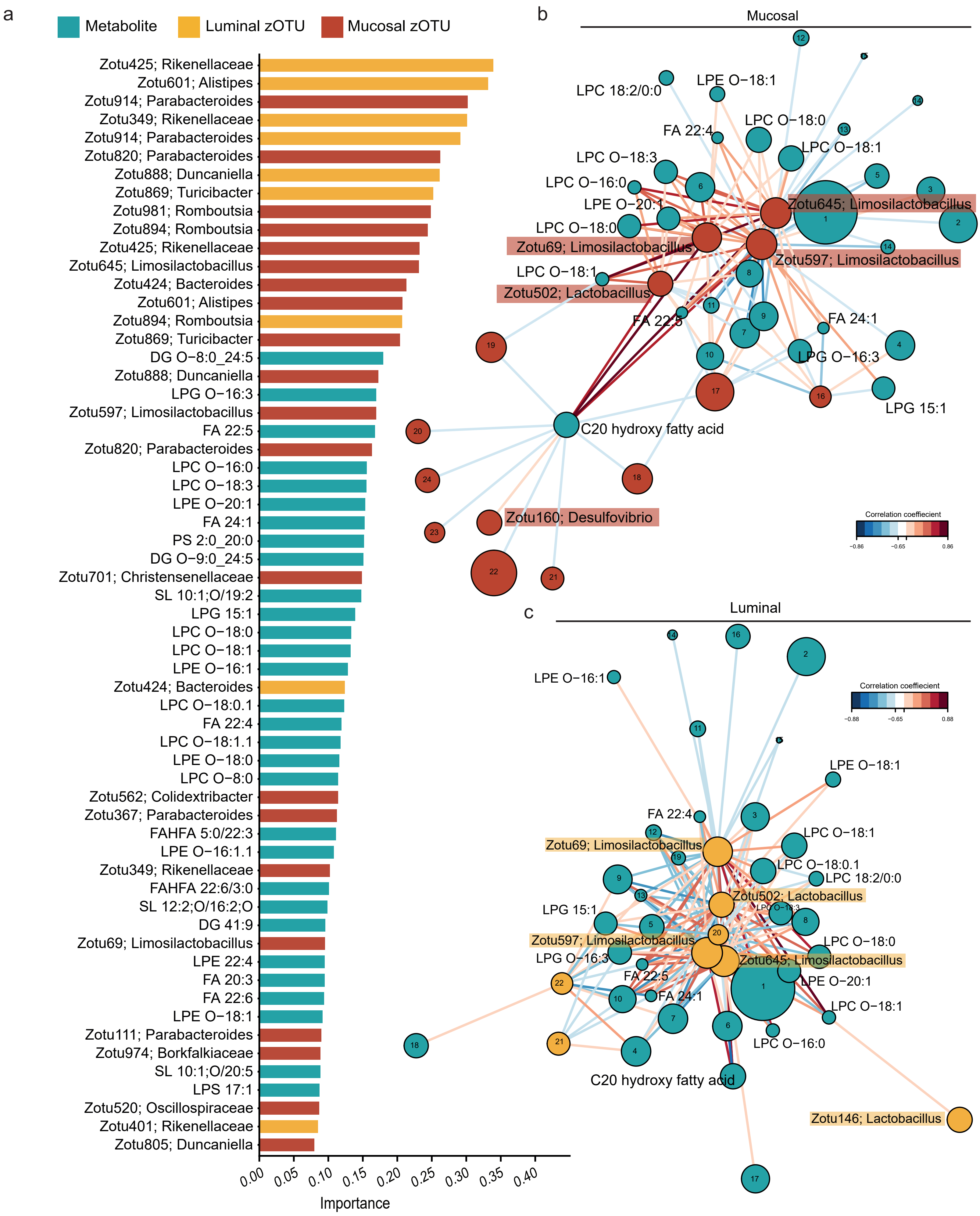

### Extended Data Fig 7

Extended Data 7.

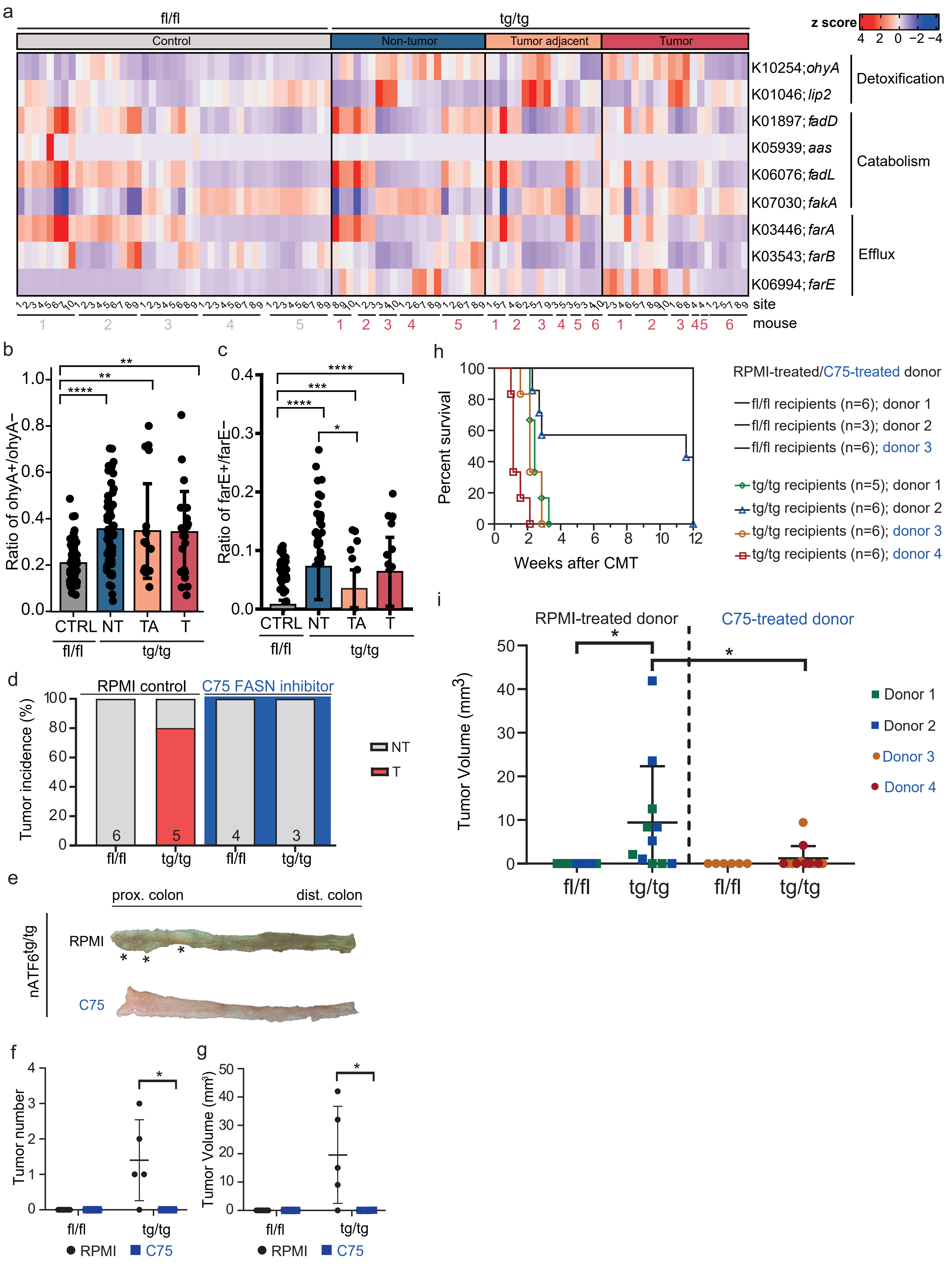

### Extended Data Fig 8

Extended Data 8.

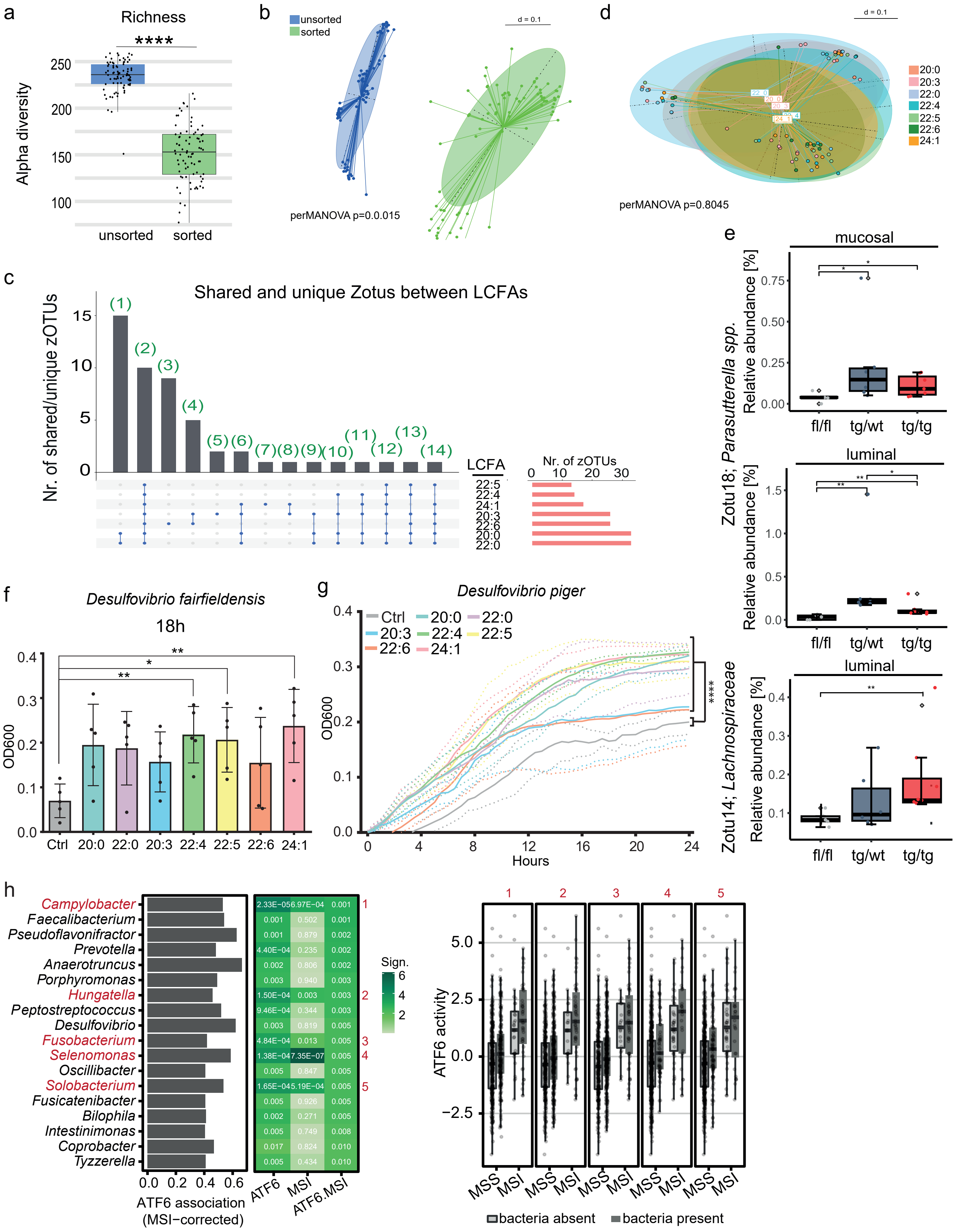
