## Supplementary material for "ATF6 activation alters colonic lipid metabolism causing tumor-associated microbial adaptation": Table 1

**Table 1.** Putative fatty acids that differ in tumor tissue compared to healthy tumor-adjacent tissue in CRC patients. *Compounds with confirmed annotations (level-1 or level-2 annotation). Duplicated compounds indicate more than one molecular feature with the same exact mass within the RT tolerance of 0.2 min.

|  | **FA Type** | **Putative Annotation** | **M-H** | **RT (min)** | **p-value** | **p-value (FDR corr)** |
| --- | --- | --- | --- | --- | --- | --- |
| 1 | C9:0 | Hydroxypelargonic acid | 173.1178 | 1.1 | 4.55E-20 | 6.83E-18 |
| 2 | C10:0 | Capric acid | 171.1385 | 1.18 | 2.91E-18 | 2.18E-16 |
| 3 | C16:1 | Palmitoleic acid, Sapienic acid | 253.2168 | 5.47 | 2.95E-17 | 1.47E-15 |
| 4 | C20:2 | Hydroxyeicosadienoid acid | 323.2587 | 5.88 | 1.90E-14 | 7.14E-13 |
| 5 | C22:2 | Docosadienoic acid* | 335.295 | 5.35 | 8.86E-13 | 2.22E-11 |
| 6 | C10:0 | Hydroxycapric acid | 187.1335 | 0.45 | 8.90E-13 | 2.22E-11 |
| 7 | C14:0 | Myristic acid | 227.2011 | 4.7 | 1.04E-12 | 2.23E-11 |
| 8 | C14:0 | Myristic acid | 227.2011 | 4.46 | 2.95E-12 | 5.52E-11 |
| 9 | C18:0 | Stearic acid | 283.2637 | 5.23 | 4.59E-12 | 7.65E-11 |
| 10 | C16:0 | Palmitic acid | 255.2324 | 4.98 | 6.51E-12 | 9.77E-11 |
| 11 | C9:0 | Hydroxypelargonic acid | 173.1178 | 2.25 | 5.93E-11 | 7.55E-10 |
| 12 | C16:1 | Palmitoleic acid, Sapienic acid | 253.2168 | 4.78 | 6.24E-11 | 7.55E-10 |
| 13 | C12:0 | Hydroxylauric acid | 215.1648 | 4.95 | 6.55E-11 | 7.55E-10 |
| 14 | C20:1 | Eicosenoic acid | 309.2794 | 5.28 | 1.54E-10 | 1.65E-09 |
| 15 | C14:1 | Myristoleic acid | 225.1854 | 4.53 | 1.92E-09 | 1.92E-08 |
| 16 | C12:0 | Lauric acid | 199.1698 | 4.3 | 5.31E-09 | 4.98E-08 |
| 17 | C20:3 | Dihomo-linolenic acid, mead acid | 305.2481 | 6.1 | 8.99E-08 | 7.93E-07 |
| 18 | C10:0 | Hydroxycapric acid | 187.1335 | 4.3 | 2.06E-07 | 1.72E-06 |
| 19 | C14:1 | Hydroxymyristoleic acid | 241.1804 | 5.23 | 4.33E-07 | 3.25E-06 |
| 20 | C12:0 | Hydroxylauric acid | 215.1648 | 5.11 | 1.05E-06 | 7.52E-06 |
| 21 | C14:0 | Hydroxymyristic acid | 243.1961 | 5.4 | 2.99E-06 | 2.00E-05 |
| 22 | C22:4 | Docosatetraenoic acid, Adrenic acid* | 331.2637 | 6.15 | 3.06E-06 | 2.00E-05 |
| 23 | C16:0 | Hydroxypalmitic acid, Hydroxypalmitic acid | 271.2274 | 5.83 | 4.49E-06 | 2.81E-05 |
| 24 | C24:6 | Herring acid | 355.2637 | 6.1 | 1.68E-05 | 0.00010074 |
| 25 | C5:0 | Hydroxyvaleric acid | 117.0552 | 1.35 | 2.43E-05 | 0.00014016 |
| 26 | C22:1 | Erucic acid | 337.3107 | 5.47 | 5.06E-05 | 0.00027089 |
| 27 | C13:0 | Hydroxytridecylic acid | 229.1804 | 4.7 | 7.17E-05 | 0.00037110 |
| 28 | C20:4 | Hydroxyeicosatetraenoic acid, Hydroxyarachidonic acid | 319.2274 | 5.5 | 0.00019 | 0.00095846 |
| 29 | C22:5 | Docosapentanoic acid, Ozubondo acid, Sardine acid* | 329.2481 | 6.02 | 0.00028 | 0.00138000 |
| 30 | C5:0 | Valeric acid | 101.0603 | 1.3 | 0.00044 | 0.00206859 |
| 31 | C20:3 | Hydroxylinolenic acid, Hydroxymead acid | 321.243 | 5.82 | 0.00054 | 0.00241711 |
| 32 | C22:6 | Hydroxydocosahexaenoic acid, Hydroxycervonic acid | 343.2274 | 5.4 | 0.00070 | 0.00300208 |
| 33 | C6:0 | Hydroxycaproic acid | 131.0709 | 2.13 | 0.00119 | 0.00485198 |
| 34 | C20:0 | Hydroxyarachidic acid | 327.29 | 6.38 | 0.00182 | 0.00701442 |
| 35 | C5:0 | Hydroxyvaleric acid | 117.0552 | 1.1 | 0.00256 | 0.00960870 |
| 36 | C20:2 | Hydroxyeicosadienoid acid | 323.2587 | 6.3 | 0.00290 | 0.01061708 |
| 37 | C19:0 | Nonadecylic acid | 297.2794 | 6.4 | 0.00309 | 0.01106635 |
| 38 | C24:5 | Tetracosapentaenoic acid | 357.2794 | 6.27 | 0.00385 | 0.01346411 |
| 39 | C26:0 | Cerotic acid | 395.3889 | 6.3 | 0.00444 | 0.01487946 |
| 40 | C14:1 | Hydroxymyristoleic acid | 241.1804 | 5.1 | 0.00446 | 0.01487946 |
| 41 | C19:0 | Nonadecylic acid | 297.2794 | 6.28 | 0.00551 | 0.01796963 |
| 42 | C6:0 | Caproic acid | 115.0759 | 1.98 | 0.00572 | 0.01813564 |
| 43 | C8:0 | Caprylic acid | 143.1072 | 3.03 | 0.00580 | 0.01813564 |
| 44 | C8:0 | Hydroxy Caprylic acid | 159.1022 | 3.7 | 0.00661 | 0.01989701 |
| 45 | C25:0 | Hydroxypentacosylic acid | 397.3682 | 6.3 | 0.00691 | 0.02034064 |
| 46 | C20:4 | Hydroxyarachidonic acid, Hydroxyeicosatetraenoic acid | 319.2274 | 5.43 | 0.01450 | 0.04184639 |
| 47 | C22:6 | Docosahexaenoic acid, Cervonic acid | 327.2324 | 5.95 | 0.01660 | 0.04612812 |
