## Supplementary Table 1 for "ATF6 activation alters colonic lipid metabolism causing tumor-associated microbial adaptation"

| ensembl_gene_id | external_gene_name | entrezgene_id | gene_biotype | baseMean | log2FoldChange | lfcSE | stat | pvalue | padj | absLog2FC |
|---|---|---|---|---|---|---|---|---|---|---|
| ENSMUSG00000100573 | 1700081H04Rik | 73477 | lncRNA | 1.277052062 | 0 | 2.58771768 | 0 | 1 | 1 | 0 |
| ENSMUSG00000113630 | 4930404H11Rik | 73815 | lncRNA | 1.60246229 | 0 | 3.03662027 | 0 | 1 | 1 | 0 |
| ENSMUSG00000031534 | Smim19 | 102032 | protein_coding | 104.3481346 | -1.24E-05 | 0.124624165 | 9.95E-05 | 0.999920582 | 0.999920582 | 1.24E-05 |
| ENSMUSG00000033712 | Ccar2 | 219158 | protein_coding | 862.6957216 | -3.23E-05 | 0.054208561 | 0.000595297 | 0.999525021 | 0.999654528 | 3.23E-05 |
| ENSMUSG00000004897 | Hdgf | 15191 | protein_coding | 1601.02326 | -7.70E-05 | 0.072677345 | 0.001059134 | 0.999154934 | 0.999457543 | 7.70E-05 |
| ENSMUSG00000044573 | Acp1 | 11431 | protein_coding | 478.8640721 | 8.73E-05 | 0.100417308 | -0.00086891 | 0.99930671 | 0.999548819 | 8.73E-05 |
| ENSMUSG00000010154 | Spire2 | 234857 | protein_coding | 707.3039452 | 9.66E-05 | 0.086247259 | -0.001120518 | 0.999105957 | 0.999457543 | 9.66E-05 |
| ENSMUSG00000004771 | Rab11a | 53869 | protein_coding | 1283.591876 | 0.00010618 | 0.079163399 | -0.001341274 | 0.998929819 | 0.999353427 | 0.00010618 |
| ENSMUSG00000020133 | 2310011J03Rik | 66374 | protein_coding | 402.9630236 | 0.000119126 | 0.077588698 | -0.001535348 | 0.99877497 | 0.999319591 | 0.000119126 |
| ENSMUSG00000026867 | Gapvd1 | 66691 | protein_coding | 1087.836205 | -0.000151106 | 0.053754663 | 0.002811026 | 0.997757128 | 0.998483602 | 0.000151106 |
| ENSMUSG00000006050 | Sra1 | 24068 | protein_coding | 841.7403856 | 0.000208183 | 0.070988323 | -0.002932641 | 0.997660094 | 0.998483602 | 0.000208183 |
| ENSMUSG00000041992 | Rapgef5 | 217944 | protein_coding | 468.4509915 | 0.000212954 | 0.078204684 | -0.002723034 | 0.997827336 | 0.998492433 | 0.000212954 |
| ENSMUSG00000028016 | Ints12 | 71793 | protein_coding | 208.5232331 | -0.00022356 | 0.077504707 | 0.00288447 | 0.997698529 | 0.998483602 | 0.00022356 |
| ENSMUSG00000018707 | Dync1h1 | 13424 | protein_coding | 6159.674525 | -0.000236844 | 0.084289837 | 0.002809874 | 0.997758048 | 0.998483602 | 0.000236844 |
| ENSMUSG00000031609 | Sap30 | 60406 | protein_coding | 86.00155943 | 0.000239647 | 0.126403354 | -0.001895894 | 0.998487296 | 0.999092293 | 0.000239647 |
| ENSMUSG00000073591 | Pcdhb22 | 93893 | protein_coding | 7.710326395 | -0.000245759 | 0.508163879 | 0.000483622 | 0.999614125 | 0.99967466 | 0.000245759 |
| ENSMUSG00000022507 | 1810013L24Rik | 69053 | protein_coding | 774.3756822 | -0.000280278 | 0.07795149 | 0.003595538 | 0.997131182 | 0.998249138 | 0.000280278 |
| ENSMUSG00000074340 | Ovgp1 | 12659 | protein_coding | 4.585169625 | 0.000378842 | 0.647902123 | -0.00058472 | 0.999533461 | 0.999654528 | 0.000378842 |
| ENSMUSG00000056612 | Ppp1r14b | 18938 | protein_coding | 380.959695 | 0.000381748 | 0.123818738 | -0.003083122 | 0.997540028 | 0.998483602 | 0.000381748 |
| ENSMUSG00000000276 | Dgke | 56077 | protein_coding | 108.0632377 | 0.000402938 | 0.109297656 | -0.003686612 | 0.997058516 | 0.998249138 | 0.000402938 |
| ENSMUSG00000033124 | Atg9a | 245860 | protein_coding | 467.9146325 | 0.000450962 | 0.073374621 | -0.006146027 | 0.995096211 | 0.997171234 | 0.000450962 |
| ENSMUSG00000019818 | Cd164 | 53599 | protein_coding | 2851.899185 | 0.00045761 | 0.110580739 | -0.004138242 | 0.99669817 | 0.99808463 | 0.00045761 |
| ENSMUSG00000022469 | Rapgef3 | 223864 | protein_coding | 271.2134217 | 0.000520859 | 0.156439441 | -0.003329464 | 0.997343477 | 0.99837123 | 0.000520859 |
| ENSMUSG00000036698 | Ago2 | 239528 | protein_coding | 1035.355603 | 0.000562776 | 0.102133777 | -0.00551018 | 0.995603535 | 0.99741548 | 0.000562776 |
| ENSMUSG00000036940 | Kdm1a | 99982 | protein_coding | 762.1568297 | -0.000564145 | 0.056022474 | 0.010069974 | 0.991965459 | 0.995389267 | 0.000564145 |
| ENSMUSG00000035829 | Ppp1r26 | 241289 | protein_coding | 84.42778239 | -0.000585851 | 0.143714117 | 0.004076505 | 0.996747428 | 0.99808463 | 0.000585851 |
| ENSMUSG00000005907 | Pex1 | 71382 | protein_coding | 186.4458389 | -0.000588582 | 0.103141592 | 0.005706548 | 0.995446858 | 0.997384323 | 0.000588582 |
| ENSMUSG00000040592 | Cd79b | 15985 | protein_coding | 2.593884132 | -0.00061115 | 0.873853767 | 0.000699374 | 0.999441981 | 1 | 0.00061115 |
| ENSMUSG00000030515 | Tarsl2 | 272396 | protein_coding | 111.4302521 | -0.000637517 | 0.170730076 | 0.003734064 | 0.997020655 | 0.998249138 | 0.000637517 |
| ENSMUSG00000033237 | Arid2 | 77044 | protein_coding | 594.575476 | 0.000672467 | 0.056968338 | -0.011804222 | 0.990581813 | 0.994433839 | 0.000672467 |
| ENSMUSG00000000134 | Tfe3 | 209446 | protein_coding | 259.9004679 | -0.000688296 | 0.078320251 | 0.008788222 | 0.992988103 | 0.995778769 | 0.000688296 |
| ENSMUSG00000022946 | Dop1b | 70028 | protein_coding | 1196.033632 | -0.000727246 | 0.10800482 | 0.006733462 | 0.994627515 | 0.996939463 | 0.000727246 |
| ENSMUSG00000053838 | Nudcd3 | 209586 | protein_coding | 567.3308542 | 0.000753768 | 0.060887952 | -0.012379584 | 0.990122773 | 0.994337599 | 0.000753768 |
| ENSMUSG00000050866 | Clrn3 | 212070 | protein_coding | 1185.907672 | -0.000761269 | 0.063606951 | 0.01196833 | 0.990450882 | 0.994433839 | 0.000761269 |
| ENSMUSG00000028811 | Yars | 107271 | protein_coding | 752.4639051 | -0.000764958 | 0.071234074 | 0.010738653 | 0.991431959 | 0.995047245 | 0.000764958 |
| ENSMUSG00000063019 | Manbal | 69161 | protein_coding | 149.7060017 | -0.00077066 | 0.100037235 | 0.007703728 | 0.993853375 | 0.996446603 | 0.00077066 |
| ENSMUSG00000035517 | Tdrd7 | 100121 | protein_coding | 770.2718511 | -0.000857793 | 0.071845862 | 0.011939353 | 0.990474001 | 0.994433839 | 0.000857793 |
| ENSMUSG00000004865 | Srpk1 | 20815 | protein_coding | 1148.339418 | 0.000922545 | 0.071661815 | -0.012873599 | 0.989728638 | 0.994183621 | 0.000922545 |
| ENSMUSG00000050174 | Nudt6 | 229228 | protein_coding | 23.39731417 | 0.000965991 | 0.211642943 | -0.004564248 | 0.99635827 | 0.998050496 | 0.000965991 |

| ensembl_gene_id | external_gene_name | entrezgene_id | gene_biotype | baseMean | log2FoldChange | lfcSE | stat | pvalue | padj | absLog2FC |
|---|---|---|---|---|---|---|---|---|---|---|
| ENSMUSG00000021892 | Sh3bp5 | 24056 | protein_coding | 69.2198607 | 0.000982382 | 0.13488677 | -0.007283011 | 0.99418905 | 0.996641735 | 0.000982382 |
| ENSMUSG00000038546 | Ranbp9 | 56705 | protein_coding | 448.7591511 | 0.001117371 | 0.062618174 | -0.017844197 | 0.985763147 | 0.99195007 | 0.001117371 |
| ENSMUSG00000069895 | Atxn1l | 52335 | protein_coding | 487.5596863 | -0.001134663 | 0.080133234 | 0.014159701 | 0.988702571 | 0.993636457 | 0.001134663 |
| ENSMUSG00000041012 | Cmtm8 | 70031 | protein_coding | 60.84010339 | -0.001139013 | 0.202496345 | 0.005624856 | 0.995512038 | 0.997384323 | 0.001139013 |
| ENSMUSG00000006740 | Kif5b | 16573 | protein_coding | 3338.247337 | -0.001154204 | 0.080724504 | 0.014298057 | 0.98859219 | 0.993636457 | 0.001154204 |
| ENSMUSG00000032508 | Myd88 | 17874 | protein_coding | 318.1733686 | -0.00117501 | 0.08901355 | 0.013200346 | 0.989467953 | 0.994042693 | 0.00117501 |
| ENSMUSG00000022792 | Yars2 | 70120 | protein_coding | 184.098991 | -0.001197818 | 0.119670116 | 0.010009334 | 0.99201384 | 0.995389267 | 0.001197818 |
| ENSMUSG00000037493 | Cib2 | 56506 | protein_coding | 22.87459745 | -0.001201816 | 0.273151742 | 0.004399813 | 0.996489469 | 0.998060837 | 0.001201816 |
| ENSMUSG00000029227 | Fip1l1 | 66899 | protein_coding | 573.6716705 | -0.001225636 | 0.056102932 | 0.021846203 | 0.982570638 | 0.990608762 | 0.001225636 |
| ENSMUSG00000022913 | Psmg1 | 56088 | protein_coding | 149.269976 | 0.001263429 | 0.105507519 | -0.011974779 | 0.990445737 | 0.994433839 | 0.001263429 |
| ENSMUSG00000028990 | Lzic | 69151 | protein_coding | 278.2900611 | -0.001291746 | 0.082913684 | 0.015579405 | 0.987569936 | 0.99304207 | 0.001291746 |
| ENSMUSG00000077714 | Snord17 | 100313519 | snoRNA | 1196.118181 | -0.001302503 | 0.212880605 | 0.006118469 | 0.995118199 | 0.997171234 | 0.001302503 |
| ENSMUSG00000024799 | Tm7sf2 | 73166 | protein_coding | 304.5683476 | 0.001329402 | 0.114056403 | -0.011655651 | 0.990700346 | 0.994433839 | 0.001329402 |
| ENSMUSG00000025047 | Pdcd11 | 18572 | protein_coding | 449.8452708 | -0.001336953 | 0.101081037 | 0.013226547 | 0.98944705 | 0.994042693 | 0.001336953 |
| ENSMUSG00000047921 | Trappc9 | 76510 | protein_coding | 548.5564266 | -0.001342385 | 0.066322407 | 0.020240298 | 0.983851681 | 0.991234621 | 0.001342385 |
| ENSMUSG00000020361 | Hspa4 | 15525 | protein_coding | 2420.20498 | 0.001356884 | 0.085419272 | -0.015884992 | 0.987326143 | 0.992978315 | 0.001356884 |
| ENSMUSG00000030291 | Med21 | 108098 | protein_coding | 185.6956275 | -0.001364518 | 0.116997893 | 0.011662759 | 0.990694675 | 0.994433839 | 0.001364518 |
| ENSMUSG00000041324 | Inhba | 16323 | protein_coding | 2.295440781 | 0.001378207 | 2.01518495 | -0.000683911 | 0.999454318 | 1 | 0.001378207 |
| ENSMUSG00000091649 | Phf11b | 236451 | protein_coding | 64.34463178 | -0.001382427 | 0.320365157 | 0.004315161 | 0.99655701 | 0.998067952 | 0.001382427 |
| ENSMUSG00000030051 | Aplf | 72103 | protein_coding | 119.6536241 | 0.001403937 | 0.101829949 | -0.013787071 | 0.988999857 | 0.993753796 | 0.001403937 |
| ENSMUSG00000028233 | Tgs1 | 116940 | protein_coding | 137.943418 | 0.001434609 | 0.112175392 | -0.01278898 | 0.989796149 | 0.994190961 | 0.001434609 |
| ENSMUSG00000024943 | Smc5 | 226026 | protein_coding | 304.0877152 | -0.001498156 | 0.063954853 | 0.023425218 | 0.981311089 | 0.98994327 | 0.001498156 |
| ENSMUSG00000033983 | Coil | 12812 | protein_coding | 96.6557696 | 0.001517499 | 0.116369794 | -0.013040313 | 0.989595631 | 0.994110484 | 0.001517499 |
| ENSMUSG00000037533 | Rapgef6 | 192786 | protein_coding | 943.0214332 | 0.001574733 | 0.07533366 | -0.02090344 | 0.983322682 | 0.990883011 | 0.001574733 |
| ENSMUSG00000024206 | Rfx2 | 19725 | protein_coding | 32.16477095 | 0.001582597 | 0.38912724 | -0.004067042 | 0.996754979 | 0.99808463 | 0.001582597 |
| ENSMUSG00000027942 | 4933434E20Rik | 99650 | protein_coding | 102.6758371 | 0.001594678 | 0.135967163 | -0.011728409 | 0.990642298 | 0.994433839 | 0.001594678 |
| ENSMUSG00000030979 | Uros | 22276 | protein_coding | 200.0758897 | 0.00163292 | 0.096274837 | -0.016961026 | 0.986467708 | 0.992356709 | 0.00163292 |
| ENSMUSG00000005417 | Mprip | 26936 | protein_coding | 1847.156355 | -0.001689492 | 0.098066393 | 0.017228039 | 0.986254694 | 0.992263312 | 0.001689492 |
| ENSMUSG00000027646 | Src | 20779 | protein_coding | 2146.887716 | 0.001730339 | 0.089822291 | -0.019264025 | 0.984630483 | 0.99157486 | 0.001730339 |
| ENSMUSG00000029826 | Zc3hav1 | 78781 | protein_coding | 1268.417383 | 0.001746693 | 0.068215835 | -0.025605393 | 0.979572084 | 0.989055615 | 0.001746693 |
| ENSMUSG00000035726 | Supt16 | 114741 | protein_coding | 996.2198014 | -0.001753737 | 0.101506634 | 0.017277067 | 0.98621558 | 0.992263312 | 0.001753737 |
| ENSMUSG00000071172 | Srsf3 | 20383 | protein_coding | 1783.902145 | -0.001799904 | 0.072637075 | 0.024779415 | 0.980230911 | 0.989337077 | 0.001799904 |
| ENSMUSG00000032020 | Ubash3b | 72828 | protein_coding | 27.54898273 | -0.001811379 | 0.224496848 | 0.008068618 | 0.993562244 | 0.99621657 | 0.001811379 |
| ENSMUSG00000026280 | Atg4b | 66615 | protein_coding | 564.5497199 | -0.001825019 | 0.054597863 | 0.033426556 | 0.973334433 | 0.98611318 | 0.001825019 |
| ENSMUSG00000039987 | Phtf2 | 68770 | protein_coding | 48.95507312 | 0.00182588 | 0.184357333 | -0.009904029 | 0.992097857 | 0.995401911 | 0.00182588 |
| ENSMUSG00000050549 | Fam241a | 70617 | protein_coding | 149.5614365 | -0.001839138 | 0.12718414 | 0.014460435 | 0.988462644 | 0.99357673 | 0.001839138 |
| ENSMUSG00000021607 | Mrpl36 | 94066 | protein_coding | 404.4919914 | -0.001846181 | 0.083942246 | 0.021993466 | 0.982453168 | 0.990550804 | 0.001846181 |
| ENSMUSG00000031364 | Grpr | 14829 | protein_coding | 30.25963417 | 0.001888063 | 0.332558804 | -0.005677381 | 0.995470129 | 0.997384323 | 0.001888063 |
| ENSMUSG00000033364 | Usp37 | 319651 | protein_coding | 462.7823755 | 0.001891098 | 0.055002314 | -0.03438215 | 0.972572417 | 0.985703995 | 0.001891098 |

Supplementary Table 1

| ensembl_gene_id | external_gene_name | entrezgene_id | gene_biotype | baseMean | log2FoldChange | lfcSE | stat | pvalue | padj | absLog2FC |
|---|---|---|---|---|---|---|---|---|---|---|
| ENSMUSG00000032185 | Carm1 | 59035 | protein_coding | 271.5097885 | -0.001898206 | 0.085777163 | 0.0221295 | 0.982344654 | 0.99050187 | 0.001898206 |
| ENSMUSG00000041390 | Mdfic | 16543 | protein_coding | 50.40902921 | 0.001907998 | 0.170086422 | -0.011217817 | 0.991049664 | 0.994724011 | 0.001907998 |
| ENSMUSG00000019978 | Epb41l2 | 13822 | protein_coding | 420.9506095 | 0.001912214 | 0.099923638 | -0.019136754 | 0.984732012 | 0.99157486 | 0.001912214 |
| ENSMUSG00000035451 | Foxa1 | 15375 | protein_coding | 566.9495321 | -0.001931471 | 0.10131365 | 0.019064274 | 0.984789832 | 0.99157486 | 0.001931471 |
| ENSMUSG00000071531 | Gprin2 | 432839 | protein_coding | 3.095350147 | -0.001993175 | 1.035042229 | 0.001925694 | 0.998463519 | 1 | 0.001993175 |
| ENSMUSG00000044350 | Lacc1 | 210808 | protein_coding | 107.8792558 | -0.0020721 | 0.115767833 | 0.017898753 | 0.985719624 | 0.99195007 | 0.0020721 |
| ENSMUSG00000022013 | Dnajc15 | 66148 | protein_coding | 331.7192118 | 0.002082908 | 0.074640486 | -0.027905876 | 0.977737222 | 0.988088396 | 0.002082908 |
| ENSMUSG00000070426 | Rnf121 | 75212 | protein_coding | 389.410576 | 0.00210336 | 0.063675807 | -0.033032318 | 0.973648815 | 0.98618968 | 0.00210336 |
| ENSMUSG00000035930 | Chst4 | 26887 | protein_coding | 755.3702886 | -0.002106814 | 0.238543778 | 0.00883198 | 0.992953191 | 0.995778769 | 0.002106814 |
| ENSMUSG00000030225 | Dera | 232449 | protein_coding | 538.3981588 | 0.002121259 | 0.082574204 | -0.025689129 | 0.979505295 | 0.989055615 | 0.002121259 |
| ENSMUSG00000089872 | Rps6kc1 | 320119 | protein_coding | 327.9857132 | -0.002123631 | 0.067276705 | 0.031565618 | 0.974818462 | 0.986890148 | 0.002123631 |
| ENSMUSG00000059475 | Zfp426 | 235028 | protein_coding | 221.8442611 | 0.002162356 | 0.094205547 | -0.0229536 | 0.981687285 | 0.990146635 | 0.002162356 |
| ENSMUSG00000096992 | Gm26788 | 108167617 | lncRNA | 17.89695839 | -0.002166188 | 0.331928649 | 0.006526066 | 0.99479299 | 0.996966345 | 0.002166188 |
| ENSMUSG00000066640 | Fbxl18 | 231863 | protein_coding | 360.0502406 | -0.00221628 | 0.085199146 | 0.026012935 | 0.979247022 | 0.989008948 | 0.00221628 |
| ENSMUSG00000028191 | Bcl10 | 12042 | protein_coding | 724.5456993 | 0.002230816 | 0.063947397 | -0.034885175 | 0.972171302 | 0.985568894 | 0.002230816 |
| ENSMUSG00000072872 | Rybp | 56353 | protein_coding | 225.3435194 | 0.002248374 | 0.105885983 | -0.021233915 | 0.98305906 | 0.990798738 | 0.002248374 |
| ENSMUSG00000024219 | Anks1 | 224650 | protein_coding | 621.6085874 | 0.002260288 | 0.059289564 | -0.038122863 | 0.969589723 | 0.984433119 | 0.002260288 |
| ENSMUSG00000026784 | Pdss1 | 56075 | protein_coding | 385.040132 | -0.002273654 | 0.071306789 | 0.03188552 | 0.974563346 | 0.986813361 | 0.002273654 |
| ENSMUSG00000031393 | Mecp2 | 17257 | protein_coding | 330.4321051 | 0.002321396 | 0.069423892 | -0.033438001 | 0.973325306 | 0.98611318 | 0.002321396 |
| ENSMUSG00000034973 | Dop1a | 320615 | protein_coding | 726.4698021 | -0.002335981 | 0.052051906 | 0.044877924 | 0.964204613 | 0.982347769 | 0.002335981 |
| ENSMUSG00000031913 | Vps4a | 116733 | protein_coding | 602.663135 | 0.002373671 | 0.08476247 | -0.028003796 | 0.977659124 | 0.988088396 | 0.002373671 |
| ENSMUSG00000043085 | Tmem82 | 213989 | protein_coding | 176.7582885 | -0.002429696 | 0.128998821 | 0.018835027 | 0.984972712 | 0.991590016 | 0.002429696 |
| ENSMUSG00000029594 | Rbm19 | 74111 | protein_coding | 204.724593 | 0.002482288 | 0.114497644 | -0.02167982 | 0.982703361 | 0.990621616 | 0.002482288 |
| ENSMUSG00000022849 | Hspbap1 | 66667 | protein_coding | 79.11399134 | -0.002490832 | 0.137500282 | 0.018115108 | 0.985547025 | 0.991913919 | 0.002490832 |
| ENSMUSG00000039714 | Cplx3 | 235415 | protein_coding | 2.511315346 | 0.002520554 | 0.965566033 | -0.002610442 | 0.997917171 | 1 | 0.002520554 |
| ENSMUSG00000039021 | Ttc16 | 338348 | protein_coding | 7.646279742 | -0.002559799 | 0.719430907 | 0.003558089 | 0.997161062 | 0.998249138 | 0.002559799 |
| ENSMUSG00000025872 | Thoc3 | 73666 | protein_coding | 282.6253644 | 0.002567986 | 0.115173649 | -0.022296642 | 0.982211327 | 0.990427908 | 0.002567986 |
| ENSMUSG00000058351 | Smim4 | 66487 | protein_coding | 70.48864374 | 0.002623413 | 0.124606275 | -0.021053618 | 0.983202884 | 0.990883011 | 0.002623413 |
| ENSMUSG00000021392 | Nol8 | 70930 | protein_coding | 209.2336982 | 0.002628763 | 0.140727023 | -0.018679875 | 0.985096483 | 0.991590016 | 0.002628763 |
| ENSMUSG00000110185 | Igip | 109169 | protein_coding | 130.994716 | 0.002638975 | 0.189326069 | -0.013938785 | 0.988878819 | 0.993692638 | 0.002638975 |
| ENSMUSG00000059796 | Eif4a1 | 13681 | protein_coding | 4015.611126 | -0.002639155 | 0.10918166 | 0.024172148 | 0.980715295 | 0.989584039 | 0.002639155 |
| ENSMUSG00000077217 | Gm23951 | 115485840 | snoRNA | 0.637227717 | -0.002661005 | 1.441072238 | 0.001846545 | 0.998526671 | 1 | 0.002661005 |
| ENSMUSG00000020124 | Usp15 | 14479 | protein_coding | 1004.507682 | 0.002662281 | 0.061563878 | -0.04324421 | 0.965506864 | 0.982706396 | 0.002662281 |
| ENSMUSG00000039063 | Echdc3 | 67856 | protein_coding | 105.7596826 | -0.00267418 | 0.125315717 | 0.021339539 | 0.982974803 | 0.990774286 | 0.00267418 |
| ENSMUSG00000020715 | Ern1 | 78943 | protein_coding | 525.2089985 | -0.002734335 | 0.072581673 | 0.037672529 | 0.969948779 | 0.984509162 | 0.002734335 |
| ENSMUSG00000021592 | Arsk | 77041 | protein_coding | 106.2054746 | -0.002758806 | 0.123492434 | 0.022339881 | 0.982176836 | 0.990427908 | 0.002758806 |
| ENSMUSG00000028224 | Nbn | 27354 | protein_coding | 333.469505 | -0.002773398 | 0.108422429 | 0.025579562 | 0.979592688 | 0.989055615 | 0.002773398 |
| ENSMUSG00000021967 | Mrpl57 | 67840 | protein_coding | 394.8801979 | -0.002825627 | 0.097413166 | 0.029006628 | 0.976859304 | 0.987811867 | 0.002825627 |
| ENSMUSG00000028597 | Gpx7 | 67305 | protein_coding | 25.12664132 | 0.002855732 | 0.204058144 | -0.013994697 | 0.988834212 | 0.993692638 | 0.002855732 |

Supplementary Table 1

| ensembl_gene_id | external_gene_name | entrezgene_id | gene_biotype | baseMean | log2FoldChange | lfcSE | stat | pvalue | padj | absLog2FC |
|---|---|---|---|---|---|---|---|---|---|---|
| ENSMUSG00000028843 | Sh3bgrl3 | 73723 | protein_coding | 692.8798783 | 0.002896364 | 0.083866948 | -0.034535222 | 0.972450356 | 0.985701275 | 0.002896364 |
| ENSMUSG00000002031 | Ift46 | 76568 | protein_coding | 289.4240698 | 0.002926048 | 0.097644162 | -0.029966443 | 0.976093815 | 0.987575865 | 0.002926048 |
| ENSMUSG00000028973 | Abcb8 | 74610 | protein_coding | 813.9932726 | -0.002956157 | 0.095507505 | 0.030952089 | 0.975307749 | 0.987234441 | 0.002956157 |
| ENSMUSG00000036398 | Ppp1r11 | 76497 | protein_coding | 483.4509429 | 0.002957935 | 0.062716144 | -0.047163856 | 0.962382634 | 0.981474584 | 0.002957935 |
| ENSMUSG00000063659 | Zbtb18 | 30928 | protein_coding | 429.3822104 | 0.002964641 | 0.057794855 | -0.051295937 | 0.959089706 | 0.980158095 | 0.002964641 |
| ENSMUSG00000002812 | Flii | 14248 | protein_coding | 2660.862153 | 0.003044211 | 0.065898347 | -0.046195565 | 0.963154378 | 0.981701728 | 0.003044211 |
| ENSMUSG00000005034 | Prkacb | 18749 | protein_coding | 1104.098423 | 0.003055641 | 0.206336045 | -0.014809053 | 0.988184517 | 0.993418099 | 0.003055641 |
| ENSMUSG00000034820 | Cpsf7 | 269061 | protein_coding | 448.0469124 | -0.003057727 | 0.060375707 | 0.050644983 | 0.959608418 | 0.980447529 | 0.003057727 |
| ENSMUSG00000030002 | Dusp11 | 72102 | protein_coding | 1283.125366 | 0.003089395 | 0.066839213 | -0.046221293 | 0.963133871 | 0.981701728 | 0.003089395 |
| ENSMUSG00000040264 | Gbp2b | 14468 | protein_coding | 0.99786071 | -0.003126751 | 2.222433082 | 0.001406905 | 0.998877453 | 1 | 0.003126751 |
| ENSMUSG00000041439 | Mfsd6 | 98682 | protein_coding | 2885.96381 | -0.003228963 | 0.094793369 | 0.034063175 | 0.972826774 | 0.985840779 | 0.003228963 |
| ENSMUSG00000061032 | Rrp1 | 18114 | protein_coding | 1134.614073 | 0.003239203 | 0.066021169 | -0.0490631 | 0.96086901 | 0.980946516 | 0.003239203 |
| ENSMUSG00000050812 | Ecpas | 230249 | protein_coding | 1760.177073 | 0.003253207 | 0.092953 | -0.034998413 | 0.972081006 | 0.985568894 | 0.003253207 |
| ENSMUSG00000041674 | BC006965 | 217294 | lncRNA | 6.70043436 | -0.003290501 | 0.707546529 | 0.004650579 | 0.996289388 | 0.998042036 | 0.003290501 |
| ENSMUSG00000039886 | Tmem120a | 215210 | protein_coding | 396.1376531 | -0.00329616 | 0.105373943 | 0.031280597 | 0.975045764 | 0.987059753 | 0.00329616 |
| ENSMUSG00000037415 | Ranbp10 | 74334 | protein_coding | 349.3132716 | 0.003303517 | 0.08652944 | -0.038177957 | 0.969545796 | 0.984433119 | 0.003303517 |
| ENSMUSG00000043671 | Dpy19l3 | 233115 | protein_coding | 68.67367529 | 0.003323706 | 0.13232717 | -0.025117335 | 0.979961373 | 0.989246416 | 0.003323706 |
| ENSMUSG00000041272 | Tox | 252838 | protein_coding | 187.0924109 | -0.003330158 | 0.113571095 | 0.029322235 | 0.976607594 | 0.987811867 | 0.003330158 |
| ENSMUSG00000025208 | Mrpl43 | 94067 | protein_coding | 320.8006508 | -0.003338138 | 0.085788234 | 0.038911374 | 0.968961048 | 0.98433949 | 0.003338138 |
| ENSMUSG00000044518 | Foxe3 | 30923 | protein_coding | 3.661887614 | -0.003344915 | 0.858920935 | 0.003894322 | 0.996892788 | 1 | 0.003344915 |
| ENSMUSG00000034601 | 2700049A03Rik | 76967 | protein_coding | 142.1892166 | 0.003347456 | 0.098834581 | -0.033869281 | 0.972981389 | 0.985876467 | 0.003347456 |
| ENSMUSG00000022089 | Bin3 | 57784 | protein_coding | 292.8845026 | -0.00340044 | 0.083020366 | 0.0409591 | 0.967328502 | 0.983635967 | 0.00340044 |
| ENSMUSG00000029462 | Vps29 | 56433 | protein_coding | 387.2708906 | 0.003438003 | 0.064694788 | -0.053141879 | 0.957618864 | 0.979202348 | 0.003438003 |
| ENSMUSG00000052419 | Smim30 | 66520 | protein_coding | 247.3876158 | -0.003464967 | 0.10393977 | 0.033336299 | 0.973406407 | 0.9861256 | 0.003464967 |
| ENSMUSG00000020056 | Washc3 | 67282 | protein_coding | 272.9817775 | 0.003543076 | 0.081970585 | -0.043223744 | 0.965523178 | 0.982706396 | 0.003543076 |
| ENSMUSG00000021339 | Mrs2 | 380836 | protein_coding | 906.7144092 | 0.003586162 | 0.07736787 | -0.046352085 | 0.963029626 | 0.981695756 | 0.003586162 |
| ENSMUSG00000024068 | Spast | 50850 | protein_coding | 444.9912384 | -0.003591132 | 0.083853356 | 0.042826335 | 0.965839971 | 0.982740683 | 0.003591132 |
| ENSMUSG00000069456 | Rdh16 | 19683 | protein_coding | 351.4552562 | -0.003654015 | 0.168491846 | 0.021686599 | 0.982697954 | 0.990621616 | 0.003654015 |
| ENSMUSG00000031781 | Ciapin1 | 109006 | protein_coding | 268.6508332 | 0.003662839 | 0.081907709 | -0.0447191 | 0.964331209 | 0.982416138 | 0.003662839 |
| ENSMUSG00000003199 | Mpnd | 68047 | protein_coding | 878.8970071 | 0.003682289 | 0.058710929 | -0.062718977 | 0.949990286 | 0.975869522 | 0.003682289 |
| ENSMUSG00000042742 | Bmt2 | 101148 | protein_coding | 280.9763168 | -0.003734192 | 0.12821368 | 0.029124752 | 0.976765095 | 0.987811867 | 0.003734192 |
| ENSMUSG00000030753 | Thap12 | 72981 | protein_coding | 564.7360381 | -0.00373743 | 0.10187173 | 0.036687611 | 0.970734087 | 0.984868386 | 0.00373743 |
| ENSMUSG00000031634 | Ufsp2 | 192169 | protein_coding | 471.5930355 | 0.003755315 | 0.11309464 | -0.033205069 | 0.973511056 | 0.986171119 | 0.003755315 |
| ENSMUSG00000049307 | Fut4 | 14345 | protein_coding | 1351.452686 | -0.003802459 | 0.245911824 | 0.015462695 | 0.987663046 | 0.993075228 | 0.003802459 |
| ENSMUSG00000056770 | Setd3 | 52690 | protein_coding | 1448.032221 | -0.003851792 | 0.074765375 | 0.051518397 | 0.958912443 | 0.980067227 | 0.003851792 |
| ENSMUSG00000099843 | Gm7160 | 635504 | lncRNA | 19.94306795 | 0.003890691 | 0.239751844 | -0.016227993 | 0.987052503 | 0.992763569 | 0.003890691 |
| ENSMUSG00000034343 | Ube2f | 67921 | protein_coding | 414.6869368 | 0.003909755 | 0.064808321 | -0.060327978 | 0.951894419 | 0.976016991 | 0.003909755 |
| ENSMUSG00000022246 | Rai14 | 75646 | protein_coding | 425.9068381 | 0.00391071 | 0.126594087 | -0.03089173 | 0.975355885 | 0.987234441 | 0.00391071 |
| ENSMUSG00000046108 | Il17c | 234836 | protein_coding | 6.85799542 | 0.003960856 | 0.469234478 | -0.008441101 | 0.993265056 | 0.995979059 | 0.003960856 |

| ensembl_gene_id | external_gene_name | entrezgene_id | gene_biotype | baseMean | log2FoldChange | lfcSE | stat | pvalue | padj | absLog2FC |
|---|---|---|---|---|---|---|---|---|---|---|
| ENSMUSG00000027868 | Tbx15 | 21384 | protein_coding | 1.7612301 | -0.003963924 | 1.570418688 | 0.002524119 | 0.997986047 | 1 | 0.003963924 |
| ENSMUSG00000031979 | Cog2 | 76332 | protein_coding | 737.7728719 | -0.00397506 | 0.094706569 | 0.041972376 | 0.96652072 | 0.983251596 | 0.00397506 |
| ENSMUSG00000114797 | Gm49336 | 71844 | protein_coding | 613.4960048 | 0.003976889 | 0.08803822 | -0.045172307 | 0.963969968 | 0.982229903 | 0.003976889 |
| ENSMUSG00000027428 | Rbbp9 | 26450 | protein_coding | 121.7880144 | 0.004015954 | 0.124369084 | -0.032290613 | 0.974240295 | 0.986657316 | 0.004015954 |
| ENSMUSG00000020721 | Helz | 78455 | protein_coding | 412.3686085 | -0.004095255 | 0.123719095 | 0.033101236 | 0.973593858 | 0.98618968 | 0.004095255 |
| ENSMUSG00000031577 | Tti2 | 234138 | protein_coding | 147.2515944 | 0.004141045 | 0.089996316 | -0.046013497 | 0.963299492 | 0.98178904 | 0.004141045 |
| ENSMUSG00000042179 | Pnliprp1 | 18946 | protein_coding | 4.730888262 | -0.004141616 | 0.617137378 | 0.006711011 | 0.994645428 | 0.996939463 | 0.004141616 |
| ENSMUSG00000026147 | Col9a1 | 12839 | protein_coding | 1.679628758 | -0.004142447 | 1.917922116 | 0.002159862 | 0.998276681 | 1 | 0.004142447 |
| ENSMUSG00000020359 | Phykpl | 72947 | protein_coding | 261.4111307 | -0.00414275 | 0.110803493 | 0.037388265 | 0.970175429 | 0.984664559 | 0.00414275 |
| ENSMUSG00000038084 | Opa1 | 74143 | protein_coding | 1410.670368 | -0.004149199 | 0.061429848 | 0.067543694 | 0.946148878 | 0.973795719 | 0.004149199 |
| ENSMUSG00000028890 | Mtf1 | 17764 | protein_coding | 328.1556375 | -0.004185849 | 0.064204123 | 0.065195946 | 0.948017989 | 0.974999631 | 0.004185849 |
| ENSMUSG00000009741 | Ubp1 | 22221 | protein_coding | 604.2559899 | 0.004215528 | 0.051913535 | -0.081202873 | 0.935280615 | 0.969552109 | 0.004215528 |
| ENSMUSG00000024841 | Eif1ad | 69860 | protein_coding | 341.2458367 | -0.004222943 | 0.102681321 | 0.041126696 | 0.967194892 | 0.983635967 | 0.004222943 |
| ENSMUSG00000055717 | Slain1 | 105439 | protein_coding | 63.57609454 | -0.004228524 | 0.151112747 | 0.027982573 | 0.97767605 | 0.988088396 | 0.004228524 |
| ENSMUSG00000056458 | Mok | 26448 | protein_coding | 8.319862078 | 0.004230274 | 0.441155846 | -0.009589069 | 0.992349147 | 0.995423402 | 0.004230274 |
| ENSMUSG00000014104 | Sval2 | 84543 | protein_coding | 1.01033778 | 0.004242161 | 3.004388395 | -0.001411988 | 0.998873397 | 1 | 0.004242161 |
| ENSMUSG00000085234 | Gm15614 | 102640951 | lncRNA | 2.917262422 | -0.004257881 | 0.925438214 | 0.004600935 | 0.996328998 | 1 | 0.004257881 |
| ENSMUSG00000095301 | Or5p55 | 258926 | protein_coding | 0.922296796 | 0.004260532 | 2.989954025 | -0.001424949 | 0.998863056 | 1 | 0.004260532 |
| ENSMUSG00000114538 | Gm31517 | 102633770 | lncRNA | 1.057898392 | 0.004278778 | 2.974199488 | -0.001438632 | 0.998852138 | 1 | 0.004278778 |
| ENSMUSG00000043727 | F830045P16Rik | 228592 | protein_coding | 1.148892156 | 0.004283669 | 2.969308392 | -0.001442649 | 0.998848933 | 1 | 0.004283669 |
| ENSMUSG00000073931 | Or52d1 | 259058 | protein_coding | 1.148892156 | 0.004283669 | 2.969308392 | -0.001442649 | 0.998848933 | 1 | 0.004283669 |
| ENSMUSG00000113056 | 6030440G07Rik | 77878 | lncRNA | 2.065391108 | -0.004292326 | 0.796202852 | 0.005390996 | 0.995698628 | 1 | 0.004292326 |
| ENSMUSG00000029595 | Lhx5 | 16873 | protein_coding | 1.244013381 | 0.004293923 | 2.960373856 | -0.001450466 | 0.998842696 | 1 | 0.004293923 |
| ENSMUSG00000011349 | Dmrtc2 | 71241 | protein_coding | 1.196452768 | 0.00429568 | 2.959386379 | -0.001451544 | 0.998841836 | 1 | 0.00429568 |
| ENSMUSG00000020168 | Or14c43 | 257929 | protein_coding | 1.196452768 | 0.00429568 | 2.959386379 | -0.001451544 | 0.998841836 | 1 | 0.00429568 |
| ENSMUSG00000050504 | Or5bh3 | 258385 | protein_coding | 1.382567757 | 0.004301634 | 2.953214332 | -0.001456594 | 0.998837807 | 1 | 0.004301634 |
| ENSMUSG00000087038 | 2900079G21Rik | 620760 | lncRNA | 1.382567757 | 0.004301634 | 2.953214332 | -0.001456594 | 0.998837807 | 1 | 0.004301634 |
| ENSMUSG00000114807 | Gm32090 | 102634531 | lncRNA | 1.382567757 | 0.004301634 | 2.953214332 | -0.001456594 | 0.998837807 | 1 | 0.004301634 |
| ENSMUSG00000086171 | Pcsk2os1 | 378776 | lncRNA | 0.867408709 | 0.004303933 | 2.956591485 | -0.001455708 | 0.998838514 | 1 | 0.004303933 |
| ENSMUSG00000092162 | Vmn2r86 | 625109 | protein_coding | 1.287446533 | 0.004304003 | 2.952051874 | -0.00145797 | 0.998836709 | 1 | 0.004304003 |
| ENSMUSG00000067522 | Or5a1 | 258677 | protein_coding | 1.335007145 | 0.004307336 | 2.949098648 | -0.00146056 | 0.998834642 | 1 | 0.004307336 |
| ENSMUSG00000056019 | Zfp709 | 236193 | protein_coding | 72.28737292 | -0.004337408 | 0.16442543 | 0.026379177 | 0.978954903 | 0.988867032 | 0.004337408 |
| ENSMUSG00000071000 | Or13c7 | 29845 | protein_coding | 1.65554905 | 0.00433769 | 2.923652944 | -0.001483654 | 0.998816216 | 1 | 0.00433769 |
| ENSMUSG00000027088 | Phospho2 | 73373 | protein_coding | 268.2992684 | 0.004337741 | 0.111131118 | -0.039032639 | 0.968864366 | 0.984301824 | 0.004337741 |
| ENSMUSG00000024790 | Sac3d1 | 66406 | protein_coding | 131.547206 | 0.004342356 | 0.146567738 | -0.02962696 | 0.976364564 | 0.98778928 | 0.004342356 |
| ENSMUSG00000001280 | Sp1 | 20683 | protein_coding | 1497.095104 | 0.004346101 | 0.079225539 | -0.054857324 | 0.956252132 | 0.978107631 | 0.004346101 |
| ENSMUSG00000037788 | Vopp1 | 232023 | protein_coding | 221.9648527 | 0.004352234 | 0.109189743 | -0.039859366 | 0.968205247 | 0.984116541 | 0.004352234 |
| ENSMUSG00000038888 | Ctu1 | 233189 | protein_coding | 162.1095651 | -0.004391502 | 0.105982214 | 0.041436212 | 0.966948144 | 0.983565239 | 0.004391502 |
| ENSMUSG00000020070 | Rufy2 | 70432 | protein_coding | 100.7477828 | -0.004422659 | 0.124998567 | 0.035381681 | 0.971775392 | 0.985501033 | 0.004422659 |

| ensembl_gene_id | external_gene_name | entrezgene_id | gene_biotype | baseMean | log2FoldChange | lfcSE | stat | pvalue | padj | absLog2FC |
|---|---|---|---|---|---|---|---|---|---|---|
| ENSMUSG00000001445 | Mrpl10 | 107732 | protein_coding | 621.5498773 | -0.004424286 | 0.093448471 | 0.047344658 | 0.962238536 | 0.981434574 | 0.004424286 |
| ENSMUSG00000035852 | Misp | 78906 | protein_coding | 7448.958347 | 0.004452934 | 0.127947274 | -0.034802879 | 0.972236925 | 0.985568894 | 0.004452934 |
| ENSMUSG00000057134 | Ado | 211488 | protein_coding | 395.1343538 | -0.004472139 | 0.113851649 | 0.039280407 | 0.968666827 | 0.984161682 | 0.004472139 |
| ENSMUSG00000051359 | Ncald | 52589 | protein_coding | 342.5184785 | -0.00447297 | 0.154245909 | 0.028998956 | 0.976865424 | 0.987811867 | 0.00447297 |
| ENSMUSG00000028101 | Pias3 | 229615 | protein_coding | 164.8290924 | 0.004481135 | 0.139042156 | -0.032228604 | 0.974289745 | 0.986657316 | 0.004481135 |
| ENSMUSG00000020491 | 2810021J22Rik | 69944 | protein_coding | 87.5773785 | -0.004513938 | 0.128978573 | 0.034997581 | 0.972081669 | 0.985568894 | 0.004513938 |
| ENSMUSG00000041769 | Ppp2r2d | 52432 | protein_coding | 358.9074714 | 0.004536676 | 0.073189828 | -0.061985062 | 0.950574728 | 0.975869522 | 0.004536676 |
| ENSMUSG00000035273 | Hpse | 15442 | protein_coding | 180.8848147 | -0.004541391 | 0.110999686 | 0.040913549 | 0.967364816 | 0.983635967 | 0.004541391 |
| ENSMUSG00000023873 | 1700010I14Rik | 66931 | protein_coding | 44.39201065 | 0.004551056 | 0.192773203 | -0.023608344 | 0.981165016 | 0.989916855 | 0.004551056 |
| ENSMUSG00000020935 | Dcakd | 68087 | protein_coding | 147.4528786 | 0.00457489 | 0.123476937 | -0.037050562 | 0.970444691 | 0.984756276 | 0.00457489 |
| ENSMUSG00000009112 | Bcl2l13 | 94044 | protein_coding | 703.1074778 | 0.00458516 | 0.05536503 | -0.082816903 | 0.933997129 | 0.968724317 | 0.00458516 |
| ENSMUSG00000041057 | Wdr43 | 72515 | protein_coding | 558.0983811 | 0.004596564 | 0.102384343 | -0.044895187 | 0.964190853 | 0.982347769 | 0.004596564 |
| ENSMUSG00000032777 | Gtf3c1 | 233863 | protein_coding | 1595.800602 | 0.004598477 | 0.039792057 | -0.115562679 | 0.907999142 | 0.955211953 | 0.004598477 |
| ENSMUSG00000020949 | Fkbp3 | 30795 | protein_coding | 182.2044994 | -0.004672805 | 0.110641409 | 0.042233778 | 0.966312336 | 0.983100167 | 0.004672805 |
| ENSMUSG00000022020 | Naa16 | 66897 | protein_coding | 124.6593066 | 0.004683871 | 0.114405066 | -0.040941114 | 0.96734284 | 0.983635967 | 0.004683871 |
| ENSMUSG00000031626 | Sorbs2 | 234214 | protein_coding | 535.9549179 | 0.004701927 | 0.152465266 | -0.030839335 | 0.975397671 | 0.987234441 | 0.004701927 |
| ENSMUSG00000024370 | Cdc23 | 52563 | protein_coding | 290.9827796 | 0.004706802 | 0.126570691 | -0.037187136 | 0.970335796 | 0.984706282 | 0.004706802 |
| ENSMUSG00000024424 | Ttc39c | 72747 | protein_coding | 17.68270885 | 0.004744183 | 0.313607601 | -0.015127767 | 0.987930248 | 0.993283417 | 0.004744183 |
| ENSMUSG00000042032 | Mat2b | 108645 | protein_coding | 993.1326032 | 0.004771693 | 0.074770689 | -0.063817701 | 0.949115384 | 0.975520722 | 0.004771693 |
| ENSMUSG00000043962 | Thrap3 | 230753 | protein_coding | 1391.323634 | 0.004776233 | 0.084724529 | -0.056373675 | 0.955044128 | 0.977538039 | 0.004776233 |
| ENSMUSG00000006304 | Arpc2 | 76709 | protein_coding | 2521.668858 | -0.004776629 | 0.051092545 | 0.09348975 | 0.925514492 | 0.964363534 | 0.004776629 |
| ENSMUSG00000115726 | Gm30083 | 102631856 | protein_coding | 1.655795596 | 0.004777584 | 2.700886196 | -0.001768895 | 0.998588627 | 1 | 0.004777584 |
| ENSMUSG00000033845 | Mrpl15 | 27395 | protein_coding | 561.2052347 | -0.004784392 | 0.06443225 | 0.074254623 | 0.940807783 | 0.971638507 | 0.004784392 |
| ENSMUSG00000029647 | Pan3 | 72587 | protein_coding | 974.8481309 | -0.004838029 | 0.049367732 | 0.097999821 | 0.921932435 | 0.962436702 | 0.004838029 |
| ENSMUSG00000097665 | Gm16853 | 330902 | lncRNA | 2.736898181 | -0.004852414 | 0.76206592 | 0.006367447 | 0.994919547 | 1 | 0.004852414 |
| ENSMUSG00000095567 | Noc2l | 57741 | protein_coding | 522.1849011 | -0.004853522 | 0.094271546 | 0.05148449 | 0.958939461 | 0.980067227 | 0.004853522 |
| ENSMUSG00000056912 | 1700017N19Rik | 66605 | protein_coding | 1.104594524 | 0.004870992 | 2.583258494 | -0.0018856 | 0.99849551 | 1 | 0.004870992 |
| ENSMUSG00000035953 | Pip4p1 | 219024 | protein_coding | 345.7412088 | 0.004871196 | 0.070596957 | -0.069000089 | 0.944989549 | 0.97354273 | 0.004871196 |
| ENSMUSG00000051190 | Or7c70 | 258305 | protein_coding | 1.242838701 | 0.004889462 | 2.572240383 | -0.001900857 | 0.998483336 | 1 | 0.004889462 |
| ENSMUSG00000029992 | Gfpt1 | 14583 | protein_coding | 11180.13193 | -0.00489726 | 0.144412537 | 0.033911596 | 0.972947647 | 0.985876467 | 0.00489726 |
| ENSMUSG00000094488 | Or1e33 | 259010 | protein_coding | 1.151844937 | 0.004897699 | 2.568206842 | -0.00190705 | 0.998478395 | 1 | 0.004897699 |
| ENSMUSG00000078190 | Dnm3os | 474332 | lncRNA | 1.151528249 | 0.004898909 | 2.567555423 | -0.001908005 | 0.998477633 | 1 | 0.004898909 |
| ENSMUSG00000048015 | Neurod4 | 11923 | protein_coding | 1.473437904 | 0.004903035 | 2.56428655 | -0.001912046 | 0.998474409 | 1 | 0.004903035 |
| ENSMUSG00000027981 | Rnpc3 | 67225 | protein_coding | 127.9954263 | 0.004907738 | 0.092932139 | -0.052809911 | 0.957883365 | 0.979351528 | 0.004907738 |
| ENSMUSG00000017740 | Slc12a5 | 57138 | protein_coding | 9.403661977 | 0.004912187 | 0.749822304 | -0.006551134 | 0.994772988 | 0.996966345 | 0.004912187 |
| ENSMUSG00000112331 | Gm33035 | 102635787 | lncRNA | 0.96265355 | 0.004913941 | 2.560772104 | -0.00191893 | 0.998468917 | 1 | 0.004913941 |
| ENSMUSG00000022791 | Tnk2 | 51789 | protein_coding | 657.4431438 | -0.004916283 | 0.068632981 | 0.07163149 | 0.942895179 | 0.972316965 | 0.004916283 |
| ENSMUSG00000074254 | Cyp2a4 | 13086 | protein_coding | 1.196329151 | 0.004923469 | 2.554489413 | -0.001927379 | 0.998462175 | 1 | 0.004923469 |
| ENSMUSG00000074628 | Tldc2 | 383766 | protein_coding | 92.10558334 | -0.004934899 | 0.219148867 | 0.022518476 | 0.982034374 | 0.990370421 | 0.004934899 |

Supplementary Table 1

| ensembl_gene_id | external_gene_name | entrezgene_id | gene_biotype | baseMean | log2FoldChange | lfcSE | stat | pvalue | padj | absLog2FC |
|---|---|---|---|---|---|---|---|---|---|---|
| ENSMUSG00000111869 | Or1j14 | 258950 | protein_coding | 1.425877292 | 0.004935419 | 2.54723809 | -0.001937557 | 0.998454054 | 1 | 0.004935419 |
| ENSMUSG00000029526 | 1700123K08Rik | 76658 | protein_coding | 1.057774775 | 0.004939593 | 2.546657721 | -0.001939638 | 0.998452394 | 1 | 0.004939593 |
| ENSMUSG00000039032 | Tsga13 | 116732 | protein_coding | 1.043535072 | 0.004940645 | 2.546149598 | -0.001940438 | 0.998451756 | 1 | 0.004940645 |
| ENSMUSG00000086022 | Rad51ap2 | 209550 | protein_coding | 0.847460503 | 0.004954383 | 2.539606907 | -0.001950846 | 0.998443451 | 1 | 0.004954383 |
| ENSMUSG00000005370 | Msh6 | 17688 | protein_coding | 435.4233794 | -0.004956253 | 0.133082432 | 0.037241979 | 0.970292068 | 0.984706282 | 0.004956253 |
| ENSMUSG00000095932 | Vmn1r49 | 24112 | protein_coding | 1.16500064 | 0.004960812 | 2.534622756 | -0.001957219 | 0.998438366 | 1 | 0.004960812 |
| ENSMUSG00000028958 | Tmub1 | 64295 | protein_coding | 259.4297212 | 0.004962895 | 0.07205475 | -0.068876717 | 0.945087752 | 0.97354273 | 0.004962895 |
| ENSMUSG00000075208 | Or5ar1 | 259017 | protein_coding | 0.874295879 | 0.004963162 | 2.535051721 | -0.001957815 | 0.998437891 | 1 | 0.004963162 |
| ENSMUSG00000080717 | B230307C23Rik | 245305 | protein_coding | 74.35730743 | -0.004967796 | 0.128932413 | 0.038530233 | 0.969264927 | 0.984403993 | 0.004967796 |
| ENSMUSG00000091468 | Vmn2r82 | 624845 | protein_coding | 1.066270333 | 0.004983316 | 2.523397126 | -0.001974844 | 0.998424304 | 1 | 0.004983316 |
| ENSMUSG00000090304 | Vmn2r99 | 665376 | protein_coding | 1.844616819 | 0.004984744 | 2.520335945 | -0.001977809 | 0.998421938 | 1 | 0.004984744 |
| ENSMUSG00000034623 | Prss55 | 71037 | protein_coding | 1.022837181 | 0.004995012 | 2.517651072 | -0.001983997 | 0.998417 | 1 | 0.004995012 |
| ENSMUSG00000060335 | Or1e25 | 193053 | protein_coding | 0.973436112 | 0.004997391 | 2.516646827 | -0.001985734 | 0.998415615 | 1 | 0.004997391 |
| ENSMUSG00000026271 | Gpr35 | 64095 | protein_coding | 361.3985963 | -0.005006522 | 0.076651362 | 0.065315505 | 0.947922797 | 0.974962449 | 0.005006522 |
| ENSMUSG00000086844 | B230206H07Rik | 320871 | lncRNA | 17.19204084 | -0.005020063 | 0.510042407 | 0.009842442 | 0.992146994 | 0.995401911 | 0.005020063 |
| ENSMUSG00000032594 | Ip6k1 | 27399 | protein_coding | 849.2192546 | 0.00502864 | 0.074594268 | -0.067413219 | 0.946252746 | 0.973795719 | 0.00502864 |
| ENSMUSG00000033972 | Zfp944 | 319615 | protein_coding | 136.5038676 | 0.005056968 | 0.127442369 | -0.039680427 | 0.968347906 | 0.984122985 | 0.005056968 |
| ENSMUSG00000050765 | Gm5084 | 100503759 | lncRNA | 1.69448389 | 0.00507117 | 2.477536774 | -0.002046859 | 0.998366844 | 1 | 0.00507117 |
| ENSMUSG00000091504 | Vmn2r118 | 383258 | protein_coding | 1.69448389 | 0.00507117 | 2.477536774 | -0.002046859 | 0.998366844 | 1 | 0.00507117 |
| ENSMUSG00000070855 | Or10ag54 | 257875 | protein_coding | 2.244326701 | 0.005094936 | 2.464912278 | -0.002066985 | 0.998350786 | 1 | 0.005094936 |
| ENSMUSG00000020950 | Foxg1 | 15228 | protein_coding | 1.040606745 | 0.005112629 | 2.458965964 | -0.002079179 | 0.998341057 | 1 | 0.005112629 |
| ENSMUSG00000029726 | Mepce | 231803 | protein_coding | 452.6238051 | 0.005114043 | 0.082631536 | -0.061889719 | 0.950650655 | 0.975869522 | 0.005114043 |
| ENSMUSG00000005893 | Nr2c2 | 22026 | protein_coding | 566.6827522 | -0.00511806 | 0.076534602 | 0.066872496 | 0.946683209 | 0.973990811 | 0.00511806 |
| ENSMUSG00000042198 | Chchd7 | 66433 | protein_coding | 322.2298501 | 0.005138394 | 0.080983426 | -0.063449949 | 0.949408214 | 0.97563953 | 0.005138394 |
| ENSMUSG00000031310 | Zmym3 | 56364 | protein_coding | 260.5321973 | -0.005163266 | 0.102853979 | 0.050199963 | 0.959963041 | 0.980506535 | 0.005163266 |
| ENSMUSG00000026675 | Hsd17b7 | 15490 | protein_coding | 306.047576 | -0.005163934 | 0.134361735 | 0.038433069 | 0.969342395 | 0.984403993 | 0.005163934 |
| ENSMUSG00000070420 | Zscan25 | 666311 | protein_coding | 139.3103244 | -0.005173196 | 0.10685669 | 0.048412465 | 0.961387525 | 0.981111951 | 0.005173196 |
| ENSMUSG00000037017 | Zscan21 | 22697 | protein_coding | 212.0206503 | -0.005181551 | 0.103095303 | 0.050259817 | 0.959915345 | 0.980506535 | 0.005181551 |
| ENSMUSG00000036339 | Tmem260 | 218989 | protein_coding | 390.399798 | 0.005212578 | 0.121172277 | -0.043017908 | 0.965687259 | 0.982706396 | 0.005212578 |
| ENSMUSG00000045180 | Shroom2 | 110380 | protein_coding | 607.6456158 | 0.005268167 | 0.075151065 | -0.070101029 | 0.944113247 | 0.973044134 | 0.005268167 |
| ENSMUSG00000035086 | Becn1 | 56208 | protein_coding | 1063.185592 | 0.005277878 | 0.045144643 | -0.116910384 | 0.906931069 | 0.954378365 | 0.005277878 |
| ENSMUSG00000033883 | Zfp267 | 241944 | protein_coding | 170.6332006 | 0.005278593 | 0.100841988 | -0.052345188 | 0.958253648 | 0.979508488 | 0.005278593 |
| ENSMUSG00000077497 | Gm23593 | 115485971 | snoRNA | 1.226548998 | 0.005310503 | 1.09477287 | -0.00485078 | 0.996129653 | 1 | 0.005310503 |
| ENSMUSG00000061559 | Wdr61 | 66317 | protein_coding | 482.4422483 | -0.005315311 | 0.09219848 | 0.057650744 | 0.954026829 | 0.977047797 | 0.005315311 |
| ENSMUSG00000045053 | Kcng3 | 225030 | protein_coding | 2.747548653 | -0.005321584 | 0.849408797 | 0.006265044 | 0.995001251 | 1 | 0.005321584 |
| ENSMUSG00000045671 | Spred2 | 114716 | protein_coding | 328.4361693 | -0.005354374 | 0.113010919 | 0.047379261 | 0.962210958 | 0.981434574 | 0.005354374 |
| ENSMUSG00000032309 | Fbxo22 | 71999 | protein_coding | 341.3702807 | -0.005363454 | 0.073891955 | 0.072585093 | 0.94213629 | 0.971883559 | 0.005363454 |
| ENSMUSG00000035439 | Haus8 | 76478 | protein_coding | 191.5829763 | -0.005395122 | 0.143277449 | 0.037655066 | 0.969962702 | 0.984509162 | 0.005395122 |
| ENSMUSG00000024985 | Tcf7l2 | 21416 | protein_coding | 558.5046978 | 0.005450128 | 0.208658745 | -0.026119818 | 0.97916177 | 0.988983332 | 0.005450128 |

Supplementary Table 1

| ensembl_gene_id | external_gene_name | entrezgene_id | gene_biotype | baseMean | log2FoldChange | lfcSE | stat | pvalue | padj | absLog2FC |
|---|---|---|---|---|---|---|---|---|---|---|
| ENSMUSG00000044224 | Dnajc21 | 78244 | protein_coding | 301.3581676 | 0.005455805 | 0.090786953 | -0.060094595 | 0.952080295 | 0.976016991 | 0.005455805 |
| ENSMUSG00000048355 | Arxes1 | 76219 | protein_coding | 0.818351343 | -0.005457427 | 1.67365432 | 0.003260785 | 0.997398275 | 1 | 0.005457427 |
| ENSMUSG00000037110 | Ralgapa2 | 241694 | protein_coding | 1902.774412 | 0.00546321 | 0.078443901 | -0.069644799 | 0.944476379 | 0.973296906 | 0.00546321 |
| ENSMUSG00000028309 | Rnf20 | 109331 | protein_coding | 802.1291369 | 0.005476486 | 0.073568068 | -0.074441077 | 0.940659424 | 0.971606814 | 0.005476486 |
| ENSMUSG00000027860 | Vangl1 | 229658 | protein_coding | 169.4605644 | -0.00552574 | 0.113142273 | 0.048838864 | 0.96104771 | 0.981007658 | 0.00552574 |
| ENSMUSG00000033306 | Lpp | 210126 | protein_coding | 1851.766542 | 0.00560402 | 0.10940236 | -0.05122394 | 0.959147076 | 0.980158095 | 0.00560402 |
| ENSMUSG00000016510 | Mtif3 | 76366 | protein_coding | 293.3454113 | 0.005610402 | 0.076495467 | -0.073342937 | 0.941533225 | 0.971840719 | 0.005610402 |
| ENSMUSG00000043911 | Or8b55 | 258777 | protein_coding | 1.272959596 | 0.005618964 | 2.243096768 | -0.002505003 | 0.998001299 | 1 | 0.005618964 |
| ENSMUSG00000094651 | Gal3st2 | 381334 | protein_coding | 159.6967969 | -0.00563163 | 0.232725273 | 0.02419862 | 0.980694179 | 0.989584039 | 0.00563163 |
| ENSMUSG00000061626 | Or52a5 | 18369 | protein_coding | 1.294086469 | 0.005639144 | 2.235361958 | -0.002522698 | 0.99798718 | 1 | 0.005639144 |
| ENSMUSG00000043314 | Or2z2 | 18329 | protein_coding | 1.234770362 | 0.005661454 | 2.227189119 | -0.002541973 | 0.997971801 | 1 | 0.005661454 |
| ENSMUSG00000027788 | Otol1 | 229389 | protein_coding | 1.566326898 | 0.005663178 | 2.225868208 | -0.002544256 | 0.99796998 | 1 | 0.005663178 |
| ENSMUSG00000020436 | Gabrg2 | 14406 | protein_coding | 1.580189068 | 0.005665099 | 2.225075121 | -0.002546026 | 0.997968567 | 1 | 0.005665099 |
| ENSMUSG00000041730 | Prrxl1 | 107751 | protein_coding | 1.913919788 | 0.005685681 | 2.217126182 | -0.002564437 | 0.997953877 | 1 | 0.005685681 |
| ENSMUSG00000033931 | Rbm34 | 52202 | protein_coding | 171.5399331 | -0.005686418 | 0.129499593 | 0.043910703 | 0.964975583 | 0.982587667 | 0.005686418 |
| ENSMUSG00000092239 | 1700031A10Rik | 73286 | lncRNA | 1.336778757 | 0.005690334 | 2.216239225 | -0.002567563 | 0.997951383 | 1 | 0.005690334 |
| ENSMUSG00000049902 | 4921517D22Rik | 70900 | protein_coding | 1.246101681 | 0.005691297 | 2.21607866 | -0.002568184 | 0.997950888 | 1 | 0.005691297 |
| ENSMUSG00000040650 | Cyp2b23 | 243881 | protein_coding | 1.245784993 | 0.005693067 | 2.215428361 | -0.002569737 | 0.997949649 | 1 | 0.005693067 |
| ENSMUSG00000021792 | Prxl2a | 70564 | protein_coding | 92.13329467 | -0.00569601 | 0.159042823 | 0.03581432 | 0.971430415 | 0.985415068 | 0.00569601 |
| ENSMUSG00000024044 | Epb41l3 | 13823 | protein_coding | 1178.556983 | 0.005703089 | 0.076003114 | -0.075037569 | 0.94018482 | 0.971417829 | 0.005703089 |
| ENSMUSG00000045381 | Or10aa1 | 258712 | protein_coding | 1.15172132 | 0.00571835 | 2.206371872 | -0.002591743 | 0.99793209 | 1 | 0.00571835 |
| ENSMUSG00000044542 | Prop1 | 19127 | protein_coding | 1.632683575 | 0.005722508 | 2.204043965 | -0.002596368 | 0.997928401 | 1 | 0.005722508 |
| ENSMUSG00000051367 | Six1 | 20471 | protein_coding | 1.202860636 | 0.005727023 | 2.203046845 | -0.002599592 | 0.997925828 | 1 | 0.005727023 |
| ENSMUSG00000031241 | Tbx22 | 245572 | protein_coding | 1.222565954 | 0.005729717 | 2.201977055 | -0.002602078 | 0.997923844 | 1 | 0.005729717 |
| ENSMUSG00000063011 | Msln | 56047 | protein_coding | 1.976206453 | -0.005733806 | 1.285055999 | 0.004461912 | 0.996439921 | 1 | 0.005733806 |
| ENSMUSG00000024593 | Megf10 | 70417 | protein_coding | 2.956303408 | 0.005749286 | 2.193130518 | -0.002621498 | 0.99790835 | 1 | 0.005749286 |
| ENSMUSG00000029685 | Asb15 | 78910 | protein_coding | 1.792681512 | 0.005755629 | 2.191874229 | -0.002625894 | 0.997904842 | 1 | 0.005755629 |
| ENSMUSG00000031358 | Msl3 | 17692 | protein_coding | 526.0892 | -0.005781457 | 0.101094143 | 0.057188843 | 0.954394765 | 0.977236803 | 0.005781457 |
| ENSMUSG00000110970 | Or13c7b | 29846 | protein_coding | 1.199158315 | 0.005781725 | 2.183562073 | -0.002647841 | 0.997887331 | 1 | 0.005781725 |
| ENSMUSG00000029387 | Gtf2h3 | 209357 | protein_coding | 377.6166657 | 0.005783371 | 0.0911347 | -0.063459593 | 0.949400534 | 0.97563953 | 0.005783371 |
| ENSMUSG00000020430 | Pes1 | 64934 | protein_coding | 575.7868989 | -0.005803929 | 0.07974034 | 0.072785349 | 0.941976929 | 0.971883559 | 0.005803929 |
| ENSMUSG00000005982 | Naa60 | 74763 | protein_coding | 332.7992821 | -0.005831284 | 0.084873637 | 0.06870548 | 0.945224056 | 0.97354273 | 0.005831284 |
| ENSMUSG00000090315 | Vmn2r104 | 22313 | protein_coding | 1.060603938 | 0.005834303 | 2.165447358 | -0.002694271 | 0.997850285 | 1 | 0.005834303 |
| ENSMUSG00000052525 | Spdya | 70891 | protein_coding | 3.810493772 | 0.00586697 | 0.577087972 | -0.01016651 | 0.991888439 | 1 | 0.00586697 |
| ENSMUSG00000028842 | Ago3 | 214150 | protein_coding | 219.0560777 | -0.005874035 | 0.128522764 | 0.045704238 | 0.963545986 | 0.981940857 | 0.005874035 |
| ENSMUSG00000024976 | Shoc2 | 56392 | protein_coding | 1044.208139 | -0.005876413 | 0.06095315 | 0.09640869 | 0.92319599 | 0.962738158 | 0.005876413 |
| ENSMUSG00000052616 | Terb1 | 320022 | protein_coding | 2.90037548 | 0.005879139 | 0.776932749 | -0.007567114 | 0.993962374 | 1 | 0.005879139 |
| ENSMUSG00000036980 | Taf6 | 21343 | protein_coding | 222.2063311 | 0.005886869 | 0.096030076 | -0.061302347 | 0.951118422 | 0.97587882 | 0.005886869 |
| ENSMUSG00000062604 | Srpk2 | 20817 | protein_coding | 219.5714223 | 0.005893837 | 0.097882297 | -0.060213514 | 0.951985582 | 0.976016991 | 0.005893837 |

| ensembl_gene_id | external_gene_name | entrezgene_id | gene_biotype | baseMean | log2FoldChange | lfcSE | stat | pvalue | padj | absLog2FC |
|---|---|---|---|---|---|---|---|---|---|---|
| ENSMUSG00000026883 | Dab2ip | 69601 | protein_coding | 1981.554412 | 0.005907888 | 0.053768054 | -0.109877283 | 0.912506699 | 0.957266079 | 0.005907888 |
| ENSMUSG00000062116 | Zfp954 | 232853 | protein_coding | 196.7074958 | -0.00592169 | 0.088978124 | 0.066552204 | 0.946938197 | 0.974163449 | 0.00592169 |
| ENSMUSG00000036781 | Rps27l | 67941 | protein_coding | 1089.435257 | -0.00595958 | 0.104555782 | 0.056999048 | 0.954545954 | 0.977331011 | 0.00595958 |
| ENSMUSG00000021913 | Ogdhl | 239017 | protein_coding | 22.24098862 | 0.005969809 | 0.22665063 | -0.026339255 | 0.978986745 | 0.988867032 | 0.005969809 |
| ENSMUSG00000003500 | Impdh1 | 23917 | protein_coding | 250.5606192 | -0.005981102 | 0.114318849 | 0.052319471 | 0.958274139 | 0.979508488 | 0.005981102 |
| ENSMUSG00000062007 | Hsh2d | 209488 | protein_coding | 39.27345323 | 0.005989992 | 0.211490261 | -0.028322781 | 0.977404711 | 0.988088396 | 0.005989992 |
| ENSMUSG00000021133 | Susd6 | 217684 | protein_coding | 1182.388743 | 0.006042569 | 0.097723774 | -0.061833149 | 0.950695705 | 0.975869522 | 0.006042569 |
| ENSMUSG00000042118 | Bhmt2 | 64918 | protein_coding | 8.154100691 | -0.006069778 | 0.386301032 | 0.015712559 | 0.987463708 | 0.992998437 | 0.006069778 |
| ENSMUSG00000046139 | Patl1 | 225929 | protein_coding | 170.0591364 | 0.006077563 | 0.098148759 | -0.061921952 | 0.950624986 | 0.975869522 | 0.006077563 |
| ENSMUSG00000038957 | Edc3 | 353190 | protein_coding | 203.5281583 | -0.006120192 | 0.09888302 | 0.061893254 | 0.95064784 | 0.975869522 | 0.006120192 |
| ENSMUSG00000024912 | Fosl1 | 14283 | protein_coding | 31.28143243 | 0.006135872 | 0.433211706 | -0.014163679 | 0.988699397 | 0.993636457 | 0.006135872 |
| ENSMUSG00000029505 | Ep400 | 75560 | protein_coding | 1086.666967 | -0.006139831 | 0.104415501 | 0.058801916 | 0.953109883 | 0.976710073 | 0.006139831 |
| ENSMUSG00000028937 | Acot7 | 70025 | protein_coding | 944.0467129 | 0.006145378 | 0.105158721 | -0.058439067 | 0.953398897 | 0.976822768 | 0.006145378 |
| ENSMUSG00000020686 | Gas2l2 | 237891 | protein_coding | 1.752291821 | 0.00621527 | 2.041670889 | -0.003044207 | 0.997571078 | 1 | 0.00621527 |
| ENSMUSG00000049482 | Ctu2 | 66965 | protein_coding | 154.7043841 | 0.006235135 | 0.110384558 | -0.056485574 | 0.954954988 | 0.977515248 | 0.006235135 |
| ENSMUSG00000004364 | Cul3 | 26554 | protein_coding | 1096.898828 | 0.006237504 | 0.08399716 | -0.074258513 | 0.940804688 | 0.971638507 | 0.006237504 |
| ENSMUSG00000030888 | Rrp8 | 101867 | protein_coding | 139.7143289 | 0.006243016 | 0.108406559 | -0.057588914 | 0.95407608 | 0.977047797 | 0.006243016 |
| ENSMUSG00000042305 | Tmem183a | 57439 | protein_coding | 1756.387409 | 0.006277025 | 0.063269957 | -0.099210194 | 0.920971381 | 0.961855464 | 0.006277025 |
| ENSMUSG00000021978 | Extl3 | 54616 | protein_coding | 525.8374457 | 0.006300884 | 0.088698153 | -0.071037374 | 0.94336801 | 0.972595566 | 0.006300884 |
| ENSMUSG00000048939 | Atp13a5 | 268878 | protein_coding | 1.987524132 | 0.006315814 | 2.011858903 | -0.003139292 | 0.997495211 | 1 | 0.006315814 |
| ENSMUSG00000031162 | Gata1 | 14460 | protein_coding | 1.809424736 | 0.006340521 | 2.004906666 | -0.003162502 | 0.997476693 | 1 | 0.006340521 |
| ENSMUSG00000020317 | Theg | 21830 | protein_coding | 1.085980126 | 0.006367339 | 1.998169094 | -0.003186587 | 0.997457476 | 1 | 0.006367339 |
| ENSMUSG00000001998 | Ap4e1 | 108011 | protein_coding | 153.6991173 | 0.006378778 | 0.091237166 | -0.069914252 | 0.94426191 | 0.973136619 | 0.006378778 |
| ENSMUSG00000085399 | Foxd2os | 100040736 | lncRNA | 386.7154176 | 0.006439132 | 0.102841241 | -0.062612354 | 0.950075192 | 0.975869522 | 0.006439132 |
| ENSMUSG00000030898 | Cckbr | 12426 | protein_coding | 1.643574589 | 0.00644893 | 1.974563613 | -0.003266003 | 0.997394111 | 1 | 0.00644893 |
| ENSMUSG00000039316 | Rftn1 | 76438 | protein_coding | 6.096606166 | 0.006475057 | 0.675262854 | -0.009588944 | 0.992349247 | 0.995423402 | 0.006475057 |
| ENSMUSG00000057541 | Pus7 | 78697 | protein_coding | 239.2222074 | 0.006486605 | 0.111375972 | -0.058240616 | 0.953556968 | 0.97686351 | 0.006486605 |
| ENSMUSG00000079625 | Tm4sf19 | 277203 | protein_coding | 1.09360816 | 0.006492409 | 1.963246436 | -0.003306976 | 0.99736142 | 1 | 0.006492409 |
| ENSMUSG00000009894 | Snap47 | 67826 | protein_coding | 217.4726997 | 0.006495774 | 0.090576475 | -0.071715907 | 0.942827997 | 0.972316965 | 0.006495774 |
| ENSMUSG00000007950 | Abhd8 | 64296 | protein_coding | 117.6948641 | -0.006511022 | 0.182283333 | 0.035719238 | 0.971506231 | 0.985415068 | 0.006511022 |
| ENSMUSG00000063549 | Or2w3 | 404346 | protein_coding | 1.150540151 | 0.006521709 | 1.955202427 | -0.003335567 | 0.997338607 | 1 | 0.006521709 |
| ENSMUSG00000036023 | Parp2 | 11546 | protein_coding | 265.9954174 | 0.006529404 | 0.08173507 | -0.079884977 | 0.936328738 | 0.969802608 | 0.006529404 |
| ENSMUSG00000002799 | Jag2 | 16450 | protein_coding | 64.74631642 | -0.006545777 | 0.222145561 | 0.029466163 | 0.976492805 | 0.987811867 | 0.006545777 |
| ENSMUSG00000049719 | Prss46 | 74306 | protein_coding | 1.025458165 | 0.006548678 | 1.948087511 | -0.003361593 | 0.997317842 | 1 | 0.006548678 |
| ENSMUSG00000033147 | Slc22a15 | 242126 | protein_coding | 174.5050995 | 0.006565754 | 0.143741831 | -0.045677406 | 0.963567372 | 0.981940857 | 0.006565754 |
| ENSMUSG00000028221 | Pip4p2 | 72519 | protein_coding | 244.1166455 | 0.006583543 | 0.090702094 | -0.072584245 | 0.942136964 | 0.971883559 | 0.006583543 |
| ENSMUSG00000032480 | Dhx30 | 72831 | protein_coding | 572.2411215 | -0.006598053 | 0.069368692 | 0.095115712 | 0.924222919 | 0.963320302 | 0.006598053 |
| ENSMUSG00000029822 | Osbpl3 | 71720 | protein_coding | 249.5851792 | -0.006621462 | 0.217397668 | 0.030457834 | 0.975701921 | 0.987300339 | 0.006621462 |
| ENSMUSG00000045232 | Rln3 | 212108 | protein_coding | 1.15530581 | 0.006624921 | 1.927763578 | -0.003436584 | 0.997258008 | 1 | 0.006624921 |

| ensembl_gene_id | external_gene_name | entrezgene_id | gene_biotype | baseMean | log2FoldChange | lfcSE | stat | pvalue | padj | absLog2FC |
|---|---|---|---|---|---|---|---|---|---|---|
| ENSMUSG00000032497 | Lrrfip2 | 71268 | protein_coding | 1199.983501 | -0.006634509 | 0.098945848 | 0.067051922 | 0.946540368 | 0.973904526 | 0.006634509 |
| ENSMUSG00000029090 | Adgra3 | 70693 | protein_coding | 1450.907578 | 0.006635848 | 0.100517312 | -0.066016966 | 0.947364318 | 0.974509428 | 0.006635848 |
| ENSMUSG00000039117 | Taf4 | 228980 | protein_coding | 436.194727 | 0.006641053 | 0.097863235 | -0.06786055 | 0.945896643 | 0.973795719 | 0.006641053 |
| ENSMUSG00000027206 | Cops2 | 12848 | protein_coding | 742.0104914 | -0.006758467 | 0.11095885 | 0.060909673 | 0.951431146 | 0.975912221 | 0.006758467 |
| ENSMUSG00000026273 | Mterf4 | 69821 | protein_coding | 153.1161409 | -0.006782039 | 0.100690994 | 0.067354969 | 0.946299117 | 0.973795719 | 0.006782039 |
| ENSMUSG00000014550 | Rbsn | 78287 | protein_coding | 404.7583959 | -0.006786905 | 0.084354431 | 0.080457006 | 0.93587379 | 0.96968875 | 0.006786905 |
| ENSMUSG00000026824 | Kcnj3 | 16519 | protein_coding | 0.944921598 | 0.006802836 | 1.882700574 | -0.003613339 | 0.997116979 | 1 | 0.006802836 |
| ENSMUSG00000000384 | Tbrg4 | 21379 | protein_coding | 906.4806778 | -0.006804982 | 0.052270479 | 0.130187861 | 0.896417798 | 0.948996349 | 0.006804982 |
| ENSMUSG00000059518 | Znhit1 | 70103 | protein_coding | 294.015806 | 0.006826791 | 0.099291014 | -0.068755372 | 0.945184342 | 0.97354273 | 0.006826791 |
| ENSMUSG00000049305 | Ccdc71 | 72454 | protein_coding | 415.9844361 | -0.006868352 | 0.070137548 | 0.097926888 | 0.921990349 | 0.962436702 | 0.006868352 |
| ENSMUSG00000034211 | Mrps17 | 66258 | protein_coding | 474.8547191 | 0.006879508 | 0.090439513 | -0.0760675 | 0.939365396 | 0.971181378 | 0.006879508 |
| ENSMUSG00000100315 | 1700031P21Rik | 73278 | lncRNA | 3.856869726 | 0.006901674 | 0.640443362 | -0.0107764 | 0.991401843 | 1 | 0.006901674 |
| ENSMUSG00000029657 | Hsph1 | 15505 | protein_coding | 754.2506714 | -0.006907372 | 0.172868423 | 0.039957395 | 0.968127093 | 0.984116541 | 0.006907372 |
| ENSMUSG00000046785 | Epm2aip1 | 77781 | protein_coding | 297.1929578 | -0.006922837 | 0.103097007 | 0.067148768 | 0.94646327 | 0.973885877 | 0.006922837 |
| ENSMUSG00000025135 | Anapc11 | 66156 | protein_coding | 245.9474447 | -0.006958912 | 0.096545918 | 0.072078781 | 0.942539213 | 0.972163488 | 0.006958912 |
| ENSMUSG00000041570 | Camsap2 | 67886 | protein_coding | 132.2481809 | -0.006963586 | 0.163984122 | 0.042465002 | 0.966128011 | 0.982973198 | 0.006963586 |
| ENSMUSG00000042520 | Ubap2l | 74383 | protein_coding | 842.7205024 | 0.006992239 | 0.061019258 | -0.114590697 | 0.908769554 | 0.955501531 | 0.006992239 |
| ENSMUSG00000038009 | Dnajc22 | 72778 | protein_coding | 779.8779283 | 0.006994416 | 0.053737597 | -0.130158708 | 0.896440863 | 0.948996349 | 0.006994416 |
| ENSMUSG00000033671 | Cep350 | 74081 | protein_coding | 1576.080099 | -0.007012866 | 0.083244174 | 0.084244524 | 0.932862019 | 0.968109605 | 0.007012866 |
| ENSMUSG00000020385 | Clk4 | 12750 | protein_coding | 330.2708626 | -0.007045781 | 0.114453522 | 0.061560196 | 0.950913076 | 0.97587882 | 0.007045781 |
| ENSMUSG00000063785 | Utp14a | 72554 | protein_coding | 289.3843583 | 0.007056744 | 0.120533074 | -0.058546122 | 0.953313625 | 0.976796005 | 0.007056744 |
| ENSMUSG00000030091 | Nup210 | 54563 | protein_coding | 382.4162076 | -0.007079155 | 0.115043475 | 0.061534607 | 0.950933454 | 0.97587882 | 0.007079155 |
| ENSMUSG00000020476 | Dbnl | 13169 | protein_coding | 1323.534683 | -0.007079984 | 0.05724227 | 0.12368455 | 0.901565045 | 0.951341404 | 0.007079984 |
| ENSMUSG00000024866 | Acy3 | 71670 | protein_coding | 184.4799706 | 0.007117378 | 0.141216887 | -0.050400331 | 0.959803373 | 0.980506535 | 0.007117378 |
| ENSMUSG00000003228 | Grk5 | 14773 | protein_coding | 119.6704512 | 0.00711781 | 0.102436804 | -0.069484894 | 0.944603657 | 0.973367327 | 0.00711781 |
| ENSMUSG00000021975 | Ints9 | 210925 | protein_coding | 182.3392182 | -0.00715615 | 0.090802087 | 0.078810415 | 0.93718342 | 0.970043107 | 0.00715615 |
| ENSMUSG00000017176 | Nt5c3b | 68106 | protein_coding | 99.27375164 | -0.007201982 | 0.127523025 | 0.056475933 | 0.954962668 | 0.977515248 | 0.007201982 |
| ENSMUSG00000078974 | Sec61g | 20335 | protein_coding | 311.5139324 | 0.007234121 | 0.167955783 | -0.04307158 | 0.965644474 | 0.982706396 | 0.007234121 |
| ENSMUSG00000061755 | Bod1l | 665775 | protein_coding | 783.475028 | 0.007269901 | 0.064729345 | -0.112312293 | 0.910575795 | 0.956641558 | 0.007269901 |
| ENSMUSG00000028793 | Rnf19b | 75234 | protein_coding | 651.4443124 | -0.007348579 | 0.136936313 | 0.053664208 | 0.9572027 | 0.978898029 | 0.007348579 |
| ENSMUSG00000027342 | Pcna | 18538 | protein_coding | 808.611401 | 0.007350089 | 0.118270375 | -0.062146495 | 0.950446171 | 0.975869522 | 0.007350089 |
| ENSMUSG00000003154 | Foxj2 | 60611 | protein_coding | 235.1554533 | 0.007352731 | 0.133835831 | -0.054938436 | 0.956187511 | 0.978107631 | 0.007352731 |
| ENSMUSG00000042558 | Adprs | 100206 | protein_coding | 253.1493821 | 0.007367214 | 0.098376482 | -0.074887957 | 0.940303859 | 0.971417829 | 0.007367214 |
| ENSMUSG00000031591 | Asah1 | 11886 | protein_coding | 1637.262148 | -0.007422949 | 0.101808308 | 0.072911037 | 0.941876911 | 0.971883559 | 0.007422949 |
| ENSMUSG00000092131 | BC050972 | 407808 | lncRNA | 0.839860736 | 0.007452894 | 1.736295048 | -0.004292412 | 0.996575161 | 1 | 0.007452894 |
| ENSMUSG00000020882 | Cacnb1 | 12295 | protein_coding | 39.43291226 | 0.007467805 | 0.172707977 | -0.043239493 | 0.965510624 | 0.982706396 | 0.007467805 |
| ENSMUSG00000030539 | Sema4b | 20352 | protein_coding | 1460.368242 | -0.007509132 | 0.120896313 | 0.062112168 | 0.950473507 | 0.975869522 | 0.007509132 |
| ENSMUSG00000044595 | Dnd1 | 213236 | protein_coding | 5.098127557 | -0.00751441 | 0.602118463 | 0.012479952 | 0.990042697 | 0.994317649 | 0.00751441 |
| ENSMUSG00000022911 | Arl13b | 68146 | protein_coding | 184.7486948 | 0.007548511 | 0.090743303 | -0.083185325 | 0.933704181 | 0.968634984 | 0.007548511 |

| ensembl_gene_id | external_gene_name | entrezgene_id | gene_biotype | baseMean | log2FoldChange | lfcSE | stat | pvalue | padj | absLog2FC |
|---|---|---|---|---|---|---|---|---|---|---|
| ENSMUSG00000040616 | Tmem51 | 214359 | protein_coding | 328.6020681 | 0.007604762 | 0.079967461 | -0.095098209 | 0.924236821 | 0.963320302 | 0.007604762 |
| ENSMUSG00000031792 | Usb1 | 101985 | protein_coding | 189.7154228 | 0.007614772 | 0.072902915 | -0.104450856 | 0.916811566 | 0.959274295 | 0.007614772 |
| ENSMUSG00000101892 | 9130401M01Rik | 75758 | protein_coding | 67.200789 | 0.007631306 | 0.158876303 | -0.048033002 | 0.961689941 | 0.981238658 | 0.007631306 |
| ENSMUSG00000045215 | Asxl3 | 211961 | protein_coding | 32.15789152 | 0.007634212 | 0.287025399 | -0.026597688 | 0.978780617 | 0.988831409 | 0.007634212 |
| ENSMUSG00000035891 | Cerk | 223753 | protein_coding | 382.239555 | 0.007645375 | 0.088378404 | -0.086507275 | 0.931063174 | 0.966840628 | 0.007645375 |
| ENSMUSG00000034875 | Nudt19 | 110959 | protein_coding | 919.0254062 | 0.007703094 | 0.104534644 | -0.073689392 | 0.94125754 | 0.971762583 | 0.007703094 |
| ENSMUSG00000037600 | Kdf1 | 69073 | protein_coding | 365.0036389 | -0.007706598 | 0.07411925 | 0.103975666 | 0.91718866 | 0.959486477 | 0.007706598 |
| ENSMUSG00000021830 | Txndc16 | 70561 | protein_coding | 334.7793098 | -0.00773743 | 0.205171412 | 0.037712026 | 0.969917288 | 0.984509162 | 0.00773743 |
| ENSMUSG00000033623 | Pcgf3 | 69587 | protein_coding | 1048.315396 | 0.007737592 | 0.08068315 | -0.09590097 | 0.923599224 | 0.962971397 | 0.007737592 |
| ENSMUSG00000047884 | Klk9 | 101533 | protein_coding | 1.659587662 | 0.007799139 | 1.667452674 | -0.004677278 | 0.996268086 | 1 | 0.007799139 |
| ENSMUSG00000022570 | Gfus | 22122 | protein_coding | 1221.227912 | 0.007799894 | 0.086939274 | -0.08971658 | 0.92851244 | 0.965886893 | 0.007799894 |
| ENSMUSG00000057181 | 5730455P16Rik | 70591 | protein_coding | 132.0121879 | -0.00780972 | 0.120637356 | 0.064737165 | 0.948383271 | 0.975173098 | 0.00780972 |
| ENSMUSG00000047671 | Dynlt4 | 242646 | protein_coding | 31.34866226 | -0.00781399 | 0.302345755 | 0.025844549 | 0.979381329 | 0.989031323 | 0.00781399 |
| ENSMUSG00000070056 | Mfhas1 | 52065 | protein_coding | 606.1597773 | 0.007835577 | 0.086931326 | -0.090135256 | 0.928179733 | 0.965818434 | 0.007835577 |
| ENSMUSG00000078572 | Ndufaf8 | 208501 | protein_coding | 256.5653784 | 0.007847581 | 0.092231638 | -0.085085562 | 0.932193368 | 0.967643553 | 0.007847581 |
| ENSMUSG00000029048 | Rer1 | 67830 | protein_coding | 1753.557343 | -0.007849546 | 0.073958246 | 0.106134829 | 0.915475378 | 0.958483509 | 0.007849546 |
| ENSMUSG00000053730 | Tmem39b | 230770 | protein_coding | 64.22735266 | -0.007885625 | 0.14288971 | 0.055186791 | 0.955989652 | 0.978020884 | 0.007885625 |
| ENSMUSG00000079487 | Med12 | 59024 | protein_coding | 535.8692453 | 0.007946554 | 0.089427209 | -0.088860589 | 0.929192705 | 0.966128972 | 0.007946554 |
| ENSMUSG00000020878 | Lrrc46 | 69297 | protein_coding | 4.212473996 | -0.007998212 | 0.715306576 | 0.011181516 | 0.991078627 | 1 | 0.007998212 |
| ENSMUSG00000021228 | Acot3 | 171281 | protein_coding | 0.990589206 | 0.008011155 | 1.628630152 | -0.004918953 | 0.996075259 | 1 | 0.008011155 |
| ENSMUSG00000023882 | Zfp54 | 22712 | protein_coding | 141.9419506 | -0.008016706 | 0.124214685 | 0.064539114 | 0.948540964 | 0.975173098 | 0.008016706 |
| ENSMUSG00000006717 | Acot13 | 66834 | protein_coding | 304.5659411 | -0.008028876 | 0.097950134 | 0.08196901 | 0.934671357 | 0.969180133 | 0.008028876 |
| ENSMUSG00000075229 | Ccdc58 | 381045 | protein_coding | 220.7892173 | -0.008032453 | 0.081848765 | 0.098137747 | 0.921822915 | 0.962383589 | 0.008032453 |
| ENSMUSG00000032743 | Katnip | 233865 | protein_coding | 602.9110334 | 0.008035023 | 0.050482642 | -0.159164075 | 0.873539607 | 0.937764615 | 0.008035023 |
| ENSMUSG00000000827 | Tpd52l2 | 66314 | protein_coding | 560.1399664 | -0.008082954 | 0.065146197 | 0.124074068 | 0.901256631 | 0.95132126 | 0.008082954 |
| ENSMUSG00000029110 | Rnf4 | 19822 | protein_coding | 1521.099862 | -0.008089478 | 0.05047531 | 0.160266044 | 0.872671506 | 0.937225143 | 0.008089478 |
| ENSMUSG00000020608 | Smc6 | 67241 | protein_coding | 693.5444421 | -0.008094068 | 0.087112076 | 0.092915567 | 0.925970639 | 0.964583514 | 0.008094068 |
| ENSMUSG00000050908 | Tvp23a | 383103 | protein_coding | 5.026772231 | 0.008104718 | 0.465001216 | -0.017429455 | 0.986094011 | 0.99222255 | 0.008104718 |
| ENSMUSG00000044026 | Slc35g1 | 240660 | protein_coding | 2181.405145 | -0.008153281 | 0.092273127 | 0.088360298 | 0.929590315 | 0.966128972 | 0.008153281 |
| ENSMUSG00000005936 | Kctd20 | 66989 | protein_coding | 490.2031704 | 0.008159648 | 0.055697441 | -0.146499505 | 0.883527081 | 0.942360409 | 0.008159648 |
| ENSMUSG00000034771 | Tle2 | 21886 | protein_coding | 43.83823658 | -0.008161934 | 0.332030892 | 0.024581851 | 0.980388496 | 0.989375191 | 0.008161934 |
| ENSMUSG00000027420 | Bfsp1 | 12075 | protein_coding | 80.36863025 | -0.008183653 | 0.183684243 | 0.044552833 | 0.964463739 | 0.982490542 | 0.008183653 |
| ENSMUSG00000031337 | Mtm1 | 17772 | protein_coding | 827.9720499 | 0.008187114 | 0.074419393 | -0.110013179 | 0.912398923 | 0.957266079 | 0.008187114 |
| ENSMUSG00000006699 | Cdc42 | 12540 | protein_coding | 4200.393197 | -0.008225363 | 0.07860573 | 0.104640758 | 0.916660872 | 0.959274295 | 0.008225363 |
| ENSMUSG00000022372 | Sla | 20491 | protein_coding | 14.26540232 | 0.008228919 | 0.429141072 | -0.019175324 | 0.984701243 | 0.99157486 | 0.008228919 |
| ENSMUSG00000006134 | Crkl | 12929 | protein_coding | 513.8858672 | 0.008260695 | 0.056261831 | -0.14682591 | 0.883269433 | 0.942211125 | 0.008260695 |
| ENSMUSG00000022476 | Polr3h | 78929 | protein_coding | 96.59048031 | 0.008282849 | 0.146638045 | -0.05648499 | 0.954955452 | 0.977515248 | 0.008282849 |
| ENSMUSG00000028180 | Zranb2 | 53861 | protein_coding | 606.6898468 | -0.008287956 | 0.092870269 | 0.089242298 | 0.928889351 | 0.966049475 | 0.008287956 |
| ENSMUSG00000027514 | Zbp1 | 58203 | protein_coding | 190.090626 | -0.008289982 | 0.293606482 | 0.02823501 | 0.977474714 | 0.988088396 | 0.008289982 |

| ensembl_gene_id | external_gene_name | entrezgene_id | gene_biotype | baseMean | log2FoldChange | lfcSE | stat | pvalue | padj | absLog2FC |
|---|---|---|---|---|---|---|---|---|---|---|
| ENSMUSG00000034006 | Slc66a2 | 66943 | protein_coding | 632.3857677 | -0.008358132 | 0.064046479 | 0.130501038 | 0.896170033 | 0.948996349 | 0.008358132 |
| ENSMUSG00000038279 | Nop2 | 110109 | protein_coding | 430.7712611 | 0.00838017 | 0.095656717 | -0.087606702 | 0.930189275 | 0.966352899 | 0.00838017 |
| ENSMUSG00000049532 | Sall2 | 50524 | protein_coding | 19.85570014 | -0.008400054 | 0.36113853 | 0.023259922 | 0.981442941 | 0.990015804 | 0.008400054 |
| ENSMUSG00000024242 | Map4k3 | 225028 | protein_coding | 1180.88525 | 0.00840102 | 0.04764649 | -0.176319807 | 0.860042702 | 0.931327553 | 0.00840102 |
| ENSMUSG00000021461 | Fancc | 14088 | protein_coding | 120.8214734 | -0.008435259 | 0.115380341 | 0.073108284 | 0.941719949 | 0.971883559 | 0.008435259 |
| ENSMUSG00000056941 | Commd7 | 99311 | protein_coding | 363.6700361 | -0.008438923 | 0.075777159 | 0.111365001 | 0.911326912 | 0.956997404 | 0.008438923 |
| ENSMUSG00000027508 | Pag1 | 94212 | protein_coding | 401.4728419 | 0.008448826 | 0.124697279 | -0.067754695 | 0.945980909 | 0.973795719 | 0.008448826 |
| ENSMUSG00000020527 | Myo19 | 66196 | protein_coding | 748.8571903 | 0.008448878 | 0.180058017 | -0.046923087 | 0.962574528 | 0.98153379 | 0.008448878 |
| ENSMUSG00000110902 | Gm33104 | 102635880 | lncRNA | 1.458210017 | 0.008449226 | 1.181774053 | -0.007149612 | 0.994295484 | 1 | 0.008449226 |
| ENSMUSG00000048731 | Ggnbp1 | 70772 | protein_coding | 17.06244312 | -0.008486421 | 0.372981713 | 0.022752915 | 0.981847367 | 0.9902423 | 0.008486421 |
| ENSMUSG00000073910 | Mob3b | 214944 | protein_coding | 356.1405204 | 0.008552435 | 0.102772228 | -0.08321737 | 0.933678701 | 0.968634984 | 0.008552435 |
| ENSMUSG00000097156 | Gm3764 | 100042277 | lncRNA | 3.722814714 | -0.00856217 | 2.485382133 | 0.003445012 | 0.997251284 | 1 | 0.00856217 |
| ENSMUSG00000034893 | Cog3 | 338337 | protein_coding | 1006.19759 | -0.008591453 | 0.08805464 | 0.097569562 | 0.922274095 | 0.962478474 | 0.008591453 |
| ENSMUSG00000040860 | Crocc | 230872 | protein_coding | 134.5599128 | -0.008597398 | 0.177071965 | 0.048553128 | 0.961275424 | 0.981095512 | 0.008597398 |
| ENSMUSG00000020841 | Cpd | 12874 | protein_coding | 4922.415684 | -0.008623041 | 0.15447904 | 0.055820138 | 0.955485092 | 0.977832194 | 0.008623041 |
| ENSMUSG00000028096 | Gpr89 | 67549 | protein_coding | 501.4979689 | -0.008633798 | 0.13800115 | 0.062563236 | 0.950114305 | 0.975869522 | 0.008633798 |
| ENSMUSG00000041605 | Inava | 67313 | protein_coding | 1456.510761 | -0.008664318 | 0.097799665 | 0.088592515 | 0.929405757 | 0.966128972 | 0.008664318 |
| ENSMUSG00000000794 | Kcnn3 | 140493 | protein_coding | 3.392877007 | 0.008668059 | 2.004069919 | -0.004325228 | 0.996548978 | 1 | 0.008668059 |
| ENSMUSG00000028397 | Kdm4c | 76804 | protein_coding | 375.5730967 | -0.008731003 | 0.066974699 | 0.130362713 | 0.896279466 | 0.948996349 | 0.008731003 |
| ENSMUSG00000001569 | Nom1 | 433864 | protein_coding | 388.9805868 | -0.008759657 | 0.096792114 | 0.090499699 | 0.927890133 | 0.965665343 | 0.008759657 |
| ENSMUSG00000021615 | Xrcc4 | 108138 | protein_coding | 99.6666059 | 0.00876372 | 0.119143263 | -0.073556153 | 0.941363562 | 0.971787076 | 0.00876372 |
| ENSMUSG00000001029 | Icam2 | 15896 | protein_coding | 1.992939436 | -0.008776091 | 1.23336237 | 0.007115582 | 0.994322635 | 1 | 0.008776091 |
| ENSMUSG00000096995 | 2810029C07Rik | 100504687 | lncRNA | 3.696688395 | 0.008788029 | 0.626796275 | -0.014020551 | 0.988813585 | 1 | 0.008788029 |
| ENSMUSG00000032666 | 1700025G04Rik | 69399 | protein_coding | 200.5153835 | -0.008807771 | 0.20016779 | 0.044001938 | 0.964902859 | 0.982587667 | 0.008807771 |
| ENSMUSG00000077416 | Gm23747 | 115485979 | snoRNA | 0.636004718 | -0.008876334 | 1.360341896 | 0.006525076 | 0.994793779 | 1 | 0.008876334 |
| ENSMUSG00000052926 | Rnaseh2a | 69724 | protein_coding | 267.2111017 | -0.008887917 | 0.099334324 | 0.089474777 | 0.928704598 | 0.966007248 | 0.008887917 |
| ENSMUSG00000005069 | Pex5 | 19305 | protein_coding | 508.6938507 | 0.00889231 | 0.069698755 | -0.127582048 | 0.898479739 | 0.950146927 | 0.00889231 |
| ENSMUSG00000075272 | Ttc30a2 | 620631 | protein_coding | 1.201385977 | 0.008898676 | 1.54841237 | -0.005746968 | 0.995414609 | 1 | 0.008898676 |
| ENSMUSG00000019470 | Xab2 | 67439 | protein_coding | 536.9825064 | -0.008914498 | 0.051245391 | 0.173957058 | 0.86189921 | 0.931842966 | 0.008914498 |
| ENSMUSG00000046380 | Jrk | 16469 | protein_coding | 68.82618931 | 0.008922725 | 0.159041539 | -0.056103112 | 0.955259665 | 0.977698054 | 0.008922725 |
| ENSMUSG00000063849 | Ppcdc | 66812 | protein_coding | 181.1365283 | -0.008925205 | 0.10909585 | 0.081810674 | 0.934797268 | 0.969249832 | 0.008925205 |
| ENSMUSG00000048379 | Socs4 | 67296 | protein_coding | 462.5769241 | -0.008933254 | 0.066968319 | 0.133395226 | 0.89388082 | 0.947705869 | 0.008933254 |
| ENSMUSG00000038347 | Tcte2 | 21646 | protein_coding | 19.58984813 | -0.008933736 | 0.380992045 | 0.023448615 | 0.981292426 | 0.98994327 | 0.008933736 |
| ENSMUSG00000095325 | Zfp870 | 240066 | protein_coding | 82.91614451 | -0.008950311 | 0.114891911 | 0.077902012 | 0.937905999 | 0.970517013 | 0.008950311 |
| ENSMUSG00000036437 | Npy1r | 18166 | protein_coding | 47.4759346 | -0.008957873 | 0.33688795 | 0.026590067 | 0.978786696 | 0.988831409 | 0.008957873 |
| ENSMUSG00000020230 | Prmt2 | 15468 | protein_coding | 168.77251 | 0.008971286 | 0.138807184 | -0.064631278 | 0.94846758 | 0.975173098 | 0.008971286 |
| ENSMUSG00000039599 | Fam149b | 105428 | protein_coding | 366.7437944 | 0.008974215 | 0.060497053 | -0.148341367 | 0.882073371 | 0.941960269 | 0.008974215 |
| ENSMUSG00000051000 | Fhip1a | 229488 | protein_coding | 354.0945245 | 0.008981453 | 0.119264705 | -0.075306878 | 0.939970549 | 0.971381329 | 0.008981453 |
| ENSMUSG00000039578 | Ccser1 | 232035 | protein_coding | 347.5532824 | 0.009062499 | 0.108994317 | -0.083146526 | 0.933735032 | 0.968634984 | 0.009062499 |

| ensembl_gene_id | external_gene_name | entrezgene_id | gene_biotype | baseMean | log2FoldChange | lfcSE | stat | pvalue | padj | absLog2FC |
|---|---|---|---|---|---|---|---|---|---|---|
| ENSMUSG00000035936 | Aldh5a1 | 214579 | protein_coding | 231.7828807 | -0.009074105 | 0.115147586 | 0.078804129 | 0.93718842 | 0.970043107 | 0.009074105 |
| ENSMUSG00000049620 | Prss33 | 353130 | protein_coding | 1.454541405 | -0.009172701 | 1.527196402 | 0.006006235 | 0.995207747 | 1 | 0.009172701 |
| ENSMUSG00000020544 | Cox11 | 69802 | protein_coding | 84.69299495 | 0.009178265 | 0.125078877 | -0.073379816 | 0.941503879 | 0.971840719 | 0.009178265 |
| ENSMUSG00000005674 | Tomm40l | 641376 | protein_coding | 130.8454661 | -0.009198506 | 0.116531896 | 0.078935517 | 0.937083913 | 0.970043107 | 0.009198506 |
| ENSMUSG00000026805 | Barhl1 | 54422 | protein_coding | 1.510755249 | -0.009268455 | 1.136153484 | 0.008157749 | 0.99349113 | 1 | 0.009268455 |
| ENSMUSG00000020386 | Sar1b | 66397 | protein_coding | 618.6740408 | 0.00933838 | 0.069215921 | -0.134916642 | 0.892677781 | 0.947202684 | 0.00933838 |
| ENSMUSG00000042814 | Mcts2 | 66405 | protein_coding | 67.81762597 | 0.009354106 | 0.15543151 | -0.06018153 | 0.952011056 | 0.976016991 | 0.009354106 |
| ENSMUSG00000047996 | Prrg1 | 546336 | protein_coding | 27.09399471 | 0.009354425 | 0.26193705 | -0.035712492 | 0.97151161 | 0.985415068 | 0.009354425 |
| ENSMUSG00000040213 | Kyat3 | 229905 | protein_coding | 92.04096636 | 0.009355696 | 0.135164194 | -0.069217265 | 0.94481668 | 0.973465352 | 0.009355696 |
| ENSMUSG00000027852 | Nras | 18176 | protein_coding | 643.3402371 | -0.009369624 | 0.068822925 | 0.13614103 | 0.891709792 | 0.947050968 | 0.009369624 |
| ENSMUSG00000022159 | Rab2b | 76338 | protein_coding | 10.7732856 | -0.009378499 | 0.369000046 | 0.025415984 | 0.979723162 | 0.989126875 | 0.009378499 |
| ENSMUSG00000058318 | Phf21a | 192285 | protein_coding | 202.5891137 | 0.009378697 | 0.128913814 | -0.072751685 | 0.942003718 | 0.971883559 | 0.009378697 |
| ENSMUSG00000018554 | Ybx2 | 53422 | protein_coding | 119.0188919 | 0.009385832 | 0.116589126 | -0.080503497 | 0.935836815 | 0.96968875 | 0.009385832 |
| ENSMUSG00000035765 | Dym | 69190 | protein_coding | 512.696312 | 0.009389227 | 0.087378563 | -0.107454578 | 0.914428358 | 0.95811623 | 0.009389227 |
| ENSMUSG00000021338 | Carmil1 | 68732 | protein_coding | 538.9739354 | -0.0094165 | 0.05498454 | 0.171257224 | 0.864021514 | 0.932456625 | 0.0094165 |
| ENSMUSG00000048787 | Dcun1d3 | 233805 | protein_coding | 144.387298 | 0.009436228 | 0.090901361 | -0.103807337 | 0.917322244 | 0.959501353 | 0.009436228 |
| ENSMUSG00000026803 | Ttf1 | 22130 | protein_coding | 342.748089 | 0.009440753 | 0.065855941 | -0.143354621 | 0.88601012 | 0.94348537 | 0.009440753 |
| ENSMUSG00000026036 | Nif3l1 | 65102 | protein_coding | 230.3399605 | 0.009479147 | 0.116630205 | -0.08127523 | 0.935223072 | 0.969552109 | 0.009479147 |
| ENSMUSG00000032897 | Nfyc | 18046 | protein_coding | 350.2025112 | -0.009483748 | 0.091382237 | 0.103781086 | 0.917343076 | 0.959501353 | 0.009483748 |
| ENSMUSG00000020453 | Patz1 | 56218 | protein_coding | 106.39046 | 0.009489831 | 0.128678664 | -0.073748285 | 0.941210677 | 0.971762583 | 0.009489831 |
| ENSMUSG00000051435 | Fhad1 | 329977 | protein_coding | 43.56735579 | 0.009495463 | 0.201404221 | -0.047146296 | 0.96239663 | 0.981474584 | 0.009495463 |
| ENSMUSG00000036109 | Mbnl3 | 171170 | protein_coding | 348.1156329 | 0.00949969 | 0.119743567 | -0.079333615 | 0.93676727 | 0.969910941 | 0.00949969 |
| ENSMUSG00000029260 | Ugt2b34 | 100727 | protein_coding | 6973.445709 | 0.009502066 | 0.085250703 | -0.111460262 | 0.911251375 | 0.956997404 | 0.009502066 |
| ENSMUSG00000028416 | Bag1 | 12017 | protein_coding | 2074.95436 | 0.009512225 | 0.080552302 | -0.11808756 | 0.90599828 | 0.953990085 | 0.009512225 |
| ENSMUSG00000005802 | Slc30a4 | 22785 | protein_coding | 1683.268306 | -0.009568772 | 0.095579894 | 0.10011281 | 0.920254765 | 0.961475844 | 0.009568772 |
| ENSMUSG00000031776 | Arl2bp | 107566 | protein_coding | 773.8108721 | -0.009589563 | 0.057376053 | 0.167135288 | 0.8672636 | 0.934002289 | 0.009589563 |
| ENSMUSG00000016833 | Mrps18c | 68735 | protein_coding | 217.917945 | 0.009603499 | 0.075536053 | -0.127137956 | 0.898831211 | 0.950352959 | 0.009603499 |
| ENSMUSG00000037112 | Sik2 | 235344 | protein_coding | 210.98679 | 0.009610915 | 0.086513717 | -0.11109123 | 0.911544003 | 0.956997404 | 0.009610915 |
| ENSMUSG00000027552 | E2f5 | 13559 | protein_coding | 330.6323299 | 0.009657573 | 0.073606918 | -0.131204698 | 0.89561338 | 0.948731562 | 0.009657573 |
| ENSMUSG00000063808 | Gpatch1 | 67471 | protein_coding | 225.8791205 | 0.009721343 | 0.090323953 | -0.107627519 | 0.914291167 | 0.95803327 | 0.009721343 |
| ENSMUSG00000020705 | Ddx42 | 72047 | protein_coding | 840.3238963 | 0.009728521 | 0.070495604 | -0.138001806 | 0.89023899 | 0.946097348 | 0.009728521 |
| ENSMUSG00000015112 | Slc25a13 | 50799 | protein_coding | 626.7551711 | 0.009741484 | 0.067620471 | -0.144061168 | 0.885452168 | 0.943195324 | 0.009741484 |
| ENSMUSG00000038374 | Rbm8a | 60365 | protein_coding | 459.2819631 | -0.009769357 | 0.072406041 | 0.134924615 | 0.892671478 | 0.947202684 | 0.009769357 |
| ENSMUSG00000021298 | Gpr132 | 56696 | protein_coding | 3.972508303 | 0.009802164 | 0.862069397 | -0.011370504 | 0.990927846 | 1 | 0.009802164 |
| ENSMUSG00000039164 | Naif1 | 71254 | protein_coding | 113.9431011 | 0.009817848 | 0.13326835 | -0.073669765 | 0.941273157 | 0.971762583 | 0.009817848 |
| ENSMUSG00000002771 | Grin2d | 14814 | protein_coding | 32.23219669 | -0.009844181 | 0.266899591 | 0.036883461 | 0.970577927 | 0.984770701 | 0.009844181 |
| ENSMUSG00000041977 | Arhgef11 | 213498 | protein_coding | 1145.033729 | 0.009862416 | 0.047147526 | -0.209182038 | 0.834306134 | 0.916804065 | 0.009862416 |
| ENSMUSG00000029014 | Dnajc2 | 22791 | protein_coding | 397.459667 | -0.009864897 | 0.089817776 | 0.109832348 | 0.912542336 | 0.957266079 | 0.009864897 |
| ENSMUSG00000045350 | Fam186a | 72277 | protein_coding | 3.76491478 | -0.009888454 | 2.379920457 | 0.004154951 | 0.996684838 | 1 | 0.009888454 |

| ensembl_gene_id | external_gene_name | entrezgene_id | gene_biotype | baseMean | log2FoldChange | lfcSE | stat | pvalue | padj | absLog2FC |
|---|---|---|---|---|---|---|---|---|---|---|
| ENSMUSG00000024137 | E4f1 | 13560 | protein_coding | 263.5657342 | -0.009944488 | 0.110175287 | 0.0902606 | 0.928080129 | 0.965802209 | 0.009944488 |
| ENSMUSG00000032030 | Cul5 | 75717 | protein_coding | 425.8412887 | -0.00995262 | 0.109427858 | 0.090951433 | 0.927531182 | 0.965445424 | 0.00995262 |
| ENSMUSG00000038456 | Dennd2a | 209773 | protein_coding | 4.693304854 | -0.00995431 | 0.991806042 | 0.010036549 | 0.991992127 | 0.995389267 | 0.00995431 |
| ENSMUSG00000026822 | Lcn2 | 16819 | protein_coding | 2.010435989 | -0.009978936 | 2.163927302 | 0.004611493 | 0.996320574 | 1 | 0.009978936 |
| ENSMUSG00000032643 | Fhl3 | 14201 | protein_coding | 5.796376185 | 0.009996746 | 0.464647701 | -0.021514678 | 0.982835095 | 0.990693936 | 0.009996746 |
| ENSMUSG00000039176 | Polg | 18975 | protein_coding | 883.165952 | -0.010000317 | 0.046617144 | 0.214520161 | 0.830141447 | 0.914175515 | 0.010000317 |
| ENSMUSG00000031585 | Gtf2e2 | 68153 | protein_coding | 230.7441799 | 0.010000963 | 0.089017762 | -0.112347949 | 0.910547525 | 0.956641558 | 0.010000963 |
| ENSMUSG00000054770 | Kctd18 | 51960 | protein_coding | 95.61795713 | -0.010049174 | 0.148775782 | 0.067545766 | 0.946147229 | 0.973795719 | 0.010049174 |
| ENSMUSG00000002455 | Prpf6 | 68879 | protein_coding | 744.3488081 | 0.010088513 | 0.082018384 | -0.123003069 | 0.902104668 | 0.951530343 | 0.010088513 |
| ENSMUSG00000090925 | 1810064F22Rik | 69862 | lncRNA | 4.760376233 | -0.010115539 | 0.50517436 | 0.020023857 | 0.984024341 | 0.991348095 | 0.010115539 |
| ENSMUSG00000032112 | Trappc4 | 60409 | protein_coding | 328.602505 | 0.010131836 | 0.075869833 | -0.133542351 | 0.893764473 | 0.947692153 | 0.010131836 |
| ENSMUSG00000021900 | Btd | 26363 | protein_coding | 140.2679393 | 0.010136004 | 0.133387966 | -0.075988896 | 0.939427933 | 0.971185231 | 0.010136004 |
| ENSMUSG00000024283 | Wac | 225131 | protein_coding | 864.3090866 | 0.010141492 | 0.058612639 | -0.173025687 | 0.862631237 | 0.932166328 | 0.010141492 |
| ENSMUSG00000047822 | Angptl8 | 624219 | protein_coding | 17.46117535 | -0.010201645 | 0.265867387 | 0.038371177 | 0.969391741 | 0.984403993 | 0.010201645 |
| ENSMUSG00000003166 | Dgcr2 | 13356 | protein_coding | 894.7944471 | -0.010207429 | 0.116114411 | 0.087908372 | 0.929949502 | 0.966286169 | 0.010207429 |
| ENSMUSG00000030330 | Ing4 | 28019 | protein_coding | 17.15423099 | 0.010230163 | 0.365547396 | -0.027985873 | 0.977673418 | 0.988088396 | 0.010230163 |
| ENSMUSG00000026275 | Ppp1r7 | 66385 | protein_coding | 400.9615479 | 0.010252637 | 0.102334536 | -0.100187461 | 0.920195501 | 0.961475844 | 0.010252637 |
| ENSMUSG00000036202 | Rif1 | 51869 | protein_coding | 594.6160475 | -0.010298591 | 0.094694966 | 0.108755419 | 0.913396485 | 0.957864571 | 0.010298591 |
| ENSMUSG00000030380 | Mzf1 | 109889 | protein_coding | 14.96875606 | 0.010300221 | 0.417009948 | -0.024700179 | 0.980294112 | 0.9893404 | 0.010300221 |
| ENSMUSG00000022895 | Ets2 | 23872 | protein_coding | 2899.056956 | 0.010307638 | 0.06601214 | -0.156147614 | 0.875916668 | 0.938630585 | 0.010307638 |
| ENSMUSG00000042492 | Tbc1d10b | 68449 | protein_coding | 672.8799773 | 0.010327065 | 0.076928509 | -0.134242364 | 0.893210927 | 0.947365784 | 0.010327065 |
| ENSMUSG00000021067 | Sav1 | 64010 | protein_coding | 267.375535 | 0.010334551 | 0.091492881 | -0.112954702 | 0.910066468 | 0.956337108 | 0.010334551 |
| ENSMUSG00000022019 | Tdrd3 | 219249 | protein_coding | 409.8904133 | 0.010356888 | 0.065434707 | -0.1582782 | 0.874237585 | 0.93820896 | 0.010356888 |
| ENSMUSG00000015994 | Fnta | 14272 | protein_coding | 543.2609857 | 0.010370724 | 0.081093775 | -0.12788558 | 0.898239523 | 0.949998977 | 0.010370724 |
| ENSMUSG00000029192 | Tbc1d14 | 100855 | protein_coding | 407.283084 | -0.010378651 | 0.077842484 | 0.133328881 | 0.893933288 | 0.947705869 | 0.010378651 |
| ENSMUSG00000058638 | Zfp110 | 65020 | protein_coding | 548.2514821 | -0.01045217 | 0.08143559 | 0.128348925 | 0.897872848 | 0.949809251 | 0.01045217 |
| ENSMUSG00000030838 | Ush1c | 72088 | protein_coding | 2531.551252 | 0.01045261 | 0.083411025 | -0.125314488 | 0.900274583 | 0.95096289 | 0.01045261 |
| ENSMUSG00000003051 | Elf3 | 13710 | protein_coding | 5854.295706 | -0.010470057 | 0.083325748 | 0.12565212 | 0.900007304 | 0.950848993 | 0.010470057 |
| ENSMUSG00000053536 | Cstf2t | 83410 | protein_coding | 750.0759245 | 0.010504451 | 0.082824764 | -0.126827418 | 0.899076995 | 0.950413359 | 0.010504451 |
| ENSMUSG00000060275 | Nrg2 | 100042150 | protein_coding | 3.471468667 | -0.010523041 | 0.852705403 | 0.012340769 | 0.990153741 | 1 | 0.010523041 |
| ENSMUSG00000034848 | Ttc21b | 73668 | protein_coding | 159.1250609 | -0.010534006 | 0.120422229 | 0.087475595 | 0.930293483 | 0.966400364 | 0.010534006 |
| ENSMUSG00000005774 | Rfx5 | 53970 | protein_coding | 124.1583742 | -0.010538784 | 0.098679119 | 0.106798519 | 0.914948822 | 0.958296749 | 0.010538784 |
| ENSMUSG00000001420 | Tmem79 | 71913 | protein_coding | 275.1308966 | -0.010570764 | 0.096619911 | 0.109405651 | 0.912880751 | 0.957419485 | 0.010570764 |
| ENSMUSG00000028653 | Trit1 | 66966 | protein_coding | 126.2777233 | 0.010574981 | 0.124563701 | -0.084896169 | 0.932343937 | 0.967739018 | 0.010574981 |
| ENSMUSG00000021028 | Mbip | 217588 | protein_coding | 92.57577092 | 0.010594604 | 0.137350237 | -0.077135679 | 0.93851561 | 0.970606637 | 0.010594604 |
| ENSMUSG00000023089 | Ndufa5 | 68202 | protein_coding | 850.4439848 | -0.01061535 | 0.085828538 | 0.123680893 | 0.901567941 | 0.951341404 | 0.01061535 |
| ENSMUSG00000021621 | Zcchc9 | 69085 | protein_coding | 287.7584964 | 0.010615756 | 0.066153636 | -0.160471243 | 0.872509872 | 0.937146539 | 0.010615756 |
| ENSMUSG00000064289 | Tank | 21353 | protein_coding | 349.1398749 | 0.010641928 | 0.090681175 | -0.117355428 | 0.906578404 | 0.954311305 | 0.010641928 |
| ENSMUSG00000016559 | H3f3b | 15081 | protein_coding | 8355.17645 | 0.010653336 | 0.103098112 | -0.103332021 | 0.917699463 | 0.959656087 | 0.010653336 |

| ensembl_gene_id | external_gene_name | entrezgene_id | gene_biotype | baseMean | log2FoldChange | lfcSE | stat | pvalue | padj | absLog2FC |
|---|---|---|---|---|---|---|---|---|---|---|
| ENSMUSG00000074305 | Peak1 | 244895 | protein_coding | 267.549827 | 0.010655563 | 0.093146567 | -0.114395655 | 0.908924158 | 0.955501531 | 0.010655563 |
| ENSMUSG00000003379 | Cd79a | 12518 | protein_coding | 2.619882015 | 0.010669654 | 1.118778056 | -0.009536882 | 0.992390785 | 1 | 0.010669654 |
| ENSMUSG00000060288 | Ppih | 66101 | protein_coding | 169.0213923 | -0.010759792 | 0.153417359 | 0.070134124 | 0.944086907 | 0.973044134 | 0.010759792 |
| ENSMUSG00000003134 | Tbc1d8 | 54610 | protein_coding | 186.2825174 | -0.010793067 | 0.128485499 | 0.084002217 | 0.933054669 | 0.968172466 | 0.010793067 |
| ENSMUSG00000008690 | Ncaph2 | 52683 | protein_coding | 1069.225707 | 0.010803882 | 0.072258561 | -0.149516983 | 0.88114571 | 0.941406719 | 0.010803882 |
| ENSMUSG00000021895 | Arhgef3 | 71704 | protein_coding | 510.03322 | -0.010829743 | 0.086882139 | 0.124648672 | 0.900801694 | 0.951246263 | 0.010829743 |
| ENSMUSG00000058624 | Gda | 14544 | protein_coding | 1153.46087 | -0.010865467 | 0.139860515 | 0.077687883 | 0.938076333 | 0.970517013 | 0.010865467 |
| ENSMUSG00000025155 | Dus1l | 68730 | protein_coding | 513.0951666 | -0.01088832 | 0.080240276 | 0.135696447 | 0.892061257 | 0.947202684 | 0.01088832 |
| ENSMUSG00000037104 | Socs5 | 56468 | protein_coding | 146.6028721 | 0.010920228 | 0.144934162 | -0.075346126 | 0.939939323 | 0.971381329 | 0.010920228 |
| ENSMUSG00000040029 | Ipo8 | 320727 | protein_coding | 879.6077946 | -0.010924571 | 0.07029391 | 0.155412762 | 0.876495924 | 0.938630585 | 0.010924571 |
| ENSMUSG00000039725 | Trp53rka | 381406 | protein_coding | 103.6473654 | 0.010928475 | 0.103850588 | -0.105232678 | 0.916191181 | 0.959151705 | 0.010928475 |
| ENSMUSG00000027079 | Clp1 | 98985 | protein_coding | 158.176021 | -0.01094104 | 0.100646173 | 0.108707955 | 0.913434132 | 0.957864571 | 0.01094104 |
| ENSMUSG00000054484 | Tmem62 | 96957 | protein_coding | 198.9971922 | -0.010951323 | 0.09088738 | 0.12049333 | 0.904092363 | 0.952605199 | 0.010951323 |
| ENSMUSG00000022412 | Mief1 | 239555 | protein_coding | 607.0434697 | 0.010952647 | 0.064127664 | -0.170794417 | 0.864385419 | 0.932569867 | 0.010952647 |
| ENSMUSG00000028792 | Ak2 | 11637 | protein_coding | 2578.085973 | 0.01096474 | 0.069687657 | -0.157341209 | 0.874975945 | 0.938415991 | 0.01096474 |
| ENSMUSG00000055065 | Ddx17 | 67040 | protein_coding | 2704.206293 | -0.010994062 | 0.074280328 | 0.148007723 | 0.882336673 | 0.941973978 | 0.010994062 |
| ENSMUSG00000078716 | Tmem8b | 242409 | protein_coding | 38.21706924 | -0.011001281 | 0.179963803 | 0.061130523 | 0.95125526 | 0.975912221 | 0.011001281 |
| ENSMUSG00000106219 | 5830416I19Rik | 74757 | lncRNA | 2.320500007 | 0.011025299 | 1.06542769 | -0.010348238 | 0.991743448 | 1 | 0.011025299 |
| ENSMUSG00000033004 | Mycbp2 | 105689 | protein_coding | 900.9162999 | 0.011048938 | 0.081811681 | -0.135053306 | 0.892569728 | 0.947202684 | 0.011048938 |
| ENSMUSG00000032839 | Trpc1 | 22063 | protein_coding | 11.23289426 | 0.011053501 | 0.383701876 | -0.028807523 | 0.977018101 | 0.987905762 | 0.011053501 |
| ENSMUSG00000048874 | Phf3 | 213109 | protein_coding | 1004.503263 | -0.01105393 | 0.062850163 | 0.175877514 | 0.860390171 | 0.931471303 | 0.01105393 |
| ENSMUSG00000028719 | Cmpk1 | 66588 | protein_coding | 5375.843685 | 0.011069918 | 0.09444353 | -0.117212032 | 0.906692033 | 0.954353007 | 0.011069918 |
| ENSMUSG00000075700 | Selenot | 69227 | protein_coding | 1443.96345 | 0.011107272 | 0.078491446 | -0.141509332 | 0.887467588 | 0.944306685 | 0.011107272 |
| ENSMUSG00000021681 | Aggf1 | 66549 | protein_coding | 618.6985321 | -0.011121304 | 0.075619904 | 0.147068474 | 0.883077973 | 0.942185661 | 0.011121304 |
| ENSMUSG00000043079 | Synpo | 104027 | protein_coding | 510.0196713 | 0.011149315 | 0.180563729 | -0.061747257 | 0.950764106 | 0.97587882 | 0.011149315 |
| ENSMUSG00000026490 | Cdc42bpa | 226751 | protein_coding | 758.5877131 | 0.011162061 | 0.132641433 | -0.084152148 | 0.932935464 | 0.968109605 | 0.011162061 |
| ENSMUSG00000048000 | Gigyf2 | 227331 | protein_coding | 1118.041789 | -0.011178753 | 0.060142121 | 0.185872276 | 0.852544924 | 0.927761944 | 0.011178753 |
| ENSMUSG00000013833 | Med16 | 216154 | protein_coding | 306.3650478 | 0.011206505 | 0.100674889 | -0.111313807 | 0.911367507 | 0.956997404 | 0.011206505 |
| ENSMUSG00000052387 | Trpm3 | 226025 | protein_coding | 6.524496426 | 0.011209403 | 2.536055899 | -0.004420014 | 0.99647335 | 0.998060837 | 0.011209403 |
| ENSMUSG00000090564 | A430057M04Rik | 319486 | lncRNA | 8.496378312 | -0.011216587 | 0.450397007 | 0.024903778 | 0.980131714 | 0.989302036 | 0.011216587 |
| ENSMUSG00000011179 | Odc1 | 18263 | protein_coding | 2462.948111 | 0.01122357 | 0.069848993 | -0.160683345 | 0.872342808 | 0.937028043 | 0.01122357 |
| ENSMUSG00000025217 | Btrc | 12234 | protein_coding | 214.110568 | -0.011224434 | 0.090426424 | 0.124127811 | 0.901214078 | 0.95132126 | 0.011224434 |
| ENSMUSG00000038181 | Chpf2 | 100910 | protein_coding | 250.8384886 | -0.011226413 | 0.101102238 | 0.111040199 | 0.91158447 | 0.956997404 | 0.011226413 |
| ENSMUSG00000024472 | Dcp2 | 70640 | protein_coding | 752.9985476 | 0.011271539 | 0.104003765 | -0.108376259 | 0.913697233 | 0.957921881 | 0.011271539 |
| ENSMUSG00000033014 | Trim33 | 94093 | protein_coding | 616.634426 | 0.011272042 | 0.052575478 | -0.214397333 | 0.830237221 | 0.914187552 | 0.011272042 |
| ENSMUSG00000041438 | Utp4 | 21771 | protein_coding | 215.804727 | -0.011292575 | 0.122946924 | 0.091849185 | 0.926817865 | 0.965096805 | 0.011292575 |
| ENSMUSG00000068742 | Cry2 | 12953 | protein_coding | 241.2815847 | -0.011304959 | 0.213060035 | 0.053059971 | 0.957684125 | 0.979208448 | 0.011304959 |
| ENSMUSG00000014426 | Map3k4 | 26407 | protein_coding | 775.7514412 | -0.011323132 | 0.056266235 | 0.20124205 | 0.840509313 | 0.920924283 | 0.011323132 |
| ENSMUSG00000112352 | Gm40617 | 105245123 | lncRNA | 1.042732304 | -0.011331091 | 1.629269024 | 0.006954709 | 0.99445099 | 1 | 0.011331091 |

| ensembl_gene_id | external_gene_name | entrezgene_id | gene_biotype | baseMean | log2FoldChange | lfcSE | stat | pvalue | padj | absLog2FC |
|---|---|---|---|---|---|---|---|---|---|---|
| ENSMUSG00000074071 | Fam169b | 434197 | protein_coding | 4.123447684 | 0.011369419 | 1.09455405 | -0.010387262 | 0.991712313 | 1 | 0.011369419 |
| ENSMUSG00000044034 | Npb | 208990 | protein_coding | 2.67898881 | -0.011374784 | 0.670709946 | 0.016959319 | 0.986469069 | 1 | 0.011374784 |
| ENSMUSG00000045294 | Insig1 | 231070 | protein_coding | 1310.999947 | 0.011391345 | 0.176441721 | -0.064561516 | 0.948523127 | 0.975173098 | 0.011391345 |
| ENSMUSG00000020925 | Ccdc43 | 52715 | protein_coding | 253.4470209 | -0.011406851 | 0.081448831 | 0.140049286 | 0.888621049 | 0.94517522 | 0.011406851 |
| ENSMUSG00000020109 | Dnajb12 | 56709 | protein_coding | 641.1710569 | 0.011419138 | 0.072669693 | -0.157137558 | 0.875136438 | 0.938415991 | 0.011419138 |
| ENSMUSG00000046168 | Kcnrg | 328424 | protein_coding | 2.776361337 | 0.011449021 | 0.709068775 | -0.016146559 | 0.98711747 | 1 | 0.011449021 |
| ENSMUSG00000029313 | Aff1 | 17355 | protein_coding | 1129.378958 | -0.011465708 | 0.092697075 | 0.12369007 | 0.901560674 | 0.951341404 | 0.011465708 |
| ENSMUSG00000032512 | Wdr48 | 67561 | protein_coding | 787.9940572 | 0.011473579 | 0.089169751 | -0.128671197 | 0.897617827 | 0.949716026 | 0.011473579 |
| ENSMUSG00000051251 | Nhlh1 | 18071 | protein_coding | 1.461124368 | -0.01147461 | 2.083152036 | 0.005508292 | 0.995605041 | 1 | 0.01147461 |
| ENSMUSG00000027167 | Elp4 | 77766 | protein_coding | 141.4407399 | 0.011493877 | 0.118396247 | -0.097079744 | 0.922663066 | 0.962530504 | 0.011493877 |
| ENSMUSG00000058921 | Slc10a5 | 241877 | protein_coding | 724.6548025 | -0.011507391 | 0.085378315 | 0.134781193 | 0.892784876 | 0.947205958 | 0.011507391 |
| ENSMUSG00000079033 | Mef2b | 17259 | protein_coding | 0.644317546 | -0.011509836 | 1.590380883 | 0.007237157 | 0.994225634 | 1 | 0.011509836 |
| ENSMUSG00000043289 | Mei4 | 75033 | protein_coding | 1.20532104 | 0.011525254 | 1.005734511 | -0.01145954 | 0.99085681 | 1 | 0.011525254 |
| ENSMUSG00000042111 | Ccdc115 | 69668 | protein_coding | 193.7637046 | 0.011535229 | 0.126083331 | -0.091488928 | 0.927104104 | 0.965151432 | 0.011535229 |
| ENSMUSG00000020143 | Dock2 | 94176 | protein_coding | 25.03219337 | 0.011563189 | 0.413276496 | -0.027979305 | 0.977678657 | 0.988088396 | 0.011563189 |
| ENSMUSG00000038375 | Trp53inp2 | 68728 | protein_coding | 1171.893606 | 0.011569362 | 0.139459849 | -0.082958374 | 0.933884638 | 0.968697092 | 0.011569362 |
| ENSMUSG00000039043 | Arpin | 70420 | protein_coding | 765.6526823 | 0.011590682 | 0.07848448 | -0.147681204 | 0.882594366 | 0.941973978 | 0.011590682 |
| ENSMUSG00000006574 | Slc4a1 | 20533 | protein_coding | 7.440871256 | -0.011594929 | 0.919993543 | 0.012603272 | 0.98994431 | 0.994279305 | 0.011594929 |
| ENSMUSG00000108030 | 9530062K07Rik | 78739 | lncRNA | 4.217707595 | -0.011620401 | 0.680284635 | 0.017081676 | 0.986371458 | 1 | 0.011620401 |
| ENSMUSG00000021646 | Mccc2 | 78038 | protein_coding | 315.6190122 | -0.011629262 | 0.129079524 | 0.090093782 | 0.92821269 | 0.965818434 | 0.011629262 |
| ENSMUSG00000046020 | Pofut1 | 140484 | protein_coding | 696.6322376 | 0.011629759 | 0.093510495 | -0.124368488 | 0.901023522 | 0.95132126 | 0.011629759 |
| ENSMUSG00000074211 | Sdhaf1 | 68332 | protein_coding | 77.86100818 | 0.011641221 | 0.157581313 | -0.073874377 | 0.941110345 | 0.971762583 | 0.011641221 |
| ENSMUSG00000021785 | Ngly1 | 59007 | protein_coding | 815.3799671 | -0.011651673 | 0.079736461 | 0.14612729 | 0.883820903 | 0.942430322 | 0.011651673 |
| ENSMUSG00000035620 | Ric8b | 237422 | protein_coding | 234.6549374 | 0.011660412 | 0.091870289 | -0.126922558 | 0.899001692 | 0.950394593 | 0.011660412 |
| ENSMUSG00000002103 | Acp2 | 11432 | protein_coding | 330.5189153 | -0.011731494 | 0.079332507 | 0.147877517 | 0.882439432 | 0.941973978 | 0.011731494 |
| ENSMUSG00000051246 | Msantd1 | 403174 | protein_coding | 18.63968727 | -0.011739617 | 0.265070368 | 0.044288682 | 0.964674293 | 0.982587667 | 0.011739617 |
| ENSMUSG00000021062 | Rab15 | 104886 | protein_coding | 1200.151933 | -0.011751116 | 0.19158962 | 0.061334826 | 0.951092556 | 0.97587882 | 0.011751116 |
| ENSMUSG00000027613 | Eif6 | 16418 | protein_coding | 2695.659702 | -0.011768204 | 0.071870484 | 0.163741822 | 0.869934393 | 0.935658236 | 0.011768204 |
| ENSMUSG00000063358 | Mapk1 | 26413 | protein_coding | 1667.886836 | 0.01176868 | 0.069978768 | -0.168175009 | 0.866445601 | 0.933791206 | 0.01176868 |
| ENSMUSG00000036769 | Wdr44 | 72404 | protein_coding | 398.2544898 | 0.011828659 | 0.099249744 | -0.11918075 | 0.905132158 | 0.953251304 | 0.011828659 |
| ENSMUSG00000007659 | Bcl2l1 | 12048 | protein_coding | 498.2058197 | 0.011847126 | 0.090134548 | -0.131438237 | 0.895428643 | 0.948680909 | 0.011847126 |
| ENSMUSG00000039765 | Cc2d2a | 231214 | protein_coding | 247.705684 | 0.011849771 | 0.106888378 | -0.110861169 | 0.911726438 | 0.956997404 | 0.011849771 |
| ENSMUSG00000025371 | Chmp6 | 208092 | protein_coding | 905.792454 | 0.01189594 | 0.082247517 | -0.144635856 | 0.884998386 | 0.943016089 | 0.01189594 |
| ENSMUSG00000032046 | Abhd12 | 76192 | protein_coding | 412.1978032 | -0.011912247 | 0.097467194 | 0.122218019 | 0.902726356 | 0.95200802 | 0.011912247 |
| ENSMUSG00000026203 | Dnajb2 | 56812 | protein_coding | 271.1336459 | -0.01193962 | 0.107231785 | 0.11134404 | 0.911343533 | 0.956997404 | 0.01193962 |
| ENSMUSG00000034940 | Synrg | 217030 | protein_coding | 581.6215687 | 0.011946745 | 0.077236928 | -0.1546766 | 0.877076279 | 0.938936708 | 0.011946745 |
| ENSMUSG00000039480 | Nt5dc1 | 319638 | protein_coding | 241.1999973 | 0.011979355 | 0.08956918 | -0.133744166 | 0.89360488 | 0.947601039 | 0.011979355 |
| ENSMUSG00000064345 | mt-Nd2 | 17717 | protein_coding | 13660.23877 | 0.011992535 | 0.419465109 | -0.028590066 | 0.977191535 | 0.988020633 | 0.011992535 |
| ENSMUSG00000029676 | Pot1a | 101185 | protein_coding | 144.8832799 | -0.01200693 | 0.104913165 | 0.114446367 | 0.90888396 | 0.955501531 | 0.01200693 |

Supplementary Table 1

| ensembl_gene_id | external_gene_name | entrezgene_id | gene_biotype | baseMean | log2FoldChange | lfcSE | stat | pvalue | padj | absLog2FC |
|---|---|---|---|---|---|---|---|---|---|---|
| ENSMUSG00000052139 | Babam2 | 107976 | protein_coding | 334.1012628 | 0.012021489 | 0.058990629 | -0.203786419 | 0.838520406 | 0.919582966 | 0.012021489 |
| ENSMUSG00000028701 | Lurap1 | 68075 | protein_coding | 29.9416668 | 0.012076745 | 0.209778181 | -0.057569119 | 0.954091848 | 0.977047797 | 0.012076745 |
| ENSMUSG00000031605 | Klhl2 | 77113 | protein_coding | 536.0524713 | 0.012077255 | 0.052922375 | -0.228206966 | 0.819485346 | 0.908985828 | 0.012077255 |
| ENSMUSG00000028622 | Mrpl37 | 56280 | protein_coding | 438.9917312 | -0.012083889 | 0.095840662 | 0.126083108 | 0.899666138 | 0.950671014 | 0.012083889 |
| ENSMUSG00000033009 | Ogfod1 | 270086 | protein_coding | 307.8644066 | 0.012098457 | 0.10914587 | -0.110846679 | 0.911737929 | 0.956997404 | 0.012098457 |
| ENSMUSG00000032890 | Rims3 | 242662 | protein_coding | 3.787188464 | -0.012122079 | 1.048114905 | 0.011565601 | 0.990772191 | 1 | 0.012122079 |
| ENSMUSG00000030327 | Necap1 | 67602 | protein_coding | 420.1771001 | 0.012134177 | 0.080344287 | -0.151027253 | 0.879954218 | 0.940676072 | 0.012134177 |
| ENSMUSG00000057133 | Chd6 | 71389 | protein_coding | 651.3093249 | 0.0121375 | 0.102275441 | -0.118674633 | 0.905533134 | 0.953575703 | 0.0121375 |
| ENSMUSG00000064994 | Gm22422 | 115486004 | snoRNA | 101.5692616 | -0.012150737 | 0.283218032 | 0.042902413 | 0.965779325 | 0.982739526 | 0.012150737 |
| ENSMUSG00000042190 | Cmklr1 | 14747 | protein_coding | 1.709466073 | -0.012170081 | 1.700768002 | 0.007155639 | 0.994290675 | 1 | 0.012170081 |
| ENSMUSG00000037149 | Ddx1 | 104721 | protein_coding | 1343.195508 | 0.012190208 | 0.057450511 | -0.212186236 | 0.831961744 | 0.915295615 | 0.012190208 |
| ENSMUSG00000043931 | Gimap7 | 231932 | protein_coding | 7.158845744 | 0.012199285 | 0.515009193 | -0.023687509 | 0.98110187 | 0.989913623 | 0.012199285 |
| ENSMUSG00000051498 | Tlr6 | 21899 | protein_coding | 12.05174064 | -0.012245484 | 0.34148159 | 0.035859867 | 0.971394097 | 0.985415068 | 0.012245484 |
| ENSMUSG00000022529 | Zfp263 | 74120 | protein_coding | 367.6305684 | -0.012253144 | 0.071682011 | 0.170937508 | 0.864272903 | 0.932569867 | 0.012253144 |
| ENSMUSG00000028559 | Osbpl9 | 100273 | protein_coding | 822.1042299 | -0.012255649 | 0.073043506 | 0.167785602 | 0.86675195 | 0.933837244 | 0.012255649 |
| ENSMUSG00000089824 | Rbm12 | 75710 | protein_coding | 139.0799931 | -0.012285136 | 0.106389741 | 0.115472937 | 0.90807027 | 0.955211953 | 0.012285136 |
| ENSMUSG00000010517 | Faf1 | 14084 | protein_coding | 786.1381523 | 0.012285373 | 0.084013692 | -0.14623061 | 0.883739342 | 0.942424291 | 0.012285373 |
| ENSMUSG00000026064 | Ptp4a1 | 19243 | protein_coding | 206.3697609 | 0.012285901 | 0.116803825 | -0.105184067 | 0.916229754 | 0.959151705 | 0.012285901 |
| ENSMUSG00000021254 | Gpatch2l | 70373 | protein_coding | 256.5887878 | 0.012289539 | 0.094252349 | -0.130389737 | 0.896258086 | 0.948996349 | 0.012289539 |
| ENSMUSG00000020307 | Cdc34 | 216150 | protein_coding | 568.2422057 | 0.012289872 | 0.086357376 | -0.142314092 | 0.886831916 | 0.94399288 | 0.012289872 |
| ENSMUSG00000022330 | Osr2 | 107587 | protein_coding | 455.5944004 | -0.01231549 | 0.312552673 | 0.039402928 | 0.968569145 | 0.984122985 | 0.01231549 |
| ENSMUSG00000060397 | Zfp128 | 243833 | protein_coding | 21.12053501 | 0.012332229 | 0.228685995 | -0.053926474 | 0.956993744 | 0.978805568 | 0.012332229 |
| ENSMUSG00000036617 | Etl4 | 208618 | protein_coding | 1473.072987 | 0.012355725 | 0.129880794 | -0.095131269 | 0.924210562 | 0.963320302 | 0.012355725 |
| ENSMUSG00000000787 | Ddx3x | 13205 | protein_coding | 4356.106278 | -0.012388067 | 0.084942991 | 0.145839777 | 0.884047874 | 0.942489774 | 0.012388067 |
| ENSMUSG00000059897 | Zfp930 | 234358 | protein_coding | 106.4358003 | -0.012420403 | 0.154181496 | 0.080557026 | 0.935794244 | 0.96968875 | 0.012420403 |
| ENSMUSG00000030551 | Nr2f2 | 11819 | protein_coding | 131.1809016 | 0.012433848 | 0.129672212 | -0.09588676 | 0.92361051 | 0.962971397 | 0.012433848 |
| ENSMUSG00000038695 | Josd2 | 66124 | protein_coding | 227.921247 | 0.012437641 | 0.104240567 | -0.119316702 | 0.905024452 | 0.953222387 | 0.012437641 |
| ENSMUSG00000041765 | Ubac2 | 68889 | protein_coding | 418.1486938 | -0.012440761 | 0.054298002 | 0.229120055 | 0.818775607 | 0.908812719 | 0.012440761 |
| ENSMUSG00000044678 | Ly6k | 76486 | protein_coding | 0.767144529 | 0.01245404 | 1.465317058 | -0.008499212 | 0.993218692 | 1 | 0.01245404 |
| ENSMUSG00000032899 | Styk1 | 243659 | protein_coding | 1483.075253 | 0.012461972 | 0.097487478 | -0.127831518 | 0.898282307 | 0.949998977 | 0.012461972 |
| ENSMUSG00000020926 | Adam11 | 11488 | protein_coding | 4.708587363 | 0.012480511 | 0.931843851 | -0.013393351 | 0.989313972 | 0.99400894 | 0.012480511 |
| ENSMUSG00000026596 | Abl2 | 11352 | protein_coding | 346.3525722 | 0.012493361 | 0.086667301 | -0.144153105 | 0.885379571 | 0.943195324 | 0.012493361 |
| ENSMUSG00000056214 | Pard6g | 93737 | protein_coding | 11.91660895 | -0.012585939 | 0.291700957 | 0.043146718 | 0.965584578 | 0.982706396 | 0.012585939 |
| ENSMUSG00000055371 | Stam2 | 56324 | protein_coding | 952.5603293 | 0.012586483 | 0.065965136 | -0.190805075 | 0.848678312 | 0.925637253 | 0.012586483 |
| ENSMUSG00000021870 | Slmap | 83997 | protein_coding | 1221.459757 | 0.012598253 | 0.069559118 | -0.181115767 | 0.856276709 | 0.930174321 | 0.012598253 |
| ENSMUSG00000025326 | Ube3a | 22215 | protein_coding | 760.263516 | 0.012635455 | 0.109701051 | -0.115180797 | 0.908301819 | 0.955333816 | 0.012635455 |
| ENSMUSG00000026812 | Tsc1 | 64930 | protein_coding | 442.0451754 | 0.012660113 | 0.068542724 | -0.184703967 | 0.853461235 | 0.92805494 | 0.012660113 |
| ENSMUSG00000027472 | Pdrg1 | 68559 | protein_coding | 211.0248283 | -0.01267436 | 0.088781966 | 0.142758268 | 0.886481097 | 0.943865173 | 0.01267436 |
| ENSMUSG00000040410 | Fbxl4 | 269514 | protein_coding | 173.1393668 | 0.012677035 | 0.114644179 | -0.110577225 | 0.911951608 | 0.957100023 | 0.012677035 |

| ensembl_gene_id | external_gene_name | entrezgene_id | gene_biotype | baseMean | log2FoldChange | lfcSE | stat | pvalue | padj | absLog2FC |
|---|---|---|---|---|---|---|---|---|---|---|
| ENSMUSG00000042133 | Ppig | 228005 | protein_coding | 563.1150185 | 0.0126772 | 0.06633441 | -0.191110467 | 0.848439047 | 0.925637253 | 0.0126772 |
| ENSMUSG00000033684 | Qsox1 | 104009 | protein_coding | 1499.281415 | 0.012688605 | 0.211503345 | -0.059992456 | 0.952161643 | 0.976016991 | 0.012688605 |
| ENSMUSG00000022200 | Golph3 | 66629 | protein_coding | 2056.407873 | 0.012702154 | 0.074065441 | -0.171499065 | 0.863831366 | 0.932373279 | 0.012702154 |
| ENSMUSG00000097690 | 4930505N22Rik | 100174921 | lncRNA | 3.853838291 | -0.012707235 | 0.615118635 | 0.020658185 | 0.983518325 | 1 | 0.012707235 |
| ENSMUSG00000031553 | Adam3 | 11497 | protein_coding | 1.634193641 | 0.012755832 | 2.126965908 | -0.005997196 | 0.995214958 | 1 | 0.012755832 |
| ENSMUSG00000022858 | Tra2b | 20462 | protein_coding | 1032.782985 | 0.012803723 | 0.084652942 | -0.151249595 | 0.879778829 | 0.940676072 | 0.012803723 |
| ENSMUSG00000039068 | Zzz3 | 108946 | protein_coding | 709.3471847 | -0.012832101 | 0.081304141 | 0.157828387 | 0.87459203 | 0.938406345 | 0.012832101 |
| ENSMUSG00000032300 | 1700017B05Rik | 74211 | protein_coding | 139.0539572 | -0.012899634 | 0.158364378 | 0.081455399 | 0.935079793 | 0.9694819 | 0.012899634 |
| ENSMUSG00000017119 | Nbr1 | 17966 | protein_coding | 2987.962052 | -0.012946497 | 0.049682337 | 0.260585519 | 0.794412158 | 0.897695523 | 0.012946497 |
| ENSMUSG00000018040 | Rrp7a | 74778 | protein_coding | 589.8311815 | 0.012998938 | 0.075241697 | -0.172762429 | 0.86283817 | 0.932166328 | 0.012998938 |
| ENSMUSG00000029684 | Wasl | 73178 | protein_coding | 821.3664904 | 0.013017536 | 0.074492075 | -0.174750615 | 0.861275595 | 0.931649419 | 0.013017536 |
| ENSMUSG00000041836 | Ptpre | 19267 | protein_coding | 151.7560572 | -0.013017718 | 0.107805548 | 0.120751837 | 0.9038876 | 0.952450218 | 0.013017718 |
| ENSMUSG00000079554 | Aox2 | 213043 | protein_coding | 9.45613553 | 0.013034423 | 0.631550621 | -0.020638762 | 0.98353382 | 0.990974832 | 0.013034423 |
| ENSMUSG00000044477 | Zfand3 | 21769 | protein_coding | 791.2260901 | -0.013047082 | 0.074668159 | 0.174734219 | 0.861288478 | 0.931649419 | 0.013047082 |
| ENSMUSG00000032905 | Atg12 | 67526 | protein_coding | 530.130965 | -0.013093908 | 0.06948659 | 0.188437906 | 0.850533387 | 0.927048271 | 0.013093908 |
| ENSMUSG00000020752 | Recql5 | 170472 | protein_coding | 596.0024789 | 0.013163109 | 0.142326614 | -0.092485224 | 0.926312531 | 0.964753099 | 0.013163109 |
| ENSMUSG00000026088 | Mitd1 | 69028 | protein_coding | 180.1034309 | 0.013199815 | 0.095276131 | -0.138542724 | 0.889811506 | 0.945840699 | 0.013199815 |
| ENSMUSG00000100017 | 2410022M11Rik | 100503842 | lncRNA | 38.13878163 | 0.013225859 | 0.220483875 | -0.059985605 | 0.9521671 | 0.976016991 | 0.013225859 |
| ENSMUSG00000025393 | Atp5b | 11947 | protein_coding | 25849.47995 | 0.013252596 | 0.074599926 | -0.177648917 | 0.858998706 | 0.931099354 | 0.013252596 |
| ENSMUSG00000030846 | Tial1 | 21843 | protein_coding | 430.7337551 | -0.01325432 | 0.085885169 | 0.154326059 | 0.877352651 | 0.939157889 | 0.01325432 |
| ENSMUSG00000062619 | 2310039H08Rik | 67101 | protein_coding | 220.96019 | -0.013260125 | 0.085294635 | 0.155462594 | 0.876456641 | 0.938630585 | 0.013260125 |
| ENSMUSG00000091896 | Ube2d2a | 56550 | protein_coding | 459.5165296 | -0.013285755 | 0.073475196 | 0.180819588 | 0.856509188 | 0.930184975 | 0.013285755 |
| ENSMUSG00000028894 | Inpp5b | 16330 | protein_coding | 345.7042876 | 0.013336644 | 0.071128728 | -0.18750011 | 0.851268538 | 0.927421073 | 0.013336644 |
| ENSMUSG00000055128 | Cgrrf1 | 68755 | protein_coding | 193.0279761 | 0.013391021 | 0.086161874 | -0.155417015 | 0.876492571 | 0.938630585 | 0.013391021 |
| ENSMUSG00000033808 | Tmem87a | 211499 | protein_coding | 865.3422952 | -0.013419428 | 0.087306592 | 0.153704636 | 0.877842629 | 0.9393553 | 0.013419428 |
| ENSMUSG00000016308 | Ube2a | 22209 | protein_coding | 398.5669924 | 0.013509571 | 0.087719428 | -0.15400888 | 0.877602734 | 0.93927028 | 0.013509571 |
| ENSMUSG00000034361 | Cpne2 | 234577 | protein_coding | 282.6125162 | -0.013582262 | 0.114006051 | 0.119136322 | 0.905167355 | 0.953251304 | 0.013582262 |
| ENSMUSG00000064030 | Pym1 | 78428 | protein_coding | 186.0286792 | 0.013583248 | 0.080338828 | -0.169074516 | 0.865738029 | 0.933333191 | 0.013583248 |
| ENSMUSG00000021457 | Syk | 20963 | protein_coding | 1050.287954 | -0.013589613 | 0.119692243 | 0.113537962 | 0.909604069 | 0.956044724 | 0.013589613 |
| ENSMUSG00000027996 | Sfrp2 | 20319 | protein_coding | 0.840895792 | 0.013612184 | 1.858973685 | -0.007322419 | 0.994157607 | 1 | 0.013612184 |
| ENSMUSG00000055239 | Kcmf1 | 74287 | protein_coding | 648.7647878 | 0.013754272 | 0.060924393 | -0.225759692 | 0.821388329 | 0.909752305 | 0.013754272 |
| ENSMUSG00000045095 | Magi1 | 14924 | protein_coding | 1030.148274 | -0.013818769 | 0.06912502 | 0.199909803 | 0.841551124 | 0.92158767 | 0.013818769 |
| ENSMUSG00000031095 | Cul4b | 72584 | protein_coding | 326.7166549 | -0.013836281 | 0.077968468 | 0.177459963 | 0.859147111 | 0.931099354 | 0.013836281 |
| ENSMUSG00000049396 | Gemin4 | 276919 | protein_coding | 8.281882771 | 0.013874443 | 0.390506081 | -0.035529391 | 0.97165761 | 0.985442104 | 0.013874443 |
| ENSMUSG00000006673 | Qrich1 | 69232 | protein_coding | 1073.185919 | -0.013875186 | 0.046583419 | 0.297856758 | 0.765812493 | 0.882651278 | 0.013875186 |
| ENSMUSG00000073079 | Srp54a | 24067 | protein_coding | 59.59895515 | -0.013947643 | 0.20499298 | 0.068039613 | 0.9457541 | 0.973795719 | 0.013947643 |
| ENSMUSG00000038459 | Abhd17c | 70178 | protein_coding | 1220.519893 | 0.013952095 | 0.109614167 | -0.127283686 | 0.898715872 | 0.950335782 | 0.013952095 |
| ENSMUSG00000024571 | Naa12 | 27366 | protein_coding | 515.9380068 | 0.013962667 | 0.094421859 | -0.147875364 | 0.882441131 | 0.941973978 | 0.013962667 |
| ENSMUSG00000025939 | Ube2w | 66799 | protein_coding | 244.1483062 | 0.013963667 | 0.1065207 | -0.131088769 | 0.895705086 | 0.948731562 | 0.013963667 |

| ensembl_gene_id | external_gene_name | entrezgene_id | gene_biotype | baseMean | log2FoldChange | lfcSE | stat | pvalue | padj | absLog2FC |
|---|---|---|---|---|---|---|---|---|---|---|
| ENSMUSG00000020694 | Tlk2 | 24086 | protein_coding | 430.3904274 | -0.013983682 | 0.058183834 | 0.240336204 | 0.810069631 | 0.906149826 | 0.013983682 |
| ENSMUSG00000042726 | Trafd1 | 231712 | protein_coding | 404.7229248 | -0.014039949 | 0.123167974 | 0.113990257 | 0.909245517 | 0.955778515 | 0.014039949 |
| ENSMUSG00000019883 | Echdc1 | 52665 | protein_coding | 175.1063838 | -0.014075545 | 0.10377942 | 0.135629448 | 0.892114224 | 0.947202684 | 0.014075545 |
| ENSMUSG00000005501 | Usp40 | 227334 | protein_coding | 397.6856995 | 0.014112116 | 0.070904043 | -0.199031188 | 0.842238348 | 0.921900136 | 0.014112116 |
| ENSMUSG00000026755 | Arpc5l | 74192 | protein_coding | 433.6306594 | -0.014123324 | 0.09070889 | 0.155699443 | 0.876269937 | 0.938630585 | 0.014123324 |
| ENSMUSG00000030774 | Pak1 | 18479 | protein_coding | 957.8264183 | -0.014132872 | 0.062206055 | 0.227194481 | 0.820272519 | 0.9092783 | 0.014132872 |
| ENSMUSG00000021552 | Gkap1 | 56278 | protein_coding | 26.73281413 | -0.014137623 | 0.233818265 | 0.060464154 | 0.951785965 | 0.976016991 | 0.014137623 |
| ENSMUSG00000019689 | Fmc1 | 66117 | protein_coding | 290.3046189 | 0.014164972 | 0.116834615 | -0.121239515 | 0.903501327 | 0.952286247 | 0.014164972 |
| ENSMUSG00000030374 | Strn4 | 97387 | protein_coding | 1084.00476 | 0.014191447 | 0.050767558 | -0.279537704 | 0.779832207 | 0.890125936 | 0.014191447 |
| ENSMUSG00000024493 | Lars | 107045 | protein_coding | 674.5675802 | 0.014281483 | 0.113030251 | -0.126350978 | 0.899454105 | 0.950621521 | 0.014281483 |
| ENSMUSG00000079022 | Col22a1 | 69700 | protein_coding | 5.079403721 | 0.014340056 | 1.879475145 | -0.007629819 | 0.993912344 | 0.996446603 | 0.014340056 |
| ENSMUSG00000027367 | Stard7 | 99138 | protein_coding | 1939.735597 | 0.014413072 | 0.076143005 | -0.189289501 | 0.849865923 | 0.926810134 | 0.014413072 |
| ENSMUSG00000027598 | Itch | 16396 | protein_coding | 1421.355536 | 0.014443171 | 0.077215552 | -0.187050031 | 0.851621406 | 0.927499565 | 0.014443171 |
| ENSMUSG00000027519 | Rab22a | 19334 | protein_coding | 770.6433578 | -0.014465097 | 0.073107691 | 0.197860123 | 0.843154505 | 0.922414938 | 0.014465097 |
| ENSMUSG00000021619 | Atg10 | 66795 | protein_coding | 45.91046229 | -0.01451792 | 0.16012536 | 0.090665965 | 0.927758015 | 0.965588697 | 0.01451792 |
| ENSMUSG00000024962 | Vegfb | 22340 | protein_coding | 85.51200619 | -0.014519983 | 0.158631036 | 0.091533056 | 0.927069042 | 0.965151432 | 0.014519983 |
| ENSMUSG00000018102 | H2bc4 | 68024 | protein_coding | 2182.446874 | -0.014543961 | 0.071015643 | 0.204799399 | 0.837728856 | 0.919158483 | 0.014543961 |
| ENSMUSG00000034308 | Sdr42e1 | 74032 | protein_coding | 550.8229917 | 0.014545815 | 0.14192852 | -0.10248691 | 0.918370203 | 0.960053524 | 0.014545815 |
| ENSMUSG00000036845 | Lin37 | 75660 | protein_coding | 123.5907519 | 0.01456435 | 0.109648948 | -0.132827081 | 0.894330136 | 0.947822351 | 0.01456435 |
| ENSMUSG00000030598 | Fbxo17 | 50760 | protein_coding | 13.75992051 | 0.014603176 | 0.306140893 | -0.047700833 | 0.96195467 | 0.981387498 | 0.014603176 |
| ENSMUSG00000014932 | Yes1 | 22612 | protein_coding | 786.8443246 | -0.014672519 | 0.099298981 | 0.147761019 | 0.882531374 | 0.941973978 | 0.014672519 |
| ENSMUSG00000000581 | C1d | 57316 | protein_coding | 392.0780266 | -0.01468401 | 0.08246343 | 0.178066932 | 0.858670411 | 0.930879221 | 0.01468401 |
| ENSMUSG00000027952 | Pmvk | 68603 | protein_coding | 307.2651352 | 0.014732778 | 0.079649511 | -0.184970105 | 0.853252484 | 0.927990748 | 0.014732778 |
| ENSMUSG00000024614 | Tmx3 | 67988 | protein_coding | 529.6782414 | -0.014736298 | 0.092153057 | 0.159911114 | 0.872951094 | 0.937394946 | 0.014736298 |
| ENSMUSG00000029415 | Sdad1 | 231452 | protein_coding | 252.004233 | -0.014736683 | 0.118671825 | 0.124180133 | 0.901172652 | 0.95132126 | 0.014736683 |
| ENSMUSG00000072692 | Rpl37rt | 100502825 | transcribed_processed_pseudogene | 90.86948823 | 0.014792855 | 0.151706788 | -0.097509514 | 0.922321779 | 0.962478474 | 0.014792855 |
| ENSMUSG00000029708 | Gcc1 | 74375 | protein_coding | 363.6904521 | -0.014806022 | 0.068164452 | 0.217210309 | 0.828044451 | 0.913297021 | 0.014806022 |
| ENSMUSG00000023104 | Rfc2 | 19718 | protein_coding | 437.7878416 | 0.014810812 | 0.10594542 | -0.13979662 | 0.888820684 | 0.945259194 | 0.014810812 |
| ENSMUSG00000070777 | Ceacam20 | 71601 | protein_coding | 1288.142015 | -0.014827299 | 0.236243253 | 0.062762846 | 0.949955352 | 0.975869522 | 0.014827299 |
| ENSMUSG00000001383 | Zmat2 | 66492 | protein_coding | 516.7085864 | 0.014834619 | 0.050359418 | -0.29457488 | 0.768318654 | 0.883975099 | 0.014834619 |
| ENSMUSG00000039849 | Pcif1 | 228866 | protein_coding | 545.2520912 | 0.014836494 | 0.055893215 | -0.265443556 | 0.790667786 | 0.895609289 | 0.014836494 |
| ENSMUSG00000038880 | Mrps34 | 79044 | protein_coding | 147.7618195 | -0.014846338 | 0.129701509 | 0.114465423 | 0.908868854 | 0.955501531 | 0.014846338 |
| ENSMUSG00000030400 | Ercc2 | 13871 | protein_coding | 245.2604769 | 0.014880085 | 0.076253902 | -0.195138669 | 0.845284391 | 0.923801556 | 0.014880085 |
| ENSMUSG00000022557 | Bop1 | 12181 | protein_coding | 559.7687859 | 0.014929909 | 0.084142868 | -0.177435226 | 0.85916654 | 0.931099354 | 0.014929909 |
| ENSMUSG00000019808 | Adat2 | 66757 | protein_coding | 45.17246423 | 0.014932681 | 0.177326103 | -0.084210279 | 0.932889246 | 0.968109605 | 0.014932681 |
| ENSMUSG00000046727 | Cystm1 | 66060 | protein_coding | 3074.971537 | 0.015085864 | 0.105906795 | -0.142444722 | 0.88672874 | 0.94399288 | 0.015085864 |
| ENSMUSG00000033083 | Tbc1d4 | 210789 | protein_coding | 365.9932362 | 0.015131838 | 0.098529916 | -0.153576077 | 0.877944001 | 0.939378465 | 0.015131838 |
| ENSMUSG00000024844 | Banf1 | 23825 | protein_coding | 586.3792076 | -0.015141907 | 0.092258262 | 0.164125216 | 0.869632572 | 0.935638301 | 0.015141907 |
| ENSMUSG00000054648 | Zfp869 | 66869 | protein_coding | 385.935471 | 0.01514961 | 0.071991935 | -0.210434818 | 0.83332832 | 0.91603434 | 0.01514961 |

| ensembl_gene_id | external_gene_name | entrezgene_id | gene_biotype | baseMean | log2FoldChange | lfcSE | stat | pvalue | padj | absLog2FC |
|---|---|---|---|---|---|---|---|---|---|---|
| ENSMUSG00000092305 | Prps1l1 | 75456 | protein_coding | 2.055605934 | 0.015155022 | 1.466906111 | -0.010331283 | 0.991756975 | 1 | 0.015155022 |
| ENSMUSG00000029617 | Ccz1 | 231874 | protein_coding | 770.9742414 | 0.015159948 | 0.089328487 | -0.16971012 | 0.865238115 | 0.932932188 | 0.015159948 |
| ENSMUSG00000027365 | Trpm7 | 58800 | protein_coding | 1008.154812 | -0.015165258 | 0.086423878 | 0.17547532 | 0.860706162 | 0.931474719 | 0.015165258 |
| ENSMUSG00000054523 | Ms4a5 | 269063 | protein_coding | 5.535614438 | -0.015166508 | 0.440895272 | 0.034399343 | 0.972558708 | 0.985703995 | 0.015166508 |
| ENSMUSG00000025784 | Clec3b | 21922 | protein_coding | 1.293241095 | -0.015179713 | 1.630025525 | 0.009312562 | 0.992569758 | 1 | 0.015179713 |
| ENSMUSG00000064302 | Clasp1 | 76707 | protein_coding | 563.8180019 | 0.015218581 | 0.079743313 | -0.190844608 | 0.848647339 | 0.925637253 | 0.015218581 |
| ENSMUSG00000087128 | Gm12655 | 115489886 | lncRNA | 3.88615255 | -0.015221604 | 0.658678118 | 0.023109322 | 0.98156307 | 1 | 0.015221604 |
| ENSMUSG00000029097 | Trmt44 | 78890 | protein_coding | 120.2870202 | 0.015258087 | 0.137018385 | -0.111357954 | 0.911332501 | 0.956997404 | 0.015258087 |
| ENSMUSG00000066839 | Ecsit | 26940 | protein_coding | 327.3597475 | -0.015286207 | 0.097942215 | 0.15607373 | 0.875974905 | 0.938630585 | 0.015286207 |
| ENSMUSG00000025283 | Sat1 | 20229 | protein_coding | 2202.827664 | -0.015318237 | 0.097553574 | 0.157023844 | 0.875226057 | 0.938415991 | 0.015318237 |
| ENSMUSG00000005148 | Klf5 | 12224 | protein_coding | 4320.015984 | -0.015324527 | 0.048258285 | 0.317552249 | 0.7508246 | 0.875049502 | 0.015324527 |
| ENSMUSG00000022234 | Cct5 | 12465 | protein_coding | 2802.304816 | 0.015347276 | 0.089345161 | -0.171775123 | 0.863614323 | 0.932314788 | 0.015347276 |
| ENSMUSG00000028986 | Klhl7 | 52323 | protein_coding | 220.0475109 | 0.015357832 | 0.131214383 | -0.117043816 | 0.906825332 | 0.954353007 | 0.015357832 |
| ENSMUSG00000042148 | Cox10 | 70383 | protein_coding | 534.0203504 | -0.015359128 | 0.073220532 | 0.20976532 | 0.833850841 | 0.916410806 | 0.015359128 |
| ENSMUSG00000018378 | Cuedc1 | 70393 | protein_coding | 69.36655545 | 0.015372014 | 0.164530233 | -0.093429721 | 0.92556218 | 0.964363534 | 0.015372014 |
| ENSMUSG00000054079 | Utp18 | 217109 | protein_coding | 225.8077873 | 0.015376197 | 0.106695733 | -0.14411258 | 0.885411571 | 0.943195324 | 0.015376197 |
| ENSMUSG00000022150 | Dab2 | 13132 | protein_coding | 7.416907053 | 0.015389577 | 0.795988958 | -0.019333908 | 0.984574735 | 0.99157486 | 0.015389577 |
| ENSMUSG00000038914 | Dido1 | 23856 | protein_coding | 1143.004386 | -0.015403445 | 0.069941211 | 0.220234183 | 0.825688776 | 0.91190529 | 0.015403445 |
| ENSMUSG00000036918 | Ttc7 | 225049 | protein_coding | 1233.867092 | 0.015457046 | 0.058994226 | -0.262009475 | 0.793314137 | 0.897130018 | 0.015457046 |
| ENSMUSG00000030397 | Mark4 | 232944 | protein_coding | 257.6057516 | -0.01549284 | 0.105302541 | 0.147126934 | 0.883031831 | 0.942185661 | 0.01549284 |
| ENSMUSG00000026417 | Pigr | 18703 | protein_coding | 52021.40112 | -0.015534351 | 0.115057928 | 0.135013303 | 0.892601356 | 0.947202684 | 0.015534351 |
| ENSMUSG00000027276 | Jag1 | 16449 | protein_coding | 327.2789961 | 0.015616142 | 0.10502873 | -0.148684479 | 0.881802609 | 0.941920329 | 0.015616142 |
| ENSMUSG00000042745 | Id1 | 15901 | protein_coding | 3257.893242 | -0.01564499 | 0.125834503 | 0.124329887 | 0.901054084 | 0.95132126 | 0.01564499 |
| ENSMUSG00000024902 | Mrpl11 | 66419 | protein_coding | 350.2820424 | -0.015655312 | 0.070003031 | 0.223637635 | 0.823039277 | 0.910297568 | 0.015655312 |
| ENSMUSG00000021711 | Trappc13 | 66975 | protein_coding | 470.5956646 | 0.015675958 | 0.071900792 | -0.218022046 | 0.827411934 | 0.913026208 | 0.015675958 |
| ENSMUSG00000046079 | Lrrc8d | 231549 | protein_coding | 372.7656508 | 0.015697787 | 0.092455024 | -0.16978836 | 0.865176581 | 0.932932188 | 0.015697787 |
| ENSMUSG00000021377 | Dek | 110052 | protein_coding | 615.1975044 | -0.015716336 | 0.147059272 | 0.10687076 | 0.914891511 | 0.958296749 | 0.015716336 |
| ENSMUSG00000034673 | Pbx2 | 18515 | protein_coding | 436.350144 | -0.015733789 | 0.085047079 | 0.185000935 | 0.853228303 | 0.927990748 | 0.015733789 |
| ENSMUSG00000038705 | Gmeb2 | 229004 | protein_coding | 373.6077898 | -0.01573431 | 0.068564225 | 0.2294828 | 0.818493687 | 0.908812719 | 0.01573431 |
| ENSMUSG00000017718 | Afmid | 71562 | protein_coding | 69.168046 | 0.015741816 | 0.170769954 | -0.092181414 | 0.926553905 | 0.964893154 | 0.015741816 |
| ENSMUSG00000047260 | Emc6 | 66048 | protein_coding | 501.5145056 | 0.015804618 | 0.113613772 | -0.139108298 | 0.889364571 | 0.945659633 | 0.015804618 |
| ENSMUSG00000046947 | Adck2 | 57869 | protein_coding | 234.9679623 | -0.015848172 | 0.086156162 | 0.183947054 | 0.854054991 | 0.928522799 | 0.015848172 |
| ENSMUSG00000055817 | Mta3 | 116871 | protein_coding | 396.3511064 | 0.015853673 | 0.061655239 | -0.25713424 | 0.797075147 | 0.899227966 | 0.015853673 |
| ENSMUSG00000019917 | Septin10 | 103080 | protein_coding | 194.9070346 | 0.015875536 | 0.144033349 | -0.11022125 | 0.91223391 | 0.957266079 | 0.015875536 |
| ENSMUSG00000011658 | Fuz | 70300 | protein_coding | 68.18011315 | -0.015893478 | 0.149390959 | 0.10638849 | 0.915274125 | 0.95833356 | 0.015893478 |
| ENSMUSG00000042997 | Nhlrc3 | 212114 | protein_coding | 847.2497336 | -0.015936325 | 0.082619907 | 0.192887225 | 0.84704729 | 0.924713357 | 0.015936325 |
| ENSMUSG00000032584 | Mst1r | 19882 | protein_coding | 849.2034151 | -0.015939615 | 0.092827827 | 0.171711604 | 0.863664262 | 0.932314788 | 0.015939615 |
| ENSMUSG00000039308 | Ndst2 | 17423 | protein_coding | 252.697605 | 0.01594823 | 0.065715993 | -0.242684146 | 0.80825009 | 0.905463808 | 0.01594823 |
| ENSMUSG00000000530 | Acvrl1 | 11482 | protein_coding | 74.4878905 | 0.015958066 | 0.169509624 | -0.094142535 | 0.924995932 | 0.964050667 | 0.015958066 |

| ensembl_gene_id | external_gene_name | entrezgene_id | gene_biotype | baseMean | log2FoldChange | lfcSE | stat | pvalue | padj | absLog2FC |
|---|---|---|---|---|---|---|---|---|---|---|
| ENSMUSG00000046070 | Igfals | 16005 | protein_coding | 21.33285162 | 0.015989234 | 0.263462459 | -0.06068885 | 0.951607012 | 0.976016284 | 0.015989234 |
| ENSMUSG00000038822 | Hace1 | 209462 | protein_coding | 229.4684836 | -0.015998138 | 0.07974938 | 0.200605166 | 0.841007318 | 0.92126689 | 0.015998138 |
| ENSMUSG00000029127 | Zbtb49 | 75079 | protein_coding | 76.97864907 | 0.016035188 | 0.119023299 | -0.134723103 | 0.892830806 | 0.947205958 | 0.016035188 |
| ENSMUSG00000037070 | Rbmxl1 | 19656 | protein_coding | 127.2006671 | -0.016068035 | 0.155981471 | 0.103012459 | 0.917953083 | 0.959738982 | 0.016068035 |
| ENSMUSG00000001419 | Mef2d | 17261 | protein_coding | 734.676898 | 0.016068125 | 0.133954342 | -0.119952252 | 0.904520972 | 0.952874415 | 0.016068125 |
| ENSMUSG00000019774 | Mtrf1l | 108853 | protein_coding | 106.7648344 | 0.016076243 | 0.109187097 | -0.147235741 | 0.882945951 | 0.942166544 | 0.016076243 |
| ENSMUSG00000027905 | Ddx20 | 53975 | protein_coding | 191.7496645 | -0.016096504 | 0.082390185 | 0.1953692 | 0.845103927 | 0.923798341 | 0.016096504 |
| ENSMUSG00000022340 | Sybu | 319613 | protein_coding | 123.8591174 | 0.016151706 | 0.189061708 | -0.085430871 | 0.931918851 | 0.967478151 | 0.016151706 |
| ENSMUSG00000030534 | Vps33b | 233405 | protein_coding | 232.5759556 | -0.016195859 | 0.079764406 | 0.203046193 | 0.839098927 | 0.919794636 | 0.016195859 |
| ENSMUSG00000030303 | Far2 | 330450 | protein_coding | 148.3282458 | -0.016215803 | 0.149992939 | 0.108110444 | 0.913908084 | 0.957979661 | 0.016215803 |
| ENSMUSG00000033577 | Myo6 | 17920 | protein_coding | 3929.935051 | 0.016218638 | 0.069797414 | -0.232367326 | 0.816252725 | 0.908409285 | 0.016218638 |
| ENSMUSG00000022353 | Mtss1 | 211401 | protein_coding | 685.0649789 | 0.01623178 | 0.067408981 | -0.240795511 | 0.809713609 | 0.905997055 | 0.01623178 |
| ENSMUSG00000020134 | Peli1 | 67245 | protein_coding | 658.0306252 | -0.016258591 | 0.073786493 | 0.22034644 | 0.825601355 | 0.91190529 | 0.016258591 |
| ENSMUSG00000024673 | Ms4a1 | 12482 | protein_coding | 4.650119365 | 0.016265968 | 0.65330297 | -0.024898048 | 0.980136285 | 0.989302036 | 0.016265968 |
| ENSMUSG00000041130 | Zfp598 | 213753 | protein_coding | 901.933276 | 0.016305221 | 0.079363855 | -0.205448954 | 0.837221376 | 0.91884587 | 0.016305221 |
| ENSMUSG00000032349 | Elovl5 | 68801 | protein_coding | 593.1708763 | -0.016305238 | 0.123050449 | 0.132508562 | 0.894582051 | 0.947906833 | 0.016305238 |
| ENSMUSG00000022801 | Lrch3 | 70144 | protein_coding | 269.49943 | -0.016309124 | 0.08826239 | 0.18477999 | 0.853401603 | 0.92805494 | 0.016309124 |
| ENSMUSG00000040747 | Cd53 | 12508 | protein_coding | 11.60564032 | 0.016344329 | 0.37751363 | -0.043294673 | 0.965466637 | 0.982706396 | 0.016344329 |
| ENSMUSG00000028975 | Pex14 | 56273 | protein_coding | 311.2471487 | -0.016366453 | 0.078329923 | 0.208942544 | 0.834493092 | 0.916826487 | 0.016366453 |
| ENSMUSG00000026074 | Map4k4 | 26921 | protein_coding | 253.0988704 | 0.016367473 | 0.093374244 | -0.175288945 | 0.860852598 | 0.931532652 | 0.016367473 |
| ENSMUSG00000066306 | Numa1 | 101706 | protein_coding | 2651.612112 | -0.016376356 | 0.078802688 | 0.20781469 | 0.835373662 | 0.917244724 | 0.016376356 |
| ENSMUSG00000078234 | Klhdc7a | 242721 | protein_coding | 228.0811618 | 0.016376416 | 0.233308227 | -0.070192191 | 0.94404069 | 0.973044134 | 0.016376416 |
| ENSMUSG00000014956 | Ppp1cb | 19046 | protein_coding | 3474.045682 | 0.016377351 | 0.07220153 | -0.226828307 | 0.820557251 | 0.9092783 | 0.016377351 |
| ENSMUSG00000029275 | Gfi1 | 14581 | protein_coding | 108.567098 | 0.016519843 | 0.190589083 | -0.086677804 | 0.93092762 | 0.966815843 | 0.016519843 |
| ENSMUSG00000044475 | Ascc1 | 69090 | protein_coding | 234.8795043 | -0.016538155 | 0.132272147 | 0.125031273 | 0.900498793 | 0.951064023 | 0.016538155 |
| ENSMUSG00000024028 | Tff2 | 21785 | protein_coding | 6.892245893 | 0.016538551 | 0.618339691 | -0.026746708 | 0.978661759 | 0.988831409 | 0.016538551 |
| ENSMUSG00000025050 | Pcgf6 | 71041 | protein_coding | 110.6307327 | 0.016549263 | 0.157973223 | -0.104759922 | 0.916566313 | 0.959274295 | 0.016549263 |
| ENSMUSG00000025041 | Nt5c2 | 76952 | protein_coding | 302.3856909 | 0.016571372 | 0.099092559 | -0.167231249 | 0.867188098 | 0.933981885 | 0.016571372 |
| ENSMUSG00000030127 | Cops7a | 26894 | protein_coding | 886.8610149 | -0.01661746 | 0.073360666 | 0.226517301 | 0.820799105 | 0.909404657 | 0.01661746 |
| ENSMUSG00000045691 | Thtpa | 105663 | protein_coding | 92.86672598 | 0.016628122 | 0.124197443 | -0.133884576 | 0.893493847 | 0.947544143 | 0.016628122 |
| ENSMUSG00000018858 | Mrpl58 | 68572 | protein_coding | 345.4457971 | -0.016651966 | 0.101387829 | 0.164240284 | 0.86954199 | 0.935632034 | 0.016651966 |
| ENSMUSG00000009073 | Nf2 | 18016 | protein_coding | 1370.0355 | 0.016671686 | 0.050009549 | -0.333370049 | 0.738854969 | 0.867566194 | 0.016671686 |
| ENSMUSG00000040407 | Akap9 | 100986 | protein_coding | 3017.535284 | -0.016739074 | 0.093325121 | 0.179363003 | 0.85765268 | 0.930374388 | 0.016739074 |
| ENSMUSG00000028967 | Errfi1 | 74155 | protein_coding | 1778.21864 | -0.01675059 | 0.202943284 | 0.082538281 | 0.934218679 | 0.968832408 | 0.01675059 |
| ENSMUSG00000079509 | Zfx | 22764 | protein_coding | 512.3693457 | 0.016752962 | 0.084770079 | -0.197628248 | 0.843335932 | 0.922491196 | 0.016752962 |
| ENSMUSG00000041278 | Ttc1 | 66827 | protein_coding | 294.5769086 | -0.016761612 | 0.072044769 | 0.232655504 | 0.816028923 | 0.908409285 | 0.016761612 |
| ENSMUSG00000074886 | Grk6 | 26385 | protein_coding | 1027.134303 | -0.016868766 | 0.095448596 | 0.176731425 | 0.859719356 | 0.931236324 | 0.016868766 |
| ENSMUSG00000077677 | Gm24468 | 115490018 | snRNA | 0.775551077 | -0.016884576 | 1.137347828 | 0.01484557 | 0.988155384 | 1 | 0.016884576 |
| ENSMUSG00000022090 | Pdlim2 | 213019 | protein_coding | 339.6611115 | 0.016902929 | 0.277015483 | -0.061017993 | 0.951344879 | 0.975912221 | 0.016902929 |

| ensembl_gene_id | external_gene_name | entrezgene_id | gene_biotype | baseMean | log2FoldChange | lfcSE | stat | pvalue | padj | absLog2FC |
|---|---|---|---|---|---|---|---|---|---|---|
| ENSMUSG00000036275 | 9530068E07Rik | 213673 | protein_coding | 1130.018705 | -0.016948443 | 0.073369458 | 0.231001338 | 0.817313762 | 0.908714429 | 0.016948443 |
| ENSMUSG00000024560 | Cxxc1 | 74322 | protein_coding | 524.1868969 | -0.016992625 | 0.058215614 | 0.291891197 | 0.770369814 | 0.884816185 | 0.016992625 |
| ENSMUSG00000041771 | Slc24a4 | 238384 | protein_coding | 2.001657045 | -0.016996014 | 2.339767042 | 0.007263977 | 0.994204236 | 1 | 0.016996014 |
| ENSMUSG00000022961 | Son | 20658 | protein_coding | 3920.002183 | 0.017121281 | 0.056630915 | -0.30233099 | 0.762399756 | 0.88093126 | 0.017121281 |
| ENSMUSG00000111375 | Btbd8 | 100503185 | protein_coding | 114.6337604 | 0.017168504 | 0.125989292 | -0.136269548 | 0.891608196 | 0.947050968 | 0.017168504 |
| ENSMUSG00000039988 | Ankrd13c | 433667 | protein_coding | 990.4967372 | -0.017232772 | 0.073251394 | 0.235255209 | 0.814010658 | 0.907710402 | 0.017232772 |
| ENSMUSG00000071014 | Ndufb6 | 230075 | protein_coding | 513.9447013 | 0.017300259 | 0.105648525 | -0.163752965 | 0.86992562 | 0.935658236 | 0.017300259 |
| ENSMUSG00000031217 | Efnb1 | 13641 | protein_coding | 1924.033483 | 0.017311088 | 0.095825152 | -0.180652866 | 0.856640058 | 0.930204755 | 0.017311088 |
| ENSMUSG00000035770 | Dync1li2 | 234663 | protein_coding | 1231.386543 | 0.017335951 | 0.082665674 | -0.209711603 | 0.833892769 | 0.916410806 | 0.017335951 |
| ENSMUSG00000012114 | Med15 | 94112 | protein_coding | 569.9550158 | 0.01735429 | 0.096992004 | -0.178924961 | 0.857996623 | 0.930446023 | 0.01735429 |
| ENSMUSG00000074746 | Pdzd8 | 107368 | protein_coding | 1115.291601 | -0.017357557 | 0.066419461 | 0.261332391 | 0.79383619 | 0.897263004 | 0.017357557 |
| ENSMUSG00000031521 | Aga | 11593 | protein_coding | 256.9650657 | -0.017371971 | 0.115393187 | 0.150545902 | 0.880333939 | 0.941010727 | 0.017371971 |
| ENSMUSG00000080797 | Gm15760 | 100042948 | transcribed_processed_pseudogene | 14.57319614 | 0.017403314 | 0.325435838 | -0.053476945 | 0.9573519 | 0.978989985 | 0.017403314 |
| ENSMUSG00000021951 | Eef1akmt1 | 68043 | protein_coding | 149.7932179 | -0.017421873 | 0.106832335 | 0.16307678 | 0.870457982 | 0.935980063 | 0.017421873 |
| ENSMUSG00000021301 | Hecw1 | 94253 | protein_coding | 4.166704329 | 0.01744059 | 1.872543386 | -0.009313851 | 0.992568729 | 1 | 0.01744059 |
| ENSMUSG00000020257 | Wdr82 | 77305 | protein_coding | 978.8072758 | -0.017467607 | 0.060091331 | 0.2906843 | 0.77129278 | 0.885199039 | 0.017467607 |
| ENSMUSG00000022761 | Lztr1 | 66863 | protein_coding | 697.2844069 | -0.017469807 | 0.053220574 | 0.328252889 | 0.742720461 | 0.870063538 | 0.017469807 |
| ENSMUSG00000016253 | Nelfcd | 57314 | protein_coding | 510.594096 | 0.017497389 | 0.066867809 | -0.261671338 | 0.79357484 | 0.897240511 | 0.017497389 |
| ENSMUSG00000009563 | Tor2a | 30933 | protein_coding | 409.6422281 | 0.017519695 | 0.085403924 | -0.20513923 | 0.837463347 | 0.918959391 | 0.017519695 |
| ENSMUSG00000053286 | Trmt1l | 98685 | protein_coding | 387.8784329 | -0.017523958 | 0.058251197 | 0.300834307 | 0.763540846 | 0.881386379 | 0.017523958 |
| ENSMUSG00000097145 | 9230114K14Rik | 414108 | lncRNA | 11.20332696 | -0.017538295 | 0.345842543 | 0.050711793 | 0.959555179 | 0.980447529 | 0.017538295 |
| ENSMUSG00000019809 | Pex3 | 56535 | protein_coding | 179.8465295 | 0.017542137 | 0.116719056 | -0.150293687 | 0.880532914 | 0.941067509 | 0.017542137 |
| ENSMUSG00000027242 | Wdr76 | 241627 | protein_coding | 132.7376792 | -0.017547219 | 0.145002383 | 0.121013316 | 0.903680488 | 0.952353515 | 0.017547219 |
| ENSMUSG00000029038 | Ssu72 | 68991 | protein_coding | 673.8437288 | -0.017556003 | 0.062304543 | 0.281777253 | 0.778114314 | 0.889135053 | 0.017556003 |
| ENSMUSG00000028522 | Mier1 | 71148 | protein_coding | 1673.265418 | -0.017564331 | 0.114111175 | 0.153922968 | 0.877670474 | 0.93927028 | 0.017564331 |
| ENSMUSG00000047731 | Wbp1l | 226178 | protein_coding | 912.2921443 | 0.017657055 | 0.05747504 | -0.307212576 | 0.758681574 | 0.878912418 | 0.017657055 |
| ENSMUSG00000022221 | Ripk3 | 56532 | protein_coding | 218.1419131 | 0.017678988 | 0.134576998 | -0.131367089 | 0.895484922 | 0.948680909 | 0.017678988 |
| ENSMUSG00000041957 | Pkp2 | 67451 | protein_coding | 745.3446082 | -0.017693643 | 0.066099772 | 0.267680843 | 0.788944997 | 0.894394019 | 0.017693643 |
| ENSMUSG00000068917 | Clk2 | 12748 | protein_coding | 272.9013151 | -0.017700546 | 0.088671553 | 0.199619213 | 0.8417784 | 0.921642147 | 0.017700546 |
| ENSMUSG00000034663 | Bmp2k | 140780 | protein_coding | 315.4500851 | -0.017712003 | 0.094511993 | 0.187404817 | 0.851343246 | 0.92744128 | 0.017712003 |
| ENSMUSG00000028496 | Mllt3 | 70122 | protein_coding | 558.007186 | 0.017717716 | 0.102643432 | -0.172614226 | 0.862954669 | 0.932166328 | 0.017717716 |
| ENSMUSG00000032469 | Dbr1 | 83703 | protein_coding | 162.8266324 | 0.017767619 | 0.113070554 | -0.157137454 | 0.87513652 | 0.938415991 | 0.017767619 |
| ENSMUSG00000039568 | Ubald1 | 207740 | protein_coding | 335.7443364 | -0.017857415 | 0.132402596 | 0.134872091 | 0.892713006 | 0.947202684 | 0.017857415 |
| ENSMUSG00000111063 | Zfp660 | 666257 | lncRNA | 1.777368946 | -0.017858784 | 1.928408944 | 0.00926089 | 0.992610984 | 1 | 0.017858784 |
| ENSMUSG00000027593 | Raly | 19383 | protein_coding | 1427.99255 | 0.017881542 | 0.063210354 | -0.282889453 | 0.77726158 | 0.888775635 | 0.017881542 |
| ENSMUSG00000028954 | Nub1 | 53312 | protein_coding | 479.6949741 | 0.01788478 | 0.059079067 | -0.302726175 | 0.762098549 | 0.880874346 | 0.01788478 |
| ENSMUSG00000020794 | Ube2g1 | 67128 | protein_coding | 497.6815876 | 0.01788768 | 0.057683536 | -0.31010027 | 0.756484707 | 0.877843331 | 0.01788768 |
| ENSMUSG00000071266 | Zfp946 | 74149 | protein_coding | 252.0376236 | -0.017935942 | 0.093980738 | 0.190846995 | 0.848645468 | 0.925637253 | 0.017935942 |
| ENSMUSG00000070934 | Rraga | 68441 | protein_coding | 336.2071035 | 0.01794139 | 0.0949977 | -0.18886131 | 0.850201518 | 0.926816617 | 0.01794139 |

| ensembl_gene_id | external_gene_name | entrezgene_id | gene_biotype | baseMean | log2FoldChange | lfcSE | stat | pvalue | padj | absLog2FC |
|---|---|---|---|---|---|---|---|---|---|---|
| ENSMUSG00000013622 | Atraid | 381629 | protein_coding | 561.7100797 | -0.017944344 | 0.087898416 | 0.204148667 | 0.838237324 | 0.919410944 | 0.017944344 |
| ENSMUSG00000037151 | Lrrc20 | 216011 | protein_coding | 113.9561715 | 0.018065019 | 0.127025042 | -0.142216201 | 0.886909235 | 0.94399288 | 0.018065019 |
| ENSMUSG00000027177 | Hipk3 | 15259 | protein_coding | 1237.763669 | -0.018085311 | 0.069559145 | 0.259999035 | 0.794864518 | 0.897960915 | 0.018085311 |
| ENSMUSG00000040331 | Nsmce4a | 67872 | protein_coding | 522.3059995 | 0.018089231 | 0.088090222 | -0.205348913 | 0.837299531 | 0.918870578 | 0.018089231 |
| ENSMUSG00000022129 | Dct | 13190 | protein_coding | 10.27179372 | 0.018116298 | 0.43888025 | -0.041278453 | 0.967073911 | 0.983632579 | 0.018116298 |
| ENSMUSG00000030469 | Zfp719 | 210105 | protein_coding | 122.687168 | -0.018121257 | 0.129945788 | 0.139452442 | 0.889092634 | 0.945487524 | 0.018121257 |
| ENSMUSG00000078920 | Ifi47 | 15953 | protein_coding | 87.03949495 | -0.018129222 | 0.292102778 | 0.06206453 | 0.950511444 | 0.975869522 | 0.018129222 |
| ENSMUSG00000039208 | Metrnl | 210029 | protein_coding | 848.5904282 | 0.018171448 | 0.138589846 | -0.131116731 | 0.895682966 | 0.948731562 | 0.018171448 |
| ENSMUSG00000062248 | Cks2 | 66197 | protein_coding | 276.9574786 | -0.018177314 | 0.157354051 | 0.115518563 | 0.908034108 | 0.955211953 | 0.018177314 |
| ENSMUSG00000029233 | Srd5a3 | 57357 | protein_coding | 170.3143399 | -0.018178256 | 0.104186503 | 0.174478031 | 0.861489795 | 0.931722347 | 0.018178256 |
| ENSMUSG00000038733 | Wdr26 | 226757 | protein_coding | 1400.837794 | 0.018179134 | 0.050836323 | -0.357601281 | 0.72064172 | 0.855240917 | 0.018179134 |
| ENSMUSG00000106380 | Gm3519 | 100041805 | lncRNA | 1.83741441 | 0.018195364 | 0.954506589 | -0.019062586 | 0.984791178 | 1 | 0.018195364 |
| ENSMUSG00000028069 | Gpatch4 | 66614 | protein_coding | 119.4556532 | 0.018211593 | 0.143926728 | -0.126533783 | 0.899309409 | 0.950598193 | 0.018211593 |
| ENSMUSG00000038533 | Cbfa2t2 | 12396 | protein_coding | 337.472033 | -0.018215001 | 0.059580215 | 0.305722312 | 0.759816084 | 0.879422681 | 0.018215001 |
| ENSMUSG00000020253 | Ppm1m | 67905 | protein_coding | 48.4403481 | -0.018310488 | 0.161290734 | 0.113524738 | 0.909614552 | 0.956044724 | 0.018310488 |
| ENSMUSG00000022300 | Dcaf13 | 223499 | protein_coding | 485.3927743 | 0.018367011 | 0.076107501 | -0.241329837 | 0.809299487 | 0.905779176 | 0.018367011 |
| ENSMUSG00000055761 | Nkain3 | 269513 | protein_coding | 1.948487245 | -0.018392149 | 2.323870799 | 0.007914445 | 0.993685252 | 1 | 0.018392149 |
| ENSMUSG00000030530 | Furin | 18550 | protein_coding | 715.3213755 | 0.018410527 | 0.082044644 | -0.224396457 | 0.822448827 | 0.910171686 | 0.018410527 |
| ENSMUSG00000114469 | C730002L08Rik | 320155 | lncRNA | 4.338382363 | 0.018421344 | 0.672139094 | -0.027407042 | 0.978135082 | 1 | 0.018421344 |
| ENSMUSG00000082101 | Slfn14 | 237890 | protein_coding | 0.862133039 | -0.018422881 | 1.726028038 | 0.01067357 | 0.991483885 | 1 | 0.018422881 |
| ENSMUSG00000001785 | Pwp1 | 103136 | protein_coding | 319.7897648 | 0.018433123 | 0.104848134 | -0.175807828 | 0.86044492 | 0.931471303 | 0.018433123 |
| ENSMUSG00000045287 | Rtn4rl1 | 237847 | protein_coding | 69.73973205 | -0.018435402 | 0.138659022 | 0.132954941 | 0.894229016 | 0.947795386 | 0.018435402 |
| ENSMUSG00000019471 | Cdc37 | 12539 | protein_coding | 1213.040388 | 0.01844402 | 0.068270385 | -0.270161364 | 0.787036115 | 0.893375436 | 0.01844402 |
| ENSMUSG00000027430 | Dtd1 | 66044 | protein_coding | 105.5882166 | 0.018446517 | 0.113476297 | -0.16255833 | 0.870866199 | 0.936355518 | 0.018446517 |
| ENSMUSG00000027303 | Ptpra | 19262 | protein_coding | 516.2957257 | 0.018453983 | 0.061993694 | -0.297675163 | 0.765951102 | 0.882687823 | 0.018453983 |
| ENSMUSG00000033318 | Gstt2 | 14872 | protein_coding | 439.13518 | 0.018488171 | 0.072028806 | -0.256677457 | 0.797427775 | 0.899441451 | 0.018488171 |
| ENSMUSG00000048047 | Zbtb33 | 56805 | protein_coding | 310.062824 | -0.018521603 | 0.088964991 | 0.208189787 | 0.835080783 | 0.917066002 | 0.018521603 |
| ENSMUSG00000038555 | Reep2 | 225362 | protein_coding | 3.84141833 | 0.018543014 | 0.768045725 | -0.024143112 | 0.980738455 | 1 | 0.018543014 |
| ENSMUSG00000025133 | Ints4 | 101861 | protein_coding | 480.1599165 | -0.018591758 | 0.090914403 | 0.204497386 | 0.837964835 | 0.919173117 | 0.018591758 |
| ENSMUSG00000028540 | Dph2 | 67728 | protein_coding | 93.69296818 | 0.018604741 | 0.131772884 | -0.141187931 | 0.88772148 | 0.94451598 | 0.018604741 |
| ENSMUSG00000038203 | Hoxa13 | 15398 | protein_coding | 470.6927732 | -0.018610167 | 0.130792439 | 0.14228779 | 0.88685269 | 0.94399288 | 0.018610167 |
| ENSMUSG00000038174 | Hycc2 | 213056 | protein_coding | 380.6835199 | 0.01862464 | 0.108052117 | -0.172367192 | 0.863148863 | 0.932166328 | 0.01862464 |
| ENSMUSG00000047379 | B4gat1 | 108902 | protein_coding | 2.800389224 | 0.018710865 | 0.642772102 | -0.029109641 | 0.976777146 | 1 | 0.018710865 |
| ENSMUSG00000101856 | 1700096K18Rik | 73571 | lncRNA | 42.2189779 | -0.018775455 | 0.190131269 | 0.098749961 | 0.9213368 | 0.962004133 | 0.018775455 |
| ENSMUSG00000098280 | Mir7687 | 102466849 | miRNA | 1.854227492 | -0.018782232 | 0.918562092 | 0.020447427 | 0.98368645 | 1 | 0.018782232 |
| ENSMUSG00000033228 | Scaf11 | 72193 | protein_coding | 1327.854413 | -0.018820556 | 0.072039952 | 0.261251648 | 0.793898451 | 0.897263004 | 0.018820556 |
| ENSMUSG00000045438 | Cox19 | 68033 | protein_coding | 282.1175846 | -0.018843576 | 0.108747442 | 0.173278341 | 0.862432647 | 0.932036168 | 0.018843576 |
| ENSMUSG00000020171 | Yeats4 | 64050 | protein_coding | 296.2940537 | 0.018865777 | 0.136877281 | -0.137829866 | 0.89037488 | 0.946160016 | 0.018865777 |
| ENSMUSG00000021958 | Pinx1 | 72400 | protein_coding | 73.40019041 | 0.018871174 | 0.190156185 | -0.099240389 | 0.920947407 | 0.961855464 | 0.018871174 |

| ensembl_gene_id | external_gene_name | entrezgene_id | gene_biotype | baseMean | log2FoldChange | lfcSE | stat | pvalue | padj | absLog2FC |
|---|---|---|---|---|---|---|---|---|---|---|
| ENSMUSG00000038022 | Mindy4 | 330323 | protein_coding | 11.13512738 | 0.018886975 | 0.453873646 | -0.041612849 | 0.96680733 | 0.983482583 | 0.018886975 |
| ENSMUSG00000049191 | Rtl5 | 331474 | protein_coding | 5.375238463 | 0.018932825 | 0.621517771 | -0.030462243 | 0.975698405 | 0.987300339 | 0.018932825 |
| ENSMUSG00000074063 | Osgin1 | 71839 | protein_coding | 159.7224408 | -0.018961722 | 0.183952979 | 0.10307918 | 0.917900129 | 0.959738982 | 0.018961722 |
| ENSMUSG00000042854 | Trp53rkb | 76367 | protein_coding | 34.08736295 | 0.018993881 | 0.191583312 | -0.099141624 | 0.921025824 | 0.961855464 | 0.018993881 |
| ENSMUSG00000027162 | Lin7c | 22343 | protein_coding | 2719.402178 | -0.018996194 | 0.096112441 | 0.197645525 | 0.843322414 | 0.922491196 | 0.018996194 |
| ENSMUSG00000054612 | Mgmt | 17314 | protein_coding | 34.00882706 | -0.018997988 | 0.222525201 | 0.085374544 | 0.93196363 | 0.967478151 | 0.018997988 |
| ENSMUSG00000029291 | Rufy3 | 52822 | protein_coding | 301.1981025 | -0.019003437 | 0.075352486 | 0.252193901 | 0.800891192 | 0.901439282 | 0.019003437 |
| ENSMUSG00000053565 | Eif3k | 73830 | protein_coding | 1326.460092 | 0.019010282 | 0.077577478 | -0.245048986 | 0.806418502 | 0.904795436 | 0.019010282 |
| ENSMUSG00000087968 | Gm25395 | 115485773 | scaRNA | 535.0798835 | 0.019094175 | 0.261327756 | -0.073066004 | 0.941753594 | 0.971883559 | 0.019094175 |
| ENSMUSG00000026175 | Vil1 | 22349 | protein_coding | 9369.642966 | 0.019108408 | 0.081984623 | -0.233073068 | 0.815704667 | 0.908392129 | 0.019108408 |
| ENSMUSG00000028121 | Bcar3 | 29815 | protein_coding | 889.8409529 | -0.019111428 | 0.109761081 | 0.174118436 | 0.861772385 | 0.931842966 | 0.019111428 |
| ENSMUSG00000025369 | Smarcc2 | 68094 | protein_coding | 1047.480611 | -0.019209235 | 0.070812914 | 0.271267404 | 0.786185374 | 0.893161222 | 0.019209235 |
| ENSMUSG00000024169 | Ift140 | 106633 | protein_coding | 266.4201372 | -0.019230488 | 0.078762987 | 0.244156407 | 0.807109687 | 0.904987729 | 0.019230488 |
| ENSMUSG00000047466 | 8030462N17Rik | 212163 | protein_coding | 332.0022407 | -0.019247588 | 0.08979248 | 0.214356343 | 0.830269184 | 0.914187552 | 0.019247588 |
| ENSMUSG00000027455 | Nsfl1c | 386649 | protein_coding | 515.4017401 | 0.019250321 | 0.060737556 | -0.316942632 | 0.751287133 | 0.875081807 | 0.019250321 |
| ENSMUSG00000007617 | Homer1 | 26556 | protein_coding | 379.7080892 | 0.019252799 | 0.089018203 | -0.21627935 | 0.828770005 | 0.913504709 | 0.019252799 |
| ENSMUSG00000026917 | Wdr5 | 140858 | protein_coding | 454.1855312 | -0.019274755 | 0.096921513 | 0.198869724 | 0.842364653 | 0.921917283 | 0.019274755 |
| ENSMUSG00000038485 | Socs7 | 192157 | protein_coding | 323.9704978 | -0.019278244 | 0.101980336 | 0.18903884 | 0.850062376 | 0.926816617 | 0.019278244 |
| ENSMUSG00000003444 | Med29 | 67224 | protein_coding | 88.61864346 | 0.019285778 | 0.131268891 | -0.146918114 | 0.883196654 | 0.942211125 | 0.019285778 |
| ENSMUSG00000028458 | Tesk1 | 21754 | protein_coding | 297.8545401 | -0.019287735 | 0.098642596 | 0.195531506 | 0.844976875 | 0.923798341 | 0.019287735 |
| ENSMUSG00000041126 | H2az2 | 77605 | protein_coding | 515.6106798 | -0.019302725 | 0.094373018 | 0.204536482 | 0.837934286 | 0.919173117 | 0.019302725 |
| ENSMUSG00000025404 | R3hdm2 | 71750 | protein_coding | 340.9762829 | -0.019337212 | 0.138224435 | 0.139897207 | 0.888741208 | 0.945235545 | 0.019337212 |
| ENSMUSG00000022552 | Sharpin | 106025 | protein_coding | 404.9786408 | -0.019346215 | 0.079961029 | 0.241945552 | 0.808822353 | 0.905678041 | 0.019346215 |
| ENSMUSG00000097471 | 5830432E09Rik | 67765 | lncRNA | 6.979710673 | -0.01936973 | 0.490058897 | 0.039525309 | 0.968471575 | 0.984122985 | 0.01936973 |
| ENSMUSG00000025809 | Itgb1 | 16412 | protein_coding | 2859.043315 | 0.019451086 | 0.064461803 | -0.301745928 | 0.762845753 | 0.881056435 | 0.019451086 |
| ENSMUSG00000030738 | Eif3c | 56347 | protein_coding | 3331.404726 | -0.019489041 | 0.061348655 | 0.317676741 | 0.750730156 | 0.87503937 | 0.019489041 |
| ENSMUSG00000031153 | Gripap1 | 54645 | protein_coding | 348.3881312 | 0.019510153 | 0.110166418 | -0.177097101 | 0.859432119 | 0.931154836 | 0.019510153 |
| ENSMUSG00000039449 | Prpf18 | 67229 | protein_coding | 470.9013564 | 0.019513658 | 0.088810763 | -0.219721773 | 0.826087847 | 0.912143267 | 0.019513658 |
| ENSMUSG00000039244 | Ints15 | 231868 | protein_coding | 201.4709446 | 0.01951705 | 0.083534705 | -0.233640023 | 0.815264453 | 0.908146859 | 0.01951705 |
| ENSMUSG00000027937 | Jtb | 23922 | protein_coding | 474.1992707 | 0.019543598 | 0.108995981 | -0.179305674 | 0.857697692 | 0.930374388 | 0.019543598 |
| ENSMUSG00000029089 | 5730480H06Rik | 66768 | protein_coding | 84.44769508 | 0.019543931 | 0.116816078 | -0.167305145 | 0.867129956 | 0.933980178 | 0.019543931 |
| ENSMUSG00000021906 | Oxnad1 | 218885 | protein_coding | 323.6506954 | -0.019579342 | 0.067038215 | 0.292062393 | 0.77023892 | 0.884771487 | 0.019579342 |
| ENSMUSG00000029389 | Ddx55 | 67848 | protein_coding | 161.5454734 | -0.019600975 | 0.105278265 | 0.186182539 | 0.852301616 | 0.927761944 | 0.019600975 |
| ENSMUSG00000022401 | Xpnpep3 | 321003 | protein_coding | 184.1290557 | 0.01960946 | 0.112426904 | -0.174419639 | 0.861535681 | 0.931722347 | 0.01960946 |
| ENSMUSG00000021668 | Polk | 27015 | protein_coding | 152.870442 | -0.019616253 | 0.158403531 | 0.123837222 | 0.90144416 | 0.951341404 | 0.019616253 |
| ENSMUSG00000078486 | Perm1 | 74183 | protein_coding | 17.06738782 | 0.019636243 | 0.374141709 | -0.052483438 | 0.958143492 | 0.979508488 | 0.019636243 |
| ENSMUSG00000059423 | Zfp933 | 242747 | protein_coding | 99.20090171 | -0.019665093 | 0.113872746 | 0.172693588 | 0.862892284 | 0.932166328 | 0.019665093 |
| ENSMUSG00000040548 | Tex2 | 21763 | protein_coding | 678.4670315 | -0.019691637 | 0.065410508 | 0.301047001 | 0.763378654 | 0.881383982 | 0.019691637 |
| ENSMUSG00000001441 | Npepps | 19155 | protein_coding | 858.043543 | 0.019720117 | 0.048637892 | -0.405447614 | 0.685148518 | 0.832151567 | 0.019720117 |

| ensembl_gene_id | external_gene_name | entrezgene_id | gene_biotype | baseMean | log2FoldChange | lfcSE | stat | pvalue | padj | absLog2FC |
|---|---|---|---|---|---|---|---|---|---|---|
| ENSMUSG00000049493 | Pls1 | 102502 | protein_coding | 3199.590583 | -0.019785213 | 0.087085864 | 0.227192013 | 0.820274438 | 0.9092783 | 0.019785213 |
| ENSMUSG00000029417 | Cxcl9 | 17329 | protein_coding | 2.534476602 | 0.019806434 | 0.819913495 | -0.024156737 | 0.980727587 | 1 | 0.019806434 |
| ENSMUSG00000030057 | Cnbp | 12785 | protein_coding | 2209.837835 | 0.019814758 | 0.10221298 | -0.193857552 | 0.846287422 | 0.924222269 | 0.019814758 |
| ENSMUSG00000050244 | Heatr1 | 217995 | protein_coding | 475.2564054 | 0.019830254 | 0.147404024 | -0.134529936 | 0.892983541 | 0.947307136 | 0.019830254 |
| ENSMUSG00000071180 | Smim15 | 75616 | protein_coding | 859.6706893 | -0.019834591 | 0.067540818 | 0.293668204 | 0.769011453 | 0.884245488 | 0.019834591 |
| ENSMUSG00000042744 | Hectd4 | 269700 | protein_coding | 1265.933277 | 0.019860354 | 0.157102629 | -0.126416431 | 0.899402296 | 0.950621521 | 0.019860354 |
| ENSMUSG00000002052 | Supt6 | 20926 | protein_coding | 1181.586085 | 0.019861008 | 0.041552381 | -0.477975207 | 0.632667852 | 0.799623213 | 0.019861008 |
| ENSMUSG00000029152 | Ociad1 | 68095 | protein_coding | 1121.666161 | 0.019960478 | 0.082566331 | -0.241750812 | 0.808973255 | 0.905720972 | 0.019960478 |
| ENSMUSG00000026587 | Astn1 | 11899 | protein_coding | 4.858387377 | 0.019960686 | 1.34357495 | -0.014856399 | 0.988146744 | 0.993418099 | 0.019960686 |
| ENSMUSG00000029004 | Kmt2e | 69188 | protein_coding | 514.3547522 | -0.019967125 | 0.157075839 | 0.127117738 | 0.898847212 | 0.950352959 | 0.019967125 |
| ENSMUSG00000046032 | Snx12 | 55988 | protein_coding | 287.850514 | -0.01998145 | 0.084337926 | 0.236921292 | 0.812717852 | 0.907147741 | 0.01998145 |
| ENSMUSG00000024941 | Scyl1 | 78891 | protein_coding | 763.9354563 | 0.019998772 | 0.064846909 | -0.308399766 | 0.75777816 | 0.87863969 | 0.019998772 |
| ENSMUSG00000074384 | AI429214 | 621080 | protein_coding | 6.465123833 | 0.019998973 | 0.440886514 | -0.045360818 | 0.963819711 | 0.982137401 | 0.019998973 |
| ENSMUSG00000038664 | Herc1 | 235439 | protein_coding | 1384.702428 | -0.020000051 | 0.070831002 | 0.282362952 | 0.77766522 | 0.888933284 | 0.020000051 |
| ENSMUSG00000036733 | Rbm42 | 68035 | protein_coding | 1.443338995 | -0.020004843 | 1.058485255 | 0.018899501 | 0.984921278 | 1 | 0.020004843 |
| ENSMUSG00000039230 | Tbcd | 108903 | protein_coding | 855.8850502 | 0.020052876 | 0.086932555 | -0.230671648 | 0.817569901 | 0.908714429 | 0.020052876 |
| ENSMUSG00000018470 | Kcnab3 | 16499 | protein_coding | 6.811425686 | -0.020054486 | 0.618093902 | 0.032445695 | 0.974116622 | 0.986602999 | 0.020054486 |
| ENSMUSG00000028468 | Rgp1 | 242406 | protein_coding | 875.9009975 | -0.020090829 | 0.08714472 | 0.230545567 | 0.81766786 | 0.908714429 | 0.020090829 |
| ENSMUSG00000112489 | 9230116L04Rik | 321017 | lncRNA | 57.50768725 | -0.020098286 | 0.186357722 | 0.107847883 | 0.914116359 | 0.958030863 | 0.020098286 |
| ENSMUSG00000031429 | Psmd10 | 53380 | protein_coding | 242.3725844 | 0.020103837 | 0.085594898 | -0.23487191 | 0.814308152 | 0.907710402 | 0.020103837 |
| ENSMUSG00000032567 | Aste1 | 66595 | protein_coding | 158.0581095 | 0.020115746 | 0.087060682 | -0.231054316 | 0.817272606 | 0.908714429 | 0.020115746 |
| ENSMUSG00000034868 | Myl12b | 67938 | protein_coding | 462.4443232 | -0.020125991 | 0.111236483 | 0.180929768 | 0.856422703 | 0.930184975 | 0.020125991 |
| ENSMUSG00000021241 | Isca2 | 74316 | protein_coding | 185.5759542 | -0.020141769 | 0.087350008 | 0.230586916 | 0.817635734 | 0.908714429 | 0.020141769 |
| ENSMUSG00000021983 | Atp8a2 | 50769 | protein_coding | 523.5551485 | 0.020191664 | 0.07184066 | -0.281061788 | 0.778663009 | 0.889454308 | 0.020191664 |
| ENSMUSG00000015533 | Itga2 | 16398 | protein_coding | 1043.140288 | 0.020216641 | 0.128151059 | -0.157756331 | 0.874648811 | 0.938406345 | 0.020216641 |
| ENSMUSG00000065637 | Gm26397 | 115487043 | snoRNA | 246.6811701 | -0.020220335 | 0.25773885 | 0.078452801 | 0.937467874 | 0.970191419 | 0.020220335 |
| ENSMUSG00000026213 | Stk11ip | 71728 | protein_coding | 258.8874745 | -0.020220892 | 0.075706609 | 0.267095462 | 0.789395662 | 0.894720656 | 0.020220892 |
| ENSMUSG00000046613 | Vwa5b2 | 328643 | protein_coding | 21.69914907 | -0.020222688 | 0.315417783 | 0.064113975 | 0.948879474 | 0.975399666 | 0.020222688 |
| ENSMUSG00000021771 | Vdac2 | 22334 | protein_coding | 4853.067908 | -0.02025008 | 0.077013559 | 0.262941749 | 0.792595477 | 0.896761253 | 0.02025008 |
| ENSMUSG00000057982 | Zfp809 | 235047 | protein_coding | 223.5133783 | -0.020292289 | 0.103804257 | 0.195486095 | 0.845012422 | 0.923798341 | 0.020292289 |
| ENSMUSG00000034928 | Rnf44 | 105239 | protein_coding | 1032.101311 | -0.020354283 | 0.085795821 | 0.237240958 | 0.812469864 | 0.907147741 | 0.020354283 |
| ENSMUSG00000034518 | Hmgxb4 | 70823 | protein_coding | 261.4082692 | -0.020366307 | 0.092118801 | 0.221087406 | 0.825024376 | 0.911456552 | 0.020366307 |
| ENSMUSG00000059689 | Zfp637 | 232337 | protein_coding | 136.0000682 | -0.020399088 | 0.095092401 | 0.214518594 | 0.830142669 | 0.914175515 | 0.020399088 |
| ENSMUSG00000027131 | Emc4 | 68032 | protein_coding | 292.5877738 | -0.020404139 | 0.071931184 | 0.283661936 | 0.776669467 | 0.888506311 | 0.020404139 |
| ENSMUSG00000020220 | Vps13d | 230895 | protein_coding | 3055.842085 | 0.020404811 | 0.107967651 | -0.18899004 | 0.850100623 | 0.926816617 | 0.020404811 |
| ENSMUSG00000003226 | Ranbp2 | 19386 | protein_coding | 2322.805956 | 0.020411815 | 0.084755328 | -0.240832231 | 0.809685148 | 0.905997055 | 0.020411815 |
| ENSMUSG00000048930 | Tada3 | 101206 | protein_coding | 297.5306447 | 0.020424733 | 0.079599374 | -0.256594138 | 0.797492101 | 0.899452571 | 0.020424733 |
| ENSMUSG00000005102 | Eif2ak4 | 27103 | protein_coding | 356.2623977 | -0.02043919 | 0.097073375 | 0.210554022 | 0.833235293 | 0.915993052 | 0.02043919 |
| ENSMUSG00000005958 | Ephb3 | 13845 | protein_coding | 356.170077 | -0.020439559 | 0.134588587 | 0.151866953 | 0.879291874 | 0.940455052 | 0.020439559 |

Supplementary Table 1

| ensembl_gene_id | external_gene_name | entrezgene_id | gene_biotype | baseMean | log2FoldChange | lfcSE | stat | pvalue | padj | absLog2FC |
|---|---|---|---|---|---|---|---|---|---|---|
| ENSMUSG00000018287 | Spag7 | 216873 | protein_coding | 408.6789057 | 0.02044203 | 0.090784649 | -0.225170561 | 0.82184659 | 0.91001573 | 0.02044203 |
| ENSMUSG00000020166 | Cnot2 | 72068 | protein_coding | 634.9309497 | 0.020475391 | 0.075226339 | -0.272183809 | 0.78548069 | 0.892918574 | 0.020475391 |
| ENSMUSG00000059273 | Zc3h4 | 330474 | protein_coding | 812.7433577 | 0.020502672 | 0.098429016 | -0.208299061 | 0.834995465 | 0.917066002 | 0.020502672 |
| ENSMUSG00000078179 | Rnf148 | 71300 | protein_coding | 2.943734535 | 0.020507883 | 0.645992793 | -0.031746303 | 0.974674369 | 1 | 0.020507883 |
| ENSMUSG00000040586 | Ofd1 | 237222 | protein_coding | 116.0143619 | -0.020518126 | 0.134850745 | 0.152154341 | 0.879065205 | 0.940340033 | 0.020518126 |
| ENSMUSG00000035772 | Mrps2 | 118451 | protein_coding | 348.6948722 | 0.02052324 | 0.101099195 | -0.203001025 | 0.839134231 | 0.919794636 | 0.02052324 |
| ENSMUSG00000085437 | 1700047A11Rik | 76604 | lncRNA | 1.672788782 | 0.020560349 | 1.050193685 | -0.019577673 | 0.984380275 | 1 | 0.020560349 |
| ENSMUSG00000039000 | Ube3c | 100763 | protein_coding | 1123.302214 | 0.02057508 | 0.07003412 | -0.293786518 | 0.768921038 | 0.884245488 | 0.02057508 |
| ENSMUSG00000035614 | Togaram1 | 328108 | protein_coding | 505.279921 | -0.020675644 | 0.082563305 | 0.250421709 | 0.802261243 | 0.90234117 | 0.020675644 |
| ENSMUSG00000037204 | Atg101 | 68118 | protein_coding | 408.4332117 | -0.020677745 | 0.080177219 | 0.257900503 | 0.796483697 | 0.898891871 | 0.020677745 |
| ENSMUSG00000040481 | Bptf | 207165 | protein_coding | 1069.330002 | 0.020706206 | 0.074872707 | -0.276552122 | 0.782124033 | 0.890758364 | 0.020706206 |
| ENSMUSG00000028920 | Fbxo42 | 213499 | protein_coding | 414.8071715 | 0.0207134 | 0.057693393 | -0.359025509 | 0.719576006 | 0.854899358 | 0.0207134 |
| ENSMUSG00000054309 | Cpsf3 | 54451 | protein_coding | 574.331252 | 0.02073004 | 0.094669916 | -0.218971778 | 0.826672032 | 0.912523774 | 0.02073004 |
| ENSMUSG00000022760 | Thap7 | 69009 | protein_coding | 163.7189817 | 0.020734925 | 0.092082094 | -0.225178688 | 0.821840269 | 0.91001573 | 0.020734925 |
| ENSMUSG00000035104 | Eva1a | 232146 | protein_coding | 27.65313407 | -0.02076387 | 0.233570966 | 0.08889748 | 0.929163386 | 0.966128972 | 0.02076387 |
| ENSMUSG00000027938 | Creb3l4 | 78284 | protein_coding | 454.1285521 | -0.020801755 | 0.237240569 | 0.087682115 | 0.930129335 | 0.966352899 | 0.020801755 |
| ENSMUSG00000010025 | Aldh3a2 | 11671 | protein_coding | 1656.789148 | 0.020830015 | 0.074645373 | -0.279053001 | 0.780204151 | 0.890125936 | 0.020830015 |
| ENSMUSG00000105387 | Gm29681 | 545809 | lncRNA | 0.877474797 | -0.020845445 | 1.731284462 | 0.01204045 | 0.990393343 | 1 | 0.020845445 |
| ENSMUSG00000034579 | Pla2g3 | 237625 | protein_coding | 231.8522878 | -0.020854421 | 0.160078799 | 0.130275974 | 0.896348088 | 0.948996349 | 0.020854421 |
| ENSMUSG00000018796 | Acsl1 | 14081 | protein_coding | 293.5978578 | 0.020936564 | 0.179648219 | -0.11654201 | 0.907222994 | 0.95457824 | 0.020936564 |
| ENSMUSG00000051185 | Fam174a | 67698 | protein_coding | 153.7962866 | -0.020940173 | 0.114680426 | 0.182595878 | 0.855115119 | 0.929465614 | 0.020940173 |
| ENSMUSG00000036257 | Pnpla8 | 67452 | protein_coding | 882.6644761 | -0.020956132 | 0.107293959 | 0.195315121 | 0.84514626 | 0.923798341 | 0.020956132 |
| ENSMUSG00000039804 | Ncoa5 | 228869 | protein_coding | 339.9128393 | 0.020961241 | 0.082964371 | -0.252653533 | 0.800535959 | 0.901189824 | 0.020961241 |
| ENSMUSG00000025870 | Arl10 | 56795 | protein_coding | 138.7139689 | 0.021029462 | 0.116863263 | -0.179949295 | 0.857192375 | 0.930374388 | 0.021029462 |
| ENSMUSG00000060240 | Cend1 | 57754 | protein_coding | 3.469343665 | 0.021061847 | 0.694291464 | -0.030335743 | 0.975799291 | 1 | 0.021061847 |
| ENSMUSG00000074582 | Arfgef2 | 99371 | protein_coding | 1319.996218 | 0.02106218 | 0.052151527 | -0.403865068 | 0.686311945 | 0.832935655 | 0.02106218 |
| ENSMUSG00000020340 | Cyfip2 | 76884 | protein_coding | 17.3379801 | -0.021073963 | 0.309311642 | 0.068131812 | 0.945680707 | 0.973795719 | 0.021073963 |
| ENSMUSG00000024668 | Sdhaf2 | 66072 | protein_coding | 426.0073436 | 0.021074307 | 0.071547334 | -0.29455056 | 0.768337235 | 0.883975099 | 0.021074307 |
| ENSMUSG00000005609 | Ctr9 | 22083 | protein_coding | 829.6526447 | -0.021169154 | 0.062127903 | 0.340735053 | 0.733303047 | 0.863385162 | 0.021169154 |
| ENSMUSG00000040435 | Ppp1r15a | 17872 | protein_coding | 549.9063937 | 0.02127365 | 0.240309692 | -0.088525975 | 0.92945864 | 0.966128972 | 0.02127365 |
| ENSMUSG00000010047 | Hyal2 | 15587 | protein_coding | 526.5150596 | -0.021279212 | 0.102119748 | 0.208375094 | 0.834936102 | 0.917066002 | 0.021279212 |
| ENSMUSG00000042364 | Snx18 | 170625 | protein_coding | 215.2708519 | -0.021315632 | 0.103429539 | 0.206088435 | 0.836721833 | 0.918419697 | 0.021315632 |
| ENSMUSG00000036112 | Metap2 | 56307 | protein_coding | 1437.723734 | 0.021326789 | 0.072509072 | -0.294125803 | 0.768661774 | 0.884145751 | 0.021326789 |
| ENSMUSG00000085876 | Gm12409 | 100040567 | lncRNA | 3.129753829 | -0.021335004 | 0.593024918 | 0.035976573 | 0.971301039 | 1 | 0.021335004 |
| ENSMUSG00000021843 | Ktn1 | 16709 | protein_coding | 746.4898419 | -0.021453397 | 0.077432342 | 0.277059897 | 0.781734114 | 0.890621398 | 0.021453397 |
| ENSMUSG00000036372 | Tmem258 | 69038 | protein_coding | 575.7129316 | 0.021566016 | 0.112637165 | -0.191464474 | 0.848161712 | 0.925501685 | 0.021566016 |
| ENSMUSG00000027787 | Nmd3 | 97112 | protein_coding | 217.9345312 | 0.021569559 | 0.093174881 | -0.231495423 | 0.816929941 | 0.908700419 | 0.021569559 |
| ENSMUSG00000035696 | Rnf38 | 73469 | protein_coding | 413.2699619 | 0.021573641 | 0.072551636 | -0.297355676 | 0.766194979 | 0.882690208 | 0.021573641 |
| ENSMUSG00000030156 | Cd69 | 12515 | protein_coding | 15.68593322 | 0.021606761 | 0.564021891 | -0.038308373 | 0.969441815 | 0.984403993 | 0.021606761 |

Supplementary Table 1

| ensembl_gene_id | external_gene_name | entrezgene_id | gene_biotype | baseMean | log2FoldChange | lfcSE | stat | pvalue | padj | absLog2FC |
|---|---|---|---|---|---|---|---|---|---|---|
| ENSMUSG00000037049 | Smpd1 | 20597 | protein_coding | 345.1177398 | -0.021641978 | 0.102475247 | 0.211192254 | 0.832737255 | 0.915689375 | 0.021641978 |
| ENSMUSG00000044150 | Bclaf3 | 382252 | protein_coding | 102.6755828 | 0.021652919 | 0.136155955 | -0.159030275 | 0.873645022 | 0.937816815 | 0.021652919 |
| ENSMUSG00000033739 | Fkbpl | 56299 | protein_coding | 65.7147216 | 0.021658001 | 0.144123268 | -0.150274144 | 0.880548332 | 0.941067509 | 0.021658001 |
| ENSMUSG00000093080 | Mir3060 | 100526466 | miRNA | 1.047837915 | -0.021662145 | 1.198550775 | 0.018073615 | 0.985580127 | 1 | 0.021662145 |
| ENSMUSG00000057363 | Uxs1 | 67883 | protein_coding | 212.5396505 | -0.021678146 | 0.075427992 | 0.287401867 | 0.773804632 | 0.88672609 | 0.021678146 |
| ENSMUSG00000025903 | Lypla1 | 18777 | protein_coding | 959.5240635 | 0.021696983 | 0.079777444 | -0.271968888 | 0.785645941 | 0.892925964 | 0.021696983 |
| ENSMUSG00000038085 | Cnbd2 | 70873 | protein_coding | 22.49029239 | 0.021736509 | 0.270302036 | -0.080415631 | 0.935906696 | 0.96968875 | 0.021736509 |
| ENSMUSG00000077744 | Gm26343 | 115486206 | snoRNA | 0.617806996 | -0.021756029 | 1.844020761 | 0.011798147 | 0.990586659 | 1 | 0.021756029 |
| ENSMUSG00000057229 | Dmac2 | 66349 | protein_coding | 527.4202969 | -0.021792468 | 0.070755181 | 0.307998203 | 0.7580837 | 0.878750865 | 0.021792468 |
| ENSMUSG00000027900 | Dram2 | 67171 | protein_coding | 1628.64894 | 0.021806856 | 0.10279053 | -0.212148494 | 0.831991188 | 0.915295615 | 0.021806856 |
| ENSMUSG00000029066 | Mrpl20 | 66448 | protein_coding | 447.6462885 | 0.021817882 | 0.071714787 | -0.304231281 | 0.760951695 | 0.880144002 | 0.021817882 |
| ENSMUSG00000036860 | Mrpl55 | 67212 | protein_coding | 266.6085474 | -0.021825514 | 0.093833487 | 0.232598353 | 0.816073306 | 0.908409285 | 0.021825514 |
| ENSMUSG00000026516 | Nvl | 67459 | protein_coding | 398.6904197 | -0.021827287 | 0.060690053 | 0.359651796 | 0.719107545 | 0.854587075 | 0.021827287 |
| ENSMUSG00000005312 | Ubqln1 | 56085 | protein_coding | 1651.432274 | 0.021852604 | 0.069779909 | -0.313164697 | 0.754155538 | 0.876856731 | 0.021852604 |
| ENSMUSG00000031379 | Pir | 69656 | protein_coding | 77.11770028 | 0.021854234 | 0.15439298 | -0.141549401 | 0.887435937 | 0.944306685 | 0.021854234 |
| ENSMUSG00000046562 | Unc119b | 106840 | protein_coding | 581.0162301 | 0.021889828 | 0.095918522 | -0.228212734 | 0.819480862 | 0.908985828 | 0.021889828 |
| ENSMUSG00000021772 | Nkiras1 | 69721 | protein_coding | 92.83251593 | 0.021902842 | 0.121195134 | -0.180723772 | 0.856584399 | 0.930204755 | 0.021902842 |
| ENSMUSG00000098274 | Rpl24 | 68193 | protein_coding | 144.2202308 | 0.022043859 | 0.151877513 | -0.145142347 | 0.884598484 | 0.942955032 | 0.022043859 |
| ENSMUSG00000023075 | Akirin1 | 68050 | protein_coding | 667.7667488 | -0.022083974 | 0.084788753 | 0.260458762 | 0.794509921 | 0.897715376 | 0.022083974 |
| ENSMUSG00000043929 | Klhl15 | 236904 | protein_coding | 124.2970006 | 0.022099092 | 0.126192061 | -0.175122684 | 0.860983234 | 0.931580268 | 0.022099092 |
| ENSMUSG00000060950 | Trmt61a | 328162 | protein_coding | 85.02265853 | -0.022143405 | 0.195843935 | 0.113066584 | 0.909977767 | 0.95630475 | 0.022143405 |
| ENSMUSG00000022035 | Ccdc25 | 67179 | protein_coding | 525.2613602 | -0.02216458 | 0.102687794 | 0.215844345 | 0.829109081 | 0.913524643 | 0.02216458 |
| ENSMUSG00000046962 | Zbtb21 | 114565 | protein_coding | 268.880694 | -0.022179271 | 0.0781331 | 0.283865242 | 0.776513654 | 0.888506311 | 0.022179271 |
| ENSMUSG00000021590 | Spata9 | 75571 | protein_coding | 3.452038104 | 0.022180583 | 0.840476937 | -0.026390472 | 0.978945894 | 1 | 0.022180583 |
| ENSMUSG00000034105 | Meak7 | 74347 | protein_coding | 253.0630914 | -0.022199311 | 0.118653643 | 0.18709338 | 0.851587419 | 0.927499565 | 0.022199311 |
| ENSMUSG00000032869 | Psmf1 | 228769 | protein_coding | 316.2964239 | 0.022283897 | 0.066534058 | -0.334924672 | 0.73768191 | 0.866866795 | 0.022283897 |
| ENSMUSG00000021025 | Nfkbia | 18035 | protein_coding | 1365.832937 | -0.022290147 | 0.23865999 | 0.093397083 | 0.925588107 | 0.964363534 | 0.022290147 |
| ENSMUSG00000042700 | Sipa1l1 | 217692 | protein_coding | 1443.128359 | 0.022308718 | 0.075384795 | -0.295931276 | 0.767282565 | 0.883364555 | 0.022308718 |
| ENSMUSG00000027651 | Rprd1b | 70470 | protein_coding | 406.2014306 | -0.022315367 | 0.055583684 | 0.401473345 | 0.688071659 | 0.833786242 | 0.022315367 |
| ENSMUSG00000001751 | Naglu | 27419 | protein_coding | 376.0621458 | 0.022328734 | 0.098364693 | -0.226999477 | 0.820424149 | 0.9092783 | 0.022328734 |
| ENSMUSG00000003346 | Abhd17a | 216169 | protein_coding | 936.1278651 | -0.022335931 | 0.098211706 | 0.227426357 | 0.820092227 | 0.909231489 | 0.022335931 |
| ENSMUSG00000021962 | Dcp1a | 75901 | protein_coding | 330.7767239 | 0.022349482 | 0.06844019 | -0.32655493 | 0.744004537 | 0.870923726 | 0.022349482 |
| ENSMUSG00000044982 | Sft2d3 | 67158 | protein_coding | 72.0861204 | -0.022354416 | 0.12057786 | 0.185394032 | 0.852919989 | 0.92788168 | 0.022354416 |
| ENSMUSG00000024036 | Slc37a1 | 224674 | protein_coding | 2369.972576 | -0.022381918 | 0.062765221 | 0.356597454 | 0.721393184 | 0.855640813 | 0.022381918 |
| ENSMUSG00000060121 | Gemin2 | 66603 | protein_coding | 72.70527017 | -0.022406353 | 0.138983713 | 0.161215671 | 0.871923538 | 0.936943344 | 0.022406353 |
| ENSMUSG00000004931 | Apba3 | 57267 | protein_coding | 451.2274431 | -0.022429974 | 0.095273669 | 0.235426789 | 0.813877496 | 0.907703989 | 0.022429974 |
| ENSMUSG00000027312 | Atrn | 11990 | protein_coding | 780.6632974 | -0.022446365 | 0.078771343 | 0.284955973 | 0.775677872 | 0.88791795 | 0.022446365 |
| ENSMUSG00000042599 | Kdm7a | 338523 | protein_coding | 1083.957843 | 0.022447067 | 0.087047049 | -0.257872808 | 0.796505071 | 0.898891871 | 0.022447067 |
| ENSMUSG00000018583 | G3bp1 | 27041 | protein_coding | 2038.528828 | 0.022477136 | 0.114540489 | -0.19623747 | 0.844424302 | 0.923559369 | 0.022477136 |

| ensembl_gene_id | external_gene_name | entrezgene_id | gene_biotype | baseMean | log2FoldChange | lfcSE | stat | pvalue | padj | absLog2FC |
|---|---|---|---|---|---|---|---|---|---|---|
| ENSMUSG00000021576 | Pdcd6 | 18570 | protein_coding | 823.4713197 | -0.022507771 | 0.056373116 | 0.399264275 | 0.689698487 | 0.834370416 | 0.022507771 |
| ENSMUSG00000001988 | Npas1 | 18142 | protein_coding | 7.849728765 | 0.022516297 | 0.510847161 | -0.044076387 | 0.964843515 | 0.982587667 | 0.022516297 |
| ENSMUSG00000060373 | Hnrnpc | 15381 | protein_coding | 1708.710343 | -0.022546667 | 0.097674299 | 0.230835207 | 0.817442828 | 0.908714429 | 0.022546667 |
| ENSMUSG00000050043 | Tmx2 | 66958 | protein_coding | 407.6303472 | 0.02255415 | 0.104000422 | -0.216865952 | 0.828312812 | 0.913317606 | 0.02255415 |
| ENSMUSG00000023987 | Pgc | 109820 | protein_coding | 1.929045266 | 0.022594638 | 1.957629537 | -0.011541835 | 0.990791152 | 1 | 0.022594638 |
| ENSMUSG00000029635 | Cdk8 | 264064 | protein_coding | 478.9562782 | 0.0226503 | 0.069437663 | -0.326196177 | 0.744275934 | 0.870990647 | 0.0226503 |
| ENSMUSG00000029384 | 2010109A12Rik | 75610 | protein_coding | 4.769351291 | 0.022757241 | 0.576617418 | -0.039466794 | 0.968518227 | 0.984122985 | 0.022757241 |
| ENSMUSG00000026014 | Raph1 | 77300 | protein_coding | 620.1006443 | 0.022789905 | 0.17117394 | -0.133138871 | 0.894083554 | 0.947786691 | 0.022789905 |
| ENSMUSG00000021494 | Ddx41 | 72935 | protein_coding | 393.822221 | 0.02279054 | 0.073753284 | -0.309010507 | 0.757313534 | 0.878311377 | 0.02279054 |
| ENSMUSG00000117648 | Gm32282 | 102634780 | lncRNA | 2.258762352 | -0.022795634 | 0.767067117 | 0.029717914 | 0.976292025 | 1 | 0.022795634 |
| ENSMUSG00000033352 | Map2k4 | 26398 | protein_coding | 590.2213792 | 0.02281479 | 0.082257008 | -0.277359827 | 0.781503825 | 0.890612766 | 0.02281479 |
| ENSMUSG00000019969 | Psen1 | 19164 | protein_coding | 2121.944964 | -0.022825967 | 0.052726637 | 0.43291149 | 0.665079072 | 0.819390913 | 0.022825967 |
| ENSMUSG00000029054 | Gabrd | 14403 | protein_coding | 9.870224868 | -0.022826165 | 0.529357605 | 0.043120501 | 0.965605477 | 0.982706396 | 0.022826165 |
| ENSMUSG00000070327 | Rnf213 | 672511 | protein_coding | 2399.813876 | 0.022838147 | 0.186993267 | -0.122133527 | 0.90279327 | 0.95200802 | 0.022838147 |
| ENSMUSG00000006205 | Htra1 | 56213 | protein_coding | 1.526988781 | -0.022848356 | 1.111123467 | 0.020563292 | 0.983594023 | 1 | 0.022848356 |
| ENSMUSG00000004591 | Pkn2 | 109333 | protein_coding | 1488.802224 | 0.022852622 | 0.056366477 | -0.405429303 | 0.685161975 | 0.832151567 | 0.022852622 |
| ENSMUSG00000029561 | Oasl2 | 23962 | protein_coding | 378.9761745 | -0.022853082 | 0.460802192 | 0.049594125 | 0.960445828 | 0.980697519 | 0.022853082 |
| ENSMUSG00000043621 | Ubxn10 | 212190 | protein_coding | 120.1361099 | 0.022899585 | 0.1468951 | -0.155890734 | 0.876119149 | 0.938630585 | 0.022899585 |
| ENSMUSG00000027502 | Rtf2 | 66404 | protein_coding | 817.1982904 | 0.022908481 | 0.05970656 | -0.383684488 | 0.701212311 | 0.842046256 | 0.022908481 |
| ENSMUSG00000004945 | Tmem242 | 70544 | protein_coding | 424.2974948 | -0.022918071 | 0.101125509 | 0.22662997 | 0.820711486 | 0.909368591 | 0.022918071 |
| ENSMUSG00000028517 | Plpp3 | 67916 | protein_coding | 65.39564912 | -0.022926999 | 0.167112857 | 0.137194704 | 0.890876897 | 0.946592528 | 0.022926999 |
| ENSMUSG00000029599 | Ddx54 | 71990 | protein_coding | 992.8838063 | -0.022938897 | 0.05107176 | 0.449150319 | 0.653323224 | 0.811872956 | 0.022938897 |
| ENSMUSG00000055408 | Hottip | 791364 | lncRNA | 188.298668 | -0.02298329 | 0.207520281 | 0.110752021 | 0.911812993 | 0.957015366 | 0.02298329 |
| ENSMUSG00000041712 | Ubr7 | 66622 | protein_coding | 311.8382255 | -0.022987788 | 0.089760269 | 0.25610204 | 0.797872047 | 0.899696756 | 0.022987788 |
| ENSMUSG00000026809 | Spaca9 | 69987 | protein_coding | 1.794774418 | 0.023038858 | 0.898219441 | -0.025649476 | 0.979536923 | 1 | 0.023038858 |
| ENSMUSG00000040842 | Szrd1 | 213491 | protein_coding | 1426.485452 | 0.023044213 | 0.064304519 | -0.358360701 | 0.720073397 | 0.855153117 | 0.023044213 |
| ENSMUSG00000010045 | Tmem115 | 56395 | protein_coding | 591.2177238 | 0.023130235 | 0.06145808 | -0.376357922 | 0.706650823 | 0.846208704 | 0.023130235 |
| ENSMUSG00000039536 | Stau1 | 20853 | protein_coding | 487.2401182 | -0.023195197 | 0.071872777 | 0.322725768 | 0.746902935 | 0.872440598 | 0.023195197 |
| ENSMUSG00000094091 | Ankrd40cl | 102639132 | protein_coding | 430.4136744 | -0.023197839 | 0.249396305 | 0.093015969 | 0.925890875 | 0.964583514 | 0.023197839 |
| ENSMUSG00000027219 | Slc28a2 | 269346 | protein_coding | 94.07853793 | -0.023255107 | 0.186645517 | 0.124595045 | 0.900844151 | 0.951246263 | 0.023255107 |
| ENSMUSG00000036376 | Abt1 | 30946 | protein_coding | 149.3202402 | -0.023271642 | 0.104197367 | 0.223341943 | 0.823269386 | 0.910297568 | 0.023271642 |
| ENSMUSG00000031497 | Tnfsf13b | 24099 | protein_coding | 51.60539284 | 0.023278894 | 0.200498436 | -0.116105116 | 0.907569235 | 0.954867376 | 0.023278894 |
| ENSMUSG00000022350 | Washc5 | 223593 | protein_coding | 560.9633071 | 0.023284061 | 0.070733802 | -0.329178692 | 0.742020628 | 0.869666246 | 0.023284061 |
| ENSMUSG00000043541 | Casc1 | 320662 | protein_coding | 10.12888402 | -0.023298723 | 0.572513563 | 0.040695496 | 0.967538653 | 0.983741738 | 0.023298723 |
| ENSMUSG00000025915 | Sgk3 | 170755 | protein_coding | 61.23617362 | -0.023300443 | 0.157142707 | 0.14827569 | 0.8821252 | 0.941960269 | 0.023300443 |
| ENSMUSG00000071019 | Sdr16c6 | 242286 | protein_coding | 8.152936471 | -0.023318047 | 0.576843466 | 0.040423525 | 0.967755475 | 0.983901614 | 0.023318047 |
| ENSMUSG00000024425 | Ndfip1 | 65113 | protein_coding | 1128.279776 | -0.023350186 | 0.081787807 | 0.285497156 | 0.775263283 | 0.887843124 | 0.023350186 |
| ENSMUSG00000039148 | Sart1 | 20227 | protein_coding | 699.9519779 | 0.023359414 | 0.052368501 | -0.446058476 | 0.655555007 | 0.81317775 | 0.023359414 |
| ENSMUSG00000042389 | Tsen2 | 381802 | protein_coding | 67.46692245 | 0.02337026 | 0.198427941 | -0.117777061 | 0.906244306 | 0.954081249 | 0.02337026 |

| ensembl_gene_id | external_gene_name | entrezgene_id | gene_biotype | baseMean | log2FoldChange | lfcSE | stat | pvalue | padj | absLog2FC |
|---|---|---|---|---|---|---|---|---|---|---|
| ENSMUSG00000090176 | Cd200r2 | 271375 | protein_coding | 67.10904905 | -0.023373302 | 0.219437612 | 0.106514568 | 0.915174098 | 0.95833356 | 0.023373302 |
| ENSMUSG00000043587 | Pxylp1 | 235534 | protein_coding | 561.0073743 | -0.023380032 | 0.149927839 | 0.1559419 | 0.876078818 | 0.938630585 | 0.023380032 |
| ENSMUSG00000100727 | 1700003P14Rik | 75890 | lncRNA | 0.767302246 | 0.02338233 | 1.702249992 | -0.013736132 | 0.989040497 | 1 | 0.02338233 |
| ENSMUSG00000027207 | Galk2 | 69976 | protein_coding | 591.7131392 | -0.023410196 | 0.102569284 | 0.228237883 | 0.819461312 | 0.908985828 | 0.023410196 |
| ENSMUSG00000048486 | Fitm2 | 228859 | protein_coding | 250.5893746 | -0.023412304 | 0.110708897 | 0.211476266 | 0.83251565 | 0.915629096 | 0.023412304 |
| ENSMUSG00000035125 | Gcfc2 | 330361 | protein_coding | 70.6160768 | 0.023415172 | 0.144942452 | -0.161548062 | 0.87166176 | 0.936825965 | 0.023415172 |
| ENSMUSG00000036885 | Arhgef26 | 622434 | protein_coding | 95.97030486 | 0.023441679 | 0.137939199 | -0.169942114 | 0.865055661 | 0.932932188 | 0.023441679 |
| ENSMUSG00000031449 | Atp4b | 11945 | protein_coding | 1.287689153 | -0.02344889 | 2.052601201 | 0.011423987 | 0.990885175 | 1 | 0.02344889 |
| ENSMUSG00000026584 | Scyl3 | 240880 | protein_coding | 327.9850113 | -0.023452992 | 0.07244851 | 0.323719459 | 0.746150437 | 0.872287153 | 0.023452992 |
| ENSMUSG00000029047 | Pex10 | 668173 | protein_coding | 102.4749671 | 0.023484124 | 0.12184457 | -0.192738372 | 0.847163871 | 0.924722861 | 0.023484124 |
| ENSMUSG00000022779 | Top3b | 21976 | protein_coding | 474.8050027 | 0.023567597 | 0.05204183 | -0.452858722 | 0.650650476 | 0.810197705 | 0.023567597 |
| ENSMUSG00000030986 | Dhx32 | 101437 | protein_coding | 1731.987154 | -0.023631275 | 0.079218769 | 0.298303989 | 0.76547116 | 0.882510888 | 0.023631275 |
| ENSMUSG00000021156 | Zmynd11 | 66505 | protein_coding | 572.5057062 | 0.023636953 | 0.04701495 | -0.502753962 | 0.615137265 | 0.787654245 | 0.023636953 |
| ENSMUSG00000019173 | Rab5c | 19345 | protein_coding | 1724.214462 | -0.023660149 | 0.068355783 | 0.346132369 | 0.729243238 | 0.86030736 | 0.023660149 |
| ENSMUSG00000046873 | Mbtps2 | 270669 | protein_coding | 150.3522418 | 0.02366344 | 0.119512173 | -0.19800025 | 0.843044868 | 0.922356748 | 0.02366344 |
| ENSMUSG00000024736 | Tmem132a | 98170 | protein_coding | 51.42025214 | -0.023671079 | 0.151263426 | 0.156489111 | 0.875647502 | 0.938630585 | 0.023671079 |
| ENSMUSG00000025218 | Poll | 56626 | protein_coding | 131.1007463 | 0.023675479 | 0.104333442 | -0.226921287 | 0.820484949 | 0.9092783 | 0.023675479 |
| ENSMUSG00000030019 | Fbxl14 | 101358 | protein_coding | 531.3006424 | 0.023705135 | 0.078607681 | -0.301562585 | 0.762985534 | 0.881056435 | 0.023705135 |
| ENSMUSG00000056153 | Socs6 | 54607 | protein_coding | 630.946538 | -0.023707059 | 0.071787304 | 0.33024028 | 0.741218413 | 0.869043726 | 0.023707059 |
| ENSMUSG00000104960 | Snhg8 | 69895 | lncRNA | 145.0502993 | -0.023738594 | 0.167215122 | 0.141964399 | 0.887108126 | 0.944106696 | 0.023738594 |
| ENSMUSG00000008429 | Herpud2 | 80517 | protein_coding | 324.1871457 | 0.023764623 | 0.095799522 | -0.248066198 | 0.804083188 | 0.903431064 | 0.023764623 |
| ENSMUSG00000086544 | Chn1os3 | 329427 | lncRNA | 2.627049784 | 0.023767254 | 0.93667094 | -0.025374177 | 0.979756508 | 1 | 0.023767254 |
| ENSMUSG00000026718 | Stam | 20844 | protein_coding | 549.9711771 | -0.023782587 | 0.077851482 | 0.305486632 | 0.759995549 | 0.879445484 | 0.023782587 |
| ENSMUSG00000031681 | Smad1 | 17125 | protein_coding | 692.3280126 | -0.023899163 | 0.066884485 | 0.357319984 | 0.720852271 | 0.855367853 | 0.023899163 |
| ENSMUSG00000092981 | Mir5125 | 100628593 | miRNA | 14.41343606 | -0.023913029 | 0.349671024 | 0.06838722 | 0.945477395 | 0.973720842 | 0.023913029 |
| ENSMUSG00000023932 | Cdc5l | 71702 | protein_coding | 797.9172009 | -0.023929167 | 0.108195761 | 0.221165476 | 0.82496359 | 0.911456552 | 0.023929167 |
| ENSMUSG00000029788 | Cpa5 | 74649 | protein_coding | 1.127913505 | -0.023936136 | 1.935120322 | 0.012369327 | 0.990130956 | 1 | 0.023936136 |
| ENSMUSG00000044308 | Ubr3 | 68795 | protein_coding | 1134.920819 | 0.023960158 | 0.075960417 | -0.315429523 | 0.752435554 | 0.875793979 | 0.023960158 |
| ENSMUSG00000030657 | Xylt1 | 233781 | protein_coding | 142.1495528 | -0.023978244 | 0.237869193 | 0.100804326 | 0.919705792 | 0.961084696 | 0.023978244 |
| ENSMUSG00000022620 | Arsa | 11883 | protein_coding | 257.9916799 | 0.023992338 | 0.068054023 | -0.352548417 | 0.724427015 | 0.857268916 | 0.023992338 |
| ENSMUSG00000025885 | Myo5b | 17919 | protein_coding | 3528.546107 | -0.024001274 | 0.083199653 | 0.288478058 | 0.772980824 | 0.886212533 | 0.024001274 |
| ENSMUSG00000027968 | Larp7 | 28036 | protein_coding | 238.6832536 | 0.024029282 | 0.106183466 | -0.226299657 | 0.820968365 | 0.909531167 | 0.024029282 |
| ENSMUSG00000060450 | Rnf14 | 56736 | protein_coding | 691.4625202 | -0.024073602 | 0.085284021 | 0.28227564 | 0.777732163 | 0.888933284 | 0.024073602 |
| ENSMUSG00000020441 | 2310033P09Rik | 67862 | protein_coding | 103.7293549 | 0.024078543 | 0.114342895 | -0.210581888 | 0.833213546 | 0.915993052 | 0.024078543 |
| ENSMUSG00000021731 | Mrps30 | 59054 | protein_coding | 244.1009014 | 0.024094087 | 0.106232345 | -0.226805565 | 0.820574936 | 0.9092783 | 0.024094087 |
| ENSMUSG00000030816 | Rnf40 | 233900 | protein_coding | 840.9840696 | 0.024135836 | 0.056402003 | -0.42792516 | 0.668705608 | 0.822256556 | 0.024135836 |
| ENSMUSG00000060317 | Acnat2 | 209186 | protein_coding | 0.77354633 | -0.024195822 | 1.94693547 | 0.012427645 | 0.990084429 | 1 | 0.024195822 |
| ENSMUSG00000022853 | Ehhadh | 74147 | protein_coding | 703.6897829 | 0.024202596 | 0.078871906 | -0.306859523 | 0.7589503 | 0.878912418 | 0.024202596 |
| ENSMUSG00000042705 | Commd10 | 69456 | protein_coding | 203.045889 | -0.024219601 | 0.091979344 | 0.263315654 | 0.792307295 | 0.896636371 | 0.024219601 |

Supplementary Table 1

| ensembl_gene_id | external_gene_name | entrezgene_id | gene_biotype | baseMean | log2FoldChange | lfcSE | stat | pvalue | padj | absLog2FC |
|---|---|---|---|---|---|---|---|---|---|---|
| ENSMUSG00000015714 | Cers2 | 76893 | protein_coding | 693.2637983 | 0.024304618 | 0.072120141 | -0.337001811 | 0.736115535 | 0.865765398 | 0.024304618 |
| ENSMUSG00000031584 | Gsr | 14782 | protein_coding | 1652.319842 | 0.024329711 | 0.097470089 | -0.249612073 | 0.802887361 | 0.902682351 | 0.024329711 |
| ENSMUSG00000038859 | Baiap2l1 | 66898 | protein_coding | 1701.702209 | -0.024338443 | 0.100031688 | 0.243307335 | 0.807767323 | 0.905233898 | 0.024338443 |
| ENSMUSG00000058748 | Zfp958 | 233987 | protein_coding | 114.5590951 | -0.024347293 | 0.150982375 | 0.161259172 | 0.871889279 | 0.936943344 | 0.024347293 |
| ENSMUSG00000029550 | Sppl3 | 74585 | protein_coding | 563.0516496 | -0.024353007 | 0.057807972 | 0.421274199 | 0.673554868 | 0.824971081 | 0.024353007 |
| ENSMUSG00000044730 | 9930104L06Rik | 194268 | protein_coding | 93.41051303 | -0.024364311 | 0.106812787 | 0.228102939 | 0.819566215 | 0.909014472 | 0.024364311 |
| ENSMUSG00000052557 | Gan | 209239 | protein_coding | 459.3403553 | -0.024397566 | 0.103149678 | 0.236525858 | 0.813024644 | 0.907217398 | 0.024397566 |
| ENSMUSG00000032263 | Bckdhb | 12040 | protein_coding | 89.94772499 | -0.024435906 | 0.150834859 | 0.162004365 | 0.871302416 | 0.936580622 | 0.024435906 |
| ENSMUSG00000021277 | Traf3 | 22031 | protein_coding | 349.6321721 | 0.024443397 | 0.109556082 | -0.223113095 | 0.823447487 | 0.910384401 | 0.024443397 |
| ENSMUSG00000027796 | Smad9 | 55994 | protein_coding | 4.826303239 | -0.024443702 | 0.515571786 | 0.04741086 | 0.962185773 | 0.981434574 | 0.024443702 |
| ENSMUSG00000027463 | Slc52a3 | 69698 | protein_coding | 493.0956375 | -0.024453735 | 0.09308978 | 0.262689791 | 0.792789686 | 0.896829747 | 0.024453735 |
| ENSMUSG00000062470 | Fbxl12os | 66662 | lncRNA | 9.547693362 | -0.024478013 | 0.390190785 | 0.062733447 | 0.949978763 | 0.975869522 | 0.024478013 |
| ENSMUSG00000054814 | Usp46 | 69727 | protein_coding | 330.4638833 | -0.024479097 | 0.106754342 | 0.229303054 | 0.81863338 | 0.908812719 | 0.024479097 |
| ENSMUSG00000047281 | Sfn | 55948 | protein_coding | 3686.64763 | -0.024486508 | 0.139501578 | 0.175528538 | 0.860664348 | 0.931474719 | 0.024486508 |
| ENSMUSG00000054894 | Dmac2l | 68055 | protein_coding | 96.13079775 | -0.024492227 | 0.125517586 | 0.195129843 | 0.845291301 | 0.923801556 | 0.024492227 |
| ENSMUSG00000048307 | Ankrd46 | 68839 | protein_coding | 255.2132908 | -0.024499205 | 0.08092599 | 0.302735928 | 0.762091115 | 0.880874346 | 0.024499205 |
| ENSMUSG00000086877 | A230072C01Rik | 320742 | lncRNA | 7.91996384 | 0.024514094 | 0.431525819 | -0.056807942 | 0.954698187 | 0.977426278 | 0.024514094 |
| ENSMUSG00000025036 | Sfxn2 | 94279 | protein_coding | 188.0910849 | 0.024598774 | 0.130759232 | -0.188122658 | 0.8507805 | 0.927198731 | 0.024598774 |
| ENSMUSG00000030909 | Anks4b | 72074 | protein_coding | 1690.25835 | -0.024638401 | 0.098365164 | 0.250478935 | 0.802216993 | 0.90234117 | 0.024638401 |
| ENSMUSG00000019826 | Zbtb24 | 268294 | protein_coding | 199.5242096 | 0.024655183 | 0.101935229 | -0.24187107 | 0.808880067 | 0.905678041 | 0.024655183 |
| ENSMUSG00000038417 | Fig4 | 103199 | protein_coding | 317.9947148 | -0.024676458 | 0.081771489 | 0.301773374 | 0.762824829 | 0.881056435 | 0.024676458 |
| ENSMUSG00000118249 | Gm36608 | 102640578 | lncRNA | 1.390703487 | 0.024717413 | 0.952711182 | -0.025944288 | 0.979301775 | 1 | 0.024717413 |
| ENSMUSG00000056310 | Tyw1 | 100929 | protein_coding | 279.783445 | 0.02476003 | 0.085170251 | -0.290712188 | 0.77127145 | 0.885199039 | 0.02476003 |
| ENSMUSG00000024381 | Bin1 | 30948 | protein_coding | 167.2717602 | 0.024769901 | 0.0985788 | -0.251270058 | 0.801605323 | 0.901751383 | 0.024769901 |
| ENSMUSG00000037351 | Actr1b | 226977 | protein_coding | 1988.520245 | -0.024774675 | 0.043999432 | 0.563068058 | 0.573388541 | 0.758644032 | 0.024774675 |
| ENSMUSG00000025340 | Rabgef1 | 56715 | protein_coding | 370.7417339 | -0.02487062 | 0.070729312 | 0.351631021 | 0.725114997 | 0.857652848 | 0.02487062 |
| ENSMUSG00000024764 | Naa40 | 70999 | protein_coding | 447.3510272 | -0.024874275 | 0.108034665 | 0.230243462 | 0.817902591 | 0.908812719 | 0.024874275 |
| ENSMUSG00000091803 | Cox16 | 66272 | protein_coding | 103.9265304 | -0.024908542 | 0.127514749 | 0.19533852 | 0.845127943 | 0.923798341 | 0.024908542 |
| ENSMUSG00000024384 | Iws1 | 73473 | protein_coding | 766.3319381 | -0.024934443 | 0.070599722 | 0.353180475 | 0.723953144 | 0.857076652 | 0.024934443 |
| ENSMUSG00000070644 | Etnk2 | 214253 | protein_coding | 15.49521301 | -0.024939959 | 0.370397797 | 0.067332904 | 0.946316683 | 0.973795719 | 0.024939959 |
| ENSMUSG00000021263 | Degs2 | 70059 | protein_coding | 2217.608908 | 0.024954827 | 0.084683484 | -0.294683518 | 0.768235655 | 0.883975099 | 0.024954827 |
| ENSMUSG00000030205 | Gprc5d | 93746 | protein_coding | 0.584102615 | -0.024958935 | 1.633164761 | 0.015282558 | 0.987806758 | 1 | 0.024958935 |
| ENSMUSG00000024218 | Taf11 | 68776 | protein_coding | 157.9914705 | -0.024964718 | 0.083743458 | 0.298109474 | 0.765619611 | 0.882510888 | 0.024964718 |
| ENSMUSG00000052428 | Tmco1 | 68944 | protein_coding | 532.4164214 | 0.024990846 | 0.113522353 | -0.220140311 | 0.825761882 | 0.91190529 | 0.024990846 |
| ENSMUSG00000040128 | Pnrc1 | 108767 | protein_coding | 1156.070224 | 0.025134395 | 0.185984149 | -0.135142671 | 0.892499073 | 0.947202684 | 0.025134395 |
| ENSMUSG00000026142 | Rhbdd1 | 76867 | protein_coding | 315.8517257 | 0.025187809 | 0.067897308 | -0.370969182 | 0.710660485 | 0.849070124 | 0.025187809 |
| ENSMUSG00000036667 | Tcaf1 | 77574 | protein_coding | 240.3176082 | 0.02519433 | 0.089672084 | -0.280960687 | 0.778740553 | 0.889481359 | 0.02519433 |
| ENSMUSG00000029621 | Arpc1a | 56443 | protein_coding | 926.287345 | -0.025243508 | 0.074585042 | 0.338452684 | 0.735022078 | 0.864910546 | 0.025243508 |
| ENSMUSG00000042535 | Gtpbp1 | 14904 | protein_coding | 519.9626541 | 0.025272472 | 0.059820242 | -0.422473591 | 0.672679373 | 0.824518194 | 0.025272472 |

| ensembl_gene_id | external_gene_name | entrezgene_id | gene_biotype | baseMean | log2FoldChange | lfcSE | stat | pvalue | padj | absLog2FC |
|---|---|---|---|---|---|---|---|---|---|---|
| ENSMUSG00000042410 | Agps | 228061 | protein_coding | 1105.740173 | -0.025283286 | 0.076524907 | 0.330392901 | 0.741103105 | 0.868970227 | 0.025283286 |
| ENSMUSG00000024400 | Wdr33 | 74320 | protein_coding | 709.0310368 | 0.025293348 | 0.051349552 | -0.492571923 | 0.622315091 | 0.792910378 | 0.025293348 |
| ENSMUSG00000028076 | Cd1d1 | 12479 | protein_coding | 57.53808719 | 0.025306402 | 0.208896807 | -0.121143079 | 0.903577708 | 0.952305972 | 0.025306402 |
| ENSMUSG00000034190 | Chmp7 | 105513 | protein_coding | 525.8498826 | 0.02533432 | 0.050162983 | -0.505040145 | 0.61353064 | 0.786448489 | 0.02533432 |
| ENSMUSG00000102923 | Gm31728 | 102634049 | lncRNA | 1.876452905 | 0.025335847 | 0.909188658 | -0.027866435 | 0.977768679 | 1 | 0.025335847 |
| ENSMUSG00000021549 | Rasa1 | 218397 | protein_coding | 571.1916931 | 0.0253641 | 0.076413315 | -0.331932988 | 0.73993986 | 0.868161055 | 0.0253641 |
| ENSMUSG00000043542 | Zc2hc1a | 67306 | protein_coding | 89.828691 | -0.025373964 | 0.144509539 | 0.175586772 | 0.860618595 | 0.931474719 | 0.025373964 |
| ENSMUSG00000045211 | Nudt18 | 213484 | protein_coding | 65.1896984 | -0.025374562 | 0.142929189 | 0.177532401 | 0.859090217 | 0.931099354 | 0.025374562 |
| ENSMUSG00000018446 | C1qbp | 12261 | protein_coding | 659.08764 | -0.025385037 | 0.101886407 | 0.249150379 | 0.803244461 | 0.902857271 | 0.025385037 |
| ENSMUSG00000029470 | P2rx4 | 18438 | protein_coding | 1492.732949 | 0.025385804 | 0.156563874 | -0.16214343 | 0.871192906 | 0.936523867 | 0.025385804 |
| ENSMUSG00000020130 | Tbc1d15 | 66687 | protein_coding | 546.4920126 | -0.02540319 | 0.146933616 | 0.172888895 | 0.862738761 | 0.932166328 | 0.02540319 |
| ENSMUSG00000032624 | Eml4 | 78798 | protein_coding | 1098.984992 | -0.025412526 | 0.109330563 | 0.232437533 | 0.8161982 | 0.908409285 | 0.025412526 |
| ENSMUSG00000024234 | Mtpap | 67440 | protein_coding | 266.1735047 | -0.025424452 | 0.094628901 | 0.268675335 | 0.788179534 | 0.894017228 | 0.025424452 |
| ENSMUSG00000022898 | Vps26c | 13185 | protein_coding | 490.1219003 | -0.02543624 | 0.055046448 | 0.462086858 | 0.644019034 | 0.80597323 | 0.02543624 |
| ENSMUSG00000046756 | Mrps7 | 50529 | protein_coding | 482.3950551 | -0.025444986 | 0.080198866 | 0.317273643 | 0.751035975 | 0.875081807 | 0.025444986 |
| ENSMUSG00000041328 | Pcf11 | 74737 | protein_coding | 1019.938773 | -0.025453101 | 0.08507407 | 0.299187528 | 0.764796965 | 0.882219689 | 0.025453101 |
| ENSMUSG00000027663 | Zmat3 | 22401 | protein_coding | 207.815639 | 0.025465434 | 0.111960651 | -0.227449858 | 0.820073955 | 0.909231489 | 0.025465434 |
| ENSMUSG00000005823 | Gpr108 | 78308 | protein_coding | 1154.099185 | 0.025467101 | 0.048101052 | -0.529449982 | 0.596493333 | 0.774662172 | 0.025467101 |
| ENSMUSG00000033382 | Trappc8 | 75964 | protein_coding | 1073.980405 | 0.025556398 | 0.065576542 | -0.389718595 | 0.696744645 | 0.839060823 | 0.025556398 |
| ENSMUSG00000098022 | Zfp82 | 330502 | protein_coding | 3.295089213 | -0.025604036 | 0.579985312 | 0.044146007 | 0.96478802 | 1 | 0.025604036 |
| ENSMUSG00000020114 | Cand1 | 71902 | protein_coding | 1318.162028 | 0.025620584 | 0.077936961 | -0.32873471 | 0.742356216 | 0.869945398 | 0.025620584 |
| ENSMUSG00000040543 | Pitpnm3 | 327958 | protein_coding | 178.9029784 | -0.025625957 | 0.145853329 | 0.175696757 | 0.860532183 | 0.931474719 | 0.025625957 |
| ENSMUSG00000094870 | Zfp131 | 72465 | protein_coding | 322.0304232 | 0.025659682 | 0.080981008 | -0.316860497 | 0.751349458 | 0.875081807 | 0.025659682 |
| ENSMUSG00000074896 | Ifit3 | 15959 | protein_coding | 48.67569434 | 0.025698077 | 0.437042705 | -0.058799922 | 0.953111471 | 0.976710073 | 0.025698077 |
| ENSMUSG00000029071 | Dvl1 | 13542 | protein_coding | 741.5323333 | -0.025735671 | 0.098510688 | 0.261247497 | 0.793901652 | 0.897263004 | 0.025735671 |
| ENSMUSG00000034863 | Ano8 | 382014 | protein_coding | 182.0174952 | 0.025772377 | 0.119081864 | -0.216425708 | 0.828655929 | 0.913504709 | 0.025772377 |
| ENSMUSG00000025358 | Cdk2 | 12566 | protein_coding | 228.5310284 | 0.025774013 | 0.141668249 | -0.18193218 | 0.85563595 | 0.929786937 | 0.025774013 |
| ENSMUSG00000052549 | Arl13a | 74448 | protein_coding | 1.62745793 | -0.025794406 | 1.121467747 | 0.023000577 | 0.981649812 | 1 | 0.025794406 |
| ENSMUSG00000041741 | Pde3a | 54611 | protein_coding | 1375.498335 | -0.025845399 | 0.090470079 | 0.285678966 | 0.775124016 | 0.88783793 | 0.025845399 |
| ENSMUSG00000036323 | Srp72 | 66661 | protein_coding | 1122.909158 | 0.025863753 | 0.10278369 | -0.251632853 | 0.801324862 | 0.901569434 | 0.025863753 |
| ENSMUSG00000029802 | Abcg2 | 26357 | protein_coding | 989.3474481 | 0.025885196 | 0.120649273 | -0.214549124 | 0.830118864 | 0.914175515 | 0.025885196 |
| ENSMUSG00000026289 | Atg16l1 | 77040 | protein_coding | 392.1685866 | -0.025970647 | 0.077351913 | 0.33574667 | 0.737061909 | 0.866569868 | 0.025970647 |
| ENSMUSG00000067365 | Tmem128 | 66309 | protein_coding | 329.7400322 | 0.02599852 | 0.087524646 | -0.297042279 | 0.766434231 | 0.882690208 | 0.02599852 |
| ENSMUSG00000025019 | Lcor | 212391 | protein_coding | 934.378249 | 0.025999358 | 0.05805942 | -0.447806027 | 0.654293193 | 0.812407823 | 0.025999358 |
| ENSMUSG00000029190 | D5Ertd579e | 320661 | protein_coding | 550.9171374 | -0.026040617 | 0.107857038 | 0.241436418 | 0.80921689 | 0.905779176 | 0.026040617 |
| ENSMUSG00000035212 | Leprot | 230514 | protein_coding | 831.3905088 | -0.0260487 | 0.07068912 | 0.368496595 | 0.712502983 | 0.850598485 | 0.0260487 |
| ENSMUSG00000009907 | Vps4b | 20479 | protein_coding | 1618.563379 | 0.026052428 | 0.052869346 | -0.492770014 | 0.6221751 | 0.792833975 | 0.026052428 |
| ENSMUSG00000035629 | Rubcn | 100502698 | protein_coding | 437.3279173 | 0.026057749 | 0.06605506 | -0.39448528 | 0.693222793 | 0.837078181 | 0.026057749 |
| ENSMUSG00000029480 | Dhx37 | 208144 | protein_coding | 186.3498753 | 0.026067189 | 0.091418504 | -0.285141277 | 0.775535907 | 0.88791795 | 0.026067189 |

| ensembl_gene_id | external_gene_name | entrezgene_id | gene_biotype | baseMean | log2FoldChange | lfcSE | stat | pvalue | padj | absLog2FC |
|---|---|---|---|---|---|---|---|---|---|---|
| ENSMUSG00000023004 | Tuba1b | 22143 | protein_coding | 395.485161 | 0.026131731 | 0.241186594 | -0.108346534 | 0.913720811 | 0.957921881 | 0.026131731 |
| ENSMUSG00000045193 | Cirbp | 12696 | protein_coding | 251.0645325 | 0.026144322 | 0.136969397 | -0.190877108 | 0.848621876 | 0.925637253 | 0.026144322 |
| ENSMUSG00000027465 | Tbc1d20 | 67231 | protein_coding | 1007.34846 | 0.026183089 | 0.067460967 | -0.388122059 | 0.697925709 | 0.839976943 | 0.026183089 |
| ENSMUSG00000100084 | 2310068J16Rik | 70281 | lncRNA | 9.689084562 | 0.026229054 | 0.371998407 | -0.070508512 | 0.943788926 | 0.97289203 | 0.026229054 |
| ENSMUSG00000026563 | Tada1 | 27878 | protein_coding | 102.7396401 | -0.026230709 | 0.109289667 | 0.240010875 | 0.810321826 | 0.906158059 | 0.026230709 |
| ENSMUSG00000042380 | Smim12 | 80284 | protein_coding | 186.844924 | -0.026254346 | 0.086357512 | 0.304019257 | 0.761113221 | 0.880245376 | 0.026254346 |
| ENSMUSG00000037296 | Lsm1 | 67207 | protein_coding | 188.9399079 | 0.026296129 | 0.073330714 | -0.35859639 | 0.719897048 | 0.855153117 | 0.026296129 |
| ENSMUSG00000050714 | Zbtb26 | 320633 | protein_coding | 87.58788839 | -0.026303387 | 0.152476899 | 0.17250736 | 0.863038675 | 0.932166328 | 0.026303387 |
| ENSMUSG00000116564 | Riok2 | 67045 | protein_coding | 229.7384512 | -0.026359277 | 0.114767794 | 0.229674859 | 0.818344433 | 0.908812719 | 0.026359277 |
| ENSMUSG00000002635 | Pdcd2l | 68079 | protein_coding | 132.1380092 | 0.02636089 | 0.107132846 | -0.246057964 | 0.805637363 | 0.904254429 | 0.02636089 |
| ENSMUSG00000029681 | Bcl7b | 12054 | protein_coding | 217.0827428 | -0.026402473 | 0.090663877 | 0.29121271 | 0.770888642 | 0.885199039 | 0.026402473 |
| ENSMUSG00000003363 | Pld3 | 18807 | protein_coding | 339.570721 | -0.026411003 | 0.08338828 | 0.316723198 | 0.751453645 | 0.875092959 | 0.026411003 |
| ENSMUSG00000026519 | Tmem63a | 208795 | protein_coding | 1442.579499 | -0.026416692 | 0.113591991 | 0.232557694 | 0.816104882 | 0.908409285 | 0.026416692 |
| ENSMUSG00000079523 | Tmsb10 | 19240 | protein_coding | 2004.191944 | 0.026449728 | 0.094469334 | -0.279982162 | 0.77949119 | 0.890072723 | 0.026449728 |
| ENSMUSG00000028959 | Fastk | 66587 | protein_coding | 816.276798 | -0.026461362 | 0.073157806 | 0.361702503 | 0.717574356 | 0.853563561 | 0.026461362 |
| ENSMUSG00000027405 | Nop56 | 67134 | protein_coding | 486.0083344 | 0.026462379 | 0.111696272 | -0.236913719 | 0.812723727 | 0.907147741 | 0.026462379 |
| ENSMUSG00000013736 | Trnt1 | 70047 | protein_coding | 286.4460552 | 0.026489297 | 0.101051022 | -0.262137845 | 0.793215171 | 0.897079532 | 0.026489297 |
| ENSMUSG00000042851 | Zc3h6 | 78751 | protein_coding | 69.59675575 | 0.02652958 | 0.195415718 | -0.135759704 | 0.892011248 | 0.947202684 | 0.02652958 |
| ENSMUSG00000019590 | Cyb561 | 13056 | protein_coding | 1222.706352 | -0.026538283 | 0.118554863 | 0.223848118 | 0.822875487 | 0.910240646 | 0.026538283 |
| ENSMUSG00000034321 | Exosc1 | 66583 | protein_coding | 108.5767924 | 0.026545764 | 0.131284365 | -0.2022005 | 0.839759982 | 0.920280009 | 0.026545764 |
| ENSMUSG00000021094 | Dhrs7 | 66375 | protein_coding | 294.3031538 | -0.02656098 | 0.140515398 | 0.189025403 | 0.850072907 | 0.926816617 | 0.02656098 |
| ENSMUSG00000033060 | Lmo7 | 380928 | protein_coding | 3213.51187 | 0.026562066 | 0.11306737 | -0.234922476 | 0.814268905 | 0.907710402 | 0.026562066 |
| ENSMUSG00000034258 | Flvcr2 | 217721 | protein_coding | 3.830669675 | 0.026574321 | 0.847378034 | -0.031360644 | 0.974981927 | 1 | 0.026574321 |
| ENSMUSG00000002210 | Smg9 | 71997 | protein_coding | 288.7587467 | 0.026580454 | 0.094184074 | -0.282218137 | 0.777776252 | 0.888933284 | 0.026580454 |
| ENSMUSG00000003380 | Rabac1 | 14470 | protein_coding | 447.3826873 | 0.026589143 | 0.069582706 | -0.382122874 | 0.702370226 | 0.84294636 | 0.026589143 |
| ENSMUSG00000045817 | Zfp36l2 | 12193 | protein_coding | 2015.553475 | 0.026604543 | 0.099917705 | -0.266264553 | 0.790035471 | 0.895242955 | 0.026604543 |
| ENSMUSG00000042429 | Adora1 | 11539 | protein_coding | 20.47639689 | -0.026621779 | 0.342411291 | 0.07774796 | 0.938028543 | 0.970517013 | 0.026621779 |
| ENSMUSG00000027281 | Slx4ip | 74243 | protein_coding | 161.5268643 | 0.026632597 | 0.113272384 | -0.235119951 | 0.814115634 | 0.907710402 | 0.026632597 |
| ENSMUSG00000004934 | Pias4 | 59004 | protein_coding | 384.084751 | 0.026680563 | 0.076756915 | -0.347598163 | 0.728141986 | 0.859565127 | 0.026680563 |
| ENSMUSG00000018509 | Cenpv | 73139 | protein_coding | 344.5544413 | -0.02670446 | 0.084266953 | 0.316903115 | 0.751317119 | 0.875081807 | 0.02670446 |
| ENSMUSG00000055204 | Ankrd17 | 81702 | protein_coding | 1775.077928 | 0.026742591 | 0.06317487 | -0.423310581 | 0.672068675 | 0.824189967 | 0.026742591 |
| ENSMUSG00000010048 | Ifrd2 | 15983 | protein_coding | 352.5350191 | -0.02675173 | 0.109541699 | 0.244215038 | 0.807064281 | 0.904987729 | 0.02675173 |
| ENSMUSG00000004846 | Plod3 | 26433 | protein_coding | 302.6770271 | 0.026757184 | 0.099440165 | -0.269078236 | 0.787869478 | 0.893788319 | 0.026757184 |
| ENSMUSG00000036172 | Cd200r3 | 74603 | protein_coding | 3.304358162 | -0.026798009 | 0.839588998 | 0.031918008 | 0.974537437 | 1 | 0.026798009 |
| ENSMUSG00000025877 | Hk3 | 212032 | protein_coding | 2.063447716 | -0.026798679 | 1.911409304 | 0.014020377 | 0.988813724 | 1 | 0.026798679 |
| ENSMUSG00000028382 | Ptbp3 | 230257 | protein_coding | 3433.702149 | -0.026840095 | 0.067325703 | 0.398660458 | 0.690143408 | 0.834763659 | 0.026840095 |
| ENSMUSG00000040463 | Mybbp1a | 18432 | protein_coding | 754.3354521 | 0.026845184 | 0.124182828 | -0.216174692 | 0.82885158 | 0.913504709 | 0.026845184 |
| ENSMUSG00000037029 | Zfp146 | 26465 | protein_coding | 348.6647308 | 0.02687483 | 0.119181357 | -0.225495252 | 0.821594019 | 0.909919095 | 0.02687483 |
| ENSMUSG00000034793 | G6pc3 | 68401 | protein_coding | 373.4205932 | -0.026883338 | 0.114913861 | 0.233943386 | 0.815028931 | 0.908068261 | 0.026883338 |

| ensembl_gene_id | external_gene_name | entrezgene_id | gene_biotype | baseMean | log2FoldChange | lfcSE | stat | pvalue | padj | absLog2FC |
|---|---|---|---|---|---|---|---|---|---|---|
| ENSMUSG00000019984 | Med23 | 70208 | protein_coding | 443.2069647 | 0.026913402 | 0.075089865 | -0.358415902 | 0.720032093 | 0.855153117 | 0.026913402 |
| ENSMUSG00000005131 | 4930550C14Rik | 75311 | protein_coding | 3.757173847 | 0.026937139 | 0.536715363 | -0.050188873 | 0.959971879 | 1 | 0.026937139 |
| ENSMUSG00000032482 | Cspg5 | 29873 | protein_coding | 14.38678507 | -0.026972572 | 0.573354154 | 0.047043475 | 0.962478578 | 0.981497544 | 0.026972572 |
| ENSMUSG00000051390 | Zbtb22 | 81630 | protein_coding | 493.1156869 | -0.026981613 | 0.070842745 | 0.380866278 | 0.703302476 | 0.843758597 | 0.026981613 |
| ENSMUSG00000029263 | Pigg | 433931 | protein_coding | 305.5888855 | 0.026988559 | 0.087085735 | -0.309907921 | 0.756630979 | 0.877889692 | 0.026988559 |
| ENSMUSG00000034926 | Dhcr24 | 74754 | protein_coding | 2077.258113 | 0.027043337 | 0.07812619 | -0.346149442 | 0.729230408 | 0.86030736 | 0.027043337 |
| ENSMUSG00000098449 | Gm7467 | 665055 | lncRNA | 18.60101864 | -0.027045576 | 0.284028017 | 0.095221509 | 0.924138886 | 0.963320302 | 0.027045576 |
| ENSMUSG00000039774 | Galnt12 | 230145 | protein_coding | 1453.212179 | -0.027061741 | 0.106646778 | 0.253751137 | 0.799687825 | 0.900821606 | 0.027061741 |
| ENSMUSG00000007035 | Msh5 | 17687 | protein_coding | 26.20037404 | 0.027072637 | 0.24366753 | -0.111104818 | 0.911533228 | 0.956997404 | 0.027072637 |
| ENSMUSG00000029670 | Ing3 | 71777 | protein_coding | 158.6952338 | -0.027091206 | 0.130651597 | 0.207354574 | 0.835732956 | 0.917517221 | 0.027091206 |
| ENSMUSG00000032312 | Csk | 12988 | protein_coding | 890.7028015 | 0.027103422 | 0.065131423 | -0.416134335 | 0.677311707 | 0.827106157 | 0.027103422 |
| ENSMUSG00000033720 | Sfxn5 | 94282 | protein_coding | 322.6708872 | -0.027123568 | 0.130225533 | 0.208281487 | 0.835009186 | 0.917066002 | 0.027123568 |
| ENSMUSG00000066829 | Zfp810 | 235050 | protein_coding | 166.6600495 | -0.027157223 | 0.123360354 | 0.220145465 | 0.825757867 | 0.91190529 | 0.027157223 |
| ENSMUSG00000024896 | Minpp1 | 17330 | protein_coding | 366.4530955 | -0.027209297 | 0.118735834 | 0.229158261 | 0.818745913 | 0.908812719 | 0.027209297 |
| ENSMUSG00000041396 | Mettl18 | 69962 | protein_coding | 43.52192248 | 0.027248547 | 0.178314638 | -0.152811609 | 0.878546843 | 0.939901695 | 0.027248547 |
| ENSMUSG00000039004 | Bmp6 | 12161 | protein_coding | 8.757105468 | -0.027262305 | 0.562025667 | 0.048507225 | 0.961312007 | 0.981095512 | 0.027262305 |
| ENSMUSG00000038777 | Sema6c | 20360 | protein_coding | 7.040356022 | 0.027274053 | 0.710684006 | -0.038377187 | 0.96938695 | 0.984403993 | 0.027274053 |
| ENSMUSG00000020572 | Nampt | 59027 | protein_coding | 862.0700147 | 0.027286382 | 0.084275868 | -0.323774556 | 0.74610872 | 0.872287153 | 0.027286382 |
| ENSMUSG00000024576 | Csnk1a1 | 93687 | protein_coding | 5000.668383 | 0.027344422 | 0.050986674 | -0.536305261 | 0.591747595 | 0.771818402 | 0.027344422 |
| ENSMUSG00000013858 | Tmem259 | 216157 | protein_coding | 1039.662488 | 0.02738981 | 0.071895604 | -0.380966404 | 0.703228178 | 0.843730756 | 0.02738981 |
| ENSMUSG00000063364 | 3300002I08Rik | 69277 | protein_coding | 14.04859083 | 0.027409734 | 0.313656674 | -0.087387697 | 0.930363348 | 0.966412148 | 0.027409734 |
| ENSMUSG00000031012 | Cask | 12361 | protein_coding | 886.6680215 | 0.027410019 | 0.076742494 | -0.357168727 | 0.720965496 | 0.855440739 | 0.027410019 |
| ENSMUSG00000038446 | Cdc40 | 71713 | protein_coding | 262.7338866 | 0.027435932 | 0.089894935 | -0.305199983 | 0.760213845 | 0.879513202 | 0.027435932 |
| ENSMUSG00000007411 | Mark3 | 17169 | protein_coding | 794.7288035 | -0.027493974 | 0.05776282 | 0.475980475 | 0.634088288 | 0.800507769 | 0.027493974 |
| ENSMUSG00000053580 | Tanc2 | 77097 | protein_coding | 107.1931513 | 0.027509125 | 0.106520842 | -0.258251104 | 0.79621312 | 0.898808084 | 0.027509125 |
| ENSMUSG00000036249 | Rbm43 | 71684 | protein_coding | 88.59528472 | -0.027532121 | 0.164185186 | 0.167689435 | 0.866827608 | 0.933837244 | 0.027532121 |
| ENSMUSG00000114828 | AI463229 | 100042802 | lncRNA | 22.77517513 | -0.027573621 | 0.300737541 | 0.091686662 | 0.926946995 | 0.965151432 | 0.027573621 |
| ENSMUSG00000027254 | Map1a | 17754 | protein_coding | 28.5173678 | 0.027575103 | 0.388284053 | -0.071017861 | 0.94338354 | 0.972595566 | 0.027575103 |
| ENSMUSG00000090751 | Vmn2r63 | 435975 | protein_coding | 2.979752546 | -0.027577402 | 1.973207236 | 0.013975928 | 0.988849186 | 1 | 0.027577402 |
| ENSMUSG00000021905 | Dph3 | 105638 | protein_coding | 327.8810703 | -0.027610296 | 0.092844337 | 0.297382657 | 0.766174383 | 0.882690208 | 0.027610296 |
| ENSMUSG00000030688 | Stard10 | 56018 | protein_coding | 6196.068577 | -0.027705043 | 0.061993557 | 0.446901975 | 0.654945838 | 0.812734264 | 0.027705043 |
| ENSMUSG00000039952 | Dag1 | 13138 | protein_coding | 2232.536582 | -0.027715113 | 0.059065059 | 0.469230268 | 0.638905052 | 0.80314212 | 0.027715113 |
| ENSMUSG00000035173 | Ccdc186 | 213993 | protein_coding | 882.7504261 | 0.027722065 | 0.084426464 | -0.328357528 | 0.742641352 | 0.870032583 | 0.027722065 |
| ENSMUSG00000040312 | Cchcr1 | 240084 | protein_coding | 175.9538727 | 0.027749132 | 0.107664065 | -0.257738107 | 0.796609035 | 0.898947765 | 0.027749132 |
| ENSMUSG00000041870 | Ankrd13a | 68420 | protein_coding | 2850.742552 | -0.027835401 | 0.040374118 | 0.689436756 | 0.49054846 | 0.69793377 | 0.027835401 |
| ENSMUSG00000020692 | Nle1 | 217011 | protein_coding | 85.90620313 | 0.027841453 | 0.130373044 | -0.213552219 | 0.830896264 | 0.914497068 | 0.027841453 |
| ENSMUSG00000057230 | Aak1 | 269774 | protein_coding | 515.0837067 | -0.0278767 | 0.088020136 | 0.316708216 | 0.751465014 | 0.875092959 | 0.0278767 |
| ENSMUSG00000069223 | 4930451E10Rik | 78111 | lncRNA | 1.515389536 | 0.027887773 | 0.834278847 | -0.033427401 | 0.973333759 | 1 | 0.027887773 |
| ENSMUSG00000017858 | Ift52 | 245866 | protein_coding | 328.5696524 | 0.027888474 | 0.076932869 | -0.362504014 | 0.716975425 | 0.853197357 | 0.027888474 |

| ensembl_gene_id | external_gene_name | entrezgene_id | gene_biotype | baseMean | log2FoldChange | lfcSE | stat | pvalue | padj | absLog2FC |
|---|---|---|---|---|---|---|---|---|---|---|
| ENSMUSG00000036208 | Nepro | 212547 | protein_coding | 174.7937779 | -0.027901499 | 0.098120941 | 0.284358246 | 0.776135853 | 0.888226437 | 0.027901499 |
| ENSMUSG00000044626 | Liph | 239759 | protein_coding | 690.688751 | -0.027920044 | 0.114779047 | 0.243250355 | 0.807811461 | 0.905233898 | 0.027920044 |
| ENSMUSG00000033444 | Specc1l | 74392 | protein_coding | 1975.60469 | -0.027944287 | 0.136245227 | 0.20510287 | 0.837491754 | 0.918959391 | 0.027944287 |
| ENSMUSG00000020097 | Sgpl1 | 20397 | protein_coding | 1672.288111 | 0.027996448 | 0.089814018 | -0.311715795 | 0.755256523 | 0.877032235 | 0.027996448 |
| ENSMUSG00000028977 | Casz1 | 69743 | protein_coding | 1151.544645 | 0.028046567 | 0.164268973 | -0.17073563 | 0.864431645 | 0.932569867 | 0.028046567 |
| ENSMUSG00000028988 | Ctnnbip1 | 67087 | protein_coding | 222.243717 | -0.028056429 | 0.103393502 | 0.27135583 | 0.786117371 | 0.893150482 | 0.028056429 |
| ENSMUSG00000027274 | Mkks | 59030 | protein_coding | 248.4332971 | 0.028107364 | 0.163181956 | -0.172245537 | 0.863244499 | 0.932166328 | 0.028107364 |
| ENSMUSG00000017781 | Pitpna | 18738 | protein_coding | 1534.071778 | 0.028121098 | 0.054505873 | -0.515927863 | 0.605904793 | 0.780796859 | 0.028121098 |
| ENSMUSG00000068270 | Shroom4 | 208431 | protein_coding | 36.5027473 | 0.028138602 | 0.292618078 | -0.096161531 | 0.923392283 | 0.962865452 | 0.028138602 |
| ENSMUSG00000037373 | Ctbp1 | 13016 | protein_coding | 1998.830201 | 0.028142367 | 0.070157989 | -0.401128482 | 0.688325532 | 0.833786242 | 0.028142367 |
| ENSMUSG00000021929 | Kpna3 | 16648 | protein_coding | 413.3639955 | 0.028148486 | 0.094589327 | -0.297586278 | 0.766018949 | 0.882690208 | 0.028148486 |
| ENSMUSG00000007837 | Prrg2 | 65116 | protein_coding | 387.9328808 | -0.028237607 | 0.099684601 | 0.283269494 | 0.77697026 | 0.888627112 | 0.028237607 |
| ENSMUSG00000042228 | Lyn | 17096 | protein_coding | 313.0568662 | -0.028258807 | 0.083480517 | 0.338507811 | 0.734980542 | 0.864910546 | 0.028258807 |
| ENSMUSG00000037119 | Fam91a1 | 210998 | protein_coding | 1315.670431 | 0.02826489 | 0.108977658 | -0.259364085 | 0.795354338 | 0.898299072 | 0.02826489 |
| ENSMUSG00000030697 | Ppp4c | 56420 | protein_coding | 1237.403828 | -0.028268208 | 0.048229985 | 0.586112715 | 0.557799772 | 0.746959571 | 0.028268208 |
| ENSMUSG00000031807 | Pgls | 66171 | protein_coding | 708.4788464 | 0.02827082 | 0.081805894 | -0.345584149 | 0.72965526 | 0.860514985 | 0.02827082 |
| ENSMUSG00000016503 | Gtf3a | 66596 | protein_coding | 152.3782087 | 0.028306623 | 0.121605214 | -0.232774744 | 0.815936325 | 0.908409285 | 0.028306623 |
| ENSMUSG00000034442 | Trmt5 | 76357 | protein_coding | 101.8643898 | 0.028382547 | 0.137315027 | -0.206696581 | 0.836246828 | 0.917959327 | 0.028382547 |
| ENSMUSG00000058715 | Fcer1g | 14127 | protein_coding | 8.645181549 | -0.028405548 | 0.396839956 | 0.071579354 | 0.942936671 | 0.972316965 | 0.028405548 |
| ENSMUSG00000022568 | Scrib | 105782 | protein_coding | 1194.248862 | 0.028451294 | 0.053470442 | -0.532093862 | 0.594660988 | 0.77336837 | 0.028451294 |
| ENSMUSG00000028850 | Gpatch3 | 242691 | protein_coding | 76.56250364 | 0.028467374 | 0.154144395 | -0.184679915 | 0.853480101 | 0.92805494 | 0.028467374 |
| ENSMUSG00000034203 | Chchd4 | 72170 | protein_coding | 322.8664127 | 0.028489091 | 0.070753979 | -0.402650025 | 0.687205703 | 0.833389267 | 0.028489091 |
| ENSMUSG00000026573 | Xcl1 | 16963 | protein_coding | 13.91269909 | -0.028496729 | 0.493814645 | 0.057707338 | 0.953981748 | 0.977047797 | 0.028496729 |
| ENSMUSG00000048755 | Mcat | 223722 | protein_coding | 192.4600739 | -0.028498825 | 0.157410882 | 0.181047365 | 0.856330398 | 0.930174321 | 0.028498825 |
| ENSMUSG00000037487 | Ubr5 | 70790 | protein_coding | 2828.794598 | 0.028516507 | 0.061917335 | -0.460557716 | 0.645115954 | 0.80668897 | 0.028516507 |
| ENSMUSG00000039456 | Morc3 | 338467 | protein_coding | 529.7794746 | -0.028518169 | 0.067095552 | 0.425038153 | 0.670808862 | 0.823378739 | 0.028518169 |
| ENSMUSG00000067577 | A430093F15Rik | 403202 | lncRNA | 1.107032597 | -0.028579537 | 2.183377368 | 0.0130896 | 0.989556308 | 1 | 0.028579537 |
| ENSMUSG00000019920 | Lims1 | 110829 | protein_coding | 1160.69259 | 0.028610992 | 0.167408724 | -0.170905022 | 0.864298448 | 0.932569867 | 0.028610992 |
| ENSMUSG00000023051 | Tarbp2 | 21357 | protein_coding | 234.0376885 | 0.028624042 | 0.121954315 | -0.234711183 | 0.814432907 | 0.907710402 | 0.028624042 |
| ENSMUSG00000013089 | Etv5 | 104156 | protein_coding | 330.2747838 | -0.028626152 | 0.205791513 | 0.139102686 | 0.889369006 | 0.945659633 | 0.028626152 |
| ENSMUSG00000036452 | Arhgap26 | 71302 | protein_coding | 418.3124529 | 0.028647924 | 0.091199782 | -0.314122721 | 0.753427834 | 0.876451623 | 0.028647924 |
| ENSMUSG00000022245 | Skor1 | 207667 | protein_coding | 2.259739784 | 0.028693105 | 1.269513002 | -0.022601663 | 0.981968017 | 1 | 0.028693105 |
| ENSMUSG00000020305 | Asb3 | 65257 | protein_coding | 111.8442959 | -0.028708403 | 0.124800424 | 0.230034495 | 0.818064965 | 0.908812719 | 0.028708403 |
| ENSMUSG00000091514 | Gm17484 | 100503003 | lncRNA | 2.496641518 | -0.028712276 | 0.682877201 | 0.042046031 | 0.966462003 | 1 | 0.028712276 |
| ENSMUSG00000097021 | 4933433G15Rik | 71274 | lncRNA | 9.197129396 | 0.028726402 | 0.416848869 | -0.06891323 | 0.945058688 | 0.97354273 | 0.028726402 |
| ENSMUSG00000029669 | Tspan12 | 269831 | protein_coding | 526.1909569 | 0.028749978 | 0.108530855 | -0.264901419 | 0.791085405 | 0.895862069 | 0.028749978 |
| ENSMUSG00000020234 | Tektip1 | 432479 | protein_coding | 10.26914474 | 0.028781523 | 0.372656433 | -0.077233399 | 0.938437872 | 0.970587024 | 0.028781523 |
| ENSMUSG00000024423 | Impact | 16210 | protein_coding | 87.98747125 | -0.028785306 | 0.154022391 | 0.186890397 | 0.851746568 | 0.927574705 | 0.028785306 |
| ENSMUSG00000033436 | Armcx2 | 67416 | protein_coding | 4.787585489 | 0.028809611 | 0.781104876 | -0.036883154 | 0.970578172 | 0.984770701 | 0.028809611 |

Supplementary Table 1

| ensembl_gene_id | external_gene_name | entrezgene_id | gene_biotype | baseMean | log2FoldChange | lfcSE | stat | pvalue | padj | absLog2FC |
|---|---|---|---|---|---|---|---|---|---|---|
| ENSMUSG00000037773 | Pced1a | 319513 | protein_coding | 94.31679933 | -0.028835112 | 0.122571265 | 0.235251811 | 0.814013295 | 0.907710402 | 0.028835112 |
| ENSMUSG00000020993 | Trappc6b | 78232 | protein_coding | 293.0651559 | 0.028892651 | 0.07579472 | -0.381196083 | 0.703057755 | 0.843587573 | 0.028892651 |
| ENSMUSG00000025027 | Xpnpep1 | 170750 | protein_coding | 905.1091454 | 0.028948817 | 0.069006745 | -0.419507068 | 0.67484559 | 0.825633433 | 0.028948817 |
| ENSMUSG00000030980 | Knop1 | 66356 | protein_coding | 395.6497464 | -0.028996154 | 0.110842012 | 0.261598951 | 0.793630653 | 0.897242186 | 0.028996154 |
| ENSMUSG00000035575 | Utp6 | 216987 | protein_coding | 472.6999581 | 0.029030912 | 0.117158376 | -0.247792035 | 0.804295318 | 0.903546454 | 0.029030912 |
| ENSMUSG00000026239 | Pde6d | 18582 | protein_coding | 105.1907647 | 0.029048925 | 0.107166994 | -0.271062239 | 0.786343164 | 0.893161222 | 0.029048925 |
| ENSMUSG00000059851 | Kmt5c | 232811 | protein_coding | 538.4007117 | 0.029063737 | 0.072444988 | -0.401183545 | 0.688284995 | 0.833786242 | 0.029063737 |
| ENSMUSG00000043391 | 2510009E07Rik | 72190 | protein_coding | 356.2078157 | -0.029074643 | 0.155975167 | 0.186405591 | 0.852126708 | 0.927743965 | 0.029074643 |
| ENSMUSG00000062234 | Gak | 231580 | protein_coding | 2480.915234 | -0.029106287 | 0.05217445 | 0.557864763 | 0.576936735 | 0.761228287 | 0.029106287 |
| ENSMUSG00000024924 | Vldlr | 22359 | protein_coding | 244.1153256 | 0.029152207 | 0.124154982 | -0.234804972 | 0.814360109 | 0.907710402 | 0.029152207 |
| ENSMUSG00000049791 | Fzd4 | 14366 | protein_coding | 85.23903084 | -0.029188957 | 0.142704269 | 0.204541584 | 0.8379303 | 0.919173117 | 0.029188957 |
| ENSMUSG00000020679 | Hnf1b | 21410 | protein_coding | 267.7460609 | -0.029199673 | 0.125794433 | 0.23212214 | 0.816443149 | 0.908418149 | 0.029199673 |
| ENSMUSG00000047767 | Atg16l2 | 73683 | protein_coding | 132.2639703 | -0.029213262 | 0.20344449 | 0.143593279 | 0.885821648 | 0.943345502 | 0.029213262 |
| ENSMUSG00000029458 | Brap | 72399 | protein_coding | 634.8668499 | -0.029296812 | 0.055342531 | 0.529372456 | 0.596547102 | 0.774662172 | 0.029296812 |
| ENSMUSG00000068039 | Tcp1 | 21454 | protein_coding | 2026.855519 | 0.029301821 | 0.106566582 | -0.274962572 | 0.783344994 | 0.891264005 | 0.029301821 |
| ENSMUSG00000078185 | Chml | 12663 | protein_coding | 255.26129 | 0.029312836 | 0.121849562 | -0.240565785 | 0.809891671 | 0.906073508 | 0.029312836 |
| ENSMUSG00000003438 | Timm50 | 66525 | protein_coding | 376.8673482 | -0.029319459 | 0.095516821 | 0.306955982 | 0.758876877 | 0.878912418 | 0.029319459 |
| ENSMUSG00000000766 | Oprm1 | 18390 | protein_coding | 2.8144158 | 0.02936544 | 1.65597877 | -0.017732981 | 0.98585187 | 1 | 0.02936544 |
| ENSMUSG00000031575 | Ash2l | 23808 | protein_coding | 582.8734968 | -0.029383797 | 0.076666118 | 0.383269665 | 0.701519829 | 0.842293038 | 0.029383797 |
| ENSMUSG00000056267 | Cep70 | 68121 | protein_coding | 109.8539129 | -0.029385487 | 0.131127825 | 0.22409803 | 0.822681026 | 0.91020732 | 0.029385487 |
| ENSMUSG00000022770 | Dlg1 | 13383 | protein_coding | 980.4266157 | -0.029389265 | 0.048621627 | 0.604448399 | 0.545545566 | 0.737366138 | 0.029389265 |
| ENSMUSG00000000804 | Usp32 | 237898 | protein_coding | 494.1788967 | -0.029405792 | 0.100398049 | 0.292892063 | 0.769604656 | 0.884510137 | 0.029405792 |
| ENSMUSG00000016252 | Atp5e | 67126 | protein_coding | 1733.34122 | 0.029424917 | 0.09112431 | -0.322909628 | 0.746763684 | 0.872440598 | 0.029424917 |
| ENSMUSG00000041351 | Rap1gap | 110351 | protein_coding | 1610.303866 | -0.029453944 | 0.120031165 | 0.245385801 | 0.806157722 | 0.904600714 | 0.029453944 |
| ENSMUSG00000024659 | Anxa1 | 16952 | protein_coding | 1826.852775 | -0.029477614 | 0.127241489 | 0.231666686 | 0.816796908 | 0.908628099 | 0.029477614 |
| ENSMUSG00000026355 | Mcm6 | 17219 | protein_coding | 671.8078104 | 0.029504568 | 0.160091535 | -0.184298115 | 0.853779593 | 0.928319475 | 0.029504568 |
| ENSMUSG00000045822 | Zswim3 | 67538 | protein_coding | 77.55914867 | -0.02951894 | 0.136947711 | 0.215548984 | 0.829339326 | 0.913717368 | 0.02951894 |
| ENSMUSG00000031901 | Dus2 | 66369 | protein_coding | 89.2006143 | 0.029537336 | 0.135049616 | -0.218714699 | 0.826872298 | 0.912640622 | 0.029537336 |
| ENSMUSG00000032077 | Bud13 | 215051 | protein_coding | 114.2296765 | 0.029551605 | 0.103389504 | -0.285827905 | 0.775009934 | 0.887804402 | 0.029551605 |
| ENSMUSG00000039968 | Rsbn1l | 242860 | protein_coding | 265.8045187 | 0.02966752 | 0.095849036 | -0.309523399 | 0.756923415 | 0.878075079 | 0.02966752 |
| ENSMUSG00000024537 | Psmg2 | 107047 | protein_coding | 120.7250568 | 0.029686351 | 0.152162016 | -0.195096985 | 0.845317023 | 0.923801556 | 0.029686351 |
| ENSMUSG00000064647 | Gm23301 | 115488997 | snoRNA | 6.45158088 | -0.029695604 | 0.458539263 | 0.064761312 | 0.948364045 | 0.975173098 | 0.029695604 |
| ENSMUSG00000041773 | Enc1 | 13803 | protein_coding | 186.3338271 | -0.029723132 | 0.118808433 | 0.250176953 | 0.802450508 | 0.902394803 | 0.029723132 |
| ENSMUSG00000085502 | Gm12320 | 102635935 | lncRNA | 34.77997628 | 0.02977079 | 0.242077625 | -0.122980346 | 0.902122662 | 0.951530343 | 0.02977079 |
| ENSMUSG00000037344 | Slc12a9 | 83704 | protein_coding | 167.1876316 | -0.029786827 | 0.122548624 | 0.243061294 | 0.807957915 | 0.905320736 | 0.029786827 |
| ENSMUSG00000115902 | D730005E14Rik | 109361 | lncRNA | 28.75777974 | 0.029852925 | 0.345290851 | -0.086457331 | 0.931102874 | 0.966840628 | 0.029852925 |
| ENSMUSG00000020590 | Snx13 | 217463 | protein_coding | 1631.237816 | 0.02985685 | 0.067605036 | -0.441636477 | 0.658752284 | 0.815856848 | 0.02985685 |
| ENSMUSG00000022765 | Snap29 | 67474 | protein_coding | 453.0733939 | 0.029869816 | 0.081626178 | -0.365934267 | 0.714414126 | 0.851829233 | 0.029869816 |
| ENSMUSG00000006390 | Elovl1 | 54325 | protein_coding | 717.9877106 | -0.029968959 | 0.071417772 | 0.41962887 | 0.674756595 | 0.825585716 | 0.029968959 |

Supplementary Table 1

| ensembl_gene_id | external_gene_name | entrezgene_id | gene_biotype | baseMean | log2FoldChange | lfcSE | stat | pvalue | padj | absLog2FC |
|---|---|---|---|---|---|---|---|---|---|---|
| ENSMUSG00000069539 | Scyl2 | 213326 | protein_coding | 810.6403533 | 0.029975812 | 0.051833343 | -0.578311375 | 0.563053916 | 0.750768863 | 0.029975812 |
| ENSMUSG00000070720 | Tmem200b | 623230 | protein_coding | 4.519332246 | 0.029986143 | 0.613033403 | -0.048914371 | 0.960987536 | 0.981006872 | 0.029986143 |
| ENSMUSG00000026927 | Entr1 | 68112 | protein_coding | 412.2007474 | 0.029986556 | 0.073382617 | -0.408632967 | 0.682809034 | 0.830576634 | 0.029986556 |
| ENSMUSG00000037085 | Trmt12 | 68260 | protein_coding | 82.62532241 | -0.029994987 | 0.155629647 | 0.192733114 | 0.847167988 | 0.924722861 | 0.029994987 |
| ENSMUSG00000048652 | Samd13 | 75015 | protein_coding | 10.51041307 | -0.030014434 | 0.339839092 | 0.088319544 | 0.929622706 | 0.966128972 | 0.030014434 |
| ENSMUSG00000034709 | Ppp1r21 | 73825 | protein_coding | 307.1307181 | 0.030052218 | 0.070916396 | -0.423769674 | 0.671733796 | 0.823993339 | 0.030052218 |
| ENSMUSG00000097119 | B230354K17Rik | 105246278 | lncRNA | 144.1508011 | 0.030054566 | 0.125989815 | -0.238547588 | 0.81145641 | 0.906704483 | 0.030054566 |
| ENSMUSG00000051469 | Zfp24 | 59057 | protein_coding | 451.040638 | 0.030060759 | 0.095321434 | -0.315362006 | 0.752486811 | 0.875793979 | 0.030060759 |
| ENSMUSG00000039981 | Zc3h12d | 237256 | protein_coding | 223.0869687 | 0.030091192 | 0.130803872 | -0.230048173 | 0.818054337 | 0.908812719 | 0.030091192 |
| ENSMUSG00000032363 | Adamts7 | 108153 | protein_coding | 8.747976147 | 0.030091325 | 0.516192896 | -0.058294729 | 0.953513866 | 0.97686351 | 0.030091325 |
| ENSMUSG00000020547 | Bzw2 | 66912 | protein_coding | 549.3890021 | -0.030096289 | 0.113716477 | 0.26466076 | 0.791270809 | 0.89588517 | 0.030096289 |
| ENSMUSG00000008976 | Gabpa | 14390 | protein_coding | 538.1918428 | 0.030113369 | 0.119166158 | -0.252700681 | 0.800499522 | 0.901189824 | 0.030113369 |
| ENSMUSG00000015757 | Ppil4 | 67418 | protein_coding | 354.3280749 | 0.030140672 | 0.077557637 | -0.388622872 | 0.697555145 | 0.839669485 | 0.030140672 |
| ENSMUSG00000047539 | Fbxo28 | 67948 | protein_coding | 383.5469783 | 0.030190042 | 0.093424549 | -0.323148916 | 0.746582466 | 0.872440598 | 0.030190042 |
| ENSMUSG00000058291 | Zfp68 | 24135 | protein_coding | 356.004324 | -0.030213846 | 0.118273752 | 0.255456904 | 0.798370225 | 0.899929685 | 0.030213846 |
| ENSMUSG00000049122 | Frmd3 | 242506 | protein_coding | 18.92785007 | 0.030251006 | 0.354394523 | -0.085359688 | 0.931975441 | 0.967478151 | 0.030251006 |
| ENSMUSG00000026321 | Tnfrsf11a | 21934 | protein_coding | 1859.811047 | 0.030256876 | 0.102934652 | -0.293942568 | 0.768801789 | 0.884245212 | 0.030256876 |
| ENSMUSG00000048012 | Zfp473 | 243963 | protein_coding | 24.92353343 | 0.030316821 | 0.263046847 | -0.115252554 | 0.908244944 | 0.955333816 | 0.030316821 |
| ENSMUSG00000024217 | Snrpc | 20630 | protein_coding | 243.9712253 | -0.030330513 | 0.126186781 | 0.240362048 | 0.810049597 | 0.906149826 | 0.030330513 |
| ENSMUSG00000027782 | Kpna4 | 16649 | protein_coding | 838.1856728 | -0.030340025 | 0.055750275 | 0.544213017 | 0.586294895 | 0.767930988 | 0.030340025 |
| ENSMUSG00000035666 | Gtf3c4 | 269252 | protein_coding | 424.1029059 | -0.030341239 | 0.082649357 | 0.367107984 | 0.713538472 | 0.851092403 | 0.030341239 |
| ENSMUSG00000018572 | Phf23 | 78246 | protein_coding | 377.3954941 | -0.030370139 | 0.094078699 | 0.322816312 | 0.746834359 | 0.872440598 | 0.030370139 |
| ENSMUSG00000015766 | Eps8 | 13860 | protein_coding | 1993.207767 | 0.030376273 | 0.13255831 | -0.229154044 | 0.81874919 | 0.908812719 | 0.030376273 |
| ENSMUSG00000038319 | Kcnh2 | 16511 | protein_coding | 39.1016121 | -0.03043773 | 0.233922232 | 0.130119012 | 0.896472269 | 0.948996349 | 0.03043773 |
| ENSMUSG00000049960 | Mrps16 | 66242 | protein_coding | 420.0186482 | -0.030440229 | 0.08200522 | 0.371198669 | 0.710489565 | 0.84898876 | 0.030440229 |
| ENSMUSG00000032860 | P2ry2 | 18442 | protein_coding | 213.9799813 | -0.030485525 | 0.193862988 | 0.15725294 | 0.875045507 | 0.938415991 | 0.030485525 |
| ENSMUSG00000074657 | Kif5a | 16572 | protein_coding | 6.316081593 | 0.030517113 | 1.150130162 | -0.026533616 | 0.978831721 | 0.988831409 | 0.030517113 |
| ENSMUSG00000006276 | Eps15l1 | 13859 | protein_coding | 627.7380668 | -0.030542375 | 0.060279674 | 0.506677843 | 0.612380879 | 0.785464687 | 0.030542375 |
| ENSMUSG00000014668 | Chfr | 231600 | protein_coding | 644.2019802 | 0.030547792 | 0.066093975 | -0.462187232 | 0.643947058 | 0.80597323 | 0.030547792 |
| ENSMUSG00000031072 | LTO1 | 72284 | protein_coding | 176.5798583 | 0.030563314 | 0.086304667 | -0.354132805 | 0.723239358 | 0.856549354 | 0.030563314 |
| ENSMUSG00000047583 | Tyw3 | 209584 | protein_coding | 29.60763766 | -0.030563584 | 0.248654533 | 0.122915851 | 0.902173733 | 0.951530343 | 0.030563584 |
| ENSMUSG00000025795 | Rassf3 | 192678 | protein_coding | 729.7293615 | -0.03059208 | 0.079271938 | 0.385913113 | 0.699561018 | 0.84101582 | 0.03059208 |
| ENSMUSG00000037461 | Ints7 | 77065 | protein_coding | 423.7242079 | 0.030598111 | 0.100248873 | -0.305221498 | 0.76019746 | 0.879513202 | 0.030598111 |
| ENSMUSG00000035206 | Sppl2b | 73218 | protein_coding | 481.4058302 | 0.030604855 | 0.0649409 | -0.471272411 | 0.637446215 | 0.802408631 | 0.030604855 |
| ENSMUSG00000025702 | Marchf8 | 71779 | protein_coding | 1468.52362 | -0.030633386 | 0.074574551 | 0.410775334 | 0.681237282 | 0.829503357 | 0.030633386 |
| ENSMUSG00000031565 | Fgfr1 | 14182 | protein_coding | 10.23204615 | -0.030644073 | 0.895245058 | 0.034229815 | 0.972693891 | 0.985766611 | 0.030644073 |
| ENSMUSG00000031021 | Tmem9b | 56786 | protein_coding | 648.4689245 | -0.030690095 | 0.093992959 | 0.326514825 | 0.744034875 | 0.870923726 | 0.030690095 |
| ENSMUSG00000037426 | Depdc5 | 277854 | protein_coding | 325.8445273 | -0.03071494 | 0.076933752 | 0.399238818 | 0.689717243 | 0.834370416 | 0.03071494 |
| ENSMUSG00000014074 | Rnf168 | 70238 | protein_coding | 329.6890739 | -0.030738698 | 0.111402539 | 0.275924569 | 0.782606003 | 0.891061468 | 0.030738698 |

| ensembl_gene_id | external_gene_name | entrezgene_id | gene_biotype | baseMean | log2FoldChange | lfcSE | stat | pvalue | padj | absLog2FC |
|---|---|---|---|---|---|---|---|---|---|---|
| ENSMUSG00000035851 | Ythdc1 | 231386 | protein_coding | 657.4636788 | 0.030777187 | 0.075409544 | -0.408133835 | 0.683175421 | 0.830769833 | 0.030777187 |
| ENSMUSG00000032580 | Rbm5 | 83486 | protein_coding | 1163.622775 | -0.030779711 | 0.060228042 | 0.511052835 | 0.609314061 | 0.783356356 | 0.030779711 |
| ENSMUSG00000020440 | Arf5 | 11844 | protein_coding | 1168.90788 | 0.030801041 | 0.074248233 | -0.414838705 | 0.678259983 | 0.82760347 | 0.030801041 |
| ENSMUSG00000021993 | Mipep | 70478 | protein_coding | 311.765382 | 0.030843846 | 0.09588494 | -0.321675609 | 0.747698458 | 0.873111243 | 0.030843846 |
| ENSMUSG00000029196 | Tada2b | 231151 | protein_coding | 201.6438262 | 0.030902571 | 0.102098009 | -0.302675546 | 0.762137135 | 0.880874346 | 0.030902571 |
| ENSMUSG00000028664 | Ephb2 | 13844 | protein_coding | 1261.735467 | -0.030921328 | 0.098717227 | 0.313231326 | 0.754104921 | 0.876856731 | 0.030921328 |
| ENSMUSG00000041261 | Car8 | 12319 | protein_coding | 697.4405106 | -0.030938537 | 0.169547481 | 0.182477122 | 0.855208307 | 0.929505724 | 0.030938537 |
| ENSMUSG00000037646 | Vps13b | 666173 | protein_coding | 1640.295709 | -0.0309433 | 0.074689308 | 0.414293564 | 0.678659127 | 0.827845828 | 0.0309433 |
| ENSMUSG00000021279 | Cdc42bpb | 217866 | protein_coding | 1627.949956 | -0.030960034 | 0.072867009 | 0.424884104 | 0.670921163 | 0.82339418 | 0.030960034 |
| ENSMUSG00000026753 | Ppp6c | 67857 | protein_coding | 759.5604352 | 0.031004551 | 0.076332116 | -0.406179637 | 0.684610615 | 0.831787794 | 0.031004551 |
| ENSMUSG00000025759 | Mfsd8 | 72175 | protein_coding | 163.5038478 | 0.031018764 | 0.112726915 | -0.275167331 | 0.783187684 | 0.891264005 | 0.031018764 |
| ENSMUSG00000051007 | Gatd1 | 213350 | protein_coding | 342.7967197 | -0.031031429 | 0.089189863 | 0.347925511 | 0.727896126 | 0.859382261 | 0.031031429 |
| ENSMUSG00000034152 | Exoc3 | 211446 | protein_coding | 848.442134 | -0.031083932 | 0.072811995 | 0.42690675 | 0.669447249 | 0.822609935 | 0.031083932 |
| ENSMUSG00000028010 | Gar1 | 68147 | protein_coding | 99.46474271 | 0.03110629 | 0.131073026 | -0.237320303 | 0.812408313 | 0.907147741 | 0.03110629 |
| ENSMUSG00000009079 | Ewsr1 | 14030 | protein_coding | 1908.382367 | 0.031107711 | 0.066229318 | -0.469696982 | 0.638571525 | 0.80301103 | 0.031107711 |
| ENSMUSG00000037643 | Prkci | 18759 | protein_coding | 1269.266364 | -0.03111156 | 0.087555455 | 0.355335479 | 0.722338279 | 0.856331252 | 0.03111156 |
| ENSMUSG00000024870 | Rab1b | 76308 | protein_coding | 1426.874461 | 0.03111494 | 0.059526397 | -0.522708278 | 0.601177277 | 0.777615016 | 0.03111494 |
| ENSMUSG00000079144 | A130010J15Rik | 319266 | protein_coding | 146.545141 | 0.031133076 | 0.117342095 | -0.265318907 | 0.7907638 | 0.895656612 | 0.031133076 |
| ENSMUSG00000036006 | Ripor2 | 193385 | protein_coding | 29.42514533 | 0.031135071 | 0.222274334 | -0.140074972 | 0.888600755 | 0.94517522 | 0.031135071 |
| ENSMUSG00000078578 | Ube2d3 | 66105 | protein_coding | 3164.330952 | 0.031144826 | 0.048966003 | -0.636049998 | 0.524743835 | 0.723223689 | 0.031144826 |
| ENSMUSG00000045205 | Dpy19l4 | 381510 | protein_coding | 332.3742949 | 0.031145823 | 0.086129701 | -0.361615366 | 0.717639481 | 0.853579543 | 0.031145823 |
| ENSMUSG00000029510 | Gpc2 | 71951 | protein_coding | 3.108183131 | 0.031159394 | 0.828036076 | -0.037630478 | 0.969982307 | 1 | 0.031159394 |
| ENSMUSG00000041238 | Rbbp8 | 225182 | protein_coding | 398.9579886 | 0.031161954 | 0.07302705 | -0.426717968 | 0.669584762 | 0.822609935 | 0.031161954 |
| ENSMUSG00000031105 | Slc25a14 | 20523 | protein_coding | 53.2358648 | 0.031180867 | 0.140347404 | -0.222169174 | 0.82418219 | 0.910891761 | 0.031180867 |
| ENSMUSG00000028581 | Laptm5 | 16792 | protein_coding | 27.46849862 | -0.031195684 | 0.388954414 | 0.080203959 | 0.936075041 | 0.969722303 | 0.031195684 |
| ENSMUSG00000032855 | Pkd1 | 18763 | protein_coding | 701.2962289 | -0.031226144 | 0.100676886 | 0.310161999 | 0.756437767 | 0.877843331 | 0.031226144 |
| ENSMUSG00000020513 | Tubd1 | 56427 | protein_coding | 50.30012727 | 0.031248598 | 0.16817374 | -0.1858114 | 0.852592665 | 0.927761944 | 0.031248598 |
| ENSMUSG00000036644 | Tbc1d9b | 76795 | protein_coding | 908.365506 | -0.031253524 | 0.070757989 | 0.441696041 | 0.658709175 | 0.815856848 | 0.031253524 |
| ENSMUSG00000042505 | Sdhaf3 | 71238 | protein_coding | 62.11396972 | 0.031255428 | 0.140463972 | -0.22251562 | 0.823912515 | 0.91065466 | 0.031255428 |
| ENSMUSG00000022672 | Prkdc | 19090 | protein_coding | 358.097704 | -0.031276672 | 0.089460539 | 0.349614165 | 0.726628279 | 0.858643248 | 0.031276672 |
| ENSMUSG00000054387 | Mdm4 | 17248 | protein_coding | 745.2782543 | -0.031294017 | 0.080406746 | 0.389196409 | 0.69713086 | 0.839403501 | 0.031294017 |
| ENSMUSG00000034636 | Zyg11b | 414872 | protein_coding | 965.9856194 | -0.031317745 | 0.067732098 | 0.462376716 | 0.643811194 | 0.805935268 | 0.031317745 |
| ENSMUSG00000088015 | Mir1932 | 100316690 | miRNA | 2.679904738 | 0.031321947 | 0.720716567 | -0.043459451 | 0.965335287 | 1 | 0.031321947 |
| ENSMUSG00000029405 | G3bp2 | 23881 | protein_coding | 1610.178834 | 0.031325158 | 0.084369257 | -0.371286402 | 0.710424225 | 0.848979902 | 0.031325158 |
| ENSMUSG00000020848 | Doc2b | 13447 | protein_coding | 3.027158117 | 0.031356704 | 1.018547892 | -0.030785694 | 0.97544045 | 1 | 0.031356704 |
| ENSMUSG00000096887 | Gm20594 | 100463512 | protein_coding | 3.622079905 | -0.03136418 | 0.714690929 | 0.043884956 | 0.964996107 | 1 | 0.03136418 |
| ENSMUSG00000097354 | 2310001H17Rik | 76432 | lncRNA | 78.00279963 | -0.031397545 | 0.182792879 | 0.171765693 | 0.863621738 | 0.932314788 | 0.031397545 |
| ENSMUSG00000032563 | Mrpl3 | 94062 | protein_coding | 385.0820845 | 0.031408059 | 0.077053308 | -0.40761468 | 0.683556585 | 0.830935108 | 0.031408059 |
| ENSMUSG00000007777 | 0610009B22Rik | 66050 | protein_coding | 170.8093752 | -0.031420495 | 0.111380025 | 0.282101709 | 0.777865523 | 0.888973789 | 0.031420495 |

| ensembl_gene_id | external_gene_name | entrezgene_id | gene_biotype | baseMean | log2FoldChange | lfcSE | stat | pvalue | padj | absLog2FC |
|---|---|---|---|---|---|---|---|---|---|---|
| ENSMUSG00000038002 | Cramp1 | 57354 | protein_coding | 492.6497258 | 0.031432357 | 0.149239286 | -0.210617173 | 0.833186011 | 0.915993052 | 0.031432357 |
| ENSMUSG00000050029 | Rap2c | 72065 | protein_coding | 498.4702012 | 0.031504161 | 0.073687595 | -0.427536835 | 0.668988362 | 0.82238042 | 0.031504161 |
| ENSMUSG00000037361 | Sf3b6 | 66055 | protein_coding | 462.7243084 | 0.031574825 | 0.075355185 | -0.419013303 | 0.67520641 | 0.825971257 | 0.031574825 |
| ENSMUSG00000036955 | Kifbp | 72320 | protein_coding | 229.913825 | 0.031606639 | 0.102945905 | -0.307021819 | 0.758826764 | 0.878912418 | 0.031606639 |
| ENSMUSG00000055835 | Zfp1 | 22640 | protein_coding | 65.26397877 | 0.031635832 | 0.131505341 | -0.240566895 | 0.809890811 | 0.906073508 | 0.031635832 |
| ENSMUSG00000035184 | Fam124a | 629059 | protein_coding | 101.0158815 | -0.031657568 | 0.181300749 | 0.174613552 | 0.861383299 | 0.931679578 | 0.031657568 |
| ENSMUSG00000005698 | Ctcf | 13018 | protein_coding | 895.236324 | 0.031658704 | 0.074667766 | -0.423994257 | 0.671570002 | 0.823884333 | 0.031658704 |
| ENSMUSG00000087424 | 5730405O15Rik | 70489 | lncRNA | 8.99928649 | -0.031712022 | 0.423850478 | 0.07481889 | 0.940358812 | 0.971417829 | 0.031712022 |
| ENSMUSG00000039194 | Rlbp1 | 19771 | protein_coding | 1.831823386 | -0.031716168 | 1.371497675 | 0.023125207 | 0.981550399 | 1 | 0.031716168 |
| ENSMUSG00000039910 | Cited2 | 17684 | protein_coding | 451.8622993 | 0.031727834 | 0.189122753 | -0.167763175 | 0.866769594 | 0.933837244 | 0.031727834 |
| ENSMUSG00000113770 | 1700019C18Rik | 75543 | lncRNA | 2.259578759 | 0.031802248 | 0.925610458 | -0.034358133 | 0.972591568 | 1 | 0.031802248 |
| ENSMUSG00000032231 | Anxa2 | 12306 | protein_coding | 6546.765741 | 0.031829773 | 0.082452243 | -0.386038898 | 0.699467861 | 0.840991064 | 0.031829773 |
| ENSMUSG00000026315 | Serpinb8 | 20725 | protein_coding | 5.549787561 | -0.031833968 | 0.571365307 | 0.055715612 | 0.955568363 | 0.977832194 | 0.031833968 |
| ENSMUSG00000087368 | BC065397 | 436230 | lncRNA | 9.592484986 | -0.031847266 | 0.372489053 | 0.085498528 | 0.931865065 | 0.967478151 | 0.031847266 |
| ENSMUSG00000028214 | Gem | 14579 | protein_coding | 12.5434369 | -0.03185669 | 0.519603248 | 0.061309644 | 0.95111261 | 0.97587882 | 0.03185669 |
| ENSMUSG00000002346 | Slc25a42 | 73095 | protein_coding | 87.8798524 | -0.031893137 | 0.247077295 | 0.129081618 | 0.897293067 | 0.949500045 | 0.031893137 |
| ENSMUSG00000027628 | Aar2 | 68295 | protein_coding | 371.3809736 | -0.031912379 | 0.072194474 | 0.442033538 | 0.658464937 | 0.815684492 | 0.031912379 |
| ENSMUSG00000037355 | Uvssa | 71101 | protein_coding | 199.5829437 | -0.03192507 | 0.105510679 | 0.302576673 | 0.762212494 | 0.880899792 | 0.03192507 |
| ENSMUSG00000088901 | Mir1943 | 100316695 | miRNA | 0.809390907 | 0.031959333 | 1.270066014 | -0.025163521 | 0.979924533 | 1 | 0.031959333 |
| ENSMUSG00000000202 | Btbd17 | 72014 | protein_coding | 1.378714198 | 0.031969009 | 1.069382416 | -0.029894834 | 0.976150926 | 1 | 0.031969009 |
| ENSMUSG00000041915 | Ammecr1l | 225339 | protein_coding | 371.2493527 | 0.031971293 | 0.075797799 | -0.421797111 | 0.673173115 | 0.824897384 | 0.031971293 |
| ENSMUSG00000016520 | Lnx2 | 140887 | protein_coding | 428.3493711 | -0.031980142 | 0.06616644 | 0.483328733 | 0.628862341 | 0.797437818 | 0.031980142 |
| ENSMUSG00000033902 | Mapkbp1 | 26390 | protein_coding | 117.7782669 | 0.031982177 | 0.138033224 | -0.23169912 | 0.816771715 | 0.908628099 | 0.031982177 |
| ENSMUSG00000031392 | Irak1 | 16179 | protein_coding | 548.3783236 | -0.032007269 | 0.07681167 | 0.416698002 | 0.676899316 | 0.826943172 | 0.032007269 |
| ENSMUSG00000001786 | Fbxo7 | 69754 | protein_coding | 172.085621 | 0.032008134 | 0.113390379 | -0.282282625 | 0.777726807 | 0.888933284 | 0.032008134 |
| ENSMUSG00000044167 | Foxo1 | 56458 | protein_coding | 200.1939317 | -0.032026609 | 0.129877254 | 0.246591359 | 0.805224492 | 0.904098264 | 0.032026609 |
| ENSMUSG00000003345 | Csnk1g2 | 103236 | protein_coding | 1319.643898 | 0.032045671 | 0.058786987 | -0.545115032 | 0.585674405 | 0.767483504 | 0.032045671 |
| ENSMUSG00000024395 | Lims2 | 225341 | protein_coding | 6.667299217 | -0.032061663 | 0.68878766 | 0.046547964 | 0.962873506 | 0.981615003 | 0.032061663 |
| ENSMUSG00000026432 | Avpr1b | 26361 | protein_coding | 7.865342352 | 0.032076551 | 0.447678293 | -0.071650896 | 0.942879735 | 0.972316965 | 0.032076551 |
| ENSMUSG00000064941 | Gm23238 | 115489085 | snRNA | 1.325466561 | -0.032102841 | 1.123883289 | 0.028564212 | 0.977212155 | 1 | 0.032102841 |
| ENSMUSG00000003458 | Ncstn | 59287 | protein_coding | 1544.545156 | -0.032104959 | 0.070750957 | 0.4537742 | 0.649991355 | 0.809917796 | 0.032104959 |
| ENSMUSG00000046722 | Cdc42se1 | 57912 | protein_coding | 1087.608143 | 0.032125114 | 0.09574914 | -0.335513344 | 0.73723788 | 0.866653357 | 0.032125114 |
| ENSMUSG00000054263 | Lifr | 16880 | protein_coding | 63.02343558 | 0.032126577 | 0.264961534 | -0.121249966 | 0.903493049 | 0.952286247 | 0.032126577 |
| ENSMUSG00000035047 | Kri1 | 215194 | protein_coding | 148.6716752 | 0.032178704 | 0.117111403 | -0.274770035 | 0.783492921 | 0.891333845 | 0.032178704 |
| ENSMUSG00000099083 | Atf7 | 223922 | protein_coding | 162.6735136 | 0.03221506 | 0.127869976 | -0.251936076 | 0.801090474 | 0.901479257 | 0.03221506 |
| ENSMUSG00000032657 | Entrep3 | 68521 | protein_coding | 24.35371786 | 0.032215498 | 0.35179179 | -0.091575469 | 0.927035343 | 0.965151432 | 0.032215498 |
| ENSMUSG00000022602 | Arc | 11838 | protein_coding | 416.6671842 | -0.032224572 | 0.406309488 | 0.079310409 | 0.936785727 | 0.969910941 | 0.032224572 |
| ENSMUSG00000035112 | Wnk4 | 69847 | protein_coding | 262.6549 | 0.032236575 | 0.165984042 | -0.194214906 | 0.846007612 | 0.924189304 | 0.032236575 |
| ENSMUSG00000039183 | Nubp2 | 26426 | protein_coding | 265.5870797 | -0.032246658 | 0.081731729 | 0.394542716 | 0.693180397 | 0.837078181 | 0.032246658 |

| ensembl_gene_id | external_gene_name | entrezgene_id | gene_biotype | baseMean | log2FoldChange | lfcSE | stat | pvalue | padj | absLog2FC |
|---|---|---|---|---|---|---|---|---|---|---|
| ENSMUSG00000036450 | Hif1an | 319594 | protein_coding | 884.8527848 | -0.032299062 | 0.065298261 | 0.494638933 | 0.620855015 | 0.791721985 | 0.032299062 |
| ENSMUSG00000031065 | Cdk16 | 18555 | protein_coding | 1454.814391 | 0.032310606 | 0.051358389 | -0.629120316 | 0.529270291 | 0.727020313 | 0.032310606 |
| ENSMUSG00000024533 | Spire1 | 68166 | protein_coding | 39.99318008 | -0.032360539 | 0.219026665 | 0.147747027 | 0.882542417 | 0.941973978 | 0.032360539 |
| ENSMUSG00000037519 | Ppfia1 | 233977 | protein_coding | 884.7975857 | 0.032373197 | 0.072440395 | -0.446894258 | 0.65495141 | 0.812734264 | 0.032373197 |
| ENSMUSG00000015733 | Capza2 | 12343 | protein_coding | 1697.129156 | -0.032480797 | 0.055884549 | 0.581212479 | 0.561097258 | 0.749612501 | 0.032480797 |
| ENSMUSG00000074218 | Cox7a1 | 12865 | protein_coding | 460.943088 | -0.032485062 | 0.12674568 | 0.256301139 | 0.797718318 | 0.899646268 | 0.032485062 |
| ENSMUSG00000003604 | Aven | 74268 | protein_coding | 268.1686161 | 0.032502468 | 0.08479752 | -0.383295031 | 0.701501023 | 0.842293038 | 0.032502468 |
| ENSMUSG00000040731 | Eif4h | 22384 | protein_coding | 3323.63161 | -0.032504999 | 0.057543229 | 0.564879649 | 0.572155627 | 0.757643976 | 0.032504999 |
| ENSMUSG00000050428 | Fbxo46 | 243867 | protein_coding | 231.2410091 | -0.032511182 | 0.072683143 | 0.447300169 | 0.654658345 | 0.812675931 | 0.032511182 |
| ENSMUSG00000026585 | Kifap3 | 16579 | protein_coding | 217.5500837 | 0.032579899 | 0.101673791 | -0.320435567 | 0.748638167 | 0.873772754 | 0.032579899 |
| ENSMUSG00000034796 | Cpne7 | 102278 | protein_coding | 3.303019233 | -0.032641091 | 1.252984267 | 0.026050679 | 0.979216916 | 1 | 0.032641091 |
| ENSMUSG00000037523 | Mavs | 228607 | protein_coding | 610.7260518 | 0.032657937 | 0.098816771 | -0.330489815 | 0.741029887 | 0.868956809 | 0.032657937 |
| ENSMUSG00000064427 | Gm22748 | 115488064 | snoRNA | 101.1570093 | 0.032662317 | 0.210144656 | -0.155427776 | 0.876484088 | 0.938630585 | 0.032662317 |
| ENSMUSG00000031819 | Emc8 | 18117 | protein_coding | 233.9503345 | 0.032667027 | 0.091306695 | -0.357772523 | 0.720513555 | 0.855204229 | 0.032667027 |
| ENSMUSG00000028629 | Exo5 | 73172 | protein_coding | 143.5116503 | -0.032683113 | 0.090356908 | 0.361711279 | 0.717567798 | 0.853563561 | 0.032683113 |
| ENSMUSG00000034254 | Agpat1 | 55979 | protein_coding | 1309.009409 | 0.032698415 | 0.079070601 | -0.413534421 | 0.679215109 | 0.827973596 | 0.032698415 |
| ENSMUSG00000027650 | Tti1 | 75425 | protein_coding | 217.4487858 | -0.032707053 | 0.123593339 | 0.264634433 | 0.791291092 | 0.89588517 | 0.032707053 |
| ENSMUSG00000029050 | Ski | 20481 | protein_coding | 788.3764415 | -0.0327459 | 0.06998082 | 0.467926783 | 0.639836949 | 0.803640659 | 0.0327459 |
| ENSMUSG00000047547 | Cltb | 74325 | protein_coding | 1780.800572 | 0.032747195 | 0.052280189 | -0.626378661 | 0.531066603 | 0.728048657 | 0.032747195 |
| ENSMUSG00000043733 | Ptpn11 | 19247 | protein_coding | 1069.936852 | 0.032754707 | 0.071415466 | -0.458650051 | 0.646485488 | 0.807324664 | 0.032754707 |
| ENSMUSG00000018923 | Med11 | 66172 | protein_coding | 144.8556266 | -0.032759723 | 0.113704754 | 0.28811217 | 0.773260877 | 0.886362513 | 0.032759723 |
| ENSMUSG00000011958 | Bnip2 | 12175 | protein_coding | 158.8131653 | 0.032786664 | 0.126428858 | -0.259328961 | 0.795381436 | 0.898299072 | 0.032786664 |
| ENSMUSG00000039048 | Foxred1 | 235169 | protein_coding | 372.3471462 | -0.032794367 | 0.077821982 | 0.421402357 | 0.673461298 | 0.824917659 | 0.032794367 |
| ENSMUSG00000039840 | Epg5 | 100502841 | protein_coding | 552.6386319 | 0.032887557 | 0.083930058 | -0.391844805 | 0.695172887 | 0.838326567 | 0.032887557 |
| ENSMUSG00000021149 | Gtpbp4 | 69237 | protein_coding | 398.9262663 | -0.032896589 | 0.105235309 | 0.312600295 | 0.754584354 | 0.876856731 | 0.032896589 |
| ENSMUSG00000051166 | Eml5 | 319670 | protein_coding | 25.97901467 | 0.032921365 | 0.317591887 | -0.103659338 | 0.917439697 | 0.959505963 | 0.032921365 |
| ENSMUSG00000037579 | Kcnh3 | 16512 | protein_coding | 257.1305457 | -0.032926179 | 0.145147649 | 0.226846105 | 0.820543412 | 0.9092783 | 0.032926179 |
| ENSMUSG00000075486 | Commd6 | 66200 | protein_coding | 301.987055 | -0.032985197 | 0.076894594 | 0.428966406 | 0.667947673 | 0.821576632 | 0.032985197 |
| ENSMUSG00000048078 | Tenm4 | 23966 | protein_coding | 156.7247463 | -0.033034674 | 0.156685861 | 0.210833792 | 0.833016969 | 0.915935963 | 0.033034674 |
| ENSMUSG00000054517 | Trim65 | 338364 | protein_coding | 177.7001396 | -0.033056967 | 0.114949302 | 0.287578666 | 0.773669277 | 0.88672609 | 0.033056967 |
| ENSMUSG00000030382 | Slc27a5 | 26459 | protein_coding | 1.122127694 | -0.033061762 | 2.540284497 | 0.013014984 | 0.989615838 | 1 | 0.033061762 |
| ENSMUSG00000038467 | Chmp4b | 75608 | protein_coding | 2798.384126 | -0.0330623 | 0.066950222 | 0.493834058 | 0.621423378 | 0.792202074 | 0.0330623 |
| ENSMUSG00000030546 | Plin1 | 103968 | protein_coding | 2.24861595 | -0.0330824 | 2.135404097 | 0.015492337 | 0.987639398 | 1 | 0.0330824 |
| ENSMUSG00000037710 | Cisd1 | 52637 | protein_coding | 807.0651137 | -0.033098591 | 0.092141833 | 0.359213505 | 0.719435374 | 0.854853631 | 0.033098591 |
| ENSMUSG00000032246 | Calml4 | 75600 | protein_coding | 1344.489793 | -0.033145132 | 0.080561177 | 0.411428098 | 0.680758654 | 0.829059617 | 0.033145132 |
| ENSMUSG00000086392 | Mccc1os | 654424 | lncRNA | 3.296867628 | 0.033160601 | 0.771177879 | -0.042999938 | 0.965701583 | 1 | 0.033160601 |
| ENSMUSG00000019782 | Rwdd1 | 66521 | protein_coding | 164.2804645 | -0.033163124 | 0.166681435 | 0.198961117 | 0.84229316 | 0.921900136 | 0.033163124 |
| ENSMUSG00000007415 | Gatad1 | 67210 | protein_coding | 717.6710778 | -0.03316914 | 0.069812834 | 0.475115218 | 0.634704852 | 0.800822518 | 0.03316914 |
| ENSMUSG00000050786 | Ccdc126 | 57895 | protein_coding | 32.45863937 | 0.033173133 | 0.213121595 | -0.155653552 | 0.876306111 | 0.938630585 | 0.033173133 |

| ensembl_gene_id | external_gene_name | entrezgene_id | gene_biotype | baseMean | log2FoldChange | lfcSE | stat | pvalue | padj | absLog2FC |
|---|---|---|---|---|---|---|---|---|---|---|
| ENSMUSG00000078905 | Gm14393 | 664987 | protein_coding | 1.469785856 | 0.03317613 | 1.154726783 | -0.028730718 | 0.977079357 | 1 | 0.03317613 |
| ENSMUSG00000025612 | Bach1 | 12013 | protein_coding | 859.3836958 | 0.03317724 | 0.095060328 | -0.349012471 | 0.727079947 | 0.858992577 | 0.03317724 |
| ENSMUSG00000004393 | Ddx56 | 52513 | protein_coding | 326.8369824 | -0.033205966 | 0.107686183 | 0.308358652 | 0.757809441 | 0.87863969 | 0.033205966 |
| ENSMUSG00000032097 | Ddx6 | 13209 | protein_coding | 1922.566536 | -0.033206768 | 0.091414417 | 0.363255268 | 0.716414205 | 0.852797822 | 0.033206768 |
| ENSMUSG00000001910 | Nacc1 | 66830 | protein_coding | 1241.63056 | -0.033276826 | 0.065758911 | 0.506042833 | 0.612826581 | 0.785880315 | 0.033276826 |
| ENSMUSG00000043190 | Rfesd | 218341 | protein_coding | 95.67391462 | -0.033297298 | 0.153012378 | 0.217611792 | 0.827731596 | 0.913225519 | 0.033297298 |
| ENSMUSG00000042532 | Golga7b | 71146 | protein_coding | 3.172997258 | -0.033320622 | 0.613868866 | 0.054279707 | 0.956712317 | 1 | 0.033320622 |
| ENSMUSG00000027394 | Ttl | 69737 | protein_coding | 95.49664183 | -0.033327432 | 0.122439747 | 0.272194552 | 0.785472431 | 0.892918574 | 0.033327432 |
| ENSMUSG00000041187 | Prkd2 | 101540 | protein_coding | 209.3480263 | -0.033332311 | 0.12027947 | 0.277123857 | 0.781685004 | 0.890621398 | 0.033332311 |
| ENSMUSG00000020228 | Helb | 117599 | protein_coding | 335.2702203 | 0.033353674 | 0.090611483 | -0.368095443 | 0.712802069 | 0.850705598 | 0.033353674 |
| ENSMUSG00000059119 | Nap1l4 | 17955 | protein_coding | 838.2488502 | -0.033367028 | 0.084761527 | 0.393657703 | 0.693833773 | 0.837509753 | 0.033367028 |
| ENSMUSG00000057969 | Sema3b | 20347 | protein_coding | 1051.546386 | -0.033399405 | 0.069166466 | 0.482884369 | 0.629177841 | 0.797446101 | 0.033399405 |
| ENSMUSG00000052031 | Tagap1 | 380608 | protein_coding | 81.4237217 | -0.033428158 | 0.154686205 | 0.216103034 | 0.828907435 | 0.913504709 | 0.033428158 |
| ENSMUSG00000031921 | Terf2 | 21750 | protein_coding | 186.6295612 | 0.033437942 | 0.084905603 | -0.393824924 | 0.693710302 | 0.837421924 | 0.033437942 |
| ENSMUSG00000046441 | Cmtr2 | 234728 | protein_coding | 131.8845981 | 0.033451135 | 0.107059297 | -0.312454277 | 0.754695306 | 0.876856731 | 0.033451135 |
| ENSMUSG00000034785 | Dio1 | 13370 | protein_coding | 682.3516825 | -0.033457262 | 0.107224522 | 0.312029947 | 0.755017765 | 0.877003824 | 0.033457262 |
| ENSMUSG00000067206 | Lrrc66 | 231296 | protein_coding | 826.7385403 | 0.033463953 | 0.095802993 | -0.349299658 | 0.726864355 | 0.85879931 | 0.033463953 |
| ENSMUSG00000030339 | Ltbr | 17000 | protein_coding | 1392.656195 | 0.033481874 | 0.074521862 | -0.449289289 | 0.653222984 | 0.811870577 | 0.033481874 |
| ENSMUSG00000079507 | H2-Q1 | 15006 | protein_coding | 214.5147542 | -0.033527393 | 0.215476202 | 0.155596733 | 0.8763509 | 0.938630585 | 0.033527393 |
| ENSMUSG00000040918 | Slc19a2 | 116914 | protein_coding | 111.0741671 | -0.033532372 | 0.124407416 | 0.269536763 | 0.787516656 | 0.893615254 | 0.033532372 |
| ENSMUSG00000042284 | Itga1 | 109700 | protein_coding | 189.760487 | 0.03353828 | 0.139820714 | -0.239866318 | 0.810433893 | 0.906179443 | 0.03353828 |
| ENSMUSG00000027843 | Ptpn22 | 19260 | protein_coding | 14.97932535 | 0.033538347 | 0.42515085 | -0.078885757 | 0.937123492 | 0.970043107 | 0.033538347 |
| ENSMUSG00000031198 | Fundc2 | 67391 | protein_coding | 183.8132229 | 0.033539784 | 0.136224434 | -0.246209751 | 0.805519868 | 0.904245469 | 0.033539784 |
| ENSMUSG00000046671 | Mtfr1l | 76824 | protein_coding | 458.1322023 | -0.033594858 | 0.076510819 | 0.43908637 | 0.660598955 | 0.81673513 | 0.033594858 |
| ENSMUSG00000022867 | Usp25 | 30940 | protein_coding | 830.0848764 | -0.033619745 | 0.058598968 | 0.573725887 | 0.566153308 | 0.752653012 | 0.033619745 |
| ENSMUSG00000098912 | 1500004A13Rik | 319830 | lncRNA | 26.65056609 | 0.033644247 | 0.225053326 | -0.149494557 | 0.881163404 | 0.941406719 | 0.033644247 |
| ENSMUSG00000028333 | Anp32b | 67628 | protein_coding | 375.038587 | 0.03366714 | 0.114273258 | -0.294619586 | 0.768284498 | 0.883975099 | 0.03366714 |
| ENSMUSG00000042182 | Bend6 | 320705 | protein_coding | 1.412359531 | -0.033668619 | 1.751520807 | 0.019222506 | 0.984663604 | 1 | 0.033668619 |
| ENSMUSG00000002319 | Ipo4 | 75751 | protein_coding | 134.3401761 | 0.033673705 | 0.130014631 | -0.258999348 | 0.795635743 | 0.898463393 | 0.033673705 |
| ENSMUSG00000029221 | Slc30a9 | 109108 | protein_coding | 753.9144217 | -0.033754858 | 0.071607097 | 0.471389836 | 0.637362373 | 0.802408631 | 0.033754858 |
| ENSMUSG00000022992 | Kansl2 | 69612 | protein_coding | 199.9678977 | -0.033778157 | 0.116526278 | 0.289875881 | 0.771911194 | 0.885478012 | 0.033778157 |
| ENSMUSG00000039461 | Tcta | 102791 | protein_coding | 423.9356025 | 0.033813907 | 0.086198109 | -0.39228131 | 0.694850372 | 0.838124245 | 0.033813907 |
| ENSMUSG00000057177 | Gsk3a | 606496 | protein_coding | 421.2772019 | 0.033844575 | 0.063088687 | -0.536460283 | 0.591640478 | 0.771785126 | 0.033844575 |
| ENSMUSG00000030868 | Dctn5 | 59288 | protein_coding | 455.2974559 | 0.033870392 | 0.073419037 | -0.461329837 | 0.644561981 | 0.806325018 | 0.033870392 |
| ENSMUSG00000024513 | Mbd2 | 17191 | protein_coding | 1295.339469 | -0.033877873 | 0.059946308 | 0.565136939 | 0.571980625 | 0.757533728 | 0.033877873 |
| ENSMUSG00000026614 | Slc30a10 | 226781 | protein_coding | 1208.93023 | -0.033882558 | 0.221717805 | 0.152818388 | 0.878541496 | 0.939901695 | 0.033882558 |
| ENSMUSG00000022471 | Xrcc6 | 14375 | protein_coding | 297.1039015 | -0.03396037 | 0.11568634 | 0.293555577 | 0.769097525 | 0.884277416 | 0.03396037 |
| ENSMUSG00000025571 | Tnrc6c | 217351 | protein_coding | 595.3098105 | 0.033991502 | 0.070816349 | -0.479995117 | 0.631230865 | 0.799065123 | 0.033991502 |
| ENSMUSG00000038861 | Pi4kb | 107650 | protein_coding | 542.8230119 | -0.033993763 | 0.048165928 | 0.705763684 | 0.48033511 | 0.691324212 | 0.033993763 |

| ensembl_gene_id | external_gene_name | entrezgene_id | gene_biotype | baseMean | log2FoldChange | lfcSE | stat | pvalue | padj | absLog2FC |
|---|---|---|---|---|---|---|---|---|---|---|
| ENSMUSG00000041945 | Mfsd9 | 211798 | protein_coding | 134.9284141 | -0.034002693 | 0.11612933 | 0.292800218 | 0.769674862 | 0.884510137 | 0.034002693 |
| ENSMUSG00000031813 | Mvb12a | 73711 | protein_coding | 751.2837275 | -0.034008544 | 0.073794653 | 0.460853776 | 0.644903517 | 0.80663006 | 0.034008544 |
| ENSMUSG00000029687 | Ezh2 | 14056 | protein_coding | 365.1070791 | -0.034010117 | 0.095791787 | 0.355042101 | 0.722558051 | 0.856468824 | 0.034010117 |
| ENSMUSG00000060149 | BC002059 | 213811 | lncRNA | 68.94001385 | -0.034045883 | 0.133967992 | 0.25413446 | 0.799391682 | 0.90054944 | 0.034045883 |
| ENSMUSG00000030757 | Zkscan2 | 210162 | protein_coding | 2.228054327 | 0.034076653 | 2.429485077 | -0.014026286 | 0.98880901 | 1 | 0.034076653 |
| ENSMUSG00000010914 | Pdhx | 27402 | protein_coding | 355.5053074 | 0.034098736 | 0.073282568 | -0.465304869 | 0.641713149 | 0.804528617 | 0.034098736 |
| ENSMUSG00000054364 | Rhob | 11852 | protein_coding | 5073.205047 | 0.034109265 | 0.137437835 | -0.248179589 | 0.803995458 | 0.903393958 | 0.034109265 |
| ENSMUSG00000029730 | Mcm7 | 17220 | protein_coding | 496.7902965 | 0.034120856 | 0.170874356 | -0.199683891 | 0.841727813 | 0.921642147 | 0.034120856 |
| ENSMUSG00000078622 | Ccdc47 | 67163 | protein_coding | 955.8667786 | -0.034138204 | 0.09192258 | 0.371379962 | 0.710354548 | 0.848979902 | 0.034138204 |
| ENSMUSG00000100594 | 2810414N06Rik | 102640772 | lncRNA | 11.23564056 | 0.034145185 | 0.310287294 | -0.110043775 | 0.912374658 | 0.957266079 | 0.034145185 |
| ENSMUSG00000018567 | Gabarap | 56486 | protein_coding | 2382.832443 | -0.03418287 | 0.046161838 | 0.740500631 | 0.458996278 | 0.673388737 | 0.03418287 |
| ENSMUSG00000049969 | Plekhf2 | 71801 | protein_coding | 843.9126404 | -0.034196606 | 0.053867963 | 0.634822691 | 0.525544063 | 0.724020576 | 0.034196606 |
| ENSMUSG00000062397 | Zfp706 | 68036 | protein_coding | 1299.842648 | -0.034277726 | 0.077184172 | 0.444103047 | 0.656968084 | 0.814465699 | 0.034277726 |
| ENSMUSG00000038780 | Smurf1 | 75788 | protein_coding | 410.4652388 | 0.034321688 | 0.090856895 | -0.377755458 | 0.705612266 | 0.845753813 | 0.034321688 |
| ENSMUSG00000093081 | Mir3075 | 100526533 | miRNA | 0.587822831 | -0.034383874 | 1.586981276 | 0.021666213 | 0.982714216 | 1 | 0.034383874 |
| ENSMUSG00000035790 | Cep19 | 66994 | protein_coding | 277.1533057 | 0.034436265 | 0.08190763 | -0.420428045 | 0.674172784 | 0.825194181 | 0.034436265 |
| ENSMUSG00000024867 | Pip5k1b | 18719 | protein_coding | 804.234879 | -0.034437815 | 0.067587449 | 0.509529737 | 0.61038096 | 0.784117555 | 0.034437815 |
| ENSMUSG00000035954 | Dock4 | 238130 | protein_coding | 23.78515086 | -0.03444417 | 0.321169017 | 0.107246242 | 0.914593631 | 0.958199105 | 0.03444417 |
| ENSMUSG00000034028 | Cd226 | 225825 | protein_coding | 7.328228876 | -0.034457624 | 0.518054365 | 0.066513529 | 0.946968987 | 0.974163449 | 0.034457624 |
| ENSMUSG00000034931 | Dhx8 | 217207 | protein_coding | 608.8919922 | 0.034514563 | 0.066784492 | -0.516805056 | 0.605292251 | 0.780311961 | 0.034514563 |
| ENSMUSG00000020064 | Herc4 | 67345 | protein_coding | 649.7763553 | 0.034548751 | 0.081065712 | -0.426182043 | 0.669975201 | 0.822722373 | 0.034548751 |
| ENSMUSG00000029474 | Rnf34 | 80751 | protein_coding | 272.0260329 | -0.034561919 | 0.106154821 | 0.325580308 | 0.744741913 | 0.8712288 | 0.034561919 |
| ENSMUSG00000034341 | Wbp2 | 22378 | protein_coding | 656.9308231 | 0.034608522 | 0.058125348 | -0.595411875 | 0.55156819 | 0.742355007 | 0.034608522 |
| ENSMUSG00000036572 | Upf3b | 68134 | protein_coding | 204.8191292 | 0.034732584 | 0.082214158 | -0.422464757 | 0.67268582 | 0.824518194 | 0.034732584 |
| ENSMUSG00000046519 | Golph3l | 229593 | protein_coding | 4145.26386 | -0.03483414 | 0.154086962 | 0.226068056 | 0.821148489 | 0.909608677 | 0.03483414 |
| ENSMUSG00000053436 | Mapk14 | 26416 | protein_coding | 765.9905329 | -0.034897639 | 0.071980634 | 0.484819829 | 0.627804156 | 0.796376973 | 0.034897639 |
| ENSMUSG00000065258 | Gm23969 | 115489598 | snoRNA | 7.174507202 | -0.034905137 | 0.465615394 | 0.074965599 | 0.940242083 | 0.971417829 | 0.034905137 |
| ENSMUSG00000056216 | Cebpg | 12611 | protein_coding | 1061.817003 | 0.0349375 | 0.071764295 | -0.486836809 | 0.626373983 | 0.795381772 | 0.0349375 |
| ENSMUSG00000083649 | Rasl2-9 | 19428 | protein_coding | 6.436345765 | -0.03504077 | 0.475710127 | 0.073659921 | 0.94128099 | 0.971762583 | 0.03504077 |
| ENSMUSG00000040701 | Ap1g2 | 11766 | protein_coding | 514.8732753 | -0.035065235 | 0.073887731 | 0.474574526 | 0.635090267 | 0.801105392 | 0.035065235 |
| ENSMUSG00000038876 | Rnf146 | 68031 | protein_coding | 143.0146489 | 0.035068555 | 0.112457226 | -0.311839052 | 0.755162843 | 0.877032235 | 0.035068555 |
| ENSMUSG00000063889 | Crem | 12916 | protein_coding | 53.10885537 | 0.035102353 | 0.174315743 | -0.201372246 | 0.840407515 | 0.920873844 | 0.035102353 |
| ENSMUSG00000023170 | Gps2 | 56310 | protein_coding | 269.8012375 | -0.035109582 | 0.073151638 | 0.479956192 | 0.631258544 | 0.799065123 | 0.035109582 |
| ENSMUSG00000070417 | Or6a2 | 18317 | protein_coding | 2.968386822 | 0.035137418 | 2.453842009 | -0.014319348 | 0.988575204 | 1 | 0.035137418 |
| ENSMUSG00000034435 | Tmem30b | 238257 | protein_coding | 2877.910642 | 0.035151261 | 0.093407234 | -0.376322686 | 0.706677016 | 0.846208704 | 0.035151261 |
| ENSMUSG00000059495 | Arhgef12 | 69632 | protein_coding | 1102.365509 | -0.035164637 | 0.055964187 | 0.62834177 | 0.529780074 | 0.72724756 | 0.035164637 |
| ENSMUSG00000005897 | Nr2c1 | 22025 | protein_coding | 146.899595 | 0.035216372 | 0.091963994 | -0.382936526 | 0.701766828 | 0.842528346 | 0.035216372 |
| ENSMUSG00000047819 | Tigd4 | 403175 | protein_coding | 4.274011072 | -0.035227161 | 0.672703615 | 0.052366541 | 0.958236634 | 1 | 0.035227161 |
| ENSMUSG00000024515 | Smad4 | 17128 | protein_coding | 1274.892756 | 0.035295516 | 0.127828697 | -0.276115743 | 0.782459169 | 0.890955714 | 0.035295516 |

| ensembl_gene_id | external_gene_name | entrezgene_id | gene_biotype | baseMean | log2FoldChange | lfcSE | stat | pvalue | padj | absLog2FC |
|---|---|---|---|---|---|---|---|---|---|---|
| ENSMUSG00000039159 | Ube2h | 22214 | protein_coding | 746.2375354 | 0.035295936 | 0.082230037 | -0.429234095 | 0.667752873 | 0.821398208 | 0.035295936 |
| ENSMUSG00000029701 | Rbm28 | 68272 | protein_coding | 352.2724992 | 0.035330781 | 0.075357727 | -0.468840845 | 0.639183402 | 0.803282676 | 0.035330781 |
| ENSMUSG00000028327 | Stra6l | 74152 | protein_coding | 269.6177593 | 0.035353514 | 0.117485024 | -0.300919319 | 0.763476018 | 0.881386379 | 0.035353514 |
| ENSMUSG00000041846 | Ppp4r3a | 68734 | protein_coding | 1067.591956 | 0.035353576 | 0.071245518 | -0.496221764 | 0.619737958 | 0.790880748 | 0.035353576 |
| ENSMUSG00000025102 | 3110040N11Rik | 67290 | protein_coding | 142.2909643 | -0.035405226 | 0.118256748 | 0.299392857 | 0.764640312 | 0.882132403 | 0.035405226 |
| ENSMUSG00000024382 | Ercc3 | 13872 | protein_coding | 421.829526 | 0.035434447 | 0.07706492 | -0.459799953 | 0.645659817 | 0.806964824 | 0.035434447 |
| ENSMUSG00000026484 | Rnf2 | 19821 | protein_coding | 288.2043309 | -0.03544895 | 0.088318603 | 0.401375803 | 0.688143462 | 0.833786242 | 0.03544895 |
| ENSMUSG00000017776 | Crk | 12928 | protein_coding | 2274.035314 | 0.035483488 | 0.07084483 | -0.50086207 | 0.616468199 | 0.788502504 | 0.035483488 |
| ENSMUSG00000040479 | Dgkz | 104418 | protein_coding | 1918.499743 | 0.035503631 | 0.056311598 | -0.630485243 | 0.528377155 | 0.726359126 | 0.035503631 |
| ENSMUSG00000036721 | Zscan12 | 22758 | protein_coding | 74.02585578 | -0.035544558 | 0.189226891 | 0.187840944 | 0.851001338 | 0.927281893 | 0.035544558 |
| ENSMUSG00000026095 | Asnsd1 | 70396 | protein_coding | 100.7332503 | 0.035551362 | 0.131947407 | -0.269435856 | 0.787594297 | 0.893615254 | 0.035551362 |
| ENSMUSG00000073599 | Ecscr | 68545 | protein_coding | 40.73092259 | 0.035553138 | 0.199330128 | -0.178363094 | 0.858437831 | 0.930749284 | 0.035553138 |
| ENSMUSG00000020128 | Vps54 | 245944 | protein_coding | 1246.139095 | -0.035569578 | 0.079129407 | 0.449511503 | 0.653062712 | 0.811818066 | 0.035569578 |
| ENSMUSG00000015120 | Ube2i | 22196 | protein_coding | 599.3902778 | 0.035613905 | 0.092391062 | -0.385469161 | 0.69988985 | 0.841086565 | 0.035613905 |
| ENSMUSG00000020246 | Hcfc2 | 67933 | protein_coding | 213.0286832 | 0.035689795 | 0.097529488 | -0.365938508 | 0.714410961 | 0.851829233 | 0.035689795 |
| ENSMUSG00000030134 | Rasgef1a | 70727 | protein_coding | 4.188339838 | 0.03571214 | 0.643904351 | -0.055461871 | 0.955770506 | 1 | 0.03571214 |
| ENSMUSG00000032217 | Rnf111 | 93836 | protein_coding | 467.8855333 | 0.035742675 | 0.064565806 | -0.553585206 | 0.579862753 | 0.763259485 | 0.035742675 |
| ENSMUSG00000015957 | Wnt11 | 22411 | protein_coding | 14.70304944 | -0.035745174 | 0.400616269 | 0.089225468 | 0.928902725 | 0.966049475 | 0.035745174 |
| ENSMUSG00000039789 | Zfp597 | 71063 | protein_coding | 123.2774803 | -0.03575064 | 0.140149774 | 0.255088816 | 0.798654502 | 0.900025962 | 0.03575064 |
| ENSMUSG00000025787 | Tgm4 | 331046 | protein_coding | 6.017114624 | -0.035782645 | 0.408129128 | 0.087674813 | 0.930135139 | 0.966352899 | 0.035782645 |
| ENSMUSG00000021585 | Cast | 12380 | protein_coding | 1835.944952 | -0.03592669 | 0.094859085 | 0.378737474 | 0.704882822 | 0.845163358 | 0.03592669 |
| ENSMUSG00000049792 | Bag5 | 70369 | protein_coding | 187.7407246 | -0.035940248 | 0.114381354 | 0.314214223 | 0.753358341 | 0.876432521 | 0.035940248 |
| ENSMUSG00000021712 | Trim23 | 81003 | protein_coding | 132.9894365 | 0.035948594 | 0.103445049 | -0.347513917 | 0.728205265 | 0.859565127 | 0.035948594 |
| ENSMUSG00000097047 | 1110020A21Rik | 68531 | lncRNA | 20.51568029 | 0.035965804 | 0.256822596 | -0.140041433 | 0.888627254 | 0.94517522 | 0.035965804 |
| ENSMUSG00000070923 | Klhl9 | 242521 | protein_coding | 1306.506231 | 0.035973012 | 0.078470556 | -0.458426876 | 0.646645787 | 0.807443873 | 0.035973012 |
| ENSMUSG00000020464 | Pnpt1 | 71701 | protein_coding | 461.9427711 | 0.035973597 | 0.103653694 | -0.347055616 | 0.728549536 | 0.859744679 | 0.035973597 |
| ENSMUSG00000024695 | Zfp91 | 109910 | protein_coding | 986.2411488 | 0.035989039 | 0.111692059 | -0.322216629 | 0.747288587 | 0.872752739 | 0.035989039 |
| ENSMUSG00000022698 | Naa50 | 72117 | protein_coding | 915.9558108 | -0.035989854 | 0.104148504 | 0.345562851 | 0.729671269 | 0.860514985 | 0.035989854 |
| ENSMUSG00000074300 | Smim31 | 407795 | protein_coding | 1243.82924 | -0.036046764 | 0.14943918 | 0.24121361 | 0.809389563 | 0.905818599 | 0.036046764 |
| ENSMUSG00000002325 | Irf9 | 16391 | protein_coding | 400.6960847 | 0.036091197 | 0.153202996 | -0.235577618 | 0.813760443 | 0.907696046 | 0.036091197 |
| ENSMUSG00000093805 | Gal3st2b | 100041596 | protein_coding | 92.00109841 | 0.036110292 | 0.469117672 | -0.076974913 | 0.938643503 | 0.970660753 | 0.036110292 |
| ENSMUSG00000030983 | Bccip | 66165 | protein_coding | 429.1934178 | -0.036146441 | 0.100158238 | 0.360893341 | 0.718179182 | 0.853791016 | 0.036146441 |
| ENSMUSG00000087366 | Junos | 230451 | lncRNA | 31.68335001 | 0.036147875 | 0.36976863 | -0.097758089 | 0.922124388 | 0.9624373 | 0.036147875 |
| ENSMUSG00000029596 | Sdsl | 257635 | protein_coding | 202.1355249 | 0.036156091 | 0.093818944 | -0.385381556 | 0.699954745 | 0.841086565 | 0.036156091 |
| ENSMUSG00000042292 | Mrtfa | 223701 | protein_coding | 328.0160912 | 0.036203629 | 0.086617994 | -0.417968915 | 0.675969845 | 0.826457838 | 0.036203629 |
| ENSMUSG00000087938 | Gm26049 | 115490020 | snRNA | 2.468360153 | 0.036218233 | 0.953608787 | -0.03798018 | 0.969703485 | 1 | 0.036218233 |
| ENSMUSG00000033618 | Map3k13 | 71751 | protein_coding | 291.5219037 | -0.036267027 | 0.131679358 | 0.275419229 | 0.782994172 | 0.891196207 | 0.036267027 |
| ENSMUSG00000063524 | Eno1 | 13806 | protein_coding | 2814.790772 | -0.036273618 | 0.117499603 | 0.308712687 | 0.757540092 | 0.878512436 | 0.036273618 |
| ENSMUSG00000005846 | Rsl1d1 | 66409 | protein_coding | 601.5452053 | 0.036341808 | 0.099935131 | -0.363653973 | 0.716116418 | 0.852650725 | 0.036341808 |

| ensembl_gene_id | external_gene_name | entrezgene_id | gene_biotype | baseMean | log2FoldChange | lfcSE | stat | pvalue | padj | absLog2FC |
|---|---|---|---|---|---|---|---|---|---|---|
| ENSMUSG00000020455 | Trim11 | 94091 | protein_coding | 296.6819996 | -0.03636857 | 0.092848978 | 0.391695961 | 0.695282874 | 0.838339763 | 0.03636857 |
| ENSMUSG00000025439 | Clns1a | 12729 | protein_coding | 469.6005279 | 0.036381845 | 0.108566478 | -0.33511122 | 0.737541189 | 0.866806221 | 0.036381845 |
| ENSMUSG00000021519 | Mterf3 | 66410 | protein_coding | 209.0082901 | 0.036443367 | 0.092834082 | -0.392564523 | 0.694641147 | 0.8379481 | 0.036443367 |
| ENSMUSG00000042015 | Wdr41 | 218460 | protein_coding | 314.3618874 | -0.036466624 | 0.099093766 | 0.368001196 | 0.712872343 | 0.850727986 | 0.036466624 |
| ENSMUSG00000039771 | Polr2j | 20022 | protein_coding | 241.1033609 | -0.036484432 | 0.072400524 | 0.503924971 | 0.614314102 | 0.78673795 | 0.036484432 |
| ENSMUSG00000041203 | Trir | 68544 | protein_coding | 859.3931811 | 0.036561438 | 0.077845306 | -0.469667851 | 0.63859234 | 0.80301103 | 0.036561438 |
| ENSMUSG00000024991 | Eif3a | 13669 | protein_coding | 2741.284954 | 0.036588535 | 0.088091533 | -0.415346781 | 0.67788806 | 0.827372024 | 0.036588535 |
| ENSMUSG00000056211 | R3hdm1 | 226412 | protein_coding | 317.0983168 | 0.036589149 | 0.106389619 | -0.343916533 | 0.730909058 | 0.861359607 | 0.036589149 |
| ENSMUSG00000004748 | Mtfp1 | 67900 | protein_coding | 150.1404937 | 0.036604208 | 0.114345428 | -0.320119556 | 0.748877701 | 0.87389673 | 0.036604208 |
| ENSMUSG00000051351 | Zfp46 | 22704 | protein_coding | 206.9641432 | -0.036656984 | 0.101433548 | 0.361389152 | 0.717808556 | 0.853596205 | 0.036656984 |
| ENSMUSG00000004980 | Hnrnpa2b1 | 53379 | protein_coding | 4108.370311 | 0.036657159 | 0.077448078 | -0.473312696 | 0.635990106 | 0.801613541 | 0.036657159 |
| ENSMUSG00000041926 | Rnpep | 215615 | protein_coding | 1024.559863 | -0.036671933 | 0.089300603 | 0.410657166 | 0.68132394 | 0.829503357 | 0.036671933 |
| ENSMUSG00000052291 | 5330438D12Rik | 102633868 | lncRNA | 16.61470006 | -0.036736391 | 0.269703756 | 0.136210159 | 0.891655145 | 0.947050968 | 0.036736391 |
| ENSMUSG00000024259 | Slc25a46 | 67453 | protein_coding | 669.841404 | 0.036748252 | 0.075788089 | -0.484881623 | 0.627760319 | 0.796376973 | 0.036748252 |
| ENSMUSG00000032281 | Acsbg1 | 94180 | protein_coding | 5.409771198 | 0.036758853 | 0.741905448 | -0.049546547 | 0.960483744 | 0.980697519 | 0.036758853 |
| ENSMUSG00000022553 | Maf1 | 68877 | protein_coding | 730.0962625 | -0.036764645 | 0.060090747 | 0.611818735 | 0.540657693 | 0.734843633 | 0.036764645 |
| ENSMUSG00000024981 | Acsl5 | 433256 | protein_coding | 1371.93623 | -0.036807802 | 0.116575334 | 0.315742628 | 0.752197868 | 0.875699372 | 0.036807802 |
| ENSMUSG00000022466 | Rpap3 | 71919 | protein_coding | 134.3974547 | 0.03684929 | 0.123597367 | -0.298139766 | 0.765596492 | 0.882510888 | 0.03684929 |
| ENSMUSG00000023967 | Mrps18a | 68565 | protein_coding | 394.1276613 | -0.036871571 | 0.073496303 | 0.501679266 | 0.615893152 | 0.788072176 | 0.036871571 |
| ENSMUSG00000015522 | Arnt | 11863 | protein_coding | 487.9069061 | -0.036881358 | 0.067194087 | 0.548878032 | 0.583089155 | 0.765756487 | 0.036881358 |
| ENSMUSG00000088958 | Scarna8 | 100217448 | scaRNA | 1.108339289 | 0.036883194 | 1.116991331 | -0.033020125 | 0.973658539 | 1 | 0.036883194 |
| ENSMUSG00000061981 | Flot2 | 14252 | protein_coding | 150.2268157 | -0.036921611 | 0.119207142 | 0.309726502 | 0.756768947 | 0.877988085 | 0.036921611 |
| ENSMUSG00000007038 | Neu1 | 18010 | protein_coding | 2008.022453 | -0.03692206 | 0.182439722 | 0.202379502 | 0.839620052 | 0.920255229 | 0.03692206 |
| ENSMUSG00000048285 | Frmd6 | 319710 | protein_coding | 39.36221791 | 0.036922643 | 0.181808045 | -0.203085858 | 0.839067925 | 0.919794636 | 0.036922643 |
| ENSMUSG00000031487 | Brf2 | 66653 | protein_coding | 81.65625327 | -0.036952361 | 0.132971384 | 0.277897094 | 0.781091355 | 0.890541867 | 0.036952361 |
| ENSMUSG00000086370 | Ftx | 78878 | lncRNA | 65.96581796 | -0.036993011 | 0.188801373 | 0.195936134 | 0.844660154 | 0.923599766 | 0.036993011 |
| ENSMUSG00000068264 | Ap5s1 | 69596 | protein_coding | 182.549863 | -0.037016436 | 0.12730924 | 0.290760006 | 0.771234875 | 0.885199039 | 0.037016436 |
| ENSMUSG00000039656 | Rxrb | 20182 | protein_coding | 554.9876103 | 0.037019186 | 0.069349601 | -0.533805324 | 0.59347623 | 0.772683055 | 0.037019186 |
| ENSMUSG00000040472 | Rabggta | 56187 | protein_coding | 238.5210526 | -0.037058488 | 0.081724993 | 0.453453549 | 0.650222185 | 0.809917796 | 0.037058488 |
| ENSMUSG00000054434 | Tmem120b | 330189 | protein_coding | 31.4089334 | 0.037134963 | 0.224037385 | -0.165753422 | 0.868351003 | 0.934807592 | 0.037134963 |
| ENSMUSG00000041293 | Adgrf1 | 77596 | protein_coding | 46.67352637 | 0.037146392 | 0.382095583 | -0.097217537 | 0.922553641 | 0.962530504 | 0.037146392 |
| ENSMUSG00000029815 | Malsu1 | 75593 | protein_coding | 164.0933818 | 0.037146804 | 0.089185976 | -0.416509469 | 0.67703724 | 0.826943172 | 0.037146804 |
| ENSMUSG00000031902 | Nfatc3 | 18021 | protein_coding | 762.2869193 | -0.037149118 | 0.050526918 | 0.735234198 | 0.462196875 | 0.675880563 | 0.037149118 |
| ENSMUSG00000025979 | Mob4 | 19070 | protein_coding | 708.3215347 | 0.037215156 | 0.087148204 | -0.427032968 | 0.669355315 | 0.822609935 | 0.037215156 |
| ENSMUSG00000044145 | 1810024B03Rik | 329509 | protein_coding | 13.55893557 | -0.037225524 | 0.349240923 | 0.106589812 | 0.915114401 | 0.95833356 | 0.037225524 |
| ENSMUSG00000035596 | Mboat7 | 77582 | protein_coding | 486.5674141 | -0.037234837 | 0.074639504 | 0.498862334 | 0.617876371 | 0.789468115 | 0.037234837 |
| ENSMUSG00000024566 | Atp9b | 50771 | protein_coding | 642.1483617 | 0.037262973 | 0.060958142 | -0.611287868 | 0.541009021 | 0.735049182 | 0.037262973 |
| ENSMUSG00000074909 | Ranbp6 | 240614 | protein_coding | 217.8213493 | -0.037305766 | 0.08613246 | 0.433120866 | 0.664926965 | 0.819267308 | 0.037305766 |
| ENSMUSG00000038618 | Rassf7 | 66985 | protein_coding | 1072.703907 | -0.037325893 | 0.096388465 | 0.387244394 | 0.698575287 | 0.840298483 | 0.037325893 |

| ensembl_gene_id | external_gene_name | entrezgene_id | gene_biotype | baseMean | log2FoldChange | lfcSE | stat | pvalue | padj | absLog2FC |
|---|---|---|---|---|---|---|---|---|---|---|
| ENSMUSG00000026792 | Lrsam1 | 227738 | protein_coding | 245.7141234 | -0.037356609 | 0.092625572 | 0.403307725 | 0.68672186 | 0.833127005 | 0.037356609 |
| ENSMUSG00000056383 | AI987944 | 233168 | protein_coding | 137.7857914 | -0.037394237 | 0.122061745 | 0.306355093 | 0.759334296 | 0.879038683 | 0.037394237 |
| ENSMUSG00000013646 | Sh3bp5l | 79566 | protein_coding | 428.9945128 | -0.03742073 | 0.063366107 | 0.590548046 | 0.554823284 | 0.744703672 | 0.03742073 |
| ENSMUSG00000061013 | Mkx | 210719 | protein_coding | 160.6123841 | -0.037429829 | 0.131996579 | 0.283566661 | 0.77674249 | 0.888506311 | 0.037429829 |
| ENSMUSG00000026065 | Slc9a4 | 110895 | protein_coding | 5.86196662 | 0.037439697 | 0.50150563 | -0.07465459 | 0.940489539 | 0.971492103 | 0.037439697 |
| ENSMUSG00000006423 | Steep1 | 77644 | protein_coding | 342.2605622 | 0.037450363 | 0.106669178 | -0.351088883 | 0.725521668 | 0.857956591 | 0.037450363 |
| ENSMUSG00000049086 | Bmyc | 107771 | protein_coding | 17.18315258 | 0.037455388 | 0.262552212 | -0.142658816 | 0.886559644 | 0.943887948 | 0.037455388 |
| ENSMUSG00000069874 | Irgm2 | 54396 | protein_coding | 175.3532193 | -0.03751705 | 0.222923218 | 0.168295839 | 0.866350547 | 0.933791206 | 0.03751705 |
| ENSMUSG00000025373 | Rnf41 | 67588 | protein_coding | 342.0705477 | -0.037532683 | 0.075241137 | 0.498831955 | 0.617897774 | 0.789468115 | 0.037532683 |
| ENSMUSG00000036537 | Rnf113a1 | 69942 | protein_coding | 26.28009886 | 0.037543065 | 0.208829483 | -0.179778567 | 0.857326411 | 0.930374388 | 0.037543065 |
| ENSMUSG00000006471 | Ndor1 | 78797 | protein_coding | 65.50198746 | 0.03757272 | 0.163798273 | -0.229384102 | 0.818570391 | 0.908812719 | 0.03757272 |
| ENSMUSG00000063801 | Ap3s2 | 11778 | protein_coding | 310.1390708 | 0.037597386 | 0.072404543 | -0.51926832 | 0.603573642 | 0.779177919 | 0.037597386 |
| ENSMUSG00000027652 | Ralgapb | 228850 | protein_coding | 1538.59055 | -0.037604375 | 0.059481961 | 0.632197971 | 0.527257523 | 0.725715181 | 0.037604375 |
| ENSMUSG00000002980 | Bcam | 57278 | protein_coding | 56.95408451 | 0.037646522 | 0.163955723 | -0.229613958 | 0.81839176 | 0.908812719 | 0.037646522 |
| ENSMUSG00000021675 | F2rl2 | 14064 | protein_coding | 19.1215894 | 0.037652298 | 0.276441214 | -0.136203635 | 0.891660302 | 0.947050968 | 0.037652298 |
| ENSMUSG00000021595 | Nsun2 | 28114 | protein_coding | 959.5752043 | 0.037660883 | 0.096628991 | -0.389747246 | 0.696723457 | 0.839060823 | 0.037660883 |
| ENSMUSG00000019578 | Ubxn6 | 66530 | protein_coding | 208.4422178 | 0.037718968 | 0.134840118 | -0.279731052 | 0.779683853 | 0.890125936 | 0.037718968 |
| ENSMUSG00000042724 | Map3k9 | 338372 | protein_coding | 252.2027264 | -0.037741239 | 0.103098376 | 0.366070159 | 0.714312724 | 0.851829233 | 0.037741239 |
| ENSMUSG00000037938 | Chchd5 | 66170 | protein_coding | 135.6819797 | -0.037770208 | 0.094510818 | 0.399638995 | 0.68942243 | 0.834319362 | 0.037770208 |
| ENSMUSG00000018924 | Alox15 | 11687 | protein_coding | 5.651957746 | 0.037782131 | 0.677812268 | -0.055741291 | 0.955547905 | 0.977832194 | 0.037782131 |
| ENSMUSG00000042306 | S100a14 | 66166 | protein_coding | 1308.369798 | -0.037799075 | 0.163984246 | 0.230504308 | 0.817699916 | 0.908714429 | 0.037799075 |
| ENSMUSG00000029432 | Nipsnap2 | 14467 | protein_coding | 983.1906584 | -0.037873196 | 0.097006606 | 0.390418736 | 0.696226935 | 0.838743187 | 0.037873196 |
| ENSMUSG00000097254 | C430042M11Rik | 320021 | lncRNA | 30.14714013 | 0.037882324 | 0.227169974 | -0.166757618 | 0.867560767 | 0.934200477 | 0.037882324 |
| ENSMUSG00000023249 | Parp3 | 235587 | protein_coding | 112.7434322 | -0.037913736 | 0.135766844 | 0.279256225 | 0.780048198 | 0.890125936 | 0.037913736 |
| ENSMUSG00000052137 | Rbm12b2 | 77604 | protein_coding | 95.44078377 | 0.037941789 | 0.15566862 | -0.243734345 | 0.807436572 | 0.90502442 | 0.037941789 |
| ENSMUSG00000028329 | Xpa | 22590 | protein_coding | 112.4650907 | -0.037942207 | 0.134583599 | 0.281922959 | 0.778002585 | 0.889068901 | 0.037942207 |
| ENSMUSG00000049739 | Zfp646 | 233905 | protein_coding | 814.9213813 | 0.037953364 | 0.038924981 | -0.975038721 | 0.329541053 | 0.551855356 | 0.037953364 |
| ENSMUSG00000026177 | Slc11a1 | 18173 | protein_coding | 4.536060121 | -0.037968171 | 1.005638461 | 0.037755289 | 0.969882793 | 0.984509162 | 0.037968171 |
| ENSMUSG00000006288 | Ttc5 | 219022 | protein_coding | 321.5649251 | -0.037987029 | 0.089371153 | 0.425047986 | 0.670801694 | 0.823378739 | 0.037987029 |
| ENSMUSG00000014767 | Tbp | 21374 | protein_coding | 150.3709788 | 0.037987586 | 0.100478066 | -0.37806845 | 0.705379745 | 0.845574994 | 0.037987586 |
| ENSMUSG00000037896 | Rcor1 | 217864 | protein_coding | 1015.378361 | 0.038002382 | 0.087586847 | -0.433882289 | 0.664373922 | 0.818950015 | 0.038002382 |
| ENSMUSG00000061759 | Armt1 | 73419 | protein_coding | 174.3880616 | 0.038006572 | 0.116748319 | -0.325542775 | 0.744770314 | 0.8712288 | 0.038006572 |
| ENSMUSG00000025917 | Cops5 | 26754 | protein_coding | 593.4415913 | 0.038014512 | 0.074776003 | -0.508378498 | 0.611187929 | 0.784646895 | 0.038014512 |
| ENSMUSG00000006599 | Gtf2h1 | 14884 | protein_coding | 574.7943877 | 0.038040133 | 0.070331387 | -0.540869942 | 0.58859723 | 0.769577776 | 0.038040133 |
| ENSMUSG00000030315 | Vgll4 | 232334 | protein_coding | 571.390717 | -0.038070931 | 0.10767631 | 0.353568306 | 0.72366243 | 0.856795141 | 0.038070931 |
| ENSMUSG00000040321 | Zfp770 | 228491 | protein_coding | 81.89772367 | 0.038073237 | 0.154880239 | -0.245823721 | 0.805818693 | 0.904335026 | 0.038073237 |
| ENSMUSG00000034751 | Mast4 | 328329 | protein_coding | 306.5120436 | 0.038076261 | 0.113779487 | -0.334649611 | 0.737889416 | 0.866982505 | 0.038076261 |
| ENSMUSG00000089235 | Gm23119 | 115486069 | scaRNA | 89.37322216 | -0.038079192 | 0.245040136 | 0.155399814 | 0.876506131 | 0.938630585 | 0.038079192 |
| ENSMUSG00000054196 | Cthrc1 | 68588 | protein_coding | 4.787686734 | 0.038091756 | 0.499491579 | -0.076261057 | 0.939211408 | 0.97108297 | 0.038091756 |

| ensembl_gene_id | external_gene_name | entrezgene_id | gene_biotype | baseMean | log2FoldChange | lfcSE | stat | pvalue | padj | absLog2FC |
|---|---|---|---|---|---|---|---|---|---|---|
| ENSMUSG00000002329 | Mdp1 | 67881 | protein_coding | 943.9921465 | -0.038114162 | 0.099001137 | 0.384987121 | 0.700246958 | 0.841315261 | 0.038114162 |
| ENSMUSG00000014349 | Ube2z | 268470 | protein_coding | 1279.828942 | 0.038120192 | 0.044865552 | -0.849653923 | 0.395517523 | 0.617841913 | 0.038120192 |
| ENSMUSG00000032966 | Fkbp1a | 14225 | protein_coding | 461.8704569 | 0.038126608 | 0.09150247 | -0.41667299 | 0.676917613 | 0.826943172 | 0.038126608 |
| ENSMUSG00000040506 | Ambra1 | 228361 | protein_coding | 601.3264084 | 0.038156116 | 0.066272316 | -0.575747439 | 0.564785905 | 0.751588072 | 0.038156116 |
| ENSMUSG00000021832 | Psmc6 | 67089 | protein_coding | 1000.992988 | 0.038157726 | 0.083149106 | -0.458907224 | 0.646300791 | 0.807226927 | 0.038157726 |
| ENSMUSG00000038708 | Golga4 | 54214 | protein_coding | 3773.399457 | 0.038165661 | 0.063391172 | -0.60206587 | 0.547130292 | 0.738779548 | 0.038165661 |
| ENSMUSG00000048921 | Zfp689 | 71131 | protein_coding | 31.65405392 | -0.038184082 | 0.182715656 | 0.208980899 | 0.83446315 | 0.916826487 | 0.038184082 |
| ENSMUSG00000028433 | Ubap2 | 68926 | protein_coding | 374.704412 | 0.038228781 | 0.090483367 | -0.422495123 | 0.67266366 | 0.824518194 | 0.038228781 |
| ENSMUSG00000009535 | Rnmt | 67897 | protein_coding | 252.7742681 | -0.038241026 | 0.112840336 | 0.338895 | 0.734688831 | 0.864656729 | 0.038241026 |
| ENSMUSG00000037661 | Gpr160 | 71862 | protein_coding | 382.7976112 | -0.038264923 | 0.134246997 | 0.285033734 | 0.775618297 | 0.88791795 | 0.038264923 |
| ENSMUSG00000038010 | Ccdc138 | 76138 | protein_coding | 50.59785374 | -0.038320542 | 0.173246458 | 0.221190912 | 0.824943785 | 0.911456552 | 0.038320542 |
| ENSMUSG00000045281 | Gpr20 | 239530 | protein_coding | 212.3629767 | -0.03833711 | 0.249467661 | 0.153675671 | 0.877865469 | 0.9393553 | 0.03833711 |
| ENSMUSG00000030095 | Tmem43 | 74122 | protein_coding | 491.6061233 | -0.038353083 | 0.110012851 | 0.348623663 | 0.727371861 | 0.859261888 | 0.038353083 |
| ENSMUSG00000033430 | Terf2ip | 57321 | protein_coding | 254.6123832 | 0.03838255 | 0.113818639 | -0.337225519 | 0.735946901 | 0.865628713 | 0.03838255 |
| ENSMUSG00000110909 | B020014A21Rik | 619314 | lncRNA | 1.092188399 | 0.038419811 | 2.29981321 | -0.016705622 | 0.986671462 | 1 | 0.038419811 |
| ENSMUSG00000025646 | Atrip | 235610 | protein_coding | 135.9673752 | 0.038484841 | 0.126622441 | -0.303933808 | 0.761178321 | 0.880259019 | 0.038484841 |
| ENSMUSG00000065864 | Gm24031 | 115488396 | snoRNA | 0.769468905 | -0.03849408 | 1.121922123 | 0.034310831 | 0.972629288 | 1 | 0.03849408 |
| ENSMUSG00000006412 | Pfdn2 | 18637 | protein_coding | 133.1162265 | -0.038542505 | 0.09409741 | 0.40960219 | 0.68209779 | 0.830017162 | 0.038542505 |
| ENSMUSG00000000605 | Clcn4 | 12727 | protein_coding | 1124.302622 | -0.038612373 | 0.083678603 | 0.461436629 | 0.644485377 | 0.806290266 | 0.038612373 |
| ENSMUSG00000001962 | Fam50a | 108160 | protein_coding | 192.5687404 | -0.038652073 | 0.104762231 | 0.368950458 | 0.712164653 | 0.85037506 | 0.038652073 |
| ENSMUSG00000009470 | Tnpo1 | 238799 | protein_coding | 1713.415294 | 0.03866323 | 0.081369408 | -0.475156832 | 0.634675193 | 0.800822518 | 0.03866323 |
| ENSMUSG00000022109 | Med4 | 67381 | protein_coding | 157.2741223 | 0.038686139 | 0.097467163 | -0.396914584 | 0.69143045 | 0.835769466 | 0.038686139 |
| ENSMUSG00000033327 | Tnxb | 81877 | protein_coding | 8.97222531 | 0.03872054 | 0.781945739 | -0.049518193 | 0.960506339 | 0.980697519 | 0.03872054 |
| ENSMUSG00000032673 | Prorsd1 | 67939 | protein_coding | 106.5104771 | -0.0387402 | 0.129874799 | 0.298288816 | 0.76548274 | 0.882510888 | 0.0387402 |
| ENSMUSG00000000938 | Hoxa10 | 15395 | protein_coding | 377.2528266 | -0.038809497 | 0.138992014 | 0.279221058 | 0.780075184 | 0.890125936 | 0.038809497 |
| ENSMUSG00000048329 | Mfsd6l | 215723 | protein_coding | 168.4053621 | -0.038823255 | 0.094903803 | 0.409080078 | 0.682480896 | 0.830405926 | 0.038823255 |
| ENSMUSG00000020121 | Srgap1 | 117600 | protein_coding | 246.2531101 | 0.038893656 | 0.138845052 | -0.280122738 | 0.77938334 | 0.890030875 | 0.038893656 |
| ENSMUSG00000024999 | Noc3l | 57753 | protein_coding | 176.7363687 | 0.03890479 | 0.130090141 | -0.299060248 | 0.764894076 | 0.882255147 | 0.03890479 |
| ENSMUSG00000015214 | Mtmr1 | 53332 | protein_coding | 391.6805975 | 0.038910263 | 0.100367764 | -0.387676889 | 0.698255161 | 0.840144679 | 0.038910263 |
| ENSMUSG00000035392 | Dennd1a | 227801 | protein_coding | 159.3726411 | -0.039021584 | 0.100126156 | 0.389724177 | 0.696740517 | 0.839060823 | 0.039021584 |
| ENSMUSG00000022661 | Cd200 | 17470 | protein_coding | 24.76322006 | -0.039034219 | 0.215057983 | 0.181505555 | 0.855970773 | 0.93002838 | 0.039034219 |
| ENSMUSG00000042541 | Sem1 | 20422 | protein_coding | 1183.951142 | 0.039040554 | 0.067809788 | -0.57573626 | 0.564793463 | 0.751588072 | 0.039040554 |
| ENSMUSG00000024309 | Pfdn6 | 14976 | protein_coding | 349.1596209 | -0.039079299 | 0.072532232 | 0.53878529 | 0.590035015 | 0.770846581 | 0.039079299 |
| ENSMUSG00000039005 | Tlr4 | 21898 | protein_coding | 385.8979073 | 0.039111478 | 0.12233283 | -0.319713671 | 0.749185395 | 0.874014884 | 0.039111478 |
| ENSMUSG00000108129 | 4930417O13Rik | 73870 | lncRNA | 2.330756465 | -0.039134584 | 0.945279735 | 0.041400003 | 0.96697701 | 1 | 0.039134584 |
| ENSMUSG00000028565 | Nfia | 18027 | protein_coding | 331.7212265 | 0.039206193 | 0.096463032 | -0.406437497 | 0.684421173 | 0.831662311 | 0.039206193 |
| ENSMUSG00000050229 | Pigm | 67556 | protein_coding | 355.7625997 | -0.039209929 | 0.080205616 | 0.488867619 | 0.624935422 | 0.794593745 | 0.039209929 |
| ENSMUSG00000036552 | Ermard | 381062 | protein_coding | 50.84057706 | -0.039283033 | 0.170249895 | 0.230737489 | 0.817518747 | 0.908714429 | 0.039283033 |
| ENSMUSG00000030591 | Psmd8 | 57296 | protein_coding | 1521.092148 | 0.039288085 | 0.061671576 | -0.637053368 | 0.524090082 | 0.722741011 | 0.039288085 |

| ensembl_gene_id | external_gene_name | entrezgene_id | gene_biotype | baseMean | log2FoldChange | lfcSE | stat | pvalue | padj | absLog2FC |
|---|---|---|---|---|---|---|---|---|---|---|
| ENSMUSG00000038965 | Ube2l3 | 22195 | protein_coding | 868.9568896 | -0.039313043 | 0.072860129 | 0.539568665 | 0.589494531 | 0.770506411 | 0.039313043 |
| ENSMUSG00000036779 | Tent4b | 214627 | protein_coding | 411.5000018 | -0.039332791 | 0.064872298 | 0.606311058 | 0.544308217 | 0.736796542 | 0.039332791 |
| ENSMUSG00000066235 | Pomgnt2 | 215494 | protein_coding | 126.833415 | -0.039357131 | 0.168599398 | 0.233435775 | 0.815423036 | 0.908262243 | 0.039357131 |
| ENSMUSG00000066798 | Zbtb6 | 241322 | protein_coding | 345.8318255 | -0.039392146 | 0.08265482 | 0.476586187 | 0.633656821 | 0.800362194 | 0.039392146 |
| ENSMUSG00000000776 | Polr3d | 67065 | protein_coding | 232.9622134 | 0.039414412 | 0.085301103 | -0.462062165 | 0.644036741 | 0.80597323 | 0.039414412 |
| ENSMUSG00000028098 | Rnf115 | 67845 | protein_coding | 541.6062068 | 0.039455656 | 0.064548029 | -0.611260438 | 0.541027178 | 0.735049182 | 0.039455656 |
| ENSMUSG00000018569 | Cldn7 | 53624 | protein_coding | 6571.834732 | -0.039494916 | 0.076344646 | 0.517323984 | 0.604930014 | 0.780088572 | 0.039494916 |
| ENSMUSG00000031365 | Zfp275 | 27081 | protein_coding | 188.358204 | 0.039502106 | 0.118161153 | -0.334307043 | 0.738147876 | 0.866982505 | 0.039502106 |
| ENSMUSG00000000441 | Raf1 | 110157 | protein_coding | 731.5476738 | 0.039522429 | 0.059289318 | -0.666602859 | 0.505025842 | 0.709123099 | 0.039522429 |
| ENSMUSG00000049225 | Pdp1 | 381511 | protein_coding | 269.0199049 | 0.039639209 | 0.162531811 | -0.243885853 | 0.807319226 | 0.904987729 | 0.039639209 |
| ENSMUSG00000015837 | Sqstm1 | 18412 | protein_coding | 2595.756012 | 0.039642848 | 0.122138378 | -0.324573232 | 0.745504089 | 0.871840133 | 0.039642848 |
| ENSMUSG00000038949 | Cnst | 226744 | protein_coding | 637.157925 | -0.039670303 | 0.075279386 | 0.526974318 | 0.598211423 | 0.775969165 | 0.039670303 |
| ENSMUSG00000107006 | Gm5553 | 433874 | processed_pseudogene | 0.607579554 | -0.039732755 | 1.860530643 | 0.021355604 | 0.982961989 | 1 | 0.039732755 |
| ENSMUSG00000020755 | Sap30bp | 57230 | protein_coding | 375.2724246 | -0.039783882 | 0.062600268 | 0.635522557 | 0.52508766 | 0.723512525 | 0.039783882 |
| ENSMUSG00000073775 | Kti12 | 100087 | protein_coding | 177.3892844 | -0.039785859 | 0.149590614 | 0.265964943 | 0.790266208 | 0.895392916 | 0.039785859 |
| ENSMUSG00000065400 | Mir200a | 387242 | miRNA | 20.66459916 | -0.039785955 | 0.309764762 | 0.128439253 | 0.897801369 | 0.949794479 | 0.039785955 |
| ENSMUSG00000040563 | Plppr2 | 235044 | protein_coding | 15.43158002 | -0.039810783 | 0.294478433 | 0.135190829 | 0.892460998 | 0.947202684 | 0.039810783 |
| ENSMUSG00000040242 | Fgfr1op2 | 67529 | protein_coding | 866.204485 | 0.039845287 | 0.071537685 | -0.556983182 | 0.57753892 | 0.761475268 | 0.039845287 |
| ENSMUSG00000040688 | Tbl3 | 213773 | protein_coding | 394.1816926 | -0.039909113 | 0.089553325 | 0.445646361 | 0.655852716 | 0.813424841 | 0.039909113 |
| ENSMUSG00000036330 | Slc18a1 | 110877 | protein_coding | 55.02963601 | -0.039968845 | 0.187152303 | 0.213563201 | 0.830887699 | 0.914497068 | 0.039968845 |
| ENSMUSG00000016028 | Celsr1 | 12614 | protein_coding | 1754.411007 | -0.04001813 | 0.09225563 | 0.433774395 | 0.664452277 | 0.818950015 | 0.04001813 |
| ENSMUSG00000040969 | Arhgef38 | 77669 | protein_coding | 794.8629897 | -0.040060477 | 0.155695225 | 0.257300612 | 0.79694672 | 0.899144506 | 0.040060477 |
| ENSMUSG00000032481 | Smarcc1 | 20588 | protein_coding | 642.3469086 | 0.040071288 | 0.084778393 | -0.472659209 | 0.636456333 | 0.801895161 | 0.040071288 |
| ENSMUSG00000010660 | Plcd1 | 18799 | protein_coding | 534.0587554 | -0.040129547 | 0.138434102 | 0.289881943 | 0.771906556 | 0.885478012 | 0.040129547 |
| ENSMUSG00000043487 | Acot6 | 217700 | protein_coding | 27.18880892 | -0.04018921 | 0.198821916 | 0.202136719 | 0.839809842 | 0.920280009 | 0.04018921 |
| ENSMUSG00000060261 | Gtf2i | 14886 | protein_coding | 760.5775839 | 0.040196434 | 0.074087295 | -0.542555017 | 0.587436218 | 0.768755187 | 0.040196434 |
| ENSMUSG00000046138 | 9930021J03Rik | 240613 | protein_coding | 576.1918315 | 0.040233696 | 0.088185961 | -0.456236979 | 0.648219577 | 0.808756278 | 0.040233696 |
| ENSMUSG00000063870 | Chd4 | 107932 | protein_coding | 3194.539178 | 0.040238538 | 0.075582233 | -0.532380916 | 0.5944622 | 0.77323163 | 0.040238538 |
| ENSMUSG00000061322 | Dnai1 | 68922 | protein_coding | 2.115843828 | -0.040247029 | 1.852426748 | 0.021726651 | 0.982666004 | 1 | 0.040247029 |
| ENSMUSG00000068854 | H2bc21 | 319190 | protein_coding | 119.8048083 | 0.040267814 | 0.134790391 | -0.298743953 | 0.765135418 | 0.882363568 | 0.040267814 |
| ENSMUSG00000020669 | Sh3yl1 | 24057 | protein_coding | 352.804777 | -0.040282664 | 0.078896819 | 0.510573999 | 0.609649386 | 0.783482488 | 0.040282664 |
| ENSMUSG00000079164 | Tlr5 | 53791 | protein_coding | 68.0372835 | 0.040285004 | 0.278299215 | -0.144754287 | 0.884904876 | 0.943016089 | 0.040285004 |
| ENSMUSG00000036061 | Smug1 | 71726 | protein_coding | 228.3211143 | -0.040295407 | 0.095591595 | 0.421537129 | 0.673362905 | 0.824897384 | 0.040295407 |
| ENSMUSG00000067825 | Pex26 | 74043 | protein_coding | 330.3867817 | 0.040304527 | 0.067723571 | -0.595132926 | 0.551754621 | 0.742476843 | 0.040304527 |
| ENSMUSG00000089847 | Timm10b | 14356 | protein_coding | 93.58132037 | -0.040310175 | 0.180079293 | 0.22384681 | 0.822876505 | 0.910240646 | 0.040310175 |
| ENSMUSG00000029518 | Rab35 | 77407 | protein_coding | 725.7809227 | 0.0403834 | 0.075341432 | -0.536005215 | 0.591954946 | 0.771976939 | 0.0403834 |
| ENSMUSG00000022528 | Hes1 | 15205 | protein_coding | 977.2623035 | 0.040505206 | 0.071309249 | -0.568021766 | 0.570020189 | 0.755664558 | 0.040505206 |
| ENSMUSG00000070565 | Rasal2 | 226525 | protein_coding | 135.1537411 | -0.04053187 | 0.10325337 | 0.392547671 | 0.694653596 | 0.8379481 | 0.04053187 |
| ENSMUSG00000045629 | Sh3tc2 | 225608 | protein_coding | 29.66374938 | -0.040557383 | 0.254053651 | 0.159641017 | 0.873163867 | 0.937483135 | 0.040557383 |

| ensembl_gene_id | external_gene_name | entrezgene_id | gene_biotype | baseMean | log2FoldChange | lfcSE | stat | pvalue | padj | absLog2FC |
|---|---|---|---|---|---|---|---|---|---|---|
| ENSMUSG00000022261 | Sdc2 | 15529 | protein_coding | 451.862126 | -0.040558437 | 0.090496484 | 0.448176937 | 0.654025506 | 0.81231093 | 0.040558437 |
| ENSMUSG00000028948 | Nol9 | 74035 | protein_coding | 332.0089935 | 0.040559146 | 0.102511423 | -0.395654897 | 0.692359633 | 0.836513744 | 0.040559146 |
| ENSMUSG00000023892 | Zfp51 | 22709 | protein_coding | 159.2934514 | -0.040561321 | 0.11810914 | 0.343422374 | 0.73128073 | 0.861613154 | 0.040561321 |
| ENSMUSG00000066149 | Cdc26 | 66440 | protein_coding | 217.8797284 | -0.040610067 | 0.086830338 | 0.467694454 | 0.640003108 | 0.8036518 | 0.040610067 |
| ENSMUSG00000025138 | Sirt7 | 209011 | protein_coding | 728.2683722 | -0.040620417 | 0.089352666 | 0.454607775 | 0.64939144 | 0.809667817 | 0.040620417 |
| ENSMUSG00000033149 | Phldb2 | 208177 | protein_coding | 106.1895724 | 0.040635919 | 0.12420191 | -0.327176284 | 0.743534558 | 0.870461483 | 0.040635919 |
| ENSMUSG00000027679 | Dnajc19 | 67713 | protein_coding | 252.0431548 | -0.040643835 | 0.077558988 | 0.524037721 | 0.600252304 | 0.77680602 | 0.040643835 |
| ENSMUSG00000031431 | Tsc22d3 | 14605 | protein_coding | 773.1679406 | -0.04064785 | 0.116759161 | 0.348134144 | 0.727739443 | 0.859322531 | 0.04064785 |
| ENSMUSG00000029191 | Rfc1 | 19687 | protein_coding | 547.3247749 | -0.040668887 | 0.106464976 | 0.381993106 | 0.702466478 | 0.843000612 | 0.040668887 |
| ENSMUSG00000030213 | Atf7ip | 54343 | protein_coding | 752.5508647 | 0.040680017 | 0.073757017 | -0.551540974 | 0.581262887 | 0.764339772 | 0.040680017 |
| ENSMUSG00000026343 | Gpr39 | 71111 | protein_coding | 106.7933073 | 0.040685779 | 0.153646187 | -0.264801748 | 0.79116219 | 0.895862069 | 0.040685779 |
| ENSMUSG00000029238 | Clock | 12753 | protein_coding | 1508.642235 | -0.040685837 | 0.11224912 | 0.362460187 | 0.71700817 | 0.853197357 | 0.040685837 |
| ENSMUSG00000028161 | Ppp3ca | 19055 | protein_coding | 886.4617117 | -0.040703809 | 0.099680378 | 0.408343249 | 0.683021692 | 0.830651757 | 0.040703809 |
| ENSMUSG00000023156 | Rpp14 | 67053 | protein_coding | 224.1607357 | -0.040734364 | 0.138323878 | 0.294485409 | 0.768387011 | 0.883975099 | 0.040734364 |
| ENSMUSG00000046822 | Slc39a3 | 106947 | protein_coding | 396.0290589 | 0.040736839 | 0.091006069 | -0.447627721 | 0.654421894 | 0.812494814 | 0.040736839 |
| ENSMUSG00000042499 | Hoxd11 | 15431 | protein_coding | 10.38919938 | -0.040739057 | 0.411826474 | 0.09892287 | 0.92119951 | 0.961976016 | 0.040739057 |
| ENSMUSG00000024422 | Dhx16 | 69192 | protein_coding | 715.3513868 | 0.04076287 | 0.058095117 | -0.701657426 | 0.48289283 | 0.693654751 | 0.04076287 |
| ENSMUSG00000041757 | Plekha6 | 240753 | protein_coding | 4665.171911 | -0.040764323 | 0.061113954 | 0.667021524 | 0.504758385 | 0.70880782 | 0.040764323 |
| ENSMUSG00000058835 | Abi1 | 11308 | protein_coding | 1498.261271 | -0.040773616 | 0.058870965 | 0.692592951 | 0.488565035 | 0.696497169 | 0.040773616 |
| ENSMUSG00000052794 | 1700030K09Rik | 72254 | protein_coding | 59.8882899 | -0.040775204 | 0.145497443 | 0.280246878 | 0.779288104 | 0.889983662 | 0.040775204 |
| ENSMUSG00000028582 | Cc2d1b | 319965 | protein_coding | 507.345905 | 0.040816689 | 0.059960271 | -0.680728899 | 0.496043047 | 0.702242166 | 0.040816689 |
| ENSMUSG00000084826 | AI847159 | 100048701 | lncRNA | 1.534471555 | -0.040858339 | 1.05403564 | 0.038763717 | 0.969078773 | 1 | 0.040858339 |
| ENSMUSG00000056537 | Rlim | 19820 | protein_coding | 1374.53376 | 0.040870562 | 0.074019324 | -0.552160703 | 0.580838255 | 0.764064128 | 0.040870562 |
| ENSMUSG00000036893 | Ehmt1 | 77683 | protein_coding | 775.0473659 | -0.040925912 | 0.070199523 | 0.582994158 | 0.559897233 | 0.748554316 | 0.040925912 |
| ENSMUSG00000093044 | Snord8 | 100217445 | snoRNA | 3.327967407 | -0.040926452 | 0.551080613 | 0.074265817 | 0.940798876 | 1 | 0.040926452 |
| ENSMUSG00000025484 | Bet1l | 54399 | protein_coding | 244.6068901 | -0.04093148 | 0.07837532 | 0.522249607 | 0.601496551 | 0.777725809 | 0.04093148 |
| ENSMUSG00000052373 | Mpp3 | 13384 | protein_coding | 106.5723709 | 0.04093945 | 0.178266424 | -0.229653177 | 0.818361282 | 0.908812719 | 0.04093945 |
| ENSMUSG00000027424 | Mgme1 | 74528 | protein_coding | 141.0405667 | -0.040941574 | 0.148616873 | 0.275484026 | 0.782944397 | 0.891196207 | 0.040941574 |
| ENSMUSG00000053470 | Kdm3a | 104263 | protein_coding | 682.1946227 | -0.040945869 | 0.073022125 | 0.560732364 | 0.574980001 | 0.760164897 | 0.040945869 |
| ENSMUSG00000025436 | Atp23 | 68876 | protein_coding | 47.58138411 | 0.040962702 | 0.171114854 | -0.239387175 | 0.810805374 | 0.906297546 | 0.040962702 |
| ENSMUSG00000016831 | Tox4 | 268741 | protein_coding | 816.5118391 | 0.04097033 | 0.057472659 | -0.71286644 | 0.475928408 | 0.687930892 | 0.04097033 |
| ENSMUSG00000050323 | Ndufaf6 | 76947 | protein_coding | 73.86264982 | -0.040994405 | 0.155595332 | 0.263468089 | 0.792189816 | 0.896636371 | 0.040994405 |
| ENSMUSG00000027522 | Stx16 | 228960 | protein_coding | 530.0125725 | -0.040995987 | 0.060272417 | 0.680178249 | 0.496391603 | 0.702494724 | 0.040995987 |
| ENSMUSG00000036282 | Naa30 | 70646 | protein_coding | 461.7903256 | 0.040996796 | 0.063282642 | -0.647836353 | 0.5170908 | 0.717711842 | 0.040996796 |
| ENSMUSG00000019726 | Lyst | 17101 | protein_coding | 298.9354274 | -0.041005409 | 0.092076427 | 0.445341013 | 0.656073333 | 0.813637356 | 0.041005409 |
| ENSMUSG00000022472 | Desi1 | 28075 | protein_coding | 214.8826824 | -0.04100962 | 0.090848444 | 0.451406959 | 0.651696269 | 0.810888501 | 0.04100962 |
| ENSMUSG00000034543 | Morc2a | 74522 | protein_coding | 416.6655941 | 0.041019985 | 0.070889303 | -0.578648447 | 0.562826408 | 0.75067522 | 0.041019985 |
| ENSMUSG00000026436 | Elk4 | 13714 | protein_coding | 553.1714474 | -0.0410376 | 0.069033311 | 0.594460839 | 0.552203927 | 0.742839333 | 0.0410376 |
| ENSMUSG00000024972 | Lgals12 | 56072 | protein_coding | 244.4386253 | 0.041070066 | 0.221289839 | -0.185594 | 0.852763159 | 0.92788168 | 0.041070066 |

| ensembl_gene_id | external_gene_name | entrezgene_id | gene_biotype | baseMean | log2FoldChange | lfcSE | stat | pvalue | padj | absLog2FC |
|---|---|---|---|---|---|---|---|---|---|---|
| ENSMUSG00000033658 | Ddx19b | 234733 | protein_coding | 429.9602003 | -0.041089899 | 0.090537261 | 0.453845176 | 0.649940266 | 0.809917796 | 0.041089899 |
| ENSMUSG00000038346 | Zfp384 | 269800 | protein_coding | 333.8183899 | -0.04111091 | 0.09465812 | 0.43430939 | 0.664063786 | 0.818817302 | 0.04111091 |
| ENSMUSG00000030788 | Rnf141 | 67150 | protein_coding | 881.4482722 | 0.041114149 | 0.099696389 | -0.412393561 | 0.680050982 | 0.828381052 | 0.041114149 |
| ENSMUSG00000086386 | Gm12108 | 104692 | lncRNA | 1.464638804 | -0.041119632 | 2.359632589 | 0.017426286 | 0.986096539 | 1 | 0.041119632 |
| ENSMUSG00000032534 | Cep63 | 28135 | protein_coding | 307.8618181 | -0.041135971 | 0.084855036 | 0.484779371 | 0.627832857 | 0.796376973 | 0.041135971 |
| ENSMUSG00000029208 | Guf1 | 231279 | protein_coding | 329.6280776 | 0.041161352 | 0.067901088 | -0.606195773 | 0.544384759 | 0.736796542 | 0.041161352 |
| ENSMUSG00000015189 | Casd1 | 213819 | protein_coding | 1151.818215 | 0.04116221 | 0.194752578 | -0.211356433 | 0.832609151 | 0.915629096 | 0.04116221 |
| ENSMUSG00000006705 | Pknox1 | 18771 | protein_coding | 263.647121 | -0.041172028 | 0.106717248 | 0.385804808 | 0.699641234 | 0.84101582 | 0.041172028 |
| ENSMUSG00000021244 | Ylpm1 | 56531 | protein_coding | 334.233419 | -0.041187607 | 0.12811465 | 0.321490222 | 0.747838921 | 0.873148468 | 0.041187607 |
| ENSMUSG00000012350 | Ehf | 13661 | protein_coding | 6098.003229 | -0.041211903 | 0.125000191 | 0.329694724 | 0.741630641 | 0.869465313 | 0.041211903 |
| ENSMUSG00000028944 | Prkag2 | 108099 | protein_coding | 426.5156573 | -0.041217747 | 0.103574322 | 0.397953335 | 0.690664584 | 0.835178936 | 0.041217747 |
| ENSMUSG00000035007 | Rundc1 | 217201 | protein_coding | 247.4381947 | 0.041230156 | 0.088128814 | -0.467839679 | 0.639899243 | 0.803650332 | 0.041230156 |
| ENSMUSG00000034957 | Cebpa | 12606 | protein_coding | 664.2369576 | 0.041243233 | 0.090105983 | -0.457719139 | 0.647154236 | 0.807915411 | 0.041243233 |
| ENSMUSG00000029075 | Tnfrsf4 | 22163 | protein_coding | 5.691792291 | 0.041252848 | 0.532884818 | -0.077414192 | 0.938294051 | 0.970559847 | 0.041252848 |
| ENSMUSG00000029249 | Rest | 19712 | protein_coding | 713.3274774 | 0.041255038 | 0.08979368 | -0.459442554 | 0.645916396 | 0.807082912 | 0.041255038 |
| ENSMUSG00000008226 | Scrn3 | 74616 | protein_coding | 147.6038897 | -0.041256468 | 0.14010886 | 0.294460091 | 0.768406355 | 0.883975099 | 0.041256468 |
| ENSMUSG00000070469 | Adamtsl3 | 269959 | protein_coding | 2.065521336 | 0.041257398 | 1.800163532 | -0.022918694 | 0.981715128 | 1 | 0.041257398 |
| ENSMUSG00000043140 | Tmem186 | 66690 | protein_coding | 279.847645 | -0.04125922 | 0.124562245 | 0.33123375 | 0.740467929 | 0.868595495 | 0.04125922 |
| ENSMUSG00000070939 | Tgfbrap1 | 73122 | protein_coding | 363.8857476 | -0.041259513 | 0.070893434 | 0.581993439 | 0.5605711 | 0.749146509 | 0.041259513 |
| ENSMUSG00000090258 | Churc1 | 211151 | protein_coding | 178.7246763 | 0.041271331 | 0.109406127 | -0.377230524 | 0.706002298 | 0.845943387 | 0.041271331 |
| ENSMUSG00000000409 | Lck | 16818 | protein_coding | 59.44306172 | 0.041288365 | 0.178477822 | -0.231336109 | 0.817053697 | 0.908714429 | 0.041288365 |
| ENSMUSG00000016918 | Sulf1 | 240725 | protein_coding | 9.009965816 | -0.041344831 | 0.821690988 | 0.050316763 | 0.959869966 | 0.980506535 | 0.041344831 |
| ENSMUSG00000047213 | Ythdf3 | 229096 | protein_coding | 1534.500626 | 0.041368567 | 0.058472491 | -0.707487684 | 0.479263464 | 0.690564248 | 0.041368567 |
| ENSMUSG00000040225 | Prrc2c | 226562 | protein_coding | 1681.050734 | 0.041389812 | 0.109081711 | -0.379438602 | 0.704362189 | 0.844600435 | 0.041389812 |
| ENSMUSG00000027763 | Mbnl1 | 56758 | protein_coding | 1423.605178 | -0.041406756 | 0.07032992 | 0.588750225 | 0.55602884 | 0.745551704 | 0.041406756 |
| ENSMUSG00000036594 | H2-Aa | 14960 | protein_coding | 21.78448967 | 0.041414017 | 0.399217022 | -0.103738105 | 0.917377187 | 0.959501353 | 0.041414017 |
| ENSMUSG00000087687 | Pet100 | 100503890 | protein_coding | 356.812572 | -0.04142131 | 0.088882737 | 0.466021991 | 0.64119976 | 0.804106371 | 0.04142131 |
| ENSMUSG00000019842 | Traf3ip2 | 103213 | protein_coding | 333.0693412 | 0.0414376 | 0.08161961 | -0.507691713 | 0.611669561 | 0.784979105 | 0.0414376 |
| ENSMUSG00000047037 | Nipa1 | 233280 | protein_coding | 47.81072901 | -0.041442386 | 0.178474637 | 0.232203223 | 0.816380174 | 0.908409285 | 0.041442386 |
| ENSMUSG00000038544 | Inip | 66209 | protein_coding | 197.2549459 | 0.041449739 | 0.08449238 | -0.490573701 | 0.623727991 | 0.793440482 | 0.041449739 |
| ENSMUSG00000036529 | Sbf1 | 77980 | protein_coding | 1475.932471 | 0.041488273 | 0.068988448 | -0.601379998 | 0.547586918 | 0.739145316 | 0.041488273 |
| ENSMUSG00000021540 | Smad5 | 17129 | protein_coding | 518.9970626 | -0.041533426 | 0.132034121 | 0.314565856 | 0.753091307 | 0.876245321 | 0.041533426 |
| ENSMUSG00000020482 | Ccdc117 | 104479 | protein_coding | 322.1895079 | -0.041536268 | 0.088971251 | 0.466850442 | 0.640606884 | 0.803690806 | 0.041536268 |
| ENSMUSG00000022390 | Zc3h7b | 20286 | protein_coding | 525.9521777 | 0.041577674 | 0.062120078 | -0.669311359 | 0.503296883 | 0.707899218 | 0.041577674 |
| ENSMUSG00000026219 | Trip12 | 14897 | protein_coding | 2751.211867 | -0.041606352 | 0.051280263 | 0.811352146 | 0.417163472 | 0.637283772 | 0.041606352 |
| ENSMUSG00000039233 | Tbce | 70430 | protein_coding | 242.0789388 | -0.0416114 | 0.108304391 | 0.38420788 | 0.700824378 | 0.841825268 | 0.0416114 |
| ENSMUSG00000103421 | Golt1a | 68338 | protein_coding | 43.41842828 | 0.041624257 | 0.169887471 | -0.245010753 | 0.806448106 | 0.904795436 | 0.041624257 |
| ENSMUSG00000020380 | Rad50 | 19360 | protein_coding | 391.5765062 | 0.041636223 | 0.099085807 | -0.4202037 | 0.674336652 | 0.825194181 | 0.041636223 |
| ENSMUSG00000041073 | Nacad | 192950 | protein_coding | 73.23832666 | -0.041679551 | 0.259036406 | 0.16090229 | 0.872170359 | 0.937028043 | 0.041679551 |

| ensembl_gene_id | external_gene_name | entrezgene_id | gene_biotype | baseMean | log2FoldChange | lfcSE | stat | pvalue | padj | absLog2FC |
|---|---|---|---|---|---|---|---|---|---|---|
| ENSMUSG00000022656 | Nectin3 | 58998 | protein_coding | 1380.410809 | -0.041702626 | 0.096189893 | 0.433544784 | 0.664619039 | 0.819068567 | 0.041702626 |
| ENSMUSG00000056035 | Cyp3a11 | 13112 | protein_coding | 6.622515776 | 0.041702782 | 0.957557381 | -0.04355121 | 0.965262144 | 0.982706396 | 0.041702782 |
| ENSMUSG00000033721 | Vav3 | 57257 | protein_coding | 218.1135615 | 0.041729755 | 0.097082261 | -0.429839132 | 0.667312665 | 0.821223739 | 0.041729755 |
| ENSMUSG00000062300 | Nectin2 | 19294 | protein_coding | 1539.913486 | -0.041733045 | 0.098488409 | 0.423735601 | 0.671758649 | 0.823993339 | 0.041733045 |
| ENSMUSG00000102869 | Norad | 347740 | lncRNA | 1233.322402 | 0.041790136 | 0.122486502 | -0.341181558 | 0.732966906 | 0.863168757 | 0.041790136 |
| ENSMUSG00000027030 | Stk39 | 53416 | protein_coding | 465.088394 | -0.0418334 | 0.130225657 | 0.321237771 | 0.748030211 | 0.873310067 | 0.0418334 |
| ENSMUSG00000034382 | AI661453 | 224833 | protein_coding | 417.5778101 | 0.04188815 | 0.144945776 | -0.288991864 | 0.772587604 | 0.885884716 | 0.04188815 |
| ENSMUSG00000040236 | Trappc5 | 66682 | protein_coding | 283.4208203 | 0.041922706 | 0.074715723 | -0.561096167 | 0.574731981 | 0.760117839 | 0.041922706 |
| ENSMUSG00000057236 | Rbbp4 | 19646 | protein_coding | 1144.388345 | 0.041942589 | 0.1027991 | -0.408005411 | 0.683269703 | 0.830769833 | 0.041942589 |
| ENSMUSG00000038369 | Ncoa6 | 56406 | protein_coding | 366.7004528 | 0.041943345 | 0.149994707 | -0.279632169 | 0.779759724 | 0.890125936 | 0.041943345 |
| ENSMUSG00000036167 | Pphln1 | 223828 | protein_coding | 227.492626 | 0.041949591 | 0.085426534 | -0.491060437 | 0.623383703 | 0.793353766 | 0.041949591 |
| ENSMUSG00000059406 | Tmprss9 | 432478 | protein_coding | 3.881028166 | -0.041978167 | 0.809757949 | 0.051840389 | 0.958655873 | 1 | 0.041978167 |
| ENSMUSG00000090110 | Cmc4 | 105886298 | protein_coding | 17.20737053 | -0.041987745 | 0.230270121 | 0.182341266 | 0.855314916 | 0.929560416 | 0.041987745 |
| ENSMUSG00000040040 | Ift88 | 21821 | protein_coding | 134.4653147 | 0.041990118 | 0.093660739 | -0.448321447 | 0.653921224 | 0.812249349 | 0.041990118 |
| ENSMUSG00000026648 | Dclre1c | 227525 | protein_coding | 95.8733748 | -0.042004155 | 0.121200869 | 0.346566452 | 0.728917053 | 0.860116914 | 0.042004155 |
| ENSMUSG00000041734 | Kirrel | 170643 | protein_coding | 6.441008468 | 0.04203288 | 0.711894991 | -0.059043652 | 0.952917339 | 0.976632343 | 0.04203288 |
| ENSMUSG00000029207 | Apbb2 | 11787 | protein_coding | 254.7053901 | -0.042037191 | 0.087738732 | 0.479117831 | 0.631854806 | 0.799444652 | 0.042037191 |
| ENSMUSG00000041702 | Btbd7 | 238386 | protein_coding | 943.8978678 | -0.042080322 | 0.06065353 | 0.693781917 | 0.487818984 | 0.696186289 | 0.042080322 |
| ENSMUSG00000017999 | Ddx27 | 228889 | protein_coding | 446.4966693 | -0.042119556 | 0.082342651 | 0.511515664 | 0.608990023 | 0.783067117 | 0.042119556 |
| ENSMUSG00000029528 | Pxn | 19303 | protein_coding | 1771.562019 | 0.0421598 | 0.070900489 | -0.594633422 | 0.552088534 | 0.742744607 | 0.0421598 |
| ENSMUSG00000024442 | Dele1 | 66839 | protein_coding | 519.3851828 | -0.042224831 | 0.096663627 | 0.436822332 | 0.662240202 | 0.81729577 | 0.042224831 |
| ENSMUSG00000049866 | Arl4c | 320982 | protein_coding | 13.03593617 | 0.042236506 | 0.350759834 | -0.120414318 | 0.90415495 | 0.952610364 | 0.042236506 |
| ENSMUSG00000050088 | 1600012H06Rik | 67912 | protein_coding | 395.6213538 | 0.042265537 | 0.083770533 | -0.50453943 | 0.613882361 | 0.78655252 | 0.042265537 |
| ENSMUSG00000049672 | Zbtb14 | 22666 | protein_coding | 169.0633892 | -0.042293498 | 0.114750091 | 0.368570502 | 0.712447886 | 0.850598485 | 0.042293498 |
| ENSMUSG00000041444 | Arhgap32 | 330914 | protein_coding | 1742.38265 | -0.042332408 | 0.068639637 | 0.616734139 | 0.5374101 | 0.732906636 | 0.042332408 |
| ENSMUSG00000000142 | Axin2 | 12006 | protein_coding | 371.478686 | 0.042348738 | 0.139211515 | -0.304204273 | 0.76097227 | 0.880144002 | 0.042348738 |
| ENSMUSG00000089874 | 9230117E06Rik | 78082 | lncRNA | 1.371531985 | 0.042350893 | 1.284481848 | -0.032971188 | 0.973697563 | 1 | 0.042350893 |
| ENSMUSG00000026618 | Iars2 | 381314 | protein_coding | 1538.983752 | -0.042393791 | 0.091409595 | 0.463778344 | 0.64280656 | 0.805165923 | 0.042393791 |
| ENSMUSG00000039756 | Dnttip2 | 99480 | protein_coding | 450.3347881 | -0.042430381 | 0.089661979 | 0.473226017 | 0.636051938 | 0.801630291 | 0.042430381 |
| ENSMUSG00000096956 | Snhg18 | 100616095 | lncRNA | 369.6357112 | -0.04243755 | 0.122634189 | 0.346049908 | 0.729305208 | 0.86030736 | 0.04243755 |
| ENSMUSG00000029843 | Slc13a4 | 243755 | protein_coding | 2.698179346 | 0.042470677 | 1.666504354 | -0.025484888 | 0.979668202 | 1 | 0.042470677 |
| ENSMUSG00000020640 | Itsn2 | 20403 | protein_coding | 1271.038871 | -0.042480054 | 0.056378513 | 0.753479501 | 0.451161825 | 0.667666133 | 0.042480054 |
| ENSMUSG00000031174 | Rpgr | 19893 | protein_coding | 77.2902948 | 0.042483263 | 0.127255828 | -0.333841397 | 0.738499243 | 0.867333511 | 0.042483263 |
| ENSMUSG00000026082 | Rev1 | 56210 | protein_coding | 102.5416283 | -0.04251491 | 0.132565784 | 0.320708018 | 0.748431671 | 0.873593484 | 0.04251491 |
| ENSMUSG00000052296 | Ppp6r1 | 243819 | protein_coding | 1827.344075 | -0.042519152 | 0.049562474 | 0.857890024 | 0.390953193 | 0.613299234 | 0.042519152 |
| ENSMUSG00000038721 | Hoxb7 | 15415 | protein_coding | 91.88997522 | 0.042522516 | 0.246069838 | -0.172806697 | 0.862803373 | 0.932166328 | 0.042522516 |
| ENSMUSG00000087138 | Gm15545 | 100502630 | lncRNA | 16.75115688 | 0.042537816 | 0.275735646 | -0.154270283 | 0.877396628 | 0.939157889 | 0.042537816 |
| ENSMUSG00000048251 | Bcl11b | 58208 | protein_coding | 294.3284053 | -0.042553678 | 0.116604193 | 0.364941236 | 0.715155272 | 0.852123921 | 0.042553678 |
| ENSMUSG00000044006 | Cilp2 | 68709 | protein_coding | 3.281935893 | 0.042557359 | 1.281894189 | -0.033198808 | 0.973516049 | 1 | 0.042557359 |

Supplementary Table 1

| ensembl_gene_id | external_gene_name | entrezgene_id | gene_biotype | baseMean | log2FoldChange | lfcSE | stat | pvalue | padj | absLog2FC |
|---|---|---|---|---|---|---|---|---|---|---|
| ENSMUSG00000028786 | Tmem54 | 66260 | protein_coding | 1597.344639 | 0.042579149 | 0.090826625 | -0.468795897 | 0.639215533 | 0.803282676 | 0.042579149 |
| ENSMUSG00000110086 | Smim32 | 115488204 | protein_coding | 11.28324883 | -0.042612387 | 0.307429303 | 0.138608735 | 0.88975934 | 0.945840699 | 0.042612387 |
| ENSMUSG00000072571 | Tmem253 | 619301 | protein_coding | 117.6058135 | -0.042621982 | 0.150816892 | 0.282607479 | 0.777477747 | 0.888933284 | 0.042621982 |
| ENSMUSG00000053684 | Kics2 | 270802 | protein_coding | 32.67400262 | -0.042626835 | 0.179389833 | 0.237621244 | 0.812174872 | 0.907147741 | 0.042626835 |
| ENSMUSG00000015971 | Actr8 | 56249 | protein_coding | 340.8433058 | -0.042635034 | 0.067919803 | 0.627726116 | 0.530183374 | 0.727438372 | 0.042635034 |
| ENSMUSG00000024853 | Sf3b2 | 319322 | protein_coding | 2138.938131 | -0.042636674 | 0.06563043 | 0.649647942 | 0.515919658 | 0.716883239 | 0.042636674 |
| ENSMUSG00000045934 | Mtmr11 | 194126 | protein_coding | 501.2978706 | -0.04265778 | 0.161908845 | 0.263467876 | 0.79218998 | 0.896636371 | 0.04265778 |
| ENSMUSG00000039936 | Pik3cd | 18707 | protein_coding | 40.04192858 | 0.042668937 | 0.34655062 | -0.123124688 | 0.902008362 | 0.951530343 | 0.042668937 |
| ENSMUSG00000036186 | Dipk1b | 56279 | protein_coding | 47.9995376 | 0.042676433 | 0.196688214 | -0.216975039 | 0.828227798 | 0.913297021 | 0.042676433 |
| ENSMUSG00000028668 | Eloa | 27224 | protein_coding | 961.0917449 | -0.042701079 | 0.061554882 | 0.693707434 | 0.487865703 | 0.696186289 | 0.042701079 |
| ENSMUSG00000020397 | Med7 | 66213 | protein_coding | 155.2412162 | 0.042702409 | 0.114625541 | -0.372538343 | 0.70949207 | 0.84834929 | 0.042702409 |
| ENSMUSG00000024072 | Yipf4 | 67864 | protein_coding | 583.2814623 | 0.042712894 | 0.111741455 | -0.382247524 | 0.702277774 | 0.84294636 | 0.042712894 |
| ENSMUSG00000020219 | Timm13 | 30055 | protein_coding | 754.5271526 | 0.042729472 | 0.087104068 | -0.490556556 | 0.62374012 | 0.793440482 | 0.042729472 |
| ENSMUSG00000030347 | D6Wsu163e | 28040 | protein_coding | 164.7560529 | 0.04273884 | 0.119428889 | -0.357860149 | 0.720447974 | 0.855204229 | 0.04273884 |
| ENSMUSG00000040990 | Sh3kbp1 | 58194 | protein_coding | 161.721298 | 0.042791168 | 0.181055362 | -0.236343002 | 0.813166521 | 0.907217398 | 0.042791168 |
| ENSMUSG00000086316 | Nbdy | 70123 | protein_coding | 60.01764469 | 0.042796584 | 0.19074394 | -0.224366675 | 0.822471999 | 0.910171686 | 0.042796584 |
| ENSMUSG00000089542 | Gm25835 | 115487914 | snoRNA | 246.1698507 | 0.042816834 | 0.187084821 | -0.228863217 | 0.81897523 | 0.90889719 | 0.042816834 |
| ENSMUSG00000054843 | Atrnl1 | 226255 | protein_coding | 1986.452605 | -0.042902399 | 0.075678726 | 0.566901703 | 0.570780967 | 0.756249943 | 0.042902399 |
| ENSMUSG00000004768 | Rab23 | 19335 | protein_coding | 70.50102242 | 0.04290368 | 0.161570207 | -0.265542026 | 0.790591939 | 0.895584805 | 0.04290368 |
| ENSMUSG00000040843 | Tiprl | 226591 | protein_coding | 296.9076699 | 0.042917766 | 0.084023803 | -0.510781047 | 0.609504382 | 0.783477444 | 0.042917766 |
| ENSMUSG00000026798 | Coq4 | 227683 | protein_coding | 151.2016363 | -0.042991592 | 0.105238011 | 0.40851772 | 0.682893624 | 0.830618347 | 0.042991592 |
| ENSMUSG00000108084 | Gm7932 | 666105 | lncRNA | 13.43988509 | 0.043024715 | 0.290527304 | -0.148091811 | 0.882270312 | 0.941973978 | 0.043024715 |
| ENSMUSG00000091827 | Speer4f2 | 100041749 | protein_coding | 0.914559436 | 0.043036239 | 2.287070191 | -0.018817192 | 0.984986939 | 1 | 0.043036239 |
| ENSMUSG00000026047 | Poglut2 | 72050 | protein_coding | 91.44322005 | 0.043079423 | 0.124530139 | -0.345935721 | 0.729391023 | 0.86030736 | 0.043079423 |
| ENSMUSG00000032109 | Nlrx1 | 270151 | protein_coding | 371.9870974 | 0.043086199 | 0.140087613 | -0.307566084 | 0.758412531 | 0.878845311 | 0.043086199 |
| ENSMUSG00000006262 | Mob1b | 68473 | protein_coding | 1096.299405 | 0.04308973 | 0.090434712 | -0.476473341 | 0.633737196 | 0.800362194 | 0.04308973 |
| ENSMUSG00000038250 | Usp38 | 74841 | protein_coding | 499.4588503 | 0.043099135 | 0.092849557 | -0.464182454 | 0.642517031 | 0.80495599 | 0.043099135 |
| ENSMUSG00000028851 | Nudc | 18221 | protein_coding | 662.4261716 | 0.043102275 | 0.103591349 | -0.416079869 | 0.67735156 | 0.827106157 | 0.043102275 |
| ENSMUSG00000020435 | Osbp2 | 74309 | protein_coding | 5.823101018 | 0.043115041 | 0.846426173 | -0.050937745 | 0.959375128 | 0.980330479 | 0.043115041 |
| ENSMUSG00000021796 | Bmpr1a | 12166 | protein_coding | 1252.688138 | -0.043149081 | 0.085336725 | 0.505633202 | 0.613114169 | 0.786099952 | 0.043149081 |
| ENSMUSG00000015092 | Edf1 | 59022 | protein_coding | 2177.94877 | -0.043169704 | 0.078913213 | 0.547052925 | 0.584342372 | 0.766711421 | 0.043169704 |
| ENSMUSG00000068580 | Zfyve19 | 72008 | protein_coding | 292.6248758 | -0.043181297 | 0.070862143 | 0.609370465 | 0.54227891 | 0.735940424 | 0.043181297 |
| ENSMUSG00000032612 | Usp4 | 22258 | protein_coding | 1340.642495 | -0.043182968 | 0.063681442 | 0.678109144 | 0.497702493 | 0.703115827 | 0.043182968 |
| ENSMUSG00000062580 | Timm17a | 21854 | protein_coding | 613.1877017 | 0.043195278 | 0.081088911 | -0.532690322 | 0.594247967 | 0.77323163 | 0.043195278 |
| ENSMUSG00000029924 | Slc37a3 | 72144 | protein_coding | 536.2576432 | 0.043218245 | 0.098550652 | -0.438538394 | 0.660996044 | 0.816884691 | 0.043218245 |
| ENSMUSG00000025423 | Pias2 | 17344 | protein_coding | 404.5180564 | 0.043219029 | 0.08433661 | -0.512458695 | 0.608330022 | 0.782626881 | 0.043219029 |
| ENSMUSG00000022974 | Paxbp1 | 67367 | protein_coding | 396.2719077 | 0.043262737 | 0.115363906 | -0.375011025 | 0.707652267 | 0.846946626 | 0.043262737 |
| ENSMUSG00000026781 | Acbd5 | 74159 | protein_coding | 928.6399199 | -0.043282933 | 0.078906258 | 0.548536124 | 0.583323832 | 0.765922697 | 0.043282933 |
| ENSMUSG00000022371 | Col14a1 | 12818 | protein_coding | 5.78303575 | -0.043482801 | 0.90011214 | 0.048308205 | 0.961470616 | 0.981136115 | 0.043482801 |

| ensembl_gene_id | external_gene_name | entrezgene_id | gene_biotype | baseMean | log2FoldChange | lfcSE | stat | pvalue | padj | absLog2FC |
|---|---|---|---|---|---|---|---|---|---|---|
| ENSMUSG00000068391 | Chrac1 | 93696 | protein_coding | 179.4573957 | -0.043499528 | 0.101811933 | 0.427253727 | 0.669194532 | 0.822558876 | 0.043499528 |
| ENSMUSG00000057691 | Zfp746 | 69228 | protein_coding | 381.3481077 | -0.043499627 | 0.062074815 | 0.700761286 | 0.483452002 | 0.693875647 | 0.043499627 |
| ENSMUSG00000038095 | Sbno1 | 243272 | protein_coding | 1507.974933 | -0.043525207 | 0.071724728 | 0.606836842 | 0.543959196 | 0.73666903 | 0.043525207 |
| ENSMUSG00000035152 | Ap2b1 | 71770 | protein_coding | 2105.976682 | -0.043526025 | 0.083057676 | 0.524045787 | 0.600246694 | 0.77680602 | 0.043526025 |
| ENSMUSG00000036114 | Rpp25l | 69961 | protein_coding | 204.9060122 | -0.043548025 | 0.094794724 | 0.459392919 | 0.645952032 | 0.807082912 | 0.043548025 |
| ENSMUSG00000014245 | Pigl | 327942 | protein_coding | 74.88668916 | -0.043643278 | 0.132490422 | 0.329407042 | 0.741848046 | 0.869560398 | 0.043643278 |
| ENSMUSG00000026749 | Nek6 | 59126 | protein_coding | 821.4336718 | 0.043647733 | 0.084563951 | -0.516150586 | 0.605749239 | 0.780657323 | 0.043647733 |
| ENSMUSG00000032475 | Nck1 | 17973 | protein_coding | 346.2868404 | -0.043667239 | 0.08964703 | 0.487101898 | 0.626186121 | 0.795381772 | 0.043667239 |
| ENSMUSG00000030431 | Tmem238 | 664968 | protein_coding | 276.5064705 | 0.043735732 | 0.151566435 | -0.288558163 | 0.772919515 | 0.886203768 | 0.043735732 |
| ENSMUSG00000033933 | Vhl | 22346 | protein_coding | 206.4146817 | -0.043774025 | 0.081411392 | 0.537689185 | 0.590791647 | 0.771232643 | 0.043774025 |
| ENSMUSG00000042229 | Rabif | 98710 | protein_coding | 332.1493859 | 0.043777291 | 0.108336158 | -0.404087533 | 0.686148353 | 0.832872319 | 0.043777291 |
| ENSMUSG00000021557 | Agtpbp1 | 67269 | protein_coding | 63.29970036 | 0.043789079 | 0.155109632 | -0.28231051 | 0.777705428 | 0.888933284 | 0.043789079 |
| ENSMUSG00000043333 | Rhbdl2 | 230726 | protein_coding | 510.3034724 | -0.043807791 | 0.105155553 | 0.416599886 | 0.676971093 | 0.826943172 | 0.043807791 |
| ENSMUSG00000022370 | Mrpl13 | 68537 | protein_coding | 311.2963923 | -0.043830918 | 0.087970069 | 0.498248078 | 0.618309199 | 0.789606655 | 0.043830918 |
| ENSMUSG00000028030 | Tbck | 271981 | protein_coding | 489.7233539 | -0.043889696 | 0.07173615 | 0.611821177 | 0.540656077 | 0.734843633 | 0.043889696 |
| ENSMUSG00000028431 | Elp1 | 230233 | protein_coding | 302.4024951 | 0.04390138 | 0.110735764 | -0.396451681 | 0.691771847 | 0.835998557 | 0.04390138 |
| ENSMUSG00000024033 | Rsph1 | 22092 | protein_coding | 15.08614931 | -0.043912336 | 0.299086791 | 0.146821383 | 0.883273006 | 0.942211125 | 0.043912336 |
| ENSMUSG00000086714 | 0610009E02Rik | 100125929 | lncRNA | 7.583474848 | 0.043932232 | 0.412831978 | -0.106416738 | 0.915251714 | 0.95833356 | 0.043932232 |
| ENSMUSG00000034101 | Ctnnd1 | 12388 | protein_coding | 831.9623403 | 0.043935492 | 0.082465173 | -0.53277632 | 0.594188428 | 0.77323163 | 0.043935492 |
| ENSMUSG00000053134 | Supt7l | 72195 | protein_coding | 201.9525074 | 0.043990668 | 0.10321372 | -0.426209503 | 0.669955194 | 0.822722373 | 0.043990668 |
| ENSMUSG00000000078 | Klf6 | 23849 | protein_coding | 7142.565263 | -0.04401657 | 0.172495988 | 0.255174456 | 0.798588359 | 0.900012841 | 0.04401657 |
| ENSMUSG00000086391 | 1700042O10Rik | 73321 | lncRNA | 2.57431512 | 0.044085524 | 0.742283079 | -0.059391795 | 0.952640048 | 1 | 0.044085524 |
| ENSMUSG00000004996 | Mri1 | 67873 | protein_coding | 198.5004799 | -0.044109369 | 0.100271775 | 0.439898154 | 0.660010873 | 0.816323585 | 0.044109369 |
| ENSMUSG00000025008 | Tctn3 | 67590 | protein_coding | 54.21534624 | 0.0441477 | 0.153239131 | -0.288096775 | 0.773272661 | 0.886362513 | 0.0441477 |
| ENSMUSG00000001482 | Def8 | 23854 | protein_coding | 244.3035699 | 0.044150999 | 0.073259978 | -0.602661925 | 0.546733616 | 0.738518372 | 0.044150999 |
| ENSMUSG00000001100 | Poldip2 | 67811 | protein_coding | 655.6451439 | 0.04419288 | 0.088672232 | -0.498384651 | 0.618212953 | 0.789572213 | 0.04419288 |
| ENSMUSG00000027907 | S100a11 | 20195 | protein_coding | 307.8448088 | 0.044243037 | 0.123472538 | -0.358322895 | 0.720101686 | 0.855153117 | 0.044243037 |
| ENSMUSG00000002608 | Ccdc97 | 52132 | protein_coding | 233.6375721 | -0.044280732 | 0.106551061 | 0.415582276 | 0.677715699 | 0.827306109 | 0.044280732 |
| ENSMUSG00000032328 | Tmem30a | 69981 | protein_coding | 1584.013196 | 0.044295052 | 0.076360104 | -0.580081078 | 0.561859943 | 0.750140815 | 0.044295052 |
| ENSMUSG00000057278 | Snrpg | 68011 | protein_coding | 220.577199 | -0.044297035 | 0.141091382 | 0.313959888 | 0.753551504 | 0.876528151 | 0.044297035 |
| ENSMUSG00000017801 | Mlx | 21428 | protein_coding | 1204.162923 | -0.044321434 | 0.110220045 | 0.402117728 | 0.687597386 | 0.833638003 | 0.044321434 |
| ENSMUSG00000034227 | Foxj1 | 15223 | protein_coding | 3.005990548 | 0.044324184 | 0.626821262 | -0.070712637 | 0.943626463 | 1 | 0.044324184 |
| ENSMUSG00000044317 | Gpr4 | 319197 | protein_coding | 8.920763787 | 0.044331964 | 0.429496065 | -0.103218558 | 0.917789512 | 0.959689482 | 0.044331964 |
| ENSMUSG00000071647 | Eml3 | 225898 | protein_coding | 283.9416586 | 0.044332856 | 0.06875219 | -0.644821004 | 0.519043189 | 0.719334577 | 0.044332856 |
| ENSMUSG00000024027 | Glp1r | 14652 | protein_coding | 14.50486651 | 0.044339225 | 0.423549649 | -0.104684835 | 0.916625896 | 0.959274295 | 0.044339225 |
| ENSMUSG00000024132 | Eci1 | 13177 | protein_coding | 618.0241444 | -0.044353019 | 0.103156111 | 0.429960172 | 0.667224613 | 0.821176574 | 0.044353019 |
| ENSMUSG00000071477 | Zfp777 | 72306 | protein_coding | 244.9605466 | -0.044381637 | 0.073443163 | 0.604299098 | 0.545644806 | 0.737439915 | 0.044381637 |
| ENSMUSG00000001552 | Jup | 16480 | protein_coding | 3801.723192 | 0.04446186 | 0.103548182 | -0.429383298 | 0.667644307 | 0.821350533 | 0.04446186 |
| ENSMUSG00000020982 | Nemf | 66244 | protein_coding | 625.5727049 | 0.044485422 | 0.06320254 | -0.703854964 | 0.481523101 | 0.692354022 | 0.044485422 |

Supplementary Table 1

| ensembl_gene_id | external_gene_name | entrezgene_id | gene_biotype | baseMean | log2FoldChange | lfcSE | stat | pvalue | padj | absLog2FC |
|---|---|---|---|---|---|---|---|---|---|---|
| ENSMUSG00000024481 | Lvrn | 74574 | protein_coding | 1.212300783 | 0.044517408 | 1.949632944 | -0.022833738 | 0.981782896 | 1 | 0.044517408 |
| ENSMUSG00000085385 | Snhg17 | 68108 | lncRNA | 116.4128262 | -0.044531457 | 0.147256267 | 0.302407891 | 0.76234114 | 0.88093126 | 0.044531457 |
| ENSMUSG00000085408 | C530005A16Rik | 654318 | lncRNA | 9.007585209 | 0.044569538 | 0.365077703 | -0.122082332 | 0.902833814 | 0.95200802 | 0.044569538 |
| ENSMUSG00000024897 | Apba1 | 319924 | protein_coding | 8.516515362 | 0.044597914 | 0.677022864 | -0.065873571 | 0.947478482 | 0.974566157 | 0.044597914 |
| ENSMUSG00000010406 | Mrpl52 | 68836 | protein_coding | 507.6556446 | 0.044666641 | 0.104759858 | -0.426371721 | 0.669837004 | 0.822722373 | 0.044666641 |
| ENSMUSG00000018651 | Tada2a | 217031 | protein_coding | 103.2184097 | 0.044670688 | 0.1175001 | -0.380175747 | 0.703814961 | 0.844104331 | 0.044670688 |
| ENSMUSG00000038132 | Rbm24 | 666794 | protein_coding | 3.029333524 | -0.04468744 | 0.816077708 | 0.054758804 | 0.95633062 | 1 | 0.04468744 |
| ENSMUSG00000029135 | Fosl2 | 14284 | protein_coding | 3196.581954 | 0.044721417 | 0.133768252 | -0.334320115 | 0.738138013 | 0.866982505 | 0.044721417 |
| ENSMUSG00000016319 | Slc25a5 | 11740 | protein_coding | 13756.28303 | 0.044776799 | 0.096891911 | -0.462131442 | 0.643987064 | 0.80597323 | 0.044776799 |
| ENSMUSG00000024293 | Esco1 | 77805 | protein_coding | 494.1642558 | 0.044784593 | 0.091165221 | -0.491246471 | 0.623252135 | 0.793308598 | 0.044784593 |
| ENSMUSG00000041303 | Gtf3c3 | 98488 | protein_coding | 311.9224918 | 0.04480071 | 0.11455063 | -0.391099635 | 0.69572359 | 0.83856979 | 0.04480071 |
| ENSMUSG00000025451 | Paip1 | 218693 | protein_coding | 566.6489282 | 0.044856845 | 0.084243832 | -0.532464437 | 0.594404366 | 0.77323163 | 0.044856845 |
| ENSMUSG00000037210 | Fam193a | 231128 | protein_coding | 441.1765218 | 0.044877696 | 0.091769601 | -0.489025726 | 0.624823484 | 0.79451259 | 0.044877696 |
| ENSMUSG00000030583 | Sipa1l3 | 74206 | protein_coding | 1161.898356 | -0.044886511 | 0.186903495 | 0.240158757 | 0.810207185 | 0.906158059 | 0.044886511 |
| ENSMUSG00000015597 | Zfp318 | 57908 | protein_coding | 232.6466497 | 0.044934582 | 0.084310587 | -0.532964886 | 0.594057888 | 0.773192935 | 0.044934582 |
| ENSMUSG00000035601 | Trmt10b | 69934 | protein_coding | 115.6779227 | 0.044939354 | 0.106796142 | -0.420795671 | 0.673904293 | 0.825096143 | 0.044939354 |
| ENSMUSG00000040322 | Slc25a24 | 229731 | protein_coding | 4125.357335 | -0.04494494 | 0.099893233 | 0.449929779 | 0.652761075 | 0.811601896 | 0.04494494 |
| ENSMUSG00000031592 | Pcm1 | 18536 | protein_coding | 897.5454437 | -0.044955719 | 0.068710822 | 0.654274207 | 0.51293516 | 0.714956306 | 0.044955719 |
| ENSMUSG00000027770 | Dhx36 | 72162 | protein_coding | 582.2647813 | -0.045010142 | 0.094558143 | 0.476004927 | 0.634070868 | 0.800507769 | 0.045010142 |
| ENSMUSG00000090330 | 9130221H12Rik | 77124 | lncRNA | 32.12441167 | 0.04502611 | 0.190402257 | -0.236478867 | 0.813061104 | 0.907217398 | 0.04502611 |
| ENSMUSG00000087259 | 2610035D17Rik | 72386 | lncRNA | 280.3508375 | 0.04504154 | 0.169417406 | -0.265861348 | 0.790345994 | 0.895392916 | 0.04504154 |
| ENSMUSG00000028114 | Mettl14 | 210529 | protein_coding | 173.0047697 | -0.045072227 | 0.112557715 | 0.400436583 | 0.688834984 | 0.833904116 | 0.045072227 |
| ENSMUSG00000025538 | Sumf2 | 67902 | protein_coding | 62.68645906 | -0.045086016 | 0.158823038 | 0.283875793 | 0.776505568 | 0.888506311 | 0.045086016 |
| ENSMUSG00000024908 | Ppp6r3 | 52036 | protein_coding | 2261.466381 | -0.04509059 | 0.066503635 | 0.678016929 | 0.49776096 | 0.703115827 | 0.04509059 |
| ENSMUSG00000044452 | Zfp507 | 668501 | protein_coding | 417.8848172 | -0.04509484 | 0.102945608 | 0.438045302 | 0.661353444 | 0.816921981 | 0.04509484 |
| ENSMUSG00000061666 | Gdpd1 | 66569 | protein_coding | 1173.649685 | -0.045173766 | 0.113353783 | 0.398520148 | 0.69024681 | 0.834766446 | 0.045173766 |
| ENSMUSG00000043648 | Pld6 | 194908 | protein_coding | 2.868979415 | -0.045173834 | 0.644638692 | 0.070076207 | 0.944133005 | 1 | 0.045173834 |
| ENSMUSG00000030849 | Fgfr2 | 14183 | protein_coding | 691.5289067 | -0.045193244 | 0.127248054 | 0.355158628 | 0.722470757 | 0.856426823 | 0.045193244 |
| ENSMUSG00000038902 | Pogz | 229584 | protein_coding | 188.9148655 | 0.045244576 | 0.112418629 | -0.402465112 | 0.687341759 | 0.833389267 | 0.045244576 |
| ENSMUSG00000033257 | Ttll4 | 67534 | protein_coding | 355.7589674 | 0.045266841 | 0.089713047 | -0.50457367 | 0.613858307 | 0.78655252 | 0.045266841 |
| ENSMUSG00000020238 | Ncln | 103425 | protein_coding | 1130.261013 | 0.045291045 | 0.062609232 | -0.72339244 | 0.469438815 | 0.683156904 | 0.045291045 |
| ENSMUSG00000021288 | Klc1 | 16593 | protein_coding | 879.0531191 | -0.045307723 | 0.103557467 | 0.437512857 | 0.661739454 | 0.816921981 | 0.045307723 |
| ENSMUSG00000028405 | Aco1 | 11428 | protein_coding | 1492.284116 | -0.045312223 | 0.088972206 | 0.509285151 | 0.610552365 | 0.784161879 | 0.045312223 |
| ENSMUSG00000059305 | Vpreb1 | 22362 | protein_coding | 2.868567947 | 0.045328285 | 0.772878665 | -0.058648644 | 0.953231965 | 1 | 0.045328285 |
| ENSMUSG00000106718 | Gm20052 | 102637176 | lncRNA | 1.121769154 | 0.04533231 | 1.99335447 | -0.02274172 | 0.981856296 | 1 | 0.04533231 |
| ENSMUSG00000030615 | Tmem126a | 66271 | protein_coding | 279.1545126 | 0.045339122 | 0.081295169 | -0.557709915 | 0.577042486 | 0.761298159 | 0.045339122 |
| ENSMUSG00000035245 | Eogt | 101351 | protein_coding | 174.9575124 | -0.045353385 | 0.093351158 | 0.48583634 | 0.627083207 | 0.795976331 | 0.045353385 |
| ENSMUSG00000060260 | Pwwp2b | 101631 | protein_coding | 219.7169153 | -0.045469352 | 0.093174016 | 0.488004636 | 0.625546557 | 0.794916857 | 0.045469352 |
| ENSMUSG00000061762 | Tac1 | 21333 | protein_coding | 39.94273795 | -0.045491951 | 0.209983565 | 0.216645292 | 0.828484786 | 0.913446239 | 0.045491951 |

| ensembl_gene_id | external_gene_name | entrezgene_id | gene_biotype | baseMean | log2FoldChange | lfcSE | stat | pvalue | padj | absLog2FC |
|---|---|---|---|---|---|---|---|---|---|---|
| ENSMUSG00000028700 | Pomgnt1 | 68273 | protein_coding | 515.3519249 | -0.045524929 | 0.069737762 | 0.652801688 | 0.513884135 | 0.715629739 | 0.045524929 |
| ENSMUSG00000022450 | Ndufa6 | 67130 | protein_coding | 2149.131824 | 0.045548285 | 0.091860804 | -0.495840258 | 0.620007119 | 0.791007228 | 0.045548285 |
| ENSMUSG00000028741 | Mrto4 | 69902 | protein_coding | 166.1042923 | -0.045605692 | 0.16443329 | 0.277350724 | 0.781510814 | 0.890612766 | 0.045605692 |
| ENSMUSG00000030335 | Mrpl51 | 66493 | protein_coding | 877.8613607 | 0.045613747 | 0.088852371 | -0.513365556 | 0.607695635 | 0.782250036 | 0.045613747 |
| ENSMUSG00000039628 | Hs3st6 | 328779 | protein_coding | 0.73548046 | -0.045708479 | 1.717482718 | 0.026613647 | 0.978767888 | 1 | 0.045708479 |
| ENSMUSG00000019731 | Slc35e1 | 270066 | protein_coding | 1512.596385 | -0.045712295 | 0.045379545 | 1.00733258 | 0.313774978 | 0.535013399 | 0.045712295 |
| ENSMUSG00000024436 | Mrps18b | 66973 | protein_coding | 184.5457018 | 0.045729328 | 0.095528892 | -0.478696306 | 0.632154694 | 0.799512805 | 0.045729328 |
| ENSMUSG00000028403 | Zdhhc21 | 68268 | protein_coding | 894.8452555 | -0.045741334 | 0.081960523 | 0.558089834 | 0.576783042 | 0.761222618 | 0.045741334 |
| ENSMUSG00000021504 | B4galt7 | 218271 | protein_coding | 199.6490538 | 0.045793305 | 0.081859307 | -0.559414766 | 0.575878686 | 0.760926598 | 0.045793305 |
| ENSMUSG00000003135 | Cnot11 | 52846 | protein_coding | 311.812095 | 0.045796223 | 0.077561074 | -0.590453701 | 0.554886517 | 0.744703672 | 0.045796223 |
| ENSMUSG00000014606 | Slc25a11 | 67863 | protein_coding | 1304.307442 | 0.045816181 | 0.089511008 | -0.511849686 | 0.608756214 | 0.782882962 | 0.045816181 |
| ENSMUSG00000024430 | Cabyr | 71132 | protein_coding | 2.532361036 | -0.045821259 | 1.026794361 | 0.044625546 | 0.96440578 | 1 | 0.045821259 |
| ENSMUSG00000022329 | Stk3 | 56274 | protein_coding | 295.9587441 | -0.045874478 | 0.072175509 | 0.635596192 | 0.525039652 | 0.723506743 | 0.045874478 |
| ENSMUSG00000058388 | Phtf1 | 18685 | protein_coding | 154.2405022 | 0.045875341 | 0.117360542 | -0.390892376 | 0.695876789 | 0.838607402 | 0.045875341 |
| ENSMUSG00000002524 | Puf60 | 67959 | protein_coding | 1285.879775 | 0.045878457 | 0.087179981 | -0.526249914 | 0.598714578 | 0.776186935 | 0.045878457 |
| ENSMUSG00000045098 | Kmt5b | 225888 | protein_coding | 805.4033878 | -0.045884171 | 0.08851597 | 0.518371669 | 0.604198978 | 0.779660446 | 0.045884171 |
| ENSMUSG00000037062 | Sh3glb1 | 54673 | protein_coding | 1353.226895 | -0.045894203 | 0.076484688 | 0.600044327 | 0.548476694 | 0.739637131 | 0.045894203 |
| ENSMUSG00000029571 | Tmem106b | 71900 | protein_coding | 830.7058683 | -0.045896755 | 0.118250775 | 0.388130691 | 0.697919321 | 0.839976943 | 0.045896755 |
| ENSMUSG00000005656 | Snx6 | 72183 | protein_coding | 613.4937303 | -0.045959911 | 0.085195705 | 0.53946277 | 0.589567579 | 0.770506411 | 0.045959911 |
| ENSMUSG00000025337 | Sbds | 66711 | protein_coding | 929.4541527 | -0.04598212 | 0.073932284 | 0.621949126 | 0.533975321 | 0.730224807 | 0.04598212 |
| ENSMUSG00000034781 | Gna11 | 14672 | protein_coding | 3849.034124 | -0.046009622 | 0.126393198 | 0.364019763 | 0.715843251 | 0.852575063 | 0.046009622 |
| ENSMUSG00000042502 | Cd2bp2 | 70233 | protein_coding | 579.6943179 | -0.046014757 | 0.064131132 | 0.717510444 | 0.473059194 | 0.685332006 | 0.046014757 |
| ENSMUSG00000024081 | Cebpz | 12607 | protein_coding | 408.8416885 | -0.046046933 | 0.126552378 | 0.363856721 | 0.715965004 | 0.852575063 | 0.046046933 |
| ENSMUSG00000005871 | Apc | 11789 | protein_coding | 1700.168744 | 0.046103368 | 0.06824224 | -0.675584031 | 0.499304783 | 0.704201299 | 0.046103368 |
| ENSMUSG00000042396 | Rbm7 | 67010 | protein_coding | 615.035543 | 0.046115593 | 0.077238111 | -0.597057488 | 0.550469 | 0.741471864 | 0.046115593 |
| ENSMUSG00000038145 | Snrk | 20623 | protein_coding | 718.945101 | -0.046117388 | 0.139152837 | 0.331415364 | 0.740330762 | 0.868496285 | 0.046117388 |
| ENSMUSG00000042189 | Tekt3 | 71062 | protein_coding | 1.834224334 | -0.046119424 | 1.30501923 | 0.035340034 | 0.971808601 | 1 | 0.046119424 |
| ENSMUSG00000089876 | Tmem102 | 380705 | protein_coding | 390.0597871 | -0.046145453 | 0.0999614 | 0.46163272 | 0.644344727 | 0.806175378 | 0.046145453 |
| ENSMUSG00000100199 | Gm20324 | 100504637 | lncRNA | 13.64636787 | 0.046161522 | 0.337085225 | -0.136943178 | 0.891075709 | 0.946708891 | 0.046161522 |
| ENSMUSG00000032178 | Ilf3 | 16201 | protein_coding | 402.3168446 | -0.046164298 | 0.104694446 | 0.440943147 | 0.659254156 | 0.816110122 | 0.046164298 |
| ENSMUSG00000085486 | Ccdc200 | 667306 | protein_coding | 0.699327368 | -0.04616548 | 1.494063146 | 0.030899283 | 0.975349862 | 1 | 0.04616548 |
| ENSMUSG00000022111 | Uchl3 | 50933 | protein_coding | 220.0731081 | -0.046190048 | 0.102422685 | 0.450974786 | 0.652007721 | 0.811153798 | 0.046190048 |
| ENSMUSG00000019951 | Uhrf1bp1l | 75089 | protein_coding | 570.3516433 | 0.046192479 | 0.087598353 | -0.527321317 | 0.597970473 | 0.77577849 | 0.046192479 |
| ENSMUSG00000071234 | Syndig1l | 627191 | protein_coding | 3.022144569 | 0.04622482 | 0.892825036 | -0.05177366 | 0.958709044 | 1 | 0.04622482 |
| ENSMUSG00000029613 | Eif2ak1 | 15467 | protein_coding | 531.2195083 | -0.046241246 | 0.071876513 | 0.643342918 | 0.520001614 | 0.720115363 | 0.046241246 |
| ENSMUSG00000049300 | Prmt6 | 99890 | protein_coding | 97.90918897 | -0.046300311 | 0.117811487 | 0.393003369 | 0.694316993 | 0.837885199 | 0.046300311 |
| ENSMUSG00000058655 | Eif4b | 75705 | protein_coding | 4476.274837 | -0.046322659 | 0.087261713 | 0.530847459 | 0.59552449 | 0.773864504 | 0.046322659 |
| ENSMUSG00000002320 | Tm9sf1 | 74140 | protein_coding | 1046.192045 | 0.046329938 | 0.096744887 | -0.478887719 | 0.632018508 | 0.799444652 | 0.046329938 |
| ENSMUSG00000039285 | Azi2 | 27215 | protein_coding | 1384.778526 | -0.046336836 | 0.062293256 | 0.743849959 | 0.456967252 | 0.671862287 | 0.046336836 |

Supplementary Table 1

| ensembl_gene_id | external_gene_name | entrezgene_id | gene_biotype | baseMean | log2FoldChange | lfcSE | stat | pvalue | padj | absLog2FC |
|---|---|---|---|---|---|---|---|---|---|---|
| ENSMUSG00000041997 | Tlk1 | 228012 | protein_coding | 1092.081191 | -0.046372767 | 0.105419718 | 0.439887039 | 0.660018924 | 0.816323585 | 0.046372767 |
| ENSMUSG00000021111 | Papola | 18789 | protein_coding | 1643.447509 | 0.046396586 | 0.078235099 | -0.593040553 | 0.553154013 | 0.743572273 | 0.046396586 |
| ENSMUSG00000036188 | Ankmy2 | 217473 | protein_coding | 385.6970401 | -0.046400416 | 0.103494931 | 0.448335161 | 0.653911329 | 0.812249349 | 0.046400416 |
| ENSMUSG00000117713 | Gm46637 | 108168397 | lncRNA | 1.814432718 | -0.046418157 | 1.097467588 | 0.042295697 | 0.966262975 | 1 | 0.046418157 |
| ENSMUSG00000020758 | Itgb4 | 192897 | protein_coding | 2307.714954 | -0.046460253 | 0.081611024 | 0.569288938 | 0.569160072 | 0.754948549 | 0.046460253 |
| ENSMUSG00000021000 | Mia2 | 338320 | protein_coding | 1257.758648 | -0.046492238 | 0.05818617 | 0.799025585 | 0.424275577 | 0.643978573 | 0.046492238 |
| ENSMUSG00000034621 | Gpatch8 | 237943 | protein_coding | 651.3406688 | 0.046502312 | 0.131326731 | -0.35409632 | 0.7232667 | 0.856549354 | 0.046502312 |
| ENSMUSG00000047638 | Nr1h4 | 20186 | protein_coding | 1471.675918 | 0.046544324 | 0.160243334 | -0.290460282 | 0.771464132 | 0.885228942 | 0.046544324 |
| ENSMUSG00000030101 | Sumf1 | 58911 | protein_coding | 395.6543987 | 0.046648536 | 0.068955496 | -0.676502071 | 0.498721932 | 0.703739933 | 0.046648536 |
| ENSMUSG00000020778 | Ten1 | 69535 | protein_coding | 205.9616663 | 0.0466692 | 0.110231586 | -0.423374111 | 0.67202233 | 0.824189967 | 0.0466692 |
| ENSMUSG00000038696 | Mapkap1 | 227743 | protein_coding | 903.0722465 | 0.046699412 | 0.071141974 | -0.65642559 | 0.51155033 | 0.71433639 | 0.046699412 |
| ENSMUSG00000029229 | Chic2 | 74277 | protein_coding | 211.9800798 | -0.046708172 | 0.112865906 | 0.413837741 | 0.678992942 | 0.827951097 | 0.046708172 |
| ENSMUSG00000097537 | 2610020C07Rik | 69918 | lncRNA | 12.66306697 | 0.046748972 | 0.299870659 | -0.155897119 | 0.876114116 | 0.938630585 | 0.046748972 |
| ENSMUSG00000033653 | Vps8 | 209018 | protein_coding | 292.6589925 | 0.046756465 | 0.07462318 | -0.626567574 | 0.530942729 | 0.728048657 | 0.046756465 |
| ENSMUSG00000028636 | Ppcs | 106564 | protein_coding | 268.9091024 | -0.046787368 | 0.09371762 | 0.499237688 | 0.617611947 | 0.789415146 | 0.046787368 |
| ENSMUSG00000020415 | Pttg1 | 30939 | protein_coding | 267.6518431 | 0.046791849 | 0.084622508 | -0.552948024 | 0.580299001 | 0.76356297 | 0.046791849 |
| ENSMUSG00000028106 | Rprd2 | 75137 | protein_coding | 496.5423049 | 0.046792797 | 0.093803073 | -0.498840766 | 0.617891566 | 0.789468115 | 0.046792797 |
| ENSMUSG00000039195 | Bbln | 73737 | protein_coding | 136.5140402 | 0.046804991 | 0.09770805 | -0.479029015 | 0.631917987 | 0.799444652 | 0.046804991 |
| ENSMUSG00000022214 | Dcaf11 | 28199 | protein_coding | 1230.547606 | 0.046843559 | 0.056608773 | -0.827496461 | 0.407955727 | 0.628624818 | 0.046843559 |
| ENSMUSG00000027668 | Mfn1 | 67414 | protein_coding | 773.8224423 | 0.046851619 | 0.065974371 | -0.710148779 | 0.47761188 | 0.68930064 | 0.046851619 |
| ENSMUSG00000033295 | Ptprf | 19268 | protein_coding | 5339.622355 | 0.046865271 | 0.051077789 | -0.9175274 | 0.358866342 | 0.581581822 | 0.046865271 |
| ENSMUSG00000022272 | Myo10 | 17909 | protein_coding | 1717.865139 | 0.04688266 | 0.061842235 | -0.758100997 | 0.448390523 | 0.665277305 | 0.04688266 |
| ENSMUSG00000019054 | Fis1 | 66437 | protein_coding | 861.1189941 | -0.046911992 | 0.072978184 | 0.642822136 | 0.520339518 | 0.720291626 | 0.046911992 |
| ENSMUSG00000020115 | Tbk1 | 56480 | protein_coding | 509.4932014 | -0.046944077 | 0.066417109 | 0.706806987 | 0.479686432 | 0.690909666 | 0.046944077 |
| ENSMUSG00000027479 | Mapre1 | 13589 | protein_coding | 1099.860583 | 0.046950346 | 0.078286647 | -0.599723551 | 0.54869049 | 0.739800355 | 0.046950346 |
| ENSMUSG00000001847 | Rac1 | 19353 | protein_coding | 3106.654676 | -0.046965088 | 0.062873424 | 0.746978369 | 0.455076618 | 0.670515281 | 0.046965088 |
| ENSMUSG00000033916 | Chmp2a | 68953 | protein_coding | 1542.473883 | -0.046969971 | 0.066981267 | 0.701240408 | 0.483152996 | 0.693738201 | 0.046969971 |
| ENSMUSG00000064370 | mt-Cytb | 17711 | protein_coding | 78985.31774 | 0.046981274 | 0.465238557 | -0.100983191 | 0.919563804 | 0.96099713 | 0.046981274 |
| ENSMUSG00000059974 | Ntm | 235106 | protein_coding | 3.659531686 | -0.046988221 | 1.023927474 | 0.045890184 | 0.963397778 | 1 | 0.046988221 |
| ENSMUSG00000054237 | Fra10ac1 | 70567 | protein_coding | 108.6580335 | 0.047016808 | 0.122438481 | -0.384003524 | 0.700975836 | 0.841852283 | 0.047016808 |
| ENSMUSG00000027575 | Arfgap1 | 228998 | protein_coding | 762.3291882 | 0.047054219 | 0.060325396 | -0.7800068 | 0.435386872 | 0.653693864 | 0.047054219 |
| ENSMUSG00000021240 | Abcd4 | 19300 | protein_coding | 130.8221782 | 0.047068275 | 0.125887222 | -0.373892399 | 0.708484372 | 0.847573958 | 0.047068275 |
| ENSMUSG00000024079 | Eif2ak2 | 19106 | protein_coding | 603.5542509 | 0.047081549 | 0.10854648 | -0.433745516 | 0.66447325 | 0.818950015 | 0.047081549 |
| ENSMUSG00000070473 | Cldn3 | 12739 | protein_coding | 7191.229362 | -0.047091336 | 0.100520267 | 0.468476035 | 0.639444205 | 0.803370142 | 0.047091336 |
| ENSMUSG00000033826 | Dnah8 | 13417 | protein_coding | 379.9825071 | -0.0471231 | 0.189340607 | 0.248880053 | 0.803453565 | 0.903013746 | 0.0471231 |
| ENSMUSG00000048756 | Foxo3 | 56484 | protein_coding | 2028.738699 | -0.047124385 | 0.140902787 | 0.334446081 | 0.738042972 | 0.866982505 | 0.047124385 |
| ENSMUSG00000030942 | Thumpd1 | 233802 | protein_coding | 347.9742012 | 0.04717634 | 0.114165543 | -0.41322748 | 0.679439956 | 0.828120806 | 0.04717634 |
| ENSMUSG00000041362 | Shtn1 | 71653 | protein_coding | 610.7565979 | -0.047202744 | 0.068916335 | 0.684928243 | 0.493389203 | 0.699749347 | 0.047202744 |
| ENSMUSG00000070332 | Trim80 | 432613 | protein_coding | 2.828269179 | 0.047217835 | 0.951188288 | -0.049640892 | 0.96040856 | 1 | 0.047217835 |

Supplementary Table 1

| ensembl_gene_id | external_gene_name | entrezgene_id | gene_biotype | baseMean | log2FoldChange | lfcSE | stat | pvalue | padj | absLog2FC |
|---|---|---|---|---|---|---|---|---|---|---|
| ENSMUSG00000106918 | Mrpl33 | 66845 | protein_coding | 276.115692 | 0.047222569 | 0.06221633 | -0.759006024 | 0.447848953 | 0.664930223 | 0.047222569 |
| ENSMUSG00000042508 | Dmtf1 | 23857 | protein_coding | 312.548522 | -0.047225954 | 0.074666458 | 0.632492217 | 0.527065293 | 0.725511065 | 0.047225954 |
| ENSMUSG00000024387 | Csnk2b | 13001 | protein_coding | 824.6334087 | -0.047242881 | 0.056245542 | 0.839939996 | 0.400942031 | 0.622737045 | 0.047242881 |
| ENSMUSG00000020152 | Actr2 | 66713 | protein_coding | 2725.701425 | 0.047256835 | 0.076886386 | -0.614632018 | 0.538797764 | 0.733468492 | 0.047256835 |
| ENSMUSG00000073616 | Cops9 | 66915 | protein_coding | 546.9661948 | -0.047266071 | 0.099600136 | 0.474558292 | 0.63510184 | 0.801105392 | 0.047266071 |
| ENSMUSG00000007817 | Zmiz1 | 328365 | protein_coding | 825.7428542 | -0.047269506 | 0.298233177 | 0.158498483 | 0.874064016 | 0.938083652 | 0.047269506 |
| ENSMUSG00000028385 | Snx30 | 209131 | protein_coding | 210.6678967 | -0.047276657 | 0.126022756 | 0.375143811 | 0.707553516 | 0.846946626 | 0.047276657 |
| ENSMUSG00000033272 | Slc35a4 | 67843 | protein_coding | 695.34763 | 0.047291452 | 0.09024819 | -0.524015513 | 0.60026775 | 0.77680602 | 0.047291452 |
| ENSMUSG00000045664 | Cdc42ep2 | 104252 | protein_coding | 161.0442645 | 0.047293877 | 0.238401883 | -0.198378789 | 0.842748711 | 0.922154269 | 0.047293877 |
| ENSMUSG00000075592 | Nynrin | 277154 | protein_coding | 34.07726207 | 0.047313562 | 0.210928168 | -0.224311253 | 0.822515121 | 0.910171686 | 0.047313562 |
| ENSMUSG00000031988 | Vps26b | 69091 | protein_coding | 608.7298483 | 0.047328472 | 0.075569781 | -0.626288333 | 0.531125837 | 0.728066081 | 0.047328472 |
| ENSMUSG00000040365 | Trim41 | 211007 | protein_coding | 539.0201825 | 0.047344629 | 0.059397091 | -0.797086663 | 0.425400702 | 0.644974953 | 0.047344629 |
| ENSMUSG00000063904 | Dpp3 | 75221 | protein_coding | 1819.678253 | -0.047368559 | 0.06888248 | 0.687672093 | 0.491659297 | 0.698491194 | 0.047368559 |
| ENSMUSG00000044715 | Gskip | 66787 | protein_coding | 175.0804953 | -0.047381531 | 0.102133535 | 0.463917464 | 0.64270688 | 0.805102133 | 0.047381531 |
| ENSMUSG00000047459 | Dynlrb1 | 67068 | protein_coding | 568.1729339 | -0.047396376 | 0.068501278 | 0.69190499 | 0.488996997 | 0.696686775 | 0.047396376 |
| ENSMUSG00000027425 | Kat14 | 228714 | protein_coding | 367.405923 | -0.047428779 | 0.063255964 | 0.749791413 | 0.453380341 | 0.669150322 | 0.047428779 |
| ENSMUSG00000086712 | Mexis | 381524 | lncRNA | 16.95192891 | -0.047478988 | 0.256187489 | 0.185329066 | 0.852970941 | 0.92788168 | 0.047478988 |
| ENSMUSG00000026705 | Klhl20 | 226541 | protein_coding | 131.6826506 | -0.047506375 | 0.091050303 | 0.521759659 | 0.601837681 | 0.77804505 | 0.047506375 |
| ENSMUSG00000038838 | Vars2 | 68915 | protein_coding | 413.874439 | 0.047511513 | 0.086573469 | -0.548799924 | 0.583142763 | 0.765756487 | 0.047511513 |
| ENSMUSG00000024240 | Epc1 | 13831 | protein_coding | 370.5920365 | -0.047516328 | 0.095964679 | 0.495143926 | 0.620498529 | 0.791450739 | 0.047516328 |
| ENSMUSG00000052151 | Plpp2 | 50784 | protein_coding | 2257.524969 | -0.047518007 | 0.102903351 | 0.461773173 | 0.644243991 | 0.806110415 | 0.047518007 |
| ENSMUSG00000017686 | Rhot1 | 59040 | protein_coding | 438.2298768 | 0.047519663 | 0.071174723 | -0.667648019 | 0.504358298 | 0.708596728 | 0.047519663 |
| ENSMUSG00000042354 | Gnl3 | 30877 | protein_coding | 284.1088853 | 0.047521426 | 0.13556444 | -0.350544923 | 0.725929782 | 0.858172551 | 0.047521426 |
| ENSMUSG00000031513 | Leprotl1 | 68192 | protein_coding | 490.885223 | -0.047525192 | 0.077375597 | 0.614214222 | 0.539073777 | 0.733662794 | 0.047525192 |
| ENSMUSG00000024858 | Grk2 | 110355 | protein_coding | 1180.37282 | 0.0475318 | 0.03937735 | -1.207084794 | 0.22739949 | 0.43599843 | 0.0475318 |
| ENSMUSG00000035142 | Nubpl | 76826 | protein_coding | 89.54622824 | -0.04754577 | 0.1396337 | 0.340503543 | 0.733477355 | 0.863523565 | 0.04754577 |
| ENSMUSG00000044894 | Uqcrq | 22272 | protein_coding | 1838.756021 | 0.047548839 | 0.109755295 | -0.433225926 | 0.664850646 | 0.819267308 | 0.047548839 |
| ENSMUSG00000024477 | Pggt1b | 225467 | protein_coding | 615.3453384 | -0.047554787 | 0.09692363 | 0.490641829 | 0.623679796 | 0.793440482 | 0.047554787 |
| ENSMUSG00000032501 | Trib1 | 211770 | protein_coding | 1772.429127 | 0.047563414 | 0.169496104 | -0.28061656 | 0.779004513 | 0.889721317 | 0.047563414 |
| ENSMUSG00000025779 | Ly96 | 17087 | protein_coding | 37.3801427 | -0.047572634 | 0.202783001 | 0.234598728 | 0.814520197 | 0.907746426 | 0.047572634 |
| ENSMUSG00000044857 | Lemd2 | 224640 | protein_coding | 512.7588254 | 0.047582818 | 0.07529911 | -0.631917404 | 0.52744085 | 0.725788994 | 0.047582818 |
| ENSMUSG00000004626 | Stxbp2 | 20911 | protein_coding | 1123.155784 | -0.047605443 | 0.05134158 | 0.927229811 | 0.353807228 | 0.576210312 | 0.047605443 |
| ENSMUSG00000024619 | Cdx1 | 12590 | protein_coding | 3646.628501 | -0.047606128 | 0.088012941 | 0.540899181 | 0.588577075 | 0.769577776 | 0.047606128 |
| ENSMUSG00000052142 | Rasal3 | 320484 | protein_coding | 12.94421837 | -0.047650937 | 0.429755421 | 0.110879199 | 0.911712141 | 0.956997404 | 0.047650937 |
| ENSMUSG00000044221 | Grsf1 | 231413 | protein_coding | 685.5788848 | 0.047695216 | 0.092614448 | -0.514986772 | 0.606562264 | 0.781400205 | 0.047695216 |
| ENSMUSG00000087475 | 4933406I18Rik | 71045 | lncRNA | 1.537798909 | 0.04771295 | 1.032737219 | -0.046200475 | 0.963150464 | 1 | 0.04771295 |
| ENSMUSG00000061288 | Taok3 | 330177 | protein_coding | 872.8916586 | 0.047738536 | 0.109351348 | -0.43656102 | 0.662429737 | 0.817346435 | 0.047738536 |
| ENSMUSG00000055413 | H2-Q5 | 15016 | protein_coding | 6.054764816 | 0.04777577 | 0.613898161 | -0.077823608 | 0.937968367 | 0.970517013 | 0.04777577 |
| ENSMUSG00000028848 | Gpn2 | 100210 | protein_coding | 165.4536528 | -0.047849465 | 0.084226503 | 0.568104613 | 0.569963936 | 0.755650645 | 0.047849465 |

| ensembl_gene_id | external_gene_name | entrezgene_id | gene_biotype | baseMean | log2FoldChange | lfcSE | stat | pvalue | padj | absLog2FC |
|---|---|---|---|---|---|---|---|---|---|---|
| ENSMUSG00000041341 | Atg2b | 76559 | protein_coding | 561.4035487 | -0.047854781 | 0.0564652 | 0.847509275 | 0.396711319 | 0.618855346 | 0.047854781 |
| ENSMUSG00000029715 | Pop7 | 74097 | protein_coding | 183.2419054 | 0.047858231 | 0.096339724 | -0.49676529 | 0.619354575 | 0.790541154 | 0.047858231 |
| ENSMUSG00000030059 | Tmf1 | 232286 | protein_coding | 1552.40419 | -0.047891953 | 0.07101713 | 0.674371836 | 0.500074944 | 0.704748047 | 0.047891953 |
| ENSMUSG00000022193 | Psmb5 | 19173 | protein_coding | 605.938988 | 0.047901874 | 0.110114455 | -0.435018943 | 0.663548678 | 0.818482437 | 0.047901874 |
| ENSMUSG00000023994 | Nfya | 18044 | protein_coding | 342.8471929 | -0.047920122 | 0.082855064 | 0.578360811 | 0.563020546 | 0.750768863 | 0.047920122 |
| ENSMUSG00000054000 | Tusc1 | 69136 | protein_coding | 46.05526213 | 0.047938046 | 0.216902991 | -0.221011458 | 0.825083512 | 0.911460908 | 0.047938046 |
| ENSMUSG00000019518 | Ap4m1 | 11781 | protein_coding | 264.9355203 | -0.047939875 | 0.075945214 | 0.631242877 | 0.527881731 | 0.726026078 | 0.047939875 |
| ENSMUSG00000030364 | Clec2h | 94071 | protein_coding | 3635.819034 | 0.047974793 | 0.129669836 | -0.369976506 | 0.711399996 | 0.849682329 | 0.047974793 |
| ENSMUSG00000031949 | Adat1 | 30947 | protein_coding | 41.73104028 | 0.047995952 | 0.186740954 | -0.257018888 | 0.797164193 | 0.899266991 | 0.047995952 |
| ENSMUSG00000094002 | Gm9866 | 636791 | lncRNA | 5.525183168 | 0.047997993 | 0.954304755 | -0.050296295 | 0.959886277 | 0.980506535 | 0.047997993 |
| ENSMUSG00000005687 | Bcas2 | 68183 | protein_coding | 377.3671205 | 0.048004974 | 0.105176401 | -0.456423428 | 0.648085522 | 0.808650118 | 0.048004974 |
| ENSMUSG00000030822 | Prr14 | 233895 | protein_coding | 389.5537016 | 0.048007443 | 0.109529247 | -0.438307067 | 0.661163704 | 0.816884691 | 0.048007443 |
| ENSMUSG00000021948 | Prkcd | 18753 | protein_coding | 2582.568836 | -0.048011112 | 0.083048691 | 0.578107991 | 0.563191212 | 0.750770075 | 0.048011112 |
| ENSMUSG00000006676 | Usp19 | 71472 | protein_coding | 1235.725008 | -0.048062252 | 0.046519464 | 1.033164347 | 0.301526995 | 0.521787361 | 0.048062252 |
| ENSMUSG00000038982 | Bloc1s5 | 17828 | protein_coding | 152.9268728 | -0.048081644 | 0.102168246 | 0.470612406 | 0.637917546 | 0.802698942 | 0.048081644 |
| ENSMUSG00000088008 | Gm25492 | 115490291 | scaRNA | 32.30073512 | -0.048086371 | 0.248124916 | 0.19379904 | 0.846333239 | 0.924222269 | 0.048086371 |
| ENSMUSG00000073608 | Gal3st2c | 619597 | protein_coding | 423.4594127 | 0.048103757 | 0.458740886 | -0.10486041 | 0.916486574 | 0.959274295 | 0.048103757 |
| ENSMUSG00000037885 | Stk35 | 67333 | protein_coding | 393.0125496 | 0.048134155 | 0.073680278 | -0.653284108 | 0.513573136 | 0.715485319 | 0.048134155 |
| ENSMUSG00000024127 | Prepl | 213760 | protein_coding | 458.21874 | 0.0481348 | 0.061503554 | -0.782634438 | 0.433841812 | 0.652382416 | 0.0481348 |
| ENSMUSG00000031995 | St14 | 19143 | protein_coding | 4321.850461 | 0.048136601 | 0.070857194 | -0.679346704 | 0.496918211 | 0.702914252 | 0.048136601 |
| ENSMUSG00000021039 | Snw1 | 66354 | protein_coding | 520.4535032 | -0.048153245 | 0.092700956 | 0.519447122 | 0.603448978 | 0.779177919 | 0.048153245 |
| ENSMUSG00000039329 | Tex19.1 | 73679 | protein_coding | 1.009463784 | 0.048201861 | 1.386177284 | -0.03477323 | 0.972260567 | 1 | 0.048201861 |
| ENSMUSG00000029269 | Sult1b1 | 56362 | protein_coding | 1841.29105 | -0.048215664 | 0.108809053 | 0.443121804 | 0.657677635 | 0.815005268 | 0.048215664 |
| ENSMUSG00000029557 | Mrm2 | 68017 | protein_coding | 84.93593561 | -0.048217584 | 0.125008559 | 0.38571426 | 0.6997083 | 0.841035219 | 0.048217584 |
| ENSMUSG00000028709 | Mob3c | 100465 | protein_coding | 207.1867882 | 0.048231634 | 0.099068864 | -0.486849571 | 0.626364938 | 0.795381772 | 0.048231634 |
| ENSMUSG00000046556 | Zfp319 | 79233 | protein_coding | 216.8318076 | 0.048247413 | 0.081178147 | -0.59433992 | 0.552284784 | 0.742887588 | 0.048247413 |
| ENSMUSG00000029600 | Rita1 | 100764 | protein_coding | 153.6895203 | -0.048301087 | 0.112496668 | 0.429355712 | 0.667664379 | 0.821350533 | 0.048301087 |
| ENSMUSG00000045598 | Zfp553 | 233887 | protein_coding | 212.0012115 | -0.048301856 | 0.088685933 | 0.544639433 | 0.586001528 | 0.767729412 | 0.048301856 |
| ENSMUSG00000029655 | N4bp2l2 | 381695 | protein_coding | 679.5250793 | -0.048331813 | 0.067066287 | 0.720657349 | 0.47112036 | 0.684193826 | 0.048331813 |
| ENSMUSG00000031701 | Dnaja2 | 56445 | protein_coding | 1887.318486 | -0.048343849 | 0.078377536 | 0.616807466 | 0.537361727 | 0.732906636 | 0.048343849 |
| ENSMUSG00000015653 | Steap2 | 74051 | protein_coding | 242.4230108 | 0.048347362 | 0.102223389 | -0.472957925 | 0.636243198 | 0.801687789 | 0.048347362 |
| ENSMUSG00000042444 | Mindy2 | 235461 | protein_coding | 319.9414456 | -0.048405449 | 0.11997355 | 0.403467668 | 0.686604215 | 0.83310074 | 0.048405449 |
| ENSMUSG00000032369 | Plscr1 | 22038 | protein_coding | 1593.878051 | -0.048432352 | 0.064310214 | 0.753105128 | 0.451386743 | 0.667939129 | 0.048432352 |
| ENSMUSG00000024983 | Vti1a | 53611 | protein_coding | 355.6216008 | 0.048452385 | 0.073365361 | -0.660425903 | 0.508980554 | 0.712495114 | 0.048452385 |
| ENSMUSG00000040865 | Ino80d | 227195 | protein_coding | 635.9744273 | 0.048466959 | 0.090076953 | -0.53806171 | 0.590534446 | 0.77103778 | 0.048466959 |
| ENSMUSG00000026162 | Nhej1 | 75570 | protein_coding | 43.41829067 | 0.048484036 | 0.152807748 | -0.317287811 | 0.751025225 | 0.875081807 | 0.048484036 |
| ENSMUSG00000107623 | Gm10415 | 100038574 | lncRNA | 0.916159086 | 0.048513294 | 2.198166724 | -0.022069888 | 0.982392206 | 1 | 0.048513294 |
| ENSMUSG00000026796 | Niban2 | 227737 | protein_coding | 1469.178001 | -0.048526086 | 0.071203843 | 0.681509364 | 0.495549244 | 0.701603242 | 0.048526086 |
| ENSMUSG00000019302 | Atp6v0a1 | 11975 | protein_coding | 665.8219351 | 0.048549246 | 0.077407679 | -0.627189012 | 0.530535345 | 0.72786082 | 0.048549246 |

Supplementary Table 1

| ensembl_gene_id | external_gene_name | entrezgene_id | gene_biotype | baseMean | log2FoldChange | lfcSE | stat | pvalue | padj | absLog2FC |
|---|---|---|---|---|---|---|---|---|---|---|
| ENSMUSG00000041440 | Gk5 | 235533 | protein_coding | 254.6984928 | -0.048567045 | 0.132868047 | 0.365528405 | 0.714717008 | 0.851944324 | 0.048567045 |
| ENSMUSG00000009681 | Bcr | 110279 | protein_coding | 506.7205755 | -0.048579325 | 0.090894454 | 0.534458629 | 0.593024265 | 0.772538316 | 0.048579325 |
| ENSMUSG00000085664 | Atxn7l1os2 | 75836 | lncRNA | 4.468521523 | -0.048607013 | 0.523158167 | 0.092910741 | 0.925974473 | 0.964583514 | 0.048607013 |
| ENSMUSG00000016940 | Kctd2 | 70382 | protein_coding | 258.0037563 | -0.048639268 | 0.085798717 | 0.566899707 | 0.570782323 | 0.756249943 | 0.048639268 |
| ENSMUSG00000028712 | Cyp4a31 | 666168 | protein_coding | 1.149834687 | 0.048650852 | 2.2086355 | -0.022027561 | 0.982425971 | 1 | 0.048650852 |
| ENSMUSG00000040188 | Scamp2 | 24044 | protein_coding | 2291.616956 | 0.048653997 | 0.066402015 | -0.732718675 | 0.463730032 | 0.677223005 | 0.048653997 |
| ENSMUSG00000015217 | Hmgb3 | 15354 | protein_coding | 203.4057761 | -0.048667171 | 0.129912579 | 0.374614774 | 0.707946984 | 0.847201146 | 0.048667171 |
| ENSMUSG00000028447 | Dctn3 | 53598 | protein_coding | 540.7918818 | -0.048694579 | 0.081784247 | 0.595402918 | 0.551574175 | 0.742355007 | 0.048694579 |
| ENSMUSG00000027739 | Rab33b | 19338 | protein_coding | 184.3075534 | 0.048704502 | 0.109258437 | -0.445773378 | 0.655760954 | 0.813372119 | 0.048704502 |
| ENSMUSG00000024290 | Rock1 | 19877 | protein_coding | 1396.35145 | 0.048708981 | 0.068869847 | -0.70726135 | 0.479404079 | 0.690650055 | 0.048708981 |
| ENSMUSG00000038253 | Hoxa5 | 15402 | protein_coding | 84.58990411 | 0.048837625 | 0.11921904 | -0.409646183 | 0.682065514 | 0.830017162 | 0.048837625 |
| ENSMUSG00000051184 | Zfp524 | 66056 | protein_coding | 61.92287201 | -0.048851594 | 0.142976796 | 0.341674977 | 0.732595505 | 0.862854445 | 0.048851594 |
| ENSMUSG00000015290 | Ubl4a | 27643 | protein_coding | 474.3453339 | 0.048876523 | 0.055385007 | -0.88248653 | 0.377513761 | 0.599649878 | 0.048876523 |
| ENSMUSG00000017132 | Cyth1 | 19157 | protein_coding | 668.8086033 | -0.048885926 | 0.080272293 | 0.609001242 | 0.542523616 | 0.736113301 | 0.048885926 |
| ENSMUSG00000038496 | Slc19a3 | 80721 | protein_coding | 2.434640418 | -0.048924273 | 0.683765647 | 0.07155123 | 0.942959054 | 1 | 0.048924273 |
| ENSMUSG00000032712 | Resf1 | 67246 | protein_coding | 569.4000688 | 0.048958114 | 0.068056224 | -0.71937747 | 0.471908374 | 0.684625749 | 0.048958114 |
| ENSMUSG00000029020 | Mfn2 | 170731 | protein_coding | 1369.714301 | 0.048967107 | 0.075091636 | -0.652098021 | 0.514337941 | 0.715988936 | 0.048967107 |
| ENSMUSG00000099045 | Mir6374 | 102465194 | miRNA | 1.129438093 | 0.048971346 | 0.994435212 | -0.049245386 | 0.960723742 | 1 | 0.048971346 |
| ENSMUSG00000021009 | Ptpn21 | 24000 | protein_coding | 133.5032855 | 0.048973918 | 0.112051114 | -0.437067655 | 0.662062284 | 0.817198338 | 0.048973918 |
| ENSMUSG00000074364 | Ehd2 | 259300 | protein_coding | 7.111448387 | 0.048993582 | 0.790049546 | -0.062013303 | 0.950552239 | 0.975869522 | 0.048993582 |
| ENSMUSG00000048497 | Mmgt2 | 216829 | protein_coding | 218.2901072 | 0.049019681 | 0.093078879 | -0.526646668 | 0.598438978 | 0.776081458 | 0.049019681 |
| ENSMUSG00000024350 | Dnajc18 | 76594 | protein_coding | 90.07851999 | 0.049033284 | 0.164065271 | -0.298864495 | 0.765043439 | 0.882319111 | 0.049033284 |
| ENSMUSG00000113450 | Zfp935 | 71508 | protein_coding | 31.02981101 | 0.04903469 | 0.198574903 | -0.246932966 | 0.804960103 | 0.903862864 | 0.04903469 |
| ENSMUSG00000035595 | Fam174c | 69770 | protein_coding | 87.33849133 | -0.049157255 | 0.125612374 | 0.391340866 | 0.695545296 | 0.838533729 | 0.049157255 |
| ENSMUSG00000018501 | Ncor1 | 20185 | protein_coding | 2917.33458 | -0.049197669 | 0.055492641 | 0.886562038 | 0.37531474 | 0.597565331 | 0.049197669 |
| ENSMUSG00000031864 | Ints10 | 70885 | protein_coding | 476.6047163 | 0.049234435 | 0.087308956 | -0.563910476 | 0.57281506 | 0.758091673 | 0.049234435 |
| ENSMUSG00000021488 | Nsd1 | 18193 | protein_coding | 2877.124776 | 0.049273017 | 0.060103803 | -0.819798656 | 0.412330897 | 0.632329199 | 0.049273017 |
| ENSMUSG00000021302 | Ggps1 | 14593 | protein_coding | 328.0285971 | -0.049291147 | 0.071653136 | 0.687913322 | 0.491507367 | 0.698491194 | 0.049291147 |
| ENSMUSG00000017707 | Serinc3 | 26943 | protein_coding | 4379.961868 | -0.049294461 | 0.144548816 | 0.341022934 | 0.733086316 | 0.86324782 | 0.049294461 |
| ENSMUSG00000039671 | Zmynd8 | 228880 | protein_coding | 1037.212499 | -0.049357596 | 0.071101014 | 0.694189759 | 0.487563214 | 0.696047947 | 0.049357596 |
| ENSMUSG00000034544 | Rsrc1 | 66880 | protein_coding | 226.4029052 | -0.049370804 | 0.083248465 | 0.593053632 | 0.55314526 | 0.743572273 | 0.049370804 |
| ENSMUSG00000027714 | Exosc9 | 50911 | protein_coding | 231.8620596 | -0.04939278 | 0.094985533 | 0.520003183 | 0.603061357 | 0.778956218 | 0.04939278 |
| ENSMUSG00000084786 | Ubl5 | 66177 | protein_coding | 982.2858235 | -0.049398381 | 0.057797433 | 0.854681222 | 0.392727649 | 0.615439779 | 0.049398381 |
| ENSMUSG00000053214 | Gm9899 | 100502829 | lncRNA | 2.710655565 | 0.049421649 | 1.282283997 | -0.03854189 | 0.969255633 | 1 | 0.049421649 |
| ENSMUSG00000021972 | Hmbox1 | 219150 | protein_coding | 262.7617553 | -0.049437431 | 0.11386734 | 0.434166914 | 0.664167237 | 0.818817302 | 0.049437431 |
| ENSMUSG00000033819 | Ppp1r16a | 73062 | protein_coding | 739.9406805 | 0.049467639 | 0.084643864 | -0.584420844 | 0.558937206 | 0.747694534 | 0.049467639 |
| ENSMUSG00000050471 | Fam118b | 109229 | protein_coding | 334.9936252 | -0.049495307 | 0.073423898 | 0.674103504 | 0.500245512 | 0.704868121 | 0.049495307 |
| ENSMUSG00000034620 | Rxylt1 | 216395 | protein_coding | 226.5558125 | -0.049503841 | 0.100677895 | 0.49170517 | 0.622927783 | 0.793281643 | 0.049503841 |
| ENSMUSG00000070814 | Zswim9 | 321008 | protein_coding | 59.71020104 | -0.049512787 | 0.161145928 | 0.307254349 | 0.758649781 | 0.878912418 | 0.049512787 |

| ensembl_gene_id | external_gene_name | entrezgene_id | gene_biotype | baseMean | log2FoldChange | lfcSE | stat | pvalue | padj | absLog2FC |
|---|---|---|---|---|---|---|---|---|---|---|
| ENSMUSG00000024830 | Rps6kb2 | 58988 | protein_coding | 189.9348894 | 0.049541332 | 0.115306751 | -0.429648149 | 0.667451607 | 0.821333519 | 0.049541332 |
| ENSMUSG00000050394 | Armcx6 | 278097 | protein_coding | 16.62218778 | 0.049614294 | 0.263969584 | -0.187954585 | 0.850912252 | 0.927277612 | 0.049614294 |
| ENSMUSG00000037996 | Slc24a2 | 76376 | protein_coding | 2.901800251 | 0.049679532 | 2.17800628 | -0.022809637 | 0.981802121 | 1 | 0.049679532 |
| ENSMUSG00000008999 | Bmp7 | 12162 | protein_coding | 28.36711592 | -0.049705675 | 0.309074267 | 0.160821136 | 0.872234278 | 0.937028043 | 0.049705675 |
| ENSMUSG00000087819 | Gm25117 | 115487522 | snoRNA | 235.3715276 | -0.049743055 | 0.210146398 | 0.236706674 | 0.812884357 | 0.907147741 | 0.049743055 |
| ENSMUSG00000010290 | AI597479 | 98404 | protein_coding | 214.3396101 | 0.049754835 | 0.122247546 | -0.407000681 | 0.684007488 | 0.831360834 | 0.049754835 |
| ENSMUSG00000025077 | Dclre1a | 55947 | protein_coding | 210.0849668 | 0.049756818 | 0.081785332 | -0.608383144 | 0.542933389 | 0.736235501 | 0.049756818 |
| ENSMUSG00000059316 | Slc27a4 | 26569 | protein_coding | 1489.337256 | 0.049761193 | 0.12624924 | -0.39415044 | 0.693469974 | 0.837254214 | 0.049761193 |
| ENSMUSG00000035305 | Ror1 | 26563 | protein_coding | 111.9186615 | -0.049783223 | 0.163842644 | 0.303847774 | 0.761243868 | 0.880273177 | 0.049783223 |
| ENSMUSG00000006906 | Stambp | 70527 | protein_coding | 132.016982 | 0.049829363 | 0.116554059 | -0.427521471 | 0.66899955 | 0.82238042 | 0.049829363 |
| ENSMUSG00000020090 | Npffr1 | 237362 | protein_coding | 4.396552229 | -0.049839244 | 0.662718111 | 0.075204289 | 0.940052172 | 0.971404891 | 0.049839244 |
| ENSMUSG00000031540 | Kat6a | 244349 | protein_coding | 968.3435318 | 0.049874132 | 0.09184231 | -0.543040912 | 0.587101634 | 0.768458962 | 0.049874132 |
| ENSMUSG00000026608 | Kctd3 | 226823 | protein_coding | 664.6157538 | 0.049878576 | 0.060575661 | -0.82340951 | 0.410275157 | 0.630552252 | 0.049878576 |
| ENSMUSG00000020413 | Hus1 | 15574 | protein_coding | 94.30768451 | -0.049886318 | 0.144966352 | 0.344123427 | 0.730753465 | 0.861267215 | 0.049886318 |
| ENSMUSG00000051506 | Wdfy4 | 545030 | protein_coding | 10.2830043 | 0.049891097 | 0.79278456 | -0.062931469 | 0.949821076 | 0.975869522 | 0.049891097 |
| ENSMUSG00000051355 | Commd1 | 17846 | protein_coding | 290.6588638 | -0.049934288 | 0.097288245 | 0.513261268 | 0.607768574 | 0.782282948 | 0.049934288 |
| ENSMUSG00000027349 | Fam98b | 68215 | protein_coding | 129.2771557 | 0.049942596 | 0.114688056 | -0.435464662 | 0.663225184 | 0.81814452 | 0.049942596 |
| ENSMUSG00000040661 | Rad54l2 | 81000 | protein_coding | 446.1354644 | -0.049953671 | 0.054580207 | 0.915234182 | 0.360068704 | 0.5825209 | 0.049953671 |
| ENSMUSG00000047180 | Neurl3 | 214854 | protein_coding | 1669.371538 | -0.050039541 | 0.11373894 | 0.439950824 | 0.659972724 | 0.816323585 | 0.050039541 |
| ENSMUSG00000029204 | Rhoh | 74734 | protein_coding | 11.16337664 | 0.050042837 | 0.686994465 | -0.072843145 | 0.941930937 | 0.971883559 | 0.050042837 |
| ENSMUSG00000024013 | Fgd2 | 26382 | protein_coding | 22.50262092 | -0.05004539 | 0.283199611 | 0.176714192 | 0.859732893 | 0.931236324 | 0.05004539 |
| ENSMUSG00000056492 | Adgrf5 | 224792 | protein_coding | 28.0662815 | -0.050063025 | 0.285375063 | 0.175428872 | 0.860742656 | 0.931474719 | 0.050063025 |
| ENSMUSG00000022841 | Ap2m1 | 11773 | protein_coding | 1383.215582 | -0.050083211 | 0.084787626 | 0.5906901 | 0.554728083 | 0.744703672 | 0.050083211 |
| ENSMUSG00000027082 | Tfpi | 21788 | protein_coding | 53.03424009 | 0.05021247 | 0.224589447 | -0.223574487 | 0.823088417 | 0.910297568 | 0.05021247 |
| ENSMUSG00000041740 | Rnf10 | 50849 | protein_coding | 1616.161243 | 0.050269601 | 0.085135906 | -0.590462988 | 0.554880293 | 0.744703672 | 0.050269601 |
| ENSMUSG00000019362 | D8Ertd738e | 101966 | protein_coding | 741.5329195 | -0.050349732 | 0.04875775 | 1.032650852 | 0.301767321 | 0.522023104 | 0.050349732 |
| ENSMUSG00000003527 | Ess2 | 27886 | protein_coding | 173.0817792 | 0.05037754 | 0.088650465 | -0.568271584 | 0.569850571 | 0.755621674 | 0.05037754 |
| ENSMUSG00000062432 | Cyp26c1 | 546726 | protein_coding | 0.899227057 | -0.050403301 | 1.25065973 | 0.04030137 | 0.967852861 | 1 | 0.050403301 |
| ENSMUSG00000030660 | Pik3c2a | 18704 | protein_coding | 1443.56607 | 0.050436341 | 0.060611807 | -0.832120736 | 0.405340799 | 0.62645534 | 0.050436341 |
| ENSMUSG00000025785 | Exosc7 | 66446 | protein_coding | 268.9503236 | 0.050447249 | 0.118353738 | -0.426241286 | 0.669932036 | 0.822722373 | 0.050447249 |
| ENSMUSG00000027439 | Gzf1 | 74533 | protein_coding | 348.1795133 | -0.050469366 | 0.078905327 | 0.639619256 | 0.522420161 | 0.721840784 | 0.050469366 |
| ENSMUSG00000055334 | Snupn | 66069 | protein_coding | 102.2881572 | -0.050512862 | 0.114321921 | 0.44184756 | 0.65859952 | 0.815790015 | 0.050512862 |
| ENSMUSG00000058301 | Upf1 | 19704 | protein_coding | 1315.148323 | 0.050519352 | 0.072441536 | -0.697381017 | 0.485564378 | 0.695380179 | 0.050519352 |
| ENSMUSG00000063800 | Prpf38a | 230596 | protein_coding | 429.3935735 | 0.050528735 | 0.078683043 | -0.642180746 | 0.520755833 | 0.720610174 | 0.050528735 |
| ENSMUSG00000016946 | Kctd5 | 69259 | protein_coding | 804.6743852 | 0.050565555 | 0.07122144 | -0.709976591 | 0.477718653 | 0.68930064 | 0.050565555 |
| ENSMUSG00000002064 | Sdf2 | 20316 | protein_coding | 590.5449718 | -0.050578465 | 0.083403298 | 0.606432432 | 0.544227638 | 0.736796542 | 0.050578465 |
| ENSMUSG00000024816 | Frmd8 | 67457 | protein_coding | 518.7423883 | -0.0505801 | 0.085560386 | 0.591162601 | 0.554411479 | 0.744656458 | 0.0505801 |
| ENSMUSG00000058239 | Usf2 | 22282 | protein_coding | 243.0113582 | -0.050624549 | 0.106165172 | 0.476847043 | 0.633471044 | 0.800270871 | 0.050624549 |
| ENSMUSG00000030313 | Dennd5b | 320560 | protein_coding | 255.8938271 | 0.050652071 | 0.102537378 | -0.493986406 | 0.62131578 | 0.792160606 | 0.050652071 |

| ensembl_gene_id | external_gene_name | entrezgene_id | gene_biotype | baseMean | log2FoldChange | lfcSE | stat | pvalue | padj | absLog2FC |
|---|---|---|---|---|---|---|---|---|---|---|
| ENSMUSG00000042672 | Dcst1 | 77772 | protein_coding | 24.96985472 | -0.050674431 | 0.319448359 | 0.158631057 | 0.873959559 | 0.938083652 | 0.050674431 |
| ENSMUSG00000071862 | Lrrtm2 | 107065 | protein_coding | 43.79545557 | 0.05072639 | 0.300963306 | -0.168546759 | 0.866153161 | 0.93371978 | 0.05072639 |
| ENSMUSG00000023845 | Lnpep | 240028 | protein_coding | 2331.272301 | -0.050753731 | 0.095486656 | 0.531526952 | 0.595053669 | 0.773528957 | 0.050753731 |
| ENSMUSG00000018537 | Pcgf2 | 22658 | protein_coding | 107.572164 | 0.050762978 | 0.226030454 | -0.224584686 | 0.82230238 | 0.910171686 | 0.050762978 |
| ENSMUSG00000021866 | Anxa11 | 11744 | protein_coding | 1329.199054 | 0.050788497 | 0.057109274 | -0.889321357 | 0.373830397 | 0.595844868 | 0.050788497 |
| ENSMUSG00000018334 | Ksr1 | 16706 | protein_coding | 356.9210958 | 0.050798661 | 0.183343821 | -0.277067753 | 0.781728082 | 0.890621398 | 0.050798661 |
| ENSMUSG00000000631 | Myo18a | 360013 | protein_coding | 3327.289707 | 0.050827413 | 0.145493298 | -0.349345391 | 0.726830025 | 0.85879931 | 0.050827413 |
| ENSMUSG00000027215 | Cd82 | 12521 | protein_coding | 1373.168635 | -0.050829774 | 0.070544599 | 0.72053389 | 0.471196341 | 0.684193826 | 0.050829774 |
| ENSMUSG00000021786 | Oxsm | 71147 | protein_coding | 355.5899188 | -0.050834415 | 0.105671892 | 0.481059003 | 0.630474565 | 0.79831751 | 0.050834415 |
| ENSMUSG00000089911 | Mfsd14a | 15247 | protein_coding | 508.559947 | 0.050849037 | 0.08650516 | -0.58781507 | 0.556656427 | 0.745972916 | 0.050849037 |
| ENSMUSG00000036977 | Anapc10 | 68999 | protein_coding | 92.0936508 | 0.050857918 | 0.121620163 | -0.418170119 | 0.675822741 | 0.826400352 | 0.050857918 |
| ENSMUSG00000011831 | Evi5 | 14020 | protein_coding | 405.7268598 | -0.050918729 | 0.076273948 | 0.667576945 | 0.504403679 | 0.708596728 | 0.050918729 |
| ENSMUSG00000005886 | Ncoa2 | 17978 | protein_coding | 837.3431736 | 0.050928396 | 0.071569439 | -0.711594176 | 0.476716115 | 0.68833522 | 0.050928396 |
| ENSMUSG00000017843 | Ppp2r5c | 26931 | protein_coding | 1406.423931 | 0.050960305 | 0.054350986 | -0.937615096 | 0.348442251 | 0.570284968 | 0.050960305 |
| ENSMUSG00000053110 | Yap1 | 22601 | protein_coding | 686.0032134 | -0.050967116 | 0.075047066 | 0.679135355 | 0.497052103 | 0.702914252 | 0.050967116 |
| ENSMUSG00000006005 | Tpr | 108989 | protein_coding | 2013.334769 | -0.051009976 | 0.092084009 | 0.553950423 | 0.579612775 | 0.762991261 | 0.051009976 |
| ENSMUSG00000026080 | Chst10 | 98388 | protein_coding | 7.787626272 | -0.051023811 | 0.527235681 | 0.096776095 | 0.922904208 | 0.96266044 | 0.051023811 |
| ENSMUSG00000018752 | Tnfsfm13 | 619441 | protein_coding | 9.20502018 | 0.051049192 | 0.317454192 | -0.160808058 | 0.872244578 | 0.937028043 | 0.051049192 |
| ENSMUSG00000072770 | Acrbp | 54137 | protein_coding | 32.18799446 | -0.051060815 | 0.307658733 | 0.16596576 | 0.868183896 | 0.934688628 | 0.051060815 |
| ENSMUSG00000058173 | Smco4 | 170748 | protein_coding | 158.824437 | 0.051110077 | 0.163553 | -0.312498559 | 0.754661658 | 0.876856731 | 0.051110077 |
| ENSMUSG00000024014 | Pim1 | 18712 | protein_coding | 181.494518 | -0.051136138 | 0.160034076 | 0.319532809 | 0.749322516 | 0.874044129 | 0.051136138 |
| ENSMUSG00000043015 | Nemp2 | 227094 | protein_coding | 49.53947012 | -0.051151355 | 0.189970695 | 0.269259189 | 0.787730235 | 0.89369175 | 0.051151355 |
| ENSMUSG00000029084 | Cd38 | 12494 | protein_coding | 2088.509137 | -0.051182396 | 0.139010393 | 0.368191149 | 0.71273071 | 0.850681912 | 0.051182396 |
| ENSMUSG00000074794 | Arrdc3 | 105171 | protein_coding | 893.8034142 | -0.05118285 | 0.137572203 | 0.372043543 | 0.70986043 | 0.84860541 | 0.05118285 |
| ENSMUSG00000029569 | Tmem168 | 101118 | protein_coding | 958.3111973 | -0.051224065 | 0.154884521 | 0.330724234 | 0.740852795 | 0.868861804 | 0.051224065 |
| ENSMUSG00000041235 | Chd7 | 320790 | protein_coding | 1233.397455 | -0.051239152 | 0.113479386 | 0.451528281 | 0.651608847 | 0.810851996 | 0.051239152 |
| ENSMUSG00000059588 | Calcrl | 54598 | protein_coding | 31.13093386 | 0.05130132 | 0.261626924 | -0.19608578 | 0.844543026 | 0.923599766 | 0.05130132 |
| ENSMUSG00000020349 | Ppp2ca | 19052 | protein_coding | 3190.806009 | 0.051309242 | 0.054699161 | -0.938026122 | 0.348230986 | 0.570165229 | 0.051309242 |
| ENSMUSG00000045538 | Ddx28 | 71986 | protein_coding | 110.7986286 | -0.05132228 | 0.109055238 | 0.47060812 | 0.637920608 | 0.802698942 | 0.05132228 |
| ENSMUSG00000091491 | Vmn2r97 | 627367 | protein_coding | 3.072694531 | 0.051360728 | 1.502732605 | -0.034178221 | 0.972735033 | 1 | 0.051360728 |
| ENSMUSG00000046605 | B3gntl1 | 210004 | protein_coding | 90.86506362 | 0.051363241 | 0.120654052 | -0.425706727 | 0.67032156 | 0.82296411 | 0.051363241 |
| ENSMUSG00000021911 | Parg | 26430 | protein_coding | 651.8892626 | 0.051424391 | 0.071517548 | -0.719045785 | 0.47211271 | 0.684706793 | 0.051424391 |
| ENSMUSG00000027677 | Ttc14 | 67120 | protein_coding | 396.3077821 | 0.051436245 | 0.091356547 | -0.563027467 | 0.57341618 | 0.758644032 | 0.051436245 |
| ENSMUSG00000059714 | Flot1 | 14251 | protein_coding | 366.4544497 | -0.05144612 | 0.101042782 | 0.509151854 | 0.610645788 | 0.784213744 | 0.05144612 |
| ENSMUSG00000019132 | BC005537 | 79555 | protein_coding | 1472.474649 | -0.051463547 | 0.06572427 | 0.783021969 | 0.433614211 | 0.652196761 | 0.051463547 |
| ENSMUSG00000096094 | A630095N17Rik | 381272 | protein_coding | 2.750910852 | 0.051543989 | 0.843127826 | -0.061134253 | 0.951252291 | 1 | 0.051543989 |
| ENSMUSG00000022696 | Sidt1 | 320007 | protein_coding | 459.499791 | -0.051640054 | 0.138015894 | 0.374160193 | 0.708285138 | 0.847451753 | 0.051640054 |
| ENSMUSG00000014791 | Elmo3 | 234683 | protein_coding | 881.8542105 | -0.051734674 | 0.064530998 | 0.80170267 | 0.422724971 | 0.642569972 | 0.051734674 |
| ENSMUSG00000049878 | Rlf | 109263 | protein_coding | 461.0060998 | 0.051741763 | 0.070202619 | -0.737034656 | 0.461101275 | 0.674995696 | 0.051741763 |

| ensembl_gene_id | external_gene_name | entrezgene_id | gene_biotype | baseMean | log2FoldChange | lfcSE | stat | pvalue | padj | absLog2FC |
|---|---|---|---|---|---|---|---|---|---|---|
| ENSMUSG00000033735 | Spr | 20751 | protein_coding | 704.6207135 | -0.051781055 | 0.079930353 | 0.647827176 | 0.517096736 | 0.717711842 | 0.051781055 |
| ENSMUSG00000003731 | Kpna6 | 16650 | protein_coding | 822.9195696 | -0.051805423 | 0.073580776 | 0.704061934 | 0.481394205 | 0.692354022 | 0.051805423 |
| ENSMUSG00000005681 | Apoa2 | 11807 | protein_coding | 6.008385744 | -0.051807693 | 0.477423468 | 0.108515179 | 0.91358704 | 0.957921881 | 0.051807693 |
| ENSMUSG00000027678 | Ncoa3 | 17979 | protein_coding | 898.9334698 | 0.051851135 | 0.060013976 | -0.863984323 | 0.387596502 | 0.610061822 | 0.051851135 |
| ENSMUSG00000024588 | Fech | 14151 | protein_coding | 569.1961052 | -0.051868586 | 0.079192628 | 0.654967354 | 0.512488773 | 0.714893745 | 0.051868586 |
| ENSMUSG00000021096 | Ppm1a | 19042 | protein_coding | 1290.19638 | -0.051908366 | 0.050408545 | 1.029753298 | 0.303125829 | 0.523653558 | 0.051908366 |
| ENSMUSG00000027004 | Frzb | 20378 | protein_coding | 1.05896454 | 0.051950003 | 2.513108405 | -0.020671613 | 0.983507614 | 1 | 0.051950003 |
| ENSMUSG00000032388 | Spg21 | 27965 | protein_coding | 383.8469005 | -0.051953249 | 0.090000048 | 0.57725802 | 0.563765168 | 0.751050176 | 0.051953249 |
| ENSMUSG00000037447 | Arid5a | 214855 | protein_coding | 90.20094355 | -0.051968271 | 0.222790545 | 0.233260667 | 0.815559 | 0.908352419 | 0.051968271 |
| ENSMUSG00000050813 | Or2av9 | 257932 | protein_coding | 1.478565516 | 0.051989979 | 2.51790858 | -0.020648081 | 0.983526386 | 1 | 0.051989979 |
| ENSMUSG00000070867 | Trabd2b | 666048 | protein_coding | 18.74676971 | -0.052010479 | 0.322049436 | 0.161498432 | 0.871700846 | 0.936825965 | 0.052010479 |
| ENSMUSG00000028141 | Oaz3 | 53814 | protein_coding | 1.717763374 | -0.052026135 | 0.858866634 | 0.060575336 | 0.951697416 | 1 | 0.052026135 |
| ENSMUSG00000027551 | Zfp64 | 22722 | protein_coding | 269.1510409 | -0.052080748 | 0.097668981 | 0.533237342 | 0.593869295 | 0.773008398 | 0.052080748 |
| ENSMUSG00000052363 | Zdhhc19 | 245308 | protein_coding | 3.924516379 | -0.052095614 | 1.097116509 | 0.047484122 | 0.962127385 | 1 | 0.052095614 |
| ENSMUSG00000058216 | Gstp3 | 225884 | protein_coding | 42.16899432 | -0.052112435 | 0.215018435 | 0.242362636 | 0.808499183 | 0.905558567 | 0.052112435 |
| ENSMUSG00000019849 | Prep | 19072 | protein_coding | 581.0961888 | 0.052114771 | 0.087357368 | -0.596569846 | 0.550794608 | 0.741668474 | 0.052114771 |
| ENSMUSG00000068130 | Zfp442 | 668923 | protein_coding | 39.59916297 | -0.052176704 | 0.164538305 | 0.317109773 | 0.751160309 | 0.875081807 | 0.052176704 |
| ENSMUSG00000004815 | Dgkq | 110524 | protein_coding | 466.345769 | -0.052187879 | 0.169750151 | 0.307439369 | 0.758508966 | 0.878895387 | 0.052187879 |
| ENSMUSG00000023027 | Atf1 | 11908 | protein_coding | 244.49034 | 0.052196441 | 0.084395253 | -0.618476033 | 0.536261592 | 0.732309926 | 0.052196441 |
| ENSMUSG00000035547 | Capn5 | 12337 | protein_coding | 1552.556781 | 0.052210473 | 0.094206873 | -0.554210868 | 0.579434542 | 0.76287826 | 0.052210473 |
| ENSMUSG00000024109 | Nrxn1 | 18189 | protein_coding | 8.614002926 | -0.05221601 | 0.691191659 | 0.075544908 | 0.939781169 | 0.97136798 | 0.05221601 |
| ENSMUSG00000099687 | Or51b6b | 259126 | protein_coding | 0.881797363 | -0.052227058 | 1.430423923 | 0.036511595 | 0.970874434 | 1 | 0.052227058 |
| ENSMUSG00000034932 | Mrpl54 | 66047 | protein_coding | 502.5721588 | -0.052246809 | 0.091264602 | 0.572476166 | 0.566999428 | 0.753307251 | 0.052246809 |
| ENSMUSG00000090035 | Galnt4 | 14426 | protein_coding | 3428.352042 | -0.0522641 | 0.087659017 | 0.59622047 | 0.551027952 | 0.741922186 | 0.0522641 |
| ENSMUSG00000035764 | Fbxo45 | 268882 | protein_coding | 264.0342665 | 0.052315278 | 0.126880526 | -0.412319206 | 0.680105473 | 0.828386324 | 0.052315278 |
| ENSMUSG00000044149 | Nkrf | 77286 | protein_coding | 79.88296582 | 0.052359824 | 0.168907223 | -0.309991622 | 0.756567328 | 0.877877519 | 0.052359824 |
| ENSMUSG00000032232 | Cgnl1 | 68178 | protein_coding | 608.7705343 | -0.052398453 | 0.116718879 | 0.448928684 | 0.653483103 | 0.812010532 | 0.052398453 |
| ENSMUSG00000054021 | Sirt5 | 68346 | protein_coding | 119.0970619 | -0.052436012 | 0.133835938 | 0.391793209 | 0.695211013 | 0.838326567 | 0.052436012 |
| ENSMUSG00000022223 | Sdr39u1 | 654795 | protein_coding | 74.95106045 | 0.052449785 | 0.155155131 | -0.338047378 | 0.735327484 | 0.865203089 | 0.052449785 |
| ENSMUSG00000025813 | Homer2 | 26557 | protein_coding | 382.7662762 | -0.052475194 | 0.102131787 | 0.513798843 | 0.607392637 | 0.781981914 | 0.052475194 |
| ENSMUSG00000078451 | Ppil6 | 73075 | protein_coding | 9.486774504 | -0.052541295 | 0.349144195 | 0.150485949 | 0.880381236 | 0.941010727 | 0.052541295 |
| ENSMUSG00000054582 | Pabpc1l | 381404 | protein_coding | 5.978271588 | 0.05255087 | 0.765171572 | -0.068678545 | 0.945245497 | 0.97354273 | 0.05255087 |
| ENSMUSG00000028563 | Tm2d1 | 94043 | protein_coding | 165.1264175 | -0.052551165 | 0.098447045 | 0.53380135 | 0.59347898 | 0.772683055 | 0.052551165 |
| ENSMUSG00000045659 | Plekha7 | 233765 | protein_coding | 1094.297527 | 0.05258218 | 0.116060367 | -0.45305888 | 0.650506344 | 0.810140405 | 0.05258218 |
| ENSMUSG00000024969 | Mark2 | 13728 | protein_coding | 1181.813416 | 0.052589553 | 0.078935438 | -0.666235022 | 0.505260891 | 0.709332513 | 0.052589553 |
| ENSMUSG00000074006 | Omp | 18378 | protein_coding | 96.66974073 | -0.052596494 | 0.167571352 | 0.313875212 | 0.753615818 | 0.876528151 | 0.052596494 |
| ENSMUSG00000033439 | Trmt13 | 229780 | protein_coding | 51.79583712 | 0.052608674 | 0.148404474 | -0.3544952 | 0.722967801 | 0.856549354 | 0.052608674 |
| ENSMUSG00000023353 | Agap3 | 213990 | protein_coding | 459.7700054 | 0.05264194 | 0.084698333 | -0.621522738 | 0.534255738 | 0.730364909 | 0.05264194 |
| ENSMUSG00000022286 | Grhl2 | 252973 | protein_coding | 858.7456566 | 0.052750351 | 0.064511071 | -0.817694543 | 0.413531627 | 0.633669972 | 0.052750351 |

| ensembl_gene_id | external_gene_name | entrezgene_id | gene_biotype | baseMean | log2FoldChange | lfcSE | stat | pvalue | padj | absLog2FC |
|---|---|---|---|---|---|---|---|---|---|---|
| ENSMUSG00000036707 | Cab39 | 12283 | protein_coding | 3787.699087 | 0.052775941 | 0.060166226 | -0.877168878 | 0.380394918 | 0.60257474 | 0.052775941 |
| ENSMUSG00000032350 | Gclc | 14629 | protein_coding | 1021.596245 | 0.052777617 | 0.100461112 | -0.525353703 | 0.599337331 | 0.77652216 | 0.052777617 |
| ENSMUSG00000038866 | Zcchc2 | 227449 | protein_coding | 408.3779774 | 0.052850138 | 0.101634139 | -0.520003797 | 0.603060929 | 0.778956218 | 0.052850138 |
| ENSMUSG00000038007 | Acer2 | 230379 | protein_coding | 88.43414376 | 0.052852237 | 0.114083069 | -0.463278532 | 0.643164732 | 0.80530915 | 0.052852237 |
| ENSMUSG00000017288 | Vps53 | 68299 | protein_coding | 687.6329507 | 0.052866334 | 0.05888638 | -0.897768443 | 0.369309012 | 0.591883639 | 0.052866334 |
| ENSMUSG00000027456 | Sdcbp2 | 228765 | protein_coding | 2405.05019 | 0.052874382 | 0.137161612 | -0.385489656 | 0.699874669 | 0.841086565 | 0.052874382 |
| ENSMUSG00000030905 | Crym | 12971 | protein_coding | 47.86465347 | 0.052881732 | 0.180660824 | -0.292712778 | 0.769741702 | 0.884525396 | 0.052881732 |
| ENSMUSG00000077167 | Gm24119 | 115487534 | snoRNA | 27.7203354 | -0.052895328 | 0.264614777 | 0.199895594 | 0.841562237 | 0.92158767 | 0.052895328 |
| ENSMUSG00000067336 | Bmpr2 | 12168 | protein_coding | 646.0065238 | -0.05291299 | 0.090864848 | 0.582326291 | 0.56034692 | 0.748973617 | 0.05291299 |
| ENSMUSG00000037111 | Setd7 | 73251 | protein_coding | 446.4293261 | -0.052971379 | 0.124651623 | 0.424955392 | 0.670869193 | 0.823391591 | 0.052971379 |
| ENSMUSG00000032766 | Gng11 | 66066 | protein_coding | 200.0483134 | 0.052986986 | 0.091568379 | -0.578660306 | 0.562818404 | 0.75067522 | 0.052986986 |
| ENSMUSG00000036867 | Smad6 | 17130 | protein_coding | 342.3636243 | -0.052997976 | 0.187278963 | 0.282989477 | 0.777184903 | 0.888773967 | 0.052997976 |
| ENSMUSG00000026096 | Osgepl1 | 72085 | protein_coding | 73.34148271 | 0.053003231 | 0.145647246 | -0.363915092 | 0.715921414 | 0.852575063 | 0.053003231 |
| ENSMUSG00000015016 | Acsf3 | 257633 | protein_coding | 295.4306026 | -0.053005182 | 0.091225864 | 0.581032395 | 0.56121862 | 0.749713986 | 0.053005182 |
| ENSMUSG00000046707 | Csnk2a2 | 13000 | protein_coding | 292.3057116 | -0.053029854 | 0.073863384 | 0.717945083 | 0.472791149 | 0.685206817 | 0.053029854 |
| ENSMUSG00000020863 | Luc7l3 | 67684 | protein_coding | 690.7410512 | 0.053031482 | 0.075227323 | -0.704949743 | 0.480841513 | 0.691751611 | 0.053031482 |
| ENSMUSG00000028436 | Dcaf12 | 68970 | protein_coding | 680.365107 | -0.053059492 | 0.061048374 | 0.869138503 | 0.384771379 | 0.607042377 | 0.053059492 |
| ENSMUSG00000038416 | Cdc16 | 69957 | protein_coding | 466.2773484 | 0.0530955 | 0.090171994 | -0.588824725 | 0.555978858 | 0.745548913 | 0.0530955 |
| ENSMUSG00000001829 | Clpb | 20480 | protein_coding | 441.7761713 | 0.053099824 | 0.070768562 | -0.750330688 | 0.453055564 | 0.669122175 | 0.053099824 |
| ENSMUSG00000027951 | Adar | 56417 | protein_coding | 763.9929948 | -0.053101282 | 0.088447672 | 0.600369466 | 0.548260032 | 0.739498445 | 0.053101282 |
| ENSMUSG00000053695 | Defb37 | 353320 | protein_coding | 51.64239128 | 0.053138109 | 0.207964142 | -0.255515726 | 0.798324799 | 0.899929685 | 0.053138109 |
| ENSMUSG00000031312 | Itgb1bp2 | 26549 | protein_coding | 4.503546455 | 0.053156053 | 0.54404091 | -0.097705986 | 0.922165762 | 0.9624373 | 0.053156053 |
| ENSMUSG00000024480 | Ap3s1 | 11777 | protein_coding | 363.8025901 | -0.053191151 | 0.096477803 | 0.551330457 | 0.581407164 | 0.764378465 | 0.053191151 |
| ENSMUSG00000014039 | Prdm15 | 114604 | protein_coding | 272.6019784 | -0.053196554 | 0.083841613 | 0.634488676 | 0.525761955 | 0.724199927 | 0.053196554 |
| ENSMUSG00000025157 | Zdhhc16 | 74168 | protein_coding | 168.080131 | 0.053215338 | 0.093613654 | -0.568457013 | 0.569724687 | 0.755515416 | 0.053215338 |
| ENSMUSG00000091476 | Catspere2 | 545391 | protein_coding | 19.83493917 | 0.053246542 | 0.29689739 | -0.179343248 | 0.857668191 | 0.930374388 | 0.053246542 |
| ENSMUSG00000041684 | Bivm | 246229 | protein_coding | 194.5308304 | 0.053251406 | 0.106373009 | -0.500610132 | 0.61664553 | 0.788668238 | 0.053251406 |
| ENSMUSG00000038605 | Samd10 | 229011 | protein_coding | 137.770031 | -0.05326527 | 0.103978971 | 0.512269646 | 0.608462306 | 0.782626881 | 0.05326527 |
| ENSMUSG00000029840 | Mtpn | 14489 | protein_coding | 1910.132463 | 0.053372672 | 0.076045192 | -0.701854652 | 0.482769812 | 0.693558997 | 0.053372672 |
| ENSMUSG00000035168 | Tanc1 | 66860 | protein_coding | 485.025639 | -0.053414515 | 0.064008153 | 0.834495487 | 0.404001829 | 0.625216588 | 0.053414515 |
| ENSMUSG00000003099 | Ppp5c | 19060 | protein_coding | 1139.887943 | 0.053429876 | 0.072553824 | -0.736417093 | 0.461476905 | 0.675186464 | 0.053429876 |
| ENSMUSG00000028744 | Slc66a1 | 212555 | protein_coding | 80.02297489 | 0.053477986 | 0.135673942 | -0.394165489 | 0.693458863 | 0.837254214 | 0.053477986 |
| ENSMUSG00000067058 | Rps15a-ps5 | 100042335 | processed_pseudogene | 2.611215804 | -0.053485821 | 0.635956718 | 0.084102926 | 0.932974599 | 1 | 0.053485821 |
| ENSMUSG00000039361 | Picalm | 233489 | protein_coding | 2691.786791 | 0.053500216 | 0.043974999 | -1.216605277 | 0.223754434 | 0.43083733 | 0.053500216 |
| ENSMUSG00000031985 | Gnpat | 14712 | protein_coding | 443.9711774 | 0.053525193 | 0.076863381 | -0.696367927 | 0.486198444 | 0.695624496 | 0.053525193 |
| ENSMUSG00000025280 | Polr3a | 218832 | protein_coding | 293.4893006 | -0.053562399 | 0.071029456 | 0.754087136 | 0.450796901 | 0.667604701 | 0.053562399 |
| ENSMUSG00000028953 | Abcf2 | 27407 | protein_coding | 502.9074116 | 0.053610205 | 0.093707174 | -0.572103529 | 0.567251839 | 0.753386941 | 0.053610205 |
| ENSMUSG00000031529 | Tnks | 21951 | protein_coding | 608.622695 | 0.053615906 | 0.080598873 | -0.665219046 | 0.5059104 | 0.709931167 | 0.053615906 |
| ENSMUSG00000030614 | Tmem126b | 68472 | protein_coding | 210.855299 | 0.053633004 | 0.096944134 | -0.553236192 | 0.580101687 | 0.763391447 | 0.053633004 |

| ensembl_gene_id | external_gene_name | entrezgene_id | gene_biotype | baseMean | log2FoldChange | lfcSE | stat | pvalue | padj | absLog2FC |
|---|---|---|---|---|---|---|---|---|---|---|
| ENSMUSG00000056148 | Rdh9 | 103142 | protein_coding | 37.94146053 | 0.053644954 | 0.310919533 | -0.172536454 | 0.863015805 | 0.932166328 | 0.053644954 |
| ENSMUSG00000034462 | Pkd2 | 18764 | protein_coding | 183.8363801 | 0.053648959 | 0.126455261 | -0.424252488 | 0.671381685 | 0.823836911 | 0.053648959 |
| ENSMUSG00000037379 | Spon2 | 100689 | protein_coding | 18.39375529 | -0.053651654 | 0.253871901 | 0.211333567 | 0.832626992 | 0.915629096 | 0.053651654 |
| ENSMUSG00000025793 | Hgs | 15239 | protein_coding | 938.9182425 | 0.053654723 | 0.079327447 | -0.676370229 | 0.498805614 | 0.703739933 | 0.053654723 |
| ENSMUSG00000001017 | Chtop | 66511 | protein_coding | 1111.814973 | 0.053663297 | 0.077241616 | -0.694745906 | 0.487214554 | 0.695767999 | 0.053663297 |
| ENSMUSG00000054509 | Parp4 | 328417 | protein_coding | 369.8639777 | 0.053776561 | 0.295414587 | -0.182037593 | 0.855553223 | 0.929758221 | 0.053776561 |
| ENSMUSG00000026319 | Relch | 227446 | protein_coding | 1355.328722 | 0.053793303 | 0.063501603 | -0.847117242 | 0.396929775 | 0.618912124 | 0.053793303 |
| ENSMUSG00000026150 | Mff | 75734 | protein_coding | 636.8211995 | -0.053820558 | 0.072379183 | 0.743591671 | 0.457123544 | 0.671972423 | 0.053820558 |
| ENSMUSG00000098132 | Rassf10 | 78748 | protein_coding | 47.26370974 | -0.053846649 | 0.254658087 | 0.211446845 | 0.832538606 | 0.915629096 | 0.053846649 |
| ENSMUSG00000009647 | Mcu | 215999 | protein_coding | 1539.40124 | 0.053853093 | 0.062858143 | -0.85674012 | 0.391588525 | 0.614062569 | 0.053853093 |
| ENSMUSG00000004843 | Chmp2b | 68942 | protein_coding | 1651.75741 | -0.053902054 | 0.078492756 | 0.686713735 | 0.49226314 | 0.698928166 | 0.053902054 |
| ENSMUSG00000021969 | Zdhhc20 | 75965 | protein_coding | 921.5298508 | 0.053927186 | 0.110354878 | -0.488670618 | 0.625074909 | 0.794593769 | 0.053927186 |
| ENSMUSG00000001323 | Srr | 27364 | protein_coding | 188.2132717 | -0.053934066 | 0.090178041 | 0.598084253 | 0.549783717 | 0.740730056 | 0.053934066 |
| ENSMUSG00000045039 | Megf8 | 269878 | protein_coding | 549.2885661 | -0.053967189 | 0.096822498 | 0.557382743 | 0.577265953 | 0.761298159 | 0.053967189 |
| ENSMUSG00000038342 | Mlxip | 208104 | protein_coding | 847.8383414 | 0.053975657 | 0.089107652 | -0.605735367 | 0.544690495 | 0.736813469 | 0.053975657 |
| ENSMUSG00000022337 | Emc2 | 66736 | protein_coding | 489.9532964 | 0.053978908 | 0.084779571 | -0.63669711 | 0.524322156 | 0.72294031 | 0.053978908 |
| ENSMUSG00000001729 | Akt1 | 11651 | protein_coding | 2120.411924 | 0.053985814 | 0.065443258 | -0.824925528 | 0.409413872 | 0.630049453 | 0.053985814 |
| ENSMUSG00000118117 | Gm30961 | 102632967 | lncRNA | 4.277422171 | 0.054036263 | 0.514605078 | -0.105005305 | 0.916371599 | 1 | 0.054036263 |
| ENSMUSG00000031112 | Stk26 | 70415 | protein_coding | 128.8569229 | 0.054066297 | 0.342506055 | -0.15785501 | 0.874571051 | 0.938406345 | 0.054066297 |
| ENSMUSG00000072969 | Armcx5 | 494468 | protein_coding | 45.65546027 | -0.054079092 | 0.181662082 | 0.297690589 | 0.765939327 | 0.882687823 | 0.054079092 |
| ENSMUSG00000026227 | 2810459M11Rik | 72792 | protein_coding | 441.2579602 | -0.054083295 | 0.106425298 | 0.508180822 | 0.611326539 | 0.784660848 | 0.054083295 |
| ENSMUSG00000056536 | Pign | 27392 | protein_coding | 554.1545413 | -0.054084415 | 0.073739355 | 0.733453866 | 0.463281656 | 0.676807614 | 0.054084415 |
| ENSMUSG00000020184 | Mdm2 | 17246 | protein_coding | 478.7893984 | 0.054105887 | 0.089125557 | -0.60707488 | 0.54380122 | 0.736636318 | 0.054105887 |
| ENSMUSG00000071203 | Naip5 | 17951 | protein_coding | 632.0577579 | -0.054182619 | 0.088260023 | 0.613897628 | 0.539282978 | 0.733766095 | 0.054182619 |
| ENSMUSG00000056481 | Cd248 | 70445 | protein_coding | 2.021936778 | -0.054192345 | 1.923882093 | 0.028168225 | 0.97752798 | 1 | 0.054192345 |
| ENSMUSG00000023307 | Marchf5 | 69104 | protein_coding | 399.8556117 | -0.054196826 | 0.067584579 | 0.801911123 | 0.422604371 | 0.642504934 | 0.054196826 |
| ENSMUSG00000021957 | Tkt | 21881 | protein_coding | 6066.757133 | 0.054241723 | 0.081151584 | -0.668400057 | 0.503878259 | 0.708294652 | 0.054241723 |
| ENSMUSG00000031652 | N4bp1 | 80750 | protein_coding | 514.173585 | 0.054316843 | 0.076756862 | -0.707648039 | 0.479163853 | 0.690541222 | 0.054316843 |
| ENSMUSG00000046516 | Cox17 | 12856 | protein_coding | 108.2471873 | 0.054344525 | 0.123561995 | -0.439815863 | 0.660070478 | 0.816326209 | 0.054344525 |
| ENSMUSG00000034194 | R3hcc1 | 71843 | protein_coding | 151.3490368 | 0.054400137 | 0.094509409 | -0.575605517 | 0.564881851 | 0.751588072 | 0.054400137 |
| ENSMUSG00000095125 | Vmn1r198 | 171254 | protein_coding | 2.393546296 | -0.054438051 | 1.932758811 | 0.028165983 | 0.977529768 | 1 | 0.054438051 |
| ENSMUSG00000003200 | Sh3gl1 | 20405 | protein_coding | 438.6154944 | 0.054452555 | 0.071419804 | -0.762429351 | 0.445803794 | 0.662983366 | 0.054452555 |
| ENSMUSG00000039747 | Orai2 | 269717 | protein_coding | 309.6367658 | -0.054505592 | 0.141253912 | 0.385869614 | 0.699593236 | 0.84101582 | 0.054505592 |
| ENSMUSG00000028059 | Arhgef2 | 16800 | protein_coding | 719.364388 | -0.054574101 | 0.117867142 | 0.463013693 | 0.643354552 | 0.805485753 | 0.054574101 |
| ENSMUSG00000035495 | Tstd2 | 272027 | protein_coding | 325.6746644 | 0.054603527 | 0.078497844 | -0.695605436 | 0.486675961 | 0.695624496 | 0.054603527 |
| ENSMUSG00000057093 | Zfp607b | 112415 | protein_coding | 11.4509136 | 0.054608505 | 0.350036181 | -0.156008172 | 0.876026579 | 0.938630585 | 0.054608505 |
| ENSMUSG00000000486 | Septin1 | 54204 | protein_coding | 17.17041654 | 0.054627264 | 0.434122024 | -0.125833892 | 0.899863413 | 0.9507578 | 0.054627264 |
| ENSMUSG00000051495 | Irf2bp2 | 270110 | protein_coding | 2286.022553 | 0.054695016 | 0.111978579 | -0.488441771 | 0.625236961 | 0.794610064 | 0.054695016 |
| ENSMUSG00000041115 | Iqsec2 | 245666 | protein_coding | 251.6417703 | 0.054701005 | 0.184922303 | -0.295805343 | 0.767378742 | 0.883408334 | 0.054701005 |

| ensembl_gene_id | external_gene_name | entrezgene_id | gene_biotype | baseMean | log2FoldChange | lfcSE | stat | pvalue | padj | absLog2FC |
|---|---|---|---|---|---|---|---|---|---|---|
| ENSMUSG00000038039 | Gcc2 | 70297 | protein_coding | 1681.6507 | -0.054766271 | 0.098116764 | 0.558174454 | 0.576725263 | 0.761222618 | 0.054766271 |
| ENSMUSG00000026739 | Bmi1 | 12151 | protein_coding | 174.840088 | -0.054832705 | 0.103898253 | 0.527753867 | 0.59767018 | 0.775625491 | 0.054832705 |
| ENSMUSG00000026494 | Kif26b | 269152 | protein_coding | 12.9990139 | -0.054861531 | 0.422591757 | 0.129821584 | 0.896707586 | 0.94900212 | 0.054861531 |
| ENSMUSG00000029465 | Arpc3 | 56378 | protein_coding | 1918.109696 | -0.054927054 | 0.076442324 | 0.718542442 | 0.472422889 | 0.684950973 | 0.054927054 |
| ENSMUSG00000027566 | Psma7 | 26444 | protein_coding | 1367.485168 | 0.05492925 | 0.07899049 | -0.695390679 | 0.4868105 | 0.695624496 | 0.05492925 |
| ENSMUSG00000055799 | Tcf7l1 | 21415 | protein_coding | 136.4244752 | -0.054929635 | 0.196444905 | 0.279618526 | 0.779770192 | 0.890125936 | 0.054929635 |
| ENSMUSG00000037541 | Shank2 | 210274 | protein_coding | 362.0286094 | -0.054941614 | 0.140484032 | 0.391087963 | 0.695732218 | 0.83856979 | 0.054941614 |
| ENSMUSG00000067219 | Nipal1 | 70701 | protein_coding | 607.5150295 | -0.054953255 | 0.103494188 | 0.530979139 | 0.595433235 | 0.773824226 | 0.054953255 |
| ENSMUSG00000073664 | Nbeal1 | 269198 | protein_coding | 2082.49584 | -0.054985646 | 0.086113614 | 0.638524432 | 0.523132355 | 0.722040793 | 0.054985646 |
| ENSMUSG00000004221 | Ikbkg | 16151 | protein_coding | 658.3508421 | -0.055000016 | 0.073406106 | 0.749256692 | 0.453702505 | 0.669388475 | 0.055000016 |
| ENSMUSG00000021340 | Gpld1 | 14756 | protein_coding | 105.3380546 | 0.055010251 | 0.155883262 | -0.352893894 | 0.724167989 | 0.857084875 | 0.055010251 |
| ENSMUSG00000033106 | Slc7a6os | 66432 | protein_coding | 214.1096019 | -0.055019765 | 0.079232657 | 0.694407674 | 0.487426582 | 0.696010599 | 0.055019765 |
| ENSMUSG00000035203 | Epn1 | 13854 | protein_coding | 2450.622276 | 0.055049933 | 0.056155948 | -0.980304576 | 0.326935797 | 0.548792208 | 0.055049933 |
| ENSMUSG00000013787 | Ehmt2 | 110147 | protein_coding | 1920.462319 | 0.055060149 | 0.033385011 | -1.649247612 | 0.099096917 | 0.245940261 | 0.055060149 |
| ENSMUSG00000036819 | Jmjd4 | 194952 | protein_coding | 162.8341734 | 0.055119834 | 0.098519752 | -0.559480036 | 0.575834152 | 0.760926598 | 0.055119834 |
| ENSMUSG00000047193 | Dync2h1 | 110350 | protein_coding | 110.9581192 | 0.05514615 | 0.131805915 | -0.418389039 | 0.675662699 | 0.826265833 | 0.05514615 |
| ENSMUSG00000042625 | Safb2 | 224902 | protein_coding | 721.0116071 | 0.055171175 | 0.144248927 | -0.382471996 | 0.702111297 | 0.842880631 | 0.055171175 |
| ENSMUSG00000079215 | Zfp664 | 269704 | protein_coding | 697.9429906 | -0.055173087 | 0.07981554 | 0.691257456 | 0.489403763 | 0.697206154 | 0.055173087 |
| ENSMUSG00000026806 | Ddx31 | 227674 | protein_coding | 103.913994 | 0.055175322 | 0.154074446 | -0.358108194 | 0.720262348 | 0.855204229 | 0.055175322 |
| ENSMUSG00000027300 | Ubox5 | 140629 | protein_coding | 133.4013872 | -0.055197944 | 0.121304872 | 0.455034844 | 0.649084172 | 0.809468057 | 0.055197944 |
| ENSMUSG00000015672 | Mrpl32 | 75398 | protein_coding | 267.4910084 | -0.055230924 | 0.086178221 | 0.640891907 | 0.521592914 | 0.721043477 | 0.055230924 |
| ENSMUSG00000026878 | Rab14 | 68365 | protein_coding | 1429.570078 | -0.055241753 | 0.049986872 | 1.105125218 | 0.269105335 | 0.484149199 | 0.055241753 |
| ENSMUSG00000030361 | Klrb1a | 17057 | protein_coding | 4.919144578 | 0.055250317 | 0.623852118 | -0.088563164 | 0.929429084 | 0.966128972 | 0.055250317 |
| ENSMUSG00000025730 | Rab40c | 224624 | protein_coding | 311.2157269 | -0.055286397 | 0.070893789 | 0.77984824 | 0.435480208 | 0.653742472 | 0.055286397 |
| ENSMUSG00000054728 | Phactr1 | 218194 | protein_coding | 4.485917552 | -0.055291844 | 2.075864657 | 0.026635573 | 0.9787504 | 0.988831409 | 0.055291844 |
| ENSMUSG00000021285 | Ppp1r13b | 21981 | protein_coding | 574.3530382 | 0.055310026 | 0.072884853 | -0.7588686 | 0.447931164 | 0.664930223 | 0.055310026 |
| ENSMUSG00000028927 | Padi2 | 18600 | protein_coding | 163.04075 | 0.055327217 | 0.188781393 | -0.293075584 | 0.769464379 | 0.884482916 | 0.055327217 |
| ENSMUSG00000037824 | Tspan14 | 52588 | protein_coding | 496.5329324 | 0.055367825 | 0.138640111 | -0.39936368 | 0.689625252 | 0.834370416 | 0.055367825 |
| ENSMUSG00000052310 | Slc39a1 | 30791 | protein_coding | 1451.271777 | -0.055391839 | 0.07151659 | 0.774531322 | 0.438616653 | 0.656764359 | 0.055391839 |
| ENSMUSG00000020898 | Ctc1 | 68964 | protein_coding | 243.5905799 | 0.055431904 | 0.118067534 | -0.469493201 | 0.638717143 | 0.80301103 | 0.055431904 |
| ENSMUSG00000032485 | Scap | 235623 | protein_coding | 776.4028261 | -0.055459263 | 0.054208612 | 1.023071075 | 0.306274242 | 0.526745764 | 0.055459263 |
| ENSMUSG00000034729 | Mrps10 | 64657 | protein_coding | 206.8942947 | 0.055466275 | 0.082189607 | -0.674857527 | 0.499766288 | 0.704451599 | 0.055466275 |
| ENSMUSG00000118537 | Shld3 | 113002583 | protein_coding | 23.23702822 | 0.055483869 | 0.285118165 | -0.194599558 | 0.845706449 | 0.924043688 | 0.055483869 |
| ENSMUSG00000037262 | Kin | 16588 | protein_coding | 154.816442 | 0.055493116 | 0.099152028 | -0.559677066 | 0.575699728 | 0.760811885 | 0.055493116 |
| ENSMUSG00000028910 | Mecr | 26922 | protein_coding | 743.187993 | -0.055533145 | 0.134140259 | 0.413993125 | 0.678879142 | 0.827931507 | 0.055533145 |
| ENSMUSG00000037410 | Tbc1d2b | 67016 | protein_coding | 387.314174 | -0.055550977 | 0.133115583 | 0.41731385 | 0.676448859 | 0.826798642 | 0.055550977 |
| ENSMUSG00000031077 | Fadd | 14082 | protein_coding | 178.2217693 | -0.055569472 | 0.106763183 | 0.520492831 | 0.602720123 | 0.778759104 | 0.055569472 |
| ENSMUSG00000046573 | Lyrm4 | 380840 | protein_coding | 84.14603203 | -0.055597655 | 0.120915525 | 0.459805765 | 0.645655645 | 0.806964824 | 0.055597655 |
| ENSMUSG00000047084 | Ngrn | 83485 | protein_coding | 236.7423489 | -0.055619525 | 0.101926737 | 0.5456814 | 0.58528496 | 0.767282764 | 0.055619525 |

| ensembl_gene_id | external_gene_name | entrezgene_id | gene_biotype | baseMean | log2FoldChange | lfcSE | stat | pvalue | padj | absLog2FC |
|---|---|---|---|---|---|---|---|---|---|---|
| ENSMUSG00000040929 | Rfx3 | 19726 | protein_coding | 105.7599651 | -0.055619553 | 0.126201552 | 0.440720039 | 0.659415688 | 0.816133095 | 0.055619553 |
| ENSMUSG00000064145 | Arih2 | 23807 | protein_coding | 723.8823985 | 0.055719893 | 0.065739122 | -0.847591083 | 0.396665742 | 0.618855346 | 0.055719893 |
| ENSMUSG00000056692 | Ilrun | 224647 | protein_coding | 1622.240605 | -0.055739553 | 0.043315378 | 1.286830577 | 0.198153347 | 0.398418975 | 0.055739553 |
| ENSMUSG00000085421 | 4732490B19Rik | 319871 | lncRNA | 20.52674784 | 0.055771267 | 0.260724839 | -0.213908531 | 0.830618387 | 0.91440345 | 0.055771267 |
| ENSMUSG00000032384 | Csnk1g1 | 214897 | protein_coding | 383.6384088 | 0.055802864 | 0.087379847 | -0.638623957 | 0.523067592 | 0.722040793 | 0.055802864 |
| ENSMUSG00000047694 | Yipf6 | 77929 | protein_coding | 1482.42088 | 0.055930669 | 0.133521102 | -0.418890108 | 0.675296447 | 0.825971257 | 0.055930669 |
| ENSMUSG00000038242 | Aox4 | 71872 | protein_coding | 3.895445327 | -0.055947985 | 1.953273305 | 0.028643193 | 0.977149163 | 1 | 0.055947985 |
| ENSMUSG00000006715 | Gmnn | 57441 | protein_coding | 182.8118387 | 0.05596374 | 0.111349589 | -0.502594943 | 0.615249086 | 0.78770474 | 0.05596374 |
| ENSMUSG00000030447 | Cyfip1 | 20430 | protein_coding | 1600.028643 | -0.055966913 | 0.070024951 | 0.799242447 | 0.424149843 | 0.643957486 | 0.055966913 |
| ENSMUSG00000025491 | Ifitm1 | 68713 | protein_coding | 1.101263219 | -0.055995054 | 0.95524994 | 0.058618223 | 0.953256195 | 1 | 0.055995054 |
| ENSMUSG00000074182 | Znhit6 | 229937 | protein_coding | 64.53388172 | -0.056159059 | 0.170500143 | 0.329378371 | 0.741869714 | 0.869560398 | 0.056159059 |
| ENSMUSG00000019320 | Noxo1 | 71893 | protein_coding | 34.67061316 | -0.056166049 | 0.187461093 | 0.299614432 | 0.764471275 | 0.882132403 | 0.056166049 |
| ENSMUSG00000037275 | Gemin5 | 216766 | protein_coding | 304.6508661 | -0.056181658 | 0.125744757 | 0.446791255 | 0.655025786 | 0.812765484 | 0.056181658 |
| ENSMUSG00000034255 | Arhgap27 | 544817 | protein_coding | 1268.875679 | -0.056229717 | 0.169587428 | 0.331567723 | 0.740215695 | 0.868422989 | 0.056229717 |
| ENSMUSG00000006728 | Cdk4 | 12567 | protein_coding | 818.5757048 | 0.056241416 | 0.117914625 | -0.476967264 | 0.633385433 | 0.800270871 | 0.056241416 |
| ENSMUSG00000002820 | Atg4d | 235040 | protein_coding | 1062.996459 | -0.056272004 | 0.062771294 | 0.896460779 | 0.370006719 | 0.59260115 | 0.056272004 |
| ENSMUSG00000069729 | Arid1b | 239985 | protein_coding | 922.9902519 | 0.056278873 | 0.056000391 | -1.004972853 | 0.314909922 | 0.535990818 | 0.056278873 |
| ENSMUSG00000034247 | Plekhm1 | 353047 | protein_coding | 561.9895551 | -0.056285209 | 0.16903237 | 0.332984794 | 0.739145761 | 0.867716116 | 0.056285209 |
| ENSMUSG00000037375 | Hhat | 226861 | protein_coding | 41.25574929 | 0.056340337 | 0.184402354 | -0.305529379 | 0.759962998 | 0.879445484 | 0.056340337 |
| ENSMUSG00000040451 | Sgms1 | 208449 | protein_coding | 523.6650687 | 0.056345394 | 0.072025247 | -0.782300596 | 0.434037937 | 0.652558493 | 0.056345394 |
| ENSMUSG00000035401 | Emsy | 233545 | protein_coding | 290.0621738 | 0.056358827 | 0.093880007 | -0.600328316 | 0.548287451 | 0.739498445 | 0.056358827 |
| ENSMUSG00000015002 | Efr3a | 76740 | protein_coding | 1075.438988 | 0.056387225 | 0.084881622 | -0.664304285 | 0.506495579 | 0.71028091 | 0.056387225 |
| ENSMUSG00000019794 | Katna1 | 23924 | protein_coding | 413.8993854 | -0.056398034 | 0.070273209 | 0.802553849 | 0.422232653 | 0.64211714 | 0.056398034 |
| ENSMUSG00000032436 | Cmtm7 | 102545 | protein_coding | 75.05868472 | -0.05640198 | 0.264395544 | 0.213324245 | 0.831074064 | 0.914530359 | 0.05640198 |
| ENSMUSG00000025034 | Trim8 | 93679 | protein_coding | 444.3672675 | -0.056436427 | 0.096069463 | 0.587454389 | 0.556898575 | 0.746166522 | 0.056436427 |
| ENSMUSG00000034681 | Rnps1 | 19826 | protein_coding | 445.1479628 | 0.056437525 | 0.08195646 | -0.688628139 | 0.491057309 | 0.69829677 | 0.056437525 |
| ENSMUSG00000021636 | Marveld2 | 218518 | protein_coding | 515.799178 | -0.056444911 | 0.064951891 | 0.869026446 | 0.384832666 | 0.607042377 | 0.056444911 |
| ENSMUSG00000032393 | Dpp8 | 74388 | protein_coding | 1666.9145 | -0.056454357 | 0.079239759 | 0.712449883 | 0.476186235 | 0.687965835 | 0.056454357 |
| ENSMUSG00000001157 | Gmcl1 | 23885 | protein_coding | 389.5082801 | -0.05646964 | 0.07290948 | 0.774517113 | 0.438625052 | 0.656764359 | 0.05646964 |
| ENSMUSG00000034164 | Emid1 | 140703 | protein_coding | 6.630223952 | 0.056476664 | 0.71688483 | -0.078780665 | 0.937207083 | 0.970043107 | 0.056476664 |
| ENSMUSG00000044502 | Bod1 | 69556 | protein_coding | 377.0111451 | -0.056506291 | 0.071663382 | 0.788496008 | 0.430406633 | 0.648871201 | 0.056506291 |
| ENSMUSG00000030016 | Zfp638 | 18139 | protein_coding | 984.4249458 | 0.056507038 | 0.091128555 | -0.620080481 | 0.535204802 | 0.731167175 | 0.056507038 |
| ENSMUSG00000004085 | Map3k20 | 65964 | protein_coding | 311.0441484 | 0.056526002 | 0.140349254 | -0.402752422 | 0.687130365 | 0.833389267 | 0.056526002 |
| ENSMUSG00000027748 | Trpc4 | 22066 | protein_coding | 2.350551478 | 0.056526052 | 1.062993028 | -0.053176315 | 0.957591427 | 1 | 0.056526052 |
| ENSMUSG00000035585 | Tsen34 | 66078 | protein_coding | 394.0749782 | 0.056544466 | 0.086363131 | -0.654729225 | 0.512642106 | 0.714893745 | 0.056544466 |
| ENSMUSG00000048039 | Isg20l2 | 229504 | protein_coding | 337.5566942 | 0.056545018 | 0.091858035 | -0.615569647 | 0.538178589 | 0.733377497 | 0.056545018 |
| ENSMUSG00000003198 | Zfp959 | 224893 | protein_coding | 59.44447192 | 0.056589668 | 0.131392987 | -0.430690175 | 0.666693664 | 0.820645435 | 0.056589668 |
| ENSMUSG00000032264 | Zw10 | 26951 | protein_coding | 345.8537793 | 0.056599846 | 0.079867209 | -0.708674402 | 0.478526554 | 0.690104577 | 0.056599846 |
| ENSMUSG00000018800 | Abca5 | 217265 | protein_coding | 21.0641141 | -0.056603231 | 0.236407281 | 0.239430998 | 0.810771396 | 0.906297546 | 0.056603231 |

| ensembl_gene_id | external_gene_name | entrezgene_id | gene_biotype | baseMean | log2FoldChange | lfcSE | stat | pvalue | padj | absLog2FC |
|---|---|---|---|---|---|---|---|---|---|---|
| ENSMUSG00000058153 | Sez6l | 56747 | protein_coding | 4.942110659 | -0.05664013 | 0.840769266 | 0.067367032 | 0.946289514 | 0.973795719 | 0.05664013 |
| ENSMUSG00000063694 | Cycs | 13063 | protein_coding | 1816.087837 | 0.056655075 | 0.109471891 | -0.517530796 | 0.604785677 | 0.779964063 | 0.056655075 |
| ENSMUSG00000117048 | Gm41584 | 105246280 | lncRNA | 3.65356735 | -0.056681818 | 0.622598126 | 0.091040778 | 0.92746019 | 1 | 0.056681818 |
| ENSMUSG00000026117 | Zap70 | 22637 | protein_coding | 13.71382398 | -0.056738393 | 0.43629812 | 0.13004501 | 0.896530816 | 0.948997494 | 0.056738393 |
| ENSMUSG00000056220 | Pla2g4a | 18783 | protein_coding | 695.5582492 | 0.056749219 | 0.167177951 | -0.339453968 | 0.734267769 | 0.864269275 | 0.056749219 |
| ENSMUSG00000028423 | Nfx1 | 74164 | protein_coding | 888.3720165 | 0.056758308 | 0.090864375 | -0.624648638 | 0.53220169 | 0.728866518 | 0.056758308 |
| ENSMUSG00000034525 | Ice1 | 218333 | protein_coding | 582.5288514 | 0.056823839 | 0.08862444 | -0.641175721 | 0.521408522 | 0.720922137 | 0.056823839 |
| ENSMUSG00000036185 | Sapcd1 | 78376 | protein_coding | 0.622570152 | -0.056823894 | 1.417370169 | 0.040091075 | 0.968020517 | 1 | 0.056823894 |
| ENSMUSG00000017307 | Acot8 | 170789 | protein_coding | 442.9423308 | 0.056837418 | 0.077346142 | -0.734844907 | 0.462433954 | 0.675987813 | 0.056837418 |
| ENSMUSG00000023977 | Ubr2 | 224826 | protein_coding | 1005.783937 | 0.056901954 | 0.055278343 | -1.029371557 | 0.30330511 | 0.523766661 | 0.056901954 |
| ENSMUSG00000089669 | Tnfsf13 | 69583 | protein_coding | 121.4931398 | 0.056916211 | 0.116461566 | -0.488712398 | 0.625045325 | 0.794593769 | 0.056916211 |
| ENSMUSG00000026558 | Uck2 | 80914 | protein_coding | 789.9579402 | -0.056949184 | 0.145640558 | 0.391025584 | 0.695778325 | 0.83856979 | 0.056949184 |
| ENSMUSG00000029624 | Ptcd1 | 71799 | protein_coding | 275.2641663 | -0.056992393 | 0.078889336 | 0.722434683 | 0.47002727 | 0.683555242 | 0.056992393 |
| ENSMUSG00000000711 | Rab5b | 19344 | protein_coding | 830.2480502 | 0.056994421 | 0.095969477 | -0.593880705 | 0.552591905 | 0.743179634 | 0.056994421 |
| ENSMUSG00000044864 | Ankrd50 | 99696 | protein_coding | 633.9345785 | -0.057001414 | 0.113109161 | 0.503950459 | 0.614296191 | 0.78673795 | 0.057001414 |
| ENSMUSG00000060591 | Ifitm2 | 80876 | protein_coding | 305.0563423 | 0.057016738 | 0.160784501 | -0.354615885 | 0.722877374 | 0.856549354 | 0.057016738 |
| ENSMUSG00000004947 | Dtx2 | 74198 | protein_coding | 102.7436778 | 0.057029675 | 0.114412735 | -0.498455654 | 0.618162918 | 0.789569374 | 0.057029675 |
| ENSMUSG00000081769 | Gm12216 | 622459 | lncRNA | 16.65513802 | -0.057039889 | 0.394000542 | 0.144771093 | 0.884891607 | 0.943016089 | 0.057039889 |
| ENSMUSG00000057346 | Apol9a | 223672 | protein_coding | 76.64459762 | 0.057048048 | 0.483625038 | -0.117959252 | 0.906099945 | 0.953990085 | 0.057048048 |
| ENSMUSG00000021112 | Pals1 | 56217 | protein_coding | 885.8978065 | -0.057054123 | 0.071140528 | 0.801991849 | 0.422557673 | 0.642504934 | 0.057054123 |
| ENSMUSG00000029486 | Mrpl1 | 94061 | protein_coding | 257.6834778 | -0.057055291 | 0.098877781 | 0.577028432 | 0.563920249 | 0.751086942 | 0.057055291 |
| ENSMUSG00000047228 | A2ml1 | 232400 | protein_coding | 10.45402249 | -0.057078456 | 0.591171272 | 0.096551472 | 0.923082596 | 0.962730128 | 0.057078456 |
| ENSMUSG00000037103 | Dcaf15 | 212123 | protein_coding | 506.2195272 | -0.057081304 | 0.076109035 | 0.749993804 | 0.453258436 | 0.669122175 | 0.057081304 |
| ENSMUSG00000062031 | Pgghg | 212974 | protein_coding | 374.4409313 | -0.05711895 | 0.087792769 | 0.650611103 | 0.515297561 | 0.716335759 | 0.05711895 |
| ENSMUSG00000021337 | Scgn | 214189 | protein_coding | 12.57146849 | 0.057136579 | 0.306989513 | -0.186118991 | 0.852351449 | 0.927761944 | 0.057136579 |
| ENSMUSG00000062691 | Cebpzos | 68554 | protein_coding | 118.6268358 | 0.05715031 | 0.118313445 | -0.483041553 | 0.629066232 | 0.797446101 | 0.05715031 |
| ENSMUSG00000031370 | Zrsr2 | 22184 | protein_coding | 242.5704049 | 0.057151045 | 0.072685253 | -0.786281159 | 0.431702788 | 0.650053783 | 0.057151045 |
| ENSMUSG00000027193 | Api5 | 11800 | protein_coding | 1363.922969 | -0.057159168 | 0.0998409 | 0.572502537 | 0.566981568 | 0.753307251 | 0.057159168 |
| ENSMUSG00000039086 | Ss18l1 | 269397 | protein_coding | 95.35596766 | -0.057199303 | 0.109127207 | 0.524152544 | 0.600172445 | 0.77680602 | 0.057199303 |
| ENSMUSG00000059995 | Atxn7l3 | 217218 | protein_coding | 729.2087119 | 0.057208424 | 0.086949212 | -0.657952177 | 0.510568862 | 0.71375046 | 0.057208424 |
| ENSMUSG00000020284 | Cfap410 | 67884 | protein_coding | 96.03045317 | -0.057211447 | 0.127487654 | 0.448760686 | 0.653604302 | 0.812100026 | 0.057211447 |
| ENSMUSG00000021546 | Hnrnpk | 15387 | protein_coding | 3988.668425 | 0.057246163 | 0.081783778 | -0.69996965 | 0.483946258 | 0.694394109 | 0.057246163 |
| ENSMUSG00000028923 | Necap2 | 66147 | protein_coding | 791.6068424 | -0.057250236 | 0.063470635 | 0.901995644 | 0.367059181 | 0.589852079 | 0.057250236 |
| ENSMUSG00000042289 | Hsd3b7 | 101502 | protein_coding | 0.54520931 | 0.057299085 | 1.865106005 | -0.030721624 | 0.975491546 | 1 | 0.057299085 |
| ENSMUSG00000035403 | Crb2 | 241324 | protein_coding | 4.000812904 | 0.057313272 | 0.797447251 | -0.071870926 | 0.942704628 | 1 | 0.057313272 |
| ENSMUSG00000024276 | Zfp397 | 69256 | protein_coding | 365.9737917 | -0.057326455 | 0.119648421 | 0.479124211 | 0.631850267 | 0.799444652 | 0.057326455 |
| ENSMUSG00000043257 | Pigv | 230801 | protein_coding | 267.3263692 | 0.057334736 | 0.078154308 | -0.733609417 | 0.46318682 | 0.676807614 | 0.057334736 |
| ENSMUSG00000060301 | 2610008E11Rik | 72128 | protein_coding | 150.3547196 | -0.057385417 | 0.124933426 | 0.459327972 | 0.645998664 | 0.807082912 | 0.057385417 |
| ENSMUSG00000042605 | Atxn2 | 20239 | protein_coding | 434.253721 | 0.057469107 | 0.088541786 | -0.649061982 | 0.516298314 | 0.717206457 | 0.057469107 |

| ensembl_gene_id | external_gene_name | entrezgene_id | gene_biotype | baseMean | log2FoldChange | lfcSE | stat | pvalue | padj | absLog2FC |
|---|---|---|---|---|---|---|---|---|---|---|
| ENSMUSG00000033970 | Rfc3 | 69263 | protein_coding | 162.3140363 | -0.057491289 | 0.138220918 | 0.415937685 | 0.677455602 | 0.827172038 | 0.057491289 |
| ENSMUSG00000037349 | Nudt22 | 68323 | protein_coding | 288.3946355 | -0.057534978 | 0.176272358 | 0.326398189 | 0.744123107 | 0.87096527 | 0.057534978 |
| ENSMUSG00000044881 | Coa4 | 68185 | protein_coding | 204.4549474 | 0.057540488 | 0.097430336 | -0.590580819 | 0.55480132 | 0.744703672 | 0.057540488 |
| ENSMUSG00000024974 | Smc3 | 13006 | protein_coding | 867.6231607 | 0.057544422 | 0.082994916 | -0.693348759 | 0.488090711 | 0.696186289 | 0.057544422 |
| ENSMUSG00000070427 | Il18bp | 16068 | protein_coding | 58.63539734 | 0.057610149 | 0.222610237 | -0.258793801 | 0.795794341 | 0.898576349 | 0.057610149 |
| ENSMUSG00000018167 | Stard3 | 59045 | protein_coding | 641.9301441 | 0.057624068 | 0.073075012 | -0.7885605 | 0.430368925 | 0.648871201 | 0.057624068 |
| ENSMUSG00000024639 | Gnaq | 14682 | protein_coding | 591.1139821 | 0.057634367 | 0.066012062 | -0.873088428 | 0.382614881 | 0.604944412 | 0.057634367 |
| ENSMUSG00000090100 | Ttbk2 | 140810 | protein_coding | 83.64902468 | -0.057683385 | 0.161866736 | 0.356363425 | 0.721568417 | 0.855725733 | 0.057683385 |
| ENSMUSG00000058440 | Nrf1 | 18181 | protein_coding | 393.713823 | 0.057683385 | 0.062880021 | -0.917356336 | 0.358955945 | 0.581669952 | 0.057683385 |
| ENSMUSG00000028478 | Clta | 12757 | protein_coding | 1651.883584 | -0.057686135 | 0.060066239 | 0.960375339 | 0.336866345 | 0.559233675 | 0.057686135 |
| ENSMUSG00000039515 | Ptpa | 110854 | protein_coding | 1054.85505 | 0.057704985 | 0.077400204 | -0.745540471 | 0.455945055 | 0.671255829 | 0.057704985 |
| ENSMUSG00000030595 | Nfkbib | 18036 | protein_coding | 223.3587594 | 0.057722597 | 0.088471235 | -0.652444796 | 0.514114274 | 0.715798257 | 0.057722597 |
| ENSMUSG00000060681 | Slc9a6 | 236794 | protein_coding | 268.9704449 | 0.057726479 | 0.075554077 | -0.764041888 | 0.444842281 | 0.662290428 | 0.057726479 |
| ENSMUSG00000015095 | Fbxw5 | 30839 | protein_coding | 510.2491016 | 0.05780426 | 0.068220821 | -0.847311109 | 0.396821736 | 0.618855346 | 0.05780426 |
| ENSMUSG00000040945 | Rcc2 | 108911 | protein_coding | 934.2887112 | 0.057819851 | 0.16604767 | -0.348212358 | 0.727680708 | 0.859322531 | 0.057819851 |
| ENSMUSG00000066232 | Ipo7 | 233726 | protein_coding | 1988.879048 | 0.057835365 | 0.114450621 | -0.505330285 | 0.613326875 | 0.786311623 | 0.057835365 |
| ENSMUSG00000021027 | Ralgapa1 | 56784 | protein_coding | 896.777353 | 0.057858639 | 0.095285808 | -0.607211512 | 0.543710553 | 0.736573919 | 0.057858639 |
| ENSMUSG00000051147 | Nat2 | 17961 | protein_coding | 212.4514293 | -0.057916479 | 0.103149428 | 0.561481342 | 0.574469446 | 0.7598549 | 0.057916479 |
| ENSMUSG00000031502 | Col4a1 | 12826 | protein_coding | 17.60117868 | -0.057926217 | 0.518633577 | 0.111690063 | 0.911069158 | 0.956997404 | 0.057926217 |
| ENSMUSG00000024914 | Drap1 | 66556 | protein_coding | 772.0048318 | 0.057944721 | 0.077420489 | -0.748441672 | 0.454193795 | 0.669812138 | 0.057944721 |
| ENSMUSG00000004994 | Ccdc130 | 67736 | protein_coding | 91.12203088 | 0.057945349 | 0.123640611 | -0.468659519 | 0.639313027 | 0.803288086 | 0.057945349 |
| ENSMUSG00000030491 | Tdrd12 | 71981 | protein_coding | 13.77284155 | 0.057948541 | 0.333612793 | -0.173699997 | 0.86210124 | 0.931908663 | 0.057948541 |
| ENSMUSG00000075307 | Klhl41 | 228003 | protein_coding | 2.864591295 | -0.057988634 | 0.726434436 | 0.079826383 | 0.93637534 | 1 | 0.057988634 |
| ENSMUSG00000020098 | Pcbd1 | 13180 | protein_coding | 307.514167 | 0.057996055 | 0.128847511 | -0.4501139 | 0.652628315 | 0.811601896 | 0.057996055 |
| ENSMUSG00000037339 | Fam53a | 74504 | protein_coding | 302.5111735 | -0.058022654 | 0.094166701 | 0.616169552 | 0.537782622 | 0.733113936 | 0.058022654 |
| ENSMUSG00000021643 | Serf1 | 20365 | protein_coding | 66.04851685 | -0.058030523 | 0.16163426 | 0.359023655 | 0.719577393 | 0.854899358 | 0.058030523 |
| ENSMUSG00000024105 | Themis3 | 74556 | protein_coding | 652.1502432 | -0.058034106 | 0.120575138 | 0.481310713 | 0.630295684 | 0.79831751 | 0.058034106 |
| ENSMUSG00000018740 | Slc25a35 | 71998 | protein_coding | 193.4874031 | 0.058050199 | 0.115038177 | -0.504616823 | 0.613827991 | 0.78655252 | 0.058050199 |
| ENSMUSG00000026048 | Ercc5 | 22592 | protein_coding | 206.8289576 | -0.058061519 | 0.090250625 | 0.643336472 | 0.520005795 | 0.720115363 | 0.058061519 |
| ENSMUSG00000021598 | Med10 | 28077 | protein_coding | 246.2721117 | 0.058080722 | 0.073934077 | -0.785574445 | 0.432116841 | 0.65055862 | 0.058080722 |
| ENSMUSG00000035299 | Mid1 | 17318 | protein_coding | 81.20401498 | 0.058095514 | 0.245201943 | -0.236929255 | 0.812711674 | 0.907147741 | 0.058095514 |
| ENSMUSG00000049288 | Lix1l | 280411 | protein_coding | 6.56769069 | 0.058098962 | 0.556007281 | -0.104493167 | 0.91677799 | 0.959274295 | 0.058098962 |
| ENSMUSG00000061461 | Smim20 | 66278 | protein_coding | 396.4474027 | 0.05812298 | 0.083792822 | -0.693651066 | 0.487901061 | 0.696186289 | 0.05812298 |
| ENSMUSG00000035382 | Pcsk7 | 18554 | protein_coding | 1871.252149 | 0.05814392 | 0.049064404 | -1.185053027 | 0.235996482 | 0.446089228 | 0.05814392 |
| ENSMUSG00000063047 | Zfp780b | 338354 | protein_coding | 128.9575993 | -0.058162163 | 0.105348586 | 0.552092484 | 0.580884991 | 0.764064128 | 0.058162163 |
| ENSMUSG00000039357 | Fut11 | 73068 | protein_coding | 117.199384 | -0.058178874 | 0.10071719 | 0.577645923 | 0.563503195 | 0.750868206 | 0.058178874 |
| ENSMUSG00000034121 | Mks1 | 380718 | protein_coding | 97.37700836 | -0.058187892 | 0.114043565 | 0.510225127 | 0.60989375 | 0.783613584 | 0.058187892 |
| ENSMUSG00000055447 | Cd47 | 16423 | protein_coding | 377.8569283 | -0.058189662 | 0.101707011 | 0.5721303 | 0.567233703 | 0.753386941 | 0.058189662 |
| ENSMUSG00000035049 | Rrp12 | 107094 | protein_coding | 195.2773882 | -0.058205101 | 0.131402494 | 0.442952789 | 0.657799883 | 0.815044063 | 0.058205101 |

| ensembl_gene_id | external_gene_name | entrezgene_id | gene_biotype | baseMean | log2FoldChange | lfcSE | stat | pvalue | padj | absLog2FC |
|---|---|---|---|---|---|---|---|---|---|---|
| ENSMUSG00000015342 | Xk | 22439 | protein_coding | 240.7235836 | -0.058304224 | 0.097167296 | 0.600039586 | 0.548479854 | 0.739637131 | 0.058304224 |
| ENSMUSG00000050213 | Snip1 | 76793 | protein_coding | 266.4318448 | -0.058379754 | 0.088357198 | 0.66072437 | 0.508789092 | 0.712306522 | 0.058379754 |
| ENSMUSG00000079243 | Xirp1 | 22437 | protein_coding | 10.75330478 | -0.058407224 | 0.499148511 | 0.117013719 | 0.906849182 | 0.954353007 | 0.058407224 |
| ENSMUSG00000085958 | A930019D19Rik | 77806 | lncRNA | 8.324259878 | -0.058425128 | 0.386606511 | 0.15112298 | 0.879878706 | 0.940676072 | 0.058425128 |
| ENSMUSG00000033688 | Inhca | 71775 | protein_coding | 2.678958123 | 0.058434591 | 1.207217104 | -0.048404376 | 0.961393972 | 1 | 0.058434591 |
| ENSMUSG00000066037 | Hnrnpr | 74326 | protein_coding | 594.531797 | 0.05846417 | 0.075681934 | -0.77249836 | 0.439819318 | 0.657836809 | 0.05846417 |
| ENSMUSG00000003299 | Mrpl4 | 66163 | protein_coding | 440.3913454 | 0.058519788 | 0.079784462 | -0.733473498 | 0.463269686 | 0.676807614 | 0.058519788 |
| ENSMUSG00000026836 | Acvr1 | 11477 | protein_coding | 127.9398337 | 0.058528695 | 0.125850258 | -0.465066153 | 0.641884084 | 0.804573016 | 0.058528695 |
| ENSMUSG00000031390 | Avpr2 | 12000 | protein_coding | 1.528653045 | -0.058586729 | 1.385351167 | 0.042290165 | 0.966267386 | 1 | 0.058586729 |
| ENSMUSG00000040511 | Pvr | 52118 | protein_coding | 159.5834414 | -0.058599884 | 0.139904794 | 0.418855443 | 0.675321783 | 0.825971257 | 0.058599884 |
| ENSMUSG00000031154 | Otud5 | 54644 | protein_coding | 878.2017388 | 0.05861439 | 0.06822845 | -0.859090156 | 0.390290778 | 0.612497263 | 0.05861439 |
| ENSMUSG00000031820 | Babam1 | 68251 | protein_coding | 614.9760069 | -0.058616945 | 0.068339135 | 0.857736127 | 0.391038186 | 0.613316042 | 0.058616945 |
| ENSMUSG00000001095 | Slc13a2 | 20500 | protein_coding | 858.574303 | 0.058629955 | 0.202762705 | -0.28915552 | 0.772462369 | 0.885864136 | 0.058629955 |
| ENSMUSG00000063522 | Ly6m | 67038 | protein_coding | 1009.146903 | 0.058696855 | 0.483281016 | -0.121454915 | 0.903330723 | 0.952286247 | 0.058696855 |
| ENSMUSG00000020781 | Tsen54 | 76265 | protein_coding | 242.553 | -0.058700559 | 0.08675417 | 0.676630982 | 0.498640117 | 0.703739933 | 0.058700559 |
| ENSMUSG00000037243 | Zfp692 | 103836 | protein_coding | 195.8978844 | -0.058715701 | 0.10699157 | 0.548788102 | 0.583150877 | 0.765756487 | 0.058715701 |
| ENSMUSG00000021103 | Mnat1 | 17420 | protein_coding | 136.7850454 | -0.058726681 | 0.121278464 | 0.484230089 | 0.628222585 | 0.796810121 | 0.058726681 |
| ENSMUSG00000041815 | Poldip3 | 73826 | protein_coding | 1117.286957 | -0.058771807 | 0.053668732 | 1.095084693 | 0.273479534 | 0.489835334 | 0.058771807 |
| ENSMUSG00000016495 | Plgrkt | 67759 | protein_coding | 815.9143735 | 0.058776888 | 0.090356171 | -0.650502203 | 0.515367879 | 0.716335759 | 0.058776888 |
| ENSMUSG00000031948 | Kars | 85305 | protein_coding | 1444.090208 | -0.058813531 | 0.078180823 | 0.752275668 | 0.451885297 | 0.668257706 | 0.058813531 |
| ENSMUSG00000024269 | Tpgs2 | 66648 | protein_coding | 83.5255598 | 0.058820115 | 0.171180539 | -0.3436145 | 0.731136218 | 0.861504353 | 0.058820115 |
| ENSMUSG00000106464 | C130083M11Rik | 414111 | lncRNA | 10.74303857 | 0.058831254 | 0.314294711 | -0.187184994 | 0.85151559 | 0.927499565 | 0.058831254 |
| ENSMUSG00000022295 | Atp6v1c1 | 66335 | protein_coding | 419.9322619 | -0.058885387 | 0.070121551 | 0.839761626 | 0.401042053 | 0.622737045 | 0.058885387 |
| ENSMUSG00000026439 | Rbbp5 | 213464 | protein_coding | 372.1020827 | 0.05890086 | 0.074717837 | -0.788310565 | 0.430515069 | 0.648975432 | 0.05890086 |
| ENSMUSG00000010376 | Nedd8 | 18002 | protein_coding | 810.615637 | -0.058937782 | 0.060602109 | 0.972536807 | 0.330783563 | 0.553114597 | 0.058937782 |
| ENSMUSG00000039047 | Pigk | 329777 | protein_coding | 505.3824965 | -0.058943045 | 0.091824573 | 0.641909278 | 0.520932089 | 0.720614094 | 0.058943045 |
| ENSMUSG00000049303 | Syt12 | 171180 | protein_coding | 12.37763451 | -0.058945688 | 0.367871996 | 0.160234235 | 0.872696562 | 0.937225143 | 0.058945688 |
| ENSMUSG00000021707 | Dhfr | 13361 | protein_coding | 184.6434921 | 0.05894921 | 0.108010815 | -0.545771371 | 0.585223105 | 0.767282764 | 0.05894921 |
| ENSMUSG00000043088 | Il17re | 57890 | protein_coding | 298.3563706 | -0.058974973 | 0.108878979 | 0.541656192 | 0.588055374 | 0.769309474 | 0.058974973 |
| ENSMUSG00000017264 | Exosc10 | 50912 | protein_coding | 446.5303307 | -0.059051944 | 0.073522457 | 0.803182408 | 0.421869312 | 0.641714691 | 0.059051944 |
| ENSMUSG00000044367 | Slc16a13 | 69309 | protein_coding | 81.84054082 | -0.059058069 | 0.120119157 | 0.491662363 | 0.622958049 | 0.793281643 | 0.059058069 |
| ENSMUSG00000011148 | Adssl1 | 11565 | protein_coding | 4.061033187 | 0.059060448 | 0.528475531 | -0.111756258 | 0.911016672 | 1 | 0.059060448 |
| ENSMUSG00000042303 | Sgsm3 | 105835 | protein_coding | 1531.056034 | 0.059096091 | 0.105140498 | -0.562067831 | 0.574069805 | 0.759387116 | 0.059096091 |
| ENSMUSG00000000194 | Gpr107 | 277463 | protein_coding | 1049.89801 | -0.05912774 | 0.052068546 | 1.135575021 | 0.256134468 | 0.468895375 | 0.05912774 |
| ENSMUSG00000038784 | Cnot4 | 53621 | protein_coding | 198.9181572 | -0.059151401 | 0.080247895 | 0.737108434 | 0.461056411 | 0.674995696 | 0.059151401 |
| ENSMUSG00000034538 | Zfp418 | 232854 | protein_coding | 39.54956756 | -0.059178904 | 0.183212918 | 0.323006176 | 0.746690565 | 0.872440598 | 0.059178904 |
| ENSMUSG00000103711 | Tstd1 | 226654 | protein_coding | 1445.58105 | -0.059182376 | 0.115701392 | 0.511509631 | 0.608994246 | 0.783067117 | 0.059182376 |
| ENSMUSG00000040848 | Sft2d2 | 108735 | protein_coding | 2626.181642 | -0.059215131 | 0.056881554 | 1.041025207 | 0.297863849 | 0.517488122 | 0.059215131 |
| ENSMUSG00000034738 | Nostrin | 329416 | protein_coding | 802.6478818 | 0.059216936 | 0.087088676 | -0.679961377 | 0.496528917 | 0.702568634 | 0.059216936 |

| ensembl_gene_id | external_gene_name | entrezgene_id | gene_biotype | baseMean | log2FoldChange | lfcSE | stat | pvalue | padj | absLog2FC |
|---|---|---|---|---|---|---|---|---|---|---|
| ENSMUSG00000091512 | Lamtor3 | 56692 | protein_coding | 315.5351406 | -0.059217509 | 0.080399566 | 0.736540157 | 0.461402038 | 0.675184655 | 0.059217509 |
| ENSMUSG00000052369 | Tmem106c | 380967 | protein_coding | 171.2943808 | -0.059222325 | 0.126758949 | 0.467204291 | 0.640353724 | 0.803678477 | 0.059222325 |
| ENSMUSG00000006638 | Abhd1 | 57742 | protein_coding | 6.178501974 | -0.059253962 | 0.614740058 | 0.096388646 | 0.923211909 | 0.962738158 | 0.059253962 |
| ENSMUSG00000035529 | Prdm4 | 72843 | protein_coding | 374.4402374 | 0.059262894 | 0.066313618 | -0.893676073 | 0.371495231 | 0.593888891 | 0.059262894 |
| ENSMUSG00000020707 | Rnf135 | 71956 | protein_coding | 78.5694516 | -0.059266943 | 0.127479031 | 0.464915227 | 0.641992166 | 0.804573016 | 0.059266943 |
| ENSMUSG00000019927 | Ube2d1 | 216080 | protein_coding | 113.3980404 | -0.059281539 | 0.125222958 | 0.473407916 | 0.635922183 | 0.801613541 | 0.059281539 |
| ENSMUSG00000001783 | Rtcb | 28088 | protein_coding | 1753.385115 | -0.059308757 | 0.102759187 | 0.577162575 | 0.563829637 | 0.751075471 | 0.059308757 |
| ENSMUSG00000055882 | Abhd16b | 241850 | protein_coding | 2.56245946 | 0.059363746 | 0.67441636 | -0.088022399 | 0.929858874 | 1 | 0.059363746 |
| ENSMUSG00000059920 | 4930453N24Rik | 67609 | protein_coding | 337.1834794 | -0.059367844 | 0.089150896 | 0.665925374 | 0.505458801 | 0.709429427 | 0.059367844 |
| ENSMUSG00000022055 | Nefl | 18039 | protein_coding | 3.368687167 | 0.059484739 | 0.664312351 | -0.089543328 | 0.928650121 | 1 | 0.059484739 |
| ENSMUSG00000030538 | Cib1 | 23991 | protein_coding | 1889.924488 | 0.059526833 | 0.065375373 | -0.910539089 | 0.362538277 | 0.584928009 | 0.059526833 |
| ENSMUSG00000024273 | 2700062C07Rik | 68046 | protein_coding | 96.32911124 | -0.059536749 | 0.124765434 | 0.477189449 | 0.633227223 | 0.800207711 | 0.059536749 |
| ENSMUSG00000014769 | Psmb1 | 19170 | protein_coding | 1661.638129 | 0.059545846 | 0.091965757 | -0.647478453 | 0.517322335 | 0.717843966 | 0.059545846 |
| ENSMUSG00000047867 | Gimap6 | 231931 | protein_coding | 26.5092813 | 0.059555951 | 0.337590962 | -0.176414532 | 0.859968289 | 0.931308041 | 0.059555951 |
| ENSMUSG00000041673 | Lrrc18 | 67580 | protein_coding | 1.367741268 | -0.059556979 | 1.282539305 | 0.046436767 | 0.962962132 | 1 | 0.059556979 |
| ENSMUSG00000031576 | Kcnu1 | 16532 | protein_coding | 4.264300142 | -0.059571653 | 0.502570972 | 0.118533812 | 0.905644705 | 1 | 0.059571653 |
| ENSMUSG00000028099 | Polr3c | 74414 | protein_coding | 448.8417663 | -0.059572485 | 0.081198667 | 0.733663328 | 0.463153954 | 0.676807614 | 0.059572485 |
| ENSMUSG00000002957 | Ap2a2 | 11772 | protein_coding | 1403.244422 | 0.059578067 | 0.057529002 | -1.035617948 | 0.300380415 | 0.520512296 | 0.059578067 |
| ENSMUSG00000057637 | Prdm2 | 110593 | protein_coding | 482.5717085 | 0.05958457 | 0.110323639 | -0.540088878 | 0.589135741 | 0.770185847 | 0.05958457 |
| ENSMUSG00000030741 | Spns1 | 73658 | protein_coding | 405.5596154 | 0.059595994 | 0.085692118 | -0.69546646 | 0.486763024 | 0.695624496 | 0.059595994 |
| ENSMUSG00000116648 | Rpl31-ps12 | 665562 | processed_pseudogene | 1.16345761 | 0.05962217 | 1.031870377 | -0.057780678 | 0.95392333 | 1 | 0.05962217 |
| ENSMUSG00000049421 | Zfp260 | 26466 | protein_coding | 428.5618219 | 0.059627729 | 0.132658126 | -0.449484183 | 0.653082415 | 0.811818066 | 0.059627729 |
| ENSMUSG00000074269 | Rec114 | 73673 | protein_coding | 2.937927397 | 0.059680115 | 0.809538556 | -0.073721152 | 0.941232268 | 1 | 0.059680115 |
| ENSMUSG00000031796 | Cfap20 | 14894 | protein_coding | 253.0089725 | 0.059707864 | 0.097129237 | -0.614725962 | 0.538735711 | 0.733468492 | 0.059707864 |
| ENSMUSG00000026988 | Wdsub1 | 72137 | protein_coding | 186.4479868 | 0.059735629 | 0.110930629 | -0.538495362 | 0.590235106 | 0.770846581 | 0.059735629 |
| ENSMUSG00000022671 | Mzt2 | 72083 | protein_coding | 144.0271302 | -0.059803919 | 0.116729032 | 0.512331152 | 0.608419266 | 0.782626881 | 0.059803919 |
| ENSMUSG00000019982 | Myb | 17863 | protein_coding | 1250.196708 | 0.059873021 | 0.146378163 | -0.409029733 | 0.682517842 | 0.830405926 | 0.059873021 |
| ENSMUSG00000025735 | Rhbdl1 | 214951 | protein_coding | 11.26657583 | -0.059940983 | 0.340977035 | 0.175791849 | 0.860457474 | 0.931471303 | 0.059940983 |
| ENSMUSG00000052917 | Senp7 | 66315 | protein_coding | 197.1584131 | -0.05996583 | 0.107379816 | 0.558446008 | 0.576539863 | 0.761222618 | 0.05996583 |
| ENSMUSG00000032812 | Arap1 | 69710 | protein_coding | 1155.941913 | 0.059988863 | 0.057848384 | -1.03700155 | 0.299735135 | 0.519697021 | 0.059988863 |
| ENSMUSG00000051786 | Tubgcp6 | 328580 | protein_coding | 295.5191081 | 0.06001989 | 0.098054065 | -0.612110164 | 0.540464873 | 0.73470834 | 0.06001989 |
| ENSMUSG00000034570 | Inpp5j | 170835 | protein_coding | 862.3470057 | 0.060069566 | 0.222961191 | -0.269417141 | 0.787608697 | 0.893615254 | 0.060069566 |
| ENSMUSG00000026578 | Ccdc181 | 74895 | protein_coding | 115.5476576 | -0.060088481 | 0.108015815 | 0.556293365 | 0.578010322 | 0.76182422 | 0.060088481 |
| ENSMUSG00000004054 | Map3k11 | 26403 | protein_coding | 1005.545006 | 0.060103976 | 0.072821742 | -0.825357564 | 0.40916862 | 0.629730716 | 0.060103976 |
| ENSMUSG00000000567 | Sox9 | 20682 | protein_coding | 520.5531139 | 0.060118921 | 0.107166314 | -0.560987116 | 0.57480632 | 0.760117839 | 0.060118921 |
| ENSMUSG00000031333 | Abcb7 | 11306 | protein_coding | 885.9039725 | -0.060120583 | 0.09279499 | 0.64788609 | 0.517058628 | 0.717711842 | 0.060120583 |
| ENSMUSG00000033615 | Cplx1 | 12889 | protein_coding | 1.071992129 | -0.060137987 | 1.814992504 | 0.033134014 | 0.973567719 | 1 | 0.060137987 |
| ENSMUSG00000032726 | Bmp8a | 12163 | protein_coding | 79.37403579 | -0.060165004 | 0.155363055 | 0.387254256 | 0.698567987 | 0.840298483 | 0.060165004 |
| ENSMUSG00000084798 | 4930533B01Rik | 75166 | lncRNA | 7.237681953 | -0.060173634 | 0.476480458 | 0.126287727 | 0.899504171 | 0.950621521 | 0.060173634 |

| ensembl_gene_id | external_gene_name | entrezgene_id | gene_biotype | baseMean | log2FoldChange | lfcSE | stat | pvalue | padj | absLog2FC |
|---|---|---|---|---|---|---|---|---|---|---|
| ENSMUSG00000051504 | Siglech | 233274 | protein_coding | 2.478526697 | -0.060247903 | 1.786072951 | 0.03373205 | 0.973090821 | 1 | 0.060247903 |
| ENSMUSG00000018849 | Wwc1 | 211652 | protein_coding | 736.3357747 | -0.06035137 | 0.079903322 | 0.755304885 | 0.450066068 | 0.666821375 | 0.06035137 |
| ENSMUSG00000006526 | Stimate | 69179 | protein_coding | 156.8328039 | 0.060355174 | 0.105867972 | -0.570098525 | 0.568610876 | 0.754583735 | 0.060355174 |
| ENSMUSG00000064090 | Vrk2 | 69922 | protein_coding | 83.47234383 | 0.060360134 | 0.118269102 | -0.510362664 | 0.609797409 | 0.783550764 | 0.060360134 |
| ENSMUSG00000025920 | Stau2 | 29819 | protein_coding | 173.0622286 | 0.060375179 | 0.092096814 | -0.655562076 | 0.512105933 | 0.714689206 | 0.060375179 |
| ENSMUSG00000028060 | Khdc4 | 74200 | protein_coding | 641.2375582 | 0.060384967 | 0.117372097 | -0.514474634 | 0.606920191 | 0.781739336 | 0.060384967 |
| ENSMUSG00000016921 | Srsf6 | 67996 | protein_coding | 1530.817171 | 0.060420672 | 0.06781649 | -0.890943657 | 0.372959398 | 0.595257447 | 0.060420672 |
| ENSMUSG00000097177 | 9330159M07Rik | 319673 | lncRNA | 19.47911415 | 0.060424378 | 0.267235111 | -0.226109429 | 0.821116311 | 0.909608677 | 0.060424378 |
| ENSMUSG00000024535 | Snx24 | 69226 | protein_coding | 241.0129987 | 0.060425642 | 0.129666375 | -0.46600857 | 0.641209366 | 0.804106371 | 0.060425642 |
| ENSMUSG00000031845 | Bco1 | 63857 | protein_coding | 27.50523309 | 0.060465097 | 0.247925522 | -0.243884116 | 0.807320571 | 0.904987729 | 0.060465097 |
| ENSMUSG00000041629 | Fam104a | 28081 | protein_coding | 176.4185183 | -0.060477996 | 0.108516768 | 0.557314761 | 0.577312392 | 0.761298159 | 0.060477996 |
| ENSMUSG00000035236 | Scai | 320271 | protein_coding | 95.66243056 | 0.060498086 | 0.137319896 | -0.440563146 | 0.659529288 | 0.8161622 | 0.060498086 |
| ENSMUSG00000074064 | Mlycd | 56690 | protein_coding | 452.9213898 | 0.060536959 | 0.063968927 | -0.94634945 | 0.343970379 | 0.56579982 | 0.060536959 |
| ENSMUSG00000042548 | Asxl1 | 228790 | protein_coding | 679.0116527 | -0.060542079 | 0.061848506 | 0.978876987 | 0.327640764 | 0.549807904 | 0.060542079 |
| ENSMUSG00000032462 | Pik3cb | 74769 | protein_coding | 172.469929 | 0.06054469 | 0.091608643 | -0.660905873 | 0.50867268 | 0.712245263 | 0.06054469 |
| ENSMUSG00000043557 | Mdga1 | 74762 | protein_coding | 13.13685766 | 0.060584822 | 0.579957088 | -0.104464319 | 0.916800883 | 0.959274295 | 0.060584822 |
| ENSMUSG00000042211 | Fbxo38 | 107035 | protein_coding | 877.4665073 | 0.060597605 | 0.052276328 | -1.159178681 | 0.246383359 | 0.458272537 | 0.060597605 |
| ENSMUSG00000037997 | Parp11 | 101187 | protein_coding | 116.4256213 | -0.06062911 | 0.131654726 | 0.460516011 | 0.645145882 | 0.80668897 | 0.06062911 |
| ENSMUSG00000035569 | Ankrd11 | 77087 | protein_coding | 2434.147604 | 0.060642478 | 0.085414731 | -0.709976809 | 0.477718517 | 0.68930064 | 0.060642478 |
| ENSMUSG00000004071 | Cdip1 | 66626 | protein_coding | 330.1607526 | -0.060657373 | 0.087653757 | 0.692011105 | 0.488930355 | 0.696651931 | 0.060657373 |
| ENSMUSG00000024776 | Stambpl1 | 76630 | protein_coding | 212.6538638 | 0.060676547 | 0.09238615 | -0.656771038 | 0.511328149 | 0.71414691 | 0.060676547 |
| ENSMUSG00000062040 | Zfp27 | 22689 | protein_coding | 124.4393726 | 0.060682221 | 0.094072654 | -0.645056971 | 0.518890267 | 0.719334577 | 0.060682221 |
| ENSMUSG00000006538 | Ihh | 16147 | protein_coding | 1115.852292 | -0.060693169 | 0.092290631 | 0.657630883 | 0.510775345 | 0.713797415 | 0.060693169 |
| ENSMUSG00000033985 | Tesk2 | 230661 | protein_coding | 377.7315813 | 0.06073987 | 0.130705104 | -0.464709244 | 0.642139688 | 0.804696828 | 0.06073987 |
| ENSMUSG00000031622 | Sin3b | 20467 | protein_coding | 825.8050992 | 0.060741916 | 0.043182706 | -1.406625985 | 0.159538324 | 0.343720271 | 0.060741916 |
| ENSMUSG00000022387 | Brd1 | 223770 | protein_coding | 1284.146688 | -0.06080027 | 0.058097237 | 1.046526006 | 0.295318244 | 0.514433539 | 0.06080027 |
| ENSMUSG00000021314 | Amph | 218038 | protein_coding | 29.39039281 | 0.060845096 | 0.255174592 | -0.23844496 | 0.811535999 | 0.906704483 | 0.060845096 |
| ENSMUSG00000030032 | Wdr54 | 75659 | protein_coding | 20.14062541 | 0.060879373 | 0.376937695 | -0.161510439 | 0.87169139 | 0.936825965 | 0.060879373 |
| ENSMUSG00000068290 | Ddrgk1 | 77006 | protein_coding | 799.0359983 | -0.060889882 | 0.066770548 | 0.911927242 | 0.36180702 | 0.584169058 | 0.060889882 |
| ENSMUSG00000044433 | Camsap3 | 69697 | protein_coding | 832.1750987 | 0.060898674 | 0.082113805 | -0.741637464 | 0.458307019 | 0.67278137 | 0.060898674 |
| ENSMUSG00000002384 | Bmp8b | 12164 | protein_coding | 13.18293562 | -0.060901687 | 0.453089911 | 0.134414132 | 0.893075108 | 0.947328174 | 0.060901687 |
| ENSMUSG00000064165 | Krt39 | 237934 | protein_coding | 0.880641899 | -0.060901777 | 1.339613039 | 0.045462216 | 0.963738891 | 1 | 0.060901777 |
| ENSMUSG00000006215 | Zbtb17 | 22642 | protein_coding | 323.9627921 | -0.060904489 | 0.065922031 | 0.923886719 | 0.355545298 | 0.578413462 | 0.060904489 |
| ENSMUSG00000010277 | Bltp2 | 72503 | protein_coding | 4626.661819 | -0.060909872 | 0.056132627 | 1.085106382 | 0.277874541 | 0.494858209 | 0.060909872 |
| ENSMUSG00000044024 | Rell2 | 225392 | protein_coding | 8.302219111 | -0.060931601 | 0.459193283 | 0.132692709 | 0.894436409 | 0.947833434 | 0.060931601 |
| ENSMUSG00000028580 | Pum1 | 80912 | protein_coding | 1063.33865 | 0.060950128 | 0.05278898 | -1.154599461 | 0.248254488 | 0.460585845 | 0.060950128 |
| ENSMUSG00000028886 | Eya3 | 14050 | protein_coding | 480.1215137 | 0.06096203 | 0.068444888 | -0.89067324 | 0.373104495 | 0.595366473 | 0.06096203 |
| ENSMUSG00000034009 | Rxfp1 | 381489 | protein_coding | 3.586262302 | 0.060966845 | 2.242687417 | -0.027184727 | 0.978312397 | 1 | 0.060966845 |
| ENSMUSG00000070661 | Rnf186 | 66825 | protein_coding | 1679.074283 | -0.061029613 | 0.118659847 | 0.514324052 | 0.607025449 | 0.781796866 | 0.061029613 |

| ensembl_gene_id | external_gene_name | entrezgene_id | gene_biotype | baseMean | log2FoldChange | lfcSE | stat | pvalue | padj | absLog2FC |
|---|---|---|---|---|---|---|---|---|---|---|
| ENSMUSG00000028771 | Ptpn12 | 19248 | protein_coding | 324.5785497 | -0.061082831 | 0.081455684 | 0.749890342 | 0.453320752 | 0.669122175 | 0.061082831 |
| ENSMUSG00000025316 | Banp | 53325 | protein_coding | 183.3974089 | 0.061128306 | 0.21471149 | -0.284699742 | 0.775874189 | 0.888050066 | 0.061128306 |
| ENSMUSG00000031016 | Wee1 | 22390 | protein_coding | 163.8924021 | 0.061129666 | 0.154545554 | -0.395544642 | 0.692440983 | 0.836513744 | 0.061129666 |
| ENSMUSG00000019370 | Calm3 | 12315 | protein_coding | 5825.809008 | -0.061129972 | 0.072255077 | 0.846030123 | 0.39753594 | 0.619446591 | 0.061129972 |
| ENSMUSG00000031145 | Prickle3 | 54630 | protein_coding | 115.4786011 | -0.061145528 | 0.117080693 | 0.522251158 | 0.601495471 | 0.777725809 | 0.061145528 |
| ENSMUSG00000098678 | Mroh6 | 223645 | protein_coding | 78.25381665 | 0.061150745 | 0.597267681 | -0.102384152 | 0.918451763 | 0.960078011 | 0.061150745 |
| ENSMUSG00000000600 | Krit1 | 79264 | protein_coding | 322.6532954 | -0.061152756 | 0.085467716 | 0.715507081 | 0.474295767 | 0.686291751 | 0.061152756 |
| ENSMUSG00000035958 | Tdp2 | 56196 | protein_coding | 273.2652868 | -0.061161461 | 0.082427405 | 0.74200397 | 0.45808493 | 0.672740084 | 0.061161461 |
| ENSMUSG00000038766 | Gabpb2 | 213054 | protein_coding | 962.1262061 | 0.061165743 | 0.078706633 | -0.777135813 | 0.437078646 | 0.655340182 | 0.061165743 |
| ENSMUSG00000026404 | Ddx59 | 67997 | protein_coding | 50.29384407 | 0.061195165 | 0.192110451 | -0.31854157 | 0.750074167 | 0.874455706 | 0.061195165 |
| ENSMUSG00000060407 | Cyp2a12 | 13085 | protein_coding | 2.384692582 | 0.061320056 | 1.378485211 | -0.044483652 | 0.964518883 | 1 | 0.061320056 |
| ENSMUSG00000005882 | Uqcc1 | 56046 | protein_coding | 642.8517209 | -0.061336181 | 0.089203118 | 0.687601313 | 0.491703881 | 0.698494442 | 0.061336181 |
| ENSMUSG00000058486 | Wdr91 | 101240 | protein_coding | 50.62106656 | -0.06134655 | 0.181742185 | 0.337547113 | 0.735704503 | 0.865466887 | 0.06134655 |
| ENSMUSG00000032565 | Nudt16 | 75686 | protein_coding | 229.6389046 | 0.061352647 | 0.083587073 | -0.733996836 | 0.462950667 | 0.676623357 | 0.061352647 |
| ENSMUSG00000039099 | Wdr93 | 626359 | protein_coding | 5.342610216 | -0.061352731 | 0.472133586 | 0.129947821 | 0.896607709 | 0.94900212 | 0.061352731 |
| ENSMUSG00000019907 | Ppp1r12a | 17931 | protein_coding | 1271.924976 | -0.06137262 | 0.068211371 | 0.899741774 | 0.368257687 | 0.5907413 | 0.06137262 |
| ENSMUSG00000042167 | Tent2 | 100715 | protein_coding | 279.4524173 | 0.061380938 | 0.092431119 | -0.664072218 | 0.50664409 | 0.710428845 | 0.061380938 |
| ENSMUSG00000022750 | Klhl22 | 224023 | protein_coding | 411.2233039 | 0.06140126 | 0.07867276 | -0.780464034 | 0.435117789 | 0.653468095 | 0.06140126 |
| ENSMUSG00000057789 | Bak1 | 12018 | protein_coding | 979.8344782 | -0.061415208 | 0.079682826 | 0.770745859 | 0.440857584 | 0.658733455 | 0.061415208 |
| ENSMUSG00000018846 | Pank3 | 211347 | protein_coding | 4424.538117 | 0.061415995 | 0.093878611 | -0.65420648 | 0.512978787 | 0.714956306 | 0.061415995 |
| ENSMUSG00000004113 | Cacna1b | 12287 | protein_coding | 96.21697865 | 0.061426303 | 0.141976312 | -0.432651771 | 0.665267772 | 0.819481407 | 0.061426303 |
| ENSMUSG00000020745 | Pafah1b1 | 18472 | protein_coding | 2167.125183 | 0.061429667 | 0.051980199 | -1.181789755 | 0.237289134 | 0.44767168 | 0.061429667 |
| ENSMUSG00000006299 | Aamp | 227290 | protein_coding | 1509.329689 | 0.061430623 | 0.051585786 | -1.190843992 | 0.233714835 | 0.443169915 | 0.061430623 |
| ENSMUSG00000062014 | Gmfb | 63985 | protein_coding | 944.1541707 | -0.061433919 | 0.094870648 | 0.647554544 | 0.517273105 | 0.717835971 | 0.061433919 |
| ENSMUSG00000026123 | Plekhb2 | 226971 | protein_coding | 4353.610701 | -0.061449204 | 0.051137646 | 1.2016432 | 0.229501797 | 0.438111017 | 0.061449204 |
| ENSMUSG00000037157 | Il22ra1 | 230828 | protein_coding | 399.7007517 | 0.061454037 | 0.169171004 | -0.363265783 | 0.716406351 | 0.852797822 | 0.061454037 |
| ENSMUSG00000071350 | Setdb2 | 239122 | protein_coding | 131.0039703 | 0.061459596 | 0.10278957 | -0.597916661 | 0.549895542 | 0.74082028 | 0.061459596 |
| ENSMUSG00000028800 | Hdac1 | 433759 | protein_coding | 925.6909707 | -0.061465602 | 0.095812491 | 0.641519715 | 0.521185076 | 0.720841738 | 0.061465602 |
| ENSMUSG00000025035 | Arl3 | 56350 | protein_coding | 63.51519192 | 0.061491482 | 0.151282512 | -0.406467879 | 0.684398854 | 0.831662311 | 0.061491482 |
| ENSMUSG00000038506 | Dcun1d2 | 102323 | protein_coding | 285.0660373 | -0.061492623 | 0.083675509 | 0.734893921 | 0.462404101 | 0.675987813 | 0.061492623 |
| ENSMUSG00000025592 | Dach2 | 93837 | protein_coding | 7.424815713 | 0.061495585 | 0.42136168 | -0.145944891 | 0.883964893 | 0.94246215 | 0.061495585 |
| ENSMUSG00000015943 | Bola1 | 69168 | protein_coding | 146.1559798 | -0.061509539 | 0.115134172 | 0.534242246 | 0.593173944 | 0.772538316 | 0.061509539 |
| ENSMUSG00000029466 | Anapc7 | 56317 | protein_coding | 304.508052 | -0.061536297 | 0.094823147 | 0.6489586 | 0.516365136 | 0.717238948 | 0.061536297 |
| ENSMUSG00000038563 | Efl1 | 101592 | protein_coding | 322.9297096 | -0.061598343 | 0.070430852 | 0.874593181 | 0.381795299 | 0.604102573 | 0.061598343 |
| ENSMUSG00000039512 | Uhrf1bp1 | 224648 | protein_coding | 208.3966934 | -0.061700116 | 0.132017059 | 0.467364725 | 0.640238956 | 0.803678477 | 0.061700116 |
| ENSMUSG00000099145 | Mir8116 | 102466887 | miRNA | 2.649642625 | 0.061735789 | 0.861004096 | -0.071702085 | 0.942838997 | 1 | 0.061735789 |
| ENSMUSG00000071856 | Mcc | 328949 | protein_coding | 329.3948469 | -0.061740899 | 0.133207244 | 0.463495055 | 0.643009558 | 0.805274537 | 0.061740899 |
| ENSMUSG00000056204 | Pgpep1 | 66522 | protein_coding | 685.0482887 | -0.061835205 | 0.089835714 | 0.688314286 | 0.491254887 | 0.698491194 | 0.061835205 |
| ENSMUSG00000035933 | Cog5 | 238123 | protein_coding | 575.6117785 | -0.061858348 | 0.073523782 | 0.841337958 | 0.400158632 | 0.622301501 | 0.061858348 |

| ensembl_gene_id | external_gene_name | entrezgene_id | gene_biotype | baseMean | log2FoldChange | lfcSE | stat | pvalue | padj | absLog2FC |
|---|---|---|---|---|---|---|---|---|---|---|
| ENSMUSG00000044156 | Hepacam2 | 101202 | protein_coding | 1616.887732 | -0.061873139 | 0.272669658 | 0.226916115 | 0.820488971 | 0.9092783 | 0.061873139 |
| ENSMUSG00000054435 | Gimap4 | 107526 | protein_coding | 32.80574215 | -0.061901439 | 0.343514482 | 0.180200376 | 0.856995263 | 0.93035707 | 0.061901439 |
| ENSMUSG00000026333 | Gin1 | 252876 | protein_coding | 68.20994423 | 0.061916074 | 0.130905009 | -0.47298476 | 0.636224052 | 0.801687789 | 0.061916074 |
| ENSMUSG00000031917 | Nip7 | 66164 | protein_coding | 203.7250508 | -0.061919432 | 0.102029963 | 0.606874976 | 0.543933886 | 0.73666903 | 0.061919432 |
| ENSMUSG00000052914 | Cyp2j6 | 13110 | protein_coding | 834.3909343 | 0.061920104 | 0.093147629 | -0.66475233 | 0.506208917 | 0.710135685 | 0.061920104 |
| ENSMUSG00000052144 | Ppp4r2 | 232314 | protein_coding | 525.6202662 | -0.06193902 | 0.100703552 | 0.615062909 | 0.538513176 | 0.733377497 | 0.06193902 |
| ENSMUSG00000017631 | Abr | 109934 | protein_coding | 2774.525899 | -0.061975022 | 0.082204708 | 0.753910855 | 0.450902752 | 0.667636772 | 0.061975022 |
| ENSMUSG00000031169 | Porcn | 53627 | protein_coding | 28.38268476 | -0.061979962 | 0.239555833 | 0.25872867 | 0.795844597 | 0.898576349 | 0.061979962 |
| ENSMUSG00000020900 | Myh10 | 77579 | protein_coding | 20.2883211 | 0.062001891 | 0.45532795 | -0.136169745 | 0.891687093 | 0.947050968 | 0.062001891 |
| ENSMUSG00000053868 | Gm5142 | 380907 | lncRNA | 1.833544745 | 0.062008962 | 0.995754509 | -0.062273343 | 0.950345156 | 1 | 0.062008962 |
| ENSMUSG00000052632 | Asap2 | 211914 | protein_coding | 461.0613836 | -0.062009271 | 0.115364505 | 0.537507365 | 0.590917199 | 0.771232643 | 0.062009271 |
| ENSMUSG00000034377 | Tulp4 | 68842 | protein_coding | 854.1876338 | -0.062024402 | 0.065497836 | 0.946968731 | 0.343654713 | 0.56547568 | 0.062024402 |
| ENSMUSG00000021820 | Camk2g | 12325 | protein_coding | 144.3587938 | 0.062061604 | 0.102098931 | -0.60785753 | 0.543281969 | 0.736456016 | 0.062061604 |
| ENSMUSG00000040717 | Il17rd | 171463 | protein_coding | 58.97202401 | -0.062083096 | 0.184850121 | 0.335856398 | 0.736979158 | 0.86653427 | 0.062083096 |
| ENSMUSG00000055493 | Epm2a | 13853 | protein_coding | 13.47981553 | -0.062097238 | 0.360304729 | 0.172346441 | 0.863165176 | 0.932166328 | 0.062097238 |
| ENSMUSG00000011837 | Snapc2 | 102209 | protein_coding | 148.1569327 | -0.062131293 | 0.093999579 | 0.660974154 | 0.508628889 | 0.712244337 | 0.062131293 |
| ENSMUSG00000063884 | Ptcd3 | 69956 | protein_coding | 520.0081868 | -0.062151163 | 0.091830734 | 0.676801327 | 0.498532016 | 0.703739933 | 0.062151163 |
| ENSMUSG00000038150 | Ormdl3 | 66612 | protein_coding | 273.7271575 | 0.06219563 | 0.095479071 | -0.651405898 | 0.514784504 | 0.716272317 | 0.06219563 |
| ENSMUSG00000022489 | Pde1b | 18574 | protein_coding | 75.35047773 | 0.062223239 | 0.124893425 | -0.498210689 | 0.618335549 | 0.789606655 | 0.062223239 |
| ENSMUSG00000055943 | Emc7 | 73024 | protein_coding | 501.0723743 | 0.062229735 | 0.095505558 | -0.65158234 | 0.514670643 | 0.716210584 | 0.062229735 |
| ENSMUSG00000022075 | Rhobtb2 | 246710 | protein_coding | 192.9243397 | -0.062271849 | 0.079162519 | 0.786632987 | 0.431496743 | 0.649802774 | 0.062271849 |
| ENSMUSG00000072825 | Cep170b | 217882 | protein_coding | 2206.006064 | -0.062341457 | 0.052783361 | 1.181081622 | 0.2375703 | 0.447960257 | 0.062341457 |
| ENSMUSG00000029790 | Cep41 | 83922 | protein_coding | 57.15421145 | 0.062348702 | 0.18083365 | -0.344784843 | 0.730256129 | 0.861143224 | 0.062348702 |
| ENSMUSG00000052957 | Gas1 | 14451 | protein_coding | 0.765117643 | -0.062364763 | 1.743161708 | 0.035776809 | 0.971460325 | 1 | 0.062364763 |
| ENSMUSG00000014554 | Dguok | 27369 | protein_coding | 202.5056706 | 0.062403652 | 0.091415846 | -0.682634956 | 0.49483754 | 0.700835947 | 0.062403652 |
| ENSMUSG00000058076 | Sdhc | 66052 | protein_coding | 2220.521394 | -0.062427277 | 0.093026025 | 0.671073245 | 0.50217387 | 0.706921771 | 0.062427277 |
| ENSMUSG00000057130 | Txnl4a | 27366 | protein_coding | 242.7498073 | 0.062490468 | 0.108085242 | -0.578159116 | 0.563156698 | 0.750770075 | 0.062490468 |
| ENSMUSG00000034021 | Pds5b | 100710 | protein_coding | 255.6994972 | 0.06252291 | 0.127405013 | -0.490741368 | 0.623609384 | 0.793440482 | 0.06252291 |
| ENSMUSG00000008859 | Rala | 56044 | protein_coding | 781.4618439 | 0.062539738 | 0.06934238 | -0.90189777 | 0.367111176 | 0.589852079 | 0.062539738 |
| ENSMUSG00000028102 | Pex11b | 18632 | protein_coding | 122.9769866 | -0.062553974 | 0.120248356 | 0.520206475 | 0.602919673 | 0.778895054 | 0.062553974 |
| ENSMUSG00000041975 | Mettl8 | 228019 | protein_coding | 98.69917926 | 0.062569794 | 0.119991431 | -0.521452183 | 0.602051808 | 0.778258299 | 0.062569794 |
| ENSMUSG00000032583 | Mon1a | 72825 | protein_coding | 503.5632156 | 0.062579735 | 0.090006032 | -0.695283786 | 0.486877473 | 0.695624496 | 0.062579735 |
| ENSMUSG00000047557 | Lxn | 17035 | protein_coding | 121.1771449 | -0.06259451 | 0.11672663 | 0.536248755 | 0.591786642 | 0.771818402 | 0.06259451 |
| ENSMUSG00000003423 | Pih1d1 | 68845 | protein_coding | 366.8195032 | 0.062607605 | 0.083465942 | -0.750097624 | 0.45319591 | 0.669122175 | 0.062607605 |
| ENSMUSG00000022808 | Snx4 | 69150 | protein_coding | 733.6582934 | -0.06266725 | 0.076948599 | 0.814404047 | 0.415413521 | 0.635668911 | 0.06266725 |
| ENSMUSG00000000531 | Tamalin | 56149 | protein_coding | 71.02418755 | 0.062735671 | 0.245182242 | -0.255873632 | 0.798048416 | 0.899834189 | 0.062735671 |
| ENSMUSG00000037594 | Clba1 | 217887 | protein_coding | 27.57640884 | 0.062800947 | 0.194237475 | -0.323320447 | 0.746452571 | 0.872440598 | 0.062800947 |
| ENSMUSG00000044636 | Csrnp2 | 207785 | protein_coding | 76.55076962 | 0.062816033 | 0.204740013 | -0.306808777 | 0.758988927 | 0.878912418 | 0.062816033 |
| ENSMUSG00000052812 | Atad2b | 320817 | protein_coding | 542.8856793 | 0.062853271 | 0.064572837 | -0.973370135 | 0.330369377 | 0.552639068 | 0.062853271 |

| ensembl_gene_id | external_gene_name | entrezgene_id | gene_biotype | baseMean | log2FoldChange | lfcSE | stat | pvalue | padj | absLog2FC |
|---|---|---|---|---|---|---|---|---|---|---|
| ENSMUSG00000024427 | Spry4 | 24066 | protein_coding | 48.32217485 | -0.062857895 | 0.259821009 | 0.241927684 | 0.808836199 | 0.905678041 | 0.062857895 |
| ENSMUSG00000058690 | Ccser2 | 72972 | protein_coding | 650.7097907 | 0.062898141 | 0.056587777 | -1.111514616 | 0.266346902 | 0.481401448 | 0.062898141 |
| ENSMUSG00000027556 | Car1 | 12346 | protein_coding | 64053.33163 | -0.062906335 | 0.178487572 | 0.35244098 | 0.724507573 | 0.857302814 | 0.062906335 |
| ENSMUSG00000071076 | Jund | 16478 | protein_coding | 2533.569084 | -0.062907297 | 0.149663282 | 0.420325522 | 0.674247668 | 0.825194181 | 0.062907297 |
| ENSMUSG00000032412 | Atp1b3 | 11933 | protein_coding | 454.2286601 | -0.062919808 | 0.118356483 | 0.531612693 | 0.594994272 | 0.773528957 | 0.062919808 |
| ENSMUSG00000021706 | Zfyve16 | 218441 | protein_coding | 415.4504947 | -0.06293298 | 0.078923585 | 0.7973913 | 0.42522381 | 0.644825161 | 0.06293298 |
| ENSMUSG00000029055 | Plch2 | 269615 | protein_coding | 847.6445335 | 0.062961947 | 0.093703582 | -0.671926793 | 0.501630302 | 0.706397443 | 0.062961947 |
| ENSMUSG00000022307 | Oxr1 | 170719 | protein_coding | 522.0198206 | 0.062972618 | 0.079839179 | -0.788743312 | 0.430262048 | 0.648871201 | 0.062972618 |
| ENSMUSG00000070643 | Sox13 | 20668 | protein_coding | 590.0932493 | 0.063016424 | 0.1249677 | -0.504261695 | 0.614077492 | 0.786724259 | 0.063016424 |
| ENSMUSG00000058407 | Txndc9 | 98258 | protein_coding | 688.3861576 | -0.063028424 | 0.074381273 | 0.847369529 | 0.396789183 | 0.618855346 | 0.063028424 |
| ENSMUSG00000021694 | Ercc8 | 71991 | protein_coding | 93.07374297 | 0.063042579 | 0.125526873 | -0.50222377 | 0.615510124 | 0.787780401 | 0.063042579 |
| ENSMUSG00000037652 | Phc3 | 241915 | protein_coding | 467.9703507 | -0.063047429 | 0.096340997 | 0.654419518 | 0.512841563 | 0.714956306 | 0.063047429 |
| ENSMUSG00000046999 | 1110032F04Rik | 68725 | protein_coding | 1.983058731 | -0.06305595 | 1.287180339 | 0.048987658 | 0.960929132 | 1 | 0.06305595 |
| ENSMUSG00000060098 | Prmt7 | 214572 | protein_coding | 276.8985201 | -0.063076619 | 0.138396747 | 0.455766634 | 0.6485578 | 0.809117134 | 0.063076619 |
| ENSMUSG00000032737 | Inppl1 | 16332 | protein_coding | 1270.805821 | -0.063077663 | 0.066469695 | 0.948968747 | 0.342636508 | 0.564644176 | 0.063077663 |
| ENSMUSG00000043279 | Trim56 | 384309 | protein_coding | 766.1801347 | -0.063087466 | 0.070705032 | 0.892262751 | 0.372252115 | 0.594533905 | 0.063087466 |
| ENSMUSG00000062944 | 9130023H24Rik | 100043133 | protein_coding | 96.30482037 | -0.063114579 | 0.150781277 | 0.418583662 | 0.675520432 | 0.826153034 | 0.063114579 |
| ENSMUSG00000047264 | Zfp358 | 140482 | protein_coding | 131.9875309 | 0.063139125 | 0.117280743 | -0.538358845 | 0.590329333 | 0.770846581 | 0.063139125 |
| ENSMUSG00000027531 | Impa1 | 55980 | protein_coding | 965.3801658 | 0.063160847 | 0.070969971 | -0.8899658 | 0.37348425 | 0.59579974 | 0.063160847 |
| ENSMUSG00000025231 | Sufu | 24069 | protein_coding | 98.91341521 | -0.063177379 | 0.133708622 | 0.472500413 | 0.636569647 | 0.801948109 | 0.063177379 |
| ENSMUSG00000024151 | Msh2 | 17685 | protein_coding | 385.5560742 | 0.063204277 | 0.143207401 | -0.441347838 | 0.658961199 | 0.815932012 | 0.063204277 |
| ENSMUSG00000022022 | Mtrf1 | 211253 | protein_coding | 66.51060607 | -0.063254893 | 0.144393908 | 0.438071756 | 0.661334268 | 0.816921981 | 0.063254893 |
| ENSMUSG00000023806 | Rsph3b | 100037282 | protein_coding | 165.1372348 | 0.063319883 | 0.127678931 | -0.495930557 | 0.619943406 | 0.790987051 | 0.063319883 |
| ENSMUSG00000028691 | Prdx1 | 18477 | protein_coding | 4199.930351 | -0.063349832 | 0.098314204 | 0.64436093 | 0.519341414 | 0.719557354 | 0.063349832 |
| ENSMUSG00000019659 | Ccdc12 | 72654 | protein_coding | 599.3843879 | -0.063366826 | 0.096378217 | 0.657480789 | 0.51087182 | 0.713811425 | 0.063366826 |
| ENSMUSG00000028182 | Lrriq3 | 74435 | protein_coding | 20.52989199 | -0.063433665 | 0.282643686 | 0.224429798 | 0.822422886 | 0.910171686 | 0.063433665 |
| ENSMUSG00000051209 | Gpr119 | 236781 | protein_coding | 4.406998855 | 0.063455466 | 0.559434421 | -0.113427889 | 0.90969133 | 0.956064572 | 0.063455466 |
| ENSMUSG00000020921 | Tmem101 | 76547 | protein_coding | 121.268534 | -0.063505803 | 0.108277979 | 0.586507094 | 0.557534795 | 0.74672584 | 0.063505803 |
| ENSMUSG00000069089 | Cdk7 | 12572 | protein_coding | 277.1983076 | 0.06350932 | 0.086606562 | -0.733308405 | 0.46337035 | 0.676855263 | 0.06350932 |
| ENSMUSG00000030246 | Ldhb | 16832 | protein_coding | 64.31383397 | -0.063523439 | 0.15852737 | 0.400709597 | 0.688633944 | 0.83386713 | 0.063523439 |
| ENSMUSG00000073411 | H2-D1 | 14964 | protein_coding | 7161.649703 | 0.063526428 | 0.084616897 | -0.750753465 | 0.45280104 | 0.669012828 | 0.063526428 |
| ENSMUSG00000038170 | Pde4dip | 83679 | protein_coding | 1856.785113 | 0.063595446 | 0.067188181 | -0.946527277 | 0.343879716 | 0.565733178 | 0.063595446 |
| ENSMUSG00000028049 | Scamp3 | 24045 | protein_coding | 145.4763278 | 0.063649089 | 0.119499359 | -0.532631215 | 0.59428889 | 0.77323163 | 0.063649089 |
| ENSMUSG00000041781 | Cpsf2 | 51786 | protein_coding | 678.6042163 | 0.063674471 | 0.093130498 | -0.683712338 | 0.494156831 | 0.700292277 | 0.063674471 |
| ENSMUSG00000031286 | Glt28d2 | 320302 | protein_coding | 104.3562161 | -0.063679242 | 0.173715661 | 0.366571679 | 0.713938537 | 0.851508089 | 0.063679242 |
| ENSMUSG00000022607 | Ptk2 | 14083 | protein_coding | 925.6641341 | 0.063683502 | 0.055973053 | -1.137752873 | 0.255223689 | 0.467839271 | 0.063683502 |
| ENSMUSG00000062190 | Lancl2 | 71835 | protein_coding | 158.5674588 | -0.063695053 | 0.140773984 | 0.45246324 | 0.650935298 | 0.810260975 | 0.063695053 |
| ENSMUSG00000032504 | Pdcd6ip | 18571 | protein_coding | 2706.283118 | 0.063722691 | 0.079138979 | -0.80519981 | 0.420704386 | 0.640678 | 0.063722691 |
| ENSMUSG00000018189 | Uchl5 | 56207 | protein_coding | 202.0796214 | 0.063769874 | 0.145638544 | -0.437863988 | 0.661484882 | 0.816921981 | 0.063769874 |

Supplementary Table 1

| ensembl_gene_id | external_gene_name | entrezgene_id | gene_biotype | baseMean | log2FoldChange | lfcSE | stat | pvalue | padj | absLog2FC |
|---|---|---|---|---|---|---|---|---|---|---|
| ENSMUSG00000041645 | Ddx24 | 27225 | protein_coding | 798.9114487 | -0.063789929 | 0.06348228 | 1.004846219 | 0.314970905 | 0.535990818 | 0.063789929 |
| ENSMUSG00000037089 | Slc35b2 | 73836 | protein_coding | 664.4692759 | 0.063797209 | 0.091440588 | -0.697690273 | 0.485370913 | 0.695380179 | 0.063797209 |
| ENSMUSG00000040025 | Ythdf2 | 213541 | protein_coding | 814.6121221 | -0.063836101 | 0.09137976 | 0.698580313 | 0.484814351 | 0.695226137 | 0.063836101 |
| ENSMUSG00000038366 | Lasp1 | 16796 | protein_coding | 4134.065352 | -0.063868814 | 0.057625947 | 1.108334312 | 0.267717462 | 0.482962166 | 0.063868814 |
| ENSMUSG00000022855 | Senp2 | 75826 | protein_coding | 460.3352808 | 0.063925753 | 0.071299849 | -0.896576269 | 0.369945066 | 0.59260115 | 0.063925753 |
| ENSMUSG00000025200 | Cwf19l1 | 72502 | protein_coding | 126.6303118 | 0.063973174 | 0.108460839 | -0.589827396 | 0.555306373 | 0.745058233 | 0.063973174 |
| ENSMUSG00000044533 | Rps2 | 16898 | protein_coding | 8938.095252 | -0.063980306 | 0.102688345 | 0.62305324 | 0.533249536 | 0.729644779 | 0.063980306 |
| ENSMUSG00000011158 | Brf1 | 72308 | protein_coding | 336.7736962 | -0.064024964 | 0.098557022 | 0.649623567 | 0.515935406 | 0.716883239 | 0.064024964 |
| ENSMUSG00000029491 | Pde6b | 18587 | protein_coding | 4.616758124 | 0.064090933 | 0.793474925 | -0.080772474 | 0.935622899 | 0.96968875 | 0.064090933 |
| ENSMUSG00000028383 | Hsdl2 | 72479 | protein_coding | 378.002759 | -0.064101488 | 0.085614494 | 0.748722378 | 0.454024553 | 0.669801811 | 0.064101488 |
| ENSMUSG00000072772 | Grcc10 | 14790 | protein_coding | 185.8288976 | -0.064109057 | 0.116257342 | 0.551440932 | 0.581331448 | 0.764339772 | 0.064109057 |
| ENSMUSG00000028992 | Nmnat1 | 66454 | protein_coding | 159.3185053 | -0.064153076 | 0.105808623 | 0.606312358 | 0.544307354 | 0.736796542 | 0.064153076 |
| ENSMUSG00000029650 | Slc46a3 | 71706 | protein_coding | 80.90939854 | 0.06415446 | 0.265741767 | -0.241416547 | 0.809232289 | 0.905779176 | 0.06415446 |
| ENSMUSG00000112980 | D430020J02Rik | 319545 | lncRNA | 27.64794032 | -0.064162828 | 0.236667772 | 0.271109273 | 0.78630699 | 0.893161222 | 0.064162828 |
| ENSMUSG00000042608 | Stk40 | 74178 | protein_coding | 544.4584298 | 0.064197434 | 0.092029766 | -0.697572503 | 0.485444582 | 0.695380179 | 0.064197434 |
| ENSMUSG00000049553 | Polr1a | 20019 | protein_coding | 380.420794 | -0.064239978 | 0.104453018 | 0.615013133 | 0.538546047 | 0.733377497 | 0.064239978 |
| ENSMUSG00000046826 | Fam187b | 76415 | protein_coding | 37.93587334 | -0.064274354 | 0.230007324 | 0.279444815 | 0.779903483 | 0.890125936 | 0.064274354 |
| ENSMUSG00000020790 | Ankfy1 | 11736 | protein_coding | 1574.870498 | 0.064281168 | 0.052269365 | -1.229805792 | 0.218769839 | 0.424931207 | 0.064281168 |
| ENSMUSG00000021264 | Yy1 | 22632 | protein_coding | 1347.484189 | 0.064360395 | 0.065991898 | -0.975277226 | 0.329422764 | 0.551796561 | 0.064360395 |
| ENSMUSG00000040043 | Rbms2 | 56516 | protein_coding | 135.3639371 | -0.064362498 | 0.08728559 | 0.737378279 | 0.460892341 | 0.674995696 | 0.064362498 |
| ENSMUSG00000035248 | Tut7 | 214290 | protein_coding | 1332.85998 | 0.064363657 | 0.052102156 | -1.235335773 | 0.216705565 | 0.422162995 | 0.064363657 |
| ENSMUSG00000031078 | Cttn | 13043 | protein_coding | 3702.805296 | -0.064378775 | 0.076537675 | 0.841138367 | 0.400270424 | 0.622315002 | 0.064378775 |
| ENSMUSG00000056666 | Retsat | 67442 | protein_coding | 1239.587024 | 0.064408483 | 0.063407628 | -1.015784466 | 0.309732035 | 0.530813078 | 0.064408483 |
| ENSMUSG00000051234 | Rnf7 | 19823 | protein_coding | 458.0844 | -0.064408548 | 0.091865138 | 0.701120675 | 0.483227709 | 0.693738201 | 0.064408548 |
| ENSMUSG00000012296 | Tjap1 | 74094 | protein_coding | 187.7393747 | 0.064433091 | 0.105154415 | -0.612747368 | 0.540043397 | 0.734487099 | 0.064433091 |
| ENSMUSG00000014198 | Zfp385c | 278304 | protein_coding | 5.025822988 | -0.064456242 | 0.819835089 | 0.078620984 | 0.937334096 | 0.970113767 | 0.064456242 |
| ENSMUSG00000026153 | Fam135a | 68187 | protein_coding | 892.8485813 | 0.06449214 | 0.07138507 | -0.903440175 | 0.366292326 | 0.588935008 | 0.06449214 |
| ENSMUSG00000074634 | Tmem267 | 633640 | protein_coding | 42.82438194 | 0.064492294 | 0.374705244 | -0.172114736 | 0.863347328 | 0.93221641 | 0.064492294 |
| ENSMUSG00000025950 | Idh1 | 15926 | protein_coding | 2420.069268 | -0.064498939 | 0.06188387 | 1.042257677 | 0.297292229 | 0.51684218 | 0.064498939 |
| ENSMUSG00000020459 | Mtif2 | 76784 | protein_coding | 384.3409984 | -0.064514127 | 0.093780752 | 0.687925033 | 0.491499992 | 0.698491194 | 0.064514127 |
| ENSMUSG00000095677 | Dynlt1f | 100040531 | protein_coding | 66.06774957 | 0.064519648 | 0.185465156 | -0.347880161 | 0.727930185 | 0.859382261 | 0.064519648 |
| ENSMUSG00000026842 | Abl1 | 11350 | protein_coding | 1188.428583 | -0.064527082 | 0.080753712 | 0.799060255 | 0.424255474 | 0.643978573 | 0.064527082 |
| ENSMUSG00000020564 | Atxn7l1 | 380753 | protein_coding | 149.1687275 | -0.064553385 | 0.160386202 | 0.402487147 | 0.687325546 | 0.833389267 | 0.064553385 |
| ENSMUSG00000008301 | Phax | 56698 | protein_coding | 291.375526 | -0.064596657 | 0.101517638 | 0.636309698 | 0.524574585 | 0.723107237 | 0.064596657 |
| ENSMUSG00000031442 | Mcf2l | 17207 | protein_coding | 1125.007219 | -0.064597491 | 0.156164491 | 0.413650314 | 0.67913022 | 0.827951097 | 0.064597491 |
| ENSMUSG00000021928 | Ebpl | 68177 | protein_coding | 166.0458532 | -0.0645994 | 0.134236605 | 0.481235351 | 0.630349239 | 0.79831751 | 0.0645994 |
| ENSMUSG00000023262 | Acy1 | 109652 | protein_coding | 470.0688601 | 0.06468491 | 0.107966192 | -0.599121904 | 0.549091596 | 0.740220295 | 0.06468491 |
| ENSMUSG00000034917 | Tjp3 | 27375 | protein_coding | 4029.363309 | -0.064691696 | 0.077025824 | 0.839870226 | 0.400981153 | 0.622737045 | 0.064691696 |
| ENSMUSG00000022354 | Ndufb9 | 66218 | protein_coding | 2185.99715 | -0.064702311 | 0.088764493 | 0.728921094 | 0.466049931 | 0.679736277 | 0.064702311 |

| ensembl_gene_id | external_gene_name | entrezgene_id | gene_biotype | baseMean | log2FoldChange | lfcSE | stat | pvalue | padj | absLog2FC |
|---|---|---|---|---|---|---|---|---|---|---|
| ENSMUSG00000085741 | 5430405H02Rik | 74487 | lncRNA | 16.29935689 | 0.064705838 | 0.296187511 | -0.218462413 | 0.827068841 | 0.912738228 | 0.064705838 |
| ENSMUSG00000028832 | Stmn1 | 16765 | protein_coding | 432.9105832 | -0.064729844 | 0.18799083 | 0.344324478 | 0.730602278 | 0.861253255 | 0.064729844 |
| ENSMUSG00000043940 | Wdfy3 | 72145 | protein_coding | 683.1700445 | 0.064741398 | 0.112056766 | -0.57775537 | 0.563429291 | 0.750868206 | 0.064741398 |
| ENSMUSG00000091575 | 2010016I18Rik | 69206 | lncRNA | 98.73580328 | 0.064744744 | 0.193579279 | -0.33446113 | 0.738031617 | 0.866982505 | 0.064744744 |
| ENSMUSG00000053929 | Cyhr1 | 54151 | protein_coding | 997.1700655 | -0.064764174 | 0.061676685 | 1.050059271 | 0.293690863 | 0.512903015 | 0.064764174 |
| ENSMUSG00000074221 | Zfp568 | 243905 | protein_coding | 262.3764332 | -0.064806081 | 0.073200182 | 0.885326773 | 0.375980414 | 0.598238321 | 0.064806081 |
| ENSMUSG00000028156 | Eif4e | 13684 | protein_coding | 573.5437982 | 0.064827018 | 0.110225275 | -0.588132065 | 0.556443652 | 0.745748293 | 0.064827018 |
| ENSMUSG00000027109 | Sp3 | 20687 | protein_coding | 1627.139229 | -0.064858456 | 0.065157376 | 0.995412333 | 0.319535763 | 0.540878801 | 0.064858456 |
| ENSMUSG00000022890 | Atp5j | 11957 | protein_coding | 1352.576686 | -0.064859737 | 0.062808652 | 1.032656087 | 0.301764871 | 0.522023104 | 0.064859737 |
| ENSMUSG00000039210 | Gpatch2 | 67769 | protein_coding | 656.0482092 | 0.064898587 | 0.100288592 | -0.647118336 | 0.517555359 | 0.717985391 | 0.064898587 |
| ENSMUSG00000026377 | Nifk | 67949 | protein_coding | 368.531189 | -0.064900897 | 0.121599477 | 0.533726782 | 0.593530577 | 0.772689314 | 0.064900897 |
| ENSMUSG00000061024 | Rrs1 | 59014 | protein_coding | 209.6325132 | 0.064911675 | 0.126581888 | -0.512803819 | 0.608088558 | 0.782450869 | 0.064911675 |
| ENSMUSG00000026159 | Agfg1 | 15463 | protein_coding | 604.5679391 | 0.064929785 | 0.067153622 | -0.966884331 | 0.333601861 | 0.555969435 | 0.064929785 |
| ENSMUSG00000074627 | Mroh8 | 629499 | protein_coding | 13.04389898 | 0.064963963 | 0.36320559 | -0.178862785 | 0.858045445 | 0.930446023 | 0.064963963 |
| ENSMUSG00000041623 | D11Wsu47e | 276852 | protein_coding | 116.3223731 | -0.064995143 | 0.122861776 | 0.529010286 | 0.596798315 | 0.774805611 | 0.064995143 |
| ENSMUSG00000044447 | Dock5 | 68813 | protein_coding | 1143.419835 | -0.064996433 | 0.095349098 | 0.681668048 | 0.495448876 | 0.701521284 | 0.064996433 |
| ENSMUSG00000038518 | Jarid2 | 16468 | protein_coding | 794.9696561 | -0.065017631 | 0.055931553 | 1.162449945 | 0.245052746 | 0.456749554 | 0.065017631 |
| ENSMUSG00000005054 | Cstb | 13014 | protein_coding | 475.1805444 | -0.065092338 | 0.082416556 | 0.789796869 | 0.429646408 | 0.648455238 | 0.065092338 |
| ENSMUSG00000071265 | 1700086L19Rik | 74284 | transcribed_unitary_pseudogene | 2.622014148 | 0.065134586 | 0.704414428 | -0.092466286 | 0.926327576 | 1 | 0.065134586 |
| ENSMUSG00000031601 | Cnot7 | 18983 | protein_coding | 461.6119699 | 0.065161182 | 0.078734064 | -0.827611062 | 0.407890801 | 0.628584238 | 0.065161182 |
| ENSMUSG00000045103 | Dmd | 13405 | protein_coding | 263.0792773 | -0.065163213 | 0.100057186 | 0.651259704 | 0.514878856 | 0.716272317 | 0.065163213 |
| ENSMUSG00000027304 | Rtf1 | 76246 | protein_coding | 344.1611472 | -0.065313944 | 0.073540356 | 0.888137459 | 0.374466815 | 0.596502845 | 0.065313944 |
| ENSMUSG00000018765 | Fxr2 | 23879 | protein_coding | 625.6009825 | 0.065347375 | 0.06148656 | -1.062791198 | 0.287876652 | 0.506822498 | 0.065347375 |
| ENSMUSG00000026073 | Il1r2 | 16178 | protein_coding | 0.505221152 | -0.065361826 | 1.901156003 | 0.034380044 | 0.972574097 | 1 | 0.065361826 |
| ENSMUSG00000003235 | Eif2b5 | 224045 | protein_coding | 717.783089 | 0.065384126 | 0.086654322 | -0.754539699 | 0.450525217 | 0.667322041 | 0.065384126 |
| ENSMUSG00000084806 | Gm15232 | 627268 | lncRNA | 1.252824341 | -0.06545449 | 1.545042961 | 0.042364188 | 0.966208377 | 1 | 0.06545449 |
| ENSMUSG00000067924 | Rtl8b | 553127 | protein_coding | 8.425670282 | -0.06549989 | 0.358205532 | 0.182855608 | 0.854911315 | 0.929305257 | 0.06549989 |
| ENSMUSG00000021763 | Cspg4b | 408066 | protein_coding | 3.019044354 | -0.065506437 | 0.762671928 | 0.085890715 | 0.931553293 | 1 | 0.065506437 |
| ENSMUSG00000000056 | Narf | 67608 | protein_coding | 690.9771543 | -0.065519835 | 0.110550248 | 0.592670177 | 0.553401904 | 0.743731716 | 0.065519835 |
| ENSMUSG00000092074 | Dynlt1a | 100310872 | protein_coding | 30.18889292 | -0.065543253 | 0.202897442 | 0.32303637 | 0.746667698 | 0.872440598 | 0.065543253 |
| ENSMUSG00000038331 | Satb2 | 212712 | protein_coding | 2427.171748 | 0.065549824 | 0.110389917 | -0.593802638 | 0.552644124 | 0.743189338 | 0.065549824 |
| ENSMUSG00000021470 | Ercc6l2 | 76251 | protein_coding | 331.6194376 | 0.065567318 | 0.119393174 | -0.549171412 | 0.582887821 | 0.765654588 | 0.065567318 |
| ENSMUSG00000063145 | Bbs5 | 72569 | protein_coding | 39.05168485 | 0.06559827 | 0.169101588 | -0.387922258 | 0.698073566 | 0.840048595 | 0.06559827 |
| ENSMUSG00000024589 | Nedd4l | 83814 | protein_coding | 1125.37471 | -0.065653972 | 0.111603802 | 0.588277198 | 0.556346248 | 0.745678268 | 0.065653972 |
| ENSMUSG00000040385 | Ppp1ca | 19045 | protein_coding | 2268.012974 | 0.065715811 | 0.063826512 | -1.029600534 | 0.303197564 | 0.523689253 | 0.065715811 |
| ENSMUSG00000022982 | Sod1 | 20655 | protein_coding | 2852.819923 | -0.065742618 | 0.065460475 | 1.004310126 | 0.315229155 | 0.536227262 | 0.065742618 |
| ENSMUSG00000031479 | Vps36 | 70160 | protein_coding | 490.2114026 | -0.065750776 | 0.073222388 | 0.897960007 | 0.369206871 | 0.591834816 | 0.065750776 |
| ENSMUSG00000002416 | Ndufb2 | 68198 | protein_coding | 407.5491078 | -0.065755677 | 0.128293075 | 0.512542688 | 0.608271253 | 0.782624968 | 0.065755677 |
| ENSMUSG00000005804 | Bloc1s6 | 18457 | protein_coding | 317.9509451 | -0.065791159 | 0.082754527 | 0.795015823 | 0.426604298 | 0.64620651 | 0.065791159 |

| ensembl_gene_id | external_gene_name | entrezgene_id | gene_biotype | baseMean | log2FoldChange | lfcSE | stat | pvalue | padj | absLog2FC |
|---|---|---|---|---|---|---|---|---|---|---|
| ENSMUSG00000070780 | Rbm47 | 245945 | protein_coding | 3598.285803 | 0.065825882 | 0.074060568 | -0.888811463 | 0.374104414 | 0.59609806 | 0.065825882 |
| ENSMUSG00000016528 | Mapkapk2 | 17164 | protein_coding | 1086.866666 | 0.065832568 | 0.066006537 | -0.997364368 | 0.31858768 | 0.540015186 | 0.065832568 |
| ENSMUSG00000055202 | Zfp811 | 240063 | protein_coding | 2.081172811 | -0.065857838 | 1.23753518 | 0.053216942 | 0.957559057 | 1 | 0.065857838 |
| ENSMUSG00000032739 | Pram1 | 378460 | protein_coding | 1.200625976 | -0.065863351 | 0.892744545 | 0.073776257 | 0.94118842 | 1 | 0.065863351 |
| ENSMUSG00000051671 | Coa6 | 67892 | protein_coding | 150.0339445 | -0.065866341 | 0.092596005 | 0.711330276 | 0.476879594 | 0.68845088 | 0.065866341 |
| ENSMUSG00000023495 | Pcbp4 | 59092 | protein_coding | 36.27696522 | 0.0658705 | 0.213561508 | -0.30843807 | 0.757749017 | 0.87863969 | 0.0658705 |
| ENSMUSG00000055003 | Lrtm2 | 211187 | protein_coding | 2.204320806 | 0.065894449 | 1.028103417 | -0.064093211 | 0.948896008 | 1 | 0.065894449 |
| ENSMUSG00000025410 | Dctn2 | 69654 | protein_coding | 1478.18125 | -0.065901413 | 0.04471788 | 1.473715042 | 0.140558331 | 0.314808425 | 0.065901413 |
| ENSMUSG00000044795 | Cyb5d1 | 327951 | protein_coding | 74.55850501 | -0.065959185 | 0.137668166 | 0.47911719 | 0.631855261 | 0.799444652 | 0.065959185 |
| ENSMUSG00000022773 | Ypel1 | 106369 | protein_coding | 13.48139131 | -0.065974512 | 0.338457407 | 0.194927074 | 0.845450038 | 0.923885782 | 0.065974512 |
| ENSMUSG00000049327 | Kmt5a | 67956 | protein_coding | 1741.01081 | 0.065986779 | 0.050648112 | -1.302847757 | 0.192626743 | 0.39045514 | 0.065986779 |
| ENSMUSG00000025020 | Slit1 | 20562 | protein_coding | 5.771477316 | 0.065987236 | 1.080253747 | -0.061084941 | 0.951291562 | 0.975912221 | 0.065987236 |
| ENSMUSG00000027375 | Mal | 17153 | protein_coding | 599.9101955 | 0.065998262 | 0.220458775 | -0.299367816 | 0.764659416 | 0.882132403 | 0.065998262 |
| ENSMUSG00000039929 | Urb1 | 207932 | protein_coding | 185.8752693 | 0.066002756 | 0.137488338 | -0.480060757 | 0.631184191 | 0.799065123 | 0.066002756 |
| ENSMUSG00000039783 | Kmo | 98256 | protein_coding | 3.576975636 | -0.066013614 | 0.798309523 | 0.082691753 | 0.934096643 | 1 | 0.066013614 |
| ENSMUSG00000028645 | Slc2a1 | 20525 | protein_coding | 1231.166972 | -0.066049192 | 0.120140826 | 0.549764756 | 0.582480735 | 0.765241595 | 0.066049192 |
| ENSMUSG00000024212 | Mllt1 | 64144 | protein_coding | 621.8322326 | 0.066053869 | 0.075197643 | -0.87840346 | 0.379724807 | 0.601975182 | 0.066053869 |
| ENSMUSG00000025766 | D3Ertd751e | 73852 | protein_coding | 109.685901 | -0.066081347 | 0.16545382 | 0.39939451 | 0.689602538 | 0.834370416 | 0.066081347 |
| ENSMUSG00000031839 | Hsbp1 | 68196 | protein_coding | 763.9332763 | -0.066107773 | 0.074060734 | 0.892615684 | 0.372063017 | 0.594336299 | 0.066107773 |
| ENSMUSG00000041241 | Mul1 | 68350 | protein_coding | 877.6765487 | 0.066108239 | 0.060045998 | -1.100959962 | 0.270914083 | 0.486449404 | 0.066108239 |
| ENSMUSG00000066042 | Med18 | 67219 | protein_coding | 75.30198101 | -0.066206484 | 0.118452877 | 0.558926774 | 0.576211696 | 0.761183901 | 0.066206484 |
| ENSMUSG00000062627 | Mysm1 | 320713 | protein_coding | 383.0374956 | -0.066255862 | 0.070089864 | 0.94529877 | 0.344506366 | 0.566265254 | 0.066255862 |
| ENSMUSG00000027509 | Rae1 | 66679 | protein_coding | 396.5610274 | 0.06628489 | 0.07497612 | -0.884080023 | 0.376653016 | 0.598887725 | 0.06628489 |
| ENSMUSG00000020530 | Ggnbp2 | 217039 | protein_coding | 1182.873977 | -0.066313716 | 0.05059649 | 1.310638666 | 0.189979869 | 0.387311023 | 0.066313716 |
| ENSMUSG00000037075 | Rnf139 | 75841 | protein_coding | 298.5705382 | -0.066386984 | 0.09691329 | 0.685014247 | 0.493334931 | 0.699749347 | 0.066386984 |
| ENSMUSG00000006335 | Tfpt | 69714 | protein_coding | 120.609035 | -0.066398877 | 0.098435625 | 0.674541121 | 0.499967352 | 0.704656555 | 0.066398877 |
| ENSMUSG00000028163 | Nfkb1 | 18033 | protein_coding | 998.2156696 | -0.066425969 | 0.046266041 | 1.435739212 | 0.151076563 | 0.331061353 | 0.066425969 |
| ENSMUSG00000036111 | Lmo1 | 109594 | protein_coding | 6.754120882 | 0.06646184 | 0.566305038 | -0.117360496 | 0.906574387 | 0.954311305 | 0.06646184 |
| ENSMUSG00000059540 | Tcea2 | 21400 | protein_coding | 17.99091363 | -0.066462149 | 0.249655536 | 0.266215403 | 0.790073321 | 0.895242955 | 0.066462149 |
| ENSMUSG00000032745 | Gpbp1 | 73274 | protein_coding | 774.9942926 | 0.066484137 | 0.063389779 | -1.048814772 | 0.294263377 | 0.513414201 | 0.066484137 |
| ENSMUSG00000089998 | Phtf1os | 100125931 | lncRNA | 2.906648543 | -0.066503815 | 0.786730839 | 0.084531852 | 0.932633579 | 1 | 0.066503815 |
| ENSMUSG00000018417 | Myo1b | 17912 | protein_coding | 824.973714 | -0.066512338 | 0.103601837 | 0.641999605 | 0.52087344 | 0.720614094 | 0.066512338 |
| ENSMUSG00000001105 | Ift20 | 55978 | protein_coding | 376.2927683 | 0.066517273 | 0.085782121 | -0.775421177 | 0.438090826 | 0.6563215 | 0.066517273 |
| ENSMUSG00000026126 | Ptpn18 | 19253 | protein_coding | 538.1380855 | -0.066524021 | 0.070868497 | 0.93869665 | 0.347886514 | 0.569827206 | 0.066524021 |
| ENSMUSG00000019804 | Snx3 | 54198 | protein_coding | 924.6592829 | 0.066530456 | 0.065985697 | -1.008255706 | 0.313331721 | 0.534596554 | 0.066530456 |
| ENSMUSG00000029366 | Dck | 13178 | protein_coding | 135.9592643 | 0.066582173 | 0.147612386 | -0.451060882 | 0.65194567 | 0.811137708 | 0.066582173 |
| ENSMUSG00000033752 | Mnd1 | 76915 | protein_coding | 16.63056592 | -0.066603199 | 0.264659418 | 0.251656258 | 0.80130677 | 0.901569434 | 0.066603199 |
| ENSMUSG00000020610 | Amz2 | 13929 | protein_coding | 363.8499462 | 0.066603924 | 0.111333663 | -0.598237062 | 0.549681765 | 0.740653122 | 0.066603924 |
| ENSMUSG00000040651 | Tasor | 218850 | protein_coding | 550.6147054 | -0.066610822 | 0.08185961 | 0.813720236 | 0.415805238 | 0.636091497 | 0.066610822 |

Supplementary Table 1

| ensembl_gene_id | external_gene_name | entrezgene_id | gene_biotype | baseMean | log2FoldChange | lfcSE | stat | pvalue | padj | absLog2FC |
|---|---|---|---|---|---|---|---|---|---|---|
| ENSMUSG00000026767 | Mindy3 | 66960 | protein_coding | 388.0117471 | 0.066611907 | 0.066160641 | -1.006820756 | 0.314020918 | 0.5351691 | 0.066611907 |
| ENSMUSG00000045690 | Wdr89 | 72338 | protein_coding | 23.33271388 | 0.066628091 | 0.319439052 | -0.208578415 | 0.834777362 | 0.917066002 | 0.066628091 |
| ENSMUSG00000073144 | 4930599N23Rik | 75379 | lncRNA | 7.478447814 | 0.066664559 | 0.446582103 | -0.149277274 | 0.881334847 | 0.941481575 | 0.066664559 |
| ENSMUSG00000086992 | Gm15941 | 100503538 | lncRNA | 7.550658364 | -0.066689411 | 0.407051275 | 0.163835407 | 0.869860717 | 0.935658236 | 0.066689411 |
| ENSMUSG00000043061 | Tmem18 | 211986 | protein_coding | 275.2259515 | 0.066754635 | 0.100792401 | -0.662298297 | 0.507780068 | 0.711603273 | 0.066754635 |
| ENSMUSG00000028868 | Wasf2 | 242687 | protein_coding | 1230.209667 | 0.066778902 | 0.067974683 | -0.982408437 | 0.325898671 | 0.54788666 | 0.066778902 |
| ENSMUSG00000012777 | Acp4 | 100503991 | protein_coding | 5.926359103 | -0.066789642 | 0.533047597 | 0.125297708 | 0.900287867 | 0.95096289 | 0.066789642 |
| ENSMUSG00000039354 | Smarcal1 | 54380 | protein_coding | 247.8665457 | 0.066822456 | 0.095305776 | -0.701137526 | 0.483217193 | 0.693738201 | 0.066822456 |
| ENSMUSG00000032352 | Lrrc1 | 214345 | protein_coding | 844.621941 | 0.066839376 | 0.073231382 | -0.91271493 | 0.36139249 | 0.583801091 | 0.066839376 |
| ENSMUSG00000033365 | Ipo13 | 230673 | protein_coding | 204.6843821 | 0.066844711 | 0.09003501 | -0.742430203 | 0.457826726 | 0.67261003 | 0.066844711 |
| ENSMUSG00000008958 | Vps72 | 21427 | protein_coding | 383.898735 | -0.066879209 | 0.067264867 | 0.994266575 | 0.320093104 | 0.541378279 | 0.066879209 |
| ENSMUSG00000002250 | Ppard | 19015 | protein_coding | 1365.93894 | 0.066900315 | 0.12712109 | -0.52627235 | 0.598698991 | 0.776186935 | 0.066900315 |
| ENSMUSG00000004530 | Coro1c | 23790 | protein_coding | 1913.291188 | -0.066905209 | 0.083080411 | 0.805306666 | 0.420642736 | 0.640643193 | 0.066905209 |
| ENSMUSG00000026626 | Ppp2r5a | 226849 | protein_coding | 1437.589895 | -0.066938276 | 0.054914744 | 1.218949073 | 0.222863522 | 0.429893421 | 0.066938276 |
| ENSMUSG00000031537 | Ikbkb | 16150 | protein_coding | 587.1183688 | 0.066949937 | 0.063416483 | -1.055718223 | 0.291096969 | 0.510000436 | 0.066949937 |
| ENSMUSG00000020868 | Xylt2 | 217119 | protein_coding | 213.1859992 | -0.066956935 | 0.116557127 | 0.574455947 | 0.565659303 | 0.752299084 | 0.066956935 |
| ENSMUSG00000037514 | Pank2 | 74450 | protein_coding | 522.2485661 | 0.066969393 | 0.064743204 | -1.034384904 | 0.300956259 | 0.521072726 | 0.066969393 |
| ENSMUSG00000047259 | Mc4r | 17202 | protein_coding | 0.931758449 | 0.066970233 | 1.715218282 | -0.039044729 | 0.968854727 | 1 | 0.066970233 |
| ENSMUSG00000022635 | Zcrb1 | 67197 | protein_coding | 266.9532603 | 0.066985873 | 0.080003681 | -0.837284897 | 0.402432446 | 0.624001556 | 0.066985873 |
| ENSMUSG00000001844 | Zdhhc4 | 72881 | protein_coding | 213.5326459 | -0.066994524 | 0.084312405 | 0.794598661 | 0.426846997 | 0.646395058 | 0.066994524 |
| ENSMUSG00000057666 | Gapdh | 14433 | protein_coding | 7129.863873 | 0.066996259 | 0.141545687 | -0.473318971 | 0.635985629 | 0.801613541 | 0.066996259 |
| ENSMUSG00000026112 | Coa5 | 76178 | protein_coding | 1311.186608 | -0.067040696 | 0.049391862 | 1.357322699 | 0.174678698 | 0.365932262 | 0.067040696 |
| ENSMUSG00000035759 | Bbs10 | 71769 | protein_coding | 24.73942644 | -0.06705613 | 0.245916897 | 0.27267801 | 0.78510074 | 0.892732467 | 0.06705613 |
| ENSMUSG00000052676 | Zmat1 | 215693 | protein_coding | 14.26073181 | 0.067100169 | 0.297847318 | -0.225283778 | 0.821758519 | 0.91001573 | 0.067100169 |
| ENSMUSG00000001472 | Tcf25 | 66855 | protein_coding | 2536.591484 | 0.067110102 | 0.033281592 | -2.016433021 | 0.043754715 | 0.134681337 | 0.067110102 |
| ENSMUSG00000005373 | Mlxipl | 58805 | protein_coding | 149.3616619 | 0.067136597 | 0.200205886 | -0.335337778 | 0.737370299 | 0.866747322 | 0.067136597 |
| ENSMUSG00000053353 | 2310001K24Rik | 69517 | lncRNA | 0.910721045 | 0.067146517 | 1.554618792 | -0.043191628 | 0.965548778 | 1 | 0.067146517 |
| ENSMUSG00000020593 | Lpin1 | 14245 | protein_coding | 126.3869156 | 0.067164075 | 0.188243928 | -0.356792784 | 0.721246939 | 0.855640813 | 0.067164075 |
| ENSMUSG00000028955 | Vamp3 | 22319 | protein_coding | 1112.859941 | -0.067212896 | 0.089622573 | 0.749954996 | 0.45328181 | 0.669122175 | 0.067212896 |
| ENSMUSG00000000339 | Rtca | 66368 | protein_coding | 422.5162335 | 0.06727258 | 0.083656357 | -0.804153837 | 0.421308135 | 0.641124452 | 0.06727258 |
| ENSMUSG00000065431 | Mir186 | 387181 | miRNA | 1.164450947 | -0.067290061 | 0.905639556 | 0.07430115 | 0.940770762 | 1 | 0.067290061 |
| ENSMUSG00000023972 | Ptk7 | 71461 | protein_coding | 277.5398839 | 0.06734848 | 0.109608623 | -0.614445088 | 0.538921249 | 0.733576122 | 0.06734848 |
| ENSMUSG00000044288 | Cnr1 | 12801 | protein_coding | 1.128127169 | 0.067350004 | 1.657543407 | -0.040632422 | 0.967588936 | 1 | 0.067350004 |
| ENSMUSG00000019295 | Tmem129 | 68366 | protein_coding | 300.5066476 | 0.067361007 | 0.089151863 | -0.755575985 | 0.449903459 | 0.666700082 | 0.067361007 |
| ENSMUSG00000054455 | Vapb | 56491 | protein_coding | 951.9244188 | 0.067370178 | 0.068677333 | -0.980966718 | 0.326609154 | 0.548411142 | 0.067370178 |
| ENSMUSG00000019461 | Plscr3 | 70310 | protein_coding | 118.0493701 | -0.067403721 | 0.12459686 | 0.540974477 | 0.588525175 | 0.769577776 | 0.067403721 |
| ENSMUSG00000016624 | Phf21b | 271305 | protein_coding | 94.70426459 | 0.067469343 | 0.148398237 | -0.454650572 | 0.649360646 | 0.809667817 | 0.067469343 |
| ENSMUSG00000008435 | Rdh13 | 108841 | protein_coding | 465.862737 | -0.067474218 | 0.10039085 | 0.672115224 | 0.501510345 | 0.706297653 | 0.067474218 |
| ENSMUSG00000032598 | Nckipsd | 80987 | protein_coding | 84.76812467 | 0.067495331 | 0.127060109 | -0.531207882 | 0.595274732 | 0.773679122 | 0.067495331 |

Supplementary Table 1

| ensembl_gene_id | external_gene_name | entrezgene_id | gene_biotype | baseMean | log2FoldChange | lfcSE | stat | pvalue | padj | absLog2FC |
|---|---|---|---|---|---|---|---|---|---|---|
| ENSMUSG00000022453 | Naga | 17939 | protein_coding | 1572.105303 | 0.067500875 | 0.05746042 | -1.174736895 | 0.240099994 | 0.450443439 | 0.067500875 |
| ENSMUSG00000039356 | Exosc2 | 227715 | protein_coding | 201.7249048 | 0.067566499 | 0.1285344 | -0.525668607 | 0.599118479 | 0.776435029 | 0.067566499 |
| ENSMUSG00000035967 | Ints6l | 236790 | protein_coding | 289.9311262 | -0.06757028 | 0.09137235 | 0.739504679 | 0.459600599 | 0.673998628 | 0.06757028 |
| ENSMUSG00000038212 | Mfsd14b | 66631 | protein_coding | 1095.09053 | -0.067581404 | 0.055596892 | 1.21556081 | 0.224152271 | 0.431277012 | 0.067581404 |
| ENSMUSG00000028455 | Stoml2 | 66592 | protein_coding | 506.2256191 | -0.067688858 | 0.085201507 | 0.794456111 | 0.426929949 | 0.646395058 | 0.067688858 |
| ENSMUSG00000036093 | Arl5a | 75423 | protein_coding | 650.6679401 | 0.06772752 | 0.083680983 | -0.809353777 | 0.418311681 | 0.638329247 | 0.06772752 |
| ENSMUSG00000020448 | Rnf185 | 193670 | protein_coding | 361.6879186 | 0.067780642 | 0.063358456 | -1.069796289 | 0.284711013 | 0.503072723 | 0.067780642 |
| ENSMUSG00000043153 | Crppa | 75847 | protein_coding | 29.91910271 | 0.067845313 | 0.230138096 | -0.294802617 | 0.768144667 | 0.883975099 | 0.067845313 |
| ENSMUSG00000027366 | Sppl2a | 66552 | protein_coding | 3568.611491 | -0.067880437 | 0.112005501 | 0.606045561 | 0.544484499 | 0.736796542 | 0.067880437 |
| ENSMUSG00000024050 | Wiz | 22404 | protein_coding | 262.7710879 | -0.067892128 | 0.091517324 | 0.741850021 | 0.45817821 | 0.672744283 | 0.067892128 |
| ENSMUSG00000020075 | Ddx21 | 56200 | protein_coding | 999.6313137 | -0.067901715 | 0.099099409 | 0.685187896 | 0.493225361 | 0.699692776 | 0.067901715 |
| ENSMUSG00000044791 | Setd2 | 235626 | protein_coding | 1121.412052 | -0.067910878 | 0.067433448 | 1.007080015 | 0.313896324 | 0.535062335 | 0.067910878 |
| ENSMUSG00000042178 | Armc5 | 233912 | protein_coding | 309.5404149 | 0.067916445 | 0.122195863 | -0.555799872 | 0.578347673 | 0.761994054 | 0.067916445 |
| ENSMUSG00000030204 | Ddx47 | 67755 | protein_coding | 471.8831773 | -0.067958032 | 0.064741156 | 1.049688266 | 0.293861461 | 0.513092425 | 0.067958032 |
| ENSMUSG00000005413 | Hmox1 | 15368 | protein_coding | 47.40455991 | 0.068022881 | 0.225382879 | -0.301810331 | 0.762796655 | 0.881056435 | 0.068022881 |
| ENSMUSG00000026426 | Arl8a | 68724 | protein_coding | 130.4229553 | -0.06804499 | 0.183272519 | 0.371277652 | 0.710430741 | 0.848979902 | 0.06804499 |
| ENSMUSG00000030236 | Slco1b2 | 28253 | protein_coding | 1.953765687 | 0.068046352 | 1.280825031 | -0.053126969 | 0.957630743 | 1 | 0.068046352 |
| ENSMUSG00000002767 | Mrpl2 | 27398 | protein_coding | 550.2904757 | -0.068054784 | 0.078883475 | 0.862725485 | 0.388288417 | 0.610859762 | 0.068054784 |
| ENSMUSG00000051278 | Zgrf1 | 71643 | protein_coding | 55.05004615 | 0.068065409 | 0.238787815 | -0.285045572 | 0.775609228 | 0.88791795 | 0.068065409 |
| ENSMUSG00000032434 | Cmtm6 | 67213 | protein_coding | 1657.734761 | -0.068092167 | 0.103319374 | 0.659045488 | 0.509866559 | 0.713191289 | 0.068092167 |
| ENSMUSG00000027007 | Itprid2 | 70599 | protein_coding | 1907.392445 | 0.068112348 | 0.150211365 | -0.453443373 | 0.650229511 | 0.809917796 | 0.068112348 |
| ENSMUSG00000061086 | Myl4 | 17896 | protein_coding | 3.145448905 | -0.068132319 | 0.879306284 | 0.077484171 | 0.938238383 | 1 | 0.068132319 |
| ENSMUSG00000018326 | Ywhab | 54401 | protein_coding | 2206.723001 | -0.068137776 | 0.082038784 | 0.830555651 | 0.406224697 | 0.627187233 | 0.068137776 |
| ENSMUSG00000117679 | Apbb3 | 225372 | protein_coding | 150.0323109 | 0.068153186 | 0.153639559 | -0.44359139 | 0.657338032 | 0.814655179 | 0.068153186 |
| ENSMUSG00000025898 | Cwf19l2 | 244672 | protein_coding | 253.5435558 | -0.068214014 | 0.079882952 | 0.853924544 | 0.393146799 | 0.615862857 | 0.068214014 |
| ENSMUSG00000051232 | Tmem199 | 195040 | protein_coding | 123.2651491 | 0.068236255 | 0.1249422 | -0.546142579 | 0.584967933 | 0.767158627 | 0.068236255 |
| ENSMUSG00000054426 | A930005H10Rik | 68161 | lncRNA | 17.20344683 | -0.068259187 | 0.271527737 | 0.251389373 | 0.801513083 | 0.901709044 | 0.068259187 |
| ENSMUSG00000037318 | Traf3ip3 | 215243 | protein_coding | 9.926112178 | 0.068284394 | 0.55092734 | -0.123944464 | 0.901359247 | 0.951341404 | 0.068284394 |
| ENSMUSG00000030682 | Cdipt | 52858 | protein_coding | 807.8695068 | 0.068298517 | 0.065265886 | -1.046465788 | 0.295346033 | 0.514433539 | 0.068298517 |
| ENSMUSG00000042992 | Borcs5 | 67774 | protein_coding | 105.3241392 | 0.068344066 | 0.123107895 | -0.555155829 | 0.578788079 | 0.762380457 | 0.068344066 |
| ENSMUSG00000016181 | Utp25 | 215193 | protein_coding | 85.07091324 | -0.068364721 | 0.150575263 | 0.454023589 | 0.649811849 | 0.809917796 | 0.068364721 |
| ENSMUSG00000078144 | Capns2 | 69543 | protein_coding | 3.315073958 | -0.06836493 | 0.533827334 | 0.128065622 | 0.898097041 | 1 | 0.06836493 |
| ENSMUSG00000050390 | C77080 | 97130 | protein_coding | 2537.273468 | -0.068366682 | 0.135404491 | 0.504907048 | 0.613624124 | 0.786448489 | 0.068366682 |
| ENSMUSG00000003546 | Klc4 | 74764 | protein_coding | 2099.753362 | -0.068373152 | 0.104775956 | 0.65256529 | 0.514036569 | 0.715750413 | 0.068373152 |
| ENSMUSG00000045140 | Pigw | 70325 | protein_coding | 89.41539518 | 0.06839503 | 0.127901283 | -0.534748589 | 0.592823718 | 0.772538316 | 0.06839503 |
| ENSMUSG00000048833 | Slc39a9 | 328133 | protein_coding | 1829.84863 | 0.068443509 | 0.071461275 | -0.957770602 | 0.338178446 | 0.559860817 | 0.068443509 |
| ENSMUSG00000009630 | Ppp2cb | 19053 | protein_coding | 925.6465058 | -0.068450587 | 0.127247403 | 0.53793308 | 0.59062325 | 0.771092762 | 0.068450587 |
| ENSMUSG00000042828 | Trim72 | 434246 | protein_coding | 83.68233834 | 0.068452154 | 0.193604526 | -0.353566912 | 0.723663475 | 0.856795141 | 0.068452154 |
| ENSMUSG00000075284 | Wipf1 | 215280 | protein_coding | 162.2899364 | -0.068455338 | 0.124108893 | 0.551574801 | 0.581239705 | 0.764339772 | 0.068455338 |

| ensembl_gene_id | external_gene_name | entrezgene_id | gene_biotype | baseMean | log2FoldChange | lfcSE | stat | pvalue | padj | absLog2FC |
|---|---|---|---|---|---|---|---|---|---|---|
| ENSMUSG00000035868 | Zfp983 | 73229 | protein_coding | 80.50408371 | -0.068461467 | 0.130259507 | 0.525577511 | 0.599181786 | 0.776435029 | 0.068461467 |
| ENSMUSG00000031833 | Mast3 | 546071 | protein_coding | 216.9694872 | 0.068471712 | 0.118871915 | -0.576012526 | 0.564606715 | 0.751588072 | 0.068471712 |
| ENSMUSG00000054556 | Gm4876 | 232599 | lncRNA | 47.3221993 | 0.068494697 | 0.229718583 | -0.298167856 | 0.765575054 | 0.882510888 | 0.068494697 |
| ENSMUSG00000022883 | Robo1 | 19876 | protein_coding | 20.81640115 | 0.068526001 | 0.336117124 | -0.203875364 | 0.838450897 | 0.919582966 | 0.068526001 |
| ENSMUSG00000046229 | Scand1 | 19018 | protein_coding | 264.5705353 | 0.068526147 | 0.152280564 | -0.449999296 | 0.652710948 | 0.811601896 | 0.068526147 |
| ENSMUSG00000031691 | Tnpo2 | 212999 | protein_coding | 973.9095529 | 0.068527928 | 0.050750055 | -1.350302541 | 0.176918958 | 0.369037513 | 0.068527928 |
| ENSMUSG00000045160 | Bola3 | 78653 | protein_coding | 274.375824 | -0.068540804 | 0.112688387 | 0.608233071 | 0.543032904 | 0.736259883 | 0.068540804 |
| ENSMUSG00000027282 | Mtch2 | 56428 | protein_coding | 2721.88699 | -0.068549875 | 0.079007824 | 0.867634004 | 0.385594725 | 0.607820554 | 0.068549875 |
| ENSMUSG00000060794 | Tssk5 | 73542 | protein_coding | 1.364849377 | 0.068603908 | 1.475939478 | -0.046481518 | 0.962926464 | 1 | 0.068603908 |
| ENSMUSG00000021065 | Fut8 | 53618 | protein_coding | 1425.17357 | 0.06865002 | 0.175898959 | -0.390280988 | 0.69632878 | 0.838747423 | 0.06865002 |
| ENSMUSG00000021428 | Riok1 | 71340 | protein_coding | 215.3313685 | 0.068686854 | 0.0912393 | -0.752820925 | 0.451557531 | 0.668047177 | 0.068686854 |
| ENSMUSG00000030189 | Ybx3 | 56449 | protein_coding | 2172.674916 | -0.068691146 | 0.099979102 | 0.687055038 | 0.492048046 | 0.698788598 | 0.068691146 |
| ENSMUSG00000025496 | Drd4 | 13491 | protein_coding | 2.658134025 | -0.068691369 | 1.21735635 | 0.056426673 | 0.955001909 | 1 | 0.068691369 |
| ENSMUSG00000016984 | Etaa1 | 68145 | protein_coding | 127.1998353 | 0.06869181 | 0.161080531 | -0.426443902 | 0.669784417 | 0.822722373 | 0.06869181 |
| ENSMUSG00000031029 | Eif3f | 66085 | protein_coding | 2033.531463 | -0.068725517 | 0.086882155 | 0.791019941 | 0.428932355 | 0.647921517 | 0.068725517 |
| ENSMUSG00000053898 | Ech1 | 51798 | protein_coding | 2546.545203 | -0.068751519 | 0.066917944 | 1.027400344 | 0.30423199 | 0.524504007 | 0.068751519 |
| ENSMUSG00000007570 | Fance | 72775 | protein_coding | 139.899566 | -0.068752066 | 0.102972701 | 0.667672744 | 0.504342512 | 0.708596728 | 0.068752066 |
| ENSMUSG00000039678 | Tbc1d13 | 70296 | protein_coding | 1223.827048 | -0.068770352 | 0.078629515 | 0.874612444 | 0.381784814 | 0.604102573 | 0.068770352 |
| ENSMUSG00000021840 | Mapk1ip1l | 218975 | protein_coding | 1007.570325 | 0.068780928 | 0.059430356 | -1.157336622 | 0.247134857 | 0.459127576 | 0.068780928 |
| ENSMUSG00000051977 | Prdm9 | 213389 | protein_coding | 74.66931531 | -0.068787541 | 0.147272295 | 0.467077265 | 0.640444599 | 0.803690806 | 0.068787541 |
| ENSMUSG00000024241 | Sos1 | 20662 | protein_coding | 399.0003076 | 0.068817842 | 0.087886208 | -0.783033472 | 0.433607457 | 0.652196761 | 0.068817842 |
| ENSMUSG00000051721 | Wdcp | 238037 | protein_coding | 37.56614496 | -0.068823127 | 0.175268716 | 0.392672056 | 0.694561712 | 0.8379481 | 0.068823127 |
| ENSMUSG00000087691 | Cd55os | 102640368 | lncRNA | 3.290089576 | 0.068866993 | 0.710284648 | -0.096956894 | 0.922760627 | 1 | 0.068866993 |
| ENSMUSG00000032621 | Srek1 | 218543 | protein_coding | 487.4957516 | 0.0689099 | 0.086933389 | -0.792674731 | 0.427967356 | 0.647057325 | 0.0689099 |
| ENSMUSG00000101268 | 2010310C07Rik | 72131 | lncRNA | 2.874707482 | -0.068911277 | 1.333757476 | 0.051667022 | 0.958794014 | 1 | 0.068911277 |
| ENSMUSG00000040302 | Rbm48 | 269623 | protein_coding | 61.32442785 | -0.06900029 | 0.149757923 | 0.460745507 | 0.644981202 | 0.806666129 | 0.06900029 |
| ENSMUSG00000070319 | Eif3g | 53356 | protein_coding | 900.2936309 | 0.069040236 | 0.086782451 | -0.795555267 | 0.426290576 | 0.646027582 | 0.069040236 |
| ENSMUSG00000031715 | Smarca5 | 93762 | protein_coding | 1363.658735 | 0.069046768 | 0.118934173 | -0.580546078 | 0.561546423 | 0.749969885 | 0.069046768 |
| ENSMUSG00000032338 | Hcn4 | 330953 | protein_coding | 5.899782784 | 0.069050852 | 0.759528523 | -0.090912783 | 0.927561893 | 0.965445424 | 0.069050852 |
| ENSMUSG00000017615 | Tnfaip1 | 21927 | protein_coding | 1422.299688 | 0.069120306 | 0.075074502 | -0.9206895 | 0.35721256 | 0.580211292 | 0.069120306 |
| ENSMUSG00000018012 | Rac3 | 170758 | protein_coding | 58.65399913 | 0.06918545 | 0.136591096 | -0.506515081 | 0.612495105 | 0.785550184 | 0.06918545 |
| ENSMUSG00000038582 | Pptc7 | 320717 | protein_coding | 677.6166279 | -0.069223161 | 0.100616631 | 0.687989261 | 0.491459544 | 0.698491194 | 0.069223161 |
| ENSMUSG00000024098 | Twsg1 | 65960 | protein_coding | 459.4848628 | -0.069225681 | 0.124799692 | 0.554694323 | 0.57910376 | 0.762564348 | 0.069225681 |
| ENSMUSG00000020077 | Srgn | 19073 | protein_coding | 16.41908765 | 0.069264202 | 0.538578256 | -0.128605642 | 0.897669702 | 0.949716026 | 0.069264202 |
| ENSMUSG00000031955 | Bcar1 | 12927 | protein_coding | 827.6346153 | -0.069264578 | 0.072449148 | 0.956044071 | 0.33904997 | 0.560532006 | 0.069264578 |
| ENSMUSG00000021759 | Plpp1 | 19012 | protein_coding | 188.2263753 | 0.069268624 | 0.258528322 | -0.267934374 | 0.788749834 | 0.894265208 | 0.069268624 |
| ENSMUSG00000002249 | Tead3 | 21678 | protein_coding | 243.3977576 | 0.069282088 | 0.132065246 | -0.524604995 | 0.599857813 | 0.7765798 | 0.069282088 |
| ENSMUSG00000032540 | Abhd5 | 67469 | protein_coding | 351.1418453 | -0.069288783 | 0.082999759 | 0.83480704 | 0.403826361 | 0.625062193 | 0.069288783 |
| ENSMUSG00000101970 | Chaserr | 69170 | lncRNA | 81.33333299 | -0.069303635 | 0.140368822 | 0.493725276 | 0.621500212 | 0.792236609 | 0.069303635 |

| ensembl_gene_id | external_gene_name | entrezgene_id | gene_biotype | baseMean | log2FoldChange | lfcSE | stat | pvalue | padj | absLog2FC |
|---|---|---|---|---|---|---|---|---|---|---|
| ENSMUSG00000032381 | Ciao2a | 68250 | protein_coding | 806.7451656 | -0.069311136 | 0.087880742 | 0.788695391 | 0.430290062 | 0.648871201 | 0.069311136 |
| ENSMUSG00000084910 | C630043F03Rik | 68285 | lncRNA | 23.65459368 | 0.069355483 | 0.206373945 | -0.33606705 | 0.736820305 | 0.86640918 | 0.069355483 |
| ENSMUSG00000031835 | Mbtps1 | 56453 | protein_coding | 1675.173016 | 0.069411262 | 0.086111121 | -0.806066176 | 0.420204693 | 0.640153165 | 0.069411262 |
| ENSMUSG00000042215 | Bag2 | 213539 | protein_coding | 31.98523335 | -0.069413042 | 0.18841 | 0.368414851 | 0.712563926 | 0.850598485 | 0.069413042 |
| ENSMUSG00000026456 | Cyb5r1 | 72017 | protein_coding | 127.2714532 | -0.069445679 | 0.13516018 | 0.513802798 | 0.607389871 | 0.781981914 | 0.069445679 |
| ENSMUSG00000037531 | Mrpl47 | 74600 | protein_coding | 245.5110236 | 0.069455857 | 0.084118474 | -0.825690882 | 0.408979466 | 0.629556944 | 0.069455857 |
| ENSMUSG00000011306 | Sugp1 | 70616 | protein_coding | 250.2585933 | 0.069502463 | 0.081454206 | -0.853270395 | 0.393509374 | 0.616138599 | 0.069502463 |
| ENSMUSG00000041671 | Pyroxd1 | 232491 | protein_coding | 221.7219353 | -0.069504034 | 0.124376884 | 0.558817943 | 0.576285976 | 0.761221133 | 0.069504034 |
| ENSMUSG00000049422 | Chchd10 | 103172 | protein_coding | 2433.51141 | 0.069525336 | 0.110035107 | -0.631846859 | 0.527486951 | 0.725788994 | 0.069525336 |
| ENSMUSG00000066324 | Bpnt2 | 242291 | protein_coding | 2087.365767 | 0.069535496 | 0.117617853 | -0.591198479 | 0.554387442 | 0.744656458 | 0.069535496 |
| ENSMUSG00000056752 | Dnah9 | 237806 | protein_coding | 17.07850251 | -0.069538975 | 0.43519727 | 0.15978725 | 0.873048668 | 0.937420397 | 0.069538975 |
| ENSMUSG00000005506 | Celf1 | 13046 | protein_coding | 1549.059469 | 0.069565635 | 0.069945508 | -0.994569019 | 0.319945922 | 0.541253166 | 0.069565635 |
| ENSMUSG00000020974 | Pole2 | 18974 | protein_coding | 47.45773765 | -0.069567315 | 0.259631548 | 0.267946307 | 0.788740649 | 0.894265208 | 0.069567315 |
| ENSMUSG00000037315 | Jade3 | 382207 | protein_coding | 132.3340814 | -0.069580816 | 0.115644125 | 0.601680511 | 0.547386825 | 0.738947603 | 0.069580816 |
| ENSMUSG00000050821 | Fam131a | 78408 | protein_coding | 5.486416865 | -0.069591608 | 0.704780641 | 0.098742224 | 0.921342943 | 0.962004133 | 0.069591608 |
| ENSMUSG00000047613 | A430005L14Rik | 97159 | protein_coding | 229.0303708 | 0.06959224 | 0.092616949 | -0.751398536 | 0.452412844 | 0.668738427 | 0.06959224 |
| ENSMUSG00000046985 | Tapt1 | 231225 | protein_coding | 456.2937628 | 0.069603202 | 0.077122917 | -0.902497014 | 0.366792907 | 0.589522317 | 0.069603202 |
| ENSMUSG00000058709 | Egln2 | 112406 | protein_coding | 922.570554 | 0.069607827 | 0.051390905 | -1.354477525 | 0.175584079 | 0.367456023 | 0.069607827 |
| ENSMUSG00000026577 | Blzf1 | 66352 | protein_coding | 234.0888566 | 0.069626746 | 0.094049652 | -0.74031902 | 0.459106443 | 0.673388737 | 0.069626746 |
| ENSMUSG00000030232 | Aebp2 | 11569 | protein_coding | 573.4951451 | 0.069649593 | 0.051956169 | -1.340545199 | 0.18006816 | 0.373667454 | 0.069649593 |
| ENSMUSG00000048429 | Timm29 | 69773 | protein_coding | 324.6561147 | 0.06967999 | 0.087780063 | -0.793802001 | 0.427310708 | 0.646801927 | 0.06967999 |
| ENSMUSG00000031231 | Cox7b | 66142 | protein_coding | 4210.71067 | -0.069681968 | 0.072157633 | 0.965690888 | 0.33419888 | 0.556515106 | 0.069681968 |
| ENSMUSG00000022771 | Ppil2 | 66053 | protein_coding | 801.0564228 | -0.06968577 | 0.043357446 | 1.607238821 | 0.108001989 | 0.261594876 | 0.06968577 |
| ENSMUSG00000042390 | Gatad2b | 229542 | protein_coding | 323.736406 | -0.069692455 | 0.112236164 | 0.620944732 | 0.534635988 | 0.730517043 | 0.069692455 |
| ENSMUSG00000084957 | Bbip1 | 100503572 | protein_coding | 240.9502092 | 0.069721217 | 0.087901356 | -0.793175676 | 0.427675477 | 0.647057325 | 0.069721217 |
| ENSMUSG00000020850 | Prpf8 | 192159 | protein_coding | 2879.78303 | 0.069759897 | 0.090332641 | -0.772255702 | 0.439962996 | 0.657874003 | 0.069759897 |
| ENSMUSG00000037503 | Fam168b | 214469 | protein_coding | 1374.744895 | -0.069818167 | 0.083485003 | 0.836295912 | 0.402988453 | 0.624327666 | 0.069818167 |
| ENSMUSG00000024926 | Kat5 | 81601 | protein_coding | 322.0692931 | -0.069826651 | 0.080445283 | 0.868001811 | 0.385393341 | 0.607636589 | 0.069826651 |
| ENSMUSG00000021555 | Naa35 | 78689 | protein_coding | 673.4273177 | 0.06984099 | 0.079338215 | -0.880294441 | 0.378699823 | 0.601173216 | 0.06984099 |
| ENSMUSG00000022952 | Runx1 | 12394 | protein_coding | 157.0660506 | -0.069900459 | 0.143463359 | 0.487235621 | 0.626091365 | 0.795381772 | 0.069900459 |
| ENSMUSG00000038756 | Ttll6 | 237930 | protein_coding | 365.3338347 | -0.069980892 | 0.123985407 | 0.564428457 | 0.572462576 | 0.757928885 | 0.069980892 |
| ENSMUSG00000029447 | Cct6a | 12466 | protein_coding | 1606.695493 | 0.070032574 | 0.086218638 | -0.812267228 | 0.416638311 | 0.636776036 | 0.070032574 |
| ENSMUSG00000078161 | Erich3 | 209601 | protein_coding | 5.577721169 | 0.070079559 | 0.876488548 | -0.079954905 | 0.936273121 | 0.969802608 | 0.070079559 |
| ENSMUSG00000027327 | 1700037H04Rik | 67326 | protein_coding | 203.744219 | -0.070095555 | 0.10259845 | 0.683202868 | 0.494478661 | 0.700667401 | 0.070095555 |
| ENSMUSG00000024664 | Fads3 | 60527 | protein_coding | 323.8414458 | 0.070109933 | 0.084769162 | -0.827068849 | 0.408198039 | 0.628822987 | 0.070109933 |
| ENSMUSG00000055024 | Ep300 | 328572 | protein_coding | 789.296829 | -0.070133057 | 0.143532235 | 0.488622341 | 0.625109093 | 0.794593769 | 0.070133057 |
| ENSMUSG00000002814 | Top3a | 21975 | protein_coding | 138.0332421 | 0.070157103 | 0.100246581 | -0.699845343 | 0.484023894 | 0.694394109 | 0.070157103 |
| ENSMUSG00000029148 | Nrbp1 | 192292 | protein_coding | 906.6638531 | -0.070174334 | 0.063658342 | 1.102358804 | 0.270305714 | 0.485621036 | 0.070174334 |
| ENSMUSG00000024913 | Lrp5 | 16973 | protein_coding | 981.8029703 | -0.070180007 | 0.077039177 | 0.910965172 | 0.362313725 | 0.584758488 | 0.070180007 |

| ensembl_gene_id | external_gene_name | entrezgene_id | gene_biotype | baseMean | log2FoldChange | lfcSE | stat | pvalue | padj | absLog2FC |
|---|---|---|---|---|---|---|---|---|---|---|
| ENSMUSG00000025878 | Uimc1 | 20184 | protein_coding | 243.6989128 | -0.07020706 | 0.092855007 | 0.756093427 | 0.449593183 | 0.666359883 | 0.07020706 |
| ENSMUSG00000034403 | Pja1 | 18744 | protein_coding | 759.8271443 | -0.070252011 | 0.07698962 | 0.912486785 | 0.361512523 | 0.583920353 | 0.070252011 |
| ENSMUSG00000025958 | Creb1 | 12912 | protein_coding | 646.6146078 | -0.070267291 | 0.104046204 | 0.675346993 | 0.499455334 | 0.704201299 | 0.070267291 |
| ENSMUSG00000019795 | Pcmt1 | 18537 | protein_coding | 457.274046 | -0.070283542 | 0.057472271 | 1.222912219 | 0.221362853 | 0.428054585 | 0.070283542 |
| ENSMUSG00000027845 | Dclre1b | 140917 | protein_coding | 119.6461029 | 0.070291785 | 0.100450395 | -0.699766134 | 0.484073368 | 0.69440476 | 0.070291785 |
| ENSMUSG00000059278 | Naa38 | 78304 | protein_coding | 227.8996355 | 0.07031446 | 0.10740591 | -0.65466099 | 0.512686046 | 0.714894653 | 0.07031446 |
| ENSMUSG00000021770 | Samd8 | 67630 | protein_coding | 475.6266307 | 0.070339615 | 0.109493849 | -0.642407003 | 0.520608954 | 0.72058807 | 0.070339615 |
| ENSMUSG00000106847 | Peg13 | 353342 | lncRNA | 128.9803648 | -0.070344017 | 0.121984846 | 0.576661933 | 0.564167853 | 0.751296404 | 0.070344017 |
| ENSMUSG00000002280 | Ciao3 | 67563 | protein_coding | 349.5834463 | -0.070355689 | 0.062669145 | 1.122652772 | 0.261584993 | 0.475244062 | 0.070355689 |
| ENSMUSG00000069227 | Gprin1 | 26913 | protein_coding | 13.39858102 | -0.070361882 | 0.430684974 | 0.163372039 | 0.870225517 | 0.935910399 | 0.070361882 |
| ENSMUSG00000039166 | Akap7 | 432442 | protein_coding | 540.7886955 | 0.070367977 | 0.089312051 | -0.787888934 | 0.430761674 | 0.649228647 | 0.070367977 |
| ENSMUSG00000022009 | Nufip1 | 27275 | protein_coding | 141.7216202 | -0.070385819 | 0.124384205 | 0.565874255 | 0.571479264 | 0.756991142 | 0.070385819 |
| ENSMUSG00000032571 | Pik3r4 | 75669 | protein_coding | 430.1078844 | -0.07038589 | 0.084222608 | 0.835712541 | 0.403316638 | 0.624624492 | 0.07038589 |
| ENSMUSG00000001632 | Brpf1 | 78783 | protein_coding | 440.5951091 | -0.070445577 | 0.095106006 | 0.740705871 | 0.458871799 | 0.673388737 | 0.070445577 |
| ENSMUSG00000004936 | Map2k1 | 26395 | protein_coding | 2706.144762 | 0.070452077 | 0.091210599 | -0.772411078 | 0.439870995 | 0.65785452 | 0.070452077 |
| ENSMUSG00000055733 | Nap1l3 | 54561 | protein_coding | 8.645070279 | -0.070470554 | 0.414177089 | 0.170145948 | 0.864895359 | 0.93291195 | 0.070470554 |
| ENSMUSG00000013160 | Atp6v0d1 | 11972 | protein_coding | 1085.36966 | 0.070551755 | 0.089598408 | -0.787421978 | 0.431034885 | 0.649494967 | 0.070551755 |
| ENSMUSG00000056124 | B4galt6 | 56386 | protein_coding | 302.622827 | -0.070572776 | 0.092020074 | 0.766928059 | 0.44312429 | 0.660786267 | 0.070572776 |
| ENSMUSG00000034292 | Traf3ip1 | 74019 | protein_coding | 118.4607734 | 0.070594894 | 0.104157336 | -0.677771692 | 0.497916462 | 0.703146268 | 0.070594894 |
| ENSMUSG00000067274 | Rplp0 | 11837 | protein_coding | 16339.61871 | -0.070613456 | 0.113472793 | 0.622294156 | 0.533748464 | 0.73008549 | 0.070613456 |
| ENSMUSG00000028952 | Zbtb48 | 100090 | protein_coding | 108.641675 | 0.070734755 | 0.11371726 | -0.622023031 | 0.533926724 | 0.730224807 | 0.070734755 |
| ENSMUSG00000025983 | Ccdc150 | 78016 | protein_coding | 2.727865709 | -0.070766411 | 1.087211555 | 0.065089826 | 0.948102481 | 1 | 0.070766411 |
| ENSMUSG00000024528 | Srfbp1 | 67222 | protein_coding | 127.6191079 | -0.070781633 | 0.102664006 | 0.689449359 | 0.490540531 | 0.69793377 | 0.070781633 |
| ENSMUSG00000047417 | Rexo1 | 66932 | protein_coding | 504.6263062 | 0.070810711 | 0.086505587 | -0.81856807 | 0.413032891 | 0.633140737 | 0.070810711 |
| ENSMUSG00000046157 | Tmem229b | 268567 | protein_coding | 25.24685237 | -0.070842518 | 0.215717657 | 0.328403891 | 0.742606301 | 0.870032583 | 0.070842518 |
| ENSMUSG00000020308 | Tpgs1 | 110012 | protein_coding | 187.8513955 | -0.070851502 | 0.139802514 | 0.506797052 | 0.612297225 | 0.785464687 | 0.070851502 |
| ENSMUSG00000022887 | Masp1 | 17174 | protein_coding | 4.283993468 | -0.070917537 | 1.794405229 | 0.039521472 | 0.968474634 | 1 | 0.070917537 |
| ENSMUSG00000030545 | Pex11a | 18631 | protein_coding | 187.3347374 | -0.071034757 | 0.089618166 | 0.792637923 | 0.427988807 | 0.647057325 | 0.071034757 |
| ENSMUSG00000024122 | Pdpk1 | 18607 | protein_coding | 1315.899722 | 0.071083007 | 0.062055281 | -1.145478782 | 0.25201088 | 0.464375257 | 0.071083007 |
| ENSMUSG00000039218 | Srrm2 | 75956 | protein_coding | 3452.68944 | 0.071088608 | 0.164365187 | -0.432504047 | 0.665375112 | 0.819511083 | 0.071088608 |
| ENSMUSG00000007610 | Gtpbp3 | 70359 | protein_coding | 241.2286097 | -0.071167132 | 0.095542128 | 0.744876984 | 0.45634609 | 0.671470886 | 0.071167132 |
| ENSMUSG00000022663 | Atg3 | 67841 | protein_coding | 373.4466004 | 0.071222665 | 0.089808822 | -0.79304754 | 0.427750126 | 0.647057325 | 0.071222665 |
| ENSMUSG00000021486 | Prelid1 | 66494 | protein_coding | 2287.833299 | -0.071228562 | 0.077084423 | 0.92403314 | 0.355469062 | 0.578346413 | 0.071228562 |
| ENSMUSG00000032249 | Anp32a | 11737 | protein_coding | 1112.254822 | 0.071245486 | 0.078083968 | -0.912421443 | 0.361546906 | 0.583920353 | 0.071245486 |
| ENSMUSG00000033799 | Tasor2 | 105203 | protein_coding | 422.3488757 | -0.07127605 | 0.090978161 | 0.783441319 | 0.433368 | 0.652026162 | 0.07127605 |
| ENSMUSG00000000562 | Adora3 | 11542 | protein_coding | 2.184868721 | 0.071293424 | 1.050683368 | -0.067854337 | 0.945901588 | 1 | 0.071293424 |
| ENSMUSG00000027314 | Dll4 | 54485 | protein_coding | 124.2143631 | 0.071298255 | 0.125172592 | -0.569599573 | 0.568949318 | 0.754850891 | 0.071298255 |
| ENSMUSG00000034371 | Tkfc | 225913 | protein_coding | 167.739711 | -0.071339385 | 0.102407997 | 0.696619282 | 0.486041086 | 0.695595302 | 0.071339385 |
| ENSMUSG00000021023 | Prorp | 66132 | protein_coding | 185.5788494 | -0.071354896 | 0.093922027 | 0.759724825 | 0.447419086 | 0.664688628 | 0.071354896 |

| ensembl_gene_id | external_gene_name | entrezgene_id | gene_biotype | baseMean | log2FoldChange | lfcSE | stat | pvalue | padj | absLog2FC |
|---|---|---|---|---|---|---|---|---|---|---|
| ENSMUSG00000026617 | Bpnt1 | 23827 | protein_coding | 1624.291107 | -0.071365493 | 0.084470758 | 0.844854417 | 0.398192128 | 0.62019321 | 0.071365493 |
| ENSMUSG00000069793 | Slfn9 | 237886 | protein_coding | 371.9677368 | 0.071370511 | 0.206051704 | -0.346371854 | 0.729063275 | 0.860227988 | 0.071370511 |
| ENSMUSG00000024146 | Cript | 56724 | protein_coding | 422.4762476 | -0.071379634 | 0.095368281 | 0.748463042 | 0.454180909 | 0.669812138 | 0.071379634 |
| ENSMUSG00000046603 | Tcaim | 382117 | protein_coding | 121.7839963 | 0.071400061 | 0.118617617 | -0.601934709 | 0.547217599 | 0.738779548 | 0.071400061 |
| ENSMUSG00000056824 | Zfp663 | 381405 | protein_coding | 1.18598668 | -0.071450831 | 1.352754965 | 0.052818753 | 0.957876319 | 1 | 0.071450831 |
| ENSMUSG00000097402 | Gm17399 | 100502695 | lncRNA | 1.429694201 | -0.071480803 | 0.855858668 | 0.0835194 | 0.933438552 | 1 | 0.071480803 |
| ENSMUSG00000022443 | Myh9 | 17886 | protein_coding | 13088.56029 | 0.07150555 | 0.065032734 | -1.099531661 | 0.271536232 | 0.487195712 | 0.07150555 |
| ENSMUSG00000020170 | Frs2 | 327826 | protein_coding | 374.043006 | 0.071624042 | 0.088230124 | -0.811786711 | 0.416914029 | 0.636961631 | 0.071624042 |
| ENSMUSG00000081534 | Slc48a1 | 67739 | protein_coding | 277.6326829 | -0.071655155 | 0.104190545 | 0.687731836 | 0.491621668 | 0.698491194 | 0.071655155 |
| ENSMUSG00000034789 | Rab24 | 19336 | protein_coding | 123.1350826 | 0.071680861 | 0.112242868 | -0.638622854 | 0.52306831 | 0.722040793 | 0.071680861 |
| ENSMUSG00000068328 | Aup1 | 11993 | protein_coding | 971.3266859 | 0.071704537 | 0.063087887 | -1.136581683 | 0.255713201 | 0.468471767 | 0.071704537 |
| ENSMUSG00000053483 | Usp21 | 30941 | protein_coding | 218.6872546 | -0.07170548 | 0.089698561 | 0.799405019 | 0.424055601 | 0.643940615 | 0.07170548 |
| ENSMUSG00000039116 | Adgrg6 | 215798 | protein_coding | 29.54077578 | 0.07170879 | 0.237133363 | -0.30239857 | 0.762348245 | 0.88093126 | 0.07170879 |
| ENSMUSG00000047434 | Xxylt1 | 268880 | protein_coding | 161.8929539 | 0.07171652 | 0.120586714 | -0.594729865 | 0.552024055 | 0.742718368 | 0.07171652 |
| ENSMUSG00000026029 | Casp8 | 12370 | protein_coding | 1302.936966 | 0.071736327 | 0.074064023 | -0.968571838 | 0.33275886 | 0.555180826 | 0.071736327 |
| ENSMUSG00000016087 | Fli1 | 14247 | protein_coding | 8.295508489 | -0.071818975 | 0.639544796 | 0.11229702 | 0.910587904 | 0.956641558 | 0.071818975 |
| ENSMUSG00000074158 | Zfp976 | 208111 | protein_coding | 63.19180948 | -0.071823046 | 0.174871666 | 0.410718602 | 0.681278885 | 0.829503357 | 0.071823046 |
| ENSMUSG00000028634 | Hivep3 | 16656 | protein_coding | 80.14312156 | -0.071841607 | 0.163305137 | 0.439922519 | 0.659993226 | 0.816323585 | 0.071841607 |
| ENSMUSG00000074445 | Sprr2a3 | 100042514 | protein_coding | 1819.801065 | -0.071866267 | 0.266745472 | 0.269418881 | 0.787607358 | 0.893615254 | 0.071866267 |
| ENSMUSG00000010097 | Nxf1 | 53319 | protein_coding | 770.0948704 | 0.071928293 | 0.061889385 | -1.162207272 | 0.245151281 | 0.456778547 | 0.071928293 |
| ENSMUSG00000012889 | Podnl1 | 244550 | protein_coding | 6.654473095 | 0.071997531 | 0.522545926 | -0.137782207 | 0.890412547 | 0.946160016 | 0.071997531 |
| ENSMUSG00000021495 | Fam193b | 212483 | protein_coding | 408.8402297 | 0.071998913 | 0.168173398 | -0.428123079 | 0.668561515 | 0.822256556 | 0.071998913 |
| ENSMUSG00000022556 | Hsf1 | 15499 | protein_coding | 287.1582588 | -0.07204896 | 0.08839347 | 0.81509369 | 0.415018683 | 0.635300198 | 0.07204896 |
| ENSMUSG00000039427 | Alg1 | 208211 | protein_coding | 144.7167449 | -0.072087646 | 0.109005137 | 0.661323384 | 0.508404948 | 0.712111524 | 0.072087646 |
| ENSMUSG00000026202 | Tuba4a | 22145 | protein_coding | 1531.051937 | 0.072095135 | 0.104638857 | -0.68899008 | 0.49082951 | 0.698213329 | 0.072095135 |
| ENSMUSG00000024925 | Rnaseh2c | 68209 | protein_coding | 191.683545 | 0.072099401 | 0.122474925 | -0.588687041 | 0.556071233 | 0.745551704 | 0.072099401 |
| ENSMUSG00000036315 | Polr1h | 66136 | protein_coding | 207.8424377 | -0.072102807 | 0.090127785 | 0.800006429 | 0.423707072 | 0.643604486 | 0.072102807 |
| ENSMUSG00000020647 | Ncoa1 | 17977 | protein_coding | 707.3254314 | -0.072103373 | 0.080932511 | 0.890907408 | 0.372978846 | 0.595257447 | 0.072103373 |
| ENSMUSG00000020802 | Ube2o | 217342 | protein_coding | 252.419927 | 0.072132168 | 0.096690617 | -0.746010002 | 0.455661373 | 0.671017649 | 0.072132168 |
| ENSMUSG00000047518 | Slfnl1 | 194219 | protein_coding | 2.394458992 | -0.072150781 | 0.877661584 | 0.082207975 | 0.934481332 | 1 | 0.072150781 |
| ENSMUSG00000003206 | Ebi3 | 50498 | protein_coding | 1.907913679 | -0.072183023 | 0.988410496 | 0.073029397 | 0.941782724 | 1 | 0.072183023 |
| ENSMUSG00000019998 | Stx7 | 53331 | protein_coding | 1048.852036 | -0.072191555 | 0.06744786 | 1.070331291 | 0.284470214 | 0.502862462 | 0.072191555 |
| ENSMUSG00000096768 | Gm47283 | 170942 | lncRNA | 9.53692367 | -0.072199419 | 0.902027698 | 0.080041244 | 0.936204454 | 0.96979553 | 0.072199419 |
| ENSMUSG00000031516 | Dctn6 | 22428 | protein_coding | 462.682918 | -0.072203482 | 0.07276957 | 0.992220803 | 0.321089826 | 0.542231046 | 0.072203482 |
| ENSMUSG00000055538 | Zcchc24 | 71918 | protein_coding | 7.561834759 | -0.072203619 | 0.50221133 | 0.143771386 | 0.885680999 | 0.943256547 | 0.072203619 |
| ENSMUSG00000053641 | Dennd4a | 102442 | protein_coding | 152.5043923 | -0.072204126 | 0.118291397 | 0.610392033 | 0.541602144 | 0.735467298 | 0.072204126 |
| ENSMUSG00000031931 | Ankrd49 | 56503 | protein_coding | 222.0559217 | -0.072231568 | 0.095155962 | 0.759086106 | 0.44780105 | 0.664930223 | 0.072231568 |
| ENSMUSG00000027324 | Rpusd2 | 271842 | protein_coding | 57.63500509 | 0.072330901 | 0.148555137 | -0.486895992 | 0.62633204 | 0.795381772 | 0.072330901 |
| ENSMUSG00000027665 | Pik3ca | 18706 | protein_coding | 774.5748466 | -0.072370836 | 0.097095344 | 0.745358455 | 0.456055052 | 0.671357918 | 0.072370836 |

Supplementary Table 1

| ensembl_gene_id | external_gene_name | entrezgene_id | gene_biotype | baseMean | log2FoldChange | lfcSE | stat | pvalue | padj | absLog2FC |
|---|---|---|---|---|---|---|---|---|---|---|
| ENSMUSG00000036270 | Edc4 | 234699 | protein_coding | 549.6862616 | 0.072372396 | 0.052743611 | -1.372154752 | 0.170015268 | 0.359584034 | 0.072372396 |
| ENSMUSG00000035561 | Aldh1b1 | 72535 | protein_coding | 3834.574957 | 0.072417252 | 0.120478592 | -0.601079833 | 0.547786815 | 0.739162904 | 0.072417252 |
| ENSMUSG00000044916 | 1700029I15Rik | 75641 | protein_coding | 10.81515042 | 0.072462769 | 0.458749334 | -0.157957219 | 0.87449051 | 0.938406345 | 0.072462769 |
| ENSMUSG00000026893 | Gca | 227960 | protein_coding | 148.4738171 | -0.072479971 | 0.102365637 | 0.708049824 | 0.478914317 | 0.690541222 | 0.072479971 |
| ENSMUSG00000091381 | Vmn2r83 | 625029 | protein_coding | 0.667830234 | -0.072485024 | 1.679189123 | 0.043166683 | 0.965568664 | 1 | 0.072485024 |
| ENSMUSG00000037279 | Ovol2 | 107586 | protein_coding | 205.0987239 | -0.072502309 | 0.145155808 | 0.499479212 | 0.617441828 | 0.789258793 | 0.072502309 |
| ENSMUSG00000041133 | Smc1a | 24061 | protein_coding | 1171.974711 | 0.07251915 | 0.106586295 | -0.680379688 | 0.496264079 | 0.702459427 | 0.07251915 |
| ENSMUSG00000029146 | Snx17 | 266781 | protein_coding | 675.0808095 | 0.072538072 | 0.066690103 | -1.087688708 | 0.276732555 | 0.493649227 | 0.072538072 |
| ENSMUSG00000024193 | Phf1 | 21652 | protein_coding | 142.8040974 | 0.072618682 | 0.16521337 | -0.439544827 | 0.660266809 | 0.816494106 | 0.072618682 |
| ENSMUSG00000029478 | Ncor2 | 20602 | protein_coding | 331.1085108 | 0.072623916 | 0.235822093 | -0.307960612 | 0.758112304 | 0.878750865 | 0.072623916 |
| ENSMUSG00000019235 | Rps6kl1 | 238323 | protein_coding | 22.05753863 | 0.072649886 | 0.232653508 | -0.312266452 | 0.754838033 | 0.876856731 | 0.072649886 |
| ENSMUSG00000037708 | Spag6 | 381350 | protein_coding | 1.305640353 | -0.072655037 | 1.181960973 | 0.061469912 | 0.950984976 | 1 | 0.072655037 |
| ENSMUSG00000038178 | Slc43a2 | 215113 | protein_coding | 154.6380287 | 0.072687287 | 0.190101289 | -0.382360831 | 0.702193739 | 0.842918326 | 0.072687287 |
| ENSMUSG00000053226 | Dand5 | 23863 | protein_coding | 27.28363634 | -0.072692751 | 0.238553042 | 0.304723637 | 0.760576646 | 0.879809661 | 0.072692751 |
| ENSMUSG00000001674 | Ddx18 | 66942 | protein_coding | 166.5279987 | 0.072702461 | 0.1185812 | -0.613102756 | 0.539808398 | 0.734239015 | 0.072702461 |
| ENSMUSG00000024871 | Doc2g | 60425 | protein_coding | 12.55454676 | 0.07272643 | 0.342386643 | -0.212410242 | 0.831786996 | 0.915204382 | 0.07272643 |
| ENSMUSG00000004096 | Cwc15 | 66070 | protein_coding | 669.1558435 | 0.072731516 | 0.066240533 | -1.0979911 | 0.272208376 | 0.488030519 | 0.072731516 |
| ENSMUSG00000060032 | H2aj | 232440 | protein_coding | 744.9012931 | -0.072734076 | 0.085220784 | 0.853478138 | 0.393394206 | 0.616016681 | 0.072734076 |
| ENSMUSG00000056394 | Lig1 | 16881 | protein_coding | 317.0370031 | 0.072735172 | 0.148202009 | -0.490783981 | 0.623579241 | 0.793440482 | 0.072735172 |
| ENSMUSG00000052997 | Uba2 | 50995 | protein_coding | 906.9369156 | 0.07273834 | 0.078853587 | -0.92244808 | 0.356294901 | 0.579347577 | 0.07273834 |
| ENSMUSG00000029250 | Polr2b | 231329 | protein_coding | 867.9206337 | 0.072742602 | 0.11955229 | -0.608458456 | 0.542883452 | 0.736235501 | 0.072742602 |
| ENSMUSG00000022536 | Glyr1 | 74022 | protein_coding | 1798.09112 | 0.072748963 | 0.054940601 | -1.324138471 | 0.185457061 | 0.380783029 | 0.072748963 |
| ENSMUSG00000033737 | Fndc3c1 | 333564 | protein_coding | 2.254231156 | 0.072761562 | 2.748693799 | -0.026471323 | 0.978881406 | 1 | 0.072761562 |
| ENSMUSG00000033208 | S100b | 20203 | protein_coding | 3.48302778 | -0.072840015 | 0.593663424 | 0.12269581 | 0.902347982 | 1 | 0.072840015 |
| ENSMUSG00000033871 | Ppargc1b | 170826 | protein_coding | 697.4297624 | -0.072841621 | 0.130423116 | 0.558502382 | 0.576501378 | 0.761222618 | 0.072841621 |
| ENSMUSG00000078908 | Mon1b | 270096 | protein_coding | 456.5177364 | -0.072863931 | 0.09206727 | 0.79142057 | 0.42869861 | 0.647656102 | 0.072863931 |
| ENSMUSG00000059482 | Cfap418 | 67157 | protein_coding | 47.49555018 | 0.072865906 | 0.14506139 | -0.502310825 | 0.615448896 | 0.787780401 | 0.072865906 |
| ENSMUSG00000059981 | Taok2 | 381921 | protein_coding | 795.949316 | 0.072873564 | 0.077951967 | -0.93485214 | 0.349864505 | 0.571706158 | 0.072873564 |
| ENSMUSG00000033769 | Exoc6b | 75914 | protein_coding | 579.7788482 | 0.072901256 | 0.093950737 | -0.775951931 | 0.437777368 | 0.656109195 | 0.072901256 |
| ENSMUSG00000057897 | Camk2b | 12323 | protein_coding | 377.720462 | -0.072901605 | 0.11898565 | 0.612692412 | 0.540079741 | 0.734487099 | 0.072901605 |
| ENSMUSG00000092335 | Zfp977 | 637776 | protein_coding | 3.441032652 | -0.072930991 | 0.68483825 | 0.106493747 | 0.915190616 | 1 | 0.072930991 |
| ENSMUSG00000039069 | Mtg2 | 52856 | protein_coding | 166.8661459 | -0.072937883 | 0.09465817 | 0.770539753 | 0.440979784 | 0.658856432 | 0.072937883 |
| ENSMUSG00000063802 | Hspbp1 | 66245 | protein_coding | 263.7071864 | 0.072962005 | 0.08925995 | -0.817410335 | 0.413693972 | 0.633801118 | 0.072962005 |
| ENSMUSG00000030970 | Ctbp2 | 13017 | protein_coding | 912.5815397 | 0.072968638 | 0.054308083 | -1.343605466 | 0.179076001 | 0.37212295 | 0.072968638 |
| ENSMUSG00000005354 | Txn2 | 56551 | protein_coding | 1103.372475 | -0.073004039 | 0.062959632 | 1.159537258 | 0.246237259 | 0.458233276 | 0.073004039 |
| ENSMUSG00000023852 | Chd1 | 12648 | protein_coding | 950.2037711 | 0.073092502 | 0.064089598 | -1.140473719 | 0.254088996 | 0.466691767 | 0.073092502 |
| ENSMUSG00000022400 | Rbx1 | 56438 | protein_coding | 541.3485084 | -0.07312623 | 0.101271862 | 0.722078456 | 0.470246243 | 0.68365582 | 0.07312623 |
| ENSMUSG00000027667 | Zfp639 | 67778 | protein_coding | 156.4773481 | -0.073156674 | 0.092564518 | 0.790331709 | 0.429334073 | 0.648260298 | 0.073156674 |
| ENSMUSG00000011960 | Ccnt1 | 12455 | protein_coding | 519.1692111 | -0.073157468 | 0.081214846 | 0.90078935 | 0.367700331 | 0.590110273 | 0.073157468 |

| ensembl_gene_id | external_gene_name | entrezgene_id | gene_biotype | baseMean | log2FoldChange | lfcSE | stat | pvalue | padj | absLog2FC |
|---|---|---|---|---|---|---|---|---|---|---|
| ENSMUSG00000032959 | Pebp1 | 23980 | protein_coding | 922.1133449 | 0.073173395 | 0.124167724 | -0.58931091 | 0.555652727 | 0.74529314 | 0.073173395 |
| ENSMUSG00000045237 | Eola1 | 68618 | protein_coding | 160.3678012 | -0.073186747 | 0.091649879 | 0.798547115 | 0.424553064 | 0.644177805 | 0.073186747 |
| ENSMUSG00000027706 | Sec62 | 69276 | protein_coding | 867.512681 | 0.073245045 | 0.123064132 | -0.595177852 | 0.551724593 | 0.742476843 | 0.073245045 |
| ENSMUSG00000021113 | Snapc1 | 75627 | protein_coding | 149.2677397 | 0.073279998 | 0.099592052 | -0.735801664 | 0.461851409 | 0.675674536 | 0.073279998 |
| ENSMUSG00000026353 | Ubxn4 | 67812 | protein_coding | 984.6877432 | 0.073323104 | 0.065265197 | -1.123464064 | 0.261240454 | 0.47491467 | 0.073323104 |
| ENSMUSG00000026201 | Stk16 | 20872 | protein_coding | 673.9597439 | 0.073332435 | 0.070718392 | -1.036964116 | 0.299752581 | 0.519697021 | 0.073332435 |
| ENSMUSG00000028243 | Ubxn2b | 68053 | protein_coding | 97.19988081 | 0.073341777 | 0.116476173 | -0.62967193 | 0.528909251 | 0.726837595 | 0.073341777 |
| ENSMUSG00000020716 | Nf1 | 18015 | protein_coding | 718.20267 | 0.073385652 | 0.107216209 | -0.684464157 | 0.493682115 | 0.699975951 | 0.073385652 |
| ENSMUSG00000003452 | Bicd1 | 12121 | protein_coding | 211.0520948 | -0.073389255 | 0.110856381 | 0.662021026 | 0.507957746 | 0.711726983 | 0.073389255 |
| ENSMUSG00000039041 | Adrm1 | 56436 | protein_coding | 503.8593163 | 0.073421736 | 0.100483275 | -0.730686137 | 0.464970885 | 0.678555072 | 0.073421736 |
| ENSMUSG00000003119 | Cdk12 | 69131 | protein_coding | 847.911418 | 0.073470601 | 0.074919903 | -0.980655319 | 0.326762744 | 0.548613253 | 0.073470601 |
| ENSMUSG00000073016 | Uprt | 331487 | protein_coding | 229.5660483 | -0.073501791 | 0.104068154 | 0.706285137 | 0.480010834 | 0.691115487 | 0.073501791 |
| ENSMUSG00000026526 | Fh1 | 14194 | protein_coding | 1550.328137 | -0.073511809 | 0.07733135 | 0.950608128 | 0.341803341 | 0.563829035 | 0.073511809 |
| ENSMUSG00000027099 | Mtx2 | 53375 | protein_coding | 603.0802267 | -0.073532494 | 0.077546356 | 0.948239193 | 0.343007699 | 0.564917636 | 0.073532494 |
| ENSMUSG00000021076 | Actr10 | 56444 | protein_coding | 606.5491003 | 0.073540235 | 0.064171578 | -1.145993868 | 0.251797689 | 0.464320478 | 0.073540235 |
| ENSMUSG00000024312 | Wdr46 | 57315 | protein_coding | 270.8799745 | 0.073558542 | 0.096084721 | -0.765559191 | 0.443938634 | 0.661243132 | 0.073558542 |
| ENSMUSG00000059288 | Cdyl | 12593 | protein_coding | 147.3746199 | 0.073575608 | 0.095850424 | -0.767608578 | 0.442719765 | 0.660497294 | 0.073575608 |
| ENSMUSG00000024078 | Ttc27 | 74196 | protein_coding | 157.4281687 | -0.073578972 | 0.156418249 | 0.470398898 | 0.638070051 | 0.802764652 | 0.073578972 |
| ENSMUSG00000047036 | Zfp445 | 235682 | protein_coding | 943.0709585 | -0.073597225 | 0.103794108 | 0.709069385 | 0.47828142 | 0.68987155 | 0.073597225 |
| ENSMUSG00000074578 | Zfas1 | 68949 | lncRNA | 322.2735656 | -0.073628468 | 0.124445015 | 0.591654623 | 0.554081888 | 0.744516542 | 0.073628468 |
| ENSMUSG00000020986 | Sec23a | 20334 | protein_coding | 1312.684775 | 0.073653502 | 0.075060895 | -0.981249987 | 0.326469479 | 0.548343875 | 0.073653502 |
| ENSMUSG00000094328 | Tdpoz9 | 668359 | protein_coding | 0.78622542 | 0.073729475 | 1.101899862 | -0.06691123 | 0.946652373 | 1 | 0.073729475 |
| ENSMUSG00000028833 | Ncdn | 26562 | protein_coding | 359.135346 | 0.073739184 | 0.061417603 | -1.200619695 | 0.229898758 | 0.438451101 | 0.073739184 |
| ENSMUSG00000020536 | Llgl1 | 16897 | protein_coding | 455.4433636 | -0.073768538 | 0.067523541 | 1.092486231 | 0.274619439 | 0.491020508 | 0.073768538 |
| ENSMUSG00000022122 | Ednrb | 13618 | protein_coding | 0.855237627 | 0.073768977 | 1.312510146 | -0.0562045 | 0.955178896 | 1 | 0.073768977 |
| ENSMUSG00000029141 | Slc4a1ap | 20534 | protein_coding | 233.4924963 | 0.073842236 | 0.073952941 | -0.998503038 | 0.318035492 | 0.539389762 | 0.073842236 |
| ENSMUSG00000020956 | Dtd2 | 328092 | protein_coding | 107.5770969 | 0.073881297 | 0.107673943 | -0.686157623 | 0.492613717 | 0.699148013 | 0.073881297 |
| ENSMUSG00000028369 | Svep1 | 64817 | protein_coding | 11.06831908 | -0.073906932 | 0.607959563 | 0.121565539 | 0.903243107 | 0.952286247 | 0.073906932 |
| ENSMUSG00000038335 | Tsr1 | 104662 | protein_coding | 444.4057602 | -0.073911507 | 0.096009817 | 0.769832809 | 0.441399075 | 0.659125086 | 0.073911507 |
| ENSMUSG00000039065 | Atpsckmt | 68073 | protein_coding | 159.4703557 | 0.07392467 | 0.107055601 | -0.690525951 | 0.489863496 | 0.697627026 | 0.07392467 |
| ENSMUSG00000018736 | Ndel1 | 83431 | protein_coding | 726.9430645 | 0.073934542 | 0.090640161 | -0.815692965 | 0.414675763 | 0.634951837 | 0.073934542 |
| ENSMUSG00000023951 | Vegfa | 22339 | protein_coding | 660.1726819 | -0.073947707 | 0.202693124 | 0.364825927 | 0.71524135 | 0.852138782 | 0.073947707 |
| ENSMUSG00000030271 | Ogg1 | 18294 | protein_coding | 83.39595566 | -0.073956186 | 0.122482911 | 0.6038082 | 0.545971167 | 0.737699873 | 0.073956186 |
| ENSMUSG00000073471 | Rsph3a | 66832 | protein_coding | 131.9807273 | 0.073966132 | 0.112721048 | -0.656187401 | 0.511703555 | 0.714354347 | 0.073966132 |
| ENSMUSG00000010609 | Psen2 | 19165 | protein_coding | 177.4624009 | -0.073974393 | 0.087972917 | 0.840876887 | 0.400416908 | 0.622382905 | 0.073974393 |
| ENSMUSG00000020706 | Ftsj3 | 56095 | protein_coding | 450.0591043 | -0.074091066 | 0.101291523 | 0.731463641 | 0.464496005 | 0.677981883 | 0.074091066 |
| ENSMUSG00000029402 | Snrnp35 | 76167 | protein_coding | 87.29959937 | 0.074103994 | 0.191104733 | -0.387766397 | 0.698188915 | 0.840126183 | 0.074103994 |
| ENSMUSG00000024696 | Lpxn | 107321 | protein_coding | 13.42599148 | -0.074105982 | 0.273559958 | 0.270894844 | 0.786471911 | 0.89324602 | 0.074105982 |
| ENSMUSG00000005580 | Adcy9 | 11515 | protein_coding | 240.8029088 | -0.074110301 | 0.091016296 | 0.814253102 | 0.41549997 | 0.635742288 | 0.074110301 |

| ensembl_gene_id | external_gene_name | entrezgene_id | gene_biotype | baseMean | log2FoldChange | lfcSE | stat | pvalue | padj | absLog2FC |
|---|---|---|---|---|---|---|---|---|---|---|
| ENSMUSG00000021901 | Bap1 | 104416 | protein_coding | 497.6567757 | 0.074132354 | 0.055463962 | -1.336585981 | 0.181357818 | 0.375447286 | 0.074132354 |
| ENSMUSG00000032288 | Imp3 | 102462 | protein_coding | 343.3363181 | -0.074144673 | 0.122004658 | 0.607720017 | 0.543373185 | 0.736456016 | 0.074144673 |
| ENSMUSG00000032053 | Pou2af1 | 18985 | protein_coding | 2.840142479 | -0.074152921 | 1.113276708 | 0.066607808 | 0.94689393 | 1 | 0.074152921 |
| ENSMUSG00000041642 | Kif21b | 16565 | protein_coding | 738.8685292 | 0.074173502 | 0.215389361 | -0.344369384 | 0.73056851 | 0.861253255 | 0.074173502 |
| ENSMUSG00000022848 | Slc49a4 | 224132 | protein_coding | 481.2973534 | -0.074214473 | 0.087141073 | 0.851658934 | 0.39440342 | 0.616875898 | 0.074214473 |
| ENSMUSG00000074569 | Gcnt7 | 654821 | protein_coding | 16.73809033 | -0.074234953 | 0.274639521 | 0.2702996 | 0.786929773 | 0.893375436 | 0.074234953 |
| ENSMUSG00000019066 | Rab3d | 19340 | protein_coding | 1976.964593 | 0.074250145 | 0.156063107 | -0.475770006 | 0.634238241 | 0.800627603 | 0.074250145 |
| ENSMUSG00000040521 | Tsfm | 66399 | protein_coding | 261.4789656 | -0.074273414 | 0.080825461 | 0.918935859 | 0.358129122 | 0.580790426 | 0.074273414 |
| ENSMUSG00000044062 | Plekhd1os | 68354 | lncRNA | 12.27190352 | -0.074288736 | 0.298056139 | 0.249244105 | 0.803171965 | 0.902837235 | 0.074288736 |
| ENSMUSG00000068250 | Amn1 | 232566 | protein_coding | 178.3719994 | 0.074306562 | 0.117853104 | -0.630501524 | 0.528366506 | 0.726359126 | 0.074306562 |
| ENSMUSG00000024238 | Zeb1 | 21417 | protein_coding | 5.488968061 | 0.074332944 | 0.801267665 | -0.09276918 | 0.926086936 | 0.964614821 | 0.074332944 |
| ENSMUSG00000040389 | Wdr47 | 99512 | protein_coding | 69.58829461 | -0.074333599 | 0.142226257 | 0.52264329 | 0.601222509 | 0.777615016 | 0.074333599 |
| ENSMUSG00000031980 | Agt | 11606 | protein_coding | 135.3727578 | -0.074359722 | 0.142351925 | 0.522365411 | 0.601415934 | 0.777725809 | 0.074359722 |
| ENSMUSG00000033557 | Fam20b | 215015 | protein_coding | 2414.706563 | 0.074404107 | 0.068432184 | -1.08726776 | 0.276918492 | 0.493794621 | 0.074404107 |
| ENSMUSG00000051910 | Sox6 | 20679 | protein_coding | 215.9482222 | 0.074469665 | 0.202453536 | -0.367835834 | 0.712995648 | 0.850752177 | 0.074469665 |
| ENSMUSG00000032806 | Slc10a3 | 214601 | protein_coding | 229.2486502 | 0.074511429 | 0.114536315 | -0.650548506 | 0.51533798 | 0.716335759 | 0.074511429 |
| ENSMUSG00000035171 | 1110059E24Rik | 66206 | protein_coding | 544.6424862 | 0.074523929 | 0.111068212 | -0.670974421 | 0.502236824 | 0.706950129 | 0.074523929 |
| ENSMUSG00000029416 | Slc15a4 | 100561 | protein_coding | 237.798877 | 0.074533763 | 0.098538216 | -0.756394482 | 0.449412715 | 0.666174493 | 0.074533763 |
| ENSMUSG00000059552 | Trp53 | 22059 | protein_coding | 286.8677191 | -0.074556251 | 0.089625467 | 0.831864575 | 0.405485389 | 0.62645577 | 0.074556251 |
| ENSMUSG00000020612 | Prkar1a | 19084 | protein_coding | 883.178799 | 0.074568781 | 0.071840486 | -1.03797713 | 0.299280703 | 0.519260509 | 0.074568781 |
| ENSMUSG00000091945 | Vmn2r114 | 666002 | protein_coding | 4.288916074 | 0.07460139 | 2.23324674 | -0.033404903 | 0.9733517 | 1 | 0.07460139 |
| ENSMUSG00000035437 | Rabgap1 | 227800 | protein_coding | 523.3318927 | -0.074629653 | 0.079079966 | 0.943723888 | 0.345310763 | 0.566901476 | 0.074629653 |
| ENSMUSG00000047040 | Prr15l | 217138 | protein_coding | 3498.917221 | 0.074636867 | 0.100953902 | -0.739316309 | 0.459714948 | 0.674101639 | 0.074636867 |
| ENSMUSG00000028274 | Rngtt | 24018 | protein_coding | 239.3114651 | -0.074652007 | 0.080375623 | 0.928789153 | 0.352998367 | 0.575403714 | 0.074652007 |
| ENSMUSG00000073460 | Pnldc1 | 240023 | protein_coding | 3.546418738 | 0.074702628 | 0.617607187 | -0.120954921 | 0.903726741 | 1 | 0.074702628 |
| ENSMUSG00000022957 | Itsn1 | 16443 | protein_coding | 467.9399724 | 0.074732163 | 0.064761853 | -1.153953436 | 0.24851926 | 0.460736668 | 0.074732163 |
| ENSMUSG00000026895 | Ndufa8 | 68375 | protein_coding | 1110.338218 | -0.074736245 | 0.084261804 | 0.886952833 | 0.375104295 | 0.597345451 | 0.074736245 |
| ENSMUSG00000069830 | Nlrp1a | 195046 | protein_coding | 1.83156864 | 0.074748784 | 1.7984264 | -0.041563438 | 0.966846721 | 1 | 0.074748784 |
| ENSMUSG00000031981 | Capn9 | 73647 | protein_coding | 429.6776005 | -0.074755613 | 0.224790723 | 0.332556488 | 0.739469093 | 0.867855348 | 0.074755613 |
| ENSMUSG00000001098 | Kctd10 | 330171 | protein_coding | 961.4025273 | -0.074801162 | 0.083412019 | 0.896767188 | 0.36984316 | 0.592567182 | 0.074801162 |
| ENSMUSG00000046792 | Zfp787 | 67109 | protein_coding | 310.8505099 | -0.074808543 | 0.064140501 | 1.166323023 | 0.243483868 | 0.454749219 | 0.074808543 |
| ENSMUSG00000032936 | Camkv | 235604 | protein_coding | 1.525313563 | -0.0748167 | 1.714758589 | 0.04363104 | 0.965198509 | 1 | 0.0748167 |
| ENSMUSG00000032370 | Lactb | 80907 | protein_coding | 154.625219 | -0.074819452 | 0.111683524 | 0.66992381 | 0.502906361 | 0.707530724 | 0.074819452 |
| ENSMUSG00000035964 | Tmem59l | 67937 | protein_coding | 1.606433954 | -0.07482141 | 1.267348308 | 0.059037764 | 0.952922029 | 1 | 0.07482141 |
| ENSMUSG00000015340 | Cybb | 13058 | protein_coding | 2.987484453 | -0.07484029 | 1.018826377 | 0.073457354 | 0.941442179 | 1 | 0.07484029 |
| ENSMUSG00000050312 | Nsun3 | 106338 | protein_coding | 191.3194327 | -0.074851784 | 0.121862036 | 0.614233824 | 0.539060825 | 0.733662794 | 0.074851784 |
| ENSMUSG00000019857 | Asf1a | 66403 | protein_coding | 106.1963009 | 0.074855708 | 0.137717163 | -0.543546689 | 0.586753454 | 0.768287864 | 0.074855708 |
| ENSMUSG00000031403 | Dkc1 | 245474 | protein_coding | 299.2094204 | 0.074898008 | 0.137098165 | -0.546309341 | 0.584853316 | 0.767077092 | 0.074898008 |
| ENSMUSG00000050188 | Lsm10 | 116748 | protein_coding | 96.27582219 | 0.074945543 | 0.122593339 | -0.611334539 | 0.54097813 | 0.735049182 | 0.074945543 |

| ensembl_gene_id | external_gene_name | entrezgene_id | gene_biotype | baseMean | log2FoldChange | lfcSE | stat | pvalue | padj | absLog2FC |
|---|---|---|---|---|---|---|---|---|---|---|
| ENSMUSG00000020671 | Rab10 | 19325 | protein_coding | 1802.655222 | 0.074947715 | 0.080021144 | -0.936598893 | 0.348964921 | 0.570683233 | 0.074947715 |
| ENSMUSG00000023805 | Synj2 | 20975 | protein_coding | 597.4651718 | -0.075001108 | 0.085892124 | 0.873201229 | 0.382553405 | 0.604944412 | 0.075001108 |
| ENSMUSG00000025528 | 2010106E10Rik | 67715 | protein_coding | 22.51025251 | 0.075008105 | 0.413956208 | -0.181198164 | 0.856212035 | 0.930174321 | 0.075008105 |
| ENSMUSG00000097080 | 1700086O06Rik | 73516 | lncRNA | 21.35540086 | -0.075022623 | 0.234965372 | 0.319292252 | 0.749504907 | 0.87416654 | 0.075022623 |
| ENSMUSG00000072501 | Phf20l1 | 239510 | protein_coding | 554.8090196 | 0.075029504 | 0.063508626 | -1.181406498 | 0.237441278 | 0.447819239 | 0.075029504 |
| ENSMUSG00000058979 | Hdhd5 | 214932 | protein_coding | 202.8695647 | -0.075035311 | 0.093434734 | 0.803077265 | 0.421930078 | 0.64171609 | 0.075035311 |
| ENSMUSG00000030402 | Ppm1n | 232941 | protein_coding | 3.115993931 | -0.075039042 | 0.947357113 | 0.079208823 | 0.936866527 | 1 | 0.075039042 |
| ENSMUSG00000049643 | 2310022A10Rik | 66367 | protein_coding | 87.5253253 | 0.075086548 | 0.135677993 | -0.553417299 | 0.579977696 | 0.763289104 | 0.075086548 |
| ENSMUSG00000022403 | St13 | 70356 | protein_coding | 1498.212399 | -0.075104917 | 0.094746517 | 0.792693172 | 0.427956609 | 0.647057325 | 0.075104917 |
| ENSMUSG00000059182 | Skap2 | 54353 | protein_coding | 668.9908224 | -0.075112319 | 0.127035598 | 0.591269847 | 0.55433963 | 0.744656458 | 0.075112319 |
| ENSMUSG00000068259 | Or9a7 | 258380 | protein_coding | 1.482554047 | -0.07511817 | 1.576488152 | 0.047649054 | 0.961995937 | 1 | 0.07511817 |
| ENSMUSG00000030064 | Frmd4b | 232288 | protein_coding | 722.9317977 | -0.075125209 | 0.051399446 | 1.461595713 | 0.14385203 | 0.320287504 | 0.075125209 |
| ENSMUSG00000020782 | Llgl2 | 217325 | protein_coding | 2854.052633 | -0.075175955 | 0.073256283 | 1.026204879 | 0.304795022 | 0.524967146 | 0.075175955 |
| ENSMUSG00000044827 | Tlr1 | 21897 | protein_coding | 872.7424452 | 0.075228301 | 0.122179372 | -0.615720152 | 0.538079235 | 0.733336674 | 0.075228301 |
| ENSMUSG00000016757 | Ttll12 | 223723 | protein_coding | 339.7872756 | 0.075245009 | 0.104987486 | -0.716704549 | 0.473556419 | 0.68565173 | 0.075245009 |
| ENSMUSG00000034757 | Tmub2 | 72053 | protein_coding | 534.4270945 | 0.075294381 | 0.052453524 | -1.435449441 | 0.151159066 | 0.33115426 | 0.075294381 |
| ENSMUSG00000086330 | 1700007J10Rik | 69320 | lncRNA | 5.386302331 | -0.075345912 | 0.442333 | 0.170337534 | 0.864744695 | 0.932810366 | 0.075345912 |
| ENSMUSG00000038968 | Klk1b16 | 16615 | protein_coding | 1.718220811 | 0.075355347 | 0.997381444 | -0.075553187 | 0.939774581 | 1 | 0.075355347 |
| ENSMUSG00000020549 | Elac2 | 68626 | protein_coding | 332.9697058 | 0.075366903 | 0.121899146 | -0.618272606 | 0.536395656 | 0.732371878 | 0.075366903 |
| ENSMUSG00000025228 | Actr1a | 54130 | protein_coding | 2086.60869 | 0.075389501 | 0.04629529 | -1.628448616 | 0.103429796 | 0.253681814 | 0.075389501 |
| ENSMUSG00000079415 | Cntf | 12803 | protein_coding | 2.207754389 | 0.07539298 | 0.670366616 | -0.112465297 | 0.910454484 | 1 | 0.07539298 |
| ENSMUSG00000032570 | Atp2c1 | 235574 | protein_coding | 969.9261378 | -0.075426461 | 0.119721435 | 0.630016345 | 0.52868389 | 0.726648836 | 0.075426461 |
| ENSMUSG00000048277 | Syngr2 | 20973 | protein_coding | 1997.156939 | 0.075517649 | 0.095544956 | -0.790388652 | 0.429300827 | 0.648260298 | 0.075517649 |
| ENSMUSG00000031755 | Bbs2 | 67378 | protein_coding | 109.4071676 | -0.075553612 | 0.105458307 | 0.716431105 | 0.473725194 | 0.685755422 | 0.075553612 |
| ENSMUSG00000028136 | Snx27 | 76742 | protein_coding | 462.2750787 | -0.075605554 | 0.094688867 | 0.798462963 | 0.424601878 | 0.644177805 | 0.075605554 |
| ENSMUSG00000045876 | Pcdhb8 | 93879 | protein_coding | 1.826113024 | 0.075607683 | 0.77753065 | -0.097240775 | 0.922535187 | 1 | 0.075607683 |
| ENSMUSG00000022827 | Rabl3 | 67657 | protein_coding | 142.6938389 | 0.075640297 | 0.106143812 | -0.712620887 | 0.476080383 | 0.687930892 | 0.075640297 |
| ENSMUSG00000021764 | Ndufs4 | 17993 | protein_coding | 945.1813423 | -0.075643591 | 0.083234125 | 0.908805025 | 0.363453054 | 0.585795796 | 0.075643591 |
| ENSMUSG00000023911 | Flywch2 | 76917 | protein_coding | 82.27139462 | 0.075695223 | 0.12494354 | -0.605835428 | 0.544624042 | 0.736796542 | 0.075695223 |
| ENSMUSG00000067148 | Polr1c | 20016 | protein_coding | 519.3510667 | -0.075700264 | 0.065902061 | 1.148678252 | 0.250688677 | 0.46291768 | 0.075700264 |
| ENSMUSG00000032435 | Dync1li1 | 235661 | protein_coding | 638.6775373 | 0.075731252 | 0.078514944 | -0.964545711 | 0.334772403 | 0.557076931 | 0.075731252 |
| ENSMUSG00000027641 | Rbl1 | 19650 | protein_coding | 403.5683313 | -0.075756947 | 0.095947136 | 0.789569649 | 0.429779139 | 0.64845799 | 0.075756947 |
| ENSMUSG00000028869 | Gnl2 | 230737 | protein_coding | 385.1463516 | 0.075791653 | 0.090201184 | -0.8402512 | 0.400767557 | 0.622662097 | 0.075791653 |
| ENSMUSG00000030866 | Ern2 | 26918 | protein_coding | 1338.548802 | -0.075816135 | 0.157072316 | 0.482682989 | 0.629320844 | 0.79746811 | 0.075816135 |
| ENSMUSG00000041660 | Bbox1 | 170442 | protein_coding | 2.231189267 | -0.075817678 | 0.984697045 | 0.076995944 | 0.938626772 | 1 | 0.075817678 |
| ENSMUSG00000035357 | Pdzrn3 | 55983 | protein_coding | 81.0640561 | -0.075825856 | 0.21608784 | 0.350902928 | 0.725661175 | 0.857991597 | 0.075825856 |
| ENSMUSG00000030527 | Crtc3 | 70461 | protein_coding | 150.7682416 | -0.075838714 | 0.1062173 | 0.713995872 | 0.475229732 | 0.687150319 | 0.075838714 |
| ENSMUSG00000024067 | Dpy30 | 66310 | protein_coding | 416.2133222 | -0.075882907 | 0.064257977 | 1.180910299 | 0.23763836 | 0.447999484 | 0.075882907 |
| ENSMUSG00000085196 | Gm14963 | 105246847 | lncRNA | 23.50274471 | 0.075908531 | 0.422926604 | -0.179483936 | 0.85755773 | 0.930374388 | 0.075908531 |

| ensembl_gene_id | external_gene_name | entrezgene_id | gene_biotype | baseMean | log2FoldChange | lfcSE | stat | pvalue | padj | absLog2FC |
|---|---|---|---|---|---|---|---|---|---|---|
| ENSMUSG00000038115 | Ano2 | 243634 | protein_coding | 2.676016662 | -0.076002207 | 3.09466081 | 0.02455914 | 0.980406611 | 1 | 0.076002207 |
| ENSMUSG00000003581 | Rnf215 | 71673 | protein_coding | 188.3011977 | -0.0760867 | 0.082703741 | 0.91999103 | 0.357577447 | 0.580632641 | 0.0760867 |
| ENSMUSG00000032316 | Clk3 | 102414 | protein_coding | 534.2491955 | -0.076103174 | 0.063678506 | 1.195115567 | 0.232041881 | 0.441111962 | 0.076103174 |
| ENSMUSG00000039770 | Ypel5 | 383295 | protein_coding | 458.3992333 | -0.076131419 | 0.104781658 | 0.726572002 | 0.467488185 | 0.681117777 | 0.076131419 |
| ENSMUSG00000008540 | Mgst1 | 56615 | protein_coding | 1665.251476 | 0.076164051 | 0.09575481 | -0.795407049 | 0.426376761 | 0.646068054 | 0.076164051 |
| ENSMUSG00000033938 | Ndufb7 | 66916 | protein_coding | 1084.944519 | -0.076176419 | 0.080387753 | 0.947612234 | 0.343326896 | 0.56521786 | 0.076176419 |
| ENSMUSG00000054931 | Zkscan4 | 544922 | protein_coding | 7.231578324 | -0.076189848 | 0.464695302 | 0.163956571 | 0.869765332 | 0.935658236 | 0.076189848 |
| ENSMUSG00000031176 | Dynlt3 | 67117 | protein_coding | 755.2625448 | 0.076247507 | 0.123915245 | -0.61531983 | 0.538343524 | 0.733377497 | 0.076247507 |
| ENSMUSG00000040687 | Madd | 228355 | protein_coding | 526.1434559 | -0.076256626 | 0.067272112 | 1.133554815 | 0.256981336 | 0.470017696 | 0.076256626 |
| ENSMUSG00000033960 | Jcad | 240185 | protein_coding | 6.191713569 | 0.076270808 | 0.69467843 | -0.10979297 | 0.912573567 | 0.957266079 | 0.076270808 |
| ENSMUSG00000039263 | Npepl1 | 228961 | protein_coding | 654.3238295 | -0.076293007 | 0.05569474 | 1.369842244 | 0.170736152 | 0.360508479 | 0.076293007 |
| ENSMUSG00000034226 | Rhov | 228543 | protein_coding | 14.61597288 | 0.076293931 | 0.332264789 | -0.229617862 | 0.818388726 | 0.908812719 | 0.076293931 |
| ENSMUSG00000079658 | Eloc | 67923 | protein_coding | 370.6795997 | 0.076307059 | 0.060812467 | -1.254793018 | 0.209553908 | 0.413054814 | 0.076307059 |
| ENSMUSG00000020594 | Pum2 | 80913 | protein_coding | 1717.042974 | 0.076308294 | 0.091086522 | -0.837756154 | 0.402167668 | 0.623781054 | 0.076308294 |
| ENSMUSG00000027499 | Pkia | 18767 | protein_coding | 72.04492766 | -0.076317538 | 0.197466639 | 0.386483198 | 0.699138844 | 0.840656682 | 0.076317538 |
| ENSMUSG00000020059 | Sycp3 | 20962 | protein_coding | 1.722972562 | 0.076317688 | 0.736914289 | -0.103563859 | 0.917515469 | 1 | 0.076317688 |
| ENSMUSG00000005615 | Pcyt1a | 13026 | protein_coding | 832.5869308 | -0.076384551 | 0.063556164 | 1.201843311 | 0.229424241 | 0.438111017 | 0.076384551 |
| ENSMUSG00000091415 | Ak9 | 633979 | protein_coding | 2.528132084 | 0.076421943 | 2.134111472 | -0.035809724 | 0.971434079 | 1 | 0.076421943 |
| ENSMUSG00000045967 | Gpr158 | 241263 | protein_coding | 1.756285756 | -0.076447155 | 1.318374878 | 0.057985901 | 0.953759859 | 1 | 0.076447155 |
| ENSMUSG00000035172 | Plekhh3 | 217198 | protein_coding | 333.3969353 | -0.076501382 | 0.134258132 | 0.569808184 | 0.568807804 | 0.754784418 | 0.076501382 |
| ENSMUSG00000028961 | Pgd | 110208 | protein_coding | 2121.539938 | 0.076550089 | 0.07980779 | -0.95918066 | 0.337467741 | 0.559438733 | 0.076550089 |
| ENSMUSG00000020843 | Timm22 | 56322 | protein_coding | 248.2184145 | 0.076580416 | 0.104116241 | -0.735528053 | 0.462017962 | 0.675841352 | 0.076580416 |
| ENSMUSG00000032463 | Faim | 23873 | protein_coding | 112.380589 | 0.076665374 | 0.118219436 | -0.648500591 | 0.516661229 | 0.717449153 | 0.076665374 |
| ENSMUSG00000028294 | Cfap206 | 69329 | protein_coding | 6.221725837 | -0.076679789 | 0.452070062 | 0.169619259 | 0.865309576 | 0.932932188 | 0.076679789 |
| ENSMUSG00000042549 | Map2k3os | 24082 | lncRNA | 16.93426453 | -0.07668874 | 0.272377585 | 0.28155305 | 0.778286245 | 0.889208458 | 0.07668874 |
| ENSMUSG00000000826 | Dnajc5 | 13002 | protein_coding | 1623.639493 | 0.076733365 | 0.054024003 | -1.420356895 | 0.155503805 | 0.337876038 | 0.076733365 |
| ENSMUSG00000037890 | Wdr19 | 213081 | protein_coding | 96.77594162 | 0.076792804 | 0.124871823 | -0.61497303 | 0.538572532 | 0.733377497 | 0.076792804 |
| ENSMUSG00000052962 | Mrpl35 | 66223 | protein_coding | 298.7019004 | -0.076840507 | 0.090221851 | 0.851684001 | 0.394389503 | 0.616875898 | 0.076840507 |
| ENSMUSG00000040936 | Ulk4 | 209012 | protein_coding | 30.66172417 | 0.076849384 | 0.311083736 | -0.247037615 | 0.804879114 | 0.90383338 | 0.076849384 |
| ENSMUSG00000059555 | Tor4a | 227612 | protein_coding | 688.1687155 | 0.076863667 | 0.088064341 | -0.872812604 | 0.382765228 | 0.605068399 | 0.076863667 |
| ENSMUSG00000036533 | Cdc42ep3 | 260409 | protein_coding | 147.9129777 | -0.076884892 | 0.16773141 | 0.458381003 | 0.646678737 | 0.807443873 | 0.076884892 |
| ENSMUSG00000042734 | Ttc9 | 69480 | protein_coding | 3.850690346 | -0.076894186 | 0.978135964 | 0.078612983 | 0.937340461 | 1 | 0.076894186 |
| ENSMUSG00000003062 | Stard3nl | 76205 | protein_coding | 328.7422013 | 0.076902322 | 0.068123392 | -1.12886808 | 0.258953488 | 0.472341459 | 0.076902322 |
| ENSMUSG00000038644 | Pold1 | 18971 | protein_coding | 305.6445021 | 0.076992794 | 0.158528845 | -0.485670564 | 0.627200757 | 0.796064353 | 0.076992794 |
| ENSMUSG00000048170 | Mcmbp | 210711 | protein_coding | 902.399064 | -0.07708786 | 0.059851214 | 1.287991581 | 0.197748894 | 0.398004294 | 0.07708786 |
| ENSMUSG00000003032 | Klf4 | 16600 | protein_coding | 8123.791838 | 0.077110614 | 0.138825982 | -0.555447998 | 0.57858827 | 0.762250234 | 0.077110614 |
| ENSMUSG00000027882 | Stxbp3 | 20912 | protein_coding | 688.1629699 | -0.077131955 | 0.069188564 | 1.114807858 | 0.264932768 | 0.479778455 | 0.077131955 |
| ENSMUSG00000040345 | Arhgap9 | 216445 | protein_coding | 64.74730205 | 0.077139304 | 0.193357758 | -0.398945999 | 0.689932995 | 0.834570281 | 0.077139304 |
| ENSMUSG00000037933 | Bicd2 | 76895 | protein_coding | 238.8591342 | -0.077159192 | 0.109280709 | 0.706064162 | 0.480148237 | 0.691115487 | 0.077159192 |

Supplementary Table 1

| ensembl_gene_id | external_gene_name | entrezgene_id | gene_biotype | baseMean | log2FoldChange | lfcSE | stat | pvalue | padj | absLog2FC |
|---|---|---|---|---|---|---|---|---|---|---|
| ENSMUSG00000000346 | Dazap2 | 23994 | protein_coding | 2093.364858 | -0.077178556 | 0.055960169 | 1.379169466 | 0.16784251 | 0.356391801 | 0.077178556 |
| ENSMUSG00000053754 | Chd8 | 67772 | protein_coding | 808.7315227 | 0.077190685 | 0.073457435 | -1.050821952 | 0.293340372 | 0.512561987 | 0.077190685 |
| ENSMUSG00000044600 | Smim7 | 66818 | protein_coding | 1180.695233 | 0.077214431 | 0.086894077 | -0.88860408 | 0.374215898 | 0.59621817 | 0.077214431 |
| ENSMUSG00000004070 | Hmox2 | 15369 | protein_coding | 303.3825702 | 0.077232424 | 0.093404901 | -0.826856226 | 0.408318557 | 0.628891312 | 0.077232424 |
| ENSMUSG00000071722 | Spin4 | 270624 | protein_coding | 12.62398837 | 0.07723982 | 0.276737812 | -0.279108298 | 0.780161715 | 0.890125936 | 0.07723982 |
| ENSMUSG00000047473 | Zfp30 | 22693 | protein_coding | 12.00107267 | 0.077240097 | 0.399610699 | -0.193288361 | 0.846733141 | 0.9245093 | 0.077240097 |
| ENSMUSG00000039253 | Fn3krp | 238024 | protein_coding | 154.0156827 | -0.077251799 | 0.118747887 | 0.650553041 | 0.515335052 | 0.716335759 | 0.077251799 |
| ENSMUSG00000034994 | Eef2 | 13629 | protein_coding | 30265.42393 | -0.077284162 | 0.073340675 | 1.053769438 | 0.291988489 | 0.510848385 | 0.077284162 |
| ENSMUSG00000000275 | Trim25 | 217069 | protein_coding | 1223.768201 | -0.077291639 | 0.123361552 | 0.626545612 | 0.530957129 | 0.728048657 | 0.077291639 |
| ENSMUSG00000031266 | Gla | 11605 | protein_coding | 246.3581704 | -0.077324265 | 0.093915201 | 0.823341313 | 0.410313926 | 0.630553153 | 0.077324265 |
| ENSMUSG00000030965 | Abraxas2 | 109359 | protein_coding | 363.7860396 | 0.077407415 | 0.080638256 | -0.959934141 | 0.337088362 | 0.559327232 | 0.077407415 |
| ENSMUSG00000020783 | Ncbp3 | 66874 | protein_coding | 457.3041882 | -0.077571713 | 0.053529282 | 1.449145414 | 0.147296978 | 0.325282468 | 0.077571713 |
| ENSMUSG00000034652 | Cd300a | 217303 | protein_coding | 4.59938374 | 0.077590407 | 1.340972645 | -0.05786129 | 0.953859118 | 0.977047797 | 0.077590407 |
| ENSMUSG00000047067 | Dusp28 | 67446 | protein_coding | 81.87106786 | -0.077617243 | 0.136471966 | 0.568741297 | 0.569531718 | 0.755320172 | 0.077617243 |
| ENSMUSG00000020585 | Laptm4a | 17775 | protein_coding | 2112.924896 | 0.077646541 | 0.091013248 | -0.853134493 | 0.393584725 | 0.616198156 | 0.077646541 |
| ENSMUSG00000031400 | G6pdx | 14381 | protein_coding | 754.2545782 | -0.077682957 | 0.066399841 | 1.169926853 | 0.242030407 | 0.45280278 | 0.077682957 |
| ENSMUSG00000003344 | Btbd2 | 208198 | protein_coding | 415.6233392 | -0.07768697 | 0.079800908 | 0.973509855 | 0.330299964 | 0.552585717 | 0.07768697 |
| ENSMUSG00000045427 | Hnrnph2 | 56258 | protein_coding | 626.0231367 | -0.077697522 | 0.124043151 | 0.626374945 | 0.53106904 | 0.728048657 | 0.077697522 |
| ENSMUSG00000019952 | Poc1b | 382406 | protein_coding | 280.7045699 | 0.077719483 | 0.089501795 | -0.8683567 | 0.385199091 | 0.607446313 | 0.077719483 |
| ENSMUSG00000042638 | Gucy2c | 14917 | protein_coding | 5198.336781 | 0.077724954 | 0.058307079 | -1.333027748 | 0.182522698 | 0.37700811 | 0.077724954 |
| ENSMUSG00000004069 | Dnaja3 | 83945 | protein_coding | 988.339349 | -0.0777377 | 0.079001527 | 0.984002495 | 0.325114286 | 0.546957754 | 0.0777377 |
| ENSMUSG00000041161 | Otud3 | 73162 | protein_coding | 238.7731989 | 0.077766676 | 0.089347386 | -0.870385578 | 0.384089726 | 0.60650772 | 0.077766676 |
| ENSMUSG00000020682 | Mmp28 | 118453 | protein_coding | 176.658549 | -0.077782535 | 0.105447715 | 0.737640781 | 0.460732767 | 0.674875015 | 0.077782535 |
| ENSMUSG00000037455 | Slc18b1 | 76306 | protein_coding | 321.0670197 | -0.077798016 | 0.094318868 | 0.824840436 | 0.409462186 | 0.630052795 | 0.077798016 |
| ENSMUSG00000054520 | Sh3bp2 | 24055 | protein_coding | 83.24620012 | -0.07781803 | 0.143863359 | 0.540916263 | 0.588565301 | 0.769577776 | 0.07781803 |
| ENSMUSG00000061393 | Acvr2b | 11481 | protein_coding | 64.3080288 | 0.077834957 | 0.173284649 | -0.449173991 | 0.653306148 | 0.811872956 | 0.077834957 |
| ENSMUSG00000031865 | Dctn1 | 13191 | protein_coding | 1687.350764 | 0.077893001 | 0.050354999 | -1.546877221 | 0.121892855 | 0.284857762 | 0.077893001 |
| ENSMUSG00000031600 | Vps37a | 52348 | protein_coding | 523.665299 | -0.077910813 | 0.062772983 | 1.241152001 | 0.214549603 | 0.419497057 | 0.077910813 |
| ENSMUSG00000039963 | Ccdc40 | 207607 | protein_coding | 4.18132282 | -0.077912531 | 0.899826743 | 0.086586147 | 0.931000477 | 1 | 0.077912531 |
| ENSMUSG00000060512 | 0610040J01Rik | 76261 | protein_coding | 854.3634461 | -0.077925905 | 0.086008001 | 0.906030883 | 0.364919506 | 0.587471313 | 0.077925905 |
| ENSMUSG00000024581 | Napg | 108123 | protein_coding | 413.4943011 | -0.077927059 | 0.083367365 | 0.934742979 | 0.349920772 | 0.571741528 | 0.077927059 |
| ENSMUSG00000058761 | Rnf169 | 108937 | protein_coding | 323.0531693 | -0.07792834 | 0.084892973 | 0.917959841 | 0.358639891 | 0.581328932 | 0.07792834 |
| ENSMUSG00000051427 | Ccdc157 | 216516 | protein_coding | 74.36784863 | 0.077995952 | 0.14282783 | -0.546083717 | 0.585008392 | 0.767158627 | 0.077995952 |
| ENSMUSG00000024829 | Mrpl21 | 353242 | protein_coding | 409.1335298 | -0.078007358 | 0.0842483 | 0.925922042 | 0.354486495 | 0.577145813 | 0.078007358 |
| ENSMUSG00000025921 | Rdh10 | 98711 | protein_coding | 268.4120197 | -0.078015998 | 0.089485849 | 0.871824969 | 0.383303871 | 0.605788125 | 0.078015998 |
| ENSMUSG00000053333 | Dis3l2 | 208718 | protein_coding | 247.5716902 | 0.078027354 | 0.083869125 | -0.930346586 | 0.352191665 | 0.574428953 | 0.078027354 |
| ENSMUSG00000039529 | Atp8b1 | 54670 | protein_coding | 7307.62984 | -0.078110375 | 0.069406015 | 1.125412179 | 0.260414413 | 0.474068167 | 0.078110375 |
| ENSMUSG00000022517 | Mgrn1 | 17237 | protein_coding | 1265.725168 | -0.078181093 | 0.059778295 | 1.307850831 | 0.190923909 | 0.388417088 | 0.078181093 |
| ENSMUSG00000042082 | Arsb | 11881 | protein_coding | 204.3937251 | -0.078195841 | 0.083726079 | 0.933948439 | 0.350330491 | 0.572354345 | 0.078195841 |

| ensembl_gene_id | external_gene_name | entrezgene_id | gene_biotype | baseMean | log2FoldChange | lfcSE | stat | pvalue | padj | absLog2FC |
|---|---|---|---|---|---|---|---|---|---|---|
| ENSMUSG00000024247 | Pkdcc | 106522 | protein_coding | 1079.410992 | 0.078213045 | 0.078691117 | -0.993924706 | 0.320259525 | 0.541493376 | 0.078213045 |
| ENSMUSG00000066442 | Mthfs | 107885 | protein_coding | 22.11071805 | 0.078214456 | 0.267340863 | -0.292564537 | 0.769855024 | 0.884532517 | 0.078214456 |
| ENSMUSG00000009549 | Srp14 | 20813 | protein_coding | 352.0434316 | 0.078226981 | 0.09788356 | -0.799184057 | 0.424183695 | 0.643957486 | 0.078226981 |
| ENSMUSG00000037991 | Rmi2 | 223970 | protein_coding | 37.88534569 | 0.078238102 | 0.285523603 | -0.274016234 | 0.784072146 | 0.891754761 | 0.078238102 |
| ENSMUSG00000039473 | Ubn1 | 170644 | protein_coding | 836.7246119 | 0.078239998 | 0.069062702 | -1.132883532 | 0.257263166 | 0.470354227 | 0.078239998 |
| ENSMUSG00000001305 | Rrp15 | 67223 | protein_coding | 131.0982258 | 0.078250199 | 0.114324102 | -0.684459331 | 0.493685162 | 0.699975951 | 0.078250199 |
| ENSMUSG00000026933 | Camsap1 | 227634 | protein_coding | 263.0049137 | 0.078264017 | 0.076469922 | -1.023461452 | 0.306089717 | 0.52648324 | 0.078264017 |
| ENSMUSG00000018585 | Atox1 | 11927 | protein_coding | 619.6648236 | 0.078272906 | 0.088170977 | -0.887740035 | 0.374680605 | 0.596728278 | 0.078272906 |
| ENSMUSG00000036561 | Ppp6r2 | 71474 | protein_coding | 459.3198868 | 0.078301509 | 0.06756412 | -1.158921464 | 0.246488199 | 0.458287111 | 0.078301509 |
| ENSMUSG00000052595 | A1cf | 69865 | protein_coding | 301.1905659 | 0.078314089 | 0.159974185 | -0.489542039 | 0.624457999 | 0.794172689 | 0.078314089 |
| ENSMUSG00000040785 | Ttc3 | 22129 | protein_coding | 912.9624596 | -0.078603092 | 0.094463425 | 0.832100806 | 0.405352047 | 0.62645534 | 0.078603092 |
| ENSMUSG00000051747 | Ttn | 22138 | protein_coding | 43.33369643 | 0.07865088 | 0.864207513 | -0.091009253 | 0.927485238 | 0.965445424 | 0.07865088 |
| ENSMUSG00000040699 | Limd2 | 67803 | protein_coding | 74.85182956 | 0.078676228 | 0.169506645 | -0.464148339 | 0.642541471 | 0.80495599 | 0.078676228 |
| ENSMUSG00000032562 | Gnai2 | 14678 | protein_coding | 1692.666622 | 0.078707682 | 0.06630682 | -1.187022423 | 0.235218776 | 0.445131871 | 0.078707682 |
| ENSMUSG00000050732 | Vamp8 | 22320 | protein_coding | 1282.993836 | -0.078742945 | 0.062490362 | 1.260081435 | 0.207639987 | 0.410708835 | 0.078742945 |
| ENSMUSG00000022811 | Zfp148 | 22661 | protein_coding | 942.3363751 | -0.078772486 | 0.076062061 | 1.035634387 | 0.300372743 | 0.520512296 | 0.078772486 |
| ENSMUSG00000026427 | Eif2d | 16865 | protein_coding | 465.7609502 | 0.078776554 | 0.061807289 | -1.274551201 | 0.202468139 | 0.404473864 | 0.078776554 |
| ENSMUSG00000024517 | Grp | 225642 | protein_coding | 1.066562172 | 0.078788717 | 2.458431847 | -0.032048363 | 0.974433482 | 1 | 0.078788717 |
| ENSMUSG00000014355 | Anapc1 | 17222 | protein_coding | 1381.470169 | 0.078825705 | 0.093635867 | -0.841832377 | 0.399881786 | 0.622046705 | 0.078825705 |
| ENSMUSG00000073982 | Rhog | 56212 | protein_coding | 241.8405984 | -0.078860687 | 0.097582762 | 0.808141572 | 0.419009088 | 0.638921152 | 0.078860687 |
| ENSMUSG00000019823 | Mical1 | 171580 | protein_coding | 149.5207038 | 0.078862671 | 0.160172921 | -0.492359575 | 0.622465172 | 0.792979237 | 0.078862671 |
| ENSMUSG00000019988 | Nedd1 | 17997 | protein_coding | 273.6782321 | 0.078870676 | 0.151380375 | -0.521009912 | 0.602359866 | 0.778438024 | 0.078870676 |
| ENSMUSG00000004264 | Phb2 | 12034 | protein_coding | 2014.644426 | -0.078879829 | 0.065412043 | 1.20589155 | 0.227859309 | 0.436477048 | 0.078879829 |
| ENSMUSG00000002578 | Ikzf4 | 22781 | protein_coding | 5.910760904 | 0.078882238 | 0.906595188 | -0.087009328 | 0.930664097 | 0.966663747 | 0.078882238 |
| ENSMUSG00000024767 | Otub1 | 107260 | protein_coding | 755.2509353 | -0.078888043 | 0.120935437 | 0.652315356 | 0.514197756 | 0.715854134 | 0.078888043 |
| ENSMUSG00000085355 | 3010003L21Rik | 109163 | lncRNA | 42.68671819 | 0.078897661 | 0.18833362 | -0.418924995 | 0.675270949 | 0.825971257 | 0.078897661 |
| ENSMUSG00000005078 | Jkamp | 104771 | protein_coding | 149.1062892 | -0.078955845 | 0.089301606 | 0.884148104 | 0.376616268 | 0.598886958 | 0.078955845 |
| ENSMUSG00000038812 | Trmt112 | 67674 | protein_coding | 81.48726205 | -0.078981785 | 0.126235284 | 0.625671223 | 0.531530612 | 0.728500002 | 0.078981785 |
| ENSMUSG00000026104 | Stat1 | 20846 | protein_coding | 236.2328057 | 0.079010309 | 0.14672699 | -0.538485176 | 0.590242137 | 0.770846581 | 0.079010309 |
| ENSMUSG00000011254 | Thg1l | 66628 | protein_coding | 98.94949089 | 0.079029534 | 0.152705222 | -0.517530001 | 0.604786232 | 0.779964063 | 0.079029534 |
| ENSMUSG00000037224 | Zfyve28 | 231125 | protein_coding | 3.076241437 | 0.079076046 | 1.821232139 | -0.043418983 | 0.965367546 | 1 | 0.079076046 |
| ENSMUSG00000026342 | Slc35f5 | 74150 | protein_coding | 1367.414973 | -0.079076132 | 0.080810807 | 0.978534129 | 0.32781022 | 0.549974786 | 0.079076132 |
| ENSMUSG00000028799 | Zfp362 | 230761 | protein_coding | 68.41644334 | 0.079111557 | 0.141740669 | -0.558142964 | 0.576746765 | 0.761222618 | 0.079111557 |
| ENSMUSG00000041974 | Spidr | 224008 | protein_coding | 147.7990195 | 0.079129305 | 0.104887232 | -0.754422659 | 0.45059547 | 0.667366241 | 0.079129305 |
| ENSMUSG00000025782 | Taf3 | 209361 | protein_coding | 84.95994785 | -0.079141408 | 0.13169648 | 0.600937915 | 0.547881339 | 0.739192192 | 0.079141408 |
| ENSMUSG00000019792 | Trmt11 | 73681 | protein_coding | 145.6598703 | 0.079152254 | 0.103364354 | -0.765759667 | 0.443819317 | 0.661197851 | 0.079152254 |
| ENSMUSG00000026017 | Carf | 241066 | protein_coding | 82.1507569 | -0.07915746 | 0.136804477 | 0.578617467 | 0.562847316 | 0.75067522 | 0.07915746 |
| ENSMUSG00000036087 | Slain2 | 75991 | protein_coding | 921.7263341 | 0.079164726 | 0.06869473 | -1.152413375 | 0.249151247 | 0.461264989 | 0.079164726 |
| ENSMUSG00000003660 | Snrnp200 | 320632 | protein_coding | 1666.581472 | 0.079173056 | 0.069120484 | -1.145435499 | 0.2520288 | 0.464375257 | 0.079173056 |

| ensembl_gene_id | external_gene_name | entrezgene_id | gene_biotype | baseMean | log2FoldChange | lfcSE | stat | pvalue | padj | absLog2FC |
|---|---|---|---|---|---|---|---|---|---|---|
| ENSMUSG00000052906 | Ubxn8 | 108159 | protein_coding | 267.9672243 | -0.079181088 | 0.135601027 | 0.583926907 | 0.55926949 | 0.747948226 | 0.079181088 |
| ENSMUSG00000078453 | Abracl | 73112 | protein_coding | 375.1839319 | 0.079209114 | 0.114175479 | -0.693748909 | 0.487839688 | 0.696186289 | 0.079209114 |
| ENSMUSG00000104011 | Gm32391 | 102634922 | lncRNA | 14.51212114 | 0.079212713 | 0.320481838 | -0.247167558 | 0.804778552 | 0.903803848 | 0.079212713 |
| ENSMUSG00000029763 | Exoc4 | 20336 | protein_coding | 660.4493929 | 0.079218182 | 0.109270822 | -0.724971046 | 0.468469793 | 0.682033868 | 0.079218182 |
| ENSMUSG00000088108 | Snora47 | 100217450 | snoRNA | 10.57154881 | 0.079247055 | 0.452824516 | -0.175006107 | 0.861074835 | 0.931590005 | 0.079247055 |
| ENSMUSG00000032244 | Fem1b | 14155 | protein_coding | 735.2895706 | -0.079279607 | 0.097727891 | 0.811228053 | 0.417234718 | 0.637333654 | 0.079279607 |
| ENSMUSG00000050912 | Tmem123 | 71929 | protein_coding | 283.885057 | 0.079299027 | 0.109811999 | -0.722134447 | 0.470211821 | 0.68365582 | 0.079299027 |
| ENSMUSG00000021327 | Zkscan3 | 72739 | protein_coding | 635.4172376 | 0.079405655 | 0.059788539 | -1.328108299 | 0.184142332 | 0.379181439 | 0.079405655 |
| ENSMUSG00000034165 | Ccnd3 | 12445 | protein_coding | 429.1339762 | 0.079419905 | 0.121988917 | -0.651041969 | 0.515019395 | 0.716272317 | 0.079419905 |
| ENSMUSG00000061718 | Ppp1r1b | 19049 | protein_coding | 2857.254842 | -0.079435355 | 0.135632819 | 0.585664706 | 0.558100856 | 0.747120423 | 0.079435355 |
| ENSMUSG00000029587 | Zfp12 | 231866 | protein_coding | 191.7825272 | -0.079471768 | 0.108732255 | 0.73089414 | 0.464843815 | 0.678429586 | 0.079471768 |
| ENSMUSG00000028029 | Aimp1 | 13722 | protein_coding | 480.0135598 | -0.079488468 | 0.091462446 | 0.869083121 | 0.384801668 | 0.607042377 | 0.079488468 |
| ENSMUSG00000042631 | Xkr7 | 228787 | protein_coding | 2.463119853 | -0.079515622 | 1.149787044 | 0.069156825 | 0.944864789 | 1 | 0.079515622 |
| ENSMUSG00000024457 | Trim26 | 22670 | protein_coding | 843.1000583 | 0.079543119 | 0.067034613 | -1.186597721 | 0.235386335 | 0.445367775 | 0.079543119 |
| ENSMUSG00000020973 | Dnaaf2 | 109065 | protein_coding | 17.8031453 | -0.079568161 | 0.251426906 | 0.316466372 | 0.751648546 | 0.875244964 | 0.079568161 |
| ENSMUSG00000045410 | Akr1e1 | 56043 | protein_coding | 431.5421654 | -0.07957032 | 0.074357986 | 1.070097851 | 0.284575266 | 0.50295958 | 0.07957032 |
| ENSMUSG00000085747 | Slc13a2os | 790911 | lncRNA | 75.72440877 | -0.079616987 | 0.203905226 | 0.39046075 | 0.696195874 | 0.838743187 | 0.079616987 |
| ENSMUSG00000056069 | Otulinl | 223433 | protein_coding | 15.97967558 | 0.079633405 | 0.32607748 | -0.244216207 | 0.807063376 | 0.904987729 | 0.079633405 |
| ENSMUSG00000048924 | Ccdc125 | 76041 | protein_coding | 85.19516631 | -0.079648607 | 0.155283899 | 0.512922506 | 0.60800553 | 0.782404996 | 0.079648607 |
| ENSMUSG00000014232 | Cluap1 | 76779 | protein_coding | 81.38410672 | 0.079669217 | 0.134735409 | -0.591301262 | 0.554318584 | 0.744656458 | 0.079669217 |
| ENSMUSG00000038740 | Mvb12b | 72543 | protein_coding | 77.69568181 | 0.079687035 | 0.233872101 | -0.340729121 | 0.733307513 | 0.863385162 | 0.079687035 |
| ENSMUSG00000042709 | Atpaf2 | 246782 | protein_coding | 304.7610393 | -0.079737567 | 0.094960884 | 0.839688551 | 0.401083035 | 0.622742125 | 0.079737567 |
| ENSMUSG00000097073 | 9430037G07Rik | 320692 | lncRNA | 11.39573895 | 0.079780151 | 0.357719607 | -0.22302426 | 0.823516625 | 0.910399889 | 0.079780151 |
| ENSMUSG00000027239 | Mdk | 17242 | protein_coding | 6.360402348 | 0.079792592 | 0.550993975 | -0.144815725 | 0.884856366 | 0.943016089 | 0.079792592 |
| ENSMUSG00000029577 | Ube3b | 117146 | protein_coding | 1554.807736 | 0.079867721 | 0.0611487 | -1.30612295 | 0.19151075 | 0.389196224 | 0.079867721 |
| ENSMUSG00000020664 | Dld | 13382 | protein_coding | 1632.108468 | -0.079869824 | 0.078185829 | 1.021538368 | 0.306999441 | 0.527501415 | 0.079869824 |
| ENSMUSG00000039239 | Tgfb2 | 21808 | protein_coding | 3.507660226 | 0.07988608 | 1.154052173 | -0.069222243 | 0.944812718 | 1 | 0.07988608 |
| ENSMUSG00000020623 | Map2k6 | 26399 | protein_coding | 237.632531 | 0.079906725 | 0.115407839 | -0.692385595 | 0.48869521 | 0.696497169 | 0.079906725 |
| ENSMUSG00000038206 | Fbxo8 | 50753 | protein_coding | 389.2928006 | 0.079922805 | 0.073641596 | -1.085294303 | 0.277791328 | 0.494763373 | 0.079922805 |
| ENSMUSG00000028796 | Phc2 | 54383 | protein_coding | 571.8248052 | -0.079952463 | 0.053234155 | 1.501901603 | 0.133122523 | 0.302752759 | 0.079952463 |
| ENSMUSG00000035545 | Leng8 | 232798 | protein_coding | 918.0719852 | 0.079985909 | 0.189371225 | -0.422376254 | 0.672750407 | 0.824536161 | 0.079985909 |
| ENSMUSG00000004661 | Arid3b | 56380 | protein_coding | 63.07989816 | -0.079988366 | 0.177824879 | 0.449815383 | 0.652843565 | 0.811643351 | 0.079988366 |
| ENSMUSG00000039483 | Asb6 | 72323 | protein_coding | 200.7363104 | 0.080011394 | 0.092710182 | -0.863027049 | 0.388122595 | 0.610657064 | 0.080011394 |
| ENSMUSG00000016619 | Nup50 | 18141 | protein_coding | 828.3949253 | 0.080014252 | 0.073267504 | -1.092083765 | 0.274796285 | 0.491230336 | 0.080014252 |
| ENSMUSG00000104945 | Gm35715 | 102639389 | lncRNA | 6.656568959 | -0.080035356 | 0.484543821 | 0.165176714 | 0.868804893 | 0.935174293 | 0.080035356 |
| ENSMUSG00000054939 | Zfp174 | 385674 | protein_coding | 24.99359651 | -0.0800538 | 0.223395249 | 0.358350505 | 0.720081027 | 0.855153117 | 0.0800538 |
| ENSMUSG00000028878 | Fam76a | 230789 | protein_coding | 267.5034866 | -0.080060024 | 0.078060019 | 1.025621374 | 0.305070089 | 0.525167043 | 0.080060024 |
| ENSMUSG00000021594 | Srd5a1 | 78925 | protein_coding | 104.0644324 | 0.080097728 | 0.152036147 | -0.526833452 | 0.59830925 | 0.776035105 | 0.080097728 |
| ENSMUSG00000004902 | Slc25a18 | 71803 | protein_coding | 8.854901957 | -0.080145895 | 0.387024411 | 0.207082274 | 0.835945606 | 0.917689672 | 0.080145895 |

| ensembl_gene_id | external_gene_name | entrezgene_id | gene_biotype | baseMean | log2FoldChange | lfcSE | stat | pvalue | padj | absLog2FC |
|---|---|---|---|---|---|---|---|---|---|---|
| ENSMUSG00000040434 | Large2 | 228366 | protein_coding | 111.7449363 | 0.080172437 | 0.107486733 | -0.745882165 | 0.4557386 | 0.671071533 | 0.080172437 |
| ENSMUSG00000066129 | Kndc1 | 76484 | protein_coding | 3.606082647 | 0.080216381 | 2.091188128 | -0.038359237 | 0.969401261 | 1 | 0.080216381 |
| ENSMUSG00000022684 | Bfar | 67118 | protein_coding | 736.4801588 | -0.080216454 | 0.085646656 | 0.93659761 | 0.348965582 | 0.570683233 | 0.080216454 |
| ENSMUSG00000005873 | Reep5 | 13476 | protein_coding | 489.1467418 | -0.080232445 | 0.122412939 | 0.655424545 | 0.512194453 | 0.714728777 | 0.080232445 |
| ENSMUSG00000026819 | Slc25a25 | 227731 | protein_coding | 785.0815469 | -0.0802749 | 0.152864852 | 0.525136413 | 0.599488364 | 0.776544903 | 0.0802749 |
| ENSMUSG00000062931 | Zfp938 | 237411 | protein_coding | 65.00570351 | -0.08029432 | 0.171041234 | 0.469444225 | 0.638752143 | 0.80301103 | 0.08029432 |
| ENSMUSG00000056493 | Foxk1 | 17425 | protein_coding | 429.2700268 | -0.080317304 | 0.092484604 | 0.868439722 | 0.385153658 | 0.607432671 | 0.080317304 |
| ENSMUSG00000024070 | Prkd3 | 75292 | protein_coding | 153.9630995 | 0.080319799 | 0.171527195 | -0.468262766 | 0.639596692 | 0.803461112 | 0.080319799 |
| ENSMUSG00000064363 | mt-Nd4 | 17719 | protein_coding | 13676.37189 | -0.08037642 | 0.414303649 | 0.19400365 | 0.846173024 | 0.924222269 | 0.08037642 |
| ENSMUSG00000007603 | Dus3l | 224907 | protein_coding | 383.9464826 | 0.080428848 | 0.06548051 | -1.228286824 | 0.219339315 | 0.425536825 | 0.080428848 |
| ENSMUSG00000002100 | Mybpc3 | 17868 | protein_coding | 3.872944523 | 0.080487706 | 1.424039478 | -0.056520699 | 0.954927007 | 1 | 0.080487706 |
| ENSMUSG00000034163 | Zfc3h1 | 216345 | protein_coding | 732.5074732 | -0.080500581 | 0.117458015 | 0.685356216 | 0.493119166 | 0.699602226 | 0.080500581 |
| ENSMUSG00000021902 | Phf7 | 71838 | protein_coding | 73.97855052 | -0.080504081 | 0.116860831 | 0.688888488 | 0.490893445 | 0.698215786 | 0.080504081 |
| ENSMUSG00000038462 | Uqcrfs1 | 66694 | protein_coding | 3233.181284 | -0.080514189 | 0.080842517 | 0.995938669 | 0.319279945 | 0.540704851 | 0.080514189 |
| ENSMUSG00000092416 | Zfp141 | 434178 | protein_coding | 115.6701353 | -0.080529887 | 0.13216181 | 0.609327966 | 0.542307074 | 0.735940424 | 0.080529887 |
| ENSMUSG00000054065 | Pkp3 | 56460 | protein_coding | 1072.214996 | -0.080568224 | 0.111875607 | 0.720158985 | 0.471427114 | 0.684288244 | 0.080568224 |
| ENSMUSG00000055932 | Fto | 26383 | protein_coding | 440.5940799 | 0.080655092 | 0.058252891 | -1.384568052 | 0.166184586 | 0.353565093 | 0.080655092 |
| ENSMUSG00000028729 | Ebna1bp2 | 69072 | protein_coding | 456.6647702 | -0.08068387 | 0.088353503 | 0.913193786 | 0.361140632 | 0.583549549 | 0.08068387 |
| ENSMUSG00000028557 | Rnf11 | 29864 | protein_coding | 907.5097372 | -0.080705496 | 0.10366212 | 0.77854376 | 0.436248519 | 0.654452039 | 0.080705496 |
| ENSMUSG00000038542 | Pcid2 | 234069 | protein_coding | 189.7837869 | 0.080759947 | 0.092500836 | -0.873072624 | 0.382623494 | 0.604944412 | 0.080759947 |
| ENSMUSG00000050675 | Gp1ba | 14723 | protein_coding | 7.845267709 | 0.080769924 | 0.540255294 | -0.149503253 | 0.881156543 | 0.941406719 | 0.080769924 |
| ENSMUSG00000028893 | Sesn2 | 230784 | protein_coding | 226.9272739 | -0.080771703 | 0.124830274 | 0.647052194 | 0.517598164 | 0.717985391 | 0.080771703 |
| ENSMUSG00000075267 | Pjvk | 381375 | protein_coding | 1.107493418 | -0.080825237 | 1.069833437 | 0.075549366 | 0.939777622 | 1 | 0.080825237 |
| ENSMUSG00000019837 | Gtf3c6 | 67371 | protein_coding | 342.9980447 | 0.080832829 | 0.067278037 | -1.201474247 | 0.22956729 | 0.438122528 | 0.080832829 |
| ENSMUSG00000037857 | Nufip2 | 68564 | protein_coding | 1244.422367 | 0.080844611 | 0.05308975 | -1.522791332 | 0.127810919 | 0.294806545 | 0.080844611 |
| ENSMUSG00000022231 | Sema5a | 20356 | protein_coding | 1266.67659 | -0.080875077 | 0.149420614 | 0.541257825 | 0.588329886 | 0.76949784 | 0.080875077 |
| ENSMUSG00000000399 | Ndufa9 | 66108 | protein_coding | 1775.628822 | -0.080890894 | 0.0924202 | 0.875251236 | 0.381437221 | 0.60376251 | 0.080890894 |
| ENSMUSG00000032547 | Ryk | 20187 | protein_coding | 648.1702737 | -0.080921488 | 0.081269167 | 0.995721879 | 0.319385297 | 0.540748207 | 0.080921488 |
| ENSMUSG00000041237 | Pklr | 18770 | protein_coding | 728.7347591 | 0.080970502 | 0.074429898 | -1.087876032 | 0.276649838 | 0.493582047 | 0.080970502 |
| ENSMUSG00000099474 | 1700097N02Rik | 67522 | lncRNA | 8.107112003 | -0.080974678 | 0.420695071 | 0.192478313 | 0.847367554 | 0.924757321 | 0.080974678 |
| ENSMUSG00000022938 | Fam3b | 52793 | protein_coding | 1221.499069 | -0.080998044 | 0.083919646 | 0.965185719 | 0.334451799 | 0.556711723 | 0.080998044 |
| ENSMUSG00000073530 | Pappa2 | 23850 | protein_coding | 66.45226844 | -0.081088709 | 0.219392182 | 0.36960619 | 0.711675938 | 0.849914408 | 0.081088709 |
| ENSMUSG00000110080 | Gm6145 | 620376 | lncRNA | 5.298191743 | 0.081107601 | 0.461175498 | -0.175871445 | 0.860394939 | 0.931471303 | 0.081107601 |
| ENSMUSG00000039477 | Tnrc18 | 231861 | protein_coding | 1524.410869 | 0.081135179 | 0.168204565 | -0.48236015 | 0.629550125 | 0.79763624 | 0.081135179 |
| ENSMUSG00000028252 | Ccnc | 51813 | protein_coding | 254.2909199 | 0.081206208 | 0.081669266 | -0.994330084 | 0.320062194 | 0.541378279 | 0.081206208 |
| ENSMUSG00000034826 | Nup54 | 269113 | protein_coding | 188.4304367 | -0.08121536 | 0.105099093 | 0.77275034 | 0.439670148 | 0.657673264 | 0.08121536 |
| ENSMUSG00000057054 | Inca1 | 103844 | protein_coding | 16.56657705 | -0.081247231 | 0.340662746 | 0.238497553 | 0.811495213 | 0.906704483 | 0.081247231 |
| ENSMUSG00000050271 | Prag1 | 244418 | protein_coding | 531.4977982 | 0.081299947 | 0.147861322 | -0.549839173 | 0.582429688 | 0.76523541 | 0.081299947 |
| ENSMUSG00000026193 | Fn1 | 14268 | protein_coding | 19.23437255 | -0.081364919 | 0.357177248 | 0.227799837 | 0.819801853 | 0.909153704 | 0.081364919 |

| ensembl_gene_id | external_gene_name | entrezgene_id | gene_biotype | baseMean | log2FoldChange | lfcSE | stat | pvalue | padj | absLog2FC |
|---|---|---|---|---|---|---|---|---|---|---|
| ENSMUSG00000055675 | Kbtbd11 | 74901 | protein_coding | 1060.439902 | -0.081374952 | 0.126063151 | 0.645509426 | 0.518597112 | 0.719129457 | 0.081374952 |
| ENSMUSG00000054874 | Pcnx3 | 104401 | protein_coding | 1057.762378 | 0.081422901 | 0.078004057 | -1.043829057 | 0.296564485 | 0.515753304 | 0.081422901 |
| ENSMUSG00000017861 | Mybl2 | 17865 | protein_coding | 82.93424163 | 0.081427824 | 0.185745411 | -0.438384045 | 0.66110791 | 0.816884691 | 0.081427824 |
| ENSMUSG00000064367 | mt-Nd5 | 17721 | protein_coding | 18023.65428 | -0.081481599 | 0.431444625 | 0.188857607 | 0.850204421 | 0.926816617 | 0.081481599 |
| ENSMUSG00000048368 | Omd | 27047 | protein_coding | 2.653625591 | 0.081484241 | 0.811325206 | -0.100433514 | 0.920000164 | 1 | 0.081484241 |
| ENSMUSG00000092607 | Scnm1 | 69269 | protein_coding | 206.0651448 | 0.081489869 | 0.076294507 | -1.068096148 | 0.285477143 | 0.503945908 | 0.081489869 |
| ENSMUSG00000028378 | Ptgr1 | 67103 | protein_coding | 248.0092734 | -0.081500764 | 0.149510409 | 0.545117659 | 0.585672598 | 0.767483504 | 0.081500764 |
| ENSMUSG00000024862 | Klc2 | 16594 | protein_coding | 157.0792134 | 0.081525814 | 0.112043486 | -0.727626535 | 0.466842233 | 0.680564322 | 0.081525814 |
| ENSMUSG00000040459 | Arglu1 | 234023 | protein_coding | 643.9211325 | -0.081531246 | 0.105096124 | 0.775777859 | 0.437880159 | 0.656184478 | 0.081531246 |
| ENSMUSG00000028414 | Fktn | 246179 | protein_coding | 259.8932964 | 0.081577299 | 0.147254164 | -0.55398976 | 0.579585854 | 0.762991261 | 0.081577299 |
| ENSMUSG00000068134 | Zfp120 | 104348 | protein_coding | 193.560182 | -0.081601253 | 0.10225911 | 0.79798517 | 0.424879096 | 0.644361593 | 0.081601253 |
| ENSMUSG00000034607 | Pof1b | 69693 | protein_coding | 2420.48784 | -0.081695219 | 0.134482793 | 0.607477109 | 0.54353433 | 0.736456016 | 0.081695219 |
| ENSMUSG00000028847 | Trappc3 | 27096 | protein_coding | 469.8684007 | -0.081713428 | 0.061770404 | 1.322857278 | 0.185882845 | 0.381372755 | 0.081713428 |
| ENSMUSG00000028809 | Srrm1 | 51796 | protein_coding | 696.9474567 | 0.081720786 | 0.08436043 | -0.968709927 | 0.332689938 | 0.55512192 | 0.081720786 |
| ENSMUSG00000029203 | Ube2k | 53323 | protein_coding | 648.0185302 | 0.081733647 | 0.060388046 | -1.353473938 | 0.175904271 | 0.367893 | 0.081733647 |
| ENSMUSG00000068876 | Cgn | 70737 | protein_coding | 2399.76795 | -0.08174684 | 0.082849778 | 0.9866875 | 0.32379586 | 0.545573394 | 0.08174684 |
| ENSMUSG00000053347 | Zfp943 | 74670 | protein_coding | 212.1605897 | -0.081747151 | 0.110633879 | 0.738897999 | 0.459968938 | 0.674354318 | 0.081747151 |
| ENSMUSG00000071653 | 1810009A15Rik | 66276 | protein_coding | 3.941485077 | 0.081748274 | 0.546003166 | -0.149721245 | 0.880984546 | 1 | 0.081748274 |
| ENSMUSG00000051306 | Usp42 | 76800 | protein_coding | 233.0502944 | 0.081773405 | 0.111777198 | -0.731575015 | 0.464428003 | 0.677942548 | 0.081773405 |
| ENSMUSG00000074030 | Exoc8 | 102058 | protein_coding | 484.5393203 | -0.081782213 | 0.068246867 | 1.198329189 | 0.230788888 | 0.439438221 | 0.081782213 |
| ENSMUSG00000036225 | Kctd1 | 106931 | protein_coding | 54.30876168 | -0.081790374 | 0.169641098 | 0.482137733 | 0.629708107 | 0.797713998 | 0.081790374 |
| ENSMUSG00000025607 | Copg2 | 54160 | protein_coding | 739.5936038 | -0.081795858 | 0.089206779 | 0.916924237 | 0.359182342 | 0.581865529 | 0.081795858 |
| ENSMUSG00000039410 | Prdm16 | 70673 | protein_coding | 95.13244527 | -0.081807102 | 0.165979957 | 0.492873374 | 0.622102062 | 0.792822461 | 0.081807102 |
| ENSMUSG00000037197 | Rbm17 | 76938 | protein_coding | 595.4068448 | 0.081811758 | 0.084055915 | -0.97330162 | 0.330403418 | 0.552639068 | 0.081811758 |
| ENSMUSG00000090019 | Gimap1 | 16205 | protein_coding | 25.46064451 | -0.08182311 | 0.262552258 | 0.311645046 | 0.755310297 | 0.877032235 | 0.08182311 |
| ENSMUSG00000029782 | Tmem209 | 72649 | protein_coding | 270.4821444 | 0.081828055 | 0.108064865 | -0.757212398 | 0.448922623 | 0.665664739 | 0.081828055 |
| ENSMUSG00000020189 | Osbpl8 | 237542 | protein_coding | 249.6747805 | 0.081844973 | 0.114384741 | -0.715523523 | 0.474285611 | 0.686291751 | 0.081844973 |
| ENSMUSG00000051343 | Rab11fip5 | 52055 | protein_coding | 58.320776 | -0.081868098 | 0.19610311 | 0.417474756 | 0.676331185 | 0.826766769 | 0.081868098 |
| ENSMUSG00000052776 | Oas1a | 246730 | protein_coding | 543.7411761 | -0.081868704 | 0.172175608 | 0.475495365 | 0.634433936 | 0.800714821 | 0.081868704 |
| ENSMUSG00000039463 | Slc9a8 | 77031 | protein_coding | 505.40761 | -0.081869707 | 0.055044567 | 1.487334939 | 0.136926371 | 0.309203076 | 0.081869707 |
| ENSMUSG00000017837 | Nkiras2 | 71966 | protein_coding | 856.689835 | -0.081876693 | 0.086628264 | 0.94514988 | 0.344582363 | 0.566268598 | 0.081876693 |
| ENSMUSG00000023905 | Tnfrsf12a | 27279 | protein_coding | 90.00946823 | 0.081905157 | 0.195911586 | -0.418072042 | 0.675894445 | 0.826426838 | 0.081905157 |
| ENSMUSG00000045248 | Med26 | 70625 | protein_coding | 135.6988479 | 0.081917136 | 0.102438936 | -0.799667966 | 0.423903197 | 0.643827591 | 0.081917136 |
| ENSMUSG00000034024 | Cct2 | 12461 | protein_coding | 2149.269614 | -0.081920156 | 0.094737375 | 0.864707896 | 0.387199134 | 0.60964446 | 0.081920156 |
| ENSMUSG00000029536 | Gatc | 384281 | protein_coding | 144.936096 | -0.081948097 | 0.113520713 | 0.721877923 | 0.470369536 | 0.683706314 | 0.081948097 |
| ENSMUSG00000035863 | Palm | 18483 | protein_coding | 29.66210753 | -0.081974743 | 0.323594137 | 0.253325797 | 0.800016463 | 0.900994567 | 0.081974743 |
| ENSMUSG00000066415 | Msl2 | 77853 | protein_coding | 835.7683303 | 0.081990475 | 0.090515064 | -0.905821322 | 0.365030434 | 0.587561035 | 0.081990475 |
| ENSMUSG00000036606 | Plxnb2 | 140570 | protein_coding | 5971.56391 | 0.082025033 | 0.0732112 | -1.120389129 | 0.262547975 | 0.476573341 | 0.082025033 |
| ENSMUSG00000021417 | Eci2 | 23986 | protein_coding | 459.6443063 | -0.082033637 | 0.072601349 | 1.129918904 | 0.258510398 | 0.471988664 | 0.082033637 |

| ensembl_gene_id | external_gene_name | entrezgene_id | gene_biotype | baseMean | log2FoldChange | lfcSE | stat | pvalue | padj | absLog2FC |
|---|---|---|---|---|---|---|---|---|---|---|
| ENSMUSG00000047090 | Tmem198b | 73827 | protein_coding | 27.81034831 | 0.082051344 | 0.209638509 | -0.391394427 | 0.695505711 | 0.838533729 | 0.082051344 |
| ENSMUSG00000036368 | Rmdn2 | 381110 | protein_coding | 55.48248884 | 0.082052654 | 0.140142474 | -0.585494545 | 0.558215234 | 0.747162726 | 0.082052654 |
| ENSMUSG00000020894 | Vamp2 | 22318 | protein_coding | 174.4743137 | 0.082094844 | 0.132644355 | -0.618909445 | 0.535976016 | 0.732041017 | 0.082094844 |
| ENSMUSG00000044847 | Lsm11 | 72290 | protein_coding | 141.2657211 | -0.082107958 | 0.135131208 | 0.607616542 | 0.543441827 | 0.736456016 | 0.082107958 |
| ENSMUSG00000030917 | Tmem159 | 233806 | protein_coding | 221.3857999 | -0.082121767 | 0.149279828 | 0.550119654 | 0.582237307 | 0.765104392 | 0.082121767 |
| ENSMUSG00000110231 | Gm6329 | 622523 | lncRNA | 6.108389897 | 0.082172411 | 0.440738727 | -0.186442457 | 0.8520978 | 0.927743965 | 0.082172411 |
| ENSMUSG00000053799 | Exoc6 | 107371 | protein_coding | 379.6808668 | -0.082176162 | 0.118557688 | 0.693132287 | 0.488226538 | 0.696271072 | 0.082176162 |
| ENSMUSG00000029706 | Pax4 | 18506 | protein_coding | 6.05739431 | -0.082180299 | 0.526544577 | 0.156074724 | 0.875974122 | 0.938630585 | 0.082180299 |
| ENSMUSG00000029634 | Rnf6 | 74132 | protein_coding | 841.7503041 | 0.082182997 | 0.058256476 | -1.410710063 | 0.158330122 | 0.342188672 | 0.082182997 |
| ENSMUSG00000018474 | Chd3 | 216848 | protein_coding | 1068.309645 | 0.082184637 | 0.158731998 | -0.517757216 | 0.604627673 | 0.779942456 | 0.082184637 |
| ENSMUSG00000029836 | Cbx3 | 12417 | protein_coding | 234.534599 | 0.082198381 | 0.144152467 | -0.570218342 | 0.568529617 | 0.754536534 | 0.082198381 |
| ENSMUSG00000023328 | Ache | 11423 | protein_coding | 35.77220164 | -0.08220532 | 0.289988879 | 0.28347749 | 0.776810835 | 0.888506311 | 0.08220532 |
| ENSMUSG00000038844 | Kif16b | 16558 | protein_coding | 1059.296738 | -0.082234385 | 0.064267063 | 1.279572794 | 0.200695423 | 0.401975042 | 0.082234385 |
| ENSMUSG00000102758 | Naaladl2 | 635702 | protein_coding | 148.4376237 | -0.082305518 | 0.183894 | 0.447570435 | 0.654463244 | 0.812494814 | 0.082305518 |
| ENSMUSG00000002205 | Vrk3 | 101568 | protein_coding | 398.7915335 | -0.082358149 | 0.073360109 | 1.122655765 | 0.261583722 | 0.475244062 | 0.082358149 |
| ENSMUSG00000087174 | 5530601H04Rik | 71445 | lncRNA | 90.67551369 | -0.082523604 | 0.138257394 | 0.596883838 | 0.550584939 | 0.741538599 | 0.082523604 |
| ENSMUSG00000023020 | Cox14 | 66379 | protein_coding | 768.9662218 | -0.082527965 | 0.072020537 | 1.145894882 | 0.251838649 | 0.464320478 | 0.082527965 |
| ENSMUSG00000025512 | Chid1 | 68038 | protein_coding | 690.5544785 | 0.082529247 | 0.129338157 | -0.638088937 | 0.523415787 | 0.722112641 | 0.082529247 |
| ENSMUSG00000039242 | B3galnt2 | 97884 | protein_coding | 343.5186353 | 0.082546348 | 0.102312354 | -0.806807239 | 0.419777548 | 0.639797547 | 0.082546348 |
| ENSMUSG00000032058 | Ppp2r1b | 73699 | protein_coding | 403.5104658 | 0.082562516 | 0.144594522 | -0.570993383 | 0.568004128 | 0.754081531 | 0.082562516 |
| ENSMUSG00000032228 | Tcf12 | 21406 | protein_coding | 839.8638007 | 0.082617352 | 0.051414572 | -1.606885933 | 0.108079393 | 0.261665898 | 0.082617352 |
| ENSMUSG00000055730 | Ces2a | 102022 | protein_coding | 2071.791551 | 0.082620146 | 0.183594449 | -0.450014401 | 0.652700057 | 0.811601896 | 0.082620146 |
| ENSMUSG00000040482 | Dxo | 112403 | protein_coding | 272.2389135 | 0.082635177 | 0.075288989 | -1.097573204 | 0.272390899 | 0.488304745 | 0.082635177 |
| ENSMUSG00000054808 | Actn4 | 60595 | protein_coding | 5271.025557 | 0.082645002 | 0.126510917 | -0.653263796 | 0.513586228 | 0.715485319 | 0.082645002 |
| ENSMUSG00000029440 | Psmd9 | 67151 | protein_coding | 241.1995837 | 0.08266454 | 0.09295957 | -0.889252604 | 0.373867338 | 0.595844868 | 0.08266454 |
| ENSMUSG00000025971 | Maip1 | 68115 | protein_coding | 117.7726876 | 0.082726463 | 0.091652877 | -0.902606282 | 0.366734891 | 0.589522317 | 0.082726463 |
| ENSMUSG00000024548 | Setbp1 | 240427 | protein_coding | 464.815649 | 0.082729268 | 0.118896969 | -0.695806362 | 0.486550104 | 0.695624496 | 0.082729268 |
| ENSMUSG00000033323 | Ctdp1 | 67655 | protein_coding | 336.0465754 | -0.082730852 | 0.082297082 | 1.005270781 | 0.314766481 | 0.535937072 | 0.082730852 |
| ENSMUSG00000022394 | L3mbtl2 | 214669 | protein_coding | 221.4514775 | 0.082752993 | 0.123494078 | -0.670096859 | 0.502796047 | 0.707478587 | 0.082752993 |
| ENSMUSG00000035754 | Wdr18 | 216156 | protein_coding | 410.2386945 | -0.082809578 | 0.088224575 | 0.93862257 | 0.347924561 | 0.569833005 | 0.082809578 |
| ENSMUSG00000026679 | Enkur | 71233 | protein_coding | 8.340947251 | 0.082816995 | 0.477991283 | -0.173260471 | 0.862446693 | 0.932036168 | 0.082816995 |
| ENSMUSG00000054611 | Kdm2a | 225876 | protein_coding | 1224.216544 | -0.082828568 | 0.059317614 | 1.396357031 | 0.162607008 | 0.348151449 | 0.082828568 |
| ENSMUSG00000039899 | Fgl2 | 14190 | protein_coding | 681.4582921 | -0.082852843 | 0.159612133 | 0.519088629 | 0.603698937 | 0.779177919 | 0.082852843 |
| ENSMUSG00000015224 | Cyp2j9 | 74519 | protein_coding | 27.80810996 | -0.082975546 | 0.231962018 | 0.357711777 | 0.720559019 | 0.855204229 | 0.082975546 |
| ENSMUSG00000045038 | Prkce | 18754 | protein_coding | 267.9625607 | -0.082993064 | 0.187708428 | 0.442138187 | 0.658389213 | 0.815651873 | 0.082993064 |
| ENSMUSG00000031993 | Snx19 | 102607 | protein_coding | 536.4723763 | 0.083014614 | 0.108473489 | -0.765298643 | 0.444093732 | 0.661295211 | 0.083014614 |
| ENSMUSG00000074579 | Lekr1 | 624866 | protein_coding | 28.91573691 | 0.083042608 | 0.259438432 | -0.320085993 | 0.748903144 | 0.87389673 | 0.083042608 |
| ENSMUSG00000032329 | Hmg20a | 66867 | protein_coding | 248.9452531 | -0.083043465 | 0.087120332 | 0.953204179 | 0.340486629 | 0.561998621 | 0.083043465 |
| ENSMUSG00000023707 | Ogfod2 | 66627 | protein_coding | 280.2170006 | 0.083045701 | 0.079906215 | -1.039289635 | 0.298670054 | 0.518527888 | 0.083045701 |

| ensembl_gene_id | external_gene_name | entrezgene_id | gene_biotype | baseMean | log2FoldChange | lfcSE | stat | pvalue | padj | absLog2FC |
|---|---|---|---|---|---|---|---|---|---|---|
| ENSMUSG00000025428 | Atp5a1 | 11946 | protein_coding | 20336.21103 | -0.083046504 | 0.091794743 | 0.904697818 | 0.365625497 | 0.588321101 | 0.083046504 |
| ENSMUSG00000015961 | Adss | 11566 | protein_coding | 828.9832117 | -0.083054571 | 0.084896454 | 0.978304357 | 0.327923816 | 0.550003442 | 0.083054571 |
| ENSMUSG00000034333 | Zbed4 | 223773 | protein_coding | 151.2548337 | -0.083075647 | 0.127945488 | 0.649305021 | 0.516141241 | 0.717108906 | 0.083075647 |
| ENSMUSG00000036737 | Oxsr1 | 108737 | protein_coding | 928.2978305 | -0.083081024 | 0.054341048 | 1.528881503 | 0.126293822 | 0.292303079 | 0.083081024 |
| ENSMUSG00000028454 | Pigo | 56703 | protein_coding | 380.9580023 | 0.083103799 | 0.080118624 | -1.037259438 | 0.299614964 | 0.519676664 | 0.083103799 |
| ENSMUSG00000021893 | Capn7 | 12339 | protein_coding | 858.5342921 | 0.083132707 | 0.075555178 | -1.100291324 | 0.271205211 | 0.486866274 | 0.083132707 |
| ENSMUSG00000017664 | Slc35c2 | 228875 | protein_coding | 239.5094485 | 0.083191077 | 0.101051387 | -0.823255173 | 0.4103629 | 0.630569734 | 0.083191077 |
| ENSMUSG00000000197 | Nalcn | 338370 | protein_coding | 3.055616801 | 0.083245831 | 1.779821974 | -0.046771999 | 0.962694946 | 1 | 0.083245831 |
| ENSMUSG00000000131 | Xpo6 | 74204 | protein_coding | 695.946306 | 0.083250422 | 0.054758206 | -1.520327767 | 0.128428619 | 0.295635135 | 0.083250422 |
| ENSMUSG00000053801 | Grwd1 | 101612 | protein_coding | 192.8949483 | -0.08326652 | 0.149257105 | 0.557873072 | 0.576931061 | 0.761228287 | 0.08326652 |
| ENSMUSG00000044393 | Dsg2 | 13511 | protein_coding | 3365.925221 | -0.083266654 | 0.119688634 | 0.695693915 | 0.486620537 | 0.695624496 | 0.083266654 |
| ENSMUSG00000030850 | Ate1 | 11907 | protein_coding | 450.2662285 | 0.083304447 | 0.070318853 | -1.184667321 | 0.236149008 | 0.446273789 | 0.083304447 |
| ENSMUSG00000044042 | Fmn1 | 14260 | protein_coding | 634.7943512 | 0.083409446 | 0.126123509 | -0.661331476 | 0.50839976 | 0.712111524 | 0.083409446 |
| ENSMUSG00000007987 | Ift22 | 67286 | protein_coding | 79.97152245 | 0.083412557 | 0.14382182 | -0.579971503 | 0.561933835 | 0.750140815 | 0.083412557 |
| ENSMUSG00000051154 | Commd3 | 12238 | protein_coding | 340.3582797 | 0.083412935 | 0.062965151 | -1.324747638 | 0.185254867 | 0.380604488 | 0.083412935 |
| ENSMUSG00000045007 | Tubg2 | 103768 | protein_coding | 10.80499974 | 0.083453624 | 0.37449224 | -0.222844734 | 0.823656351 | 0.910493405 | 0.083453624 |
| ENSMUSG00000015671 | Psma2 | 19166 | protein_coding | 1270.740536 | 0.083459507 | 0.080404767 | -1.037992031 | 0.299273765 | 0.519260509 | 0.083459507 |
| ENSMUSG00000031125 | 3830403N18Rik | 70691 | protein_coding | 2.378487937 | 0.08345999 | 1.397174329 | -0.059734844 | 0.95236682 | 1 | 0.08345999 |
| ENSMUSG00000031101 | Sash3 | 74131 | protein_coding | 11.79917071 | -0.08346793 | 0.417676195 | 0.199838849 | 0.841606617 | 0.92158767 | 0.08346793 |
| ENSMUSG00000040123 | Zmym5 | 219105 | protein_coding | 332.9204503 | -0.083484068 | 0.081316382 | 1.026657432 | 0.3045818 | 0.524764095 | 0.083484068 |
| ENSMUSG00000030231 | Plekha5 | 109135 | protein_coding | 1295.800449 | 0.083499433 | 0.057286098 | -1.457586334 | 0.144954599 | 0.322004337 | 0.083499433 |
| ENSMUSG00000087020 | Gm16283 | 102635853 | lncRNA | 3.533157523 | 0.083502453 | 0.755776669 | -0.110485618 | 0.912024255 | 1 | 0.083502453 |
| ENSMUSG00000040383 | Aqr | 11834 | protein_coding | 702.3621942 | 0.083503748 | 0.075285356 | -1.109163229 | 0.267359772 | 0.482533255 | 0.083503748 |
| ENSMUSG00000085704 | 4921531C22Rik | 66730 | lncRNA | 57.8816305 | 0.083504658 | 0.254831906 | -0.327685255 | 0.743149653 | 0.870239999 | 0.083504658 |
| ENSMUSG00000005371 | Fbxo11 | 225055 | protein_coding | 401.7987009 | 0.083509159 | 0.095354688 | -0.875774027 | 0.381152892 | 0.603601732 | 0.083509159 |
| ENSMUSG00000002396 | Ocel1 | 77090 | protein_coding | 319.060215 | -0.083528305 | 0.087042277 | 0.95962913 | 0.337241903 | 0.559393767 | 0.083528305 |
| ENSMUSG00000039901 | Armh3 | 71617 | protein_coding | 215.0153421 | -0.083558086 | 0.107592982 | 0.776612798 | 0.437387249 | 0.65562431 | 0.083558086 |
| ENSMUSG00000033128 | Gga1 | 106039 | protein_coding | 744.9021485 | 0.083584228 | 0.058749046 | -1.422733383 | 0.154813459 | 0.336792184 | 0.083584228 |
| ENSMUSG00000048271 | Rbm33 | 381626 | protein_coding | 585.1870221 | 0.083615209 | 0.102605527 | -0.814919146 | 0.415118593 | 0.635394239 | 0.083615209 |
| ENSMUSG00000036693 | Nop14 | 75416 | protein_coding | 188.052456 | 0.08365503 | 0.112988532 | -0.74038514 | 0.459066333 | 0.673388737 | 0.08365503 |
| ENSMUSG00000029032 | Arhgef16 | 230972 | protein_coding | 1654.107241 | -0.083680106 | 0.062461544 | 1.33970601 | 0.180340943 | 0.374092492 | 0.083680106 |
| ENSMUSG00000020659 | Cbll1 | 104836 | protein_coding | 83.25620086 | -0.083722358 | 0.156338743 | 0.535518944 | 0.592291062 | 0.772293296 | 0.083722358 |
| ENSMUSG00000027091 | Zc3h15 | 69082 | protein_coding | 851.5654738 | 0.083729551 | 0.068355598 | -1.224911401 | 0.220608604 | 0.427096189 | 0.083729551 |
| ENSMUSG00000071757 | Zhx2 | 387609 | protein_coding | 188.4805382 | -0.083739055 | 0.082776127 | 1.011632913 | 0.311713603 | 0.532495959 | 0.083739055 |
| ENSMUSG00000052298 | Cdc42se2 | 72729 | protein_coding | 883.2458195 | 0.083741491 | 0.095581333 | -0.876128097 | 0.3809604 | 0.603354756 | 0.083741491 |
| ENSMUSG00000040697 | Dnajc16 | 214063 | protein_coding | 353.9336687 | -0.083749131 | 0.06485853 | 1.291258544 | 0.196614041 | 0.396311693 | 0.083749131 |
| ENSMUSG00000020287 | Mpg | 268395 | protein_coding | 68.82915167 | 0.083755843 | 0.127327559 | -0.657798227 | 0.510667794 | 0.713786099 | 0.083755843 |
| ENSMUSG00000030102 | Itpr1 | 16438 | protein_coding | 414.8143605 | -0.083771402 | 0.190609644 | 0.439491939 | 0.660305122 | 0.816494106 | 0.083771402 |
| ENSMUSG00000055341 | Zfp457 | 431706 | protein_coding | 3.771408617 | -0.083806607 | 0.583444773 | 0.143641027 | 0.885783941 | 1 | 0.083806607 |

Supplementary Table 1

| ensembl_gene_id | external_gene_name | entrezgene_id | gene_biotype | baseMean | log2FoldChange | lfcSE | stat | pvalue | padj | absLog2FC |
|---|---|---|---|---|---|---|---|---|---|---|
| ENSMUSG00000038214 | Bend3 | 331623 | protein_coding | 135.1912546 | 0.083839938 | 0.103442595 | -0.810497244 | 0.417654448 | 0.63773884 | 0.083839938 |
| ENSMUSG00000066233 | Tmem42 | 66079 | protein_coding | 60.68036362 | 0.083841264 | 0.154711543 | -0.541919903 | 0.587873685 | 0.769205771 | 0.083841264 |
| ENSMUSG00000031555 | Adam9 | 11502 | protein_coding | 1108.191737 | 0.083876763 | 0.080659423 | -1.039887959 | 0.298391957 | 0.51821109 | 0.083876763 |
| ENSMUSG00000047117 | Ankdd1b | 271144 | protein_coding | 33.63362269 | -0.083911854 | 0.236925672 | 0.354169529 | 0.723211837 | 0.856549354 | 0.083911854 |
| ENSMUSG00000010607 | Pigyl | 66268 | protein_coding | 111.2358106 | -0.083969951 | 0.133514035 | 0.628922276 | 0.529399941 | 0.727088376 | 0.083969951 |
| ENSMUSG00000068184 | Ndufaf2 | 75597 | protein_coding | 131.6499938 | -0.083997547 | 0.097140019 | 0.864705892 | 0.387200235 | 0.60964446 | 0.083997547 |
| ENSMUSG00000045210 | Vcpip1 | 70675 | protein_coding | 614.1568775 | 0.083999471 | 0.084547712 | -0.993515605 | 0.32045875 | 0.541719295 | 0.083999471 |
| ENSMUSG00000035226 | Rims4 | 241770 | protein_coding | 194.7520098 | 0.084020043 | 0.310498861 | -0.270596943 | 0.786701047 | 0.893302015 | 0.084020043 |
| ENSMUSG00000047797 | Gjb1 | 14618 | protein_coding | 574.0310504 | 0.084031207 | 0.123872036 | -0.678371076 | 0.497536443 | 0.703090606 | 0.084031207 |
| ENSMUSG00000021483 | Cdk20 | 105278 | protein_coding | 30.65604135 | 0.084040926 | 0.208685697 | -0.402715314 | 0.687157667 | 0.833389267 | 0.084040926 |
| ENSMUSG00000036948 | Trappc14 | 231807 | protein_coding | 243.5670798 | -0.084047818 | 0.084639899 | 0.993004708 | 0.32070766 | 0.541897747 | 0.084047818 |
| ENSMUSG00000089661 | Mia | 12587 | protein_coding | 5.744699605 | -0.08409649 | 0.422663872 | 0.198967775 | 0.842287953 | 0.921900136 | 0.08409649 |
| ENSMUSG00000021413 | Prpf4b | 19134 | protein_coding | 1282.578097 | 0.084101066 | 0.056388032 | -1.491470161 | 0.1358381 | 0.307249743 | 0.084101066 |
| ENSMUSG00000072704 | Smim10l1 | 381820 | protein_coding | 1136.870505 | 0.084108821 | 0.091190584 | -0.922341076 | 0.356350695 | 0.579378227 | 0.084108821 |
| ENSMUSG00000089781 | Gm15756 | 102631993 | lncRNA | 0.776972506 | 0.084141501 | 1.282152531 | -0.065625188 | 0.947676235 | 1 | 0.084141501 |
| ENSMUSG00000073131 | Vma21 | 67048 | protein_coding | 186.4280695 | 0.084174027 | 0.126703409 | -0.664339092 | 0.506473306 | 0.71028091 | 0.084174027 |
| ENSMUSG00000026457 | Adipor1 | 72674 | protein_coding | 2638.029858 | -0.084234605 | 0.063976688 | 1.316645296 | 0.187957548 | 0.38424498 | 0.084234605 |
| ENSMUSG00000022521 | Crebbp | 12914 | protein_coding | 1011.501531 | 0.084267748 | 0.12161295 | -0.692917558 | 0.48836129 | 0.696363286 | 0.084267748 |
| ENSMUSG00000022949 | Clic6 | 209195 | protein_coding | 710.4623773 | -0.08427539 | 0.185687055 | 0.453857111 | 0.649931675 | 0.809917796 | 0.08427539 |
| ENSMUSG00000048920 | Fkrp | 243853 | protein_coding | 209.4919459 | 0.084340561 | 0.093569633 | -0.901366804 | 0.367393325 | 0.590040437 | 0.084340561 |
| ENSMUSG00000002496 | Tsc2 | 22084 | protein_coding | 753.4543215 | -0.084403674 | 0.053901398 | 1.565890267 | 0.117374309 | 0.277378268 | 0.084403674 |
| ENSMUSG00000097405 | D630044L22Rik | 240054 | lncRNA | 1.607990219 | 0.084439837 | 1.051730712 | -0.080286556 | 0.93600935 | 1 | 0.084439837 |
| ENSMUSG00000020224 | Llph | 66225 | protein_coding | 121.0579614 | -0.084471881 | 0.133365532 | 0.633386152 | 0.526481506 | 0.724949186 | 0.084471881 |
| ENSMUSG00000021704 | Mtx3 | 382793 | protein_coding | 239.981147 | -0.084503919 | 0.137419285 | 0.614934933 | 0.538597692 | 0.733377497 | 0.084503919 |
| ENSMUSG00000029265 | Dr1 | 13486 | protein_coding | 413.3455001 | 0.084562741 | 0.089801005 | -0.94166809 | 0.346362597 | 0.568119977 | 0.084562741 |
| ENSMUSG00000025858 | Get4 | 67604 | protein_coding | 289.025544 | 0.084564145 | 0.096688711 | -0.874602051 | 0.381790471 | 0.604102573 | 0.084564145 |
| ENSMUSG00000036180 | Gatad2a | 234366 | protein_coding | 1130.68463 | 0.084615897 | 0.055358125 | -1.528518121 | 0.126383948 | 0.292373241 | 0.084615897 |
| ENSMUSG00000027956 | Tmem144 | 70652 | protein_coding | 103.2534975 | -0.084641179 | 0.162374869 | 0.521270194 | 0.602178562 | 0.778302948 | 0.084641179 |
| ENSMUSG00000066861 | Oas1g | 23960 | protein_coding | 199.9957288 | -0.084642546 | 0.195783546 | 0.43232717 | 0.665503643 | 0.819563591 | 0.084642546 |
| ENSMUSG00000021534 | 1700001L19Rik | 69315 | protein_coding | 4.122505672 | 0.084654693 | 0.560043654 | -0.151157312 | 0.879851623 | 1 | 0.084654693 |
| ENSMUSG00000016257 | Prelid3b | 66390 | protein_coding | 1700.464768 | 0.084655725 | 0.080784489 | -1.047920542 | 0.294675217 | 0.513807046 | 0.084655725 |
| ENSMUSG00000027014 | Cwc22 | 80744 | protein_coding | 332.7712011 | 0.084673044 | 0.0923135 | -0.917233605 | 0.359020241 | 0.581717059 | 0.084673044 |
| ENSMUSG00000022756 | Slc7a4 | 224022 | protein_coding | 213.8582661 | 0.084681737 | 0.145244085 | -0.583030537 | 0.559872743 | 0.748554316 | 0.084681737 |
| ENSMUSG00000018171 | Vmp1 | 75909 | protein_coding | 1363.21093 | 0.084705934 | 0.097229943 | -0.871191851 | 0.383649408 | 0.606155692 | 0.084705934 |
| ENSMUSG00000035671 | Zswim4 | 212168 | protein_coding | 228.9599354 | 0.084786377 | 0.162616833 | -0.521387454 | 0.602096889 | 0.778258299 | 0.084786377 |
| ENSMUSG00000018882 | Mrpl45 | 67036 | protein_coding | 430.3129331 | -0.084827987 | 0.092182184 | 0.920221052 | 0.357457255 | 0.580534441 | 0.084827987 |
| ENSMUSG00000049858 | Suox | 211389 | protein_coding | 240.1947429 | -0.084856108 | 0.143932669 | 0.589554185 | 0.555489575 | 0.745195357 | 0.084856108 |
| ENSMUSG00000078201 | Tmem203 | 227615 | protein_coding | 39.68713246 | -0.084863328 | 0.177451474 | 0.478233997 | 0.632483668 | 0.799512805 | 0.084863328 |
| ENSMUSG00000031731 | Ap1g1 | 11765 | protein_coding | 1290.882232 | 0.084877314 | 0.06303284 | -1.346557048 | 0.178122935 | 0.370562126 | 0.084877314 |

Supplementary Table 1

| ensembl_gene_id | external_gene_name | entrezgene_id | gene_biotype | baseMean | log2FoldChange | lfcSE | stat | pvalue | padj | absLog2FC |
|---|---|---|---|---|---|---|---|---|---|---|
| ENSMUSG00000097290 | 1300002E11Rik | 100043489 | lncRNA | 99.24224419 | -0.08488835 | 0.12409893 | 0.684037725 | 0.493951344 | 0.700180271 | 0.08488835 |
| ENSMUSG00000056648 | Hoxb8 | 15416 | protein_coding | 49.44547246 | -0.084901887 | 0.341205718 | 0.248829029 | 0.803493036 | 0.903013746 | 0.084901887 |
| ENSMUSG00000018293 | Pfn1 | 18643 | protein_coding | 5870.0814 | -0.084922422 | 0.089605576 | 0.947735905 | 0.343263917 | 0.565170522 | 0.084922422 |
| ENSMUSG00000027475 | Kif3b | 16569 | protein_coding | 744.0651421 | 0.084925827 | 0.068315697 | -1.243137835 | 0.213817046 | 0.418758859 | 0.084925827 |
| ENSMUSG00000043243 | Niban3 | 100037278 | protein_coding | 10.75696494 | 0.084958069 | 0.387157528 | -0.219440571 | 0.826306869 | 0.912263113 | 0.084958069 |
| ENSMUSG00000047126 | Cltc | 67300 | protein_coding | 6589.824786 | 0.084985675 | 0.093587158 | -0.908091198 | 0.36383004 | 0.586288962 | 0.084985675 |
| ENSMUSG00000024181 | Mrpl28 | 68611 | protein_coding | 627.8144038 | 0.085064893 | 0.076684585 | -1.109282827 | 0.267308191 | 0.482492892 | 0.085064893 |
| ENSMUSG00000046056 | Sbsn | 282619 | protein_coding | 4.410968919 | -0.085099447 | 0.50080992 | 0.169923646 | 0.865070185 | 0.932932188 | 0.085099447 |
| ENSMUSG00000022008 | Gpalpp1 | 67467 | protein_coding | 493.0278922 | 0.085163916 | 0.103689563 | -0.821335472 | 0.411455208 | 0.631719162 | 0.085163916 |
| ENSMUSG00000058799 | Nap1l1 | 53605 | protein_coding | 541.6312878 | 0.08517561 | 0.144747433 | -0.588442974 | 0.556235 | 0.745631404 | 0.08517561 |
| ENSMUSG00000051890 | Klhdc1 | 271005 | protein_coding | 37.82838666 | -0.085276132 | 0.204675672 | 0.416640294 | 0.676941532 | 0.826943172 | 0.085276132 |
| ENSMUSG00000022377 | Asap1 | 13196 | protein_coding | 251.9615957 | 0.085279526 | 0.258049261 | -0.330477697 | 0.741039042 | 0.868956809 | 0.085279526 |
| ENSMUSG00000007041 | Clic1 | 114584 | protein_coding | 2243.990554 | -0.08533874 | 0.064011978 | 1.333168307 | 0.182476577 | 0.377007156 | 0.08533874 |
| ENSMUSG00000020697 | Lig3 | 16882 | protein_coding | 441.532217 | 0.085377459 | 0.096031695 | -0.889055012 | 0.373973515 | 0.595946987 | 0.085377459 |
| ENSMUSG00000024958 | Gpr137 | 107173 | protein_coding | 282.3076436 | -0.085378271 | 0.11135125 | 0.766747306 | 0.443231772 | 0.660786267 | 0.085378271 |
| ENSMUSG00000025728 | Pigq | 14755 | protein_coding | 716.5103422 | -0.085411872 | 0.061932264 | 1.379117564 | 0.167858509 | 0.356391801 | 0.085411872 |
| ENSMUSG00000072568 | Lratd2 | 399603 | protein_coding | 517.5846693 | 0.085441291 | 0.096776458 | -0.88287268 | 0.377305066 | 0.599520433 | 0.085441291 |
| ENSMUSG00000033590 | Myo5c | 208943 | protein_coding | 2854.646692 | 0.085446586 | 0.096894096 | -0.881855443 | 0.377854985 | 0.600047814 | 0.085446586 |
| ENSMUSG00000057777 | Mab21l2 | 23937 | protein_coding | 6.449792549 | -0.085455187 | 0.483103397 | 0.176887987 | 0.859596375 | 0.93121061 | 0.085455187 |
| ENSMUSG00000027380 | Acoxl | 74121 | protein_coding | 2.745458749 | 0.085509496 | 0.726821222 | -0.117648595 | 0.906346099 | 1 | 0.085509496 |
| ENSMUSG00000032301 | Psma4 | 26441 | protein_coding | 1142.949208 | 0.08552117 | 0.083442307 | -1.02491378 | 0.305403872 | 0.525632052 | 0.08552117 |
| ENSMUSG00000025235 | Bbs4 | 102774 | protein_coding | 58.48496626 | -0.08553112 | 0.170191668 | 0.502557623 | 0.615275329 | 0.78770474 | 0.08553112 |
| ENSMUSG00000059263 | Usp47 | 74996 | protein_coding | 2184.258744 | 0.085533281 | 0.062372775 | -1.371323957 | 0.170273991 | 0.359946836 | 0.085533281 |
| ENSMUSG00000054764 | Mtnr1a | 17773 | protein_coding | 1.365097336 | -0.085537984 | 1.17626515 | 0.072719985 | 0.942028945 | 1 | 0.085537984 |
| ENSMUSG00000030042 | Pole4 | 66979 | protein_coding | 254.4463464 | 0.085538382 | 0.103767602 | -0.824326474 | 0.40975408 | 0.630279329 | 0.085538382 |
| ENSMUSG00000036160 | Surf6 | 20935 | protein_coding | 374.0021679 | -0.08554724 | 0.07649802 | 1.118293516 | 0.263441655 | 0.477601877 | 0.08554724 |
| ENSMUSG00000035623 | Rsf1 | 233532 | protein_coding | 493.5247481 | -0.085573049 | 0.076514612 | 1.118388322 | 0.26340118 | 0.477601877 | 0.085573049 |
| ENSMUSG00000041415 | Dicer1 | 192119 | protein_coding | 901.7598496 | 0.085588031 | 0.074024271 | -1.156215794 | 0.247592901 | 0.459757614 | 0.085588031 |
| ENSMUSG00000003528 | Slc25a1 | 13358 | protein_coding | 988.037439 | -0.085595312 | 0.075912034 | 1.127559197 | 0.259506127 | 0.472954995 | 0.085595312 |
| ENSMUSG00000033581 | Igf2bp2 | 319765 | protein_coding | 635.7329994 | 0.085611314 | 0.153536892 | -0.557594419 | 0.577121368 | 0.761298159 | 0.085611314 |
| ENSMUSG00000057531 | Dtnbp1 | 94245 | protein_coding | 164.6894005 | -0.085645151 | 0.110707492 | 0.773616576 | 0.439157567 | 0.657120605 | 0.085645151 |
| ENSMUSG00000042699 | Dhx9 | 13211 | protein_coding | 1002.806736 | 0.085650851 | 0.142504955 | -0.601037708 | 0.547814871 | 0.739162904 | 0.085650851 |
| ENSMUSG00000041638 | Gcn1 | 231659 | protein_coding | 1224.23084 | 0.085754776 | 0.07842269 | -1.093494451 | 0.274176764 | 0.490619315 | 0.085754776 |
| ENSMUSG00000043702 | Pde12 | 211948 | protein_coding | 401.3737513 | 0.085769598 | 0.07115307 | -1.205423713 | 0.228039772 | 0.436620149 | 0.085769598 |
| ENSMUSG00000042272 | Sestd1 | 228071 | protein_coding | 509.4756121 | 0.085779607 | 0.073709495 | -1.163752468 | 0.24452434 | 0.456022015 | 0.085779607 |
| ENSMUSG00000025266 | Gnl3l | 237107 | protein_coding | 665.2886205 | 0.085797205 | 0.087138456 | -0.984607818 | 0.324816748 | 0.546769252 | 0.085797205 |
| ENSMUSG00000022884 | Eif4a2 | 13682 | protein_coding | 3468.785449 | -0.085814688 | 0.078882664 | 1.087877662 | 0.276649119 | 0.493582047 | 0.085814688 |
| ENSMUSG00000047108 | Dnajb7 | 57755 | protein_coding | 4.580369568 | -0.085836778 | 0.494794829 | 0.173479537 | 0.862274511 | 0.932001143 | 0.085836778 |
| ENSMUSG00000021221 | Dpf3 | 70127 | protein_coding | 41.26573558 | -0.085850713 | 0.293382162 | 0.292624174 | 0.769809434 | 0.884532517 | 0.085850713 |

| ensembl_gene_id | external_gene_name | entrezgene_id | gene_biotype | baseMean | log2FoldChange | lfcSE | stat | pvalue | padj | absLog2FC |
|---|---|---|---|---|---|---|---|---|---|---|
| ENSMUSG00000070697 | Utp3 | 65961 | protein_coding | 289.7086351 | 0.085877723 | 0.091841843 | -0.935060973 | 0.349756878 | 0.571586846 | 0.085877723 |
| ENSMUSG00000030798 | Cd37 | 12493 | protein_coding | 10.85166666 | 0.085897477 | 0.35571215 | -0.2414803 | 0.809182883 | 0.905779176 | 0.085897477 |
| ENSMUSG00000032560 | Dnajc13 | 235567 | protein_coding | 1568.426671 | 0.085971647 | 0.065919855 | -1.304184403 | 0.192170716 | 0.38986575 | 0.085971647 |
| ENSMUSG00000027810 | Eif2a | 229317 | protein_coding | 643.1805375 | 0.086003748 | 0.108571807 | -0.792137027 | 0.428280783 | 0.647320963 | 0.086003748 |
| ENSMUSG00000067370 | B3galt4 | 54218 | protein_coding | 104.7652715 | 0.086039437 | 0.102694981 | -0.837815402 | 0.402134387 | 0.623781054 | 0.086039437 |
| ENSMUSG00000026918 | Brd3 | 67382 | protein_coding | 429.5603543 | -0.086125284 | 0.09499418 | 0.906637483 | 0.364598533 | 0.587183556 | 0.086125284 |
| ENSMUSG00000006941 | Eif1b | 68969 | protein_coding | 132.9535484 | 0.0861439 | 0.124220385 | -0.693476357 | 0.488010658 | 0.696186289 | 0.0861439 |
| ENSMUSG00000030521 | Mphosph10 | 67973 | protein_coding | 210.0242118 | 0.086168725 | 0.160063325 | -0.538341465 | 0.590341329 | 0.770846581 | 0.086168725 |
| ENSMUSG00000026341 | Actr3 | 74117 | protein_coding | 3437.310809 | -0.086171998 | 0.083827759 | 1.027964942 | 0.303966319 | 0.524360158 | 0.086171998 |
| ENSMUSG00000024955 | Esrra | 26379 | protein_coding | 2221.328652 | 0.086206116 | 0.09083301 | -0.94906154 | 0.342589314 | 0.564622747 | 0.086206116 |
| ENSMUSG00000061898 | Rbak | 57782 | protein_coding | 104.3612038 | -0.08621808 | 0.121820828 | 0.707744985 | 0.479103636 | 0.690541222 | 0.08621808 |
| ENSMUSG00000021203 | Otub2 | 68149 | protein_coding | 27.37900818 | -0.086270681 | 0.226803805 | 0.38037581 | 0.703666469 | 0.844104331 | 0.086270681 |
| ENSMUSG00000032468 | Armc8 | 74125 | protein_coding | 500.1575359 | -0.086273863 | 0.080381047 | 1.073311015 | 0.283131595 | 0.501193606 | 0.086273863 |
| ENSMUSG00000031644 | Nek1 | 18004 | protein_coding | 259.9087613 | 0.086334605 | 0.09135708 | -0.945023689 | 0.344646782 | 0.566318105 | 0.086334605 |
| ENSMUSG00000090659 | Zfp493 | 72958 | protein_coding | 34.01283736 | -0.086348796 | 0.214289495 | 0.402953939 | 0.68698211 | 0.8333203 | 0.086348796 |
| ENSMUSG00000036686 | Cc2d1a | 212139 | protein_coding | 682.6734724 | -0.086355597 | 0.093002682 | 0.928528037 | 0.353133731 | 0.57545396 | 0.086355597 |
| ENSMUSG00000034674 | Tdg | 21665 | protein_coding | 264.728122 | -0.0863714 | 0.102837004 | 0.839886391 | 0.400972089 | 0.622737045 | 0.0863714 |
| ENSMUSG00000028398 | Dmac1 | 66928 | protein_coding | 210.3406246 | 0.086382785 | 0.125242525 | -0.689724083 | 0.490367718 | 0.69793377 | 0.086382785 |
| ENSMUSG00000026083 | Eif5b | 226982 | protein_coding | 540.6758949 | 0.086399418 | 0.078757621 | -1.097029304 | 0.272628582 | 0.488518708 | 0.086399418 |
| ENSMUSG00000042185 | Nfrkb | 235134 | protein_coding | 284.5119275 | 0.086419484 | 0.07205065 | -1.199426849 | 0.230362013 | 0.439030384 | 0.086419484 |
| ENSMUSG00000094396 | Vmn2r124 | 637021 | protein_coding | 1.553447458 | -0.086488948 | 2.631079336 | 0.032872041 | 0.973776629 | 1 | 0.086488948 |
| ENSMUSG00000017670 | Elmo2 | 140579 | protein_coding | 194.7409318 | 0.086501559 | 0.08942775 | -0.967278708 | 0.333404725 | 0.555809168 | 0.086501559 |
| ENSMUSG00000034645 | Zyg11a | 230590 | protein_coding | 5.359491047 | -0.086502189 | 0.703925806 | 0.122885378 | 0.902197865 | 0.951530343 | 0.086502189 |
| ENSMUSG00000000216 | Scnn1g | 20278 | protein_coding | 25.60705508 | -0.086551059 | 0.288819456 | 0.299671843 | 0.764427478 | 0.882132403 | 0.086551059 |
| ENSMUSG00000021171 | Esyt2 | 52635 | protein_coding | 2047.840396 | -0.086655095 | 0.066837817 | 1.296497992 | 0.194803972 | 0.393901406 | 0.086655095 |
| ENSMUSG00000026641 | Usf1 | 22278 | protein_coding | 440.7036777 | -0.086778545 | 0.060394881 | 1.436852653 | 0.150759866 | 0.330823349 | 0.086778545 |
| ENSMUSG00000011877 | Git1 | 216963 | protein_coding | 715.4736689 | -0.086779416 | 0.095731437 | 0.906488182 | 0.364677517 | 0.587211164 | 0.086779416 |
| ENSMUSG00000024610 | Cd74 | 16149 | protein_coding | 61.81853981 | 0.08679968 | 0.324880106 | -0.267174501 | 0.789334808 | 0.894720656 | 0.08679968 |
| ENSMUSG00000030672 | Mylpf | 17907 | protein_coding | 15.38439179 | 0.086801538 | 0.466541409 | -0.186053235 | 0.852403015 | 0.927761944 | 0.086801538 |
| ENSMUSG00000033411 | Ctdspl2 | 329506 | protein_coding | 340.966244 | -0.086807802 | 0.098705664 | 0.879461196 | 0.379151264 | 0.60160505 | 0.086807802 |
| ENSMUSG00000024491 | Rbm27 | 225432 | protein_coding | 587.4703764 | 0.08681126 | 0.056029123 | -1.549395284 | 0.121286727 | 0.284063113 | 0.08681126 |
| ENSMUSG00000033033 | Calhm2 | 72691 | protein_coding | 20.55098041 | 0.086813004 | 0.233096729 | -0.372433385 | 0.709570201 | 0.848381285 | 0.086813004 |
| ENSMUSG00000002372 | Ranbp3 | 71810 | protein_coding | 676.5916014 | 0.086835264 | 0.053212145 | -1.631869273 | 0.102707028 | 0.252397465 | 0.086835264 |
| ENSMUSG00000028483 | Snapc3 | 77634 | protein_coding | 158.4577113 | -0.086838266 | 0.132608275 | 0.654848017 | 0.512565612 | 0.714893745 | 0.086838266 |
| ENSMUSG00000004864 | Mapk13 | 26415 | protein_coding | 2087.058654 | 0.086852496 | 0.077035874 | -1.127429231 | 0.259561046 | 0.472954995 | 0.086852496 |
| ENSMUSG00000042050 | Dync2i1 | 217935 | protein_coding | 117.8703249 | -0.086867136 | 0.126666134 | 0.685796065 | 0.492841718 | 0.699388858 | 0.086867136 |
| ENSMUSG00000042298 | Ttc19 | 72795 | protein_coding | 270.4744083 | -0.086867349 | 0.084502541 | 1.027985045 | 0.303956862 | 0.524360158 | 0.086867349 |
| ENSMUSG00000018677 | Slc25a39 | 68066 | protein_coding | 1969.12372 | -0.086885052 | 0.056224514 | 1.545323309 | 0.12226808 | 0.285349785 | 0.086885052 |
| ENSMUSG00000029504 | Ddx51 | 69663 | protein_coding | 138.2042885 | -0.08692926 | 0.129338753 | 0.672105289 | 0.501516669 | 0.706297653 | 0.08692926 |

| ensembl_gene_id | external_gene_name | entrezgene_id | gene_biotype | baseMean | log2FoldChange | lfcSE | stat | pvalue | padj | absLog2FC |
|---|---|---|---|---|---|---|---|---|---|---|
| ENSMUSG00000038510 | Rpf2 | 67239 | protein_coding | 118.770027 | -0.086935342 | 0.135970577 | 0.639368784 | 0.522583051 | 0.721840784 | 0.086935342 |
| ENSMUSG00000021891 | Mettl6 | 67011 | protein_coding | 169.4492327 | 0.086938513 | 0.080234296 | -1.083558008 | 0.278560818 | 0.495777145 | 0.086938513 |
| ENSMUSG00000021400 | Wrnip1 | 78903 | protein_coding | 377.3311514 | 0.086948757 | 0.089215571 | -0.974591721 | 0.329762822 | 0.552030063 | 0.086948757 |
| ENSMUSG00000058488 | Kl | 16591 | protein_coding | 4.224595511 | 0.086975651 | 1.139615011 | -0.076320205 | 0.939164352 | 1 | 0.086975651 |
| ENSMUSG00000030401 | Rtn2 | 20167 | protein_coding | 29.56129584 | 0.086982771 | 0.308815512 | -0.28166581 | 0.778199773 | 0.889171179 | 0.086982771 |
| ENSMUSG00000025162 | Csnk1d | 104318 | protein_coding | 1635.65125 | 0.08701654 | 0.057472363 | -1.514058871 | 0.130010909 | 0.297863506 | 0.08701654 |
| ENSMUSG00000046404 | Yod1 | 226418 | protein_coding | 156.6423495 | 0.087052276 | 0.104938282 | -0.829556897 | 0.406789355 | 0.627629974 | 0.087052276 |
| ENSMUSG00000019494 | Cops6 | 26893 | protein_coding | 500.6867127 | 0.087103422 | 0.076087247 | -1.14478347 | 0.252298864 | 0.464617944 | 0.087103422 |
| ENSMUSG00000001558 | Klhl10 | 66720 | protein_coding | 2.140100407 | 0.087126778 | 0.741480773 | -0.117503758 | 0.906460866 | 1 | 0.087126778 |
| ENSMUSG00000061904 | Slc25a3 | 18674 | protein_coding | 7224.770198 | -0.087135293 | 0.070469285 | 1.236500306 | 0.216272652 | 0.421530084 | 0.087135293 |
| ENSMUSG00000021395 | Spin1 | 20729 | protein_coding | 718.7436703 | -0.087139987 | 0.075676762 | 1.151476148 | 0.249536402 | 0.461770971 | 0.087139987 |
| ENSMUSG00000004099 | Dnmt1 | 13433 | protein_coding | 954.3335969 | 0.087229584 | 0.138046505 | -0.631885499 | 0.527461699 | 0.725788994 | 0.087229584 |
| ENSMUSG00000063108 | Zfp26 | 22688 | protein_coding | 370.8039221 | -0.087239471 | 0.070491092 | 1.237595687 | 0.215866014 | 0.421123611 | 0.087239471 |
| ENSMUSG00000044674 | Fzd1 | 14362 | protein_coding | 368.0297195 | 0.087243268 | 0.104868234 | -0.831932269 | 0.405447176 | 0.62645534 | 0.087243268 |
| ENSMUSG00000007036 | Abhd16a | 193742 | protein_coding | 790.063873 | -0.087253526 | 0.080327388 | 1.086223867 | 0.27737996 | 0.494297255 | 0.087253526 |
| ENSMUSG00000045314 | Sowahb | 78088 | protein_coding | 519.1512499 | -0.087280416 | 0.121289345 | 0.719604975 | 0.471768248 | 0.684542733 | 0.087280416 |
| ENSMUSG00000038806 | Sde2 | 208768 | protein_coding | 269.8663414 | -0.087304798 | 0.071070541 | 1.228424551 | 0.219287636 | 0.425536548 | 0.087304798 |
| ENSMUSG00000034274 | Thoc5 | 107829 | protein_coding | 380.2156107 | -0.087321949 | 0.101388497 | 0.861260903 | 0.389094363 | 0.611719756 | 0.087321949 |
| ENSMUSG00000020859 | Spag9 | 70834 | protein_coding | 1172.069214 | -0.08732851 | 0.092346497 | 0.945661316 | 0.344321359 | 0.56607413 | 0.08732851 |
| ENSMUSG00000035686 | Thrsp | 21835 | protein_coding | 1.204472807 | -0.087365452 | 2.016817188 | 0.043318479 | 0.965447661 | 1 | 0.087365452 |
| ENSMUSG00000019897 | Ccdc59 | 52713 | protein_coding | 226.7257678 | 0.087384268 | 0.071047589 | -1.22993995 | 0.218719593 | 0.424883585 | 0.087384268 |
| ENSMUSG00000019082 | Slc25a22 | 68267 | protein_coding | 209.9047824 | 0.087390606 | 0.161988984 | -0.539484873 | 0.589552332 | 0.770506411 | 0.087390606 |
| ENSMUSG00000035834 | Polr3g | 67486 | protein_coding | 126.8418686 | 0.087397785 | 0.121583342 | -0.718830258 | 0.472245512 | 0.684753919 | 0.087397785 |
| ENSMUSG00000028217 | Cdh17 | 12557 | protein_coding | 9368.265391 | -0.087428268 | 0.086196269 | 1.014292956 | 0.310442984 | 0.53139398 | 0.087428268 |
| ENSMUSG00000051022 | Hs3st1 | 15476 | protein_coding | 135.8372902 | 0.087460763 | 0.105073137 | -0.832379857 | 0.405194568 | 0.626416691 | 0.087460763 |
| ENSMUSG00000037720 | Tmem33 | 67878 | protein_coding | 732.3518865 | -0.087518655 | 0.086317847 | 1.013911469 | 0.310624998 | 0.531626202 | 0.087518655 |
| ENSMUSG00000010122 | Slc47a1 | 67473 | protein_coding | 22.98406562 | 0.087561559 | 0.367254457 | -0.238422046 | 0.81155377 | 0.906704483 | 0.087561559 |
| ENSMUSG00000030752 | Kdm8 | 77035 | protein_coding | 71.06086522 | 0.087563344 | 0.183753877 | -0.476525153 | 0.633700292 | 0.800362194 | 0.087563344 |
| ENSMUSG00000090210 | Itga10 | 213119 | protein_coding | 14.11981055 | 0.08756749 | 0.469171803 | -0.186642696 | 0.851940789 | 0.927725037 | 0.08756749 |
| ENSMUSG00000043881 | Kbtbd7 | 211255 | protein_coding | 173.139623 | -0.087578661 | 0.101907953 | 0.859389854 | 0.390125465 | 0.612475705 | 0.087578661 |
| ENSMUSG00000037640 | Zfp60 | 22718 | protein_coding | 119.0250904 | -0.087625557 | 0.136948689 | 0.639842248 | 0.522275163 | 0.721805342 | 0.087625557 |
| ENSMUSG00000052299 | Ltn1 | 78913 | protein_coding | 811.381982 | 0.087634669 | 0.092749417 | -0.944854122 | 0.344733356 | 0.566362983 | 0.087634669 |
| ENSMUSG00000027864 | Ptgfrn | 19221 | protein_coding | 2156.407059 | 0.087660114 | 0.055872758 | -1.568924065 | 0.116665634 | 0.276098635 | 0.087660114 |
| ENSMUSG00000039485 | Tspyl4 | 72480 | protein_coding | 9.727491583 | -0.087685452 | 0.333139137 | 0.2632097 | 0.792388955 | 0.896636371 | 0.087685452 |
| ENSMUSG00000039414 | Heatr5b | 320473 | protein_coding | 397.9568725 | 0.087735563 | 0.074881817 | -1.171653762 | 0.241336094 | 0.451964647 | 0.087735563 |
| ENSMUSG00000021123 | Rdh12 | 77974 | protein_coding | 8.261797029 | -0.087743166 | 0.410240825 | 0.213882092 | 0.830639005 | 0.91440345 | 0.087743166 |
| ENSMUSG00000024759 | Atl3 | 109168 | protein_coding | 877.5037634 | -0.087790499 | 0.09626023 | 0.912012148 | 0.361762322 | 0.584163582 | 0.087790499 |
| ENSMUSG00000087630 | Gm11513 | 102640343 | lncRNA | 1.930360315 | -0.087819811 | 0.811508045 | 0.108218041 | 0.913822735 | 1 | 0.087819811 |
| ENSMUSG00000041236 | Vps41 | 218035 | protein_coding | 833.2065943 | -0.087832254 | 0.073226221 | 1.199464522 | 0.230347372 | 0.439030384 | 0.087832254 |

| ensembl_gene_id | external_gene_name | entrezgene_id | gene_biotype | baseMean | log2FoldChange | lfcSE | stat | pvalue | padj | absLog2FC |
|---|---|---|---|---|---|---|---|---|---|---|
| ENSMUSG00000061360 | Phf5a | 68479 | protein_coding | 443.7434928 | 0.087844644 | 0.108444828 | -0.810039963 | 0.417917208 | 0.637944042 | 0.087844644 |
| ENSMUSG00000026127 | Imp4 | 27993 | protein_coding | 515.7899613 | -0.087866425 | 0.085936468 | 1.022457953 | 0.306564203 | 0.527079777 | 0.087866425 |
| ENSMUSG00000046782 | Ttc6 | 70846 | protein_coding | 73.4206902 | 0.087931677 | 0.166006492 | -0.529688185 | 0.596328141 | 0.774560557 | 0.087931677 |
| ENSMUSG00000024251 | Thada | 240174 | protein_coding | 249.6851263 | 0.087949953 | 0.121871332 | -0.721662357 | 0.470502091 | 0.683786987 | 0.087949953 |
| ENSMUSG00000067150 | Xpo5 | 72322 | protein_coding | 511.8374872 | 0.088017173 | 0.083990016 | -1.047948043 | 0.294662546 | 0.513807046 | 0.088017173 |
| ENSMUSG00000035284 | Vps13c | 320528 | protein_coding | 1045.858842 | -0.088034521 | 0.144841662 | 0.607798335 | 0.543321234 | 0.736456016 | 0.088034521 |
| ENSMUSG00000019718 | L3hypdh | 67217 | protein_coding | 9.409085421 | -0.088071462 | 0.356009668 | 0.247385028 | 0.80461026 | 0.903777298 | 0.088071462 |
| ENSMUSG00000026941 | Mamdc4 | 381352 | protein_coding | 53.96107224 | 0.088094764 | 0.23944918 | -0.367905893 | 0.712943406 | 0.850751312 | 0.088094764 |
| ENSMUSG00000044566 | Cage1 | 71213 | protein_coding | 28.0292417 | 0.08810175 | 0.221038492 | -0.398581031 | 0.690201941 | 0.834766446 | 0.08810175 |
| ENSMUSG00000030649 | Anapc15 | 75430 | protein_coding | 61.5840391 | 0.088116759 | 0.139606456 | -0.631179684 | 0.527923044 | 0.726026078 | 0.088116759 |
| ENSMUSG00000073077 | Cfap47 | 636104 | protein_coding | 30.39217275 | -0.088121306 | 0.256919015 | 0.342992544 | 0.731604069 | 0.861932625 | 0.088121306 |
| ENSMUSG00000036913 | Trim67 | 330863 | protein_coding | 4.263515847 | -0.088143735 | 1.266021426 | 0.069622625 | 0.944494028 | 1 | 0.088143735 |
| ENSMUSG00000085241 | Snhg3 | 399101 | lncRNA | 2.233972557 | 0.088154168 | 0.76924297 | -0.1145986 | 0.908763289 | 1 | 0.088154168 |
| ENSMUSG00000038291 | Snx25 | 102141 | protein_coding | 346.6310805 | -0.088155482 | 0.093877785 | 0.93904518 | 0.347707548 | 0.569590562 | 0.088155482 |
| ENSMUSG00000020888 | Dvl2 | 13543 | protein_coding | 102.610294 | -0.088250779 | 0.127961164 | 0.689668458 | 0.490402706 | 0.69793377 | 0.088250779 |
| ENSMUSG00000041565 | L3mbtl4 | 320858 | protein_coding | 4.886827072 | 0.088286864 | 0.718523377 | -0.12287264 | 0.902207952 | 0.951530343 | 0.088286864 |
| ENSMUSG00000027620 | Rbm39 | 170791 | protein_coding | 2632.99219 | 0.088299276 | 0.049347989 | -1.789318625 | 0.073563516 | 0.197983688 | 0.088299276 |
| ENSMUSG00000032101 | Ddx25 | 30959 | protein_coding | 6.502323357 | -0.088300684 | 0.603596625 | 0.146290884 | 0.883691762 | 0.942424291 | 0.088300684 |
| ENSMUSG00000094936 | Rbm4 | 19653 | protein_coding | 53.76943733 | 0.088365408 | 0.145078987 | -0.609084812 | 0.542468224 | 0.736098624 | 0.088365408 |
| ENSMUSG00000050447 | Lypd6 | 320343 | protein_coding | 2.119288815 | 0.088386276 | 1.177679207 | -0.075051232 | 0.940173949 | 1 | 0.088386276 |
| ENSMUSG00000027001 | Dusp19 | 68082 | protein_coding | 112.1927233 | 0.088399733 | 0.122301571 | -0.722801288 | 0.469801976 | 0.683370901 | 0.088399733 |
| ENSMUSG00000061544 | Zfp229 | 381067 | protein_coding | 65.43624497 | -0.088543631 | 0.165050475 | 0.536463955 | 0.591637941 | 0.771785126 | 0.088543631 |
| ENSMUSG00000065847 | Gm25188 | 115489822 | snoRNA | 0.979078393 | 0.088566162 | 1.186229521 | -0.07466191 | 0.940483714 | 1 | 0.088566162 |
| ENSMUSG00000043284 | Tmem11 | 216821 | protein_coding | 365.1345699 | -0.088596841 | 0.096925215 | 0.914074226 | 0.360677845 | 0.583005439 | 0.088596841 |
| ENSMUSG00000025937 | Lactb2 | 212442 | protein_coding | 284.34044 | 0.08864314 | 0.099045463 | -0.894974262 | 0.370800847 | 0.593065878 | 0.08864314 |
| ENSMUSG00000029330 | Cds1 | 74596 | protein_coding | 2220.100154 | -0.088647554 | 0.08833409 | 1.00354862 | 0.31559623 | 0.53655334 | 0.088647554 |
| ENSMUSG00000021079 | Timm9 | 30056 | protein_coding | 115.6158279 | -0.088660999 | 0.144556253 | 0.613332159 | 0.539656733 | 0.734194205 | 0.088660999 |
| ENSMUSG00000060475 | Wtap | 60532 | protein_coding | 627.6446884 | -0.088680469 | 0.085802543 | 1.033541274 | 0.301350666 | 0.521536878 | 0.088680469 |
| ENSMUSG00000009210 | Prr29 | 75573 | protein_coding | 2.353087113 | 0.08870595 | 0.873998602 | -0.101494384 | 0.919158015 | 1 | 0.08870595 |
| ENSMUSG00000027108 | Ola1 | 67059 | protein_coding | 842.1984811 | 0.088728563 | 0.101821795 | -0.871410313 | 0.383530157 | 0.606087752 | 0.088728563 |
| ENSMUSG00000019877 | Serinc1 | 56442 | protein_coding | 1061.985152 | -0.088761611 | 0.113136247 | 0.784555032 | 0.432714504 | 0.651280288 | 0.088761611 |
| ENSMUSG00000042225 | Ammecr1 | 56068 | protein_coding | 276.4674192 | 0.088773214 | 0.166744663 | -0.532390138 | 0.594455814 | 0.77323163 | 0.088773214 |
| ENSMUSG00000026966 | Ssna1 | 68475 | protein_coding | 298.8328443 | -0.088775865 | 0.11528665 | 0.770044626 | 0.441273422 | 0.659125086 | 0.088775865 |
| ENSMUSG00000038299 | Wdr36 | 225348 | protein_coding | 534.6067043 | -0.088793859 | 0.089505226 | 0.99205224 | 0.321172043 | 0.542303804 | 0.088793859 |
| ENSMUSG00000048696 | Mex3d | 237400 | protein_coding | 84.45859521 | -0.088824355 | 0.134326617 | 0.661256551 | 0.5084478 | 0.712111524 | 0.088824355 |
| ENSMUSG00000047388 | Atmin | 234776 | protein_coding | 405.8792573 | 0.088864952 | 0.096373832 | -0.922085901 | 0.35648377 | 0.579378227 | 0.088864952 |
| ENSMUSG00000039960 | Rhou | 69581 | protein_coding | 487.9694294 | -0.088935174 | 0.107122786 | 0.830217155 | 0.406416018 | 0.627248048 | 0.088935174 |
| ENSMUSG00000030079 | Ruvbl1 | 56505 | protein_coding | 167.2010427 | 0.088954121 | 0.157358822 | -0.565294785 | 0.571873275 | 0.757452299 | 0.088954121 |
| ENSMUSG00000037366 | Pafah2 | 100163 | protein_coding | 878.904703 | -0.08896003 | 0.070562809 | 1.260721209 | 0.20740931 | 0.410345914 | 0.08896003 |

| ensembl_gene_id | external_gene_name | entrezgene_id | gene_biotype | baseMean | log2FoldChange | lfcSE | stat | pvalue | padj | absLog2FC |
|---|---|---|---|---|---|---|---|---|---|---|
| ENSMUSG00000001767 | Crnkl1 | 66877 | protein_coding | 378.3996272 | 0.088969499 | 0.074674511 | -1.191430618 | 0.233484579 | 0.442885865 | 0.088969499 |
| ENSMUSG00000032718 | Mansc1 | 67729 | protein_coding | 759.3207405 | 0.089016089 | 0.142615624 | -0.624167862 | 0.532517351 | 0.729065788 | 0.089016089 |
| ENSMUSG00000021501 | Caml | 12328 | protein_coding | 250.4509871 | 0.089037896 | 0.084269999 | -1.056578817 | 0.290703852 | 0.50962669 | 0.089037896 |
| ENSMUSG00000030990 | Pgap2 | 233575 | protein_coding | 622.2495058 | -0.089074036 | 0.117602775 | 0.757414409 | 0.448801626 | 0.665664739 | 0.089074036 |
| ENSMUSG00000001285 | Myg1 | 60315 | protein_coding | 231.067875 | 0.089087044 | 0.095874198 | -0.929207715 | 0.35278145 | 0.575163675 | 0.089087044 |
| ENSMUSG00000076271 | Mir697 | 735277 | miRNA | 0.905942072 | 0.089104948 | 1.081789121 | -0.082368131 | 0.934353978 | 1 | 0.089104948 |
| ENSMUSG00000026276 | Septin2 | 18000 | protein_coding | 37.88971125 | -0.08914291 | 0.264658406 | 0.336822516 | 0.736250699 | 0.8658627 | 0.08914291 |
| ENSMUSG00000026205 | Slc23a3 | 22626 | protein_coding | 50.42628245 | 0.089146135 | 0.155537325 | -0.573149466 | 0.566543497 | 0.753111109 | 0.089146135 |
| ENSMUSG00000040396 | Abhd13 | 68904 | protein_coding | 1025.655518 | 0.089164857 | 0.09606676 | -0.928155133 | 0.353327104 | 0.575712264 | 0.089164857 |
| ENSMUSG00000021693 | Kif2a | 16563 | protein_coding | 317.0311523 | -0.089172253 | 0.094297269 | 0.945650431 | 0.344326913 | 0.56607413 | 0.089172253 |
| ENSMUSG00000021481 | Zfp346 | 26919 | protein_coding | 95.02506379 | 0.08917513 | 0.152433922 | -0.585008436 | 0.558542044 | 0.747347538 | 0.08917513 |
| ENSMUSG00000025025 | Mxi1 | 17859 | protein_coding | 637.6971302 | -0.089228796 | 0.139167449 | 0.641161397 | 0.521417827 | 0.720922137 | 0.089228796 |
| ENSMUSG00000026483 | Niban1 | 63913 | protein_coding | 1164.926466 | -0.08924032 | 0.086665188 | 1.029713568 | 0.303144484 | 0.523653558 | 0.08924032 |
| ENSMUSG00000005204 | Senp3 | 80886 | protein_coding | 542.2316589 | -0.089329804 | 0.068072383 | 1.312276721 | 0.189426784 | 0.386626363 | 0.089329804 |
| ENSMUSG00000025938 | Slco5a1 | 240726 | protein_coding | 2.271420429 | -0.089336811 | 1.762437285 | 0.050689356 | 0.959573059 | 1 | 0.089336811 |
| ENSMUSG00000022945 | Chaf1b | 110749 | protein_coding | 114.6352875 | 0.089386956 | 0.178695695 | -0.500218857 | 0.616920982 | 0.788881899 | 0.089386956 |
| ENSMUSG00000055660 | Mettl4 | 76781 | protein_coding | 272.1301617 | -0.089420819 | 0.134434133 | 0.665164548 | 0.505945253 | 0.709931167 | 0.089420819 |
| ENSMUSG00000034880 | Mrpl34 | 94065 | protein_coding | 409.3700315 | -0.089450532 | 0.101906672 | 0.877769139 | 0.380069015 | 0.60223179 | 0.089450532 |
| ENSMUSG00000102976 | Zc3h11a | 70579 | protein_coding | 111.7819893 | 0.089470755 | 0.110538045 | -0.80941141 | 0.418278541 | 0.638329247 | 0.089470755 |
| ENSMUSG00000002059 | Rab34 | 19376 | protein_coding | 2.946569567 | -0.089472058 | 0.798383738 | 0.112066484 | 0.910770692 | 1 | 0.089472058 |
| ENSMUSG00000071456 | 1110002L01Rik | 100043040 | lncRNA | 16.97612085 | -0.089473638 | 0.261327408 | 0.342381379 | 0.7320639 | 0.862351327 | 0.089473638 |
| ENSMUSG00000043782 | Bicdl2 | 212733 | protein_coding | 274.8778627 | -0.089499341 | 0.089688192 | 0.997894353 | 0.318330591 | 0.539723961 | 0.089499341 |
| ENSMUSG00000027341 | Tmem230 | 70612 | protein_coding | 146.5393624 | 0.089529875 | 0.091759265 | -0.975703922 | 0.329211208 | 0.551602465 | 0.089529875 |
| ENSMUSG00000032913 | Lrig2 | 269473 | protein_coding | 351.5835528 | -0.089539329 | 0.085047399 | 1.052816784 | 0.292424971 | 0.511503652 | 0.089539329 |
| ENSMUSG00000024084 | Qpct | 70536 | protein_coding | 7.144603626 | 0.089550597 | 0.457113634 | -0.195904455 | 0.84468495 | 0.923599766 | 0.089550597 |
| ENSMUSG00000003072 | Atp5d | 66043 | protein_coding | 4002.816721 | -0.089569095 | 0.083806568 | 1.068759843 | 0.285177898 | 0.503574403 | 0.089569095 |
| ENSMUSG00000022833 | Ccdc14 | 239839 | protein_coding | 69.48795703 | -0.089607611 | 0.154795443 | 0.578877582 | 0.562671778 | 0.75067522 | 0.089607611 |
| ENSMUSG00000034218 | Atm | 11920 | protein_coding | 576.2834829 | -0.089615418 | 0.156681992 | 0.571957353 | 0.567350867 | 0.753395965 | 0.089615418 |
| ENSMUSG00000024319 | Vps52 | 224705 | protein_coding | 497.7404191 | 0.089617978 | 0.076773712 | -1.167300307 | 0.243089115 | 0.454166041 | 0.089617978 |
| ENSMUSG00000033446 | Lpar6 | 67168 | protein_coding | 171.2750344 | 0.089635477 | 0.099000041 | -0.905408478 | 0.365249026 | 0.587772599 | 0.089635477 |
| ENSMUSG00000026176 | Ctdsp1 | 227292 | protein_coding | 712.9638435 | -0.089636795 | 0.07126693 | 1.257761412 | 0.208478055 | 0.411572816 | 0.089636795 |
| ENSMUSG00000028696 | Ipp | 16351 | protein_coding | 130.3827786 | -0.089662617 | 0.12295285 | 0.729243912 | 0.465852474 | 0.679601393 | 0.089662617 |
| ENSMUSG00000001247 | Lsr | 54135 | protein_coding | 621.9465722 | 0.089748263 | 0.216910682 | -0.413756771 | 0.679052246 | 0.827951097 | 0.089748263 |
| ENSMUSG00000071648 | Rom1 | 19881 | protein_coding | 32.79686737 | -0.089785096 | 0.283260027 | 0.316970584 | 0.751265923 | 0.875081807 | 0.089785096 |
| ENSMUSG00000020978 | Klhdc2 | 69554 | protein_coding | 295.3317119 | -0.089788555 | 0.095853435 | 0.936727565 | 0.348898713 | 0.570683233 | 0.089788555 |
| ENSMUSG00000029623 | Pdap1 | 231887 | protein_coding | 680.8168822 | 0.089830196 | 0.086248878 | -1.041523063 | 0.297632855 | 0.517311687 | 0.089830196 |
| ENSMUSG00000043323 | Fbrsl1 | 381668 | protein_coding | 368.286246 | -0.089875403 | 0.141233895 | 0.636358598 | 0.524542719 | 0.723107237 | 0.089875403 |
| ENSMUSG00000030034 | Ino80b | 70020 | protein_coding | 172.2200786 | -0.089888508 | 0.105390158 | 0.852911785 | 0.393708226 | 0.616333078 | 0.089888508 |
| ENSMUSG00000052572 | Dlg2 | 23859 | protein_coding | 10.15676131 | -0.089922424 | 0.580675886 | 0.154858203 | 0.876933105 | 0.938919382 | 0.089922424 |

| ensembl_gene_id | external_gene_name | entrezgene_id | gene_biotype | baseMean | log2FoldChange | lfcSE | stat | pvalue | padj | absLog2FC |
|---|---|---|---|---|---|---|---|---|---|---|
| ENSMUSG00000043909 | Trp53bp1 | 27223 | protein_coding | 412.0176718 | 0.089979206 | 0.08505024 | -1.057953588 | 0.290076602 | 0.50901339 | 0.089979206 |
| ENSMUSG00000077394 | Gm24339 | 115487283 | snoRNA | 183.1854907 | -0.089984546 | 0.148666121 | 0.60527944 | 0.54499334 | 0.736964526 | 0.089984546 |
| ENSMUSG00000009035 | Tmem184b | 223693 | protein_coding | 301.7265981 | 0.090010238 | 0.106234451 | -0.847279175 | 0.396839532 | 0.618855346 | 0.090010238 |
| ENSMUSG00000018362 | Kpna2 | 16647 | protein_coding | 1084.955951 | -0.090030363 | 0.144053438 | 0.624978923 | 0.531984891 | 0.728820184 | 0.090030363 |
| ENSMUSG00000045435 | Tmem60 | 212090 | protein_coding | 114.3807168 | 0.090071035 | 0.114682294 | -0.78539618 | 0.43222132 | 0.650656598 | 0.090071035 |
| ENSMUSG00000037525 | Bcdin3d | 75284 | protein_coding | 60.27020649 | -0.090088264 | 0.137795193 | 0.653783789 | 0.513251112 | 0.715139121 | 0.090088264 |
| ENSMUSG00000060657 | Marf1 | 223989 | protein_coding | 930.3898782 | -0.090098275 | 0.106725547 | 0.844205328 | 0.398554678 | 0.620508339 | 0.090098275 |
| ENSMUSG00000077505 | Gm24233 | 115485895 | snoRNA | 13.31865458 | -0.090106975 | 0.506469241 | 0.177912037 | 0.858792057 | 0.930949982 | 0.090106975 |
| ENSMUSG00000045019 | Acer1 | 171168 | protein_coding | 587.555246 | -0.0901324 | 0.145816112 | 0.618123738 | 0.536493775 | 0.732400135 | 0.0901324 |
| ENSMUSG00000079466 | Prdm12 | 381359 | protein_coding | 0.978957204 | -0.090160963 | 2.274192978 | 0.039645256 | 0.968375947 | 1 | 0.090160963 |
| ENSMUSG00000000708 | Kat2b | 18519 | protein_coding | 559.2131352 | -0.090176727 | 0.08986772 | 1.003438464 | 0.315649354 | 0.53655334 | 0.090176727 |
| ENSMUSG00000013663 | Pten | 19211 | protein_coding | 1440.493439 | -0.090188496 | 0.072354975 | 1.246472638 | 0.212590928 | 0.417180016 | 0.090188496 |
| ENSMUSG00000032440 | Tgfbr2 | 21813 | protein_coding | 786.8752464 | -0.090217474 | 0.103563752 | 0.871129835 | 0.383683266 | 0.606155692 | 0.090217474 |
| ENSMUSG00000034297 | Med13 | 327987 | protein_coding | 1207.899547 | 0.090245199 | 0.068081342 | -1.325549655 | 0.184988912 | 0.380389352 | 0.090245199 |
| ENSMUSG00000029179 | Zcchc4 | 78796 | protein_coding | 135.0003293 | 0.090271276 | 0.094115101 | -0.959158254 | 0.337479027 | 0.559438733 | 0.090271276 |
| ENSMUSG00000062373 | Tmem65 | 74868 | protein_coding | 421.1174626 | 0.090287147 | 0.084738793 | -1.065475957 | 0.2866606 | 0.50522539 | 0.090287147 |
| ENSMUSG00000085595 | Gm16090 | 102633478 | lncRNA | 5.94240911 | 0.09030837 | 0.573844958 | -0.157374164 | 0.874949974 | 0.938415991 | 0.09030837 |
| ENSMUSG00000029500 | Pgam5 | 72542 | protein_coding | 361.8155275 | 0.09032613 | 0.105720462 | -0.854386452 | 0.3928909 | 0.615578778 | 0.09032613 |
| ENSMUSG00000043262 | Uevld | 54122 | protein_coding | 236.0878077 | -0.090335706 | 0.115862122 | 0.779682816 | 0.435577596 | 0.653742472 | 0.090335706 |
| ENSMUSG00000021147 | Wdr37 | 207615 | protein_coding | 232.1176158 | -0.090413519 | 0.082060211 | 1.101794868 | 0.270550862 | 0.485955725 | 0.090413519 |
| ENSMUSG00000110012 | Or51a43 | 259125 | protein_coding | 1.367230395 | 0.090433702 | 2.345153977 | -0.038561946 | 0.969239642 | 1 | 0.090433702 |
| ENSMUSG00000034714 | Ttyh2 | 117160 | protein_coding | 328.4724258 | -0.090448272 | 0.147318556 | 0.613963874 | 0.5392392 | 0.733766095 | 0.090448272 |
| ENSMUSG00000040102 | Klhl42 | 232539 | protein_coding | 295.4695803 | -0.090454789 | 0.110689625 | 0.817193019 | 0.413818131 | 0.633873724 | 0.090454789 |
| ENSMUSG00000050608 | Micos10 | 433771 | protein_coding | 543.0706089 | -0.090474771 | 0.098829056 | 0.915467325 | 0.359946349 | 0.582491838 | 0.090474771 |
| ENSMUSG00000024026 | Glo1 | 109801 | protein_coding | 868.3804269 | -0.090515557 | 0.081734662 | 1.107431711 | 0.268107322 | 0.48334534 | 0.090515557 |
| ENSMUSG00000058498 | Rnf207 | 433809 | protein_coding | 3.705894836 | -0.090531864 | 1.073822301 | 0.08430805 | 0.932811512 | 1 | 0.090531864 |
| ENSMUSG00000003279 | Dlgap1 | 224997 | protein_coding | 9.311617682 | 0.090538825 | 0.838683794 | -0.107953469 | 0.914032603 | 0.958005484 | 0.090538825 |
| ENSMUSG00000040888 | Gfer | 11692 | protein_coding | 463.1890007 | 0.090594195 | 0.088184233 | -1.027328713 | 0.304265706 | 0.524504007 | 0.090594195 |
| ENSMUSG00000020156 | Pwwp3a | 68114 | protein_coding | 327.2983054 | -0.0906138 | 0.09362241 | 0.967864428 | 0.333112082 | 0.55548954 | 0.0906138 |
| ENSMUSG00000001774 | Chordc1 | 66917 | protein_coding | 601.1358465 | 0.090616373 | 0.115900739 | -0.78184465 | 0.434305877 | 0.652783039 | 0.090616373 |
| ENSMUSG00000038683 | Pak1ip1 | 68083 | protein_coding | 473.449776 | 0.090632618 | 0.081902982 | -1.106585087 | 0.268473358 | 0.483542311 | 0.090632618 |
| ENSMUSG00000057506 | Bloc1s2 | 73689 | protein_coding | 165.9683531 | -0.090662602 | 0.143535902 | 0.631637108 | 0.527624033 | 0.725856654 | 0.090662602 |
| ENSMUSG00000047514 | Tspyl1 | 22110 | protein_coding | 483.5082896 | 0.090697981 | 0.072056496 | -1.258706516 | 0.208136357 | 0.41124734 | 0.090697981 |
| ENSMUSG00000022003 | Slc25a30 | 67554 | protein_coding | 55.29017026 | -0.09070042 | 0.175496899 | 0.516820641 | 0.60528137 | 0.780311961 | 0.09070042 |
| ENSMUSG00000024359 | Hspa9 | 15526 | protein_coding | 4096.984852 | -0.090750127 | 0.090763694 | 0.999850521 | 0.317382852 | 0.538827241 | 0.090750127 |
| ENSMUSG00000022100 | Xpo7 | 65246 | protein_coding | 747.9754155 | 0.090763885 | 0.0729948 | -1.24342946 | 0.21370962 | 0.418744149 | 0.090763885 |
| ENSMUSG00000032285 | Dnaja4 | 58233 | protein_coding | 39.77389029 | -0.090769039 | 0.224659675 | 0.404029066 | 0.686191345 | 0.832872319 | 0.090769039 |
| ENSMUSG00000005718 | Tfap4 | 83383 | protein_coding | 187.9396536 | -0.090814079 | 0.121333673 | 0.748465591 | 0.454179372 | 0.669812138 | 0.090814079 |
| ENSMUSG00000037902 | Sirpa | 19261 | protein_coding | 31.0083744 | 0.090818599 | 0.28037495 | -0.323918378 | 0.74599983 | 0.872287153 | 0.090818599 |

Supplementary Table 1

| ensembl_gene_id | external_gene_name | entrezgene_id | gene_biotype | baseMean | log2FoldChange | lfcSE | stat | pvalue | padj | absLog2FC |
|---|---|---|---|---|---|---|---|---|---|---|
| ENSMUSG00000025764 | Jade1 | 269424 | protein_coding | 538.5897295 | 0.09081907 | 0.072541791 | -1.251955162 | 0.210586203 | 0.414644158 | 0.09081907 |
| ENSMUSG00000037822 | Smim14 | 68552 | protein_coding | 1559.01755 | 0.090836138 | 0.126647329 | -0.717236908 | 0.47322793 | 0.685335967 | 0.090836138 |
| ENSMUSG00000028394 | Pole3 | 59001 | protein_coding | 306.8486804 | 0.09084653 | 0.123238012 | -0.737163225 | 0.461023094 | 0.674995696 | 0.09084653 |
| ENSMUSG00000114531 | A530065N20Rik | 328263 | lncRNA | 0.761390086 | -0.090873239 | 2.21583847 | 0.041010769 | 0.967287311 | 1 | 0.090873239 |
| ENSMUSG00000068115 | Ninl | 78177 | protein_coding | 24.25192258 | 0.090913909 | 0.226133051 | -0.402037245 | 0.687656616 | 0.833648609 | 0.090913909 |
| ENSMUSG00000046062 | Ppp1r15b | 108954 | protein_coding | 2453.875502 | -0.090927435 | 0.064880889 | 1.401451739 | 0.16107903 | 0.345901869 | 0.090927435 |
| ENSMUSG00000070798 | Psg25 | 114868 | protein_coding | 2.077192107 | -0.090938768 | 1.101029255 | 0.082594325 | 0.934174114 | 1 | 0.090938768 |
| ENSMUSG00000040652 | Oaz2 | 18247 | protein_coding | 232.2889813 | -0.090973852 | 0.127109991 | 0.715709683 | 0.47417063 | 0.686291751 | 0.090973852 |
| ENSMUSG00000029994 | Anxa4 | 11746 | protein_coding | 3760.299753 | -0.091081481 | 0.053875138 | 1.690603212 | 0.090912611 | 0.230406822 | 0.091081481 |
| ENSMUSG00000037685 | Atp8a1 | 11980 | protein_coding | 1415.028919 | 0.091148908 | 0.114345872 | -0.797133344 | 0.425373593 | 0.644974953 | 0.091148908 |
| ENSMUSG00000028437 | Ubap1 | 67123 | protein_coding | 730.8447152 | -0.091158433 | 0.059862768 | 1.522790141 | 0.127811217 | 0.294806545 | 0.091158433 |
| ENSMUSG00000048310 | Pskh1 | 244631 | protein_coding | 352.614403 | -0.091181207 | 0.075592461 | 1.206220918 | 0.22773232 | 0.43633502 | 0.091181207 |
| ENSMUSG00000021131 | Erh | 13877 | protein_coding | 229.2491588 | -0.091196889 | 0.157841496 | 0.577775118 | 0.563415956 | 0.750868206 | 0.091196889 |
| ENSMUSG00000027361 | Gabpb1 | 14391 | protein_coding | 200.2355533 | 0.091200822 | 0.095164058 | -0.95835364 | 0.337884464 | 0.559633454 | 0.091200822 |
| ENSMUSG00000026502 | Desi2 | 78825 | protein_coding | 401.7540063 | 0.091201844 | 0.071735032 | -1.271371067 | 0.203596671 | 0.405906838 | 0.091201844 |
| ENSMUSG00000000420 | Galnt1 | 14423 | protein_coding | 1177.144389 | 0.091246691 | 0.086314966 | -1.057136377 | 0.290449352 | 0.509468423 | 0.091246691 |
| ENSMUSG00000074604 | Mgst2 | 211666 | protein_coding | 976.5929977 | -0.091247846 | 0.072703903 | 1.25506118 | 0.209456551 | 0.412914333 | 0.091247846 |
| ENSMUSG00000057788 | Ddx49 | 234374 | protein_coding | 294.5102007 | 0.091265471 | 0.087659796 | -1.041132602 | 0.297814011 | 0.517476912 | 0.091265471 |
| ENSMUSG00000041594 | Tmtc4 | 70551 | protein_coding | 325.802073 | 0.091282593 | 0.095807281 | -0.95277302 | 0.340705087 | 0.562302999 | 0.091282593 |
| ENSMUSG00000038072 | Galnt11 | 231050 | protein_coding | 236.8636733 | -0.091291434 | 0.088288429 | 1.034013568 | 0.30112982 | 0.521287512 | 0.091291434 |
| ENSMUSG00000048087 | Ahcyl | 11615 | protein_coding | 9.003623966 | -0.091298584 | 0.35099371 | 0.260114587 | 0.794775386 | 0.897921648 | 0.091298584 |
| ENSMUSG00000026192 | Atic | 108147 | protein_coding | 869.1200869 | -0.091298591 | 0.113875957 | 0.801737201 | 0.422704992 | 0.642569972 | 0.091298591 |
| ENSMUSG00000028111 | Ctsk | 13038 | protein_coding | 2.186025295 | 0.091327564 | 0.766117817 | -0.11920825 | 0.905110371 | 1 | 0.091327564 |
| ENSMUSG00000041058 | Wwp1 | 107568 | protein_coding | 930.4823602 | 0.09141366 | 0.068761669 | -1.329427595 | 0.183706939 | 0.378649075 | 0.09141366 |
| ENSMUSG00000019889 | Ptprk | 19272 | protein_coding | 2077.456133 | -0.091447142 | 0.088918487 | 1.028437899 | 0.303743889 | 0.524250269 | 0.091447142 |
| ENSMUSG00000036523 | Greb1 | 268527 | protein_coding | 143.0634574 | -0.091479106 | 0.110155427 | 0.830454826 | 0.406281678 | 0.62720573 | 0.091479106 |
| ENSMUSG00000028982 | Slc25a33 | 70556 | protein_coding | 74.75868366 | -0.091493942 | 0.150264477 | 0.608886036 | 0.54259998 | 0.736156431 | 0.091493942 |
| ENSMUSG00000044018 | Mrpl50 | 28028 | protein_coding | 447.4224522 | 0.091591616 | 0.103669255 | -0.883498351 | 0.376967073 | 0.599156327 | 0.091591616 |
| ENSMUSG00000097162 | 2310010J17Rik | 78329 | lncRNA | 22.5970299 | 0.091604329 | 0.220470521 | -0.415494681 | 0.677779808 | 0.827323214 | 0.091604329 |
| ENSMUSG00000026940 | Ccdc183 | 77058 | protein_coding | 2.158202852 | -0.091623253 | 0.797940101 | 0.114824726 | 0.90858405 | 1 | 0.091623253 |
| ENSMUSG00000032187 | Smarca4 | 20586 | protein_coding | 1864.270814 | 0.091664664 | 0.06450989 | -1.420939699 | 0.15533429 | 0.337697097 | 0.091664664 |
| ENSMUSG00000033075 | Senp1 | 223870 | protein_coding | 356.9392813 | 0.091683102 | 0.074133004 | -1.236737976 | 0.216184375 | 0.421495722 | 0.091683102 |
| ENSMUSG00000031983 | 2310022B05Rik | 69551 | protein_coding | 224.4264115 | -0.091694937 | 0.093661531 | 0.979003181 | 0.327578408 | 0.549759128 | 0.091694937 |
| ENSMUSG00000037475 | Thoc2 | 331401 | protein_coding | 815.4182992 | 0.091772245 | 0.065231564 | -1.406868685 | 0.159466331 | 0.343609993 | 0.091772245 |
| ENSMUSG00000023452 | Pisd | 320951 | protein_coding | 585.7610711 | -0.091793861 | 0.087364871 | 1.050695318 | 0.293398547 | 0.512561987 | 0.091793861 |
| ENSMUSG00000048581 | E130311K13Rik | 329659 | protein_coding | 37.54466638 | 0.091811691 | 0.237170428 | -0.387112726 | 0.698672757 | 0.840340998 | 0.091811691 |
| ENSMUSG00000020267 | Hint1 | 15254 | protein_coding | 3289.977407 | -0.091818325 | 0.095914638 | 0.957292097 | 0.338419843 | 0.560042619 | 0.091818325 |
| ENSMUSG00000085105 | Gm12758 | 100126229 | lncRNA | 4.895105878 | -0.091860709 | 0.622390842 | 0.147593285 | 0.882663755 | 0.941987155 | 0.091860709 |
| ENSMUSG00000026499 | Acbd3 | 170760 | protein_coding | 1130.368925 | 0.091889589 | 0.063146465 | -1.455181834 | 0.145618929 | 0.322785367 | 0.091889589 |

Supplementary Table 1

| ensembl_gene_id | external_gene_name | entrezgene_id | gene_biotype | baseMean | log2FoldChange | lfcSE | stat | pvalue | padj | absLog2FC |
|---|---|---|---|---|---|---|---|---|---|---|
| ENSMUSG00000030265 | Kras | 16653 | protein_coding | 1260.398754 | -0.092026371 | 0.058107507 | 1.583726019 | 0.113256081 | 0.270127107 | 0.092026371 |
| ENSMUSG00000047909 | Ankrd16 | 320816 | protein_coding | 203.0675355 | -0.092043833 | 0.152080416 | 0.605231336 | 0.545025297 | 0.736964526 | 0.092043833 |
| ENSMUSG00000022048 | Dpysl2 | 12934 | protein_coding | 908.7598952 | 0.092049697 | 0.162021885 | -0.568131255 | 0.569945846 | 0.755650645 | 0.092049697 |
| ENSMUSG00000028134 | Ptbp2 | 56195 | protein_coding | 117.1926193 | -0.092135697 | 0.156948627 | 0.587043666 | 0.55717438 | 0.746364189 | 0.092135697 |
| ENSMUSG00000005836 | Gata6 | 14465 | protein_coding | 626.9015689 | -0.092158831 | 0.063452024 | 1.45241752 | 0.146385548 | 0.32396287 | 0.092158831 |
| ENSMUSG00000028690 | Mmachc | 67096 | protein_coding | 164.9900424 | -0.092184279 | 0.121198185 | 0.760607747 | 0.446891391 | 0.664203441 | 0.092184279 |
| ENSMUSG00000027012 | Dync1i2 | 13427 | protein_coding | 1110.268729 | 0.092256529 | 0.064093351 | -1.439408722 | 0.150034755 | 0.329724247 | 0.092256529 |
| ENSMUSG00000047415 | Gpr68 | 238377 | protein_coding | 14.92524598 | -0.092273549 | 0.448083085 | 0.205929552 | 0.836845941 | 0.918494874 | 0.092273549 |
| ENSMUSG00000060708 | Bloc1s4 | 117197 | protein_coding | 87.68975658 | -0.092306725 | 0.148016345 | 0.623625215 | 0.532873747 | 0.72936091 | 0.092306725 |
| ENSMUSG00000057738 | Sptan1 | 20740 | protein_coding | 6224.069403 | -0.092311379 | 0.05088899 | 1.81397544 | 0.069681507 | 0.190422043 | 0.092311379 |
| ENSMUSG00000046010 | Zfp830 | 66983 | protein_coding | 158.9921388 | 0.09231494 | 0.100096392 | -0.922260418 | 0.356392755 | 0.579378227 | 0.09231494 |
| ENSMUSG00000097292 | A230107N01Rik | 320919 | lncRNA | 3.586617157 | 0.092365976 | 0.683781069 | -0.135081213 | 0.892547664 | 1 | 0.092365976 |
| ENSMUSG00000026358 | Rgs1 | 50778 | protein_coding | 56.10332297 | -0.092376724 | 0.265046668 | 0.348530033 | 0.727442163 | 0.859261888 | 0.092376724 |
| ENSMUSG00000021782 | Dlg5 | 71228 | protein_coding | 375.8630646 | 0.092386395 | 0.121040584 | -0.763267924 | 0.445303627 | 0.662536791 | 0.092386395 |
| ENSMUSG00000059013 | Sh2d3c | 27387 | protein_coding | 45.25823103 | -0.092462208 | 0.159526334 | 0.579604671 | 0.562181242 | 0.750196731 | 0.092462208 |
| ENSMUSG00000042446 | Zmym4 | 67785 | protein_coding | 247.137144 | -0.092505817 | 0.079586569 | 1.16232951 | 0.245101644 | 0.456768986 | 0.092505817 |
| ENSMUSG00000034379 | Wdr5b | 69544 | protein_coding | 53.79782624 | -0.092587041 | 0.182973725 | 0.506012768 | 0.612847687 | 0.785880315 | 0.092587041 |
| ENSMUSG00000057841 | Rpl32 | 19951 | protein_coding | 5424.702469 | -0.092603847 | 0.097672678 | 0.948103899 | 0.343076564 | 0.564974708 | 0.092603847 |
| ENSMUSG00000026356 | Dars | 226414 | protein_coding | 715.4848051 | -0.092606183 | 0.097094265 | 0.953776034 | 0.340197023 | 0.561689026 | 0.092606183 |
| ENSMUSG00000030726 | Pold3 | 67967 | protein_coding | 373.2768737 | -0.092667172 | 0.096475954 | 0.960520922 | 0.336793107 | 0.559233675 | 0.092667172 |
| ENSMUSG00000027751 | Supt20 | 56790 | protein_coding | 481.3795144 | 0.092693033 | 0.067509084 | -1.373045333 | 0.169738254 | 0.359143126 | 0.092693033 |
| ENSMUSG00000026116 | Tmem131 | 56030 | protein_coding | 1670.119934 | 0.092719154 | 0.086923984 | -1.066669403 | 0.286121149 | 0.50453867 | 0.092719154 |
| ENSMUSG00000007097 | Atp1a2 | 98660 | protein_coding | 5.598462455 | -0.092819696 | 0.79312944 | 0.117029694 | 0.906836523 | 0.954353007 | 0.092819696 |
| ENSMUSG00000049401 | Ogfr | 72075 | protein_coding | 720.9925613 | -0.092835198 | 0.070051219 | 1.325247421 | 0.185089102 | 0.380500613 | 0.092835198 |
| ENSMUSG00000009356 | Lpo | 76113 | protein_coding | 243.532443 | 0.09284753 | 0.22980771 | -0.404022692 | 0.686196032 | 0.832872319 | 0.09284753 |
| ENSMUSG00000079109 | Pms2 | 18861 | protein_coding | 100.8493115 | 0.092871516 | 0.120637872 | -0.76983715 | 0.441396499 | 0.659125086 | 0.092871516 |
| ENSMUSG00000059325 | Hopx | 74318 | protein_coding | 204.2778739 | 0.092887444 | 0.176081542 | -0.527525164 | 0.597828946 | 0.775716761 | 0.092887444 |
| ENSMUSG00000022297 | Fzd6 | 14368 | protein_coding | 130.0004297 | -0.092931966 | 0.178839735 | 0.519638245 | 0.603315736 | 0.779139367 | 0.092931966 |
| ENSMUSG00000025066 | Sfr1 | 67788 | protein_coding | 394.4284715 | 0.092932261 | 0.087156282 | -1.066271512 | 0.286300923 | 0.504747886 | 0.092932261 |
| ENSMUSG00000028607 | Cpt2 | 12896 | protein_coding | 1138.57578 | -0.092945958 | 0.06978565 | 1.331877809 | 0.182900344 | 0.377599236 | 0.092945958 |
| ENSMUSG00000085683 | Tmem238l | 71576 | protein_coding | 899.7247968 | 0.092950106 | 0.08818128 | -1.054079804 | 0.291846382 | 0.510653862 | 0.092950106 |
| ENSMUSG00000025981 | Coq10b | 67876 | protein_coding | 390.7919251 | -0.092952642 | 0.108896144 | 0.853589837 | 0.393332291 | 0.615978138 | 0.092952642 |
| ENSMUSG00000025470 | Zfp511 | 69752 | protein_coding | 112.0648217 | -0.092978395 | 0.122097996 | 0.761506316 | 0.446354708 | 0.663584952 | 0.092978395 |
| ENSMUSG00000079283 | 2310009B15Rik | 69549 | protein_coding | 74.65546005 | 0.093005324 | 0.115344525 | -0.8063263 | 0.42005473 | 0.63998633 | 0.093005324 |
| ENSMUSG00000015937 | Macroh2a1 | 26914 | protein_coding | 1279.524277 | 0.093015369 | 0.053052409 | -1.75327323 | 0.079555118 | 0.208999876 | 0.093015369 |
| ENSMUSG00000030842 | Lamtor1 | 66508 | protein_coding | 758.8787466 | -0.093060685 | 0.070432103 | 1.321282222 | 0.186407279 | 0.3822113 | 0.093060685 |
| ENSMUSG00000028782 | Adgrb2 | 230775 | protein_coding | 4.879714499 | -0.093094372 | 1.30857393 | 0.071141851 | 0.94328486 | 0.972595566 | 0.093094372 |
| ENSMUSG00000005107 | Slc2a9 | 117591 | protein_coding | 83.5984349 | 0.093112136 | 0.166038714 | -0.560785695 | 0.57494364 | 0.760164897 | 0.093112136 |
| ENSMUSG00000038827 | Abitram | 230234 | protein_coding | 101.6828616 | 0.093116051 | 0.123562813 | -0.753592838 | 0.451093746 | 0.667666133 | 0.093116051 |

| ensembl_gene_id | external_gene_name | entrezgene_id | gene_biotype | baseMean | log2FoldChange | lfcSE | stat | pvalue | padj | absLog2FC |
|---|---|---|---|---|---|---|---|---|---|---|
| ENSMUSG00000052544 | St6galnac3 | 20447 | protein_coding | 2.121003391 | 0.093176513 | 1.033264382 | -0.090176837 | 0.928146691 | 1 | 0.093176513 |
| ENSMUSG00000052593 | Adam17 | 11491 | protein_coding | 558.0451558 | 0.093181482 | 0.085175514 | -1.093993766 | 0.273957712 | 0.490474594 | 0.093181482 |
| ENSMUSG00000025060 | Slk | 20874 | protein_coding | 2173.74461 | -0.09325568 | 0.0724723 | 1.286776891 | 0.198172064 | 0.398418975 | 0.09325568 |
| ENSMUSG00000023079 | Gtf2ird1 | 57080 | protein_coding | 900.9864121 | 0.093257303 | 0.099925819 | -0.933265328 | 0.350682992 | 0.572816907 | 0.093257303 |
| ENSMUSG00000078762 | Haus5 | 71909 | protein_coding | 12.40532544 | -0.093270804 | 0.384583813 | 0.242523998 | 0.808374164 | 0.905485545 | 0.093270804 |
| ENSMUSG00000020875 | Hoxb9 | 15417 | protein_coding | 499.254042 | 0.093278128 | 0.080460467 | -1.159303836 | 0.246332359 | 0.458272537 | 0.093278128 |
| ENSMUSG00000067194 | Eif1ax | 66235 | protein_coding | 815.8694392 | -0.093296601 | 0.125933041 | 0.740842911 | 0.458788693 | 0.673341315 | 0.093296601 |
| ENSMUSG00000024052 | Lpin2 | 64898 | protein_coding | 724.2800779 | -0.093349617 | 0.232522968 | 0.401464069 | 0.688078487 | 0.833786242 | 0.093349617 |
| ENSMUSG00000071078 | Nr2c2ap | 75692 | protein_coding | 27.36724317 | -0.093359538 | 0.212870721 | 0.438573882 | 0.660970326 | 0.816884691 | 0.093359538 |
| ENSMUSG00000060227 | Golm2 | 319996 | protein_coding | 233.903554 | -0.093363359 | 0.198385357 | 0.470616182 | 0.63791485 | 0.802698942 | 0.093363359 |
| ENSMUSG00000036435 | Exoc1 | 69940 | protein_coding | 441.0878136 | 0.093364045 | 0.07343903 | -1.27131371 | 0.203617067 | 0.405906838 | 0.093364045 |
| ENSMUSG00000050890 | Pdik1l | 230809 | protein_coding | 363.3990395 | 0.09340545 | 0.097603786 | -0.95698593 | 0.338574357 | 0.560242177 | 0.09340545 |
| ENSMUSG00000039428 | Tmem135 | 72759 | protein_coding | 552.8010373 | 0.09340854 | 0.104851186 | -0.890867748 | 0.373000125 | 0.595257447 | 0.09340854 |
| ENSMUSG00000003360 | Ddx23 | 74351 | protein_coding | 966.0473989 | 0.093433642 | 0.044532193 | -2.098114574 | 0.035895025 | 0.115594862 | 0.093433642 |
| ENSMUSG00000089900 | Tbc1d22bos | 74092 | lncRNA | 4.248032864 | 0.093466233 | 0.5417496 | -0.17252663 | 0.863023527 | 1 | 0.093466233 |
| ENSMUSG00000037992 | Rara | 19401 | protein_coding | 302.0788982 | -0.093472237 | 0.1539054 | 0.607335654 | 0.543628181 | 0.736522749 | 0.093472237 |
| ENSMUSG00000026260 | Ndufa10 | 67273 | protein_coding | 2077.738484 | -0.0935175 | 0.08311199 | 1.125198673 | 0.260504855 | 0.474068167 | 0.0935175 |
| ENSMUSG00000026603 | Smyd2 | 226830 | protein_coding | 193.3625396 | -0.093523672 | 0.104899856 | 0.891551957 | 0.372633131 | 0.594901732 | 0.093523672 |
| ENSMUSG00000019966 | Kitl | 17311 | protein_coding | 1467.224304 | -0.093557679 | 0.118164919 | 0.791755111 | 0.428503481 | 0.647539027 | 0.093557679 |
| ENSMUSG00000030811 | Fbxl19 | 233902 | protein_coding | 606.5415955 | 0.093584246 | 0.074934938 | -1.248873328 | 0.211711408 | 0.416065952 | 0.093584246 |
| ENSMUSG00000006585 | Cdt1 | 67177 | protein_coding | 199.7967066 | -0.093601819 | 0.140547484 | 0.665980041 | 0.505423858 | 0.709429427 | 0.093601819 |
| ENSMUSG00000026240 | Cops7b | 26895 | protein_coding | 145.7033779 | 0.09370928 | 0.096153229 | -0.974582772 | 0.329767263 | 0.552030063 | 0.09370928 |
| ENSMUSG00000038535 | Zfp280d | 235469 | protein_coding | 244.4694919 | -0.093720495 | 0.096552804 | 0.970665694 | 0.33171478 | 0.554166705 | 0.093720495 |
| ENSMUSG00000066637 | Ttc32 | 75516 | protein_coding | 89.60241738 | 0.093771418 | 0.131548139 | -0.71282968 | 0.475951157 | 0.687930892 | 0.093771418 |
| ENSMUSG00000079560 | Hoxa3 | 15400 | protein_coding | 76.1928106 | -0.093797113 | 0.16025992 | 0.585281165 | 0.558358678 | 0.747162726 | 0.093797113 |
| ENSMUSG00000074766 | Ism1 | 319909 | protein_coding | 21.46593545 | 0.093813026 | 0.751873405 | -0.124772369 | 0.900703763 | 0.951219667 | 0.093813026 |
| ENSMUSG00000063787 | Chchd1 | 66121 | protein_coding | 258.7221668 | -0.093852755 | 0.092942914 | 1.009789246 | 0.312596274 | 0.533617414 | 0.093852755 |
| ENSMUSG00000036442 | Thap11 | 59016 | protein_coding | 282.5939577 | -0.093899262 | 0.095465589 | 0.983592756 | 0.325315788 | 0.547073822 | 0.093899262 |
| ENSMUSG00000031851 | Ntpcr | 66566 | protein_coding | 156.442917 | -0.093915214 | 0.111203024 | 0.844538306 | 0.398368668 | 0.620277218 | 0.093915214 |
| ENSMUSG00000028328 | Tmod1 | 21916 | protein_coding | 2.319459546 | -0.093948985 | 1.044177952 | 0.089974112 | 0.928307787 | 1 | 0.093948985 |
| ENSMUSG00000097487 | Ptges3l | 73635 | protein_coding | 19.66054709 | -0.09395027 | 0.230389846 | 0.40778824 | 0.683429148 | 0.830902522 | 0.09395027 |
| ENSMUSG00000057342 | Sphk2 | 56632 | protein_coding | 1009.191439 | -0.093965992 | 0.098371545 | 0.955215172 | 0.339468895 | 0.560935494 | 0.093965992 |
| ENSMUSG00000047248 | C2cd3 | 277939 | protein_coding | 392.0152112 | 0.093967873 | 0.090750565 | -1.035452204 | 0.300457776 | 0.520523758 | 0.093967873 |
| ENSMUSG00000040370 | Etfrf1 | 67636 | protein_coding | 252.453004 | 0.093979141 | 0.098458888 | -0.954501348 | 0.339829927 | 0.561307404 | 0.093979141 |
| ENSMUSG00000046034 | Otulin | 432940 | protein_coding | 211.2440725 | -0.09399677 | 0.097893978 | 0.960189501 | 0.33695985 | 0.559251755 | 0.09399677 |
| ENSMUSG00000038848 | Ythdf1 | 228994 | protein_coding | 448.1380149 | 0.094055118 | 0.068718274 | -1.368706052 | 0.171091178 | 0.361027307 | 0.094055118 |
| ENSMUSG00000071072 | Ptges3 | 56351 | protein_coding | 974.5639108 | -0.094095038 | 0.122566506 | 0.767705969 | 0.442661889 | 0.660473296 | 0.094095038 |
| ENSMUSG00000071302 | 2610044O15Rik8 | 72139 | protein_coding | 93.91449155 | -0.094100813 | 0.155314056 | 0.605874414 | 0.544598151 | 0.736796542 | 0.094100813 |
| ENSMUSG00000031976 | Urb2 | 382038 | protein_coding | 283.1867295 | -0.094116681 | 0.113460483 | 0.829510664 | 0.406815505 | 0.627629974 | 0.094116681 |

| ensembl_gene_id | external_gene_name | entrezgene_id | gene_biotype | baseMean | log2FoldChange | lfcSE | stat | pvalue | padj | absLog2FC |
|---|---|---|---|---|---|---|---|---|---|---|
| ENSMUSG00000018372 | Cep95 | 320162 | protein_coding | 142.0176079 | -0.094126294 | 0.099507632 | 0.945920348 | 0.344189214 | 0.566016798 | 0.094126294 |
| ENSMUSG00000020734 | Grin2c | 14813 | protein_coding | 2.565877376 | -0.094153306 | 1.06226862 | 0.088634178 | 0.929372644 | 1 | 0.094153306 |
| ENSMUSG00000028901 | Gmeb1 | 56809 | protein_coding | 216.8798913 | -0.094200846 | 0.078568783 | 1.198960226 | 0.230543411 | 0.43912271 | 0.094200846 |
| ENSMUSG00000046201 | Scaf8 | 106583 | protein_coding | 661.0821937 | -0.094216556 | 0.052754668 | 1.785937809 | 0.074109328 | 0.198998609 | 0.094216556 |
| ENSMUSG00000026896 | Ifih1 | 71586 | protein_coding | 896.1111371 | -0.094228756 | 0.107343093 | 0.877827844 | 0.380037152 | 0.60223179 | 0.094228756 |
| ENSMUSG00000074890 | Lcmt2 | 329504 | protein_coding | 104.8531656 | -0.094234284 | 0.130644159 | 0.721304991 | 0.470721887 | 0.683906563 | 0.094234284 |
| ENSMUSG00000031483 | Erlin2 | 244373 | protein_coding | 1600.535497 | -0.094255673 | 0.078895972 | 1.194682962 | 0.232210922 | 0.441179379 | 0.094255673 |
| ENSMUSG00000030652 | Coq7 | 12850 | protein_coding | 259.5809379 | -0.094261788 | 0.135536441 | 0.695471916 | 0.486759605 | 0.695624496 | 0.094261788 |
| ENSMUSG00000089290 | Gm23207 | 115488981 | snRNA | 1.198806553 | 0.094273203 | 1.507247034 | -0.062546617 | 0.950127539 | 1 | 0.094273203 |
| ENSMUSG00000050944 | Efcab5 | 319634 | protein_coding | 54.79870693 | 0.094335114 | 0.163782904 | -0.575976558 | 0.564631027 | 0.751588072 | 0.094335114 |
| ENSMUSG00000021120 | Pigh | 110417 | protein_coding | 127.3475748 | -0.094360595 | 0.111690934 | 0.844836652 | 0.398202047 | 0.62019321 | 0.094360595 |
| ENSMUSG00000039458 | Mtmr12 | 268783 | protein_coding | 460.6496001 | -0.094366858 | 0.059590661 | 1.583584686 | 0.113288261 | 0.270127107 | 0.094366858 |
| ENSMUSG00000039740 | Alg2 | 56737 | protein_coding | 210.7378678 | 0.09442524 | 0.112723623 | -0.837670379 | 0.402215854 | 0.623797203 | 0.09442524 |
| ENSMUSG00000001524 | Gtf2h4 | 14885 | protein_coding | 171.4065912 | 0.094434992 | 0.119130597 | -0.792701413 | 0.427951807 | 0.647057325 | 0.094434992 |
| ENSMUSG00000030200 | Bcl2l14 | 66813 | protein_coding | 1019.285578 | 0.094493735 | 0.091319699 | -1.034757409 | 0.300782218 | 0.520880615 | 0.094493735 |
| ENSMUSG00000015647 | Lama5 | 16776 | protein_coding | 220.6118158 | 0.094496581 | 0.263514974 | -0.358600422 | 0.719894031 | 0.855153117 | 0.094496581 |
| ENSMUSG00000020079 | Supv3l1 | 338359 | protein_coding | 299.6652534 | -0.094522555 | 0.101894275 | 0.927653245 | 0.353587469 | 0.575966015 | 0.094522555 |
| ENSMUSG00000020650 | Bcap29 | 12033 | protein_coding | 80.12767736 | -0.094526413 | 0.14406658 | 0.656130054 | 0.511740448 | 0.714354347 | 0.094526413 |
| ENSMUSG00000033707 | Lrrc24 | 378937 | protein_coding | 2.749347438 | -0.094536151 | 0.87339646 | 0.108239677 | 0.913805572 | 1 | 0.094536151 |
| ENSMUSG00000015474 | Ppt2 | 54397 | protein_coding | 124.7190533 | -0.09454113 | 0.12628686 | 0.748622064 | 0.454085029 | 0.669812138 | 0.09454113 |
| ENSMUSG00000045631 | Tmprss12 | 75002 | protein_coding | 1.606814593 | 0.094644394 | 0.885553427 | -0.106875984 | 0.914887366 | 1 | 0.094644394 |
| ENSMUSG00000086359 | 9630013K17Rik | 78883 | lncRNA | 4.45759765 | -0.094722778 | 0.602462963 | 0.157225895 | 0.875066821 | 0.938415991 | 0.094722778 |
| ENSMUSG00000037703 | Lzts3 | 241638 | protein_coding | 1213.483171 | 0.094725775 | 0.066025833 | -1.434677475 | 0.151379026 | 0.331436869 | 0.094725775 |
| ENSMUSG00000020801 | Med31 | 67279 | protein_coding | 59.58406693 | 0.094729703 | 0.147566252 | -0.641946936 | 0.520907637 | 0.720614094 | 0.094729703 |
| ENSMUSG00000027291 | Vps39 | 269338 | protein_coding | 380.6155154 | -0.094756948 | 0.076156203 | 1.244244654 | 0.213409534 | 0.418308017 | 0.094756948 |
| ENSMUSG00000039050 | Osbpl2 | 228983 | protein_coding | 1095.904788 | -0.094768403 | 0.105747138 | 0.896179363 | 0.370156976 | 0.592724939 | 0.094768403 |
| ENSMUSG00000042810 | Krba1 | 77827 | protein_coding | 326.9760194 | 0.094772853 | 0.121873209 | -0.77763484 | 0.436784316 | 0.654986223 | 0.094772853 |
| ENSMUSG00000036990 | Otud4 | 73945 | protein_coding | 1042.666477 | -0.094788875 | 0.086992724 | 1.089618425 | 0.275881263 | 0.492718348 | 0.094788875 |
| ENSMUSG00000019878 | Hsf2 | 15500 | protein_coding | 87.67201104 | -0.094824152 | 0.133285458 | 0.711436596 | 0.476813728 | 0.688415973 | 0.094824152 |
| ENSMUSG00000028453 | Fancg | 60534 | protein_coding | 102.4198065 | 0.094853144 | 0.119548445 | -0.793428505 | 0.427528209 | 0.647057325 | 0.094853144 |
| ENSMUSG00000029920 | Smarcad1 | 13990 | protein_coding | 357.4596729 | -0.094916435 | 0.085252347 | 1.113358622 | 0.265554439 | 0.480535393 | 0.094916435 |
| ENSMUSG00000032615 | Nt5m | 103850 | protein_coding | 146.5455911 | -0.094950132 | 0.10028333 | 0.946818697 | 0.343731172 | 0.565545141 | 0.094950132 |
| ENSMUSG00000025078 | Nhlrc2 | 66866 | protein_coding | 1005.867775 | -0.094979856 | 0.073495954 | 1.292314068 | 0.196248402 | 0.395804361 | 0.094979856 |
| ENSMUSG00000085492 | Trmt61b | 68789 | lncRNA | 59.4461778 | 0.094988042 | 0.132872523 | -0.714881002 | 0.474682577 | 0.686659782 | 0.094988042 |
| ENSMUSG00000022304 | Dpys | 64705 | protein_coding | 1.987987616 | 0.094995913 | 1.337272027 | -0.07103709 | 0.943368237 | 1 | 0.094995913 |
| ENSMUSG00000001518 | Itfg2 | 101142 | protein_coding | 135.5512881 | 0.095096243 | 0.107848678 | -0.881756221 | 0.377908651 | 0.600075333 | 0.095096243 |
| ENSMUSG00000047003 | Zfp41 | 22701 | protein_coding | 80.52509336 | 0.095096447 | 0.198729502 | -0.478522043 | 0.632278689 | 0.799512805 | 0.095096447 |
| ENSMUSG00000041220 | Elovl6 | 170439 | protein_coding | 1115.9085 | -0.095098631 | 0.099314342 | 0.957551834 | 0.338288797 | 0.55988186 | 0.095098631 |
| ENSMUSG00000044700 | Tmem201 | 230917 | protein_coding | 182.8831733 | 0.095151444 | 0.095450409 | -0.99686785 | 0.318828659 | 0.540266681 | 0.095151444 |

| ensembl_gene_id | external_gene_name | entrezgene_id | gene_biotype | baseMean | log2FoldChange | lfcSE | stat | pvalue | padj | absLog2FC |
|---|---|---|---|---|---|---|---|---|---|---|
| ENSMUSG00000086728 | Man2c1os | 76794 | lncRNA | 12.83534159 | -0.095283461 | 0.298970188 | 0.318705558 | 0.7499498 | 0.874376659 | 0.095283461 |
| ENSMUSG00000022437 | Samm50 | 68653 | protein_coding | 1575.537244 | 0.095306404 | 0.074729334 | -1.275354648 | 0.202183743 | 0.404116133 | 0.095306404 |
| ENSMUSG00000030609 | Aen | 68048 | protein_coding | 222.1027474 | -0.095308512 | 0.152126056 | 0.626510108 | 0.530980409 | 0.728048657 | 0.095308512 |
| ENSMUSG00000031309 | Rps6ka3 | 110651 | protein_coding | 1731.852806 | -0.09531119 | 0.115035119 | 0.828539934 | 0.407364789 | 0.628183969 | 0.09531119 |
| ENSMUSG00000029482 | Aacs | 78894 | protein_coding | 1255.705574 | -0.09534827 | 0.135475258 | 0.703805786 | 0.48155373 | 0.692354022 | 0.09534827 |
| ENSMUSG00000024073 | Birc6 | 12211 | protein_coding | 3018.825341 | 0.09535494 | 0.07209273 | -1.322670676 | 0.185944919 | 0.38145272 | 0.09535494 |
| ENSMUSG00000038112 | AW551984 | 244810 | protein_coding | 19.94353953 | -0.095396961 | 0.263617273 | 0.361876746 | 0.717444138 | 0.853563561 | 0.095396961 |
| ENSMUSG00000002844 | Adprh | 11544 | protein_coding | 310.8894472 | 0.095411885 | 0.091198856 | -1.046196082 | 0.295470511 | 0.514582498 | 0.095411885 |
| ENSMUSG00000045854 | Lyrm2 | 108755 | protein_coding | 115.6380785 | -0.095429241 | 0.135025459 | 0.706749983 | 0.479721862 | 0.690909666 | 0.095429241 |
| ENSMUSG00000027165 | Iftap | 68170 | protein_coding | 55.17319935 | -0.095456102 | 0.173450188 | 0.550337263 | 0.58208807 | 0.765030035 | 0.095456102 |
| ENSMUSG00000022141 | Nipbl | 71175 | protein_coding | 1174.539931 | -0.095466018 | 0.086796819 | 1.099879235 | 0.271384743 | 0.48708267 | 0.095466018 |
| ENSMUSG00000037958 | Nsrp1 | 237859 | protein_coding | 235.9718144 | 0.095488452 | 0.094782942 | -1.007443425 | 0.313721732 | 0.534985614 | 0.095488452 |
| ENSMUSG00000097357 | Gm16793 | 100504714 | lncRNA | 3.801329604 | -0.095504279 | 0.640101629 | 0.149201744 | 0.881394444 | 1 | 0.095504279 |
| ENSMUSG00000030061 | Uba3 | 22200 | protein_coding | 579.4614206 | -0.095504751 | 0.06269104 | 1.523419462 | 0.127653796 | 0.294670784 | 0.095504751 |
| ENSMUSG00000043421 | Hilpda | 69573 | protein_coding | 89.08040033 | -0.095511281 | 0.110100634 | 0.867490747 | 0.38567318 | 0.607820554 | 0.095511281 |
| ENSMUSG00000010067 | Rassf1 | 56289 | protein_coding | 198.7444279 | 0.095527919 | 0.142457249 | -0.670572536 | 0.502492884 | 0.707250275 | 0.095527919 |
| ENSMUSG00000024426 | Atat1 | 73242 | protein_coding | 51.74323032 | -0.095569594 | 0.204524244 | 0.467277579 | 0.640301296 | 0.803678477 | 0.095569594 |
| ENSMUSG00000001507 | Itga3 | 16400 | protein_coding | 1774.84768 | 0.095588798 | 0.07835121 | -1.220004106 | 0.222463318 | 0.429533455 | 0.095588798 |
| ENSMUSG00000002968 | Med25 | 75613 | protein_coding | 862.2584561 | 0.095599878 | 0.073801827 | -1.295359237 | 0.195196333 | 0.394331932 | 0.095599878 |
| ENSMUSG00000063235 | Ptpmt1 | 66461 | protein_coding | 292.5469103 | -0.095626877 | 0.091500146 | 1.04510081 | 0.29597638 | 0.515205432 | 0.095626877 |
| ENSMUSG00000046806 | Cyren | 78412 | protein_coding | 23.53843467 | 0.095670293 | 0.237013145 | -0.403649736 | 0.686470306 | 0.833066625 | 0.095670293 |
| ENSMUSG00000028295 | Smim8 | 66291 | protein_coding | 139.1077029 | 0.095670924 | 0.133044861 | -0.719087709 | 0.47208688 | 0.684706793 | 0.095670924 |
| ENSMUSG00000027411 | Vps16 | 80743 | protein_coding | 418.046946 | -0.095694264 | 0.06148658 | 1.556343902 | 0.119626337 | 0.281411586 | 0.095694264 |
| ENSMUSG00000033790 | Tubgcp5 | 233276 | protein_coding | 152.6557495 | 0.095804183 | 0.108051768 | -0.886650769 | 0.375266951 | 0.597546855 | 0.095804183 |
| ENSMUSG00000015357 | Clpx | 270166 | protein_coding | 603.6437515 | -0.095831487 | 0.067953853 | 1.410243624 | 0.158467759 | 0.342262172 | 0.095831487 |
| ENSMUSG00000028647 | Mycbp | 56309 | protein_coding | 177.5250763 | -0.09584154 | 0.099123047 | 0.966894607 | 0.333596723 | 0.555969435 | 0.09584154 |
| ENSMUSG00000090326 | Dthd1 | 100322896 | protein_coding | 3.279251972 | 0.095876308 | 0.589041374 | -0.162766679 | 0.870702144 | 1 | 0.095876308 |
| ENSMUSG00000035274 | Tpbg | 21983 | protein_coding | 45.49741922 | -0.095879838 | 0.179101948 | 0.535336655 | 0.592417084 | 0.772328749 | 0.095879838 |
| ENSMUSG00000048899 | Rimkla | 194237 | protein_coding | 24.89882453 | 0.095920559 | 0.344082041 | -0.278772349 | 0.780419536 | 0.890186044 | 0.095920559 |
| ENSMUSG00000028086 | Fbxw7 | 50754 | protein_coding | 247.4532085 | -0.095984172 | 0.073576858 | 1.304542949 | 0.192048526 | 0.389807821 | 0.095984172 |
| ENSMUSG00000017734 | Dbndd2 | 52840 | protein_coding | 36.28633515 | -0.095993311 | 0.203263947 | 0.47225941 | 0.636741639 | 0.801948109 | 0.095993311 |
| ENSMUSG00000028751 | Pla2g2e | 26970 | protein_coding | 2.279100459 | -0.096019963 | 1.30839098 | 0.073387821 | 0.941497509 | 1 | 0.096019963 |
| ENSMUSG00000028089 | Chd1l | 68058 | protein_coding | 213.4654384 | -0.096070814 | 0.11345236 | 0.846794327 | 0.397109771 | 0.619075876 | 0.096070814 |
| ENSMUSG00000082315 | Gm16523 | 100042584 | transcribed_processed_pseudogene | 3.656768762 | -0.09608863 | 0.77732303 | 0.123614798 | 0.901620275 | 1 | 0.09608863 |
| ENSMUSG00000001467 | Cyp51 | 13121 | protein_coding | 1493.539929 | 0.096155637 | 0.13556247 | -0.709308679 | 0.478132944 | 0.689777883 | 0.096155637 |
| ENSMUSG00000020962 | Gtf2a1 | 83602 | protein_coding | 444.1433746 | -0.096171082 | 0.086450088 | 1.112446313 | 0.265946301 | 0.480980968 | 0.096171082 |
| ENSMUSG00000038056 | Kmt2c | 231051 | protein_coding | 1692.979842 | 0.096185689 | 0.139230925 | -0.690835667 | 0.489668819 | 0.697463418 | 0.096185689 |
| ENSMUSG00000073926 | Or52d3 | 57250 | protein_coding | 3.352536062 | 0.096345071 | 0.711279421 | -0.135453197 | 0.892253567 | 1 | 0.096345071 |
| ENSMUSG00000033454 | Zbtb1 | 268564 | protein_coding | 318.1200781 | 0.096347787 | 0.099401122 | -0.969282688 | 0.332404164 | 0.554813258 | 0.096347787 |

| ensembl_gene_id | external_gene_name | entrezgene_id | gene_biotype | baseMean | log2FoldChange | lfcSE | stat | pvalue | padj | absLog2FC |
|---|---|---|---|---|---|---|---|---|---|---|
| ENSMUSG00000022956 | Atp5o | 28080 | protein_coding | 2770.003752 | -0.096354634 | 0.08696735 | 1.107940323 | 0.26788759 | 0.483116267 | 0.096354634 |
| ENSMUSG00000037553 | Zdhhc18 | 503610 | protein_coding | 296.3816779 | 0.096372468 | 0.102697495 | -0.938411086 | 0.348033191 | 0.569954395 | 0.096372468 |
| ENSMUSG00000022680 | Pdxdc1 | 94184 | protein_coding | 7144.321256 | -0.096390217 | 0.138250513 | 0.697214171 | 0.485668772 | 0.695380179 | 0.096390217 |
| ENSMUSG00000090069 | E430024P14Rik | 100040822 | lncRNA | 12.23048543 | 0.096412519 | 0.313355241 | -0.307678015 | 0.75832735 | 0.878808271 | 0.096412519 |
| ENSMUSG00000044501 | Zfp758 | 224598 | protein_coding | 160.5172214 | 0.096438162 | 0.146674356 | -0.657498449 | 0.510860468 | 0.713811425 | 0.096438162 |
| ENSMUSG00000000563 | Atp5pb | 11950 | protein_coding | 3963.396068 | -0.096477089 | 0.08231809 | 1.172003487 | 0.241195656 | 0.451804114 | 0.096477089 |
| ENSMUSG00000029804 | Herc3 | 73998 | protein_coding | 233.9247998 | -0.096480381 | 0.097919291 | 0.985305142 | 0.324474208 | 0.546493327 | 0.096480381 |
| ENSMUSG00000035314 | Gdpd5 | 233552 | protein_coding | 117.5064908 | -0.096488854 | 0.197762327 | 0.487903109 | 0.625618472 | 0.794916857 | 0.096488854 |
| ENSMUSG00000022515 | Anks3 | 72615 | protein_coding | 259.4382235 | 0.096536076 | 0.084801416 | -1.138378121 | 0.254962626 | 0.467516411 | 0.096536076 |
| ENSMUSG00000074457 | S100a16 | 67860 | protein_coding | 1428.508977 | -0.096565076 | 0.149766047 | 0.644772816 | 0.519074421 | 0.719334577 | 0.096565076 |
| ENSMUSG00000000282 | Mnt | 17428 | protein_coding | 192.266028 | 0.096594361 | 0.126461004 | -0.763827244 | 0.444970199 | 0.662372535 | 0.096594361 |
| ENSMUSG00000020515 | Cnot8 | 69125 | protein_coding | 417.2630813 | -0.096635826 | 0.079075373 | 1.222072337 | 0.221680273 | 0.42856802 | 0.096635826 |
| ENSMUSG00000033728 | Lrrc14 | 223664 | protein_coding | 203.4043011 | 0.096638218 | 0.105323179 | -0.917539895 | 0.358859797 | 0.581581822 | 0.096638218 |
| ENSMUSG00000039081 | Zfp503 | 218820 | protein_coding | 24.52005863 | -0.096642767 | 0.287282993 | 0.336402674 | 0.736567233 | 0.866173274 | 0.096642767 |
| ENSMUSG00000028530 | Jak1 | 16451 | protein_coding | 1421.880104 | -0.09668242 | 0.077131855 | 1.253469405 | 0.210034927 | 0.413854764 | 0.09668242 |
| ENSMUSG00000031511 | Arhgef7 | 54126 | protein_coding | 482.7505443 | 0.096690459 | 0.055493767 | -1.742366131 | 0.081444398 | 0.212471132 | 0.096690459 |
| ENSMUSG00000028048 | Gba | 14466 | protein_coding | 666.3890191 | 0.096696508 | 0.075591146 | -1.27920416 | 0.200825171 | 0.402137369 | 0.096696508 |
| ENSMUSG00000073563 | Csnk1g3 | 70425 | protein_coding | 565.5615583 | -0.096742499 | 0.071273486 | 1.357342035 | 0.174672557 | 0.365932262 | 0.096742499 |
| ENSMUSG00000074797 | Itpa | 16434 | protein_coding | 369.3641328 | -0.096768417 | 0.130457333 | 0.741762954 | 0.458230969 | 0.672761934 | 0.096768417 |
| ENSMUSG00000032002 | Dcun1d5 | 76863 | protein_coding | 401.9225579 | 0.096775869 | 0.071888572 | -1.346192675 | 0.178240387 | 0.37057407 | 0.096775869 |
| ENSMUSG00000023048 | Prr13 | 66151 | protein_coding | 1560.919312 | -0.096780977 | 0.105422509 | 0.91802954 | 0.358603401 | 0.581328932 | 0.096780977 |
| ENSMUSG00000008035 | Mid1ip1 | 68041 | protein_coding | 816.4118149 | 0.096857153 | 0.110203353 | -0.878894794 | 0.379458321 | 0.601768559 | 0.096857153 |
| ENSMUSG00000026254 | Eif4e2 | 26987 | protein_coding | 798.0777801 | -0.096864931 | 0.076028443 | 1.274061755 | 0.202641531 | 0.404694914 | 0.096864931 |
| ENSMUSG00000027610 | Gss | 14854 | protein_coding | 1873.027529 | -0.096879107 | 0.079123185 | 1.224408588 | 0.220798131 | 0.427312825 | 0.096879107 |
| ENSMUSG00000028017 | Egf | 13645 | protein_coding | 84.14889033 | -0.096909638 | 0.216051279 | 0.448549245 | 0.653756854 | 0.812167358 | 0.096909638 |
| ENSMUSG00000051978 | Erich1 | 234086 | protein_coding | 88.4623656 | -0.096919013 | 0.12511101 | 0.774664137 | 0.438538148 | 0.656764359 | 0.096919013 |
| ENSMUSG00000072582 | Ptrh2 | 217057 | protein_coding | 163.9294927 | 0.096932877 | 0.120418911 | -0.804963912 | 0.420840505 | 0.640714813 | 0.096932877 |
| ENSMUSG00000038227 | Hoxa9 | 15405 | protein_coding | 306.3192661 | -0.096970766 | 0.101481211 | 0.955553884 | 0.339297669 | 0.560708667 | 0.096970766 |
| ENSMUSG00000035266 | Helq | 191578 | protein_coding | 96.99089434 | -0.096977854 | 0.125965548 | 0.769876012 | 0.441373444 | 0.659125086 | 0.096977854 |
| ENSMUSG00000068823 | Csde1 | 229663 | protein_coding | 4033.627671 | -0.0969835 | 0.088375014 | 1.097408592 | 0.272462819 | 0.488337435 | 0.0969835 |
| ENSMUSG00000030924 | Rexo5 | 434234 | protein_coding | 74.84881116 | -0.097022951 | 0.148171975 | 0.654799606 | 0.512596784 | 0.714893745 | 0.097022951 |
| ENSMUSG00000097585 | E230029C05Rik | 319711 | lncRNA | 3.0196279 | -0.097033289 | 0.685583262 | 0.141533924 | 0.887448162 | 1 | 0.097033289 |
| ENSMUSG00000020812 | Snhg16 | 66293 | lncRNA | 65.36258322 | -0.097119863 | 0.129935563 | 0.747446352 | 0.454794175 | 0.670218722 | 0.097119863 |
| ENSMUSG00000004268 | Emg1 | 14791 | protein_coding | 664.1098304 | 0.097154246 | 0.099634548 | -0.975105997 | 0.329507684 | 0.551855356 | 0.097154246 |
| ENSMUSG00000023118 | Sympk | 68188 | protein_coding | 1257.885784 | 0.097160802 | 0.055382998 | -1.754343489 | 0.079371668 | 0.208654479 | 0.097160802 |
| ENSMUSG00000038495 | Otud7b | 229603 | protein_coding | 561.1072242 | 0.097195866 | 0.08388111 | -1.158733666 | 0.246564764 | 0.458326261 | 0.097195866 |
| ENSMUSG00000033216 | Eefsec | 65967 | protein_coding | 210.293974 | 0.097199558 | 0.091695461 | -1.060025837 | 0.289132845 | 0.508005086 | 0.097199558 |
| ENSMUSG00000030201 | Lrp6 | 16974 | protein_coding | 2094.149072 | -0.097238143 | 0.061273232 | 1.586959589 | 0.112521797 | 0.269146142 | 0.097238143 |
| ENSMUSG00000020840 | Blmh | 104184 | protein_coding | 754.7798093 | 0.097270548 | 0.064177705 | -1.515643907 | 0.129609418 | 0.297320449 | 0.097270548 |

| ensembl_gene_id | external_gene_name | entrezgene_id | gene_biotype | baseMean | log2FoldChange | lfcSE | stat | pvalue | padj | absLog2FC |
|---|---|---|---|---|---|---|---|---|---|---|
| ENSMUSG00000022361 | Zhx1 | 22770 | protein_coding | 403.1193151 | -0.097273246 | 0.086189616 | 1.128595888 | 0.259068346 | 0.472369953 | 0.097273246 |
| ENSMUSG00000041164 | Zmiz2 | 52915 | protein_coding | 482.1686083 | -0.097286875 | 0.122229754 | 0.795934474 | 0.426070122 | 0.645752753 | 0.097286875 |
| ENSMUSG00000015882 | Lcorl | 209707 | protein_coding | 568.9023919 | -0.0973149 | 0.088158513 | 1.103862771 | 0.26965267 | 0.484817006 | 0.0973149 |
| ENSMUSG00000025262 | Fam120c | 207375 | protein_coding | 106.1704831 | -0.097327135 | 0.142207664 | 0.684401477 | 0.493721684 | 0.699975951 | 0.097327135 |
| ENSMUSG00000033396 | Spg11 | 214585 | protein_coding | 668.4212716 | -0.09733751 | 0.051478999 | 1.890819789 | 0.058648405 | 0.167999957 | 0.09733751 |
| ENSMUSG00000055313 | Pgbd1 | 319207 | protein_coding | 14.23247548 | 0.097408769 | 0.31758949 | -0.306712822 | 0.75906197 | 0.878912418 | 0.097408769 |
| ENSMUSG00000021670 | Hmgcr | 15357 | protein_coding | 2649.025496 | 0.097416251 | 0.13532286 | -0.719880226 | 0.471598744 | 0.684381431 | 0.097416251 |
| ENSMUSG00000113621 | 2900060N12Rik | 102641621 | lncRNA | 1.900169643 | -0.097416817 | 2.165715112 | 0.044981363 | 0.964122164 | 1 | 0.097416817 |
| ENSMUSG00000057551 | Zfp317 | 244713 | protein_coding | 340.2104578 | -0.09743063 | 0.099825718 | 0.976007313 | 0.329060841 | 0.551494032 | 0.09743063 |
| ENSMUSG00000064068 | Mtx1 | 17827 | protein_coding | 354.4106156 | 0.097479148 | 0.071787419 | -1.357886238 | 0.174499785 | 0.365696631 | 0.097479148 |
| ENSMUSG00000028911 | Srsf4 | 57317 | protein_coding | 132.3113178 | -0.097487416 | 0.09556569 | 1.020108957 | 0.307676789 | 0.52811293 | 0.097487416 |
| ENSMUSG00000022052 | Ppp2r2a | 71978 | protein_coding | 229.8579573 | 0.097543934 | 0.195969005 | -0.497751848 | 0.618658959 | 0.789897483 | 0.097543934 |
| ENSMUSG00000020226 | Slc5a4b | 64454 | protein_coding | 9.386331724 | 0.097555095 | 0.600427605 | -0.162476032 | 0.870931001 | 0.936356422 | 0.097555095 |
| ENSMUSG00000032750 | Gab3 | 210710 | protein_coding | 1.64261965 | 0.097568554 | 1.068805422 | -0.091287481 | 0.927264165 | 1 | 0.097568554 |
| ENSMUSG00000061950 | Ppp4r1 | 70351 | protein_coding | 854.2728103 | 0.097569114 | 0.072194445 | -1.351476752 | 0.176542762 | 0.36866808 | 0.097569114 |
| ENSMUSG00000024286 | Ccny | 67974 | protein_coding | 1097.207222 | -0.097604421 | 0.097774685 | 0.998258611 | 0.318153972 | 0.539479894 | 0.097604421 |
| ENSMUSG00000023078 | Cxcl13 | 55985 | protein_coding | 3.725476048 | -0.097625161 | 0.518688028 | 0.188215565 | 0.850707671 | 1 | 0.097625161 |
| ENSMUSG00000034390 | Cmip | 74440 | protein_coding | 787.2449189 | -0.09765765 | 0.14089806 | 0.69310855 | 0.488241432 | 0.696271072 | 0.09765765 |
| ENSMUSG00000030281 | Il17rc | 171095 | protein_coding | 1245.03069 | -0.097676161 | 0.105958606 | 0.921833204 | 0.356615585 | 0.579526645 | 0.097676161 |
| ENSMUSG00000104368 | 2010103J01Rik | 69889 | lncRNA | 6.384529396 | 0.097677562 | 0.542305604 | -0.180115347 | 0.857062014 | 0.93035707 | 0.097677562 |
| ENSMUSG00000054589 | Gm9949 | 225609 | lncRNA | 3.597948128 | -0.097691717 | 0.531229097 | 0.183897527 | 0.854093845 | 1 | 0.097691717 |
| ENSMUSG00000030619 | Eed | 13626 | protein_coding | 363.6695245 | 0.097725079 | 0.080194599 | -1.21859926 | 0.22299633 | 0.429954629 | 0.097725079 |
| ENSMUSG00000098702 | 1500015A07Rik | 68982 | lncRNA | 27.82996088 | 0.097854321 | 0.293601173 | -0.333289951 | 0.738915424 | 0.867575493 | 0.097854321 |
| ENSMUSG00000044322 | Dsc1 | 13505 | protein_coding | 1.598105152 | 0.097860256 | 2.666357085 | -0.036701857 | 0.970722728 | 1 | 0.097860256 |
| ENSMUSG00000032290 | Ptpn9 | 56294 | protein_coding | 654.0680618 | 0.097919177 | 0.061528283 | -1.591449838 | 0.111508376 | 0.267419303 | 0.097919177 |
| ENSMUSG00000086427 | Hoxa11os | 15397 | lncRNA | 275.465696 | -0.097953495 | 0.160461162 | 0.610449865 | 0.541563844 | 0.735467298 | 0.097953495 |
| ENSMUSG00000074151 | Nlrc5 | 434341 | protein_coding | 30.26315127 | -0.097955839 | 0.374076222 | 0.261860641 | 0.793428885 | 0.897198344 | 0.097955839 |
| ENSMUSG00000043592 | Unc5cl | 76589 | protein_coding | 133.0830562 | 0.097984731 | 0.136288604 | -0.71895029 | 0.472171549 | 0.684706793 | 0.097984731 |
| ENSMUSG00000038658 | Ric1 | 226089 | protein_coding | 365.4118696 | -0.098016529 | 0.091768028 | 1.068090183 | 0.285479834 | 0.503945908 | 0.098016529 |
| ENSMUSG00000071253 | Slc25a16 | 73132 | protein_coding | 386.3097794 | 0.098027916 | 0.095691257 | -1.024418724 | 0.305637542 | 0.525979404 | 0.098027916 |
| ENSMUSG00000036026 | Tmem63b | 224807 | protein_coding | 913.2597842 | -0.098043743 | 0.076879274 | 1.275294868 | 0.202204893 | 0.404116133 | 0.098043743 |
| ENSMUSG00000028358 | Zfp618 | 72701 | protein_coding | 429.2888367 | -0.098065834 | 0.124523871 | 0.787526386 | 0.430973788 | 0.649489062 | 0.098065834 |
| ENSMUSG00000020477 | Mrps24 | 64660 | protein_coding | 546.3330773 | -0.09810415 | 0.068264319 | 1.437121926 | 0.150683352 | 0.330726326 | 0.09810415 |
| ENSMUSG00000030714 | Sgf29 | 75565 | protein_coding | 310.7734134 | -0.098129227 | 0.073707511 | 1.331332788 | 0.183079534 | 0.377733062 | 0.098129227 |
| ENSMUSG00000061207 | Stk19 | 54402 | protein_coding | 179.1444467 | -0.098192667 | 0.106892558 | 0.918610885 | 0.358299137 | 0.580947662 | 0.098192667 |
| ENSMUSG00000024331 | Dsc2 | 13506 | protein_coding | 3077.280873 | -0.098262823 | 0.100682294 | 0.975969253 | 0.329079701 | 0.551494032 | 0.098262823 |
| ENSMUSG00000017286 | Glod4 | 67201 | protein_coding | 572.8588186 | -0.098297037 | 0.109217957 | 0.900008023 | 0.368115981 | 0.59060209 | 0.098297037 |
| ENSMUSG00000021806 | Nid2 | 18074 | protein_coding | 94.11891529 | -0.098309335 | 0.138867017 | 0.7079387 | 0.478983325 | 0.690541222 | 0.098309335 |
| ENSMUSG00000027427 | Polr3f | 70408 | protein_coding | 212.5351216 | 0.098326987 | 0.077389422 | -1.270548143 | 0.203889445 | 0.406017465 | 0.098326987 |

Supplementary Table 1

| ensembl_gene_id | external_gene_name | entrezgene_id | gene_biotype | baseMean | log2FoldChange | lfcSE | stat | pvalue | padj | absLog2FC |
|---|---|---|---|---|---|---|---|---|---|---|
| ENSMUSG00000022551 | Cyc1 | 66445 | protein_coding | 3374.269143 | -0.098339805 | 0.070076005 | 1.403330652 | 0.160518262 | 0.345021291 | 0.098339805 |
| ENSMUSG00000044134 | Pheta1 | 231717 | protein_coding | 856.6406299 | 0.098348062 | 0.104889027 | -0.937639193 | 0.348429863 | 0.570284968 | 0.098348062 |
| ENSMUSG00000037098 | Rab11fip3 | 215445 | protein_coding | 272.3149248 | 0.098381466 | 0.097238135 | -1.011758051 | 0.311653752 | 0.532495959 | 0.098381466 |
| ENSMUSG00000071037 | Camkmt | 73582 | protein_coding | 88.32981505 | -0.098408935 | 0.133606899 | 0.736555787 | 0.46139253 | 0.675184655 | 0.098408935 |
| ENSMUSG00000015747 | Vps45 | 22365 | protein_coding | 219.6935938 | 0.098420604 | 0.082637347 | -1.190994237 | 0.233655847 | 0.443108941 | 0.098420604 |
| ENSMUSG00000040124 | Gorab | 98376 | protein_coding | 188.1480892 | -0.098427428 | 0.117588527 | 0.837049585 | 0.402564697 | 0.624030976 | 0.098427428 |
| ENSMUSG00000051224 | Tceanc | 245695 | protein_coding | 65.83440949 | -0.098448193 | 0.17148889 | 0.574079131 | 0.565914255 | 0.752577549 | 0.098448193 |
| ENSMUSG00000026005 | Rpe | 66646 | protein_coding | 535.2417745 | 0.098472815 | 0.111345937 | -0.884386244 | 0.376487746 | 0.598740239 | 0.098472815 |
| ENSMUSG00000102748 | Pcdhgb2 | 93700 | protein_coding | 7.892418808 | 0.098495431 | 0.403786388 | -0.243929549 | 0.807285384 | 0.904987729 | 0.098495431 |
| ENSMUSG00000004056 | Akt2 | 11652 | protein_coding | 430.2851353 | -0.098506512 | 0.065099062 | 1.513178671 | 0.130234281 | 0.298143983 | 0.098506512 |
| ENSMUSG00000068329 | Htra2 | 64704 | protein_coding | 179.9089845 | -0.09855077 | 0.095954862 | 1.027053436 | 0.304395303 | 0.524552231 | 0.09855077 |
| ENSMUSG00000037740 | Mrps26 | 99045 | protein_coding | 369.6257949 | 0.098620534 | 0.090245619 | -1.092801343 | 0.274481032 | 0.490866888 | 0.098620534 |
| ENSMUSG00000031631 | Cfap97 | 66756 | protein_coding | 203.928897 | -0.098671033 | 0.127968286 | 0.771058486 | 0.440672266 | 0.658605063 | 0.098671033 |
| ENSMUSG00000022538 | Lsg1 | 224092 | protein_coding | 265.3562693 | 0.098736864 | 0.072294575 | -1.365757575 | 0.172015066 | 0.362441146 | 0.098736864 |
| ENSMUSG00000001416 | Cct3 | 12462 | protein_coding | 1826.016615 | 0.098758567 | 0.113038541 | -0.873671631 | 0.382297104 | 0.604659935 | 0.098758567 |
| ENSMUSG00000030744 | Rps3 | 27050 | protein_coding | 8881.17455 | -0.098769374 | 0.102231453 | 0.96613489 | 0.333976688 | 0.556313398 | 0.098769374 |
| ENSMUSG00000028080 | Lrba | 80877 | protein_coding | 2828.716916 | -0.098777694 | 0.094499123 | 1.045276303 | 0.295895287 | 0.515123206 | 0.098777694 |
| ENSMUSG00000059923 | Grb2 | 14784 | protein_coding | 613.4558992 | -0.098781129 | 0.067872906 | 1.455383811 | 0.145563036 | 0.322749862 | 0.098781129 |
| ENSMUSG00000030663 | 1110004F10Rik | 56372 | protein_coding | 477.0475237 | 0.098810296 | 0.069615231 | -1.419377564 | 0.15578897 | 0.338423974 | 0.098810296 |
| ENSMUSG00000061313 | Ddhd2 | 72108 | protein_coding | 206.6101282 | -0.098815645 | 0.077462803 | 1.275652846 | 0.202078264 | 0.404039964 | 0.098815645 |
| ENSMUSG00000035769 | Xylb | 102448 | protein_coding | 60.77354562 | -0.098827827 | 0.141652954 | 0.697675721 | 0.485380015 | 0.695380179 | 0.098827827 |
| ENSMUSG00000022335 | Zfat | 380993 | protein_coding | 83.57723578 | 0.098828226 | 0.154845509 | -0.638237596 | 0.523319027 | 0.722098569 | 0.098828226 |
| ENSMUSG00000078671 | Chd2 | 244059 | protein_coding | 964.7212595 | 0.098837354 | 0.102600798 | -0.963319544 | 0.335387188 | 0.557762742 | 0.098837354 |
| ENSMUSG00000021550 | 2210016F16Rik | 70153 | protein_coding | 896.8275867 | -0.098867361 | 0.100701459 | 0.981786779 | 0.326204903 | 0.548178261 | 0.098867361 |
| ENSMUSG00000116114 | Gm35853 | 102639573 | lncRNA | 4.03156485 | -0.098877487 | 0.603194123 | 0.16392316 | 0.869791635 | 1 | 0.098877487 |
| ENSMUSG00000027649 | Ctnnbl1 | 66642 | protein_coding | 342.8706471 | 0.098925557 | 0.073470652 | -1.346463581 | 0.178153058 | 0.37057407 | 0.098925557 |
| ENSMUSG00000073733 | Cplane2 | 76166 | protein_coding | 6.872035166 | 0.098935843 | 0.402231534 | -0.245967395 | 0.805707472 | 0.904271659 | 0.098935843 |
| ENSMUSG00000051716 | Apon | 28194 | protein_coding | 1.317633634 | -0.098936224 | 1.162926148 | 0.085075242 | 0.932201573 | 1 | 0.098936224 |
| ENSMUSG00000097727 | F630040K05Rik | 433111 | lncRNA | 5.42108129 | -0.098940547 | 0.918992986 | 0.107661917 | 0.91426388 | 0.95803327 | 0.098940547 |
| ENSMUSG00000007833 | Aldh16a1 | 69748 | protein_coding | 1073.283288 | -0.09894586 | 0.063939259 | 1.547497763 | 0.121743264 | 0.28456734 | 0.09894586 |
| ENSMUSG00000005683 | Cs | 12974 | protein_coding | 7001.289517 | -0.099126372 | 0.078770859 | 1.258414259 | 0.208241978 | 0.41130117 | 0.099126372 |
| ENSMUSG00000024399 | Ltb | 16994 | protein_coding | 10.49663326 | 0.09917413 | 0.551156303 | -0.179938302 | 0.857201005 | 0.930374388 | 0.09917413 |
| ENSMUSG00000020955 | Ap4s1 | 11782 | protein_coding | 110.1639707 | -0.099208014 | 0.108067254 | 0.918021046 | 0.358607847 | 0.581328932 | 0.099208014 |
| ENSMUSG00000028470 | Hint2 | 68917 | protein_coding | 324.3859735 | -0.099208085 | 0.063107702 | 1.572043998 | 0.115940348 | 0.274894315 | 0.099208085 |
| ENSMUSG00000026455 | Klhl12 | 240756 | protein_coding | 255.0302668 | -0.099216367 | 0.078541877 | 1.263228868 | 0.206506941 | 0.409247045 | 0.099216367 |
| ENSMUSG00000067199 | Frat1 | 14296 | protein_coding | 253.5171458 | -0.0992342 | 0.084051636 | 1.180633774 | 0.237748241 | 0.447999484 | 0.0992342 |
| ENSMUSG00000019814 | Ltv1 | 353258 | protein_coding | 281.4381197 | 0.099236674 | 0.106083286 | -0.93546003 | 0.349551273 | 0.571420483 | 0.099236674 |
| ENSMUSG00000038895 | Zfp653 | 319601 | protein_coding | 47.33391911 | 0.099307603 | 0.16964384 | -0.585388792 | 0.558286323 | 0.747162726 | 0.099307603 |
| ENSMUSG00000024750 | Zfand5 | 22682 | protein_coding | 1548.089782 | 0.099331477 | 0.141111992 | -0.703919459 | 0.481482933 | 0.692354022 | 0.099331477 |

Supplementary Table 1

| ensembl_gene_id | external_gene_name | entrezgene_id | gene_biotype | baseMean | log2FoldChange | lfcSE | stat | pvalue | padj | absLog2FC |
|---|---|---|---|---|---|---|---|---|---|---|
| ENSMUSG00000038569 | Rad9b | 231724 | protein_coding | 30.16659814 | -0.099349938 | 0.239308456 | 0.415154312 | 0.678028942 | 0.827422396 | 0.099349938 |
| ENSMUSG00000031982 | Arv1 | 68865 | protein_coding | 28.50481231 | 0.099370991 | 0.208486877 | -0.476629477 | 0.633625989 | 0.800362194 | 0.099370991 |
| ENSMUSG00000036748 | Cuedc2 | 67116 | protein_coding | 228.7380023 | -0.099377048 | 0.076794914 | 1.294057688 | 0.195645495 | 0.394964884 | 0.099377048 |
| ENSMUSG00000015149 | Sirt2 | 64383 | protein_coding | 395.4592805 | -0.099390893 | 0.066649914 | 1.491238129 | 0.135898986 | 0.307285909 | 0.099390893 |
| ENSMUSG00000050711 | Scg2 | 20254 | protein_coding | 15.0753977 | 0.099424998 | 0.309108091 | -0.321651231 | 0.747716928 | 0.873111243 | 0.099424998 |
| ENSMUSG00000054256 | Msi1 | 17690 | protein_coding | 17.537195 | 0.099439838 | 0.36682872 | -0.271079751 | 0.786329695 | 0.893161222 | 0.099439838 |
| ENSMUSG00000035285 | Nat14 | 269854 | protein_coding | 15.86673525 | 0.099525931 | 0.366046721 | -0.271894065 | 0.785703475 | 0.892925964 | 0.099525931 |
| ENSMUSG00000035085 | 1700020L24Rik | 66330 | protein_coding | 25.89045284 | -0.09953166 | 0.255985853 | 0.388817033 | 0.6974115 | 0.839586229 | 0.09953166 |
| ENSMUSG00000074715 | Ccl28 | 56838 | protein_coding | 1687.535385 | -0.099565611 | 0.178633009 | 0.557375211 | 0.577271099 | 0.761298159 | 0.099565611 |
| ENSMUSG00000038241 | Cep250 | 16328 | protein_coding | 251.9515672 | 0.099576157 | 0.120174873 | -0.828593821 | 0.407334285 | 0.628183969 | 0.099576157 |
| ENSMUSG00000029575 | Mmab | 77697 | protein_coding | 206.2814438 | -0.099655163 | 0.133637404 | 0.745713101 | 0.455840743 | 0.671162093 | 0.099655163 |
| ENSMUSG00000028037 | Ifi44 | 99899 | protein_coding | 41.73573182 | -0.099655356 | 0.557131804 | 0.178872137 | 0.858038101 | 0.930446023 | 0.099655356 |
| ENSMUSG00000041429 | Nthl1 | 18207 | protein_coding | 26.5466328 | -0.099678024 | 0.1872879 | 0.53221817 | 0.5945749 | 0.773317311 | 0.099678024 |
| ENSMUSG00000037608 | Bclaf1 | 72567 | protein_coding | 1574.430206 | 0.099711735 | 0.082145798 | -1.213838544 | 0.224809385 | 0.432138537 | 0.099711735 |
| ENSMUSG00000046798 | Cldn12 | 64945 | protein_coding | 942.0426725 | 0.099776418 | 0.13395902 | -0.744827917 | 0.456375756 | 0.671470886 | 0.099776418 |
| ENSMUSG00000038872 | Zfhx3 | 11906 | protein_coding | 336.9483217 | 0.099804867 | 0.122126317 | -0.817226539 | 0.413798979 | 0.633873724 | 0.099804867 |
| ENSMUSG00000044229 | Nxpe4 | 244853 | protein_coding | 625.9074155 | 0.099812006 | 0.299259978 | -0.333529417 | 0.738734688 | 0.867548331 | 0.099812006 |
| ENSMUSG00000024188 | Luc7l | 66978 | protein_coding | 477.6278202 | 0.099822191 | 0.08664082 | -1.152138109 | 0.249264325 | 0.461422606 | 0.099822191 |
| ENSMUSG00000022420 | Dnal4 | 54152 | protein_coding | 174.4436091 | 0.099833088 | 0.08326396 | -1.198995192 | 0.230529815 | 0.43912271 | 0.099833088 |
| ENSMUSG00000071669 | Snx29 | 74478 | protein_coding | 89.13755821 | -0.099860303 | 0.242044834 | 0.412569447 | 0.67992209 | 0.828297243 | 0.099860303 |
| ENSMUSG00000043866 | Taf10 | 24075 | protein_coding | 734.0956585 | 0.099874568 | 0.092698174 | -1.077416778 | 0.281294112 | 0.499023496 | 0.099874568 |
| ENSMUSG00000026709 | Dars2 | 226539 | protein_coding | 310.5499348 | -0.09991662 | 0.124137146 | 0.804888973 | 0.420883752 | 0.640714813 | 0.09991662 |
| ENSMUSG00000005625 | Psmd4 | 19185 | protein_coding | 1084.441408 | 0.099933448 | 0.052462458 | -1.904856368 | 0.056798744 | 0.164036653 | 0.099933448 |
| ENSMUSG00000032216 | Nedd4 | 17999 | protein_coding | 4438.321901 | 0.099940143 | 0.10720429 | -0.93224015 | 0.35121243 | 0.573211901 | 0.099940143 |
| ENSMUSG00000033373 | Fntb | 110606 | protein_coding | 192.4461214 | 0.099987792 | 0.081034118 | -1.233897452 | 0.217241119 | 0.422906972 | 0.099987792 |
| ENSMUSG00000024740 | Ddb1 | 13194 | protein_coding | 4802.556638 | 0.099989581 | 0.107413286 | -0.930886529 | 0.351912264 | 0.574143363 | 0.099989581 |
| ENSMUSG00000112449 | Srp54b | 665155 | protein_coding | 0.559318665 | -0.099995834 | 1.765006635 | 0.05665465 | 0.9548203 | 1 | 0.099995834 |
| ENSMUSG00000027198 | Ext2 | 14043 | protein_coding | 580.1368562 | 0.100006854 | 0.063734646 | -1.569112874 | 0.116621641 | 0.276057789 | 0.100006854 |
| ENSMUSG00000054199 | Gon4l | 76022 | protein_coding | 335.3622922 | 0.100007581 | 0.06967341 | -1.435376577 | 0.151179817 | 0.33115579 | 0.100007581 |
| ENSMUSG00000063077 | Kif1b | 16561 | protein_coding | 1115.183368 | -0.100018059 | 0.063165619 | 1.583425616 | 0.113324488 | 0.270127107 | 0.100018059 |
| ENSMUSG00000057191 | AB124611 | 382062 | protein_coding | 2.161476846 | 0.100065706 | 0.74274581 | -0.134724026 | 0.892830077 | 1 | 0.100065706 |
| ENSMUSG00000022244 | Amacr | 17117 | protein_coding | 136.4631411 | -0.100125107 | 0.107830388 | 0.928542576 | 0.353126193 | 0.57545396 | 0.100125107 |
| ENSMUSG00000014470 | Rnf166 | 68718 | protein_coding | 238.2362781 | 0.100151062 | 0.085958222 | -1.165113237 | 0.243973159 | 0.4551483 | 0.100151062 |
| ENSMUSG00000028964 | Park7 | 57320 | protein_coding | 803.6597066 | 0.100154769 | 0.084747819 | -1.181797594 | 0.237286023 | 0.44767168 | 0.100154769 |
| ENSMUSG00000053897 | Slc39a8 | 67547 | protein_coding | 528.3273163 | 0.100159145 | 0.138066891 | -0.725439272 | 0.468182588 | 0.681791678 | 0.100159145 |
| ENSMUSG00000058812 | 0610039K10Rik | 68386 | lncRNA | 19.83594179 | 0.10017729 | 0.3476896 | -0.288122767 | 0.773252766 | 0.886362513 | 0.10017729 |
| ENSMUSG00000097248 | Gm2694 | 100040294 | lncRNA | 2.890023421 | -0.100267185 | 1.322454346 | 0.075819015 | 0.939563088 | 1 | 0.100267185 |
| ENSMUSG00000050856 | Atp5k | 11958 | protein_coding | 1964.017933 | -0.100293981 | 0.10672752 | 0.939719957 | 0.347361226 | 0.569192626 | 0.100293981 |
| ENSMUSG00000019428 | Fkbp8 | 14232 | protein_coding | 3293.1903 | 0.100312008 | 0.057978734 | -1.730151736 | 0.083603169 | 0.216806333 | 0.100312008 |

| ensembl_gene_id | external_gene_name | entrezgene_id | gene_biotype | baseMean | log2FoldChange | lfcSE | stat | pvalue | padj | absLog2FC |
|---|---|---|---|---|---|---|---|---|---|---|
| ENSMUSG00000028621 | Cyb5rl | 230582 | protein_coding | 66.67876503 | -0.100325673 | 0.158832464 | 0.631644631 | 0.527619116 | 0.725856654 | 0.100325673 |
| ENSMUSG00000028830 | AU040320 | 100317 | protein_coding | 1386.402599 | -0.100353293 | 0.044627792 | 2.248672623 | 0.024533332 | 0.086109128 | 0.100353293 |
| ENSMUSG00000046442 | Ppm1e | 320472 | protein_coding | 9.441679398 | 0.100369596 | 0.464697827 | -0.21598895 | 0.828996361 | 0.913504709 | 0.100369596 |
| ENSMUSG00000026885 | Ttll11 | 74410 | protein_coding | 14.6043498 | 0.100379667 | 0.341077898 | -0.294301296 | 0.768527682 | 0.884053089 | 0.100379667 |
| ENSMUSG00000038481 | Cdk19 | 78334 | protein_coding | 375.2294911 | -0.100414931 | 0.103549038 | 0.969733107 | 0.332179543 | 0.554606509 | 0.100414931 |
| ENSMUSG00000031220 | Awat2 | 245532 | protein_coding | 6.83322058 | 0.100445412 | 0.396711748 | -0.253194951 | 0.800117568 | 0.900998399 | 0.100445412 |
| ENSMUSG00000050621 | Rps27rt | 100043813 | protein_coding | 20.93454735 | 0.100493595 | 0.372504166 | -0.269778447 | 0.787330705 | 0.893606822 | 0.100493595 |
| ENSMUSG00000027259 | Adal | 75894 | protein_coding | 93.69636391 | 0.100546845 | 0.130234094 | -0.772047025 | 0.440086576 | 0.657938595 | 0.100546845 |
| ENSMUSG00000053553 | 3110082I17Rik | 73212 | protein_coding | 53.97363731 | 0.100549361 | 0.182945903 | -0.549612533 | 0.582585161 | 0.765317902 | 0.100549361 |
| ENSMUSG00000035268 | Pkig | 18769 | protein_coding | 294.0699519 | 0.100553072 | 0.118253644 | -0.850316902 | 0.395148922 | 0.617417854 | 0.100553072 |
| ENSMUSG00000025485 | Ric8a | 101489 | protein_coding | 671.8936772 | -0.100565581 | 0.062738096 | 1.602942835 | 0.108947276 | 0.26324771 | 0.100565581 |
| ENSMUSG00000030762 | Aqp8 | 11833 | protein_coding | 2018.274923 | -0.100575707 | 0.406865031 | 0.247196735 | 0.804755972 | 0.903803848 | 0.100575707 |
| ENSMUSG00000032551 | 1110059G10Rik | 66202 | protein_coding | 128.9935714 | -0.100580158 | 0.101015059 | 0.99569469 | 0.319398511 | 0.540748207 | 0.100580158 |
| ENSMUSG00000024975 | Pdcd4 | 18569 | protein_coding | 2012.089842 | 0.100585091 | 0.154607226 | -0.650584668 | 0.51531463 | 0.716335759 | 0.100585091 |
| ENSMUSG00000078923 | Ube2v1 | 66589 | protein_coding | 14.30578298 | -0.100588826 | 0.331620894 | 0.303324753 | 0.761642384 | 0.880610679 | 0.100588826 |
| ENSMUSG00000002846 | Timmdc1 | 76916 | protein_coding | 281.8916255 | -0.100661874 | 0.067504098 | 1.491196502 | 0.135909912 | 0.307285909 | 0.100661874 |
| ENSMUSG00000023965 | Fbxl17 | 50758 | protein_coding | 270.7417162 | -0.100688418 | 0.074547483 | 1.350661528 | 0.176803882 | 0.368933447 | 0.100688418 |
| ENSMUSG00000079043 | Fastkd5 | 380601 | protein_coding | 62.55686403 | -0.10071539 | 0.168288385 | 0.598469048 | 0.549527006 | 0.740505016 | 0.10071539 |
| ENSMUSG00000054051 | Ercc6 | 319955 | protein_coding | 336.5859435 | -0.10076437 | 0.073520667 | 1.370558428 | 0.17051265 | 0.360359086 | 0.10076437 |
| ENSMUSG00000051341 | Zfp52 | 22710 | protein_coding | 58.83800224 | -0.100781574 | 0.171281712 | 0.588396582 | 0.556266132 | 0.745631404 | 0.100781574 |
| ENSMUSG00000055013 | Agap1 | 347722 | protein_coding | 826.5808074 | -0.100786087 | 0.078059071 | 1.291151501 | 0.196651149 | 0.396326223 | 0.100786087 |
| ENSMUSG00000062198 | 2700097O09Rik | 72658 | protein_coding | 61.85161149 | 0.100823035 | 0.153451661 | -0.657034495 | 0.511158736 | 0.714031075 | 0.100823035 |
| ENSMUSG00000054951 | 9130008F23Rik | 71583 | protein_coding | 200.3400827 | 0.100845033 | 0.14151809 | -0.71259464 | 0.476096629 | 0.687930892 | 0.100845033 |
| ENSMUSG00000033981 | Gria2 | 14800 | protein_coding | 5.711257716 | -0.100856517 | 1.61436547 | 0.062474402 | 0.950185046 | 0.975869522 | 0.100856517 |
| ENSMUSG00000071645 | Tut1 | 70044 | protein_coding | 249.5397104 | 0.100872213 | 0.082103802 | -1.228593689 | 0.219224183 | 0.425500655 | 0.100872213 |
| ENSMUSG00000025869 | Nop16 | 28126 | protein_coding | 252.9412923 | 0.100878603 | 0.097758133 | -1.031920313 | 0.302109449 | 0.522421155 | 0.100878603 |
| ENSMUSG00000020628 | Trappc12 | 217449 | protein_coding | 497.8466927 | -0.100933818 | 0.067894043 | 1.486637302 | 0.13711063 | 0.309427758 | 0.100933818 |
| ENSMUSG00000067656 | Slc22a27 | 171405 | protein_coding | 2.222976911 | -0.10099841 | 0.719400543 | 0.140392458 | 0.888349917 | 1 | 0.10099841 |
| ENSMUSG00000062202 | Btbd9 | 224671 | protein_coding | 459.8879082 | -0.101010594 | 0.065096798 | 1.551698342 | 0.12073442 | 0.283211393 | 0.101010594 |
| ENSMUSG00000078584 | AU022252 | 230696 | protein_coding | 209.6983516 | 0.101110758 | 0.085233671 | -1.186277166 | 0.235512861 | 0.445505084 | 0.101110758 |
| ENSMUSG00000031667 | Aktip | 14339 | protein_coding | 508.1421673 | -0.101125659 | 0.060311293 | 1.676728429 | 0.093595599 | 0.235436057 | 0.101125659 |
| ENSMUSG00000052684 | Jun | 16476 | protein_coding | 10893.12234 | -0.101203975 | 0.115077918 | 0.879438696 | 0.379163459 | 0.60160505 | 0.101203975 |
| ENSMUSG00000030887 | Pdzd9 | 67983 | protein_coding | 7.046305552 | -0.101210839 | 0.462309012 | 0.218924651 | 0.826708743 | 0.912523774 | 0.101210839 |
| ENSMUSG00000054474 | Thnsl2 | 232078 | protein_coding | 269.2060895 | 0.101218754 | 0.133396762 | -0.758779691 | 0.447984356 | 0.664930223 | 0.101218754 |
| ENSMUSG00000031446 | Cul4a | 99375 | protein_coding | 1068.338099 | 0.101256667 | 0.077301888 | -1.309886073 | 0.190234379 | 0.387604015 | 0.101256667 |
| ENSMUSG00000002342 | Tmem161a | 234371 | protein_coding | 343.8457682 | -0.101295018 | 0.07706008 | 1.314494055 | 0.188679997 | 0.385300595 | 0.101295018 |
| ENSMUSG00000066720 | Cldn9 | 56863 | protein_coding | 3.692596347 | -0.101316685 | 0.71910321 | 0.140893107 | 0.887954388 | 1 | 0.101316685 |
| ENSMUSG00000056167 | Cnot10 | 78893 | protein_coding | 481.3768611 | 0.101367657 | 0.053766672 | -1.885325107 | 0.059385958 | 0.169583213 | 0.101367657 |
| ENSMUSG00000025899 | Alkbh8 | 67667 | protein_coding | 204.1690826 | 0.101444402 | 0.128137957 | -0.791681127 | 0.42854663 | 0.647544976 | 0.101444402 |

| ensembl_gene_id | external_gene_name | entrezgene_id | gene_biotype | baseMean | log2FoldChange | lfcSE | stat | pvalue | padj | absLog2FC |
|---|---|---|---|---|---|---|---|---|---|---|
| ENSMUSG00000024095 | Hnrnpll | 72692 | protein_coding | 747.8826423 | 0.101453637 | 0.060849297 | -1.667293497 | 0.095456059 | 0.238987471 | 0.101453637 |
| ENSMUSG00000024773 | Atg2a | 329015 | protein_coding | 705.5939272 | -0.101460874 | 0.121681066 | 0.83382631 | 0.404378867 | 0.62546825 | 0.101460874 |
| ENSMUSG00000019797 | Mtres1 | 67851 | protein_coding | 295.271451 | -0.101565692 | 0.085753437 | 1.184392084 | 0.236257892 | 0.44640307 | 0.101565692 |
| ENSMUSG00000102428 | Pcdhga12 | 93724 | protein_coding | 6.017960096 | -0.101596661 | 0.414898887 | 0.244870893 | 0.806556399 | 0.904855461 | 0.101596661 |
| ENSMUSG00000014748 | Tex261 | 21766 | protein_coding | 877.6581805 | -0.101597236 | 0.066159871 | 1.535632319 | 0.124628583 | 0.289400593 | 0.101597236 |
| ENSMUSG00000037580 | Gch1 | 14528 | protein_coding | 264.9180874 | -0.101646069 | 0.140528039 | 0.72331522 | 0.469486244 | 0.683156904 | 0.101646069 |
| ENSMUSG00000018841 | Rad51d | 19364 | protein_coding | 263.2998261 | -0.101662923 | 0.095961413 | 1.05941461 | 0.289410999 | 0.50827749 | 0.101662923 |
| ENSMUSG00000037960 | Card19 | 68480 | protein_coding | 387.1994243 | -0.101672773 | 0.116321773 | 0.87406485 | 0.382082938 | 0.604440039 | 0.101672773 |
| ENSMUSG00000058325 | Dock1 | 330662 | protein_coding | 1451.907829 | -0.101691332 | 0.045831768 | 2.21879573 | 0.026500625 | 0.091363531 | 0.101691332 |
| ENSMUSG00000020289 | Nprl3 | 17168 | protein_coding | 141.3475538 | -0.101727108 | 0.119401351 | 0.851976185 | 0.39422731 | 0.616875898 | 0.101727108 |
| ENSMUSG00000013973 | Dedd | 21945 | protein_coding | 103.0458816 | -0.10179726 | 0.124704086 | 0.816310536 | 0.41432255 | 0.634528665 | 0.10179726 |
| ENSMUSG00000049299 | Trappc1 | 245828 | protein_coding | 332.487863 | 0.101828932 | 0.085984565 | -1.1842699 | 0.23630624 | 0.446443342 | 0.101828932 |
| ENSMUSG00000092417 | Gpank1 | 81845 | protein_coding | 117.8317273 | -0.101833678 | 0.10622241 | 0.958683554 | 0.337718186 | 0.559610488 | 0.101833678 |
| ENSMUSG00000098766 | Mir6386 | 102465203 | miRNA | 7.980301981 | -0.101841474 | 0.402365684 | 0.25310676 | 0.800185715 | 0.901013699 | 0.101841474 |
| ENSMUSG00000050106 | Tmc8 | 217356 | protein_coding | 142.5382406 | 0.10184174 | 0.139896984 | -0.727976665 | 0.466627871 | 0.68031188 | 0.10184174 |
| ENSMUSG00000024997 | Prdx3 | 11757 | protein_coding | 1370.966002 | -0.10185792 | 0.067257074 | 1.514456598 | 0.129910074 | 0.29771794 | 0.10185792 |
| ENSMUSG00000048027 | Rgmb | 68799 | protein_coding | 795.9202801 | 0.101865072 | 0.106241009 | -0.958811224 | 0.337653854 | 0.559563944 | 0.101865072 |
| ENSMUSG00000068860 | Gm128 | 229588 | protein_coding | 3.57693077 | -0.101869059 | 0.606077677 | 0.168079214 | 0.866520962 | 1 | 0.101869059 |
| ENSMUSG00000043411 | Usp48 | 170707 | protein_coding | 798.9166092 | 0.101886064 | 0.049336128 | -2.065141057 | 0.038909656 | 0.123307245 | 0.101886064 |
| ENSMUSG00000026171 | Rnf25 | 57751 | protein_coding | 212.0772359 | -0.101923733 | 0.080018425 | 1.273753302 | 0.20275086 | 0.404864292 | 0.101923733 |
| ENSMUSG00000026848 | Tor1b | 30934 | protein_coding | 461.8046635 | -0.10201792 | 0.067789699 | 1.504917744 | 0.132345226 | 0.301713259 | 0.10201792 |
| ENSMUSG00000025264 | Tsr2 | 69499 | protein_coding | 115.3163958 | 0.102020548 | 0.155528224 | -0.655961631 | 0.511848812 | 0.714390744 | 0.102020548 |
| ENSMUSG00000046897 | Zfp740 | 68744 | protein_coding | 853.0021571 | -0.102040634 | 0.077494054 | 1.316754366 | 0.187920974 | 0.384217774 | 0.102040634 |
| ENSMUSG00000003923 | Tfam | 21780 | protein_coding | 453.3928952 | -0.102042716 | 0.087742669 | 1.16297711 | 0.24483879 | 0.456505337 | 0.102042716 |
| ENSMUSG00000051832 | E230016K23Rik | 100504464 | lncRNA | 2.122951362 | -0.10204974 | 0.906372257 | 0.112591421 | 0.910354487 | 1 | 0.10204974 |
| ENSMUSG00000026672 | Optn | 71648 | protein_coding | 493.352055 | 0.102084671 | 0.153883989 | -0.663387219 | 0.50708259 | 0.710854454 | 0.102084671 |
| ENSMUSG00000075419 | Dolk | 227697 | protein_coding | 194.7713663 | 0.102119786 | 0.077531649 | -1.317136785 | 0.187792779 | 0.384098334 | 0.102119786 |
| ENSMUSG00000028567 | Txndc12 | 66073 | protein_coding | 356.2975786 | -0.102127446 | 0.107075444 | 0.953789614 | 0.340190148 | 0.561689026 | 0.102127446 |
| ENSMUSG00000064351 | mt-Co1 | 17708 | protein_coding | 639120.1894 | -0.102183676 | 0.436549577 | 0.234071183 | 0.814929717 | 0.908068261 | 0.102183676 |
| ENSMUSG00000028187 | Rpf1 | 70285 | protein_coding | 317.2765411 | -0.102185402 | 0.096396667 | 1.060051195 | 0.289121309 | 0.508005086 | 0.102185402 |
| ENSMUSG00000024392 | Bag6 | 224727 | protein_coding | 1428.854144 | -0.102197466 | 0.081621205 | 1.252094546 | 0.210535415 | 0.414593589 | 0.102197466 |
| ENSMUSG00000039046 | Usp6nl | 98910 | protein_coding | 584.9452705 | -0.102202882 | 0.068343461 | 1.495430283 | 0.134802182 | 0.3056602 | 0.102202882 |
| ENSMUSG00000026383 | Epb41l5 | 226352 | protein_coding | 220.010906 | -0.102219515 | 0.101660236 | 1.005501448 | 0.314655453 | 0.535858528 | 0.102219515 |
| ENSMUSG00000037957 | Wdr20 | 69641 | protein_coding | 324.1455388 | 0.10224128 | 0.075614148 | -1.352144831 | 0.176328988 | 0.368268232 | 0.10224128 |
| ENSMUSG00000029176 | Anapc4 | 52206 | protein_coding | 412.4802661 | 0.102258674 | 0.094138488 | -1.086257878 | 0.277364917 | 0.494297255 | 0.102258674 |
| ENSMUSG00000021066 | Atl1 | 73991 | protein_coding | 24.87899126 | -0.102269651 | 0.27372099 | 0.37362736 | 0.708681575 | 0.847687059 | 0.102269651 |
| ENSMUSG00000026857 | Ntmt1 | 66617 | protein_coding | 207.367212 | 0.102344118 | 0.073015583 | -1.401675006 | 0.161012317 | 0.345858144 | 0.102344118 |
| ENSMUSG00000030172 | Erc1 | 111173 | protein_coding | 391.8836755 | 0.102368436 | 0.075932412 | -1.348152033 | 0.177609489 | 0.370029503 | 0.102368436 |
| ENSMUSG00000056185 | Snx32 | 225861 | protein_coding | 39.63488868 | -0.102368693 | 0.270733269 | 0.378116413 | 0.705344116 | 0.845574994 | 0.102368693 |

| ensembl_gene_id | external_gene_name | entrezgene_id | gene_biotype | baseMean | log2FoldChange | lfcSE | stat | pvalue | padj | absLog2FC |
|---|---|---|---|---|---|---|---|---|---|---|
| ENSMUSG00000035027 | Map2k2 | 26396 | protein_coding | 1352.38048 | 0.102404195 | 0.074364454 | -1.377058384 | 0.168494198 | 0.357236254 | 0.102404195 |
| ENSMUSG00000056851 | Pcbp2 | 18521 | protein_coding | 755.7165134 | 0.102406927 | 0.150921871 | -0.678542655 | 0.497427688 | 0.702997077 | 0.102406927 |
| ENSMUSG00000022323 | Rida | 15473 | protein_coding | 198.9566846 | -0.102408367 | 0.111672067 | 0.917045498 | 0.359118799 | 0.581819665 | 0.102408367 |
| ENSMUSG00000000278 | Scpep1 | 74617 | protein_coding | 361.358767 | -0.1024139 | 0.083650213 | 1.22431129 | 0.22083482 | 0.427333749 | 0.1024139 |
| ENSMUSG00000046791 | Riox1 | 71952 | protein_coding | 148.5930353 | -0.10247479 | 0.100049266 | 1.024243297 | 0.305720373 | 0.526067129 | 0.10247479 |
| ENSMUSG00000003039 | Fam32a | 67922 | protein_coding | 1145.541053 | -0.102547448 | 0.092952545 | 1.103223672 | 0.269930044 | 0.485210052 | 0.102547448 |
| ENSMUSG00000040167 | Ikzf5 | 67143 | protein_coding | 31.0312011 | -0.102550694 | 0.197096216 | 0.520307776 | 0.602849077 | 0.778864783 | 0.102550694 |
| ENSMUSG00000040618 | Pck2 | 74551 | protein_coding | 447.598437 | 0.102563503 | 0.108137299 | -0.948456301 | 0.342897209 | 0.564917636 | 0.102563503 |
| ENSMUSG00000056632 | Dsg3 | 13512 | protein_coding | 8.400734732 | -0.102571717 | 0.467217662 | 0.219537327 | 0.826231506 | 0.912240897 | 0.102571717 |
| ENSMUSG00000109186 | Gm34821 | 102638204 | lncRNA | 8.573993317 | 0.102600288 | 0.389729649 | -0.263260157 | 0.792350067 | 0.896636371 | 0.102600288 |
| ENSMUSG00000036281 | Snapc4 | 227644 | protein_coding | 155.0726039 | 0.102614407 | 0.099670683 | -1.02953451 | 0.303228572 | 0.523689253 | 0.102614407 |
| ENSMUSG00000031858 | Mau2 | 74549 | protein_coding | 1226.329803 | -0.102641446 | 0.051305204 | 2.000604997 | 0.045434975 | 0.138818348 | 0.102641446 |
| ENSMUSG00000019528 | Gyg | 27357 | protein_coding | 234.6494006 | -0.102658406 | 0.121945579 | 0.841837861 | 0.399878716 | 0.622046705 | 0.102658406 |
| ENSMUSG00000036298 | Slc2a13 | 239606 | protein_coding | 568.8871784 | 0.102665484 | 0.101423012 | -1.012250391 | 0.311418348 | 0.53237708 | 0.102665484 |
| ENSMUSG00000041515 | Irf8 | 15900 | protein_coding | 1077.247636 | 0.102710968 | 0.101865175 | -1.00830307 | 0.31330899 | 0.534596554 | 0.102710968 |
| ENSMUSG00000034290 | Nek9 | 217718 | protein_coding | 781.7891733 | -0.102722942 | 0.051902531 | 1.979150907 | 0.047799018 | 0.144095106 | 0.102722942 |
| ENSMUSG00000025658 | Cnksr2 | 245684 | protein_coding | 2.280681024 | 0.102733846 | 2.097870843 | -0.04897053 | 0.960942782 | 1 | 0.102733846 |
| ENSMUSG00000028677 | Rnf220 | 66743 | protein_coding | 500.2995915 | 0.102746836 | 0.079457004 | -1.293112387 | 0.195972192 | 0.395392153 | 0.102746836 |
| ENSMUSG00000024012 | Mtch1 | 56462 | protein_coding | 1693.823553 | -0.102757997 | 0.076858965 | 1.336968262 | 0.181232997 | 0.375235922 | 0.102757997 |
| ENSMUSG00000058006 | Mdn1 | 100019 | protein_coding | 496.0830249 | 0.102797169 | 0.12609278 | -0.815250239 | 0.414929086 | 0.635253445 | 0.102797169 |
| ENSMUSG00000083394 | Gm11703 | 108167871 | processed_pseudogene | 0.641152516 | -0.102873761 | 1.443491784 | 0.0712673 | 0.94318502 | 1 | 0.102873761 |
| ENSMUSG00000030801 | Kat8 | 67773 | protein_coding | 208.0402352 | 0.102874521 | 0.085596169 | -1.201858938 | 0.229418186 | 0.438111017 | 0.102874521 |
| ENSMUSG00000027710 | Acad9 | 229211 | protein_coding | 516.6957296 | -0.102935359 | 0.104300378 | 0.986912619 | 0.323685478 | 0.545498722 | 0.102935359 |
| ENSMUSG00000038637 | Lrrc56 | 70552 | protein_coding | 101.5752832 | 0.102938429 | 0.122007381 | -0.843706571 | 0.398833395 | 0.620766699 | 0.102938429 |
| ENSMUSG00000034832 | Tet3 | 194388 | protein_coding | 1396.022975 | -0.102941168 | 0.111999569 | 0.919121112 | 0.358032228 | 0.580790426 | 0.102941168 |
| ENSMUSG00000029434 | Vps33a | 77573 | protein_coding | 388.3946719 | 0.102972337 | 0.073419588 | -1.402518595 | 0.160760442 | 0.345496868 | 0.102972337 |
| ENSMUSG00000033467 | Crlf2 | 57914 | protein_coding | 38.62118195 | -0.102987395 | 0.209601024 | 0.491349673 | 0.623179153 | 0.793304622 | 0.102987395 |
| ENSMUSG00000042644 | Itpr3 | 16440 | protein_coding | 2871.248831 | -0.103016924 | 0.057286745 | 1.798268061 | 0.072134539 | 0.195389167 | 0.103016924 |
| ENSMUSG00000022433 | Csnk1e | 27373 | protein_coding | 164.3559332 | 0.103046206 | 0.135935468 | -0.758052382 | 0.448419625 | 0.665277305 | 0.103046206 |
| ENSMUSG00000036813 | Entpd8 | 72090 | protein_coding | 1379.528626 | -0.103135126 | 0.097724602 | 1.05536501 | 0.291258419 | 0.510066755 | 0.103135126 |
| ENSMUSG00000097404 | Gm10814 | 100502985 | lncRNA | 7.960126162 | -0.103177514 | 0.570580657 | 0.180828972 | 0.856501822 | 0.930184975 | 0.103177514 |
| ENSMUSG00000073406 | H2-Bl | 14963 | protein_coding | 11.23352825 | 0.10319093 | 0.55106271 | -0.187258053 | 0.85145831 | 0.927499565 | 0.10319093 |
| ENSMUSG00000022500 | Litaf | 56722 | protein_coding | 1050.220678 | -0.103205542 | 0.065650418 | 1.572046982 | 0.115939656 | 0.274894315 | 0.103205542 |
| ENSMUSG00000043822 | Adamtsl5 | 66548 | protein_coding | 70.51789367 | -0.103216041 | 0.278466809 | 0.370658327 | 0.710892033 | 0.849223889 | 0.103216041 |
| ENSMUSG00000064138 | Fam172a | 68675 | protein_coding | 174.0886253 | -0.103222116 | 0.087655523 | 1.177588268 | 0.238960789 | 0.449146195 | 0.103222116 |
| ENSMUSG00000030610 | Det1 | 76375 | protein_coding | 150.1344388 | 0.103246041 | 0.100214318 | -1.030252391 | 0.302891542 | 0.523380864 | 0.103246041 |
| ENSMUSG00000041408 | Wapl | 218914 | protein_coding | 2069.195697 | 0.103252645 | 0.063066182 | -1.637210955 | 0.101586399 | 0.250425106 | 0.103252645 |
| ENSMUSG00000039232 | Stx11 | 74732 | protein_coding | 3.709874668 | -0.103268329 | 0.815885045 | 0.126572156 | 0.899279036 | 1 | 0.103268329 |
| ENSMUSG00000024899 | Papss2 | 23972 | protein_coding | 6410.762028 | -0.103269891 | 0.135663573 | 0.761220484 | 0.446525385 | 0.663778937 | 0.103269891 |

Supplementary Table 1

| ensembl_gene_id | external_gene_name | entrezgene_id | gene_biotype | baseMean | log2FoldChange | lfcSE | stat | pvalue | padj | absLog2FC |
|---|---|---|---|---|---|---|---|---|---|---|
| ENSMUSG00000075232 | Amd1 | 11702 | protein_coding | 402.6814976 | -0.103296923 | 0.153069422 | 0.674837088 | 0.499779274 | 0.704451599 | 0.103296923 |
| ENSMUSG00000020454 | Eif4enif1 | 74203 | protein_coding | 413.038094 | 0.103353276 | 0.058175032 | -1.77659164 | 0.075635452 | 0.20160769 | 0.103353276 |
| ENSMUSG00000030879 | Mrpl17 | 27397 | protein_coding | 471.2054793 | 0.10337485 | 0.097959103 | -1.055285802 | 0.291294633 | 0.510066755 | 0.10337485 |
| ENSMUSG00000030911 | Zp2 | 22787 | protein_coding | 5.418931895 | -0.103404622 | 0.658441335 | 0.157044548 | 0.875209739 | 0.938415991 | 0.103404622 |
| ENSMUSG00000046456 | Tmem150b | 330460 | protein_coding | 199.6146188 | -0.103410617 | 0.123680629 | 0.836110053 | 0.403092994 | 0.624336681 | 0.103410617 |
| ENSMUSG00000039997 | Ifi203 | 15950 | protein_coding | 6.048377488 | 0.103437671 | 0.735304645 | -0.140673218 | 0.888128104 | 0.94482687 | 0.103437671 |
| ENSMUSG00000040048 | Ndufb10 | 68342 | protein_coding | 1503.821282 | -0.103524619 | 0.083658642 | 1.237464733 | 0.215914599 | 0.421168638 | 0.103524619 |
| ENSMUSG00000057058 | Skap1 | 78473 | protein_coding | 9.003785644 | 0.103527451 | 0.533825431 | -0.19393503 | 0.846226754 | 0.924222269 | 0.103527451 |
| ENSMUSG00000052459 | Atp6v1a | 11964 | protein_coding | 991.831285 | -0.103531983 | 0.089905987 | 1.151558262 | 0.249502641 | 0.461760239 | 0.103531983 |
| ENSMUSG00000015461 | Atf6b | 12915 | protein_coding | 455.1214415 | 0.103588122 | 0.080246683 | -1.290871077 | 0.196748386 | 0.396473807 | 0.103588122 |
| ENSMUSG00000030403 | Vasp | 22323 | protein_coding | 4728.198015 | -0.103645905 | 0.079460269 | 1.304373957 | 0.19210611 | 0.389807821 | 0.103645905 |
| ENSMUSG00000080365 | Gm25776 | 115488078 | snoRNA | 122.5279008 | -0.103657309 | 0.287670418 | 0.360333572 | 0.718597697 | 0.854165577 | 0.103657309 |
| ENSMUSG00000031823 | Zdhhc7 | 102193 | protein_coding | 933.3012635 | 0.103805042 | 0.069577009 | -1.491944599 | 0.13571367 | 0.30705241 | 0.103805042 |
| ENSMUSG00000038522 | Mfsd4b1 | 215929 | protein_coding | 1.340139256 | 0.103830259 | 1.100828384 | -0.094320114 | 0.924854872 | 1 | 0.103830259 |
| ENSMUSG00000057554 | Lgals8 | 56048 | protein_coding | 644.1824923 | 0.103838861 | 0.082433046 | -1.259675172 | 0.207786565 | 0.410708835 | 0.103838861 |
| ENSMUSG00000050796 | B3galt6 | 117592 | protein_coding | 92.95029415 | 0.103858599 | 0.143944842 | -0.721516644 | 0.470591704 | 0.683821668 | 0.103858599 |
| ENSMUSG00000024789 | Jak2 | 16452 | protein_coding | 916.3442459 | -0.103872877 | 0.069926072 | 1.485467065 | 0.137420142 | 0.309858682 | 0.103872877 |
| ENSMUSG00000037408 | Cnnm4 | 94220 | protein_coding | 6137.144141 | -0.103887441 | 0.088901785 | 1.168564175 | 0.242579269 | 0.45367543 | 0.103887441 |
| ENSMUSG00000038975 | Rabggtb | 19352 | protein_coding | 440.8189274 | -0.103933569 | 0.089181431 | 1.165417141 | 0.243850182 | 0.455021685 | 0.103933569 |
| ENSMUSG00000038324 | Trpc4ap | 56407 | protein_coding | 1159.503051 | 0.103950863 | 0.056693883 | -1.833546384 | 0.066721357 | 0.184376923 | 0.103950863 |
| ENSMUSG00000098112 | Bin2 | 668218 | protein_coding | 16.29595073 | 0.104052263 | 0.358430312 | -0.290299843 | 0.771586859 | 0.885228942 | 0.104052263 |
| ENSMUSG00000029551 | Psmg3 | 66506 | protein_coding | 98.5049566 | 0.104070987 | 0.117949524 | -0.882334951 | 0.377595701 | 0.599693731 | 0.104070987 |
| ENSMUSG00000029580 | Actb | 11461 | protein_coding | 54482.32294 | 0.104087843 | 0.109089955 | -0.954146903 | 0.340009287 | 0.561491337 | 0.104087843 |
| ENSMUSG00000085049 | Parvaos | 102636183 | lncRNA | 2.01205628 | 0.104097177 | 1.091989858 | -0.095327971 | 0.924054326 | 1 | 0.104097177 |
| ENSMUSG00000033767 | Tmem131l | 229473 | protein_coding | 196.2909425 | -0.104100214 | 0.101835227 | 1.02224168 | 0.306666528 | 0.527150555 | 0.104100214 |
| ENSMUSG00000041308 | Sntb2 | 20650 | protein_coding | 169.8832226 | 0.10412869 | 0.115171589 | -0.904117854 | 0.365932912 | 0.588586355 | 0.10412869 |
| ENSMUSG00000090150 | Acad11 | 102632 | protein_coding | 1469.791732 | -0.104148292 | 0.088212286 | 1.180655176 | 0.237739735 | 0.447999484 | 0.104148292 |
| ENSMUSG00000011096 | Akt1s1 | 67605 | protein_coding | 545.6327809 | 0.104157795 | 0.081076974 | -1.284677868 | 0.198904875 | 0.39947392 | 0.104157795 |
| ENSMUSG00000029605 | Oas1b | 23961 | protein_coding | 19.87747968 | 0.104181109 | 0.40149335 | -0.25948402 | 0.79526181 | 0.898286835 | 0.104181109 |
| ENSMUSG00000038871 | Bpgm | 12183 | protein_coding | 575.2319686 | 0.104217551 | 0.073977611 | -1.408771526 | 0.158902737 | 0.342932541 | 0.104217551 |
| ENSMUSG00000032705 | Exd2 | 97827 | protein_coding | 246.3547716 | 0.104264937 | 0.099610447 | -1.046726929 | 0.29522554 | 0.514332163 | 0.104264937 |
| ENSMUSG00000030244 | Gys2 | 232493 | protein_coding | 1.698799735 | 0.104299873 | 1.286980721 | -0.081042296 | 0.935408316 | 1 | 0.104299873 |
| ENSMUSG00000042572 | Ube2q1 | 70093 | protein_coding | 828.7091037 | -0.10432357 | 0.054946596 | 1.898635727 | 0.057612388 | 0.16577019 | 0.10432357 |
| ENSMUSG00000028256 | Odf2l | 52184 | protein_coding | 157.840942 | 0.104334504 | 0.087883062 | -1.187196957 | 0.235149941 | 0.445073481 | 0.104334504 |
| ENSMUSG00000021810 | Ecd | 70601 | protein_coding | 492.5904637 | 0.104360966 | 0.077060007 | -1.354281817 | 0.175646485 | 0.367540047 | 0.104360966 |
| ENSMUSG00000025139 | Tollip | 54473 | protein_coding | 685.0697994 | -0.104375461 | 0.052468478 | 1.989298434 | 0.046668272 | 0.141461059 | 0.104375461 |
| ENSMUSG00000014980 | Tsen15 | 66637 | protein_coding | 59.83425618 | 0.104392778 | 0.177758638 | -0.587272604 | 0.557020637 | 0.746218772 | 0.104392778 |
| ENSMUSG00000051934 | Spats2 | 72572 | protein_coding | 154.4600214 | 0.104433152 | 0.108679578 | -0.960927104 | 0.336588821 | 0.559203284 | 0.104433152 |
| ENSMUSG00000039221 | Rpl22l1 | 68028 | protein_coding | 565.2474978 | 0.104483562 | 0.128822268 | -0.811067552 | 0.417326878 | 0.637415471 | 0.104483562 |

| ensembl_gene_id | external_gene_name | entrezgene_id | gene_biotype | baseMean | log2FoldChange | lfcSE | stat | pvalue | padj | absLog2FC |
|---|---|---|---|---|---|---|---|---|---|---|
| ENSMUSG00000020023 | Tmcc3 | 319880 | protein_coding | 1496.282863 | 0.104556564 | 0.173950968 | -0.601069173 | 0.547793914 | 0.739162904 | 0.104556564 |
| ENSMUSG00000029145 | Eif2b4 | 13667 | protein_coding | 358.870762 | -0.104578941 | 0.091736034 | 1.139998502 | 0.254286925 | 0.466951438 | 0.104578941 |
| ENSMUSG00000028587 | Orc1 | 18392 | protein_coding | 129.3687287 | 0.104580809 | 0.115627016 | -0.904466903 | 0.365747876 | 0.588390676 | 0.104580809 |
| ENSMUSG00000037217 | Syn1 | 20964 | protein_coding | 5.550659735 | -0.104615605 | 0.521213663 | 0.200715393 | 0.840921123 | 0.921253246 | 0.104615605 |
| ENSMUSG00000047719 | Ubiad1 | 71707 | protein_coding | 229.8652062 | 0.104618749 | 0.103551095 | -1.010310408 | 0.312346596 | 0.533456622 | 0.104618749 |
| ENSMUSG00000019088 | Dnase1l1 | 69537 | protein_coding | 153.1493468 | -0.104649825 | 0.119207939 | 0.877876302 | 0.380010851 | 0.60223179 | 0.104649825 |
| ENSMUSG00000103409 | Lsmem2 | 434436 | protein_coding | 9.973412545 | -0.104708918 | 0.38702323 | 0.27054944 | 0.786737587 | 0.893302015 | 0.104708918 |
| ENSMUSG00000009640 | Dmap1 | 66233 | protein_coding | 209.3706332 | -0.104754458 | 0.096312179 | 1.087655369 | 0.276747277 | 0.493649227 | 0.104754458 |
| ENSMUSG00000045827 | Serpinb9 | 20723 | protein_coding | 36.60481177 | -0.104772317 | 0.177744906 | 0.589453277 | 0.555557246 | 0.7452256 | 0.104772317 |
| ENSMUSG00000084803 | 5830444B04Rik | 641454 | lncRNA | 44.05052523 | -0.104915572 | 0.358304693 | 0.292811047 | 0.769666584 | 0.884510137 | 0.104915572 |
| ENSMUSG00000051295 | 9630028B13Rik | 319562 | lncRNA | 30.65846421 | 0.104929082 | 0.321710081 | -0.326160379 | 0.744303017 | 0.870990647 | 0.104929082 |
| ENSMUSG00000088951 | Gm24940 | 115487682 | snRNA | 0.760793376 | 0.104947673 | 1.61978264 | -0.064791208 | 0.948340242 | 1 | 0.104947673 |
| ENSMUSG00000045767 | B230219D22Rik | 78521 | protein_coding | 587.3948086 | 0.10499056 | 0.094623896 | -1.109556504 | 0.267190183 | 0.482454733 | 0.10499056 |
| ENSMUSG00000020137 | Thap2 | 66816 | protein_coding | 228.274071 | 0.104995771 | 0.127351816 | -0.824454449 | 0.409681388 | 0.630279329 | 0.104995771 |
| ENSMUSG00000026765 | Lypd6b | 71897 | protein_coding | 58.5044114 | -0.10512821 | 0.276828551 | 0.379759275 | 0.704124115 | 0.844376272 | 0.10512821 |
| ENSMUSG00000003970 | Rpl8 | 26961 | protein_coding | 8930.751957 | -0.105173886 | 0.088247823 | 1.191801487 | 0.233339093 | 0.442718029 | 0.105173886 |
| ENSMUSG00000018196 | Glrx2 | 69367 | protein_coding | 207.7533573 | 0.105193699 | 0.096489788 | -1.090205512 | 0.275622626 | 0.492426209 | 0.105193699 |
| ENSMUSG00000062960 | Kdr | 16542 | protein_coding | 3.086564233 | -0.105236747 | 2.428591127 | 0.043332427 | 0.965436543 | 1 | 0.105236747 |
| ENSMUSG00000049940 | Pgrmc2 | 70804 | protein_coding | 375.9280373 | 0.105258194 | 0.081814893 | -1.286540745 | 0.198254408 | 0.3984875 | 0.105258194 |
| ENSMUSG00000024431 | Nr3c1 | 14815 | protein_coding | 1489.699065 | -0.105259843 | 0.05767575 | 1.825027735 | 0.067996844 | 0.186963018 | 0.105259843 |
| ENSMUSG00000038803 | Ost4 | 67695 | protein_coding | 393.2972045 | 0.105290383 | 0.05706754 | -1.845013534 | 0.06503557 | 0.181112546 | 0.105290383 |
| ENSMUSG00000028127 | Abcd3 | 19299 | protein_coding | 2944.723536 | 0.105321402 | 0.064859821 | -1.62383122 | 0.104411835 | 0.255483337 | 0.105321402 |
| ENSMUSG00000025332 | Kdm5c | 20591 | protein_coding | 2456.467971 | 0.105344485 | 0.064919844 | -1.622685436 | 0.104656666 | 0.255796319 | 0.105344485 |
| ENSMUSG00000044496 | 2510039O18Rik | 77034 | protein_coding | 1362.549504 | 0.105377236 | 0.065278769 | -1.614265064 | 0.106469944 | 0.259060655 | 0.105377236 |
| ENSMUSG00000042978 | Sbk1 | 104175 | protein_coding | 263.7722422 | -0.105403075 | 0.107033121 | 0.984770639 | 0.324736746 | 0.546768212 | 0.105403075 |
| ENSMUSG00000098332 | Pigbos1 | 66364 | protein_coding | 85.91415225 | -0.105426074 | 0.128492733 | 0.820482774 | 0.411940945 | 0.632229811 | 0.105426074 |
| ENSMUSG00000102440 | Pcdhga9 | 93717 | protein_coding | 11.45871078 | -0.105464497 | 0.3029474 | 0.348128082 | 0.727743996 | 0.859322531 | 0.105464497 |
| ENSMUSG00000056820 | Tsnax | 53424 | protein_coding | 585.5018824 | -0.105471187 | 0.095239921 | 1.107426225 | 0.268109692 | 0.48334534 | 0.105471187 |
| ENSMUSG00000058297 | Spock2 | 94214 | protein_coding | 14.09051112 | -0.10548779 | 0.455786338 | 0.231441316 | 0.816971971 | 0.908700419 | 0.10548779 |
| ENSMUSG00000021282 | Eif5 | 217869 | protein_coding | 2591.574461 | 0.105506508 | 0.06805499 | -1.550312591 | 0.121066507 | 0.283788828 | 0.105506508 |
| ENSMUSG00000036504 | Phpt1 | 75454 | protein_coding | 17.78360501 | 0.105522907 | 0.262108622 | -0.402592278 | 0.687248191 | 0.833389267 | 0.105522907 |
| ENSMUSG00000030650 | Tmc5 | 74424 | protein_coding | 1653.523475 | -0.105545448 | 0.115245646 | 0.915830245 | 0.359755938 | 0.582349713 | 0.105545448 |
| ENSMUSG00000071291 | Zfp58 | 238693 | protein_coding | 33.00017572 | -0.105581036 | 0.177281881 | 0.59555458 | 0.551472827 | 0.742339604 | 0.105581036 |
| ENSMUSG00000003882 | Il7r | 16197 | protein_coding | 2.494114433 | -0.105611954 | 0.818417381 | 0.129044124 | 0.897322735 | 1 | 0.105611954 |
| ENSMUSG00000042203 | Tbc1d22b | 381085 | protein_coding | 282.2710126 | 0.105614135 | 0.084339871 | -1.252244435 | 0.210480809 | 0.414545055 | 0.105614135 |
| ENSMUSG00000043183 | Simc1 | 319719 | protein_coding | 55.46419195 | 0.105654242 | 0.19984997 | -0.52866779 | 0.597035925 | 0.774998778 | 0.105654242 |
| ENSMUSG00000071424 | Grid2 | 14804 | protein_coding | 27.74865513 | 0.105707909 | 0.298401897 | -0.354246773 | 0.723153953 | 0.856549354 | 0.105707909 |
| ENSMUSG00000048779 | P2ry6 | 233571 | protein_coding | 173.6679412 | -0.105777513 | 0.10669613 | 0.991390344 | 0.321495014 | 0.542471509 | 0.105777513 |
| ENSMUSG00000002033 | Cd3g | 12502 | protein_coding | 21.8240925 | -0.105789813 | 0.373147012 | 0.283507063 | 0.776788168 | 0.888506311 | 0.105789813 |

| ensembl_gene_id | external_gene_name | entrezgene_id | gene_biotype | baseMean | log2FoldChange | lfcSE | stat | pvalue | padj | absLog2FC |
|---|---|---|---|---|---|---|---|---|---|---|
| ENSMUSG00000089857 | Zfp882 | 382019 | protein_coding | 24.14027483 | 0.105800009 | 0.267535322 | -0.395461835 | 0.692502082 | 0.836513744 | 0.105800009 |
| ENSMUSG00000073147 | 5031425E22Rik | 269630 | lncRNA | 205.1424808 | -0.105862805 | 0.227183466 | 0.465979356 | 0.641230277 | 0.804106371 | 0.105862805 |
| ENSMUSG00000062761 | Zfp512 | 269639 | protein_coding | 182.843183 | -0.105891186 | 0.086508382 | 1.224056945 | 0.220930746 | 0.427441699 | 0.105891186 |
| ENSMUSG00000044465 | Fhip1b | 74349 | protein_coding | 732.3457401 | -0.105906149 | 0.08187546 | 1.293502953 | 0.195837163 | 0.395168001 | 0.105906149 |
| ENSMUSG00000028683 | Eif2b3 | 108067 | protein_coding | 105.8057144 | -0.10592928 | 0.15418689 | 0.687018717 | 0.492070933 | 0.698788598 | 0.10592928 |
| ENSMUSG00000058997 | Vwa8 | 219189 | protein_coding | 2526.719602 | -0.105933378 | 0.084668489 | 1.251154695 | 0.210878044 | 0.415010021 | 0.105933378 |
| ENSMUSG00000029064 | Gnb1 | 14688 | protein_coding | 3513.604006 | 0.105957494 | 0.060762593 | -1.743794816 | 0.081194875 | 0.211992437 | 0.105957494 |
| ENSMUSG00000022704 | Qtrt2 | 106248 | protein_coding | 166.0286267 | -0.105962013 | 0.102809905 | 1.030659578 | 0.302700487 | 0.523269741 | 0.105962013 |
| ENSMUSG00000087177 | E130307A14Rik | 327744 | lncRNA | 31.78932449 | 0.105969401 | 0.250520506 | -0.422996914 | 0.672297513 | 0.824348168 | 0.105969401 |
| ENSMUSG00000031803 | B3gnt3 | 72297 | protein_coding | 3052.958989 | 0.105995821 | 0.068163055 | -1.555033305 | 0.119938137 | 0.281984397 | 0.105995821 |
| ENSMUSG00000055302 | Mrfap1 | 67568 | protein_coding | 1484.797484 | -0.105999115 | 0.100010432 | 1.05988058 | 0.289198932 | 0.508065109 | 0.105999115 |
| ENSMUSG00000038530 | Rgs4 | 19736 | protein_coding | 6.238012696 | 0.10600377 | 0.597616281 | -0.177377648 | 0.859211763 | 0.931099354 | 0.10600377 |
| ENSMUSG00000031023 | Akip1 | 57373 | protein_coding | 98.42193533 | -0.106011516 | 0.165938378 | 0.638860746 | 0.522913526 | 0.722022907 | 0.106011516 |
| ENSMUSG00000021037 | Ahsa1 | 217737 | protein_coding | 848.1617248 | -0.106036018 | 0.117418201 | 0.90306287 | 0.366492529 | 0.589199534 | 0.106036018 |
| ENSMUSG00000047878 | A4galt | 239559 | protein_coding | 170.5537055 | -0.10606364 | 0.256362298 | 0.413725578 | 0.679075093 | 0.827951097 | 0.10606364 |
| ENSMUSG00000030228 | Pik3c2g | 18705 | protein_coding | 18.78719684 | -0.106086921 | 0.304692726 | 0.348176741 | 0.727707454 | 0.859322531 | 0.106086921 |
| ENSMUSG00000052033 | Pfdn4 | 109054 | protein_coding | 95.68201487 | 0.10610949 | 0.123306393 | -0.860535187 | 0.389494096 | 0.612056856 | 0.10610949 |
| ENSMUSG00000030780 | Rusf1 | 233913 | protein_coding | 334.9807159 | 0.106135561 | 0.072808209 | -1.457741672 | 0.144911761 | 0.321952485 | 0.106135561 |
| ENSMUSG00000022255 | Mtdh | 67154 | protein_coding | 1453.468859 | 0.106193626 | 0.08519825 | -1.246429659 | 0.212606698 | 0.417180016 | 0.106193626 |
| ENSMUSG00000030126 | Tmcc1 | 330401 | protein_coding | 426.3760262 | -0.106256142 | 0.068991812 | 1.540126852 | 0.123529436 | 0.287399986 | 0.106256142 |
| ENSMUSG00000092341 | Malat1 | 72289 | lncRNA | 45152.54829 | -0.106279632 | 0.145800247 | 0.728939998 | 0.466038367 | 0.679736277 | 0.106279632 |
| ENSMUSG00000022545 | Ercc4 | 50505 | protein_coding | 164.3916374 | -0.10628205 | 0.082396592 | 1.289884053 | 0.197090918 | 0.397106786 | 0.10628205 |
| ENSMUSG00000037257 | Aagab | 66939 | protein_coding | 761.6564489 | -0.106290111 | 0.10505401 | 1.011766333 | 0.311649791 | 0.532495959 | 0.106290111 |
| ENSMUSG00000019916 | P4ha1 | 18451 | protein_coding | 222.5658689 | 0.106296831 | 0.098635779 | -1.07767011 | 0.281181002 | 0.498917274 | 0.106296831 |
| ENSMUSG00000024556 | Me2 | 107029 | protein_coding | 2078.302351 | -0.106304017 | 0.100134054 | 1.061617023 | 0.288409583 | 0.50724071 | 0.106304017 |
| ENSMUSG00000056014 | A430033K04Rik | 243308 | protein_coding | 140.2643715 | 0.106329086 | 0.090683941 | -1.172523885 | 0.240986788 | 0.45166903 | 0.106329086 |
| ENSMUSG00000022175 | Lrp10 | 65107 | protein_coding | 3466.077545 | 0.106347724 | 0.07092091 | -1.499525646 | 0.133737321 | 0.303828141 | 0.106347724 |
| ENSMUSG00000008036 | Ap2s1 | 232910 | protein_coding | 1052.019843 | 0.10636781 | 0.065547712 | -1.622753973 | 0.104642009 | 0.255796319 | 0.10636781 |
| ENSMUSG00000043683 | Fem1a | 14154 | protein_coding | 789.0453671 | -0.106408165 | 0.08471132 | 1.256126862 | 0.209069979 | 0.412396276 | 0.106408165 |
| ENSMUSG00000040390 | Map3k10 | 269881 | protein_coding | 134.3249288 | 0.106420932 | 0.116131292 | -0.916384637 | 0.35946519 | 0.582152413 | 0.106420932 |
| ENSMUSG00000002550 | Uck1 | 22245 | protein_coding | 341.0699447 | -0.106424583 | 0.105221061 | 1.011438032 | 0.311806826 | 0.532600117 | 0.106424583 |
| ENSMUSG00000021456 | Fbp2 | 14120 | protein_coding | 1616.048715 | -0.106529392 | 0.076237213 | 1.397341109 | 0.162311019 | 0.347734431 | 0.106529392 |
| ENSMUSG00000001707 | Eef1e1 | 66143 | protein_coding | 146.4284245 | 0.106536372 | 0.11979448 | -0.88932622 | 0.373827784 | 0.595844868 | 0.106536372 |
| ENSMUSG00000040464 | Gtpbp10 | 207704 | protein_coding | 167.3405463 | -0.106546148 | 0.128389089 | 0.829869183 | 0.40661275 | 0.627434401 | 0.106546148 |
| ENSMUSG00000042308 | Setd1a | 233904 | protein_coding | 100.1811584 | 0.106581194 | 0.139762326 | -0.762588868 | 0.445708626 | 0.662922836 | 0.106581194 |
| ENSMUSG00000085169 | Gm10785 | 100038392 | lncRNA | 13.51222303 | -0.106586909 | 0.34748776 | 0.306735722 | 0.759044537 | 0.878912418 | 0.106586909 |
| ENSMUSG00000078954 | Arhgap8 | 73167 | protein_coding | 67.08843854 | -0.106590161 | 0.159208234 | 0.669501558 | 0.503175587 | 0.707788896 | 0.106590161 |
| ENSMUSG00000020231 | Dip2a | 64451 | protein_coding | 1124.506352 | -0.106610291 | 0.049821494 | 2.139845304 | 0.032367271 | 0.10695305 | 0.106610291 |
| ENSMUSG00000024725 | Ostf1 | 20409 | protein_coding | 948.4176055 | 0.106620729 | 0.081860944 | -1.302461512 | 0.192758667 | 0.390530809 | 0.106620729 |

| ensembl_gene_id | external_gene_name | entrezgene_id | gene_biotype | baseMean | log2FoldChange | lfcSE | stat | pvalue | padj | absLog2FC |
|---|---|---|---|---|---|---|---|---|---|---|
| ENSMUSG00000019297 | Nop9 | 67842 | protein_coding | 240.2118745 | 0.106631506 | 0.121813043 | -0.875370192 | 0.381372513 | 0.603717953 | 0.106631506 |
| ENSMUSG00000003721 | Insig2 | 72999 | protein_coding | 305.4759099 | 0.106635889 | 0.08967483 | -1.189139575 | 0.234384748 | 0.443879555 | 0.106635889 |
| ENSMUSG00000025940 | Tmem70 | 70397 | protein_coding | 405.2832558 | -0.106641258 | 0.11015074 | 0.96813928 | 0.332974815 | 0.555316713 | 0.106641258 |
| ENSMUSG00000042719 | Naa25 | 231713 | protein_coding | 358.7958188 | 0.10666058 | 0.084152415 | -1.267469032 | 0.20498763 | 0.407409522 | 0.10666058 |
| ENSMUSG00000006273 | Atp6v1b2 | 11966 | protein_coding | 775.5482221 | -0.106660774 | 0.087389729 | 1.220518423 | 0.22226841 | 0.429396526 | 0.106660774 |
| ENSMUSG00000100627 | A830008E24Rik | 108167736 | lncRNA | 2.678573221 | -0.106679485 | 1.159010551 | 0.092043584 | 0.926663412 | 1 | 0.106679485 |
| ENSMUSG00000018217 | Pmp22 | 18858 | protein_coding | 2285.728431 | 0.106688928 | 0.183292986 | -0.582067707 | 0.560521076 | 0.749145763 | 0.106688928 |
| ENSMUSG00000028902 | Sf3a3 | 75062 | protein_coding | 427.5666805 | 0.106737627 | 0.11946297 | -0.893478763 | 0.37160084 | 0.593942729 | 0.106737627 |
| ENSMUSG00000061458 | Nol10 | 217431 | protein_coding | 79.50673311 | 0.106803025 | 0.146902398 | -0.727033911 | 0.467205182 | 0.680853016 | 0.106803025 |
| ENSMUSG00000039768 | Dnajc11 | 230935 | protein_coding | 972.1653897 | -0.106820343 | 0.073949143 | 1.444510912 | 0.148595313 | 0.327361658 | 0.106820343 |
| ENSMUSG00000033900 | Map9 | 213582 | protein_coding | 23.23895291 | 0.106856114 | 0.262335914 | -0.407325527 | 0.683768917 | 0.83113204 | 0.106856114 |
| ENSMUSG00000008167 | Fbxw9 | 68628 | protein_coding | 218.3596666 | 0.106874896 | 0.091952161 | -1.162288022 | 0.24511849 | 0.456768986 | 0.106874896 |
| ENSMUSG00000039201 | Tbc1d25 | 209815 | protein_coding | 96.9241324 | -0.106900275 | 0.118271201 | 0.903857188 | 0.366071133 | 0.588694 | 0.106900275 |
| ENSMUSG00000038023 | Atp6v0a2 | 21871 | protein_coding | 2174.584519 | 0.10690178 | 0.069206898 | -1.544669437 | 0.122426241 | 0.285476835 | 0.10690178 |
| ENSMUSG00000023723 | Mrps23 | 64656 | protein_coding | 592.9324733 | -0.106948879 | 0.08498686 | 1.258416643 | 0.208241116 | 0.41130117 | 0.106948879 |
| ENSMUSG00000005447 | Pafah1b3 | 18476 | protein_coding | 690.1722408 | 0.107040492 | 0.100463222 | -1.06546943 | 0.286663552 | 0.50522539 | 0.107040492 |
| ENSMUSG00000118559 | A930007A09Rik | 432999 | protein_coding | 6.520009181 | -0.107052009 | 0.621359675 | 0.172286702 | 0.863212139 | 0.932166328 | 0.107052009 |
| ENSMUSG00000060038 | Dhps | 330817 | protein_coding | 190.9566499 | 0.107056447 | 0.091710129 | -1.167335047 | 0.24307509 | 0.454166041 | 0.107056447 |
| ENSMUSG00000021650 | Ptcd2 | 68927 | protein_coding | 569.7018585 | 0.107060871 | 0.098867061 | -1.082877045 | 0.278863002 | 0.496167516 | 0.107060871 |
| ENSMUSG00000071662 | Polr2g | 67710 | protein_coding | 251.0951382 | 0.10707354 | 0.092903683 | -1.152522019 | 0.249106627 | 0.461264989 | 0.10707354 |
| ENSMUSG00000020419 | Hormad2 | 75828 | protein_coding | 1.475333014 | 0.107088192 | 1.154310206 | -0.092772456 | 0.926084334 | 1 | 0.107088192 |
| ENSMUSG00000057894 | Zfp329 | 67230 | protein_coding | 141.4898355 | -0.10710373 | 0.137506397 | 0.778899982 | 0.436038635 | 0.654196603 | 0.10710373 |
| ENSMUSG00000029310 | Nudt9 | 74167 | protein_coding | 391.2299778 | -0.10710905 | 0.073058756 | 1.466067257 | 0.142629965 | 0.318339132 | 0.10710905 |
| ENSMUSG00000090000 | Ier3ip1 | 66191 | protein_coding | 351.8918202 | 0.107175299 | 0.092918214 | -1.153436927 | 0.248731092 | 0.460883095 | 0.107175299 |
| ENSMUSG00000032403 | 2300009A05Rik | 69478 | protein_coding | 41.41350372 | -0.107217269 | 0.204115365 | 0.525277794 | 0.599390092 | 0.77652216 | 0.107217269 |
| ENSMUSG00000006932 | Ctnnb1 | 12387 | protein_coding | 10595.83715 | 0.107219554 | 0.042576335 | -2.518289919 | 0.011792621 | 0.04843157 | 0.107219554 |
| ENSMUSG00000049657 | Zbtb5 | 230119 | protein_coding | 122.3670956 | -0.107222751 | 0.131527079 | 0.815214266 | 0.414949674 | 0.635253445 | 0.107222751 |
| ENSMUSG00000031766 | Slc12a3 | 20497 | protein_coding | 8.452439679 | 0.107230702 | 0.689801239 | -0.155451594 | 0.876465313 | 0.938630585 | 0.107230702 |
| ENSMUSG00000007656 | Arpp19 | 59046 | protein_coding | 867.517463 | -0.107238624 | 0.060192718 | 1.781587985 | 0.074816445 | 0.19999509 | 0.107238624 |
| ENSMUSG00000033632 | AW554918 | 225289 | protein_coding | 91.50225169 | 0.107242396 | 0.112148085 | -0.956257034 | 0.338942392 | 0.560514185 | 0.107242396 |
| ENSMUSG00000024104 | Washc2 | 28006 | protein_coding | 569.1278381 | -0.107255342 | 0.081965993 | 1.308534653 | 0.190692029 | 0.388133006 | 0.107255342 |
| ENSMUSG00000054514 | Atad3aos | 70448 | lncRNA | 5.010153085 | -0.107310818 | 0.506015231 | 0.212070331 | 0.832052166 | 0.915301723 | 0.107310818 |
| ENSMUSG00000069306 | H4c17 | 100041230 | protein_coding | 0.773698078 | -0.107329467 | 1.658733238 | 0.064705683 | 0.948408338 | 1 | 0.107329467 |
| ENSMUSG00000021396 | Nxnl2 | 75124 | protein_coding | 3.068097215 | 0.107383007 | 0.725716643 | -0.14796823 | 0.882367841 | 1 | 0.107383007 |
| ENSMUSG00000057396 | Zfp759 | 268670 | protein_coding | 77.42661503 | -0.107390658 | 0.134160646 | 0.800463187 | 0.423442484 | 0.643305352 | 0.107390658 |
| ENSMUSG00000041729 | Coro2b | 235431 | protein_coding | 13.00555245 | 0.10739154 | 0.592872865 | -0.181137552 | 0.85625961 | 0.930174321 | 0.10739154 |
| ENSMUSG00000000958 | Slc7a7 | 20540 | protein_coding | 420.9106072 | 0.10740681 | 0.078393103 | -1.370105342 | 0.17065402 | 0.360508479 | 0.10740681 |
| ENSMUSG00000032998 | Foxj3 | 230700 | protein_coding | 401.53314 | -0.107414385 | 0.094905823 | 1.13179973 | 0.257718641 | 0.47089684 | 0.107414385 |
| ENSMUSG00000021819 | Zswim8 | 268721 | protein_coding | 558.7977277 | 0.10743396 | 0.09313172 | -1.153570021 | 0.248676495 | 0.460883095 | 0.10743396 |

| ensembl_gene_id | external_gene_name | entrezgene_id | gene_biotype | baseMean | log2FoldChange | lfcSE | stat | pvalue | padj | absLog2FC |
|---|---|---|---|---|---|---|---|---|---|---|
| ENSMUSG00000018821 | Avpi1 | 69534 | protein_coding | 123.5903826 | 0.107502194 | 0.1723511 | -0.623739528 | 0.532798659 | 0.72936091 | 0.107502194 |
| ENSMUSG00000031696 | Vps35 | 65114 | protein_coding | 1407.355784 | 0.107551381 | 0.083214303 | -1.292462668 | 0.196196966 | 0.395770858 | 0.107551381 |
| ENSMUSG00000032846 | Zswim6 | 67263 | protein_coding | 290.8133656 | -0.107552582 | 0.127934594 | 0.840684123 | 0.400524917 | 0.622394781 | 0.107552582 |
| ENSMUSG00000018199 | Ro60 | 20822 | protein_coding | 301.8138463 | 0.107559139 | 0.108788349 | -0.988700907 | 0.322809499 | 0.544244621 | 0.107559139 |
| ENSMUSG00000041459 | Tardbp | 230908 | protein_coding | 1216.332788 | 0.107621515 | 0.088615786 | -1.214473412 | 0.224566997 | 0.431773126 | 0.107621515 |
| ENSMUSG00000001583 | Tnk1 | 83813 | protein_coding | 355.7738152 | -0.10767586 | 0.095110205 | 1.132116788 | 0.257585338 | 0.470794255 | 0.10767586 |
| ENSMUSG00000031328 | Flna | 192176 | protein_coding | 339.285902 | 0.107757987 | 0.117612347 | -0.916213221 | 0.359555073 | 0.582240877 | 0.107757987 |
| ENSMUSG00000049728 | Zfp668 | 244219 | protein_coding | 108.7536669 | 0.107789184 | 0.126640369 | -0.851143949 | 0.394689394 | 0.617166997 | 0.107789184 |
| ENSMUSG00000001739 | Cldn15 | 60363 | protein_coding | 2021.985921 | -0.107833568 | 0.109920414 | 0.981014939 | 0.326585374 | 0.548411142 | 0.107833568 |
| ENSMUSG00000032489 | Kif9 | 16578 | protein_coding | 24.33026664 | -0.107841425 | 0.249173241 | 0.432796976 | 0.66516227 | 0.819432282 | 0.107841425 |
| ENSMUSG00000025821 | Zfp282 | 101095 | protein_coding | 317.2746568 | -0.107874902 | 0.073342631 | 1.470834913 | 0.141335768 | 0.316005805 | 0.107874902 |
| ENSMUSG00000027478 | Dnmt3b | 13436 | protein_coding | 43.96226413 | -0.107899985 | 0.224938306 | 0.479687014 | 0.631449963 | 0.799123664 | 0.107899985 |
| ENSMUSG00000038697 | Taf5l | 102162 | protein_coding | 425.0357075 | -0.107961682 | 0.057107924 | 1.890485148 | 0.058693105 | 0.168040558 | 0.107961682 |
| ENSMUSG00000041360 | Pum3 | 52874 | protein_coding | 420.7868047 | -0.108066435 | 0.102249262 | 1.056892083 | 0.290560842 | 0.509484153 | 0.108066435 |
| ENSMUSG00000042590 | Ipo11 | 76582 | protein_coding | 283.5944447 | 0.108071343 | 0.118378976 | -0.912926833 | 0.361281024 | 0.583719287 | 0.108071343 |
| ENSMUSG00000024982 | Zdhhc6 | 66980 | protein_coding | 440.2541734 | 0.108071392 | 0.085250704 | -1.26768915 | 0.204908981 | 0.407353559 | 0.108071392 |
| ENSMUSG00000020250 | Txnrd1 | 50493 | protein_coding | 3215.801385 | -0.108097001 | 0.067394137 | 1.603952595 | 0.108724502 | 0.262880883 | 0.108097001 |
| ENSMUSG00000030795 | Fus | 233908 | protein_coding | 431.9428224 | 0.108128774 | 0.164128858 | -0.658804158 | 0.510021537 | 0.71331253 | 0.108128774 |
| ENSMUSG00000031438 | Rnf128 | 66889 | protein_coding | 2348.199402 | -0.108143726 | 0.061765899 | 1.750864592 | 0.079969237 | 0.209820779 | 0.108143726 |
| ENSMUSG00000019732 | Calr3 | 73316 | protein_coding | 7.354353238 | -0.10820249 | 0.414215417 | 0.26122275 | 0.793920735 | 0.897263004 | 0.10820249 |
| ENSMUSG00000054312 | Mrps21 | 66292 | protein_coding | 458.5677327 | -0.108214728 | 0.084236718 | 1.284650339 | 0.198914499 | 0.39947392 | 0.108214728 |
| ENSMUSG00000018377 | Vezf1 | 22344 | protein_coding | 470.5724864 | -0.108240815 | 0.088187104 | 1.227399586 | 0.219672442 | 0.425947586 | 0.108240815 |
| ENSMUSG00000027983 | Cyp2u1 | 71519 | protein_coding | 18.67955595 | -0.108287819 | 0.363233879 | 0.298121474 | 0.765610453 | 0.882510888 | 0.108287819 |
| ENSMUSG00000023939 | Mrpl14 | 68463 | protein_coding | 316.0296976 | -0.108294409 | 0.083230873 | 1.301132677 | 0.193213044 | 0.391355356 | 0.108294409 |
| ENSMUSG00000039841 | Zfp800 | 627049 | protein_coding | 314.7046692 | 0.108318894 | 0.086577084 | -1.251126612 | 0.210888288 | 0.415010021 | 0.108318894 |
| ENSMUSG00000032423 | Syncrip | 56403 | protein_coding | 903.8417648 | 0.108338535 | 0.103559723 | -1.046145472 | 0.295493874 | 0.514582498 | 0.108338535 |
| ENSMUSG00000031974 | Abcb10 | 56199 | protein_coding | 376.1863104 | -0.108349771 | 0.085104389 | 1.27313964 | 0.202968495 | 0.405213026 | 0.108349771 |
| ENSMUSG00000007207 | Stx1a | 20907 | protein_coding | 21.91743188 | 0.108349855 | 0.41605528 | -0.260421775 | 0.794538448 | 0.897715376 | 0.108349855 |
| ENSMUSG00000024683 | Mrpl16 | 94063 | protein_coding | 250.4964966 | 0.108351037 | 0.090089164 | -1.202708876 | 0.229088998 | 0.438019649 | 0.108351037 |
| ENSMUSG00000031897 | Psmb10 | 19171 | protein_coding | 1309.538983 | -0.108428406 | 0.0651897 | 1.66327511 | 0.096257379 | 0.240265168 | 0.108428406 |
| ENSMUSG00000022571 | Pycrl | 66194 | protein_coding | 644.3923307 | -0.108440908 | 0.086403091 | 1.255058197 | 0.209457634 | 0.412914333 | 0.108440908 |
| ENSMUSG00000025995 | Wdr75 | 73674 | protein_coding | 314.5104785 | 0.108446246 | 0.101553154 | -1.067876693 | 0.285576137 | 0.504062027 | 0.108446246 |
| ENSMUSG00000026235 | Epha4 | 13838 | protein_coding | 260.7468617 | 0.108463443 | 0.147271495 | -0.736486331 | 0.461434783 | 0.675184655 | 0.108463443 |
| ENSMUSG00000024645 | Timm21 | 67105 | protein_coding | 188.6113742 | -0.108523541 | 0.127540953 | 0.850891718 | 0.394829506 | 0.617210759 | 0.108523541 |
| ENSMUSG00000004565 | Pnpla6 | 50767 | protein_coding | 706.1092591 | -0.108554729 | 0.10004872 | 1.085018669 | 0.277913387 | 0.494874021 | 0.108554729 |
| ENSMUSG00000057594 | Arl16 | 70317 | protein_coding | 78.36348307 | 0.108565151 | 0.143425114 | -0.756946591 | 0.449081861 | 0.665841071 | 0.108565151 |
| ENSMUSG00000033419 | Snap91 | 20616 | protein_coding | 3.251829716 | -0.108598233 | 1.34161614 | 0.080945831 | 0.935485032 | 1 | 0.108598233 |
| ENSMUSG00000061607 | Mdc1 | 240087 | protein_coding | 248.3376121 | 0.108639744 | 0.123041776 | -0.882950063 | 0.377263253 | 0.599511678 | 0.108639744 |
| ENSMUSG00000091370 | 5730435O14Rik | 70538 | lncRNA | 0.780683872 | -0.108695826 | 1.5339423 | 0.07086044 | 0.943508829 | 1 | 0.108695826 |

Supplementary Table 1

| ensembl_gene_id | external_gene_name | entrezgene_id | gene_biotype | baseMean | log2FoldChange | lfcSE | stat | pvalue | padj | absLog2FC |
|---|---|---|---|---|---|---|---|---|---|---|
| ENSMUSG00000024570 | Rbfa | 68731 | protein_coding | 110.2766775 | -0.108698347 | 0.132641313 | 0.819490888 | 0.412506399 | 0.632451088 | 0.108698347 |
| ENSMUSG00000049612 | Omg | 18377 | protein_coding | 5.007887367 | 0.108714692 | 0.513591721 | -0.21167532 | 0.832360344 | 0.915579745 | 0.108714692 |
| ENSMUSG00000024646 | Cyb5a | 109672 | protein_coding | 999.7410295 | 0.108781453 | 0.069075889 | -1.574810759 | 0.115300132 | 0.273824144 | 0.108781453 |
| ENSMUSG00000084880 | Tomm6os | 100415787 | lncRNA | 5.93748596 | -0.108824458 | 0.531589342 | 0.204715274 | 0.837794586 | 0.919169532 | 0.108824458 |
| ENSMUSG00000092260 | Zfp963 | 620419 | protein_coding | 29.98041749 | 0.108825834 | 0.210165282 | -0.517810712 | 0.604590344 | 0.779942456 | 0.108825834 |
| ENSMUSG00000087260 | Lamtor5 | 68576 | protein_coding | 360.4939305 | -0.108916173 | 0.11904269 | 0.914933738 | 0.360226417 | 0.582642414 | 0.108916173 |
| ENSMUSG00000090164 | BC035044 | 232406 | protein_coding | 2.675241534 | -0.108941103 | 0.741355171 | 0.146948599 | 0.883172592 | 1 | 0.108941103 |
| ENSMUSG00000029178 | Klf3 | 16599 | protein_coding | 2954.586206 | -0.109008839 | 0.084683284 | 1.287253325 | 0.198006006 | 0.398279073 | 0.109008839 |
| ENSMUSG00000004630 | Pcp2 | 18545 | protein_coding | 1.498441324 | -0.10901701 | 1.333665544 | 0.081742391 | 0.934851568 | 1 | 0.10901701 |
| ENSMUSG00000046733 | Gprc5a | 232431 | protein_coding | 1239.746564 | -0.109088388 | 0.22494434 | 0.484957249 | 0.627706671 | 0.796376973 | 0.109088388 |
| ENSMUSG00000021738 | Atxn7 | 246103 | protein_coding | 463.1648533 | 0.109133206 | 0.058195336 | -1.875291266 | 0.060752664 | 0.172412699 | 0.109133206 |
| ENSMUSG00000029153 | Ociad2 | 433904 | protein_coding | 2149.281875 | -0.109161253 | 0.072499908 | 1.505674357 | 0.13215079 | 0.3013667 | 0.109161253 |
| ENSMUSG00000088835 | Gm23547 | 115490496 | snoRNA | 4.742595552 | -0.109174543 | 0.544357266 | 0.200556785 | 0.841045152 | 0.92126689 | 0.109174543 |
| ENSMUSG00000020520 | Galnt10 | 171212 | protein_coding | 2016.380478 | -0.109202698 | 0.132508004 | 0.824121525 | 0.409870511 | 0.630340996 | 0.109202698 |
| ENSMUSG00000021097 | Clmn | 94040 | protein_coding | 2578.882791 | -0.109223692 | 0.125152169 | 0.872727115 | 0.382811834 | 0.605068399 | 0.109223692 |
| ENSMUSG00000021803 | Cdhr1 | 170677 | protein_coding | 87.45868768 | 0.109234846 | 0.213907263 | -0.510664504 | 0.609586 | 0.783477444 | 0.109234846 |
| ENSMUSG00000022283 | Pabpc1 | 18458 | protein_coding | 3903.83026 | -0.109339784 | 0.098733328 | 1.10742528 | 0.268110101 | 0.48334534 | 0.109339784 |
| ENSMUSG00000022962 | Gart | 14450 | protein_coding | 533.733064 | -0.109344088 | 0.130428147 | 0.838347318 | 0.401835668 | 0.623441772 | 0.109344088 |
| ENSMUSG00000026657 | Frmd4a | 209630 | protein_coding | 155.0714926 | 0.109368854 | 0.091811809 | -1.191228619 | 0.233563847 | 0.442985342 | 0.109368854 |
| ENSMUSG00000020483 | Dynll2 | 68097 | protein_coding | 866.2964104 | 0.109407329 | 0.091058864 | -1.201501135 | 0.229556867 | 0.438122528 | 0.109407329 |
| ENSMUSG00000025651 | Uqcrc1 | 22273 | protein_coding | 5157.735676 | -0.10941094 | 0.080408507 | 1.360688608 | 0.173612116 | 0.364391266 | 0.10941094 |
| ENSMUSG00000038970 | Lmtk2 | 231876 | protein_coding | 1503.861004 | 0.109506812 | 0.0719232 | -1.522551995 | 0.127870828 | 0.294844608 | 0.109506812 |
| ENSMUSG00000075010 | AW112010 | 107350 | lncRNA | 367.0131301 | 0.109514569 | 0.146089336 | -0.749641091 | 0.453470896 | 0.669164362 | 0.109514569 |
| ENSMUSG00000024947 | Men1 | 17283 | protein_coding | 405.627589 | 0.109526443 | 0.11012627 | -0.994553281 | 0.319953579 | 0.541253166 | 0.109526443 |
| ENSMUSG00000032405 | Pias1 | 56469 | protein_coding | 282.0144843 | -0.109526903 | 0.071856311 | 1.5242489 | 0.127446547 | 0.294356962 | 0.109526903 |
| ENSMUSG00000045725 | Prr15 | 78004 | protein_coding | 503.3092539 | 0.109540237 | 0.176817838 | -0.619508971 | 0.535581114 | 0.73156216 | 0.109540237 |
| ENSMUSG00000086995 | Gm13544 | 100502932 | lncRNA | 1.751811137 | -0.109568855 | 0.952497181 | 0.115033259 | 0.90841876 | 1 | 0.109568855 |
| ENSMUSG00000028862 | Map3k6 | 53608 | protein_coding | 129.5734996 | -0.109629865 | 0.167946423 | 0.652766895 | 0.513906569 | 0.715629739 | 0.109629865 |
| ENSMUSG00000029919 | Hpgds | 54486 | protein_coding | 76.46245632 | 0.109630084 | 0.148744458 | -0.737036428 | 0.461100197 | 0.674995696 | 0.109630084 |
| ENSMUSG00000039809 | Gabbr2 | 242425 | protein_coding | 4.523670617 | 0.109660769 | 1.017481438 | -0.107776677 | 0.914172844 | 0.958030863 | 0.109660769 |
| ENSMUSG00000048799 | Cep120 | 225523 | protein_coding | 196.4794581 | -0.109663195 | 0.075418915 | 1.454054261 | 0.145931263 | 0.323295567 | 0.109663195 |
| ENSMUSG00000032299 | Commd4 | 66199 | protein_coding | 230.7066956 | -0.109664079 | 0.07521555 | 1.45799743 | 0.144841251 | 0.321916639 | 0.109664079 |
| ENSMUSG00000027433 | Xrn2 | 24128 | protein_coding | 907.6336118 | -0.10967475 | 0.053097663 | 2.065528783 | 0.038872995 | 0.123253915 | 0.10967475 |
| ENSMUSG00000070501 | Ifi214 | 545384 | protein_coding | 1.413395245 | -0.109693985 | 2.054609328 | 0.053389218 | 0.957421796 | 1 | 0.109693985 |
| ENSMUSG00000075271 | Ttc30a1 | 78802 | protein_coding | 50.97136333 | -0.109730398 | 0.19325422 | 0.567803373 | 0.570168489 | 0.755800484 | 0.109730398 |
| ENSMUSG00000025613 | Cct8 | 12469 | protein_coding | 1583.966873 | 0.109736869 | 0.09753411 | -1.125112733 | 0.260541266 | 0.474068167 | 0.109736869 |
| ENSMUSG00000041949 | Tango6 | 272538 | protein_coding | 64.50478636 | -0.109740156 | 0.143074437 | 0.767014415 | 0.443072945 | 0.660786267 | 0.109740156 |
| ENSMUSG00000027374 | Mrps5 | 77721 | protein_coding | 434.82836 | -0.109779779 | 0.097109061 | 1.130479259 | 0.258274333 | 0.471755595 | 0.109779779 |
| ENSMUSG00000028115 | Bnipl | 171388 | protein_coding | 89.38591824 | -0.109797578 | 0.161707084 | 0.67899053 | 0.497143862 | 0.702914252 | 0.109797578 |

Supplementary Table 1

| ensembl_gene_id | external_gene_name | entrezgene_id | gene_biotype | baseMean | log2FoldChange | lfcSE | stat | pvalue | padj | absLog2FC |
|---|---|---|---|---|---|---|---|---|---|---|
| ENSMUSG00000022438 | Parvb | 170736 | protein_coding | 3.454062976 | -0.109804593 | 1.60840352 | 0.068269306 | 0.945571257 | 1 | 0.109804593 |
| ENSMUSG00000033809 | Alg3 | 208624 | protein_coding | 90.47625297 | 0.109832544 | 0.164563317 | -0.667418144 | 0.50450508 | 0.708596728 | 0.109832544 |
| ENSMUSG00000022706 | Mrpl40 | 18100 | protein_coding | 340.0230489 | -0.109858762 | 0.098311207 | 1.117459196 | 0.263798037 | 0.47814299 | 0.109858762 |
| ENSMUSG00000075000 | Nrbf2 | 641340 | protein_coding | 104.7938198 | -0.109867695 | 0.106878505 | 1.027968108 | 0.30396483 | 0.524360158 | 0.109867695 |
| ENSMUSG00000061306 | Slc38a10 | 72055 | protein_coding | 1589.40058 | 0.10988507 | 0.077181891 | -1.423715706 | 0.154528785 | 0.336447148 | 0.10988507 |
| ENSMUSG00000053119 | Chmp3 | 66700 | protein_coding | 1810.308388 | -0.109923846 | 0.072181709 | 1.522876729 | 0.127789549 | 0.294806545 | 0.109923846 |
| ENSMUSG00000017291 | Taok1 | 216965 | protein_coding | 1669.588264 | 0.109935622 | 0.055289042 | -1.988379928 | 0.046769687 | 0.141690444 | 0.109935622 |
| ENSMUSG00000033088 | Triobp | 110253 | protein_coding | 805.3844684 | 0.10994406 | 0.048246836 | -2.278782809 | 0.022679979 | 0.08105111 | 0.10994406 |
| ENSMUSG00000068663 | Clec16a | 74374 | protein_coding | 411.9323834 | 0.110018766 | 0.069233373 | -1.589100171 | 0.112037779 | 0.268218365 | 0.110018766 |
| ENSMUSG00000032324 | Tspan3 | 56434 | protein_coding | 3441.092879 | 0.110024268 | 0.120211068 | -0.915259056 | 0.360055648 | 0.5825209 | 0.110024268 |
| ENSMUSG00000037234 | Hook3 | 320191 | protein_coding | 362.8608812 | -0.110108377 | 0.080548941 | 1.36697487 | 0.171633183 | 0.361939775 | 0.110108377 |
| ENSMUSG00000029320 | 1700016H13Rik | 74218 | protein_coding | 7.02443844 | 0.110113476 | 0.412903926 | -0.266680623 | 0.789715074 | 0.894959836 | 0.110113476 |
| ENSMUSG00000066368 | Actl11 | 67722 | protein_coding | 3.028851954 | 0.11015461 | 1.475571974 | -0.074652143 | 0.940491486 | 1 | 0.11015461 |
| ENSMUSG00000074166 | AW146154 | 101835 | protein_coding | 94.40368696 | -0.110188149 | 0.111640499 | 0.986990835 | 0.323647131 | 0.545489766 | 0.110188149 |
| ENSMUSG00000032373 | Car12 | 76459 | protein_coding | 845.9142005 | 0.110210495 | 0.129716347 | -0.849626883 | 0.395532561 | 0.617841913 | 0.110210495 |
| ENSMUSG00000040532 | Abhd11 | 68758 | protein_coding | 530.1723571 | 0.110235222 | 0.072178884 | -1.527250307 | 0.126698781 | 0.292957669 | 0.110235222 |
| ENSMUSG00000006567 | Atp7b | 11979 | protein_coding | 25.50514424 | 0.110256476 | 0.322428473 | -0.341956389 | 0.732383713 | 0.862666522 | 0.110256476 |
| ENSMUSG00000054191 | Klf1 | 16596 | protein_coding | 9.522377488 | -0.110290175 | 0.412818857 | 0.267163608 | 0.789343195 | 0.894720656 | 0.110290175 |
| ENSMUSG00000037458 | Azin1 | 54375 | protein_coding | 2332.013389 | -0.110290323 | 0.067205144 | 1.641099409 | 0.100776782 | 0.249173195 | 0.110290323 |
| ENSMUSG00000047371 | Zfp768 | 233890 | protein_coding | 429.6306211 | -0.11032699 | 0.069010999 | 1.598687057 | 0.109890155 | 0.264768897 | 0.11032699 |
| ENSMUSG00000072640 | Lyrm9 | 66274 | protein_coding | 108.9574634 | -0.110337859 | 0.139860371 | 0.788914391 | 0.430162044 | 0.648857873 | 0.110337859 |
| ENSMUSG00000020021 | Fgd6 | 13998 | protein_coding | 368.7037482 | -0.110361324 | 0.201400477 | 0.547969525 | 0.583712828 | 0.766128886 | 0.110361324 |
| ENSMUSG00000085078 | C030013C21Rik | 77417 | lncRNA | 9.287337574 | 0.110377278 | 0.387088727 | -0.285147228 | 0.775531348 | 0.88791795 | 0.110377278 |
| ENSMUSG00000019850 | Tnfaip3 | 21929 | protein_coding | 538.0938343 | -0.11044017 | 0.227288196 | 0.485903678 | 0.62703546 | 0.795976331 | 0.11044017 |
| ENSMUSG00000067851 | Arfgef1 | 211673 | protein_coding | 2759.553327 | 0.110486128 | 0.08998446 | -1.227835648 | 0.219508671 | 0.42576535 | 0.110486128 |
| ENSMUSG00000028268 | Gbp3 | 55932 | protein_coding | 41.46117599 | 0.110497383 | 0.420261025 | -0.262925601 | 0.792607924 | 0.896761253 | 0.110497383 |
| ENSMUSG00000022674 | Ube2v2 | 70620 | protein_coding | 92.32951449 | -0.110530718 | 0.164111592 | 0.67350951 | 0.500623201 | 0.705219785 | 0.110530718 |
| ENSMUSG00000025010 | Ccnj | 240665 | protein_coding | 63.33964729 | -0.110564745 | 0.177837442 | 0.621718037 | 0.534127289 | 0.730361684 | 0.110564745 |
| ENSMUSG00000034235 | Usp54 | 78787 | protein_coding | 107.8276544 | 0.110567062 | 0.157736616 | -0.700960026 | 0.483327962 | 0.693817626 | 0.110567062 |
| ENSMUSG00000035621 | Midn | 59090 | protein_coding | 2123.847966 | 0.110570008 | 0.141871268 | -0.779368578 | 0.435762628 | 0.653940181 | 0.110570008 |
| ENSMUSG00000022463 | Srebf2 | 20788 | protein_coding | 1725.959458 | -0.1106203 | 0.057907537 | 1.9102919 | 0.05609564 | 0.162548412 | 0.1106203 |
| ENSMUSG00000021807 | Rtraf | 68045 | protein_coding | 1074.166309 | -0.110621133 | 0.101375154 | 1.091205574 | 0.275182436 | 0.49170772 | 0.110621133 |
| ENSMUSG00000046840 | Hnf4aos | 68314 | lncRNA | 5.77961545 | -0.110642867 | 0.473211634 | 0.233812651 | 0.815130428 | 0.908120076 | 0.110642867 |
| ENSMUSG00000052675 | Zfp112 | 57745 | protein_coding | 26.77296208 | -0.110661266 | 0.253558033 | 0.43643368 | 0.662522107 | 0.817399333 | 0.110661266 |
| ENSMUSG00000022788 | Fgd4 | 224014 | protein_coding | 1029.285362 | -0.11066316 | 0.081078099 | 1.364895841 | 0.172285789 | 0.36280925 | 0.11066316 |
| ENSMUSG00000031156 | Slc35a2 | 22232 | protein_coding | 1187.094006 | 0.110669591 | 0.141307662 | -0.783181818 | 0.43352035 | 0.652195961 | 0.110669591 |
| ENSMUSG00000032602 | Slc25a20 | 57279 | protein_coding | 1778.61799 | -0.110675044 | 0.136935252 | 0.80822902 | 0.418958755 | 0.638903396 | 0.110675044 |
| ENSMUSG00000017478 | Zc3h18 | 76014 | protein_coding | 1198.153698 | 0.110710125 | 0.06822933 | -1.622617786 | 0.104671136 | 0.255796319 | 0.110710125 |
| ENSMUSG00000025364 | Pa2g4 | 18813 | protein_coding | 854.5705535 | 0.110723341 | 0.087348671 | -1.267601902 | 0.204940153 | 0.407364189 | 0.110723341 |

Supplementary Table 1

| ensembl_gene_id | external_gene_name | entrezgene_id | gene_biotype | baseMean | log2FoldChange | lfcSE | stat | pvalue | padj | absLog2FC |
|---|---|---|---|---|---|---|---|---|---|---|
| ENSMUSG00000028536 | 2610528J11Rik | 66451 | protein_coding | 482.0731302 | -0.110730221 | 0.08455737 | 1.309527727 | 0.190355652 | 0.387766717 | 0.110730221 |
| ENSMUSG00000053253 | Ndfip2 | 76273 | protein_coding | 1879.43666 | 0.110736408 | 0.091996237 | -1.203705839 | 0.228703295 | 0.437687589 | 0.110736408 |
| ENSMUSG00000027893 | Ahcyl1 | 229709 | protein_coding | 1030.850838 | 0.11074014 | 0.102157754 | -1.084011103 | 0.278359876 | 0.495562203 | 0.11074014 |
| ENSMUSG00000032320 | Rcn2 | 26611 | protein_coding | 268.4426736 | 0.110764216 | 0.082271518 | -1.346325181 | 0.178197668 | 0.37057407 | 0.110764216 |
| ENSMUSG00000100200 | H2al1m | 76383 | protein_coding | 1.282989417 | -0.110773252 | 0.98042542 | 0.112984883 | 0.91004254 | 1 | 0.110773252 |
| ENSMUSG00000030745 | Il21r | 60504 | protein_coding | 8.121029649 | -0.110777557 | 0.503631599 | 0.219957519 | 0.82590424 | 0.912001512 | 0.110777557 |
| ENSMUSG00000043091 | Tuba1c | 22146 | protein_coding | 1515.453147 | 0.11080861 | 0.116046441 | -0.954864349 | 0.339646301 | 0.561116348 | 0.11080861 |
| ENSMUSG00000020083 | Fam241b | 69894 | protein_coding | 166.9779528 | -0.110812508 | 0.089566563 | 1.237208444 | 0.216009708 | 0.421254642 | 0.110812508 |
| ENSMUSG00000014601 | Strip1 | 229707 | protein_coding | 495.038811 | 0.110830979 | 0.066599066 | -1.664152141 | 0.096082029 | 0.239900005 | 0.110830979 |
| ENSMUSG00000074024 | 4632427E13Rik | 666737 | lncRNA | 135.8535309 | -0.110839329 | 0.176917735 | 0.626502078 | 0.530985674 | 0.728048657 | 0.110839329 |
| ENSMUSG00000037139 | Myom3 | 242702 | protein_coding | 39.73891553 | -0.110858157 | 0.354633172 | 0.312599514 | 0.754584947 | 0.876856731 | 0.110858157 |
| ENSMUSG00000025239 | Limd1 | 29806 | protein_coding | 1066.455482 | -0.110878435 | 0.069291811 | 1.600166508 | 0.10956165 | 0.264291499 | 0.110878435 |
| ENSMUSG00000025888 | Casp1 | 12362 | protein_coding | 1676.690301 | -0.110879105 | 0.083875311 | 1.32195164 | 0.186184255 | 0.38181259 | 0.110879105 |
| ENSMUSG00000032186 | Tmod2 | 50876 | protein_coding | 9.389283513 | 0.110890363 | 0.617728866 | -0.179513002 | 0.857534909 | 0.930374388 | 0.110890363 |
| ENSMUSG00000063663 | Brwd3 | 382236 | protein_coding | 425.2912314 | 0.110921861 | 0.090252659 | -1.229014872 | 0.21906623 | 0.42530681 | 0.110921861 |
| ENSMUSG00000116903 | Gm19522 | 100503040 | lncRNA | 10.3700997 | 0.110923618 | 0.383513372 | -0.289230118 | 0.772405285 | 0.885864136 | 0.110923618 |
| ENSMUSG00000038236 | Hoxa7 | 15404 | protein_coding | 277.9025816 | 0.110932344 | 0.089999549 | -1.232587781 | 0.217729598 | 0.423608221 | 0.110932344 |
| ENSMUSG00000024927 | Rela | 19697 | protein_coding | 389.0372732 | -0.110959475 | 0.064012628 | 1.733399787 | 0.083024635 | 0.215780426 | 0.110959475 |
| ENSMUSG00000026604 | Ptpn14 | 19250 | protein_coding | 60.64017323 | -0.110959643 | 0.14642227 | 0.757805781 | 0.448567261 | 0.665436557 | 0.110959643 |
| ENSMUSG00000109813 | Or4g16 | 258388 | protein_coding | 1.020055681 | -0.110988143 | 1.759389925 | 0.063083311 | 0.949700163 | 1 | 0.110988143 |
| ENSMUSG00000035367 | Rmi1 | 74386 | protein_coding | 114.8636548 | -0.111048941 | 0.108982884 | 1.018957629 | 0.308223082 | 0.528838189 | 0.111048941 |
| ENSMUSG00000027579 | Srms | 20811 | protein_coding | 5.487685274 | -0.111063911 | 0.502207342 | 0.221151507 | 0.824974466 | 0.911456552 | 0.111063911 |
| ENSMUSG00000058258 | Idi1 | 319554 | protein_coding | 209.9063587 | -0.111099338 | 0.202668915 | 0.548181443 | 0.583567323 | 0.766020418 | 0.111099338 |
| ENSMUSG00000041112 | Elmo1 | 140580 | protein_coding | 208.6714144 | 0.111160457 | 0.164865251 | -0.674250374 | 0.500152149 | 0.704796705 | 0.111160457 |
| ENSMUSG00000026087 | Mrpl30 | 107734 | protein_coding | 647.5118251 | -0.111164362 | 0.066671373 | 1.667347717 | 0.095445284 | 0.238987471 | 0.111164362 |
| ENSMUSG00000022905 | Kpna1 | 16646 | protein_coding | 761.7622851 | 0.111182047 | 0.073356761 | -1.515634618 | 0.129611768 | 0.297320449 | 0.111182047 |
| ENSMUSG00000044017 | Adgrd1 | 243277 | protein_coding | 7.094397557 | 0.11123815 | 0.830388685 | -0.133959136 | 0.893434888 | 0.947542466 | 0.11123815 |
| ENSMUSG00000069114 | Zbtb10 | 229055 | protein_coding | 551.8484185 | 0.111323167 | 0.086475084 | -1.287343846 | 0.197974467 | 0.398279073 | 0.111323167 |
| ENSMUSG00000021518 | Ptdss1 | 19210 | protein_coding | 1228.74279 | -0.111348059 | 0.065257345 | 1.70629159 | 0.087953803 | 0.22515016 | 0.111348059 |
| ENSMUSG00000033554 | Dph5 | 69740 | protein_coding | 98.4881183 | -0.111358207 | 0.169294517 | 0.657777986 | 0.510680802 | 0.713786099 | 0.111358207 |
| ENSMUSG00000032018 | Sc5d | 235293 | protein_coding | 739.1845923 | 0.111361626 | 0.074107688 | -1.502700044 | 0.132916411 | 0.30270362 | 0.111361626 |
| ENSMUSG00000025949 | Pikfyve | 18711 | protein_coding | 517.9017231 | -0.111374776 | 0.054831235 | 2.031228676 | 0.042231803 | 0.131176532 | 0.111374776 |
| ENSMUSG00000025737 | Wdr24 | 268933 | protein_coding | 400.7469501 | 0.111380828 | 0.06727042 | -1.655717742 | 0.097778993 | 0.243290987 | 0.111380828 |
| ENSMUSG00000001436 | Slc19a1 | 20509 | protein_coding | 111.996956 | 0.111449673 | 0.13631875 | -0.81756672 | 0.413604637 | 0.633723045 | 0.111449673 |
| ENSMUSG00000018861 | Fdxr | 14149 | protein_coding | 199.5812501 | -0.111454278 | 0.11161919 | 0.998522549 | 0.318026036 | 0.539389762 | 0.111454278 |
| ENSMUSG00000090877 | Hspa1b | 15511 | protein_coding | 1185.438871 | 0.11147638 | 0.470115059 | -0.237125737 | 0.812559246 | 0.907147741 | 0.11147638 |
| ENSMUSG00000014496 | Ankrd28 | 105522 | protein_coding | 584.4076719 | 0.111489575 | 0.101810176 | -1.095073009 | 0.273484652 | 0.489835334 | 0.111489575 |
| ENSMUSG00000028139 | Riiad1 | 66353 | protein_coding | 14.46614412 | -0.111519458 | 0.496893935 | 0.224433124 | 0.822420299 | 0.910171686 | 0.111519458 |
| ENSMUSG00000058503 | Fam133b | 68152 | protein_coding | 194.6558142 | 0.111528484 | 0.09493285 | -1.174814447 | 0.240068959 | 0.450443439 | 0.111528484 |

| ensembl_gene_id | external_gene_name | entrezgene_id | gene_biotype | baseMean | log2FoldChange | lfcSE | stat | pvalue | padj | absLog2FC |
|---|---|---|---|---|---|---|---|---|---|---|
| ENSMUSG00000097781 | 9330136K24Rik | 319870 | lncRNA | 1.622827424 | -0.111581537 | 0.886344028 | 0.125889647 | 0.899819278 | 1 | 0.111581537 |
| ENSMUSG00000000881 | Dlg3 | 53310 | protein_coding | 751.5365116 | -0.111650023 | 0.072516354 | 1.539653013 | 0.123644956 | 0.28762823 | 0.111650023 |
| ENSMUSG00000031327 | Chic1 | 12212 | protein_coding | 239.0571319 | 0.111661969 | 0.129101336 | -0.864917222 | 0.387084224 | 0.609604127 | 0.111661969 |
| ENSMUSG00000061540 | Orm2 | 18406 | protein_coding | 1.171780014 | -0.111686962 | 1.592154687 | 0.070148311 | 0.944075615 | 1 | 0.111686962 |
| ENSMUSG00000041035 | Dpcd | 226162 | protein_coding | 185.9118463 | 0.1117364 | 0.088007453 | -1.269624288 | 0.204218493 | 0.406467904 | 0.1117364 |
| ENSMUSG00000062797 | Hikeshi | 67669 | protein_coding | 176.7357568 | -0.111769038 | 0.123099703 | 0.90795538 | 0.363901796 | 0.586347377 | 0.111769038 |
| ENSMUSG00000032596 | Uba7 | 74153 | protein_coding | 510.5974075 | -0.111817972 | 0.115848202 | 0.965211121 | 0.334439078 | 0.556711723 | 0.111817972 |
| ENSMUSG00000036636 | Clcn7 | 26373 | protein_coding | 347.8130011 | -0.111843962 | 0.069443774 | 1.610568607 | 0.107273781 | 0.260339278 | 0.111843962 |
| ENSMUSG00000036104 | Rab3gap1 | 226407 | protein_coding | 955.8959493 | -0.111849565 | 0.087995454 | 1.271083443 | 0.203698965 | 0.405918699 | 0.111849565 |
| ENSMUSG00000054693 | Adam10 | 11487 | protein_coding | 1382.500117 | 0.111941509 | 0.079143832 | -1.414405982 | 0.157242735 | 0.340507018 | 0.111941509 |
| ENSMUSG00000061143 | Maml3 | 433586 | protein_coding | 314.9984307 | 0.111960158 | 0.088451362 | -1.265782187 | 0.205591074 | 0.408255959 | 0.111960158 |
| ENSMUSG00000026020 | Nop58 | 55989 | protein_coding | 734.2137714 | 0.111967589 | 0.089161315 | -1.25578666 | 0.209193329 | 0.412590307 | 0.111967589 |
| ENSMUSG00000026971 | Itgb6 | 16420 | protein_coding | 528.3669595 | -0.111975807 | 0.119037592 | 0.94067601 | 0.346870917 | 0.568558443 | 0.111975807 |
| ENSMUSG00000000759 | Tubgcp3 | 259279 | protein_coding | 391.1456918 | -0.111976306 | 0.070328601 | 1.592187318 | 0.111342622 | 0.267099369 | 0.111976306 |
| ENSMUSG00000049354 | Dcaf7 | 71833 | protein_coding | 1278.278334 | 0.112015514 | 0.10011094 | -1.118913819 | 0.263176907 | 0.477488842 | 0.112015514 |
| ENSMUSG00000021998 | Lcp1 | 18826 | protein_coding | 146.6785962 | 0.112036296 | 0.169829544 | -0.659698506 | 0.509447326 | 0.712785998 | 0.112036296 |
| ENSMUSG00000020865 | Abcc3 | 76408 | protein_coding | 4603.498921 | -0.112038452 | 0.137889067 | 0.812526 | 0.416489874 | 0.636725956 | 0.112038452 |
| ENSMUSG00000029199 | Lias | 79464 | protein_coding | 326.2443609 | -0.112064557 | 0.076745208 | 1.460215676 | 0.144230808 | 0.320998127 | 0.112064557 |
| ENSMUSG00000028882 | Ppp1r8 | 100336 | protein_coding | 347.2255198 | 0.112100002 | 0.085751601 | -1.307264239 | 0.191122985 | 0.388695194 | 0.112100002 |
| ENSMUSG00000020922 | Lsm12 | 268490 | protein_coding | 519.4260763 | 0.112109648 | 0.08897811 | -1.259968856 | 0.207680598 | 0.410708835 | 0.112109648 |
| ENSMUSG00000037787 | Coa8 | 68020 | protein_coding | 196.0186938 | -0.11212835 | 0.085496498 | 1.311496407 | 0.189690107 | 0.386972505 | 0.11212835 |
| ENSMUSG00000022755 | Adgrg7 | 239853 | protein_coding | 3829.312265 | -0.112155717 | 0.112435174 | 0.997514508 | 0.318514835 | 0.539980904 | 0.112155717 |
| ENSMUSG00000036099 | Vezt | 215008 | protein_coding | 222.7275296 | -0.112164204 | 0.08861523 | 1.265744086 | 0.205604719 | 0.408255959 | 0.112164204 |
| ENSMUSG00000024604 | Rbm22 | 66810 | protein_coding | 417.5412993 | -0.112166691 | 0.063861483 | 1.756405983 | 0.079019112 | 0.208221097 | 0.112166691 |
| ENSMUSG00000047139 | Cd24a | 12484 | protein_coding | 3877.486936 | 0.112223957 | 0.153790722 | -0.729718642 | 0.465562184 | 0.67923791 | 0.112223957 |
| ENSMUSG00000089682 | Bcl2l2 | 12050 | protein_coding | 598.7159045 | 0.11223478 | 0.097520481 | -1.150884193 | 0.249779881 | 0.462014445 | 0.11223478 |
| ENSMUSG00000024160 | Spsb3 | 79043 | protein_coding | 84.0507282 | 0.112294181 | 0.116542839 | -0.963544234 | 0.335274477 | 0.557631454 | 0.112294181 |
| ENSMUSG00000038619 | Ensa | 56205 | protein_coding | 1173.359601 | 0.112298853 | 0.055025163 | -2.040863636 | 0.041264383 | 0.128987322 | 0.112298853 |
| ENSMUSG00000027680 | Fxr1 | 14359 | protein_coding | 507.537419 | 0.112305695 | 0.083235809 | -1.349247365 | 0.177257526 | 0.369553185 | 0.112305695 |
| ENSMUSG00000050914 | Ankrd37 | 654824 | protein_coding | 67.20030449 | -0.112325719 | 0.30052778 | 0.373761516 | 0.708581754 | 0.847629053 | 0.112325719 |
| ENSMUSG00000024732 | Ccdc86 | 108673 | protein_coding | 121.3690289 | -0.112338445 | 0.151742087 | 0.740324897 | 0.459102878 | 0.673388737 | 0.112338445 |
| ENSMUSG00000071632 | 2510002D24Rik | 72307 | protein_coding | 177.8050013 | -0.112352711 | 0.083985149 | 1.337768786 | 0.18097182 | 0.374977244 | 0.112352711 |
| ENSMUSG00000042447 | Mios | 252875 | protein_coding | 194.3466528 | -0.112391783 | 0.087832895 | 1.279609231 | 0.200682602 | 0.401975042 | 0.112391783 |
| ENSMUSG00000022817 | Itgb5 | 16419 | protein_coding | 464.9284882 | 0.112427706 | 0.101746726 | -1.104976157 | 0.26916992 | 0.484159903 | 0.112427706 |
| ENSMUSG00000011008 | Mcoln2 | 68279 | protein_coding | 373.1536352 | 0.112486818 | 0.145229872 | -0.774543257 | 0.438609598 | 0.656764359 | 0.112486818 |
| ENSMUSG00000022565 | Plec | 18810 | protein_coding | 11731.88422 | -0.112577065 | 0.180113983 | 0.625032343 | 0.531949831 | 0.728820184 | 0.112577065 |
| ENSMUSG00000090553 | Snrpe | 20643 | protein_coding | 383.5655275 | -0.112602412 | 0.1284625 | 0.876539167 | 0.380736994 | 0.603058768 | 0.112602412 |
| ENSMUSG00000020537 | Drg2 | 13495 | protein_coding | 410.4384943 | 0.112639271 | 0.088564496 | -1.271833251 | 0.203432372 | 0.405734564 | 0.112639271 |
| ENSMUSG00000003184 | Irf3 | 54131 | protein_coding | 759.0073551 | -0.112646669 | 0.054398522 | 2.070767108 | 0.038380565 | 0.122005131 | 0.112646669 |

| ensembl_gene_id | external_gene_name | entrezgene_id | gene_biotype | baseMean | log2FoldChange | lfcSE | stat | pvalue | padj | absLog2FC |
|---|---|---|---|---|---|---|---|---|---|---|
| ENSMUSG00000042323 | Pbrm1 | 66923 | protein_coding | 1280.358423 | 0.112711464 | 0.087253494 | -1.291770207 | 0.196436736 | 0.396067528 | 0.112711464 |
| ENSMUSG00000059975 | Zfp74 | 72723 | protein_coding | 49.78500075 | -0.112748615 | 0.163928442 | 0.68779166 | 0.491583989 | 0.698491194 | 0.112748615 |
| ENSMUSG00000027573 | Gid8 | 76425 | protein_coding | 748.4144508 | -0.112760915 | 0.070031743 | 1.610140067 | 0.107367282 | 0.260514736 | 0.112760915 |
| ENSMUSG00000017950 | Hnf4a | 15378 | protein_coding | 9439.864415 | -0.112770085 | 0.073720744 | 1.529692711 | 0.126092809 | 0.292047215 | 0.112770085 |
| ENSMUSG00000039844 | Rapgef1 | 107746 | protein_coding | 546.81881 | 0.112780096 | 0.062107102 | -1.815896932 | 0.069386184 | 0.18970918 | 0.112780096 |
| ENSMUSG00000063065 | Mapk3 | 26417 | protein_coding | 2997.508925 | -0.112801281 | 0.068168173 | 1.654749949 | 0.09797523 | 0.243645564 | 0.112801281 |
| ENSMUSG00000067369 | Trmt2b | 215201 | protein_coding | 511.9423622 | -0.112843963 | 0.074140402 | 1.522030628 | 0.128001407 | 0.295101944 | 0.112843963 |
| ENSMUSG00000074250 | Gm10653 | 677044 | transcribed_processed_pseudogene | 2.216509913 | -0.112845829 | 0.915111964 | 0.123313686 | 0.901858704 | 1 | 0.112845829 |
| ENSMUSG00000008200 | Fnbp4 | 55935 | protein_coding | 448.5136126 | 0.11284666 | 0.111020671 | -1.016447287 | 0.309416438 | 0.53032725 | 0.11284666 |
| ENSMUSG00000044033 | Ccdc141 | 545428 | protein_coding | 18.76409479 | -0.112855343 | 0.354056488 | 0.31874954 | 0.749916445 | 0.874376659 | 0.112855343 |
| ENSMUSG00000070394 | Tmem256 | 69186 | protein_coding | 367.18172 | -0.112882847 | 0.088015403 | 1.282535136 | 0.199654988 | 0.40075936 | 0.112882847 |
| ENSMUSG00000005813 | Metap1 | 75624 | protein_coding | 778.1805512 | 0.112915958 | 0.098776573 | -1.143145132 | 0.252978338 | 0.465324602 | 0.112915958 |
| ENSMUSG00000096199 | Ptrhd1 | 69709 | protein_coding | 65.48513862 | 0.112947564 | 0.158621587 | -0.712056702 | 0.476429664 | 0.688162291 | 0.112947564 |
| ENSMUSG00000019254 | Ppp1r12c | 232807 | protein_coding | 403.9937018 | 0.112948118 | 0.0856636 | -1.318507724 | 0.187333741 | 0.383501006 | 0.112948118 |
| ENSMUSG00000091971 | Hspa1a | 193740 | protein_coding | 429.081421 | 0.112958486 | 0.565285547 | -0.199825533 | 0.841617031 | 0.92158767 | 0.112958486 |
| ENSMUSG00000026466 | Tor1aip1 | 208263 | protein_coding | 266.9270102 | -0.112962648 | 0.095794804 | 1.179214778 | 0.238312663 | 0.448589299 | 0.112962648 |
| ENSMUSG00000023341 | Mx2 | 17858 | protein_coding | 20.72591541 | 0.112965071 | 0.463219401 | -0.243869472 | 0.807331914 | 0.904987729 | 0.112965071 |
| ENSMUSG00000027245 | Hypk | 67693 | protein_coding | 19.06586084 | 0.112978758 | 0.34676582 | -0.325807075 | 0.744570325 | 0.871118268 | 0.112978758 |
| ENSMUSG00000018412 | Kansl1 | 76719 | protein_coding | 687.5247721 | -0.113013012 | 0.07556417 | 1.495589931 | 0.134760549 | 0.305611042 | 0.113013012 |
| ENSMUSG00000035078 | Mtmr9 | 210376 | protein_coding | 352.403288 | -0.113052217 | 0.100485853 | 1.125056045 | 0.260565286 | 0.474068167 | 0.113052217 |
| ENSMUSG00000069259 | Prl6a1 | 19111 | protein_coding | 3.035506202 | 0.113110437 | 0.597776014 | -0.189218761 | 0.849921364 | 1 | 0.113110437 |
| ENSMUSG00000078812 | Eif5a | 276770 | protein_coding | 3379.974848 | 0.113114946 | 0.078007577 | -1.450050756 | 0.147044366 | 0.324941878 | 0.113114946 |
| ENSMUSG00000015341 | Golga7 | 57437 | protein_coding | 418.4278076 | -0.113116825 | 0.087963419 | 1.285953025 | 0.198459456 | 0.398802563 | 0.113116825 |
| ENSMUSG00000092134 | Gm17089 | 118027456 | lncRNA | 4.527573319 | 0.113116873 | 0.693345718 | -0.163146422 | 0.87040315 | 0.935980063 | 0.113116873 |
| ENSMUSG00000024644 | Cndp2 | 66054 | protein_coding | 654.2474763 | -0.113125767 | 0.085042993 | 1.330218556 | 0.183446272 | 0.378300667 | 0.113125767 |
| ENSMUSG00000002307 | Daxx | 13163 | protein_coding | 317.9125459 | -0.113130192 | 0.073334742 | 1.542654816 | 0.12291455 | 0.286211349 | 0.113130192 |
| ENSMUSG00000097167 | Gm16740 | 100504029 | lncRNA | 4.191643076 | 0.113140319 | 0.503380278 | -0.224761128 | 0.822165108 | 1 | 0.113140319 |
| ENSMUSG00000050114 | Prdx6b | 320769 | protein_coding | 0.920862446 | 0.11316769 | 1.567166285 | -0.072211667 | 0.942433461 | 1 | 0.11316769 |
| ENSMUSG00000020648 | Dus4l | 71916 | protein_coding | 41.17680244 | 0.113169026 | 0.18324657 | -0.617577867 | 0.536853638 | 0.732676764 | 0.113169026 |
| ENSMUSG00000019179 | Mdh2 | 17448 | protein_coding | 2884.943669 | 0.113184491 | 0.07757605 | -1.459013319 | 0.144561442 | 0.321507788 | 0.113184491 |
| ENSMUSG00000022838 | Eaf2 | 106389 | protein_coding | 6.326626064 | 0.113232521 | 0.488117448 | -0.231978023 | 0.816555083 | 0.908481483 | 0.113232521 |
| ENSMUSG00000073434 | Wdr90 | 106618 | protein_coding | 171.0514665 | 0.113232958 | 0.138966915 | -0.814819539 | 0.415175615 | 0.635422624 | 0.113232958 |
| ENSMUSG00000021196 | Pfkp | 56421 | protein_coding | 2353.111196 | -0.113283151 | 0.074848514 | 1.513498995 | 0.130152956 | 0.298087027 | 0.113283151 |
| ENSMUSG00000087370 | Tmem170b | 621976 | protein_coding | 293.114318 | 0.113322522 | 0.121151919 | -0.935375381 | 0.34959488 | 0.571435202 | 0.113322522 |
| ENSMUSG00000041361 | Myzap | 102371 | protein_coding | 191.6829207 | -0.113376109 | 0.247834175 | 0.457467616 | 0.647334974 | 0.808079957 | 0.113376109 |
| ENSMUSG00000037868 | Egr2 | 13654 | protein_coding | 76.93892788 | -0.11343386 | 0.333746568 | 0.339880228 | 0.733946727 | 0.86401456 | 0.11343386 |
| ENSMUSG00000079562 | Maea | 59003 | protein_coding | 1286.222438 | 0.113439989 | 0.049037974 | -2.313309028 | 0.020705655 | 0.075334803 | 0.113439989 |
| ENSMUSG00000038984 | Tspyl5 | 239364 | protein_coding | 7.828824251 | -0.113474161 | 0.422278605 | 0.268718708 | 0.788146154 | 0.894017228 | 0.113474161 |
| ENSMUSG00000027347 | Rasgrp1 | 19419 | protein_coding | 13.88361056 | -0.113506371 | 0.4072895 | 0.278687201 | 0.780484885 | 0.890186044 | 0.113506371 |

Supplementary Table 1

| ensembl_gene_id | external_gene_name | entrezgene_id | gene_biotype | baseMean | log2FoldChange | lfcSE | stat | pvalue | padj | absLog2FC |
|---|---|---|---|---|---|---|---|---|---|---|
| ENSMUSG00000033434 | Gtpbp6 | 107999 | protein_coding | 94.45075298 | -0.113547024 | 0.145309668 | 0.781414104 | 0.434558979 | 0.652884901 | 0.113547024 |
| ENSMUSG00000018565 | Elp5 | 54351 | protein_coding | 415.1402663 | 0.113549337 | 0.07221332 | -1.572415401 | 0.115854245 | 0.274864513 | 0.113549337 |
| ENSMUSG00000067714 | Lpar5 | 381810 | protein_coding | 93.26879754 | 0.113568519 | 0.107892112 | -1.052611881 | 0.29251891 | 0.511613776 | 0.113568519 |
| ENSMUSG00000021749 | Fam3d | 18300 | protein_coding | 10609.73214 | -0.113572816 | 0.181703794 | 0.625043724 | 0.531942361 | 0.728820184 | 0.113572816 |
| ENSMUSG00000021794 | Glud1 | 14661 | protein_coding | 2313.637158 | -0.113577449 | 0.080185083 | 1.416441122 | 0.156646391 | 0.339527301 | 0.113577449 |
| ENSMUSG00000002105 | Slc39a13 | 68427 | protein_coding | 75.74068633 | 0.113584677 | 0.135044328 | -0.841091801 | 0.400296508 | 0.622315002 | 0.113584677 |
| ENSMUSG00000032333 | Stoml1 | 69106 | protein_coding | 128.4566268 | -0.113669354 | 0.118321317 | 0.960683641 | 0.336711259 | 0.559233675 | 0.113669354 |
| ENSMUSG00000097403 | 9230116N13Rik | 320054 | lncRNA | 3.565625583 | -0.113767407 | 0.672198221 | 0.169246813 | 0.865602509 | 1 | 0.113767407 |
| ENSMUSG00000013275 | Slc41a1 | 98396 | protein_coding | 180.3469236 | -0.113774259 | 0.134910327 | 0.843332465 | 0.399042531 | 0.621033677 | 0.113774259 |
| ENSMUSG00000039100 | Marchf6 | 223455 | protein_coding | 1420.184248 | -0.113801951 | 0.064073343 | 1.776120087 | 0.075713126 | 0.201763202 | 0.113801951 |
| ENSMUSG00000028861 | Mrps15 | 66407 | protein_coding | 347.6677283 | 0.113838129 | 0.077458057 | -1.469674472 | 0.14164994 | 0.316622511 | 0.113838129 |
| ENSMUSG00000092558 | Med20 | 56771 | protein_coding | 228.1130456 | 0.113863998 | 0.088287955 | -1.289688926 | 0.197158686 | 0.397106786 | 0.113863998 |
| ENSMUSG00000033307 | Mif | 17319 | protein_coding | 564.7395936 | 0.113891886 | 0.13887761 | -0.820088173 | 0.412165844 | 0.632329199 | 0.113891886 |
| ENSMUSG00000093661 | Eif4e3 | 66892 | protein_coding | 66.25397919 | 0.113908239 | 0.250994496 | -0.453827639 | 0.649952889 | 0.809917796 | 0.113908239 |
| ENSMUSG00000080465 | Snord94 | 115490493 | snoRNA | 145.4550709 | 0.113909207 | 0.266158246 | -0.427975496 | 0.668668961 | 0.822256556 | 0.113909207 |
| ENSMUSG00000025059 | Gk | 14933 | protein_coding | 736.5370709 | -0.113919281 | 0.226923748 | 0.502015685 | 0.615656488 | 0.787858751 | 0.113919281 |
| ENSMUSG00000023953 | Polh | 80905 | protein_coding | 164.1775412 | 0.114039097 | 0.130547561 | -0.873544448 | 0.38236639 | 0.604711604 | 0.114039097 |
| ENSMUSG00000030465 | Psd3 | 234353 | protein_coding | 871.129074 | -0.114063772 | 0.073543556 | 1.550968946 | 0.120909126 | 0.283500399 | 0.114063772 |
| ENSMUSG00000029270 | Dipk1a | 67266 | protein_coding | 87.67492513 | -0.114074408 | 0.241183012 | 0.47297862 | 0.636228433 | 0.801687789 | 0.114074408 |
| ENSMUSG00000031708 | Tecr | 106529 | protein_coding | 1163.358896 | 0.114090558 | 0.062320764 | -1.830698962 | 0.067145488 | 0.185177119 | 0.114090558 |
| ENSMUSG00000046731 | Kctd11 | 216858 | protein_coding | 95.40016898 | -0.114094265 | 0.142822649 | 0.798852744 | 0.424375803 | 0.644012315 | 0.114094265 |
| ENSMUSG00000025004 | Cyp2c40 | 13099 | protein_coding | 2.154628473 | 0.114111188 | 0.948761118 | -0.120273888 | 0.904266189 | 1 | 0.114111188 |
| ENSMUSG00000030264 | Thumpd3 | 14911 | protein_coding | 611.7116579 | -0.114112816 | 0.075757161 | 1.506297407 | 0.131990843 | 0.301146282 | 0.114112816 |
| ENSMUSG00000029642 | Polr1d | 20018 | protein_coding | 910.7901649 | -0.114120605 | 0.083673596 | 1.363878334 | 0.172605861 | 0.363248781 | 0.114120605 |
| ENSMUSG00000063382 | Bcl9l | 80288 | protein_coding | 173.8979565 | -0.114123842 | 0.181530801 | 0.628674813 | 0.529561971 | 0.727166502 | 0.114123842 |
| ENSMUSG00000039662 | Icmt | 57295 | protein_coding | 761.7155222 | 0.114265722 | 0.086300429 | -1.324045814 | 0.18548783 | 0.380798859 | 0.114265722 |
| ENSMUSG00000039686 | Zer1 | 227693 | protein_coding | 255.2272495 | -0.114276384 | 0.071704317 | 1.593716946 | 0.110999447 | 0.266624708 | 0.114276384 |
| ENSMUSG00000031433 | Rbm41 | 237073 | protein_coding | 70.60691746 | -0.114287233 | 0.144389183 | 0.791522126 | 0.42863937 | 0.647625851 | 0.114287233 |
| ENSMUSG00000053560 | Ier2 | 15936 | protein_coding | 4317.336684 | -0.114318763 | 0.113493143 | 1.007274624 | 0.313802821 | 0.535013399 | 0.114318763 |
| ENSMUSG00000024797 | Vps51 | 68505 | protein_coding | 441.7293096 | -0.11433075 | 0.065131497 | 1.755383429 | 0.079193745 | 0.208527092 | 0.11433075 |
| ENSMUSG00000048603 | Gm9828 | 105246056 | lncRNA | 10.18873096 | 0.114351377 | 0.374142253 | -0.305636094 | 0.759881735 | 0.87943703 | 0.114351377 |
| ENSMUSG00000022824 | Muc13 | 17063 | protein_coding | 15820.20597 | -0.114360371 | 0.102898627 | 1.111388697 | 0.266401076 | 0.481401448 | 0.114360371 |
| ENSMUSG00000034675 | Dbn1 | 56320 | protein_coding | 12.64149407 | -0.114365012 | 0.467219243 | 0.244778044 | 0.806628295 | 0.904874646 | 0.114365012 |
| ENSMUSG00000021222 | Dcaf4 | 73828 | protein_coding | 150.676184 | -0.114392342 | 0.122143923 | 0.93653732 | 0.348996607 | 0.570683233 | 0.114392342 |
| ENSMUSG00000021678 | F2rl1 | 14063 | protein_coding | 1109.857099 | 0.114428297 | 0.074569262 | -1.534523665 | 0.124900876 | 0.289855686 | 0.114428297 |
| ENSMUSG00000086894 | Gm15708 | 100504231 | lncRNA | 26.08976304 | -0.114433063 | 0.265225117 | 0.431456359 | 0.666136578 | 0.820143104 | 0.114433063 |
| ENSMUSG00000028514 | Usp24 | 329908 | protein_coding | 1116.858959 | 0.114468715 | 0.098307058 | -1.164399767 | 0.244262042 | 0.455584296 | 0.114468715 |
| ENSMUSG00000009090 | Ap1b1 | 11764 | protein_coding | 1548.938373 | -0.114494244 | 0.063981463 | 1.789490886 | 0.073535794 | 0.197941337 | 0.114494244 |
| ENSMUSG00000022667 | Cd200r1 | 57781 | protein_coding | 5.961290507 | -0.114557637 | 0.618171983 | 0.185316772 | 0.852980584 | 0.92788168 | 0.114557637 |

Supplementary Table 1

| ensembl_gene_id | external_gene_name | entrezgene_id | gene_biotype | baseMean | log2FoldChange | lfcSE | stat | pvalue | padj | absLog2FC |
|---|---|---|---|---|---|---|---|---|---|---|
| ENSMUSG00000031059 | Ndufb11 | 104130 | protein_coding | 1318.24369 | 0.114569112 | 0.085071082 | -1.346745675 | 0.178062156 | 0.370529039 | 0.114569112 |
| ENSMUSG00000040774 | Cept1 | 99712 | protein_coding | 1387.327697 | 0.114645513 | 0.075016602 | -1.528268535 | 0.126445879 | 0.292413842 | 0.114645513 |
| ENSMUSG00000044469 | Tnfaip8l1 | 66443 | protein_coding | 194.3668154 | 0.114711712 | 0.120766973 | -0.949859958 | 0.342183415 | 0.564082441 | 0.114711712 |
| ENSMUSG00000035506 | Slc12a8 | 171286 | protein_coding | 1245.649984 | 0.114712342 | 0.206328342 | -0.555969873 | 0.57823145 | 0.76196251 | 0.114712342 |
| ENSMUSG00000108383 | Gm6567 | 625243 | lncRNA | 1.861736789 | -0.114761332 | 1.223286696 | 0.09381393 | 0.925256966 | 1 | 0.114761332 |
| ENSMUSG00000004896 | Rrnad1 | 229503 | protein_coding | 115.0814237 | -0.114775717 | 0.130804386 | 0.877460765 | 0.380236421 | 0.602439245 | 0.114775717 |
| ENSMUSG00000034216 | Vps18 | 228545 | protein_coding | 297.1567281 | -0.114780349 | 0.082350426 | 1.393803957 | 0.163376813 | 0.349482473 | 0.114780349 |
| ENSMUSG00000073139 | Tmem185a | 236848 | protein_coding | 147.9911379 | 0.114795161 | 0.107107476 | -1.071775432 | 0.28382091 | 0.502090896 | 0.114795161 |
| ENSMUSG00000022805 | Cfap91 | 320214 | protein_coding | 9.867286605 | -0.114816954 | 0.458700333 | 0.250309289 | 0.802348173 | 0.90234117 | 0.114816954 |
| ENSMUSG00000067931 | Zfp948 | 381066 | protein_coding | 192.4283584 | -0.114825162 | 0.098952685 | 1.160404709 | 0.24588407 | 0.457679163 | 0.114825162 |
| ENSMUSG00000031314 | Taf1 | 270627 | protein_coding | 834.5065612 | 0.114833907 | 0.094636799 | -1.213417066 | 0.224970405 | 0.432209358 | 0.114833907 |
| ENSMUSG00000047632 | Fgfbp3 | 72514 | protein_coding | 10.58154737 | -0.114985079 | 0.327546903 | 0.351049204 | 0.725551435 | 0.857956591 | 0.114985079 |
| ENSMUSG00000027346 | Gpcpd1 | 74182 | protein_coding | 438.2336428 | -0.114987279 | 0.10159592 | 1.131810017 | 0.257714315 | 0.47089684 | 0.114987279 |
| ENSMUSG00000014177 | Tvp23b | 67510 | protein_coding | 671.4679294 | 0.114997891 | 0.114328306 | -1.005856682 | 0.314484516 | 0.535752227 | 0.114997891 |
| ENSMUSG00000036863 | Syde2 | 214804 | protein_coding | 201.0733423 | 0.11502852 | 0.083154984 | -1.383302777 | 0.166572047 | 0.354161295 | 0.11502852 |
| ENSMUSG00000001763 | Tspan33 | 232670 | protein_coding | 202.2864137 | 0.115114444 | 0.183741297 | -0.626502836 | 0.530985177 | 0.728048657 | 0.115114444 |
| ENSMUSG00000000976 | Heatr6 | 217026 | protein_coding | 295.0485245 | 0.115139903 | 0.061606986 | -1.868942329 | 0.061630838 | 0.174007807 | 0.115139903 |
| ENSMUSG00000024383 | Map3k2 | 26405 | protein_coding | 828.4402793 | -0.11516469 | 0.093399924 | 1.233027667 | 0.217565442 | 0.423388606 | 0.11516469 |
| ENSMUSG00000031574 | Star | 20845 | protein_coding | 25.8581114 | -0.115178679 | 0.218468482 | 0.527209591 | 0.598048049 | 0.775818184 | 0.115178679 |
| ENSMUSG00000014195 | Dnajc7 | 56354 | protein_coding | 882.2527227 | -0.115263675 | 0.072151583 | 1.597521075 | 0.110149604 | 0.265200548 | 0.115263675 |
| ENSMUSG00000020003 | Pex7 | 18634 | protein_coding | 472.6262791 | 0.115365128 | 0.096394583 | -1.196800938 | 0.231384153 | 0.440188363 | 0.115365128 |
| ENSMUSG00000022636 | Alcam | 11658 | protein_coding | 571.9139148 | 0.115392704 | 0.165982849 | -0.695208598 | 0.486924585 | 0.695624496 | 0.115392704 |
| ENSMUSG00000034252 | Senp6 | 215351 | protein_coding | 1198.325844 | 0.115410063 | 0.070870313 | -1.628468374 | 0.10342561 | 0.253681814 | 0.115410063 |
| ENSMUSG00000078429 | Ctdsp2 | 52468 | protein_coding | 1552.978946 | -0.115413094 | 0.07038142 | 1.639823335 | 0.101041905 | 0.249608668 | 0.115413094 |
| ENSMUSG00000016942 | Tmprss6 | 71753 | protein_coding | 4.911360699 | 0.115431761 | 0.717999424 | -0.160768598 | 0.872275659 | 0.937028043 | 0.115431761 |
| ENSMUSG00000004508 | Gab2 | 14389 | protein_coding | 240.6721597 | 0.115462023 | 0.104616621 | -1.103668062 | 0.269737155 | 0.484916108 | 0.115462023 |
| ENSMUSG00000054823 | Nsd3 | 234135 | protein_coding | 631.986976 | 0.115487159 | 0.096651878 | -1.19487755 | 0.232134876 | 0.441179379 | 0.115487159 |
| ENSMUSG00000031066 | Usp11 | 236733 | protein_coding | 30.04660988 | -0.115547236 | 0.21639164 | 0.533972737 | 0.593360396 | 0.772650496 | 0.115547236 |
| ENSMUSG00000039294 | Cybc1 | 217370 | protein_coding | 225.0308874 | -0.115552216 | 0.078034219 | 1.480789035 | 0.138662799 | 0.311717339 | 0.115552216 |
| ENSMUSG00000026270 | Capn10 | 23830 | protein_coding | 297.0444992 | 0.115554644 | 0.072214275 | -1.600163464 | 0.109562325 | 0.264291499 | 0.115554644 |
| ENSMUSG00000027247 | Arhgap1 | 228359 | protein_coding | 1469.622678 | -0.115578096 | 0.09858117 | 1.172415538 | 0.241030264 | 0.451699249 | 0.115578096 |
| ENSMUSG00000039114 | Nrn1 | 68404 | protein_coding | 62.00897819 | -0.115594917 | 0.190634555 | 0.60636917 | 0.544269636 | 0.736796542 | 0.115594917 |
| ENSMUSG00000028779 | Pef1 | 67898 | protein_coding | 890.0870871 | -0.115597723 | 0.048745547 | 2.371451981 | 0.017718349 | 0.066683451 | 0.115597723 |
| ENSMUSG00000074212 | Dnajb14 | 70604 | protein_coding | 318.946203 | 0.115640608 | 0.092557623 | -1.249390424 | 0.211522309 | 0.415941821 | 0.115640608 |
| ENSMUSG00000025792 | Slc25a10 | 27376 | protein_coding | 1972.588701 | 0.11565207 | 0.086708242 | -1.333807111 | 0.18226708 | 0.376762869 | 0.11565207 |
| ENSMUSG00000069744 | Psmb3 | 26446 | protein_coding | 856.9587278 | 0.115730626 | 0.116874197 | -0.99021537 | 0.322068861 | 0.543328754 | 0.115730626 |
| ENSMUSG00000025616 | Usp16 | 74112 | protein_coding | 583.9991483 | 0.115733642 | 0.074063138 | -1.562634863 | 0.118138503 | 0.278825103 | 0.115733642 |
| ENSMUSG00000049957 | Ccdc137 | 67291 | protein_coding | 182.4960553 | -0.115779276 | 0.088520239 | 1.307941289 | 0.190893224 | 0.388417088 | 0.115779276 |
| ENSMUSG00000027387 | Zc3h8 | 57432 | protein_coding | 30.61782197 | -0.115784445 | 0.227937155 | 0.507966528 | 0.611476817 | 0.784792738 | 0.115784445 |

| ensembl_gene_id | external_gene_name | entrezgene_id | gene_biotype | baseMean | log2FoldChange | lfcSE | stat | pvalue | padj | absLog2FC |
|---|---|---|---|---|---|---|---|---|---|---|
| ENSMUSG00000037916 | Ndufv1 | 17995 | protein_coding | 2627.668385 | -0.115785827 | 0.082568441 | 1.402301236 | 0.160825312 | 0.345591307 | 0.115785827 |
| ENSMUSG00000027884 | Clcc1 | 229725 | protein_coding | 363.1078127 | 0.115804554 | 0.139086531 | -0.832607966 | 0.405065865 | 0.62627635 | 0.115804554 |
| ENSMUSG00000028189 | Ctbs | 74245 | protein_coding | 105.8112336 | -0.11581193 | 0.151220231 | 0.765849448 | 0.443765888 | 0.661197851 | 0.11581193 |
| ENSMUSG00000039130 | Zc3hc1 | 232679 | protein_coding | 112.1481715 | -0.115835523 | 0.10178907 | 1.137995692 | 0.255122281 | 0.467705301 | 0.115835523 |
| ENSMUSG00000013662 | Atad1 | 67979 | protein_coding | 1148.638371 | -0.1158555 | 0.091956841 | 1.259889952 | 0.207709064 | 0.410708835 | 0.1158555 |
| ENSMUSG00000045730 | Adrb2 | 11555 | protein_coding | 14.5949692 | 0.115864518 | 0.591396649 | -0.195916765 | 0.844675315 | 0.923599766 | 0.115864518 |
| ENSMUSG00000055917 | Zfp277 | 246196 | protein_coding | 255.7402109 | -0.115885444 | 0.083086998 | 1.394748233 | 0.163091774 | 0.349008366 | 0.115885444 |
| ENSMUSG00000029169 | Dhx15 | 13204 | protein_coding | 1713.059324 | 0.115947462 | 0.088372366 | -1.312033026 | 0.189508991 | 0.386709513 | 0.115947462 |
| ENSMUSG00000021945 | Zmym2 | 76007 | protein_coding | 777.057143 | -0.1159501 | 0.077328042 | 1.499457328 | 0.133755031 | 0.303828141 | 0.1159501 |
| ENSMUSG00000015944 | Castor2 | 80909 | protein_coding | 47.70126298 | -0.115963604 | 0.368391492 | 0.31478361 | 0.752925957 | 0.876176397 | 0.115963604 |
| ENSMUSG00000039735 | Fnbp1l | 214459 | protein_coding | 829.5247202 | -0.115982013 | 0.069959091 | 1.657854783 | 0.097346781 | 0.242325103 | 0.115982013 |
| ENSMUSG00000074344 | Tmigd3 | 69296 | protein_coding | 1.278748165 | -0.115986423 | 1.078347004 | 0.107559462 | 0.914345155 | 1 | 0.115986423 |
| ENSMUSG00000050982 | Apol10a | 245282 | protein_coding | 2324.087725 | -0.116012518 | 0.139307327 | 0.83278116 | 0.404968162 | 0.626183916 | 0.116012518 |
| ENSMUSG00000021779 | Thrb | 21834 | protein_coding | 421.219062 | -0.116073158 | 0.068133653 | 1.703609794 | 0.088454011 | 0.22586956 | 0.116073158 |
| ENSMUSG00000034269 | Setd5 | 72895 | protein_coding | 1717.700806 | 0.116073737 | 0.118632446 | -0.978431624 | 0.327860894 | 0.549974786 | 0.116073737 |
| ENSMUSG00000033192 | Lpcat2 | 270084 | protein_coding | 186.8488295 | 0.116113613 | 0.186958493 | -0.621066265 | 0.534556024 | 0.730517043 | 0.116113613 |
| ENSMUSG00000045409 | Trim39 | 79263 | protein_coding | 253.3546579 | 0.116114806 | 0.107075037 | -1.08442461 | 0.278176577 | 0.495289271 | 0.116114806 |
| ENSMUSG00000047098 | Rnf31 | 268749 | protein_coding | 402.5241713 | -0.11617086 | 0.148524927 | 0.78216406 | 0.434118163 | 0.652619695 | 0.11617086 |
| ENSMUSG00000024575 | Pde6a | 225600 | protein_coding | 18.60275242 | -0.116186329 | 0.396094897 | 0.293329526 | 0.769270287 | 0.884414475 | 0.116186329 |
| ENSMUSG00000031730 | Dhodh | 56749 | protein_coding | 142.6438489 | 0.116202641 | 0.139303912 | -0.834166388 | 0.404187229 | 0.625349723 | 0.116202641 |
| ENSMUSG00000008384 | Sertad1 | 55942 | protein_coding | 666.6342118 | 0.11621412 | 0.160226929 | -0.725309541 | 0.468262154 | 0.681791678 | 0.11621412 |
| ENSMUSG00000004535 | Tax1bp1 | 52440 | protein_coding | 5914.090463 | 0.116229797 | 0.104051961 | -1.117036104 | 0.263978889 | 0.47841828 | 0.116229797 |
| ENSMUSG00000003038 | Hmgn2 | 15331 | protein_coding | 542.1518034 | 0.116261581 | 0.112955261 | -1.029271058 | 0.30335232 | 0.523793413 | 0.116261581 |
| ENSMUSG00000045005 | Fzd5 | 14367 | protein_coding | 2191.467994 | -0.116268209 | 0.069748408 | 1.66696577 | 0.095521212 | 0.239041868 | 0.116268209 |
| ENSMUSG00000027792 | Bche | 12038 | protein_coding | 553.0046021 | -0.116289601 | 0.098613918 | 1.179241261 | 0.23830212 | 0.448589299 | 0.116289601 |
| ENSMUSG00000036019 | Tmtc2 | 278279 | protein_coding | 579.4166189 | -0.11629577 | 0.085016099 | 1.367926436 | 0.171335104 | 0.361449656 | 0.11629577 |
| ENSMUSG00000027284 | Cdan1 | 68968 | protein_coding | 105.1097377 | 0.116305635 | 0.121301905 | -0.958811285 | 0.337653823 | 0.559563944 | 0.116305635 |
| ENSMUSG00000038593 | Tctn1 | 654470 | protein_coding | 63.84356858 | -0.116310211 | 0.184200523 | 0.631432581 | 0.527757719 | 0.725980089 | 0.116310211 |
| ENSMUSG00000035234 | Abraxas1 | 70681 | protein_coding | 91.51164593 | 0.116317973 | 0.108807399 | -1.069026315 | 0.285057812 | 0.503470022 | 0.116317973 |
| ENSMUSG00000018415 | Gid4 | 66771 | protein_coding | 397.5020583 | -0.116318888 | 0.098852018 | 1.176697154 | 0.239316404 | 0.449590928 | 0.116318888 |
| ENSMUSG00000030689 | Ino80e | 233875 | protein_coding | 189.0051074 | 0.116375259 | 0.100714526 | -1.155496274 | 0.247887258 | 0.460059583 | 0.116375259 |
| ENSMUSG00000021748 | Pdhb | 68263 | protein_coding | 1143.439619 | 0.116428499 | 0.06214429 | -1.873518841 | 0.060996774 | 0.172956854 | 0.116428499 |
| ENSMUSG00000022024 | Sugt1 | 67955 | protein_coding | 663.990884 | -0.116444223 | 0.048639518 | 2.394025006 | 0.016664613 | 0.063600511 | 0.116444223 |
| ENSMUSG00000094724 | Rnaset2b | 68195 | protein_coding | 8.024548272 | 0.116447158 | 0.450568929 | -0.258444713 | 0.796063713 | 0.89873561 | 0.116447158 |
| ENSMUSG00000037730 | Mynn | 80732 | protein_coding | 214.4702668 | -0.116451578 | 0.082466151 | 1.412113649 | 0.157916501 | 0.341473496 | 0.116451578 |
| ENSMUSG00000051375 | Pcdh1 | 75599 | protein_coding | 1412.087919 | -0.116452518 | 0.121819896 | 0.955940053 | 0.339102522 | 0.560532006 | 0.116452518 |
| ENSMUSG00000024414 | Mrpl27 | 94064 | protein_coding | 294.8358312 | -0.116455745 | 0.087473034 | 1.331333085 | 0.183079437 | 0.377733062 | 0.116455745 |
| ENSMUSG00000026965 | Anapc2 | 99152 | protein_coding | 617.3986561 | -0.116471856 | 0.062606122 | 1.860390855 | 0.062830248 | 0.176389198 | 0.116471856 |
| ENSMUSG00000028670 | Lypla2 | 26394 | protein_coding | 861.0476234 | -0.11649178 | 0.082600136 | 1.410309787 | 0.15844823 | 0.342262172 | 0.11649178 |

| ensembl_gene_id | external_gene_name | entrezgene_id | gene_biotype | baseMean | log2FoldChange | lfcSE | stat | pvalue | padj | absLog2FC |
|---|---|---|---|---|---|---|---|---|---|---|
| ENSMUSG00000008730 | Hipk1 | 15257 | protein_coding | 1809.544795 | -0.116502989 | 0.048688817 | 2.392807944 | 0.016719991 | 0.06376765 | 0.116502989 |
| ENSMUSG00000031388 | Naa10 | 56292 | protein_coding | 278.6029068 | 0.116512663 | 0.105989298 | -1.09928705 | 0.271642879 | 0.487334112 | 0.116512663 |
| ENSMUSG00000038013 | Wipf2 | 68524 | protein_coding | 516.1103071 | -0.116592734 | 0.130611111 | 0.892670875 | 0.372033452 | 0.594336299 | 0.116592734 |
| ENSMUSG00000038607 | Gng10 | 14700 | protein_coding | 7.810376033 | 0.116613596 | 0.365583681 | -0.318979216 | 0.749742274 | 0.874258149 | 0.116613596 |
| ENSMUSG00000024603 | Dctn4 | 67665 | protein_coding | 898.4175777 | -0.116671036 | 0.06915132 | 1.687184523 | 0.091567876 | 0.231605439 | 0.116671036 |
| ENSMUSG00000022120 | Obi1 | 72486 | protein_coding | 132.9310276 | -0.116676029 | 0.123781641 | 0.942595589 | 0.345887797 | 0.567566681 | 0.116676029 |
| ENSMUSG00000026730 | Pter | 19212 | protein_coding | 122.3086332 | -0.116738343 | 0.179846349 | 0.649100432 | 0.516273462 | 0.717206457 | 0.116738343 |
| ENSMUSG00000003131 | Pafah1b2 | 18475 | protein_coding | 799.2895445 | 0.116760408 | 0.071109657 | -1.641976817 | 0.100594809 | 0.248835031 | 0.116760408 |
| ENSMUSG00000010392 | Gosr1 | 53334 | protein_coding | 701.7600756 | 0.116766091 | 0.093911528 | -1.243362705 | 0.213734207 | 0.418744149 | 0.116766091 |
| ENSMUSG00000041921 | Metap1d | 66559 | protein_coding | 147.4012318 | -0.116780087 | 0.099402294 | 1.174822854 | 0.240065595 | 0.450443439 | 0.116780087 |
| ENSMUSG00000038622 | Med30 | 69790 | protein_coding | 152.6073145 | 0.116797053 | 0.127647638 | -0.914995802 | 0.360193834 | 0.582642414 | 0.116797053 |
| ENSMUSG00000030982 | Vps35l | 71517 | protein_coding | 583.9463777 | -0.116845196 | 0.054724801 | 2.135141565 | 0.032749448 | 0.108013659 | 0.116845196 |
| ENSMUSG00000021686 | Ap3b1 | 11774 | protein_coding | 1803.215302 | -0.116868314 | 0.046646786 | 2.505388343 | 0.012231701 | 0.049808611 | 0.116868314 |
| ENSMUSG00000033478 | Fhip2a | 226252 | protein_coding | 317.6267656 | 0.116869665 | 0.081985676 | -1.425488832 | 0.154015946 | 0.335676301 | 0.116869665 |
| ENSMUSG00000038152 | 5033430I15Rik | 76000 | lncRNA | 19.58418481 | 0.116875952 | 0.282922738 | -0.41310201 | 0.679531877 | 0.828120806 | 0.116875952 |
| ENSMUSG00000080440 | Gm25848 | 115488713 | snoRNA | 15.651527 | 0.116877118 | 0.388442298 | -0.300886692 | 0.763500898 | 0.881386379 | 0.116877118 |
| ENSMUSG00000032410 | Xrn1 | 24127 | protein_coding | 587.0815179 | -0.11691987 | 0.094088007 | 1.24266497 | 0.213991317 | 0.419000785 | 0.11691987 |
| ENSMUSG00000029703 | Lrwd1 | 71735 | protein_coding | 212.4697788 | 0.116938982 | 0.101325088 | -1.154097012 | 0.248460398 | 0.460735958 | 0.116938982 |
| ENSMUSG00000038650 | Rnh1 | 107702 | protein_coding | 1399.213453 | 0.117007712 | 0.075252244 | -1.554873378 | 0.119976229 | 0.2820338 | 0.117007712 |
| ENSMUSG00000020456 | Ogdh | 18293 | protein_coding | 4470.084894 | -0.117035974 | 0.059302786 | 1.973532488 | 0.048434927 | 0.145560397 | 0.117035974 |
| ENSMUSG00000042680 | Garem1 | 381126 | protein_coding | 340.7383349 | -0.117049234 | 0.136944702 | 0.854718969 | 0.392706747 | 0.615439779 | 0.117049234 |
| ENSMUSG00000015846 | Rxra | 20181 | protein_coding | 1391.706035 | -0.117063044 | 0.083393063 | 1.403750391 | 0.160393191 | 0.344842229 | 0.117063044 |
| ENSMUSG00000043131 | Mob1a | 232157 | protein_coding | 1566.708822 | -0.117065781 | 0.109047234 | 1.073532784 | 0.283032138 | 0.501169049 | 0.117065781 |
| ENSMUSG00000040146 | Rgl3 | 71746 | protein_coding | 69.96124208 | 0.117066525 | 0.220930668 | -0.529879016 | 0.596195816 | 0.774449596 | 0.117066525 |
| ENSMUSG00000024002 | Brd4 | 57261 | protein_coding | 440.0022091 | 0.117094102 | 0.189661309 | -0.617385291 | 0.536980621 | 0.732746487 | 0.117094102 |
| ENSMUSG00000032253 | Phip | 83946 | protein_coding | 837.0753281 | 0.117166738 | 0.084074491 | -1.393606271 | 0.163436534 | 0.349519674 | 0.117166738 |
| ENSMUSG00000063236 | 1110038F14Rik | 117171 | protein_coding | 156.3978304 | -0.117219621 | 0.08886809 | 1.319029375 | 0.187159292 | 0.383324885 | 0.117219621 |
| ENSMUSG00000004100 | Ppan | 235036 | protein_coding | 239.966713 | 0.117230704 | 0.111712962 | -1.049392135 | 0.293997677 | 0.513159947 | 0.117230704 |
| ENSMUSG00000029128 | Rab28 | 100972 | protein_coding | 301.8024362 | 0.117268377 | 0.075459412 | -1.554058984 | 0.120170348 | 0.282329367 | 0.117268377 |
| ENSMUSG00000028755 | Cda | 72269 | protein_coding | 190.8067967 | 0.117318272 | 0.169006493 | -0.69416429 | 0.487579184 | 0.696047947 | 0.117318272 |
| ENSMUSG00000026458 | Ppfia4 | 68507 | protein_coding | 20.72674013 | -0.11734245 | 0.437802158 | 0.268026202 | 0.78867915 | 0.894265208 | 0.11734245 |
| ENSMUSG00000054400 | Cklf | 75458 | protein_coding | 59.49395383 | 0.117375929 | 0.145151217 | -0.808645852 | 0.418718883 | 0.638714542 | 0.117375929 |
| ENSMUSG00000031904 | Slc7a6 | 330836 | protein_coding | 145.5871566 | 0.117412127 | 0.156613545 | -0.749693314 | 0.453439435 | 0.669164362 | 0.117412127 |
| ENSMUSG00000066191 | Anks6 | 75691 | protein_coding | 121.596989 | 0.117446658 | 0.147013228 | -0.79888497 | 0.424357115 | 0.644012315 | 0.117446658 |
| ENSMUSG00000029781 | Fkbp9 | 27055 | protein_coding | 888.5527658 | 0.117466263 | 0.15735523 | -0.746503712 | 0.45536319 | 0.67085817 | 0.117466263 |
| ENSMUSG00000031907 | Zfp90 | 22751 | protein_coding | 59.4683702 | -0.117482848 | 0.138761871 | 0.846650794 | 0.397189793 | 0.619083741 | 0.117482848 |
| ENSMUSG00000038215 | Cep44 | 382010 | protein_coding | 84.42305707 | 0.11749998 | 0.158961382 | -0.739173117 | 0.459801883 | 0.674169253 | 0.11749998 |
| ENSMUSG00000090038 | Gm16573 | 102635191 | lncRNA | 4.696953068 | -0.117531255 | 0.705392107 | 0.166618331 | 0.86767037 | 0.934257579 | 0.117531255 |
| ENSMUSG00000068740 | Celsr2 | 53883 | protein_coding | 58.76810813 | -0.117587558 | 0.220090369 | 0.534269439 | 0.593155133 | 0.772538316 | 0.117587558 |

| ensembl_gene_id | external_gene_name | entrezgene_id | gene_biotype | baseMean | log2FoldChange | lfcSE | stat | pvalue | padj | absLog2FC |
|---|---|---|---|---|---|---|---|---|---|---|
| ENSMUSG00000046230 | Vps13a | 271564 | protein_coding | 964.1414764 | 0.117614369 | 0.073939066 | -1.590693195 | 0.111678639 | 0.26767214 | 0.117614369 |
| ENSMUSG00000056763 | Cspp1 | 211660 | protein_coding | 328.6503539 | -0.11761941 | 0.108836965 | 1.080693588 | 0.279833436 | 0.49722072 | 0.11761941 |
| ENSMUSG00000024780 | Cdc37l1 | 67072 | protein_coding | 448.6978157 | -0.117622086 | 0.09784758 | 1.202094992 | 0.229326726 | 0.438111017 | 0.117622086 |
| ENSMUSG00000031827 | Cotl1 | 72042 | protein_coding | 708.4557323 | 0.117632084 | 0.117335733 | -1.002525664 | 0.316089778 | 0.536972183 | 0.117632084 |
| ENSMUSG00000002660 | Clpp | 53895 | protein_coding | 356.6302607 | -0.11763216 | 0.082946252 | 1.418173299 | 0.156140175 | 0.338874866 | 0.11763216 |
| ENSMUSG00000058444 | Map2k5 | 23938 | protein_coding | 161.1708463 | 0.117770691 | 0.098817107 | -1.191804676 | 0.233337842 | 0.442718029 | 0.117770691 |
| ENSMUSG00000033423 | Eri3 | 140546 | protein_coding | 371.5135347 | 0.117799406 | 0.077235534 | -1.525197021 | 0.127209963 | 0.293933865 | 0.117799406 |
| ENSMUSG00000064341 | mt-Nd1 | 17716 | protein_coding | 63758.75734 | 0.117803819 | 0.485753778 | -0.242517555 | 0.808379156 | 0.905485545 | 0.117803819 |
| ENSMUSG00000029198 | Grpel1 | 17713 | protein_coding | 1025.287478 | 0.117816197 | 0.057687428 | -2.042320151 | 0.041119785 | 0.128705861 | 0.117816197 |
| ENSMUSG00000032582 | Rbm6 | 19654 | protein_coding | 919.6861948 | -0.117823004 | 0.054524689 | 2.160911061 | 0.03070221 | 0.102718051 | 0.117823004 |
| ENSMUSG00000037949 | Ano10 | 102566 | protein_coding | 869.1009443 | -0.117844759 | 0.07504178 | 1.570388646 | 0.116324722 | 0.275528753 | 0.117844759 |
| ENSMUSG00000041747 | Utp15 | 105372 | protein_coding | 184.6651461 | 0.117862443 | 0.124773197 | -0.944613475 | 0.344856245 | 0.566380511 | 0.117862443 |
| ENSMUSG00000074405 | Zfp865 | 319748 | protein_coding | 160.8252661 | 0.11789385 | 0.132393836 | -0.890478388 | 0.373209068 | 0.595475802 | 0.11789385 |
| ENSMUSG00000022174 | Dad1 | 13135 | protein_coding | 828.5965287 | 0.117921429 | 0.123018612 | -0.958565763 | 0.337777547 | 0.559613498 | 0.117921429 |
| ENSMUSG00000055491 | Pprc1 | 226169 | protein_coding | 220.3131817 | 0.117936481 | 0.1592947 | -0.740366636 | 0.459077558 | 0.673388737 | 0.117936481 |
| ENSMUSG00000027384 | Ndufaf5 | 69487 | protein_coding | 145.3228924 | 0.117948536 | 0.09822441 | -1.200806762 | 0.229826168 | 0.438404222 | 0.117948536 |
| ENSMUSG00000024446 | Rpp21 | 67676 | protein_coding | 205.3452701 | -0.117961078 | 0.086538206 | 1.363109808 | 0.172847906 | 0.363480239 | 0.117961078 |
| ENSMUSG00000018405 | Mrm1 | 217038 | protein_coding | 140.4971331 | -0.117980817 | 0.129030617 | 0.914362963 | 0.360526156 | 0.582866455 | 0.117980817 |
| ENSMUSG00000020904 | Cfap52 | 71860 | protein_coding | 17.76975557 | -0.117987381 | 0.252994461 | 0.466363494 | 0.640955338 | 0.804026642 | 0.117987381 |
| ENSMUSG00000054008 | Ndst1 | 15531 | protein_coding | 715.4484963 | -0.118005254 | 0.069785218 | 1.690977798 | 0.090841043 | 0.230331489 | 0.118005254 |
| ENSMUSG00000028899 | Taf12 | 66464 | protein_coding | 270.9071581 | 0.118007568 | 0.079725973 | -1.480164659 | 0.138829309 | 0.31192207 | 0.118007568 |
| ENSMUSG00000041347 | Bdkrb1 | 12061 | protein_coding | 0.687309041 | 0.118021865 | 1.279262972 | -0.092257704 | 0.926493292 | 1 | 0.118021865 |
| ENSMUSG00000025521 | Tmem192 | 73067 | protein_coding | 113.1960046 | -0.118032662 | 0.15818115 | 0.746186651 | 0.455554671 | 0.670920346 | 0.118032662 |
| ENSMUSG00000018809 | Smyd4 | 319822 | protein_coding | 72.41245556 | -0.118057828 | 0.166923474 | 0.707257197 | 0.47940666 | 0.690650055 | 0.118057828 |
| ENSMUSG00000107002 | 0610012G03Rik | 106264 | protein_coding | 416.943223 | -0.118099356 | 0.078043327 | 1.513253727 | 0.130215222 | 0.298143983 | 0.118099356 |
| ENSMUSG00000030522 | Mtmr10 | 233315 | protein_coding | 206.1602176 | -0.118101172 | 0.118304029 | 0.998285291 | 0.318141038 | 0.539479894 | 0.118101172 |
| ENSMUSG00000097383 | 1500026H17Rik | 69002 | lncRNA | 12.32945406 | -0.118116826 | 0.400481309 | 0.294937176 | 0.768041872 | 0.883975099 | 0.118116826 |
| ENSMUSG00000046523 | Kctd4 | 67516 | protein_coding | 9.493713567 | 0.118120261 | 0.370891584 | -0.318476519 | 0.750123503 | 0.874455706 | 0.118120261 |
| ENSMUSG00000045302 | Preb | 50907 | protein_coding | 946.7595375 | 0.118122092 | 0.079783348 | -1.480535667 | 0.13873035 | 0.311742141 | 0.118122092 |
| ENSMUSG00000045625 | Pigz | 239827 | protein_coding | 2.52924997 | -0.118132246 | 0.832262803 | 0.141941038 | 0.887126579 | 1 | 0.118132246 |
| ENSMUSG00000005362 | Crbn | 58799 | protein_coding | 275.0400213 | -0.11815933 | 0.091010512 | 1.298304193 | 0.194182826 | 0.392885958 | 0.11815933 |
| ENSMUSG00000047907 | Tshz2 | 228911 | protein_coding | 166.4024931 | -0.118189928 | 0.096965327 | 1.21888857 | 0.222886489 | 0.429893421 | 0.118189928 |
| ENSMUSG00000030067 | Foxp1 | 108655 | protein_coding | 1295.992018 | -0.118234223 | 0.101520213 | 1.164637268 | 0.244165852 | 0.455456329 | 0.118234223 |
| ENSMUSG00000038260 | Trpm4 | 68667 | protein_coding | 2943.158723 | -0.118248885 | 0.12260455 | 0.964473873 | 0.334808401 | 0.557080699 | 0.118248885 |
| ENSMUSG00000024963 | Dnajc4 | 57431 | protein_coding | 247.0073784 | -0.118255223 | 0.085018477 | 1.390935561 | 0.164244968 | 0.350538094 | 0.118255223 |
| ENSMUSG00000032215 | Rsl24d1 | 225215 | protein_coding | 163.2196902 | -0.118272961 | 0.126262379 | 0.936723687 | 0.348900709 | 0.570683233 | 0.118272961 |
| ENSMUSG00000028572 | Hook1 | 77963 | protein_coding | 2309.312772 | 0.118286999 | 0.088847431 | -1.331349689 | 0.183073976 | 0.377733062 | 0.118286999 |
| ENSMUSG00000029490 | Mfsd7a | 243197 | protein_coding | 174.0801337 | 0.118345312 | 0.154312425 | -0.766920173 | 0.443128979 | 0.660786267 | 0.118345312 |
| ENSMUSG00000024177 | Nme4 | 56520 | protein_coding | 52.31643567 | 0.118352001 | 0.15855138 | -0.746458349 | 0.455390583 | 0.67085817 | 0.118352001 |

| ensembl_gene_id | external_gene_name | entrezgene_id | gene_biotype | baseMean | log2FoldChange | lfcSE | stat | pvalue | padj | absLog2FC |
|---|---|---|---|---|---|---|---|---|---|---|
| ENSMUSG00000021218 | Gdi2 | 14569 | protein_coding | 4521.497369 | 0.118373127 | 0.084091579 | -1.407669222 | 0.159229039 | 0.343330529 | 0.118373127 |
| ENSMUSG00000064368 | mt-Nd6 | 17722 | protein_coding | 3380.479065 | -0.118377272 | 0.377206165 | 0.313826452 | 0.753652853 | 0.876528151 | 0.118377272 |
| ENSMUSG00000015787 | Abo | 80908 | protein_coding | 125.4478922 | -0.118389905 | 0.292899417 | 0.404199866 | 0.686065753 | 0.832872319 | 0.118389905 |
| ENSMUSG00000052533 | Nup188 | 227699 | protein_coding | 343.9027707 | 0.118395703 | 0.085576255 | -1.383511147 | 0.166508192 | 0.354116714 | 0.118395703 |
| ENSMUSG00000033377 | Palmd | 114301 | protein_coding | 31.41619875 | 0.118444569 | 0.249348842 | -0.475015517 | 0.634775913 | 0.800822518 | 0.118444569 |
| ENSMUSG00000027630 | Tbl1xr1 | 81004 | protein_coding | 1493.81903 | 0.118455264 | 0.077914494 | -1.52032385 | 0.128429603 | 0.295635135 | 0.118455264 |
| ENSMUSG00000020828 | Pld2 | 18806 | protein_coding | 27.54689787 | 0.118481364 | 0.32583115 | -0.363628105 | 0.716135737 | 0.852650725 | 0.118481364 |
| ENSMUSG00000040621 | Gemin8 | 237221 | protein_coding | 22.62203591 | -0.118486233 | 0.244210855 | 0.485180045 | 0.627548636 | 0.796376973 | 0.118486233 |
| ENSMUSG00000062082 | Cd200r4 | 239849 | protein_coding | 4.777698601 | -0.11850523 | 0.718501347 | 0.1649339 | 0.868996009 | 0.935258087 | 0.11850523 |
| ENSMUSG00000062963 | Ufc1 | 66155 | protein_coding | 851.9250555 | 0.118519353 | 0.097151374 | -1.219945202 | 0.222485649 | 0.429533455 | 0.118519353 |
| ENSMUSG00000051111 | Sv2c | 75209 | protein_coding | 61.14187876 | 0.118528846 | 0.178984264 | -0.662230539 | 0.507823485 | 0.711603273 | 0.118528846 |
| ENSMUSG00000037791 | Phf12 | 268448 | protein_coding | 395.8222313 | 0.118560042 | 0.111980859 | -1.058752747 | 0.289712399 | 0.508545065 | 0.118560042 |
| ENSMUSG00000040351 | Ankib1 | 70797 | protein_coding | 1219.921176 | -0.118560419 | 0.078901033 | 1.502647225 | 0.132930038 | 0.30270362 | 0.118560419 |
| ENSMUSG00000063765 | Chadl | 214685 | protein_coding | 6.697538576 | 0.118563965 | 0.398877232 | -0.297244252 | 0.766280039 | 0.882690208 | 0.118563965 |
| ENSMUSG00000039676 | Capsl | 75568 | protein_coding | 55.54902213 | 0.11857262 | 0.20712821 | -0.572460022 | 0.567010363 | 0.753307251 | 0.11857262 |
| ENSMUSG00000054892 | Txk | 22165 | protein_coding | 12.12931953 | 0.118604285 | 0.631785023 | -0.187728865 | 0.851089202 | 0.927286869 | 0.118604285 |
| ENSMUSG00000052656 | Rnf103 | 22644 | protein_coding | 1274.858828 | 0.118628188 | 0.077747266 | -1.52581812 | 0.127055167 | 0.293636687 | 0.118628188 |
| ENSMUSG00000067942 | Zfp160 | 224585 | protein_coding | 218.3979525 | -0.118650409 | 0.091991179 | 1.289802025 | 0.197119405 | 0.397106786 | 0.118650409 |
| ENSMUSG00000100632 | 9430060I03Rik | 100037260 | lncRNA | 12.32560271 | 0.118676941 | 0.362389479 | -0.32748451 | 0.743301457 | 0.870311987 | 0.118676941 |
| ENSMUSG00000092203 | 1110038B12Rik | 68763 | lncRNA | 572.1586308 | -0.118687339 | 0.095018248 | 1.249100471 | 0.211628328 | 0.416065952 | 0.118687339 |
| ENSMUSG00000042557 | Sin3a | 20466 | protein_coding | 658.1158104 | 0.118700745 | 0.072192442 | -1.644226749 | 0.100129374 | 0.247943691 | 0.118700745 |
| ENSMUSG00000001065 | Zfp276 | 57247 | protein_coding | 291.906634 | 0.118799048 | 0.102571628 | -1.158205731 | 0.24678009 | 0.458623273 | 0.118799048 |
| ENSMUSG00000006732 | Mettl1 | 17299 | protein_coding | 149.1692129 | 0.118805461 | 0.144968979 | -0.819523332 | 0.412487896 | 0.632451088 | 0.118805461 |
| ENSMUSG00000052512 | Nav2 | 78286 | protein_coding | 241.3770995 | -0.118814883 | 0.181621636 | 0.654189036 | 0.512990024 | 0.714956306 | 0.118814883 |
| ENSMUSG00000048772 | Tmem53 | 68777 | protein_coding | 56.47752859 | -0.11883062 | 0.147373407 | 0.806323356 | 0.420056427 | 0.63998633 | 0.11883062 |
| ENSMUSG00000031740 | Mmp2 | 17390 | protein_coding | 7.797256383 | -0.118846273 | 0.459922817 | 0.258404821 | 0.796094496 | 0.89873561 | 0.118846273 |
| ENSMUSG00000037536 | Fbxo34 | 78938 | protein_coding | 448.8205931 | 0.118860047 | 0.104505453 | -1.137357362 | 0.255388924 | 0.468075728 | 0.118860047 |
| ENSMUSG00000028789 | Azin2 | 242669 | protein_coding | 11.0550222 | -0.118968129 | 0.623489119 | 0.190810272 | 0.84867424 | 0.925637253 | 0.118968129 |
| ENSMUSG00000023286 | Ube2j2 | 140499 | protein_coding | 580.4621 | -0.119096763 | 0.074014502 | 1.609100372 | 0.107594396 | 0.26091246 | 0.119096763 |
| ENSMUSG00000024451 | Arap3 | 106952 | protein_coding | 12.45013857 | -0.119106285 | 0.634197434 | 0.187806318 | 0.851028483 | 0.927281893 | 0.119106285 |
| ENSMUSG00000027848 | Olfml3 | 99543 | protein_coding | 4.033768734 | 0.11915205 | 0.641218864 | -0.185821186 | 0.85258499 | 1 | 0.11915205 |
| ENSMUSG00000047714 | Ppp1r2 | 66849 | protein_coding | 601.1592578 | -0.119153807 | 0.085496128 | 1.393674901 | 0.163415799 | 0.349519674 | 0.119153807 |
| ENSMUSG00000030835 | Nomo1 | 211548 | protein_coding | 1342.110227 | 0.119157169 | 0.070876324 | -1.681198503 | 0.092724364 | 0.233671624 | 0.119157169 |
| ENSMUSG00000027438 | Napb | 17957 | protein_coding | 25.46144702 | -0.119175787 | 0.203479344 | 0.585689852 | 0.558083955 | 0.747120423 | 0.119175787 |
| ENSMUSG00000031353 | Rbbp7 | 245688 | protein_coding | 1492.146113 | 0.11922535 | 0.097694419 | -1.220390597 | 0.22231684 | 0.429396526 | 0.11922535 |
| ENSMUSG00000043051 | Disc1 | 244667 | protein_coding | 1.647001954 | -0.119284033 | 1.171645512 | 0.101808979 | 0.918908299 | 1 | 0.119284033 |
| ENSMUSG00000067713 | Prkag1 | 19082 | protein_coding | 355.1301635 | 0.119293576 | 0.074649018 | -1.598059541 | 0.110029727 | 0.265066507 | 0.119293576 |
| ENSMUSG00000030881 | Arfip2 | 76932 | protein_coding | 712.2689597 | 0.119326817 | 0.073651736 | -1.620149417 | 0.105200184 | 0.256689693 | 0.119326817 |
| ENSMUSG00000041471 | Shld2 | 75698 | protein_coding | 67.45092635 | 0.119389968 | 0.135083199 | -0.883825444 | 0.376790449 | 0.598990901 | 0.119389968 |

| ensembl_gene_id | external_gene_name | entrezgene_id | gene_biotype | baseMean | log2FoldChange | lfcSE | stat | pvalue | padj | absLog2FC |
|---|---|---|---|---|---|---|---|---|---|---|
| ENSMUSG00000040952 | Rps19 | 20085 | protein_coding | 5195.67392 | 0.119461886 | 0.113095763 | -1.056289661 | 0.290835898 | 0.509749949 | 0.119461886 |
| ENSMUSG00000010751 | Tnfrsf22 | 79202 | protein_coding | 5.914618913 | -0.119498811 | 0.662440323 | 0.180391814 | 0.856844981 | 0.93035707 | 0.119498811 |
| ENSMUSG00000001794 | Capns1 | 12336 | protein_coding | 2228.855101 | -0.119510108 | 0.076405804 | 1.564149613 | 0.117782437 | 0.278183519 | 0.119510108 |
| ENSMUSG00000027377 | Mall | 228576 | protein_coding | 1223.183997 | -0.119540049 | 0.123089029 | 0.971167375 | 0.331464937 | 0.553861375 | 0.119540049 |
| ENSMUSG00000021134 | Srsf5 | 20384 | protein_coding | 3080.480499 | -0.119557357 | 0.081651806 | 1.464234084 | 0.14313 | 0.319067066 | 0.119557357 |
| ENSMUSG00000048200 | Cracr2b | 213573 | protein_coding | 192.2015774 | -0.11961301 | 0.186498613 | 0.641361393 | 0.52128791 | 0.720922137 | 0.11961301 |
| ENSMUSG00000022544 | Eef2kmt | 70511 | protein_coding | 167.7158871 | -0.119614751 | 0.115550138 | 1.035176181 | 0.30058664 | 0.52065112 | 0.119614751 |
| ENSMUSG00000049775 | Tmsb4x | 19241 | protein_coding | 3622.360051 | -0.119621311 | 0.074000686 | 1.616489226 | 0.105988581 | 0.258193749 | 0.119621311 |
| ENSMUSG00000029592 | Usp30 | 100756 | protein_coding | 224.949513 | 0.119633671 | 0.085461433 | -1.399855667 | 0.161556544 | 0.346538122 | 0.119633671 |
| ENSMUSG00000048905 | Bnip5 | 207819 | protein_coding | 1137.465037 | -0.119815796 | 0.347984154 | 0.344313941 | 0.730610201 | 0.861253255 | 0.119815796 |
| ENSMUSG00000037025 | Foxa2 | 15376 | protein_coding | 325.9542937 | 0.119831409 | 0.233220712 | -0.513811182 | 0.60738401 | 0.781981914 | 0.119831409 |
| ENSMUSG00000021583 | Erap1 | 80898 | protein_coding | 1249.550482 | -0.119865215 | 0.09419163 | 1.272567577 | 0.20317153 | 0.40541018 | 0.119865215 |
| ENSMUSG00000071550 | Cfap44 | 212517 | protein_coding | 10.96760611 | 0.119942121 | 0.53478481 | -0.224281092 | 0.822538588 | 0.910171686 | 0.119942121 |
| ENSMUSG00000008348 | Ubc | 22190 | protein_coding | 1413.173933 | -0.119945677 | 0.089448994 | 1.340939364 | 0.17994014 | 0.373495661 | 0.119945677 |
| ENSMUSG00000032422 | Snx14 | 244962 | protein_coding | 534.4327796 | -0.119972533 | 0.08493576 | 1.412509094 | 0.157800115 | 0.341333371 | 0.119972533 |
| ENSMUSG00000021559 | Dapk1 | 69635 | protein_coding | 203.2765934 | -0.11999038 | 0.085698218 | 1.400150225 | 0.161468338 | 0.346538122 | 0.11999038 |
| ENSMUSG00000038507 | Parp12 | 243771 | protein_coding | 874.2743617 | -0.120017362 | 0.131216636 | 0.914650503 | 0.360375137 | 0.582711741 | 0.120017362 |
| ENSMUSG00000046541 | Zfp526 | 210172 | protein_coding | 159.9761532 | 0.120044788 | 0.098771458 | -1.215379318 | 0.224221452 | 0.431309617 | 0.120044788 |
| ENSMUSG00000039852 | Rere | 68703 | protein_coding | 748.2248829 | -0.12011419 | 0.158937564 | 0.755731921 | 0.449809942 | 0.666621321 | 0.12011419 |
| ENSMUSG00000040987 | Mill2 | 243864 | protein_coding | 56.37309173 | -0.120201425 | 0.198365487 | 0.605959369 | 0.544541734 | 0.736796542 | 0.120201425 |
| ENSMUSG00000039115 | Itga9 | 104099 | protein_coding | 4.771444485 | 0.12026515 | 0.976585302 | -0.123148638 | 0.901989397 | 0.951530343 | 0.12026515 |
| ENSMUSG00000057110 | Cntrl | 26920 | protein_coding | 817.6802486 | -0.120297732 | 0.096940803 | 1.240940123 | 0.214627869 | 0.419600407 | 0.120297732 |
| ENSMUSG00000004667 | Polr2e | 66420 | protein_coding | 464.7758358 | -0.120314898 | 0.063756869 | 1.887089187 | 0.059148331 | 0.169109338 | 0.120314898 |
| ENSMUSG00000056735 | A930024E05Rik | 109202 | lncRNA | 6.847511813 | 0.120328276 | 0.40499567 | -0.297110032 | 0.766382506 | 0.882690208 | 0.120328276 |
| ENSMUSG00000027896 | Slc16a4 | 229699 | protein_coding | 11.60240637 | 0.120340998 | 0.333182965 | -0.361185927 | 0.717960461 | 0.853653903 | 0.120340998 |
| ENSMUSG00000014551 | Mrps25 | 64658 | protein_coding | 272.3355759 | -0.120344483 | 0.084053036 | 1.431768428 | 0.152210106 | 0.332794609 | 0.120344483 |
| ENSMUSG00000055301 | Adh7 | 11529 | protein_coding | 4.681506268 | -0.120349597 | 0.512480856 | 0.234837254 | 0.814335051 | 0.907710402 | 0.120349597 |
| ENSMUSG00000031399 | Fam3a | 66294 | protein_coding | 367.3257733 | 0.120371699 | 0.071480857 | -1.683971116 | 0.092187251 | 0.232815456 | 0.120371699 |
| ENSMUSG00000022744 | Cldnd1 | 224250 | protein_coding | 274.2034426 | -0.120434094 | 0.098015418 | 1.228726014 | 0.219174549 | 0.425467087 | 0.120434094 |
| ENSMUSG00000032376 | Usp3 | 235441 | protein_coding | 1010.131598 | -0.120475591 | 0.071296141 | 1.689791198 | 0.091067908 | 0.23065881 | 0.120475591 |
| ENSMUSG00000033596 | Rfwd3 | 234736 | protein_coding | 563.8760748 | 0.120490053 | 0.104498237 | -1.153034307 | 0.248896303 | 0.461051436 | 0.120490053 |
| ENSMUSG00000028962 | Slc4a2 | 20535 | protein_coding | 2677.040284 | -0.120491158 | 0.132016723 | 0.912696169 | 0.361402359 | 0.583801091 | 0.120491158 |
| ENSMUSG00000022000 | Zc3h13 | 67302 | protein_coding | 316.7294425 | 0.120506834 | 0.079191311 | -1.52171789 | 0.128079783 | 0.295198531 | 0.120506834 |
| ENSMUSG00000052658 | 5830454E08Rik | 76100 | lncRNA | 6.960215435 | 0.120569639 | 0.494775201 | -0.243685696 | 0.807474253 | 0.90502442 | 0.120569639 |
| ENSMUSG00000071337 | Tia1 | 21841 | protein_coding | 335.1727928 | 0.120621319 | 0.089203226 | -1.352208038 | 0.176308773 | 0.368268232 | 0.120621319 |
| ENSMUSG00000003031 | Cdkn1b | 12576 | protein_coding | 780.1322636 | -0.120642609 | 0.073560686 | 1.640041913 | 0.100996453 | 0.249566874 | 0.120642609 |
| ENSMUSG00000032340 | Neo1 | 18007 | protein_coding | 1167.688291 | -0.120660439 | 0.069605616 | 1.733487128 | 0.083009123 | 0.215774069 | 0.120660439 |
| ENSMUSG00000024957 | Kcnk4 | 16528 | protein_coding | 6.670262305 | 0.120662689 | 0.509744232 | -0.236712221 | 0.812880054 | 0.907147741 | 0.120662689 |
| ENSMUSG00000040363 | Bcor | 71458 | protein_coding | 600.4378349 | 0.12071086 | 0.068800668 | -1.754501284 | 0.07934465 | 0.208654479 | 0.12071086 |

| ensembl_gene_id | external_gene_name | entrezgene_id | gene_biotype | baseMean | log2FoldChange | lfcSE | stat | pvalue | padj | absLog2FC |
|---|---|---|---|---|---|---|---|---|---|---|
| ENSMUSG00000051950 | B3glct | 381694 | protein_coding | 181.5499657 | -0.120711225 | 0.105587731 | 1.143231542 | 0.252942469 | 0.465324602 | 0.120711225 |
| ENSMUSG00000024125 | Sbpl | 638345 | protein_coding | 2.266870592 | 0.120754529 | 0.919508761 | -0.131325045 | 0.89551818 | 1 | 0.120754529 |
| ENSMUSG00000027102 | Hoxd8 | 15437 | protein_coding | 28.29488974 | -0.120769856 | 0.337600645 | 0.357729932 | 0.720545431 | 0.855204229 | 0.120769856 |
| ENSMUSG00000056737 | Capg | 12332 | protein_coding | 499.9260518 | 0.12082473 | 0.16149399 | -0.748168586 | 0.454358477 | 0.669875536 | 0.12082473 |
| ENSMUSG00000006731 | B4galnt1 | 14421 | protein_coding | 1147.660956 | 0.12084814 | 0.097823891 | -1.235364278 | 0.216694961 | 0.422162995 | 0.12084814 |
| ENSMUSG00000037808 | Fam76b | 72826 | protein_coding | 289.3586737 | 0.120955736 | 0.086025492 | -1.406045273 | 0.159710682 | 0.343912139 | 0.120955736 |
| ENSMUSG00000021116 | Eif2s1 | 13665 | protein_coding | 710.1420045 | 0.120968618 | 0.082551804 | -1.465366136 | 0.142821051 | 0.318636429 | 0.120968618 |
| ENSMUSG00000020300 | Cpeb4 | 67579 | protein_coding | 1115.494207 | -0.120981286 | 0.080521586 | 1.50247024 | 0.132975708 | 0.302752759 | 0.120981286 |
| ENSMUSG00000023266 | Frs3 | 107971 | protein_coding | 24.9503182 | -0.120983524 | 0.219997699 | 0.549930864 | 0.582366794 | 0.765213657 | 0.120983524 |
| ENSMUSG00000028293 | Slc35a1 | 24060 | protein_coding | 2225.243424 | 0.120984115 | 0.174801669 | -0.692122196 | 0.488860593 | 0.696638145 | 0.120984115 |
| ENSMUSG00000004677 | Myo9b | 17925 | protein_coding | 616.6139027 | 0.120989033 | 0.107309334 | -1.127479109 | 0.259539968 | 0.472954995 | 0.120989033 |
| ENSMUSG00000042659 | Arrdc4 | 66412 | protein_coding | 988.0380608 | 0.12106909 | 0.101043745 | -1.19818491 | 0.230845039 | 0.439494463 | 0.12106909 |
| ENSMUSG00000038797 | Zscan2 | 22691 | protein_coding | 33.10194188 | -0.12115792 | 0.210436665 | 0.575745295 | 0.564787355 | 0.751588072 | 0.12115792 |
| ENSMUSG00000033400 | Agl | 77559 | protein_coding | 357.8835194 | 0.121166637 | 0.109025228 | -1.111363297 | 0.266412004 | 0.481401448 | 0.121166637 |
| ENSMUSG00000051346 | Spryd4 | 66701 | protein_coding | 196.3296636 | -0.121172183 | 0.1528622 | 0.792688987 | 0.427959048 | 0.647057325 | 0.121172183 |
| ENSMUSG00000024277 | Mapre2 | 212307 | protein_coding | 832.7077104 | -0.121181381 | 0.056827586 | 2.132439351 | 0.032970745 | 0.108526787 | 0.121181381 |
| ENSMUSG00000013878 | Rnf170 | 77733 | protein_coding | 192.6777183 | 0.121181835 | 0.102553572 | -1.181644207 | 0.237346905 | 0.44769238 | 0.121181835 |
| ENSMUSG00000078440 | Dohh | 102115 | protein_coding | 98.27667242 | 0.121196264 | 0.115987414 | -1.044908746 | 0.296065147 | 0.515232502 | 0.121196264 |
| ENSMUSG00000020271 | Fbxw11 | 103583 | protein_coding | 874.9049676 | -0.121203179 | 0.071157489 | 1.703308837 | 0.088510288 | 0.225948198 | 0.121203179 |
| ENSMUSG00000036568 | Bicral | 210982 | protein_coding | 323.4636603 | -0.121214157 | 0.079255765 | 1.529404911 | 0.126164096 | 0.292089426 | 0.121214157 |
| ENSMUSG00000040325 | Dcaf1 | 321006 | protein_coding | 700.3522082 | 0.121218801 | 0.071038747 | -1.706375827 | 0.087938128 | 0.225148876 | 0.121218801 |
| ENSMUSG00000048410 | Zfp407 | 240476 | protein_coding | 400.5667338 | 0.121220315 | 0.099039643 | -1.223957511 | 0.220968256 | 0.427441699 | 0.121220315 |
| ENSMUSG00000049588 | Ccdc69 | 52570 | protein_coding | 4.935166174 | -0.121250271 | 0.488219569 | 0.248351929 | 0.803862124 | 0.903305601 | 0.121250271 |
| ENSMUSG00000022889 | Mrpl39 | 27393 | protein_coding | 404.7960814 | -0.121257537 | 0.080214881 | 1.51165888 | 0.130620664 | 0.298680373 | 0.121257537 |
| ENSMUSG00000022477 | Aco2 | 11429 | protein_coding | 6383.475493 | -0.121270551 | 0.080588691 | 1.504808542 | 0.132373307 | 0.301713259 | 0.121270551 |
| ENSMUSG00000072572 | Slc39a2 | 214922 | protein_coding | 5.398220294 | 0.121299615 | 0.609390043 | -0.199050865 | 0.842222956 | 0.921900136 | 0.121299615 |
| ENSMUSG00000039193 | Nlrc4 | 268973 | protein_coding | 696.4504825 | -0.121349191 | 0.074322985 | 1.632727643 | 0.102526291 | 0.252139861 | 0.121349191 |
| ENSMUSG00000022404 | Slc25a17 | 20524 | protein_coding | 528.7877429 | -0.121354717 | 0.098325957 | 1.234208351 | 0.217125276 | 0.422731291 | 0.121354717 |
| ENSMUSG00000021606 | Ndufs6 | 407785 | protein_coding | 945.0829601 | 0.121360166 | 0.091709288 | -1.323313795 | 0.185731046 | 0.381203392 | 0.121360166 |
| ENSMUSG00000028798 | Eif3i | 54709 | protein_coding | 1555.300578 | -0.121479156 | 0.106985611 | 1.135471902 | 0.256177648 | 0.468911293 | 0.121479156 |
| ENSMUSG00000049658 | Bdp1 | 544971 | protein_coding | 828.0313123 | 0.121528755 | 0.073012668 | -1.66448861 | 0.096014825 | 0.239840996 | 0.121528755 |
| ENSMUSG00000030876 | Mettl9 | 59052 | protein_coding | 761.3448473 | 0.121541636 | 0.090112731 | -1.348773205 | 0.177409824 | 0.36973067 | 0.121541636 |
| ENSMUSG00000037337 | Map4k1 | 26411 | protein_coding | 16.09114518 | 0.121541913 | 0.447374033 | -0.271678515 | 0.785869223 | 0.892991423 | 0.121541913 |
| ENSMUSG00000026790 | Odf2 | 18286 | protein_coding | 396.2724986 | 0.121543239 | 0.088395194 | -1.374998274 | 0.16913198 | 0.358358419 | 0.121543239 |
| ENSMUSG00000021361 | Tmem14c | 66154 | protein_coding | 1033.824755 | -0.121589799 | 0.090224873 | 1.347630589 | 0.177777227 | 0.370076028 | 0.121589799 |
| ENSMUSG00000022440 | C1qtnf6 | 72709 | protein_coding | 226.592101 | 0.121669318 | 0.131494071 | -0.925283684 | 0.354818364 | 0.577401503 | 0.121669318 |
| ENSMUSG00000014633 | Cmc2 | 66531 | protein_coding | 155.4183076 | 0.121703439 | 0.109183334 | -1.114670476 | 0.264991657 | 0.479832481 | 0.121703439 |
| ENSMUSG00000014763 | Fam120b | 67544 | protein_coding | 639.8427545 | 0.121739307 | 0.047223197 | -2.577955631 | 0.009938676 | 0.042377303 | 0.121739307 |
| ENSMUSG00000043090 | Zfp866 | 330788 | protein_coding | 169.2176307 | -0.121813501 | 0.148089851 | 0.822564819 | 0.410755512 | 0.630971948 | 0.121813501 |

| ensembl_gene_id | external_gene_name | entrezgene_id | gene_biotype | baseMean | log2FoldChange | lfcSE | stat | pvalue | padj | absLog2FC |
|---|---|---|---|---|---|---|---|---|---|---|
| ENSMUSG00000031378 | Abcd1 | 11666 | protein_coding | 317.6248089 | -0.121829149 | 0.071404649 | 1.706179499 | 0.087974665 | 0.22515016 | 0.121829149 |
| ENSMUSG00000085112 | A530072M11Rik | 100415915 | lncRNA | 20.57573978 | 0.12187632 | 0.234746851 | -0.519181917 | 0.603633888 | 0.779177919 | 0.12187632 |
| ENSMUSG00000031467 | Agpat5 | 52123 | protein_coding | 584.2269825 | -0.121884186 | 0.108951483 | 1.11870148 | 0.263267514 | 0.477583041 | 0.121884186 |
| ENSMUSG00000004568 | Arhgef18 | 102098 | protein_coding | 721.9985317 | -0.121912206 | 0.071083211 | 1.715063282 | 0.086333625 | 0.222142956 | 0.121912206 |
| ENSMUSG00000008763 | Man1a2 | 17156 | protein_coding | 1972.564438 | 0.121932418 | 0.081245985 | -1.500780835 | 0.133412257 | 0.303216352 | 0.121932418 |
| ENSMUSG00000019777 | Hdac2 | 15182 | protein_coding | 531.090065 | 0.121951241 | 0.081180434 | -1.502224554 | 0.133039126 | 0.302752759 | 0.121951241 |
| ENSMUSG00000020463 | Ppp4r3b | 104570 | protein_coding | 1234.375528 | 0.121953317 | 0.06233527 | -1.956409548 | 0.050416928 | 0.149799416 | 0.121953317 |
| ENSMUSG00000079173 | Zan | 22635 | protein_coding | 1169.427704 | -0.121984913 | 0.137997327 | 0.883965769 | 0.376714692 | 0.598928123 | 0.121984913 |
| ENSMUSG00000040147 | Maob | 109731 | protein_coding | 356.5946709 | 0.121998628 | 0.151364656 | -0.805991513 | 0.420247743 | 0.640159692 | 0.121998628 |
| ENSMUSG00000068587 | Mgam | 232714 | protein_coding | 6147.07189 | 0.122116725 | 0.096327713 | -1.26772163 | 0.204897378 | 0.407353559 | 0.122116725 |
| ENSMUSG00000051124 | Gimap9 | 317758 | protein_coding | 73.78849664 | 0.122201742 | 0.149759904 | -0.815984378 | 0.41450907 | 0.634755452 | 0.122201742 |
| ENSMUSG00000024404 | Riok3 | 66878 | protein_coding | 1534.01495 | 0.122375271 | 0.049236253 | -2.485470863 | 0.012938018 | 0.05194233 | 0.122375271 |
| ENSMUSG00000039813 | Tbc1d2 | 381605 | protein_coding | 809.961464 | 0.12237575 | 0.107423841 | -1.139186132 | 0.254625529 | 0.467157647 | 0.12237575 |
| ENSMUSG00000079610 | Ankrd39 | 109346 | protein_coding | 47.64952645 | 0.122409833 | 0.169803273 | -0.720892069 | 0.470975923 | 0.684066582 | 0.122409833 |
| ENSMUSG00000010057 | Nprl2 | 56032 | protein_coding | 189.844649 | -0.122435425 | 0.101844889 | 1.202175446 | 0.22929556 | 0.438111017 | 0.122435425 |
| ENSMUSG00000019987 | Arg1 | 11846 | protein_coding | 10.52819663 | 0.122545771 | 0.324613173 | -0.377513241 | 0.705792226 | 0.845823862 | 0.122545771 |
| ENSMUSG00000022299 | Slc25a32 | 69906 | protein_coding | 285.7645602 | 0.122573297 | 0.080812844 | -1.516755148 | 0.129328515 | 0.29700057 | 0.122573297 |
| ENSMUSG00000037683 | Armc3 | 70882 | protein_coding | 10.16162001 | -0.122600446 | 0.451201594 | 0.271719887 | 0.785837409 | 0.892991423 | 0.122600446 |
| ENSMUSG00000044676 | Zfp612 | 234725 | protein_coding | 135.1671635 | 0.122614267 | 0.130246586 | -0.941401008 | 0.346499397 | 0.568175062 | 0.122614267 |
| ENSMUSG00000028347 | Tmeff1 | 230157 | protein_coding | 123.5380524 | -0.122641723 | 0.169984215 | 0.721488893 | 0.470608772 | 0.683821668 | 0.122641723 |
| ENSMUSG00000022561 | Gpaa1 | 14731 | protein_coding | 538.2218699 | -0.122758173 | 0.084663592 | 1.44995234 | 0.14707181 | 0.324959041 | 0.122758173 |
| ENSMUSG00000031891 | Hsd11b2 | 15484 | protein_coding | 2275.465186 | -0.122818186 | 0.202389382 | 0.606841052 | 0.543956402 | 0.73666903 | 0.122818186 |
| ENSMUSG00000028797 | Tmem234 | 76799 | protein_coding | 940.7174323 | -0.122849592 | 0.06986518 | 1.758380817 | 0.078682735 | 0.207566563 | 0.122849592 |
| ENSMUSG00000022641 | Bbx | 70508 | protein_coding | 467.6579809 | -0.122868724 | 0.103645338 | 1.185472756 | 0.23583058 | 0.445952845 | 0.122868724 |
| ENSMUSG00000024054 | Smchd1 | 74355 | protein_coding | 816.9455969 | 0.122886226 | 0.077729797 | -1.580941033 | 0.113891521 | 0.271186338 | 0.122886226 |
| ENSMUSG00000032293 | Ireb2 | 64602 | protein_coding | 598.023779 | -0.122889577 | 0.219537187 | 0.559766562 | 0.575638674 | 0.760792082 | 0.122889577 |
| ENSMUSG00000000838 | Fmr1 | 14265 | protein_coding | 690.0949544 | 0.122891754 | 0.081573146 | -1.506522192 | 0.131933175 | 0.30108314 | 0.122891754 |
| ENSMUSG00000022809 | Nr1i2 | 18171 | protein_coding | 973.9203917 | -0.122893928 | 0.073960425 | 1.661617394 | 0.096589513 | 0.240948524 | 0.122893928 |
| ENSMUSG00000007815 | Rhoa | 11848 | protein_coding | 2563.817113 | -0.122896041 | 0.08986591 | 1.367549064 | 0.17145327 | 0.361606551 | 0.122896041 |
| ENSMUSG00000025199 | Chuk | 12675 | protein_coding | 1064.376508 | 0.122947246 | 0.068373493 | -1.798171203 | 0.072149882 | 0.195389167 | 0.122947246 |
| ENSMUSG00000033644 | Piwil2 | 57746 | protein_coding | 8.089036415 | 0.122973564 | 0.453094625 | -0.271408127 | 0.786077152 | 0.893150482 | 0.122973564 |
| ENSMUSG00000056130 | Ticam2 | 225471 | protein_coding | 4.09464322 | -0.123072428 | 0.675032683 | 0.182320696 | 0.855331058 | 1 | 0.123072428 |
| ENSMUSG00000018076 | Med13l | 76199 | protein_coding | 685.8742953 | 0.12311756 | 0.131385594 | -0.937070463 | 0.348722314 | 0.570573694 | 0.12311756 |
| ENSMUSG00000024413 | Npc1 | 18145 | protein_coding | 451.3867505 | -0.123126854 | 0.123578938 | 0.996341739 | 0.31908413 | 0.540502136 | 0.123126854 |
| ENSMUSG00000028360 | Slc44a5 | 242259 | protein_coding | 3.988389921 | 0.123137285 | 1.028455698 | -0.119730276 | 0.904696816 | 1 | 0.123137285 |
| ENSMUSG00000026858 | Miga2 | 108958 | protein_coding | 655.4814443 | 0.123140639 | 0.08095351 | -1.521127848 | 0.128227757 | 0.295417576 | 0.123140639 |
| ENSMUSG00000099759 | 1700030C10Rik | 69513 | lncRNA | 16.87164499 | 0.123267501 | 0.295222678 | -0.417540758 | 0.676282918 | 0.826766769 | 0.123267501 |
| ENSMUSG00000078515 | Ddi2 | 68817 | protein_coding | 1399.41296 | 0.123297721 | 0.059252699 | -2.08087941 | 0.037444945 | 0.119675987 | 0.123297721 |
| ENSMUSG00000018547 | Pip4k2b | 108083 | protein_coding | 124.8194004 | 0.123300656 | 0.114344537 | -1.078325729 | 0.280888419 | 0.498558829 | 0.123300656 |

Supplementary Table 1

| ensembl_gene_id | external_gene_name | entrezgene_id | gene_biotype | baseMean | log2FoldChange | lfcSE | stat | pvalue | padj | absLog2FC |
|---|---|---|---|---|---|---|---|---|---|---|
| ENSMUSG00000019373 | Cops3 | 26572 | protein_coding | 574.9497688 | 0.123320868 | 0.064724864 | -1.905309041 | 0.056739911 | 0.163972315 | 0.123320868 |
| ENSMUSG00000049625 | Tifab | 212937 | protein_coding | 2.632548469 | -0.123353022 | 1.22815353 | 0.100437786 | 0.919996772 | 1 | 0.123353022 |
| ENSMUSG00000030967 | Zranb1 | 360216 | protein_coding | 628.0893306 | -0.123402115 | 0.08902304 | 1.386181765 | 0.165691408 | 0.352762472 | 0.123402115 |
| ENSMUSG00000013150 | Gfod2 | 70575 | protein_coding | 210.927637 | -0.123412521 | 0.119122444 | 1.036014012 | 0.300195606 | 0.520301242 | 0.123412521 |
| ENSMUSG00000061758 | Akr1b10 | 67861 | protein_coding | 66.39966856 | -0.12348973 | 0.124115995 | 0.994954198 | 0.319758541 | 0.541034072 | 0.12348973 |
| ENSMUSG00000024335 | Brd2 | 14312 | protein_coding | 2456.713081 | 0.123490018 | 0.104950805 | -1.1766467 | 0.23933655 | 0.449590928 | 0.123490018 |
| ENSMUSG00000021973 | Micu2 | 68514 | protein_coding | 767.5458014 | -0.123499749 | 0.06884398 | 1.793907751 | 0.072827894 | 0.19654843 | 0.123499749 |
| ENSMUSG00000028149 | Rap1gds1 | 229877 | protein_coding | 607.6853407 | -0.123585188 | 0.07356977 | 1.679836533 | 0.092989125 | 0.234124471 | 0.123585188 |
| ENSMUSG00000015143 | Actn1 | 109711 | protein_coding | 1269.54438 | -0.123598939 | 0.108109828 | 1.143271996 | 0.252925678 | 0.465324602 | 0.123598939 |
| ENSMUSG00000026632 | Tatdn3 | 68972 | protein_coding | 58.87668535 | 0.123685226 | 0.150037208 | -0.824363689 | 0.40973294 | 0.630279329 | 0.123685226 |
| ENSMUSG00000090935 | Synj2bp | 24071 | protein_coding | 744.1883865 | -0.123750917 | 0.103647982 | 1.193953954 | 0.232495981 | 0.441568559 | 0.123750917 |
| ENSMUSG00000002617 | Zfp40 | 22700 | protein_coding | 58.81674738 | 0.123761023 | 0.154703459 | -0.799988726 | 0.423717329 | 0.643604486 | 0.123761023 |
| ENSMUSG00000030549 | Rhcg | 56315 | protein_coding | 1.423525057 | 0.123811418 | 2.325279831 | -0.053245814 | 0.957536053 | 1 | 0.123811418 |
| ENSMUSG00000089862 | Umad1 | 100036521 | protein_coding | 259.4510169 | 0.123929279 | 0.161240185 | -0.768600447 | 0.44213054 | 0.659918993 | 0.123929279 |
| ENSMUSG00000033499 | Larp4b | 217980 | protein_coding | 1915.027239 | -0.123935258 | 0.066747456 | 1.856778744 | 0.063342639 | 0.177475457 | 0.123935258 |
| ENSMUSG00000030557 | Mef2a | 17258 | protein_coding | 344.3861875 | 0.123936123 | 0.074205153 | -1.670182157 | 0.094883329 | 0.237733773 | 0.123936123 |
| ENSMUSG00000053332 | Gas5 | 14455 | lncRNA | 969.2952358 | -0.123945791 | 0.079958964 | 1.550117515 | 0.121113313 | 0.283858253 | 0.123945791 |
| ENSMUSG00000057963 | Itpk1 | 217837 | protein_coding | 2356.349752 | 0.123958487 | 0.12083528 | -1.025846809 | 0.304963798 | 0.525093542 | 0.123958487 |
| ENSMUSG00000024805 | Pcgf5 | 76073 | protein_coding | 333.5506832 | -0.123974174 | 0.149050204 | 0.831761184 | 0.405543757 | 0.626487335 | 0.123974174 |
| ENSMUSG00000020899 | Pfas | 237823 | protein_coding | 440.8823239 | -0.123984889 | 0.116921248 | 1.060413661 | 0.28895645 | 0.507895701 | 0.123984889 |
| ENSMUSG00000091091 | Kcnmb3 | 100502876 | protein_coding | 2.515263353 | 0.124025873 | 0.929261951 | -0.133467073 | 0.893824003 | 1 | 0.124025873 |
| ENSMUSG00000015501 | Hivep2 | 15273 | protein_coding | 406.3929505 | 0.124051037 | 0.130760552 | -0.948688536 | 0.342779046 | 0.564822707 | 0.124051037 |
| ENSMUSG00000058174 | Gm5148 | 381438 | protein_coding | 13.91959111 | -0.124057468 | 0.324603421 | 0.382181642 | 0.702326638 | 0.84294636 | 0.124057468 |
| ENSMUSG00000022897 | Dyrk1a | 13548 | protein_coding | 898.2351066 | 0.124065643 | 0.043496662 | -2.852302602 | 0.004340376 | 0.021615492 | 0.124065643 |
| ENSMUSG00000026926 | Pmpca | 66865 | protein_coding | 707.5510641 | 0.124067114 | 0.061925632 | -2.003485619 | 0.04512519 | 0.137999515 | 0.124067114 |
| ENSMUSG00000042063 | Zfp386 | 56220 | protein_coding | 305.9727509 | -0.12408917 | 0.106449519 | 1.16570907 | 0.243732092 | 0.454904132 | 0.12408917 |
| ENSMUSG00000021597 | Slf1 | 105377 | protein_coding | 179.8872853 | 0.124133808 | 0.096433017 | -1.28725422 | 0.198005695 | 0.398279073 | 0.124133808 |
| ENSMUSG00000001948 | Spa17 | 20686 | protein_coding | 8.922361452 | 0.12415305 | 0.315941853 | -0.392961709 | 0.694347763 | 0.837885199 | 0.12415305 |
| ENSMUSG00000047617 | Paxx | 227622 | protein_coding | 109.7005956 | -0.124189245 | 0.100371753 | 1.237292774 | 0.215978409 | 0.421243351 | 0.124189245 |
| ENSMUSG00000027166 | Dnajc24 | 99349 | protein_coding | 83.8747143 | 0.124214846 | 0.146609468 | -0.847249825 | 0.396855887 | 0.618855346 | 0.124214846 |
| ENSMUSG00000018507 | Trpv2 | 22368 | protein_coding | 4.520845049 | 0.124246983 | 0.899544461 | -0.138122115 | 0.890143908 | 0.946057182 | 0.124246983 |
| ENSMUSG00000062115 | Rai1 | 19377 | protein_coding | 269.6462434 | 0.124276318 | 0.143860218 | -0.863868551 | 0.387660105 | 0.610103781 | 0.124276318 |
| ENSMUSG00000024826 | Dpf2 | 19708 | protein_coding | 439.9365561 | -0.124304533 | 0.058902062 | 2.110359616 | 0.034827392 | 0.113038434 | 0.124304533 |
| ENSMUSG00000013465 | Nelfb | 58202 | protein_coding | 802.7123969 | -0.124437344 | 0.047231856 | 2.634606244 | 0.008423494 | 0.037065168 | 0.124437344 |
| ENSMUSG00000029406 | Pitpnm2 | 19679 | protein_coding | 58.36391149 | -0.124438768 | 0.285363136 | 0.436071631 | 0.662784759 | 0.817662297 | 0.124438768 |
| ENSMUSG00000028552 | Eps15 | 13858 | protein_coding | 573.1905474 | -0.124482976 | 0.09550139 | 1.303467685 | 0.192415142 | 0.390165244 | 0.124482976 |
| ENSMUSG00000014498 | Ankrd52 | 237615 | protein_coding | 596.7664929 | 0.124538692 | 0.102703429 | -1.212604995 | 0.225280878 | 0.432685604 | 0.124538692 |
| ENSMUSG00000022014 | Epsti1 | 108670 | protein_coding | 498.0470268 | -0.124540569 | 0.085653905 | 1.453997565 | 0.145946981 | 0.323295567 | 0.124540569 |
| ENSMUSG00000067028 | Cntnap5b | 241175 | protein_coding | 2.620401536 | 0.124562745 | 2.358344032 | -0.052817885 | 0.957877011 | 1 | 0.124562745 |

| ensembl_gene_id | external_gene_name | entrezgene_id | gene_biotype | baseMean | log2FoldChange | lfcSE | stat | pvalue | padj | absLog2FC |
|---|---|---|---|---|---|---|---|---|---|---|
| ENSMUSG00000007216 | Zfp775 | 243372 | protein_coding | 90.80748979 | 0.12457221 | 0.128987261 | -0.965771422 | 0.334158572 | 0.5565041 | 0.12457221 |
| ENSMUSG00000056919 | Cep162 | 382090 | protein_coding | 200.064871 | 0.124579968 | 0.082016952 | -1.518953892 | 0.128774102 | 0.29613919 | 0.124579968 |
| ENSMUSG00000021144 | Mta1 | 116870 | protein_coding | 226.0357485 | -0.124600296 | 0.100511754 | 1.239658961 | 0.215101563 | 0.420139706 | 0.124600296 |
| ENSMUSG00000036815 | Dpp10 | 269109 | protein_coding | 3.714102894 | -0.124615777 | 2.175237052 | 0.057288366 | 0.954315487 | 1 | 0.124615777 |
| ENSMUSG00000043207 | Zmpste24 | 230709 | protein_coding | 588.4330586 | 0.124663252 | 0.087288949 | -1.42816763 | 0.153243615 | 0.334272057 | 0.124663252 |
| ENSMUSG00000021635 | Rad17 | 19356 | protein_coding | 246.7175769 | 0.124759208 | 0.080274055 | -1.554166009 | 0.120144823 | 0.28232699 | 0.124759208 |
| ENSMUSG00000038141 | Tmem181a | 77106 | protein_coding | 397.3048453 | -0.124970532 | 0.12888988 | 0.969591495 | 0.332250154 | 0.554612255 | 0.124970532 |
| ENSMUSG00000073771 | Btbd19 | 78611 | protein_coding | 63.80694196 | -0.124986058 | 0.179176695 | 0.697557558 | 0.485453932 | 0.695380179 | 0.124986058 |
| ENSMUSG00000028392 | Bspry | 192120 | protein_coding | 2842.800909 | -0.125005849 | 0.046997821 | 2.659822247 | 0.00781819 | 0.034960626 | 0.125005849 |
| ENSMUSG00000026667 | Uhmk1 | 16589 | protein_coding | 1453.529275 | -0.125022563 | 0.079386131 | 1.574866572 | 0.115287246 | 0.273824144 | 0.125022563 |
| ENSMUSG00000020132 | Rab21 | 216344 | protein_coding | 568.8758004 | 0.125052294 | 0.074604706 | -1.676198467 | 0.093699324 | 0.235661079 | 0.125052294 |
| ENSMUSG00000024811 | Tnks2 | 74493 | protein_coding | 1822.920889 | 0.125065092 | 0.075022173 | -1.667041722 | 0.095506109 | 0.239040298 | 0.125065092 |
| ENSMUSG00000047446 | Arl4a | 11861 | protein_coding | 1355.774172 | -0.125096295 | 0.174114905 | 0.718469764 | 0.472467685 | 0.684955785 | 0.125096295 |
| ENSMUSG00000026821 | Ralgds | 19730 | protein_coding | 1279.053183 | 0.125135883 | 0.127525313 | -0.98126309 | 0.326463019 | 0.548343875 | 0.125135883 |
| ENSMUSG00000042594 | Sh2b3 | 16923 | protein_coding | 165.3305508 | -0.125153364 | 0.088684994 | 1.411212413 | 0.158181991 | 0.341996258 | 0.125153364 |
| ENSMUSG00000055296 | Tmem245 | 242474 | protein_coding | 557.524107 | -0.125200643 | 0.116733615 | 1.072532904 | 0.283480743 | 0.501585257 | 0.125200643 |
| ENSMUSG00000037815 | Ctnna1 | 12385 | protein_coding | 6087.065458 | -0.125241774 | 0.082341782 | 1.520999073 | 0.12826007 | 0.295450802 | 0.125241774 |
| ENSMUSG00000015176 | Nolc1 | 70769 | protein_coding | 534.0002211 | 0.125328134 | 0.128637201 | -0.974275974 | 0.329919531 | 0.552117058 | 0.125328134 |
| ENSMUSG00000071451 | Psmg4 | 69666 | protein_coding | 64.16319132 | -0.125345998 | 0.145320779 | 0.862546975 | 0.388386595 | 0.610900332 | 0.125345998 |
| ENSMUSG00000066036 | Ubr4 | 69116 | protein_coding | 3298.129105 | 0.125358345 | 0.137927294 | -0.908872647 | 0.363417354 | 0.585795796 | 0.125358345 |
| ENSMUSG00000022634 | Yaf2 | 67057 | protein_coding | 105.2669957 | 0.125359905 | 0.111009171 | -1.129275211 | 0.258781755 | 0.472212364 | 0.125359905 |
| ENSMUSG00000026166 | Ccl20 | 20297 | protein_coding | 11.93398347 | -0.125360955 | 0.409000391 | 0.306505708 | 0.759219634 | 0.87897876 | 0.125360955 |
| ENSMUSG00000041040 | Fam117b | 72750 | protein_coding | 735.3023302 | 0.125385018 | 0.065988713 | -1.900097952 | 0.057420266 | 0.165344954 | 0.125385018 |
| ENSMUSG00000040423 | Rc3h1 | 381305 | protein_coding | 1081.954256 | 0.125420112 | 0.084710769 | -1.480568674 | 0.138721548 | 0.311742141 | 0.125420112 |
| ENSMUSG00000078597 | Cyp4a12b | 13118 | protein_coding | 1.222943214 | 0.125460556 | 2.64261903 | -0.047475839 | 0.962133986 | 1 | 0.125460556 |
| ENSMUSG00000024485 | Slc4a9 | 240215 | protein_coding | 5.911253614 | -0.125462538 | 0.583849004 | 0.214888673 | 0.829854116 | 0.914162556 | 0.125462538 |
| ENSMUSG00000069631 | Strada | 72149 | protein_coding | 68.73738379 | -0.125506281 | 0.129846642 | 0.966573174 | 0.33375745 | 0.556110732 | 0.125506281 |
| ENSMUSG00000026849 | Tor1a | 30931 | protein_coding | 325.6520304 | 0.125522773 | 0.07190841 | -1.745592373 | 0.080881809 | 0.211442486 | 0.125522773 |
| ENSMUSG00000044066 | Cep68 | 216543 | protein_coding | 293.3842999 | 0.12553171 | 0.103071241 | -1.217912083 | 0.223257384 | 0.430343308 | 0.12553171 |
| ENSMUSG00000026303 | Mlph | 171531 | protein_coding | 801.1275227 | 0.125531824 | 0.106033687 | -1.183886241 | 0.236458097 | 0.446602599 | 0.125531824 |
| ENSMUSG00000020150 | Gamt | 14431 | protein_coding | 17.04155812 | -0.125547897 | 0.284642275 | 0.441072559 | 0.659160469 | 0.816056375 | 0.125547897 |
| ENSMUSG00000005103 | Wdr1 | 22388 | protein_coding | 4534.050505 | -0.125565441 | 0.061403719 | 2.044915912 | 0.040863149 | 0.128048205 | 0.125565441 |
| ENSMUSG00000028776 | Tinagl1 | 94242 | protein_coding | 799.9261926 | 0.125566324 | 0.160489753 | -0.78239465 | 0.433982677 | 0.65253482 | 0.125566324 |
| ENSMUSG00000031918 | Mtmr2 | 77116 | protein_coding | 645.5744003 | 0.125622335 | 0.087948977 | -1.428354706 | 0.153189789 | 0.334228588 | 0.125622335 |
| ENSMUSG00000026435 | Slc45a3 | 212980 | protein_coding | 428.4746985 | -0.125626928 | 0.15254239 | 0.823554211 | 0.410192902 | 0.630552252 | 0.125626928 |
| ENSMUSG00000061979 | Rcc1l | 94254 | protein_coding | 199.4339522 | -0.12568036 | 0.141793857 | 0.886359691 | 0.375423733 | 0.597623629 | 0.12568036 |
| ENSMUSG00000049470 | Aff4 | 93736 | protein_coding | 1724.622974 | -0.125715761 | 0.074694443 | 1.683067116 | 0.0923621 | 0.233078807 | 0.125715761 |
| ENSMUSG00000054452 | Tle5 | 14797 | protein_coding | 5086.158442 | -0.125763056 | 0.082165027 | 1.530615412 | 0.12586447 | 0.291600149 | 0.125763056 |
| ENSMUSG00000029328 | Hnrnpdl | 50926 | protein_coding | 1275.635719 | 0.125774845 | 0.058881669 | -2.136061147 | 0.03267443 | 0.107787762 | 0.125774845 |

| ensembl_gene_id | external_gene_name | entrezgene_id | gene_biotype | baseMean | log2FoldChange | lfcSE | stat | pvalue | padj | absLog2FC |
|---|---|---|---|---|---|---|---|---|---|---|
| ENSMUSG00000020867 | Spata20 | 217116 | protein_coding | 2.447455494 | 0.125827252 | 1.013083393 | -0.124202265 | 0.901155129 | 1 | 0.125827252 |
| ENSMUSG00000060601 | Nr1h2 | 22260 | protein_coding | 626.3491183 | -0.125849079 | 0.085886628 | 1.465293058 | 0.14284098 | 0.318637841 | 0.125849079 |
| ENSMUSG00000001855 | Nup214 | 227720 | protein_coding | 314.9213827 | 0.125870121 | 0.090222927 | -1.395101277 | 0.162985301 | 0.348916161 | 0.125870121 |
| ENSMUSG00000025437 | Usp33 | 170822 | protein_coding | 919.6892473 | -0.125877043 | 0.073400185 | 1.714941761 | 0.086355905 | 0.222165667 | 0.125877043 |
| ENSMUSG00000039917 | Rhbdd2 | 215160 | protein_coding | 144.0658447 | -0.125881057 | 0.108322711 | 1.162092941 | 0.245197714 | 0.456813522 | 0.125881057 |
| ENSMUSG00000029501 | Ankle2 | 71782 | protein_coding | 534.2463162 | 0.125896564 | 0.076686174 | -1.641711385 | 0.100649831 | 0.248933849 | 0.125896564 |
| ENSMUSG00000027395 | Polr1b | 20017 | protein_coding | 301.8966296 | -0.125896865 | 0.125922793 | 0.999794101 | 0.317410161 | 0.538827241 | 0.125896865 |
| ENSMUSG00000042439 | Zfp532 | 328977 | protein_coding | 132.7755007 | -0.125897705 | 0.105760433 | 1.190404595 | 0.233887408 | 0.443446229 | 0.125897705 |
| ENSMUSG00000020248 | Nfyb | 18045 | protein_coding | 305.8483908 | 0.12591422 | 0.080387145 | -1.56634771 | 0.117267237 | 0.27724426 | 0.12591422 |
| ENSMUSG00000034586 | Hid1 | 217310 | protein_coding | 998.6323538 | 0.125931446 | 0.197306348 | -0.638253395 | 0.523308744 | 0.722098569 | 0.125931446 |
| ENSMUSG00000020196 | Cabin1 | 104248 | protein_coding | 917.2522083 | 0.125938486 | 0.080697648 | -1.560621513 | 0.11861308 | 0.279625468 | 0.125938486 |
| ENSMUSG00000064262 | Gimap8 | 243374 | protein_coding | 27.02826674 | -0.125962717 | 0.218843735 | 0.575582923 | 0.564897127 | 0.751588072 | 0.125962717 |
| ENSMUSG00000042121 | Ssh1 | 231637 | protein_coding | 141.3231193 | 0.125996664 | 0.110569472 | -1.139524877 | 0.254484298 | 0.467106114 | 0.125996664 |
| ENSMUSG00000023025 | Larp4 | 207214 | protein_coding | 1537.026427 | -0.126058108 | 0.087203539 | 1.445561844 | 0.148300135 | 0.326885802 | 0.126058108 |
| ENSMUSG00000018126 | Baiap2l2 | 207495 | protein_coding | 638.6510066 | -0.126087469 | 0.167546565 | 0.7525518 | 0.451719291 | 0.668072037 | 0.126087469 |
| ENSMUSG00000101225 | 1700008J07Rik | 629159 | lncRNA | 25.11920886 | -0.126146865 | 0.253248655 | 0.49811465 | 0.618403235 | 0.789632028 | 0.126146865 |
| ENSMUSG00000097180 | 2700038G22Rik | 67194 | lncRNA | 8.414676053 | -0.126147937 | 0.405606653 | 0.311010522 | 0.755792623 | 0.877286805 | 0.126147937 |
| ENSMUSG00000115219 | Eef1akmt4 | 110599566 | protein_coding | 51.9799717 | -0.126151102 | 0.191208538 | 0.659756638 | 0.509410014 | 0.712785998 | 0.126151102 |
| ENSMUSG00000000861 | Bcl11a | 14025 | protein_coding | 55.53108243 | -0.126154601 | 0.169957868 | 0.742269847 | 0.457923857 | 0.67261003 | 0.126154601 |
| ENSMUSG00000024077 | Strn | 268980 | protein_coding | 840.5082035 | -0.126234263 | 0.06627215 | 1.904785976 | 0.056807898 | 0.164036653 | 0.126234263 |
| ENSMUSG00000026615 | Eprs | 107508 | protein_coding | 2193.020555 | 0.126237336 | 0.097039224 | -1.300889784 | 0.193296183 | 0.391427734 | 0.126237336 |
| ENSMUSG00000003316 | Glg1 | 20340 | protein_coding | 1708.542241 | -0.126244425 | 0.080223294 | 1.57366295 | 0.115565392 | 0.274320379 | 0.126244425 |
| ENSMUSG00000022637 | Cblb | 208650 | protein_coding | 254.7499099 | -0.126296637 | 0.112268398 | 1.124952691 | 0.260609082 | 0.474080016 | 0.126296637 |
| ENSMUSG00000022023 | Wbp4 | 22380 | protein_coding | 326.3596072 | 0.126298247 | 0.085174169 | -1.482823365 | 0.138121347 | 0.310726208 | 0.126298247 |
| ENSMUSG00000060427 | Zfp868 | 234362 | protein_coding | 145.4520578 | -0.126304039 | 0.103857156 | 1.216132274 | 0.223934538 | 0.431058976 | 0.126304039 |
| ENSMUSG00000044726 | Erich5 | 239368 | protein_coding | 7.615997893 | 0.126308052 | 0.406088176 | -0.311036024 | 0.755773236 | 0.877286805 | 0.126308052 |
| ENSMUSG00000072066 | 6720489N17Rik | 211378 | protein_coding | 15.42648362 | 0.126350312 | 0.267985923 | -0.471481154 | 0.637297174 | 0.802403594 | 0.126350312 |
| ENSMUSG00000032114 | Slc37a4 | 14385 | protein_coding | 412.1042178 | -0.126358306 | 0.103106744 | 1.225509607 | 0.220383273 | 0.426788709 | 0.126358306 |
| ENSMUSG00000043991 | Pura | 19290 | protein_coding | 748.8876873 | -0.12643976 | 0.074131893 | 1.705605441 | 0.088081566 | 0.225215631 | 0.12643976 |
| ENSMUSG00000019977 | Hbs1l | 56422 | protein_coding | 562.9253552 | 0.126487096 | 0.067548739 | -1.872530821 | 0.061133202 | 0.1731653 | 0.126487096 |
| ENSMUSG00000006289 | Osgep | 66246 | protein_coding | 313.3150836 | 0.126523678 | 0.107065566 | -1.181740151 | 0.237308822 | 0.44767168 | 0.126523678 |
| ENSMUSG00000074867 | Zfp808 | 630579 | protein_coding | 21.06093209 | 0.126527544 | 0.253512811 | -0.49909724 | 0.617710882 | 0.789468115 | 0.126527544 |
| ENSMUSG00000026932 | Nacc2 | 67991 | protein_coding | 400.3580114 | -0.126534121 | 0.084797246 | 1.492196105 | 0.135647743 | 0.306987369 | 0.126534121 |
| ENSMUSG00000027540 | Ptpn1 | 19246 | protein_coding | 347.8962339 | 0.126554552 | 0.085129996 | -1.48660353 | 0.137119555 | 0.309427758 | 0.126554552 |
| ENSMUSG00000002129 | Sf3a1 | 67465 | protein_coding | 834.0778207 | 0.126583057 | 0.078295023 | -1.616744625 | 0.105933417 | 0.258135524 | 0.126583057 |
| ENSMUSG00000037306 | Man1c1 | 230815 | protein_coding | 18.05065698 | -0.126588247 | 0.335561086 | 0.377243525 | 0.705992638 | 0.845943387 | 0.126588247 |
| ENSMUSG00000035109 | Shc4 | 271849 | protein_coding | 3.630532206 | 0.12659767 | 0.695432155 | -0.182041727 | 0.85554998 | 1 | 0.12659767 |
| ENSMUSG00000075576 | Gm12359 | 100037262 | lncRNA | 8.389093063 | -0.126608568 | 0.372684989 | 0.339720063 | 0.734067351 | 0.864094963 | 0.126608568 |
| ENSMUSG00000021468 | Sptlc1 | 268656 | protein_coding | 761.1247772 | 0.126697493 | 0.082506431 | -1.535607495 | 0.124634675 | 0.289400593 | 0.126697493 |

| ensembl_gene_id | external_gene_name | entrezgene_id | gene_biotype | baseMean | log2FoldChange | lfcSE | stat | pvalue | padj | absLog2FC |
|---|---|---|---|---|---|---|---|---|---|---|
| ENSMUSG00000058587 | Tmod3 | 50875 | protein_coding | 1297.800695 | -0.126703089 | 0.068116122 | 1.86010427 | 0.062870775 | 0.17645275 | 0.126703089 |
| ENSMUSG00000031822 | Gse1 | 382034 | protein_coding | 584.5893379 | 0.126704782 | 0.175454209 | -0.72215299 | 0.470200422 | 0.68365582 | 0.126704782 |
| ENSMUSG00000073008 | Gpr174 | 213439 | protein_coding | 2.573285249 | 0.12674758 | 1.072801217 | -0.11814638 | 0.905951675 | 1 | 0.12674758 |
| ENSMUSG00000026754 | Golga1 | 76899 | protein_coding | 505.8567937 | 0.12675344 | 0.073513061 | -1.724230202 | 0.084666294 | 0.219013029 | 0.12675344 |
| ENSMUSG00000033460 | Armcx1 | 78248 | protein_coding | 32.71425468 | 0.126768776 | 0.215754226 | -0.587561033 | 0.556826973 | 0.746140914 | 0.126768776 |
| ENSMUSG00000062743 | Zfp677 | 210503 | protein_coding | 44.0197094 | -0.126797659 | 0.203984196 | 0.621605307 | 0.534201431 | 0.730364909 | 0.126797659 |
| ENSMUSG00000018541 | Cwc25 | 67480 | protein_coding | 209.5030988 | -0.126868689 | 0.090769539 | 1.397701155 | 0.162202827 | 0.347646657 | 0.126868689 |
| ENSMUSG00000023094 | Msrb2 | 76467 | protein_coding | 66.92287338 | -0.126896403 | 0.176764034 | 0.717885878 | 0.472827656 | 0.685206817 | 0.126896403 |
| ENSMUSG00000024037 | Wdr4 | 57773 | protein_coding | 207.2091162 | 0.126898505 | 0.084040758 | -1.509963832 | 0.131052653 | 0.299543739 | 0.126898505 |
| ENSMUSG00000025246 | Tbl1x | 21372 | protein_coding | 1012.300908 | -0.126901547 | 0.073178082 | 1.73414693 | 0.082892016 | 0.215537515 | 0.126901547 |
| ENSMUSG00000045576 | St7l | 229681 | protein_coding | 311.0057431 | -0.126927442 | 0.062430289 | 2.033106748 | 0.042041739 | 0.130773645 | 0.126927442 |
| ENSMUSG00000022054 | Nefm | 18040 | protein_coding | 8.286032868 | 0.126939464 | 0.54959192 | -0.230970397 | 0.8173378 | 0.908714429 | 0.126939464 |
| ENSMUSG00000028249 | Sdcbp | 53378 | protein_coding | 2651.544183 | -0.12695233 | 0.084732907 | 1.498264776 | 0.134064472 | 0.304481573 | 0.12695233 |
| ENSMUSG00000034903 | Cobll1 | 319876 | protein_coding | 289.511799 | 0.126956501 | 0.08017816 | -1.583429968 | 0.113323497 | 0.270127107 | 0.126956501 |
| ENSMUSG00000027171 | Prrg4 | 228413 | protein_coding | 15.53347481 | 0.127003099 | 0.38558391 | -0.329378626 | 0.741869521 | 0.869560398 | 0.127003099 |
| ENSMUSG00000021033 | Gstz1 | 14874 | protein_coding | 359.9660706 | -0.127028155 | 0.078253107 | 1.623298555 | 0.104525598 | 0.255685933 | 0.127028155 |
| ENSMUSG00000003234 | Abcf3 | 27406 | protein_coding | 589.0444762 | 0.127036762 | 0.066355077 | -1.914499518 | 0.055556361 | 0.161179189 | 0.127036762 |
| ENSMUSG00000040483 | Xaf1 | 327959 | protein_coding | 77.46434433 | -0.127053613 | 0.283726056 | 0.447803824 | 0.654294783 | 0.812407823 | 0.127053613 |
| ENSMUSG00000066057 | Gm1976 | 100038746 | lncRNA | 28.01576986 | -0.127072234 | 0.20830415 | 0.610032174 | 0.541840495 | 0.735609498 | 0.127072234 |
| ENSMUSG00000049717 | Lig4 | 319583 | protein_coding | 189.1916665 | 0.127078846 | 0.084131114 | -1.510485718 | 0.130919531 | 0.299280888 | 0.127078846 |
| ENSMUSG00000036687 | Tmem184a | 231832 | protein_coding | 430.3316956 | -0.127214842 | 0.071310178 | 1.783964717 | 0.074429399 | 0.199468856 | 0.127214842 |
| ENSMUSG00000064125 | Prr36 | 73072 | protein_coding | 12.99490529 | -0.127260315 | 0.624638937 | 0.203734201 | 0.838561214 | 0.919582966 | 0.127260315 |
| ENSMUSG00000022177 | Haus4 | 219072 | protein_coding | 155.4182726 | 0.127267392 | 0.142802326 | -0.891213714 | 0.372814529 | 0.595133797 | 0.127267392 |
| ENSMUSG00000019478 | Rab4a | 19341 | protein_coding | 386.7084748 | 0.127271171 | 0.115247109 | -1.104332872 | 0.269448768 | 0.484503153 | 0.127271171 |
| ENSMUSG00000030766 | Arhgap17 | 70497 | protein_coding | 407.5697076 | 0.127332759 | 0.179917418 | -0.707728915 | 0.479113617 | 0.690541222 | 0.127332759 |
| ENSMUSG00000070705 | Eid2b | 434156 | protein_coding | 28.0679257 | -0.127345766 | 0.210395715 | 0.605267871 | 0.545001026 | 0.736964526 | 0.127345766 |
| ENSMUSG00000015889 | Lta4h | 16993 | protein_coding | 1209.27889 | -0.127349993 | 0.070896803 | 1.796272732 | 0.072451152 | 0.195920934 | 0.127349993 |
| ENSMUSG00000026510 | Trp53bp2 | 209456 | protein_coding | 584.314862 | -0.127366449 | 0.053973572 | 2.359792851 | 0.018285142 | 0.068347858 | 0.127366449 |
| ENSMUSG00000014226 | Cacybp | 12301 | protein_coding | 454.0041175 | 0.127372805 | 0.094269292 | -1.351159028 | 0.176644496 | 0.36877688 | 0.127372805 |
| ENSMUSG00000016933 | Plcg1 | 18803 | protein_coding | 267.7134409 | -0.127388738 | 0.094463755 | 1.348546205 | 0.17748277 | 0.36983602 | 0.127388738 |
| ENSMUSG00000039831 | Arhgap29 | 214137 | protein_coding | 166.7729657 | -0.127390372 | 0.117235402 | 1.086620341 | 0.277204631 | 0.494198131 | 0.127390372 |
| ENSMUSG00000047515 | BC049715 | 320135 | protein_coding | 33.06041037 | 0.127398317 | 0.228516537 | -0.557501522 | 0.577184819 | 0.761298159 | 0.127398317 |
| ENSMUSG00000036461 | Elf1 | 13709 | protein_coding | 1315.932882 | -0.127399497 | 0.067494666 | 1.887549113 | 0.059086507 | 0.168985435 | 0.127399497 |
| ENSMUSG00000044320 | 1700001O22Rik | 73598 | protein_coding | 29.81158916 | -0.127412708 | 0.218221157 | 0.583869641 | 0.55930802 | 0.747948226 | 0.127412708 |
| ENSMUSG00000059326 | Csf2ra | 12982 | protein_coding | 173.0963835 | 0.127493769 | 0.206457591 | -0.617530061 | 0.536885159 | 0.732676764 | 0.127493769 |
| ENSMUSG00000035064 | Eef2k | 13631 | protein_coding | 1066.491226 | -0.127532578 | 0.088223248 | 1.445566564 | 0.148298811 | 0.326885802 | 0.127532578 |
| ENSMUSG00000063253 | Scoc | 56367 | protein_coding | 49.11445205 | 0.127534489 | 0.207377215 | -0.61498795 | 0.538562678 | 0.733377497 | 0.127534489 |
| ENSMUSG00000024101 | Washc1 | 68767 | protein_coding | 279.9942724 | -0.127546581 | 0.06616407 | 1.927731809 | 0.053888493 | 0.157814254 | 0.127546581 |
| ENSMUSG00000020831 | 0610010K14Rik | 104457 | protein_coding | 5.887441764 | 0.127571036 | 0.489431162 | -0.260651642 | 0.794361161 | 0.897695523 | 0.127571036 |

| ensembl_gene_id | external_gene_name | entrezgene_id | gene_biotype | baseMean | log2FoldChange | lfcSE | stat | pvalue | padj | absLog2FC |
|---|---|---|---|---|---|---|---|---|---|---|
| ENSMUSG00000084834 | 4930565N06Rik | 68306 | lncRNA | 36.35492176 | -0.127643205 | 0.209786955 | 0.608442049 | 0.54289433 | 0.736235501 | 0.127643205 |
| ENSMUSG00000035835 | Plppr3 | 216152 | protein_coding | 50.40882313 | 0.127722705 | 0.16986438 | -0.751909879 | 0.452105257 | 0.66852312 | 0.127722705 |
| ENSMUSG00000057092 | Fxyd3 | 17178 | protein_coding | 259.2705776 | -0.127730214 | 0.169718823 | 0.75259898 | 0.45169093 | 0.668072037 | 0.127730214 |
| ENSMUSG00000028672 | Hmgcl | 15356 | protein_coding | 998.1519118 | -0.127759666 | 0.065148566 | 1.961051083 | 0.049873064 | 0.148611292 | 0.127759666 |
| ENSMUSG00000006850 | Tmco6 | 71983 | protein_coding | 179.9565955 | -0.127804343 | 0.087717891 | 1.456992884 | 0.145118344 | 0.322298818 | 0.127804343 |
| ENSMUSG00000061132 | Blnk | 17060 | protein_coding | 1026.459518 | -0.127804627 | 0.08694939 | 1.469873769 | 0.141595945 | 0.316544664 | 0.127804627 |
| ENSMUSG00000115970 | 8430426J06Rik | 78102 | lncRNA | 2.242083502 | -0.127825736 | 1.091691392 | 0.117089626 | 0.906789031 | 1 | 0.127825736 |
| ENSMUSG00000028248 | Pnisr | 66625 | protein_coding | 710.7415833 | -0.127856644 | 0.092619274 | 1.380453963 | 0.167446915 | 0.355746604 | 0.127856644 |
| ENSMUSG00000024769 | Cdc42bpg | 240505 | protein_coding | 1780.072349 | 0.127891327 | 0.099739176 | -1.282257699 | 0.199752262 | 0.40082018 | 0.127891327 |
| ENSMUSG00000036264 | Fstl4 | 320027 | protein_coding | 36.67332891 | -0.127953805 | 0.203260952 | 0.629505095 | 0.529018434 | 0.726903567 | 0.127953805 |
| ENSMUSG00000057469 | E2f6 | 50496 | protein_coding | 154.4987255 | 0.127975903 | 0.116988045 | -1.093922917 | 0.273988787 | 0.490477055 | 0.127975903 |
| ENSMUSG00000049339 | Retreg2 | 227298 | protein_coding | 866.7240549 | -0.128003453 | 0.064596089 | 1.981597554 | 0.047524301 | 0.143502708 | 0.128003453 |
| ENSMUSG00000029427 | Zcchc8 | 70650 | protein_coding | 441.3032106 | -0.12806405 | 0.05293532 | 2.419255252 | 0.015552323 | 0.060119629 | 0.12806405 |
| ENSMUSG00000025580 | Eif4a3 | 192170 | protein_coding | 957.5209637 | 0.128105252 | 0.073589523 | -1.740808285 | 0.081717189 | 0.213086636 | 0.128105252 |
| ENSMUSG00000022983 | Scaf4 | 224432 | protein_coding | 274.1670147 | 0.128129495 | 0.133429523 | -0.960278451 | 0.336915092 | 0.559233675 | 0.128129495 |
| ENSMUSG00000047879 | Usp14 | 59025 | protein_coding | 589.9173727 | 0.128185122 | 0.09379351 | -1.366673691 | 0.171727608 | 0.36209266 | 0.128185122 |
| ENSMUSG00000022119 | Rbm26 | 74213 | protein_coding | 482.0974359 | 0.1281971 | 0.074596174 | -1.718547935 | 0.085696719 | 0.220951697 | 0.1281971 |
| ENSMUSG00000031109 | Enox2 | 209224 | protein_coding | 278.5351999 | -0.128205667 | 0.090652331 | 1.414256703 | 0.157286544 | 0.34055723 | 0.128205667 |
| ENSMUSG00000021385 | Ippk | 75678 | protein_coding | 454.0991197 | 0.128256942 | 0.13026176 | -0.984609317 | 0.324816012 | 0.546769252 | 0.128256942 |
| ENSMUSG00000038014 | Fam120a | 218236 | protein_coding | 6539.024078 | 0.128281167 | 0.048334351 | -2.654037231 | 0.007953505 | 0.035440956 | 0.128281167 |
| ENSMUSG00000051065 | Mb21d2 | 239796 | protein_coding | 164.5820633 | 0.128306367 | 0.089371121 | -1.435658029 | 0.151099674 | 0.331068065 | 0.128306367 |
| ENSMUSG00000108504 | Gm4265 | 100043150 | lncRNA | 1.703767149 | 0.128326409 | 0.86670071 | -0.148063117 | 0.882292957 | 1 | 0.128326409 |
| ENSMUSG00000037211 | Spry1 | 24063 | protein_coding | 136.9784133 | -0.128338896 | 0.121300496 | 1.058024497 | 0.290044274 | 0.509010748 | 0.128338896 |
| ENSMUSG00000087362 | Gm13710 | 672763 | lncRNA | 0.873141777 | 0.128342114 | 1.455359131 | -0.088185872 | 0.929728946 | 1 | 0.128342114 |
| ENSMUSG00000028532 | Cachd1 | 320508 | protein_coding | 872.2975185 | 0.128364967 | 0.068522509 | -1.873325556 | 0.061023443 | 0.173002023 | 0.128364967 |
| ENSMUSG00000090626 | Tex9 | 21778 | protein_coding | 107.8371029 | 0.128461956 | 0.186490577 | -0.688838859 | 0.490924679 | 0.698215786 | 0.128461956 |
| ENSMUSG00000021234 | Fam161b | 217705 | protein_coding | 35.04438247 | 0.12848565 | 0.200162234 | -0.641907551 | 0.520933211 | 0.720614094 | 0.12848565 |
| ENSMUSG00000020329 | Polrmt | 216151 | protein_coding | 359.6032937 | -0.12857194 | 0.068766035 | 1.869701243 | 0.061525317 | 0.173816344 | 0.12857194 |
| ENSMUSG00000059669 | Taf1b | 21340 | protein_coding | 57.56724226 | 0.128609813 | 0.170852278 | -0.752754451 | 0.451597483 | 0.668047177 | 0.128609813 |
| ENSMUSG00000030814 | Bcl7c | 12055 | protein_coding | 97.05556954 | -0.128611203 | 0.146336993 | 0.878870068 | 0.379471729 | 0.601768559 | 0.128611203 |
| ENSMUSG00000021276 | Cinp | 67236 | protein_coding | 160.2961646 | 0.128636043 | 0.144181193 | -0.892183235 | 0.372294726 | 0.594533905 | 0.128636043 |
| ENSMUSG00000024142 | Mlst8 | 56716 | protein_coding | 273.5018638 | 0.128639658 | 0.071253971 | -1.805368263 | 0.071017074 | 0.192986007 | 0.128639658 |
| ENSMUSG00000034795 | Ccdc122 | 108811 | protein_coding | 98.32966307 | 0.128658723 | 0.148302015 | -0.867545347 | 0.385643277 | 0.607820554 | 0.128658723 |
| ENSMUSG00000005161 | Prdx2 | 21672 | protein_coding | 1643.461573 | -0.128664774 | 0.087379851 | 1.472476473 | 0.140892256 | 0.315313011 | 0.128664774 |
| ENSMUSG00000022359 | Wdyhv1 | 76773 | protein_coding | 128.1128278 | -0.128677131 | 0.108727507 | 1.183482768 | 0.236617872 | 0.446793087 | 0.128677131 |
| ENSMUSG00000005779 | Psmb4 | 19172 | protein_coding | 1677.892811 | 0.128678614 | 0.076079511 | -1.691370152 | 0.090766129 | 0.230247597 | 0.128678614 |
| ENSMUSG00000034055 | Phka1 | 18679 | protein_coding | 214.558948 | 0.128682725 | 0.087991065 | -1.462452183 | 0.143617338 | 0.319894352 | 0.128682725 |
| ENSMUSG00000073609 | D2hgdh | 98314 | protein_coding | 211.2524378 | -0.128692712 | 0.08261827 | 1.55767861 | 0.119309454 | 0.280906233 | 0.128692712 |
| ENSMUSG00000058833 | Rex1bd | 66462 | protein_coding | 288.4281024 | 0.128718247 | 0.111257766 | -1.156937196 | 0.24729802 | 0.459379022 | 0.128718247 |

| ensembl_gene_id | external_gene_name | entrezgene_id | gene_biotype | baseMean | log2FoldChange | lfcSE | stat | pvalue | padj | absLog2FC |
|---|---|---|---|---|---|---|---|---|---|---|
| ENSMUSG00000034574 | Daam1 | 208846 | protein_coding | 900.2476254 | -0.128737188 | 0.067198254 | 1.915781735 | 0.055392884 | 0.160822447 | 0.128737188 |
| ENSMUSG00000068394 | Cep152 | 99100 | protein_coding | 157.8494312 | 0.128743451 | 0.112610806 | -1.143260188 | 0.252930579 | 0.465324602 | 0.128743451 |
| ENSMUSG00000046711 | Hmga1 | 15361 | protein_coding | 333.8735469 | -0.128801104 | 0.142247731 | 0.905470358 | 0.365216257 | 0.587772599 | 0.128801104 |
| ENSMUSG00000028749 | Pla2g2f | 26971 | protein_coding | 517.2429252 | 0.128842355 | 0.398004634 | -0.323720741 | 0.746149467 | 0.872287153 | 0.128842355 |
| ENSMUSG00000046082 | Tmem174 | 68344 | protein_coding | 23.90126583 | -0.128850439 | 0.528261395 | 0.243914168 | 0.807297296 | 0.904987729 | 0.128850439 |
| ENSMUSG00000017421 | Zfp207 | 22680 | protein_coding | 1412.451618 | 0.128864973 | 0.04497139 | -2.865487915 | 0.004163672 | 0.020854981 | 0.128864973 |
| ENSMUSG00000037946 | Fgd3 | 30938 | protein_coding | 197.7264452 | -0.128869497 | 0.121922438 | 1.056979331 | 0.290521021 | 0.509468423 | 0.128869497 |
| ENSMUSG00000025812 | Pard3 | 93742 | protein_coding | 949.5614785 | -0.128903026 | 0.046182798 | 2.791148042 | 0.005252144 | 0.025397924 | 0.128903026 |
| ENSMUSG00000032399 | Rpl4 | 67891 | protein_coding | 12120.75198 | -0.128908991 | 0.090222589 | 1.428788425 | 0.153065053 | 0.334133019 | 0.128908991 |
| ENSMUSG00000023018 | Smarcd1 | 83797 | protein_coding | 150.45321 | -0.128933374 | 0.09928391 | 1.298633127 | 0.194069864 | 0.392753643 | 0.128933374 |
| ENSMUSG00000097915 | A330009N23Rik | 380977 | lncRNA | 6.768139305 | -0.129040413 | 0.604568086 | 0.213442317 | 0.830981977 | 0.914497068 | 0.129040413 |
| ENSMUSG00000021910 | Nisch | 64652 | protein_coding | 1484.991774 | -0.129050616 | 0.069201093 | 1.864863848 | 0.062200491 | 0.17504753 | 0.129050616 |
| ENSMUSG00000028107 | Tars2 | 71807 | protein_coding | 365.5726028 | -0.129056064 | 0.120052201 | 1.074999569 | 0.282374921 | 0.500444242 | 0.129056064 |
| ENSMUSG00000031967 | Afg3l1 | 114896 | protein_coding | 435.7180204 | -0.129072279 | 0.063116736 | 2.044977094 | 0.040857116 | 0.128048205 | 0.129072279 |
| ENSMUSG00000036879 | Phkb | 102093 | protein_coding | 598.4234219 | -0.12909908 | 0.104070319 | 1.240498555 | 0.214791048 | 0.419775423 | 0.12909908 |
| ENSMUSG00000044617 | Zbtb39 | 320080 | protein_coding | 224.2881725 | 0.129122739 | 0.098654695 | -1.30883522 | 0.190590174 | 0.388089535 | 0.129122739 |
| ENSMUSG00000024833 | Pola2 | 18969 | protein_coding | 233.8658238 | 0.129130681 | 0.119157766 | -1.083695049 | 0.278500032 | 0.495758276 | 0.129130681 |
| ENSMUSG00000036501 | Fam13b | 225358 | protein_coding | 598.3037126 | -0.129149796 | 0.086952123 | 1.485297784 | 0.137464959 | 0.309868458 | 0.129149796 |
| ENSMUSG00000051844 | B230319C09Rik | 320775 | lncRNA | 28.36220451 | -0.129167085 | 0.246877046 | 0.523204111 | 0.600832221 | 0.777353752 | 0.129167085 |
| ENSMUSG00000038074 | Fkbp14 | 231997 | protein_coding | 43.35868784 | 0.129199114 | 0.167904734 | -0.769478683 | 0.441609195 | 0.659260011 | 0.129199114 |
| ENSMUSG00000052253 | Zfp622 | 52521 | protein_coding | 682.2767959 | -0.12921977 | 0.075670038 | 1.707674175 | 0.087696817 | 0.224854857 | 0.12921977 |
| ENSMUSG00000046096 | Mosmo | 233812 | protein_coding | 78.90351952 | -0.129271852 | 0.137407497 | 0.940791846 | 0.34681154 | 0.568538439 | 0.129271852 |
| ENSMUSG00000074829 | 2010315B03Rik | 630836 | protein_coding | 385.6370174 | -0.129275634 | 0.085312002 | 1.515327621 | 0.129689457 | 0.297380166 | 0.129275634 |
| ENSMUSG00000035378 | Shq1 | 72171 | protein_coding | 96.68676732 | -0.129292577 | 0.12115368 | 1.06717829 | 0.285891334 | 0.50434884 | 0.129292577 |
| ENSMUSG00000076432 | Ywhaq | 22630 | protein_coding | 1117.109388 | 0.129305359 | 0.110894454 | -1.166021867 | 0.243605604 | 0.454822267 | 0.129305359 |
| ENSMUSG00000026305 | Lrrfip1 | 16978 | protein_coding | 1378.652832 | -0.129414219 | 0.078003225 | 1.659088065 | 0.097098048 | 0.241894213 | 0.129414219 |
| ENSMUSG00000030612 | Mrpl46 | 67308 | protein_coding | 339.3754319 | -0.129533347 | 0.091003179 | 1.423393649 | 0.154622073 | 0.336553171 | 0.129533347 |
| ENSMUSG00000021188 | Trip11 | 109181 | protein_coding | 1188.868932 | 0.129547622 | 0.071119054 | -1.821559975 | 0.068521779 | 0.188061933 | 0.129547622 |
| ENSMUSG00000003208 | Yju2 | 72886 | protein_coding | 111.3243278 | 0.129575606 | 0.108072378 | -1.198970623 | 0.230539369 | 0.43912271 | 0.129575606 |
| ENSMUSG00000085184 | 4933439K11Rik | 71319 | lncRNA | 35.42683183 | 0.129616667 | 0.213032537 | -0.608436009 | 0.542898335 | 0.736235501 | 0.129616667 |
| ENSMUSG00000052026 | Slc6a7 | 240332 | protein_coding | 54.92117695 | -0.129634926 | 0.276457168 | 0.468915047 | 0.63913036 | 0.803282676 | 0.129634926 |
| ENSMUSG00000076434 | Wfdc3 | 71856 | protein_coding | 4.597358712 | 0.129683498 | 0.54869269 | -0.236349964 | 0.813161119 | 0.907217398 | 0.129683498 |
| ENSMUSG00000057440 | Mpp7 | 75739 | protein_coding | 274.0268437 | -0.129696672 | 0.095199618 | 1.362365477 | 0.173082573 | 0.363688435 | 0.129696672 |
| ENSMUSG00000007721 | Ccdc124 | 234388 | protein_coding | 480.7330138 | 0.129768613 | 0.098399943 | -1.318787484 | 0.18724017 | 0.383442977 | 0.129768613 |
| ENSMUSG00000024817 | Uhrf2 | 109113 | protein_coding | 1267.936366 | 0.129792672 | 0.085391836 | -1.519965822 | 0.128519566 | 0.29575977 | 0.129792672 |
| ENSMUSG00000042873 | Lhfpl4 | 269788 | protein_coding | 4.589798181 | -0.129823941 | 0.837972825 | 0.154926195 | 0.876879503 | 0.938919382 | 0.129823941 |
| ENSMUSG00000040669 | Phc1 | 13619 | protein_coding | 107.796515 | -0.129880725 | 0.132279289 | 0.98186743 | 0.326165163 | 0.548167261 | 0.129880725 |
| ENSMUSG00000042417 | Ccno | 218630 | protein_coding | 5.188191461 | 0.129887485 | 0.755703582 | -0.171876234 | 0.863534831 | 0.932314788 | 0.129887485 |
| ENSMUSG00000020457 | Drg1 | 13494 | protein_coding | 484.9991154 | 0.129891196 | 0.094215257 | -1.378664137 | 0.167998332 | 0.356505326 | 0.129891196 |

| ensembl_gene_id | external_gene_name | entrezgene_id | gene_biotype | baseMean | log2FoldChange | lfcSE | stat | pvalue | padj | absLog2FC |
|---|---|---|---|---|---|---|---|---|---|---|
| ENSMUSG00000035673 | Sbno2 | 216161 | protein_coding | 1092.612557 | 0.12990104 | 0.113376199 | -1.145752293 | 0.25189766 | 0.464371283 | 0.12990104 |
| ENSMUSG00000028807 | Zbtb8a | 73680 | protein_coding | 200.9695165 | -0.129907262 | 0.111942496 | 1.160482094 | 0.24585258 | 0.457679163 | 0.129907262 |
| ENSMUSG00000015377 | Dennd6b | 69440 | protein_coding | 93.00451493 | -0.129936162 | 0.14280142 | 0.909908052 | 0.362871003 | 0.585225632 | 0.129936162 |
| ENSMUSG00000035949 | Fbxw2 | 30050 | protein_coding | 491.8534557 | 0.129984703 | 0.060772581 | -2.138870861 | 0.032446129 | 0.107141645 | 0.129984703 |
| ENSMUSG00000030611 | Mrps11 | 67994 | protein_coding | 178.0638643 | 0.130009236 | 0.087318933 | -1.488900881 | 0.136513472 | 0.308481591 | 0.130009236 |
| ENSMUSG00000020526 | Znhit3 | 448850 | protein_coding | 63.69480418 | 0.130014742 | 0.151918568 | -0.855818633 | 0.392098107 | 0.614686552 | 0.130014742 |
| ENSMUSG00000022414 | Tab1 | 66513 | protein_coding | 358.657703 | -0.130031592 | 0.087409481 | 1.487614284 | 0.136852644 | 0.309078852 | 0.130031592 |
| ENSMUSG00000055839 | Elob | 67673 | protein_coding | 1378.699015 | -0.130036941 | 0.075815386 | 1.715178783 | 0.086312454 | 0.222126524 | 0.130036941 |
| ENSMUSG00000095845 | Gng14 | 100503710 | protein_coding | 1.260157496 | 0.130064266 | 1.156542885 | -0.112459527 | 0.910459059 | 1 | 0.130064266 |
| ENSMUSG00000022828 | Gtf2e1 | 74197 | protein_coding | 135.6752349 | 0.13008191 | 0.096684052 | -1.345432956 | 0.178485458 | 0.370989158 | 0.13008191 |
| ENSMUSG00000063273 | Naa15 | 74838 | protein_coding | 592.7965227 | 0.130108747 | 0.10496353 | -1.239561462 | 0.215137642 | 0.420139706 | 0.130108747 |
| ENSMUSG00000042694 | Stn1 | 108689 | protein_coding | 106.601017 | 0.130119279 | 0.128482197 | -1.012741704 | 0.311183552 | 0.532085855 | 0.130119279 |
| ENSMUSG00000031278 | Acsl4 | 50790 | protein_coding | 207.2236614 | 0.130213002 | 0.108988225 | -1.194743763 | 0.232187158 | 0.441179379 | 0.130213002 |
| ENSMUSG00000006058 | Snf8 | 27681 | protein_coding | 492.4095448 | -0.130230157 | 0.065078343 | 2.001128962 | 0.045378494 | 0.138710092 | 0.130230157 |
| ENSMUSG00000068566 | Myadm | 50918 | protein_coding | 729.5271106 | 0.130238842 | 0.063921883 | -2.037468791 | 0.041603086 | 0.129751344 | 0.130238842 |
| ENSMUSG00000032741 | Tpcn1 | 252972 | protein_coding | 1179.441028 | 0.130255164 | 0.060456676 | -2.154520777 | 0.031199354 | 0.104013109 | 0.130255164 |
| ENSMUSG00000043004 | Gng2 | 14702 | protein_coding | 10.29380901 | -0.130264722 | 0.346520124 | 0.375922532 | 0.706974489 | 0.846503532 | 0.130264722 |
| ENSMUSG00000035504 | Reep6 | 70335 | protein_coding | 174.5240926 | -0.130285768 | 0.137065761 | 0.950534745 | 0.341840608 | 0.563829035 | 0.130285768 |
| ENSMUSG00000020740 | Gga3 | 260302 | protein_coding | 168.3214584 | 0.130308353 | 0.125119331 | -1.041472585 | 0.29765627 | 0.517311687 | 0.130308353 |
| ENSMUSG00000022095 | Fhip2b | 239170 | protein_coding | 269.6380338 | -0.130329577 | 0.109588035 | 1.189268308 | 0.234334102 | 0.443834541 | 0.130329577 |
| ENSMUSG00000027185 | Nat10 | 98956 | protein_coding | 310.8481827 | -0.130344504 | 0.07205049 | 1.809071722 | 0.070439862 | 0.191827816 | 0.130344504 |
| ENSMUSG00000042650 | Alkbh5 | 268420 | protein_coding | 2030.404804 | 0.130392134 | 0.056199314 | -2.320173047 | 0.020331518 | 0.074331346 | 0.130392134 |
| ENSMUSG00000031166 | Wdr13 | 73447 | protein_coding | 825.0941752 | -0.130473986 | 0.098241945 | 1.328088377 | 0.184148913 | 0.379181439 | 0.130473986 |
| ENSMUSG00000022247 | Brix1 | 67832 | protein_coding | 269.7309277 | 0.13048472 | 0.107374698 | -1.21522782 | 0.224279213 | 0.431320242 | 0.13048472 |
| ENSMUSG00000032067 | Pts | 19286 | protein_coding | 469.0749963 | 0.13054717 | 0.087903989 | -1.485110888 | 0.137514452 | 0.309937719 | 0.13054717 |
| ENSMUSG00000031916 | Cog8 | 97484 | protein_coding | 428.497522 | -0.130552071 | 0.076528965 | 1.705917116 | 0.088023512 | 0.22515016 | 0.130552071 |
| ENSMUSG00000001918 | Slc1a5 | 20514 | protein_coding | 1456.892046 | -0.13055567 | 0.086297732 | 1.512851682 | 0.130317338 | 0.298275885 | 0.13055567 |
| ENSMUSG00000025278 | Flnb | 286940 | protein_coding | 9435.182382 | -0.130571323 | 0.093134382 | 1.401966923 | 0.160925124 | 0.345760799 | 0.130571323 |
| ENSMUSG00000027122 | Arl14ep | 212772 | protein_coding | 168.1962635 | 0.130601548 | 0.122358936 | -1.0673642 | 0.285807408 | 0.504312604 | 0.130601548 |
| ENSMUSG00000033545 | Znrf1 | 170737 | protein_coding | 190.064011 | 0.130642706 | 0.086368456 | -1.512620606 | 0.130376057 | 0.298368931 | 0.130642706 |
| ENSMUSG00000074282 | Zfp94 | 22756 | protein_coding | 26.32235995 | -0.130668179 | 0.216809147 | 0.602687576 | 0.546716548 | 0.738518372 | 0.130668179 |
| ENSMUSG00000037605 | Adgrl3 | 319387 | protein_coding | 3.679928652 | 0.130685933 | 1.394834872 | -0.093692763 | 0.925353219 | 1 | 0.130685933 |
| ENSMUSG00000033751 | Gadd45gip1 | 102060 | protein_coding | 209.5034419 | 0.130734679 | 0.121537145 | -1.075676734 | 0.282071856 | 0.499960785 | 0.130734679 |
| ENSMUSG00000080316 | Spaca6 | 75202 | protein_coding | 20.37794132 | -0.130750905 | 0.457867753 | 0.285564782 | 0.775211481 | 0.887843124 | 0.130750905 |
| ENSMUSG00000071652 | Ints5 | 109077 | protein_coding | 260.3633168 | 0.13078074 | 0.107804105 | -1.213133211 | 0.225078894 | 0.432354642 | 0.13078074 |
| ENSMUSG00000042404 | Dennd4b | 229541 | protein_coding | 28.94609477 | 0.130787054 | 0.279673333 | -0.467642203 | 0.640040479 | 0.8036518 | 0.130787054 |
| ENSMUSG00000031299 | Pdha1 | 18597 | protein_coding | 3288.160209 | -0.130800858 | 0.082203896 | 1.591175909 | 0.111569993 | 0.267528222 | 0.130800858 |
| ENSMUSG00000032336 | Nptn | 20320 | protein_coding | 1042.911922 | -0.130805097 | 0.04058792 | 3.222759295 | 0.001269622 | 0.00775103 | 0.130805097 |
| ENSMUSG00000046865 | Fbl | 14113 | protein_coding | 280.533028 | -0.130829399 | 0.131347529 | 0.996055279 | 0.319223287 | 0.540682396 | 0.130829399 |

Supplementary Table 1

| ensembl_gene_id | external_gene_name | entrezgene_id | gene_biotype | baseMean | log2FoldChange | lfcSE | stat | pvalue | padj | absLog2FC |
|---|---|---|---|---|---|---|---|---|---|---|
| ENSMUSG00000035200 | Chrnb4 | 108015 | protein_coding | 52.8690522 | -0.130858009 | 0.185489231 | 0.705474964 | 0.480514707 | 0.691465456 | 0.130858009 |
| ENSMUSG00000001833 | Septin7 | 235072 | protein_coding | 1085.006817 | 0.130860135 | 0.080521323 | -1.625161277 | 0.104128199 | 0.254940412 | 0.130860135 |
| ENSMUSG00000038406 | Scaf1 | 233208 | protein_coding | 826.7846388 | 0.130872242 | 0.122267727 | -1.070374379 | 0.284450827 | 0.502862462 | 0.130872242 |
| ENSMUSG00000027823 | Gmps | 229363 | protein_coding | 670.9014105 | 0.130975801 | 0.111159765 | -1.17826626 | 0.238690474 | 0.448874722 | 0.130975801 |
| ENSMUSG00000006010 | Odr4 | 226499 | protein_coding | 445.6288598 | 0.130994098 | 0.111264427 | -1.177322367 | 0.239066863 | 0.449192192 | 0.130994098 |
| ENSMUSG00000039202 | Abhd2 | 54608 | protein_coding | 6667.120591 | -0.131000946 | 0.075010201 | 1.746441733 | 0.080734224 | 0.211290803 | 0.131000946 |
| ENSMUSG00000022822 | Abcc5 | 27416 | protein_coding | 132.6428053 | -0.131023627 | 0.235425545 | 0.556539551 | 0.577842065 | 0.761753322 | 0.131023627 |
| ENSMUSG00000101031 | Ms4a12 | 381213 | protein_coding | 257.8282058 | -0.13104117 | 0.180577731 | 0.725677354 | 0.468036588 | 0.68170367 | 0.13104117 |
| ENSMUSG00000051864 | Tbc1d22a | 223754 | protein_coding | 729.0061965 | -0.131111521 | 0.093036536 | 1.409247662 | 0.158761949 | 0.342763083 | 0.131111521 |
| ENSMUSG00000036898 | Zfp157 | 72154 | protein_coding | 119.1959828 | 0.131161602 | 0.121356188 | -1.080798631 | 0.279786697 | 0.49722072 | 0.131161602 |
| ENSMUSG00000072487 | Mroh5 | 268816 | protein_coding | 5.75787416 | 0.131186957 | 1.808197934 | -0.072551215 | 0.942163249 | 0.971883559 | 0.131186957 |
| ENSMUSG00000023460 | Rab12 | 19328 | protein_coding | 85.78545312 | 0.131192375 | 0.138536138 | -0.946990271 | 0.343643736 | 0.56547568 | 0.131192375 |
| ENSMUSG00000031546 | Gins4 | 109145 | protein_coding | 195.2876209 | 0.131208021 | 0.091994952 | -1.426252407 | 0.153795497 | 0.335284335 | 0.131208021 |
| ENSMUSG00000013653 | 1810065E05Rik | 69864 | protein_coding | 10784.0731 | 0.131210633 | 0.190295456 | -0.689510069 | 0.490502339 | 0.69793377 | 0.131210633 |
| ENSMUSG00000029016 | Clcn6 | 26372 | protein_coding | 35.62719059 | -0.131222619 | 0.216556895 | 0.605949854 | 0.544548053 | 0.736796542 | 0.131222619 |
| ENSMUSG00000025962 | Fastkd2 | 75619 | protein_coding | 259.4936421 | 0.131250477 | 0.135201245 | -0.970778612 | 0.331658535 | 0.554128799 | 0.131250477 |
| ENSMUSG00000040177 | 2310057M21Rik | 68277 | protein_coding | 76.79736757 | 0.131277836 | 0.14659906 | -0.895488935 | 0.370525778 | 0.593022866 | 0.131277836 |
| ENSMUSG00000020091 | Eif4ebp2 | 13688 | protein_coding | 1257.811302 | -0.131321922 | 0.089177572 | 1.472589124 | 0.140861859 | 0.315287712 | 0.131321922 |
| ENSMUSG00000062646 | Ganc | 76051 | protein_coding | 203.7773796 | -0.131331587 | 0.091993614 | 1.427616351 | 0.153402315 | 0.334559671 | 0.131331587 |
| ENSMUSG00000046949 | Nqo2 | 18105 | protein_coding | 359.4065381 | -0.131371773 | 0.103398718 | 1.270535804 | 0.203893838 | 0.406017465 | 0.131371773 |
| ENSMUSG00000036499 | Eea1 | 216238 | protein_coding | 1455.90765 | 0.131389135 | 0.078433269 | -1.675170974 | 0.093900691 | 0.235970255 | 0.131389135 |
| ENSMUSG00000068798 | Rap1a | 109905 | protein_coding | 527.3019075 | -0.131421529 | 0.107078339 | 1.227340002 | 0.219694827 | 0.425947586 | 0.131421529 |
| ENSMUSG00000106943 | Dancr | 70036 | lncRNA | 32.6243551 | -0.131470466 | 0.328162342 | 0.400626305 | 0.688695276 | 0.83386713 | 0.131470466 |
| ENSMUSG00000028617 | Lrrc42 | 77809 | protein_coding | 732.9766904 | -0.131517116 | 0.105387429 | 1.247939324 | 0.212053278 | 0.41642117 | 0.131517116 |
| ENSMUSG00000026078 | Pdcl3 | 68833 | protein_coding | 442.2466726 | -0.131574421 | 0.077275839 | 1.702659233 | 0.088631858 | 0.226153663 | 0.131574421 |
| ENSMUSG00000102418 | Sh2d1b1 | 26904 | protein_coding | 4.355743129 | 0.131588434 | 0.938796146 | -0.140167207 | 0.888527881 | 1 | 0.131588434 |
| ENSMUSG00000018666 | Cbx1 | 12412 | protein_coding | 610.4485483 | 0.131589435 | 0.114671624 | -1.147532668 | 0.25116154 | 0.463479905 | 0.131589435 |
| ENSMUSG00000035011 | Zbtb7a | 16969 | protein_coding | 1965.712822 | 0.131691343 | 0.124589812 | -1.056999295 | 0.290511909 | 0.509468423 | 0.131691343 |
| ENSMUSG00000038534 | Osbpl7 | 71240 | protein_coding | 775.9324625 | 0.131732455 | 0.143228741 | -0.919734785 | 0.35771137 | 0.580678813 | 0.131732455 |
| ENSMUSG00000026977 | Marchf7 | 57438 | protein_coding | 1126.926087 | -0.131813583 | 0.067292818 | 1.9588061 | 0.050135499 | 0.149231729 | 0.131813583 |
| ENSMUSG00000057246 | BC051142 | 407788 | protein_coding | 18.01994418 | 0.131840314 | 0.337812507 | -0.390276591 | 0.696332032 | 0.838747423 | 0.131840314 |
| ENSMUSG00000056999 | Ide | 15925 | protein_coding | 704.2452246 | 0.131911824 | 0.100815189 | -1.308451891 | 0.190720082 | 0.388133006 | 0.131911824 |
| ENSMUSG00000051517 | Arhgef39 | 230098 | protein_coding | 209.448188 | 0.131977983 | 0.095761101 | -1.378200343 | 0.168141441 | 0.356671517 | 0.131977983 |
| ENSMUSG00000033020 | Polr2f | 69833 | protein_coding | 406.0119582 | -0.132002401 | 0.075199774 | 1.755356356 | 0.079198372 | 0.208527092 | 0.132002401 |
| ENSMUSG00000073728 | Tmem51os1 | 100038693 | lncRNA | 20.74899355 | -0.132024817 | 0.301112918 | 0.438456172 | 0.661055634 | 0.816884691 | 0.132024817 |
| ENSMUSG00000025558 | Dock9 | 105445 | protein_coding | 535.9716094 | -0.132028323 | 0.081954856 | 1.610988406 | 0.107182249 | 0.26018046 | 0.132028323 |
| ENSMUSG00000020280 | Pus10 | 74467 | protein_coding | 328.3127441 | 0.132059834 | 0.066307011 | -1.991642103 | 0.04641034 | 0.140963833 | 0.132059834 |
| ENSMUSG00000000561 | Wdr77 | 70465 | protein_coding | 143.0322137 | 0.132107956 | 0.145696545 | -0.906733624 | 0.364547678 | 0.587158915 | 0.132107956 |
| ENSMUSG00000028082 | Sh3d19 | 27059 | protein_coding | 1482.28052 | -0.132115188 | 0.053863892 | 2.452759757 | 0.014176498 | 0.05583573 | 0.132115188 |

| ensembl_gene_id | external_gene_name | entrezgene_id | gene_biotype | baseMean | log2FoldChange | lfcSE | stat | pvalue | padj | absLog2FC |
|---|---|---|---|---|---|---|---|---|---|---|
| ENSMUSG00000085944 | 1700003D09Rik | 69338 | lncRNA | 24.88163404 | -0.132186574 | 0.291434109 | 0.453572763 | 0.650136361 | 0.809917796 | 0.132186574 |
| ENSMUSG00000019810 | Fuca2 | 66848 | protein_coding | 983.1473107 | -0.132201924 | 0.09282046 | 1.424275679 | 0.154366685 | 0.336218865 | 0.132201924 |
| ENSMUSG00000028601 | Echdc2 | 52430 | protein_coding | 149.8874021 | -0.132203556 | 0.089790332 | 1.472358476 | 0.1409241 | 0.315341543 | 0.132203556 |
| ENSMUSG00000020611 | Gna13 | 14674 | protein_coding | 2096.710386 | -0.132206396 | 0.059410753 | 2.225294081 | 0.026061503 | 0.090207431 | 0.132206396 |
| ENSMUSG00000059734 | Ndufs8 | 225887 | protein_coding | 1216.933323 | -0.132211397 | 0.076464916 | 1.729046517 | 0.08380077 | 0.217114201 | 0.132211397 |
| ENSMUSG00000035824 | Tk2 | 57813 | protein_coding | 158.1685014 | -0.132220228 | 0.087429645 | 1.512304294 | 0.130456469 | 0.298470232 | 0.132220228 |
| ENSMUSG00000102805 | Gm37240 | 99889 | protein_coding | 10.19236841 | 0.132223698 | 0.424284408 | -0.311639304 | 0.75531466 | 0.877032235 | 0.132223698 |
| ENSMUSG00000059883 | Irak4 | 266632 | protein_coding | 513.8082731 | 0.132247188 | 0.088391748 | -1.49614857 | 0.134614943 | 0.305377683 | 0.132247188 |
| ENSMUSG00000036661 | Dennd3 | 105841 | protein_coding | 189.292885 | -0.132251671 | 0.093047587 | 1.421333703 | 0.15521977 | 0.337542702 | 0.132251671 |
| ENSMUSG00000029601 | Iqcd | 75732 | protein_coding | 17.40210234 | 0.132299014 | 0.341655561 | -0.387229214 | 0.698586525 | 0.840298483 | 0.132299014 |
| ENSMUSG00000039953 | Clstn1 | 65945 | protein_coding | 3047.543092 | -0.132369401 | 0.057647335 | 2.296192891 | 0.021664855 | 0.078201839 | 0.132369401 |
| ENSMUSG00000066440 | Zfyve26 | 211978 | protein_coding | 333.4367724 | 0.132450996 | 0.114100482 | -1.160827658 | 0.245711993 | 0.45758404 | 0.132450996 |
| ENSMUSG00000041117 | Ccdc8 | 434130 | protein_coding | 1.967294293 | 0.132473234 | 1.109585018 | -0.119389891 | 0.90496647 | 1 | 0.132473234 |
| ENSMUSG00000056260 | Lrif1 | 321000 | protein_coding | 161.2748825 | 0.132546374 | 0.105447813 | -1.256985524 | 0.208758877 | 0.412028696 | 0.132546374 |
| ENSMUSG00000062012 | Zfp13 | 22654 | protein_coding | 40.79942229 | 0.132588839 | 0.174883347 | -0.758155887 | 0.448357666 | 0.665277305 | 0.132588839 |
| ENSMUSG00000037202 | Prf1 | 18646 | protein_coding | 1.09479915 | -0.132594253 | 1.377040576 | 0.096289286 | 0.92329082 | 1 | 0.132594253 |
| ENSMUSG00000018899 | Irf1 | 16362 | protein_coding | 1328.362785 | -0.132602695 | 0.115317015 | 1.149897048 | 0.250186277 | 0.462507128 | 0.132602695 |
| ENSMUSG00000100121 | 1700025N23Rik | 69986 | lncRNA | 0.983285879 | -0.132605229 | 1.352632698 | 0.098034913 | 0.92190457 | 1 | 0.132605229 |
| ENSMUSG00000022722 | Arl6 | 56297 | protein_coding | 22.28030652 | 0.132667147 | 0.227781328 | -0.582432053 | 0.560275696 | 0.748973617 | 0.132667147 |
| ENSMUSG00000036873 | 2410004B18Rik | 66421 | protein_coding | 165.9535922 | 0.132677766 | 0.090205077 | -1.470845875 | 0.141332803 | 0.316005805 | 0.132677766 |
| ENSMUSG00000044453 | Ffar1 | 233081 | protein_coding | 5.774044218 | -0.132709041 | 0.637517297 | 0.208165396 | 0.835099827 | 0.917066002 | 0.132709041 |
| ENSMUSG00000025867 | Cplx2 | 12890 | protein_coding | 56.33984595 | -0.132720122 | 0.179711297 | 0.738518522 | 0.460199416 | 0.674460666 | 0.132720122 |
| ENSMUSG00000032609 | Klhdc8b | 78267 | protein_coding | 36.72062243 | -0.132740124 | 0.219481916 | 0.604788434 | 0.54531958 | 0.737121023 | 0.132740124 |
| ENSMUSG00000002870 | Mcm2 | 17216 | protein_coding | 446.4823686 | 0.132745919 | 0.126611628 | -1.048449662 | 0.294431483 | 0.513653234 | 0.132745919 |
| ENSMUSG00000021143 | Pacs2 | 217893 | protein_coding | 525.1120661 | -0.132749499 | 0.053254091 | 2.492756845 | 0.012675562 | 0.051167007 | 0.132749499 |
| ENSMUSG00000002010 | Idh3g | 15929 | protein_coding | 1547.092056 | -0.132808792 | 0.078124674 | 1.699959635 | 0.089138518 | 0.227130611 | 0.132808792 |
| ENSMUSG00000034473 | Sec22a | 317717 | protein_coding | 332.2584714 | 0.132829618 | 0.090685106 | -1.464734657 | 0.142993326 | 0.318805424 | 0.132829618 |
| ENSMUSG00000001036 | Epn2 | 13855 | protein_coding | 190.6138576 | -0.132857634 | 0.106532061 | 1.247114086 | 0.212355669 | 0.416836029 | 0.132857634 |
| ENSMUSG00000005947 | Itgae | 16407 | protein_coding | 34.96667659 | -0.13286126 | 0.331838647 | 0.400379103 | 0.688877313 | 0.833904116 | 0.13286126 |
| ENSMUSG00000071074 | Yipf3 | 28064 | protein_coding | 505.7984822 | -0.132884631 | 0.064827335 | 2.049824051 | 0.040381604 | 0.126876294 | 0.132884631 |
| ENSMUSG00000028232 | Tmem68 | 72098 | protein_coding | 155.1992167 | 0.13288994 | 0.125242259 | -1.061063101 | 0.288661225 | 0.507554459 | 0.13288994 |
| ENSMUSG00000079036 | Alkbh1 | 211064 | protein_coding | 179.0594048 | -0.132893461 | 0.117150485 | 1.134382509 | 0.256634133 | 0.4695907 | 0.132893461 |
| ENSMUSG00000029714 | Gigyf1 | 57330 | protein_coding | 230.299878 | -0.132923701 | 0.175072396 | 0.759249912 | 0.447703074 | 0.664871274 | 0.132923701 |
| ENSMUSG00000009566 | Fpgs | 14287 | protein_coding | 225.7609451 | 0.132926294 | 0.10204319 | -1.302647381 | 0.192695174 | 0.390530809 | 0.132926294 |
| ENSMUSG00000050240 | Hic2 | 58180 | protein_coding | 153.3479668 | -0.132929216 | 0.101069063 | 1.315231512 | 0.188432108 | 0.384976843 | 0.132929216 |
| ENSMUSG00000053007 | Creb5 | 231991 | protein_coding | 7.872798789 | 0.132947593 | 2.300962454 | -0.057779123 | 0.953924568 | 0.977047797 | 0.132947593 |
| ENSMUSG00000033099 | Nol12 | 97961 | protein_coding | 160.8394 | 0.132968882 | 0.103255287 | -1.287768263 | 0.197826643 | 0.398112257 | 0.132968882 |
| ENSMUSG00000024366 | Gfra3 | 14587 | protein_coding | 18.94717527 | 0.132974263 | 0.27526514 | -0.483077017 | 0.629041052 | 0.797446101 | 0.132974263 |
| ENSMUSG00000046909 | Tefm | 68550 | protein_coding | 90.02804128 | -0.13297915 | 0.151482545 | 0.877851303 | 0.380024419 | 0.60223179 | 0.13297915 |

| ensembl_gene_id | external_gene_name | entrezgene_id | gene_biotype | baseMean | log2FoldChange | lfcSE | stat | pvalue | padj | absLog2FC |
|---|---|---|---|---|---|---|---|---|---|---|
| ENSMUSG00000097392 | Thoc2l | 100042165 | protein_coding | 75.90756144 | -0.13299648 | 0.151399485 | 0.878447371 | 0.379700987 | 0.601975182 | 0.13299648 |
| ENSMUSG00000030505 | Prmt3 | 71974 | protein_coding | 170.584704 | -0.133001107 | 0.14256682 | 0.932903657 | 0.350869714 | 0.572963936 | 0.133001107 |
| ENSMUSG00000022338 | Eny2 | 223527 | protein_coding | 478.1035033 | 0.133002856 | 0.078130473 | -1.702317304 | 0.088695903 | 0.226282116 | 0.133002856 |
| ENSMUSG00000029686 | Cul1 | 26965 | protein_coding | 1250.234759 | -0.133037951 | 0.059534471 | 2.234637322 | 0.025441169 | 0.088524117 | 0.133037951 |
| ENSMUSG00000078348 | Sf3b5 | 66125 | protein_coding | 336.3953393 | 0.133054808 | 0.108458627 | -1.226779382 | 0.219905522 | 0.426134686 | 0.133054808 |
| ENSMUSG00000047789 | Slc38a9 | 268706 | protein_coding | 104.9599605 | -0.133058807 | 0.142783359 | 0.93189296 | 0.351391846 | 0.573407604 | 0.133058807 |
| ENSMUSG00000003575 | Crtc1 | 382056 | protein_coding | 131.7731293 | 0.133076042 | 0.14217968 | -0.935970891 | 0.349288175 | 0.571046919 | 0.133076042 |
| ENSMUSG00000050668 | Gpatch11 | 53951 | protein_coding | 159.7813347 | -0.133135751 | 0.095196362 | 1.398538217 | 0.161951505 | 0.347172272 | 0.133135751 |
| ENSMUSG00000024851 | Pitpnm1 | 18739 | protein_coding | 366.6858031 | -0.13313691 | 0.179244857 | 0.74276558 | 0.457623618 | 0.672408259 | 0.13313691 |
| ENSMUSG00000061808 | Ttr | 22139 | protein_coding | 152.4335955 | -0.133172329 | 0.400872001 | 0.332206611 | 0.739733252 | 0.867980313 | 0.133172329 |
| ENSMUSG00000044811 | Cd300c2 | 140497 | protein_coding | 0.796804571 | -0.133173146 | 1.563577982 | 0.085172053 | 0.932124609 | 1 | 0.133173146 |
| ENSMUSG00000024029 | Tff3 | 21786 | protein_coding | 5983.949861 | -0.133194113 | 0.316954814 | 0.420230605 | 0.674316999 | 0.825194181 | 0.133194113 |
| ENSMUSG00000026374 | Tsn | 22099 | protein_coding | 607.516533 | 0.133207702 | 0.097807041 | -1.361943885 | 0.173215595 | 0.363760214 | 0.133207702 |
| ENSMUSG00000024735 | Prpf19 | 28000 | protein_coding | 1138.866871 | 0.133235357 | 0.10949932 | -1.216768804 | 0.223692192 | 0.43083733 | 0.133235357 |
| ENSMUSG00000028553 | Angptl3 | 30924 | protein_coding | 7.010377194 | 0.133238586 | 0.558550751 | -0.238543383 | 0.811459672 | 0.906704483 | 0.133238586 |
| ENSMUSG00000015980 | Lrrc27 | 76612 | protein_coding | 7.061607266 | -0.133385571 | 0.533807308 | 0.24987588 | 0.802683336 | 0.902595167 | 0.133385571 |
| ENSMUSG00000029780 | Nt5c3 | 107569 | protein_coding | 716.8673324 | -0.13339822 | 0.093995473 | 1.419198354 | 0.155841196 | 0.338448383 | 0.13339822 |
| ENSMUSG00000028676 | Srsf10 | 14105 | protein_coding | 756.7466317 | 0.133427899 | 0.127784306 | -1.044164992 | 0.296409061 | 0.515610058 | 0.133427899 |
| ENSMUSG00000006191 | Cdkal1 | 68916 | protein_coding | 200.6936062 | -0.133461028 | 0.088459073 | 1.508731928 | 0.131367303 | 0.300075862 | 0.133461028 |
| ENSMUSG00000046668 | Cxxc5 | 67393 | protein_coding | 123.2444396 | 0.133481403 | 0.156695882 | -0.851850096 | 0.394297297 | 0.616875898 | 0.133481403 |
| ENSMUSG00000024771 | Lipk | 240633 | protein_coding | 4.681873987 | 0.13349536 | 0.563623417 | -0.236852047 | 0.812771572 | 0.907147741 | 0.13349536 |
| ENSMUSG00000072762 | 4930522L14Rik | 100041734 | protein_coding | 38.25845462 | -0.133566613 | 0.213202854 | 0.626476665 | 0.531002338 | 0.728048657 | 0.133566613 |
| ENSMUSG00000006024 | Napa | 108124 | protein_coding | 1070.605921 | 0.133610619 | 0.047190118 | -2.831326233 | 0.004635541 | 0.022844324 | 0.133610619 |
| ENSMUSG00000002222 | Rmnd5a | 68477 | protein_coding | 2406.283831 | 0.133649867 | 0.066412105 | -2.012432326 | 0.044174384 | 0.135669662 | 0.133649867 |
| ENSMUSG00000065799 | Gm25517 | 115488411 | snoRNA | 2.464038408 | 0.13368368 | 0.822414892 | -0.162550169 | 0.870872624 | 1 | 0.13368368 |
| ENSMUSG00000020018 | Snrpf | 69878 | protein_coding | 129.7247619 | 0.133690852 | 0.130133735 | -1.027334323 | 0.304263066 | 0.524504007 | 0.133690852 |
| ENSMUSG00000015023 | Ddx19a | 13680 | protein_coding | 283.9410931 | 0.133698864 | 0.109686419 | -1.218919032 | 0.222874925 | 0.429893421 | 0.133698864 |
| ENSMUSG00000052707 | Tnrc6a | 233833 | protein_coding | 649.5973494 | -0.13370557 | 0.10135461 | 1.319185881 | 0.187106977 | 0.383265272 | 0.13370557 |
| ENSMUSG00000071847 | Apcdd1 | 494504 | protein_coding | 3.994677581 | 0.133765702 | 0.925520496 | -0.144530243 | 0.885081777 | 1 | 0.133765702 |
| ENSMUSG00000092212 | Slc22a13b | 109280 | protein_coding | 4.043778091 | 0.13378055 | 0.794116862 | -0.168464562 | 0.86621782 | 1 | 0.13378055 |
| ENSMUSG00000020523 | Fam114a2 | 67726 | protein_coding | 828.4750121 | -0.133804799 | 0.071082049 | 1.882399298 | 0.059781821 | 0.170333437 | 0.133804799 |
| ENSMUSG00000074918 | Inafm2 | 100043272 | protein_coding | 265.6418298 | -0.133805619 | 0.106208963 | 1.259833584 | 0.207729402 | 0.410708835 | 0.133805619 |
| ENSMUSG00000028525 | Pde4b | 18578 | protein_coding | 28.29420545 | -0.133833435 | 0.299674782 | 0.446595585 | 0.655167084 | 0.812818663 | 0.133833435 |
| ENSMUSG00000035443 | Thyn1 | 77862 | protein_coding | 226.9185197 | 0.1338362 | 0.11367569 | -1.177351113 | 0.239055394 | 0.449192192 | 0.1338362 |
| ENSMUSG00000046027 | Stard5 | 170460 | protein_coding | 513.6841835 | -0.133863187 | 0.120890695 | 1.107307611 | 0.268160955 | 0.48334534 | 0.133863187 |
| ENSMUSG00000034032 | Rpap1 | 68925 | protein_coding | 368.0669027 | 0.133942796 | 0.0915112 | -1.463676534 | 0.14328235 | 0.319339495 | 0.133942796 |
| ENSMUSG00000039623 | Ap5z1 | 231855 | protein_coding | 215.0415307 | -0.134015087 | 0.11106222 | 1.206666738 | 0.227560514 | 0.436092673 | 0.134015087 |
| ENSMUSG00000055015 | Gm9961 | 791359 | lncRNA | 2.457962739 | -0.134069603 | 0.637706234 | 0.210237246 | 0.833482511 | 1 | 0.134069603 |
| ENSMUSG00000024360 | Etf1 | 225363 | protein_coding | 1648.873343 | 0.134092839 | 0.07487208 | -1.790959184 | 0.073299845 | 0.197442677 | 0.134092839 |

| ensembl_gene_id | external_gene_name | entrezgene_id | gene_biotype | baseMean | log2FoldChange | lfcSE | stat | pvalue | padj | absLog2FC |
|---|---|---|---|---|---|---|---|---|---|---|
| ENSMUSG00000022503 | Nubp1 | 26425 | protein_coding | 247.5933791 | 0.134110327 | 0.104360399 | -1.285069131 | 0.198768127 | 0.399325652 | 0.134110327 |
| ENSMUSG00000058594 | Fbh1 | 50755 | protein_coding | 1368.485815 | -0.134117749 | 0.060734615 | 2.208258824 | 0.027226236 | 0.093397187 | 0.134117749 |
| ENSMUSG00000022781 | Pak2 | 224105 | protein_coding | 1611.781312 | -0.13412644 | 0.068228524 | 1.965841153 | 0.049316962 | 0.147459771 | 0.13412644 |
| ENSMUSG00000086158 | Pierce2 | 546143 | protein_coding | 54.81141277 | -0.134128737 | 0.156036588 | 0.85959799 | 0.390010684 | 0.612460672 | 0.134128737 |
| ENSMUSG00000022617 | Chkb | 12651 | protein_coding | 102.7680814 | -0.134147132 | 0.202274787 | 0.66319255 | 0.507207242 | 0.710916686 | 0.134147132 |
| ENSMUSG00000026740 | Dnajc1 | 13418 | protein_coding | 384.4692246 | 0.134198479 | 0.089128608 | -1.505672338 | 0.132151308 | 0.3013667 | 0.134198479 |
| ENSMUSG00000015879 | Fam184b | 58227 | protein_coding | 12.07817338 | 0.134198893 | 0.454204965 | -0.295458886 | 0.767643356 | 0.883589766 | 0.134198893 |
| ENSMUSG00000053644 | Aldh7a1 | 110695 | protein_coding | 253.5989402 | 0.134212973 | 0.077875088 | -1.72343912 | 0.084809146 | 0.219348196 | 0.134212973 |
| ENSMUSG00000000326 | Comt | 12846 | protein_coding | 698.2828375 | -0.134255242 | 0.061496478 | 2.183137088 | 0.029025716 | 0.098364596 | 0.134255242 |
| ENSMUSG00000000915 | Hip1r | 29816 | protein_coding | 1364.168767 | 0.134256759 | 0.050718345 | -2.64710448 | 0.008118427 | 0.036020338 | 0.134256759 |
| ENSMUSG00000059791 | Nrm | 106582 | protein_coding | 76.95679048 | -0.134276027 | 0.19802358 | 0.678080999 | 0.497720338 | 0.703115827 | 0.134276027 |
| ENSMUSG00000066406 | Akap13 | 75547 | protein_coding | 2864.148041 | -0.134281163 | 0.137368137 | 0.977527727 | 0.328307958 | 0.55048001 | 0.134281163 |
| ENSMUSG00000034041 | Lyl1 | 17095 | protein_coding | 5.155064776 | -0.134357609 | 0.436680981 | 0.307679095 | 0.758326529 | 0.878808271 | 0.134357609 |
| ENSMUSG00000028195 | Ccn1 | 16007 | protein_coding | 429.4117269 | -0.134371613 | 0.287253912 | 0.467779922 | 0.63994198 | 0.803650332 | 0.134371613 |
| ENSMUSG00000032342 | Mto1 | 68291 | protein_coding | 192.7281982 | 0.13439791 | 0.09022412 | -1.489600681 | 0.136329263 | 0.308134243 | 0.13439791 |
| ENSMUSG00000030922 | Lyrm1 | 73919 | protein_coding | 22.58693813 | 0.134490871 | 0.233856595 | -0.575099759 | 0.565223831 | 0.751901591 | 0.134490871 |
| ENSMUSG00000091764 | Zfp964 | 636741 | protein_coding | 25.93050097 | 0.134527291 | 0.251322911 | -0.535276671 | 0.592458556 | 0.772328749 | 0.134527291 |
| ENSMUSG00000050730 | Arhgap42 | 71544 | protein_coding | 414.6566278 | 0.134532476 | 0.084803178 | -1.58640842 | 0.112646692 | 0.269245209 | 0.134532476 |
| ENSMUSG00000049882 | Vcpkmt | 207965 | protein_coding | 44.90295989 | -0.134539136 | 0.15640765 | 0.860182584 | 0.389688403 | 0.612245675 | 0.134539136 |
| ENSMUSG00000021500 | Ddx46 | 212880 | protein_coding | 845.5158993 | 0.134544057 | 0.0605358 | -2.22255357 | 0.026245918 | 0.09071266 | 0.134544057 |
| ENSMUSG00000024287 | Thoc1 | 225160 | protein_coding | 224.9615936 | 0.134555609 | 0.094698892 | -1.420878391 | 0.155352116 | 0.337697097 | 0.134555609 |
| ENSMUSG00000010080 | Epn3 | 71889 | protein_coding | 618.0992612 | 0.134587517 | 0.083317277 | -1.615361448 | 0.106232444 | 0.258635202 | 0.134587517 |
| ENSMUSG00000037935 | Smarce1 | 57376 | protein_coding | 649.2246245 | -0.134591281 | 0.072933418 | 1.845399328 | 0.064979471 | 0.181047913 | 0.134591281 |
| ENSMUSG00000044468 | Tent5c | 74645 | protein_coding | 144.4420188 | -0.134603939 | 0.271591821 | 0.495611164 | 0.620168775 | 0.791152348 | 0.134603939 |
| ENSMUSG00000055991 | Zkscan5 | 22757 | protein_coding | 178.1272042 | -0.134609502 | 0.095247317 | 1.41326292 | 0.157578432 | 0.340965701 | 0.134609502 |
| ENSMUSG00000086549 | Gm13648 | 100415901 | lncRNA | 122.0450129 | 0.134615101 | 0.238686427 | -0.563983059 | 0.572765662 | 0.758091673 | 0.134615101 |
| ENSMUSG00000003662 | Ciao1 | 26371 | protein_coding | 437.1498273 | -0.134626227 | 0.070660182 | 1.905262948 | 0.056745899 | 0.163972315 | 0.134626227 |
| ENSMUSG00000033540 | Idua | 15932 | protein_coding | 114.5553171 | -0.134633597 | 0.11094973 | 1.213464848 | 0.224952146 | 0.432209358 | 0.134633597 |
| ENSMUSG00000026643 | Nmt2 | 18108 | protein_coding | 437.032003 | -0.134644982 | 0.088005492 | 1.529961133 | 0.12602635 | 0.291934232 | 0.134644982 |
| ENSMUSG00000019432 | Ddx39b | 53817 | protein_coding | 1665.63617 | 0.134662622 | 0.069607315 | -1.934604471 | 0.053038851 | 0.155998286 | 0.134662622 |
| ENSMUSG00000031134 | Rbmx | 19655 | protein_coding | 367.9470669 | 0.134833744 | 0.09648789 | -1.397416228 | 0.162288442 | 0.347734431 | 0.134833744 |
| ENSMUSG00000007888 | Crlf1 | 12931 | protein_coding | 9.123960002 | -0.134841214 | 0.485103451 | 0.277963832 | 0.781040123 | 0.890541867 | 0.134841214 |
| ENSMUSG00000025085 | Ablim1 | 226251 | protein_coding | 1378.235025 | -0.134864807 | 0.122737434 | 1.09880745 | 0.271852062 | 0.487629303 | 0.134864807 |
| ENSMUSG00000079108 | Srp54c | 100101806 | protein_coding | 82.88430199 | 0.134867704 | 0.155238562 | -0.868777076 | 0.384969074 | 0.60719955 | 0.134867704 |
| ENSMUSG00000028857 | Tmem222 | 52174 | protein_coding | 313.7052145 | 0.134868298 | 0.091507614 | -1.47384782 | 0.14052257 | 0.314808425 | 0.134868298 |
| ENSMUSG00000001056 | Nhp2 | 52530 | protein_coding | 302.7219409 | -0.134897981 | 0.103569126 | 1.302492216 | 0.192748177 | 0.390530809 | 0.134897981 |
| ENSMUSG00000007021 | Syngr3 | 20974 | protein_coding | 2.728499952 | -0.134914549 | 0.715705783 | 0.188505601 | 0.850480325 | 1 | 0.134914549 |
| ENSMUSG00000026028 | Trak2 | 70827 | protein_coding | 786.8992732 | -0.134983299 | 0.099346933 | 1.358706249 | 0.174239691 | 0.365337006 | 0.134983299 |
| ENSMUSG00000004018 | Fancl | 67030 | protein_coding | 93.71346347 | 0.134988823 | 0.130205782 | -1.036734473 | 0.299859622 | 0.519828028 | 0.134988823 |

Supplementary Table 1

| ensembl_gene_id | external_gene_name | entrezgene_id | gene_biotype | baseMean | log2FoldChange | lfcSE | stat | pvalue | padj | absLog2FC |
|---|---|---|---|---|---|---|---|---|---|---|
| ENSMUSG00000048521 | Cxcr6 | 80901 | protein_coding | 9.174290609 | -0.134994136 | 0.40517402 | 0.333175695 | 0.739001664 | 0.867615063 | 0.134994136 |
| ENSMUSG00000069495 | Epc2 | 227867 | protein_coding | 400.1733828 | 0.135020704 | 0.102539792 | -1.316763971 | 0.187917753 | 0.384217774 | 0.135020704 |
| ENSMUSG00000020776 | Fbf1 | 217335 | protein_coding | 202.5922164 | 0.135027057 | 0.114558461 | -1.178673805 | 0.23852809 | 0.448702897 | 0.135027057 |
| ENSMUSG00000059336 | Slc14a1 | 108052 | protein_coding | 1.912013781 | 0.135057803 | 1.190384361 | -0.113457306 | 0.909668009 | 1 | 0.135057803 |
| ENSMUSG00000037236 | Matr3 | 17184 | protein_coding | 748.5534765 | -0.135060695 | 0.112242791 | 1.203290602 | 0.228863885 | 0.437741278 | 0.135060695 |
| ENSMUSG00000054150 | Syne3 | 212073 | protein_coding | 140.1294119 | -0.135067676 | 0.18534626 | 0.728731595 | 0.466165862 | 0.679754841 | 0.135067676 |
| ENSMUSG00000042369 | Rbm45 | 241490 | protein_coding | 424.0458552 | -0.135070311 | 0.090932861 | 1.485385032 | 0.137441858 | 0.309858682 | 0.135070311 |
| ENSMUSG00000047284 | Neurl4 | 216860 | protein_coding | 627.4043111 | -0.135330493 | 0.060447123 | 2.238824405 | 0.025167344 | 0.087793307 | 0.135330493 |
| ENSMUSG00000055471 | Alk | 11682 | protein_coding | 4.145192173 | 0.135331144 | 2.26314147 | -0.059797916 | 0.952316585 | 1 | 0.135331144 |
| ENSMUSG00000075420 | Smim6 | 68528 | protein_coding | 897.3642135 | -0.13539381 | 0.176838885 | 0.765633704 | 0.443894284 | 0.661236714 | 0.13539381 |
| ENSMUSG00000053477 | Tcf4 | 21413 | protein_coding | 346.35157 | 0.135397243 | 0.094424301 | -1.433923702 | 0.151594037 | 0.331843045 | 0.135397243 |
| ENSMUSG00000026354 | Lct | 226413 | protein_coding | 30.95146894 | 0.135429858 | 0.874644071 | -0.154839966 | 0.876947483 | 0.938919382 | 0.135429858 |
| ENSMUSG00000033319 | Fem1c | 240263 | protein_coding | 459.4611467 | -0.135564924 | 0.077638362 | 1.746107468 | 0.080792279 | 0.21130879 | 0.135564924 |
| ENSMUSG00000038217 | Tlcd2 | 380712 | protein_coding | 58.41941336 | -0.135611992 | 0.173214885 | 0.782911887 | 0.433678857 | 0.652196761 | 0.135611992 |
| ENSMUSG00000101348 | Gm7135 | 634346 | transcribed_unprocessed_pseudogene | 0.681628359 | -0.135661762 | 1.584297475 | 0.085628971 | 0.931761368 | 1 | 0.135661762 |
| ENSMUSG00000027840 | Wnt2b | 22414 | protein_coding | 9.642866508 | -0.135675439 | 0.4719957 | 0.287450583 | 0.773767335 | 0.88672609 | 0.135675439 |
| ENSMUSG00000055723 | Rras2 | 66922 | protein_coding | 626.0289645 | -0.135691158 | 0.070235914 | 1.931934123 | 0.053367637 | 0.156761502 | 0.135691158 |
| ENSMUSG00000040723 | Rcsd1 | 226594 | protein_coding | 12.88445651 | -0.135699935 | 0.63462454 | 0.213827115 | 0.830681879 | 0.91440345 | 0.135699935 |
| ENSMUSG00000040187 | Arntl2 | 272322 | protein_coding | 14.58403305 | -0.135761938 | 0.301432612 | 0.450389016 | 0.652429963 | 0.8114957 | 0.135761938 |
| ENSMUSG00000004612 | Nkg7 | 72310 | protein_coding | 12.57757806 | 0.135763359 | 0.466840979 | -0.290812859 | 0.77119445 | 0.885199039 | 0.135763359 |
| ENSMUSG00000027329 | Spef1 | 70997 | protein_coding | 105.0135703 | -0.135802938 | 0.166209923 | 0.817056741 | 0.413896003 | 0.633934205 | 0.135802938 |
| ENSMUSG00000032239 | Rp9 | 55934 | protein_coding | 407.267924 | -0.135811735 | 0.078197322 | 1.736782445 | 0.082425579 | 0.214493541 | 0.135811735 |
| ENSMUSG00000068267 | Cenpb | 12616 | protein_coding | 2721.108475 | -0.135814784 | 0.049445492 | 2.746757661 | 0.006018759 | 0.02823687 | 0.135814784 |
| ENSMUSG00000070738 | Dgkd | 227333 | protein_coding | 1608.338521 | -0.135831722 | 0.060393033 | 2.249128989 | 0.024504291 | 0.08602548 | 0.135831722 |
| ENSMUSG00000026384 | Ptpn4 | 19258 | protein_coding | 300.3082407 | -0.135850484 | 0.104795908 | 1.296333863 | 0.194860487 | 0.393919217 | 0.135850484 |
| ENSMUSG00000008575 | Nfib | 18028 | protein_coding | 674.7439564 | 0.135911586 | 0.120840526 | -1.124718588 | 0.260708303 | 0.474208273 | 0.135911586 |
| ENSMUSG00000014859 | E2f4 | 104394 | protein_coding | 451.2474567 | 0.135946588 | 0.062365487 | -2.179836862 | 0.029269556 | 0.09897928 | 0.135946588 |
| ENSMUSG00000033964 | Zbtb41 | 226470 | protein_coding | 440.1438559 | 0.135949316 | 0.102652757 | -1.324361078 | 0.185383155 | 0.380725956 | 0.135949316 |
| ENSMUSG00000027363 | Usp8 | 84092 | protein_coding | 769.9885913 | 0.136204868 | 0.0468612 | -2.906559578 | 0.003654273 | 0.018770347 | 0.136204868 |
| ENSMUSG00000025630 | Hprt | 15452 | protein_coding | 740.159001 | 0.136244781 | 0.104285905 | -1.306454415 | 0.191398071 | 0.389062992 | 0.136244781 |
| ENSMUSG00000033066 | Gas7 | 14457 | protein_coding | 109.6808095 | -0.136260659 | 0.127453218 | 1.069103329 | 0.285023112 | 0.503462581 | 0.136260659 |
| ENSMUSG00000060206 | Zfp462 | 242466 | protein_coding | 253.2242154 | -0.136429204 | 0.113879927 | 1.198009239 | 0.23091342 | 0.439526427 | 0.136429204 |
| ENSMUSG00000001366 | Fbxo9 | 71538 | protein_coding | 540.0865011 | 0.136448193 | 0.07205285 | -1.893723748 | 0.058261687 | 0.16718219 | 0.136448193 |
| ENSMUSG00000056665 | Them6 | 223626 | protein_coding | 18.83623671 | 0.136504276 | 0.270562512 | -0.504520285 | 0.613895811 | 0.78655252 | 0.136504276 |
| ENSMUSG00000050373 | Snx21 | 101113 | protein_coding | 190.1166748 | 0.136517893 | 0.099904619 | -1.366482295 | 0.171787634 | 0.362126754 | 0.136517893 |
| ENSMUSG00000042766 | Trim46 | 360213 | protein_coding | 157.5687987 | -0.136527749 | 0.188764481 | 0.723270333 | 0.469513815 | 0.683156904 | 0.136527749 |
| ENSMUSG00000071054 | Safb | 224903 | protein_coding | 802.6656485 | 0.136591378 | 0.112265645 | -1.21668012 | 0.223725946 | 0.43083733 | 0.136591378 |
| ENSMUSG00000086796 | 4932702P03Rik | 73326 | lncRNA | 4.432403258 | 0.13660625 | 0.854379666 | -0.159889397 | 0.872968201 | 0.937394946 | 0.13660625 |
| ENSMUSG00000064390 | Snord73b | 19871 | snoRNA | 6.303939264 | 0.136606542 | 0.445992448 | -0.306297881 | 0.759377852 | 0.879038683 | 0.136606542 |

| ensembl_gene_id | external_gene_name | entrezgene_id | gene_biotype | baseMean | log2FoldChange | lfcSE | stat | pvalue | padj | absLog2FC |
|---|---|---|---|---|---|---|---|---|---|---|
| ENSMUSG00000024048 | Myl12a | 67268 | protein_coding | 565.4570145 | -0.13660876 | 0.083467831 | 1.636663606 | 0.101700777 | 0.250669647 | 0.13660876 |
| ENSMUSG00000037316 | Bag4 | 67384 | protein_coding | 275.353017 | -0.136661497 | 0.094750751 | 1.442326267 | 0.149210353 | 0.32840994 | 0.136661497 |
| ENSMUSG00000031848 | Lsm4 | 50783 | protein_coding | 554.9203043 | 0.136666636 | 0.094798094 | -1.441660164 | 0.149398267 | 0.328779714 | 0.136666636 |
| ENSMUSG00000019775 | Rgs17 | 56533 | protein_coding | 167.8923628 | -0.136684318 | 0.207820515 | 0.657703684 | 0.510728554 | 0.71379243 | 0.136684318 |
| ENSMUSG00000024845 | Tmem134 | 66990 | protein_coding | 708.0906157 | -0.136696277 | 0.064981544 | 2.103616927 | 0.035411876 | 0.114552737 | 0.136696277 |
| ENSMUSG00000030770 | Parva | 57342 | protein_coding | 1049.893741 | -0.136703295 | 0.114415754 | 1.194794336 | 0.232167394 | 0.441179379 | 0.136703295 |
| ENSMUSG00000028331 | Trmo | 74753 | protein_coding | 61.27883652 | -0.136730783 | 0.169846654 | 0.805024884 | 0.42080532 | 0.640714813 | 0.136730783 |
| ENSMUSG00000022248 | Rad1 | 19355 | protein_coding | 76.49876496 | 0.136753108 | 0.128520629 | -1.0640557 | 0.287303468 | 0.506201691 | 0.136753108 |
| ENSMUSG00000029213 | Commd8 | 27784 | protein_coding | 305.8896121 | 0.136764172 | 0.084867669 | -1.611499095 | 0.107070984 | 0.260075004 | 0.136764172 |
| ENSMUSG00000047824 | Pygo2 | 68911 | protein_coding | 554.6380191 | 0.136764349 | 0.055759672 | -2.452746664 | 0.014177014 | 0.05583573 | 0.136764349 |
| ENSMUSG00000062929 | Cfl2 | 12632 | protein_coding | 262.5608052 | -0.136767271 | 0.199130008 | 0.68682401 | 0.492193638 | 0.698889573 | 0.136767271 |
| ENSMUSG00000001150 | Mcm3ap | 54387 | protein_coding | 570.4286609 | 0.136812948 | 0.072208648 | -1.89468924 | 0.058133584 | 0.166901601 | 0.136812948 |
| ENSMUSG00000087305 | A430035B10Rik | 320312 | lncRNA | 22.21264256 | -0.136828443 | 0.24721133 | 0.55348775 | 0.579929466 | 0.76328646 | 0.136828443 |
| ENSMUSG00000020785 | Camkk1 | 55984 | protein_coding | 34.18290483 | 0.136838146 | 0.185326576 | -0.73836224 | 0.460294354 | 0.674531987 | 0.136838146 |
| ENSMUSG00000087598 | Zfp111 | 56707 | protein_coding | 138.9887755 | -0.136846968 | 0.12011895 | 1.13926211 | 0.254593847 | 0.467151421 | 0.136846968 |
| ENSMUSG00000026889 | Rbm18 | 67889 | protein_coding | 227.3988285 | 0.136850286 | 0.090681378 | -1.509133285 | 0.131264725 | 0.299903938 | 0.136850286 |
| ENSMUSG00000023043 | Krt18 | 16668 | protein_coding | 6615.610191 | -0.136909412 | 0.083078967 | 1.647943125 | 0.099364343 | 0.246352764 | 0.136909412 |
| ENSMUSG00000008393 | Carhsp1 | 52502 | protein_coding | 1302.71217 | -0.136929174 | 0.092884934 | 1.474180667 | 0.140432954 | 0.314711603 | 0.136929174 |
| ENSMUSG00000027534 | Snx16 | 74718 | protein_coding | 88.10294702 | -0.136944335 | 0.104490705 | 1.310588679 | 0.189996766 | 0.387311023 | 0.136944335 |
| ENSMUSG00000015932 | Dstn | 56431 | protein_coding | 12626.54385 | 0.136957725 | 0.099673521 | -1.37406327 | 0.169422041 | 0.358767397 | 0.136957725 |
| ENSMUSG00000058135 | Gstm1 | 14862 | protein_coding | 2427.491274 | 0.136958051 | 0.142527938 | -0.96092074 | 0.336592022 | 0.559203284 | 0.136958051 |
| ENSMUSG00000036078 | Sigmar1 | 18391 | protein_coding | 486.1848236 | 0.136969181 | 0.104507177 | -1.310619846 | 0.189986231 | 0.387311023 | 0.136969181 |
| ENSMUSG00000032094 | Cd3d | 12500 | protein_coding | 9.210008843 | -0.136987282 | 0.494940188 | 0.276775427 | 0.781952551 | 0.890747356 | 0.136987282 |
| ENSMUSG00000063450 | Syne2 | 319565 | protein_coding | 3017.1959 | -0.137023255 | 0.113067252 | 1.211873935 | 0.225560641 | 0.433049501 | 0.137023255 |
| ENSMUSG00000010453 | Kansl3 | 226976 | protein_coding | 818.009483 | -0.13706659 | 0.079444673 | 1.725308756 | 0.084471845 | 0.218595521 | 0.13706659 |
| ENSMUSG00000034930 | Rtkn | 20166 | protein_coding | 167.1817196 | -0.137103837 | 0.114971116 | 1.192506799 | 0.233062586 | 0.442542894 | 0.137103837 |
| ENSMUSG00000046982 | Tshz1 | 110796 | protein_coding | 865.1781386 | -0.13714312 | 0.064462708 | 2.127479948 | 0.033380226 | 0.109497021 | 0.13714312 |
| ENSMUSG00000036513 | Commd2 | 52245 | protein_coding | 165.1172451 | 0.137153678 | 0.104611961 | -1.311070717 | 0.189833873 | 0.387122324 | 0.137153678 |
| ENSMUSG00000028126 | Pip5k1a | 18720 | protein_coding | 233.2506158 | -0.137156237 | 0.080645566 | 1.700728799 | 0.088993925 | 0.226867192 | 0.137156237 |
| ENSMUSG00000052056 | Zfp217 | 228913 | protein_coding | 814.976122 | -0.137178636 | 0.067932743 | 2.019330145 | 0.043452917 | 0.133877139 | 0.137178636 |
| ENSMUSG00000058392 | Rrp1b | 72462 | protein_coding | 135.8044974 | 0.137192223 | 0.1619028 | -0.847373999 | 0.396786693 | 0.618855346 | 0.137192223 |
| ENSMUSG00000049411 | Tmem241 | 338363 | protein_coding | 115.713243 | 0.137204007 | 0.111164394 | -1.234244182 | 0.217111929 | 0.422731291 | 0.137204007 |
| ENSMUSG00000037686 | Aspg | 104816 | protein_coding | 670.927295 | 0.137206121 | 0.151034644 | -0.908441382 | 0.36364507 | 0.586048082 | 0.137206121 |
| ENSMUSG00000014850 | Msh3 | 17686 | protein_coding | 292.8901036 | -0.137212072 | 0.085726878 | 1.600572369 | 0.109471666 | 0.264184582 | 0.137212072 |
| ENSMUSG00000044814 | Or55b3 | 257947 | protein_coding | 1.131997778 | 0.137231877 | 0.967554186 | -0.141833789 | 0.887211294 | 1 | 0.137231877 |
| ENSMUSG00000034610 | Tut4 | 230594 | protein_coding | 239.6838012 | -0.137246971 | 0.09762278 | 1.405890824 | 0.159756547 | 0.343942789 | 0.137246971 |
| ENSMUSG00000055833 | 1700034H15Rik | 98736 | lncRNA | 6.846739016 | -0.137255876 | 0.496440319 | 0.276480114 | 0.782179331 | 0.890759912 | 0.137255876 |
| ENSMUSG00000017978 | Cadps2 | 320405 | protein_coding | 905.3223004 | -0.137264916 | 0.078123461 | 1.757025529 | 0.078913458 | 0.208009073 | 0.137264916 |
| ENSMUSG00000025158 | Rfng | 19719 | protein_coding | 184.0891524 | 0.137269106 | 0.107913819 | -1.272025287 | 0.203364135 | 0.405696463 | 0.137269106 |

| ensembl_gene_id | external_gene_name | entrezgene_id | gene_biotype | baseMean | log2FoldChange | lfcSE | stat | pvalue | padj | absLog2FC |
|---|---|---|---|---|---|---|---|---|---|---|
| ENSMUSG00000026565 | Pou2f1 | 18986 | protein_coding | 234.2035066 | 0.137285784 | 0.121156756 | -1.13312529 | 0.257161642 | 0.470243313 | 0.137285784 |
| ENSMUSG00000036864 | Proser3 | 333193 | protein_coding | 38.53986395 | 0.137302194 | 0.213574329 | -0.642877798 | 0.520303397 | 0.720291626 | 0.137302194 |
| ENSMUSG00000029015 | Slc26a5 | 80979 | protein_coding | 1.510887764 | 0.137307388 | 2.671289263 | -0.051401168 | 0.959005854 | 1 | 0.137307388 |
| ENSMUSG00000025147 | Mob2 | 101513 | protein_coding | 654.1347338 | 0.137319925 | 0.051385521 | -2.672346659 | 0.007532279 | 0.033883971 | 0.137319925 |
| ENSMUSG00000002778 | Kdelr1 | 68137 | protein_coding | 1267.817866 | 0.137326653 | 0.086747958 | -1.583053447 | 0.113409283 | 0.270251212 | 0.137326653 |
| ENSMUSG00000021932 | Rnaseh2b | 67153 | protein_coding | 148.5686196 | 0.13734676 | 0.120613375 | -1.138735733 | 0.254813394 | 0.467346556 | 0.13734676 |
| ENSMUSG00000024064 | Galnt14 | 71685 | protein_coding | 3.496291461 | -0.137396105 | 1.99844696 | 0.06875144 | 0.945187472 | 1 | 0.137396105 |
| ENSMUSG00000072494 | Ppp1r3e | 105651 | protein_coding | 48.43480083 | -0.137399029 | 0.252154015 | 0.544901211 | 0.585821463 | 0.767554399 | 0.137399029 |
| ENSMUSG00000066043 | Phactr4 | 100169 | protein_coding | 422.7290478 | -0.137401817 | 0.12753324 | 1.077380431 | 0.281310343 | 0.499023496 | 0.137401817 |
| ENSMUSG00000054792 | Klhl18 | 270201 | protein_coding | 249.6503138 | -0.137468516 | 0.076547095 | 1.795868483 | 0.072515436 | 0.196025521 | 0.137468516 |
| ENSMUSG00000028430 | Nol6 | 230082 | protein_coding | 364.8617309 | 0.137489578 | 0.106860645 | -1.286625 | 0.198225026 | 0.398476941 | 0.137489578 |
| ENSMUSG00000028657 | Ppt1 | 19063 | protein_coding | 383.6484916 | -0.137584844 | 0.083626782 | 1.645224653 | 0.099923491 | 0.247508105 | 0.137584844 |
| ENSMUSG00000022560 | Slc52a2 | 52710 | protein_coding | 239.673431 | -0.13760615 | 0.141389217 | 0.973243595 | 0.330432249 | 0.552639068 | 0.13760615 |
| ENSMUSG00000034109 | Golim4 | 73124 | protein_coding | 1055.517088 | 0.137688874 | 0.060355836 | -2.28128517 | 0.022531578 | 0.080642929 | 0.137688874 |
| ENSMUSG00000030722 | Nfatc2ip | 18020 | protein_coding | 70.32055125 | 0.137715295 | 0.152725192 | -0.901719573 | 0.367205853 | 0.589852079 | 0.137715295 |
| ENSMUSG00000028007 | Snx7 | 76561 | protein_coding | 694.0637143 | -0.13772874 | 0.081441881 | 1.691129165 | 0.090812136 | 0.230328923 | 0.13772874 |
| ENSMUSG00000021270 | Hsp90aa1 | 15519 | protein_coding | 3121.300132 | -0.137835439 | 0.159862453 | 0.86221271 | 0.388570477 | 0.611070646 | 0.137835439 |
| ENSMUSG00000021022 | Ppp2r3c | 59032 | protein_coding | 119.8297512 | -0.137842148 | 0.11689681 | 1.179178012 | 0.2383273 | 0.448589299 | 0.137842148 |
| ENSMUSG00000028639 | Ybx1 | 22608 | protein_coding | 5081.886756 | -0.137842537 | 0.076303266 | 1.806509014 | 0.070838867 | 0.192691988 | 0.137842537 |
| ENSMUSG00000032407 | U2surp | 67958 | protein_coding | 1078.919948 | -0.137873643 | 0.067478801 | 2.043214164 | 0.041031242 | 0.128477424 | 0.137873643 |
| ENSMUSG00000071064 | Zfp827 | 622675 | protein_coding | 316.157415 | -0.137894724 | 0.101371387 | 1.360292369 | 0.173737422 | 0.364468978 | 0.137894724 |
| ENSMUSG00000038418 | Egr1 | 13653 | protein_coding | 3255.422974 | -0.137911946 | 0.198397089 | 0.695130896 | 0.486973274 | 0.695624496 | 0.137911946 |
| ENSMUSG00000089695 | Gm15946 | 108168333 | processed_pseudogene | 0.614287737 | -0.137912922 | 1.441593439 | 0.095667002 | 0.92378505 | 1 | 0.137912922 |
| ENSMUSG00000024847 | Aip | 11632 | protein_coding | 276.093547 | -0.137932931 | 0.073161101 | 1.885331538 | 0.059385091 | 0.169583213 | 0.137932931 |
| ENSMUSG00000031596 | Slc7a2 | 11988 | protein_coding | 13.13805403 | 0.137970071 | 0.343909075 | -0.401181826 | 0.68828626 | 0.833786242 | 0.137970071 |
| ENSMUSG00000041245 | Wnk3 | 279561 | protein_coding | 4.449226845 | -0.138005888 | 0.94390916 | 0.146206747 | 0.883758179 | 0.942424291 | 0.138005888 |
| ENSMUSG00000020775 | Mrpl38 | 60441 | protein_coding | 432.3968237 | -0.13806635 | 0.114333395 | 1.20757676 | 0.227210103 | 0.435789505 | 0.13806635 |
| ENSMUSG00000033417 | Cacul1 | 78832 | protein_coding | 691.8688026 | 0.138071963 | 0.052370547 | -2.636443043 | 0.008378026 | 0.036904435 | 0.138071963 |
| ENSMUSG00000028245 | Nsmaf | 18201 | protein_coding | 194.2933861 | -0.138076355 | 0.083106001 | 1.661448669 | 0.09662337 | 0.240996576 | 0.138076355 |
| ENSMUSG00000038290 | Smg6 | 103677 | protein_coding | 459.5467225 | -0.13808157 | 0.08781662 | 1.572385388 | 0.115861201 | 0.274864513 | 0.13808157 |
| ENSMUSG00000103168 | Gm30948 | 102633020 | lncRNA | 0.952893882 | 0.138082061 | 3.078699063 | -0.044850782 | 0.964226248 | 1 | 0.138082061 |
| ENSMUSG00000021666 | Gfm2 | 320806 | protein_coding | 661.1070842 | -0.138100581 | 0.07007648 | 1.970712306 | 0.048756792 | 0.14626152 | 0.138100581 |
| ENSMUSG00000020630 | Rnaseh1 | 19819 | protein_coding | 127.2281801 | -0.138105601 | 0.107803143 | 1.281090672 | 0.20016182 | 0.40124694 | 0.138105601 |
| ENSMUSG00000076431 | Sox4 | 20677 | protein_coding | 582.0213123 | 0.138116618 | 0.084201914 | -1.640302607 | 0.100942264 | 0.2494703 | 0.138116618 |
| ENSMUSG00000004642 | Slbp | 20492 | protein_coding | 448.0132009 | 0.13812042 | 0.076667157 | -1.801559182 | 0.071614788 | 0.194226738 | 0.13812042 |
| ENSMUSG00000023960 | Enpp5 | 83965 | protein_coding | 549.7231653 | 0.138218184 | 0.149332194 | -0.925575261 | 0.354666755 | 0.577325459 | 0.138218184 |
| ENSMUSG00000084799 | Ino80dos | 100380944 | lncRNA | 20.37349578 | 0.138259297 | 0.294963496 | -0.468733584 | 0.639260078 | 0.803282676 | 0.138259297 |
| ENSMUSG00000031527 | Eri1 | 67276 | protein_coding | 377.5312863 | 0.138298828 | 0.106211361 | -1.302109555 | 0.192878937 | 0.390726542 | 0.138298828 |
| ENSMUSG00000040170 | Fmo2 | 55990 | protein_coding | 6.682885309 | -0.138316445 | 0.462045561 | 0.299356724 | 0.764667878 | 0.882132403 | 0.138316445 |

Supplementary Table 1

| ensembl_gene_id | external_gene_name | entrezgene_id | gene_biotype | baseMean | log2FoldChange | lfcSE | stat | pvalue | padj | absLog2FC |
|---|---|---|---|---|---|---|---|---|---|---|
| ENSMUSG00000052738 | Suclg1 | 56451 | protein_coding | 2791.888596 | -0.138325348 | 0.087823903 | 1.575030757 | 0.115249345 | 0.273806314 | 0.138325348 |
| ENSMUSG00000038954 | Supt3 | 109115 | protein_coding | 35.09588863 | 0.138392882 | 0.216440197 | -0.639404715 | 0.522559682 | 0.721840784 | 0.138392882 |
| ENSMUSG00000062981 | Mrpl42 | 67270 | protein_coding | 657.2712602 | -0.138406779 | 0.077016354 | 1.797108942 | 0.072318326 | 0.195652825 | 0.138406779 |
| ENSMUSG00000097099 | Gm9917 | 100038755 | lncRNA | 75.87248311 | -0.138419664 | 0.165329696 | 0.83723413 | 0.402460977 | 0.624001556 | 0.138419664 |
| ENSMUSG00000019791 | Hint3 | 66847 | protein_coding | 291.8754322 | -0.138459229 | 0.071525465 | 1.935803252 | 0.052891803 | 0.155751645 | 0.138459229 |
| ENSMUSG00000055320 | Tead1 | 21676 | protein_coding | 200.8670039 | -0.138488099 | 0.17432305 | 0.794433664 | 0.426943013 | 0.646395058 | 0.138488099 |
| ENSMUSG00000085772 | D630024D03Rik | 414116 | lncRNA | 9.88362935 | -0.138505122 | 0.395318865 | 0.350363048 | 0.726066254 | 0.858172551 | 0.138505122 |
| ENSMUSG00000034487 | Poglut3 | 68304 | protein_coding | 180.6178559 | 0.1385304 | 0.127307587 | -1.0881551 | 0.276526642 | 0.493575548 | 0.1385304 |
| ENSMUSG00000038267 | Slc22a23 | 73102 | protein_coding | 1624.623001 | -0.138575492 | 0.098996975 | 1.399795217 | 0.161574651 | 0.346538122 | 0.138575492 |
| ENSMUSG00000031311 | Nono | 53610 | protein_coding | 1686.546673 | -0.138575724 | 0.100694693 | 1.376196889 | 0.168760685 | 0.357709402 | 0.138575724 |
| ENSMUSG00000024580 | Grpel2 | 17714 | protein_coding | 699.9215511 | -0.138581711 | 0.079937925 | 1.733616568 | 0.082986139 | 0.215748283 | 0.138581711 |
| ENSMUSG00000118423 | Lrrc70 | 113523645 | transcribed_unitary_pseudogene | 4.531903047 | -0.138591338 | 0.544336168 | 0.254606153 | 0.799027306 | 0.900323208 | 0.138591338 |
| ENSMUSG00000040712 | Camta2 | 216874 | protein_coding | 665.2457163 | 0.138613189 | 0.158332295 | -0.875457459 | 0.381325047 | 0.603700684 | 0.138613189 |
| ENSMUSG00000030045 | Mrpl19 | 56284 | protein_coding | 293.3041964 | -0.138653223 | 0.08908313 | 1.55644759 | 0.119601696 | 0.281393704 | 0.138653223 |
| ENSMUSG00000023169 | Slc38a1 | 105727 | protein_coding | 673.7516263 | -0.13872558 | 0.130561157 | 1.062533324 | 0.287993638 | 0.506866348 | 0.13872558 |
| ENSMUSG00000026032 | Ndufb3 | 66495 | protein_coding | 501.4098443 | -0.138749685 | 0.094441535 | 1.469159574 | 0.141789512 | 0.316830918 | 0.138749685 |
| ENSMUSG00000098789 | Jmjd7 | 433466 | protein_coding | 16.17866178 | -0.138777224 | 0.258709554 | 0.536420947 | 0.591667658 | 0.771785126 | 0.138777224 |
| ENSMUSG00000057409 | Zfp53 | 24132 | protein_coding | 189.6915033 | 0.138877252 | 0.103439057 | -1.342599755 | 0.17940161 | 0.372658891 | 0.138877252 |
| ENSMUSG00000028879 | Stx12 | 100226 | protein_coding | 834.8270405 | -0.138881809 | 0.073389597 | 1.892390937 | 0.058438913 | 0.167508599 | 0.138881809 |
| ENSMUSG00000030802 | Bckdk | 12041 | protein_coding | 617.3560972 | 0.138898965 | 0.087700884 | -1.583780668 | 0.11324364 | 0.270127107 | 0.138898965 |
| ENSMUSG00000028053 | Ash1l | 192195 | protein_coding | 1157.755275 | -0.138904572 | 0.080672662 | 1.721829544 | 0.085100401 | 0.219864981 | 0.138904572 |
| ENSMUSG00000030386 | Zfp606 | 67370 | protein_coding | 140.434808 | -0.139003951 | 0.10858894 | 1.280093077 | 0.200512403 | 0.401803401 | 0.139003951 |
| ENSMUSG00000032802 | Srxn1 | 76650 | protein_coding | 1647.499408 | -0.139009811 | 0.123827387 | 1.122609585 | 0.261603343 | 0.475244062 | 0.139009811 |
| ENSMUSG00000017210 | Med24 | 23989 | protein_coding | 483.6462172 | 0.139040917 | 0.069383904 | -2.003936203 | 0.045076895 | 0.137877355 | 0.139040917 |
| ENSMUSG00000029188 | Slc34a2 | 20531 | protein_coding | 615.1368902 | 0.139080201 | 0.315579328 | -0.440713915 | 0.659420121 | 0.816133095 | 0.139080201 |
| ENSMUSG00000063239 | Grm4 | 268934 | protein_coding | 7.165093042 | -0.139110934 | 0.571926703 | 0.243232102 | 0.8078256 | 0.905233898 | 0.139110934 |
| ENSMUSG00000030733 | Sh2b1 | 20399 | protein_coding | 472.4766372 | 0.139134272 | 0.108870207 | -1.277982981 | 0.201255427 | 0.402657151 | 0.139134272 |
| ENSMUSG00000030536 | Iqgap1 | 29875 | protein_coding | 5398.671723 | -0.139148227 | 0.096991431 | 1.434644538 | 0.151388416 | 0.331436869 | 0.139148227 |
| ENSMUSG00000032220 | Myo1e | 71602 | protein_coding | 1337.304645 | -0.139149131 | 0.06905746 | 2.014976102 | 0.043907153 | 0.135006119 | 0.139149131 |
| ENSMUSG00000026365 | Cfh | 12628 | protein_coding | 6.117805254 | -0.139161465 | 0.509732365 | 0.273008886 | 0.784846385 | 0.892504697 | 0.139161465 |
| ENSMUSG00000113836 | Gm3325 | 100041420 | transcribed_processed_pseudogene | 36.59338682 | -0.139190354 | 0.200189402 | 0.695293321 | 0.486871499 | 0.695624496 | 0.139190354 |
| ENSMUSG00000086529 | Acss2os | 74945 | lncRNA | 2.143250408 | -0.139267811 | 0.680522574 | 0.204648334 | 0.837846889 | 1 | 0.139267811 |
| ENSMUSG00000052539 | Magi3 | 99470 | protein_coding | 1464.067504 | 0.139281 | 0.046367349 | -3.003859497 | 0.002665784 | 0.01440064 | 0.139281 |
| ENSMUSG00000026313 | Hdac4 | 208727 | protein_coding | 684.8505781 | -0.139325935 | 0.119835654 | 1.162641755 | 0.244974882 | 0.456655966 | 0.139325935 |
| ENSMUSG00000027067 | Ssrp1 | 20833 | protein_coding | 1194.231544 | 0.13935266 | 0.101537873 | -1.372420513 | 0.169932568 | 0.359501207 | 0.13935266 |
| ENSMUSG00000026390 | Marco | 17167 | protein_coding | 1.485611066 | -0.139363042 | 1.771317194 | 0.078677632 | 0.937289038 | 1 | 0.139363042 |
| ENSMUSG00000028322 | Exosc3 | 66362 | protein_coding | 232.892623 | 0.139393426 | 0.077992904 | -1.78725781 | 0.07389583 | 0.198651429 | 0.139393426 |
| ENSMUSG00000021715 | Cwc27 | 67285 | protein_coding | 133.2553665 | 0.139412604 | 0.104343956 | -1.336087005 | 0.181520837 | 0.375643497 | 0.139412604 |
| ENSMUSG00000027582 | Zgpat | 229007 | protein_coding | 388.260569 | -0.139426343 | 0.068280005 | 2.041979118 | 0.041153603 | 0.128762902 | 0.139426343 |

| ensembl_gene_id | external_gene_name | entrezgene_id | gene_biotype | baseMean | log2FoldChange | lfcSE | stat | pvalue | padj | absLog2FC |
|---|---|---|---|---|---|---|---|---|---|---|
| ENSMUSG00000032042 | Srpr | 67398 | protein_coding | 1393.289148 | 0.139427072 | 0.091726791 | -1.520025609 | 0.128504539 | 0.29575977 | 0.139427072 |
| ENSMUSG00000043987 | Cep164 | 214552 | protein_coding | 476.364828 | -0.139454471 | 0.079937016 | 1.74455437 | 0.08106247 | 0.211747172 | 0.139454471 |
| ENSMUSG00000056602 | Fry | 320365 | protein_coding | 229.3975717 | -0.139492179 | 0.115911016 | 1.203441949 | 0.228805343 | 0.437741278 | 0.139492179 |
| ENSMUSG00000022555 | Dgat1 | 13350 | protein_coding | 415.4187623 | -0.139520578 | 0.110631602 | 1.261127701 | 0.207262842 | 0.410105269 | 0.139520578 |
| ENSMUSG00000024792 | Zfpl1 | 81909 | protein_coding | 262.3323067 | 0.139533741 | 0.081925369 | -1.703181112 | 0.08853418 | 0.225974259 | 0.139533741 |
| ENSMUSG00000022056 | Adam7 | 11500 | protein_coding | 2.606074123 | 0.139548937 | 0.787496613 | -0.177205762 | 0.85934677 | 1 | 0.139548937 |
| ENSMUSG00000024993 | Dennd10 | 67894 | protein_coding | 269.248535 | 0.139564349 | 0.070992365 | -1.965906446 | 0.049309418 | 0.147459771 | 0.139564349 |
| ENSMUSG00000043036 | Ccdc63 | 330188 | protein_coding | 3.495792971 | -0.139580788 | 1.265736687 | 0.110276323 | 0.912190234 | 1 | 0.139580788 |
| ENSMUSG00000025132 | Arhgdia | 192662 | protein_coding | 2931.661644 | 0.139584217 | 0.070137561 | -1.99014929 | 0.046574492 | 0.141280523 | 0.139584217 |
| ENSMUSG00000088183 | Gm22215 | 115488890 | snRNA | 0.698573286 | -0.139586877 | 1.393595094 | 0.100163152 | 0.9202148 | 1 | 0.139586877 |
| ENSMUSG00000002395 | Use1 | 67023 | protein_coding | 815.0658561 | -0.139595577 | 0.065938042 | 2.117071922 | 0.034253744 | 0.111571269 | 0.139595577 |
| ENSMUSG00000033732 | Sf3b3 | 101943 | protein_coding | 1324.648794 | 0.139614548 | 0.092188182 | -1.514451697 | 0.129911316 | 0.29771794 | 0.139614548 |
| ENSMUSG00000022856 | Tmem41a | 66664 | protein_coding | 186.1213655 | -0.139657132 | 0.095784601 | 1.458033244 | 0.14483138 | 0.321916639 | 0.139657132 |
| ENSMUSG00000066148 | Prpf4 | 70052 | protein_coding | 280.9187884 | 0.139659853 | 0.108118498 | -1.291729502 | 0.196450837 | 0.396067528 | 0.139659853 |
| ENSMUSG00000041037 | Irgq | 210146 | protein_coding | 457.8695003 | 0.139671534 | 0.085198763 | -1.639361068 | 0.101138085 | 0.24962192 | 0.139671534 |
| ENSMUSG00000017760 | Ctsa | 19025 | protein_coding | 1597.910186 | -0.139713124 | 0.05096273 | 2.741476458 | 0.006116374 | 0.028605439 | 0.139713124 |
| ENSMUSG00000040018 | Cox15 | 226139 | protein_coding | 666.4446293 | -0.139739064 | 0.094966871 | 1.471450658 | 0.141169282 | 0.315761821 | 0.139739064 |
| ENSMUSG00000015755 | Tab2 | 68652 | protein_coding | 961.9808755 | 0.139853493 | 0.059489126 | -2.350908527 | 0.018727637 | 0.069749255 | 0.139853493 |
| ENSMUSG00000022142 | Nup155 | 170762 | protein_coding | 270.1099852 | 0.139879991 | 0.247121018 | -0.566038423 | 0.571367661 | 0.756904024 | 0.139879991 |
| ENSMUSG00000029649 | Pomp | 66537 | protein_coding | 713.8378287 | 0.139885155 | 0.073819138 | -1.89497139 | 0.058096192 | 0.166823251 | 0.139885155 |
| ENSMUSG00000078652 | Psme3 | 19192 | protein_coding | 751.575389 | 0.139915814 | 0.070000509 | -1.998782797 | 0.04563186 | 0.13918813 | 0.139915814 |
| ENSMUSG00000028788 | Ptp4a2 | 19244 | protein_coding | 5043.244296 | 0.139943567 | 0.088841205 | -1.575210141 | 0.115207947 | 0.273747344 | 0.139943567 |
| ENSMUSG00000031659 | Adcy7 | 11513 | protein_coding | 25.02913141 | 0.139949517 | 0.308342939 | -0.453876187 | 0.649917944 | 0.809917796 | 0.139949517 |
| ENSMUSG00000030921 | Trim30a | 20128 | protein_coding | 80.66718771 | -0.139957002 | 0.394690906 | 0.354599005 | 0.722890022 | 0.856549354 | 0.139957002 |
| ENSMUSG00000058656 | Samd12 | 320679 | protein_coding | 275.3088588 | 0.139988387 | 0.136367646 | -1.026551321 | 0.304631786 | 0.524795464 | 0.139988387 |
| ENSMUSG00000030685 | Kctd13 | 233877 | protein_coding | 56.45497762 | 0.140077824 | 0.162001706 | -0.864668825 | 0.387220585 | 0.60964446 | 0.140077824 |
| ENSMUSG00000031445 | Proz | 66901 | protein_coding | 20.70397448 | 0.140105451 | 0.266263353 | -0.526191268 | 0.598755321 | 0.776186935 | 0.140105451 |
| ENSMUSG00000024182 | Axin1 | 12005 | protein_coding | 710.4561102 | 0.140119047 | 0.064693516 | -2.165890129 | 0.030319584 | 0.101747126 | 0.140119047 |
| ENSMUSG00000039878 | Slc39a5 | 72002 | protein_coding | 1179.049675 | -0.140179494 | 0.080578096 | 1.739672452 | 0.081916549 | 0.213539052 | 0.140179494 |
| ENSMUSG00000030509 | Asb7 | 117589 | protein_coding | 390.0790262 | 0.140209834 | 0.087472401 | -1.602903676 | 0.108955922 | 0.26324771 | 0.140209834 |
| ENSMUSG00000038525 | Armc10 | 67211 | protein_coding | 375.7459975 | -0.140214552 | 0.117614965 | 1.19214891 | 0.233202862 | 0.442685693 | 0.140214552 |
| ENSMUSG00000022160 | Mettl3 | 56335 | protein_coding | 207.5742565 | -0.14022034 | 0.125386668 | 1.118303425 | 0.263437425 | 0.477601877 | 0.14022034 |
| ENSMUSG00000000753 | Serpinf1 | 20317 | protein_coding | 3.142259092 | 0.140223366 | 1.343423424 | -0.10437764 | 0.916869667 | 1 | 0.140223366 |
| ENSMUSG00000017747 | Ghdc | 80860 | protein_coding | 148.1797926 | -0.140225178 | 0.08639833 | 1.62300797 | 0.104587701 | 0.255796319 | 0.140225178 |
| ENSMUSG00000042377 | Fam83g | 69640 | protein_coding | 565.1230476 | -0.140242123 | 0.094410761 | 1.485446377 | 0.137425618 | 0.309858682 | 0.140242123 |
| ENSMUSG00000038280 | Ostm1 | 14628 | protein_coding | 162.591141 | -0.140289796 | 0.114013794 | 1.230463357 | 0.21852364 | 0.424602824 | 0.140289796 |
| ENSMUSG00000025420 | Katnal2 | 71206 | protein_coding | 5.264833715 | -0.140294461 | 0.660529759 | 0.212396881 | 0.831797419 | 0.915204382 | 0.140294461 |
| ENSMUSG00000032786 | Alas1 | 11655 | protein_coding | 2578.379321 | -0.140343419 | 0.127750151 | 1.098577323 | 0.271952474 | 0.487730576 | 0.140343419 |
| ENSMUSG00000090247 | Bloc1s1 | 14533 | protein_coding | 96.52255526 | -0.14034462 | 0.12746756 | 1.101022258 | 0.27088697 | 0.486449404 | 0.14034462 |

| ensembl_gene_id | external_gene_name | entrezgene_id | gene_biotype | baseMean | log2FoldChange | lfcSE | stat | pvalue | padj | absLog2FC |
|---|---|---|---|---|---|---|---|---|---|---|
| ENSMUSG00000041837 | Pdcd7 | 50996 | protein_coding | 156.8995055 | 0.140385396 | 0.095177596 | -1.474983625 | 0.140216946 | 0.314312833 | 0.140385396 |
| ENSMUSG00000075595 | Zfp652 | 268469 | protein_coding | 839.4536825 | 0.140478469 | 0.087186861 | -1.61123439 | 0.107128645 | 0.260126811 | 0.140478469 |
| ENSMUSG00000046719 | Nxph3 | 104079 | protein_coding | 4.356429966 | 0.140489111 | 0.774573933 | -0.181375987 | 0.856072466 | 1 | 0.140489111 |
| ENSMUSG00000034303 | Ccdc15 | 245902 | protein_coding | 39.47211795 | 0.140493183 | 0.197122465 | -0.712720303 | 0.47601885 | 0.687930892 | 0.140493183 |
| ENSMUSG00000037022 | Mmaa | 109136 | protein_coding | 257.2953813 | -0.140513448 | 0.076795734 | 1.829703822 | 0.067294239 | 0.185483473 | 0.140513448 |
| ENSMUSG00000031723 | Txnl4b | 234723 | protein_coding | 94.67669286 | 0.140557058 | 0.128533668 | -1.093542728 | 0.274155579 | 0.490619315 | 0.140557058 |
| ENSMUSG00000022185 | Acin1 | 56215 | protein_coding | 1511.198482 | 0.140602625 | 0.037841797 | -3.715537716 | 0.000202772 | 0.001599893 | 0.140602625 |
| ENSMUSG00000031935 | Med17 | 234959 | protein_coding | 348.382282 | -0.140626553 | 0.085391848 | 1.64683815 | 0.099591318 | 0.246796371 | 0.140626553 |
| ENSMUSG00000097277 | 2900076A07Rik | 100504421 | lncRNA | 32.10222618 | -0.140641719 | 0.223916459 | 0.628099066 | 0.529939045 | 0.727284417 | 0.140641719 |
| ENSMUSG00000036371 | Serbp1 | 66870 | protein_coding | 3247.029824 | 0.140683366 | 0.083360671 | -1.687646755 | 0.091479058 | 0.231451688 | 0.140683366 |
| ENSMUSG00000021243 | Fcf1 | 73736 | protein_coding | 253.1261959 | -0.140703634 | 0.123557202 | 1.13877323 | 0.25479775 | 0.467346556 | 0.140703634 |
| ENSMUSG00000029570 | Lfng | 16848 | protein_coding | 171.058869 | 0.140714232 | 0.164682145 | -0.854459551 | 0.392850412 | 0.615573746 | 0.140714232 |
| ENSMUSG00000000171 | Sdhd | 66925 | protein_coding | 3460.198097 | -0.140730134 | 0.086742184 | 1.622395552 | 0.104718681 | 0.255833124 | 0.140730134 |
| ENSMUSG00000009995 | Tafazzin | 66826 | protein_coding | 589.9714738 | -0.140733863 | 0.061332508 | 2.294604713 | 0.021755788 | 0.078410101 | 0.140733863 |
| ENSMUSG00000034083 | Ccdc174 | 232236 | protein_coding | 215.554625 | -0.140756955 | 0.075068015 | 1.875058969 | 0.060784611 | 0.172473724 | 0.140756955 |
| ENSMUSG00000013495 | Tmem175 | 72392 | protein_coding | 161.006676 | -0.140793103 | 0.098933786 | 1.423104359 | 0.154705905 | 0.336646899 | 0.140793103 |
| ENSMUSG00000022125 | Cln5 | 211286 | protein_coding | 318.2476164 | -0.140903338 | 0.078970991 | 1.78424173 | 0.074384394 | 0.19938276 | 0.140903338 |
| ENSMUSG00000017831 | Rab5a | 271457 | protein_coding | 849.3327004 | 0.140944967 | 0.067329922 | -2.093348147 | 0.036318088 | 0.116638837 | 0.140944967 |
| ENSMUSG00000006392 | Med8 | 80509 | protein_coding | 413.1369198 | -0.140959726 | 0.070661628 | 1.994855346 | 0.046058659 | 0.140153435 | 0.140959726 |
| ENSMUSG00000038412 | Higd1a | 56295 | protein_coding | 2563.934196 | 0.140982549 | 0.126545035 | -1.114089929 | 0.265240609 | 0.480125333 | 0.140982549 |
| ENSMUSG00000043602 | Zfp3 | 193043 | protein_coding | 43.78835612 | 0.140985199 | 0.197850465 | -0.712584619 | 0.476102832 | 0.687930892 | 0.140985199 |
| ENSMUSG00000017299 | Dnttip1 | 76233 | protein_coding | 230.3419712 | 0.140996855 | 0.078684754 | -1.791920896 | 0.073145638 | 0.197148208 | 0.140996855 |
| ENSMUSG00000038240 | Pdss2 | 71365 | protein_coding | 407.2387427 | -0.141035191 | 0.073309791 | 1.923824763 | 0.054376552 | 0.158736855 | 0.141035191 |
| ENSMUSG00000055612 | Cdca7 | 66953 | protein_coding | 432.6264585 | -0.141036577 | 0.123178121 | 1.144980747 | 0.252217133 | 0.464617944 | 0.141036577 |
| ENSMUSG00000044709 | Gemin7 | 69731 | protein_coding | 227.4774152 | -0.141037557 | 0.078928373 | 1.786905655 | 0.073952738 | 0.198739711 | 0.141037557 |
| ENSMUSG00000026817 | Ak1 | 11636 | protein_coding | 13.00955309 | -0.141043282 | 0.369419328 | 0.381797246 | 0.702611761 | 0.843113692 | 0.141043282 |
| ENSMUSG00000035847 | Ids | 15931 | protein_coding | 324.9353823 | -0.141055641 | 0.113566731 | 1.242050732 | 0.214217845 | 0.419245495 | 0.141055641 |
| ENSMUSG00000005699 | Pard6a | 56513 | protein_coding | 44.79860983 | -0.141056481 | 0.164174225 | 0.859187734 | 0.39023695 | 0.612497263 | 0.141056481 |
| ENSMUSG00000039238 | Zfp750 | 319530 | protein_coding | 101.8290454 | -0.141065027 | 0.170217859 | 0.828732236 | 0.407255941 | 0.628133427 | 0.141065027 |
| ENSMUSG00000059149 | Mfsd4a | 213006 | protein_coding | 1756.262415 | -0.141133389 | 0.180633196 | 0.78132587 | 0.434610858 | 0.652884901 | 0.141133389 |
| ENSMUSG00000045078 | Rnf216 | 108086 | protein_coding | 469.7915932 | -0.141149551 | 0.072379179 | 1.95014026 | 0.051159404 | 0.151705224 | 0.141149551 |
| ENSMUSG00000073725 | Lmbrd1 | 68421 | protein_coding | 645.0474803 | -0.141151239 | 0.076643389 | 1.841662281 | 0.065524565 | 0.181983293 | 0.141151239 |
| ENSMUSG00000073062 | Zxdb | 668166 | protein_coding | 49.24936466 | -0.141200252 | 0.187390343 | 0.753508688 | 0.451144292 | 0.667666133 | 0.141200252 |
| ENSMUSG00000040725 | Hnrnpul1 | 232989 | protein_coding | 2018.114763 | 0.141207473 | 0.044886277 | -3.145894092 | 0.0016558 | 0.009689537 | 0.141207473 |
| ENSMUSG00000065581 | Mir194-1 | 387189 | miRNA | 2.322930858 | 0.141232963 | 0.693096651 | -0.20377095 | 0.838532495 | 1 | 0.141232963 |
| ENSMUSG00000022210 | Dhrs4 | 28200 | protein_coding | 402.4074178 | -0.141263478 | 0.097334115 | 1.45132545 | 0.146689258 | 0.324504542 | 0.141263478 |
| ENSMUSG00000061371 | Zfp873 | 408062 | protein_coding | 64.44908187 | -0.141345295 | 0.136284883 | 1.037131136 | 0.299674746 | 0.519697021 | 0.141345295 |
| ENSMUSG00000025786 | Zdhhc3 | 69035 | protein_coding | 1483.44268 | 0.141354793 | 0.064588791 | -2.188534423 | 0.028630697 | 0.097285458 | 0.141354793 |
| ENSMUSG00000021572 | Cep72 | 74470 | protein_coding | 77.22189387 | 0.141376907 | 0.134537834 | -1.050833823 | 0.293334918 | 0.512561987 | 0.141376907 |

| ensembl_gene_id | external_gene_name | entrezgene_id | gene_biotype | baseMean | log2FoldChange | lfcSE | stat | pvalue | padj | absLog2FC |
|---|---|---|---|---|---|---|---|---|---|---|
| ENSMUSG00000030796 | Tead2 | 21677 | protein_coding | 45.57470188 | 0.141407164 | 0.182734916 | -0.77383768 | 0.439026787 | 0.65700846 | 0.141407164 |
| ENSMUSG00000042772 | Smg7 | 226517 | protein_coding | 870.4462135 | 0.141418787 | 0.064592241 | -2.189408261 | 0.02856718 | 0.097089608 | 0.141418787 |
| ENSMUSG00000118522 | Gm31160 | 102633301 | protein_coding | 20.51492307 | 0.141441557 | 0.316404943 | -0.447027015 | 0.654855555 | 0.812734264 | 0.141441557 |
| ENSMUSG00000024479 | Mal2 | 105853 | protein_coding | 3444.332555 | -0.141463849 | 0.077119061 | 1.834356479 | 0.066601095 | 0.184167865 | 0.141463849 |
| ENSMUSG00000047443 | Erfe | 227358 | protein_coding | 92.94328302 | 0.141465272 | 0.128251508 | -1.103030089 | 0.270014099 | 0.485271256 | 0.141465272 |
| ENSMUSG00000032816 | Igdcc4 | 56741 | protein_coding | 899.2614112 | 0.141467818 | 0.089213672 | -1.585719045 | 0.112803057 | 0.269506609 | 0.141467818 |
| ENSMUSG00000032782 | Cntrob | 216846 | protein_coding | 162.6974529 | 0.141473446 | 0.110445825 | -1.280930682 | 0.200218015 | 0.401310875 | 0.141473446 |
| ENSMUSG00000018347 | Zkscan6 | 52712 | protein_coding | 224.7153379 | -0.141489984 | 0.085149977 | 1.66165616 | 0.096581736 | 0.240948524 | 0.141489984 |
| ENSMUSG00000050919 | Zfp366 | 238803 | protein_coding | 2.465416625 | -0.141519751 | 1.742673467 | 0.081208416 | 0.935276207 | 1 | 0.141519751 |
| ENSMUSG00000029797 | Sspo | 243369 | protein_coding | 83.30183518 | -0.141532377 | 0.243862574 | 0.580377605 | 0.561660004 | 0.750060917 | 0.141532377 |
| ENSMUSG00000053158 | Fes | 14159 | protein_coding | 112.9389638 | -0.141604108 | 0.111680895 | 1.267934935 | 0.204821187 | 0.40732383 | 0.141604108 |
| ENSMUSG00000024621 | Csf1r | 12978 | protein_coding | 4.959883036 | -0.141634131 | 0.615831419 | 0.229988477 | 0.818100723 | 0.908812719 | 0.141634131 |
| ENSMUSG00000062028 | Irgc1 | 210145 | protein_coding | 5.620827915 | 0.141659904 | 0.556567578 | -0.254524175 | 0.79909063 | 0.900333128 | 0.141659904 |
| ENSMUSG00000064181 | Rab3ip | 216363 | protein_coding | 393.05894 | -0.1416793 | 0.08838432 | 1.60299134 | 0.108936566 | 0.26324771 | 0.1416793 |
| ENSMUSG00000029513 | Prkab1 | 19079 | protein_coding | 1376.859034 | -0.141688938 | 0.088804556 | 1.595514279 | 0.11059728 | 0.265929452 | 0.141688938 |
| ENSMUSG00000021068 | Nin | 18080 | protein_coding | 83.06623531 | -0.141710199 | 0.149286281 | 0.949251317 | 0.342492807 | 0.564520034 | 0.141710199 |
| ENSMUSG00000019852 | Arfgef3 | 215821 | protein_coding | 948.7429017 | 0.141728017 | 0.176869343 | -0.801314771 | 0.422949442 | 0.64285201 | 0.141728017 |
| ENSMUSG00000040721 | Zfhx2 | 239102 | protein_coding | 71.6515451 | 0.141730664 | 0.323616113 | -0.437959231 | 0.661415838 | 0.816921981 | 0.141730664 |
| ENSMUSG00000077042 | Mir574 | 100124451 | miRNA | 1.675876616 | 0.141751176 | 1.112935752 | -0.127366899 | 0.898650014 | 1 | 0.141751176 |
| ENSMUSG00000020522 | Mfap3 | 216760 | protein_coding | 288.7674094 | 0.141790006 | 0.076611873 | -1.850757601 | 0.064204433 | 0.179403047 | 0.141790006 |
| ENSMUSG00000038591 | Colec10 | 239447 | protein_coding | 1.008547246 | -0.141851686 | 2.073177533 | 0.068422354 | 0.945449428 | 1 | 0.141851686 |
| ENSMUSG00000031683 | Lsm6 | 78651 | protein_coding | 270.7683395 | 0.141894095 | 0.099761042 | -1.422339742 | 0.154927647 | 0.336996201 | 0.141894095 |
| ENSMUSG00000045624 | Esf1 | 66580 | protein_coding | 244.5976547 | 0.141947683 | 0.135216016 | -1.049784535 | 0.293817187 | 0.513069369 | 0.141947683 |
| ENSMUSG00000033763 | Mtss2 | 244654 | protein_coding | 10.02930454 | 0.141971894 | 0.86435432 | -0.164251963 | 0.869532796 | 0.935632034 | 0.141971894 |
| ENSMUSG00000030393 | Zik1 | 22775 | protein_coding | 4.115317907 | 0.141982838 | 0.754099517 | -0.188281301 | 0.850656143 | 1 | 0.141982838 |
| ENSMUSG00000101942 | Gm19582 | 100503175 | lncRNA | 3.09434699 | 0.141992706 | 0.5601784 | -0.253477653 | 0.799899128 | 1 | 0.141992706 |
| ENSMUSG00000068856 | Sf3b4 | 107701 | protein_coding | 163.2557641 | 0.142005456 | 0.138020289 | -1.028873776 | 0.303538993 | 0.52400616 | 0.142005456 |
| ENSMUSG00000078815 | Cacng6 | 54378 | protein_coding | 0.61496024 | 0.142065916 | 1.416521024 | -0.100292134 | 0.920112402 | 1 | 0.142065916 |
| ENSMUSG00000027998 | Plrg1 | 53317 | protein_coding | 468.0823841 | 0.142101763 | 0.076195547 | -1.864961507 | 0.0621868 | 0.17503883 | 0.142101763 |
| ENSMUSG00000042258 | Isl1 | 16392 | protein_coding | 8.982573981 | 0.142105515 | 0.48864653 | -0.290814538 | 0.771193166 | 0.885199039 | 0.142105515 |
| ENSMUSG00000006641 | Slc5a6 | 330064 | protein_coding | 358.654102 | -0.142106661 | 0.18346632 | 0.774565386 | 0.438596517 | 0.656764359 | 0.142106661 |
| ENSMUSG00000036352 | Ubac1 | 98766 | protein_coding | 432.8361834 | -0.142113898 | 0.084908188 | 1.673736087 | 0.094182478 | 0.236336338 | 0.142113898 |
| ENSMUSG00000036751 | Cox6b1 | 110323 | protein_coding | 4449.02711 | -0.142115531 | 0.066601503 | 2.133818674 | 0.032857627 | 0.10826234 | 0.142115531 |
| ENSMUSG00000000402 | Egfl6 | 54156 | protein_coding | 2.687023648 | -0.142166392 | 0.775743431 | 0.183264706 | 0.854590326 | 1 | 0.142166392 |
| ENSMUSG00000057219 | Armc7 | 276905 | protein_coding | 72.83403963 | 0.14216663 | 0.140772545 | -1.009903109 | 0.312541713 | 0.533579433 | 0.14216663 |
| ENSMUSG00000047412 | Zbtb44 | 235132 | protein_coding | 482.601323 | -0.142193071 | 0.135777302 | 1.047252145 | 0.294983301 | 0.514127096 | 0.142193071 |
| ENSMUSG00000031207 | Msn | 17698 | protein_coding | 27.04623497 | 0.142201588 | 0.322835685 | -0.440476671 | 0.659591905 | 0.8161622 | 0.142201588 |
| ENSMUSG00000041215 | Yeats2 | 208146 | protein_coding | 421.0575941 | 0.142222496 | 0.081534814 | -1.744316184 | 0.081103971 | 0.21178858 | 0.142222496 |
| ENSMUSG00000030779 | Rbbp6 | 19647 | protein_coding | 633.7332516 | 0.142265703 | 0.0678487 | -2.096808082 | 0.036010568 | 0.115831422 | 0.142265703 |

Supplementary Table 1

| ensembl_gene_id | external_gene_name | entrezgene_id | gene_biotype | baseMean | log2FoldChange | lfcSE | stat | pvalue | padj | absLog2FC |
|---|---|---|---|---|---|---|---|---|---|---|
| ENSMUSG00000045917 | Tmem268 | 230279 | protein_coding | 379.6886918 | 0.142276882 | 0.10371607 | -1.371792077 | 0.170128174 | 0.359776754 | 0.142276882 |
| ENSMUSG00000072945 | Ripply1 | 622473 | protein_coding | 8.667513059 | -0.142283558 | 0.497804714 | 0.285822038 | 0.775014428 | 0.887804402 | 0.142283558 |
| ENSMUSG00000039385 | Cdh6 | 12563 | protein_coding | 2.304618159 | 0.142284816 | 1.118623878 | -0.127196298 | 0.898785035 | 1 | 0.142284816 |
| ENSMUSG00000028576 | Ift74 | 67694 | protein_coding | 100.6256802 | 0.142297601 | 0.1331843 | -1.068426235 | 0.285328288 | 0.503786095 | 0.142297601 |
| ENSMUSG00000020780 | Srp68 | 217337 | protein_coding | 640.884532 | 0.142314847 | 0.067712848 | -2.101740677 | 0.035575998 | 0.114836207 | 0.142314847 |
| ENSMUSG00000027318 | Adam33 | 110751 | protein_coding | 2.199383616 | -0.142347903 | 1.48833061 | 0.095642663 | 0.92380438 | 1 | 0.142347903 |
| ENSMUSG00000020409 | Slu7 | 193116 | protein_coding | 690.6729968 | -0.142355965 | 0.073605159 | 1.934048728 | 0.053107137 | 0.156162974 | 0.142355965 |
| ENSMUSG00000027164 | Traf6 | 22034 | protein_coding | 399.9330407 | 0.142434844 | 0.079328628 | -1.795503697 | 0.072573484 | 0.196079582 | 0.142434844 |
| ENSMUSG00000036916 | Zfp280c | 208968 | protein_coding | 95.05042106 | -0.142443001 | 0.13878245 | 1.026376177 | 0.304714303 | 0.524882862 | 0.142443001 |
| ENSMUSG00000040822 | 1700123O20Rik | 58248 | protein_coding | 319.2678475 | 0.142486438 | 0.082300405 | -1.731296923 | 0.08339882 | 0.216412337 | 0.142486438 |
| ENSMUSG00000026554 | Dcaf8 | 98193 | protein_coding | 1267.788483 | -0.1425366 | 0.06185844 | 2.30423852 | 0.021209258 | 0.076884085 | 0.1425366 |
| ENSMUSG00000047786 | Lix1 | 66643 | protein_coding | 7.89025562 | 0.14256569 | 0.474019563 | -0.300759084 | 0.763598211 | 0.881390987 | 0.14256569 |
| ENSMUSG00000058298 | Mcm9 | 71567 | protein_coding | 175.086336 | -0.142595973 | 0.09728278 | 1.46578843 | 0.142705934 | 0.318422618 | 0.142595973 |
| ENSMUSG00000025956 | Mettl21a | 67099 | protein_coding | 92.11944108 | 0.142637372 | 0.142150783 | -1.003423047 | 0.315656789 | 0.53655334 | 0.142637372 |
| ENSMUSG00000055639 | Dach1 | 13134 | protein_coding | 31.74633781 | -0.142707267 | 0.225449822 | 0.632989044 | 0.526740798 | 0.725185288 | 0.142707267 |
| ENSMUSG00000040044 | Orc3 | 50793 | protein_coding | 310.357224 | -0.142856973 | 0.075627205 | 1.888962756 | 0.058896819 | 0.168536142 | 0.142856973 |
| ENSMUSG00000024082 | Ndufaf7 | 73694 | protein_coding | 222.7307749 | -0.142949252 | 0.101564498 | 1.407472649 | 0.159287282 | 0.343403417 | 0.142949252 |
| ENSMUSG00000070371 | Prss36 | 77613 | protein_coding | 26.06666559 | 0.142994366 | 0.205609389 | -0.69546613 | 0.48676323 | 0.695624496 | 0.142994366 |
| ENSMUSG00000030272 | Camk1 | 52163 | protein_coding | 238.3051859 | -0.143037327 | 0.107500965 | 1.330567847 | 0.183331248 | 0.37815791 | 0.143037327 |
| ENSMUSG00000028599 | Tnfrsf1b | 21938 | protein_coding | 292.2581752 | -0.143042196 | 0.13941355 | 1.026027933 | 0.304878418 | 0.525029915 | 0.143042196 |
| ENSMUSG00000022752 | Tomm70a | 28185 | protein_coding | 1477.672194 | 0.143057634 | 0.081543007 | -1.754382627 | 0.079364966 | 0.208654479 | 0.143057634 |
| ENSMUSG00000026589 | Sec16b | 89867 | protein_coding | 458.8487622 | -0.143068665 | 0.07352421 | 1.9458715 | 0.051670177 | 0.152917797 | 0.143068665 |
| ENSMUSG00000032842 | Abcc10 | 224814 | protein_coding | 352.9141666 | 0.143115349 | 0.133299885 | -1.073634453 | 0.28298655 | 0.50115173 | 0.143115349 |
| ENSMUSG00000036198 | Arhgap36 | 75404 | protein_coding | 2.230688868 | 0.143120635 | 1.376718714 | -0.10395779 | 0.917202846 | 1 | 0.143120635 |
| ENSMUSG00000026131 | Dst | 13518 | protein_coding | 3941.365959 | -0.143124414 | 0.160136193 | 0.893766806 | 0.371446673 | 0.593868753 | 0.143124414 |
| ENSMUSG00000086186 | Ccdc34os | 75517 | lncRNA | 1.163777638 | 0.143220052 | 1.995832425 | -0.071759558 | 0.942793258 | 1 | 0.143220052 |
| ENSMUSG00000029735 | Tpk1 | 29807 | protein_coding | 359.0778597 | -0.143236148 | 0.074583186 | 1.920488469 | 0.054796228 | 0.15967971 | 0.143236148 |
| ENSMUSG00000033773 | Rpap2 | 231571 | protein_coding | 146.7359965 | -0.143280586 | 0.097651847 | 1.467259367 | 0.142305513 | 0.31779121 | 0.143280586 |
| ENSMUSG00000025324 | Atp10a | 11982 | protein_coding | 19.57432217 | 0.143440679 | 0.361937386 | -0.39631352 | 0.691873755 | 0.836060529 | 0.143440679 |
| ENSMUSG00000021374 | Nup153 | 218210 | protein_coding | 708.6936044 | 0.143447551 | 0.061661759 | -2.326361649 | 0.019999266 | 0.073392861 | 0.143447551 |
| ENSMUSG00000000148 | Brat1 | 231841 | protein_coding | 218.5106924 | 0.143462782 | 0.117505476 | -1.220902953 | 0.222122766 | 0.429222486 | 0.143462782 |
| ENSMUSG00000030805 | Stx4a | 20909 | protein_coding | 294.3675415 | -0.14346539 | 0.117002914 | 1.226169374 | 0.220134944 | 0.426485613 | 0.14346539 |
| ENSMUSG00000060166 | Zdhhc8 | 27801 | protein_coding | 77.79322777 | -0.143472067 | 0.158974831 | 0.902482902 | 0.3668004 | 0.589522317 | 0.143472067 |
| ENSMUSG00000040009 | Gnaz | 14687 | protein_coding | 9.521500254 | 0.143576367 | 0.367412279 | -0.39077727 | 0.695961878 | 0.838607402 | 0.143576367 |
| ENSMUSG00000028693 | Nasp | 50927 | protein_coding | 428.0915598 | -0.143610408 | 0.115335472 | 1.245153858 | 0.2130752 | 0.417801455 | 0.143610408 |
| ENSMUSG00000039545 | Flicr | 78185 | lncRNA | 2.190747627 | 0.143624926 | 1.233363121 | -0.11644983 | 0.907296045 | 1 | 0.143624926 |
| ENSMUSG00000039156 | Stim2 | 116873 | protein_coding | 548.971765 | -0.143645678 | 0.126076021 | 1.139357643 | 0.254554015 | 0.467148794 | 0.143645678 |
| ENSMUSG00000041797 | Abca9 | 217262 | protein_coding | 3.530862016 | -0.143669806 | 1.55725913 | 0.092258124 | 0.926492958 | 1 | 0.143669806 |
| ENSMUSG00000026072 | Il1r1 | 16177 | protein_coding | 304.532062 | 0.143679373 | 0.136405702 | -1.053323809 | 0.292192611 | 0.511151353 | 0.143679373 |

| ensembl_gene_id | external_gene_name | entrezgene_id | gene_biotype | baseMean | log2FoldChange | lfcSE | stat | pvalue | padj | absLog2FC |
|---|---|---|---|---|---|---|---|---|---|---|
| ENSMUSG00000034871 | Fam151a | 230579 | protein_coding | 0.787522863 | -0.143795386 | 1.95826814 | 0.073429876 | 0.941464044 | 1 | 0.143795386 |
| ENSMUSG00000038646 | Ramac | 67148 | protein_coding | 295.1300081 | -0.143809191 | 0.118673416 | 1.211806279 | 0.225586544 | 0.433049501 | 0.143809191 |
| ENSMUSG00000003555 | Cyp17a1 | 13074 | protein_coding | 2.035742069 | -0.14384171 | 1.219562726 | 0.117945315 | 0.906110987 | 1 | 0.14384171 |
| ENSMUSG00000018604 | Tbx3 | 21386 | protein_coding | 162.7735875 | 0.143844022 | 0.27359172 | -0.525761605 | 0.599053854 | 0.776435029 | 0.143844022 |
| ENSMUSG00000097006 | 9530082P21Rik | 638247 | lncRNA | 101.3756101 | 0.143875865 | 0.175474822 | -0.819923127 | 0.412259932 | 0.632329199 | 0.143875865 |
| ENSMUSG00000038235 | F11r | 16456 | protein_coding | 2532.069226 | -0.143913059 | 0.059369127 | 2.424038644 | 0.015348972 | 0.059503296 | 0.143913059 |
| ENSMUSG00000038909 | Kat7 | 217127 | protein_coding | 892.1738583 | 0.143916953 | 0.061274025 | -2.34874326 | 0.01883689 | 0.070067307 | 0.143916953 |
| ENSMUSG00000028479 | Gne | 50798 | protein_coding | 6891.926151 | -0.143961874 | 0.137726902 | 1.045270541 | 0.295897949 | 0.515123206 | 0.143961874 |
| ENSMUSG00000038034 | Igsf8 | 140559 | protein_coding | 340.3466292 | -0.144008091 | 0.096678554 | 1.489555694 | 0.136341099 | 0.308134243 | 0.144008091 |
| ENSMUSG00000028409 | Smu1 | 74255 | protein_coding | 969.6134858 | 0.144013782 | 0.082398229 | -1.747777632 | 0.08050254 | 0.210915627 | 0.144013782 |
| ENSMUSG00000020283 | Pex13 | 72129 | protein_coding | 411.1283689 | -0.14403403 | 0.061437597 | 2.344395567 | 0.019057947 | 0.070692483 | 0.14403403 |
| ENSMUSG00000027550 | Lrrcc1 | 71710 | protein_coding | 206.3337199 | -0.144054457 | 0.15035799 | 0.958076499 | 0.338024185 | 0.559780524 | 0.144054457 |
| ENSMUSG00000000154 | Slc22a18 | 18400 | protein_coding | 934.0731162 | -0.14407841 | 0.102138226 | 1.410621811 | 0.158356157 | 0.342200154 | 0.14407841 |
| ENSMUSG00000047013 | Fbxo41 | 330369 | protein_coding | 4.583202195 | 0.144108044 | 1.072553622 | -0.134359757 | 0.893118102 | 0.947328174 | 0.144108044 |
| ENSMUSG00000002379 | Ndufa11 | 69875 | protein_coding | 868.9266312 | 0.144117667 | 0.090984844 | -1.583974444 | 0.113199535 | 0.270127107 | 0.144117667 |
| ENSMUSG00000036938 | Try5 | 103964 | protein_coding | 0.688408279 | 0.144117814 | 2.165730331 | -0.066544672 | 0.946944194 | 1 | 0.144117814 |
| ENSMUSG00000003068 | Stk11 | 20869 | protein_coding | 917.3606012 | 0.144135979 | 0.068535762 | -2.103076922 | 0.035459046 | 0.114637957 | 0.144135979 |
| ENSMUSG00000101174 | Hoxd4 | 15436 | protein_coding | 24.44118061 | -0.144141378 | 0.351213407 | 0.410409669 | 0.681505455 | 0.829602026 | 0.144141378 |
| ENSMUSG00000031887 | Tradd | 71609 | protein_coding | 875.0650637 | -0.144168096 | 0.084967061 | 1.69675278 | 0.089743409 | 0.228322063 | 0.144168096 |
| ENSMUSG00000073043 | Atoh1 | 11921 | protein_coding | 975.0809339 | 0.144182762 | 0.222347895 | -0.648455708 | 0.516690249 | 0.717449153 | 0.144182762 |
| ENSMUSG00000029559 | 2210016L21Rik | 72357 | protein_coding | 279.1918441 | -0.144208112 | 0.076123348 | 1.894400553 | 0.058171863 | 0.166982469 | 0.144208112 |
| ENSMUSG00000002825 | Qtrt1 | 60507 | protein_coding | 141.0349038 | -0.14422675 | 0.123971931 | 1.163382305 | 0.244674425 | 0.456250391 | 0.14422675 |
| ENSMUSG00000025354 | Dnajc14 | 74330 | protein_coding | 1087.44961 | -0.144247371 | 0.043248785 | 3.33529305 | 0.000852096 | 0.00556186 | 0.144247371 |
| ENSMUSG00000023456 | Tpi1 | 21991 | protein_coding | 5783.675939 | -0.144261699 | 0.057728197 | 2.498981538 | 0.01245508 | 0.050511589 | 0.144261699 |
| ENSMUSG00000020474 | Polm | 54125 | protein_coding | 123.1791818 | -0.144300109 | 0.107111362 | 1.347197039 | 0.177916782 | 0.370319855 | 0.144300109 |
| ENSMUSG00000031657 | Heatr3 | 234549 | protein_coding | 235.2878471 | -0.144328026 | 0.150973927 | 0.955979809 | 0.339082435 | 0.560532006 | 0.144328026 |
| ENSMUSG00000034312 | Iqsec1 | 232227 | protein_coding | 281.1373398 | 0.144330854 | 0.140075692 | -1.030377593 | 0.302832788 | 0.5233341 | 0.144330854 |
| ENSMUSG00000053574 | 4930563E22Rik | 75304 | protein_coding | 4.053925356 | -0.144348937 | 0.685159021 | 0.210679466 | 0.833137399 | 1 | 0.144348937 |
| ENSMUSG00000005378 | Bud23 | 66138 | protein_coding | 341.3918117 | 0.144365477 | 0.085893721 | -1.680745404 | 0.092812377 | 0.233822059 | 0.144365477 |
| ENSMUSG00000051391 | Ywhag | 22628 | protein_coding | 766.9862369 | -0.144399607 | 0.08757981 | 1.648777355 | 0.099193256 | 0.246105382 | 0.144399607 |
| ENSMUSG00000032080 | Apoa4 | 11808 | protein_coding | 19.15411222 | 0.144449811 | 1.106416613 | -0.130556437 | 0.896126206 | 0.948996349 | 0.144449811 |
| ENSMUSG00000054720 | Lrrc8c | 100604 | protein_coding | 9.6039644 | -0.144470543 | 0.463549431 | 0.311661569 | 0.755297738 | 0.877032235 | 0.144470543 |
| ENSMUSG00000043059 | Zfp513 | 101023 | protein_coding | 219.4775546 | 0.144512445 | 0.110764579 | -1.304681028 | 0.192001484 | 0.389807821 | 0.144512445 |
| ENSMUSG00000055652 | Klhl25 | 207952 | protein_coding | 211.2225385 | 0.144607202 | 0.089739845 | -1.611404623 | 0.10709156 | 0.260075004 | 0.144607202 |
| ENSMUSG00000063646 | Jakmip1 | 76071 | protein_coding | 117.2701958 | 0.14462592 | 0.107902402 | -1.340340138 | 0.180134788 | 0.37375875 | 0.14462592 |
| ENSMUSG00000033161 | Atp1a1 | 11928 | protein_coding | 30040.49394 | -0.144645557 | 0.128485172 | 1.125776271 | 0.26026023 | 0.473946662 | 0.144645557 |
| ENSMUSG00000079477 | Rab7 | 19349 | protein_coding | 1138.312815 | -0.14464811 | 0.057066041 | 2.534749354 | 0.011252781 | 0.046796074 | 0.14464811 |
| ENSMUSG00000007379 | Dennd2c | 329727 | protein_coding | 24.28135563 | 0.144681679 | 0.231642197 | -0.624591205 | 0.532239393 | 0.728866518 | 0.144681679 |
| ENSMUSG00000026743 | Mllt10 | 17354 | protein_coding | 494.2928466 | -0.144725416 | 0.057847741 | 2.501833499 | 0.012355202 | 0.050180472 | 0.144725416 |

| ensembl_gene_id | external_gene_name | entrezgene_id | gene_biotype | baseMean | log2FoldChange | lfcSE | stat | pvalue | padj | absLog2FC |
|---|---|---|---|---|---|---|---|---|---|---|
| ENSMUSG00000041886 | Macc1 | 238455 | protein_coding | 83.5106868 | -0.144772081 | 0.204508427 | 0.707902765 | 0.479005642 | 0.690541222 | 0.144772081 |
| ENSMUSG00000018160 | Med1 | 19014 | protein_coding | 839.5223917 | 0.144795428 | 0.045905495 | -3.154206894 | 0.001609349 | 0.009447848 | 0.144795428 |
| ENSMUSG00000075543 | Urad | 231903 | protein_coding | 16.55649068 | 0.144860183 | 0.29932529 | -0.483955709 | 0.628417303 | 0.796934675 | 0.144860183 |
| ENSMUSG00000075040 | Zfp408 | 381410 | protein_coding | 176.5478492 | -0.144883171 | 0.118768815 | 1.219875528 | 0.222512064 | 0.429533455 | 0.144883171 |
| ENSMUSG00000038425 | Poli | 26447 | protein_coding | 67.58444083 | -0.144899832 | 0.14575136 | 0.994157672 | 0.320146112 | 0.541384622 | 0.144899832 |
| ENSMUSG00000024006 | Stk38 | 106504 | protein_coding | 956.1633139 | -0.14490986 | 0.05486962 | 2.640985289 | 0.008266531 | 0.036559585 | 0.14490986 |
| ENSMUSG00000025968 | Ndufs1 | 227197 | protein_coding | 2111.139465 | -0.145002567 | 0.075210849 | 1.927947477 | 0.05386166 | 0.157763648 | 0.145002567 |
| ENSMUSG00000028991 | Mtor | 56717 | protein_coding | 963.0921367 | 0.145045602 | 0.053801653 | -2.695932074 | 0.007019198 | 0.031967742 | 0.145045602 |
| ENSMUSG00000078599 | Skint8 | 639774 | protein_coding | 1.809730397 | -0.14509077 | 1.423772282 | 0.101905882 | 0.918831381 | 1 | 0.14509077 |
| ENSMUSG00000037486 | Asxl2 | 75302 | protein_coding | 642.0545228 | -0.145105092 | 0.060221892 | 2.409507356 | 0.015974074 | 0.061433595 | 0.145105092 |
| ENSMUSG00000022789 | Dnm1l | 74006 | protein_coding | 1131.267453 | 0.145106225 | 0.065511489 | -2.214973691 | 0.02676187 | 0.092148775 | 0.145106225 |
| ENSMUSG00000035278 | Plekhj1 | 78670 | protein_coding | 215.0580731 | -0.1451095 | 0.07491124 | 1.937085803 | 0.052734856 | 0.155400323 | 0.1451095 |
| ENSMUSG00000070733 | Fryl | 72313 | protein_coding | 3009.082645 | 0.145122104 | 0.128981337 | -1.125140331 | 0.260529573 | 0.474068167 | 0.145122104 |
| ENSMUSG00000015467 | Egfl8 | 81701 | protein_coding | 10.17254917 | -0.145140361 | 0.414269331 | 0.350352658 | 0.726074051 | 0.858172551 | 0.145140361 |
| ENSMUSG00000001665 | Gstt3 | 103140 | protein_coding | 587.8363315 | 0.14516053 | 0.114663595 | -1.265968771 | 0.205524263 | 0.408230416 | 0.14516053 |
| ENSMUSG00000032034 | Kcnj5 | 16521 | protein_coding | 3.725660928 | 0.145207187 | 1.301002893 | -0.111611733 | 0.911131269 | 1 | 0.145207187 |
| ENSMUSG00000024561 | Mbd1 | 17190 | protein_coding | 758.3660209 | -0.145259016 | 0.070837271 | 2.050601515 | 0.040305769 | 0.126662125 | 0.145259016 |
| ENSMUSG00000057193 | Slc44a2 | 68682 | protein_coding | 345.6675785 | -0.145434454 | 0.108034029 | 1.346191156 | 0.178240877 | 0.37057407 | 0.145434454 |
| ENSMUSG00000047414 | Flrt2 | 399558 | protein_coding | 22.19431058 | 0.145454923 | 0.24006527 | -0.605897403 | 0.544582884 | 0.736796542 | 0.145454923 |
| ENSMUSG00000026987 | Baz2b | 407823 | protein_coding | 728.7810802 | -0.145466887 | 0.100935726 | 1.441183341 | 0.149532894 | 0.328900667 | 0.145466887 |
| ENSMUSG00000039318 | Rab3gap2 | 98732 | protein_coding | 700.2376524 | 0.145483117 | 0.059477557 | -2.446017053 | 0.014444417 | 0.056578534 | 0.145483117 |
| ENSMUSG00000096917 | 2500002B13Rik | 78353 | lncRNA | 14.80205201 | -0.145522167 | 0.49644003 | 0.293131412 | 0.769421707 | 0.884482916 | 0.145522167 |
| ENSMUSG00000024672 | Ms4a7 | 109225 | protein_coding | 11.5790626 | -0.145547021 | 0.366630641 | 0.396985425 | 0.691378209 | 0.835769466 | 0.145547021 |
| ENSMUSG00000021548 | Ccnh | 66671 | protein_coding | 154.3866665 | 0.145652888 | 0.116520996 | -1.250014106 | 0.211294394 | 0.415642124 | 0.145652888 |
| ENSMUSG00000049680 | Urgcp | 72046 | protein_coding | 997.8816612 | 0.145671134 | 0.077984032 | -1.86796104 | 0.061767502 | 0.174214948 | 0.145671134 |
| ENSMUSG00000001173 | Ocrl | 320634 | protein_coding | 280.1648003 | -0.145677095 | 0.098986379 | 1.47168829 | 0.141105071 | 0.315703719 | 0.145677095 |
| ENSMUSG00000020521 | Rnft1 | 76892 | protein_coding | 500.9125185 | 0.145684672 | 0.066518939 | -2.190123203 | 0.028515303 | 0.096986852 | 0.145684672 |
| ENSMUSG00000040525 | Cblc | 80794 | protein_coding | 902.5919015 | -0.145736187 | 0.079461944 | 1.834037535 | 0.066648423 | 0.184267881 | 0.145736187 |
| ENSMUSG00000029538 | Srsf9 | 108014 | protein_coding | 332.6248162 | 0.145772993 | 0.089717068 | -1.624807817 | 0.104203515 | 0.255086991 | 0.145772993 |
| ENSMUSG00000039481 | Nrtn | 18188 | protein_coding | 15.69840363 | 0.145798939 | 0.304865423 | -0.478240325 | 0.632479164 | 0.799512805 | 0.145798939 |
| ENSMUSG00000117011 | E130008D07Rik | 545207 | lncRNA | 5.663770056 | -0.145818592 | 0.592449192 | 0.246128434 | 0.805582813 | 0.904254429 | 0.145818592 |
| ENSMUSG00000009145 | Dqx1 | 93838 | protein_coding | 512.9304682 | -0.145964937 | 0.10257782 | 1.422967814 | 0.154745486 | 0.336688663 | 0.145964937 |
| ENSMUSG00000066551 | Hmgb1 | 15289 | protein_coding | 1435.588607 | 0.145968861 | 0.143890358 | -1.01444505 | 0.310370437 | 0.53139398 | 0.145968861 |
| ENSMUSG00000073927 | Or52h7 | 259050 | protein_coding | 2.055307622 | -0.146031743 | 1.298845839 | 0.112431929 | 0.91048094 | 1 | 0.146031743 |
| ENSMUSG00000020708 | Psmc5 | 19184 | protein_coding | 797.6732475 | 0.14603488 | 0.090325639 | -1.616760006 | 0.105930096 | 0.258135524 | 0.14603488 |
| ENSMUSG00000063281 | Zfp35 | 22694 | protein_coding | 141.1866167 | 0.146047842 | 0.120572967 | -1.211281812 | 0.225787417 | 0.433362785 | 0.146047842 |
| ENSMUSG00000032199 | Polr2m | 28015 | protein_coding | 1141.664073 | 0.14604897 | 0.073344328 | -1.991278318 | 0.046450298 | 0.141033318 | 0.14604897 |
| ENSMUSG00000022774 | Ncbp2 | 68092 | protein_coding | 178.869323 | 0.146093921 | 0.151537956 | -0.964074776 | 0.335008438 | 0.557245099 | 0.146093921 |
| ENSMUSG00000111356 | Rpl32l | 621697 | processed_pseudogene | 12.21977783 | 0.146134235 | 0.338335155 | -0.431921521 | 0.665798452 | 0.819849053 | 0.146134235 |

Supplementary Table 1

| ensembl_gene_id | external_gene_name | entrezgene_id | gene_biotype | baseMean | log2FoldChange | lfcSE | stat | pvalue | padj | absLog2FC |
|---|---|---|---|---|---|---|---|---|---|---|
| ENSMUSG00000024118 | Tedc2 | 72016 | protein_coding | 55.25510727 | -0.146147613 | 0.158062395 | 0.924619759 | 0.355163731 | 0.577906578 | 0.146147613 |
| ENSMUSG00000014907 | Naf1 | 234344 | protein_coding | 92.9100874 | 0.146149947 | 0.139735408 | -1.045904899 | 0.295604942 | 0.514721638 | 0.146149947 |
| ENSMUSG00000034602 | Mon2 | 67074 | protein_coding | 1641.660205 | 0.146179206 | 0.037729176 | -3.874434123 | 0.000106873 | 0.000913981 | 0.146179206 |
| ENSMUSG00000024317 | Rnf138 | 56515 | protein_coding | 193.4523752 | -0.146196555 | 0.105332674 | 1.387950665 | 0.165152068 | 0.352113109 | 0.146196555 |
| ENSMUSG00000059183 | Mtfmt | 69606 | protein_coding | 170.560391 | -0.146245656 | 0.102722063 | 1.423702479 | 0.154532616 | 0.336447148 | 0.146245656 |
| ENSMUSG00000024325 | Ring1 | 19763 | protein_coding | 252.9407056 | -0.146263334 | 0.08444657 | 1.732022197 | 0.08326961 | 0.216212948 | 0.146263334 |
| ENSMUSG00000034430 | Zxdc | 80292 | protein_coding | 279.5139735 | -0.146363094 | 0.091915664 | 1.592362904 | 0.111303186 | 0.26704356 | 0.146363094 |
| ENSMUSG00000040761 | Spen | 56381 | protein_coding | 788.2303577 | 0.146443952 | 0.182818199 | -0.801035964 | 0.423110826 | 0.642978943 | 0.146443952 |
| ENSMUSG00000001865 | Cpa3 | 12873 | protein_coding | 1.531213025 | -0.14645552 | 1.316487145 | 0.111247208 | 0.911420317 | 1 | 0.14645552 |
| ENSMUSG00000032330 | Cox7a2 | 12866 | protein_coding | 2799.102643 | -0.146467257 | 0.083531747 | 1.753432228 | 0.079527843 | 0.208994717 | 0.146467257 |
| ENSMUSG00000050705 | 2310061I04Rik | 69662 | protein_coding | 215.5701841 | -0.146476616 | 0.124955405 | 1.172231137 | 0.241104271 | 0.451758428 | 0.146476616 |
| ENSMUSG00000080574 | Mir466i | 100316665 | miRNA | 1.551329845 | -0.146570712 | 0.839599343 | 0.174572209 | 0.861415787 | 1 | 0.146570712 |
| ENSMUSG00000057497 | Fam136a | 66488 | protein_coding | 425.446747 | -0.146623179 | 0.12230049 | 1.198876458 | 0.230575987 | 0.439134108 | 0.146623179 |
| ENSMUSG00000048439 | Nupl2 | 231042 | protein_coding | 109.9223268 | 0.146625546 | 0.121493592 | -1.206858268 | 0.227486732 | 0.436052025 | 0.146625546 |
| ENSMUSG00000033016 | Nfatc1 | 18018 | protein_coding | 134.4118108 | -0.146631426 | 0.094960895 | 1.54412431 | 0.122558221 | 0.285669361 | 0.146631426 |
| ENSMUSG00000036957 | Lrfn3 | 233067 | protein_coding | 146.7846539 | 0.146638729 | 0.21146133 | -0.693454113 | 0.488024613 | 0.696186289 | 0.146638729 |
| ENSMUSG00000074826 | Gm10767 | 100038538 | protein_coding | 25.06724779 | -0.146669899 | 0.198960214 | 0.737182054 | 0.461011645 | 0.674995696 | 0.146669899 |
| ENSMUSG00000036941 | Elac1 | 114615 | protein_coding | 150.1587693 | 0.146672168 | 0.127524571 | -1.150148294 | 0.250082798 | 0.462419363 | 0.146672168 |
| ENSMUSG00000025997 | Ikzf2 | 22779 | protein_coding | 202.1234934 | 0.146679679 | 0.146856353 | -0.998796961 | 0.317893059 | 0.539328021 | 0.146679679 |
| ENSMUSG00000014075 | Dynlt2b | 66061 | protein_coding | 22.99217873 | 0.146697733 | 0.240284404 | -0.610517083 | 0.541519331 | 0.735467298 | 0.146697733 |
| ENSMUSG00000056342 | Usp34 | 17847 | protein_coding | 1494.162943 | 0.146824655 | 0.128970796 | -1.138433345 | 0.254939577 | 0.467516411 | 0.146824655 |
| ENSMUSG00000031952 | Chst5 | 56773 | protein_coding | 1.52594337 | 0.146826344 | 0.828065128 | -0.177312556 | 0.859262889 | 1 | 0.146826344 |
| ENSMUSG00000038175 | Mylip | 218203 | protein_coding | 218.6844744 | -0.146904962 | 0.089967542 | 1.632866245 | 0.102497131 | 0.252105692 | 0.146904962 |
| ENSMUSG00000002345 | Borcs8 | 72368 | protein_coding | 190.4113525 | -0.146945391 | 0.083115968 | 1.767956204 | 0.077068226 | 0.204680714 | 0.146945391 |
| ENSMUSG00000028960 | Ube4b | 63958 | protein_coding | 641.0509984 | 0.146958218 | 0.079613883 | -1.845886826 | 0.06490864 | 0.180947387 | 0.146958218 |
| ENSMUSG00000044348 | Slc25a53 | 67062 | protein_coding | 25.35956333 | -0.146977991 | 0.205081269 | 0.716681694 | 0.473570524 | 0.68565173 | 0.146977991 |
| ENSMUSG00000087714 | Gm25827 | 115490026 | snRNA | 0.6877759 | 0.146979359 | 1.669689937 | -0.088027936 | 0.929854473 | 1 | 0.146979359 |
| ENSMUSG00000042790 | Rnf214 | 235315 | protein_coding | 261.7405882 | 0.146983789 | 0.085643802 | -1.716222131 | 0.086121394 | 0.221735064 | 0.146983789 |
| ENSMUSG00000024960 | Plcb3 | 18797 | protein_coding | 4777.978611 | -0.147009188 | 0.056076533 | 2.621581249 | 0.00875229 | 0.038236855 | 0.147009188 |
| ENSMUSG00000061613 | U2af1 | 108121 | protein_coding | 426.0997156 | -0.147076656 | 0.132288456 | 1.111787536 | 0.266229512 | 0.481307968 | 0.147076656 |
| ENSMUSG00000020570 | Sypl | 19027 | protein_coding | 1930.627441 | 0.147107011 | 0.140173812 | -1.049461443 | 0.293965792 | 0.513159947 | 0.147107011 |
| ENSMUSG00000085227 | 6330418K02Rik | 69004 | lncRNA | 8.501587164 | -0.147155345 | 0.403262822 | 0.364911758 | 0.715177277 | 0.852123921 | 0.147155345 |
| ENSMUSG00000037331 | Larp1 | 73158 | protein_coding | 1958.681898 | 0.147158716 | 0.073576396 | -2.000080513 | 0.045491571 | 0.138914164 | 0.147158716 |
| ENSMUSG00000031505 | Naxd | 69225 | protein_coding | 530.0679456 | -0.147194002 | 0.068322567 | 2.154398022 | 0.031208972 | 0.104013109 | 0.147194002 |
| ENSMUSG00000090191 | 9230105E05Rik | 320626 | lncRNA | 80.41281884 | -0.147226743 | 0.146542973 | 1.004666009 | 0.315057701 | 0.535990818 | 0.147226743 |
| ENSMUSG00000074406 | Zfp628 | 232816 | protein_coding | 126.3487006 | 0.147244921 | 0.139519751 | -1.055369722 | 0.291256265 | 0.510066755 | 0.147244921 |
| ENSMUSG00000021890 | Eaf1 | 74427 | protein_coding | 466.6377235 | 0.147253102 | 0.058423506 | -2.520442764 | 0.01172073 | 0.048232278 | 0.147253102 |
| ENSMUSG00000045519 | Zfp560 | 434377 | protein_coding | 51.67393339 | -0.147281636 | 0.164889125 | 0.893216185 | 0.371741412 | 0.594020826 | 0.147281636 |
| ENSMUSG00000002910 | Arrdc2 | 70807 | protein_coding | 220.7885124 | 0.147286896 | 0.181836916 | -0.809994465 | 0.417943357 | 0.637944042 | 0.147286896 |

| ensembl_gene_id | external_gene_name | entrezgene_id | gene_biotype | baseMean | log2FoldChange | lfcSE | stat | pvalue | padj | absLog2FC |
|---|---|---|---|---|---|---|---|---|---|---|
| ENSMUSG00000029352 | Crybb3 | 12962 | protein_coding | 9.396343387 | 0.147288446 | 0.567267393 | -0.259645535 | 0.795137209 | 0.898207529 | 0.147288446 |
| ENSMUSG00000021639 | Gtf2h2 | 23894 | protein_coding | 183.3134357 | 0.147357818 | 0.080124079 | -1.839120267 | 0.065897499 | 0.182803846 | 0.147357818 |
| ENSMUSG00000030880 | Polr3e | 26939 | protein_coding | 334.7993247 | -0.147410012 | 0.109168599 | 1.350296814 | 0.176920794 | 0.369037513 | 0.147410012 |
| ENSMUSG00000068921 | Dap3 | 65111 | protein_coding | 683.7040482 | -0.147483353 | 0.071072672 | 2.075106369 | 0.037976678 | 0.120977402 | 0.147483353 |
| ENSMUSG00000038668 | Lpar1 | 14745 | protein_coding | 489.6090911 | -0.147490881 | 0.074683245 | 1.974885818 | 0.048281107 | 0.1452832 | 0.147490881 |
| ENSMUSG00000003778 | Brd8 | 78656 | protein_coding | 690.7837688 | 0.147545936 | 0.080012611 | -1.844033508 | 0.065178257 | 0.181269925 | 0.147545936 |
| ENSMUSG00000027580 | Helz2 | 229003 | protein_coding | 1644.243965 | 0.147618105 | 0.18908173 | -0.780710572 | 0.434972741 | 0.653309672 | 0.147618105 |
| ENSMUSG00000041079 | Rwdd2b | 53858 | protein_coding | 196.2004387 | -0.147633572 | 0.145775377 | 1.01274697 | 0.311181037 | 0.532085855 | 0.147633572 |
| ENSMUSG00000038855 | Itpkb | 320404 | protein_coding | 157.3725608 | 0.147644813 | 0.128009245 | -1.153391794 | 0.248749608 | 0.460883095 | 0.147644813 |
| ENSMUSG00000035944 | Ttc38 | 239570 | protein_coding | 1139.337446 | 0.147685263 | 0.09577314 | -1.542032175 | 0.123065775 | 0.286482691 | 0.147685263 |
| ENSMUSG00000034501 | Pcnx4 | 67708 | protein_coding | 63.1862134 | 0.147739789 | 0.143775885 | -1.027570026 | 0.30415213 | 0.524504007 | 0.147739789 |
| ENSMUSG00000021209 | Ppp4r4 | 74521 | protein_coding | 7.848079816 | 0.147783642 | 0.413345142 | -0.357530855 | 0.720694432 | 0.855242013 | 0.147783642 |
| ENSMUSG00000040850 | Psme4 | 103554 | protein_coding | 2104.015171 | 0.147793898 | 0.083576979 | -1.768356536 | 0.077001318 | 0.204535912 | 0.147793898 |
| ENSMUSG00000024354 | Slc23a1 | 20522 | protein_coding | 9.420136467 | 0.147885247 | 0.498072044 | -0.296915374 | 0.766531118 | 0.882740229 | 0.147885247 |
| ENSMUSG00000033542 | Arhgef5 | 54324 | protein_coding | 1822.326907 | -0.147912609 | 0.076131836 | 1.942848298 | 0.05203449 | 0.153913232 | 0.147912609 |
| ENSMUSG00000021669 | Cert1 | 68018 | protein_coding | 820.7787678 | 0.147919782 | 0.110784093 | -1.335207767 | 0.181808356 | 0.376049998 | 0.147919782 |
| ENSMUSG00000055963 | Triqk | 208820 | protein_coding | 75.60360982 | 0.147957296 | 0.17402477 | -0.850208256 | 0.395209312 | 0.617453792 | 0.147957296 |
| ENSMUSG00000001376 | Vps50 | 73288 | protein_coding | 488.8429866 | 0.148030648 | 0.091045669 | -1.625894455 | 0.10397211 | 0.254671525 | 0.148030648 |
| ENSMUSG00000036817 | Sun1 | 77053 | protein_coding | 1066.772875 | -0.148034804 | 0.090836199 | 1.629689557 | 0.103167126 | 0.253263255 | 0.148034804 |
| ENSMUSG00000035498 | Cdcp1 | 109332 | protein_coding | 1515.50675 | 0.148036815 | 0.06919615 | -2.139379357 | 0.032404957 | 0.107027093 | 0.148036815 |
| ENSMUSG00000061959 | Ces1e | 13897 | protein_coding | 215.2919061 | -0.14806101 | 0.116170705 | 1.274512459 | 0.20248186 | 0.404473864 | 0.14806101 |
| ENSMUSG00000068252 | Apol7b | 278679 | protein_coding | 1.310323023 | -0.148064162 | 1.31642155 | 0.112474733 | 0.910447003 | 1 | 0.148064162 |
| ENSMUSG00000038736 | Nudcd1 | 67429 | protein_coding | 418.0448302 | 0.148073233 | 0.114282302 | -1.295679466 | 0.195085939 | 0.394262295 | 0.148073233 |
| ENSMUSG00000000916 | Nsun5 | 100609 | protein_coding | 103.7024453 | 0.14808885 | 0.138744096 | -1.067352448 | 0.285812713 | 0.504312604 | 0.14808885 |
| ENSMUSG00000040446 | Rprd1a | 225283 | protein_coding | 96.37218463 | -0.148119612 | 0.118628695 | 1.248598508 | 0.211811958 | 0.416114999 | 0.148119612 |
| ENSMUSG00000073678 | Pgap1 | 241062 | protein_coding | 572.4578457 | 0.148131484 | 0.078553253 | -1.885746014 | 0.059329189 | 0.169479714 | 0.148131484 |
| ENSMUSG00000027010 | Slc25a12 | 78830 | protein_coding | 326.8098605 | 0.148236071 | 0.174765385 | -0.848200406 | 0.396326371 | 0.618730733 | 0.148236071 |
| ENSMUSG00000024300 | Myo1f | 17916 | protein_coding | 17.11563977 | 0.148313366 | 0.395058252 | -0.375421513 | 0.707347008 | 0.846814623 | 0.148313366 |
| ENSMUSG00000079418 | Atg4a | 666468 | protein_coding | 173.4152957 | -0.148319481 | 0.108859493 | 1.362485498 | 0.173044718 | 0.363688435 | 0.148319481 |
| ENSMUSG00000030452 | Nipa2 | 93790 | protein_coding | 536.1609971 | 0.148346943 | 0.066406469 | -2.233923075 | 0.025488135 | 0.088668856 | 0.148346943 |
| ENSMUSG00000034616 | Ssh3 | 245857 | protein_coding | 1175.850803 | -0.148401398 | 0.059436087 | 2.496823131 | 0.012531144 | 0.050745295 | 0.148401398 |
| ENSMUSG00000047728 | Ly6g2 | 223631 | protein_coding | 142.8527222 | 0.148406258 | 0.189870474 | -0.781618411 | 0.434438864 | 0.652884901 | 0.148406258 |
| ENSMUSG00000087230 | Mroh3 | 76422 | protein_coding | 78.80372371 | -0.148411194 | 0.209415167 | 0.708693628 | 0.47851462 | 0.690104577 | 0.148411194 |
| ENSMUSG00000021811 | Dnajc9 | 108671 | protein_coding | 193.2649202 | 0.148414171 | 0.184432935 | -0.804705361 | 0.420989725 | 0.640805153 | 0.148414171 |
| ENSMUSG00000037573 | Tob1 | 22057 | protein_coding | 2138.375132 | -0.148443941 | 0.115335874 | 1.287057841 | 0.198074128 | 0.398335908 | 0.148443941 |
| ENSMUSG00000001741 | Il16 | 16170 | protein_coding | 14.08138229 | 0.14845343 | 0.418693003 | -0.354563914 | 0.722916314 | 0.856549354 | 0.14845343 |
| ENSMUSG00000031878 | Nae1 | 234664 | protein_coding | 277.1434762 | -0.148489336 | 0.095459801 | 1.555516934 | 0.119823005 | 0.281753823 | 0.148489336 |
| ENSMUSG00000028656 | Cap1 | 12331 | protein_coding | 3002.012387 | -0.148506912 | 0.060569013 | 2.451862829 | 0.014211883 | 0.055893077 | 0.148506912 |
| ENSMUSG00000071252 | 2210408I21Rik | 72371 | protein_coding | 42.66694025 | -0.148524097 | 0.154795742 | 0.959484383 | 0.337314783 | 0.559393767 | 0.148524097 |

| ensembl_gene_id | external_gene_name | entrezgene_id | gene_biotype | baseMean | log2FoldChange | lfcSE | stat | pvalue | padj | absLog2FC |
|---|---|---|---|---|---|---|---|---|---|---|
| ENSMUSG00000026727 | Rsu1 | 20163 | protein_coding | 186.0108377 | 0.148538461 | 0.110217255 | -1.34768791 | 0.177758782 | 0.370076028 | 0.148538461 |
| ENSMUSG00000024053 | Emilin2 | 246707 | protein_coding | 4.570408644 | -0.148554024 | 0.887112512 | 0.167457929 | 0.867009748 | 0.933936484 | 0.148554024 |
| ENSMUSG00000002748 | Baz1b | 22385 | protein_coding | 1482.053825 | 0.148559454 | 0.072306204 | -2.054587929 | 0.039918823 | 0.125757237 | 0.148559454 |
| ENSMUSG00000035093 | Secisbp2l | 70354 | protein_coding | 1257.879835 | 0.148561483 | 0.094085993 | -1.578996815 | 0.114336788 | 0.272029637 | 0.148561483 |
| ENSMUSG00000074922 | Fam122a | 68034 | protein_coding | 78.0497284 | 0.14860527 | 0.155135028 | -0.957909197 | 0.338108548 | 0.559860817 | 0.14860527 |
| ENSMUSG00000025185 | Loxl4 | 67573 | protein_coding | 5.879074114 | -0.148609345 | 0.748678474 | 0.198495549 | 0.842657367 | 0.922115416 | 0.148609345 |
| ENSMUSG00000108659 | Gm34121 | 102637256 | lncRNA | 9.349209097 | -0.148611826 | 0.415156719 | 0.357965605 | 0.720369054 | 0.855204229 | 0.148611826 |
| ENSMUSG00000033909 | Usp36 | 72344 | protein_coding | 339.5713787 | 0.148639279 | 0.115047477 | -1.291982089 | 0.196363347 | 0.395987827 | 0.148639279 |
| ENSMUSG00000015659 | Serac1 | 321007 | protein_coding | 75.06439237 | -0.148646848 | 0.15606579 | 0.952462726 | 0.340862361 | 0.562506348 | 0.148646848 |
| ENSMUSG00000001815 | Evx2 | 14029 | protein_coding | 8.993358845 | 0.1486565 | 0.540726032 | -0.274920184 | 0.78337756 | 0.891264005 | 0.1486565 |
| ENSMUSG00000028898 | Trnau1ap | 71787 | protein_coding | 243.1865703 | 0.148677489 | 0.093496931 | -1.590185779 | 0.111792935 | 0.26786833 | 0.148677489 |
| ENSMUSG00000031583 | Wrn | 22427 | protein_coding | 330.3244311 | -0.148680992 | 0.080551044 | 1.845798443 | 0.064921477 | 0.180947387 | 0.148680992 |
| ENSMUSG00000090550 | Prlh | 623503 | protein_coding | 0.880501033 | -0.14873706 | 1.344428498 | 0.110632184 | 0.911908025 | 1 | 0.14873706 |
| ENSMUSG00000023348 | Trip6 | 22051 | protein_coding | 115.055607 | 0.148747681 | 0.136547006 | -1.089351466 | 0.275998925 | 0.49283969 | 0.148747681 |
| ENSMUSG00000022971 | Ifnar2 | 15976 | protein_coding | 35.12856088 | -0.148803814 | 0.237981955 | 0.625273518 | 0.531791557 | 0.728736684 | 0.148803814 |
| ENSMUSG00000111080 | Gm20300 | 100504586 | lncRNA | 172.1510412 | -0.148813043 | 0.122641417 | 1.213399576 | 0.224977089 | 0.432209358 | 0.148813043 |
| ENSMUSG00000063568 | Jazf1 | 231986 | protein_coding | 11.20033409 | 0.148829377 | 0.340643807 | -0.43690616 | 0.662179405 | 0.817281815 | 0.148829377 |
| ENSMUSG00000032601 | Prkar2a | 19087 | protein_coding | 1726.122523 | 0.148832258 | 0.080090756 | -1.858295079 | 0.063127122 | 0.176991731 | 0.148832258 |
| ENSMUSG00000066952 | Myo1h | 231646 | protein_coding | 10.88546173 | 0.148859218 | 0.395443174 | -0.376436434 | 0.706592464 | 0.846208704 | 0.148859218 |
| ENSMUSG00000020142 | Slc1a4 | 55963 | protein_coding | 545.5931425 | 0.148875054 | 0.113609599 | -1.310409122 | 0.190057469 | 0.387358014 | 0.148875054 |
| ENSMUSG00000032417 | Rwdd2a | 69519 | protein_coding | 5.648116711 | -0.148916697 | 0.454398979 | 0.32772234 | 0.74312161 | 0.870239999 | 0.148916697 |
| ENSMUSG00000021275 | Tecpr2 | 104859 | protein_coding | 264.1933498 | -0.148921648 | 0.076303881 | 1.951691662 | 0.050974823 | 0.151293715 | 0.148921648 |
| ENSMUSG00000025350 | Rdh5 | 19682 | protein_coding | 33.56962929 | -0.148934597 | 0.200445136 | 0.743019261 | 0.457470019 | 0.672242383 | 0.148934597 |
| ENSMUSG00000008450 | Nutf2 | 68051 | protein_coding | 109.9091208 | -0.148939903 | 0.158934715 | 0.937113726 | 0.348700062 | 0.570573694 | 0.148939903 |
| ENSMUSG00000079469 | Pigb | 55981 | protein_coding | 120.9704376 | 0.149014322 | 0.129476714 | -1.150896689 | 0.24977474 | 0.462014445 | 0.149014322 |
| ENSMUSG00000020116 | Pno1 | 66249 | protein_coding | 289.4476729 | 0.149037793 | 0.115992711 | -1.284889297 | 0.198830972 | 0.399403317 | 0.149037793 |
| ENSMUSG00000029338 | Antxr2 | 71914 | protein_coding | 529.3485051 | -0.149124561 | 0.078786687 | 1.892763459 | 0.058389333 | 0.167403029 | 0.149124561 |
| ENSMUSG00000023030 | Slc11a2 | 18174 | protein_coding | 441.9655707 | 0.149263784 | 0.112443058 | -1.327461085 | 0.184356205 | 0.379371759 | 0.149263784 |
| ENSMUSG00000034748 | Sirt6 | 50721 | protein_coding | 171.7348456 | 0.149288964 | 0.087989154 | -1.696674613 | 0.089758195 | 0.228322063 | 0.149288964 |
| ENSMUSG00000045362 | Tnfrsf26 | 244237 | protein_coding | 2.036740981 | -0.149297217 | 1.109195574 | 0.134599543 | 0.892928503 | 1 | 0.149297217 |
| ENSMUSG00000045071 | E130308A19Rik | 230259 | protein_coding | 100.9316689 | 0.149300633 | 0.102925219 | -1.450573874 | 0.146898555 | 0.324750031 | 0.149300633 |
| ENSMUSG00000079426 | Arpc4 | 68089 | protein_coding | 1155.549884 | 0.14931355 | 0.061219603 | -2.438982662 | 0.014728675 | 0.057501025 | 0.14931355 |
| ENSMUSG00000040565 | Btaf1 | 107182 | protein_coding | 812.3548347 | 0.149398896 | 0.072468411 | -2.061572671 | 0.039248438 | 0.124214011 | 0.149398896 |
| ENSMUSG00000023791 | Pigx | 72084 | protein_coding | 251.8388068 | -0.149408463 | 0.07438824 | 2.008495743 | 0.044590638 | 0.136567098 | 0.149408463 |
| ENSMUSG00000026283 | Ing5 | 66262 | protein_coding | 298.4357207 | -0.149433585 | 0.101716592 | 1.469117099 | 0.141801031 | 0.316830918 | 0.149433585 |
| ENSMUSG00000025982 | Sf3b1 | 81898 | protein_coding | 3699.483714 | 0.149452312 | 0.037429668 | -3.992883676 | 6.53E-05 | 0.000592929 | 0.149452312 |
| ENSMUSG00000015291 | Gdi1 | 14567 | protein_coding | 661.8729403 | -0.149454096 | 0.061657846 | 2.423926628 | 0.015353707 | 0.059503296 | 0.149454096 |
| ENSMUSG00000031970 | Dbndd1 | 72185 | protein_coding | 12.80893886 | 0.149487867 | 0.360116395 | -0.415109863 | 0.67806148 | 0.827422396 | 0.149487867 |
| ENSMUSG00000024410 | Rmc1 | 76482 | protein_coding | 439.8526301 | -0.149489104 | 0.059678921 | 2.504889509 | 0.012248965 | 0.049834789 | 0.149489104 |

| ensembl_gene_id | external_gene_name | entrezgene_id | gene_biotype | baseMean | log2FoldChange | lfcSE | stat | pvalue | padj | absLog2FC |
|---|---|---|---|---|---|---|---|---|---|---|
| ENSMUSG00000025880 | Smad7 | 17131 | protein_coding | 901.5879665 | -0.149518484 | 0.182633915 | 0.818678631 | 0.412969792 | 0.63310278 | 0.149518484 |
| ENSMUSG00000107283 | Mpv17 | 17527 | protein_coding | 137.1643973 | -0.14953302 | 0.099041499 | 1.509801658 | 0.131094042 | 0.299596873 | 0.14953302 |
| ENSMUSG00000020849 | Ywhae | 22627 | protein_coding | 1785.667618 | 0.149572391 | 0.067452273 | -2.217455159 | 0.026592004 | 0.091602076 | 0.149572391 |
| ENSMUSG00000115667 | Gm10389 | 100038608 | lncRNA | 9.42329016 | -0.149668163 | 0.599490516 | 0.249658934 | 0.802851119 | 0.902682351 | 0.149668163 |
| ENSMUSG00000032443 | Zcwpw2 | 100039681 | protein_coding | 2.760267322 | -0.149694484 | 1.119209881 | 0.133750145 | 0.893600151 | 1 | 0.149694484 |
| ENSMUSG00000033508 | Asprv1 | 67855 | protein_coding | 6.408574903 | 0.149764411 | 0.64481047 | -0.232261134 | 0.816335197 | 0.908409285 | 0.149764411 |
| ENSMUSG00000003348 | Mob3a | 208228 | protein_coding | 364.9557869 | 0.149782392 | 0.059078729 | -2.535301554 | 0.011235056 | 0.046746214 | 0.149782392 |
| ENSMUSG00000022179 | 4931414P19Rik | 74359 | protein_coding | 98.4914423 | -0.149794724 | 0.137793693 | 1.087094197 | 0.276995182 | 0.493878043 | 0.149794724 |
| ENSMUSG00000013418 | B4galnt2 | 14422 | protein_coding | 4895.674482 | -0.149824261 | 0.114434488 | 1.309257939 | 0.190446992 | 0.387893639 | 0.149824261 |
| ENSMUSG00000087625 | 4930419G24Rik | 73874 | lncRNA | 0.986827362 | 0.14983293 | 1.294788939 | -0.115719964 | 0.907874484 | 1 | 0.14983293 |
| ENSMUSG00000014402 | Tsg101 | 22088 | protein_coding | 524.5910612 | -0.149878845 | 0.0725885 | 2.064773966 | 0.038944392 | 0.123393648 | 0.149878845 |
| ENSMUSG00000020917 | Acly | 104112 | protein_coding | 5124.893006 | -0.149914517 | 0.179483646 | 0.835254465 | 0.40357445 | 0.62490656 | 0.149914517 |
| ENSMUSG00000021451 | Sema4d | 20354 | protein_coding | 101.7713482 | 0.149950164 | 0.103595113 | -1.447463679 | 0.147767102 | 0.326059049 | 0.149950164 |
| ENSMUSG00000074480 | Mex3a | 72640 | protein_coding | 34.62614591 | 0.149998997 | 0.204859319 | -0.732204898 | 0.464043515 | 0.677560974 | 0.149998997 |
| ENSMUSG00000041827 | Oasl1 | 231655 | protein_coding | 233.3759016 | -0.150077717 | 0.293043338 | 0.512134889 | 0.608556608 | 0.782687214 | 0.150077717 |
| ENSMUSG00000026872 | Zeb2 | 24136 | protein_coding | 7.378744722 | 0.150087513 | 1.256374754 | -0.119460784 | 0.904910308 | 0.953204491 | 0.150087513 |
| ENSMUSG00000022051 | Bnip3l | 12177 | protein_coding | 617.3414146 | -0.150125895 | 0.094085174 | 1.595638168 | 0.110569601 | 0.265929452 | 0.150125895 |
| ENSMUSG00000030782 | Tgfb1i1 | 21804 | protein_coding | 4.714040252 | -0.150128869 | 0.99286946 | 0.151207057 | 0.879812384 | 0.940676072 | 0.150128869 |
| ENSMUSG00000074892 | B3galt5 | 93961 | protein_coding | 7152.506842 | 0.150210324 | 0.141504478 | -1.061523472 | 0.288452072 | 0.50724071 | 0.150210324 |
| ENSMUSG00000020639 | Pfn4 | 382562 | protein_coding | 1.761904997 | 0.150221459 | 0.976113311 | -0.153897562 | 0.877690506 | 1 | 0.150221459 |
| ENSMUSG00000065778 | Gm22154 | 115490050 | snoRNA | 292.5540673 | 0.150253357 | 0.209449086 | -0.717374134 | 0.473143276 | 0.685335967 | 0.150253357 |
| ENSMUSG00000038677 | Scube3 | 268935 | protein_coding | 8.172535952 | 0.150277433 | 0.773375841 | -0.194313586 | 0.845930349 | 0.924166034 | 0.150277433 |
| ENSMUSG00000038528 | Mfsd4b5 | 215928 | protein_coding | 11.38620434 | 0.150292162 | 0.504189989 | -0.298086367 | 0.765637247 | 0.882510888 | 0.150292162 |
| ENSMUSG00000026187 | Xrcc5 | 22596 | protein_coding | 289.7646833 | -0.150302657 | 0.116868923 | 1.286078911 | 0.198415523 | 0.398762803 | 0.150302657 |
| ENSMUSG00000029185 | Fam114a1 | 68303 | protein_coding | 1048.498491 | 0.150315233 | 0.092294513 | -1.628647559 | 0.10338765 | 0.253653789 | 0.150315233 |
| ENSMUSG00000028261 | Ndufaf4 | 68493 | protein_coding | 353.1687524 | 0.150352954 | 0.095572163 | -1.57318773 | 0.115675356 | 0.274502489 | 0.150352954 |
| ENSMUSG00000039865 | Slc44a3 | 213603 | protein_coding | 951.0497586 | -0.150355741 | 0.130174916 | 1.15502852 | 0.248078747 | 0.460363236 | 0.150355741 |
| ENSMUSG00000020107 | Anapc16 | 52717 | protein_coding | 431.051953 | 0.150363189 | 0.066814528 | -2.250456503 | 0.024419982 | 0.085802463 | 0.150363189 |
| ENSMUSG00000024782 | Ak3 | 56248 | protein_coding | 2644.987741 | -0.150388703 | 0.106373399 | 1.41378112 | 0.157426179 | 0.340684159 | 0.150388703 |
| ENSMUSG00000033813 | Tcea1 | 21399 | protein_coding | 514.9973281 | 0.150453218 | 0.094057524 | -1.599587275 | 0.109690173 | 0.264518622 | 0.150453218 |
| ENSMUSG00000035476 | Tab3 | 66724 | protein_coding | 162.2963182 | 0.15048345 | 0.095525319 | -1.575325287 | 0.11518138 | 0.273724003 | 0.15048345 |
| ENSMUSG00000029102 | Hgfac | 54426 | protein_coding | 2123.576893 | -0.150526403 | 0.172821428 | 0.870993861 | 0.383757505 | 0.606204104 | 0.150526403 |
| ENSMUSG00000004105 | Angptl2 | 26360 | protein_coding | 16.92717173 | 0.150528804 | 0.435061328 | -0.345994448 | 0.729346887 | 0.86030736 | 0.150528804 |
| ENSMUSG00000047821 | Trim16 | 94092 | protein_coding | 1508.441922 | -0.150585904 | 0.095308388 | 1.579985843 | 0.114110109 | 0.271607716 | 0.150585904 |
| ENSMUSG00000052302 | Tbc1d30 | 74694 | protein_coding | 275.7739104 | -0.150668286 | 0.082728752 | 1.821232424 | 0.068571534 | 0.188104702 | 0.150668286 |
| ENSMUSG00000029775 | Klhdc10 | 76788 | protein_coding | 692.962787 | 0.150762931 | 0.068324036 | -2.206587006 | 0.027342926 | 0.093703879 | 0.150762931 |
| ENSMUSG00000055818 | A230083G16Rik | 442825 | lncRNA | 15.22140655 | 0.150766756 | 0.319112256 | -0.472456803 | 0.636600768 | 0.801948109 | 0.150766756 |
| ENSMUSG00000022800 | Fyttd1 | 69823 | protein_coding | 496.6851887 | 0.150790318 | 0.115501383 | -1.305528251 | 0.191713034 | 0.389463531 | 0.150790318 |
| ENSMUSG00000068749 | Psma5 | 26442 | protein_coding | 585.155524 | -0.150823881 | 0.148111196 | 1.018315193 | 0.30852819 | 0.529189296 | 0.150823881 |

Supplementary Table 1

| ensembl_gene_id | external_gene_name | entrezgene_id | gene_biotype | baseMean | log2FoldChange | lfcSE | stat | pvalue | padj | absLog2FC |
|---|---|---|---|---|---|---|---|---|---|---|
| ENSMUSG00000078202 | Nrarp | 67122 | protein_coding | 79.01685135 | 0.150862568 | 0.158102738 | -0.954205915 | 0.339979421 | 0.561491337 | 0.150862568 |
| ENSMUSG00000050965 | Prkca | 18750 | protein_coding | 1625.190949 | -0.150887234 | 0.064611772 | 2.335290127 | 0.01952827 | 0.07201649 | 0.150887234 |
| ENSMUSG00000046836 | Brox | 71678 | protein_coding | 628.4200648 | -0.150893215 | 0.09309345 | 1.620878974 | 0.105043597 | 0.256383382 | 0.150893215 |
| ENSMUSG00000057375 | Yipf1 | 230584 | protein_coding | 1238.380621 | -0.150906221 | 0.116472933 | 1.295633386 | 0.195101821 | 0.394262295 | 0.150906221 |
| ENSMUSG00000021987 | Mtmr6 | 219135 | protein_coding | 1455.636617 | -0.150946486 | 0.075694892 | 1.994143631 | 0.04613636 | 0.140312312 | 0.150946486 |
| ENSMUSG00000024168 | Tmem204 | 407831 | protein_coding | 5.094998447 | -0.151014151 | 0.479631756 | 0.314854363 | 0.752872233 | 0.876175621 | 0.151014151 |
| ENSMUSG00000026181 | Ppm1f | 68606 | protein_coding | 84.02915785 | 0.151021222 | 0.149167563 | -1.012426687 | 0.311334084 | 0.532288131 | 0.151021222 |
| ENSMUSG00000063714 | Sp3os | 100503849 | lncRNA | 24.37508724 | 0.151040136 | 0.210316034 | -0.71815797 | 0.47265989 | 0.685174282 | 0.151040136 |
| ENSMUSG00000061436 | Hipk2 | 15258 | protein_coding | 445.6101467 | -0.151172858 | 0.210962009 | 0.716588065 | 0.473628311 | 0.685675281 | 0.151172858 |
| ENSMUSG00000025474 | Tubgcp2 | 74237 | protein_coding | 199.3752751 | 0.151210126 | 0.105130931 | -1.438302926 | 0.150348122 | 0.330166076 | 0.151210126 |
| ENSMUSG00000040749 | Siah1b | 20438 | protein_coding | 55.34559573 | 0.151219682 | 0.195815858 | -0.772254519 | 0.439963697 | 0.657874003 | 0.151219682 |
| ENSMUSG00000002948 | Map2k7 | 26400 | protein_coding | 213.9767761 | -0.15122196 | 0.075002573 | 2.016223641 | 0.043776595 | 0.134723574 | 0.15122196 |
| ENSMUSG00000032078 | Zpr1 | 22687 | protein_coding | 323.6301268 | -0.151262874 | 0.109213036 | 1.385025813 | 0.166044575 | 0.353403803 | 0.151262874 |
| ENSMUSG00000002043 | Trappc6a | 67091 | protein_coding | 834.0377947 | 0.151315055 | 0.088531839 | -1.709159733 | 0.087421365 | 0.224312683 | 0.151315055 |
| ENSMUSG00000059146 | Ntrk3 | 18213 | protein_coding | 10.71417191 | 0.151337418 | 2.00608059 | -0.075439351 | 0.939865151 | 0.971381329 | 0.151337418 |
| ENSMUSG00000023944 | Hsp90ab1 | 15516 | protein_coding | 10969.34108 | -0.151384748 | 0.109212236 | 1.386151896 | 0.165700527 | 0.352762472 | 0.151384748 |
| ENSMUSG00000019647 | Sema6a | 20358 | protein_coding | 239.4680644 | -0.15142593 | 0.217458411 | 0.69634432 | 0.486213224 | 0.695624496 | 0.15142593 |
| ENSMUSG00000022723 | Crybg3 | 224273 | protein_coding | 346.7037727 | -0.151427048 | 0.074182059 | 2.041289365 | 0.041222074 | 0.128879463 | 0.151427048 |
| ENSMUSG00000028413 | B4galt1 | 14595 | protein_coding | 1284.480197 | -0.151459495 | 0.13137012 | 1.152921957 | 0.248942419 | 0.46108514 | 0.151459495 |
| ENSMUSG00000025159 | Mms19 | 72199 | protein_coding | 325.073273 | -0.151462642 | 0.077176827 | 1.962540398 | 0.049699602 | 0.148254918 | 0.151462642 |
| ENSMUSG00000029062 | Cdk11b | 12537 | protein_coding | 566.8403893 | 0.151512323 | 0.097051534 | -1.561153309 | 0.118487583 | 0.279437874 | 0.151512323 |
| ENSMUSG00000021432 | Slc35b3 | 108652 | protein_coding | 587.4095805 | 0.151524198 | 0.060764947 | -2.493611945 | 0.01264507 | 0.051106386 | 0.151524198 |
| ENSMUSG00000003421 | Nosip | 66394 | protein_coding | 387.375426 | -0.151538811 | 0.06495597 | 2.332946618 | 0.019650947 | 0.072388075 | 0.151538811 |
| ENSMUSG00000014444 | Piezo1 | 234839 | protein_coding | 237.2644011 | -0.151574627 | 0.102124742 | 1.484210624 | 0.137753052 | 0.310179152 | 0.151574627 |
| ENSMUSG00000022766 | Serpind1 | 15160 | protein_coding | 2.309397462 | 0.151582558 | 0.88112562 | -0.172032857 | 0.863411698 | 1 | 0.151582558 |
| ENSMUSG00000054200 | Ffar4 | 107221 | protein_coding | 283.180685 | 0.151641968 | 0.23734619 | -0.638906264 | 0.522883913 | 0.722022907 | 0.151641968 |
| ENSMUSG00000004038 | Gstm3 | 14864 | protein_coding | 578.5111532 | -0.151696725 | 0.139435986 | 1.087930953 | 0.27662559 | 0.493582047 | 0.151696725 |
| ENSMUSG00000045284 | Dcaf12l1 | 245404 | protein_coding | 11.72648822 | -0.151706967 | 0.310502524 | 0.488585295 | 0.625135326 | 0.794593769 | 0.151706967 |
| ENSMUSG00000004043 | Stat5a | 20850 | protein_coding | 143.2243409 | -0.151716931 | 0.097690182 | 1.553041741 | 0.120413163 | 0.282675077 | 0.151716931 |
| ENSMUSG00000006763 | Saal1 | 78935 | protein_coding | 149.2256565 | 0.151766876 | 0.121586084 | -1.248225715 | 0.211948409 | 0.416333535 | 0.151766876 |
| ENSMUSG00000038764 | Ptpn3 | 545622 | protein_coding | 1230.480631 | -0.151782832 | 0.087757164 | 1.729577678 | 0.083705757 | 0.216936107 | 0.151782832 |
| ENSMUSG00000025374 | Nabp2 | 69917 | protein_coding | 319.2694171 | 0.151787312 | 0.088353256 | -1.717959456 | 0.085804011 | 0.22115998 | 0.151787312 |
| ENSMUSG00000024228 | Nudt12 | 67993 | protein_coding | 195.5921657 | -0.151828367 | 0.102799143 | 1.476941959 | 0.139691196 | 0.313346973 | 0.151828367 |
| ENSMUSG00000034007 | Scaper | 244891 | protein_coding | 276.4869789 | -0.151873941 | 0.097714758 | 1.554257962 | 0.120122897 | 0.28232699 | 0.151873941 |
| ENSMUSG00000040985 | Sun3 | 194974 | protein_coding | 4.789634065 | 0.151915543 | 0.619112263 | -0.245376408 | 0.806164994 | 0.904600714 | 0.151915543 |
| ENSMUSG00000009569 | Mrtfb | 239719 | protein_coding | 694.243977 | -0.151917429 | 0.091161508 | 1.666464626 | 0.095620908 | 0.239182623 | 0.151917429 |
| ENSMUSG00000070699 | Sars2 | 71984 | protein_coding | 162.9186859 | 0.151932978 | 0.126494548 | -1.201102975 | 0.229711259 | 0.438295983 | 0.151932978 |
| ENSMUSG00000028389 | Zfp37 | 22696 | protein_coding | 95.9516319 | -0.151933846 | 0.129826607 | 1.170282805 | 0.241887179 | 0.452586095 | 0.151933846 |
| ENSMUSG00000025171 | Ubtd1 | 226122 | protein_coding | 58.20730642 | -0.151936894 | 0.158130419 | 0.960832805 | 0.336636241 | 0.55922049 | 0.151936894 |

| ensembl_gene_id | external_gene_name | entrezgene_id | gene_biotype | baseMean | log2FoldChange | lfcSE | stat | pvalue | padj | absLog2FC |
|---|---|---|---|---|---|---|---|---|---|---|
| ENSMUSG00000002580 | Mien1 | 103742 | protein_coding | 481.9702706 | 0.151942232 | 0.088927503 | -1.708607885 | 0.087523607 | 0.224470391 | 0.151942232 |
| ENSMUSG00000028229 | Rmdn1 | 66302 | protein_coding | 112.418735 | -0.151956063 | 0.108050921 | 1.406337514 | 0.159623926 | 0.343804463 | 0.151956063 |
| ENSMUSG00000018428 | Akap1 | 11640 | protein_coding | 1449.432005 | -0.151956125 | 0.091173032 | 1.666678426 | 0.095578365 | 0.239112425 | 0.151956125 |
| ENSMUSG00000027255 | Arfgap2 | 77038 | protein_coding | 406.2449256 | 0.151990283 | 0.06217943 | -2.444382044 | 0.014510052 | 0.056795212 | 0.151990283 |
| ENSMUSG00000024085 | Man2a1 | 17158 | protein_coding | 1649.70066 | 0.151994039 | 0.091327068 | -1.664282475 | 0.096055993 | 0.239895137 | 0.151994039 |
| ENSMUSG00000035133 | Arhgap5 | 11855 | protein_coding | 3151.211937 | -0.152000852 | 0.086976771 | 1.747602841 | 0.080532823 | 0.210915627 | 0.152000852 |
| ENSMUSG00000029673 | Auts2 | 319974 | protein_coding | 388.0644268 | -0.152047087 | 0.147311486 | 1.03214685 | 0.302003329 | 0.522337974 | 0.152047087 |
| ENSMUSG00000044876 | Zfp444 | 72667 | protein_coding | 215.3105253 | 0.152064424 | 0.086200158 | -1.764085207 | 0.077717635 | 0.205842665 | 0.152064424 |
| ENSMUSG00000091530 | Cldn20 | 621628 | processed_pseudogene | 2.439288402 | 0.152084204 | 0.672847236 | -0.226030807 | 0.82117746 | 1 | 0.152084204 |
| ENSMUSG00000003948 | Mmd | 67468 | protein_coding | 85.18995449 | -0.152095721 | 0.160998704 | 0.944701527 | 0.344811277 | 0.566362983 | 0.152095721 |
| ENSMUSG00000013822 | Elof1 | 66126 | protein_coding | 671.9700388 | 0.152101069 | 0.07572672 | -2.008552185 | 0.044584646 | 0.136567098 | 0.152101069 |
| ENSMUSG00000024998 | Plce1 | 74055 | protein_coding | 1575.495755 | -0.152109621 | 0.082795517 | 1.837172177 | 0.066184482 | 0.183322799 | 0.152109621 |
| ENSMUSG00000079104 | Prps1l3 | 328099 | protein_coding | 289.1564668 | 0.15212059 | 0.13315461 | -1.142435776 | 0.253272927 | 0.465710848 | 0.15212059 |
| ENSMUSG00000022707 | Gbe1 | 74185 | protein_coding | 295.3048401 | -0.1521273 | 0.107227277 | 1.418736947 | 0.155975721 | 0.338606949 | 0.1521273 |
| ENSMUSG00000043003 | Rasef | 242505 | protein_coding | 1739.600568 | 0.152181014 | 0.08458226 | -1.799207224 | 0.071985907 | 0.195041062 | 0.152181014 |
| ENSMUSG00000015536 | Mocs2 | 17434 | protein_coding | 364.3859448 | -0.152211721 | 0.077142045 | 1.973135674 | 0.048480107 | 0.145669666 | 0.152211721 |
| ENSMUSG00000001158 | Snrnp27 | 66618 | protein_coding | 250.8513462 | 0.152219365 | 0.083602918 | -1.820742238 | 0.068646048 | 0.188215315 | 0.152219365 |
| ENSMUSG00000033047 | Eif3l | 223691 | protein_coding | 1842.775235 | -0.152230701 | 0.111599742 | 1.364077538 | 0.172543164 | 0.363163115 | 0.152230701 |
| ENSMUSG00000027615 | Hps3 | 12807 | protein_coding | 141.0155408 | -0.152320069 | 0.125338812 | 1.215266575 | 0.224264436 | 0.431320242 | 0.152320069 |
| ENSMUSG00000029106 | Add1 | 11518 | protein_coding | 3152.639566 | -0.152387581 | 0.061764464 | 2.467237167 | 0.013616014 | 0.054116692 | 0.152387581 |
| ENSMUSG00000005251 | Ripk4 | 72388 | protein_coding | 644.7540681 | -0.152418298 | 0.103493324 | 1.472735546 | 0.140822358 | 0.315242025 | 0.152418298 |
| ENSMUSG00000026111 | Unc50 | 67387 | protein_coding | 494.8076963 | 0.152526495 | 0.090173244 | -1.691482845 | 0.090744622 | 0.230228404 | 0.152526495 |
| ENSMUSG00000044345 | Marveld1 | 277010 | protein_coding | 54.87199256 | -0.152629213 | 0.184700059 | 0.826362555 | 0.408598458 | 0.629263727 | 0.152629213 |
| ENSMUSG00000020844 | Nxn | 18230 | protein_coding | 30.78198882 | 0.152635009 | 0.178772158 | -0.85379631 | 0.393217859 | 0.615915748 | 0.152635009 |
| ENSMUSG00000002428 | Hltf | 20585 | protein_coding | 566.4980161 | 0.152642701 | 0.100921637 | -1.512487372 | 0.130409923 | 0.29840508 | 0.152642701 |
| ENSMUSG00000022203 | Efs | 13644 | protein_coding | 4.591170666 | -0.152643317 | 0.652434631 | 0.233959557 | 0.815016377 | 0.908068261 | 0.152643317 |
| ENSMUSG00000047187 | Rab2a | 59021 | protein_coding | 1956.898274 | 0.152695905 | 0.103610312 | -1.473752006 | 0.140548375 | 0.314808425 | 0.152695905 |
| ENSMUSG00000068206 | Pick1 | 18693 | protein_coding | 201.0547693 | 0.152727987 | 0.087901661 | -1.73748692 | 0.082301261 | 0.214305112 | 0.152727987 |
| ENSMUSG00000031879 | Ciao2b | 68523 | protein_coding | 281.1495088 | -0.152738017 | 0.105089789 | 1.453404925 | 0.14611136 | 0.323442761 | 0.152738017 |
| ENSMUSG00000026915 | Strbp | 20744 | protein_coding | 1857.814973 | -0.152770843 | 0.061544404 | 2.482286497 | 0.013054227 | 0.05235005 | 0.152770843 |
| ENSMUSG00000041488 | Stx3 | 20908 | protein_coding | 700.6841598 | -0.15280746 | 0.07706032 | 1.982959059 | 0.047372003 | 0.143173729 | 0.15280746 |
| ENSMUSG00000040488 | Ltbp4 | 108075 | protein_coding | 16.44956176 | 0.152813908 | 0.461826843 | -0.330890052 | 0.740727536 | 0.868781842 | 0.152813908 |
| ENSMUSG00000013766 | Ly6g6e | 70274 | protein_coding | 107.674528 | -0.152819857 | 0.216620943 | 0.705471294 | 0.48051699 | 0.691465456 | 0.152819857 |
| ENSMUSG00000070003 | Ssbp4 | 76900 | protein_coding | 444.0025175 | 0.152833773 | 0.136303994 | -1.121271422 | 0.262172346 | 0.47603234 | 0.152833773 |
| ENSMUSG00000047368 | Abhd17b | 226016 | protein_coding | 695.9202747 | 0.152863935 | 0.092753189 | -1.648072006 | 0.099337896 | 0.246352764 | 0.152863935 |
| ENSMUSG00000056870 | Gulp1 | 70676 | protein_coding | 64.70572429 | 0.15288093 | 0.184108929 | -0.830383029 | 0.406322257 | 0.62720573 | 0.15288093 |
| ENSMUSG00000031425 | Plp1 | 18823 | protein_coding | 6.920297362 | -0.15304047 | 0.570889455 | 0.268073737 | 0.788642561 | 0.894265208 | 0.15304047 |
| ENSMUSG00000043290 | Zfp784 | 654801 | protein_coding | 5.786559283 | 0.15307731 | 0.429476599 | -0.356427591 | 0.72152037 | 0.855725733 | 0.15307731 |
| ENSMUSG00000085826 | Gm15638 | 100503378 | lncRNA | 14.09439051 | -0.153081281 | 0.35860518 | 0.426879725 | 0.669466933 | 0.822609935 | 0.153081281 |

Supplementary Table 1

| ensembl_gene_id | external_gene_name | entrezgene_id | gene_biotype | baseMean | log2FoldChange | lfcSE | stat | pvalue | padj | absLog2FC |
|---|---|---|---|---|---|---|---|---|---|---|
| ENSMUSG00000065715 | Snord7 | 100302731 | snoRNA | 1.233024909 | 0.153174558 | 1.175925136 | -0.130258767 | 0.896361701 | 1 | 0.153174558 |
| ENSMUSG00000038611 | Phrf1 | 101471 | protein_coding | 848.9383617 | 0.153264582 | 0.092542602 | -1.656151632 | 0.097691116 | 0.243108963 | 0.153264582 |
| ENSMUSG00000054280 | Prr14l | 215476 | protein_coding | 950.7701099 | -0.153287546 | 0.065302526 | 2.347344814 | 0.018907748 | 0.07028111 | 0.153287546 |
| ENSMUSG00000062352 | Itgb1bp1 | 16413 | protein_coding | 86.64239956 | 0.153297333 | 0.108303269 | -1.415445116 | 0.15693803 | 0.340070151 | 0.153297333 |
| ENSMUSG00000031256 | Cstf2 | 108062 | protein_coding | 795.8820411 | 0.153305313 | 0.083998989 | -1.825085219 | 0.06798817 | 0.186963018 | 0.153305313 |
| ENSMUSG00000002897 | Il17ra | 16172 | protein_coding | 266.9436012 | -0.153343982 | 0.066479575 | 2.306633018 | 0.021075287 | 0.076491713 | 0.153343982 |
| ENSMUSG00000006575 | Rundc3a | 51799 | protein_coding | 87.71933739 | -0.153369473 | 0.110175728 | 1.392044116 | 0.163909039 | 0.350167383 | 0.153369473 |
| ENSMUSG00000042102 | Dmgdh | 74129 | protein_coding | 10.22489634 | 0.153378031 | 0.362138719 | -0.423533921 | 0.671905755 | 0.82411257 | 0.153378031 |
| ENSMUSG00000032459 | Mrps22 | 64655 | protein_coding | 253.8975596 | -0.153424741 | 0.088865164 | 1.726489147 | 0.08425945 | 0.218131456 | 0.153424741 |
| ENSMUSG00000002949 | Timm44 | 21856 | protein_coding | 710.2846818 | -0.153425596 | 0.060112905 | 2.552290482 | 0.010701725 | 0.044980477 | 0.153425596 |
| ENSMUSG00000058230 | Arhgap35 | 232906 | protein_coding | 987.2526262 | -0.153438831 | 0.099241527 | 1.546115179 | 0.122076753 | 0.285167413 | 0.153438831 |
| ENSMUSG00000088625 | Gm23318 | 115489368 | snRNA | 0.753951854 | 0.153466152 | 1.440795007 | -0.106514911 | 0.915173825 | 1 | 0.153466152 |
| ENSMUSG00000026021 | Sumo1 | 22218 | protein_coding | 542.0052041 | -0.153510933 | 0.104947161 | 1.462744977 | 0.143537174 | 0.319845216 | 0.153510933 |
| ENSMUSG00000026922 | Agpat2 | 67512 | protein_coding | 1089.124154 | -0.153541647 | 0.10226455 | 1.501416147 | 0.13324796 | 0.302968031 | 0.153541647 |
| ENSMUSG00000025086 | Trub1 | 72133 | protein_coding | 99.1890106 | -0.153545974 | 0.134472868 | 1.141836091 | 0.253522157 | 0.465961592 | 0.153545974 |
| ENSMUSG00000091455 | Otogl | 628870 | protein_coding | 2.768052669 | 0.153650272 | 2.025336005 | -0.075864089 | 0.939527227 | 1 | 0.153650272 |
| ENSMUSG00000062906 | Hdac10 | 170787 | protein_coding | 207.3683616 | -0.153659663 | 0.071787142 | 2.140490051 | 0.032315184 | 0.10685821 | 0.153659663 |
| ENSMUSG00000032431 | Crtap | 56693 | protein_coding | 85.02002527 | -0.15368164 | 0.12123006 | 1.267685913 | 0.204910138 | 0.407353559 | 0.15368164 |
| ENSMUSG00000035735 | Dagla | 269060 | protein_coding | 175.7919717 | -0.153711871 | 0.128168068 | 1.199299267 | 0.2304116 | 0.439074217 | 0.153711871 |
| ENSMUSG00000028689 | Ccdc163 | 68394 | protein_coding | 67.37163982 | -0.153792249 | 0.131213296 | 1.172078236 | 0.241165647 | 0.451799148 | 0.153792249 |
| ENSMUSG00000020032 | Nuak1 | 77976 | protein_coding | 13.17647467 | 0.153832017 | 0.376198608 | -0.408911711 | 0.682604454 | 0.830450122 | 0.153832017 |
| ENSMUSG00000033855 | Ston1 | 77057 | protein_coding | 57.27315809 | -0.15384153 | 0.205089367 | 0.750119486 | 0.453182745 | 0.669122175 | 0.15384153 |
| ENSMUSG00000015702 | Anxa9 | 71790 | protein_coding | 22.18715301 | 0.153871454 | 0.328141171 | -0.468918463 | 0.639127919 | 0.803282676 | 0.153871454 |
| ENSMUSG00000040591 | Cstpp1 | 228356 | protein_coding | 41.36288346 | 0.153939345 | 0.16926879 | -0.909437264 | 0.363119359 | 0.585429376 | 0.153939345 |
| ENSMUSG00000027806 | Tsc22d2 | 72033 | protein_coding | 406.90775 | 0.153979681 | 0.147100991 | -1.04676168 | 0.295209508 | 0.514332163 | 0.153979681 |
| ENSMUSG00000032346 | Ooep | 67968 | protein_coding | 1.165099727 | -0.154027658 | 0.999164295 | 0.154156488 | 0.87748635 | 1 | 0.154027658 |
| ENSMUSG00000060548 | Tnfrsf19 | 29820 | protein_coding | 33.41747318 | 0.154040064 | 0.251462382 | -0.61257697 | 0.54015609 | 0.734530441 | 0.154040064 |
| ENSMUSG00000059363 | Fxn | 14297 | protein_coding | 75.09941831 | 0.154062457 | 0.122623233 | -1.256388806 | 0.208975038 | 0.412258246 | 0.154062457 |
| ENSMUSG00000031503 | Col4a2 | 12827 | protein_coding | 16.77000391 | -0.154158132 | 0.579907342 | 0.265832351 | 0.790368326 | 0.895392916 | 0.154158132 |
| ENSMUSG00000023473 | Celsr3 | 107934 | protein_coding | 17.8054604 | -0.154172415 | 0.543643359 | 0.283591093 | 0.776723763 | 0.888506311 | 0.154172415 |
| ENSMUSG00000003309 | Ap1m2 | 11768 | protein_coding | 1791.21643 | 0.154194122 | 0.081692783 | -1.887487684 | 0.059094761 | 0.168985435 | 0.154194122 |
| ENSMUSG00000025584 | Pde8a | 18584 | protein_coding | 93.91882526 | -0.154196457 | 0.163905047 | 0.940766983 | 0.346824284 | 0.568538439 | 0.154196457 |
| ENSMUSG00000055415 | Atp10b | 319767 | protein_coding | 3229.405372 | 0.154197449 | 0.076909475 | -2.004921366 | 0.044971454 | 0.137605817 | 0.154197449 |
| ENSMUSG00000001127 | Araf | 11836 | protein_coding | 904.9185017 | 0.154226666 | 0.067361428 | -2.289539745 | 0.022048011 | 0.079188614 | 0.154226666 |
| ENSMUSG00000079316 | Rab9 | 56382 | protein_coding | 298.7499233 | -0.154255467 | 0.078550484 | 1.963774884 | 0.049556204 | 0.14798755 | 0.154255467 |
| ENSMUSG00000022948 | Setd4 | 224440 | protein_coding | 108.4114842 | -0.154258443 | 0.143306749 | 1.07642134 | 0.281738863 | 0.499562412 | 0.154258443 |
| ENSMUSG00000051735 | Rinl | 320435 | protein_coding | 21.55371536 | 0.154276229 | 0.338549176 | -0.455698139 | 0.648607061 | 0.809117465 | 0.154276229 |
| ENSMUSG00000091586 | Cyp4f17 | 208285 | protein_coding | 37.3747132 | -0.154294215 | 0.186045109 | 0.829337657 | 0.406913368 | 0.627713038 | 0.154294215 |
| ENSMUSG00000029502 | Golga3 | 269682 | protein_coding | 1300.471166 | -0.15429926 | 0.099568277 | 1.549682938 | 0.121217635 | 0.283960728 | 0.15429926 |

| ensembl_gene_id | external_gene_name | entrezgene_id | gene_biotype | baseMean | log2FoldChange | lfcSE | stat | pvalue | padj | absLog2FC |
|---|---|---|---|---|---|---|---|---|---|---|
| ENSMUSG00000085498 | Gm14023 | 100503468 | lncRNA | 5.630308155 | -0.154339955 | 0.526683081 | 0.293041415 | 0.769490496 | 0.884482916 | 0.154339955 |
| ENSMUSG00000009555 | Cdk9 | 107951 | protein_coding | 428.490346 | -0.15434096 | 0.064480013 | 2.393624825 | 0.016682804 | 0.063640522 | 0.15434096 |
| ENSMUSG00000042419 | Nfkbil1 | 18038 | protein_coding | 166.0682744 | 0.15434413 | 0.09213839 | -1.675133773 | 0.093907988 | 0.235970255 | 0.15434413 |
| ENSMUSG00000025453 | Nnt | 18115 | protein_coding | 8.919812186 | -0.15437655 | 0.385834805 | 0.400110481 | 0.689075145 | 0.833960201 | 0.15437655 |
| ENSMUSG00000075296 | Aldh3b2 | 621603 | protein_coding | 94.73288813 | 0.154437771 | 0.125422208 | -1.231343107 | 0.218194566 | 0.42426293 | 0.154437771 |
| ENSMUSG00000020911 | Krt19 | 16669 | protein_coding | 39271.83275 | -0.154497562 | 0.11614446 | 1.330218953 | 0.183446141 | 0.378300667 | 0.154497562 |
| ENSMUSG00000021716 | Srek1ip1 | 67288 | protein_coding | 276.9479802 | -0.154500631 | 0.094695688 | 1.631548739 | 0.102774584 | 0.25251889 | 0.154500631 |
| ENSMUSG00000006154 | Eps8l1 | 67425 | protein_coding | 66.3002116 | 0.15453759 | 0.227227101 | -0.680101929 | 0.496439923 | 0.702502904 | 0.15453759 |
| ENSMUSG00000042496 | Prdm10 | 382066 | protein_coding | 245.7047692 | 0.15455135 | 0.079635148 | -1.940742927 | 0.052289467 | 0.154446119 | 0.15455135 |
| ENSMUSG00000067424 | Zfp563 | 240068 | protein_coding | 97.85510103 | -0.154586599 | 0.119609142 | 1.292431296 | 0.196207824 | 0.395770858 | 0.154586599 |
| ENSMUSG00000027799 | Nbea | 26422 | protein_coding | 229.4425089 | 0.154617537 | 0.154150998 | -1.003026502 | 0.315848074 | 0.536652594 | 0.154617537 |
| ENSMUSG00000073448 | Gm10509 | 100310809 | protein_coding | 25.46150153 | -0.154620429 | 0.219652548 | 0.703931872 | 0.481475202 | 0.692354022 | 0.154620429 |
| ENSMUSG00000061079 | Zfp143 | 20841 | protein_coding | 233.345022 | 0.154727552 | 0.088459486 | -1.74913466 | 0.080267745 | 0.21050366 | 0.154727552 |
| ENSMUSG00000023143 | Nagpa | 27426 | protein_coding | 210.1540503 | -0.154773928 | 0.105485575 | 1.467252071 | 0.142307497 | 0.31779121 | 0.154773928 |
| ENSMUSG00000048763 | Hoxb3 | 15410 | protein_coding | 45.84458906 | -0.154816658 | 0.39893854 | 0.38807145 | 0.697963159 | 0.839976943 | 0.154816658 |
| ENSMUSG00000099881 | 2810013P06Rik | 100503178 | lncRNA | 159.4390802 | -0.154842553 | 0.129250337 | 1.19800503 | 0.230915059 | 0.439526427 | 0.154842553 |
| ENSMUSG00000003955 | Fam162a | 70186 | protein_coding | 1099.613046 | -0.154857898 | 0.121701793 | 1.272437282 | 0.203217795 | 0.405453505 | 0.154857898 |
| ENSMUSG00000002504 | Slc9a3r2 | 65962 | protein_coding | 111.7672352 | -0.154919656 | 0.143681027 | 1.078219294 | 0.280935904 | 0.498589523 | 0.154919656 |
| ENSMUSG00000060152 | Pop5 | 117109 | protein_coding | 223.7617528 | -0.154920333 | 0.102210292 | 1.515701893 | 0.129594749 | 0.297320449 | 0.154920333 |
| ENSMUSG00000025024 | Smndc1 | 76479 | protein_coding | 269.6054969 | -0.154920355 | 0.106586852 | 1.453465899 | 0.146094441 | 0.323442761 | 0.154920355 |
| ENSMUSG00000088242 | Gm25913 | 115485866 | scaRNA | 0.727869722 | -0.154931699 | 1.556620252 | 0.099530826 | 0.920716814 | 1 | 0.154931699 |
| ENSMUSG00000023039 | Krt7 | 110310 | protein_coding | 2104.849289 | 0.15499578 | 0.088774204 | -1.745955173 | 0.080818741 | 0.211344528 | 0.15499578 |
| ENSMUSG00000027498 | Cstf1 | 67337 | protein_coding | 311.233901 | 0.155044453 | 0.11352519 | -1.365727317 | 0.172024567 | 0.362441146 | 0.155044453 |
| ENSMUSG00000015083 | C8g | 69379 | protein_coding | 24.06824754 | -0.15508757 | 0.207457481 | 0.747563157 | 0.454723695 | 0.670174662 | 0.15508757 |
| ENSMUSG00000057778 | Cyb5d2 | 192986 | protein_coding | 98.09467096 | -0.155109308 | 0.103838961 | 1.49374865 | 0.135241328 | 0.306361493 | 0.155109308 |
| ENSMUSG00000033068 | Entpd6 | 12497 | protein_coding | 777.6490143 | 0.155121424 | 0.078163828 | -1.984567904 | 0.047192566 | 0.142788208 | 0.155121424 |
| ENSMUSG00000021366 | Hivep1 | 110521 | protein_coding | 683.6867437 | -0.155195792 | 0.10502362 | 1.477722754 | 0.139482002 | 0.313005269 | 0.155195792 |
| ENSMUSG00000060216 | Arrb2 | 216869 | protein_coding | 35.11956412 | 0.15525835 | 0.229871072 | -0.675414911 | 0.499412194 | 0.704201299 | 0.15525835 |
| ENSMUSG00000030768 | Disp1 | 68897 | protein_coding | 113.2067798 | 0.155263497 | 0.108423633 | -1.432007883 | 0.152141565 | 0.332776929 | 0.155263497 |
| ENSMUSG00000028345 | Tex10 | 269536 | protein_coding | 179.7706571 | -0.155329035 | 0.111914859 | 1.387921458 | 0.165160963 | 0.352113109 | 0.155329035 |
| ENSMUSG00000025907 | Rb1cc1 | 12421 | protein_coding | 994.8854898 | -0.155329564 | 0.090320246 | 1.719764624 | 0.085475235 | 0.220621763 | 0.155329564 |
| ENSMUSG00000040524 | Zfp609 | 214812 | protein_coding | 287.5600983 | -0.155368447 | 0.11975672 | 1.297367254 | 0.194504854 | 0.393392916 | 0.155368447 |
| ENSMUSG00000034807 | Colgalt1 | 234407 | protein_coding | 610.3715325 | -0.155427521 | 0.069221678 | 2.245359049 | 0.024745092 | 0.086686561 | 0.155427521 |
| ENSMUSG00000064120 | Mocs1 | 56738 | protein_coding | 300.4099617 | -0.155465815 | 0.088742832 | 1.751868982 | 0.079796339 | 0.209433685 | 0.155465815 |
| ENSMUSG00000020159 | Gabrp | 216643 | protein_coding | 3.185763747 | 0.155480949 | 1.123853342 | -0.138346298 | 0.889966737 | 1 | 0.155480949 |
| ENSMUSG00000036931 | Nfkbid | 243910 | protein_coding | 54.12428243 | 0.155487116 | 0.272366735 | -0.570874102 | 0.568084987 | 0.754128254 | 0.155487116 |
| ENSMUSG00000026035 | Ppil3 | 70225 | protein_coding | 111.8224253 | 0.155530895 | 0.123059077 | -1.263871781 | 0.206276052 | 0.408887615 | 0.155530895 |
| ENSMUSG00000074283 | Zfp109 | 56869 | protein_coding | 60.41686287 | -0.155604184 | 0.168839833 | 0.921608258 | 0.356732949 | 0.579660329 | 0.155604184 |
| ENSMUSG00000041258 | Zfp236 | 329002 | protein_coding | 582.9771293 | 0.155611926 | 0.0694226 | -2.24151683 | 0.024992617 | 0.087325531 | 0.155611926 |

| ensembl_gene_id | external_gene_name | entrezgene_id | gene_biotype | baseMean | log2FoldChange | lfcSE | stat | pvalue | padj | absLog2FC |
|---|---|---|---|---|---|---|---|---|---|---|
| ENSMUSG00000085427 | 6430710C18Rik | 76218 | lncRNA | 5.413965063 | 0.155634789 | 0.465524612 | -0.334321291 | 0.738137126 | 0.866982505 | 0.155634789 |
| ENSMUSG00000050855 | Zfp940 | 233057 | protein_coding | 21.99464052 | 0.15563629 | 0.21558927 | -0.721911115 | 0.470349127 | 0.683706314 | 0.15563629 |
| ENSMUSG00000027524 | Edn3 | 13616 | protein_coding | 80.22262962 | 0.155649413 | 0.285524838 | -0.545134406 | 0.585661081 | 0.767483504 | 0.155649413 |
| ENSMUSG00000037465 | Klf10 | 21847 | protein_coding | 1369.147879 | 0.155675479 | 0.127524655 | -1.220748082 | 0.222181417 | 0.429285587 | 0.155675479 |
| ENSMUSG00000024812 | Tjp2 | 21873 | protein_coding | 2205.954975 | 0.155704573 | 0.063052907 | -2.469427351 | 0.013532949 | 0.053877321 | 0.155704573 |
| ENSMUSG00000029787 | Avl9 | 78937 | protein_coding | 1317.433956 | -0.155790786 | 0.092708193 | 1.68044248 | 0.092871257 | 0.233934706 | 0.155790786 |
| ENSMUSG00000039330 | Tsga10ip | 78306 | protein_coding | 2.800449909 | -0.155817219 | 1.181200122 | 0.131914327 | 0.895052057 | 1 | 0.155817219 |
| ENSMUSG00000027570 | Col9a3 | 12841 | protein_coding | 9.50997761 | -0.155841883 | 0.524394339 | 0.297184527 | 0.766325635 | 0.882690208 | 0.155841883 |
| ENSMUSG00000039634 | Zfp189 | 230162 | protein_coding | 23.24689143 | -0.155915278 | 0.23022212 | 0.677238477 | 0.498254659 | 0.703443399 | 0.155915278 |
| ENSMUSG00000048534 | Jaml | 270152 | protein_coding | 17.69351534 | 0.155980962 | 0.296653513 | -0.525801838 | 0.599025897 | 0.776435029 | 0.155980962 |
| ENSMUSG00000007039 | Ddah2 | 51793 | protein_coding | 15.70547817 | -0.156019503 | 0.376865745 | 0.413992266 | 0.678879771 | 0.827931507 | 0.156019503 |
| ENSMUSG00000038305 | Spats2l | 67198 | protein_coding | 141.0291859 | -0.156145043 | 0.223608518 | 0.698296489 | 0.484991794 | 0.695300529 | 0.156145043 |
| ENSMUSG00000046314 | Stxbp6 | 217517 | protein_coding | 271.2728788 | -0.1561621 | 0.121278417 | 1.287633069 | 0.197873722 | 0.39815848 | 0.1561621 |
| ENSMUSG00000027163 | Commd9 | 76501 | protein_coding | 275.6485141 | 0.156172208 | 0.087117247 | -1.792666921 | 0.073026199 | 0.196922705 | 0.156172208 |
| ENSMUSG00000013593 | Ndufs2 | 226646 | protein_coding | 3334.428724 | -0.156221235 | 0.078420824 | 1.992088687 | 0.046361328 | 0.14084087 | 0.156221235 |
| ENSMUSG00000073002 | Vamp5 | 53620 | protein_coding | 10.28557819 | 0.156239383 | 0.413684869 | -0.377677295 | 0.705670336 | 0.845753813 | 0.156239383 |
| ENSMUSG00000040195 | Nemp1 | 72243 | protein_coding | 122.6003624 | 0.156241582 | 0.157741451 | -0.990491598 | 0.321933894 | 0.543156551 | 0.156241582 |
| ENSMUSG00000033701 | Acbd6 | 72482 | protein_coding | 211.0248237 | -0.15628415 | 0.131353633 | 1.189796938 | 0.234126213 | 0.443634979 | 0.15628415 |
| ENSMUSG00000004895 | Prcc | 94315 | protein_coding | 533.6840781 | -0.156295462 | 0.057828923 | 2.70272129 | 0.006877439 | 0.031443528 | 0.156295462 |
| ENSMUSG00000060073 | Psma3 | 19167 | protein_coding | 810.4499901 | -0.156337342 | 0.111260673 | 1.405144666 | 0.159978264 | 0.344285818 | 0.156337342 |
| ENSMUSG00000038722 | Bud31 | 231889 | protein_coding | 335.7754621 | 0.156416099 | 0.079699281 | -1.962578547 | 0.049695166 | 0.148254918 | 0.156416099 |
| ENSMUSG00000036292 | Gramd1c | 207798 | protein_coding | 129.1398443 | -0.156451806 | 0.114847303 | 1.362259296 | 0.173116069 | 0.363688435 | 0.156451806 |
| ENSMUSG00000046679 | C87436 | 232196 | protein_coding | 142.7843888 | -0.156473941 | 0.118682469 | 1.318425058 | 0.187361397 | 0.383501006 | 0.156473941 |
| ENSMUSG00000003435 | Supt5 | 20924 | protein_coding | 1100.792404 | 0.156500161 | 0.05171416 | -3.026253585 | 0.002476045 | 0.01350377 | 0.156500161 |
| ENSMUSG00000030621 | Me3 | 109264 | protein_coding | 19.70898627 | -0.15654281 | 0.449724386 | 0.348086105 | 0.727775519 | 0.859322531 | 0.15654281 |
| ENSMUSG00000073787 | Gm10575 | 100126795 | lncRNA | 14.11834696 | -0.15656965 | 0.390479095 | 0.400968073 | 0.68844363 | 0.83386713 | 0.15656965 |
| ENSMUSG00000003534 | Ddr1 | 12305 | protein_coding | 1570.915281 | -0.156590423 | 0.091780474 | 1.706140926 | 0.087981844 | 0.22515016 | 0.156590423 |
| ENSMUSG00000029471 | Camkk2 | 207565 | protein_coding | 53.09128985 | 0.156638143 | 0.209392487 | -0.748059996 | 0.45442397 | 0.669912287 | 0.156638143 |
| ENSMUSG00000030421 | Uri1 | 19777 | protein_coding | 398.7418389 | 0.15667653 | 0.105361645 | -1.487035719 | 0.137005377 | 0.309296897 | 0.15667653 |
| ENSMUSG00000031984 | 2810004N23Rik | 66523 | protein_coding | 253.3436657 | -0.156677062 | 0.108562518 | 1.443196649 | 0.148965083 | 0.328001251 | 0.156677062 |
| ENSMUSG00000042670 | Immp1l | 66541 | protein_coding | 174.4201784 | -0.156679447 | 0.13272415 | 1.180489357 | 0.237805641 | 0.448045905 | 0.156679447 |
| ENSMUSG00000023909 | Paqr4 | 76498 | protein_coding | 2166.79389 | 0.156730201 | 0.059036683 | -2.654793483 | 0.007935698 | 0.035388872 | 0.156730201 |
| ENSMUSG00000051339 | 2900026A02Rik | 243219 | protein_coding | 3618.16354 | -0.156778446 | 0.176221497 | 0.889666974 | 0.373644732 | 0.595844868 | 0.156778446 |
| ENSMUSG00000032590 | Apeh | 235606 | protein_coding | 806.4781105 | -0.15677922 | 0.076116887 | 2.059716667 | 0.039425635 | 0.12467923 | 0.15677922 |
| ENSMUSG00000003033 | Ap1m1 | 11767 | protein_coding | 132.3115318 | 0.156787116 | 0.115840479 | -1.353474341 | 0.175904142 | 0.367893 | 0.156787116 |
| ENSMUSG00000019832 | Rab32 | 67844 | protein_coding | 178.8260189 | -0.156803125 | 0.227369878 | 0.689638953 | 0.490421265 | 0.69793377 | 0.156803125 |
| ENSMUSG00000040600 | Eps8l3 | 99662 | protein_coding | 2039.571743 | -0.156825862 | 0.123991511 | 1.264811284 | 0.205938985 | 0.408576247 | 0.156825862 |
| ENSMUSG00000029029 | Wrap73 | 59002 | protein_coding | 155.0830297 | 0.156841378 | 0.107584023 | -1.457850089 | 0.144881869 | 0.321929383 | 0.156841378 |
| ENSMUSG00000000627 | Sema4f | 20355 | protein_coding | 3.814325651 | 0.156850784 | 0.856214303 | -0.183191035 | 0.854648129 | 1 | 0.156850784 |

| ensembl_gene_id | external_gene_name | entrezgene_id | gene_biotype | baseMean | log2FoldChange | lfcSE | stat | pvalue | padj | absLog2FC |
|---|---|---|---|---|---|---|---|---|---|---|
| ENSMUSG00000021054 | Sgpp1 | 81535 | protein_coding | 411.2425374 | -0.156853183 | 0.08955951 | 1.751385018 | 0.079879612 | 0.209618927 | 0.156853183 |
| ENSMUSG00000029022 | Miip | 28010 | protein_coding | 199.2010524 | 0.156938057 | 0.092270918 | -1.700839886 | 0.088973057 | 0.226849015 | 0.156938057 |
| ENSMUSG00000032977 | Fam207a | 108707 | protein_coding | 339.8313313 | 0.156954947 | 0.069611326 | -2.254732907 | 0.024150099 | 0.085053259 | 0.156954947 |
| ENSMUSG00000034432 | Cops8 | 108679 | protein_coding | 334.0243595 | 0.156975723 | 0.072739184 | -2.158062744 | 0.030922954 | 0.103308737 | 0.156975723 |
| ENSMUSG00000025413 | Ttc4 | 72354 | protein_coding | 536.1758669 | 0.156981598 | 0.061452142 | -2.554534191 | 0.010632999 | 0.044782795 | 0.156981598 |
| ENSMUSG00000090231 | Cfb | 14962 | protein_coding | 103.6827595 | -0.157026186 | 0.136547157 | 1.14997771 | 0.250153052 | 0.462497481 | 0.157026186 |
| ENSMUSG00000022201 | Zfr | 22763 | protein_coding | 961.7960191 | 0.157028384 | 0.056305086 | -2.788884555 | 0.005288991 | 0.02553125 | 0.157028384 |
| ENSMUSG00000035311 | Gnptab | 432486 | protein_coding | 536.944865 | -0.157036493 | 0.128951576 | 1.217794286 | 0.223302156 | 0.430343308 | 0.157036493 |
| ENSMUSG00000021018 | Polr2h | 245841 | protein_coding | 233.6282871 | -0.15707163 | 0.086261518 | 1.820877179 | 0.068625529 | 0.1881903 | 0.15707163 |
| ENSMUSG00000040818 | Dennd6a | 211922 | protein_coding | 555.5204888 | -0.157111962 | 0.115461874 | 1.360725894 | 0.173600329 | 0.364391266 | 0.157111962 |
| ENSMUSG00000037400 | Atp11b | 76295 | protein_coding | 1942.866814 | 0.15712495 | 0.075172463 | -2.090192927 | 0.036600473 | 0.11738594 | 0.15712495 |
| ENSMUSG00000042350 | Arel1 | 68497 | protein_coding | 724.3174308 | -0.15719216 | 0.064693179 | 2.429810408 | 0.015106723 | 0.058719535 | 0.15719216 |
| ENSMUSG00000045268 | Zfp691 | 195522 | protein_coding | 78.58897589 | -0.157236975 | 0.122978398 | 1.278573943 | 0.201047131 | 0.402386658 | 0.157236975 |
| ENSMUSG00000090534 | Gm4675 | 105245869 | lncRNA | 9.848144355 | -0.15724788 | 0.358975946 | 0.43804573 | 0.661353133 | 0.816921981 | 0.15724788 |
| ENSMUSG00000086859 | Snhg20 | 76972 | lncRNA | 35.5765524 | -0.157261829 | 0.18790288 | 0.836931444 | 0.402631104 | 0.624030976 | 0.157261829 |
| ENSMUSG00000032942 | Ucp3 | 22229 | protein_coding | 1.496369095 | 0.157333059 | 1.074386859 | -0.146439858 | 0.883574164 | 1 | 0.157333059 |
| ENSMUSG00000031621 | Isx | 71597 | protein_coding | 1237.892139 | -0.157380836 | 0.206052815 | 0.763788815 | 0.444993103 | 0.662372535 | 0.157380836 |
| ENSMUSG00000098905 | Zfp953 | 629016 | protein_coding | 97.28190839 | -0.157395496 | 0.129057674 | 1.219574866 | 0.222626079 | 0.429692271 | 0.157395496 |
| ENSMUSG00000024777 | Ppp2r5b | 225849 | protein_coding | 412.7646232 | -0.157397483 | 0.101835097 | 1.545611362 | 0.122198456 | 0.285308256 | 0.157397483 |
| ENSMUSG00000060639 | H4c9 | 319158 | protein_coding | 160.1689433 | 0.15740548 | 0.165439514 | -0.95143824 | 0.341381956 | 0.563251236 | 0.15740548 |
| ENSMUSG00000074748 | Atxn7l3b | 382423 | protein_coding | 1244.986204 | -0.157415013 | 0.105812464 | 1.487679305 | 0.136835488 | 0.309078852 | 0.157415013 |
| ENSMUSG00000015869 | Prpsap1 | 67763 | protein_coding | 1118.333419 | -0.157454797 | 0.119610711 | 1.316393788 | 0.188041906 | 0.384369853 | 0.157454797 |
| ENSMUSG00000039828 | Wdr70 | 545085 | protein_coding | 122.2934343 | 0.157462231 | 0.212930882 | -0.739499267 | 0.459603885 | 0.673998628 | 0.157462231 |
| ENSMUSG00000041842 | Fhdc1 | 229474 | protein_coding | 250.7260641 | 0.157509124 | 0.235071002 | -0.670049146 | 0.502826461 | 0.707478587 | 0.157509124 |
| ENSMUSG00000023074 | Mospd1 | 70380 | protein_coding | 169.6334941 | -0.157534991 | 0.120777789 | 1.304337433 | 0.192118558 | 0.389807821 | 0.157534991 |
| ENSMUSG00000029165 | Agbl5 | 231093 | protein_coding | 131.7318227 | 0.157544674 | 0.100235193 | -1.571750093 | 0.11600852 | 0.274998498 | 0.157544674 |
| ENSMUSG00000029499 | Pxmp2 | 19301 | protein_coding | 42.65069727 | -0.157556578 | 0.175119179 | 0.899710577 | 0.368274293 | 0.5907413 | 0.157556578 |
| ENSMUSG00000036989 | Trim3 | 55992 | protein_coding | 78.227704 | -0.157559259 | 0.17529618 | 0.898817415 | 0.368749923 | 0.591274514 | 0.157559259 |
| ENSMUSG00000026811 | St6galnac6 | 50935 | protein_coding | 7907.948593 | 0.157564674 | 0.090131697 | -1.748160526 | 0.080436235 | 0.210845076 | 0.157564674 |
| ENSMUSG00000059890 | Ube4a | 140630 | protein_coding | 583.7229615 | 0.157655864 | 0.074719339 | -2.109974012 | 0.034860594 | 0.11310174 | 0.157655864 |
| ENSMUSG00000026975 | Dph7 | 67228 | protein_coding | 83.50354579 | 0.157685076 | 0.134076557 | -1.176082384 | 0.239561957 | 0.449849479 | 0.157685076 |
| ENSMUSG00000042763 | Maneal | 215090 | protein_coding | 5.648226952 | -0.157834368 | 0.533359036 | 0.295925179 | 0.767287222 | 0.883364555 | 0.157834368 |
| ENSMUSG00000032251 | Irak1bp1 | 65099 | protein_coding | 2.927674926 | 0.157897695 | 0.773946867 | -0.204016196 | 0.838340843 | 1 | 0.157897695 |
| ENSMUSG00000039254 | Pomt1 | 99011 | protein_coding | 322.9467453 | 0.157918568 | 0.087102852 | -1.813012591 | 0.069829879 | 0.190674731 | 0.157918568 |
| ENSMUSG00000037697 | Ddhd1 | 114874 | protein_coding | 202.3166707 | 0.158004082 | 0.076124451 | -2.075602249 | 0.037930753 | 0.120901073 | 0.158004082 |
| ENSMUSG00000021661 | Ankra2 | 68558 | protein_coding | 200.4555883 | -0.158010356 | 0.095196153 | 1.659839716 | 0.096946702 | 0.24162056 | 0.158010356 |
| ENSMUSG00000024346 | Pfdn1 | 67199 | protein_coding | 480.4818061 | 0.158037351 | 0.06790117 | -2.327461366 | 0.019940723 | 0.07322684 | 0.158037351 |
| ENSMUSG00000028790 | Khdrbs1 | 20218 | protein_coding | 842.778024 | 0.158088623 | 0.072387137 | -2.183932521 | 0.028967207 | 0.098226787 | 0.158088623 |
| ENSMUSG00000026349 | Ccnt2 | 72949 | protein_coding | 716.2202884 | -0.15810535 | 0.057389477 | 2.754953665 | 0.005870045 | 0.027715628 | 0.15810535 |

| ensembl_gene_id | external_gene_name | entrezgene_id | gene_biotype | baseMean | log2FoldChange | lfcSE | stat | pvalue | padj | absLog2FC |
|---|---|---|---|---|---|---|---|---|---|---|
| ENSMUSG00000068923 | Syt11 | 229521 | protein_coding | 32.70692022 | -0.158125923 | 0.196710953 | 0.803849105 | 0.421484126 | 0.641215003 | 0.158125923 |
| ENSMUSG00000032245 | Cln6 | 76524 | protein_coding | 245.9019106 | -0.158146758 | 0.127306944 | 1.24224769 | 0.214145189 | 0.419152975 | 0.158146758 |
| ENSMUSG00000029640 | Usp12 | 22217 | protein_coding | 959.0244582 | 0.158209676 | 0.075250122 | -2.10245075 | 0.03551381 | 0.114725166 | 0.158209676 |
| ENSMUSG00000041319 | Thoc6 | 386612 | protein_coding | 138.859101 | 0.158395807 | 0.148490465 | -1.066706923 | 0.2861042 | 0.50453867 | 0.158395807 |
| ENSMUSG00000052406 | Rexo4 | 227656 | protein_coding | 388.681795 | 0.158520777 | 0.092716777 | -1.709731302 | 0.087315572 | 0.224110872 | 0.158520777 |
| ENSMUSG00000024911 | Fibp | 58249 | protein_coding | 427.7657657 | -0.158520963 | 0.061934155 | 2.559507955 | 0.010482046 | 0.044259909 | 0.158520963 |
| ENSMUSG00000000374 | Trappc10 | 216131 | protein_coding | 2033.573954 | 0.158524524 | 0.066362506 | -2.388766383 | 0.016905048 | 0.064265646 | 0.158524524 |
| ENSMUSG00000029202 | Pds5a | 71521 | protein_coding | 1575.933899 | 0.158525468 | 0.069652026 | -2.275963477 | 0.022848196 | 0.081458356 | 0.158525468 |
| ENSMUSG00000020372 | Rack1 | 14694 | protein_coding | 11895.48733 | -0.158530305 | 0.102615797 | 1.544891817 | 0.122372433 | 0.285472293 | 0.158530305 |
| ENSMUSG00000024340 | Btnl2 | 547431 | protein_coding | 36.40047298 | -0.158570466 | 0.395006246 | 0.40143787 | 0.688097773 | 0.833786242 | 0.158570466 |
| ENSMUSG00000053263 | Gm12592 | 791413 | lncRNA | 3.418988032 | 0.158648128 | 0.815765754 | -0.194477553 | 0.84580197 | 1 | 0.158648128 |
| ENSMUSG00000026069 | Il1rl1 | 17082 | protein_coding | 3.475580188 | -0.15870246 | 1.023984497 | 0.154985218 | 0.876832971 | 1 | 0.15870246 |
| ENSMUSG00000042460 | C1galt1 | 94192 | protein_coding | 3304.404108 | 0.158770538 | 0.150044339 | -1.058157474 | 0.289983655 | 0.508958453 | 0.158770538 |
| ENSMUSG00000107167 | B3galt9 | 102637973 | protein_coding | 0.902170874 | -0.158824566 | 1.553972013 | 0.102205551 | 0.918593522 | 1 | 0.158824566 |
| ENSMUSG00000029713 | Gnb2 | 14693 | protein_coding | 3181.631548 | 0.158826515 | 0.049648944 | -3.198990804 | 0.001379096 | 0.00832093 | 0.158826515 |
| ENSMUSG00000024527 | Afg3l2 | 69597 | protein_coding | 1571.543025 | -0.1588558 | 0.05510094 | 2.882996181 | 0.003939122 | 0.01996653 | 0.1588558 |
| ENSMUSG00000000823 | Zfp512b | 269401 | protein_coding | 244.5038747 | -0.159002906 | 0.090112431 | 1.764494685 | 0.077648729 | 0.205759165 | 0.159002906 |
| ENSMUSG00000030029 | Lrig1 | 16206 | protein_coding | 1960.805077 | 0.159007014 | 0.144603707 | -1.09960538 | 0.271504097 | 0.487190988 | 0.159007014 |
| ENSMUSG00000015804 | Med28 | 66999 | protein_coding | 493.0832596 | -0.159020027 | 0.087687675 | 1.813482076 | 0.069757501 | 0.190566644 | 0.159020027 |
| ENSMUSG00000020961 | Ston2 | 108800 | protein_coding | 401.0276201 | -0.15908998 | 0.155835463 | 1.020884313 | 0.307309252 | 0.527754626 | 0.15908998 |
| ENSMUSG00000024544 | Ldlrad4 | 52662 | protein_coding | 35.10476049 | -0.15909421 | 0.226246143 | 0.703190816 | 0.481936842 | 0.692663621 | 0.15909421 |
| ENSMUSG00000049504 | Proser1 | 212127 | protein_coding | 85.42094046 | 0.159094406 | 0.221705519 | -0.71759335 | 0.473008059 | 0.685318046 | 0.159094406 |
| ENSMUSG00000025500 | Lmntd2 | 72000 | protein_coding | 59.7741332 | -0.159104902 | 0.218755556 | 0.727318222 | 0.467031038 | 0.680659303 | 0.159104902 |
| ENSMUSG00000085028 | Slc2a4rg-ps | 329584 | transcribed_unitary_pseudogene | 50.7606642 | -0.159129009 | 0.194089288 | 0.819875279 | 0.412287211 | 0.632329199 | 0.159129009 |
| ENSMUSG00000042626 | Shc1 | 20416 | protein_coding | 941.049274 | 0.159132153 | 0.059786981 | -2.661652257 | 0.007775816 | 0.034799413 | 0.159132153 |
| ENSMUSG00000025220 | Oga | 76055 | protein_coding | 1615.433346 | -0.159182519 | 0.048414739 | 3.287893798 | 0.001009399 | 0.006402953 | 0.159182519 |
| ENSMUSG00000029629 | Phf14 | 75725 | protein_coding | 388.4857469 | -0.15919458 | 0.094156977 | 1.690735887 | 0.090887257 | 0.230377922 | 0.15919458 |
| ENSMUSG00000030729 | Pgm2l1 | 70974 | protein_coding | 239.2441964 | -0.159203345 | 0.135163793 | 1.177854966 | 0.238854431 | 0.449052273 | 0.159203345 |
| ENSMUSG00000039616 | Mocos | 68591 | protein_coding | 138.9512822 | -0.159224397 | 0.191844824 | 0.829964517 | 0.406558845 | 0.627409847 | 0.159224397 |
| ENSMUSG00000091618 | H60c | 670558 | protein_coding | 34.13455698 | 0.159247408 | 0.286923381 | -0.555017189 | 0.578882904 | 0.762395109 | 0.159247408 |
| ENSMUSG00000036632 | Alg5 | 66248 | protein_coding | 374.1002987 | 0.159304801 | 0.108747103 | -1.464910751 | 0.142945269 | 0.318770942 | 0.159304801 |
| ENSMUSG00000033392 | Clasp2 | 76499 | protein_coding | 315.7409898 | 0.159335574 | 0.082577153 | -1.929535816 | 0.053664378 | 0.157297398 | 0.159335574 |
| ENSMUSG00000047832 | Cdca4 | 71963 | protein_coding | 275.3491748 | 0.159339116 | 0.11024835 | -1.445274393 | 0.148380828 | 0.326976379 | 0.159339116 |
| ENSMUSG00000021175 | Cdca7l | 217946 | protein_coding | 125.7894856 | 0.159344422 | 0.137821383 | -1.156166182 | 0.24761319 | 0.459757614 | 0.159344422 |
| ENSMUSG00000032307 | Ube2q2 | 109161 | protein_coding | 278.7101655 | -0.159354973 | 0.117833347 | 1.352375843 | 0.176255113 | 0.368253661 | 0.159354973 |
| ENSMUSG00000046618 | Olfml2a | 241327 | protein_coding | 2.356296725 | -0.159395148 | 0.686807087 | 0.232081396 | 0.816474794 | 1 | 0.159395148 |
| ENSMUSG00000037742 | Eef1a1 | 13627 | protein_coding | 66955.30617 | -0.159397411 | 0.110622645 | 1.440911224 | 0.149609765 | 0.328982111 | 0.159397411 |
| ENSMUSG00000062075 | Lmnb2 | 16907 | protein_coding | 860.6863236 | -0.15948353 | 0.076367928 | 2.088357429 | 0.036765605 | 0.117801163 | 0.15948353 |
| ENSMUSG00000020024 | Cep83 | 77048 | protein_coding | 276.7792167 | 0.159512584 | 0.089960322 | -1.773143768 | 0.076204887 | 0.202844537 | 0.159512584 |

Supplementary Table 1

| ensembl_gene_id | external_gene_name | entrezgene_id | gene_biotype | baseMean | log2FoldChange | lfcSE | stat | pvalue | padj | absLog2FC |
|---|---|---|---|---|---|---|---|---|---|---|
| ENSMUSG00000031352 | Hccs | 15159 | protein_coding | 313.0555181 | 0.159664382 | 0.064467087 | -2.476680565 | 0.013261053 | 0.053024211 | 0.159664382 |
| ENSMUSG00000031200 | Mtcp1 | 17763 | protein_coding | 36.33218065 | 0.159685863 | 0.21445454 | -0.744614049 | 0.456505073 | 0.671491432 | 0.159685863 |
| ENSMUSG00000052974 | Cyp2f2 | 13107 | protein_coding | 152.3939504 | -0.159703978 | 0.276439555 | 0.577717535 | 0.563454839 | 0.750868206 | 0.159703978 |
| ENSMUSG00000052681 | Rap1b | 215449 | protein_coding | 1097.10265 | 0.159746657 | 0.075113827 | -2.126727719 | 0.033442713 | 0.109621469 | 0.159746657 |
| ENSMUSG00000032261 | Sh3bgrl2 | 212531 | protein_coding | 2571.618076 | -0.159759555 | 0.077234943 | 2.06848804 | 0.038594154 | 0.122566127 | 0.159759555 |
| ENSMUSG00000041143 | Tmco4 | 77056 | protein_coding | 849.9724127 | -0.159765634 | 0.0957339 | 1.668851199 | 0.095146873 | 0.238311216 | 0.159765634 |
| ENSMUSG00000106629 | Gm40155 | 105244562 | lncRNA | 1.331891768 | 0.159777445 | 1.09231206 | -0.146274541 | 0.883704663 | 1 | 0.159777445 |
| ENSMUSG00000058240 | Cryzl1 | 66609 | protein_coding | 285.1971739 | -0.159841578 | 0.076765435 | 2.082207671 | 0.037323505 | 0.119334048 | 0.159841578 |
| ENSMUSG00000033282 | Rpgrip1l | 244585 | protein_coding | 122.9498009 | 0.159861674 | 0.109483075 | -1.460149662 | 0.144248946 | 0.320998127 | 0.159861674 |
| ENSMUSG00000032314 | Etfa | 110842 | protein_coding | 1179.25782 | -0.159964614 | 0.069130247 | 2.313959819 | 0.020669926 | 0.075235434 | 0.159964614 |
| ENSMUSG00000002804 | Nudt14 | 66174 | protein_coding | 276.8796533 | -0.159965961 | 0.082635109 | 1.935811097 | 0.052890841 | 0.155751645 | 0.159965961 |
| ENSMUSG00000041355 | Ssr2 | 66256 | protein_coding | 1548.889367 | 0.16001605 | 0.100751257 | -1.588228821 | 0.112234606 | 0.268614824 | 0.16001605 |
| ENSMUSG00000079614 | Seh1l | 72124 | protein_coding | 504.4110429 | 0.160023847 | 0.107639795 | -1.486660653 | 0.13710446 | 0.309427758 | 0.160023847 |
| ENSMUSG00000045795 | Whamm | 434204 | protein_coding | 257.1419507 | 0.160037877 | 0.082887494 | -1.930784358 | 0.053509726 | 0.157015607 | 0.160037877 |
| ENSMUSG00000085154 | C130046K22Rik | 399609 | lncRNA | 6.315412206 | -0.160043341 | 0.405131123 | 0.395040846 | 0.692812744 | 0.836797094 | 0.160043341 |
| ENSMUSG00000072618 | Gm10384 | 100038735 | lncRNA | 29.42477144 | -0.160051253 | 0.246243092 | 0.649972562 | 0.515709945 | 0.716728293 | 0.160051253 |
| ENSMUSG00000031758 | Cdyl2 | 75796 | protein_coding | 28.44988265 | 0.160052384 | 0.289905594 | -0.552084495 | 0.580890464 | 0.764064128 | 0.160052384 |
| ENSMUSG00000064264 | Zfp428 | 232969 | protein_coding | 21.03887605 | 0.160063876 | 0.254313138 | -0.629396803 | 0.529089311 | 0.726903567 | 0.160063876 |
| ENSMUSG00000008140 | Emc10 | 69683 | protein_coding | 648.4352733 | -0.160072055 | 0.064990048 | 2.463024122 | 0.013777065 | 0.054559819 | 0.160072055 |
| ENSMUSG00000025064 | Col17a1 | 12821 | protein_coding | 14.92945471 | 0.160136552 | 0.426195617 | -0.375734864 | 0.707114016 | 0.846609212 | 0.160136552 |
| ENSMUSG00000024511 | Rab27b | 80718 | protein_coding | 918.5472493 | 0.160137711 | 0.181918241 | -0.880272975 | 0.378711449 | 0.601173216 | 0.160137711 |
| ENSMUSG00000040738 | Ints8 | 72656 | protein_coding | 390.6399349 | 0.160158267 | 0.089107672 | -1.797356656 | 0.072279017 | 0.19557897 | 0.160158267 |
| ENSMUSG00000041720 | Pi4ka | 224020 | protein_coding | 834.5102092 | -0.16016862 | 0.067909514 | 2.358559373 | 0.018346025 | 0.068544401 | 0.16016862 |
| ENSMUSG00000020741 | Cluh | 74148 | protein_coding | 2974.017835 | -0.160190289 | 0.070440509 | 2.27412168 | 0.022958672 | 0.081727701 | 0.160190289 |
| ENSMUSG00000048445 | Ccdc57 | 71276 | protein_coding | 45.00227416 | -0.160201647 | 0.208338734 | 0.768947971 | 0.441924198 | 0.659670632 | 0.160201647 |
| ENSMUSG00000026301 | Iqca | 74918 | protein_coding | 2.41105814 | 0.160232371 | 2.184985957 | -0.073333364 | 0.941540843 | 1 | 0.160232371 |
| ENSMUSG00000025531 | Chm | 12662 | protein_coding | 297.9115274 | 0.16025776 | 0.071421349 | -2.243835531 | 0.024842985 | 0.086937288 | 0.16025776 |
| ENSMUSG00000041263 | Rusc1 | 72296 | protein_coding | 95.96319436 | -0.160342808 | 0.1023962 | 1.565905851 | 0.11737066 | 0.277378268 | 0.160342808 |
| ENSMUSG00000035649 | Zcchc7 | 319885 | protein_coding | 430.1043593 | 0.160343405 | 0.081207922 | -1.974479839 | 0.048327208 | 0.145342471 | 0.160343405 |
| ENSMUSG00000052605 | Isoc2b | 67441 | protein_coding | 157.8760174 | -0.160359781 | 0.122852342 | 1.305305033 | 0.191789001 | 0.389522022 | 0.160359781 |
| ENSMUSG00000072258 | Taf1a | 21339 | protein_coding | 87.24456913 | -0.160384564 | 0.115248564 | 1.39164046 | 0.1640313 | 0.350247335 | 0.160384564 |
| ENSMUSG00000047804 | Akap10 | 56697 | protein_coding | 415.5328382 | -0.160391194 | 0.060158435 | 2.666146396 | 0.007672627 | 0.034419117 | 0.160391194 |
| ENSMUSG00000020720 | Psmd12 | 66997 | protein_coding | 576.924462 | 0.160408311 | 0.08668766 | -1.850416907 | 0.064253484 | 0.179449017 | 0.160408311 |
| ENSMUSG00000022010 | Tsc22d1 | 21807 | protein_coding | 4251.953711 | -0.160414272 | 0.066738962 | 2.403607531 | 0.016234192 | 0.062245517 | 0.160414272 |
| ENSMUSG00000040820 | Hlcs | 110948 | protein_coding | 175.1739812 | -0.16042608 | 0.14279538 | 1.123468279 | 0.261238665 | 0.47491467 | 0.16042608 |
| ENSMUSG00000024292 | Cyp4f14 | 64385 | protein_coding | 1529.276432 | 0.160426508 | 0.184511347 | -0.869466892 | 0.384591809 | 0.607010336 | 0.160426508 |
| ENSMUSG00000033054 | Npat | 244879 | protein_coding | 302.0202261 | 0.160458388 | 0.097493467 | -1.645837325 | 0.099797256 | 0.247269599 | 0.160458388 |
| ENSMUSG00000048668 | Rhno1 | 72440 | protein_coding | 120.8270483 | 0.160484055 | 0.140445617 | -1.142677557 | 0.253172491 | 0.465629861 | 0.160484055 |
| ENSMUSG00000022836 | Mylk | 107589 | protein_coding | 629.219247 | -0.160506778 | 0.133076105 | 1.206127716 | 0.22776825 | 0.436353234 | 0.160506778 |

Supplementary Table 1

| ensembl_gene_id | external_gene_name | entrezgene_id | gene_biotype | baseMean | log2FoldChange | lfcSE | stat | pvalue | padj | absLog2FC |
|---|---|---|---|---|---|---|---|---|---|---|
| ENSMUSG00000020644 | Id2 | 15902 | protein_coding | 188.6474569 | 0.160518974 | 0.211332188 | -0.759557621 | 0.447519059 | 0.664777345 | 0.160518974 |
| ENSMUSG00000033862 | Cdk10 | 234854 | protein_coding | 222.5807347 | -0.16052879 | 0.079318035 | 2.023862413 | 0.04298431 | 0.132820779 | 0.16052879 |
| ENSMUSG00000059486 | Kbtbd2 | 210973 | protein_coding | 489.7357074 | -0.160570809 | 0.086048685 | 1.866046052 | 0.062034923 | 0.174819917 | 0.160570809 |
| ENSMUSG00000020832 | Eral1 | 57837 | protein_coding | 184.0202816 | -0.160608647 | 0.105990062 | 1.515317988 | 0.129691895 | 0.297380166 | 0.160608647 |
| ENSMUSG00000040021 | Lats1 | 16798 | protein_coding | 1105.37841 | 0.160613071 | 0.061544234 | -2.609717598 | 0.009061699 | 0.039359523 | 0.160613071 |
| ENSMUSG00000032026 | Rexo2 | 104444 | protein_coding | 213.5023952 | -0.16063478 | 0.116615846 | 1.377469582 | 0.168367113 | 0.357012648 | 0.16063478 |
| ENSMUSG00000001018 | Snapin | 20615 | protein_coding | 185.5619404 | -0.16065561 | 0.103971412 | 1.54519023 | 0.122300257 | 0.285383751 | 0.16065561 |
| ENSMUSG00000066568 | Lsm14a | 67070 | protein_coding | 642.5193262 | 0.160683743 | 0.076987969 | -2.087127964 | 0.036876569 | 0.118133785 | 0.160683743 |
| ENSMUSG00000042302 | Ehbp1 | 216565 | protein_coding | 139.8924183 | 0.160686382 | 0.128318085 | -1.252250471 | 0.21047861 | 0.414545055 | 0.160686382 |
| ENSMUSG00000024038 | Ndufv3 | 78330 | protein_coding | 1468.095508 | 0.160761979 | 0.063943501 | -2.514125383 | 0.0119328 | 0.048873577 | 0.160761979 |
| ENSMUSG00000087679 | Tmem250-ps | 109299 | protein_coding | 674.5217882 | -0.160792589 | 0.075928962 | 2.117671357 | 0.03420291 | 0.111471652 | 0.160792589 |
| ENSMUSG00000062203 | Gspt1 | 14852 | protein_coding | 950.1009512 | 0.160817617 | 0.069823749 | -2.303193679 | 0.021267948 | 0.077038583 | 0.160817617 |
| ENSMUSG00000034248 | Slc25a37 | 67712 | protein_coding | 310.6465352 | -0.160838064 | 0.131191319 | 1.225980994 | 0.220205827 | 0.426566456 | 0.160838064 |
| ENSMUSG00000086213 | A330040F15Rik | 74333 | lncRNA | 8.306844335 | -0.160864149 | 0.428552568 | 0.3753662 | 0.707388139 | 0.846814623 | 0.160864149 |
| ENSMUSG00000026775 | Yme1l1 | 27377 | protein_coding | 2053.474203 | 0.160885625 | 0.071712374 | -2.243484853 | 0.024865566 | 0.086997871 | 0.160885625 |
| ENSMUSG00000002625 | Akap8l | 54194 | protein_coding | 399.2981533 | 0.161001466 | 0.148897585 | -1.081289975 | 0.279568146 | 0.496963225 | 0.161001466 |
| ENSMUSG00000021476 | Habp4 | 56541 | protein_coding | 153.820358 | 0.161058902 | 0.092298145 | -1.74498526 | 0.080987435 | 0.211618117 | 0.161058902 |
| ENSMUSG00000040891 | Foxa3 | 15377 | protein_coding | 703.9279152 | 0.161096529 | 0.166791323 | -0.965856775 | 0.334115854 | 0.556489079 | 0.161096529 |
| ENSMUSG00000073436 | Eme2 | 193838 | protein_coding | 103.9156813 | 0.161170053 | 0.130217429 | -1.237699546 | 0.215827487 | 0.421098196 | 0.161170053 |
| ENSMUSG00000039967 | Zfp292 | 30046 | protein_coding | 958.8566734 | -0.161193712 | 0.070786576 | 2.277179114 | 0.022775532 | 0.081322191 | 0.161193712 |
| ENSMUSG00000054321 | Taf4b | 72504 | protein_coding | 128.3238018 | 0.161264525 | 0.156775516 | -1.028633357 | 0.303651997 | 0.524146449 | 0.161264525 |
| ENSMUSG00000026991 | Pkp4 | 227937 | protein_coding | 1002.183333 | 0.161337103 | 0.061130904 | -2.63920687 | 0.008310025 | 0.036683176 | 0.161337103 |
| ENSMUSG00000020948 | Klhl28 | 66689 | protein_coding | 152.2844911 | -0.1613404 | 0.127096444 | 1.26943284 | 0.20428673 | 0.406554718 | 0.1613404 |
| ENSMUSG00000028651 | Ppie | 56031 | protein_coding | 195.6697184 | -0.161444766 | 0.14393565 | 1.121645443 | 0.262013222 | 0.475826228 | 0.161444766 |
| ENSMUSG00000051034 | Zfp11 | 22648 | protein_coding | 26.95791606 | -0.161548138 | 0.204549719 | 0.789774431 | 0.429659514 | 0.648455238 | 0.161548138 |
| ENSMUSG00000022427 | Tomm22 | 223696 | protein_coding | 1057.55404 | -0.161576683 | 0.089747938 | 1.800338677 | 0.071807177 | 0.194652614 | 0.161576683 |
| ENSMUSG00000051285 | Pcmtd1 | 319263 | protein_coding | 529.7252526 | -0.16161984 | 0.111425345 | 1.450476456 | 0.1469257 | 0.324759105 | 0.16161984 |
| ENSMUSG00000105552 | Gm19710 | 100503469 | lncRNA | 1.797380846 | -0.16164054 | 0.865593043 | 0.186739648 | 0.851864769 | 1 | 0.16164054 |
| ENSMUSG00000074925 | Ptar1 | 72351 | protein_coding | 447.0602808 | 0.161648619 | 0.11886474 | -1.359937515 | 0.173849698 | 0.364658188 | 0.161648619 |
| ENSMUSG00000087396 | 4933407K13Rik | 74396 | lncRNA | 36.065512 | 0.16165878 | 0.236718646 | -0.682915278 | 0.494660379 | 0.700763122 | 0.16165878 |
| ENSMUSG00000058704 | Memo1 | 76890 | protein_coding | 325.5931556 | -0.161667423 | 0.080808021 | 2.000635853 | 0.045431647 | 0.138818348 | 0.161667423 |
| ENSMUSG00000068036 | Afdn | 17356 | protein_coding | 2354.289793 | -0.161791765 | 0.069072 | 2.342363965 | 0.019162018 | 0.070982855 | 0.161791765 |
| ENSMUSG00000070002 | Ell | 13716 | protein_coding | 329.420306 | 0.161797163 | 0.159353995 | -1.015331705 | 0.309947736 | 0.530962336 | 0.161797163 |
| ENSMUSG00000022462 | Slc38a2 | 67760 | protein_coding | 2294.750585 | 0.161838996 | 0.064422206 | -2.512161657 | 0.011999411 | 0.049101931 | 0.161838996 |
| ENSMUSG00000037242 | Clic4 | 29876 | protein_coding | 651.1385301 | -0.161839735 | 0.109108819 | 1.483287393 | 0.13799807 | 0.31052626 | 0.161839735 |
| ENSMUSG00000038564 | Ift172 | 67661 | protein_coding | 333.9554467 | 0.161843658 | 0.065017864 | -2.489218333 | 0.012802432 | 0.051565697 | 0.161843658 |
| ENSMUSG00000064179 | Tnnt1 | 21955 | protein_coding | 22.64541217 | -0.161913455 | 0.351756682 | 0.460299586 | 0.645301198 | 0.806814922 | 0.161913455 |
| ENSMUSG00000027936 | Crtc2 | 74343 | protein_coding | 132.298355 | -0.161967883 | 0.125150933 | 1.294180381 | 0.195603122 | 0.394941012 | 0.161967883 |
| ENSMUSG00000058192 | Zfp846 | 244721 | protein_coding | 102.2629385 | 0.161972551 | 0.105424391 | -1.536385927 | 0.124443756 | 0.289079222 | 0.161972551 |

Supplementary Table 1

| ensembl_gene_id | external_gene_name | entrezgene_id | gene_biotype | baseMean | log2FoldChange | lfcSE | stat | pvalue | padj | absLog2FC |
|---|---|---|---|---|---|---|---|---|---|---|
| ENSMUSG00000002733 | Plekha3 | 83435 | protein_coding | 460.7360407 | -0.161996021 | 0.10966964 | 1.477127314 | 0.139641513 | 0.31327808 | 0.161996021 |
| ENSMUSG00000097327 | E030030I06Rik | 319887 | transcribed_unprocessed_pseudogene | 60.82592792 | 0.162017959 | 0.163223106 | -0.992616564 | 0.32089685 | 0.542047573 | 0.162017959 |
| ENSMUSG00000053164 | Gpr21 | 338346 | protein_coding | 11.69089603 | -0.16202883 | 0.346977147 | 0.466972629 | 0.640519461 | 0.803690806 | 0.16202883 |
| ENSMUSG00000020677 | Ddx52 | 78394 | protein_coding | 346.8247014 | 0.162033713 | 0.086151102 | -1.880808357 | 0.059997993 | 0.170740454 | 0.162033713 |
| ENSMUSG00000034833 | Tespa1 | 67596 | protein_coding | 3.540361985 | 0.16204845 | 0.885961639 | -0.182906847 | 0.85487111 | 1 | 0.16204845 |
| ENSMUSG00000021258 | Ccnk | 12454 | protein_coding | 130.6706647 | 0.162063427 | 0.152942512 | -1.059636233 | 0.289310123 | 0.508154368 | 0.162063427 |
| ENSMUSG00000022718 | Dgcr8 | 94223 | protein_coding | 261.661673 | 0.162100157 | 0.071699865 | -2.260815351 | 0.023770694 | 0.084039656 | 0.162100157 |
| ENSMUSG00000045667 | Smtnl2 | 276829 | protein_coding | 9.655294015 | -0.162189666 | 0.369909499 | 0.438457694 | 0.661054532 | 0.816884691 | 0.162189666 |
| ENSMUSG00000029534 | St7 | 64213 | protein_coding | 119.0924106 | 0.162210006 | 0.131723091 | -1.231447011 | 0.218155724 | 0.42426293 | 0.162210006 |
| ENSMUSG00000040028 | Elavl1 | 15568 | protein_coding | 834.5032088 | 0.162226244 | 0.084653439 | -1.916357399 | 0.055319619 | 0.160666232 | 0.162226244 |
| ENSMUSG00000033943 | Mga | 29808 | protein_coding | 756.0824431 | -0.162263617 | 0.085105283 | 1.906622141 | 0.056569536 | 0.163634492 | 0.162263617 |
| ENSMUSG00000052423 | B4galt3 | 57370 | protein_coding | 227.8352097 | 0.162264635 | 0.090920062 | -1.784695613 | 0.074310703 | 0.199366082 | 0.162264635 |
| ENSMUSG00000003429 | Rps11 | 27207 | protein_coding | 4184.715633 | -0.162334347 | 0.082828072 | 1.959895266 | 0.050008033 | 0.148948255 | 0.162334347 |
| ENSMUSG00000019790 | Stxbp5 | 78808 | protein_coding | 474.7146937 | 0.162385172 | 0.057244799 | -2.836679929 | 0.004558528 | 0.022511824 | 0.162385172 |
| ENSMUSG00000022407 | Adsl | 11564 | protein_coding | 562.2937834 | -0.162436534 | 0.086936277 | 1.868455145 | 0.061698657 | 0.174109982 | 0.162436534 |
| ENSMUSG00000048186 | Bend7 | 209645 | protein_coding | 90.55242041 | 0.162444936 | 0.122985515 | -1.320846083 | 0.186552691 | 0.382367026 | 0.162444936 |
| ENSMUSG00000026976 | Pax8 | 18510 | protein_coding | 47.78504022 | -0.162475332 | 0.186913348 | 0.869254837 | 0.384707759 | 0.607019295 | 0.162475332 |
| ENSMUSG00000040720 | Virma | 66185 | protein_coding | 670.4472199 | 0.162479068 | 0.059720244 | -2.720669864 | 0.006514979 | 0.030010701 | 0.162479068 |
| ENSMUSG00000073889 | Il11ra1 | 16157 | protein_coding | 99.82988062 | -0.162502395 | 0.137559537 | 1.181324086 | 0.237474002 | 0.447829813 | 0.162502395 |
| ENSMUSG00000113800 | 2210039B01Rik | 73644 | lncRNA | 15.08915867 | -0.162512005 | 0.297215216 | 0.54678225 | 0.584528339 | 0.766833571 | 0.162512005 |
| ENSMUSG00000031551 | Ido1 | 15930 | protein_coding | 9.16238824 | -0.162518991 | 0.681204054 | 0.238576078 | 0.811434317 | 0.906704483 | 0.162518991 |
| ENSMUSG00000023147 | Get1 | 71446 | protein_coding | 99.65362463 | 0.162559758 | 0.144811243 | -1.122563102 | 0.261623093 | 0.475244062 | 0.162559758 |
| ENSMUSG00000021408 | Ripk1 | 19766 | protein_coding | 637.6770698 | -0.162564276 | 0.064351972 | 2.526173949 | 0.011531237 | 0.04769017 | 0.162564276 |
| ENSMUSG00000038671 | Arfrp1 | 76688 | protein_coding | 273.9720005 | -0.162612948 | 0.075148436 | 2.163889996 | 0.030472793 | 0.102074585 | 0.162612948 |
| ENSMUSG00000108948 | Or2ag13 | 258591 | protein_coding | 1.548386771 | -0.162619868 | 2.361640836 | 0.068858848 | 0.945101975 | 1 | 0.162619868 |
| ENSMUSG00000021458 | Aopep | 72061 | protein_coding | 829.5946266 | -0.162634152 | 0.079370135 | 2.049059795 | 0.040456268 | 0.127038375 | 0.162634152 |
| ENSMUSG00000034063 | 4930590J08Rik | 381798 | protein_coding | 6.112190554 | 0.162668215 | 0.646482295 | -0.251620525 | 0.801334392 | 0.901569434 | 0.162668215 |
| ENSMUSG00000026480 | Ncf2 | 17970 | protein_coding | 325.3550459 | 0.162678084 | 0.098994957 | -1.643296685 | 0.100321564 | 0.248345121 | 0.162678084 |
| ENSMUSG00000019710 | Mrpl24 | 67707 | protein_coding | 593.934937 | -0.162773681 | 0.051843611 | 3.139705702 | 0.001691176 | 0.009868582 | 0.162773681 |
| ENSMUSG00000032033 | Barx2 | 12023 | protein_coding | 18.35335699 | -0.162774023 | 0.249682756 | 0.651923366 | 0.514450611 | 0.716025064 | 0.162774023 |
| ENSMUSG00000035578 | Iqcg | 69707 | protein_coding | 12.3706547 | -0.16288473 | 0.382691791 | 0.425629015 | 0.670378194 | 0.822972458 | 0.16288473 |
| ENSMUSG00000037419 | Endod1 | 71946 | protein_coding | 4368.752447 | -0.162888583 | 0.090437279 | 1.801122119 | 0.071683634 | 0.194381533 | 0.162888583 |
| ENSMUSG00000026470 | Stx6 | 58244 | protein_coding | 360.621276 | -0.162888758 | 0.067215106 | 2.423395102 | 0.015376193 | 0.059550294 | 0.162888758 |
| ENSMUSG00000002831 | Plin4 | 57435 | protein_coding | 6.29293453 | 0.162894302 | 0.607460351 | -0.268156271 | 0.788579034 | 0.894265208 | 0.162894302 |
| ENSMUSG00000036792 | Mbd5 | 109241 | protein_coding | 164.401558 | -0.162907097 | 0.161027122 | 1.011674899 | 0.311693521 | 0.532495959 | 0.162907097 |
| ENSMUSG00000016018 | Mtrex | 72198 | protein_coding | 554.9383702 | 0.162912161 | 0.087045589 | -1.871572845 | 0.061265723 | 0.173362261 | 0.162912161 |
| ENSMUSG00000026828 | Galnt5 | 241391 | protein_coding | 624.0795487 | 0.162953911 | 0.23696064 | -0.687683454 | 0.491652142 | 0.698491194 | 0.162953911 |
| ENSMUSG00000021051 | Ppp2r5e | 26932 | protein_coding | 455.2702314 | 0.163016336 | 0.087431912 | -1.864494692 | 0.062252266 | 0.175163388 | 0.163016336 |
| ENSMUSG00000069208 | Zfp825 | 235956 | protein_coding | 46.3728346 | -0.163033519 | 0.188751037 | 0.863748994 | 0.387725793 | 0.610149013 | 0.163033519 |

| ensembl_gene_id | external_gene_name | entrezgene_id | gene_biotype | baseMean | log2FoldChange | lfcSE | stat | pvalue | padj | absLog2FC |
|---|---|---|---|---|---|---|---|---|---|---|
| ENSMUSG00000029279 | Brdt | 114642 | protein_coding | 11.96621103 | -0.163048896 | 0.435390096 | 0.374489217 | 0.708040377 | 0.847226925 | 0.163048896 |
| ENSMUSG00000038732 | Mboat1 | 218121 | protein_coding | 3081.649454 | -0.163068279 | 0.11932381 | 1.366603017 | 0.171749771 | 0.362093159 | 0.163068279 |
| ENSMUSG00000040265 | Dnm3 | 103967 | protein_coding | 8.258163436 | 0.163119614 | 0.839293567 | -0.194353467 | 0.845899123 | 0.924166034 | 0.163119614 |
| ENSMUSG00000020634 | Ubxn2a | 217379 | protein_coding | 874.4050732 | -0.163128163 | 0.072165027 | 2.260487804 | 0.023790993 | 0.084093418 | 0.163128163 |
| ENSMUSG00000042901 | Aida | 108909 | protein_coding | 184.8736614 | -0.163202695 | 0.115832824 | 1.408950335 | 0.158849854 | 0.34286322 | 0.163202695 |
| ENSMUSG00000062127 | Cttnbp2nl | 80281 | protein_coding | 841.0084442 | -0.163278354 | 0.085805662 | 1.902885561 | 0.057055479 | 0.164593053 | 0.163278354 |
| ENSMUSG00000049950 | Rpp38 | 227522 | protein_coding | 46.11038147 | 0.163293647 | 0.199167965 | -0.819879074 | 0.412285047 | 0.632329199 | 0.163293647 |
| ENSMUSG00000036054 | Sugp2 | 234373 | protein_coding | 263.7561372 | -0.163319813 | 0.096364108 | 1.694819951 | 0.090109582 | 0.228911821 | 0.163319813 |
| ENSMUSG00000032041 | Tirap | 117149 | protein_coding | 226.842317 | 0.163330649 | 0.08474879 | -1.927232818 | 0.053950622 | 0.157940182 | 0.163330649 |
| ENSMUSG00000024754 | Cemip2 | 83921 | protein_coding | 648.0135713 | -0.163379362 | 0.091194767 | 1.79154317 | 0.073206173 | 0.197279169 | 0.163379362 |
| ENSMUSG00000044627 | Swi5 | 72931 | protein_coding | 832.2462993 | 0.163404725 | 0.086639158 | -1.886037783 | 0.059289863 | 0.169396679 | 0.163404725 |
| ENSMUSG00000036565 | Ttyh3 | 78339 | protein_coding | 397.9657913 | 0.163405228 | 0.131269759 | -1.244804814 | 0.213203506 | 0.417979787 | 0.163405228 |
| ENSMUSG00000049800 | Sertad2 | 58172 | protein_coding | 628.7468077 | -0.163426084 | 0.062606988 | 2.610348933 | 0.009044991 | 0.039297286 | 0.163426084 |
| ENSMUSG00000069769 | Msi2 | 76626 | protein_coding | 1156.507155 | 0.163473931 | 0.136225368 | -1.200025617 | 0.230129391 | 0.438743376 | 0.163473931 |
| ENSMUSG00000040034 | Nup43 | 69912 | protein_coding | 84.06453145 | 0.163557894 | 0.179023814 | -0.913609703 | 0.360921966 | 0.583310369 | 0.163557894 |
| ENSMUSG00000018433 | Nol11 | 68979 | protein_coding | 266.0209135 | 0.163660072 | 0.138152512 | -1.184633347 | 0.236162446 | 0.446273789 | 0.163660072 |
| ENSMUSG00000070520 | Nsmce3 | 66647 | protein_coding | 220.8028734 | 0.163662119 | 0.108956726 | -1.502083669 | 0.133075502 | 0.302752759 | 0.163662119 |
| ENSMUSG00000026946 | Nmi | 64685 | protein_coding | 278.1947461 | -0.163685435 | 0.098727858 | 1.657945773 | 0.097328412 | 0.242315905 | 0.163685435 |
| ENSMUSG00000060586 | H2-Eb1 | 14969 | protein_coding | 21.48435743 | 0.163705279 | 0.304688299 | -0.537287711 | 0.591068893 | 0.771308708 | 0.163705279 |
| ENSMUSG00000022941 | Ripply3 | 170765 | protein_coding | 7.430426374 | -0.163713693 | 0.404113752 | 0.405117846 | 0.685390889 | 0.832307165 | 0.163713693 |
| ENSMUSG00000021608 | Lpcat1 | 210992 | protein_coding | 169.198609 | -0.163725881 | 0.159662673 | 1.025448708 | 0.305151515 | 0.525252463 | 0.163725881 |
| ENSMUSG00000044312 | Neurog3 | 11925 | protein_coding | 8.649443589 | 0.163773686 | 0.420773802 | -0.389220254 | 0.697113222 | 0.839403501 | 0.163773686 |
| ENSMUSG00000033906 | Zdhhc15 | 108672 | protein_coding | 83.1685708 | -0.163786325 | 0.172394621 | 0.950066327 | 0.342078552 | 0.564006111 | 0.163786325 |
| ENSMUSG00000038121 | Fam210a | 108654 | protein_coding | 569.088552 | -0.163869172 | 0.105889031 | 1.547555681 | 0.121729309 | 0.28456734 | 0.163869172 |
| ENSMUSG00000040498 | Igsf23 | 70080 | protein_coding | 104.2160978 | 0.163905892 | 0.143082989 | -1.145530248 | 0.251989573 | 0.464375257 | 0.163905892 |
| ENSMUSG00000042099 | Kank3 | 80880 | protein_coding | 30.70987712 | 0.163912064 | 0.195919623 | -0.836629133 | 0.402801065 | 0.624177234 | 0.163912064 |
| ENSMUSG00000034509 | Mad2l1bp | 66591 | protein_coding | 223.4193403 | 0.163928727 | 0.113494158 | -1.444380309 | 0.148632026 | 0.327398864 | 0.163928727 |
| ENSMUSG00000036309 | Skp1 | 21402 | protein_coding | 1852.957205 | -0.163957196 | 0.089616865 | 1.829535067 | 0.067319491 | 0.185502098 | 0.163957196 |
| ENSMUSG00000022601 | Zbtb11 | 271377 | protein_coding | 287.4767542 | -0.164005067 | 0.07551742 | 2.171751462 | 0.029874415 | 0.100633238 | 0.164005067 |
| ENSMUSG00000001249 | Hpn | 15451 | protein_coding | 78.97831495 | -0.164014115 | 0.260941717 | 0.628546929 | 0.529645714 | 0.727184014 | 0.164014115 |
| ENSMUSG00000031549 | Ido2 | 209176 | protein_coding | 4.101948238 | 0.164039524 | 0.873708753 | -0.187750808 | 0.851071999 | 1 | 0.164039524 |
| ENSMUSG00000036775 | Decr2 | 26378 | protein_coding | 655.9835127 | -0.164182188 | 0.066506653 | 2.468658091 | 0.013562073 | 0.053943227 | 0.164182188 |
| ENSMUSG00000027669 | Gnb4 | 14696 | protein_coding | 16.22140488 | 0.164194263 | 0.349750521 | -0.469461096 | 0.638740087 | 0.80301103 | 0.164194263 |
| ENSMUSG00000030703 | Gdpd3 | 68616 | protein_coding | 23.68018604 | 0.164217332 | 0.28325543 | -0.579749987 | 0.562083229 | 0.750140815 | 0.164217332 |
| ENSMUSG00000065551 | Mir210 | 387206 | miRNA | 2.840240944 | 0.16422262 | 0.668081134 | -0.245812389 | 0.805827466 | 1 | 0.16422262 |
| ENSMUSG00000031024 | Denn2b | 76954 | protein_coding | 586.4428695 | 0.164227303 | 0.087293496 | -1.88132347 | 0.05992793 | 0.1706035 | 0.164227303 |
| ENSMUSG00000035597 | Prpf39 | 328110 | protein_coding | 314.6916448 | 0.164261422 | 0.108661099 | -1.511685626 | 0.130613857 | 0.298680373 | 0.164261422 |
| ENSMUSG00000040913 | Fbxw4 | 30838 | protein_coding | 105.6788354 | 0.164273921 | 0.169389846 | -0.969797927 | 0.332147226 | 0.554606509 | 0.164273921 |
| ENSMUSG00000006333 | Rps9 | 76846 | protein_coding | 7216.915476 | -0.164284977 | 0.090728327 | 1.81073521 | 0.07018185 | 0.191366503 | 0.164284977 |

Supplementary Table 1

| ensembl_gene_id | external_gene_name | entrezgene_id | gene_biotype | baseMean | log2FoldChange | lfcSE | stat | pvalue | padj | absLog2FC |
|---|---|---|---|---|---|---|---|---|---|---|
| ENSMUSG00000028266 | Lmo4 | 16911 | protein_coding | 272.8171628 | -0.164320079 | 0.081236121 | 2.02274651 | 0.043099289 | 0.133085576 | 0.164320079 |
| ENSMUSG00000042642 | Flad1 | 319945 | protein_coding | 435.5378725 | -0.164321052 | 0.088349254 | 1.859903117 | 0.062899234 | 0.176502626 | 0.164321052 |
| ENSMUSG00000002107 | Celf2 | 14007 | protein_coding | 44.27859336 | 0.164329055 | 0.290035671 | -0.56658222 | 0.570998058 | 0.756475084 | 0.164329055 |
| ENSMUSG00000041297 | Cdk13 | 69562 | protein_coding | 1133.106161 | 0.164347696 | 0.055328378 | -2.970405112 | 0.002974073 | 0.015751711 | 0.164347696 |
| ENSMUSG00000042729 | Wdr74 | 107071 | protein_coding | 250.5090517 | 0.164426664 | 0.093343109 | -1.761529751 | 0.078148782 | 0.206686257 | 0.164426664 |
| ENSMUSG00000028330 | Ncbp1 | 433702 | protein_coding | 590.2276357 | 0.164431194 | 0.109110049 | -1.50702154 | 0.131805136 | 0.300890309 | 0.164431194 |
| ENSMUSG00000036430 | Tbcc | 72726 | protein_coding | 162.4293287 | 0.164448104 | 0.09819136 | -1.674771627 | 0.093979048 | 0.236076969 | 0.164448104 |
| ENSMUSG00000044328 | Trp53i13 | 216964 | protein_coding | 56.19827722 | -0.16445423 | 0.197153965 | 0.834141127 | 0.404201462 | 0.625349723 | 0.16445423 |
| ENSMUSG00000030674 | Qprt | 67375 | protein_coding | 1.686399452 | 0.164470625 | 1.746121812 | -0.094191954 | 0.924956676 | 1 | 0.164470625 |
| ENSMUSG00000028649 | Macf1 | 11426 | protein_coding | 2476.787166 | 0.164499244 | 0.103149465 | -1.594765848 | 0.110764607 | 0.266215502 | 0.164499244 |
| ENSMUSG00000075327 | Zbtb2 | 381990 | protein_coding | 179.5453621 | -0.164529238 | 0.108047002 | 1.522756166 | 0.12781972 | 0.294806545 | 0.164529238 |
| ENSMUSG00000039745 | Htatip2 | 53415 | protein_coding | 270.7881585 | -0.16453561 | 0.09154316 | 1.7973556 | 0.072279185 | 0.19557897 | 0.16453561 |
| ENSMUSG00000022861 | Dgkg | 110197 | protein_coding | 86.22307417 | -0.164604129 | 0.219195543 | 0.750946513 | 0.452684846 | 0.668926246 | 0.164604129 |
| ENSMUSG00000021713 | Ppwd1 | 238831 | protein_coding | 166.0299303 | 0.164609604 | 0.111580598 | -1.475252922 | 0.140144558 | 0.314193217 | 0.164609604 |
| ENSMUSG00000032096 | Arcn1 | 213827 | protein_coding | 3064.99834 | 0.164631389 | 0.088602435 | -1.858091008 | 0.063156092 | 0.177042895 | 0.164631389 |
| ENSMUSG00000085776 | 5430402O13Rik | 73183 | lncRNA | 1.794236721 | 0.164647841 | 0.997758978 | -0.165017649 | 0.86893009 | 1 | 0.164647841 |
| ENSMUSG00000114755 | Galr3 | 14429 | protein_coding | 2.472506609 | 0.164656311 | 0.700842951 | -0.234940383 | 0.814255005 | 1 | 0.164656311 |
| ENSMUSG00000003873 | Bax | 12028 | protein_coding | 266.6889685 | 0.164710168 | 0.086904308 | -1.895304988 | 0.058052008 | 0.166725366 | 0.164710168 |
| ENSMUSG00000029027 | Dffb | 13368 | protein_coding | 92.07855077 | -0.164720326 | 0.12449581 | 1.323099363 | 0.185802336 | 0.381254943 | 0.164720326 |
| ENSMUSG00000029865 | Sval1 | 71578 | protein_coding | 1757.576689 | 0.164766271 | 0.532383246 | -0.309488085 | 0.756950274 | 0.878075079 | 0.164766271 |
| ENSMUSG00000003484 | Cyp4f18 | 72054 | protein_coding | 1.294035413 | 0.164771898 | 1.217563119 | -0.135329246 | 0.892351563 | 1 | 0.164771898 |
| ENSMUSG00000001700 | Gramd3 | 107022 | protein_coding | 853.8991258 | -0.164779161 | 0.124128568 | 1.327487808 | 0.18434737 | 0.379371759 | 0.164779161 |
| ENSMUSG00000078588 | Ccdc24 | 381546 | protein_coding | 2.996058309 | 0.164782867 | 0.680513911 | -0.242144745 | 0.808668008 | 1 | 0.164782867 |
| ENSMUSG00000001062 | Vps9d1 | 72325 | protein_coding | 209.6321519 | -0.164787235 | 0.085843901 | 1.919614933 | 0.054906556 | 0.15977566 | 0.164787235 |
| ENSMUSG00000054659 | Pm20d2 | 242377 | protein_coding | 23.26128489 | -0.164824177 | 0.214199434 | 0.76948932 | 0.441602882 | 0.659260011 | 0.164824177 |
| ENSMUSG00000025645 | Ccdc51 | 66658 | protein_coding | 206.0809152 | -0.164825682 | 0.103059422 | 1.599326663 | 0.109748038 | 0.264619521 | 0.164825682 |
| ENSMUSG00000036199 | Ndufa13 | 67184 | protein_coding | 1742.216452 | -0.16482818 | 0.073965214 | 2.228455387 | 0.025850163 | 0.089682688 | 0.16482818 |
| ENSMUSG00000079429 | Mroh2a | 100040766 | protein_coding | 112.012606 | 0.164861989 | 0.255654876 | -0.644861509 | 0.519016937 | 0.719334577 | 0.164861989 |
| ENSMUSG00000036613 | Eipr1 | 380752 | protein_coding | 289.9350197 | 0.164882076 | 0.081104469 | -2.032959193 | 0.042056646 | 0.130795377 | 0.164882076 |
| ENSMUSG00000061780 | Cfd | 11537 | protein_coding | 1.654985867 | 0.164946992 | 1.466594165 | -0.112469418 | 0.910451217 | 1 | 0.164946992 |
| ENSMUSG00000039264 | Gimap3 | 83408 | protein_coding | 30.42232825 | 0.164951606 | 0.302326177 | -0.545608082 | 0.585335367 | 0.767282764 | 0.164951606 |
| ENSMUSG00000001138 | Cnnm3 | 94218 | protein_coding | 279.2170946 | 0.164971308 | 0.081264286 | -2.030059142 | 0.042350528 | 0.131338332 | 0.164971308 |
| ENSMUSG00000039810 | Zc3h10 | 103284 | protein_coding | 195.8625891 | -0.164983355 | 0.087838711 | 1.878253359 | 0.060346514 | 0.171466333 | 0.164983355 |
| ENSMUSG00000060716 | Plekhh1 | 211945 | protein_coding | 1467.468619 | -0.165005736 | 0.0836772 | 1.971931857 | 0.048617386 | 0.145896332 | 0.165005736 |
| ENSMUSG00000069237 | Fam8a1 | 97863 | protein_coding | 826.6915626 | 0.165026026 | 0.08702888 | -1.896221415 | 0.057930773 | 0.166521981 | 0.165026026 |
| ENSMUSG00000021930 | Spryd7 | 66674 | protein_coding | 267.4386052 | -0.165036259 | 0.096956798 | 1.702162837 | 0.088724847 | 0.226286043 | 0.165036259 |
| ENSMUSG00000018848 | Rars | 104458 | protein_coding | 963.5441283 | 0.165102278 | 0.06688193 | -2.468563326 | 0.013565665 | 0.053943227 | 0.165102278 |
| ENSMUSG00000035139 | Secisbp2 | 75420 | protein_coding | 262.447109 | -0.165103024 | 0.079426525 | 2.078688761 | 0.037645967 | 0.120138597 | 0.165103024 |
| ENSMUSG00000002028 | Kmt2a | 214162 | protein_coding | 595.769029 | 0.165118514 | 0.176043625 | -0.937940893 | 0.348274787 | 0.570180413 | 0.165118514 |

Supplementary Table 1

| ensembl_gene_id | external_gene_name | entrezgene_id | gene_biotype | baseMean | log2FoldChange | lfcSE | stat | pvalue | padj | absLog2FC |
|---|---|---|---|---|---|---|---|---|---|---|
| ENSMUSG00000031557 | Plekha2 | 83436 | protein_coding | 852.5406775 | -0.165133198 | 0.097402387 | 1.695371168 | 0.090005032 | 0.228807636 | 0.165133198 |
| ENSMUSG00000020346 | Mgat1 | 17308 | protein_coding | 970.1125763 | -0.165244858 | 0.063135112 | 2.617321054 | 0.008862294 | 0.038640192 | 0.165244858 |
| ENSMUSG00000055137 | Sugct | 192136 | protein_coding | 21.55472587 | -0.165250186 | 0.237216115 | 0.696622932 | 0.486038801 | 0.695595302 | 0.165250186 |
| ENSMUSG00000005150 | Wdr83 | 67836 | protein_coding | 152.2971956 | -0.165250811 | 0.105684911 | 1.563617817 | 0.117907348 | 0.278398906 | 0.165250811 |
| ENSMUSG00000075704 | Txnrd2 | 26462 | protein_coding | 128.4815588 | -0.165251588 | 0.100786445 | 1.639621164 | 0.10108396 | 0.249608668 | 0.165251588 |
| ENSMUSG00000042043 | Tbca | 21371 | protein_coding | 526.8622185 | -0.165274441 | 0.097985087 | 1.686730567 | 0.091655172 | 0.231775376 | 0.165274441 |
| ENSMUSG00000032845 | Alpk2 | 225638 | protein_coding | 1.551064803 | -0.165343922 | 2.136455917 | 0.077391685 | 0.938311956 | 1 | 0.165343922 |
| ENSMUSG00000020803 | Txndc17 | 52700 | protein_coding | 1040.514391 | -0.165386368 | 0.067681222 | 2.443607869 | 0.014541222 | 0.056890248 | 0.165386368 |
| ENSMUSG00000049323 | Smcr8 | 237782 | protein_coding | 561.3732788 | 0.165390786 | 0.080062483 | -2.065771386 | 0.038850071 | 0.123236664 | 0.165390786 |
| ENSMUSG00000048826 | Dact2 | 240025 | protein_coding | 320.0927784 | -0.165416307 | 0.140920508 | 1.173827071 | 0.240464298 | 0.450945654 | 0.165416307 |
| ENSMUSG00000034707 | Gns | 75612 | protein_coding | 777.9486858 | -0.165467197 | 0.050073324 | 3.304497989 | 0.000951467 | 0.00610432 | 0.165467197 |
| ENSMUSG00000021996 | Esd | 13885 | protein_coding | 1755.685558 | -0.165486692 | 0.077261548 | 2.141902366 | 0.032201339 | 0.106567719 | 0.165486692 |
| ENSMUSG00000022969 | Il10rb | 16155 | protein_coding | 484.1918311 | -0.165515571 | 0.07070035 | 2.341085604 | 0.019227758 | 0.071194438 | 0.165515571 |
| ENSMUSG00000054342 | Kcnn4 | 16534 | protein_coding | 628.1862516 | -0.165637367 | 0.153534663 | 1.07882718 | 0.280664776 | 0.498301136 | 0.165637367 |
| ENSMUSG00000029472 | Anapc5 | 59008 | protein_coding | 1808.196903 | 0.165695677 | 0.087393287 | -1.895977162 | 0.057963064 | 0.166571051 | 0.165695677 |
| ENSMUSG00000101013 | A630072M18Rik | 320770 | lncRNA | 2.912692288 | 0.165749891 | 0.822768775 | -0.201453794 | 0.840343756 | 1 | 0.165749891 |
| ENSMUSG00000026970 | Rbms1 | 56878 | protein_coding | 214.5223176 | -0.165763687 | 0.103663655 | 1.59905308 | 0.109808809 | 0.264727397 | 0.165763687 |
| ENSMUSG00000102629 | Gm29999 | 102631733 | lncRNA | 1.10648684 | 0.16577657 | 2.644859176 | -0.062678789 | 0.950022288 | 1 | 0.16577657 |
| ENSMUSG00000024298 | Zfp871 | 208292 | protein_coding | 677.253412 | -0.165835918 | 0.102275384 | 1.621464635 | 0.104918029 | 0.256152621 | 0.165835918 |
| ENSMUSG00000036285 | Noa1 | 56412 | protein_coding | 185.7815432 | -0.165840565 | 0.119632864 | 1.386245879 | 0.165671837 | 0.352762472 | 0.165840565 |
| ENSMUSG00000019900 | Rfx6 | 320995 | protein_coding | 11.36528297 | -0.165873451 | 0.39414067 | 0.420848352 | 0.673865821 | 0.825096143 | 0.165873451 |
| ENSMUSG00000020621 | Rdh14 | 105014 | protein_coding | 337.4347965 | 0.165881628 | 0.099871418 | -1.66095197 | 0.096723093 | 0.241136045 | 0.165881628 |
| ENSMUSG00000012211 | Tex22 | 75671 | protein_coding | 1.772839404 | 0.165917784 | 1.113741863 | -0.148973285 | 0.881574713 | 1 | 0.165917784 |
| ENSMUSG00000051853 | Arf3 | 11842 | protein_coding | 1469.931698 | -0.165964079 | 0.04125137 | 4.023238013 | 5.74E-05 | 0.000528406 | 0.165964079 |
| ENSMUSG00000005505 | Kbtbd4 | 67136 | protein_coding | 74.85362703 | -0.166016781 | 0.138720673 | 1.196770296 | 0.231396099 | 0.440188363 | 0.166016781 |
| ENSMUSG00000027661 | Slc2a10 | 170441 | protein_coding | 141.578363 | -0.166050882 | 0.146442464 | 1.133898443 | 0.25683715 | 0.469910115 | 0.166050882 |
| ENSMUSG00000068479 | Mfap1a | 67532 | protein_coding | 188.0579281 | -0.166251927 | 0.125920371 | 1.320294134 | 0.186736833 | 0.382601993 | 0.166251927 |
| ENSMUSG00000065273 | Gm25128 | 115488831 | snoRNA | 4.719228525 | -0.166263377 | 0.483197167 | 0.344090132 | 0.730778504 | 0.861267215 | 0.166263377 |
| ENSMUSG00000072049 | Vmn2r121 | 100038941 | protein_coding | 1.976995173 | 0.16630737 | 2.655015217 | -0.062638952 | 0.950054011 | 1 | 0.16630737 |
| ENSMUSG00000021809 | Nudt13 | 67725 | protein_coding | 145.8437399 | -0.166322551 | 0.092879191 | 1.790740742 | 0.073334908 | 0.197497175 | 0.166322551 |
| ENSMUSG00000025533 | Asl | 109900 | protein_coding | 361.3716769 | -0.1663472 | 0.084025347 | 1.979726412 | 0.047734279 | 0.143952499 | 0.1663472 |
| ENSMUSG00000068959 | Zfp619 | 70227 | protein_coding | 33.49358062 | -0.166352933 | 0.210653029 | 0.789701119 | 0.429702337 | 0.64845799 | 0.166352933 |
| ENSMUSG00000038398 | Upf3a | 67031 | protein_coding | 381.4934391 | -0.166376949 | 0.083350307 | 1.996116815 | 0.045921209 | 0.139760938 | 0.166376949 |
| ENSMUSG00000025198 | Erlin1 | 226144 | protein_coding | 633.4024936 | 0.166389202 | 0.08664647 | -1.920322923 | 0.054817123 | 0.159701725 | 0.166389202 |
| ENSMUSG00000068794 | Col28a1 | 213945 | protein_coding | 19.81307103 | 0.166441673 | 0.252264318 | -0.659790785 | 0.509388098 | 0.712785998 | 0.166441673 |
| ENSMUSG00000039703 | Nploc4 | 217365 | protein_coding | 885.3897604 | 0.166452471 | 0.066892886 | -2.488343419 | 0.012833974 | 0.051654946 | 0.166452471 |
| ENSMUSG00000039640 | Mrpl12 | 56282 | protein_coding | 884.0264894 | -0.166458093 | 0.076018181 | 2.189714246 | 0.028544967 | 0.097034086 | 0.166458093 |
| ENSMUSG00000047143 | Dmrta2 | 242620 | protein_coding | 4.909099811 | 0.1664749 | 0.623877742 | -0.26683898 | 0.78959314 | 0.894883064 | 0.1664749 |
| ENSMUSG00000071659 | Hnrnpul2 | 68693 | protein_coding | 2042.047083 | 0.166475988 | 0.05133705 | -3.242803954 | 0.001183596 | 0.00731937 | 0.166475988 |

### Supplementary Table 1

| ensembl_gene_id | external_gene_name | entrezgene_id | gene_biotype | baseMean | log2FoldChange | lfcSE | stat | pvalue | padj | absLog2FC |
|---|---|---|---|---|---|---|---|---|---|---|
| ENSMUSG00000022682 | Rrn3 | 106298 | protein_coding | 479.4440879 | 0.166506312 | 0.09898916 | -1.682066111 | 0.092556021 | 0.233354218 | 0.166506312 |
| ENSMUSG00000021211 | Akr1c12 | 622402 | protein_coding | 1355.208157 | -0.166607961 | 0.073795543 | 2.257696812 | 0.023964568 | 0.084507982 | 0.166607961 |
| ENSMUSG00000024180 | Pgap6 | 60455 | protein_coding | 1398.940373 | -0.166614852 | 0.059034657 | 2.822322701 | 0.004767718 | 0.023369313 | 0.166614852 |
| ENSMUSG00000034320 | Slc26a2 | 13521 | protein_coding | 13764.11954 | 0.166637559 | 0.116545882 | -1.429802197 | 0.152773798 | 0.33362953 | 0.166637559 |
| ENSMUSG00000034259 | Exosc4 | 109075 | protein_coding | 197.5294121 | 0.166658817 | 0.0934882 | -1.78267222 | 0.074639676 | 0.199902629 | 0.166658817 |
| ENSMUSG00000021497 | Txndc15 | 69672 | protein_coding | 420.0435906 | -0.166673964 | 0.073431965 | 2.269773992 | 0.023221299 | 0.082503558 | 0.166673964 |
| ENSMUSG00000071112 | Spx | 319552 | protein_coding | 2.455548208 | 0.166723789 | 0.808225378 | -0.206283783 | 0.836569245 | 1 | 0.166723789 |
| ENSMUSG00000040455 | Usp45 | 77593 | protein_coding | 323.547897 | -0.166725135 | 0.093336267 | 1.786284582 | 0.074053192 | 0.198945226 | 0.166725135 |
| ENSMUSG00000010021 | Kif19a | 286942 | protein_coding | 12.68471478 | -0.16672846 | 0.545303421 | 0.305753556 | 0.759792293 | 0.879422681 | 0.16672846 |
| ENSMUSG00000062762 | Ei24 | 13663 | protein_coding | 1031.47018 | -0.16675286 | 0.089109762 | 1.871319777 | 0.061300771 | 0.173431717 | 0.16675286 |
| ENSMUSG00000021192 | Golga5 | 27277 | protein_coding | 734.3687518 | 0.166800903 | 0.078134721 | -2.134785919 | 0.032778501 | 0.108084245 | 0.166800903 |
| ENSMUSG00000021012 | Zc3h14 | 75553 | protein_coding | 787.8053597 | 0.166804714 | 0.069211501 | -2.410072195 | 0.015949364 | 0.061381449 | 0.166804714 |
| ENSMUSG00000000544 | Gpa33 | 59290 | protein_coding | 6357.056546 | -0.166807686 | 0.052328496 | 3.18770267 | 0.001434079 | 0.008569101 | 0.166807686 |
| ENSMUSG00000047423 | AI837181 | 107242 | protein_coding | 308.2879792 | -0.166821661 | 0.068086265 | 2.450151442 | 0.014279614 | 0.056092661 | 0.166821661 |
| ENSMUSG00000038167 | Plekhg6 | 213522 | protein_coding | 317.2407175 | -0.166865058 | 0.279903205 | 0.596152724 | 0.551073205 | 0.741922624 | 0.166865058 |
| ENSMUSG00000044361 | BC024139 | 271278 | protein_coding | 27.36639905 | 0.166866626 | 0.232532286 | -0.717606269 | 0.473000091 | 0.685318046 | 0.166866626 |
| ENSMUSG00000029364 | Wsb2 | 59043 | protein_coding | 800.7578525 | -0.16687334 | 0.086687956 | 1.924988734 | 0.054230768 | 0.158479366 | 0.16687334 |
| ENSMUSG00000021357 | Exoc2 | 66482 | protein_coding | 325.6037746 | -0.166901082 | 0.110974634 | 1.503957041 | 0.132592428 | 0.302143144 | 0.166901082 |
| ENSMUSG00000028292 | Rars2 | 109093 | protein_coding | 441.1938409 | -0.166949868 | 0.097765687 | 1.707653006 | 0.087700747 | 0.224854857 | 0.166949868 |
| ENSMUSG00000029017 | Pmpcb | 73078 | protein_coding | 871.6992277 | -0.166960047 | 0.084233507 | 1.982109657 | 0.047466969 | 0.143382024 | 0.166960047 |
| ENSMUSG00000002963 | Pnkp | 59047 | protein_coding | 338.8132175 | -0.167080175 | 0.072571193 | 2.302293356 | 0.021318634 | 0.077202358 | 0.167080175 |
| ENSMUSG00000041849 | Card6 | 239319 | protein_coding | 44.92230794 | 0.167096959 | 0.262112377 | -0.637501214 | 0.523798419 | 0.722399123 | 0.167096959 |
| ENSMUSG00000044456 | Rin3 | 217835 | protein_coding | 52.8999575 | -0.167127467 | 0.256651555 | 0.651184313 | 0.514927516 | 0.716272317 | 0.167127467 |
| ENSMUSG00000013523 | Bcas1 | 76960 | protein_coding | 1057.469802 | 0.167165719 | 0.188643344 | -0.886146921 | 0.375538362 | 0.597748482 | 0.167165719 |
| ENSMUSG00000063796 | Slc22a8 | 19879 | protein_coding | 2.62233006 | 0.167173104 | 2.672967045 | -0.062542149 | 0.950131097 | 1 | 0.167173104 |
| ENSMUSG00000031293 | Rs1 | 20147 | protein_coding | 2.9890196 | 0.167290115 | 1.386910386 | -0.12062071 | 0.903991465 | 1 | 0.167290115 |
| ENSMUSG00000021981 | Cab39l | 69008 | protein_coding | 465.6122326 | -0.167341678 | 0.108182978 | 1.546839259 | 0.121902011 | 0.284857762 | 0.167341678 |
| ENSMUSG00000034453 | Polr3b | 70428 | protein_coding | 351.5863137 | -0.167405446 | 0.106475255 | 1.572247431 | 0.115893179 | 0.274894315 | 0.167405446 |
| ENSMUSG00000041762 | Gpr155 | 68526 | protein_coding | 8.743753689 | 0.167409017 | 0.392647423 | -0.426359647 | 0.669845801 | 0.822722373 | 0.167409017 |
| ENSMUSG00000032305 | Fam219b | 78323 | protein_coding | 152.1632095 | -0.167428877 | 0.109537518 | 1.52850713 | 0.126386675 | 0.292373241 | 0.167428877 |
| ENSMUSG00000026113 | Inpp4a | 269180 | protein_coding | 153.6491237 | -0.167433332 | 0.136386829 | 1.227635642 | 0.219583776 | 0.425861007 | 0.167433332 |
| ENSMUSG00000070985 | Acnat1 | 230161 | protein_coding | 358.0610235 | -0.167437031 | 0.138765608 | 1.206617649 | 0.227579427 | 0.436092673 | 0.167437031 |
| ENSMUSG00000026810 | Dpm2 | 13481 | protein_coding | 241.979861 | 0.167492718 | 0.104606786 | -1.601164936 | 0.109340393 | 0.263984696 | 0.167492718 |
| ENSMUSG00000032733 | Snx33 | 235406 | protein_coding | 318.865961 | -0.1675095 | 0.077401666 | 2.164158833 | 0.030452161 | 0.102046874 | 0.1675095 |
| ENSMUSG00000055670 | Zzef1 | 195018 | protein_coding | 2593.609642 | 0.167539874 | 0.090851951 | -1.8440977 | 0.065168903 | 0.181269925 | 0.167539874 |
| ENSMUSG00000008333 | Snrpb2 | 20639 | protein_coding | 250.4870873 | 0.167545761 | 0.103466939 | -1.619316879 | 0.105379101 | 0.257088266 | 0.167545761 |
| ENSMUSG00000029125 | Stx18 | 71116 | protein_coding | 231.6783941 | 0.167558 | 0.089925816 | -1.863291414 | 0.062421278 | 0.175579114 | 0.167558 |
| ENSMUSG00000039630 | Hnrnpu | 51810 | protein_coding | 4573.946112 | 0.167562522 | 0.052716113 | -3.17858264 | 0.00147997 | 0.008791449 | 0.167562522 |
| ENSMUSG00000041685 | Fcho2 | 218503 | protein_coding | 633.9611959 | -0.16756448 | 0.105754348 | 1.58446894 | 0.113087044 | 0.270067889 | 0.16756448 |

Supplementary Table 1

| ensembl_gene_id | external_gene_name | entrezgene_id | gene_biotype | baseMean | log2FoldChange | lfcSE | stat | pvalue | padj | absLog2FC |
|---|---|---|---|---|---|---|---|---|---|---|
| ENSMUSG00000030243 | Recql | 19691 | protein_coding | 164.8688465 | 0.16765257 | 0.09381347 | -1.787084198 | 0.073923881 | 0.198694494 | 0.16765257 |
| ENSMUSG00000024507 | Hsd17b4 | 15488 | protein_coding | 1030.408261 | -0.167678014 | 0.067358481 | 2.489337819 | 0.012798129 | 0.051560944 | 0.167678014 |
| ENSMUSG00000047746 | Fbxo40 | 207215 | protein_coding | 1.064085804 | -0.167710837 | 2.026577229 | 0.08275571 | 0.934045787 | 1 | 0.167710837 |
| ENSMUSG00000028646 | Rragc | 54170 | protein_coding | 565.012456 | 0.167724267 | 0.094145414 | -1.781544737 | 0.074823503 | 0.19999509 | 0.167724267 |
| ENSMUSG00000053091 | Lins1 | 72635 | protein_coding | 98.42505994 | 0.167724894 | 0.116795044 | -1.43606174 | 0.150984774 | 0.331035921 | 0.167724894 |
| ENSMUSG00000028270 | Gbp2 | 14469 | protein_coding | 61.95497333 | -0.167739672 | 0.366091102 | 0.458191064 | 0.646815179 | 0.807553177 | 0.167739672 |
| ENSMUSG00000063929 | Cyp4a32 | 100040843 | protein_coding | 1.644414415 | -0.167749584 | 2.693693896 | 0.062274924 | 0.950343897 | 1 | 0.167749584 |
| ENSMUSG00000024622 | Hmgxb3 | 106894 | protein_coding | 530.3086374 | -0.167755075 | 0.060668577 | 2.765106488 | 0.005690422 | 0.027104593 | 0.167755075 |
| ENSMUSG00000020272 | Stk10 | 20868 | protein_coding | 477.6252499 | -0.167846688 | 0.254972233 | 0.658293987 | 0.510349241 | 0.71356425 | 0.167846688 |
| ENSMUSG00000041644 | Slc5a12 | 241612 | protein_coding | 4.957144405 | 0.167899258 | 1.706013127 | -0.098416158 | 0.921601845 | 0.962213622 | 0.167899258 |
| ENSMUSG00000071716 | Apol7e | 666348 | protein_coding | 4.857197178 | 0.167900568 | 0.664625671 | -0.252624259 | 0.800558582 | 0.901189824 | 0.167900568 |
| ENSMUSG00000022677 | Cep20 | 66086 | protein_coding | 446.8079597 | 0.167934176 | 0.079031157 | -2.124911007 | 0.033594039 | 0.109921135 | 0.167934176 |
| ENSMUSG00000024726 | Carnmt1 | 67383 | protein_coding | 92.29481802 | -0.167934854 | 0.145900541 | 1.151022832 | 0.249722843 | 0.462014445 | 0.167934854 |
| ENSMUSG00000036197 | Gxylt1 | 223827 | protein_coding | 458.0819575 | 0.167976544 | 0.083422017 | -2.013575661 | 0.044054104 | 0.1353506 | 0.167976544 |
| ENSMUSG00000074652 | Myh7b | 668940 | protein_coding | 6.613884007 | 0.168046712 | 0.733457708 | -0.229115749 | 0.818778953 | 0.908812719 | 0.168046712 |
| ENSMUSG00000024747 | Aldh1a7 | 26358 | protein_coding | 238.9203625 | -0.16809526 | 0.116616975 | 1.441430466 | 0.149463108 | 0.328844638 | 0.16809526 |
| ENSMUSG00000057113 | Npm1 | 18148 | protein_coding | 2400.795357 | -0.168248369 | 0.128890918 | 1.305354732 | 0.191772085 | 0.389522022 | 0.168248369 |
| ENSMUSG00000061130 | Ppm1b | 19043 | protein_coding | 1080.938619 | 0.168252461 | 0.061381831 | -2.741079205 | 0.006123774 | 0.028631939 | 0.168252461 |
| ENSMUSG00000023991 | Foxp4 | 74123 | protein_coding | 585.1177393 | 0.168391377 | 0.129090091 | -1.304448513 | 0.192080704 | 0.389807821 | 0.168391377 |
| ENSMUSG00000063851 | Rnf183 | 76072 | protein_coding | 8.183677877 | 0.168398387 | 0.538407395 | -0.312771312 | 0.754454412 | 0.876856731 | 0.168398387 |
| ENSMUSG00000051998 | Lax1 | 240754 | protein_coding | 4.405173518 | -0.168415654 | 0.664209991 | 0.253557846 | 0.799837167 | 0.900866933 | 0.168415654 |
| ENSMUSG00000015750 | Aph1a | 226548 | protein_coding | 740.4180466 | -0.168437137 | 0.108172647 | 1.557113947 | 0.119443434 | 0.28110145 | 0.168437137 |
| ENSMUSG00000036334 | Igsf10 | 242050 | protein_coding | 2.216569948 | -0.168458909 | 1.461314054 | 0.115279059 | 0.908223936 | 1 | 0.168458909 |
| ENSMUSG00000075908 | Gm23940 | 115489070 | snRNA | 1.682673142 | -0.168485179 | 0.86624076 | 0.19450156 | 0.845783174 | 1 | 0.168485179 |
| ENSMUSG00000031103 | Elf4 | 56501 | protein_coding | 643.8904753 | -0.168549501 | 0.132286996 | 1.27411995 | 0.202620909 | 0.404694914 | 0.168549501 |
| ENSMUSG00000078765 | U2af1l4 | 233073 | protein_coding | 4.85003259 | -0.168661187 | 0.522285797 | 0.322928918 | 0.746749075 | 0.872440598 | 0.168661187 |
| ENSMUSG00000095315 | Gm10130 | 102633750 | protein_coding | 36.53737851 | -0.168708817 | 0.226467613 | 0.74495781 | 0.456297225 | 0.671470886 | 0.168708817 |
| ENSMUSG00000032854 | Ugt8a | 22239 | protein_coding | 115.0030754 | 0.168712269 | 0.223813831 | -0.753806272 | 0.450965557 | 0.667636772 | 0.168712269 |
| ENSMUSG00000052446 | Zfp961 | 234413 | protein_coding | 173.4539545 | 0.168754977 | 0.106197593 | -1.589065934 | 0.112045508 | 0.268218365 | 0.168754977 |
| ENSMUSG00000061410 | Zcchc14 | 142682 | protein_coding | 545.1655815 | 0.168756357 | 0.054826335 | -3.078016359 | 0.002083835 | 0.01173204 | 0.168756357 |
| ENSMUSG00000051730 | Mettl5 | 75422 | protein_coding | 80.94092471 | -0.168940978 | 0.119160032 | 1.417765469 | 0.156259249 | 0.338979766 | 0.168940978 |
| ENSMUSG00000031570 | Plpp5 | 71910 | protein_coding | 553.9112411 | 0.168945861 | 0.07081883 | -2.385606489 | 0.017050985 | 0.06471615 | 0.168945861 |
| ENSMUSG00000034263 | Ints14 | 69882 | protein_coding | 358.0111263 | 0.168971093 | 0.089748282 | -1.882722313 | 0.05973801 | 0.17026467 | 0.168971093 |
| ENSMUSG00000026556 | Vangl2 | 93840 | protein_coding | 30.59334461 | 0.168972015 | 0.231287971 | -0.730569838 | 0.465041941 | 0.6785988 | 0.168972015 |
| ENSMUSG00000062861 | Zfp28 | 22690 | protein_coding | 28.12978736 | -0.169053516 | 0.263954661 | 0.640464218 | 0.521870845 | 0.721306924 | 0.169053516 |
| ENSMUSG00000027805 | Pfn2 | 18645 | protein_coding | 13.13998437 | 0.169057923 | 0.33804728 | -0.500101414 | 0.61700367 | 0.788881899 | 0.169057923 |
| ENSMUSG00000027673 | Ndufb5 | 66046 | protein_coding | 1201.668883 | -0.169109173 | 0.120173344 | 1.407210346 | 0.159365025 | 0.343481339 | 0.169109173 |
| ENSMUSG00000039633 | Lonrf1 | 244421 | protein_coding | 59.50320072 | 0.169196489 | 0.176827313 | -0.956845896 | 0.338645043 | 0.560303 | 0.169196489 |
| ENSMUSG00000025127 | Gcgr | 14527 | protein_coding | 2.038236806 | 0.169203707 | 2.174010987 | -0.077830198 | 0.937963124 | 1 | 0.169203707 |

| ensembl_gene_id | external_gene_name | entrezgene_id | gene_biotype | baseMean | log2FoldChange | lfcSE | stat | pvalue | padj | absLog2FC |
|---|---|---|---|---|---|---|---|---|---|---|
| ENSMUSG00000056201 | Cfl1 | 12631 | protein_coding | 3177.317836 | -0.169244777 | 0.07783061 | 2.174527181 | 0.029665566 | 0.10006477 | 0.169244777 |
| ENSMUSG00000117231 | Gm41609 | 105246306 | lncRNA | 2.783411581 | -0.169264519 | 0.842677912 | 0.200865024 | 0.840804117 | 1 | 0.169264519 |
| ENSMUSG00000061577 | Adgrg5 | 382045 | protein_coding | 6.589052167 | -0.169297381 | 0.581688948 | 0.291044521 | 0.77101727 | 0.885199039 | 0.169297381 |
| ENSMUSG00000004788 | Eif2b2 | 217715 | protein_coding | 335.7621947 | 0.169346511 | 0.074274163 | -2.28001912 | 0.022606555 | 0.080865411 | 0.169346511 |
| ENSMUSG00000037971 | 1110032A03Rik | 68721 | protein_coding | 211.5008061 | -0.169369603 | 0.093896558 | 1.803789252 | 0.071264351 | 0.193498767 | 0.169369603 |
| ENSMUSG00000059554 | Ccdc28a | 215814 | protein_coding | 161.9738904 | -0.169433923 | 0.129021685 | 1.31322051 | 0.189108655 | 0.386024762 | 0.169433923 |
| ENSMUSG00000030532 | Hddc3 | 68695 | protein_coding | 24.7222715 | -0.169440409 | 0.237840825 | 0.712410954 | 0.476210334 | 0.687965835 | 0.169440409 |
| ENSMUSG00000027180 | Fbxo3 | 57443 | protein_coding | 638.4196973 | -0.169492921 | 0.101538651 | 1.669245344 | 0.095068767 | 0.23816226 | 0.169492921 |
| ENSMUSG00000036611 | Eepd1 | 67484 | protein_coding | 67.36514369 | -0.169550877 | 0.188225527 | 0.90078577 | 0.367702235 | 0.590110273 | 0.169550877 |
| ENSMUSG00000021532 | Fastkd3 | 69577 | protein_coding | 228.9781356 | 0.169560687 | 0.081132264 | -2.089929186 | 0.036624161 | 0.117439107 | 0.169560687 |
| ENSMUSG00000038976 | Ppp1r9b | 217124 | protein_coding | 703.8971078 | -0.169606394 | 0.105882374 | 1.601837849 | 0.109191471 | 0.263739645 | 0.169606394 |
| ENSMUSG00000022474 | Pmm1 | 29858 | protein_coding | 300.4413957 | -0.1696416 | 0.125722719 | 1.349331296 | 0.177230578 | 0.369543657 | 0.1696416 |
| ENSMUSG00000010476 | Ebf3 | 13593 | protein_coding | 8.512643667 | -0.169677051 | 0.546051933 | 0.310734274 | 0.75600264 | 0.877468906 | 0.169677051 |
| ENSMUSG00000020864 | Ankrd40 | 71452 | protein_coding | 591.0432317 | -0.169695984 | 0.0640579 | 2.649103148 | 0.00807057 | 0.035875474 | 0.169695984 |
| ENSMUSG00000042328 | Hps4 | 192232 | protein_coding | 336.985424 | 0.169700712 | 0.060796886 | -2.791273077 | 0.005250116 | 0.025395551 | 0.169700712 |
| ENSMUSG00000000983 | Wfdc18 | 14038 | protein_coding | 61.86872149 | 0.169779619 | 0.221107892 | -0.767858702 | 0.442571135 | 0.660397553 | 0.169779619 |
| ENSMUSG00000038910 | Plcl2 | 224860 | protein_coding | 1060.775626 | -0.16980924 | 0.091176828 | 1.862416629 | 0.062544387 | 0.175865487 | 0.16980924 |
| ENSMUSG00000022325 | Pop1 | 67724 | protein_coding | 102.3439424 | 0.169889177 | 0.120689334 | -1.407656924 | 0.159232683 | 0.343330529 | 0.169889177 |
| ENSMUSG00000028780 | Sema3c | 20348 | protein_coding | 1664.906088 | -0.169891974 | 0.128039538 | 1.326871191 | 0.184551296 | 0.379678597 | 0.169891974 |
| ENSMUSG00000049892 | Rasd1 | 19416 | protein_coding | 411.3524952 | 0.169918052 | 0.180761121 | -0.940014373 | 0.347210188 | 0.569001592 | 0.169918052 |
| ENSMUSG00000056515 | Rab31 | 106572 | protein_coding | 92.85626726 | -0.169948066 | 0.144631822 | 1.175039241 | 0.239979017 | 0.450393623 | 0.169948066 |
| ENSMUSG00000005262 | Ufd1 | 22230 | protein_coding | 396.216614 | 0.169952975 | 0.058527706 | -2.903803784 | 0.003686591 | 0.018918697 | 0.169952975 |
| ENSMUSG00000024069 | Slc30a6 | 210148 | protein_coding | 245.3608576 | 0.169962353 | 0.09578267 | -1.774458292 | 0.075987375 | 0.20239605 | 0.169962353 |
| ENSMUSG00000029433 | Diablo | 66593 | protein_coding | 65.1761842 | -0.170000975 | 0.154718724 | 1.098774411 | 0.271866476 | 0.487629303 | 0.170000975 |
| ENSMUSG00000066877 | Nck2 | 17974 | protein_coding | 853.0925249 | -0.170137254 | 0.078946269 | 2.155101903 | 0.031153861 | 0.103913322 | 0.170137254 |
| ENSMUSG00000030678 | Maz | 17188 | protein_coding | 677.0743007 | 0.17013984 | 0.113286576 | -1.501853499 | 0.133134948 | 0.302752759 | 0.17013984 |
| ENSMUSG00000009863 | Sdhb | 67680 | protein_coding | 2465.321882 | -0.170195002 | 0.083677173 | 2.033947808 | 0.041956858 | 0.130534204 | 0.170195002 |
| ENSMUSG00000035258 | Abi3bp | 320712 | protein_coding | 8.075400673 | 0.170225729 | 0.611631218 | -0.278314324 | 0.780771081 | 0.890384202 | 0.170225729 |
| ENSMUSG00000053441 | Adamts19 | 240322 | protein_coding | 2.363738757 | -0.170290293 | 2.653636139 | 0.064172435 | 0.948832926 | 1 | 0.170290293 |
| ENSMUSG00000030216 | Wbp11 | 60321 | protein_coding | 560.6834918 | 0.170303702 | 0.080641149 | -2.111870983 | 0.034697517 | 0.112749861 | 0.170303702 |
| ENSMUSG00000032898 | Fbxo21 | 231670 | protein_coding | 257.7404179 | 0.170309681 | 0.152725582 | -1.115135259 | 0.264792464 | 0.479629566 | 0.170309681 |
| ENSMUSG00000049148 | Plcxd3 | 239318 | protein_coding | 2.655909859 | -0.170329931 | 0.812474232 | 0.209643486 | 0.833945936 | 1 | 0.170329931 |
| ENSMUSG00000022346 | Myc | 17869 | protein_coding | 267.7016921 | 0.170352867 | 0.13754758 | -1.23850138 | 0.215530211 | 0.420617573 | 0.170352867 |
| ENSMUSG00000031751 | Amfr | 23802 | protein_coding | 1974.139966 | -0.170374538 | 0.078460778 | 2.171461225 | 0.029896325 | 0.100654009 | 0.170374538 |
| ENSMUSG00000036850 | Mrpl41 | 107733 | protein_coding | 238.8957244 | -0.170399382 | 0.094681196 | 1.799717253 | 0.071905295 | 0.194854619 | 0.170399382 |
| ENSMUSG00000029049 | Morn1 | 76866 | protein_coding | 39.02538386 | -0.170458897 | 0.16953786 | 1.005432632 | 0.314688574 | 0.535859673 | 0.170458897 |
| ENSMUSG00000078784 | Rbis | 69126 | protein_coding | 238.5433821 | -0.170471728 | 0.110047096 | 1.549079748 | 0.121362551 | 0.284132491 | 0.170471728 |
| ENSMUSG00000026827 | Gpd2 | 14571 | protein_coding | 685.0719151 | -0.170491181 | 0.089117929 | 1.91309631 | 0.055735724 | 0.161590545 | 0.170491181 |
| ENSMUSG00000041354 | Rgl2 | 19732 | protein_coding | 394.967256 | -0.170539866 | 0.073781456 | 2.311419081 | 0.020809718 | 0.075644218 | 0.170539866 |

| ensembl_gene_id | external_gene_name | entrezgene_id | gene_biotype | baseMean | log2FoldChange | lfcSE | stat | pvalue | padj | absLog2FC |
|---|---|---|---|---|---|---|---|---|---|---|
| ENSMUSG00000026000 | Lancl1 | 14768 | protein_coding | 241.1745976 | 0.170554406 | 0.122483777 | -1.392465274 | 0.16378155 | 0.349988901 | 0.170554406 |
| ENSMUSG00000071533 | Pcnp | 76302 | protein_coding | 450.8615899 | -0.170590964 | 0.108607489 | 1.570710882 | 0.11624982 | 0.275430349 | 0.170590964 |
| ENSMUSG00000031158 | Timm17b | 21855 | protein_coding | 315.7352061 | 0.170622167 | 0.11112171 | -1.535453036 | 0.124672585 | 0.289447921 | 0.170622167 |
| ENSMUSG00000020153 | Ndufs7 | 75406 | protein_coding | 981.0932989 | -0.170691191 | 0.075576235 | 2.258529948 | 0.02391264 | 0.08439695 | 0.170691191 |
| ENSMUSG00000032599 | Ip6k2 | 76500 | protein_coding | 197.2940681 | -0.17071889 | 0.082880479 | 2.059820277 | 0.039415725 | 0.124671766 | 0.17071889 |
| ENSMUSG00000045679 | Pqlc3 | 217430 | protein_coding | 288.3912373 | 0.170729261 | 0.107160403 | -1.59321219 | 0.111112598 | 0.26676599 | 0.170729261 |
| ENSMUSG00000008682 | Rpl10 | 110954 | protein_coding | 5582.379804 | -0.170755077 | 0.139736081 | 1.221982723 | 0.22171416 | 0.42858336 | 0.170755077 |
| ENSMUSG00000046318 | Ccbe1 | 320924 | protein_coding | 4.398483674 | 0.170767771 | 1.902813687 | -0.089744872 | 0.928489957 | 0.965886893 | 0.170767771 |
| ENSMUSG00000024191 | Bnip1 | 224630 | protein_coding | 218.323894 | -0.170774652 | 0.096245484 | 1.774365352 | 0.076002737 | 0.202404322 | 0.170774652 |
| ENSMUSG00000026098 | Pms1 | 227099 | protein_coding | 61.22562543 | 0.170795797 | 0.151912662 | -1.124302579 | 0.260884685 | 0.474424588 | 0.170795797 |
| ENSMUSG00000084346 | Cyp4a30b | 435802 | protein_coding | 1.160799633 | 0.170815812 | 1.636217724 | -0.10439675 | 0.916854502 | 1 | 0.170815812 |
| ENSMUSG00000051113 | Garin5a | 75538 | protein_coding | 8.502407951 | 0.17090064 | 0.363896356 | -0.469640978 | 0.638611543 | 0.80301103 | 0.17090064 |
| ENSMUSG00000043154 | Ppp2r3a | 235542 | protein_coding | 201.0436454 | 0.171008971 | 0.094124972 | -1.816828921 | 0.069243313 | 0.189412634 | 0.171008971 |
| ENSMUSG00000023971 | Rrp36 | 224823 | protein_coding | 296.5645507 | -0.171013795 | 0.103026051 | 1.659908271 | 0.096932907 | 0.24162056 | 0.171013795 |
| ENSMUSG00000020451 | Limk2 | 16886 | protein_coding | 1933.541994 | -0.17108866 | 0.094011356 | 1.819872263 | 0.06877846 | 0.188515765 | 0.17108866 |
| ENSMUSG00000012640 | Zfp715 | 69930 | protein_coding | 248.6556988 | -0.17109319 | 0.088807972 | 1.92655216 | 0.054035466 | 0.158048473 | 0.17109319 |
| ENSMUSG00000030154 | Klrb1f | 232408 | protein_coding | 1.919863929 | -0.171172317 | 1.066216123 | 0.160541858 | 0.872454252 | 1 | 0.171172317 |
| ENSMUSG00000037826 | Ppm1k | 243382 | protein_coding | 176.9474208 | -0.171200662 | 0.15394875 | 1.112062693 | 0.266111196 | 0.481173797 | 0.171200662 |
| ENSMUSG00000052915 | Msl1 | 74026 | protein_coding | 687.8258428 | -0.171252102 | 0.055707142 | 3.074149836 | 0.002111034 | 0.011857691 | 0.171252102 |
| ENSMUSG00000036890 | Gtdc1 | 227835 | protein_coding | 53.80791273 | -0.171264319 | 0.149006115 | 1.14937779 | 0.250400234 | 0.462725114 | 0.171264319 |
| ENSMUSG00000053291 | Rab4b | 19342 | protein_coding | 44.92491362 | 0.171328702 | 0.201335049 | -0.850963126 | 0.394789836 | 0.617207172 | 0.171328702 |
| ENSMUSG00000031906 | Smpd3 | 58994 | protein_coding | 2464.821688 | 0.17136173 | 0.148186864 | -1.156389475 | 0.247521885 | 0.459691453 | 0.17136173 |
| ENSMUSG00000026659 | Dusp12 | 80915 | protein_coding | 61.63445272 | 0.171532738 | 0.166851776 | -1.028054613 | 0.303924139 | 0.524360158 | 0.171532738 |
| ENSMUSG00000025894 | Aasdhppt | 67618 | protein_coding | 207.1647365 | -0.171651483 | 0.094464014 | 1.817109778 | 0.069200306 | 0.189357722 | 0.171651483 |
| ENSMUSG00000008496 | Pou2f2 | 18987 | protein_coding | 7.88182033 | -0.171683851 | 1.271015986 | 0.135076076 | 0.892551726 | 0.947202684 | 0.171683851 |
| ENSMUSG00000037818 | Abhd18 | 269423 | protein_coding | 102.6564891 | -0.171703444 | 0.139051312 | 1.234820741 | 0.216897226 | 0.42248653 | 0.171703444 |
| ENSMUSG00000062210 | Tnfaip8 | 106869 | protein_coding | 429.1067703 | -0.171705298 | 0.186819036 | 0.919099584 | 0.358043487 | 0.580790426 | 0.171705298 |
| ENSMUSG00000024597 | Slc12a2 | 20496 | protein_coding | 5013.733157 | -0.171719424 | 0.168251959 | 1.020608767 | 0.307439834 | 0.527870807 | 0.171719424 |
| ENSMUSG00000041225 | Arhgap12 | 75415 | protein_coding | 1227.356828 | -0.171733077 | 0.084715335 | 2.027178152 | 0.042644197 | 0.132001174 | 0.171733077 |
| ENSMUSG00000022772 | Senp5 | 320213 | protein_coding | 434.6488476 | 0.171735071 | 0.070936162 | -2.420980576 | 0.015478704 | 0.059891125 | 0.171735071 |
| ENSMUSG00000027536 | Chmp4c | 66371 | protein_coding | 2119.13127 | -0.171736476 | 0.067389507 | 2.548415667 | 0.010821345 | 0.045275826 | 0.171736476 |
| ENSMUSG00000028909 | Ptpru | 19273 | protein_coding | 69.21857558 | -0.171761272 | 0.172469566 | 0.995893222 | 0.319302029 | 0.540704851 | 0.171761272 |
| ENSMUSG00000033253 | Szt2 | 230676 | protein_coding | 643.4119986 | 0.171821339 | 0.13306677 | -1.291241523 | 0.196619941 | 0.396311693 | 0.171821339 |
| ENSMUSG00000028708 | Mknk1 | 17346 | protein_coding | 130.5007595 | 0.171824492 | 0.13138629 | -1.307780993 | 0.190947603 | 0.388417088 | 0.171824492 |
| ENSMUSG00000033157 | Abhd10 | 213012 | protein_coding | 116.2981329 | 0.171858112 | 0.126844375 | -1.354873733 | 0.17545779 | 0.36728482 | 0.171858112 |
| ENSMUSG00000020282 | Rhbdf1 | 13650 | protein_coding | 305.1985913 | -0.171867201 | 0.162471134 | 1.057832225 | 0.290131938 | 0.509056399 | 0.171867201 |
| ENSMUSG00000032279 | Idh3a | 67834 | protein_coding | 1747.102829 | -0.171940485 | 0.090332934 | 1.903408617 | 0.056987248 | 0.164439526 | 0.171940485 |
| ENSMUSG00000030541 | Idh2 | 269951 | protein_coding | 1539.143062 | -0.172061091 | 0.089631924 | 1.919640717 | 0.054903297 | 0.15977566 | 0.172061091 |
| ENSMUSG00000079363 | Gbp4 | 17472 | protein_coding | 7.214883495 | 0.172082969 | 0.651609091 | -0.26408927 | 0.791711132 | 0.896267126 | 0.172082969 |

Supplementary Table 1

| ensembl_gene_id | external_gene_name | entrezgene_id | gene_biotype | baseMean | log2FoldChange | lfcSE | stat | pvalue | padj | absLog2FC |
|---|---|---|---|---|---|---|---|---|---|---|
| ENSMUSG00000085282 | Gm15663 | 327812 | lncRNA | 21.293689 | -0.172102161 | 0.234630181 | 0.733503934 | 0.463251129 | 0.676807614 | 0.172102161 |
| ENSMUSG00000069020 | Urm1 | 68205 | protein_coding | 281.6857773 | 0.172140047 | 0.100467913 | -1.713383323 | 0.086642042 | 0.222728314 | 0.172140047 |
| ENSMUSG00000025575 | Cant1 | 76025 | protein_coding | 3681.528215 | 0.172140087 | 0.075210636 | -2.288773175 | 0.022092535 | 0.0793122 | 0.172140087 |
| ENSMUSG00000033502 | Cdc14a | 229776 | protein_coding | 535.68606 | -0.172175971 | 0.105572431 | 1.630880047 | 0.102915633 | 0.252796186 | 0.172175971 |
| ENSMUSG00000025204 | Ndufb8 | 67264 | protein_coding | 2001.561507 | -0.172178342 | 0.088985338 | 1.934906874 | 0.053001724 | 0.155964091 | 0.172178342 |
| ENSMUSG00000087963 | Gm25394 | 115488237 | snoRNA | 33.30204228 | 0.172212435 | 0.247773179 | -0.695040663 | 0.487029819 | 0.695624496 | 0.172212435 |
| ENSMUSG00000037395 | Rcor3 | 214742 | protein_coding | 211.9110384 | -0.172223505 | 0.108110282 | 1.593035387 | 0.111152253 | 0.26676599 | 0.172223505 |
| ENSMUSG00000064844 | Gm26202 | 115488998 | snoRNA | 8.735059899 | 0.172226306 | 0.36857548 | -0.467275537 | 0.640302756 | 0.803678477 | 0.172226306 |
| ENSMUSG00000050854 | Tmem125 | 230678 | protein_coding | 256.680974 | 0.172234233 | 0.110471484 | -1.55908319 | 0.118976692 | 0.280242631 | 0.172234233 |
| ENSMUSG00000026869 | Psmd5 | 66998 | protein_coding | 659.7948577 | 0.17223714 | 0.055188689 | -3.120877523 | 0.00180313 | 0.010407861 | 0.17223714 |
| ENSMUSG00000038451 | Spsb2 | 14794 | protein_coding | 174.5364503 | -0.172239348 | 0.118616639 | 1.452067346 | 0.14648288 | 0.32409141 | 0.172239348 |
| ENSMUSG00000018750 | Zbtb4 | 75580 | protein_coding | 284.4891581 | -0.172246549 | 0.119821755 | 1.437523179 | 0.150569391 | 0.330520129 | 0.172246549 |
| ENSMUSG00000049091 | Sephs2 | 20768 | protein_coding | 754.6929956 | -0.172298804 | 0.082209972 | 2.095838255 | 0.036096542 | 0.11606275 | 0.172298804 |
| ENSMUSG00000020736 | Nt5c | 50773 | protein_coding | 856.4839924 | -0.172349181 | 0.133103642 | 1.294849475 | 0.195372161 | 0.394567184 | 0.172349181 |
| ENSMUSG00000009828 | Cilk1 | 56542 | protein_coding | 1327.982695 | 0.172358345 | 0.087948077 | -1.959773884 | 0.050022225 | 0.148948255 | 0.172358345 |
| ENSMUSG00000034889 | Cactin | 70312 | protein_coding | 353.1525777 | 0.172369045 | 0.08566289 | -2.012178739 | 0.044201099 | 0.135701235 | 0.172369045 |
| ENSMUSG00000046417 | Lrrc75a | 192976 | protein_coding | 651.9882889 | 0.172385865 | 0.083845277 | -2.055999708 | 0.039782544 | 0.125530417 | 0.172385865 |
| ENSMUSG00000020923 | Ubtf | 21429 | protein_coding | 1662.608608 | 0.17239468 | 0.052891124 | -3.259425565 | 0.001116381 | 0.0069833 | 0.17239468 |
| ENSMUSG00000022351 | Sqle | 20775 | protein_coding | 1321.294961 | 0.172412637 | 0.150028017 | -1.149202927 | 0.250472314 | 0.462725114 | 0.172412637 |
| ENSMUSG00000009076 | Zmat5 | 67178 | protein_coding | 111.8835088 | -0.172424996 | 0.115509578 | 1.49273332 | 0.135507008 | 0.30685209 | 0.172424996 |
| ENSMUSG00000002221 | Paxip1 | 55982 | protein_coding | 316.7921165 | 0.172452817 | 0.081634205 | -2.112506848 | 0.034642999 | 0.112617025 | 0.172452817 |
| ENSMUSG00000043384 | Gprasp1 | 67298 | protein_coding | 107.6814591 | -0.172508263 | 0.109043962 | 1.582006561 | 0.113648073 | 0.270742105 | 0.172508263 |
| ENSMUSG00000030180 | Kdm5a | 214899 | protein_coding | 823.7274639 | -0.172566555 | 0.058182384 | 2.965958846 | 0.003017409 | 0.015940335 | 0.172566555 |
| ENSMUSG00000047747 | Rnf150 | 330812 | protein_coding | 120.4593266 | 0.172581647 | 0.168521874 | -1.024090482 | 0.305792539 | 0.526136486 | 0.172581647 |
| ENSMUSG00000084946 | Dlx1as | 111970 | lncRNA | 2.05402942 | 0.172653526 | 0.997350318 | -0.173112219 | 0.862563221 | 1 | 0.172653526 |
| ENSMUSG00000032589 | Bsn | 12217 | protein_coding | 192.5502294 | -0.172689663 | 0.115466346 | 1.495584379 | 0.134761996 | 0.305611042 | 0.172689663 |
| ENSMUSG00000036943 | Rab8b | 235442 | protein_coding | 153.2613813 | 0.172700427 | 0.107015304 | -1.613791864 | 0.10657258 | 0.259233994 | 0.172700427 |
| ENSMUSG00000025579 | Gaa | 14387 | protein_coding | 541.2897674 | -0.172721401 | 0.099854116 | 1.729737418 | 0.0836772 | 0.216896136 | 0.172721401 |
| ENSMUSG00000059811 | Atl2 | 56298 | protein_coding | 1039.986973 | 0.172760603 | 0.075080009 | -2.301020006 | 0.0213905 | 0.077346775 | 0.172760603 |
| ENSMUSG00000056268 | Dennd1b | 329260 | protein_coding | 989.3114626 | -0.172816984 | 0.10269773 | 1.682773164 | 0.092419012 | 0.233151171 | 0.172816984 |
| ENSMUSG00000022280 | Rnf19a | 30945 | protein_coding | 417.8323338 | -0.172851916 | 0.1127859 | 1.532566708 | 0.12538265 | 0.290646979 | 0.172851916 |
| ENSMUSG00000019505 | Ubb | 22187 | protein_coding | 3635.678153 | 0.172890592 | 0.088282694 | -1.95837468 | 0.050186064 | 0.149272747 | 0.172890592 |
| ENSMUSG00000027506 | Tpd52 | 21985 | protein_coding | 4673.407281 | 0.172925983 | 0.099190481 | -1.74337276 | 0.081268523 | 0.212151185 | 0.172925983 |
| ENSMUSG00000030699 | Tbx6 | 21389 | protein_coding | 12.06950303 | -0.172933351 | 0.491770297 | 0.351654729 | 0.725097215 | 0.857652848 | 0.172933351 |
| ENSMUSG00000050891 | Tatdn1 | 69694 | protein_coding | 104.6384511 | 0.172936226 | 0.103235219 | -1.675166935 | 0.093901483 | 0.235970255 | 0.172936226 |
| ENSMUSG00000003476 | Crhr2 | 12922 | protein_coding | 2.160563904 | 0.172974823 | 1.945892549 | -0.088892279 | 0.92916752 | 1 | 0.172974823 |
| ENSMUSG00000053113 | Socs3 | 12702 | protein_coding | 283.7209276 | -0.172990601 | 0.273391347 | 0.632758144 | 0.526891594 | 0.725332426 | 0.172990601 |
| ENSMUSG00000048264 | Dip2c | 208440 | protein_coding | 239.8391302 | -0.172997052 | 0.075560039 | 2.289530994 | 0.022048519 | 0.079188614 | 0.172997052 |
| ENSMUSG00000027940 | Tpm3 | 59069 | protein_coding | 3357.576382 | 0.173027899 | 0.093885989 | -1.842957621 | 0.065335198 | 0.181579513 | 0.173027899 |

| ensembl_gene_id | external_gene_name | entrezgene_id | gene_biotype | baseMean | log2FoldChange | lfcSE | stat | pvalue | padj | absLog2FC |
|---|---|---|---|---|---|---|---|---|---|---|
| ENSMUSG00000004328 | Hif3a | 53417 | protein_coding | 78.65091293 | 0.173199069 | 0.178754884 | -0.968919368 | 0.332585421 | 0.555003602 | 0.173199069 |
| ENSMUSG00000033940 | Brk1 | 101314 | protein_coding | 786.2218737 | -0.173205111 | 0.08538197 | 2.028591174 | 0.042499948 | 0.131725022 | 0.173205111 |
| ENSMUSG00000036528 | Ppfibp2 | 19024 | protein_coding | 361.0320146 | -0.173324512 | 0.098381582 | 1.761757729 | 0.07811024 | 0.206650512 | 0.173324512 |
| ENSMUSG00000033991 | Ttc37 | 218343 | protein_coding | 511.8681114 | 0.173333367 | 0.105196352 | -1.647712711 | 0.099411638 | 0.246388083 | 0.173333367 |
| ENSMUSG00000071713 | Csf2rb | 12983 | protein_coding | 6.008248577 | 0.173376172 | 1.01070774 | -0.171539373 | 0.863799674 | 0.932373279 | 0.173376172 |
| ENSMUSG00000048120 | Entpd1 | 12495 | protein_coding | 4.759266183 | -0.173439466 | 0.756823361 | 0.229167696 | 0.81873858 | 0.908812719 | 0.173439466 |
| ENSMUSG00000035151 | Elmod2 | 244548 | protein_coding | 363.1655186 | 0.173510182 | 0.090851337 | -1.909825303 | 0.05615571 | 0.162665392 | 0.173510182 |
| ENSMUSG00000044155 | Lsm8 | 76522 | protein_coding | 268.0913019 | 0.173554796 | 0.099312487 | -1.747562687 | 0.080539781 | 0.210915627 | 0.173554796 |
| ENSMUSG00000031007 | Atp6ap2 | 70495 | protein_coding | 1766.570917 | 0.173647109 | 0.14545848 | -1.193791585 | 0.232559504 | 0.441638415 | 0.173647109 |
| ENSMUSG00000027317 | Ppp1r14d | 72112 | protein_coding | 1478.858046 | 0.173665569 | 0.078563777 | -2.210504323 | 0.027070181 | 0.092985996 | 0.173665569 |
| ENSMUSG00000021055 | Esr2 | 13983 | protein_coding | 18.32718754 | -0.173678646 | 0.240821673 | 0.721191927 | 0.470791438 | 0.683906563 | 0.173678646 |
| ENSMUSG00000024091 | Vapa | 30960 | protein_coding | 1721.675562 | 0.173684274 | 0.059942849 | -2.897497829 | 0.003761523 | 0.01920773 | 0.173684274 |
| ENSMUSG00000114362 | 4933433G19Rik | 71226 | lncRNA | 10.43578278 | 0.173695632 | 0.35369934 | -0.491082714 | 0.623367947 | 0.793353766 | 0.173695632 |
| ENSMUSG00000039382 | Wdr45 | 54636 | protein_coding | 116.9800346 | -0.173725973 | 0.130420935 | 1.332040543 | 0.182846866 | 0.377583238 | 0.173725973 |
| ENSMUSG00000028854 | Slc9a1 | 20544 | protein_coding | 610.5255289 | 0.173775387 | 0.15156205 | -1.146562658 | 0.251562417 | 0.464012259 | 0.173775387 |
| ENSMUSG00000037722 | Gnpnat1 | 54342 | protein_coding | 82.27110402 | 0.173812327 | 0.189073566 | -0.919284122 | 0.357946981 | 0.580790426 | 0.173812327 |
| ENSMUSG00000033096 | Apmap | 71881 | protein_coding | 436.6915348 | 0.173820212 | 0.106441701 | -1.633008579 | 0.102467192 | 0.252075246 | 0.173820212 |
| ENSMUSG00000021510 | Zfp729a | 212281 | protein_coding | 168.8757126 | -0.173845623 | 0.110404714 | 1.574621378 | 0.115343865 | 0.27387327 | 0.173845623 |
| ENSMUSG00000049382 | Krt8 | 16691 | protein_coding | 59862.67777 | -0.173874505 | 0.125855999 | 1.381535294 | 0.167114434 | 0.355177317 | 0.173874505 |
| ENSMUSG00000039660 | Spout1 | 227695 | protein_coding | 214.2834508 | 0.173887909 | 0.102846509 | -1.690751688 | 0.090884238 | 0.230377922 | 0.173887909 |
| ENSMUSG00000034484 | Snx2 | 67804 | protein_coding | 779.2736883 | -0.17401915 | 0.078996711 | 2.202865765 | 0.027604211 | 0.09441921 | 0.17401915 |
| ENSMUSG00000020638 | Cmpk2 | 22169 | protein_coding | 139.6178052 | 0.174040396 | 0.233790292 | -0.744429526 | 0.456616662 | 0.671509918 | 0.174040396 |
| ENSMUSG00000030055 | Rab43 | 69834 | protein_coding | 102.1552357 | -0.174077476 | 0.197793284 | 0.880098011 | 0.378806217 | 0.601173216 | 0.174077476 |
| ENSMUSG00000097766 | 5730420D15Rik | 70523 | lncRNA | 5.08579696 | 0.17409334 | 0.643017031 | -0.270744523 | 0.786587531 | 0.893302015 | 0.17409334 |
| ENSMUSG00000005686 | Ampd3 | 11717 | protein_coding | 628.5802577 | -0.174133955 | 0.070770184 | 2.460555348 | 0.013872217 | 0.05487085 | 0.174133955 |
| ENSMUSG00000025234 | Arih1 | 23806 | protein_coding | 1216.622711 | 0.174156726 | 0.04751826 | -3.665048464 | 0.000247292 | 0.001911596 | 0.174156726 |
| ENSMUSG00000030616 | Sytl2 | 83671 | protein_coding | 1121.749075 | -0.17417268 | 0.188891985 | 0.922075548 | 0.35648917 | 0.579378227 | 0.17417268 |
| ENSMUSG00000007589 | Tinf2 | 28113 | protein_coding | 243.1878222 | 0.174223691 | 0.102064727 | -1.706992175 | 0.087823507 | 0.224960042 | 0.174223691 |
| ENSMUSG00000039983 | Ccdc32 | 269336 | protein_coding | 449.0769836 | -0.174238427 | 0.137381517 | 1.268281427 | 0.204697467 | 0.407126819 | 0.174238427 |
| ENSMUSG00000026155 | Smap1 | 98366 | protein_coding | 307.8278901 | 0.174260439 | 0.111308657 | -1.565560518 | 0.117451539 | 0.277481361 | 0.174260439 |
| ENSMUSG00000043219 | Hoxa6 | 15403 | protein_coding | 23.88076203 | 0.174316355 | 0.24283036 | -0.717852394 | 0.472848304 | 0.685206817 | 0.174316355 |
| ENSMUSG00000024176 | Sox8 | 20681 | protein_coding | 1.275494418 | -0.17431704 | 1.716071656 | 0.101579115 | 0.919090758 | 1 | 0.17431704 |
| ENSMUSG00000019851 | Perp | 64058 | protein_coding | 2729.870833 | -0.17433226 | 0.102104959 | 1.707382881 | 0.08775091 | 0.224907418 | 0.17433226 |
| ENSMUSG00000074342 | I830077J02Rik | 433638 | protein_coding | 1.909749086 | 0.174367958 | 1.369073078 | -0.127362053 | 0.898653849 | 1 | 0.174367958 |
| ENSMUSG00000035818 | Plekhs1 | 226245 | protein_coding | 13.21684127 | 0.174394487 | 0.371005043 | -0.470059614 | 0.638312427 | 0.802947244 | 0.174394487 |
| ENSMUSG00000038486 | Sv2a | 64051 | protein_coding | 5.212951589 | -0.1744047 | 1.018340905 | 0.171263571 | 0.864016523 | 0.932456625 | 0.1744047 |
| ENSMUSG00000052942 | Glis3 | 226075 | protein_coding | 161.4940293 | -0.174481752 | 0.12956759 | 1.346646579 | 0.178094085 | 0.370548786 | 0.174481752 |
| ENSMUSG00000001380 | Hars | 15115 | protein_coding | 495.6326869 | 0.174504142 | 0.084836514 | -2.056946161 | 0.039691405 | 0.12530374 | 0.174504142 |
| ENSMUSG00000027288 | Zfp106 | 20402 | protein_coding | 2039.282062 | 0.174504857 | 0.073562149 | -2.372209893 | 0.017682043 | 0.066621322 | 0.174504857 |

| ensembl_gene_id | external_gene_name | entrezgene_id | gene_biotype | baseMean | log2FoldChange | lfcSE | stat | pvalue | padj | absLog2FC |
|---|---|---|---|---|---|---|---|---|---|---|
| ENSMUSG00000019856 | Fam184a | 75906 | protein_coding | 2.874458202 | -0.174595324 | 0.800094593 | 0.218218352 | 0.827258987 | 1 | 0.174595324 |
| ENSMUSG00000038903 | Ccdc68 | 381175 | protein_coding | 311.4916174 | -0.174607454 | 0.175725769 | 0.993636022 | 0.3204001 | 0.5416756 | 0.174607454 |
| ENSMUSG00000008475 | Arpc5 | 67771 | protein_coding | 2619.45069 | -0.174626115 | 0.0953489 | 1.83144342 | 0.067034385 | 0.184994292 | 0.174626115 |
| ENSMUSG00000035376 | Hacd2 | 70757 | protein_coding | 202.325567 | -0.174645406 | 0.108771827 | 1.605612517 | 0.108359074 | 0.262189266 | 0.174645406 |
| ENSMUSG00000064542 | Gm22980 | 115490044 | snoRNA | 3.830785792 | 0.174670336 | 0.573881101 | -0.304366768 | 0.760848483 | 1 | 0.174670336 |
| ENSMUSG00000031637 | Lrp2bp | 67620 | protein_coding | 5.122365411 | -0.174683051 | 0.835776667 | 0.209006854 | 0.834442888 | 0.916826487 | 0.174683051 |
| ENSMUSG00000096188 | Cmtm4 | 97487 | protein_coding | 3384.260396 | -0.174697363 | 0.085045423 | 2.054165371 | 0.039959689 | 0.125813975 | 0.174697363 |
| ENSMUSG00000049232 | Tigd2 | 68140 | protein_coding | 168.7531409 | -0.17472415 | 0.108955718 | 1.603625339 | 0.108796662 | 0.263016846 | 0.17472415 |
| ENSMUSG00000020315 | Sptbn1 | 20742 | protein_coding | 9189.480466 | -0.174854227 | 0.060696866 | 2.880778519 | 0.003966943 | 0.020064348 | 0.174854227 |
| ENSMUSG00000091017 | Garin4 | 619288 | protein_coding | 24.70148789 | 0.174859788 | 0.709341185 | -0.24651013 | 0.805287364 | 0.904107385 | 0.174859788 |
| ENSMUSG00000021213 | Akr1c13 | 27384 | protein_coding | 2180.60445 | -0.174915562 | 0.067080407 | 2.607550713 | 0.009119256 | 0.039567892 | 0.174915562 |
| ENSMUSG00000029610 | Aimp2 | 231872 | protein_coding | 139.1958718 | 0.174974742 | 0.101941942 | -1.716415627 | 0.086085998 | 0.221678493 | 0.174974742 |
| ENSMUSG00000030166 | Rad52 | 19365 | protein_coding | 99.62807433 | -0.175027508 | 0.122773876 | 1.425608715 | 0.153981319 | 0.33564513 | 0.175027508 |
| ENSMUSG00000039715 | Dync2i2 | 71820 | protein_coding | 97.84335668 | 0.175033866 | 0.126931668 | -1.37896137 | 0.167906665 | 0.356436387 | 0.175033866 |
| ENSMUSG00000054161 | Fam83e | 73813 | protein_coding | 4718.969309 | -0.175057959 | 0.057945285 | 3.021090647 | 0.002518659 | 0.013704495 | 0.175057959 |
| ENSMUSG00000031402 | Mpp1 | 17524 | protein_coding | 269.9622816 | -0.175122649 | 0.141292724 | 1.239431472 | 0.215185752 | 0.420143948 | 0.175122649 |
| ENSMUSG00000092118 | Fancf | 100040608 | protein_coding | 31.0877655 | 0.175255043 | 0.246232335 | -0.711746661 | 0.476621667 | 0.68833522 | 0.175255043 |
| ENSMUSG00000071646 | Mta2 | 23942 | protein_coding | 1403.700857 | 0.175316295 | 0.06685436 | -2.622361438 | 0.008732277 | 0.038193919 | 0.175316295 |
| ENSMUSG00000047641 | Krt87 | 406219 | protein_coding | 43.40028126 | 0.175320934 | 0.184455273 | -0.950479382 | 0.341868725 | 0.563829035 | 0.175320934 |
| ENSMUSG00000000355 | Mcts1 | 68995 | protein_coding | 188.2292551 | 0.175364306 | 0.094345287 | -1.858750042 | 0.063062577 | 0.176840786 | 0.175364306 |
| ENSMUSG00000024096 | Ralbp1 | 19765 | protein_coding | 1173.059871 | -0.175364417 | 0.067029018 | 2.61624623 | 0.008890242 | 0.038706422 | 0.175364417 |
| ENSMUSG00000035215 | Lsm7 | 66094 | protein_coding | 92.66807274 | -0.175438969 | 0.139067402 | 1.261539134 | 0.20711467 | 0.409951417 | 0.175438969 |
| ENSMUSG00000044147 | Arf6 | 11845 | protein_coding | 2538.103893 | 0.175471746 | 0.092217963 | -1.902793554 | 0.057067489 | 0.164593053 | 0.175471746 |
| ENSMUSG00000026779 | Mastl | 67121 | protein_coding | 173.1964412 | 0.175477532 | 0.194743903 | -0.901068171 | 0.367552075 | 0.590046157 | 0.175477532 |
| ENSMUSG00000069743 | Zfp820 | 75424 | protein_coding | 79.36585352 | -0.175481835 | 0.130629054 | 1.343359917 | 0.179155459 | 0.372241225 | 0.175481835 |
| ENSMUSG00000047045 | Tmem164 | 209497 | protein_coding | 1808.606806 | -0.175497181 | 0.082215237 | 2.13460652 | 0.032793164 | 0.108093077 | 0.175497181 |
| ENSMUSG00000040287 | Stac3 | 237611 | protein_coding | 8.352956818 | 0.17551007 | 0.55009147 | -0.319056157 | 0.749683929 | 0.874258149 | 0.17551007 |
| ENSMUSG00000069184 | Zfp72 | 238722 | protein_coding | 40.58875079 | -0.175520964 | 0.236215665 | 0.743053868 | 0.457449068 | 0.672242383 | 0.175520964 |
| ENSMUSG00000024174 | Pot1b | 72836 | protein_coding | 57.92408862 | -0.17554468 | 0.140034249 | 1.253583897 | 0.209993287 | 0.413822094 | 0.17554468 |
| ENSMUSG00000019822 | Smpd2 | 20598 | protein_coding | 230.9495383 | -0.175555228 | 0.097905695 | 1.793105376 | 0.072956076 | 0.196830035 | 0.175555228 |
| ENSMUSG00000036835 | Psenen | 66340 | protein_coding | 245.6149462 | -0.175641356 | 0.121713298 | 1.443074501 | 0.148999485 | 0.328033263 | 0.175641356 |
| ENSMUSG00000028467 | Gba2 | 230101 | protein_coding | 167.5303187 | -0.175653476 | 0.115928616 | 1.515186515 | 0.129725177 | 0.297415185 | 0.175653476 |
| ENSMUSG00000050989 | Selenon | 74777 | protein_coding | 35.40701628 | 0.175703725 | 0.231130551 | -0.76019256 | 0.447139492 | 0.664452584 | 0.175703725 |
| ENSMUSG00000069835 | Sat2 | 69215 | protein_coding | 21.08351167 | -0.175730876 | 0.277203709 | 0.633941285 | 0.526119141 | 0.724510632 | 0.175730876 |
| ENSMUSG00000035473 | Galm | 319625 | protein_coding | 674.3127596 | -0.175734473 | 0.058819463 | 2.987692576 | 0.002810921 | 0.015076179 | 0.175734473 |
| ENSMUSG00000028745 | Capzb | 12345 | protein_coding | 2498.718381 | 0.175741607 | 0.056278962 | -3.122687447 | 0.001792079 | 0.010353962 | 0.175741607 |
| ENSMUSG00000005763 | Cd247 | 12503 | protein_coding | 6.68586058 | -0.175791099 | 0.452154951 | 0.38878508 | 0.697435139 | 0.839586229 | 0.175791099 |
| ENSMUSG00000031660 | Brd7 | 26992 | protein_coding | 426.1900261 | -0.175792955 | 0.069477879 | 2.530200353 | 0.011399741 | 0.0473047 | 0.175792955 |
| ENSMUSG00000030727 | Rabep2 | 70314 | protein_coding | 338.1891267 | -0.17581703 | 0.097219867 | 1.808447547 | 0.070536873 | 0.192028673 | 0.17581703 |

| ensembl_gene_id | external_gene_name | entrezgene_id | gene_biotype | baseMean | log2FoldChange | lfcSE | stat | pvalue | padj | absLog2FC |
|---|---|---|---|---|---|---|---|---|---|---|
| ENSMUSG00000097267 | 1700109K24Rik | 74303 | lncRNA | 6.624655469 | 0.175827774 | 0.46441947 | -0.378596904 | 0.704987221 | 0.845227165 | 0.175827774 |
| ENSMUSG00000074576 | Mocs3 | 69372 | protein_coding | 122.8511559 | 0.175842133 | 0.11339825 | -1.550660031 | 0.120983178 | 0.283633759 | 0.175842133 |
| ENSMUSG00000047735 | Samd9l | 209086 | protein_coding | 1545.603148 | -0.175849829 | 0.139593481 | 1.259728085 | 0.20776747 | 0.410708835 | 0.175849829 |
| ENSMUSG00000022305 | Lrp12 | 239393 | protein_coding | 166.4805285 | -0.175887926 | 0.148420756 | 1.185062867 | 0.235992592 | 0.446089228 | 0.175887926 |
| ENSMUSG00000027820 | Mme | 17380 | protein_coding | 15.65207168 | 0.175899131 | 0.757405157 | -0.232239151 | 0.816352271 | 0.908409285 | 0.175899131 |
| ENSMUSG00000020516 | Rps6kb1 | 72508 | protein_coding | 493.8829122 | 0.175919614 | 0.07479144 | -2.352135692 | 0.018665964 | 0.069550933 | 0.175919614 |
| ENSMUSG00000043162 | Pyurf | 66459 | protein_coding | 159.2522166 | -0.17599588 | 0.093701377 | 1.878263546 | 0.060345121 | 0.171466333 | 0.17599588 |
| ENSMUSG00000030307 | Slc6a11 | 243616 | protein_coding | 2.04742851 | 0.176008836 | 2.687610432 | -0.065488969 | 0.947784689 | 1 | 0.176008836 |
| ENSMUSG00000026890 | Lhx6 | 16874 | protein_coding | 1.882689748 | 0.176009083 | 2.697732885 | -0.065243332 | 0.94798026 | 1 | 0.176009083 |
| ENSMUSG00000094950 | Vmn2r66 | 233437 | protein_coding | 1.650431378 | 0.176009164 | 2.725787284 | -0.064571863 | 0.948514888 | 1 | 0.176009164 |
| ENSMUSG00000086118 | Gm14169 | 100504350 | lncRNA | 1.407836067 | 0.176009277 | 2.802444811 | -0.062805618 | 0.949921293 | 1 | 0.176009277 |
| ENSMUSG00000057447 | Or4c11c | 258898 | protein_coding | 1.589823595 | 0.176009793 | 2.734047731 | -0.064377001 | 0.948670043 | 1 | 0.176009793 |
| ENSMUSG00000100459 | Gm5833 | 545363 | lncRNA | 1.589823595 | 0.176009793 | 2.734047731 | -0.064377001 | 0.948670043 | 1 | 0.176009793 |
| ENSMUSG00000023433 | Cela3b | 67868 | protein_coding | 1.050223206 | 0.176009995 | 2.834549745 | -0.062094516 | 0.950487564 | 1 | 0.176009995 |
| ENSMUSG00000031441 | Atp11a | 50770 | protein_coding | 806.0789567 | 0.176010103 | 0.178910495 | -0.983788586 | 0.325219472 | 0.547052145 | 0.176010103 |
| ENSMUSG00000087469 | Gm16142 | 102633324 | lncRNA | 1.585696135 | 0.176010272 | 2.711261486 | -0.064918221 | 0.948239113 | 1 | 0.176010272 |
| ENSMUSG00000039717 | Ralyl | 76897 | protein_coding | 1.451269219 | 0.176010352 | 2.739785544 | -0.064242383 | 0.94877723 | 1 | 0.176010352 |
| ENSMUSG00000025776 | Crispld1 | 83691 | protein_coding | 1.494702371 | 0.176010528 | 2.719971762 | -0.064710425 | 0.948404563 | 1 | 0.176010528 |
| ENSMUSG00000116210 | Spn-ps | 20738 | transcribed_processed_pseudogene | 1.494702371 | 0.176010528 | 2.719971762 | -0.064710425 | 0.948404563 | 1 | 0.176010528 |
| ENSMUSG00000045871 | Slitrk6 | 239250 | protein_coding | 1.443014299 | 0.17601055 | 2.728879874 | -0.064499193 | 0.948572749 | 1 | 0.17601055 |
| ENSMUSG00000008658 | Rbfox1 | 268859 | protein_coding | 1.399581147 | 0.176010904 | 2.723944528 | -0.064616185 | 0.948479598 | 1 | 0.176010904 |
| ENSMUSG00000073010 | Gm5127 | 331493 | protein_coding | 1.076674987 | 0.17601115 | 2.781198248 | -0.063286086 | 0.949538696 | 1 | 0.17601115 |
| ENSMUSG00000053228 | Ceacam3 | 384557 | protein_coding | 1.26515423 | 0.176011399 | 2.739116354 | -0.06425846 | 0.948764429 | 1 | 0.176011399 |
| ENSMUSG00000101874 | Or2y1e | 258460 | protein_coding | 1.217593618 | 0.176011683 | 2.738801253 | -0.064265957 | 0.94875846 | 1 | 0.176011683 |
| ENSMUSG00000063881 | Or1e1c | 258924 | protein_coding | 1.122472394 | 0.17601179 | 2.76309771 | -0.063700892 | 0.949208394 | 1 | 0.17601179 |
| ENSMUSG00000059361 | Nrsn2 | 228777 | protein_coding | 1.126599854 | 0.176012105 | 2.749163318 | -0.064023881 | 0.948951212 | 1 | 0.176012105 |
| ENSMUSG00000022112 | Gpc5 | 103978 | protein_coding | 1.079039242 | 0.176012346 | 2.753149854 | -0.063931263 | 0.949024959 | 1 | 0.176012346 |
| ENSMUSG00000041907 | Gpr45 | 93690 | protein_coding | 1.079039242 | 0.176012346 | 2.753149854 | -0.063931263 | 0.949024959 | 1 | 0.176012346 |
| ENSMUSG00000095941 | Tle7 | 102638837 | protein_coding | 1.079039242 | 0.176012346 | 2.753149854 | -0.063931263 | 0.949024959 | 1 | 0.176012346 |
| ENSMUSG00000097547 | B230110C06Rik | 77981 | lncRNA | 1.589699978 | 0.176016293 | 2.415464129 | -0.072870589 | 0.941909098 | 1 | 0.176016293 |
| ENSMUSG00000118259 | Gm5095 | 328953 | lncRNA | 1.276122551 | 0.176017408 | 2.415767062 | -0.072861912 | 0.941916003 | 1 | 0.176017408 |
| ENSMUSG00000085820 | 4933406K04Rik | 71033 | lncRNA | 1.672438822 | 0.176017572 | 2.329068894 | -0.075574223 | 0.939757845 | 1 | 0.176017572 |
| ENSMUSG00000066189 | Cacng3 | 54376 | protein_coding | 1.451811383 | 0.176017891 | 2.339733829 | -0.075229878 | 0.940031812 | 1 | 0.176017891 |
| ENSMUSG00000004296 | Il12b | 16160 | protein_coding | 1.041341862 | 0.176018703 | 2.385074045 | -0.0738001 | 0.941169447 | 1 | 0.176018703 |
| ENSMUSG00000109528 | Or4f15 | 258439 | protein_coding | 1.256775693 | 0.176018739 | 2.349692258 | -0.074911401 | 0.940285205 | 1 | 0.176018739 |
| ENSMUSG00000015365 | Mov10l1 | 83456 | protein_coding | 1.449847305 | 0.176018776 | 2.298166862 | -0.076590947 | 0.938948961 | 1 | 0.176018776 |
| ENSMUSG00000086277 | 4930558K02Rik | 75368 | protein_coding | 1.038952644 | 0.176019069 | 2.358529837 | -0.074630843 | 0.940508434 | 1 | 0.176019069 |
| ENSMUSG00000004894 | Hapln2 | 73940 | protein_coding | 1.084310157 | 0.176019157 | 2.367716487 | -0.074341315 | 0.940738803 | 1 | 0.176019157 |
| ENSMUSG00000047976 | Kcna1 | 16485 | protein_coding | 1.169909389 | 0.176019553 | 2.330297273 | -0.075535235 | 0.939788864 | 1 | 0.176019553 |

Supplementary Table 1

| ensembl_gene_id | external_gene_name | entrezgene_id | gene_biotype | baseMean | log2FoldChange | lfcSE | stat | pvalue | padj | absLog2FC |
|---|---|---|---|---|---|---|---|---|---|---|
| ENSMUSG00000053268 | Dspp | 666279 | protein_coding | 1.169909389 | 0.176019553 | 2.330297273 | -0.075535235 | 0.939788864 | 1 | 0.176019553 |
| ENSMUSG00000100798 | Gm19589 | 100503197 | lncRNA | 1.112236534 | 0.176019694 | 2.340384841 | -0.075209722 | 0.940047849 | 1 | 0.176019694 |
| ENSMUSG00000110874 | D030045P18Rik | 320363 | lncRNA | 1.395082835 | 0.176019727 | 2.262947058 | -0.077783405 | 0.938000348 | 1 | 0.176019727 |
| ENSMUSG00000039323 | Igfbp2 | 16008 | protein_coding | 1.098676867 | 0.176019762 | 2.319360953 | -0.075891491 | 0.939505426 | 1 | 0.176019762 |
| ENSMUSG00000004344 | Gpx5 | 14780 | protein_coding | 1.031355012 | 0.176020317 | 2.327788968 | -0.075616956 | 0.939723846 | 1 | 0.176020317 |
| ENSMUSG00000052417 | Or2t6 | 258387 | protein_coding | 1.031355012 | 0.176020317 | 2.327788968 | -0.075616956 | 0.939723846 | 1 | 0.176020317 |
| ENSMUSG00000036218 | Pdzrn4 | 239618 | protein_coding | 0.9837944 | 0.176020424 | 2.336098839 | -0.075348021 | 0.939937815 | 1 | 0.176020424 |
| ENSMUSG00000066772 | Obox3 | 246791 | protein_coding | 0.940191321 | 0.176021222 | 2.306916041 | -0.076301529 | 0.939179209 | 1 | 0.176021222 |
| ENSMUSG00000062314 | Or9i1b | 258151 | protein_coding | 1.410036582 | 0.17602362 | 2.019225106 | -0.087173847 | 0.930533327 | 1 | 0.17602362 |
| ENSMUSG00000050343 | Or1ad6 | 258912 | protein_coding | 1.115065698 | 0.176024266 | 2.038717679 | -0.086340678 | 0.931195603 | 1 | 0.176024266 |
| ENSMUSG00000070460 | Or5ae2 | 258410 | protein_coding | 1.080687237 | 0.176024513 | 2.031040429 | -0.086667163 | 0.930936078 | 1 | 0.176024513 |
| ENSMUSG00000035861 | Tmprss11b | 319875 | protein_coding | 1.126553544 | 0.176024601 | 2.023587208 | -0.086986417 | 0.930682309 | 1 | 0.176024601 |
| ENSMUSG00000069441 | Dsg1a | 13510 | protein_coding | 1.930068697 | 0.176025634 | 1.816178701 | -0.096920878 | 0.922789229 | 1 | 0.176025634 |
| ENSMUSG00000012396 | Nanog | 71950 | protein_coding | 0.93524569 | 0.176026289 | 1.959774512 | -0.089819664 | 0.928430521 | 1 | 0.176026289 |
| ENSMUSG00000031872 | Bean1 | 65115 | protein_coding | 0.74251537 | 0.176026823 | 1.949037062 | -0.090314764 | 0.928037088 | 1 | 0.176026823 |
| ENSMUSG00000092734 | Mir5100 | 100628625 | miRNA | 0.706042491 | 0.176029121 | 1.75855527 | -0.100098714 | 0.920265956 | 1 | 0.176029121 |
| ENSMUSG00000040151 | Hs2st1 | 23908 | protein_coding | 251.7515126 | 0.176068944 | 0.08322577 | -2.115558004 | 0.034382417 | 0.11194538 | 0.176068944 |
| ENSMUSG00000011034 | Slc5a1 | 20537 | protein_coding | 2802.657482 | 0.176083509 | 0.066054419 | -2.665733985 | 0.007682045 | 0.034445098 | 0.176083509 |
| ENSMUSG00000041958 | Pigs | 276846 | protein_coding | 577.6311717 | -0.176091603 | 0.081694357 | 2.155492851 | 0.031123287 | 0.103865546 | 0.176091603 |
| ENSMUSG00000052534 | Pbx1 | 18514 | protein_coding | 504.7017705 | -0.176102333 | 0.093205018 | 1.889408283 | 0.058837142 | 0.168423741 | 0.176102333 |
| ENSMUSG00000056055 | Sag | 20215 | protein_coding | 4.432524784 | -0.176151823 | 0.784185124 | 0.224630406 | 0.822266808 | 0.910171686 | 0.176151823 |
| ENSMUSG00000015542 | Nat9 | 66176 | protein_coding | 215.3127448 | -0.176209193 | 0.076176809 | 2.313160585 | 0.020713812 | 0.07534535 | 0.176209193 |
| ENSMUSG00000056888 | Glipr1 | 73690 | protein_coding | 8.691910961 | 0.176218212 | 0.487029086 | -0.361822768 | 0.717484477 | 0.853563561 | 0.176218212 |
| ENSMUSG00000109737 | Spem3 | 105243794 | protein_coding | 2.179110047 | -0.176233663 | 1.939169271 | 0.090881011 | 0.927587139 | 1 | 0.176233663 |
| ENSMUSG00000026627 | Pacc1 | 66950 | protein_coding | 63.86419866 | -0.176243354 | 0.136653615 | 1.289708685 | 0.197151823 | 0.397106786 | 0.176243354 |
| ENSMUSG00000026960 | Arl6ip6 | 65103 | protein_coding | 187.6321521 | 0.176281864 | 0.102328712 | -1.722701879 | 0.08494245 | 0.219589797 | 0.176281864 |
| ENSMUSG00000113634 | Gm36756 | 102640764 | lncRNA | 7.91542274 | 0.176289553 | 0.418946771 | -0.420792248 | 0.673906793 | 0.825096143 | 0.176289553 |
| ENSMUSG00000041566 | Tssk1 | 22114 | protein_coding | 5.999294809 | -0.176367481 | 0.750284268 | 0.235067546 | 0.814156308 | 0.907710402 | 0.176367481 |
| ENSMUSG00000029999 | Tgfa | 21802 | protein_coding | 485.9824924 | -0.176390707 | 0.078979372 | 2.23337693 | 0.025524098 | 0.088763622 | 0.176390707 |
| ENSMUSG00000025609 | Mkln1 | 27418 | protein_coding | 909.5875918 | -0.176537717 | 0.048250194 | 3.65879809 | 0.000253401 | 0.001953624 | 0.176537717 |
| ENSMUSG00000040250 | Ints13 | 71177 | protein_coding | 338.1239509 | 0.176621724 | 0.071626576 | -2.465868593 | 0.013668147 | 0.0542586 | 0.176621724 |
| ENSMUSG00000020890 | Gucy2e | 14919 | protein_coding | 2.712523321 | -0.176643049 | 1.925958506 | 0.091716955 | 0.926922926 | 1 | 0.176643049 |
| ENSMUSG00000040054 | Baz2a | 116848 | protein_coding | 1175.888387 | -0.176648288 | 0.101658593 | 1.737662132 | 0.082270365 | 0.214258446 | 0.176648288 |
| ENSMUSG00000023979 | Guca1b | 107477 | protein_coding | 3.974569906 | -0.176707197 | 0.62150492 | 0.284321477 | 0.776164029 | 1 | 0.176707197 |
| ENSMUSG00000022721 | Trmt2a | 15547 | protein_coding | 334.3868072 | 0.176715969 | 0.082390759 | -2.144851828 | 0.031964696 | 0.106039571 | 0.176715969 |
| ENSMUSG00000059820 | Nkapd1 | 270156 | protein_coding | 187.6684085 | -0.176817636 | 0.143393597 | 1.233092969 | 0.21754108 | 0.423388606 | 0.176817636 |
| ENSMUSG00000007908 | Hmgcll1 | 208982 | protein_coding | 5.474922205 | 0.176855391 | 0.496531792 | -0.356181404 | 0.721704718 | 0.855825916 | 0.176855391 |
| ENSMUSG00000115987 | Vps28 | 66914 | protein_coding | 1209.643518 | -0.176871092 | 0.06002995 | 2.946380786 | 0.003215163 | 0.016858279 | 0.176871092 |
| ENSMUSG00000032908 | Sgpp2 | 433323 | protein_coding | 1654.866879 | 0.176931631 | 0.057901389 | -3.055740716 | 0.002245053 | 0.012437037 | 0.176931631 |

| ensembl_gene_id | external_gene_name | entrezgene_id | gene_biotype | baseMean | log2FoldChange | lfcSE | stat | pvalue | padj | absLog2FC |
|---|---|---|---|---|---|---|---|---|---|---|
| ENSMUSG00000026259 | Ngef | 53972 | protein_coding | 581.7892318 | -0.176961836 | 0.085376581 | 2.072721056 | 0.038198247 | 0.12161285 | 0.176961836 |
| ENSMUSG00000031604 | Msmo1 | 66234 | protein_coding | 1324.568999 | -0.176986244 | 0.170933922 | 1.035407375 | 0.300478703 | 0.520523758 | 0.176986244 |
| ENSMUSG00000025857 | Dnaaf5 | 433956 | protein_coding | 149.7114021 | 0.177050252 | 0.118684135 | -1.491776916 | 0.135757637 | 0.307109811 | 0.177050252 |
| ENSMUSG00000020733 | Slc9a3r1 | 26941 | protein_coding | 2277.753961 | 0.177061485 | 0.163569296 | -1.082486076 | 0.279036599 | 0.496338905 | 0.177061485 |
| ENSMUSG00000027963 | Extl2 | 58193 | protein_coding | 107.8183022 | 0.177165959 | 0.124944052 | -1.417962332 | 0.156201763 | 0.338919446 | 0.177165959 |
| ENSMUSG00000039782 | Cpeb2 | 231207 | protein_coding | 254.2731588 | 0.17720516 | 0.11259324 | -1.573852568 | 0.115521538 | 0.274255704 | 0.17720516 |
| ENSMUSG00000055720 | Ubl7 | 69459 | protein_coding | 242.1836611 | -0.177261468 | 0.096159755 | 1.843405988 | 0.065269756 | 0.18142817 | 0.177261468 |
| ENSMUSG00000034867 | Ankrd27 | 245886 | protein_coding | 429.6197494 | -0.177370369 | 0.074619467 | 2.376998597 | 0.017454154 | 0.065958331 | 0.177370369 |
| ENSMUSG00000037993 | Dhx38 | 64340 | protein_coding | 647.649623 | 0.177393589 | 0.065749695 | -2.698013876 | 0.006975453 | 0.031812382 | 0.177393589 |
| ENSMUSG00000066892 | Fbxl12 | 30843 | protein_coding | 77.113051 | 0.177416184 | 0.132289366 | -1.341122044 | 0.179880831 | 0.37341949 | 0.177416184 |
| ENSMUSG00000002393 | Nr2f6 | 13864 | protein_coding | 770.7801627 | 0.177418388 | 0.101495268 | -1.748045916 | 0.080456077 | 0.210863618 | 0.177418388 |
| ENSMUSG00000028466 | Creb3 | 12913 | protein_coding | 593.3087554 | 0.177456754 | 0.069704091 | -2.54585852 | 0.010900936 | 0.045517455 | 0.177456754 |
| ENSMUSG00000040648 | Ppip5k2 | 227399 | protein_coding | 1272.718426 | 0.177458431 | 0.082622943 | -2.147810586 | 0.031728806 | 0.10540525 | 0.177458431 |
| ENSMUSG00000030533 | Unc45a | 101869 | protein_coding | 612.1239803 | -0.177466907 | 0.081650738 | 2.173488095 | 0.029743601 | 0.100242005 | 0.177466907 |
| ENSMUSG00000044030 | Irf2bp1 | 272359 | protein_coding | 524.817632 | 0.177498107 | 0.078866795 | -2.250606306 | 0.024410484 | 0.085787344 | 0.177498107 |
| ENSMUSG00000025277 | Abhd6 | 66082 | protein_coding | 530.0203297 | 0.177519614 | 0.096352151 | -1.842404267 | 0.065416038 | 0.181743011 | 0.177519614 |
| ENSMUSG00000037190 | Cyb561d2 | 56368 | protein_coding | 110.509105 | -0.177587151 | 0.110119792 | 1.612672422 | 0.106815695 | 0.259672367 | 0.177587151 |
| ENSMUSG00000020827 | Mink1 | 50932 | protein_coding | 812.6688476 | -0.177741446 | 0.103614665 | 1.715408198 | 0.086270413 | 0.222084117 | 0.177741446 |
| ENSMUSG00000068284 | Usf3 | 207806 | protein_coding | 2385.964938 | 0.177766438 | 0.094923319 | -1.872737272 | 0.061104674 | 0.173114185 | 0.177766438 |
| ENSMUSG00000067629 | Syngap1 | 240057 | protein_coding | 14.18256161 | 0.177793202 | 0.545599469 | -0.325867623 | 0.744524513 | 0.871118268 | 0.177793202 |
| ENSMUSG00000069601 | Ank3 | 11735 | protein_coding | 1334.653517 | -0.177828747 | 0.165945113 | 1.071611834 | 0.283894416 | 0.502167136 | 0.177828747 |
| ENSMUSG00000041890 | Git2 | 26431 | protein_coding | 240.6842119 | -0.177893981 | 0.118749899 | 1.498055847 | 0.134118741 | 0.304528653 | 0.177893981 |
| ENSMUSG00000030314 | Atg7 | 74244 | protein_coding | 385.8506075 | -0.177924221 | 0.090561059 | 1.964687941 | 0.049450366 | 0.147805132 | 0.177924221 |
| ENSMUSG00000052566 | Hook2 | 170833 | protein_coding | 1074.607049 | -0.178026227 | 0.067377801 | 2.642208917 | 0.008236723 | 0.036437516 | 0.178026227 |
| ENSMUSG00000034189 | Hsdl1 | 72552 | protein_coding | 321.0364023 | -0.178037972 | 0.110214533 | 1.615376551 | 0.106229175 | 0.258635202 | 0.178037972 |
| ENSMUSG00000026698 | Pigc | 67292 | protein_coding | 270.8289083 | 0.178061439 | 0.10417096 | -1.709319368 | 0.087391808 | 0.224271688 | 0.178061439 |
| ENSMUSG00000036046 | 5031439G07Rik | 223739 | protein_coding | 563.7955456 | -0.178071831 | 0.058061658 | 3.066943622 | 0.002162596 | 0.01206932 | 0.178071831 |
| ENSMUSG00000006019 | Dhx34 | 71723 | protein_coding | 191.0802932 | -0.17808645 | 0.098474559 | 1.808451347 | 0.070536283 | 0.192028673 | 0.17808645 |
| ENSMUSG00000003378 | Grik5 | 14809 | protein_coding | 42.42152807 | -0.178097172 | 0.256820157 | 0.69347038 | 0.488014408 | 0.696186289 | 0.178097172 |
| ENSMUSG00000057649 | Brd9 | 105246 | protein_coding | 442.6543068 | 0.178158296 | 0.069785679 | -2.552934899 | 0.010681946 | 0.044908772 | 0.178158296 |
| ENSMUSG00000020100 | Slc29a3 | 71279 | protein_coding | 96.12917847 | 0.178240058 | 0.118738325 | -1.501116494 | 0.133325433 | 0.303102451 | 0.178240058 |
| ENSMUSG00000030854 | Ptpn5 | 19259 | protein_coding | 7.035586618 | -0.178256312 | 0.947574355 | 0.188118549 | 0.850783721 | 0.927198731 | 0.178256312 |
| ENSMUSG00000027784 | Ppm1l | 242083 | protein_coding | 169.6123724 | -0.178304405 | 0.11519384 | 1.547864063 | 0.121655029 | 0.284460467 | 0.178304405 |
| ENSMUSG00000057388 | Mrpl18 | 67681 | protein_coding | 503.8007814 | 0.178445449 | 0.079041874 | -2.257606494 | 0.023970203 | 0.084509808 | 0.178445449 |
| ENSMUSG00000039450 | Dcxr | 67880 | protein_coding | 646.5563522 | -0.178453314 | 0.113540794 | 1.571710986 | 0.116017593 | 0.274998498 | 0.178453314 |
| ENSMUSG00000020182 | Ddc | 13195 | protein_coding | 823.9024085 | -0.178487246 | 0.128082374 | 1.393534798 | 0.16345813 | 0.349520595 | 0.178487246 |
| ENSMUSG00000021375 | Kif13a | 16553 | protein_coding | 1239.219767 | -0.178578875 | 0.101432623 | 1.760566473 | 0.078311807 | 0.20691859 | 0.178578875 |
| ENSMUSG00000040714 | Klc3 | 232943 | protein_coding | 13.50388459 | 0.178600657 | 0.281557843 | -0.634330249 | 0.52586532 | 0.724255209 | 0.178600657 |
| ENSMUSG00000020770 | Unk | 217331 | protein_coding | 222.3298838 | 0.178602962 | 0.098175517 | -1.819220993 | 0.068877721 | 0.188725185 | 0.178602962 |

Supplementary Table 1

| ensembl_gene_id | external_gene_name | entrezgene_id | gene_biotype | baseMean | log2FoldChange | lfcSE | stat | pvalue | padj | absLog2FC |
|---|---|---|---|---|---|---|---|---|---|---|
| ENSMUSG00000085568 | Gm14858 | 434825 | lncRNA | 1.276331499 | -0.178627024 | 1.296808544 | 0.137743559 | 0.890443092 | 1 | 0.178627024 |
| ENSMUSG00000053094 | Tmem248 | 71667 | protein_coding | 652.640138 | 0.178727561 | 0.09310019 | -1.919733589 | 0.054891559 | 0.15977566 | 0.178727561 |
| ENSMUSG00000025163 | Cd7 | 12516 | protein_coding | 49.3415542 | 0.178760942 | 0.320312873 | -0.558082289 | 0.576788194 | 0.761222618 | 0.178760942 |
| ENSMUSG00000046697 | Enpp7 | 238011 | protein_coding | 13.01578833 | 0.1788072 | 1.175221264 | -0.15214769 | 0.879070451 | 0.940340033 | 0.1788072 |
| ENSMUSG00000031453 | Rasa3 | 19414 | protein_coding | 402.5957482 | 0.178837624 | 0.128035113 | -1.396785771 | 0.162478002 | 0.347924739 | 0.178837624 |
| ENSMUSG00000036478 | Btg1 | 12226 | protein_coding | 2195.753303 | -0.178855532 | 0.111236736 | 1.60788187 | 0.107861053 | 0.261367195 | 0.178855532 |
| ENSMUSG00000043964 | Orai3 | 269999 | protein_coding | 166.6032916 | -0.178856878 | 0.103534643 | 1.727507553 | 0.08407655 | 0.217760374 | 0.178856878 |
| ENSMUSG00000023026 | Dip2b | 239667 | protein_coding | 618.9014202 | -0.178912128 | 0.089961713 | 1.988758575 | 0.046727857 | 0.141589694 | 0.178912128 |
| ENSMUSG00000026923 | Notch1 | 18128 | protein_coding | 497.4295688 | 0.178926165 | 0.101032518 | -1.770976005 | 0.076564692 | 0.203670961 | 0.178926165 |
| ENSMUSG00000029439 | Sfswap | 231769 | protein_coding | 559.6159748 | 0.179010479 | 0.115509838 | -1.549742277 | 0.121203387 | 0.283960728 | 0.179010479 |
| ENSMUSG00000020817 | Rabep1 | 54189 | protein_coding | 417.9536927 | -0.179059982 | 0.100821353 | 1.776012498 | 0.075730857 | 0.201777892 | 0.179059982 |
| ENSMUSG00000022664 | Slc35a5 | 74102 | protein_coding | 273.0526953 | 0.179075475 | 0.091261387 | -1.962226105 | 0.049736166 | 0.148337194 | 0.179075475 |
| ENSMUSG00000000088 | Cox5a | 12858 | protein_coding | 2338.513058 | -0.179095835 | 0.074500742 | 2.403946983 | 0.016219125 | 0.06221406 | 0.179095835 |
| ENSMUSG00000011263 | Exoc3l2 | 74463 | protein_coding | 3.34495287 | 0.179111933 | 0.782680426 | -0.228844272 | 0.818989955 | 1 | 0.179111933 |
| ENSMUSG00000073565 | Prr16 | 71373 | protein_coding | 6.019642926 | 0.179212321 | 0.496134619 | -0.361217125 | 0.717937141 | 0.853653903 | 0.179212321 |
| ENSMUSG00000038160 | Atg5 | 11793 | protein_coding | 508.4919772 | 0.179233654 | 0.078208081 | -2.291753627 | 0.021919864 | 0.07888094 | 0.179233654 |
| ENSMUSG00000020069 | Hnrnph3 | 432467 | protein_coding | 167.0932923 | -0.179254643 | 0.110462775 | 1.622760643 | 0.104640582 | 0.255796319 | 0.179254643 |
| ENSMUSG00000089832 | Shkbp1 | 192192 | protein_coding | 288.267888 | 0.179256691 | 0.09299245 | -1.927647804 | 0.053898949 | 0.157816886 | 0.179256691 |
| ENSMUSG00000030772 | Dkk3 | 50781 | protein_coding | 1.169304703 | 0.179285966 | 1.3915633 | -0.128837809 | 0.897485988 | 1 | 0.179285966 |
| ENSMUSG00000028407 | Smim27 | 100504309 | protein_coding | 52.88781394 | -0.17930853 | 0.139784654 | 1.282748325 | 0.199580263 | 0.400664779 | 0.17930853 |
| ENSMUSG00000029718 | Pcolce | 18542 | protein_coding | 43.02328535 | 0.179319592 | 0.242771058 | -0.738636615 | 0.460127684 | 0.674460666 | 0.179319592 |
| ENSMUSG00000023923 | Tbc1d5 | 72238 | protein_coding | 265.4259748 | -0.179325013 | 0.079112366 | 2.266712803 | 0.023407776 | 0.083023198 | 0.179325013 |
| ENSMUSG00000071649 | B3gat3 | 72727 | protein_coding | 354.4497509 | -0.179335563 | 0.09890002 | 1.813301575 | 0.06978532 | 0.190611112 | 0.179335563 |
| ENSMUSG00000027835 | Pdcd10 | 56426 | protein_coding | 439.9407687 | -0.179367932 | 0.101662218 | 1.764351946 | 0.077672743 | 0.205789777 | 0.179367932 |
| ENSMUSG00000074781 | Ube2n | 93765 | protein_coding | 466.1934498 | -0.179387276 | 0.109822013 | 1.633436417 | 0.102377243 | 0.251954286 | 0.179387276 |
| ENSMUSG00000020154 | Ptprb | 19263 | protein_coding | 7.561740164 | -0.17940021 | 0.801374404 | 0.22386566 | 0.822861837 | 0.910240646 | 0.17940021 |
| ENSMUSG00000036932 | Aifm1 | 26926 | protein_coding | 871.0550483 | -0.179428751 | 0.093365406 | 1.921790506 | 0.054632123 | 0.159370233 | 0.179428751 |
| ENSMUSG00000029370 | Rassf6 | 73246 | protein_coding | 691.7237114 | -0.179450175 | 0.12106309 | 1.482286431 | 0.138264097 | 0.310948291 | 0.179450175 |
| ENSMUSG00000027774 | Gfm1 | 28030 | protein_coding | 719.6378551 | -0.179458292 | 0.08845762 | 2.028748827 | 0.04248388 | 0.13170242 | 0.179458292 |
| ENSMUSG00000038119 | Cdon | 57810 | protein_coding | 297.7249513 | 0.17951289 | 0.098631654 | -1.820033257 | 0.068753941 | 0.188479843 | 0.17951289 |
| ENSMUSG00000032641 | Gpr19 | 14760 | protein_coding | 45.98129182 | 0.179520976 | 0.208152438 | -0.862449546 | 0.388440186 | 0.610923926 | 0.179520976 |
| ENSMUSG00000030592 | Ryr1 | 20190 | protein_coding | 16.45818891 | 0.179545151 | 1.632586463 | -0.109975891 | 0.912428495 | 0.957266079 | 0.179545151 |
| ENSMUSG00000028015 | Ctso | 229445 | protein_coding | 253.4276358 | -0.179568632 | 0.099585596 | 1.803158678 | 0.071363298 | 0.193640076 | 0.179568632 |
| ENSMUSG00000046449 | Nexmif | 245555 | protein_coding | 32.63272915 | -0.179604336 | 0.309553663 | 0.580204201 | 0.561776921 | 0.750095737 | 0.179604336 |
| ENSMUSG00000056445 | Hoxaas2 | 105242657 | lncRNA | 13.63957841 | -0.179620915 | 0.321249853 | 0.55913151 | 0.576071972 | 0.761121093 | 0.179620915 |
| ENSMUSG00000070047 | Fat1 | 14107 | protein_coding | 5200.872292 | -0.179665939 | 0.129013605 | 1.392612343 | 0.163737048 | 0.349981053 | 0.179665939 |
| ENSMUSG00000028463 | Car9 | 230099 | protein_coding | 581.7876409 | -0.179674971 | 0.066507664 | 2.701567931 | 0.006901338 | 0.031544062 | 0.179674971 |
| ENSMUSG00000006464 | Bbs1 | 52028 | protein_coding | 37.38226508 | -0.179697031 | 0.185593987 | 0.968226575 | 0.332931225 | 0.555300096 | 0.179697031 |
| ENSMUSG00000059921 | Unc5c | 22253 | protein_coding | 4.138213019 | 0.179822962 | 2.416937868 | -0.074401152 | 0.940691192 | 1 | 0.179822962 |

Supplementary Table 1

| ensembl_gene_id | external_gene_name | entrezgene_id | gene_biotype | baseMean | log2FoldChange | lfcSE | stat | pvalue | padj | absLog2FC |
|---|---|---|---|---|---|---|---|---|---|---|
| ENSMUSG00000024246 | Thumpd2 | 72167 | protein_coding | 33.75852895 | -0.179831394 | 0.213678278 | 0.841598855 | 0.40001253 | 0.622191478 | 0.179831394 |
| ENSMUSG00000023980 | Taf8 | 63856 | protein_coding | 236.5783648 | 0.179860769 | 0.089983744 | -1.998814009 | 0.045628481 | 0.13918813 | 0.179860769 |
| ENSMUSG00000026174 | Cnot9 | 58184 | protein_coding | 442.4016042 | 0.179881314 | 0.091681479 | -1.962024566 | 0.049759624 | 0.14838036 | 0.179881314 |
| ENSMUSG00000049516 | Spty2d1 | 101685 | protein_coding | 605.8779915 | 0.179886373 | 0.077761923 | -2.313296363 | 0.020706351 | 0.075334803 | 0.179886373 |
| ENSMUSG00000002326 | Gmpr2 | 105446 | protein_coding | 46.96642887 | -0.179953792 | 0.185009213 | 0.972674761 | 0.330714973 | 0.553055905 | 0.179953792 |
| ENSMUSG00000084507 | Gm25321 | 115487006 | snRNA | 1.0140295 | -0.179979246 | 1.22276789 | 0.147190033 | 0.882982027 | 1 | 0.179979246 |
| ENSMUSG00000001672 | Marveld3 | 73608 | protein_coding | 707.9073144 | -0.180056657 | 0.082976245 | 2.169978371 | 0.030008485 | 0.100990445 | 0.180056657 |
| ENSMUSG00000008373 | Prpf31 | 68988 | protein_coding | 382.5358855 | 0.180060287 | 0.085478213 | -2.106505061 | 0.035160504 | 0.113873419 | 0.180060287 |
| ENSMUSG00000039182 | AW209491 | 105351 | protein_coding | 123.9720337 | -0.180095485 | 0.105592142 | 1.705576586 | 0.088086942 | 0.225215631 | 0.180095485 |
| ENSMUSG00000044734 | Serpinb1a | 66222 | protein_coding | 4021.194329 | -0.180123596 | 0.115349254 | 1.56154972 | 0.118394102 | 0.279268705 | 0.180123596 |
| ENSMUSG00000049755 | Zfp672 | 319475 | protein_coding | 447.5346529 | -0.180139095 | 0.078105021 | 2.306370242 | 0.021089953 | 0.076528123 | 0.180139095 |
| ENSMUSG00000031684 | Slc10a7 | 76775 | protein_coding | 396.4327941 | 0.180181697 | 0.082703095 | -2.178657246 | 0.02935714 | 0.099121614 | 0.180181697 |
| ENSMUSG00000022708 | Zbtb20 | 56490 | protein_coding | 246.5632567 | -0.180237396 | 0.156195123 | 1.153924609 | 0.248531079 | 0.460736668 | 0.180237396 |
| ENSMUSG00000021838 | Samd4 | 74480 | protein_coding | 8.802200767 | 0.180267765 | 0.585204844 | -0.308042161 | 0.758050251 | 0.878750865 | 0.180267765 |
| ENSMUSG00000031834 | Pik3r2 | 18709 | protein_coding | 683.1170899 | 0.180341867 | 0.084649701 | -2.130448953 | 0.033134566 | 0.10878414 | 0.180341867 |
| ENSMUSG00000020884 | Asgr1 | 11889 | protein_coding | 2.445511912 | -0.180346818 | 1.055577157 | 0.170851384 | 0.864340625 | 1 | 0.180346818 |
| ENSMUSG00000060904 | Arl1 | 104303 | protein_coding | 2655.052022 | 0.180371935 | 0.09547337 | -1.88923817 | 0.058859922 | 0.16845975 | 0.180371935 |
| ENSMUSG00000024231 | Cul2 | 71745 | protein_coding | 442.6391054 | 0.180377797 | 0.093210033 | -1.935175767 | 0.05296873 | 0.155922569 | 0.180377797 |
| ENSMUSG00000062376 | Borcs7 | 66439 | protein_coding | 64.91878825 | -0.180392135 | 0.168044669 | 1.073477284 | 0.283057026 | 0.501169049 | 0.180392135 |
| ENSMUSG00000003752 | Itpkc | 233011 | protein_coding | 396.2151726 | -0.180418862 | 0.160560022 | 1.123684835 | 0.261146752 | 0.474848873 | 0.180418862 |
| ENSMUSG00000030423 | Pop4 | 66161 | protein_coding | 169.8511522 | 0.18044093 | 0.113672976 | -1.587368749 | 0.112429153 | 0.268968865 | 0.18044093 |
| ENSMUSG00000036377 | Cracd | 320827 | protein_coding | 919.1218993 | -0.180449318 | 0.113195888 | 1.594133158 | 0.110906214 | 0.266439513 | 0.180449318 |
| ENSMUSG00000088254 | Gm24289 | 115487307 | scaRNA | 3.429383626 | 0.180451529 | 0.723449655 | -0.249432047 | 0.803026598 | 1 | 0.180451529 |
| ENSMUSG00000021492 | F12 | 58992 | protein_coding | 1.842344972 | 0.180456164 | 1.250173021 | -0.144344951 | 0.885228084 | 1 | 0.180456164 |
| ENSMUSG00000034424 | Gcsh | 68133 | protein_coding | 459.7913758 | 0.180474439 | 0.076134438 | -2.370470496 | 0.017765462 | 0.066769512 | 0.180474439 |
| ENSMUSG00000054920 | Klhl5 | 71778 | protein_coding | 238.9642914 | 0.180503304 | 0.146745443 | -1.230043675 | 0.21868075 | 0.424858107 | 0.180503304 |
| ENSMUSG00000057751 | Megf6 | 230971 | protein_coding | 10.81094352 | -0.180511378 | 0.598593874 | 0.301559013 | 0.762988257 | 0.881056435 | 0.180511378 |
| ENSMUSG00000020903 | Stx8 | 55943 | protein_coding | 269.723962 | -0.18055801 | 0.085137545 | 2.12078009 | 0.033940313 | 0.11076884 | 0.18055801 |
| ENSMUSG00000034639 | Setmar | 74729 | protein_coding | 41.61054378 | -0.180572251 | 0.211993507 | 0.851781988 | 0.394335105 | 0.616875898 | 0.180572251 |
| ENSMUSG00000045973 | Slc25a51 | 230125 | protein_coding | 906.7155624 | -0.180649557 | 0.064610313 | 2.795986419 | 0.005174157 | 0.025072194 | 0.180649557 |
| ENSMUSG00000047454 | Gphn | 268566 | protein_coding | 263.1613846 | -0.180659188 | 0.108056756 | 1.671891645 | 0.094545691 | 0.236959711 | 0.180659188 |
| ENSMUSG00000025016 | Tm9sf3 | 107358 | protein_coding | 6794.225974 | 0.180676347 | 0.134044552 | -1.347882803 | 0.17769608 | 0.370029503 | 0.180676347 |
| ENSMUSG00000026843 | Fubp3 | 320267 | protein_coding | 526.5798514 | 0.180704805 | 0.061674122 | -2.929993971 | 0.003389686 | 0.017619538 | 0.180704805 |
| ENSMUSG00000019929 | Dcn | 13179 | protein_coding | 4.849207902 | -0.180725958 | 0.716848431 | 0.252111813 | 0.800954639 | 0.901449255 | 0.180725958 |
| ENSMUSG00000052748 | Swt1 | 66875 | protein_coding | 252.7922563 | -0.180742139 | 0.080942481 | 2.232970081 | 0.025550917 | 0.08879374 | 0.180742139 |
| ENSMUSG00000029469 | Ift81 | 12589 | protein_coding | 48.80113871 | -0.180814868 | 0.173444259 | 1.042495553 | 0.297181986 | 0.516704919 | 0.180814868 |
| ENSMUSG00000005973 | Rcn1 | 19672 | protein_coding | 81.38356448 | 0.180977815 | 0.146364824 | -1.23648436 | 0.216278576 | 0.421530084 | 0.180977815 |
| ENSMUSG00000041840 | Haus1 | 225745 | protein_coding | 76.01081472 | 0.18108326 | 0.145118714 | -1.247828449 | 0.212093888 | 0.41642117 | 0.18108326 |
| ENSMUSG00000024617 | Camk2a | 12322 | protein_coding | 8.64747986 | -0.181152585 | 0.566986888 | 0.319500483 | 0.749347025 | 0.874044129 | 0.181152585 |

| ensembl_gene_id | external_gene_name | entrezgene_id | gene_biotype | baseMean | log2FoldChange | lfcSE | stat | pvalue | padj | absLog2FC |
|---|---|---|---|---|---|---|---|---|---|---|
| ENSMUSG00000026322 | Htr4 | 15562 | protein_coding | 310.7280628 | -0.181221918 | 0.191822399 | 0.944738046 | 0.344792628 | 0.566362983 | 0.181221918 |
| ENSMUSG00000076333 | Mir599 | 100316736 | miRNA | 0.532854639 | 0.181223808 | 2.079475567 | -0.087148804 | 0.930553233 | 1 | 0.181223808 |
| ENSMUSG00000021265 | Slc25a29 | 214663 | protein_coding | 56.78914337 | 0.181243293 | 0.171616584 | -1.056094279 | 0.290925144 | 0.509852258 | 0.181243293 |
| ENSMUSG00000021140 | Pcnx | 54604 | protein_coding | 549.1348683 | -0.181272851 | 0.107311735 | 1.689217409 | 0.091177774 | 0.230830869 | 0.181272851 |
| ENSMUSG00000028837 | Psmb2 | 26445 | protein_coding | 940.3981802 | 0.181310364 | 0.070415277 | -2.574872534 | 0.010027703 | 0.042679767 | 0.181310364 |
| ENSMUSG00000018770 | Atp5g3 | 228033 | protein_coding | 4721.173918 | 0.181316147 | 0.083490253 | -2.171704378 | 0.029877968 | 0.100633238 | 0.181316147 |
| ENSMUSG00000034801 | Sos2 | 20663 | protein_coding | 441.716427 | -0.181338418 | 0.082032332 | 2.210572506 | 0.027065454 | 0.092985996 | 0.181338418 |
| ENSMUSG00000002881 | Nab1 | 17936 | protein_coding | 700.4551356 | -0.181343356 | 0.101737473 | 1.782463729 | 0.074673641 | 0.199909758 | 0.181343356 |
| ENSMUSG00000032024 | Clmp | 71566 | protein_coding | 62.2060426 | 0.181351451 | 0.276385218 | -0.656154668 | 0.511724613 | 0.714354347 | 0.181351451 |
| ENSMUSG00000018425 | Dhx40 | 67487 | protein_coding | 405.7652694 | 0.181393691 | 0.095219848 | -1.904998744 | 0.056780234 | 0.164036653 | 0.181393691 |
| ENSMUSG00000014503 | Pkd2l2 | 53871 | protein_coding | 74.42289108 | -0.181425422 | 0.13378887 | 1.356057654 | 0.175080825 | 0.366681641 | 0.181425422 |
| ENSMUSG00000048175 | Asb8 | 78541 | protein_coding | 332.4202544 | -0.181527179 | 0.065170937 | 2.785400807 | 0.005346159 | 0.025753241 | 0.181527179 |
| ENSMUSG00000025860 | Xiap | 11798 | protein_coding | 1491.39162 | -0.181686392 | 0.08520091 | 2.132446608 | 0.032970149 | 0.108526787 | 0.181686392 |
| ENSMUSG00000022419 | Deptor | 97998 | protein_coding | 1478.966416 | -0.181686584 | 0.137789308 | 1.318582598 | 0.187308694 | 0.383501006 | 0.181686584 |
| ENSMUSG00000063317 | Usp31 | 76179 | protein_coding | 194.499862 | 0.18168734 | 0.172119765 | -1.05558673 | 0.291157066 | 0.510042197 | 0.18168734 |
| ENSMUSG00000026102 | Inpp1 | 16329 | protein_coding | 274.099415 | 0.181690787 | 0.094979222 | -1.912952995 | 0.05575407 | 0.161615362 | 0.181690787 |
| ENSMUSG00000030413 | Pglyrp1 | 21946 | protein_coding | 1161.635649 | -0.181753777 | 0.082570381 | 2.201198255 | 0.027721992 | 0.094754639 | 0.181753777 |
| ENSMUSG00000043065 | Spice1 | 212514 | protein_coding | 158.7177065 | -0.181772024 | 0.093926433 | 1.935259523 | 0.052958457 | 0.15592012 | 0.181772024 |
| ENSMUSG00000038344 | Txlng | 353170 | protein_coding | 230.7691194 | -0.181808272 | 0.076492447 | 2.376813394 | 0.01746292 | 0.065976358 | 0.181808272 |
| ENSMUSG00000040003 | Magi2 | 50791 | protein_coding | 7.714947144 | -0.181876693 | 0.638341734 | 0.284920574 | 0.775704993 | 0.88791795 | 0.181876693 |
| ENSMUSG00000028692 | Akr1a1 | 58810 | protein_coding | 5507.489697 | 0.181899603 | 0.062334999 | -2.918097482 | 0.003521742 | 0.018197138 | 0.181899603 |
| ENSMUSG00000040174 | Alkbh3 | 69113 | protein_coding | 164.1506408 | 0.181911439 | 0.092740709 | -1.9615058 | 0.049820049 | 0.14848011 | 0.181911439 |
| ENSMUSG00000024592 | C330018D20Rik | 77422 | protein_coding | 167.2154287 | -0.181911462 | 0.097188945 | 1.871729986 | 0.061243969 | 0.173330403 | 0.181911462 |
| ENSMUSG00000017188 | Coa3 | 52469 | protein_coding | 945.7892459 | -0.181947395 | 0.07399172 | 2.459023729 | 0.013931541 | 0.055052756 | 0.181947395 |
| ENSMUSG00000051669 | AU021092 | 239691 | protein_coding | 1.627256968 | -0.181971562 | 1.629419264 | 0.111678784 | 0.911078102 | 1 | 0.181971562 |
| ENSMUSG00000018481 | Appbp2 | 66884 | protein_coding | 768.7052454 | 0.181977521 | 0.090457685 | -2.011741967 | 0.044247144 | 0.135792109 | 0.181977521 |
| ENSMUSG00000034160 | Ogt | 108155 | protein_coding | 1068.10919 | -0.181980338 | 0.100548201 | 1.809881579 | 0.070314153 | 0.191611869 | 0.181980338 |
| ENSMUSG00000033335 | Dnm2 | 13430 | protein_coding | 2864.847894 | 0.182026993 | 0.096016331 | -1.89579201 | 0.057987553 | 0.166571051 | 0.182026993 |
| ENSMUSG00000027351 | Spred1 | 114715 | protein_coding | 456.7733831 | 0.182070272 | 0.127180411 | -1.431590538 | 0.152261039 | 0.332817843 | 0.182070272 |
| ENSMUSG00000044906 | 4930503L19Rik | 269033 | protein_coding | 3.550454355 | 0.182093066 | 0.657380274 | -0.276998069 | 0.781781589 | 1 | 0.182093066 |
| ENSMUSG00000024099 | Ndufv2 | 72900 | protein_coding | 1733.994081 | -0.182096143 | 0.075129039 | 2.423778422 | 0.015359974 | 0.059503296 | 0.182096143 |
| ENSMUSG00000020074 | Ccar1 | 67500 | protein_coding | 878.7161724 | 0.182135619 | 0.047667168 | -3.820986827 | 0.000132919 | 0.001108596 | 0.182135619 |
| ENSMUSG00000040771 | Oard1 | 106821 | protein_coding | 279.1676283 | 0.182139262 | 0.089789202 | -2.028520787 | 0.042507124 | 0.131725022 | 0.182139262 |
| ENSMUSG00000061887 | Ssbp3 | 72475 | protein_coding | 901.4786314 | 0.182139263 | 0.098230255 | -1.854207366 | 0.063709499 | 0.178321807 | 0.182139263 |
| ENSMUSG00000027243 | Harbi1 | 241547 | protein_coding | 151.0920785 | 0.182170754 | 0.115041124 | -1.583527249 | 0.113301341 | 0.270127107 | 0.182170754 |
| ENSMUSG00000078676 | Casc3 | 192160 | protein_coding | 465.3674467 | -0.18220477 | 0.052446734 | 3.474091853 | 0.000512586 | 0.003624129 | 0.18220477 |
| ENSMUSG00000026317 | Cln8 | 26889 | protein_coding | 207.7092447 | 0.182325815 | 0.080608133 | -2.261878653 | 0.023704902 | 0.083878883 | 0.182325815 |
| ENSMUSG00000026109 | Tmeff2 | 56363 | protein_coding | 3.742430174 | -0.182360696 | 0.996661534 | 0.18297154 | 0.854820349 | 1 | 0.182360696 |
| ENSMUSG00000042208 | Sanbr | 71675 | protein_coding | 270.0824699 | 0.182438581 | 0.097245821 | -1.87605574 | 0.060647626 | 0.172144191 | 0.182438581 |

| ensembl_gene_id | external_gene_name | entrezgene_id | gene_biotype | baseMean | log2FoldChange | lfcSE | stat | pvalue | padj | absLog2FC |
|---|---|---|---|---|---|---|---|---|---|---|
| ENSMUSG00000028465 | Tln1 | 21894 | protein_coding | 2012.792541 | 0.182438884 | 0.066643987 | -2.737514561 | 0.006190538 | 0.028864834 | 0.182438884 |
| ENSMUSG00000114245 | Percc1 | 102643076 | protein_coding | 1.227521128 | -0.182488975 | 0.99527473 | 0.183355378 | 0.854519185 | 1 | 0.182488975 |
| ENSMUSG00000091537 | Tma7 | 66167 | protein_coding | 681.0476596 | -0.182515858 | 0.100212394 | 1.82129026 | 0.068562746 | 0.188104702 | 0.182515858 |
| ENSMUSG00000000439 | Mkrn2 | 67027 | protein_coding | 193.1944081 | -0.182567695 | 0.085816311 | 2.127424175 | 0.033384855 | 0.109497021 | 0.182567695 |
| ENSMUSG00000013698 | Pea15a | 18611 | protein_coding | 170.1746743 | -0.18263175 | 0.090192694 | 2.024906254 | 0.042876991 | 0.132607819 | 0.18263175 |
| ENSMUSG00000087861 | Gm24440 | 115488686 | snRNA | 0.91515639 | 0.182675591 | 1.347566906 | -0.13555957 | 0.892169469 | 1 | 0.182675591 |
| ENSMUSG00000055436 | Srsf11 | 69207 | protein_coding | 914.306711 | 0.182698537 | 0.063797759 | -2.863714012 | 0.004187059 | 0.020927689 | 0.182698537 |
| ENSMUSG00000024388 | Myo7b | 17922 | protein_coding | 2929.274068 | -0.18276335 | 0.161970662 | 1.128373175 | 0.259162352 | 0.472489189 | 0.18276335 |
| ENSMUSG00000054717 | Hmgb2 | 97165 | protein_coding | 939.3975758 | 0.182776853 | 0.152218724 | -1.20075145 | 0.22984763 | 0.438404222 | 0.182776853 |
| ENSMUSG00000036503 | Rnf13 | 24017 | protein_coding | 491.9927761 | -0.1828391 | 0.093913847 | 1.946881172 | 0.051548982 | 0.152668561 | 0.1828391 |
| ENSMUSG00000008090 | Fgfrl1 | 116701 | protein_coding | 234.1431804 | 0.182856205 | 0.13582576 | -1.34625571 | 0.178220064 | 0.37057407 | 0.182856205 |
| ENSMUSG00000039206 | Daglb | 231871 | protein_coding | 603.2413283 | 0.182872779 | 0.133455865 | -1.37028656 | 0.170597467 | 0.360481579 | 0.182872779 |
| ENSMUSG00000018395 | Kif3a | 16568 | protein_coding | 93.52070602 | 0.182899557 | 0.13212344 | -1.384308162 | 0.166264116 | 0.35368873 | 0.182899557 |
| ENSMUSG00000002981 | Clptm1 | 56457 | protein_coding | 2254.661046 | -0.182915722 | 0.063640536 | 2.874201475 | 0.004050505 | 0.020374669 | 0.182915722 |
| ENSMUSG00000020910 | Adprm | 66358 | protein_coding | 130.3003949 | -0.182920612 | 0.115642275 | 1.581779772 | 0.113699854 | 0.270826395 | 0.182920612 |
| ENSMUSG00000063626 | Unc5d | 210801 | protein_coding | 2.717044933 | -0.182922171 | 1.586952384 | 0.115266326 | 0.908234028 | 1 | 0.182922171 |
| ENSMUSG00000030168 | Adipor2 | 68465 | protein_coding | 2421.492345 | -0.182937003 | 0.074438022 | 2.457574733 | 0.01398787 | 0.055222494 | 0.182937003 |
| ENSMUSG00000027935 | Rab13 | 68328 | protein_coding | 109.1300243 | -0.182980407 | 0.101094931 | 1.809985977 | 0.070297962 | 0.191599363 | 0.182980407 |
| ENSMUSG00000042472 | Zfp410 | 52708 | protein_coding | 351.0208031 | 0.182990857 | 0.071203589 | -2.569966772 | 0.010170827 | 0.043155457 | 0.182990857 |
| ENSMUSG00000046991 | Wdr27 | 71682 | protein_coding | 14.43255473 | -0.183024963 | 0.394040478 | 0.464482644 | 0.642301991 | 0.804778079 | 0.183024963 |
| ENSMUSG00000016554 | Eif3d | 55944 | protein_coding | 945.4496071 | 0.183095674 | 0.08874507 | -2.063164449 | 0.039097007 | 0.123758475 | 0.183095674 |
| ENSMUSG00000027170 | Eif3m | 98221 | protein_coding | 875.4602911 | -0.183104839 | 0.117682474 | 1.555922749 | 0.119726463 | 0.281607009 | 0.183104839 |
| ENSMUSG00000086682 | Gm16023 | 100502734 | lncRNA | 18.64020479 | -0.183190557 | 0.256609243 | 0.713889161 | 0.47529572 | 0.687185565 | 0.183190557 |
| ENSMUSG00000027977 | Ndst3 | 83398 | protein_coding | 2.301911773 | 0.183203815 | 3.05671988 | -0.059934774 | 0.952207584 | 1 | 0.183203815 |
| ENSMUSG00000040112 | Mrps35 | 232536 | protein_coding | 534.0297774 | -0.183206939 | 0.081738483 | 2.241379231 | 0.025001521 | 0.087325531 | 0.183206939 |
| ENSMUSG00000026004 | Kansl1l | 68691 | protein_coding | 127.0605047 | -0.183283717 | 0.130440796 | 1.405110386 | 0.159988456 | 0.344285818 | 0.183283717 |
| ENSMUSG00000025236 | Adpgk | 72141 | protein_coding | 207.0232119 | 0.183322101 | 0.101134874 | -1.812649723 | 0.069885864 | 0.190791065 | 0.183322101 |
| ENSMUSG00000037263 | Aldh3b3 | 73458 | protein_coding | 3.420348808 | -0.183329782 | 0.767923681 | 0.238734378 | 0.811311558 | 1 | 0.183329782 |
| ENSMUSG00000029326 | Enoph1 | 67870 | protein_coding | 366.0294546 | -0.183332984 | 0.089076157 | 2.058160005 | 0.039574776 | 0.125007242 | 0.183332984 |
| ENSMUSG00000029784 | Ssmem1 | 75647 | protein_coding | 1.137673775 | 0.183333759 | 1.343011946 | -0.136509403 | 0.891418591 | 1 | 0.183333759 |
| ENSMUSG00000040007 | Bahd1 | 228536 | protein_coding | 428.4519866 | -0.183361829 | 0.067806555 | 2.704190299 | 0.006847106 | 0.031346224 | 0.183361829 |
| ENSMUSG00000006169 | Clint1 | 216705 | protein_coding | 3157.748375 | -0.183368135 | 0.04585679 | 3.998712857 | 6.37E-05 | 0.000580752 | 0.183368135 |
| ENSMUSG00000034758 | Tle6 | 114606 | protein_coding | 53.9456524 | -0.183413905 | 0.200046537 | 0.916856187 | 0.359218005 | 0.581866222 | 0.183413905 |
| ENSMUSG00000029056 | Pank4 | 269614 | protein_coding | 252.4608979 | -0.183495459 | 0.08749781 | 2.097143439 | 0.03598088 | 0.115758474 | 0.183495459 |
| ENSMUSG00000042712 | Tceal9 | 22381 | protein_coding | 918.7143962 | 0.183540381 | 0.132064701 | -1.38977622 | 0.164596841 | 0.351137093 | 0.183540381 |
| ENSMUSG00000045665 | Mfsd5 | 106073 | protein_coding | 270.658823 | 0.183575234 | 0.081084545 | -2.263997838 | 0.023574248 | 0.083506033 | 0.183575234 |
| ENSMUSG00000022386 | Trmu | 72026 | protein_coding | 133.1568378 | -0.183610443 | 0.105626589 | 1.738297573 | 0.082158394 | 0.214101896 | 0.183610443 |
| ENSMUSG00000026999 | Nup35 | 69482 | protein_coding | 61.32594081 | -0.183613766 | 0.186949342 | 0.982157859 | 0.326022085 | 0.548038346 | 0.183613766 |
| ENSMUSG00000004394 | Tmed4 | 103694 | protein_coding | 772.9101958 | 0.18362889 | 0.07971689 | -2.303512966 | 0.021249998 | 0.076990451 | 0.18362889 |

Supplementary Table 1

| ensembl_gene_id | external_gene_name | entrezgene_id | gene_biotype | baseMean | log2FoldChange | lfcSE | stat | pvalue | padj | absLog2FC |
|---|---|---|---|---|---|---|---|---|---|---|
| ENSMUSG00000021248 | Tmed10 | 68581 | protein_coding | 1760.584824 | 0.183718112 | 0.100197214 | -1.83356507 | 0.066718581 | 0.184376923 | 0.183718112 |
| ENSMUSG00000017667 | Zfp334 | 228876 | protein_coding | 48.04106586 | 0.183720105 | 0.174252135 | -1.054334886 | 0.291729622 | 0.510557756 | 0.183720105 |
| ENSMUSG00000020604 | Arsg | 74008 | protein_coding | 74.29437661 | -0.183733766 | 0.160082468 | 1.147744458 | 0.251074072 | 0.463422066 | 0.183733766 |
| ENSMUSG00000057101 | Zfp180 | 210135 | protein_coding | 361.6833312 | -0.183733857 | 0.075276007 | 2.440802391 | 0.014654671 | 0.057293381 | 0.183733857 |
| ENSMUSG00000097251 | 5033417F24Rik | 75995 | lncRNA | 7.891876247 | -0.183758554 | 0.373857624 | 0.491520145 | 0.623058608 | 0.793281643 | 0.183758554 |
| ENSMUSG00000041775 | Mapk1ip1 | 69546 | protein_coding | 77.61654925 | -0.183758932 | 0.145328057 | 1.264442227 | 0.206071345 | 0.408658758 | 0.183758932 |
| ENSMUSG00000042320 | Prox2 | 73422 | protein_coding | 11.99104817 | -0.183759201 | 0.38025108 | 0.48325754 | 0.628912884 | 0.797440677 | 0.183759201 |
| ENSMUSG00000024910 | Ctsw | 13041 | protein_coding | 27.82554063 | 0.183914416 | 0.364872434 | -0.504051275 | 0.614225346 | 0.78673795 | 0.183914416 |
| ENSMUSG00000022408 | Fam83f | 213956 | protein_coding | 416.4902136 | 0.183937295 | 0.070493794 | -2.609269324 | 0.00907358 | 0.039390404 | 0.183937295 |
| ENSMUSG00000001630 | Stk38l | 232533 | protein_coding | 769.8535882 | -0.183941685 | 0.137573416 | 1.337043811 | 0.181208337 | 0.375235922 | 0.183941685 |
| ENSMUSG00000025745 | Hadha | 97212 | protein_coding | 3844.630925 | -0.183964884 | 0.062466902 | 2.944997724 | 0.00322957 | 0.016920408 | 0.183964884 |
| ENSMUSG00000031465 | Angpt2 | 11601 | protein_coding | 24.28725615 | -0.184020724 | 0.248026717 | 0.74193912 | 0.458124223 | 0.672740084 | 0.184020724 |
| ENSMUSG00000025359 | Pmel | 20431 | protein_coding | 32.6022608 | -0.184035397 | 0.200225063 | 0.919142657 | 0.35802096 | 0.580790426 | 0.184035397 |
| ENSMUSG00000036934 | 4921524J17Rik | 66714 | protein_coding | 62.61869877 | -0.184059112 | 0.137045335 | 1.343052736 | 0.179254898 | 0.37240098 | 0.184059112 |
| ENSMUSG00000112654 | 4930455C13Rik | 73990 | lncRNA | 3.587447036 | -0.184100068 | 0.829299757 | 0.2219946 | 0.824318087 | 1 | 0.184100068 |
| ENSMUSG00000045962 | Wnk1 | 232341 | protein_coding | 1151.013674 | 0.184155949 | 0.111369839 | -1.653553162 | 0.098218336 | 0.244200181 | 0.184155949 |
| ENSMUSG00000022508 | Bcl6 | 12053 | protein_coding | 66.67620448 | -0.184164335 | 0.180110425 | 1.022507915 | 0.306540569 | 0.527079777 | 0.184164335 |
| ENSMUSG00000037999 | Arap2 | 212285 | protein_coding | 1180.123084 | -0.184181658 | 0.099890881 | 1.843828539 | 0.065208132 | 0.181293038 | 0.184181658 |
| ENSMUSG00000020620 | Abca8b | 27404 | protein_coding | 7.568748282 | -0.184256913 | 0.754741789 | 0.244132385 | 0.807128292 | 0.904987729 | 0.184256913 |
| ENSMUSG00000003541 | Ier3 | 15937 | protein_coding | 209.1625551 | 0.184308874 | 0.235923337 | -0.78122358 | 0.434671007 | 0.652915864 | 0.184308874 |
| ENSMUSG00000029507 | Pus1 | 56361 | protein_coding | 296.4712922 | -0.184321559 | 0.123543182 | 1.491960598 | 0.135709475 | 0.30705241 | 0.184321559 |
| ENSMUSG00000032380 | Dapk2 | 13143 | protein_coding | 431.5321326 | -0.184321716 | 0.071082365 | 2.593072349 | 0.009512276 | 0.040902139 | 0.184321716 |
| ENSMUSG00000078919 | Dpm1 | 13480 | protein_coding | 46.37977861 | -0.184326674 | 0.16390169 | 1.124617289 | 0.260751244 | 0.474234146 | 0.184326674 |
| ENSMUSG00000015488 | Cacfd1 | 381356 | protein_coding | 406.3464197 | 0.184340247 | 0.078855763 | -2.337688958 | 0.019403389 | 0.071716108 | 0.184340247 |
| ENSMUSG00000041417 | Pik3r1 | 18708 | protein_coding | 1878.950061 | -0.184474289 | 0.059238869 | 3.114075136 | 0.001845225 | 0.010614744 | 0.184474289 |
| ENSMUSG00000024642 | Tle4 | 21888 | protein_coding | 786.2792254 | -0.184505952 | 0.129525427 | 1.424476691 | 0.154308528 | 0.336218865 | 0.184505952 |
| ENSMUSG00000029475 | Kdm2b | 30841 | protein_coding | 182.0046918 | -0.184530422 | 0.092405265 | 1.996968691 | 0.045828584 | 0.139530465 | 0.184530422 |
| ENSMUSG00000026447 | Pik3c2b | 240752 | protein_coding | 1164.099362 | -0.184539354 | 0.113638993 | 1.623908738 | 0.104395287 | 0.255480702 | 0.184539354 |
| ENSMUSG00000053070 | Cfap300 | 234912 | protein_coding | 12.2928017 | -0.184569957 | 0.301442122 | 0.612289868 | 0.540345992 | 0.734651013 | 0.184569957 |
| ENSMUSG00000060678 | H4c3 | 319155 | protein_coding | 2895.116891 | 0.18458529 | 0.113282319 | -1.629427184 | 0.103222618 | 0.253324166 | 0.18458529 |
| ENSMUSG00000026395 | Ptprc | 19264 | protein_coding | 142.514823 | -0.184586818 | 0.177520574 | 1.039805212 | 0.298430407 | 0.518220794 | 0.184586818 |
| ENSMUSG00000031875 | Cmtm3 | 68119 | protein_coding | 9.840673046 | -0.184614238 | 0.456052163 | 0.404809478 | 0.685617561 | 0.832521206 | 0.184614238 |
| ENSMUSG00000024948 | Map4k2 | 26412 | protein_coding | 135.571432 | 0.184618634 | 0.151670444 | -1.217235404 | 0.223514663 | 0.430702585 | 0.184618634 |
| ENSMUSG00000026348 | Acmsd | 266645 | protein_coding | 1.401756215 | 0.184639395 | 1.251163208 | -0.147574188 | 0.882678827 | 1 | 0.184639395 |
| ENSMUSG00000098286 | Mir8091 | 102465884 | miRNA | 4.49435316 | -0.184648679 | 0.524737761 | 0.351887538 | 0.724922605 | 0.857548131 | 0.184648679 |
| ENSMUSG00000026269 | Rnpepl1 | 108657 | protein_coding | 1077.073099 | -0.184672665 | 0.060648875 | 3.044947888 | 0.002327206 | 0.012806226 | 0.184672665 |
| ENSMUSG00000002679 | Med6 | 69792 | protein_coding | 168.411221 | 0.184705161 | 0.109788652 | -1.682370249 | 0.092497067 | 0.233277344 | 0.184705161 |
| ENSMUSG00000001687 | Ubl3 | 24109 | protein_coding | 1267.387206 | -0.184735152 | 0.046314395 | 3.988719976 | 6.64E-05 | 0.000602437 | 0.184735152 |
| ENSMUSG00000041119 | Pde9a | 18585 | protein_coding | 1788.703457 | 0.184802742 | 0.131580433 | -1.404484974 | 0.160174483 | 0.344596328 | 0.184802742 |

Supplementary Table 1

| ensembl_gene_id | external_gene_name | entrezgene_id | gene_biotype | baseMean | log2FoldChange | lfcSE | stat | pvalue | padj | absLog2FC |
|---|---|---|---|---|---|---|---|---|---|---|
| ENSMUSG00000117869 | Snhg4 | 100503380 | lncRNA | 135.0134672 | -0.18483802 | 0.382936166 | 0.482686244 | 0.629318532 | 0.79746811 | 0.18483802 |
| ENSMUSG00000041716 | Gm20604 | 100859931 | protein_coding | 49.8223088 | 0.184880188 | 0.156901581 | -1.178319475 | 0.238669267 | 0.448874722 | 0.184880188 |
| ENSMUSG00000026928 | Card9 | 332579 | protein_coding | 3.633058164 | -0.184908364 | 0.700085901 | 0.264122393 | 0.791685609 | 1 | 0.184908364 |
| ENSMUSG00000031832 | Taf1c | 21341 | protein_coding | 208.4102031 | 0.184940097 | 0.075360982 | -2.454056359 | 0.014125484 | 0.055659328 | 0.184940097 |
| ENSMUSG00000037253 | Mex3c | 240396 | protein_coding | 685.2613902 | 0.18494024 | 0.068725817 | -2.690986399 | 0.00712411 | 0.032409794 | 0.18494024 |
| ENSMUSG00000004561 | Mettl17 | 52535 | protein_coding | 196.8463147 | -0.1849698 | 0.091315319 | 2.025616324 | 0.042804117 | 0.132446542 | 0.1849698 |
| ENSMUSG00000020680 | Taf15 | 70439 | protein_coding | 320.6616031 | 0.185038753 | 0.07877424 | -2.348975402 | 0.01882515 | 0.070049242 | 0.185038753 |
| ENSMUSG00000065399 | Mir133a-1 | 387151 | miRNA | 0.557688373 | -0.185079915 | 1.419747568 | 0.130361143 | 0.896280708 | 1 | 0.185079915 |
| ENSMUSG00000090112 | Shprh | 268281 | protein_coding | 372.7825435 | -0.185108346 | 0.080758148 | 2.292132121 | 0.02189802 | 0.078853883 | 0.185108346 |
| ENSMUSG00000048732 | Klhl11 | 217194 | protein_coding | 143.4568342 | 0.185147526 | 0.098748015 | -1.874949335 | 0.060799694 | 0.172486883 | 0.185147526 |
| ENSMUSG00000032698 | Lmo2 | 16909 | protein_coding | 8.289993688 | -0.185189524 | 0.423241925 | 0.437550048 | 0.661712489 | 0.816921981 | 0.185189524 |
| ENSMUSG00000086914 | Gm16124 | 102636154 | lncRNA | 5.016178277 | -0.185302015 | 0.468544773 | 0.395484116 | 0.692485642 | 0.836513744 | 0.185302015 |
| ENSMUSG00000060733 | Ipmk | 69718 | protein_coding | 2554.942005 | -0.185342078 | 0.103222253 | 1.795563199 | 0.072564013 | 0.196079582 | 0.185342078 |
| ENSMUSG00000044986 | Tst | 22117 | protein_coding | 2241.347036 | -0.185352884 | 0.095115974 | 1.948704042 | 0.051330779 | 0.152129476 | 0.185352884 |
| ENSMUSG00000030981 | Mmp21 | 214766 | protein_coding | 2.821747315 | 0.18535423 | 0.712242278 | -0.260240421 | 0.794678327 | 1 | 0.18535423 |
| ENSMUSG00000028042 | Zbtb7b | 22724 | protein_coding | 3334.971975 | 0.185366685 | 0.10744199 | -1.725272257 | 0.084478419 | 0.218595521 | 0.185366685 |
| ENSMUSG00000029556 | Hnf1a | 21405 | protein_coding | 335.3414884 | 0.185506413 | 0.149424711 | -1.241470782 | 0.214431885 | 0.419407924 | 0.185506413 |
| ENSMUSG00000026405 | C4bp | 12269 | protein_coding | 98.58385734 | -0.185603197 | 0.294387209 | 0.630473036 | 0.528385139 | 0.726359126 | 0.185603197 |
| ENSMUSG00000010608 | Rbm25 | 67039 | protein_coding | 1048.274841 | 0.185623132 | 0.066817884 | -2.77804567 | 0.005468693 | 0.026243183 | 0.185623132 |
| ENSMUSG00000060131 | Atp8b4 | 241633 | protein_coding | 4.59029099 | 0.185715921 | 1.260224836 | -0.147367292 | 0.882842121 | 0.942116626 | 0.185715921 |
| ENSMUSG00000072647 | Adam1a | 280668 | protein_coding | 8.458561603 | -0.185731514 | 0.427321219 | 0.43464145 | 0.663822703 | 0.818698142 | 0.185731514 |
| ENSMUSG00000031862 | Atp13a1 | 170759 | protein_coding | 1056.698855 | 0.185763476 | 0.059313999 | -3.131865665 | 0.001736993 | 0.010082498 | 0.185763476 |
| ENSMUSG00000026816 | Gtf3c5 | 70239 | protein_coding | 256.8159073 | 0.185785782 | 0.087930839 | -2.112862611 | 0.034612529 | 0.112562288 | 0.185785782 |
| ENSMUSG00000025223 | Ldb1 | 16825 | protein_coding | 405.1099837 | -0.185808453 | 0.091940428 | 2.020965723 | 0.043283314 | 0.133504042 | 0.185808453 |
| ENSMUSG00000020747 | Tmem94 | 71947 | protein_coding | 1790.974787 | -0.185816154 | 0.083624258 | 2.2220365 | 0.026280839 | 0.090782827 | 0.185816154 |
| ENSMUSG00000002413 | Braf | 109880 | protein_coding | 384.4369258 | 0.185841084 | 0.105243009 | -1.765828303 | 0.077424659 | 0.205363124 | 0.185841084 |
| ENSMUSG00000032667 | Pon2 | 330260 | protein_coding | 1995.336025 | -0.185996689 | 0.081730077 | 2.275743462 | 0.022861368 | 0.081487727 | 0.185996689 |
| ENSMUSG00000018733 | Pex12 | 103737 | protein_coding | 89.96585362 | 0.18601982 | 0.116840128 | -1.592088464 | 0.111364829 | 0.267113839 | 0.18601982 |
| ENSMUSG00000021190 | Lgmn | 19141 | protein_coding | 394.593738 | -0.186028111 | 0.152727478 | 1.218039572 | 0.223208935 | 0.430314308 | 0.186028111 |
| ENSMUSG00000042055 | Wdr11 | 207425 | protein_coding | 623.1331384 | -0.186120503 | 0.08679278 | 2.14442381 | 0.031998944 | 0.106094619 | 0.186120503 |
| ENSMUSG00000019087 | Atp6ap1 | 54411 | protein_coding | 1134.369132 | -0.186121318 | 0.073357592 | 2.53717869 | 0.01117499 | 0.046543198 | 0.186121318 |
| ENSMUSG00000110444 | Gm10033 | 378466 | protein_coding | 70.23930316 | -0.186121636 | 0.147235167 | 1.264111283 | 0.206190087 | 0.408815356 | 0.186121636 |
| ENSMUSG00000031816 | Mthfsd | 234814 | protein_coding | 247.5170257 | 0.186123325 | 0.086823949 | -2.143686475 | 0.032058017 | 0.106242441 | 0.186123325 |
| ENSMUSG00000023830 | Igf2r | 16004 | protein_coding | 2570.864696 | -0.186127996 | 0.121249915 | 1.535077337 | 0.124764832 | 0.289621372 | 0.186127996 |
| ENSMUSG00000055707 | Klhl26 | 234378 | protein_coding | 164.0265144 | -0.186132205 | 0.097769619 | 1.903783672 | 0.056938364 | 0.164355907 | 0.186132205 |
| ENSMUSG00000036905 | C1qb | 12260 | protein_coding | 2.699318208 | -0.186175041 | 0.669580984 | 0.278047085 | 0.780976215 | 1 | 0.186175041 |
| ENSMUSG00000021737 | Psmd6 | 66413 | protein_coding | 675.7804153 | 0.186264494 | 0.084358088 | -2.208021753 | 0.027242757 | 0.093415051 | 0.186264494 |
| ENSMUSG00000039501 | Znfx1 | 98999 | protein_coding | 778.126088 | -0.18629565 | 0.112694952 | 1.65309667 | 0.098311191 | 0.244320693 | 0.18629565 |
| ENSMUSG00000029512 | Ulk1 | 22241 | protein_coding | 256.1727649 | -0.186322652 | 0.104019913 | 1.79122099 | 0.073257839 | 0.197386189 | 0.186322652 |

| ensembl_gene_id | external_gene_name | entrezgene_id | gene_biotype | baseMean | log2FoldChange | lfcSE | stat | pvalue | padj | absLog2FC |
|---|---|---|---|---|---|---|---|---|---|---|
| ENSMUSG00000075376 | Rc3h2 | 319817 | protein_coding | 840.6026989 | 0.186479356 | 0.068126745 | -2.73724153 | 0.006195679 | 0.028870044 | 0.186479356 |
| ENSMUSG00000078317 | F8a | 14070 | protein_coding | 216.0620502 | 0.186490528 | 0.107300215 | -1.738025672 | 0.08220629 | 0.214159123 | 0.186490528 |
| ENSMUSG00000083307 | AA414768 | 245350 | processed_pseudogene | 14.36870771 | -0.186502172 | 0.304135912 | 0.613219829 | 0.539730995 | 0.734194205 | 0.186502172 |
| ENSMUSG00000109644 | 0610005C13Rik | 71661 | lncRNA | 209.945329 | 0.186539086 | 0.230519809 | -0.809210657 | 0.418393986 | 0.63839585 | 0.186539086 |
| ENSMUSG00000027487 | Cdk5rap1 | 66971 | protein_coding | 191.4592255 | -0.186539481 | 0.260062569 | 0.717286927 | 0.473197072 | 0.685335967 | 0.186539481 |
| ENSMUSG00000040006 | Ginm1 | 215751 | protein_coding | 682.17493 | -0.186622981 | 0.084750011 | 2.202040787 | 0.027662427 | 0.094598742 | 0.186622981 |
| ENSMUSG00000039191 | Rbpj | 19664 | protein_coding | 211.9850773 | -0.186624033 | 0.109032289 | 1.711640052 | 0.086963023 | 0.223449098 | 0.186624033 |
| ENSMUSG00000029777 | Gars | 353172 | protein_coding | 2084.438496 | 0.186627435 | 0.067810204 | -2.752202847 | 0.005919584 | 0.027890446 | 0.186627435 |
| ENSMUSG00000041852 | Tcf20 | 21411 | protein_coding | 2493.85197 | -0.186628557 | 0.039288299 | 4.750232498 | 2.03E-06 | 2.65E-05 | 0.186628557 |
| ENSMUSG00000022191 | Drosha | 14000 | protein_coding | 546.3963208 | 0.186696205 | 0.059470331 | -3.139316736 | 0.001693423 | 0.009870396 | 0.186696205 |
| ENSMUSG00000062382 | Ftl1-ps1 | 434624 | protein_coding | 5.32222936 | -0.186720451 | 0.572247057 | 0.326293423 | 0.744202364 | 0.870990647 | 0.186720451 |
| ENSMUSG00000019806 | Aig1 | 66253 | protein_coding | 614.0977316 | -0.186724873 | 0.075101043 | 2.486315319 | 0.012907354 | 0.051861813 | 0.186724873 |
| ENSMUSG00000038500 | Prr3 | 75210 | protein_coding | 33.11220749 | -0.186756041 | 0.198417452 | 0.941227897 | 0.346588083 | 0.56826406 | 0.186756041 |
| ENSMUSG00000066900 | Suds3 | 71954 | protein_coding | 1170.516927 | -0.186783791 | 0.059536129 | 3.137318382 | 0.001705009 | 0.009910776 | 0.186783791 |
| ENSMUSG00000027708 | Dcun1d1 | 114893 | protein_coding | 381.3928971 | -0.186902974 | 0.090311168 | 2.069544415 | 0.038495028 | 0.122282111 | 0.186902974 |
| ENSMUSG00000001964 | Emd | 13726 | protein_coding | 272.3423534 | 0.186977924 | 0.095416675 | -1.959593798 | 0.050043287 | 0.148984107 | 0.186977924 |
| ENSMUSG00000050846 | Zfp623 | 78834 | protein_coding | 187.0898312 | -0.18700087 | 0.12317254 | 1.51820259 | 0.128963335 | 0.296522686 | 0.18700087 |
| ENSMUSG00000028556 | Dock7 | 67299 | protein_coding | 829.6434261 | 0.187003865 | 0.054277561 | -3.445325531 | 0.000570372 | 0.003959279 | 0.187003865 |
| ENSMUSG00000045948 | Mrps12 | 24030 | protein_coding | 389.3644304 | 0.187004908 | 0.107978153 | -1.731877258 | 0.083295419 | 0.216245959 | 0.187004908 |
| ENSMUSG00000050945 | Zfp438 | 240186 | protein_coding | 36.74386659 | -0.187064701 | 0.208071271 | 0.899041467 | 0.368630574 | 0.591140542 | 0.187064701 |
| ENSMUSG00000070000 | Fcho1 | 74015 | protein_coding | 76.40664514 | -0.187071256 | 0.219748081 | 0.851298701 | 0.394603446 | 0.617091033 | 0.187071256 |
| ENSMUSG00000109771 | Gm35315 | 102638847 | protein_coding | 22.3020258 | 0.18709842 | 0.322706232 | -0.579779383 | 0.562063402 | 0.750140815 | 0.18709842 |
| ENSMUSG00000037953 | A4gnt | 333424 | protein_coding | 173.2133307 | -0.187126369 | 0.234121605 | 0.799269971 | 0.424133887 | 0.643957486 | 0.187126369 |
| ENSMUSG00000015745 | Plekho1 | 67220 | protein_coding | 16.23690695 | 0.187157055 | 0.283269502 | -0.66070316 | 0.508802697 | 0.712306522 | 0.187157055 |
| ENSMUSG00000027263 | Tubgcp4 | 51885 | protein_coding | 321.4791305 | -0.187165731 | 0.082015212 | 2.282085568 | 0.022484289 | 0.080560978 | 0.187165731 |
| ENSMUSG00000020015 | Cdk17 | 237459 | protein_coding | 156.3773392 | -0.187181625 | 0.149875806 | 1.24891155 | 0.211697427 | 0.416065952 | 0.187181625 |
| ENSMUSG00000007739 | Cct4 | 12464 | protein_coding | 1730.542364 | -0.187230009 | 0.090398411 | 2.071164831 | 0.038343395 | 0.12191044 | 0.187230009 |
| ENSMUSG00000058886 | Deaf1 | 54006 | protein_coding | 228.0635256 | -0.187243465 | 0.109424005 | 1.711173573 | 0.087049076 | 0.223580206 | 0.187243465 |
| ENSMUSG00000010797 | Wnt2 | 22413 | protein_coding | 0.968317744 | -0.187247226 | 2.272048037 | 0.08241341 | 0.934317973 | 1 | 0.187247226 |
| ENSMUSG00000030730 | Atp2a1 | 11937 | protein_coding | 6.891946206 | 0.187247558 | 0.668897827 | -0.279934469 | 0.779527782 | 0.890072723 | 0.187247558 |
| ENSMUSG00000014418 | Hps5 | 246694 | protein_coding | 846.0389007 | 0.18727808 | 0.079765922 | -2.347845742 | 0.01888234 | 0.070214581 | 0.18727808 |
| ENSMUSG00000029290 | Zfp326 | 54367 | protein_coding | 281.1072066 | 0.187318491 | 0.075080575 | -2.494899542 | 0.012599279 | 0.050983705 | 0.187318491 |
| ENSMUSG00000063334 | Krr1 | 52705 | protein_coding | 344.0576366 | 0.187393365 | 0.105015073 | -1.784442553 | 0.074351782 | 0.19938276 | 0.187393365 |
| ENSMUSG00000037860 | Aim2 | 383619 | protein_coding | 534.0264468 | 0.187437247 | 0.086667047 | -2.162727971 | 0.030562108 | 0.102353003 | 0.187437247 |
| ENSMUSG00000022514 | Il1rap | 16180 | protein_coding | 116.7767418 | 0.187461559 | 0.141965758 | -1.32047024 | 0.186678066 | 0.382529046 | 0.187461559 |
| ENSMUSG00000054676 | 1600014C10Rik | 72244 | protein_coding | 114.3424494 | 0.187466507 | 0.116649527 | -1.607091869 | 0.108034217 | 0.261594876 | 0.187466507 |
| ENSMUSG00000036764 | Dnajc12 | 30045 | protein_coding | 47.30278606 | -0.187471245 | 0.177764844 | 1.054602476 | 0.291607171 | 0.510451645 | 0.187471245 |
| ENSMUSG00000092103 | Gm17039 | 102636903 | lncRNA | 4.362966152 | -0.187484117 | 0.533535746 | 0.351399355 | 0.725288767 | 1 | 0.187484117 |
| ENSMUSG00000035441 | Myo1d | 338367 | protein_coding | 4271.874136 | 0.187488349 | 0.063911373 | -2.93356783 | 0.003350904 | 0.017451579 | 0.187488349 |

Supplementary Table 1

| ensembl_gene_id | external_gene_name | entrezgene_id | gene_biotype | baseMean | log2FoldChange | lfcSE | stat | pvalue | padj | absLog2FC |
|---|---|---|---|---|---|---|---|---|---|---|
| ENSMUSG00000031778 | Cx3cl1 | 20312 | protein_coding | 948.5127061 | -0.187521295 | 0.079321854 | 2.36405587 | 0.018076086 | 0.067704352 | 0.187521295 |
| ENSMUSG00000029430 | Ran | 19384 | protein_coding | 1403.930419 | 0.187539409 | 0.144939164 | -1.293918112 | 0.195693708 | 0.394975053 | 0.187539409 |
| ENSMUSG00000045868 | Gvin1 | 74558 | protein_coding | 2.851248343 | -0.187554569 | 0.848886321 | 0.220941915 | 0.82513766 | 1 | 0.187554569 |
| ENSMUSG00000056592 | Zfp658 | 210104 | protein_coding | 30.75279394 | -0.187578712 | 0.206176921 | 0.909794906 | 0.362930682 | 0.585225632 | 0.187578712 |
| ENSMUSG00000022312 | Eif3h | 68135 | protein_coding | 1724.331813 | -0.187589884 | 0.090510133 | 2.072584331 | 0.038210981 | 0.121629941 | 0.187589884 |
| ENSMUSG00000056228 | Cars2 | 71941 | protein_coding | 259.8898063 | 0.187606873 | 0.070357102 | -2.666495165 | 0.007664671 | 0.03440456 | 0.187606873 |
| ENSMUSG00000097431 | Gm26782 | 102640338 | lncRNA | 23.93572516 | -0.187607246 | 0.208968872 | 0.897776037 | 0.369304963 | 0.591883639 | 0.187607246 |
| ENSMUSG00000032834 | Pwp2 | 110816 | protein_coding | 193.3288858 | -0.187656143 | 0.110684252 | 1.695418636 | 0.089996033 | 0.228807636 | 0.187656143 |
| ENSMUSG00000020235 | Fzr1 | 56371 | protein_coding | 668.375646 | 0.187680457 | 0.06633203 | -2.829409206 | 0.004663403 | 0.022933722 | 0.187680457 |
| ENSMUSG00000038415 | Foxq1 | 15220 | protein_coding | 44.28055085 | -0.187766252 | 0.305065087 | 0.615495708 | 0.538227403 | 0.733377497 | 0.187766252 |
| ENSMUSG00000004791 | Pgf | 18654 | protein_coding | 1.058952313 | -0.187806878 | 1.782718398 | 0.105348594 | 0.916099205 | 1 | 0.187806878 |
| ENSMUSG00000017485 | Top2b | 21974 | protein_coding | 1621.776954 | -0.18782445 | 0.102655648 | 1.829655294 | 0.0673015 | 0.185483473 | 0.18782445 |
| ENSMUSG00000026944 | Abca2 | 11305 | protein_coding | 116.5822681 | -0.18788942 | 0.102307219 | 1.836521625 | 0.066280547 | 0.1835581 | 0.18788942 |
| ENSMUSG00000025873 | Faf2 | 76577 | protein_coding | 898.1574473 | 0.187928364 | 0.081644846 | -2.301778659 | 0.021347658 | 0.077259526 | 0.187928364 |
| ENSMUSG00000015133 | Lrrk1 | 233328 | protein_coding | 669.2644442 | 0.187937915 | 0.067558597 | -2.781850488 | 0.005404993 | 0.025977317 | 0.187937915 |
| ENSMUSG00000001995 | Sipa1l2 | 244668 | protein_coding | 308.4540463 | 0.187938501 | 0.074714199 | -2.515432169 | 0.011888655 | 0.048729637 | 0.187938501 |
| ENSMUSG00000030357 | Fkbp4 | 14228 | protein_coding | 3688.442181 | 0.18793986 | 0.117240869 | -1.603023429 | 0.108929482 | 0.26324771 | 0.18793986 |
| ENSMUSG00000032303 | Chrna3 | 110834 | protein_coding | 2.478116299 | 0.188036499 | 1.30002003 | -0.144641232 | 0.884994141 | 1 | 0.188036499 |
| ENSMUSG00000022526 | Zfp251 | 71591 | protein_coding | 133.4246832 | -0.188040799 | 0.122298025 | 1.537562017 | 0.124155741 | 0.288572541 | 0.188040799 |
| ENSMUSG00000035919 | Bbs9 | 319845 | protein_coding | 59.77272503 | 0.188041025 | 0.141033784 | -1.333304825 | 0.182431791 | 0.377007156 | 0.188041025 |
| ENSMUSG00000074238 | Ap1ar | 211556 | protein_coding | 1105.642848 | -0.188062766 | 0.07303533 | 2.574956068 | 0.010025282 | 0.042679767 | 0.188062766 |
| ENSMUSG00000005043 | Sgsh | 27029 | protein_coding | 180.1868489 | -0.188114615 | 0.134231279 | 1.401421608 | 0.161088034 | 0.345901869 | 0.188114615 |
| ENSMUSG00000014773 | Dll1 | 13388 | protein_coding | 154.8938433 | -0.188128956 | 0.156771565 | 1.200019637 | 0.230131714 | 0.438743376 | 0.188128956 |
| ENSMUSG00000001864 | Aif1l | 108897 | protein_coding | 1.638956795 | 0.188246416 | 1.521612894 | -0.123715051 | 0.901540894 | 1 | 0.188246416 |
| ENSMUSG00000046658 | Zfp316 | 54201 | protein_coding | 124.7983666 | -0.188254171 | 0.124042656 | 1.517656723 | 0.129100959 | 0.296643 | 0.188254171 |
| ENSMUSG00000063838 | Cdc42ep5 | 58804 | protein_coding | 1538.642253 | 0.188353623 | 0.120617997 | -1.561571475 | 0.118388974 | 0.279268705 | 0.188353623 |
| ENSMUSG00000039219 | Arid4b | 94246 | protein_coding | 605.0346967 | -0.188426651 | 0.08131068 | 2.317366555 | 0.020483773 | 0.074722558 | 0.188426651 |
| ENSMUSG00000042569 | Dhrs7b | 216820 | protein_coding | 515.7550502 | -0.188452186 | 0.066246496 | 2.844711755 | 0.004445163 | 0.022069064 | 0.188452186 |
| ENSMUSG00000028804 | Csmd2 | 329942 | protein_coding | 7.580897611 | 0.188477542 | 2.293762919 | -0.082169583 | 0.934511861 | 0.969075596 | 0.188477542 |
| ENSMUSG00000031561 | Tenm3 | 23965 | protein_coding | 5.592257253 | -0.188518534 | 1.010978684 | 0.186471325 | 0.852075164 | 0.927743965 | 0.188518534 |
| ENSMUSG00000036968 | Cnpy4 | 66455 | protein_coding | 130.8694483 | -0.18857758 | 0.120150191 | 1.569515448 | 0.116527883 | 0.275891248 | 0.18857758 |
| ENSMUSG00000059534 | Uqcr10 | 66152 | protein_coding | 2293.749251 | -0.188600188 | 0.089702442 | 2.10250896 | 0.035508716 | 0.114725166 | 0.188600188 |
| ENSMUSG00000030199 | Etv6 | 14011 | protein_coding | 469.2241528 | 0.188609481 | 0.095531113 | -1.974325175 | 0.04834478 | 0.145350463 | 0.188609481 |
| ENSMUSG00000024697 | Gna14 | 14675 | protein_coding | 1063.559601 | 0.188691319 | 0.190579342 | -0.990093247 | 0.322128544 | 0.54337393 | 0.188691319 |
| ENSMUSG00000015165 | Hnrnpl | 15388 | protein_coding | 2599.261171 | 0.188758882 | 0.054735807 | -3.448544805 | 0.000563616 | 0.003917321 | 0.188758882 |
| ENSMUSG00000034459 | Ifit1 | 15957 | protein_coding | 217.7121121 | 0.188761527 | 0.403239312 | -0.468112907 | 0.63970385 | 0.803534599 | 0.188761527 |
| ENSMUSG00000033416 | Gucd1 | 68778 | protein_coding | 1917.921619 | -0.188828874 | 0.089519094 | 2.109369812 | 0.034912673 | 0.113248455 | 0.188828874 |
| ENSMUSG00000047648 | Fbxo30 | 71865 | protein_coding | 177.5646882 | -0.188832511 | 0.121186864 | 1.558192894 | 0.119187529 | 0.280659184 | 0.188832511 |
| ENSMUSG00000067610 | Klri1 | 503550 | protein_coding | 0.857917052 | 0.188851216 | 1.192080529 | -0.158421526 | 0.874124653 | 1 | 0.188851216 |

| ensembl_gene_id | external_gene_name | entrezgene_id | gene_biotype | baseMean | log2FoldChange | lfcSE | stat | pvalue | padj | absLog2FC |
|---|---|---|---|---|---|---|---|---|---|---|
| ENSMUSG00000059493 | Nhs | 195727 | protein_coding | 18.62653022 | -0.188858939 | 0.337158463 | 0.560148889 | 0.575377886 | 0.760569151 | 0.188858939 |
| ENSMUSG00000033793 | Atp6v1h | 108664 | protein_coding | 609.229472 | -0.188864273 | 0.116457955 | 1.621737841 | 0.104859493 | 0.256047562 | 0.188864273 |
| ENSMUSG00000034640 | Tiparp | 99929 | protein_coding | 255.2296017 | -0.188942668 | 0.1341922 | 1.408000372 | 0.15913096 | 0.343290486 | 0.188942668 |
| ENSMUSG00000031729 | Ist1 | 71955 | protein_coding | 1570.242977 | -0.188976946 | 0.054249065 | 3.483506052 | 0.000494892 | 0.003519658 | 0.188976946 |
| ENSMUSG00000022831 | Hcls1 | 15163 | protein_coding | 19.4352479 | -0.188991813 | 0.359969267 | 0.525022078 | 0.599567843 | 0.776544903 | 0.188991813 |
| ENSMUSG00000034656 | Cacna1a | 12286 | protein_coding | 82.03336297 | -0.189037328 | 0.219563935 | 0.860967117 | 0.389256154 | 0.611857618 | 0.189037328 |
| ENSMUSG00000039103 | Nexn | 68810 | protein_coding | 13.66460469 | 0.189045893 | 0.309094166 | -0.611612622 | 0.540794085 | 0.734913884 | 0.189045893 |
| ENSMUSG00000056952 | Tatdn2 | 381801 | protein_coding | 723.4105804 | 0.189079595 | 0.063360235 | -2.984199698 | 0.002843211 | 0.015214774 | 0.189079595 |
| ENSMUSG00000019979 | Apaf1 | 11783 | protein_coding | 559.676502 | 0.189144672 | 0.10192296 | -1.855761171 | 0.063487607 | 0.177826758 | 0.189144672 |
| ENSMUSG00000029625 | Cpsf4 | 54188 | protein_coding | 194.3935628 | 0.189225734 | 0.09124196 | -2.073889409 | 0.038089583 | 0.121313669 | 0.189225734 |
| ENSMUSG00000020402 | Vdac1 | 22333 | protein_coding | 3343.433747 | -0.189324535 | 0.082259117 | 2.30156294 | 0.021359832 | 0.077273602 | 0.189324535 |
| ENSMUSG00000002083 | Bbc3 | 170770 | protein_coding | 90.00110663 | -0.189342567 | 0.216493564 | 0.874587509 | 0.381798387 | 0.604102573 | 0.189342567 |
| ENSMUSG00000000682 | Cd52 | 23833 | protein_coding | 21.02868964 | -0.189347919 | 0.366581148 | 0.516523886 | 0.605488561 | 0.780450414 | 0.189347919 |
| ENSMUSG00000045731 | Pnoc | 18155 | protein_coding | 1.042114838 | 0.189415146 | 1.804789568 | -0.104951375 | 0.916414393 | 1 | 0.189415146 |
| ENSMUSG00000070533 | Wfdc8 | 277343 | protein_coding | 2.080660926 | -0.189469803 | 1.148962646 | 0.164905103 | 0.869018676 | 1 | 0.189469803 |
| ENSMUSG00000029998 | Pcyox1 | 66881 | protein_coding | 946.4617921 | -0.189487827 | 0.094179799 | 2.011979523 | 0.044222096 | 0.135740462 | 0.189487827 |
| ENSMUSG00000058446 | Znrf2 | 387524 | protein_coding | 962.0838925 | -0.189580124 | 0.070595156 | 2.685455139 | 0.00724311 | 0.032806559 | 0.189580124 |
| ENSMUSG00000002728 | Naa20 | 67877 | protein_coding | 433.3014968 | -0.189585912 | 0.083379775 | 2.273763777 | 0.022980193 | 0.081770074 | 0.189585912 |
| ENSMUSG00000086541 | Has2os | 594843 | lncRNA | 7.161959022 | -0.189594184 | 0.42309136 | 0.448116417 | 0.65406918 | 0.81231093 | 0.189594184 |
| ENSMUSG00000054499 | Dedd2 | 67379 | protein_coding | 570.1075613 | -0.189698026 | 0.153526528 | 1.235604222 | 0.216605715 | 0.422068056 | 0.189698026 |
| ENSMUSG00000020823 | Sec14l1 | 74136 | protein_coding | 415.1477947 | -0.189703806 | 0.090406793 | 2.098335758 | 0.035875496 | 0.115554504 | 0.189703806 |
| ENSMUSG00000048280 | Zfp738 | 408068 | protein_coding | 135.0589432 | -0.189716893 | 0.098379829 | 1.928412505 | 0.053803838 | 0.157622241 | 0.189716893 |
| ENSMUSG00000005534 | Insr | 16337 | protein_coding | 1538.706535 | -0.189775812 | 0.074044684 | 2.562990349 | 0.010377493 | 0.043863303 | 0.189775812 |
| ENSMUSG00000024045 | Akap8 | 56399 | protein_coding | 871.8097951 | 0.189804282 | 0.049813337 | -3.810310494 | 0.000138792 | 0.001150611 | 0.189804282 |
| ENSMUSG00000033579 | Fa2h | 338521 | protein_coding | 1092.551901 | 0.189889826 | 0.210573025 | -0.9017766 | 0.367175553 | 0.589852079 | 0.189889826 |
| ENSMUSG00000059474 | Mbtd1 | 103537 | protein_coding | 280.7835957 | -0.189937069 | 0.082216839 | 2.310196674 | 0.020877268 | 0.075856371 | 0.189937069 |
| ENSMUSG00000031171 | Ftsj1 | 54632 | protein_coding | 166.0142181 | 0.189937435 | 0.093514517 | -2.031101048 | 0.042244745 | 0.131182723 | 0.189937435 |
| ENSMUSG00000022843 | Clcn2 | 12724 | protein_coding | 3105.045655 | -0.189943672 | 0.171857371 | 1.105240179 | 0.269055531 | 0.484112338 | 0.189943672 |
| ENSMUSG00000029710 | Ephb4 | 13846 | protein_coding | 1331.420506 | -0.189944455 | 0.10182123 | 1.865470043 | 0.062115547 | 0.17492772 | 0.189944455 |
| ENSMUSG00000010054 | Tusc2 | 80385 | protein_coding | 423.6079197 | 0.189947267 | 0.082266444 | -2.308927633 | 0.020947597 | 0.07607843 | 0.189947267 |
| ENSMUSG00000045316 | Fahd1 | 68636 | protein_coding | 539.5082848 | -0.190106298 | 0.106299873 | 1.788396292 | 0.073712094 | 0.198286614 | 0.190106298 |
| ENSMUSG00000022110 | Sucla2 | 20916 | protein_coding | 1141.389884 | -0.190163996 | 0.069266211 | 2.745407812 | 0.006043574 | 0.028321899 | 0.190163996 |
| ENSMUSG00000020744 | Slc25a19 | 67283 | protein_coding | 202.6596348 | 0.190168097 | 0.092154772 | -2.063572973 | 0.039058222 | 0.12365941 | 0.190168097 |
| ENSMUSG00000015697 | Setdb1 | 84505 | protein_coding | 468.7945429 | -0.19019734 | 0.084449308 | 2.252207203 | 0.024309181 | 0.08552233 | 0.19019734 |
| ENSMUSG00000024665 | Fads2 | 56473 | protein_coding | 2791.062787 | -0.190202777 | 0.229778332 | 0.827766376 | 0.407802821 | 0.628507412 | 0.190202777 |
| ENSMUSG00000024007 | Ppil1 | 68816 | protein_coding | 351.6669474 | 0.190254354 | 0.128927436 | -1.47567003 | 0.140032495 | 0.313984606 | 0.190254354 |
| ENSMUSG00000017428 | Psmd11 | 69077 | protein_coding | 606.4387418 | 0.1902565 | 0.073091408 | -2.602994046 | 0.009241356 | 0.039961181 | 0.1902565 |
| ENSMUSG00000073433 | Arhgdig | 14570 | protein_coding | 14.42961814 | -0.190309147 | 0.269473816 | 0.706225005 | 0.480048223 | 0.691115487 | 0.190309147 |
| ENSMUSG00000002332 | Dhrs1 | 52585 | protein_coding | 370.8718201 | -0.190312078 | 0.090498963 | 2.102919981 | 0.035472765 | 0.114659863 | 0.190312078 |

| ensembl_gene_id | external_gene_name | entrezgene_id | gene_biotype | baseMean | log2FoldChange | lfcSE | stat | pvalue | padj | absLog2FC |
|---|---|---|---|---|---|---|---|---|---|---|
| ENSMUSG00000043668 | Tox3 | 244579 | protein_coding | 189.0585882 | -0.190359934 | 0.088635164 | 2.14767959 | 0.031739218 | 0.105418633 | 0.190359934 |
| ENSMUSG00000061894 | Zscan20 | 269585 | protein_coding | 93.18035012 | -0.190382776 | 0.120212052 | 1.583724534 | 0.113256419 | 0.270127107 | 0.190382776 |
| ENSMUSG00000094018 | S100a2 | 628324 | protein_coding | 3.870576199 | 0.190383578 | 1.669073507 | -0.114065425 | 0.90918593 | 1 | 0.190383578 |
| ENSMUSG00000022614 | Lmf2 | 105847 | protein_coding | 242.494835 | 0.190394311 | 0.102009294 | -1.866440825 | 0.061979716 | 0.174694151 | 0.190394311 |
| ENSMUSG00000084128 | Esrp2 | 77411 | protein_coding | 884.4901349 | -0.19040145 | 0.065175005 | 2.921387594 | 0.00348476 | 0.018045571 | 0.19040145 |
| ENSMUSG00000052819 | Best2 | 212989 | protein_coding | 341.6619233 | 0.190428732 | 0.515698106 | -0.369263975 | 0.711930975 | 0.850157504 | 0.190428732 |
| ENSMUSG00000036009 | Mettl25 | 216292 | protein_coding | 58.43224581 | 0.190499946 | 0.160684409 | -1.185553393 | 0.235798717 | 0.445943657 | 0.190499946 |
| ENSMUSG00000036875 | Dna2 | 327762 | protein_coding | 72.24268012 | 0.190515312 | 0.18348814 | -1.03829769 | 0.299131484 | 0.51911069 | 0.190515312 |
| ENSMUSG00000085637 | 1700048M11Rik | 100503166 | lncRNA | 1.216277845 | -0.190532002 | 1.056903872 | 0.180273728 | 0.856937679 | 1 | 0.190532002 |
| ENSMUSG00000013921 | Clip3 | 76686 | protein_coding | 4.350059724 | 0.190537368 | 1.03876095 | -0.183427542 | 0.854462567 | 1 | 0.190537368 |
| ENSMUSG00000046101 | Mcmdc2 | 240697 | protein_coding | 4.659237809 | -0.190560289 | 0.745644402 | 0.255564567 | 0.798287081 | 0.899929685 | 0.190560289 |
| ENSMUSG00000001755 | Coasy | 71743 | protein_coding | 394.6218828 | -0.190562338 | 0.100243494 | 1.900994577 | 0.057302723 | 0.165147848 | 0.190562338 |
| ENSMUSG00000052920 | Prkg1 | 19091 | protein_coding | 3.586128684 | -0.190699139 | 1.230130667 | 0.155023482 | 0.876802805 | 1 | 0.190699139 |
| ENSMUSG00000020777 | Acox1 | 11430 | protein_coding | 3744.49281 | -0.190702452 | 0.125620882 | 1.518079237 | 0.128994425 | 0.296522686 | 0.190702452 |
| ENSMUSG00000026594 | Ralgps2 | 78255 | protein_coding | 744.9811165 | 0.190710583 | 0.053488712 | -3.565436085 | 0.000363252 | 0.002700918 | 0.190710583 |
| ENSMUSG00000028282 | Casp8ap2 | 26885 | protein_coding | 350.3451636 | 0.190731283 | 0.077917975 | -2.447847028 | 0.014371266 | 0.056385621 | 0.190731283 |
| ENSMUSG00000025781 | Atp5c1 | 11949 | protein_coding | 4427.719366 | -0.190761343 | 0.092095713 | 2.071337918 | 0.038327228 | 0.12190868 | 0.190761343 |
| ENSMUSG00000020641 | Rsad2 | 58185 | protein_coding | 87.61965689 | -0.190771432 | 0.330945787 | 0.576443149 | 0.564315686 | 0.751419871 | 0.190771432 |
| ENSMUSG00000025815 | Dhtkd1 | 209692 | protein_coding | 3.520276031 | -0.190784915 | 1.291852297 | 0.147683227 | 0.88259277 | 1 | 0.190784915 |
| ENSMUSG00000020183 | Cpm | 70574 | protein_coding | 638.5859354 | -0.190815389 | 0.119530264 | 1.596377205 | 0.110404602 | 0.265659566 | 0.190815389 |
| ENSMUSG00000031986 | Sprtn | 244666 | protein_coding | 221.3593752 | -0.190867477 | 0.090715188 | 2.104030011 | 0.035375829 | 0.114480981 | 0.190867477 |
| ENSMUSG00000021114 | Atp6v1d | 73834 | protein_coding | 438.6902015 | 0.190984481 | 0.063100362 | -3.026678048 | 0.002472572 | 0.013498197 | 0.190984481 |
| ENSMUSG00000025477 | Inpp5a | 212111 | protein_coding | 400.2037013 | -0.191026144 | 0.089868572 | 2.125616766 | 0.033535182 | 0.109773894 | 0.191026144 |
| ENSMUSG00000097534 | Gm16675 | 100503498 | lncRNA | 16.92282244 | -0.191090598 | 0.281441487 | 0.678970965 | 0.497156259 | 0.702914252 | 0.191090598 |
| ENSMUSG00000004270 | Lpcat3 | 14792 | protein_coding | 897.3946128 | 0.191096024 | 0.071953325 | -2.655833111 | 0.007911276 | 0.035329044 | 0.191096024 |
| ENSMUSG00000016128 | Stard13 | 243362 | protein_coding | 352.2275489 | 0.191110107 | 0.071181961 | -2.684810944 | 0.007257084 | 0.032851834 | 0.191110107 |
| ENSMUSG00000028057 | Rit1 | 19769 | protein_coding | 397.7155226 | -0.191267786 | 0.066263168 | 2.88648718 | 0.003895686 | 0.019776625 | 0.191267786 |
| ENSMUSG00000006678 | Pola1 | 18968 | protein_coding | 255.8829336 | 0.191282757 | 0.176984891 | -1.0807858 | 0.279792406 | 0.49722072 | 0.191282757 |
| ENSMUSG00000016477 | E2f3 | 13557 | protein_coding | 224.892631 | 0.191383164 | 0.090278825 | -2.119911986 | 0.034013468 | 0.110941817 | 0.191383164 |
| ENSMUSG00000001823 | Hoxd12 | 15432 | protein_coding | 10.22649461 | -0.191406363 | 0.481016998 | 0.397920165 | 0.690689035 | 0.835178936 | 0.191406363 |
| ENSMUSG00000000085 | Scmh1 | 29871 | protein_coding | 298.5073388 | 0.191411215 | 0.075486479 | -2.535701994 | 0.011222219 | 0.046704567 | 0.191411215 |
| ENSMUSG00000050122 | Vwa3b | 70853 | protein_coding | 26.85601329 | -0.191426123 | 0.310639202 | 0.616232986 | 0.537740761 | 0.733113936 | 0.191426123 |
| ENSMUSG00000036986 | Pml | 18854 | protein_coding | 372.188564 | -0.191435803 | 0.10572552 | 1.810686789 | 0.070189349 | 0.191366503 | 0.191435803 |
| ENSMUSG00000032872 | Cyb5r4 | 266690 | protein_coding | 748.5624766 | -0.191438113 | 0.071238893 | 2.68726963 | 0.007203878 | 0.032709605 | 0.191438113 |
| ENSMUSG00000030282 | Cmas | 12764 | protein_coding | 2483.538607 | 0.191449857 | 0.119857228 | -1.597315915 | 0.110195305 | 0.265233241 | 0.191449857 |
| ENSMUSG00000031295 | Phka2 | 110094 | protein_coding | 197.6154133 | 0.191468665 | 0.108348921 | -1.767148786 | 0.077203314 | 0.204907671 | 0.191468665 |
| ENSMUSG00000024873 | Cnih2 | 12794 | protein_coding | 13.44018519 | 0.191557042 | 0.319808978 | -0.59897331 | 0.549190683 | 0.740233019 | 0.191557042 |
| ENSMUSG00000073490 | Ifi207 | 226691 | protein_coding | 1.292580119 | 0.191583003 | 1.476612139 | -0.129744974 | 0.896768199 | 1 | 0.191583003 |
| ENSMUSG00000097333 | Zfp87 | 170763 | protein_coding | 104.7743611 | -0.191598915 | 0.14563136 | 1.315643245 | 0.188293812 | 0.384741896 | 0.191598915 |

| ensembl_gene_id | external_gene_name | entrezgene_id | gene_biotype | baseMean | log2FoldChange | lfcSE | stat | pvalue | padj | absLog2FC |
|---|---|---|---|---|---|---|---|---|---|---|
| ENSMUSG00000028917 | Plekhm2 | 69582 | protein_coding | 238.7318635 | -0.191657862 | 0.097175387 | 1.972288141 | 0.048576723 | 0.145821162 | 0.191657862 |
| ENSMUSG00000035674 | Ndufa3 | 66091 | protein_coding | 1093.480778 | -0.191737499 | 0.071628408 | 2.676835982 | 0.007432101 | 0.033524642 | 0.191737499 |
| ENSMUSG00000032207 | Lipc | 15450 | protein_coding | 6.567225643 | 0.191752716 | 0.528274303 | -0.362979451 | 0.716620235 | 0.852920113 | 0.191752716 |
| ENSMUSG00000051331 | Cacna1c | 12288 | protein_coding | 39.91222212 | 0.191757959 | 0.343298723 | -0.558574636 | 0.576452054 | 0.761222618 | 0.191757959 |
| ENSMUSG00000037174 | Elf2 | 69257 | protein_coding | 706.4038061 | -0.19177291 | 0.069792962 | 2.747739967 | 0.006000758 | 0.028168424 | 0.19177291 |
| ENSMUSG00000030149 | Klrk1 | 27007 | protein_coding | 5.244775589 | -0.19180404 | 0.759847258 | 0.252424468 | 0.80071299 | 0.901300137 | 0.19180404 |
| ENSMUSG00000028974 | Dffa | 13347 | protein_coding | 264.6029564 | -0.191814187 | 0.075501326 | 2.540540637 | 0.011068123 | 0.046167968 | 0.191814187 |
| ENSMUSG00000049971 | Glt1d1 | 319804 | protein_coding | 210.6982253 | -0.191863439 | 0.135244176 | 1.418644745 | 0.156002614 | 0.338620816 | 0.191863439 |
| ENSMUSG00000021687 | Scamp1 | 107767 | protein_coding | 1374.936727 | -0.191938781 | 0.165419145 | 1.16031782 | 0.245919432 | 0.457693395 | 0.191938781 |
| ENSMUSG00000033885 | Pxk | 218699 | protein_coding | 624.4310511 | -0.191946335 | 0.070040153 | 2.7405185 | 0.006134233 | 0.028656498 | 0.191946335 |
| ENSMUSG00000053768 | Chchd3 | 66075 | protein_coding | 1832.004477 | -0.191972113 | 0.054459614 | 3.525036238 | 0.000423425 | 0.003084446 | 0.191972113 |
| ENSMUSG00000028064 | Sema4a | 20351 | protein_coding | 2099.128769 | 0.191979675 | 0.113214693 | -1.695713424 | 0.089940165 | 0.228714488 | 0.191979675 |
| ENSMUSG00000008604 | Ubqln4 | 94232 | protein_coding | 562.8310387 | -0.192071241 | 0.066626594 | 2.882801452 | 0.003941558 | 0.01996653 | 0.192071241 |
| ENSMUSG00000038025 | Phf2 | 18676 | protein_coding | 384.9751733 | 0.192078361 | 0.077027209 | -2.493642997 | 0.012643964 | 0.051106386 | 0.192078361 |
| ENSMUSG00000118379 | Gm41790 | 105246509 | lncRNA | 1.497901698 | -0.192081739 | 2.353914626 | 0.081600979 | 0.934964023 | 1 | 0.192081739 |
| ENSMUSG00000025583 | Rptor | 74370 | protein_coding | 660.771738 | 0.192110089 | 0.076564349 | -2.509132407 | 0.012102811 | 0.049435028 | 0.192110089 |
| ENSMUSG00000034422 | Parp14 | 547253 | protein_coding | 759.6205408 | -0.192124859 | 0.187009877 | 1.027351401 | 0.304255027 | 0.524504007 | 0.192124859 |
| ENSMUSG00000043923 | Ccdc84 | 382073 | protein_coding | 119.4097834 | 0.192134521 | 0.150943734 | -1.272888354 | 0.203057662 | 0.405231932 | 0.192134521 |
| ENSMUSG00000104986 | 1600023N17Rik | 69788 | TEC | 5.529444828 | 0.192147661 | 0.606386796 | -0.316873095 | 0.751339898 | 0.875081807 | 0.192147661 |
| ENSMUSG00000062168 | Ppef1 | 237178 | protein_coding | 1.923626493 | 0.192174345 | 1.086906263 | -0.176808573 | 0.859658755 | 1 | 0.192174345 |
| ENSMUSG00000047507 | Baiap3 | 545192 | protein_coding | 35.65977367 | -0.192223821 | 0.225162165 | 0.853712798 | 0.393264141 | 0.615929821 | 0.192223821 |
| ENSMUSG00000047656 | Trpt1 | 107328 | protein_coding | 87.904714 | -0.192234852 | 0.13199558 | 1.456373407 | 0.145289423 | 0.3224445 | 0.192234852 |
| ENSMUSG00000020646 | Mboat2 | 67216 | protein_coding | 105.7056611 | 0.192281998 | 0.195518895 | -0.983444586 | 0.325388676 | 0.547140677 | 0.192281998 |
| ENSMUSG00000065738 | Gm24494 | 115489817 | snoRNA | 7.866320771 | -0.192317492 | 0.366473279 | 0.52477903 | 0.599736811 | 0.776544903 | 0.192317492 |
| ENSMUSG00000018595 | Glra4 | 14657 | protein_coding | 1.553709676 | 0.192332033 | 1.169090136 | -0.16451429 | 0.869326298 | 1 | 0.192332033 |
| ENSMUSG00000047227 | Gm527 | 217648 | protein_coding | 16.81453139 | -0.192365418 | 0.290794053 | 0.661517715 | 0.508280357 | 0.71205818 | 0.192365418 |
| ENSMUSG00000021108 | Prkch | 18755 | protein_coding | 120.2662438 | 0.192365602 | 0.142588648 | -1.349094792 | 0.177306521 | 0.36956202 | 0.192365602 |
| ENSMUSG00000052488 | Cherp | 27967 | protein_coding | 199.4633712 | 0.192384422 | 0.150886998 | -1.275023194 | 0.202301032 | 0.404218578 | 0.192384422 |
| ENSMUSG00000029632 | Ndufa4 | 17992 | protein_coding | 1763.759497 | 0.192418305 | 0.078860644 | -2.439978867 | 0.014688121 | 0.0573834 | 0.192418305 |
| ENSMUSG00000031783 | Polr2c | 20021 | protein_coding | 398.5322317 | -0.192422867 | 0.078224365 | 2.459884053 | 0.013898191 | 0.054947264 | 0.192422867 |
| ENSMUSG00000032065 | Tex12 | 66654 | protein_coding | 17.48556049 | -0.192424504 | 0.336386056 | 0.572034723 | 0.567298451 | 0.753386941 | 0.192424504 |
| ENSMUSG00000030603 | Psmc4 | 23996 | protein_coding | 999.4915012 | 0.192471022 | 0.064116898 | -3.001876707 | 0.002683208 | 0.014480556 | 0.192471022 |
| ENSMUSG00000034117 | Ptgdr2 | 14764 | protein_coding | 1.88932073 | 0.1925089 | 0.924521621 | -0.208225417 | 0.835052964 | 1 | 0.1925089 |
| ENSMUSG00000030157 | Clec2d | 93694 | protein_coding | 1093.154734 | -0.192533784 | 0.166777723 | 1.154433461 | 0.248322503 | 0.460660281 | 0.192533784 |
| ENSMUSG00000019143 | Hars2 | 70791 | protein_coding | 284.2598119 | -0.192603783 | 0.087082921 | 2.211728552 | 0.026985426 | 0.092763808 | 0.192603783 |
| ENSMUSG00000033032 | Afap1l1 | 106877 | protein_coding | 183.5250928 | 0.192613384 | 0.104605161 | -1.841337302 | 0.065572145 | 0.1820236 | 0.192613384 |
| ENSMUSG00000003644 | Rps6ka1 | 20111 | protein_coding | 3461.438561 | 0.192634951 | 0.059175914 | -3.255293226 | 0.001132753 | 0.007066978 | 0.192634951 |
| ENSMUSG00000049721 | Gal3st1 | 53897 | protein_coding | 569.4297558 | -0.192647031 | 0.106308589 | 1.812149267 | 0.069963136 | 0.190901365 | 0.192647031 |
| ENSMUSG00000021177 | Tdp1 | 104884 | protein_coding | 64.55294037 | 0.192686022 | 0.164379743 | -1.172200532 | 0.241116555 | 0.451758428 | 0.192686022 |

| ensembl_gene_id | external_gene_name | entrezgene_id | gene_biotype | baseMean | log2FoldChange | lfcSE | stat | pvalue | padj | absLog2FC |
|---|---|---|---|---|---|---|---|---|---|---|
| ENSMUSG00000026495 | Efcab2 | 68226 | protein_coding | 38.94513098 | -0.192699685 | 0.195728122 | 0.984527326 | 0.324856302 | 0.546769252 | 0.192699685 |
| ENSMUSG00000022594 | Lynx1 | 23936 | protein_coding | 104.1813425 | -0.192730023 | 0.154117127 | 1.250542535 | 0.211101427 | 0.415328565 | 0.192730023 |
| ENSMUSG00000027184 | Caprin1 | 53872 | protein_coding | 4318.415462 | 0.192759987 | 0.072497283 | -2.658858102 | 0.007840598 | 0.035051335 | 0.192759987 |
| ENSMUSG00000047181 | Samd14 | 217125 | protein_coding | 29.98517964 | -0.192843225 | 0.296143699 | 0.651181255 | 0.514929489 | 0.716272317 | 0.192843225 |
| ENSMUSG00000005621 | Zfp592 | 233410 | protein_coding | 529.6606298 | -0.192936324 | 0.072996022 | 2.64310738 | 0.008214898 | 0.036352985 | 0.192936324 |
| ENSMUSG00000042359 | Osbpl6 | 99031 | protein_coding | 167.2964436 | -0.192960778 | 0.110690831 | 1.743240853 | 0.081291552 | 0.212177761 | 0.192960778 |
| ENSMUSG00000104301 | Wdr49 | 213248 | protein_coding | 1.8590389 | 0.192962241 | 1.602042445 | -0.120447646 | 0.90412855 | 1 | 0.192962241 |
| ENSMUSG00000042498 | Radx | 102871 | protein_coding | 62.27479171 | -0.193004684 | 0.168189794 | 1.147540995 | 0.2511581 | 0.463479905 | 0.193004684 |
| ENSMUSG00000025006 | Sorbs1 | 20411 | protein_coding | 209.8932557 | 0.193045869 | 0.265413004 | -0.727341411 | 0.467016836 | 0.680659303 | 0.193045869 |
| ENSMUSG00000018820 | Zfyve27 | 319740 | protein_coding | 366.1668607 | -0.193054713 | 0.060598969 | 3.185775544 | 0.001443666 | 0.008606748 | 0.193054713 |
| ENSMUSG00000022228 | Zscan26 | 432731 | protein_coding | 414.0563171 | -0.193072506 | 0.084277391 | 2.290916975 | 0.021968216 | 0.079002436 | 0.193072506 |
| ENSMUSG00000024208 | Uqcc2 | 67267 | protein_coding | 348.4102897 | -0.193097982 | 0.103341817 | 1.868536736 | 0.061687295 | 0.174107671 | 0.193097982 |
| ENSMUSG00000018486 | Wnt9b | 22412 | protein_coding | 1.359347322 | -0.193115726 | 2.346964842 | 0.082283178 | 0.934421531 | 1 | 0.193115726 |
| ENSMUSG00000040653 | Ppp1r14c | 76142 | protein_coding | 18.84796126 | -0.193120914 | 0.291138909 | 0.663329111 | 0.507119796 | 0.710854454 | 0.193120914 |
| ENSMUSG00000078619 | Smarcd2 | 83796 | protein_coding | 1069.429312 | -0.193132644 | 0.081406468 | 2.372448394 | 0.017670631 | 0.066608721 | 0.193132644 |
| ENSMUSG00000109511 | Nup62 | 18226 | protein_coding | 372.9055563 | 0.193184584 | 0.072336609 | -2.670633683 | 0.007570822 | 0.034020286 | 0.193184584 |
| ENSMUSG00000029911 | Ssbp1 | 381760 | protein_coding | 224.9552479 | 0.193253449 | 0.112552641 | -1.717005007 | 0.085978257 | 0.22143558 | 0.193253449 |
| ENSMUSG00000099481 | Xndc1 | 102443350 | protein_coding | 34.74330764 | -0.193320809 | 0.165031989 | 1.171414165 | 0.241432341 | 0.452093626 | 0.193320809 |
| ENSMUSG00000025083 | Afap1l2 | 226250 | protein_coding | 34.87886495 | -0.193325161 | 0.348242089 | 0.55514588 | 0.578794883 | 0.762380457 | 0.193325161 |
| ENSMUSG00000038383 | Pigu | 228812 | protein_coding | 326.6759704 | 0.193375873 | 0.084080992 | -2.299876201 | 0.021455235 | 0.077538047 | 0.193375873 |
| ENSMUSG00000034427 | Myo15b | 217328 | protein_coding | 8292.18235 | 0.193388031 | 0.227938809 | -0.848420819 | 0.396203653 | 0.618597629 | 0.193388031 |
| ENSMUSG00000020528 | Prpsap2 | 212627 | protein_coding | 234.8889403 | -0.193409315 | 0.112009464 | 1.726722978 | 0.084217427 | 0.21805685 | 0.193409315 |
| ENSMUSG00000056131 | Pgm3 | 109785 | protein_coding | 578.962675 | 0.193415723 | 0.111660101 | -1.732182951 | 0.083240993 | 0.216206632 | 0.193415723 |
| ENSMUSG00000022619 | Mapk8ip2 | 60597 | protein_coding | 8.627449759 | 0.193441616 | 0.620983165 | -0.311508631 | 0.755413982 | 0.877032235 | 0.193441616 |
| ENSMUSG00000029009 | Mthfr | 17769 | protein_coding | 246.2972323 | -0.193450985 | 0.089753218 | 2.15536545 | 0.031133248 | 0.103865546 | 0.193450985 |
| ENSMUSG00000036555 | Iqce | 74239 | protein_coding | 136.8278952 | 0.193457944 | 0.11001291 | -1.758502204 | 0.078662097 | 0.207566563 | 0.193457944 |
| ENSMUSG00000018286 | Psmb6 | 19175 | protein_coding | 1358.8076 | 0.193478242 | 0.070183504 | -2.756748125 | 0.00583793 | 0.027610883 | 0.193478242 |
| ENSMUSG00000034528 | Hsd17b13 | 243168 | protein_coding | 197.3560681 | 0.193518789 | 0.16138439 | -1.199117143 | 0.230482399 | 0.43912271 | 0.193518789 |
| ENSMUSG00000008136 | Fhl2 | 14200 | protein_coding | 168.2223578 | -0.193584625 | 0.104636473 | 1.850068335 | 0.064303701 | 0.179528541 | 0.193584625 |
| ENSMUSG00000000303 | Cdh1 | 12550 | protein_coding | 11260.16188 | -0.193595812 | 0.080099678 | 2.416936193 | 0.015651761 | 0.060376822 | 0.193595812 |
| ENSMUSG00000037552 | Plekhg2 | 101497 | protein_coding | 175.9535712 | 0.19364533 | 0.247837408 | -0.781340199 | 0.434602433 | 0.652884901 | 0.19364533 |
| ENSMUSG00000078606 | Gvin2 | 100042856 | protein_coding | 4.896840877 | -0.193682646 | 0.550244674 | 0.351993677 | 0.724843004 | 0.857515393 | 0.193682646 |
| ENSMUSG00000047495 | Dlgap2 | 244310 | protein_coding | 3.060931308 | 0.193720031 | 1.429644488 | -0.135502241 | 0.892214793 | 1 | 0.193720031 |
| ENSMUSG00000034800 | Zfp661 | 72180 | protein_coding | 56.20341954 | -0.193876613 | 0.142473134 | 1.360794184 | 0.17357874 | 0.364391266 | 0.193876613 |
| ENSMUSG00000073643 | Wdfy1 | 69368 | protein_coding | 794.0306957 | -0.19389187 | 0.070550167 | 2.748283651 | 0.005990816 | 0.02813775 | 0.19389187 |
| ENSMUSG00000020390 | Ube2b | 22210 | protein_coding | 613.771816 | -0.193907755 | 0.073990478 | 2.620712289 | 0.008774628 | 0.038324307 | 0.193907755 |
| ENSMUSG00000035199 | Arl6ip5 | 65106 | protein_coding | 378.5860503 | -0.193922194 | 0.070045751 | 2.768507597 | 0.005631367 | 0.026854288 | 0.193922194 |
| ENSMUSG00000009291 | Pttg1ip | 108705 | protein_coding | 2348.08714 | -0.193923967 | 0.086848984 | 2.232887 | 0.025556397 | 0.088794097 | 0.193923967 |
| ENSMUSG00000037750 | Fam222b | 216971 | protein_coding | 101.658057 | 0.193948209 | 0.134700635 | -1.439846278 | 0.149910895 | 0.329600388 | 0.193948209 |

| ensembl_gene_id | external_gene_name | entrezgene_id | gene_biotype | baseMean | log2FoldChange | lfcSE | stat | pvalue | padj | absLog2FC |
|---|---|---|---|---|---|---|---|---|---|---|
| ENSMUSG00000031889 | Phaf1 | 234678 | protein_coding | 177.7361938 | -0.19396128 | 0.096217114 | 2.015870891 | 0.043813478 | 0.134811958 | 0.19396128 |
| ENSMUSG00000038894 | Irs2 | 384783 | protein_coding | 908.2452513 | 0.193978431 | 0.205954907 | -0.941849036 | 0.346269936 | 0.568024408 | 0.193978431 |
| ENSMUSG00000070304 | Scn2b | 72821 | protein_coding | 33.70106633 | -0.194032456 | 0.248975144 | 0.779324604 | 0.435788525 | 0.653940181 | 0.194032456 |
| ENSMUSG00000021109 | Hif1a | 15251 | protein_coding | 1298.655051 | -0.194085906 | 0.170297866 | 1.139684902 | 0.254417599 | 0.467050404 | 0.194085906 |
| ENSMUSG00000064330 | Pde6h | 78600 | protein_coding | 3.002431167 | 0.194109036 | 1.18538007 | -0.163752573 | 0.869925929 | 1 | 0.194109036 |
| ENSMUSG00000026245 | Farsb | 23874 | protein_coding | 693.3992454 | 0.194157525 | 0.104322003 | -1.861136865 | 0.062724851 | 0.176210159 | 0.194157525 |
| ENSMUSG00000038332 | Sesn1 | 140742 | protein_coding | 131.7678533 | -0.19428507 | 0.110074156 | 1.765038022 | 0.077557376 | 0.205616071 | 0.19428507 |
| ENSMUSG00000036181 | H1f2 | 50708 | protein_coding | 7024.998537 | 0.194288914 | 0.092726819 | -2.095282862 | 0.036145855 | 0.116153464 | 0.194288914 |
| ENSMUSG00000057234 | Mettl15 | 76894 | protein_coding | 46.56581644 | -0.194344923 | 0.165382686 | 1.175122546 | 0.239945692 | 0.450382263 | 0.194344923 |
| ENSMUSG00000001924 | Uba1 | 22201 | protein_coding | 4139.393078 | 0.194351526 | 0.070435884 | -2.759268629 | 0.005793089 | 0.027443223 | 0.194351526 |
| ENSMUSG00000018168 | Ikzf3 | 22780 | protein_coding | 17.14691942 | -0.194372978 | 0.429298942 | 0.452768361 | 0.650715548 | 0.81021764 | 0.194372978 |
| ENSMUSG00000044950 | Pwwp2a | 70802 | protein_coding | 243.8137758 | 0.194429103 | 0.083696907 | -2.323014186 | 0.02017839 | 0.073869638 | 0.194429103 |
| ENSMUSG00000062154 | Tex33 | 73376 | protein_coding | 1.878796304 | -0.194459857 | 1.219563996 | 0.15945031 | 0.873314105 | 1 | 0.194459857 |
| ENSMUSG00000025156 | Gps1 | 209318 | protein_coding | 597.5506498 | -0.194503214 | 0.134169466 | 1.449683147 | 0.147146897 | 0.32503797 | 0.194503214 |
| ENSMUSG00000060510 | Zfp266 | 77519 | protein_coding | 621.9637289 | -0.194531265 | 0.134494313 | 1.446390266 | 0.14806777 | 0.326547965 | 0.194531265 |
| ENSMUSG00000015850 | Adamtsl4 | 229595 | protein_coding | 67.07242817 | 0.194568137 | 0.161879798 | -1.201929697 | 0.229390767 | 0.438111017 | 0.194568137 |
| ENSMUSG00000030706 | Mrpl48 | 52443 | protein_coding | 318.1567863 | -0.194570263 | 0.096917491 | 2.007586679 | 0.044687231 | 0.136786828 | 0.194570263 |
| ENSMUSG00000030629 | Zfand6 | 65098 | protein_coding | 898.159949 | 0.19457895 | 0.076676651 | -2.537655836 | 0.011159767 | 0.046491522 | 0.19457895 |
| ENSMUSG00000038742 | Angptl6 | 70726 | protein_coding | 14.15671115 | 0.194579052 | 0.390201396 | -0.498663137 | 0.618016718 | 0.789504764 | 0.194579052 |
| ENSMUSG00000028826 | Maco1 | 66146 | protein_coding | 555.8738028 | 0.194590276 | 0.119216002 | -1.63224964 | 0.102626907 | 0.252289573 | 0.194590276 |
| ENSMUSG00000057858 | Fam204a | 76539 | protein_coding | 162.0763627 | 0.194592319 | 0.10026662 | -1.940748763 | 0.052288759 | 0.154446119 | 0.194592319 |
| ENSMUSG00000028488 | Sh3gl2 | 20404 | protein_coding | 159.573154 | -0.194611615 | 0.149046845 | 1.305707716 | 0.191651973 | 0.3894353 | 0.194611615 |
| ENSMUSG00000031896 | Ctrl | 109660 | protein_coding | 7.224315802 | 0.194619958 | 0.574306904 | -0.338877971 | 0.73470166 | 0.864656729 | 0.194619958 |
| ENSMUSG00000035390 | Brsk1 | 381979 | protein_coding | 18.16368006 | -0.194649664 | 0.314811239 | 0.618305957 | 0.536373676 | 0.732371878 | 0.194649664 |
| ENSMUSG00000022847 | Thpo | 21832 | protein_coding | 13.50957582 | 0.194673451 | 0.318677514 | -0.610879157 | 0.541279584 | 0.735271124 | 0.194673451 |
| ENSMUSG00000026649 | Cfap126 | 75472 | protein_coding | 9.379494128 | -0.19472588 | 0.549294203 | 0.354501975 | 0.722962725 | 0.856549354 | 0.19472588 |
| ENSMUSG00000024039 | Cbs | 12411 | protein_coding | 773.6441896 | -0.194737851 | 0.125303054 | 1.55413491 | 0.12015224 | 0.28232699 | 0.194737851 |
| ENSMUSG00000030793 | Pycard | 66824 | protein_coding | 1925.760933 | 0.194780653 | 0.10391052 | -1.874503683 | 0.060861035 | 0.172631249 | 0.194780653 |
| ENSMUSG00000069565 | Dazap1 | 70248 | protein_coding | 901.1296809 | 0.194790937 | 0.083141687 | -2.342879299 | 0.019135573 | 0.070900795 | 0.194790937 |
| ENSMUSG00000079502 | Cfap77 | 329375 | protein_coding | 4.211051705 | -0.194792546 | 0.587224215 | 0.331717497 | 0.740102588 | 1 | 0.194792546 |
| ENSMUSG00000020872 | Tac4 | 93670 | protein_coding | 2.718418975 | -0.194850801 | 0.771038285 | 0.252712226 | 0.800490599 | 1 | 0.194850801 |
| ENSMUSG00000022016 | Akap11 | 219181 | protein_coding | 797.8371958 | -0.194913311 | 0.079270702 | 2.458831635 | 0.013938997 | 0.055069042 | 0.194913311 |
| ENSMUSG00000045106 | Ccdc73 | 211936 | protein_coding | 11.03500545 | -0.194918901 | 0.357019063 | 0.545962166 | 0.585091944 | 0.767207271 | 0.194918901 |
| ENSMUSG00000025269 | Apex2 | 77622 | protein_coding | 80.45622748 | 0.194958757 | 0.133336569 | -1.462155193 | 0.143698687 | 0.320032382 | 0.194958757 |
| ENSMUSG00000004356 | Utp20 | 70683 | protein_coding | 1602.983358 | 0.194962854 | 0.065097266 | -2.994946888 | 0.002744926 | 0.014765379 | 0.194962854 |
| ENSMUSG00000025893 | Kbtbd3 | 69149 | protein_coding | 47.41092475 | 0.194964953 | 0.150648099 | -1.294174667 | 0.195605095 | 0.394941012 | 0.194964953 |
| ENSMUSG00000078153 | Psme2b | 621823 | protein_coding | 25.58499774 | -0.195164037 | 0.198890939 | 0.981261576 | 0.326463766 | 0.548343875 | 0.195164037 |
| ENSMUSG00000024889 | Rce1 | 19671 | protein_coding | 315.7941362 | -0.195168156 | 0.086258185 | 2.262604476 | 0.023660082 | 0.083756183 | 0.195168156 |
| ENSMUSG00000040229 | Gpr34 | 23890 | protein_coding | 5.042864139 | 0.195184478 | 0.656614153 | -0.297259018 | 0.766268767 | 0.882690208 | 0.195184478 |

Supplementary Table 1

| ensembl_gene_id | external_gene_name | entrezgene_id | gene_biotype | baseMean | log2FoldChange | lfcSE | stat | pvalue | padj | absLog2FC |
|---|---|---|---|---|---|---|---|---|---|---|
| ENSMUSG00000006782 | Cnp | 12799 | protein_coding | 412.7897524 | 0.19520105 | 0.119055735 | -1.639577045 | 0.101093139 | 0.249608668 | 0.19520105 |
| ENSMUSG00000037152 | Ndufc1 | 66377 | protein_coding | 637.7277788 | -0.195205326 | 0.082422357 | 2.368354082 | 0.017867427 | 0.067059702 | 0.195205326 |
| ENSMUSG00000031246 | Sh3bgrl | 56726 | protein_coding | 1425.480609 | -0.195221113 | 0.132705053 | 1.471090273 | 0.141266705 | 0.315936941 | 0.195221113 |
| ENSMUSG00000018363 | Smurf2 | 66313 | protein_coding | 758.3995272 | -0.195260785 | 0.074464017 | 2.622216637 | 0.008735988 | 0.038193919 | 0.195260785 |
| ENSMUSG00000036766 | Dner | 227325 | protein_coding | 4.500829553 | -0.195273738 | 0.798125129 | 0.244665568 | 0.80671539 | 0.904906279 | 0.195273738 |
| ENSMUSG00000024758 | Rtn3 | 20168 | protein_coding | 1578.916992 | -0.195281465 | 0.063763914 | 3.062570234 | 0.00219445 | 0.01219993 | 0.195281465 |
| ENSMUSG00000029723 | Spacdr | 78829 | protein_coding | 940.9655193 | -0.195376404 | 0.105780576 | 1.846996975 | 0.064747579 | 0.180584617 | 0.195376404 |
| ENSMUSG00000096145 | Vkorc1 | 27973 | protein_coding | 97.62373016 | 0.195388126 | 0.168377907 | -1.160414271 | 0.245880179 | 0.457679163 | 0.195388126 |
| ENSMUSG00000054909 | Wbscr25 | 71304 | lncRNA | 15.82449563 | -0.195397004 | 0.492394442 | 0.396830239 | 0.69149265 | 0.835783476 | 0.195397004 |
| ENSMUSG00000018761 | Mpdu1 | 24070 | protein_coding | 486.9664778 | -0.195410741 | 0.08353078 | 2.339386043 | 0.019315463 | 0.071455098 | 0.195410741 |
| ENSMUSG00000002546 | Golga2 | 99412 | protein_coding | 1278.53553 | 0.195443386 | 0.046224763 | -4.228110077 | 2.36E-05 | 0.000238903 | 0.195443386 |
| ENSMUSG00000036822 | Topors | 106021 | protein_coding | 469.7404297 | -0.195504927 | 0.060513117 | 3.230785924 | 0.001234504 | 0.007575843 | 0.195504927 |
| ENSMUSG00000048485 | Zbtb8b | 215627 | protein_coding | 1.009513459 | -0.195537689 | 1.614358968 | 0.121124045 | 0.903592784 | 1 | 0.195537689 |
| ENSMUSG00000073791 | Efcab7 | 230500 | protein_coding | 34.82606105 | 0.195567907 | 0.221545433 | -0.88274402 | 0.377374592 | 0.599541641 | 0.195567907 |
| ENSMUSG00000027804 | Ppid | 67738 | protein_coding | 302.1327699 | -0.195615964 | 0.106014953 | 1.845173336 | 0.065012328 | 0.181090121 | 0.195615964 |
| ENSMUSG00000035048 | Anapc13 | 69010 | protein_coding | 382.3010268 | -0.195633979 | 0.094031319 | 2.080519348 | 0.037477923 | 0.11971188 | 0.195633979 |
| ENSMUSG00000007646 | Rad51c | 114714 | protein_coding | 23.02806926 | 0.195677826 | 0.306494801 | -0.638437668 | 0.523188817 | 0.722040793 | 0.195677826 |
| ENSMUSG00000002475 | Abhd3 | 106861 | protein_coding | 89.38957869 | -0.195707423 | 0.175862452 | 1.112843708 | 0.26577556 | 0.480830149 | 0.195707423 |
| ENSMUSG00000000538 | Tom1l2 | 216810 | protein_coding | 888.8988228 | 0.195770464 | 0.1336443 | -1.464862059 | 0.142958556 | 0.318770942 | 0.195770464 |
| ENSMUSG00000014303 | Glis2 | 83396 | protein_coding | 17.53368092 | 0.195811174 | 0.360163812 | -0.54367254 | 0.586666832 | 0.768287864 | 0.195811174 |
| ENSMUSG00000037007 | Zfp113 | 56314 | protein_coding | 168.0004753 | -0.195834674 | 0.092117391 | 2.125925109 | 0.033509496 | 0.109753236 | 0.195834674 |
| ENSMUSG00000034354 | Mtmr3 | 74302 | protein_coding | 1072.548974 | -0.195898851 | 0.066489062 | 2.946332011 | 0.00321567 | 0.016858279 | 0.195898851 |
| ENSMUSG00000031831 | Dnaaf1 | 68270 | protein_coding | 22.26647925 | 0.195906286 | 0.304768214 | -0.642804193 | 0.520351162 | 0.720291626 | 0.195906286 |
| ENSMUSG00000018822 | Sfrp5 | 54612 | protein_coding | 3.040096984 | -0.196012997 | 0.687497269 | 0.285110947 | 0.775559144 | 1 | 0.196012997 |
| ENSMUSG00000050010 | Shisa3 | 330096 | protein_coding | 4.052201081 | 0.19602882 | 0.620167103 | -0.316090323 | 0.751933952 | 1 | 0.19602882 |
| ENSMUSG00000012483 | Rpa3 | 68240 | protein_coding | 159.3919843 | 0.196037525 | 0.138275978 | -1.417726548 | 0.156270616 | 0.338979766 | 0.196037525 |
| ENSMUSG00000033114 | Slc35d2 | 70484 | protein_coding | 64.7439584 | -0.196060195 | 0.141419652 | 1.386371636 | 0.165633453 | 0.352762472 | 0.196060195 |
| ENSMUSG00000034064 | Poglut1 | 224143 | protein_coding | 176.0436635 | -0.196072645 | 0.135365794 | 1.448465223 | 0.147486985 | 0.325571458 | 0.196072645 |
| ENSMUSG00000046312 | Myorg | 329828 | protein_coding | 239.9469705 | -0.196114635 | 0.150058695 | 1.306919503 | 0.191240052 | 0.388837505 | 0.196114635 |
| ENSMUSG00000022685 | Parn | 74108 | protein_coding | 253.5333977 | 0.196175208 | 0.098611562 | -1.989373298 | 0.046660015 | 0.141461059 | 0.196175208 |
| ENSMUSG00000041703 | Zic5 | 65100 | protein_coding | 1.093696463 | 0.196198178 | 1.843358759 | -0.106435156 | 0.915237101 | 1 | 0.196198178 |
| ENSMUSG00000104585 | 4921511C10Rik | 70924 | lncRNA | 6.866514195 | 0.196222695 | 0.466586269 | -0.420549655 | 0.674083963 | 0.825190702 | 0.196222695 |
| ENSMUSG00000024170 | Telo2 | 71718 | protein_coding | 145.884298 | 0.196286987 | 0.130289578 | -1.506544042 | 0.13192757 | 0.30108314 | 0.196286987 |
| ENSMUSG00000001228 | Uhrf1 | 18140 | protein_coding | 433.5344856 | 0.196287684 | 0.163124897 | -1.203296903 | 0.228861447 | 0.437741278 | 0.196287684 |
| ENSMUSG00000042462 | Dctpp1 | 66422 | protein_coding | 223.1751078 | -0.19629983 | 0.107157712 | 1.831877768 | 0.066969634 | 0.184932165 | 0.19629983 |
| ENSMUSG00000026870 | Cutal | 77996 | protein_coding | 883.9593632 | -0.196301507 | 0.128929637 | 1.522547584 | 0.127871932 | 0.294844608 | 0.196301507 |
| ENSMUSG00000036214 | Polr1has | 76416 | protein_coding | 65.24610163 | 0.196308254 | 0.13635654 | -1.439668781 | 0.14996113 | 0.329666946 | 0.196308254 |
| ENSMUSG00000098602 | Mir6946 | 102465571 | miRNA | 1.459373607 | 0.196329335 | 0.929913088 | -0.211126543 | 0.832788529 | 1 | 0.196329335 |
| ENSMUSG00000052698 | Tln2 | 70549 | protein_coding | 856.7780176 | -0.19636358 | 0.145687676 | 1.347839335 | 0.177710064 | 0.370029503 | 0.19636358 |

| ensembl_gene_id | external_gene_name | entrezgene_id | gene_biotype | baseMean | log2FoldChange | lfcSE | stat | pvalue | padj | absLog2FC |
|---|---|---|---|---|---|---|---|---|---|---|
| ENSMUSG00000074649 | BC029722 | 613262 | protein_coding | 57.02699517 | -0.196394691 | 0.146823117 | 1.337627851 | 0.181017781 | 0.375025422 | 0.196394691 |
| ENSMUSG00000029173 | Sepsecs | 211006 | protein_coding | 557.0187626 | -0.196404464 | 0.083179401 | 2.361215162 | 0.018215159 | 0.068117105 | 0.196404464 |
| ENSMUSG00000085633 | Bloc1s6os | 75915 | lncRNA | 2.241721581 | 0.196519823 | 0.819373448 | -0.239841581 | 0.810453071 | 1 | 0.196519823 |
| ENSMUSG00000061232 | H2-K1 | 14972 | protein_coding | 6213.373324 | -0.196530637 | 0.095923711 | 2.048822295 | 0.040479495 | 0.127087144 | 0.196530637 |
| ENSMUSG00000053453 | Thoc7 | 66231 | protein_coding | 333.9483129 | -0.196636839 | 0.070087335 | 2.805597306 | 0.005022341 | 0.024436932 | 0.196636839 |
| ENSMUSG00000032578 | Cish | 12700 | protein_coding | 90.64616293 | -0.196682269 | 0.161085249 | 1.220982496 | 0.222092647 | 0.429214508 | 0.196682269 |
| ENSMUSG00000027080 | Med19 | 381379 | protein_coding | 252.6959852 | 0.196692533 | 0.08225912 | -2.39113344 | 0.016796447 | 0.063995287 | 0.196692533 |
| ENSMUSG00000037012 | Hk1 | 15275 | protein_coding | 2447.478373 | -0.196722697 | 0.070879392 | 2.775456896 | 0.005512421 | 0.026363195 | 0.196722697 |
| ENSMUSG00000067722 | BC003965 | 214489 | protein_coding | 292.6760521 | -0.19675496 | 0.084011092 | 2.342011702 | 0.019180114 | 0.071033953 | 0.19675496 |
| ENSMUSG00000025261 | Huwe1 | 59026 | protein_coding | 3430.262596 | 0.196891648 | 0.082089074 | -2.398512243 | 0.016461826 | 0.06297211 | 0.196891648 |
| ENSMUSG00000022533 | Atp13a3 | 224088 | protein_coding | 2257.360283 | 0.196902184 | 0.112178623 | -1.755255854 | 0.079215554 | 0.208539082 | 0.196902184 |
| ENSMUSG00000020652 | Cenpo | 52504 | protein_coding | 107.5297666 | 0.196912148 | 0.121728711 | -1.617631092 | 0.105742124 | 0.25785531 | 0.196912148 |
| ENSMUSG00000030353 | Tead4 | 21679 | protein_coding | 47.00008369 | 0.196920407 | 0.162283794 | -1.21343236 | 0.224964561 | 0.432209358 | 0.196920407 |
| ENSMUSG00000082769 | Gm15793 | 102640707 | processed_pseudogene | 4.956336054 | 0.196937996 | 0.509364562 | -0.386634662 | 0.699026693 | 0.840583034 | 0.196937996 |
| ENSMUSG00000024503 | Spink1 | 20730 | protein_coding | 1764.852772 | -0.196978643 | 0.370510721 | 0.531640873 | 0.594974751 | 0.773528957 | 0.196978643 |
| ENSMUSG00000022235 | Cmbl | 69574 | protein_coding | 473.3796486 | -0.197050982 | 0.097808043 | 2.014670529 | 0.043939182 | 0.135072906 | 0.197050982 |
| ENSMUSG00000034850 | Tmem127 | 69470 | protein_coding | 1269.330798 | -0.197062849 | 0.057746398 | 3.412556529 | 0.000643566 | 0.004377202 | 0.197062849 |
| ENSMUSG00000068205 | Macrod2 | 72899 | protein_coding | 81.3838651 | 0.197100373 | 0.116189911 | -1.696363914 | 0.089816983 | 0.228436418 | 0.197100373 |
| ENSMUSG00000020599 | Rgs9 | 19739 | protein_coding | 7.810350848 | 0.197148146 | 0.719499266 | -0.274007431 | 0.784078911 | 0.891754761 | 0.197148146 |
| ENSMUSG00000106864 | Gtf3c2 | 71752 | protein_coding | 215.9801971 | -0.197174105 | 0.077360721 | 2.548762503 | 0.010810589 | 0.045254367 | 0.197174105 |
| ENSMUSG00000028876 | Epha10 | 230735 | protein_coding | 3.266303825 | 0.19719824 | 1.41530966 | -0.139332222 | 0.889187628 | 1 | 0.19719824 |
| ENSMUSG00000038057 | Dbil5 | 13168 | protein_coding | 2.730616878 | -0.197261975 | 0.625043194 | 0.315597349 | 0.75230815 | 1 | 0.197261975 |
| ENSMUSG00000032119 | Hinfp | 102423 | protein_coding | 306.9718023 | 0.19731082 | 0.092558322 | -2.131745871 | 0.033027744 | 0.108584543 | 0.19731082 |
| ENSMUSG00000056917 | Sipa1 | 20469 | protein_coding | 52.73060474 | -0.197311384 | 0.262876077 | 0.75058707 | 0.452901205 | 0.669100957 | 0.197311384 |
| ENSMUSG00000079557 | Marchf2 | 224703 | protein_coding | 543.5442313 | -0.19731383 | 0.075755782 | 2.604604234 | 0.009198044 | 0.039805161 | 0.19731383 |
| ENSMUSG00000030435 | U2af2 | 22185 | protein_coding | 1400.742857 | 0.197333245 | 0.074103733 | -2.662932582 | 0.007746293 | 0.034686086 | 0.197333245 |
| ENSMUSG00000047242 | Taf9b | 407786 | protein_coding | 64.50351298 | 0.197398864 | 0.171746621 | -1.149360978 | 0.250407164 | 0.462725114 | 0.197398864 |
| ENSMUSG00000026637 | Traf5 | 22033 | protein_coding | 30.98674677 | -0.197412391 | 0.19492619 | 1.012754577 | 0.311177402 | 0.532085855 | 0.197412391 |
| ENSMUSG00000012126 | Ubxn11 | 67586 | protein_coding | 22.92696816 | 0.197454805 | 0.283251467 | -0.697100731 | 0.485739757 | 0.695380179 | 0.197454805 |
| ENSMUSG00000017679 | Ttpal | 76080 | protein_coding | 313.9083776 | 0.197481588 | 0.066253747 | -2.980685563 | 0.002876039 | 0.01536058 | 0.197481588 |
| ENSMUSG00000030966 | Trim21 | 20821 | protein_coding | 150.1656992 | -0.19751841 | 0.106614449 | 1.852642028 | 0.063933684 | 0.178888658 | 0.19751841 |
| ENSMUSG00000042564 | Fam227a | 75729 | protein_coding | 13.61292002 | -0.197538548 | 0.327921487 | 0.602395867 | 0.54691066 | 0.738667101 | 0.197538548 |
| ENSMUSG00000048706 | Lurap1l | 52829 | protein_coding | 1670.505441 | -0.197558065 | 0.070073743 | 2.81928803 | 0.004813031 | 0.023543362 | 0.197558065 |
| ENSMUSG00000004937 | Sgta | 52551 | protein_coding | 936.5555878 | 0.197577559 | 0.061681431 | -3.203193501 | 0.001359126 | 0.008215451 | 0.197577559 |
| ENSMUSG00000061665 | Cd2ap | 12488 | protein_coding | 4697.422268 | 0.197604011 | 0.069684844 | -2.835681341 | 0.004572805 | 0.022575574 | 0.197604011 |
| ENSMUSG00000061118 | Dnajc30 | 66114 | protein_coding | 125.03056 | -0.197632001 | 0.100631963 | 1.96390883 | 0.049540666 | 0.147967906 | 0.197632001 |
| ENSMUSG00000027775 | Mfsd1 | 66868 | protein_coding | 711.259561 | -0.197705104 | 0.090541677 | 2.183581215 | 0.028993035 | 0.098274011 | 0.197705104 |
| ENSMUSG00000048720 | Tbc1d12 | 209478 | protein_coding | 143.6882113 | 0.197749349 | 0.133319364 | -1.483275516 | 0.138001225 | 0.31052626 | 0.197749349 |
| ENSMUSG00000044719 | E230025N22Rik | 240216 | protein_coding | 61.15496917 | 0.197785107 | 0.158501125 | -1.24784671 | 0.212087199 | 0.41642117 | 0.197785107 |

Supplementary Table 1

| ensembl_gene_id | external_gene_name | entrezgene_id | gene_biotype | baseMean | log2FoldChange | lfcSE | stat | pvalue | padj | absLog2FC |
|---|---|---|---|---|---|---|---|---|---|---|
| ENSMUSG00000023764 | Sfi1 | 78887 | protein_coding | 437.1792321 | -0.197837409 | 0.270222646 | 0.732127423 | 0.464090798 | 0.677570103 | 0.197837409 |
| ENSMUSG00000002781 | Tmem143 | 70209 | protein_coding | 240.0918913 | -0.197885238 | 0.109733683 | 1.803322673 | 0.071337554 | 0.193614755 | 0.197885238 |
| ENSMUSG00000029422 | Rsrc2 | 208606 | protein_coding | 851.8249423 | -0.197909709 | 0.060120833 | 3.291865722 | 0.000995251 | 0.006340887 | 0.197909709 |
| ENSMUSG00000053930 | Shisa6 | 380702 | protein_coding | 3.31131204 | -0.197927954 | 1.548913125 | 0.127785058 | 0.898319075 | 1 | 0.197927954 |
| ENSMUSG00000087354 | 4930404I05Rik | 67394 | lncRNA | 4.095154259 | -0.197951695 | 0.60954229 | 0.324754653 | 0.745366767 | 1 | 0.197951695 |
| ENSMUSG00000042197 | Zfp451 | 98403 | protein_coding | 310.4047953 | 0.198000369 | 0.078580864 | -2.519702098 | 0.011745419 | 0.048307661 | 0.198000369 |
| ENSMUSG00000038473 | Nos1ap | 70729 | protein_coding | 65.64001598 | -0.198054579 | 0.239083073 | 0.82839231 | 0.40744836 | 0.628254175 | 0.198054579 |
| ENSMUSG00000027636 | Sla2 | 77799 | protein_coding | 8.808779535 | 0.198112766 | 0.531224895 | -0.372935771 | 0.709196248 | 0.84811839 | 0.198112766 |
| ENSMUSG00000031242 | Chmp1b2 | 67028 | protein_coding | 404.8938859 | 0.198115143 | 0.10005738 | -1.980015299 | 0.04770181 | 0.143880855 | 0.198115143 |
| ENSMUSG00000025372 | Baiap2 | 108100 | protein_coding | 301.2198203 | 0.198193977 | 0.102522112 | -1.933182746 | 0.053213689 | 0.156392751 | 0.198193977 |
| ENSMUSG00000032023 | Jhy | 70989 | protein_coding | 2.07486104 | 0.198197081 | 1.106486704 | -0.179122876 | 0.85784122 | 1 | 0.198197081 |
| ENSMUSG00000040859 | Bsdc1 | 100383 | protein_coding | 986.8975794 | -0.198341418 | 0.080453709 | 2.46528619 | 0.013690385 | 0.054307717 | 0.198341418 |
| ENSMUSG00000042363 | Lgalsl | 216551 | protein_coding | 566.7422814 | 0.19835771 | 0.088556161 | -2.239908633 | 0.025096855 | 0.087584418 | 0.19835771 |
| ENSMUSG00000041231 | Ublcp1 | 79560 | protein_coding | 271.4871247 | -0.198412402 | 0.093121803 | 2.130676129 | 0.033115833 | 0.108765884 | 0.198412402 |
| ENSMUSG00000099689 | Zfp383 | 73729 | protein_coding | 24.37610379 | -0.198467244 | 0.241410954 | 0.822113665 | 0.411012209 | 0.631215067 | 0.198467244 |
| ENSMUSG00000025508 | Rplp2 | 67186 | protein_coding | 3825.499404 | -0.198513168 | 0.112518573 | 1.764270227 | 0.077686494 | 0.205793193 | 0.198513168 |
| ENSMUSG00000020541 | Tom1l1 | 71943 | protein_coding | 856.1949994 | -0.198516232 | 0.063236859 | 3.139248761 | 0.001693816 | 0.009870396 | 0.198516232 |
| ENSMUSG00000057421 | Las1l | 76130 | protein_coding | 355.9559262 | 0.198523252 | 0.068103923 | -2.915004665 | 0.003556832 | 0.018344012 | 0.198523252 |
| ENSMUSG00000020163 | Uqcr11 | 66594 | protein_coding | 2359.194875 | -0.198526172 | 0.092135448 | 2.154720863 | 0.031183684 | 0.103991793 | 0.198526172 |
| ENSMUSG00000038781 | Stap2 | 106766 | protein_coding | 877.8323372 | -0.198559053 | 0.097917655 | 2.027816656 | 0.042578964 | 0.131848679 | 0.198559053 |
| ENSMUSG00000069516 | Lyz2 | 17105 | protein_coding | 13.27462715 | -0.198579455 | 0.545770669 | 0.363851459 | 0.715968934 | 0.852575063 | 0.198579455 |
| ENSMUSG00000098923 | Tmem185b | 226351 | protein_coding | 151.3188015 | -0.198619151 | 0.112801574 | 1.760783511 | 0.078275052 | 0.20691859 | 0.198619151 |
| ENSMUSG00000117684 | 4930449E18Rik | 78120 | lncRNA | 2.075727916 | 0.198660024 | 1.096271159 | -0.181214312 | 0.856199361 | 1 | 0.198660024 |
| ENSMUSG00000049606 | Zfp644 | 52397 | protein_coding | 984.7164987 | -0.198712684 | 0.080948132 | 2.454814947 | 0.014095713 | 0.055555275 | 0.198712684 |
| ENSMUSG00000048897 | Zfp710 | 209225 | protein_coding | 641.2033248 | 0.198715087 | 0.1006183 | -1.974939815 | 0.048274978 | 0.1452832 | 0.198715087 |
| ENSMUSG00000020251 | Glt8d2 | 74782 | protein_coding | 4.840622786 | 0.198831088 | 0.523055783 | -0.380133621 | 0.70384623 | 0.844104331 | 0.198831088 |
| ENSMUSG00000021024 | Psma6 | 26443 | protein_coding | 1067.799567 | 0.198852342 | 0.070547107 | -2.818717158 | 0.004821598 | 0.023571307 | 0.198852342 |
| ENSMUSG00000037656 | Slc20a2 | 20516 | protein_coding | 232.252308 | -0.198884607 | 0.112099304 | 1.774182353 | 0.076032992 | 0.202452247 | 0.198884607 |
| ENSMUSG00000041199 | Rpusd1 | 106707 | protein_coding | 60.83445706 | 0.198895912 | 0.139990027 | -1.420786297 | 0.155378896 | 0.337710856 | 0.198895912 |
| ENSMUSG00000059857 | Ntng1 | 80883 | protein_coding | 2.561819837 | -0.198904473 | 1.350431268 | 0.147289594 | 0.882903446 | 1 | 0.198904473 |
| ENSMUSG00000029725 | Ppp1r35 | 69871 | protein_coding | 98.05067474 | -0.19891838 | 0.14115209 | 1.409248562 | 0.158761683 | 0.342763083 | 0.19891838 |
| ENSMUSG00000060477 | Irak2 | 108960 | protein_coding | 435.3691408 | -0.198923691 | 0.168260769 | 1.182234525 | 0.237112657 | 0.447506105 | 0.198923691 |
| ENSMUSG00000030869 | Ndufab1 | 70316 | protein_coding | 704.447861 | -0.198982872 | 0.101186715 | 1.966492062 | 0.049241798 | 0.147315047 | 0.198982872 |
| ENSMUSG00000029404 | Arl6ip4 | 65105 | protein_coding | 296.5077556 | 0.199044086 | 0.075432646 | -2.638699488 | 0.008322472 | 0.036718488 | 0.199044086 |
| ENSMUSG00000014243 | Zswim7 | 69747 | protein_coding | 57.47589174 | -0.199054215 | 0.144788669 | 1.374791382 | 0.169196131 | 0.358402373 | 0.199054215 |
| ENSMUSG00000056718 | Gm13199 | 791411 | protein_coding | 4.150639168 | 0.199122558 | 0.504533473 | -0.394666694 | 0.693088886 | 1 | 0.199122558 |
| ENSMUSG00000046879 | Irgm1 | 15944 | protein_coding | 535.3159919 | 0.199169953 | 0.134320228 | -1.4827994 | 0.138127716 | 0.310726208 | 0.199169953 |
| ENSMUSG00000071359 | Tbpl1 | 237336 | protein_coding | 222.3835769 | 0.199172937 | 0.115675826 | -1.721819873 | 0.085102153 | 0.219864981 | 0.199172937 |
| ENSMUSG00000075602 | Ly6a | 110454 | protein_coding | 828.0159047 | -0.19919659 | 0.207860232 | 0.958319869 | 0.337901488 | 0.559633454 | 0.19919659 |

| ensembl_gene_id | external_gene_name | entrezgene_id | gene_biotype | baseMean | log2FoldChange | lfcSE | stat | pvalue | padj | absLog2FC |
|---|---|---|---|---|---|---|---|---|---|---|
| ENSMUSG00000020375 | Rufy1 | 216724 | protein_coding | 296.9477303 | -0.199242355 | 0.078309298 | 2.544300085 | 0.010949697 | 0.045697068 | 0.199242355 |
| ENSMUSG00000032396 | Dis3l | 213550 | protein_coding | 281.0596975 | -0.19924656 | 0.107653327 | 1.850816555 | 0.064195949 | 0.179403047 | 0.19924656 |
| ENSMUSG00000061353 | Cxcl12 | 20315 | protein_coding | 2.454987812 | 0.199276268 | 1.758689324 | -0.113309534 | 0.909785159 | 1 | 0.199276268 |
| ENSMUSG00000025825 | Iscu | 66383 | protein_coding | 317.7030967 | -0.199311961 | 0.079423941 | 2.509469546 | 0.012091264 | 0.049412308 | 0.199311961 |
| ENSMUSG00000038354 | Ankrd35 | 213121 | protein_coding | 27.10780235 | 0.19932657 | 0.250933355 | -0.794340674 | 0.426997131 | 0.646395058 | 0.19932657 |
| ENSMUSG00000071644 | Eef1g | 67160 | protein_coding | 4154.50887 | -0.199441905 | 0.104663936 | 1.90554561 | 0.056709184 | 0.163952289 | 0.199441905 |
| ENSMUSG00000058267 | Mrps14 | 64659 | protein_coding | 380.3598086 | 0.199465172 | 0.121454869 | -1.64229869 | 0.100528118 | 0.248744583 | 0.199465172 |
| ENSMUSG00000023915 | Tnfrsf21 | 94185 | protein_coding | 968.4059235 | -0.199497964 | 0.085846749 | 2.323884915 | 0.020131663 | 0.073733764 | 0.199497964 |
| ENSMUSG00000040249 | Lrp1 | 16971 | protein_coding | 3759.472412 | 0.199530824 | 0.217885049 | -0.91576189 | 0.359791796 | 0.582349713 | 0.199530824 |
| ENSMUSG00000035239 | Neu3 | 50877 | protein_coding | 50.201232 | -0.199631346 | 0.261774812 | 0.762607161 | 0.445697713 | 0.662922836 | 0.199631346 |
| ENSMUSG00000025794 | Rpl14 | 67115 | protein_coding | 4128.125655 | -0.199632438 | 0.120174145 | 1.661192911 | 0.096674709 | 0.241063164 | 0.199632438 |
| ENSMUSG00000015575 | Atp6v0e | 11974 | protein_coding | 463.1379784 | -0.199640393 | 0.074714139 | 2.672056388 | 0.007538798 | 0.03389483 | 0.199640393 |
| ENSMUSG00000052763 | Zfp212 | 232784 | protein_coding | 165.4047057 | -0.199653195 | 0.10606354 | 1.882392332 | 0.059782766 | 0.170333437 | 0.199653195 |
| ENSMUSG00000022575 | Gsdmd | 69146 | protein_coding | 1054.782998 | -0.199658384 | 0.083392061 | 2.394213328 | 0.016656058 | 0.063582557 | 0.199658384 |
| ENSMUSG00000060029 | 4930473A02Rik | 321014 | lncRNA | 2.987681182 | -0.199750992 | 0.583636816 | 0.342252214 | 0.732161094 | 1 | 0.199750992 |
| ENSMUSG00000032415 | Ube2cbp | 70348 | protein_coding | 30.70457409 | -0.19975783 | 0.224558706 | 0.889557269 | 0.373703659 | 0.595844868 | 0.19975783 |
| ENSMUSG00000001143 | Lman2l | 214895 | protein_coding | 193.5999613 | 0.199794765 | 0.117897613 | -1.694646398 | 0.09014252 | 0.228911821 | 0.199794765 |
| ENSMUSG00000035640 | Cbarp | 100503659 | protein_coding | 29.96439395 | -0.199842875 | 0.266010469 | 0.751259437 | 0.452496536 | 0.668802273 | 0.199842875 |
| ENSMUSG00000028550 | Atg4c | 242557 | protein_coding | 164.1856513 | 0.199893855 | 0.104301184 | -1.916506095 | 0.055300708 | 0.160639559 | 0.199893855 |
| ENSMUSG00000025184 | R3hcc1l | 52013 | protein_coding | 263.5271315 | -0.199963284 | 0.06907181 | 2.895005703 | 0.003791516 | 0.019336968 | 0.199963284 |
| ENSMUSG00000066278 | Vps37b | 330192 | protein_coding | 287.6005844 | 0.200015287 | 0.151807455 | -1.31755906 | 0.187651298 | 0.383856501 | 0.200015287 |
| ENSMUSG00000066151 | Fkbp15 | 338355 | protein_coding | 679.3133015 | -0.200036453 | 0.063949844 | 3.128020954 | 0.001759876 | 0.010183111 | 0.200036453 |
| ENSMUSG00000054519 | Zfp867 | 237775 | protein_coding | 57.21362027 | -0.200054711 | 0.1376547 | 1.453308253 | 0.146138187 | 0.323458788 | 0.200054711 |
| ENSMUSG00000063952 | Brpf3 | 268936 | protein_coding | 587.9393225 | 0.200108723 | 0.229036744 | -0.873697031 | 0.382283268 | 0.604659935 | 0.200108723 |
| ENSMUSG00000021326 | Trim27 | 19720 | protein_coding | 564.9555826 | 0.200130512 | 0.078520804 | -2.548757801 | 0.010810735 | 0.045254367 | 0.200130512 |
| ENSMUSG00000024505 | Dtwd2 | 68857 | protein_coding | 35.4864083 | 0.200135044 | 0.234263737 | -0.854315081 | 0.392930433 | 0.615582315 | 0.200135044 |
| ENSMUSG00000029076 | Sdf4 | 20318 | protein_coding | 1957.333345 | -0.200221638 | 0.067081963 | 2.98473136 | 0.002838275 | 0.015198206 | 0.200221638 |
| ENSMUSG00000104768 | Gm38412 | 329730 | lncRNA | 2.196713347 | -0.200233358 | 2.118043488 | 0.094536944 | 0.924682637 | 1 | 0.200233358 |
| ENSMUSG00000067786 | Nnat | 18111 | protein_coding | 62.92630572 | 0.200255877 | 0.43072697 | -0.464925326 | 0.641984933 | 0.804573016 | 0.200255877 |
| ENSMUSG00000050335 | Lgals3 | 16854 | protein_coding | 15954.24501 | -0.200282251 | 0.147350211 | 1.35922609 | 0.174074956 | 0.365037946 | 0.200282251 |
| ENSMUSG00000092680 | Snord55 | 100216533 | snoRNA | 14.03553992 | 0.200286192 | 0.346420725 | -0.578158805 | 0.563156909 | 0.750770075 | 0.200286192 |
| ENSMUSG00000026478 | Lamc1 | 226519 | protein_coding | 141.173497 | 0.200293831 | 0.129916437 | -1.541712781 | 0.123143405 | 0.286623 | 0.200293831 |
| ENSMUSG00000000568 | Hnrnpd | 11991 | protein_coding | 590.0392508 | 0.200349854 | 0.058140263 | -3.445974319 | 0.000569004 | 0.003951446 | 0.200349854 |
| ENSMUSG00000044703 | Phf11a | 219131 | protein_coding | 1.942545018 | 0.200356946 | 1.046244087 | -0.19150115 | 0.848132979 | 1 | 0.200356946 |
| ENSMUSG00000041141 | Pnma8a | 71691 | protein_coding | 5.862582738 | -0.200393557 | 0.443520477 | 0.451824814 | 0.651395192 | 0.81075823 | 0.200393557 |
| ENSMUSG00000071691 | Gm960 | 381196 | protein_coding | 5.3350233 | 0.200475922 | 0.776716942 | -0.258106797 | 0.796324486 | 0.898810919 | 0.200475922 |
| ENSMUSG00000022650 | Retnlb | 57263 | protein_coding | 1123.237453 | 0.200509549 | 0.248429862 | -0.807107276 | 0.419604681 | 0.639652146 | 0.200509549 |
| ENSMUSG00000030120 | Mlf2 | 30853 | protein_coding | 2125.669826 | 0.200535082 | 0.065114518 | -3.079729206 | 0.002071889 | 0.01168152 | 0.200535082 |
| ENSMUSG00000031373 | Car5b | 56078 | protein_coding | 9.238668342 | 0.200616561 | 0.338382562 | -0.592869088 | 0.553268767 | 0.743665995 | 0.200616561 |

### Supplementary Table 1

| ensembl_gene_id | external_gene_name | entrezgene_id | gene_biotype | baseMean | log2FoldChange | lfcSE | stat | pvalue | padj | absLog2FC |
|---|---|---|---|---|---|---|---|---|---|---|
| ENSMUSG00000035678 | Tnfsf9 | 21950 | protein_coding | 4.07814195 | -0.200619626 | 0.576230804 | 0.348158454 | 0.727721187 | 1 | 0.200619626 |
| ENSMUSG00000062098 | Btbd3 | 228662 | protein_coding | 259.4957261 | -0.200627796 | 0.094095131 | 2.132180423 | 0.032992017 | 0.108571968 | 0.200627796 |
| ENSMUSG00000078517 | Emc1 | 230866 | protein_coding | 792.8036004 | 0.200644265 | 0.073528204 | -2.728806815 | 0.006356393 | 0.02939498 | 0.200644265 |
| ENSMUSG00000035914 | Cd276 | 102657 | protein_coding | 56.98418587 | 0.20069789 | 0.170777661 | -1.175199902 | 0.23991475 | 0.450375375 | 0.20069789 |
| ENSMUSG00000002108 | Nr1h3 | 22259 | protein_coding | 218.0999375 | -0.200752426 | 0.135507427 | 1.481486517 | 0.138476975 | 0.311384636 | 0.200752426 |
| ENSMUSG00000035878 | Hykk | 235386 | protein_coding | 97.48062837 | -0.200756996 | 0.13662186 | 1.469435391 | 0.141714734 | 0.316724471 | 0.200756996 |
| ENSMUSG00000025001 | Hells | 15201 | protein_coding | 223.7573138 | 0.200789495 | 0.192884389 | -1.040983653 | 0.297883135 | 0.517488122 | 0.200789495 |
| ENSMUSG00000004709 | Cd244a | 18106 | protein_coding | 12.94387094 | -0.200797063 | 0.434797411 | 0.461817523 | 0.644212184 | 0.806110415 | 0.200797063 |
| ENSMUSG00000026034 | Clk1 | 12747 | protein_coding | 1186.570251 | -0.200799445 | 0.131342906 | 1.528818353 | 0.126309481 | 0.292303079 | 0.200799445 |
| ENSMUSG00000032059 | Alg9 | 102580 | protein_coding | 533.0050753 | 0.200972237 | 0.089225932 | -2.252397185 | 0.024297183 | 0.085498335 | 0.200972237 |
| ENSMUSG00000047369 | Dnah14 | 240960 | protein_coding | 10.13073476 | -0.201009636 | 0.721335392 | 0.27866321 | 0.780503299 | 0.890186044 | 0.201009636 |
| ENSMUSG00000044243 | Bhlha9 | 320522 | protein_coding | 1.330308751 | 0.201121741 | 1.099606038 | -0.182903453 | 0.854873773 | 1 | 0.201121741 |
| ENSMUSG00000044098 | Rsbn1 | 229675 | protein_coding | 256.6177166 | -0.201151475 | 0.073503736 | 2.736615643 | 0.006207477 | 0.028892413 | 0.201151475 |
| ENSMUSG00000072707 | Or2t1 | 18330 | protein_coding | 1.793092102 | 0.201205015 | 2.135180054 | -0.094233278 | 0.924923851 | 1 | 0.201205015 |
| ENSMUSG00000026961 | Lrrc26 | 227618 | protein_coding | 337.1660421 | -0.201206019 | 0.158058537 | 1.272984192 | 0.203023651 | 0.405213026 | 0.201206019 |
| ENSMUSG00000029314 | Gpat3 | 231510 | protein_coding | 188.5665395 | 0.201242566 | 0.13794416 | -1.458869773 | 0.144600954 | 0.321521497 | 0.201242566 |
| ENSMUSG00000046410 | Kcnk6 | 52150 | protein_coding | 725.0816454 | 0.201257376 | 0.106973356 | -1.881378536 | 0.059920445 | 0.1706035 | 0.201257376 |
| ENSMUSG00000086013 | Gm15706 | 100039495 | lncRNA | 35.97571296 | -0.20133178 | 0.177024785 | 1.137308427 | 0.255409373 | 0.468075728 | 0.20133178 |
| ENSMUSG00000050954 | Zfp169 | 67911 | protein_coding | 106.7852293 | -0.201365141 | 0.153532716 | 1.311545486 | 0.189673537 | 0.386972505 | 0.201365141 |
| ENSMUSG00000026785 | Pkn3 | 263803 | protein_coding | 96.19740908 | 0.201393666 | 0.169275196 | -1.189741152 | 0.234148146 | 0.443634979 | 0.201393666 |
| ENSMUSG00000040620 | Dhx33 | 216877 | protein_coding | 299.8533508 | 0.20141471 | 0.094496967 | -2.131441001 | 0.033052828 | 0.108625296 | 0.20141471 |
| ENSMUSG00000070877 | Ldlrad1 | 546840 | protein_coding | 4.236338044 | 0.201435053 | 0.644377768 | -0.312603977 | 0.754581556 | 1 | 0.201435053 |
| ENSMUSG00000033918 | Parl | 381038 | protein_coding | 379.9914651 | -0.201471274 | 0.082406686 | 2.444841355 | 0.014491587 | 0.056736385 | 0.201471274 |
| ENSMUSG00000021290 | Atp5mpl | 70257 | protein_coding | 1582.998141 | -0.201532603 | 0.069960412 | 2.880666348 | 0.003968355 | 0.020065343 | 0.201532603 |
| ENSMUSG00000034334 | Fam151b | 73942 | protein_coding | 38.77052933 | 0.201544158 | 0.189571978 | -1.063153745 | 0.287712235 | 0.506641059 | 0.201544158 |
| ENSMUSG00000074643 | Cpne1 | 266692 | protein_coding | 2.208897915 | -0.201573836 | 0.779111198 | 0.258722806 | 0.795849122 | 1 | 0.201573836 |
| ENSMUSG00000036928 | Stag3 | 50878 | protein_coding | 4.12582339 | 0.201640609 | 1.235012796 | -0.163270057 | 0.870305809 | 1 | 0.201640609 |
| ENSMUSG00000039137 | Whrn | 73750 | protein_coding | 112.3632648 | -0.201892257 | 0.204009312 | 0.989622756 | 0.322358543 | 0.54358318 | 0.201892257 |
| ENSMUSG00000032308 | Ulk3 | 71742 | protein_coding | 86.87394365 | -0.20189601 | 0.154536154 | 1.30646457 | 0.19139462 | 0.389062992 | 0.20189601 |
| ENSMUSG00000037761 | Actr5 | 109275 | protein_coding | 128.2185048 | -0.201962493 | 0.1117608 | 1.807095987 | 0.070747314 | 0.192474654 | 0.201962493 |
| ENSMUSG00000053178 | Mterf1b | 208595 | protein_coding | 10.49258331 | -0.202011851 | 0.340855383 | 0.592661467 | 0.553407734 | 0.743731716 | 0.202011851 |
| ENSMUSG00000027296 | Itpka | 228550 | protein_coding | 147.4346566 | -0.202044025 | 0.138832262 | 1.455310335 | 0.145583367 | 0.322749862 | 0.202044025 |
| ENSMUSG00000022840 | Adcy5 | 224129 | protein_coding | 59.20751456 | -0.202046577 | 0.216742849 | 0.932194892 | 0.351235815 | 0.573211901 | 0.202046577 |
| ENSMUSG00000035235 | Trim13 | 66597 | protein_coding | 45.57385387 | -0.202054945 | 0.173282462 | 1.166043828 | 0.243596726 | 0.454822267 | 0.202054945 |
| ENSMUSG00000002763 | Pex6 | 224824 | protein_coding | 720.4521626 | -0.202078852 | 0.05326876 | 3.793571569 | 0.000148496 | 0.001218211 | 0.202078852 |
| ENSMUSG00000029388 | Eif2b1 | 209354 | protein_coding | 812.7800603 | 0.202082711 | 0.090049512 | -2.244128894 | 0.024824109 | 0.086889644 | 0.202082711 |
| ENSMUSG00000047293 | Gpr15 | 71223 | protein_coding | 1.829130367 | 0.202108632 | 1.113161499 | -0.181562722 | 0.855925906 | 1 | 0.202108632 |
| ENSMUSG00000023022 | Lima1 | 65970 | protein_coding | 4427.735788 | -0.202111738 | 0.094372794 | 2.141631388 | 0.032223156 | 0.106618553 | 0.202111738 |
| ENSMUSG00000006241 | Ccdc159 | 67119 | protein_coding | 20.8570792 | -0.2021179 | 0.296228182 | 0.682304767 | 0.49504626 | 0.701011313 | 0.2021179 |

| ensembl_gene_id | external_gene_name | entrezgene_id | gene_biotype | baseMean | log2FoldChange | lfcSE | stat | pvalue | padj | absLog2FC |
|---|---|---|---|---|---|---|---|---|---|---|
| ENSMUSG00000020381 | Mrnip | 68067 | protein_coding | 19.23696217 | 0.202155634 | 0.286800972 | -0.704863838 | 0.480894976 | 0.691768261 | 0.202155634 |
| ENSMUSG00000016382 | Pls3 | 102866 | protein_coding | 168.870552 | -0.202167731 | 0.095110453 | 2.125610015 | 0.033535745 | 0.109773894 | 0.202167731 |
| ENSMUSG00000089281 | Scarna6 | 100217415 | scaRNA | 509.0355506 | -0.202326323 | 0.165241769 | 1.224426027 | 0.220791556 | 0.427312825 | 0.202326323 |
| ENSMUSG00000075054 | Yae1d1 | 67008 | protein_coding | 144.0469453 | -0.202396138 | 0.09288543 | 2.178986915 | 0.02933264 | 0.099059146 | 0.202396138 |
| ENSMUSG00000036459 | Wtip | 101543 | protein_coding | 157.2762602 | 0.202427497 | 0.151742475 | -1.334020006 | 0.1821973 | 0.376665775 | 0.202427497 |
| ENSMUSG00000110344 | Smim36 | 121811722 | protein_coding | 12.54430808 | -0.202471634 | 0.324069583 | 0.624778271 | 0.532116594 | 0.728866518 | 0.202471634 |
| ENSMUSG00000029147 | Ppm1g | 14208 | protein_coding | 994.6693396 | 0.202479927 | 0.087052615 | -2.32594881 | 0.020021282 | 0.073420411 | 0.202479927 |
| ENSMUSG00000058056 | Palld | 72333 | protein_coding | 235.5317194 | 0.202510826 | 0.170140494 | -1.190256482 | 0.2339456 | 0.44350564 | 0.202510826 |
| ENSMUSG00000107456 | Gm10400 | 100093700 | TEC | 16.75350537 | -0.202520675 | 0.32912214 | 0.61533592 | 0.5383329 | 0.733377497 | 0.202520675 |
| ENSMUSG00000036155 | Mgat5 | 107895 | protein_coding | 1894.759269 | -0.202537969 | 0.075961378 | 2.666328274 | 0.007668477 | 0.034412292 | 0.202537969 |
| ENSMUSG00000032458 | Copb2 | 50797 | protein_coding | 3276.41685 | 0.202612087 | 0.097001844 | -2.088744693 | 0.036730711 | 0.1177122 | 0.202612087 |
| ENSMUSG00000043099 | Hic1 | 15248 | protein_coding | 12.28360544 | 0.20262558 | 0.477495218 | -0.424351015 | 0.67130984 | 0.823809965 | 0.20262558 |
| ENSMUSG00000044912 | Syt16 | 238266 | protein_coding | 3.122624443 | -0.202640894 | 1.159310261 | 0.174794359 | 0.861241221 | 1 | 0.202640894 |
| ENSMUSG00000080364 | Gm25777 | 115488712 | snoRNA | 3.282207993 | 0.202643272 | 0.588156422 | -0.344539759 | 0.730440401 | 1 | 0.202643272 |
| ENSMUSG00000024294 | Mib1 | 225164 | protein_coding | 1557.703235 | 0.202725377 | 0.076883533 | -2.636785397 | 0.008369576 | 0.036886891 | 0.202725377 |
| ENSMUSG00000023150 | Ivns1abp | 117198 | protein_coding | 4510.427662 | -0.20276088 | 0.133246821 | 1.521693936 | 0.128085788 | 0.295198531 | 0.20276088 |
| ENSMUSG00000033624 | Pdpr | 319518 | protein_coding | 550.70092 | -0.202818708 | 0.084429014 | 2.402239449 | 0.016295038 | 0.062415953 | 0.202818708 |
| ENSMUSG00000040562 | Gstm2 | 14863 | protein_coding | 979.5791652 | 0.202850356 | 0.131370059 | -1.544114067 | 0.122560701 | 0.285669361 | 0.202850356 |
| ENSMUSG00000044763 | Trmt10c | 52575 | protein_coding | 228.6717956 | -0.202917557 | 0.113853164 | 1.782274198 | 0.074704528 | 0.199909758 | 0.202917557 |
| ENSMUSG00000077565 | Gm23201 | 115486177 | snoRNA | 8.143552718 | -0.202936087 | 0.398016655 | 0.509868329 | 0.610143711 | 0.783873755 | 0.202936087 |
| ENSMUSG00000078862 | Gm14326 | 628147 | protein_coding | 43.29275658 | -0.202950656 | 0.228418457 | 0.888503751 | 0.374269839 | 0.596246587 | 0.202950656 |
| ENSMUSG00000050270 | Tmem220 | 338369 | protein_coding | 60.50280818 | 0.202993545 | 0.192466335 | -1.054696372 | 0.291564212 | 0.510430551 | 0.202993545 |
| ENSMUSG00000025577 | Cbx2 | 12416 | protein_coding | 79.10411912 | -0.203022728 | 0.149445008 | 1.358511271 | 0.174301509 | 0.365420225 | 0.203022728 |
| ENSMUSG00000030701 | Plekhb1 | 27276 | protein_coding | 18.75973865 | 0.203028532 | 0.365750646 | -0.555100951 | 0.578825613 | 0.762380457 | 0.203028532 |
| ENSMUSG00000029003 | Mad2l2 | 71890 | protein_coding | 57.44301246 | -0.203044015 | 0.139168849 | 1.458976023 | 0.144571707 | 0.321507788 | 0.203044015 |
| ENSMUSG00000087516 | Tbx3os1 | 100039542 | lncRNA | 2.116243267 | 0.203096321 | 1.902048111 | -0.106777699 | 0.91496534 | 1 | 0.203096321 |
| ENSMUSG00000078607 | 1810010H24Rik | 69066 | protein_coding | 14.89573155 | 0.203162423 | 0.284330703 | -0.714528612 | 0.47490037 | 0.686795739 | 0.203162423 |
| ENSMUSG00000079334 | Naa80 | 56441 | protein_coding | 115.4106227 | -0.203232771 | 0.113678976 | 1.78777798 | 0.073811835 | 0.198490252 | 0.203232771 |
| ENSMUSG00000013997 | Nit1 | 27045 | protein_coding | 523.401006 | -0.203279563 | 0.073924809 | 2.74981521 | 0.005962888 | 0.028062449 | 0.203279563 |
| ENSMUSG00000042129 | Rassf4 | 213391 | protein_coding | 219.1180433 | 0.203301273 | 0.178180461 | -1.140985224 | 0.253876072 | 0.466508229 | 0.203301273 |
| ENSMUSG00000006127 | Inpp5k | 19062 | protein_coding | 277.5798478 | -0.203310899 | 0.084077018 | 2.418150703 | 0.015599615 | 0.06023867 | 0.203310899 |
| ENSMUSG00000027272 | Ubr1 | 22222 | protein_coding | 363.8184511 | -0.20338824 | 0.08404987 | 2.419851933 | 0.015526828 | 0.060049189 | 0.20338824 |
| ENSMUSG00000025225 | Nfkb2 | 18034 | protein_coding | 615.784582 | 0.203444516 | 0.081542207 | -2.494959643 | 0.012597146 | 0.050983705 | 0.203444516 |
| ENSMUSG00000018339 | Gpx3 | 14778 | protein_coding | 16.98333351 | 0.203462073 | 0.301866724 | -0.67401292 | 0.500303099 | 0.704889121 | 0.203462073 |
| ENSMUSG00000031958 | Ldhd | 52815 | protein_coding | 81.7865209 | -0.203502518 | 0.117683694 | 1.729232926 | 0.083767416 | 0.217061839 | 0.203502518 |
| ENSMUSG00000036805 | Noxa1 | 241275 | protein_coding | 941.2347946 | 0.203528044 | 0.105661473 | -1.926227575 | 0.054075964 | 0.158138918 | 0.203528044 |
| ENSMUSG00000029111 | Nelfa | 24116 | protein_coding | 283.3223238 | 0.203579743 | 0.076111433 | -2.674759028 | 0.007478298 | 0.033677833 | 0.203579743 |
| ENSMUSG00000034126 | Pomt2 | 217734 | protein_coding | 221.9372219 | 0.203694917 | 0.092763794 | -2.195845037 | 0.028103037 | 0.095748618 | 0.203694917 |
| ENSMUSG00000042506 | Usp22 | 216825 | protein_coding | 963.9994378 | 0.203705068 | 0.073479484 | -2.772271338 | 0.005566661 | 0.026568742 | 0.203705068 |

Supplementary Table 1

| ensembl_gene_id | external_gene_name | entrezgene_id | gene_biotype | baseMean | log2FoldChange | lfcSE | stat | pvalue | padj | absLog2FC |
|---|---|---|---|---|---|---|---|---|---|---|
| ENSMUSG00000003848 | Nob1 | 67619 | protein_coding | 149.9183417 | -0.203709448 | 0.105621793 | 1.928668721 | 0.053772002 | 0.157556926 | 0.203709448 |
| ENSMUSG00000027709 | Mccc1 | 72039 | protein_coding | 270.0785606 | -0.203724572 | 0.097715497 | 2.084874752 | 0.037080671 | 0.118695523 | 0.203724572 |
| ENSMUSG00000028291 | Akirin2 | 433693 | protein_coding | 272.7017353 | 0.20378194 | 0.089250417 | -2.283260382 | 0.022415035 | 0.080365152 | 0.20378194 |
| ENSMUSG00000074093 | Svip | 75744 | protein_coding | 222.007557 | -0.203819874 | 0.091814363 | 2.219912745 | 0.026424691 | 0.091158836 | 0.203819874 |
| ENSMUSG00000039521 | Foxp3 | 20371 | protein_coding | 4.861232003 | -0.203826683 | 0.777181058 | 0.262264091 | 0.793117845 | 0.897030894 | 0.203826683 |
| ENSMUSG00000048807 | Slc35e4 | 103710 | protein_coding | 149.4091779 | -0.203865924 | 0.118595444 | 1.719002992 | 0.085613827 | 0.220841415 | 0.203865924 |
| ENSMUSG00000038429 | Usp5 | 22225 | protein_coding | 1226.077587 | 0.203903161 | 0.07582818 | -2.689015624 | 0.007166306 | 0.032583807 | 0.203903161 |
| ENSMUSG00000001415 | Smg5 | 229512 | protein_coding | 939.8515114 | 0.203957351 | 0.054982425 | -3.709500806 | 0.000207668 | 0.001633842 | 0.203957351 |
| ENSMUSG00000001034 | Mapk7 | 23939 | protein_coding | 179.2333901 | 0.203983351 | 0.107420847 | -1.898917736 | 0.057575293 | 0.165730938 | 0.203983351 |
| ENSMUSG00000061689 | Dlgap4 | 228836 | protein_coding | 301.6139486 | 0.203990609 | 0.096651401 | -2.110580989 | 0.034808343 | 0.112998816 | 0.203990609 |
| ENSMUSG00000025049 | Taf5 | 226182 | protein_coding | 105.172549 | 0.204040164 | 0.142106457 | -1.435826135 | 0.151051821 | 0.331061353 | 0.204040164 |
| ENSMUSG00000022208 | Jph4 | 319984 | protein_coding | 7.959427254 | 0.204239736 | 1.006164699 | -0.202988373 | 0.83914412 | 0.919794636 | 0.204239736 |
| ENSMUSG00000027173 | Depdc7 | 211896 | protein_coding | 7.89618765 | -0.204311489 | 0.438370661 | 0.46607017 | 0.641165274 | 0.804106371 | 0.204311489 |
| ENSMUSG00000034154 | Ino80 | 68142 | protein_coding | 520.8437612 | -0.204328061 | 0.078431857 | 2.605166675 | 0.009182958 | 0.039779225 | 0.204328061 |
| ENSMUSG00000020577 | Tspan13 | 66109 | protein_coding | 1332.714959 | 0.204359956 | 0.175940526 | -1.161528615 | 0.245426993 | 0.457189099 | 0.204359956 |
| ENSMUSG00000001052 | Sec24b | 99683 | protein_coding | 368.63899 | 0.204368377 | 0.114036679 | -1.792128458 | 0.073112391 | 0.197090766 | 0.204368377 |
| ENSMUSG00000047866 | Lonp2 | 66887 | protein_coding | 1022.882594 | -0.204405321 | 0.059563558 | 3.431717788 | 0.000599771 | 0.004126525 | 0.204405321 |
| ENSMUSG00000084651 | Mir1904 | 100316683 | miRNA | 0.675602507 | 0.204456635 | 1.435162427 | -0.142462366 | 0.886714804 | 1 | 0.204456635 |
| ENSMUSG00000044340 | Phlpp1 | 98432 | protein_coding | 477.7477236 | -0.20448194 | 0.077307457 | 2.645048065 | 0.008167931 | 0.036230249 | 0.20448194 |
| ENSMUSG00000042473 | Tbc1d8b | 245638 | protein_coding | 560.6046263 | 0.204548559 | 0.108869242 | -1.878846183 | 0.060265499 | 0.171354072 | 0.204548559 |
| ENSMUSG00000010095 | Slc3a2 | 17254 | protein_coding | 546.909011 | 0.204563725 | 0.070591325 | -2.89785927 | 0.003757191 | 0.019203423 | 0.204563725 |
| ENSMUSG00000004233 | Wars2 | 70560 | protein_coding | 194.3922823 | -0.204580529 | 0.10414034 | 1.964469577 | 0.049475661 | 0.147853975 | 0.204580529 |
| ENSMUSG00000097397 | Gm16861 | 100503704 | lncRNA | 11.07403314 | -0.204589542 | 0.35407428 | 0.577815315 | 0.563388814 | 0.750868206 | 0.204589542 |
| ENSMUSG00000022762 | Ncam2 | 17968 | protein_coding | 2.610747217 | 0.20459884 | 1.017737255 | -0.201033065 | 0.84067272 | 1 | 0.20459884 |
| ENSMUSG00000034998 | Foxn2 | 14236 | protein_coding | 295.7868371 | -0.204606586 | 0.125810224 | 1.626311284 | 0.103883453 | 0.254529872 | 0.204606586 |
| ENSMUSG00000004789 | Dlst | 78920 | protein_coding | 2117.436162 | -0.204672331 | 0.046726582 | 4.380211867 | 1.19E-05 | 0.000130357 | 0.204672331 |
| ENSMUSG00000017548 | Suz12 | 52615 | protein_coding | 665.0752688 | 0.204687387 | 0.107501831 | -1.904036287 | 0.056905458 | 0.16428964 | 0.204687387 |
| ENSMUSG00000029093 | Sorcs2 | 81840 | protein_coding | 212.3508389 | 0.204757325 | 0.17119009 | -1.19608165 | 0.231664698 | 0.440496352 | 0.204757325 |
| ENSMUSG00000034022 | Cpsf1 | 94230 | protein_coding | 917.256618 | 0.204784486 | 0.070532388 | -2.903410663 | 0.003691223 | 0.01893658 | 0.204784486 |
| ENSMUSG00000026568 | Mpc2 | 70456 | protein_coding | 804.5309891 | -0.204788994 | 0.056848509 | 3.602363492 | 0.000315337 | 0.002381578 | 0.204788994 |
| ENSMUSG00000039328 | Rnf122 | 68867 | protein_coding | 32.63764557 | -0.204811839 | 0.233212589 | 0.878219482 | 0.379824622 | 0.602075619 | 0.204811839 |
| ENSMUSG00000061374 | Fiz1 | 23877 | protein_coding | 290.306065 | 0.204891219 | 0.071454964 | -2.86741755 | 0.004138366 | 0.020759714 | 0.204891219 |
| ENSMUSG00000032360 | Hcrtr2 | 387285 | protein_coding | 1.9351828 | -0.204989019 | 1.574825439 | 0.130166185 | 0.896434948 | 1 | 0.204989019 |
| ENSMUSG00000091264 | Smim13 | 108934 | protein_coding | 144.036121 | -0.204991241 | 0.107142709 | 1.913254233 | 0.055715514 | 0.161560315 | 0.204991241 |
| ENSMUSG00000009670 | Tex11 | 83558 | protein_coding | 1.238471737 | 0.20506807 | 1.441110905 | -0.142298604 | 0.886844149 | 1 | 0.20506807 |
| ENSMUSG00000028234 | Rps20 | 67427 | protein_coding | 4198.758247 | -0.205123907 | 0.102689612 | 1.997513696 | 0.045769408 | 0.139401698 | 0.205123907 |
| ENSMUSG00000028228 | Cpne3 | 70568 | protein_coding | 1555.022533 | -0.205168325 | 0.10024275 | 2.046714856 | 0.04068609 | 0.127638694 | 0.205168325 |
| ENSMUSG00000021038 | Vipas39 | 104799 | protein_coding | 701.7430237 | -0.20538029 | 0.068912603 | 2.980300845 | 0.002879654 | 0.015369945 | 0.20538029 |
| ENSMUSG00000040841 | Six5 | 20475 | protein_coding | 58.5653385 | 0.205409627 | 0.225229637 | -0.91200088 | 0.361768254 | 0.584163582 | 0.205409627 |

Supplementary Table 1

| ensembl_gene_id | external_gene_name | entrezgene_id | gene_biotype | baseMean | log2FoldChange | lfcSE | stat | pvalue | padj | absLog2FC |
|---|---|---|---|---|---|---|---|---|---|---|
| ENSMUSG00000005087 | Cd44 | 12505 | protein_coding | 496.5265804 | 0.205437056 | 0.197758354 | -1.038828712 | 0.298884405 | 0.518790947 | 0.205437056 |
| ENSMUSG00000114649 | 3110006O06Rik | 73093 | lncRNA | 2.548999809 | 0.205472176 | 1.544164987 | -0.133063615 | 0.89414307 | 1 | 0.205472176 |
| ENSMUSG00000059022 | Kcp | 333088 | protein_coding | 3.291207224 | 0.205546605 | 1.18587215 | -0.173329481 | 0.862392452 | 1 | 0.205546605 |
| ENSMUSG00000038816 | Ctnnal1 | 54366 | protein_coding | 203.9524693 | -0.205562702 | 0.153548315 | 1.338749313 | 0.180652297 | 0.374503142 | 0.205562702 |
| ENSMUSG00000020525 | Ppm1d | 53892 | protein_coding | 147.3267132 | -0.205594074 | 0.09706466 | 2.118114604 | 0.034165363 | 0.11137126 | 0.205594074 |
| ENSMUSG00000020561 | Polr1f | 28071 | protein_coding | 196.3199756 | 0.20560435 | 0.132960488 | -1.546356756 | 0.122018432 | 0.285089471 | 0.20560435 |
| ENSMUSG00000024601 | Isoc1 | 66307 | protein_coding | 531.4508305 | -0.205626265 | 0.123465587 | 1.665454072 | 0.095822199 | 0.239504737 | 0.205626265 |
| ENSMUSG00000005057 | Sh2b2 | 23921 | protein_coding | 6.22385315 | -0.205670149 | 0.900414004 | 0.228417314 | 0.81932183 | 0.908985828 | 0.205670149 |
| ENSMUSG00000026398 | Nr5a2 | 26424 | protein_coding | 1303.449784 | -0.205762785 | 0.066631589 | 3.088066609 | 0.002014633 | 0.011401366 | 0.205762785 |
| ENSMUSG00000032411 | Tfdp2 | 211586 | protein_coding | 676.1181821 | 0.205773809 | 0.10066949 | -2.044053365 | 0.040948275 | 0.128287829 | 0.205773809 |
| ENSMUSG00000071256 | Zfp213 | 449521 | protein_coding | 147.0426323 | 0.20578974 | 0.142522097 | -1.443914622 | 0.148762993 | 0.327599956 | 0.20578974 |
| ENSMUSG00000055900 | Tmem69 | 230657 | protein_coding | 106.4279622 | -0.205835852 | 0.118944097 | 1.730526006 | 0.083536339 | 0.216667049 | 0.205835852 |
| ENSMUSG00000034345 | Gtf2h5 | 66467 | protein_coding | 288.7923274 | 0.20588449 | 0.104112325 | -1.977522732 | 0.047982574 | 0.144595661 | 0.20588449 |
| ENSMUSG00000035873 | Pawr | 114774 | protein_coding | 543.7583012 | 0.205887907 | 0.079094431 | -2.603064523 | 0.009239456 | 0.039961181 | 0.205887907 |
| ENSMUSG00000025510 | Cd151 | 12476 | protein_coding | 1394.274901 | -0.205911825 | 0.054361451 | 3.787827968 | 0.00015197 | 0.001242392 | 0.205911825 |
| ENSMUSG00000002803 | Btbd6 | 399566 | protein_coding | 249.165376 | -0.205913948 | 0.088075229 | 2.337932582 | 0.019390746 | 0.071685421 | 0.205913948 |
| ENSMUSG00000023235 | Ccl25 | 20300 | protein_coding | 71.47910292 | -0.205960895 | 0.19122051 | 1.077085794 | 0.281441939 | 0.499165737 | 0.205960895 |
| ENSMUSG00000031148 | Gpkow | 209416 | protein_coding | 243.2497155 | -0.206048286 | 0.08312496 | 2.478777563 | 0.013183349 | 0.052790938 | 0.206048286 |
| ENSMUSG00000049489 | Ccnq | 69109 | protein_coding | 126.4359471 | -0.206116429 | 0.147150995 | 1.400713799 | 0.161299674 | 0.346267391 | 0.206116429 |
| ENSMUSG00000030681 | Mvp | 78388 | protein_coding | 5262.323786 | -0.206183586 | 0.084261715 | 2.446942676 | 0.014407375 | 0.056473628 | 0.206183586 |
| ENSMUSG00000029312 | Klhl8 | 246293 | protein_coding | 54.18650627 | 0.206305236 | 0.203276291 | -1.01490063 | 0.310153198 | 0.531259196 | 0.206305236 |
| ENSMUSG00000000690 | Hoxb6 | 15414 | protein_coding | 195.9673662 | -0.20637576 | 0.267268359 | 0.772166825 | 0.440015628 | 0.657892085 | 0.20637576 |
| ENSMUSG00000028339 | Col15a1 | 12819 | protein_coding | 25.42196414 | -0.206398129 | 0.402884517 | 0.51230097 | 0.608440387 | 0.782626881 | 0.206398129 |
| ENSMUSG00000034613 | Ppm1h | 319468 | protein_coding | 731.8127045 | -0.20649176 | 0.04793075 | 4.308127074 | 1.65E-05 | 0.000173843 | 0.20649176 |
| ENSMUSG00000041028 | Ghitm | 66092 | protein_coding | 3596.526255 | -0.206507975 | 0.052841777 | 3.908043715 | 9.30E-05 | 0.000809147 | 0.206507975 |
| ENSMUSG00000049285 | Mblac1 | 330216 | protein_coding | 32.11125278 | -0.206511157 | 0.197843177 | 1.043812378 | 0.296572204 | 0.515753304 | 0.206511157 |
| ENSMUSG00000028619 | Tceanc2 | 66526 | protein_coding | 202.2862231 | -0.206523212 | 0.089578648 | 2.305495964 | 0.021138813 | 0.076654886 | 0.206523212 |
| ENSMUSG00000072763 | 5430403G16Rik | 77200 | protein_coding | 23.04182159 | -0.206556857 | 0.233927067 | 0.882996823 | 0.377237988 | 0.599511678 | 0.206556857 |
| ENSMUSG00000041530 | Ago1 | 236511 | protein_coding | 454.6063499 | -0.206617782 | 0.099950114 | 2.06720907 | 0.038714458 | 0.122877294 | 0.206617782 |
| ENSMUSG00000004798 | Ulk2 | 29869 | protein_coding | 240.2942813 | 0.206631235 | 0.091097359 | -2.26824618 | 0.023314206 | 0.082744638 | 0.206631235 |
| ENSMUSG00000112503 | Gm18409 | 100417125 | transcribed_processed_pseudogene | 1.048005245 | -0.206662656 | 1.473628462 | 0.140240679 | 0.888469832 | 1 | 0.206662656 |
| ENSMUSG00000086896 | Cyp4x1os | 105244662 | lncRNA | 0.889061718 | 0.206677864 | 1.759107218 | -0.117490203 | 0.906471607 | 1 | 0.206677864 |
| ENSMUSG00000048490 | Nrip1 | 268903 | protein_coding | 1634.877353 | -0.206714916 | 0.065911164 | 3.136265597 | 0.001711142 | 0.009942925 | 0.206714916 |
| ENSMUSG00000000308 | Ckmt1 | 12716 | protein_coding | 20666.60126 | -0.206748307 | 0.16326732 | 1.266317759 | 0.205399341 | 0.408031364 | 0.206748307 |
| ENSMUSG00000038268 | Ovca2 | 246257 | protein_coding | 99.58783479 | -0.206751445 | 0.130381207 | 1.585745743 | 0.112796998 | 0.269506609 | 0.206751445 |
| ENSMUSG00000051451 | Crebzf | 233490 | protein_coding | 554.0778256 | 0.206758492 | 0.08440822 | -2.449506602 | 0.014305209 | 0.056179837 | 0.206758492 |
| ENSMUSG00000093046 | Mir3081 | 100526478 | miRNA | 3.10805031 | 0.206824022 | 0.738887574 | -0.279912709 | 0.779544477 | 1 | 0.206824022 |
| ENSMUSG00000050697 | Prkaa1 | 105787 | protein_coding | 880.2944289 | -0.206853265 | 0.079157611 | 2.613182284 | 0.008970344 | 0.039003757 | 0.206853265 |
| ENSMUSG00000032382 | Snx1 | 56440 | protein_coding | 1394.478593 | -0.206881826 | 0.064282319 | 3.218331707 | 0.001289386 | 0.007860067 | 0.206881826 |

| ensembl_gene_id | external_gene_name | entrezgene_id | gene_biotype | baseMean | log2FoldChange | lfcSE | stat | pvalue | padj | absLog2FC |
|---|---|---|---|---|---|---|---|---|---|---|
| ENSMUSG00000042428 | Mgat3 | 17309 | protein_coding | 1908.2898 | 0.206885104 | 0.055161023 | -3.750566847 | 0.000176435 | 0.001419217 | 0.206885104 |
| ENSMUSG00000002109 | Ddb2 | 107986 | protein_coding | 72.44921646 | -0.206984326 | 0.125384914 | 1.650791314 | 0.098781194 | 0.24534105 | 0.206984326 |
| ENSMUSG00000005677 | Nr1i3 | 12355 | protein_coding | 10.5371726 | 0.206984762 | 0.37747649 | -0.548338155 | 0.583459733 | 0.766020418 | 0.206984762 |
| ENSMUSG00000031093 | Dock11 | 75974 | protein_coding | 138.9536522 | -0.207000826 | 0.139898449 | 1.479650614 | 0.138966512 | 0.312060508 | 0.207000826 |
| ENSMUSG00000043311 | D17H6S53E | 114585 | protein_coding | 401.9062859 | -0.207029614 | 0.060879493 | 3.400646187 | 0.000672268 | 0.00453877 | 0.207029614 |
| ENSMUSG00000063488 | Zkscan7 | 382118 | protein_coding | 170.4805137 | -0.207092189 | 0.088349776 | 2.344003541 | 0.01907799 | 0.070735054 | 0.207092189 |
| ENSMUSG00000049076 | Acap2 | 78618 | protein_coding | 1072.740486 | -0.207100257 | 0.096232255 | 2.152087738 | 0.031390445 | 0.104512461 | 0.207100257 |
| ENSMUSG00000027601 | Mtfr1 | 67472 | protein_coding | 519.3396278 | -0.207294188 | 0.07238311 | 2.863847501 | 0.004185295 | 0.020925206 | 0.207294188 |
| ENSMUSG00000117042 | 2700054A10Rik | 72578 | lncRNA | 14.77129315 | 0.207315338 | 0.308781755 | -0.671397628 | 0.501967256 | 0.706691156 | 0.207315338 |
| ENSMUSG00000032028 | Nxpe2 | 78252 | protein_coding | 1115.157147 | 0.207419392 | 0.401614609 | -0.516463763 | 0.605530542 | 0.780450414 | 0.207419392 |
| ENSMUSG00000000942 | Hoxa4 | 15401 | protein_coding | 8.188936131 | -0.207424078 | 0.371064091 | 0.558997983 | 0.576163097 | 0.761180591 | 0.207424078 |
| ENSMUSG00000025503 | Taldo1 | 21351 | protein_coding | 3564.379783 | -0.207428904 | 0.05969615 | 3.474745087 | 0.000511339 | 0.003617932 | 0.207428904 |
| ENSMUSG00000021278 | Amn | 93835 | protein_coding | 101.6828334 | 0.20743774 | 0.251504523 | -0.824787316 | 0.409492349 | 0.630052795 | 0.20743774 |
| ENSMUSG00000090641 | Zfp712 | 78251 | protein_coding | 44.95270168 | -0.207454343 | 0.159183107 | 1.303243458 | 0.192491658 | 0.390229222 | 0.207454343 |
| ENSMUSG00000048373 | Fgfbp1 | 14181 | protein_coding | 1346.403658 | 0.207457459 | 0.110272633 | -1.881314088 | 0.059929206 | 0.1706035 | 0.207457459 |
| ENSMUSG00000039850 | Endov | 338371 | protein_coding | 221.1012313 | -0.207492062 | 0.099191462 | 2.09183388 | 0.036453378 | 0.117005071 | 0.207492062 |
| ENSMUSG00000027722 | Spata5 | 57815 | protein_coding | 204.3676331 | -0.207697153 | 0.13012507 | 1.596134807 | 0.110458699 | 0.265751013 | 0.207697153 |
| ENSMUSG00000006642 | Tcf23 | 69852 | protein_coding | 51.30864262 | 0.207715901 | 0.18788101 | -1.105571557 | 0.268912006 | 0.484040442 | 0.207715901 |
| ENSMUSG00000025862 | Stag2 | 20843 | protein_coding | 1553.464545 | -0.207737079 | 0.082104067 | 2.530167959 | 0.011400794 | 0.0473047 | 0.207737079 |
| ENSMUSG00000053985 | Zfp14 | 243906 | protein_coding | 24.20739645 | -0.207805155 | 0.211754851 | 0.98134779 | 0.326421263 | 0.548343875 | 0.207805155 |
| ENSMUSG00000031226 | Pbdc1 | 67683 | protein_coding | 241.2586578 | 0.207911922 | 0.131291648 | -1.58358833 | 0.113287431 | 0.270127107 | 0.207911922 |
| ENSMUSG00000019158 | Tmem160 | 69094 | protein_coding | 147.6660645 | -0.207931635 | 0.129004047 | 1.611822579 | 0.107000553 | 0.260075004 | 0.207931635 |
| ENSMUSG00000035242 | Oaz1 | 18245 | protein_coding | 1681.83226 | -0.207967974 | 0.117436371 | 1.770899189 | 0.076577467 | 0.203672136 | 0.207967974 |
| ENSMUSG00000032530 | Lyzl4 | 69032 | protein_coding | 20.7651797 | 0.20800613 | 0.323121998 | -0.643738683 | 0.519744901 | 0.719935187 | 0.20800613 |
| ENSMUSG00000065304 | Gm23245 | 115490273 | snoRNA | 6.466674523 | 0.208082124 | 0.437635612 | -0.475468902 | 0.634452793 | 0.800714821 | 0.208082124 |
| ENSMUSG00000039157 | Fam102a | 98952 | protein_coding | 1615.432752 | -0.208098 | 0.124322102 | 1.673861666 | 0.09415779 | 0.236336338 | 0.208098 |
| ENSMUSG00000021061 | Sptb | 20741 | protein_coding | 41.58560082 | -0.208103374 | 0.306283905 | 0.679445999 | 0.496855312 | 0.702910017 | 0.208103374 |
| ENSMUSG00000033187 | BC016579 | 212998 | protein_coding | 416.7133934 | 0.208201138 | 0.090826076 | -2.292305764 | 0.021888005 | 0.07883501 | 0.208201138 |
| ENSMUSG00000017144 | Rnd3 | 74194 | protein_coding | 708.3211325 | -0.208203945 | 0.166694968 | 1.249011573 | 0.211660841 | 0.416065952 | 0.208203945 |
| ENSMUSG00000031608 | Galnt7 | 108150 | protein_coding | 3285.135978 | 0.208262201 | 0.163888461 | -1.270755731 | 0.203815562 | 0.406008467 | 0.208262201 |
| ENSMUSG00000030956 | Fam53b | 77938 | protein_coding | 330.0124419 | -0.208314742 | 0.093804874 | 2.220724079 | 0.026369655 | 0.090987984 | 0.208314742 |
| ENSMUSG00000000532 | Acvr1b | 11479 | protein_coding | 1416.073448 | -0.208322326 | 0.077532493 | 2.686903489 | 0.007211779 | 0.03273648 | 0.208322326 |
| ENSMUSG00000021072 | Tmx1 | 72736 | protein_coding | 853.8944756 | 0.208391728 | 0.11998267 | -1.736848554 | 0.082413906 | 0.214493541 | 0.208391728 |
| ENSMUSG00000023284 | Zfp605 | 675812 | protein_coding | 38.9507602 | -0.208490169 | 0.172568346 | 1.208159976 | 0.226985734 | 0.435409735 | 0.208490169 |
| ENSMUSG00000074165 | Zfp788 | 67607 | protein_coding | 87.99735588 | -0.208493813 | 0.173314167 | 1.202981939 | 0.22898331 | 0.437918979 | 0.208493813 |
| ENSMUSG00000052131 | Akr1b7 | 11997 | protein_coding | 40.79996727 | -0.208541884 | 0.215640617 | 0.967080723 | 0.333503681 | 0.555918017 | 0.208541884 |
| ENSMUSG00000098184 | Gm27011 | 115486886 | lncRNA | 3.560601229 | 0.208577349 | 0.55378526 | -0.376639402 | 0.706441602 | 1 | 0.208577349 |
| ENSMUSG00000114613 | Gm10732 | 100038690 | lncRNA | 11.55635613 | 0.208581517 | 0.325555715 | -0.640693766 | 0.521721665 | 0.721161093 | 0.208581517 |
| ENSMUSG00000042207 | Kdm5b | 75605 | protein_coding | 986.3470608 | -0.20861463 | 0.0674074 | 3.094832747 | 0.001969239 | 0.011209933 | 0.20861463 |

| ensembl_gene_id | external_gene_name | entrezgene_id | gene_biotype | baseMean | log2FoldChange | lfcSE | stat | pvalue | padj | absLog2FC |
|---|---|---|---|---|---|---|---|---|---|---|
| ENSMUSG00000034042 | Gpbp1l1 | 77110 | protein_coding | 1823.904987 | -0.20878117 | 0.065310306 | 3.196756853 | 0.00138982 | 0.008373401 | 0.20878117 |
| ENSMUSG00000042807 | Hecw2 | 329152 | protein_coding | 6.586202508 | -0.208793391 | 0.871761039 | 0.239507596 | 0.810712007 | 0.906297546 | 0.208793391 |
| ENSMUSG00000031808 | Slc27a1 | 26457 | protein_coding | 329.5812887 | -0.208807856 | 0.11456553 | 1.822606289 | 0.068363042 | 0.187813556 | 0.208807856 |
| ENSMUSG00000032477 | Cdc25a | 12530 | protein_coding | 658.6607289 | 0.208944955 | 0.101475055 | -2.059077034 | 0.039486859 | 0.124778744 | 0.208944955 |
| ENSMUSG00000078681 | Tm2d3 | 68634 | protein_coding | 278.0047087 | -0.208957782 | 0.120441904 | 1.734925929 | 0.082753926 | 0.215212336 | 0.208957782 |
| ENSMUSG00000072844 | G530011O06Rik | 654820 | lncRNA | 3.095660523 | -0.20897905 | 0.739162714 | 0.282724014 | 0.777388406 | 1 | 0.20897905 |
| ENSMUSG00000089214 | Mir1931 | 100316689 | miRNA | 1.157888644 | -0.208989211 | 1.184711852 | 0.176405098 | 0.8599757 | 1 | 0.208989211 |
| ENSMUSG00000047989 | Ino80c | 225280 | protein_coding | 286.6083973 | 0.209001466 | 0.150165216 | -1.39181011 | 0.163979908 | 0.350228168 | 0.209001466 |
| ENSMUSG00000001761 | Smo | 319757 | protein_coding | 337.6317393 | 0.209003491 | 0.118642168 | -1.761629067 | 0.07813199 | 0.206674944 | 0.209003491 |
| ENSMUSG00000083674 | Zfp133-ps | 668917 | transcribed_unitary_pseudogene | 6.779029773 | -0.209009054 | 0.464483465 | 0.449981688 | 0.652723644 | 0.811601896 | 0.209009054 |
| ENSMUSG00000035929 | H2-Q4 | 15015 | protein_coding | 209.9076726 | 0.20906416 | 0.135937674 | -1.537941272 | 0.124062975 | 0.288478734 | 0.20906416 |
| ENSMUSG00000072946 | Ptgr2 | 77219 | protein_coding | 505.755306 | -0.209076631 | 0.080199359 | 2.606961378 | 0.009134966 | 0.03960484 | 0.209076631 |
| ENSMUSG00000058248 | Kcnh1 | 16510 | protein_coding | 16.9023302 | -0.209118626 | 0.46308753 | 0.451574729 | 0.651575379 | 0.810851996 | 0.209118626 |
| ENSMUSG00000074466 | Gm15417 | 545539 | lncRNA | 63.45001478 | 0.209146019 | 0.185254487 | -1.128966015 | 0.25891217 | 0.472341459 | 0.209146019 |
| ENSMUSG00000060862 | Zbtb40 | 230848 | protein_coding | 294.9972238 | 0.209167938 | 0.122153017 | -1.712343599 | 0.086833366 | 0.223150671 | 0.209167938 |
| ENSMUSG00000002997 | Prkar2b | 19088 | protein_coding | 44.73563027 | 0.209205883 | 0.251444164 | -0.832017253 | 0.405399206 | 0.62645534 | 0.209205883 |
| ENSMUSG00000085208 | Brip1os | 74038 | lncRNA | 765.0448961 | 0.209234744 | 0.105427237 | -1.984636513 | 0.047184927 | 0.142788208 | 0.209234744 |
| ENSMUSG00000055762 | Eef1d | 66656 | protein_coding | 1426.872211 | 0.209293048 | 0.084879215 | -2.465775017 | 0.013671718 | 0.054259732 | 0.209293048 |
| ENSMUSG00000034853 | Acot11 | 329910 | protein_coding | 931.1586209 | -0.209320032 | 0.085198067 | 2.456863641 | 0.014015587 | 0.055305475 | 0.209320032 |
| ENSMUSG00000089239 | Gm22200 | 115489001 | snoRNA | 1.124106392 | 0.209359243 | 1.041889755 | -0.200941839 | 0.840744052 | 1 | 0.209359243 |
| ENSMUSG00000019868 | Vta1 | 66201 | protein_coding | 361.4492776 | 0.209502614 | 0.085932254 | -2.437997427 | 0.014768879 | 0.057630735 | 0.209502614 |
| ENSMUSG00000022543 | Dnaaf8 | 74684 | transcribed_unitary_pseudogene | 12.4420275 | 0.209532359 | 0.306993503 | -0.682530273 | 0.494903707 | 0.70086955 | 0.209532359 |
| ENSMUSG00000029276 | Glmn | 170823 | protein_coding | 46.26622113 | 0.209542592 | 0.203167292 | -1.031379556 | 0.302362864 | 0.522795554 | 0.209542592 |
| ENSMUSG00000038463 | Olfml2b | 320078 | protein_coding | 2.180191761 | 0.209544901 | 0.797627352 | -0.262710275 | 0.792773897 | 1 | 0.209544901 |
| ENSMUSG00000005225 | Plekha8 | 231999 | protein_coding | 1176.084052 | -0.209579068 | 0.053157538 | 3.942602978 | 8.06E-05 | 0.000714089 | 0.209579068 |
| ENSMUSG00000038069 | Cdkn2aip | 70925 | protein_coding | 206.6587952 | -0.209609606 | 0.094333383 | 2.222008794 | 0.026282711 | 0.090782827 | 0.209609606 |
| ENSMUSG00000026277 | Stk25 | 59041 | protein_coding | 2623.700028 | -0.209617363 | 0.095933384 | 2.185030435 | 0.028886615 | 0.097993747 | 0.209617363 |
| ENSMUSG00000057367 | Birc2 | 11797 | protein_coding | 485.0603058 | 0.209677081 | 0.085089599 | -2.464191667 | 0.013732266 | 0.054421559 | 0.209677081 |
| ENSMUSG00000037001 | Zfp39 | 22698 | protein_coding | 52.30052857 | -0.209687226 | 0.175123286 | 1.197369181 | 0.231162691 | 0.439947064 | 0.209687226 |
| ENSMUSG00000028825 | Rhd | 19746 | protein_coding | 1.008669881 | 0.209703173 | 1.25284208 | -0.167381968 | 0.867069512 | 1 | 0.209703173 |
| ENSMUSG00000006818 | Sod2 | 20656 | protein_coding | 1145.691561 | -0.209705163 | 0.070904511 | 2.957571525 | 0.003100728 | 0.01632826 | 0.209705163 |
| ENSMUSG00000116220 | A430088P11Rik | 106059 | lncRNA | 21.26649393 | -0.209774966 | 0.252639608 | 0.830332853 | 0.406350618 | 0.62720573 | 0.209774966 |
| ENSMUSG00000029068 | Ccnl2 | 56036 | protein_coding | 1269.798626 | 0.209822827 | 0.120409602 | -1.742575545 | 0.081407785 | 0.212413992 | 0.209822827 |
| ENSMUSG00000026600 | Soat1 | 20652 | protein_coding | 760.0751491 | -0.209833715 | 0.18154282 | 1.155835932 | 0.247748273 | 0.45990501 | 0.209833715 |
| ENSMUSG00000030105 | Arl8b | 67166 | protein_coding | 956.0467507 | 0.209863442 | 0.059777538 | -3.510740829 | 0.00044686 | 0.003226691 | 0.209863442 |
| ENSMUSG00000039853 | Trim14 | 74735 | protein_coding | 188.529233 | -0.2099016 | 0.086238801 | 2.433957784 | 0.014934737 | 0.058209168 | 0.2099016 |
| ENSMUSG00000029101 | Rgs12 | 71729 | protein_coding | 68.43575291 | 0.209919485 | 0.166022954 | -1.264400372 | 0.206086359 | 0.408658758 | 0.209919485 |
| ENSMUSG00000037845 | Fdxacb1 | 382137 | protein_coding | 49.29478893 | -0.20993285 | 0.161689031 | 1.298374098 | 0.194158815 | 0.392885514 | 0.20993285 |
| ENSMUSG00000049119 | Fam110b | 242297 | protein_coding | 15.78287163 | -0.209956646 | 0.315047274 | 0.666429021 | 0.505136918 | 0.709218761 | 0.209956646 |

| ensembl_gene_id | external_gene_name | entrezgene_id | gene_biotype | baseMean | log2FoldChange | lfcSE | stat | pvalue | padj | absLog2FC |
|---|---|---|---|---|---|---|---|---|---|---|
| ENSMUSG00000029104 | Htt | 15194 | protein_coding | 1133.20288 | 0.209984163 | 0.0617342 | -3.401423551 | 0.000670359 | 0.004529584 | 0.209984163 |
| ENSMUSG00000019779 | Frk | 14302 | protein_coding | 1425.054743 | -0.210016851 | 0.082300245 | 2.551837499 | 0.010715648 | 0.045027536 | 0.210016851 |
| ENSMUSG00000063052 | Lrrc40 | 67144 | protein_coding | 315.6628322 | 0.210094154 | 0.11643566 | -1.804379807 | 0.071171786 | 0.1933428 | 0.210094154 |
| ENSMUSG00000100334 | C230024C17Rik | 329271 | lncRNA | 1.39172506 | -0.210115282 | 1.514351122 | 0.138749382 | 0.889648194 | 1 | 0.210115282 |
| ENSMUSG00000062949 | Atp11c | 320940 | protein_coding | 295.9548987 | 0.210178365 | 0.090030579 | -2.334521983 | 0.019568406 | 0.072116193 | 0.210178365 |
| ENSMUSG00000002365 | Snx9 | 66616 | protein_coding | 973.715712 | -0.210181756 | 0.076055473 | 2.763532281 | 0.005717944 | 0.027204299 | 0.210181756 |
| ENSMUSG00000025188 | Hps1 | 192236 | protein_coding | 224.0090569 | -0.210219103 | 0.103884343 | 2.023587938 | 0.043012567 | 0.132867477 | 0.210219103 |
| ENSMUSG00000030337 | Vamp1 | 22317 | protein_coding | 124.9562162 | 0.210257245 | 0.186478231 | -1.1275163 | 0.259524253 | 0.472954995 | 0.210257245 |
| ENSMUSG00000030096 | Slc6a6 | 21366 | protein_coding | 1820.535582 | -0.210258143 | 0.081647583 | 2.575191273 | 0.010018467 | 0.042673448 | 0.210258143 |
| ENSMUSG00000020122 | Egfr | 13649 | protein_coding | 657.5154683 | -0.210287316 | 0.071400866 | 2.945164792 | 0.003227827 | 0.01691664 | 0.210287316 |
| ENSMUSG00000071267 | Zfp942 | 73233 | protein_coding | 227.9632678 | -0.210371738 | 0.084936916 | 2.476799805 | 0.013256624 | 0.053020073 | 0.210371738 |
| ENSMUSG00000060176 | Kif27 | 75050 | protein_coding | 31.74037844 | -0.210378264 | 0.295607036 | 0.711682195 | 0.476661596 | 0.68833522 | 0.210378264 |
| ENSMUSG00000043639 | Rbm20 | 73713 | protein_coding | 53.07914798 | -0.210387907 | 0.152517045 | 1.379438652 | 0.167759549 | 0.356273302 | 0.210387907 |
| ENSMUSG00000013539 | Tango2 | 27883 | protein_coding | 198.6100624 | 0.210400062 | 0.088909099 | -2.366462658 | 0.017958986 | 0.067342119 | 0.210400062 |
| ENSMUSG00000040746 | Rnf167 | 70510 | protein_coding | 583.0663972 | -0.210534138 | 0.068537992 | 3.071787382 | 0.002127812 | 0.011930684 | 0.210534138 |
| ENSMUSG00000075184 | F930017D23Rik | 414125 | lncRNA | 2.898767311 | 0.210604882 | 0.69374657 | -0.303576106 | 0.761450858 | 1 | 0.210604882 |
| ENSMUSG00000022584 | Ly6c2 | 100041546 | protein_coding | 3.649709887 | 0.210619312 | 1.008171375 | -0.208912212 | 0.83451677 | 1 | 0.210619312 |
| ENSMUSG00000027744 | Stoml3 | 229277 | protein_coding | 1.461318199 | 0.210626998 | 2.071065045 | -0.101699847 | 0.918994924 | 1 | 0.210626998 |
| ENSMUSG00000005533 | Igf1r | 16001 | protein_coding | 234.7836478 | -0.210690577 | 0.103300397 | 2.039591158 | 0.041391063 | 0.129309879 | 0.210690577 |
| ENSMUSG00000027624 | Epb41l1 | 13821 | protein_coding | 977.6629875 | -0.210715183 | 0.101028099 | 2.085708688 | 0.037005019 | 0.118476325 | 0.210715183 |
| ENSMUSG00000002996 | Hbp1 | 73389 | protein_coding | 471.840677 | -0.210794757 | 0.140845612 | 1.49663702 | 0.134487732 | 0.305198626 | 0.210794757 |
| ENSMUSG00000020877 | Scrn2 | 217140 | protein_coding | 197.1588123 | -0.210806363 | 0.08109543 | 2.599485117 | 0.009336373 | 0.040329809 | 0.210806363 |
| ENSMUSG00000027589 | Pcmtd2 | 245867 | protein_coding | 534.0745173 | -0.210903127 | 0.120194446 | 1.754682803 | 0.079313579 | 0.208654479 | 0.210903127 |
| ENSMUSG00000030861 | Acadsb | 66885 | protein_coding | 272.1895655 | -0.21090754 | 0.132400578 | 1.592950294 | 0.111171342 | 0.26676599 | 0.21090754 |
| ENSMUSG00000028706 | Nsun4 | 72181 | protein_coding | 140.4189279 | -0.210923407 | 0.106275067 | 1.984693244 | 0.047178611 | 0.142788208 | 0.210923407 |
| ENSMUSG00000037343 | Taf2 | 319944 | protein_coding | 449.6341745 | 0.210957105 | 0.063976885 | -3.29739569 | 0.000975859 | 0.006239 | 0.210957105 |
| ENSMUSG00000023017 | Asic1 | 11419 | protein_coding | 47.06770427 | -0.211041736 | 0.15937937 | 1.324147136 | 0.185454184 | 0.380783029 | 0.211041736 |
| ENSMUSG00000052504 | Epha3 | 13837 | protein_coding | 3.18078286 | -0.211102461 | 0.959396272 | 0.220036774 | 0.825842515 | 1 | 0.211102461 |
| ENSMUSG00000052293 | Taf9 | 108143 | protein_coding | 1.378840637 | -0.211122395 | 0.942960294 | 0.223893197 | 0.82284041 | 1 | 0.211122395 |
| ENSMUSG00000028838 | Extl1 | 56219 | protein_coding | 13.56566019 | 0.211144467 | 0.336976024 | -0.626586025 | 0.530930631 | 0.728048657 | 0.211144467 |
| ENSMUSG00000038302 | Afg1l | 215951 | protein_coding | 359.9376698 | -0.211282593 | 0.082886691 | 2.5490533 | 0.010801579 | 0.045254367 | 0.211282593 |
| ENSMUSG00000028145 | Them4 | 75778 | protein_coding | 92.23787718 | -0.211301966 | 0.131469807 | 1.607228078 | 0.108004345 | 0.261594876 | 0.211301966 |
| ENSMUSG00000064247 | Plcxd1 | 403178 | protein_coding | 13.3442995 | -0.211312571 | 0.358294205 | 0.589773901 | 0.555342242 | 0.745058233 | 0.211312571 |
| ENSMUSG00000023912 | Slc25a27 | 74011 | protein_coding | 12.82463405 | -0.211342791 | 0.328599704 | 0.643161843 | 0.52011909 | 0.72021186 | 0.211342791 |
| ENSMUSG00000052889 | Prkcb | 18751 | protein_coding | 7.335619186 | -0.211397181 | 0.719865681 | 0.293661979 | 0.76901621 | 0.884245488 | 0.211397181 |
| ENSMUSG00000030555 | Ttc23 | 67009 | protein_coding | 144.7492363 | 0.211430048 | 0.083982618 | -2.517545334 | 0.011817576 | 0.048509931 | 0.211430048 |
| ENSMUSG00000028699 | Tspan1 | 66805 | protein_coding | 13305.48047 | -0.211514293 | 0.085898349 | 2.462379016 | 0.013801873 | 0.05461877 | 0.211514293 |
| ENSMUSG00000000959 | Oxa1l | 69089 | protein_coding | 630.3510781 | -0.211525832 | 0.086292794 | 2.451257187 | 0.01423582 | 0.055973887 | 0.211525832 |
| ENSMUSG00000036241 | Ube2r2 | 67615 | protein_coding | 890.0589716 | -0.211556805 | 0.08643015 | 2.447719983 | 0.014376334 | 0.056392107 | 0.211556805 |

| ensembl_gene_id | external_gene_name | entrezgene_id | gene_biotype | baseMean | log2FoldChange | lfcSE | stat | pvalue | padj | absLog2FC |
|---|---|---|---|---|---|---|---|---|---|---|
| ENSMUSG00000003868 | Ruvbl2 | 20174 | protein_coding | 598.1396622 | 0.211559989 | 0.098702512 | -2.143410384 | 0.03208016 | 0.106287366 | 0.211559989 |
| ENSMUSG00000035051 | Dhx57 | 106794 | protein_coding | 287.1984503 | 0.211724449 | 0.06602234 | -3.206860721 | 0.001341919 | 0.008129295 | 0.211724449 |
| ENSMUSG00000030089 | Slc41a3 | 71699 | protein_coding | 25.89017749 | 0.211791925 | 0.224817672 | -0.942060843 | 0.346161491 | 0.567902926 | 0.211791925 |
| ENSMUSG00000016496 | Cd274 | 60533 | protein_coding | 22.9978959 | -0.211807789 | 0.277357862 | 0.763662465 | 0.445068414 | 0.662372535 | 0.211807789 |
| ENSMUSG00000045374 | Wdr81 | 192652 | protein_coding | 406.8684636 | -0.211839846 | 0.071659304 | 2.956208524 | 0.003114464 | 0.016390142 | 0.211839846 |
| ENSMUSG00000003271 | Sult2b1 | 54200 | protein_coding | 226.6600151 | -0.211897405 | 0.175930304 | 1.204439485 | 0.228419758 | 0.437296996 | 0.211897405 |
| ENSMUSG00000035778 | Ggta1 | 14594 | protein_coding | 4.873030973 | 0.211898521 | 0.768892726 | -0.275589187 | 0.782863616 | 0.891186777 | 0.211898521 |
| ENSMUSG00000012535 | Tnpo3 | 320938 | protein_coding | 983.6304492 | 0.211917761 | 0.080051234 | -2.647276638 | 0.008114294 | 0.036011679 | 0.211917761 |
| ENSMUSG00000037363 | Letm2 | 270035 | protein_coding | 19.45939686 | 0.21192054 | 0.303186776 | -0.698976858 | 0.484566493 | 0.695051774 | 0.21192054 |
| ENSMUSG00000031430 | Vsig1 | 78789 | protein_coding | 86.60410586 | 0.211923331 | 0.31028291 | -0.683000334 | 0.494606631 | 0.700749306 | 0.211923331 |
| ENSMUSG00000067873 | Htatsf1 | 72459 | protein_coding | 417.1381847 | 0.211956075 | 0.090866115 | -2.332619547 | 0.019668122 | 0.07241317 | 0.211956075 |
| ENSMUSG00000064128 | Cenpj | 219103 | protein_coding | 67.83136536 | 0.211956891 | 0.132667062 | -1.597660255 | 0.110118609 | 0.265192684 | 0.211956891 |
| ENSMUSG00000071192 | Wfikkn1 | 215001 | protein_coding | 17.86020514 | 0.212049342 | 0.312435699 | -0.678697546 | 0.497329521 | 0.702918497 | 0.212049342 |
| ENSMUSG00000039737 | Prkrip1 | 66801 | protein_coding | 176.9186959 | -0.212052646 | 0.096614089 | 2.194841842 | 0.028174945 | 0.095954019 | 0.212052646 |
| ENSMUSG00000046329 | Slc25a23 | 66972 | protein_coding | 125.473777 | 0.212075495 | 0.160730649 | -1.319446516 | 0.187019879 | 0.383134386 | 0.212075495 |
| ENSMUSG00000003929 | Zfp81 | 224694 | protein_coding | 108.6522589 | -0.212085531 | 0.143222999 | 1.480806383 | 0.138658175 | 0.311717339 | 0.212085531 |
| ENSMUSG00000038538 | Ubn2 | 320538 | protein_coding | 627.1263135 | 0.212090516 | 0.109233534 | -1.941624587 | 0.052182564 | 0.154268325 | 0.212090516 |
| ENSMUSG00000067787 | Blcap | 53619 | protein_coding | 342.739281 | -0.21211707 | 0.098875176 | 2.145301567 | 0.031928743 | 0.105941585 | 0.21211707 |
| ENSMUSG00000021185 | Dglucy | 217830 | protein_coding | 359.8030972 | 0.212248403 | 0.097953865 | -2.166820087 | 0.030248575 | 0.101632751 | 0.212248403 |
| ENSMUSG00000043008 | Klhl6 | 239743 | protein_coding | 3.949150141 | 0.212325548 | 0.672263047 | -0.315837007 | 0.752126228 | 1 | 0.212325548 |
| ENSMUSG00000084883 | Ccdc85c | 668158 | protein_coding | 1333.350434 | 0.212380865 | 0.089594698 | -2.370462431 | 0.017765849 | 0.066769512 | 0.212380865 |
| ENSMUSG00000084845 | Tmem240 | 381582 | protein_coding | 5.388108631 | -0.212399623 | 1.043064255 | 0.20363043 | 0.838642311 | 0.919610832 | 0.212399623 |
| ENSMUSG00000032235 | Ice2 | 93697 | protein_coding | 179.3292818 | -0.21243441 | 0.095013247 | 2.235839906 | 0.02536226 | 0.08834262 | 0.21243441 |
| ENSMUSG00000021752 | Kctd6 | 71393 | protein_coding | 83.97150941 | -0.212458934 | 0.115639064 | 1.837259197 | 0.06617164 | 0.183322799 | 0.212458934 |
| ENSMUSG00000031885 | Cbfb | 12400 | protein_coding | 534.0730638 | 0.212476646 | 0.077839125 | -2.729689536 | 0.0063394 | 0.029324606 | 0.212476646 |
| ENSMUSG00000019338 | Zfp687 | 78266 | protein_coding | 474.6390035 | 0.21251164 | 0.066671121 | -3.187461615 | 0.001435275 | 0.008569101 | 0.21251164 |
| ENSMUSG00000030588 | Yif1b | 77254 | protein_coding | 294.9339213 | 0.212587654 | 0.121747496 | -1.746135734 | 0.080787369 | 0.21130879 | 0.212587654 |
| ENSMUSG00000021245 | Mlh3 | 217716 | protein_coding | 168.8800542 | 0.212711077 | 0.115951642 | -1.834480936 | 0.066582635 | 0.184147653 | 0.212711077 |
| ENSMUSG00000051790 | Nlgn2 | 216856 | protein_coding | 28.66968994 | 0.212713872 | 0.432255364 | -0.492102329 | 0.622647006 | 0.793088519 | 0.212713872 |
| ENSMUSG00000032295 | Man2c1 | 73744 | protein_coding | 458.7438721 | -0.212745211 | 0.061334851 | 3.468586085 | 0.000523205 | 0.003686094 | 0.212745211 |
| ENSMUSG00000049439 | Cyp20a1 | 77951 | protein_coding | 245.4383426 | 0.212746581 | 0.158775004 | -1.339924899 | 0.180269762 | 0.373991816 | 0.212746581 |
| ENSMUSG00000027253 | Lrp4 | 228357 | protein_coding | 738.5799724 | -0.212780098 | 0.079636378 | 2.671895739 | 0.007542408 | 0.03390183 | 0.212780098 |
| ENSMUSG00000025742 | Prps2 | 110639 | protein_coding | 498.215185 | -0.212953285 | 0.149224826 | 1.427063381 | 0.153561626 | 0.334862894 | 0.212953285 |
| ENSMUSG00000038604 | Ripor1 | 75687 | protein_coding | 68.96785528 | -0.212969173 | 0.21786414 | 0.977532021 | 0.328305833 | 0.55048001 | 0.212969173 |
| ENSMUSG00000097091 | 4930526L06Rik | 75138 | lncRNA | 0.603910236 | -0.212971876 | 1.461890278 | 0.145682531 | 0.884172012 | 1 | 0.212971876 |
| ENSMUSG00000051502 | Ufsp1 | 70240 | protein_coding | 50.43454267 | -0.21304229 | 0.216564089 | 0.983737848 | 0.325244426 | 0.547052145 | 0.21304229 |
| ENSMUSG00000025487 | Psmd13 | 23997 | protein_coding | 889.6202986 | 0.213076246 | 0.064213816 | -3.318230573 | 0.000905897 | 0.005852887 | 0.213076246 |
| ENSMUSG00000027376 | Prom2 | 192212 | protein_coding | 3.749898875 | 0.213096806 | 1.591851693 | -0.133867248 | 0.893507549 | 1 | 0.213096806 |
| ENSMUSG00000057411 | Antkmt | 214917 | protein_coding | 188.3063755 | -0.213123578 | 0.096661479 | 2.204845005 | 0.027464973 | 0.09404034 | 0.213123578 |

Supplementary Table 1

| ensembl_gene_id | external_gene_name | entrezgene_id | gene_biotype | baseMean | log2FoldChange | lfcSE | stat | pvalue | padj | absLog2FC |
|---|---|---|---|---|---|---|---|---|---|---|
| ENSMUSG00000031517 | Gpm6a | 234267 | protein_coding | 0.805656837 | 0.213204618 | 1.395955444 | -0.152730246 | 0.878611008 | 1 | 0.213204618 |
| ENSMUSG00000055923 | Aasdh | 231326 | protein_coding | 105.6313234 | -0.213207738 | 0.120369786 | 1.771272882 | 0.076515335 | 0.203572456 | 0.213207738 |
| ENSMUSG00000020839 | Tmigd1 | 66601 | protein_coding | 708.8029619 | 0.213235384 | 0.214829799 | -0.99257824 | 0.320915533 | 0.542047573 | 0.213235384 |
| ENSMUSG00000021918 | Nek4 | 23955 | protein_coding | 218.3299937 | 0.213292478 | 0.103269435 | -2.065397936 | 0.038885364 | 0.123253915 | 0.213292478 |
| ENSMUSG00000029918 | Mrps33 | 14548 | protein_coding | 77.91356926 | -0.213295131 | 0.152516166 | 1.398508344 | 0.161960469 | 0.347172272 | 0.213295131 |
| ENSMUSG00000020514 | Mrpl22 | 216767 | protein_coding | 150.0328645 | -0.213305022 | 0.112260936 | 1.900082344 | 0.057422314 | 0.165344954 | 0.213305022 |
| ENSMUSG00000024875 | Yif1a | 68090 | protein_coding | 835.8305966 | 0.213337242 | 0.074227625 | -2.874094938 | 0.004051871 | 0.020375337 | 0.213337242 |
| ENSMUSG00000079470 | Utp14b | 195434 | protein_coding | 1360.485252 | -0.213374858 | 0.148504379 | 1.436825359 | 0.150767623 | 0.330823349 | 0.213374858 |
| ENSMUSG00000048109 | Rbm15 | 229700 | protein_coding | 651.6564679 | 0.21341483 | 0.065857501 | -3.240554637 | 0.001192974 | 0.00736753 | 0.21341483 |
| ENSMUSG00000074656 | Eif2s2 | 67204 | protein_coding | 2202.84368 | -0.213416593 | 0.127505647 | 1.673781507 | 0.094173548 | 0.236336338 | 0.213416593 |
| ENSMUSG00000038239 | Hrc | 15464 | protein_coding | 25.76938739 | -0.213504228 | 0.445864944 | 0.478854036 | 0.632042472 | 0.799444652 | 0.213504228 |
| ENSMUSG00000040471 | Ggt6 | 71522 | protein_coding | 652.5119818 | -0.213516817 | 0.098871835 | 2.159531249 | 0.030808976 | 0.102970942 | 0.213516817 |
| ENSMUSG00000023961 | Enpp4 | 224794 | protein_coding | 151.5026769 | -0.213765396 | 0.128866477 | 1.658813075 | 0.097153465 | 0.241953298 | 0.213765396 |
| ENSMUSG00000022749 | Tbc1d23 | 67581 | protein_coding | 413.6418274 | 0.213775139 | 0.090985248 | -2.34955823 | 0.018795704 | 0.069955433 | 0.213775139 |
| ENSMUSG00000086424 | Gm15569 | 102638208 | lncRNA | 2.105176772 | -0.213777194 | 0.975651621 | 0.219112221 | 0.826562631 | 1 | 0.213777194 |
| ENSMUSG00000024730 | Ms4a8a | 64381 | protein_coding | 4049.477522 | -0.213784696 | 0.073199596 | 2.920572064 | 0.003493894 | 0.018083738 | 0.213784696 |
| ENSMUSG00000041592 | Sdk2 | 237979 | protein_coding | 7.301054416 | -0.213800064 | 1.228968708 | 0.173967053 | 0.861891355 | 0.931842966 | 0.213800064 |
| ENSMUSG00000002006 | Pdzd4 | 245469 | protein_coding | 18.38897037 | 0.213813944 | 0.296602028 | -0.720878228 | 0.47098444 | 0.684066582 | 0.213813944 |
| ENSMUSG00000032252 | Glce | 93683 | protein_coding | 560.5823529 | 0.213895075 | 0.116489593 | -1.836173247 | 0.066332038 | 0.183669899 | 0.213895075 |
| ENSMUSG00000093806 | Asmt | 107626 | protein_coding | 1.859854988 | 0.213912542 | 1.460277445 | -0.146487603 | 0.883536476 | 1 | 0.213912542 |
| ENSMUSG00000005299 | Letm1 | 56384 | protein_coding | 1739.600115 | -0.213915551 | 0.074306798 | 2.878815353 | 0.00399172 | 0.020134166 | 0.213915551 |
| ENSMUSG00000036073 | Galt | 14430 | protein_coding | 220.1748724 | -0.213932089 | 0.073348426 | 2.916655474 | 0.003538063 | 0.018266571 | 0.213932089 |
| ENSMUSG00000097814 | Panct2 | 329217 | lncRNA | 24.59897379 | 0.213961224 | 0.265990882 | -0.804393076 | 0.421169998 | 0.640973307 | 0.213961224 |
| ENSMUSG00000025103 | Btbd1 | 83962 | protein_coding | 652.2586915 | -0.213985996 | 0.091028078 | 2.35076915 | 0.018734653 | 0.069759652 | 0.213985996 |
| ENSMUSG00000027111 | Itga6 | 16403 | protein_coding | 3278.155289 | -0.214051926 | 0.071521013 | 2.99285364 | 0.002763823 | 0.014862184 | 0.214051926 |
| ENSMUSG00000026389 | Steap3 | 68428 | protein_coding | 343.2584034 | -0.214164316 | 0.081311681 | 2.633869001 | 0.008441805 | 0.037106194 | 0.214164316 |
| ENSMUSG00000020661 | Dnmt3a | 13435 | protein_coding | 667.1065112 | -0.214170615 | 0.088050162 | 2.432370466 | 0.015000356 | 0.05840978 | 0.214170615 |
| ENSMUSG00000041831 | Sytl3 | 83672 | protein_coding | 41.66489984 | -0.214174434 | 0.349820381 | 0.612241156 | 0.540378216 | 0.734651013 | 0.214174434 |
| ENSMUSG00000036752 | Tubb4b | 227613 | protein_coding | 7018.308984 | 0.214218861 | 0.096797943 | -2.213051798 | 0.026894075 | 0.09253023 | 0.214218861 |
| ENSMUSG00000038482 | Tfdp1 | 21781 | protein_coding | 658.5221146 | 0.214222661 | 0.121869238 | -1.757807504 | 0.078780268 | 0.207790664 | 0.214222661 |
| ENSMUSG00000037966 | Ninj1 | 18081 | protein_coding | 194.1435797 | -0.214267032 | 0.079806279 | 2.684839264 | 0.007256469 | 0.032851834 | 0.214267032 |
| ENSMUSG00000040022 | Rab11fip2 | 74998 | protein_coding | 76.04458807 | -0.214303435 | 0.149259967 | 1.435773033 | 0.151066936 | 0.331061353 | 0.214303435 |
| ENSMUSG00000059112 | Or4c52 | 258790 | protein_coding | 1.084298549 | -0.214345386 | 2.046639544 | 0.104730404 | 0.916589736 | 1 | 0.214345386 |
| ENSMUSG00000064967 | Gm24405 | 115487650 | snRNA | 1.174392592 | 0.214389654 | 1.45459771 | -0.147387592 | 0.882826099 | 1 | 0.214389654 |
| ENSMUSG00000066842 | Hmcn1 | 545370 | protein_coding | 421.9518776 | 0.214412657 | 0.175013385 | -1.225121477 | 0.220529454 | 0.426993013 | 0.214412657 |
| ENSMUSG00000038524 | Fchsd1 | 319262 | protein_coding | 94.75259267 | -0.214466197 | 0.136054364 | 1.576327215 | 0.114950413 | 0.273253364 | 0.214466197 |
| ENSMUSG00000010936 | Vac14 | 234729 | protein_coding | 330.893306 | 0.21448811 | 0.068943544 | -3.111068802 | 0.001864115 | 0.01071493 | 0.21448811 |
| ENSMUSG00000040356 | Skiv2l | 108077 | protein_coding | 1132.095621 | 0.214503202 | 0.065785003 | -3.26067022 | 0.001111492 | 0.006960631 | 0.214503202 |
| ENSMUSG00000037847 | Nmrk1 | 225994 | protein_coding | 48.1001834 | 0.214557823 | 0.168686233 | -1.271934407 | 0.203396426 | 0.40571187 | 0.214557823 |

Supplementary Table 1

| ensembl_gene_id | external_gene_name | entrezgene_id | gene_biotype | baseMean | log2FoldChange | lfcSE | stat | pvalue | padj | absLog2FC |
|---|---|---|---|---|---|---|---|---|---|---|
| ENSMUSG00000018476 | Kdm6b | 216850 | protein_coding | 397.2410629 | 0.214704055 | 0.275597517 | -0.779049308 | 0.435950669 | 0.654124055 | 0.214704055 |
| ENSMUSG00000095620 | Csta2 | 76770 | protein_coding | 5.714403864 | 0.214851854 | 0.788240914 | -0.272571304 | 0.785182773 | 0.892764273 | 0.214851854 |
| ENSMUSG00000020918 | Kat2a | 14534 | protein_coding | 279.0384518 | -0.214922585 | 0.107509367 | 1.999105666 | 0.045596922 | 0.139132959 | 0.214922585 |
| ENSMUSG00000086382 | Chrna1os | 102633437 | lncRNA | 6.592769829 | -0.214923332 | 0.375169999 | 0.572869187 | 0.566733269 | 0.753213832 | 0.214923332 |
| ENSMUSG00000041961 | Znrf3 | 407821 | protein_coding | 161.9108852 | -0.214977172 | 0.128558501 | 1.672212803 | 0.094482367 | 0.236872902 | 0.214977172 |
| ENSMUSG00000006576 | Slc4a3 | 20536 | protein_coding | 13.22576273 | -0.21498554 | 0.517232477 | 0.415645865 | 0.67766916 | 0.827306109 | 0.21498554 |
| ENSMUSG00000055210 | Foxd2 | 17301 | protein_coding | 714.9593691 | 0.214993114 | 0.100747611 | -2.133977288 | 0.03284464 | 0.108241146 | 0.214993114 |
| ENSMUSG00000039615 | Stub1 | 56424 | protein_coding | 408.9604448 | 0.215018155 | 0.075987494 | -2.82965187 | 0.004659868 | 0.022923163 | 0.215018155 |
| ENSMUSG00000006342 | Susd2 | 71733 | protein_coding | 5.764627995 | 0.215060266 | 0.756227432 | -0.284385698 | 0.776114818 | 0.888226437 | 0.215060266 |
| ENSMUSG00000026860 | Sh3glb2 | 227700 | protein_coding | 726.4911253 | -0.215090895 | 0.080765101 | 2.663166315 | 0.007740914 | 0.034680809 | 0.215090895 |
| ENSMUSG00000029409 | U90926 | 57425 | lncRNA | 0.856508635 | 0.215179619 | 1.46875409 | -0.146504865 | 0.88352285 | 1 | 0.215179619 |
| ENSMUSG00000018999 | Slc35b4 | 58246 | protein_coding | 246.3249837 | 0.215243237 | 0.104770683 | -2.054422395 | 0.039934827 | 0.125783662 | 0.215243237 |
| ENSMUSG00000027993 | Trim2 | 80890 | protein_coding | 4173.692062 | -0.215274243 | 0.077343895 | 2.783338535 | 0.005380263 | 0.025873521 | 0.215274243 |
| ENSMUSG00000070574 | 2310016G11Rik | 69578 | lncRNA | 60.31334871 | -0.215311655 | 0.255569014 | 0.842479498 | 0.399519611 | 0.621711048 | 0.215311655 |
| ENSMUSG00000038641 | Akr1d1 | 208665 | protein_coding | 0.955054165 | -0.215330907 | 1.286974435 | 0.167315606 | 0.867121726 | 1 | 0.215330907 |
| ENSMUSG00000085696 | Hoxaas3 | 72628 | lncRNA | 128.6348398 | 0.215336826 | 0.127706324 | -1.686187653 | 0.091759661 | 0.231983933 | 0.215336826 |
| ENSMUSG00000007850 | Hnrnph1 | 59013 | protein_coding | 1972.792708 | 0.215338231 | 0.049783095 | -4.325529238 | 1.52E-05 | 0.000161703 | 0.215338231 |
| ENSMUSG00000034573 | Ptpn13 | 19249 | protein_coding | 329.5704706 | -0.215384847 | 0.123225613 | 1.747890251 | 0.080483033 | 0.210900795 | 0.215384847 |
| ENSMUSG00000006589 | Aprt | 11821 | protein_coding | 894.4897775 | -0.215392005 | 0.073873065 | 2.915704204 | 0.003548868 | 0.018308655 | 0.215392005 |
| ENSMUSG00000006281 | Tep1 | 21745 | protein_coding | 2204.324474 | -0.215507074 | 0.070136695 | 3.072672218 | 0.002121514 | 0.011900366 | 0.215507074 |
| ENSMUSG00000027901 | Dennd2d | 72121 | protein_coding | 891.9642785 | 0.215604586 | 0.093316189 | -2.310473557 | 0.020861951 | 0.075817398 | 0.215604586 |
| ENSMUSG00000027655 | Dhx35 | 71715 | protein_coding | 261.2147952 | -0.215632743 | 0.111682108 | 1.9307725 | 0.053511193 | 0.157015607 | 0.215632743 |
| ENSMUSG00000006611 | Hfe | 15216 | protein_coding | 105.6602328 | -0.215655529 | 0.108352034 | 1.990322853 | 0.046555382 | 0.141278128 | 0.215655529 |
| ENSMUSG00000043467 | Zbtb37 | 240869 | protein_coding | 111.5949704 | -0.21566214 | 0.104840557 | 2.057048783 | 0.039681534 | 0.12530374 | 0.21566214 |
| ENSMUSG00000118346 | Tmem179b | 67706 | protein_coding | 176.9657056 | -0.215694279 | 0.093151111 | 2.315530927 | 0.020583894 | 0.075004948 | 0.215694279 |
| ENSMUSG00000110424 | Zbed5-ps | 75479 | lncRNA | 13.89530209 | 0.215717013 | 0.376528881 | -0.572909606 | 0.5667059 | 0.753213832 | 0.215717013 |
| ENSMUSG00000028337 | Coro2a | 107684 | protein_coding | 3492.778217 | -0.215733137 | 0.125830513 | 1.714473953 | 0.086441717 | 0.222351793 | 0.215733137 |
| ENSMUSG00000054662 | Ano9 | 71345 | protein_coding | 835.1249309 | -0.215758157 | 0.132585052 | 1.627318869 | 0.103669392 | 0.254141273 | 0.215758157 |
| ENSMUSG00000063229 | Ldha | 16828 | protein_coding | 9330.667281 | -0.215785753 | 0.074195024 | 2.908358816 | 0.003633312 | 0.018683911 | 0.215785753 |
| ENSMUSG00000059208 | Hnrnpm | 76936 | protein_coding | 2163.964709 | 0.215788923 | 0.070333765 | -3.06807012 | 0.002154461 | 0.012036117 | 0.215788923 |
| ENSMUSG00000024937 | Ehbp1l1 | 114601 | protein_coding | 727.847952 | 0.215798718 | 0.143644966 | -1.502306165 | 0.133018057 | 0.302752759 | 0.215798718 |
| ENSMUSG00000033039 | Micall1 | 27008 | protein_coding | 145.5588945 | -0.215816589 | 0.100339471 | 2.150864338 | 0.031486909 | 0.104749157 | 0.215816589 |
| ENSMUSG00000046971 | Pla2g4f | 271844 | protein_coding | 124.4601281 | 0.215826651 | 0.199902969 | -1.079657055 | 0.280294924 | 0.497933561 | 0.215826651 |
| ENSMUSG00000027189 | Trim44 | 80985 | protein_coding | 499.0788136 | -0.215903311 | 0.131470251 | 1.642221783 | 0.10054405 | 0.248746733 | 0.215903311 |
| ENSMUSG00000030203 | Dusp16 | 70686 | protein_coding | 407.2800466 | -0.215936183 | 0.118847906 | 1.816911976 | 0.069230593 | 0.189409212 | 0.215936183 |
| ENSMUSG00000066150 | Slc31a1 | 20529 | protein_coding | 1213.844122 | -0.215956803 | 0.081811809 | 2.639677651 | 0.008298492 | 0.036642767 | 0.215956803 |
| ENSMUSG00000029036 | Atad3a | 108888 | protein_coding | 685.3869675 | -0.215973112 | 0.07922641 | 2.726024224 | 0.006410229 | 0.02958595 | 0.215973112 |
| ENSMUSG00000034659 | Tmem109 | 68539 | protein_coding | 329.1040426 | -0.216030968 | 0.130840555 | 1.651100969 | 0.098717959 | 0.245220876 | 0.216030968 |
| ENSMUSG00000001827 | Folr1 | 14275 | protein_coding | 26.07124678 | 0.216040871 | 0.257832947 | -0.837910259 | 0.402081106 | 0.623763963 | 0.216040871 |

Supplementary Table 1

| ensembl_gene_id | external_gene_name | entrezgene_id | gene_biotype | baseMean | log2FoldChange | lfcSE | stat | pvalue | padj | absLog2FC |
|---|---|---|---|---|---|---|---|---|---|---|
| ENSMUSG00000028464 | Tpm2 | 22004 | protein_coding | 43.54387531 | -0.216059168 | 0.348126304 | 0.620634424 | 0.534840185 | 0.730731431 | 0.216059168 |
| ENSMUSG00000007564 | Ppp2r1a | 51792 | protein_coding | 3157.046018 | 0.216107751 | 0.062370661 | -3.464894326 | 0.00053044 | 0.00373044 | 0.216107751 |
| ENSMUSG00000040505 | Abcg5 | 27409 | protein_coding | 31.56531548 | -0.216167954 | 0.282100645 | 0.766279546 | 0.443509986 | 0.660955754 | 0.216167954 |
| ENSMUSG00000086401 | Gm15559 | 100504049 | lncRNA | 4.930663608 | -0.216214251 | 0.5587316 | 0.386973372 | 0.698775921 | 0.840342672 | 0.216214251 |
| ENSMUSG00000028578 | Caap1 | 67770 | protein_coding | 128.4473468 | 0.216230368 | 0.095287174 | -2.269249456 | 0.02325316 | 0.082564172 | 0.216230368 |
| ENSMUSG00000028284 | Map3k7 | 26409 | protein_coding | 634.418284 | 0.216332754 | 0.063213145 | -3.422274834 | 0.000620995 | 0.004253469 | 0.216332754 |
| ENSMUSG00000041264 | Uspl1 | 231915 | protein_coding | 303.0697174 | 0.216355551 | 0.102272489 | -2.115481415 | 0.034388937 | 0.11194538 | 0.216355551 |
| ENSMUSG00000009075 | Cabp7 | 192650 | protein_coding | 1.646629071 | 0.21637376 | 1.426541108 | -0.151677199 | 0.879441541 | 1 | 0.21637376 |
| ENSMUSG00000044783 | Hjurp | 212427 | protein_coding | 1736.373403 | 0.216395705 | 0.073666933 | -2.937487635 | 0.003308834 | 0.017269937 | 0.216395705 |
| ENSMUSG00000028164 | Manba | 110173 | protein_coding | 357.5683004 | -0.216431642 | 0.103871252 | 2.083652959 | 0.037191746 | 0.118981886 | 0.216431642 |
| ENSMUSG00000041718 | Alg13 | 67574 | protein_coding | 58.81036003 | 0.216474217 | 0.163069538 | -1.327496348 | 0.184344547 | 0.379371759 | 0.216474217 |
| ENSMUSG00000017802 | Retreg3 | 67998 | protein_coding | 1053.464675 | -0.216533853 | 0.087037998 | 2.487808279 | 0.0128533 | 0.051707527 | 0.216533853 |
| ENSMUSG00000061411 | Nol4l | 329540 | protein_coding | 749.0827583 | 0.216547925 | 0.139526345 | -1.552021772 | 0.120657014 | 0.283150481 | 0.216547925 |
| ENSMUSG00000028140 | Mrpl9 | 78523 | protein_coding | 697.381977 | -0.21658563 | 0.091605863 | 2.364320622 | 0.018063172 | 0.06767935 | 0.21658563 |
| ENSMUSG00000028434 | Epb41l4b | 54357 | protein_coding | 2776.812878 | -0.216604453 | 0.093135928 | 2.325680942 | 0.020035578 | 0.073420411 | 0.216604453 |
| ENSMUSG00000029246 | Ppat | 231327 | protein_coding | 250.3846128 | -0.216638819 | 0.164733232 | 1.31508874 | 0.18848008 | 0.38502722 | 0.216638819 |
| ENSMUSG00000038608 | Dock10 | 210293 | protein_coding | 31.99631213 | -0.216642107 | 0.482153888 | 0.449321496 | 0.653199753 | 0.811870577 | 0.216642107 |
| ENSMUSG00000028907 | Utp11 | 67205 | protein_coding | 333.9096094 | 0.216672073 | 0.110686065 | -1.957537048 | 0.05028436 | 0.14945931 | 0.216672073 |
| ENSMUSG00000003190 | Bcl2l12 | 75736 | protein_coding | 82.0519419 | 0.216683866 | 0.135329072 | -1.601162727 | 0.109340882 | 0.263984696 | 0.216683866 |
| ENSMUSG00000026839 | Upp2 | 76654 | protein_coding | 41.83729662 | -0.216686317 | 0.238000018 | 0.910446642 | 0.36258701 | 0.584928009 | 0.216686317 |
| ENSMUSG00000005951 | Shpk | 74637 | protein_coding | 231.206665 | -0.216707482 | 0.118309948 | 1.83169282 | 0.066997199 | 0.184936371 | 0.216707482 |
| ENSMUSG00000031925 | Maml2 | 270118 | protein_coding | 4.791131102 | -0.216729065 | 0.651353286 | 0.332736581 | 0.739333134 | 0.867757454 | 0.216729065 |
| ENSMUSG00000042831 | Alkbh6 | 233065 | protein_coding | 243.1301651 | 0.216907619 | 0.071414473 | -3.037306149 | 0.002387029 | 0.013089272 | 0.216907619 |
| ENSMUSG00000056476 | Med12l | 329650 | protein_coding | 27.39616749 | -0.216913132 | 0.341033893 | 0.636045673 | 0.524746654 | 0.723223689 | 0.216913132 |
| ENSMUSG00000074811 | Hps6 | 20170 | protein_coding | 98.41573524 | -0.217042573 | 0.142880716 | 1.519047352 | 0.128750577 | 0.29613919 | 0.217042573 |
| ENSMUSG00000040659 | Efhd2 | 27984 | protein_coding | 1267.412956 | -0.21715897 | 0.082346826 | 2.637126174 | 0.008361173 | 0.036859691 | 0.21715897 |
| ENSMUSG00000110008 | Or51f23b | 259117 | protein_coding | 1.810633158 | 0.217195352 | 1.09065503 | -0.199142117 | 0.842151576 | 1 | 0.217195352 |
| ENSMUSG00000049562 | Ap5b1 | 381201 | protein_coding | 196.5784772 | -0.21737057 | 0.104213253 | 2.08582464 | 0.036994511 | 0.118465648 | 0.21737057 |
| ENSMUSG00000014077 | Chp1 | 56398 | protein_coding | 3496.50742 | -0.217386496 | 0.114499209 | 1.898585139 | 0.057619044 | 0.16577019 | 0.217386496 |
| ENSMUSG00000037134 | Prmt9 | 102182 | protein_coding | 211.7503919 | 0.217392491 | 0.097568921 | -2.228091578 | 0.025874409 | 0.089747949 | 0.217392491 |
| ENSMUSG00000019854 | Reps1 | 19707 | protein_coding | 201.5044184 | 0.217424082 | 0.078988633 | -2.752599618 | 0.005912416 | 0.027880534 | 0.217424082 |
| ENSMUSG00000039835 | Nhsl1 | 215819 | protein_coding | 1006.218923 | 0.217431567 | 0.107795218 | -2.017079895 | 0.043687176 | 0.134498514 | 0.217431567 |
| ENSMUSG00000020311 | Erlec1 | 66753 | protein_coding | 259.4756912 | -0.217433186 | 0.138822377 | 1.566269005 | 0.117285654 | 0.277248109 | 0.217433186 |
| ENSMUSG00000020614 | Fam20a | 208659 | protein_coding | 4.145936921 | -0.21744222 | 1.219529919 | 0.178300029 | 0.858487356 | 1 | 0.21744222 |
| ENSMUSG00000115548 | 5430440P10Rik | 71362 | lncRNA | 3.561693523 | -0.217569946 | 0.637977079 | 0.341030976 | 0.733080263 | 1 | 0.217569946 |
| ENSMUSG00000053693 | Mast1 | 56527 | protein_coding | 12.05954678 | 0.21770641 | 0.446213831 | -0.487897045 | 0.625622768 | 0.794916857 | 0.21770641 |
| ENSMUSG00000029578 | Wipi2 | 74781 | protein_coding | 957.8094773 | -0.217728199 | 0.05042604 | 4.317773133 | 1.58E-05 | 0.000167167 | 0.217728199 |
| ENSMUSG00000028410 | Dnaja1 | 15502 | protein_coding | 1066.128986 | -0.217816695 | 0.136983872 | 1.590090073 | 0.111814503 | 0.267881142 | 0.217816695 |
| ENSMUSG00000003161 | Sri | 109552 | protein_coding | 4026.608217 | -0.217829092 | 0.088053893 | 2.473815558 | 0.013367869 | 0.053374514 | 0.217829092 |

| ensembl_gene_id | external_gene_name | entrezgene_id | gene_biotype | baseMean | log2FoldChange | lfcSE | stat | pvalue | padj | absLog2FC |
|---|---|---|---|---|---|---|---|---|---|---|
| ENSMUSG00000036438 | Calm2 | 12314 | protein_coding | 4263.96488 | -0.217836947 | 0.082771404 | 2.631789929 | 0.008493636 | 0.037314155 | 0.217836947 |
| ENSMUSG00000027404 | Snrpb | 20638 | protein_coding | 1362.467341 | 0.217990473 | 0.086977584 | -2.506283383 | 0.012200779 | 0.049728326 | 0.217990473 |
| ENSMUSG00000041782 | Lad1 | 16763 | protein_coding | 4728.11922 | -0.218014625 | 0.094175617 | 2.314979514 | 0.020614053 | 0.07508171 | 0.218014625 |
| ENSMUSG00000034342 | Cbl | 12402 | protein_coding | 792.8974228 | -0.218044978 | 0.12933036 | 1.685953536 | 0.091804749 | 0.232062395 | 0.218044978 |
| ENSMUSG00000043366 | Or51e2 | 170639 | protein_coding | 12.00240214 | -0.218159245 | 0.391910371 | 0.556655964 | 0.577762509 | 0.761709251 | 0.218159245 |
| ENSMUSG00000059586 | Nsmce2 | 68501 | protein_coding | 148.6638496 | 0.21832304 | 0.094519768 | -2.309813532 | 0.02089848 | 0.075916739 | 0.21832304 |
| ENSMUSG00000050565 | Tor1aip2 | 240832 | protein_coding | 2805.636559 | 0.218441622 | 0.070592825 | -3.094388459 | 0.001972191 | 0.011213534 | 0.218441622 |
| ENSMUSG00000026154 | Sdhaf4 | 68002 | protein_coding | 93.83073687 | -0.218474652 | 0.168459887 | 1.296894211 | 0.194667589 | 0.393673838 | 0.218474652 |
| ENSMUSG00000033379 | Atp6v0b | 114143 | protein_coding | 984.1342883 | -0.218491762 | 0.069825413 | 3.129115227 | 0.001753335 | 0.010159502 | 0.218491762 |
| ENSMUSG00000092618 | Btnl6 | 624681 | protein_coding | 1292.519803 | -0.218548376 | 0.166782221 | 1.310381732 | 0.190066731 | 0.387358014 | 0.218548376 |
| ENSMUSG00000032409 | Atr | 245000 | protein_coding | 304.2579594 | 0.218553107 | 0.126126766 | -1.732805128 | 0.083130311 | 0.215987092 | 0.218553107 |
| ENSMUSG00000027944 | Hax1 | 23897 | protein_coding | 213.1775652 | -0.21860328 | 0.141447712 | 1.545470592 | 0.122232477 | 0.285320309 | 0.21860328 |
| ENSMUSG00000033326 | Kdm4a | 230674 | protein_coding | 1061.135728 | 0.218674428 | 0.069699624 | -3.137383187 | 0.001704632 | 0.009910776 | 0.218674428 |
| ENSMUSG00000025967 | Eef1b2 | 55949 | protein_coding | 3862.142575 | -0.218693747 | 0.097295593 | 2.247725097 | 0.024593725 | 0.086284421 | 0.218693747 |
| ENSMUSG00000025810 | Nrp1 | 18186 | protein_coding | 92.72404439 | 0.218713719 | 0.181720158 | -1.20357434 | 0.228754142 | 0.437734172 | 0.218713719 |
| ENSMUSG00000039611 | Pgap4 | 67063 | protein_coding | 752.4011917 | -0.218721751 | 0.101349879 | 2.15808596 | 0.030921149 | 0.103308737 | 0.218721751 |
| ENSMUSG00000021640 | Naip1 | 17940 | protein_coding | 2834.582642 | -0.218745696 | 0.109448184 | 1.998623352 | 0.045649122 | 0.139215069 | 0.218745696 |
| ENSMUSG00000027016 | Zfp385b | 241494 | protein_coding | 33.96168312 | 0.218757334 | 0.31687151 | -0.690366054 | 0.489964019 | 0.697703355 | 0.218757334 |
| ENSMUSG00000039304 | Tnfsf10 | 22035 | protein_coding | 111.9336686 | -0.218770409 | 0.159592011 | 1.370810528 | 0.170434028 | 0.36023903 | 0.218770409 |
| ENSMUSG00000018143 | Mafk | 17135 | protein_coding | 1161.232265 | -0.218823186 | 0.164661919 | 1.328924056 | 0.183873026 | 0.37880229 | 0.218823186 |
| ENSMUSG00000080538 | Gm25541 | 115489638 | snRNA | 24.84958873 | 0.218858431 | 0.323571339 | -0.676383861 | 0.498796961 | 0.703739933 | 0.218858431 |
| ENSMUSG00000027203 | Dut | 110074 | protein_coding | 229.5167819 | 0.218862716 | 0.169797604 | -1.28896233 | 0.197411183 | 0.397518385 | 0.218862716 |
| ENSMUSG00000021271 | Zfp839 | 72805 | protein_coding | 147.9405459 | -0.218887605 | 0.098330809 | 2.226032777 | 0.026011986 | 0.090073507 | 0.218887605 |
| ENSMUSG00000112627 | 4933412E12Rik | 71086 | lncRNA | 6.971958774 | 0.218926199 | 0.453324301 | -0.482935061 | 0.629141846 | 0.797446101 | 0.218926199 |
| ENSMUSG00000027002 | Nckap1 | 50884 | protein_coding | 2816.744636 | 0.218966373 | 0.094697636 | -2.312268641 | 0.020762884 | 0.07549059 | 0.218966373 |
| ENSMUSG00000036206 | Sh3bp4 | 98402 | protein_coding | 225.6560132 | -0.219025198 | 0.121937802 | 1.796204242 | 0.07246204 | 0.195920934 | 0.219025198 |
| ENSMUSG00000097577 | 6230400D17Rik | 76133 | lncRNA | 15.00562911 | -0.219128842 | 0.339403231 | 0.645629805 | 0.51851913 | 0.719081703 | 0.219128842 |
| ENSMUSG00000005566 | Trim28 | 21849 | protein_coding | 1988.247209 | 0.219274124 | 0.09638145 | -2.27506563 | 0.022901993 | 0.081579713 | 0.219274124 |
| ENSMUSG00000045482 | Trrap | 100683 | protein_coding | 1001.348926 | 0.219275372 | 0.079273434 | -2.766063755 | 0.005673744 | 0.027040753 | 0.219275372 |
| ENSMUSG00000026504 | Sdccag8 | 76816 | protein_coding | 111.8290881 | 0.219290487 | 0.11454545 | -1.91444084 | 0.055563851 | 0.161179189 | 0.219290487 |
| ENSMUSG00000046637 | Ttc34 | 242800 | protein_coding | 1.751341168 | 0.219302825 | 1.469057886 | -0.149281268 | 0.881331695 | 1 | 0.219302825 |
| ENSMUSG00000040690 | Col16a1 | 107581 | protein_coding | 315.9584279 | 0.219310753 | 0.112428648 | -1.950666101 | 0.051096778 | 0.151546731 | 0.219310753 |
| ENSMUSG00000090264 | Eif4ebp3 | 108112 | protein_coding | 252.495357 | -0.219422088 | 0.154828143 | 1.417197699 | 0.156425134 | 0.339181285 | 0.219422088 |
| ENSMUSG00000002768 | Mea1 | 17256 | protein_coding | 640.8441848 | -0.219540193 | 0.068614035 | 3.199639713 | 0.001375995 | 0.008308292 | 0.219540193 |
| ENSMUSG00000052713 | Zfp608 | 269023 | protein_coding | 300.1532992 | 0.219622711 | 0.124421953 | -1.765144388 | 0.077539502 | 0.205601692 | 0.219622711 |
| ENSMUSG00000006315 | Tmem147 | 69804 | protein_coding | 454.4500834 | -0.219638218 | 0.110899012 | 1.980524577 | 0.047644615 | 0.143734593 | 0.219638218 |
| ENSMUSG00000025572 | Tmc6 | 217353 | protein_coding | 884.5412072 | 0.219645712 | 0.055003687 | -3.993290678 | 6.52E-05 | 0.000592563 | 0.219645712 |
| ENSMUSG00000065547 | Mir199a-1 | 387194 | miRNA | 3.142845306 | -0.219656101 | 0.617791972 | 0.355550268 | 0.722177394 | 1 | 0.219656101 |
| ENSMUSG00000028137 | Celf3 | 78784 | protein_coding | 47.78239214 | 0.219679176 | 0.319939786 | -0.686626628 | 0.492318044 | 0.698946027 | 0.219679176 |

| ensembl_gene_id | external_gene_name | entrezgene_id | gene_biotype | baseMean | log2FoldChange | lfcSE | stat | pvalue | padj | absLog2FC |
|---|---|---|---|---|---|---|---|---|---|---|
| ENSMUSG00000057672 | Pkn1 | 320795 | protein_coding | 848.4685309 | -0.219807619 | 0.057340961 | 3.833343837 | 0.000126413 | 0.001061844 | 0.219807619 |
| ENSMUSG00000040247 | Tbc1d10c | 108995 | protein_coding | 5.877846684 | -0.220036688 | 0.516547624 | 0.425975608 | 0.67012562 | 0.822845898 | 0.220036688 |
| ENSMUSG00000040998 | Npnt | 114249 | protein_coding | 7.76835469 | -0.220121672 | 0.714790075 | 0.307952893 | 0.758118177 | 0.878750865 | 0.220121672 |
| ENSMUSG00000025504 | Eps8l2 | 98845 | protein_coding | 1666.166582 | -0.220130961 | 0.183395062 | 1.200310185 | 0.230018896 | 0.438629566 | 0.220130961 |
| ENSMUSG00000074824 | Rslcan18 | 432770 | protein_coding | 6.257869338 | -0.220226288 | 0.706866813 | 0.311552734 | 0.75538046 | 0.877032235 | 0.220226288 |
| ENSMUSG00000022205 | Sub1 | 20024 | protein_coding | 699.4548439 | 0.220230803 | 0.095795815 | -2.298960591 | 0.021507177 | 0.077700618 | 0.220230803 |
| ENSMUSG00000039262 | Prrc2b | 227723 | protein_coding | 993.692088 | 0.220245947 | 0.071831553 | -3.066144846 | 0.002168382 | 0.012093437 | 0.220245947 |
| ENSMUSG00000020788 | Atp2a3 | 53313 | protein_coding | 12336.37218 | -0.220284116 | 0.116360365 | 1.893119843 | 0.058341934 | 0.167325234 | 0.220284116 |
| ENSMUSG00000051457 | Spn | 20737 | protein_coding | 14.04256165 | -0.220292071 | 0.399449067 | 0.551489762 | 0.581297983 | 0.764339772 | 0.220292071 |
| ENSMUSG00000026491 | Ahctf1 | 226747 | protein_coding | 931.7488091 | 0.220346822 | 0.081302912 | -2.71019594 | 0.006724347 | 0.030863221 | 0.220346822 |
| ENSMUSG00000037260 | Hgsnat | 52120 | protein_coding | 347.3907778 | -0.220409555 | 0.112033809 | 1.967348586 | 0.049143037 | 0.147109787 | 0.220409555 |
| ENSMUSG00000027676 | Ccdc39 | 51938 | protein_coding | 4.635969341 | 0.220425022 | 0.700892959 | -0.314491706 | 0.753147615 | 0.876249099 | 0.220425022 |
| ENSMUSG00000038286 | Bphl | 68021 | protein_coding | 273.2504621 | -0.220533272 | 0.096725649 | 2.279987507 | 0.022608429 | 0.080865411 | 0.220533272 |
| ENSMUSG00000023070 | Rgn | 19733 | protein_coding | 18.46717262 | -0.220579854 | 0.260743674 | 0.845964353 | 0.397572631 | 0.619446591 | 0.220579854 |
| ENSMUSG00000053390 | Zfp952 | 240067 | protein_coding | 120.6461288 | -0.220580786 | 0.10346824 | 2.131869519 | 0.033017575 | 0.108572726 | 0.220580786 |
| ENSMUSG00000026581 | Sell | 20343 | protein_coding | 4.805919455 | 0.220596102 | 0.954159002 | -0.231194279 | 0.817163875 | 0.908714429 | 0.220596102 |
| ENSMUSG00000002007 | Srpk3 | 56504 | protein_coding | 7.043315339 | 0.220679545 | 0.450777867 | -0.489552752 | 0.624450417 | 0.794172689 | 0.220679545 |
| ENSMUSG00000038244 | Mical2 | 320878 | protein_coding | 1316.770004 | -0.22071757 | 0.085545757 | 2.580111233 | 0.00987685 | 0.042179026 | 0.22071757 |
| ENSMUSG00000003808 | Farsa | 66590 | protein_coding | 374.6676012 | 0.220791324 | 0.090105896 | -2.450353804 | 0.01427159 | 0.056074481 | 0.220791324 |
| ENSMUSG00000034902 | Pip5k1c | 18717 | protein_coding | 511.4238602 | 0.220828368 | 0.061479035 | -3.591929608 | 0.000328239 | 0.00246612 | 0.220828368 |
| ENSMUSG00000030494 | Rhpn2 | 52428 | protein_coding | 1541.871169 | -0.220836277 | 0.059686703 | 3.699924213 | 0.000215664 | 0.001688702 | 0.220836277 |
| ENSMUSG00000028273 | Pdlim5 | 56376 | protein_coding | 1183.24845 | -0.220838988 | 0.08085237 | 2.731385461 | 0.006306866 | 0.029215031 | 0.220838988 |
| ENSMUSG00000011114 | Tbrg1 | 21376 | protein_coding | 1495.700411 | 0.220848082 | 0.081098699 | -2.723201284 | 0.006465265 | 0.029831627 | 0.220848082 |
| ENSMUSG00000040943 | Tet2 | 214133 | protein_coding | 486.3317079 | -0.220848987 | 0.124218335 | 1.777909737 | 0.075418681 | 0.201270862 | 0.220848987 |
| ENSMUSG00000058897 | Col25a1 | 77018 | protein_coding | 12.03261193 | 0.220972527 | 0.497956698 | -0.443758519 | 0.657217182 | 0.814655179 | 0.220972527 |
| ENSMUSG00000003531 | Dgcr6 | 13353 | protein_coding | 199.6056077 | -0.220990277 | 0.080529211 | 2.744225043 | 0.006065394 | 0.028407451 | 0.220990277 |
| ENSMUSG00000022216 | Psme1 | 19186 | protein_coding | 1057.636013 | -0.221013526 | 0.07589412 | 2.91212977 | 0.003589735 | 0.018490605 | 0.221013526 |
| ENSMUSG00000036992 | Nxt1 | 56488 | protein_coding | 119.091934 | 0.221015324 | 0.142232767 | -1.553898789 | 0.120208561 | 0.282356022 | 0.221015324 |
| ENSMUSG00000037058 | Paip2 | 67869 | protein_coding | 779.1739832 | -0.221062455 | 0.112267301 | 1.969072502 | 0.048944766 | 0.146772084 | 0.221062455 |
| ENSMUSG00000041658 | Rragb | 245670 | protein_coding | 6.288948453 | 0.221143236 | 0.452672756 | -0.488527824 | 0.625176023 | 0.794593769 | 0.221143236 |
| ENSMUSG00000036469 | Marchf1 | 72925 | protein_coding | 3.779489773 | 0.221182356 | 0.732218138 | -0.30207167 | 0.762597428 | 1 | 0.221182356 |
| ENSMUSG00000024369 | Nelfe | 27632 | protein_coding | 355.1625546 | 0.221210374 | 0.085633258 | -2.583229672 | 0.009788014 | 0.041864611 | 0.221210374 |
| ENSMUSG00000047205 | Dusp18 | 75219 | protein_coding | 172.1605755 | -0.22121872 | 0.105361551 | 2.099615263 | 0.035762699 | 0.115258628 | 0.22121872 |
| ENSMUSG00000034168 | Irf2bpl | 238330 | protein_coding | 601.3125042 | -0.221279471 | 0.110160237 | 2.008705466 | 0.044568379 | 0.136567098 | 0.221279471 |
| ENSMUSG00000022096 | Hr | 15460 | protein_coding | 396.8626151 | 0.221279592 | 0.097306641 | -2.274044084 | 0.022963336 | 0.081727701 | 0.221279592 |
| ENSMUSG00000024787 | Snx15 | 69024 | protein_coding | 325.4406259 | -0.221301584 | 0.077484647 | 2.856070102 | 0.004289204 | 0.021386449 | 0.221301584 |
| ENSMUSG00000017386 | Traf4 | 22032 | protein_coding | 687.8439043 | -0.221346285 | 0.079991985 | 2.767105799 | 0.00565564 | 0.02696225 | 0.221346285 |
| ENSMUSG00000055560 | Zfp459 | 328274 | protein_coding | 32.5358627 | -0.221360056 | 0.194083668 | 1.140539326 | 0.254061679 | 0.466691767 | 0.221360056 |
| ENSMUSG00000097610 | A930012L18Rik | 626275 | lncRNA | 3.483185116 | -0.221377447 | 0.590141677 | 0.375125932 | 0.707566812 | 1 | 0.221377447 |

| ensembl_gene_id | external_gene_name | entrezgene_id | gene_biotype | baseMean | log2FoldChange | lfcSE | stat | pvalue | padj | absLog2FC |
|---|---|---|---|---|---|---|---|---|---|---|
| ENSMUSG00000026019 | Wdr12 | 57750 | protein_coding | 125.3659619 | -0.221380281 | 0.148965262 | 1.486120175 | 0.137247337 | 0.309673797 | 0.221380281 |
| ENSMUSG00000073437 | D330041H03Rik | 654822 | lncRNA | 19.76405829 | -0.22138433 | 0.327822358 | 0.675317971 | 0.499473768 | 0.704201299 | 0.22138433 |
| ENSMUSG00000062867 | Impdh2 | 23918 | protein_coding | 933.9784893 | -0.221384731 | 0.152581638 | 1.450926427 | 0.146800348 | 0.324596932 | 0.221384731 |
| ENSMUSG00000037169 | Mycn | 18109 | protein_coding | 34.25547326 | 0.221398125 | 0.28378553 | -0.78016002 | 0.435296692 | 0.653617891 | 0.221398125 |
| ENSMUSG00000027599 | Armc1 | 74252 | protein_coding | 403.9909623 | 0.221441095 | 0.094409252 | -2.345544425 | 0.018999314 | 0.070538371 | 0.221441095 |
| ENSMUSG00000040560 | Wdr7 | 104082 | protein_coding | 393.0683621 | 0.221444842 | 0.059590539 | -3.716107395 | 0.000202316 | 0.001597055 | 0.221444842 |
| ENSMUSG00000075703 | Selenoi | 28042 | protein_coding | 1039.9155 | 0.221505477 | 0.062987669 | -3.516648281 | 0.000437032 | 0.003173763 | 0.221505477 |
| ENSMUSG00000053012 | Krcc1 | 57896 | protein_coding | 1091.830338 | 0.221526346 | 0.080505161 | -2.751703666 | 0.005928614 | 0.027917061 | 0.221526346 |
| ENSMUSG00000022032 | Scara5 | 71145 | protein_coding | 50.2361419 | 0.221602307 | 0.263159176 | -0.842084666 | 0.399740564 | 0.621944194 | 0.221602307 |
| ENSMUSG00000029168 | Dpysl5 | 65254 | protein_coding | 4.672932231 | 0.221608535 | 1.192166816 | -0.185887186 | 0.852533231 | 0.927761944 | 0.221608535 |
| ENSMUSG00000041798 | Gck | 103988 | protein_coding | 7.057151893 | -0.221623936 | 0.465624594 | 0.475971284 | 0.634094836 | 0.800507769 | 0.221623936 |
| ENSMUSG00000048222 | Mfap1b | 100034361 | protein_coding | 112.2865801 | -0.221626809 | 0.135937234 | 1.630361331 | 0.103025152 | 0.252989943 | 0.221626809 |
| ENSMUSG00000032449 | Slc25a36 | 192287 | protein_coding | 588.1384348 | -0.221667028 | 0.124192365 | 1.784868397 | 0.074282666 | 0.199334407 | 0.221667028 |
| ENSMUSG00000035279 | Ssc5d | 269855 | protein_coding | 7.049063205 | 0.221682627 | 0.961489464 | -0.230561681 | 0.81765534 | 0.908714429 | 0.221682627 |
| ENSMUSG00000058355 | Abce1 | 24015 | protein_coding | 1267.479482 | 0.221760713 | 0.115670466 | -1.91717662 | 0.055215497 | 0.160458367 | 0.221760713 |
| ENSMUSG00000040121 | Rep15 | 66532 | protein_coding | 1122.441361 | 0.221836125 | 0.345927463 | -0.64127931 | 0.521341229 | 0.720922137 | 0.221836125 |
| ENSMUSG00000109408 | A930037H05Rik | 78439 | lncRNA | 30.10461172 | -0.221850432 | 0.275074455 | 0.806510485 | 0.419948565 | 0.63998633 | 0.221850432 |
| ENSMUSG00000069844 | Sco1 | 52892 | protein_coding | 679.2118065 | -0.221886201 | 0.082199431 | 2.699364184 | 0.006947211 | 0.031711558 | 0.221886201 |
| ENSMUSG00000020876 | Snx11 | 74479 | protein_coding | 215.539809 | 0.221908695 | 0.093581306 | -2.371293004 | 0.017725973 | 0.066683451 | 0.221908695 |
| ENSMUSG00000032911 | Cspg4 | 121021 | protein_coding | 28.10052929 | 0.221914528 | 0.414464957 | -0.535424104 | 0.592356626 | 0.772317806 | 0.221914528 |
| ENSMUSG00000039994 | Timeless | 21853 | protein_coding | 104.2854225 | 0.221951017 | 0.173521321 | -1.279099397 | 0.200862056 | 0.402162462 | 0.221951017 |
| ENSMUSG00000018697 | Aatf | 56321 | protein_coding | 260.0394028 | -0.222078377 | 0.099280507 | 2.23687794 | 0.025294319 | 0.088198982 | 0.222078377 |
| ENSMUSG00000016150 | Tenm1 | 23963 | protein_coding | 3.780300254 | -0.22214601 | 2.442525307 | 0.090949318 | 0.927532863 | 1 | 0.22214601 |
| ENSMUSG00000022428 | Cby1 | 73739 | protein_coding | 139.7407449 | 0.222237512 | 0.132097947 | -1.682369158 | 0.092497278 | 0.233277344 | 0.222237512 |
| ENSMUSG00000020687 | Cdc27 | 217232 | protein_coding | 771.8710917 | 0.22226015 | 0.088145629 | -2.521510726 | 0.011685211 | 0.048151079 | 0.22226015 |
| ENSMUSG00000031578 | Mak16 | 67920 | protein_coding | 149.6415882 | 0.222305524 | 0.116584503 | -1.906818817 | 0.056544054 | 0.163589437 | 0.222305524 |
| ENSMUSG00000022452 | Smdt1 | 69029 | protein_coding | 1228.525411 | 0.22233244 | 0.085666732 | -2.59531834 | 0.009450334 | 0.040715581 | 0.22233244 |
| ENSMUSG00000068601 | Gm10244 | 621427 | lncRNA | 2.768496189 | -0.222342903 | 0.720244807 | 0.308704626 | 0.757546225 | 1 | 0.222342903 |
| ENSMUSG00000057842 | Zfp595 | 218314 | protein_coding | 81.36903636 | -0.222379133 | 0.137070955 | 1.622365097 | 0.104725198 | 0.255833124 | 0.222379133 |
| ENSMUSG00000020387 | Jade2 | 76901 | protein_coding | 209.4295142 | -0.222497193 | 0.135477269 | 1.642321215 | 0.100523453 | 0.248744583 | 0.222497193 |
| ENSMUSG00000066880 | Zfp617 | 170938 | protein_coding | 155.4006519 | -0.222511243 | 0.099893956 | 2.227474527 | 0.025915576 | 0.089852998 | 0.222511243 |
| ENSMUSG00000025743 | Sdc3 | 20970 | protein_coding | 8.17590166 | -0.222540145 | 0.522583052 | 0.425846465 | 0.670219727 | 0.822900265 | 0.222540145 |
| ENSMUSG00000026707 | Nsun6 | 74455 | protein_coding | 116.4349971 | 0.222618592 | 0.109586243 | -2.031446514 | 0.04220972 | 0.131147943 | 0.222618592 |
| ENSMUSG00000054893 | Zfp667 | 384763 | protein_coding | 46.28666482 | -0.222738361 | 0.196039179 | 1.136193094 | 0.25587576 | 0.468566456 | 0.222738361 |
| ENSMUSG00000068011 | Mkrn2os | 70291 | protein_coding | 311.9619695 | -0.222814579 | 0.098986862 | 2.250951031 | 0.024388639 | 0.085735413 | 0.222814579 |
| ENSMUSG00000025722 | Wdr73 | 71968 | protein_coding | 146.7822528 | -0.222820253 | 0.146892608 | 1.516892208 | 0.129293901 | 0.296962376 | 0.222820253 |
| ENSMUSG00000041323 | Ak7 | 78801 | protein_coding | 8.392695435 | -0.222882263 | 0.579966198 | 0.384302161 | 0.700754506 | 0.841825268 | 0.222882263 |
| ENSMUSG00000039834 | Zfp335 | 329559 | protein_coding | 392.2488115 | 0.222898704 | 0.112993995 | -1.972659731 | 0.048534343 | 0.145806101 | 0.222898704 |
| ENSMUSG00000032182 | Yipf2 | 74766 | protein_coding | 200.8399887 | 0.222941913 | 0.1004413 | -2.219623929 | 0.026444307 | 0.091207452 | 0.222941913 |

| ensembl_gene_id | external_gene_name | entrezgene_id | gene_biotype | baseMean | log2FoldChange | lfcSE | stat | pvalue | padj | absLog2FC |
|---|---|---|---|---|---|---|---|---|---|---|
| ENSMUSG00000037570 | Mcrs1 | 51812 | protein_coding | 478.1159824 | -0.222960333 | 0.092554291 | 2.408968081 | 0.015997697 | 0.061495803 | 0.222960333 |
| ENSMUSG00000029461 | Fam168a | 319604 | protein_coding | 176.4999734 | -0.223162447 | 0.111063373 | 2.009325314 | 0.044502645 | 0.13639879 | 0.223162447 |
| ENSMUSG00000018921 | Pelp1 | 75273 | protein_coding | 229.9136434 | 0.223169938 | 0.102251628 | -2.182556335 | 0.029068498 | 0.098469165 | 0.223169938 |
| ENSMUSG00000033488 | Cryzl2 | 226527 | protein_coding | 135.6442445 | -0.223260569 | 0.156193133 | 1.42938786 | 0.152892786 | 0.333845229 | 0.223260569 |
| ENSMUSG00000025082 | Vwa2 | 240675 | protein_coding | 10.35239946 | 0.223264401 | 0.425229381 | -0.525044627 | 0.599552168 | 0.776544903 | 0.223264401 |
| ENSMUSG00000093674 | Rpl41 | 67945 | protein_coding | 6863.15504 | -0.223286823 | 0.103593275 | 2.15541814 | 0.031129128 | 0.103865546 | 0.223286823 |
| ENSMUSG00000030751 | Psma1 | 26440 | protein_coding | 1219.994108 | 0.223306131 | 0.095532104 | -2.337498298 | 0.019413289 | 0.071736644 | 0.223306131 |
| ENSMUSG00000020225 | Tmbim4 | 68212 | protein_coding | 687.014301 | -0.22334698 | 0.062869426 | 3.552553184 | 0.000381512 | 0.002813884 | 0.22334698 |
| ENSMUSG00000034159 | Mab21l4 | 71874 | protein_coding | 163.6459061 | -0.223357072 | 0.184205088 | 1.212545618 | 0.225303592 | 0.432685604 | 0.223357072 |
| ENSMUSG00000034724 | Cnot6l | 231464 | protein_coding | 478.9499914 | -0.223361095 | 0.12976644 | 1.721254709 | 0.085204614 | 0.220060838 | 0.223361095 |
| ENSMUSG00000031987 | Egln1 | 112405 | protein_coding | 472.5763813 | -0.223366085 | 0.070190438 | 3.18228654 | 0.001461172 | 0.008692083 | 0.223366085 |
| ENSMUSG00000024145 | Pigf | 18701 | protein_coding | 160.1128867 | -0.223394815 | 0.101648574 | 2.19771715 | 0.027969269 | 0.095391264 | 0.223394815 |
| ENSMUSG00000034394 | Lif | 16878 | protein_coding | 56.94524545 | 0.223402994 | 0.210894745 | -1.059310389 | 0.289458445 | 0.508294925 | 0.223402994 |
| ENSMUSG00000034171 | Faah | 14073 | protein_coding | 315.9281172 | -0.223426399 | 0.088674288 | 2.519630028 | 0.011747824 | 0.048307661 | 0.223426399 |
| ENSMUSG00000094840 | Muc3a | 619309 | protein_coding | 380.8969121 | 0.223449498 | 0.325969611 | -0.685491808 | 0.493033628 | 0.699601077 | 0.223449498 |
| ENSMUSG00000037145 | Lypd8l | 78354 | protein_coding | 370.8912982 | 0.223458023 | 0.310323594 | -0.720080673 | 0.471475326 | 0.684298079 | 0.223458023 |
| ENSMUSG00000022184 | Fbxo4 | 106052 | protein_coding | 149.7388075 | 0.223496202 | 0.139894282 | -1.597607847 | 0.110130279 | 0.265192684 | 0.223496202 |
| ENSMUSG00000027569 | Mrgbp | 73247 | protein_coding | 209.662514 | -0.223547447 | 0.085420411 | 2.617026113 | 0.008869956 | 0.038648666 | 0.223547447 |
| ENSMUSG00000032386 | Trip4 | 56404 | protein_coding | 339.9809949 | 0.223567771 | 0.098361338 | -2.272923237 | 0.023030806 | 0.081931007 | 0.223567771 |
| ENSMUSG00000074129 | Rpl13a | 22121 | protein_coding | 332.2726482 | 0.223584741 | 0.082789186 | -2.700651531 | 0.00692038 | 0.031622347 | 0.223584741 |
| ENSMUSG00000029130 | Rnf32 | 56874 | protein_coding | 97.16527528 | -0.223588568 | 0.114584493 | 1.951298664 | 0.051021528 | 0.151405123 | 0.223588568 |
| ENSMUSG00000046185 | Zfp84 | 74352 | protein_coding | 311.2110946 | -0.223644412 | 0.088913016 | 2.515316893 | 0.011892544 | 0.048732869 | 0.223644412 |
| ENSMUSG00000115145 | Gm35166 | 102638653 | lncRNA | 3.051455361 | -0.223702538 | 0.620293002 | 0.360640113 | 0.718368499 | 1 | 0.223702538 |
| ENSMUSG00000026413 | Pkp1 | 18772 | protein_coding | 465.8069533 | -0.22370453 | 0.146102073 | 1.531152338 | 0.125731747 | 0.291415309 | 0.22370453 |
| ENSMUSG00000024456 | Diaph1 | 13367 | protein_coding | 1261.55468 | -0.223706951 | 0.078764137 | 2.840213323 | 0.004508337 | 0.022343376 | 0.223706951 |
| ENSMUSG00000028969 | Cdk5 | 12568 | protein_coding | 281.3475556 | -0.223736017 | 0.088572268 | 2.526027875 | 0.011536033 | 0.047698058 | 0.223736017 |
| ENSMUSG00000039680 | Mrps6 | 121022 | protein_coding | 347.0422005 | -0.223746531 | 0.116286561 | 1.924096206 | 0.054342526 | 0.158736855 | 0.223746531 |
| ENSMUSG00000020472 | Zkscan17 | 268417 | protein_coding | 185.4149093 | 0.223837065 | 0.143723609 | -1.557413332 | 0.119372383 | 0.280974277 | 0.223837065 |
| ENSMUSG00000043241 | Upf2 | 326622 | protein_coding | 355.2781338 | 0.223855619 | 0.064266794 | -3.483223693 | 0.000495414 | 0.003520597 | 0.223855619 |
| ENSMUSG00000014609 | Chrne | 11448 | protein_coding | 1.793716851 | 0.223855923 | 1.342067757 | -0.166799271 | 0.867527993 | 1 | 0.223855923 |
| ENSMUSG00000052783 | Grk4 | 14772 | protein_coding | 7.506368899 | -0.223985171 | 0.634595736 | 0.352957258 | 0.724120484 | 0.857084875 | 0.223985171 |
| ENSMUSG00000005510 | Ndufs3 | 68349 | protein_coding | 918.9246752 | -0.223989478 | 0.082126136 | 2.727383629 | 0.006383877 | 0.029489048 | 0.223989478 |
| ENSMUSG00000004707 | Ly9 | 17085 | protein_coding | 6.066840388 | -0.224028974 | 0.673822875 | 0.332474574 | 0.739530936 | 0.867862447 | 0.224028974 |
| ENSMUSG00000029817 | Tra2a | 101214 | protein_coding | 939.77074 | 0.224106756 | 0.063232651 | -3.544161933 | 0.000393863 | 0.002893353 | 0.224106756 |
| ENSMUSG00000029722 | Agfg2 | 231801 | protein_coding | 319.7216833 | -0.224152755 | 0.090699367 | 2.471381703 | 0.013459207 | 0.053648407 | 0.224152755 |
| ENSMUSG00000070817 | Vmn1r85 | 252909 | protein_coding | 2.27187974 | 0.224163717 | 1.946325357 | -0.115172788 | 0.908308168 | 1 | 0.224163717 |
| ENSMUSG00000097726 | 9530036O11Rik | 654796 | lncRNA | 10.70244573 | -0.224218508 | 0.32360781 | 0.692871127 | 0.488390431 | 0.696363286 | 0.224218508 |
| ENSMUSG00000024906 | Mus81 | 71711 | protein_coding | 154.9660529 | 0.224330484 | 0.096179671 | -2.332410603 | 0.019679101 | 0.07241317 | 0.224330484 |
| ENSMUSG00000074754 | Smim26 | 228715 | protein_coding | 61.25202312 | -0.224378272 | 0.137234517 | 1.634998806 | 0.102049298 | 0.251303624 | 0.224378272 |

| ensembl_gene_id | external_gene_name | entrezgene_id | gene_biotype | baseMean | log2FoldChange | lfcSE | stat | pvalue | padj | absLog2FC |
|---|---|---|---|---|---|---|---|---|---|---|
| ENSMUSG00000017688 | Hnf4g | 30942 | protein_coding | 410.8536428 | -0.224418672 | 0.099162653 | 2.263137045 | 0.023627243 | 0.083675806 | 0.224418672 |
| ENSMUSG00000057835 | Zfp119a | 104349 | protein_coding | 49.92334411 | -0.224433701 | 0.141466441 | 1.586480151 | 0.112630431 | 0.269245209 | 0.224433701 |
| ENSMUSG00000042507 | Mideas | 238317 | protein_coding | 799.5124373 | 0.224464725 | 0.164987672 | -1.360493925 | 0.173673674 | 0.364427834 | 0.224464725 |
| ENSMUSG00000073424 | Cyp4f15 | 106648 | protein_coding | 3.060354926 | -0.224498508 | 1.002283105 | 0.223987122 | 0.822767324 | 1 | 0.224498508 |
| ENSMUSG00000053977 | Cd8a | 12525 | protein_coding | 23.68407837 | -0.224533596 | 0.38873887 | 0.57759492 | 0.563537637 | 0.750868206 | 0.224533596 |
| ENSMUSG00000023904 | Hcfc1r1 | 353502 | protein_coding | 442.2787874 | -0.224546995 | 0.103536313 | 2.168775262 | 0.03009975 | 0.101276951 | 0.224546995 |
| ENSMUSG00000051169 | Rpusd3 | 101122 | protein_coding | 48.3547032 | 0.224566174 | 0.180339052 | -1.245244282 | 0.21304197 | 0.417785903 | 0.224566174 |
| ENSMUSG00000054836 | Elp6 | 72341 | protein_coding | 70.45007565 | 0.224568303 | 0.13680469 | -1.641524877 | 0.100688507 | 0.248992215 | 0.224568303 |
| ENSMUSG00000074884 | Serf2 | 378702 | protein_coding | 1164.597632 | 0.224570317 | 0.081833857 | -2.744222572 | 0.00606544 | 0.028407451 | 0.224570317 |
| ENSMUSG00000025279 | Dnase1l3 | 13421 | protein_coding | 3.160079395 | 0.224572311 | 0.910608949 | -0.246617729 | 0.805204082 | 1 | 0.224572311 |
| ENSMUSG00000042312 | S100a13 | 20196 | protein_coding | 324.5390215 | -0.22461974 | 0.096073099 | 2.338008675 | 0.019386798 | 0.071685421 | 0.22461974 |
| ENSMUSG00000032267 | Usp28 | 235323 | protein_coding | 194.5454755 | -0.224634691 | 0.118227869 | 1.900014712 | 0.057431189 | 0.165344954 | 0.224634691 |
| ENSMUSG00000034212 | Ankmy1 | 241158 | protein_coding | 3.868101144 | 0.224643501 | 0.78193047 | -0.287293448 | 0.773887639 | 1 | 0.224643501 |
| ENSMUSG00000016995 | Matn4 | 17183 | protein_coding | 10.61125571 | -0.224719481 | 0.420327739 | 0.534629195 | 0.592906291 | 0.772538316 | 0.224719481 |
| ENSMUSG00000001983 | Taco1 | 70207 | protein_coding | 154.9256379 | -0.224732268 | 0.105018416 | 2.139931996 | 0.032360263 | 0.10695305 | 0.224732268 |
| ENSMUSG00000050377 | Il31ra | 218624 | protein_coding | 1.865747881 | -0.224764564 | 1.837923369 | 0.122292675 | 0.902667233 | 1 | 0.224764564 |
| ENSMUSG00000020475 | Pgam2 | 56012 | protein_coding | 21.83916482 | -0.224779337 | 0.300437394 | 0.748173634 | 0.454355432 | 0.669875536 | 0.224779337 |
| ENSMUSG00000054027 | Nt5dc3 | 103466 | protein_coding | 78.47696497 | -0.224800948 | 0.105307026 | 2.134719367 | 0.03278394 | 0.108084245 | 0.224800948 |
| ENSMUSG00000062818 | Vmn1r51 | 22296 | protein_coding | 2.514230485 | 0.224826849 | 0.992074245 | -0.226623007 | 0.8207169 | 1 | 0.224826849 |
| ENSMUSG00000027175 | Tcp11l1 | 320554 | protein_coding | 5.274779665 | 0.224839263 | 0.61719339 | -0.36429305 | 0.715639189 | 0.852489762 | 0.224839263 |
| ENSMUSG00000020580 | Rock2 | 19878 | protein_coding | 1847.285505 | 0.22484787 | 0.090742839 | -2.477858006 | 0.013217373 | 0.05288871 | 0.22484787 |
| ENSMUSG00000020605 | Hs1bp3 | 58240 | protein_coding | 198.4345727 | -0.224892319 | 0.079673856 | 2.82266142 | 0.004762684 | 0.023359359 | 0.224892319 |
| ENSMUSG00000087586 | Gm6938 | 629024 | lncRNA | 0.673731784 | 0.224912598 | 1.667038992 | -0.134917419 | 0.892677167 | 1 | 0.224912598 |
| ENSMUSG00000035900 | Gramd4 | 223752 | protein_coding | 484.5163238 | -0.224973736 | 0.075792629 | 2.968279888 | 0.002994715 | 0.015840719 | 0.224973736 |
| ENSMUSG00000054469 | Lclat1 | 225010 | protein_coding | 994.1073221 | 0.225036891 | 0.084104693 | -2.675675798 | 0.007457875 | 0.0336207 | 0.225036891 |
| ENSMUSG00000037214 | Thap1 | 73754 | protein_coding | 94.68272896 | 0.225046941 | 0.120926805 | -1.861017837 | 0.062741658 | 0.176210159 | 0.225046941 |
| ENSMUSG00000029136 | Rbks | 71336 | protein_coding | 113.0762204 | 0.225052316 | 0.125547297 | -1.792569992 | 0.073041708 | 0.19693237 | 0.225052316 |
| ENSMUSG00000034275 | Igsf9b | 235086 | protein_coding | 330.2080785 | 0.225058802 | 0.120929227 | -1.861078642 | 0.062733072 | 0.176210159 | 0.225058802 |
| ENSMUSG00000078768 | Zfp566 | 72556 | protein_coding | 11.79619746 | -0.225104958 | 0.30109144 | 0.747629882 | 0.454683435 | 0.670174662 | 0.225104958 |
| ENSMUSG00000003810 | Mast2 | 17776 | protein_coding | 1239.948857 | -0.225215945 | 0.065913335 | 3.41684947 | 0.000633503 | 0.004319434 | 0.225215945 |
| ENSMUSG00000063116 | Or3a1 | 258702 | protein_coding | 1.525866186 | -0.225319151 | 2.105714513 | 0.107003656 | 0.91478608 | 1 | 0.225319151 |
| ENSMUSG00000036560 | Lgi4 | 243914 | protein_coding | 2.294198655 | 0.225349054 | 1.591081068 | -0.141632666 | 0.887370163 | 1 | 0.225349054 |
| ENSMUSG00000045252 | Zfp574 | 232976 | protein_coding | 238.6608123 | 0.225399399 | 0.104619448 | -2.1544694 | 0.031203379 | 0.104013109 | 0.225399399 |
| ENSMUSG00000051989 | Smim11 | 68936 | protein_coding | 172.4953979 | -0.225466172 | 0.103086381 | 2.187157696 | 0.028731014 | 0.097546045 | 0.225466172 |
| ENSMUSG00000056962 | Jmjd6 | 107817 | protein_coding | 417.0530269 | 0.225515674 | 0.100821518 | -2.236781193 | 0.025300644 | 0.088202415 | 0.225515674 |
| ENSMUSG00000033356 | Pus7l | 78895 | protein_coding | 51.49750604 | 0.22551766 | 0.213893372 | -1.054346181 | 0.291724453 | 0.510557756 | 0.22551766 |
| ENSMUSG00000078612 | Fyb2 | 242594 | protein_coding | 78.81859586 | -0.225604322 | 0.222570269 | 1.013631886 | 0.310758437 | 0.531744361 | 0.225604322 |
| ENSMUSG00000000724 | Cryba1 | 12957 | protein_coding | 2.787825654 | -0.225729281 | 1.200644918 | 0.188006694 | 0.850871404 | 1 | 0.225729281 |
| ENSMUSG00000042453 | Reln | 19699 | protein_coding | 6.368356984 | 0.225747796 | 2.524471777 | -0.089423775 | 0.92874513 | 0.966007248 | 0.225747796 |

Supplementary Table 1

| ensembl_gene_id | external_gene_name | entrezgene_id | gene_biotype | baseMean | log2FoldChange | lfcSE | stat | pvalue | padj | absLog2FC |
|---|---|---|---|---|---|---|---|---|---|---|
| ENSMUSG00000021124 | Vti1b | 53612 | protein_coding | 653.885327 | -0.225748829 | 0.068820748 | 3.280243736 | 0.001037174 | 0.0065599 | 0.225748829 |
| ENSMUSG00000021709 | Erbin | 59079 | protein_coding | 5974.937996 | -0.225750849 | 0.097169439 | 2.32327006 | 0.020164649 | 0.073835702 | 0.225750849 |
| ENSMUSG00000032803 | Cdv3 | 321022 | protein_coding | 1667.766374 | 0.225772383 | 0.090144424 | -2.504562941 | 0.012260279 | 0.049857012 | 0.225772383 |
| ENSMUSG00000035372 | 1810055G02Rik | 72056 | protein_coding | 183.0355441 | -0.225786796 | 0.120750496 | 1.869862268 | 0.061502947 | 0.173816344 | 0.225786796 |
| ENSMUSG00000000168 | Dlat | 235339 | protein_coding | 1207.408457 | -0.225787931 | 0.096900569 | 2.330099114 | 0.019800913 | 0.07274578 | 0.225787931 |
| ENSMUSG00000026888 | Grb14 | 50915 | protein_coding | 23.1997682 | -0.225796331 | 0.216193003 | 1.04442016 | 0.29629104 | 0.515481484 | 0.225796331 |
| ENSMUSG00000041153 | Osgin2 | 209212 | protein_coding | 87.10794107 | 0.225968671 | 0.112898341 | -2.001523406 | 0.045336014 | 0.138618577 | 0.225968671 |
| ENSMUSG00000039001 | Rps21 | 66481 | protein_coding | 2352.891104 | -0.225974219 | 0.126818969 | 1.781864499 | 0.074771331 | 0.19999509 | 0.225974219 |
| ENSMUSG00000041287 | Sox15 | 20670 | protein_coding | 2.717478564 | -0.22599297 | 0.900971707 | 0.250832483 | 0.801943626 | 1 | 0.22599297 |
| ENSMUSG00000030560 | Ctsc | 13032 | protein_coding | 898.6581489 | -0.22602808 | 0.09266861 | 2.439100786 | 0.014723861 | 0.057495825 | 0.22602808 |
| ENSMUSG00000012429 | Mplkip | 66308 | protein_coding | 128.6959589 | -0.226064428 | 0.09954788 | 2.270911521 | 0.023152335 | 0.082276233 | 0.226064428 |
| ENSMUSG00000102106 | 2310043O21Rik | 69679 | lncRNA | 4.06074612 | 0.226070465 | 0.508952242 | -0.444187974 | 0.656906686 | 1 | 0.226070465 |
| ENSMUSG00000002658 | Gtf2f1 | 98053 | protein_coding | 692.6946203 | 0.22613214 | 0.050069119 | -4.516399438 | 6.29E-06 | 7.37E-05 | 0.22613214 |
| ENSMUSG00000024942 | Capn1 | 12333 | protein_coding | 1613.044064 | -0.226140874 | 0.064854904 | 3.486873924 | 0.000488702 | 0.003481631 | 0.226140874 |
| ENSMUSG00000026385 | Dbi | 13167 | protein_coding | 2844.518478 | 0.226212963 | 0.074418115 | -3.03975668 | 0.002367694 | 0.0129987 | 0.226212963 |
| ENSMUSG00000039205 | Ciz1 | 68379 | protein_coding | 442.6914575 | -0.226227331 | 0.093233282 | 2.426465387 | 0.015246705 | 0.059173697 | 0.226227331 |
| ENSMUSG00000042589 | Cux2 | 13048 | protein_coding | 30.60182346 | 0.226316672 | 0.433719374 | -0.521804386 | 0.601806536 | 0.77804505 | 0.226316672 |
| ENSMUSG00000020946 | Gosr2 | 56494 | protein_coding | 785.9501357 | 0.226318794 | 0.062104157 | -3.644181064 | 0.000268245 | 0.002060368 | 0.226318794 |
| ENSMUSG00000032456 | Nmnat3 | 74080 | protein_coding | 136.0641211 | -0.226351082 | 0.183695613 | 1.232207335 | 0.217871644 | 0.423784727 | 0.226351082 |
| ENSMUSG00000034075 | Zdhhc5 | 228136 | protein_coding | 2297.73727 | 0.226441342 | 0.044299928 | -5.11155099 | 3.20E-07 | 4.87E-06 | 0.226441342 |
| ENSMUSG00000030417 | Pdcd5 | 56330 | protein_coding | 289.4570775 | -0.226554546 | 0.094901754 | 2.387253505 | 0.016974782 | 0.064474124 | 0.226554546 |
| ENSMUSG00000007880 | Arid1a | 93760 | protein_coding | 760.7269786 | 0.226572572 | 0.15048071 | -1.505658579 | 0.132154842 | 0.3013667 | 0.226572572 |
| ENSMUSG00000028399 | Ptprd | 19266 | protein_coding | 1362.932505 | -0.226648828 | 0.070978489 | 3.193204489 | 0.001407033 | 0.00846073 | 0.226648828 |
| ENSMUSG00000022378 | Cyrib | 223601 | protein_coding | 398.2074156 | 0.226658909 | 0.072586316 | -3.122612092 | 0.001792538 | 0.010353962 | 0.226658909 |
| ENSMUSG00000072964 | Bhlhb9 | 70237 | protein_coding | 23.93651225 | 0.226666937 | 0.260558863 | -0.869926027 | 0.384340831 | 0.606725963 | 0.226666937 |
| ENSMUSG00000029012 | Orc5 | 26429 | protein_coding | 150.6407756 | -0.226684321 | 0.109006017 | 2.079557877 | 0.037566104 | 0.119923959 | 0.226684321 |
| ENSMUSG00000024667 | Tmem216 | 68642 | protein_coding | 44.27243977 | 0.226736725 | 0.164440265 | -1.378839447 | 0.167944261 | 0.356436387 | 0.226736725 |
| ENSMUSG00000037013 | Ss18 | 268996 | protein_coding | 730.5029599 | 0.226801209 | 0.086290518 | -2.628344508 | 0.008580157 | 0.037634187 | 0.226801209 |
| ENSMUSG00000003308 | Keap1 | 50868 | protein_coding | 613.2364637 | 0.226855078 | 0.068845719 | -3.295122484 | 0.000983788 | 0.00627754 | 0.226855078 |
| ENSMUSG00000002477 | Snrpd1 | 20641 | protein_coding | 296.7412377 | 0.226856357 | 0.139418199 | -1.627164593 | 0.103702145 | 0.254161061 | 0.226856357 |
| ENSMUSG00000033966 | Cdkl4 | 381113 | protein_coding | 3.291153594 | -0.226887055 | 0.822916438 | 0.275710929 | 0.782770101 | 1 | 0.226887055 |
| ENSMUSG00000063286 | Gvin-ps7 | 668139 | transcribed_unprocessed_pseudogene | 20.01980945 | 0.22691042 | 0.324050713 | -0.700231202 | 0.483782929 | 0.694229344 | 0.22691042 |
| ENSMUSG00000026851 | BC005624 | 227707 | protein_coding | 299.0272797 | -0.226963208 | 0.063715938 | 3.562110415 | 0.000367886 | 0.002731683 | 0.226963208 |
| ENSMUSG00000004837 | Grap | 71520 | protein_coding | 4.578400126 | 0.226975633 | 0.570853475 | -0.397607518 | 0.690919518 | 0.835396465 | 0.226975633 |
| ENSMUSG00000042595 | Fam199x | 245622 | protein_coding | 417.9875504 | 0.227059536 | 0.126834659 | -1.790201016 | 0.073421602 | 0.197666177 | 0.227059536 |
| ENSMUSG00000090093 | Gm14399 | 100043761 | protein_coding | 24.67886116 | -0.227065639 | 0.2886287 | 0.786704991 | 0.431454582 | 0.649798537 | 0.227065639 |
| ENSMUSG00000060923 | Acyp2 | 75572 | protein_coding | 37.0417996 | -0.227130817 | 0.17143161 | 1.324906284 | 0.185202236 | 0.380591057 | 0.227130817 |
| ENSMUSG00000058600 | Rpl30 | 19946 | protein_coding | 1757.1322 | -0.227138977 | 0.123396991 | 1.840717296 | 0.065662998 | 0.182183929 | 0.227138977 |
| ENSMUSG00000025808 | Ccdc7a | 74703 | protein_coding | 4.431189896 | -0.227145412 | 0.917023379 | 0.247698606 | 0.804367611 | 0.903566201 | 0.227145412 |

Supplementary Table 1

| ensembl_gene_id | external_gene_name | entrezgene_id | gene_biotype | baseMean | log2FoldChange | lfcSE | stat | pvalue | padj | absLog2FC |
|---|---|---|---|---|---|---|---|---|---|---|
| ENSMUSG00000054428 | Atpif1 | 11983 | protein_coding | 1706.029637 | -0.227151539 | 0.083731448 | 2.712858123 | 0.006670566 | 0.030641927 | 0.227151539 |
| ENSMUSG00000028633 | Ctps | 51797 | protein_coding | 198.1828158 | 0.227239417 | 0.145672182 | -1.559936937 | 0.118774785 | 0.279846882 | 0.227239417 |
| ENSMUSG00000027947 | Il6ra | 16194 | protein_coding | 33.64422472 | -0.227370779 | 0.351879707 | 0.646160533 | 0.518175396 | 0.718665364 | 0.227370779 |
| ENSMUSG00000012848 | Rps5 | 20103 | protein_coding | 6368.247198 | -0.227419325 | 0.089329366 | 2.545851777 | 0.010901147 | 0.045517455 | 0.227419325 |
| ENSMUSG00000062312 | Erbb2 | 13866 | protein_coding | 2558.146127 | -0.227425714 | 0.115880438 | 1.962589353 | 0.049693909 | 0.148254918 | 0.227425714 |
| ENSMUSG00000026509 | Capn2 | 12334 | protein_coding | 921.9104565 | -0.227436121 | 0.132280201 | 1.719351192 | 0.085550444 | 0.220716112 | 0.227436121 |
| ENSMUSG00000023277 | Twf2 | 23999 | protein_coding | 218.51959 | 0.22744504 | 0.095873969 | -2.37233361 | 0.017676123 | 0.066614215 | 0.22744504 |
| ENSMUSG00000053964 | Lgals4 | 16855 | protein_coding | 42014.74337 | -0.227481825 | 0.086724212 | 2.62304862 | 0.008714684 | 0.038133091 | 0.227481825 |
| ENSMUSG00000110195 | Pde2a | 207728 | protein_coding | 24.30640153 | -0.227492908 | 0.314584066 | 0.723154577 | 0.469584922 | 0.683166089 | 0.227492908 |
| ENSMUSG00000026694 | Eef1aknmt | 71449 | protein_coding | 128.9449848 | -0.227507447 | 0.131427828 | 1.731044711 | 0.08344379 | 0.216495012 | 0.227507447 |
| ENSMUSG00000034192 | Lsm3 | 67678 | protein_coding | 179.6825719 | 0.227690123 | 0.114939763 | -1.980951741 | 0.047596686 | 0.143668739 | 0.227690123 |
| ENSMUSG00000024892 | Pcx | 18563 | protein_coding | 1016.520746 | -0.227728568 | 0.191595454 | 1.188590662 | 0.234600784 | 0.444186809 | 0.227728568 |
| ENSMUSG00000029832 | Nfe2l3 | 18025 | protein_coding | 36.71294383 | -0.227731792 | 0.227901519 | 0.999255261 | 0.317671052 | 0.539052565 | 0.227731792 |
| ENSMUSG00000030839 | Sergef | 27414 | protein_coding | 65.08503692 | -0.227788337 | 0.132525156 | 1.718830932 | 0.085645161 | 0.220887739 | 0.227788337 |
| ENSMUSG00000048546 | Tob2 | 57259 | protein_coding | 415.3398682 | -0.227833951 | 0.096002831 | 2.373200326 | 0.017634697 | 0.066534014 | 0.227833951 |
| ENSMUSG00000031749 | St3gal2 | 20444 | protein_coding | 13.27869836 | -0.227901892 | 0.45444056 | 0.501499892 | 0.616019354 | 0.788172589 | 0.227901892 |
| ENSMUSG00000118391 | Gm19519 | 100503033 | lncRNA | 1.414797724 | -0.227903792 | 1.105079269 | 0.206232981 | 0.836608926 | 1 | 0.227903792 |
| ENSMUSG00000028357 | Kif12 | 16552 | protein_coding | 275.7928326 | 0.227919154 | 0.121377951 | -1.877764062 | 0.060413449 | 0.17162699 | 0.227919154 |
| ENSMUSG00000022114 | Spry2 | 24064 | protein_coding | 391.298366 | -0.227944023 | 0.129775289 | 1.756451663 | 0.079011318 | 0.208221097 | 0.227944023 |
| ENSMUSG00000041135 | Ripk2 | 192656 | protein_coding | 154.047095 | 0.228061108 | 0.108864733 | -2.094903475 | 0.036179574 | 0.116216589 | 0.228061108 |
| ENSMUSG00000022518 | 4930562C15Rik | 78809 | protein_coding | 10.73904534 | 0.228097867 | 0.546875522 | -0.417092844 | 0.676610499 | 0.826873809 | 0.228097867 |
| ENSMUSG00000030707 | Coro1a | 12721 | protein_coding | 49.78359922 | 0.228135668 | 0.324196964 | -0.703694647 | 0.481622955 | 0.692385331 | 0.228135668 |
| ENSMUSG00000032606 | Nicn1 | 66257 | protein_coding | 90.31346324 | -0.228141558 | 0.125597705 | 1.816446876 | 0.06930185 | 0.189541363 | 0.228141558 |
| ENSMUSG00000020589 | Cyria | 76820 | protein_coding | 203.8355814 | -0.228145019 | 0.118748494 | 1.921245581 | 0.054700754 | 0.159457768 | 0.228145019 |
| ENSMUSG00000038304 | Cd160 | 54215 | protein_coding | 19.22000425 | 0.228151048 | 0.341572247 | -0.667943752 | 0.504169498 | 0.708596728 | 0.228151048 |
| ENSMUSG00000038122 | Tbc1d32 | 544696 | protein_coding | 325.5681595 | -0.228263217 | 0.063965538 | 3.568534294 | 0.000358984 | 0.002672795 | 0.228263217 |
| ENSMUSG00000031482 | Slc25a15 | 18408 | protein_coding | 384.3245897 | -0.228280049 | 0.101306974 | 2.253349806 | 0.024237102 | 0.085323278 | 0.228280049 |
| ENSMUSG00000055567 | Unc80 | 329178 | protein_coding | 11.29465575 | 0.228329761 | 0.637509202 | -0.35815916 | 0.720224208 | 0.855204229 | 0.228329761 |
| ENSMUSG00000039087 | Rreb1 | 68750 | protein_coding | 1219.66146 | 0.228469704 | 0.087086843 | -2.623469813 | 0.008703916 | 0.038096068 | 0.228469704 |
| ENSMUSG00000038991 | Txndc5 | 105245 | protein_coding | 4055.139002 | 0.228551931 | 0.137218973 | -1.665600077 | 0.095793096 | 0.239468239 | 0.228551931 |
| ENSMUSG00000021952 | Xpo4 | 57258 | protein_coding | 342.4705488 | 0.228610613 | 0.096478358 | -2.369553321 | 0.017809587 | 0.066903439 | 0.228610613 |
| ENSMUSG00000024563 | Smad2 | 17126 | protein_coding | 670.0783359 | 0.228635455 | 0.060365461 | -3.787521099 | 0.000152158 | 0.001243312 | 0.228635455 |
| ENSMUSG00000113722 | Snhg10 | 69434 | lncRNA | 5.614346775 | -0.228644344 | 0.581845446 | 0.392964053 | 0.694346032 | 0.837885199 | 0.228644344 |
| ENSMUSG00000022336 | Eif3e | 16341 | protein_coding | 1335.356268 | -0.228775714 | 0.11877771 | 1.92608288 | 0.054094026 | 0.158163729 | 0.228775714 |
| ENSMUSG00000077733 | Gm23608 | 115486143 | snoRNA | 1.305305294 | -0.228812627 | 0.833708038 | 0.274451747 | 0.783737481 | 1 | 0.228812627 |
| ENSMUSG00000015627 | Gata5 | 14464 | protein_coding | 12.84395841 | 0.228823025 | 0.445902582 | -0.513168201 | 0.607833668 | 0.782288135 | 0.228823025 |
| ENSMUSG00000061589 | Dot1l | 208266 | protein_coding | 450.7040136 | 0.228836537 | 0.174415816 | -1.312017123 | 0.189514357 | 0.386709513 | 0.228836537 |
| ENSMUSG00000110916 | 2310003N18Rik | 71873 | lncRNA | 0.678458349 | -0.22892818 | 1.512890836 | 0.151318373 | 0.879724577 | 1 | 0.22892818 |
| ENSMUSG00000050552 | Lamtor4 | 66096 | protein_coding | 421.2642511 | -0.228957998 | 0.087736057 | 2.609622612 | 0.009064216 | 0.039360099 | 0.228957998 |

| ensembl_gene_id | external_gene_name | entrezgene_id | gene_biotype | baseMean | log2FoldChange | lfcSE | stat | pvalue | padj | absLog2FC |
|---|---|---|---|---|---|---|---|---|---|---|
| ENSMUSG00000030094 | Xpc | 22591 | protein_coding | 264.8076829 | -0.228977192 | 0.105414167 | 2.172167153 | 0.029843057 | 0.100556672 | 0.228977192 |
| ENSMUSG00000044279 | Crb3 | 224912 | protein_coding | 837.9752525 | -0.228990498 | 0.101362686 | 2.259120265 | 0.023875905 | 0.084321363 | 0.228990498 |
| ENSMUSG00000032226 | Gcnt3 | 72077 | protein_coding | 3813.256002 | -0.229009389 | 0.143362309 | 1.597417001 | 0.110172785 | 0.265217693 | 0.229009389 |
| ENSMUSG00000053062 | Jam2 | 67374 | protein_coding | 4.589631191 | 0.229142376 | 0.7204438 | -0.318057253 | 0.750441508 | 0.874764669 | 0.229142376 |
| ENSMUSG00000028132 | Tlcd4 | 99887 | protein_coding | 473.4181673 | 0.22916987 | 0.103049794 | -2.223875103 | 0.026156849 | 0.090461612 | 0.22916987 |
| ENSMUSG00000016256 | Ctsz | 64138 | protein_coding | 994.3734663 | -0.22922222 | 0.054334432 | 4.21872858 | 2.46E-05 | 0.0002483 | 0.22922222 |
| ENSMUSG00000038276 | Asic3 | 171209 | protein_coding | 6.974158078 | 0.229247531 | 0.508995377 | -0.450392168 | 0.652427691 | 0.8114957 | 0.229247531 |
| ENSMUSG00000020180 | Snrpd3 | 67332 | protein_coding | 232.3243346 | 0.229273077 | 0.102213443 | -2.243081431 | 0.024891564 | 0.087070385 | 0.229273077 |
| ENSMUSG00000045975 | C2cd2 | 207781 | protein_coding | 211.7460594 | -0.229414959 | 0.113223804 | 2.02620784 | 0.042743491 | 0.132283734 | 0.229414959 |
| ENSMUSG00000034471 | Caskin2 | 140721 | protein_coding | 1047.197357 | -0.229423106 | 0.119052484 | 1.927075344 | 0.053970241 | 0.157957314 | 0.229423106 |
| ENSMUSG00000064927 | Gm22043 | 115489855 | snRNA | 2.181553871 | 0.229463926 | 0.783875279 | -0.292730148 | 0.769728424 | 1 | 0.229463926 |
| ENSMUSG00000052062 | Pard3b | 72823 | protein_coding | 496.2263229 | -0.229512675 | 0.076124596 | 3.014960842 | 0.002570124 | 0.013943178 | 0.229512675 |
| ENSMUSG00000089844 | A530032D15Rik | 381287 | protein_coding | 6.100045147 | -0.229616486 | 0.789415049 | 0.290869153 | 0.771151394 | 0.885199039 | 0.229616486 |
| ENSMUSG00000041959 | S100a10 | 20194 | protein_coding | 3326.055497 | -0.229622442 | 0.081427981 | 2.819945182 | 0.004803185 | 0.023502823 | 0.229622442 |
| ENSMUSG00000045838 | Ccdc9b | 214239 | protein_coding | 208.7233072 | 0.229829549 | 0.252381474 | -0.910643502 | 0.362483242 | 0.58491775 | 0.229829549 |
| ENSMUSG00000020601 | Trib2 | 217410 | protein_coding | 41.31984004 | 0.229835835 | 0.186498048 | -1.232376624 | 0.217808429 | 0.423711673 | 0.229835835 |
| ENSMUSG00000040430 | Pitpnc1 | 71795 | protein_coding | 35.91477986 | -0.229870922 | 0.230601906 | 0.996830106 | 0.318846982 | 0.540266681 | 0.229870922 |
| ENSMUSG00000031954 | Cfdp1 | 23837 | protein_coding | 608.0543459 | 0.229876885 | 0.100346181 | -2.290838398 | 0.021972762 | 0.079002436 | 0.229876885 |
| ENSMUSG00000008683 | Rps15a | 267019 | protein_coding | 2789.327745 | -0.229899931 | 0.137257447 | 1.674954153 | 0.093943227 | 0.23602289 | 0.229899931 |
| ENSMUSG00000050619 | Zscan29 | 99334 | protein_coding | 334.144663 | -0.230043323 | 0.080968017 | 2.841162869 | 0.004494935 | 0.022291098 | 0.230043323 |
| ENSMUSG00000022002 | Erich6b | 75272 | protein_coding | 1.597328555 | 0.230063495 | 1.443578464 | -0.159370274 | 0.873377158 | 1 | 0.230063495 |
| ENSMUSG00000045259 | Klhdc9 | 68874 | protein_coding | 7.866343854 | -0.23019347 | 0.410426565 | 0.560863963 | 0.574890278 | 0.760164897 | 0.23019347 |
| ENSMUSG00000040387 | Klhl32 | 212390 | protein_coding | 2.265121357 | -0.23024918 | 0.915902921 | 0.251390376 | 0.801512307 | 1 | 0.23024918 |
| ENSMUSG00000027562 | Car2 | 12349 | protein_coding | 12114.95619 | -0.230348782 | 0.127117836 | 1.812088611 | 0.069972506 | 0.190901365 | 0.230348782 |
| ENSMUSG00000044148 | 1810030O07Rik | 69155 | protein_coding | 360.3167868 | 0.230442729 | 0.107462387 | -2.144403587 | 0.032000563 | 0.106094619 | 0.230442729 |
| ENSMUSG00000015776 | Med22 | 20933 | protein_coding | 444.9595013 | -0.230503783 | 0.121699059 | 1.894047378 | 0.058218721 | 0.167087933 | 0.230503783 |
| ENSMUSG00000027957 | Slc35a3 | 229782 | protein_coding | 4145.818085 | 0.230547823 | 0.140505518 | -1.640845327 | 0.100829526 | 0.249266287 | 0.230547823 |
| ENSMUSG00000022426 | Josd1 | 74158 | protein_coding | 487.9580867 | -0.230575542 | 0.073224764 | 3.148873793 | 0.00163901 | 0.009604899 | 0.230575542 |
| ENSMUSG00000039178 | Tbc1d19 | 67249 | protein_coding | 71.06338958 | 0.230634623 | 0.189260064 | -1.218612198 | 0.222991417 | 0.429954629 | 0.230634623 |
| ENSMUSG00000111243 | Gm4668 | 100043817 | lncRNA | 2.238561892 | 0.230667468 | 0.806766536 | -0.285916009 | 0.774942452 | 1 | 0.230667468 |
| ENSMUSG00000029821 | Gsdme | 54722 | protein_coding | 267.2471653 | -0.230765165 | 0.088169804 | 2.617281131 | 0.008863331 | 0.038640192 | 0.230765165 |
| ENSMUSG00000048070 | Pirt | 193003 | protein_coding | 2.990424314 | -0.230798898 | 1.189717029 | 0.193994784 | 0.846179966 | 1 | 0.230798898 |
| ENSMUSG00000038717 | Atp5l | 27425 | protein_coding | 265.0681505 | -0.230800596 | 0.109345613 | 2.110743998 | 0.034794322 | 0.112975508 | 0.230800596 |
| ENSMUSG00000025747 | Tyms | 22171 | protein_coding | 367.597761 | 0.230809853 | 0.159133334 | -1.450418 | 0.146941991 | 0.324759105 | 0.230809853 |
| ENSMUSG00000001089 | Luzp1 | 269593 | protein_coding | 1839.660607 | -0.230859779 | 0.063045539 | 3.661794055 | 0.000250455 | 0.001933622 | 0.230859779 |
| ENSMUSG00000029847 | Slc23a4 | 243753 | protein_coding | 56.2727512 | -0.230953648 | 0.167408781 | 1.379579057 | 0.167716289 | 0.356246496 | 0.230953648 |
| ENSMUSG00000042138 | Msantd2 | 235184 | protein_coding | 75.63600806 | -0.231012867 | 0.128694262 | 1.795051807 | 0.072645446 | 0.196184284 | 0.231012867 |
| ENSMUSG00000030316 | Tamm41 | 68971 | protein_coding | 139.6288062 | 0.231052467 | 0.110270295 | -2.095328284 | 0.03614182 | 0.116153464 | 0.231052467 |
| ENSMUSG00000056698 | Elmod3 | 232089 | protein_coding | 143.2504337 | -0.23119745 | 0.104548945 | 2.211380037 | 0.027009531 | 0.092827346 | 0.23119745 |

Supplementary Table 1

| ensembl_gene_id | external_gene_name | entrezgene_id | gene_biotype | baseMean | log2FoldChange | lfcSE | stat | pvalue | padj | absLog2FC |
|---|---|---|---|---|---|---|---|---|---|---|
| ENSMUSG00000037940 | Inpp4b | 234515 | protein_coding | 221.384742 | 0.231197738 | 0.236300343 | -0.978406274 | 0.327873426 | 0.549974786 | 0.231197738 |
| ENSMUSG00000096423 | Gm24978 | 115489655 | snRNA | 1.217576751 | -0.23121461 | 1.185887791 | 0.194971743 | 0.845415069 | 1 | 0.23121461 |
| ENSMUSG00000041301 | Cftr | 12638 | protein_coding | 2781.028435 | -0.231367937 | 0.128546728 | 1.799874184 | 0.071880507 | 0.194819414 | 0.231367937 |
| ENSMUSG00000079497 | Gm13420 | 433415 | protein_coding | 5.037716649 | 0.231388363 | 0.560588266 | -0.412759911 | 0.679782526 | 0.828297243 | 0.231388363 |
| ENSMUSG00000035354 | Uvrag | 78610 | protein_coding | 229.1592827 | 0.231406436 | 0.090547074 | -2.555647857 | 0.010599032 | 0.044651126 | 0.231406436 |
| ENSMUSG00000055862 | Izumo4 | 71564 | protein_coding | 95.12427262 | -0.231442821 | 0.127888403 | 1.809724859 | 0.070338466 | 0.1916312 | 0.231442821 |
| ENSMUSG00000053870 | Fpgt | 75540 | protein_coding | 598.148702 | -0.231500117 | 0.125707358 | 1.841579685 | 0.065536655 | 0.181986265 | 0.231500117 |
| ENSMUSG00000069873 | 4930438A08Rik | 73988 | protein_coding | 1.923551896 | 0.231526289 | 1.283721098 | -0.1803556 | 0.856873409 | 1 | 0.231526289 |
| ENSMUSG00000000317 | Bcl6b | 12029 | protein_coding | 5.791817074 | 0.231535838 | 0.539100476 | -0.429485501 | 0.667569943 | 0.821350533 | 0.231535838 |
| ENSMUSG00000041923 | Nol4 | 319211 | protein_coding | 2.914386858 | 0.231618473 | 1.103053309 | -0.209979401 | 0.83368375 | 1 | 0.231618473 |
| ENSMUSG00000026634 | Angel2 | 52477 | protein_coding | 567.0722012 | -0.231708141 | 0.076800916 | 3.016997087 | 0.002552923 | 0.013872644 | 0.231708141 |
| ENSMUSG00000024959 | Bad | 12015 | protein_coding | 446.6765485 | 0.231876788 | 0.075647375 | -3.065232441 | 0.002175009 | 0.012122206 | 0.231876788 |
| ENSMUSG00000061080 | Lsamp | 268890 | protein_coding | 6.458188463 | -0.231944788 | 3.607792987 | 0.064289938 | 0.948739365 | 0.975316352 | 0.231944788 |
| ENSMUSG00000050677 | Ccdc96 | 66717 | protein_coding | 6.975297706 | 0.231984455 | 0.459439957 | -0.504928776 | 0.613608862 | 0.786448489 | 0.231984455 |
| ENSMUSG00000100860 | 2010009K17Rik | 72340 | lncRNA | 1.386261216 | 0.232039283 | 1.030355683 | -0.22520309 | 0.821821286 | 1 | 0.232039283 |
| ENSMUSG00000040282 | Cdin1 | 399568 | protein_coding | 56.24825096 | 0.232068636 | 0.196146753 | -1.183137791 | 0.236754543 | 0.446932387 | 0.232068636 |
| ENSMUSG00000054716 | Zfp771 | 244216 | protein_coding | 116.0597467 | 0.232085628 | 0.138194085 | -1.679417957 | 0.093070616 | 0.234222517 | 0.232085628 |
| ENSMUSG00000110277 | Gm45871 | 108168395 | protein_coding | 233.7345473 | -0.232119309 | 0.134485016 | 1.725986405 | 0.084349859 | 0.21833128 | 0.232119309 |
| ENSMUSG00000002017 | Fam98a | 72722 | protein_coding | 382.5689866 | 0.232142081 | 0.114855537 | -2.021165778 | 0.043262607 | 0.133465102 | 0.232142081 |
| ENSMUSG00000002885 | Adgre5 | 26364 | protein_coding | 384.6948306 | -0.232216876 | 0.20726835 | 1.120368233 | 0.262556876 | 0.476573341 | 0.232216876 |
| ENSMUSG00000059355 | Wdr83os | 414077 | protein_coding | 155.5010708 | -0.232254705 | 0.117202825 | 1.981647668 | 0.047518688 | 0.143502708 | 0.232254705 |
| ENSMUSG00000112532 | Gm36283 | 102640148 | lncRNA | 2.820260988 | -0.232289089 | 1.043424806 | 0.222621781 | 0.823829883 | 1 | 0.232289089 |
| ENSMUSG00000045411 | 2410002F23Rik | 668661 | protein_coding | 320.7675763 | -0.232308211 | 0.118144199 | 1.966310779 | 0.049262722 | 0.147350951 | 0.232308211 |
| ENSMUSG00000047109 | Cldn14 | 56173 | protein_coding | 24.45098626 | 0.232336736 | 0.22112845 | -1.050686769 | 0.293402475 | 0.512561987 | 0.232336736 |
| ENSMUSG00000022681 | Ntan1 | 18203 | protein_coding | 270.3831182 | -0.232338195 | 0.068901561 | 3.372030935 | 0.000746161 | 0.004948633 | 0.232338195 |
| ENSMUSG00000032076 | Cadm1 | 54725 | protein_coding | 15.9009785 | -0.232359512 | 0.281904609 | 0.824248716 | 0.409798252 | 0.630288565 | 0.232359512 |
| ENSMUSG00000021007 | Spata7 | 104871 | protein_coding | 57.71625872 | -0.232383858 | 0.143034575 | 1.624669127 | 0.104233079 | 0.255121545 | 0.232383858 |
| ENSMUSG00000037108 | Zcwpw1 | 381678 | protein_coding | 12.32763039 | -0.232432788 | 0.328747552 | 0.70702515 | 0.479550849 | 0.690797515 | 0.232432788 |
| ENSMUSG00000046862 | Pramel12 | 242736 | protein_coding | 235.5849336 | -0.232497429 | 0.097308392 | 2.389284467 | 0.016881226 | 0.064232353 | 0.232497429 |
| ENSMUSG00000047678 | Gpr82 | 319200 | protein_coding | 3.200362035 | -0.232519271 | 0.572169093 | 0.406382088 | 0.684461879 | 1 | 0.232519271 |
| ENSMUSG00000030725 | Lipt2 | 67164 | protein_coding | 105.0548147 | 0.232612012 | 0.17070334 | -1.362668192 | 0.172987107 | 0.363680341 | 0.232612012 |
| ENSMUSG00000078942 | Naip6 | 17952 | protein_coding | 1314.518957 | -0.232613997 | 0.086751498 | 2.681383072 | 0.007331853 | 0.033126737 | 0.232613997 |
| ENSMUSG00000039108 | Lsm14b | 241846 | protein_coding | 320.8934685 | -0.232634373 | 0.079783209 | 2.915831235 | 0.003547423 | 0.018306921 | 0.232634373 |
| ENSMUSG00000038515 | Grtp1 | 66790 | protein_coding | 893.6135834 | -0.232640647 | 0.063379391 | 3.670604008 | 0.000241978 | 0.001872551 | 0.232640647 |
| ENSMUSG00000028381 | Ugcg | 22234 | protein_coding | 838.6370373 | -0.232648656 | 0.110508464 | 2.105256436 | 0.035268992 | 0.114179991 | 0.232648656 |
| ENSMUSG00000030263 | Irag2 | 16970 | protein_coding | 284.0956152 | -0.232653945 | 0.189373945 | 1.228542531 | 0.219243374 | 0.425500655 | 0.232653945 |
| ENSMUSG00000024831 | Ighmbp2 | 20589 | protein_coding | 159.7298818 | 0.232730988 | 0.099779684 | -2.332448623 | 0.019677103 | 0.07241317 | 0.232730988 |
| ENSMUSG00000024213 | Nudt3 | 56409 | protein_coding | 564.2392665 | -0.232733993 | 0.099954816 | 2.328391986 | 0.019891299 | 0.07306159 | 0.232733993 |
| ENSMUSG00000115111 | Gm4632 | 100043765 | lncRNA | 1.88818489 | 0.232779832 | 0.83499639 | -0.278779447 | 0.780414089 | 1 | 0.232779832 |

| ensembl_gene_id | external_gene_name | entrezgene_id | gene_biotype | baseMean | log2FoldChange | lfcSE | stat | pvalue | padj | absLog2FC |
|---|---|---|---|---|---|---|---|---|---|---|
| ENSMUSG00000020905 | Usp43 | 216835 | protein_coding | 470.3969182 | 0.232810021 | 0.071208714 | -3.269403531 | 0.001077745 | 0.006772404 | 0.232810021 |
| ENSMUSG00000041528 | Rnf123 | 84585 | protein_coding | 339.7735242 | -0.232823964 | 0.073532275 | 3.166282605 | 0.001544007 | 0.009122624 | 0.232823964 |
| ENSMUSG00000028555 | Ttc39a | 230603 | protein_coding | 1728.851015 | 0.232845173 | 0.153021555 | -1.521649497 | 0.128096928 | 0.295198531 | 0.232845173 |
| ENSMUSG00000095687 | Rnaset2a | 100037283 | protein_coding | 5.574443753 | -0.232874097 | 0.530002816 | 0.439382755 | 0.66038422 | 0.816530773 | 0.232874097 |
| ENSMUSG00000025195 | Dnmbp | 71972 | protein_coding | 2249.445197 | -0.232977146 | 0.083955992 | 2.774991273 | 0.005520319 | 0.026393326 | 0.232977146 |
| ENSMUSG00000066979 | Bub3 | 12237 | protein_coding | 630.621801 | 0.233088273 | 0.104448244 | -2.231615046 | 0.025640416 | 0.089020995 | 0.233088273 |
| ENSMUSG00000030929 | Eri2 | 71151 | protein_coding | 89.33175073 | 0.233097028 | 0.153950887 | -1.514099938 | 0.130000494 | 0.297863506 | 0.233097028 |
| ENSMUSG00000021506 | Pitx1 | 18740 | protein_coding | 2.34029626 | 0.233122813 | 1.282403927 | -0.181785791 | 0.855750836 | 1 | 0.233122813 |
| ENSMUSG00000034401 | Spata6 | 67946 | protein_coding | 66.68340662 | -0.233148978 | 0.195205177 | 1.194379075 | 0.232329718 | 0.441303539 | 0.233148978 |
| ENSMUSG00000033628 | Pik3c3 | 225326 | protein_coding | 560.0652115 | -0.233160622 | 0.113598983 | 2.05248865 | 0.040122197 | 0.126157266 | 0.233160622 |
| ENSMUSG00000086968 | 4933431E20Rik | 329735 | lncRNA | 274.3853539 | -0.233198748 | 0.120602095 | 1.933621045 | 0.053159738 | 0.156261999 | 0.233198748 |
| ENSMUSG00000035798 | Zdhhc17 | 320150 | protein_coding | 281.9068463 | -0.233233221 | 0.089914015 | 2.593958479 | 0.009487794 | 0.040813086 | 0.233233221 |
| ENSMUSG00000018844 | Fndc8 | 78919 | protein_coding | 4.011348827 | 0.233242093 | 0.531521742 | -0.438819478 | 0.660792345 | 1 | 0.233242093 |
| ENSMUSG00000040811 | Eml2 | 72205 | protein_coding | 212.8136152 | -0.23333828 | 0.108865803 | 2.143356981 | 0.032084445 | 0.106287366 | 0.23333828 |
| ENSMUSG00000073802 | Cdkn2b | 12579 | protein_coding | 635.0130847 | -0.233392585 | 0.12336317 | 1.891914614 | 0.058502358 | 0.167639761 | 0.233392585 |
| ENSMUSG00000029442 | Wdr66 | 269701 | protein_coding | 8.130333841 | 0.233397386 | 1.067462688 | -0.21864688 | 0.826925131 | 0.912640622 | 0.233397386 |
| ENSMUSG00000038422 | Hdhd3 | 72748 | protein_coding | 232.0661039 | 0.23343732 | 0.13090396 | -1.783271646 | 0.074542095 | 0.199738463 | 0.23343732 |
| ENSMUSG00000061531 | Tmem236 | 625286 | protein_coding | 1118.486781 | -0.233612531 | 0.175641449 | 1.330053538 | 0.183500632 | 0.378318282 | 0.233612531 |
| ENSMUSG00000001901 | Kcnh6 | 192775 | protein_coding | 44.99816516 | 0.233619655 | 0.200379186 | -1.165887831 | 0.2436598 | 0.454872025 | 0.233619655 |
| ENSMUSG00000034855 | Cxcl10 | 15945 | protein_coding | 9.926883353 | 0.2336812 | 0.659381701 | -0.354394428 | 0.72304331 | 0.856549354 | 0.2336812 |
| ENSMUSG00000020211 | Sf3a2 | 20222 | protein_coding | 227.7736423 | 0.233720914 | 0.137121933 | -1.704475055 | 0.088292372 | 0.22556625 | 0.233720914 |
| ENSMUSG00000110507 | 8030455M16Rik | 77171 | lncRNA | 1.410516161 | -0.233724331 | 1.770689959 | 0.131996192 | 0.894987304 | 1 | 0.233724331 |
| ENSMUSG00000026873 | Phf19 | 74016 | protein_coding | 73.61687591 | -0.233734691 | 0.192159728 | 1.216356277 | 0.223849232 | 0.430945001 | 0.233734691 |
| ENSMUSG00000029535 | Triap1 | 69076 | protein_coding | 258.9212069 | -0.233787832 | 0.097976739 | 2.386156488 | 0.017025505 | 0.064634295 | 0.233787832 |
| ENSMUSG00000020496 | Rnf187 | 108660 | protein_coding | 828.8957294 | 0.23380682 | 0.079836688 | -2.928563607 | 0.003405321 | 0.017689676 | 0.23380682 |
| ENSMUSG00000019863 | Qrsl1 | 76563 | protein_coding | 209.8659142 | -0.233816149 | 0.100502106 | 2.326480092 | 0.019992953 | 0.073386004 | 0.233816149 |
| ENSMUSG00000032298 | Neil1 | 72774 | protein_coding | 99.8489168 | -0.233821428 | 0.102617355 | 2.278575868 | 0.02269229 | 0.081077558 | 0.233821428 |
| ENSMUSG00000025402 | Nab2 | 17937 | protein_coding | 14.34941495 | 0.233872208 | 0.331594795 | -0.705295172 | 0.480626564 | 0.691562872 | 0.233872208 |
| ENSMUSG00000020178 | Adora2a | 11540 | protein_coding | 7.409289904 | 0.234010139 | 0.685187938 | -0.341526939 | 0.732706928 | 0.862924134 | 0.234010139 |
| ENSMUSG00000050530 | Fam171a1 | 269233 | protein_coding | 241.499486 | -0.234081723 | 0.103513366 | 2.26136712 | 0.023736533 | 0.083972816 | 0.234081723 |
| ENSMUSG00000022554 | Hgh1 | 59053 | protein_coding | 84.6234852 | 0.234111935 | 0.126155059 | -1.855747491 | 0.063489558 | 0.177826758 | 0.234111935 |
| ENSMUSG00000063888 | Rpl7l1 | 66229 | protein_coding | 790.3374468 | 0.234200637 | 0.094109302 | -2.488602417 | 0.012824629 | 0.051642509 | 0.234200637 |
| ENSMUSG00000079197 | Psme2 | 19188 | protein_coding | 1151.241627 | -0.234254911 | 0.10178902 | 2.301377005 | 0.021370331 | 0.077290766 | 0.234254911 |
| ENSMUSG00000041688 | Amot | 27494 | protein_coding | 546.022296 | 0.234256784 | 0.113153428 | -2.070257954 | 0.038428195 | 0.122133027 | 0.234256784 |
| ENSMUSG00000069206 | Zfp874a | 238692 | protein_coding | 118.1293685 | -0.234266194 | 0.187610117 | 1.248686359 | 0.211779812 | 0.416114999 | 0.234266194 |
| ENSMUSG00000041168 | Lonp1 | 74142 | protein_coding | 983.0723049 | -0.234269364 | 0.06612828 | 3.542650212 | 0.000396128 | 0.002907402 | 0.234269364 |
| ENSMUSG00000046311 | Zfp62 | 22720 | protein_coding | 343.9804255 | -0.234280574 | 0.086278656 | 2.715394332 | 0.00661969 | 0.030425148 | 0.234280574 |
| ENSMUSG00000053749 | Gm9920 | 100503794 | lncRNA | 33.26333551 | 0.234288352 | 0.202430828 | -1.157374864 | 0.247119239 | 0.459127576 | 0.234288352 |
| ENSMUSG00000018909 | Arrb1 | 109689 | protein_coding | 1246.253235 | -0.234311667 | 0.093104304 | 2.516657726 | 0.011847386 | 0.048608132 | 0.234311667 |

| ensembl_gene_id | external_gene_name | entrezgene_id | gene_biotype | baseMean | log2FoldChange | lfcSE | stat | pvalue | padj | absLog2FC |
|---|---|---|---|---|---|---|---|---|---|---|
| ENSMUSG00000005575 | Ube2m | 22192 | protein_coding | 798.3029637 | 0.234340819 | 0.104338106 | -2.245975403 | 0.024705583 | 0.086566519 | 0.234340819 |
| ENSMUSG00000030428 | Ttyh1 | 57776 | protein_coding | 63.19619484 | -0.234370351 | 0.214461727 | 1.092830664 | 0.274468156 | 0.490866888 | 0.234370351 |
| ENSMUSG00000021368 | Tbc1d7 | 67046 | protein_coding | 77.74681006 | 0.234372434 | 0.117914182 | -1.987652627 | 0.046850122 | 0.141882067 | 0.234372434 |
| ENSMUSG00000030325 | Klrb1c | 17059 | protein_coding | 4.579012026 | 0.234475128 | 0.585144761 | -0.400713027 | 0.688631419 | 0.83386713 | 0.234475128 |
| ENSMUSG00000036138 | Acaa1a | 113868 | protein_coding | 1023.735748 | -0.234479419 | 0.084854751 | 2.763303348 | 0.005721956 | 0.027207712 | 0.234479419 |
| ENSMUSG00000021411 | Pxdc1 | 66895 | protein_coding | 26.7556612 | 0.234608125 | 0.307561494 | -0.762800708 | 0.445582259 | 0.662854285 | 0.234608125 |
| ENSMUSG00000107499 | Ccdc142 | 243510 | protein_coding | 4.115128326 | 0.234636888 | 0.597148058 | -0.392929166 | 0.694371799 | 1 | 0.234636888 |
| ENSMUSG00000118401 | Gpr52 | 620246 | protein_coding | 7.355030286 | -0.234822031 | 0.44381556 | 0.529098239 | 0.596737303 | 0.774787312 | 0.234822031 |
| ENSMUSG00000117391 | 4930500A05Rik | 74972 | lncRNA | 0.816735893 | 0.234830986 | 1.925621837 | -0.121950728 | 0.90293804 | 1 | 0.234830986 |
| ENSMUSG00000059436 | Max | 17187 | protein_coding | 1020.829914 | -0.234913992 | 0.107263346 | 2.190067714 | 0.028519326 | 0.096986852 | 0.234913992 |
| ENSMUSG00000022523 | Fgf12 | 14167 | protein_coding | 15.6365556 | -0.235057345 | 0.373205876 | 0.629832914 | 0.528803909 | 0.726753308 | 0.235057345 |
| ENSMUSG00000024043 | Arhgap28 | 268970 | protein_coding | 3.377703861 | 0.235133483 | 0.895777832 | -0.262490848 | 0.79294304 | 1 | 0.235133483 |
| ENSMUSG00000041559 | Fmod | 14264 | protein_coding | 1.842474253 | 0.23518286 | 1.481824879 | -0.158711642 | 0.873896065 | 1 | 0.23518286 |
| ENSMUSG00000063268 | Parp10 | 671535 | protein_coding | 403.0783102 | -0.235210081 | 0.086042881 | 2.733637894 | 0.006263889 | 0.029064867 | 0.235210081 |
| ENSMUSG00000045165 | AI467606 | 101602 | protein_coding | 35.02286716 | 0.235264256 | 0.213002579 | -1.104513651 | 0.269370385 | 0.484414956 | 0.235264256 |
| ENSMUSG00000006586 | Runx1t1 | 12395 | protein_coding | 9.716754044 | -0.235306503 | 0.485323657 | 0.484844494 | 0.627786658 | 0.796376973 | 0.235306503 |
| ENSMUSG00000020691 | Mettl2 | 52686 | protein_coding | 152.3381851 | -0.235331816 | 0.127981079 | 1.838801631 | 0.065944368 | 0.182841722 | 0.235331816 |
| ENSMUSG00000031818 | Cox4i1 | 12857 | protein_coding | 8281.545296 | -0.235336612 | 0.087072262 | 2.702773615 | 0.006876356 | 0.031443528 | 0.235336612 |
| ENSMUSG00000064901 | Snora21 | 100302498 | snoRNA | 32.03401835 | 0.235401744 | 0.267122279 | -0.881250879 | 0.378182046 | 0.600451717 | 0.235401744 |
| ENSMUSG00000077575 | Gm24888 | 115485611 | snoRNA | 2.840927562 | 0.235418516 | 0.630064163 | -0.373642129 | 0.708670586 | 1 | 0.235418516 |
| ENSMUSG00000020902 | Ntn1 | 18208 | protein_coding | 5.303619582 | 0.23544405 | 0.81890679 | -0.287510194 | 0.773721698 | 0.88672609 | 0.23544405 |
| ENSMUSG00000118134 | 4930549G23Rik | 75286 | lncRNA | 3.109585598 | -0.235539032 | 0.62550717 | 0.376556885 | 0.706502934 | 1 | 0.235539032 |
| ENSMUSG00000031642 | Sh3rf1 | 59009 | protein_coding | 515.7326429 | -0.235564031 | 0.094476447 | 2.493362511 | 0.012653958 | 0.051129793 | 0.235564031 |
| ENSMUSG00000055692 | Tmem191 | 224019 | protein_coding | 44.05203251 | 0.235586613 | 0.21004401 | -1.121605957 | 0.262030018 | 0.475826228 | 0.235586613 |
| ENSMUSG00000049047 | Armcx3 | 71703 | protein_coding | 124.1914394 | -0.235595253 | 0.19590508 | 1.202599001 | 0.229131534 | 0.438050261 | 0.235595253 |
| ENSMUSG00000016637 | Ift27 | 67042 | protein_coding | 92.1451884 | 0.235597043 | 0.133281883 | -1.767659921 | 0.077117774 | 0.204779373 | 0.235597043 |
| ENSMUSG00000116505 | Gm10865 | 100038602 | lncRNA | 3.662639595 | 0.235641122 | 0.635656597 | -0.370705068 | 0.710857215 | 1 | 0.235641122 |
| ENSMUSG00000017418 | Arl5b | 75869 | protein_coding | 446.5651473 | -0.235659297 | 0.112363025 | 2.097302896 | 0.035966771 | 0.11573563 | 0.235659297 |
| ENSMUSG00000026972 | Arrdc1 | 215705 | protein_coding | 738.0458655 | -0.235693881 | 0.061816529 | 3.812797081 | 0.000137403 | 0.00114081 | 0.235693881 |
| ENSMUSG00000020684 | Rasl10b | 276952 | protein_coding | 2.231998047 | 0.235705705 | 1.7805403 | -0.132378753 | 0.894684719 | 1 | 0.235705705 |
| ENSMUSG00000113902 | Ndufb1 | 102631912 | protein_coding | 697.9415576 | -0.235723633 | 0.097918702 | 2.407340255 | 0.016069191 | 0.061727521 | 0.235723633 |
| ENSMUSG00000031376 | Atp2b3 | 320707 | protein_coding | 4.479601901 | 0.235750183 | 1.439757319 | -0.163743 | 0.869933465 | 0.935658236 | 0.235750183 |
| ENSMUSG00000051316 | Taf7 | 24074 | protein_coding | 176.5163238 | -0.23582262 | 0.107465768 | 2.194397571 | 0.02820684 | 0.096042838 | 0.23582262 |
| ENSMUSG00000038365 | Fbxo25 | 66822 | protein_coding | 1012.350395 | -0.235829337 | 0.102729456 | 2.295635011 | 0.021696759 | 0.078282779 | 0.235829337 |
| ENSMUSG00000030800 | Prss8 | 76560 | protein_coding | 736.1082687 | 0.235835327 | 0.068256983 | -3.455109152 | 0.00055007 | 0.003837987 | 0.235835327 |
| ENSMUSG00000030604 | Zfp626 | 71163 | protein_coding | 150.4785 | -0.235846754 | 0.104898516 | 2.248332606 | 0.024554989 | 0.086166827 | 0.235846754 |
| ENSMUSG00000002343 | Armc6 | 76813 | protein_coding | 109.92289 | 0.235861173 | 0.146971326 | -1.604810809 | 0.108535446 | 0.262539089 | 0.235861173 |
| ENSMUSG00000045282 | Tmem86b | 68255 | protein_coding | 29.79116338 | -0.235874776 | 0.239701993 | 0.98403344 | 0.325099071 | 0.546957754 | 0.235874776 |
| ENSMUSG00000036526 | Card11 | 108723 | protein_coding | 40.71905824 | 0.235899471 | 0.21018941 | -1.122318536 | 0.261727028 | 0.475380569 | 0.235899471 |

| ensembl_gene_id | external_gene_name | entrezgene_id | gene_biotype | baseMean | log2FoldChange | lfcSE | stat | pvalue | padj | absLog2FC |
|---|---|---|---|---|---|---|---|---|---|---|
| ENSMUSG00000030763 | Lcmt1 | 30949 | protein_coding | 217.115282 | 0.235939358 | 0.125194831 | -1.884577472 | 0.059486906 | 0.169754064 | 0.235939358 |
| ENSMUSG00000035283 | Adrb1 | 11554 | protein_coding | 2.918723112 | -0.236030828 | 1.323695363 | 0.178312046 | 0.858477919 | 1 | 0.236030828 |
| ENSMUSG00000078129 | Actl10 | 70362 | protein_coding | 0.827606995 | 0.23604865 | 1.336594099 | -0.176604588 | 0.85981899 | 1 | 0.23604865 |
| ENSMUSG00000035004 | Igsf6 | 80719 | protein_coding | 7.223716171 | -0.236072785 | 0.421537885 | 0.560027446 | 0.575460717 | 0.760617759 | 0.236072785 |
| ENSMUSG00000015337 | Endog | 13804 | protein_coding | 139.4899574 | 0.236127773 | 0.13519711 | -1.746544535 | 0.080716376 | 0.211290803 | 0.236127773 |
| ENSMUSG00000050666 | Vstm4 | 320736 | protein_coding | 1.121168604 | -0.236172321 | 1.991821193 | 0.118571045 | 0.905615206 | 1 | 0.236172321 |
| ENSMUSG00000015027 | Galns | 50917 | protein_coding | 320.9466154 | -0.236173256 | 0.062046973 | 3.806362265 | 0.000141026 | 0.001167368 | 0.236173256 |
| ENSMUSG00000038257 | Glra3 | 110304 | protein_coding | 0.801418616 | -0.236173773 | 2.199936324 | 0.107354822 | 0.914507494 | 1 | 0.236173773 |
| ENSMUSG00000038024 | Dennd4c | 329877 | protein_coding | 1522.476698 | -0.236175374 | 0.066126961 | 3.57154434 | 0.000354882 | 0.002648228 | 0.236175374 |
| ENSMUSG00000019787 | Trdn | 76757 | protein_coding | 1.753672952 | -0.236177031 | 2.186911291 | 0.107995707 | 0.913999098 | 1 | 0.236177031 |
| ENSMUSG00000096029 | Or56a3 | 259046 | protein_coding | 0.897527938 | -0.236181457 | 2.67509266 | 0.088289075 | 0.929646922 | 1 | 0.236181457 |
| ENSMUSG00000031645 | F11 | 109821 | protein_coding | 0.992649162 | -0.236181783 | 2.658179121 | 0.088850966 | 0.929200353 | 1 | 0.236181783 |
| ENSMUSG00000069421 | Or6c69b | 258543 | protein_coding | 0.945212167 | -0.236182559 | 2.719997668 | 0.086831897 | 0.930805133 | 1 | 0.236182559 |
| ENSMUSG00000096069 | Or13a28 | 18362 | protein_coding | 1.135454616 | -0.236183625 | 2.714333134 | 0.0870135 | 0.930660782 | 1 | 0.236183625 |
| ENSMUSG00000030647 | Ndufc2 | 68197 | protein_coding | 977.234188 | -0.236191772 | 0.08778758 | 2.690491885 | 0.007134677 | 0.032448928 | 0.236191772 |
| ENSMUSG00000036480 | Prss56 | 69453 | protein_coding | 1.088017621 | -0.2361943 | 3.271757651 | 0.072191869 | 0.942449216 | 1 | 0.2361943 |
| ENSMUSG00000096584 | Or51aa5 | 258722 | protein_coding | 1.088017621 | -0.2361943 | 3.271757651 | 0.072191869 | 0.942449216 | 1 | 0.2361943 |
| ENSMUSG00000070547 | Mrgprb1 | 233231 | protein_coding | 1.365126374 | -0.23619505 | 3.207033373 | 0.073649078 | 0.941289619 | 1 | 0.23619505 |
| ENSMUSG00000075478 | Slitrk1 | 76965 | protein_coding | 1.365126374 | -0.23619505 | 3.207033373 | 0.073649078 | 0.941289619 | 1 | 0.23619505 |
| ENSMUSG00000058692 | Or7g28 | 258279 | protein_coding | 1.229524778 | -0.236195156 | 3.241237349 | 0.07287191 | 0.941908047 | 1 | 0.236195156 |
| ENSMUSG00000075174 | Or8k3b | 258843 | protein_coding | 1.230575841 | -0.236196306 | 3.310240875 | 0.071353208 | 0.943116649 | 1 | 0.236196306 |
| ENSMUSG00000030871 | Ears2 | 67417 | protein_coding | 242.9588444 | 0.23622927 | 0.131636108 | -1.794562851 | 0.072723376 | 0.196362629 | 0.23622927 |
| ENSMUSG00000032050 | Rdx | 19684 | protein_coding | 252.0668247 | 0.236292504 | 0.366139676 | -0.645361645 | 0.518692853 | 0.719201828 | 0.236292504 |
| ENSMUSG00000040260 | Daam2 | 76441 | protein_coding | 10.98173043 | 0.236349465 | 0.501586989 | -0.471203341 | 0.637495533 | 0.802408631 | 0.236349465 |
| ENSMUSG00000037326 | Capn15 | 50817 | protein_coding | 305.6705679 | 0.236378888 | 0.09625002 | -2.455884032 | 0.01405385 | 0.055403505 | 0.236378888 |
| ENSMUSG00000036534 | Slc38a7 | 234595 | protein_coding | 141.3194117 | -0.236455066 | 0.094663735 | 2.497842142 | 0.012495182 | 0.050636917 | 0.236455066 |
| ENSMUSG00000023921 | Mmut | 17850 | protein_coding | 1024.766578 | 0.236487191 | 0.093506186 | -2.52910743 | 0.011435302 | 0.047391714 | 0.236487191 |
| ENSMUSG00000050473 | Slc35d3 | 76157 | protein_coding | 4.196902302 | 0.236591341 | 0.775132965 | -0.305226783 | 0.760193435 | 1 | 0.236591341 |
| ENSMUSG00000038776 | Ephx1 | 13849 | protein_coding | 190.471193 | 0.236594649 | 0.199187959 | -1.187795936 | 0.234913816 | 0.444677528 | 0.236594649 |
| ENSMUSG00000051550 | Zfp579 | 68490 | protein_coding | 66.64285499 | -0.236656787 | 0.189389311 | 1.249578374 | 0.211453607 | 0.415856243 | 0.236656787 |
| ENSMUSG00000006587 | Snai3 | 30927 | protein_coding | 10.15148274 | 0.236709171 | 0.391329645 | -0.604884333 | 0.545255853 | 0.737095216 | 0.236709171 |
| ENSMUSG00000115518 | Gm10791 | 100038614 | lncRNA | 10.7694913 | -0.23673454 | 0.361938631 | 0.654073701 | 0.513064324 | 0.714999514 | 0.23673454 |
| ENSMUSG00000047197 | Gjd3 | 353155 | protein_coding | 0.880193528 | -0.236784909 | 1.871208629 | 0.126541159 | 0.899303571 | 1 | 0.236784909 |
| ENSMUSG00000024371 | C2 | 12263 | protein_coding | 17.94925614 | -0.23678542 | 0.272209795 | 0.869863699 | 0.384374895 | 0.606725963 | 0.23678542 |
| ENSMUSG00000033487 | Fndc3a | 319448 | protein_coding | 2135.944928 | 0.236790242 | 0.062866691 | -3.766545341 | 0.000165522 | 0.001338605 | 0.236790242 |
| ENSMUSG00000048232 | Fbxo10 | 269529 | protein_coding | 49.58875382 | 0.236814524 | 0.186028046 | -1.273004419 | 0.203016474 | 0.405213026 | 0.236814524 |
| ENSMUSG00000042271 | Nxt2 | 237082 | protein_coding | 187.7217339 | -0.236894626 | 0.122250568 | 1.937779349 | 0.052650148 | 0.155233806 | 0.236894626 |
| ENSMUSG00000020053 | Igf1 | 16000 | protein_coding | 3.250390382 | -0.236938311 | 1.645909347 | 0.14395587 | 0.885535318 | 1 | 0.236938311 |
| ENSMUSG00000039395 | Mreg | 381269 | protein_coding | 10.68253615 | -0.236941855 | 0.424310189 | 0.558416604 | 0.576559937 | 0.761222618 | 0.236941855 |

| ensembl_gene_id | external_gene_name | entrezgene_id | gene_biotype | baseMean | log2FoldChange | lfcSE | stat | pvalue | padj | absLog2FC |
|---|---|---|---|---|---|---|---|---|---|---|
| ENSMUSG00000023827 | Agpat4 | 68262 | protein_coding | 796.2489715 | -0.236959181 | 0.109381724 | 2.166350764 | 0.030284393 | 0.101670354 | 0.236959181 |
| ENSMUSG00000034768 | Asb16 | 217217 | protein_coding | 11.72894317 | 0.237007156 | 0.342222159 | -0.692553507 | 0.488589796 | 0.696497169 | 0.237007156 |
| ENSMUSG00000089804 | Gm16136 | 105245924 | lncRNA | 68.88767446 | -0.237023563 | 0.138517659 | 1.711143292 | 0.087054664 | 0.223580206 | 0.237023563 |
| ENSMUSG00000041936 | Agrn | 11603 | protein_coding | 850.8461615 | -0.237027958 | 0.133976377 | 1.769177253 | 0.076864299 | 0.204314673 | 0.237027958 |
| ENSMUSG00000021553 | Slc28a3 | 114304 | protein_coding | 59.39375905 | -0.237043566 | 0.24949127 | 0.950107659 | 0.342057552 | 0.564006111 | 0.237043566 |
| ENSMUSG00000028078 | Dclk2 | 70762 | protein_coding | 23.04028338 | 0.237078445 | 0.240427143 | -0.986071881 | 0.324097841 | 0.545970799 | 0.237078445 |
| ENSMUSG00000068246 | Apol9b | 71898 | protein_coding | 37.57521582 | -0.237096316 | 0.49568827 | 0.478317383 | 0.632424326 | 0.799512805 | 0.237096316 |
| ENSMUSG00000036427 | Gpi1 | 14751 | protein_coding | 5563.1764 | -0.23710081 | 0.072469142 | 3.271748538 | 0.001068846 | 0.006724161 | 0.23710081 |
| ENSMUSG00000028179 | Cth | 107869 | protein_coding | 1076.870089 | -0.237203918 | 0.083677301 | 2.834746298 | 0.004586209 | 0.022621462 | 0.237203918 |
| ENSMUSG00000026103 | Gls | 14660 | protein_coding | 1160.410907 | -0.237292381 | 0.056811821 | 4.176813507 | 2.96E-05 | 0.000292855 | 0.237292381 |
| ENSMUSG00000002102 | Psmc3 | 19182 | protein_coding | 1721.501858 | 0.237298826 | 0.076575208 | -3.098898869 | 0.001942413 | 0.011080142 | 0.237298826 |
| ENSMUSG00000047502 | Mroh7 | 381538 | protein_coding | 9.26069389 | 0.237349209 | 0.689774207 | -0.344096961 | 0.730773368 | 0.861267215 | 0.237349209 |
| ENSMUSG00000118516 | Cyp2d13 | 68444 | protein_coding | 102.2889808 | -0.237349922 | 0.258223662 | 0.919164108 | 0.358009741 | 0.580790426 | 0.237349922 |
| ENSMUSG00000000001 | Gnai3 | 14679 | protein_coding | 2306.328636 | 0.237352357 | 0.069655391 | -3.407523132 | 0.000655554 | 0.004448039 | 0.237352357 |
| ENSMUSG00000019828 | Grm1 | 14816 | protein_coding | 2.14702026 | 0.237379294 | 1.743389937 | -0.13615961 | 0.891695104 | 1 | 0.237379294 |
| ENSMUSG00000038520 | Tbc1d17 | 233204 | protein_coding | 385.7286094 | -0.237386142 | 0.070004971 | 3.390989812 | 0.000696407 | 0.00466172 | 0.237386142 |
| ENSMUSG00000038665 | Dgki | 320127 | protein_coding | 21.67724023 | -0.237398506 | 0.407995966 | 0.581864837 | 0.560657727 | 0.749146509 | 0.237398506 |
| ENSMUSG00000003233 | Dvl3 | 13544 | protein_coding | 271.0858033 | 0.237422529 | 0.169917088 | -1.397284588 | 0.162328008 | 0.347734431 | 0.237422529 |
| ENSMUSG00000033107 | Rnf125 | 67664 | protein_coding | 65.34931151 | -0.237530626 | 0.202808066 | 1.171208971 | 0.24151479 | 0.452145476 | 0.237530626 |
| ENSMUSG00000062961 | Ccdc177 | 380768 | protein_coding | 8.838276383 | 0.237598968 | 0.378468606 | -0.627790427 | 0.530141238 | 0.727438372 | 0.237598968 |
| ENSMUSG00000032355 | Mlip | 69642 | protein_coding | 5.825166649 | -0.237604966 | 0.940555745 | 0.252621886 | 0.800560416 | 0.901189824 | 0.237604966 |
| ENSMUSG00000031355 | Arhgap6 | 11856 | protein_coding | 26.90622287 | 0.237610217 | 0.237225541 | -1.00162156 | 0.316526404 | 0.537548029 | 0.237610217 |
| ENSMUSG00000031774 | Psme3ip1 | 102122 | protein_coding | 512.7235777 | 0.237611184 | 0.059796413 | -3.973669497 | 7.08E-05 | 0.000636513 | 0.237611184 |
| ENSMUSG00000025959 | Klf7 | 93691 | protein_coding | 260.7094764 | 0.237652152 | 0.109691631 | -2.166547705 | 0.030269359 | 0.101661214 | 0.237652152 |
| ENSMUSG00000030654 | Arl6ip1 | 54208 | protein_coding | 1892.634373 | 0.237727575 | 0.071951608 | -3.303992531 | 0.000953184 | 0.006112962 | 0.237727575 |
| ENSMUSG00000026655 | Fam107b | 66540 | protein_coding | 1637.815533 | -0.237833673 | 0.132291527 | 1.797799735 | 0.072208749 | 0.195501402 | 0.237833673 |
| ENSMUSG00000062526 | Mppe1 | 225651 | protein_coding | 177.9951539 | -0.237848838 | 0.132572745 | 1.794100578 | 0.072797117 | 0.196529604 | 0.237848838 |
| ENSMUSG00000029644 | Pdx1 | 18609 | protein_coding | 6.555628951 | -0.237866903 | 0.400762879 | 0.59353527 | 0.552822986 | 0.743248294 | 0.237866903 |
| ENSMUSG00000014725 | Adam28 | 13522 | protein_coding | 3.115529418 | -0.237939496 | 1.57082402 | 0.151474317 | 0.879601569 | 1 | 0.237939496 |
| ENSMUSG00000090083 | Rnf8 | 58230 | protein_coding | 97.79898766 | -0.238007866 | 0.13876622 | 1.715171498 | 0.086313789 | 0.222126524 | 0.238007866 |
| ENSMUSG00000053617 | Sh3pxd2a | 14218 | protein_coding | 232.1460335 | -0.238033201 | 0.122862836 | 1.937389761 | 0.052697717 | 0.155346323 | 0.238033201 |
| ENSMUSG00000087331 | 1810021B22Rik | 69120 | lncRNA | 24.32364556 | 0.23804411 | 0.206131743 | -1.154815395 | 0.248166031 | 0.460473465 | 0.23804411 |
| ENSMUSG00000025733 | Rhot2 | 214952 | protein_coding | 574.5055749 | -0.238129694 | 0.107179043 | 2.22179344 | 0.026297268 | 0.090795126 | 0.238129694 |
| ENSMUSG00000087579 | Hectd2os | 668215 | lncRNA | 1.219167261 | 0.238188536 | 1.47460953 | -0.161526513 | 0.871678731 | 1 | 0.238188536 |
| ENSMUSG00000018209 | Stk4 | 58231 | protein_coding | 702.0451212 | -0.238222266 | 0.08238673 | 2.891512572 | 0.003833923 | 0.019529117 | 0.238222266 |
| ENSMUSG00000005397 | Nid1 | 18073 | protein_coding | 6.687141421 | 0.238227582 | 0.442415979 | -0.538469659 | 0.590252846 | 0.770846581 | 0.238227582 |
| ENSMUSG00000049687 | Pheta2 | 338368 | protein_coding | 9.243610834 | 0.238236169 | 0.388069215 | -0.613901231 | 0.539280597 | 0.733766095 | 0.238236169 |
| ENSMUSG00000027597 | Ahcy | 269378 | protein_coding | 1096.492587 | -0.2382436 | 0.102042162 | 2.3347565 | 0.019556145 | 0.072087093 | 0.2382436 |
| ENSMUSG00000064669 | Gm24461 | 115489568 | snRNA | 1.314840088 | 0.238332654 | 1.074878906 | -0.221729772 | 0.824524253 | 1 | 0.238332654 |

| ensembl_gene_id | external_gene_name | entrezgene_id | gene_biotype | baseMean | log2FoldChange | lfcSE | stat | pvalue | padj | absLog2FC |
|---|---|---|---|---|---|---|---|---|---|---|
| ENSMUSG00000032883 | Acsl3 | 74205 | protein_coding | 1871.102864 | -0.238352532 | 0.097039812 | 2.45623449 | 0.014040151 | 0.055375938 | 0.238352532 |
| ENSMUSG00000020737 | Jpt1 | 15374 | protein_coding | 1983.96074 | -0.238362701 | 0.075079003 | 3.174825077 | 0.001499269 | 0.008896486 | 0.238362701 |
| ENSMUSG00000038172 | Ttc39b | 69863 | protein_coding | 904.7816593 | -0.238384944 | 0.091748128 | 2.598254026 | 0.009369915 | 0.040464115 | 0.238384944 |
| ENSMUSG00000037706 | Cd81 | 12520 | protein_coding | 896.2942862 | -0.238476279 | 0.176924706 | 1.347896988 | 0.177691517 | 0.370029503 | 0.238476279 |
| ENSMUSG00000029426 | Scarb2 | 12492 | protein_coding | 1905.683768 | -0.23871833 | 0.055582477 | 4.294848729 | 1.75E-05 | 0.000182944 | 0.23871833 |
| ENSMUSG00000031558 | Slit2 | 20563 | protein_coding | 69.61443798 | 0.238760466 | 0.233891093 | -1.020818979 | 0.307340211 | 0.527754626 | 0.238760466 |
| ENSMUSG00000004567 | Mcoln1 | 94178 | protein_coding | 317.3939204 | -0.238786506 | 0.098955496 | 2.413069656 | 0.015818796 | 0.060935759 | 0.238786506 |
| ENSMUSG00000006998 | Psmd2 | 21762 | protein_coding | 2369.167352 | 0.238840555 | 0.083064431 | -2.875364978 | 0.004035607 | 0.020312105 | 0.238840555 |
| ENSMUSG00000049287 | Iba57 | 216792 | protein_coding | 241.437919 | -0.238882402 | 0.096504115 | 2.475359765 | 0.013310202 | 0.053169976 | 0.238882402 |
| ENSMUSG00000089842 | Pitpnm2os2 | 100047482 | lncRNA | 1.397514702 | 0.23893431 | 1.341022527 | -0.178173226 | 0.858586936 | 1 | 0.23893431 |
| ENSMUSG00000018995 | Nars2 | 244141 | protein_coding | 189.8799629 | -0.238945893 | 0.130590577 | 1.829733042 | 0.067289867 | 0.185483473 | 0.238945893 |
| ENSMUSG00000031799 | Tpm4 | 326618 | protein_coding | 1422.721371 | -0.239012743 | 0.116277336 | 2.055540231 | 0.039826854 | 0.125611281 | 0.239012743 |
| ENSMUSG00000074500 | Zfp558 | 72230 | protein_coding | 21.25250594 | 0.239015042 | 0.268755716 | -0.889339379 | 0.373820714 | 0.595844868 | 0.239015042 |
| ENSMUSG00000052301 | Doc2a | 13446 | protein_coding | 1.387111745 | 0.239025907 | 1.855973745 | -0.128787332 | 0.89752593 | 1 | 0.239025907 |
| ENSMUSG00000070493 | Chchd2 | 14004 | protein_coding | 2816.881232 | -0.239091055 | 0.08563278 | 2.792050622 | 0.005237516 | 0.025356887 | 0.239091055 |
| ENSMUSG00000097779 | 4833407H14Rik | 73899 | lncRNA | 19.50218618 | 0.23915415 | 0.423896318 | -0.564180768 | 0.572631115 | 0.758030479 | 0.23915415 |
| ENSMUSG00000086379 | 1700026D11Rik | 70371 | lncRNA | 1.484035425 | 0.239168176 | 2.077808118 | -0.115105998 | 0.908361106 | 1 | 0.239168176 |
| ENSMUSG00000035021 | Baz1a | 217578 | protein_coding | 421.7778791 | 0.239177704 | 0.097670614 | -2.448819507 | 0.014332525 | 0.056246986 | 0.239177704 |
| ENSMUSG00000028673 | Fuca1 | 71665 | protein_coding | 3072.602607 | -0.239184501 | 0.060468702 | 3.955509097 | 7.64E-05 | 0.000680996 | 0.239184501 |
| ENSMUSG00000022993 | Tex49 | 73863 | protein_coding | 21.05659747 | -0.239267745 | 0.267673001 | 0.89388076 | 0.371385693 | 0.593868753 | 0.239267745 |
| ENSMUSG00000001240 | Ramp2 | 54409 | protein_coding | 1.922389149 | 0.239293354 | 0.875039139 | -0.273465886 | 0.784495112 | 1 | 0.239293354 |
| ENSMUSG00000106583 | 4930447N08Rik | 71647 | lncRNA | 9.541758434 | -0.239353748 | 0.329764413 | 0.72583256 | 0.467941424 | 0.681625181 | 0.239353748 |
| ENSMUSG00000021609 | Slc6a3 | 13162 | protein_coding | 1.204093892 | -0.239410528 | 1.809667784 | 0.132295292 | 0.894750731 | 1 | 0.239410528 |
| ENSMUSG00000031775 | Pllp | 67801 | protein_coding | 248.3067963 | -0.239637798 | 0.124716505 | 1.921460178 | 0.054673718 | 0.159435243 | 0.239637798 |
| ENSMUSG00000031349 | Nsdhl | 18194 | protein_coding | 385.614839 | 0.239649309 | 0.096317395 | -2.488120745 | 0.012842012 | 0.051674706 | 0.239649309 |
| ENSMUSG00000028419 | Chmp5 | 76959 | protein_coding | 1351.553118 | 0.239679837 | 0.070377334 | -3.40563964 | 0.000660092 | 0.004471192 | 0.239679837 |
| ENSMUSG00000049823 | Zbtb12 | 193736 | protein_coding | 36.77641783 | 0.239684847 | 0.188704359 | -1.270160628 | 0.204027419 | 0.406185509 | 0.239684847 |
| ENSMUSG00000072620 | Slfn2 | 20556 | protein_coding | 16.46149834 | 0.239705966 | 0.324750378 | -0.738123749 | 0.460439253 | 0.674684457 | 0.239705966 |
| ENSMUSG00000019820 | Utrn | 22288 | protein_coding | 1940.570233 | 0.239819647 | 0.06630065 | -3.617153781 | 0.00029786 | 0.002264671 | 0.239819647 |
| ENSMUSG00000036646 | Man1b1 | 227619 | protein_coding | 1612.718236 | 0.239836018 | 0.078122091 | -3.07001533 | 0.002140478 | 0.011982322 | 0.239836018 |
| ENSMUSG00000015942 | Gtf2ird2 | 114674 | protein_coding | 106.4166042 | 0.239934506 | 0.128525978 | -1.866817205 | 0.061927119 | 0.174605505 | 0.239934506 |
| ENSMUSG00000026656 | Fcgr2b | 14130 | protein_coding | 2.208400069 | 0.239950506 | 1.282658142 | -0.187072843 | 0.85160352 | 1 | 0.239950506 |
| ENSMUSG00000003559 | As3mt | 57344 | protein_coding | 547.2438961 | 0.239962467 | 0.12050437 | -1.991317556 | 0.046445987 | 0.141033318 | 0.239962467 |
| ENSMUSG00000029828 | 4921507P07Rik | 70821 | protein_coding | 8.6366949 | -0.2399992 | 0.379013582 | 0.633220578 | 0.526589609 | 0.725037586 | 0.2399992 |
| ENSMUSG00000040824 | Snrpd2 | 107686 | protein_coding | 424.1903831 | -0.240000183 | 0.099320449 | 2.416422657 | 0.015673856 | 0.060447934 | 0.240000183 |
| ENSMUSG00000077450 | Rab11b | 19326 | protein_coding | 627.2813281 | -0.240007918 | 0.085633353 | 2.802738752 | 0.00506707 | 0.024640043 | 0.240007918 |
| ENSMUSG00000053819 | Camk2d | 108058 | protein_coding | 1244.302031 | 0.24003223 | 0.051854279 | -4.628976276 | 3.67E-06 | 4.53E-05 | 0.24003223 |
| ENSMUSG00000031826 | Usp10 | 22224 | protein_coding | 503.1833669 | 0.24003568 | 0.090667998 | -2.64741347 | 0.008111011 | 0.036011679 | 0.24003568 |
| ENSMUSG00000035429 | Ptprh | 545902 | protein_coding | 743.855371 | -0.240053651 | 0.301926944 | 0.79507197 | 0.426571638 | 0.64620651 | 0.240053651 |

| ensembl_gene_id | external_gene_name | entrezgene_id | gene_biotype | baseMean | log2FoldChange | lfcSE | stat | pvalue | padj | absLog2FC |
|---|---|---|---|---|---|---|---|---|---|---|
| ENSMUSG00000020929 | Eftud2 | 20624 | protein_coding | 685.7322169 | 0.240107607 | 0.095672619 | -2.509679452 | 0.01208408 | 0.049395172 | 0.240107607 |
| ENSMUSG00000027742 | Cog6 | 67542 | protein_coding | 824.3721224 | 0.240121072 | 0.083396905 | -2.879256381 | 0.003986141 | 0.020124469 | 0.240121072 |
| ENSMUSG00000076123 | Mir700 | 735285 | miRNA | 0.604723788 | 0.240182063 | 1.875238775 | -0.12808079 | 0.898085038 | 1 | 0.240182063 |
| ENSMUSG00000028104 | Polr3gl | 69870 | protein_coding | 106.0592122 | -0.240248018 | 0.102390262 | 2.346395196 | 0.018955996 | 0.070425045 | 0.240248018 |
| ENSMUSG00000022710 | Usp7 | 252870 | protein_coding | 2146.651576 | 0.240249991 | 0.05470695 | -4.391580763 | 1.13E-05 | 0.000124365 | 0.240249991 |
| ENSMUSG00000054099 | Slc25a40 | 319653 | protein_coding | 240.3767146 | -0.24025578 | 0.0982439 | 2.445503278 | 0.014465013 | 0.056645774 | 0.24025578 |
| ENSMUSG00000061517 | Sox21 | 223227 | protein_coding | 61.43197519 | 0.240271925 | 0.179960825 | -1.335134604 | 0.181832296 | 0.376052415 | 0.240271925 |
| ENSMUSG00000033589 | Reep4 | 72549 | protein_coding | 444.2415006 | 0.240281594 | 0.076836856 | -3.127165873 | 0.001765003 | 0.010209199 | 0.240281594 |
| ENSMUSG00000025505 | Tmem80 | 71448 | protein_coding | 91.90616517 | -0.240356148 | 0.124071977 | 1.937231542 | 0.052717046 | 0.155375567 | 0.240356148 |
| ENSMUSG00000071657 | Bscl2 | 14705 | protein_coding | 435.5047249 | -0.240393397 | 0.087852322 | 2.736335167 | 0.006212771 | 0.028908905 | 0.240393397 |
| ENSMUSG00000091931 | Gon7 | 100233175 | protein_coding | 42.69745708 | 0.240432196 | 0.189056975 | -1.271744646 | 0.203463862 | 0.405748366 | 0.240432196 |
| ENSMUSG00000074738 | Fndc10 | 230991 | protein_coding | 24.42888891 | -0.240483407 | 0.21010249 | 1.14460046 | 0.252374702 | 0.464617944 | 0.240483407 |
| ENSMUSG00000021474 | Sfxn1 | 14057 | protein_coding | 632.0734894 | -0.24048732 | 0.112160226 | 2.144140828 | 0.032021605 | 0.10614307 | 0.24048732 |
| ENSMUSG00000051768 | Xrcc1 | 22594 | protein_coding | 283.7842762 | 0.24057364 | 0.106605577 | -2.256670315 | 0.024028683 | 0.084679828 | 0.24057364 |
| ENSMUSG00000070803 | Cited4 | 56222 | protein_coding | 105.8485021 | 0.240655552 | 0.127294749 | -1.89053793 | 0.058686053 | 0.168040558 | 0.240655552 |
| ENSMUSG00000054383 | Pnma1 | 70481 | protein_coding | 5.437831497 | 0.240697832 | 0.800983479 | -0.300502867 | 0.76379361 | 0.881554912 | 0.240697832 |
| ENSMUSG00000039089 | L3mbtl3 | 237339 | protein_coding | 269.3276332 | 0.240742129 | 0.086374825 | -2.787179344 | 0.005316904 | 0.025643505 | 0.240742129 |
| ENSMUSG00000076448 | Mir763 | 791076 | miRNA | 0.741381929 | -0.240744467 | 1.507491673 | 0.159698704 | 0.873118422 | 1 | 0.240744467 |
| ENSMUSG00000021699 | Pde4d | 238871 | protein_coding | 306.7545952 | -0.240803983 | 0.085600046 | 2.813129118 | 0.004906195 | 0.023928204 | 0.240803983 |
| ENSMUSG00000021696 | Elovl7 | 74559 | protein_coding | 1065.020211 | -0.240859919 | 0.122657412 | 1.963680103 | 0.049567201 | 0.14799363 | 0.240859919 |
| ENSMUSG00000012117 | Dhdds | 67422 | protein_coding | 339.3509756 | 0.240868698 | 0.080788645 | -2.981467224 | 0.002868708 | 0.015327126 | 0.240868698 |
| ENSMUSG00000033904 | Ccp110 | 101565 | protein_coding | 57.23302247 | 0.240905139 | 0.173633617 | -1.387433742 | 0.165309542 | 0.352338897 | 0.240905139 |
| ENSMUSG00000054737 | Zfp182 | 319535 | protein_coding | 112.1191758 | -0.240942543 | 0.118091724 | 2.040299987 | 0.041320456 | 0.129138155 | 0.240942543 |
| ENSMUSG00000030107 | Usp18 | 24110 | protein_coding | 45.18966116 | 0.240998068 | 0.441855464 | -0.545422854 | 0.585462726 | 0.767388797 | 0.240998068 |
| ENSMUSG00000022199 | Slc22a17 | 59049 | protein_coding | 16.48130905 | 0.241039502 | 0.296731238 | -0.812315897 | 0.416610391 | 0.636776036 | 0.241039502 |
| ENSMUSG00000037172 | Dennd11 | 243780 | protein_coding | 721.412624 | -0.241064563 | 0.204437997 | 1.179157331 | 0.238335533 | 0.448589299 | 0.241064563 |
| ENSMUSG00000046567 | 4930430F08Rik | 68281 | protein_coding | 35.73222637 | 0.241135254 | 0.210669059 | -1.144616371 | 0.252368108 | 0.464617944 | 0.241135254 |
| ENSMUSG00000063457 | Rps15 | 20054 | protein_coding | 5145.310868 | -0.241257334 | 0.091563215 | 2.634871799 | 0.008416907 | 0.037046055 | 0.241257334 |
| ENSMUSG00000038745 | Nlrp6 | 101613 | protein_coding | 3094.516356 | -0.241266475 | 0.113114546 | 2.132939438 | 0.032929695 | 0.108456517 | 0.241266475 |
| ENSMUSG00000074793 | Hspa12b | 72630 | protein_coding | 2.987102186 | 0.241436457 | 0.778400885 | -0.310169813 | 0.756431825 | 1 | 0.241436457 |
| ENSMUSG00000016194 | Hsd11b1 | 15483 | protein_coding | 2.518097602 | 0.241461699 | 0.854051905 | -0.282724853 | 0.777387764 | 1 | 0.241461699 |
| ENSMUSG00000004849 | Ap1s1 | 11769 | protein_coding | 764.783446 | -0.241536477 | 0.079071653 | 3.054653184 | 0.002253209 | 0.012461318 | 0.241536477 |
| ENSMUSG00000118087 | 4833438C02Rik | 100503392 | lncRNA | 81.7076119 | -0.241596158 | 0.172596417 | 1.39977505 | 0.161580692 | 0.346538122 | 0.241596158 |
| ENSMUSG00000040857 | Erf | 13875 | protein_coding | 400.2094692 | -0.241689696 | 0.095669602 | 2.526295622 | 0.011527244 | 0.047685597 | 0.241689696 |
| ENSMUSG00000057147 | Dph6 | 66632 | protein_coding | 61.75814032 | 0.241763558 | 0.15459733 | -1.563827516 | 0.117858081 | 0.278322371 | 0.241763558 |
| ENSMUSG00000031641 | Cbr4 | 234309 | protein_coding | 218.2895902 | -0.241766209 | 0.122265596 | 1.977385433 | 0.047998079 | 0.144615997 | 0.241766209 |
| ENSMUSG00000026571 | Dcaf6 | 74106 | protein_coding | 331.0316023 | -0.241769659 | 0.079406853 | 3.044695145 | 0.002329163 | 0.012812723 | 0.241769659 |
| ENSMUSG00000047749 | Zc3hav1l | 209032 | protein_coding | 23.11025002 | 0.241799636 | 0.236820955 | -1.021022977 | 0.307243553 | 0.527698391 | 0.241799636 |
| ENSMUSG00000030220 | Arhgdib | 11857 | protein_coding | 15.07776002 | 0.241825111 | 0.369817015 | -0.653904771 | 0.51317316 | 0.715090842 | 0.241825111 |

| ensembl_gene_id | external_gene_name | entrezgene_id | gene_biotype | baseMean | log2FoldChange | lfcSE | stat | pvalue | padj | absLog2FC |
|---|---|---|---|---|---|---|---|---|---|---|
| ENSMUSG00000038368 | Focad | 230393 | protein_coding | 230.8519469 | 0.24196797 | 0.094079071 | -2.571963837 | 0.010112345 | 0.042953833 | 0.24196797 |
| ENSMUSG00000041679 | Fbxl9 | 234684 | protein_coding | 43.35937392 | -0.241980938 | 0.156270415 | 1.548475685 | 0.121507811 | 0.284375933 | 0.241980938 |
| ENSMUSG00000002486 | Tchp | 77832 | protein_coding | 87.65235191 | 0.241984867 | 0.158215648 | -1.529462295 | 0.126149879 | 0.292089426 | 0.241984867 |
| ENSMUSG00000004266 | Ptpn6 | 15170 | protein_coding | 326.55576 | -0.242190736 | 0.090211426 | 2.684701328 | 0.007259465 | 0.032853604 | 0.242190736 |
| ENSMUSG00000033998 | Kcnk1 | 16525 | protein_coding | 1080.223591 | 0.242280872 | 0.096596158 | -2.508183315 | 0.012135369 | 0.049543507 | 0.242280872 |
| ENSMUSG00000020022 | Ndufa12 | 66414 | protein_coding | 879.6929161 | -0.242329024 | 0.084272066 | 2.875555748 | 0.004033169 | 0.020306023 | 0.242329024 |
| ENSMUSG00000025498 | Irf7 | 54123 | protein_coding | 985.4363221 | -0.242350224 | 0.397370524 | 0.609884753 | 0.541938153 | 0.7356816 | 0.242350224 |
| ENSMUSG00000025421 | Hdhd2 | 76987 | protein_coding | 117.6664947 | -0.242431836 | 0.128654571 | 1.884362394 | 0.059515973 | 0.169807667 | 0.242431836 |
| ENSMUSG00000028675 | Pnrc2 | 52830 | protein_coding | 1129.71745 | 0.242479789 | 0.086329773 | -2.808762042 | 0.004973239 | 0.024219425 | 0.242479789 |
| ENSMUSG00000078816 | Prkcg | 18752 | protein_coding | 10.68703957 | 0.242566707 | 0.407803194 | -0.594813161 | 0.551968369 | 0.742703956 | 0.242566707 |
| ENSMUSG00000045983 | Eif4g1 | 208643 | protein_coding | 6219.577783 | 0.242584003 | 0.061604052 | -3.937792966 | 8.22E-05 | 0.000726214 | 0.242584003 |
| ENSMUSG00000027223 | Mapk8ip1 | 19099 | protein_coding | 26.68077339 | -0.24260367 | 0.326393322 | 0.743286254 | 0.457308392 | 0.672184316 | 0.24260367 |
| ENSMUSG00000021102 | Glrx5 | 73046 | protein_coding | 614.0312147 | 0.242606887 | 0.089600733 | -2.707643998 | 0.006776267 | 0.031092878 | 0.242606887 |
| ENSMUSG00000023050 | Map3k12 | 26404 | protein_coding | 32.96156865 | -0.242652785 | 0.270168367 | 0.898153947 | 0.369103482 | 0.591783971 | 0.242652785 |
| ENSMUSG00000003410 | Elavl3 | 15571 | protein_coding | 5.832940472 | -0.242670161 | 1.067000965 | 0.227431997 | 0.820087842 | 0.909231489 | 0.242670161 |
| ENSMUSG00000026344 | Lypd1 | 72585 | protein_coding | 3.760208786 | -0.242850518 | 0.744509438 | 0.326188636 | 0.744281639 | 1 | 0.242850518 |
| ENSMUSG00000024258 | Polr2d | 69241 | protein_coding | 137.2645973 | 0.242852057 | 0.109875185 | -2.210253898 | 0.027087546 | 0.092992945 | 0.242852057 |
| ENSMUSG00000025058 | Tasl | 71398 | protein_coding | 3.212031609 | 0.242938041 | 0.691143003 | -0.351501846 | 0.725211888 | 1 | 0.242938041 |
| ENSMUSG00000027035 | Cers6 | 241447 | protein_coding | 2359.595539 | -0.242953649 | 0.06252775 | 3.885533234 | 0.000102106 | 0.000875933 | 0.242953649 |
| ENSMUSG00000012405 | Rpl15 | 66480 | protein_coding | 2405.234837 | -0.242956348 | 0.123357083 | 1.969537074 | 0.04889145 | 0.14663883 | 0.242956348 |
| ENSMUSG00000038256 | Bcl9 | 77578 | protein_coding | 182.4540923 | 0.243009828 | 0.191177308 | -1.271122765 | 0.203684978 | 0.405918699 | 0.243009828 |
| ENSMUSG00000021311 | Mtr | 238505 | protein_coding | 123.4979709 | 0.24301863 | 0.139714181 | -1.739398458 | 0.081964699 | 0.213630847 | 0.24301863 |
| ENSMUSG00000060771 | Tsga10 | 211484 | protein_coding | 64.5137722 | -0.243057407 | 0.159958678 | 1.519501229 | 0.128636379 | 0.295987343 | 0.243057407 |
| ENSMUSG00000059772 | Slx1b | 75764 | protein_coding | 147.4670254 | -0.243061274 | 0.099323796 | 2.447160542 | 0.014398669 | 0.0564529 | 0.243061274 |
| ENSMUSG00000026761 | Orc4 | 26428 | protein_coding | 331.7678693 | 0.243087854 | 0.067971426 | -3.576324143 | 0.00034846 | 0.002603828 | 0.243087854 |
| ENSMUSG00000021224 | Numb | 18222 | protein_coding | 736.5076042 | 0.243103679 | 0.074739489 | -3.252680494 | 0.00114322 | 0.007110783 | 0.243103679 |
| ENSMUSG00000033948 | Zswim5 | 74464 | protein_coding | 165.9298045 | -0.243145437 | 0.094730573 | 2.566705014 | 0.010266991 | 0.043464619 | 0.243145437 |
| ENSMUSG00000020063 | Sirt1 | 93759 | protein_coding | 244.4208574 | -0.243166647 | 0.11435142 | 2.126485587 | 0.033462848 | 0.109658699 | 0.243166647 |
| ENSMUSG00000021044 | Adck1 | 72113 | protein_coding | 162.2766267 | -0.243167255 | 0.096522188 | 2.519288676 | 0.01175922 | 0.048327356 | 0.243167255 |
| ENSMUSG00000078435 | AU041133 | 216177 | protein_coding | 51.34367716 | -0.243200906 | 0.169816583 | 1.432138737 | 0.15210412 | 0.332739097 | 0.243200906 |
| ENSMUSG00000050721 | Plekho2 | 102595 | protein_coding | 13.89762513 | -0.243238413 | 0.430528002 | 0.564976987 | 0.572089417 | 0.757617052 | 0.243238413 |
| ENSMUSG00000070509 | Rgma | 244058 | protein_coding | 20.70429734 | 0.243266584 | 0.32232676 | -0.754720409 | 0.450416758 | 0.667221236 | 0.243266584 |
| ENSMUSG00000024074 | Crim1 | 50766 | protein_coding | 698.7661528 | -0.243320595 | 0.070400575 | 3.456230198 | 0.000547787 | 0.003829873 | 0.243320595 |
| ENSMUSG00000022787 | Wdr53 | 68980 | protein_coding | 89.41475805 | -0.243326463 | 0.134744629 | 1.805834221 | 0.070944238 | 0.192819808 | 0.243326463 |
| ENSMUSG00000024197 | Plin3 | 66905 | protein_coding | 1359.701036 | 0.243331455 | 0.059351183 | -4.099858549 | 4.13E-05 | 0.000396224 | 0.243331455 |
| ENSMUSG00000085990 | Gm16731 | 105247126 | lncRNA | 10.71589802 | 0.243424062 | 0.38560898 | -0.631271768 | 0.527862843 | 0.726026078 | 0.243424062 |
| ENSMUSG00000031536 | Polb | 18970 | protein_coding | 198.5747644 | -0.243426479 | 0.091332648 | 2.665273423 | 0.007692575 | 0.034482949 | 0.243426479 |
| ENSMUSG00000045867 | Cradd | 12905 | protein_coding | 76.37058878 | -0.243428454 | 0.141278575 | 1.723038719 | 0.084881523 | 0.219498448 | 0.243428454 |
| ENSMUSG00000034161 | Scx | 20289 | protein_coding | 6.346028324 | 0.243435777 | 0.522589169 | -0.465826296 | 0.641339841 | 0.804182698 | 0.243435777 |

| ensembl_gene_id | external_gene_name | entrezgene_id | gene_biotype | baseMean | log2FoldChange | lfcSE | stat | pvalue | padj | absLog2FC |
|---|---|---|---|---|---|---|---|---|---|---|
| ENSMUSG00000026766 | Mmadhc | 109129 | protein_coding | 635.5301556 | 0.24356335 | 0.103457044 | -2.354246167 | 0.018560315 | 0.069204119 | 0.24356335 |
| ENSMUSG00000020467 | Efemp1 | 216616 | protein_coding | 0.715411271 | -0.243586078 | 1.89008975 | 0.128875403 | 0.89745624 | 1 | 0.243586078 |
| ENSMUSG00000064358 | mt-Co3 | 17710 | protein_coding | 15.8555211 | 0.243654134 | 0.669048226 | -0.364180226 | 0.715723432 | 0.852528618 | 0.243654134 |
| ENSMUSG00000073639 | Rab18 | 19330 | protein_coding | 816.7695485 | 0.243661458 | 0.071043913 | -3.429730228 | 0.000604182 | 0.004152083 | 0.243661458 |
| ENSMUSG00000065591 | Gm22753 | 115487847 | snRNA | 1.910692349 | -0.243669145 | 0.732450794 | 0.332676471 | 0.739378512 | 1 | 0.243669145 |
| ENSMUSG00000017561 | Crlf3 | 54394 | protein_coding | 125.2709568 | -0.243676696 | 0.108656041 | 2.242642871 | 0.024919853 | 0.087113984 | 0.243676696 |
| ENSMUSG00000069892 | 9930111J21Rik2 | 245240 | protein_coding | 2.618218965 | -0.243742361 | 0.84666315 | 0.287885874 | 0.7734341 | 1 | 0.243742361 |
| ENSMUSG00000020277 | Pfkl | 18641 | protein_coding | 3188.534636 | -0.243780343 | 0.098268277 | 2.48076339 | 0.013110137 | 0.052535985 | 0.243780343 |
| ENSMUSG00000070421 | Or52n4 | 259051 | protein_coding | 1.341653401 | -0.243790244 | 1.427930403 | 0.17072978 | 0.864436245 | 1 | 0.243790244 |
| ENSMUSG00000037995 | Igsf9 | 93842 | protein_coding | 1379.981352 | 0.243829632 | 0.154034453 | -1.58295516 | 0.113431685 | 0.270265596 | 0.243829632 |
| ENSMUSG00000019861 | Gopc | 94221 | protein_coding | 519.9334188 | 0.243930569 | 0.131526121 | -1.854616915 | 0.063650951 | 0.178188134 | 0.243930569 |
| ENSMUSG00000038967 | Pdk2 | 18604 | protein_coding | 501.7473971 | 0.243998832 | 0.114868646 | -2.124155199 | 0.033657168 | 0.110042513 | 0.243998832 |
| ENSMUSG00000026430 | Rassf5 | 54354 | protein_coding | 13.41573901 | 0.244073354 | 0.428956701 | -0.568992985 | 0.5693609 | 0.755154277 | 0.244073354 |
| ENSMUSG00000052407 | Ccdc171 | 320226 | protein_coding | 58.80388543 | 0.244119833 | 0.178288432 | -1.369241015 | 0.170923949 | 0.360812744 | 0.244119833 |
| ENSMUSG00000038879 | Nipal2 | 223473 | protein_coding | 428.927085 | 0.244148158 | 0.082100168 | -2.973783908 | 0.002941521 | 0.015594313 | 0.244148158 |
| ENSMUSG00000025495 | Ptdss2 | 27388 | protein_coding | 270.9568346 | -0.244170228 | 0.091441213 | 2.670242667 | 0.007579645 | 0.034050667 | 0.244170228 |
| ENSMUSG00000022906 | Parp9 | 80285 | protein_coding | 605.8744435 | -0.244278254 | 0.133941542 | 1.823767673 | 0.068187203 | 0.187361642 | 0.244278254 |
| ENSMUSG00000063200 | Nol7 | 70078 | protein_coding | 512.561897 | 0.244279695 | 0.088081287 | -2.773343834 | 0.005548346 | 0.026496643 | 0.244279695 |
| ENSMUSG00000019715 | Gle1 | 74412 | protein_coding | 295.1931402 | 0.244310141 | 0.124108939 | -1.968513649 | 0.049008967 | 0.146868346 | 0.244310141 |
| ENSMUSG00000025757 | Hspa4l | 18415 | protein_coding | 825.986033 | -0.244330243 | 0.138449588 | 1.764759621 | 0.077604174 | 0.205674101 | 0.244330243 |
| ENSMUSG00000081984 | Dnajb3 | 15504 | protein_coding | 19.85369476 | 0.244415257 | 0.338341117 | -0.722393008 | 0.470052884 | 0.683555242 | 0.244415257 |
| ENSMUSG00000085004 | 5430427M07Rik | 71349 | lncRNA | 49.53493919 | -0.244464994 | 0.532751078 | 0.458872828 | 0.646325492 | 0.807226927 | 0.244464994 |
| ENSMUSG00000020105 | Lrig3 | 320398 | protein_coding | 1248.363228 | -0.244646108 | 0.105506059 | 2.318787278 | 0.020406573 | 0.074572727 | 0.244646108 |
| ENSMUSG00000006021 | Kptn | 70394 | protein_coding | 165.2797366 | -0.244983635 | 0.096071807 | 2.55000548 | 0.010772123 | 0.045218819 | 0.244983635 |
| ENSMUSG00000003437 | Paf1 | 54624 | protein_coding | 445.4813817 | 0.244992368 | 0.079020291 | -3.10037287 | 0.001932772 | 0.011044218 | 0.244992368 |
| ENSMUSG00000025534 | Gusb | 110006 | protein_coding | 471.0964945 | 0.245089897 | 0.093151388 | -2.631092264 | 0.008511093 | 0.037373936 | 0.245089897 |
| ENSMUSG00000099254 | Mir7030 | 102465623 | miRNA | 0.659985141 | -0.245204787 | 1.592158797 | 0.154007746 | 0.877603628 | 1 | 0.245204787 |
| ENSMUSG00000030286 | Emc3 | 66087 | protein_coding | 780.1422895 | 0.245284029 | 0.076515745 | -3.205667407 | 0.001347496 | 0.008157095 | 0.245284029 |
| ENSMUSG00000030224 | Strap | 20901 | protein_coding | 1041.658248 | 0.245400228 | 0.076564459 | -3.205145446 | 0.001349942 | 0.008168908 | 0.245400228 |
| ENSMUSG00000049502 | Dtx3l | 209200 | protein_coding | 684.6200442 | -0.245512516 | 0.149951522 | 1.637279259 | 0.101572133 | 0.250425106 | 0.245512516 |
| ENSMUSG00000052456 | Get3 | 56495 | protein_coding | 652.7423513 | 0.245551989 | 0.073484026 | -3.341569623 | 0.000833061 | 0.005454866 | 0.245551989 |
| ENSMUSG00000033059 | Pygb | 110078 | protein_coding | 3121.316241 | 0.245619338 | 0.111116359 | -2.210469643 | 0.027072585 | 0.092985996 | 0.245619338 |
| ENSMUSG00000016494 | Cd34 | 12490 | protein_coding | 4.557053465 | 0.245630213 | 0.767993325 | -0.319833786 | 0.749094334 | 0.874014884 | 0.245630213 |
| ENSMUSG00000099706 | Gm29536 | 102636764 | lncRNA | 1.081133225 | -0.245695568 | 2.318270806 | 0.105982255 | 0.915596431 | 1 | 0.245695568 |
| ENSMUSG00000019437 | Tlcd1 | 68385 | protein_coding | 113.8579512 | -0.245814151 | 0.138055797 | 1.780542043 | 0.074987295 | 0.200249061 | 0.245814151 |
| ENSMUSG00000032656 | Marchf3 | 320253 | protein_coding | 78.80990822 | 0.245864775 | 0.38669118 | -0.635816866 | 0.524895793 | 0.723368866 | 0.245864775 |
| ENSMUSG00000112925 | Gm31763 | 102634101 | lncRNA | 4.773041758 | -0.245868815 | 0.609450792 | 0.403426853 | 0.686634236 | 0.83310074 | 0.245868815 |
| ENSMUSG00000030498 | Gas2 | 14453 | protein_coding | 35.57208848 | 0.245869253 | 0.181708102 | -1.353100112 | 0.17602365 | 0.368018849 | 0.245869253 |
| ENSMUSG00000024824 | Rad9a | 19367 | protein_coding | 88.55185857 | 0.245919877 | 0.161165764 | -1.525881619 | 0.127039349 | 0.293636687 | 0.245919877 |

| ensembl_gene_id | external_gene_name | entrezgene_id | gene_biotype | baseMean | log2FoldChange | lfcSE | stat | pvalue | padj | absLog2FC |
|---|---|---|---|---|---|---|---|---|---|---|
| ENSMUSG00000028885 | Smpdl3b | 100340 | protein_coding | 71.3783588 | 0.24596875 | 0.170663737 | -1.441247884 | 0.149514665 | 0.328900667 | 0.24596875 |
| ENSMUSG00000087620 | 5330434G04Rik | 100043213 | lncRNA | 1.951641558 | -0.246013033 | 2.32173845 | 0.105960701 | 0.915613533 | 1 | 0.246013033 |
| ENSMUSG00000057530 | Ece1 | 230857 | protein_coding | 1815.649389 | -0.246221209 | 0.061937563 | 3.975313171 | 7.03E-05 | 0.000632887 | 0.246221209 |
| ENSMUSG00000016409 | Nkap | 67050 | protein_coding | 176.8909683 | -0.246231382 | 0.116424777 | 2.114939686 | 0.034435088 | 0.112051438 | 0.246231382 |
| ENSMUSG00000028737 | Aldh4a1 | 212647 | protein_coding | 239.9479563 | 0.24623306 | 0.090014287 | -2.735488629 | 0.006228774 | 0.028942592 | 0.24623306 |
| ENSMUSG00000047409 | Ctdspl | 69274 | protein_coding | 1185.281631 | 0.246316386 | 0.082724495 | -2.977550805 | 0.002905615 | 0.015463527 | 0.246316386 |
| ENSMUSG00000051319 | Mtln | 67885 | protein_coding | 149.9519705 | -0.246458558 | 0.107371665 | 2.295378015 | 0.02171147 | 0.078315179 | 0.246458558 |
| ENSMUSG00000089671 | Gm16537 | 102638434 | lncRNA | 1.234546564 | 0.246567514 | 1.094063827 | -0.225368491 | 0.821692622 | 1 | 0.246567514 |
| ENSMUSG00000025494 | Sigirr | 24058 | protein_coding | 350.0996862 | -0.246650413 | 0.070886494 | 3.479512077 | 0.000502328 | 0.003561803 | 0.246650413 |
| ENSMUSG00000027457 | Snph | 241727 | protein_coding | 77.98691047 | -0.246688606 | 0.191672203 | 1.287033808 | 0.198082505 | 0.398335908 | 0.246688606 |
| ENSMUSG00000020444 | Guk1 | 14923 | protein_coding | 762.4364379 | 0.246729482 | 0.108250343 | -2.279248953 | 0.022652271 | 0.08100219 | 0.246729482 |
| ENSMUSG00000048707 | Tprn | 97031 | protein_coding | 1616.994224 | 0.246818892 | 0.115532697 | -2.136355327 | 0.032650462 | 0.107730217 | 0.246818892 |
| ENSMUSG00000022559 | Fbxl6 | 30840 | protein_coding | 172.693566 | -0.247000629 | 0.091306149 | 2.705191615 | 0.0068265 | 0.03127124 | 0.247000629 |
| ENSMUSG00000003746 | Man1a | 17155 | protein_coding | 1001.314946 | -0.247025764 | 0.066094094 | 3.737486192 | 0.000185869 | 0.001488577 | 0.247025764 |
| ENSMUSG00000039126 | Prune2 | 353211 | protein_coding | 26.25242973 | 0.247037023 | 0.383288263 | -0.644520187 | 0.519238171 | 0.719474674 | 0.247037023 |
| ENSMUSG00000025173 | Wdr45b | 66840 | protein_coding | 479.0900109 | 0.247043644 | 0.049718117 | -4.96888582 | 6.73E-07 | 9.64E-06 | 0.247043644 |
| ENSMUSG00000033703 | Fcsk | 234730 | protein_coding | 763.5356677 | 0.247056246 | 0.06308783 | -3.916068196 | 9.00E-05 | 0.000786839 | 0.247056246 |
| ENSMUSG00000046352 | Gjb2 | 14619 | protein_coding | 152.3780895 | 0.247062026 | 0.134459487 | -1.837445847 | 0.066144104 | 0.183317181 | 0.247062026 |
| ENSMUSG00000076011 | Mir652 | 723976 | miRNA | 3.178487793 | -0.247075957 | 0.674056176 | 0.366550988 | 0.713953973 | 1 | 0.247075957 |
| ENSMUSG00000065822 | Snord15a | 449630 | snoRNA | 103.7713765 | -0.247107515 | 0.1686556 | 1.465160448 | 0.142877148 | 0.318675475 | 0.247107515 |
| ENSMUSG00000071708 | Sms | 20603 | protein_coding | 101.7950746 | -0.247142637 | 0.141033669 | 1.752366222 | 0.079710856 | 0.209275846 | 0.247142637 |
| ENSMUSG00000026156 | B3gat2 | 280645 | protein_coding | 18.16490968 | -0.247168251 | 0.261636213 | 0.944701988 | 0.344811042 | 0.566362983 | 0.247168251 |
| ENSMUSG00000035885 | Cox8a | 12868 | protein_coding | 5437.324101 | -0.247177347 | 0.077097353 | 3.206041935 | 0.001345744 | 0.008149473 | 0.247177347 |
| ENSMUSG00000020009 | Ifngr1 | 15979 | protein_coding | 2390.429537 | -0.247185188 | 0.089101855 | 2.774186763 | 0.00553399 | 0.026443376 | 0.247185188 |
| ENSMUSG00000022580 | Rhpn1 | 14787 | protein_coding | 1.353787863 | 0.247323032 | 1.813675407 | -0.136365654 | 0.891532225 | 1 | 0.247323032 |
| ENSMUSG00000048076 | Arf1 | 11840 | protein_coding | 4975.350239 | 0.247394042 | 0.079806105 | -3.099938802 | 0.001935606 | 0.011045129 | 0.247394042 |
| ENSMUSG00000031728 | Zfp821 | 75871 | protein_coding | 151.7814857 | -0.247452553 | 0.140319855 | 1.763489228 | 0.077818012 | 0.205991262 | 0.247452553 |
| ENSMUSG00000045327 | 6330549D23Rik | 229613 | transcribed_processed_pseudogene | 11.52152403 | -0.24760125 | 0.34914579 | 0.709162926 | 0.478223377 | 0.689848083 | 0.24760125 |
| ENSMUSG00000025817 | Nudt5 | 53893 | protein_coding | 319.054395 | 0.247611719 | 0.143077831 | -1.730608565 | 0.083521603 | 0.216662858 | 0.247611719 |
| ENSMUSG00000060679 | Mrps9 | 69527 | protein_coding | 422.7652655 | -0.247619051 | 0.109673547 | 2.25778282 | 0.023959203 | 0.084507107 | 0.247619051 |
| ENSMUSG00000089984 | Fbxo24 | 71176 | protein_coding | 17.26199983 | -0.247817723 | 0.354580077 | 0.698904814 | 0.484611519 | 0.695055986 | 0.247817723 |
| ENSMUSG00000037989 | Wnk2 | 75607 | protein_coding | 256.1705839 | 0.247827227 | 0.205581431 | -1.205494223 | 0.228012567 | 0.436620149 | 0.247827227 |
| ENSMUSG00000034243 | Golgb1 | 224139 | protein_coding | 3597.660359 | 0.247843936 | 0.061099857 | -4.056375072 | 4.98E-05 | 0.000465269 | 0.247843936 |
| ENSMUSG00000010554 | Mettl16 | 67493 | protein_coding | 271.3846053 | 0.247894613 | 0.092488993 | -2.680260701 | 0.007356484 | 0.033210765 | 0.247894613 |
| ENSMUSG00000043614 | Vps37d | 194309 | protein_coding | 12.45044718 | -0.247907228 | 0.362883657 | 0.683158976 | 0.494506392 | 0.700667401 | 0.247907228 |
| ENSMUSG00000020792 | Exoc7 | 53413 | protein_coding | 588.5834499 | 0.247980608 | 0.059079162 | -4.197429316 | 2.70E-05 | 0.000269538 | 0.247980608 |
| ENSMUSG00000068747 | Sort1 | 20661 | protein_coding | 5526.846597 | -0.248021799 | 0.08454647 | 2.933555917 | 0.003351033 | 0.017451579 | 0.248021799 |
| ENSMUSG00000028184 | Adgrl2 | 99633 | protein_coding | 321.675023 | -0.248041887 | 0.096638744 | 2.566691954 | 0.010267378 | 0.043464619 | 0.248041887 |
| ENSMUSG00000118100 | Gm50340 | 115489003 | lncRNA | 9.651272823 | -0.248159575 | 0.342517846 | 0.724515753 | 0.46874916 | 0.682320284 | 0.248159575 |

Supplementary Table 1

| ensembl_gene_id | external_gene_name | entrezgene_id | gene_biotype | baseMean | log2FoldChange | lfcSE | stat | pvalue | padj | absLog2FC |
|---|---|---|---|---|---|---|---|---|---|---|
| ENSMUSG00000068882 | Ssb | 20823 | protein_coding | 937.8280708 | 0.248247857 | 0.095484526 | -2.599875275 | 0.009325765 | 0.040294527 | 0.248247857 |
| ENSMUSG00000024220 | Zfp523 | 224656 | protein_coding | 226.19673 | 0.248274647 | 0.08090479 | -3.068726189 | 0.002149735 | 0.012021919 | 0.248274647 |
| ENSMUSG00000021660 | Btf3 | 218490 | protein_coding | 1718.639353 | -0.248312568 | 0.077373534 | 3.209270071 | 0.001330724 | 0.008079258 | 0.248312568 |
| ENSMUSG00000006931 | P3h4 | 66180 | protein_coding | 39.28415494 | -0.248356801 | 0.189764237 | 1.308765049 | 0.190613949 | 0.388090095 | 0.248356801 |
| ENSMUSG00000073423 | Zfp414 | 328801 | protein_coding | 72.7554379 | -0.24837521 | 0.134220816 | 1.850496937 | 0.064241959 | 0.179449017 | 0.24837521 |
| ENSMUSG00000017057 | Il13ra1 | 16164 | protein_coding | 1703.244687 | 0.248392818 | 0.176197739 | -1.409738963 | 0.158616775 | 0.342539221 | 0.248392818 |
| ENSMUSG00000079499 | 6530402F18Rik | 76220 | lncRNA | 4.483674929 | 0.248432237 | 0.919328737 | -0.270232211 | 0.786981613 | 0.893375436 | 0.248432237 |
| ENSMUSG00000020434 | 4921536K21Rik | 67430 | protein_coding | 32.17336813 | 0.24844281 | 0.204722762 | -1.213557338 | 0.224916806 | 0.432209358 | 0.24844281 |
| ENSMUSG00000028411 | Aptx | 66408 | protein_coding | 131.5270672 | 0.248454833 | 0.135430483 | -1.834556205 | 0.066571473 | 0.184147622 | 0.248454833 |
| ENSMUSG00000028035 | Dnajb4 | 67035 | protein_coding | 167.629072 | -0.248472362 | 0.126890362 | 1.958165758 | 0.050210566 | 0.149293713 | 0.248472362 |
| ENSMUSG00000068327 | Tlx2 | 21909 | protein_coding | 0.922438628 | -0.24861591 | 1.586021037 | 0.156754484 | 0.875438346 | 1 | 0.24861591 |
| ENSMUSG00000006784 | Odad4 | 74407 | protein_coding | 19.41512155 | -0.248659147 | 0.31146916 | 0.798342753 | 0.424671615 | 0.644224421 | 0.248659147 |
| ENSMUSG00000097331 | F420014N23Rik | 100038591 | lncRNA | 9.366613986 | -0.248663811 | 0.428022149 | 0.580960148 | 0.561267313 | 0.749718386 | 0.248663811 |
| ENSMUSG00000036291 | Ap5m1 | 74385 | protein_coding | 505.2603806 | -0.248731664 | 0.107154012 | 2.321253855 | 0.020273147 | 0.074167202 | 0.248731664 |
| ENSMUSG00000031167 | Rbm3 | 19652 | protein_coding | 1182.615487 | 0.248781557 | 0.145705091 | -1.707432154 | 0.087741758 | 0.224907418 | 0.248781557 |
| ENSMUSG00000097743 | Gm16973 | 100503823 | lncRNA | 19.28364219 | -0.24879377 | 0.238610656 | 1.042676694 | 0.297098055 | 0.516613382 | 0.24879377 |
| ENSMUSG00000045502 | Hcar2 | 80885 | protein_coding | 5.368212514 | -0.248815788 | 0.631702472 | 0.393881296 | 0.69366868 | 0.837421924 | 0.248815788 |
| ENSMUSG00000022285 | Ywhaz | 22631 | protein_coding | 3328.027835 | 0.248837314 | 0.085268197 | -2.918289849 | 0.00351957 | 0.018191607 | 0.248837314 |
| ENSMUSG00000023952 | Gtpbp2 | 56055 | protein_coding | 1661.783552 | 0.248852786 | 0.188851321 | -1.317718005 | 0.187598065 | 0.383842701 | 0.248852786 |
| ENSMUSG00000025068 | Gsto1 | 14873 | protein_coding | 3875.776188 | -0.248922437 | 0.076518839 | 3.253086957 | 0.001141586 | 0.007103295 | 0.248922437 |
| ENSMUSG00000034744 | Nagk | 56174 | protein_coding | 456.1158945 | -0.249019696 | 0.066510305 | 3.744076879 | 0.000181058 | 0.001453571 | 0.249019696 |
| ENSMUSG00000020814 | Mxra7 | 67622 | protein_coding | 26.94068279 | 0.249186016 | 0.209596704 | -1.188883278 | 0.234485601 | 0.444019633 | 0.249186016 |
| ENSMUSG00000027605 | Acss2 | 60525 | protein_coding | 1611.045378 | -0.249188588 | 0.179663659 | 1.386972689 | 0.165450091 | 0.352592954 | 0.249188588 |
| ENSMUSG00000096948 | Gm4221 | 100043089 | lncRNA | 3.317957483 | 0.249219268 | 0.743418377 | -0.335234204 | 0.737448422 | 1 | 0.249219268 |
| ENSMUSG00000035725 | Prkx | 19108 | protein_coding | 370.5476149 | -0.249269734 | 0.111335099 | 2.238914199 | 0.025161499 | 0.087791464 | 0.249269734 |
| ENSMUSG00000044052 | Ccr10 | 12777 | protein_coding | 12.61034892 | -0.2492891 | 0.447550391 | 0.557007892 | 0.577522037 | 0.761475268 | 0.2492891 |
| ENSMUSG00000049807 | Arhgap23 | 58996 | protein_coding | 12.31250878 | 0.249346818 | 0.566460572 | -0.44018389 | 0.659803927 | 0.816323585 | 0.249346818 |
| ENSMUSG00000030309 | Caprin2 | 232560 | protein_coding | 74.70183851 | -0.249437277 | 0.19937423 | 1.251100897 | 0.210897668 | 0.415010021 | 0.249437277 |
| ENSMUSG00000033222 | Ttf2 | 74044 | protein_coding | 192.6705253 | 0.249560693 | 0.136163871 | -1.832796698 | 0.06683281 | 0.184592245 | 0.249560693 |
| ENSMUSG00000025094 | Slc18a2 | 214084 | protein_coding | 34.56931683 | -0.249566938 | 0.223526464 | 1.116498391 | 0.264208858 | 0.478777869 | 0.249566938 |
| ENSMUSG00000021620 | Acot12 | 74156 | protein_coding | 2.701134488 | -0.249603534 | 1.401988247 | 0.178035397 | 0.858695177 | 1 | 0.249603534 |
| ENSMUSG00000096938 | 9530052E02Rik | 619321 | lncRNA | 1.871774418 | 0.249615533 | 1.201820333 | -0.207697878 | 0.835464875 | 1 | 0.249615533 |
| ENSMUSG00000005983 | 1700037C18Rik | 73261 | protein_coding | 40.11056603 | -0.249616005 | 0.16694201 | 1.495225828 | 0.134855515 | 0.305727036 | 0.249616005 |
| ENSMUSG00000031255 | Sytl4 | 27359 | protein_coding | 284.36999 | -0.249616128 | 0.123115079 | 2.02750248 | 0.042611051 | 0.131923303 | 0.249616128 |
| ENSMUSG00000021816 | Ppp3cb | 19056 | protein_coding | 514.1811904 | 0.249753759 | 0.097245747 | -2.568274369 | 0.010220622 | 0.043322217 | 0.249753759 |
| ENSMUSG00000002731 | Prkra | 23992 | protein_coding | 346.1960507 | -0.249812214 | 0.091152365 | 2.740600476 | 0.006132703 | 0.028656498 | 0.249812214 |
| ENSMUSG00000032387 | Rbpms2 | 71973 | protein_coding | 5.143771636 | 0.249823359 | 0.572189385 | -0.436609565 | 0.662394525 | 0.817346435 | 0.249823359 |
| ENSMUSG00000052609 | Plekhg3 | 263406 | protein_coding | 699.7721007 | -0.249850456 | 0.091269119 | 2.737513629 | 0.006190555 | 0.028864834 | 0.249850456 |
| ENSMUSG00000028028 | Alpk1 | 71481 | protein_coding | 428.1280956 | 0.249872765 | 0.077675713 | -3.216871202 | 0.001295968 | 0.007888281 | 0.249872765 |

| ensembl_gene_id | external_gene_name | entrezgene_id | gene_biotype | baseMean | log2FoldChange | lfcSE | stat | pvalue | padj | absLog2FC |
|---|---|---|---|---|---|---|---|---|---|---|
| ENSMUSG00000021460 | Auh | 11992 | protein_coding | 251.5255861 | -0.249912331 | 0.098034413 | 2.549230664 | 0.010796087 | 0.045254367 | 0.249912331 |
| ENSMUSG00000033543 | Gtf2a2 | 235459 | protein_coding | 294.9137802 | 0.249974304 | 0.103425874 | -2.416941665 | 0.015651525 | 0.060376822 | 0.249974304 |
| ENSMUSG00000029121 | Crmp1 | 12933 | protein_coding | 6.654970246 | 0.249984945 | 0.913484157 | -0.273660953 | 0.784345188 | 0.891996173 | 0.249984945 |
| ENSMUSG00000022742 | Cpox | 12892 | protein_coding | 324.4683896 | 0.250062777 | 0.109567839 | -2.282264379 | 0.022473736 | 0.080540642 | 0.250062777 |
| ENSMUSG00000034947 | Tmem106a | 217203 | protein_coding | 187.479996 | -0.25019233 | 0.163698432 | 1.528373405 | 0.126419854 | 0.292394604 | 0.25019233 |
| ENSMUSG00000051590 | Map3k19 | 22625 | protein_coding | 3.338909151 | -0.250205433 | 1.399154711 | 0.178826138 | 0.858074221 | 1 | 0.250205433 |
| ENSMUSG00000046834 | Krt1 | 16678 | protein_coding | 1.615613748 | -0.250252317 | 1.719776255 | 0.145514462 | 0.884304698 | 1 | 0.250252317 |
| ENSMUSG00000036466 | Megf11 | 214058 | protein_coding | 40.56098549 | -0.250300596 | 0.231871256 | 1.079480919 | 0.280373395 | 0.498019388 | 0.250300596 |
| ENSMUSG00000028643 | Svbp | 69216 | protein_coding | 153.5661911 | -0.250322238 | 0.130112432 | 1.923891786 | 0.054368149 | 0.158736855 | 0.250322238 |
| ENSMUSG00000062794 | Zfp599 | 235048 | protein_coding | 68.44548954 | -0.250405266 | 0.157749628 | 1.587358839 | 0.112431396 | 0.268968865 | 0.250405266 |
| ENSMUSG00000038005 | Hpf1 | 72612 | protein_coding | 138.8724749 | 0.250466121 | 0.138441843 | -1.809179329 | 0.070423148 | 0.191813931 | 0.250466121 |
| ENSMUSG00000074655 | Gm1527 | 385263 | protein_coding | 0.836582822 | 0.250471944 | 1.452043343 | -0.17249619 | 0.863047457 | 1 | 0.250471944 |
| ENSMUSG00000003623 | Crot | 74114 | protein_coding | 733.9077185 | 0.250493667 | 0.070186798 | -3.568957042 | 0.000358405 | 0.002670749 | 0.250493667 |
| ENSMUSG00000031562 | Dctd | 320685 | protein_coding | 60.59424156 | 0.25050746 | 0.267243273 | -0.937376109 | 0.348565126 | 0.57042954 | 0.25050746 |
| ENSMUSG00000020412 | Ascc2 | 75452 | protein_coding | 689.2759561 | -0.250608369 | 0.057379731 | 4.367541738 | 1.26E-05 | 0.000137056 | 0.250608369 |
| ENSMUSG00000086290 | Snhg12 | 100039864 | lncRNA | 126.9600314 | -0.250630292 | 0.155874174 | 1.607901334 | 0.10785679 | 0.261367195 | 0.250630292 |
| ENSMUSG00000040675 | Mthfd1l | 270685 | protein_coding | 199.5872377 | 0.250642946 | 0.128803258 | -1.945936377 | 0.051662383 | 0.152917797 | 0.250642946 |
| ENSMUSG00000027236 | Eif3j1 | 78655 | protein_coding | 213.0253005 | -0.250683139 | 0.117008466 | 2.142435915 | 0.03215842 | 0.106468355 | 0.250683139 |
| ENSMUSG00000035842 | Ddx11 | 320209 | protein_coding | 135.0800011 | 0.250841204 | 0.127053138 | -1.974301518 | 0.048347468 | 0.145350463 | 0.250841204 |
| ENSMUSG00000026799 | Med27 | 68975 | protein_coding | 111.2291501 | 0.250842986 | 0.104752947 | -2.394615083 | 0.016637821 | 0.063542317 | 0.250842986 |
| ENSMUSG00000034158 | Lrrc58 | 320184 | protein_coding | 913.0095054 | 0.250850083 | 0.100643865 | -2.492452791 | 0.012686419 | 0.051198321 | 0.250850083 |
| ENSMUSG00000068154 | Insm1 | 53626 | protein_coding | 25.9646222 | 0.250862861 | 0.272850095 | -0.919416434 | 0.357877797 | 0.580790426 | 0.250862861 |
| ENSMUSG00000020458 | Rtn4 | 68585 | protein_coding | 862.0291734 | -0.250927107 | 0.071737119 | 3.497869861 | 0.00046899 | 0.003361502 | 0.250927107 |
| ENSMUSG00000022702 | Hira | 15260 | protein_coding | 114.2780495 | 0.250933978 | 0.099556316 | -2.520522948 | 0.01171806 | 0.048232278 | 0.250933978 |
| ENSMUSG00000047888 | Tnrc6b | 213988 | protein_coding | 1188.607526 | 0.250983823 | 0.085022068 | -2.951984446 | 0.003157388 | 0.016594879 | 0.250983823 |
| ENSMUSG00000067071 | Hes6 | 55927 | protein_coding | 593.9466114 | -0.251026462 | 0.132003707 | 1.901662221 | 0.057215328 | 0.164953548 | 0.251026462 |
| ENSMUSG00000032127 | Vps11 | 71732 | protein_coding | 413.7603677 | 0.251036541 | 0.072359362 | -3.469302854 | 0.000521811 | 0.003679413 | 0.251036541 |
| ENSMUSG00000032044 | Rpusd4 | 71989 | protein_coding | 158.3902666 | -0.251072953 | 0.097417424 | 2.577290009 | 0.009957837 | 0.042442029 | 0.251072953 |
| ENSMUSG00000026434 | Nucks1 | 98415 | protein_coding | 1700.768538 | 0.251089478 | 0.143248294 | -1.752827003 | 0.079631707 | 0.209167807 | 0.251089478 |
| ENSMUSG00000035247 | Hectd1 | 207304 | protein_coding | 3509.654419 | 0.251130333 | 0.058365032 | -4.302753317 | 1.69E-05 | 0.000177661 | 0.251130333 |
| ENSMUSG00000030884 | Uqcrc2 | 67003 | protein_coding | 3910.210541 | -0.251367085 | 0.082576743 | 3.044042135 | 0.002334224 | 0.012832018 | 0.251367085 |
| ENSMUSG00000038526 | Car14 | 23831 | protein_coding | 6.612092239 | 0.251478914 | 0.493751357 | -0.509322984 | 0.61052585 | 0.784161879 | 0.251478914 |
| ENSMUSG00000035530 | Eif1 | 20918 | protein_coding | 1898.356646 | -0.251489454 | 0.104322921 | 2.410682635 | 0.015922697 | 0.061293105 | 0.251489454 |
| ENSMUSG00000079020 | Slc45a4 | 106068 | protein_coding | 350.0184222 | 0.251495018 | 0.090143571 | -2.789938495 | 0.005271806 | 0.025463175 | 0.251495018 |
| ENSMUSG00000034300 | Fam53c | 66306 | protein_coding | 287.2285929 | 0.251501755 | 0.089492469 | -2.810311954 | 0.00494935 | 0.024110197 | 0.251501755 |
| ENSMUSG00000018559 | Ctdnep1 | 67181 | protein_coding | 373.5761126 | 0.251516192 | 0.120272767 | -2.091214813 | 0.036508812 | 0.117137463 | 0.251516192 |
| ENSMUSG00000033022 | Cdo1 | 12583 | protein_coding | 333.6885399 | 0.251526659 | 0.160890951 | -1.563336275 | 0.11797352 | 0.278515327 | 0.251526659 |
| ENSMUSG00000020265 | Sumo3 | 20610 | protein_coding | 937.6505378 | 0.251624722 | 0.087147094 | -2.887356441 | 0.003884938 | 0.019745882 | 0.251624722 |
| ENSMUSG00000026207 | Speg | 11790 | protein_coding | 10.00927154 | -0.251661373 | 1.50015844 | 0.167756529 | 0.866774823 | 0.933837244 | 0.251661373 |

| ensembl_gene_id | external_gene_name | entrezgene_id | gene_biotype | baseMean | log2FoldChange | lfcSE | stat | pvalue | padj | absLog2FC |
|---|---|---|---|---|---|---|---|---|---|---|
| ENSMUSG00000084453 | Gm24596 | 115486854 | snRNA | 0.602479715 | 0.251757321 | 1.681821878 | -0.149693213 | 0.881006663 | 1 | 0.251757321 |
| ENSMUSG00000050567 | Maml1 | 103806 | protein_coding | 211.9123572 | -0.251779992 | 0.121130257 | 2.07858877 | 0.037655164 | 0.120138597 | 0.251779992 |
| ENSMUSG00000117698 | BC037704 | 100502982 | lncRNA | 10.31888751 | 0.251818163 | 0.335022099 | -0.751646426 | 0.452263716 | 0.668637691 | 0.251818163 |
| ENSMUSG00000055053 | Nfic | 18029 | protein_coding | 140.0744978 | -0.251842605 | 0.194324872 | 1.295987499 | 0.194979792 | 0.394112152 | 0.251842605 |
| ENSMUSG00000021692 | Dimt1 | 66254 | protein_coding | 52.25722425 | -0.251879036 | 0.167380214 | 1.504831609 | 0.132367375 | 0.301713259 | 0.251879036 |
| ENSMUSG00000015759 | Cnih1 | 12793 | protein_coding | 539.9719113 | 0.251894288 | 0.084697798 | -2.974035859 | 0.002939107 | 0.015586518 | 0.251894288 |
| ENSMUSG00000020821 | Kif1c | 16562 | protein_coding | 3928.726624 | -0.251947473 | 0.060799676 | 4.143894978 | 3.41E-05 | 0.000334053 | 0.251947473 |
| ENSMUSG00000023883 | Phf10 | 72057 | protein_coding | 350.7423977 | 0.251949314 | 0.082671741 | -3.047586899 | 0.002306868 | 0.012715494 | 0.251949314 |
| ENSMUSG00000022724 | Riox2 | 67014 | protein_coding | 225.9234368 | -0.251950313 | 0.096731279 | 2.604641609 | 0.009197041 | 0.039805161 | 0.251950313 |
| ENSMUSG00000026037 | Orc2 | 18393 | protein_coding | 175.949398 | 0.252050682 | 0.11295752 | -2.231375851 | 0.025656243 | 0.08904733 | 0.252050682 |
| ENSMUSG00000032177 | Pde4a | 18577 | protein_coding | 27.90905933 | -0.25214508 | 0.231414456 | 1.089582234 | 0.275897212 | 0.492718348 | 0.25214508 |
| ENSMUSG00000033955 | Tnks1bp1 | 228140 | protein_coding | 1887.823527 | -0.252188643 | 0.066116459 | 3.814309557 | 0.000136564 | 0.001134417 | 0.252188643 |
| ENSMUSG00000074749 | Kiz | 228730 | protein_coding | 212.1644345 | -0.252256733 | 0.116753663 | 2.160589445 | 0.030727067 | 0.102738771 | 0.252256733 |
| ENSMUSG00000109398 | Gm3854 | 100416477 | protein_coding | 1.691607249 | -0.252267148 | 1.330659974 | 0.189580474 | 0.849637889 | 1 | 0.252267148 |
| ENSMUSG00000025142 | Aspscr1 | 68938 | protein_coding | 511.5567168 | 0.252295865 | 0.071594976 | -3.523932546 | 0.000425192 | 0.003094592 | 0.252295865 |
| ENSMUSG00000038187 | Btbd10 | 68815 | protein_coding | 163.5340108 | -0.252325877 | 0.104367264 | 2.41767262 | 0.015620123 | 0.060282943 | 0.252325877 |
| ENSMUSG00000098557 | Kctd12 | 239217 | protein_coding | 650.0495667 | 0.252332132 | 0.196788472 | -1.282250579 | 0.199754759 | 0.40082018 | 0.252332132 |
| ENSMUSG00000000876 | Pxmp4 | 59038 | protein_coding | 403.3845128 | 0.252365948 | 0.07788878 | -3.240080911 | 0.001194958 | 0.007373206 | 0.252365948 |
| ENSMUSG00000030655 | Smg1 | 233789 | protein_coding | 2521.196632 | 0.252378721 | 0.128856335 | -1.958605457 | 0.05015901 | 0.149272747 | 0.252378721 |
| ENSMUSG00000032744 | Heyl | 56198 | protein_coding | 2.761000628 | 0.252409205 | 1.270981508 | -0.198593924 | 0.842580407 | 1 | 0.252409205 |
| ENSMUSG00000075706 | Gpx4 | 625249 | protein_coding | 812.5209727 | -0.252477931 | 0.112964205 | 2.23502596 | 0.025415645 | 0.088476845 | 0.252477931 |
| ENSMUSG00000096010 | H4c16 | 320332 | protein_coding | 787.5414501 | 0.252478734 | 0.09996556 | -2.525657186 | 0.011548211 | 0.047736459 | 0.252478734 |
| ENSMUSG00000052331 | Ankrd44 | 329154 | protein_coding | 204.3607902 | -0.252516896 | 0.149789728 | 1.685809157 | 0.091832563 | 0.232097177 | 0.252516896 |
| ENSMUSG00000054733 | Msra | 110265 | protein_coding | 429.1323605 | -0.252537831 | 0.063160261 | 3.998365827 | 6.38E-05 | 0.000581283 | 0.252537831 |
| ENSMUSG00000090266 | Mettl23 | 74319 | protein_coding | 149.2489083 | -0.252551919 | 0.106718616 | 2.366521685 | 0.017956122 | 0.067342119 | 0.252551919 |
| ENSMUSG00000022106 | Rcbtb2 | 105670 | protein_coding | 193.366014 | 0.252600335 | 0.105940494 | -2.384360549 | 0.017108831 | 0.06492078 | 0.252600335 |
| ENSMUSG00000026579 | F5 | 14067 | protein_coding | 8.697328289 | -0.252624748 | 0.577204903 | 0.437669095 | 0.661626176 | 0.816921981 | 0.252624748 |
| ENSMUSG00000034560 | Washc4 | 319277 | protein_coding | 441.4067678 | 0.252771906 | 0.080542719 | -3.138358235 | 0.001698971 | 0.009886119 | 0.252771906 |
| ENSMUSG00000036834 | Plch1 | 269437 | protein_coding | 646.6568773 | 0.252789409 | 0.145672316 | -1.735329098 | 0.082682531 | 0.215060531 | 0.252789409 |
| ENSMUSG00000039646 | Vasn | 246154 | protein_coding | 76.91545062 | -0.252852353 | 0.157257172 | 1.607890757 | 0.107859107 | 0.261367195 | 0.252852353 |
| ENSMUSG00000055681 | Cope | 59042 | protein_coding | 1556.51221 | 0.252854648 | 0.083303393 | -3.035346335 | 0.002402596 | 0.013164059 | 0.252854648 |
| ENSMUSG00000025934 | Gsta3 | 14859 | protein_coding | 189.8485891 | -0.252868645 | 0.153829498 | 1.64382416 | 0.100212529 | 0.2481124 | 0.252868645 |
| ENSMUSG00000025362 | Rps26 | 27370 | protein_coding | 3506.955216 | -0.252881204 | 0.097697445 | 2.588411646 | 0.009641968 | 0.041336308 | 0.252881204 |
| ENSMUSG00000053111 | Fank1 | 66930 | protein_coding | 7.633535255 | -0.252881425 | 0.455751398 | 0.554867031 | 0.578985614 | 0.762469572 | 0.252881425 |
| ENSMUSG00000022885 | St6gal1 | 20440 | protein_coding | 1193.89072 | -0.25290226 | 0.374397729 | 0.675490904 | 0.499363928 | 0.704201299 | 0.25290226 |
| ENSMUSG00000021420 | Fars2 | 69955 | protein_coding | 485.2328471 | -0.252955912 | 0.066493129 | 3.804241375 | 0.000142239 | 0.001176235 | 0.252955912 |
| ENSMUSG00000048701 | Ccdc6 | 76551 | protein_coding | 1165.142705 | 0.253030164 | 0.081110473 | -3.1195745 | 0.001811125 | 0.010450353 | 0.253030164 |
| ENSMUSG00000001804 | Dsg4 | 16769 | protein_coding | 3.1744412 | -0.253049098 | 0.786703525 | 0.321657511 | 0.74771217 | 1 | 0.253049098 |
| ENSMUSG00000061028 | Clasrp | 53609 | protein_coding | 401.1485904 | 0.253051485 | 0.155847095 | -1.623716401 | 0.104436349 | 0.255505462 | 0.253051485 |

| ensembl_gene_id | external_gene_name | entrezgene_id | gene_biotype | baseMean | log2FoldChange | lfcSE | stat | pvalue | padj | absLog2FC |
|---|---|---|---|---|---|---|---|---|---|---|
| ENSMUSG00000003382 | Etv3 | 27049 | protein_coding | 857.9848066 | 0.253097732 | 0.097550754 | -2.59452359 | 0.009472211 | 0.040777919 | 0.253097732 |
| ENSMUSG00000064655 | Gm25788 | 115490049 | snoRNA | 184.7197399 | 0.253147144 | 0.294571408 | -0.859374457 | 0.390133957 | 0.612475705 | 0.253147144 |
| ENSMUSG00000028495 | Rps6 | 20104 | protein_coding | 2270.231933 | -0.253173975 | 0.148848245 | 1.700886534 | 0.088964296 | 0.226849015 | 0.253173975 |
| ENSMUSG00000024271 | Elp2 | 58523 | protein_coding | 340.4827967 | 0.253220658 | 0.075653929 | -3.347091983 | 0.000816641 | 0.00535797 | 0.253220658 |
| ENSMUSG00000065990 | Aurkaip1 | 66077 | protein_coding | 678.2502808 | -0.253221126 | 0.066363437 | 3.815672273 | 0.000135813 | 0.001128742 | 0.253221126 |
| ENSMUSG00000045414 | Dipk2a | 68861 | protein_coding | 338.3034945 | 0.253263853 | 0.119263364 | -2.123567917 | 0.033706291 | 0.110135672 | 0.253263853 |
| ENSMUSG00000090622 | A930033H14Rik | 320700 | lncRNA | 5.978900149 | 0.253347587 | 0.69273455 | -0.365721022 | 0.714573258 | 0.851895956 | 0.253347587 |
| ENSMUSG00000100475 | Gm29543 | 105245595 | lncRNA | 0.786870164 | 0.253375255 | 1.20184436 | -0.21082202 | 0.833026156 | 1 | 0.253375255 |
| ENSMUSG00000028307 | Aldob | 230163 | protein_coding | 2097.32192 | -0.253386872 | 0.079394035 | 3.191510196 | 0.001415311 | 0.008495983 | 0.253386872 |
| ENSMUSG00000017639 | Rab11fip4 | 268451 | protein_coding | 569.3544164 | -0.253449716 | 0.136894776 | 1.851419931 | 0.064109164 | 0.179258167 | 0.253449716 |
| ENSMUSG00000025380 | Fscn2 | 238021 | protein_coding | 4.373327456 | 0.253543611 | 0.586810052 | -0.432071009 | 0.665689804 | 1 | 0.253543611 |
| ENSMUSG00000024891 | Slc29a2 | 13340 | protein_coding | 17.8861044 | -0.253548807 | 0.303886244 | 0.83435434 | 0.404081339 | 0.625281037 | 0.253548807 |
| ENSMUSG00000039307 | Hexdc | 238023 | protein_coding | 160.6752596 | -0.253560193 | 0.117269733 | 2.162196386 | 0.030603042 | 0.102448537 | 0.253560193 |
| ENSMUSG00000004040 | Stat3 | 20848 | protein_coding | 1727.539255 | 0.253605191 | 0.088931607 | -2.851687955 | 0.004348777 | 0.021650799 | 0.253605191 |
| ENSMUSG00000031285 | Dcx | 13193 | protein_coding | 4.36715505 | -0.253638579 | 1.168565526 | 0.217051225 | 0.828168424 | 1 | 0.253638579 |
| ENSMUSG00000021936 | Mapk8 | 26419 | protein_coding | 316.8993019 | 0.253680061 | 0.085850857 | -2.954892581 | 0.003127779 | 0.016452754 | 0.253680061 |
| ENSMUSG00000026279 | Thap4 | 67026 | protein_coding | 748.5009064 | -0.25381308 | 0.068020636 | 3.731412926 | 0.000190409 | 0.001515379 | 0.25381308 |
| ENSMUSG00000087060 | Eldr | 72735 | lncRNA | 39.8785474 | 0.253877432 | 0.233226252 | -1.088545696 | 0.276354275 | 0.493321208 | 0.253877432 |
| ENSMUSG00000021135 | Slc10a1 | 20493 | protein_coding | 18.92924193 | 0.253935273 | 0.406514512 | -0.624664718 | 0.532191134 | 0.728866518 | 0.253935273 |
| ENSMUSG00000029401 | Rilpl2 | 80291 | protein_coding | 134.6422837 | 0.25398574 | 0.132495404 | -1.91694 | 0.055245555 | 0.160507581 | 0.25398574 |
| ENSMUSG00000064811 | Gm22358 | 115488216 | snoRNA | 3.280849832 | -0.254010285 | 0.621996724 | 0.408378815 | 0.682995585 | 1 | 0.254010285 |
| ENSMUSG00000031340 | Gabre | 14404 | protein_coding | 25.5460836 | 0.254027744 | 0.430006893 | -0.590752725 | 0.554686115 | 0.744703672 | 0.254027744 |
| ENSMUSG00000090115 | Usp49 | 224836 | protein_coding | 114.43641 | -0.254193928 | 0.142695203 | 1.781376827 | 0.074850911 | 0.200014231 | 0.254193928 |
| ENSMUSG00000090290 | Tarbp1 | 212728 | protein_coding | 179.320561 | -0.254197095 | 0.135936575 | 1.869968368 | 0.061488211 | 0.173816344 | 0.254197095 |
| ENSMUSG00000031703 | Itfg1 | 71927 | protein_coding | 1079.121204 | -0.254221983 | 0.090875221 | 2.797484081 | 0.005150231 | 0.024970906 | 0.254221983 |
| ENSMUSG00000032180 | Tmed1 | 17083 | protein_coding | 88.40913091 | -0.254269882 | 0.143330083 | 1.774016149 | 0.076060479 | 0.202492786 | 0.254269882 |
| ENSMUSG00000087281 | Gm16015 | 115490183 | lncRNA | 12.26252466 | -0.254339719 | 0.468122976 | 0.54331817 | 0.586910756 | 0.768390968 | 0.254339719 |
| ENSMUSG00000034278 | Dnajc17 | 69408 | protein_coding | 86.78381278 | -0.254347653 | 0.115803726 | 2.196368472 | 0.02806558 | 0.095640734 | 0.254347653 |
| ENSMUSG00000001103 | Sebox | 18292 | protein_coding | 0.670790068 | -0.254355966 | 1.541807372 | 0.164972597 | 0.868965551 | 1 | 0.254355966 |
| ENSMUSG00000023232 | Serinc2 | 230779 | protein_coding | 1458.525435 | -0.254431116 | 0.06261987 | 4.063105158 | 4.84E-05 | 0.000454104 | 0.254431116 |
| ENSMUSG00000042106 | Inka1 | 68176 | protein_coding | 3.382475087 | -0.254445992 | 0.741089359 | 0.343340501 | 0.731342315 | 1 | 0.254445992 |
| ENSMUSG00000015468 | Notch4 | 18132 | protein_coding | 126.9360663 | 0.254500311 | 0.160160392 | -1.589034017 | 0.112052713 | 0.268218365 | 0.254500311 |
| ENSMUSG00000096433 | Zfp994 | 240038 | protein_coding | 172.3467726 | -0.254569546 | 0.10991919 | 2.315969999 | 0.020559907 | 0.074950618 | 0.254569546 |
| ENSMUSG00000030613 | Ccdc90b | 66365 | protein_coding | 186.0625569 | -0.254635434 | 0.148100405 | 1.719343272 | 0.085551885 | 0.220716112 | 0.254635434 |
| ENSMUSG00000050520 | Cldn8 | 54420 | protein_coding | 1534.067741 | 0.254750674 | 0.186831397 | -1.363532457 | 0.172714763 | 0.363275924 | 0.254750674 |
| ENSMUSG00000078945 | Naip2 | 17948 | protein_coding | 1593.315335 | -0.254757567 | 0.090973029 | 2.800363686 | 0.005104506 | 0.024790788 | 0.254757567 |
| ENSMUSG00000036975 | Tmem177 | 66343 | protein_coding | 113.0761886 | -0.25484957 | 0.11594054 | 2.198105775 | 0.027941569 | 0.095316478 | 0.25484957 |
| ENSMUSG00000058454 | Dhcr7 | 13360 | protein_coding | 696.3686814 | -0.254984669 | 0.087901179 | 2.900810551 | 0.003721988 | 0.0190648 | 0.254984669 |
| ENSMUSG00000021611 | Tert | 21752 | protein_coding | 49.58289233 | 0.255003196 | 0.186153709 | -1.369852883 | 0.17073283 | 0.360508479 | 0.255003196 |

Supplementary Table 1

| ensembl_gene_id | external_gene_name | entrezgene_id | gene_biotype | baseMean | log2FoldChange | lfcSE | stat | pvalue | padj | absLog2FC |
|---|---|---|---|---|---|---|---|---|---|---|
| ENSMUSG00000037692 | Ahdc1 | 230793 | protein_coding | 385.3053635 | 0.255020697 | 0.18501217 | -1.378399577 | 0.168079954 | 0.356592544 | 0.255020697 |
| ENSMUSG00000029729 | Zkscan1 | 74570 | protein_coding | 1425.653857 | -0.255064076 | 0.100852267 | 2.529086204 | 0.011435994 | 0.047391714 | 0.255064076 |
| ENSMUSG00000037813 | D630003M21Rik | 228846 | protein_coding | 6.693502661 | 0.25515146 | 0.642736619 | -0.396976697 | 0.691384645 | 0.835769466 | 0.25515146 |
| ENSMUSG00000049600 | Zbtb45 | 232879 | protein_coding | 132.1783258 | 0.255243938 | 0.111854963 | -2.281918761 | 0.022494137 | 0.080578781 | 0.255243938 |
| ENSMUSG00000027018 | Hat1 | 107435 | protein_coding | 222.7543857 | 0.255247622 | 0.150262779 | -1.698674969 | 0.089380442 | 0.227592128 | 0.255247622 |
| ENSMUSG00000051703 | Tmem198 | 319998 | protein_coding | 14.5698324 | -0.255263435 | 0.300036042 | 0.850775905 | 0.394893849 | 0.617252913 | 0.255263435 |
| ENSMUSG00000032724 | Abtb2 | 99382 | protein_coding | 87.29706344 | -0.255284593 | 0.217358261 | 1.174487651 | 0.240199755 | 0.450500711 | 0.255284593 |
| ENSMUSG00000027305 | Ndufaf1 | 69702 | protein_coding | 158.5594265 | -0.255291736 | 0.115109329 | 2.217819695 | 0.026567129 | 0.091535482 | 0.255291736 |
| ENSMUSG00000079317 | Trappc2 | 66226 | protein_coding | 30.37040877 | -0.255306066 | 0.175349559 | 1.45598351 | 0.145397178 | 0.322533618 | 0.255306066 |
| ENSMUSG00000046204 | Pnma2 | 239157 | protein_coding | 2.717920323 | -0.255338328 | 1.382775038 | 0.184656449 | 0.853498507 | 1 | 0.255338328 |
| ENSMUSG00000077274 | Gm22786 | 115485822 | snoRNA | 12.57717166 | 0.255374521 | 0.506658556 | -0.504036729 | 0.614235567 | 0.78673795 | 0.255374521 |
| ENSMUSG00000030207 | Fam234b | 74525 | protein_coding | 4.864155826 | 0.255391215 | 0.877587817 | -0.291014995 | 0.771039851 | 0.885199039 | 0.255391215 |
| ENSMUSG00000022793 | B4galt4 | 56375 | protein_coding | 711.5027562 | 0.25551756 | 0.11585446 | -2.205504736 | 0.027418696 | 0.093901357 | 0.25551756 |
| ENSMUSG00000028051 | Hcn3 | 15168 | protein_coding | 13.98544794 | 0.255559828 | 0.40124681 | -0.636914293 | 0.524180672 | 0.72280558 | 0.255559828 |
| ENSMUSG00000031245 | Hmgn5 | 50887 | protein_coding | 162.6336161 | -0.255563637 | 0.142458116 | 1.793956322 | 0.072820141 | 0.19654843 | 0.255563637 |
| ENSMUSG00000037266 | Rsrp1 | 27981 | protein_coding | 1540.201249 | -0.255588353 | 0.109983825 | 2.323872196 | 0.020132345 | 0.073733764 | 0.255588353 |
| ENSMUSG00000053604 | Rpia | 19895 | protein_coding | 325.416866 | -0.255635863 | 0.10285147 | 2.485485746 | 0.012937477 | 0.05194233 | 0.255635863 |
| ENSMUSG00000028871 | Rspo1 | 192199 | protein_coding | 1.815758265 | 0.255671986 | 1.554332181 | -0.164489926 | 0.869345476 | 1 | 0.255671986 |
| ENSMUSG00000022313 | Utp23 | 78581 | protein_coding | 86.56827302 | 0.255690646 | 0.137206063 | -1.8635521 | 0.06238463 | 0.175505924 | 0.255690646 |
| ENSMUSG00000031201 | Brcc3 | 210766 | protein_coding | 357.9422538 | 0.255694053 | 0.080216082 | -3.187565968 | 0.001434757 | 0.008569101 | 0.255694053 |
| ENSMUSG00000022241 | Tars | 110960 | protein_coding | 1260.137458 | 0.255698025 | 0.099536497 | -2.568887133 | 0.010202568 | 0.043279016 | 0.255698025 |
| ENSMUSG00000053716 | Dusp7 | 235584 | protein_coding | 89.54766608 | 0.255776407 | 0.159330162 | -1.605323209 | 0.108422694 | 0.262304772 | 0.255776407 |
| ENSMUSG00000039031 | Arhgap18 | 73910 | protein_coding | 571.3383387 | -0.255776764 | 0.10995216 | 2.32625501 | 0.020004951 | 0.073397413 | 0.255776764 |
| ENSMUSG00000022263 | Trio | 223435 | protein_coding | 1418.049079 | 0.255867475 | 0.165859156 | -1.542679229 | 0.122908624 | 0.286211349 | 0.255867475 |
| ENSMUSG00000015247 | Nipsnap3b | 66536 | protein_coding | 678.6299117 | 0.255923476 | 0.086687174 | -2.952264609 | 0.003154525 | 0.016585107 | 0.255923476 |
| ENSMUSG00000036333 | Kidins220 | 77480 | protein_coding | 1448.29918 | -0.255963493 | 0.064787509 | 3.950815491 | 7.79E-05 | 0.000692995 | 0.255963493 |
| ENSMUSG00000064919 | Gm22572 | 115489570 | snRNA | 1.032727485 | -0.255965213 | 1.247502146 | 0.205182182 | 0.83742979 | 1 | 0.255965213 |
| ENSMUSG00000031264 | Btk | 12229 | protein_coding | 9.019640469 | 0.25606685 | 0.489868078 | -0.522726141 | 0.601164844 | 0.777615016 | 0.25606685 |
| ENSMUSG00000036158 | Prickle1 | 106042 | protein_coding | 16.73154585 | -0.256081676 | 0.254138377 | 1.007646618 | 0.313624141 | 0.534874425 | 0.256081676 |
| ENSMUSG00000086938 | 4930481A15Rik | 74931 | lncRNA | 77.19805723 | 0.256211543 | 0.192620793 | -1.3301344 | 0.183473993 | 0.378310591 | 0.256211543 |
| ENSMUSG00000045257 | Morn2 | 378462 | protein_coding | 40.18825948 | 0.256393602 | 0.189765885 | -1.351104824 | 0.176661856 | 0.36877688 | 0.256393602 |
| ENSMUSG00000047675 | Rps8 | 20116 | protein_coding | 6731.473885 | -0.256416246 | 0.115285524 | 2.224184252 | 0.02613605 | 0.090408616 | 0.256416246 |
| ENSMUSG00000022814 | Umps | 22247 | protein_coding | 334.1984266 | -0.256447647 | 0.102430506 | 2.503625689 | 0.012292801 | 0.049963897 | 0.256447647 |
| ENSMUSG00000033350 | Chst2 | 54371 | protein_coding | 2.288815945 | 0.256486385 | 1.819366736 | -0.140975637 | 0.887889189 | 1 | 0.256486385 |
| ENSMUSG00000039523 | Cep104 | 230967 | protein_coding | 235.4705219 | 0.25672109 | 0.099087047 | -2.590864269 | 0.009573524 | 0.041117602 | 0.25672109 |
| ENSMUSG00000040234 | Tm7sf3 | 67623 | protein_coding | 673.8590881 | -0.256804676 | 0.067872483 | 3.783634585 | 0.000154555 | 0.001258539 | 0.256804676 |
| ENSMUSG00000044551 | 9930012K11Rik | 268759 | protein_coding | 23.43677691 | -0.25680999 | 0.234863908 | 1.093441698 | 0.274199914 | 0.490619315 | 0.25680999 |
| ENSMUSG00000097316 | Gm10516 | 100038712 | lncRNA | 98.89545416 | -0.25685421 | 0.131550383 | 1.952515871 | 0.050876989 | 0.151084803 | 0.25685421 |
| ENSMUSG00000041939 | Mvk | 17855 | protein_coding | 369.0059602 | 0.256875626 | 0.122190612 | -2.102253367 | 0.035531087 | 0.114758532 | 0.256875626 |

| ensembl_gene_id | external_gene_name | entrezgene_id | gene_biotype | baseMean | log2FoldChange | lfcSE | stat | pvalue | padj | absLog2FC |
|---|---|---|---|---|---|---|---|---|---|---|
| ENSMUSG00000071655 | Ubxn1 | 225896 | protein_coding | 888.1224897 | -0.256979498 | 0.07230569 | 3.55407018 | 0.000379318 | 0.00280271 | 0.256979498 |
| ENSMUSG00000034485 | Uaca | 72565 | protein_coding | 1233.980279 | 0.257203664 | 0.067735812 | -3.797159238 | 0.000146364 | 0.001204311 | 0.257203664 |
| ENSMUSG00000027878 | Notch2 | 18129 | protein_coding | 194.3909633 | 0.257209039 | 0.095945254 | -2.680789582 | 0.007344868 | 0.033167393 | 0.257209039 |
| ENSMUSG00000027488 | Snta1 | 20648 | protein_coding | 68.52561231 | 0.257212276 | 0.208881618 | -1.231378226 | 0.218181437 | 0.42426293 | 0.257212276 |
| ENSMUSG00000024120 | Lrpprc | 72416 | protein_coding | 996.1747949 | -0.257255983 | 0.088662907 | 2.901506291 | 0.003713733 | 0.019034324 | 0.257255983 |
| ENSMUSG00000037572 | Wdhd1 | 218973 | protein_coding | 156.962421 | 0.257263831 | 0.134031033 | -1.919434812 | 0.054929329 | 0.159785615 | 0.257263831 |
| ENSMUSG00000038127 | Ccdc50 | 67501 | protein_coding | 526.0414112 | -0.257350533 | 0.068227537 | 3.771945258 | 0.00016198 | 0.001312529 | 0.257350533 |
| ENSMUSG00000050777 | Tmem37 | 170706 | protein_coding | 181.5634369 | -0.257389828 | 0.195810775 | 1.314482454 | 0.188683899 | 0.385300595 | 0.257389828 |
| ENSMUSG00000003037 | Rab8a | 17274 | protein_coding | 1489.743873 | 0.257437932 | 0.046007329 | -5.59558525 | 2.20E-08 | 4.01E-07 | 0.257437932 |
| ENSMUSG00000021575 | Ahrr | 11624 | protein_coding | 4.727294468 | 0.257478073 | 0.881576736 | -0.292065412 | 0.770236611 | 0.884771487 | 0.257478073 |
| ENSMUSG00000015314 | Slamf6 | 30925 | protein_coding | 3.751085303 | 0.257506042 | 1.42367839 | -0.180873745 | 0.856466677 | 1 | 0.257506042 |
| ENSMUSG00000019892 | Lrriq1 | 74978 | protein_coding | 1.927943457 | -0.257543301 | 1.767529175 | 0.145708091 | 0.884151834 | 1 | 0.257543301 |
| ENSMUSG00000051510 | Mafg | 17134 | protein_coding | 343.8226761 | -0.257581208 | 0.109025414 | 2.362579496 | 0.018148249 | 0.067913025 | 0.257581208 |
| ENSMUSG00000024887 | Asah2 | 54447 | protein_coding | 288.8512722 | 0.257744841 | 0.145864039 | -1.76702114 | 0.077224688 | 0.204931463 | 0.257744841 |
| ENSMUSG00000071650 | Ganab | 14376 | protein_coding | 2872.642326 | 0.257787275 | 0.094062429 | -2.740597674 | 0.006132755 | 0.028656498 | 0.257787275 |
| ENSMUSG00000023336 | Wfdc1 | 67866 | protein_coding | 1.182022123 | 0.257896853 | 1.41275044 | -0.182549476 | 0.85515153 | 1 | 0.257896853 |
| ENSMUSG00000086843 | E030013I19Rik | 338535 | lncRNA | 1.991781034 | 0.258010087 | 0.832555709 | -0.309901288 | 0.756636023 | 1 | 0.258010087 |
| ENSMUSG00000020309 | Chac2 | 68044 | protein_coding | 110.5356076 | 0.258079314 | 0.115451263 | -2.235396202 | 0.025391349 | 0.088425293 | 0.258079314 |
| ENSMUSG00000003436 | Dll3 | 13389 | protein_coding | 4.481916371 | 0.258346906 | 0.605313149 | -0.426798768 | 0.669525904 | 0.822609935 | 0.258346906 |
| ENSMUSG00000020268 | Lyrm7 | 75530 | protein_coding | 61.6628164 | 0.258639057 | 0.168313714 | -1.536648744 | 0.124379349 | 0.288970255 | 0.258639057 |
| ENSMUSG00000047030 | Spata2 | 263876 | protein_coding | 624.9711988 | -0.258643938 | 0.061503977 | 4.205320519 | 2.61E-05 | 0.000261409 | 0.258643938 |
| ENSMUSG00000023990 | Tfeb | 21425 | protein_coding | 235.6683713 | -0.258696578 | 0.118629037 | 2.180718865 | 0.029204217 | 0.098824478 | 0.258696578 |
| ENSMUSG00000039367 | Sec24c | 218811 | protein_coding | 1755.822346 | 0.258748702 | 0.051693129 | -5.005475753 | 5.57E-07 | 8.14E-06 | 0.258748702 |
| ENSMUSG00000061882 | Ccdc62 | 208908 | protein_coding | 22.13431048 | 0.258752684 | 0.347504182 | -0.74460308 | 0.456511706 | 0.671491432 | 0.258752684 |
| ENSMUSG00000094410 | Zbed6 | 667118 | protein_coding | 64.56103782 | 0.258761987 | 0.200813401 | -1.288569315 | 0.197547857 | 0.397745099 | 0.258761987 |
| ENSMUSG00000047603 | Zfp235 | 56525 | protein_coding | 117.132934 | -0.258769747 | 0.103096718 | 2.509970759 | 0.012074116 | 0.049378888 | 0.258769747 |
| ENSMUSG00000021235 | Coq6 | 217707 | protein_coding | 276.4942108 | -0.258800349 | 0.077134527 | 3.355181657 | 0.000793129 | 0.005227463 | 0.258800349 |
| ENSMUSG00000065430 | Mir26a-2 | 723962 | miRNA | 0.861081691 | 0.258802322 | 1.135238467 | -0.227971769 | 0.819668187 | 1 | 0.258802322 |
| ENSMUSG00000017466 | Timp2 | 21858 | protein_coding | 60.34412327 | -0.258949718 | 0.202027208 | 1.281756659 | 0.199928022 | 0.400972961 | 0.258949718 |
| ENSMUSG00000019558 | Slc6a8 | 102857 | protein_coding | 3072.635941 | -0.258954601 | 0.16785704 | 1.542709207 | 0.122901347 | 0.286211349 | 0.258954601 |
| ENSMUSG00000092827 | Mir3091 | 100526556 | miRNA | 2.286772159 | -0.258984816 | 0.729480014 | 0.355026609 | 0.722569657 | 1 | 0.258984816 |
| ENSMUSG00000027230 | Creb3l1 | 26427 | protein_coding | 1394.048887 | 0.259004983 | 0.128947863 | -2.008602362 | 0.044579321 | 0.136567098 | 0.259004983 |
| ENSMUSG00000007892 | Rplp1 | 56040 | protein_coding | 8039.370389 | -0.259050386 | 0.096080903 | 2.696169358 | 0.007014199 | 0.031962608 | 0.259050386 |
| ENSMUSG00000025777 | Gdap1 | 14545 | protein_coding | 2.092761502 | 0.259092501 | 1.391288791 | -0.186224818 | 0.852268463 | 1 | 0.259092501 |
| ENSMUSG00000030037 | Mrpl53 | 68499 | protein_coding | 0.731297528 | -0.259346446 | 1.299964776 | 0.199502672 | 0.841869554 | 1 | 0.259346446 |
| ENSMUSG00000043336 | Filip1l | 78749 | protein_coding | 368.3515593 | -0.259406357 | 0.17658554 | 1.469012457 | 0.141829411 | 0.316830918 | 0.259406357 |
| ENSMUSG00000020260 | Pofut2 | 80294 | protein_coding | 460.8748154 | 0.25940684 | 0.072234985 | -3.591152419 | 0.000329219 | 0.002472362 | 0.25940684 |
| ENSMUSG00000059674 | Cdh24 | 239096 | protein_coding | 30.03725878 | 0.25944757 | 0.201194552 | -1.289535763 | 0.197211892 | 0.397165509 | 0.25944757 |
| ENSMUSG00000032091 | Tmprss4 | 214523 | protein_coding | 2320.444617 | 0.259533107 | 0.049101476 | -5.285647811 | 1.25E-07 | 2.03E-06 | 0.259533107 |

| ensembl_gene_id | external_gene_name | entrezgene_id | gene_biotype | baseMean | log2FoldChange | lfcSE | stat | pvalue | padj | absLog2FC |
|---|---|---|---|---|---|---|---|---|---|---|
| ENSMUSG00000030754 | Copb1 | 70349 | protein_coding | 2438.932301 | 0.259569979 | 0.114013141 | -2.276667207 | 0.022806106 | 0.081360993 | 0.259569979 |
| ENSMUSG00000025241 | Fyco1 | 17281 | protein_coding | 536.4247948 | -0.259571049 | 0.10059322 | 2.580403031 | 0.009868507 | 0.042154299 | 0.259571049 |
| ENSMUSG00000030208 | Emp1 | 13730 | protein_coding | 7661.737637 | -0.259580256 | 0.208905982 | 1.24256976 | 0.214026419 | 0.419019832 | 0.259580256 |
| ENSMUSG00000029060 | Mib2 | 76580 | protein_coding | 420.4859231 | -0.259604599 | 0.104213355 | 2.491087621 | 0.012735271 | 0.051357818 | 0.259604599 |
| ENSMUSG00000024827 | Gldc | 104174 | protein_coding | 5.387436725 | 0.25964698 | 0.653389937 | -0.397384418 | 0.691084004 | 0.835534172 | 0.25964698 |
| ENSMUSG00000052397 | Ezr | 22350 | protein_coding | 7506.532088 | -0.259688605 | 0.103590056 | 2.50688739 | 0.012179951 | 0.049685066 | 0.259688605 |
| ENSMUSG00000015355 | Cd48 | 12506 | protein_coding | 5.659539645 | -0.259689033 | 0.548007178 | 0.47387889 | 0.635586274 | 0.80133583 | 0.259689033 |
| ENSMUSG00000039542 | Ncam1 | 17967 | protein_coding | 4.960362913 | 0.259696345 | 1.196352399 | -0.217073452 | 0.828151103 | 0.913297021 | 0.259696345 |
| ENSMUSG00000075227 | Znhit2 | 29805 | protein_coding | 194.5982238 | 0.259735632 | 0.131947076 | -1.968483426 | 0.049012441 | 0.146868346 | 0.259735632 |
| ENSMUSG00000032531 | Amotl2 | 56332 | protein_coding | 89.01379768 | -0.259789731 | 0.225418522 | 1.152477303 | 0.249124991 | 0.461264989 | 0.259789731 |
| ENSMUSG00000075590 | Nrbp2 | 223649 | protein_coding | 264.3590152 | -0.259830427 | 0.099089597 | 2.622176637 | 0.008737014 | 0.038193919 | 0.259830427 |
| ENSMUSG00000079559 | Pou2af3 | 100502940 | protein_coding | 11.53395845 | -0.259866375 | 0.373570535 | 0.695628672 | 0.486661406 | 0.695624496 | 0.259866375 |
| ENSMUSG00000047910 | Pcdhb16 | 93887 | protein_coding | 4.047278109 | 0.259885989 | 0.659539472 | -0.3940416 | 0.693550326 | 1 | 0.259885989 |
| ENSMUSG00000002741 | Ykt6 | 56418 | protein_coding | 1279.16101 | 0.259913361 | 0.065640401 | -3.95965529 | 7.51E-05 | 0.000670367 | 0.259913361 |
| ENSMUSG00000020078 | Vps26a | 30930 | protein_coding | 626.502784 | -0.259915418 | 0.057191473 | 4.544653324 | 5.50E-06 | 6.52E-05 | 0.259915418 |
| ENSMUSG00000030882 | Dnhd1 | 77505 | protein_coding | 13.24027291 | 0.259944861 | 0.538362611 | -0.482843451 | 0.629206897 | 0.797446101 | 0.259944861 |
| ENSMUSG00000032375 | Aph1b | 208117 | protein_coding | 182.8007955 | 0.260026006 | 0.108167879 | -2.403911469 | 0.016220701 | 0.06221406 | 0.260026006 |
| ENSMUSG00000022881 | Rfc4 | 106344 | protein_coding | 117.3732331 | 0.260069865 | 0.166110899 | -1.565639982 | 0.117432924 | 0.27747708 | 0.260069865 |
| ENSMUSG00000003812 | Dnase2a | 13423 | protein_coding | 87.59658385 | -0.260077381 | 0.166973851 | 1.557593474 | 0.119329647 | 0.280913726 | 0.260077381 |
| ENSMUSG00000024135 | Srbd1 | 78586 | protein_coding | 240.5399569 | 0.260098263 | 0.107562622 | -2.418110089 | 0.015601356 | 0.06023867 | 0.260098263 |
| ENSMUSG00000048832 | Vps37c | 107305 | protein_coding | 590.565986 | 0.260129223 | 0.104103309 | -2.498760364 | 0.012462855 | 0.050530713 | 0.260129223 |
| ENSMUSG00000026831 | Pierce1 | 69327 | protein_coding | 0.536437832 | 0.260141527 | 1.289019517 | -0.20181349 | 0.840062535 | 1 | 0.260141527 |
| ENSMUSG00000021587 | Pcsk1 | 18548 | protein_coding | 252.5357674 | -0.260163536 | 0.171674065 | 1.515450431 | 0.129658374 | 0.297380166 | 0.260163536 |
| ENSMUSG00000051329 | Nup160 | 59015 | protein_coding | 366.4279562 | 0.260181646 | 0.106343377 | -2.446618242 | 0.014420349 | 0.056508103 | 0.260181646 |
| ENSMUSG00000054226 | Tprkb | 69786 | protein_coding | 157.9202734 | -0.260202647 | 0.101312093 | 2.568327625 | 0.010219052 | 0.043322217 | 0.260202647 |
| ENSMUSG00000045555 | Mettl24 | 327747 | protein_coding | 15.40316303 | 0.260257892 | 0.285646743 | -0.911118012 | 0.362233197 | 0.584685663 | 0.260257892 |
| ENSMUSG00000098661 | Mir7052 | 102465637 | miRNA | 0.75556368 | 0.260285738 | 1.291148032 | -0.201592483 | 0.840235322 | 1 | 0.260285738 |
| ENSMUSG00000056608 | Chd9 | 109151 | protein_coding | 992.9852421 | 0.260324238 | 0.101715946 | -2.559325741 | 0.010487542 | 0.044271797 | 0.260324238 |
| ENSMUSG00000061536 | Sec22c | 215474 | protein_coding | 93.37995713 | 0.260350338 | 0.187203958 | -1.390730945 | 0.16430703 | 0.35060942 | 0.260350338 |
| ENSMUSG00000060636 | Rpl35a | 57808 | protein_coding | 2022.019922 | -0.260361813 | 0.102250664 | 2.546309261 | 0.010886869 | 0.045492347 | 0.260361813 |
| ENSMUSG00000027765 | P2ry1 | 18441 | protein_coding | 287.0346013 | -0.260364969 | 0.113912239 | 2.285662819 | 0.022273992 | 0.079911515 | 0.260364969 |
| ENSMUSG00000084925 | 1810062O18Rik | 75602 | lncRNA | 7.985126977 | -0.2604238 | 0.414523323 | 0.628248847 | 0.529840935 | 0.727270652 | 0.2604238 |
| ENSMUSG00000028703 | Lrrc41 | 230654 | protein_coding | 418.144449 | 0.260442749 | 0.090093179 | -2.890815396 | 0.003842438 | 0.019566455 | 0.260442749 |
| ENSMUSG00000007030 | Vwa7 | 27762 | protein_coding | 8.388714714 | -0.260451487 | 0.554601526 | 0.469619132 | 0.638627153 | 0.80301103 | 0.260451487 |
| ENSMUSG00000025069 | Gsto2 | 68214 | protein_coding | 11.75948833 | 0.260488366 | 0.311421116 | -0.836450558 | 0.40290148 | 0.624274258 | 0.260488366 |
| ENSMUSG00000039345 | Mettl22 | 239706 | protein_coding | 52.89631017 | -0.26050645 | 0.185244574 | 1.406283834 | 0.159639859 | 0.343804463 | 0.26050645 |
| ENSMUSG00000021250 | Fos | 14281 | protein_coding | 19884.91458 | -0.260515864 | 0.162831376 | 1.599911953 | 0.109618117 | 0.264383465 | 0.260515864 |
| ENSMUSG00000000305 | Cdh4 | 12561 | protein_coding | 6.699020939 | -0.26063473 | 0.859697978 | 0.30317011 | 0.761760227 | 0.880656876 | 0.26063473 |
| ENSMUSG00000070287 | Slc35g2 | 245020 | protein_coding | 68.03923766 | -0.260637041 | 0.258181872 | 1.009509453 | 0.31273037 | 0.533791146 | 0.260637041 |

Supplementary Table 1

| ensembl_gene_id | external_gene_name | entrezgene_id | gene_biotype | baseMean | log2FoldChange | lfcSE | stat | pvalue | padj | absLog2FC |
|---|---|---|---|---|---|---|---|---|---|---|
| ENSMUSG00000050212 | Eva1b | 230752 | protein_coding | 21.24225523 | -0.260685031 | 0.2782236 | 0.936962324 | 0.348777938 | 0.570608171 | 0.260685031 |
| ENSMUSG00000027809 | Etfdh | 66841 | protein_coding | 1270.387727 | -0.260719489 | 0.069484232 | 3.752210854 | 0.000175282 | 0.001412003 | 0.260719489 |
| ENSMUSG00000107388 | Gm42788 | 105242579 | lncRNA | 0.768561446 | -0.26072896 | 1.551990769 | 0.167996463 | 0.866586061 | 1 | 0.26072896 |
| ENSMUSG00000023959 | Clic5 | 224796 | protein_coding | 1981.182429 | -0.260753968 | 0.17298097 | 1.507414185 | 0.131704525 | 0.30074233 | 0.260753968 |
| ENSMUSG00000023942 | Slc29a1 | 63959 | protein_coding | 192.4175495 | 0.260776268 | 0.163917486 | -1.590899627 | 0.111632167 | 0.267599593 | 0.260776268 |
| ENSMUSG00000022807 | Osbpl11 | 106326 | protein_coding | 308.7592968 | -0.260816368 | 0.115960091 | 2.249190779 | 0.024500361 | 0.08602548 | 0.260816368 |
| ENSMUSG00000069806 | Cacng7 | 81904 | protein_coding | 1.732613599 | 0.260864737 | 2.083864862 | -0.125183135 | 0.900378568 | 1 | 0.260864737 |
| ENSMUSG00000035992 | Fnip1 | 216742 | protein_coding | 420.6990944 | 0.260892133 | 0.113926804 | -2.289997814 | 0.022021443 | 0.079125787 | 0.260892133 |
| ENSMUSG00000055370 | Gm9968 | 791329 | lncRNA | 6.194056111 | 0.260895364 | 0.498033482 | -0.523851052 | 0.600382142 | 0.776893175 | 0.260895364 |
| ENSMUSG00000061859 | Patj | 12695 | protein_coding | 1448.16689 | -0.260943477 | 0.066644199 | 3.915471716 | 9.02E-05 | 0.000787953 | 0.260943477 |
| ENSMUSG00000065049 | Gm24041 | 115488605 | snRNA | 1.122783556 | -0.261046832 | 1.143269787 | 0.228333535 | 0.819386956 | 1 | 0.261046832 |
| ENSMUSG00000027423 | Snx5 | 69178 | protein_coding | 1241.541243 | -0.261062845 | 0.090520746 | 2.884011181 | 0.003926448 | 0.019914424 | 0.261062845 |
| ENSMUSG00000030007 | Cct7 | 12468 | protein_coding | 1984.552697 | 0.261070971 | 0.088165404 | -2.961149834 | 0.003064928 | 0.016170677 | 0.261070971 |
| ENSMUSG00000087480 | Gm15910 | 100504616 | lncRNA | 3.393473218 | -0.261151365 | 0.823452958 | 0.317141813 | 0.751135999 | 1 | 0.261151365 |
| ENSMUSG00000106617 | Gm36266 | 108169076 | lncRNA | 30.17615316 | -0.2611736 | 0.243927922 | 1.07069989 | 0.284304392 | 0.502676953 | 0.2611736 |
| ENSMUSG00000040220 | Gas8 | 104346 | protein_coding | 99.16321582 | 0.261236759 | 0.124916261 | -2.09129506 | 0.036501622 | 0.117137153 | 0.261236759 |
| ENSMUSG00000024130 | Abca3 | 27410 | protein_coding | 944.5200439 | -0.261263943 | 0.087824703 | 2.974834361 | 0.002931468 | 0.015558323 | 0.261263943 |
| ENSMUSG00000057315 | Arhgap24 | 231532 | protein_coding | 172.8827879 | -0.261300781 | 0.174227958 | 1.499763776 | 0.133675604 | 0.303731277 | 0.261300781 |
| ENSMUSG00000021963 | Sap18 | 20220 | protein_coding | 447.7547265 | -0.261338514 | 0.102525023 | 2.549021748 | 0.010802556 | 0.045254367 | 0.261338514 |
| ENSMUSG00000014776 | Nol3 | 78688 | protein_coding | 33.99063604 | 0.261357411 | 0.186135103 | -1.404127468 | 0.160280896 | 0.34473544 | 0.261357411 |
| ENSMUSG00000028070 | Naxe | 246703 | protein_coding | 592.0439133 | -0.261362402 | 0.082039366 | 3.185816923 | 0.001443459 | 0.008606748 | 0.261362402 |
| ENSMUSG00000037638 | Zbtb42 | 382639 | protein_coding | 283.202064 | -0.261364675 | 0.090542266 | 2.88665932 | 0.003893556 | 0.019774484 | 0.261364675 |
| ENSMUSG00000000295 | Hddc2 | 69692 | protein_coding | 137.1426641 | 0.261376926 | 0.124756153 | -2.09510248 | 0.036161884 | 0.116182363 | 0.261376926 |
| ENSMUSG00000054630 | Ugt2b5 | 22238 | protein_coding | 504.7003075 | 0.261403209 | 0.147195617 | -1.775889894 | 0.075751067 | 0.201799181 | 0.261403209 |
| ENSMUSG00000041560 | Nop53 | 68077 | protein_coding | 1627.534098 | -0.261471086 | 0.078768226 | 3.319499494 | 0.00090179 | 0.005835484 | 0.261471086 |
| ENSMUSG00000045763 | Basp1 | 70350 | protein_coding | 3.034096066 | -0.261471636 | 0.800154926 | 0.326776262 | 0.743837115 | 1 | 0.261471636 |
| ENSMUSG00000018678 | Sp2 | 78912 | protein_coding | 105.1233526 | 0.261572026 | 0.219836655 | -1.18984719 | 0.234106458 | 0.443634979 | 0.261572026 |
| ENSMUSG00000026566 | Mpzl1 | 68481 | protein_coding | 243.5235323 | 0.26158711 | 0.116386556 | -2.247571523 | 0.024603525 | 0.08629118 | 0.26158711 |
| ENSMUSG00000029063 | Nadk | 192185 | protein_coding | 2370.977621 | -0.261607421 | 0.051898485 | 5.040752602 | 4.64E-07 | 6.90E-06 | 0.261607421 |
| ENSMUSG00000019189 | Rnf145 | 74315 | protein_coding | 999.0539551 | 0.261634785 | 0.111395023 | -2.348711612 | 0.018838491 | 0.070067307 | 0.261634785 |
| ENSMUSG00000089712 | Gm15889 | 115486954 | lncRNA | 9.739251029 | -0.26169447 | 0.538155745 | 0.486280175 | 0.626768533 | 0.795699228 | 0.26169447 |
| ENSMUSG00000055555 | Ct55 | 75013 | protein_coding | 0.776276256 | 0.261695311 | 1.551387458 | -0.168684689 | 0.866044663 | 1 | 0.261695311 |
| ENSMUSG00000027466 | Rbck1 | 24105 | protein_coding | 1055.365863 | 0.261733392 | 0.053042994 | -4.934363142 | 8.04E-07 | 1.13E-05 | 0.261733392 |
| ENSMUSG00000041000 | Trim62 | 67525 | protein_coding | 70.56497415 | -0.261775564 | 0.203615491 | 1.285636781 | 0.198569854 | 0.398975857 | 0.261775564 |
| ENSMUSG00000037031 | Tspan15 | 70423 | protein_coding | 1429.986491 | -0.261794868 | 0.087214207 | 3.001745657 | 0.002684364 | 0.014482058 | 0.261794868 |
| ENSMUSG00000029614 | Rpl6 | 19988 | protein_coding | 3733.03617 | -0.261878671 | 0.122023846 | 2.146127006 | 0.031862847 | 0.105765439 | 0.261878671 |
| ENSMUSG00000028919 | Arhgef19 | 213649 | protein_coding | 317.6013869 | -0.261987632 | 0.148373318 | 1.76573279 | 0.077440689 | 0.205372657 | 0.261987632 |
| ENSMUSG00000025245 | Lztfl1 | 93730 | protein_coding | 228.6916818 | 0.262079705 | 0.092357158 | -2.837676146 | 0.004544326 | 0.022468565 | 0.262079705 |
| ENSMUSG00000062980 | Cped1 | 214642 | protein_coding | 10.07539681 | 0.262126885 | 0.41049658 | -0.638560461 | 0.52310891 | 0.722040793 | 0.262126885 |

| ensembl_gene_id | external_gene_name | entrezgene_id | gene_biotype | baseMean | log2FoldChange | lfcSE | stat | pvalue | padj | absLog2FC |
|---|---|---|---|---|---|---|---|---|---|---|
| ENSMUSG00000006546 | Cryba2 | 12958 | protein_coding | 3.815871008 | 0.262361024 | 0.691512268 | -0.379401836 | 0.704389487 | 1 | 0.262361024 |
| ENSMUSG00000074415 | Mir100hg | 73144 | lncRNA | 7.137061925 | 0.262373999 | 1.443822281 | -0.18172181 | 0.855801049 | 0.929905153 | 0.262373999 |
| ENSMUSG00000029661 | Col1a2 | 12843 | protein_coding | 10.94207406 | 0.2624602 | 0.622488327 | -0.421630718 | 0.673294581 | 0.824897384 | 0.2624602 |
| ENSMUSG00000032086 | Bace1 | 23821 | protein_coding | 253.2294733 | -0.262601166 | 0.104804016 | 2.505640299 | 0.01222299 | 0.049790441 | 0.262601166 |
| ENSMUSG00000026024 | Als2 | 74018 | protein_coding | 197.8154299 | 0.262620974 | 0.145845134 | -1.800683825 | 0.071752729 | 0.194536949 | 0.262620974 |
| ENSMUSG00000032212 | Sltm | 66660 | protein_coding | 807.7801081 | 0.262658188 | 0.060259657 | -4.358773354 | 1.31E-05 | 0.000142007 | 0.262658188 |
| ENSMUSG00000045394 | Epcam | 17075 | protein_coding | 16955.67628 | -0.262666805 | 0.069394517 | 3.78512331 | 0.000153632 | 0.001253501 | 0.262666805 |
| ENSMUSG00000025395 | Prim1 | 19075 | protein_coding | 133.539156 | 0.262696864 | 0.213492199 | -1.230475236 | 0.218519195 | 0.424602824 | 0.262696864 |
| ENSMUSG00000023572 | Ccndbp1 | 17151 | protein_coding | 317.0115465 | -0.262711944 | 0.092809665 | 2.830652874 | 0.004645311 | 0.022878813 | 0.262711944 |
| ENSMUSG00000063810 | Alms1 | 236266 | protein_coding | 176.1286211 | -0.262768658 | 0.106026827 | 2.478322365 | 0.013200182 | 0.052845529 | 0.262768658 |
| ENSMUSG00000071893 | Vmn1r4 | 171194 | protein_coding | 3.164694368 | -0.262808786 | 0.697330773 | 0.376878228 | 0.706264102 | 1 | 0.262808786 |
| ENSMUSG00000114213 | A430090L17Rik | 319803 | lncRNA | 4.695104736 | -0.262835114 | 0.523360255 | 0.502206868 | 0.615522012 | 0.787780401 | 0.262835114 |
| ENSMUSG00000021189 | Atxn3 | 110616 | protein_coding | 465.270507 | -0.262869782 | 0.136700194 | 1.922965677 | 0.05448436 | 0.159023459 | 0.262869782 |
| ENSMUSG00000024268 | Celf4 | 108013 | protein_coding | 14.81588526 | -0.262903676 | 0.567201105 | 0.463510514 | 0.64299848 | 0.805274537 | 0.262903676 |
| ENSMUSG00000028040 | Efna4 | 13639 | protein_coding | 90.08655391 | 0.262945115 | 0.142456005 | -1.845798747 | 0.064921433 | 0.180947387 | 0.262945115 |
| ENSMUSG00000029186 | Pi4k2b | 67073 | protein_coding | 1724.302847 | 0.263014818 | 0.065096458 | -4.040385973 | 5.34E-05 | 0.000494524 | 0.263014818 |
| ENSMUSG00000020485 | Supt4a | 20922 | protein_coding | 222.5152332 | -0.263100487 | 0.08515012 | 3.08984281 | 0.002002625 | 0.011349122 | 0.263100487 |
| ENSMUSG00000022151 | Ttc33 | 67515 | protein_coding | 209.5925292 | -0.263103331 | 0.108277999 | 2.429887259 | 0.01510352 | 0.058719535 | 0.263103331 |
| ENSMUSG00000041607 | Mbp | 17196 | protein_coding | 862.2582654 | -0.263165105 | 0.068299513 | 3.853103672 | 0.00011663 | 0.000987961 | 0.263165105 |
| ENSMUSG00000031787 | Katnb1 | 74187 | protein_coding | 253.710702 | 0.263184848 | 0.085940353 | -3.062412943 | 0.002195603 | 0.01219993 | 0.263184848 |
| ENSMUSG00000047407 | Tgif1 | 21815 | protein_coding | 435.2934613 | -0.263247305 | 0.100619414 | 2.61626753 | 0.008889688 | 0.038706422 | 0.263247305 |
| ENSMUSG00000025002 | Cyp2c55 | 72082 | protein_coding | 18703.26013 | -0.263253524 | 0.171990115 | 1.530631708 | 0.12586044 | 0.291600149 | 0.263253524 |
| ENSMUSG00000086003 | B230206L02Rik | 100039440 | lncRNA | 8.601336583 | -0.263268727 | 0.441811352 | 0.595884932 | 0.5512521 | 0.742102974 | 0.263268727 |
| ENSMUSG00000004317 | Clcn5 | 12728 | protein_coding | 449.7042287 | -0.263382488 | 0.102960636 | 2.55808917 | 0.01052491 | 0.044395495 | 0.263382488 |
| ENSMUSG00000035934 | Pknox2 | 208076 | protein_coding | 9.259520741 | -0.263383919 | 0.959712111 | 0.274440549 | 0.783746085 | 0.891560436 | 0.263383919 |
| ENSMUSG00000026963 | Tmem210 | 78217 | protein_coding | 2.621367255 | 0.263396723 | 1.055450873 | -0.249558487 | 0.802928805 | 1 | 0.263396723 |
| ENSMUSG00000047246 | H2bc6 | 319179 | protein_coding | 712.2944283 | 0.263444962 | 0.183028582 | -1.439365145 | 0.150047094 | 0.329724247 | 0.263444962 |
| ENSMUSG00000058793 | Cds2 | 110911 | protein_coding | 1176.890767 | -0.263544963 | 0.099277292 | 2.654634886 | 0.007939429 | 0.035388872 | 0.263544963 |
| ENSMUSG00000011382 | Dhdh | 71755 | protein_coding | 17.14731111 | -0.263562419 | 0.340706367 | 0.773576442 | 0.439181308 | 0.657120605 | 0.263562419 |
| ENSMUSG00000025417 | Pip4k2c | 117150 | protein_coding | 962.4817936 | -0.263571795 | 0.062565583 | 4.21272816 | 2.52E-05 | 0.00025468 | 0.263571795 |
| ENSMUSG00000051223 | Bzw1 | 66882 | protein_coding | 3147.795137 | 0.263647282 | 0.075787199 | -3.478783814 | 0.000503695 | 0.003569964 | 0.263647282 |
| ENSMUSG00000031119 | Gpc4 | 14735 | protein_coding | 283.4190775 | -0.263666824 | 0.140773504 | 1.872986159 | 0.061070296 | 0.173046478 | 0.263666824 |
| ENSMUSG00000050379 | Septin6 | 56526 | protein_coding | 74.54523466 | 0.263671249 | 0.14582667 | -1.808114034 | 0.070588754 | 0.192129489 | 0.263671249 |
| ENSMUSG00000097101 | 1810034E14Rik | 66503 | lncRNA | 75.91789605 | -0.263673776 | 0.174695273 | 1.509335496 | 0.131213068 | 0.299827398 | 0.263673776 |
| ENSMUSG00000063953 | Amd2 | 100041585 | protein_coding | 1.172139577 | -0.263685449 | 0.970198381 | 0.271785084 | 0.785787275 | 1 | 0.263685449 |
| ENSMUSG00000032306 | Mpi | 110119 | protein_coding | 1075.836843 | 0.263807306 | 0.071157688 | -3.707361962 | 0.00020943 | 0.00164613 | 0.263807306 |
| ENSMUSG00000087231 | E230016M11Rik | 320172 | lncRNA | 13.21269008 | -0.263807569 | 0.375281231 | 0.702959667 | 0.482080885 | 0.69281035 | 0.263807569 |
| ENSMUSG00000036810 | Cnep1r1 | 382030 | protein_coding | 200.2360959 | 0.263808343 | 0.102832259 | -2.565423966 | 0.01030498 | 0.043612621 | 0.263808343 |
| ENSMUSG00000042423 | Fbrs | 14123 | protein_coding | 401.3916878 | 0.263815366 | 0.196583827 | -1.341999339 | 0.17959621 | 0.372922396 | 0.263815366 |

Supplementary Table 1

| ensembl_gene_id | external_gene_name | entrezgene_id | gene_biotype | baseMean | log2FoldChange | lfcSE | stat | pvalue | padj | absLog2FC |
|---|---|---|---|---|---|---|---|---|---|---|
| ENSMUSG00000006418 | Rnf114 | 81018 | protein_coding | 1139.45537 | 0.263866543 | 0.059183197 | -4.458470604 | 8.25E-06 | 9.45E-05 | 0.263866543 |
| ENSMUSG00000040350 | Trim7 | 94089 | protein_coding | 36.19661002 | -0.264021182 | 0.26615097 | 0.991997819 | 0.32119859 | 0.542303804 | 0.264021182 |
| ENSMUSG00000030086 | Chchd6 | 66098 | protein_coding | 45.06712271 | 0.264055874 | 0.187387578 | -1.409142893 | 0.15879292 | 0.342785135 | 0.264055874 |
| ENSMUSG00000044792 | Isca1 | 69046 | protein_coding | 521.4084149 | -0.264072423 | 0.112122569 | 2.355212022 | 0.01851214 | 0.069055676 | 0.264072423 |
| ENSMUSG00000037259 | Dzank1 | 241688 | protein_coding | 3.197509579 | -0.264109551 | 1.404283351 | 0.188074259 | 0.850818439 | 1 | 0.264109551 |
| ENSMUSG00000071369 | Map3k5 | 26408 | protein_coding | 695.4748417 | 0.264135492 | 0.057190462 | -4.618523511 | 3.86E-06 | 4.75E-05 | 0.264135492 |
| ENSMUSG00000059070 | Rpl18 | 19899 | protein_coding | 4929.390872 | -0.26416624 | 0.102589588 | 2.574980989 | 0.01002456 | 0.042679767 | 0.26416624 |
| ENSMUSG00000096914 | Galntl6 | 270049 | protein_coding | 91.41345196 | 0.26417453 | 0.116230621 | -2.272847959 | 0.023035344 | 0.081931007 | 0.26417453 |
| ENSMUSG00000028932 | Psmc2 | 19181 | protein_coding | 791.196072 | 0.264309364 | 0.066273585 | -3.988155534 | 6.66E-05 | 0.000602545 | 0.264309364 |
| ENSMUSG00000087413 | Gm11266 | 102635783 | lncRNA | 1.689110431 | 0.264322206 | 0.983360061 | -0.268794937 | 0.78808749 | 1 | 0.264322206 |
| ENSMUSG00000022994 | Adcy6 | 11512 | protein_coding | 3021.129186 | -0.26436138 | 0.059358279 | 4.453656444 | 8.44E-06 | 9.64E-05 | 0.26436138 |
| ENSMUSG00000026309 | Ilkap | 67444 | protein_coding | 422.4847328 | -0.264480017 | 0.066637821 | 3.968917564 | 7.22E-05 | 0.000647282 | 0.264480017 |
| ENSMUSG00000049624 | Slc17a5 | 235504 | protein_coding | 278.5013415 | -0.264490006 | 0.096193902 | 2.749550642 | 0.005967704 | 0.028073652 | 0.264490006 |
| ENSMUSG00000042275 | Pelo | 105083 | protein_coding | 47.63686693 | 0.264584206 | 0.177557333 | -1.490133929 | 0.136189025 | 0.307874819 | 0.264584206 |
| ENSMUSG00000035890 | Rnf126 | 70294 | protein_coding | 363.4571646 | 0.26474536 | 0.091945776 | -2.879364031 | 0.003984781 | 0.020123752 | 0.26474536 |
| ENSMUSG00000079465 | Col4a3 | 12828 | protein_coding | 7.222849078 | 0.264750771 | 0.742403423 | -0.35661308 | 0.721381484 | 0.855640813 | 0.264750771 |
| ENSMUSG00000026771 | Spopl | 76857 | protein_coding | 237.6256255 | -0.264894251 | 0.090653885 | 2.922039697 | 0.003477472 | 0.01801348 | 0.264894251 |
| ENSMUSG00000023088 | Abcc1 | 17250 | protein_coding | 517.2947491 | -0.264899968 | 0.09167938 | 2.8894171 | 0.003859568 | 0.019641571 | 0.264899968 |
| ENSMUSG00000014361 | Mertk | 17289 | protein_coding | 554.3284877 | -0.26490581 | 0.127891722 | 2.071328825 | 0.038328077 | 0.12190868 | 0.26490581 |
| ENSMUSG00000113679 | Gm10432 | 100038713 | lncRNA | 4.514224801 | 0.264907021 | 0.724572782 | -0.365604433 | 0.714660266 | 0.851938182 | 0.264907021 |
| ENSMUSG00000025762 | Larp1b | 214048 | protein_coding | 774.5338933 | 0.264943341 | 0.064123139 | -4.131789971 | 3.60E-05 | 0.000350484 | 0.264943341 |
| ENSMUSG00000029445 | Hpd | 15445 | protein_coding | 582.5544579 | -0.264970043 | 0.271254802 | 0.976830791 | 0.328652929 | 0.550946551 | 0.264970043 |
| ENSMUSG00000074358 | Ccdc61 | 232933 | protein_coding | 232.0258526 | 0.264992986 | 0.07759522 | -3.415068429 | 0.00063766 | 0.004340608 | 0.264992986 |
| ENSMUSG00000058626 | Capn11 | 268958 | protein_coding | 7.396333509 | -0.265055527 | 1.376404334 | 0.192570977 | 0.847294977 | 0.924739227 | 0.265055527 |
| ENSMUSG00000026547 | Tagln2 | 21346 | protein_coding | 2280.863053 | -0.265071697 | 0.066461407 | 3.988355192 | 6.65E-05 | 0.000602545 | 0.265071697 |
| ENSMUSG00000031357 | Syap1 | 67043 | protein_coding | 823.9303119 | -0.265077278 | 0.069559782 | 3.810783631 | 0.000138527 | 0.001148988 | 0.265077278 |
| ENSMUSG00000018427 | Ypel2 | 77864 | protein_coding | 262.1144766 | -0.265093265 | 0.179152625 | 1.479706286 | 0.138951647 | 0.312060508 | 0.265093265 |
| ENSMUSG00000032867 | Fbxw8 | 231672 | protein_coding | 681.6777526 | 0.265118971 | 0.067943424 | -3.902054937 | 9.54E-05 | 0.000825955 | 0.265118971 |
| ENSMUSG00000032344 | Cgas | 214763 | protein_coding | 47.42962671 | 0.265137297 | 0.217910429 | -1.216726058 | 0.223708461 | 0.43083733 | 0.265137297 |
| ENSMUSG00000004668 | Abca13 | 268379 | protein_coding | 15.13685905 | -0.265192401 | 0.552728005 | 0.479788248 | 0.63137797 | 0.799093792 | 0.265192401 |
| ENSMUSG00000063760 | Rnf217 | 268291 | protein_coding | 137.0850299 | 0.26525939 | 0.111842592 | -2.371720704 | 0.017705469 | 0.066679159 | 0.26525939 |
| ENSMUSG00000026662 | Sephs1 | 109079 | protein_coding | 428.7277544 | 0.265279174 | 0.093782691 | -2.828658165 | 0.00467436 | 0.022973921 | 0.265279174 |
| ENSMUSG00000031849 | Comp | 12845 | protein_coding | 5.605842113 | 0.265283484 | 0.847434002 | -0.313043238 | 0.754247813 | 0.876856731 | 0.265283484 |
| ENSMUSG00000087380 | 2210408F21Rik | 73652 | lncRNA | 3.195610584 | -0.265341472 | 0.964286772 | 0.275168632 | 0.783186685 | 1 | 0.265341472 |
| ENSMUSG00000068962 | Zfp114 | 232966 | protein_coding | 2.436509382 | -0.265394098 | 0.774630555 | 0.34260732 | 0.731893894 | 1 | 0.265394098 |
| ENSMUSG00000050222 | Il17d | 239114 | protein_coding | 23.89819608 | -0.265415575 | 0.268517268 | 0.988448812 | 0.322932892 | 0.544366934 | 0.265415575 |
| ENSMUSG00000074272 | Ceacam1 | 26365 | protein_coding | 22651.19201 | -0.265434745 | 0.124723713 | 2.128181877 | 0.033322007 | 0.109351285 | 0.265434745 |
| ENSMUSG00000020709 | Adap2 | 216991 | protein_coding | 33.03879242 | -0.265454152 | 0.252924238 | 1.049540185 | 0.29392957 | 0.513157091 | 0.265454152 |
| ENSMUSG00000025165 | Sectm1a | 209588 | protein_coding | 51.33279591 | -0.265479796 | 0.362193041 | 0.73297873 | 0.463571402 | 0.677051219 | 0.265479796 |

Supplementary Table 1

| ensembl_gene_id | external_gene_name | entrezgene_id | gene_biotype | baseMean | log2FoldChange | lfcSE | stat | pvalue | padj | absLog2FC |
|---|---|---|---|---|---|---|---|---|---|---|
| ENSMUSG00000024772 | Ehd1 | 13660 | protein_coding | 835.3329967 | 0.265508081 | 0.082512819 | -3.217779784 | 0.00129187 | 0.007872301 | 0.265508081 |
| ENSMUSG00000025555 | Farp1 | 223254 | protein_coding | 604.8358411 | 0.265597705 | 0.078403953 | -3.387555012 | 0.000705186 | 0.004714752 | 0.265597705 |
| ENSMUSG00000079659 | Tmem243 | 652925 | protein_coding | 193.5276001 | -0.265605478 | 0.123466003 | 2.15124384 | 0.031456958 | 0.104670605 | 0.265605478 |
| ENSMUSG00000091243 | Vgll3 | 73569 | protein_coding | 2.785309681 | 0.265686819 | 1.260564125 | -0.21076819 | 0.833068162 | 1 | 0.265686819 |
| ENSMUSG00000097619 | 4833422M21Rik | 75768 | lncRNA | 2.191453282 | -0.265714533 | 1.66437426 | 0.159648307 | 0.873158123 | 1 | 0.265714533 |
| ENSMUSG00000042816 | Gpr151 | 240239 | protein_coding | 2.4240118 | 0.26575842 | 0.691737181 | -0.384189873 | 0.700837723 | 1 | 0.26575842 |
| ENSMUSG00000003824 | Syce2 | 71846 | protein_coding | 37.6998944 | -0.265847962 | 0.192236568 | 1.38292087 | 0.16668913 | 0.354364611 | 0.265847962 |
| ENSMUSG00000020405 | Fabp6 | 16204 | protein_coding | 45.72191873 | 0.265855022 | 0.332531703 | -0.799487748 | 0.424007648 | 0.64392701 | 0.265855022 |
| ENSMUSG00000029376 | Mthfd2l | 665563 | protein_coding | 97.0674133 | 0.265857032 | 0.112855681 | -2.355725724 | 0.018486562 | 0.068975843 | 0.265857032 |
| ENSMUSG00000031666 | Rbl2 | 19651 | protein_coding | 432.6219393 | -0.265960896 | 0.078218271 | 3.400240049 | 0.000673267 | 0.004543659 | 0.265960896 |
| ENSMUSG00000045348 | Nyap1 | 243300 | protein_coding | 102.4867385 | -0.265979928 | 0.346183835 | 0.768319896 | 0.442297157 | 0.660108021 | 0.265979928 |
| ENSMUSG00000047841 | Fndc11 | 332713 | protein_coding | 1.608105901 | 0.266048492 | 1.136492655 | -0.234096094 | 0.814910378 | 1 | 0.266048492 |
| ENSMUSG00000029420 | Rimbp2 | 231760 | protein_coding | 45.48543567 | 0.266077247 | 0.288027305 | -0.923791745 | 0.355594754 | 0.578423233 | 0.266077247 |
| ENSMUSG00000023908 | Pkmyt1 | 268930 | protein_coding | 151.5662956 | 0.266111782 | 0.133169279 | -1.998297079 | 0.045684462 | 0.139280674 | 0.266111782 |
| ENSMUSG00000063320 | 1190007I07Rik | 544717 | protein_coding | 77.31736265 | -0.266304661 | 0.129598144 | 2.05484934 | 0.039893559 | 0.125701628 | 0.266304661 |
| ENSMUSG00000090700 | Cyp4f40 | 631304 | protein_coding | 1028.11333 | 0.266322418 | 0.081073543 | -3.284948551 | 0.00102001 | 0.006461237 | 0.266322418 |
| ENSMUSG00000020297 | Nsg2 | 18197 | protein_coding | 6.354313957 | 0.266338809 | 0.565764245 | -0.470759352 | 0.637812595 | 0.802698942 | 0.266338809 |
| ENSMUSG00000029860 | Zyx | 22793 | protein_coding | 523.582034 | -0.266370601 | 0.122243214 | 2.179021577 | 0.029330065 | 0.099059146 | 0.266370601 |
| ENSMUSG00000079260 | Tmppe | 100504715 | protein_coding | 123.5686371 | -0.266479309 | 0.112810042 | 2.362194915 | 0.018167088 | 0.067952726 | 0.266479309 |
| ENSMUSG00000058435 | Btnl4 | 632126 | protein_coding | 926.0097216 | -0.266506715 | 0.113691532 | 2.344121071 | 0.019071979 | 0.070728647 | 0.266506715 |
| ENSMUSG00000034723 | Tmx4 | 52837 | protein_coding | 173.0669407 | 0.266527235 | 0.183051826 | -1.456020628 | 0.145386917 | 0.322533618 | 0.266527235 |
| ENSMUSG00000078552 | Dcdc2b | 100504491 | protein_coding | 10.43441428 | -0.266568397 | 0.40282096 | 0.66175404 | 0.508128865 | 0.711906345 | 0.266568397 |
| ENSMUSG00000022892 | App | 11820 | protein_coding | 3986.685218 | -0.266707999 | 0.036856101 | 7.236468105 | 4.61E-13 | 1.44E-11 | 0.266707999 |
| ENSMUSG00000024921 | Smarca2 | 67155 | protein_coding | 449.4488855 | -0.266761734 | 0.073003949 | 3.654072675 | 0.000258113 | 0.001988097 | 0.266761734 |
| ENSMUSG00000005986 | Ankrd13d | 68423 | protein_coding | 8.670700268 | -0.266779481 | 0.470576833 | 0.566920132 | 0.570768446 | 0.756249943 | 0.266779481 |
| ENSMUSG00000028341 | Nr4a3 | 18124 | protein_coding | 10.41121024 | -0.266866655 | 0.597924767 | 0.446321461 | 0.655365056 | 0.813003196 | 0.266866655 |
| ENSMUSG00000002985 | Apoe | 11816 | protein_coding | 35.46939338 | 0.266898909 | 0.338396645 | -0.788716179 | 0.430277909 | 0.648871201 | 0.266898909 |
| ENSMUSG00000014856 | Tmem208 | 66320 | protein_coding | 425.2527982 | -0.266920151 | 0.076868313 | 3.472434105 | 0.000515762 | 0.00364298 | 0.266920151 |
| ENSMUSG00000099632 | 2900093K20Rik | 108168734 | lncRNA | 11.31669735 | 0.266920753 | 0.328244774 | -0.813175942 | 0.416117191 | 0.636395319 | 0.266920753 |
| ENSMUSG00000102543 | Pcdhgc5 | 93708 | protein_coding | 16.83475698 | 0.266933356 | 0.318924583 | -0.836979558 | 0.402604059 | 0.624030976 | 0.266933356 |
| ENSMUSG00000074785 | Plxnc1 | 54712 | protein_coding | 6.289313702 | -0.26706617 | 0.548224687 | 0.487147289 | 0.626153956 | 0.795381772 | 0.26706617 |
| ENSMUSG00000047021 | Cfap65 | 241116 | protein_coding | 8.537294073 | 0.267195197 | 0.726960902 | -0.367550987 | 0.713208067 | 0.850882677 | 0.267195197 |
| ENSMUSG00000025237 | Parp6 | 67287 | protein_coding | 291.7437838 | -0.267210669 | 0.092240714 | 2.896884221 | 0.003768888 | 0.019227501 | 0.267210669 |
| ENSMUSG00000087006 | Gm13889 | 620695 | protein_coding | 2.280246643 | -0.267261024 | 1.069891093 | 0.249802084 | 0.802740408 | 1 | 0.267261024 |
| ENSMUSG00000104346 | Pcdhga3 | 93711 | protein_coding | 3.043964228 | 0.267338922 | 0.628034542 | -0.425675507 | 0.670344312 | 1 | 0.267338922 |
| ENSMUSG00000025780 | Itih5 | 209378 | protein_coding | 2887.531755 | -0.267475664 | 0.190001974 | 1.407752031 | 0.159204509 | 0.343330529 | 0.267475664 |
| ENSMUSG00000089417 | Gm22009 | 115489854 | scaRNA | 1584.437012 | -0.267648297 | 0.092001192 | 2.909182933 | 0.003623748 | 0.018648354 | 0.267648297 |
| ENSMUSG00000037270 | Bltp1 | 229227 | protein_coding | 1384.478442 | 0.267672642 | 0.134688183 | -1.987350609 | 0.046883557 | 0.14195729 | 0.267672642 |
| ENSMUSG00000050605 | Zfp61 | 22719 | protein_coding | 52.09649368 | 0.267673627 | 0.160755898 | -1.66509366 | 0.095894071 | 0.239648108 | 0.267673627 |

Supplementary Table 1

| ensembl_gene_id | external_gene_name | entrezgene_id | gene_biotype | baseMean | log2FoldChange | lfcSE | stat | pvalue | padj | absLog2FC |
|---|---|---|---|---|---|---|---|---|---|---|
| ENSMUSG00000015013 | Trappc2l | 59005 | protein_coding | 237.5010327 | -0.267710744 | 0.088858216 | 3.012785509 | 0.002588618 | 0.014029681 | 0.267710744 |
| ENSMUSG00000087543 | Gm16576 | 100504191 | lncRNA | 21.12522806 | -0.267773004 | 0.336662679 | 0.795374779 | 0.426395526 | 0.646068054 | 0.267773004 |
| ENSMUSG00000041872 | Il17f | 257630 | protein_coding | 0.950061646 | 0.267850176 | 1.448789142 | -0.184878647 | 0.85332422 | 1 | 0.267850176 |
| ENSMUSG00000006057 | Atp5g1 | 11951 | protein_coding | 3021.225711 | 0.267901041 | 0.10654568 | -2.514424255 | 0.011922691 | 0.048844287 | 0.267901041 |
| ENSMUSG00000033576 | Apol6 | 71939 | protein_coding | 146.5409936 | -0.267918649 | 0.218177923 | 1.227982398 | 0.219453576 | 0.425708487 | 0.267918649 |
| ENSMUSG00000031530 | Dusp4 | 319520 | protein_coding | 80.1937705 | -0.267940005 | 0.177957933 | 1.505636753 | 0.132160448 | 0.3013667 | 0.267940005 |
| ENSMUSG00000057572 | Zbtb8os | 67106 | protein_coding | 165.8178384 | -0.267971371 | 0.10551565 | 2.539636262 | 0.011096781 | 0.046264135 | 0.267971371 |
| ENSMUSG00000071660 | Ttc9c | 70387 | protein_coding | 485.0010832 | 0.26802445 | 0.082688907 | -3.24135919 | 0.001189612 | 0.007349514 | 0.26802445 |
| ENSMUSG00000031753 | Cog4 | 102339 | protein_coding | 800.3621095 | -0.268034234 | 0.060732061 | 4.413389428 | 1.02E-05 | 0.000113627 | 0.268034234 |
| ENSMUSG00000062175 | Tgif2 | 228839 | protein_coding | 4.888765697 | -0.268092956 | 0.510789328 | 0.524860135 | 0.599680424 | 0.776544903 | 0.268092956 |
| ENSMUSG00000046364 | Rpl27a | 26451 | protein_coding | 2556.229272 | -0.268105641 | 0.114526258 | 2.340997129 | 0.019232315 | 0.071195349 | 0.268105641 |
| ENSMUSG00000040557 | Mettl27 | 79565 | protein_coding | 132.4037764 | -0.268131828 | 0.110654717 | 2.423139611 | 0.015387012 | 0.059578222 | 0.268131828 |
| ENSMUSG00000061518 | Cox5b | 12859 | protein_coding | 4041.931803 | -0.268136749 | 0.096679269 | 2.773466856 | 0.005546249 | 0.026494289 | 0.268136749 |
| ENSMUSG00000007653 | Gabrb2 | 14401 | protein_coding | 2.216204121 | -0.268155215 | 1.957931288 | 0.13695844 | 0.891063646 | 1 | 0.268155215 |
| ENSMUSG00000041147 | Brca2 | 12190 | protein_coding | 81.26192107 | 0.26816452 | 0.161079257 | -1.664798592 | 0.095952945 | 0.239722683 | 0.26816452 |
| ENSMUSG00000028577 | Plaa | 18786 | protein_coding | 735.5395576 | 0.268179658 | 0.076327031 | -3.513560735 | 0.000442143 | 0.003202436 | 0.268179658 |
| ENSMUSG00000041598 | Cdc42ep4 | 56699 | protein_coding | 942.0719346 | -0.26821291 | 0.054891147 | 4.886268981 | 1.03E-06 | 1.41E-05 | 0.26821291 |
| ENSMUSG00000005949 | Ctns | 83429 | protein_coding | 185.7954293 | -0.268215736 | 0.127557184 | 2.102709764 | 0.035491148 | 0.114696833 | 0.268215736 |
| ENSMUSG00000027510 | Rbm38 | 56190 | protein_coding | 106.1387205 | -0.268231023 | 0.216069128 | 1.241412992 | 0.214453222 | 0.419407924 | 0.268231023 |
| ENSMUSG00000047730 | Fcgbp | 215384 | protein_coding | 68572.87662 | -0.268258989 | 0.28466448 | 0.942369026 | 0.346003739 | 0.567700522 | 0.268258989 |
| ENSMUSG00000034173 | Chchd2l | 71970 | protein_coding | 7.410225192 | -0.268472016 | 0.440341731 | 0.609690149 | 0.542067081 | 0.735735661 | 0.268472016 |
| ENSMUSG00000018661 | Cog1 | 16834 | protein_coding | 867.935164 | 0.268568629 | 0.056873216 | -4.722233869 | 2.33E-06 | 2.99E-05 | 0.268568629 |
| ENSMUSG00000071281 | Zfp65 | 235907 | protein_coding | 158.954708 | -0.268593678 | 0.153062248 | 1.754800293 | 0.079293473 | 0.208654479 | 0.268593678 |
| ENSMUSG00000039826 | Trub2 | 227682 | protein_coding | 135.8706827 | -0.268647349 | 0.103199351 | 2.603188351 | 0.00923612 | 0.039959466 | 0.268647349 |
| ENSMUSG00000039552 | Rsph4a | 212892 | protein_coding | 1.612324807 | 0.26865254 | 1.428357573 | -0.188084934 | 0.850810071 | 1 | 0.26865254 |
| ENSMUSG00000027546 | Atp9a | 11981 | protein_coding | 3528.267587 | -0.268777541 | 0.094738126 | 2.837057818 | 0.004553137 | 0.022498653 | 0.268777541 |
| ENSMUSG00000020092 | Pald1 | 27355 | protein_coding | 19.59007254 | 0.26885565 | 0.292574456 | -0.918930698 | 0.358131822 | 0.580790426 | 0.26885565 |
| ENSMUSG00000040528 | Milr1 | 380732 | protein_coding | 5.94045404 | 0.268882492 | 0.529012942 | -0.508272049 | 0.611262569 | 0.784646895 | 0.268882492 |
| ENSMUSG00000049659 | Aftph | 216549 | protein_coding | 1921.60378 | 0.268937693 | 0.074487594 | -3.610503128 | 0.000305604 | 0.002315017 | 0.268937693 |
| ENSMUSG00000007812 | Zfp655 | 72611 | protein_coding | 413.3374332 | -0.269107521 | 0.092439513 | 2.911174157 | 0.003600733 | 0.018535691 | 0.269107521 |
| ENSMUSG00000045503 | Sys1 | 66460 | protein_coding | 290.9878546 | 0.269108395 | 0.068191596 | -3.946357158 | 7.93E-05 | 0.000704123 | 0.269108395 |
| ENSMUSG00000037887 | Dusp8 | 18218 | protein_coding | 253.5845112 | 0.269138306 | 0.201893988 | -1.333067461 | 0.182509667 | 0.37700811 | 0.269138306 |
| ENSMUSG00000026939 | Tmem141 | 51875 | protein_coding | 83.16784697 | -0.269155431 | 0.109300943 | 2.462517002 | 0.013796563 | 0.054614131 | 0.269155431 |
| ENSMUSG00000032773 | Chrm1 | 12669 | protein_coding | 84.04993598 | -0.269243859 | 0.233399798 | 1.153573661 | 0.248675002 | 0.460883095 | 0.269243859 |
| ENSMUSG00000017721 | Pigt | 78928 | protein_coding | 1145.179823 | 0.269246706 | 0.076270885 | -3.530137427 | 0.000415344 | 0.003030927 | 0.269246706 |
| ENSMUSG00000028469 | Npr2 | 230103 | protein_coding | 3.451989686 | -0.269251144 | 1.26395485 | 0.213022755 | 0.831309214 | 1 | 0.269251144 |
| ENSMUSG00000023106 | Denr | 68184 | protein_coding | 384.1782947 | 0.269257583 | 0.077245904 | -3.485719893 | 0.000490815 | 0.003492164 | 0.269257583 |
| ENSMUSG00000038497 | Tmco3 | 234076 | protein_coding | 986.1891214 | -0.269259672 | 0.103691858 | 2.596729155 | 0.00941161 | 0.040601707 | 0.269259672 |
| ENSMUSG00000028760 | Eif4g3 | 230861 | protein_coding | 795.4265474 | 0.269472181 | 0.065082787 | -4.140452407 | 3.47E-05 | 0.000338705 | 0.269472181 |

| ensembl_gene_id | external_gene_name | entrezgene_id | gene_biotype | baseMean | log2FoldChange | lfcSE | stat | pvalue | padj | absLog2FC |
|---|---|---|---|---|---|---|---|---|---|---|
| ENSMUSG00000039518 | Cdsn | 386463 | protein_coding | 10.40607751 | 0.269495136 | 0.366524788 | -0.735271242 | 0.462174319 | 0.675880563 | 0.269495136 |
| ENSMUSG00000043448 | Gjc2 | 118454 | protein_coding | 2.334076099 | 0.269525444 | 0.794067258 | -0.339423948 | 0.73429038 | 1 | 0.269525444 |
| ENSMUSG00000026640 | Plxna2 | 18845 | protein_coding | 733.726721 | -0.269610799 | 0.104878568 | 2.570694878 | 0.010149471 | 0.043086943 | 0.269610799 |
| ENSMUSG00000001134 | Uxt | 22294 | protein_coding | 116.0806418 | -0.269723869 | 0.123854135 | 2.177754246 | 0.029424338 | 0.099328192 | 0.269723869 |
| ENSMUSG00000069135 | Cep43 | 75296 | protein_coding | 183.3879932 | 0.26974494 | 0.110837894 | -2.433688794 | 0.014945839 | 0.058238694 | 0.26974494 |
| ENSMUSG00000035898 | Uba6 | 231380 | protein_coding | 345.4347032 | 0.2697567 | 0.099776871 | -2.703599512 | 0.006859291 | 0.031386624 | 0.2697567 |
| ENSMUSG00000078630 | Tomt | 791260 | protein_coding | 6.135913503 | -0.269841857 | 0.524067688 | 0.514898863 | 0.606623695 | 0.781418386 | 0.269841857 |
| ENSMUSG00000089789 | Rdh1 | 107605 | protein_coding | 31.85633339 | 0.269895299 | 0.212363719 | -1.270910584 | 0.203760461 | 0.405947672 | 0.269895299 |
| ENSMUSG00000062585 | Cnr2 | 12802 | protein_coding | 6.75457184 | 0.269955169 | 0.532731652 | -0.50673762 | 0.61233893 | 0.785464687 | 0.269955169 |
| ENSMUSG00000035551 | Igfbpl1 | 75426 | protein_coding | 1.739899678 | 0.270057249 | 1.12608331 | -0.239819955 | 0.810469836 | 1 | 0.270057249 |
| ENSMUSG00000020275 | Rel | 19696 | protein_coding | 74.92110834 | -0.270230229 | 0.203210816 | 1.329802394 | 0.183583387 | 0.378441649 | 0.270230229 |
| ENSMUSG00000045594 | Glb1 | 12091 | protein_coding | 988.9602574 | -0.27024638 | 0.102450057 | 2.637835334 | 0.008343709 | 0.036802352 | 0.27024638 |
| ENSMUSG00000040296 | Rigi | 230073 | protein_coding | 246.5890796 | -0.270275284 | 0.170368961 | 1.586411524 | 0.112645988 | 0.269245209 | 0.270275284 |
| ENSMUSG00000019261 | Map1s | 270058 | protein_coding | 135.2637085 | 0.270300741 | 0.111379637 | -2.426841653 | 0.015230902 | 0.059126261 | 0.270300741 |
| ENSMUSG00000025227 | Mfsd13a | 75146 | protein_coding | 148.6336079 | -0.27037843 | 0.10701931 | 2.526445281 | 0.011522334 | 0.047677228 | 0.27037843 |
| ENSMUSG00000030084 | Plxna1 | 18844 | protein_coding | 498.5922761 | 0.270395221 | 0.08820088 | -3.065674881 | 0.002171793 | 0.012108371 | 0.270395221 |
| ENSMUSG00000032193 | Ldlr | 16835 | protein_coding | 3066.898394 | -0.270399951 | 0.144777728 | 1.86769025 | 0.061805259 | 0.174291673 | 0.270399951 |
| ENSMUSG00000063882 | Uqcrh | 66576 | protein_coding | 3833.611837 | -0.270433893 | 0.075799904 | 3.567733966 | 0.000360082 | 0.002678555 | 0.270433893 |
| ENSMUSG00000030451 | Herc2 | 15204 | protein_coding | 2191.557104 | 0.270520376 | 0.067722937 | -3.994516333 | 6.48E-05 | 0.000589831 | 0.270520376 |
| ENSMUSG00000021215 | Net1 | 56349 | protein_coding | 2401.676144 | -0.270537662 | 0.071263767 | 3.796286291 | 0.00014688 | 0.001207955 | 0.270537662 |
| ENSMUSG00000015451 | C4a | 625018 | lncRNA | 8.486682681 | -0.270547007 | 0.67528489 | 0.400641286 | 0.688684244 | 0.83386713 | 0.270547007 |
| ENSMUSG00000015289 | Lage3 | 66192 | protein_coding | 134.2868906 | -0.270589834 | 0.113566016 | 2.382665555 | 0.017187802 | 0.06516055 | 0.270589834 |
| ENSMUSG00000047759 | Hs3st3a1 | 15478 | protein_coding | 12.77610813 | 0.270611999 | 0.319802337 | -0.846185183 | 0.397449446 | 0.619371535 | 0.270611999 |
| ENSMUSG00000032128 | Robo3 | 19649 | protein_coding | 4.115989278 | 0.270623254 | 1.916879387 | -0.141179072 | 0.887728478 | 1 | 0.270623254 |
| ENSMUSG00000045251 | Zfp688 | 69234 | protein_coding | 85.65081839 | -0.270692285 | 0.108118509 | 2.503662761 | 0.012291513 | 0.049963897 | 0.270692285 |
| ENSMUSG00000040681 | Hmgn1 | 15312 | protein_coding | 707.7778371 | 0.270745173 | 0.123830694 | -2.186414082 | 0.028785324 | 0.097670196 | 0.270745173 |
| ENSMUSG00000026253 | Chrng | 11449 | protein_coding | 2.318033037 | -0.270894699 | 1.632050941 | 0.165984218 | 0.86816937 | 1 | 0.270894699 |
| ENSMUSG00000022039 | Adam2 | 11495 | protein_coding | 1.57713395 | -0.270985641 | 1.836832216 | 0.147528793 | 0.882714654 | 1 | 0.270985641 |
| ENSMUSG00000023938 | Aars2 | 224805 | protein_coding | 271.9829597 | -0.27101562 | 0.085112212 | 3.184215424 | 0.001451469 | 0.008643912 | 0.27101562 |
| ENSMUSG00000027834 | Serpini1 | 20713 | protein_coding | 5.89140624 | -0.27104583 | 0.582882419 | 0.465009445 | 0.641924693 | 0.804573016 | 0.27104583 |
| ENSMUSG00000042770 | Hebp1 | 15199 | protein_coding | 4.489521221 | -0.271063955 | 0.561258309 | 0.482957581 | 0.629125855 | 0.797446101 | 0.271063955 |
| ENSMUSG00000024402 | Lta | 16992 | protein_coding | 2.009876145 | 0.271370248 | 1.218240328 | -0.222755922 | 0.823725475 | 1 | 0.271370248 |
| ENSMUSG00000022863 | Btg3 | 12228 | protein_coding | 75.74599489 | -0.271385934 | 0.232278772 | 1.168363047 | 0.242660354 | 0.453724307 | 0.271385934 |
| ENSMUSG00000009739 | Pou6f1 | 19009 | protein_coding | 27.73103172 | -0.271471073 | 0.267896795 | 1.013341997 | 0.310896834 | 0.531926058 | 0.271471073 |
| ENSMUSG00000036120 | Rfxank | 19727 | protein_coding | 186.5110694 | -0.271474988 | 0.084176716 | 3.225060318 | 0.001259461 | 0.007703947 | 0.271474988 |
| ENSMUSG00000055531 | Cpsf6 | 432508 | protein_coding | 925.5996912 | 0.271602087 | 0.064575567 | -4.205957476 | 2.60E-05 | 0.000260832 | 0.271602087 |
| ENSMUSG00000112481 | Gm29684 | 628062 | lncRNA | 1.107287656 | -0.271615637 | 1.318152018 | 0.2060579 | 0.836745684 | 1 | 0.271615637 |
| ENSMUSG00000030761 | Myo7a | 17921 | protein_coding | 290.7957043 | 0.271646505 | 0.159159411 | -1.706757415 | 0.087867151 | 0.225018236 | 0.271646505 |
| ENSMUSG00000022999 | Lmbr1l | 74775 | protein_coding | 208.2188384 | -0.271720106 | 0.14058925 | 1.932723204 | 0.053270305 | 0.156531285 | 0.271720106 |

| ensembl_gene_id | external_gene_name | entrezgene_id | gene_biotype | baseMean | log2FoldChange | lfcSE | stat | pvalue | padj | absLog2FC |
|---|---|---|---|---|---|---|---|---|---|---|
| ENSMUSG00000051314 | Ffar2 | 233079 | protein_coding | 249.7548097 | 0.271798773 | 0.192046834 | -1.415273387 | 0.156988355 | 0.340132673 | 0.271798773 |
| ENSMUSG00000038400 | Pmepa1 | 65112 | protein_coding | 98.37356302 | -0.271823642 | 0.166378383 | 1.633767787 | 0.102307619 | 0.251864642 | 0.271823642 |
| ENSMUSG00000050957 | Insl6 | 27356 | protein_coding | 35.66475697 | -0.271833052 | 0.214776288 | 1.265656719 | 0.20563601 | 0.408255959 | 0.271833052 |
| ENSMUSG00000066152 | Slc31a2 | 20530 | protein_coding | 143.044918 | -0.271947285 | 0.122161078 | 2.226136916 | 0.026005012 | 0.090068533 | 0.271947285 |
| ENSMUSG00000032492 | Pth1r | 19228 | protein_coding | 1.780535617 | 0.272001368 | 1.692807558 | -0.16068062 | 0.872344954 | 1 | 0.272001368 |
| ENSMUSG00000022130 | Tgds | 76355 | protein_coding | 178.4023564 | 0.272027242 | 0.106143782 | -2.562818441 | 0.010382633 | 0.043873796 | 0.272027242 |
| ENSMUSG00000031367 | Ap1s2 | 108012 | protein_coding | 23.04839775 | 0.272061674 | 0.283750331 | -0.958806542 | 0.337656213 | 0.559563944 | 0.272061674 |
| ENSMUSG00000034765 | Dusp5 | 240672 | protein_coding | 309.9442989 | -0.272105109 | 0.37331392 | 0.728890872 | 0.466068419 | 0.679736277 | 0.272105109 |
| ENSMUSG00000036123 | Slc9a3 | 105243 | protein_coding | 4235.618702 | -0.272136944 | 0.201440574 | 1.350953976 | 0.176710176 | 0.368831124 | 0.272136944 |
| ENSMUSG00000052155 | Acvr2a | 11480 | protein_coding | 556.1834559 | 0.272170651 | 0.076700461 | -3.548487815 | 0.00038745 | 0.002851314 | 0.272170651 |
| ENSMUSG00000018001 | Cyth3 | 19159 | protein_coding | 195.2166562 | -0.272185123 | 0.088666939 | 3.069747606 | 0.002142397 | 0.011989004 | 0.272185123 |
| ENSMUSG00000060012 | Kif13b | 16554 | protein_coding | 2807.153215 | -0.272240509 | 0.088896276 | 3.062451212 | 0.002195323 | 0.01219993 | 0.272240509 |
| ENSMUSG00000047655 | Vmn1r74 | 171240 | protein_coding | 0.990263078 | -0.272243953 | 2.653927822 | 0.102581521 | 0.918295109 | 1 | 0.272243953 |
| ENSMUSG00000051615 | Rap2a | 76108 | protein_coding | 930.4860802 | 0.272261631 | 0.072667284 | -3.746687884 | 0.000179185 | 0.001439931 | 0.272261631 |
| ENSMUSG00000038351 | Sgsm2 | 97761 | protein_coding | 370.3515703 | -0.2723031 | 0.08888544 | 3.063528741 | 0.002187432 | 0.012175009 | 0.2723031 |
| ENSMUSG00000020941 | Map3k14 | 53859 | protein_coding | 88.13626945 | -0.272340137 | 0.136838598 | 1.99022894 | 0.046565722 | 0.141279869 | 0.272340137 |
| ENSMUSG00000020532 | Acaca | 107476 | protein_coding | 919.7559771 | 0.272408031 | 0.098951601 | -2.752942121 | 0.005906234 | 0.027859338 | 0.272408031 |
| ENSMUSG00000001123 | Lgals9 | 16859 | protein_coding | 821.7128348 | 0.272414692 | 0.185675278 | -1.467156505 | 0.142333486 | 0.317806272 | 0.272414692 |
| ENSMUSG00000035456 | Prdm8 | 77630 | protein_coding | 1.496919971 | -0.272415437 | 2.656245923 | 0.102556557 | 0.918314923 | 1 | 0.272415437 |
| ENSMUSG00000041733 | Coq5 | 52064 | protein_coding | 557.3082598 | -0.272448239 | 0.063701891 | 4.276925449 | 1.89E-05 | 0.000196193 | 0.272448239 |
| ENSMUSG00000042613 | Pbxip1 | 229534 | protein_coding | 981.2944105 | -0.272464731 | 0.085332107 | 3.192991953 | 0.001408069 | 0.008461735 | 0.272464731 |
| ENSMUSG00000036904 | Fzd8 | 14370 | protein_coding | 291.9464418 | 0.272491763 | 0.116642906 | -2.336119458 | 0.019485016 | 0.071937304 | 0.272491763 |
| ENSMUSG00000025591 | Tma16 | 66282 | protein_coding | 82.92596611 | -0.272534271 | 0.159250028 | 1.711360905 | 0.08701451 | 0.22354661 | 0.272534271 |
| ENSMUSG00000026925 | Inpp5e | 64436 | protein_coding | 230.521519 | -0.272545729 | 0.110285278 | 2.471279344 | 0.013463061 | 0.053650817 | 0.272545729 |
| ENSMUSG00000086433 | Rsf1os1 | 75448 | lncRNA | 1.353482348 | -0.272557988 | 1.264771391 | 0.215499805 | 0.829377664 | 1 | 0.272557988 |
| ENSMUSG00000022040 | Ephx2 | 13850 | protein_coding | 300.8501988 | -0.272598269 | 0.10154382 | 2.684538259 | 0.007263007 | 0.03286063 | 0.272598269 |
| ENSMUSG00000056313 | Tcim | 69068 | protein_coding | 1698.250943 | -0.272622025 | 0.125404713 | 2.173937632 | 0.029709819 | 0.10016904 | 0.272622025 |
| ENSMUSG00000052229 | Gpr17 | 574402 | protein_coding | 7.245093141 | -0.272645162 | 0.693539802 | 0.393121146 | 0.694230007 | 0.837885199 | 0.272645162 |
| ENSMUSG00000021127 | Zfp36l1 | 12192 | protein_coding | 297.4017026 | -0.272718646 | 0.127033564 | 2.146823537 | 0.031807333 | 0.105623626 | 0.272718646 |
| ENSMUSG00000026238 | Ptma | 19231 | protein_coding | 4956.985153 | 0.272740592 | 0.155347687 | -1.755678481 | 0.079143323 | 0.208481869 | 0.272740592 |
| ENSMUSG00000032702 | Kank1 | 107351 | protein_coding | 534.3326967 | 0.272754803 | 0.129198444 | -2.111130702 | 0.034761079 | 0.112911971 | 0.272754803 |
| ENSMUSG00000074647 | Fam83c | 71405 | protein_coding | 164.3508341 | -0.272812866 | 0.122079728 | 2.234710632 | 0.025436352 | 0.088524117 | 0.272812866 |
| ENSMUSG00000053774 | Ubxn7 | 224111 | protein_coding | 402.4362548 | -0.272814798 | 0.098006388 | 2.783643033 | 0.005375216 | 0.025856776 | 0.272814798 |
| ENSMUSG00000019948 | Actr6 | 67019 | protein_coding | 44.43404449 | 0.272896673 | 0.193864771 | -1.407665103 | 0.15923026 | 0.343330529 | 0.272896673 |
| ENSMUSG00000055976 | Cldn23 | 71908 | protein_coding | 1184.808521 | -0.272932919 | 0.218255632 | 1.250519478 | 0.211109845 | 0.415328565 | 0.272932919 |
| ENSMUSG00000024766 | Lipo3 | 381236 | protein_coding | 208.5524654 | -0.27300293 | 0.170966085 | 1.596825063 | 0.110304707 | 0.265457875 | 0.27300293 |
| ENSMUSG00000026548 | Slamf9 | 98365 | protein_coding | 6.239719076 | 0.273004265 | 0.521396277 | -0.523602253 | 0.600555214 | 0.777056241 | 0.273004265 |
| ENSMUSG00000026489 | Coq8a | 67426 | protein_coding | 566.0013298 | -0.2730084 | 0.165126563 | 1.653328183 | 0.09826409 | 0.244240395 | 0.2730084 |
| ENSMUSG00000073468 | Sft2d1 | 106489 | protein_coding | 205.0871782 | -0.273123548 | 0.125778299 | 2.171467976 | 0.029895815 | 0.100654009 | 0.273123548 |

Supplementary Table 1

| ensembl_gene_id | external_gene_name | entrezgene_id | gene_biotype | baseMean | log2FoldChange | lfcSE | stat | pvalue | padj | absLog2FC |
|---|---|---|---|---|---|---|---|---|---|---|
| ENSMUSG00000096370 | Gm21992 | 102902673 | protein_coding | 49.54391665 | 0.273166941 | 0.160233811 | -1.704802121 | 0.088231336 | 0.225445669 | 0.273166941 |
| ENSMUSG00000030651 | Art2b | 11872 | protein_coding | 3.093952423 | -0.273349773 | 0.814348482 | 0.33566683 | 0.737122122 | 1 | 0.273349773 |
| ENSMUSG00000055198 | Gm12830 | 433746 | lncRNA | 7.278838361 | -0.273391961 | 0.387135327 | 0.706192234 | 0.480068599 | 0.691115487 | 0.273391961 |
| ENSMUSG00000013701 | Timm23 | 53600 | protein_coding | 357.4352556 | -0.273500589 | 0.168598265 | 1.622202865 | 0.104759918 | 0.255880091 | 0.273500589 |
| ENSMUSG00000103749 | Pcdhgb5 | 93702 | protein_coding | 11.52110366 | -0.273523263 | 0.323047538 | 0.846696635 | 0.397164235 | 0.619083741 | 0.273523263 |
| ENSMUSG00000024558 | Mapk4 | 225724 | protein_coding | 3.433521055 | 0.27354868 | 0.867809541 | -0.315217414 | 0.752596584 | 1 | 0.27354868 |
| ENSMUSG00000001349 | Cnn1 | 12797 | protein_coding | 1.730209806 | -0.273727896 | 1.201032647 | 0.227910454 | 0.819715855 | 1 | 0.273727896 |
| ENSMUSG00000078653 | Cntd1 | 68107 | protein_coding | 18.59462968 | 0.273752144 | 0.271899201 | -1.006814817 | 0.314023773 | 0.5351691 | 0.273752144 |
| ENSMUSG00000030987 | Stim1 | 20866 | protein_coding | 1706.451417 | -0.273772599 | 0.120327016 | 2.275238004 | 0.022891656 | 0.081578077 | 0.273772599 |
| ENSMUSG00000029175 | Slc35f6 | 74919 | protein_coding | 586.7981472 | 0.273821121 | 0.084910108 | -3.224835372 | 0.001260451 | 0.007703947 | 0.273821121 |
| ENSMUSG00000022972 | Cfap298 | 68001 | protein_coding | 69.82455986 | -0.273910794 | 0.164890261 | 1.661170233 | 0.096679262 | 0.241063164 | 0.273910794 |
| ENSMUSG00000022865 | Cxadr | 13052 | protein_coding | 2541.696885 | -0.274014119 | 0.076120023 | 3.59976401 | 0.000318506 | 0.002397361 | 0.274014119 |
| ENSMUSG00000001627 | Ifrd1 | 15982 | protein_coding | 1018.895549 | -0.274226544 | 0.23326798 | 1.17558588 | 0.239760401 | 0.450136796 | 0.274226544 |
| ENSMUSG00000035342 | Lzts2 | 226154 | protein_coding | 857.8108083 | -0.274238837 | 0.067865663 | 4.040907069 | 5.32E-05 | 0.000493703 | 0.274238837 |
| ENSMUSG00000007476 | Lrrc8a | 241296 | protein_coding | 170.6372748 | -0.274350294 | 0.150951381 | 1.817474555 | 0.069144481 | 0.189251307 | 0.274350294 |
| ENSMUSG00000054302 | Eapp | 66266 | protein_coding | 331.9961671 | -0.274406334 | 0.084219187 | 3.258240119 | 0.001121055 | 0.007009884 | 0.274406334 |
| ENSMUSG00000027876 | Reg4 | 67709 | protein_coding | 1841.309659 | 0.27444871 | 0.514558916 | -0.533366932 | 0.593779604 | 0.772952576 | 0.27444871 |
| ENSMUSG00000030695 | Aldoa | 11674 | protein_coding | 12023.19891 | -0.27447424 | 0.073567265 | 3.730928994 | 0.000190775 | 0.001517562 | 0.27447424 |
| ENSMUSG00000031434 | Morc4 | 75746 | protein_coding | 103.8819292 | 0.274486581 | 0.172529689 | -1.590952741 | 0.111620212 | 0.267599593 | 0.274486581 |
| ENSMUSG00000016179 | Camk1g | 215303 | protein_coding | 7.676924393 | 0.274591236 | 0.605268814 | -0.453668237 | 0.650067631 | 0.809917796 | 0.274591236 |
| ENSMUSG00000075467 | Dnlz | 52838 | protein_coding | 263.0263576 | -0.274625287 | 0.090150371 | 3.046302367 | 0.002316747 | 0.012761429 | 0.274625287 |
| ENSMUSG00000071262 | Zfp957 | 105590 | protein_coding | 1.695652921 | -0.274639868 | 1.510760892 | 0.181789104 | 0.855748236 | 1 | 0.274639868 |
| ENSMUSG00000030031 | Kbtbd8 | 243574 | protein_coding | 139.0976541 | 0.274713722 | 0.140175064 | -1.95979024 | 0.050020312 | 0.148948255 | 0.274713722 |
| ENSMUSG00000047635 | Mtrfr | 72650 | protein_coding | 76.02140049 | -0.274846038 | 0.1443833 | 1.903586061 | 0.056964116 | 0.164401504 | 0.274846038 |
| ENSMUSG00000020546 | Stxbp4 | 20913 | protein_coding | 35.31941154 | 0.274848767 | 0.173347473 | -1.585536622 | 0.112844463 | 0.269566536 | 0.274848767 |
| ENSMUSG00000052712 | BC004004 | 80748 | protein_coding | 505.3211445 | -0.2748639 | 0.063078876 | 4.357463493 | 1.32E-05 | 0.000142578 | 0.2748639 |
| ENSMUSG00000030691 | Fchsd2 | 207278 | protein_coding | 675.5212477 | -0.27488335 | 0.098330428 | 2.795506487 | 0.005181846 | 0.025095735 | 0.27488335 |
| ENSMUSG00000028609 | Magoh | 17149 | protein_coding | 349.6860809 | 0.27492277 | 0.10991512 | -2.501227944 | 0.012376349 | 0.050254003 | 0.27492277 |
| ENSMUSG00000065725 | Gm26165 | 115486130 | snoRNA | 49.02067933 | 0.274930224 | 0.235660161 | -1.166638534 | 0.243356375 | 0.454613934 | 0.274930224 |
| ENSMUSG00000041654 | Slc39a11 | 69806 | protein_coding | 1622.606275 | 0.274952073 | 0.100512099 | -2.735512202 | 0.006228328 | 0.028942592 | 0.274952073 |
| ENSMUSG00000040680 | Kremen2 | 73016 | protein_coding | 118.2973199 | -0.274967325 | 0.15014895 | 1.831297017 | 0.067056223 | 0.185023636 | 0.274967325 |
| ENSMUSG00000021768 | Dusp13 | 27389 | protein_coding | 5.501553555 | -0.275016412 | 0.535452523 | 0.513614933 | 0.607521237 | 0.782086507 | 0.275016412 |
| ENSMUSG00000043572 | Pars2 | 230577 | protein_coding | 97.60868005 | -0.275026553 | 0.145272995 | 1.893170534 | 0.058335195 | 0.167325234 | 0.275026553 |
| ENSMUSG00000031706 | Rfx1 | 19724 | protein_coding | 378.1262583 | 0.275064467 | 0.124847758 | -2.203199086 | 0.02758072 | 0.094358402 | 0.275064467 |
| ENSMUSG00000026031 | Cflar | 12633 | protein_coding | 924.7004247 | 0.275200917 | 0.084076959 | -3.273202568 | 0.001063362 | 0.006694122 | 0.275200917 |
| ENSMUSG00000026335 | Pam | 18484 | protein_coding | 2699.569679 | -0.275263022 | 0.164458158 | 1.673757178 | 0.094178331 | 0.236336338 | 0.275263022 |
| ENSMUSG00000005980 | Dnase1 | 13419 | protein_coding | 12.87564026 | 0.275285559 | 0.348028812 | -0.790984967 | 0.428952764 | 0.647921517 | 0.275285559 |
| ENSMUSG00000026339 | Ccdc93 | 70829 | protein_coding | 1048.404304 | -0.27529202 | 0.101803289 | 2.70415644 | 0.006847804 | 0.031346224 | 0.27529202 |
| ENSMUSG00000035069 | Oma1 | 67013 | protein_coding | 555.8675535 | -0.27533947 | 0.091183332 | 3.019625018 | 0.002530878 | 0.013766443 | 0.27533947 |

| ensembl_gene_id | external_gene_name | entrezgene_id | gene_biotype | baseMean | log2FoldChange | lfcSE | stat | pvalue | padj | absLog2FC |
|---|---|---|---|---|---|---|---|---|---|---|
| ENSMUSG00000062519 | Zfp398 | 272347 | protein_coding | 207.320157 | -0.27536192 | 0.087230874 | 3.156702513 | 0.00159564 | 0.009377369 | 0.27536192 |
| ENSMUSG00000034135 | Sik3 | 70661 | protein_coding | 287.7969799 | -0.275365548 | 0.103681707 | 2.655873985 | 0.007910317 | 0.035329044 | 0.275365548 |
| ENSMUSG00000031410 | Nxf7 | 170722 | protein_coding | 7.918472339 | -0.275393423 | 0.463591955 | 0.594042713 | 0.552483545 | 0.743094418 | 0.275393423 |
| ENSMUSG00000038388 | Pals2 | 56524 | protein_coding | 725.2380489 | 0.275434678 | 0.135665557 | -2.030247649 | 0.042331373 | 0.131303585 | 0.275434678 |
| ENSMUSG00000028568 | Btf3l4 | 70533 | protein_coding | 148.7525809 | -0.275445414 | 0.101548839 | 2.712442769 | 0.006678931 | 0.030671822 | 0.275445414 |
| ENSMUSG00000022105 | Rb1 | 19645 | protein_coding | 559.4823068 | -0.275463893 | 0.080875434 | 3.406026783 | 0.000659157 | 0.004469372 | 0.275463893 |
| ENSMUSG00000037126 | Psd | 73728 | protein_coding | 81.2862406 | 0.275475078 | 0.164027495 | -1.679444521 | 0.093065443 | 0.234222517 | 0.275475078 |
| ENSMUSG00000024539 | Ptpn2 | 19255 | protein_coding | 417.2228819 | 0.275575619 | 0.07437968 | -3.704985279 | 0.000211403 | 0.001657697 | 0.275575619 |
| ENSMUSG00000035773 | Kiss1r | 114229 | protein_coding | 18.25819984 | -0.275686987 | 0.293545727 | 0.939161981 | 0.347647586 | 0.569548832 | 0.275686987 |
| ENSMUSG00000038156 | Spon1 | 233744 | protein_coding | 16.94585536 | 0.27578355 | 0.412218097 | -0.669023394 | 0.503480555 | 0.708014489 | 0.27578355 |
| ENSMUSG00000037280 | Galnt6 | 207839 | protein_coding | 1620.472235 | -0.275804738 | 0.092414377 | 2.984435418 | 0.002841022 | 0.015207984 | 0.275804738 |
| ENSMUSG00000027176 | Cstf3 | 228410 | protein_coding | 297.3904968 | 0.275812519 | 0.100206156 | -2.752450854 | 0.005915102 | 0.027885242 | 0.275812519 |
| ENSMUSG00000022973 | Synj1 | 104015 | protein_coding | 316.4321615 | -0.27582191 | 0.147757654 | 1.866718258 | 0.061940943 | 0.174614668 | 0.27582191 |
| ENSMUSG00000096740 | Lbhd1 | 102308570 | protein_coding | 58.36055564 | 0.275878877 | 0.151780956 | -1.81761193 | 0.069123467 | 0.189251307 | 0.275878877 |
| ENSMUSG00000015087 | Rabl6 | 227624 | protein_coding | 763.9880864 | 0.275901557 | 0.066893308 | -4.124501633 | 3.72E-05 | 0.000359858 | 0.275901557 |
| ENSMUSG00000037221 | Mospd3 | 68929 | protein_coding | 62.60449672 | 0.275952199 | 0.246518876 | -1.119395819 | 0.262971314 | 0.477220689 | 0.275952199 |
| ENSMUSG00000028945 | Rheb | 19744 | protein_coding | 584.45135 | 0.276014289 | 0.054666249 | -5.049080407 | 4.44E-07 | 6.62E-06 | 0.276014289 |
| ENSMUSG00000030739 | Myh14 | 71960 | protein_coding | 11567.04495 | 0.276036414 | 0.165080265 | -1.672134551 | 0.094497793 | 0.236875615 | 0.276036414 |
| ENSMUSG00000025507 | Pidd1 | 57913 | protein_coding | 56.29588947 | 0.276041931 | 0.173558396 | -1.590484449 | 0.111725648 | 0.267745952 | 0.276041931 |
| ENSMUSG00000038930 | Rccd1 | 269955 | protein_coding | 68.7241451 | -0.276067487 | 0.177679236 | 1.553740853 | 0.120246244 | 0.282387155 | 0.276067487 |
| ENSMUSG00000041078 | Grid1 | 14803 | protein_coding | 3.455580334 | 0.276184852 | 1.337119771 | -0.206552067 | 0.836359698 | 1 | 0.276184852 |
| ENSMUSG00000022687 | Boc | 117606 | protein_coding | 881.4940742 | -0.276228171 | 0.121926139 | 2.265536939 | 0.02347975 | 0.083216624 | 0.276228171 |
| ENSMUSG00000108171 | Gm43915 | 215458 | lncRNA | 9.634940761 | 0.276235261 | 0.610535459 | -0.452447531 | 0.650946612 | 0.810260975 | 0.276235261 |
| ENSMUSG00000085929 | Gm13421 | 105244102 | lncRNA | 16.19569016 | 0.276239497 | 0.304515816 | -0.90714335 | 0.364330996 | 0.586981666 | 0.276239497 |
| ENSMUSG00000031170 | Slc38a5 | 209837 | protein_coding | 2.979517742 | 0.276288938 | 0.72733978 | -0.379862268 | 0.704047658 | 1 | 0.276288938 |
| ENSMUSG00000021676 | Iqgap2 | 544963 | protein_coding | 6303.936767 | 0.276303098 | 0.093727826 | -2.947930298 | 0.003199092 | 0.016787354 | 0.276303098 |
| ENSMUSG00000040359 | Ufl1 | 67490 | protein_coding | 382.3544299 | 0.27631276 | 0.121238866 | -2.279077416 | 0.022662464 | 0.081006045 | 0.27631276 |
| ENSMUSG00000041426 | Hibch | 227095 | protein_coding | 324.855701 | -0.276357358 | 0.095973328 | 2.879522516 | 0.003982779 | 0.020119794 | 0.276357358 |
| ENSMUSG00000085499 | Gm15713 | 626693 | lncRNA | 2.878286837 | 0.276430578 | 0.677241503 | -0.40817135 | 0.68314788 | 1 | 0.276430578 |
| ENSMUSG00000100622 | Gm20379 | 115488195 | protein_coding | 2.475493123 | -0.276435203 | 0.816168811 | 0.338698562 | 0.734836824 | 1 | 0.276435203 |
| ENSMUSG00000021577 | Sdha | 66945 | protein_coding | 5115.202895 | -0.276543195 | 0.069316252 | 3.989586674 | 6.62E-05 | 0.00060057 | 0.276543195 |
| ENSMUSG00000025486 | Sirt3 | 64384 | protein_coding | 172.8727581 | -0.276546223 | 0.130071809 | 2.126104218 | 0.033494583 | 0.109726154 | 0.276546223 |
| ENSMUSG00000050144 | Slc25a44 | 229517 | protein_coding | 567.9009961 | 0.276599484 | 0.095107087 | -2.908295203 | 0.003634051 | 0.018683911 | 0.276599484 |
| ENSMUSG00000072612 | Gm10382 | 639281 | lncRNA | 238.8920579 | 0.276737404 | 0.234352979 | -1.180857205 | 0.237659455 | 0.447999484 | 0.276737404 |
| ENSMUSG00000022711 | Pmm2 | 54128 | protein_coding | 2269.755702 | 0.276747727 | 0.050922905 | -5.434641426 | 5.49E-08 | 9.36E-07 | 0.276747727 |
| ENSMUSG00000040412 | Elapor1 | 229722 | protein_coding | 3228.096386 | -0.27679775 | 0.173757495 | 1.593011857 | 0.111157531 | 0.26676599 | 0.27679775 |
| ENSMUSG00000022037 | Clu | 12759 | protein_coding | 46.70961116 | -0.27685908 | 0.30879098 | 0.896590567 | 0.369937434 | 0.59260115 | 0.27685908 |
| ENSMUSG00000035329 | Fbxo33 | 70611 | protein_coding | 329.3806769 | 0.276874185 | 0.077868148 | -3.555679584 | 0.000377003 | 0.002788738 | 0.276874185 |
| ENSMUSG00000027309 | Dnaaf9 | 228602 | protein_coding | 208.9466692 | -0.277009716 | 0.158366785 | 1.749165499 | 0.080262416 | 0.21050366 | 0.277009716 |

| ensembl_gene_id | external_gene_name | entrezgene_id | gene_biotype | baseMean | log2FoldChange | lfcSE | stat | pvalue | padj | absLog2FC |
|---|---|---|---|---|---|---|---|---|---|---|
| ENSMUSG00000098176 | Ccdc166 | 223648 | protein_coding | 74.12393305 | -0.277208705 | 0.164229931 | 1.68793047 | 0.091424577 | 0.231384745 | 0.277208705 |
| ENSMUSG00000093803 | Ppp2r3d | 19054 | protein_coding | 295.97457 | -0.277265644 | 0.122669521 | 2.260265158 | 0.023804799 | 0.084106216 | 0.277265644 |
| ENSMUSG00000033159 | Cnppd1 | 69171 | protein_coding | 673.9541981 | -0.277268821 | 0.082814081 | 3.348087895 | 0.000813712 | 0.005342998 | 0.277268821 |
| ENSMUSG00000060002 | Chpt1 | 212862 | protein_coding | 1068.314852 | -0.277308752 | 0.173792168 | 1.595634346 | 0.110570455 | 0.265929452 | 0.277308752 |
| ENSMUSG00000027333 | Smox | 228608 | protein_coding | 246.4206691 | -0.277338309 | 0.120225671 | 2.306814391 | 0.02106517 | 0.076471798 | 0.277338309 |
| ENSMUSG00000032661 | Oas3 | 246727 | protein_coding | 131.7518807 | -0.277440447 | 0.524076319 | 0.529389397 | 0.596535352 | 0.774662172 | 0.277440447 |
| ENSMUSG00000029646 | Cdx2 | 12591 | protein_coding | 3237.993453 | 0.277472726 | 0.1369935 | -2.025444465 | 0.042821746 | 0.132476266 | 0.277472726 |
| ENSMUSG00000048878 | Hexim1 | 192231 | protein_coding | 862.9329806 | 0.27747629 | 0.190897605 | -1.453534679 | 0.146075358 | 0.323442761 | 0.27747629 |
| ENSMUSG00000056234 | Ncoa4 | 27057 | protein_coding | 639.7812737 | -0.277497911 | 0.083837279 | 3.309958448 | 0.000933098 | 0.006002799 | 0.277497911 |
| ENSMUSG00000073779 | Lrp8os2 | 619295 | lncRNA | 4.06116636 | 0.277540126 | 0.848443327 | -0.32711687 | 0.743579493 | 1 | 0.277540126 |
| ENSMUSG00000031939 | Taf1d | 75316 | protein_coding | 1162.213486 | -0.277595716 | 0.107062421 | 2.592839895 | 0.009518707 | 0.040914089 | 0.277595716 |
| ENSMUSG00000029521 | Chek2 | 50883 | protein_coding | 184.2800168 | 0.277651595 | 0.17351533 | -1.600155992 | 0.109563982 | 0.264291499 | 0.277651595 |
| ENSMUSG00000066571 | Garre1 | 233103 | protein_coding | 215.5443069 | -0.277687105 | 0.09582988 | 2.897708983 | 0.003758992 | 0.019206682 | 0.277687105 |
| ENSMUSG00000054128 | H2-T3 | 15043 | protein_coding | 556.5987613 | -0.277804604 | 0.180604062 | 1.53819688 | 0.124000484 | 0.28837403 | 0.277804604 |
| ENSMUSG00000042351 | Grap2 | 17444 | protein_coding | 24.59776655 | 0.277958622 | 0.228821519 | -1.214739867 | 0.224465322 | 0.431627891 | 0.277958622 |
| ENSMUSG00000061136 | Prpf40a | 56194 | protein_coding | 1314.840574 | 0.278095914 | 0.071051163 | -3.914023388 | 9.08E-05 | 0.000792278 | 0.278095914 |
| ENSMUSG00000035297 | Cops4 | 26891 | protein_coding | 846.6458988 | 0.278097043 | 0.090274244 | -3.080580144 | 0.002065977 | 0.01165615 | 0.278097043 |
| ENSMUSG00000015748 | Prpf3 | 70767 | protein_coding | 196.1107853 | 0.278151339 | 0.09707284 | -2.865387873 | 0.004164987 | 0.020855246 | 0.278151339 |
| ENSMUSG00000019210 | Atp6v1e1 | 11973 | protein_coding | 1086.52366 | -0.278196959 | 0.062968898 | 4.418005854 | 9.96E-06 | 0.000111756 | 0.278196959 |
| ENSMUSG00000024608 | Rps14 | 20044 | protein_coding | 6383.655202 | -0.278269296 | 0.091094027 | 3.054747973 | 0.002252497 | 0.012461318 | 0.278269296 |
| ENSMUSG00000025207 | Sema4g | 26456 | protein_coding | 3469.686647 | 0.278343287 | 0.112868055 | -2.466094484 | 0.01365953 | 0.054237431 | 0.278343287 |
| ENSMUSG00000039197 | Adk | 11534 | protein_coding | 1766.617388 | -0.278367492 | 0.044937996 | 6.194479459 | 5.85E-10 | 1.32E-08 | 0.278367492 |
| ENSMUSG00000118376 | Gm9926 | 100503565 | lncRNA | 118.3821918 | -0.278392762 | 0.117221425 | 2.374930702 | 0.017552245 | 0.066267555 | 0.278392762 |
| ENSMUSG00000080059 | Rps19-ps3 | 277692 | processed_pseudogene | 7.008002651 | -0.278420033 | 0.399966346 | 0.696108649 | 0.486360791 | 0.695624496 | 0.278420033 |
| ENSMUSG00000029253 | Cenpc1 | 12617 | protein_coding | 254.4039697 | 0.278451186 | 0.103687314 | -2.685489441 | 0.007242366 | 0.032806559 | 0.278451186 |
| ENSMUSG00000047654 | Tssk6 | 83984 | protein_coding | 15.09213172 | -0.278521165 | 0.269707467 | 1.032678734 | 0.301754269 | 0.522023104 | 0.278521165 |
| ENSMUSG00000031590 | Frg1 | 14300 | protein_coding | 512.2861752 | -0.278632783 | 0.117950197 | 2.362291792 | 0.01816234 | 0.06795036 | 0.278632783 |
| ENSMUSG00000072109 | A530040E14Rik | 621875 | lncRNA | 3.832151378 | 0.278647845 | 1.825451062 | -0.152646023 | 0.878677429 | 1 | 0.278647845 |
| ENSMUSG00000078861 | Zfp931 | 353208 | protein_coding | 46.57902727 | -0.278672379 | 0.144100304 | 1.933877805 | 0.053128154 | 0.156196961 | 0.278672379 |
| ENSMUSG00000022701 | Ccdc191 | 212153 | protein_coding | 92.31026356 | -0.27871388 | 0.132626129 | 2.101500522 | 0.035597052 | 0.114868009 | 0.27871388 |
| ENSMUSG00000030047 | Arhgap25 | 232201 | protein_coding | 7.560596836 | 0.278733219 | 0.536975262 | -0.519080187 | 0.603704825 | 0.779177919 | 0.278733219 |
| ENSMUSG00000056145 | AI504432 | 229694 | lncRNA | 4.115888 | 0.278775442 | 1.000013494 | -0.278771681 | 0.780420049 | 1 | 0.278775442 |
| ENSMUSG00000001911 | Nfix | 18032 | protein_coding | 252.0123995 | -0.278852007 | 0.08478384 | 3.288975898 | 0.001005526 | 0.006384236 | 0.278852007 |
| ENSMUSG00000027695 | Pld1 | 18805 | protein_coding | 1605.020211 | -0.278904581 | 0.073786418 | 3.779890506 | 0.000156897 | 0.00127605 | 0.278904581 |
| ENSMUSG00000025221 | Kcnip2 | 80906 | protein_coding | 5.615970765 | -0.279023329 | 0.797089209 | 0.350052824 | 0.726299054 | 0.858315628 | 0.279023329 |
| ENSMUSG00000012519 | Mlkl | 74568 | protein_coding | 228.5532404 | -0.279087006 | 0.092746572 | 3.00913554 | 0.002619922 | 0.014180725 | 0.279087006 |
| ENSMUSG00000021217 | Tshz3 | 243931 | protein_coding | 1.301668583 | -0.279093086 | 1.885054265 | 0.14805573 | 0.882298787 | 1 | 0.279093086 |
| ENSMUSG00000024421 | Lama3 | 16774 | protein_coding | 624.249901 | -0.279122031 | 0.428743946 | 0.651022675 | 0.51503185 | 0.716272317 | 0.279122031 |
| ENSMUSG00000060336 | Zfp937 | 245174 | protein_coding | 33.18158986 | -0.279157085 | 0.206328575 | 1.352973452 | 0.176064112 | 0.368018849 | 0.279157085 |

Supplementary Table 1

| ensembl_gene_id | external_gene_name | entrezgene_id | gene_biotype | baseMean | log2FoldChange | lfcSE | stat | pvalue | padj | absLog2FC |
|---|---|---|---|---|---|---|---|---|---|---|
| ENSMUSG00000060703 | Cd302 | 66205 | protein_coding | 34.81159356 | 0.279381909 | 0.184636009 | -1.51314963 | 0.130241655 | 0.298143983 | 0.279381909 |
| ENSMUSG00000038729 | Pakap | 677884 | protein_coding | 25.52320714 | -0.279435872 | 0.30704444 | 0.910082827 | 0.36277883 | 0.585166009 | 0.279435872 |
| ENSMUSG00000098490 | Mir7089 | 102465660 | miRNA | 0.684017244 | -0.27945767 | 1.520190873 | 0.183830646 | 0.854146314 | 1 | 0.27945767 |
| ENSMUSG00000064293 | Cntn4 | 269784 | protein_coding | 3.540275142 | 0.279475958 | 0.958951346 | -0.291439142 | 0.770715482 | 1 | 0.279475958 |
| ENSMUSG00000000385 | Tmprss2 | 50528 | protein_coding | 5690.980347 | -0.279479413 | 0.152569275 | 1.831819763 | 0.066978278 | 0.184932165 | 0.279479413 |
| ENSMUSG00000014704 | Hoxa2 | 15399 | protein_coding | 19.06172562 | 0.279495057 | 0.232493424 | -1.202163283 | 0.229300272 | 0.438111017 | 0.279495057 |
| ENSMUSG00000070283 | Ndufaf3 | 66706 | protein_coding | 255.9193059 | 0.279528977 | 0.090598004 | -3.085376767 | 0.002032944 | 0.011497276 | 0.279528977 |
| ENSMUSG00000086596 | Susd5 | 382111 | protein_coding | 1.505291913 | -0.279559256 | 2.295075609 | 0.121808299 | 0.903050841 | 1 | 0.279559256 |
| ENSMUSG00000034647 | Ankrd12 | 106585 | protein_coding | 279.9576617 | -0.279579555 | 0.098556843 | 2.836734074 | 0.004557756 | 0.022511824 | 0.279579555 |
| ENSMUSG00000053182 | Cd200l1 | 208166 | protein_coding | 117.0043917 | -0.279580111 | 0.162537291 | 1.720098257 | 0.085414583 | 0.220499674 | 0.279580111 |
| ENSMUSG00000026158 | Ogfrl1 | 70155 | protein_coding | 35.20765603 | -0.27963571 | 0.189070271 | 1.479004117 | 0.139139214 | 0.312363375 | 0.27963571 |
| ENSMUSG00000031994 | Adamts8 | 30806 | protein_coding | 2.933012214 | 0.279682305 | 1.408612659 | -0.198551605 | 0.842613513 | 1 | 0.279682305 |
| ENSMUSG00000026576 | Atp1b1 | 11931 | protein_coding | 9681.290842 | -0.279694768 | 0.097385517 | 2.872036576 | 0.004078358 | 0.020502282 | 0.279694768 |
| ENSMUSG00000045639 | Zfp629 | 320683 | protein_coding | 268.9488206 | -0.279701255 | 0.108750882 | 2.5719447 | 0.010112904 | 0.042953833 | 0.279701255 |
| ENSMUSG00000033166 | Dis3 | 72662 | protein_coding | 232.1353528 | 0.2797226 | 0.146112329 | -1.914435299 | 0.055564559 | 0.161179189 | 0.2797226 |
| ENSMUSG00000049350 | Zg16 | 69036 | protein_coding | 28919.21281 | 0.279760853 | 0.345492335 | -0.809745469 | 0.418086479 | 0.638103523 | 0.279760853 |
| ENSMUSG00000005089 | Slc1a2 | 20511 | protein_coding | 12.82068931 | 0.279765404 | 0.559349707 | -0.500161886 | 0.616961093 | 0.788881899 | 0.279765404 |
| ENSMUSG00000026721 | Rabgap1l | 29809 | protein_coding | 987.4648096 | -0.27986703 | 0.078873284 | 3.548312125 | 0.000387708 | 0.002851946 | 0.27986703 |
| ENSMUSG00000080626 | Mir1192 | 100316672 | miRNA | 3.899898349 | -0.279976511 | 0.576581712 | 0.485579936 | 0.627265025 | 1 | 0.279976511 |
| ENSMUSG00000031221 | Igbp1 | 18518 | protein_coding | 783.1075684 | -0.279996079 | 0.09131013 | 3.06642951 | 0.002166319 | 0.01208601 | 0.279996079 |
| ENSMUSG00000027133 | Nop10 | 66181 | protein_coding | 331.2885307 | -0.280081829 | 0.1124728 | 2.490218329 | 0.012766464 | 0.05145848 | 0.280081829 |
| ENSMUSG00000044244 | Il20rb | 213208 | protein_coding | 15.23688356 | -0.280188043 | 0.340102817 | 0.823833348 | 0.410034256 | 0.630476965 | 0.280188043 |
| ENSMUSG00000001289 | Pfdn5 | 56612 | protein_coding | 932.5534379 | -0.280222629 | 0.069851679 | 4.011680661 | 6.03E-05 | 0.000552495 | 0.280222629 |
| ENSMUSG00000035572 | Dcaf10 | 242418 | protein_coding | 398.5828702 | 0.280249385 | 0.094101393 | -2.978164028 | 0.002899808 | 0.015447555 | 0.280249385 |
| ENSMUSG00000042202 | Slc35e2 | 320541 | protein_coding | 378.1969699 | 0.28028161 | 0.061870255 | -4.530151171 | 5.89E-06 | 6.93E-05 | 0.28028161 |
| ENSMUSG00000061315 | Naca | 17938 | protein_coding | 2724.967511 | -0.280372855 | 0.094192597 | 2.976591187 | 0.002914724 | 0.015492034 | 0.280372855 |
| ENSMUSG00000046994 | Mars2 | 212679 | protein_coding | 265.0592007 | -0.280389402 | 0.124291903 | 2.255894354 | 0.024077248 | 0.084814777 | 0.280389402 |
| ENSMUSG00000020288 | Ahsa2 | 268390 | protein_coding | 161.0010325 | -0.280466921 | 0.10575009 | 2.652167211 | 0.007997692 | 0.035589839 | 0.280466921 |
| ENSMUSG00000026553 | Copa | 12847 | protein_coding | 4823.750438 | 0.280497255 | 0.087727116 | -3.19738376 | 0.001386803 | 0.008358271 | 0.280497255 |
| ENSMUSG00000025224 | Gbf1 | 107338 | protein_coding | 1315.261492 | 0.280540841 | 0.06878846 | -4.078312566 | 4.54E-05 | 0.000429797 | 0.280540841 |
| ENSMUSG00000046330 | Rpl37a | 19981 | protein_coding | 3648.62809 | -0.280688302 | 0.107285466 | 2.616275176 | 0.008889488 | 0.038706422 | 0.280688302 |
| ENSMUSG00000060470 | Adgrg3 | 54672 | protein_coding | 4.949994743 | -0.280719598 | 0.869884707 | 0.322708971 | 0.746915658 | 0.872440598 | 0.280719598 |
| ENSMUSG00000025209 | Twnk | 226153 | protein_coding | 212.5054192 | -0.28074595 | 0.08285856 | 3.388255231 | 0.000703388 | 0.004704636 | 0.28074595 |
| ENSMUSG00000096458 | Moap1 | 64113 | protein_coding | 4.181102908 | 0.280776623 | 0.542389368 | -0.517666162 | 0.604691211 | 1 | 0.280776623 |
| ENSMUSG00000042903 | Foxo4 | 54601 | protein_coding | 386.5439698 | -0.28081082 | 0.083126326 | 3.378121404 | 0.000729829 | 0.004853962 | 0.28081082 |
| ENSMUSG00000021418 | Rpp40 | 208366 | protein_coding | 36.85646396 | -0.280876633 | 0.253488675 | 1.108044108 | 0.267842767 | 0.483088189 | 0.280876633 |
| ENSMUSG00000032184 | Lysmd2 | 70082 | protein_coding | 22.47833881 | 0.280890067 | 0.247546381 | -1.13469672 | 0.256502412 | 0.469401688 | 0.280890067 |
| ENSMUSG00000030789 | Itgax | 16411 | protein_coding | 11.1584588 | 0.280930756 | 0.646069412 | -0.434830609 | 0.663685385 | 0.818589921 | 0.280930756 |
| ENSMUSG00000026248 | Mrpl44 | 69163 | protein_coding | 364.33407 | 0.280955266 | 0.097487729 | -2.881955177 | 0.00395216 | 0.020007959 | 0.280955266 |

Supplementary Table 1

| ensembl_gene_id | external_gene_name | entrezgene_id | gene_biotype | baseMean | log2FoldChange | lfcSE | stat | pvalue | padj | absLog2FC |
|---|---|---|---|---|---|---|---|---|---|---|
| ENSMUSG00000063275 | Hacd1 | 30963 | protein_coding | 231.4904676 | -0.280984393 | 0.131016376 | 2.144650924 | 0.031980767 | 0.10607158 | 0.280984393 |
| ENSMUSG00000061778 | Mospd2 | 76763 | protein_coding | 253.2571572 | 0.281019103 | 0.093796674 | -2.99604552 | 0.002735056 | 0.014731479 | 0.281019103 |
| ENSMUSG00000024019 | Cmtr1 | 74157 | protein_coding | 72.71332969 | 0.281108642 | 0.18788546 | -1.496170285 | 0.134609286 | 0.305377683 | 0.281108642 |
| ENSMUSG00000071506 | Tmem139 | 109218 | protein_coding | 371.9740942 | -0.28112956 | 0.145564757 | 1.931302369 | 0.053445671 | 0.156906987 | 0.28112956 |
| ENSMUSG00000039754 | Alkbh4 | 72041 | protein_coding | 99.0284082 | -0.281198232 | 0.125455289 | 2.241421901 | 0.024998759 | 0.087325531 | 0.281198232 |
| ENSMUSG00000021710 | Nln | 75805 | protein_coding | 538.6498595 | -0.281269231 | 0.071377114 | 3.940608076 | 8.13E-05 | 0.000718512 | 0.281269231 |
| ENSMUSG00000040263 | Klhdc4 | 234825 | protein_coding | 329.5586398 | 0.281296752 | 0.102869934 | -2.734489481 | 0.006247709 | 0.02899794 | 0.281296752 |
| ENSMUSG00000007867 | Ift43 | 76411 | protein_coding | 184.4509084 | -0.281369883 | 0.112708795 | 2.496432364 | 0.012544959 | 0.050788785 | 0.281369883 |
| ENSMUSG00000032202 | Rab27a | 11891 | protein_coding | 560.1514198 | 0.281396592 | 0.159216726 | -1.767380848 | 0.077164468 | 0.20483749 | 0.281396592 |
| ENSMUSG00000041881 | Ndufa7 | 66416 | protein_coding | 1366.383817 | -0.281451382 | 0.091714487 | 3.068777808 | 0.002149364 | 0.012021919 | 0.281451382 |
| ENSMUSG00000034813 | Grip1 | 74053 | protein_coding | 31.94376817 | -0.281466573 | 0.242867987 | 1.158928257 | 0.24648543 | 0.458287111 | 0.281466573 |
| ENSMUSG00000048007 | Timm8a1 | 30058 | protein_coding | 43.08266745 | -0.281470582 | 0.183534927 | 1.533607725 | 0.125126187 | 0.290256194 | 0.281470582 |
| ENSMUSG00000020916 | Krt36 | 16673 | protein_coding | 30.28762411 | 0.281476524 | 0.429585316 | -0.655228458 | 0.512320674 | 0.714747285 | 0.281476524 |
| ENSMUSG00000046111 | Cep295 | 319675 | protein_coding | 197.6715514 | 0.281560125 | 0.117943893 | -2.387237841 | 0.016975505 | 0.064474124 | 0.281560125 |
| ENSMUSG00000033917 | Gde1 | 56209 | protein_coding | 529.3451247 | -0.281578786 | 0.150974159 | 1.865079345 | 0.062170283 | 0.175022172 | 0.281578786 |
| ENSMUSG00000048440 | Cyp4f16 | 70101 | protein_coding | 440.5954303 | -0.28158371 | 0.093043282 | 3.026373355 | 0.002475065 | 0.013502881 | 0.28158371 |
| ENSMUSG00000079445 | B3gnt7 | 227327 | protein_coding | 4128.456134 | -0.281607624 | 0.421168303 | 0.66863442 | 0.50372871 | 0.708204999 | 0.281607624 |
| ENSMUSG00000055407 | Map6 | 17760 | protein_coding | 10.37842152 | 0.281686991 | 0.511951446 | -0.550222083 | 0.582167059 | 0.76507296 | 0.281686991 |
| ENSMUSG00000053886 | Sh2d4a | 72281 | protein_coding | 1138.950122 | -0.281759647 | 0.123846823 | 2.275065593 | 0.022901995 | 0.081579713 | 0.281759647 |
| ENSMUSG00000023988 | Bysl | 53414 | protein_coding | 222.8126808 | 0.28176839 | 0.121643382 | -2.316347872 | 0.020539282 | 0.074891965 | 0.28176839 |
| ENSMUSG00000056050 | Mia3 | 338366 | protein_coding | 3166.743659 | 0.281786686 | 0.148092753 | -1.902771605 | 0.057070354 | 0.164593053 | 0.281786686 |
| ENSMUSG00000030341 | Tnfrsf1a | 21937 | protein_coding | 424.2590791 | 0.281813301 | 0.091663913 | -3.074419264 | 0.002109128 | 0.011855051 | 0.281813301 |
| ENSMUSG00000022876 | Samsn1 | 67742 | protein_coding | 3.382096871 | -0.281832727 | 0.727270683 | 0.387521088 | 0.698370476 | 1 | 0.281832727 |
| ENSMUSG00000046861 | Hectd3 | 76608 | protein_coding | 1273.321477 | -0.281874681 | 0.094624752 | 2.978868378 | 0.00289315 | 0.015417065 | 0.281874681 |
| ENSMUSG00000039470 | Zdhhc2 | 70546 | protein_coding | 216.7699618 | 0.281966631 | 0.188930674 | -1.492434369 | 0.135585311 | 0.306972281 | 0.281966631 |
| ENSMUSG00000026638 | Irf6 | 54139 | protein_coding | 2224.719441 | -0.282009491 | 0.0645995 | 4.365505748 | 1.27E-05 | 0.000137975 | 0.282009491 |
| ENSMUSG00000073607 | Gm10548 | 633395 | transcribed_processed_pseudogene | 9.560768231 | 0.282034638 | 0.422367364 | -0.667747233 | 0.504294954 | 0.708596728 | 0.282034638 |
| ENSMUSG00000073557 | Ppp1r12b | 329251 | protein_coding | 144.1516595 | 0.282114342 | 0.135592508 | -2.080604209 | 0.037470148 | 0.119710201 | 0.282114342 |
| ENSMUSG00000025427 | Rnf165 | 225743 | protein_coding | 3.741195933 | -0.282264374 | 2.012589419 | 0.140249358 | 0.888462975 | 1 | 0.282264374 |
| ENSMUSG00000020954 | Strn3 | 94186 | protein_coding | 672.8073073 | 0.282326006 | 0.055455563 | -5.09103125 | 3.56E-07 | 5.40E-06 | 0.282326006 |
| ENSMUSG00000041697 | Cox6a1 | 12861 | protein_coding | 5145.288174 | -0.282346546 | 0.080470331 | 3.508703697 | 0.000450296 | 0.003244412 | 0.282346546 |
| ENSMUSG00000018927 | Ccl6 | 20305 | protein_coding | 2732.501412 | -0.282361431 | 0.369957898 | 0.763225851 | 0.445328714 | 0.662536791 | 0.282361431 |
| ENSMUSG00000031545 | Gpat4 | 102247 | protein_coding | 1143.704641 | 0.282416461 | 0.072356274 | -3.903137139 | 9.50E-05 | 0.000823133 | 0.282416461 |
| ENSMUSG00000005338 | Cadm3 | 94332 | protein_coding | 5.235101377 | 0.282420525 | 3.175992912 | -0.088923538 | 0.929142677 | 0.966128972 | 0.282420525 |
| ENSMUSG00000068735 | Trp53i11 | 277414 | protein_coding | 2342.180673 | 0.28242601 | 0.120700587 | -2.339889282 | 0.019289457 | 0.071374882 | 0.28242601 |
| ENSMUSG00000047517 | Dmbt1 | 12945 | protein_coding | 91353.67156 | -0.282472945 | 0.11062084 | 2.553523783 | 0.0106639 | 0.044864238 | 0.282472945 |
| ENSMUSG00000049550 | Clip1 | 56430 | protein_coding | 670.1303144 | -0.2825768 | 0.073663868 | 3.836029876 | 0.000125039 | 0.001052445 | 0.2825768 |
| ENSMUSG00000026854 | Usp20 | 74270 | protein_coding | 214.7805317 | 0.282617256 | 0.098466714 | -2.870180622 | 0.004102374 | 0.020610466 | 0.282617256 |
| ENSMUSG00000022540 | Rogdi | 66049 | protein_coding | 89.18914941 | 0.282622183 | 0.146673416 | -1.926880758 | 0.053994492 | 0.157984593 | 0.282622183 |

| ensembl_gene_id | external_gene_name | entrezgene_id | gene_biotype | baseMean | log2FoldChange | lfcSE | stat | pvalue | padj | absLog2FC |
|---|---|---|---|---|---|---|---|---|---|---|
| ENSMUSG00000023057 | Fabp2 | 14079 | protein_coding | 3502.633321 | 0.282729487 | 0.124798239 | -2.265492601 | 0.023482468 | 0.083216624 | 0.282729487 |
| ENSMUSG00000074733 | Zfp950 | 414758 | protein_coding | 205.47405 | -0.282847741 | 0.132202129 | 2.139509725 | 0.032394409 | 0.107013657 | 0.282847741 |
| ENSMUSG00000039741 | Bahcc1 | 268515 | protein_coding | 246.3683421 | 0.282853021 | 0.167370312 | -1.689983231 | 0.091031163 | 0.230625736 | 0.282853021 |
| ENSMUSG00000037225 | Fgf2 | 14173 | protein_coding | 9.458392223 | 0.282893298 | 0.448374648 | -0.630930627 | 0.528085885 | 0.72618955 | 0.282893298 |
| ENSMUSG00000028085 | Gatb | 229487 | protein_coding | 183.5377499 | -0.282902604 | 0.089681469 | 3.154526881 | 0.001607586 | 0.009444208 | 0.282902604 |
| ENSMUSG00000027959 | Sass6 | 72776 | protein_coding | 62.01024492 | 0.282904583 | 0.197447065 | -1.4328123 | 0.151911485 | 0.33244981 | 0.282904583 |
| ENSMUSG00000030317 | Timp4 | 110595 | protein_coding | 32.10263439 | 0.282948793 | 0.330848128 | -0.85522259 | 0.392427932 | 0.615086833 | 0.282948793 |
| ENSMUSG00000073427 | Gm4924 | 237412 | protein_coding | 36.67678443 | -0.282994823 | 0.197798337 | 1.430723977 | 0.152509339 | 0.333254034 | 0.282994823 |
| ENSMUSG00000059401 | Mamld1 | 333639 | protein_coding | 13.40253352 | -0.283044729 | 0.299872341 | 0.943884079 | 0.345228888 | 0.566884219 | 0.283044729 |
| ENSMUSG00000097032 | 4930539J05Rik | 319587 | lncRNA | 12.7014376 | 0.283100423 | 0.546780447 | -0.517758864 | 0.604626523 | 0.779942456 | 0.283100423 |
| ENSMUSG00000005981 | Trap1 | 68015 | protein_coding | 1169.531432 | -0.283101573 | 0.088802173 | 3.188002754 | 0.001432592 | 0.008569101 | 0.283101573 |
| ENSMUSG00000070604 | Vsig10l | 75690 | protein_coding | 26.74741685 | 0.28318488 | 0.329497938 | -0.859443559 | 0.390095847 | 0.612475705 | 0.28318488 |
| ENSMUSG00000078656 | Vps25 | 28084 | protein_coding | 9.242983358 | -0.28321466 | 0.359720769 | 0.787318066 | 0.431095697 | 0.649494967 | 0.28321466 |
| ENSMUSG00000051403 | Ppp1r37 | 232947 | protein_coding | 1312.181862 | -0.283306873 | 0.048578057 | 5.831992643 | 5.48E-09 | 1.10E-07 | 0.283306873 |
| ENSMUSG00000073676 | Hspe1 | 15528 | protein_coding | 809.5418139 | -0.283415942 | 0.131874781 | 2.149129187 | 0.031624161 | 0.105142217 | 0.283415942 |
| ENSMUSG00000027883 | Gpsm2 | 76123 | protein_coding | 730.145371 | 0.283680531 | 0.099151244 | -2.861088974 | 0.004221885 | 0.021082618 | 0.283680531 |
| ENSMUSG00000022257 | Laptm4b | 114128 | protein_coding | 1023.171683 | 0.283792302 | 0.086845295 | -3.267791318 | 0.001083903 | 0.006808508 | 0.283792302 |
| ENSMUSG00000056121 | Fez2 | 225020 | protein_coding | 272.8114918 | -0.283814893 | 0.104281639 | 2.72161902 | 0.006496299 | 0.029941356 | 0.283814893 |
| ENSMUSG00000057522 | Spop | 20747 | protein_coding | 723.7392424 | -0.283830657 | 0.101477096 | 2.79699232 | 0.005158076 | 0.025001604 | 0.283830657 |
| ENSMUSG00000040111 | Gramd1b | 235283 | protein_coding | 167.6937489 | -0.283870707 | 0.183388799 | 1.547917369 | 0.121642192 | 0.284460467 | 0.283870707 |
| ENSMUSG00000085862 | Gm13483 | 100502624 | lncRNA | 1.41208787 | -0.283888703 | 1.069404888 | 0.265464191 | 0.790651892 | 1 | 0.283888703 |
| ENSMUSG00000109865 | Hspa14 | 50497 | protein_coding | 342.3548531 | 0.283899858 | 0.103401832 | -2.745597935 | 0.006040074 | 0.028320777 | 0.283899858 |
| ENSMUSG00000001016 | Ilf2 | 67781 | protein_coding | 599.0691827 | 0.283957736 | 0.101753554 | -2.79064196 | 0.005260363 | 0.025430219 | 0.283957736 |
| ENSMUSG00000030218 | Mgp | 17313 | protein_coding | 1.922316784 | -0.284041886 | 1.324702686 | 0.214419348 | 0.830220055 | 1 | 0.284041886 |
| ENSMUSG00000020804 | Aanat | 11298 | protein_coding | 3.012966882 | -0.284077796 | 0.955967431 | 0.29716263 | 0.766342351 | 1 | 0.284077796 |
| ENSMUSG00000022288 | 4930447A16Rik | 74892 | lncRNA | 2.073562015 | -0.284105303 | 0.82794337 | 0.343145815 | 0.731488766 | 1 | 0.284105303 |
| ENSMUSG00000042606 | Hirip3 | 233876 | protein_coding | 123.1190782 | 0.284166499 | 0.168678105 | -1.684667365 | 0.092052766 | 0.232532135 | 0.284166499 |
| ENSMUSG00000086607 | 4930511M06Rik | 75084 | lncRNA | 2.971304886 | -0.284180066 | 0.876040502 | 0.32439147 | 0.745641677 | 1 | 0.284180066 |
| ENSMUSG00000065395 | Mir193a | 387188 | miRNA | 1.018051733 | -0.28424178 | 1.5345271 | 0.185230863 | 0.853047962 | 1 | 0.28424178 |
| ENSMUSG00000025137 | Pcyt2 | 68671 | protein_coding | 743.2073929 | -0.284246121 | 0.063339605 | 4.487652261 | 7.20E-06 | 8.35E-05 | 0.284246121 |
| ENSMUSG00000079003 | Samd1 | 666704 | protein_coding | 188.4374237 | -0.284322042 | 0.140445293 | 2.024432681 | 0.042925652 | 0.132673445 | 0.284322042 |
| ENSMUSG00000019841 | Rev3l | 19714 | protein_coding | 621.4562572 | 0.284323627 | 0.078840862 | -3.606297811 | 0.000310597 | 0.002350684 | 0.284323627 |
| ENSMUSG00000108034 | 4930557K07Rik | 78202 | lncRNA | 4.469464205 | 0.284410908 | 0.560939788 | -0.507025734 | 0.612136763 | 0.785395579 | 0.284410908 |
| ENSMUSG00000059810 | Rgs3 | 50780 | protein_coding | 280.2856093 | 0.284539229 | 0.077835603 | -3.655643654 | 0.000256538 | 0.001976883 | 0.284539229 |
| ENSMUSG00000058331 | Zfp85 | 22746 | protein_coding | 26.8020056 | -0.284588989 | 0.234051785 | 1.215923176 | 0.224014188 | 0.431111794 | 0.284588989 |
| ENSMUSG00000029088 | Kcnip4 | 80334 | protein_coding | 5.908667535 | -0.284707939 | 0.519274575 | 0.548280144 | 0.583499559 | 0.766020418 | 0.284707939 |
| ENSMUSG00000021040 | Slirp | 380773 | protein_coding | 119.7636619 | 0.284732985 | 0.144903311 | -1.964986055 | 0.049415852 | 0.147728706 | 0.284732985 |
| ENSMUSG00000026475 | Rgs16 | 19734 | protein_coding | 21.46153405 | -0.284755502 | 0.315012217 | 0.903950659 | 0.366021565 | 0.588671613 | 0.284755502 |
| ENSMUSG00000030138 | Bms1 | 213895 | protein_coding | 521.3168353 | 0.284760069 | 0.069649483 | -4.088473529 | 4.34E-05 | 0.000413537 | 0.284760069 |

| ensembl_gene_id | external_gene_name | entrezgene_id | gene_biotype | baseMean | log2FoldChange | lfcSE | stat | pvalue | padj | absLog2FC |
|---|---|---|---|---|---|---|---|---|---|---|
| ENSMUSG00000021371 | Mcur1 | 76137 | protein_coding | 608.4284643 | -0.284784087 | 0.071755874 | 3.968791257 | 7.22E-05 | 0.000647282 | 0.284784087 |
| ENSMUSG00000005610 | Eif4g2 | 13690 | protein_coding | 7380.913851 | 0.284799863 | 0.073981729 | -3.849597309 | 0.000118312 | 0.001000926 | 0.284799863 |
| ENSMUSG00000024835 | Coro1b | 23789 | protein_coding | 2944.85137 | -0.284825042 | 0.060709439 | 4.69161055 | 2.71E-06 | 3.41E-05 | 0.284825042 |
| ENSMUSG00000045382 | Cxcr4 | 12767 | protein_coding | 10.65614432 | -0.284864223 | 0.376620189 | 0.756370029 | 0.449427372 | 0.666174493 | 0.284864223 |
| ENSMUSG00000021363 | Mak | 17152 | protein_coding | 2.626865201 | 0.284915268 | 1.141935749 | -0.249502013 | 0.802972484 | 1 | 0.284915268 |
| ENSMUSG00000055633 | Zfp580 | 68992 | protein_coding | 1.908987423 | -0.284976369 | 1.152651964 | 0.2472354 | 0.80472605 | 1 | 0.284976369 |
| ENSMUSG00000026393 | Nek7 | 59125 | protein_coding | 658.2849535 | 0.285002625 | 0.103872562 | -2.74377199 | 0.006073771 | 0.028422287 | 0.285002625 |
| ENSMUSG00000075600 | Zc3h3 | 223642 | protein_coding | 79.22584204 | -0.285047502 | 0.194905016 | 1.462494435 | 0.143605768 | 0.319894352 | 0.285047502 |
| ENSMUSG00000018008 | Cyth4 | 72318 | protein_coding | 15.43700199 | 0.285123282 | 0.524878167 | -0.543218027 | 0.586979696 | 0.768401324 | 0.285123282 |
| ENSMUSG00000040728 | Esrp1 | 207920 | protein_coding | 3237.534697 | 0.285134628 | 0.055160869 | -5.169146753 | 2.35E-07 | 3.65E-06 | 0.285134628 |
| ENSMUSG00000030407 | Qpctl | 67369 | protein_coding | 166.8552056 | 0.285163315 | 0.103477033 | -2.755812627 | 0.005854653 | 0.02767136 | 0.285163315 |
| ENSMUSG00000028199 | Cryz | 12972 | protein_coding | 172.5381559 | -0.285283978 | 0.117140355 | 2.435403058 | 0.01487521 | 0.058004537 | 0.285283978 |
| ENSMUSG00000037278 | Tmem97 | 69071 | protein_coding | 569.0764841 | 0.285292363 | 0.152697101 | -1.86835481 | 0.061712632 | 0.174119665 | 0.285292363 |
| ENSMUSG00000005355 | Casp14 | 12365 | protein_coding | 34.26789104 | -0.285329079 | 0.367964504 | 0.775425552 | 0.438088241 | 0.6563215 | 0.285329079 |
| ENSMUSG00000020085 | Aifm2 | 71361 | protein_coding | 141.5467454 | -0.285332756 | 0.140083632 | 2.036874341 | 0.041662636 | 0.12986349 | 0.285332756 |
| ENSMUSG00000069305 | H4c18 | 319161 | protein_coding | 875.018648 | 0.28536644 | 0.159162559 | -1.792924424 | 0.072985009 | 0.196850041 | 0.28536644 |
| ENSMUSG00000047721 | Bola2 | 66162 | protein_coding | 299.6729182 | -0.285371928 | 0.077532447 | 3.680677422 | 0.000232615 | 0.001809424 | 0.285371928 |
| ENSMUSG00000029319 | Coq2 | 71883 | protein_coding | 502.7051177 | -0.285377935 | 0.112543108 | 2.535721109 | 0.011221606 | 0.046704567 | 0.285377935 |
| ENSMUSG00000034981 | Parm1 | 231440 | protein_coding | 1887.908261 | -0.285379801 | 0.082192936 | 3.472072137 | 0.000516458 | 0.003646336 | 0.285379801 |
| ENSMUSG00000031068 | Glrx3 | 30926 | protein_coding | 565.4202544 | -0.285526646 | 0.116449725 | 2.451930608 | 0.014209206 | 0.055893077 | 0.285526646 |
| ENSMUSG00000097011 | Gm4651 | 100043793 | lncRNA | 13.96596615 | -0.285588798 | 0.296490137 | 0.963232036 | 0.335431091 | 0.557777472 | 0.285588798 |
| ENSMUSG00000046748 | Tmem45a2 | 69457 | protein_coding | 0.728947028 | 0.285604521 | 1.875190965 | -0.152306899 | 0.878944884 | 1 | 0.285604521 |
| ENSMUSG00000063439 | B9d2 | 232987 | protein_coding | 18.67206116 | -0.285615074 | 0.241021936 | 1.185016928 | 0.236010754 | 0.446089228 | 0.285615074 |
| ENSMUSG00000028931 | Kcnab2 | 16498 | protein_coding | 12.20204014 | 0.285639032 | 0.657864542 | -0.434191256 | 0.664149562 | 0.818817302 | 0.285639032 |
| ENSMUSG00000084745 | Gm25290 | 115490269 | snoRNA | 1.18978069 | -0.285658439 | 1.042505645 | 0.274011408 | 0.784075855 | 1 | 0.285658439 |
| ENSMUSG00000025578 | Cbx8 | 30951 | protein_coding | 119.7410823 | 0.285718204 | 0.118556155 | -2.409982044 | 0.015953306 | 0.061382313 | 0.285718204 |
| ENSMUSG00000028426 | Rad23b | 19359 | protein_coding | 1907.320276 | 0.285798149 | 0.07685457 | -3.718687765 | 0.00020026 | 0.00158386 | 0.285798149 |
| ENSMUSG00000053835 | H2-T24 | 15042 | protein_coding | 18.68947736 | 0.285833508 | 0.306567616 | -0.932366935 | 0.351146927 | 0.573211901 | 0.285833508 |
| ENSMUSG00000033184 | Tmed7 | 66676 | protein_coding | 2271.935811 | 0.285926541 | 0.091183477 | -3.135727557 | 0.001714284 | 0.009957679 | 0.285926541 |
| ENSMUSG00000062822 | 4833420G17Rik | 67392 | protein_coding | 279.3079547 | -0.285964596 | 0.092184173 | 3.102100786 | 0.001921525 | 0.010995173 | 0.285964596 |
| ENSMUSG00000033405 | Nudt15 | 214254 | protein_coding | 21.08043286 | -0.285999816 | 0.295349353 | 0.968344142 | 0.332872526 | 0.555258272 | 0.285999816 |
| ENSMUSG00000070737 | Tmem35b | 100039968 | protein_coding | 92.9752356 | -0.286090244 | 0.148461571 | 1.927032306 | 0.053975604 | 0.157957314 | 0.286090244 |
| ENSMUSG00000035295 | Wdr38 | 76646 | protein_coding | 5.034171088 | 0.286175955 | 0.690399908 | -0.414507522 | 0.67850246 | 0.827715862 | 0.286175955 |
| ENSMUSG00000038371 | Sbf2 | 319934 | protein_coding | 953.5920327 | 0.286197659 | 0.068528793 | -4.176312559 | 2.96E-05 | 0.000293324 | 0.286197659 |
| ENSMUSG00000041189 | Chrnb1 | 11443 | protein_coding | 18.34427008 | -0.286301706 | 0.294769554 | 0.97127299 | 0.331412355 | 0.553829552 | 0.286301706 |
| ENSMUSG00000056468 | 5730596B20Rik | 77580 | TEC | 19.35143286 | -0.286303893 | 0.287703966 | 0.995133635 | 0.319671274 | 0.54094184 | 0.286303893 |
| ENSMUSG00000039713 | Plekhg5 | 269608 | protein_coding | 121.7499048 | -0.286316801 | 0.108777763 | 2.632126208 | 0.008485234 | 0.03728716 | 0.286316801 |
| ENSMUSG00000018398 | Septin8 | 20362 | protein_coding | 503.7809586 | -0.286377749 | 0.075289369 | 3.80369437 | 0.000142554 | 0.001178247 | 0.286377749 |
| ENSMUSG00000006527 | Sfmbt1 | 54650 | protein_coding | 174.2451667 | 0.286395276 | 0.119607583 | -2.394457525 | 0.016644971 | 0.063554926 | 0.286395276 |

| ensembl_gene_id | external_gene_name | entrezgene_id | gene_biotype | baseMean | log2FoldChange | lfcSE | stat | pvalue | padj | absLog2FC |
|---|---|---|---|---|---|---|---|---|---|---|
| ENSMUSG00000028124 | Gclm | 14630 | protein_coding | 872.249796 | -0.28639703 | 0.081993863 | 3.492908126 | 0.000477791 | 0.003410008 | 0.28639703 |
| ENSMUSG00000062991 | Nrg1 | 211323 | protein_coding | 5.382176543 | 0.286410199 | 1.220141509 | -0.234735231 | 0.814414241 | 0.907710402 | 0.286410199 |
| ENSMUSG00000032727 | Mier3 | 218613 | protein_coding | 948.3770307 | -0.286453238 | 0.124148783 | 2.30733827 | 0.02103597 | 0.076382586 | 0.286453238 |
| ENSMUSG00000027999 | Pla2g12a | 66350 | protein_coding | 90.50164269 | -0.286505298 | 0.155628341 | 1.840958374 | 0.065627659 | 0.182116478 | 0.286505298 |
| ENSMUSG00000042331 | Specc1 | 432572 | protein_coding | 650.3511003 | -0.286519446 | 0.097603638 | 2.935540637 | 0.00332967 | 0.017356745 | 0.286519446 |
| ENSMUSG00000070372 | Capza1 | 12340 | protein_coding | 789.169861 | -0.286551824 | 0.108097008 | 2.650876553 | 0.008028318 | 0.035697265 | 0.286551824 |
| ENSMUSG00000093989 | Rnasek | 52898 | protein_coding | 4.596643476 | -0.286568966 | 0.488679953 | 0.58641441 | 0.557597063 | 0.746748674 | 0.286568966 |
| ENSMUSG00000021814 | Anxa7 | 11750 | protein_coding | 2004.027689 | -0.286596447 | 0.117277576 | 2.443744645 | 0.014535711 | 0.056882163 | 0.286596447 |
| ENSMUSG00000097852 | 4933405D12Rik | 74057 | lncRNA | 7.163167024 | 0.286672409 | 0.405096245 | -0.707664938 | 0.479153356 | 0.690541222 | 0.286672409 |
| ENSMUSG00000029397 | Rchy1 | 68098 | protein_coding | 400.1650067 | 0.286672591 | 0.07296797 | -3.928745608 | 8.54E-05 | 0.000751269 | 0.286672591 |
| ENSMUSG00000093445 | Lrch4 | 231798 | protein_coding | 5.323629597 | 0.286712022 | 0.698135636 | -0.410682404 | 0.681305431 | 0.829503357 | 0.286712022 |
| ENSMUSG00000036923 | Stox1 | 216021 | protein_coding | 5.85741463 | 0.286718592 | 0.448465678 | -0.639332296 | 0.522606783 | 0.721840784 | 0.286718592 |
| ENSMUSG00000087029 | Gm14133 | 100503289 | lncRNA | 0.785192368 | -0.286739242 | 1.304156744 | 0.219865628 | 0.825975806 | 1 | 0.286739242 |
| ENSMUSG00000039246 | Lyplal1 | 226791 | protein_coding | 93.09813448 | 0.286750292 | 0.136015601 | -2.108216186 | 0.035012294 | 0.113460169 | 0.286750292 |
| ENSMUSG00000020657 | Dnajc27 | 217378 | protein_coding | 19.95841801 | 0.286769818 | 0.277715429 | -1.032603119 | 0.301789668 | 0.522023104 | 0.286769818 |
| ENSMUSG00000022236 | Ropn1l | 252967 | protein_coding | 2.435551349 | 0.286805303 | 0.740249285 | -0.387444214 | 0.698427376 | 1 | 0.286805303 |
| ENSMUSG00000017550 | Atad5 | 237877 | protein_coding | 118.6480393 | 0.286812775 | 0.127262411 | -2.253711636 | 0.024214315 | 0.085261236 | 0.286812775 |
| ENSMUSG00000078995 | Zfp456 | 408065 | protein_coding | 35.85525306 | -0.286878508 | 0.211006589 | 1.359571328 | 0.173965616 | 0.36485499 | 0.286878508 |
| ENSMUSG00000018965 | Ywhah | 22629 | protein_coding | 1404.141201 | -0.286909327 | 0.107189898 | 2.676645218 | 0.007436333 | 0.033534573 | 0.286909327 |
| ENSMUSG00000085024 | C230035I16Rik | 320842 | lncRNA | 3.163590988 | -0.286911267 | 0.615739785 | 0.465961878 | 0.641242788 | 1 | 0.286911267 |
| ENSMUSG00000031480 | Thsd1 | 56229 | protein_coding | 3.374912589 | 0.286961766 | 0.853984513 | -0.336026897 | 0.736850584 | 1 | 0.286961766 |
| ENSMUSG00000040268 | Plekha1 | 101476 | protein_coding | 634.9130397 | -0.286997142 | 0.099838477 | 2.874614581 | 0.00404521 | 0.020354233 | 0.286997142 |
| ENSMUSG00000071654 | Uqcc3 | 107197 | protein_coding | 209.430851 | -0.287059784 | 0.110298485 | 2.602572328 | 0.00925273 | 0.039999889 | 0.287059784 |
| ENSMUSG00000021679 | S100z | 268686 | protein_coding | 2.623236859 | -0.287136009 | 0.610466687 | 0.470354919 | 0.638101467 | 1 | 0.287136009 |
| ENSMUSG00000032528 | Vipr1 | 22354 | protein_coding | 3656.004552 | -0.287141306 | 0.092171698 | 3.115287139 | 0.001837659 | 0.010584969 | 0.287141306 |
| ENSMUSG00000033458 | Fan1 | 330554 | protein_coding | 102.4799543 | -0.287151679 | 0.119651284 | 2.399904706 | 0.01639934 | 0.062762157 | 0.287151679 |
| ENSMUSG00000033313 | Fbxl8 | 50788 | protein_coding | 140.8687522 | -0.287241085 | 0.112335759 | 2.556987085 | 0.010558314 | 0.044513657 | 0.287241085 |
| ENSMUSG00000050295 | Foxc1 | 17300 | protein_coding | 24.96779171 | 0.287296977 | 0.345484607 | -0.831576782 | 0.405647872 | 0.626589558 | 0.287296977 |
| ENSMUSG00000019996 | Map7 | 17761 | protein_coding | 2168.028734 | 0.287382664 | 0.069031923 | -4.163040132 | 3.14E-05 | 0.000309798 | 0.287382664 |
| ENSMUSG00000044641 | Pard6b | 58220 | protein_coding | 457.4460441 | -0.287388739 | 0.185198708 | 1.551785875 | 0.120713467 | 0.283202471 | 0.287388739 |
| ENSMUSG00000064672 | Gm22806 | 115488214 | snoRNA | 3.327123334 | -0.287395315 | 0.623290344 | 0.461093803 | 0.644731307 | 1 | 0.287395315 |
| ENSMUSG00000039033 | Tasp1 | 75812 | protein_coding | 160.6486602 | 0.287493419 | 0.095389201 | -3.013899006 | 0.002579136 | 0.013987474 | 0.287493419 |
| ENSMUSG00000032763 | Ilvbl | 216136 | protein_coding | 616.2683379 | -0.28755931 | 0.064869504 | 4.432888968 | 9.30E-06 | 0.000105168 | 0.28755931 |
| ENSMUSG00000021256 | Vash1 | 238328 | protein_coding | 2.886258572 | -0.28766464 | 0.958627913 | 0.300079557 | 0.764116472 | 1 | 0.28766464 |
| ENSMUSG00000059602 | Syn3 | 27204 | protein_coding | 25.02034235 | -0.287697579 | 0.2792033 | 1.030423279 | 0.30281135 | 0.5233341 | 0.287697579 |
| ENSMUSG00000028300 | C9orf72 | 73205 | protein_coding | 158.0226828 | 0.28771223 | 0.12004144 | -2.396774239 | 0.016540111 | 0.063213005 | 0.28771223 |
| ENSMUSG00000020131 | Pcsk4 | 18551 | protein_coding | 30.82095813 | -0.287755825 | 0.283355301 | 1.015530059 | 0.309853225 | 0.530888475 | 0.287755825 |
| ENSMUSG00000086690 | 4933406J10Rik | 77800 | lncRNA | 2.568998568 | -0.287779256 | 0.749735614 | 0.383840984 | 0.701096309 | 1 | 0.287779256 |
| ENSMUSG00000039914 | Coq10a | 210582 | protein_coding | 130.2269776 | -0.287802162 | 0.115380521 | 2.494373924 | 0.012617954 | 0.051034263 | 0.287802162 |

| ensembl_gene_id | external_gene_name | entrezgene_id | gene_biotype | baseMean | log2FoldChange | lfcSE | stat | pvalue | padj | absLog2FC |
|---|---|---|---|---|---|---|---|---|---|---|
| ENSMUSG00000022217 | Emc9 | 85308 | protein_coding | 49.39440565 | -0.287824563 | 0.172029351 | 1.673113113 | 0.094305031 | 0.236536038 | 0.287824563 |
| ENSMUSG00000024403 | Atp6v1g2 | 66237 | protein_coding | 14.33366429 | 0.287888831 | 0.351960471 | -0.817957851 | 0.413381255 | 0.633557127 | 0.287888831 |
| ENSMUSG00000071855 | Ccdc112 | 240261 | protein_coding | 33.21168901 | 0.287904226 | 0.206038507 | -1.397332132 | 0.162313718 | 0.347734431 | 0.287904226 |
| ENSMUSG00000029545 | Acads | 11409 | protein_coding | 3593.541056 | -0.288006537 | 0.068498856 | 4.204545209 | 2.62E-05 | 0.000261952 | 0.288006537 |
| ENSMUSG00000026750 | Psmb7 | 19177 | protein_coding | 742.9782924 | 0.28808445 | 0.078007105 | -3.693053962 | 0.000221577 | 0.001731719 | 0.28808445 |
| ENSMUSG00000030516 | Tjp1 | 21872 | protein_coding | 1349.560291 | -0.288189771 | 0.093130557 | 3.094470607 | 0.001971645 | 0.011213534 | 0.288189771 |
| ENSMUSG00000062328 | Rpl17 | 319195 | protein_coding | 5286.330961 | -0.28821999 | 0.111329176 | 2.588898984 | 0.009628333 | 0.041288574 | 0.28821999 |
| ENSMUSG00000019970 | Sgk1 | 20393 | protein_coding | 2154.152609 | -0.288281038 | 0.238878604 | 1.206809791 | 0.227505405 | 0.436052025 | 0.288281038 |
| ENSMUSG00000028393 | Alad | 17025 | protein_coding | 471.1127707 | -0.288543872 | 0.102399401 | 2.817827724 | 0.004834974 | 0.023622712 | 0.288543872 |
| ENSMUSG00000038712 | Mindy1 | 75007 | protein_coding | 1077.118146 | -0.288582934 | 0.077334547 | 3.731617319 | 0.000190254 | 0.00151488 | 0.288582934 |
| ENSMUSG00000032579 | Hemk1 | 69536 | protein_coding | 97.12007879 | -0.288632816 | 0.112382161 | 2.568315228 | 0.010219418 | 0.043322217 | 0.288632816 |
| ENSMUSG00000045467 | Ttll13 | 269954 | protein_coding | 9.734603502 | 0.288635389 | 0.488318569 | -0.591080102 | 0.554466752 | 0.744666516 | 0.288635389 |
| ENSMUSG00000076430 | Hus1b | 210554 | protein_coding | 1.022839979 | 0.28870106 | 1.236236422 | -0.233532239 | 0.815348138 | 1 | 0.28870106 |
| ENSMUSG00000022131 | Gpr180 | 58245 | protein_coding | 344.6795673 | 0.288715599 | 0.121836781 | -2.36969161 | 0.017802928 | 0.06689364 | 0.288715599 |
| ENSMUSG00000116665 | E130310I04Rik | 278725 | lncRNA | 4.695095975 | 0.288730807 | 0.738772736 | -0.390824936 | 0.695926642 | 0.838607402 | 0.288730807 |
| ENSMUSG00000002944 | Cd36 | 12491 | protein_coding | 2.337678092 | -0.288752939 | 1.31348983 | 0.219836448 | 0.825998532 | 1 | 0.288752939 |
| ENSMUSG00000021178 | Psmc1 | 19179 | protein_coding | 1068.429891 | 0.288825077 | 0.083256487 | -3.469099999 | 0.000522205 | 0.003680621 | 0.288825077 |
| ENSMUSG00000025481 | Urah | 76974 | protein_coding | 141.0284378 | -0.288881082 | 0.118383968 | 2.440204421 | 0.014678953 | 0.057361152 | 0.288881082 |
| ENSMUSG00000039018 | Mtg1 | 212508 | protein_coding | 106.0851846 | -0.288900697 | 0.134230235 | 2.152277364 | 0.031375516 | 0.104504894 | 0.288900697 |
| ENSMUSG00000074865 | Zfp934 | 77117 | protein_coding | 28.0031055 | -0.289105972 | 0.261677657 | 1.104817185 | 0.269238813 | 0.484231077 | 0.289105972 |
| ENSMUSG00000096169 | Olfr1564 | 100043474 | protein_coding | 1.993834474 | -0.289116031 | 2.136508922 | 0.135321705 | 0.892357524 | 1 | 0.289116031 |
| ENSMUSG00000031938 | 4931406C07Rik | 70984 | protein_coding | 1752.950469 | -0.289151623 | 0.153164314 | 1.887852432 | 0.059045764 | 0.168933082 | 0.289151623 |
| ENSMUSG00000049106 | Dcaf5 | 320808 | protein_coding | 606.9068177 | 0.289167671 | 0.059121973 | -4.89103552 | 1.00E-06 | 1.39E-05 | 0.289167671 |
| ENSMUSG00000029699 | Ssc4d | 109267 | protein_coding | 48.95522193 | 0.289184993 | 0.254447754 | -1.136520126 | 0.255738947 | 0.468471767 | 0.289184993 |
| ENSMUSG00000015711 | Prune1 | 229589 | protein_coding | 218.8802271 | -0.289216849 | 0.086204948 | 3.354991274 | 0.000793675 | 0.005227463 | 0.289216849 |
| ENSMUSG00000049922 | Slc35c1 | 228368 | protein_coding | 2384.218455 | 0.289253496 | 0.081023874 | -3.569978581 | 0.00035701 | 0.002661702 | 0.289253496 |
| ENSMUSG00000001909 | Trmt1 | 212528 | protein_coding | 468.3596643 | -0.289310082 | 0.09134763 | 3.167132889 | 0.0015395 | 0.009099247 | 0.289310082 |
| ENSMUSG00000023330 | Dtwd1 | 69185 | protein_coding | 49.72229384 | 0.289313759 | 0.168188929 | -1.720171247 | 0.085401318 | 0.220499674 | 0.289313759 |
| ENSMUSG00000038213 | Tapbpl | 213233 | protein_coding | 416.4343254 | 0.289407673 | 0.093465775 | -3.096402651 | 0.001958842 | 0.011158438 | 0.289407673 |
| ENSMUSG00000009772 | Nuak2 | 74137 | protein_coding | 523.9410574 | 0.289517925 | 0.117853552 | -2.456590571 | 0.014026244 | 0.055334304 | 0.289517925 |
| ENSMUSG00000032012 | Nectin1 | 58235 | protein_coding | 143.238199 | 0.289543878 | 0.117260186 | -2.469242879 | 0.013539928 | 0.053892113 | 0.289543878 |
| ENSMUSG00000097712 | Gm10532 | 100038353 | lncRNA | 2.843628676 | 0.289559025 | 0.855949992 | -0.338289653 | 0.735144921 | 1 | 0.289559025 |
| ENSMUSG00000055373 | Fut9 | 14348 | protein_coding | 196.7253644 | 0.289703156 | 0.415579334 | -0.697106744 | 0.485735994 | 0.695380179 | 0.289703156 |
| ENSMUSG00000030103 | Bhlhe40 | 20893 | protein_coding | 1843.336453 | -0.289909527 | 0.114980197 | 2.521386591 | 0.011689335 | 0.048151079 | 0.289909527 |
| ENSMUSG00000048938 | Nr1h5 | 381463 | protein_coding | 1.913324202 | -0.289913592 | 2.11424801 | 0.137123739 | 0.890932989 | 1 | 0.289913592 |
| ENSMUSG00000035509 | Fbxl21 | 213311 | protein_coding | 5.516442851 | -0.290007858 | 0.49680251 | 0.583748777 | 0.559389345 | 0.747996408 | 0.290007858 |
| ENSMUSG00000022562 | Oplah | 75475 | protein_coding | 420.6415778 | -0.290026714 | 0.086211749 | 3.364120535 | 0.00076788 | 0.005084511 | 0.290026714 |
| ENSMUSG00000024215 | Spdef | 30051 | protein_coding | 1556.002787 | 0.290029529 | 0.172297199 | -1.683309602 | 0.092315173 | 0.23299599 | 0.290029529 |
| ENSMUSG00000043020 | Dnai3 | 242253 | protein_coding | 8.097700296 | -0.29010248 | 0.386019621 | 0.751522627 | 0.452338189 | 0.668687928 | 0.29010248 |

Supplementary Table 1

| ensembl_gene_id | external_gene_name | entrezgene_id | gene_biotype | baseMean | log2FoldChange | lfcSE | stat | pvalue | padj | absLog2FC |
|---|---|---|---|---|---|---|---|---|---|---|
| ENSMUSG00000048661 | Lemd3 | 380664 | protein_coding | 340.302 | 0.290225818 | 0.080940003 | -3.585690743 | 0.000336187 | 0.00251811 | 0.290225818 |
| ENSMUSG00000026979 | Psd4 | 215632 | protein_coding | 1069.882507 | -0.290240298 | 0.089207567 | 3.253538996 | 0.001139771 | 0.007094676 | 0.290240298 |
| ENSMUSG00000039158 | Akna | 100182 | protein_coding | 91.87337162 | 0.290265388 | 0.124724732 | -2.327248049 | 0.019952067 | 0.073252209 | 0.290265388 |
| ENSMUSG00000095098 | Ccdc85b | 240514 | protein_coding | 164.3266939 | -0.290324241 | 0.131901325 | 2.201071454 | 0.027730966 | 0.094754639 | 0.290324241 |
| ENSMUSG00000073096 | Lrrc61 | 243371 | protein_coding | 60.0999971 | -0.290378518 | 0.162409034 | 1.787945596 | 0.073784786 | 0.198449829 | 0.290378518 |
| ENSMUSG00000033294 | Noc4l | 100608 | protein_coding | 181.419763 | 0.290382613 | 0.10692519 | -2.715754945 | 0.006612484 | 0.030408957 | 0.290382613 |
| ENSMUSG00000014504 | Srp19 | 66384 | protein_coding | 293.9973561 | 0.290424732 | 0.0915403 | -3.172643441 | 0.001510579 | 0.008943393 | 0.290424732 |
| ENSMUSG00000091625 | Lsm5 | 66373 | protein_coding | 73.59632009 | 0.290448374 | 0.162962783 | -1.782298805 | 0.074700518 | 0.199909758 | 0.290448374 |
| ENSMUSG00000034575 | Tent4a | 210106 | protein_coding | 293.2991915 | 0.290473902 | 0.072002822 | -4.03420161 | 5.48E-05 | 0.000506309 | 0.290473902 |
| ENSMUSG00000021886 | Gpr65 | 14744 | protein_coding | 2.342899374 | -0.290563612 | 0.845215072 | 0.343774764 | 0.731015681 | 1 | 0.290563612 |
| ENSMUSG00000071379 | Hpcal1 | 53602 | protein_coding | 770.7716787 | -0.290656035 | 0.071328745 | 4.074879427 | 4.60E-05 | 0.000434691 | 0.290656035 |
| ENSMUSG00000036622 | Atp13a2 | 74772 | protein_coding | 447.2218301 | 0.290707555 | 0.075448871 | -3.853040504 | 0.00011666 | 0.000987961 | 0.290707555 |
| ENSMUSG00000097069 | Gm16998 | 100502920 | lncRNA | 1.57763972 | 0.290797749 | 1.065457172 | -0.272932368 | 0.784905205 | 1 | 0.290797749 |
| ENSMUSG00000037601 | Nme1 | 18102 | protein_coding | 304.499851 | 0.290819314 | 0.137739897 | -2.111365836 | 0.034740879 | 0.112868558 | 0.290819314 |
| ENSMUSG00000111329 | A830035O19Rik | 100101457 | lncRNA | 9.89619027 | -0.290867214 | 0.36047287 | 0.806904589 | 0.419721455 | 0.6397711 | 0.290867214 |
| ENSMUSG00000055026 | Gabrg3 | 14407 | protein_coding | 3.321163435 | -0.290931807 | 1.099226336 | 0.264669611 | 0.791263989 | 1 | 0.290931807 |
| ENSMUSG00000066613 | Zfp932 | 69504 | protein_coding | 70.80768165 | -0.290937998 | 0.162984124 | 1.785069553 | 0.074250036 | 0.199311621 | 0.290937998 |
| ENSMUSG00000038298 | Pdzk1 | 59020 | protein_coding | 3.874629485 | 0.290971077 | 1.039543364 | -0.279902779 | 0.779552095 | 1 | 0.290971077 |
| ENSMUSG00000038102 | Trappc11 | 320714 | protein_coding | 870.6608411 | 0.291088313 | 0.068044505 | -4.27791068 | 1.89E-05 | 0.000195572 | 0.291088313 |
| ENSMUSG00000033880 | Lgals3bp | 19039 | protein_coding | 2070.14766 | 0.291101277 | 0.173909062 | -1.673870665 | 0.094156021 | 0.236336338 | 0.291101277 |
| ENSMUSG00000062488 | Ifit3b | 667370 | protein_coding | 19.52741256 | -0.291152679 | 0.518977063 | 0.561012615 | 0.574788937 | 0.760117839 | 0.291152679 |
| ENSMUSG00000114203 | Gm35279 | 102638802 | lncRNA | 2.119244396 | 0.291226016 | 0.859251629 | -0.338929839 | 0.734662585 | 1 | 0.291226016 |
| ENSMUSG00000030077 | Chl1 | 12661 | protein_coding | 3.355168515 | -0.291337049 | 1.181158164 | 0.246653715 | 0.80517623 | 1 | 0.291337049 |
| ENSMUSG00000050503 | Fbxl22 | 74165 | protein_coding | 23.12027733 | 0.291363506 | 0.254860078 | -1.143229287 | 0.252943405 | 0.465324602 | 0.291363506 |
| ENSMUSG00000079293 | Clec7a | 56644 | protein_coding | 1.192262421 | -0.291376147 | 1.447254141 | 0.201330326 | 0.840440292 | 1 | 0.291376147 |
| ENSMUSG00000050840 | Cdh20 | 23836 | protein_coding | 1.807692416 | 0.291403192 | 1.76176707 | -0.165403927 | 0.868626063 | 1 | 0.291403192 |
| ENSMUSG00000041453 | Rpl21 | 19933 | protein_coding | 4333.555043 | -0.291555507 | 0.116663847 | 2.499107606 | 0.01245065 | 0.050506026 | 0.291555507 |
| ENSMUSG00000031374 | Zfp92 | 22754 | protein_coding | 4.108695213 | 0.291564351 | 0.787878207 | -0.370062719 | 0.711335759 | 1 | 0.291564351 |
| ENSMUSG00000046675 | Tmem251 | 320351 | protein_coding | 113.3849813 | 0.291622427 | 0.102334245 | -2.849705184 | 0.004375977 | 0.021779652 | 0.291622427 |
| ENSMUSG00000070563 | Spaca4 | 69363 | protein_coding | 116.5352562 | -0.291677068 | 0.19660808 | 1.483545675 | 0.137929491 | 0.310491768 | 0.291677068 |
| ENSMUSG00000029823 | Luc7l2 | 192196 | protein_coding | 1391.947578 | 0.291707299 | 0.088245763 | -3.30562385 | 0.000947652 | 0.00608931 | 0.291707299 |
| ENSMUSG00000021824 | Ap3m1 | 55946 | protein_coding | 793.6076594 | 0.291772179 | 0.08554874 | -3.410595856 | 0.000648211 | 0.004405167 | 0.291772179 |
| ENSMUSG00000020392 | Cdkn2aipnl | 52626 | protein_coding | 412.7546711 | 0.29181261 | 0.118556843 | -2.461372976 | 0.01384064 | 0.054759062 | 0.29181261 |
| ENSMUSG00000118462 | Lhb | 100040019 | protein_coding | 2.032214403 | 0.29181549 | 1.621095076 | -0.180011336 | 0.857143669 | 1 | 0.29181549 |
| ENSMUSG00000060063 | Alox5ap | 11690 | protein_coding | 30.17882555 | -0.291859223 | 0.228799712 | 1.275610102 | 0.202093381 | 0.404039964 | 0.291859223 |
| ENSMUSG00000058729 | Lin9 | 72568 | protein_coding | 106.8479264 | 0.29187458 | 0.136943569 | -2.131349293 | 0.033060377 | 0.108626953 | 0.29187458 |
| ENSMUSG00000055980 | Irs1 | 16367 | protein_coding | 92.9877376 | 0.29199639 | 0.235127894 | -1.241861975 | 0.214287492 | 0.419287439 | 0.29199639 |
| ENSMUSG00000030235 | Slco1c1 | 58807 | protein_coding | 3.800668815 | -0.292004448 | 1.021944274 | 0.285734218 | 0.775081695 | 1 | 0.292004448 |
| ENSMUSG00000030664 | Sox6os | 68469 | lncRNA | 1.598836572 | 0.292023816 | 0.901504125 | -0.323929539 | 0.745991379 | 1 | 0.292023816 |

| ensembl_gene_id | external_gene_name | entrezgene_id | gene_biotype | baseMean | log2FoldChange | lfcSE | stat | pvalue | padj | absLog2FC |
|---|---|---|---|---|---|---|---|---|---|---|
| ENSMUSG00000050947 | Amigo1 | 229715 | protein_coding | 141.1691325 | -0.29209991 | 0.125595748 | 2.325714958 | 0.020033762 | 0.073420411 | 0.29209991 |
| ENSMUSG00000042688 | Mapk6 | 50772 | protein_coding | 1132.244422 | -0.292107672 | 0.096586065 | 3.02432523 | 0.002491884 | 0.013581179 | 0.292107672 |
| ENSMUSG00000078937 | Cpt1b | 12895 | protein_coding | 6.768835035 | 0.292127614 | 0.473279037 | -0.617241821 | 0.537075235 | 0.732754497 | 0.292127614 |
| ENSMUSG00000073402 | Gm8909 | 667977 | protein_coding | 49.97551737 | -0.292162539 | 0.377322226 | 0.774305138 | 0.438750366 | 0.656832884 | 0.292162539 |
| ENSMUSG00000031347 | Cetn2 | 26370 | protein_coding | 170.2792819 | 0.292231568 | 0.141164995 | -2.070141882 | 0.03843906 | 0.122144051 | 0.292231568 |
| ENSMUSG00000043460 | Elfn2 | 207393 | protein_coding | 26.88350363 | 0.292239761 | 0.284521671 | -1.027126545 | 0.30436088 | 0.52454765 | 0.292239761 |
| ENSMUSG00000028194 | Ddah1 | 69219 | protein_coding | 759.2474552 | 0.292301735 | 0.093117873 | -3.13905082 | 0.00169496 | 0.009870825 | 0.292301735 |
| ENSMUSG00000028630 | Dyrk2 | 69181 | protein_coding | 1274.801428 | -0.29240713 | 0.1160697 | 2.5192374 | 0.011760933 | 0.048327356 | 0.29240713 |
| ENSMUSG00000031934 | Panx1 | 55991 | protein_coding | 266.6679967 | -0.292412637 | 0.098220107 | 2.977115838 | 0.00290974 | 0.015480494 | 0.292412637 |
| ENSMUSG00000059323 | Tonsl | 72749 | protein_coding | 268.3818435 | 0.292495762 | 0.117978681 | -2.479225575 | 0.013166801 | 0.05273746 | 0.292495762 |
| ENSMUSG00000008601 | Rab25 | 53868 | protein_coding | 1413.978257 | -0.292548452 | 0.078802909 | 3.71240675 | 0.000205298 | 0.001618275 | 0.292548452 |
| ENSMUSG00000029193 | Cckar | 12425 | protein_coding | 2.051617315 | -0.292593854 | 2.156048651 | 0.135708373 | 0.892051828 | 1 | 0.292593854 |
| ENSMUSG00000040729 | Cep126 | 234915 | protein_coding | 37.94135907 | 0.292751161 | 0.190386621 | -1.537666667 | 0.124130138 | 0.288572541 | 0.292751161 |
| ENSMUSG00000046338 | Gpat2 | 215456 | protein_coding | 10.11522933 | -0.292755171 | 0.34682433 | 0.844102174 | 0.398612313 | 0.620539568 | 0.292755171 |
| ENSMUSG00000061244 | Exoc5 | 105504 | protein_coding | 872.4276925 | 0.292799482 | 0.059882301 | -4.889583014 | 1.01E-06 | 1.39E-05 | 0.292799482 |
| ENSMUSG00000020553 | Pctp | 18559 | protein_coding | 238.7040467 | -0.292808614 | 0.116229703 | 2.519223636 | 0.011761393 | 0.048327356 | 0.292808614 |
| ENSMUSG00000073481 | Mtarc2 | 67247 | protein_coding | 940.4328762 | -0.292849032 | 0.072890863 | 4.017637036 | 5.88E-05 | 0.000540217 | 0.292849032 |
| ENSMUSG00000053877 | Srcap | 100043597 | protein_coding | 127.2978072 | 0.292921331 | 0.194155218 | -1.508696674 | 0.131376316 | 0.300075862 | 0.292921331 |
| ENSMUSG00000028391 | Wdr31 | 71354 | protein_coding | 44.82824573 | 0.29299872 | 0.165764894 | -1.76755592 | 0.077135173 | 0.204792643 | 0.29299872 |
| ENSMUSG00000020739 | Nup85 | 445007 | protein_coding | 347.8979032 | 0.29314291 | 0.142656614 | -2.054884811 | 0.039890132 | 0.125701628 | 0.29314291 |
| ENSMUSG00000037907 | Ankrd13b | 268445 | protein_coding | 88.36566004 | -0.29322344 | 0.156645277 | 1.871894545 | 0.061221194 | 0.17331619 | 0.29322344 |
| ENSMUSG00000021057 | Akap5 | 238276 | protein_coding | 5.422027834 | 0.293337524 | 0.812429298 | -0.361062218 | 0.718052936 | 0.85370239 | 0.293337524 |
| ENSMUSG00000048495 | Tyw5 | 68736 | protein_coding | 51.57329318 | 0.2933697 | 0.146890163 | -1.997204541 | 0.045802968 | 0.139478188 | 0.2933697 |
| ENSMUSG00000005410 | Mcm5 | 17218 | protein_coding | 524.3027787 | 0.2934684 | 0.178967057 | -1.639790054 | 0.101048827 | 0.249608668 | 0.2934684 |
| ENSMUSG00000036306 | Lzts1 | 211134 | protein_coding | 5.077591883 | 0.293670084 | 0.558075198 | -0.526219558 | 0.598735667 | 0.776186935 | 0.293670084 |
| ENSMUSG00000055061 | 4931431C16Rik | 74364 | lncRNA | 2.502415758 | -0.293816311 | 0.964381228 | 0.304668219 | 0.760618857 | 1 | 0.293816311 |
| ENSMUSG00000016534 | Lamp2 | 16784 | protein_coding | 1377.733359 | -0.293822425 | 0.079195002 | 3.710113235 | 0.000207167 | 0.001631449 | 0.293822425 |
| ENSMUSG00000078580 | E430018J23Rik | 101604 | protein_coding | 123.2827398 | -0.293836313 | 0.099551918 | 2.951588676 | 0.003161438 | 0.016610875 | 0.293836313 |
| ENSMUSG00000041879 | Ipo9 | 226432 | protein_coding | 445.2200558 | 0.293867801 | 0.133507564 | -2.201132229 | 0.027726664 | 0.094754639 | 0.293867801 |
| ENSMUSG00000074623 | Gm826 | 329554 | protein_coding | 3.220125009 | -0.293874231 | 0.819649188 | 0.358536597 | 0.719941785 | 1 | 0.293874231 |
| ENSMUSG00000031594 | Fgl1 | 234199 | protein_coding | 12.74139141 | -0.293897 | 0.280358461 | 1.048290102 | 0.294504968 | 0.513672903 | 0.293897 |
| ENSMUSG00000019194 | Scn1b | 20266 | protein_coding | 4.66818688 | 0.294104738 | 0.733696701 | -0.400853293 | 0.68852814 | 0.83386713 | 0.294104738 |
| ENSMUSG00000016526 | Dyrk3 | 226419 | protein_coding | 12.89876351 | 0.294104743 | 0.293616188 | -1.001663925 | 0.316505936 | 0.537548029 | 0.294104743 |
| ENSMUSG00000058546 | Rpl23a | 268449 | protein_coding | 1833.197884 | -0.294126466 | 0.117294751 | 2.50758421 | 0.012155961 | 0.04960305 | 0.294126466 |
| ENSMUSG00000038690 | Atp5j2 | 57423 | protein_coding | 2721.202412 | -0.294175284 | 0.08132283 | 3.61737636 | 0.000297604 | 0.002263768 | 0.294175284 |
| ENSMUSG00000056459 | Zbtb25 | 109929 | protein_coding | 25.48307226 | 0.294242402 | 0.215801233 | -1.363488048 | 0.172728749 | 0.363275924 | 0.294242402 |
| ENSMUSG00000063316 | Rpl27 | 19942 | protein_coding | 1155.504046 | -0.294252395 | 0.124191205 | 2.369349713 | 0.017819395 | 0.066917689 | 0.294252395 |
| ENSMUSG00000037390 | Muc3 | 666339 | protein_coding | 8415.002334 | -0.294300607 | 0.223582281 | 1.31629665 | 0.188074495 | 0.384388887 | 0.294300607 |
| ENSMUSG00000059439 | Bcas3 | 192197 | protein_coding | 218.4451014 | -0.294337255 | 0.145144835 | 2.02788652 | 0.042571831 | 0.131848679 | 0.294337255 |

Supplementary Table 1

| ensembl_gene_id | external_gene_name | entrezgene_id | gene_biotype | baseMean | log2FoldChange | lfcSE | stat | pvalue | padj | absLog2FC |
|---|---|---|---|---|---|---|---|---|---|---|
| ENSMUSG00000021262 | Evl | 14026 | protein_coding | 80.31801567 | -0.294346368 | 0.207994146 | 1.415166598 | 0.157019656 | 0.340132673 | 0.294346368 |
| ENSMUSG00000031682 | 1700011L22Rik | 67687 | protein_coding | 0.682712175 | -0.294356578 | 1.472861608 | 0.199853521 | 0.841595142 | 1 | 0.294356578 |
| ENSMUSG00000096472 | Cdkn2d | 12581 | protein_coding | 78.13233696 | 0.29436986 | 0.177592134 | -1.657561365 | 0.097406033 | 0.242436056 | 0.29436986 |
| ENSMUSG00000034071 | Zfp551 | 619331 | protein_coding | 32.42119753 | -0.294387204 | 0.211315704 | 1.393115601 | 0.163584836 | 0.349700969 | 0.294387204 |
| ENSMUSG00000032068 | Plet1 | 76509 | protein_coding | 802.1542491 | -0.294412439 | 0.2425028 | 1.214057897 | 0.224725616 | 0.432027803 | 0.294412439 |
| ENSMUSG00000031711 | Zfp330 | 30932 | protein_coding | 425.6325122 | 0.294574089 | 0.087968969 | -3.348613632 | 0.00081217 | 0.005334992 | 0.294574089 |
| ENSMUSG00000107476 | Zfp862-ps | 58894 | transcribed_unitary_pseudogene | 88.99500903 | -0.294682865 | 0.145757822 | 2.021729343 | 0.043204321 | 0.133360029 | 0.294682865 |
| ENSMUSG00000090291 | Lrrc10b | 278795 | protein_coding | 2.293005535 | 0.294798275 | 0.830208707 | -0.355089356 | 0.72252265 | 1 | 0.294798275 |
| ENSMUSG00000001942 | Siae | 22619 | protein_coding | 317.4246897 | -0.294850006 | 0.098432459 | 2.995455029 | 0.002740357 | 0.014751683 | 0.294850006 |
| ENSMUSG00000034353 | Ramp1 | 51801 | protein_coding | 567.6754566 | -0.29497883 | 0.283324294 | 1.041134967 | 0.297812913 | 0.517476912 | 0.29497883 |
| ENSMUSG00000030155 | Clec2e | 232409 | protein_coding | 297.2433641 | -0.295085333 | 0.291339203 | 1.012858312 | 0.311127843 | 0.532085855 | 0.295085333 |
| ENSMUSG00000043716 | Rpl7 | 19989 | protein_coding | 8169.21816 | -0.295241172 | 0.11942709 | 2.472145754 | 0.013430475 | 0.053546803 | 0.295241172 |
| ENSMUSG00000004267 | Eno2 | 13807 | protein_coding | 4.609587994 | 0.295268979 | 1.320899878 | -0.22353623 | 0.823118189 | 0.910297568 | 0.295268979 |
| ENSMUSG00000027257 | Pacsin3 | 80708 | protein_coding | 77.98923128 | -0.295277783 | 0.135020916 | 2.186904014 | 0.028749532 | 0.097588853 | 0.295277783 |
| ENSMUSG00000018727 | Cpsf4l | 52670 | protein_coding | 4.995080223 | 0.295406404 | 0.558284132 | -0.529132724 | 0.596713382 | 0.774787312 | 0.295406404 |
| ENSMUSG00000021798 | Ldb3 | 24131 | protein_coding | 146.4849421 | 0.295446893 | 0.171622154 | -1.72149624 | 0.085160814 | 0.219982118 | 0.295446893 |
| ENSMUSG00000001281 | Itgb7 | 16421 | protein_coding | 23.40675992 | 0.295506007 | 0.419969874 | -0.703636201 | 0.481659361 | 0.692385331 | 0.295506007 |
| ENSMUSG00000042073 | Abhd14b | 76491 | protein_coding | 323.1566715 | -0.295575166 | 0.089867447 | 3.289012599 | 0.001005395 | 0.006384236 | 0.295575166 |
| ENSMUSG00000062157 | Ifnlr1 | 242700 | protein_coding | 166.4272197 | -0.295675139 | 0.1004065 | 2.944780864 | 0.003231835 | 0.016926901 | 0.295675139 |
| ENSMUSG00000010492 | Uckl1os | 100271841 | lncRNA | 23.56098448 | -0.295680621 | 0.302546531 | 0.97730627 | 0.32841755 | 0.55060786 | 0.295680621 |
| ENSMUSG00000014294 | Ndufa2 | 17991 | protein_coding | 1087.156071 | 0.295704783 | 0.091163448 | -3.243677034 | 0.001179975 | 0.007309115 | 0.295704783 |
| ENSMUSG00000088252 | Snord13 | 100217422 | snoRNA | 122.0335719 | 0.295717737 | 0.237385065 | -1.245730172 | 0.212863473 | 0.417634239 | 0.295717737 |
| ENSMUSG00000028251 | Tstd3 | 77032 | protein_coding | 78.52046811 | -0.295888761 | 0.164546773 | 1.798204579 | 0.072144594 | 0.195389167 | 0.295888761 |
| ENSMUSG00000044548 | Dact1 | 59036 | protein_coding | 2.2809594 | -0.295941968 | 0.958334127 | 0.308808754 | 0.75746701 | 1 | 0.295941968 |
| ENSMUSG00000049281 | Scn3b | 235281 | protein_coding | 2.765669134 | 0.296056832 | 0.788995611 | -0.375232546 | 0.707487527 | 1 | 0.296056832 |
| ENSMUSG00000085272 | Sbk3 | 381835 | protein_coding | 1.997001449 | 0.296063347 | 1.971555758 | -0.150167372 | 0.880632567 | 1 | 0.296063347 |
| ENSMUSG00000015127 | Unkl | 74154 | protein_coding | 58.65172907 | -0.296071246 | 0.217341947 | 1.362237024 | 0.173123095 | 0.363688435 | 0.296071246 |
| ENSMUSG00000027829 | Ccnl1 | 56706 | protein_coding | 1116.24025 | -0.296099288 | 0.135707426 | 2.181894518 | 0.029117319 | 0.098614315 | 0.296099288 |
| ENSMUSG00000031639 | Tlr3 | 142980 | protein_coding | 314.0775447 | 0.296175953 | 0.07562197 | -3.916533167 | 8.98E-05 | 0.00078574 | 0.296175953 |
| ENSMUSG00000067367 | Lyar | 17089 | protein_coding | 194.2987943 | 0.296220836 | 0.113361364 | -2.613066961 | 0.008973372 | 0.03900665 | 0.296220836 |
| ENSMUSG00000047141 | Zfp654 | 72020 | protein_coding | 341.9919037 | 0.296238497 | 0.114124541 | -2.59574755 | 0.009438538 | 0.040685989 | 0.296238497 |
| ENSMUSG00000055214 | Pld5 | 319455 | protein_coding | 4.237035004 | 0.296267717 | 0.916480482 | -0.323266805 | 0.746493192 | 1 | 0.296267717 |
| ENSMUSG00000030409 | Dmpk | 13400 | protein_coding | 55.95799308 | 0.296340018 | 0.195965945 | -1.51220161 | 0.130482582 | 0.298488621 | 0.296340018 |
| ENSMUSG00000028367 | Txn1 | 22166 | protein_coding | 6987.792315 | -0.296362763 | 0.087306858 | 3.394495825 | 0.000687551 | 0.004614708 | 0.296362763 |
| ENSMUSG00000009376 | Met | 17295 | protein_coding | 848.4251323 | -0.29638196 | 0.087755395 | 3.377364539 | 0.00073184 | 0.004863751 | 0.29638196 |
| ENSMUSG00000026062 | Slc9a2 | 226999 | protein_coding | 3888.855354 | -0.296567909 | 0.127273248 | 2.330166887 | 0.019797332 | 0.07274578 | 0.296567909 |
| ENSMUSG00000063931 | Pepd | 18624 | protein_coding | 554.9297218 | 0.296618873 | 0.069849743 | -4.246527783 | 2.17E-05 | 0.00022214 | 0.296618873 |
| ENSMUSG00000031754 | Nudt21 | 68219 | protein_coding | 227.7820049 | -0.296673119 | 0.145517902 | 2.038739673 | 0.041476016 | 0.129452831 | 0.296673119 |
| ENSMUSG00000099988 | 1700054A03Rik | 78457 | lncRNA | 6.004438827 | -0.29683787 | 0.617033201 | 0.481072768 | 0.630464782 | 0.79831751 | 0.29683787 |

| ensembl_gene_id | external_gene_name | entrezgene_id | gene_biotype | baseMean | log2FoldChange | lfcSE | stat | pvalue | padj | absLog2FC |
|---|---|---|---|---|---|---|---|---|---|---|
| ENSMUSG00000032047 | Acat1 | 110446 | protein_coding | 1973.464263 | -0.296849264 | 0.117120776 | 2.534556831 | 0.011258966 | 0.046798532 | 0.296849264 |
| ENSMUSG00000063767 | S100a7a | 381493 | protein_coding | 2.069222869 | 0.296859154 | 0.847567951 | -0.350248206 | 0.726152432 | 1 | 0.296859154 |
| ENSMUSG00000047642 | D930020B18Rik | 216393 | protein_coding | 21.42500458 | 0.296861095 | 0.359427841 | -0.825926825 | 0.408845603 | 0.629409554 | 0.296861095 |
| ENSMUSG00000039637 | Coro7 | 78885 | protein_coding | 291.0859764 | 0.296941956 | 0.124671296 | -2.381798908 | 0.017228303 | 0.065269142 | 0.296941956 |
| ENSMUSG00000047428 | Dlk2 | 106565 | protein_coding | 2.994804949 | 0.297077112 | 1.074290021 | -0.276533437 | 0.782138381 | 1 | 0.297077112 |
| ENSMUSG00000086358 | Gm13270 | 105244084 | lncRNA | 6.265431712 | 0.297078903 | 0.519607023 | -0.571737658 | 0.567499718 | 0.753472451 | 0.297078903 |
| ENSMUSG00000098973 | Mir6236 | 102465959 | miRNA | 33850.15109 | 0.297110487 | 0.720734547 | -0.412232892 | 0.680168731 | 0.828402273 | 0.297110487 |
| ENSMUSG00000021257 | Angel1 | 68737 | protein_coding | 59.47326982 | -0.297158992 | 0.143473595 | 2.071175478 | 0.0383424 | 0.12191044 | 0.297158992 |
| ENSMUSG00000085396 | Firre | 103012 | lncRNA | 687.0072759 | 0.297161967 | 0.146224942 | -2.032224884 | 0.042130895 | 0.130976957 | 0.297161967 |
| ENSMUSG00000026433 | Rab29 | 226422 | protein_coding | 120.5681597 | -0.297217402 | 0.137740596 | 2.157805395 | 0.030942965 | 0.103335112 | 0.297217402 |
| ENSMUSG00000042210 | Abhd14a | 68644 | protein_coding | 114.0873029 | -0.297398973 | 0.174323508 | 1.706017599 | 0.088004803 | 0.22515016 | 0.297398973 |
| ENSMUSG00000022031 | Elp3 | 74195 | protein_coding | 600.7050092 | 0.297469239 | 0.107081239 | -2.777977185 | 0.005469846 | 0.026243183 | 0.297469239 |
| ENSMUSG00000031782 | Coq9 | 67914 | protein_coding | 1052.411883 | -0.297584653 | 0.069496561 | 4.282005447 | 1.85E-05 | 0.000192981 | 0.297584653 |
| ENSMUSG00000028980 | H6pd | 100198 | protein_coding | 441.7330111 | 0.29761894 | 0.123528875 | -2.40930666 | 0.015982862 | 0.061453081 | 0.29761894 |
| ENSMUSG00000036390 | Gadd45a | 13197 | protein_coding | 395.2938373 | 0.297635228 | 0.196495949 | -1.514714326 | 0.129844765 | 0.297648036 | 0.297635228 |
| ENSMUSG00000031386 | Hcfc1 | 15161 | protein_coding | 1346.196222 | 0.297638565 | 0.076506413 | -3.890374038 | 0.00010009 | 0.000861774 | 0.297638565 |
| ENSMUSG00000027800 | Tm4sf1 | 17112 | protein_coding | 23.38318779 | -0.2976762 | 0.252460884 | 1.179098304 | 0.238359034 | 0.448589299 | 0.2976762 |
| ENSMUSG00000040711 | Sh3pxd2b | 268396 | protein_coding | 98.90978726 | -0.297692568 | 0.134756503 | 2.209114666 | 0.027166666 | 0.093231573 | 0.297692568 |
| ENSMUSG00000026134 | Prim2 | 19076 | protein_coding | 96.62734088 | 0.297815765 | 0.17469238 | -1.704801122 | 0.088231522 | 0.225445669 | 0.297815765 |
| ENSMUSG00000087406 | E130215H24Rik | 109344 | lncRNA | 8.711785259 | 0.297899637 | 0.368355923 | -0.808727694 | 0.418671795 | 0.638701711 | 0.297899637 |
| ENSMUSG00000021645 | Smn1 | 20595 | protein_coding | 86.90196801 | 0.297911938 | 0.133835706 | -2.2259526 | 0.026017357 | 0.090073507 | 0.297911938 |
| ENSMUSG00000034285 | Nipsnap1 | 18082 | protein_coding | 592.7549703 | -0.297979209 | 0.092632225 | 3.216798573 | 0.001296296 | 0.007888281 | 0.297979209 |
| ENSMUSG00000073764 | Gm12888 | 545677 | protein_coding | 0.631497094 | 0.297991747 | 1.50908882 | -0.197464684 | 0.843463915 | 1 | 0.297991747 |
| ENSMUSG00000039155 | Cdh26 | 381409 | protein_coding | 21.34531719 | 0.298036901 | 0.242209107 | -1.2304942 | 0.218512098 | 0.424602824 | 0.298036901 |
| ENSMUSG00000110754 | 4930405J17Rik | 73811 | lncRNA | 12.87500499 | -0.298121852 | 0.43216107 | 0.689839674 | 0.490295016 | 0.69793377 | 0.298121852 |
| ENSMUSG00000028998 | Tomm7 | 66169 | protein_coding | 1052.543416 | -0.298167841 | 0.070484895 | 4.230237409 | 2.33E-05 | 0.000236946 | 0.298167841 |
| ENSMUSG00000070570 | Slc17a7 | 72961 | protein_coding | 2.930631261 | -0.298210033 | 1.981825654 | 0.150472385 | 0.880391936 | 1 | 0.298210033 |
| ENSMUSG00000002808 | Epdr1 | 105298 | protein_coding | 12.35386077 | 0.298334027 | 0.46006166 | -0.64846531 | 0.51668404 | 0.717449153 | 0.298334027 |
| ENSMUSG00000114953 | 4930519K11Rik | 78019 | lncRNA | 3.677895781 | -0.298419904 | 0.54923255 | 0.5433398 | 0.586895866 | 1 | 0.298419904 |
| ENSMUSG00000059866 | Tnip2 | 231130 | protein_coding | 149.1086606 | 0.298459575 | 0.100612111 | -2.966437851 | 0.003012712 | 0.015925715 | 0.298459575 |
| ENSMUSG00000045930 | Clec14a | 66864 | protein_coding | 296.1161789 | -0.298587518 | 0.163936103 | 1.821365227 | 0.068551357 | 0.188104702 | 0.298587518 |
| ENSMUSG00000072244 | Trim6 | 94088 | protein_coding | 102.3608969 | -0.298646566 | 0.137850172 | 2.166457693 | 0.030276229 | 0.101663614 | 0.298646566 |
| ENSMUSG00000048612 | Myof | 226101 | protein_coding | 692.9328978 | 0.298748849 | 0.093881553 | -3.182189045 | 0.001461664 | 0.008692083 | 0.298748849 |
| ENSMUSG00000041625 | Ggact | 223267 | protein_coding | 470.4285392 | -0.298770069 | 0.06702156 | 4.457820288 | 8.28E-06 | 9.48E-05 | 0.298770069 |
| ENSMUSG00000041540 | Sox5 | 20678 | protein_coding | 8.254653456 | 0.298798081 | 0.545689065 | -0.547561057 | 0.583993335 | 0.766436139 | 0.298798081 |
| ENSMUSG00000027217 | Tspan18 | 241556 | protein_coding | 7.936200751 | -0.298806777 | 0.668470534 | 0.44700067 | 0.654874576 | 0.812734264 | 0.298806777 |
| ENSMUSG00000058254 | Tspan7 | 21912 | protein_coding | 613.4151732 | -0.298894797 | 0.070841535 | 4.219202732 | 2.45E-05 | 0.00024793 | 0.298894797 |
| ENSMUSG00000033287 | Kctd17 | 72844 | protein_coding | 247.4441389 | -0.298910656 | 0.15532482 | 1.924423003 | 0.054301583 | 0.158658236 | 0.298910656 |
| ENSMUSG00000040896 | Kcnd3 | 56543 | protein_coding | 10.70243785 | -0.298957849 | 0.61602265 | 0.485303339 | 0.627461187 | 0.796333696 | 0.298957849 |

Supplementary Table 1

| ensembl_gene_id | external_gene_name | entrezgene_id | gene_biotype | baseMean | log2FoldChange | lfcSE | stat | pvalue | padj | absLog2FC |
|---|---|---|---|---|---|---|---|---|---|---|
| ENSMUSG00000033706 | Smyd5 | 232187 | protein_coding | 156.2788817 | 0.298996428 | 0.116162198 | -2.573956357 | 0.010054295 | 0.042770901 | 0.298996428 |
| ENSMUSG00000038255 | Neurod2 | 18013 | protein_coding | 1.524772059 | 0.299073516 | 1.214382313 | -0.246276245 | 0.805468397 | 1 | 0.299073516 |
| ENSMUSG00000005237 | Dnah2 | 327954 | protein_coding | 27.61773084 | 0.299090581 | 0.611290544 | -0.489277291 | 0.624645396 | 0.794347303 | 0.299090581 |
| ENSMUSG00000020674 | Pxdn | 69675 | protein_coding | 7.880933473 | -0.299132342 | 0.660816195 | 0.452671022 | 0.650785649 | 0.810243834 | 0.299132342 |
| ENSMUSG00000050310 | Rictor | 78757 | protein_coding | 605.4360134 | -0.299151228 | 0.068120978 | 4.391469948 | 1.13E-05 | 0.000124365 | 0.299151228 |
| ENSMUSG00000027168 | Pax6 | 18508 | protein_coding | 12.89517148 | -0.299231118 | 0.634986985 | 0.471239765 | 0.637469525 | 0.802408631 | 0.299231118 |
| ENSMUSG00000050931 | Sgms2 | 74442 | protein_coding | 835.6117263 | -0.299281368 | 0.110084147 | 2.718660011 | 0.006554694 | 0.030176812 | 0.299281368 |
| ENSMUSG00000078307 | AI593442 | 330941 | protein_coding | 2.288712613 | 0.299323804 | 2.412708601 | -0.124061316 | 0.901266727 | 1 | 0.299323804 |
| ENSMUSG00000007613 | Tgfbr1 | 21812 | protein_coding | 569.0541805 | 0.299333795 | 0.105058445 | -2.849212131 | 0.004382765 | 0.021806863 | 0.299333795 |
| ENSMUSG00000072774 | Zfp951 | 626391 | protein_coding | 19.36301331 | -0.29940898 | 0.356708408 | 0.839366197 | 0.401263847 | 0.622900809 | 0.29940898 |
| ENSMUSG00000097191 | Mm2pr | 433966 | lncRNA | 0.991943658 | -0.299410194 | 1.289060567 | 0.232270075 | 0.816328254 | 1 | 0.299410194 |
| ENSMUSG00000036672 | Cenpt | 320394 | protein_coding | 91.97220197 | 0.299419714 | 0.126079381 | -2.374850766 | 0.017556047 | 0.066267555 | 0.299419714 |
| ENSMUSG00000024124 | Prss30 | 30943 | protein_coding | 1663.590572 | -0.299606637 | 0.213399445 | 1.403971022 | 0.160327479 | 0.344790726 | 0.299606637 |
| ENSMUSG00000038000 | Acd | 497652 | protein_coding | 120.2061509 | -0.299636472 | 0.101212664 | 2.960464231 | 0.003071758 | 0.016201536 | 0.299636472 |
| ENSMUSG00000006494 | Pdk1 | 228026 | protein_coding | 713.1242666 | -0.299776199 | 0.134654272 | 2.226265792 | 0.025996383 | 0.090057536 | 0.299776199 |
| ENSMUSG00000014313 | Cox6c | 12864 | protein_coding | 3353.840418 | -0.299830428 | 0.109262551 | 2.744128031 | 0.006067187 | 0.028407578 | 0.299830428 |
| ENSMUSG00000061455 | Stx17 | 67727 | protein_coding | 454.674596 | -0.299840374 | 0.090978996 | 3.295709848 | 0.000981734 | 0.006266853 | 0.299840374 |
| ENSMUSG00000039492 | Ccdc27 | 381580 | protein_coding | 1.224253683 | -0.29986656 | 1.684759944 | 0.177987708 | 0.858732629 | 1 | 0.29986656 |
| ENSMUSG00000036411 | 9530077C05Rik | 68283 | protein_coding | 6.861403604 | -0.299962366 | 0.433412948 | 0.692093689 | 0.488878495 | 0.696638145 | 0.299962366 |
| ENSMUSG00000035399 | Oser1 | 66680 | protein_coding | 128.8691533 | -0.300040219 | 0.151235422 | 1.983928197 | 0.047263845 | 0.142911457 | 0.300040219 |
| ENSMUSG00000047875 | Gpr157 | 269604 | protein_coding | 22.81360963 | -0.30009567 | 0.224117706 | 1.339009201 | 0.180567678 | 0.374374718 | 0.30009567 |
| ENSMUSG00000004610 | Etfb | 110826 | protein_coding | 463.9506483 | 0.3001368 | 0.078917353 | -3.803178759 | 0.000142851 | 0.001180112 | 0.3001368 |
| ENSMUSG00000091183 | Gm5141 | 380850 | protein_coding | 16.82076323 | -0.3001414 | 0.302239664 | 0.993057614 | 0.320681878 | 0.541897747 | 0.3001414 |
| ENSMUSG00000015126 | Tsr3 | 68327 | protein_coding | 89.01849926 | 0.300167798 | 0.124066893 | -2.419402889 | 0.015546011 | 0.060109302 | 0.300167798 |
| ENSMUSG00000028412 | Slc44a1 | 100434 | protein_coding | 4697.03714 | -0.300187531 | 0.097648495 | 3.074164446 | 0.00211093 | 0.011857691 | 0.300187531 |
| ENSMUSG00000041817 | Fam169a | 320557 | protein_coding | 5.941815034 | 0.30026026 | 0.451015131 | -0.665743207 | 0.505575251 | 0.709532566 | 0.30026026 |
| ENSMUSG00000041445 | Mmrn2 | 105450 | protein_coding | 3.396282476 | 0.300269039 | 0.713657791 | -0.420746529 | 0.673940181 | 1 | 0.300269039 |
| ENSMUSG00000060938 | Rpl26 | 19941 | protein_coding | 4155.911559 | -0.300292945 | 0.112381813 | 2.672077767 | 0.007538318 | 0.03389483 | 0.300292945 |
| ENSMUSG00000070709 | Zfp974 | 73430 | protein_coding | 78.22390138 | -0.300516691 | 0.141024737 | 2.130950197 | 0.033093245 | 0.108713319 | 0.300516691 |
| ENSMUSG00000031452 | 1700029H14Rik | 66501 | protein_coding | 3.810759855 | 0.300557464 | 0.570440484 | -0.52688663 | 0.598272319 | 1 | 0.300557464 |
| ENSMUSG00000004383 | Large1 | 16795 | protein_coding | 159.1880848 | -0.300581186 | 0.125341295 | 2.398101805 | 0.016480284 | 0.063028118 | 0.300581186 |
| ENSMUSG00000032413 | Rasa2 | 114713 | protein_coding | 609.0438241 | -0.300635324 | 0.061337536 | 4.901327075 | 9.52E-07 | 1.32E-05 | 0.300635324 |
| ENSMUSG00000053329 | Gatd3a | 28295 | protein_coding | 538.2661955 | -0.300643179 | 0.083057 | 3.61972116 | 0.000294921 | 0.002244387 | 0.300643179 |
| ENSMUSG00000107355 | AI839979 | 100740 | lncRNA | 3.431780723 | 0.300773371 | 0.582703218 | -0.516169058 | 0.605736339 | 1 | 0.300773371 |
| ENSMUSG00000074227 | Spint2 | 20733 | protein_coding | 6804.875222 | -0.300778371 | 0.078528459 | 3.830183029 | 0.000128048 | 0.001074484 | 0.300778371 |
| ENSMUSG00000039879 | Heca | 380629 | protein_coding | 736.4708239 | -0.300806345 | 0.090609119 | 3.319824187 | 0.000900742 | 0.005830986 | 0.300806345 |
| ENSMUSG00000051527 | Usp29 | 57775 | protein_coding | 5.201539669 | 0.301110256 | 1.44786313 | -0.20796873 | 0.835253384 | 0.91717364 | 0.301110256 |
| ENSMUSG00000035875 | AI182371 | 98870 | protein_coding | 8.157668239 | -0.301134933 | 0.486688943 | 0.618742088 | 0.536086278 | 0.732131062 | 0.301134933 |
| ENSMUSG00000026196 | Bard1 | 12021 | protein_coding | 67.6023253 | 0.301161035 | 0.163308558 | -1.844122798 | 0.065165246 | 0.181269925 | 0.301161035 |

Supplementary Table 1

| ensembl_gene_id | external_gene_name | entrezgene_id | gene_biotype | baseMean | log2FoldChange | lfcSE | stat | pvalue | padj | absLog2FC |
|---|---|---|---|---|---|---|---|---|---|---|
| ENSMUSG00000024525 | Impa2 | 114663 | protein_coding | 243.7186004 | -0.301170544 | 0.116435345 | 2.586590382 | 0.009693075 | 0.041512301 | 0.301170544 |
| ENSMUSG00000027953 | Slc50a1 | 19729 | protein_coding | 417.2787955 | -0.301172524 | 0.161442068 | 1.865514532 | 0.062109317 | 0.17492772 | 0.301172524 |
| ENSMUSG00000038046 | Mrm3 | 67390 | protein_coding | 86.39727467 | -0.301197364 | 0.124731888 | 2.41475832 | 0.015745653 | 0.060682336 | 0.301197364 |
| ENSMUSG00000054889 | Dsp | 109620 | protein_coding | 8770.832075 | -0.301353385 | 0.100354313 | 3.002894196 | 0.002674254 | 0.014436949 | 0.301353385 |
| ENSMUSG00000023007 | Prpf40b | 54614 | protein_coding | 130.2856517 | -0.301362929 | 0.200398552 | 1.503817899 | 0.132628261 | 0.302183099 | 0.301362929 |
| ENSMUSG00000000743 | Chmp1a | 234852 | protein_coding | 2211.975879 | -0.301485374 | 0.084884122 | 3.55172873 | 0.000382709 | 0.00281989 | 0.301485374 |
| ENSMUSG00000026042 | Col5a2 | 12832 | protein_coding | 4.684123938 | 0.301595154 | 1.03789276 | -0.290584119 | 0.771369407 | 0.885225461 | 0.301595154 |
| ENSMUSG00000078713 | Tomm5 | 68512 | protein_coding | 380.7257603 | -0.301605186 | 0.109758332 | 2.747902414 | 0.005997786 | 0.028162478 | 0.301605186 |
| ENSMUSG00000050824 | Sstr5 | 20609 | protein_coding | 6.727346483 | -0.301664632 | 0.464149034 | 0.64993054 | 0.51573709 | 0.716728293 | 0.301664632 |
| ENSMUSG00000040624 | Plekhg1 | 213783 | protein_coding | 242.8810111 | -0.301791238 | 0.10294432 | 2.931596786 | 0.003372243 | 0.01753991 | 0.301791238 |
| ENSMUSG00000054850 | Smim10l2a | 320237 | protein_coding | 31.80226361 | 0.301878233 | 0.279095736 | -1.081629684 | 0.279417109 | 0.496825672 | 0.301878233 |
| ENSMUSG00000068922 | Msto1 | 229524 | protein_coding | 111.980296 | -0.301880571 | 0.111984127 | 2.695744296 | 0.007023156 | 0.031968135 | 0.301880571 |
| ENSMUSG00000032358 | Fam83b | 208994 | protein_coding | 483.0777262 | -0.301958417 | 0.081276832 | 3.71518436 | 0.000203056 | 0.001601366 | 0.301958417 |
| ENSMUSG00000022178 | Ajuba | 16475 | protein_coding | 61.63960021 | 0.301972293 | 0.153797898 | -1.96343576 | 0.049595562 | 0.148051539 | 0.301972293 |
| ENSMUSG00000024302 | Dtna | 13527 | protein_coding | 5.066544218 | 0.302033553 | 1.322524846 | -0.228376468 | 0.819353582 | 0.908985828 | 0.302033553 |
| ENSMUSG00000032120 | C2cd2l | 71764 | protein_coding | 709.8284277 | 0.30217044 | 0.120730196 | -2.502857194 | 0.012319524 | 0.050060192 | 0.30217044 |
| ENSMUSG00000031990 | Jam3 | 83964 | protein_coding | 1.664061895 | 0.302182297 | 0.930924879 | -0.324604384 | 0.745480508 | 1 | 0.302182297 |
| ENSMUSG00000040658 | Dnph1 | 381101 | protein_coding | 35.51102689 | -0.302354623 | 0.211196947 | 1.431624026 | 0.15225145 | 0.332817843 | 0.302354623 |
| ENSMUSG00000027452 | Acss1 | 68738 | protein_coding | 1861.529848 | 0.302446943 | 0.088547376 | -3.415651104 | 0.000636297 | 0.004334906 | 0.302446943 |
| ENSMUSG00000027660 | Skil | 20482 | protein_coding | 1389.429241 | 0.302487539 | 0.115647151 | -2.61560735 | 0.008906892 | 0.038748264 | 0.302487539 |
| ENSMUSG00000023087 | Noct | 12457 | protein_coding | 411.9515873 | -0.302504131 | 0.221853448 | 1.363531349 | 0.172715111 | 0.363275924 | 0.302504131 |
| ENSMUSG00000039990 | Edrf1 | 214764 | protein_coding | 346.7716783 | 0.30260664 | 0.08634719 | -3.504533745 | 0.000457408 | 0.003288476 | 0.30260664 |
| ENSMUSG00000049521 | Cdc42ep1 | 104445 | protein_coding | 263.8802214 | -0.302608899 | 0.128511652 | 2.354719549 | 0.01853669 | 0.069131639 | 0.302608899 |
| ENSMUSG00000030750 | Nsmce1 | 67711 | protein_coding | 218.4240825 | 0.302643268 | 0.098221732 | -3.081225124 | 0.002061507 | 0.011638882 | 0.302643268 |
| ENSMUSG00000001666 | Ddt | 13202 | protein_coding | 237.2367162 | 0.302655211 | 0.09780965 | -3.094328739 | 0.001972588 | 0.011213534 | 0.302655211 |
| ENSMUSG00000022829 | Stxbp5l | 207227 | protein_coding | 5.531905107 | -0.302663943 | 0.772536509 | 0.391779469 | 0.695221166 | 0.838326567 | 0.302663943 |
| ENSMUSG00000001054 | Rmnd5b | 66089 | protein_coding | 220.2758414 | -0.302707093 | 0.107229871 | 2.822973585 | 0.004758049 | 0.02334356 | 0.302707093 |
| ENSMUSG00000037251 | Pomk | 74653 | protein_coding | 147.5273132 | -0.302711636 | 0.123447295 | 2.452152843 | 0.014200433 | 0.05587466 | 0.302711636 |
| ENSMUSG00000021823 | Vcl | 22330 | protein_coding | 768.1632426 | -0.302733326 | 0.078682503 | 3.847530435 | 0.000119314 | 0.001007326 | 0.302733326 |
| ENSMUSG00000074513 | Arfip1 | 99889 | protein_coding | 285.2762268 | 0.302735177 | 0.092759423 | -3.263659559 | 0.001099832 | 0.006900696 | 0.302735177 |
| ENSMUSG00000091705 | H2-Q2 | 15013 | protein_coding | 1602.710067 | 0.302765899 | 0.148818892 | -2.034458762 | 0.041905362 | 0.130398559 | 0.302765899 |
| ENSMUSG00000039270 | Megf9 | 230316 | protein_coding | 196.9933914 | 0.302775524 | 0.148549572 | -2.038212027 | 0.041528733 | 0.129592876 | 0.302775524 |
| ENSMUSG00000038195 | Rilp | 280408 | protein_coding | 156.3817357 | -0.302840535 | 0.134540697 | 2.25092141 | 0.024390515 | 0.085735413 | 0.302840535 |
| ENSMUSG00000026411 | Tmem9 | 66241 | protein_coding | 113.0458281 | -0.302877609 | 0.166187235 | 1.822508265 | 0.068377901 | 0.18782313 | 0.302877609 |
| ENSMUSG00000048473 | Sult6b2 | 330440 | protein_coding | 1.645984505 | 0.302994137 | 1.423901236 | -0.21279154 | 0.831489563 | 1 | 0.302994137 |
| ENSMUSG00000030279 | C2cd5 | 74741 | protein_coding | 371.2575392 | 0.303018298 | 0.068969054 | -4.393540005 | 1.12E-05 | 0.000123434 | 0.303018298 |
| ENSMUSG00000026611 | Spata17 | 74717 | protein_coding | 15.664591 | -0.303048782 | 0.355402023 | 0.852692899 | 0.39382963 | 0.616464692 | 0.303048782 |
| ENSMUSG00000010086 | Rnf112 | 22671 | protein_coding | 2.874758228 | 0.303102066 | 0.844882606 | -0.35875051 | 0.719781739 | 1 | 0.303102066 |
| ENSMUSG00000025529 | Zfp711 | 245595 | protein_coding | 28.45476812 | 0.303189728 | 0.236978775 | -1.279396132 | 0.200757595 | 0.402050804 | 0.303189728 |

Supplementary Table 1

| ensembl_gene_id | external_gene_name | entrezgene_id | gene_biotype | baseMean | log2FoldChange | lfcSE | stat | pvalue | padj | absLog2FC |
|---|---|---|---|---|---|---|---|---|---|---|
| ENSMUSG00000050199 | Lgr4 | 107515 | protein_coding | 3417.034626 | -0.303244413 | 0.113599952 | 2.669406167 | 0.00759855 | 0.034126314 | 0.303244413 |
| ENSMUSG00000064141 | Zfp69 | 381549 | protein_coding | 22.26014415 | 0.303302672 | 0.271147593 | -1.118588842 | 0.263315586 | 0.477583041 | 0.303302672 |
| ENSMUSG00000080932 | Gm10224 | 100043805 | processed_pseudogene | 2.305757843 | -0.303326585 | 0.841593908 | 0.360419179 | 0.718533686 | 1 | 0.303326585 |
| ENSMUSG00000031779 | Ccl22 | 20299 | protein_coding | 2.328077985 | 0.303356684 | 0.978022146 | -0.310173635 | 0.756428919 | 1 | 0.303356684 |
| ENSMUSG00000020273 | Papolg | 216578 | protein_coding | 204.2903423 | -0.303362071 | 0.075223518 | 4.032808878 | 5.51E-05 | 0.000509035 | 0.303362071 |
| ENSMUSG00000039585 | Myo9a | 270163 | protein_coding | 228.7526898 | -0.303402546 | 0.126911956 | 2.390653773 | 0.016818405 | 0.064054228 | 0.303402546 |
| ENSMUSG00000027011 | Ube2e3 | 22193 | protein_coding | 341.2589165 | 0.303438357 | 0.095613694 | -3.173586796 | 0.001505679 | 0.0089185 | 0.303438357 |
| ENSMUSG00000085236 | 2610206C17Rik | 72495 | lncRNA | 4.203006931 | 0.303483748 | 0.697651177 | -0.435007864 | 0.663556719 | 1 | 0.303483748 |
| ENSMUSG00000039067 | Psmd7 | 17463 | protein_coding | 696.3359626 | 0.303489312 | 0.066440436 | -4.567840488 | 4.93E-06 | 5.89E-05 | 0.303489312 |
| ENSMUSG00000018907 | Alox12e | 11685 | protein_coding | 1.522378619 | -0.303511924 | 2.073163792 | 0.146400359 | 0.883605344 | 1 | 0.303511924 |
| ENSMUSG00000025980 | Hspd1 | 15510 | protein_coding | 2851.908288 | -0.303515744 | 0.141621772 | 2.143143241 | 0.032101598 | 0.106322863 | 0.303515744 |
| ENSMUSG00000034936 | Arl4d | 80981 | protein_coding | 47.03801556 | 0.303662853 | 0.313165677 | -0.969655602 | 0.332218188 | 0.554612255 | 0.303662853 |
| ENSMUSG00000074622 | Mafb | 16658 | protein_coding | 154.7516673 | 0.303673614 | 0.286824964 | -1.058741924 | 0.289717329 | 0.508545065 | 0.303673614 |
| ENSMUSG00000063362 | Alg11 | 207958 | protein_coding | 363.3534138 | 0.303740767 | 0.116594799 | -2.605097045 | 0.009184824 | 0.039779225 | 0.303740767 |
| ENSMUSG00000027412 | Lpin3 | 64899 | protein_coding | 670.2651064 | 0.303753467 | 0.110186731 | -2.756715478 | 0.005838513 | 0.027610883 | 0.303753467 |
| ENSMUSG00000002015 | Bcap31 | 27061 | protein_coding | 1447.602573 | -0.303757459 | 0.090750836 | 3.347158805 | 0.000816444 | 0.00535797 | 0.303757459 |
| ENSMUSG00000067567 | Hdac8 | 70315 | protein_coding | 86.36797598 | -0.303783708 | 0.125055822 | 2.429184844 | 0.015132815 | 0.058800778 | 0.303783708 |
| ENSMUSG00000054052 | Rdh19 | 216453 | protein_coding | 6.291819702 | -0.303788079 | 0.501650618 | 0.605577005 | 0.544795676 | 0.736895388 | 0.303788079 |
| ENSMUSG00000078350 | Smim1 | 68859 | protein_coding | 8.871264425 | 0.303887818 | 0.367296832 | -0.827363023 | 0.408031332 | 0.628624818 | 0.303887818 |
| ENSMUSG00000009293 | Ube2g2 | 22213 | protein_coding | 345.7386962 | 0.30391006 | 0.095414914 | -3.185142098 | 0.001446829 | 0.008622498 | 0.30391006 |
| ENSMUSG00000045062 | Pcdhb7 | 93878 | protein_coding | 1.848662838 | 0.303946872 | 0.820171449 | -0.370589432 | 0.710943355 | 1 | 0.303946872 |
| ENSMUSG00000022314 | Rad21 | 19357 | protein_coding | 1818.859854 | 0.303948725 | 0.099194453 | -3.06417059 | 0.002182744 | 0.012153013 | 0.303948725 |
| ENSMUSG00000036273 | Lrrk2 | 66725 | protein_coding | 272.1380098 | 0.30395797 | 0.268459254 | -1.132231299 | 0.257537205 | 0.470773677 | 0.30395797 |
| ENSMUSG00000020087 | Tysnd1 | 71767 | protein_coding | 389.3471629 | 0.303997092 | 0.09332747 | -3.257316335 | 0.00112471 | 0.007027417 | 0.303997092 |
| ENSMUSG00000092080 | Vmn2r16 | 384220 | protein_coding | 1.882770371 | -0.304010144 | 2.046527197 | 0.148549281 | 0.881909297 | 1 | 0.304010144 |
| ENSMUSG00000035469 | Rcbtb1 | 71330 | protein_coding | 680.5054487 | 0.304323442 | 0.117303543 | -2.594324396 | 0.009477702 | 0.040790921 | 0.304323442 |
| ENSMUSG00000042340 | Ctf1 | 13019 | protein_coding | 20.75261484 | -0.304457004 | 0.281407525 | 1.081907826 | 0.279293487 | 0.496688848 | 0.304457004 |
| ENSMUSG00000051579 | Tceal8 | 66684 | protein_coding | 233.4139916 | 0.304484462 | 0.146367508 | -2.080273593 | 0.037500445 | 0.119750527 | 0.304484462 |
| ENSMUSG00000006711 | D130043K22Rik | 210108 | protein_coding | 21.99730057 | -0.30456319 | 0.298110294 | 1.021646003 | 0.306948476 | 0.527501415 | 0.30456319 |
| ENSMUSG00000000440 | Pparg | 19016 | protein_coding | 1086.922247 | -0.304584627 | 0.130342265 | 2.336806311 | 0.019449258 | 0.071853476 | 0.304584627 |
| ENSMUSG00000026650 | Meig1 | 104362 | protein_coding | 1.002649023 | -0.304593136 | 1.156025873 | 0.263482975 | 0.792178344 | 1 | 0.304593136 |
| ENSMUSG00000070315 | 4930581F22Rik | 78934 | lncRNA | 49.22519084 | -0.304772382 | 0.211963262 | 1.437854746 | 0.150475272 | 0.330401362 | 0.304772382 |
| ENSMUSG00000022683 | Pla2g10 | 26565 | protein_coding | 683.6162985 | -0.304825594 | 0.149477248 | 2.03927753 | 0.041422337 | 0.129358637 | 0.304825594 |
| ENSMUSG00000025357 | Dgka | 13139 | protein_coding | 1389.302797 | -0.304870513 | 0.118196788 | 2.579346848 | 0.009898734 | 0.042239714 | 0.304870513 |
| ENSMUSG00000027954 | Efna1 | 13636 | protein_coding | 921.8314469 | -0.304931311 | 0.137245557 | 2.221793689 | 0.026297251 | 0.090795126 | 0.304931311 |
| ENSMUSG00000040957 | Cables1 | 63955 | protein_coding | 375.6738166 | -0.304936894 | 0.163068394 | 1.869993854 | 0.061484671 | 0.173816344 | 0.304936894 |
| ENSMUSG00000039903 | Eva1c | 70967 | protein_coding | 10.21877705 | -0.304980647 | 0.567343575 | 0.537559004 | 0.590881539 | 0.771232643 | 0.304980647 |
| ENSMUSG00000033595 | Lgi3 | 213469 | protein_coding | 1.733234576 | -0.305007438 | 1.834323986 | 0.166277844 | 0.867938301 | 1 | 0.305007438 |
| ENSMUSG00000073295 | Nudt11 | 58242 | protein_coding | 17.38907589 | 0.305079298 | 0.291252086 | -1.047475065 | 0.294880527 | 0.514056479 | 0.305079298 |

| ensembl_gene_id | external_gene_name | entrezgene_id | gene_biotype | baseMean | log2FoldChange | lfcSE | stat | pvalue | padj | absLog2FC |
|---|---|---|---|---|---|---|---|---|---|---|
| ENSMUSG00000034761 | Map4k5 | 399510 | protein_coding | 325.3416142 | -0.305091141 | 0.076518043 | 3.987179107 | 6.69E-05 | 0.000603709 | 0.305091141 |
| ENSMUSG00000071753 | Cdr1os | 331424 | lncRNA | 4.466318064 | -0.305096388 | 3.14706773 | 0.096946241 | 0.922769086 | 0.962580298 | 0.305096388 |
| ENSMUSG00000034993 | Vat1 | 26949 | protein_coding | 533.601369 | -0.305172211 | 0.100845574 | 3.026133896 | 0.002477026 | 0.013504657 | 0.305172211 |
| ENSMUSG00000033174 | Mgll | 23945 | protein_coding | 788.4843876 | 0.305269977 | 0.213994853 | -1.426529532 | 0.153715549 | 0.335154287 | 0.305269977 |
| ENSMUSG00000022102 | Dok2 | 13449 | protein_coding | 6.723452598 | -0.305369322 | 0.58172254 | 0.524939814 | 0.599625031 | 0.776544903 | 0.305369322 |
| ENSMUSG00000028125 | Abca4 | 11304 | protein_coding | 4.108984253 | 0.305404623 | 1.701354723 | -0.17950673 | 0.857539833 | 1 | 0.305404623 |
| ENSMUSG00000079049 | Serpinb1c | 380839 | protein_coding | 2.149348383 | -0.305508397 | 0.833859729 | 0.366378644 | 0.714082553 | 1 | 0.305508397 |
| ENSMUSG00000027087 | Itgav | 16410 | protein_coding | 468.3685802 | 0.305510796 | 0.115701727 | -2.640503336 | 0.008278298 | 0.036592026 | 0.305510796 |
| ENSMUSG00000022226 | Mcpt2 | 17225 | protein_coding | 3.958337632 | -0.305526885 | 0.637949286 | 0.478920334 | 0.631995304 | 1 | 0.305526885 |
| ENSMUSG00000038080 | Kdm1b | 218214 | protein_coding | 380.6362165 | -0.3057676 | 0.111696474 | 2.737486577 | 0.006191065 | 0.028864834 | 0.3057676 |
| ENSMUSG00000020805 | Slc13a5 | 237831 | protein_coding | 1.940004603 | 0.305779722 | 1.522286502 | -0.200868707 | 0.840801237 | 1 | 0.305779722 |
| ENSMUSG00000025525 | Apool | 68117 | protein_coding | 317.6691265 | -0.305893605 | 0.094164448 | 3.248504223 | 0.001160135 | 0.007207851 | 0.305893605 |
| ENSMUSG00000040209 | Zfp704 | 170753 | protein_coding | 823.5977352 | 0.30593439 | 0.124087545 | -2.465472182 | 0.01368328 | 0.054292572 | 0.30593439 |
| ENSMUSG00000057182 | Scn3a | 20269 | protein_coding | 28.12957521 | -0.305998471 | 0.29306679 | 1.044125372 | 0.296427389 | 0.515610058 | 0.305998471 |
| ENSMUSG00000118506 | Cfap141 | 73545 | protein_coding | 4.221437255 | 0.306024165 | 0.57119495 | -0.535761329 | 0.592123512 | 1 | 0.306024165 |
| ENSMUSG00000031488 | Rab11fip1 | 75767 | protein_coding | 2327.12458 | -0.306073352 | 0.071518793 | 4.279621321 | 1.87E-05 | 0.000194686 | 0.306073352 |
| ENSMUSG00000037434 | Slc30a1 | 22782 | protein_coding | 764.8918148 | 0.306142437 | 0.108049927 | -2.833342384 | 0.004606402 | 0.022714279 | 0.306142437 |
| ENSMUSG00000023755 | Rhebl1 | 69159 | protein_coding | 12.40027194 | -0.306151179 | 0.356000323 | 0.859974444 | 0.38980313 | 0.612367665 | 0.306151179 |
| ENSMUSG00000032479 | Map4 | 17758 | protein_coding | 1117.13714 | 0.306155515 | 0.098840187 | -3.097480132 | 0.001951735 | 0.011125629 | 0.306155515 |
| ENSMUSG00000026773 | Pfkfb3 | 170768 | protein_coding | 34.22727905 | 0.306242944 | 0.2180027 | -1.404766753 | 0.160090649 | 0.344460843 | 0.306242944 |
| ENSMUSG00000041491 | Cep78 | 208518 | protein_coding | 90.23261706 | 0.30624331 | 0.180767061 | -1.69413226 | 0.090240153 | 0.229089299 | 0.30624331 |
| ENSMUSG00000024949 | Sf1 | 22668 | protein_coding | 1365.569028 | 0.306321978 | 0.058692444 | -5.219104148 | 1.80E-07 | 2.84E-06 | 0.306321978 |
| ENSMUSG00000030226 | Lmo3 | 109593 | protein_coding | 1.722639271 | -0.30632302 | 2.391605225 | 0.128082602 | 0.898083604 | 1 | 0.30632302 |
| ENSMUSG00000027297 | Ltk | 17005 | protein_coding | 14.15298958 | -0.306331579 | 0.356600676 | 0.859032526 | 0.390322571 | 0.612497263 | 0.306331579 |
| ENSMUSG00000061397 | Krt79 | 223917 | protein_coding | 2.77448124 | -0.306562632 | 0.811386716 | 0.37782555 | 0.705560192 | 1 | 0.306562632 |
| ENSMUSG00000115338 | Pnp | 18950 | protein_coding | 271.3796399 | -0.306579144 | 0.1049789 | 2.920388224 | 0.003495956 | 0.018083738 | 0.306579144 |
| ENSMUSG00000026094 | Stk17b | 98267 | protein_coding | 774.4068002 | -0.306580081 | 0.1621503 | 1.890715472 | 0.058662336 | 0.168010721 | 0.306580081 |
| ENSMUSG00000028821 | Syf2 | 68592 | protein_coding | 485.1075575 | -0.306714575 | 0.084605615 | 3.625227194 | 0.000288708 | 0.002203196 | 0.306714575 |
| ENSMUSG00000030020 | Prickle2 | 243548 | protein_coding | 4.074078195 | -0.306764784 | 1.027283496 | 0.298617456 | 0.765231945 | 1 | 0.306764784 |
| ENSMUSG00000067995 | Gtf2f2 | 68705 | protein_coding | 239.0662523 | 0.306771585 | 0.089102415 | -3.442909887 | 0.000575491 | 0.003986434 | 0.306771585 |
| ENSMUSG00000035762 | Tmem161b | 72745 | protein_coding | 250.5641697 | 0.306867181 | 0.087371329 | -3.512218309 | 0.000444383 | 0.003214427 | 0.306867181 |
| ENSMUSG00000074923 | Pak6 | 214230 | protein_coding | 27.54455496 | -0.306907482 | 0.223982013 | 1.370232719 | 0.170614268 | 0.360481579 | 0.306907482 |
| ENSMUSG00000004151 | Etv1 | 14009 | protein_coding | 10.7289061 | -0.306928895 | 0.424755427 | 0.722601466 | 0.469924767 | 0.683489308 | 0.306928895 |
| ENSMUSG00000028551 | Cdkn2c | 12580 | protein_coding | 138.779042 | 0.306931123 | 0.17800503 | -1.724283427 | 0.08465669 | 0.219013029 | 0.306931123 |
| ENSMUSG00000039478 | Micu3 | 78506 | protein_coding | 16.50012273 | 0.306968967 | 0.305494536 | -1.004826374 | 0.314980462 | 0.535990818 | 0.306968967 |
| ENSMUSG00000025377 | Tepsin | 78777 | protein_coding | 376.1950276 | 0.306980166 | 0.087863032 | -3.493849 | 0.00047611 | 0.003403673 | 0.306980166 |
| ENSMUSG00000059713 | Rcan3 | 53902 | protein_coding | 459.7019171 | 0.30703113 | 0.107079838 | -2.867310383 | 0.004139768 | 0.02076044 | 0.30703113 |
| ENSMUSG00000028004 | Npy2r | 18167 | protein_coding | 5.856195652 | -0.307039448 | 0.51280471 | 0.598745373 | 0.549342695 | 0.740342018 | 0.307039448 |
| ENSMUSG00000038059 | Smim3 | 106878 | protein_coding | 55.97877006 | 0.30705683 | 0.255527499 | -1.201658649 | 0.229495808 | 0.438111017 | 0.30705683 |

Supplementary Table 1

| ensembl_gene_id | external_gene_name | entrezgene_id | gene_biotype | baseMean | log2FoldChange | lfcSE | stat | pvalue | padj | absLog2FC |
|---|---|---|---|---|---|---|---|---|---|---|
| ENSMUSG00000038301 | Snx10 | 71982 | protein_coding | 122.4360142 | -0.307094572 | 0.13997321 | 2.19395248 | 0.028238826 | 0.096124511 | 0.307094572 |
| ENSMUSG00000001555 | Fkbp10 | 14230 | protein_coding | 2.103093283 | -0.307135473 | 1.063052967 | 0.288918315 | 0.772643887 | 1 | 0.307135473 |
| ENSMUSG00000110605 | Gm32856 | 102635552 | lncRNA | 25.11593868 | 0.307195774 | 0.220613695 | -1.392460131 | 0.163783107 | 0.349988901 | 0.307195774 |
| ENSMUSG00000003518 | Dusp3 | 72349 | protein_coding | 322.3325746 | -0.307201474 | 0.101144142 | 3.037264143 | 0.002387362 | 0.013089272 | 0.307201474 |
| ENSMUSG00000031432 | Prps1 | 19139 | protein_coding | 169.877847 | 0.307344335 | 0.1628163 | -1.887675468 | 0.059069532 | 0.168971808 | 0.307344335 |
| ENSMUSG00000028271 | Gtf2b | 229906 | protein_coding | 440.8867106 | 0.307405453 | 0.09300424 | -3.305284296 | 0.000948801 | 0.006091952 | 0.307405453 |
| ENSMUSG00000078763 | Slfn1 | 20555 | protein_coding | 1.283975828 | -0.30743757 | 1.479256954 | 0.207832432 | 0.835359809 | 1 | 0.30743757 |
| ENSMUSG00000023345 | Poc1a | 70235 | protein_coding | 177.5815267 | 0.307441905 | 0.109812229 | -2.799705535 | 0.005114924 | 0.024814295 | 0.307441905 |
| ENSMUSG00000097368 | Gm10390 | 100503427 | lncRNA | 12.5643057 | -0.30761217 | 0.489236991 | 0.628759019 | 0.529506833 | 0.727166502 | 0.30761217 |
| ENSMUSG00000113204 | Gm46430 | 108168101 | lncRNA | 76.64037619 | -0.307653642 | 0.186483675 | 1.649761794 | 0.098991666 | 0.245826823 | 0.307653642 |
| ENSMUSG00000032198 | Dock6 | 319899 | protein_coding | 356.1381211 | 0.307697431 | 0.198559452 | -1.549648869 | 0.121225817 | 0.283960728 | 0.307697431 |
| ENSMUSG00000022673 | Mcm4 | 17217 | protein_coding | 334.979159 | 0.307698572 | 0.158725789 | -1.938554366 | 0.052555625 | 0.154982783 | 0.307698572 |
| ENSMUSG00000028936 | Rpl22 | 19934 | protein_coding | 2181.722287 | -0.30771121 | 0.108740271 | 2.829781523 | 0.00465798 | 0.022923163 | 0.30771121 |
| ENSMUSG00000078921 | Tgtp2 | 100039796 | protein_coding | 4.428257961 | 0.307740669 | 0.60482844 | -0.508806546 | 0.610887831 | 0.784402584 | 0.307740669 |
| ENSMUSG00000002458 | Rgs19 | 56470 | protein_coding | 35.29536985 | 0.307855181 | 0.27593296 | -1.115688323 | 0.264555571 | 0.479304239 | 0.307855181 |
| ENSMUSG00000041779 | Tram2 | 170829 | protein_coding | 22.89650644 | 0.307991391 | 0.315613695 | -0.975849259 | 0.32913917 | 0.551537722 | 0.307991391 |
| ENSMUSG00000027985 | Lef1 | 16842 | protein_coding | 4.714976867 | -0.308015273 | 0.97882764 | 0.314677744 | 0.753006344 | 0.8762082 | 0.308015273 |
| ENSMUSG00000037300 | Ttc13 | 234875 | protein_coding | 607.8674656 | 0.308060764 | 0.077733917 | -3.963016084 | 7.40E-05 | 0.000661879 | 0.308060764 |
| ENSMUSG00000084771 | A230072E10Rik | 331547 | lncRNA | 7.412120672 | -0.308074459 | 0.403421799 | 0.763653475 | 0.445073773 | 0.662372535 | 0.308074459 |
| ENSMUSG00000059970 | Hspa2 | 15512 | protein_coding | 25.19448088 | 0.308259224 | 0.277656646 | -1.110217343 | 0.266905381 | 0.48202925 | 0.308259224 |
| ENSMUSG00000034557 | Zfyve9 | 230597 | protein_coding | 415.1646789 | -0.308385659 | 0.070958448 | 4.346003435 | 1.39E-05 | 0.000149154 | 0.308385659 |
| ENSMUSG00000025648 | Pfkfb4 | 270198 | protein_coding | 355.1117951 | -0.308385675 | 0.13935447 | 2.212958614 | 0.026900499 | 0.09253023 | 0.308385675 |
| ENSMUSG00000009585 | Apobec3 | 80287 | protein_coding | 157.4644568 | -0.308439635 | 0.205648923 | 1.499835884 | 0.133656919 | 0.303730613 | 0.308439635 |
| ENSMUSG00000020258 | Glyctk | 235582 | protein_coding | 13.20524941 | -0.308471618 | 0.278877598 | 1.106118315 | 0.268675312 | 0.483691716 | 0.308471618 |
| ENSMUSG00000041859 | Mcm3 | 17215 | protein_coding | 342.6520783 | 0.308704048 | 0.176161689 | -1.75239038 | 0.079706705 | 0.209275846 | 0.308704048 |
| ENSMUSG00000042046 | Dstyk | 213452 | protein_coding | 180.4671145 | -0.3087111 | 0.091180658 | 3.385708194 | 0.000709948 | 0.004741185 | 0.3087111 |
| ENSMUSG00000034591 | Slc41a2 | 338365 | protein_coding | 283.8871807 | 0.30876327 | 0.165821388 | -1.862023195 | 0.062599821 | 0.175901555 | 0.30876327 |
| ENSMUSG00000085286 | Ube4bos3 | 74310 | lncRNA | 3.963826637 | 0.308788257 | 0.814985005 | -0.378888268 | 0.704770836 | 1 | 0.308788257 |
| ENSMUSG00000036550 | Cnot1 | 234594 | protein_coding | 3622.800164 | 0.308936937 | 0.064629434 | -4.780127549 | 1.75E-06 | 2.29E-05 | 0.308936937 |
| ENSMUSG00000023966 | Rsph9 | 75564 | protein_coding | 10.98454688 | -0.308983274 | 0.365850768 | 0.844560955 | 0.398356017 | 0.620277218 | 0.308983274 |
| ENSMUSG00000024800 | Rpp30 | 54364 | protein_coding | 74.03605533 | 0.309020721 | 0.14520619 | -2.128151159 | 0.033324553 | 0.109351285 | 0.309020721 |
| ENSMUSG00000043430 | Psapl1 | 76943 | protein_coding | 3.595908902 | -0.309243197 | 0.847804703 | 0.364757587 | 0.715292368 | 1 | 0.309243197 |
| ENSMUSG00000024429 | Gnl1 | 14670 | protein_coding | 397.2212928 | -0.309436149 | 0.090853967 | 3.405862824 | 0.000659553 | 0.004469372 | 0.309436149 |
| ENSMUSG00000031493 | Ggn | 243897 | protein_coding | 6.607679439 | 0.30944396 | 0.692874319 | -0.446609077 | 0.655157341 | 0.812818663 | 0.30944396 |
| ENSMUSG00000117284 | Gm7072 | 631624 | protein_coding | 285.2379513 | -0.309444036 | 0.126739554 | 2.441574296 | 0.014623379 | 0.05718458 | 0.309444036 |
| ENSMUSG00000035776 | Cd99l2 | 171486 | protein_coding | 284.7507908 | -0.309623051 | 0.105490749 | 2.93507301 | 0.003334692 | 0.01737744 | 0.309623051 |
| ENSMUSG00000047844 | Bex4 | 406217 | protein_coding | 25.95833606 | 0.309644738 | 0.274337332 | -1.128700695 | 0.259024116 | 0.472341459 | 0.309644738 |
| ENSMUSG00000059878 | Zfp422 | 67255 | protein_coding | 276.6760462 | -0.309660704 | 0.091735778 | 3.375571804 | 0.000736625 | 0.004889317 | 0.309660704 |
| ENSMUSG00000030867 | Plk1 | 18817 | protein_coding | 655.5858751 | 0.309719941 | 0.14209232 | -2.17970922 | 0.029279023 | 0.09897928 | 0.309719941 |

| ensembl_gene_id | external_gene_name | entrezgene_id | gene_biotype | baseMean | log2FoldChange | lfcSE | stat | pvalue | padj | absLog2FC |
|---|---|---|---|---|---|---|---|---|---|---|
| ENSMUSG00000028874 | Fgr | 14191 | protein_coding | 19.58799669 | 0.309742593 | 0.307381439 | -1.007681511 | 0.313607384 | 0.534874425 | 0.309742593 |
| ENSMUSG00000027533 | Fabp5 | 16592 | protein_coding | 41.15700856 | -0.309765689 | 0.228650551 | 1.354755924 | 0.175495334 | 0.367316849 | 0.309765689 |
| ENSMUSG00000050064 | Zfp697 | 242109 | protein_coding | 57.85552946 | -0.309785995 | 0.170030115 | 1.821947802 | 0.068462906 | 0.187931588 | 0.309785995 |
| ENSMUSG00000088712 | Gm22728 | 115490275 | scaRNA | 2.013300495 | -0.309796186 | 0.81337812 | 0.380875977 | 0.703295279 | 1 | 0.309796186 |
| ENSMUSG00000034210 | Efcab14 | 230648 | protein_coding | 763.9784164 | -0.309869821 | 0.085869889 | 3.608596949 | 0.000307857 | 0.002331022 | 0.309869821 |
| ENSMUSG00000032688 | Malt1 | 240354 | protein_coding | 364.5250924 | 0.309882321 | 0.080016037 | -3.872752672 | 0.000107613 | 0.000919359 | 0.309882321 |
| ENSMUSG00000025366 | Esyt1 | 23943 | protein_coding | 584.9950417 | 0.309896286 | 0.103941387 | -2.981452299 | 0.002868847 | 0.015327126 | 0.309896286 |
| ENSMUSG00000034614 | Pik3ip1 | 216505 | protein_coding | 27.30474844 | -0.309987369 | 0.222710534 | 1.39188463 | 0.163957337 | 0.350225257 | 0.309987369 |
| ENSMUSG00000051499 | Zfp786 | 330301 | protein_coding | 21.28906884 | -0.309988077 | 0.246636995 | 1.256859609 | 0.208804477 | 0.412069446 | 0.309988077 |
| ENSMUSG00000003657 | Calb2 | 12308 | protein_coding | 1.966699971 | -0.310003909 | 2.088874513 | 0.148407148 | 0.882021459 | 1 | 0.310003909 |
| ENSMUSG00000078963 | Hsbp1l1 | 66255 | protein_coding | 33.39149787 | 0.310073751 | 0.208836437 | -1.484768441 | 0.137605175 | 0.31001526 | 0.310073751 |
| ENSMUSG00000037418 | Best1 | 24115 | protein_coding | 17.48140812 | 0.310283159 | 0.667368472 | -0.464935298 | 0.641977792 | 0.804573016 | 0.310283159 |
| ENSMUSG00000002012 | Pnck | 93843 | protein_coding | 11.04187126 | 0.310589711 | 0.632155131 | -0.491318817 | 0.623200973 | 0.793304622 | 0.310589711 |
| ENSMUSG00000045100 | Slc25a26 | 67582 | protein_coding | 74.84497139 | -0.310610865 | 0.157196675 | 1.975937885 | 0.048161813 | 0.145029937 | 0.310610865 |
| ENSMUSG00000048617 | Rtbdn | 234542 | protein_coding | 3.096937161 | 0.310700085 | 0.891771311 | -0.348407805 | 0.727533942 | 1 | 0.310700085 |
| ENSMUSG00000085333 | 1700030A11Rik | 74179 | lncRNA | 20.83685856 | -0.310736049 | 0.256752886 | 1.210253383 | 0.22618168 | 0.433917781 | 0.310736049 |
| ENSMUSG00000021373 | Cap2 | 67252 | protein_coding | 10.77192565 | 0.310901922 | 0.413950107 | -0.751061339 | 0.452615741 | 0.668918592 | 0.310901922 |
| ENSMUSG00000014353 | Tmem87b | 72477 | protein_coding | 1112.306228 | -0.310902744 | 0.078397156 | 3.965740105 | 7.32E-05 | 0.000654908 | 0.310902744 |
| ENSMUSG00000095432 | Zfp748 | 212276 | protein_coding | 175.2934566 | -0.310935421 | 0.135198922 | 2.299836531 | 0.021457483 | 0.077538047 | 0.310935421 |
| ENSMUSG00000061286 | Exosc5 | 27998 | protein_coding | 200.2961834 | -0.31094203 | 0.108104173 | 2.876318474 | 0.004023435 | 0.020269375 | 0.31094203 |
| ENSMUSG00000033249 | Hsf4 | 26386 | protein_coding | 9.314780543 | -0.310948473 | 0.366173751 | 0.849182858 | 0.39577955 | 0.618169251 | 0.310948473 |
| ENSMUSG00000028062 | Lamtor2 | 83409 | protein_coding | 490.8462374 | -0.311051292 | 0.079975253 | 3.889344262 | 0.000100515 | 0.000864538 | 0.311051292 |
| ENSMUSG00000026198 | Abcb6 | 74104 | protein_coding | 65.67808401 | -0.311077401 | 0.150156632 | 2.07168606 | 0.038294728 | 0.121849545 | 0.311077401 |
| ENSMUSG00000052117 | D630039A03Rik | 242484 | protein_coding | 493.4986947 | -0.311266161 | 0.204253152 | 1.523923415 | 0.127527844 | 0.294503539 | 0.311266161 |
| ENSMUSG00000112343 | Sfta3-ps | 104798 | lncRNA | 2.400106364 | 0.311361547 | 2.37939655 | -0.130857358 | 0.895888148 | 1 | 0.311361547 |
| ENSMUSG00000047230 | Cldn2 | 12738 | protein_coding | 2145.946905 | 0.311386101 | 0.213743712 | -1.456819939 | 0.14516609 | 0.322338142 | 0.311386101 |
| ENSMUSG00000040272 | Accs | 329470 | protein_coding | 48.1644694 | -0.311444374 | 0.202805163 | 1.535682669 | 0.124616228 | 0.289400593 | 0.311444374 |
| ENSMUSG00000030301 | Ccdc91 | 67015 | protein_coding | 418.351977 | -0.311548838 | 0.099638191 | 3.126801427 | 0.001767192 | 0.010218283 | 0.311548838 |
| ENSMUSG00000097170 | Gm16982 | 100036523 | lncRNA | 9.653436583 | -0.311560839 | 0.378363101 | 0.823444035 | 0.410255531 | 0.630552252 | 0.311560839 |
| ENSMUSG00000021198 | Unc79 | 217843 | protein_coding | 24.84916131 | -0.311658921 | 0.321194246 | 0.970312901 | 0.331890548 | 0.554292124 | 0.311658921 |
| ENSMUSG00000050332 | Amer1 | 72345 | protein_coding | 146.4391598 | -0.311739941 | 0.130164329 | 2.394972143 | 0.016621628 | 0.063495157 | 0.311739941 |
| ENSMUSG00000030446 | Zfp273 | 212569 | protein_coding | 47.29763679 | -0.311783975 | 0.16080382 | 1.938909005 | 0.052512419 | 0.154934252 | 0.311783975 |
| ENSMUSG00000011257 | Pabpc4 | 230721 | protein_coding | 720.5894427 | -0.311887456 | 0.101329003 | 3.077968277 | 0.002084171 | 0.01173204 | 0.311887456 |
| ENSMUSG00000095440 | Fignl2 | 668225 | protein_coding | 39.49388611 | -0.312150931 | 0.27696934 | 1.127023414 | 0.259732582 | 0.473163139 | 0.312150931 |
| ENSMUSG00000072972 | Adam4 | 11498 | protein_coding | 12.0033713 | 0.312326449 | 0.322493431 | -0.968473831 | 0.332807782 | 0.555206356 | 0.312326449 |
| ENSMUSG00000019188 | H13 | 14950 | protein_coding | 1292.964796 | 0.312398462 | 0.085042653 | -3.673432703 | 0.000239314 | 0.001854541 | 0.312398462 |
| ENSMUSG00000023110 | Prmt5 | 27374 | protein_coding | 485.3655788 | 0.312504328 | 0.124398094 | -2.512131164 | 0.012000448 | 0.049101931 | 0.312504328 |
| ENSMUSG00000026380 | Tfcp2l1 | 81879 | protein_coding | 6841.680259 | -0.312557686 | 0.102254369 | 3.056668265 | 0.002238118 | 0.01240278 | 0.312557686 |
| ENSMUSG00000036502 | Tmem255a | 245386 | protein_coding | 4.159201623 | 0.312567196 | 0.805662118 | -0.387963129 | 0.698043319 | 1 | 0.312567196 |

| ensembl_gene_id | external_gene_name | entrezgene_id | gene_biotype | baseMean | log2FoldChange | lfcSE | stat | pvalue | padj | absLog2FC |
|---|---|---|---|---|---|---|---|---|---|---|
| ENSMUSG00000031360 | Ctps2 | 55936 | protein_coding | 278.3985948 | -0.312668638 | 0.119473295 | 2.617058791 | 0.008869106 | 0.038648666 | 0.312668638 |
| ENSMUSG00000028318 | Polr1e | 64424 | protein_coding | 102.2777721 | 0.312674405 | 0.126724215 | -2.467361154 | 0.0136113 | 0.054110978 | 0.312674405 |
| ENSMUSG00000019080 | Mfsd3 | 69572 | protein_coding | 50.29048862 | -0.312790291 | 0.160309386 | 1.951166417 | 0.051037252 | 0.151424575 | 0.312790291 |
| ENSMUSG00000028999 | Rint1 | 72772 | protein_coding | 328.9620044 | 0.312852097 | 0.093853571 | -3.333406422 | 0.000857895 | 0.005597504 | 0.312852097 |
| ENSMUSG00000079110 | Capn3 | 12335 | protein_coding | 8.826591099 | 0.312889127 | 0.574192926 | -0.544919858 | 0.585808638 | 0.767554399 | 0.312889127 |
| ENSMUSG00000059142 | Zfp945 | 240041 | protein_coding | 109.4464929 | -0.312918986 | 0.165060185 | 1.895787192 | 0.05798819 | 0.166571051 | 0.312918986 |
| ENSMUSG00000054978 | Kbtbd13 | 74492 | protein_coding | 32.69638902 | 0.312956657 | 0.214351098 | -1.460018912 | 0.144284876 | 0.321034822 | 0.312956657 |
| ENSMUSG00000000184 | Ccnd2 | 12444 | protein_coding | 2225.177164 | -0.313018104 | 0.168398539 | 1.858793468 | 0.063056419 | 0.176840786 | 0.313018104 |
| ENSMUSG00000029830 | Svopl | 320590 | protein_coding | 3.701174892 | 0.313177278 | 1.102798485 | -0.283984139 | 0.776422536 | 1 | 0.313177278 |
| ENSMUSG00000021708 | Rasgrf2 | 19418 | protein_coding | 122.4727549 | 0.313194295 | 0.140496722 | -2.229192898 | 0.025801073 | 0.089531187 | 0.313194295 |
| ENSMUSG00000017376 | Nlk | 18099 | protein_coding | 206.0390343 | 0.313197746 | 0.105116092 | -2.97954139 | 0.002886802 | 0.015398144 | 0.313197746 |
| ENSMUSG00000018570 | 2810408A11Rik | 70419 | protein_coding | 59.26404825 | -0.313259287 | 0.206362243 | 1.518006793 | 0.129012686 | 0.296522686 | 0.313259287 |
| ENSMUSG00000020629 | Adi1 | 104923 | protein_coding | 170.5638355 | -0.313278699 | 0.137324888 | 2.281295859 | 0.022530946 | 0.080642929 | 0.313278699 |
| ENSMUSG00000042826 | Fgf11 | 14166 | protein_coding | 38.68027627 | 0.313298631 | 0.296870283 | -1.055338473 | 0.291270552 | 0.510066755 | 0.313298631 |
| ENSMUSG00000020495 | Smg8 | 74133 | protein_coding | 189.2580013 | 0.313318037 | 0.097005057 | -3.229914485 | 0.001238272 | 0.007596149 | 0.313318037 |
| ENSMUSG00000027684 | Mecom | 14013 | protein_coding | 1993.493236 | -0.313331381 | 0.098807697 | 3.171123215 | 0.001518507 | 0.0089816 | 0.313331381 |
| ENSMUSG00000031490 | Eif4ebp1 | 13685 | protein_coding | 172.3207177 | 0.313514057 | 0.113568312 | -2.760576884 | 0.005769938 | 0.027372809 | 0.313514057 |
| ENSMUSG00000029547 | Ints1 | 68510 | protein_coding | 966.4975283 | 0.313551424 | 0.078254774 | -4.006802483 | 6.15E-05 | 0.000562777 | 0.313551424 |
| ENSMUSG00000026525 | Opn3 | 13603 | protein_coding | 12.31463748 | 0.313553905 | 0.289627381 | -1.0826114 | 0.278980945 | 0.496293367 | 0.313553905 |
| ENSMUSG00000021916 | Glt8d1 | 76485 | protein_coding | 96.95076436 | 0.313578353 | 0.133718409 | -2.345064943 | 0.019023766 | 0.070613276 | 0.313578353 |
| ENSMUSG00000039639 | Kcne1 | 16509 | protein_coding | 1.678192659 | 0.313603331 | 1.819368007 | -0.172369378 | 0.863147145 | 1 | 0.313603331 |
| ENSMUSG00000117602 | 9130209A04Rik | 77668 | lncRNA | 30.89964646 | -0.313612729 | 0.360731001 | 0.869381141 | 0.384638695 | 0.607019295 | 0.313612729 |
| ENSMUSG00000031520 | Vegfc | 22341 | protein_coding | 46.43083456 | 0.31363143 | 0.175893803 | -1.783072652 | 0.074574477 | 0.199760409 | 0.31363143 |
| ENSMUSG00000042737 | Dpm3 | 68563 | protein_coding | 227.3283986 | -0.313758526 | 0.096868278 | 3.239022451 | 0.001199401 | 0.007393398 | 0.313758526 |
| ENSMUSG00000034522 | Zfp395 | 380912 | protein_coding | 117.6602315 | -0.313822958 | 0.136693479 | 2.295815132 | 0.021686454 | 0.078262696 | 0.313822958 |
| ENSMUSG00000052337 | Immt | 76614 | protein_coding | 3455.618749 | 0.313946739 | 0.064640777 | -4.856790959 | 1.19E-06 | 1.62E-05 | 0.313946739 |
| ENSMUSG00000026003 | Acadl | 11363 | protein_coding | 1224.472796 | -0.314209249 | 0.083201437 | 3.776488255 | 0.000159055 | 0.00129073 | 0.314209249 |
| ENSMUSG00000098269 | Mir8094 | 102465886 | miRNA | 1.4427443 | 0.31428815 | 0.985263064 | -0.318989071 | 0.7497348 | 1 | 0.31428815 |
| ENSMUSG00000050973 | Gdpgp1 | 269952 | protein_coding | 154.439607 | -0.314341631 | 0.119083636 | 2.63967108 | 0.008298653 | 0.036642767 | 0.314341631 |
| ENSMUSG00000043432 | Leng9 | 243813 | protein_coding | 662.0572498 | 0.314348943 | 0.104220977 | -3.016177276 | 0.002559836 | 0.013891924 | 0.314348943 |
| ENSMUSG00000110949 | Nudt8 | 66387 | protein_coding | 2.264023428 | 0.314357973 | 0.654203804 | -0.48051994 | 0.630857728 | 1 | 0.314357973 |
| ENSMUSG00000023073 | Slc10a2 | 20494 | protein_coding | 884.0612118 | -0.314369667 | 0.289385225 | 1.086336275 | 0.277330244 | 0.494297255 | 0.314369667 |
| ENSMUSG00000002661 | Alkbh7 | 66400 | protein_coding | 180.1184416 | -0.31439002 | 0.114957934 | 2.734826624 | 0.006241314 | 0.028986639 | 0.31439002 |
| ENSMUSG00000037679 | Inf2 | 70435 | protein_coding | 199.1891956 | -0.314456481 | 0.16779994 | 1.873996379 | 0.060930924 | 0.172799808 | 0.314456481 |
| ENSMUSG00000021360 | Gcnt2 | 14538 | protein_coding | 118.27705 | -0.314471705 | 0.299356521 | 1.050492251 | 0.293491852 | 0.512663892 | 0.314471705 |
| ENSMUSG00000034686 | Prr7 | 432763 | protein_coding | 6.982176276 | -0.314512755 | 0.455711336 | 0.690157848 | 0.490094928 | 0.697829595 | 0.314512755 |
| ENSMUSG00000047155 | Cyp4x1 | 81906 | protein_coding | 5.900181616 | -0.314601322 | 0.741687331 | 0.424169739 | 0.671442028 | 0.823849409 | 0.314601322 |
| ENSMUSG00000064823 | Snord82 | 80828 | snoRNA | 9.073406557 | 0.314654047 | 0.408154369 | -0.770919217 | 0.440754816 | 0.658639492 | 0.314654047 |
| ENSMUSG00000017868 | Sgk2 | 27219 | protein_coding | 623.6600271 | 0.314714149 | 0.25421722 | -1.23797337 | 0.215725935 | 0.420949793 | 0.314714149 |

| ensembl_gene_id | external_gene_name | entrezgene_id | gene_biotype | baseMean | log2FoldChange | lfcSE | stat | pvalue | padj | absLog2FC |
|---|---|---|---|---|---|---|---|---|---|---|
| ENSMUSG00000030960 | Eef1akmt2 | 72096 | protein_coding | 136.4301531 | 0.314774741 | 0.097070494 | -3.242743781 | 0.001183846 | 0.00731937 | 0.314774741 |
| ENSMUSG00000044430 | Klk12 | 69511 | protein_coding | 1.742873645 | 0.314797527 | 2.719977863 | -0.115735327 | 0.907862308 | 1 | 0.314797527 |
| ENSMUSG00000028803 | Nipal3 | 74552 | protein_coding | 1283.69388 | 0.314799236 | 0.054895266 | -5.734542482 | 9.78E-09 | 1.88E-07 | 0.314799236 |
| ENSMUSG00000037432 | Fer1l5 | 100534273 | protein_coding | 40.33816458 | 0.314838993 | 0.246930281 | -1.275011681 | 0.202305108 | 0.404218578 | 0.314838993 |
| ENSMUSG00000047044 | D030056L22Rik | 225995 | protein_coding | 166.9608603 | 0.314871008 | 0.146328908 | -2.151803177 | 0.03141286 | 0.104544934 | 0.314871008 |
| ENSMUSG00000031107 | Rbmx2 | 209003 | protein_coding | 42.03037357 | 0.314873842 | 0.200356923 | -1.571564571 | 0.116051568 | 0.275039552 | 0.314873842 |
| ENSMUSG00000117406 | Ntn3 | 18209 | protein_coding | 3.962712511 | -0.314894761 | 0.861978746 | 0.365316155 | 0.714875421 | 1 | 0.314894761 |
| ENSMUSG00000074825 | Itpripl1 | 73338 | protein_coding | 50.56958869 | 0.314913388 | 0.149688473 | -2.103791842 | 0.035396609 | 0.114525783 | 0.314913388 |
| ENSMUSG00000075028 | Prdm11 | 100042784 | protein_coding | 23.62410191 | 0.314915263 | 0.294088508 | -1.070817982 | 0.28425128 | 0.502676953 | 0.314915263 |
| ENSMUSG00000086598 | Btbd18 | 100270744 | protein_coding | 2.442160673 | -0.314996763 | 0.91753819 | 0.343306433 | 0.731367942 | 1 | 0.314996763 |
| ENSMUSG00000020431 | Adcy1 | 432530 | protein_coding | 7.674253062 | -0.315069784 | 0.694159196 | 0.453886927 | 0.649910213 | 0.809917796 | 0.315069784 |
| ENSMUSG00000024248 | Cox7a2l | 20463 | protein_coding | 1933.402661 | -0.315219251 | 0.108787199 | 2.897576677 | 0.003760578 | 0.01920773 | 0.315219251 |
| ENSMUSG00000071068 | Treml2 | 328833 | protein_coding | 5.409347774 | 0.315226323 | 1.07792604 | -0.292437803 | 0.769951908 | 0.884582288 | 0.315226323 |
| ENSMUSG00000079083 | Jrkl | 77532 | protein_coding | 131.1730597 | -0.315356696 | 0.107762854 | 2.926395182 | 0.00342915 | 0.017779903 | 0.315356696 |
| ENSMUSG00000066704 | Cyp2b19 | 13090 | protein_coding | 3.382391211 | -0.31541726 | 1.812520559 | 0.174021342 | 0.86184869 | 1 | 0.31541726 |
| ENSMUSG00000024682 | Cblif | 14603 | protein_coding | 0.769826449 | -0.315448324 | 1.689041225 | 0.186761767 | 0.851847425 | 1 | 0.315448324 |
| ENSMUSG00000048647 | Exd1 | 241624 | protein_coding | 26.95258218 | 0.315472773 | 0.238187404 | -1.324472946 | 0.185346022 | 0.380697042 | 0.315472773 |
| ENSMUSG00000061650 | Med9 | 192191 | protein_coding | 231.0019456 | 0.315497312 | 0.110776675 | -2.848048214 | 0.004398826 | 0.021880184 | 0.315497312 |
| ENSMUSG00000073155 | 1810058I24Rik | 67705 | protein_coding | 117.1544906 | -0.3155147 | 0.131520963 | 2.398968905 | 0.016441311 | 0.062908204 | 0.3155147 |
| ENSMUSG00000025289 | Prdx4 | 53381 | protein_coding | 241.571296 | 0.315604037 | 0.173179624 | -1.822408605 | 0.06839301 | 0.187833389 | 0.315604037 |
| ENSMUSG00000086446 | Prkag2os1 | 71161 | lncRNA | 5.27118103 | 0.315642962 | 0.536397215 | -0.588450038 | 0.55623026 | 0.745631404 | 0.315642962 |
| ENSMUSG00000027076 | Timm10 | 30059 | protein_coding | 166.0803308 | -0.315644232 | 0.116129736 | 2.718031075 | 0.006567167 | 0.030217385 | 0.315644232 |
| ENSMUSG00000038751 | Ptk6 | 20459 | protein_coding | 345.569062 | 0.315779698 | 0.227786422 | -1.386297285 | 0.165656145 | 0.352762472 | 0.315779698 |
| ENSMUSG00000027071 | P2rx3 | 228139 | protein_coding | 6.661052192 | 0.315866511 | 0.505051049 | -0.625415019 | 0.531698708 | 0.728669914 | 0.315866511 |
| ENSMUSG00000027186 | Elf5 | 13711 | protein_coding | 3.146927862 | -0.315896703 | 1.563945062 | 0.201987084 | 0.839926821 | 1 | 0.315896703 |
| ENSMUSG00000021226 | Acot2 | 171210 | protein_coding | 85.51475551 | -0.316007831 | 0.132337438 | 2.387894424 | 0.016945209 | 0.064388674 | 0.316007831 |
| ENSMUSG00000033538 | Casp4 | 12363 | protein_coding | 271.2059602 | 0.316025709 | 0.199216237 | -1.586345136 | 0.112661039 | 0.269245209 | 0.316025709 |
| ENSMUSG00000041124 | Msantd4 | 78100 | protein_coding | 518.0671926 | 0.316058857 | 0.058261324 | -5.424848515 | 5.80E-08 | 9.81E-07 | 0.316058857 |
| ENSMUSG00000047150 | 1700001C19Rik | 75462 | protein_coding | 5.270014231 | -0.316074301 | 0.61046837 | 0.517757047 | 0.60462779 | 0.779942456 | 0.316074301 |
| ENSMUSG00000048550 | Thnsl1 | 208967 | protein_coding | 97.15261463 | -0.316326043 | 0.114704765 | 2.757741084 | 0.005820228 | 0.027548077 | 0.316326043 |
| ENSMUSG00000061983 | Rps12 | 20042 | protein_coding | 5463.873793 | -0.316378833 | 0.105269572 | 3.005415786 | 0.00265218 | 0.014342673 | 0.316378833 |
| ENSMUSG00000025644 | Gm7628 | 115487215 | lncRNA | 3.168498646 | 0.316423756 | 0.82794387 | -0.382180202 | 0.702327706 | 1 | 0.316423756 |
| ENSMUSG00000028865 | Cd164l2 | 69655 | protein_coding | 7.834879444 | -0.316493183 | 0.514402495 | 0.615263701 | 0.538380585 | 0.733377497 | 0.316493183 |
| ENSMUSG00000094483 | Purb | 19291 | protein_coding | 2923.262862 | 0.316513189 | 0.110871746 | -2.854768705 | 0.004306818 | 0.021461314 | 0.316513189 |
| ENSMUSG00000019845 | Tube1 | 71924 | protein_coding | 45.88777051 | 0.316607861 | 0.193836406 | -1.633376657 | 0.102389803 | 0.251954286 | 0.316607861 |
| ENSMUSG00000026278 | Bok | 51800 | protein_coding | 162.0690791 | -0.316684385 | 0.110024969 | 2.8782956 | 0.003998303 | 0.020161214 | 0.316684385 |
| ENSMUSG00000022455 | Wbp2nl | 74716 | protein_coding | 4.600310878 | -0.316758321 | 0.541070792 | 0.585428609 | 0.558259557 | 0.747162726 | 0.316758321 |
| ENSMUSG00000085795 | Zfp703 | 353310 | protein_coding | 169.6751107 | 0.316840867 | 0.226203897 | -1.400687038 | 0.16130768 | 0.346267391 | 0.316840867 |
| ENSMUSG00000036944 | Tmem71 | 213068 | protein_coding | 5.249241061 | 0.316843357 | 0.539401509 | -0.587397979 | 0.556936451 | 0.746166522 | 0.316843357 |

| ensembl_gene_id | external_gene_name | entrezgene_id | gene_biotype | baseMean | log2FoldChange | lfcSE | stat | pvalue | padj | absLog2FC |
|---|---|---|---|---|---|---|---|---|---|---|
| ENSMUSG00000022240 | Ctnnd2 | 18163 | protein_coding | 6.590599064 | 0.316860209 | 0.749718085 | -0.422639143 | 0.672558563 | 0.824518194 | 0.316860209 |
| ENSMUSG00000088695 | Gm22334 | 115486131 | snoRNA | 0.958428519 | 0.316869955 | 1.604605319 | -0.197475324 | 0.84345559 | 1 | 0.316869955 |
| ENSMUSG00000075470 | Alg10b | 380959 | protein_coding | 798.3295329 | 0.316904107 | 0.100480258 | -3.153894235 | 0.001611075 | 0.009451256 | 0.316904107 |
| ENSMUSG00000049090 | Zadh2 | 225791 | protein_coding | 435.9917053 | -0.316944001 | 0.084800174 | 3.737539502 | 0.00018583 | 0.001488577 | 0.316944001 |
| ENSMUSG00000022550 | Adck5 | 268822 | protein_coding | 775.8290033 | -0.317158277 | 0.095654076 | 3.315679677 | 0.000914206 | 0.00589734 | 0.317158277 |
| ENSMUSG00000033781 | Asb13 | 142688 | protein_coding | 167.0950045 | -0.31717792 | 0.114871452 | 2.761155307 | 0.005759728 | 0.027332226 | 0.31717792 |
| ENSMUSG00000037894 | H2az1 | 51788 | protein_coding | 1235.160633 | 0.317482039 | 0.152579953 | -2.080758532 | 0.037456014 | 0.119688198 | 0.317482039 |
| ENSMUSG00000037977 | 6430571L13Rik | 235599 | protein_coding | 3.575455706 | 0.317483112 | 0.6580804 | -0.482438182 | 0.629494703 | 1 | 0.317483112 |
| ENSMUSG00000020042 | Btbd11 | 74007 | protein_coding | 19.4818469 | 0.317565868 | 0.350383244 | -0.906338626 | 0.364756647 | 0.587266381 | 0.317565868 |
| ENSMUSG00000025329 | Padi1 | 18599 | protein_coding | 0.80691154 | 0.317630035 | 1.445676598 | -0.219710297 | 0.826096784 | 1 | 0.317630035 |
| ENSMUSG00000021850 | ccdc198 | 67082 | protein_coding | 82.96260562 | -0.317645901 | 0.152697557 | 2.080229093 | 0.037504525 | 0.119750527 | 0.317645901 |
| ENSMUSG00000055737 | Ghr | 14600 | protein_coding | 185.503506 | -0.317715381 | 0.144467207 | 2.199221462 | 0.027862179 | 0.095110269 | 0.317715381 |
| ENSMUSG00000097842 | 9330104G04Rik | 320950 | lncRNA | 6.875550342 | 0.317761913 | 0.486625351 | -0.652990873 | 0.513762163 | 0.715609679 | 0.317761913 |
| ENSMUSG00000051695 | Pcbp1 | 23983 | protein_coding | 1361.272616 | 0.317810273 | 0.083967136 | -3.784936444 | 0.000153748 | 0.001253823 | 0.317810273 |
| ENSMUSG00000042540 | Acot5 | 217698 | protein_coding | 2.77379508 | -0.317817493 | 0.743642841 | 0.427379214 | 0.669103144 | 1 | 0.317817493 |
| ENSMUSG00000029394 | Cdk2ap1 | 13445 | protein_coding | 146.016263 | -0.317892845 | 0.188964572 | 1.68228807 | 0.092512993 | 0.233281352 | 0.317892845 |
| ENSMUSG00000044442 | N6amt1 | 67768 | protein_coding | 100.5326031 | -0.317973672 | 0.115126597 | 2.761947985 | 0.005745763 | 0.027285164 | 0.317973672 |
| ENSMUSG00000060036 | Rpl3 | 27367 | protein_coding | 6307.52096 | -0.318056981 | 0.128615927 | 2.472920643 | 0.01340139 | 0.053482492 | 0.318056981 |
| ENSMUSG00000052214 | Opa3 | 403187 | protein_coding | 954.6977226 | 0.318188124 | 0.056636111 | -5.618113876 | 1.93E-08 | 3.55E-07 | 0.318188124 |
| ENSMUSG00000023781 | Hes7 | 84653 | protein_coding | 2.332024242 | 0.318205469 | 0.798904598 | -0.398302213 | 0.69040743 | 1 | 0.318205469 |
| ENSMUSG00000039942 | Ptger4 | 19219 | protein_coding | 439.0006206 | -0.318260173 | 0.115774951 | 2.748955365 | 0.005978553 | 0.028104133 | 0.318260173 |
| ENSMUSG00000100733 | 4932411K12Rik | 71174 | lncRNA | 1.005729404 | -0.318431978 | 1.731783574 | 0.183875158 | 0.854111394 | 1 | 0.318431978 |
| ENSMUSG00000040432 | Ltb4r2 | 57260 | protein_coding | 10.93280527 | 0.318565136 | 0.479228019 | -0.664746474 | 0.506212664 | 0.710135685 | 0.318565136 |
| ENSMUSG00000028420 | Tmem38b | 52076 | protein_coding | 117.0611175 | -0.318597051 | 0.137437484 | 2.31812343 | 0.020442614 | 0.074638365 | 0.318597051 |
| ENSMUSG00000026788 | Zbtb43 | 71834 | protein_coding | 449.4241456 | -0.318631918 | 0.072078343 | 4.420633263 | 9.84E-06 | 0.000110481 | 0.318631918 |
| ENSMUSG00000031563 | Wwc2 | 52357 | protein_coding | 272.4394587 | -0.318651869 | 0.112288443 | 2.837797545 | 0.004542599 | 0.022466748 | 0.318651869 |
| ENSMUSG00000070034 | Sp110 | 109032 | protein_coding | 41.33612642 | -0.318664831 | 0.729721555 | 0.436693734 | 0.662333474 | 0.817346435 | 0.318664831 |
| ENSMUSG00000045045 | Lrfn4 | 225875 | protein_coding | 323.2355971 | -0.318739351 | 0.106941141 | 2.980511989 | 0.00287767 | 0.015364319 | 0.318739351 |
| ENSMUSG00000028223 | Decr1 | 67460 | protein_coding | 1011.458846 | -0.318755893 | 0.103481447 | 3.080319237 | 0.002067788 | 0.011662382 | 0.318755893 |
| ENSMUSG00000029098 | Acox3 | 80911 | protein_coding | 438.1112908 | -0.318775112 | 0.074871636 | 4.257621863 | 2.07E-05 | 0.00021219 | 0.318775112 |
| ENSMUSG00000032366 | Tpm1 | 22003 | protein_coding | 2775.032069 | -0.31878597 | 0.063740356 | 5.001320793 | 5.69E-07 | 8.30E-06 | 0.31878597 |
| ENSMUSG00000089917 | Uckl1 | 68556 | protein_coding | 405.8772084 | -0.318791195 | 0.066727396 | 4.777515913 | 1.77E-06 | 2.32E-05 | 0.318791195 |
| ENSMUSG00000041632 | Mrps27 | 218506 | protein_coding | 277.6087768 | 0.318859296 | 0.099888147 | -3.192163479 | 0.001412114 | 0.008482957 | 0.318859296 |
| ENSMUSG00000014599 | Csf1 | 12977 | protein_coding | 11.97825786 | 0.318879481 | 0.58567435 | -0.544465505 | 0.58612118 | 0.767773712 | 0.318879481 |
| ENSMUSG00000025429 | Pstpip2 | 19201 | protein_coding | 130.8540548 | -0.318972698 | 0.112716017 | 2.829879073 | 0.00465656 | 0.022923163 | 0.318972698 |
| ENSMUSG00000009687 | Fxyd5 | 18301 | protein_coding | 10.3317659 | 0.318983918 | 0.703318245 | -0.453541367 | 0.650158963 | 0.809917796 | 0.318983918 |
| ENSMUSG00000071604 | Entrep1 | 381217 | protein_coding | 542.1517923 | 0.319008233 | 0.114928701 | -2.775705541 | 0.005508207 | 0.026350675 | 0.319008233 |
| ENSMUSG00000060244 | Alyref2 | 56009 | protein_coding | 26.67509324 | -0.319019698 | 0.197553066 | 1.614855713 | 0.106341945 | 0.258825479 | 0.319019698 |
| ENSMUSG00000075524 | 4930407I10Rik | 328573 | protein_coding | 2.919992514 | 0.31932635 | 2.371393359 | -0.134657689 | 0.892882528 | 1 | 0.31932635 |

| ensembl_gene_id | external_gene_name | entrezgene_id | gene_biotype | baseMean | log2FoldChange | lfcSE | stat | pvalue | padj | absLog2FC |
|---|---|---|---|---|---|---|---|---|---|---|
| ENSMUSG00000045404 | Kcnk13 | 217826 | protein_coding | 1.011936034 | 0.319385866 | 1.620814049 | -0.19705275 | 0.843786258 | 1 | 0.319385866 |
| ENSMUSG00000039163 | Cmc1 | 67899 | protein_coding | 202.4417651 | -0.319493567 | 0.110770996 | 2.884270958 | 0.00392321 | 0.019904115 | 0.319493567 |
| ENSMUSG00000028520 | 4921539E11Rik | 70941 | protein_coding | 1.392657679 | -0.319545924 | 1.646079373 | 0.194125465 | 0.846077642 | 1 | 0.319545924 |
| ENSMUSG00000057137 | Tmem140 | 68487 | protein_coding | 42.23697594 | -0.319803166 | 0.405531766 | 0.78860201 | 0.430344656 | 0.648871201 | 0.319803166 |
| ENSMUSG00000052751 | Repin1 | 58887 | protein_coding | 226.623741 | -0.319923343 | 0.180044016 | 1.776917385 | 0.075581834 | 0.20154342 | 0.319923343 |
| ENSMUSG00000079677 | Fdx2 | 68165 | protein_coding | 1.145932549 | -0.319990589 | 1.147122146 | 0.278950755 | 0.780282617 | 1 | 0.319990589 |
| ENSMUSG00000094500 | Smim18 | 72632 | protein_coding | 1.07568703 | -0.320127289 | 1.138746055 | 0.281122633 | 0.778616342 | 1 | 0.320127289 |
| ENSMUSG00000007338 | Mrpl49 | 18120 | protein_coding | 371.4628082 | -0.320212182 | 0.077614738 | 4.125662086 | 3.70E-05 | 0.000358259 | 0.320212182 |
| ENSMUSG00000031320 | Rps4x | 20102 | protein_coding | 6713.270678 | -0.3202655 | 0.128575431 | 2.490876341 | 0.012742846 | 0.051375821 | 0.3202655 |
| ENSMUSG00000029530 | Ccr9 | 12769 | protein_coding | 10.49062579 | -0.320306317 | 0.575861349 | 0.556221245 | 0.578059617 | 0.76182422 | 0.320306317 |
| ENSMUSG00000029456 | Acad10 | 71985 | protein_coding | 195.5872024 | -0.320425756 | 0.106856204 | 2.998663106 | 0.00271167 | 0.014624596 | 0.320425756 |
| ENSMUSG00000012422 | Tmem167 | 66074 | protein_coding | 680.2606148 | 0.320463659 | 0.102914454 | -3.113883867 | 0.001846422 | 0.010616924 | 0.320463659 |
| ENSMUSG00000025977 | Boll | 75388 | protein_coding | 2.980627632 | -0.320578684 | 0.738419087 | 0.434141925 | 0.664185382 | 1 | 0.320578684 |
| ENSMUSG00000039128 | Cdc123 | 98828 | protein_coding | 632.4841308 | 0.320629917 | 0.093316214 | -3.435950772 | 0.000590478 | 0.004076571 | 0.320629917 |
| ENSMUSG00000093606 | B130034C11Rik | 414095 | lncRNA | 9.188327546 | 0.32063232 | 0.340120781 | -0.942701354 | 0.345833681 | 0.567534275 | 0.32063232 |
| ENSMUSG00000017715 | Pgs1 | 74451 | protein_coding | 551.9821958 | 0.320663934 | 0.093428374 | -3.432190023 | 0.000598728 | 0.004121541 | 0.320663934 |
| ENSMUSG00000035811 | Ugt2b35 | 243085 | protein_coding | 498.452295 | 0.320715253 | 0.125032455 | -2.565056041 | 0.010315914 | 0.043636528 | 0.320715253 |
| ENSMUSG00000085810 | Gm16325 | 100286843 | lncRNA | 1.354338746 | 0.320731476 | 0.935968905 | -0.342673218 | 0.731844313 | 1 | 0.320731476 |
| ENSMUSG00000006777 | Krt23 | 94179 | protein_coding | 116.0193685 | 0.320807655 | 0.157552603 | -2.036193937 | 0.041730885 | 0.130002609 | 0.320807655 |
| ENSMUSG00000021208 | Ifi27l2b | 217845 | protein_coding | 1312.821643 | -0.320873157 | 0.175733944 | 1.82590312 | 0.067864856 | 0.18675558 | 0.320873157 |
| ENSMUSG00000052221 | Ppp1r36 | 210762 | protein_coding | 1.650178958 | -0.321008244 | 1.524124651 | 0.210618104 | 0.833185284 | 1 | 0.321008244 |
| ENSMUSG00000040354 | Mars1 | 216443 | protein_coding | 479.9886973 | 0.321042605 | 0.123170311 | -2.606493422 | 0.009147458 | 0.039648589 | 0.321042605 |
| ENSMUSG00000033429 | Mcee | 73724 | protein_coding | 311.2222834 | -0.32104658 | 0.102276102 | 3.139018544 | 0.001695147 | 0.009870825 | 0.32104658 |
| ENSMUSG00000040570 | Rundc3b | 242819 | protein_coding | 314.9085561 | -0.321051475 | 0.084623797 | 3.793867525 | 0.000148319 | 0.001217364 | 0.321051475 |
| ENSMUSG00000087175 | Gm15133 | 622070 | lncRNA | 14.67817255 | -0.321075954 | 0.389002375 | 0.825383018 | 0.409154173 | 0.629730716 | 0.321075954 |
| ENSMUSG00000116848 | 4930542D17Rik | 75211 | lncRNA | 1.924505704 | 0.321126796 | 2.372065424 | -0.135378558 | 0.892312577 | 1 | 0.321126796 |
| ENSMUSG00000077689 | Gm22499 | 115485942 | snoRNA | 0.519056765 | 0.32125799 | 1.52620921 | -0.210494071 | 0.833282079 | 1 | 0.32125799 |
| ENSMUSG00000051705 | Senp8 | 71599 | protein_coding | 75.12571387 | -0.32128484 | 0.161121171 | 1.994057254 | 0.046145798 | 0.140315174 | 0.32128484 |
| ENSMUSG00000038893 | Fam117a | 215512 | protein_coding | 79.03766668 | -0.321491147 | 0.175189819 | 1.835101771 | 0.066490612 | 0.184012672 | 0.321491147 |
| ENSMUSG00000029438 | Bcl7a | 77045 | protein_coding | 102.4724472 | -0.321494965 | 0.142327741 | 2.258835572 | 0.023893615 | 0.084361596 | 0.321494965 |
| ENSMUSG00000090862 | Rps13 | 68052 | protein_coding | 2231.644036 | -0.321575563 | 0.131667913 | 2.442322923 | 0.014593087 | 0.057079639 | 0.321575563 |
| ENSMUSG00000003464 | Pex19 | 19298 | protein_coding | 809.3313464 | -0.32159642 | 0.057380563 | 5.604622951 | 2.09E-08 | 3.83E-07 | 0.32159642 |
| ENSMUSG00000051212 | Gpr183 | 321019 | protein_coding | 13.7658368 | 0.321644595 | 0.367302079 | -0.875695002 | 0.381195863 | 0.603611898 | 0.321644595 |
| ENSMUSG00000035559 | Mpv17l2 | 234384 | protein_coding | 391.3319125 | 0.321776233 | 0.091684304 | -3.509610896 | 0.000448763 | 0.003236188 | 0.321776233 |
| ENSMUSG00000025816 | Sec61a2 | 57743 | protein_coding | 115.167796 | 0.321779516 | 0.13339201 | -2.412284777 | 0.015852894 | 0.061038631 | 0.321779516 |
| ENSMUSG00000036880 | Acaa2 | 52538 | protein_coding | 1533.572137 | -0.321901941 | 0.067863225 | 4.74339293 | 2.10E-06 | 2.73E-05 | 0.321901941 |
| ENSMUSG00000027160 | Ccdc34 | 68201 | protein_coding | 238.2078044 | 0.322103617 | 0.142349176 | -2.262771204 | 0.023649797 | 0.083737724 | 0.322103617 |
| ENSMUSG00000000093 | Tbx2 | 21385 | protein_coding | 2.79133405 | 0.322110222 | 0.827415446 | -0.389296844 | 0.697056571 | 1 | 0.322110222 |
| ENSMUSG00000010051 | Hyal1 | 15586 | protein_coding | 24.0130484 | 0.322510783 | 0.276498455 | -1.166410797 | 0.243448395 | 0.454734397 | 0.322510783 |

| ensembl_gene_id | external_gene_name | entrezgene_id | gene_biotype | baseMean | log2FoldChange | lfcSE | stat | pvalue | padj | absLog2FC |
|---|---|---|---|---|---|---|---|---|---|---|
| ENSMUSG00000021520 | Uqcrb | 67530 | protein_coding | 2463.672785 | -0.322614932 | 0.092620161 | 3.48320417 | 0.00049545 | 0.003520597 | 0.322614932 |
| ENSMUSG00000052749 | Trim30b | 244183 | protein_coding | 3.599797518 | 0.322614971 | 0.906399401 | -0.355930257 | 0.721892796 | 1 | 0.322614971 |
| ENSMUSG00000037341 | Slc9a7 | 236727 | protein_coding | 11.5650912 | 0.322690791 | 0.452345266 | -0.71337276 | 0.475615125 | 0.687587163 | 0.322690791 |
| ENSMUSG00000115852 | Gm52969 | 102633938 | transcribed_processed_pseudogene | 2.145470673 | 0.322707696 | 1.40427879 | -0.229803154 | 0.818244733 | 1 | 0.322707696 |
| ENSMUSG00000020120 | Plek | 56193 | protein_coding | 37.29012428 | 0.322841297 | 0.215664374 | -1.496961649 | 0.134403237 | 0.305048799 | 0.322841297 |
| ENSMUSG00000051455 | Meioc | 268491 | protein_coding | 2.684008742 | 0.322939255 | 0.732457581 | -0.440898236 | 0.659286671 | 1 | 0.322939255 |
| ENSMUSG00000042079 | Hnrnpf | 98758 | protein_coding | 1023.73977 | -0.323087454 | 0.135227702 | 2.38921057 | 0.016884622 | 0.064232353 | 0.323087454 |
| ENSMUSG00000007458 | M6pr | 17113 | protein_coding | 2615.400493 | 0.32317392 | 0.097880359 | -3.301723877 | 0.000960926 | 0.006157833 | 0.32317392 |
| ENSMUSG00000088408 | Gm25881 | 115489577 | snRNA | 2.223350512 | -0.323191781 | 0.852775576 | 0.378988083 | 0.704696713 | 1 | 0.323191781 |
| ENSMUSG00000022197 | Pdzd2 | 68070 | protein_coding | 38.6756161 | -0.323250083 | 0.255475197 | 1.265289495 | 0.205767571 | 0.408369867 | 0.323250083 |
| ENSMUSG00000029086 | Prom1 | 19126 | protein_coding | 4922.35339 | -0.323409648 | 0.112854723 | 2.865716557 | 0.004160666 | 0.020846249 | 0.323409648 |
| ENSMUSG00000038188 | Scarf1 | 380713 | protein_coding | 44.7877686 | -0.323531022 | 0.215387212 | 1.502090207 | 0.133073814 | 0.302752759 | 0.323531022 |
| ENSMUSG00000088208 | Gm23751 | 115490492 | scaRNA | 2.297367509 | -0.323640972 | 0.772148666 | 0.419143341 | 0.675111377 | 1 | 0.323640972 |
| ENSMUSG00000005503 | Evx1 | 14028 | protein_coding | 27.69509638 | 0.323648812 | 0.23908886 | -1.353675832 | 0.175839822 | 0.367865096 | 0.323648812 |
| ENSMUSG00000003282 | Plag1 | 56711 | protein_coding | 35.53715859 | -0.323655344 | 0.185014924 | 1.749347224 | 0.080231018 | 0.21047419 | 0.323655344 |
| ENSMUSG00000008398 | Elk3 | 13713 | protein_coding | 23.61902344 | -0.323663063 | 0.216829458 | 1.492707985 | 0.135513642 | 0.30685209 | 0.323663063 |
| ENSMUSG00000001819 | Hoxd13 | 15433 | protein_coding | 129.721534 | -0.323667915 | 0.427426394 | 0.757248311 | 0.448901112 | 0.665664739 | 0.323667915 |
| ENSMUSG00000038700 | Hoxb5 | 15413 | protein_coding | 46.14178143 | -0.323697613 | 0.364563208 | 0.887905322 | 0.374591682 | 0.596644197 | 0.323697613 |
| ENSMUSG00000034685 | Fam171a2 | 217219 | protein_coding | 8.583221428 | 0.323716638 | 0.493014382 | -0.656606885 | 0.51143372 | 0.71423395 | 0.323716638 |
| ENSMUSG00000029363 | Rfc5 | 72151 | protein_coding | 268.1711963 | 0.324005288 | 0.156350729 | -2.072297905 | 0.038237668 | 0.121691433 | 0.324005288 |
| ENSMUSG00000037669 | Ldah | 68832 | protein_coding | 666.2664129 | 0.324012713 | 0.083058863 | -3.901001077 | 9.58E-05 | 0.000829125 | 0.324012713 |
| ENSMUSG00000019866 | Crybg1 | 11630 | protein_coding | 1724.301617 | 0.324050898 | 0.094809447 | -3.417917798 | 0.000631022 | 0.0043132 | 0.324050898 |
| ENSMUSG00000074183 | Gsta1 | 14857 | protein_coding | 88.64165165 | 0.32407895 | 0.213745341 | -1.516191882 | 0.12947084 | 0.297203427 | 0.32407895 |
| ENSMUSG00000070639 | Lrrc8b | 433926 | protein_coding | 318.6946558 | 0.324090969 | 0.081689877 | -3.967333291 | 7.27E-05 | 0.0006509 | 0.324090969 |
| ENSMUSG00000083041 | Gmfg-ps | 100040018 | processed_pseudogene | 1.089915481 | 0.324116066 | 1.259546478 | -0.257327595 | 0.796925892 | 1 | 0.324116066 |
| ENSMUSG00000094622 | Gm3055 | 100040944 | protein_coding | 5.085343695 | -0.324257985 | 0.476953063 | 0.67985303 | 0.496597525 | 0.702605511 | 0.324257985 |
| ENSMUSG00000032085 | Tagln | 21345 | protein_coding | 16.70482724 | 0.324298969 | 0.292976583 | -1.106910887 | 0.268332458 | 0.483440174 | 0.324298969 |
| ENSMUSG00000027107 | Chrna1 | 11435 | protein_coding | 20.41056025 | -0.324317872 | 0.299507175 | 1.082838406 | 0.278880155 | 0.496167516 | 0.324317872 |
| ENSMUSG00000029598 | Plbd2 | 71772 | protein_coding | 261.1084495 | -0.324324551 | 0.07545721 | 4.298125402 | 1.72E-05 | 0.000180604 | 0.324324551 |
| ENSMUSG00000076437 | Selenoh | 72657 | protein_coding | 169.8483574 | 0.324374005 | 0.154787214 | -2.095612401 | 0.036116588 | 0.116104602 | 0.324374005 |
| ENSMUSG00000029368 | Alb | 11657 | protein_coding | 13.29974312 | 0.324413195 | 0.39894782 | -0.813171995 | 0.416119453 | 0.636395319 | 0.324413195 |
| ENSMUSG00000098347 | Mir6339 | 102466628 | miRNA | 3.41881392 | 0.324463581 | 0.70447594 | -0.460574396 | 0.645103985 | 1 | 0.324463581 |
| ENSMUSG00000068129 | Cst7 | 13011 | protein_coding | 3.316451659 | 0.324560635 | 0.66698986 | -0.486605051 | 0.626538244 | 1 | 0.324560635 |
| ENSMUSG00000027618 | Nfs1 | 18041 | protein_coding | 472.7081416 | -0.324572356 | 0.079273142 | 4.094354614 | 4.23E-05 | 0.000404346 | 0.324572356 |
| ENSMUSG00000010601 | Apol7a | 75761 | protein_coding | 116.0037701 | -0.324595026 | 0.362472479 | 0.895502541 | 0.370518508 | 0.593022866 | 0.324595026 |
| ENSMUSG00000021982 | Cdadc1 | 71891 | protein_coding | 210.291062 | -0.324596698 | 0.100891099 | 3.217297672 | 0.001294043 | 0.007882634 | 0.324596698 |
| ENSMUSG00000018166 | Erbb3 | 13867 | protein_coding | 4674.942508 | -0.324708628 | 0.110593873 | 2.936045359 | 0.003324257 | 0.017334001 | 0.324708628 |
| ENSMUSG00000006219 | Fblim1 | 74202 | protein_coding | 674.5872437 | -0.324754597 | 0.073899448 | 4.394546981 | 1.11E-05 | 0.000122946 | 0.324754597 |
| ENSMUSG00000053714 | 4732471J01Rik | 654804 | lncRNA | 51.94766484 | 0.324823822 | 0.23822289 | -1.363529013 | 0.172715847 | 0.363275924 | 0.324823822 |

| ensembl_gene_id | external_gene_name | entrezgene_id | gene_biotype | baseMean | log2FoldChange | lfcSE | stat | pvalue | padj | absLog2FC |
|---|---|---|---|---|---|---|---|---|---|---|
| ENSMUSG00000071415 | Rpl23 | 65019 | protein_coding | 4564.948076 | -0.324891988 | 0.123531273 | 2.63003836 | 0.008537523 | 0.03747705 | 0.324891988 |
| ENSMUSG00000000751 | Rpa1 | 68275 | protein_coding | 536.3567917 | 0.324892458 | 0.145487395 | -2.233131322 | 0.025540286 | 0.088775474 | 0.324892458 |
| ENSMUSG00000038225 | Primpol | 408022 | protein_coding | 101.3645654 | -0.324924498 | 0.167787573 | 1.936523027 | 0.052803675 | 0.155547608 | 0.324924498 |
| ENSMUSG00000049488 | Tmem67 | 329795 | protein_coding | 62.76575439 | 0.324970124 | 0.162949731 | -1.994296777 | 0.046119631 | 0.140291806 | 0.324970124 |
| ENSMUSG00000028073 | Pear1 | 73182 | protein_coding | 56.42051547 | 0.324986066 | 0.188119561 | -1.72755063 | 0.084068821 | 0.217760374 | 0.324986066 |
| ENSMUSG00000027955 | Gask1b | 68659 | protein_coding | 11.59797536 | 0.325040201 | 0.39115002 | -0.830986028 | 0.405981523 | 0.626929013 | 0.325040201 |
| ENSMUSG00000004730 | Adgre1 | 13733 | protein_coding | 2.567413694 | -0.325065846 | 0.953161254 | 0.341039719 | 0.733073681 | 1 | 0.325065846 |
| ENSMUSG00000018239 | Zcchc10 | 67966 | protein_coding | 17.82974131 | -0.325203919 | 0.310321896 | 1.047956728 | 0.294658544 | 0.513807046 | 0.325203919 |
| ENSMUSG00000004446 | Bid | 12122 | protein_coding | 130.8285758 | -0.32523774 | 0.111432126 | 2.918707127 | 0.003514863 | 0.018172963 | 0.32523774 |
| ENSMUSG00000032265 | Tent5a | 212943 | protein_coding | 1498.066844 | -0.325252827 | 0.093534066 | 3.477372918 | 0.000506353 | 0.003584189 | 0.325252827 |
| ENSMUSG00000062006 | Rpl34 | 68436 | protein_coding | 2381.690994 | -0.325253138 | 0.133832097 | 2.430307414 | 0.015086021 | 0.058674176 | 0.325253138 |
| ENSMUSG00000065476 | Mir30b | 387226 | miRNA | 1.570088225 | -0.325306091 | 1.196960565 | 0.271776783 | 0.785793658 | 1 | 0.325306091 |
| ENSMUSG00000040938 | Slc16a11 | 216867 | protein_coding | 33.00610827 | -0.325417119 | 0.21937323 | 1.483394848 | 0.137969535 | 0.31052626 | 0.325417119 |
| ENSMUSG00000029235 | Pdcl2 | 79455 | protein_coding | 0.70998615 | 0.325422102 | 1.431476783 | -0.227333133 | 0.820164711 | 1 | 0.325422102 |
| ENSMUSG00000091996 | Smim35 | 408059 | protein_coding | 9.909708889 | 0.325534315 | 0.435073604 | -0.748228145 | 0.454322557 | 0.669875536 | 0.325534315 |
| ENSMUSG00000010066 | Cacna2d2 | 56808 | protein_coding | 87.11161187 | -0.325569244 | 0.171937957 | 1.893527461 | 0.05828776 | 0.167227948 | 0.325569244 |
| ENSMUSG00000092032 | Vmn2r59 | 628444 | protein_coding | 3.14997079 | -0.325572464 | 1.972285767 | 0.165073677 | 0.868885991 | 1 | 0.325572464 |
| ENSMUSG00000001418 | Glmp | 56700 | protein_coding | 664.7783026 | -0.325575765 | 0.062999502 | 5.167910155 | 2.37E-07 | 3.66E-06 | 0.325575765 |
| ENSMUSG00000031389 | Arhgap4 | 171207 | protein_coding | 41.4516654 | -0.325671029 | 0.221302509 | 1.4716102 | 0.141126169 | 0.31570815 | 0.325671029 |
| ENSMUSG00000029030 | Tprgl | 67808 | protein_coding | 835.8385533 | -0.325689518 | 0.069481319 | 4.687440092 | 2.77E-06 | 3.47E-05 | 0.325689518 |
| ENSMUSG00000004637 | Wwox | 80707 | protein_coding | 128.5253158 | 0.325700174 | 0.098070851 | -3.321070119 | 0.00089673 | 0.005807294 | 0.325700174 |
| ENSMUSG00000063296 | Tmem117 | 320709 | protein_coding | 295.5126244 | -0.325766143 | 0.151490187 | 2.150410859 | 0.03152273 | 0.104847204 | 0.325766143 |
| ENSMUSG00000031303 | Map3k15 | 270672 | protein_coding | 37.62904205 | 0.325822745 | 0.213512227 | -1.526014455 | 0.127006265 | 0.293627531 | 0.325822745 |
| ENSMUSG00000023995 | Tspo2 | 70026 | protein_coding | 4.02871342 | -0.325849092 | 0.655032594 | 0.497454775 | 0.618868387 | 1 | 0.325849092 |
| ENSMUSG00000037705 | Tecta | 21683 | protein_coding | 2.324789347 | -0.325928564 | 1.869870719 | 0.1743054 | 0.861625455 | 1 | 0.325928564 |
| ENSMUSG00000073542 | Cep76 | 225659 | protein_coding | 90.12638527 | 0.326024802 | 0.146340183 | -2.227855642 | 0.025890143 | 0.089783667 | 0.326024802 |
| ENSMUSG00000026172 | Bcs1l | 66821 | protein_coding | 138.7936149 | -0.326025129 | 0.114238599 | 2.853896423 | 0.004318661 | 0.021513836 | 0.326025129 |
| ENSMUSG00000088990 | Gm22767 | 115490051 | scaRNA | 10.02024492 | 0.32603211 | 0.405552629 | -0.803920593 | 0.421442836 | 0.641211258 | 0.32603211 |
| ENSMUSG00000040549 | Ckap5 | 75786 | protein_coding | 1113.770644 | 0.326096937 | 0.105139847 | -3.101554239 | 0.001925076 | 0.011004052 | 0.326096937 |
| ENSMUSG00000025257 | Ribc1 | 66611 | protein_coding | 26.55848979 | -0.326099701 | 0.230654636 | 1.413800764 | 0.157420409 | 0.340684159 | 0.326099701 |
| ENSMUSG00000047793 | Sned1 | 208777 | protein_coding | 234.9429766 | -0.326181896 | 0.1194318 | 2.731114302 | 0.006312058 | 0.029222685 | 0.326181896 |
| ENSMUSG00000029651 | Mtus2 | 77521 | protein_coding | 44.12555704 | 0.326498441 | 0.350858994 | -0.930568824 | 0.352076648 | 0.574354812 | 0.326498441 |
| ENSMUSG00000008822 | Acyp1 | 66204 | protein_coding | 74.6327845 | -0.326530646 | 0.139320234 | 2.343741725 | 0.019091387 | 0.070752953 | 0.326530646 |
| ENSMUSG00000022705 | Drd3 | 13490 | protein_coding | 7.844231049 | -0.326577474 | 0.472497573 | 0.691172806 | 0.489456951 | 0.697221779 | 0.326577474 |
| ENSMUSG00000035165 | Kcne3 | 57442 | protein_coding | 1342.14672 | -0.326633126 | 0.20204281 | 1.616653051 | 0.105953193 | 0.258145624 | 0.326633126 |
| ENSMUSG00000042622 | Maff | 17133 | protein_coding | 420.9130001 | -0.32672142 | 0.241501175 | 1.352877147 | 0.176094881 | 0.368018849 | 0.32672142 |
| ENSMUSG00000084950 | Gm5577 | 434064 | lncRNA | 1.896004816 | 0.326917305 | 0.989876564 | -0.330260678 | 0.741203002 | 1 | 0.326917305 |
| ENSMUSG00000026361 | Cdc73 | 214498 | protein_coding | 440.7482187 | 0.326925199 | 0.094122327 | -3.473407517 | 0.000513894 | 0.003631345 | 0.326925199 |
| ENSMUSG00000030188 | Magohb | 66441 | protein_coding | 101.0553448 | 0.326968244 | 0.166087307 | -1.96865281 | 0.048992974 | 0.146863309 | 0.326968244 |

| ensembl_gene_id | external_gene_name | entrezgene_id | gene_biotype | baseMean | log2FoldChange | lfcSE | stat | pvalue | padj | absLog2FC |
|---|---|---|---|---|---|---|---|---|---|---|
| ENSMUSG00000024587 | Nars | 70223 | protein_coding | 2300.34225 | 0.326983013 | 0.076362224 | -4.28199959 | 1.85E-05 | 0.000192981 | 0.326983013 |
| ENSMUSG00000028549 | Itgb3bp | 67733 | protein_coding | 56.14732233 | 0.326999157 | 0.16104839 | -2.030440403 | 0.042311793 | 0.131292175 | 0.326999157 |
| ENSMUSG00000020719 | Ddx5 | 13207 | protein_coding | 7846.135244 | -0.327018404 | 0.05674902 | 5.762538348 | 8.29E-09 | 1.62E-07 | 0.327018404 |
| ENSMUSG00000085763 | Smc2os | 72648 | lncRNA | 1.406289038 | -0.327097333 | 0.959652674 | 0.340849707 | 0.733216728 | 1 | 0.327097333 |
| ENSMUSG00000036057 | Ptpn23 | 104831 | protein_coding | 289.5156694 | 0.327184347 | 0.185615477 | -1.762699708 | 0.07795115 | 0.206262665 | 0.327184347 |
| ENSMUSG00000078789 | Dph1 | 116905 | protein_coding | 115.3273093 | -0.32731663 | 0.118587363 | 2.760130786 | 0.005777823 | 0.027402344 | 0.32731663 |
| ENSMUSG00000006235 | Epor | 13857 | protein_coding | 2.289641013 | -0.327321048 | 1.196851489 | 0.273485099 | 0.784480345 | 1 | 0.327321048 |
| ENSMUSG00000031512 | Tex29 | 75528 | protein_coding | 2.126293241 | 0.327326124 | 1.363299762 | -0.240098424 | 0.810253956 | 1 | 0.327326124 |
| ENSMUSG00000098188 | Sowahc | 268301 | protein_coding | 840.2308738 | -0.327467688 | 0.093488639 | 3.502753829 | 0.000460475 | 0.003307101 | 0.327467688 |
| ENSMUSG00000095961 | Vmn2r102 | 224572 | protein_coding | 4.263653931 | 0.327555553 | 1.774015776 | -0.184640722 | 0.853510844 | 1 | 0.327555553 |
| ENSMUSG00000037321 | Tap1 | 21354 | protein_coding | 242.7449895 | -0.327598158 | 0.21044266 | 1.556709832 | 0.119539393 | 0.281287196 | 0.327598158 |
| ENSMUSG00000049881 | 2810025M15Rik | 69953 | transcribed_processed_pseudogene | 74.3116588 | 0.327619451 | 0.15575536 | -2.103423287 | 0.035428785 | 0.114584988 | 0.327619451 |
| ENSMUSG00000108621 | Gm33989 | 102637087 | lncRNA | 232.8901115 | -0.327623624 | 0.130692624 | 2.50682567 | 0.012182078 | 0.049685066 | 0.327623624 |
| ENSMUSG00000021578 | Ccdc127 | 67433 | protein_coding | 356.8246997 | 0.327624074 | 0.101377532 | -3.23172272 | 0.001230464 | 0.007553859 | 0.327624074 |
| ENSMUSG00000069678 | Pcgf1 | 69837 | protein_coding | 62.65418776 | -0.327637655 | 0.133051072 | 2.46249542 | 0.013797394 | 0.054614131 | 0.327637655 |
| ENSMUSG00000029131 | Dnajb6 | 23950 | protein_coding | 368.91685 | -0.327659776 | 0.083130289 | 3.941520944 | 8.10E-05 | 0.000716166 | 0.327659776 |
| ENSMUSG00000084678 | Gm26020 | 115489260 | snRNA | 2.543665007 | 0.327672183 | 0.7873799 | -0.416155129 | 0.677296492 | 1 | 0.327672183 |
| ENSMUSG00000060279 | Ap2a1 | 11771 | protein_coding | 788.8169379 | 0.327674585 | 0.074205576 | -4.415767714 | 1.01E-05 | 0.000112689 | 0.327674585 |
| ENSMUSG00000079103 | Tgm7 | 640543 | protein_coding | 4.428075409 | 0.32771691 | 0.779631635 | -0.420348399 | 0.674230958 | 0.825194181 | 0.32771691 |
| ENSMUSG00000021701 | Plk2 | 20620 | protein_coding | 435.9368009 | -0.327721486 | 0.169725922 | 1.930886462 | 0.053497095 | 0.157015607 | 0.327721486 |
| ENSMUSG00000025856 | Pdgfa | 18590 | protein_coding | 439.1293052 | -0.327748176 | 0.109629352 | 2.989602414 | 0.002793408 | 0.014996859 | 0.327748176 |
| ENSMUSG00000045662 | Henmt1 | 66715 | protein_coding | 2.175776258 | 0.327789703 | 0.939544492 | -0.348881512 | 0.727178266 | 1 | 0.327789703 |
| ENSMUSG00000038886 | Man2a2 | 140481 | protein_coding | 95.40982354 | 0.327855301 | 0.146635243 | -2.235856095 | 0.025361199 | 0.08834262 | 0.327855301 |
| ENSMUSG00000020017 | Hal | 15109 | protein_coding | 3.016966196 | 0.327868004 | 1.266596139 | -0.258857574 | 0.795745134 | 1 | 0.327868004 |
| ENSMUSG00000019865 | Nmbr | 18101 | protein_coding | 1.104007993 | 0.327877305 | 1.466522755 | -0.223574645 | 0.823088294 | 1 | 0.327877305 |
| ENSMUSG00000043531 | Sorcs1 | 58178 | protein_coding | 2.456876349 | -0.327877411 | 1.823736955 | 0.179783279 | 0.857322711 | 1 | 0.327877411 |
| ENSMUSG00000021003 | Galc | 14420 | protein_coding | 61.20589836 | 0.327963839 | 0.146854 | -2.233264597 | 0.025531501 | 0.088763622 | 0.327963839 |
| ENSMUSG00000003847 | Nfat5 | 54446 | protein_coding | 364.0062262 | -0.328119322 | 0.181871537 | 1.804126849 | 0.071211424 | 0.193418659 | 0.328119322 |
| ENSMUSG00000091086 | Rpl6l | 432502 | processed_pseudogene | 4.569947312 | 0.328164506 | 0.51919138 | -0.63206848 | 0.527342131 | 0.725771143 | 0.328164506 |
| ENSMUSG00000028759 | Hp1bp3 | 15441 | protein_coding | 1682.846115 | 0.328268874 | 0.056297725 | -5.830943903 | 5.51E-09 | 1.10E-07 | 0.328268874 |
| ENSMUSG00000042035 | Igsf3 | 78908 | protein_coding | 477.6689561 | -0.328278663 | 0.131318429 | 2.499867419 | 0.012423979 | 0.050422608 | 0.328278663 |
| ENSMUSG00000062064 | Slc2a7 | 435818 | protein_coding | 2.815142809 | 0.328343916 | 1.166249306 | -0.281538359 | 0.778297511 | 1 | 0.328343916 |
| ENSMUSG00000021702 | Thbs4 | 21828 | protein_coding | 2318.485996 | -0.328465371 | 0.236151228 | 1.390911128 | 0.164252378 | 0.350538094 | 0.328465371 |
| ENSMUSG00000098866 | Mir6539 | 102465310 | miRNA | 1.029048646 | -0.32847835 | 1.304722171 | 0.251761147 | 0.80122569 | 1 | 0.32847835 |
| ENSMUSG00000091754 | Gm3636 | 100041151 | protein_coding | 2.323818863 | -0.328495342 | 0.751823312 | 0.436931573 | 0.662160974 | 1 | 0.328495342 |
| ENSMUSG00000008028 | 1700008O03Rik | 69349 | protein_coding | 3.494263005 | 0.328557138 | 1.668194871 | -0.196953692 | 0.843863777 | 1 | 0.328557138 |
| ENSMUSG00000065750 | Gm23346 | 115490470 | snoRNA | 5.467598206 | 0.328608324 | 0.499962992 | -0.657265297 | 0.511010346 | 0.713884177 | 0.328608324 |
| ENSMUSG00000042747 | Krtcap2 | 66059 | protein_coding | 597.6373589 | 0.328621987 | 0.119399964 | -2.75227878 | 0.005918212 | 0.027890446 | 0.328621987 |
| ENSMUSG00000028484 | Psip1 | 101739 | protein_coding | 215.3461363 | 0.328638088 | 0.149550121 | -2.197511354 | 0.027983947 | 0.095392863 | 0.328638088 |

| ensembl_gene_id | external_gene_name | entrezgene_id | gene_biotype | baseMean | log2FoldChange | lfcSE | stat | pvalue | padj | absLog2FC |
|---|---|---|---|---|---|---|---|---|---|---|
| ENSMUSG00000033285 | Wdr3 | 269470 | protein_coding | 328.9247399 | 0.328640846 | 0.095191602 | -3.452414284 | 0.000555594 | 0.003866447 | 0.328640846 |
| ENSMUSG00000040283 | Btnl9 | 237754 | protein_coding | 5.49571421 | 0.328688612 | 0.726325428 | -0.452536287 | 0.650882686 | 0.810260975 | 0.328688612 |
| ENSMUSG00000065669 | Gm22581 | 115489818 | snoRNA | 2.830239605 | -0.328745788 | 0.647565576 | 0.507664089 | 0.611688937 | 1 | 0.328745788 |
| ENSMUSG00000007034 | Slc44a4 | 70129 | protein_coding | 4978.067283 | -0.32878964 | 0.06454245 | 5.094161109 | 3.50E-07 | 5.32E-06 | 0.32878964 |
| ENSMUSG00000060803 | Gstp1 | 14870 | protein_coding | 1513.103542 | -0.328816551 | 0.102464998 | 3.209062203 | 0.001331687 | 0.00808213 | 0.328816551 |
| ENSMUSG00000022195 | 6030458C11Rik | 77877 | protein_coding | 410.9536593 | 0.328846767 | 0.080276491 | -4.096426766 | 4.20E-05 | 0.000401209 | 0.328846767 |
| ENSMUSG00000090015 | Gm15446 | 100642166 | protein_coding | 68.55705781 | -0.328891661 | 0.173473447 | 1.895919326 | 0.057970713 | 0.166571051 | 0.328891661 |
| ENSMUSG00000019813 | Cep57l1 | 103268 | protein_coding | 42.6276725 | 0.328901725 | 0.18992797 | -1.731718213 | 0.083323746 | 0.216285499 | 0.328901725 |
| ENSMUSG00000055553 | Kxd1 | 75620 | protein_coding | 467.1456214 | -0.328984117 | 0.07107467 | 4.628711166 | 3.68E-06 | 4.53E-05 | 0.328984117 |
| ENSMUSG00000105352 | C030018K13Rik | 77382 | lncRNA | 1.22926799 | 0.329042794 | 1.490185899 | -0.220806541 | 0.82524307 | 1 | 0.329042794 |
| ENSMUSG00000022668 | Gtpbp8 | 66067 | protein_coding | 246.2040834 | -0.329058442 | 0.099409999 | 3.310114133 | 0.000932579 | 0.006001799 | 0.329058442 |
| ENSMUSG00000029833 | Trim24 | 21848 | protein_coding | 775.2969861 | 0.329125603 | 0.053321555 | -6.172468192 | 6.72E-10 | 1.51E-08 | 0.329125603 |
| ENSMUSG00000021831 | Ero1a | 50527 | protein_coding | 565.6525988 | 0.329188136 | 0.077612936 | -4.241408123 | 2.22E-05 | 0.000226707 | 0.329188136 |
| ENSMUSG00000075254 | Heg1 | 77446 | protein_coding | 116.7881646 | 0.329208603 | 0.171118918 | -1.923858602 | 0.05437231 | 0.158736855 | 0.329208603 |
| ENSMUSG00000039062 | Anpep | 16790 | protein_coding | 146.7071497 | -0.329218342 | 0.598181182 | 0.550365594 | 0.582068642 | 0.765030035 | 0.329218342 |
| ENSMUSG00000015305 | Sash1 | 70097 | protein_coding | 949.8679935 | -0.329252812 | 0.13037675 | 2.525395158 | 0.011556826 | 0.047749183 | 0.329252812 |
| ENSMUSG00000038201 | Kcna7 | 16495 | protein_coding | 1.221273544 | -0.329275573 | 1.153266835 | 0.285515514 | 0.77524922 | 1 | 0.329275573 |
| ENSMUSG00000020151 | Ptprr | 19279 | protein_coding | 247.6263121 | -0.329284976 | 0.264378012 | 1.245508178 | 0.212945011 | 0.417696153 | 0.329284976 |
| ENSMUSG00000063972 | Nr6a1 | 14536 | protein_coding | 126.0547618 | -0.329288658 | 0.201267982 | 1.636070748 | 0.101824781 | 0.250862963 | 0.329288658 |
| ENSMUSG00000035228 | Ccdc106 | 232821 | protein_coding | 3.381885486 | 0.329518308 | 0.739039501 | -0.445873743 | 0.65568845 | 1 | 0.329518308 |
| ENSMUSG00000028763 | Hspg2 | 15530 | protein_coding | 494.711838 | -0.329543414 | 0.113415268 | 2.905635376 | 0.003665083 | 0.018819941 | 0.329543414 |
| ENSMUSG00000024121 | Atp6v0c | 11984 | protein_coding | 601.5625673 | -0.329553712 | 0.103875967 | 3.172569393 | 0.001510964 | 0.008943393 | 0.329553712 |
| ENSMUSG00000027612 | Mmp24 | 17391 | protein_coding | 7.278261621 | -0.329554093 | 0.531525683 | 0.620015371 | 0.535247667 | 0.731167175 | 0.329554093 |
| ENSMUSG00000031881 | Cdh16 | 12556 | protein_coding | 2.1285986 | 0.329555822 | 1.535221205 | -0.214663412 | 0.830029751 | 1 | 0.329555822 |
| ENSMUSG00000001542 | Ell2 | 192657 | protein_coding | 698.8523013 | 0.329599082 | 0.077084209 | -4.275831432 | 1.90E-05 | 0.000197035 | 0.329599082 |
| ENSMUSG00000031959 | Wdr59 | 319481 | protein_coding | 231.2036618 | 0.329600991 | 0.127306751 | -2.589029959 | 0.009624672 | 0.041288574 | 0.329600991 |
| ENSMUSG00000027086 | Fastkd1 | 320720 | protein_coding | 316.3305267 | -0.329615412 | 0.098733692 | 3.338428918 | 0.000842536 | 0.005507949 | 0.329615412 |
| ENSMUSG00000059040 | Eno1b | 433182 | protein_coding | 2.408980379 | 0.329946079 | 0.703218397 | -0.469194322 | 0.638930744 | 1 | 0.329946079 |
| ENSMUSG00000099301 | Mir3473e | 102466677 | miRNA | 0.594424915 | 0.329994827 | 1.242514481 | -0.265586303 | 0.790557836 | 1 | 0.329994827 |
| ENSMUSG00000046793 | Gpr61 | 229714 | protein_coding | 2.296110606 | -0.330123715 | 0.875349322 | 0.377133685 | 0.706074259 | 1 | 0.330123715 |
| ENSMUSG00000034035 | Ccdc17 | 622665 | protein_coding | 46.84070774 | -0.330149738 | 0.168294744 | 1.96173529 | 0.049793311 | 0.148427209 | 0.330149738 |
| ENSMUSG00000021556 | Golm1 | 105348 | protein_coding | 8577.456403 | 0.330271326 | 0.104551668 | -3.158929276 | 0.001583499 | 0.009312644 | 0.330271326 |
| ENSMUSG00000040522 | Tlr8 | 170744 | protein_coding | 1.775747267 | -0.330409503 | 1.258193437 | 0.262606284 | 0.792854056 | 1 | 0.330409503 |
| ENSMUSG00000033233 | Trim45 | 229644 | protein_coding | 45.29145739 | 0.33043343 | 0.170428112 | -1.938843452 | 0.052520403 | 0.154934252 | 0.33043343 |
| ENSMUSG00000020362 | Cnot6 | 104625 | protein_coding | 755.08091 | 0.330454269 | 0.094030407 | -3.51433413 | 0.000440858 | 0.003195929 | 0.330454269 |
| ENSMUSG00000010358 | Ifi35 | 70110 | protein_coding | 265.0714238 | -0.330489449 | 0.124837479 | 2.647357596 | 0.008112352 | 0.036011679 | 0.330489449 |
| ENSMUSG00000090236 | Car15 | 80733 | protein_coding | 7.294955998 | -0.330494451 | 0.46712957 | 0.7075006 | 0.47925544 | 0.690564248 | 0.330494451 |
| ENSMUSG00000032841 | Prr5l | 72446 | protein_coding | 55.0448305 | -0.330690124 | 0.23621456 | 1.399956562 | 0.161526326 | 0.346538122 | 0.330690124 |
| ENSMUSG00000095115 | Itpripl2 | 319622 | protein_coding | 94.39265214 | -0.330692082 | 0.213201596 | 1.551076957 | 0.120883243 | 0.283485892 | 0.330692082 |

| ensembl_gene_id | external_gene_name | entrezgene_id | gene_biotype | baseMean | log2FoldChange | lfcSE | stat | pvalue | padj | absLog2FC |
|---|---|---|---|---|---|---|---|---|---|---|
| ENSMUSG00000028571 | Cyp2j13 | 230459 | protein_coding | 8.492776229 | 0.330788697 | 0.472129075 | -0.700631913 | 0.483532757 | 0.693930647 | 0.330788697 |
| ENSMUSG00000042487 | Leo1 | 235497 | protein_coding | 394.0237129 | 0.330835398 | 0.097273956 | -3.401068585 | 0.00067123 | 0.004533615 | 0.330835398 |
| ENSMUSG00000086837 | Gm16618 | 100504291 | lncRNA | 3.673615392 | -0.330850672 | 0.561497349 | 0.589229268 | 0.555707486 | 1 | 0.330850672 |
| ENSMUSG00000029622 | Arpc1b | 11867 | protein_coding | 3650.446721 | -0.330978232 | 0.078687442 | 4.206239555 | 2.60E-05 | 0.000260665 | 0.330978232 |
| ENSMUSG00000084804 | 1700095J12Rik | 73563 | lncRNA | 1.564889482 | -0.331038669 | 1.175832962 | 0.281535456 | 0.778299737 | 1 | 0.331038669 |
| ENSMUSG00000003153 | Slc2a3 | 20527 | protein_coding | 4.796960898 | -0.331108635 | 0.825275573 | 0.401209784 | 0.688265678 | 0.833786242 | 0.331108635 |
| ENSMUSG00000053600 | Zfp472 | 224691 | protein_coding | 102.1422363 | -0.331236943 | 0.146742278 | 2.257270004 | 0.023991209 | 0.084565809 | 0.331236943 |
| ENSMUSG00000085645 | Hoxb5os | 75395 | lncRNA | 21.29178218 | -0.331280186 | 0.419317511 | 0.790046152 | 0.429500816 | 0.64833423 | 0.331280186 |
| ENSMUSG00000062101 | Zfp119b | 240120 | protein_coding | 35.81965475 | -0.331328164 | 0.189470677 | 1.748704179 | 0.080342167 | 0.210665378 | 0.331328164 |
| ENSMUSG00000040520 | Manea | 242362 | protein_coding | 284.3061845 | 0.331373519 | 0.088156516 | -3.758922583 | 0.000170647 | 0.001378698 | 0.331373519 |
| ENSMUSG00000041845 | Rhod | 11854 | protein_coding | 109.1867582 | 0.331375315 | 0.24222359 | -1.3680555 | 0.171294705 | 0.361410598 | 0.331375315 |
| ENSMUSG00000052395 | Rft1 | 328370 | protein_coding | 229.0351121 | 0.331404405 | 0.103691954 | -3.196047448 | 0.001393242 | 0.008390955 | 0.331404405 |
| ENSMUSG00000033111 | 3830406C13Rik | 218734 | protein_coding | 177.0094226 | -0.331464336 | 0.117407019 | 2.823207167 | 0.004754584 | 0.023333491 | 0.331464336 |
| ENSMUSG00000000957 | Mmp14 | 17387 | protein_coding | 630.9506752 | 0.331557817 | 0.090794655 | -3.651732771 | 0.000260477 | 0.002004434 | 0.331557817 |
| ENSMUSG00000061731 | Ext1 | 14042 | protein_coding | 473.9098162 | 0.331589994 | 0.070072002 | -4.732132424 | 2.22E-06 | 2.87E-05 | 0.331589994 |
| ENSMUSG00000067818 | Myl9 | 98932 | protein_coding | 1.156438844 | -0.331605523 | 1.276788566 | 0.259718431 | 0.795080975 | 1 | 0.331605523 |
| ENSMUSG00000046269 | Usp27x | 54651 | protein_coding | 25.25459994 | -0.331697895 | 0.227477789 | 1.458155086 | 0.144797801 | 0.321915843 | 0.331697895 |
| ENSMUSG00000017677 | Wsb1 | 78889 | protein_coding | 1261.365348 | -0.331841446 | 0.140290405 | 2.36538947 | 0.018011119 | 0.067522273 | 0.331841446 |
| ENSMUSG00000035198 | Tubg1 | 103733 | protein_coding | 189.6115995 | 0.33189879 | 0.1420511 | -2.336474626 | 0.019466519 | 0.07190116 | 0.33189879 |
| ENSMUSG00000030123 | Plxnd1 | 67784 | protein_coding | 25.74337841 | -0.331986626 | 0.263819154 | 1.258387122 | 0.208251787 | 0.41130117 | 0.331986626 |
| ENSMUSG00000074277 | Phldb3 | 232970 | protein_coding | 78.7771785 | -0.33203871 | 0.188247675 | 1.763839631 | 0.077758983 | 0.205917797 | 0.33203871 |
| ENSMUSG00000002699 | Lcp2 | 16822 | protein_coding | 10.50200992 | 0.332057008 | 0.56150781 | -0.591366678 | 0.554274762 | 0.744656458 | 0.332057008 |
| ENSMUSG00000025076 | Casp7 | 12369 | protein_coding | 1841.89003 | -0.332100012 | 0.105295755 | 3.153973414 | 0.001610638 | 0.009451256 | 0.332100012 |
| ENSMUSG00000039084 | Chad | 12643 | protein_coding | 28.69203514 | -0.332171844 | 0.280751351 | 1.183153147 | 0.236748458 | 0.446932387 | 0.332171844 |
| ENSMUSG00000020743 | Mif4gd | 69674 | protein_coding | 414.5996211 | -0.332206987 | 0.07963419 | 4.171662774 | 3.02E-05 | 0.000298838 | 0.332206987 |
| ENSMUSG00000095789 | Nupr1l | 69034 | protein_coding | 6.9537229 | 0.33221243 | 0.430870945 | -0.77102537 | 0.440691894 | 0.658605063 | 0.33221243 |
| ENSMUSG00000068335 | Dok1 | 13448 | protein_coding | 152.5507721 | -0.332319378 | 0.143496066 | 2.315877971 | 0.020564932 | 0.074952393 | 0.332319378 |
| ENSMUSG00000019699 | Akt3 | 23797 | protein_coding | 8.768642076 | -0.332320684 | 0.418065038 | 0.794901879 | 0.426670581 | 0.646247636 | 0.332320684 |
| ENSMUSG00000074639 | Rdh16f2 | 216454 | protein_coding | 5.810058366 | -0.332348626 | 0.516836231 | 0.643044364 | 0.520195314 | 0.720257015 | 0.332348626 |
| ENSMUSG00000071724 | Smpd5 | 100503915 | protein_coding | 4.69529745 | -0.332413846 | 0.481890411 | 0.68981212 | 0.490312346 | 0.69793377 | 0.332413846 |
| ENSMUSG00000021193 | Pitrm1 | 69617 | protein_coding | 558.3400146 | 0.332451084 | 0.088027051 | -3.776692301 | 0.000158925 | 0.001290307 | 0.332451084 |
| ENSMUSG00000003269 | Cyth2 | 19158 | protein_coding | 703.7946589 | 0.33249873 | 0.077350241 | -4.298612744 | 1.72E-05 | 0.000180437 | 0.33249873 |
| ENSMUSG00000039851 | 4932438H23Rik | 74387 | protein_coding | 12.17160751 | 0.332536195 | 0.367609448 | -0.904590991 | 0.36568211 | 0.588354867 | 0.332536195 |
| ENSMUSG00000052516 | Robo2 | 268902 | protein_coding | 350.9332882 | 0.332566963 | 0.250443849 | -1.327910288 | 0.184207746 | 0.379255294 | 0.332566963 |
| ENSMUSG00000032418 | Me1 | 17436 | protein_coding | 222.9460377 | 0.332633958 | 0.195071809 | -1.705187238 | 0.088159509 | 0.225331394 | 0.332633958 |
| ENSMUSG00000026189 | Pecr | 111175 | protein_coding | 83.54268441 | 0.332637573 | 0.174550312 | -1.905683059 | 0.056691339 | 0.163933457 | 0.332637573 |
| ENSMUSG00000046854 | Pip5kl1 | 227733 | protein_coding | 3.133264272 | 0.332773483 | 0.815490132 | -0.408065616 | 0.683225503 | 1 | 0.332773483 |
| ENSMUSG00000053317 | Sec61b | 66212 | protein_coding | 1085.581679 | 0.33286972 | 0.110346065 | -3.016598002 | 0.002556286 | 0.013886349 | 0.33286972 |
| ENSMUSG00000109228 | Fam81b | 238726 | protein_coding | 1.329190379 | -0.332893795 | 1.626703972 | 0.204643131 | 0.837850954 | 1 | 0.332893795 |

| ensembl_gene_id | external_gene_name | entrezgene_id | gene_biotype | baseMean | log2FoldChange | lfcSE | stat | pvalue | padj | absLog2FC |
|---|---|---|---|---|---|---|---|---|---|---|
| ENSMUSG00000052560 | Cpne8 | 66871 | protein_coding | 241.8838955 | 0.332953162 | 0.086142333 | -3.865151417 | 0.00011102 | 0.000944559 | 0.332953162 |
| ENSMUSG00000056116 | H2-T22 | 15039 | protein_coding | 394.3735863 | -0.332971258 | 0.093440939 | 3.563440856 | 0.000366025 | 0.002719091 | 0.332971258 |
| ENSMUSG00000055235 | Wdr86 | 269633 | protein_coding | 4.348801381 | -0.333074247 | 0.953645163 | 0.349264338 | 0.726890869 | 1 | 0.333074247 |
| ENSMUSG00000029630 | Cyp3a25 | 56388 | protein_coding | 29.63697323 | 0.333080698 | 0.404778226 | -0.822872074 | 0.410580745 | 0.630787089 | 0.333080698 |
| ENSMUSG00000028698 | Pik3r3 | 18710 | protein_coding | 291.1591737 | 0.333096947 | 0.125134828 | -2.661904373 | 0.007769995 | 0.034782786 | 0.333096947 |
| ENSMUSG00000031936 | Hephl1 | 244698 | protein_coding | 3.481866073 | 0.333149048 | 1.286888984 | -0.2588794 | 0.795728293 | 1 | 0.333149048 |
| ENSMUSG00000049604 | Hoxb13 | 15408 | protein_coding | 2359.510884 | -0.333371281 | 0.332142507 | 1.00369954 | 0.315523459 | 0.53655334 | 0.333371281 |
| ENSMUSG00000020393 | Kremen1 | 84035 | protein_coding | 301.2583722 | -0.333388009 | 0.096809142 | 3.443765769 | 0.000573672 | 0.003977173 | 0.333388009 |
| ENSMUSG00000046667 | Rbm12b1 | 72397 | protein_coding | 15.19406535 | 0.333403821 | 0.29034956 | -1.148284231 | 0.250851247 | 0.463099872 | 0.333403821 |
| ENSMUSG00000006373 | Pgrmc1 | 53328 | protein_coding | 1302.994746 | -0.333444916 | 0.067663507 | 4.927987509 | 8.31E-07 | 1.17E-05 | 0.333444916 |
| ENSMUSG00000056209 | Npm3 | 18150 | protein_coding | 237.1279803 | -0.333495105 | 0.148390072 | 2.247421948 | 0.024613074 | 0.086297303 | 0.333495105 |
| ENSMUSG00000035298 | Klhl35 | 72184 | protein_coding | 5.938956 | 0.333499315 | 0.575046726 | -0.57995168 | 0.561947203 | 0.750140815 | 0.333499315 |
| ENSMUSG00000092738 | Mir5135 | 100628604 | miRNA | 2.482308048 | -0.333547335 | 0.729598012 | 0.457165905 | 0.647551803 | 1 | 0.333547335 |
| ENSMUSG00000022940 | Pigp | 56176 | protein_coding | 184.2591153 | -0.333577769 | 0.096473053 | 3.457729994 | 0.000544747 | 0.00381346 | 0.333577769 |
| ENSMUSG00000038762 | Abcf1 | 224742 | protein_coding | 1382.801041 | 0.333644807 | 0.059848058 | -5.574864394 | 2.48E-08 | 4.48E-07 | 0.333644807 |
| ENSMUSG00000032507 | Fbxl2 | 72179 | protein_coding | 7.835455882 | -0.333708396 | 0.485744478 | 0.687003995 | 0.492080211 | 0.698788598 | 0.333708396 |
| ENSMUSG00000036091 | Hyal3 | 109685 | protein_coding | 7.110948342 | -0.333784804 | 0.438149638 | 0.761805501 | 0.446176096 | 0.663438867 | 0.333784804 |
| ENSMUSG00000042414 | Prdm14 | 383491 | protein_coding | 14.70294844 | 0.333819272 | 0.339355411 | -0.983686312 | 0.325269772 | 0.547052145 | 0.333819272 |
| ENSMUSG00000056972 | Magel2 | 27385 | protein_coding | 1.515053078 | 0.333895151 | 2.110792113 | -0.158184763 | 0.87431121 | 1 | 0.333895151 |
| ENSMUSG00000039753 | Fbxl5 | 242960 | protein_coding | 1205.800414 | 0.334136804 | 0.06637875 | -5.033791781 | 4.81E-07 | 7.13E-06 | 0.334136804 |
| ENSMUSG00000060376 | Bckdha | 12039 | protein_coding | 169.4368071 | -0.334150474 | 0.127069132 | 2.62967463 | 0.008546662 | 0.037507197 | 0.334150474 |
| ENSMUSG00000070815 | Vmn1r87 | 171261 | protein_coding | 1.173573317 | -0.334180396 | 2.3171638 | 0.144219583 | 0.885327078 | 1 | 0.334180396 |
| ENSMUSG00000026149 | Tm4sf20 | 66261 | protein_coding | 1618.159119 | -0.334186696 | 0.092870939 | 3.598399022 | 0.000320182 | 0.002408878 | 0.334186696 |
| ENSMUSG00000033792 | Atp7a | 11977 | protein_coding | 432.2208069 | -0.334192883 | 0.084758759 | 3.942871333 | 8.05E-05 | 0.000713673 | 0.334192883 |
| ENSMUSG00000044968 | Napepld | 242864 | protein_coding | 137.9597204 | -0.334211493 | 0.141820793 | 2.356576119 | 0.018444287 | 0.068863768 | 0.334211493 |
| ENSMUSG00000066438 | Plekhd1 | 217682 | protein_coding | 48.08359622 | -0.334251802 | 0.22594843 | 1.479327832 | 0.139052718 | 0.312211636 | 0.334251802 |
| ENSMUSG00000022893 | Adamts1 | 11504 | protein_coding | 2.318913282 | -0.334274532 | 1.477382879 | 0.226261274 | 0.820998217 | 1 | 0.334274532 |
| ENSMUSG00000026492 | Tfb2m | 15278 | protein_coding | 135.2763403 | -0.334314597 | 0.112543333 | 2.970541106 | 0.002972756 | 0.015750587 | 0.334314597 |
| ENSMUSG00000041954 | Tnfrsf18 | 21936 | protein_coding | 21.85718948 | 0.334393219 | 0.22035625 | -1.517511846 | 0.129137505 | 0.296685693 | 0.334393219 |
| ENSMUSG00000018920 | Cxcl16 | 66102 | protein_coding | 40.22026556 | -0.334500512 | 0.498202785 | 0.671414377 | 0.501956589 | 0.706691156 | 0.334500512 |
| ENSMUSG00000077391 | Gm24336 | 115489055 | snoRNA | 363.4043518 | -0.334516564 | 0.095524588 | 3.501889629 | 0.000461971 | 0.00331551 | 0.334516564 |
| ENSMUSG00000018341 | Il12rb2 | 16162 | protein_coding | 3.664138448 | -0.33455936 | 0.687675117 | 0.486507874 | 0.626607124 | 1 | 0.33455936 |
| ENSMUSG00000037295 | Ldlrap1 | 100017 | protein_coding | 231.033187 | -0.33460773 | 0.095077057 | 3.519332011 | 0.000432635 | 0.003147371 | 0.33460773 |
| ENSMUSG00000029381 | Shroom3 | 27428 | protein_coding | 3014.014056 | -0.334654093 | 0.102650173 | 3.260141534 | 0.001113566 | 0.006968334 | 0.334654093 |
| ENSMUSG00000068739 | Sars | 20226 | protein_coding | 1417.324866 | 0.334904228 | 0.068358184 | -4.899255816 | 9.62E-07 | 1.33E-05 | 0.334904228 |
| ENSMUSG00000023008 | Fmnl3 | 22379 | protein_coding | 31.43141714 | 0.334971772 | 0.296771187 | -1.128720664 | 0.259015689 | 0.472341459 | 0.334971772 |
| ENSMUSG00000040782 | Cop1 | 26374 | protein_coding | 873.1296702 | -0.335020882 | 0.085413128 | 3.922358171 | 8.77E-05 | 0.000768199 | 0.335020882 |
| ENSMUSG00000026199 | Ankzf1 | 52231 | protein_coding | 297.8663641 | -0.335042185 | 0.063587768 | 5.268972217 | 1.37E-07 | 2.21E-06 | 0.335042185 |
| ENSMUSG00000096146 | Kcnj11 | 16514 | protein_coding | 6.783093094 | -0.335156614 | 0.489942534 | 0.684073315 | 0.493928871 | 0.700180271 | 0.335156614 |

| ensembl_gene_id | external_gene_name | entrezgene_id | gene_biotype | baseMean | log2FoldChange | lfcSE | stat | pvalue | padj | absLog2FC |
|---|---|---|---|---|---|---|---|---|---|---|
| ENSMUSG00000032553 | Srprb | 20818 | protein_coding | 501.809713 | 0.33524281 | 0.105611619 | -3.174298558 | 0.001501991 | 0.00890944 | 0.33524281 |
| ENSMUSG00000037188 | Grhl3 | 230824 | protein_coding | 13.73789308 | -0.335277211 | 0.298005188 | 1.125071724 | 0.260558642 | 0.474068167 | 0.335277211 |
| ENSMUSG00000029705 | Cux1 | 13047 | protein_coding | 1411.515856 | -0.335426486 | 0.052946516 | 6.335194682 | 2.37E-10 | 5.67E-09 | 0.335426486 |
| ENSMUSG00000075225 | Ccdc162 | 75973 | protein_coding | 11.62163564 | -0.335436347 | 0.566802627 | 0.591804503 | 0.553981508 | 0.744442234 | 0.335436347 |
| ENSMUSG00000026197 | Zfand2b | 68818 | protein_coding | 288.1580147 | -0.335504095 | 0.07492246 | 4.478017638 | 7.53E-06 | 8.70E-05 | 0.335504095 |
| ENSMUSG00000054702 | Ap1s3 | 252903 | protein_coding | 142.206841 | -0.335531869 | 0.164364158 | 2.04139317 | 0.041211763 | 0.128871625 | 0.335531869 |
| ENSMUSG00000089313 | Gm23182 | 115485729 | snoRNA | 2.782626561 | -0.33561454 | 0.637488919 | 0.526463331 | 0.598566323 | 1 | 0.33561454 |
| ENSMUSG00000063480 | Snu13 | 20826 | protein_coding | 298.3020006 | -0.335847375 | 0.142524792 | 2.356413714 | 0.018452354 | 0.068863768 | 0.335847375 |
| ENSMUSG00000021774 | Ube2e1 | 22194 | protein_coding | 216.2063816 | 0.335850655 | 0.094869491 | -3.540133413 | 0.000399925 | 0.002930061 | 0.335850655 |
| ENSMUSG00000040631 | Dok4 | 114255 | protein_coding | 1217.418812 | -0.335872293 | 0.100494971 | 3.342180101 | 0.000831231 | 0.005445042 | 0.335872293 |
| ENSMUSG00000030450 | Oca2 | 18431 | protein_coding | 2.882194114 | -0.335976907 | 2.416464055 | 0.139036584 | 0.88942124 | 1 | 0.335976907 |
| ENSMUSG00000055170 | Ifng | 15978 | protein_coding | 0.949112482 | -0.336066571 | 1.515460347 | 0.221758736 | 0.824501704 | 1 | 0.336066571 |
| ENSMUSG00000018974 | Sart3 | 53890 | protein_coding | 533.4700765 | 0.336166906 | 0.068037858 | -4.9408802 | 7.78E-07 | 1.10E-05 | 0.336166906 |
| ENSMUSG00000035757 | Selenoo | 223776 | protein_coding | 286.3331158 | -0.336194924 | 0.071056889 | 4.731348735 | 2.23E-06 | 2.88E-05 | 0.336194924 |
| ENSMUSG00000031969 | Acad8 | 66948 | protein_coding | 485.7879196 | -0.336297867 | 0.097153954 | 3.461494379 | 0.000537185 | 0.003770114 | 0.336297867 |
| ENSMUSG00000026469 | Xpr1 | 19775 | protein_coding | 1297.393655 | -0.336364815 | 0.091784998 | 3.664703637 | 0.000247625 | 0.001912667 | 0.336364815 |
| ENSMUSG00000067724 | Gbx1 | 231044 | protein_coding | 1.699974173 | 0.336367624 | 1.220512587 | -0.275595375 | 0.782858863 | 1 | 0.336367624 |
| ENSMUSG00000032280 | Tle3 | 21887 | protein_coding | 582.368779 | -0.336389075 | 0.152959881 | 2.19919807 | 0.027863842 | 0.095110269 | 0.336389075 |
| ENSMUSG00000028684 | Urod | 22275 | protein_coding | 294.0694349 | -0.336395793 | 0.09576872 | 3.512585257 | 0.00044377 | 0.003211398 | 0.336395793 |
| ENSMUSG00000026674 | Ddr2 | 18214 | protein_coding | 3.979259346 | 0.336472226 | 1.256766259 | -0.267728565 | 0.788908261 | 1 | 0.336472226 |
| ENSMUSG00000073414 | Mpig6b | 106722 | protein_coding | 7.692916513 | -0.336646781 | 0.713745897 | 0.471661949 | 0.637168101 | 0.802306417 | 0.336646781 |
| ENSMUSG00000064558 | Gm24620 | 115488217 | snoRNA | 11.14152052 | 0.336675332 | 0.58854619 | -0.572045725 | 0.567290997 | 0.753386941 | 0.336675332 |
| ENSMUSG00000035697 | Arhgap45 | 70719 | protein_coding | 41.68005492 | 0.336769938 | 0.366366412 | -0.919216191 | 0.357982504 | 0.580790426 | 0.336769938 |
| ENSMUSG00000007080 | Pole | 18973 | protein_coding | 212.2240194 | 0.336781049 | 0.175190842 | -1.922366743 | 0.054559627 | 0.15918687 | 0.336781049 |
| ENSMUSG00000021139 | Gm20498 | 105940408 | protein_coding | 7.797719984 | 0.336817056 | 0.366196517 | -0.919771325 | 0.357692271 | 0.580678813 | 0.336817056 |
| ENSMUSG00000034653 | Ythdc2 | 240255 | protein_coding | 433.0618497 | 0.336823707 | 0.080499545 | -4.184169097 | 2.86E-05 | 0.000284557 | 0.336823707 |
| ENSMUSG00000031791 | Tmem38a | 74166 | protein_coding | 48.41541246 | -0.336916754 | 0.166501975 | 2.02350004 | 0.043021619 | 0.132870586 | 0.336916754 |
| ENSMUSG00000048677 | Tpcn2 | 233979 | protein_coding | 166.4817481 | 0.336971253 | 0.148725677 | -2.265723439 | 0.023468322 | 0.083204714 | 0.336971253 |
| ENSMUSG00000053289 | Ddx10 | 77591 | protein_coding | 382.8725877 | 0.33702096 | 0.102966209 | -3.273121944 | 0.001063666 | 0.006694122 | 0.33702096 |
| ENSMUSG00000068547 | Clca4a | 99663 | protein_coding | 1035.136583 | 0.337174935 | 0.351081757 | -0.960388653 | 0.336859647 | 0.559233675 | 0.337174935 |
| ENSMUSG00000009030 | Pdcl | 67466 | protein_coding | 650.2719161 | 0.337183529 | 0.112220075 | -3.004663185 | 0.002658751 | 0.014369954 | 0.337183529 |
| ENSMUSG00000060981 | H4c8 | 69386 | protein_coding | 3119.732364 | 0.3372335 | 0.115819263 | -2.911722029 | 0.003594424 | 0.018508983 | 0.3372335 |
| ENSMUSG00000021690 | Jmy | 57748 | protein_coding | 300.8625783 | -0.337502282 | 0.127628913 | 2.644403007 | 0.008183516 | 0.036261245 | 0.337502282 |
| ENSMUSG00000063179 | Pstk | 214580 | protein_coding | 177.878734 | 0.337633179 | 0.100995103 | -3.343064843 | 0.000828585 | 0.005429865 | 0.337633179 |
| ENSMUSG00000097281 | Gm26685 | 102635083 | lncRNA | 11.31956123 | -0.337792839 | 0.310594766 | 1.087567712 | 0.276785991 | 0.493664959 | 0.337792839 |
| ENSMUSG00000002147 | Stat6 | 20852 | protein_coding | 1081.217133 | -0.337793239 | 0.051786505 | 6.522804276 | 6.90E-11 | 1.77E-09 | 0.337793239 |
| ENSMUSG00000030245 | Golt1b | 66964 | protein_coding | 376.7172749 | 0.337998042 | 0.127298249 | -2.655166469 | 0.007926928 | 0.035366953 | 0.337998042 |
| ENSMUSG00000020592 | Sdc1 | 20969 | protein_coding | 3487.253821 | -0.338069504 | 0.073237891 | 4.61604638 | 3.91E-06 | 4.80E-05 | 0.338069504 |
| ENSMUSG00000092985 | Mir378b | 100628571 | miRNA | 1.796434961 | 0.338174944 | 0.902565447 | -0.374681908 | 0.707897049 | 1 | 0.338174944 |

| ensembl_gene_id | external_gene_name | entrezgene_id | gene_biotype | baseMean | log2FoldChange | lfcSE | stat | pvalue | padj | absLog2FC |
|---|---|---|---|---|---|---|---|---|---|---|
| ENSMUSG00000048234 | Rnf149 | 67702 | protein_coding | 507.0911776 | -0.338211148 | 0.09999565 | 3.382258611 | 0.000718924 | 0.004789552 | 0.338211148 |
| ENSMUSG00000000884 | Gnb1l | 13972 | protein_coding | 35.88478301 | 0.338238164 | 0.175245938 | -1.930077052 | 0.053597291 | 0.157212375 | 0.338238164 |
| ENSMUSG00000028334 | Nans | 94181 | protein_coding | 6530.111198 | 0.338246564 | 0.125471955 | -2.695794158 | 0.007022105 | 0.031968135 | 0.338246564 |
| ENSMUSG00000051098 | Mblac2 | 72852 | protein_coding | 43.7218868 | -0.338477617 | 0.164548563 | 2.057007432 | 0.039685511 | 0.12530374 | 0.338477617 |
| ENSMUSG00000074452 | Pate2 | 330921 | protein_coding | 3.708818975 | 0.338608699 | 0.942236876 | -0.359366851 | 0.71932067 | 1 | 0.338608699 |
| ENSMUSG00000047710 | Champ1 | 101994 | protein_coding | 246.4927137 | -0.338611481 | 0.110889835 | 3.053584502 | 0.00226125 | 0.012501603 | 0.338611481 |
| ENSMUSG00000037649 | H2-DMa | 14998 | protein_coding | 32.232851 | 0.33865056 | 0.239306045 | -1.415135837 | 0.157028672 | 0.340132673 | 0.33865056 |
| ENSMUSG00000096210 | H1f0 | 14958 | protein_coding | 4148.075524 | 0.338662798 | 0.120348859 | -2.814009225 | 0.004892782 | 0.023883952 | 0.338662798 |
| ENSMUSG00000021917 | Spcs1 | 69019 | protein_coding | 807.8661316 | 0.338678296 | 0.099783902 | -3.394117585 | 0.000688501 | 0.00461816 | 0.338678296 |
| ENSMUSG00000017221 | Psmd3 | 22123 | protein_coding | 1250.797615 | 0.338721871 | 0.06070996 | -5.579345953 | 2.41E-08 | 4.37E-07 | 0.338721871 |
| ENSMUSG00000047828 | A730018C14Rik | 100504733 | lncRNA | 1.447388305 | -0.338774122 | 2.675537406 | 0.126619094 | 0.899241884 | 1 | 0.338774122 |
| ENSMUSG00000033186 | Mzt1 | 76789 | protein_coding | 476.2058652 | 0.338796292 | 0.103338296 | -3.278516329 | 0.001043543 | 0.00659513 | 0.338796292 |
| ENSMUSG00000033760 | Rbm4b | 66704 | protein_coding | 107.5134477 | 0.338826785 | 0.134101767 | -2.526639231 | 0.011515973 | 0.047662852 | 0.338826785 |
| ENSMUSG00000040269 | Mrps28 | 66230 | protein_coding | 260.2369146 | 0.339056804 | 0.087302327 | -3.88370868 | 0.000102875 | 0.000882077 | 0.339056804 |
| ENSMUSG00000000869 | Il4 | 16189 | protein_coding | 3.584584434 | 0.339143864 | 0.674329082 | -0.502935248 | 0.615009798 | 1 | 0.339143864 |
| ENSMUSG00000066838 | Zfp772 | 232855 | protein_coding | 61.82415854 | -0.339146786 | 0.180518134 | 1.878740808 | 0.060279893 | 0.171365493 | 0.339146786 |
| ENSMUSG00000072974 | Gm4787 | 214321 | protein_coding | 8.275664938 | -0.339154334 | 0.377123311 | 0.899319465 | 0.368482522 | 0.591017907 | 0.339154334 |
| ENSMUSG00000036825 | Ssx2ip | 99167 | protein_coding | 923.5946227 | 0.339158903 | 0.068004568 | -4.987295884 | 6.12E-07 | 8.86E-06 | 0.339158903 |
| ENSMUSG00000051177 | Plcb1 | 18795 | protein_coding | 80.02445606 | -0.339167147 | 0.180776698 | 1.876166286 | 0.06063245 | 0.172130699 | 0.339167147 |
| ENSMUSG00000042523 | Dnal1 | 105000 | protein_coding | 18.91950664 | 0.339192921 | 0.271850569 | -1.24771827 | 0.212134248 | 0.4164509 | 0.339192921 |
| ENSMUSG00000052146 | Rps10 | 67097 | protein_coding | 130.037676 | 0.339197626 | 0.210275326 | -1.613111877 | 0.106720204 | 0.259543427 | 0.339197626 |
| ENSMUSG00000023963 | Cyp39a1 | 56050 | protein_coding | 170.4036972 | -0.33929377 | 0.138053123 | 2.457704423 | 0.013982821 | 0.055215758 | 0.33929377 |
| ENSMUSG00000039220 | Ppp1r10 | 52040 | protein_coding | 1053.556411 | 0.339313513 | 0.108591239 | -3.124685894 | 0.00177995 | 0.010288448 | 0.339313513 |
| ENSMUSG00000058183 | Mmel1 | 27390 | protein_coding | 2.424476123 | 0.3393232 | 1.052176063 | -0.322496597 | 0.747076515 | 1 | 0.3393232 |
| ENSMUSG00000061751 | Kalrn | 545156 | protein_coding | 530.7705397 | -0.339402219 | 0.15015521 | 2.260342612 | 0.023799995 | 0.084106216 | 0.339402219 |
| ENSMUSG00000068957 | 4930589L23Rik | 75879 | lncRNA | 2.403820046 | -0.339570143 | 1.397830447 | 0.24292656 | 0.808062289 | 1 | 0.339570143 |
| ENSMUSG00000026737 | Pip4k2a | 18718 | protein_coding | 23.60369808 | 0.339574524 | 0.282749417 | -1.20097338 | 0.229761528 | 0.438341251 | 0.339574524 |
| ENSMUSG00000043110 | Lrrn4 | 320974 | protein_coding | 2.419293682 | 0.339620203 | 1.313829126 | -0.258496479 | 0.796023766 | 1 | 0.339620203 |
| ENSMUSG00000113733 | Gm34785 | 102638153 | lncRNA | 1.591771574 | -0.33984417 | 2.377925107 | 0.142916263 | 0.886356314 | 1 | 0.33984417 |
| ENSMUSG00000011832 | Evi5l | 213027 | protein_coding | 201.4260437 | 0.339960502 | 0.101795091 | -3.339655184 | 0.000838825 | 0.005488253 | 0.339960502 |
| ENSMUSG00000044505 | Lingo4 | 320747 | protein_coding | 6.151755563 | 0.340146623 | 0.632686701 | -0.537622527 | 0.590837675 | 0.771232643 | 0.340146623 |
| ENSMUSG00000042041 | 2010003K11Rik | 69861 | protein_coding | 229.4280833 | -0.340261596 | 0.219657216 | 1.549057218 | 0.121367966 | 0.284132491 | 0.340261596 |
| ENSMUSG00000051557 | Pusl1 | 433813 | protein_coding | 64.19289146 | -0.340270623 | 0.175873199 | 1.934749726 | 0.053021015 | 0.15599306 | 0.340270623 |
| ENSMUSG00000000214 | Th | 21823 | protein_coding | 1.329020868 | -0.340409452 | 1.221681322 | 0.278640138 | 0.780521007 | 1 | 0.340409452 |
| ENSMUSG00000039145 | Camk1d | 227541 | protein_coding | 1227.512325 | -0.340423449 | 0.147032131 | 2.315299697 | 0.020596536 | 0.075034459 | 0.340423449 |
| ENSMUSG00000002409 | Dyrk1b | 13549 | protein_coding | 140.5312619 | -0.340468081 | 0.20146846 | 1.689932417 | 0.091040885 | 0.230625736 | 0.340468081 |
| ENSMUSG00000064612 | Gm26457 | 115487641 | snoRNA | 0.934973304 | -0.340483625 | 1.557490309 | 0.218610429 | 0.826953528 | 1 | 0.340483625 |
| ENSMUSG00000079652 | Garin1a | 245884 | protein_coding | 2.770867153 | 0.340578967 | 0.745923202 | -0.456587174 | 0.6479678 | 1 | 0.340578967 |
| ENSMUSG00000052392 | Acot4 | 171282 | protein_coding | 101.6846364 | 0.340652983 | 0.197252849 | -1.726986381 | 0.084170109 | 0.21796851 | 0.340652983 |

| ensembl_gene_id | external_gene_name | entrezgene_id | gene_biotype | baseMean | log2FoldChange | lfcSE | stat | pvalue | padj | absLog2FC |
|---|---|---|---|---|---|---|---|---|---|---|
| ENSMUSG00000034858 | Atosa | 235493 | protein_coding | 161.3943623 | -0.340728366 | 0.115580763 | 2.947967774 | 0.003198704 | 0.016787354 | 0.340728366 |
| ENSMUSG00000073684 | Faap20 | 67513 | protein_coding | 140.0149796 | -0.340909081 | 0.097074283 | 3.511837242 | 0.00044502 | 0.003216222 | 0.340909081 |
| ENSMUSG00000029597 | Sds | 231691 | protein_coding | 1.006901766 | 0.341000126 | 1.200162472 | -0.284128303 | 0.776312058 | 1 | 0.341000126 |
| ENSMUSG00000026414 | Tnnt2 | 21956 | protein_coding | 41.98883802 | -0.341055711 | 0.360028994 | 0.947300682 | 0.343485583 | 0.565422739 | 0.341055711 |
| ENSMUSG00000085933 | Tmem61 | 329909 | protein_coding | 2.067779304 | 0.341075155 | 1.638752617 | -0.20813096 | 0.835126714 | 1 | 0.341075155 |
| ENSMUSG00000105814 | Mir703 | 735265 | miRNA | 13.81022801 | -0.34109014 | 0.495670164 | 0.68813934 | 0.491365039 | 0.698491194 | 0.34109014 |
| ENSMUSG00000063354 | Slc39a4 | 72027 | protein_coding | 2083.852109 | 0.341123246 | 0.139934015 | -2.437743585 | 0.014779253 | 0.057657592 | 0.341123246 |
| ENSMUSG00000087166 | L1td1 | 381591 | protein_coding | 2.178468644 | 0.341128873 | 2.39350521 | -0.14252272 | 0.886667134 | 1 | 0.341128873 |
| ENSMUSG00000104158 | Becn2 | 226720 | protein_coding | 1.225701694 | -0.341282537 | 2.338677206 | 0.145929731 | 0.883976861 | 1 | 0.341282537 |
| ENSMUSG00000046794 | Ppp1r3b | 244416 | protein_coding | 451.9678914 | -0.341336207 | 0.107541917 | 3.17398293 | 0.001503625 | 0.008912731 | 0.341336207 |
| ENSMUSG00000040705 | A930016O22Rik | 100503495 | lncRNA | 1.77064334 | 0.341357266 | 1.464428698 | -0.233099274 | 0.815684319 | 1 | 0.341357266 |
| ENSMUSG00000029311 | Hsd17b11 | 114664 | protein_coding | 830.6145914 | -0.341410768 | 0.086219807 | 3.959771884 | 7.50E-05 | 0.000670367 | 0.341410768 |
| ENSMUSG00000074194 | Zfp791 | 244556 | protein_coding | 47.43435594 | -0.341474808 | 0.263592543 | 1.295464596 | 0.195160007 | 0.394331623 | 0.341474808 |
| ENSMUSG00000067529 | Or4d11 | 258675 | protein_coding | 1.654415108 | -0.341483741 | 1.627978525 | 0.209759364 | 0.83385549 | 1 | 0.341483741 |
| ENSMUSG00000003549 | Ercc1 | 13870 | protein_coding | 136.5481908 | 0.341558297 | 0.123775763 | -2.759492556 | 0.005789121 | 0.027440166 | 0.341558297 |
| ENSMUSG00000050368 | Hoxd10 | 15430 | protein_coding | 6.855629936 | -0.341565452 | 0.432779325 | 0.789236991 | 0.429973506 | 0.648691985 | 0.341565452 |
| ENSMUSG00000038900 | Rpl12 | 269261 | protein_coding | 4875.85955 | -0.341605305 | 0.13552416 | 2.520622931 | 0.011714731 | 0.04823163 | 0.341605305 |
| ENSMUSG00000036103 | Colec12 | 140792 | protein_coding | 3.103964521 | 0.341821474 | 1.479239283 | -0.23107923 | 0.817253251 | 1 | 0.341821474 |
| ENSMUSG00000094649 | Gm7102 | 633057 | protein_coding | 0.582982676 | 0.341900338 | 1.478778636 | -0.231204542 | 0.817155901 | 1 | 0.341900338 |
| ENSMUSG00000050860 | Phospho1 | 237928 | protein_coding | 31.20995311 | 0.341914631 | 0.443124604 | -0.771599293 | 0.440351793 | 0.658275506 | 0.341914631 |
| ENSMUSG00000037166 | Ppp1r14a | 68458 | protein_coding | 13.99927368 | -0.341939703 | 0.511142778 | 0.668971015 | 0.503513967 | 0.708014489 | 0.341939703 |
| ENSMUSG00000045678 | Or5b124 | 258628 | protein_coding | 3.236212819 | -0.341953918 | 2.649750935 | 0.129051343 | 0.897317022 | 1 | 0.341953918 |
| ENSMUSG00000030748 | Il4ra | 16190 | protein_coding | 541.6680457 | 0.341956837 | 0.118771324 | -2.879119514 | 0.003987872 | 0.020127052 | 0.341956837 |
| ENSMUSG00000028247 | Coq3 | 230027 | protein_coding | 126.4632877 | -0.342101144 | 0.119189481 | 2.870229317 | 0.004101742 | 0.020610466 | 0.342101144 |
| ENSMUSG00000009281 | Rarres2 | 71660 | protein_coding | 0.849920138 | 0.342162104 | 1.428486129 | -0.239527775 | 0.810696362 | 1 | 0.342162104 |
| ENSMUSG00000078877 | Gm14295 | 628147 | protein_coding | 27.23764193 | -0.342200973 | 0.301511054 | 1.134953323 | 0.256394876 | 0.469256898 | 0.342200973 |
| ENSMUSG00000046761 | Fam83h | 105732 | protein_coding | 1594.695384 | -0.342203447 | 0.095002121 | 3.60206113 | 0.000315704 | 0.002381698 | 0.342203447 |
| ENSMUSG00000022912 | Pros1 | 19128 | protein_coding | 97.90920003 | 0.342300552 | 0.237308523 | -1.442428397 | 0.149181558 | 0.328390328 | 0.342300552 |
| ENSMUSG00000041841 | Rpl37 | 67281 | protein_coding | 2977.514515 | -0.342301202 | 0.109921648 | 3.114047205 | 0.0018454 | 0.010614744 | 0.342301202 |
| ENSMUSG00000044122 | Proca1 | 216974 | protein_coding | 6.958356059 | -0.342376118 | 0.631867613 | 0.541847865 | 0.587923314 | 0.769209762 | 0.342376118 |
| ENSMUSG00000025352 | Gdf11 | 14561 | protein_coding | 15.09307297 | 0.342422059 | 0.351008435 | -0.975537978 | 0.329293473 | 0.551684327 | 0.342422059 |
| ENSMUSG00000027890 | Gstm4 | 14865 | protein_coding | 134.3776161 | 0.342614347 | 0.132481623 | -2.586127344 | 0.009706106 | 0.041557335 | 0.342614347 |
| ENSMUSG00000058022 | Adtrp | 109254 | protein_coding | 345.6876245 | -0.342629448 | 0.111994395 | 3.059344615 | 0.002218218 | 0.01230489 | 0.342629448 |
| ENSMUSG00000018322 | Tomm34 | 67145 | protein_coding | 264.8394164 | -0.342639135 | 0.083791626 | 4.089181116 | 4.33E-05 | 0.000412515 | 0.342639135 |
| ENSMUSG00000018500 | Adora2b | 11541 | protein_coding | 15.76454196 | -0.342686723 | 0.359490704 | 0.953256146 | 0.340460305 | 0.561998621 | 0.342686723 |
| ENSMUSG00000057132 | Rpgrip1 | 77945 | protein_coding | 13.17024536 | -0.342701509 | 0.512287199 | 0.66896364 | 0.503518673 | 0.708014489 | 0.342701509 |
| ENSMUSG00000087385 | Frg2f1 | 433752 | protein_coding | 39.3366598 | -0.342762545 | 0.17953134 | 1.909207295 | 0.056235356 | 0.162838973 | 0.342762545 |
| ENSMUSG00000030122 | Ptms | 69202 | protein_coding | 1857.836468 | 0.342824453 | 0.061814801 | -5.545992946 | 2.92E-08 | 5.22E-07 | 0.342824453 |
| ENSMUSG00000055541 | Lair1 | 52855 | protein_coding | 2.705883775 | -0.342862773 | 1.28821576 | 0.26615322 | 0.790121209 | 1 | 0.342862773 |

| ensembl_gene_id | external_gene_name | entrezgene_id | gene_biotype | baseMean | log2FoldChange | lfcSE | stat | pvalue | padj | absLog2FC |
|---|---|---|---|---|---|---|---|---|---|---|
| ENSMUSG00000023206 | Il15ra | 16169 | protein_coding | 92.138871 | -0.342916562 | 0.107476298 | 3.190624989 | 0.001419654 | 0.008514609 | 0.342916562 |
| ENSMUSG00000029195 | Klb | 83379 | protein_coding | 104.4256924 | -0.342947575 | 0.244287997 | 1.403865842 | 0.160358803 | 0.344813186 | 0.342947575 |
| ENSMUSG00000000411 | Tssk3 | 58864 | protein_coding | 2.158301263 | -0.343101692 | 0.770106421 | 0.445525038 | 0.65594037 | 1 | 0.343101692 |
| ENSMUSG00000054604 | Cggbp1 | 106143 | protein_coding | 1354.661379 | 0.343139345 | 0.090175499 | -3.805239217 | 0.000141667 | 0.00117209 | 0.343139345 |
| ENSMUSG00000051413 | Plagl2 | 54711 | protein_coding | 456.363815 | 0.343181875 | 0.067508084 | -5.083567126 | 3.70E-07 | 5.60E-06 | 0.343181875 |
| ENSMUSG00000023186 | Vwa5a | 67776 | protein_coding | 844.5287409 | -0.343184019 | 0.110030997 | 3.118975817 | 0.001814809 | 0.010467953 | 0.343184019 |
| ENSMUSG00000022799 | Arhgap31 | 12549 | protein_coding | 124.0547977 | 0.343203079 | 0.113600645 | -3.021136717 | 0.002518276 | 0.013704495 | 0.343203079 |
| ENSMUSG00000022415 | Syngr1 | 20972 | protein_coding | 24.14276917 | -0.343314016 | 0.321821272 | 1.066784721 | 0.286069059 | 0.50453867 | 0.343314016 |
| ENSMUSG00000054277 | Arfgap3 | 66251 | protein_coding | 1678.476631 | -0.343491191 | 0.079742615 | 4.307498427 | 1.65E-05 | 0.000174227 | 0.343491191 |
| ENSMUSG00000028238 | Atp6v0d2 | 242341 | protein_coding | 1.057275941 | -0.343551436 | 1.799260992 | 0.190940302 | 0.848572365 | 1 | 0.343551436 |
| ENSMUSG00000026601 | Axdnd1 | 77352 | protein_coding | 5.687297155 | 0.343584538 | 0.723297344 | -0.4750253 | 0.634768941 | 0.800822518 | 0.343584538 |
| ENSMUSG00000020227 | Irak3 | 73914 | protein_coding | 54.87642455 | 0.343635098 | 0.214929384 | -1.598827908 | 0.109858846 | 0.264732086 | 0.343635098 |
| ENSMUSG00000015605 | Srf | 20807 | protein_coding | 603.7535988 | 0.343718029 | 0.113525273 | -3.027678514 | 0.002464401 | 0.013458042 | 0.343718029 |
| ENSMUSG00000018932 | Map2k3 | 26397 | protein_coding | 664.2537028 | 0.343745927 | 0.110618968 | -3.107477262 | 0.001886915 | 0.010834671 | 0.343745927 |
| ENSMUSG00000035150 | Eif2s3x | 26905 | protein_coding | 673.3378471 | -0.34378638 | 0.123793281 | 2.777100473 | 0.005484622 | 0.026283536 | 0.34378638 |
| ENSMUSG00000000126 | Wnt9a | 216795 | protein_coding | 2.57348249 | -0.343805938 | 0.86540762 | 0.3972763 | 0.691163722 | 1 | 0.343805938 |
| ENSMUSG00000026887 | Mrrf | 67871 | protein_coding | 166.884311 | -0.343927164 | 0.126826136 | 2.711800392 | 0.006691888 | 0.030722778 | 0.343927164 |
| ENSMUSG00000031757 | Mt4 | 17752 | protein_coding | 1.146383389 | 0.343940545 | 1.425877599 | -0.241213233 | 0.809389855 | 1 | 0.343940545 |
| ENSMUSG00000078779 | Zfp59 | 22717 | protein_coding | 63.68863395 | -0.344005867 | 0.205123802 | 1.677064598 | 0.093529851 | 0.235306513 | 0.344005867 |
| ENSMUSG00000062270 | Morf4l1 | 21761 | protein_coding | 707.8509806 | -0.344036446 | 0.120047212 | 2.865842855 | 0.004159006 | 0.020844259 | 0.344036446 |
| ENSMUSG00000065546 | Mir196a-1 | 387191 | miRNA | 38.601088 | -0.34404645 | 0.245149561 | 1.403414505 | 0.16049327 | 0.345012479 | 0.34404645 |
| ENSMUSG00000054385 | Ceacam2 | 26367 | protein_coding | 114.0129266 | -0.344088807 | 0.251936562 | 1.365775592 | 0.172009409 | 0.362441146 | 0.344088807 |
| ENSMUSG00000053004 | Hrh1 | 15465 | protein_coding | 103.4545315 | -0.344127542 | 0.178311567 | 1.929922708 | 0.053616415 | 0.157212621 | 0.344127542 |
| ENSMUSG00000021624 | Cd180 | 17079 | protein_coding | 2.095844052 | 0.344131581 | 0.838290692 | -0.410515809 | 0.681427609 | 1 | 0.344131581 |
| ENSMUSG00000065616 | Mir375 | 723900 | miRNA | 1.286860372 | -0.344142716 | 1.073068562 | 0.320708972 | 0.748430947 | 1 | 0.344142716 |
| ENSMUSG00000022219 | Cideb | 12684 | protein_coding | 73.52983364 | -0.34416287 | 0.195163869 | 1.763455872 | 0.077823634 | 0.205991262 | 0.34416287 |
| ENSMUSG00000001768 | Rin2 | 74030 | protein_coding | 1003.603423 | -0.344221389 | 0.07060838 | 4.875078402 | 1.09E-06 | 1.49E-05 | 0.344221389 |
| ENSMUSG00000032402 | Smad3 | 17127 | protein_coding | 827.9745455 | 0.344354164 | 0.07050594 | -4.884044747 | 1.04E-06 | 1.43E-05 | 0.344354164 |
| ENSMUSG00000066735 | Vkorc1l1 | 69568 | protein_coding | 504.7301261 | 0.344433126 | 0.083297413 | -4.134979863 | 3.55E-05 | 0.000346059 | 0.344433126 |
| ENSMUSG00000054046 | Klk13 | 626834 | protein_coding | 2.100461105 | -0.344554053 | 2.368073076 | 0.145499755 | 0.884316309 | 1 | 0.344554053 |
| ENSMUSG00000016427 | Ndufa1 | 54405 | protein_coding | 1017.648649 | -0.344610636 | 0.101259369 | 3.403246933 | 0.000665901 | 0.004503147 | 0.344610636 |
| ENSMUSG00000014547 | Wdfy2 | 268752 | protein_coding | 210.5125414 | 0.344699425 | 0.135910461 | -2.536224378 | 0.011205491 | 0.046658467 | 0.344699425 |
| ENSMUSG00000061798 | Or4c106 | 258455 | protein_coding | 1.312567999 | -0.344702227 | 2.369486238 | 0.145475513 | 0.884335448 | 1 | 0.344702227 |
| ENSMUSG00000089133 | Gm24771 | 115485992 | snoRNA | 1.343199634 | 0.344713418 | 0.971710887 | -0.354748951 | 0.722777675 | 1 | 0.344713418 |
| ENSMUSG00000052833 | Sae1 | 56459 | protein_coding | 886.8062332 | 0.344821331 | 0.112385442 | -3.068202828 | 0.002153504 | 0.012034844 | 0.344821331 |
| ENSMUSG00000069520 | Tmem19 | 67226 | protein_coding | 484.8328188 | 0.344831042 | 0.078280476 | -4.405070851 | 1.06E-05 | 0.000117654 | 0.344831042 |
| ENSMUSG00000030433 | Sbk2 | 381836 | protein_coding | 1.363890202 | -0.344845787 | 1.516577459 | 0.227384223 | 0.820124987 | 1 | 0.344845787 |
| ENSMUSG00000097583 | 6430590A07Rik | 619719 | lncRNA | 22.56253102 | 0.345129368 | 0.32468834 | -1.062955842 | 0.287801977 | 0.506745053 | 0.345129368 |
| ENSMUSG00000066009 | Zfp987 | 626316 | protein_coding | 1.805231828 | -0.345208357 | 0.752500752 | 0.458748189 | 0.646415005 | 1 | 0.345208357 |

| ensembl_gene_id | external_gene_name | entrezgene_id | gene_biotype | baseMean | log2FoldChange | lfcSE | stat | pvalue | padj | absLog2FC |
|---|---|---|---|---|---|---|---|---|---|---|
| ENSMUSG00000020538 | Srebf1 | 20787 | protein_coding | 1439.499973 | 0.345332054 | 0.103770711 | -3.327837415 | 0.000875229 | 0.005694853 | 0.345332054 |
| ENSMUSG00000078776 | 9530053A07Rik | 319482 | protein_coding | 11.59892865 | 0.345348693 | 0.66031766 | -0.523003872 | 0.600971559 | 0.777473116 | 0.345348693 |
| ENSMUSG00000029878 | Dbpht2 | 386753 | lncRNA | 5.929207797 | -0.345383519 | 0.589611443 | 0.585781573 | 0.558022308 | 0.747120423 | 0.345383519 |
| ENSMUSG00000029859 | Epha1 | 13835 | protein_coding | 388.2805688 | -0.345421699 | 0.136924834 | 2.522710366 | 0.011645427 | 0.048018124 | 0.345421699 |
| ENSMUSG00000010721 | Lmbr1 | 56873 | protein_coding | 225.8401383 | -0.345495763 | 0.104599381 | 3.303038323 | 0.000956433 | 0.00613142 | 0.345495763 |
| ENSMUSG00000009013 | Dynll1 | 56455 | protein_coding | 1292.205892 | 0.345499564 | 0.077532941 | -4.456164801 | 8.34E-06 | 9.54E-05 | 0.345499564 |
| ENSMUSG00000022758 | P2rx6 | 18440 | protein_coding | 4.838238797 | -0.345541711 | 0.55434741 | 0.623330613 | 0.533067284 | 0.729516295 | 0.345541711 |
| ENSMUSG00000027603 | Ggt7 | 207182 | protein_coding | 3.844706428 | 0.345580011 | 0.782031211 | -0.441900536 | 0.658561183 | 1 | 0.345580011 |
| ENSMUSG00000027406 | Idh3b | 170718 | protein_coding | 2601.791602 | -0.345681295 | 0.065753554 | 5.257226054 | 1.46E-07 | 2.34E-06 | 0.345681295 |
| ENSMUSG00000049154 | Fam183b | 75429 | protein_coding | 4.200643105 | 0.345803344 | 0.545384042 | -0.634054752 | 0.526045091 | 1 | 0.345803344 |
| ENSMUSG00000042105 | Inpp5f | 101490 | protein_coding | 134.7144716 | -0.345844839 | 0.127935821 | 2.703268214 | 0.006866132 | 0.031409224 | 0.345844839 |
| ENSMUSG00000074653 | Lrrc31 | 320352 | protein_coding | 185.2377497 | -0.345849376 | 0.200857326 | 1.721865876 | 0.085093818 | 0.219864981 | 0.345849376 |
| ENSMUSG00000057948 | Unc13d | 70450 | protein_coding | 291.6363164 | -0.345943387 | 0.13648276 | 2.534703921 | 0.01125424 | 0.046796074 | 0.345943387 |
| ENSMUSG00000068893 | Sprr2a2 | 100303744 | protein_coding | 182.1677889 | -0.345943638 | 0.409571839 | 0.844647032 | 0.398307941 | 0.620277218 | 0.345943638 |
| ENSMUSG00000017314 | Mpp2 | 50997 | protein_coding | 17.23928318 | 0.346097785 | 0.371762617 | -0.930964464 | 0.351871947 | 0.574134308 | 0.346097785 |
| ENSMUSG00000020668 | Kif3c | 16570 | protein_coding | 22.92728246 | -0.346237137 | 0.272494041 | 1.270622784 | 0.203862878 | 0.406017465 | 0.346237137 |
| ENSMUSG00000038807 | Rap1gap2 | 380711 | protein_coding | 1551.29128 | -0.346263225 | 0.066217791 | 5.229157023 | 1.70E-07 | 2.70E-06 | 0.346263225 |
| ENSMUSG00000096935 | 1700113A16Rik | 76642 | lncRNA | 10.59052926 | 0.346336151 | 0.342892634 | -1.01004255 | 0.312474905 | 0.533520531 | 0.346336151 |
| ENSMUSG00000003123 | Lipe | 16890 | protein_coding | 79.3060354 | -0.346352363 | 0.154880557 | 2.236254636 | 0.025335096 | 0.088285245 | 0.346352363 |
| ENSMUSG00000058925 | Ccdc192 | 75444 | protein_coding | 1.70323952 | 0.346384551 | 0.892298343 | -0.38819365 | 0.697872732 | 1 | 0.346384551 |
| ENSMUSG00000030499 | Kctd15 | 233107 | protein_coding | 18.03618501 | -0.346411048 | 0.326697327 | 1.06034246 | 0.288988829 | 0.507895701 | 0.346411048 |
| ENSMUSG00000096856 | Gm10778 | 100233208 | protein_coding | 3.883453264 | 0.346414964 | 0.686712695 | -0.504453997 | 0.613942382 | 1 | 0.346414964 |
| ENSMUSG00000020460 | Rps27a | 78294 | protein_coding | 3153.000897 | -0.346554033 | 0.145650413 | 2.379354967 | 0.017342966 | 0.065598201 | 0.346554033 |
| ENSMUSG00000030056 | Isy1 | 57905 | protein_coding | 168.3930389 | 0.346894072 | 0.131014055 | -2.647762267 | 0.008102648 | 0.036008378 | 0.346894072 |
| ENSMUSG00000052105 | Mtcl1 | 68617 | protein_coding | 87.22911153 | 0.346912171 | 0.165198912 | -2.099966437 | 0.035731794 | 0.115181503 | 0.346912171 |
| ENSMUSG00000051627 | H1f4 | 50709 | protein_coding | 4849.101536 | 0.346923182 | 0.093491578 | -3.710742612 | 0.000206652 | 0.001628174 | 0.346923182 |
| ENSMUSG00000078521 | Aunip | 69885 | protein_coding | 28.67691774 | 0.346931407 | 0.26583519 | -1.305062011 | 0.191871733 | 0.38964213 | 0.346931407 |
| ENSMUSG00000025544 | Tm9sf2 | 68059 | protein_coding | 7385.485686 | 0.346963685 | 0.108843963 | -3.187716383 | 0.001434011 | 0.008569101 | 0.346963685 |
| ENSMUSG00000020354 | Sgcd | 24052 | protein_coding | 2.776965027 | 0.34705364 | 1.400212559 | -0.247857825 | 0.804244412 | 1 | 0.34705364 |
| ENSMUSG00000050017 | Pitpnb | 56305 | protein_coding | 335.9596429 | 0.347152647 | 0.085699774 | -4.050800007 | 5.10E-05 | 0.000475421 | 0.347152647 |
| ENSMUSG00000038685 | Rtel1 | 269400 | protein_coding | 278.208045 | 0.347158925 | 0.100513188 | -3.453864429 | 0.000552615 | 0.003848961 | 0.347158925 |
| ENSMUSG00000042333 | Tnfrsf14 | 230979 | protein_coding | 138.008373 | -0.347184124 | 0.132099183 | 2.628207962 | 0.008583602 | 0.037639301 | 0.347184124 |
| ENSMUSG00000014852 | Adamts13 | 279028 | protein_coding | 7.838655263 | -0.347194811 | 0.551921806 | 0.629065219 | 0.52930636 | 0.727020313 | 0.347194811 |
| ENSMUSG00000025260 | Hsd17b10 | 15108 | protein_coding | 612.3846672 | -0.347278546 | 0.085996502 | 4.038286885 | 5.38E-05 | 0.00049841 | 0.347278546 |
| ENSMUSG00000024978 | Gpam | 14732 | protein_coding | 437.2591239 | -0.347281619 | 0.141891633 | 2.447513009 | 0.014384593 | 0.056411107 | 0.347281619 |
| ENSMUSG00000028159 | Dapp1 | 26377 | protein_coding | 116.7102726 | 0.347509934 | 0.189820633 | -1.830727929 | 0.067141162 | 0.185177119 | 0.347509934 |
| ENSMUSG00000050921 | P2ry10 | 78826 | protein_coding | 13.35735642 | -0.34766872 | 0.534073041 | 0.650975979 | 0.515061993 | 0.716272317 | 0.34766872 |
| ENSMUSG00000058402 | Zfp420 | 233058 | protein_coding | 35.18450881 | -0.347673551 | 0.201889331 | 1.722099673 | 0.085051464 | 0.219837202 | 0.347673551 |
| ENSMUSG00000020205 | Phlda1 | 21664 | protein_coding | 110.58274 | 0.347692886 | 0.189220351 | -1.837502592 | 0.066135734 | 0.183317181 | 0.347692886 |

| ensembl_gene_id | external_gene_name | entrezgene_id | gene_biotype | baseMean | log2FoldChange | lfcSE | stat | pvalue | padj | absLog2FC |
|---|---|---|---|---|---|---|---|---|---|---|
| ENSMUSG00000029378 | Areg | 11839 | protein_coding | 1037.869365 | -0.34775534 | 0.3442481 | 1.010188118 | 0.312405171 | 0.533456622 | 0.34775534 |
| ENSMUSG00000039509 | Nup133 | 234865 | protein_coding | 286.7905266 | 0.347759456 | 0.14382998 | -2.417850966 | 0.01561247 | 0.060267491 | 0.347759456 |
| ENSMUSG00000054932 | Afp | 11576 | protein_coding | 1.534981418 | 0.347832427 | 1.154261437 | -0.301346312 | 0.763150429 | 1 | 0.347832427 |
| ENSMUSG00000049517 | Rps23 | 66475 | protein_coding | 3336.722903 | -0.347859962 | 0.120790424 | 2.879863745 | 0.003978471 | 0.020104181 | 0.347859962 |
| ENSMUSG00000024260 | Sap130 | 269003 | protein_coding | 416.7302454 | 0.347871366 | 0.102401646 | -3.397126702 | 0.000680974 | 0.004584431 | 0.347871366 |
| ENSMUSG00000051228 | Nyx | 236690 | protein_coding | 5.662617772 | -0.347889985 | 0.639880631 | 0.543679505 | 0.586662038 | 0.768287864 | 0.347889985 |
| ENSMUSG00000020400 | Tnip1 | 57783 | protein_coding | 988.7857202 | -0.347907761 | 0.136490116 | 2.548959375 | 0.010804489 | 0.045254367 | 0.347907761 |
| ENSMUSG00000095193 | Gm20939 | 100044193 | lncRNA | 25.21277573 | -0.347960265 | 0.229580616 | 1.515634339 | 0.129611839 | 0.297320449 | 0.347960265 |
| ENSMUSG00000059900 | Tmem40 | 94346 | protein_coding | 14.40508409 | 0.347977558 | 0.320349197 | -1.086244516 | 0.277370827 | 0.494297255 | 0.347977558 |
| ENSMUSG00000023947 | Nfkbie | 18037 | protein_coding | 32.46302274 | -0.348082347 | 0.199107674 | 1.74821161 | 0.080427392 | 0.210845076 | 0.348082347 |
| ENSMUSG00000026980 | Ly75 | 17076 | protein_coding | 425.4134548 | -0.348131063 | 0.084297797 | 4.129776534 | 3.63E-05 | 0.000352943 | 0.348131063 |
| ENSMUSG00000005514 | Por | 18984 | protein_coding | 1218.840006 | -0.348132454 | 0.140071893 | 2.485384096 | 0.012941172 | 0.05194233 | 0.348132454 |
| ENSMUSG00000047344 | Lancl3 | 236285 | protein_coding | 57.65016083 | -0.3481577 | 0.18805857 | 1.851325895 | 0.064122683 | 0.179265615 | 0.3481577 |
| ENSMUSG00000038437 | Mllt6 | 246198 | protein_coding | 810.1890366 | -0.348178642 | 0.09775389 | 3.5617881 | 0.000368338 | 0.00273381 | 0.348178642 |
| ENSMUSG00000027894 | Slc6a17 | 229706 | protein_coding | 3.647026552 | -0.348197811 | 2.328118831 | 0.149561872 | 0.881110291 | 1 | 0.348197811 |
| ENSMUSG00000015839 | Nfe2l2 | 18024 | protein_coding | 2656.824541 | -0.348219031 | 0.074199649 | 4.693001049 | 2.69E-06 | 3.39E-05 | 0.348219031 |
| ENSMUSG00000019734 | Tmc4 | 353499 | protein_coding | 2303.699576 | -0.34823279 | 0.055662743 | 6.25611985 | 3.95E-10 | 9.15E-09 | 0.34823279 |
| ENSMUSG00000026163 | Sphkap | 77629 | protein_coding | 1.966832869 | -0.348241529 | 2.058507315 | 0.169171868 | 0.865661457 | 1 | 0.348241529 |
| ENSMUSG00000041617 | Ccdc74a | 72315 | protein_coding | 11.9913792 | 0.348495088 | 0.393349812 | -0.885967343 | 0.375635125 | 0.597844879 | 0.348495088 |
| ENSMUSG00000040950 | Mgl2 | 216864 | protein_coding | 1.160401234 | 0.348574296 | 2.294876222 | -0.151892417 | 0.879271789 | 1 | 0.348574296 |
| ENSMUSG00000024613 | Tcof1 | 21453 | protein_coding | 468.9502343 | 0.34861853 | 0.108087224 | -3.225344456 | 0.001258212 | 0.007701301 | 0.34861853 |
| ENSMUSG00000025925 | Terf1 | 21749 | protein_coding | 136.6388503 | 0.348717967 | 0.153001631 | -2.279178103 | 0.02265648 | 0.08100219 | 0.348717967 |
| ENSMUSG00000079037 | Prnp | 19122 | protein_coding | 1078.07028 | -0.348750755 | 0.11397079 | 3.060001213 | 0.002213361 | 0.012282071 | 0.348750755 |
| ENSMUSG00000020358 | Hnrnpab | 15384 | protein_coding | 2395.196954 | 0.348964589 | 0.067385854 | -5.178603034 | 2.24E-07 | 3.48E-06 | 0.348964589 |
| ENSMUSG00000029298 | Gbp9 | 236573 | protein_coding | 8.819260975 | 0.349006463 | 0.440495341 | -0.79230455 | 0.428183119 | 0.647232589 | 0.349006463 |
| ENSMUSG00000063049 | Ing2 | 69260 | protein_coding | 99.71774672 | -0.349014454 | 0.10402774 | 3.355013338 | 0.000793612 | 0.005227463 | 0.349014454 |
| ENSMUSG00000039795 | Zfand1 | 66361 | protein_coding | 206.4562184 | -0.349057424 | 0.120327373 | 2.9008979 | 0.003720951 | 0.0190648 | 0.349057424 |
| ENSMUSG00000028914 | Casp9 | 12371 | protein_coding | 234.146135 | -0.34910763 | 0.082585003 | 4.227252149 | 2.37E-05 | 0.000239668 | 0.34910763 |
| ENSMUSG00000040429 | Mterf1a | 545725 | protein_coding | 21.44467532 | -0.349142498 | 0.266841716 | 1.30842547 | 0.190729038 | 0.388133006 | 0.349142498 |
| ENSMUSG00000022416 | Cacna1i | 239556 | protein_coding | 4.06962951 | -0.349183109 | 1.646788165 | 0.212038874 | 0.832076707 | 1 | 0.349183109 |
| ENSMUSG00000026090 | Cracdl | 72097 | protein_coding | 761.7755874 | -0.349353152 | 0.091466819 | 3.819452317 | 0.000133748 | 0.001112705 | 0.349353152 |
| ENSMUSG00000050708 | Ftl1 | 14325 | protein_coding | 5272.320298 | -0.349361102 | 0.105276705 | 3.318503395 | 0.000905012 | 0.005849461 | 0.349361102 |
| ENSMUSG00000028081 | Rps3a1 | 20091 | protein_coding | 4645.301009 | -0.349629833 | 0.144933349 | 2.412349089 | 0.015850098 | 0.061038631 | 0.349629833 |
| ENSMUSG00000002718 | Cse1l | 110750 | protein_coding | 626.2203008 | 0.349680453 | 0.143269797 | -2.44071298 | 0.0146583 | 0.057294003 | 0.349680453 |
| ENSMUSG00000022763 | Aifm3 | 72168 | protein_coding | 20.69217114 | -0.349786271 | 0.282863205 | 1.236591629 | 0.216238729 | 0.421530084 | 0.349786271 |
| ENSMUSG00000085007 | Smim43 | 100503068 | protein_coding | 8.810297549 | 0.349816756 | 0.517282506 | -0.676258624 | 0.498876458 | 0.703779756 | 0.349816756 |
| ENSMUSG00000095217 | H2bc15 | 319187 | protein_coding | 847.0901938 | 0.349882429 | 0.196051351 | -1.784646865 | 0.074318615 | 0.199366082 | 0.349882429 |
| ENSMUSG00000049164 | Zfp518a | 72672 | protein_coding | 479.3053875 | -0.349947335 | 0.156816512 | 2.231571987 | 0.025643265 | 0.089020995 | 0.349947335 |
| ENSMUSG00000025381 | Cnpy2 | 56530 | protein_coding | 706.6510287 | -0.349978797 | 0.105061963 | 3.331165594 | 0.000864831 | 0.005638305 | 0.349978797 |

Supplementary Table 1

| ensembl_gene_id | external_gene_name | entrezgene_id | gene_biotype | baseMean | log2FoldChange | lfcSE | stat | pvalue | padj | absLog2FC |
|---|---|---|---|---|---|---|---|---|---|---|
| ENSMUSG00000066441 | Rdh11 | 17252 | protein_coding | 395.9058985 | 0.350066971 | 0.147545642 | -2.372601227 | 0.017663322 | 0.066596371 | 0.350066971 |
| ENSMUSG00000070691 | Runx3 | 12399 | protein_coding | 7.421216286 | 0.350216221 | 0.593055036 | -0.590529039 | 0.554836023 | 0.744703672 | 0.350216221 |
| ENSMUSG00000033256 | Shf | 435684 | protein_coding | 137.9026778 | 0.350254661 | 0.123415594 | -2.838009763 | 0.004539579 | 0.022458542 | 0.350254661 |
| ENSMUSG00000039462 | Col10a1 | 12813 | protein_coding | 7.345775253 | -0.350292002 | 0.482140291 | 0.726535427 | 0.467510598 | 0.681117777 | 0.350292002 |
| ENSMUSG00000063430 | Wscd2 | 320916 | protein_coding | 5.152776439 | -0.350323427 | 1.260560326 | 0.277910878 | 0.781080773 | 0.890541867 | 0.350323427 |
| ENSMUSG00000017774 | Myo1c | 17913 | protein_coding | 1170.482851 | -0.350337015 | 0.051734747 | 6.771793387 | 1.27E-11 | 3.52E-10 | 0.350337015 |
| ENSMUSG00000019891 | Dcbld1 | 66686 | protein_coding | 415.0273793 | 0.350342809 | 0.082767185 | -4.232870908 | 2.31E-05 | 0.000234331 | 0.350342809 |
| ENSMUSG00000090693 | Trim43a | 547109 | protein_coding | 1.779807132 | 0.350361804 | 0.894704007 | -0.39159521 | 0.695357328 | 1 | 0.350361804 |
| ENSMUSG00000079481 | Nhsl2 | 100042480 | protein_coding | 6.267534532 | 0.350535327 | 1.635786648 | -0.214291593 | 0.830319674 | 0.914187552 | 0.350535327 |
| ENSMUSG00000049676 | Catsperg1 | 320225 | protein_coding | 7.624021209 | 0.350574887 | 0.712358375 | -0.492132751 | 0.622625501 | 0.793088519 | 0.350574887 |
| ENSMUSG00000028518 | Prkaa2 | 108079 | protein_coding | 1740.606832 | -0.350623494 | 0.085763116 | 4.088278408 | 4.35E-05 | 0.000413646 | 0.350623494 |
| ENSMUSG00000037474 | Dtl | 76843 | protein_coding | 135.947748 | 0.35063343 | 0.178248191 | -1.967107925 | 0.04917077 | 0.147129207 | 0.35063343 |
| ENSMUSG00000019890 | Nts | 67405 | protein_coding | 89.28142259 | -0.350852711 | 0.334099712 | 1.05014371 | 0.293652045 | 0.512889463 | 0.350852711 |
| ENSMUSG00000020108 | Ddit4 | 74747 | protein_coding | 752.6388956 | 0.35097501 | 0.130022226 | -2.699346261 | 0.006947585 | 0.031711558 | 0.35097501 |
| ENSMUSG00000085175 | Gm11423 | 100503272 | lncRNA | 18.27416697 | -0.350985805 | 0.275127432 | 1.275720865 | 0.20205421 | 0.404039964 | 0.350985805 |
| ENSMUSG00000087497 | 2810001G20Rik | 66456 | lncRNA | 41.41646092 | -0.351042011 | 0.18320395 | 1.916126865 | 0.05534895 | 0.16072315 | 0.351042011 |
| ENSMUSG00000018574 | Acadvl | 11370 | protein_coding | 1406.102522 | -0.351134618 | 0.070334929 | 4.992322045 | 5.97E-07 | 8.65E-06 | 0.351134618 |
| ENSMUSG00000031788 | Kifc3 | 16582 | protein_coding | 367.6169216 | 0.351166145 | 0.116153124 | -3.023303485 | 0.002500314 | 0.013622626 | 0.351166145 |
| ENSMUSG00000022604 | Cep97 | 74201 | protein_coding | 56.35596095 | -0.351197761 | 0.155331166 | 2.260961342 | 0.023761651 | 0.084025676 | 0.351197761 |
| ENSMUSG00000020072 | Pbld2 | 67307 | protein_coding | 210.9804743 | -0.351264371 | 0.102775506 | 3.417782948 | 0.000631334 | 0.004313552 | 0.351264371 |
| ENSMUSG00000034177 | Rnf43 | 207742 | protein_coding | 463.4665767 | -0.351375734 | 0.08621873 | 4.075399108 | 4.59E-05 | 0.000434218 | 0.351375734 |
| ENSMUSG00000024454 | Hdac3 | 15183 | protein_coding | 785.6234261 | 0.351510112 | 0.063302053 | -5.552902234 | 2.81E-08 | 5.04E-07 | 0.351510112 |
| ENSMUSG00000084466 | Gm22960 | 115488256 | snRNA | 0.599327296 | 0.35151934 | 1.59696346 | -0.220117334 | 0.825779775 | 1 | 0.35151934 |
| ENSMUSG00000031903 | Pla2g15 | 192654 | protein_coding | 93.47732099 | 0.351539495 | 0.113150362 | -3.106834915 | 0.00189102 | 0.010850695 | 0.351539495 |
| ENSMUSG00000059291 | Rpl11 | 67025 | protein_coding | 3009.443102 | -0.351586576 | 0.13374213 | 2.628839364 | 0.008567682 | 0.037589453 | 0.351586576 |
| ENSMUSG00000106514 | Mir6976 | 102466778 | miRNA | 3.210492038 | -0.351664531 | 0.641923075 | 0.547829709 | 0.583808837 | 1 | 0.351664531 |
| ENSMUSG00000029344 | Tpst2 | 22022 | protein_coding | 192.7116517 | 0.351725731 | 0.119787993 | -2.936235274 | 0.003322222 | 0.017328863 | 0.351725731 |
| ENSMUSG00000074259 | Gramd2 | 546134 | protein_coding | 38.52623928 | 0.351761653 | 0.21859483 | -1.609194752 | 0.107573764 | 0.260900739 | 0.351761653 |
| ENSMUSG00000058975 | Kcnc1 | 16502 | protein_coding | 7.782633571 | -0.351774051 | 0.730682535 | 0.481432132 | 0.630209404 | 0.798287804 | 0.351774051 |
| ENSMUSG00000021496 | Pcbd2 | 72562 | protein_coding | 113.7070219 | -0.351825851 | 0.153340523 | 2.29440884 | 0.021767026 | 0.078433486 | 0.351825851 |
| ENSMUSG00000039579 | Grin3a | 242443 | protein_coding | 141.1328394 | -0.351916155 | 0.320002265 | 1.099730201 | 0.271449692 | 0.487146296 | 0.351916155 |
| ENSMUSG00000018042 | Cyb5r3 | 109754 | protein_coding | 1270.979313 | 0.35195601 | 0.046046088 | -7.643559485 | 2.11E-14 | 7.33E-13 | 0.35195601 |
| ENSMUSG00000028165 | Cisd2 | 67006 | protein_coding | 659.9207995 | 0.352007401 | 0.061350241 | -5.737669422 | 9.60E-09 | 1.85E-07 | 0.352007401 |
| ENSMUSG00000036985 | Zdhhc9 | 208884 | protein_coding | 364.3501193 | 0.352015358 | 0.102761314 | -3.425563041 | 0.000613527 | 0.004211046 | 0.352015358 |
| ENSMUSG00000024654 | Asrgl1 | 66514 | protein_coding | 66.06263678 | 0.35203249 | 0.187502427 | -1.877482309 | 0.06045202 | 0.171677499 | 0.35203249 |
| ENSMUSG00000024944 | Arl2 | 56327 | protein_coding | 153.5096912 | -0.352060368 | 0.096878099 | 3.634055298 | 0.000279001 | 0.002138015 | 0.352060368 |
| ENSMUSG00000026942 | Traf2 | 22030 | protein_coding | 312.8508368 | 0.352063563 | 0.090414923 | -3.89386565 | 9.87E-05 | 0.000850344 | 0.352063563 |
| ENSMUSG00000044254 | Pcsk9 | 100102 | protein_coding | 265.1200426 | -0.352114598 | 0.227839966 | 1.545447027 | 0.122238173 | 0.285320309 | 0.352114598 |
| ENSMUSG00000028822 | Tmem50a | 71817 | protein_coding | 488.2967619 | -0.35235745 | 0.09286083 | 3.794468032 | 0.00014796 | 0.00121563 | 0.35235745 |

| ensembl_gene_id | external_gene_name | entrezgene_id | gene_biotype | baseMean | log2FoldChange | lfcSE | stat | pvalue | padj | absLog2FC |
|---|---|---|---|---|---|---|---|---|---|---|
| ENSMUSG00000086629 | 2810403D21Rik | 69964 | lncRNA | 28.32632591 | -0.352359755 | 0.238714332 | 1.476072892 | 0.139924325 | 0.313784669 | 0.352359755 |
| ENSMUSG00000085887 | Arhgap27os3 | 653030 | lncRNA | 24.75717895 | -0.352467833 | 0.363237073 | 0.970352036 | 0.331871048 | 0.554292124 | 0.352467833 |
| ENSMUSG00000078485 | Plekhn1 | 231002 | protein_coding | 185.1140361 | -0.352485367 | 0.131424862 | 2.682029576 | 0.007317699 | 0.033071834 | 0.352485367 |
| ENSMUSG00000038570 | Saxo2 | 330577 | protein_coding | 6.896763813 | 0.352494171 | 0.43275679 | -0.814531809 | 0.415340357 | 0.635615852 | 0.352494171 |
| ENSMUSG00000042677 | Zc3h12a | 230738 | protein_coding | 402.8412792 | 0.35254461 | 0.210748398 | -1.672822256 | 0.094362292 | 0.236643719 | 0.35254461 |
| ENSMUSG00000041995 | Zbed3 | 72114 | protein_coding | 115.1686903 | 0.352558459 | 0.161145305 | -2.187829542 | 0.028682021 | 0.097419765 | 0.352558459 |
| ENSMUSG00000027246 | Ell3 | 269344 | protein_coding | 5.357924613 | -0.352696046 | 0.4282429 | 0.823588777 | 0.410173255 | 0.630552252 | 0.352696046 |
| ENSMUSG00000063146 | Clip2 | 269713 | protein_coding | 1318.705423 | 0.352747826 | 0.076573963 | -4.606628829 | 4.09E-06 | 5.00E-05 | 0.352747826 |
| ENSMUSG00000053907 | Mat2a | 232087 | protein_coding | 2633.791637 | -0.352889003 | 0.108744626 | 3.245116693 | 0.001174025 | 0.0072831 | 0.352889003 |
| ENSMUSG00000020806 | Rhbdf2 | 217344 | protein_coding | 209.2401804 | -0.353076302 | 0.102183144 | 3.455328237 | 0.000549623 | 0.003837834 | 0.353076302 |
| ENSMUSG00000034799 | Unc13a | 382018 | protein_coding | 33.43981098 | -0.353127858 | 0.417929059 | 0.84494689 | 0.398140493 | 0.62019321 | 0.353127858 |
| ENSMUSG00000091475 | Cerox1 | 72834 | lncRNA | 1.385414542 | -0.353138341 | 1.154471662 | 0.305887406 | 0.759690376 | 1 | 0.353138341 |
| ENSMUSG00000055401 | Fbxo6 | 50762 | protein_coding | 187.1528825 | -0.353158595 | 0.115446101 | 3.059077727 | 0.002220195 | 0.012311721 | 0.353158595 |
| ENSMUSG00000047787 | Flrt1 | 396184 | protein_coding | 11.9978969 | -0.353208784 | 0.469246768 | 0.752714367 | 0.451621575 | 0.668047177 | 0.353208784 |
| ENSMUSG00000038224 | Serpinf2 | 18816 | protein_coding | 11.66815033 | 0.353320148 | 0.395291081 | -0.893822716 | 0.371416753 | 0.593868753 | 0.353320148 |
| ENSMUSG00000027195 | Hsd17b12 | 56348 | protein_coding | 1137.909397 | 0.353370118 | 0.102855859 | -3.43558571 | 0.000591274 | 0.004080362 | 0.353370118 |
| ENSMUSG00000034472 | Rasd2 | 75141 | protein_coding | 261.8393969 | -0.353387711 | 0.283730115 | 1.24550653 | 0.212945617 | 0.417696153 | 0.353387711 |
| ENSMUSG00000047631 | Apof | 103161 | protein_coding | 1.956177607 | -0.353400241 | 1.429002684 | 0.247305512 | 0.804671793 | 1 | 0.353400241 |
| ENSMUSG00000034584 | Exph5 | 320051 | protein_coding | 896.2302551 | -0.35349995 | 0.104516646 | 3.382235876 | 0.000718984 | 0.004789552 | 0.35349995 |
| ENSMUSG00000048758 | Rpl29 | 19944 | protein_coding | 2303.426833 | -0.353580718 | 0.142267081 | 2.485330517 | 0.01294312 | 0.05194233 | 0.353580718 |
| ENSMUSG00000030486 | Zfp108 | 54678 | protein_coding | 29.63454191 | -0.353808633 | 0.225532773 | 1.568768159 | 0.11670197 | 0.276145055 | 0.353808633 |
| ENSMUSG00000075569 | Rsph10b | 18861 | protein_coding | 2.161217055 | 0.353951842 | 0.776347504 | -0.455919341 | 0.648447981 | 1 | 0.353951842 |
| ENSMUSG00000036327 | Qsox2 | 227638 | protein_coding | 374.595413 | 0.353977425 | 0.172259551 | -2.054907392 | 0.03988795 | 0.125701628 | 0.353977425 |
| ENSMUSG00000032359 | Ctsh | 13036 | protein_coding | 479.914797 | -0.35403182 | 0.086190528 | 4.107549018 | 4.00E-05 | 0.000384378 | 0.35403182 |
| ENSMUSG00000030720 | Cln3 | 12752 | protein_coding | 261.8821675 | -0.354088317 | 0.089558796 | 3.953696697 | 7.70E-05 | 0.000685437 | 0.354088317 |
| ENSMUSG00000024583 | Txnl1 | 53382 | protein_coding | 874.4860714 | 0.354098756 | 0.117148031 | -3.022660764 | 0.00250563 | 0.013647086 | 0.354098756 |
| ENSMUSG00000029267 | Mtf2 | 17765 | protein_coding | 393.7188048 | 0.354109098 | 0.08146585 | -4.346718279 | 1.38E-05 | 0.00014896 | 0.354109098 |
| ENSMUSG00000021990 | Spata13 | 219140 | protein_coding | 698.7758846 | -0.35418316 | 0.105488405 | 3.357555359 | 0.00078635 | 0.005188088 | 0.35418316 |
| ENSMUSG00000033105 | Lss | 16987 | protein_coding | 755.4421813 | 0.354211154 | 0.103449781 | -3.423991347 | 0.000617086 | 0.004230202 | 0.354211154 |
| ENSMUSG00000029410 | Ppef2 | 19023 | protein_coding | 3.116932454 | -0.354363248 | 0.831069037 | 0.426394478 | 0.669820425 | 1 | 0.354363248 |
| ENSMUSG00000025499 | Hras | 15461 | protein_coding | 760.4029002 | 0.354954551 | 0.05916076 | -5.999830802 | 1.98E-09 | 4.21E-08 | 0.354954551 |
| ENSMUSG00000069808 | Tlcd3a | 116972 | protein_coding | 494.2554124 | 0.355033847 | 0.114156666 | -3.110057966 | 0.001870506 | 0.010747927 | 0.355033847 |
| ENSMUSG00000085436 | Zfp335os | 629777 | lncRNA | 14.15310467 | 0.355035605 | 0.291761199 | -1.216870528 | 0.223653481 | 0.43083733 | 0.355035605 |
| ENSMUSG00000000740 | Rpl13 | 270106 | protein_coding | 4237.277656 | -0.355193453 | 0.121144275 | 2.931987106 | 0.003368007 | 0.017528923 | 0.355193453 |
| ENSMUSG00000029028 | Lrrc47 | 72946 | protein_coding | 381.8416015 | 0.355201264 | 0.067641278 | -5.251250057 | 1.51E-07 | 2.42E-06 | 0.355201264 |
| ENSMUSG00000031732 | Phlpp2 | 244650 | protein_coding | 585.2810748 | -0.355212281 | 0.174469735 | 2.035953583 | 0.041755016 | 0.130053252 | 0.355212281 |
| ENSMUSG00000016239 | Lonrf3 | 74365 | protein_coding | 93.15501068 | 0.355292817 | 0.154705366 | -2.296577207 | 0.0216429 | 0.078139671 | 0.355292817 |
| ENSMUSG00000052837 | Junb | 16477 | protein_coding | 5504.789745 | -0.355331694 | 0.163468503 | 2.173701276 | 0.029727577 | 0.100208451 | 0.355331694 |
| ENSMUSG00000099825 | 1810012K16Rik | 69108 | lncRNA | 3.483418035 | -0.355476897 | 0.606229919 | 0.586373067 | 0.557624839 | 1 | 0.355476897 |

| ensembl_gene_id | external_gene_name | entrezgene_id | gene_biotype | baseMean | log2FoldChange | lfcSE | stat | pvalue | padj | absLog2FC |
|---|---|---|---|---|---|---|---|---|---|---|
| ENSMUSG00000087408 | Cers1 | 14559 | protein_coding | 2.17285123 | 0.355499576 | 1.390554731 | -0.255653063 | 0.798218741 | 1 | 0.355499576 |
| ENSMUSG00000087440 | 4930577N17Rik | 67746 | TEC | 14.82732485 | 0.355535808 | 0.274756783 | -1.294001935 | 0.195664752 | 0.394964884 | 0.355535808 |
| ENSMUSG00000029761 | Cald1 | 109624 | protein_coding | 19.29350427 | 0.355634978 | 0.315841282 | -1.1259927 | 0.260168609 | 0.473905185 | 0.355634978 |
| ENSMUSG00000112963 | Gm6093 | 619715 | lncRNA | 2.169034596 | -0.355759482 | 2.368965223 | 0.150175055 | 0.880626506 | 1 | 0.355759482 |
| ENSMUSG00000028573 | Fggy | 75578 | protein_coding | 101.5934836 | -0.355782765 | 0.131570647 | 2.704119606 | 0.006848563 | 0.031346224 | 0.355782765 |
| ENSMUSG00000050022 | Amz1 | 231842 | protein_coding | 5.562665818 | 0.355813514 | 0.635018079 | -0.560320289 | 0.575260991 | 0.760475505 | 0.355813514 |
| ENSMUSG00000029769 | Ccdc136 | 232664 | protein_coding | 5.522273823 | -0.356143431 | 0.658511382 | 0.540831093 | 0.588624009 | 0.769577776 | 0.356143431 |
| ENSMUSG00000043644 | 0610009L18Rik | 66838 | lncRNA | 27.30046995 | -0.356237644 | 0.217930367 | 1.63463977 | 0.102124586 | 0.251451529 | 0.356237644 |
| ENSMUSG00000031161 | Hdac6 | 15185 | protein_coding | 229.6286234 | 0.35627635 | 0.091972354 | -3.87373308 | 0.000107181 | 0.000916141 | 0.35627635 |
| ENSMUSG00000058706 | 0610030E20Rik | 68364 | protein_coding | 218.4395863 | -0.356318573 | 0.095664175 | 3.724681413 | 0.000195562 | 0.001548948 | 0.356318573 |
| ENSMUSG00000116632 | Magef1 | 76222 | transcribed_processed_pseudogene | 25.90277464 | -0.356357595 | 0.203913255 | 1.747594071 | 0.080534343 | 0.210915627 | 0.356357595 |
| ENSMUSG00000027224 | Duoxa1 | 213696 | protein_coding | 14.95224597 | 0.356549314 | 0.420518885 | -0.847879434 | 0.396505119 | 0.618855346 | 0.356549314 |
| ENSMUSG00000001576 | Ergic1 | 67458 | protein_coding | 1411.913263 | 0.356610932 | 0.122734849 | -2.90553935 | 0.003666207 | 0.018819941 | 0.356610932 |
| ENSMUSG00000028457 | Atp8b5 | 320571 | protein_coding | 24.79013147 | -0.356622989 | 0.279555212 | 1.275679987 | 0.202068666 | 0.404039964 | 0.356622989 |
| ENSMUSG00000005899 | Smpd4 | 77626 | protein_coding | 305.4861794 | 0.356660698 | 0.068279876 | -5.223511175 | 1.76E-07 | 2.77E-06 | 0.356660698 |
| ENSMUSG00000039105 | Atp6v1g1 | 66290 | protein_coding | 704.216962 | -0.356669561 | 0.105904786 | 3.36783233 | 0.000757617 | 0.005018564 | 0.356669561 |
| ENSMUSG00000023072 | Cep89 | 72140 | protein_coding | 248.1485723 | 0.356809937 | 0.079150913 | -4.507969921 | 6.55E-06 | 7.64E-05 | 0.356809937 |
| ENSMUSG00000026049 | Tex30 | 75623 | protein_coding | 66.76533749 | 0.356877088 | 0.170538859 | -2.092643808 | 0.036380963 | 0.116818048 | 0.356877088 |
| ENSMUSG00000031508 | Ankrd10 | 102334 | protein_coding | 614.3969833 | -0.356986066 | 0.103725324 | 3.441648131 | 0.000578182 | 0.004001262 | 0.356986066 |
| ENSMUSG00000049897 | Stkld1 | 279029 | protein_coding | 5.319311173 | -0.357102687 | 0.562094579 | 0.635307118 | 0.525228132 | 0.723645701 | 0.357102687 |
| ENSMUSG00000024232 | Bambi | 68010 | protein_coding | 65.10629957 | 0.35714635 | 0.209557901 | -1.704284824 | 0.088327889 | 0.225622081 | 0.35714635 |
| ENSMUSG00000041966 | Dcaf17 | 75763 | protein_coding | 223.9542386 | -0.357287771 | 0.099695053 | 3.583806418 | 0.000338623 | 0.002533766 | 0.357287771 |
| ENSMUSG00000115816 | Gm34589 | 102637893 | lncRNA | 3.243340397 | 0.357325267 | 0.982931235 | -0.36353028 | 0.716208798 | 1 | 0.357325267 |
| ENSMUSG00000034480 | Diaph2 | 54004 | protein_coding | 190.4627504 | -0.357378827 | 0.090291432 | 3.958059128 | 7.56E-05 | 0.000674132 | 0.357378827 |
| ENSMUSG00000043398 | Gpr135 | 238252 | protein_coding | 2.617281806 | 0.357475585 | 0.638747313 | -0.559651019 | 0.575717498 | 1 | 0.357475585 |
| ENSMUSG00000033610 | Pank1 | 75735 | protein_coding | 409.5749029 | -0.357483776 | 0.08536185 | 4.187863494 | 2.82E-05 | 0.000280811 | 0.357483776 |
| ENSMUSG00000026921 | Egfl7 | 353156 | protein_coding | 19.85732057 | 0.357536595 | 0.407527653 | -0.877330882 | 0.380306943 | 0.602493175 | 0.357536595 |
| ENSMUSG00000006498 | Ptbp1 | 19205 | protein_coding | 2524.212974 | 0.357607024 | 0.062889004 | -5.686320392 | 1.30E-08 | 2.45E-07 | 0.357607024 |
| ENSMUSG00000109450 | Gm39043 | 105242985 | lncRNA | 1.275980282 | 0.357821229 | 2.319062956 | -0.154295608 | 0.87737666 | 1 | 0.357821229 |
| ENSMUSG00000006920 | Ezh1 | 14055 | protein_coding | 570.4435737 | -0.357921085 | 0.066691086 | 5.366850469 | 8.01E-08 | 1.34E-06 | 0.357921085 |
| ENSMUSG00000051985 | Igfn1 | 226438 | protein_coding | 7.095364019 | 0.357977556 | 0.830585848 | -0.430994047 | 0.666472699 | 0.820495762 | 0.357977556 |
| ENSMUSG00000067430 | Zfp763 | 73451 | protein_coding | 67.86384842 | -0.358057571 | 0.149898926 | 2.388660024 | 0.016909942 | 0.064269456 | 0.358057571 |
| ENSMUSG00000073164 | 2410018L13Rik | 69732 | lncRNA | 3.10327556 | 0.358068425 | 0.651225324 | -0.549837993 | 0.582430497 | 1 | 0.358068425 |
| ENSMUSG00000046324 | Ermp1 | 226090 | protein_coding | 1489.593293 | -0.358088825 | 0.080755928 | 4.434211008 | 9.24E-06 | 0.000104668 | 0.358088825 |
| ENSMUSG00000026288 | Inpp5d | 16331 | protein_coding | 73.40425278 | -0.358100587 | 0.210805483 | 1.698725199 | 0.089370973 | 0.227592128 | 0.358100587 |
| ENSMUSG00000034919 | Ttc22 | 230576 | protein_coding | 1043.399703 | 0.358113362 | 0.146034134 | -2.452257921 | 0.014196287 | 0.055871658 | 0.358113362 |
| ENSMUSG00000010911 | Apip | 56369 | protein_coding | 83.67163471 | 0.35824535 | 0.129711695 | -2.761858512 | 0.005747338 | 0.027285164 | 0.35824535 |
| ENSMUSG00000027078 | Ube2l6 | 56791 | protein_coding | 74.12809324 | -0.358318851 | 0.23088581 | 1.551931021 | 0.120678729 | 0.283161202 | 0.358318851 |
| ENSMUSG00000029627 | Zkscan14 | 67235 | protein_coding | 95.98420759 | -0.358433428 | 0.141985361 | 2.524439318 | 0.011588301 | 0.047830343 | 0.358433428 |

| ensembl_gene_id | external_gene_name | entrezgene_id | gene_biotype | baseMean | log2FoldChange | lfcSE | stat | pvalue | padj | absLog2FC |
|---|---|---|---|---|---|---|---|---|---|---|
| ENSMUSG00000087648 | E130018N17Rik | 77017 | lncRNA | 0.844759267 | -0.358477698 | 1.342347163 | 0.267052897 | 0.789428434 | 1 | 0.358477698 |
| ENSMUSG00000040502 | Marchf9 | 216438 | protein_coding | 51.29466239 | -0.35859327 | 0.185640807 | 1.931651104 | 0.053402584 | 0.156808369 | 0.35859327 |
| ENSMUSG00000070524 | Fcrlb | 435653 | protein_coding | 0.994003205 | 0.358622093 | 2.264973191 | -0.158333924 | 0.874193678 | 1 | 0.358622093 |
| ENSMUSG00000019055 | Plod1 | 18822 | protein_coding | 165.1631335 | 0.35875691 | 0.147993586 | -2.424138234 | 0.015344763 | 0.059503296 | 0.35875691 |
| ENSMUSG00000036580 | Spg20 | 229285 | protein_coding | 158.4447611 | 0.358771191 | 0.120418547 | -2.979368206 | 0.002888435 | 0.015401876 | 0.358771191 |
| ENSMUSG00000097494 | 4933406C10Rik | 74076 | lncRNA | 30.36306031 | 0.358776799 | 0.207411249 | -1.729784676 | 0.083668753 | 0.216896136 | 0.358776799 |
| ENSMUSG00000026114 | Cnga3 | 12790 | protein_coding | 3.137280409 | -0.358820897 | 0.665247915 | 0.539379212 | 0.589625222 | 1 | 0.358820897 |
| ENSMUSG00000025348 | Itga7 | 16404 | protein_coding | 377.5038996 | -0.358826737 | 0.148693962 | 2.413189701 | 0.015813587 | 0.060929904 | 0.358826737 |
| ENSMUSG00000057863 | Rpl36 | 54217 | protein_coding | 2298.916531 | -0.358833261 | 0.112037063 | 3.202808538 | 0.001360944 | 0.00822343 | 0.358833261 |
| ENSMUSG00000030058 | Copg1 | 54161 | protein_coding | 2091.569461 | 0.358847273 | 0.078979336 | -4.543559025 | 5.53E-06 | 6.55E-05 | 0.358847273 |
| ENSMUSG00000018566 | Slc2a4 | 20528 | protein_coding | 15.29549454 | 0.358858365 | 0.31230306 | -1.149070921 | 0.250526738 | 0.462773887 | 0.358858365 |
| ENSMUSG00000025532 | Crcp | 12909 | protein_coding | 197.0150354 | 0.35887156 | 0.079219909 | -4.53006783 | 5.90E-06 | 6.93E-05 | 0.35887156 |
| ENSMUSG00000036002 | Atosb | 230088 | protein_coding | 600.7783592 | 0.358925418 | 0.135114687 | -2.656450057 | 0.007896815 | 0.035293098 | 0.358925418 |
| ENSMUSG00000064778 | Snord59a | 115487514 | snoRNA | 2.021526508 | 0.358975826 | 0.751793938 | -0.477492312 | 0.633011594 | 1 | 0.358975826 |
| ENSMUSG00000041020 | Map7d2 | 78283 | protein_coding | 2.588392455 | 0.358995413 | 1.280999718 | -0.280246286 | 0.779288558 | 1 | 0.358995413 |
| ENSMUSG00000020034 | Tcp11l2 | 216198 | protein_coding | 78.71791407 | -0.359002722 | 0.204298233 | 1.757248294 | 0.078875497 | 0.207979566 | 0.359002722 |
| ENSMUSG00000074059 | Fbxw18 | 546161 | protein_coding | 1.252245074 | 0.35907496 | 2.667387635 | -0.134616715 | 0.892914925 | 1 | 0.35907496 |
| ENSMUSG00000035615 | Frmpd1 | 666060 | protein_coding | 1.906504513 | -0.359207925 | 1.608180963 | 0.223362876 | 0.823253095 | 1 | 0.359207925 |
| ENSMUSG00000031697 | Orc6 | 56452 | protein_coding | 119.7567386 | 0.35952199 | 0.114319286 | -3.144893598 | 0.001661472 | 0.009712409 | 0.35952199 |
| ENSMUSG00000017929 | B4galt5 | 56336 | protein_coding | 714.5709613 | -0.359573837 | 0.088716675 | 4.053058099 | 5.06E-05 | 0.000471385 | 0.359573837 |
| ENSMUSG00000046085 | 4931422A03Rik | 70986 | lncRNA | 5.853868128 | 0.359586301 | 0.676458076 | -0.53157219 | 0.59502233 | 0.773528957 | 0.359586301 |
| ENSMUSG00000014747 | Ankrd53 | 75305 | protein_coding | 2.561874909 | -0.359625845 | 0.874077946 | 0.411434525 | 0.680753942 | 1 | 0.359625845 |
| ENSMUSG00000019256 | Ahr | 11622 | protein_coding | 181.1430776 | -0.359646522 | 0.099316273 | 3.621224524 | 0.000293212 | 0.002234472 | 0.359646522 |
| ENSMUSG00000036925 | Mucl2 | 20770 | protein_coding | 1.481793215 | 0.359735414 | 2.680059852 | -0.134226634 | 0.893223365 | 1 | 0.359735414 |
| ENSMUSG00000075334 | Rprm | 67874 | protein_coding | 19.60200025 | 0.359977983 | 0.357690542 | -1.006395028 | 0.314225584 | 0.535446743 | 0.359977983 |
| ENSMUSG00000091144 | Phf11c | 628705 | protein_coding | 20.37515397 | 0.36014316 | 0.296258774 | -1.215637111 | 0.224123191 | 0.431271309 | 0.36014316 |
| ENSMUSG00000020462 | Cfap36 | 216618 | protein_coding | 146.5046556 | 0.360215577 | 0.108256678 | -3.32742132 | 0.000876537 | 0.005701118 | 0.360215577 |
| ENSMUSG00000038692 | Hoxb4 | 15412 | protein_coding | 45.96910916 | -0.360280872 | 0.323781259 | 1.11272923 | 0.265824738 | 0.480866438 | 0.360280872 |
| ENSMUSG00000034951 | Cog7 | 233824 | protein_coding | 530.4384339 | 0.360318651 | 0.081349318 | -4.429276876 | 9.45E-06 | 0.000106652 | 0.360318651 |
| ENSMUSG00000030177 | Ccdc77 | 67200 | protein_coding | 117.1861639 | 0.3603478 | 0.130866366 | -2.753555476 | 0.005895178 | 0.027823083 | 0.3603478 |
| ENSMUSG00000028603 | Scp2 | 20280 | protein_coding | 1506.66314 | -0.360442928 | 0.114031884 | 3.160896022 | 0.001572846 | 0.009273111 | 0.360442928 |
| ENSMUSG00000079071 | Slc28a2b | 381417 | protein_coding | 0.822665944 | 0.360457426 | 2.153715791 | -0.167365363 | 0.867082577 | 1 | 0.360457426 |
| ENSMUSG00000022391 | Rangap1 | 19387 | protein_coding | 1534.704299 | 0.360585148 | 0.118204151 | -3.050528626 | 0.002284389 | 0.012608423 | 0.360585148 |
| ENSMUSG00000022877 | Hrg | 94175 | protein_coding | 4.617059125 | -0.360585956 | 0.732289728 | 0.492408868 | 0.622430332 | 0.792979237 | 0.360585956 |
| ENSMUSG00000073805 | Insyn2a | 627214 | protein_coding | 25.00545805 | -0.360720444 | 0.247623526 | 1.456729297 | 0.145191119 | 0.322338142 | 0.360720444 |
| ENSMUSG00000091337 | Eid1 | 58521 | protein_coding | 154.0341746 | -0.360838848 | 0.2057783 | 1.753532069 | 0.07951072 | 0.208982974 | 0.360838848 |
| ENSMUSG00000032849 | Abcc4 | 239273 | protein_coding | 191.7466898 | 0.361022037 | 0.175332388 | -2.059072154 | 0.039487327 | 0.124778744 | 0.361022037 |
| ENSMUSG00000031853 | Map3k21 | 234878 | protein_coding | 999.7690053 | -0.361139119 | 0.047159683 | 7.657793648 | 1.89E-14 | 6.60E-13 | 0.361139119 |
| ENSMUSG00000079339 | Ifit1bl1 | 667373 | protein_coding | 893.4063939 | -0.361249044 | 0.341061612 | 1.059189984 | 0.289513266 | 0.508294925 | 0.361249044 |

| ensembl_gene_id | external_gene_name | entrezgene_id | gene_biotype | baseMean | log2FoldChange | lfcSE | stat | pvalue | padj | absLog2FC |
|---|---|---|---|---|---|---|---|---|---|---|
| ENSMUSG00000068966 | Zbtb34 | 241311 | protein_coding | 69.53147174 | -0.361287847 | 0.189373077 | 1.907809986 | 0.05641578 | 0.163304151 | 0.361287847 |
| ENSMUSG00000022686 | B3gnt5 | 108105 | protein_coding | 1168.683129 | 0.361301907 | 0.349214656 | -1.034612666 | 0.300849836 | 0.520943084 | 0.361301907 |
| ENSMUSG00000053965 | Pde5a | 242202 | protein_coding | 1301.162266 | -0.361306066 | 0.065472405 | 5.518448062 | 3.42E-08 | 6.04E-07 | 0.361306066 |
| ENSMUSG00000074240 | Cib3 | 234421 | protein_coding | 0.741128 | -0.361309263 | 1.51163196 | 0.239019333 | 0.811090592 | 1 | 0.361309263 |
| ENSMUSG00000031533 | Mrps31 | 57312 | protein_coding | 380.45752 | -0.361377994 | 0.080727024 | 4.476543004 | 7.59E-06 | 8.75E-05 | 0.361377994 |
| ENSMUSG00000028041 | Adam15 | 11490 | protein_coding | 595.1730466 | -0.361492351 | 0.091641555 | 3.944633518 | 7.99E-05 | 0.000708826 | 0.361492351 |
| ENSMUSG00000032561 | Acpp | 56318 | protein_coding | 391.2246115 | -0.361533542 | 0.273779712 | 1.320527148 | 0.186659078 | 0.382529046 | 0.361533542 |
| ENSMUSG00000046574 | Prr12 | 233210 | protein_coding | 120.5012842 | 0.361542477 | 0.294027806 | -1.229620021 | 0.218839429 | 0.425016386 | 0.361542477 |
| ENSMUSG00000047810 | Ccdc88b | 78317 | protein_coding | 35.3234608 | 0.361701309 | 0.308681275 | -1.171763037 | 0.241292207 | 0.451933708 | 0.361701309 |
| ENSMUSG00000085347 | Hk1os | 70162 | lncRNA | 6.535562295 | 0.361765089 | 0.473754574 | -0.763612868 | 0.445097979 | 0.662372535 | 0.361765089 |
| ENSMUSG00000020290 | Xpo1 | 103573 | protein_coding | 1209.672737 | 0.361798521 | 0.108846106 | -3.323945472 | 0.000887535 | 0.005758032 | 0.361798521 |
| ENSMUSG00000057098 | Ebf1 | 13591 | protein_coding | 5.203127784 | 0.36191929 | 1.976591579 | -0.183102718 | 0.854717423 | 0.929155653 | 0.36191929 |
| ENSMUSG00000006362 | Cbfa2t3 | 12398 | protein_coding | 181.5269668 | -0.361963822 | 0.104721596 | 3.456439117 | 0.000547363 | 0.003828526 | 0.361963822 |
| ENSMUSG00000038679 | Trps1 | 83925 | protein_coding | 4.203120742 | 0.362109116 | 1.003784831 | -0.360743762 | 0.718291007 | 1 | 0.362109116 |
| ENSMUSG00000034429 | Zfp707 | 69020 | protein_coding | 127.8979956 | -0.362190521 | 0.132394388 | 2.735693915 | 0.00622489 | 0.02894083 | 0.362190521 |
| ENSMUSG00000037851 | Iars | 105148 | protein_coding | 678.9028413 | 0.362209272 | 0.132181722 | -2.740237209 | 0.006139486 | 0.028672926 | 0.362209272 |
| ENSMUSG00000043372 | Hexim2 | 71059 | protein_coding | 47.16847377 | 0.362209931 | 0.204808444 | -1.768530264 | 0.076972297 | 0.204491718 | 0.362209931 |
| ENSMUSG00000039016 | Timm8b | 30057 | protein_coding | 441.1677435 | -0.362225916 | 0.113637357 | 3.187560189 | 0.001434786 | 0.008569101 | 0.362225916 |
| ENSMUSG00000044768 | Macir | 52392 | protein_coding | 647.2872067 | -0.362357196 | 0.097165884 | 3.729263614 | 0.00019204 | 0.001525421 | 0.362357196 |
| ENSMUSG00000021835 | Bmp4 | 12159 | protein_coding | 1.105873731 | 0.362360113 | 1.751121076 | -0.206930359 | 0.836064247 | 1 | 0.362360113 |
| ENSMUSG00000025226 | Fbxl15 | 68431 | protein_coding | 64.48056532 | -0.362447163 | 0.181380565 | 1.99826902 | 0.045687502 | 0.139280674 | 0.362447163 |
| ENSMUSG00000023868 | Pde10a | 23984 | protein_coding | 6.331258319 | -0.362535489 | 1.181699505 | 0.306791606 | 0.759001998 | 0.878912418 | 0.362535489 |
| ENSMUSG00000043313 | Pcdhb19 | 93890 | protein_coding | 4.481621139 | 0.362702124 | 0.628095351 | -0.577463475 | 0.563626405 | 0.750925894 | 0.362702124 |
| ENSMUSG00000031785 | Adgrg1 | 14766 | protein_coding | 622.7651661 | 0.36274867 | 0.084794794 | -4.277959216 | 1.89E-05 | 0.000195572 | 0.36274867 |
| ENSMUSG00000040097 | Flywch1 | 224613 | protein_coding | 316.3528538 | 0.362769133 | 0.079472286 | -4.564725029 | 5.00E-06 | 5.97E-05 | 0.362769133 |
| ENSMUSG00000008438 | Adam21 | 56622 | protein_coding | 1.630940061 | 0.362818828 | 1.352317505 | -0.268294115 | 0.788472936 | 1 | 0.362818828 |
| ENSMUSG00000025314 | Ptprj | 19271 | protein_coding | 1476.531566 | -0.363008651 | 0.07595705 | 4.779130475 | 1.76E-06 | 2.30E-05 | 0.363008651 |
| ENSMUSG00000091311 | Spata31d1b | 238662 | protein_coding | 1.756207656 | -0.363032201 | 2.251319296 | 0.161253093 | 0.871894066 | 1 | 0.363032201 |
| ENSMUSG00000029096 | Htra3 | 78558 | protein_coding | 2.119687911 | 0.363145202 | 1.102119217 | -0.329497205 | 0.741779906 | 1 | 0.363145202 |
| ENSMUSG00000024878 | Cbwd1 | 226043 | protein_coding | 143.1361766 | 0.363233685 | 0.112501389 | -3.228704013 | 0.001243525 | 0.007622708 | 0.363233685 |
| ENSMUSG00000027330 | Cdc25b | 12531 | protein_coding | 261.2853911 | 0.363292342 | 0.132090113 | -2.750337124 | 0.005953398 | 0.028025774 | 0.363292342 |
| ENSMUSG00000005267 | Zfp287 | 170740 | protein_coding | 34.20938363 | 0.36330209 | 0.216813243 | -1.675645292 | 0.093807692 | 0.235825882 | 0.36330209 |
| ENSMUSG00000022797 | Tfrc | 22042 | protein_coding | 3689.141464 | 0.363321044 | 0.196994821 | -1.844317743 | 0.065136847 | 0.181269925 | 0.363321044 |
| ENSMUSG00000057378 | Ryr3 | 20192 | protein_coding | 15.04678787 | -0.363326489 | 0.738881634 | 0.491724889 | 0.622913841 | 0.793281643 | 0.363326489 |
| ENSMUSG00000060572 | Mfap2 | 17150 | protein_coding | 2.426237999 | 0.363339231 | 0.941275789 | -0.386007199 | 0.699491337 | 1 | 0.363339231 |
| ENSMUSG00000006301 | Tmbim1 | 69660 | protein_coding | 1959.161183 | -0.363354537 | 0.059147964 | 6.143145311 | 8.09E-10 | 1.80E-08 | 0.363354537 |
| ENSMUSG00000038623 | Tm6sf1 | 107769 | protein_coding | 3.430445202 | -0.36339921 | 1.009453447 | 0.359996006 | 0.71885012 | 1 | 0.36339921 |
| ENSMUSG00000102252 | Snrpn | 20646 | protein_coding | 90.11721924 | -0.363446474 | 0.148189803 | 2.452574112 | 0.014183816 | 0.055849197 | 0.363446474 |
| ENSMUSG00000021868 | Ppif | 105675 | protein_coding | 1024.298652 | -0.36377097 | 0.140157994 | 2.595435038 | 0.009447126 | 0.040712379 | 0.36377097 |

| ensembl_gene_id | external_gene_name | entrezgene_id | gene_biotype | baseMean | log2FoldChange | lfcSE | stat | pvalue | padj | absLog2FC |
|---|---|---|---|---|---|---|---|---|---|---|
| ENSMUSG00000110682 | A530010L16Rik | 382036 | lncRNA | 1.051893037 | -0.363817815 | 2.267273745 | 0.160464882 | 0.872514883 | 1 | 0.363817815 |
| ENSMUSG00000037583 | Nr0b2 | 23957 | protein_coding | 0.740836427 | 0.363824673 | 1.485811428 | -0.244865981 | 0.806560203 | 1 | 0.363824673 |
| ENSMUSG00000017009 | Sdc4 | 20971 | protein_coding | 4041.24277 | -0.363882613 | 0.093014715 | 3.912097293 | 9.15E-05 | 0.000797335 | 0.363882613 |
| ENSMUSG00000086315 | Arl4aos | 100043606 | lncRNA | 4.968866543 | -0.363956028 | 0.561746993 | 0.64790027 | 0.517049456 | 0.717711842 | 0.363956028 |
| ENSMUSG00000026012 | Cd28 | 12487 | protein_coding | 3.38776774 | 0.36396969 | 0.848792151 | -0.428808972 | 0.668062249 | 1 | 0.36396969 |
| ENSMUSG00000015335 | Zdhhc12 | 66220 | protein_coding | 197.5562389 | 0.364046426 | 0.1306517 | -2.786388746 | 0.005329891 | 0.025698634 | 0.364046426 |
| ENSMUSG00000035161 | Ints6 | 18130 | protein_coding | 595.3029205 | 0.364096697 | 0.102687873 | -3.545664014 | 0.000391625 | 0.002879422 | 0.364096697 |
| ENSMUSG00000020723 | Cacng4 | 54377 | protein_coding | 3.811869888 | -0.364151483 | 1.016625298 | 0.358196361 | 0.72019637 | 1 | 0.364151483 |
| ENSMUSG00000036620 | Mgat4b | 103534 | protein_coding | 3186.738883 | 0.364188809 | 0.059358472 | -6.135414147 | 8.49E-10 | 1.89E-08 | 0.364188809 |
| ENSMUSG00000026009 | Icos | 54167 | protein_coding | 1.315116046 | 0.364274993 | 1.966191913 | -0.185269297 | 0.853017819 | 1 | 0.364274993 |
| ENSMUSG00000032398 | Snapc5 | 330959 | protein_coding | 324.9787641 | -0.364277342 | 0.083580947 | 4.358377779 | 1.31E-05 | 0.00014217 | 0.364277342 |
| ENSMUSG00000062647 | Rpl7a | 27176 | protein_coding | 7851.891397 | -0.364298193 | 0.10429048 | 3.493110704 | 0.000477429 | 0.003410008 | 0.364298193 |
| ENSMUSG00000111748 | Gm34885 | 102638284 | lncRNA | 2.608768835 | 0.364340214 | 0.762232615 | -0.477990848 | 0.632656719 | 1 | 0.364340214 |
| ENSMUSG00000000125 | Wnt3 | 22415 | protein_coding | 1.172382201 | -0.364404492 | 1.143027307 | 0.318806463 | 0.749873277 | 1 | 0.364404492 |
| ENSMUSG00000030523 | Trpm1 | 17364 | protein_coding | 5.896013806 | 0.364507488 | 1.67771845 | -0.217263802 | 0.828002765 | 0.913297021 | 0.364507488 |
| ENSMUSG00000031010 | Usp9x | 22284 | protein_coding | 3161.038092 | 0.364525237 | 0.078142459 | -4.664880523 | 3.09E-06 | 3.85E-05 | 0.364525237 |
| ENSMUSG00000097133 | Gm26628 | 102632638 | lncRNA | 5.672486284 | 0.364569232 | 0.548442973 | -0.664734986 | 0.506220013 | 0.710135685 | 0.364569232 |
| ENSMUSG00000031139 | Mcf2 | 109904 | protein_coding | 0.968284731 | -0.364627328 | 2.621964826 | 0.139066445 | 0.889397643 | 1 | 0.364627328 |
| ENSMUSG00000042857 | Gm9776 | 328309 | lncRNA | 27.59171516 | -0.364629863 | 0.216445742 | 1.684624794 | 0.092060985 | 0.232532135 | 0.364629863 |
| ENSMUSG00000033498 | Strc | 140476 | protein_coding | 28.88549158 | 0.364666112 | 0.331178106 | -1.101117815 | 0.270845385 | 0.48643183 | 0.364666112 |
| ENSMUSG00000059898 | Dsc3 | 13507 | protein_coding | 1.427791439 | -0.364774476 | 1.228632038 | 0.296894811 | 0.766546818 | 1 | 0.364774476 |
| ENSMUSG00000062209 | Erbb4 | 13869 | protein_coding | 4.333702883 | 0.364785995 | 1.432303337 | -0.254684874 | 0.7989665 | 1 | 0.364785995 |
| ENSMUSG00000023067 | Cdkn1a | 12575 | protein_coding | 1579.839772 | 0.364809767 | 0.18528591 | -1.968901828 | 0.048964366 | 0.1468042 | 0.364809767 |
| ENSMUSG00000044934 | Zfp367 | 238673 | protein_coding | 177.0728838 | 0.364902485 | 0.195170644 | -1.869658666 | 0.061531233 | 0.173816344 | 0.364902485 |
| ENSMUSG00000034120 | Srsf2 | 20382 | protein_coding | 1667.331754 | 0.365132936 | 0.061188521 | -5.967343666 | 2.41E-09 | 5.09E-08 | 0.365132936 |
| ENSMUSG00000002257 | Def6 | 23853 | protein_coding | 14.67321342 | 0.3652394 | 0.343825125 | -1.062282461 | 0.288107474 | 0.506958639 | 0.3652394 |
| ENSMUSG00000022339 | Ebag9 | 55960 | protein_coding | 287.3136681 | 0.365378522 | 0.097733376 | -3.738523504 | 0.000185104 | 0.001483889 | 0.365378522 |
| ENSMUSG00000047221 | Fam185a | 330050 | protein_coding | 168.8650543 | 0.365414807 | 0.111247814 | -3.284692019 | 0.001020939 | 0.006462165 | 0.365414807 |
| ENSMUSG00000029560 | Snx8 | 231834 | protein_coding | 73.45353405 | 0.365497185 | 0.168543682 | -2.168560587 | 0.03011606 | 0.101311187 | 0.365497185 |
| ENSMUSG00000052934 | Fbxo31 | 76454 | protein_coding | 214.671038 | -0.365511835 | 0.107045586 | 3.41454372 | 0.000638889 | 0.004347186 | 0.365511835 |
| ENSMUSG00000109179 | Gm35339 | 102638882 | protein_coding | 28.11538796 | 0.365572437 | 0.313089902 | -1.167627682 | 0.242956979 | 0.454124681 | 0.365572437 |
| ENSMUSG00000064853 | Gm23442 | 115488838 | snoRNA | 350.4057424 | 0.365572869 | 0.231760576 | -1.577372975 | 0.11470973 | 0.272838324 | 0.365572869 |
| ENSMUSG00000049686 | Orai1 | 109305 | protein_coding | 286.5680725 | 0.365691698 | 0.092889116 | -3.936862727 | 8.26E-05 | 0.000728644 | 0.365691698 |
| ENSMUSG00000040495 | Chrm4 | 12672 | protein_coding | 2.673958511 | -0.365734776 | 0.655603209 | 0.557859954 | 0.576940019 | 1 | 0.365734776 |
| ENSMUSG00000103472 | Pcdhga7 | 93715 | protein_coding | 11.91914712 | -0.365748765 | 0.324318727 | 1.127744822 | 0.259427702 | 0.47292075 | 0.365748765 |
| ENSMUSG00000022914 | Brwd1 | 93871 | protein_coding | 1106.817086 | 0.365872768 | 0.097925156 | -3.736249029 | 0.000186786 | 0.00149302 | 0.365872768 |
| ENSMUSG00000048478 | Spata33 | 320869 | protein_coding | 5.392142092 | -0.36587796 | 0.570220392 | 0.641643064 | 0.521104965 | 0.720791305 | 0.36587796 |
| ENSMUSG00000034327 | Kctd9 | 105440 | protein_coding | 228.9717852 | 0.366007939 | 0.106464764 | -3.437831702 | 0.000586392 | 0.004051748 | 0.366007939 |
| ENSMUSG00000071866 | Ppia | 268373 | protein_coding | 3724.795969 | -0.366009675 | 0.163663011 | 2.236361618 | 0.025328093 | 0.08827947 | 0.366009675 |

| ensembl_gene_id | external_gene_name | entrezgene_id | gene_biotype | baseMean | log2FoldChange | lfcSE | stat | pvalue | padj | absLog2FC |
|---|---|---|---|---|---|---|---|---|---|---|
| ENSMUSG00000023010 | Tmbim6 | 110213 | protein_coding | 14146.59156 | 0.366027588 | 0.06307834 | -5.802746031 | 6.52E-09 | 1.29E-07 | 0.366027588 |
| ENSMUSG00000037926 | Ssh2 | 237860 | protein_coding | 711.7469507 | 0.366342818 | 0.075216096 | -4.87053755 | 1.11E-06 | 1.52E-05 | 0.366342818 |
| ENSMUSG00000001661 | Hoxc6 | 15425 | protein_coding | 1.269656671 | -0.366378233 | 2.029044672 | 0.180566864 | 0.856707567 | 1 | 0.366378233 |
| ENSMUSG00000116802 | Gm5165 | 102640673 | lncRNA | 71.85713028 | -0.366557633 | 0.172566918 | 2.124147766 | 0.033657789 | 0.110042513 | 0.366557633 |
| ENSMUSG00000075829 | Gm23692 | 115488709 | snRNA | 0.519997961 | 0.366672309 | 1.61681408 | -0.226786935 | 0.820589424 | 1 | 0.366672309 |
| ENSMUSG00000027315 | Spint1 | 20732 | protein_coding | 3021.392747 | -0.366739034 | 0.087580548 | 4.187448474 | 2.82E-05 | 0.000281155 | 0.366739034 |
| ENSMUSG00000025323 | Sp4 | 20688 | protein_coding | 103.5391442 | -0.366818087 | 0.114674161 | 3.198785883 | 0.001380076 | 0.008323805 | 0.366818087 |
| ENSMUSG00000074698 | Csnk2a1 | 12995 | protein_coding | 1229.347599 | -0.366927986 | 0.073689068 | 4.979408701 | 6.38E-07 | 9.17E-06 | 0.366927986 |
| ENSMUSG00000043251 | Exoc3l | 277978 | protein_coding | 6.966016804 | 0.367049993 | 0.417452993 | -0.879260659 | 0.379259962 | 0.601649715 | 0.367049993 |
| ENSMUSG00000024055 | Cyp4f13 | 170716 | protein_coding | 413.6088856 | -0.367096703 | 0.107482161 | 3.415419812 | 0.000636838 | 0.0043368 | 0.367096703 |
| ENSMUSG00000109293 | Dcst2 | 329702 | protein_coding | 25.33108856 | 0.367150158 | 0.224149456 | -1.637970328 | 0.101427884 | 0.25018373 | 0.367150158 |
| ENSMUSG00000027298 | Tyro3 | 22174 | protein_coding | 170.4327221 | -0.367174751 | 0.144073774 | 2.548519001 | 0.010818139 | 0.045273886 | 0.367174751 |
| ENSMUSG00000023122 | Sult1c2 | 69083 | protein_coding | 471.216222 | -0.367192166 | 0.192683799 | 1.905672235 | 0.056692744 | 0.163933457 | 0.367192166 |
| ENSMUSG00000037752 | Xkr8 | 381560 | protein_coding | 16.67130374 | -0.367334546 | 0.287229144 | 1.278890229 | 0.200935714 | 0.402261169 | 0.367334546 |
| ENSMUSG00000029716 | Tfr2 | 50765 | protein_coding | 2.559339327 | -0.367345285 | 1.66136062 | 0.221111107 | 0.825005922 | 1 | 0.367345285 |
| ENSMUSG00000025134 | Alyref | 21681 | protein_coding | 697.1903988 | 0.367359453 | 0.098313218 | -3.736623197 | 0.000186508 | 0.001492245 | 0.367359453 |
| ENSMUSG00000036403 | Cep135 | 381644 | protein_coding | 124.1916632 | 0.367392087 | 0.120449351 | -3.050179053 | 0.00228705 | 0.012618891 | 0.367392087 |
| ENSMUSG00000035948 | Acss3 | 380660 | protein_coding | 3.727997968 | -0.367394129 | 0.644600509 | 0.569956312 | 0.56870733 | 1 | 0.367394129 |
| ENSMUSG00000025384 | Faap100 | 71885 | protein_coding | 291.8385743 | -0.36743241 | 0.09789374 | 3.753380051 | 0.000174466 | 0.001406803 | 0.36743241 |
| ENSMUSG00000001156 | Mxd1 | 17119 | protein_coding | 5861.726798 | 0.367444627 | 0.19670543 | -1.867994323 | 0.061762862 | 0.174214948 | 0.367444627 |
| ENSMUSG00000034445 | Cyb561a3 | 225912 | protein_coding | 249.1868227 | -0.367473755 | 0.072721326 | 5.05317733 | 4.35E-07 | 6.49E-06 | 0.367473755 |
| ENSMUSG00000039199 | Zdhhc1 | 70796 | protein_coding | 130.9788502 | -0.367497545 | 0.122601793 | 2.99748915 | 0.002722135 | 0.014666671 | 0.367497545 |
| ENSMUSG00000093904 | Tomm20 | 67952 | protein_coding | 558.8886389 | -0.367567977 | 0.122207752 | 3.007730464 | 0.002632065 | 0.014241781 | 0.367567977 |
| ENSMUSG00000044296 | Zfp879 | 214779 | protein_coding | 7.462595675 | -0.367741001 | 0.464991331 | 0.790855607 | 0.429028256 | 0.647976278 | 0.367741001 |
| ENSMUSG00000103677 | Pcdhga4 | 93712 | protein_coding | 2.704708252 | -0.367741167 | 0.778095429 | 0.472617052 | 0.636486415 | 1 | 0.367741167 |
| ENSMUSG00000000738 | Spg7 | 234847 | protein_coding | 1010.419472 | -0.367742173 | 0.073770032 | 4.984980517 | 6.20E-07 | 8.95E-06 | 0.367742173 |
| ENSMUSG00000031458 | Coprs | 66423 | protein_coding | 121.3252242 | -0.367783028 | 0.120801231 | 3.044530466 | 0.002330438 | 0.01281547 | 0.367783028 |
| ENSMUSG00000068299 | Nat8f4 | 75541 | protein_coding | 160.5825172 | -0.367813191 | 0.152723422 | 2.408361371 | 0.016024311 | 0.061569443 | 0.367813191 |
| ENSMUSG00000030350 | Prmt8 | 381813 | protein_coding | 1.273069236 | -0.367817315 | 2.658809964 | 0.138339077 | 0.889972444 | 1 | 0.367817315 |
| ENSMUSG00000096773 | Or52e8b | 258147 | protein_coding | 1.502617377 | -0.367846978 | 2.659121938 | 0.138334002 | 0.889976454 | 1 | 0.367846978 |
| ENSMUSG00000020062 | Slc5a8 | 216225 | protein_coding | 4559.823663 | -0.367856439 | 0.140637397 | 2.615637421 | 0.008906107 | 0.038748264 | 0.367856439 |
| ENSMUSG00000067847 | Romo1 | 67067 | protein_coding | 309.8356556 | -0.367896222 | 0.119314367 | 3.083419295 | 0.002046366 | 0.01156526 | 0.367896222 |
| ENSMUSG00000036743 | Psma8 | 73677 | protein_coding | 1.471404427 | 0.367904381 | 2.33176238 | -0.157779534 | 0.874630526 | 1 | 0.367904381 |
| ENSMUSG00000023044 | Csad | 246277 | protein_coding | 326.2438401 | -0.367923171 | 0.15318852 | 2.401767254 | 0.016316086 | 0.062458007 | 0.367923171 |
| ENSMUSG00000030304 | Ergic2 | 67456 | protein_coding | 364.9244456 | 0.367950831 | 0.071863809 | -5.120113114 | 3.05E-07 | 4.67E-06 | 0.367950831 |
| ENSMUSG00000030471 | Zdhhc13 | 243983 | protein_coding | 1632.912806 | 0.367966118 | 0.090495239 | -4.06613789 | 4.78E-05 | 0.000448493 | 0.367966118 |
| ENSMUSG00000071528 | Atp5md | 66477 | protein_coding | 1154.171185 | -0.367979712 | 0.112337175 | 3.275671755 | 0.00105411 | 0.006649187 | 0.367979712 |
| ENSMUSG00000028012 | Rrh | 20132 | protein_coding | 4.132266736 | 0.367981946 | 0.574903239 | -0.640076314 | 0.522122987 | 1 | 0.367981946 |
| ENSMUSG00000026546 | Cfap45 | 71870 | protein_coding | 40.70866607 | -0.368028588 | 0.257640177 | 1.428459615 | 0.153159611 | 0.334228588 | 0.368028588 |

Supplementary Table 1

| ensembl_gene_id | external_gene_name | entrezgene_id | gene_biotype | baseMean | log2FoldChange | lfcSE | stat | pvalue | padj | absLog2FC |
|---|---|---|---|---|---|---|---|---|---|---|
| ENSMUSG00000041229 | Phf8 | 320595 | protein_coding | 467.4915633 | 0.368073409 | 0.06611664 | -5.567031386 | 2.59E-08 | 4.67E-07 | 0.368073409 |
| ENSMUSG00000069805 | Fbp1 | 14121 | protein_coding | 26.62847414 | -0.368086939 | 0.217223304 | 1.694509441 | 0.090168519 | 0.228942639 | 0.368086939 |
| ENSMUSG00000034115 | Scn11a | 24046 | protein_coding | 3.837939344 | -0.368125398 | 1.047068046 | 0.35157734 | 0.725155262 | 1 | 0.368125398 |
| ENSMUSG00000009927 | Rps25 | 75617 | protein_coding | 3272.024089 | -0.368139364 | 0.115065479 | 3.199390192 | 0.001377186 | 0.008312447 | 0.368139364 |
| ENSMUSG00000033629 | Hacd3 | 57874 | protein_coding | 759.2491393 | -0.368146147 | 0.123933001 | 2.970525565 | 0.002972907 | 0.015750587 | 0.368146147 |
| ENSMUSG00000001504 | Irx2 | 16372 | protein_coding | 1.441664084 | 0.368177229 | 1.88620038 | -0.195195183 | 0.84524015 | 1 | 0.368177229 |
| ENSMUSG00000026200 | Glb1l | 74577 | protein_coding | 25.44663093 | -0.368204191 | 0.209882474 | 1.754335105 | 0.079373104 | 0.208654479 | 0.368204191 |
| ENSMUSG00000067928 | Zfp760 | 240034 | protein_coding | 62.40520674 | -0.368220635 | 0.222695626 | 1.653470443 | 0.098235157 | 0.244205235 | 0.368220635 |
| ENSMUSG00000078787 | Cyp2t4 | 384724 | protein_coding | 3.678384508 | -0.368228148 | 1.111441589 | 0.331306793 | 0.740412761 | 1 | 0.368228148 |
| ENSMUSG00000040274 | Cdk6 | 12571 | protein_coding | 535.6265 | -0.368289224 | 0.116145483 | 3.17093024 | 0.001519516 | 0.008984351 | 0.368289224 |
| ENSMUSG00000022180 | Slc7a8 | 50934 | protein_coding | 55.9297678 | -0.368383034 | 0.258779413 | 1.423540726 | 0.154579465 | 0.336504783 | 0.368383034 |
| ENSMUSG00000089204 | Mir1927 | 100316784 | miRNA | 0.808328913 | -0.36877076 | 1.748635983 | 0.210890525 | 0.832972698 | 1 | 0.36877076 |
| ENSMUSG00000050100 | Hmx2 | 15372 | protein_coding | 10.75714585 | -0.368930291 | 0.316634168 | 1.165162601 | 0.243953181 | 0.4551483 | 0.368930291 |
| ENSMUSG00000040148 | Hmx3 | 15373 | protein_coding | 12.40317648 | -0.368933831 | 0.302540631 | 1.219452178 | 0.222672616 | 0.429717907 | 0.368933831 |
| ENSMUSG00000018379 | Srsf1 | 110809 | protein_coding | 2515.858567 | 0.368957906 | 0.108774255 | -3.39195984 | 0.000693946 | 0.004647131 | 0.368957906 |
| ENSMUSG00000062937 | Mtap | 66902 | protein_coding | 82.54539445 | 0.36896313 | 0.182225751 | -2.024758456 | 0.042892173 | 0.132619611 | 0.36896313 |
| ENSMUSG00000054766 | Set | 56086 | protein_coding | 1779.18401 | 0.369019146 | 0.114717119 | -3.21677488 | 0.001296403 | 0.007888281 | 0.369019146 |
| ENSMUSG00000024749 | Tmc1 | 13409 | protein_coding | 14.0006371 | -0.369036731 | 0.438890114 | 0.840840838 | 0.400437106 | 0.622382905 | 0.369036731 |
| ENSMUSG00000032892 | Rangrf | 57785 | protein_coding | 2.125793818 | 0.369043629 | 0.813242203 | -0.453793012 | 0.649977814 | 1 | 0.369043629 |
| ENSMUSG00000074361 | C5ar2 | 319430 | protein_coding | 1.769504278 | -0.369051302 | 1.869579998 | 0.197397973 | 0.843516116 | 1 | 0.369051302 |
| ENSMUSG00000025626 | Phf6 | 70998 | protein_coding | 296.0560183 | 0.36911082 | 0.135928285 | -2.715482065 | 0.006617936 | 0.030425148 | 0.36911082 |
| ENSMUSG00000045273 | Cenph | 26886 | protein_coding | 39.56665013 | 0.369132197 | 0.246489262 | -1.497558939 | 0.134247882 | 0.304738078 | 0.369132197 |
| ENSMUSG00000052040 | Klf13 | 50794 | protein_coding | 917.6883872 | -0.369223238 | 0.056918221 | 6.486907586 | 8.76E-11 | 2.22E-09 | 0.369223238 |
| ENSMUSG00000034187 | Nsf | 18195 | protein_coding | 783.9279586 | 0.369241946 | 0.081275126 | -4.543111353 | 5.54E-06 | 6.56E-05 | 0.369241946 |
| ENSMUSG00000005628 | Tmod4 | 50874 | protein_coding | 5.86270183 | -0.369326391 | 0.531367993 | 0.695048245 | 0.487025067 | 0.695624496 | 0.369326391 |
| ENSMUSG00000054381 | Zfp747 | 269997 | protein_coding | 147.4618541 | -0.369334696 | 0.129835267 | 2.844640781 | 0.004446153 | 0.022069064 | 0.369334696 |
| ENSMUSG00000024401 | Tnf | 21926 | protein_coding | 47.22344395 | 0.36948065 | 0.420448166 | -0.878778124 | 0.37952159 | 0.601768559 | 0.36948065 |
| ENSMUSG00000015652 | Steap1 | 70358 | protein_coding | 51.08661632 | 0.369644266 | 0.193773656 | -1.907608465 | 0.05644184 | 0.163322332 | 0.369644266 |
| ENSMUSG00000033825 | Tpsb2 | 17229 | protein_coding | 3.467266505 | 0.369668109 | 0.753357757 | -0.490693971 | 0.623642911 | 1 | 0.369668109 |
| ENSMUSG00000062257 | Opcml | 330908 | protein_coding | 4.802744365 | 0.36974358 | 1.565622677 | -0.236163914 | 0.81330548 | 0.907311132 | 0.36974358 |
| ENSMUSG00000049321 | Zfp2 | 22678 | protein_coding | 42.21486268 | -0.369762654 | 0.188267796 | 1.964024975 | 0.049527196 | 0.147954434 | 0.369762654 |
| ENSMUSG00000028756 | Pink1 | 68943 | protein_coding | 1489.90727 | 0.369800761 | 0.099017631 | -3.7346961 | 0.000187942 | 0.00150081 | 0.369800761 |
| ENSMUSG00000096714 | Or2ag17 | 258180 | protein_coding | 0.872725862 | 0.369830886 | 1.764694139 | -0.20957223 | 0.834001556 | 1 | 0.369830886 |
| ENSMUSG00000045092 | S1pr1 | 13609 | protein_coding | 6.011860627 | 0.369972398 | 0.605371024 | -0.611149828 | 0.541100395 | 0.73508818 | 0.369972398 |
| ENSMUSG00000030683 | Sez6l2 | 233878 | protein_coding | 27.99294349 | -0.369985957 | 0.264012314 | 1.401396591 | 0.161095511 | 0.345901869 | 0.369985957 |
| ENSMUSG00000079434 | Neu2 | 23956 | protein_coding | 16.31354835 | 0.370063176 | 0.250968392 | -1.474540973 | 0.140335994 | 0.314536999 | 0.370063176 |
| ENSMUSG00000059248 | Septin9 | 53860 | protein_coding | 740.4849849 | 0.370223421 | 0.083736222 | -4.421305512 | 9.81E-06 | 0.000110288 | 0.370223421 |
| ENSMUSG00000029385 | Ccng2 | 12452 | protein_coding | 1000.418281 | -0.370365461 | 0.193335658 | 1.915660384 | 0.055408338 | 0.16083904 | 0.370365461 |
| ENSMUSG00000024158 | Hagh | 14651 | protein_coding | 797.0307647 | 0.370381466 | 0.072889247 | -5.081428058 | 3.75E-07 | 5.65E-06 | 0.370381466 |

| ensembl_gene_id | external_gene_name | entrezgene_id | gene_biotype | baseMean | log2FoldChange | lfcSE | stat | pvalue | padj | absLog2FC |
|---|---|---|---|---|---|---|---|---|---|---|
| ENSMUSG00000036591 | Arhgap21 | 71435 | protein_coding | 1686.590094 | 0.370448132 | 0.113511831 | -3.263520012 | 0.001100374 | 0.006901473 | 0.370448132 |
| ENSMUSG00000042116 | Vwa1 | 246228 | protein_coding | 10.96385472 | -0.37065897 | 0.483683886 | 0.766324826 | 0.44348305 | 0.660955754 | 0.37065897 |
| ENSMUSG00000023036 | Pcdhgc4 | 93707 | protein_coding | 86.62277068 | 0.370757756 | 0.204137766 | -1.816213448 | 0.069337636 | 0.189607836 | 0.370757756 |
| ENSMUSG00000005483 | Dnajb1 | 81489 | protein_coding | 1720.22231 | -0.370763676 | 0.159091259 | 2.330509409 | 0.019779243 | 0.072698514 | 0.370763676 |
| ENSMUSG00000040940 | Arhgef1 | 16801 | protein_coding | 593.8326909 | 0.370837352 | 0.097544805 | -3.801712998 | 0.000143699 | 0.001185931 | 0.370837352 |
| ENSMUSG00000003762 | Coq8b | 76889 | protein_coding | 314.1990708 | 0.370866081 | 0.070687338 | -5.246570182 | 1.55E-07 | 2.47E-06 | 0.370866081 |
| ENSMUSG00000034205 | Loxl2 | 94352 | protein_coding | 1.062377492 | -0.37109947 | 1.201372248 | 0.308896323 | 0.757400394 | 1 | 0.37109947 |
| ENSMUSG00000020437 | Myo1g | 246177 | protein_coding | 17.74860056 | 0.371154811 | 0.487840951 | -0.760811101 | 0.446769903 | 0.664082645 | 0.371154811 |
| ENSMUSG00000035033 | Tbr1 | 21375 | protein_coding | 2.546540648 | 0.371169709 | 1.200389528 | -0.30920772 | 0.757163522 | 1 | 0.371169709 |
| ENSMUSG00000039699 | Batf2 | 74481 | protein_coding | 205.5196637 | -0.37119916 | 0.147987152 | 2.508320191 | 0.012130669 | 0.049536564 | 0.37119916 |
| ENSMUSG00000022818 | Cyp2ab1 | 224044 | protein_coding | 2.028871797 | -0.37122336 | 1.571428895 | 0.236232999 | 0.813251875 | 1 | 0.37122336 |
| ENSMUSG00000025855 | Prkar1b | 19085 | protein_coding | 7.065192715 | -0.371254677 | 0.53210083 | 0.6977149 | 0.485355508 | 0.695380179 | 0.371254677 |
| ENSMUSG00000000416 | Cttnbp2 | 30785 | protein_coding | 160.6270681 | 0.371307988 | 0.21622968 | -1.717192512 | 0.085944003 | 0.221416423 | 0.371307988 |
| ENSMUSG00000032397 | Tipin | 66131 | protein_coding | 107.4275607 | 0.371349067 | 0.135537224 | -2.739830851 | 0.006147081 | 0.028700282 | 0.371349067 |
| ENSMUSG00000011884 | Gltp | 56356 | protein_coding | 674.0650229 | 0.371393315 | 0.12548038 | -2.959772008 | 0.003078668 | 0.016227616 | 0.371393315 |
| ENSMUSG00000046876 | Atxn1 | 20238 | protein_coding | 640.277287 | -0.371395024 | 0.131639611 | 2.821301439 | 0.004782924 | 0.023423844 | 0.371395024 |
| ENSMUSG00000043831 | Lysmd4 | 75099 | protein_coding | 185.3594674 | 0.371468471 | 0.098139069 | -3.785123256 | 0.000153632 | 0.001253501 | 0.371468471 |
| ENSMUSG00000037921 | Ddx60 | 234311 | protein_coding | 430.6013194 | 0.371573418 | 0.460017507 | -0.807737557 | 0.419241678 | 0.639216792 | 0.371573418 |
| ENSMUSG00000019301 | Hsd17b1 | 15485 | protein_coding | 1.49276165 | 0.371606949 | 1.117658864 | -0.332486916 | 0.739521618 | 1 | 0.371606949 |
| ENSMUSG00000030275 | Etnk1 | 75320 | protein_coding | 1162.066988 | 0.371642322 | 0.093481773 | -3.97555919 | 7.02E-05 | 0.000632578 | 0.371642322 |
| ENSMUSG00000039943 | Plcb4 | 18798 | protein_coding | 331.2181571 | 0.371680148 | 0.155444693 | -2.391076472 | 0.016799053 | 0.063995287 | 0.371680148 |
| ENSMUSG00000047649 | Cd3eap | 70333 | protein_coding | 76.4126091 | -0.371685767 | 0.15842803 | 2.346085896 | 0.018971735 | 0.070451816 | 0.371685767 |
| ENSMUSG00000000555 | Itga5 | 16402 | protein_coding | 100.4401499 | -0.37179649 | 0.145559821 | 2.554252186 | 0.010641615 | 0.044806045 | 0.37179649 |
| ENSMUSG00000045636 | Mtus1 | 102103 | protein_coding | 2496.660997 | -0.371801121 | 0.074644841 | 4.980935277 | 6.33E-07 | 9.12E-06 | 0.371801121 |
| ENSMUSG00000029798 | Herc6 | 67138 | protein_coding | 40.89739116 | -0.371965456 | 0.328571401 | 1.132068874 | 0.25760548 | 0.470794255 | 0.371965456 |
| ENSMUSG00000022422 | Dscc1 | 72107 | protein_coding | 22.69344562 | 0.371988643 | 0.272961513 | -1.362787883 | 0.172949371 | 0.363647302 | 0.371988643 |
| ENSMUSG00000111952 | Gm4489 | 100043513 | lncRNA | 1.161039685 | -0.372010254 | 1.225124108 | 0.303651076 | 0.761393735 | 1 | 0.372010254 |
| ENSMUSG00000020481 | Ankrd36 | 76389 | protein_coding | 2.461475965 | -0.372021377 | 1.760917887 | 0.211265602 | 0.832680023 | 1 | 0.372021377 |
| ENSMUSG00000054905 | Stfa3 | 20863 | protein_coding | 1.043837783 | -0.372038226 | 2.649742664 | 0.140405418 | 0.888339678 | 1 | 0.372038226 |
| ENSMUSG00000075273 | Ttc30b | 72421 | protein_coding | 127.8890125 | -0.372067206 | 0.133757295 | 2.781659158 | 0.005408181 | 0.025985072 | 0.372067206 |
| ENSMUSG00000049690 | Nckap5 | 210356 | protein_coding | 58.39141168 | 0.372072339 | 0.406741877 | -0.914762804 | 0.360316166 | 0.582711741 | 0.372072339 |
| ENSMUSG00000024140 | Epas1 | 13819 | protein_coding | 1165.044084 | -0.372131699 | 0.140558481 | 2.64752219 | 0.008108404 | 0.036011679 | 0.372131699 |
| ENSMUSG00000092183 | 4930515G01Rik | 67642 | lncRNA | 12.02847996 | -0.372178165 | 0.324520184 | 1.146856755 | 0.251440828 | 0.463839794 | 0.372178165 |
| ENSMUSG00000062169 | Cnih4 | 98417 | protein_coding | 456.8286678 | -0.37219464 | 0.095291967 | 3.905834376 | 9.39E-05 | 0.00081486 | 0.37219464 |
| ENSMUSG00000068732 | Tmem167b | 67495 | protein_coding | 821.5397099 | 0.372219332 | 0.102494734 | -3.631594678 | 0.000281675 | 0.002154013 | 0.372219332 |
| ENSMUSG00000034175 | Rhbdd3 | 279766 | protein_coding | 98.31341374 | -0.37223745 | 0.114345363 | 3.255378604 | 0.001132413 | 0.007066978 | 0.37223745 |
| ENSMUSG00000051037 | Zfp455 | 218311 | protein_coding | 15.90003902 | -0.372242241 | 0.317887843 | 1.170986086 | 0.24160437 | 0.45221065 | 0.372242241 |
| ENSMUSG00000028654 | Mycl | 16918 | protein_coding | 167.6538569 | -0.372279038 | 0.125151748 | 2.974621165 | 0.002933506 | 0.01556181 | 0.372279038 |
| ENSMUSG00000079184 | Mphosph8 | 75339 | protein_coding | 164.9043928 | 0.372326902 | 0.114707607 | -3.24587804 | 0.00117089 | 0.00726647 | 0.372326902 |

Supplementary Table 1

| ensembl_gene_id | external_gene_name | entrezgene_id | gene_biotype | baseMean | log2FoldChange | lfcSE | stat | pvalue | padj | absLog2FC |
|---|---|---|---|---|---|---|---|---|---|---|
| ENSMUSG00000026820 | Ptges2 | 96979 | protein_coding | 458.0624826 | -0.372578599 | 0.09200872 | 4.049383576 | 5.14E-05 | 0.000477499 | 0.372578599 |
| ENSMUSG00000057329 | Bcl2 | 12043 | protein_coding | 209.3342526 | 0.372820247 | 0.151302415 | -2.464073347 | 0.0137368 | 0.054426466 | 0.372820247 |
| ENSMUSG00000029618 | Ocm | 18261 | protein_coding | 1.786016222 | 0.372875438 | 1.453064465 | -0.256613142 | 0.797477429 | 1 | 0.372875438 |
| ENSMUSG00000078650 | G6pc | 14377 | protein_coding | 1.75634793 | 0.372897139 | 1.678752245 | -0.22212756 | 0.824214584 | 1 | 0.372897139 |
| ENSMUSG00000048001 | Hes5 | 15208 | protein_coding | 1.411274838 | -0.373139596 | 1.118157902 | 0.333709215 | 0.738598995 | 1 | 0.373139596 |
| ENSMUSG00000065536 | Mir98 | 723947 | miRNA | 0.753702534 | -0.373143504 | 1.231512038 | 0.302996229 | 0.761892735 | 1 | 0.373143504 |
| ENSMUSG00000055760 | Gemin6 | 67242 | protein_coding | 57.89797195 | 0.373210074 | 0.22553855 | -1.654750705 | 0.097975077 | 0.243645564 | 0.373210074 |
| ENSMUSG00000035067 | Xkr6 | 219149 | protein_coding | 32.65467725 | -0.373239946 | 0.233156626 | 1.600812093 | 0.109418544 | 0.264120665 | 0.373239946 |
| ENSMUSG00000021048 | Mthfd1 | 108156 | protein_coding | 858.9911121 | -0.373247267 | 0.139095285 | 2.683392672 | 0.007287936 | 0.032955362 | 0.373247267 |
| ENSMUSG00000014778 | Fhod1 | 234686 | protein_coding | 328.4245009 | 0.373247786 | 0.136618582 | -2.732042599 | 0.0062943 | 0.029171357 | 0.373247786 |
| ENSMUSG00000030781 | Slc5a2 | 246787 | protein_coding | 2.995941002 | 0.373301133 | 0.931483376 | -0.400759845 | 0.688596946 | 1 | 0.373301133 |
| ENSMUSG00000055254 | Ntrk2 | 18212 | protein_coding | 114.625146 | 0.373409077 | 0.200158367 | -1.865568164 | 0.062101807 | 0.17492772 | 0.373409077 |
| ENSMUSG00000027429 | Sec23b | 27054 | protein_coding | 1005.87877 | 0.373494526 | 0.098526936 | -3.79078596 | 0.000150171 | 0.001229514 | 0.373494526 |
| ENSMUSG00000028138 | Adh5 | 11532 | protein_coding | 1220.414968 | -0.373553052 | 0.105015628 | 3.557118665 | 0.000374945 | 0.002777855 | 0.373553052 |
| ENSMUSG00000039531 | Zup1 | 72580 | protein_coding | 167.9050686 | -0.373556329 | 0.088066233 | 4.241765726 | 2.22E-05 | 0.000226486 | 0.373556329 |
| ENSMUSG00000085418 | Grip1os1 | 78105 | lncRNA | 1.268973061 | -0.373557935 | 1.289362977 | 0.289722865 | 0.772028263 | 1 | 0.373557935 |
| ENSMUSG00000097328 | Tnfsf12 | 21944 | protein_coding | 4.70156422 | -0.373626051 | 0.58904357 | 0.634292726 | 0.525889803 | 0.724255209 | 0.373626051 |
| ENSMUSG00000055430 | Nap1l5 | 58243 | protein_coding | 2.125323568 | -0.373632923 | 0.94842236 | 0.393952039 | 0.693616449 | 1 | 0.373632923 |
| ENSMUSG00000021466 | Ptch1 | 19206 | protein_coding | 229.2515766 | -0.373668009 | 0.079028668 | 4.728258962 | 2.26E-06 | 2.91E-05 | 0.373668009 |
| ENSMUSG00000049760 | Micos13 | 224904 | protein_coding | 902.386781 | -0.373741273 | 0.093710245 | 3.988264829 | 6.66E-05 | 0.000602545 | 0.373741273 |
| ENSMUSG00000041346 | Wrap53 | 216853 | protein_coding | 187.9854117 | 0.373764104 | 0.084173784 | -4.440386123 | 8.98E-06 | 0.000102059 | 0.373764104 |
| ENSMUSG00000055865 | Tafa3 | 329731 | protein_coding | 6.648731599 | -0.373855918 | 0.605056635 | 0.617885825 | 0.536650604 | 0.732538277 | 0.373855918 |
| ENSMUSG00000025511 | Tspan4 | 64540 | protein_coding | 18.81446361 | 0.373866554 | 0.305452837 | -1.223974731 | 0.22096176 | 0.427441699 | 0.373866554 |
| ENSMUSG00000001229 | Dpp9 | 224897 | protein_coding | 685.4643441 | 0.373874438 | 0.088654674 | -4.217199374 | 2.47E-05 | 0.000249836 | 0.373874438 |
| ENSMUSG00000029273 | Sult1d1 | 53315 | protein_coding | 3732.462429 | -0.373921239 | 0.089222347 | 4.190892197 | 2.78E-05 | 0.000277255 | 0.373921239 |
| ENSMUSG00000074398 | Gm15441 | 100038464 | lncRNA | 8.893832201 | -0.373956043 | 0.422734998 | 0.884611031 | 0.376366455 | 0.598604992 | 0.373956043 |
| ENSMUSG00000074796 | Slc4a11 | 269356 | protein_coding | 5.207805302 | 0.374024715 | 0.748385872 | -0.499775221 | 0.61723336 | 0.789053391 | 0.374024715 |
| ENSMUSG00000021493 | Pdlim7 | 67399 | protein_coding | 64.55576322 | 0.374051371 | 0.204926157 | -1.825298325 | 0.067956023 | 0.186913018 | 0.374051371 |
| ENSMUSG00000059939 | 9430015G10Rik | 230996 | protein_coding | 109.4381657 | 0.374077802 | 0.12204811 | -3.065002819 | 0.002176679 | 0.012123334 | 0.374077802 |
| ENSMUSG00000037286 | Stag1 | 20842 | protein_coding | 690.5960594 | 0.374090899 | 0.08821937 | -4.240462129 | 2.23E-05 | 0.000227524 | 0.374090899 |
| ENSMUSG00000026564 | Dusp27 | 240892 | protein_coding | 5.17840358 | -0.374105603 | 0.670383597 | 0.558047071 | 0.576812242 | 0.761222618 | 0.374105603 |
| ENSMUSG00000038208 | Pgap3 | 320655 | protein_coding | 369.6452711 | 0.374125464 | 0.091903572 | -4.07084792 | 4.68E-05 | 0.000440521 | 0.374125464 |
| ENSMUSG00000013091 | Tmem190 | 78052 | protein_coding | 10.58430065 | -0.374299028 | 0.3229984 | 1.158826261 | 0.246527011 | 0.458307673 | 0.374299028 |
| ENSMUSG00000059834 | Sclt1 | 67161 | protein_coding | 145.8573364 | 0.374385841 | 0.112931937 | -3.315145827 | 0.000915953 | 0.005906307 | 0.374385841 |
| ENSMUSG00000021700 | Rab3c | 67295 | protein_coding | 22.98476268 | -0.374487517 | 0.281713851 | 1.329318797 | 0.183742815 | 0.378653073 | 0.374487517 |
| ENSMUSG00000001348 | Acp5 | 11433 | protein_coding | 1653.527593 | -0.374886724 | 0.104908207 | 3.573473751 | 0.000352277 | 0.002629971 | 0.374886724 |
| ENSMUSG00000051444 | Bbs12 | 241950 | protein_coding | 8.872638281 | 0.374953451 | 0.341696653 | -1.097328428 | 0.272497848 | 0.488337435 | 0.374953451 |
| ENSMUSG00000065490 | Mir30c-1 | 387227 | miRNA | 1.398686512 | 0.375042826 | 1.246356438 | -0.300911373 | 0.763482078 | 1 | 0.375042826 |
| ENSMUSG00000065436 | Mir342 | 723909 | miRNA | 3.395535395 | -0.375045051 | 0.569903321 | 0.658085394 | 0.510483261 | 1 | 0.375045051 |

Supplementary Table 1

| ensembl_gene_id | external_gene_name | entrezgene_id | gene_biotype | baseMean | log2FoldChange | lfcSE | stat | pvalue | padj | absLog2FC |
|---|---|---|---|---|---|---|---|---|---|---|
| ENSMUSG00000022018 | Rgcc | 66214 | protein_coding | 79.70284167 | 0.37511982 | 0.185658518 | -2.020482682 | 0.043333345 | 0.133608449 | 0.37511982 |
| ENSMUSG00000040139 | 9430038I01Rik | 77252 | protein_coding | 109.2211386 | -0.375357306 | 0.151262263 | 2.481499998 | 0.013083071 | 0.052452983 | 0.375357306 |
| ENSMUSG00000027285 | Haus2 | 66296 | protein_coding | 138.7119214 | -0.375386336 | 0.129980104 | 2.888029201 | 0.003876639 | 0.019710226 | 0.375386336 |
| ENSMUSG00000057322 | Rpl38 | 67671 | protein_coding | 2246.914901 | -0.37548697 | 0.126006205 | 2.979908574 | 0.002883344 | 0.015384668 | 0.37548697 |
| ENSMUSG00000021638 | Ocln | 18260 | protein_coding | 1379.949565 | -0.3755925 | 0.074200392 | 5.06186681 | 4.15E-07 | 6.21E-06 | 0.3755925 |
| ENSMUSG00000026229 | Psmd1 | 70247 | protein_coding | 1326.404854 | 0.375603187 | 0.068911414 | -5.45052213 | 5.02E-08 | 8.60E-07 | 0.375603187 |
| ENSMUSG00000030054 | Gp9 | 54368 | protein_coding | 2.530686409 | -0.375622641 | 0.764433732 | 0.491373713 | 0.623162153 | 1 | 0.375622641 |
| ENSMUSG00000089707 | Slain1os | 70898 | lncRNA | 3.171450805 | 0.375676415 | 0.888393677 | -0.422871554 | 0.672388978 | 1 | 0.375676415 |
| ENSMUSG00000097325 | Gm16897 | 320400 | lncRNA | 7.538915236 | 0.375705714 | 0.454097139 | -0.8273686 | 0.408028172 | 0.628624818 | 0.375705714 |
| ENSMUSG00000038615 | Nfe2l1 | 18023 | protein_coding | 2849.939657 | 0.375839699 | 0.057943929 | -6.486265376 | 8.80E-11 | 2.23E-09 | 0.375839699 |
| ENSMUSG00000035397 | Klf16 | 118445 | protein_coding | 202.8614781 | -0.376046997 | 0.118442715 | 3.174927191 | 0.001498741 | 0.008896486 | 0.376046997 |
| ENSMUSG00000053559 | Smagp | 207818 | protein_coding | 1223.022636 | -0.376066808 | 0.097039943 | 3.875381596 | 0.000106458 | 0.000910903 | 0.376066808 |
| ENSMUSG00000027671 | Actl6a | 56456 | protein_coding | 452.6510068 | 0.376070265 | 0.073918632 | -5.087624757 | 3.63E-07 | 5.49E-06 | 0.376070265 |
| ENSMUSG00000061787 | Rps17 | 20068 | protein_coding | 3196.137274 | -0.376092653 | 0.116262273 | 3.234864097 | 0.001217006 | 0.007482994 | 0.376092653 |
| ENSMUSG00000025255 | Zfhx4 | 80892 | protein_coding | 4.582601074 | 0.376118448 | 1.886350043 | -0.19938953 | 0.84195805 | 0.921777727 | 0.376118448 |
| ENSMUSG00000021427 | Ssr1 | 107513 | protein_coding | 2266.144613 | 0.376163407 | 0.130572857 | -2.88086985 | 0.003965794 | 0.020064348 | 0.376163407 |
| ENSMUSG00000045996 | Polr2k | 17749 | protein_coding | 77.79481917 | -0.376330824 | 0.148558332 | 2.533219234 | 0.011302024 | 0.046953868 | 0.376330824 |
| ENSMUSG00000019945 | Cabcoco1 | 73287 | protein_coding | 37.44826747 | 0.37634742 | 0.175693988 | -2.142062024 | 0.032188491 | 0.106546551 | 0.37634742 |
| ENSMUSG00000041935 | Rimoc1 | 106064 | protein_coding | 292.1502941 | -0.376486691 | 0.103431028 | 3.639978254 | 0.000272661 | 0.002092344 | 0.376486691 |
| ENSMUSG00000035840 | Lysmd3 | 80289 | protein_coding | 459.2087043 | 0.376538697 | 0.07956471 | -4.732483762 | 2.22E-06 | 2.87E-05 | 0.376538697 |
| ENSMUSG00000075707 | Dio3 | 107585 | protein_coding | 3.051772979 | -0.37657571 | 0.712201224 | 0.528749035 | 0.596979556 | 1 | 0.37657571 |
| ENSMUSG00000060093 | H4c1 | 326619 | protein_coding | 637.2264932 | 0.376576473 | 0.153121543 | -2.45933045 | 0.013919643 | 0.055018905 | 0.376576473 |
| ENSMUSG00000031843 | Mphosph6 | 68533 | protein_coding | 160.6538595 | 0.376626745 | 0.105813463 | -3.55934619 | 0.000371779 | 0.002758114 | 0.376626745 |
| ENSMUSG00000024900 | Cpt1a | 12894 | protein_coding | 4499.121888 | -0.376751668 | 0.084946953 | 4.435140436 | 9.20E-06 | 0.000104289 | 0.376751668 |
| ENSMUSG00000025927 | Tfap2b | 21419 | protein_coding | 3.62961883 | 0.376800034 | 2.059913728 | -0.182920298 | 0.854860556 | 1 | 0.376800034 |
| ENSMUSG00000078664 | Sprr2a1 | 20755 | protein_coding | 294.2020933 | 0.37693391 | 0.804671417 | -0.468432085 | 0.639475628 | 0.803370142 | 0.37693391 |
| ENSMUSG00000025287 | Acot9 | 56360 | protein_coding | 188.7606751 | -0.376979036 | 0.123375271 | 3.055547778 | 0.002246498 | 0.012440868 | 0.376979036 |
| ENSMUSG00000037563 | Rps16 | 20055 | protein_coding | 1139.973889 | -0.377137828 | 0.164682046 | 2.290096816 | 0.022015705 | 0.079122383 | 0.377137828 |
| ENSMUSG00000025290 | Rps24 | 20088 | protein_coding | 5629.019648 | -0.377192592 | 0.116965399 | 3.224821997 | 0.00126051 | 0.007703947 | 0.377192592 |
| ENSMUSG00000061099 | Gapdhs | 14447 | protein_coding | 5.151040313 | -0.377192899 | 0.491891332 | 0.766821601 | 0.443187592 | 0.660786267 | 0.377192899 |
| ENSMUSG00000038774 | Ascc3 | 77987 | protein_coding | 1103.257027 | 0.377199131 | 0.074879672 | -5.037403603 | 4.72E-07 | 7.00E-06 | 0.377199131 |
| ENSMUSG00000071723 | Gspt2 | 14853 | protein_coding | 43.09054626 | 0.377217138 | 0.200809982 | -1.878478024 | 0.060315801 | 0.171438061 | 0.377217138 |
| ENSMUSG00000069922 | Ces3a | 382053 | protein_coding | 2.176648266 | -0.377287152 | 1.077019666 | 0.350306651 | 0.726108574 | 1 | 0.377287152 |
| ENSMUSG00000087739 | Gm24644 | 115489635 | scaRNA | 1.932033912 | 0.377386716 | 1.145756963 | -0.329377633 | 0.741870272 | 1 | 0.377386716 |
| ENSMUSG00000070298 | Trcg1 | 541610 | protein_coding | 3.866956192 | -0.377466739 | 0.973235512 | 0.387847273 | 0.69812906 | 1 | 0.377466739 |
| ENSMUSG00000006930 | Hap1 | 15114 | protein_coding | 38.76364469 | -0.377551306 | 0.216389066 | 1.744779962 | 0.081023178 | 0.21167802 | 0.377551306 |
| ENSMUSG00000051076 | Vtcn1 | 242122 | protein_coding | 13.35244912 | 0.377622921 | 0.400567203 | -0.94272052 | 0.345823875 | 0.567534275 | 0.377622921 |
| ENSMUSG00000015968 | Cacna1d | 12289 | protein_coding | 28.85793307 | 0.377686796 | 0.321026924 | -1.176495702 | 0.239396849 | 0.449607593 | 0.377686796 |
| ENSMUSG00000035877 | Zhx3 | 320799 | protein_coding | 215.7609004 | 0.377713306 | 0.119553424 | -3.15936837 | 0.001581115 | 0.009308568 | 0.377713306 |

| ensembl_gene_id | external_gene_name | entrezgene_id | gene_biotype | baseMean | log2FoldChange | lfcSE | stat | pvalue | padj | absLog2FC |
|---|---|---|---|---|---|---|---|---|---|---|
| ENSMUSG00000031842 | Pde4c | 110385 | protein_coding | 9.776557292 | 0.377980687 | 0.586249824 | -0.64474337 | 0.519093506 | 0.719334577 | 0.377980687 |
| ENSMUSG00000097787 | 2700046G09Rik | 67188 | lncRNA | 17.5155948 | -0.378028814 | 0.240925417 | 1.569069877 | 0.116631658 | 0.276057789 | 0.378028814 |
| ENSMUSG00000032313 | Tmem266 | 244886 | protein_coding | 14.33247724 | 0.378029251 | 0.368535563 | -1.025760575 | 0.305004454 | 0.525108794 | 0.378029251 |
| ENSMUSG00000037816 | Fbxw17 | 109082 | protein_coding | 75.89895135 | -0.378182632 | 0.157429512 | 2.402234675 | 0.016295251 | 0.062415953 | 0.378182632 |
| ENSMUSG00000036639 | Nudt1 | 17766 | protein_coding | 160.5664415 | -0.378430372 | 0.098696265 | 3.834292738 | 0.000125926 | 0.001058292 | 0.378430372 |
| ENSMUSG00000077438 | Gm22595 | 115485710 | snoRNA | 0.730776855 | -0.378459916 | 1.489460187 | 0.254091999 | 0.799424485 | 1 | 0.378459916 |
| ENSMUSG00000092622 | Khdc3 | 66991 | protein_coding | 3.129958707 | -0.378473653 | 0.766704677 | 0.493636813 | 0.621562697 | 1 | 0.378473653 |
| ENSMUSG00000015966 | Il17rb | 50905 | protein_coding | 36.0296325 | -0.378481184 | 0.249323386 | 1.518033228 | 0.129006022 | 0.296522686 | 0.378481184 |
| ENSMUSG00000022332 | Khdrbs3 | 13992 | protein_coding | 268.632847 | 0.378545026 | 0.139813057 | -2.7075084 | 0.006779036 | 0.031096942 | 0.378545026 |
| ENSMUSG00000031647 | Mfap3l | 71306 | protein_coding | 590.2475571 | -0.378614591 | 0.164197892 | 2.305843192 | 0.021119396 | 0.076601296 | 0.378614591 |
| ENSMUSG00000039824 | Myl6b | 216459 | protein_coding | 16.93961804 | 0.378683869 | 0.299186978 | -1.265709731 | 0.205617023 | 0.408255959 | 0.378683869 |
| ENSMUSG00000063576 | Klhdc3 | 71765 | protein_coding | 267.4209456 | -0.378738331 | 0.097553446 | 3.882367527 | 0.000103444 | 0.000886497 | 0.378738331 |
| ENSMUSG00000028310 | Ppp3r2 | 19059 | protein_coding | 1.752935216 | -0.378748481 | 1.085534825 | 0.348904957 | 0.727160664 | 1 | 0.378748481 |
| ENSMUSG00000036242 | Armh4 | 67419 | protein_coding | 84.72228682 | -0.378771392 | 0.150029639 | 2.524643757 | 0.011581563 | 0.047814481 | 0.378771392 |
| ENSMUSG00000038489 | Polr2l | 66491 | protein_coding | 86.75773656 | -0.378845599 | 0.140021463 | 2.705625196 | 0.006817594 | 0.031247781 | 0.378845599 |
| ENSMUSG00000052911 | Lamb2 | 16779 | protein_coding | 375.1405986 | 0.37886112 | 0.105326576 | -3.597013544 | 0.000321892 | 0.002420638 | 0.37886112 |
| ENSMUSG00000004952 | Rasa4 | 54153 | protein_coding | 375.3183499 | -0.378967558 | 0.141730493 | 2.673860437 | 0.007498365 | 0.033749795 | 0.378967558 |
| ENSMUSG00000097276 | 4930525G20Rik | 320163 | lncRNA | 6.211962058 | 0.378981075 | 0.422978338 | -0.895982232 | 0.370262253 | 0.592811812 | 0.378981075 |
| ENSMUSG00000040596 | Pogk | 71592 | protein_coding | 257.8635811 | -0.37902932 | 0.122024047 | 3.106185446 | 0.001895178 | 0.010863233 | 0.37902932 |
| ENSMUSG00000043102 | Qrfp | 227717 | protein_coding | 5.230202856 | -0.379042677 | 0.685540395 | 0.552910784 | 0.580324503 | 0.76356297 | 0.379042677 |
| ENSMUSG00000055546 | Timd4 | 276891 | protein_coding | 1.338644369 | 0.379136957 | 1.839733768 | -0.206082513 | 0.836726459 | 1 | 0.379136957 |
| ENSMUSG00000090799 | Klhl33 | 546611 | protein_coding | 4.15706893 | -0.379301387 | 0.711252039 | 0.533286889 | 0.593835002 | 1 | 0.379301387 |
| ENSMUSG00000021662 | Arhgef28 | 110596 | protein_coding | 1121.941498 | -0.379374636 | 0.072565351 | 5.228041064 | 1.71E-07 | 2.72E-06 | 0.379374636 |
| ENSMUSG00000055866 | Per2 | 18627 | protein_coding | 345.4918205 | 0.379523539 | 0.286779907 | -1.323396548 | 0.185703539 | 0.381194313 | 0.379523539 |
| ENSMUSG00000042628 | Zfyve1 | 217695 | protein_coding | 642.7604009 | -0.379644665 | 0.131932786 | 2.877561197 | 0.004007622 | 0.02019882 | 0.379644665 |
| ENSMUSG00000037005 | Xpnpep2 | 170745 | protein_coding | 74.70777283 | -0.379691094 | 0.135822566 | 2.795493456 | 0.005182055 | 0.025095735 | 0.379691094 |
| ENSMUSG00000067586 | S1pr3 | 13610 | protein_coding | 1.599838458 | -0.379699641 | 1.374355979 | 0.276274594 | 0.782337168 | 1 | 0.379699641 |
| ENSMUSG00000027879 | Sec22b | 20333 | protein_coding | 645.962207 | 0.379798282 | 0.105808163 | -3.58949886 | 0.000331314 | 0.002485836 | 0.379798282 |
| ENSMUSG00000049353 | Rd3 | 74023 | protein_coding | 3.396721554 | -0.379881399 | 1.011779904 | 0.375458533 | 0.70731948 | 1 | 0.379881399 |
| ENSMUSG00000037239 | Spred3 | 101809 | protein_coding | 11.74235666 | 0.379994896 | 0.575545498 | -0.660234329 | 0.509103466 | 0.712606767 | 0.379994896 |
| ENSMUSG00000064380 | Gm26448 | 115487256 | snoRNA | 825.4893378 | 0.380136693 | 0.201515819 | -1.886386363 | 0.059242909 | 0.169321114 | 0.380136693 |
| ENSMUSG00000027220 | Syt13 | 80976 | protein_coding | 18.72321956 | -0.380168546 | 0.357973222 | 1.06200275 | 0.288234437 | 0.507128009 | 0.380168546 |
| ENSMUSG00000024966 | Stip1 | 20867 | protein_coding | 1193.273994 | 0.380276057 | 0.123343578 | -3.083063295 | 0.002048816 | 0.011575143 | 0.380276057 |
| ENSMUSG00000015950 | Ncf1 | 17969 | protein_coding | 11.45428723 | -0.380322031 | 0.39141973 | 0.971647573 | 0.331225907 | 0.553630732 | 0.380322031 |
| ENSMUSG00000049848 | Ceacam19 | 319930 | protein_coding | 2.070724396 | 0.380383504 | 1.286223431 | -0.295736724 | 0.767431149 | 1 | 0.380383504 |
| ENSMUSG00000043872 | Zmym1 | 68310 | protein_coding | 57.05730672 | 0.380697276 | 0.192995832 | -1.972567347 | 0.048544876 | 0.14581122 | 0.380697276 |
| ENSMUSG00000030823 | 9130019O22Rik | 78921 | protein_coding | 63.45172627 | -0.380721855 | 0.187437 | 2.031199037 | 0.042234808 | 0.131176532 | 0.380721855 |
| ENSMUSG00000063458 | Lrmda | 76633 | protein_coding | 2.528547714 | -0.380737312 | 1.18224839 | 0.322045109 | 0.747418521 | 1 | 0.380737312 |
| ENSMUSG00000074881 | Mageb3 | 17147 | protein_coding | 1.566394407 | 0.38075342 | 1.504223655 | -0.253122877 | 0.800173261 | 1 | 0.38075342 |

| ensembl_gene_id | external_gene_name | entrezgene_id | gene_biotype | baseMean | log2FoldChange | lfcSE | stat | pvalue | padj | absLog2FC |
|---|---|---|---|---|---|---|---|---|---|---|
| ENSMUSG00000020111 | Micu1 | 216001 | protein_coding | 1306.66792 | -0.380986705 | 0.039482782 | 9.649439313 | 4.94E-22 | 2.80E-20 | 0.380986705 |
| ENSMUSG00000020952 | Scfd1 | 76983 | protein_coding | 709.6162205 | 0.381004577 | 0.08297506 | -4.591796361 | 4.39E-06 | 5.32E-05 | 0.381004577 |
| ENSMUSG00000117232 | 1600002D24Rik | 69776 | lncRNA | 3.217013469 | 0.381136985 | 0.759816226 | -0.501617327 | 0.615936728 | 1 | 0.381136985 |
| ENSMUSG00000034566 | Atp5h | 71679 | protein_coding | 2426.365602 | -0.381146027 | 0.09980534 | 3.818894117 | 0.000134051 | 0.001114664 | 0.381146027 |
| ENSMUSG00000044286 | Or5au1 | 258420 | protein_coding | 1.249069549 | 0.381223364 | 1.585650225 | -0.240420843 | 0.810004022 | 1 | 0.381223364 |
| ENSMUSG00000064442 | Gm26225 | 115488410 | snoRNA | 9.978634917 | 0.381321403 | 0.459008415 | -0.830750353 | 0.406114674 | 0.627075992 | 0.381321403 |
| ENSMUSG00000022139 | Mbnl2 | 105559 | protein_coding | 2544.778113 | 0.381326426 | 0.06728056 | -5.667705918 | 1.45E-08 | 2.71E-07 | 0.381326426 |
| ENSMUSG00000037979 | Ccdc92 | 215707 | protein_coding | 19.1628266 | 0.381332092 | 0.303208832 | -1.257654961 | 0.208516567 | 0.411599641 | 0.381332092 |
| ENSMUSG00000044991 | Shld1 | 73747 | protein_coding | 53.74655213 | -0.381415277 | 0.187198374 | 2.037492465 | 0.041600716 | 0.129751344 | 0.381415277 |
| ENSMUSG00000034958 | Atcay | 16467 | protein_coding | 2.941082851 | -0.381451324 | 1.27844374 | 0.298371616 | 0.76541955 | 1 | 0.381451324 |
| ENSMUSG00000060519 | Tor3a | 30935 | protein_coding | 348.1300018 | -0.38152458 | 0.115040379 | 3.316440567 | 0.00091172 | 0.005884119 | 0.38152458 |
| ENSMUSG00000048351 | Coa7 | 69893 | protein_coding | 186.763013 | -0.381601728 | 0.113015962 | 3.376529483 | 0.000734065 | 0.004876247 | 0.381601728 |
| ENSMUSG00000041506 | Rrp9 | 27966 | protein_coding | 244.9337626 | 0.381768866 | 0.127028269 | -3.005385098 | 0.002652448 | 0.014342673 | 0.381768866 |
| ENSMUSG00000028527 | Ak4 | 11639 | protein_coding | 91.52010506 | 0.381909911 | 0.32044991 | -1.191792849 | 0.233342481 | 0.442718029 | 0.381909911 |
| ENSMUSG00000036596 | Cpz | 242939 | protein_coding | 1.717281649 | -0.382021156 | 2.490834039 | 0.153370778 | 0.878105888 | 1 | 0.382021156 |
| ENSMUSG00000025754 | Agbl1 | 244071 | protein_coding | 1.122583772 | -0.3820576 | 2.479767014 | 0.154069958 | 0.877554575 | 1 | 0.3820576 |
| ENSMUSG00000031633 | Slc25a4 | 11739 | protein_coding | 263.6299243 | 0.382132798 | 0.078610014 | -4.861121099 | 1.17E-06 | 1.59E-05 | 0.382132798 |
| ENSMUSG00000037060 | Cavin3 | 109042 | protein_coding | 10.27469055 | -0.38214037 | 0.349541294 | 1.093262445 | 0.274278587 | 0.490623547 | 0.38214037 |
| ENSMUSG00000066258 | Trim12a | 76681 | protein_coding | 159.5066008 | -0.382146648 | 0.1619307 | 2.359939457 | 0.018277918 | 0.068336322 | 0.382146648 |
| ENSMUSG00000016541 | Atxn10 | 54138 | protein_coding | 258.6014449 | 0.382164347 | 0.110874003 | -3.44683456 | 0.000567196 | 0.003940543 | 0.382164347 |
| ENSMUSG00000036168 | Ccdc38 | 237465 | protein_coding | 4.491768956 | 0.382238788 | 0.691698164 | -0.552609227 | 0.580531023 | 0.763773846 | 0.382238788 |
| ENSMUSG00000038587 | Akap12 | 83397 | protein_coding | 5.595848448 | 0.382244655 | 0.677381119 | -0.564297769 | 0.572551499 | 0.757985847 | 0.382244655 |
| ENSMUSG00000032116 | Stt3a | 16430 | protein_coding | 2485.978908 | 0.382411951 | 0.11908674 | -3.21120514 | 0.001321795 | 0.008036866 | 0.382411951 |
| ENSMUSG00000037443 | Cep85 | 70012 | protein_coding | 408.4721543 | 0.38245962 | 0.076474148 | -5.001162217 | 5.70E-07 | 8.30E-06 | 0.38245962 |
| ENSMUSG00000022546 | Gpt | 76282 | protein_coding | 680.4671163 | -0.38247666 | 0.215237893 | 1.776994998 | 0.075569063 | 0.201541909 | 0.38247666 |
| ENSMUSG00000065904 | Gm26109 | 115488999 | snoRNA | 0.754857076 | -0.382581941 | 1.353214105 | 0.282720923 | 0.777390776 | 1 | 0.382581941 |
| ENSMUSG00000027871 | Hsd3b1 | 15492 | protein_coding | 2.257820628 | -0.382698444 | 0.953672425 | 0.4012892 | 0.688207214 | 1 | 0.382698444 |
| ENSMUSG00000024235 | Map3k8 | 26410 | protein_coding | 15.29321221 | 0.382719754 | 0.324956805 | -1.177755775 | 0.238893984 | 0.449071742 | 0.382719754 |
| ENSMUSG00000075415 | Fnbp1 | 14269 | protein_coding | 183.9259945 | -0.382803717 | 0.097963238 | 3.907626215 | 9.32E-05 | 0.000810119 | 0.382803717 |
| ENSMUSG00000105834 | 4930592C13Rik | 381498 | lncRNA | 1.056006653 | -0.382912941 | 1.121910207 | 0.341304445 | 0.732874402 | 1 | 0.382912941 |
| ENSMUSG00000005732 | Ranbp1 | 19385 | protein_coding | 413.205177 | 0.382948762 | 0.109038081 | -3.512064406 | 0.00044464 | 0.003214881 | 0.382948762 |
| ENSMUSG00000004665 | Cnn2 | 12798 | protein_coding | 138.0961622 | 0.3829495 | 0.159979219 | -2.393745268 | 0.016677327 | 0.063634329 | 0.3829495 |
| ENSMUSG00000019359 | Gdpd2 | 71584 | protein_coding | 29.60188966 | -0.38295664 | 0.373006462 | 1.026675616 | 0.304573235 | 0.524764095 | 0.38295664 |
| ENSMUSG00000038058 | Nod1 | 107607 | protein_coding | 196.9956085 | 0.382983384 | 0.123294192 | -3.106256478 | 0.001894723 | 0.010863233 | 0.382983384 |
| ENSMUSG00000059895 | Ptp4a3 | 19245 | protein_coding | 117.9206965 | -0.383030525 | 0.149003971 | 2.57060616 | 0.010152071 | 0.043086943 | 0.383030525 |
| ENSMUSG00000032840 | Spring1 | 76792 | protein_coding | 177.4431317 | 0.383093724 | 0.112709222 | -3.398956347 | 0.000676435 | 0.004563174 | 0.383093724 |
| ENSMUSG00000026956 | Uap1l1 | 227620 | protein_coding | 212.6560169 | -0.383141486 | 0.118161482 | 3.242524387 | 0.001184758 | 0.007322266 | 0.383141486 |
| ENSMUSG00000025871 | 4833439L19Rik | 97820 | protein_coding | 2349.102617 | 0.383177946 | 0.059558701 | -6.433618276 | 1.25E-10 | 3.05E-09 | 0.383177946 |
| ENSMUSG00000073457 | Smok2b | 236574 | protein_coding | 3.695090602 | 0.383184039 | 2.057832128 | -0.186207628 | 0.852281942 | 1 | 0.383184039 |

| ensembl_gene_id | external_gene_name | entrezgene_id | gene_biotype | baseMean | log2FoldChange | lfcSE | stat | pvalue | padj | absLog2FC |
|---|---|---|---|---|---|---|---|---|---|---|
| ENSMUSG00000066516 | Klk1b21 | 16616 | protein_coding | 0.786071127 | 0.383197788 | 1.637126203 | -0.234067348 | 0.814932695 | 1 | 0.383197788 |
| ENSMUSG00000031504 | Rab20 | 19332 | protein_coding | 253.5450897 | 0.383266046 | 0.095983222 | -3.993052511 | 6.52E-05 | 0.000592833 | 0.383266046 |
| ENSMUSG00000047342 | Zfp286 | 192651 | protein_coding | 14.03382548 | 0.383312771 | 0.365356504 | -1.049147248 | 0.294110353 | 0.513201434 | 0.383312771 |
| ENSMUSG00000047153 | Khnyn | 219094 | protein_coding | 1081.231787 | -0.383377027 | 0.094174934 | 4.07090308 | 4.68E-05 | 0.000440521 | 0.383377027 |
| ENSMUSG00000015094 | Npdc1 | 18146 | protein_coding | 398.5012754 | -0.38344183 | 0.076411788 | 5.018097868 | 5.22E-07 | 7.68E-06 | 0.38344183 |
| ENSMUSG00000105975 | Gm9831 | 101056032 | lncRNA | 5.394376818 | -0.383445196 | 0.716892354 | 0.534871371 | 0.592738806 | 0.772538316 | 0.383445196 |
| ENSMUSG00000004654 | Ghrhr | 14602 | protein_coding | 4.029737241 | 0.383483404 | 1.136266867 | -0.337494135 | 0.735744432 | 1 | 0.383483404 |
| ENSMUSG00000007107 | Atp1a4 | 27222 | protein_coding | 4.296479189 | 0.38360083 | 1.229868679 | -0.311903894 | 0.755113563 | 1 | 0.38360083 |
| ENSMUSG00000023066 | Rttn | 246102 | protein_coding | 180.7592191 | 0.383631875 | 0.114310613 | -3.356047741 | 0.000790649 | 0.005214369 | 0.383631875 |
| ENSMUSG00000089069 | Gm22840 | 115486335 | snoRNA | 0.727132154 | 0.383646685 | 1.214429351 | -0.315906961 | 0.752073128 | 1 | 0.383646685 |
| ENSMUSG00000032089 | Il10ra | 16154 | protein_coding | 1.919824091 | -0.383728671 | 0.983784269 | 0.390053676 | 0.696496856 | 1 | 0.383728671 |
| ENSMUSG00000027637 | Rab5if | 67388 | protein_coding | 367.1865554 | 0.38374015 | 0.112233984 | -3.419108345 | 0.000628267 | 0.00429793 | 0.38374015 |
| ENSMUSG00000024393 | Prrc2a | 53761 | protein_coding | 2315.334918 | 0.383742871 | 0.129746363 | -2.957638765 | 0.003100052 | 0.01632826 | 0.383742871 |
| ENSMUSG00000039652 | Cpeb3 | 208922 | protein_coding | 51.67375434 | -0.383744492 | 0.144249513 | 2.660282755 | 0.007807508 | 0.034924193 | 0.383744492 |
| ENSMUSG00000026696 | Vamp4 | 53330 | protein_coding | 211.4178536 | -0.383776367 | 0.106536133 | 3.602311768 | 0.0003154 | 0.002381578 | 0.383776367 |
| ENSMUSG00000036022 | Fam122b | 78755 | protein_coding | 46.83030279 | 0.383779122 | 0.190100648 | -2.018820689 | 0.04350586 | 0.134003711 | 0.383779122 |
| ENSMUSG00000067103 | AY702103 | 446210 | lncRNA | 1.111252454 | -0.383808552 | 2.100607098 | 0.182713156 | 0.855023093 | 1 | 0.383808552 |
| ENSMUSG00000031919 | Tmed6 | 66269 | protein_coding | 21.0134008 | -0.383887951 | 0.326796522 | 1.174700233 | 0.240114666 | 0.450443439 | 0.383887951 |
| ENSMUSG00000051956 | Rnf133 | 386611 | protein_coding | 4.363465835 | 0.383896985 | 0.576066805 | -0.666410531 | 0.505148733 | 1 | 0.383896985 |
| ENSMUSG00000070730 | Rmdn3 | 67809 | protein_coding | 664.779333 | -0.383922202 | 0.096524104 | 3.977474926 | 6.97E-05 | 0.000627847 | 0.383922202 |
| ENSMUSG00000026628 | Atf3 | 11910 | protein_coding | 6886.389736 | -0.384005668 | 0.353594844 | 1.086004716 | 0.277476906 | 0.494416662 | 0.384005668 |
| ENSMUSG00000031556 | Tm2d2 | 69742 | protein_coding | 1019.523434 | 0.384084225 | 0.10297649 | -3.729824388 | 0.000191613 | 0.001522763 | 0.384084225 |
| ENSMUSG00000029863 | Casp2 | 12366 | protein_coding | 565.4762914 | -0.384119068 | 0.091784759 | 4.184998359 | 2.85E-05 | 0.000283691 | 0.384119068 |
| ENSMUSG00000072591 | Fzd10os | 319616 | lncRNA | 1.560035394 | 0.384365898 | 1.729498521 | -0.222241241 | 0.824126091 | 1 | 0.384365898 |
| ENSMUSG00000030785 | Cox6a2 | 12862 | protein_coding | 1.847168346 | -0.384401541 | 0.713507419 | 0.538749185 | 0.59005993 | 1 | 0.384401541 |
| ENSMUSG00000006476 | Nsmf | 56876 | protein_coding | 599.9405758 | -0.384559589 | 0.09144909 | 4.205176785 | 2.61E-05 | 0.000261416 | 0.384559589 |
| ENSMUSG00000069307 | H2bc23 | 665596 | protein_coding | 0.801861309 | -0.384562364 | 1.394339497 | 0.275802532 | 0.782699739 | 1 | 0.384562364 |
| ENSMUSG00000028277 | Ube2j1 | 56228 | protein_coding | 834.3883723 | 0.384676223 | 0.064944956 | -5.923111588 | 3.16E-09 | 6.55E-08 | 0.384676223 |
| ENSMUSG00000036718 | Micall2 | 231830 | protein_coding | 427.634514 | 0.38471319 | 0.138150112 | -2.784747584 | 0.005356941 | 0.025783888 | 0.38471319 |
| ENSMUSG00000037499 | Nenf | 66208 | protein_coding | 203.0310228 | -0.384776578 | 0.097404523 | 3.950294758 | 7.81E-05 | 0.000694131 | 0.384776578 |
| ENSMUSG00000068040 | Tm9sf4 | 99237 | protein_coding | 2496.352728 | 0.384829086 | 0.116422094 | -3.305464391 | 0.000948192 | 0.006090406 | 0.384829086 |
| ENSMUSG00000039372 | Marchf4 | 381270 | protein_coding | 4.129041232 | 0.384868275 | 0.914648693 | -0.420782621 | 0.673913823 | 1 | 0.384868275 |
| ENSMUSG00000022292 | Rrm2b | 382985 | protein_coding | 66.94917364 | -0.384871865 | 0.178121258 | 2.160729543 | 0.030716237 | 0.102723358 | 0.384871865 |
| ENSMUSG00000073705 | Cenps | 69928 | protein_coding | 26.49058505 | 0.384907365 | 0.256152404 | -1.502649825 | 0.132929368 | 0.30270362 | 0.384907365 |
| ENSMUSG00000031147 | Magix | 54634 | protein_coding | 3.321987036 | -0.384947457 | 0.80545818 | 0.47792358 | 0.632704598 | 1 | 0.384947457 |
| ENSMUSG00000036402 | Gng12 | 14701 | protein_coding | 2481.492076 | -0.384948004 | 0.081296882 | 4.735089424 | 2.19E-06 | 2.83E-05 | 0.384948004 |
| ENSMUSG00000079555 | Haus3 | 231123 | protein_coding | 89.54648382 | 0.385115226 | 0.136627244 | -2.818729381 | 0.004821415 | 0.023571307 | 0.385115226 |
| ENSMUSG00000023052 | Npff | 54615 | protein_coding | 2.818483738 | 0.385130425 | 0.629986915 | -0.611330832 | 0.540980584 | 1 | 0.385130425 |
| ENSMUSG00000065130 | Gm24457 | 115489766 | snRNA | 1.779350454 | -0.38521708 | 0.872111233 | 0.441706362 | 0.658701705 | 1 | 0.38521708 |

| ensembl_gene_id | external_gene_name | entrezgene_id | gene_biotype | baseMean | log2FoldChange | lfcSE | stat | pvalue | padj | absLog2FC |
|---|---|---|---|---|---|---|---|---|---|---|
| ENSMUSG00000038633 | Degs1 | 13244 | protein_coding | 215.0721421 | 0.385305024 | 0.134819272 | -2.857937278 | 0.004264047 | 0.021273857 | 0.385305024 |
| ENSMUSG00000028600 | Podn | 242608 | protein_coding | 5.028462797 | -0.385359027 | 1.06049421 | 0.363376833 | 0.716323405 | 0.852797822 | 0.385359027 |
| ENSMUSG00000025876 | Unc5a | 107448 | protein_coding | 63.18040595 | 0.385385077 | 0.141003066 | -2.733168074 | 0.006272831 | 0.029098184 | 0.385385077 |
| ENSMUSG00000085451 | Gm40614 | 105245120 | lncRNA | 9.086184668 | -0.385425974 | 0.351516555 | 1.096466068 | 0.272874865 | 0.488906968 | 0.385425974 |
| ENSMUSG00000101026 | Ly6g6g | 78725 | protein_coding | 1.003024077 | 0.38554842 | 1.748097254 | -0.220553187 | 0.825440355 | 1 | 0.38554842 |
| ENSMUSG00000033126 | Ybey | 216119 | protein_coding | 69.69656848 | -0.385696573 | 0.150350121 | 2.565322664 | 0.01030799 | 0.04361418 | 0.385696573 |
| ENSMUSG00000042808 | Gpx2 | 14776 | protein_coding | 4073.146887 | 0.385701532 | 0.160508827 | -2.402992657 | 0.016261514 | 0.062321337 | 0.385701532 |
| ENSMUSG00000030041 | M1ap | 110958 | protein_coding | 6.640345389 | 0.385757696 | 0.435150148 | -0.886493314 | 0.375351756 | 0.597566653 | 0.385757696 |
| ENSMUSG00000025403 | Shmt2 | 108037 | protein_coding | 215.8646248 | 0.385823059 | 0.114835092 | -3.359801015 | 0.000779986 | 0.005158468 | 0.385823059 |
| ENSMUSG00000073633 | Fbxo36 | 66153 | protein_coding | 66.24697476 | 0.385850604 | 0.143006086 | -2.698141137 | 0.006972787 | 0.031809007 | 0.385850604 |
| ENSMUSG00000021236 | Entpd5 | 12499 | protein_coding | 3252.950939 | -0.385866626 | 0.088590697 | 4.355611124 | 1.33E-05 | 0.000143449 | 0.385866626 |
| ENSMUSG00000039891 | Txlnb | 378431 | protein_coding | 2.799877452 | 0.385879457 | 0.900421089 | -0.428554441 | 0.668247507 | 1 | 0.385879457 |
| ENSMUSG00000070306 | Ccdc153 | 270150 | protein_coding | 1.184555884 | -0.385949568 | 1.333362693 | 0.289455802 | 0.772232598 | 1 | 0.385949568 |
| ENSMUSG00000048163 | Selplg | 20345 | protein_coding | 23.53220069 | 0.386026488 | 0.384422123 | -1.004173446 | 0.315295019 | 0.53622883 | 0.386026488 |
| ENSMUSG00000070564 | Ntn5 | 243967 | protein_coding | 9.69480135 | 0.386193364 | 0.449216201 | -0.859704889 | 0.389951739 | 0.612426345 | 0.386193364 |
| ENSMUSG00000087026 | A230103J11Rik | 320466 | lncRNA | 10.9929108 | -0.386234676 | 0.325718992 | 1.185791082 | 0.235704814 | 0.445817123 | 0.386234676 |
| ENSMUSG00000024855 | Pacs1 | 107975 | protein_coding | 89.41355998 | -0.386273991 | 0.120925402 | 3.194316374 | 0.001401624 | 0.008435285 | 0.386273991 |
| ENSMUSG00000024483 | Ankhd1 | 108857 | protein_coding | 1052.071487 | 0.386290581 | 0.061523395 | -6.278759139 | 3.41E-10 | 8.01E-09 | 0.386290581 |
| ENSMUSG00000036995 | Asap3 | 230837 | protein_coding | 29.04400561 | 0.386404717 | 0.327560533 | -1.179643694 | 0.238141957 | 0.448564665 | 0.386404717 |
| ENSMUSG00000072889 | Nfxl1 | 100978 | protein_coding | 225.8046805 | 0.386626195 | 0.089599404 | -4.315053221 | 1.60E-05 | 0.000169022 | 0.386626195 |
| ENSMUSG00000001473 | Tubb6 | 67951 | protein_coding | 53.71543349 | -0.386666957 | 0.146655624 | 2.636564125 | 0.008375037 | 0.03690111 | 0.386666957 |
| ENSMUSG00000104529 | Rbakdn | 100042605 | lncRNA | 2.236369447 | -0.386701882 | 0.919339361 | 0.420630181 | 0.674025151 | 1 | 0.386701882 |
| ENSMUSG00000026479 | Lamc2 | 16782 | protein_coding | 704.8006561 | -0.386721083 | 0.078790702 | 4.908207105 | 9.19E-07 | 1.28E-05 | 0.386721083 |
| ENSMUSG00000021947 | Cryl1 | 68631 | protein_coding | 1152.351243 | -0.386799961 | 0.07759668 | 4.984748825 | 6.20E-07 | 8.96E-06 | 0.386799961 |
| ENSMUSG00000087022 | 9130024F11Rik | 329160 | lncRNA | 170.759575 | 0.386843586 | 0.127823108 | -3.026397909 | 0.002474864 | 0.013502881 | 0.386843586 |
| ENSMUSG00000040258 | Nxph4 | 104080 | protein_coding | 1.087769733 | -0.386998962 | 1.721435343 | 0.224811791 | 0.822125693 | 1 | 0.386998962 |
| ENSMUSG00000067455 | H4c11 | 319159 | protein_coding | 705.9345883 | 0.387032223 | 0.195813455 | -1.976535386 | 0.048094172 | 0.144852664 | 0.387032223 |
| ENSMUSG00000056687 | Gm11696 | 100036768 | lncRNA | 26.90192985 | 0.387050916 | 0.282066164 | -1.372199028 | 0.170001488 | 0.359584034 | 0.387050916 |
| ENSMUSG00000054598 | 9130230L23Rik | 231253 | lncRNA | 53.79332513 | -0.387092689 | 0.22275769 | 1.737729857 | 0.082258425 | 0.214258446 | 0.387092689 |
| ENSMUSG00000079435 | Rpl36a | 19982 | protein_coding | 1793.777072 | -0.38710609 | 0.117315328 | 3.299705984 | 0.000967862 | 0.006195065 | 0.38710609 |
| ENSMUSG00000070808 | Bicra | 243842 | protein_coding | 128.2067668 | 0.387114297 | 0.287208862 | -1.347849418 | 0.17770682 | 0.370029503 | 0.387114297 |
| ENSMUSG00000031950 | Gabarapl2 | 93739 | protein_coding | 425.2433339 | -0.387235582 | 0.103245286 | 3.750636919 | 0.000176386 | 0.001419217 | 0.387235582 |
| ENSMUSG00000109901 | Chmp1b | 67064 | protein_coding | 681.1050592 | -0.387320338 | 0.08024207 | 4.826898649 | 1.39E-06 | 1.86E-05 | 0.387320338 |
| ENSMUSG00000018698 | Lhx1 | 16869 | protein_coding | 1.331511612 | 0.387521988 | 2.118247508 | -0.182944621 | 0.854841471 | 1 | 0.387521988 |
| ENSMUSG00000020027 | Socs2 | 216233 | protein_coding | 321.5386418 | -0.387527001 | 0.097642312 | 3.96884293 | 7.22E-05 | 0.000647282 | 0.387527001 |
| ENSMUSG00000024922 | Ovol1 | 18426 | protein_coding | 306.2852745 | -0.387611879 | 0.137341476 | 2.822249271 | 0.004768809 | 0.023369313 | 0.387611879 |
| ENSMUSG00000022421 | Nptxr | 73340 | protein_coding | 8.411736894 | 0.387679682 | 0.620891342 | -0.624392153 | 0.532370077 | 0.72898503 | 0.387679682 |
| ENSMUSG00000017756 | Slc12a7 | 20499 | protein_coding | 1183.374339 | -0.387753409 | 0.073716462 | 5.260065404 | 1.44E-07 | 2.32E-06 | 0.387753409 |
| ENSMUSG00000002668 | Dennd1c | 70785 | protein_coding | 272.7072676 | -0.387759958 | 0.099024807 | 3.915786043 | 9.01E-05 | 0.000787343 | 0.387759958 |

| ensembl_gene_id | external_gene_name | entrezgene_id | gene_biotype | baseMean | log2FoldChange | lfcSE | stat | pvalue | padj | absLog2FC |
|---|---|---|---|---|---|---|---|---|---|---|
| ENSMUSG00000052727 | Map1b | 17755 | protein_coding | 42.71415155 | -0.387782735 | 0.311907188 | 1.243263219 | 0.213770853 | 0.418744149 | 0.387782735 |
| ENSMUSG00000032392 | Parp16 | 214424 | protein_coding | 186.73551 | 0.387835003 | 0.109076537 | -3.555622632 | 0.000377085 | 0.002788738 | 0.387835003 |
| ENSMUSG00000027692 | Tnik | 665113 | protein_coding | 365.3945849 | -0.387879843 | 0.100209806 | 3.870677533 | 0.000108533 | 0.000926263 | 0.387879843 |
| ENSMUSG00000060739 | Nsa2 | 59050 | protein_coding | 566.5537046 | -0.387886614 | 0.131181242 | 2.956875608 | 0.003107735 | 0.016359939 | 0.387886614 |
| ENSMUSG00000021877 | Arf4 | 11843 | protein_coding | 2789.773457 | 0.388066535 | 0.073110356 | -5.307955775 | 1.11E-07 | 1.82E-06 | 0.388066535 |
| ENSMUSG00000078546 | Zfp995 | 70081 | protein_coding | 111.1440899 | -0.388099301 | 0.17631079 | 2.201222631 | 0.027720267 | 0.094754639 | 0.388099301 |
| ENSMUSG00000026623 | Lpgat1 | 226856 | protein_coding | 800.0147537 | 0.388119087 | 0.088891826 | -4.366195476 | 1.26E-05 | 0.000137721 | 0.388119087 |
| ENSMUSG00000034667 | Xpot | 73192 | protein_coding | 638.5925855 | 0.388129961 | 0.114846955 | -3.379540724 | 0.00072607 | 0.004832861 | 0.388129961 |
| ENSMUSG00000066894 | Vsig10 | 231668 | protein_coding | 1899.947731 | -0.388271857 | 0.066448301 | 5.843217237 | 5.12E-09 | 1.03E-07 | 0.388271857 |
| ENSMUSG00000063564 | Col23a1 | 237759 | protein_coding | 9.456731132 | 0.388336566 | 0.523162132 | -0.742287223 | 0.457913331 | 0.67261003 | 0.388336566 |
| ENSMUSG00000020088 | Sar1a | 20224 | protein_coding | 1404.553693 | 0.388365974 | 0.112391803 | -3.455465275 | 0.000549344 | 0.003837834 | 0.388365974 |
| ENSMUSG00000032526 | Ss18l2 | 26901 | protein_coding | 187.3489857 | 0.388395202 | 0.111592726 | -3.480470605 | 0.000500534 | 0.003550608 | 0.388395202 |
| ENSMUSG00000033717 | Adra2a | 11551 | protein_coding | 552.388221 | 0.388700367 | 0.242821773 | -1.60076406 | 0.109429187 | 0.264120665 | 0.388700367 |
| ENSMUSG00000018931 | Natd1 | 24083 | protein_coding | 106.8937824 | -0.388812583 | 0.144723462 | 2.686589844 | 0.007218553 | 0.032744159 | 0.388812583 |
| ENSMUSG00000034066 | Farp2 | 227377 | protein_coding | 476.1827756 | -0.38895656 | 0.083576804 | 4.653881704 | 3.26E-06 | 4.04E-05 | 0.38895656 |
| ENSMUSG00000022812 | Gsk3b | 56637 | protein_coding | 1126.072129 | 0.388999268 | 0.084913198 | -4.581140241 | 4.62E-06 | 5.57E-05 | 0.388999268 |
| ENSMUSG00000030323 | Ift122 | 81896 | protein_coding | 105.5201337 | 0.389173285 | 0.106798539 | -3.64399448 | 0.000268439 | 0.002060904 | 0.389173285 |
| ENSMUSG00000091558 | Gm17227 | 102639888 | lncRNA | 4.062402064 | -0.389228912 | 0.554646721 | 0.701760052 | 0.482828816 | 1 | 0.389228912 |
| ENSMUSG00000018659 | Pnpo | 103711 | protein_coding | 1138.468085 | 0.389248968 | 0.11808125 | -3.296450271 | 0.000979149 | 0.006255193 | 0.389248968 |
| ENSMUSG00000031196 | F8 | 14069 | protein_coding | 10.72845721 | 0.389275851 | 0.390314855 | -0.997338036 | 0.318600457 | 0.540015186 | 0.389275851 |
| ENSMUSG00000024553 | Galr1 | 14427 | protein_coding | 3.610916737 | -0.38935857 | 0.733400229 | 0.530895075 | 0.595491491 | 1 | 0.38935857 |
| ENSMUSG00000044258 | Ctla2a | 13024 | protein_coding | 1.448729426 | -0.389518394 | 1.160548834 | 0.33563292 | 0.737147696 | 1 | 0.389518394 |
| ENSMUSG00000036246 | Gmip | 78816 | protein_coding | 172.2782648 | -0.389721487 | 0.110566495 | 3.524770213 | 0.00042385 | 0.003086183 | 0.389721487 |
| ENSMUSG00000016356 | Col20a1 | 73368 | protein_coding | 62.19146546 | 0.389751308 | 0.293204314 | -1.329282309 | 0.183754848 | 0.378653073 | 0.389751308 |
| ENSMUSG00000037353 | Letmd1 | 68614 | protein_coding | 244.4812048 | -0.389822192 | 0.080541564 | 4.840012713 | 1.30E-06 | 1.75E-05 | 0.389822192 |
| ENSMUSG00000020395 | Itk | 16428 | protein_coding | 12.31732638 | -0.389887912 | 0.545323213 | 0.714966652 | 0.47462965 | 0.686643367 | 0.389887912 |
| ENSMUSG00000044707 | Ccnjl | 380694 | protein_coding | 65.62020252 | 0.389902964 | 0.19819781 | -1.967241536 | 0.049155372 | 0.147109787 | 0.389902964 |
| ENSMUSG00000078994 | Zfp429 | 72807 | protein_coding | 74.66629445 | -0.389923986 | 0.151562938 | 2.5726869 | 0.010091245 | 0.042894936 | 0.389923986 |
| ENSMUSG00000030787 | Lyve1 | 114332 | protein_coding | 28.7049331 | 0.389983214 | 0.231644889 | -1.683539043 | 0.092270788 | 0.232919564 | 0.389983214 |
| ENSMUSG00000004880 | Lbr | 98386 | protein_coding | 1027.576695 | 0.390024479 | 0.117361731 | -3.323267957 | 0.000889694 | 0.005763989 | 0.390024479 |
| ENSMUSG00000084780 | Gm15350 | 100504249 | lncRNA | 1.221605614 | 0.390062999 | 1.141318143 | -0.341765354 | 0.732527485 | 1 | 0.390062999 |
| ENSMUSG00000009207 | Lnpk | 69605 | protein_coding | 193.8150024 | 0.390262478 | 0.12347151 | -3.160749202 | 0.001573639 | 0.009274475 | 0.390262478 |
| ENSMUSG00000040666 | Sh3bgr | 50795 | protein_coding | 40.73821556 | -0.390281466 | 0.307251356 | 1.270235127 | 0.204000889 | 0.406181658 | 0.390281466 |
| ENSMUSG00000022564 | Grina | 66168 | protein_coding | 402.5585633 | -0.39028532 | 0.130580149 | 2.988856455 | 0.002800237 | 0.015028634 | 0.39028532 |
| ENSMUSG00000026463 | Atp2b4 | 381290 | protein_coding | 17.70396357 | 0.390564948 | 0.642557059 | -0.607829208 | 0.543300755 | 0.736456016 | 0.390564948 |
| ENSMUSG00000054206 | Gzmm | 16904 | protein_coding | 7.588896483 | 0.390635187 | 0.585312655 | -0.667395764 | 0.504519371 | 0.708596728 | 0.390635187 |
| ENSMUSG00000046691 | Chtf8 | 214987 | protein_coding | 62.69309374 | 0.390639056 | 0.144350303 | -2.706187996 | 0.00680605 | 0.03120353 | 0.390639056 |
| ENSMUSG00000031447 | Lamp1 | 16783 | protein_coding | 3846.132265 | -0.390796402 | 0.080582994 | 4.849613819 | 1.24E-06 | 1.67E-05 | 0.390796402 |
| ENSMUSG00000065523 | Mir192 | 387187 | miRNA | 27.09477451 | 0.390934888 | 0.447262552 | -0.8740613 | 0.382084871 | 0.604440039 | 0.390934888 |

Supplementary Table 1

| ensembl_gene_id | external_gene_name | entrezgene_id | gene_biotype | baseMean | log2FoldChange | lfcSE | stat | pvalue | padj | absLog2FC |
|---|---|---|---|---|---|---|---|---|---|---|
| ENSMUSG00000025969 | Nrp2 | 18187 | protein_coding | 1215.760381 | -0.390953957 | 0.148151475 | 2.638879949 | 0.008318043 | 0.036708756 | 0.390953957 |
| ENSMUSG00000018672 | Copz2 | 56358 | protein_coding | 73.90551393 | -0.391064135 | 0.185454896 | 2.108675175 | 0.034972629 | 0.113383099 | 0.391064135 |
| ENSMUSG00000030890 | Ilk | 16202 | protein_coding | 15.38558968 | 0.391098554 | 0.300905588 | -1.29973842 | 0.193690638 | 0.392130341 | 0.391098554 |
| ENSMUSG00000022074 | Tnfrsf10b | 21933 | protein_coding | 89.99753697 | -0.391308826 | 0.178856209 | 2.187840318 | 0.028681236 | 0.097419765 | 0.391308826 |
| ENSMUSG00000028995 | Hycc1 | 84652 | protein_coding | 269.7259587 | 0.391327861 | 0.120963796 | -3.235082527 | 0.001216076 | 0.007482219 | 0.391327861 |
| ENSMUSG00000051518 | Rps19bp1 | 66538 | protein_coding | 152.5398663 | 0.391424511 | 0.110166533 | -3.553025608 | 0.000380827 | 0.002810091 | 0.391424511 |
| ENSMUSG00000065118 | Gm23297 | 115486827 | snoRNA | 12.66185364 | -0.391507903 | 0.502112682 | 0.779721199 | 0.435554998 | 0.653742472 | 0.391507903 |
| ENSMUSG00000026228 | Htr2b | 15559 | protein_coding | 8.606103259 | -0.391574078 | 0.452305553 | 0.865729097 | 0.38663874 | 0.609134912 | 0.391574078 |
| ENSMUSG00000005493 | Msh4 | 55993 | protein_coding | 3.241025313 | 0.391689404 | 0.756716222 | -0.517617295 | 0.604725312 | 1 | 0.391689404 |
| ENSMUSG00000029072 | Tas1r3 | 83771 | protein_coding | 49.30427049 | 0.391801048 | 0.250809429 | -1.562146404 | 0.118253503 | 0.27902128 | 0.391801048 |
| ENSMUSG00000007891 | Ctsd | 13033 | protein_coding | 1540.757913 | -0.391853446 | 0.079157037 | 4.950329892 | 7.41E-07 | 1.05E-05 | 0.391853446 |
| ENSMUSG00000055319 | Sec23ip | 207352 | protein_coding | 645.6758765 | 0.391986317 | 0.081512624 | -4.808903169 | 1.52E-06 | 2.01E-05 | 0.391986317 |
| ENSMUSG00000021431 | Snrnp48 | 67797 | protein_coding | 322.6115111 | 0.392071961 | 0.099540368 | -3.938823718 | 8.19E-05 | 0.000723489 | 0.392071961 |
| ENSMUSG00000022802 | Lmln | 239833 | protein_coding | 83.15391701 | -0.392090767 | 0.141866302 | 2.763804802 | 0.005713171 | 0.027189423 | 0.392090767 |
| ENSMUSG00000028753 | Vwa5b1 | 75718 | protein_coding | 4.544532348 | 0.392115453 | 1.522195644 | -0.257598591 | 0.796716717 | 0.899007849 | 0.392115453 |
| ENSMUSG00000075324 | Fign | 60344 | protein_coding | 5.923438168 | 0.392175428 | 0.782841842 | -0.500963806 | 0.616396596 | 0.788502504 | 0.392175428 |
| ENSMUSG00000026687 | Aldh9a1 | 56752 | protein_coding | 836.1810411 | 0.392184019 | 0.063027514 | -6.222425624 | 4.90E-10 | 1.12E-08 | 0.392184019 |
| ENSMUSG00000042345 | Ubash3a | 328795 | protein_coding | 4.77214624 | 0.392543432 | 1.263957414 | -0.310566976 | 0.756129836 | 0.87755486 | 0.392543432 |
| ENSMUSG00000048865 | Arhgap30 | 226652 | protein_coding | 18.79270464 | 0.392554268 | 0.438601153 | -0.895014218 | 0.370779488 | 0.593065878 | 0.392554268 |
| ENSMUSG00000032536 | Trak1 | 67095 | protein_coding | 3127.364489 | -0.3927503 | 0.10601994 | 3.704494656 | 0.000211813 | 0.00166012 | 0.3927503 |
| ENSMUSG00000001588 | Acap1 | 216859 | protein_coding | 16.76449914 | -0.392755286 | 0.409298644 | 0.959581205 | 0.337266032 | 0.559393767 | 0.392755286 |
| ENSMUSG00000034201 | Gas2l1 | 78926 | protein_coding | 506.8465825 | -0.392831179 | 0.102747744 | 3.823258427 | 0.0001317 | 0.001100095 | 0.392831179 |
| ENSMUSG00000025790 | Slco3a1 | 108116 | protein_coding | 13.99898509 | -0.392969933 | 0.470714021 | 0.834837959 | 0.40380895 | 0.625062193 | 0.392969933 |
| ENSMUSG00000025091 | Pnliprp2 | 18947 | protein_coding | 13.2403378 | 0.393024111 | 0.372285408 | -1.055706462 | 0.291102344 | 0.510000436 | 0.393024111 |
| ENSMUSG00000033102 | Cdc14b | 218294 | protein_coding | 218.8864918 | -0.393038146 | 0.13545179 | 2.901682916 | 0.00371164 | 0.019029502 | 0.393038146 |
| ENSMUSG00000049871 | Nlrc3 | 268857 | protein_coding | 12.70991135 | 0.393284643 | 0.545784599 | -0.72058582 | 0.471164381 | 0.684193826 | 0.393284643 |
| ENSMUSG00000041216 | Clvs1 | 74438 | protein_coding | 1.752887874 | 0.39347572 | 2.687419417 | -0.14641396 | 0.883594607 | 1 | 0.39347572 |
| ENSMUSG00000027074 | Slc43a3 | 58207 | protein_coding | 14.50789535 | -0.393513263 | 0.444729071 | 0.884838183 | 0.376243912 | 0.59846773 | 0.393513263 |
| ENSMUSG00000029503 | P2rx2 | 231602 | protein_coding | 5.218769846 | -0.393575688 | 0.502701632 | 0.782921047 | 0.433673477 | 0.652196761 | 0.393575688 |
| ENSMUSG00000024487 | Yipf5 | 67180 | protein_coding | 475.6935054 | 0.393592081 | 0.086035433 | -4.574767245 | 4.77E-06 | 5.72E-05 | 0.393592081 |
| ENSMUSG00000030336 | Cd27 | 21940 | protein_coding | 8.417064767 | 0.393661995 | 0.539395972 | -0.72982005 | 0.465500187 | 0.679207466 | 0.393661995 |
| ENSMUSG00000100714 | 2010308F09Rik | 72122 | lncRNA | 1.331520012 | -0.393686544 | 1.048813567 | 0.375363703 | 0.707389996 | 1 | 0.393686544 |
| ENSMUSG00000003418 | St8sia6 | 241230 | protein_coding | 1.585898502 | -0.393803154 | 1.689778673 | 0.233050139 | 0.815722473 | 1 | 0.393803154 |
| ENSMUSG00000068263 | Efcc1 | 58229 | protein_coding | 2.602393823 | -0.393824995 | 1.636575174 | 0.240639722 | 0.809834361 | 1 | 0.393824995 |
| ENSMUSG00000030859 | Fam24a | 68223 | protein_coding | 0.931062152 | 0.393942932 | 1.701076255 | -0.231584523 | 0.81686073 | 1 | 0.393942932 |
| ENSMUSG00000032518 | Rpsa | 16785 | protein_coding | 8056.599601 | -0.393985027 | 0.13897556 | 2.834923111 | 0.004583672 | 0.022615701 | 0.393985027 |
| ENSMUSG00000053054 | Adh6a | 69117 | protein_coding | 6.280655339 | 0.394074675 | 0.851996872 | -0.462530659 | 0.643700821 | 0.805858188 | 0.394074675 |
| ENSMUSG00000021939 | Ctsb | 13030 | protein_coding | 3572.910923 | -0.394125291 | 0.054920568 | 7.17627844 | 7.16E-13 | 2.22E-11 | 0.394125291 |
| ENSMUSG00000021259 | Cyp46a1 | 13116 | protein_coding | 2.350557374 | -0.394156911 | 1.927824999 | 0.20445679 | 0.837996555 | 1 | 0.394156911 |

| ensembl_gene_id | external_gene_name | entrezgene_id | gene_biotype | baseMean | log2FoldChange | lfcSE | stat | pvalue | padj | absLog2FC |
|---|---|---|---|---|---|---|---|---|---|---|
| ENSMUSG00000009633 | G0s2 | 14373 | protein_coding | 3.144475123 | -0.394168435 | 0.644114379 | 0.611954101 | 0.540568126 | 1 | 0.394168435 |
| ENSMUSG00000065687 | Gm23734 | 115487646 | snoRNA | 11.81589892 | -0.394285306 | 0.323498734 | 1.218815608 | 0.222914186 | 0.429896633 | 0.394285306 |
| ENSMUSG00000011427 | Zfp790 | 233056 | protein_coding | 225.1513943 | 0.394346085 | 0.08115215 | -4.85934243 | 1.18E-06 | 1.60E-05 | 0.394346085 |
| ENSMUSG00000023175 | Bsg | 12215 | protein_coding | 7166.135774 | -0.394371247 | 0.066694741 | 5.913078628 | 3.36E-09 | 6.93E-08 | 0.394371247 |
| ENSMUSG00000029674 | Limk1 | 16885 | protein_coding | 46.89637463 | 0.394482481 | 0.176731714 | -2.232097867 | 0.025608495 | 0.088956393 | 0.394482481 |
| ENSMUSG00000024968 | Rcor2 | 104383 | protein_coding | 12.40995403 | 0.394508453 | 0.3986003 | -0.98973446 | 0.322303927 | 0.54358318 | 0.394508453 |
| ENSMUSG00000036615 | Rfxap | 170767 | protein_coding | 126.9790956 | 0.394555323 | 0.101156461 | -3.900446087 | 9.60E-05 | 0.000830593 | 0.394555323 |
| ENSMUSG00000021733 | Slc4a7 | 218756 | protein_coding | 2889.599175 | 0.394831665 | 0.158388243 | -2.492809175 | 0.012673694 | 0.051167007 | 0.394831665 |
| ENSMUSG00000028648 | Ndufs5 | 595136 | protein_coding | 243.5734784 | -0.394890355 | 0.160788832 | 2.455956355 | 0.014051022 | 0.055403505 | 0.394890355 |
| ENSMUSG00000012123 | Crybg2 | 230806 | protein_coding | 680.9024377 | -0.394969245 | 0.20026544 | 1.97222868 | 0.048583507 | 0.145821162 | 0.394969245 |
| ENSMUSG00000093314 | Mir5136 | 100628605 | miRNA | 0.565156851 | 0.395151452 | 1.294375891 | -0.305283384 | 0.76015033 | 1 | 0.395151452 |
| ENSMUSG00000059562 | Ccdc154 | 207209 | protein_coding | 3.770811276 | 0.39518159 | 0.949198899 | -0.416331699 | 0.6771673 | 1 | 0.39518159 |
| ENSMUSG00000027454 | Gins1 | 69270 | protein_coding | 34.82824187 | 0.395348533 | 0.216258375 | -1.828130505 | 0.067529968 | 0.186051032 | 0.395348533 |
| ENSMUSG00000028896 | Rcc1 | 100088 | protein_coding | 487.3965379 | 0.395533607 | 0.087614832 | -4.514459472 | 6.35E-06 | 7.43E-05 | 0.395533607 |
| ENSMUSG00000038079 | Tmem237 | 381259 | protein_coding | 16.12294454 | 0.395606089 | 0.284716584 | -1.389473289 | 0.164688878 | 0.351288056 | 0.395606089 |
| ENSMUSG00000060560 | Ces4a | 234677 | protein_coding | 1.678301082 | -0.395697268 | 2.386219263 | 0.16582603 | 0.868293861 | 1 | 0.395697268 |
| ENSMUSG00000024097 | Srsf7 | 225027 | protein_coding | 1167.810585 | 0.395706161 | 0.080913553 | -4.890480621 | 1.01E-06 | 1.39E-05 | 0.395706161 |
| ENSMUSG00000059456 | Ptk2b | 19229 | protein_coding | 609.1502107 | -0.395746131 | 0.123374805 | 3.207673804 | 0.001338132 | 0.00811528 | 0.395746131 |
| ENSMUSG00000021388 | Aspn | 66695 | protein_coding | 2.907484934 | 0.39575539 | 0.598512719 | -0.661231378 | 0.508463941 | 1 | 0.39575539 |
| ENSMUSG00000020420 | Zfp607a | 545938 | protein_coding | 58.66899731 | -0.395795275 | 0.143248429 | 2.762999067 | 0.005727293 | 0.027217415 | 0.395795275 |
| ENSMUSG00000007836 | Hnrnpa0 | 77134 | protein_coding | 1506.71077 | 0.395797622 | 0.086070405 | -4.5985333 | 4.25E-06 | 5.17E-05 | 0.395797622 |
| ENSMUSG00000001175 | Calm1 | 12313 | protein_coding | 15731.1525 | 0.395861048 | 0.065376796 | -6.055069588 | 1.40E-09 | 3.04E-08 | 0.395861048 |
| ENSMUSG00000026311 | Asb1 | 65247 | protein_coding | 58.04103891 | 0.39592655 | 0.172493776 | -2.295309189 | 0.021715411 | 0.078315179 | 0.39592655 |
| ENSMUSG00000073158 | 9030624G23Rik | 66808 | protein_coding | 5.512831898 | -0.395966082 | 0.526778571 | 0.751674619 | 0.452246757 | 0.668637691 | 0.395966082 |
| ENSMUSG00000014873 | Surf2 | 20931 | protein_coding | 137.0924415 | -0.396054205 | 0.092460816 | 4.283481621 | 1.84E-05 | 0.000192306 | 0.396054205 |
| ENSMUSG00000068227 | Il2rb | 16185 | protein_coding | 62.39576062 | 0.396071968 | 0.36170581 | -1.095011351 | 0.273511663 | 0.489835334 | 0.396071968 |
| ENSMUSG00000050079 | Rspry1 | 67610 | protein_coding | 540.8335613 | 0.396072281 | 0.057802138 | -6.852208159 | 7.27E-12 | 2.06E-10 | 0.396072281 |
| ENSMUSG00000071984 | Fndc1 | 68655 | protein_coding | 5.816281746 | 0.396148735 | 1.313975189 | -0.301488748 | 0.763041829 | 0.881056689 | 0.396148735 |
| ENSMUSG00000018411 | Mapt | 17762 | protein_coding | 85.62385355 | 0.396353214 | 0.144871169 | -2.735901265 | 0.006220969 | 0.028930746 | 0.396353214 |
| ENSMUSG00000096377 | Mir5123 | 100628614 | miRNA | 0.995281493 | -0.396396717 | 1.121037778 | 0.353598001 | 0.723640173 | 1 | 0.396396717 |
| ENSMUSG00000062444 | Ap3b2 | 11775 | protein_coding | 7.511840001 | 0.396420627 | 0.762939063 | -0.519596711 | 0.603344691 | 0.779139367 | 0.396420627 |
| ENSMUSG00000065874 | Gm25683 | 115486588 | snoRNA | 0.974049656 | -0.396427707 | 1.117231813 | 0.354830307 | 0.722716722 | 1 | 0.396427707 |
| ENSMUSG00000034949 | Zfr2 | 103406 | protein_coding | 76.864209 | 0.396459958 | 0.283278589 | -1.399540852 | 0.161650857 | 0.346643585 | 0.396459958 |
| ENSMUSG00000042579 | 4632404H12Rik | 74034 | lncRNA | 51.42270263 | -0.396497277 | 0.176968657 | 2.240494358 | 0.025058847 | 0.087507252 | 0.396497277 |
| ENSMUSG00000030972 | Acsm5 | 272428 | protein_coding | 1.725627493 | -0.396527332 | 1.859917811 | 0.213196158 | 0.831173965 | 1 | 0.396527332 |
| ENSMUSG00000017311 | Pyy | 217212 | protein_coding | 345.769881 | -0.396566613 | 0.204263447 | 1.941446785 | 0.052204108 | 0.154304393 | 0.396566613 |
| ENSMUSG00000020364 | Zfp354a | 21408 | protein_coding | 12.25051881 | -0.396613925 | 0.336137231 | 1.179916677 | 0.238033356 | 0.448423779 | 0.396613925 |
| ENSMUSG00000028863 | Meaf6 | 70088 | protein_coding | 150.1787806 | -0.396618848 | 0.092651128 | 4.280777349 | 1.86E-05 | 0.000193921 | 0.396618848 |
| ENSMUSG00000033953 | Ppp3r1 | 19058 | protein_coding | 749.7106257 | 0.396721043 | 0.06642645 | -5.972335476 | 2.34E-09 | 4.95E-08 | 0.396721043 |

| ensembl_gene_id | external_gene_name | entrezgene_id | gene_biotype | baseMean | log2FoldChange | lfcSE | stat | pvalue | padj | absLog2FC |
|---|---|---|---|---|---|---|---|---|---|---|
| ENSMUSG00000060798 | Intu | 380614 | protein_coding | 112.3812884 | 0.396808192 | 0.136836407 | -2.899872935 | 0.00373314 | 0.019107412 | 0.396808192 |
| ENSMUSG00000111769 | Gm36278 | 102640141 | lncRNA | 0.768378386 | 0.396939462 | 1.472507019 | -0.26956711 | 0.787493306 | 1 | 0.396939462 |
| ENSMUSG00000049233 | Apoo-ps | 621156 | processed_pseudogene | 6.901797808 | 0.397031003 | 0.478765289 | -0.829281094 | 0.406945366 | 0.627713038 | 0.397031003 |
| ENSMUSG00000092274 | Neat1 | 66961 | lncRNA | 13206.44079 | 0.397126026 | 0.22271522 | -1.783111302 | 0.074568187 | 0.199760409 | 0.397126026 |
| ENSMUSG00000085315 | A430018G15Rik | 115489452 | lncRNA | 7.886794312 | -0.397192569 | 0.400034878 | 0.992894847 | 0.320761201 | 0.541897747 | 0.397192569 |
| ENSMUSG00000024524 | Gnal | 14680 | protein_coding | 47.16907468 | -0.397322709 | 0.291119363 | 1.364810314 | 0.172312676 | 0.36280925 | 0.397322709 |
| ENSMUSG00000027902 | Chil6 | 229688 | protein_coding | 8.715010541 | 0.397339685 | 0.407415855 | -0.975268097 | 0.329427291 | 0.551796561 | 0.397339685 |
| ENSMUSG00000035208 | Slfn8 | 276950 | protein_coding | 3.124181101 | 0.397544269 | 0.952009709 | -0.417584259 | 0.676251107 | 1 | 0.397544269 |
| ENSMUSG00000032577 | Mapkapk3 | 102626 | protein_coding | 987.4317593 | -0.39763091 | 0.115487936 | 3.4430515 | 0.00057519 | 0.003986019 | 0.39763091 |
| ENSMUSG00000027750 | Postn | 50706 | protein_coding | 3.863682583 | 0.397649441 | 0.920341255 | -0.432067387 | 0.665692437 | 1 | 0.397649441 |
| ENSMUSG00000001166 | Oas1c | 114643 | protein_coding | 34.15534306 | -0.397880752 | 0.228307651 | 1.742739462 | 0.081379135 | 0.212372794 | 0.397880752 |
| ENSMUSG00000052798 | Nup107 | 103468 | protein_coding | 214.6455589 | 0.397887859 | 0.148399479 | -2.681194456 | 0.007335987 | 0.03313635 | 0.397887859 |
| ENSMUSG00000024222 | Fkbp5 | 14229 | protein_coding | 2708.065445 | -0.398096238 | 0.146194793 | 2.723053473 | 0.006468159 | 0.02983307 | 0.398096238 |
| ENSMUSG00000109946 | Brd3os | 100504116 | protein_coding | 2.39458305 | 0.398122754 | 0.804121596 | -0.495102676 | 0.620527645 | 1 | 0.398122754 |
| ENSMUSG00000003949 | Hlf | 217082 | protein_coding | 2.558689088 | 0.398200479 | 1.559909591 | -0.255271511 | 0.798513402 | 1 | 0.398200479 |
| ENSMUSG00000028949 | Smarcd3 | 66993 | protein_coding | 8.194857215 | 0.398521749 | 0.420480794 | -0.947776343 | 0.343243326 | 0.565170522 | 0.398521749 |
| ENSMUSG00000037216 | Lipt1 | 623661 | protein_coding | 31.60448138 | -0.398561132 | 0.217195303 | 1.835035692 | 0.066500401 | 0.184012672 | 0.398561132 |
| ENSMUSG00000026135 | Zfp142 | 77264 | protein_coding | 156.6495373 | 0.39861677 | 0.114516158 | -3.480877972 | 0.000499773 | 0.00354826 | 0.39861677 |
| ENSMUSG00000020186 | Csrp2 | 13008 | protein_coding | 1896.143937 | -0.39862706 | 0.096284991 | 4.140074749 | 3.47E-05 | 0.000339062 | 0.39862706 |
| ENSMUSG00000034825 | Nrip3 | 78593 | protein_coding | 7.838650674 | -0.398691668 | 0.575733752 | 0.69249313 | 0.488627699 | 0.696497169 | 0.398691668 |
| ENSMUSG00000030259 | Rassf8 | 71323 | protein_coding | 20.17374288 | 0.398712208 | 0.286622098 | -1.391072812 | 0.164203348 | 0.350538094 | 0.398712208 |
| ENSMUSG00000060538 | Tmem219 | 68742 | protein_coding | 186.8231772 | -0.399006382 | 0.114608411 | 3.48147556 | 0.000498659 | 0.003541875 | 0.399006382 |
| ENSMUSG00000039958 | Etfbkmt | 320204 | protein_coding | 176.4759036 | -0.399071296 | 0.128862899 | 3.096867293 | 0.001955774 | 0.011144808 | 0.399071296 |
| ENSMUSG00000038598 | Shoc1 | 100155 | protein_coding | 12.30177602 | -0.399128144 | 0.393732486 | 1.013703868 | 0.310724078 | 0.531740666 | 0.399128144 |
| ENSMUSG00000024594 | Prrc1 | 73137 | protein_coding | 686.8998891 | 0.399186394 | 0.074909565 | -5.328910847 | 9.88E-08 | 1.63E-06 | 0.399186394 |
| ENSMUSG00000028860 | Sytl1 | 269589 | protein_coding | 706.0781379 | -0.399213284 | 0.160230023 | 2.491501137 | 0.012720456 | 0.05132314 | 0.399213284 |
| ENSMUSG00000048537 | Phldb1 | 102693 | protein_coding | 1117.032136 | -0.399220091 | 0.080169139 | 4.979722848 | 6.37E-07 | 9.17E-06 | 0.399220091 |
| ENSMUSG00000026097 | Ormdl1 | 227102 | protein_coding | 188.8633996 | 0.399332938 | 0.148641555 | -2.686549784 | 0.007219419 | 0.032744159 | 0.399332938 |
| ENSMUSG00000035781 | R3hdm4 | 109284 | protein_coding | 1349.272026 | -0.399492336 | 0.071942555 | 5.552935037 | 2.81E-08 | 5.04E-07 | 0.399492336 |
| ENSMUSG00000022516 | Nudt16l1 | 66911 | protein_coding | 148.0807693 | 0.399513685 | 0.114358981 | -3.49350512 | 0.000476724 | 0.003406585 | 0.399513685 |
| ENSMUSG00000029524 | Sirt4 | 75387 | protein_coding | 58.31240879 | -0.39954807 | 0.154085408 | 2.593029895 | 0.00951345 | 0.040902139 | 0.39954807 |
| ENSMUSG00000030834 | Abcc6 | 27421 | protein_coding | 142.6910011 | 0.399563256 | 0.183795 | -2.173961511 | 0.029708026 | 0.10016904 | 0.399563256 |
| ENSMUSG00000086915 | Gm16364 | 100504626 | lncRNA | 5.700380337 | 0.399576048 | 0.48242543 | -0.828264894 | 0.4075205 | 0.628274069 | 0.399576048 |
| ENSMUSG00000025089 | Gfra1 | 14585 | protein_coding | 2.936660048 | -0.399578926 | 0.949262761 | 0.42093606 | 0.673801773 | 1 | 0.399578926 |
| ENSMUSG00000031712 | Il15 | 16168 | protein_coding | 142.1420713 | -0.399597964 | 0.142675484 | 2.800747209 | 0.005098444 | 0.024770728 | 0.399597964 |
| ENSMUSG00000018819 | Lsp1 | 16985 | protein_coding | 19.08222659 | 0.399657582 | 0.446441339 | -0.895207381 | 0.37067624 | 0.593065878 | 0.399657582 |
| ENSMUSG00000058396 | Gpr182 | 11536 | protein_coding | 7.586931324 | 0.399749373 | 0.535648915 | -0.746289895 | 0.455492315 | 0.670888341 | 0.399749373 |
| ENSMUSG00000019768 | Esr1 | 13982 | protein_coding | 34.42371914 | -0.399840883 | 0.313326483 | 1.276115826 | 0.201914578 | 0.403926995 | 0.399840883 |
| ENSMUSG00000041375 | Ccdc9 | 243846 | protein_coding | 676.9065328 | -0.399862762 | 0.07069329 | 5.656304296 | 1.55E-08 | 2.89E-07 | 0.399862762 |

Supplementary Table 1

| ensembl_gene_id | external_gene_name | entrezgene_id | gene_biotype | baseMean | log2FoldChange | lfcSE | stat | pvalue | padj | absLog2FC |
|---|---|---|---|---|---|---|---|---|---|---|
| ENSMUSG00000112023 | Lilrb4b | 14727 | protein_coding | 2.329151448 | 0.399868314 | 1.047460365 | -0.381750305 | 0.702646583 | 1 | 0.399868314 |
| ENSMUSG00000026179 | Pnkd | 56695 | protein_coding | 226.3418164 | 0.399989846 | 0.114731487 | -3.486312741 | 0.000489728 | 0.003485935 | 0.399989846 |
| ENSMUSG00000053846 | Lipg | 16891 | protein_coding | 700.2904616 | -0.400148083 | 0.179962587 | 2.223507063 | 0.026181628 | 0.09052835 | 0.400148083 |
| ENSMUSG00000050304 | Slc25a2 | 83885 | protein_coding | 4.025802316 | -0.400246036 | 0.571937745 | 0.699806998 | 0.484047844 | 1 | 0.400246036 |
| ENSMUSG00000043760 | Pkhd1 | 241035 | protein_coding | 160.5985171 | 0.400502636 | 0.112221665 | -3.568853104 | 0.000358547 | 0.002670749 | 0.400502636 |
| ENSMUSG00000022227 | Mcpt1 | 17224 | protein_coding | 9.685281371 | -0.400502756 | 0.546720843 | 0.732554394 | 0.463830256 | 0.677309474 | 0.400502756 |
| ENSMUSG00000048992 | Prss32 | 69814 | protein_coding | 4928.401481 | -0.400520895 | 0.087428884 | 4.581104977 | 4.63E-06 | 5.57E-05 | 0.400520895 |
| ENSMUSG00000025497 | Cdhr5 | 72040 | protein_coding | 6413.323795 | -0.400697645 | 0.131953302 | 3.036662503 | 0.002392131 | 0.013111071 | 0.400697645 |
| ENSMUSG00000020681 | Ace | 11421 | protein_coding | 249.1719677 | -0.400844034 | 0.216991635 | 1.847278742 | 0.064706752 | 0.18053173 | 0.400844034 |
| ENSMUSG00000084861 | 1700095J07Rik | 115490402 | lncRNA | 1.412887162 | 0.400988745 | 1.69079589 | -0.237159758 | 0.812532854 | 1 | 0.400988745 |
| ENSMUSG00000023832 | Acat2 | 110460 | protein_coding | 335.9198837 | 0.401062355 | 0.137008272 | -2.927285695 | 0.003419346 | 0.017740208 | 0.401062355 |
| ENSMUSG00000049928 | Glp2r | 93896 | protein_coding | 5.298122 | 0.401155393 | 0.951747635 | -0.421493448 | 0.673394794 | 0.824897384 | 0.401155393 |
| ENSMUSG00000020846 | Rflnb | 76566 | protein_coding | 1347.030903 | -0.401277487 | 0.116579285 | 3.442099409 | 0.000577218 | 0.003996721 | 0.401277487 |
| ENSMUSG00000054405 | Dnajc8 | 68598 | protein_coding | 449.7354076 | -0.401302135 | 0.072999804 | 5.49730426 | 3.86E-08 | 6.75E-07 | 0.401302135 |
| ENSMUSG00000063698 | Sfxn4 | 94281 | protein_coding | 4.221702561 | 0.401390554 | 0.642797416 | -0.624443322 | 0.532336481 | 1 | 0.401390554 |
| ENSMUSG00000044786 | Zfp36 | 22695 | protein_coding | 3799.777969 | -0.401545645 | 0.195745304 | 2.051367958 | 0.040231127 | 0.126475696 | 0.401545645 |
| ENSMUSG00000022748 | Cmss1 | 66497 | protein_coding | 63.71296895 | 0.40157833 | 0.174808521 | -2.297246885 | 0.02160469 | 0.078035837 | 0.40157833 |
| ENSMUSG00000068551 | Zfp467 | 68910 | protein_coding | 323.7323645 | -0.401751524 | 0.103751791 | 3.872237022 | 0.000107841 | 0.000920831 | 0.401751524 |
| ENSMUSG00000002233 | Rhoc | 11853 | protein_coding | 1016.064078 | -0.401960489 | 0.187011002 | 2.149394876 | 0.031603111 | 0.105093391 | 0.401960489 |
| ENSMUSG00000028680 | Plk3 | 12795 | protein_coding | 1078.890588 | -0.401992116 | 0.202011085 | 1.989950775 | 0.046596358 | 0.141320893 | 0.401992116 |
| ENSMUSG00000051396 | Gm45902 | 119065974 | protein_coding | 51.59804981 | 0.402198045 | 0.175880354 | -2.286770723 | 0.022209209 | 0.07969641 | 0.402198045 |
| ENSMUSG00000022098 | Bmp1 | 12153 | protein_coding | 53.69668115 | 0.402348706 | 0.29627527 | -1.35802325 | 0.174456307 | 0.365696631 | 0.402348706 |
| ENSMUSG00000079355 | Ackr4 | 252837 | protein_coding | 45.57932078 | -0.40239054 | 0.191480541 | 2.10146962 | 0.035599762 | 0.114868009 | 0.40239054 |
| ENSMUSG00000097587 | 4930578M01Rik | 75951 | lncRNA | 4.731075024 | 0.402695269 | 0.62702278 | -0.642233874 | 0.520721342 | 0.720610174 | 0.402695269 |
| ENSMUSG00000032113 | Chek1 | 12649 | protein_coding | 63.50856503 | 0.402730711 | 0.259647628 | -1.551066398 | 0.120885773 | 0.283485892 | 0.402730711 |
| ENSMUSG00000094995 | Pate3 | 100312956 | protein_coding | 0.952377064 | 0.402746905 | 1.990264018 | -0.202358532 | 0.839636444 | 1 | 0.402746905 |
| ENSMUSG00000030366 | Ceacam12 | 67315 | protein_coding | 18.90920295 | -0.402780086 | 0.324916651 | 1.239641258 | 0.215108113 | 0.420139706 | 0.402780086 |
| ENSMUSG00000040613 | Apobec1 | 11810 | protein_coding | 369.0642877 | 0.402832415 | 0.205086996 | -1.964202616 | 0.0495066 | 0.147919664 | 0.402832415 |
| ENSMUSG00000021822 | Plau | 18792 | protein_coding | 1.567206475 | 0.402841882 | 2.041983928 | -0.197279654 | 0.8436087 | 1 | 0.402841882 |
| ENSMUSG00000087172 | Gm12153 | 102638460 | lncRNA | 1.257814786 | 0.402967328 | 1.826519097 | -0.220620375 | 0.825388035 | 1 | 0.402967328 |
| ENSMUSG00000062054 | Iah1 | 67732 | protein_coding | 297.1767974 | -0.403023626 | 0.091980062 | 4.381641159 | 1.18E-05 | 0.000129591 | 0.403023626 |
| ENSMUSG00000020190 | Mknk2 | 17347 | protein_coding | 2461.109709 | -0.403256502 | 0.075297813 | 5.355487573 | 8.53E-08 | 1.42E-06 | 0.403256502 |
| ENSMUSG00000032292 | Nr2e3 | 23958 | protein_coding | 50.96211015 | 0.403281118 | 0.214150567 | -1.883166236 | 0.059677843 | 0.170151917 | 0.403281118 |
| ENSMUSG00000002058 | Unc119 | 22248 | protein_coding | 267.3049594 | -0.403285452 | 0.199680738 | 2.019651253 | 0.043419575 | 0.133824348 | 0.403285452 |
| ENSMUSG00000086503 | Xist | 213742 | lncRNA | 4057.971602 | -0.403302884 | 0.143454529 | 2.811363898 | 0.004933196 | 0.024045689 | 0.403302884 |
| ENSMUSG00000036853 | Mcoln3 | 171166 | protein_coding | 9.452455448 | 0.403349639 | 0.5053395 | -0.798175561 | 0.424768618 | 0.64425321 | 0.403349639 |
| ENSMUSG00000064280 | Ccdc146 | 75172 | protein_coding | 3.487828517 | 0.403355433 | 0.917983417 | -0.43939294 | 0.660376841 | 1 | 0.403355433 |
| ENSMUSG00000027752 | Exosc8 | 69639 | protein_coding | 113.4935711 | 0.403362635 | 0.173966607 | -2.318621033 | 0.020415594 | 0.074589184 | 0.403362635 |
| ENSMUSG00000028188 | Spata1 | 70951 | protein_coding | 15.60441947 | -0.403372461 | 0.302367068 | 1.334048921 | 0.182187824 | 0.376665775 | 0.403372461 |

| ensembl_gene_id | external_gene_name | entrezgene_id | gene_biotype | baseMean | log2FoldChange | lfcSE | stat | pvalue | padj | absLog2FC |
|---|---|---|---|---|---|---|---|---|---|---|
| ENSMUSG00000093289 | Snord92 | 100217462 | snoRNA | 0.752580739 | 0.403406466 | 1.668387279 | -0.241794259 | 0.808939588 | 1 | 0.403406466 |
| ENSMUSG00000051650 | B3gnt2 | 53625 | protein_coding | 77.41885872 | 0.403989351 | 0.263517403 | -1.533065162 | 0.1252598 | 0.290432101 | 0.403989351 |
| ENSMUSG00000041481 | Serpina3g | 20715 | protein_coding | 5.025363542 | -0.404039213 | 0.625249616 | 0.646204657 | 0.518146824 | 0.718665364 | 0.404039213 |
| ENSMUSG00000004285 | Atp6v1f | 66144 | protein_coding | 524.0618 | -0.404148447 | 0.065963659 | 6.126834846 | 8.96E-10 | 1.98E-08 | 0.404148447 |
| ENSMUSG00000019487 | Trip10 | 106628 | protein_coding | 568.5023952 | -0.404301997 | 0.10760129 | 3.757408476 | 0.000171682 | 0.001385242 | 0.404301997 |
| ENSMUSG00000027997 | Casp6 | 12368 | protein_coding | 250.4280422 | -0.404337168 | 0.091093162 | 4.438721411 | 9.05E-06 | 0.00010271 | 0.404337168 |
| ENSMUSG00000061482 | H4c4 | 319156 | protein_coding | 3470.831971 | 0.404400776 | 0.129853507 | -3.11428459 | 0.001843915 | 0.010614744 | 0.404400776 |
| ENSMUSG00000000127 | Fer | 14158 | protein_coding | 219.4842831 | 0.404584724 | 0.093407587 | -4.331390403 | 1.48E-05 | 0.000157966 | 0.404584724 |
| ENSMUSG00000084941 | Gm11944 | 100302648 | lncRNA | 3.453327496 | -0.404594709 | 0.56658222 | 0.714097081 | 0.475167151 | 1 | 0.404594709 |
| ENSMUSG00000001168 | Oas1h | 246729 | protein_coding | 5.133349268 | 0.404649722 | 0.699438257 | -0.5785353 | 0.562902772 | 0.750688555 | 0.404649722 |
| ENSMUSG00000043019 | Edem3 | 66967 | protein_coding | 2164.056912 | 0.404739527 | 0.103494596 | -3.910731031 | 9.20E-05 | 0.00080104 | 0.404739527 |
| ENSMUSG00000054091 | 1810037I17Rik | 67704 | protein_coding | 844.9011446 | -0.40483404 | 0.086108759 | 4.701426967 | 2.58E-06 | 3.27E-05 | 0.40483404 |
| ENSMUSG00000033177 | Tmprss7 | 208171 | protein_coding | 20.49299184 | 0.40487886 | 0.301953397 | -1.340865395 | 0.179964159 | 0.373498571 | 0.40487886 |
| ENSMUSG00000020649 | Rrm2 | 20135 | protein_coding | 183.589983 | 0.404879921 | 0.21057928 | -1.922695912 | 0.05451825 | 0.159094255 | 0.404879921 |
| ENSMUSG00000061576 | Dpp6 | 13483 | protein_coding | 4.914617198 | 0.404882578 | 2.088390358 | -0.193873035 | 0.846275298 | 0.924222269 | 0.404882578 |
| ENSMUSG00000028013 | Ppa2 | 74776 | protein_coding | 868.1982255 | -0.405004667 | 0.074379137 | 5.44513804 | 5.18E-08 | 8.85E-07 | 0.405004667 |
| ENSMUSG00000046753 | Ccdc66 | 320234 | protein_coding | 151.0990289 | 0.405024683 | 0.090948786 | -4.453326978 | 8.45E-06 | 9.65E-05 | 0.405024683 |
| ENSMUSG00000063652 | Slc22a21 | 56517 | protein_coding | 119.7352017 | -0.405131841 | 0.165308836 | 2.450757318 | 0.014255603 | 0.056038331 | 0.405131841 |
| ENSMUSG00000031428 | Zcchc18 | 66995 | protein_coding | 3.211898331 | -0.405268059 | 0.717345937 | 0.564954839 | 0.572104482 | 1 | 0.405268059 |
| ENSMUSG00000036304 | Zdhhc23 | 332175 | protein_coding | 234.1547085 | -0.405300215 | 0.12323 | 3.288973598 | 0.001005535 | 0.006384236 | 0.405300215 |
| ENSMUSG00000021773 | Comtd1 | 69156 | protein_coding | 76.67169163 | -0.405307043 | 0.153474049 | 2.640883233 | 0.008269021 | 0.036560808 | 0.405307043 |
| ENSMUSG00000058470 | Gm8369 | 666926 | protein_coding | 3.362067935 | 0.405436363 | 1.133674183 | -0.35763041 | 0.720619917 | 1 | 0.405436363 |
| ENSMUSG00000058740 | Kcnt1 | 227632 | protein_coding | 6.90951613 | 0.405521822 | 1.095738023 | -0.370090125 | 0.711315339 | 0.849668102 | 0.405521822 |
| ENSMUSG00000061477 | Rps7 | 20115 | protein_coding | 3952.441881 | -0.405628197 | 0.133526485 | 3.037810799 | 0.002383035 | 0.013074234 | 0.405628197 |
| ENSMUSG00000054715 | Zscan22 | 232878 | protein_coding | 84.03098568 | 0.40564216 | 0.145803556 | -2.78211431 | 0.005400601 | 0.025963765 | 0.40564216 |
| ENSMUSG00000028167 | Bdh2 | 69772 | protein_coding | 2.589970441 | 0.405653735 | 1.05613847 | -0.38409143 | 0.700910683 | 1 | 0.405653735 |
| ENSMUSG00000029334 | Prkg2 | 19092 | protein_coding | 608.8810466 | -0.405721418 | 0.184087739 | 2.203956769 | 0.027527385 | 0.0942345 | 0.405721418 |
| ENSMUSG00000021567 | Nkd2 | 72293 | protein_coding | 5.314085686 | 0.405792032 | 0.773224867 | -0.524804684 | 0.599718975 | 0.776544903 | 0.405792032 |
| ENSMUSG00000045980 | Tmem104 | 320534 | protein_coding | 101.6701797 | 0.405795093 | 0.15950395 | -2.544106866 | 0.010955756 | 0.045710802 | 0.405795093 |
| ENSMUSG00000034282 | Evpl | 14027 | protein_coding | 920.6237005 | -0.405824624 | 0.146163571 | 2.776510051 | 0.005494594 | 0.026309923 | 0.405824624 |
| ENSMUSG00000032498 | Mlh1 | 17350 | protein_coding | 255.2824509 | 0.405876855 | 0.080714162 | -5.028570535 | 4.94E-07 | 7.31E-06 | 0.405876855 |
| ENSMUSG00000078444 | Gm10941 | 100169878 | lncRNA | 1.812385002 | -0.405931714 | 0.754178395 | 0.538243626 | 0.590408865 | 1 | 0.405931714 |
| ENSMUSG00000090213 | Peds1 | 407243 | protein_coding | 553.679367 | 0.406011041 | 0.114949217 | -3.532090524 | 0.000412288 | 0.00301129 | 0.406011041 |
| ENSMUSG00000025129 | Ppp1r27 | 68701 | protein_coding | 3.31974573 | 0.406077082 | 0.893836284 | -0.45430812 | 0.649607073 | 1 | 0.406077082 |
| ENSMUSG00000063015 | Ccni | 12453 | protein_coding | 2077.841335 | -0.406089434 | 0.08931016 | 4.546956725 | 5.44E-06 | 6.46E-05 | 0.406089434 |
| ENSMUSG00000027200 | Sema6d | 214968 | protein_coding | 690.2369689 | 0.406164491 | 0.148026232 | -2.743868337 | 0.006071989 | 0.028422001 | 0.406164491 |
| ENSMUSG00000027746 | Ufm1 | 67890 | protein_coding | 733.3893813 | 0.406175366 | 0.084569504 | -4.80285858 | 1.56E-06 | 2.07E-05 | 0.406175366 |
| ENSMUSG00000029171 | Pgm2 | 66681 | protein_coding | 897.7788634 | -0.40619376 | 0.058245407 | 6.973833322 | 3.08E-12 | 9.01E-11 | 0.40619376 |
| ENSMUSG00000037016 | Frem2 | 242022 | protein_coding | 25.77171125 | 0.406228867 | 0.30508991 | -1.331505414 | 0.183022765 | 0.377733062 | 0.406228867 |

| ensembl_gene_id | external_gene_name | entrezgene_id | gene_biotype | baseMean | log2FoldChange | lfcSE | stat | pvalue | padj | absLog2FC |
|---|---|---|---|---|---|---|---|---|---|---|
| ENSMUSG00000037593 | Rskr | 216976 | protein_coding | 7.20426931 | 0.40631249 | 0.561792683 | -0.723242759 | 0.469530753 | 0.683156904 | 0.40631249 |
| ENSMUSG00000109032 | Gm7972 | 666190 | transcribed_unprocessed_pseudogene | 5.979304134 | 0.406331812 | 0.55075975 | -0.737765988 | 0.460656665 | 0.674823398 | 0.406331812 |
| ENSMUSG00000051256 | Jagn1 | 67767 | protein_coding | 314.6824037 | 0.406474007 | 0.087309228 | -4.655567513 | 3.23E-06 | 4.01E-05 | 0.406474007 |
| ENSMUSG00000031996 | Aplp2 | 11804 | protein_coding | 2189.793415 | -0.406533732 | 0.106169768 | 3.829091287 | 0.000128617 | 0.001078713 | 0.406533732 |
| ENSMUSG00000038020 | Rapgefl1 | 268480 | protein_coding | 875.0866052 | -0.406588354 | 0.123029918 | 3.30479253 | 0.000950467 | 0.006100279 | 0.406588354 |
| ENSMUSG00000027356 | Fermt1 | 241639 | protein_coding | 2353.9896 | 0.406625197 | 0.078035554 | -5.210768346 | 1.88E-07 | 2.96E-06 | 0.406625197 |
| ENSMUSG00000017607 | Tns4 | 217169 | protein_coding | 1735.160878 | -0.406723963 | 0.198294357 | 2.051112136 | 0.040256028 | 0.12652989 | 0.406723963 |
| ENSMUSG00000036098 | Myrf | 225908 | protein_coding | 8.283474487 | 0.406730402 | 0.900810977 | -0.451515815 | 0.65161783 | 0.810851996 | 0.406730402 |
| ENSMUSG00000037681 | Esyt3 | 272636 | protein_coding | 6.672380396 | -0.406733526 | 0.813012391 | 0.500279615 | 0.616878206 | 0.788881899 | 0.406733526 |
| ENSMUSG00000069814 | Ccdc92b | 432582 | protein_coding | 20.71704413 | 0.407043986 | 0.306561016 | -1.327774784 | 0.18425252 | 0.379300188 | 0.407043986 |
| ENSMUSG00000053205 | Styx | 56291 | protein_coding | 232.9961024 | -0.407172245 | 0.15788648 | 2.578892409 | 0.009911765 | 0.042284395 | 0.407172245 |
| ENSMUSG00000005320 | Fgfr4 | 14186 | protein_coding | 273.682636 | 0.407225331 | 0.081289309 | -5.009580394 | 5.45E-07 | 7.99E-06 | 0.407225331 |
| ENSMUSG00000009621 | Vav2 | 22325 | protein_coding | 243.4076303 | -0.407274156 | 0.199697567 | 2.039454768 | 0.041404661 | 0.129327894 | 0.407274156 |
| ENSMUSG00000046242 | Nme9 | 69797 | protein_coding | 4.869880917 | -0.407314113 | 0.660506806 | 0.616669062 | 0.537453031 | 0.732906636 | 0.407314113 |
| ENSMUSG00000030806 | Stx1b | 56216 | protein_coding | 4.475847153 | -0.407314907 | 1.372814384 | 0.29670064 | 0.766695068 | 0.882867468 | 0.407314907 |
| ENSMUSG00000056076 | Eif3b | 27979 | protein_coding | 1597.762711 | 0.407328055 | 0.072653084 | -5.60647988 | 2.06E-08 | 3.79E-07 | 0.407328055 |
| ENSMUSG00000041343 | Ankrd42 | 73845 | protein_coding | 30.36852708 | 0.40734954 | 0.196270962 | -2.075444762 | 0.037945333 | 0.120924206 | 0.40734954 |
| ENSMUSG00000079396 | Gm3411 | 100861615 | protein_coding | 2.839144112 | 0.407412464 | 0.826911838 | -0.492691537 | 0.622230559 | 1 | 0.407412464 |
| ENSMUSG00000032870 | Smap2 | 69780 | protein_coding | 298.0693265 | 0.407493644 | 0.118567541 | -3.436806058 | 0.000588617 | 0.00406542 | 0.407493644 |
| ENSMUSG00000033669 | Zfp7 | 223669 | protein_coding | 255.1219643 | -0.407567321 | 0.123962596 | 3.287825 | 0.001009646 | 0.006402953 | 0.407567321 |
| ENSMUSG00000056598 | Drc3 | 74665 | protein_coding | 26.71722158 | 0.407597803 | 0.221643544 | -1.838978914 | 0.065918288 | 0.182830803 | 0.407597803 |
| ENSMUSG00000029413 | Naaa | 67111 | protein_coding | 193.3425136 | -0.407670322 | 0.152513273 | 2.673015371 | 0.007517281 | 0.033825714 | 0.407670322 |
| ENSMUSG00000058756 | Thra | 21833 | protein_coding | 528.8281767 | -0.407737595 | 0.110869363 | 3.677639923 | 0.000235402 | 0.001828517 | 0.407737595 |
| ENSMUSG00000026728 | Vim | 22352 | protein_coding | 72.31679523 | 0.407864367 | 0.157756916 | -2.585397691 | 0.009726673 | 0.0416346 | 0.407864367 |
| ENSMUSG00000040997 | Abhd4 | 105501 | protein_coding | 591.9989339 | -0.407917768 | 0.063376493 | 6.436420676 | 1.22E-10 | 3.00E-09 | 0.407917768 |
| ENSMUSG00000044252 | Osbpl1a | 64291 | protein_coding | 121.5594616 | -0.407951259 | 0.091662485 | 4.450580411 | 8.56E-06 | 9.76E-05 | 0.407951259 |
| ENSMUSG00000048206 | Dnajb8 | 56691 | protein_coding | 1.294606934 | -0.407956891 | 2.296413759 | 0.177649559 | 0.858998201 | 1 | 0.407956891 |
| ENSMUSG00000044712 | Slc38a6 | 625098 | protein_coding | 57.12879755 | 0.407959031 | 0.149298562 | -2.732504764 | 0.006285476 | 0.029148653 | 0.407959031 |
| ENSMUSG00000111709 | Gm3776 | 100042295 | protein_coding | 7.982764519 | 0.407966089 | 0.480516653 | -0.849015505 | 0.395872665 | 0.618256212 | 0.407966089 |
| ENSMUSG00000038578 | Susd1 | 634731 | protein_coding | 19.89475593 | 0.408038979 | 0.375816737 | -1.085739242 | 0.277594374 | 0.494519255 | 0.408038979 |
| ENSMUSG00000059277 | R74862 | 97423 | lncRNA | 7.770301938 | 0.408302088 | 0.619909911 | -0.658647459 | 0.51012218 | 0.713367573 | 0.408302088 |
| ENSMUSG00000079299 | Klrb1 | 100043861 | protein_coding | 5.286382301 | -0.408377706 | 0.57141929 | 0.714672594 | 0.474811376 | 0.686785939 | 0.408377706 |
| ENSMUSG00000020738 | Sumo2 | 170930 | protein_coding | 126.5173315 | -0.408389726 | 0.22888763 | 1.784236773 | 0.074385199 | 0.19938276 | 0.408389726 |
| ENSMUSG00000077526 | Gm25379 | 115485641 | snoRNA | 2.571234655 | 0.408473279 | 0.620636136 | -0.658152588 | 0.510440087 | 1 | 0.408473279 |
| ENSMUSG00000029228 | Lnx1 | 16924 | protein_coding | 310.3765878 | -0.408532995 | 0.159151975 | 2.566936378 | 0.010260144 | 0.043456274 | 0.408532995 |
| ENSMUSG00000063410 | Stk24 | 223255 | protein_coding | 866.4352967 | 0.408569297 | 0.063328102 | -6.45162709 | 1.11E-10 | 2.72E-09 | 0.408569297 |
| ENSMUSG00000055480 | Zfp458 | 238690 | protein_coding | 36.92975604 | -0.408588786 | 0.180840306 | 2.259390033 | 0.023859134 | 0.084280157 | 0.408588786 |
| ENSMUSG00000097207 | 6030443J06Rik | 320719 | lncRNA | 16.13184403 | -0.408634798 | 0.37558665 | 1.087990741 | 0.276599195 | 0.493582047 | 0.408634798 |
| ENSMUSG00000032939 | Nup93 | 71805 | protein_coding | 312.7117105 | 0.408779329 | 0.108404648 | -3.770865313 | 0.000162682 | 0.00131693 | 0.408779329 |

| ensembl_gene_id | external_gene_name | entrezgene_id | gene_biotype | baseMean | log2FoldChange | lfcSE | stat | pvalue | padj | absLog2FC |
|---|---|---|---|---|---|---|---|---|---|---|
| ENSMUSG00000073758 | Sh3d21 | 66938 | protein_coding | 538.27075 | -0.408800062 | 0.204380557 | 2.000190565 | 0.04547969 | 0.13890357 | 0.408800062 |
| ENSMUSG00000025466 | Fuom | 69064 | protein_coding | 289.2664188 | -0.408809043 | 0.101795932 | 4.015966385 | 5.92E-05 | 0.000543756 | 0.408809043 |
| ENSMUSG00000022149 | C9 | 12279 | protein_coding | 6.463641515 | 0.408835041 | 0.461864913 | -0.885183155 | 0.376057856 | 0.598287036 | 0.408835041 |
| ENSMUSG00000097571 | Jpx | 70252 | lncRNA | 26.25287773 | -0.408847483 | 0.2877082 | 1.421049114 | 0.155302482 | 0.337678102 | 0.408847483 |
| ENSMUSG00000025272 | Tro | 56191 | protein_coding | 2.658784496 | -0.408967407 | 1.125940048 | 0.363223075 | 0.716438251 | 1 | 0.408967407 |
| ENSMUSG00000002871 | Tpra1 | 24100 | protein_coding | 320.8771257 | -0.409121075 | 0.057208934 | 7.151349418 | 8.59E-13 | 2.62E-11 | 0.409121075 |
| ENSMUSG00000026959 | Grin1 | 14810 | protein_coding | 35.56391231 | 0.409139368 | 0.243016723 | -1.683585236 | 0.092261854 | 0.232919564 | 0.409139368 |
| ENSMUSG00000055485 | Soga1 | 320706 | protein_coding | 107.1175943 | 0.40920219 | 0.288661801 | -1.417583444 | 0.156312417 | 0.339025906 | 0.40920219 |
| ENSMUSG00000051483 | Cbr1 | 12408 | protein_coding | 2514.504006 | -0.409425473 | 0.107979608 | 3.791692528 | 0.000149624 | 0.001225641 | 0.409425473 |
| ENSMUSG00000034593 | Myo5a | 17918 | protein_coding | 22.87853839 | 0.409445697 | 0.281208895 | -1.456019719 | 0.145387168 | 0.322533618 | 0.409445697 |
| ENSMUSG00000058317 | Ube2e2 | 218793 | protein_coding | 82.19895282 | 0.409561494 | 0.124686859 | -3.284720607 | 0.001020835 | 0.006462165 | 0.409561494 |
| ENSMUSG00000019864 | Rtn4ip1 | 170728 | protein_coding | 226.3461736 | -0.409615093 | 0.09171209 | 4.466315114 | 7.96E-06 | 9.14E-05 | 0.409615093 |
| ENSMUSG00000022479 | Vdr | 22337 | protein_coding | 3216.802767 | -0.409628891 | 0.086778244 | 4.720410019 | 2.35E-06 | 3.01E-05 | 0.409628891 |
| ENSMUSG00000024712 | Rfk | 54391 | protein_coding | 4689.976775 | -0.409658522 | 0.085233761 | 4.806294112 | 1.54E-06 | 2.04E-05 | 0.409658522 |
| ENSMUSG00000078570 | 1110065P20Rik | 68920 | protein_coding | 88.30916753 | 0.40975405 | 0.147609837 | -2.775926453 | 0.005504466 | 0.026340408 | 0.40975405 |
| ENSMUSG00000049908 | Gja8 | 14616 | protein_coding | 3.122585891 | 0.409802034 | 1.00832342 | -0.406419236 | 0.684434588 | 1 | 0.409802034 |
| ENSMUSG00000042246 | Tmc7 | 209760 | protein_coding | 107.6433223 | -0.409809071 | 0.211170953 | 1.940650765 | 0.052300652 | 0.154451532 | 0.409809071 |
| ENSMUSG00000036585 | Fgf1 | 14164 | protein_coding | 433.1556149 | 0.409908226 | 0.165732343 | -2.473314615 | 0.013386624 | 0.053436476 | 0.409908226 |
| ENSMUSG00000031133 | Arhgef6 | 73341 | protein_coding | 9.141096346 | 0.410173008 | 0.573271993 | -0.715494586 | 0.474303485 | 0.686291751 | 0.410173008 |
| ENSMUSG00000107121 | 1110006O24Rik | 66123 | lncRNA | 6.582179506 | 0.410185365 | 0.566136321 | -0.724534621 | 0.468737581 | 0.682320284 | 0.410185365 |
| ENSMUSG00000084989 | Crocc2 | 381284 | protein_coding | 6.341726365 | 0.410272984 | 0.566464978 | -0.72426893 | 0.468900648 | 0.682480636 | 0.410272984 |
| ENSMUSG00000041548 | Hspb8 | 80888 | protein_coding | 3.511719659 | -0.410420414 | 1.065871536 | 0.385056173 | 0.700195798 | 1 | 0.410420414 |
| ENSMUSG00000025736 | Jmjd8 | 72106 | protein_coding | 247.7908566 | 0.410494182 | 0.108652279 | -3.778054022 | 0.000158059 | 0.001283905 | 0.410494182 |
| ENSMUSG00000097637 | 4933417D19Rik | 71186 | lncRNA | 4.587657589 | -0.41064561 | 0.522979733 | 0.785203678 | 0.43233416 | 0.650767142 | 0.41064561 |
| ENSMUSG00000075553 | Gm5464 | 432870 | lncRNA | 3.988332135 | -0.410709652 | 0.59286303 | 0.69275639 | 0.488462445 | 1 | 0.410709652 |
| ENSMUSG00000036095 | Dgkb | 217480 | protein_coding | 3.538448214 | 0.410747998 | 2.041793829 | -0.201170163 | 0.840565521 | 1 | 0.410747998 |
| ENSMUSG00000015599 | Ttbk1 | 106763 | protein_coding | 7.701822479 | 0.410928984 | 1.335143342 | -0.307778926 | 0.758250558 | 0.878808271 | 0.410928984 |
| ENSMUSG00000061878 | Sphk1 | 20698 | protein_coding | 3.218030528 | -0.410966841 | 0.87838688 | 0.467865414 | 0.639880838 | 1 | 0.410966841 |
| ENSMUSG00000001288 | Rarg | 19411 | protein_coding | 135.8272374 | -0.411080453 | 0.155729658 | 2.639705617 | 0.008297807 | 0.036642767 | 0.411080453 |
| ENSMUSG00000041774 | Ydjc | 69101 | protein_coding | 343.0547874 | 0.411269552 | 0.09573773 | -4.295793867 | 1.74E-05 | 0.000182282 | 0.411269552 |
| ENSMUSG00000036412 | Arsi | 545260 | protein_coding | 1.347292927 | 0.411292248 | 1.663868634 | -0.247190337 | 0.804760923 | 1 | 0.411292248 |
| ENSMUSG00000029103 | Lrpap1 | 16976 | protein_coding | 567.94931 | -0.411619941 | 0.082169266 | 5.009414835 | 5.46E-07 | 7.99E-06 | 0.411619941 |
| ENSMUSG00000063172 | Hspb11 | 72938 | protein_coding | 108.0959343 | -0.411689647 | 0.140572439 | 2.928665463 | 0.003404206 | 0.017689445 | 0.411689647 |
| ENSMUSG00000037805 | Rpl10a | 19896 | protein_coding | 2970.473524 | -0.411722483 | 0.126527478 | 3.254016349 | 0.001137857 | 0.007085436 | 0.411722483 |
| ENSMUSG00000068748 | Ptprz1 | 19283 | protein_coding | 3.162340113 | 0.411786625 | 2.415064787 | -0.170507486 | 0.864611048 | 1 | 0.411786625 |
| ENSMUSG00000100164 | 2610306M01Rik | 67170 | lncRNA | 31.91827091 | -0.411824729 | 0.216635835 | 1.901000028 | 0.057302009 | 0.165147848 | 0.411824729 |
| ENSMUSG00000049764 | Zfp280b | 64453 | protein_coding | 228.0917877 | 0.411904259 | 0.091822578 | -4.485871232 | 7.26E-06 | 8.40E-05 | 0.411904259 |
| ENSMUSG00000062590 | Armc9 | 78795 | protein_coding | 82.94303056 | 0.412267579 | 0.165268941 | -2.494525445 | 0.012612568 | 0.051024976 | 0.412267579 |
| ENSMUSG00000031026 | Trim66 | 330627 | protein_coding | 5.312342899 | -0.412327019 | 1.490681665 | 0.276602999 | 0.782084962 | 0.890758364 | 0.412327019 |

| ensembl_gene_id | external_gene_name | entrezgene_id | gene_biotype | baseMean | log2FoldChange | lfcSE | stat | pvalue | padj | absLog2FC |
|---|---|---|---|---|---|---|---|---|---|---|
| ENSMUSG00000030602 | Pak4 | 70584 | protein_coding | 1462.60188 | 0.412332765 | 0.062335032 | -6.614783918 | 3.72E-11 | 9.85E-10 | 0.412332765 |
| ENSMUSG00000064127 | Med14 | 26896 | protein_coding | 562.5313857 | 0.412353565 | 0.101655414 | -4.056385674 | 4.98E-05 | 0.000465269 | 0.412353565 |
| ENSMUSG00000039168 | Dap | 223453 | protein_coding | 490.0122428 | 0.412401096 | 0.079331371 | -5.198461722 | 2.01E-07 | 3.15E-06 | 0.412401096 |
| ENSMUSG00000034918 | Cdhr2 | 268663 | protein_coding | 7592.048941 | -0.412417106 | 0.126379394 | 3.26332556 | 0.001101129 | 0.006903589 | 0.412417106 |
| ENSMUSG00000037072 | Selenof | 93684 | protein_coding | 2022.303617 | 0.412440072 | 0.121187083 | -3.403333597 | 0.000665689 | 0.004503147 | 0.412440072 |
| ENSMUSG00000021180 | Rps6ka5 | 73086 | protein_coding | 44.9060221 | 0.412441705 | 0.198025491 | -2.082770772 | 0.037272123 | 0.119192842 | 0.412441705 |
| ENSMUSG00000020422 | Tns3 | 319939 | protein_coding | 1689.31868 | -0.412545758 | 0.070033731 | 5.890672302 | 3.85E-09 | 7.89E-08 | 0.412545758 |
| ENSMUSG00000031767 | Nudt7 | 67528 | protein_coding | 217.7972619 | -0.412576771 | 0.088569625 | 4.658219702 | 3.19E-06 | 3.96E-05 | 0.412576771 |
| ENSMUSG00000089715 | Cbx6 | 494448 | protein_coding | 82.99351455 | 0.412682315 | 0.219098283 | -1.88354883 | 0.059626029 | 0.170033542 | 0.412682315 |
| ENSMUSG00000031828 | Klhl36 | 234796 | protein_coding | 68.32190722 | -0.41268764 | 0.126764307 | 3.255550793 | 0.001131727 | 0.00706591 | 0.41268764 |
| ENSMUSG00000029162 | Khk | 16548 | protein_coding | 134.4899403 | -0.412748484 | 0.127711538 | 3.231880938 | 0.001229783 | 0.007552486 | 0.412748484 |
| ENSMUSG00000033565 | Rbfox2 | 93686 | protein_coding | 361.2651183 | 0.412781419 | 0.071794693 | -5.749469794 | 8.95E-09 | 1.74E-07 | 0.412781419 |
| ENSMUSG00000030704 | Rab6a | 19346 | protein_coding | 1432.979333 | 0.412851889 | 0.06305605 | -6.547379499 | 5.86E-11 | 1.52E-09 | 0.412851889 |
| ENSMUSG00000062833 | Sval3 | 387564 | protein_coding | 4.790788896 | 0.4128569 | 0.687174612 | -0.600803483 | 0.547970884 | 0.739252608 | 0.4128569 |
| ENSMUSG00000082361 | Btc | 12223 | protein_coding | 63.08619396 | -0.412942233 | 0.186326413 | 2.216230258 | 0.026675737 | 0.091871348 | 0.412942233 |
| ENSMUSG00000014301 | Pam16 | 66449 | protein_coding | 87.09081804 | -0.413026042 | 0.137173338 | 3.010979014 | 0.002604068 | 0.014104161 | 0.413026042 |
| ENSMUSG00000024304 | Cdh2 | 12558 | protein_coding | 6.464159667 | 0.413066099 | 0.719828952 | -0.573839241 | 0.566076592 | 0.752646336 | 0.413066099 |
| ENSMUSG00000099354 | 1700124L16Rik | 67934 | lncRNA | 0.685650798 | -0.413123186 | 1.39877487 | 0.295346445 | 0.76772924 | 1 | 0.413123186 |
| ENSMUSG00000054753 | AU018091 | 245128 | protein_coding | 2.392429008 | 0.413282614 | 2.1753595 | -0.189983593 | 0.849321987 | 1 | 0.413282614 |
| ENSMUSG00000029736 | Nobox | 18291 | protein_coding | 0.833255486 | 0.413340268 | 1.692946202 | -0.244154402 | 0.80711124 | 1 | 0.413340268 |
| ENSMUSG00000000296 | Tpd52l1 | 21987 | protein_coding | 78.33771218 | 0.413547683 | 0.190720977 | -2.168338735 | 0.030132923 | 0.101319853 | 0.413547683 |
| ENSMUSG00000042426 | Dhx29 | 218629 | protein_coding | 249.0460981 | 0.413667927 | 0.120111257 | -3.444039626 | 0.000573092 | 0.003974816 | 0.413667927 |
| ENSMUSG00000097113 | Gm19705 | 100503460 | lncRNA | 2.616431859 | 0.413701185 | 1.006424355 | -0.411060387 | 0.681028256 | 1 | 0.413701185 |
| ENSMUSG00000019773 | Fbxo5 | 67141 | protein_coding | 97.9753439 | 0.413797233 | 0.241405237 | -1.714118708 | 0.086506927 | 0.222484875 | 0.413797233 |
| ENSMUSG00000027854 | Sike1 | 66641 | protein_coding | 585.5447606 | -0.413898966 | 0.130931177 | 3.161194878 | 0.001571233 | 0.00926691 | 0.413898966 |
| ENSMUSG00000005374 | Tbl2 | 27368 | protein_coding | 379.9095726 | 0.413921531 | 0.112253463 | -3.687383161 | 0.000226572 | 0.001766577 | 0.413921531 |
| ENSMUSG00000020101 | Vsir | 74048 | protein_coding | 16.64102361 | 0.414086236 | 0.437578318 | -0.946313424 | 0.343988749 | 0.56579982 | 0.414086236 |
| ENSMUSG00000031805 | Jak3 | 16453 | protein_coding | 181.3281232 | 0.414112329 | 0.12827471 | -3.228324037 | 0.001245178 | 0.00763001 | 0.414112329 |
| ENSMUSG00000061039 | Or8b53 | 258783 | protein_coding | 3.838933201 | -0.414137856 | 0.800215186 | 0.517533112 | 0.60478406 | 1 | 0.414137856 |
| ENSMUSG00000117797 | 2900057B20Rik | 73009 | lncRNA | 5.78138608 | 0.414138901 | 0.43057094 | -0.961836628 | 0.336131676 | 0.558663295 | 0.414138901 |
| ENSMUSG00000032555 | Topbp1 | 235559 | protein_coding | 694.566745 | 0.414152072 | 0.146765096 | -2.821870341 | 0.004774448 | 0.02338927 | 0.414152072 |
| ENSMUSG00000086287 | Gm15972 | 100502834 | lncRNA | 13.3700495 | -0.414267303 | 0.317792167 | 1.303579336 | 0.19237705 | 0.390140563 | 0.414267303 |
| ENSMUSG00000021482 | Prxl2c | 66129 | protein_coding | 448.7733581 | -0.414276826 | 0.129844445 | 3.190562569 | 0.001419961 | 0.008514609 | 0.414276826 |
| ENSMUSG00000024339 | Tap2 | 21355 | protein_coding | 217.4976814 | -0.414388914 | 0.092837232 | 4.463606945 | 8.06E-06 | 9.24E-05 | 0.414388914 |
| ENSMUSG00000002324 | Rec8 | 56739 | protein_coding | 48.38913692 | 0.414504439 | 0.333245067 | -1.243842684 | 0.213557467 | 0.418548305 | 0.414504439 |
| ENSMUSG00000004032 | Gstm5 | 14866 | protein_coding | 346.8949509 | 0.414606999 | 0.122686725 | -3.379395768 | 0.000726453 | 0.004833462 | 0.414606999 |
| ENSMUSG00000041477 | Dcp1b | 319618 | protein_coding | 148.6414247 | -0.41463763 | 0.163986875 | 2.528480592 | 0.011455742 | 0.047449241 | 0.41463763 |
| ENSMUSG00000029541 | Cyp2w1 | 545817 | protein_coding | 86.99935248 | 0.414683309 | 0.288060409 | -1.439570644 | 0.14998891 | 0.329684129 | 0.414683309 |
| ENSMUSG00000038831 | Ralgps1 | 241308 | protein_coding | 161.0108101 | 0.414733361 | 0.105608655 | -3.92707737 | 8.60E-05 | 0.000754888 | 0.414733361 |

| ensembl_gene_id | external_gene_name | entrezgene_id | gene_biotype | baseMean | log2FoldChange | lfcSE | stat | pvalue | padj | absLog2FC |
|---|---|---|---|---|---|---|---|---|---|---|
| ENSMUSG00000036887 | C1qa | 12259 | protein_coding | 5.829170395 | 0.414858017 | 0.515574654 | -0.804651691 | 0.421020704 | 0.640805153 | 0.414858017 |
| ENSMUSG00000022483 | Col2a1 | 12824 | protein_coding | 8.296944246 | 0.414867241 | 1.33311189 | -0.311202116 | 0.755646975 | 0.877241064 | 0.414867241 |
| ENSMUSG00000006720 | Zfp184 | 193452 | protein_coding | 14.12924868 | 0.415103248 | 0.311607123 | -1.332136583 | 0.182815311 | 0.377565289 | 0.415103248 |
| ENSMUSG00000106193 | Mir7090 | 102465661 | miRNA | 0.821431154 | -0.415127297 | 1.133423718 | 0.366259582 | 0.714171386 | 1 | 0.415127297 |
| ENSMUSG00000026708 | Cenpl | 70454 | protein_coding | 90.95224322 | 0.415274284 | 0.176226256 | -2.356483614 | 0.018448882 | 0.068863768 | 0.415274284 |
| ENSMUSG00000032637 | Atxn2l | 233871 | protein_coding | 676.6525257 | 0.415279316 | 0.161402603 | -2.57294064 | 0.01008385 | 0.042874537 | 0.415279316 |
| ENSMUSG00000055067 | Smyd3 | 69726 | protein_coding | 100.4989981 | -0.415305598 | 0.132629476 | 3.131322012 | 0.001740212 | 0.010094213 | 0.415305598 |
| ENSMUSG00000016481 | Cr1l | 12946 | protein_coding | 753.9549658 | -0.41540833 | 0.095546314 | 4.347716977 | 1.38E-05 | 0.00014838 | 0.41540833 |
| ENSMUSG00000029163 | Emilin1 | 100952 | protein_coding | 32.71842405 | 0.415574175 | 0.220588394 | -1.883934905 | 0.059573781 | 0.169937908 | 0.415574175 |
| ENSMUSG00000030083 | Abtb1 | 80283 | protein_coding | 260.8181473 | -0.415717512 | 0.11156283 | 3.726308427 | 0.000194305 | 0.001541925 | 0.415717512 |
| ENSMUSG00000038612 | Mcl1 | 17210 | protein_coding | 4154.089562 | -0.415719009 | 0.083076759 | 5.004034981 | 5.61E-07 | 8.20E-06 | 0.415719009 |
| ENSMUSG00000034209 | Rasl10a | 75668 | protein_coding | 3.444711318 | -0.415752372 | 0.583815028 | 0.712130302 | 0.476384092 | 1 | 0.415752372 |
| ENSMUSG00000063730 | Hsd3b2 | 15493 | protein_coding | 341.619233 | -0.415919087 | 0.169175627 | 2.458504773 | 0.013951692 | 0.055106014 | 0.415919087 |
| ENSMUSG00000078970 | Dnaaf10 | 103784 | protein_coding | 241.0445353 | 0.41601399 | 0.126122728 | -3.298485501 | 0.000972079 | 0.00621724 | 0.41601399 |
| ENSMUSG00000042262 | Ccr8 | 12776 | protein_coding | 0.835391382 | -0.416086163 | 1.616693917 | 0.257368546 | 0.796894282 | 1 | 0.416086163 |
| ENSMUSG00000003863 | Ppfia3 | 76787 | protein_coding | 353.5353585 | -0.416127215 | 0.237463575 | 1.752383352 | 0.079707912 | 0.209275846 | 0.416127215 |
| ENSMUSG00000090667 | Mdfic2 | 330390 | protein_coding | 1.486721491 | 0.416245099 | 1.575926929 | -0.264127157 | 0.791681939 | 1 | 0.416245099 |
| ENSMUSG00000071573 | Rnls | 67795 | protein_coding | 35.01362822 | -0.416341212 | 0.199323735 | 2.088768864 | 0.036728534 | 0.1177122 | 0.416341212 |
| ENSMUSG00000040795 | Iqcc | 230767 | protein_coding | 93.52777409 | -0.416386795 | 0.112882264 | 3.688682179 | 0.000225419 | 0.001759245 | 0.416386795 |
| ENSMUSG00000100826 | Snhg14 | 16353 | lncRNA | 30.58017426 | -0.416559237 | 0.288474313 | 1.444008073 | 0.148736704 | 0.327585748 | 0.416559237 |
| ENSMUSG00000008855 | Hdac5 | 15184 | protein_coding | 507.4996003 | -0.416716365 | 0.112610238 | 3.700519353 | 0.000215159 | 0.001685546 | 0.416716365 |
| ENSMUSG00000034438 | Gbp8 | 76074 | protein_coding | 7.810724718 | -0.416906976 | 0.525741826 | 0.792988031 | 0.427784796 | 0.647057325 | 0.416906976 |
| ENSMUSG00000061451 | Tmem151a | 381199 | protein_coding | 15.93672208 | 0.417005729 | 0.321931193 | -1.295325642 | 0.195207917 | 0.394331932 | 0.417005729 |
| ENSMUSG00000034112 | Atp2c2 | 69047 | protein_coding | 745.9400414 | -0.417013785 | 0.257750979 | 1.617894092 | 0.105685424 | 0.257759428 | 0.417013785 |
| ENSMUSG00000040433 | Zbtb38 | 245007 | protein_coding | 689.081318 | -0.417106968 | 0.089030744 | 4.684976781 | 2.80E-06 | 3.51E-05 | 0.417106968 |
| ENSMUSG00000025933 | Tmem14a | 75712 | protein_coding | 29.27309587 | -0.417110704 | 0.244784543 | 1.703991188 | 0.088382734 | 0.22572726 | 0.417110704 |
| ENSMUSG00000055305 | Zfp93 | 22755 | protein_coding | 70.12331837 | -0.417169818 | 0.151596063 | 2.751851261 | 0.005925943 | 0.027912442 | 0.417169818 |
| ENSMUSG00000029156 | Sgcb | 24051 | protein_coding | 28.94618125 | -0.417222293 | 0.181778015 | 2.295229665 | 0.021719966 | 0.078315179 | 0.417222293 |
| ENSMUSG00000053931 | Cnn3 | 71994 | protein_coding | 189.0305939 | -0.417245677 | 0.148287236 | 2.813766644 | 0.004896476 | 0.023887859 | 0.417245677 |
| ENSMUSG00000060675 | Plaat3 | 225845 | protein_coding | 503.1832793 | -0.417444726 | 0.107384193 | 3.887394532 | 0.000101326 | 0.000870602 | 0.417444726 |
| ENSMUSG00000048498 | Cd300e | 217306 | protein_coding | 0.750245638 | -0.41753209 | 1.675542718 | 0.249192149 | 0.803212152 | 1 | 0.41753209 |
| ENSMUSG00000048481 | Mypop | 232934 | protein_coding | 20.6380703 | -0.417565205 | 0.306592419 | 1.361955416 | 0.173211956 | 0.363760214 | 0.417565205 |
| ENSMUSG00000050994 | Adgb | 215772 | protein_coding | 6.186712567 | 0.417919196 | 0.955372219 | -0.437441228 | 0.66179139 | 0.816925028 | 0.417919196 |
| ENSMUSG00000062204 | Or2y8 | 211472 | protein_coding | 1.526547279 | -0.418026196 | 1.494334917 | 0.279740633 | 0.779676502 | 1 | 0.418026196 |
| ENSMUSG00000057880 | Abat | 268860 | protein_coding | 3251.617225 | -0.418034489 | 0.142943452 | 2.924474557 | 0.003450383 | 0.017884376 | 0.418034489 |
| ENSMUSG00000041650 | Pcca | 110821 | protein_coding | 479.6104837 | -0.418037183 | 0.091030756 | 4.592263099 | 4.38E-06 | 5.31E-05 | 0.418037183 |
| ENSMUSG00000030411 | Nova2 | 384569 | protein_coding | 5.986011185 | 0.418197266 | 0.874027329 | -0.478471613 | 0.632314574 | 0.799512805 | 0.418197266 |
| ENSMUSG00000021303 | Gng4 | 14706 | protein_coding | 9.516874143 | -0.418210736 | 0.42917396 | 0.974455058 | 0.329830643 | 0.552046615 | 0.418210736 |
| ENSMUSG00000056643 | Chst13 | 71797 | protein_coding | 2.458583394 | 0.418272136 | 0.771890281 | -0.541880299 | 0.587900969 | 1 | 0.418272136 |

| ensembl_gene_id | external_gene_name | entrezgene_id | gene_biotype | baseMean | log2FoldChange | lfcSE | stat | pvalue | padj | absLog2FC |
|---|---|---|---|---|---|---|---|---|---|---|
| ENSMUSG00000020423 | Btg2 | 12227 | protein_coding | 6560.98182 | -0.418378903 | 0.16128749 | 2.593994755 | 0.009486793 | 0.040813086 | 0.418378903 |
| ENSMUSG00000010142 | Tnfrsf13b | 57916 | protein_coding | 2.288464675 | 0.418415659 | 1.051021896 | -0.398103655 | 0.69055378 | 1 | 0.418415659 |
| ENSMUSG00000043556 | Fbxl7 | 448987 | protein_coding | 2.043911232 | -0.41852882 | 3.094800825 | 0.135236108 | 0.892425199 | 1 | 0.41852882 |
| ENSMUSG00000039431 | Mtmr7 | 54384 | protein_coding | 255.3294178 | 0.418618779 | 0.096109949 | -4.355623752 | 1.33E-05 | 0.000143449 | 0.418618779 |
| ENSMUSG00000036086 | Zranb3 | 226409 | protein_coding | 67.09382733 | 0.418656907 | 0.17788128 | -2.353574845 | 0.018593864 | 0.069313561 | 0.418656907 |
| ENSMUSG00000068015 | Lrch1 | 380916 | protein_coding | 319.0378765 | -0.418659108 | 0.088729375 | 4.718382276 | 2.38E-06 | 3.04E-05 | 0.418659108 |
| ENSMUSG00000024854 | Pold4 | 69745 | protein_coding | 156.599412 | 0.418883845 | 0.118253425 | -3.542255504 | 0.000396721 | 0.002910462 | 0.418883845 |
| ENSMUSG00000029468 | P2rx7 | 18439 | protein_coding | 41.83642883 | -0.419010578 | 0.234605307 | 1.786023442 | 0.074095463 | 0.198998609 | 0.419010578 |
| ENSMUSG00000021767 | Kat6b | 54169 | protein_coding | 592.6017423 | 0.41901964 | 0.092616078 | -4.524264573 | 6.06E-06 | 7.11E-05 | 0.41901964 |
| ENSMUSG00000027875 | Hmgcs2 | 15360 | protein_coding | 13878.88844 | -0.41916398 | 0.126489755 | 3.313817638 | 0.000920315 | 0.005929802 | 0.41916398 |
| ENSMUSG00000030209 | Grin2b | 14812 | protein_coding | 12.16565868 | -0.419194825 | 0.778328852 | 0.538583176 | 0.590174499 | 0.770846581 | 0.419194825 |
| ENSMUSG00000026853 | Crat | 12908 | protein_coding | 636.1319498 | -0.419260572 | 0.06375544 | 6.576075248 | 4.83E-11 | 1.27E-09 | 0.419260572 |
| ENSMUSG00000100210 | H3c7 | 260423 | protein_coding | 1190.312132 | 0.419485542 | 0.196292553 | -2.137042575 | 0.032594529 | 0.107588657 | 0.419485542 |
| ENSMUSG00000020471 | Pold2 | 18972 | protein_coding | 445.3870034 | 0.419496795 | 0.115349389 | -3.636749166 | 0.000276101 | 0.002116771 | 0.419496795 |
| ENSMUSG00000020886 | Dlg4 | 13385 | protein_coding | 14.10521162 | 0.419656632 | 0.764481736 | -0.548942653 | 0.583044806 | 0.765756487 | 0.419656632 |
| ENSMUSG00000021754 | Map3k1 | 26401 | protein_coding | 544.2573616 | -0.419675227 | 0.065505315 | 6.40673546 | 1.49E-10 | 3.61E-09 | 0.419675227 |
| ENSMUSG00000051439 | Cd14 | 12475 | protein_coding | 378.8724501 | 0.419793558 | 0.218200702 | -1.923887292 | 0.054368712 | 0.158736855 | 0.419793558 |
| ENSMUSG00000024313 | Mep1b | 17288 | protein_coding | 1092.926499 | -0.419936155 | 0.159042706 | 2.640398708 | 0.008280854 | 0.036593531 | 0.419936155 |
| ENSMUSG00000010755 | Cars | 27267 | protein_coding | 358.041771 | 0.419942097 | 0.074504199 | -5.636488994 | 1.74E-08 | 3.22E-07 | 0.419942097 |
| ENSMUSG00000026027 | Stradb | 227154 | protein_coding | 173.9445148 | -0.419942398 | 0.112745963 | 3.724677912 | 0.000195565 | 0.001548948 | 0.419942398 |
| ENSMUSG00000027950 | Chrnb2 | 11444 | protein_coding | 35.67048712 | -0.419950478 | 0.172708636 | 2.431554602 | 0.015034182 | 0.058500114 | 0.419950478 |
| ENSMUSG00000038980 | Rbbp8nl | 271887 | protein_coding | 53.67869189 | 0.419971073 | 0.181439858 | -2.314657204 | 0.0206317 | 0.075129413 | 0.419971073 |
| ENSMUSG00000028772 | Zcchc17 | 619605 | protein_coding | 380.6274014 | 0.420082396 | 0.077705023 | -5.406116337 | 6.44E-08 | 1.09E-06 | 0.420082396 |
| ENSMUSG00000032289 | Thsd4 | 207596 | protein_coding | 414.198872 | -0.420332611 | 0.150984236 | 2.783950317 | 0.005370126 | 0.025839819 | 0.420332611 |
| ENSMUSG00000037526 | Atg14 | 100504663 | protein_coding | 152.2864395 | -0.42045472 | 0.121457427 | 3.461745643 | 0.000536684 | 0.003768198 | 0.42045472 |
| ENSMUSG00000047880 | Cxcr5 | 12145 | protein_coding | 5.137772515 | 0.420498878 | 0.646141822 | -0.650784184 | 0.515185811 | 0.716335759 | 0.420498878 |
| ENSMUSG00000005198 | Polr2a | 20020 | protein_coding | 583.2022804 | 0.420697282 | 0.3025741 | -1.390394227 | 0.164409198 | 0.350782106 | 0.420697282 |
| ENSMUSG00000050002 | Idnk | 75731 | protein_coding | 257.5339971 | -0.420714233 | 0.07420308 | 5.669767767 | 1.43E-08 | 2.69E-07 | 0.420714233 |
| ENSMUSG00000087273 | Gm13203 | 115489366 | lncRNA | 5.95722193 | -0.42073824 | 0.548983427 | 0.766395158 | 0.443441213 | 0.660955754 | 0.42073824 |
| ENSMUSG00000020461 | Clhc1 | 73324 | protein_coding | 1.993614873 | 0.42111257 | 1.002016435 | -0.420265133 | 0.674291778 | 1 | 0.42111257 |
| ENSMUSG00000045094 | Arhgef37 | 328967 | protein_coding | 117.6867656 | -0.421130106 | 0.135843806 | 3.100105327 | 0.001934518 | 0.011044625 | 0.421130106 |
| ENSMUSG00000100583 | Gm29443 | 102634559 | lncRNA | 1.214411167 | -0.421171803 | 1.853718199 | 0.227203791 | 0.82026528 | 1 | 0.421171803 |
| ENSMUSG00000015316 | Slamf1 | 27218 | protein_coding | 0.858326258 | -0.421269889 | 1.859354727 | 0.226567789 | 0.820759842 | 1 | 0.421269889 |
| ENSMUSG00000037325 | Bbs7 | 71492 | protein_coding | 78.30475049 | 0.42131402 | 0.135778432 | -3.102952467 | 0.001916004 | 0.01096738 | 0.42131402 |
| ENSMUSG00000045246 | Kcng4 | 66733 | protein_coding | 1.00490653 | -0.421410452 | 1.867371819 | 0.22567035 | 0.821457821 | 1 | 0.421410452 |
| ENSMUSG00000089125 | Gm26403 | 115488986 | snRNA | 1.693491278 | -0.421417963 | 0.899822807 | 0.468334387 | 0.639545481 | 1 | 0.421417963 |
| ENSMUSG00000112941 | Gm48623 | 12890 | lncRNA | 3.777080459 | -0.421612106 | 0.809882957 | 0.520583996 | 0.6026566 | 1 | 0.421612106 |
| ENSMUSG00000027286 | Lrrc57 | 66606 | protein_coding | 84.87838517 | -0.421615767 | 0.128323947 | 3.285557976 | 0.001017806 | 0.00644975 | 0.421615767 |
| ENSMUSG00000019873 | Reep3 | 28193 | protein_coding | 876.4242171 | -0.421705063 | 0.071666058 | 5.88430666 | 4.00E-09 | 8.18E-08 | 0.421705063 |

| ensembl_gene_id | external_gene_name | entrezgene_id | gene_biotype | baseMean | log2FoldChange | lfcSE | stat | pvalue | padj | absLog2FC |
|---|---|---|---|---|---|---|---|---|---|---|
| ENSMUSG00000033713 | Foxn3 | 71375 | protein_coding | 850.1754611 | -0.421755796 | 0.091750823 | 4.596752191 | 4.29E-06 | 5.21E-05 | 0.421755796 |
| ENSMUSG00000063142 | Kcnma1 | 16531 | protein_coding | 198.847557 | -0.421793642 | 0.216038398 | 1.952401272 | 0.050890582 | 0.151098 | 0.421793642 |
| ENSMUSG00000064326 | Siva1 | 30954 | protein_coding | 92.09133519 | -0.421875945 | 0.137342372 | 3.071710058 | 0.002128364 | 0.011930684 | 0.421875945 |
| ENSMUSG00000027962 | Vcam1 | 22329 | protein_coding | 6.427941917 | -0.421985447 | 0.686537449 | 0.614657581 | 0.538780879 | 0.733468492 | 0.421985447 |
| ENSMUSG00000055912 | Tmem150a | 232086 | protein_coding | 8.42677998 | -0.42212035 | 0.49168472 | 0.858518341 | 0.390606308 | 0.612813278 | 0.42212035 |
| ENSMUSG00000071711 | Mpst | 246221 | protein_coding | 2582.223104 | 0.422374913 | 0.17906956 | -2.358719788 | 0.018338097 | 0.068530286 | 0.422374913 |
| ENSMUSG00000056738 | A730036I17Rik | 329513 | lncRNA | 2.437069142 | -0.422516346 | 0.99650065 | 0.424000071 | 0.671565762 | 1 | 0.422516346 |
| ENSMUSG00000020135 | Apc2 | 23805 | protein_coding | 15.61390409 | 0.422626105 | 0.532864901 | -0.793120552 | 0.42770759 | 0.647057325 | 0.422626105 |
| ENSMUSG00000035024 | Ncapd3 | 78658 | protein_coding | 389.2009247 | 0.422683285 | 0.10983479 | -3.848355189 | 0.000118914 | 0.001004984 | 0.422683285 |
| ENSMUSG00000047881 | Rell1 | 100532 | protein_coding | 653.5286486 | 0.422717325 | 0.073243184 | -5.771422032 | 7.86E-09 | 1.54E-07 | 0.422717325 |
| ENSMUSG00000027861 | Casq2 | 12373 | protein_coding | 1.614685553 | -0.422827309 | 1.72494433 | 0.245125192 | 0.806359498 | 1 | 0.422827309 |
| ENSMUSG00000021703 | Serinc5 | 218442 | protein_coding | 1040.890113 | -0.423007546 | 0.136715717 | 3.094066697 | 0.001974331 | 0.011219581 | 0.423007546 |
| ENSMUSG00000032294 | Pkm | 18746 | protein_coding | 7768.212112 | -0.423037532 | 0.103991276 | 4.06800983 | 4.74E-05 | 0.000445412 | 0.423037532 |
| ENSMUSG00000062997 | Rpl35 | 66489 | protein_coding | 1045.800546 | -0.423060183 | 0.137574484 | 3.075135526 | 0.002104069 | 0.011831403 | 0.423060183 |
| ENSMUSG00000042371 | Slc5a10 | 109342 | protein_coding | 5.357167305 | 0.423119598 | 0.797136874 | -0.530799178 | 0.59555795 | 0.773864504 | 0.423119598 |
| ENSMUSG00000086241 | 4930483K19Rik | 74953 | lncRNA | 3.193818796 | -0.423302653 | 0.673170819 | 0.628819076 | 0.52946751 | 1 | 0.423302653 |
| ENSMUSG00000000934 | Top1mt | 72960 | protein_coding | 69.74926484 | -0.423332295 | 0.153976646 | 2.749327936 | 0.005971761 | 0.028080198 | 0.423332295 |
| ENSMUSG00000021036 | Sptlc2 | 20773 | protein_coding | 3490.325014 | 0.423421325 | 0.053607084 | -7.898607685 | 2.82E-15 | 1.05E-13 | 0.423421325 |
| ENSMUSG00000030852 | Tacc2 | 57752 | protein_coding | 652.5990578 | -0.423441562 | 0.166071035 | 2.549761692 | 0.010779658 | 0.045227455 | 0.423441562 |
| ENSMUSG00000024498 | Tcerg1 | 56070 | protein_coding | 511.0436432 | 0.423452835 | 0.087398022 | -4.845107787 | 1.27E-06 | 1.71E-05 | 0.423452835 |
| ENSMUSG00000034108 | Ccs | 12460 | protein_coding | 440.7266292 | -0.423458229 | 0.08565375 | 4.943837606 | 7.66E-07 | 1.09E-05 | 0.423458229 |
| ENSMUSG00000038541 | Srd5a2 | 94224 | protein_coding | 33.3534861 | -0.423570324 | 0.341457435 | 1.24047767 | 0.214798769 | 0.419775423 | 0.423570324 |
| ENSMUSG00000064147 | Rab44 | 442827 | protein_coding | 3.886678246 | -0.423579577 | 1.195623944 | 0.35427492 | 0.723132862 | 1 | 0.423579577 |
| ENSMUSG00000069833 | Ahnak | 66395 | protein_coding | 15650.19897 | -0.423686964 | 0.139804322 | 3.030571292 | 0.002440916 | 0.013347443 | 0.423686964 |
| ENSMUSG00000060579 | Fhit | 14198 | protein_coding | 69.54354836 | 0.423700373 | 0.147897082 | -2.864832538 | 0.004172298 | 0.020879192 | 0.423700373 |
| ENSMUSG00000036136 | Fam110c | 104943 | protein_coding | 435.6957207 | -0.423701253 | 0.097742115 | 4.334889358 | 1.46E-05 | 0.000155877 | 0.423701253 |
| ENSMUSG00000068744 | Psrc1 | 56742 | protein_coding | 44.07955277 | 0.423807377 | 0.205278413 | -2.064549167 | 0.038965677 | 0.123437405 | 0.423807377 |
| ENSMUSG00000025978 | Rftn2 | 74013 | protein_coding | 49.10049141 | -0.423819683 | 0.175884957 | 2.40964145 | 0.015968205 | 0.061425328 | 0.423819683 |
| ENSMUSG00000050410 | Tcf19 | 106795 | protein_coding | 143.3826344 | 0.424011127 | 0.208655389 | -2.032112041 | 0.042142315 | 0.130987801 | 0.424011127 |
| ENSMUSG00000034107 | Ano7 | 404545 | protein_coding | 430.9897189 | -0.424018368 | 0.130709414 | 3.243977265 | 0.001178732 | 0.007304155 | 0.424018368 |
| ENSMUSG00000020286 | 1700093K21Rik | 67358 | protein_coding | 0.778796039 | -0.424199072 | 2.049897089 | 0.206936765 | 0.836059244 | 1 | 0.424199072 |
| ENSMUSG00000050612 | Txndc2 | 213272 | protein_coding | 0.807100321 | -0.424310931 | 2.058222134 | 0.2061541 | 0.836670541 | 1 | 0.424310931 |
| ENSMUSG00000001520 | Nrip2 | 60345 | protein_coding | 8.123189256 | 0.424324004 | 0.41643322 | -1.018948499 | 0.308227417 | 0.528838189 | 0.424324004 |
| ENSMUSG00000017830 | Dhx58 | 80861 | protein_coding | 104.0843335 | -0.424373158 | 0.206649329 | 2.053590782 | 0.040015315 | 0.125929956 | 0.424373158 |
| ENSMUSG00000087141 | Plcxd2 | 433022 | protein_coding | 213.4427945 | 0.424377379 | 0.11143597 | -3.808262075 | 0.000139947 | 0.001159019 | 0.424377379 |
| ENSMUSG00000045903 | Npas4 | 225872 | protein_coding | 1.95850209 | 0.424507006 | 1.468043082 | -0.289165224 | 0.772454943 | 1 | 0.424507006 |
| ENSMUSG00000039842 | Mcph1 | 244329 | protein_coding | 162.6017771 | 0.424533018 | 0.108972375 | -3.895785692 | 9.79E-05 | 0.000844958 | 0.424533018 |
| ENSMUSG00000070489 | 4930527J03Rik | 75164 | pseudogene | 1.638540011 | -0.424564057 | 0.870362873 | 0.487801204 | 0.625690659 | 1 | 0.424564057 |
| ENSMUSG00000035849 | Krt222 | 268481 | protein_coding | 8.810325165 | -0.424577185 | 0.439287723 | 0.966512751 | 0.33378767 | 0.556110732 | 0.424577185 |

| ensembl_gene_id | external_gene_name | entrezgene_id | gene_biotype | baseMean | log2FoldChange | lfcSE | stat | pvalue | padj | absLog2FC |
|---|---|---|---|---|---|---|---|---|---|---|
| ENSMUSG00000046408 | 1700067K01Rik | 73453 | protein_coding | 3.970932652 | -0.42467318 | 0.733686831 | 0.578820775 | 0.562710112 | 1 | 0.42467318 |
| ENSMUSG00000024187 | Fam234a | 106581 | protein_coding | 3133.495001 | -0.424675033 | 0.061051344 | 6.95603086 | 3.50E-12 | 1.02E-10 | 0.424675033 |
| ENSMUSG00000024565 | Sall3 | 20689 | protein_coding | 1.97652087 | -0.42467932 | 2.086225832 | 0.203563446 | 0.838694661 | 1 | 0.42467932 |
| ENSMUSG00000027863 | Cd2 | 12481 | protein_coding | 2.814823708 | -0.424758546 | 0.841185783 | 0.504952121 | 0.613592465 | 1 | 0.424758546 |
| ENSMUSG00000054793 | Cadm4 | 260299 | protein_coding | 8.976264229 | 0.424916982 | 0.474719057 | -0.895091477 | 0.37073819 | 0.593065878 | 0.424916982 |
| ENSMUSG00000013643 | Lypd8 | 70163 | protein_coding | 50651.70691 | -0.424951336 | 0.101626552 | 4.181499091 | 2.90E-05 | 0.000287573 | 0.424951336 |
| ENSMUSG00000038259 | Gdf5 | 14563 | protein_coding | 1.079146846 | -0.425057561 | 2.116749004 | 0.200806784 | 0.840849658 | 1 | 0.425057561 |
| ENSMUSG00000032604 | Qars | 97541 | protein_coding | 2147.676628 | 0.425082825 | 0.070418913 | -6.036486621 | 1.58E-09 | 3.40E-08 | 0.425082825 |
| ENSMUSG00000036678 | Aaas | 223921 | protein_coding | 167.7605063 | 0.425184094 | 0.140347426 | -3.029511173 | 0.002449498 | 0.013385512 | 0.425184094 |
| ENSMUSG00000036882 | Arhgap33 | 233071 | protein_coding | 29.67500558 | 0.425186007 | 0.353699448 | -1.202111029 | 0.229320514 | 0.438111017 | 0.425186007 |
| ENSMUSG00000044667 | Plppr4 | 229791 | protein_coding | 1.619479293 | -0.42524228 | 2.131793987 | 0.199476254 | 0.841890216 | 1 | 0.42524228 |
| ENSMUSG00000046049 | Rp1l1 | 271209 | protein_coding | 1.710789745 | -0.425289985 | 2.135870801 | 0.199117842 | 0.842170565 | 1 | 0.425289985 |
| ENSMUSG00000069041 | Slc25a31 | 73333 | protein_coding | 0.823246668 | 0.425301347 | 1.372281726 | -0.309922765 | 0.75661969 | 1 | 0.425301347 |
| ENSMUSG00000059114 | Lrrc74a | 627607 | protein_coding | 1.429863731 | -0.425304125 | 2.137037686 | 0.199015735 | 0.842250436 | 1 | 0.425304125 |
| ENSMUSG00000029074 | Ttll10 | 330010 | protein_coding | 186.7450912 | -0.425335724 | 0.209286215 | 2.032316003 | 0.042121675 | 0.130972952 | 0.425335724 |
| ENSMUSG00000018593 | Sparc | 20692 | protein_coding | 6.325948919 | 0.425387635 | 0.676686072 | -0.628633649 | 0.529588926 | 0.727166502 | 0.425387635 |
| ENSMUSG00000022432 | Smc1b | 140557 | protein_coding | 1.771050683 | -0.425391833 | 2.144296372 | 0.198382947 | 0.842745459 | 1 | 0.425391833 |
| ENSMUSG00000043259 | Fam13c | 71721 | protein_coding | 2.035142422 | -0.425416316 | 2.146331835 | 0.198206218 | 0.842883723 | 1 | 0.425416316 |
| ENSMUSG00000096916 | Zfp850 | 100043772 | protein_coding | 58.13774274 | 0.425442114 | 0.142285798 | -2.990053252 | 0.002789288 | 0.014979611 | 0.425442114 |
| ENSMUSG00000038770 | Kpna7 | 381686 | protein_coding | 1.520058982 | -0.42545685 | 2.149902108 | 0.197895917 | 0.843126499 | 1 | 0.42545685 |
| ENSMUSG00000040767 | Snrnp25 | 78372 | protein_coding | 147.0721174 | -0.425458636 | 0.137170018 | 3.101688272 | 0.001924205 | 0.01100288 | 0.425458636 |
| ENSMUSG00000054708 | Ankrd24 | 70615 | protein_coding | 112.410448 | -0.42546107 | 0.145874844 | 2.91661713 | 0.003538498 | 0.018266571 | 0.42546107 |
| ENSMUSG00000022269 | Marchf11 | 211147 | protein_coding | 0.978956374 | -0.425475504 | 2.151729647 | 0.197736507 | 0.843251225 | 1 | 0.425475504 |
| ENSMUSG00000031928 | Mre11a | 17535 | protein_coding | 274.1701846 | 0.42552679 | 0.098397801 | -4.32455589 | 1.53E-05 | 0.000162315 | 0.42552679 |
| ENSMUSG00000024973 | Plaat5 | 66727 | protein_coding | 3.856383712 | 0.425595983 | 0.628307128 | -0.677369337 | 0.498171648 | 1 | 0.425595983 |
| ENSMUSG00000034220 | Gpc1 | 14733 | protein_coding | 35.96399926 | -0.425637148 | 0.197303714 | 2.157268804 | 0.030984726 | 0.103453653 | 0.425637148 |
| ENSMUSG00000042961 | Egflam | 268780 | protein_coding | 5.89378942 | 0.425700475 | 1.05603688 | -0.403111371 | 0.686866296 | 0.83324102 | 0.425700475 |
| ENSMUSG00000070883 | Ccdc173 | 75051 | protein_coding | 8.128672288 | 0.425730796 | 0.370452829 | -1.149217288 | 0.250466394 | 0.462725114 | 0.425730796 |
| ENSMUSG00000060126 | Tpt1 | 22070 | protein_coding | 8051.526135 | -0.425796484 | 0.097175452 | 4.381728856 | 1.18E-05 | 0.000129591 | 0.425796484 |
| ENSMUSG00000062093 | Gm10110 | 328451 | transcribed_processed_pseudogene | 6.210441477 | 0.425796703 | 0.495509066 | -0.85931163 | 0.39016861 | 0.612475705 | 0.425796703 |
| ENSMUSG00000061533 | Cep128 | 75216 | protein_coding | 106.2754082 | 0.425902809 | 0.178287828 | -2.388849614 | 0.016901219 | 0.064265646 | 0.425902809 |
| ENSMUSG00000002057 | Foxn1 | 15218 | protein_coding | 21.24176094 | 0.426083349 | 0.36793535 | -1.158038632 | 0.246848272 | 0.458698364 | 0.426083349 |
| ENSMUSG00000021182 | Ccdc88c | 68339 | protein_coding | 1296.866558 | 0.426123328 | 0.077437168 | -5.502826879 | 3.74E-08 | 6.56E-07 | 0.426123328 |
| ENSMUSG00000029922 | Mkrn1 | 54484 | protein_coding | 2460.244198 | -0.42612796 | 0.070361348 | 6.0562791 | 1.39E-09 | 3.02E-08 | 0.42612796 |
| ENSMUSG00000087165 | 2010001A14Rik | 76516 | lncRNA | 54.9024863 | -0.426299491 | 0.153033997 | 2.785652209 | 0.005342015 | 0.025742061 | 0.426299491 |
| ENSMUSG00000020919 | Stat5b | 20851 | protein_coding | 281.305574 | -0.426376201 | 0.071972895 | 5.924121889 | 3.14E-09 | 6.51E-08 | 0.426376201 |
| ENSMUSG00000005824 | Tnfsf14 | 50930 | protein_coding | 2.183785361 | 0.426423076 | 0.991325072 | -0.430154637 | 0.667083158 | 1 | 0.426423076 |
| ENSMUSG00000016206 | H2-M3 | 14991 | protein_coding | 60.70927599 | 0.426503751 | 0.164276453 | -2.596256149 | 0.009424578 | 0.040647029 | 0.426503751 |
| ENSMUSG00000048148 | Nwd1 | 319555 | protein_coding | 4.261356306 | 0.426548366 | 1.56029956 | -0.273375944 | 0.784564242 | 1 | 0.426548366 |

Supplementary Table 1

| ensembl_gene_id | external_gene_name | entrezgene_id | gene_biotype | baseMean | log2FoldChange | lfcSE | stat | pvalue | padj | absLog2FC |
|---|---|---|---|---|---|---|---|---|---|---|
| ENSMUSG00000024194 | Cuta | 67675 | protein_coding | 415.1820319 | 0.426591251 | 0.095328637 | -4.474953853 | 7.64E-06 | 8.81E-05 | 0.426591251 |
| ENSMUSG00000095253 | Zfp799 | 240064 | protein_coding | 161.8967245 | -0.426699732 | 0.148015478 | 2.8828048 | 0.003941516 | 0.01996653 | 0.426699732 |
| ENSMUSG00000066113 | Adamtsl1 | 77739 | protein_coding | 4.451857525 | 0.426709736 | 1.609615805 | -0.265100364 | 0.790932147 | 0.89578585 | 0.426709736 |
| ENSMUSG00000033276 | Stk36 | 269209 | protein_coding | 9.827059859 | 0.426763775 | 0.396129862 | -1.077333006 | 0.281331522 | 0.499023496 | 0.426763775 |
| ENSMUSG00000067889 | Sptbn2 | 20743 | protein_coding | 223.3785657 | -0.42676438 | 0.16364717 | 2.607832323 | 0.009111758 | 0.039545746 | 0.42676438 |
| ENSMUSG00000019278 | Dpep1 | 13479 | protein_coding | 3476.834646 | -0.426998924 | 0.147296142 | 2.898914509 | 0.00374457 | 0.019150767 | 0.426998924 |
| ENSMUSG00000025911 | Adhfe1 | 76187 | protein_coding | 17.42564255 | -0.427057615 | 0.308193512 | 1.385680098 | 0.165844608 | 0.353023699 | 0.427057615 |
| ENSMUSG00000038884 | Shfl | 319278 | protein_coding | 25.32259633 | -0.427097759 | 0.225698884 | 1.892334387 | 0.058446442 | 0.167508599 | 0.427097759 |
| ENSMUSG00000029851 | Tcaf2 | 232748 | protein_coding | 295.52622 | -0.427202432 | 0.163622684 | 2.610899792 | 0.009030435 | 0.03924437 | 0.427202432 |
| ENSMUSG00000048583 | Igf2 | 16002 | protein_coding | 1.886294065 | -0.427325 | 1.752262815 | 0.243870381 | 0.807331209 | 1 | 0.427325 |
| ENSMUSG00000115149 | 9330188P03Rik | 380930 | lncRNA | 15.26499233 | -0.427361405 | 0.263774404 | 1.620177693 | 0.105194112 | 0.256689693 | 0.427361405 |
| ENSMUSG00000044770 | Scml4 | 268297 | protein_coding | 607.8272101 | -0.427429495 | 0.16901251 | 2.528981411 | 0.011439409 | 0.047393475 | 0.427429495 |
| ENSMUSG00000040329 | Il7 | 16196 | protein_coding | 104.7828703 | 0.427540654 | 0.167388881 | -2.554175954 | 0.010643945 | 0.044806045 | 0.427540654 |
| ENSMUSG00000074261 | Erich4 | 632778 | protein_coding | 54.29682545 | 0.427615698 | 0.240700271 | -1.776548469 | 0.075642561 | 0.20160769 | 0.427615698 |
| ENSMUSG00000041313 | Slc7a1 | 11987 | protein_coding | 485.5017483 | 0.427665761 | 0.104250346 | -4.10229584 | 4.09E-05 | 0.000392527 | 0.427665761 |
| ENSMUSG00000032999 | Nlrp4f | 97895 | protein_coding | 1.118845089 | -0.427671646 | 2.366524379 | 0.180717194 | 0.856589562 | 1 | 0.427671646 |
| ENSMUSG00000016493 | Cd46 | 17221 | protein_coding | 23.22949885 | -0.42768496 | 0.251055612 | 1.703546704 | 0.088465806 | 0.22586956 | 0.42768496 |
| ENSMUSG00000028938 | Galntl5 | 67909 | protein_coding | 0.937598424 | -0.427686945 | 2.368391335 | 0.180581198 | 0.856696315 | 1 | 0.427686945 |
| ENSMUSG00000083193 | 4930595D18Rik | 78062 | lncRNA | 2.917893146 | -0.42780435 | 0.781597905 | 0.547345824 | 0.584141167 | 1 | 0.42780435 |
| ENSMUSG00000073198 | Bnip3l-ps | 100043324 | transcribed_processed_pseudogene | 3.019667564 | 0.427812265 | 0.60398664 | -0.708314119 | 0.478750211 | 1 | 0.427812265 |
| ENSMUSG00000069727 | Zfp975 | 434179 | protein_coding | 54.56090511 | -0.427890543 | 0.173924457 | 2.460209165 | 0.013885606 | 0.054910657 | 0.427890543 |
| ENSMUSG00000099564 | Gm28729 | 102635744 | protein_coding | 1.383043982 | -0.4278909 | 2.391412243 | 0.178928121 | 0.857994141 | 1 | 0.4278909 |
| ENSMUSG00000020811 | Wscd1 | 216881 | protein_coding | 3.935572492 | 0.427904967 | 0.620930011 | -0.689135586 | 0.490737948 | 1 | 0.427904967 |
| ENSMUSG00000026914 | Psmd14 | 59029 | protein_coding | 783.175018 | 0.427909313 | 0.075439083 | -5.672249672 | 1.41E-08 | 2.65E-07 | 0.427909313 |
| ENSMUSG00000035805 | Mlc1 | 170790 | protein_coding | 1.013459738 | -0.427944429 | 2.39786038 | 0.178469286 | 0.85835444 | 1 | 0.427944429 |
| ENSMUSG00000026473 | Glul | 14645 | protein_coding | 6370.506637 | -0.427968029 | 0.173724128 | 2.463492174 | 0.01375909 | 0.054501706 | 0.427968029 |
| ENSMUSG00000045942 | BC049762 | 193286 | protein_coding | 2.677712198 | -0.427971386 | 0.742624386 | 0.576295897 | 0.564415196 | 1 | 0.427971386 |
| ENSMUSG00000116976 | 4930500H12Rik | 74991 | lncRNA | 1.515106644 | -0.427974407 | 2.400990927 | 0.178249073 | 0.858527372 | 1 | 0.427974407 |
| ENSMUSG00000050808 | Muc15 | 269328 | protein_coding | 1.196928993 | -0.428002642 | 2.404511493 | 0.177999832 | 0.858723108 | 1 | 0.428002642 |
| ENSMUSG00000038055 | Dexi | 58239 | protein_coding | 157.884108 | 0.428048998 | 0.126100357 | -3.394510587 | 0.000687514 | 0.004614708 | 0.428048998 |
| ENSMUSG00000058093 | Zfp729b | 100416706 | protein_coding | 116.107383 | -0.428051879 | 0.194277208 | 2.20330467 | 0.027573282 | 0.094352503 | 0.428051879 |
| ENSMUSG00000114961 | A930002C04Rik | 319467 | lncRNA | 0.871395148 | -0.428080566 | 2.414007611 | 0.177331904 | 0.859247692 | 1 | 0.428080566 |
| ENSMUSG00000040536 | Necab1 | 69352 | protein_coding | 1.715040822 | -0.428129123 | 2.41916303 | 0.176974068 | 0.859528759 | 1 | 0.428129123 |
| ENSMUSG00000087410 | 2310065F04Rik | 74184 | lncRNA | 1.485175994 | -0.428142042 | 2.420830284 | 0.176857521 | 0.859620306 | 1 | 0.428142042 |
| ENSMUSG00000022820 | Ndufb4 | 68194 | protein_coding | 279.5903331 | -0.4281476 | 0.110670151 | 3.868681799 | 0.000109425 | 0.000932912 | 0.4281476 |
| ENSMUSG00000075090 | Or4a15 | 258975 | protein_coding | 1.0263964 | -0.42816378 | 2.423775049 | 0.176651616 | 0.859782048 | 1 | 0.42816378 |
| ENSMUSG00000064010 | Dppa1 | 347708 | protein_coding | 1.117073477 | -0.428165315 | 2.42387617 | 0.17664488 | 0.859787339 | 1 | 0.428165315 |
| ENSMUSG00000032035 | Ets1 | 23871 | protein_coding | 58.69123534 | 0.42819345 | 0.261317699 | -1.638593373 | 0.101297973 | 0.249900617 | 0.42819345 |
| ENSMUSG00000074965 | Or4k35 | 258391 | protein_coding | 1.073640325 | -0.428204989 | 2.428702309 | 0.176310199 | 0.86005025 | 1 | 0.428204989 |

| ensembl_gene_id | external_gene_name | entrezgene_id | gene_biotype | baseMean | log2FoldChange | lfcSE | stat | pvalue | padj | absLog2FC |
|---|---|---|---|---|---|---|---|---|---|---|
| ENSMUSG00000099696 | 2900052N01Rik | 73040 | lncRNA | 1.580613906 | -0.428229571 | 2.43132314 | 0.176130258 | 0.86019161 | 1 | 0.428229571 |
| ENSMUSG00000057564 | Or4e1 | 57270 | protein_coding | 1.703709504 | -0.428252262 | 2.434070474 | 0.175940782 | 0.860340466 | 1 | 0.428252262 |
| ENSMUSG00000071322 | Tcp10a | 21461 | protein_coding | 1.430728211 | -0.428252751 | 2.434244458 | 0.175928408 | 0.860350187 | 1 | 0.428252751 |
| ENSMUSG00000103567 | Pcdhga5 | 93713 | protein_coding | 8.765933606 | 0.428260669 | 0.383596042 | -1.116436621 | 0.264235285 | 0.478777869 | 0.428260669 |
| ENSMUSG00000073971 | Or52m1 | 258322 | protein_coding | 1.200439207 | -0.42827397 | 2.436996049 | 0.175738475 | 0.860499407 | 1 | 0.42827397 |
| ENSMUSG00000043331 | Or10d5 | 258825 | protein_coding | 1.201180071 | -0.428274842 | 2.437101932 | 0.175731198 | 0.860505124 | 1 | 0.428274842 |
| ENSMUSG00000024306 | Ccdc178 | 70950 | protein_coding | 2.452871333 | -0.428279948 | 2.077035203 | 0.206197732 | 0.836636459 | 1 | 0.428279948 |
| ENSMUSG00000025932 | Eya1 | 14048 | protein_coding | 1.194688356 | -0.4284176 | 2.454632218 | 0.174534334 | 0.86144555 | 1 | 0.4284176 |
| ENSMUSG00000050334 | C130071C03Rik | 320203 | lncRNA | 1.343861907 | -0.42850014 | 2.464949766 | 0.173837271 | 0.861993352 | 1 | 0.42850014 |
| ENSMUSG00000016386 | Mpped2 | 77015 | protein_coding | 0.924071318 | -0.428553829 | 2.472131722 | 0.173353962 | 0.86237321 | 1 | 0.428553829 |
| ENSMUSG00000085336 | Gm11732 | 102637448 | lncRNA | 0.984131332 | -0.428556401 | 2.472317665 | 0.173341965 | 0.86238264 | 1 | 0.428556401 |
| ENSMUSG00000067780 | Pi15 | 94227 | protein_coding | 1.258658764 | -0.428599756 | 2.477603638 | 0.172989638 | 0.862659573 | 1 | 0.428599756 |
| ENSMUSG00000038760 | Trhr | 22045 | protein_coding | 1.295250233 | -0.428634943 | 2.482104673 | 0.172690116 | 0.862895014 | 1 | 0.428634943 |
| ENSMUSG00000015668 | Pdzd11 | 72621 | protein_coding | 383.3642497 | 0.428648805 | 0.085459225 | -5.01582839 | 5.28E-07 | 7.76E-06 | 0.428648805 |
| ENSMUSG00000037033 | Clca3b | 229927 | protein_coding | 3844.558357 | 0.428701723 | 0.188300123 | -2.276693805 | 0.022804517 | 0.081360993 | 0.428701723 |
| ENSMUSG00000115727 | 4930432J09Rik | 73877 | lncRNA | 1.44618699 | -0.428755724 | 2.497692154 | 0.171660756 | 0.86370424 | 1 | 0.428755724 |
| ENSMUSG00000063687 | Pcdhb5 | 93876 | protein_coding | 3.031933483 | -0.42875702 | 0.67002971 | 0.639907475 | 0.522232754 | 1 | 0.42875702 |
| ENSMUSG00000069804 | Gm10277 | 791303 | protein_coding | 32.06453063 | -0.428995458 | 0.303425016 | 1.413843403 | 0.157407887 | 0.340684159 | 0.428995458 |
| ENSMUSG00000025040 | Fundc1 | 72018 | protein_coding | 256.9252873 | 0.429016933 | 0.08759206 | -4.897897534 | 9.69E-07 | 1.34E-05 | 0.429016933 |
| ENSMUSG00000045087 | S1pr5 | 94226 | protein_coding | 1.278137616 | -0.429027311 | 1.834587051 | 0.233854976 | 0.815097568 | 1 | 0.429027311 |
| ENSMUSG00000064600 | Gm25636 | 115486823 | snoRNA | 8.03350012 | -0.429177642 | 0.413844441 | 1.037050639 | 0.299712258 | 0.519697021 | 0.429177642 |
| ENSMUSG00000028459 | Cd72 | 12517 | protein_coding | 6.847993452 | 0.429537533 | 0.603152148 | -0.712154527 | 0.476369092 | 0.688134988 | 0.429537533 |
| ENSMUSG00000033685 | Ucp2 | 22228 | protein_coding | 2733.747347 | -0.429541233 | 0.06947815 | 6.182393075 | 6.31E-10 | 1.42E-08 | 0.429541233 |
| ENSMUSG00000036553 | Sh3tc1 | 231147 | protein_coding | 567.6742219 | 0.42975634 | 0.144466183 | -2.974788491 | 0.002931906 | 0.015558323 | 0.42975634 |
| ENSMUSG00000047843 | Bri3 | 55950 | protein_coding | 228.2942448 | -0.429840708 | 0.102056171 | 4.211805147 | 2.53E-05 | 0.000254944 | 0.429840708 |
| ENSMUSG00000029073 | Cptp | 79554 | protein_coding | 400.4141575 | 0.430032689 | 0.074798244 | -5.74923511 | 8.96E-09 | 1.74E-07 | 0.430032689 |
| ENSMUSG00000041219 | Arhgap11a | 228482 | protein_coding | 429.601323 | 0.430087556 | 0.128259667 | -3.353256462 | 0.000798667 | 0.005252561 | 0.430087556 |
| ENSMUSG00000041567 | Serpina12 | 68054 | protein_coding | 1.158044699 | -0.430202891 | 2.623980958 | 0.163950462 | 0.869770141 | 1 | 0.430202891 |
| ENSMUSG00000025318 | Jph3 | 57340 | protein_coding | 5.334700026 | 0.430262557 | 0.615855485 | -0.698642081 | 0.484775739 | 0.695226137 | 0.430262557 |
| ENSMUSG00000000579 | Dynlt1c | 100040563 | protein_coding | 19.2245882 | -0.430344196 | 0.313222424 | 1.373925246 | 0.169464891 | 0.358787591 | 0.430344196 |
| ENSMUSG00000033863 | Klf9 | 16601 | protein_coding | 999.2982016 | -0.430382996 | 0.132462776 | 3.24908634 | 0.001157763 | 0.007195824 | 0.430382996 |
| ENSMUSG00000065728 | Gm26175 | 115490468 | snoRNA | 5.111862217 | 0.430388218 | 0.477563669 | -0.901216416 | 0.367473265 | 0.590046157 | 0.430388218 |
| ENSMUSG00000022124 | Fbxl3 | 50789 | protein_coding | 1137.411194 | -0.430407229 | 0.097510202 | 4.413971242 | 1.01E-05 | 0.000113436 | 0.430407229 |
| ENSMUSG00000094338 | H2bc13 | 319185 | protein_coding | 398.5385631 | 0.430525053 | 0.204839225 | -2.101770567 | 0.035573379 | 0.114836207 | 0.430525053 |
| ENSMUSG00000073900 | Or2ag20 | 257902 | protein_coding | 1.616966817 | -0.430808918 | 2.816833125 | 0.152940873 | 0.878444903 | 1 | 0.430808918 |
| ENSMUSG00000022586 | Ly6i | 57248 | protein_coding | 0.791931008 | 0.430878747 | 1.933419247 | -0.222858414 | 0.823645703 | 1 | 0.430878747 |
| ENSMUSG00000040935 | Padi6 | 242726 | protein_coding | 1.387418677 | -0.430905498 | 2.835192463 | 0.151984567 | 0.879199108 | 1 | 0.430905498 |
| ENSMUSG00000060303 | Or5b97 | 258698 | protein_coding | 1.387418677 | -0.430905498 | 2.835192463 | 0.151984567 | 0.879199108 | 1 | 0.430905498 |
| ENSMUSG00000061584 | Lyg2 | 332427 | protein_coding | 1.530100513 | -0.430943803 | 2.842269904 | 0.151619592 | 0.87948698 | 1 | 0.430943803 |

Supplementary Table 1

| ensembl_gene_id | external_gene_name | entrezgene_id | gene_biotype | baseMean | log2FoldChange | lfcSE | stat | pvalue | padj | absLog2FC |
|---|---|---|---|---|---|---|---|---|---|---|
| ENSMUSG00000035560 | Wdr20rt | 70948 | protein_coding | 1.343985524 | -0.43094792 | 2.843308224 | 0.151565671 | 0.879529511 | 1 | 0.43094792 |
| ENSMUSG00000040133 | Gpr176 | 381413 | protein_coding | 1.25299176 | -0.431021368 | 2.857547532 | 0.150836115 | 0.880104996 | 1 | 0.431021368 |
| ENSMUSG00000073209 | Klf14 | 619665 | protein_coding | 1.25299176 | -0.431021368 | 2.857547532 | 0.150836115 | 0.880104996 | 1 | 0.431021368 |
| ENSMUSG00000030638 | Sh3gl3 | 20408 | protein_coding | 1.066876772 | -0.431065384 | 2.866438772 | 0.150383601 | 0.880461979 | 1 | 0.431065384 |
| ENSMUSG00000050926 | Dcaf12l2 | 245403 | protein_coding | 1.173979019 | -0.431072745 | 2.867518758 | 0.150329529 | 0.880504637 | 1 | 0.431072745 |
| ENSMUSG00000052085 | Dock8 | 76088 | protein_coding | 321.690361 | -0.431159217 | 0.089137987 | 4.836986248 | 1.32E-06 | 1.77E-05 | 0.431159217 |
| ENSMUSG00000034387 | Ssu2 | 243612 | protein_coding | 1.016951905 | -0.431220179 | 2.897002335 | 0.148850477 | 0.881671619 | 1 | 0.431220179 |
| ENSMUSG00000021179 | Nrde2 | 217827 | protein_coding | 307.5056019 | 0.431224004 | 0.102378769 | -4.21204522 | 2.53E-05 | 0.000254828 | 0.431224004 |
| ENSMUSG00000010538 | Tsacc | 76927 | protein_coding | 12.34153434 | -0.431236851 | 0.28543273 | 1.510817805 | 0.130834877 | 0.299128777 | 0.431236851 |
| ENSMUSG00000038042 | Ptpdc1 | 218232 | protein_coding | 8.971125172 | 0.431246988 | 0.514009428 | -0.838986532 | 0.401476867 | 0.62300216 | 0.431246988 |
| ENSMUSG00000049097 | Ankrd34a | 545554 | protein_coding | 1.204794997 | 0.431293603 | 1.79916423 | -0.239718863 | 0.810548211 | 1 | 0.431293603 |
| ENSMUSG00000075093 | Or4c1 | 258446 | protein_coding | 0.971755547 | -0.431308886 | 2.915317934 | 0.147945746 | 0.882385586 | 1 | 0.431308886 |
| ENSMUSG00000009905 | Kdsr | 70750 | protein_coding | 350.0899859 | -0.4314308 | 0.125096911 | 3.448772618 | 0.000563141 | 0.003915665 | 0.4314308 |
| ENSMUSG00000061111 | Mcrip1 | 192173 | protein_coding | 85.78828509 | -0.431594617 | 0.152832882 | 2.823964395 | 0.004743365 | 0.023285356 | 0.431594617 |
| ENSMUSG00000072761 | Gm6712 | 100862203 | lncRNA | 32.81463177 | -0.43160334 | 0.189044061 | 2.283083313 | 0.022425461 | 0.080385081 | 0.43160334 |
| ENSMUSG00000056215 | Lrguk | 74354 | protein_coding | 5.444960781 | 0.43168399 | 0.979999046 | -0.440494296 | 0.659579142 | 0.8161622 | 0.43168399 |
| ENSMUSG00000026459 | Myog | 17928 | protein_coding | 1.357843179 | -0.431800025 | 1.502308173 | 0.2874244 | 0.773787381 | 1 | 0.431800025 |
| ENSMUSG00000031930 | Wwp2 | 66894 | protein_coding | 1498.28454 | -0.431805071 | 0.111101598 | 3.886578391 | 0.000101667 | 0.000872625 | 0.431805071 |
| ENSMUSG00000026443 | Lrrn2 | 16980 | protein_coding | 0.840252914 | 0.431950242 | 1.902827205 | -0.22700445 | 0.820420282 | 1 | 0.431950242 |
| ENSMUSG00000014329 | Bicc1 | 83675 | protein_coding | 9.452400303 | -0.43205503 | 0.460504835 | 0.9382204 | 0.348131157 | 0.570058297 | 0.43205503 |
| ENSMUSG00000035202 | Lars2 | 102436 | protein_coding | 11475.44353 | -0.432136069 | 0.563011242 | 0.767544299 | 0.442757965 | 0.660497294 | 0.432136069 |
| ENSMUSG00000031886 | Ces2e | 234673 | protein_coding | 2622.98925 | -0.432214419 | 0.132957883 | 3.250761891 | 0.001150962 | 0.007156245 | 0.432214419 |
| ENSMUSG00000118292 | Gm10549 | 433171 | lncRNA | 3.413631735 | -0.432515945 | 0.6577194 | 0.657599494 | 0.51079552 | 1 | 0.432515945 |
| ENSMUSG00000020519 | Sap30l | 50724 | protein_coding | 356.9772235 | 0.432530532 | 0.134854993 | -3.207374998 | 0.001339523 | 0.008117753 | 0.432530532 |
| ENSMUSG00000085440 | Sorbs2os | 319940 | lncRNA | 6.670762508 | 0.432574454 | 0.490195349 | -0.882453199 | 0.377531778 | 0.599649878 | 0.432574454 |
| ENSMUSG00000019467 | Arhgef25 | 52666 | protein_coding | 3.310672309 | 0.432704668 | 1.114051032 | -0.388406506 | 0.697715231 | 1 | 0.432704668 |
| ENSMUSG00000078253 | Krtap16-1 | 100504183 | protein_coding | 2.227888009 | 0.432831223 | 2.211732655 | -0.195697804 | 0.844846704 | 1 | 0.432831223 |
| ENSMUSG00000019986 | Ahi1 | 52906 | protein_coding | 22.2274704 | 0.432934872 | 0.289768905 | -1.494069461 | 0.135157465 | 0.30625554 | 0.432934872 |
| ENSMUSG00000035783 | Acta2 | 11475 | protein_coding | 2.37141596 | -0.43305208 | 0.811992069 | 0.533320578 | 0.593811685 | 1 | 0.43305208 |
| ENSMUSG00000016756 | Cmah | 12763 | protein_coding | 608.2699785 | -0.433267392 | 0.252111382 | 1.718555463 | 0.085695347 | 0.220951697 | 0.433267392 |
| ENSMUSG00000058101 | 1700084J12Rik | 73486 | transcribed_processed_pseudogene | 2.911161007 | 0.43329874 | 0.663160237 | -0.653384681 | 0.513508312 | 1 | 0.43329874 |
| ENSMUSG00000015656 | Hspa8 | 15481 | protein_coding | 14797.55801 | -0.433498722 | 0.136699526 | 3.171179403 | 0.001518213 | 0.0089816 | 0.433498722 |
| ENSMUSG00000038393 | Txnip | 56338 | protein_coding | 9188.776409 | -0.43362476 | 0.110952658 | 3.908196226 | 9.30E-05 | 0.000809062 | 0.43362476 |
| ENSMUSG00000024729 | Ms4a20 | 69369 | protein_coding | 1.967640148 | -0.43374342 | 1.708121792 | 0.253930031 | 0.799549613 | 1 | 0.43374342 |
| ENSMUSG00000036840 | Siah1a | 20437 | protein_coding | 153.7875753 | -0.433743598 | 0.102148392 | 4.24621072 | 2.17E-05 | 0.000222179 | 0.433743598 |
| ENSMUSG00000033170 | Card10 | 105844 | protein_coding | 420.8433466 | -0.433993238 | 0.076626476 | 5.66375043 | 1.48E-08 | 2.77E-07 | 0.433993238 |
| ENSMUSG00000006445 | Epha2 | 13836 | protein_coding | 738.358647 | 0.434031298 | 0.282321519 | -1.537365269 | 0.124203886 | 0.288643819 | 0.434031298 |
| ENSMUSG00000096606 | Tpbgl | 100503386 | protein_coding | 3.205554922 | -0.434164969 | 0.671061847 | 0.646982049 | 0.517643562 | 1 | 0.434164969 |
| ENSMUSG00000035325 | Sec31a | 69162 | protein_coding | 2489.776038 | 0.434269707 | 0.050256479 | -8.641069117 | 5.57E-18 | 2.57E-16 | 0.434269707 |

| ensembl_gene_id | external_gene_name | entrezgene_id | gene_biotype | baseMean | log2FoldChange | lfcSE | stat | pvalue | padj | absLog2FC |
|---|---|---|---|---|---|---|---|---|---|---|
| ENSMUSG00000028933 | Xrcc2 | 57434 | protein_coding | 40.86995964 | 0.434435715 | 0.241653861 | -1.797760292 | 0.072215002 | 0.195501402 | 0.434435715 |
| ENSMUSG00000038759 | Nup205 | 70699 | protein_coding | 702.0150011 | 0.434540331 | 0.121096983 | -3.588366293 | 0.000332757 | 0.00249439 | 0.434540331 |
| ENSMUSG00000043913 | Ccdc60 | 269693 | protein_coding | 3.338921294 | 0.434549439 | 1.971247295 | -0.220443899 | 0.82552546 | 1 | 0.434549439 |
| ENSMUSG00000036500 | Akp3 | 11648 | protein_coding | 4.015933001 | 0.434551839 | 1.330754564 | -0.326545443 | 0.744011713 | 1 | 0.434551839 |
| ENSMUSG00000032714 | Syde1 | 71709 | protein_coding | 10.83176034 | 0.434587946 | 0.402958339 | -1.078493492 | 0.280813585 | 0.498479581 | 0.434587946 |
| ENSMUSG00000113918 | Gm6566 | 625237 | lncRNA | 6.227928959 | -0.434599514 | 0.471798395 | 0.921155136 | 0.356969438 | 0.579930477 | 0.434599514 |
| ENSMUSG00000038763 | Alpk3 | 116904 | protein_coding | 4.048740204 | 0.434669836 | 1.311641531 | -0.331393773 | 0.740347068 | 1 | 0.434669836 |
| ENSMUSG00000093176 | Mir5119 | 100628613 | miRNA | 0.63535501 | 0.43472486 | 1.623565259 | -0.2677594 | 0.788884524 | 1 | 0.43472486 |
| ENSMUSG00000038845 | Phb | 18673 | protein_coding | 662.893273 | -0.434729246 | 0.117277171 | 3.706853107 | 0.000209851 | 0.001648655 | 0.434729246 |
| ENSMUSG00000049670 | Morn4 | 226123 | protein_coding | 2.968266116 | -0.434804829 | 0.6251333 | 0.695539382 | 0.48671734 | 1 | 0.434804829 |
| ENSMUSG00000096299 | Gm21814 | 436100 | lncRNA | 8.893010522 | -0.434812763 | 0.432295132 | 1.00582387 | 0.314500303 | 0.535752227 | 0.434812763 |
| ENSMUSG00000062464 | Cyp4f37 | 677156 | protein_coding | 1.919440296 | -0.434832899 | 1.412725199 | 0.307797227 | 0.758236631 | 1 | 0.434832899 |
| ENSMUSG00000034990 | Otoa | 246190 | protein_coding | 3.970591941 | 0.434836182 | 0.953111106 | -0.456228218 | 0.648225876 | 1 | 0.434836182 |
| ENSMUSG00000049314 | Fbxw13 | 211305 | protein_coding | 1.485909581 | -0.434890619 | 1.718351247 | 0.253085986 | 0.800201768 | 1 | 0.434890619 |
| ENSMUSG00000006800 | Sulf2 | 72043 | protein_coding | 285.7922779 | 0.43490427 | 0.191443313 | -2.271713036 | 0.023103848 | 0.082121599 | 0.43490427 |
| ENSMUSG00000024669 | Cd5 | 12507 | protein_coding | 4.4947527 | 0.434960568 | 0.757904145 | -0.57389918 | 0.566036028 | 0.752646336 | 0.434960568 |
| ENSMUSG00000111422 | 1700027J07Rik | 69443 | lncRNA | 1.70197375 | 0.435032375 | 1.799182593 | -0.241794455 | 0.808939436 | 1 | 0.435032375 |
| ENSMUSG00000098134 | Rnf113a2 | 66381 | protein_coding | 152.886509 | 0.43508864 | 0.114434352 | -3.802080687 | 0.000143486 | 0.001184764 | 0.43508864 |
| ENSMUSG00000060508 | Nlrp9b | 243874 | protein_coding | 192.4738956 | 0.435134402 | 0.37166245 | -1.170778491 | 0.241687826 | 0.452315589 | 0.435134402 |
| ENSMUSG00000015962 | 1700016C15Rik | 69428 | protein_coding | 1.380578869 | -0.435212109 | 1.017793982 | 0.427603343 | 0.668939932 | 1 | 0.435212109 |
| ENSMUSG00000035637 | Grhpr | 76238 | protein_coding | 283.1358386 | 0.435256048 | 0.102377062 | -4.251499695 | 2.12E-05 | 0.000217399 | 0.435256048 |
| ENSMUSG00000092009 | Myh15 | 667772 | protein_coding | 1.841525791 | 0.435301111 | 2.134261025 | -0.203958703 | 0.838385771 | 1 | 0.435301111 |
| ENSMUSG00000050627 | Gpd1l | 333433 | protein_coding | 2079.998984 | 0.435313399 | 0.063196944 | -6.888203371 | 5.65E-12 | 1.61E-10 | 0.435313399 |
| ENSMUSG00000031910 | Has3 | 15118 | protein_coding | 13.89680433 | -0.43553238 | 0.379711967 | 1.147007251 | 0.251378624 | 0.463828671 | 0.43553238 |
| ENSMUSG00000032014 | Oaf | 102644 | protein_coding | 173.248456 | -0.43557327 | 0.144320757 | 3.018091641 | 0.00254372 | 0.013831738 | 0.43557327 |
| ENSMUSG00000047798 | Cd300lf | 246746 | protein_coding | 32.18037812 | -0.435617234 | 0.345206928 | 1.261901772 | 0.206984134 | 0.409799304 | 0.435617234 |
| ENSMUSG00000037053 | Azgp1 | 12007 | protein_coding | 53.43271107 | -0.435776376 | 0.222591111 | 1.957743839 | 0.050260078 | 0.14941403 | 0.435776376 |
| ENSMUSG00000009575 | Cbx5 | 12419 | protein_coding | 1496.984328 | 0.435816628 | 0.149086248 | -2.9232517 | 0.003463964 | 0.017949136 | 0.435816628 |
| ENSMUSG00000002274 | Metrn | 70083 | protein_coding | 27.78141983 | 0.435866018 | 0.259649131 | -1.678673124 | 0.093215767 | 0.23455206 | 0.435866018 |
| ENSMUSG00000031812 | Map1lc3b | 67443 | protein_coding | 994.8523245 | -0.435938881 | 0.083256852 | 5.236072136 | 1.64E-07 | 2.61E-06 | 0.435938881 |
| ENSMUSG00000040478 | Prdm13 | 230025 | protein_coding | 1.413184018 | 0.435947839 | 1.879743994 | -0.23191873 | 0.816601137 | 1 | 0.435947839 |
| ENSMUSG00000071658 | Gng3 | 14704 | protein_coding | 3.229612148 | 0.435993152 | 0.747085521 | -0.583592025 | 0.559494827 | 1 | 0.435993152 |
| ENSMUSG00000027340 | Slc23a2 | 54338 | protein_coding | 801.0425191 | -0.43599524 | 0.103508569 | 4.21216566 | 2.53E-05 | 0.000254828 | 0.43599524 |
| ENSMUSG00000064360 | mt-Nd3 | 17718 | protein_coding | 4.026843207 | 0.436025201 | 0.680983224 | -0.640287727 | 0.521985558 | 1 | 0.436025201 |
| ENSMUSG00000024953 | Prdx5 | 54683 | protein_coding | 1732.303799 | -0.436094008 | 0.078502753 | 5.555142834 | 2.77E-08 | 4.98E-07 | 0.436094008 |
| ENSMUSG00000023861 | Mpc1 | 55951 | protein_coding | 434.7616214 | -0.436137889 | 0.095262088 | 4.578294435 | 4.69E-06 | 5.63E-05 | 0.436137889 |
| ENSMUSG00000013155 | Enkd1 | 102124 | protein_coding | 27.1461322 | 0.436191956 | 0.234903666 | -1.856897184 | 0.063325783 | 0.177458338 | 0.436191956 |
| ENSMUSG00000044461 | Shisa2 | 219134 | protein_coding | 610.8043837 | -0.436254972 | 0.120226475 | 3.628609862 | 0.000284951 | 0.002177552 | 0.436254972 |
| ENSMUSG00000055945 | Prr18 | 320111 | protein_coding | 4.268634207 | -0.436286151 | 0.70953394 | 0.614891164 | 0.538626598 | 1 | 0.436286151 |

| ensembl_gene_id | external_gene_name | entrezgene_id | gene_biotype | baseMean | log2FoldChange | lfcSE | stat | pvalue | padj | absLog2FC |
|---|---|---|---|---|---|---|---|---|---|---|
| ENSMUSG00000070305 | Mpzl3 | 319742 | protein_coding | 357.2052248 | -0.436384536 | 0.140839161 | 3.098460206 | 0.001945291 | 0.011092726 | 0.436384536 |
| ENSMUSG00000030161 | Gabarapl1 | 57436 | protein_coding | 255.7253235 | -0.436466443 | 0.106863653 | 4.084330193 | 4.42E-05 | 0.000420016 | 0.436466443 |
| ENSMUSG00000028186 | Uox | 22262 | protein_coding | 19.31647302 | 0.436497435 | 0.228130313 | -1.91336885 | 0.055700849 | 0.161546158 | 0.436497435 |
| ENSMUSG00000060212 | Pcnx2 | 270109 | protein_coding | 5.294769774 | 0.436674395 | 0.796622211 | -0.548157444 | 0.5835838 | 0.766020418 | 0.436674395 |
| ENSMUSG00000037020 | Wdr62 | 233064 | protein_coding | 79.06598951 | 0.436692213 | 0.19410204 | -2.249807431 | 0.024461172 | 0.085928908 | 0.436692213 |
| ENSMUSG00000086468 | Etaa1os | 102633673 | lncRNA | 6.248461861 | -0.437075845 | 0.446326264 | 0.979274313 | 0.327444459 | 0.549590181 | 0.437075845 |
| ENSMUSG00000002984 | Tomm40 | 53333 | protein_coding | 811.1435228 | 0.437179665 | 0.091219634 | -4.792604889 | 1.65E-06 | 2.17E-05 | 0.437179665 |
| ENSMUSG00000039298 | Cdk5rap2 | 214444 | protein_coding | 316.7777503 | 0.437215006 | 0.094078302 | -4.647352241 | 3.36E-06 | 4.16E-05 | 0.437215006 |
| ENSMUSG00000035435 | Abca17 | 381072 | protein_coding | 3.488875326 | 0.437312016 | 1.727339182 | -0.2531709 | 0.800136152 | 1 | 0.437312016 |
| ENSMUSG00000073755 | 5730409E04Rik | 230757 | protein_coding | 8.76238218 | -0.437312687 | 0.422939338 | 1.033984422 | 0.301143446 | 0.521287512 | 0.437312687 |
| ENSMUSG00000097615 | Gm2061 | 100039138 | lncRNA | 113.1899271 | 0.437466598 | 0.259603616 | -1.685132913 | 0.091962931 | 0.232355545 | 0.437466598 |
| ENSMUSG00000030894 | Tpp1 | 12751 | protein_coding | 337.2842054 | -0.437643545 | 0.06663193 | 6.568075457 | 5.10E-11 | 1.34E-09 | 0.437643545 |
| ENSMUSG00000089960 | Ugt1a1 | 394436 | protein_coding | 285.5276743 | -0.437672443 | 0.188781348 | 2.318409355 | 0.020427084 | 0.07460396 | 0.437672443 |
| ENSMUSG00000078496 | Zfp982 | 195531 | protein_coding | 3.38353899 | -0.437790437 | 0.745000556 | 0.587637732 | 0.556775479 | 1 | 0.437790437 |
| ENSMUSG00000066621 | Tecpr1 | 70381 | protein_coding | 865.4918131 | 0.437856356 | 0.057842989 | -7.569739483 | 3.74E-14 | 1.29E-12 | 0.437856356 |
| ENSMUSG00000019303 | Psmc3ip | 19183 | protein_coding | 21.58298918 | 0.437865057 | 0.24938446 | -1.755783248 | 0.079125426 | 0.208467978 | 0.437865057 |
| ENSMUSG00000025991 | Cps1 | 227231 | protein_coding | 338.1852598 | 0.437874025 | 0.198474906 | -2.20619339 | 0.027370462 | 0.093755613 | 0.437874025 |
| ENSMUSG00000030173 | Klra5 | 16636 | protein_coding | 5.718169354 | -0.437938073 | 0.624446932 | 0.701321523 | 0.483102385 | 0.693738201 | 0.437938073 |
| ENSMUSG00000083012 | Fam220a | 67238 | protein_coding | 5.207960021 | 0.438174238 | 0.507103159 | -0.864073179 | 0.387547692 | 0.61004314 | 0.438174238 |
| ENSMUSG00000056222 | Spock1 | 20745 | protein_coding | 4.680521767 | -0.438238364 | 0.832422168 | 0.526461669 | 0.598567478 | 0.776186935 | 0.438238364 |
| ENSMUSG00000031627 | Irf2 | 16363 | protein_coding | 289.3605141 | -0.438265742 | 0.149676324 | 2.928089965 | 0.003410513 | 0.017711075 | 0.438265742 |
| ENSMUSG00000030060 | Hmces | 232210 | protein_coding | 112.9710063 | -0.438433935 | 0.12174911 | 3.601126416 | 0.000316841 | 0.002388097 | 0.438433935 |
| ENSMUSG00000075318 | Scn2a | 110876 | protein_coding | 3.251113992 | -0.438456361 | 2.442427295 | 0.179516648 | 0.857532047 | 1 | 0.438456361 |
| ENSMUSG00000017007 | Rbpjl | 19668 | protein_coding | 3.570849102 | 0.438554823 | 0.99317598 | -0.441568093 | 0.658801777 | 1 | 0.438554823 |
| ENSMUSG00000053769 | Lysmd1 | 217779 | protein_coding | 17.57136594 | -0.43875895 | 0.246934016 | 1.776826691 | 0.075596759 | 0.201550674 | 0.43875895 |
| ENSMUSG00000005124 | Ccn4 | 22402 | protein_coding | 2.07521022 | 0.438863489 | 1.564212618 | -0.280565112 | 0.779043978 | 1 | 0.438863489 |
| ENSMUSG00000057103 | Nat8f1 | 66116 | protein_coding | 14.9047493 | -0.438998381 | 0.322660175 | 1.360559543 | 0.173652924 | 0.364427834 | 0.438998381 |
| ENSMUSG00000000706 | Btn1a1 | 12231 | protein_coding | 64.98481613 | 0.439057655 | 0.193261291 | -2.271834425 | 0.023096513 | 0.0821132 | 0.439057655 |
| ENSMUSG00000017453 | Pipox | 19193 | protein_coding | 56.69456944 | 0.439060457 | 0.184522758 | -2.379437973 | 0.017339061 | 0.065598201 | 0.439060457 |
| ENSMUSG00000092819 | Gm23639 | 115489002 | snoRNA | 70.69377676 | 0.439076428 | 0.297948351 | -1.473666249 | 0.140571474 | 0.314808425 | 0.439076428 |
| ENSMUSG00000017832 | Hspb9 | 75482 | protein_coding | 1.614354407 | 0.439175598 | 0.861465125 | -0.509800786 | 0.610191035 | 1 | 0.439175598 |
| ENSMUSG00000023931 | Efhb | 211482 | protein_coding | 18.16969602 | -0.439327912 | 0.289669768 | 1.516650893 | 0.129354849 | 0.297019741 | 0.439327912 |
| ENSMUSG00000026388 | 3110009E18Rik | 73103 | protein_coding | 11.91757192 | -0.439583656 | 0.301298722 | 1.458962897 | 0.14457532 | 0.321507788 | 0.439583656 |
| ENSMUSG00000024143 | Rhoq | 104215 | protein_coding | 399.7375403 | 0.439630616 | 0.122468651 | -3.589739999 | 0.000331008 | 0.002484666 | 0.439630616 |
| ENSMUSG00000053192 | Mllt11 | 56772 | protein_coding | 69.20473161 | -0.439631178 | 0.153456726 | 2.864854412 | 0.00417201 | 0.020879192 | 0.439631178 |
| ENSMUSG00000044724 | Gpr152 | 269053 | protein_coding | 1.913782085 | 0.439992976 | 1.21291006 | -0.362758122 | 0.716785578 | 1 | 0.439992976 |
| ENSMUSG00000028702 | Rad54l | 19366 | protein_coding | 84.62725531 | 0.440134448 | 0.215172015 | -2.045500421 | 0.040805548 | 0.127940538 | 0.440134448 |
| ENSMUSG00000027984 | Hadh | 15107 | protein_coding | 3226.117383 | -0.440178053 | 0.091731581 | 4.798544293 | 1.60E-06 | 2.11E-05 | 0.440178053 |
| ENSMUSG00000032353 | Tmed3 | 66111 | protein_coding | 1299.249991 | 0.440179002 | 0.170020115 | -2.588981909 | 0.009626015 | 0.041288574 | 0.440179002 |

Supplementary Table 1

| ensembl_gene_id | external_gene_name | entrezgene_id | gene_biotype | baseMean | log2FoldChange | lfcSE | stat | pvalue | padj | absLog2FC |
|---|---|---|---|---|---|---|---|---|---|---|
| ENSMUSG00000026748 | Plxdc2 | 67448 | protein_coding | 1.330671848 | 0.440264514 | 2.128354374 | -0.206856771 | 0.836121718 | 1 | 0.440264514 |
| ENSMUSG00000070495 | Ctcfl | 664799 | protein_coding | 11.14434052 | -0.440272828 | 0.360911906 | 1.219890007 | 0.222506575 | 0.429533455 | 0.440272828 |
| ENSMUSG00000060791 | Gmfg | 63986 | protein_coding | 179.9138322 | 0.440410638 | 0.11158623 | -3.946818849 | 7.92E-05 | 0.000703145 | 0.440410638 |
| ENSMUSG00000026825 | Dnm1 | 13429 | protein_coding | 47.49953848 | 0.440519662 | 0.366054853 | -1.203425273 | 0.228811793 | 0.437741278 | 0.440519662 |
| ENSMUSG00000042010 | Acacb | 100705 | protein_coding | 418.6456114 | 0.440976235 | 0.207942412 | -2.120665187 | 0.033949988 | 0.110778523 | 0.440976235 |
| ENSMUSG00000028211 | Trp53inp1 | 60599 | protein_coding | 782.0176475 | -0.441026455 | 0.146209856 | 3.01639347 | 0.002558011 | 0.013886585 | 0.441026455 |
| ENSMUSG00000031451 | Gas6 | 14456 | protein_coding | 4193.708382 | -0.441076375 | 0.117964695 | 3.739054097 | 0.000184714 | 0.0014822 | 0.441076375 |
| ENSMUSG00000031150 | Ccdc120 | 54648 | protein_coding | 136.7964492 | -0.441077938 | 0.187099231 | 2.357454578 | 0.018400706 | 0.068717608 | 0.441077938 |
| ENSMUSG00000070081 | Mir505 | 751545 | miRNA | 0.507643072 | -0.441096104 | 1.491859967 | 0.295668571 | 0.767483201 | 1 | 0.441096104 |
| ENSMUSG00000028521 | Slc35d1 | 242585 | protein_coding | 503.6798964 | -0.441140831 | 0.226348543 | 1.948944863 | 0.05130201 | 0.152100788 | 0.441140831 |
| ENSMUSG00000065653 | Gm23105 | 115487521 | snoRNA | 138.5365793 | 0.441162266 | 0.25358685 | -1.739689048 | 0.081913633 | 0.213539052 | 0.441162266 |
| ENSMUSG00000022265 | Ank | 11732 | protein_coding | 835.0839124 | -0.441193401 | 0.099220599 | 4.446590773 | 8.72E-06 | 9.92E-05 | 0.441193401 |
| ENSMUSG00000038323 | 1700066M21Rik | 73467 | protein_coding | 94.34043391 | 0.441218886 | 0.199824006 | -2.208037435 | 0.027241664 | 0.093415051 | 0.441218886 |
| ENSMUSG00000027555 | Car13 | 71934 | protein_coding | 541.9713108 | 0.441239847 | 0.080026914 | -5.513643185 | 3.51E-08 | 6.19E-07 | 0.441239847 |
| ENSMUSG00000049551 | Fzd9 | 14371 | protein_coding | 26.4801668 | -0.441241007 | 0.260937219 | 1.69098532 | 0.090839606 | 0.230331489 | 0.441241007 |
| ENSMUSG00000000340 | Dbt | 13171 | protein_coding | 396.8491352 | -0.441389494 | 0.115535458 | 3.820381206 | 0.000133246 | 0.0011102 | 0.441389494 |
| ENSMUSG00000079084 | Ccdc82 | 66396 | protein_coding | 146.1379082 | -0.441393526 | 0.130739204 | 3.376137467 | 0.000735112 | 0.004881237 | 0.441393526 |
| ENSMUSG00000007591 | Tssk4 | 71099 | protein_coding | 21.97588492 | 0.441470647 | 0.249425015 | -1.769953372 | 0.076734909 | 0.204058016 | 0.441470647 |
| ENSMUSG00000038836 | Agbl3 | 76223 | protein_coding | 15.74774053 | 0.441682726 | 0.332498874 | -1.328373598 | 0.184054717 | 0.379085971 | 0.441682726 |
| ENSMUSG00000033544 | Angptl1 | 72713 | protein_coding | 7.539300965 | 0.441696484 | 0.453450058 | -0.974079672 | 0.330016982 | 0.552168231 | 0.441696484 |
| ENSMUSG00000056073 | Grik2 | 14806 | protein_coding | 3.108925784 | -0.441698759 | 2.400522119 | 0.18400112 | 0.854012576 | 1 | 0.441698759 |
| ENSMUSG00000008668 | Rps18 | 20084 | protein_coding | 5419.960139 | -0.441755199 | 0.118829387 | 3.717558516 | 0.000201157 | 0.001589432 | 0.441755199 |
| ENSMUSG00000022900 | Ildr1 | 106347 | protein_coding | 3769.495805 | 0.441977427 | 0.103217514 | -4.282000328 | 1.85E-05 | 0.000192981 | 0.441977427 |
| ENSMUSG00000102047 | 2010010A06Rik | 100503117 | transcribed_processed_pseudogene | 18.57494253 | -0.441984319 | 0.485773084 | 0.909857573 | 0.362897627 | 0.585225632 | 0.441984319 |
| ENSMUSG00000045775 | Slc16a5 | 217316 | protein_coding | 1424.148791 | 0.442114644 | 0.182758047 | -2.419125459 | 0.015557873 | 0.060127011 | 0.442114644 |
| ENSMUSG00000010044 | Zmynd10 | 114602 | protein_coding | 3.300740004 | 0.44218702 | 0.954294101 | -0.46336556 | 0.643102361 | 1 | 0.44218702 |
| ENSMUSG00000054414 | Slc30a7 | 66500 | protein_coding | 930.3959601 | 0.442439938 | 0.102665212 | -4.309540969 | 1.64E-05 | 0.000172846 | 0.442439938 |
| ENSMUSG00000039058 | Ak5 | 229949 | protein_coding | 3.849746196 | -0.442447584 | 0.740218428 | 0.597725707 | 0.550022969 | 1 | 0.442447584 |
| ENSMUSG00000040710 | St8sia4 | 20452 | protein_coding | 14.39143101 | 0.442553511 | 0.600320585 | -0.737195296 | 0.461003594 | 0.674995696 | 0.442553511 |
| ENSMUSG00000049751 | Rpl36al | 66483 | protein_coding | 241.3721747 | 0.442724948 | 0.167496693 | -2.643186196 | 0.008212986 | 0.036352985 | 0.442724948 |
| ENSMUSG00000067235 | H2-Q10 | 15007 | protein_coding | 50.19584864 | -0.442751685 | 0.222238756 | 1.992234353 | 0.046345351 | 0.140818237 | 0.442751685 |
| ENSMUSG00000052821 | Cysltr1 | 58861 | protein_coding | 4.094267078 | 0.442780331 | 0.808485926 | -0.547666096 | 0.583921196 | 1 | 0.442780331 |
| ENSMUSG00000023935 | Spats1 | 71020 | protein_coding | 7.424574327 | 0.44279073 | 0.399080864 | -1.109526341 | 0.267203188 | 0.482454733 | 0.44279073 |
| ENSMUSG00000045128 | Rpl18a | 76808 | protein_coding | 4357.431313 | -0.442819122 | 0.125361688 | 3.532332161 | 0.000411912 | 0.003009871 | 0.442819122 |
| ENSMUSG00000022783 | Spag6l | 50525 | protein_coding | 3.941934199 | -0.442895326 | 0.655219782 | 0.675949258 | 0.499072862 | 1 | 0.442895326 |
| ENSMUSG00000027634 | Ndrg3 | 29812 | protein_coding | 621.5140633 | -0.442987737 | 0.092051589 | 4.812385559 | 1.49E-06 | 1.98E-05 | 0.442987737 |
| ENSMUSG00000021095 | Gsc | 14836 | protein_coding | 0.733220376 | 0.442989588 | 1.563733826 | -0.283289637 | 0.776954821 | 1 | 0.442989588 |
| ENSMUSG00000040813 | Tex264 | 21767 | protein_coding | 602.1598606 | -0.443043409 | 0.064680794 | 6.849690319 | 7.40E-12 | 2.09E-10 | 0.443043409 |
| ENSMUSG00000027358 | Bmp2 | 12156 | protein_coding | 291.5161615 | -0.44310701 | 0.239584812 | 1.849478713 | 0.064388719 | 0.179735514 | 0.44310701 |

| ensembl_gene_id | external_gene_name | entrezgene_id | gene_biotype | baseMean | log2FoldChange | lfcSE | stat | pvalue | padj | absLog2FC |
|---|---|---|---|---|---|---|---|---|---|---|
| ENSMUSG00000028778 | Hcrtr1 | 230777 | protein_coding | 2.785054448 | 0.443131966 | 0.876812312 | -0.505389762 | 0.613285108 | 1 | 0.443131966 |
| ENSMUSG00000022475 | Hdac7 | 56233 | protein_coding | 83.55447829 | 0.44317491 | 0.179430888 | -2.469891975 | 0.013515386 | 0.053833353 | 0.44317491 |
| ENSMUSG00000024462 | Gabbr1 | 54393 | protein_coding | 82.78885852 | 0.443238017 | 0.22071991 | -2.008146961 | 0.044627677 | 0.136655194 | 0.443238017 |
| ENSMUSG00000059991 | Nptx2 | 53324 | protein_coding | 13.61438182 | 0.443274809 | 0.434132844 | -1.021057988 | 0.307226966 | 0.527698391 | 0.443274809 |
| ENSMUSG00000112280 | A830082N09Rik | 414093 | lncRNA | 5.301018894 | 0.443357974 | 0.582161879 | -0.761571636 | 0.446315709 | 0.663584952 | 0.443357974 |
| ENSMUSG00000032062 | 2310030G06Rik | 66952 | protein_coding | 209.0092851 | 0.44338908 | 0.098652938 | -4.49443362 | 6.98E-06 | 8.11E-05 | 0.44338908 |
| ENSMUSG00000017404 | Rpl19 | 19921 | protein_coding | 4196.104562 | -0.443413852 | 0.128105608 | 3.461314907 | 0.000537544 | 0.003771026 | 0.443413852 |
| ENSMUSG00000070544 | Top1 | 21969 | protein_coding | 1458.307412 | 0.443452829 | 0.059058362 | -7.50872205 | 5.97E-14 | 2.01E-12 | 0.443452829 |
| ENSMUSG00000063445 | Nmral1 | 67824 | protein_coding | 75.90380393 | 0.443454409 | 0.190133355 | -2.332333582 | 0.019683149 | 0.07241317 | 0.443454409 |
| ENSMUSG00000055421 | Pcdh9 | 211712 | protein_coding | 1.523347713 | 0.443608746 | 2.37420918 | -0.186844845 | 0.851782285 | 1 | 0.443608746 |
| ENSMUSG00000044628 | Rnf208 | 68846 | protein_coding | 8.11183613 | -0.443641531 | 0.515058537 | 0.861341963 | 0.389049731 | 0.611707822 | 0.443641531 |
| ENSMUSG00000095562 | Gm55594 | 170942 | lncRNA | 29.69464996 | -0.443715301 | 0.874819145 | 0.507208037 | 0.612008856 | 0.785292483 | 0.443715301 |
| ENSMUSG00000030775 | Trat1 | 77647 | protein_coding | 4.158100137 | -0.444044077 | 0.814595254 | 0.545110071 | 0.585677816 | 1 | 0.444044077 |
| ENSMUSG00000045912 | C2cd4c | 237397 | protein_coding | 29.14695195 | -0.444098878 | 0.223834838 | 1.984047172 | 0.047250581 | 0.142911457 | 0.444098878 |
| ENSMUSG00000063873 | Slc24a3 | 94249 | protein_coding | 41.99309544 | -0.444154527 | 0.226805831 | 1.958302946 | 0.050194475 | 0.149272747 | 0.444154527 |
| ENSMUSG00000006456 | Rbm14 | 56275 | protein_coding | 494.4200081 | 0.444162972 | 0.066475768 | -6.6815771 | 2.36E-11 | 6.36E-10 | 0.444162972 |
| ENSMUSG00000049804 | Armcx4 | 100503043 | protein_coding | 12.02141982 | 0.444340431 | 0.532326883 | -0.834713491 | 0.403879044 | 0.625085148 | 0.444340431 |
| ENSMUSG00000031391 | L1cam | 16728 | protein_coding | 1325.043731 | -0.444510066 | 0.093620934 | 4.747977262 | 2.05E-06 | 2.67E-05 | 0.444510066 |
| ENSMUSG00000042942 | Greb1l | 381157 | protein_coding | 103.9857606 | -0.444671902 | 0.139473255 | 3.188223443 | 0.001431499 | 0.008568238 | 0.444671902 |
| ENSMUSG00000031538 | Plat | 18791 | protein_coding | 10.20670937 | 0.444688975 | 0.639750493 | -0.69509751 | 0.486994195 | 0.695624496 | 0.444688975 |
| ENSMUSG00000028820 | Sfpq | 71514 | protein_coding | 2146.021868 | 0.445087799 | 0.062204855 | -7.155193876 | 8.36E-13 | 2.56E-11 | 0.445087799 |
| ENSMUSG00000029464 | Gpn3 | 68080 | protein_coding | 199.6008751 | 0.445115208 | 0.113765429 | -3.912570038 | 9.13E-05 | 0.000796221 | 0.445115208 |
| ENSMUSG00000091813 | Ces2h | 436059 | protein_coding | 3.747487908 | 0.445177391 | 0.690452576 | -0.64476172 | 0.519081613 | 1 | 0.445177391 |
| ENSMUSG00000099146 | 0610031O16Rik | 68369 | lncRNA | 2.883680877 | 0.445215205 | 0.846881922 | -0.525711074 | 0.599088968 | 1 | 0.445215205 |
| ENSMUSG00000030298 | Sec13 | 110379 | protein_coding | 1710.738724 | 0.44522746 | 0.070077479 | -6.353360135 | 2.11E-10 | 5.06E-09 | 0.44522746 |
| ENSMUSG00000110062 | B430319F04Rik | 619306 | TEC | 20.31040183 | 0.445266052 | 0.28928253 | -1.53920823 | 0.123753469 | 0.287840111 | 0.445266052 |
| ENSMUSG00000020838 | Slc6a4 | 15567 | protein_coding | 157.7763391 | -0.445366262 | 0.311801098 | 1.428366562 | 0.153186378 | 0.334228588 | 0.445366262 |
| ENSMUSG00000024992 | Pde6c | 110855 | protein_coding | 1.60711077 | -0.445516611 | 1.564678537 | 0.284733637 | 0.775848218 | 1 | 0.445516611 |
| ENSMUSG00000050910 | Cdr2l | 237988 | protein_coding | 119.5561089 | 0.445594217 | 0.181362677 | -2.456923468 | 0.014013254 | 0.055305475 | 0.445594217 |
| ENSMUSG00000085287 | 4833418N02Rik | 74597 | lncRNA | 51.11624493 | 0.445836499 | 0.227943841 | -1.955905001 | 0.050476346 | 0.14994898 | 0.445836499 |
| ENSMUSG00000028066 | Pmf1 | 67037 | protein_coding | 216.2148066 | 0.445849232 | 0.101891739 | -4.375715196 | 1.21E-05 | 0.000132912 | 0.445849232 |
| ENSMUSG00000028688 | Toe1 | 68276 | protein_coding | 165.311386 | 0.445860824 | 0.105582605 | -4.222862512 | 2.41E-05 | 0.000244236 | 0.445860824 |
| ENSMUSG00000058145 | Adamts17 | 233332 | protein_coding | 70.90786864 | 0.445894914 | 0.179188384 | -2.488414167 | 0.012831421 | 0.051654946 | 0.445894914 |
| ENSMUSG00000035828 | Pim3 | 223775 | protein_coding | 1138.718354 | -0.445952152 | 0.130779458 | 3.409955654 | 0.000649734 | 0.004413704 | 0.445952152 |
| ENSMUSG00000028018 | Gstcd | 67553 | protein_coding | 134.8455573 | 0.44597737 | 0.146585819 | -3.042431886 | 0.002346749 | 0.012896578 | 0.44597737 |
| ENSMUSG00000029209 | Gnpda2 | 67980 | protein_coding | 61.99082544 | -0.446021702 | 0.144624574 | 3.083996663 | 0.002042399 | 0.01154679 | 0.446021702 |
| ENSMUSG00000104667 | Gm4961 | 242914 | lncRNA | 6.723290657 | 0.446026964 | 0.526367663 | -0.847367716 | 0.396790193 | 0.618855346 | 0.446026964 |
| ENSMUSG00000105496 | Mir7004 | 102466784 | miRNA | 0.710300434 | -0.446045312 | 1.184992124 | 0.376412048 | 0.70661059 | 1 | 0.446045312 |
| ENSMUSG00000034111 | Tmed8 | 382620 | protein_coding | 71.76214383 | -0.446089143 | 0.154760981 | 2.882439353 | 0.003946091 | 0.019983363 | 0.446089143 |

| ensembl_gene_id | external_gene_name | entrezgene_id | gene_biotype | baseMean | log2FoldChange | lfcSE | stat | pvalue | padj | absLog2FC |
|---|---|---|---|---|---|---|---|---|---|---|
| ENSMUSG00000116396 | Gm20556 | 328576 | lncRNA | 1.743750709 | -0.446138843 | 0.948192549 | 0.470515027 | 0.637987101 | 1 | 0.446138843 |
| ENSMUSG00000061838 | Suclg2 | 20917 | protein_coding | 2478.300946 | -0.44636377 | 0.104558569 | 4.269030965 | 1.96E-05 | 0.000202378 | 0.44636377 |
| ENSMUSG00000020834 | Dhrs13 | 70451 | protein_coding | 63.80426074 | -0.4464199 | 0.176448125 | 2.530034824 | 0.011405121 | 0.047310767 | 0.4464199 |
| ENSMUSG00000019975 | Ikbip | 67454 | protein_coding | 168.2562871 | 0.446698242 | 0.129359894 | -3.453143222 | 0.000554095 | 0.003857639 | 0.446698242 |
| ENSMUSG00000029189 | Sel1l3 | 231238 | protein_coding | 2624.965694 | 0.446737562 | 0.081715918 | -5.466958867 | 4.58E-08 | 7.90E-07 | 0.446737562 |
| ENSMUSG00000059409 | Ppp2r5d | 21770 | protein_coding | 838.4114419 | 0.446748508 | 0.072817242 | -6.135202236 | 8.51E-10 | 1.89E-08 | 0.446748508 |
| ENSMUSG00000064065 | Ipcef1 | 320495 | protein_coding | 15.15853716 | 0.447123242 | 0.352874744 | -1.267087684 | 0.205123939 | 0.407579707 | 0.447123242 |
| ENSMUSG00000018865 | Sult4a1 | 29859 | protein_coding | 9.336353813 | 0.447445005 | 0.442221385 | -1.01181223 | 0.311627841 | 0.532495959 | 0.447445005 |
| ENSMUSG00000026787 | Gad2 | 14417 | protein_coding | 25.80302935 | -0.447506512 | 0.27156914 | 1.647854801 | 0.09938247 | 0.246352764 | 0.447506512 |
| ENSMUSG00000024691 | Fam111a | 107373 | protein_coding | 143.2875634 | 0.447590662 | 0.251582861 | -1.779098385 | 0.075223633 | 0.200815241 | 0.447590662 |
| ENSMUSG00000024411 | Aqp4 | 11829 | protein_coding | 14478.62701 | 0.447625109 | 0.190695821 | -2.347325217 | 0.018908742 | 0.07028111 | 0.447625109 |
| ENSMUSG00000054942 | Miga1 | 215708 | protein_coding | 125.807203 | -0.447827378 | 0.109682022 | 4.082960623 | 4.45E-05 | 0.000421772 | 0.447827378 |
| ENSMUSG00000046808 | Atp10d | 231287 | protein_coding | 9.185001778 | -0.4478678 | 0.463515 | 0.966242302 | 0.33392295 | 0.556279996 | 0.4478678 |
| ENSMUSG00000042155 | Klhl23 | 277396 | protein_coding | 54.73108486 | 0.44787949 | 0.254513182 | -1.759749679 | 0.078450258 | 0.207184962 | 0.44787949 |
| ENSMUSG00000023926 | Rhag | 19743 | protein_coding | 0.947043583 | 0.448018642 | 1.114554425 | -0.401971077 | 0.687705312 | 1 | 0.448018642 |
| ENSMUSG00000074646 | 6430550D23Rik | 320095 | protein_coding | 12.34203345 | -0.448086357 | 0.430162653 | 1.041667272 | 0.297565968 | 0.51726362 | 0.448086357 |
| ENSMUSG00000038047 | Haus6 | 230376 | protein_coding | 141.6992519 | 0.448176509 | 0.14513634 | -3.087968934 | 0.002015295 | 0.011401366 | 0.448176509 |
| ENSMUSG00000022451 | Twf1 | 19230 | protein_coding | 1843.717767 | 0.448198056 | 0.072213454 | -6.206572744 | 5.42E-10 | 1.23E-08 | 0.448198056 |
| ENSMUSG00000030978 | Rrm1 | 20133 | protein_coding | 673.3457049 | 0.448203667 | 0.191504613 | -2.340432742 | 0.019261407 | 0.071287064 | 0.448203667 |
| ENSMUSG00000000394 | Gcg | 14526 | protein_coding | 415.3484431 | -0.448330828 | 0.131236124 | 3.416215096 | 0.000634981 | 0.004327722 | 0.448330828 |
| ENSMUSG00000014164 | Klhl3 | 100503085 | protein_coding | 26.06305303 | -0.448341885 | 0.251666107 | 1.781494899 | 0.074831637 | 0.19999509 | 0.448341885 |
| ENSMUSG00000034708 | Grn | 14824 | protein_coding | 871.9073227 | -0.448419247 | 0.061425098 | 7.300260974 | 2.87E-13 | 9.10E-12 | 0.448419247 |
| ENSMUSG00000035000 | Dpp4 | 13482 | protein_coding | 758.5165014 | 0.448495547 | 0.101393308 | -4.423324948 | 9.72E-06 | 0.000109485 | 0.448495547 |
| ENSMUSG00000050623 | Catsperz | 67077 | protein_coding | 5.305472626 | 0.448574867 | 0.44054907 | -1.018217713 | 0.308574504 | 0.529213766 | 0.448574867 |
| ENSMUSG00000049115 | Agtr1a | 11607 | protein_coding | 4.486803755 | 0.448584026 | 0.624800589 | -0.717963513 | 0.472779785 | 0.685206817 | 0.448584026 |
| ENSMUSG00000003545 | Fosb | 14282 | protein_coding | 5081.997123 | -0.448713739 | 0.117286808 | 3.82578184 | 0.000130358 | 0.001090541 | 0.448713739 |
| ENSMUSG00000021390 | Ogn | 18295 | protein_coding | 4.382227197 | 0.448714766 | 0.527533824 | -0.850589565 | 0.394997388 | 1 | 0.448714766 |
| ENSMUSG00000029217 | Tec | 21682 | protein_coding | 168.2900672 | -0.448841359 | 0.100872712 | 4.449581547 | 8.60E-06 | 9.80E-05 | 0.448841359 |
| ENSMUSG00000021118 | Plek2 | 27260 | protein_coding | 406.2310013 | -0.448957168 | 0.092625318 | 4.847024334 | 1.25E-06 | 1.69E-05 | 0.448957168 |
| ENSMUSG00000012609 | Ttll5 | 320244 | protein_coding | 248.2222762 | 0.449024949 | 0.084451559 | -5.316952728 | 1.06E-07 | 1.74E-06 | 0.449024949 |
| ENSMUSG00000036782 | Klhl13 | 67455 | protein_coding | 36.68910002 | 0.449052102 | 0.283476292 | -1.584090502 | 0.113173125 | 0.270127107 | 0.449052102 |
| ENSMUSG00000012076 | Brms1l | 52592 | protein_coding | 83.69881485 | 0.449066429 | 0.180541502 | -2.487330743 | 0.012870568 | 0.051764384 | 0.449066429 |
| ENSMUSG00000043895 | S1pr2 | 14739 | protein_coding | 27.16675731 | 0.449113823 | 0.206535542 | -2.174511078 | 0.029666774 | 0.10006477 | 0.449113823 |
| ENSMUSG00000030493 | Faap24 | 101831 | protein_coding | 83.12339132 | 0.449223967 | 0.128538453 | -3.494860538 | 0.00047431 | 0.003393738 | 0.449223967 |
| ENSMUSG00000055197 | Fev | 260298 | protein_coding | 3.814250231 | -0.44954904 | 0.719535603 | 0.624776645 | 0.532117661 | 1 | 0.44954904 |
| ENSMUSG00000055240 | Zfp101 | 22643 | protein_coding | 121.6451827 | 0.449612786 | 0.165410746 | -2.71815947 | 0.006564619 | 0.03021408 | 0.449612786 |
| ENSMUSG00000091648 | C2cd4d | 271944 | protein_coding | 4.150391142 | 0.449623403 | 0.611000114 | -0.735881047 | 0.461803092 | 1 | 0.449623403 |
| ENSMUSG00000018068 | Ints2 | 70422 | protein_coding | 325.6084509 | 0.449655412 | 0.121819539 | -3.691160011 | 0.000223234 | 0.001743841 | 0.449655412 |
| ENSMUSG00000040964 | Arhgef10l | 72754 | protein_coding | 799.219105 | 0.449780034 | 0.066299006 | -6.784114296 | 1.17E-11 | 3.25E-10 | 0.449780034 |

| ensembl_gene_id | external_gene_name | entrezgene_id | gene_biotype | baseMean | log2FoldChange | lfcSE | stat | pvalue | padj | absLog2FC |
|---|---|---|---|---|---|---|---|---|---|---|
| ENSMUSG00000079427 | Mthfsl | 100039707 | protein_coding | 55.76990498 | -0.449799404 | 0.145608533 | 3.089100578 | 0.002007635 | 0.011373612 | 0.449799404 |
| ENSMUSG00000022780 | Meltf | 30060 | protein_coding | 1.886135024 | -0.449908038 | 1.199274405 | 0.375150205 | 0.707548761 | 1 | 0.449908038 |
| ENSMUSG00000078941 | Ak6 | 108143 | protein_coding | 131.639024 | 0.449935957 | 0.110430286 | -4.074389114 | 4.61E-05 | 0.00043511 | 0.449935957 |
| ENSMUSG00000040812 | Agbl2 | 271813 | protein_coding | 35.95918759 | -0.450271445 | 0.354880552 | 1.268797184 | 0.20451341 | 0.406858746 | 0.450271445 |
| ENSMUSG00000051736 | Fam229b | 66337 | protein_coding | 5.091200518 | 0.450307274 | 0.482642716 | -0.933003356 | 0.350818235 | 0.572963936 | 0.450307274 |
| ENSMUSG00000044037 | Als2cl | 235633 | protein_coding | 455.3478948 | -0.450449635 | 0.171204284 | 2.631065207 | 0.00851177 | 0.037373936 | 0.450449635 |
| ENSMUSG00000079580 | Tmem217 | 71138 | protein_coding | 3.863646088 | 0.450702474 | 0.706020037 | -0.638370655 | 0.523232428 | 1 | 0.450702474 |
| ENSMUSG00000032000 | Birc3 | 11796 | protein_coding | 485.6661365 | -0.450965353 | 0.103404968 | 4.361157504 | 1.29E-05 | 0.000140653 | 0.450965353 |
| ENSMUSG00000038128 | Camk4 | 12326 | protein_coding | 4.076186762 | 0.451089194 | 0.664461584 | -0.678879268 | 0.497214363 | 1 | 0.451089194 |
| ENSMUSG00000073650 | Catip | 241112 | protein_coding | 2.263683746 | 0.451112105 | 1.508792966 | -0.298988738 | 0.764948639 | 1 | 0.451112105 |
| ENSMUSG00000032527 | Pccb | 66904 | protein_coding | 2502.654338 | 0.451131835 | 0.107833508 | -4.183596013 | 2.87E-05 | 0.000285104 | 0.451131835 |
| ENSMUSG00000039131 | Gipc2 | 54120 | protein_coding | 1710.843042 | -0.451146109 | 0.061112426 | 7.382231994 | 1.56E-13 | 5.08E-12 | 0.451146109 |
| ENSMUSG00000021376 | Tpmt | 22017 | protein_coding | 145.7373507 | -0.451275875 | 0.095740994 | 4.713507327 | 2.43E-06 | 3.10E-05 | 0.451275875 |
| ENSMUSG00000041460 | Cacna2d4 | 319734 | protein_coding | 12.89654523 | 0.451567643 | 0.647614113 | -0.69727888 | 0.485628283 | 0.695380179 | 0.451567643 |
| ENSMUSG00000005142 | Man2b1 | 17159 | protein_coding | 1359.027933 | -0.451616444 | 0.128334779 | 3.519049537 | 0.000433096 | 0.003149337 | 0.451616444 |
| ENSMUSG00000090565 | 4930405O22Rik | 414079 | lncRNA | 6.020608153 | -0.451755639 | 0.479820388 | 0.941509885 | 0.346443626 | 0.568175062 | 0.451755639 |
| ENSMUSG00000064307 | Lrrc51 | 69358 | protein_coding | 37.54127648 | -0.45176234 | 0.19156642 | 2.358254336 | 0.018361108 | 0.068585239 | 0.45176234 |
| ENSMUSG00000100680 | 1810044D09Rik | 69798 | lncRNA | 12.67821067 | -0.451855516 | 0.320399448 | 1.410288062 | 0.158454642 | 0.342262172 | 0.451855516 |
| ENSMUSG00000029607 | Ankrd61 | 66729 | protein_coding | 11.12867942 | -0.451867836 | 0.455774973 | 0.991427486 | 0.321476884 | 0.542471509 | 0.451867836 |
| ENSMUSG00000031385 | Plxnb3 | 140571 | protein_coding | 4.225033661 | 0.451929709 | 1.514466372 | -0.298408547 | 0.765391367 | 1 | 0.451929709 |
| ENSMUSG00000070498 | Tmem132b | 208151 | protein_coding | 4.920391046 | 0.451948092 | 1.007307355 | -0.448669505 | 0.653670087 | 0.81212066 | 0.451948092 |
| ENSMUSG00000026639 | Lamb3 | 16780 | protein_coding | 3251.876467 | 0.452145529 | 0.153752807 | -2.940730252 | 0.003274395 | 0.01711111 | 0.452145529 |
| ENSMUSG00000038630 | Zkscan16 | 100041581 | protein_coding | 1.072622222 | 0.452167249 | 1.784841166 | -0.253337528 | 0.800007399 | 1 | 0.452167249 |
| ENSMUSG00000070568 | Slc6a21 | 76713 | protein_coding | 2.3357246 | -0.45224324 | 1.772390476 | 0.255160048 | 0.798599487 | 1 | 0.45224324 |
| ENSMUSG00000027654 | Fam83d | 71878 | protein_coding | 164.8221123 | 0.452283151 | 0.159749782 | -2.831197291 | 0.004637411 | 0.022846718 | 0.452283151 |
| ENSMUSG00000011589 | Fsd1 | 240121 | protein_coding | 7.001741867 | -0.452358353 | 0.61540664 | 0.735056017 | 0.46230538 | 0.675942878 | 0.452358353 |
| ENSMUSG00000005672 | Kit | 16590 | protein_coding | 617.1475563 | -0.452503741 | 0.232208998 | 1.948691673 | 0.051332257 | 0.152129476 | 0.452503741 |
| ENSMUSG00000033705 | Stard9 | 668880 | protein_coding | 11.87644145 | 0.452526152 | 0.70831004 | -0.638881459 | 0.52290005 | 0.722022907 | 0.452526152 |
| ENSMUSG00000084669 | Gm23206 | 115488234 | snRNA | 2.878228843 | -0.452575428 | 0.741123946 | 0.610660917 | 0.541424085 | 1 | 0.452575428 |
| ENSMUSG00000114590 | 5930438M14Rik | 328324 | lncRNA | 1.053439962 | -0.452618558 | 1.378899454 | 0.328246238 | 0.74272549 | 1 | 0.452618558 |
| ENSMUSG00000044813 | Shb | 230126 | protein_coding | 340.6828314 | 0.452790123 | 0.107863966 | -4.197788564 | 2.70E-05 | 0.000269273 | 0.452790123 |
| ENSMUSG00000018925 | Heatr9 | 629303 | protein_coding | 2.647235069 | 0.452901766 | 1.073988543 | -0.421700742 | 0.673243462 | 1 | 0.452901766 |
| ENSMUSG00000022012 | Enox1 | 239188 | protein_coding | 6.268478808 | -0.453112849 | 0.747931416 | 0.605821389 | 0.544633365 | 0.736796542 | 0.453112849 |
| ENSMUSG00000056155 | Nanos3 | 244551 | protein_coding | 0.995244453 | 0.453308281 | 1.663730324 | -0.272464999 | 0.7852645 | 1 | 0.453308281 |
| ENSMUSG00000008153 | Clstn3 | 232370 | protein_coding | 5.733117623 | 0.453366472 | 1.390823417 | -0.32596983 | 0.744447182 | 0.871097624 | 0.453366472 |
| ENSMUSG00000019961 | Tmpo | 21917 | protein_coding | 1693.729055 | 0.453574005 | 0.140215271 | -3.234840275 | 0.001217108 | 0.007482994 | 0.453574005 |
| ENSMUSG00000080727 | C920021L13Rik | 100042889 | lncRNA | 7.519670238 | -0.453741391 | 0.473741766 | 0.957782116 | 0.338172639 | 0.559860817 | 0.453741391 |
| ENSMUSG00000026815 | Gfi1b | 14582 | protein_coding | 8.037330404 | -0.453853575 | 0.414504525 | 1.094930329 | 0.273547161 | 0.489845783 | 0.453853575 |
| ENSMUSG00000027698 | Nceh1 | 320024 | protein_coding | 730.4143298 | -0.45390039 | 0.119461343 | 3.79955875 | 0.000144954 | 0.001195092 | 0.45390039 |

| ensembl_gene_id | external_gene_name | entrezgene_id | gene_biotype | baseMean | log2FoldChange | lfcSE | stat | pvalue | padj | absLog2FC |
|---|---|---|---|---|---|---|---|---|---|---|
| ENSMUSG00000014786 | Slc9a5 | 277973 | protein_coding | 56.98383599 | 0.454128697 | 0.203965719 | -2.226495215 | 0.025981029 | 0.09002323 | 0.454128697 |
| ENSMUSG00000030760 | Acer3 | 66190 | protein_coding | 274.052669 | -0.454192757 | 0.092869284 | 4.890667173 | 1.00E-06 | 1.39E-05 | 0.454192757 |
| ENSMUSG00000038274 | Fau | 14109 | protein_coding | 3570.908476 | -0.454229761 | 0.093544207 | 4.855776508 | 1.20E-06 | 1.62E-05 | 0.454229761 |
| ENSMUSG00000047419 | Cmya5 | 76469 | protein_coding | 3.455408347 | 0.45430362 | 1.18958425 | -0.381901172 | 0.702534671 | 1 | 0.45430362 |
| ENSMUSG00000017390 | Aldoc | 11676 | protein_coding | 153.143405 | -0.45437845 | 0.183628293 | 2.474446839 | 0.013344268 | 0.053293166 | 0.45437845 |
| ENSMUSG00000019843 | Fyn | 14360 | protein_coding | 31.69116201 | 0.454397269 | 0.275647626 | -1.64847155 | 0.099255944 | 0.246223924 | 0.454397269 |
| ENSMUSG00000020866 | Cacna1g | 12291 | protein_coding | 7.075889621 | 0.45463428 | 1.135955808 | -0.400221801 | 0.688993158 | 0.833930977 | 0.45463428 |
| ENSMUSG00000043439 | Epop | 103551 | protein_coding | 38.94792219 | 0.45475179 | 0.428890573 | -1.060297937 | 0.289009078 | 0.507895701 | 0.45475179 |
| ENSMUSG00000028943 | Espn | 56226 | protein_coding | 362.5170049 | -0.454883364 | 0.23358103 | 1.947432821 | 0.051482866 | 0.1525001 | 0.454883364 |
| ENSMUSG00000094584 | Ms4a18 | 76002 | protein_coding | 9.715614168 | -0.454909834 | 0.395992945 | 1.148782673 | 0.250645606 | 0.462889905 | 0.454909834 |
| ENSMUSG00000004347 | Pde1c | 18575 | protein_coding | 17.56158493 | 0.454936704 | 0.672609406 | -0.676375768 | 0.498802098 | 0.703739933 | 0.454936704 |
| ENSMUSG00000040270 | Bach2 | 12014 | protein_coding | 37.29151923 | -0.455048061 | 0.255939996 | 1.777948222 | 0.075412359 | 0.201270862 | 0.455048061 |
| ENSMUSG00000063894 | Zkscan8 | 93681 | protein_coding | 187.2335421 | -0.455062337 | 0.153283676 | 2.968759297 | 0.002990047 | 0.015825683 | 0.455062337 |
| ENSMUSG00000056501 | Cebpb | 12608 | protein_coding | 151.7238841 | 0.455458921 | 0.251218274 | -1.813000755 | 0.069831705 | 0.190674731 | 0.455458921 |
| ENSMUSG00000026712 | Mrc1 | 17533 | protein_coding | 11.60092243 | 0.455507691 | 0.409924963 | -1.111197735 | 0.266483246 | 0.481477497 | 0.455507691 |
| ENSMUSG00000062545 | Tlr12 | 384059 | protein_coding | 2.555266575 | 0.455574448 | 1.016191082 | -0.448315731 | 0.653925349 | 1 | 0.455574448 |
| ENSMUSG00000046215 | Rprml | 104582 | protein_coding | 2.0241664 | 0.455747102 | 0.799778587 | -0.56984159 | 0.568785144 | 1 | 0.455747102 |
| ENSMUSG00000033805 | Ephx4 | 384214 | protein_coding | 53.59836069 | -0.455749266 | 0.169902638 | 2.6824143 | 0.007309288 | 0.033042863 | 0.455749266 |
| ENSMUSG00000031618 | Nr3c2 | 110784 | protein_coding | 2264.670624 | -0.455786415 | 0.094962865 | 4.799627877 | 1.59E-06 | 2.10E-05 | 0.455786415 |
| ENSMUSG00000032717 | Mdfi | 17240 | protein_coding | 7.98383472 | 0.455927113 | 0.501159805 | -0.909743975 | 0.362957547 | 0.585225632 | 0.455927113 |
| ENSMUSG00000073680 | Tmem88b | 320587 | protein_coding | 2.585082203 | 0.45595923 | 1.057909174 | -0.431000355 | 0.666468112 | 1 | 0.45595923 |
| ENSMUSG00000037852 | Cpe | 12876 | protein_coding | 60.54527734 | -0.456128968 | 0.193868004 | 2.352781058 | 0.018633602 | 0.069446017 | 0.456128968 |
| ENSMUSG00000020149 | Rab1a | 19324 | protein_coding | 3547.676157 | 0.456192869 | 0.068623614 | -6.64775345 | 2.98E-11 | 7.94E-10 | 0.456192869 |
| ENSMUSG00000038670 | Mybpc2 | 233199 | protein_coding | 4.359735534 | -0.456204822 | 1.13155128 | 0.403167607 | 0.686824928 | 1 | 0.456204822 |
| ENSMUSG00000044921 | Rassf9 | 237504 | protein_coding | 87.23444602 | -0.456358219 | 0.211071144 | 2.162106153 | 0.030609995 | 0.102451045 | 0.456358219 |
| ENSMUSG00000028063 | Lmna | 16905 | protein_coding | 5558.575528 | -0.456558786 | 0.083707878 | 5.454191342 | 4.92E-08 | 8.45E-07 | 0.456558786 |
| ENSMUSG00000026255 | Efhd1 | 98363 | protein_coding | 4.653546524 | 0.456594202 | 0.55159799 | -0.827766253 | 0.40780289 | 0.628507412 | 0.456594202 |
| ENSMUSG00000076436 | Oxct2a | 64059 | protein_coding | 6.396688263 | -0.456607455 | 0.532618174 | 0.857288537 | 0.391285442 | 0.613645564 | 0.456607455 |
| ENSMUSG00000027824 | Vmn2r1 | 56544 | protein_coding | 1.151493856 | -0.456743966 | 1.985518595 | 0.230037617 | 0.818062539 | 1 | 0.456743966 |
| ENSMUSG00000050555 | Hyls1 | 76832 | protein_coding | 80.23776213 | 0.456807046 | 0.138744845 | -3.292425352 | 0.000993273 | 0.006330723 | 0.456807046 |
| ENSMUSG00000029553 | Tfec | 21426 | protein_coding | 0.883982192 | 0.456878508 | 1.953914082 | -0.233827328 | 0.815119033 | 1 | 0.456878508 |
| ENSMUSG00000031146 | Plp2 | 18824 | protein_coding | 816.3280395 | -0.456933864 | 0.119953858 | 3.809246901 | 0.000139391 | 0.001154992 | 0.456933864 |
| ENSMUSG00000087593 | Gm16174 | 102638408 | lncRNA | 14.16951093 | 0.456981812 | 0.292861043 | -1.56040492 | 0.118664223 | 0.279706106 | 0.456981812 |
| ENSMUSG00000073529 | F830208F22Rik | 619308 | lncRNA | 5.545066673 | -0.456991537 | 0.695795003 | 0.656790483 | 0.511315644 | 0.71414691 | 0.456991537 |
| ENSMUSG00000069270 | H2ac6 | 319164 | protein_coding | 287.5560934 | 0.457078711 | 0.14936921 | -3.060059783 | 0.002212928 | 0.012282071 | 0.457078711 |
| ENSMUSG00000074673 | Ttll9 | 74711 | protein_coding | 2.64675703 | -0.457094242 | 1.32198849 | 0.345762649 | 0.729521098 | 1 | 0.457094242 |
| ENSMUSG00000030554 | Synm | 233335 | protein_coding | 6.048099546 | 0.457195294 | 0.664317764 | -0.688217776 | 0.491315652 | 0.698491194 | 0.457195294 |
| ENSMUSG00000029581 | Fscn1 | 14086 | protein_coding | 3.301436769 | 0.457604946 | 0.801353136 | -0.571040313 | 0.567972317 | 1 | 0.457604946 |
| ENSMUSG00000062760 | Shisal1 | 72301 | protein_coding | 6.666743207 | 0.457634652 | 0.691055048 | -0.662226047 | 0.507826364 | 0.711603273 | 0.457634652 |

| ensembl_gene_id | external_gene_name | entrezgene_id | gene_biotype | baseMean | log2FoldChange | lfcSE | stat | pvalue | padj | absLog2FC |
|---|---|---|---|---|---|---|---|---|---|---|
| ENSMUSG00000005881 | Ergic3 | 66366 | protein_coding | 1201.49326 | -0.457682286 | 0.072376529 | 6.323628566 | 2.55E-10 | 6.08E-09 | 0.457682286 |
| ENSMUSG00000056305 | Usp39 | 28035 | protein_coding | 310.9536195 | 0.458124786 | 0.087779968 | -5.219012877 | 1.80E-07 | 2.84E-06 | 0.458124786 |
| ENSMUSG00000044250 | Pced1b | 239647 | protein_coding | 2.731902687 | 0.458130668 | 0.870285682 | -0.526414116 | 0.59860051 | 1 | 0.458130668 |
| ENSMUSG00000035699 | Slc51a | 106407 | protein_coding | 562.9237994 | -0.458183881 | 0.276894457 | 1.654723919 | 0.097980513 | 0.243645564 | 0.458183881 |
| ENSMUSG00000030432 | Rpl28 | 19943 | protein_coding | 3355.339182 | -0.458245854 | 0.123128253 | 3.721695422 | 0.00019789 | 0.00156603 | 0.458245854 |
| ENSMUSG00000032311 | Nrg4 | 83961 | protein_coding | 14.44962026 | -0.458523111 | 0.335769405 | 1.365589313 | 0.172067903 | 0.362486203 | 0.458523111 |
| ENSMUSG00000037428 | Vgf | 381677 | protein_coding | 6.244944824 | -0.458907835 | 0.878519106 | 0.522365229 | 0.601416061 | 0.777725809 | 0.458907835 |
| ENSMUSG00000004187 | Kifc2 | 16581 | protein_coding | 135.2386902 | -0.459052278 | 0.261011333 | 1.758744623 | 0.078620894 | 0.207507849 | 0.459052278 |
| ENSMUSG00000005952 | Trpv1 | 193034 | protein_coding | 7.557979413 | -0.459110636 | 0.477997125 | 0.960488278 | 0.336809528 | 0.559233675 | 0.459110636 |
| ENSMUSG00000042115 | Klhdc8a | 213417 | protein_coding | 102.3951251 | -0.45916776 | 0.148493182 | 3.092180751 | 0.001986918 | 0.011271717 | 0.45916776 |
| ENSMUSG00000058794 | Nfe2 | 18022 | protein_coding | 7.285971883 | 0.459223805 | 0.645341388 | -0.711598254 | 0.476713588 | 0.68833522 | 0.459223805 |
| ENSMUSG00000017778 | Cox7c | 12867 | protein_coding | 1621.738254 | -0.459314199 | 0.111264576 | 4.128126096 | 3.66E-05 | 0.00035465 | 0.459314199 |
| ENSMUSG00000101581 | C430002N11Rik | 319707 | lncRNA | 5.035726409 | -0.45938012 | 0.589126154 | 0.779765278 | 0.435529047 | 0.653742472 | 0.45938012 |
| ENSMUSG00000031227 | Magee1 | 107528 | protein_coding | 11.24022182 | 0.459395617 | 0.315385199 | -1.456617553 | 0.14522198 | 0.322338142 | 0.459395617 |
| ENSMUSG00000055884 | Fancm | 104806 | protein_coding | 119.961835 | 0.459563448 | 0.118972804 | -3.862760516 | 0.000112113 | 0.000952873 | 0.459563448 |
| ENSMUSG00000025409 | Mbd6 | 110962 | protein_coding | 197.1092124 | 0.45957216 | 0.25338527 | -1.813728792 | 0.06971949 | 0.190494318 | 0.45957216 |
| ENSMUSG00000022369 | Mtbp | 105837 | protein_coding | 78.31923301 | 0.459655699 | 0.155361934 | -2.958612103 | 0.003090278 | 0.016283616 | 0.459655699 |
| ENSMUSG00000041571 | Selenow | 20364 | protein_coding | 464.0144297 | -0.459776682 | 0.064884179 | 7.086113908 | 1.38E-12 | 4.11E-11 | 0.459776682 |
| ENSMUSG00000025791 | Pgm1 | 72157 | protein_coding | 1251.022536 | 0.459862032 | 0.111805004 | -4.11307202 | 3.90E-05 | 0.000376608 | 0.459862032 |
| ENSMUSG00000042784 | Muc1 | 17829 | protein_coding | 89.57783023 | -0.459865791 | 0.222649925 | 2.06542082 | 0.0388832 | 0.123253915 | 0.459865791 |
| ENSMUSG00000035478 | Mbd3 | 17192 | protein_coding | 457.1232174 | 0.460005642 | 0.091659642 | -5.018627944 | 5.20E-07 | 7.67E-06 | 0.460005642 |
| ENSMUSG00000024810 | Il33 | 77125 | protein_coding | 8.907093825 | 0.460083997 | 0.71444519 | -0.643973819 | 0.519592411 | 0.719844721 | 0.460083997 |
| ENSMUSG00000044788 | Fads6 | 328035 | protein_coding | 15.90813112 | -0.460118197 | 0.347140987 | 1.325450507 | 0.185021775 | 0.38040956 | 0.460118197 |
| ENSMUSG00000063558 | Aox1 | 11761 | protein_coding | 2.739077259 | -0.460212891 | 1.407880861 | 0.326883406 | 0.743756072 | 1 | 0.460212891 |
| ENSMUSG00000035580 | Kcnh8 | 211468 | protein_coding | 10.37192118 | -0.460373834 | 0.450862559 | 1.02109573 | 0.307209086 | 0.527698391 | 0.460373834 |
| ENSMUSG00000057914 | Cacnb2 | 12296 | protein_coding | 5.299847693 | 0.460428863 | 0.889443723 | -0.517659354 | 0.604695962 | 0.779964063 | 0.460428863 |
| ENSMUSG00000056429 | Tgoln1 | 22134 | protein_coding | 11969.85452 | -0.460470368 | 0.102624595 | 4.486939704 | 7.23E-06 | 8.37E-05 | 0.460470368 |
| ENSMUSG00000025597 | Klhl4 | 237010 | protein_coding | 0.9043219 | -0.460517273 | 2.160109019 | 0.213191681 | 0.831177456 | 1 | 0.460517273 |
| ENSMUSG00000031075 | Ano1 | 101772 | protein_coding | 1838.864703 | -0.460563075 | 0.189410359 | 2.431562225 | 0.015033866 | 0.058500114 | 0.460563075 |
| ENSMUSG00000062078 | Qki | 19317 | protein_coding | 9.394283861 | 0.461382129 | 0.435341929 | -1.059815511 | 0.289228539 | 0.508065109 | 0.461382129 |
| ENSMUSG00000061273 | Mmgt1 | 236792 | protein_coding | 503.5561078 | 0.461414286 | 0.120625825 | -3.825169986 | 0.000130682 | 0.001092699 | 0.461414286 |
| ENSMUSG00000074519 | Zfp971 | 626848 | protein_coding | 77.87987359 | -0.461612222 | 0.158435208 | 2.913570969 | 0.003573206 | 0.018411209 | 0.461612222 |
| ENSMUSG00000020818 | Mfsd11 | 69900 | protein_coding | 264.7616294 | 0.461795869 | 0.077378901 | -5.967981731 | 2.40E-09 | 5.07E-08 | 0.461795869 |
| ENSMUSG00000070871 | Ccnyl1 | 227210 | protein_coding | 134.6604215 | 0.46199229 | 0.10737518 | -4.302598524 | 1.69E-05 | 0.000177672 | 0.46199229 |
| ENSMUSG00000074136 | 4930513N10Rik | 319960 | lncRNA | 26.40119587 | 0.462029547 | 0.322264546 | -1.433696484 | 0.151658896 | 0.331941022 | 0.462029547 |
| ENSMUSG00000117916 | 9630028I04Rik | 319720 | lncRNA | 1.85538576 | 0.462083447 | 1.746348493 | -0.264599791 | 0.791317781 | 1 | 0.462083447 |
| ENSMUSG00000021565 | Slc6a19 | 74338 | protein_coding | 24.93142059 | -0.462087883 | 0.515134121 | 0.897024414 | 0.36970589 | 0.592404722 | 0.462087883 |
| ENSMUSG00000052135 | Foxo6 | 329934 | protein_coding | 20.1092518 | -0.462381178 | 0.363201816 | 1.273069564 | 0.202993358 | 0.405213026 | 0.462381178 |
| ENSMUSG00000047053 | Teddm1a | 240819 | protein_coding | 1.054587016 | 0.462645232 | 2.266044493 | -0.204164231 | 0.838225162 | 1 | 0.462645232 |

Supplementary Table 1

| ensembl_gene_id | external_gene_name | entrezgene_id | gene_biotype | baseMean | log2FoldChange | lfcSE | stat | pvalue | padj | absLog2FC |
|---|---|---|---|---|---|---|---|---|---|---|
| ENSMUSG00000029033 | Acap3 | 140500 | protein_coding | 133.9596917 | -0.462733605 | 0.120594794 | 3.837094377 | 0.000124499 | 0.00104843 | 0.462733605 |
| ENSMUSG00000067860 | Zic3 | 22773 | protein_coding | 1.206198641 | 0.46280127 | 2.277154727 | -0.203236637 | 0.838950078 | 1 | 0.46280127 |
| ENSMUSG00000099826 | Scgb2b10 | 624584 | unprocessed_pseudogene | 1.292615626 | 0.462857951 | 1.224906896 | -0.377871945 | 0.705525725 | 1 | 0.462857951 |
| ENSMUSG00000049013 | Prss48 | 368202 | protein_coding | 10.35613046 | -0.462909157 | 0.44116386 | 1.049290749 | 0.294044322 | 0.513159947 | 0.462909157 |
| ENSMUSG00000032952 | Ap4b1 | 67489 | protein_coding | 46.85776379 | 0.46291648 | 0.157864793 | -2.932360471 | 0.00336396 | 0.01751338 | 0.46291648 |
| ENSMUSG00000031714 | Gab1 | 14388 | protein_coding | 332.626056 | -0.462958407 | 0.09330694 | 4.961671736 | 6.99E-07 | 9.98E-06 | 0.462958407 |
| ENSMUSG00000031398 | Plxna3 | 18846 | protein_coding | 55.62183707 | -0.462968882 | 0.164177078 | 2.819936184 | 0.00480332 | 0.023502823 | 0.462968882 |
| ENSMUSG00000025351 | Cd63 | 12512 | protein_coding | 2203.851664 | -0.463254931 | 0.074101251 | 6.251647928 | 4.06E-10 | 9.39E-09 | 0.463254931 |
| ENSMUSG00000048029 | Eno4 | 226265 | protein_coding | 3.25085477 | 0.463692611 | 0.848533594 | -0.546463468 | 0.584747393 | 1 | 0.463692611 |
| ENSMUSG00000020689 | Itgb3 | 16416 | protein_coding | 3.206184111 | 0.463708688 | 1.024566992 | -0.452589915 | 0.650844062 | 1 | 0.463708688 |
| ENSMUSG00000100153 | Ppp1ccb | 434233 | transcribed_processed_pseudogene | 9.960299267 | 0.463773596 | 0.366476395 | -1.265493774 | 0.205694379 | 0.408322752 | 0.463773596 |
| ENSMUSG00000028889 | Yrdc | 230734 | protein_coding | 278.6296466 | 0.463784071 | 0.098563978 | -4.705411476 | 2.53E-06 | 3.22E-05 | 0.463784071 |
| ENSMUSG00000091670 | Vmn2r105 | 627743 | protein_coding | 2.683204595 | 0.463887774 | 2.730137059 | -0.169913731 | 0.865077982 | 1 | 0.463887774 |
| ENSMUSG00000109714 | Gm35021 | 102638463 | lncRNA | 1.473126396 | 0.463900576 | 1.826772404 | -0.25394547 | 0.799537686 | 1 | 0.463900576 |
| ENSMUSG00000029759 | Pon3 | 269823 | protein_coding | 647.7494364 | -0.464141254 | 0.076556625 | 6.062718323 | 1.34E-09 | 2.91E-08 | 0.464141254 |
| ENSMUSG00000020181 | Nav3 | 260315 | protein_coding | 4.621563981 | 0.464176267 | 1.935627895 | -0.23980656 | 0.810480221 | 0.906179443 | 0.464176267 |
| ENSMUSG00000051486 | Pcdhb11 | 93882 | protein_coding | 2.100825372 | -0.46423241 | 0.916891424 | 0.506311214 | 0.612638192 | 1 | 0.46423241 |
| ENSMUSG00000022935 | Grik1 | 14805 | protein_coding | 2.310974618 | 0.464255028 | 2.745473404 | -0.169098352 | 0.865719281 | 1 | 0.464255028 |
| ENSMUSG00000057719 | Sh3rf2 | 269016 | protein_coding | 218.0298426 | -0.464331038 | 0.163966377 | 2.8318674 | 0.004627704 | 0.022812506 | 0.464331038 |
| ENSMUSG00000024299 | Adamts10 | 224697 | protein_coding | 36.67201618 | 0.464415301 | 0.196430179 | -2.36427673 | 0.018065313 | 0.06767935 | 0.464415301 |
| ENSMUSG00000000732 | Icosl | 50723 | protein_coding | 64.11463815 | -0.464489028 | 0.188488994 | 2.464276663 | 0.013729009 | 0.054421559 | 0.464489028 |
| ENSMUSG00000027881 | Prpf38b | 66921 | protein_coding | 921.0098777 | 0.464541548 | 0.063574881 | -7.306998265 | 2.73E-13 | 8.69E-12 | 0.464541548 |
| ENSMUSG00000024502 | Jakmip2 | 76217 | protein_coding | 4.085529913 | 0.464637386 | 1.341658748 | -0.346315624 | 0.729105528 | 1 | 0.464637386 |
| ENSMUSG00000044807 | Zfp354c | 30944 | protein_coding | 3.627428433 | 0.464730089 | 0.628778843 | -0.739099437 | 0.459846619 | 1 | 0.464730089 |
| ENSMUSG00000026274 | Pask | 269224 | protein_coding | 88.13422769 | 0.46487801 | 0.145695569 | -3.190749127 | 0.001419044 | 0.008514609 | 0.46487801 |
| ENSMUSG00000047238 | Mageh1 | 75625 | protein_coding | 64.92819832 | 0.465195928 | 0.178418155 | -2.607335162 | 0.009124999 | 0.039582411 | 0.465195928 |
| ENSMUSG00000109372 | Gm19410 | 100502846 | protein_coding | 14.21064954 | 0.465267349 | 0.51643945 | -0.900913647 | 0.367634235 | 0.590110273 | 0.465267349 |
| ENSMUSG00000069633 | Pex11g | 69129 | protein_coding | 132.0350097 | 0.465341194 | 0.10193752 | -4.564964834 | 5.00E-06 | 5.97E-05 | 0.465341194 |
| ENSMUSG00000049538 | Adamts16 | 271127 | protein_coding | 12.24699685 | -0.465456359 | 0.370440254 | 1.256495085 | 0.208936527 | 0.412258246 | 0.465456359 |
| ENSMUSG00000043456 | Zfp536 | 243937 | protein_coding | 4.945321792 | 0.465475275 | 2.143822795 | -0.217123951 | 0.828111749 | 0.913297021 | 0.465475275 |
| ENSMUSG00000073408 | Mucl3 | 268949 | protein_coding | 1.03076368 | -0.465534602 | 2.170611155 | 0.214471671 | 0.830179256 | 1 | 0.465534602 |
| ENSMUSG00000029166 | Mapre3 | 100732 | protein_coding | 117.5871664 | 0.465640739 | 0.373410057 | -1.24699571 | 0.212399071 | 0.416871672 | 0.465640739 |
| ENSMUSG00000048827 | Pkd1l3 | 244646 | protein_coding | 17.51327116 | -0.465668024 | 0.34230304 | 1.360396987 | 0.173704332 | 0.364445856 | 0.465668024 |
| ENSMUSG00000029149 | Krtcap3 | 69815 | protein_coding | 623.5015703 | -0.465836225 | 0.085802141 | 5.429191128 | 5.66E-08 | 9.61E-07 | 0.465836225 |
| ENSMUSG00000075520 | Malrd1 | 102635496 | protein_coding | 307.4158627 | -0.465886769 | 0.220056926 | 2.117119323 | 0.034249722 | 0.111571269 | 0.465886769 |
| ENSMUSG00000012428 | Steap4 | 117167 | protein_coding | 65.51503263 | 0.465989625 | 0.264957494 | -1.758733519 | 0.078622781 | 0.207507849 | 0.465989625 |
| ENSMUSG00000034842 | Art3 | 109979 | protein_coding | 3.547614589 | 0.466068923 | 0.870394929 | -0.535468334 | 0.592326049 | 1 | 0.466068923 |
| ENSMUSG00000022194 | Pabpn1 | 12050 | protein_coding | 285.5957835 | 0.466081556 | 0.185736517 | -2.509369533 | 0.012094688 | 0.049414074 | 0.466081556 |
| ENSMUSG00000044646 | Zbtb7c | 207259 | protein_coding | 378.7141027 | 0.466165452 | 0.128933521 | -3.615548922 | 0.000299712 | 0.002276086 | 0.466165452 |

| ensembl_gene_id | external_gene_name | entrezgene_id | gene_biotype | baseMean | log2FoldChange | lfcSE | stat | pvalue | padj | absLog2FC |
|---|---|---|---|---|---|---|---|---|---|---|
| ENSMUSG00000028614 | Ndc1 | 72787 | protein_coding | 267.2450282 | 0.466266324 | 0.182619342 | -2.553214353 | 0.010673379 | 0.04488418 | 0.466266324 |
| ENSMUSG00000090220 | Pitpnm2os1 | 75927 | lncRNA | 0.711202118 | -0.466445507 | 1.653849914 | 0.282036177 | 0.777915771 | 1 | 0.466445507 |
| ENSMUSG00000035183 | Slc24a5 | 317750 | protein_coding | 5.438212479 | 0.466547101 | 0.537835822 | -0.867452636 | 0.385694053 | 0.607820554 | 0.466547101 |
| ENSMUSG00000096795 | Zfp433 | 73610 | protein_coding | 7.878656976 | -0.466646119 | 0.422292054 | 1.105031729 | 0.269145841 | 0.484159903 | 0.466646119 |
| ENSMUSG00000020328 | Nudcd2 | 52653 | protein_coding | 307.0368417 | 0.466846025 | 0.113821322 | -4.101569185 | 4.10E-05 | 0.000393534 | 0.466846025 |
| ENSMUSG00000047938 | 4930483J18Rik | 67638 | lncRNA | 0.927086421 | 0.46693282 | 1.609942913 | -0.290030669 | 0.771792775 | 1 | 0.46693282 |
| ENSMUSG00000028841 | Cnksr1 | 194231 | protein_coding | 537.1333268 | 0.466984204 | 0.133391962 | -3.500842158 | 0.00046379 | 0.003327122 | 0.466984204 |
| ENSMUSG00000039934 | Gsap | 212167 | protein_coding | 41.62402427 | 0.467064606 | 0.285894761 | -1.633694175 | 0.102323082 | 0.251865163 | 0.467064606 |
| ENSMUSG00000026407 | Cacna1s | 12292 | protein_coding | 3.050140602 | 0.467157861 | 2.21281001 | -0.211115215 | 0.832797368 | 1 | 0.467157861 |
| ENSMUSG00000021591 | Glrx | 93692 | protein_coding | 693.7739106 | -0.467243866 | 0.083393457 | 5.602884027 | 2.11E-08 | 3.86E-07 | 0.467243866 |
| ENSMUSG00000032532 | Cck | 12424 | protein_coding | 14.94979161 | -0.467267735 | 0.378579261 | 1.234266592 | 0.21710358 | 0.422731291 | 0.467267735 |
| ENSMUSG00000004263 | Atn1 | 13498 | protein_coding | 20.89237574 | 0.467294103 | 0.483409504 | -0.966663044 | 0.333712508 | 0.556097714 | 0.467294103 |
| ENSMUSG00000037944 | Ccr7 | 12775 | protein_coding | 2.80737558 | 0.467307482 | 0.919880942 | -0.508008657 | 0.611447272 | 1 | 0.467307482 |
| ENSMUSG00000033021 | Gmppa | 69080 | protein_coding | 877.7133015 | 0.467421653 | 0.073473519 | -6.361770371 | 1.99E-10 | 4.80E-09 | 0.467421653 |
| ENSMUSG00000022070 | Bora | 77744 | protein_coding | 192.3632293 | 0.467734146 | 0.107111422 | -4.366799896 | 1.26E-05 | 0.000137432 | 0.467734146 |
| ENSMUSG00000048997 | Atxn7l2 | 72522 | protein_coding | 45.31718395 | -0.467747712 | 0.207150235 | 2.258011976 | 0.023944913 | 0.084474747 | 0.467747712 |
| ENSMUSG00000074088 | Snrnp40 | 66585 | protein_coding | 469.0111552 | 0.467791684 | 0.071604935 | -6.532953143 | 6.45E-11 | 1.66E-09 | 0.467791684 |
| ENSMUSG00000026068 | Il18rap | 16174 | protein_coding | 4.677070469 | 0.467890195 | 0.804333389 | -0.581711765 | 0.560760846 | 0.749223673 | 0.467890195 |
| ENSMUSG00000061769 | Klra6 | 16637 | protein_coding | 1.447507332 | -0.467966747 | 1.239059719 | 0.377678929 | 0.705669123 | 1 | 0.467966747 |
| ENSMUSG00000045757 | Zfp764 | 233893 | protein_coding | 79.89149714 | -0.46798189 | 0.156434754 | 2.991546815 | 0.00277568 | 0.014921086 | 0.46798189 |
| ENSMUSG00000093626 | Ly6l | 102637705 | protein_coding | 0.841356297 | -0.468603076 | 2.138869838 | 0.219089104 | 0.826580639 | 1 | 0.468603076 |
| ENSMUSG00000039682 | Lap3 | 66988 | protein_coding | 956.4867207 | 0.468610868 | 0.087853181 | -5.334022767 | 9.61E-08 | 1.59E-06 | 0.468610868 |
| ENSMUSG00000025075 | Habp2 | 226243 | protein_coding | 2.916101105 | -0.468630298 | 0.633609673 | 0.739619861 | 0.459530687 | 1 | 0.468630298 |
| ENSMUSG00000036216 | Leap2 | 259301 | protein_coding | 2.078990679 | -0.468677653 | 0.771463923 | 0.607517266 | 0.543507688 | 1 | 0.468677653 |
| ENSMUSG00000038146 | Notch3 | 18131 | protein_coding | 6.442227996 | -0.469118693 | 0.910107261 | 0.515454291 | 0.606235603 | 0.781162199 | 0.469118693 |
| ENSMUSG00000037364 | Srrt | 83701 | protein_coding | 1172.271533 | 0.469244548 | 0.065796176 | -7.131790527 | 9.91E-13 | 3.00E-11 | 0.469244548 |
| ENSMUSG00000066224 | Arid3c | 550619 | protein_coding | 1.317453713 | -0.469258782 | 1.84893786 | 0.25379911 | 0.799650761 | 1 | 0.469258782 |
| ENSMUSG00000035045 | Zc3h12b | 547176 | protein_coding | 5.290560113 | 0.469294466 | 0.606149426 | -0.774222405 | 0.438799281 | 0.656846567 | 0.469294466 |
| ENSMUSG00000017764 | Zswim1 | 71971 | protein_coding | 117.9526382 | 0.469334349 | 0.133521737 | -3.51504077 | 0.000439686 | 0.003190048 | 0.469334349 |
| ENSMUSG00000107877 | 4933427D14Rik | 74477 | protein_coding | 6.482535297 | -0.469439722 | 0.704879763 | 0.66598553 | 0.50542035 | 0.709429427 | 0.469439722 |
| ENSMUSG00000058952 | Cfi | 12630 | protein_coding | 3.165295345 | 0.469450627 | 0.634859873 | -0.739455504 | 0.459630449 | 1 | 0.469450627 |
| ENSMUSG00000101930 | Gm5441 | 432713 | lncRNA | 2.836536298 | 0.469637098 | 1.01447057 | -0.46293812 | 0.643408723 | 1 | 0.469637098 |
| ENSMUSG00000026688 | Mgst3 | 66447 | protein_coding | 5649.409416 | -0.469680091 | 0.093948616 | 4.999329508 | 5.75E-07 | 8.36E-06 | 0.469680091 |
| ENSMUSG00000029648 | Flt1 | 14254 | protein_coding | 6.389910573 | 0.469821249 | 1.617721542 | -0.290421582 | 0.771493735 | 0.885228942 | 0.469821249 |
| ENSMUSG00000025193 | Cutc | 66388 | protein_coding | 117.4187564 | -0.469842636 | 0.172302668 | 2.726844814 | 0.00639431 | 0.029520727 | 0.469842636 |
| ENSMUSG00000037674 | Rfx7 | 319758 | protein_coding | 460.230867 | 0.469885054 | 0.099976746 | -4.699943455 | 2.60E-06 | 3.29E-05 | 0.469885054 |
| ENSMUSG00000027544 | Nfatc2 | 18019 | protein_coding | 59.31645313 | -0.469996099 | 0.200521914 | 2.343864023 | 0.019085128 | 0.070745635 | 0.469996099 |
| ENSMUSG00000037490 | Slc2a12 | 353169 | protein_coding | 4.662177629 | -0.470034754 | 0.762728199 | 0.616254591 | 0.537726504 | 0.733113936 | 0.470034754 |
| ENSMUSG00000025347 | Mettl7b | 71664 | protein_coding | 3232.229152 | -0.470090749 | 0.156798386 | 2.998058593 | 0.002717054 | 0.014644071 | 0.470090749 |

| ensembl_gene_id | external_gene_name | entrezgene_id | gene_biotype | baseMean | log2FoldChange | lfcSE | stat | pvalue | padj | absLog2FC |
|---|---|---|---|---|---|---|---|---|---|---|
| ENSMUSG00000097768 | 2310043M15Rik | 102633000 | lncRNA | 6.888303282 | -0.47015899 | 0.490108588 | 0.959295555 | 0.337409874 | 0.559438733 | 0.47015899 |
| ENSMUSG00000041476 | Smpx | 66106 | protein_coding | 74.92847309 | 0.470258404 | 0.209492738 | -2.244748003 | 0.024784314 | 0.086787144 | 0.470258404 |
| ENSMUSG00000050108 | Bpifc | 270757 | protein_coding | 1.892381843 | 0.470303085 | 1.541031407 | -0.305187216 | 0.760223568 | 1 | 0.470303085 |
| ENSMUSG00000057286 | St6galnac2 | 20446 | protein_coding | 490.9551484 | -0.470468499 | 0.118226555 | 3.979380922 | 6.91E-05 | 0.000623516 | 0.470468499 |
| ENSMUSG00000020847 | Rph3al | 380714 | protein_coding | 335.1317162 | 0.470589922 | 0.166743455 | -2.822239239 | 0.004768959 | 0.023369313 | 0.470589922 |
| ENSMUSG00000038126 | Mphosph9 | 269702 | protein_coding | 158.4800113 | 0.470719327 | 0.147298306 | -3.19568731 | 0.001394982 | 0.008398372 | 0.470719327 |
| ENSMUSG00000025480 | Syce1 | 74075 | protein_coding | 1.05413698 | 0.470792025 | 1.479181207 | -0.318278803 | 0.750273461 | 1 | 0.470792025 |
| ENSMUSG00000022519 | Srl | 106393 | protein_coding | 101.2181997 | 0.470860619 | 0.149357371 | -3.15257705 | 0.001618361 | 0.00949063 | 0.470860619 |
| ENSMUSG00000007029 | Vars | 22321 | protein_coding | 1309.767714 | 0.470933205 | 0.124573818 | -3.780354593 | 0.000156605 | 0.001274607 | 0.470933205 |
| ENSMUSG00000021874 | 4933413J09Rik | 71106 | lncRNA | 2.500832805 | -0.471011559 | 1.012885108 | 0.465019729 | 0.641917328 | 1 | 0.471011559 |
| ENSMUSG00000094282 | Cfap73 | 546886 | protein_coding | 0.697544318 | -0.47102371 | 2.209119698 | 0.213217831 | 0.831157061 | 1 | 0.47102371 |
| ENSMUSG00000047945 | Marcksl1 | 17357 | protein_coding | 186.1303169 | 0.471033818 | 0.159859421 | -2.946550261 | 0.003213402 | 0.01685709 | 0.471033818 |
| ENSMUSG00000066964 | Tmem211 | 333048 | protein_coding | 2.046128449 | 0.471132947 | 1.44435147 | -0.326189959 | 0.744280638 | 1 | 0.471132947 |
| ENSMUSG00000029776 | Hibadh | 58875 | protein_coding | 742.1563521 | -0.471201004 | 0.105207247 | 4.478788478 | 7.51E-06 | 8.68E-05 | 0.471201004 |
| ENSMUSG00000090841 | Myl6 | 17904 | protein_coding | 5529.678099 | -0.471235529 | 0.083466523 | 5.645802787 | 1.64E-08 | 3.06E-07 | 0.471235529 |
| ENSMUSG00000037492 | Zmat4 | 320158 | protein_coding | 13.51560238 | -0.471481987 | 0.505673073 | 0.932384998 | 0.351137595 | 0.573211901 | 0.471481987 |
| ENSMUSG00000102752 | Gm7694 | 665574 | protein_coding | 69.77448562 | -0.471555955 | 0.146337205 | 3.222392791 | 0.001271247 | 0.007756249 | 0.471555955 |
| ENSMUSG00000032348 | Gsta4 | 14860 | protein_coding | 473.1813114 | -0.471573847 | 0.121957312 | 3.866712369 | 0.000110312 | 0.00093902 | 0.471573847 |
| ENSMUSG00000070732 | Rbm44 | 329207 | protein_coding | 1.538714407 | -0.471634295 | 1.062702971 | 0.44380632 | 0.657182619 | 1 | 0.471634295 |
| ENSMUSG00000024818 | Slc25a45 | 107375 | protein_coding | 23.5128208 | -0.471782667 | 0.217916582 | 2.164969103 | 0.030390051 | 0.101900774 | 0.471782667 |
| ENSMUSG00000058586 | Serhl | 68607 | protein_coding | 448.8863745 | -0.471869945 | 0.101530985 | 4.647546225 | 3.36E-06 | 4.16E-05 | 0.471869945 |
| ENSMUSG00000031662 | Snx20 | 71607 | protein_coding | 170.0368053 | 0.471883486 | 0.149375038 | -3.159051816 | 0.001582833 | 0.009312046 | 0.471883486 |
| ENSMUSG00000071719 | Nalf2 | 620592 | protein_coding | 2.786751286 | -0.471954244 | 1.248217088 | 0.378102694 | 0.705354307 | 1 | 0.471954244 |
| ENSMUSG00000033065 | Pfkm | 18642 | protein_coding | 127.3495707 | -0.471978446 | 0.128443704 | 3.674593862 | 0.000238228 | 0.001847862 | 0.471978446 |
| ENSMUSG00000041353 | Tmem29 | 382245 | protein_coding | 99.35241978 | 0.47227642 | 0.131754316 | -3.58452333 | 0.000337694 | 0.002527962 | 0.47227642 |
| ENSMUSG00000004609 | Cd33 | 12489 | protein_coding | 5.682651899 | -0.472446099 | 0.979825699 | 0.482173614 | 0.629682619 | 0.797713998 | 0.472446099 |
| ENSMUSG00000020376 | Rnf130 | 59044 | protein_coding | 648.0970419 | -0.472623409 | 0.116038754 | 4.072978997 | 4.64E-05 | 0.000437255 | 0.472623409 |
| ENSMUSG00000061046 | Haghl | 68977 | protein_coding | 72.90591381 | -0.472633771 | 0.151683567 | 3.115919413 | 0.001833724 | 0.010565985 | 0.472633771 |
| ENSMUSG00000029436 | Mmp17 | 23948 | protein_coding | 7.614402339 | 0.472811361 | 0.508517527 | -0.929783804 | 0.352483034 | 0.574847405 | 0.472811361 |
| ENSMUSG00000044139 | Prss53 | 330657 | protein_coding | 3.736389824 | 0.472823971 | 0.5551043 | -0.85177501 | 0.394338979 | 1 | 0.472823971 |
| ENSMUSG00000067599 | Klra7 | 16638 | protein_coding | 4.907183715 | 0.472829546 | 0.67336036 | -0.702193913 | 0.482558241 | 0.693375689 | 0.472829546 |
| ENSMUSG00000029864 | Gstk1 | 76263 | protein_coding | 567.3508664 | -0.472879579 | 0.098720541 | 4.790082917 | 1.67E-06 | 2.19E-05 | 0.472879579 |
| ENSMUSG00000034362 | Csta1 | 209294 | protein_coding | 20.03341865 | 0.473400164 | 0.320401895 | -1.477519861 | 0.139536338 | 0.313084659 | 0.473400164 |
| ENSMUSG00000047115 | Fam221a | 231946 | protein_coding | 55.49319286 | -0.473422634 | 0.157899937 | 2.998244598 | 0.002715396 | 0.014639914 | 0.473422634 |
| ENSMUSG00000039873 | Neurl2 | 415115 | protein_coding | 3.637728304 | -0.473489351 | 0.579770401 | 0.816684242 | 0.414108899 | 1 | 0.473489351 |
| ENSMUSG00000020534 | Shmt1 | 20425 | protein_coding | 1066.360831 | 0.473524223 | 0.130357801 | -3.632496245 | 0.000280693 | 0.002147988 | 0.473524223 |
| ENSMUSG00000022132 | Cldn10 | 58187 | protein_coding | 3.200519201 | 0.47352596 | 0.685545931 | -0.690728277 | 0.489736316 | 1 | 0.47352596 |
| ENSMUSG00000044702 | Palb2 | 233826 | protein_coding | 56.87086493 | 0.473754177 | 0.149892972 | -3.160616345 | 0.001574357 | 0.009275396 | 0.473754177 |
| ENSMUSG00000034324 | Tmem132c | 208213 | protein_coding | 25.26704998 | -0.473793114 | 0.290137044 | 1.632997664 | 0.102469488 | 0.252075246 | 0.473793114 |

| ensembl_gene_id | external_gene_name | entrezgene_id | gene_biotype | baseMean | log2FoldChange | lfcSE | stat | pvalue | padj | absLog2FC |
|---|---|---|---|---|---|---|---|---|---|---|
| ENSMUSG00000061119 | Prcp | 72461 | protein_coding | 563.4440398 | -0.473807839 | 0.089804116 | 5.276014738 | 1.32E-07 | 2.14E-06 | 0.473807839 |
| ENSMUSG00000067613 | Krt83 | 100126226 | protein_coding | 3.543466219 | 0.473813813 | 0.623047273 | -0.76047811 | 0.446968849 | 1 | 0.473813813 |
| ENSMUSG00000049555 | Tmie | 20776 | protein_coding | 15.95387722 | -0.473873471 | 0.381283613 | 1.242837232 | 0.213927819 | 0.418926125 | 0.473873471 |
| ENSMUSG00000027761 | Aadac | 67758 | protein_coding | 752.610648 | -0.473954241 | 0.163453103 | 2.899634395 | 0.003735981 | 0.019112763 | 0.473954241 |
| ENSMUSG00000003680 | Taf6l | 225895 | protein_coding | 108.4813491 | 0.474185589 | 0.151566757 | -3.128559317 | 0.001756656 | 0.010174535 | 0.474185589 |
| ENSMUSG00000038523 | 1700003F12Rik | 75480 | protein_coding | 0.699402081 | -0.474188315 | 1.524629972 | 0.311018623 | 0.755786464 | 1 | 0.474188315 |
| ENSMUSG00000086693 | A730081D07Rik | 102633880 | lncRNA | 1.405513142 | -0.474390274 | 1.140741681 | 0.415861261 | 0.677511528 | 1 | 0.474390274 |
| ENSMUSG00000013974 | Mcemp1 | 69189 | protein_coding | 2.406692044 | 0.474411821 | 1.919164666 | -0.247197038 | 0.804755738 | 1 | 0.474411821 |
| ENSMUSG00000039887 | Alg14 | 66789 | protein_coding | 201.2565418 | 0.474511873 | 0.124229614 | -3.81963572 | 0.000133649 | 0.001112439 | 0.474511873 |
| ENSMUSG00000031157 | Pqbp1 | 54633 | protein_coding | 332.8393723 | 0.474591074 | 0.084986586 | -5.584305647 | 2.35E-08 | 4.27E-07 | 0.474591074 |
| ENSMUSG00000022148 | Fyb | 23880 | protein_coding | 70.13702095 | -0.474597277 | 0.213346109 | 2.224541521 | 0.026112032 | 0.090344458 | 0.474597277 |
| ENSMUSG00000027132 | Katnbl1 | 72425 | protein_coding | 134.382643 | 0.474629343 | 0.117830272 | -4.028076451 | 5.62E-05 | 0.000518229 | 0.474629343 |
| ENSMUSG00000026308 | Klhl30 | 70788 | protein_coding | 1.4756976 | 0.474833706 | 1.431423844 | -0.331721249 | 0.740099754 | 1 | 0.474833706 |
| ENSMUSG00000047766 | Lrrc49 | 102747 | protein_coding | 100.5995998 | 0.475097122 | 0.144883597 | -3.279164321 | 0.00104115 | 0.006582523 | 0.475097122 |
| ENSMUSG00000035513 | Ntng2 | 171171 | protein_coding | 2.852251825 | 0.475135774 | 1.8115756 | -0.262277641 | 0.793107399 | 1 | 0.475135774 |
| ENSMUSG00000059842 | Zfp341 | 228807 | protein_coding | 42.38202689 | -0.475230041 | 0.160081791 | 2.968670198 | 0.002990914 | 0.015825683 | 0.475230041 |
| ENSMUSG00000026173 | Plcd4 | 18802 | protein_coding | 7.50826548 | -0.475356803 | 0.578577424 | 0.821595837 | 0.41130696 | 0.631550268 | 0.475356803 |
| ENSMUSG00000013033 | Adgrl1 | 330814 | protein_coding | 50.08510284 | 0.475357012 | 0.26616321 | -1.785960622 | 0.074105634 | 0.198998609 | 0.475357012 |
| ENSMUSG00000029161 | Cgref1 | 68567 | protein_coding | 1315.163731 | 0.475403327 | 0.085140966 | -5.583720112 | 2.35E-08 | 4.28E-07 | 0.475403327 |
| ENSMUSG00000037617 | Spag1 | 26942 | protein_coding | 35.07199557 | 0.475652166 | 0.252730611 | -1.882052053 | 0.059828948 | 0.170406218 | 0.475652166 |
| ENSMUSG00000044055 | Otos | 260301 | protein_coding | 0.922942345 | 0.475764025 | 2.268129417 | -0.209760528 | 0.833854581 | 1 | 0.475764025 |
| ENSMUSG00000016344 | Ppdpf | 66496 | protein_coding | 517.0560215 | -0.475829602 | 0.114483547 | 4.156314301 | 3.23E-05 | 0.000318106 | 0.475829602 |
| ENSMUSG00000118061 | Rbfaos | 105246506 | lncRNA | 2.20174845 | 0.475835412 | 1.076903266 | -0.441855297 | 0.658593921 | 1 | 0.475835412 |
| ENSMUSG00000042510 | AA986860 | 212439 | protein_coding | 454.4262282 | 0.475896206 | 0.159858881 | -2.976976972 | 0.002911059 | 0.01548252 | 0.475896206 |
| ENSMUSG00000042842 | Serpinb6b | 20708 | protein_coding | 5.428352434 | 0.47606318 | 0.549101548 | -0.866985682 | 0.385949855 | 0.608165639 | 0.47606318 |
| ENSMUSG00000028133 | Rwdd3 | 66568 | protein_coding | 25.72703948 | 0.476073202 | 0.255906568 | -1.860339913 | 0.06283745 | 0.176389198 | 0.476073202 |
| ENSMUSG00000116810 | Gm32624 | 102635233 | lncRNA | 1.002291532 | 0.476172055 | 2.290609601 | -0.207880057 | 0.835322621 | 1 | 0.476172055 |
| ENSMUSG00000056758 | Hmga2 | 15364 | protein_coding | 105.837822 | -0.476212359 | 0.123093659 | 3.868699354 | 0.000109417 | 0.000932912 | 0.476212359 |
| ENSMUSG00000029369 | Afm | 280662 | protein_coding | 39.12336394 | 0.476386138 | 0.234885162 | -2.028166162 | 0.042543292 | 0.131812369 | 0.476386138 |
| ENSMUSG00000033055 | Ankrd54 | 223690 | protein_coding | 209.9735642 | 0.476415029 | 0.082080035 | -5.804274222 | 6.46E-09 | 1.28E-07 | 0.476415029 |
| ENSMUSG00000040537 | Adam22 | 11496 | protein_coding | 38.13222131 | -0.476431975 | 0.263267721 | 1.809686254 | 0.070344455 | 0.1916312 | 0.476431975 |
| ENSMUSG00000070425 | Xntrpc | 102443351 | protein_coding | 1.896769625 | 0.476760364 | 0.969013883 | -0.492005711 | 0.622715307 | 1 | 0.476760364 |
| ENSMUSG00000031104 | Rab33a | 19337 | protein_coding | 3.338804102 | -0.476841013 | 1.029349749 | 0.463244892 | 0.643188842 | 1 | 0.476841013 |
| ENSMUSG00000106445 | Gm21190 | 100861755 | protein_coding | 2.574209659 | -0.477079216 | 0.989027572 | 0.482372009 | 0.629541702 | 1 | 0.477079216 |
| ENSMUSG00000093772 | 4931403E22Rik | 70942 | lncRNA | 1.654459326 | -0.477217294 | 0.950420129 | 0.502111939 | 0.615588782 | 1 | 0.477217294 |
| ENSMUSG00000041420 | Meis3 | 17537 | protein_coding | 98.54348924 | -0.47727088 | 0.175273871 | 2.72300074 | 0.006469191 | 0.02983307 | 0.47727088 |
| ENSMUSG00000022488 | Nckap1l | 105855 | protein_coding | 15.47486199 | 0.477491643 | 0.357525411 | -1.335546028 | 0.181697701 | 0.375915289 | 0.477491643 |
| ENSMUSG00000086638 | 4930405A21Rik | 74846 | lncRNA | 2.991033092 | 0.477511936 | 0.605997196 | -0.787977138 | 0.430710079 | 1 | 0.477511936 |
| ENSMUSG00000117226 | Gm31235 | 102633401 | lncRNA | 27.35772577 | -0.477528797 | 0.352422669 | 1.35498888 | 0.1754211 | 0.36725457 | 0.477528797 |

| ensembl_gene_id | external_gene_name | entrezgene_id | gene_biotype | baseMean | log2FoldChange | lfcSE | stat | pvalue | padj | absLog2FC |
|---|---|---|---|---|---|---|---|---|---|---|
| ENSMUSG00000070044 | Fam149a | 212326 | protein_coding | 107.2583727 | 0.477555724 | 0.136750706 | -3.492162769 | 0.000479126 | 0.003417675 | 0.477555724 |
| ENSMUSG00000048960 | Prex2 | 109294 | protein_coding | 5.246609321 | 0.477672308 | 1.076826871 | -0.443592485 | 0.65733724 | 0.814655179 | 0.477672308 |
| ENSMUSG00000020696 | Rffl | 67338 | protein_coding | 291.271079 | -0.477713772 | 0.100889704 | 4.735010161 | 2.19E-06 | 2.83E-05 | 0.477713772 |
| ENSMUSG00000004933 | Matk | 17179 | protein_coding | 101.7852543 | -0.47775195 | 0.220470668 | 2.16696377 | 0.030237617 | 0.101616606 | 0.47775195 |
| ENSMUSG00000020609 | Apob | 238055 | protein_coding | 954.8512561 | -0.477753101 | 0.216150705 | 2.21027778 | 0.02708589 | 0.092992945 | 0.477753101 |
| ENSMUSG00000028750 | Pla2g2c | 18781 | protein_coding | 48.28758472 | 0.477786207 | 0.174571124 | -2.736914303 | 0.006201845 | 0.028888694 | 0.477786207 |
| ENSMUSG00000071042 | Rasgrp3 | 240168 | protein_coding | 3.002817608 | -0.477833185 | 1.482943929 | 0.322219321 | 0.747286549 | 1 | 0.477833185 |
| ENSMUSG00000027523 | Gnas | 14683 | protein_coding | 257.8998902 | 0.478133337 | 0.23056712 | -2.073727331 | 0.038104641 | 0.121338228 | 0.478133337 |
| ENSMUSG00000031197 | Vbp1 | 22327 | protein_coding | 393.2624129 | 0.47823885 | 0.132074961 | -3.620965283 | 0.000293506 | 0.002235681 | 0.47823885 |
| ENSMUSG00000101304 | Plet1os | 69631 | lncRNA | 22.52912879 | -0.478255816 | 0.368241909 | 1.298754445 | 0.194028213 | 0.392717478 | 0.478255816 |
| ENSMUSG00000026575 | Nme7 | 171567 | protein_coding | 126.9604432 | 0.478366609 | 0.125008356 | -3.826677062 | 0.000129885 | 0.001087135 | 0.478366609 |
| ENSMUSG00000034786 | Gpsm3 | 106512 | protein_coding | 6.403999712 | 0.478485687 | 0.579144155 | -0.826194451 | 0.408693796 | 0.629272248 | 0.478485687 |
| ENSMUSG00000000792 | Slc5a5 | 114479 | protein_coding | 6.212422916 | 0.478489045 | 0.529059428 | -0.90441455 | 0.365775626 | 0.588390676 | 0.478489045 |
| ENSMUSG00000091712 | Sec14l5 | 665119 | protein_coding | 1.645349379 | 0.478491789 | 1.629939457 | -0.293564149 | 0.769090974 | 1 | 0.478491789 |
| ENSMUSG00000020988 | L2hgdh | 217666 | protein_coding | 346.3587161 | 0.478767778 | 0.097826875 | -4.89403118 | 9.88E-07 | 1.37E-05 | 0.478767778 |
| ENSMUSG00000028743 | Akr7a5 | 110198 | protein_coding | 512.7153375 | -0.47877837 | 0.107518716 | 4.452977017 | 8.47E-06 | 9.66E-05 | 0.47877837 |
| ENSMUSG00000031073 | Fgf15 | 14170 | protein_coding | 1.388515282 | 0.47895882 | 2.053376696 | -0.23325424 | 0.81556399 | 1 | 0.47895882 |
| ENSMUSG00000039701 | Usp53 | 99526 | protein_coding | 1356.181103 | -0.478967936 | 0.118930232 | 4.027301777 | 5.64E-05 | 0.000519649 | 0.478967936 |
| ENSMUSG00000113186 | A330076C08Rik | 100503592 | lncRNA | 2.605007127 | -0.479451038 | 1.954438008 | 0.245314016 | 0.8062133 | 1 | 0.479451038 |
| ENSMUSG00000039738 | Slx4 | 52864 | protein_coding | 145.4503025 | 0.479459139 | 0.105653697 | -4.538025214 | 5.68E-06 | 6.71E-05 | 0.479459139 |
| ENSMUSG00000022844 | Pdia5 | 72599 | protein_coding | 805.3038839 | 0.479466259 | 0.186996644 | -2.564036706 | 0.01034626 | 0.043750335 | 0.479466259 |
| ENSMUSG00000049313 | Sorl1 | 20660 | protein_coding | 2222.236355 | -0.479479594 | 0.094714411 | 5.062372128 | 4.14E-07 | 6.20E-06 | 0.479479594 |
| ENSMUSG00000031628 | Casp3 | 12367 | protein_coding | 719.6075695 | -0.479510148 | 0.075739043 | 6.331082714 | 2.43E-10 | 5.80E-09 | 0.479510148 |
| ENSMUSG00000014959 | Gorasp2 | 70231 | protein_coding | 1426.67588 | 0.479547706 | 0.090302778 | -5.310442434 | 1.09E-07 | 1.79E-06 | 0.479547706 |
| ENSMUSG00000057766 | Ankrd29 | 225187 | protein_coding | 7.423794955 | -0.479647614 | 0.456377999 | 1.050987591 | 0.293264289 | 0.512561987 | 0.479647614 |
| ENSMUSG00000061474 | Mrps36 | 66128 | protein_coding | 234.5395077 | 0.479652838 | 0.135871477 | -3.530195215 | 0.000415253 | 0.003030927 | 0.479652838 |
| ENSMUSG00000024590 | Lmnb1 | 16906 | protein_coding | 653.4535611 | 0.479803974 | 0.120608372 | -3.978197918 | 6.94E-05 | 0.000626283 | 0.479803974 |
| ENSMUSG00000024190 | Dusp1 | 19252 | protein_coding | 2915.3634 | -0.479814061 | 0.182811725 | 2.624635055 | 0.008674188 | 0.037986091 | 0.479814061 |
| ENSMUSG00000052396 | Mogat2 | 233549 | protein_coding | 54.99598574 | 0.47997555 | 0.334143303 | -1.436436243 | 0.150878246 | 0.330978129 | 0.47997555 |
| ENSMUSG00000033213 | AA467197 | 433470 | protein_coding | 1523.986115 | -0.48004957 | 0.118577752 | 4.048394928 | 5.16E-05 | 0.000479251 | 0.48004957 |
| ENSMUSG00000024349 | Sting1 | 72512 | protein_coding | 74.88788078 | 0.48014115 | 0.212636612 | -2.258036117 | 0.023943408 | 0.084474747 | 0.48014115 |
| ENSMUSG00000065560 | Mir148b | 724064 | miRNA | 0.492069034 | -0.480228289 | 1.808031506 | 0.265608363 | 0.790540844 | 1 | 0.480228289 |
| ENSMUSG00000000365 | Rnf17 | 30054 | protein_coding | 3.619223913 | 0.480237868 | 1.193597199 | -0.402345002 | 0.687430139 | 1 | 0.480237868 |
| ENSMUSG00000040852 | Plekhh2 | 213556 | protein_coding | 37.65888765 | 0.48027151 | 0.203173325 | -2.363851214 | 0.018086075 | 0.067726402 | 0.48027151 |
| ENSMUSG00000020718 | Polg2 | 50776 | protein_coding | 139.369675 | -0.480325534 | 0.132208122 | 3.633101557 | 0.000280035 | 0.002143947 | 0.480325534 |
| ENSMUSG00000035351 | Nup37 | 69736 | protein_coding | 137.4359806 | 0.480403289 | 0.172320748 | -2.787843573 | 0.005306016 | 0.025598465 | 0.480403289 |
| ENSMUSG00000002265 | Peg3 | 18616 | protein_coding | 45.93140291 | -0.480404892 | 0.232770656 | 2.063855038 | 0.039031463 | 0.123598385 | 0.480404892 |
| ENSMUSG00000069867 | Pabpn1l | 382035 | protein_coding | 0.653230925 | -0.480472962 | 1.611291574 | 0.298191197 | 0.76555724 | 1 | 0.480472962 |
| ENSMUSG00000029554 | Mad1l1 | 17120 | protein_coding | 243.9778441 | 0.480524586 | 0.093599069 | -5.133860772 | 2.84E-07 | 4.36E-06 | 0.480524586 |

| ensembl_gene_id | external_gene_name | entrezgene_id | gene_biotype | baseMean | log2FoldChange | lfcSE | stat | pvalue | padj | absLog2FC |
|---|---|---|---|---|---|---|---|---|---|---|
| ENSMUSG00000025582 | Nptx1 | 18164 | protein_coding | 6.89709185 | -0.480659273 | 0.629062846 | 0.764087843 | 0.444814896 | 0.662290428 | 0.480659273 |
| ENSMUSG00000070167 | Snora57 | 118027464 | snoRNA | 328.7843409 | 0.480731066 | 0.338042465 | -1.422102593 | 0.15499647 | 0.337101502 | 0.480731066 |
| ENSMUSG00000027332 | Ivd | 56357 | protein_coding | 1044.549836 | 0.48106977 | 0.084556211 | -5.689348709 | 1.28E-08 | 2.42E-07 | 0.48106977 |
| ENSMUSG00000050966 | Lin28a | 83557 | protein_coding | 1.768529121 | -0.481120496 | 1.924648259 | 0.249978402 | 0.802604051 | 1 | 0.481120496 |
| ENSMUSG00000065315 | Gm26064 | 115488251 | snRNA | 0.732790944 | 0.481223191 | 1.60003383 | -0.300758135 | 0.763598934 | 1 | 0.481223191 |
| ENSMUSG00000071202 | Ccdc78 | 381077 | protein_coding | 3.802585954 | 0.481294859 | 0.634230414 | -0.758864362 | 0.447933699 | 1 | 0.481294859 |
| ENSMUSG00000023216 | Epb42 | 13828 | protein_coding | 2.858335945 | 0.481298027 | 1.09869857 | -0.43806194 | 0.661341383 | 1 | 0.481298027 |
| ENSMUSG00000020895 | Tmem107 | 66910 | protein_coding | 82.35769612 | 0.481310817 | 0.210307009 | -2.288610445 | 0.022101997 | 0.079328922 | 0.481310817 |
| ENSMUSG00000049001 | Ndnf | 68169 | protein_coding | 28.73696684 | 0.481339897 | 0.272142614 | -1.768704612 | 0.076943182 | 0.204447258 | 0.481339897 |
| ENSMUSG00000027364 | Usp50 | 75083 | protein_coding | 3.753573894 | -0.481346922 | 0.760428959 | 0.632993939 | 0.526737601 | 1 | 0.481346922 |
| ENSMUSG00000062124 | Defb45 | 433490 | protein_coding | 17.4900985 | -0.481370283 | 0.451125503 | 1.067042941 | 0.285952446 | 0.504402765 | 0.481370283 |
| ENSMUSG00000102742 | Pcdhga11 | 93723 | protein_coding | 2.676218197 | 0.481396978 | 0.740827576 | -0.649809744 | 0.515815124 | 1 | 0.481396978 |
| ENSMUSG00000049109 | Themis | 210757 | protein_coding | 4.37431913 | 0.481545056 | 0.889796094 | -0.541185851 | 0.588379489 | 1 | 0.481545056 |
| ENSMUSG00000078502 | Zfp268 | 433801 | protein_coding | 23.52489724 | -0.481557533 | 0.302265882 | 1.593158744 | 0.111124584 | 0.26676599 | 0.481557533 |
| ENSMUSG00000021514 | Zfp369 | 170936 | protein_coding | 189.8545434 | -0.481573082 | 0.098764872 | 4.875955115 | 1.08E-06 | 1.48E-05 | 0.481573082 |
| ENSMUSG00000029446 | Psph | 100678 | protein_coding | 80.84162608 | 0.481605765 | 0.138805234 | -3.469651334 | 0.000521134 | 0.003676212 | 0.481605765 |
| ENSMUSG00000027359 | Slc27a2 | 26458 | protein_coding | 62.16010087 | 0.481636847 | 0.155970455 | -3.088000537 | 0.002015081 | 0.011401366 | 0.481636847 |
| ENSMUSG00000028212 | Ccne2 | 12448 | protein_coding | 98.33362052 | 0.481776503 | 0.159655153 | -3.01760697 | 0.002547791 | 0.013849316 | 0.481776503 |
| ENSMUSG00000042660 | Wdr55 | 67936 | protein_coding | 183.8828718 | -0.48195423 | 0.120649732 | 3.994656454 | 6.48E-05 | 0.000589808 | 0.48195423 |
| ENSMUSG00000026880 | Stom | 13830 | protein_coding | 409.2167108 | -0.482105248 | 0.205640535 | 2.344407671 | 0.019057328 | 0.070692483 | 0.482105248 |
| ENSMUSG00000091184 | Gm17059 | 102640356 | lncRNA | 2.299271622 | -0.482209316 | 0.79072597 | 0.609831135 | 0.541973675 | 1 | 0.482209316 |
| ENSMUSG00000027339 | Rassf2 | 215653 | protein_coding | 12.39395348 | 0.482290007 | 0.483892961 | -0.996687379 | 0.318916278 | 0.5402916 | 0.482290007 |
| ENSMUSG00000057123 | Gja5 | 14613 | protein_coding | 4.18170667 | 0.482520714 | 0.971187189 | -0.496835954 | 0.619304739 | 1 | 0.482520714 |
| ENSMUSG00000046434 | Hnrnpa1 | 15382 | protein_coding | 1534.742962 | -0.48266035 | 0.103080275 | 4.682373537 | 2.84E-06 | 3.55E-05 | 0.48266035 |
| ENSMUSG00000038521 | C1s1 | 50908 | protein_coding | 1.781254396 | 0.482696432 | 1.120191405 | -0.430905317 | 0.666537217 | 1 | 0.482696432 |
| ENSMUSG00000022621 | Rabl2 | 68708 | protein_coding | 57.10279319 | 0.482708549 | 0.133085941 | -3.627043882 | 0.000286685 | 0.00218877 | 0.482708549 |
| ENSMUSG00000050157 | Gm867 | 333670 | protein_coding | 3.275827834 | -0.483007423 | 0.794946766 | 0.607597193 | 0.543454663 | 1 | 0.483007423 |
| ENSMUSG00000100738 | 2010106C02Rik | 72104 | lncRNA | 6.433518297 | -0.483116358 | 0.523936706 | 0.922089161 | 0.35648207 | 0.579378227 | 0.483116358 |
| ENSMUSG00000048385 | Scrt1 | 170729 | protein_coding | 1.876992989 | -0.483231204 | 1.51272964 | 0.319443205 | 0.749390453 | 1 | 0.483231204 |
| ENSMUSG00000064210 | Ano6 | 105722 | protein_coding | 366.868342 | 0.483262743 | 0.094502495 | -5.113756427 | 3.16E-07 | 4.82E-06 | 0.483262743 |
| ENSMUSG00000060224 | Pyroxd2 | 74580 | protein_coding | 93.14470477 | -0.48340776 | 0.12043487 | 4.013852151 | 5.97E-05 | 0.000548043 | 0.48340776 |
| ENSMUSG00000037772 | Mrpl23 | 19935 | protein_coding | 82.66838866 | -0.483635253 | 0.169971732 | 2.845386387 | 0.004435758 | 0.022037337 | 0.483635253 |
| ENSMUSG00000047854 | Stx19 | 68159 | protein_coding | 26.75244992 | 0.483800063 | 0.255575921 | -1.89297983 | 0.058360552 | 0.167349566 | 0.483800063 |
| ENSMUSG00000017697 | Ada | 11486 | protein_coding | 97.23261115 | 0.483890981 | 0.133396241 | -3.62747088 | 0.000286211 | 0.002186165 | 0.483890981 |
| ENSMUSG00000087881 | Gm22442 | 115487906 | scaRNA | 559.6408806 | -0.4839547 | 0.122259443 | 3.958423901 | 7.54E-05 | 0.000673467 | 0.4839547 |
| ENSMUSG00000089371 | Mir1938 | 100316693 | miRNA | 44.54097687 | -0.483955685 | 0.301242915 | 1.606529684 | 0.108157578 | 0.261778433 | 0.483955685 |
| ENSMUSG00000109324 | Prmt1 | 15469 | protein_coding | 320.0149496 | 0.484223374 | 0.091352192 | -5.300621309 | 1.15E-07 | 1.88E-06 | 0.484223374 |
| ENSMUSG00000087113 | Gm11714 | 108167930 | lncRNA | 0.587287972 | -0.484460529 | 1.798650965 | 0.269346604 | 0.787662972 | 1 | 0.484460529 |
| ENSMUSG00000022753 | Tmem30c | 71027 | protein_coding | 2.492230077 | 0.484464144 | 1.511035571 | -0.320617299 | 0.748500426 | 1 | 0.484464144 |

| ensembl_gene_id | external_gene_name | entrezgene_id | gene_biotype | baseMean | log2FoldChange | lfcSE | stat | pvalue | padj | absLog2FC |
|---|---|---|---|---|---|---|---|---|---|---|
| ENSMUSG00000025396 | Hsd17b6 | 27400 | protein_coding | 0.961759712 | 0.484485058 | 1.816476746 | -0.266716906 | 0.789687136 | 1 | 0.484485058 |
| ENSMUSG00000117328 | 4930474M22Rik | 74917 | lncRNA | 1.54601673 | 0.484557718 | 1.105204393 | -0.438432675 | 0.661072665 | 1 | 0.484557718 |
| ENSMUSG00000086847 | Tbx3os2 | 433944 | lncRNA | 1.960580594 | -0.48465604 | 1.493028233 | 0.324612776 | 0.745474156 | 1 | 0.48465604 |
| ENSMUSG00000028497 | Hacd4 | 66775 | protein_coding | 31.23029906 | -0.484771825 | 0.194948443 | 2.486666818 | 0.01289461 | 0.05182755 | 0.484771825 |
| ENSMUSG00000108622 | Gm36864 | 102640913 | protein_coding | 5.782901445 | 0.484884374 | 0.49056611 | -0.988418002 | 0.322947974 | 0.544366934 | 0.484884374 |
| ENSMUSG00000090946 | Ccdc71l | 72123 | protein_coding | 1240.427268 | 0.484978735 | 0.149819092 | -3.237095681 | 0.001207529 | 0.007435174 | 0.484978735 |
| ENSMUSG00000049184 | Purg | 75029 | protein_coding | 16.54499121 | -0.485040841 | 0.346739066 | 1.398864126 | 0.161853732 | 0.347033567 | 0.485040841 |
| ENSMUSG00000063856 | Gpx1 | 14775 | protein_coding | 2230.788943 | -0.485087889 | 0.096727535 | 5.014992769 | 5.30E-07 | 7.79E-06 | 0.485087889 |
| ENSMUSG00000014837 | 4931428F04Rik | 74356 | protein_coding | 30.51017272 | 0.485110742 | 0.244524813 | -1.98389168 | 0.047267917 | 0.142911457 | 0.485110742 |
| ENSMUSG00000074028 | Slc22a13 | 102570 | protein_coding | 1.088063369 | 0.485131108 | 1.825286708 | -0.26578351 | 0.790405944 | 1 | 0.485131108 |
| ENSMUSG00000042485 | Mustn1 | 66175 | protein_coding | 3.311059811 | 0.485344954 | 0.657576593 | -0.738081251 | 0.460465076 | 1 | 0.485344954 |
| ENSMUSG00000028093 | Acp6 | 66659 | protein_coding | 369.3779006 | -0.485462568 | 0.080826034 | 6.006264876 | 1.90E-09 | 4.05E-08 | 0.485462568 |
| ENSMUSG00000048216 | Gpr85 | 64450 | protein_coding | 6.226617011 | 0.485671811 | 0.440665407 | -1.102132828 | 0.27040393 | 0.485744642 | 0.485671811 |
| ENSMUSG00000071656 | Lrrn4cl | 68852 | protein_coding | 1.568358496 | 0.485921008 | 1.08083425 | -0.44957958 | 0.653013615 | 1 | 0.485921008 |
| ENSMUSG00000030718 | Ppme1 | 72590 | protein_coding | 597.5768195 | 0.485979681 | 0.063275271 | -7.680404565 | 1.59E-14 | 5.57E-13 | 0.485979681 |
| ENSMUSG00000025665 | Rps6ka6 | 67071 | protein_coding | 48.88601925 | 0.486045934 | 0.195527473 | -2.485819137 | 0.012925364 | 0.051921541 | 0.486045934 |
| ENSMUSG00000024308 | Tapbp | 21356 | protein_coding | 1700.502227 | -0.486059838 | 0.060563259 | 8.025655301 | 1.01E-15 | 3.93E-14 | 0.486059838 |
| ENSMUSG00000048911 | Rnf24 | 51902 | protein_coding | 50.98819505 | 0.486061153 | 0.266053546 | -1.826929807 | 0.067710324 | 0.186423523 | 0.486061153 |
| ENSMUSG00000027459 | Fam110a | 73847 | protein_coding | 126.560225 | 0.486156588 | 0.110142205 | -4.413899178 | 1.02E-05 | 0.000113436 | 0.486156588 |
| ENSMUSG00000001521 | Tulp3 | 22158 | protein_coding | 27.00002826 | 0.486286036 | 0.266252338 | -1.826410386 | 0.067788469 | 0.186607563 | 0.486286036 |
| ENSMUSG00000021260 | Hhipl1 | 214305 | protein_coding | 2.820235089 | 0.486330355 | 2.125476706 | -0.228810014 | 0.819016583 | 1 | 0.486330355 |
| ENSMUSG00000021200 | Asb2 | 65256 | protein_coding | 12.23886726 | 0.486525386 | 0.4842377 | -1.004724302 | 0.315029623 | 0.535990818 | 0.486525386 |
| ENSMUSG00000095362 | Gm14325 | 329575 | protein_coding | 45.55378092 | -0.486725087 | 0.272479136 | 1.786283874 | 0.074053307 | 0.198945226 | 0.486725087 |
| ENSMUSG00000048796 | Cyb561d1 | 72023 | protein_coding | 275.3524968 | -0.486933506 | 0.105552564 | 4.613184996 | 3.97E-06 | 4.86E-05 | 0.486933506 |
| ENSMUSG00000020432 | Tcn2 | 21452 | protein_coding | 1024.033092 | -0.486957032 | 0.084343689 | 5.773485095 | 7.76E-09 | 1.52E-07 | 0.486957032 |
| ENSMUSG00000031342 | Gpm6b | 14758 | protein_coding | 6.914623785 | -0.486996575 | 0.632443708 | 0.770023591 | 0.441285899 | 0.659125086 | 0.486996575 |
| ENSMUSG00000019899 | Lama2 | 16773 | protein_coding | 3.657374594 | -0.487244144 | 1.290928544 | 0.377436959 | 0.705848905 | 1 | 0.487244144 |
| ENSMUSG00000050822 | Slc29a4 | 243328 | protein_coding | 13.26677078 | -0.487267894 | 0.373573129 | 1.304344064 | 0.192116298 | 0.389807821 | 0.487267894 |
| ENSMUSG00000032754 | Slc8b1 | 170756 | protein_coding | 522.0735575 | -0.487322681 | 0.137148921 | 3.553237447 | 0.000380521 | 0.002809084 | 0.487322681 |
| ENSMUSG00000079018 | Ly6c1 | 17067 | protein_coding | 13.48075454 | 0.48746065 | 0.486478428 | -1.002019046 | 0.316334394 | 0.537332461 | 0.48746065 |
| ENSMUSG00000019763 | Rmnd1 | 66084 | protein_coding | 213.6411049 | -0.487500825 | 0.093907547 | 5.191284858 | 2.09E-07 | 3.25E-06 | 0.487500825 |
| ENSMUSG00000063455 | D630045J12Rik | 330286 | protein_coding | 20.76930276 | 0.487558809 | 0.386132985 | -1.262670705 | 0.206707546 | 0.409464624 | 0.487558809 |
| ENSMUSG00000029765 | Plxna4 | 243743 | protein_coding | 5.089643079 | 0.487563046 | 1.557665067 | -0.313008911 | 0.754273892 | 0.876856731 | 0.487563046 |
| ENSMUSG00000074912 | Gm14207 | 100043609 | lncRNA | 32.50199648 | -0.487587963 | 0.210683402 | 2.314315981 | 0.020650396 | 0.075180916 | 0.487587963 |
| ENSMUSG00000023032 | Slc4a8 | 59033 | protein_coding | 31.15778632 | 0.487591769 | 0.336037729 | -1.451003047 | 0.146779012 | 0.324596932 | 0.487591769 |
| ENSMUSG00000055850 | Rnf181 | 66510 | protein_coding | 833.3842525 | 0.487697626 | 0.074239391 | -6.569256809 | 5.06E-11 | 1.33E-09 | 0.487697626 |
| ENSMUSG00000117924 | Tmem223 | 66836 | protein_coding | 127.5750152 | -0.487772079 | 0.240202118 | 2.030673515 | 0.042288125 | 0.13128129 | 0.487772079 |
| ENSMUSG00000028047 | Thbs3 | 21827 | protein_coding | 13.71837464 | -0.487785561 | 0.351896311 | 1.386162758 | 0.165697211 | 0.352762472 | 0.487785561 |
| ENSMUSG00000103897 | Pcdhga8 | 93716 | protein_coding | 4.337395447 | -0.487802845 | 0.552950631 | 0.882181551 | 0.377678637 | 1 | 0.487802845 |

| ensembl_gene_id | external_gene_name | entrezgene_id | gene_biotype | baseMean | log2FoldChange | lfcSE | stat | pvalue | padj | absLog2FC |
|---|---|---|---|---|---|---|---|---|---|---|
| ENSMUSG00000028671 | Gale | 74246 | protein_coding | 3837.840522 | 0.487860425 | 0.082262863 | -5.930506265 | 3.02E-09 | 6.28E-08 | 0.487860425 |
| ENSMUSG00000113035 | 5830428M24Rik | 76062 | lncRNA | 19.02143589 | -0.487921157 | 0.255449173 | 1.910051816 | 0.056126542 | 0.162609424 | 0.487921157 |
| ENSMUSG00000030352 | Tspan9 | 109246 | protein_coding | 109.7059206 | 0.487955868 | 0.16868713 | -2.892668032 | 0.003819848 | 0.019469435 | 0.487955868 |
| ENSMUSG00000026023 | Cdk15 | 271697 | protein_coding | 1.255966403 | 0.488097818 | 1.700515199 | -0.287029377 | 0.774089827 | 1 | 0.488097818 |
| ENSMUSG00000032009 | Sesn3 | 75747 | protein_coding | 389.4746678 | 0.48840629 | 0.093971356 | -5.197395387 | 2.02E-07 | 3.16E-06 | 0.48840629 |
| ENSMUSG00000028289 | Epha7 | 13841 | protein_coding | 18.53793495 | 0.488552136 | 0.354313763 | -1.378868637 | 0.16793526 | 0.356436387 | 0.488552136 |
| ENSMUSG00000040339 | Fam102b | 329739 | protein_coding | 916.8670228 | 0.488600574 | 0.089521805 | -5.457894601 | 4.82E-08 | 8.28E-07 | 0.488600574 |
| ENSMUSG00000089774 | Slc5a3 | 53881 | protein_coding | 1060.367175 | -0.488684188 | 0.133902133 | 3.649562395 | 0.000262687 | 0.002018623 | 0.488684188 |
| ENSMUSG00000032013 | Trim29 | 72169 | protein_coding | 7.933615581 | -0.488965175 | 0.428908465 | 1.140022207 | 0.25427705 | 0.466951438 | 0.488965175 |
| ENSMUSG00000030512 | Snrpa1 | 68981 | protein_coding | 237.7426247 | 0.48901604 | 0.11865661 | -4.12127096 | 3.77E-05 | 0.000364515 | 0.48901604 |
| ENSMUSG00000026616 | Cr2 | 12902 | protein_coding | 1.96562298 | -0.489086351 | 1.093897848 | 0.447104226 | 0.654799808 | 1 | 0.489086351 |
| ENSMUSG00000033967 | Rnf225 | 381845 | protein_coding | 7.824438053 | 0.489095891 | 0.57714751 | -0.847436543 | 0.396751843 | 0.618855346 | 0.489095891 |
| ENSMUSG00000097115 | 1810019N24Rik | 69086 | lncRNA | 6.072506162 | 0.489097189 | 0.520031533 | -0.940514485 | 0.346953723 | 0.568637732 | 0.489097189 |
| ENSMUSG00000027811 | 4930579G24Rik | 75939 | protein_coding | 20.06394072 | 0.48915868 | 0.302992847 | -1.614423194 | 0.106435664 | 0.259015408 | 0.48915868 |
| ENSMUSG00000026409 | Pfkfb2 | 18640 | protein_coding | 370.7094317 | 0.489187412 | 0.060204988 | -8.125363506 | 4.46E-16 | 1.79E-14 | 0.489187412 |
| ENSMUSG00000079235 | Ccdc13 | 100502861 | protein_coding | 3.5687713 | 0.489531188 | 0.820206968 | -0.596838612 | 0.550615136 | 1 | 0.489531188 |
| ENSMUSG00000032902 | Slc16a1 | 20501 | protein_coding | 4504.265076 | -0.489836068 | 0.141781663 | 3.454861911 | 0.000550575 | 0.003837987 | 0.489836068 |
| ENSMUSG00000042750 | Bex2 | 12069 | protein_coding | 26.82692195 | 0.489982418 | 0.224695553 | -2.180650271 | 0.029209294 | 0.098824478 | 0.489982418 |
| ENSMUSG00000050105 | Fam110d | 72690 | protein_coding | 1.072019769 | -0.490009715 | 1.214099302 | 0.403599371 | 0.686507348 | 1 | 0.490009715 |
| ENSMUSG00000033029 | 1700088E04Rik | 27660 | protein_coding | 11.493256 | -0.490215107 | 0.429200072 | 1.142159891 | 0.253387564 | 0.465869766 | 0.490215107 |
| ENSMUSG00000043753 | Dmrta1 | 242523 | protein_coding | 8.914909272 | 0.490367274 | 0.461830074 | -1.061791559 | 0.288330323 | 0.50718864 | 0.490367274 |
| ENSMUSG00000030329 | Pianp | 319352 | protein_coding | 3.96666673 | -0.490419903 | 0.989710114 | 0.495518735 | 0.620234001 | 1 | 0.490419903 |
| ENSMUSG00000090840 | Tex54 | 74307 | protein_coding | 2.966724422 | 0.490433924 | 0.767002937 | -0.639415966 | 0.522552365 | 1 | 0.490433924 |
| ENSMUSG00000031506 | Ptpn7 | 320139 | protein_coding | 8.531459135 | 0.490440165 | 0.768926293 | -0.637824677 | 0.523587814 | 0.722169631 | 0.490440165 |
| ENSMUSG00000029810 | Tmem176b | 65963 | protein_coding | 636.9505517 | 0.490539803 | 0.107296192 | -4.571828639 | 4.83E-06 | 5.79E-05 | 0.490539803 |
| ENSMUSG00000021464 | Ror2 | 26564 | protein_coding | 100.8057168 | 0.4905476 | 0.151402864 | -3.240015333 | 0.001195233 | 0.007373206 | 0.4905476 |
| ENSMUSG00000005803 | Sqor | 59010 | protein_coding | 5734.612306 | -0.490602079 | 0.082239021 | 5.96556325 | 2.44E-09 | 5.14E-08 | 0.490602079 |
| ENSMUSG00000098922 | Mir8106 | 102465896 | miRNA | 0.82024253 | 0.490629303 | 1.231069304 | -0.398539141 | 0.690232812 | 1 | 0.490629303 |
| ENSMUSG00000051984 | Sec31b | 240667 | protein_coding | 14.85243918 | -0.490653356 | 0.376285026 | 1.303940691 | 0.192253805 | 0.389986406 | 0.490653356 |
| ENSMUSG00000049811 | Fam161a | 73873 | protein_coding | 83.1265319 | -0.490754942 | 0.182653534 | 2.686807803 | 0.007213845 | 0.03273686 | 0.490754942 |
| ENSMUSG00000022747 | St3gal6 | 54613 | protein_coding | 1031.774233 | -0.490825459 | 0.261068993 | 1.880060338 | 0.060099855 | 0.170971405 | 0.490825459 |
| ENSMUSG00000024254 | Abcg8 | 67470 | protein_coding | 21.95050198 | -0.490842949 | 0.391853799 | 1.252617557 | 0.210344922 | 0.414416136 | 0.490842949 |
| ENSMUSG00000073293 | Nudt10 | 102954 | protein_coding | 17.08424415 | 0.49095158 | 0.268911452 | -1.825699781 | 0.067895496 | 0.186808768 | 0.49095158 |
| ENSMUSG00000062826 | Ces2f | 71903 | protein_coding | 4.964370409 | -0.490990165 | 0.62113169 | 0.790476759 | 0.42924939 | 0.648250976 | 0.490990165 |
| ENSMUSG00000042757 | Tmem108 | 81907 | protein_coding | 7.748625609 | 0.49102474 | 0.891932155 | -0.550518038 | 0.581964108 | 0.765030035 | 0.49102474 |
| ENSMUSG00000031273 | Col4a6 | 94216 | protein_coding | 4.291506031 | 0.491208113 | 1.301083334 | -0.37753778 | 0.705773994 | 1 | 0.491208113 |
| ENSMUSG00000002289 | Angptl4 | 57875 | protein_coding | 24.40689371 | -0.49129389 | 0.425166142 | 1.1555339 | 0.247871859 | 0.460059583 | 0.49129389 |
| ENSMUSG00000030693 | Klk10 | 69540 | protein_coding | 1.574461976 | 0.491312364 | 1.364086994 | -0.360176709 | 0.718714991 | 1 | 0.491312364 |
| ENSMUSG00000044162 | Tnip3 | 414084 | protein_coding | 335.0642948 | 0.49133867 | 0.241406994 | -2.035312492 | 0.04181944 | 0.130180251 | 0.49133867 |

| ensembl_gene_id | external_gene_name | entrezgene_id | gene_biotype | baseMean | log2FoldChange | lfcSE | stat | pvalue | padj | absLog2FC |
|---|---|---|---|---|---|---|---|---|---|---|
| ENSMUSG00000026511 | Srp9 | 27058 | protein_coding | 595.5305796 | 0.491421245 | 0.081637236 | -6.019572286 | 1.75E-09 | 3.74E-08 | 0.491421245 |
| ENSMUSG00000056531 | Ccdc18 | 73254 | protein_coding | 74.67031896 | 0.491464478 | 0.178383444 | -2.755101406 | 0.005867395 | 0.027715628 | 0.491464478 |
| ENSMUSG00000053318 | Slamf8 | 74748 | protein_coding | 1.667832329 | 0.491643796 | 1.460769454 | -0.336564949 | 0.736444883 | 1 | 0.491643796 |
| ENSMUSG00000062901 | Klhl24 | 75785 | protein_coding | 777.3567233 | -0.491702114 | 0.218608381 | 2.249237248 | 0.024497406 | 0.08602548 | 0.491702114 |
| ENSMUSG00000070972 | Dnajc25 | 72429 | protein_coding | 110.4155126 | 0.491775293 | 0.118271064 | -4.158035585 | 3.21E-05 | 0.000316096 | 0.491775293 |
| ENSMUSG00000078813 | Leng1 | 69757 | protein_coding | 211.3720974 | 0.491847093 | 0.10414543 | -4.722694908 | 2.33E-06 | 2.98E-05 | 0.491847093 |
| ENSMUSG00000083011 | Gm12816 | 433745 | processed_pseudogene | 0.733314262 | 0.492109242 | 1.87727104 | -0.262140752 | 0.793212929 | 1 | 0.492109242 |
| ENSMUSG00000033730 | Egr3 | 13655 | protein_coding | 7.584895595 | 0.492161909 | 0.625336452 | -0.787035375 | 0.431261158 | 0.649597228 | 0.492161909 |
| ENSMUSG00000049598 | Vsig8 | 240916 | protein_coding | 70.22733458 | 0.492267962 | 0.616727892 | -0.798193122 | 0.424758428 | 0.64425321 | 0.492267962 |
| ENSMUSG00000006307 | Kmt2b | 75410 | protein_coding | 673.4690644 | 0.49232009 | 0.199119716 | -2.472482878 | 0.013417814 | 0.05352217 | 0.49232009 |
| ENSMUSG00000096687 | Mfsd4b4 | 213332 | protein_coding | 26.24642192 | 0.492484066 | 0.232843189 | -2.115088995 | 0.034422363 | 0.112032105 | 0.492484066 |
| ENSMUSG00000111761 | 9230112J17Rik | 100504561 | lncRNA | 8.442196576 | -0.49250319 | 0.419734514 | 1.173368341 | 0.240648127 | 0.451187917 | 0.49250319 |
| ENSMUSG00000021884 | Hacl1 | 56794 | protein_coding | 15.14609076 | -0.492670701 | 0.361638386 | 1.362329664 | 0.17309387 | 0.363688435 | 0.492670701 |
| ENSMUSG00000032827 | Ppp1r9a | 243725 | protein_coding | 539.3770863 | -0.492674421 | 0.097992935 | 5.027652464 | 4.97E-07 | 7.34E-06 | 0.492674421 |
| ENSMUSG00000089308 | Gm24837 | 115490071 | snoRNA | 1.428546032 | 0.492951877 | 1.14968795 | -0.428770152 | 0.668090503 | 1 | 0.492951877 |
| ENSMUSG00000037621 | Atoh8 | 71093 | protein_coding | 25.32528985 | 0.493048074 | 0.260674218 | -1.89143398 | 0.058566436 | 0.167794262 | 0.493048074 |
| ENSMUSG00000062591 | Tubb4a | 22153 | protein_coding | 2.866452807 | 0.493257067 | 1.109467845 | -0.44458888 | 0.656616885 | 1 | 0.493257067 |
| ENSMUSG00000037849 | Ifi206 | 102639543 | protein_coding | 2.404894845 | -0.493258781 | 0.750573158 | 0.657176154 | 0.511067657 | 1 | 0.493258781 |
| ENSMUSG00000058063 | Trim31 | 224762 | protein_coding | 1046.417811 | 0.493320339 | 0.075622932 | -6.523422521 | 6.87E-11 | 1.76E-09 | 0.493320339 |
| ENSMUSG00000015790 | Surf1 | 20930 | protein_coding | 608.4397266 | -0.493373869 | 0.061400355 | 8.035358559 | 9.33E-16 | 3.66E-14 | 0.493373869 |
| ENSMUSG00000002835 | Chaf1a | 27221 | protein_coding | 170.3902762 | 0.493400382 | 0.140680216 | -3.507247831 | 0.000452767 | 0.003259372 | 0.493400382 |
| ENSMUSG00000026830 | Ermn | 77767 | protein_coding | 8.039979152 | 0.493474007 | 0.38478242 | -1.282475449 | 0.199675912 | 0.40075936 | 0.493474007 |
| ENSMUSG00000090272 | Mndal | 100040462 | protein_coding | 6.047281846 | -0.493502402 | 0.613860073 | 0.803933051 | 0.421435641 | 0.641211258 | 0.493502402 |
| ENSMUSG00000021115 | Vrk1 | 22367 | protein_coding | 203.4709154 | 0.493533219 | 0.14599914 | -3.380384431 | 0.000723845 | 0.004819991 | 0.493533219 |
| ENSMUSG00000057143 | Trim12c | 319236 | protein_coding | 217.9759074 | -0.493748641 | 0.126781963 | 3.894470715 | 9.84E-05 | 0.000848668 | 0.493748641 |
| ENSMUSG00000038037 | Socs1 | 12703 | protein_coding | 42.65635143 | 0.493836298 | 0.330552534 | -1.493972206 | 0.135182884 | 0.306271113 | 0.493836298 |
| ENSMUSG00000029602 | Rasal1 | 19415 | protein_coding | 5.245952773 | -0.493846566 | 0.70967343 | 0.695878619 | 0.486504848 | 0.695624496 | 0.493846566 |
| ENSMUSG00000042848 | Vmn1r226 | 171225 | protein_coding | 0.89118345 | 0.493953082 | 1.956466402 | -0.252472049 | 0.800676216 | 1 | 0.493953082 |
| ENSMUSG00000025145 | Lrrc45 | 217366 | protein_coding | 471.9088699 | 0.494027343 | 0.089415061 | -5.525102127 | 3.29E-08 | 5.82E-07 | 0.494027343 |
| ENSMUSG00000027713 | 1810062G17Rik | 72282 | protein_coding | 10.34280748 | 0.494039467 | 0.351601065 | -1.405113679 | 0.159987477 | 0.344285818 | 0.494039467 |
| ENSMUSG00000002489 | Tiam1 | 21844 | protein_coding | 219.6785982 | 0.494135786 | 0.154320695 | -3.202005962 | 0.001364742 | 0.00824336 | 0.494135786 |
| ENSMUSG00000049649 | Gpr3 | 14748 | protein_coding | 3.915725966 | -0.494400093 | 0.830151434 | 0.595554103 | 0.551473145 | 1 | 0.494400093 |
| ENSMUSG00000036231 | Agr3 | 403205 | protein_coding | 13.4058293 | -0.494478272 | 0.29406428 | 1.681531234 | 0.092659775 | 0.233580144 | 0.494478272 |
| ENSMUSG00000053367 | Epp13 | 627821 | protein_coding | 2.23436834 | 0.494518344 | 2.409977282 | -0.205196268 | 0.837418785 | 1 | 0.494518344 |
| ENSMUSG00000057596 | Trim30d | 209387 | protein_coding | 228.6143693 | -0.494571577 | 0.17055255 | 2.899819298 | 0.003733778 | 0.019107412 | 0.494571577 |
| ENSMUSG00000097084 | Foxl1 | 14241 | protein_coding | 0.980516654 | 0.49463165 | 1.967522297 | -0.251398244 | 0.801506226 | 1 | 0.49463165 |
| ENSMUSG00000019312 | Grb7 | 14786 | protein_coding | 750.9292366 | -0.494795458 | 0.105278029 | 4.699892893 | 2.60E-06 | 3.29E-05 | 0.494795458 |
| ENSMUSG00000029094 | Afap1 | 70292 | protein_coding | 161.0428446 | 0.494842644 | 0.11216687 | -4.411664918 | 1.03E-05 | 0.000114459 | 0.494842644 |
| ENSMUSG00000002020 | Ltbp2 | 16997 | protein_coding | 6.976178037 | -0.494914792 | 0.817507374 | 0.605394897 | 0.544916641 | 0.736964526 | 0.494914792 |

Supplementary Table 1

| ensembl_gene_id | external_gene_name | entrezgene_id | gene_biotype | baseMean | log2FoldChange | lfcSE | stat | pvalue | padj | absLog2FC |
|---|---|---|---|---|---|---|---|---|---|---|
| ENSMUSG00000030577 | Cd22 | 12483 | protein_coding | 3.064228491 | 0.494931743 | 1.329231282 | -0.372344339 | 0.70963649 | 1 | 0.494931743 |
| ENSMUSG00000074671 | Tspyl3 | 241732 | protein_coding | 6.001366393 | 0.495405745 | 0.46706647 | -1.060675036 | 0.288837609 | 0.507809662 | 0.495405745 |
| ENSMUSG00000085457 | 1110046J04Rik | 68808 | lncRNA | 5.467816483 | -0.495462571 | 0.50068463 | 0.989570164 | 0.322384259 | 0.54358318 | 0.495462571 |
| ENSMUSG00000097429 | Gm26520 | 105244630 | lncRNA | 1.309208534 | -0.495588692 | 1.147208415 | 0.431995342 | 0.665744799 | 1 | 0.495588692 |
| ENSMUSG00000027399 | Il1a | 16175 | protein_coding | 2.196080359 | 0.495667151 | 1.0403055 | -0.476463068 | 0.633744512 | 1 | 0.495667151 |
| ENSMUSG00000059060 | Rad51b | 19363 | protein_coding | 25.72174688 | 0.495773749 | 0.258415262 | -1.91851575 | 0.055045647 | 0.160039404 | 0.495773749 |
| ENSMUSG00000030472 | Ceacam18 | 72431 | protein_coding | 72.69657171 | -0.495887862 | 0.16165436 | 3.067581119 | 0.002157989 | 0.012051751 | 0.495887862 |
| ENSMUSG00000064288 | H4c12 | 319160 | protein_coding | 969.0656754 | 0.495966229 | 0.178587736 | -2.77715726 | 0.005483664 | 0.026283536 | 0.495966229 |
| ENSMUSG00000061414 | Cracr2a | 381812 | protein_coding | 733.2993923 | -0.496050193 | 0.178892727 | 2.772891897 | 0.005556057 | 0.026525797 | 0.496050193 |
| ENSMUSG00000025647 | Shisa5 | 66940 | protein_coding | 1952.372672 | -0.496146153 | 0.076101757 | 6.519509874 | 7.05E-11 | 1.80E-09 | 0.496146153 |
| ENSMUSG00000042632 | Pla2g6 | 53357 | protein_coding | 153.4175473 | 0.49619889 | 0.149508036 | -3.318877736 | 0.0009038 | 0.005843912 | 0.49619889 |
| ENSMUSG00000114284 | 9630015K15Rik | 78569 | lncRNA | 1.849747 | -0.496377971 | 1.427331044 | 0.347766535 | 0.728015524 | 1 | 0.496377971 |
| ENSMUSG00000030287 | Itpr2 | 16439 | protein_coding | 743.7250978 | -0.496423785 | 0.094180808 | 5.270965433 | 1.36E-07 | 2.19E-06 | 0.496423785 |
| ENSMUSG00000032607 | Amt | 434437 | protein_coding | 213.9268687 | -0.496436176 | 0.119198579 | 4.164782681 | 3.12E-05 | 0.000307626 | 0.496436176 |
| ENSMUSG00000040606 | Kazn | 71529 | protein_coding | 121.9965725 | -0.496478459 | 0.14235934 | 3.487501841 | 0.000487556 | 0.003474964 | 0.496478459 |
| ENSMUSG00000034336 | Ina | 226180 | protein_coding | 1.602556247 | -0.496494529 | 1.184372224 | 0.419204806 | 0.67506646 | 1 | 0.496494529 |
| ENSMUSG00000000632 | Sez6 | 20370 | protein_coding | 2.45980291 | 0.496646017 | 1.517420108 | -0.32729632 | 0.743443776 | 1 | 0.496646017 |
| ENSMUSG00000023243 | Kcnk5 | 16529 | protein_coding | 1110.066698 | -0.496678342 | 0.11260101 | 4.410958157 | 1.03E-05 | 0.000114755 | 0.496678342 |
| ENSMUSG00000010110 | Stx5a | 56389 | protein_coding | 993.4831042 | 0.496848208 | 0.05557355 | -8.940371921 | 3.88E-19 | 1.89E-17 | 0.496848208 |
| ENSMUSG00000037254 | Itih2 | 16425 | protein_coding | 109.3908525 | 0.496957881 | 0.416872588 | -1.19210976 | 0.233218211 | 0.442685693 | 0.496957881 |
| ENSMUSG00000074220 | Zfp382 | 233060 | protein_coding | 3.172095675 | 0.496961002 | 0.577999326 | -0.859795125 | 0.389901987 | 1 | 0.496961002 |
| ENSMUSG00000033533 | Acsm1 | 117147 | protein_coding | 1.026291417 | 0.4970555 | 2.008279129 | -0.247503195 | 0.804518819 | 1 | 0.4970555 |
| ENSMUSG00000026879 | Gsn | 227753 | protein_coding | 2169.49228 | -0.497172228 | 0.197087669 | 2.522594293 | 0.011649271 | 0.048021984 | 0.497172228 |
| ENSMUSG00000073792 | Alg6 | 320438 | protein_coding | 248.7222711 | 0.497206612 | 0.1495832 | -3.323946882 | 0.000887531 | 0.005758032 | 0.497206612 |
| ENSMUSG00000051855 | Mest | 17294 | protein_coding | 8.163757029 | -0.497244424 | 0.422616268 | 1.176586095 | 0.239360751 | 0.449590928 | 0.497244424 |
| ENSMUSG00000078157 | Fem1al | 216622 | protein_coding | 25.28884062 | -0.497317549 | 0.305442701 | 1.628186061 | 0.103485439 | 0.253780598 | 0.497317549 |
| ENSMUSG00000054679 | Srsf12 | 272009 | protein_coding | 1.45040971 | 0.49745976 | 1.298308031 | -0.383160043 | 0.701601103 | 1 | 0.49745976 |
| ENSMUSG00000097415 | AU020206 | 108167440 | lncRNA | 81.84781693 | 0.497592225 | 0.178336217 | -2.790191657 | 0.005267685 | 0.025458165 | 0.497592225 |
| ENSMUSG00000028024 | Enpep | 13809 | protein_coding | 77.36476061 | -0.497725747 | 0.407823647 | 1.220443568 | 0.22229677 | 0.429396526 | 0.497725747 |
| ENSMUSG00000031461 | Myom2 | 17930 | protein_coding | 4.692608166 | 0.497745364 | 1.797217612 | -0.276953309 | 0.781815959 | 0.890653197 | 0.497745364 |
| ENSMUSG00000049744 | Arhgap15 | 76117 | protein_coding | 5.454381744 | 0.497782971 | 0.790888131 | -0.629397448 | 0.529088888 | 0.726903567 | 0.497782971 |
| ENSMUSG00000112366 | C730027H18Rik | 319572 | lncRNA | 1.115101584 | 0.497794706 | 2.02113682 | -0.246294413 | 0.805454335 | 1 | 0.497794706 |
| ENSMUSG00000117322 | 6330415G19Rik | 320020 | lncRNA | 6.163317022 | 0.497820361 | 0.498193865 | -0.999250283 | 0.317673463 | 0.539052565 | 0.497820361 |
| ENSMUSG00000029092 | D5Ertd615e | 52401 | lncRNA | 1.138702293 | 0.49787755 | 2.022597619 | -0.246157489 | 0.805560322 | 1 | 0.49787755 |
| ENSMUSG00000029993 | Nfu1 | 56748 | protein_coding | 252.8494328 | -0.497880726 | 0.095976973 | 5.18750186 | 2.13E-07 | 3.32E-06 | 0.497880726 |
| ENSMUSG00000089665 | Fcor | 100503924 | protein_coding | 1.728574944 | 0.497883308 | 0.926792021 | -0.537211474 | 0.591121547 | 1 | 0.497883308 |
| ENSMUSG00000031354 | Amelx | 11704 | protein_coding | 1.205089304 | 0.497897697 | 1.735718188 | -0.286853995 | 0.774224117 | 1 | 0.497897697 |
| ENSMUSG00000066072 | Cyp4a10 | 13117 | protein_coding | 1.16223802 | 0.497988766 | 2.024547027 | -0.245975401 | 0.805701275 | 1 | 0.497988766 |
| ENSMUSG00000037736 | Limch1 | 77569 | protein_coding | 78.62669851 | 0.498252014 | 0.205002079 | -2.430472978 | 0.01507913 | 0.058661192 | 0.498252014 |

Supplementary Table 1

| ensembl_gene_id | external_gene_name | entrezgene_id | gene_biotype | baseMean | log2FoldChange | lfcSE | stat | pvalue | padj | absLog2FC |
|---|---|---|---|---|---|---|---|---|---|---|
| ENSMUSG00000087050 | Dhrs13os | 100504178 | lncRNA | 1.605931618 | -0.498380352 | 0.890640603 | 0.559575154 | 0.575769256 | 1 | 0.498380352 |
| ENSMUSG00000024987 | Cyp26a1 | 13082 | protein_coding | 1.56517835 | 0.498589624 | 1.937573898 | -0.257326766 | 0.796926532 | 1 | 0.498589624 |
| ENSMUSG00000032192 | Gnb5 | 14697 | protein_coding | 92.75091182 | -0.498836276 | 0.130356147 | 3.826718483 | 0.000129863 | 0.001087135 | 0.498836276 |
| ENSMUSG00000090966 | Vmn2r116 | 619697 | protein_coding | 2.101601676 | -0.498982911 | 3.126517541 | 0.159597029 | 0.87319852 | 1 | 0.498982911 |
| ENSMUSG00000030082 | Sec61a1 | 53421 | protein_coding | 3156.508255 | 0.499039766 | 0.085764466 | -5.818724119 | 5.93E-09 | 1.18E-07 | 0.499039766 |
| ENSMUSG00000023927 | Satb1 | 20230 | protein_coding | 6.934883477 | 0.499278753 | 0.716261529 | -0.697062083 | 0.485763942 | 0.695380179 | 0.499278753 |
| ENSMUSG00000000301 | Pemt | 18618 | protein_coding | 50.38014578 | 0.499304506 | 0.155646152 | -3.207946344 | 0.001336864 | 0.008110572 | 0.499304506 |
| ENSMUSG00000063383 | Zfp947 | 210853 | protein_coding | 48.93611044 | -0.499436719 | 0.177902641 | 2.807359774 | 0.004994942 | 0.024310779 | 0.499436719 |
| ENSMUSG00000022751 | Nit2 | 52633 | protein_coding | 155.2646978 | -0.499440768 | 0.120400715 | 4.148154496 | 3.35E-05 | 0.000328678 | 0.499440768 |
| ENSMUSG00000039883 | Lrrc17 | 74511 | protein_coding | 3.272838309 | 0.499449224 | 0.597822239 | -0.835447716 | 0.403465673 | 1 | 0.499449224 |
| ENSMUSG00000052479 | A330008L17Rik | 234624 | lncRNA | 0.933043422 | -0.499587708 | 1.715794059 | 0.291169972 | 0.770921327 | 1 | 0.499587708 |
| ENSMUSG00000054752 | Fsd1l | 319636 | protein_coding | 148.0965399 | -0.499593933 | 0.149333341 | 3.345494912 | 0.000821359 | 0.005386782 | 0.499593933 |
| ENSMUSG00000050600 | Zfp831 | 100043757 | protein_coding | 8.056869683 | 0.499660112 | 0.787386048 | -0.634580856 | 0.525701817 | 0.724177495 | 0.499660112 |
| ENSMUSG00000096215 | Smim22 | 432995 | protein_coding | 247.3666497 | -0.499669921 | 0.077770959 | 6.424890869 | 1.32E-10 | 3.22E-09 | 0.499669921 |
| ENSMUSG00000004455 | Ppp1cc | 19047 | protein_coding | 580.0025146 | -0.499672757 | 0.139930715 | 3.570858324 | 0.000355813 | 0.002653975 | 0.499672757 |
| ENSMUSG00000026818 | Cel | 12613 | protein_coding | 3.915205761 | -0.499883509 | 1.016634551 | 0.49170423 | 0.622928448 | 1 | 0.499883509 |
| ENSMUSG00000021671 | Poc5 | 67463 | protein_coding | 200.3242605 | 0.499897239 | 0.083835298 | -5.962849177 | 2.48E-09 | 5.21E-08 | 0.499897239 |
| ENSMUSG00000087621 | 1700003G18Rik | 69350 | lncRNA | 1.086228262 | 0.499954999 | 1.239758513 | -0.40326805 | 0.686751043 | 1 | 0.499954999 |
| ENSMUSG00000031015 | Swap70 | 20947 | protein_coding | 391.1514585 | -0.499991448 | 0.085414795 | 5.853686711 | 4.81E-09 | 9.72E-08 | 0.499991448 |
| ENSMUSG00000001131 | Timp1 | 21857 | protein_coding | 1.594885996 | -0.500072711 | 1.244423891 | 0.40185078 | 0.687793848 | 1 | 0.500072711 |
| ENSMUSG00000026986 | Hnmt | 140483 | protein_coding | 9.559748509 | -0.500072962 | 0.398305257 | 1.255501788 | 0.20929666 | 0.412695527 | 0.500072962 |
| ENSMUSG00000030872 | Gga2 | 74105 | protein_coding | 139.6403424 | 0.500352856 | 0.161393024 | -3.100213647 | 0.001933811 | 0.011044625 | 0.500352856 |
| ENSMUSG00000048154 | Kmt2d | 381022 | protein_coding | 1624.447867 | 0.500422312 | 0.266152246 | -1.88021074 | 0.060079363 | 0.170942556 | 0.500422312 |
| ENSMUSG00000045008 | 9030612E09Rik | 74530 | lncRNA | 5.117111018 | -0.500529123 | 0.460291923 | 1.087416699 | 0.276852695 | 0.493730605 | 0.500529123 |
| ENSMUSG00000035642 | Aamdc | 66273 | protein_coding | 296.596199 | -0.500570725 | 0.091122263 | 5.493396544 | 3.94E-08 | 6.88E-07 | 0.500570725 |
| ENSMUSG00000027381 | Bcl2l11 | 12125 | protein_coding | 106.9442639 | -0.50058175 | 0.15706056 | 3.187189394 | 0.001436627 | 0.008574071 | 0.50058175 |
| ENSMUSG00000085162 | Gm12295 | 327946 | lncRNA | 1.52606114 | 0.500712722 | 2.073940733 | -0.241430584 | 0.809221411 | 1 | 0.500712722 |
| ENSMUSG00000037798 | Mat1a | 11720 | protein_coding | 1.350361076 | 0.500885236 | 1.589911057 | -0.315039784 | 0.752731447 | 1 | 0.500885236 |
| ENSMUSG00000043068 | Fam89a | 69627 | protein_coding | 4.781976366 | 0.500922036 | 0.49461621 | -1.012748926 | 0.311180102 | 0.532085855 | 0.500922036 |
| ENSMUSG00000019464 | Ptger1 | 19216 | protein_coding | 14.35152911 | -0.501025959 | 0.350403696 | 1.429853524 | 0.152759063 | 0.33362953 | 0.501025959 |
| ENSMUSG00000027403 | Tgm6 | 241636 | protein_coding | 1.489831846 | 0.501207938 | 2.083239831 | -0.240590608 | 0.80987243 | 1 | 0.501207938 |
| ENSMUSG00000108714 | Gm21057 | 100859962 | lncRNA | 4.363724774 | 0.501399488 | 0.509334547 | -0.984420734 | 0.324908688 | 1 | 0.501399488 |
| ENSMUSG00000020883 | Fbxl20 | 72194 | protein_coding | 350.5993513 | -0.501437948 | 0.129866379 | 3.861183725 | 0.000112839 | 0.000958551 | 0.501437948 |
| ENSMUSG00000035356 | Nfkbiz | 80859 | protein_coding | 1312.777144 | 0.50153355 | 0.146056596 | -3.433830209 | 0.000595117 | 0.004103449 | 0.50153355 |
| ENSMUSG00000092518 | Garin5b | 243822 | protein_coding | 2.581261561 | 0.501900509 | 1.655113594 | -0.303242334 | 0.761705189 | 1 | 0.501900509 |
| ENSMUSG00000027460 | Angpt4 | 11602 | protein_coding | 3.131264802 | 0.501902499 | 0.864467982 | -0.580591196 | 0.561516007 | 1 | 0.501902499 |
| ENSMUSG00000040163 | 1700034J05Rik | 73344 | protein_coding | 5.041759544 | 0.501941412 | 0.564084359 | -0.889833948 | 0.373555055 | 0.595844868 | 0.501941412 |
| ENSMUSG00000046179 | E2f8 | 108961 | protein_coding | 174.1595085 | 0.502016328 | 0.211696859 | -2.371392425 | 0.017721205 | 0.066683451 | 0.502016328 |
| ENSMUSG00000031229 | Atrx | 22589 | protein_coding | 1650.406056 | 0.502099735 | 0.072525492 | -6.923079331 | 4.42E-12 | 1.26E-10 | 0.502099735 |

| ensembl_gene_id | external_gene_name | entrezgene_id | gene_biotype | baseMean | log2FoldChange | lfcSE | stat | pvalue | padj | absLog2FC |
|---|---|---|---|---|---|---|---|---|---|---|
| ENSMUSG00000064595 | Gm22300 | 115486242 | snoRNA | 1.489634103 | 0.502121167 | 0.983225863 | -0.510687509 | 0.609569889 | 1 | 0.502121167 |
| ENSMUSG00000104600 | Dmrt1i | 100579137 | lncRNA | 2.552912214 | 0.502545266 | 0.846710718 | -0.59352652 | 0.55282884 | 1 | 0.502545266 |
| ENSMUSG00000051243 | Islr2 | 320563 | protein_coding | 4.056208884 | -0.502606809 | 0.930362842 | 0.540226658 | 0.589040731 | 1 | 0.502606809 |
| ENSMUSG00000020389 | Cdkl3 | 213084 | protein_coding | 32.27928053 | 0.502618879 | 0.194817738 | -2.579944124 | 0.00988163 | 0.042188532 | 0.502618879 |
| ENSMUSG00000008307 | 1700109H08Rik | 77036 | protein_coding | 14.66873373 | -0.502737841 | 0.338066455 | 1.487097679 | 0.136989014 | 0.309296897 | 0.502737841 |
| ENSMUSG00000038312 | Edem2 | 108687 | protein_coding | 804.8077526 | 0.503062774 | 0.082170248 | -6.122200969 | 9.23E-10 | 2.03E-08 | 0.503062774 |
| ENSMUSG00000024842 | Cabp4 | 73660 | protein_coding | 4.699903503 | 0.503197912 | 0.869273765 | -0.578871619 | 0.562675801 | 0.75067522 | 0.503197912 |
| ENSMUSG00000045107 | Saysd1 | 67509 | protein_coding | 130.3578529 | -0.503397237 | 0.115134493 | 4.372253922 | 1.23E-05 | 0.000134753 | 0.503397237 |
| ENSMUSG00000029772 | Ahcyl2 | 74340 | protein_coding | 8720.013898 | -0.503589482 | 0.120310189 | 4.185759204 | 2.84E-05 | 0.000282913 | 0.503589482 |
| ENSMUSG00000049526 | Tmem202 | 73893 | protein_coding | 4.447425263 | -0.50362081 | 0.531044835 | 0.948358362 | 0.342947049 | 0.564917636 | 0.50362081 |
| ENSMUSG00000097216 | 4932441J04Rik | 319216 | lncRNA | 1.134497324 | 0.503646779 | 2.050204037 | -0.245656905 | 0.805947835 | 1 | 0.503646779 |
| ENSMUSG00000021959 | Lats2 | 50523 | protein_coding | 44.09607795 | -0.503685764 | 0.284657476 | 1.769445057 | 0.076819632 | 0.204250427 | 0.503685764 |
| ENSMUSG00000024598 | Fbn2 | 14119 | protein_coding | 4.981467514 | 0.503744998 | 1.179643326 | -0.427031618 | 0.669356298 | 0.822609935 | 0.503744998 |
| ENSMUSG00000028218 | Cibar1 | 68099 | protein_coding | 65.5488406 | 0.503752284 | 0.214343568 | -2.350209473 | 0.018762849 | 0.069848891 | 0.503752284 |
| ENSMUSG00000089407 | Gm24878 | 115490251 | snRNA | 0.60775806 | -0.503827488 | 1.833127623 | 0.274845833 | 0.783434684 | 1 | 0.503827488 |
| ENSMUSG00000025931 | Paqr8 | 74229 | protein_coding | 420.1311123 | 0.503905725 | 0.15032067 | -3.352205155 | 0.000801706 | 0.005268353 | 0.503905725 |
| ENSMUSG00000063260 | Syt10 | 54526 | protein_coding | 4.460012711 | 0.503914994 | 0.503588392 | -1.000648549 | 0.31699675 | 0.538291447 | 0.503914994 |
| ENSMUSG00000004371 | Il11 | 16156 | protein_coding | 3.080454138 | 0.503922781 | 0.825264335 | -0.610619845 | 0.541451282 | 1 | 0.503922781 |
| ENSMUSG00000032548 | Slco2a1 | 24059 | protein_coding | 397.2595052 | -0.504116465 | 0.287288883 | 1.75473711 | 0.079304285 | 0.208654479 | 0.504116465 |
| ENSMUSG00000045968 | Teddm2 | 240817 | protein_coding | 18.5937443 | 0.504208949 | 0.364473349 | -1.383390446 | 0.166545178 | 0.354149765 | 0.504208949 |
| ENSMUSG00000079508 | Apoo | 68316 | protein_coding | 109.9759766 | -0.504362398 | 0.182469342 | 2.764093918 | 0.005708111 | 0.027173174 | 0.504362398 |
| ENSMUSG00000051965 | Nanos2 | 378430 | protein_coding | 1.753838352 | 0.504365099 | 2.145100433 | -0.235124235 | 0.814112309 | 1 | 0.504365099 |
| ENSMUSG00000068523 | Gng5 | 14707 | protein_coding | 682.6391533 | -0.504510291 | 0.115831439 | 4.355555756 | 1.33E-05 | 0.000143449 | 0.504510291 |
| ENSMUSG00000041577 | Prelp | 116847 | protein_coding | 84.15823308 | 0.504534135 | 0.258917022 | -1.948632547 | 0.051339322 | 0.152129476 | 0.504534135 |
| ENSMUSG00000030406 | Gipr | 381853 | protein_coding | 7.381684959 | -0.50454266 | 0.606805212 | 0.831473841 | 0.405706 | 0.626602117 | 0.50454266 |
| ENSMUSG00000025509 | Pnpla2 | 66853 | protein_coding | 712.5582861 | -0.504721614 | 0.087232388 | 5.785942858 | 7.21E-09 | 1.42E-07 | 0.504721614 |
| ENSMUSG00000021477 | Ctsl | 13039 | protein_coding | 862.0390301 | -0.504894193 | 0.100217151 | 5.038001883 | 4.70E-07 | 6.99E-06 | 0.504894193 |
| ENSMUSG00000026994 | Galnt3 | 14425 | protein_coding | 934.347507 | 0.504932721 | 0.124421439 | -4.058245293 | 4.94E-05 | 0.000462865 | 0.504932721 |
| ENSMUSG00000029322 | Plac8 | 231507 | protein_coding | 5827.885613 | -0.505043288 | 0.121498387 | 4.156790074 | 3.23E-05 | 0.000317634 | 0.505043288 |
| ENSMUSG00000023055 | Calcoco1 | 67488 | protein_coding | 179.1582805 | -0.505092921 | 0.131958143 | 3.827675271 | 0.000129359 | 0.001083835 | 0.505092921 |
| ENSMUSG00000017737 | Mmp9 | 17395 | protein_coding | 1.595089041 | 0.505237597 | 1.96188002 | -0.257527265 | 0.79677177 | 1 | 0.505237597 |
| ENSMUSG00000000204 | Slfn4 | 20558 | protein_coding | 503.4769073 | -0.505368349 | 0.237049269 | 2.131912707 | 0.033014023 | 0.108572726 | 0.505368349 |
| ENSMUSG00000026234 | Ncl | 17975 | protein_coding | 2369.46601 | 0.505389193 | 0.08568482 | -5.8982349 | 3.67E-09 | 7.57E-08 | 0.505389193 |
| ENSMUSG00000036655 | Colec11 | 71693 | protein_coding | 1.500195684 | -0.505409218 | 1.283557401 | 0.393756616 | 0.693760738 | 1 | 0.505409218 |
| ENSMUSG00000030844 | Rgs10 | 67865 | protein_coding | 8.387356946 | 0.505431459 | 0.447463573 | -1.129547721 | 0.25866685 | 0.472054853 | 0.505431459 |
| ENSMUSG00000069308 | H2bc22 | 319188 | protein_coding | 39.83875093 | 0.505440086 | 0.25036924 | -2.018778685 | 0.043510227 | 0.134003711 | 0.505440086 |
| ENSMUSG00000035246 | Pcyt1b | 236899 | protein_coding | 5.016587553 | -0.505672678 | 0.718944869 | 0.703353901 | 0.481835228 | 0.692577853 | 0.505672678 |
| ENSMUSG00000078922 | Tgtp1 | 21822 | protein_coding | 3.544248653 | 0.505678027 | 0.778576661 | -0.649490349 | 0.516021483 | 1 | 0.505678027 |
| ENSMUSG00000085852 | Gm13807 | 639658 | lncRNA | 15.67093967 | 0.505754873 | 0.317358827 | -1.593637328 | 0.111017289 | 0.266628782 | 0.505754873 |

| ensembl_gene_id | external_gene_name | entrezgene_id | gene_biotype | baseMean | log2FoldChange | lfcSE | stat | pvalue | padj | absLog2FC |
|---|---|---|---|---|---|---|---|---|---|---|
| ENSMUSG00000022987 | Zfp641 | 239652 | protein_coding | 8.183868113 | 0.50595657 | 0.601301816 | -0.841435294 | 0.40010412 | 0.622275328 | 0.50595657 |
| ENSMUSG00000038116 | Phf20 | 228829 | protein_coding | 215.6712539 | 0.506004464 | 0.12857853 | -3.935372916 | 8.31E-05 | 0.000732396 | 0.506004464 |
| ENSMUSG00000073413 | Ly6g6d | 114654 | protein_coding | 31.13353512 | -0.506035273 | 0.183225828 | 2.761811904 | 0.005748158 | 0.027285164 | 0.506035273 |
| ENSMUSG00000071637 | Cebpd | 12609 | protein_coding | 398.4816021 | 0.50604147 | 0.195554014 | -2.587732463 | 0.009660998 | 0.041407144 | 0.50604147 |
| ENSMUSG00000091994 | E130317F20Rik | 414101 | lncRNA | 31.11296209 | 0.506302105 | 0.30590829 | -1.655078076 | 0.097908662 | 0.243576926 | 0.506302105 |
| ENSMUSG00000019997 | Ccn2 | 14219 | protein_coding | 1.266879813 | -0.506338118 | 1.325671801 | 0.381948321 | 0.702499698 | 1 | 0.506338118 |
| ENSMUSG00000030889 | Vwa3a | 233813 | protein_coding | 2.150081629 | 0.506369295 | 2.375869598 | -0.213130088 | 0.831225497 | 1 | 0.506369295 |
| ENSMUSG00000105044 | Gm10416 | 667213 | TEC | 4.049690312 | -0.506374562 | 0.53686506 | 0.943206401 | 0.34557534 | 1 | 0.506374562 |
| ENSMUSG00000022837 | Iqcb1 | 320299 | protein_coding | 24.12882195 | 0.506403595 | 0.206653469 | -2.450496465 | 0.014265936 | 0.056065605 | 0.506403595 |
| ENSMUSG00000042581 | Thsd7b | 210417 | protein_coding | 2.741922176 | 0.506436241 | 2.376770433 | -0.213077474 | 0.831266534 | 1 | 0.506436241 |
| ENSMUSG00000034145 | Tmem63c | 217733 | protein_coding | 2.482427863 | -0.506459492 | 1.46321939 | 0.346126832 | 0.729247399 | 1 | 0.506459492 |
| ENSMUSG00000027130 | Slc12a6 | 107723 | protein_coding | 361.8327922 | -0.506606195 | 0.117674186 | 4.305159951 | 1.67E-05 | 0.000175853 | 0.506606195 |
| ENSMUSG00000030946 | Lhpp | 76429 | protein_coding | 157.5787001 | -0.50683217 | 0.119528299 | 4.240269245 | 2.23E-05 | 0.000227579 | 0.50683217 |
| ENSMUSG00000036395 | Glb1l2 | 244757 | protein_coding | 3.934803969 | -0.506915755 | 1.065922151 | 0.475565457 | 0.634383989 | 1 | 0.506915755 |
| ENSMUSG00000017417 | Plxdc1 | 72324 | protein_coding | 1.334465089 | 0.506927872 | 1.625804457 | -0.311801257 | 0.755191569 | 1 | 0.506927872 |
| ENSMUSG00000056144 | Trim34a | 94094 | protein_coding | 76.75558173 | -0.507121452 | 0.186464046 | 2.719674183 | 0.006534627 | 0.030092813 | 0.507121452 |
| ENSMUSG00000050953 | Gja1 | 14609 | protein_coding | 70.93052808 | -0.507256316 | 0.249683156 | 2.031600062 | 0.04219416 | 0.131124269 | 0.507256316 |
| ENSMUSG00000069911 | Insyn2b | 574403 | protein_coding | 2.2769706 | -0.507463879 | 1.409522937 | 0.360025272 | 0.718828235 | 1 | 0.507463879 |
| ENSMUSG00000045319 | Proser2 | 227545 | protein_coding | 141.599712 | -0.507470304 | 0.115160264 | 4.406644131 | 1.05E-05 | 0.000116906 | 0.507470304 |
| ENSMUSG00000039231 | Suv39h1 | 20937 | protein_coding | 150.6723411 | 0.507584554 | 0.165055656 | -3.075232714 | 0.002103384 | 0.011831403 | 0.507584554 |
| ENSMUSG00000042759 | Apobr | 171504 | protein_coding | 10.61399909 | 0.5076056 | 0.4587434 | -1.106513139 | 0.26850448 | 0.483542311 | 0.5076056 |
| ENSMUSG00000021125 | Arg2 | 11847 | protein_coding | 123.1205895 | -0.507644369 | 0.251340109 | 2.01975073 | 0.04340925 | 0.133817502 | 0.507644369 |
| ENSMUSG00000030257 | Srgap3 | 259302 | protein_coding | 145.7170457 | -0.507682509 | 0.184282434 | 2.754915363 | 0.005870732 | 0.027715628 | 0.507682509 |
| ENSMUSG00000079038 | D130040H23Rik | 211135 | protein_coding | 23.51296037 | -0.507739662 | 0.26444579 | 1.920014159 | 0.054856111 | 0.159769633 | 0.507739662 |
| ENSMUSG00000019947 | Arid5b | 71371 | protein_coding | 301.8476431 | -0.507751662 | 0.126551911 | 4.012200679 | 6.02E-05 | 0.000551585 | 0.507751662 |
| ENSMUSG00000097651 | 4930461G14Rik | 74879 | lncRNA | 1.163263742 | 0.507917825 | 2.063214883 | -0.24617786 | 0.805544553 | 1 | 0.507917825 |
| ENSMUSG00000068699 | Flnc | 68794 | protein_coding | 6.067758367 | 0.507925078 | 1.348554732 | -0.376644022 | 0.706438168 | 0.846158956 | 0.507925078 |
| ENSMUSG00000109864 | Eid3 | 66341 | protein_coding | 32.49999974 | -0.507993576 | 0.229585613 | 2.212654223 | 0.026921494 | 0.092582579 | 0.507993576 |
| ENSMUSG00000030513 | Pcsk6 | 18553 | protein_coding | 13.22141694 | 0.508014691 | 0.325001493 | -1.563114945 | 0.118025561 | 0.278598358 | 0.508014691 |
| ENSMUSG00000064023 | Klk8 | 259277 | protein_coding | 1.749793244 | 0.508051854 | 1.522836503 | -0.333622062 | 0.738664768 | 1 | 0.508051854 |
| ENSMUSG00000007655 | Cav1 | 12389 | protein_coding | 2.905999782 | 0.508178023 | 0.826518256 | -0.614841861 | 0.538659161 | 1 | 0.508178023 |
| ENSMUSG00000095930 | Nim1k | 245269 | protein_coding | 46.54474582 | 0.508281003 | 0.501023295 | -1.014485769 | 0.310351017 | 0.53139398 | 0.508281003 |
| ENSMUSG00000029223 | Uchl1 | 22223 | protein_coding | 4.689119246 | -0.50830404 | 0.525374093 | 0.967508764 | 0.333289762 | 0.555673613 | 0.50830404 |
| ENSMUSG00000049134 | Nrap | 18175 | protein_coding | 235.5054168 | -0.508316611 | 0.271558285 | 1.871850865 | 0.061227239 | 0.17331619 | 0.508316611 |
| ENSMUSG00000045216 | Hs6st1 | 50785 | protein_coding | 1315.702261 | 0.508625012 | 0.086829573 | -5.857739411 | 4.69E-09 | 9.51E-08 | 0.508625012 |
| ENSMUSG00000072949 | Acot1 | 26897 | protein_coding | 79.15264391 | 0.508825661 | 0.155714508 | -3.267683061 | 0.001084317 | 0.006808522 | 0.508825661 |
| ENSMUSG00000031568 | Rwdd4a | 192174 | protein_coding | 249.7340397 | 0.508845717 | 0.091186024 | -5.580303792 | 2.40E-08 | 4.35E-07 | 0.508845717 |
| ENSMUSG00000038990 | Cables2 | 252966 | protein_coding | 337.7807084 | 0.509220928 | 0.087668997 | -5.808449362 | 6.31E-09 | 1.25E-07 | 0.509220928 |
| ENSMUSG00000074529 | Zfp972 | 78703 | protein_coding | 6.218561936 | -0.509268665 | 0.413817696 | 1.230659466 | 0.218450254 | 0.424602824 | 0.509268665 |

| ensembl_gene_id | external_gene_name | entrezgene_id | gene_biotype | baseMean | log2FoldChange | lfcSE | stat | pvalue | padj | absLog2FC |
|---|---|---|---|---|---|---|---|---|---|---|
| ENSMUSG00000086010 | Gm15318 | 102631757 | lncRNA | 1.287453336 | -0.509270857 | 1.348230509 | 0.377732779 | 0.705629115 | 1 | 0.509270857 |
| ENSMUSG00000085551 | 1810063I02Rik | 69854 | lncRNA | 7.84748168 | -0.509334934 | 0.448071291 | 1.136727445 | 0.255652242 | 0.468416857 | 0.509334934 |
| ENSMUSG00000029671 | Wnt16 | 93735 | protein_coding | 1.290351472 | -0.509361761 | 1.704648485 | 0.298807505 | 0.765086924 | 1 | 0.509361761 |
| ENSMUSG00000031284 | Pak3 | 18481 | protein_coding | 4.504699415 | 0.509472072 | 1.291370307 | -0.39452051 | 0.693196788 | 0.837078181 | 0.509472072 |
| ENSMUSG00000063160 | Numbl | 18223 | protein_coding | 11.94047696 | 0.509499162 | 0.507104461 | -1.004722304 | 0.315030585 | 0.535990818 | 0.509499162 |
| ENSMUSG00000024734 | Zp1 | 22786 | protein_coding | 3.889098907 | -0.50950679 | 0.885199012 | 0.575584454 | 0.564896091 | 1 | 0.50950679 |
| ENSMUSG00000019888 | Mgat4c | 67569 | protein_coding | 7858.549368 | 0.509526724 | 0.119261809 | -4.272337698 | 1.93E-05 | 0.000199648 | 0.509526724 |
| ENSMUSG00000027222 | Pex16 | 18633 | protein_coding | 327.2194497 | -0.50961397 | 0.088986661 | 5.72685798 | 1.02E-08 | 1.97E-07 | 0.50961397 |
| ENSMUSG00000044177 | Wfikkn2 | 278507 | protein_coding | 16.44897817 | -0.509765926 | 0.302035224 | 1.68776979 | 0.091455429 | 0.23142736 | 0.509765926 |
| ENSMUSG00000110622 | Iqcn | 637079 | protein_coding | 29.45513317 | 0.509780792 | 0.483616932 | -1.054100381 | 0.291836962 | 0.510653862 | 0.509780792 |
| ENSMUSG00000027712 | Anxa5 | 11747 | protein_coding | 541.2115075 | 0.509826575 | 0.103414999 | -4.929909418 | 8.23E-07 | 1.16E-05 | 0.509826575 |
| ENSMUSG00000044952 | Kctd21 | 622320 | protein_coding | 96.70780767 | -0.50988532 | 0.143809879 | 3.545551419 | 0.000391793 | 0.002879422 | 0.50988532 |
| ENSMUSG00000026018 | Ica1l | 70375 | protein_coding | 9.901296416 | -0.509951478 | 0.403007278 | 1.265365431 | 0.205740361 | 0.408364943 | 0.509951478 |
| ENSMUSG00000044072 | Eml6 | 237711 | protein_coding | 57.12016769 | 0.510011702 | 0.19903284 | -2.562450005 | 0.010393655 | 0.043909138 | 0.510011702 |
| ENSMUSG00000005442 | Cic | 71722 | protein_coding | 816.0206427 | 0.510088394 | 0.15499884 | -3.290917501 | 0.000998612 | 0.006350049 | 0.510088394 |
| ENSMUSG00000002500 | Rpl3l | 66211 | protein_coding | 2.520674722 | -0.510303875 | 1.067098308 | 0.478216366 | 0.632496215 | 1 | 0.510303875 |
| ENSMUSG00000039697 | Ncoa7 | 211329 | protein_coding | 599.7248279 | 0.510311412 | 0.242324052 | -2.105904913 | 0.035212613 | 0.114019821 | 0.510311412 |
| ENSMUSG00000000058 | Cav2 | 12390 | protein_coding | 78.72912934 | -0.510403562 | 0.258778563 | 1.972356428 | 0.048568932 | 0.145821162 | 0.510403562 |
| ENSMUSG00000019905 | Gprc6a | 210198 | protein_coding | 0.972439092 | -0.510424769 | 1.481036894 | 0.344640144 | 0.730364922 | 1 | 0.510424769 |
| ENSMUSG00000035031 | C8a | 230558 | protein_coding | 1.067705722 | 0.510427675 | 2.277989343 | -0.224069387 | 0.822703313 | 1 | 0.510427675 |
| ENSMUSG00000073739 | Gm16287 | 100038595 | lncRNA | 9.810216747 | -0.510568392 | 0.461927722 | 1.10529931 | 0.269029917 | 0.484112338 | 0.510568392 |
| ENSMUSG00000102737 | 1700016A09Rik | 74227 | TEC | 12.02173426 | 0.510669738 | 0.551842867 | -0.925389758 | 0.354763204 | 0.577368637 | 0.510669738 |
| ENSMUSG00000043998 | Mgat2 | 217664 | protein_coding | 674.5210581 | 0.51069531 | 0.085921732 | -5.943726921 | 2.79E-09 | 5.82E-08 | 0.51069531 |
| ENSMUSG00000024530 | Prelid3a | 225655 | protein_coding | 0.968465792 | -0.510726449 | 2.157971355 | 0.236669707 | 0.812913037 | 1 | 0.510726449 |
| ENSMUSG00000026281 | Dtymk | 21915 | protein_coding | 220.9684691 | 0.510810865 | 0.134481453 | -3.798374087 | 0.000145648 | 0.001199022 | 0.510810865 |
| ENSMUSG00000048574 | Ccnb1-ps | 434175 | processed_pseudogene | 1.46824283 | 0.510816437 | 0.902735756 | -0.565853777 | 0.571493186 | 1 | 0.510816437 |
| ENSMUSG00000032021 | Crtam | 54698 | protein_coding | 3.791085823 | 0.510897655 | 1.206687713 | -0.423388462 | 0.672011862 | 1 | 0.510897655 |
| ENSMUSG00000074527 | Gm14296 | 102639598 | protein_coding | 0.85474171 | 0.510921632 | 1.502026334 | -0.340154909 | 0.733739873 | 1 | 0.510921632 |
| ENSMUSG00000044375 | Pcare | 225004 | protein_coding | 1.171006337 | 0.511083409 | 2.293689768 | -0.222821506 | 0.823674429 | 1 | 0.511083409 |
| ENSMUSG00000091350 | Vmn2r92 | 627111 | protein_coding | 1.408730805 | 0.511093042 | 2.293965353 | -0.222798937 | 0.823691995 | 1 | 0.511093042 |
| ENSMUSG00000051379 | Flrt3 | 71436 | protein_coding | 637.4973351 | 0.511167322 | 0.113459674 | -4.505277553 | 6.63E-06 | 7.73E-05 | 0.511167322 |
| ENSMUSG00000031669 | Gins3 | 78833 | protein_coding | 38.94982231 | 0.511262708 | 0.24864147 | -2.056224606 | 0.039760872 | 0.125499051 | 0.511262708 |
| ENSMUSG00000085851 | 4921518K17Rik | 78758 | lncRNA | 3.483799646 | -0.511320371 | 0.670622289 | 0.762456571 | 0.445787554 | 1 | 0.511320371 |
| ENSMUSG00000066705 | Fxyd6 | 59095 | protein_coding | 9.245328294 | -0.511478235 | 0.371287122 | 1.377581404 | 0.168332566 | 0.356993365 | 0.511478235 |
| ENSMUSG00000056553 | Ptprn2 | 19276 | protein_coding | 501.9940044 | -0.511481213 | 0.197092697 | 2.595130218 | 0.009455509 | 0.040727249 | 0.511481213 |
| ENSMUSG00000102018 | Iqank1 | 432964 | protein_coding | 34.79404344 | -0.511484417 | 0.213907973 | 2.391142372 | 0.016796038 | 0.063995287 | 0.511484417 |
| ENSMUSG00000005705 | Agrp | 11604 | protein_coding | 2.963673552 | -0.511579138 | 0.894116536 | 0.572161589 | 0.567212507 | 1 | 0.511579138 |
| ENSMUSG00000021391 | Cenpp | 66336 | protein_coding | 41.55312148 | 0.511673202 | 0.198022118 | -2.583919444 | 0.009768461 | 0.041791803 | 0.511673202 |
| ENSMUSG00000068220 | Lgals1 | 16852 | protein_coding | 10.22465687 | 0.511848104 | 0.382126193 | -1.339474012 | 0.180416409 | 0.374186529 | 0.511848104 |

| ensembl_gene_id | external_gene_name | entrezgene_id | gene_biotype | baseMean | log2FoldChange | lfcSE | stat | pvalue | padj | absLog2FC |
|---|---|---|---|---|---|---|---|---|---|---|
| ENSMUSG00000000730 | Dnmt3l | 54427 | protein_coding | 2.192145368 | 0.511849205 | 2.312474698 | -0.221342618 | 0.824825668 | 1 | 0.511849205 |
| ENSMUSG00000053783 | 1700016K19Rik | 74230 | protein_coding | 15.43039778 | -0.511864115 | 0.269621189 | 1.898456563 | 0.057635965 | 0.165789988 | 0.511864115 |
| ENSMUSG00000065634 | Gm24252 | 115486082 | snoRNA | 0.525772766 | -0.511873304 | 1.75593078 | 0.291511095 | 0.77066046 | 1 | 0.511873304 |
| ENSMUSG00000037905 | Bri3bp | 76809 | protein_coding | 259.5550527 | -0.51215617 | 0.113077354 | 4.529254995 | 5.92E-06 | 6.95E-05 | 0.51215617 |
| ENSMUSG00000038271 | Iffo1 | 320678 | protein_coding | 18.84333219 | 0.512216825 | 0.610336047 | -0.83923738 | 0.401336115 | 0.622900809 | 0.512216825 |
| ENSMUSG00000042408 | Zmym6 | 100177 | protein_coding | 134.3919918 | -0.512217309 | 0.124842858 | 4.102896364 | 4.08E-05 | 0.000391737 | 0.512217309 |
| ENSMUSG00000022652 | Morc1 | 17450 | protein_coding | 4.660858369 | 0.512262703 | 0.857623213 | -0.597305081 | 0.550303713 | 0.741309692 | 0.512262703 |
| ENSMUSG00000039982 | Dtx4 | 207521 | protein_coding | 301.0553691 | 0.512270416 | 0.112715327 | -4.544815958 | 5.50E-06 | 6.52E-05 | 0.512270416 |
| ENSMUSG00000047146 | Tet1 | 52463 | protein_coding | 25.06290022 | -0.512342663 | 0.252652216 | 2.02785739 | 0.042574805 | 0.131848679 | 0.512342663 |
| ENSMUSG00000026429 | Ube2t | 67196 | protein_coding | 54.56170215 | 0.512374784 | 0.213295459 | -2.402183275 | 0.016297541 | 0.062415953 | 0.512374784 |
| ENSMUSG00000074247 | Dda1 | 66498 | protein_coding | 534.250737 | 0.512412498 | 0.071914799 | -7.125271915 | 1.04E-12 | 3.14E-11 | 0.512412498 |
| ENSMUSG00000097736 | 9530059O14Rik | 319626 | lncRNA | 1.793922131 | 0.512642239 | 2.332182355 | -0.219812245 | 0.826017383 | 1 | 0.512642239 |
| ENSMUSG00000024666 | Tmem138 | 72982 | protein_coding | 26.09673211 | 0.512665914 | 0.205892573 | -2.489967985 | 0.01277546 | 0.051482174 | 0.512665914 |
| ENSMUSG00000066097 | Cyp2j11 | 100066 | protein_coding | 1.054819085 | 0.512838238 | 2.337002811 | -0.219442713 | 0.8263052 | 1 | 0.512838238 |
| ENSMUSG00000057265 | Bbof1 | 72873 | protein_coding | 18.01130676 | 0.512903277 | 0.287786586 | -1.782234832 | 0.074710945 | 0.199909758 | 0.512903277 |
| ENSMUSG00000030276 | Ttll3 | 101100 | protein_coding | 92.94946798 | 0.51296032 | 0.306307924 | -1.674655727 | 0.094001799 | 0.236098206 | 0.51296032 |
| ENSMUSG00000097908 | 4933404O12Rik | 66752 | lncRNA | 16.94339911 | -0.512972965 | 0.296089557 | 1.732492599 | 0.083185894 | 0.216097506 | 0.512972965 |
| ENSMUSG00000025271 | Pfkfb1 | 18639 | protein_coding | 19.28043606 | 0.513141449 | 0.281328152 | -1.823996092 | 0.068152663 | 0.187297899 | 0.513141449 |
| ENSMUSG00000053460 | Ggcx | 56316 | protein_coding | 354.3743782 | 0.51314176 | 0.085021686 | -6.035422106 | 1.59E-09 | 3.41E-08 | 0.51314176 |
| ENSMUSG00000048279 | Sacs | 50720 | protein_coding | 13.2431464 | 0.513204728 | 0.515686545 | -0.995187354 | 0.319645151 | 0.54094184 | 0.513204728 |
| ENSMUSG00000035713 | Usp35 | 244144 | protein_coding | 26.21957457 | -0.513235693 | 0.247005204 | 2.077833519 | 0.037724695 | 0.120313948 | 0.513235693 |
| ENSMUSG00000086496 | Gm14204 | 100045469 | lncRNA | 1.808272886 | 0.513276255 | 2.348291028 | -0.21857438 | 0.826981612 | 1 | 0.513276255 |
| ENSMUSG00000020263 | Appl2 | 216190 | protein_coding | 1446.882347 | -0.513311183 | 0.103246846 | 4.971688726 | 6.64E-07 | 9.51E-06 | 0.513311183 |
| ENSMUSG00000017412 | Cacnb4 | 12298 | protein_coding | 5.117057194 | -0.513354527 | 0.690224816 | 0.74374974 | 0.457027891 | 0.671891623 | 0.513354527 |
| ENSMUSG00000113666 | Gm35725 | 102639399 | lncRNA | 1.775882028 | 0.513577127 | 2.356027562 | -0.217984346 | 0.827441308 | 1 | 0.513577127 |
| ENSMUSG00000072235 | Tuba1a | 22142 | protein_coding | 74.51632354 | -0.513972688 | 0.184495616 | 2.785826025 | 0.005339152 | 0.025735772 | 0.513972688 |
| ENSMUSG00000032036 | Kirrel3 | 67703 | protein_coding | 20.07396661 | -0.514034302 | 0.345985462 | 1.485710697 | 0.13735566 | 0.309791228 | 0.514034302 |
| ENSMUSG00000103622 | B430319G15Rik | 78575 | TEC | 2.008856583 | 0.514130511 | 1.230930149 | -0.41767643 | 0.676183707 | 1 | 0.514130511 |
| ENSMUSG00000020698 | Cct6b | 12467 | protein_coding | 5.854963438 | -0.514141832 | 0.507961091 | 1.012167744 | 0.311457856 | 0.532389508 | 0.514141832 |
| ENSMUSG00000087590 | Epb41l4aos | 69749 | lncRNA | 112.7907359 | -0.514164761 | 0.130672121 | 3.934770158 | 8.33E-05 | 0.000733845 | 0.514164761 |
| ENSMUSG00000085984 | 1700001G11Rik | 69303 | lncRNA | 0.891793862 | -0.514264336 | 1.25643509 | 0.409304341 | 0.68231633 | 1 | 0.514264336 |
| ENSMUSG00000047534 | Mis18bp1 | 217653 | protein_coding | 131.1768324 | 0.514441893 | 0.182054606 | -2.825755984 | 0.004716918 | 0.023162411 | 0.514441893 |
| ENSMUSG00000085975 | Gm13572 | 102636022 | lncRNA | 4.280342473 | 0.514543232 | 0.557196855 | -0.923449633 | 0.355772937 | 1 | 0.514543232 |
| ENSMUSG00000115386 | Gm35909 | 102639647 | lncRNA | 1.311541828 | 0.514569867 | 2.381987073 | -0.216025466 | 0.828967897 | 1 | 0.514569867 |
| ENSMUSG00000018900 | Slc22a5 | 20520 | protein_coding | 327.7862008 | -0.51461134 | 0.0937723 | 5.487882244 | 4.07E-08 | 7.09E-07 | 0.51461134 |
| ENSMUSG00000000037 | Scml2 | 107815 | protein_coding | 25.16642504 | 0.514633019 | 0.235953742 | -2.181075897 | 0.029177803 | 0.098758402 | 0.514633019 |
| ENSMUSG00000043885 | Slc36a4 | 234967 | protein_coding | 204.2228724 | 0.51475725 | 0.141031121 | -3.649955039 | 0.000262286 | 0.002016478 | 0.51475725 |
| ENSMUSG00000030131 | Mug2 | 17837 | protein_coding | 1.471539765 | 0.514766746 | 2.387275816 | -0.215629356 | 0.829276672 | 1 | 0.514766746 |
| ENSMUSG00000026670 | Uap1 | 107652 | protein_coding | 3320.072722 | 0.514886443 | 0.064022397 | -8.042286267 | 8.82E-16 | 3.47E-14 | 0.514886443 |

| ensembl_gene_id | external_gene_name | entrezgene_id | gene_biotype | baseMean | log2FoldChange | lfcSE | stat | pvalue | padj | absLog2FC |
|---|---|---|---|---|---|---|---|---|---|---|
| ENSMUSG00000065852 | Gm22353 | 115488556 | snoRNA | 0.468825176 | -0.514968673 | 1.607478376 | 0.320358072 | 0.748696906 | 1 | 0.514968673 |
| ENSMUSG00000065036 | Gm22362 | 115486329 | snoRNA | 9.795350042 | -0.515082102 | 0.399805359 | 1.288332161 | 0.197630362 | 0.397814213 | 0.515082102 |
| ENSMUSG00000024376 | Epb41l4a | 13824 | protein_coding | 116.8235645 | -0.515180725 | 0.173728965 | 2.965427926 | 0.003022622 | 0.015962767 | 0.515180725 |
| ENSMUSG00000001313 | Rnd2 | 11858 | protein_coding | 16.67319024 | -0.515281634 | 0.358786422 | 1.436179303 | 0.150951327 | 0.331006534 | 0.515281634 |
| ENSMUSG00000090171 | Ugt1a2 | 22236 | protein_coding | 9.341590375 | -0.515379353 | 0.519719499 | 0.99164906 | 0.321368748 | 0.54236929 | 0.515379353 |
| ENSMUSG00000053199 | Arhgap20 | 244867 | protein_coding | 367.0029925 | -0.515505259 | 0.117971501 | 4.369744017 | 1.24E-05 | 0.000136041 | 0.515505259 |
| ENSMUSG00000089736 | Tgfbr3l | 100044509 | protein_coding | 9.819729671 | -0.515704215 | 0.481985032 | 1.069958984 | 0.284637771 | 0.502997128 | 0.515704215 |
| ENSMUSG00000071103 | 1700029J07Rik | 69479 | protein_coding | 8.643542161 | -0.515784415 | 0.363148149 | 1.420314042 | 0.155516275 | 0.337876038 | 0.515784415 |
| ENSMUSG00000032652 | Crebl2 | 232430 | protein_coding | 214.0079459 | -0.51579826 | 0.093938974 | 5.490780202 | 4.00E-08 | 6.98E-07 | 0.51579826 |
| ENSMUSG00000027961 | Lrrc39 | 109245 | protein_coding | 5.051831921 | 0.515809416 | 0.53158442 | -0.970324557 | 0.331884741 | 0.554292124 | 0.515809416 |
| ENSMUSG00000045441 | Gprin3 | 243385 | protein_coding | 420.6592377 | -0.515851111 | 0.110025784 | 4.688456596 | 2.75E-06 | 3.46E-05 | 0.515851111 |
| ENSMUSG00000007050 | Lsm2 | 27756 | protein_coding | 116.8283073 | 0.516023764 | 0.151834575 | -3.398591949 | 0.000677337 | 0.004567391 | 0.516023764 |
| ENSMUSG00000024743 | Syt7 | 54525 | protein_coding | 575.1188144 | -0.51610166 | 0.127893656 | 4.035396877 | 5.45E-05 | 0.00050402 | 0.51610166 |
| ENSMUSG00000053841 | Txlna | 109658 | protein_coding | 1082.327675 | 0.5161024 | 0.06648207 | -7.763031434 | 8.29E-15 | 3.00E-13 | 0.5161024 |
| ENSMUSG00000005968 | Tuft1 | 22156 | protein_coding | 407.4177592 | -0.51642958 | 0.127282963 | 4.057334687 | 4.96E-05 | 0.000464149 | 0.51642958 |
| ENSMUSG00000096981 | Gm16845 | 100503652 | lncRNA | 8.17429894 | -0.516486036 | 0.353865634 | 1.459554099 | 0.144412662 | 0.321275859 | 0.516486036 |
| ENSMUSG00000071181 | 3830408C21Rik | 100040322 | lncRNA | 5.2796161 | 0.516543546 | 0.682924981 | -0.756369382 | 0.44942776 | 0.666174493 | 0.516543546 |
| ENSMUSG00000016487 | Ppfibp1 | 67533 | protein_coding | 343.475849 | 0.516766696 | 0.079832231 | -6.473158612 | 9.60E-11 | 2.40E-09 | 0.516766696 |
| ENSMUSG00000053101 | Gpr141 | 353346 | protein_coding | 1.330544162 | -0.516860945 | 1.748520988 | 0.295598937 | 0.767536386 | 1 | 0.516860945 |
| ENSMUSG00000032549 | Rab6b | 270192 | protein_coding | 13.65694364 | 0.516862324 | 0.371008237 | -1.39312897 | 0.163580794 | 0.349700969 | 0.516862324 |
| ENSMUSG00000028950 | Tas1r1 | 110326 | protein_coding | 6.580227007 | 0.517132158 | 0.534432864 | -0.967627916 | 0.33323023 | 0.555630454 | 0.517132158 |
| ENSMUSG00000019913 | Sim1 | 20464 | protein_coding | 0.946463493 | -0.517281878 | 2.278676742 | 0.227009768 | 0.820416147 | 1 | 0.517281878 |
| ENSMUSG00000042515 | Pwwp3b | 245631 | protein_coding | 6.336708305 | 0.51754662 | 0.445135184 | -1.162672911 | 0.244962236 | 0.456655966 | 0.51754662 |
| ENSMUSG00000089687 | Rab42 | 242681 | protein_coding | 4.691587371 | 0.517588778 | 0.639816991 | -0.80896379 | 0.418535975 | 0.6385535 | 0.517588778 |
| ENSMUSG00000096727 | Psmb9 | 16912 | protein_coding | 143.0375545 | -0.51761319 | 0.157925939 | 3.277569175 | 0.001047051 | 0.006612235 | 0.51761319 |
| ENSMUSG00000029005 | Draxin | 70433 | protein_coding | 1.879706988 | -0.517626611 | 1.87974542 | 0.275370593 | 0.783031534 | 1 | 0.517626611 |
| ENSMUSG00000019796 | Lrp11 | 237253 | protein_coding | 18.39600049 | -0.517819641 | 0.256925984 | 2.015442865 | 0.043858267 | 0.134899501 | 0.517819641 |
| ENSMUSG00000031214 | Ophn1 | 94190 | protein_coding | 243.8496035 | -0.517827964 | 0.093505318 | 5.53795202 | 3.06E-08 | 5.44E-07 | 0.517827964 |
| ENSMUSG00000053950 | Adnp2 | 240442 | protein_coding | 404.2030438 | 0.517890174 | 0.113000352 | -4.583084615 | 4.58E-06 | 5.53E-05 | 0.517890174 |
| ENSMUSG00000081234 | Or12e8 | 259031 | protein_coding | 1.647598506 | 0.518113827 | 2.704273395 | -0.191590772 | 0.848062772 | 1 | 0.518113827 |
| ENSMUSG00000031210 | Gpr165 | 76206 | protein_coding | 2.64432076 | 0.518417111 | 2.425820334 | -0.213707959 | 0.830774805 | 1 | 0.518417111 |
| ENSMUSG00000062518 | Zfp534 | 100503584 | protein_coding | 0.804521636 | -0.518563762 | 1.70765573 | 0.303669969 | 0.76137934 | 1 | 0.518563762 |
| ENSMUSG00000020014 | Cfap54 | 380654 | protein_coding | 6.596073363 | 0.518575689 | 1.616494021 | -0.320802726 | 0.748359893 | 0.873591524 | 0.518575689 |
| ENSMUSG00000044117 | Bmerb1 | 67254 | protein_coding | 1.217817684 | -0.518795814 | 1.443298099 | 0.359451602 | 0.719257277 | 1 | 0.518795814 |
| ENSMUSG00000090523 | Gypc | 71683 | protein_coding | 1.193252061 | -0.518871199 | 1.34987327 | 0.38438512 | 0.700693027 | 1 | 0.518871199 |
| ENSMUSG00000097990 | Gm19557 | 100503104 | lncRNA | 6.593066357 | -0.518939625 | 0.450875629 | 1.150959583 | 0.249748863 | 0.462014445 | 0.518939625 |
| ENSMUSG00000040253 | Gbp7 | 229900 | protein_coding | 71.96623213 | -0.519233951 | 0.268792945 | 1.931724627 | 0.053393503 | 0.156808369 | 0.519233951 |
| ENSMUSG00000028461 | Ccdc107 | 622404 | protein_coding | 288.4509246 | -0.519251682 | 0.089324923 | 5.813066085 | 6.13E-09 | 1.22E-07 | 0.519251682 |
| ENSMUSG00000029603 | Dtx1 | 14357 | protein_coding | 8.640209245 | 0.519269587 | 0.425839313 | -1.219402652 | 0.222691404 | 0.429717907 | 0.519269587 |

| ensembl_gene_id | external_gene_name | entrezgene_id | gene_biotype | baseMean | log2FoldChange | lfcSE | stat | pvalue | padj | absLog2FC |
|---|---|---|---|---|---|---|---|---|---|---|
| ENSMUSG00000024114 | Prss41 | 71003 | protein_coding | 7.674035785 | -0.519334748 | 0.395214783 | 1.314056989 | 0.188827027 | 0.38549753 | 0.519334748 |
| ENSMUSG00000018387 | Shroom1 | 71774 | protein_coding | 23.88982113 | 0.519348264 | 0.270529969 | -1.919743923 | 0.054890253 | 0.15977566 | 0.519348264 |
| ENSMUSG00000076435 | Acsf2 | 264895 | protein_coding | 1290.933426 | -0.519602816 | 0.154137717 | 3.371029658 | 0.000748878 | 0.004964661 | 0.519602816 |
| ENSMUSG00000039653 | Baat | 12012 | protein_coding | 11.09724891 | -0.519744587 | 0.371282363 | 1.399863392 | 0.16155423 | 0.346538122 | 0.519744587 |
| ENSMUSG00000035951 | Ascl3 | 56787 | protein_coding | 1.599933037 | -0.51986389 | 1.316421448 | 0.394906882 | 0.69291161 | 1 | 0.51986389 |
| ENSMUSG00000032557 | Uba5 | 66663 | protein_coding | 522.2286642 | 0.520026901 | 0.092327399 | -5.632422269 | 1.78E-08 | 3.29E-07 | 0.520026901 |
| ENSMUSG00000017405 | Nek8 | 140859 | protein_coding | 121.8248556 | -0.520046707 | 0.098018093 | 5.305619517 | 1.12E-07 | 1.84E-06 | 0.520046707 |
| ENSMUSG00000001604 | Tcea3 | 21401 | protein_coding | 508.4731912 | -0.520275318 | 0.071637308 | 7.262630744 | 3.80E-13 | 1.20E-11 | 0.520275318 |
| ENSMUSG00000063550 | Nup98 | 269966 | protein_coding | 1351.289708 | 0.520492194 | 0.05605013 | -9.286190703 | 1.60E-20 | 8.33E-19 | 0.520492194 |
| ENSMUSG00000037660 | Gdf7 | 238057 | protein_coding | 3.783470322 | 0.520553101 | 0.5900337 | -0.882243 | 0.377645413 | 1 | 0.520553101 |
| ENSMUSG00000029247 | Paics | 67054 | protein_coding | 793.8303592 | -0.520580928 | 0.176798001 | 2.944495561 | 0.003234816 | 0.016937143 | 0.520580928 |
| ENSMUSG00000032327 | Stra6 | 20897 | protein_coding | 4.350551065 | -0.520630548 | 0.966775573 | 0.538522655 | 0.590216269 | 1 | 0.520630548 |
| ENSMUSG00000042708 | Shcbp1l | 71836 | protein_coding | 2.587959184 | 0.520844859 | 1.022807435 | -0.509230615 | 0.610590586 | 1 | 0.520844859 |
| ENSMUSG00000036966 | Spryd3 | 223918 | protein_coding | 545.3156925 | 0.520896648 | 0.064450771 | -8.082085591 | 6.37E-16 | 2.53E-14 | 0.520896648 |
| ENSMUSG00000022123 | Scel | 64929 | protein_coding | 32.72433805 | -0.520963941 | 0.288246055 | 1.80735844 | 0.070706409 | 0.192395064 | 0.520963941 |
| ENSMUSG00000098470 | C1rb | 667277 | protein_coding | 1.736324631 | 0.521055642 | 2.572982882 | -0.202510342 | 0.839517775 | 1 | 0.521055642 |
| ENSMUSG00000004366 | Sst | 20604 | protein_coding | 52.27491203 | -0.521098357 | 0.178807717 | 2.91429456 | 0.003564934 | 0.018374318 | 0.521098357 |
| ENSMUSG00000046192 | Iqub | 214704 | protein_coding | 1.26926997 | -0.521117247 | 1.143891988 | 0.455565081 | 0.648702758 | 1 | 0.521117247 |
| ENSMUSG00000001891 | Ugp2 | 216558 | protein_coding | 2776.149567 | 0.521121239 | 0.079828098 | -6.528042791 | 6.66E-11 | 1.72E-09 | 0.521121239 |
| ENSMUSG00000061104 | Sap18b | 100041953 | protein_coding | 7.884524017 | 0.521239451 | 0.346686729 | -1.503488332 | 0.132713164 | 0.30233483 | 0.521239451 |
| ENSMUSG00000059005 | Hnrnpa3 | 229279 | protein_coding | 1529.29859 | 0.521270281 | 0.110798493 | -4.704669428 | 2.54E-06 | 3.23E-05 | 0.521270281 |
| ENSMUSG00000032123 | Dpagt1 | 13478 | protein_coding | 399.9553705 | 0.521307119 | 0.073948005 | -7.049644165 | 1.79E-12 | 5.32E-11 | 0.521307119 |
| ENSMUSG00000034413 | Neurl1b | 240055 | protein_coding | 836.2344903 | -0.521324639 | 0.097433954 | 5.350543818 | 8.77E-08 | 1.46E-06 | 0.521324639 |
| ENSMUSG00000029034 | Ints11 | 71957 | protein_coding | 387.5706442 | 0.521367499 | 0.080554027 | -6.472271069 | 9.65E-11 | 2.41E-09 | 0.521367499 |
| ENSMUSG00000046432 | Bex3 | 12070 | protein_coding | 216.2672744 | 0.521588167 | 0.150138095 | -3.474056122 | 0.000512654 | 0.003624129 | 0.521588167 |
| ENSMUSG00000039496 | Cdnf | 227526 | protein_coding | 15.61664035 | 0.521651741 | 0.282388937 | -1.84728108 | 0.064706414 | 0.18053173 | 0.521651741 |
| ENSMUSG00000059854 | Hydin | 244653 | protein_coding | 12.42055934 | 0.521677697 | 0.919956381 | -0.567067861 | 0.570668077 | 0.756249943 | 0.521677697 |
| ENSMUSG00000031953 | Tmem170 | 66817 | protein_coding | 428.7236117 | 0.521822596 | 0.087670296 | -5.952102611 | 2.65E-09 | 5.55E-08 | 0.521822596 |
| ENSMUSG00000069581 | Tspear | 252974 | protein_coding | 5.722300621 | -0.521827552 | 0.456729627 | 1.14253055 | 0.253233555 | 0.465690304 | 0.521827552 |
| ENSMUSG00000040158 | Tax1bp3 | 76281 | protein_coding | 556.5618103 | -0.521853896 | 0.120330735 | 4.336829623 | 1.45E-05 | 0.000154732 | 0.521853896 |
| ENSMUSG00000027520 | Zdbf2 | 73884 | protein_coding | 1.915585744 | 0.521945372 | 2.365757108 | -0.220625089 | 0.825384364 | 1 | 0.521945372 |
| ENSMUSG00000028174 | Rpe65 | 19892 | protein_coding | 1.254067945 | 0.522237567 | 2.612194443 | -0.19992293 | 0.841540857 | 1 | 0.522237567 |
| ENSMUSG00000090919 | Pabpc4l | 241989 | protein_coding | 1.382757918 | 0.522248131 | 2.612591336 | -0.199896602 | 0.841561448 | 1 | 0.522248131 |
| ENSMUSG00000058620 | Adra2b | 11552 | protein_coding | 1.069155956 | 0.52228199 | 2.613636431 | -0.199829626 | 0.841613831 | 1 | 0.52228199 |
| ENSMUSG00000073961 | Or52r1b | 259055 | protein_coding | 1.298704097 | 0.522284398 | 2.613799216 | -0.199818102 | 0.841622844 | 1 | 0.522284398 |
| ENSMUSG00000001260 | Gabrg1 | 14405 | protein_coding | 1.780750352 | 0.522289866 | 2.614084375 | -0.199798396 | 0.841638256 | 1 | 0.522289866 |
| ENSMUSG00000032051 | Fdx1 | 14148 | protein_coding | 448.1968523 | 0.522340887 | 0.067513686 | -7.73681482 | 1.02E-14 | 3.64E-13 | 0.522340887 |
| ENSMUSG00000024253 | Dync2li1 | 213575 | protein_coding | 17.00117746 | 0.522341618 | 0.278842217 | -1.873251559 | 0.061033656 | 0.173002023 | 0.522341618 |
| ENSMUSG00000040632 | Nrl | 18185 | protein_coding | 3.712380319 | 0.522351535 | 2.616411617 | -0.19964425 | 0.841758818 | 1 | 0.522351535 |

Supplementary Table 1

| ensembl_gene_id | external_gene_name | entrezgene_id | gene_biotype | baseMean | log2FoldChange | lfcSE | stat | pvalue | padj | absLog2FC |
|---|---|---|---|---|---|---|---|---|---|---|
| ENSMUSG00000036557 | Stpg4 | 75467 | protein_coding | 1.26586186 | -0.522400073 | 1.257307445 | 0.415491115 | 0.677782418 | 1 | 0.522400073 |
| ENSMUSG00000074359 | Psg23 | 56868 | protein_coding | 0.996861088 | 0.52245806 | 2.619583783 | -0.199443157 | 0.841916104 | 1 | 0.52245806 |
| ENSMUSG00000059246 | Foxb1 | 64290 | protein_coding | 1.178848617 | 0.522701262 | 2.628024402 | -0.198895133 | 0.842344777 | 1 | 0.522701262 |
| ENSMUSG00000042804 | Gpr153 | 100129 | protein_coding | 5.793462483 | 0.522786078 | 0.724880469 | -0.721203151 | 0.470784533 | 0.683906563 | 0.522786078 |
| ENSMUSG00000035095 | Fam167a | 219148 | protein_coding | 14.82319738 | -0.522835259 | 0.47249258 | 1.106547024 | 0.268489823 | 0.483542311 | 0.522835259 |
| ENSMUSG00000020303 | Stc2 | 20856 | protein_coding | 315.3387758 | 0.522853157 | 0.131110001 | -3.987896815 | 6.67E-05 | 0.000602545 | 0.522853157 |
| ENSMUSG00000091624 | Vmn2r9 | 435864 | protein_coding | 1.883199052 | 0.522883946 | 2.634539305 | -0.19847263 | 0.842675297 | 1 | 0.522883946 |
| ENSMUSG00000043461 | Sptssb | 66183 | protein_coding | 1079.122323 | -0.522907799 | 0.206713361 | 2.529627488 | 0.011418368 | 0.047353826 | 0.522907799 |
| ENSMUSG00000115699 | Gm6961 | 629300 | lncRNA | 1.055683565 | 0.522911611 | 2.635268847 | -0.198428184 | 0.842710068 | 1 | 0.522911611 |
| ENSMUSG00000026924 | Sec16a | 227648 | protein_coding | 2558.789121 | 0.522916488 | 0.068366296 | -7.648746855 | 2.03E-14 | 7.06E-13 | 0.522916488 |
| ENSMUSG00000023009 | Nckap5l | 380969 | protein_coding | 5.882912622 | 0.522960601 | 0.46981871 | -1.113111483 | 0.265660553 | 0.480674742 | 0.522960601 |
| ENSMUSG00000060860 | Ube2s | 77891 | protein_coding | 394.5601662 | 0.52299207 | 0.158670652 | -3.296085722 | 0.000980421 | 0.006260894 | 0.52299207 |
| ENSMUSG00000056899 | Immp2l | 93757 | protein_coding | 57.99159141 | -0.523197476 | 0.148320435 | 3.527480734 | 0.000419534 | 0.003057453 | 0.523197476 |
| ENSMUSG00000020096 | Tbata | 65971 | protein_coding | 1.87112687 | 0.523248649 | 2.647286425 | -0.197654717 | 0.843315221 | 1 | 0.523248649 |
| ENSMUSG00000075046 | Duxf3 | 74399 | protein_coding | 2.291658323 | 0.523251474 | 2.647454607 | -0.197643228 | 0.843324211 | 1 | 0.523251474 |
| ENSMUSG00000070983 | Or13d1 | 258600 | protein_coding | 1.116716568 | 0.523284755 | 2.648366116 | -0.19758777 | 0.843367604 | 1 | 0.523284755 |
| ENSMUSG00000033774 | Npbwr1 | 226304 | protein_coding | 1.689139342 | 0.523295795 | 2.648909098 | -0.197551436 | 0.843396035 | 1 | 0.523295795 |
| ENSMUSG00000067144 | Slc22a7 | 108114 | protein_coding | 1.290660921 | 0.52331583 | 2.649515788 | -0.197513762 | 0.843425513 | 1 | 0.52331583 |
| ENSMUSG00000024365 | Cyp21a1 | 13079 | protein_coding | 1.139790159 | 0.52340196 | 2.652489123 | -0.197324828 | 0.843573351 | 1 | 0.52340196 |
| ENSMUSG00000059366 | Or10d1 | 258501 | protein_coding | 1.241798554 | 0.523403065 | 2.652580406 | -0.197318454 | 0.843578339 | 1 | 0.523403065 |
| ENSMUSG00000117863 | Vmn1r61 | 636731 | protein_coding | 1.551325829 | 0.523413292 | 2.65302805 | -0.197289016 | 0.843601375 | 1 | 0.523413292 |
| ENSMUSG00000053093 | Myh7 | 140781 | protein_coding | 5.715906189 | 0.523426532 | 1.155181526 | -0.453111931 | 0.650468145 | 0.810140405 | 0.523426532 |
| ENSMUSG00000030148 | Clec4a2 | 26888 | protein_coding | 1.277603672 | 0.52345303 | 2.654351397 | -0.197205626 | 0.843666628 | 1 | 0.52345303 |
| ENSMUSG00000038372 | Gmds | 218138 | protein_coding | 4057.675968 | 0.523495274 | 0.092174144 | -5.67941566 | 1.35E-08 | 2.55E-07 | 0.523495274 |
| ENSMUSG00000022442 | Ttll1 | 319953 | protein_coding | 43.01323659 | -0.523509333 | 0.175543843 | 2.982214149 | 0.002861717 | 0.015303887 | 0.523509333 |
| ENSMUSG00000050587 | Lrrc4c | 241568 | protein_coding | 1.321036824 | 0.523570038 | 2.658525909 | -0.19693998 | 0.843874507 | 1 | 0.523570038 |
| ENSMUSG00000078651 | Aoc2 | 237940 | protein_coding | 22.13929508 | 0.523586482 | 0.269965539 | -1.939456735 | 0.052445748 | 0.154769314 | 0.523586482 |
| ENSMUSG00000031212 | Pgr15l | 245526 | protein_coding | 1.325164285 | 0.523792341 | 2.666469626 | -0.196436643 | 0.844268418 | 1 | 0.523792341 |
| ENSMUSG00000017943 | Gdap1l1 | 228858 | protein_coding | 2.971097214 | -0.523842901 | 0.800563922 | 0.654342379 | 0.512891249 | 1 | 0.523842901 |
| ENSMUSG00000029454 | Mapkapk5 | 17165 | protein_coding | 34.27272828 | 0.524091661 | 0.238304505 | -2.199252009 | 0.027860008 | 0.095110269 | 0.524091661 |
| ENSMUSG00000058046 | 4933430I17Rik | 214106 | protein_coding | 6.416298174 | 0.524252336 | 0.460614332 | -1.138158974 | 0.255054106 | 0.467632232 | 0.524252336 |
| ENSMUSG00000026989 | Dapl1 | 76747 | protein_coding | 0.858948479 | -0.524279594 | 2.164809197 | 0.242182819 | 0.808638507 | 1 | 0.524279594 |
| ENSMUSG00000117334 | A730006G06Rik | 320638 | lncRNA | 1.186486291 | 0.524287384 | 2.684341596 | -0.195313214 | 0.845147753 | 1 | 0.524287384 |
| ENSMUSG00000085148 | Mir22hg | 100042498 | lncRNA | 205.334199 | -0.524665042 | 0.131574386 | 3.987592561 | 6.67E-05 | 0.000602988 | 0.524665042 |
| ENSMUSG00000004633 | Chn2 | 69993 | protein_coding | 259.7737334 | -0.524709094 | 0.148105053 | 3.542816969 | 0.000395877 | 0.002906856 | 0.524709094 |
| ENSMUSG00000020793 | Galr2 | 14428 | protein_coding | 7.489831201 | 0.524761597 | 0.442097331 | -1.186982052 | 0.2352347 | 0.445131871 | 0.524761597 |
| ENSMUSG00000003872 | Lin7b | 22342 | protein_coding | 2.688329188 | 0.524773528 | 0.68143926 | -0.770095823 | 0.441243054 | 1 | 0.524773528 |
| ENSMUSG00000050350 | Gpr18 | 110168 | protein_coding | 10.51317967 | -0.524943897 | 0.408898565 | 1.283799803 | 0.199212013 | 0.40002276 | 0.524943897 |
| ENSMUSG00000055044 | Pdlim1 | 54132 | protein_coding | 2502.535434 | -0.52498427 | 0.071370044 | 7.355806964 | 1.90E-13 | 6.16E-12 | 0.52498427 |

| ensembl_gene_id | external_gene_name | entrezgene_id | gene_biotype | baseMean | log2FoldChange | lfcSE | stat | pvalue | padj | absLog2FC |
|---|---|---|---|---|---|---|---|---|---|---|
| ENSMUSG00000052485 | Tmem171 | 380863 | protein_coding | 1227.585639 | -0.525082814 | 0.105956332 | 4.955652983 | 7.21E-07 | 1.03E-05 | 0.525082814 |
| ENSMUSG00000028116 | Myoz2 | 59006 | protein_coding | 2.109090717 | -0.525161794 | 1.015426779 | 0.517183322 | 0.605028194 | 1 | 0.525161794 |
| ENSMUSG00000097709 | 2810429I04Rik | 76937 | lncRNA | 7.433537569 | 0.525206505 | 0.674911854 | -0.778185332 | 0.436459763 | 0.654709467 | 0.525206505 |
| ENSMUSG00000097039 | Pvt1 | 19296 | lncRNA | 16.76289498 | 0.525929299 | 0.303498477 | -1.732889419 | 0.083115325 | 0.215982137 | 0.525929299 |
| ENSMUSG00000045802 | Hsf3 | 245525 | protein_coding | 16.76327919 | 0.525978534 | 0.479319481 | -1.09734437 | 0.272490882 | 0.488337435 | 0.525978534 |
| ENSMUSG00000045589 | Frrs1l | 230235 | protein_coding | 2.669089585 | -0.526001228 | 0.972687915 | 0.540770806 | 0.588665568 | 1 | 0.526001228 |
| ENSMUSG00000044957 | Pp2d1 | 110332 | protein_coding | 3.34565897 | 0.526002022 | 0.717911953 | -0.732683193 | 0.463751677 | 1 | 0.526002022 |
| ENSMUSG00000037235 | Mxd4 | 17122 | protein_coding | 1578.967971 | 0.526087807 | 0.08707494 | -6.041781994 | 1.52E-09 | 3.29E-08 | 0.526087807 |
| ENSMUSG00000024660 | Incenp | 16319 | protein_coding | 433.8553312 | 0.526097888 | 0.141096382 | -3.728641955 | 0.000192514 | 0.001528454 | 0.526097888 |
| ENSMUSG00000029283 | Cdc7 | 12545 | protein_coding | 69.91657441 | 0.526131385 | 0.182152285 | -2.888414964 | 0.003871887 | 0.019692129 | 0.526131385 |
| ENSMUSG00000033409 | Syce1l | 668110 | protein_coding | 1.556443344 | 0.526192666 | 1.215924493 | -0.432751104 | 0.665195599 | 1 | 0.526192666 |
| ENSMUSG00000032377 | Plscr4 | 235527 | protein_coding | 21.55035763 | -0.526208512 | 0.321586752 | 1.636287903 | 0.101779347 | 0.250788441 | 0.526208512 |
| ENSMUSG00000075389 | Reno1 | 100042332 | lncRNA | 24.68024499 | -0.526297273 | 0.220902816 | 2.382483314 | 0.017196311 | 0.065177848 | 0.526297273 |
| ENSMUSG00000073598 | Smim33 | 73449 | protein_coding | 281.1122334 | 0.526494829 | 0.181138481 | -2.906587415 | 0.003653948 | 0.018770347 | 0.526494829 |
| ENSMUSG00000052273 | Dnah3 | 381917 | protein_coding | 5.635976471 | -0.526749853 | 1.680440936 | 0.313459308 | 0.753931732 | 0.876790748 | 0.526749853 |
| ENSMUSG00000036745 | Ttll7 | 70892 | protein_coding | 116.3306082 | 0.526987429 | 0.139456442 | -3.778867604 | 0.000157543 | 0.001280348 | 0.526987429 |
| ENSMUSG00000066800 | Rnasel | 24014 | protein_coding | 4244.73001 | -0.52715815 | 0.117958546 | 4.469011915 | 7.86E-06 | 9.04E-05 | 0.52715815 |
| ENSMUSG00000041372 | B4galnt3 | 330406 | protein_coding | 678.3087274 | -0.5271946 | 0.189836347 | 2.777100422 | 0.005484623 | 0.026283536 | 0.5271946 |
| ENSMUSG00000020198 | Ap3d1 | 11776 | protein_coding | 2302.872104 | 0.527215526 | 0.052578134 | -10.02727725 | 1.16E-23 | 7.02E-22 | 0.527215526 |
| ENSMUSG00000039716 | Dock3 | 208869 | protein_coding | 13.5469117 | 0.527468382 | 0.552449744 | -0.954780751 | 0.339688584 | 0.561130066 | 0.527468382 |
| ENSMUSG00000022615 | Tymp | 72962 | protein_coding | 38.95195373 | -0.527562445 | 0.20543422 | 2.56803587 | 0.010227657 | 0.043340911 | 0.527562445 |
| ENSMUSG00000031629 | Cenpu | 71876 | protein_coding | 50.0315795 | 0.528008918 | 0.211411422 | -2.497542058 | 0.012505763 | 0.050667362 | 0.528008918 |
| ENSMUSG00000070462 | Tlnrd1 | 80889 | protein_coding | 1165.577723 | 0.528025642 | 0.078841084 | -6.697341226 | 2.12E-11 | 5.74E-10 | 0.528025642 |
| ENSMUSG00000020169 | Best3 | 382427 | protein_coding | 1.582097142 | -0.528102515 | 1.31330371 | 0.402117584 | 0.687597492 | 1 | 0.528102515 |
| ENSMUSG00000053293 | Pom121 | 107939 | protein_coding | 217.4586468 | 0.528411049 | 0.130327038 | -4.054500576 | 5.02E-05 | 0.000468751 | 0.528411049 |
| ENSMUSG00000110151 | Gm38416 | 330850 | lncRNA | 33.4522553 | 0.528455548 | 0.290769383 | -1.817438764 | 0.069149957 | 0.189251307 | 0.528455548 |
| ENSMUSG00000066090 | Insl5 | 23919 | protein_coding | 240.7028363 | -0.52854416 | 0.274483614 | 1.92559458 | 0.054155016 | 0.158286006 | 0.52854416 |
| ENSMUSG00000071551 | Akr1c19 | 432720 | protein_coding | 2592.620475 | -0.528600033 | 0.120000866 | 4.404968469 | 1.06E-05 | 0.000117654 | 0.528600033 |
| ENSMUSG00000066643 | Wdr35 | 74682 | protein_coding | 32.69709805 | 0.528624841 | 0.197276484 | -2.679614062 | 0.007370709 | 0.033256799 | 0.528624841 |
| ENSMUSG00000053175 | Bcl3 | 12051 | protein_coding | 373.8011499 | 0.52882196 | 0.162487095 | -3.254547438 | 0.001135732 | 0.007074869 | 0.52882196 |
| ENSMUSG00000069094 | Pde7a | 18583 | protein_coding | 765.085364 | -0.528879216 | 0.17858197 | 2.961548791 | 0.00306096 | 0.016154903 | 0.528879216 |
| ENSMUSG00000086404 | Gm6787 | 627782 | lncRNA | 2.649810308 | 0.528975272 | 0.868106388 | -0.60934383 | 0.54229656 | 1 | 0.528975272 |
| ENSMUSG00000055188 | Rbm3os | 70226 | lncRNA | 12.15584071 | -0.528980787 | 0.337386584 | 1.567877361 | 0.116909756 | 0.276478264 | 0.528980787 |
| ENSMUSG00000054723 | Vmac | 106639 | protein_coding | 263.0757089 | 0.529192328 | 0.109867649 | -4.816634679 | 1.46E-06 | 1.95E-05 | 0.529192328 |
| ENSMUSG00000075112 | Or4c15 | 258025 | protein_coding | 1.855822837 | 0.529241695 | 2.397627983 | -0.220735535 | 0.825298361 | 1 | 0.529241695 |
| ENSMUSG00000100572 | 2410021H03Rik | 69750 | lncRNA | 2.734459572 | -0.529249725 | 0.76077021 | 0.695676196 | 0.486631636 | 1 | 0.529249725 |
| ENSMUSG00000073242 | Dnmt3aos | 100038522 | lncRNA | 2.461950367 | -0.529259691 | 0.812430465 | 0.651452296 | 0.514754562 | 1 | 0.529259691 |
| ENSMUSG00000064043 | Trerf1 | 224829 | protein_coding | 8.539017445 | 0.529344127 | 0.867666322 | -0.610077992 | 0.541810145 | 0.735609498 | 0.529344127 |
| ENSMUSG00000114212 | Gm47985 | 118567800 | protein_coding | 1.165741945 | -0.529372561 | 1.328921309 | 0.39834756 | 0.690374007 | 1 | 0.529372561 |

| ensembl_gene_id | external_gene_name | entrezgene_id | gene_biotype | baseMean | log2FoldChange | lfcSE | stat | pvalue | padj | absLog2FC |
|---|---|---|---|---|---|---|---|---|---|---|
| ENSMUSG00000027104 | Atf2 | 11909 | protein_coding | 665.2724847 | 0.529709807 | 0.07851867 | -6.746291106 | 1.52E-11 | 4.15E-10 | 0.529709807 |
| ENSMUSG00000021604 | Irx4 | 50916 | protein_coding | 22.27319942 | 0.529853126 | 0.354528638 | -1.494528423 | 0.135037559 | 0.306025833 | 0.529853126 |
| ENSMUSG00000045521 | Tssk2 | 22115 | protein_coding | 1.593402834 | -0.529883082 | 0.835976974 | 0.633848896 | 0.526179439 | 1 | 0.529883082 |
| ENSMUSG00000036391 | Sec24a | 77371 | protein_coding | 691.4178169 | 0.529931236 | 0.131469167 | -4.030840457 | 5.56E-05 | 0.000512457 | 0.529931236 |
| ENSMUSG00000086782 | E130102H24Rik | 77866 | lncRNA | 12.47199838 | -0.530033995 | 0.435905147 | 1.215938831 | 0.224008225 | 0.431111794 | 0.530033995 |
| ENSMUSG00000063535 | Zfp773 | 76373 | protein_coding | 13.60061688 | -0.530055361 | 0.311400852 | 1.702164135 | 0.088724604 | 0.226286043 | 0.530055361 |
| ENSMUSG00000072694 | 1500011B03Rik | 66236 | protein_coding | 66.4830767 | 0.530106007 | 0.150914561 | -3.51262333 | 0.000443706 | 0.003211398 | 0.530106007 |
| ENSMUSG00000026107 | Nabp1 | 109019 | protein_coding | 239.4180103 | 0.530220168 | 0.178747815 | -2.966302939 | 0.003014034 | 0.015927605 | 0.530220168 |
| ENSMUSG00000023885 | Thbs2 | 21826 | protein_coding | 1.446538311 | 0.530563254 | 1.693643906 | -0.3132673 | 0.754077591 | 1 | 0.530563254 |
| ENSMUSG00000029923 | Rab19 | 19331 | protein_coding | 119.6142982 | -0.530564488 | 0.131004046 | 4.049985504 | 5.12E-05 | 0.000476541 | 0.530564488 |
| ENSMUSG00000027273 | Snap25 | 20614 | protein_coding | 19.59283174 | 0.530573616 | 0.312442789 | -1.698146458 | 0.089480123 | 0.227755048 | 0.530573616 |
| ENSMUSG00000006958 | Chrd | 12667 | protein_coding | 22.65079349 | 0.530584817 | 0.332536064 | -1.595570747 | 0.110584663 | 0.265929452 | 0.530584817 |
| ENSMUSG00000039672 | Kcne2 | 246133 | protein_coding | 4.94021569 | -0.530592737 | 0.495846838 | 1.070073853 | 0.284586067 | 0.50295958 | 0.530592737 |
| ENSMUSG00000019761 | Krt10 | 16661 | protein_coding | 18.56513741 | -0.530771881 | 0.243141375 | 2.182976388 | 0.029037549 | 0.098384506 | 0.530771881 |
| ENSMUSG00000025169 | Ogfod3 | 66179 | protein_coding | 103.391647 | -0.530935979 | 0.12216198 | 4.346163834 | 1.39E-05 | 0.000149142 | 0.530935979 |
| ENSMUSG00000019923 | Zwint | 52696 | protein_coding | 386.914665 | 0.53111505 | 0.076462237 | -6.946109265 | 3.75E-12 | 1.09E-10 | 0.53111505 |
| ENSMUSG00000030030 | 1700003E16Rik | 71837 | protein_coding | 41.18127142 | -0.531151576 | 0.194673824 | 2.728418053 | 0.00636389 | 0.029421411 | 0.531151576 |
| ENSMUSG00000078853 | Igtp | 16145 | protein_coding | 106.2849146 | 0.531183499 | 0.310872997 | -1.708683302 | 0.087509629 | 0.224469402 | 0.531183499 |
| ENSMUSG00000020037 | Rfx4 | 71137 | protein_coding | 2.552077194 | 0.531197576 | 1.701336491 | -0.312223701 | 0.754870521 | 1 | 0.531197576 |
| ENSMUSG00000064918 | Gm22571 | 115487260 | snoRNA | 2.697983584 | -0.531382501 | 0.824589039 | 0.64442101 | 0.519302464 | 1 | 0.531382501 |
| ENSMUSG00000047492 | Inhbe | 16326 | protein_coding | 2.306730932 | 0.531428888 | 1.676473356 | -0.316992147 | 0.751249561 | 1 | 0.531428888 |
| ENSMUSG00000069085 | Dytn | 241073 | protein_coding | 2.616451305 | 0.531585655 | 2.993824229 | -0.177560743 | 0.859067957 | 1 | 0.531585655 |
| ENSMUSG00000050511 | Oprd1 | 18386 | protein_coding | 16.01861015 | 0.531627519 | 0.344149228 | -1.544758713 | 0.122404637 | 0.285476835 | 0.531627519 |
| ENSMUSG00000006269 | Atp6v1b1 | 110935 | protein_coding | 1.489515604 | -0.53166271 | 2.360060704 | 0.225275015 | 0.821765335 | 1 | 0.53166271 |
| ENSMUSG00000040016 | Ptger3 | 19218 | protein_coding | 58.18208393 | -0.531663768 | 0.260967654 | 2.037278412 | 0.04162215 | 0.129774424 | 0.531663768 |
| ENSMUSG00000037712 | Fermt2 | 218952 | protein_coding | 2.731988644 | 0.53175057 | 1.146577225 | -0.463772138 | 0.642811007 | 1 | 0.53175057 |
| ENSMUSG00000024791 | Cdca5 | 67849 | protein_coding | 100.4654962 | 0.531836714 | 0.239267833 | -2.222767296 | 0.026231495 | 0.090681791 | 0.531836714 |
| ENSMUSG00000060147 | Serpinb6a | 20719 | protein_coding | 4266.271499 | -0.531956275 | 0.091479833 | 5.815011422 | 6.06E-09 | 1.21E-07 | 0.531956275 |
| ENSMUSG00000059460 | Or10ad1b | 629524 | protein_coding | 1.923679423 | 0.53219246 | 3.02392225 | -0.175994095 | 0.860298582 | 1 | 0.53219246 |
| ENSMUSG00000094910 | D430019H16Rik | 268595 | protein_coding | 1.971240035 | 0.532209608 | 3.024806138 | -0.175948336 | 0.860334531 | 1 | 0.532209608 |
| ENSMUSG00000034457 | Eda2r | 245527 | protein_coding | 24.20326561 | 0.532216894 | 0.450794561 | -1.1806196 | 0.237753874 | 0.447999484 | 0.532216894 |
| ENSMUSG00000020524 | Gria1 | 14799 | protein_coding | 3.479732554 | 0.532280245 | 3.42857188 | -0.155248384 | 0.876625505 | 1 | 0.532280245 |
| ENSMUSG00000056328 | Myh1 | 17879 | protein_coding | 1.694131282 | 0.532488595 | 3.038922046 | -0.175222854 | 0.860904527 | 1 | 0.532488595 |
| ENSMUSG00000066262 | Or51i1 | 258819 | protein_coding | 1.694131282 | 0.532488595 | 3.038922046 | -0.175222854 | 0.860904527 | 1 | 0.532488595 |
| ENSMUSG00000029059 | Prxl2b | 66469 | protein_coding | 97.09202127 | -0.53260639 | 0.142978471 | 3.725081025 | 0.000195252 | 0.00154796 | 0.53260639 |
| ENSMUSG00000111079 | Gm47112 | 115487203 | lncRNA | 4.668719157 | 0.532637789 | 0.6115671 | -0.870939246 | 0.383787326 | 0.606204104 | 0.532637789 |
| ENSMUSG00000029119 | Man2b2 | 17160 | protein_coding | 547.5665742 | -0.532700995 | 0.089819093 | 5.930821377 | 3.01E-09 | 6.28E-08 | 0.532700995 |
| ENSMUSG00000067768 | Xlr4b | 27083 | protein_coding | 1.788347881 | -0.532750844 | 1.32067933 | 0.403391521 | 0.686660223 | 1 | 0.532750844 |
| ENSMUSG00000091734 | Vmn1r29 | 113859 | protein_coding | 1.65069813 | 0.532759063 | 3.052864501 | -0.174511205 | 0.861463726 | 1 | 0.532759063 |

| ensembl_gene_id | external_gene_name | entrezgene_id | gene_biotype | baseMean | log2FoldChange | lfcSE | stat | pvalue | padj | absLog2FC |
|---|---|---|---|---|---|---|---|---|---|---|
| ENSMUSG00000045176 | Borcs6 | 71923 | protein_coding | 182.3381629 | 0.532798775 | 0.101596458 | -5.244265276 | 1.57E-07 | 2.50E-06 | 0.532798775 |
| ENSMUSG00000030587 | 2200002D01Rik | 72275 | protein_coding | 3228.790195 | -0.532823754 | 0.181188519 | 2.940714767 | 0.003274559 | 0.01711111 | 0.532823754 |
| ENSMUSG00000030092 | Cntn6 | 53870 | protein_coding | 1.41702253 | 0.532854418 | 3.057711595 | -0.174265754 | 0.861656611 | 1 | 0.532854418 |
| ENSMUSG00000045608 | Dbx2 | 223843 | protein_coding | 1.41702253 | 0.532854418 | 3.057711595 | -0.174265754 | 0.861656611 | 1 | 0.532854418 |
| ENSMUSG00000000606 | Vmn2r88 | 669149 | protein_coding | 1.512143754 | 0.532875078 | 3.058840141 | -0.174208214 | 0.86170183 | 1 | 0.532875078 |
| ENSMUSG00000049799 | Lrrc19 | 100061 | protein_coding | 745.8959293 | -0.532925227 | 0.129881906 | 4.103152192 | 4.08E-05 | 0.000391532 | 0.532925227 |
| ENSMUSG00000084866 | A930006K02Rik | 100503120 | lncRNA | 3.080098471 | -0.532934316 | 0.818138808 | 0.651398406 | 0.514789339 | 1 | 0.532934316 |
| ENSMUSG00000064495 | Gm24313 | 115487881 | snoRNA | 2.198320747 | 0.532987298 | 0.849618832 | -0.627325193 | 0.530446092 | 1 | 0.532987298 |
| ENSMUSG00000071909 | Nutm2 | 328250 | protein_coding | 1.698258743 | 0.532996705 | 3.065301719 | -0.173880666 | 0.861959247 | 1 | 0.532996705 |
| ENSMUSG00000091601 | Or2h1b | 258051 | protein_coding | 1.373589377 | 0.532998869 | 3.065245442 | -0.173884565 | 0.861956184 | 1 | 0.532998869 |
| ENSMUSG00000027635 | Dsn1 | 66934 | protein_coding | 84.02113394 | 0.532999956 | 0.189325867 | -2.815251628 | 0.004873906 | 0.02379884 | 0.532999956 |
| ENSMUSG00000044716 | Dok7 | 231134 | protein_coding | 52.67664409 | -0.533036659 | 0.194609041 | 2.739012821 | 0.006162397 | 0.028755531 | 0.533036659 |
| ENSMUSG00000091635 | Vmn2r13 | 231589 | protein_coding | 1.559704366 | 0.5331027 | 3.070818344 | -0.173602812 | 0.862177622 | 1 | 0.5331027 |
| ENSMUSG00000027859 | Ngf | 18049 | protein_coding | 1.235035001 | 0.533133096 | 3.072224138 | -0.173533269 | 0.86223228 | 1 | 0.533133096 |
| ENSMUSG00000109801 | Or4f14d | 258737 | protein_coding | 1.235035001 | 0.533133096 | 3.072224138 | -0.173533269 | 0.86223228 | 1 | 0.533133096 |
| ENSMUSG00000090733 | Rps27 | 57294 | protein_coding | 4361.529613 | -0.533207997 | 0.104654405 | 5.094940816 | 3.49E-07 | 5.30E-06 | 0.533207997 |
| ENSMUSG00000050201 | Otop2 | 237987 | protein_coding | 1.187474389 | 0.533213713 | 3.076459404 | -0.173320575 | 0.862399452 | 1 | 0.533213713 |
| ENSMUSG00000034127 | Tspan8 | 216350 | protein_coding | 5442.5878 | -0.533258525 | 0.088022725 | 6.058191487 | 1.38E-09 | 2.99E-08 | 0.533258525 |
| ENSMUSG00000061984 | Or1e32 | 259008 | protein_coding | 1.096480625 | 0.533283132 | 3.080065938 | -0.173140167 | 0.862541253 | 1 | 0.533283132 |
| ENSMUSG00000033491 | Prss35 | 244954 | protein_coding | 1.282595613 | 0.533297156 | 3.080964434 | -0.173094227 | 0.862577363 | 1 | 0.533297156 |
| ENSMUSG00000033383 | Rtp1 | 239766 | protein_coding | 1.144041237 | 0.533355542 | 3.083968093 | -0.172944572 | 0.862694996 | 1 | 0.533355542 |
| ENSMUSG00000048982 | Gphb5 | 217674 | protein_coding | 1.144041237 | 0.533355542 | 3.083968093 | -0.172944572 | 0.862694996 | 1 | 0.533355542 |
| ENSMUSG00000020123 | Avpr1a | 54140 | protein_coding | 1.344202367 | -0.533398855 | 1.025833997 | 0.519966054 | 0.603087235 | 1 | 0.533398855 |
| ENSMUSG00000062046 | 5730460C07Rik | 70594 | lncRNA | 0.953675171 | 0.533448186 | 3.0887437 | -0.172707171 | 0.862881607 | 1 | 0.533448186 |
| ENSMUSG00000005553 | Atp4a | 11944 | protein_coding | 6.219905057 | -0.533476572 | 0.603794446 | 0.883540046 | 0.376944556 | 0.599156327 | 0.533476572 |
| ENSMUSG00000011256 | Adam19 | 11492 | protein_coding | 8.440402058 | -0.533547491 | 0.725657568 | 0.735260699 | 0.462180739 | 0.675880563 | 0.533547491 |
| ENSMUSG00000042406 | Atf4 | 11911 | protein_coding | 3016.844137 | 0.533551041 | 0.097966525 | -5.446258749 | 5.14E-08 | 8.80E-07 | 0.533551041 |
| ENSMUSG00000055960 | Skint4 | 320640 | protein_coding | 1.048920012 | 0.533689598 | 3.101873172 | -0.172053971 | 0.863395099 | 1 | 0.533689598 |
| ENSMUSG00000038352 | Arl5c | 217151 | protein_coding | 27.44063405 | 0.533726961 | 0.24543696 | -2.174598968 | 0.029660182 | 0.10006477 | 0.533726961 |
| ENSMUSG00000049404 | Rarres1 | 109222 | protein_coding | 5.87296663 | 0.533830748 | 0.52221869 | -1.022236006 | 0.306669213 | 0.527150555 | 0.533830748 |
| ENSMUSG00000085252 | 8430437L04Rik | 74541 | lncRNA | 1.330156225 | 0.533832433 | 3.109890985 | -0.171656317 | 0.86370773 | 1 | 0.533832433 |
| ENSMUSG00000020773 | Trim47 | 217333 | protein_coding | 42.60660643 | 0.53410827 | 0.24944207 | -2.141211667 | 0.032256973 | 0.106687692 | 0.53410827 |
| ENSMUSG00000062729 | Ppox | 19044 | protein_coding | 233.7963566 | 0.534249813 | 0.087708679 | -6.091185272 | 1.12E-09 | 2.46E-08 | 0.534249813 |
| ENSMUSG00000030470 | Csrp3 | 13009 | protein_coding | 1.180626026 | 0.534451239 | 1.98297875 | -0.269519398 | 0.787530017 | 1 | 0.534451239 |
| ENSMUSG00000075702 | Selenom | 114679 | protein_coding | 1105.112626 | 0.534515268 | 0.097455697 | -5.484700076 | 4.14E-08 | 7.21E-07 | 0.534515268 |
| ENSMUSG00000047501 | Cldn4 | 12740 | protein_coding | 491.78253 | -0.534639453 | 0.264500195 | 2.021319694 | 0.043246682 | 0.133440901 | 0.534639453 |
| ENSMUSG00000056296 | Synpr | 72003 | protein_coding | 1.809170622 | 0.534660783 | 1.756724131 | -0.304351021 | 0.760860478 | 1 | 0.534660783 |
| ENSMUSG00000090655 | Vmn2r120 | 224916 | protein_coding | 5.039463333 | 0.535088639 | 2.720487882 | -0.196688485 | 0.844071322 | 0.923234456 | 0.535088639 |
| ENSMUSG00000039543 | Cfap70 | 76670 | protein_coding | 2.029929628 | -0.535184724 | 0.921386882 | 0.580846911 | 0.561343635 | 1 | 0.535184724 |

| ensembl_gene_id | external_gene_name | entrezgene_id | gene_biotype | baseMean | log2FoldChange | lfcSE | stat | pvalue | padj | absLog2FC |
|---|---|---|---|---|---|---|---|---|---|---|
| ENSMUSG00000037148 | Arhgap10 | 78514 | protein_coding | 34.61732018 | 0.535239542 | 0.225717964 | -2.371275784 | 0.017726799 | 0.066683451 | 0.535239542 |
| ENSMUSG00000057895 | Zfp105 | 22646 | protein_coding | 11.29314475 | 0.535418196 | 0.311378437 | -1.719509546 | 0.085521631 | 0.220707019 | 0.535418196 |
| ENSMUSG00000097767 | Miat | 330166 | lncRNA | 5.568825288 | -0.535831858 | 0.721835091 | 0.742318937 | 0.457894121 | 0.67261003 | 0.535831858 |
| ENSMUSG00000075012 | Fjx1 | 14221 | protein_coding | 3.774155887 | -0.53587768 | 0.551400454 | 0.971848457 | 0.331125945 | 1 | 0.53587768 |
| ENSMUSG00000071636 | Rimbp3 | 239731 | protein_coding | 12.85928688 | -0.53602458 | 0.351544085 | 1.524772009 | 0.127315974 | 0.29413766 | 0.53602458 |
| ENSMUSG00000042029 | Ncapg2 | 76044 | protein_coding | 293.8525411 | 0.536084343 | 0.195266484 | -2.745398662 | 0.006043743 | 0.028321899 | 0.536084343 |
| ENSMUSG00000055725 | Paqr3 | 231474 | protein_coding | 10.43407428 | 0.536135521 | 0.474976938 | -1.128761164 | 0.2589986 | 0.472341459 | 0.536135521 |
| ENSMUSG00000065405 | Mir30a | 387225 | miRNA | 0.778119863 | 0.536189771 | 1.16318439 | -0.460967131 | 0.644822187 | 1 | 0.536189771 |
| ENSMUSG00000032011 | Thy1 | 21838 | protein_coding | 6.791415958 | -0.536355159 | 0.683842418 | 0.784325664 | 0.432849044 | 0.651423414 | 0.536355159 |
| ENSMUSG00000033450 | Tagap | 72536 | protein_coding | 40.69315477 | -0.536442181 | 0.241467339 | 2.221593127 | 0.026310815 | 0.090822909 | 0.536442181 |
| ENSMUSG00000011171 | Vipr2 | 22355 | protein_coding | 42.98949957 | -0.53657793 | 0.250580286 | 2.14134136 | 0.03224652 | 0.106674486 | 0.53657793 |
| ENSMUSG00000034595 | Ppp1r18 | 76448 | protein_coding | 9.360378259 | 0.536609906 | 0.55603316 | -0.965068174 | 0.334510666 | 0.556753592 | 0.536609906 |
| ENSMUSG00000026874 | Hc | 15139 | protein_coding | 16.80967271 | -0.536689919 | 0.36908063 | 1.454126486 | 0.145911242 | 0.323295567 | 0.536689919 |
| ENSMUSG00000101972 | H3c11 | 319153 | protein_coding | 420.9484567 | 0.536716404 | 0.19379295 | -2.769535233 | 0.005613633 | 0.026785189 | 0.536716404 |
| ENSMUSG00000003849 | Nqo1 | 18104 | protein_coding | 759.4173881 | -0.536760072 | 0.140459719 | 3.821451994 | 0.000132668 | 0.001107066 | 0.536760072 |
| ENSMUSG00000117292 | E330032C10Rik | 328855 | lncRNA | 1.705820219 | -0.536901519 | 1.295928616 | 0.414298683 | 0.678655378 | 1 | 0.536901519 |
| ENSMUSG00000027357 | Crls1 | 66586 | protein_coding | 512.5044076 | 0.536962787 | 0.097977374 | -5.480477459 | 4.24E-08 | 7.37E-07 | 0.536962787 |
| ENSMUSG00000032425 | Zfp949 | 71640 | protein_coding | 104.4703508 | -0.536994076 | 0.108282914 | 4.959176447 | 7.08E-07 | 1.01E-05 | 0.536994076 |
| ENSMUSG00000028039 | Efna3 | 13638 | protein_coding | 49.80659441 | -0.537145893 | 0.444862672 | 1.207442043 | 0.227261952 | 0.435838332 | 0.537145893 |
| ENSMUSG00000056174 | Col8a2 | 329941 | protein_coding | 3.697207687 | 0.537163376 | 0.799760731 | -0.671655102 | 0.50180329 | 1 | 0.537163376 |
| ENSMUSG00000087651 | 1500009L16Rik | 69784 | protein_coding | 48.49928563 | -0.537236273 | 0.175273502 | 3.065131157 | 0.002175746 | 0.012122221 | 0.537236273 |
| ENSMUSG00000044359 | P2ry4 | 57385 | protein_coding | 52.25663513 | 0.537571471 | 0.182807322 | -2.940645181 | 0.003275295 | 0.01711111 | 0.537571471 |
| ENSMUSG00000050107 | Haspin | 14841 | protein_coding | 109.8725713 | 0.537757385 | 0.171373839 | -3.137919922 | 0.001701514 | 0.009897427 | 0.537757385 |
| ENSMUSG00000112640 | Gm32687 | 102635315 | protein_coding | 10.08481324 | -0.537825537 | 0.403905331 | 1.331563355 | 0.183003713 | 0.377733062 | 0.537825537 |
| ENSMUSG00000053414 | Hunk | 26559 | protein_coding | 343.8340092 | -0.537986083 | 0.094091925 | 5.717664746 | 1.08E-08 | 2.07E-07 | 0.537986083 |
| ENSMUSG00000079440 | Alpi | 76768 | protein_coding | 282.4458554 | -0.538150862 | 0.273446089 | 1.968032766 | 0.049064268 | 0.146943656 | 0.538150862 |
| ENSMUSG00000031478 | Nek3 | 23954 | protein_coding | 104.0331997 | -0.538188865 | 0.155534291 | 3.460258581 | 0.000539657 | 0.003784244 | 0.538188865 |
| ENSMUSG00000084479 | Gm26424 | 115490012 | snRNA | 1.937409418 | -0.538221675 | 0.948341226 | 0.567540101 | 0.57034729 | 1 | 0.538221675 |
| ENSMUSG00000056413 | Adap1 | 231821 | protein_coding | 565.8801632 | -0.538344282 | 0.132353542 | 4.067471661 | 4.75E-05 | 0.000446188 | 0.538344282 |
| ENSMUSG00000034908 | Sidt2 | 214597 | protein_coding | 689.8967094 | -0.538344729 | 0.0677289 | 7.94852314 | 1.89E-15 | 7.15E-14 | 0.538344729 |
| ENSMUSG00000028415 | Spink4 | 20731 | protein_coding | 6251.19093 | 0.538533378 | 0.268346505 | -2.006858182 | 0.044764766 | 0.136998766 | 0.538533378 |
| ENSMUSG00000022364 | Tbc1d31 | 210544 | protein_coding | 95.26284631 | 0.538566674 | 0.164474694 | -3.274465275 | 0.001058621 | 0.006667458 | 0.538566674 |
| ENSMUSG00000074217 | Misp3 | 70134 | protein_coding | 32.77151717 | -0.538655253 | 0.22178128 | 2.428767901 | 0.015150228 | 0.058854589 | 0.538655253 |
| ENSMUSG00000020167 | Tcf3 | 21423 | protein_coding | 547.9289732 | 0.53868721 | 0.079762085 | -6.753675158 | 1.44E-11 | 3.96E-10 | 0.53868721 |
| ENSMUSG00000047330 | Kcne4 | 57814 | protein_coding | 2.234560571 | -0.538873187 | 0.703183168 | 0.766334024 | 0.443477578 | 1 | 0.538873187 |
| ENSMUSG00000096910 | Zfp955b | 100043468 | protein_coding | 132.9402198 | -0.539142591 | 0.185772376 | 2.902167708 | 0.003705901 | 0.019005977 | 0.539142591 |
| ENSMUSG00000026602 | Nphs2 | 170484 | protein_coding | 1.037106996 | -0.539196761 | 1.753871212 | 0.307432357 | 0.758514303 | 1 | 0.539196761 |
| ENSMUSG00000002565 | Scin | 20259 | protein_coding | 2170.492442 | -0.539353162 | 0.203324259 | 2.652674918 | 0.007985674 | 0.035574288 | 0.539353162 |
| ENSMUSG00000064061 | Dzip3 | 224170 | protein_coding | 102.713584 | 0.539469407 | 0.123392884 | -4.371965314 | 1.23E-05 | 0.000134753 | 0.539469407 |

| ensembl_gene_id | external_gene_name | entrezgene_id | gene_biotype | baseMean | log2FoldChange | lfcSE | stat | pvalue | padj | absLog2FC |
|---|---|---|---|---|---|---|---|---|---|---|
| ENSMUSG00000031971 | Ccsap | 73420 | protein_coding | 37.5940563 | 0.539494968 | 0.215629747 | -2.501950571 | 0.012351117 | 0.050176222 | 0.539494968 |
| ENSMUSG00000026497 | Mixl1 | 27217 | protein_coding | 1.076976271 | -0.539502203 | 1.302797623 | 0.414110521 | 0.678793168 | 1 | 0.539502203 |
| ENSMUSG00000021596 | Mctp1 | 78771 | protein_coding | 34.85645579 | -0.539664233 | 0.200107374 | 2.696873303 | 0.00699939 | 0.031903925 | 0.539664233 |
| ENSMUSG00000079685 | Ulbp1 | 77777 | protein_coding | 52.7370977 | 0.539785564 | 0.356656429 | -1.513460912 | 0.130162623 | 0.298087027 | 0.539785564 |
| ENSMUSG00000037731 | Themis2 | 230787 | protein_coding | 1.90571781 | 0.539902022 | 1.00994274 | -0.534586764 | 0.592935638 | 1 | 0.539902022 |
| ENSMUSG00000027009 | Itga4 | 16401 | protein_coding | 15.38525955 | -0.53991356 | 0.339596573 | 1.589867515 | 0.111864671 | 0.267923594 | 0.53991356 |
| ENSMUSG00000051238 | Swsap1 | 66962 | protein_coding | 158.6590075 | 0.539935531 | 0.130060207 | -4.151427574 | 3.30E-05 | 0.00032459 | 0.539935531 |
| ENSMUSG00000065176 | Rnu12 | 104307 | snRNA | 46.70530613 | 0.540385968 | 0.440457001 | -1.226875646 | 0.219869333 | 0.426114561 | 0.540385968 |
| ENSMUSG00000086103 | Gm11832 | 102639547 | lncRNA | 1.840835384 | 0.540584131 | 0.809426422 | -0.667860743 | 0.504222489 | 1 | 0.540584131 |
| ENSMUSG00000064357 | mt-Atp6 | 17705 | protein_coding | 4.663179079 | 0.540662769 | 0.888431746 | -0.608558588 | 0.542817061 | 0.736235501 | 0.540662769 |
| ENSMUSG00000040550 | Otud6b | 72201 | protein_coding | 501.7091278 | 0.540954873 | 0.097641726 | -5.540201881 | 3.02E-08 | 5.38E-07 | 0.540954873 |
| ENSMUSG00000036959 | Bcorl1 | 320376 | protein_coding | 142.4598504 | -0.5410071 | 0.227737578 | 2.375572376 | 0.017521756 | 0.066168368 | 0.5410071 |
| ENSMUSG00000031844 | Hsd17b2 | 15486 | protein_coding | 1624.072572 | -0.541161357 | 0.15396034 | 3.514939992 | 0.000439853 | 0.003190048 | 0.541161357 |
| ENSMUSG00000096141 | Dnah7a | 627872 | protein_coding | 4.364639932 | 0.541622678 | 0.805494974 | -0.672409755 | 0.501322874 | 1 | 0.541622678 |
| ENSMUSG00000040734 | Ppp1r13l | 333654 | protein_coding | 44.47141235 | -0.54172722 | 0.278355876 | 1.946167721 | 0.051634596 | 0.152894696 | 0.54172722 |
| ENSMUSG00000054083 | Capn12 | 60594 | protein_coding | 11.5484825 | 0.541754855 | 0.534136313 | -1.014263292 | 0.310457135 | 0.53139398 | 0.541754855 |
| ENSMUSG00000067925 | Rtl8a | 66158 | protein_coding | 11.95112403 | -0.541806209 | 0.351009378 | 1.543566191 | 0.122693461 | 0.285938444 | 0.541806209 |
| ENSMUSG00000028173 | Wls | 68151 | protein_coding | 1298.625597 | -0.541842829 | 0.104369443 | 5.191584949 | 2.09E-07 | 3.25E-06 | 0.541842829 |
| ENSMUSG00000040093 | Bmf | 171543 | protein_coding | 141.8311132 | -0.54187613 | 0.156606253 | 3.460118101 | 0.000539939 | 0.003784612 | 0.54187613 |
| ENSMUSG00000066071 | Cyp4a12a | 277753 | protein_coding | 1.183346929 | 0.541957066 | 3.668259146 | -0.147742306 | 0.882546142 | 1 | 0.541957066 |
| ENSMUSG00000062372 | Otof | 83762 | protein_coding | 5.923892525 | 0.542097017 | 2.5083557 | -0.216116485 | 0.82889695 | 0.913504709 | 0.542097017 |
| ENSMUSG00000024131 | Slc3a1 | 20532 | protein_coding | 428.6073139 | 0.542286752 | 0.221653509 | -2.446551625 | 0.014423014 | 0.056508103 | 0.542286752 |
| ENSMUSG00000046572 | Zfp518b | 100515 | protein_coding | 62.4614864 | 0.542342411 | 0.189337018 | -2.864428819 | 0.00417762 | 0.020899492 | 0.542342411 |
| ENSMUSG00000053862 | Slc51b | 330962 | protein_coding | 500.6281086 | 0.542537598 | 0.185144008 | -2.930354609 | 0.003385754 | 0.017604641 | 0.542537598 |
| ENSMUSG00000041046 | Ramp3 | 56089 | protein_coding | 1.894304613 | -0.542625815 | 0.854951417 | 0.63468614 | 0.525633135 | 1 | 0.542625815 |
| ENSMUSG00000021815 | Mss51 | 74843 | protein_coding | 4.953597143 | 0.542688142 | 0.522643447 | -1.038352523 | 0.299105964 | 0.51911069 | 0.542688142 |
| ENSMUSG00000004207 | Psap | 19156 | protein_coding | 7201.131386 | -0.542890657 | 0.05760924 | 9.42367323 | 4.36E-21 | 2.37E-19 | 0.542890657 |
| ENSMUSG00000051839 | Gypa | 14934 | protein_coding | 8.226264766 | 0.542924421 | 0.395194917 | -1.373814281 | 0.169499346 | 0.358814538 | 0.542924421 |
| ENSMUSG00000048965 | Mrgpre | 244238 | protein_coding | 8.931064174 | -0.542932448 | 0.38942812 | 1.394178847 | 0.163263604 | 0.349330805 | 0.542932448 |
| ENSMUSG00000044471 | Lncpint | 232685 | lncRNA | 145.2280116 | 0.543038536 | 0.224265063 | -2.421413883 | 0.015460264 | 0.059833793 | 0.543038536 |
| ENSMUSG00000042293 | Gm5617 | 434402 | protein_coding | 63.96623822 | -0.543058475 | 0.158159785 | 3.433606554 | 0.000595608 | 0.004105122 | 0.543058475 |
| ENSMUSG00000111119 | Gm34005 | 102637110 | processed_pseudogene | 0.825113516 | 0.543076625 | 1.481303846 | -0.366620681 | 0.71390198 | 1 | 0.543076625 |
| ENSMUSG00000005465 | Il27ra | 50931 | protein_coding | 5.128138065 | 0.54342507 | 0.910640169 | -0.596750603 | 0.550673902 | 0.741566405 | 0.54342507 |
| ENSMUSG00000043252 | Tmem64 | 100201 | protein_coding | 137.8085561 | 0.543518839 | 0.17720906 | -3.067105247 | 0.002161427 | 0.012066874 | 0.543518839 |
| ENSMUSG00000034265 | Zdhhc14 | 224454 | protein_coding | 67.96391091 | -0.543590933 | 0.162059631 | 3.354264913 | 0.000795761 | 0.005237626 | 0.543590933 |
| ENSMUSG00000073491 | Ifi213 | 623121 | protein_coding | 2.753632518 | 0.543656685 | 1.12122325 | -0.484878177 | 0.627762764 | 1 | 0.543656685 |
| ENSMUSG00000070796 | Psg21 | 72242 | protein_coding | 1.510606819 | 0.543755776 | 2.106624066 | -0.258117139 | 0.796316505 | 1 | 0.543755776 |
| ENSMUSG00000044231 | Nhlrc1 | 105193 | protein_coding | 38.17180403 | -0.543798367 | 0.233256952 | 2.33132759 | 0.019736092 | 0.072556062 | 0.543798367 |
| ENSMUSG00000068245 | Phf11d | 219132 | protein_coding | 102.2539057 | -0.54383583 | 0.278744392 | 1.951019807 | 0.05105469 | 0.151449101 | 0.54383583 |

| ensembl_gene_id | external_gene_name | entrezgene_id | gene_biotype | baseMean | log2FoldChange | lfcSE | stat | pvalue | padj | absLog2FC |
|---|---|---|---|---|---|---|---|---|---|---|
| ENSMUSG00000036983 | Tfb1m | 224481 | protein_coding | 107.4024185 | 0.543874835 | 0.147617252 | -3.684358218 | 0.00022928 | 0.001786002 | 0.543874835 |
| ENSMUSG00000047215 | Rpl9 | 20005 | protein_coding | 5242.709024 | -0.543976227 | 0.143557322 | 3.789261464 | 0.000151096 | 0.00123647 | 0.543976227 |
| ENSMUSG00000097633 | Gm16617 | 100502764 | lncRNA | 1.658414959 | 0.544087708 | 0.815738436 | -0.666987952 | 0.504779829 | 1 | 0.544087708 |
| ENSMUSG00000025161 | Slc16a3 | 80879 | protein_coding | 45.11597159 | -0.544246657 | 0.301807147 | 1.803292805 | 0.071342242 | 0.193614755 | 0.544246657 |
| ENSMUSG00000021797 | 9230112D13Rik | 78243 | protein_coding | 0.967340173 | 0.544270779 | 1.046271819 | -0.520200171 | 0.602924066 | 1 | 0.544270779 |
| ENSMUSG00000020029 | Nudt4 | 71207 | protein_coding | 3705.74115 | -0.544301673 | 0.110583022 | 4.922108891 | 8.56E-07 | 1.20E-05 | 0.544301673 |
| ENSMUSG00000030321 | Efcab12 | 212516 | protein_coding | 7.161011241 | 0.544336616 | 0.459801253 | -1.183851964 | 0.236471668 | 0.446602599 | 0.544336616 |
| ENSMUSG00000103793 | Pcdhga6 | 93714 | protein_coding | 6.364885912 | 0.544439093 | 0.462231286 | -1.177849941 | 0.238856435 | 0.449052273 | 0.544439093 |
| ENSMUSG00000032278 | Paqr5 | 74090 | protein_coding | 751.3433826 | -0.544481413 | 0.132942605 | 4.095612629 | 4.21E-05 | 0.000402389 | 0.544481413 |
| ENSMUSG00000086189 | Gm15462 | 108168861 | lncRNA | 22.38448967 | -0.544691053 | 0.262341722 | 2.076265447 | 0.037869407 | 0.12072884 | 0.544691053 |
| ENSMUSG00000032514 | Ttc21a | 74052 | protein_coding | 8.846208462 | 0.544729276 | 0.577182083 | -0.943773709 | 0.345285298 | 0.566901476 | 0.544729276 |
| ENSMUSG00000048442 | Smim5 | 66528 | protein_coding | 69.69313443 | -0.544754562 | 0.193830534 | 2.810468255 | 0.004946947 | 0.024105601 | 0.544754562 |
| ENSMUSG00000034450 | Gulo | 268756 | protein_coding | 2.11578078 | 0.544766265 | 1.469337705 | -0.370756337 | 0.710819025 | 1 | 0.544766265 |
| ENSMUSG00000029449 | Rhof | 23912 | protein_coding | 344.5064235 | 0.544968471 | 0.215006247 | -2.534663432 | 0.011255541 | 0.046796074 | 0.544968471 |
| ENSMUSG00000008489 | Elavl2 | 15569 | protein_coding | 3.730497464 | -0.544999166 | 0.905408963 | 0.601937013 | 0.547216066 | 1 | 0.544999166 |
| ENSMUSG00000025144 | Cenpx | 20892 | protein_coding | 295.9511757 | 0.545023139 | 0.095768693 | -5.691036617 | 1.26E-08 | 2.40E-07 | 0.545023139 |
| ENSMUSG00000040966 | Slc22a2 | 20518 | protein_coding | 1.200024373 | -0.545196172 | 2.151909492 | 0.253354602 | 0.799994206 | 1 | 0.545196172 |
| ENSMUSG00000066196 | Spag8 | 433700 | protein_coding | 2.689563305 | -0.545366518 | 0.832531946 | 0.655069779 | 0.512422828 | 1 | 0.545366518 |
| ENSMUSG00000072966 | Gprasp2 | 245607 | protein_coding | 4.390389265 | 0.545410123 | 0.554032574 | -0.984436925 | 0.32490073 | 0.546769252 | 0.545410123 |
| ENSMUSG00000032258 | Lca5 | 75782 | protein_coding | 24.59047822 | 0.545597411 | 0.225566832 | -2.418783855 | 0.01557249 | 0.060155346 | 0.545597411 |
| ENSMUSG00000032010 | Usp2 | 53376 | protein_coding | 186.51999 | 0.545725005 | 0.23260382 | -2.346156682 | 0.018968132 | 0.070451816 | 0.545725005 |
| ENSMUSG00000003573 | Homer3 | 26558 | protein_coding | 16.14789052 | 0.545734606 | 0.305859917 | -1.784263241 | 0.0743809 | 0.19938276 | 0.545734606 |
| ENSMUSG00000079414 | Gm11110 | 100169874 | lncRNA | 4.031247988 | -0.545862139 | 0.73812304 | 0.739527301 | 0.459586868 | 1 | 0.545862139 |
| ENSMUSG00000044948 | Cfap43 | 100048534 | protein_coding | 7.669018232 | 0.546068857 | 0.805979255 | -0.677522223 | 0.498074675 | 0.703249417 | 0.546068857 |
| ENSMUSG00000015970 | Chdh | 218865 | protein_coding | 1446.539444 | -0.546239051 | 0.135150896 | 4.04169759 | 5.31E-05 | 0.000492318 | 0.546239051 |
| ENSMUSG00000035521 | Gnptg | 214505 | protein_coding | 191.7101904 | -0.546261873 | 0.100549325 | 5.432775161 | 5.55E-08 | 9.45E-07 | 0.546261873 |
| ENSMUSG00000049357 | Brd8dc | 271508 | protein_coding | 5.058716614 | 0.546279411 | 0.635934922 | -0.859017791 | 0.390330701 | 0.612497263 | 0.546279411 |
| ENSMUSG00000100599 | 1700120C14Rik | 73600 | lncRNA | 5.972595888 | -0.54636972 | 0.448110077 | 1.219275683 | 0.222739575 | 0.429760642 | 0.54636972 |
| ENSMUSG00000000782 | Tcf7 | 21414 | protein_coding | 26.81986434 | -0.546414731 | 0.225898501 | 2.418850631 | 0.015569632 | 0.060155346 | 0.546414731 |
| ENSMUSG00000025868 | Higd2a | 67044 | protein_coding | 1290.740793 | -0.546504134 | 0.094455325 | 5.785847814 | 7.21E-09 | 1.42E-07 | 0.546504134 |
| ENSMUSG00000024990 | Rbp4 | 19662 | protein_coding | 70.36693696 | -0.54658956 | 0.22979101 | 2.378637702 | 0.017376745 | 0.065695873 | 0.54658956 |
| ENSMUSG00000032493 | Prss44 | 73336 | protein_coding | 2.946252739 | 0.546730673 | 2.467016433 | -0.221616146 | 0.824612713 | 1 | 0.546730673 |
| ENSMUSG00000037754 | Ppp1r16b | 228852 | protein_coding | 14.55511459 | 0.546759104 | 0.474030791 | -1.153425293 | 0.248735865 | 0.460883095 | 0.546759104 |
| ENSMUSG00000079298 | Klrb1b | 80782 | protein_coding | 2.913299162 | 0.546950286 | 0.780058633 | -0.701165609 | 0.48319967 | 1 | 0.546950286 |
| ENSMUSG00000045751 | Mms22l | 212377 | protein_coding | 135.7928098 | 0.54695205 | 0.161155377 | -3.3939423 | 0.000688942 | 0.00461924 | 0.54695205 |
| ENSMUSG00000101172 | Gm28535 | 105246665 | lncRNA | 0.588140355 | 0.547230313 | 1.675298822 | -0.32664639 | 0.743935352 | 1 | 0.547230313 |
| ENSMUSG00000054619 | Mettl7a1 | 70152 | protein_coding | 1436.589299 | -0.547496873 | 0.121013109 | 4.524277391 | 6.06E-06 | 7.11E-05 | 0.547496873 |
| ENSMUSG00000032194 | Kank2 | 235041 | protein_coding | 278.8201617 | -0.54753588 | 0.174287306 | 3.141570616 | 0.001680443 | 0.009812883 | 0.54753588 |
| ENSMUSG00000031860 | Pbx4 | 80720 | protein_coding | 2.579733967 | -0.547685546 | 0.836403835 | 0.654809942 | 0.512590129 | 1 | 0.547685546 |

Supplementary Table 1

| ensembl_gene_id | external_gene_name | entrezgene_id | gene_biotype | baseMean | log2FoldChange | lfcSE | stat | pvalue | padj | absLog2FC |
|---|---|---|---|---|---|---|---|---|---|---|
| ENSMUSG00000090071 | Cdk5r2 | 12570 | protein_coding | 2.991550296 | -0.547772562 | 0.870351986 | 0.629369003 | 0.529107506 | 1 | 0.547772562 |
| ENSMUSG00000027778 | Ift80 | 68259 | protein_coding | 96.97896627 | 0.547831556 | 0.148189164 | -3.696839503 | 0.0002183 | 0.001708536 | 0.547831556 |
| ENSMUSG00000078247 | Airn | 104103 | lncRNA | 60.67719207 | -0.548011557 | 0.172109509 | 3.184086462 | 0.001452116 | 0.008644646 | 0.548011557 |
| ENSMUSG00000001300 | Efnb2 | 13642 | protein_coding | 1375.811314 | 0.5481742 | 0.161414122 | -3.39607336 | 0.0006836 | 0.004596489 | 0.5481742 |
| ENSMUSG00000029916 | Agk | 69923 | protein_coding | 83.24762216 | -0.548647068 | 0.146975048 | 3.732926614 | 0.000189268 | 0.001509936 | 0.548647068 |
| ENSMUSG00000039037 | St6galnac5 | 26938 | protein_coding | 1.290821616 | -0.548788999 | 1.555380397 | 0.352832658 | 0.724213899 | 1 | 0.548788999 |
| ENSMUSG00000021091 | Serpina3n | 20716 | protein_coding | 1.175826134 | -0.548816421 | 1.784099221 | 0.307615414 | 0.75837499 | 1 | 0.548816421 |
| ENSMUSG00000070880 | Gad1 | 14415 | protein_coding | 3.707415272 | 0.549011203 | 1.574128155 | -0.348771605 | 0.727260783 | 1 | 0.549011203 |
| ENSMUSG00000091594 | Gm17067 | 115486519 | protein_coding | 1.283827425 | -0.549169421 | 1.624303311 | 0.338095365 | 0.735291323 | 1 | 0.549169421 |
| ENSMUSG00000096001 | 2610528A11Rik | 70045 | protein_coding | 3520.145776 | -0.549188157 | 0.08906305 | 6.166285092 | 6.99E-10 | 1.57E-08 | 0.549188157 |
| ENSMUSG00000061825 | Ces2c | 234671 | protein_coding | 1342.433451 | -0.549224499 | 0.193157506 | 2.843402314 | 0.004463469 | 0.022141702 | 0.549224499 |
| ENSMUSG00000043510 | Hscb | 100900 | protein_coding | 114.6580696 | -0.549263114 | 0.115565888 | 4.752813504 | 2.01E-06 | 2.61E-05 | 0.549263114 |
| ENSMUSG00000086550 | Pla2g10os | 76684 | lncRNA | 65.9912417 | -0.549292782 | 0.178380547 | 3.0793312 | 0.002074659 | 0.011690393 | 0.549292782 |
| ENSMUSG00000035459 | Stab2 | 192188 | protein_coding | 5.104264511 | 0.549458023 | 1.142114212 | -0.481088509 | 0.630453595 | 0.79831751 | 0.549458023 |
| ENSMUSG00000021313 | Ryr2 | 20191 | protein_coding | 9.736620849 | 0.549796845 | 2.546080458 | -0.21593852 | 0.829035671 | 0.913504709 | 0.549796845 |
| ENSMUSG00000013846 | St3gal1 | 20442 | protein_coding | 180.2627919 | 0.549809583 | 0.376106739 | -1.461844542 | 0.143783815 | 0.320178792 | 0.549809583 |
| ENSMUSG00000025401 | Myo1a | 432516 | protein_coding | 6150.778071 | -0.549824885 | 0.126177154 | 4.357562908 | 1.32E-05 | 0.000142578 | 0.549824885 |
| ENSMUSG00000041025 | Iffo2 | 212632 | protein_coding | 231.5425336 | -0.550072931 | 0.122070198 | 4.506201685 | 6.60E-06 | 7.70E-05 | 0.550072931 |
| ENSMUSG00000096957 | E230013L22Rik | 102132 | lncRNA | 32.7861578 | 0.550082355 | 0.260248231 | -2.113683363 | 0.03454232 | 0.112356092 | 0.550082355 |
| ENSMUSG00000019943 | Atp2b1 | 67972 | protein_coding | 5960.995826 | 0.550112654 | 0.09121022 | -6.031261136 | 1.63E-09 | 3.50E-08 | 0.550112654 |
| ENSMUSG00000022629 | Kif21a | 16564 | protein_coding | 1183.9298 | 0.550300244 | 0.101878275 | -5.401546542 | 6.61E-08 | 1.11E-06 | 0.550300244 |
| ENSMUSG00000097061 | 9330151L19Rik | 414085 | lncRNA | 17.58013462 | -0.550331165 | 0.267229242 | 2.059397242 | 0.0394562 | 0.124751997 | 0.550331165 |
| ENSMUSG00000031700 | Gpt2 | 108682 | protein_coding | 397.3234362 | -0.550332062 | 0.148029319 | 3.717723386 | 0.000201026 | 0.001589156 | 0.550332062 |
| ENSMUSG00000041707 | Tmem273 | 69069 | protein_coding | 1.985334551 | -0.550500249 | 0.853648259 | 0.644879484 | 0.519005288 | 1 | 0.550500249 |
| ENSMUSG00000018548 | Trim37 | 68729 | protein_coding | 372.773454 | 0.55058312 | 0.136980522 | -4.019426345 | 5.83E-05 | 0.000536429 | 0.55058312 |
| ENSMUSG00000063087 | Gm10125 | 791318 | lncRNA | 2.358748414 | 0.55070705 | 2.35734884 | -0.233612879 | 0.815285528 | 1 | 0.55070705 |
| ENSMUSG00000064037 | Gpn1 | 74254 | protein_coding | 236.6301738 | 0.550713098 | 0.103325552 | -5.329882972 | 9.83E-08 | 1.62E-06 | 0.550713098 |
| ENSMUSG00000007989 | Fzd3 | 14365 | protein_coding | 150.2022341 | 0.55077718 | 0.157443116 | -3.498261419 | 0.000468302 | 0.003358027 | 0.55077718 |
| ENSMUSG00000078786 | BC024978 | 414069 | protein_coding | 108.0000885 | 0.550946056 | 0.217083033 | -2.537950794 | 0.011150366 | 0.046464079 | 0.550946056 |
| ENSMUSG00000114865 | Cirop | 101056084 | protein_coding | 2.248387847 | 0.55124056 | 1.045823486 | -0.52708757 | 0.598132778 | 1 | 0.55124056 |
| ENSMUSG00000053581 | Zfand2a | 100494 | protein_coding | 284.5641921 | -0.551304422 | 0.138816137 | 3.971472148 | 7.14E-05 | 0.000641781 | 0.551304422 |
| ENSMUSG00000048970 | C1galt1c1 | 59048 | protein_coding | 813.7999444 | 0.551378957 | 0.149198241 | -3.695612996 | 0.000219357 | 0.00171518 | 0.551378957 |
| ENSMUSG00000028801 | Stpg1 | 78806 | protein_coding | 11.46335694 | -0.551389485 | 0.380004833 | 1.451006508 | 0.146778048 | 0.324596932 | 0.551389485 |
| ENSMUSG00000018919 | Tm4sf5 | 75604 | protein_coding | 26.40032588 | 0.551638575 | 0.276150423 | -1.997601773 | 0.045759851 | 0.139398299 | 0.551638575 |
| ENSMUSG00000007122 | Casq1 | 12372 | protein_coding | 2.914259742 | 0.551687212 | 0.941577427 | -0.585918052 | 0.557930586 | 1 | 0.551687212 |
| ENSMUSG00000067149 | Jchain | 16069 | protein_coding | 1.112912085 | 0.551696973 | 1.462678917 | -0.377182557 | 0.706037942 | 1 | 0.551696973 |
| ENSMUSG00000101062 | Gm28821 | 115486482 | lncRNA | 2.159740414 | -0.551740533 | 0.823697233 | 0.669834147 | 0.502963523 | 1 | 0.551740533 |
| ENSMUSG00000019838 | Slc16a10 | 72472 | protein_coding | 108.6126366 | 0.551786608 | 0.141255857 | -3.906291885 | 9.37E-05 | 0.000813747 | 0.551786608 |
| ENSMUSG00000002297 | Dbf4 | 27214 | protein_coding | 269.8232827 | 0.551844212 | 0.129017858 | -4.277269975 | 1.89E-05 | 0.000196012 | 0.551844212 |

| ensembl_gene_id | external_gene_name | entrezgene_id | gene_biotype | baseMean | log2FoldChange | lfcSE | stat | pvalue | padj | absLog2FC |
|---|---|---|---|---|---|---|---|---|---|---|
| ENSMUSG00000022043 | Trim35 | 66854 | protein_coding | 278.1452918 | 0.551869803 | 0.133510519 | -4.133530505 | 3.57E-05 | 0.000348044 | 0.551869803 |
| ENSMUSG00000097527 | 1700112J16Rik | 76628 | lncRNA | 7.849402203 | 0.552096082 | 0.421753608 | -1.309048868 | 0.190517798 | 0.387990001 | 0.552096082 |
| ENSMUSG00000052353 | Cemip | 80982 | protein_coding | 7.647713366 | 0.552173621 | 0.945589939 | -0.583946167 | 0.559256531 | 0.747948226 | 0.552173621 |
| ENSMUSG00000039985 | Sinhcaf | 56306 | protein_coding | 112.6868426 | -0.552217253 | 0.167380633 | 3.299170528 | 0.00096971 | 0.00620449 | 0.552217253 |
| ENSMUSG00000047298 | Kcnv2 | 240595 | protein_coding | 4.99854139 | 0.552219762 | 0.513060934 | -1.076323931 | 0.28178241 | 0.499562412 | 0.552219762 |
| ENSMUSG00000002014 | Ssr4 | 20832 | protein_coding | 1895.29294 | 0.552283634 | 0.083520377 | -6.612561583 | 3.78E-11 | 9.98E-10 | 0.552283634 |
| ENSMUSG00000035293 | G2e3 | 217558 | protein_coding | 351.4091722 | 0.552301792 | 0.147963572 | -3.732687618 | 0.000189447 | 0.00151064 | 0.552301792 |
| ENSMUSG00000008845 | Cd163 | 93671 | protein_coding | 3.210958519 | 0.552313536 | 1.97660741 | -0.279425005 | 0.779918684 | 1 | 0.552313536 |
| ENSMUSG00000090215 | Trim34b | 434218 | protein_coding | 5.365755737 | -0.552372834 | 0.462411556 | 1.194548074 | 0.232263647 | 0.44122879 | 0.552372834 |
| ENSMUSG00000117545 | Gm30794 | 102632818 | lncRNA | 3.669354123 | 0.552833786 | 0.615309019 | -0.898465273 | 0.368937551 | 1 | 0.552833786 |
| ENSMUSG00000024327 | Slc39a7 | 14977 | protein_coding | 229.5242044 | 0.553030599 | 0.128643051 | -4.298954311 | 1.72E-05 | 0.000180273 | 0.553030599 |
| ENSMUSG00000028618 | Tmem59 | 56374 | protein_coding | 3417.720235 | -0.553035475 | 0.11311907 | 4.888967668 | 1.01E-06 | 1.39E-05 | 0.553035475 |
| ENSMUSG00000101360 | 4933417O13Rik | 71153 | lncRNA | 10.42820399 | -0.553108978 | 0.553496914 | 0.999299118 | 0.317649813 | 0.539052565 | 0.553108978 |
| ENSMUSG00000024713 | Pcsk5 | 18552 | protein_coding | 654.3597682 | -0.553678905 | 0.224292371 | 2.468558798 | 0.013565836 | 0.053943227 | 0.553678905 |
| ENSMUSG00000087444 | Gm5475 | 432982 | lncRNA | 7.017491925 | -0.553704743 | 0.6369725 | 0.869275743 | 0.384696327 | 0.607019295 | 0.553704743 |
| ENSMUSG00000031789 | Cngb1 | 333329 | protein_coding | 7.107627212 | -0.553734208 | 1.773273347 | 0.312266695 | 0.754837849 | 0.876856731 | 0.553734208 |
| ENSMUSG00000061048 | Cdh3 | 12560 | protein_coding | 6.972327754 | 0.553759129 | 0.520637688 | -1.063617064 | 0.287502209 | 0.50637921 | 0.553759129 |
| ENSMUSG00000038702 | Dsel | 319901 | protein_coding | 94.11444825 | 0.553876144 | 0.140134951 | -3.95244828 | 7.74E-05 | 0.000688653 | 0.553876144 |
| ENSMUSG00000062181 | Ces3b | 13909 | protein_coding | 1.160852324 | -0.553898484 | 1.540215971 | 0.359623906 | 0.719128403 | 1 | 0.553898484 |
| ENSMUSG00000021414 | Fam217a | 71864 | protein_coding | 3.647364968 | 0.553925582 | 0.865725723 | -0.639839579 | 0.522276898 | 1 | 0.553925582 |
| ENSMUSG00000033278 | Ptprm | 19274 | protein_coding | 64.79417742 | 0.554053145 | 0.206112384 | -2.688111862 | 0.007185732 | 0.032645166 | 0.554053145 |
| ENSMUSG00000015202 | Cnksr3 | 215748 | protein_coding | 353.1122566 | -0.554244482 | 0.141139741 | 3.926920084 | 8.60E-05 | 0.000754981 | 0.554244482 |
| ENSMUSG00000022860 | Chodl | 246048 | protein_coding | 11.07354859 | -0.554365257 | 0.483903348 | 1.145611534 | 0.251955923 | 0.464375257 | 0.554365257 |
| ENSMUSG00000034151 | Zbbx | 213234 | protein_coding | 1.643552009 | 0.554500283 | 2.27780939 | -0.24343577 | 0.807667836 | 1 | 0.554500283 |
| ENSMUSG00000019971 | Cep290 | 216274 | protein_coding | 239.5572713 | 0.554570361 | 0.094512302 | -5.867705537 | 4.42E-09 | 8.98E-08 | 0.554570361 |
| ENSMUSG00000024661 | Fth1 | 14319 | protein_coding | 10812.79316 | -0.554955622 | 0.138390486 | 4.010070616 | 6.07E-05 | 0.000555968 | 0.554955622 |
| ENSMUSG00000026258 | Snorc | 73234 | protein_coding | 4.805119132 | 0.555035832 | 0.68489499 | -0.81039552 | 0.417712892 | 0.63776911 | 0.555035832 |
| ENSMUSG00000031022 | BC051019 | 57355 | protein_coding | 2.330533305 | -0.555131892 | 0.843384297 | 0.658219384 | 0.510397171 | 1 | 0.555131892 |
| ENSMUSG00000036299 | BC031181 | 407819 | protein_coding | 974.693346 | 0.55534349 | 0.073371016 | -7.56897641 | 3.76E-14 | 1.29E-12 | 0.55534349 |
| ENSMUSG00000005481 | Ddx39a | 68278 | protein_coding | 799.4887701 | 0.555343722 | 0.096083021 | -5.779832034 | 7.48E-09 | 1.47E-07 | 0.555343722 |
| ENSMUSG00000042265 | Trem1 | 58217 | protein_coding | 2.157325546 | 0.555368883 | 1.465857336 | -0.378869669 | 0.704784649 | 1 | 0.555368883 |
| ENSMUSG00000045377 | Tmem88 | 67020 | protein_coding | 3.22537194 | -0.555381719 | 0.697660434 | 0.796063087 | 0.425995368 | 1 | 0.555381719 |
| ENSMUSG00000050232 | Cxcr3 | 12766 | protein_coding | 4.817952944 | 0.555482437 | 0.584816372 | -0.949840777 | 0.342193163 | 0.564082441 | 0.555482437 |
| ENSMUSG00000042476 | Abcb4 | 18670 | protein_coding | 4.442506335 | -0.555492136 | 0.840269023 | 0.661088438 | 0.5085556 | 0.712202101 | 0.555492136 |
| ENSMUSG00000037664 | Cdkn1c | 12577 | protein_coding | 118.6317483 | 0.555515277 | 0.218715306 | -2.539901236 | 0.011088378 | 0.046240775 | 0.555515277 |
| ENSMUSG00000036062 | Phf24 | 230085 | protein_coding | 2.19554064 | 0.555593914 | 1.336077232 | -0.415839669 | 0.677527329 | 1 | 0.555593914 |
| ENSMUSG00000028480 | Glipr2 | 384009 | protein_coding | 88.65112482 | -0.555836272 | 0.242626622 | 2.290912129 | 0.021968496 | 0.079002436 | 0.555836272 |
| ENSMUSG00000030865 | Chp2 | 70261 | protein_coding | 1471.909993 | -0.555846431 | 0.172832134 | 3.216105811 | 0.001299429 | 0.007903784 | 0.555846431 |
| ENSMUSG00000034959 | Rubcnl | 271221 | protein_coding | 2.064831693 | -0.555936055 | 1.293907742 | 0.429656641 | 0.667445428 | 1 | 0.555936055 |

| ensembl_gene_id | external_gene_name | entrezgene_id | gene_biotype | baseMean | log2FoldChange | lfcSE | stat | pvalue | padj | absLog2FC |
|---|---|---|---|---|---|---|---|---|---|---|
| ENSMUSG00000090394 | 4930523C07Rik | 67647 | protein_coding | 86.8960717 | -0.556252925 | 0.179245608 | 3.103300165 | 0.001913754 | 0.0109583 | 0.556252925 |
| ENSMUSG00000001750 | Tcirg1 | 27060 | protein_coding | 324.2696236 | -0.556270107 | 0.155246918 | 3.58313141 | 0.0003395 | 0.002539175 | 0.556270107 |
| ENSMUSG00000037348 | Paqr7 | 71904 | protein_coding | 34.37628223 | -0.556272197 | 0.180090031 | 3.088856129 | 0.002009287 | 0.01137907 | 0.556272197 |
| ENSMUSG00000049904 | Tmem17 | 103765 | protein_coding | 12.96498757 | 0.556371535 | 0.300500468 | -1.851483088 | 0.064100085 | 0.179258167 | 0.556371535 |
| ENSMUSG00000034245 | Hdac11 | 232232 | protein_coding | 323.1930195 | -0.556693217 | 0.113485248 | 4.905423636 | 9.32E-07 | 1.30E-05 | 0.556693217 |
| ENSMUSG00000026686 | Lmx1a | 110648 | protein_coding | 15.8347407 | -0.556879319 | 0.312644308 | 1.781191291 | 0.074881206 | 0.200044477 | 0.556879319 |
| ENSMUSG00000051235 | Gen1 | 209334 | protein_coding | 87.51029223 | 0.556944416 | 0.156627937 | -3.555843405 | 0.000376768 | 0.002788738 | 0.556944416 |
| ENSMUSG00000046245 | Pilra | 231805 | protein_coding | 0.954515715 | -0.557049385 | 1.454891548 | 0.38288035 | 0.701808481 | 1 | 0.557049385 |
| ENSMUSG00000020173 | Cobl | 12808 | protein_coding | 641.8647108 | -0.557096983 | 0.168398678 | 3.308202832 | 0.000938968 | 0.006038206 | 0.557096983 |
| ENSMUSG00000037376 | Trmt6 | 66926 | protein_coding | 234.5410527 | 0.557458166 | 0.098597411 | -5.653882387 | 1.57E-08 | 2.93E-07 | 0.557458166 |
| ENSMUSG00000105402 | Gm3716 | 100042198 | lncRNA | 9.742741721 | -0.557510072 | 0.395899391 | 1.408211491 | 0.159068455 | 0.343200478 | 0.557510072 |
| ENSMUSG00000104794 | Mir28c | 100628567 | miRNA | 0.577322777 | 0.557609492 | 1.476062017 | -0.377768336 | 0.705602698 | 1 | 0.557609492 |
| ENSMUSG00000009248 | Ascl2 | 17173 | protein_coding | 67.36472926 | 0.557635643 | 0.235176631 | -2.371135434 | 0.017733532 | 0.066693587 | 0.557635643 |
| ENSMUSG00000033594 | Spata2l | 78779 | protein_coding | 92.4907854 | -0.557777911 | 0.132827139 | 4.19927671 | 2.68E-05 | 0.000267672 | 0.557777911 |
| ENSMUSG00000025608 | Podxl | 27205 | protein_coding | 8.484783945 | 0.557885901 | 0.606283025 | -0.920174041 | 0.357481817 | 0.580534441 | 0.557885901 |
| ENSMUSG00000063887 | Nlgn1 | 192167 | protein_coding | 2.747693292 | 0.557997456 | 2.342020293 | -0.238254749 | 0.811683516 | 1 | 0.557997456 |
| ENSMUSG00000051344 | Plekhm3 | 241075 | protein_coding | 34.91699769 | -0.558260776 | 0.262325861 | 2.128119487 | 0.033327178 | 0.109351285 | 0.558260776 |
| ENSMUSG00000084821 | Gm15880 | 100504375 | lncRNA | 4.340207157 | -0.558399375 | 0.683132571 | 0.817409971 | 0.41369418 | 1 | 0.558399375 |
| ENSMUSG00000058900 | Rsl1 | 380855 | protein_coding | 58.24140801 | -0.558516054 | 0.168677536 | 3.311146626 | 0.000929145 | 0.005982028 | 0.558516054 |
| ENSMUSG00000024338 | Psmb8 | 16913 | protein_coding | 178.4772943 | -0.558530861 | 0.180744166 | 3.090173655 | 0.002000395 | 0.011340379 | 0.558530861 |
| ENSMUSG00000042787 | Exog | 208194 | protein_coding | 103.832786 | 0.558622832 | 0.117977601 | -4.734990601 | 2.19E-06 | 2.83E-05 | 0.558622832 |
| ENSMUSG00000040713 | Creg1 | 433375 | protein_coding | 542.4008661 | 0.558802346 | 0.110367691 | -5.063097205 | 4.12E-07 | 6.18E-06 | 0.558802346 |
| ENSMUSG00000067872 | Ccdc87 | 399599 | protein_coding | 1.476029626 | 0.559110095 | 1.370996664 | -0.407812878 | 0.683411058 | 1 | 0.559110095 |
| ENSMUSG00000039748 | Exo1 | 26909 | protein_coding | 91.70268717 | 0.559179316 | 0.177755575 | -3.145776528 | 0.001656465 | 0.009689999 | 0.559179316 |
| ENSMUSG00000034912 | Mdga2 | 320772 | protein_coding | 2.774717349 | -0.559315516 | 2.701181676 | 0.207063272 | 0.835960445 | 1 | 0.559315516 |
| ENSMUSG00000078492 | 1700045H11Rik | 73348 | lncRNA | 9.62801239 | -0.559353966 | 0.355194364 | 1.574782773 | 0.115306594 | 0.273824144 | 0.559353966 |
| ENSMUSG00000025545 | Clybl | 69634 | protein_coding | 332.6510199 | -0.559641152 | 0.082747578 | 6.763233041 | 1.35E-11 | 3.71E-10 | 0.559641152 |
| ENSMUSG00000019817 | Plagl1 | 22634 | protein_coding | 32.42381062 | 0.55972028 | 0.254372546 | -2.200395791 | 0.027778826 | 0.094898539 | 0.55972028 |
| ENSMUSG00000020366 | Mapk9 | 26420 | protein_coding | 694.0597389 | 0.559728718 | 0.082644325 | -6.772742299 | 1.26E-11 | 3.51E-10 | 0.559728718 |
| ENSMUSG00000003809 | Gcdh | 270076 | protein_coding | 693.7451715 | 0.559825317 | 0.102167738 | -5.479472589 | 4.27E-08 | 7.39E-07 | 0.559825317 |
| ENSMUSG00000040877 | Wdr25 | 212198 | protein_coding | 61.7115518 | -0.559893055 | 0.136766978 | 4.093773699 | 4.24E-05 | 0.000405126 | 0.559893055 |
| ENSMUSG00000062908 | Acadm | 11364 | protein_coding | 1460.239015 | -0.559956657 | 0.077553361 | 7.220275775 | 5.19E-13 | 1.62E-11 | 0.559956657 |
| ENSMUSG00000002076 | Hsf2bp | 74377 | protein_coding | 12.14003849 | 0.560003179 | 0.358951385 | -1.560108702 | 0.118734196 | 0.279831098 | 0.560003179 |
| ENSMUSG00000030340 | Scnn1a | 20276 | protein_coding | 622.610839 | -0.56012119 | 0.142716752 | 3.924705286 | 8.68E-05 | 0.000761151 | 0.56012119 |
| ENSMUSG00000032093 | Cd3e | 12501 | protein_coding | 9.881757897 | -0.560137202 | 0.422808252 | 1.324801962 | 0.185236844 | 0.380604488 | 0.560137202 |
| ENSMUSG00000071454 | Dtnb | 13528 | protein_coding | 78.19295699 | 0.560275799 | 0.149331424 | -3.751894828 | 0.000175503 | 0.001413095 | 0.560275799 |
| ENSMUSG00000022220 | Adcy4 | 104110 | protein_coding | 5.176028415 | -0.56054962 | 1.199761337 | 0.467217606 | 0.640344198 | 0.803678477 | 0.56054962 |
| ENSMUSG00000042797 | Aqp11 | 66333 | protein_coding | 198.6880231 | -0.560682123 | 0.097547442 | 5.747789087 | 9.04E-09 | 1.75E-07 | 0.560682123 |
| ENSMUSG00000022899 | Slc15a2 | 57738 | protein_coding | 65.06354121 | 0.560739804 | 0.625865249 | -0.895943344 | 0.370283023 | 0.592811812 | 0.560739804 |

| ensembl_gene_id | external_gene_name | entrezgene_id | gene_biotype | baseMean | log2FoldChange | lfcSE | stat | pvalue | padj | absLog2FC |
|---|---|---|---|---|---|---|---|---|---|---|
| ENSMUSG00000039396 | Neil3 | 234258 | protein_coding | 59.16425014 | 0.560784771 | 0.256790509 | -2.183822026 | 0.028975328 | 0.098234156 | 0.560784771 |
| ENSMUSG00000022311 | Csmd3 | 239420 | protein_coding | 4.174836806 | 0.560991442 | 2.031278405 | -0.27617654 | 0.782412475 | 1 | 0.560991442 |
| ENSMUSG00000048330 | Ric3 | 320360 | protein_coding | 12.90229425 | -0.561225558 | 0.332595061 | 1.687413987 | 0.091523776 | 0.231529357 | 0.561225558 |
| ENSMUSG00000054423 | Cadps | 27062 | protein_coding | 20.67589415 | -0.561476645 | 0.417116493 | 1.346090731 | 0.178273258 | 0.370594735 | 0.561476645 |
| ENSMUSG00000061740 | Cyp2d22 | 56448 | protein_coding | 150.0555051 | -0.562030869 | 0.184212338 | 3.050994713 | 0.002280846 | 0.012597287 | 0.562030869 |
| ENSMUSG00000036030 | Prtg | 235472 | protein_coding | 2.591025502 | -0.562097334 | 1.059488781 | 0.530536372 | 0.595740098 | 1 | 0.562097334 |
| ENSMUSG00000022967 | Ifnar1 | 15975 | protein_coding | 640.0862127 | 0.562231304 | 0.059289352 | -9.482837807 | 2.47E-21 | 1.36E-19 | 0.562231304 |
| ENSMUSG00000059003 | Grin2a | 14811 | protein_coding | 5.582863867 | 0.562239646 | 2.098701654 | -0.267898796 | 0.788777221 | 0.894265208 | 0.562239646 |
| ENSMUSG00000066357 | Wdr6 | 83669 | protein_coding | 249.8663839 | 0.562252352 | 0.102041544 | -5.510033742 | 3.59E-08 | 6.31E-07 | 0.562252352 |
| ENSMUSG00000079641 | Rpl39 | 67248 | protein_coding | 1808.358923 | -0.562380217 | 0.17880422 | 3.145229001 | 0.001659569 | 0.009704716 | 0.562380217 |
| ENSMUSG00000069792 | Wfdc17 | 100034251 | protein_coding | 27.47337603 | 0.562392989 | 0.268806598 | -2.092184471 | 0.036422017 | 0.116927137 | 0.562392989 |
| ENSMUSG00000019738 | Polr2i | 69920 | protein_coding | 270.6146966 | 0.562399853 | 0.0997955 | -5.635523182 | 1.75E-08 | 3.24E-07 | 0.562399853 |
| ENSMUSG00000002228 | Ppm1j | 71887 | protein_coding | 79.85912106 | -0.562407619 | 0.355750885 | 1.580902938 | 0.113900233 | 0.271186338 | 0.562407619 |
| ENSMUSG00000031672 | Got2 | 14719 | protein_coding | 885.6093272 | 0.562417364 | 0.101568916 | -5.537298071 | 3.07E-08 | 5.45E-07 | 0.562417364 |
| ENSMUSG00000031790 | Mmp15 | 17388 | protein_coding | 1595.959359 | -0.562535022 | 0.103594764 | 5.430149156 | 5.63E-08 | 9.57E-07 | 0.562535022 |
| ENSMUSG00000025465 | Echs1 | 93747 | protein_coding | 1475.0035 | -0.562855336 | 0.072699543 | 7.742212891 | 9.77E-15 | 3.50E-13 | 0.562855336 |
| ENSMUSG00000048349 | Pou4f1 | 18996 | protein_coding | 7.857423422 | 0.562859809 | 0.489181463 | -1.150615573 | 0.249890423 | 0.462129017 | 0.562859809 |
| ENSMUSG00000001211 | Agpat3 | 28169 | protein_coding | 718.1640128 | 0.562901964 | 0.049733369 | -11.31839602 | 1.06E-29 | 8.17E-28 | 0.562901964 |
| ENSMUSG00000020573 | Pik3cg | 30955 | protein_coding | 123.5724372 | -0.562926674 | 0.162912109 | 3.455401068 | 0.000549475 | 0.003837834 | 0.562926674 |
| ENSMUSG00000070802 | Pnma8b | 434128 | protein_coding | 33.46259719 | -0.563002711 | 0.20105795 | 2.800201192 | 0.005107077 | 0.024790788 | 0.563002711 |
| ENSMUSG00000027208 | Fgf7 | 14178 | protein_coding | 1.168695729 | 0.563054826 | 1.456781096 | -0.386506131 | 0.699121863 | 1 | 0.563054826 |
| ENSMUSG00000071113 | Mboat4 | 234155 | protein_coding | 2.685481267 | 0.563284862 | 0.714502459 | -0.788359585 | 0.430486403 | 1 | 0.563284862 |
| ENSMUSG00000009731 | Kcnd1 | 16506 | protein_coding | 7.689911548 | -0.56331929 | 0.57365511 | 0.981982521 | 0.32610846 | 0.548127746 | 0.56331929 |
| ENSMUSG00000073888 | Ccl27a | 20301 | protein_coding | 14.44288015 | -0.563348988 | 0.309868743 | 1.818024571 | 0.069060377 | 0.189172946 | 0.563348988 |
| ENSMUSG00000069170 | Adgrv1 | 110789 | protein_coding | 13.30078587 | 0.563400664 | 0.693587969 | -0.812298784 | 0.416620208 | 0.636776036 | 0.563400664 |
| ENSMUSG00000024727 | Trpm6 | 225997 | protein_coding | 1954.262229 | 0.563489158 | 0.213068132 | -2.644643071 | 0.008177713 | 0.036254161 | 0.563489158 |
| ENSMUSG00000024909 | Efemp2 | 58859 | protein_coding | 20.52269929 | 0.563523196 | 0.283133115 | -1.990311856 | 0.046556593 | 0.141278128 | 0.563523196 |
| ENSMUSG00000092130 | D030025P21Rik | 100303738 | lncRNA | 1.583223529 | -0.563556867 | 1.637844103 | 0.344084559 | 0.730782694 | 1 | 0.563556867 |
| ENSMUSG00000046058 | Eid2 | 386655 | protein_coding | 11.96390511 | 0.563710593 | 0.350012595 | -1.610543739 | 0.107279205 | 0.260339278 | 0.563710593 |
| ENSMUSG00000037369 | Kdm6a | 22289 | protein_coding | 1074.801903 | 0.564064706 | 0.063165556 | -8.929941265 | 4.26E-19 | 2.07E-17 | 0.564064706 |
| ENSMUSG00000031383 | Dusp9 | 75590 | protein_coding | 23.0779186 | 0.564103837 | 0.368813862 | -1.529508231 | 0.1261385 | 0.292089426 | 0.564103837 |
| ENSMUSG00000033268 | Duox1 | 99439 | protein_coding | 11.65184292 | 0.564164703 | 0.34423399 | -1.638898887 | 0.10123432 | 0.249818224 | 0.564164703 |
| ENSMUSG00000038987 | Cfap157 | 227736 | protein_coding | 67.45051899 | 0.564253254 | 0.221341779 | -2.549239719 | 0.010795806 | 0.045254367 | 0.564253254 |
| ENSMUSG00000107455 | Gm43868 | 115490363 | TEC | 10.08060048 | -0.564437136 | 0.377649761 | 1.494604776 | 0.135017619 | 0.306022641 | 0.564437136 |
| ENSMUSG00000030518 | Entrep2 | 70638 | protein_coding | 39.05247276 | -0.564496836 | 0.247417085 | 2.281559642 | 0.022515352 | 0.080637284 | 0.564496836 |
| ENSMUSG00000012819 | Cdh23 | 22295 | protein_coding | 15.07462475 | 0.564561708 | 0.774785683 | -0.728668225 | 0.466204634 | 0.679754841 | 0.564561708 |
| ENSMUSG00000046818 | Ddit4l | 73284 | protein_coding | 4.231561489 | 0.564700978 | 0.496797602 | -1.136682173 | 0.255671175 | 1 | 0.564700978 |
| ENSMUSG00000039801 | Cplane1 | 73692 | protein_coding | 92.34640547 | 0.564715605 | 0.209256264 | -2.698679571 | 0.006961517 | 0.031766369 | 0.564715605 |
| ENSMUSG00000078931 | Pdf | 68023 | protein_coding | 3.251515795 | -0.564898346 | 0.571382314 | 0.988652138 | 0.322833368 | 1 | 0.564898346 |

| ensembl_gene_id | external_gene_name | entrezgene_id | gene_biotype | baseMean | log2FoldChange | lfcSE | stat | pvalue | padj | absLog2FC |
|---|---|---|---|---|---|---|---|---|---|---|
| ENSMUSG00000046093 | Hpcal4 | 170638 | protein_coding | 5.085120462 | -0.565128959 | 0.83641931 | 0.675652693 | 0.499261178 | 0.704201299 | 0.565128959 |
| ENSMUSG00000090235 | Gm16244 | 102637720 | lncRNA | 2.06572292 | 0.565195897 | 0.972328905 | -0.581280567 | 0.561051375 | 1 | 0.565195897 |
| ENSMUSG00000070577 | Gm572 | 230909 | protein_coding | 1.498739165 | -0.565232441 | 1.183530234 | 0.477581751 | 0.632947922 | 1 | 0.565232441 |
| ENSMUSG00000034116 | Vav1 | 22324 | protein_coding | 105.7930336 | -0.565368244 | 0.130095083 | 4.345807937 | 1.39E-05 | 0.000149189 | 0.565368244 |
| ENSMUSG00000098065 | Gm5177 | 382450 | transcribed_processed_pseudogene | 2.25863705 | 0.565429753 | 1.142945208 | -0.494712913 | 0.620802785 | 1 | 0.565429753 |
| ENSMUSG00000092713 | Gm22858 | 115488835 | snoRNA | 1.466232724 | -0.565698695 | 0.933101552 | 0.606256301 | 0.544344572 | 1 | 0.565698695 |
| ENSMUSG00000099378 | 1700067G17Rik | 73475 | lncRNA | 0.79119059 | -0.566052803 | 1.638736962 | 0.345420172 | 0.729778514 | 1 | 0.566052803 |
| ENSMUSG00000060402 | Chst8 | 68947 | protein_coding | 22.34058676 | 0.566166302 | 0.347989446 | -1.626964001 | 0.103744744 | 0.254227734 | 0.566166302 |
| ENSMUSG00000020798 | Spns3 | 77577 | protein_coding | 3.837419839 | -0.566211581 | 0.984246545 | 0.575274136 | 0.565105911 | 1 | 0.566211581 |
| ENSMUSG00000017146 | Brca1 | 12189 | protein_coding | 177.7352296 | 0.566276424 | 0.170854022 | -3.314387441 | 0.000918441 | 0.00592004 | 0.566276424 |
| ENSMUSG00000086126 | Evx1os | 70593 | lncRNA | 7.667362019 | 0.566318329 | 0.361323508 | -1.567344269 | 0.117034243 | 0.276733031 | 0.566318329 |
| ENSMUSG00000034773 | Hrob | 217216 | protein_coding | 29.88968162 | 0.566360625 | 0.224922062 | -2.518030553 | 0.011801308 | 0.048455198 | 0.566360625 |
| ENSMUSG00000028364 | Tnc | 21923 | protein_coding | 4.694568534 | 0.566471238 | 0.763843246 | -0.741606659 | 0.458325688 | 0.67278137 | 0.566471238 |
| ENSMUSG00000026664 | Phyh | 16922 | protein_coding | 2156.069756 | 0.566557303 | 0.061530209 | -9.20779098 | 3.33E-20 | 1.69E-18 | 0.566557303 |
| ENSMUSG00000042251 | Pm20d1 | 212933 | protein_coding | 191.075585 | -0.566689389 | 0.140271102 | 4.039958211 | 5.35E-05 | 0.000495149 | 0.566689389 |
| ENSMUSG00000060743 | H3f3a | 15078 | protein_coding | 1511.43576 | -0.566694806 | 0.128724687 | 4.402378576 | 1.07E-05 | 0.000118988 | 0.566694806 |
| ENSMUSG00000102101 | Zbtb11os1 | 66391 | lncRNA | 11.37648257 | 0.566837705 | 0.320400824 | -1.769151836 | 0.076868539 | 0.204314673 | 0.566837705 |
| ENSMUSG00000042348 | Arl15 | 218639 | protein_coding | 108.6586598 | 0.566854012 | 0.121895815 | -4.650315616 | 3.31E-06 | 4.11E-05 | 0.566854012 |
| ENSMUSG00000031838 | Ifi30 | 65972 | protein_coding | 909.3986556 | -0.566961178 | 0.108216683 | 5.239129132 | 1.61E-07 | 2.57E-06 | 0.566961178 |
| ENSMUSG00000079436 | Kcnj13 | 100040591 | protein_coding | 6.218126507 | -0.566974342 | 0.450570194 | 1.258348531 | 0.208265737 | 0.41130117 | 0.566974342 |
| ENSMUSG00000095061 | E030018B13Rik | 381994 | protein_coding | 12.34472938 | -0.566976216 | 0.316130304 | 1.793488977 | 0.072894772 | 0.196696774 | 0.566976216 |
| ENSMUSG00000078490 | Cfap74 | 544678 | protein_coding | 30.15859224 | 0.567348782 | 0.539123813 | -1.052353408 | 0.292637437 | 0.511766878 | 0.567348782 |
| ENSMUSG00000085833 | Gm13003 | 100503595 | lncRNA | 1.537995978 | -0.567356641 | 1.544205751 | 0.367410004 | 0.713313211 | 1 | 0.567356641 |
| ENSMUSG00000031129 | Slc9a9 | 331004 | protein_coding | 16.48853191 | -0.567504468 | 0.303591911 | 1.86930036 | 0.061581038 | 0.173926674 | 0.567504468 |
| ENSMUSG00000062859 | Tcp11 | 21463 | protein_coding | 9.602118481 | -0.567563051 | 0.614418175 | 0.923740661 | 0.355621357 | 0.578423233 | 0.567563051 |
| ENSMUSG00000062683 | Atp5g2 | 67942 | protein_coding | 465.5268885 | -0.567568289 | 0.179323032 | 3.165060738 | 0.001550506 | 0.009157744 | 0.567568289 |
| ENSMUSG00000024542 | Cep192 | 70799 | protein_coding | 427.1563159 | 0.567637928 | 0.12396205 | -4.579126653 | 4.67E-06 | 5.62E-05 | 0.567637928 |
| ENSMUSG00000031972 | Acta1 | 11459 | protein_coding | 232.96575 | 0.567640258 | 0.309168695 | -1.836021136 | 0.066354531 | 0.183701378 | 0.567640258 |
| ENSMUSG00000029201 | Ugdh | 22235 | protein_coding | 6006.739695 | -0.567940653 | 0.120660303 | 4.706938715 | 2.51E-06 | 3.20E-05 | 0.567940653 |
| ENSMUSG00000097639 | Platr4 | 78706 | lncRNA | 2.340459414 | -0.56801265 | 1.003688599 | 0.565925179 | 0.571444644 | 1 | 0.56801265 |
| ENSMUSG00000050545 | Fam228b | 207921 | protein_coding | 2.119335428 | -0.568205562 | 0.910804543 | 0.623850163 | 0.532725993 | 1 | 0.568205562 |
| ENSMUSG00000073856 | Iqck | 434232 | protein_coding | 5.458315499 | -0.568273778 | 0.586443747 | 0.969016689 | 0.332536862 | 0.55497865 | 0.568273778 |
| ENSMUSG00000041920 | Slc16a6 | 104681 | protein_coding | 89.89744198 | 0.568850848 | 0.170935131 | -3.327875582 | 0.000875109 | 0.005694853 | 0.568850848 |
| ENSMUSG00000004127 | Trmt10a | 108943 | protein_coding | 59.4298527 | 0.568862593 | 0.182672496 | -3.114111894 | 0.001844995 | 0.010614744 | 0.568862593 |
| ENSMUSG00000092060 | Bend4 | 666938 | protein_coding | 3.750550234 | -0.569086141 | 1.052464246 | 0.54071779 | 0.588702115 | 1 | 0.569086141 |
| ENSMUSG00000028545 | Bend5 | 67621 | protein_coding | 3.741493506 | -0.569104155 | 0.760397647 | 0.748429664 | 0.454201036 | 1 | 0.569104155 |
| ENSMUSG00000065434 | Mir7-1 | 723902 | miRNA | 5.887883357 | -0.569246782 | 0.495295079 | 1.149308373 | 0.250428847 | 0.462725114 | 0.569246782 |
| ENSMUSG00000087330 | Bloodlinc | 105463053 | lncRNA | 7.612848531 | 0.569251038 | 0.469765725 | -1.211776441 | 0.225597969 | 0.433049501 | 0.569251038 |
| ENSMUSG00000024172 | St6gal2 | 240119 | protein_coding | 3.366691636 | -0.569471219 | 0.758004376 | 0.751276954 | 0.452485996 | 1 | 0.569471219 |

| ensembl_gene_id | external_gene_name | entrezgene_id | gene_biotype | baseMean | log2FoldChange | lfcSE | stat | pvalue | padj | absLog2FC |
|---|---|---|---|---|---|---|---|---|---|---|
| ENSMUSG00000040722 | Scamp5 | 56807 | protein_coding | 527.6834062 | -0.56951309 | 0.088054636 | 6.467724061 | 9.95E-11 | 2.47E-09 | 0.56951309 |
| ENSMUSG00000059839 | Zfp874b | 408067 | protein_coding | 54.15107147 | -0.569898157 | 0.171712415 | 3.318910618 | 0.000903694 | 0.005843912 | 0.569898157 |
| ENSMUSG00000021242 | Npc2 | 67963 | protein_coding | 1966.272651 | -0.570144575 | 0.151618698 | 3.760384333 | 0.000169653 | 0.001371337 | 0.570144575 |
| ENSMUSG00000097482 | Gm17634 | 100504568 | lncRNA | 6.919700811 | -0.570239822 | 0.528593737 | 1.078786565 | 0.280682885 | 0.498301136 | 0.570239822 |
| ENSMUSG00000093042 | Mir3109 | 100526560 | miRNA | 18.60094354 | -0.570251625 | 0.285664497 | 1.99622855 | 0.045909051 | 0.13974969 | 0.570251625 |
| ENSMUSG00000086962 | Gm12248 | 676527 | protein_coding | 5.038033725 | -0.570325805 | 0.592135794 | 0.96316725 | 0.335463597 | 0.557777472 | 0.570325805 |
| ENSMUSG00000026974 | Zmynd19 | 67187 | protein_coding | 235.5425989 | 0.570366321 | 0.123696051 | -4.611030971 | 4.01E-06 | 4.90E-05 | 0.570366321 |
| ENSMUSG00000086228 | Ubap1l | 208080 | protein_coding | 3.535532308 | 0.5703913 | 0.684958707 | -0.832738229 | 0.404992379 | 1 | 0.5703913 |
| ENSMUSG00000038843 | Gcnt1 | 14537 | protein_coding | 325.0870474 | -0.570754986 | 0.102477608 | 5.569558052 | 2.55E-08 | 4.60E-07 | 0.570754986 |
| ENSMUSG00000099139 | 1110028F18Rik | 68692 | lncRNA | 0.789063942 | -0.570780949 | 1.455694878 | 0.392102052 | 0.694982811 | 1 | 0.570780949 |
| ENSMUSG00000010175 | Prox1 | 19130 | protein_coding | 25.16933408 | 0.570917658 | 0.24630406 | -2.317938475 | 0.020452665 | 0.074658557 | 0.570917658 |
| ENSMUSG00000001930 | Vwf | 22371 | protein_coding | 45.23177969 | 0.570943865 | 0.269664842 | -2.117235083 | 0.034239901 | 0.111570191 | 0.570943865 |
| ENSMUSG00000096883 | Shisa8 | 435145 | protein_coding | 6.608708716 | 0.570947653 | 0.578802006 | -0.986429985 | 0.323922157 | 0.545730514 | 0.570947653 |
| ENSMUSG00000067297 | Ifit1bl2 | 112419 | protein_coding | 93.28629514 | -0.57099506 | 0.376724103 | 1.515684966 | 0.129599031 | 0.297320449 | 0.57099506 |
| ENSMUSG00000087547 | Platr27 | 78316 | lncRNA | 2.097617941 | -0.571127052 | 0.766240375 | 0.745362774 | 0.456052441 | 1 | 0.571127052 |
| ENSMUSG00000075377 | Or1b1 | 259053 | protein_coding | 1.46447461 | 0.571474164 | 2.644047923 | -0.216136084 | 0.828881673 | 1 | 0.571474164 |
| ENSMUSG00000054057 | A930004D18Rik | 77940 | lncRNA | 10.36926843 | -0.571531775 | 0.341012805 | 1.675983327 | 0.093741458 | 0.235723563 | 0.571531775 |
| ENSMUSG00000035042 | Ccl5 | 20304 | protein_coding | 27.46207979 | 0.57179397 | 0.370561935 | -1.543045619 | 0.122819708 | 0.286151898 | 0.57179397 |
| ENSMUSG00000052102 | Gnpda1 | 26384 | protein_coding | 181.1823804 | -0.571897071 | 0.152167905 | 3.758329133 | 0.000171052 | 0.001381295 | 0.571897071 |
| ENSMUSG00000092035 | Peg10 | 170676 | protein_coding | 5.23423079 | 0.571933738 | 1.16075717 | -0.492724709 | 0.622207117 | 0.792833975 | 0.571933738 |
| ENSMUSG00000064620 | Gm22303 | 115488066 | snoRNA | 1.300159392 | 0.572079206 | 1.251475642 | -0.457123724 | 0.647582119 | 1 | 0.572079206 |
| ENSMUSG00000048578 | Mlec | 109154 | protein_coding | 8055.14432 | 0.572127174 | 0.103273061 | -5.539945928 | 3.03E-08 | 5.38E-07 | 0.572127174 |
| ENSMUSG00000061972 | Or1o4 | 258508 | protein_coding | 2.889299291 | -0.572352393 | 0.92299527 | 0.620103278 | 0.535189794 | 1 | 0.572352393 |
| ENSMUSG00000052372 | Il1rapl1 | 331461 | protein_coding | 2.437061672 | 0.572487634 | 1.957927443 | -0.292394713 | 0.76998485 | 1 | 0.572487634 |
| ENSMUSG00000051682 | Treml4 | 224840 | protein_coding | 1.020135995 | 0.57251034 | 2.391110192 | -0.239432855 | 0.810769956 | 1 | 0.57251034 |
| ENSMUSG00000056812 | St8sia3 | 20451 | protein_coding | 2.808816487 | 0.572585409 | 2.462795888 | -0.232494058 | 0.816154302 | 1 | 0.572585409 |
| ENSMUSG00000041483 | Zfp281 | 226442 | protein_coding | 116.2628982 | 0.572593861 | 0.129715261 | -4.414236658 | 1.01E-05 | 0.000113413 | 0.572593861 |
| ENSMUSG00000070576 | Mn1 | 433938 | protein_coding | 8.939216877 | -0.572598878 | 0.576674379 | 0.992932751 | 0.320742728 | 0.541897747 | 0.572598878 |
| ENSMUSG00000100002 | Gm29642 | 102631685 | lncRNA | 1.900137887 | -0.5729566 | 0.79064498 | 0.724669876 | 0.46865458 | 1 | 0.5729566 |
| ENSMUSG00000058558 | Rpl5 | 100503670 | protein_coding | 3587.320565 | -0.573032684 | 0.167469089 | 3.421722097 | 0.000622259 | 0.004258592 | 0.573032684 |
| ENSMUSG00000020086 | Macroh2a2 | 404634 | protein_coding | 15.81137237 | 0.573274029 | 0.298994663 | -1.917338668 | 0.05519492 | 0.160445154 | 0.573274029 |
| ENSMUSG00000091794 | Vmn2r18 | 632671 | protein_coding | 2.12421253 | 0.573337695 | 1.809104947 | -0.316917875 | 0.751305918 | 1 | 0.573337695 |
| ENSMUSG00000020057 | Dram1 | 71712 | protein_coding | 1.411066823 | -0.573362416 | 1.274208884 | 0.449975214 | 0.652728312 | 1 | 0.573362416 |
| ENSMUSG00000069274 | H4c6 | 319157 | protein_coding | 1443.234425 | 0.573575679 | 0.208696036 | -2.748378407 | 0.005989084 | 0.028137622 | 0.573575679 |
| ENSMUSG00000023914 | Mep1a | 17287 | protein_coding | 4138.69259 | -0.57363312 | 0.115889652 | 4.949821741 | 7.43E-07 | 1.06E-05 | 0.57363312 |
| ENSMUSG00000074345 | Tnfaip8l3 | 244882 | protein_coding | 1.751349089 | 0.573664545 | 2.111751358 | -0.271653451 | 0.785888496 | 1 | 0.573664545 |
| ENSMUSG00000058244 | Or5p78 | 258215 | protein_coding | 1.474240336 | 0.573716614 | 2.101638486 | -0.272985396 | 0.784864442 | 1 | 0.573716614 |
| ENSMUSG00000027316 | Gfra4 | 14588 | protein_coding | 0.894158328 | 0.573749941 | 2.095552437 | -0.273794123 | 0.784242841 | 1 | 0.573749941 |
| ENSMUSG00000024903 | Lao1 | 100470 | protein_coding | 1.015884918 | 0.573823015 | 2.080078992 | -0.275865973 | 0.78265101 | 1 | 0.573823015 |

| ensembl_gene_id | external_gene_name | entrezgene_id | gene_biotype | baseMean | log2FoldChange | lfcSE | stat | pvalue | padj | absLog2FC |
|---|---|---|---|---|---|---|---|---|---|---|
| ENSMUSG00000069255 | Dusp22 | 105352 | protein_coding | 47.35275688 | 0.573922108 | 0.207688946 | -2.763373403 | 0.005720728 | 0.027207712 | 0.573922108 |
| ENSMUSG00000028608 | Czib | 74098 | protein_coding | 299.0997567 | 0.573999054 | 0.068103137 | -8.428379026 | 3.50E-17 | 1.50E-15 | 0.573999054 |
| ENSMUSG00000097744 | D030040B21Rik | 329101 | lncRNA | 0.858716145 | 0.574051576 | 2.039713674 | -0.281437333 | 0.778374987 | 1 | 0.574051576 |
| ENSMUSG00000020411 | Nipal4 | 214112 | protein_coding | 1.1500646 | 0.574209913 | 2.01201312 | -0.28539074 | 0.775344801 | 1 | 0.574209913 |
| ENSMUSG00000038963 | Slco4a1 | 108115 | protein_coding | 76.2438406 | -0.574474184 | 0.193635307 | 2.966784279 | 0.00300932 | 0.015912877 | 0.574474184 |
| ENSMUSG00000052565 | H1f3 | 14957 | protein_coding | 3036.405989 | 0.574496049 | 0.147108124 | -3.905263923 | 9.41E-05 | 0.000816355 | 0.574496049 |
| ENSMUSG00000050195 | Scd4 | 329065 | protein_coding | 4.379974526 | 0.574519014 | 0.857408618 | -0.670064426 | 0.502816721 | 1 | 0.574519014 |
| ENSMUSG00000064177 | Ghrl | 58991 | protein_coding | 10.96325217 | 0.574555746 | 0.344292821 | -1.668799669 | 0.095157088 | 0.238311216 | 0.574555746 |
| ENSMUSG00000057561 | Eif1a | 13664 | protein_coding | 1137.613111 | 0.574569815 | 0.084877032 | -6.769438103 | 1.29E-11 | 3.58E-10 | 0.574569815 |
| ENSMUSG00000005125 | Ndrg1 | 17988 | protein_coding | 1258.190956 | -0.574661273 | 0.110222644 | 5.213640811 | 1.85E-07 | 2.92E-06 | 0.574661273 |
| ENSMUSG00000041891 | Lman1 | 70361 | protein_coding | 1199.054393 | 0.574678953 | 0.151470154 | -3.794007851 | 0.000148235 | 0.001217281 | 0.574678953 |
| ENSMUSG00000109776 | Gm30400 | 102632284 | lncRNA | 1.213284915 | 0.574709928 | 1.932499711 | -0.297391987 | 0.766167261 | 1 | 0.574709928 |
| ENSMUSG00000097855 | A930007I19Rik | 77779 | lncRNA | 9.109035024 | 0.574746681 | 0.527967357 | -1.088602682 | 0.276329134 | 0.493321208 | 0.574746681 |
| ENSMUSG00000054967 | Zfp647 | 239546 | protein_coding | 9.153234316 | -0.574765446 | 0.440972201 | 1.303405169 | 0.192436473 | 0.390165244 | 0.574765446 |
| ENSMUSG00000031756 | Cenpn | 72155 | protein_coding | 44.64151667 | 0.574765665 | 0.225505976 | -2.548782408 | 0.010809972 | 0.045254367 | 0.574765665 |
| ENSMUSG00000085886 | D030047H15Rik | 100037396 | lncRNA | 1.401581125 | -0.57510283 | 1.027417794 | 0.559755567 | 0.575646175 | 1 | 0.57510283 |
| ENSMUSG00000078532 | Nkain1 | 67149 | protein_coding | 95.95467469 | 0.575226853 | 0.154042379 | -3.734211704 | 0.000188304 | 0.001502974 | 0.575226853 |
| ENSMUSG00000041268 | Dmxl2 | 235380 | protein_coding | 32.48155368 | -0.575338649 | 0.263016977 | 2.187458218 | 0.02870909 | 0.097491654 | 0.575338649 |
| ENSMUSG00000001985 | Grik3 | 14807 | protein_coding | 7.552280681 | 0.575373878 | 1.348257085 | -0.426753833 | 0.669558636 | 0.822609935 | 0.575373878 |
| ENSMUSG00000059213 | Ddn | 13199 | protein_coding | 0.814631172 | 0.575495109 | 1.826218589 | -0.315129368 | 0.752663431 | 1 | 0.575495109 |
| ENSMUSG00000026077 | Npas2 | 18143 | protein_coding | 292.8365715 | 0.575570935 | 0.223764602 | -2.572216202 | 0.010104977 | 0.04294225 | 0.575570935 |
| ENSMUSG00000042269 | Cibar2 | 436062 | protein_coding | 2.022611009 | 0.575623305 | 0.885260443 | -0.65023046 | 0.515543368 | 1 | 0.575623305 |
| ENSMUSG00000097748 | Gm26533 | 115488895 | lncRNA | 1.927125016 | -0.575632906 | 1.204659981 | 0.477838489 | 0.632765164 | 1 | 0.575632906 |
| ENSMUSG00000034892 | Rps29 | 20090 | protein_coding | 2946.534189 | -0.575773312 | 0.163642071 | 3.518491963 | 0.000434007 | 0.003154573 | 0.575773312 |
| ENSMUSG00000038502 | Ptov1 | 84113 | protein_coding | 683.1220831 | 0.57598415 | 0.07348344 | -7.838285047 | 4.57E-15 | 1.67E-13 | 0.57598415 |
| ENSMUSG00000003273 | Car11 | 12348 | protein_coding | 12.82622922 | -0.576051022 | 0.363779017 | 1.583519101 | 0.113303197 | 0.270127107 | 0.576051022 |
| ENSMUSG00000087030 | Gm16143 | 102632251 | lncRNA | 5.411854269 | -0.576060354 | 0.502520899 | 1.146341088 | 0.251654048 | 0.464129434 | 0.576060354 |
| ENSMUSG00000038205 | Prkab2 | 108097 | protein_coding | 39.15043045 | -0.576131795 | 0.193626604 | 2.975478487 | 0.002925319 | 0.015543344 | 0.576131795 |
| ENSMUSG00000050493 | Fam167b | 230766 | protein_coding | 2.046811566 | -0.576253798 | 1.094055264 | 0.52671361 | 0.598392483 | 1 | 0.576253798 |
| ENSMUSG00000032796 | Lama1 | 16772 | protein_coding | 2.960819993 | 0.576362611 | 2.183547339 | -0.263957003 | 0.791813051 | 1 | 0.576362611 |
| ENSMUSG00000034452 | Slc24a1 | 214111 | protein_coding | 3.5209167 | 0.576558323 | 1.508816504 | -0.382126204 | 0.702367756 | 1 | 0.576558323 |
| ENSMUSG00000069072 | Slc7a14 | 241919 | protein_coding | 1.640543333 | 0.576691348 | 1.4896724 | -0.387126289 | 0.698662716 | 1 | 0.576691348 |
| ENSMUSG00000065954 | Tacc1 | 320165 | protein_coding | 121.256068 | 0.576863487 | 0.159299672 | -3.621247178 | 0.000293186 | 0.002234472 | 0.576863487 |
| ENSMUSG00000050296 | Abca12 | 74591 | protein_coding | 765.1311478 | 0.576938512 | 0.257754714 | -2.238323799 | 0.025199947 | 0.087888475 | 0.576938512 |
| ENSMUSG00000029861 | Fam131b | 76156 | protein_coding | 1.456091127 | -0.577063309 | 1.860929976 | 0.310094048 | 0.756489438 | 1 | 0.577063309 |
| ENSMUSG00000024066 | Xdh | 22436 | protein_coding | 1007.905707 | 0.577077496 | 0.162505682 | -3.55112197 | 0.000383593 | 0.002824186 | 0.577077496 |
| ENSMUSG00000090891 | D6Ertd527e | 52372 | lncRNA | 4.802405877 | -0.577182737 | 0.727743203 | 0.793113195 | 0.427711876 | 0.647057325 | 0.577182737 |
| ENSMUSG00000048118 | Arid4a | 238247 | protein_coding | 720.3428085 | 0.57723544 | 0.102684195 | -5.62146336 | 1.89E-08 | 3.49E-07 | 0.57723544 |
| ENSMUSG00000026834 | Acvr1c | 269275 | protein_coding | 170.5386175 | -0.577390317 | 0.15479721 | 3.729978844 | 0.000191496 | 0.001522563 | 0.577390317 |

| ensembl_gene_id | external_gene_name | entrezgene_id | gene_biotype | baseMean | log2FoldChange | lfcSE | stat | pvalue | padj | absLog2FC |
|---|---|---|---|---|---|---|---|---|---|---|
| ENSMUSG00000030935 | Acsm3 | 20216 | protein_coding | 1775.311301 | -0.57749439 | 0.242676331 | 2.379689802 | 0.017327217 | 0.065583696 | 0.57749439 |
| ENSMUSG00000030346 | Rad51ap1 | 19362 | protein_coding | 62.2111568 | 0.577606627 | 0.243065888 | -2.376337671 | 0.017485453 | 0.06604638 | 0.577606627 |
| ENSMUSG00000028524 | Sgip1 | 73094 | protein_coding | 3.65099241 | 0.577611265 | 2.227377866 | -0.259323428 | 0.795385704 | 1 | 0.577611265 |
| ENSMUSG00000024270 | Slc39a6 | 106957 | protein_coding | 44.524232 | 0.577963723 | 0.182951109 | -3.159115706 | 0.001582486 | 0.009312046 | 0.577963723 |
| ENSMUSG00000027249 | F2 | 14061 | protein_coding | 10.07411915 | -0.577990849 | 0.516684296 | 1.118653796 | 0.263287864 | 0.477583041 | 0.577990849 |
| ENSMUSG00000001802 | Lrp3 | 435965 | protein_coding | 8.347857374 | 0.578009842 | 0.542010054 | -1.066419042 | 0.286234258 | 0.504684233 | 0.578009842 |
| ENSMUSG00000055632 | Hmcn2 | 665700 | protein_coding | 14.31865255 | 0.578022003 | 1.076633533 | -0.536879064 | 0.591351154 | 0.771555104 | 0.578022003 |
| ENSMUSG00000075025 | Gm10804 | 100038525 | lncRNA | 12.17597003 | -0.578146532 | 0.483006287 | 1.196975169 | 0.231316233 | 0.440150633 | 0.578146532 |
| ENSMUSG00000022108 | Itm2b | 16432 | protein_coding | 6243.894978 | -0.57818649 | 0.062395765 | 9.266438039 | 1.92E-20 | 9.93E-19 | 0.57818649 |
| ENSMUSG00000047123 | Ticam1 | 106759 | protein_coding | 487.509111 | 0.578237568 | 0.119656749 | -4.832469309 | 1.35E-06 | 1.81E-05 | 0.578237568 |
| ENSMUSG00000031960 | Aars | 234734 | protein_coding | 1425.639303 | 0.578521617 | 0.074964995 | -7.717223472 | 1.19E-14 | 4.21E-13 | 0.578521617 |
| ENSMUSG00000050248 | Evc2 | 68525 | protein_coding | 2.457771558 | 0.578791019 | 2.137005454 | -0.270842088 | 0.786512488 | 1 | 0.578791019 |
| ENSMUSG00000067377 | Tspan6 | 56496 | protein_coding | 111.6847432 | -0.578804103 | 0.175993051 | 3.288789529 | 0.001006192 | 0.006385957 | 0.578804103 |
| ENSMUSG00000077561 | Gm24607 | 115486260 | snoRNA | 0.507797213 | 0.5788114 | 1.614256998 | -0.358562113 | 0.719922694 | 1 | 0.5788114 |
| ENSMUSG00000023224 | Serping1 | 12258 | protein_coding | 3.199114966 | 0.578989801 | 0.724581115 | -0.799068302 | 0.424250809 | 1 | 0.578989801 |
| ENSMUSG00000029925 | Tbxas1 | 21391 | protein_coding | 11.24965375 | -0.578994238 | 0.494330219 | 1.171270165 | 0.2414902 | 0.452145476 | 0.578994238 |
| ENSMUSG00000086898 | Itpr3os | 105246244 | lncRNA | 4.303813424 | 0.579026056 | 0.641772105 | -0.902230015 | 0.366934693 | 1 | 0.579026056 |
| ENSMUSG00000002908 | Kcnn1 | 84036 | protein_coding | 8.299787355 | -0.57921561 | 0.463873318 | 1.248650414 | 0.211792965 | 0.416114999 | 0.57921561 |
| ENSMUSG00000029484 | Anxa3 | 11745 | protein_coding | 1151.657986 | -0.579354 | 0.188535595 | 3.072915753 | 0.002119783 | 0.011898742 | 0.579354 |
| ENSMUSG00000002797 | Ggct | 110175 | protein_coding | 51.08451844 | 0.579543722 | 0.167102861 | -3.46818552 | 0.000523985 | 0.003689141 | 0.579543722 |
| ENSMUSG00000036949 | Slc39a12 | 277468 | protein_coding | 1.509730784 | -0.579581062 | 2.081280083 | 0.278473362 | 0.780649011 | 1 | 0.579581062 |
| ENSMUSG00000053198 | Prx | 19153 | protein_coding | 51.76450744 | 0.579711663 | 0.16369549 | -3.541402778 | 0.000398006 | 0.002918524 | 0.579711663 |
| ENSMUSG00000001227 | Sema6b | 20359 | protein_coding | 46.41257593 | 0.579725687 | 0.247269322 | -2.34451117 | 0.01905204 | 0.070692483 | 0.579725687 |
| ENSMUSG00000037405 | Icam1 | 15894 | protein_coding | 21.3681052 | 0.579762745 | 0.373346324 | -1.552881891 | 0.120451354 | 0.282708024 | 0.579762745 |
| ENSMUSG00000065422 | Mir221 | 723827 | miRNA | 0.414436208 | -0.579847218 | 2.330744654 | 0.248781958 | 0.803529448 | 1 | 0.579847218 |
| ENSMUSG00000042671 | Rgs8 | 67792 | protein_coding | 10.70662031 | -0.580010957 | 0.580762414 | 0.998706086 | 0.317937092 | 0.539333655 | 0.580010957 |
| ENSMUSG00000046814 | Gchfr | 320415 | protein_coding | 44.21936161 | 0.580029602 | 0.180001784 | -3.222354744 | 0.001271416 | 0.007756249 | 0.580029602 |
| ENSMUSG00000065979 | Cpped1 | 223978 | protein_coding | 1080.030046 | 0.580059138 | 0.06862441 | -8.452664814 | 2.85E-17 | 1.23E-15 | 0.580059138 |
| ENSMUSG00000055805 | Fmnl1 | 57778 | protein_coding | 13.64551596 | 0.580371865 | 0.570783837 | -1.016798002 | 0.309249533 | 0.530096199 | 0.580371865 |
| ENSMUSG00000061292 | Cyp3a59 | 100041449 | protein_coding | 0.938880794 | -0.58071659 | 1.332122887 | 0.435933198 | 0.662885198 | 1 | 0.58071659 |
| ENSMUSG00000021994 | Wnt5a | 22418 | protein_coding | 25.01681272 | -0.580994703 | 0.41595322 | 1.396778955 | 0.162480053 | 0.347924739 | 0.580994703 |
| ENSMUSG00000056708 | Ier5 | 15939 | protein_coding | 788.1053497 | 0.581227142 | 0.222046752 | -2.617589027 | 0.008855339 | 0.038625742 | 0.581227142 |
| ENSMUSG00000027244 | Atg13 | 51897 | protein_coding | 378.3280077 | 0.581261302 | 0.081332637 | -7.146716515 | 8.89E-13 | 2.71E-11 | 0.581261302 |
| ENSMUSG00000000167 | Pih1d2 | 72614 | protein_coding | 15.25363993 | 0.581261847 | 0.273980374 | -2.12154556 | 0.033875918 | 0.110624266 | 0.581261847 |
| ENSMUSG00000020102 | Slc16a7 | 20503 | protein_coding | 67.31749607 | -0.581645132 | 0.209166497 | 2.780775791 | 0.005422918 | 0.026043699 | 0.581645132 |
| ENSMUSG00000017897 | Eya2 | 14049 | protein_coding | 23.78281259 | 0.581768439 | 0.298956483 | -1.945997067 | 0.051655092 | 0.152917797 | 0.581768439 |
| ENSMUSG00000029001 | Fbxo44 | 230903 | protein_coding | 27.96264951 | -0.58187091 | 0.218726702 | 2.660264633 | 0.007807928 | 0.034924193 | 0.58187091 |
| ENSMUSG00000002845 | Tmem39a | 67846 | protein_coding | 320.3764542 | 0.581981554 | 0.090614943 | -6.422578174 | 1.34E-10 | 3.26E-09 | 0.581981554 |
| ENSMUSG00000071537 | Klrg2 | 74253 | protein_coding | 5.310946075 | 0.581999818 | 0.522354878 | -1.11418471 | 0.265199953 | 0.480125333 | 0.581999818 |

| ensembl_gene_id | external_gene_name | entrezgene_id | gene_biotype | baseMean | log2FoldChange | lfcSE | stat | pvalue | padj | absLog2FC |
|---|---|---|---|---|---|---|---|---|---|---|
| ENSMUSG00000028546 | Elavl4 | 15572 | protein_coding | 2.085747077 | 0.582309075 | 1.402949835 | -0.415060511 | 0.678097607 | 1 | 0.582309075 |
| ENSMUSG00000117079 | Gm41611 | 105246309 | lncRNA | 1.515898279 | 0.582362571 | 1.726658414 | -0.337277232 | 0.735907921 | 1 | 0.582362571 |
| ENSMUSG00000031840 | Rab3a | 19339 | protein_coding | 110.8308061 | 0.582522966 | 0.127236716 | -4.578261565 | 4.69E-06 | 5.63E-05 | 0.582522966 |
| ENSMUSG00000020829 | Slc46a1 | 52466 | protein_coding | 164.98649 | -0.582589573 | 0.190204775 | 3.062959769 | 0.002191595 | 0.012189964 | 0.582589573 |
| ENSMUSG00000028538 | St3gal3 | 20441 | protein_coding | 986.5207203 | -0.582591085 | 0.124452735 | 4.681223622 | 2.85E-06 | 3.57E-05 | 0.582591085 |
| ENSMUSG00000098488 | Pla2g4b | 211429 | protein_coding | 6.871397003 | 0.582673604 | 0.526344337 | -1.107019802 | 0.268285367 | 0.483440174 | 0.582673604 |
| ENSMUSG00000029771 | Irf5 | 27056 | protein_coding | 8.4367831 | -0.582689614 | 0.37694609 | 1.545816838 | 0.122148809 | 0.285273007 | 0.582689614 |
| ENSMUSG00000049939 | Lrrc4 | 192198 | protein_coding | 12.59474646 | 0.582920693 | 0.357293953 | -1.631487707 | 0.102787452 | 0.25251889 | 0.582920693 |
| ENSMUSG00000046610 | Oacyl | 319888 | protein_coding | 3.642891903 | 0.583004837 | 0.753270599 | -0.773964678 | 0.43895168 | 1 | 0.583004837 |
| ENSMUSG00000027194 | Ttc17 | 74569 | protein_coding | 1018.855946 | 0.583396273 | 0.055066566 | -10.59438265 | 3.16E-26 | 2.13E-24 | 0.583396273 |
| ENSMUSG00000020261 | Slc36a1 | 215335 | protein_coding | 763.6859031 | -0.583405989 | 0.257594794 | 2.264820574 | 0.023523692 | 0.083344831 | 0.583405989 |
| ENSMUSG00000098552 | Gm27217 | 108168350 | lncRNA | 2.14182386 | -0.583617893 | 0.913926846 | 0.638582722 | 0.523094424 | 1 | 0.583617893 |
| ENSMUSG00000041482 | Piezo2 | 667742 | protein_coding | 31.06101085 | -0.583706057 | 0.407658345 | 1.431851117 | 0.152186434 | 0.332794609 | 0.583706057 |
| ENSMUSG00000043155 | Hpdl | 242642 | protein_coding | 102.1343195 | -0.584138069 | 0.203531858 | 2.870008036 | 0.004104614 | 0.020615447 | 0.584138069 |
| ENSMUSG00000048644 | Ctxn1 | 330695 | protein_coding | 5.276256059 | 0.584447873 | 0.574711367 | -1.016941558 | 0.309181232 | 0.53003414 | 0.584447873 |
| ENSMUSG00000086015 | 4833417C18Rik | 73906 | lncRNA | 2.280989674 | -0.584625111 | 0.723765698 | 0.807754654 | 0.419231834 | 1 | 0.584625111 |
| ENSMUSG00000051675 | Trim32 | 69807 | protein_coding | 285.4194785 | 0.585274623 | 0.094153968 | -6.216143983 | 5.10E-10 | 1.16E-08 | 0.585274623 |
| ENSMUSG00000037466 | Tedc1 | 104732 | protein_coding | 200.3549866 | 0.585293518 | 0.157779706 | -3.709561465 | 0.000207619 | 0.001633842 | 0.585293518 |
| ENSMUSG00000099267 | Mir6955 | 102466210 | miRNA | 0.998861144 | -0.585396729 | 1.305002401 | 0.448579044 | 0.653735354 | 1 | 0.585396729 |
| ENSMUSG00000051149 | Adnp | 11538 | protein_coding | 31.34265925 | 0.585588904 | 0.247230673 | -2.368593249 | 0.017855879 | 0.067031593 | 0.585588904 |
| ENSMUSG00000062661 | Ncs1 | 14299 | protein_coding | 9.984198302 | 0.585676899 | 0.404387016 | -1.448307873 | 0.147530967 | 0.325625018 | 0.585676899 |
| ENSMUSG00000019960 | Dusp6 | 67603 | protein_coding | 1063.857534 | 0.585678529 | 0.145872239 | -4.0150102 | 5.94E-05 | 0.000545662 | 0.585678529 |
| ENSMUSG00000053675 | Tgm5 | 74176 | protein_coding | 4.43781756 | 0.585748457 | 0.749133126 | -0.78190169 | 0.434272352 | 0.652783039 | 0.585748457 |
| ENSMUSG00000033308 | Dpyd | 99586 | protein_coding | 369.0815404 | -0.586024374 | 0.097795378 | 5.992352453 | 2.07E-09 | 4.40E-08 | 0.586024374 |
| ENSMUSG00000049265 | Kcnk3 | 16527 | protein_coding | 9.630624239 | -0.586067569 | 0.512334008 | 1.143916976 | 0.25265807 | 0.465046296 | 0.586067569 |
| ENSMUSG00000086273 | Rbm46os | 78212 | lncRNA | 0.893032899 | -0.586158512 | 2.159517602 | 0.271430301 | 0.7860601 | 1 | 0.586158512 |
| ENSMUSG00000033722 | BC034090 | 207792 | protein_coding | 3.757394429 | 0.58616817 | 1.049837202 | -0.558341969 | 0.576610891 | 1 | 0.58616817 |
| ENSMUSG00000033952 | Aspm | 12316 | protein_coding | 373.5260561 | 0.586278765 | 0.190257733 | -3.081497688 | 0.002059621 | 0.011632208 | 0.586278765 |
| ENSMUSG00000002602 | Axl | 26362 | protein_coding | 4.506941285 | -0.586506061 | 0.764890195 | 0.766784651 | 0.443209564 | 0.660786267 | 0.586506061 |
| ENSMUSG00000039670 | Oxld1 | 66431 | protein_coding | 87.46277316 | -0.586512891 | 0.127962285 | 4.583482464 | 4.57E-06 | 5.53E-05 | 0.586512891 |
| ENSMUSG00000040035 | Disp2 | 214240 | protein_coding | 58.24088396 | 0.586623481 | 0.285425355 | -2.055260584 | 0.039853842 | 0.125648406 | 0.586623481 |
| ENSMUSG00000064232 | Gm5414 | 406223 | protein_coding | 1.656019503 | -0.586630135 | 1.406015354 | 0.417228826 | 0.676511043 | 1 | 0.586630135 |
| ENSMUSG00000028494 | Plin2 | 11520 | protein_coding | 1175.799828 | -0.586742052 | 0.119090315 | 4.92686623 | 8.36E-07 | 1.17E-05 | 0.586742052 |
| ENSMUSG00000020600 | Slc7a15 | 328059 | protein_coding | 4.868926363 | -0.586795041 | 0.69539141 | 0.84383418 | 0.398762073 | 0.620714192 | 0.586795041 |
| ENSMUSG00000027966 | Col11a1 | 12814 | protein_coding | 3.304522555 | 0.587014181 | 2.337682706 | -0.251109434 | 0.801729502 | 1 | 0.587014181 |
| ENSMUSG00000058163 | Gm5431 | 432555 | protein_coding | 67.7411378 | -0.587054697 | 0.355884441 | 1.649565502 | 0.099031836 | 0.245852636 | 0.587054697 |
| ENSMUSG00000028071 | Sh2d2a | 27371 | protein_coding | 13.52799918 | 0.587226714 | 0.565234385 | -1.038908336 | 0.298847369 | 0.518781189 | 0.587226714 |
| ENSMUSG00000041777 | Cir1 | 66935 | protein_coding | 65.08668441 | -0.587283857 | 0.173553139 | 3.383884952 | 0.000714679 | 0.004768572 | 0.587283857 |
| ENSMUSG00000070336 | Fbxo47 | 72973 | protein_coding | 7.487278102 | 0.58738718 | 0.473246252 | -1.241187179 | 0.21453661 | 0.419497057 | 0.58738718 |

| ensembl_gene_id | external_gene_name | entrezgene_id | gene_biotype | baseMean | log2FoldChange | lfcSE | stat | pvalue | padj | absLog2FC |
|---|---|---|---|---|---|---|---|---|---|---|
| ENSMUSG00000034057 | Myrfl | 237558 | protein_coding | 8.292656042 | -0.587516585 | 0.61439121 | 0.956258122 | 0.338941842 | 0.560514185 | 0.587516585 |
| ENSMUSG00000039110 | Mycbpap | 104601 | protein_coding | 73.37937439 | -0.587676186 | 0.173018484 | 3.396609257 | 0.000682263 | 0.004591236 | 0.587676186 |
| ENSMUSG00000030342 | Cd9 | 12527 | protein_coding | 1943.335883 | -0.587787499 | 0.147053312 | 3.997104788 | 6.41E-05 | 0.000584065 | 0.587787499 |
| ENSMUSG00000015806 | Qdpr | 110391 | protein_coding | 1081.672604 | 0.587883296 | 0.073339628 | -8.015902358 | 1.09E-15 | 4.24E-14 | 0.587883296 |
| ENSMUSG00000026938 | Fcna | 14133 | protein_coding | 53.95250645 | -0.587894866 | 0.180306552 | 3.260529687 | 0.001112043 | 0.006961441 | 0.587894866 |
| ENSMUSG00000035960 | Apex1 | 11792 | protein_coding | 149.0747826 | 0.58794394 | 0.109936417 | -5.348036221 | 8.89E-08 | 1.48E-06 | 0.58794394 |
| ENSMUSG00000040424 | Hipk4 | 233020 | protein_coding | 8.948047804 | 0.587968261 | 0.467341126 | -1.258113673 | 0.208350649 | 0.411419661 | 0.587968261 |
| ENSMUSG00000029174 | Tbc1d1 | 57915 | protein_coding | 610.8230925 | 0.588132746 | 0.090343571 | -6.509956832 | 7.52E-11 | 1.91E-09 | 0.588132746 |
| ENSMUSG00000040594 | Ranbp17 | 66011 | protein_coding | 20.78222147 | 0.588147826 | 0.363248304 | -1.619134406 | 0.105418347 | 0.257146025 | 0.588147826 |
| ENSMUSG00000015085 | Entpd2 | 12496 | protein_coding | 92.72657571 | -0.588167864 | 0.132782423 | 4.42956118 | 9.44E-06 | 0.000106585 | 0.588167864 |
| ENSMUSG00000022679 | Mpv17l | 93734 | protein_coding | 107.9125638 | 0.588269552 | 0.122727982 | -4.793279775 | 1.64E-06 | 2.16E-05 | 0.588269552 |
| ENSMUSG00000064158 | Izumo1 | 73456 | protein_coding | 5.362582277 | 0.588356086 | 0.634121929 | -0.927828007 | 0.353496794 | 0.575875116 | 0.588356086 |
| ENSMUSG00000042489 | Clspn | 269582 | protein_coding | 155.2730327 | 0.588582817 | 0.189595251 | -3.104417512 | 0.001906541 | 0.010922096 | 0.588582817 |
| ENSMUSG00000003420 | Fcgrt | 14132 | protein_coding | 666.0896721 | -0.588596695 | 0.109029515 | 5.398507885 | 6.72E-08 | 1.13E-06 | 0.588596695 |
| ENSMUSG00000022657 | Cd96 | 84544 | protein_coding | 10.76292824 | -0.588853677 | 0.407113176 | 1.446412721 | 0.148061476 | 0.326547965 | 0.588853677 |
| ENSMUSG00000066108 | Muc5b | 74180 | protein_coding | 13.31629033 | 0.588996325 | 1.030028195 | -0.571825439 | 0.567440241 | 0.75345406 | 0.588996325 |
| ENSMUSG00000020869 | Lrrc59 | 98238 | protein_coding | 2397.954138 | 0.588996746 | 0.078392002 | -7.513480065 | 5.76E-14 | 1.94E-12 | 0.588996746 |
| ENSMUSG00000005469 | Prkaca | 18747 | protein_coding | 861.921895 | 0.589032676 | 0.06273498 | -9.389222375 | 6.04E-21 | 3.25E-19 | 0.589032676 |
| ENSMUSG00000024979 | Tectb | 21684 | protein_coding | 1.121678095 | 0.589064346 | 1.779417361 | -0.331043385 | 0.740611715 | 1 | 0.589064346 |
| ENSMUSG00000060962 | Dmkn | 73712 | protein_coding | 6.904323125 | -0.589143245 | 0.702701148 | 0.838398012 | 0.401807206 | 0.623441772 | 0.589143245 |
| ENSMUSG00000032496 | Ltf | 17002 | protein_coding | 5.495797411 | 0.589356295 | 0.581038484 | -1.014315422 | 0.310432268 | 0.53139398 | 0.589356295 |
| ENSMUSG00000025178 | Pi4k2a | 84095 | protein_coding | 374.4114285 | 0.58943222 | 0.077275583 | -7.627664499 | 2.39E-14 | 8.28E-13 | 0.58943222 |
| ENSMUSG00000057614 | Gnai1 | 14677 | protein_coding | 147.3246835 | -0.589613932 | 0.173304543 | 3.402183939 | 0.000668496 | 0.004518849 | 0.589613932 |
| ENSMUSG00000048388 | Fam171b | 241520 | protein_coding | 2.050043001 | 0.58962249 | 1.961463866 | -0.300603289 | 0.763717023 | 1 | 0.58962249 |
| ENSMUSG00000021509 | Slc25a48 | 328258 | protein_coding | 11.21186885 | 0.589629989 | 0.39139678 | -1.506476339 | 0.131944937 | 0.30108314 | 0.589629989 |
| ENSMUSG00000043705 | Capn13 | 381122 | protein_coding | 625.0681224 | -0.589659632 | 0.375506458 | 1.570304903 | 0.116344194 | 0.275535353 | 0.589659632 |
| ENSMUSG00000027422 | Rrbp1 | 81910 | protein_coding | 8251.310301 | 0.589794179 | 0.044530059 | -13.24485506 | 4.83E-40 | 5.02E-38 | 0.589794179 |
| ENSMUSG00000054252 | Fgfr3 | 14184 | protein_coding | 415.8503676 | -0.589893449 | 0.222139158 | 2.655513122 | 0.007918785 | 0.035343465 | 0.589893449 |
| ENSMUSG00000081683 | Fzd10 | 93897 | protein_coding | 7.56082684 | -0.589953978 | 0.427251192 | 1.380812948 | 0.167336481 | 0.355557727 | 0.589953978 |
| ENSMUSG00000021879 | Dnah12 | 110083 | protein_coding | 3.754372978 | 0.589992711 | 1.273633765 | -0.463235765 | 0.643195383 | 1 | 0.589992711 |
| ENSMUSG00000032491 | Nradd | 67169 | protein_coding | 17.37153657 | -0.589992978 | 0.28864857 | 2.043983723 | 0.040955155 | 0.128287829 | 0.589992978 |
| ENSMUSG00000032092 | Mpzl2 | 14012 | protein_coding | 207.4289205 | -0.590132845 | 0.140193215 | 4.209425155 | 2.56E-05 | 0.000257174 | 0.590132845 |
| ENSMUSG00000040010 | Slc7a5 | 20539 | protein_coding | 184.966728 | 0.590161834 | 0.130065815 | -4.537409257 | 5.69E-06 | 6.72E-05 | 0.590161834 |
| ENSMUSG00000070385 | Ampd1 | 229665 | protein_coding | 1.788888638 | -0.590198579 | 1.858439001 | 0.31757759 | 0.750805375 | 1 | 0.590198579 |
| ENSMUSG00000069662 | Marcks | 17118 | protein_coding | 891.4750581 | 0.590294814 | 0.060966997 | -9.682202591 | 3.59E-22 | 2.04E-20 | 0.590294814 |
| ENSMUSG00000029345 | Tfip11 | 54723 | protein_coding | 575.1755518 | 0.590470796 | 0.061904142 | -9.538469874 | 1.45E-21 | 8.09E-20 | 0.590470796 |
| ENSMUSG00000027378 | Nphp1 | 53885 | protein_coding | 63.38938725 | 0.590596561 | 0.180831072 | -3.266012613 | 0.001090734 | 0.00684621 | 0.590596561 |
| ENSMUSG00000024486 | Hbegf | 15200 | protein_coding | 3581.620554 | -0.590601325 | 0.185038389 | 3.191777283 | 0.001414003 | 0.008491218 | 0.590601325 |
| ENSMUSG00000030269 | Mtmr14 | 97287 | protein_coding | 508.8711985 | 0.590609869 | 0.069132546 | -8.543152324 | 1.31E-17 | 5.80E-16 | 0.590609869 |

Supplementary Table 1

| ensembl_gene_id | external_gene_name | entrezgene_id | gene_biotype | baseMean | log2FoldChange | lfcSE | stat | pvalue | padj | absLog2FC |
|---|---|---|---|---|---|---|---|---|---|---|
| ENSMUSG00000073418 | C4b | 12268 | protein_coding | 6.3727011 | 0.590666306 | 0.888832838 | -0.664541499 | 0.506343797 | 0.710248999 | 0.590666306 |
| ENSMUSG00000092570 | Plut | 70147 | lncRNA | 5.017004524 | -0.590796393 | 0.593563306 | 0.995338471 | 0.319571673 | 0.540884146 | 0.590796393 |
| ENSMUSG00000020256 | Aldh1l2 | 216188 | protein_coding | 30.4245904 | 0.590806957 | 0.256794926 | -2.300695602 | 0.021408843 | 0.077396154 | 0.590806957 |
| ENSMUSG00000050930 | Map10 | 74393 | protein_coding | 44.92837692 | -0.590928767 | 0.164131222 | 3.600343428 | 0.000317797 | 0.002393317 | 0.590928767 |
| ENSMUSG00000000811 | Txnrd3 | 232223 | protein_coding | 109.8992262 | -0.590978359 | 0.149933372 | 3.941606528 | 8.09E-05 | 0.000716166 | 0.590978359 |
| ENSMUSG00000034684 | Sema3f | 20350 | protein_coding | 30.866551 | -0.591032177 | 0.279562334 | 2.11413379 | 0.034503842 | 0.112253043 | 0.591032177 |
| ENSMUSG00000043664 | Tmem221 | 434325 | protein_coding | 1.526443448 | 0.591214622 | 1.13721067 | -0.519881353 | 0.603146273 | 1 | 0.591214622 |
| ENSMUSG00000050315 | Synpo2 | 118449 | protein_coding | 19.012666 | -0.591605858 | 0.430282944 | 1.374922864 | 0.16915536 | 0.358361978 | 0.591605858 |
| ENSMUSG00000080478 | Snord23 | 100217441 | snoRNA | 1.678757321 | 0.591675737 | 1.103910708 | -0.535981518 | 0.591971324 | 1 | 0.591675737 |
| ENSMUSG00000051452 | Gm11437 | 628813 | protein_coding | 68.4532107 | -0.5919031 | 0.205812825 | 2.875929129 | 0.004028401 | 0.020288204 | 0.5919031 |
| ENSMUSG00000032470 | Mras | 17532 | protein_coding | 14.66640195 | -0.591931532 | 0.320733422 | 1.845556128 | 0.064956682 | 0.181014958 | 0.591931532 |
| ENSMUSG00000005950 | P2rx5 | 94045 | protein_coding | 3.167042324 | 0.591941223 | 0.702015123 | -0.843202951 | 0.399114948 | 1 | 0.591941223 |
| ENSMUSG00000094559 | Cyp2d34 | 223706 | protein_coding | 1038.947016 | -0.591985018 | 0.209875166 | 2.820653006 | 0.004792601 | 0.02346428 | 0.591985018 |
| ENSMUSG00000021187 | Tc2n | 74413 | protein_coding | 595.4972909 | 0.592089927 | 0.101920395 | -5.809337023 | 6.27E-09 | 1.24E-07 | 0.592089927 |
| ENSMUSG00000031099 | Smarca1 | 93761 | protein_coding | 4.87958971 | 0.592131297 | 1.061155678 | -0.558006058 | 0.576840247 | 0.761222618 | 0.592131297 |
| ENSMUSG00000050092 | Sprr2b | 20756 | protein_coding | 2.173006469 | 0.592196747 | 1.247762457 | -0.47460696 | 0.635067144 | 1 | 0.592196747 |
| ENSMUSG00000078252 | Krtap17-1 | 77914 | protein_coding | 0.524748482 | 0.592272377 | 1.763643398 | -0.335823204 | 0.737004191 | 1 | 0.592272377 |
| ENSMUSG00000028678 | Kif2c | 73804 | protein_coding | 195.1594968 | 0.59260171 | 0.139335434 | -4.253058216 | 2.11E-05 | 0.000216159 | 0.59260171 |
| ENSMUSG00000038545 | Cul7 | 66515 | protein_coding | 217.374334 | 0.59277508 | 0.116279264 | -5.097857189 | 3.44E-07 | 5.22E-06 | 0.59277508 |
| ENSMUSG00000018986 | Slfn3 | 20557 | protein_coding | 2.219657269 | 0.593079597 | 1.290084043 | -0.459721675 | 0.64571601 | 1 | 0.593079597 |
| ENSMUSG00000117431 | 8430430B14Rik | 78148 | lncRNA | 14.27239228 | 0.593145948 | 0.320094124 | -1.853036041 | 0.063877193 | 0.178760882 | 0.593145948 |
| ENSMUSG00000085601 | Meiosin | 243866 | protein_coding | 3.809621314 | 0.593304769 | 1.027247303 | -0.577567609 | 0.563556081 | 1 | 0.593304769 |
| ENSMUSG00000037833 | Sh2d4b | 328381 | protein_coding | 5.688520315 | 0.593351874 | 0.529584245 | -1.120410736 | 0.262538772 | 0.476573341 | 0.593351874 |
| ENSMUSG00000017713 | Tha1 | 71776 | protein_coding | 106.1900603 | -0.593429597 | 0.119097283 | 4.982729925 | 6.27E-07 | 9.04E-06 | 0.593429597 |
| ENSMUSG00000058717 | Lypd11 | 210155 | protein_coding | 1.013252438 | -0.593449832 | 3.019106223 | 0.196564741 | 0.844168165 | 1 | 0.593449832 |
| ENSMUSG00000033209 | Ttc28 | 209683 | protein_coding | 5.930642552 | 0.593665151 | 0.875722875 | -0.677914403 | 0.497825967 | 0.703138729 | 0.593665151 |
| ENSMUSG00000029862 | Clcn1 | 12723 | protein_coding | 9.619799519 | 0.593807223 | 0.609078137 | -0.974927825 | 0.329596063 | 0.551855356 | 0.593807223 |
| ENSMUSG00000059498 | Fcgr3 | 14131 | protein_coding | 2.666901021 | 0.59395473 | 0.761466687 | -0.78001407 | 0.435382593 | 1 | 0.59395473 |
| ENSMUSG00000032356 | Rasgrf1 | 19417 | protein_coding | 4.856652444 | 0.594201213 | 1.901301503 | -0.312523401 | 0.754642781 | 0.876856731 | 0.594201213 |
| ENSMUSG00000073838 | Tufm | 233870 | protein_coding | 1106.94598 | -0.594394325 | 0.131684462 | 4.513777214 | 6.37E-06 | 7.45E-05 | 0.594394325 |
| ENSMUSG00000097591 | A330032B11Rik | 319931 | lncRNA | 1.657019918 | -0.594587674 | 1.132656842 | 0.524949527 | 0.599618278 | 1 | 0.594587674 |
| ENSMUSG00000031257 | Nox1 | 237038 | protein_coding | 459.6538583 | 0.594670989 | 0.238452023 | -2.493881079 | 0.012635487 | 0.051092662 | 0.594670989 |
| ENSMUSG00000040584 | Abcb1a | 18671 | protein_coding | 7518.659721 | -0.594954129 | 0.120791881 | 4.925448022 | 8.42E-07 | 1.18E-05 | 0.594954129 |
| ENSMUSG00000033152 | Podxl2 | 319655 | protein_coding | 5.056367279 | 0.595181463 | 0.809242338 | -0.735479886 | 0.462047285 | 0.675841352 | 0.595181463 |
| ENSMUSG00000056290 | Ms4a4b | 60361 | protein_coding | 9.017681769 | 0.595374995 | 0.593311927 | -1.003477207 | 0.315630669 | 0.53655334 | 0.595374995 |
| ENSMUSG00000028068 | Iqgap3 | 404710 | protein_coding | 359.356754 | 0.595464345 | 0.15670263 | -3.799963964 | 0.000144717 | 0.001193736 | 0.595464345 |
| ENSMUSG00000034040 | Galnt17 | 212996 | protein_coding | 6.803508153 | -0.595562351 | 0.581898599 | 1.023481328 | 0.306080324 | 0.52648324 | 0.595562351 |
| ENSMUSG00000048153 | Or6e1 | 18348 | protein_coding | 1.777989764 | -0.595741371 | 1.553471352 | 0.383490413 | 0.701356177 | 1 | 0.595741371 |
| ENSMUSG00000029380 | Cxcl1 | 14825 | protein_coding | 36.22891872 | 0.595836128 | 0.464836884 | -1.281817662 | 0.199906617 | 0.400972961 | 0.595836128 |

| ensembl_gene_id | external_gene_name | entrezgene_id | gene_biotype | baseMean | log2FoldChange | lfcSE | stat | pvalue | padj | absLog2FC |
|---|---|---|---|---|---|---|---|---|---|---|
| ENSMUSG00000012187 | Mogat1 | 68393 | protein_coding | 1.042848859 | -0.59598025 | 1.219740285 | 0.488612418 | 0.625116119 | 1 | 0.59598025 |
| ENSMUSG00000006218 | Fam131c | 277743 | protein_coding | 5.947916945 | 0.596232356 | 0.621548717 | -0.959268904 | 0.337423297 | 0.559438733 | 0.596232356 |
| ENSMUSG00000051354 | Samd3 | 268288 | protein_coding | 3.905155872 | 0.596244478 | 1.166773927 | -0.511019713 | 0.609337253 | 1 | 0.596244478 |
| ENSMUSG00000020232 | Hmg20b | 15353 | protein_coding | 722.6653145 | 0.596362682 | 0.112724681 | -5.290435767 | 1.22E-07 | 1.99E-06 | 0.596362682 |
| ENSMUSG00000000149 | Gna12 | 14673 | protein_coding | 361.5355622 | 0.596628062 | 0.0814317 | -7.326729805 | 2.36E-13 | 7.57E-12 | 0.596628062 |
| ENSMUSG00000062825 | Actg1 | 11465 | protein_coding | 11280.97666 | -0.596757894 | 0.12943785 | 4.610381679 | 4.02E-06 | 4.92E-05 | 0.596757894 |
| ENSMUSG00000051198 | 4930548G14Rik | 75281 | lncRNA | 0.529598418 | -0.597022121 | 1.477198744 | 0.404158292 | 0.686096322 | 1 | 0.597022121 |
| ENSMUSG00000035413 | Tmem98 | 103743 | protein_coding | 1663.458539 | 0.597201824 | 0.058307189 | -10.24233605 | 1.28E-24 | 8.11E-23 | 0.597201824 |
| ENSMUSG00000030170 | Wnt5b | 22419 | protein_coding | 6.252459498 | 0.597316244 | 0.561517294 | -1.063753958 | 0.287440174 | 0.50636895 | 0.597316244 |
| ENSMUSG00000032757 | Bet1 | 12068 | protein_coding | 412.9124013 | 0.597419435 | 0.129456191 | -4.614838673 | 3.93E-06 | 4.83E-05 | 0.597419435 |
| ENSMUSG00000050640 | Tmem150c | 231503 | protein_coding | 3.976362083 | 0.597471716 | 0.633431626 | -0.943230005 | 0.345563269 | 1 | 0.597471716 |
| ENSMUSG00000025232 | Hexa | 15211 | protein_coding | 1420.229302 | -0.597486339 | 0.117815962 | 5.071353058 | 3.95E-07 | 5.94E-06 | 0.597486339 |
| ENSMUSG00000003526 | Prodh | 19125 | protein_coding | 179.9389747 | -0.59761599 | 0.129332015 | 4.62078928 | 3.82E-06 | 4.70E-05 | 0.59761599 |
| ENSMUSG00000022487 | Gtsf1 | 74174 | protein_coding | 1.265052598 | -0.597630618 | 1.39947252 | 0.427039909 | 0.66935026 | 1 | 0.597630618 |
| ENSMUSG00000068452 | Duox2 | 214593 | protein_coding | 1782.509454 | 0.598252058 | 0.250387478 | -2.389305019 | 0.016880281 | 0.064232353 | 0.598252058 |
| ENSMUSG00000109564 | Muc16 | 73732 | protein_coding | 11.47326436 | 0.598254479 | 1.222076712 | -0.489539219 | 0.624459996 | 0.794172689 | 0.598254479 |
| ENSMUSG00000027994 | Mcub | 66815 | protein_coding | 37.24514732 | -0.59834082 | 0.212285965 | 2.818560421 | 0.004823953 | 0.023575839 | 0.59834082 |
| ENSMUSG00000051146 | Camk2n2 | 73047 | protein_coding | 8.433447872 | -0.598445731 | 0.403190592 | 1.484275038 | 0.13773597 | 0.310179152 | 0.598445731 |
| ENSMUSG00000037613 | Tnfrsf23 | 79201 | protein_coding | 17.24687922 | -0.598609813 | 0.321399566 | 1.862509714 | 0.062531278 | 0.175858569 | 0.598609813 |
| ENSMUSG00000036894 | Rap2b | 74012 | protein_coding | 32.52834932 | 0.598932904 | 0.2347578 | -2.5512801 | 0.010732803 | 0.045076681 | 0.598932904 |
| ENSMUSG00000095990 | Zfp97 | 22759 | protein_coding | 44.0756932 | -0.598940483 | 0.231103198 | 2.591658135 | 0.009551463 | 0.041033524 | 0.598940483 |
| ENSMUSG00000027353 | Mcm8 | 66634 | protein_coding | 79.49338628 | 0.59899831 | 0.209418682 | -2.860290703 | 0.004232528 | 0.021129375 | 0.59899831 |
| ENSMUSG00000028710 | Atpaf1 | 230649 | protein_coding | 112.1992641 | -0.599116347 | 0.136386905 | 4.392770321 | 1.12E-05 | 0.000123789 | 0.599116347 |
| ENSMUSG00000032633 | Flcn | 216805 | protein_coding | 266.2019216 | -0.599253067 | 0.117823706 | 5.086014413 | 3.66E-07 | 5.53E-06 | 0.599253067 |
| ENSMUSG00000069310 | H3c3 | 319148 | protein_coding | 751.4548042 | 0.599528388 | 0.237400536 | -2.52538768 | 0.011557072 | 0.047749183 | 0.599528388 |
| ENSMUSG00000032040 | Dcps | 69305 | protein_coding | 257.9162321 | 0.599714219 | 0.097833705 | -6.129934636 | 8.79E-10 | 1.95E-08 | 0.599714219 |
| ENSMUSG00000021536 | Adcy2 | 210044 | protein_coding | 38.92830595 | 0.599871758 | 0.271628434 | -2.208427701 | 0.027214473 | 0.093376231 | 0.599871758 |
| ENSMUSG00000026930 | Gpsm1 | 67839 | protein_coding | 64.84433339 | -0.600084292 | 0.179465419 | 3.343732161 | 0.000826595 | 0.005418972 | 0.600084292 |
| ENSMUSG00000025789 | St8sia2 | 20450 | protein_coding | 2.359634158 | -0.600112377 | 2.165985575 | 0.277062038 | 0.781732471 | 1 | 0.600112377 |
| ENSMUSG00000049536 | Tceal1 | 237052 | protein_coding | 12.52778383 | -0.600144985 | 0.311664925 | 1.925609641 | 0.054153134 | 0.158286006 | 0.600144985 |
| ENSMUSG00000025574 | Tk1 | 21877 | protein_coding | 365.2548891 | 0.60026401 | 0.135773176 | -4.421079519 | 9.82E-06 | 0.000110328 | 0.60026401 |
| ENSMUSG00000058513 | Or6c211 | 258282 | protein_coding | 1.199548508 | 0.600398854 | 2.630211917 | -0.228270145 | 0.819436233 | 1 | 0.600398854 |
| ENSMUSG00000018401 | Mtmr4 | 170749 | protein_coding | 1539.056134 | 0.600549347 | 0.056412297 | -10.64571696 | 1.83E-26 | 1.25E-24 | 0.600549347 |
| ENSMUSG00000020331 | Hcn2 | 15166 | protein_coding | 118.9320728 | -0.600678683 | 0.171113362 | 3.51041365 | 0.00044741 | 0.003229253 | 0.600678683 |
| ENSMUSG00000038543 | BC028528 | 229600 | protein_coding | 2.455654548 | 0.600784028 | 0.863671287 | -0.695616535 | 0.486669008 | 1 | 0.600784028 |
| ENSMUSG00000094493 | Or2ag18 | 258593 | protein_coding | 1.068737194 | 0.600814137 | 1.080828675 | -0.555882862 | 0.578290935 | 1 | 0.600814137 |
| ENSMUSG00000031375 | Bgn | 12111 | protein_coding | 4.841613463 | 0.60082056 | 1.446703611 | -0.415303146 | 0.677919999 | 0.827372024 | 0.60082056 |
| ENSMUSG00000019539 | Rcn3 | 52377 | protein_coding | 1.898094235 | -0.600874202 | 1.301504486 | 0.461676628 | 0.644313234 | 1 | 0.600874202 |
| ENSMUSG00000063415 | Cyp26b1 | 232174 | protein_coding | 4.214163805 | 0.601011698 | 0.815998411 | -0.736535379 | 0.461404945 | 1 | 0.601011698 |

| ensembl_gene_id | external_gene_name | entrezgene_id | gene_biotype | baseMean | log2FoldChange | lfcSE | stat | pvalue | padj | absLog2FC |
|---|---|---|---|---|---|---|---|---|---|---|
| ENSMUSG00000038725 | Pkhd1l1 | 192190 | protein_coding | 5.872596386 | 0.6010388 | 1.106846937 | -0.543018894 | 0.587116794 | 0.768458962 | 0.6010388 |
| ENSMUSG00000090066 | 1110002E22Rik | 102634333 | protein_coding | 2.186469535 | 0.601075598 | 2.709200505 | -0.221864567 | 0.824419314 | 1 | 0.601075598 |
| ENSMUSG00000096054 | Syne1 | 64009 | protein_coding | 212.6149553 | -0.60114991 | 0.231484497 | 2.596933775 | 0.009406006 | 0.040588131 | 0.60114991 |
| ENSMUSG00000030165 | Klrd1 | 16643 | protein_coding | 13.39378766 | 0.601223253 | 0.353585309 | -1.700362648 | 0.089062733 | 0.227007559 | 0.601223253 |
| ENSMUSG00000057156 | Homez | 239099 | protein_coding | 121.5122953 | 0.601231002 | 0.114147498 | -5.267141295 | 1.39E-07 | 2.23E-06 | 0.601231002 |
| ENSMUSG00000078866 | Zfp970 | 628308 | protein_coding | 29.8949896 | -0.601634035 | 0.255792065 | 2.352043389 | 0.018670597 | 0.069552501 | 0.601634035 |
| ENSMUSG00000042675 | Ypel3 | 66090 | protein_coding | 240.258328 | -0.601644833 | 0.169307413 | 3.553564608 | 0.000380048 | 0.002806847 | 0.601644833 |
| ENSMUSG00000026646 | Suv39h2 | 64707 | protein_coding | 49.52783296 | 0.601931173 | 0.192229313 | -3.131318347 | 0.001740234 | 0.010094213 | 0.601931173 |
| ENSMUSG00000018862 | Otop3 | 69602 | protein_coding | 1.664230554 | 0.601985824 | 1.714916088 | -0.351029318 | 0.725566354 | 1 | 0.601985824 |
| ENSMUSG00000033569 | Adgrb3 | 210933 | protein_coding | 11.49346979 | -0.602157759 | 0.558671663 | 1.077838378 | 0.281105889 | 0.498837595 | 0.602157759 |
| ENSMUSG00000104117 | Gm20743 | 433374 | lncRNA | 4.134966227 | -0.602226083 | 0.554477766 | 1.086114034 | 0.277428544 | 1 | 0.602226083 |
| ENSMUSG00000045009 | Prrt3 | 210673 | protein_coding | 11.10247406 | 0.602368027 | 0.399568647 | -1.507545779 | 0.131670819 | 0.300706943 | 0.602368027 |
| ENSMUSG00000029053 | Prkcz | 18762 | protein_coding | 1414.748949 | 0.602926382 | 0.056824821 | -10.61026448 | 2.67E-26 | 1.81E-24 | 0.602926382 |
| ENSMUSG00000055403 | 4933427D06Rik | 232217 | lncRNA | 1.380438309 | -0.603215399 | 2.52836691 | 0.238579059 | 0.811432005 | 1 | 0.603215399 |
| ENSMUSG00000019230 | Lhx9 | 16876 | protein_coding | 2.773220092 | 0.603343659 | 1.789641652 | -0.337130988 | 0.736018158 | 1 | 0.603343659 |
| ENSMUSG00000114279 | H2bc14 | 319186 | protein_coding | 546.0688645 | 0.603353663 | 0.233027479 | -2.589195342 | 0.009620051 | 0.041285217 | 0.603353663 |
| ENSMUSG00000062638 | Btnl1 | 100038862 | protein_coding | 754.7528686 | -0.603370987 | 0.17965059 | 3.358580597 | 0.000783439 | 0.005170946 | 0.603370987 |
| ENSMUSG00000036492 | Rnf39 | 386454 | protein_coding | 87.19305318 | -0.603445817 | 0.128522098 | 4.695268957 | 2.66E-06 | 3.36E-05 | 0.603445817 |
| ENSMUSG00000073987 | Ggh | 14590 | protein_coding | 1821.956859 | -0.603473472 | 0.191806717 | 3.146258287 | 0.001653739 | 0.009680909 | 0.603473472 |
| ENSMUSG00000067813 | Xkr9 | 381246 | protein_coding | 25.63024441 | -0.603513012 | 0.244365172 | 2.469717786 | 0.013521968 | 0.053846584 | 0.603513012 |
| ENSMUSG00000019214 | Chtf18 | 214901 | protein_coding | 92.00403667 | 0.603565347 | 0.167560954 | -3.602064392 | 0.0003157 | 0.002381698 | 0.603565347 |
| ENSMUSG00000021071 | Trim9 | 94090 | protein_coding | 9.887189484 | -0.603726686 | 0.593057229 | 1.017990603 | 0.308682422 | 0.529343875 | 0.603726686 |
| ENSMUSG00000075120 | Or4a66 | 258456 | protein_coding | 1.011471311 | -0.603728828 | 2.551213339 | 0.236643803 | 0.812933135 | 1 | 0.603728828 |
| ENSMUSG00000007033 | Hspa1l | 15482 | protein_coding | 3.909212129 | 0.603784355 | 0.746752568 | -0.808546741 | 0.41877591 | 1 | 0.603784355 |
| ENSMUSG00000111443 | Gm46123 | 118567674 | lncRNA | 1.34705256 | 0.603896801 | 1.838909387 | -0.328399433 | 0.742609671 | 1 | 0.603896801 |
| ENSMUSG00000064984 | Snord73a | 19870 | snoRNA | 2.691649099 | 0.603899616 | 0.705016069 | -0.856575676 | 0.391679433 | 1 | 0.603899616 |
| ENSMUSG00000020484 | Xbp1 | 22433 | protein_coding | 3261.561833 | 0.603947618 | 0.090289047 | -6.689046351 | 2.25E-11 | 6.05E-10 | 0.603947618 |
| ENSMUSG00000005682 | Pan2 | 103135 | protein_coding | 560.5673078 | 0.60403047 | 0.095913098 | -6.297684942 | 3.02E-10 | 7.13E-09 | 0.60403047 |
| ENSMUSG00000027639 | Samhd1 | 56045 | protein_coding | 454.3063465 | -0.604057825 | 0.098792725 | 6.114395822 | 9.69E-10 | 2.13E-08 | 0.604057825 |
| ENSMUSG00000089953 | Rnf224 | 329360 | protein_coding | 1.282003345 | 0.604092215 | 1.853104861 | -0.325989224 | 0.744432508 | 1 | 0.604092215 |
| ENSMUSG00000113387 | Gm31508 | 102633758 | lncRNA | 1.449484205 | 0.604125724 | 1.85562748 | -0.325564118 | 0.744754164 | 1 | 0.604125724 |
| ENSMUSG00000049249 | Gm5316 | 102634348 | transcribed_processed_pseudogene | 1.020713639 | 0.604203923 | 1.861301337 | -0.324613705 | 0.745473453 | 1 | 0.604203923 |
| ENSMUSG00000074758 | Gm5535 | 433481 | lncRNA | 4.410444759 | 0.604257666 | 0.698391956 | -0.865212808 | 0.386921997 | 0.609455738 | 0.604257666 |
| ENSMUSG00000038936 | Sccpdh | 109232 | protein_coding | 191.6394947 | 0.604258588 | 0.085996843 | -7.02652057 | 2.12E-12 | 6.24E-11 | 0.604258588 |
| ENSMUSG00000028444 | Cntfr | 12804 | protein_coding | 3.579396452 | -0.604313151 | 1.784470608 | 0.338651222 | 0.734872491 | 1 | 0.604313151 |
| ENSMUSG00000024694 | Keg1 | 64697 | protein_coding | 0.933453316 | 0.604372832 | 1.873988215 | -0.322506207 | 0.747069236 | 1 | 0.604372832 |
| ENSMUSG00000081402 | Gm15455 | 433287 | processed_pseudogene | 2.606411933 | 0.604386275 | 0.783375097 | -0.771515813 | 0.440401253 | 1 | 0.604386275 |
| ENSMUSG00000040928 | S100pbp | 74648 | protein_coding | 154.7620912 | -0.604595806 | 0.162683264 | 3.716398303 | 0.000202083 | 0.001595982 | 0.604595806 |
| ENSMUSG00000040140 | Tdrd6 | 210510 | protein_coding | 1.396146154 | 0.604606687 | 1.892126109 | -0.319538262 | 0.749318382 | 1 | 0.604606687 |

| ensembl_gene_id | external_gene_name | entrezgene_id | gene_biotype | baseMean | log2FoldChange | lfcSE | stat | pvalue | padj | absLog2FC |
|---|---|---|---|---|---|---|---|---|---|---|
| ENSMUSG00000040883 | Tmem205 | 235043 | protein_coding | 294.4643376 | -0.604652278 | 0.101099793 | 5.980746951 | 2.22E-09 | 4.71E-08 | 0.604652278 |
| ENSMUSG00000103088 | Pcdhgb6 | 93703 | protein_coding | 24.68370083 | 0.604734102 | 0.249831057 | -2.420572167 | 0.015496103 | 0.059944401 | 0.604734102 |
| ENSMUSG00000089296 | Gm23205 | 115488985 | snRNA | 1.802963842 | -0.604763763 | 0.792833632 | 0.762787726 | 0.445590002 | 1 | 0.604763763 |
| ENSMUSG00000035653 | Lrfn5 | 238205 | protein_coding | 1.557115261 | 0.604778438 | 1.905709788 | -0.317350754 | 0.75097747 | 1 | 0.604778438 |
| ENSMUSG00000072419 | Dppa2 | 73703 | protein_coding | 1.34865431 | 0.604796954 | 1.90717178 | -0.317117189 | 0.751154682 | 1 | 0.604796954 |
| ENSMUSG00000086376 | 1700071G01Rik | 73485 | lncRNA | 1.451852676 | 0.60484691 | 1.023028672 | -0.591231631 | 0.554365232 | 1 | 0.60484691 |
| ENSMUSG00000109238 | Tex51 | 102638514 | protein_coding | 1.244300333 | 0.604891481 | 1.914767132 | -0.31590864 | 0.752071854 | 1 | 0.604891481 |
| ENSMUSG00000106230 | Gm29811 | 101243624 | lncRNA | 1.113329767 | -0.605068704 | 1.02770477 | 0.588757318 | 0.556024082 | 1 | 0.605068704 |
| ENSMUSG00000032343 | Impg1 | 63859 | protein_coding | 2.081647146 | 0.605248273 | 1.263317574 | -0.479094319 | 0.631871531 | 1 | 0.605248273 |
| ENSMUSG00000040183 | Ankrd6 | 140577 | protein_coding | 5.634927152 | 0.605254995 | 0.780037926 | -0.775930214 | 0.437790192 | 0.656109195 | 0.605254995 |
| ENSMUSG00000057163 | Prss2 | 22072 | protein_coding | 0.714371033 | 0.60581203 | 2.126784355 | -0.284848827 | 0.775759962 | 1 | 0.60581203 |
| ENSMUSG00000023019 | Gpd1 | 14555 | protein_coding | 303.5486617 | -0.605845886 | 0.097957298 | 6.184795809 | 6.22E-10 | 1.40E-08 | 0.605845886 |
| ENSMUSG00000091405 | H4c14 | 97122 | protein_coding | 757.3587616 | 0.605855018 | 0.168174296 | -3.602542321 | 0.00031512 | 0.002381578 | 0.605855018 |
| ENSMUSG00000040794 | C1qtnf4 | 67445 | protein_coding | 17.66535332 | -0.605886954 | 0.309383628 | 1.95836786 | 0.050186863 | 0.149272747 | 0.605886954 |
| ENSMUSG00000065468 | Mir26b | 387219 | miRNA | 2.815497489 | 0.605896151 | 0.690953352 | -0.876898779 | 0.38054162 | 1 | 0.605896151 |
| ENSMUSG00000035032 | Nek11 | 208583 | protein_coding | 4.812387641 | 0.605916899 | 0.53001576 | -1.143205438 | 0.252953305 | 0.465324602 | 0.605916899 |
| ENSMUSG00000082316 | Vmn1r221 | 100312485 | protein_coding | 1.486751827 | 0.606159305 | 2.674508112 | -0.226643285 | 0.820701131 | 1 | 0.606159305 |
| ENSMUSG00000034911 | Ushbp1 | 234395 | protein_coding | 3.983828385 | 0.60616496 | 1.022855236 | -0.592620479 | 0.553435171 | 1 | 0.60616496 |
| ENSMUSG00000031786 | Drc7 | 330830 | protein_coding | 1.902414956 | 0.606312709 | 2.68171774 | -0.226091173 | 0.82113051 | 1 | 0.606312709 |
| ENSMUSG00000059408 | Mrgprh | 80978 | protein_coding | 0.936309009 | 0.606360805 | 2.045600732 | -0.296421875 | 0.766907922 | 1 | 0.606360805 |
| ENSMUSG00000065464 | Mir185 | 387180 | miRNA | 0.947310143 | 0.606501033 | 2.059533245 | -0.294484701 | 0.768387552 | 1 | 0.606501033 |
| ENSMUSG00000074947 | Or4f59 | 258359 | protein_coding | 1.300636838 | 0.606530338 | 2.687539546 | -0.225682386 | 0.821448458 | 1 | 0.606530338 |
| ENSMUSG00000005547 | Cyp2a5 | 13087 | protein_coding | 1.311082323 | 0.606578239 | 2.067495975 | -0.293387869 | 0.769225697 | 1 | 0.606578239 |
| ENSMUSG00000097350 | 4732491K20Rik | 224523 | lncRNA | 19.97849917 | 0.606616605 | 0.331326784 | -1.830871014 | 0.067119798 | 0.185168116 | 0.606616605 |
| ENSMUSG00000074003 | Gucy2d | 14918 | protein_coding | 1.810680329 | 0.606749035 | 2.693689456 | -0.225248324 | 0.821786098 | 1 | 0.606749035 |
| ENSMUSG00000045932 | Ifit2 | 15958 | protein_coding | 86.44172036 | 0.606780511 | 0.454672238 | -1.334544888 | 0.182025346 | 0.376404528 | 0.606780511 |
| ENSMUSG00000052012 | Or10p1 | 258933 | protein_coding | 0.984734102 | 0.606783977 | 2.08851565 | -0.290533603 | 0.771408047 | 1 | 0.606783977 |
| ENSMUSG00000038602 | Slc35f1 | 215085 | protein_coding | 1.342872228 | 0.606789438 | 2.089222344 | -0.290437942 | 0.77148122 | 1 | 0.606789438 |
| ENSMUSG00000115020 | Vmn1r218 | 171256 | protein_coding | 1.949975568 | 0.606819589 | 2.695669049 | -0.225109084 | 0.821894415 | 1 | 0.606819589 |
| ENSMUSG00000032640 | Chsy1 | 269941 | protein_coding | 94.07946088 | 0.606845219 | 0.162009726 | -3.745733265 | 0.000179868 | 0.001444715 | 0.606845219 |
| ENSMUSG00000040855 | Reps2 | 194590 | protein_coding | 568.9060431 | 0.606960586 | 0.098888628 | -6.137819864 | 8.37E-10 | 1.86E-08 | 0.606960586 |
| ENSMUSG00000094441 | Zfp955a | 77652 | protein_coding | 112.8963671 | -0.607027612 | 0.170128778 | 3.568047808 | 0.000359651 | 0.002676555 | 0.607027612 |
| ENSMUSG00000061802 | Odad2 | 74934 | protein_coding | 1.57147858 | 0.607036172 | 2.115599193 | -0.286933448 | 0.774163279 | 1 | 0.607036172 |
| ENSMUSG00000024223 | Armc12 | 67645 | protein_coding | 0.933154481 | 0.607046636 | 2.116458262 | -0.286821926 | 0.774248673 | 1 | 0.607046636 |
| ENSMUSG00000073774 | Kncn | 654462 | protein_coding | 1.021512152 | 0.607060289 | 2.117968482 | -0.286623854 | 0.774400348 | 1 | 0.607060289 |
| ENSMUSG00000053166 | Cdh22 | 104010 | protein_coding | 1.139483582 | 0.607091558 | 2.121512073 | -0.286159841 | 0.7747557 | 1 | 0.607091558 |
| ENSMUSG00000059816 | Shisal2a | 545667 | protein_coding | 0.90391929 | 0.607092563 | 2.121544292 | -0.286155969 | 0.774758666 | 1 | 0.607092563 |
| ENSMUSG00000054667 | Irs4 | 16370 | protein_coding | 1.414692394 | 0.607149811 | 2.128017131 | -0.285312464 | 0.775404765 | 1 | 0.607149811 |
| ENSMUSG00000024164 | C3 | 12266 | protein_coding | 69.49180941 | -0.60718107 | 0.248875475 | 2.439698284 | 0.014699533 | 0.057414402 | 0.60718107 |

| ensembl_gene_id | external_gene_name | entrezgene_id | gene_biotype | baseMean | log2FoldChange | lfcSE | stat | pvalue | padj | absLog2FC |
|---|---|---|---|---|---|---|---|---|---|---|
| ENSMUSG00000059934 | Prh1 | 19131 | protein_coding | 0.931460176 | 0.607182593 | 2.131366805 | -0.284879445 | 0.775736504 | 1 | 0.607182593 |
| ENSMUSG00000085037 | 4933421O10Rik | 100504166 | lncRNA | 69.40746991 | 0.607193072 | 0.147330393 | -4.121302183 | 3.77E-05 | 0.000364515 | 0.607193072 |
| ENSMUSG00000099370 | Platr10 | 69193 | lncRNA | 1.257264713 | 0.607206888 | 2.134302511 | -0.28449898 | 0.776028015 | 1 | 0.607206888 |
| ENSMUSG00000068463 | B630019A10Rik | 105246089 | lncRNA | 1.480608433 | 0.607287872 | 2.143429374 | -0.283325347 | 0.776927449 | 1 | 0.607287872 |
| ENSMUSG00000036422 | Pcdh8 | 18530 | protein_coding | 1.342995845 | 0.6073392 | 2.149194889 | -0.28258917 | 0.777491783 | 1 | 0.6073392 |
| ENSMUSG00000073897 | Or10a4 | 18314 | protein_coding | 1.080596189 | 0.607389671 | 2.154849126 | -0.281871089 | 0.778042358 | 1 | 0.607389671 |
| ENSMUSG00000079480 | Pin4 | 69713 | protein_coding | 101.1980024 | -0.607398007 | 0.18401763 | 3.300759858 | 0.000964234 | 0.006174237 | 0.607398007 |
| ENSMUSG00000036117 | Il5 | 16191 | protein_coding | 1.082436646 | 0.607410622 | 2.157254754 | -0.281566477 | 0.778275948 | 1 | 0.607410622 |
| ENSMUSG00000054672 | Scart2 | 244234 | protein_coding | 1.815914896 | 0.607425542 | 2.159302808 | -0.281306327 | 0.778475458 | 1 | 0.607425542 |
| ENSMUSG00000029586 | Spdye4b | 330228 | protein_coding | 2.031640267 | 0.607430715 | 2.159920582 | -0.281228264 | 0.778535328 | 1 | 0.607430715 |
| ENSMUSG00000028610 | Dmrtb1 | 56296 | protein_coding | 1.089432325 | 0.607470915 | 2.164221885 | -0.280687909 | 0.778949784 | 1 | 0.607470915 |
| ENSMUSG00000091721 | Gimd1 | 433653 | protein_coding | 1.55838159 | 0.607476643 | 2.165095281 | -0.280577325 | 0.779034609 | 1 | 0.607476643 |
| ENSMUSG00000047963 | Stbd1 | 52331 | protein_coding | 4.689402735 | -0.607747782 | 0.571921459 | 1.062642033 | 0.287944318 | 0.50683356 | 0.607747782 |
| ENSMUSG00000036196 | Slc26a8 | 224661 | protein_coding | 4.351475399 | 0.607960158 | 0.819438124 | -0.741923204 | 0.458133866 | 1 | 0.607960158 |
| ENSMUSG00000000489 | Pdgfb | 18591 | protein_coding | 38.93633764 | 0.607973579 | 0.23841904 | -2.550021089 | 0.01077164 | 0.045218819 | 0.607973579 |
| ENSMUSG00000079111 | Kdelr2 | 66913 | protein_coding | 2590.911418 | 0.60805151 | 0.097560841 | -6.23253657 | 4.59E-10 | 1.05E-08 | 0.60805151 |
| ENSMUSG00000015437 | Gzmb | 14939 | protein_coding | 56.69086977 | 0.608181754 | 0.3652847 | -1.664952717 | 0.09592219 | 0.239682106 | 0.608181754 |
| ENSMUSG00000073702 | Rpl31 | 114641 | protein_coding | 3091.079943 | -0.608260287 | 0.143569565 | 4.236693822 | 2.27E-05 | 0.000231089 | 0.608260287 |
| ENSMUSG00000041298 | Katnal1 | 231912 | protein_coding | 51.24583016 | 0.608407839 | 0.160772597 | -3.784275735 | 0.000154157 | 0.001255918 | 0.608407839 |
| ENSMUSG00000021453 | Gadd45g | 23882 | protein_coding | 198.36364 | 0.608431062 | 0.209796902 | -2.900095555 | 0.003730489 | 0.019102418 | 0.608431062 |
| ENSMUSG00000068122 | Agtr2 | 11609 | protein_coding | 3.461451196 | 0.608612596 | 0.597839985 | -1.018019221 | 0.308668822 | 1 | 0.608612596 |
| ENSMUSG00000031144 | Syp | 20977 | protein_coding | 12.23458653 | -0.608626497 | 0.355760591 | 1.710775482 | 0.087122567 | 0.223685024 | 0.608626497 |
| ENSMUSG00000052125 | F730043M19Rik | 320046 | lncRNA | 3.651666744 | -0.60881501 | 0.936591951 | 0.65003229 | 0.515671364 | 1 | 0.60881501 |
| ENSMUSG00000040703 | Cyp2s1 | 74134 | protein_coding | 958.270963 | -0.60903742 | 0.145724883 | 4.179364609 | 2.92E-05 | 0.000290111 | 0.60903742 |
| ENSMUSG00000103948 | 4930594C11Rik | 77633 | TEC | 15.59208567 | -0.609041168 | 0.336847889 | 1.808059925 | 0.070597175 | 0.192129489 | 0.609041168 |
| ENSMUSG00000028795 | Ccdc28b | 66264 | protein_coding | 123.8253798 | -0.609058719 | 0.115408413 | 5.27742045 | 1.31E-07 | 2.13E-06 | 0.609058719 |
| ENSMUSG00000026141 | Col19a1 | 12823 | protein_coding | 1.837261565 | 0.609176246 | 1.565701479 | -0.389075603 | 0.697220221 | 1 | 0.609176246 |
| ENSMUSG00000025955 | Akr1cl | 70861 | protein_coding | 1.065686167 | 0.609211532 | 2.399346828 | -0.253907241 | 0.799567221 | 1 | 0.609211532 |
| ENSMUSG00000039092 | Sptlc3 | 228677 | protein_coding | 1.160807392 | 0.609300338 | 2.413558415 | -0.252448971 | 0.800694052 | 1 | 0.609300338 |
| ENSMUSG00000034731 | Dgkh | 380921 | protein_coding | 936.9362139 | -0.609312179 | 0.139354835 | 4.372379175 | 1.23E-05 | 0.000134753 | 0.609312179 |
| ENSMUSG00000099878 | Gm28178 | 102634661 | lncRNA | 0.975758551 | 0.609336272 | 2.419195608 | -0.25187557 | 0.801137244 | 1 | 0.609336272 |
| ENSMUSG00000099724 | 4930554C24Rik | 75253 | lncRNA | 0.927131791 | 0.609350104 | 2.421329709 | -0.251659286 | 0.80130443 | 1 | 0.609350104 |
| ENSMUSG00000085139 | A730046J19Rik | 319764 | protein_coding | 1.22111464 | 0.609377624 | 2.426185685 | -0.251166936 | 0.801685047 | 1 | 0.609377624 |
| ENSMUSG00000086338 | Or8h6 | 257901 | protein_coding | 1.394280297 | -0.609382284 | 2.084357827 | 0.292359726 | 0.770011597 | 1 | 0.609382284 |
| ENSMUSG00000057402 | Cldn34b2 | 73347 | protein_coding | 1.090296805 | 0.60939749 | 2.429300002 | -0.250853122 | 0.801927669 | 1 | 0.60939749 |
| ENSMUSG00000087194 | Skint6 | 230622 | protein_coding | 1.459114386 | 0.609416498 | 2.432764594 | -0.250503686 | 0.802197854 | 1 | 0.609416498 |
| ENSMUSG00000037124 | Trim58 | 216781 | protein_coding | 1.231775643 | 0.609434461 | 2.435503319 | -0.25022937 | 0.802409974 | 1 | 0.609434461 |
| ENSMUSG00000100147 | 1700047M11Rik | 67330 | lncRNA | 1.055235595 | 0.609451405 | 2.438212569 | -0.249958274 | 0.802619617 | 1 | 0.609451405 |
| ENSMUSG00000021943 | Gdf10 | 14560 | protein_coding | 1.572466678 | 0.609456085 | 2.439298095 | -0.249848957 | 0.802704158 | 1 | 0.609456085 |

| ensembl_gene_id | external_gene_name | entrezgene_id | gene_biotype | baseMean | log2FoldChange | lfcSE | stat | pvalue | padj | absLog2FC |
|---|---|---|---|---|---|---|---|---|---|---|
| ENSMUSG00000095239 | Or5p72 | 258733 | protein_coding | 1.160931009 | 0.609456397 | 2.439000375 | -0.249879583 | 0.802680473 | 1 | 0.609456397 |
| ENSMUSG00000042863 | Or5m13b | 258577 | protein_coding | 1.069196381 | 0.609460502 | 2.439575069 | -0.249822401 | 0.802724695 | 1 | 0.609460502 |
| ENSMUSG00000087115 | Pcsk2os2 | 378775 | lncRNA | 1.299485386 | 0.609465581 | 2.440653856 | -0.249714059 | 0.802808485 | 1 | 0.609465581 |
| ENSMUSG00000074991 | Gabrr3 | 328699 | protein_coding | 1.342918538 | 0.609470935 | 2.441586654 | -0.249620849 | 0.802880573 | 1 | 0.609470935 |
| ENSMUSG00000062753 | Smim29 | 106672 | protein_coding | 71.58332275 | -0.60947472 | 0.15575958 | 3.912919635 | 9.12E-05 | 0.000795489 | 0.60947472 |
| ENSMUSG00000085171 | D830026I12Rik | 319682 | lncRNA | 1.208491621 | 0.609491162 | 2.44481998 | -0.249298994 | 0.803129509 | 1 | 0.609491162 |
| ENSMUSG00000023192 | Grm2 | 108068 | protein_coding | 1.576594138 | 0.609496082 | 2.44594764 | -0.249186071 | 0.803216853 | 1 | 0.609496082 |
| ENSMUSG00000086823 | 1700028P15Rik | 75655 | lncRNA | 1.346305134 | 0.609519167 | 2.449630832 | -0.248820826 | 0.803499381 | 1 | 0.609519167 |
| ENSMUSG00000091450 | Vmn2r11 | 384219 | protein_coding | 1.204364161 | 0.609523642 | 2.45026279 | -0.248758478 | 0.803547611 | 1 | 0.609523642 |
| ENSMUSG00000049044 | Rapgef4 | 56508 | protein_coding | 58.27156965 | -0.609527777 | 0.257705348 | 2.365211978 | 0.018019754 | 0.067539312 | 0.609527777 |
| ENSMUSG00000020617 | 1700012B07Rik | 69324 | protein_coding | 0.986267925 | 0.609533344 | 2.451680797 | -0.248618558 | 0.803655852 | 1 | 0.609533344 |
| ENSMUSG00000020702 | Ccl1 | 20290 | protein_coding | 1.485600374 | 0.609544515 | 2.454007498 | -0.248387389 | 0.80383469 | 1 | 0.609544515 |
| ENSMUSG00000043969 | Emx2 | 13797 | protein_coding | 1.414966171 | 0.609569473 | 2.458225656 | -0.247971325 | 0.804156593 | 1 | 0.609569473 |
| ENSMUSG00000085271 | E130006D01Rik | 269683 | lncRNA | 1.076700798 | 0.609618502 | 2.466227808 | -0.247186614 | 0.804763804 | 1 | 0.609618502 |
| ENSMUSG00000099517 | H3c8 | 97908 | protein_coding | 230.456894 | 0.609648058 | 0.214852845 | -2.837514477 | 0.004546628 | 0.022473218 | 0.609648058 |
| ENSMUSG00000086807 | Platr21 | 78526 | lncRNA | 1.326896867 | 0.609787913 | 2.496025301 | -0.244303578 | 0.806995712 | 1 | 0.609787913 |
| ENSMUSG00000006522 | Itih3 | 16426 | protein_coding | 1.165058469 | 0.609791749 | 2.49646756 | -0.244261836 | 0.807028039 | 1 | 0.609791749 |
| ENSMUSG00000109270 | Gm10584 | 100043682 | lncRNA | 1.326130299 | -0.609823365 | 0.898887263 | 0.678420299 | 0.497505242 | 1 | 0.609823365 |
| ENSMUSG00000040660 | Cyp2b9 | 13094 | protein_coding | 1.441426359 | 0.609860286 | 2.509005737 | -0.24306851 | 0.807952325 | 1 | 0.609860286 |
| ENSMUSG00000071494 | Or10ac1 | 546896 | protein_coding | 1.580721599 | 0.609882319 | 2.513103476 | -0.242680942 | 0.808252572 | 1 | 0.609882319 |
| ENSMUSG00000040447 | Spns2 | 216892 | protein_coding | 40.30875939 | -0.610026426 | 0.241937037 | 2.521426374 | 0.011688013 | 0.048151079 | 0.610026426 |
| ENSMUSG00000111489 | 1700019J19Rik | 76412 | lncRNA | 3.425902722 | 0.610037767 | 2.214647626 | -0.275455905 | 0.782965998 | 1 | 0.610037767 |
| ENSMUSG00000024663 | Rab3il1 | 74760 | protein_coding | 5.026965649 | 0.610058429 | 0.883478746 | -0.690518512 | 0.489868172 | 0.697627026 | 0.610058429 |
| ENSMUSG00000060935 | Tmem263 | 103266 | protein_coding | 398.5472191 | 0.610063138 | 0.147009139 | -4.14983139 | 3.33E-05 | 0.000326473 | 0.610063138 |
| ENSMUSG00000061601 | Pclo | 26875 | protein_coding | 23.474097 | 0.610218476 | 0.471742052 | -1.293542674 | 0.195823435 | 0.395168001 | 0.610218476 |
| ENSMUSG00000036473 | Tbc1d24 | 224617 | protein_coding | 186.0344888 | -0.610226133 | 0.225551208 | 2.705488199 | 0.006820407 | 0.031251999 | 0.610226133 |
| ENSMUSG00000041523 | Upk2 | 22269 | protein_coding | 0.879121542 | 0.610355913 | 1.303368263 | -0.46829122 | 0.639576347 | 1 | 0.610355913 |
| ENSMUSG00000092517 | Art2a | 11871 | protein_coding | 62.13498513 | 0.610583107 | 0.251555833 | -2.427226984 | 0.015214733 | 0.059077383 | 0.610583107 |
| ENSMUSG00000030825 | Hsd17b14 | 66065 | protein_coding | 10.53473488 | -0.610693973 | 0.424466511 | 1.438732989 | 0.150226189 | 0.330029969 | 0.610693973 |
| ENSMUSG00000053647 | Gper1 | 76854 | protein_coding | 8.066882027 | -0.610774758 | 0.37462378 | 1.63036836 | 0.103023668 | 0.252989943 | 0.610774758 |
| ENSMUSG00000061175 | Fnip2 | 329679 | protein_coding | 345.4606018 | -0.610884435 | 0.129988847 | 4.699514233 | 2.61E-06 | 3.29E-05 | 0.610884435 |
| ENSMUSG00000029762 | Akr1b8 | 14187 | protein_coding | 940.0807141 | -0.610910261 | 0.097444738 | 6.269299629 | 3.63E-10 | 8.42E-09 | 0.610910261 |
| ENSMUSG00000002308 | Cd320 | 54219 | protein_coding | 230.2488439 | -0.610940542 | 0.147982321 | 4.128469778 | 3.65E-05 | 0.000354329 | 0.610940542 |
| ENSMUSG00000020787 | P2rx1 | 18436 | protein_coding | 20.38864022 | -0.610990228 | 0.308462485 | 1.98076025 | 0.047618167 | 0.143681053 | 0.610990228 |
| ENSMUSG00000022218 | Tgm1 | 21816 | protein_coding | 2.385979295 | -0.611037511 | 1.500660276 | 0.407179107 | 0.683876445 | 1 | 0.611037511 |
| ENSMUSG00000033355 | Rtp4 | 67775 | protein_coding | 130.6307312 | 0.611064181 | 0.335887055 | -1.819254929 | 0.068872546 | 0.188725185 | 0.611064181 |
| ENSMUSG00000046490 | Rnf222 | 320040 | protein_coding | 1.085177761 | 0.611169847 | 2.233872989 | -0.27359203 | 0.784398159 | 1 | 0.611169847 |
| ENSMUSG00000025422 | Agap2 | 216439 | protein_coding | 15.93739319 | 0.611227851 | 0.565268322 | -1.081305688 | 0.279561158 | 0.496963225 | 0.611227851 |
| ENSMUSG00000042888 | Prr30 | 76627 | protein_coding | 1.853826509 | 0.611278724 | 2.812008013 | -0.217381573 | 0.827910991 | 1 | 0.611278724 |

| ensembl_gene_id | external_gene_name | entrezgene_id | gene_biotype | baseMean | log2FoldChange | lfcSE | stat | pvalue | padj | absLog2FC |
|---|---|---|---|---|---|---|---|---|---|---|
| ENSMUSG00000108827 | Or4f6 | 258441 | protein_coding | 1.620150908 | 0.611331128 | 2.82509923 | -0.216392798 | 0.828681579 | 1 | 0.611331128 |
| ENSMUSG00000049010 | Or10s1 | 258853 | protein_coding | 1.624278368 | 0.611337663 | 2.82677698 | -0.216266677 | 0.828779883 | 1 | 0.611337663 |
| ENSMUSG00000096405 | 4930474N05Rik | 218921 | protein_coding | 1.902453268 | 0.611342111 | 2.8283178 | -0.216150431 | 0.828870491 | 1 | 0.611342111 |
| ENSMUSG00000010760 | Phlda2 | 22113 | protein_coding | 17.05450127 | -0.611364172 | 0.272721944 | 2.241712432 | 0.024979964 | 0.087305635 | 0.611364172 |
| ENSMUSG00000094858 | Or4k47 | 258890 | protein_coding | 1.485723991 | 0.611367532 | 2.834294878 | -0.215703573 | 0.829218816 | 1 | 0.611367532 |
| ENSMUSG00000037101 | Ttc29 | 73301 | protein_coding | 1.438163379 | 0.611375929 | 2.836404576 | -0.215546095 | 0.829341578 | 1 | 0.611375929 |
| ENSMUSG00000101480 | Or4c115 | 258902 | protein_coding | 1.438163379 | 0.611375929 | 2.836404576 | -0.215546095 | 0.829341578 | 1 | 0.611375929 |
| ENSMUSG00000045013 | Or6b6 | 259037 | protein_coding | 1.394730227 | 0.611405462 | 2.844034045 | -0.21497825 | 0.829784276 | 1 | 0.611405462 |
| ENSMUSG00000029414 | Kntc1 | 208628 | protein_coding | 190.5636383 | 0.611412041 | 0.208847944 | -2.92754637 | 0.003416481 | 0.017730914 | 0.611412041 |
| ENSMUSG00000059832 | Kprp | 433619 | protein_coding | 1.299609003 | 0.611426782 | 2.849476522 | -0.214575125 | 0.830098589 | 1 | 0.611426782 |
| ENSMUSG00000075069 | Or4c3 | 258206 | protein_coding | 1.442290839 | 0.611430152 | 2.85060517 | -0.21449135 | 0.830163912 | 1 | 0.611430152 |
| ENSMUSG00000036131 | Frmd7 | 385354 | protein_coding | 1.117621474 | 0.611480109 | 2.863233927 | -0.213562749 | 0.830888051 | 1 | 0.611480109 |
| ENSMUSG00000050397 | Foxl2 | 26927 | protein_coding | 1.117621474 | 0.611480109 | 2.863233927 | -0.213562749 | 0.830888051 | 1 | 0.611480109 |
| ENSMUSG00000046248 | Krtap5-3 | 77215 | protein_coding | 1.165182086 | 0.611484357 | 2.864474543 | -0.213471737 | 0.830959031 | 1 | 0.611484357 |
| ENSMUSG00000109145 | Gm20744 | 434205 | lncRNA | 1.165182086 | 0.611484357 | 2.864474543 | -0.213471737 | 0.830959031 | 1 | 0.611484357 |
| ENSMUSG00000044248 | Vmn1r45 | 22297 | protein_coding | 1.167818179 | 0.611500265 | 2.868854114 | -0.213151398 | 0.831208876 | 1 | 0.611500265 |
| ENSMUSG00000045792 | Or51g1 | 259119 | protein_coding | 1.351297075 | 0.611509086 | 2.87150292 | -0.212957849 | 0.83135984 | 1 | 0.611509086 |
| ENSMUSG00000023443 | Esx1 | 13984 | protein_coding | 1.070060862 | 0.611520734 | 2.874047098 | -0.212773386 | 0.831503723 | 1 | 0.611520734 |
| ENSMUSG00000034087 | Nlrp4b | 210045 | protein_coding | 1.070060862 | 0.611520734 | 2.874047098 | -0.212773386 | 0.831503723 | 1 | 0.611520734 |
| ENSMUSG00000098769 | Mir8103 | 102465893 | miRNA | 1.629023243 | -0.611545011 | 0.885011065 | 0.691002672 | 0.489563862 | 1 | 0.611545011 |
| ENSMUSG00000042501 | Cpa6 | 329093 | protein_coding | 1.31592923 | 0.611545215 | 2.881300069 | -0.212246278 | 0.831914905 | 1 | 0.611545215 |
| ENSMUSG00000050030 | Or4k1 | 258039 | protein_coding | 1.172262327 | 0.611578976 | 2.890253376 | -0.211600471 | 0.832418742 | 1 | 0.611578976 |
| ENSMUSG00000079330 | Lemd1 | 213409 | protein_coding | 1.260303311 | 0.611660012 | 2.912908344 | -0.209982581 | 0.833681268 | 1 | 0.611660012 |
| ENSMUSG00000115177 | 4930523O13Rik | 74726 | lncRNA | 1.260303311 | 0.611660012 | 2.912908344 | -0.209982581 | 0.833681268 | 1 | 0.611660012 |
| ENSMUSG00000085057 | Gm13415 | 102638285 | lncRNA | 10.32841357 | -0.611806577 | 0.427417984 | 1.431401111 | 0.15231529 | 0.332892364 | 0.611806577 |
| ENSMUSG00000030167 | Klrc1 | 16641 | protein_coding | 2.125506891 | 0.611923538 | 0.931541606 | -0.656893405 | 0.511249459 | 1 | 0.611923538 |
| ENSMUSG00000032446 | Eomes | 13813 | protein_coding | 3.508564499 | -0.61196717 | 0.922442133 | 0.663420661 | 0.507061177 | 1 | 0.61196717 |
| ENSMUSG00000079553 | Kifc1 | 100502766 | protein_coding | 183.1352002 | 0.612047629 | 0.182195042 | -3.359299042 | 0.000781405 | 0.005165779 | 0.612047629 |
| ENSMUSG00000025738 | Fbxl16 | 214931 | protein_coding | 9.394924731 | -0.612309859 | 0.425697001 | 1.438370148 | 0.150329058 | 0.330166076 | 0.612309859 |
| ENSMUSG00000000673 | Haao | 107766 | protein_coding | 4.520634495 | 0.612378877 | 0.861528667 | -0.710804991 | 0.477205087 | 0.68880035 | 0.612378877 |
| ENSMUSG00000044022 | Pcdhb21 | 93892 | protein_coding | 1.141719804 | -0.612629817 | 1.168258592 | 0.52439573 | 0.600003326 | 1 | 0.612629817 |
| ENSMUSG00000049539 | H1f1 | 80838 | protein_coding | 1208.252444 | 0.612771002 | 0.188935331 | -3.243284349 | 0.001181602 | 0.007316453 | 0.612771002 |
| ENSMUSG00000034617 | Mtrr | 210009 | protein_coding | 156.308789 | 0.612812836 | 0.116286819 | -5.269839198 | 1.37E-07 | 2.21E-06 | 0.612812836 |
| ENSMUSG00000058159 | T2 | 21331 | protein_coding | 1.24870035 | -0.612971421 | 1.352016358 | 0.453375743 | 0.6502782 | 1 | 0.612971421 |
| ENSMUSG00000064284 | Cdpf1 | 72355 | protein_coding | 310.5604786 | 0.613017293 | 0.090283957 | -6.789880633 | 1.12E-11 | 3.14E-10 | 0.613017293 |
| ENSMUSG00000021214 | Akr1c18 | 105349 | protein_coding | 19.54115467 | -0.613045359 | 0.358337194 | 1.710805827 | 0.087116964 | 0.223685024 | 0.613045359 |
| ENSMUSG00000019326 | Aoc3 | 11754 | protein_coding | 3.577226937 | 0.613246445 | 1.861191679 | -0.329491289 | 0.741784377 | 1 | 0.613246445 |
| ENSMUSG00000114288 | Gm32123 | 102634576 | lncRNA | 0.730749059 | -0.61327051 | 1.798285648 | 0.341030643 | 0.733080514 | 1 | 0.61327051 |
| ENSMUSG00000025961 | 4933402D24Rik | 74426 | lncRNA | 1.152047539 | 0.613585324 | 1.078274527 | -0.569043698 | 0.569326485 | 1 | 0.613585324 |

Supplementary Table 1

| ensembl_gene_id | external_gene_name | entrezgene_id | gene_biotype | baseMean | log2FoldChange | lfcSE | stat | pvalue | padj | absLog2FC |
|---|---|---|---|---|---|---|---|---|---|---|
| ENSMUSG00000037336 | Mfsd2b | 432628 | protein_coding | 15.30449836 | 0.613827405 | 0.425850699 | -1.441414577 | 0.149467594 | 0.328844638 | 0.613827405 |
| ENSMUSG00000028175 | Depdc1a | 76131 | protein_coding | 56.46356996 | 0.614533295 | 0.223013313 | -2.755590175 | 0.005858635 | 0.027682261 | 0.614533295 |
| ENSMUSG00000039217 | Il18 | 16173 | protein_coding | 935.9836996 | -0.614745025 | 0.222286813 | 2.765548781 | 0.00568271 | 0.027075673 | 0.614745025 |
| ENSMUSG00000086040 | Wipf3 | 330319 | protein_coding | 16.43272754 | -0.614846692 | 0.366086083 | 1.679513971 | 0.093051918 | 0.234222517 | 0.614846692 |
| ENSMUSG00000058070 | Eml1 | 68519 | protein_coding | 38.34061234 | 0.614906687 | 0.192779657 | -3.189686592 | 0.001424272 | 0.008537358 | 0.614906687 |
| ENSMUSG00000037962 | Rflna | 73121 | protein_coding | 9.499126829 | -0.614968663 | 0.367370665 | 1.673973242 | 0.094135858 | 0.236336338 | 0.614968663 |
| ENSMUSG00000027344 | Fsip1 | 71313 | protein_coding | 2.481844465 | 0.614987733 | 1.25151563 | -0.491394369 | 0.623147546 | 1 | 0.614987733 |
| ENSMUSG00000090843 | Heatr4 | 101055670 | protein_coding | 2.194075772 | 0.615389379 | 1.289299475 | -0.477305227 | 0.633144789 | 1 | 0.615389379 |
| ENSMUSG00000065999 | Zfp985 | 433804 | protein_coding | 2.978723775 | -0.615504892 | 0.986355955 | 0.624019036 | 0.532615084 | 1 | 0.615504892 |
| ENSMUSG00000027447 | Cst3 | 13010 | protein_coding | 770.2207571 | -0.615528668 | 0.111240105 | 5.533334112 | 3.14E-08 | 5.57E-07 | 0.615528668 |
| ENSMUSG00000078695 | Cisd3 | 217149 | protein_coding | 970.7512999 | -0.615834858 | 0.098183707 | 6.272271421 | 3.56E-10 | 8.30E-09 | 0.615834858 |
| ENSMUSG00000003779 | Kif20a | 19348 | protein_coding | 318.577639 | 0.615902921 | 0.175279906 | -3.513825031 | 0.000441704 | 0.003200656 | 0.615902921 |
| ENSMUSG00000067594 | Krt77 | 406220 | protein_coding | 2.634616148 | 0.615942425 | 3.022434925 | -0.203790136 | 0.838517501 | 1 | 0.615942425 |
| ENSMUSG00000029392 | Rilpl1 | 75695 | protein_coding | 14.45366454 | 0.61597314 | 0.289782318 | -2.125640874 | 0.033533174 | 0.109773894 | 0.61597314 |
| ENSMUSG00000029286 | Enam | 13801 | protein_coding | 2.452628619 | 0.616165291 | 3.032264381 | -0.203203024 | 0.83897635 | 1 | 0.616165291 |
| ENSMUSG00000016262 | Sertad4 | 214791 | protein_coding | 4.37488616 | -0.616195021 | 0.70517701 | 0.873816094 | 0.382218415 | 1 | 0.616195021 |
| ENSMUSG00000043424 | Eif3j2 | 100042807 | protein_coding | 6.878788824 | 0.616203422 | 0.514807926 | -1.196957914 | 0.231322959 | 0.440150633 | 0.616203422 |
| ENSMUSG00000059654 | Reg1 | 19692 | protein_coding | 9.201342679 | -0.616744231 | 0.471880001 | 1.306993791 | 0.19121482 | 0.388834077 | 0.616744231 |
| ENSMUSG00000007946 | Phox2a | 11859 | protein_coding | 2.395748284 | 0.616759946 | 1.008977571 | -0.611272206 | 0.541019388 | 1 | 0.616759946 |
| ENSMUSG00000021365 | Nedd9 | 18003 | protein_coding | 1474.37095 | -0.616871972 | 0.077674756 | 7.941730409 | 1.99E-15 | 7.52E-14 | 0.616871972 |
| ENSMUSG00000044125 | 9530080O11Rik | 319247 | lncRNA | 11.65270883 | 0.616975399 | 0.374053988 | -1.649428742 | 0.09905983 | 0.24588517 | 0.616975399 |
| ENSMUSG00000056755 | Grm7 | 108073 | protein_coding | 1.807417349 | 0.617017344 | 3.070855911 | -0.20092683 | 0.840755788 | 1 | 0.617017344 |
| ENSMUSG00000037936 | Scarb1 | 20778 | protein_coding | 156.1443636 | -0.617074463 | 0.109579552 | 5.631292063 | 1.79E-08 | 3.31E-07 | 0.617074463 |
| ENSMUSG00000097075 | Cdiptos | 381922 | protein_coding | 7.164284197 | -0.617211159 | 0.534768354 | 1.154165452 | 0.248432343 | 0.460735958 | 0.617211159 |
| ENSMUSG00000048226 | Or4c107 | 258241 | protein_coding | 1.581996668 | 0.617332913 | 3.085554819 | -0.200071932 | 0.841424325 | 1 | 0.617332913 |
| ENSMUSG00000057457 | Phex | 18675 | protein_coding | 1.581996668 | 0.617332913 | 3.085554819 | -0.200071932 | 0.841424325 | 1 | 0.617332913 |
| ENSMUSG00000069294 | Vmn1r197 | 171278 | protein_coding | 1.486875444 | 0.617443583 | 3.09075588 | -0.199771061 | 0.841659635 | 1 | 0.617443583 |
| ENSMUSG00000021286 | Zfyve21 | 68520 | protein_coding | 390.8456675 | 0.617462567 | 0.084203566 | -7.332974058 | 2.25E-13 | 7.26E-12 | 0.617462567 |
| ENSMUSG00000035455 | Fignl1 | 60530 | protein_coding | 114.5499219 | 0.617537287 | 0.245715902 | -2.513216617 | 0.011963585 | 0.048978652 | 0.617537287 |
| ENSMUSG00000051220 | Ercc6l | 236930 | protein_coding | 134.8392815 | 0.61764252 | 0.19970817 | -3.092725346 | 0.001983276 | 0.011262661 | 0.61764252 |
| ENSMUSG00000061298 | Agbl4 | 78933 | protein_coding | 2.433172551 | 0.617701676 | 2.155107472 | -0.286622214 | 0.774401603 | 1 | 0.617701676 |
| ENSMUSG00000044550 | Tceal3 | 594844 | protein_coding | 1.715857564 | -0.61780601 | 1.22132478 | 0.505849075 | 0.612962604 | 1 | 0.61780601 |
| ENSMUSG00000040061 | Plcb2 | 18796 | protein_coding | 58.31037085 | -0.617837852 | 0.180012983 | 3.432184945 | 0.000598739 | 0.004121541 | 0.617837852 |
| ENSMUSG00000029638 | Glcci1 | 170772 | protein_coding | 27.82757817 | 0.617840667 | 0.295708651 | -2.089356075 | 0.036675682 | 0.117581482 | 0.617840667 |
| ENSMUSG00000027547 | Sall4 | 99377 | protein_coding | 1.3773909 | 0.617893207 | 3.112334075 | -0.198530489 | 0.842630032 | 1 | 0.617893207 |
| ENSMUSG00000024136 | Dnase1l2 | 66705 | protein_coding | 75.92547408 | -0.617902131 | 0.322724006 | 1.914645704 | 0.055537702 | 0.161179189 | 0.617902131 |
| ENSMUSG00000031344 | Gabrq | 57249 | protein_coding | 1.953605236 | -0.618004071 | 1.907974652 | 0.323905808 | 0.746009347 | 1 | 0.618004071 |
| ENSMUSG00000029869 | Ephb6 | 13848 | protein_coding | 110.3489401 | -0.618074239 | 0.305297075 | 2.024501016 | 0.042918628 | 0.132673445 | 0.618074239 |
| ENSMUSG00000028978 | Nos3 | 18127 | protein_coding | 2.829694754 | -0.618133974 | 1.057476851 | 0.584536648 | 0.558859316 | 1 | 0.618133974 |

| ensembl_gene_id | external_gene_name | entrezgene_id | gene_biotype | baseMean | log2FoldChange | lfcSE | stat | pvalue | padj | absLog2FC |
|---|---|---|---|---|---|---|---|---|---|---|
| ENSMUSG00000036833 | Pnpla7 | 241274 | protein_coding | 131.8662624 | -0.618161654 | 0.124919606 | 4.948475855 | 7.48E-07 | 1.06E-05 | 0.618161654 |
| ENSMUSG00000048776 | Pthlh | 19227 | protein_coding | 3.366283624 | 0.6182015 | 0.626906696 | -0.986114048 | 0.324077151 | 1 | 0.6182015 |
| ENSMUSG00000037801 | Iqch | 78250 | protein_coding | 11.66647211 | -0.61820706 | 0.462349644 | 1.337098596 | 0.181190455 | 0.375235922 | 0.61820706 |
| ENSMUSG00000024784 | Gpha2 | 170458 | protein_coding | 1.518002497 | -0.61828507 | 1.116305323 | 0.55386735 | 0.579669632 | 1 | 0.61828507 |
| ENSMUSG00000078498 | Zfp988 | 115489950 | protein_coding | 2.190326761 | -0.618488683 | 1.003240795 | 0.616490763 | 0.537570667 | 1 | 0.618488683 |
| ENSMUSG00000029070 | Mxra8 | 74761 | protein_coding | 3.991846256 | -0.618600201 | 0.785210181 | 0.787814799 | 0.430805044 | 1 | 0.618600201 |
| ENSMUSG00000030528 | Blm | 12144 | protein_coding | 83.26498655 | 0.618719317 | 0.193452396 | -3.198302683 | 0.001382391 | 0.008334723 | 0.618719317 |
| ENSMUSG00000101731 | 3830432H09Rik | 320069 | lncRNA | 1.528035428 | 0.618772462 | 1.262654189 | -0.490056951 | 0.6240936 | 1 | 0.618772462 |
| ENSMUSG00000033762 | Recql4 | 79456 | protein_coding | 75.42847927 | 0.618975119 | 0.160837495 | -3.848450377 | 0.000118867 | 0.001004984 | 0.618975119 |
| ENSMUSG00000083820 | Ndufs6b | 623286 | processed_pseudogene | 0.692718363 | 0.618978987 | 1.150926171 | -0.537809464 | 0.590708598 | 1 | 0.618978987 |
| ENSMUSG00000063704 | Mapk15 | 332110 | protein_coding | 39.17615213 | -0.619000568 | 0.231941859 | 2.668774706 | 0.00761285 | 0.034181239 | 0.619000568 |
| ENSMUSG00000075217 | Fads2b | 228151 | protein_coding | 1.485897172 | -0.619345334 | 1.145567141 | 0.540645163 | 0.588752182 | 1 | 0.619345334 |
| ENSMUSG00000032815 | Fanca | 14087 | protein_coding | 137.0552758 | 0.619353788 | 0.212495716 | -2.914664827 | 0.003560707 | 0.018358264 | 0.619353788 |
| ENSMUSG00000021720 | Rnf180 | 71816 | protein_coding | 51.81762407 | -0.619559902 | 0.174413659 | 3.552244154 | 0.00038196 | 0.002815934 | 0.619559902 |
| ENSMUSG00000046743 | Fat4 | 329628 | protein_coding | 9.389221367 | 0.619711252 | 0.656862195 | -0.943441801 | 0.345454971 | 0.567081848 | 0.619711252 |
| ENSMUSG00000002833 | Hdgfl2 | 15193 | protein_coding | 422.559622 | 0.619734325 | 0.100869951 | -6.143894371 | 8.05E-10 | 1.80E-08 | 0.619734325 |
| ENSMUSG00000026950 | Neb | 17996 | protein_coding | 12.77598847 | 0.619746829 | 0.964902799 | -0.642289389 | 0.520685303 | 0.720610174 | 0.619746829 |
| ENSMUSG00000023829 | Slc22a1 | 20517 | protein_coding | 333.641529 | -0.619845144 | 0.082093192 | 7.550506079 | 4.34E-14 | 1.47E-12 | 0.619845144 |
| ENSMUSG00000001020 | S100a4 | 20198 | protein_coding | 1.471666417 | 0.619983919 | 0.854787903 | -0.725307315 | 0.468263519 | 1 | 0.619983919 |
| ENSMUSG00000062480 | Acat3 | 224530 | protein_coding | 3.043661468 | 0.619988018 | 0.816812165 | -0.759033771 | 0.447832355 | 1 | 0.619988018 |
| ENSMUSG00000037904 | Ankrd9 | 74251 | protein_coding | 325.9430341 | -0.620065562 | 0.134721228 | 4.602582456 | 4.17E-06 | 5.09E-05 | 0.620065562 |
| ENSMUSG00000001424 | Snd1 | 56463 | protein_coding | 2218.053816 | 0.620253078 | 0.100514686 | -6.170770664 | 6.80E-10 | 1.53E-08 | 0.620253078 |
| ENSMUSG00000071226 | Cecr2 | 330409 | protein_coding | 9.400156062 | 0.620318356 | 0.925461108 | -0.670280307 | 0.502679119 | 0.707452102 | 0.620318356 |
| ENSMUSG00000038390 | Gpr162 | 14788 | protein_coding | 3.748183603 | 0.620343885 | 1.245099169 | -0.498228495 | 0.618323 | 1 | 0.620343885 |
| ENSMUSG00000037876 | Jmjd1c | 108829 | protein_coding | 1523.040346 | 0.620370298 | 0.077206549 | -8.035203004 | 9.34E-16 | 3.66E-14 | 0.620370298 |
| ENSMUSG00000031673 | Cdh11 | 12552 | protein_coding | 6.23948563 | 0.62048485 | 0.647364573 | -0.958478229 | 0.337821664 | 0.559613498 | 0.62048485 |
| ENSMUSG00000039621 | Prex1 | 277360 | protein_coding | 91.73980435 | -0.620560337 | 0.234786272 | 2.643086118 | 0.008215414 | 0.036352985 | 0.620560337 |
| ENSMUSG00000059327 | Eda | 13607 | protein_coding | 5.356794397 | 0.62061009 | 0.950536904 | -0.652904782 | 0.513817666 | 0.715626628 | 0.62061009 |
| ENSMUSG00000026415 | Fcamr | 64435 | protein_coding | 61.65693116 | 0.620654336 | 0.214248204 | -2.89689399 | 0.003768771 | 0.019227501 | 0.620654336 |
| ENSMUSG00000103332 | Pcdhga2 | 93710 | protein_coding | 2.967082388 | 0.620874308 | 0.736011345 | -0.843566219 | 0.398911848 | 1 | 0.620874308 |
| ENSMUSG00000027115 | Kif18a | 228421 | protein_coding | 117.4282059 | 0.620899531 | 0.188666196 | -3.29099512 | 0.000998336 | 0.006350049 | 0.620899531 |
| ENSMUSG00000045545 | Krt14 | 16664 | protein_coding | 17.00490827 | -0.621421376 | 0.355812124 | 1.746487356 | 0.080726302 | 0.211290803 | 0.621421376 |
| ENSMUSG00000075265 | Tnk2os | 675921 | lncRNA | 2.977480838 | -0.621755725 | 0.727120914 | 0.855092617 | 0.392499876 | 1 | 0.621755725 |
| ENSMUSG00000032988 | Slc16a8 | 57274 | protein_coding | 17.77396449 | 0.621802695 | 0.314839631 | -1.974982291 | 0.048270158 | 0.1452832 | 0.621802695 |
| ENSMUSG00000021629 | Slc30a5 | 69048 | protein_coding | 996.8522968 | 0.621806905 | 0.095032181 | -6.543119368 | 6.02E-11 | 1.56E-09 | 0.621806905 |
| ENSMUSG00000020936 | Nmt1 | 18107 | protein_coding | 1434.760509 | 0.621851991 | 0.068590891 | -9.066101704 | 1.23E-19 | 6.12E-18 | 0.621851991 |
| ENSMUSG00000025375 | Aatk | 11302 | protein_coding | 5.101388652 | 0.621940972 | 1.127474092 | -0.551623293 | 0.581206474 | 0.764339772 | 0.621940972 |
| ENSMUSG00000022253 | Nadk2 | 68646 | protein_coding | 464.5988984 | 0.622028917 | 0.096979909 | -6.413997729 | 1.42E-10 | 3.45E-09 | 0.622028917 |
| ENSMUSG00000036902 | Neto2 | 74513 | protein_coding | 30.87917158 | 0.622118272 | 0.27626307 | -2.2519053 | 0.024328257 | 0.085571212 | 0.622118272 |

| ensembl_gene_id | external_gene_name | entrezgene_id | gene_biotype | baseMean | log2FoldChange | lfcSE | stat | pvalue | padj | absLog2FC |
|---|---|---|---|---|---|---|---|---|---|---|
| ENSMUSG00000037622 | Wdtc1 | 230796 | protein_coding | 496.374024 | -0.622139035 | 0.104702095 | 5.941992206 | 2.82E-09 | 5.88E-08 | 0.622139035 |
| ENSMUSG00000021697 | Depdc1b | 218581 | protein_coding | 54.58608888 | 0.622139345 | 0.211602521 | -2.940132005 | 0.003280724 | 0.017134056 | 0.622139345 |
| ENSMUSG00000021253 | Tgfb3 | 21809 | protein_coding | 4.64917903 | 0.622333711 | 0.580073034 | -1.072854062 | 0.2833366 | 0.501395265 | 0.622333711 |
| ENSMUSG00000078593 | P3r3urf | 67323 | protein_coding | 1.340765365 | -0.622480635 | 1.091570688 | 0.570261406 | 0.568500413 | 1 | 0.622480635 |
| ENSMUSG00000065191 | Gm23458 | 115490288 | snRNA | 0.832032919 | -0.622491999 | 1.196445993 | 0.520284244 | 0.602865476 | 1 | 0.622491999 |
| ENSMUSG00000054146 | Krt15 | 16665 | protein_coding | 15.47004257 | 0.62257051 | 0.52252074 | -1.191475213 | 0.233467082 | 0.442885865 | 0.62257051 |
| ENSMUSG00000014602 | Kif1a | 16560 | protein_coding | 22.54427299 | -0.622793874 | 0.341977746 | 1.821153223 | 0.068583569 | 0.18810647 | 0.622793874 |
| ENSMUSG00000004814 | Ccl24 | 56221 | protein_coding | 0.675005381 | -0.622873649 | 2.016564542 | 0.308878608 | 0.75741387 | 1 | 0.622873649 |
| ENSMUSG00000044005 | Gls2 | 216456 | protein_coding | 11.20235964 | 0.622893965 | 0.333159466 | -1.869657111 | 0.061531449 | 0.173816344 | 0.622893965 |
| ENSMUSG00000021010 | Npas3 | 27386 | protein_coding | 5.965773354 | -0.62293962 | 0.610443382 | 1.020470758 | 0.307505251 | 0.527928237 | 0.62293962 |
| ENSMUSG00000038580 | Sct | 20287 | protein_coding | 107.2990433 | -0.62302567 | 0.271852626 | 2.291777275 | 0.021918499 | 0.07888094 | 0.62302567 |
| ENSMUSG00000108660 | Gm21284 | 100861870 | lncRNA | 2.093057074 | 0.623059425 | 0.984691999 | -0.632745493 | 0.526899856 | 1 | 0.623059425 |
| ENSMUSG00000071540 | 3425401B19Rik | 100504518 | protein_coding | 11.22652524 | -0.623155065 | 0.443737611 | 1.404332311 | 0.160219917 | 0.344649174 | 0.623155065 |
| ENSMUSG00000087088 | Gm16638 | 102631889 | lncRNA | 11.15179848 | 0.623434119 | 0.447231017 | -1.393986764 | 0.163321602 | 0.349409629 | 0.623434119 |
| ENSMUSG00000022505 | Emp2 | 13731 | protein_coding | 331.3285398 | -0.6234834 | 0.132908622 | 4.691068119 | 2.72E-06 | 3.42E-05 | 0.6234834 |
| ENSMUSG00000090137 | Uba52 | 22186 | protein_coding | 114.6114229 | 0.623509656 | 0.144143551 | -4.325616025 | 1.52E-05 | 0.000161703 | 0.623509656 |
| ENSMUSG00000031258 | Xkrx | 331524 | protein_coding | 4.252390739 | 0.623552579 | 0.865973663 | -0.720059519 | 0.47148835 | 1 | 0.623552579 |
| ENSMUSG00000027323 | Rad51 | 19361 | protein_coding | 54.59433559 | 0.623652281 | 0.217471082 | -2.867748094 | 0.004134045 | 0.020744341 | 0.623652281 |
| ENSMUSG00000085631 | 9630028H03Rik | 320684 | lncRNA | 2.968707919 | 0.623723543 | 0.847238929 | -0.736183764 | 0.461618872 | 1 | 0.623723543 |
| ENSMUSG00000027690 | Slc2a2 | 20526 | protein_coding | 10.22241635 | 0.623756782 | 0.897736931 | -0.694810206 | 0.487174251 | 0.695767999 | 0.623756782 |
| ENSMUSG00000022587 | Ly6e | 17069 | protein_coding | 1155.2834 | -0.623876107 | 0.273170499 | 2.283834119 | 0.022381282 | 0.08026156 | 0.623876107 |
| ENSMUSG00000025701 | Alox5 | 11689 | protein_coding | 81.16670742 | -0.62423902 | 0.238753175 | 2.61457893 | 0.008933752 | 0.038854879 | 0.62423902 |
| ENSMUSG00000086537 | Gnasas1 | 56802 | lncRNA | 0.959690653 | -0.62440712 | 1.350741992 | 0.462269718 | 0.643887912 | 1 | 0.62440712 |
| ENSMUSG00000022947 | Cbr3 | 109857 | protein_coding | 518.6061095 | -0.624685924 | 0.1931227 | 3.234658193 | 0.001217884 | 0.00748498 | 0.624685924 |
| ENSMUSG00000089706 | B230216N24Rik | 78603 | lncRNA | 10.59602312 | -0.624738192 | 0.342453065 | 1.824303112 | 0.06810626 | 0.187201528 | 0.624738192 |
| ENSMUSG00000085980 | Gm12408 | 100040557 | lncRNA | 27.7906184 | 0.62478894 | 0.259749042 | -2.405356089 | 0.016156714 | 0.06203487 | 0.62478894 |
| ENSMUSG00000034329 | Brip1 | 237911 | protein_coding | 114.4674328 | 0.625359714 | 0.194806456 | -3.210159078 | 0.001326615 | 0.008063204 | 0.625359714 |
| ENSMUSG00000031861 | Lpar2 | 53978 | protein_coding | 17.93358588 | -0.625675204 | 0.291562405 | 2.145939235 | 0.031877827 | 0.105793898 | 0.625675204 |
| ENSMUSG00000074207 | Adh1 | 11522 | protein_coding | 5892.910036 | -0.625904759 | 0.316967253 | 1.974666952 | 0.048305956 | 0.145305019 | 0.625904759 |
| ENSMUSG00000015134 | Aldh1a3 | 56847 | protein_coding | 27.632863 | 0.625965398 | 0.282739283 | -2.213931471 | 0.026833493 | 0.09235688 | 0.625965398 |
| ENSMUSG00000104713 | Gbp6 | 100702 | protein_coding | 7.51884915 | -0.62610467 | 0.48926317 | 1.279688946 | 0.200654554 | 0.401975042 | 0.62610467 |
| ENSMUSG00000039007 | Cpq | 54381 | protein_coding | 574.3656056 | -0.626106737 | 0.178539399 | 3.506826728 | 0.000453484 | 0.003263111 | 0.626106737 |
| ENSMUSG00000051008 | 4930412M03Rik | 100504140 | lncRNA | 12.2103283 | -0.626437708 | 0.356525839 | 1.757061171 | 0.078907383 | 0.208009073 | 0.626437708 |
| ENSMUSG00000047343 | Mettl21c | 433294 | protein_coding | 7.413718735 | 0.626539115 | 0.531434036 | -1.17895933 | 0.238414371 | 0.448589299 | 0.626539115 |
| ENSMUSG00000047441 | Antxrl | 239029 | protein_coding | 2.824858596 | 0.626642972 | 3.676585424 | -0.170441565 | 0.864662887 | 1 | 0.626642972 |
| ENSMUSG00000028158 | Mttp | 17777 | protein_coding | 73.44296495 | 0.626916371 | 0.369931025 | -1.694684492 | 0.090135289 | 0.228911821 | 0.626916371 |
| ENSMUSG00000025140 | Pycr1 | 209027 | protein_coding | 95.42489065 | 0.626967498 | 0.148046415 | -4.234938727 | 2.29E-05 | 0.000232612 | 0.626967498 |
| ENSMUSG00000020953 | Coch | 12810 | protein_coding | 4.450683667 | 0.626973613 | 0.672077133 | -0.93288937 | 0.350877092 | 0.572963936 | 0.626973613 |
| ENSMUSG00000020994 | Pnn | 18949 | protein_coding | 685.6694221 | 0.626985182 | 0.096143302 | -6.521361041 | 6.97E-11 | 1.78E-09 | 0.626985182 |

Supplementary Table 1

| ensembl_gene_id | external_gene_name | entrezgene_id | gene_biotype | baseMean | log2FoldChange | lfcSE | stat | pvalue | padj | absLog2FC |
|---|---|---|---|---|---|---|---|---|---|---|
| ENSMUSG00000037747 | Phyhipl | 70911 | protein_coding | 2.984335632 | 0.62701474 | 1.994169127 | -0.314424053 | 0.75319899 | 1 | 0.62701474 |
| ENSMUSG00000038943 | Prc1 | 233406 | protein_coding | 367.4350801 | 0.627473099 | 0.166231396 | -3.774696689 | 0.000160202 | 0.001299403 | 0.627473099 |
| ENSMUSG00000086494 | 2210417A02Rik | 70138 | lncRNA | 3.656803106 | -0.627623562 | 0.608111768 | 1.032085869 | 0.302031893 | 1 | 0.627623562 |
| ENSMUSG00000064833 | Gm25926 | 115488083 | snRNA | 2.146683097 | -0.627638781 | 0.753283365 | 0.833204091 | 0.404729636 | 1 | 0.627638781 |
| ENSMUSG00000002633 | Shh | 20423 | protein_coding | 15.24165918 | -0.627863609 | 0.341445525 | 1.838839766 | 0.065938757 | 0.182841722 | 0.627863609 |
| ENSMUSG00000036641 | Ccdc148 | 227933 | protein_coding | 10.30476007 | -0.627978064 | 0.676180167 | 0.928714113 | 0.353037265 | 0.575410323 | 0.627978064 |
| ENSMUSG00000036964 | Trim17 | 56631 | protein_coding | 6.421505727 | 0.628256228 | 0.566543098 | -1.108929278 | 0.267460692 | 0.482609908 | 0.628256228 |
| ENSMUSG00000031090 | Nadsyn1 | 78914 | protein_coding | 493.4867996 | -0.628277989 | 0.078641172 | 7.989173754 | 1.36E-15 | 5.19E-14 | 0.628277989 |
| ENSMUSG00000037709 | Fam13a | 58909 | protein_coding | 643.5168173 | 0.628312422 | 0.112554321 | -5.582303865 | 2.37E-08 | 4.31E-07 | 0.628312422 |
| ENSMUSG00000023800 | Tiam2 | 24001 | protein_coding | 379.2341237 | 0.628365322 | 0.127177134 | -4.940867146 | 7.78E-07 | 1.10E-05 | 0.628365322 |
| ENSMUSG00000096971 | 4930556M19Rik | 75259 | lncRNA | 14.58900378 | -0.628513935 | 0.297489307 | 2.112727818 | 0.034624071 | 0.112577655 | 0.628513935 |
| ENSMUSG00000031922 | Cep57 | 74360 | protein_coding | 270.2494935 | 0.62877666 | 0.109683163 | -5.732663465 | 9.89E-09 | 1.90E-07 | 0.62877666 |
| ENSMUSG00000002983 | Relb | 19698 | protein_coding | 432.3079431 | -0.62891054 | 0.092871808 | 6.771813248 | 1.27E-11 | 3.52E-10 | 0.62891054 |
| ENSMUSG00000040907 | Atp1a3 | 232975 | protein_coding | 3.98767334 | 0.628939636 | 1.463210528 | -0.429835368 | 0.667315403 | 1 | 0.628939636 |
| ENSMUSG00000094942 | Gm3604 | 100041979 | protein_coding | 23.68189041 | -0.628958075 | 0.327638963 | 1.91966813 | 0.054899832 | 0.15977566 | 0.628958075 |
| ENSMUSG00000077349 | Gm24867 | 115488880 | snRNA | 0.811469629 | 0.629322304 | 1.314297677 | -0.478827829 | 0.632061117 | 1 | 0.629322304 |
| ENSMUSG00000118362 | Gm19466 | 100502945 | lncRNA | 2.284909832 | 0.629326914 | 0.793836141 | -0.79276677 | 0.42791372 | 1 | 0.629326914 |
| ENSMUSG00000080268 | Brms1 | 107392 | protein_coding | 6.995831406 | 0.629538821 | 0.56056483 | -1.123043736 | 0.26141892 | 0.47508221 | 0.629538821 |
| ENSMUSG00000070704 | Ugt2b36 | 231396 | protein_coding | 66.12903984 | 0.62960802 | 0.171106506 | -3.679626428 | 0.000233576 | 0.001816041 | 0.62960802 |
| ENSMUSG00000057604 | Lmcd1 | 30937 | protein_coding | 5.652788741 | 0.629680333 | 0.552057512 | -1.140606404 | 0.254033751 | 0.466691767 | 0.629680333 |
| ENSMUSG00000024474 | Ik | 24010 | protein_coding | 1207.837766 | 0.629751281 | 0.04653226 | -13.53364907 | 9.90E-42 | 1.09E-39 | 0.629751281 |
| ENSMUSG00000052496 | Pkdrej | 18766 | protein_coding | 51.86884049 | 0.629765167 | 0.187504225 | -3.358671878 | 0.00078318 | 0.005170946 | 0.629765167 |
| ENSMUSG00000034587 | 8430429K09Rik | 71523 | lncRNA | 31.11698513 | -0.629837565 | 0.185602832 | 3.393469587 | 0.000690132 | 0.004625342 | 0.629837565 |
| ENSMUSG00000022965 | Ifngr2 | 15980 | protein_coding | 1304.201965 | -0.629989054 | 0.093718273 | 6.722158164 | 1.79E-11 | 4.87E-10 | 0.629989054 |
| ENSMUSG00000035062 | Zc4h2 | 245522 | protein_coding | 13.07284153 | 0.630001704 | 0.311750978 | -2.020849168 | 0.043295381 | 0.133516326 | 0.630001704 |
| ENSMUSG00000024677 | Ms4a6b | 69774 | protein_coding | 5.231814383 | 0.630040929 | 0.634351115 | -0.993205361 | 0.320609886 | 0.541863848 | 0.630040929 |
| ENSMUSG00000019942 | Cdk1 | 12534 | protein_coding | 460.7715529 | 0.630445207 | 0.163031063 | -3.867025055 | 0.000110171 | 0.000938301 | 0.630445207 |
| ENSMUSG00000013236 | Ptprs | 19280 | protein_coding | 329.1258264 | -0.630476369 | 0.148879743 | 4.234802899 | 2.29E-05 | 0.000232612 | 0.630476369 |
| ENSMUSG00000100585 | 1700108J01Rik | 68245 | lncRNA | 2.96920173 | 0.630564452 | 0.8620956 | -0.731432166 | 0.464515224 | 1 | 0.630564452 |
| ENSMUSG00000024386 | Proc | 19123 | protein_coding | 34.46660144 | -0.630938839 | 0.203058722 | 3.107174287 | 0.00188885 | 0.010842012 | 0.630938839 |
| ENSMUSG00000035864 | Syt1 | 20979 | protein_coding | 3.341945087 | 0.630967764 | 1.622859629 | -0.388799963 | 0.697424128 | 1 | 0.630967764 |
| ENSMUSG00000064917 | Snord138 | 115488275 | TEC | 2.218097754 | -0.631051825 | 0.698623476 | 0.903278872 | 0.366377907 | 1 | 0.631051825 |
| ENSMUSG00000030954 | Gp2 | 67133 | protein_coding | 2.318788903 | -0.631427425 | 1.350500953 | 0.467550522 | 0.640106055 | 1 | 0.631427425 |
| ENSMUSG00000027577 | Chrna4 | 11438 | protein_coding | 2.737351681 | 0.631733653 | 1.313207202 | -0.481061673 | 0.630472667 | 1 | 0.631733653 |
| ENSMUSG00000039908 | Slc26a11 | 268512 | protein_coding | 51.52468246 | -0.631964588 | 0.167509603 | 3.772706609 | 0.000161486 | 0.001309171 | 0.631964588 |
| ENSMUSG00000025964 | Adam23 | 23792 | protein_coding | 4.478118283 | 0.632050875 | 2.293569826 | -0.275575161 | 0.78287439 | 0.891186777 | 0.632050875 |
| ENSMUSG00000046169 | Adamts6 | 108154 | protein_coding | 2.168313461 | 0.63207025 | 1.23583656 | -0.511451329 | 0.609035061 | 1 | 0.63207025 |
| ENSMUSG00000039236 | Isg20 | 57444 | protein_coding | 221.1949788 | 0.632108231 | 0.121355653 | -5.20872507 | 1.90E-07 | 2.99E-06 | 0.632108231 |
| ENSMUSG00000054640 | Slc8a1 | 20541 | protein_coding | 1048.212444 | -0.632623181 | 0.158158707 | 3.999926349 | 6.34E-05 | 0.000578421 | 0.632623181 |

| ensembl_gene_id | external_gene_name | entrezgene_id | gene_biotype | baseMean | log2FoldChange | lfcSE | stat | pvalue | padj | absLog2FC |
|---|---|---|---|---|---|---|---|---|---|---|
| ENSMUSG00000074768 | Bhmt | 12116 | protein_coding | 2.612891025 | 0.632719358 | 0.9858264 | -0.641816204 | 0.520992527 | 1 | 0.632719358 |
| ENSMUSG00000032181 | Scg3 | 20255 | protein_coding | 13.37975785 | -0.633129393 | 0.371231973 | 1.705481853 | 0.088104594 | 0.225225892 | 0.633129393 |
| ENSMUSG00000009418 | Nav1 | 215690 | protein_coding | 67.53671949 | -0.633138012 | 0.190156327 | 3.329565847 | 0.000869815 | 0.005666321 | 0.633138012 |
| ENSMUSG00000024516 | Sec11c | 66286 | protein_coding | 1554.519082 | 0.633188462 | 0.11216212 | -5.645296867 | 1.65E-08 | 3.07E-07 | 0.633188462 |
| ENSMUSG00000028640 | Tfap2c | 21420 | protein_coding | 1.263104942 | -0.633207558 | 1.986227309 | 0.31879914 | 0.749878831 | 1 | 0.633207558 |
| ENSMUSG00000030873 | Scnn1b | 20277 | protein_coding | 147.8803138 | 0.63371326 | 0.180188081 | -3.516954378 | 0.000436529 | 0.003171508 | 0.63371326 |
| ENSMUSG00000026425 | Srgap2 | 14270 | protein_coding | 173.6670466 | 0.633715224 | 0.106860264 | -5.930316833 | 3.02E-09 | 6.28E-08 | 0.633715224 |
| ENSMUSG00000024397 | Aif1 | 11629 | protein_coding | 1.903083418 | 0.633844835 | 1.313413158 | -0.482593639 | 0.629384297 | 1 | 0.633844835 |
| ENSMUSG00000036564 | Ndrg4 | 234593 | protein_coding | 6.285609188 | 0.634593663 | 0.667536363 | -0.950650329 | 0.341781911 | 0.563829035 | 0.634593663 |
| ENSMUSG00000025330 | Padi4 | 18602 | protein_coding | 2.762245287 | 0.635110209 | 1.848086484 | -0.343658273 | 0.731103296 | 1 | 0.635110209 |
| ENSMUSG00000032452 | Clstn2 | 64085 | protein_coding | 3.34128718 | -0.63515752 | 0.934778702 | 0.679473675 | 0.496837782 | 1 | 0.63515752 |
| ENSMUSG00000031004 | Mki67 | 17345 | protein_coding | 3257.468974 | 0.635531071 | 0.156838281 | -4.052142552 | 5.08E-05 | 0.000472967 | 0.635531071 |
| ENSMUSG00000025732 | Mcrip2 | 68241 | protein_coding | 267.4228459 | -0.635847541 | 0.117096585 | 5.43011175 | 5.63E-08 | 9.57E-07 | 0.635847541 |
| ENSMUSG00000019802 | Sec63 | 140740 | protein_coding | 1124.060621 | 0.635887004 | 0.077294752 | -8.226781062 | 1.92E-16 | 7.94E-15 | 0.635887004 |
| ENSMUSG00000034845 | Plvap | 84094 | protein_coding | 5.635113407 | 0.63603861 | 0.712170569 | -0.893098701 | 0.371804319 | 0.594037976 | 0.63603861 |
| ENSMUSG00000104445 | Rhbg | 58176 | protein_coding | 17.55100453 | -0.636272287 | 0.261345677 | 2.434600393 | 0.014908244 | 0.058119626 | 0.636272287 |
| ENSMUSG00000029126 | Nsg1 | 18196 | protein_coding | 3.047989579 | -0.636386418 | 0.870051695 | 0.73143518 | 0.464513384 | 1 | 0.636386418 |
| ENSMUSG00000008892 | Vdac3 | 22335 | protein_coding | 1803.216358 | -0.636598621 | 0.104276501 | 6.104909673 | 1.03E-09 | 2.26E-08 | 0.636598621 |
| ENSMUSG00000020660 | Pomc | 18976 | protein_coding | 4.053953675 | 0.637027201 | 0.662397949 | -0.961698632 | 0.33620101 | 1 | 0.637027201 |
| ENSMUSG00000063681 | Crb1 | 170788 | protein_coding | 4.131321026 | 0.637131204 | 1.490315328 | -0.42751436 | 0.669004728 | 1 | 0.637131204 |
| ENSMUSG00000021944 | Gata4 | 14463 | protein_coding | 6.865347558 | -0.637158516 | 0.617871788 | 1.031214773 | 0.302440114 | 0.522874377 | 0.637158516 |
| ENSMUSG00000075408 | Smim41 | 77717 | protein_coding | 2.444119182 | 0.637463961 | 1.033847893 | -0.616593568 | 0.537502838 | 1 | 0.637463961 |
| ENSMUSG00000044338 | Aplnr | 23796 | protein_coding | 1.79248052 | 0.637723598 | 1.769109495 | -0.360477178 | 0.718490321 | 1 | 0.637723598 |
| ENSMUSG00000047733 | Tmem262 | 433215 | protein_coding | 1.914393561 | -0.637778654 | 1.02807353 | 0.620362878 | 0.535018907 | 1 | 0.637778654 |
| ENSMUSG00000060621 | Nkpd1 | 69547 | protein_coding | 10.51847832 | -0.637824635 | 0.343034218 | 1.859361552 | 0.062975908 | 0.176657744 | 0.637824635 |
| ENSMUSG00000021448 | Shc3 | 20418 | protein_coding | 1.702635949 | -0.63784734 | 1.666456946 | 0.382756567 | 0.701900268 | 1 | 0.63784734 |
| ENSMUSG00000074354 | Arhgap20os | 408028 | lncRNA | 7.252326901 | -0.638172094 | 0.450052086 | 1.41799608 | 0.156191909 | 0.338919446 | 0.638172094 |
| ENSMUSG00000021335 | Slc17a1 | 20504 | protein_coding | 15.1472516 | -0.638223752 | 0.388986419 | 1.640735309 | 0.100852372 | 0.249285447 | 0.638223752 |
| ENSMUSG00000030895 | Hpx | 15458 | protein_coding | 1.883057144 | 0.638340145 | 0.910016326 | -0.701460103 | 0.483015925 | 1 | 0.638340145 |
| ENSMUSG00000079588 | Tmem182 | 381339 | protein_coding | 31.31951679 | 0.638519506 | 0.215382878 | -2.96457876 | 0.003030976 | 0.016001773 | 0.638519506 |
| ENSMUSG00000053646 | Plxnb1 | 235611 | protein_coding | 171.1542602 | 0.638639882 | 0.147260779 | -4.33679549 | 1.45E-05 | 0.000154732 | 0.638639882 |
| ENSMUSG00000028965 | Tnfrsf9 | 21942 | protein_coding | 4.645446846 | 0.638700319 | 0.631188774 | -1.011900632 | 0.311585567 | 0.532495959 | 0.638700319 |
| ENSMUSG00000025912 | Mybl1 | 17864 | protein_coding | 27.45023566 | 0.638818225 | 0.258934301 | -2.467105446 | 0.013621024 | 0.054123578 | 0.638818225 |
| ENSMUSG00000006736 | Tspan31 | 67125 | protein_coding | 514.9297115 | 0.639019193 | 0.102735419 | -6.220047589 | 4.97E-10 | 1.13E-08 | 0.639019193 |
| ENSMUSG00000021186 | Fbln5 | 23876 | protein_coding | 3.807202719 | 0.639394526 | 1.010154783 | -0.632966885 | 0.526755268 | 1 | 0.639394526 |
| ENSMUSG00000025270 | Alas2 | 11656 | protein_coding | 21.36039889 | -0.639439873 | 0.350963715 | 1.821954368 | 0.06846191 | 0.187931588 | 0.639439873 |
| ENSMUSG00000089929 | Bcl2a1b | 12045 | protein_coding | 0.773773846 | 0.639586945 | 1.311182865 | -0.487793856 | 0.625695864 | 1 | 0.639586945 |
| ENSMUSG00000032322 | Pstpip1 | 19200 | protein_coding | 21.53156374 | -0.639639407 | 0.283886586 | 2.253151218 | 0.024249616 | 0.085349139 | 0.639639407 |
| ENSMUSG00000116056 | Gm4544 | 100043600 | lncRNA | 13.27213223 | 0.639765289 | 0.350153037 | -1.827101929 | 0.067684445 | 0.186383347 | 0.639765289 |

| ensembl_gene_id | external_gene_name | entrezgene_id | gene_biotype | baseMean | log2FoldChange | lfcSE | stat | pvalue | padj | absLog2FC |
|---|---|---|---|---|---|---|---|---|---|---|
| ENSMUSG00000031748 | Gnao1 | 14681 | protein_coding | 22.11421869 | -0.639866091 | 0.355134844 | 1.801755309 | 0.071583912 | 0.194206788 | 0.639866091 |
| ENSMUSG00000086390 | 1810019D21Rik | 69771 | lncRNA | 126.3623567 | 0.640013837 | 0.178672233 | -3.582055395 | 0.000340901 | 0.002548505 | 0.640013837 |
| ENSMUSG00000067212 | H2-T23 | 15040 | protein_coding | 1799.202273 | -0.640256264 | 0.124182795 | 5.15575659 | 2.53E-07 | 3.90E-06 | 0.640256264 |
| ENSMUSG00000051335 | Gfod1 | 328232 | protein_coding | 70.13294484 | -0.640330516 | 0.225483813 | 2.839807023 | 0.004514083 | 0.022359199 | 0.640330516 |
| ENSMUSG00000028128 | F3 | 14066 | protein_coding | 190.9629364 | -0.640420256 | 0.244603203 | 2.618200611 | 0.008839482 | 0.038576958 | 0.640420256 |
| ENSMUSG00000022504 | Ciita | 12265 | protein_coding | 16.15598159 | 0.640447137 | 0.452649504 | -1.414885319 | 0.157102123 | 0.340247143 | 0.640447137 |
| ENSMUSG00000068686 | Cd59b | 333883 | protein_coding | 2.722840912 | -0.64047218 | 0.881635086 | 0.726459496 | 0.467557129 | 1 | 0.64047218 |
| ENSMUSG00000024209 | Acsbg3 | 78625 | protein_coding | 4.500531478 | 0.640776847 | 0.832573109 | -0.76963433 | 0.441516835 | 0.659241321 | 0.640776847 |
| ENSMUSG00000035121 | Neil2 | 382913 | protein_coding | 3.234872154 | -0.640796509 | 0.831879236 | 0.770299919 | 0.441122004 | 1 | 0.640796509 |
| ENSMUSG00000029359 | Tesc | 57816 | protein_coding | 31.86312359 | 0.640951883 | 0.267381428 | -2.397144364 | 0.016523412 | 0.063163803 | 0.640951883 |
| ENSMUSG00000117286 | Gm1043 | 381634 | protein_coding | 11.90059293 | 0.64106431 | 0.637389127 | -1.005765996 | 0.314528148 | 0.535752227 | 0.64106431 |
| ENSMUSG00000008305 | Tle1 | 21885 | protein_coding | 710.2704032 | 0.641174236 | 0.092081097 | -6.96314725 | 3.33E-12 | 9.71E-11 | 0.641174236 |
| ENSMUSG00000028976 | Slc2a5 | 56485 | protein_coding | 49.72873704 | 0.641175575 | 0.30801552 | -2.081633984 | 0.037375915 | 0.119478486 | 0.641175575 |
| ENSMUSG00000062110 | Scfd2 | 212986 | protein_coding | 226.8258665 | 0.641349532 | 0.093606412 | -6.851555547 | 7.31E-12 | 2.06E-10 | 0.641349532 |
| ENSMUSG00000091844 | Gm8251 | 102636082 | protein_coding | 10.10064716 | 0.641796044 | 0.534884177 | -1.199878537 | 0.230186517 | 0.438797199 | 0.641796044 |
| ENSMUSG00000032513 | Gorasp1 | 74498 | protein_coding | 580.4119549 | 0.641893662 | 0.088520176 | -7.251382558 | 4.13E-13 | 1.30E-11 | 0.641893662 |
| ENSMUSG00000034098 | Fstl5 | 213262 | protein_coding | 1.553113234 | 0.641953013 | 1.888354661 | -0.339953625 | 0.733891452 | 1 | 0.641953013 |
| ENSMUSG00000048240 | Gng7 | 14708 | protein_coding | 5.349445982 | -0.64228231 | 0.480619235 | 1.33636414 | 0.181430281 | 0.375503154 | 0.64228231 |
| ENSMUSG00000024008 | Cpne5 | 240058 | protein_coding | 8.233761038 | -0.642308601 | 0.825708158 | 0.777888163 | 0.436634947 | 0.654912763 | 0.642308601 |
| ENSMUSG00000027602 | Map1lc3a | 66734 | protein_coding | 261.8027012 | -0.642346015 | 0.126449471 | 5.079863194 | 3.78E-07 | 5.69E-06 | 0.642346015 |
| ENSMUSG00000028443 | Nudt2 | 66401 | protein_coding | 213.1032817 | 0.642456651 | 0.08956748 | -7.172878533 | 7.34E-13 | 2.27E-11 | 0.642456651 |
| ENSMUSG00000090336 | Cfap97d2 | 403185 | protein_coding | 2.209736822 | -0.642567281 | 1.950850984 | 0.329377941 | 0.741870039 | 1 | 0.642567281 |
| ENSMUSG00000015476 | Prrt1 | 260297 | protein_coding | 4.02484217 | -0.642589663 | 0.851026117 | 0.755076313 | 0.450203194 | 1 | 0.642589663 |
| ENSMUSG00000035020 | Epgn | 71920 | protein_coding | 1.891221232 | 0.64294922 | 1.247762206 | -0.515281851 | 0.606356079 | 1 | 0.64294922 |
| ENSMUSG00000074580 | 4931440P22Rik | 71004 | transcribed_processed_pseudogene | 21.96091139 | 0.642968723 | 0.397491642 | -1.617565391 | 0.105756293 | 0.25785531 | 0.642968723 |
| ENSMUSG00000040752 | Myh6 | 17888 | protein_coding | 3.866606692 | 0.643021398 | 2.209451271 | -0.29103217 | 0.771026716 | 1 | 0.643021398 |
| ENSMUSG00000069540 | Gm4925 | 237433 | transcribed_processed_pseudogene | 8.937401087 | -0.643033905 | 0.33504519 | 1.919245297 | 0.054953297 | 0.159827185 | 0.643033905 |
| ENSMUSG00000021938 | Pspc1 | 66645 | protein_coding | 307.9779822 | 0.643123294 | 0.088690677 | -7.251306634 | 4.13E-13 | 1.30E-11 | 0.643123294 |
| ENSMUSG00000106799 | Gm20756 | 626904 | lncRNA | 1.938160249 | -0.643126742 | 1.295209571 | 0.496542611 | 0.619511632 | 1 | 0.643126742 |
| ENSMUSG00000023387 | Kcnk16 | 74571 | protein_coding | 4.464729399 | 0.643236407 | 0.710137647 | -0.905791165 | 0.365046398 | 0.587561035 | 0.643236407 |
| ENSMUSG00000038357 | Camp | 12796 | protein_coding | 1.071798451 | 0.64350361 | 1.387875721 | -0.463660831 | 0.642890764 | 1 | 0.64350361 |
| ENSMUSG00000001506 | Col1a1 | 12842 | protein_coding | 18.70443679 | 0.643760423 | 0.56653316 | -1.136315522 | 0.255824537 | 0.46852461 | 0.643760423 |
| ENSMUSG00000068877 | Selenbp2 | 20342 | protein_coding | 68.95000377 | -0.644180786 | 0.15449269 | 4.169652214 | 3.05E-05 | 0.000301306 | 0.644180786 |
| ENSMUSG00000024742 | Fen1 | 14156 | protein_coding | 30.10726278 | 0.64444907 | 0.231412194 | -2.784853546 | 0.00535519 | 0.025782978 | 0.64444907 |
| ENSMUSG00000022270 | Retreg1 | 66270 | protein_coding | 1155.760462 | -0.644461846 | 0.161604344 | 3.987899269 | 6.67E-05 | 0.000602545 | 0.644461846 |
| ENSMUSG00000086799 | 4930502E09Rik | 74995 | lncRNA | 2.482588506 | 0.644653523 | 0.83024023 | -0.776466256 | 0.437473738 | 1 | 0.644653523 |
| ENSMUSG00000065582 | Mir194-2 | 723957 | miRNA | 15.28487087 | 0.64503219 | 0.395272469 | -1.631867233 | 0.102707458 | 0.252397465 | 0.64503219 |
| ENSMUSG00000074635 | 3110070M22Rik | 67304 | lncRNA | 5.132709625 | -0.645053728 | 0.574339351 | 1.123122987 | 0.261385265 | 0.475073328 | 0.645053728 |
| ENSMUSG00000039976 | Tbc1d16 | 207592 | protein_coding | 172.9818568 | -0.645180207 | 0.182283997 | 3.539423188 | 0.000401002 | 0.002935352 | 0.645180207 |

| ensembl_gene_id | external_gene_name | entrezgene_id | gene_biotype | baseMean | log2FoldChange | lfcSE | stat | pvalue | padj | absLog2FC |
|---|---|---|---|---|---|---|---|---|---|---|
| ENSMUSG00000004552 | Ctse | 13034 | protein_coding | 483.5272528 | -0.645210824 | 0.261347037 | 2.468789518 | 0.013557093 | 0.053943227 | 0.645210824 |
| ENSMUSG00000039763 | Dnajc28 | 246738 | protein_coding | 64.00971274 | -0.645275281 | 0.197233181 | 3.271636534 | 0.001069269 | 0.006724263 | 0.645275281 |
| ENSMUSG00000044294 | Krt84 | 16680 | protein_coding | 55.8504908 | 0.645300408 | 0.21690048 | -2.975099029 | 0.00292894 | 0.015552576 | 0.645300408 |
| ENSMUSG00000028946 | Hes3 | 15207 | protein_coding | 1.153618548 | 0.6453065 | 1.386298208 | -0.465488952 | 0.641581347 | 1 | 0.6453065 |
| ENSMUSG00000038642 | Ctss | 13040 | protein_coding | 1908.375619 | -0.64554174 | 0.099445043 | 6.491442087 | 8.50E-11 | 2.16E-09 | 0.64554174 |
| ENSMUSG00000024056 | Ndc80 | 67052 | protein_coding | 126.7551507 | 0.645670261 | 0.201555664 | -3.203433968 | 0.001357992 | 0.008211599 | 0.645670261 |
| ENSMUSG00000020635 | Fkbp1b | 14226 | protein_coding | 13.10192325 | -0.646088512 | 0.342957635 | 1.88387266 | 0.059582202 | 0.169937908 | 0.646088512 |
| ENSMUSG00000043633 | Fam221b | 242408 | protein_coding | 15.96143919 | 0.646121746 | 0.349045839 | -1.851108577 | 0.064153934 | 0.179322626 | 0.646121746 |
| ENSMUSG00000056271 | Lman1l | 235416 | protein_coding | 4.677546701 | 0.646271909 | 0.8029138 | -0.80490821 | 0.42087265 | 0.640714813 | 0.646271909 |
| ENSMUSG00000024201 | Kdm4b | 193796 | protein_coding | 580.6059666 | 0.646384095 | 0.071013073 | -9.102325348 | 8.84E-20 | 4.45E-18 | 0.646384095 |
| ENSMUSG00000046598 | Bdh1 | 71911 | protein_coding | 1493.842978 | -0.646448708 | 0.112881521 | 5.726789509 | 1.02E-08 | 1.97E-07 | 0.646448708 |
| ENSMUSG00000027006 | Dnajc10 | 66861 | protein_coding | 2031.637272 | 0.646553531 | 0.175936024 | -3.674935443 | 0.00023791 | 0.001846439 | 0.646553531 |
| ENSMUSG00000046295 | Ankle1 | 234396 | protein_coding | 62.74805471 | 0.646631014 | 0.208972164 | -3.094340413 | 0.00197251 | 0.011213534 | 0.646631014 |
| ENSMUSG00000040428 | Plekha4 | 69217 | protein_coding | 10.2212386 | -0.646638811 | 0.567558905 | 1.139333389 | 0.254564127 | 0.467148794 | 0.646638811 |
| ENSMUSG00000023826 | Prkn | 50873 | protein_coding | 16.63563602 | -0.646672762 | 0.352673574 | 1.833629766 | 0.066708971 | 0.184376923 | 0.646672762 |
| ENSMUSG00000027287 | Snap23 | 20619 | protein_coding | 818.1725065 | 0.646703301 | 0.064851169 | -9.972114732 | 2.02E-23 | 1.21E-21 | 0.646703301 |
| ENSMUSG00000113986 | Gm31333 | 102633526 | lncRNA | 1.656854123 | -0.646713089 | 1.72000608 | 0.375994653 | 0.706920871 | 1 | 0.646713089 |
| ENSMUSG00000032776 | Mctp2 | 244049 | protein_coding | 823.4108827 | 0.647163144 | 0.075684987 | -8.550746616 | 1.22E-17 | 5.46E-16 | 0.647163144 |
| ENSMUSG00000004044 | Cavin1 | 19285 | protein_coding | 47.99883801 | -0.647184841 | 0.227304923 | 2.847209955 | 0.004410426 | 0.02193128 | 0.647184841 |
| ENSMUSG00000040605 | Bace2 | 56175 | protein_coding | 593.4202777 | -0.64736694 | 0.238535872 | 2.713918596 | 0.00664925 | 0.03055251 | 0.64736694 |
| ENSMUSG00000035704 | Alg8 | 381903 | protein_coding | 146.6383167 | 0.647408299 | 0.173770946 | -3.725641795 | 0.000194819 | 0.001545264 | 0.647408299 |
| ENSMUSG00000060802 | B2m | 12010 | protein_coding | 2813.149375 | -0.647593839 | 0.12181391 | 5.316255255 | 1.06E-07 | 1.74E-06 | 0.647593839 |
| ENSMUSG00000022033 | Pbk | 52033 | protein_coding | 236.4539884 | 0.647630097 | 0.205317663 | -3.154283403 | 0.001608928 | 0.009447848 | 0.647630097 |
| ENSMUSG00000042353 | Frem3 | 333315 | protein_coding | 2.181345866 | 0.647912871 | 1.69190355 | -0.382949058 | 0.701757536 | 1 | 0.647912871 |
| ENSMUSG00000064945 | Rny3 | 19874 | misc_RNA | 356.2866081 | 0.647916849 | 0.301550719 | -2.148616493 | 0.031664813 | 0.105235002 | 0.647916849 |
| ENSMUSG00000105960 | 6330410L21Rik | 319224 | lncRNA | 1.049803358 | -0.64854794 | 1.790864039 | 0.362142478 | 0.717245562 | 1 | 0.64854794 |
| ENSMUSG00000035126 | Dnai4 | 242584 | protein_coding | 33.70390144 | 0.648887577 | 0.259034931 | -2.50501959 | 0.012244461 | 0.049834789 | 0.648887577 |
| ENSMUSG00000035395 | Dcaf8l | 18630 | protein_coding | 1.290204042 | 0.649177342 | 2.099795899 | -0.309162116 | 0.757198211 | 1 | 0.649177342 |
| ENSMUSG00000055602 | Tcp10b | 21462 | protein_coding | 1.974897736 | -0.649266125 | 1.046444521 | 0.620449639 | 0.534961801 | 1 | 0.649266125 |
| ENSMUSG00000033200 | Tpsg1 | 26945 | protein_coding | 326.8627857 | -0.64927211 | 0.241469376 | 2.688838318 | 0.007170114 | 0.032592144 | 0.64927211 |
| ENSMUSG00000042249 | Grk3 | 320129 | protein_coding | 93.17452922 | -0.649403898 | 0.18671318 | 3.478082789 | 0.000505014 | 0.003576244 | 0.649403898 |
| ENSMUSG00000031060 | Rbm10 | 236732 | protein_coding | 1110.591322 | 0.649415596 | 0.073262457 | -8.864234401 | 7.70E-19 | 3.71E-17 | 0.649415596 |
| ENSMUSG00000020493 | Prr11 | 270906 | protein_coding | 78.15972353 | 0.649520891 | 0.206838765 | -3.140228041 | 0.001688164 | 0.009854484 | 0.649520891 |
| ENSMUSG00000040797 | Iqsec3 | 243621 | protein_coding | 3.751167707 | 0.64952353 | 1.842490163 | -0.352524829 | 0.724444701 | 1 | 0.64952353 |
| ENSMUSG00000022324 | Matn2 | 17181 | protein_coding | 74.54029764 | 0.649527625 | 0.262265954 | -2.476599095 | 0.01326408 | 0.053024211 | 0.649527625 |
| ENSMUSG00000067642 | Adgrf3 | 381628 | protein_coding | 7.17784765 | -0.649554456 | 0.611457351 | 1.062305418 | 0.288097056 | 0.506958639 | 0.649554456 |
| ENSMUSG00000001151 | Pcnt | 18541 | protein_coding | 991.1720672 | 0.649581368 | 0.086511486 | -7.508614142 | 5.98E-14 | 2.01E-12 | 0.649581368 |
| ENSMUSG00000000817 | Fasl | 14103 | protein_coding | 4.742267606 | -0.649880907 | 0.614776116 | 1.057101748 | 0.290465153 | 0.509468423 | 0.649880907 |
| ENSMUSG00000038773 | Kdm3b | 277250 | protein_coding | 1445.595182 | 0.650945843 | 0.054518623 | -11.93988047 | 7.33E-33 | 6.21E-31 | 0.650945843 |

Supplementary Table 1

| ensembl_gene_id | external_gene_name | entrezgene_id | gene_biotype | baseMean | log2FoldChange | lfcSE | stat | pvalue | padj | absLog2FC |
|---|---|---|---|---|---|---|---|---|---|---|
| ENSMUSG00000065549 | Mir200b | 387243 | miRNA | 3.603083579 | 0.650993501 | 0.609568947 | -1.067957127 | 0.285539852 | 1 | 0.650993501 |
| ENSMUSG00000088151 | Gm26347 | 115487535 | scaRNA | 0.925172757 | 0.651192431 | 1.436380167 | -0.453356602 | 0.650291981 | 1 | 0.651192431 |
| ENSMUSG00000058773 | H1f5 | 56702 | protein_coding | 1312.537761 | 0.651194211 | 0.186279229 | -3.495796142 | 0.00047265 | 0.003383327 | 0.651194211 |
| ENSMUSG00000048100 | Taf13 | 99730 | protein_coding | 245.4623431 | 0.651204042 | 0.087660238 | -7.428727766 | 1.10E-13 | 3.61E-12 | 0.651204042 |
| ENSMUSG00000032254 | Kif23 | 71819 | protein_coding | 460.9356929 | 0.651446664 | 0.110973828 | -5.87027298 | 4.35E-09 | 8.87E-08 | 0.651446664 |
| ENSMUSG00000030088 | Aldh1l1 | 107747 | protein_coding | 1402.494451 | -0.65183092 | 0.236232472 | 2.759277392 | 0.005792934 | 0.027443223 | 0.65183092 |
| ENSMUSG00000026778 | Prkcq | 18761 | protein_coding | 6.402147587 | 0.652086935 | 0.627076148 | -1.039884769 | 0.298393439 | 0.51821109 | 0.652086935 |
| ENSMUSG00000034349 | Smc4 | 70099 | protein_coding | 966.9200741 | 0.652310704 | 0.167326177 | -3.898437873 | 9.68E-05 | 0.000837072 | 0.652310704 |
| ENSMUSG00000022262 | Dnah5 | 110082 | protein_coding | 165.8156137 | -0.652361845 | 0.249001763 | 2.619908538 | 0.008795335 | 0.038404592 | 0.652361845 |
| ENSMUSG00000028519 | Dab1 | 13131 | protein_coding | 22.55843907 | -0.652376679 | 0.479319587 | 1.361047403 | 0.17349871 | 0.364292142 | 0.652376679 |
| ENSMUSG00000087519 | AV039307 | 100502700 | lncRNA | 2.530678867 | -0.652395656 | 0.889292217 | 0.733612241 | 0.463185098 | 1 | 0.652395656 |
| ENSMUSG00000032357 | Tinag | 26944 | protein_coding | 326.41022 | -0.652526777 | 0.125614333 | 5.194684087 | 2.05E-07 | 3.20E-06 | 0.652526777 |
| ENSMUSG00000022960 | Donson | 60364 | protein_coding | 118.0375706 | 0.652821656 | 0.159717821 | -4.087343859 | 4.36E-05 | 0.000415076 | 0.652821656 |
| ENSMUSG00000089798 | 1700028K03Rik | 76421 | protein_coding | 2.700651123 | -0.652858256 | 0.834237658 | 0.782580659 | 0.433873403 | 1 | 0.652858256 |
| ENSMUSG00000050071 | Bex1 | 19716 | protein_coding | 31.54436924 | 0.652908525 | 0.248997914 | -2.622144553 | 0.008737836 | 0.038193919 | 0.652908525 |
| ENSMUSG00000043505 | Gimap5 | 317757 | protein_coding | 11.25104343 | 0.653095381 | 0.49165253 | -1.328367781 | 0.184056638 | 0.379085971 | 0.653095381 |
| ENSMUSG00000085628 | Appbp2os | 654810 | lncRNA | 10.32905212 | -0.653468682 | 0.330327028 | 1.978247695 | 0.047900771 | 0.144375493 | 0.653468682 |
| ENSMUSG00000040276 | Pacsin1 | 23969 | protein_coding | 64.46011696 | -0.653652825 | 0.315847552 | 2.069519994 | 0.038497317 | 0.122282111 | 0.653652825 |
| ENSMUSG00000030670 | Cyp2r1 | 244209 | protein_coding | 29.71576668 | 0.653684921 | 0.202561034 | -3.227101031 | 0.001250513 | 0.007657016 | 0.653684921 |
| ENSMUSG00000058385 | H2bc8 | 319181 | protein_coding | 905.7388384 | 0.653836796 | 0.172762319 | -3.784603035 | 0.000153954 | 0.001254886 | 0.653836796 |
| ENSMUSG00000030322 | Mbd4 | 17193 | protein_coding | 46.83320735 | 0.653849274 | 0.190772875 | -3.427370235 | 0.000609458 | 0.004186598 | 0.653849274 |
| ENSMUSG00000088891 | Gm22516 | 115489634 | snoRNA | 2.128510728 | 0.653947575 | 0.904170462 | -0.723256955 | 0.469522033 | 1 | 0.653947575 |
| ENSMUSG00000022186 | Oxct1 | 67041 | protein_coding | 1759.194847 | -0.654144017 | 0.15300748 | 4.275242086 | 1.91E-05 | 0.000197434 | 0.654144017 |
| ENSMUSG00000026121 | Sema4c | 20353 | protein_coding | 63.85309506 | 0.654377704 | 0.152100888 | -4.302260906 | 1.69E-05 | 0.00017783 | 0.654377704 |
| ENSMUSG00000040751 | Lat2 | 56743 | protein_coding | 7.286513789 | 0.654588086 | 0.53016454 | -1.234688547 | 0.21694644 | 0.422532552 | 0.654588086 |
| ENSMUSG00000084592 | Mir1898 | 100316684 | miRNA | 1.881781164 | -0.654632623 | 0.802799143 | 0.81543762 | 0.414821858 | 1 | 0.654632623 |
| ENSMUSG00000022665 | Ccdc80 | 67896 | protein_coding | 2.083119214 | -0.654696908 | 1.788230519 | 0.36611438 | 0.714279727 | 1 | 0.654696908 |
| ENSMUSG00000040972 | Igsf21 | 230868 | protein_coding | 1.238976682 | 0.655295163 | 1.154580332 | -0.567561342 | 0.570332863 | 1 | 0.655295163 |
| ENSMUSG00000098325 | Mir7048 | 102466794 | miRNA | 1.982475488 | -0.655373555 | 0.854090962 | 0.76733461 | 0.442882596 | 1 | 0.655373555 |
| ENSMUSG00000038718 | Pbx3 | 18516 | protein_coding | 52.46388659 | 0.655393352 | 0.197025367 | -3.326441479 | 0.000879625 | 0.005716696 | 0.655393352 |
| ENSMUSG00000022382 | Wnt7b | 22422 | protein_coding | 3.80946628 | 0.655407157 | 1.011196834 | -0.64814993 | 0.516887982 | 1 | 0.655407157 |
| ENSMUSG00000029678 | Hyal5 | 74468 | protein_coding | 1.293492555 | -0.655460365 | 1.420839393 | 0.46131911 | 0.644569676 | 1 | 0.655460365 |
| ENSMUSG00000037129 | Tmprss13 | 214531 | protein_coding | 38.0874085 | -0.655587529 | 0.404195151 | 1.621957929 | 0.104812357 | 0.255970313 | 0.655587529 |
| ENSMUSG00000028344 | Invs | 16348 | protein_coding | 83.21851963 | 0.655704547 | 0.130693058 | -5.017133695 | 5.24E-07 | 7.71E-06 | 0.655704547 |
| ENSMUSG00000062070 | Pgk1 | 18655 | protein_coding | 2232.519938 | 0.65590269 | 0.146169331 | -4.487279815 | 7.21E-06 | 8.36E-05 | 0.65590269 |
| ENSMUSG00000021756 | Il6st | 16195 | protein_coding | 724.5265795 | 0.655943921 | 0.07792793 | -8.417314855 | 3.85E-17 | 1.64E-15 | 0.655943921 |
| ENSMUSG00000010825 | Grid2ip | 170935 | protein_coding | 4.793659575 | 0.656068166 | 0.926309151 | -0.708260482 | 0.478783513 | 0.690414856 | 0.656068166 |
| ENSMUSG00000028032 | Papss1 | 23971 | protein_coding | 3020.41073 | 0.656151447 | 0.062169462 | -10.55424041 | 4.86E-26 | 3.26E-24 | 0.656151447 |
| ENSMUSG00000025467 | Prap1 | 22264 | protein_coding | 188.1866966 | -0.656766067 | 0.260925071 | 2.517067692 | 0.011833609 | 0.048563673 | 0.656766067 |

| ensembl_gene_id | external_gene_name | entrezgene_id | gene_biotype | baseMean | log2FoldChange | lfcSE | stat | pvalue | padj | absLog2FC |
|---|---|---|---|---|---|---|---|---|---|---|
| ENSMUSG00000117975 | Itprip | 414801 | protein_coding | 36.25048772 | 0.656787274 | 0.281674236 | -2.331726474 | 0.019715085 | 0.072511117 | 0.656787274 |
| ENSMUSG00000042570 | Mier2 | 70427 | protein_coding | 440.5752587 | 0.657373216 | 0.12694549 | -5.178389676 | 2.24E-07 | 3.48E-06 | 0.657373216 |
| ENSMUSG00000038599 | Capn8 | 170725 | protein_coding | 327.8151305 | 0.657451209 | 0.101813851 | -6.457384771 | 1.07E-10 | 2.63E-09 | 0.657451209 |
| ENSMUSG00000027188 | Pamr1 | 210622 | protein_coding | 16.67503782 | -0.657741935 | 0.335250673 | 1.961940684 | 0.04976939 | 0.148382689 | 0.657741935 |
| ENSMUSG00000031290 | Lrch2 | 210297 | protein_coding | 1.782851897 | -0.658123968 | 1.837333042 | 0.35819525 | 0.720197202 | 1 | 0.658123968 |
| ENSMUSG00000041062 | Mslnl | 328783 | protein_coding | 1.119068541 | 0.658175512 | 1.72295224 | -0.382004502 | 0.702458025 | 1 | 0.658175512 |
| ENSMUSG00000003665 | Has1 | 15116 | protein_coding | 3.417774592 | 0.658199533 | 1.73248416 | -0.379916624 | 0.704007307 | 1 | 0.658199533 |
| ENSMUSG00000009487 | Otog | 18419 | protein_coding | 8.198626486 | 0.658603471 | 1.483088333 | -0.444075687 | 0.656987864 | 0.814465699 | 0.658603471 |
| ENSMUSG00000035189 | Ano4 | 320091 | protein_coding | 2.763162046 | 0.658650289 | 2.103392271 | -0.313137163 | 0.754176456 | 1 | 0.658650289 |
| ENSMUSG00000029167 | Ppargc1a | 19017 | protein_coding | 365.4287647 | -0.658740408 | 0.117603797 | 5.601353229 | 2.13E-08 | 3.89E-07 | 0.658740408 |
| ENSMUSG00000034271 | Jdp2 | 81703 | protein_coding | 4.324290884 | 0.658837429 | 0.77787487 | -0.846970965 | 0.397011305 | 1 | 0.658837429 |
| ENSMUSG00000041144 | Dnah7b | 227058 | protein_coding | 16.54243519 | 0.65915375 | 0.437389125 | -1.507019064 | 0.131805771 | 0.300890309 | 0.65915375 |
| ENSMUSG00000037606 | Osbpl5 | 79196 | protein_coding | 334.582524 | -0.659189444 | 0.117230387 | 5.623025409 | 1.88E-08 | 3.46E-07 | 0.659189444 |
| ENSMUSG00000059447 | Hadhb | 231086 | protein_coding | 1370.998084 | -0.659285547 | 0.109391691 | 6.026833874 | 1.67E-09 | 3.59E-08 | 0.659285547 |
| ENSMUSG00000028836 | Slc30a2 | 230810 | protein_coding | 8.664474408 | 0.65950395 | 0.646782946 | -1.01966812 | 0.307885886 | 0.528416912 | 0.65950395 |
| ENSMUSG00000070695 | Cntnap5a | 636808 | protein_coding | 4.181834291 | 0.659518501 | 1.49982136 | -0.43973137 | 0.660131679 | 1 | 0.659518501 |
| ENSMUSG00000118245 | Gm30082 | 102631855 | lncRNA | 1.146219678 | -0.659556872 | 1.056945471 | 0.624021664 | 0.532613358 | 1 | 0.659556872 |
| ENSMUSG00000023034 | Nr4a1 | 15370 | protein_coding | 1358.619906 | -0.659590838 | 0.238194794 | 2.769123648 | 0.00562073 | 0.026811304 | 0.659590838 |
| ENSMUSG00000023151 | Lrrc69 | 73314 | protein_coding | 0.625640322 | -0.659672398 | 1.67681855 | 0.393407144 | 0.694018794 | 1 | 0.659672398 |
| ENSMUSG00000034906 | Ncaph | 215387 | protein_coding | 241.7185018 | 0.65983178 | 0.167762609 | -3.933127793 | 8.38E-05 | 0.000738092 | 0.65983178 |
| ENSMUSG00000058569 | Tmed9 | 67511 | protein_coding | 1438.947976 | 0.660203402 | 0.082458392 | -8.006503499 | 1.18E-15 | 4.55E-14 | 0.660203402 |
| ENSMUSG00000079019 | Insl3 | 16336 | protein_coding | 2.059116676 | 0.660478101 | 1.03933844 | -0.635479335 | 0.52511584 | 1 | 0.660478101 |
| ENSMUSG00000035105 | Egln3 | 112407 | protein_coding | 323.0141889 | -0.660510356 | 0.223158689 | 2.959823605 | 0.003078152 | 0.016227616 | 0.660510356 |
| ENSMUSG00000044562 | Rasip1 | 69903 | protein_coding | 24.88255034 | 0.660565929 | 0.255636694 | -2.584002782 | 0.009766101 | 0.041791803 | 0.660565929 |
| ENSMUSG00000027495 | Fam210b | 67017 | protein_coding | 281.0702813 | -0.660733136 | 0.073328903 | 9.010541716 | 2.05E-19 | 1.01E-17 | 0.660733136 |
| ENSMUSG00000030254 | Rad18 | 58186 | protein_coding | 83.89608356 | 0.661076796 | 0.154513186 | -4.278449064 | 1.88E-05 | 0.000195468 | 0.661076796 |
| ENSMUSG00000028312 | Smc2 | 14211 | protein_coding | 470.8973308 | 0.661373369 | 0.175362067 | -3.771473397 | 0.000162286 | 0.001314369 | 0.661373369 |
| ENSMUSG00000004655 | Aqp1 | 11826 | protein_coding | 817.0294749 | -0.661444779 | 0.170040325 | 3.889928937 | 0.000100274 | 0.000862907 | 0.661444779 |
| ENSMUSG00000107504 | Gm35549 | 102639178 | protein_coding | 2.28302113 | 0.661463994 | 0.78130223 | -0.846617312 | 0.397208461 | 1 | 0.661463994 |
| ENSMUSG00000023393 | Slc17a9 | 228993 | protein_coding | 889.9881792 | 0.661614046 | 0.071030659 | -9.314485492 | 1.23E-20 | 6.47E-19 | 0.661614046 |
| ENSMUSG00000020766 | Galk1 | 14635 | protein_coding | 381.454463 | 0.661642074 | 0.135307958 | -4.889897693 | 1.01E-06 | 1.39E-05 | 0.661642074 |
| ENSMUSG00000037738 | Nek5 | 330721 | protein_coding | 10.15696603 | -0.661665687 | 0.389675491 | 1.697991542 | 0.089509358 | 0.227794351 | 0.661665687 |
| ENSMUSG00000030036 | Mogs | 57377 | protein_coding | 1365.857313 | 0.661877126 | 0.11382856 | -5.814684182 | 6.07E-09 | 1.21E-07 | 0.661877126 |
| ENSMUSG00000077426 | Gm26387 | 115490267 | snoRNA | 2.669848869 | 0.661947139 | 0.772402411 | -0.856997764 | 0.391446121 | 1 | 0.661947139 |
| ENSMUSG00000032297 | Celf6 | 76183 | protein_coding | 21.28248602 | 0.66201581 | 0.325053242 | -2.036638075 | 0.041686324 | 0.129891785 | 0.66201581 |
| ENSMUSG00000052212 | Cd177 | 68891 | protein_coding | 3002.072234 | 0.662546294 | 0.25294368 | -2.619343149 | 0.008809928 | 0.038458141 | 0.662546294 |
| ENSMUSG00000032788 | Pdxk | 216134 | protein_coding | 460.8769031 | 0.662634973 | 0.106043475 | -6.248710457 | 4.14E-10 | 9.55E-09 | 0.662634973 |
| ENSMUSG00000022598 | Psca | 72373 | protein_coding | 1.185774993 | -0.6626908 | 1.76708368 | 0.375019478 | 0.70764598 | 1 | 0.6626908 |
| ENSMUSG00000030278 | Cidec | 14311 | protein_coding | 87.73009866 | 0.662755433 | 0.367876075 | -1.801572538 | 0.071612685 | 0.194226738 | 0.662755433 |

| ensembl_gene_id | external_gene_name | entrezgene_id | gene_biotype | baseMean | log2FoldChange | lfcSE | stat | pvalue | padj | absLog2FC |
|---|---|---|---|---|---|---|---|---|---|---|
| ENSMUSG00000021336 | Slc17a4 | 319848 | protein_coding | 664.5024284 | -0.662940141 | 0.128943446 | 5.141324843 | 2.73E-07 | 4.19E-06 | 0.662940141 |
| ENSMUSG00000011752 | Pgam1 | 18648 | protein_coding | 445.6113916 | -0.663293676 | 0.157115636 | 4.221691067 | 2.42E-05 | 0.000245359 | 0.663293676 |
| ENSMUSG00000062044 | Lmtk3 | 381983 | protein_coding | 10.59437076 | -0.663379117 | 0.673342037 | 0.985203776 | 0.324523986 | 0.546521427 | 0.663379117 |
| ENSMUSG00000024846 | Cst6 | 73720 | protein_coding | 18.06181963 | -0.663417721 | 0.369180809 | 1.796999476 | 0.072335703 | 0.19566778 | 0.663417721 |
| ENSMUSG00000099419 | 1700001D01Rik | 69279 | lncRNA | 3.639582623 | -0.663468425 | 0.614283914 | 1.080068044 | 0.280111881 | 1 | 0.663468425 |
| ENSMUSG00000092399 | 4930556N13Rik | 75285 | lncRNA | 3.944348191 | -0.663472849 | 0.579688974 | 1.144532464 | 0.252402883 | 1 | 0.663472849 |
| ENSMUSG00000031292 | Cdkl5 | 382253 | protein_coding | 60.89600919 | 0.663709184 | 0.15280841 | -4.343407444 | 1.40E-05 | 0.000150633 | 0.663709184 |
| ENSMUSG00000099590 | C330022C24Rik | 78520 | lncRNA | 1.077410819 | 0.663750616 | 1.231109832 | -0.539148172 | 0.589784619 | 1 | 0.663750616 |
| ENSMUSG00000029095 | Ablim2 | 231148 | protein_coding | 108.4462024 | -0.663828869 | 0.218842219 | 3.033367476 | 0.002418409 | 0.013237523 | 0.663828869 |
| ENSMUSG00000039055 | Eme1 | 268465 | protein_coding | 56.30456099 | 0.663876478 | 0.204889711 | -3.24016503 | 0.001194605 | 0.007373206 | 0.663876478 |
| ENSMUSG00000091523 | AU040972 | 104522 | lncRNA | 5.235283962 | -0.663966723 | 0.606342839 | 1.095035151 | 0.273501237 | 0.489835334 | 0.663966723 |
| ENSMUSG00000019853 | Hebp2 | 56016 | protein_coding | 100.1440457 | 0.664026119 | 0.10994285 | -6.039739007 | 1.54E-09 | 3.33E-08 | 0.664026119 |
| ENSMUSG00000048249 | Crebrf | 77128 | protein_coding | 451.7508565 | -0.664046989 | 0.158611821 | 4.186617278 | 2.83E-05 | 0.000282016 | 0.664046989 |
| ENSMUSG00000022034 | Esco2 | 71988 | protein_coding | 140.6699307 | 0.664150522 | 0.222044247 | -2.991072872 | 0.002779991 | 0.014939401 | 0.664150522 |
| ENSMUSG00000038828 | Tmem214 | 68796 | protein_coding | 928.6617058 | 0.664339172 | 0.070742721 | -9.390919187 | 5.95E-21 | 3.21E-19 | 0.664339172 |
| ENSMUSG00000009733 | Tfcp2 | 21422 | protein_coding | 156.2401976 | 0.664405729 | 0.092906355 | -7.151348597 | 8.59E-13 | 2.62E-11 | 0.664405729 |
| ENSMUSG00000073412 | Lst1 | 16988 | protein_coding | 1.468612103 | 0.664522581 | 1.059826969 | -0.627010447 | 0.530652387 | 1 | 0.664522581 |
| ENSMUSG00000078495 | Zfp984 | 100041677 | protein_coding | 77.27809986 | -0.664580647 | 0.193663192 | 3.431631182 | 0.000599963 | 0.004126525 | 0.664580647 |
| ENSMUSG00000107709 | 9430018G01Rik | 319812 | lncRNA | 4.970645048 | -0.664679326 | 0.580299464 | 1.145407445 | 0.252040416 | 0.464375257 | 0.664679326 |
| ENSMUSG00000000378 | Ccm2 | 216527 | protein_coding | 274.2462586 | 0.66481035 | 0.083656626 | -7.946894151 | 1.91E-15 | 7.23E-14 | 0.66481035 |
| ENSMUSG00000031165 | Was | 22376 | protein_coding | 8.263453428 | 0.664997785 | 0.650134659 | -1.022861612 | 0.306373282 | 0.526861229 | 0.664997785 |
| ENSMUSG00000022365 | Derl1 | 67819 | protein_coding | 1392.2759 | 0.665069758 | 0.099314637 | -6.696593532 | 2.13E-11 | 5.76E-10 | 0.665069758 |
| ENSMUSG00000092752 | Mir3473a | 100499528 | miRNA | 0.708083812 | -0.6653201 | 1.340327249 | 0.496386312 | 0.619621881 | 1 | 0.6653201 |
| ENSMUSG00000050050 | Ccdc158 | 320696 | protein_coding | 4.648665409 | -0.665419524 | 0.578325109 | 1.150597672 | 0.249897791 | 0.462129017 | 0.665419524 |
| ENSMUSG00000086741 | Gm15816 | 101056408 | lncRNA | 1.517509045 | 0.665426062 | 1.634074146 | -0.407219013 | 0.683847138 | 1 | 0.665426062 |
| ENSMUSG00000025432 | Avil | 11567 | protein_coding | 156.4711401 | -0.665484301 | 0.14634507 | 4.547364 | 5.43E-06 | 6.46E-05 | 0.665484301 |
| ENSMUSG00000020447 | Npc1l1 | 237636 | protein_coding | 15.72134773 | -0.66562591 | 0.775269689 | 0.85857337 | 0.390575935 | 0.612813278 | 0.66562591 |
| ENSMUSG00000084575 | Gm23656 | 115489188 | snRNA | 0.690794652 | 0.665754391 | 1.20875696 | -0.550776056 | 0.5817872 | 1 | 0.665754391 |
| ENSMUSG00000064254 | Ethe1 | 66071 | protein_coding | 5170.799991 | -0.665764782 | 0.113935971 | 5.84332391 | 5.12E-09 | 1.03E-07 | 0.665764782 |
| ENSMUSG00000109311 | AI314278 | 101521 | lncRNA | 0.7934529 | 0.665809835 | 1.579666179 | -0.421487681 | 0.673399005 | 1 | 0.665809835 |
| ENSMUSG00000070348 | Ccnd1 | 12443 | protein_coding | 1171.354373 | 0.665820005 | 0.126239138 | -5.274275579 | 1.33E-07 | 2.16E-06 | 0.665820005 |
| ENSMUSG00000041431 | Ccnb1 | 268697 | protein_coding | 350.3668427 | 0.66603387 | 0.167591496 | -3.974150763 | 7.06E-05 | 0.000635639 | 0.66603387 |
| ENSMUSG00000060961 | Slc4a4 | 54403 | protein_coding | 1349.922103 | -0.666046466 | 0.082639428 | 8.05966934 | 7.65E-16 | 3.02E-14 | 0.666046466 |
| ENSMUSG00000029675 | Eln | 13717 | protein_coding | 6.487385712 | 0.66626444 | 1.450886227 | -0.459212051 | 0.646081898 | 0.807125838 | 0.66626444 |
| ENSMUSG00000031137 | Fgf13 | 14168 | protein_coding | 1.209523277 | -0.666400007 | 1.473748175 | 0.452180378 | 0.651139043 | 1 | 0.666400007 |
| ENSMUSG00000030824 | Nucb1 | 18220 | protein_coding | 6016.377918 | 0.666561587 | 0.042039425 | -15.85563048 | 1.29E-56 | 2.02E-54 | 0.666561587 |
| ENSMUSG00000031377 | Bmx | 12169 | protein_coding | 37.18954421 | -0.667076207 | 0.195935406 | 3.404572048 | 0.000662678 | 0.004486866 | 0.667076207 |
| ENSMUSG00000040441 | Slc26a10 | 216441 | protein_coding | 23.60309676 | 0.667310087 | 0.430719494 | -1.549291583 | 0.121311642 | 0.284081177 | 0.667310087 |
| ENSMUSG00000024793 | Tnfrsf25 | 85030 | protein_coding | 7.697971426 | -0.66810726 | 0.472921612 | 1.412723045 | 0.157737173 | 0.341264466 | 0.66810726 |

| ensembl_gene_id | external_gene_name | entrezgene_id | gene_biotype | baseMean | log2FoldChange | lfcSE | stat | pvalue | padj | absLog2FC |
|---|---|---|---|---|---|---|---|---|---|---|
| ENSMUSG00000063727 | Tnfrsf11b | 18383 | protein_coding | 10.203781 | 0.668129811 | 0.368275273 | -1.814213063 | 0.06964493 | 0.190353587 | 0.668129811 |
| ENSMUSG00000024501 | Dpysl3 | 22240 | protein_coding | 138.2796414 | -0.668799942 | 0.172291727 | 3.881787915 | 0.000103691 | 0.000888152 | 0.668799942 |
| ENSMUSG00000005357 | Slc1a6 | 20513 | protein_coding | 1.245792686 | -0.668906048 | 2.00840731 | 0.333052984 | 0.739094288 | 1 | 0.668906048 |
| ENSMUSG00000051851 | Rtl8c | 72865 | protein_coding | 1.059355334 | -0.668937546 | 1.282918109 | 0.521418741 | 0.602075099 | 1 | 0.668937546 |
| ENSMUSG00000031028 | Tub | 22141 | protein_coding | 3.251949007 | 0.668985978 | 1.550869617 | -0.431361844 | 0.666205289 | 1 | 0.668985978 |
| ENSMUSG00000044933 | Sstr3 | 20607 | protein_coding | 14.65845974 | 0.669615663 | 0.366978248 | -1.824673991 | 0.06805024 | 0.187078687 | 0.669615663 |
| ENSMUSG00000095026 | Gm3336 | 100502950 | protein_coding | 633.5629222 | -0.669694441 | 0.087366723 | 7.665326318 | 1.78E-14 | 6.25E-13 | 0.669694441 |
| ENSMUSG00000012443 | Kif11 | 16551 | protein_coding | 697.1691314 | 0.66976925 | 0.190939433 | -3.507757622 | 0.000451901 | 0.003254551 | 0.66976925 |
| ENSMUSG00000110140 | 5430421F17Rik | 74489 | lncRNA | 1.871010492 | -0.669777044 | 1.600868794 | 0.418383472 | 0.675666769 | 1 | 0.669777044 |
| ENSMUSG00000091780 | Sco2 | 100126824 | protein_coding | 0.889701953 | 0.66984408 | 1.348333701 | -0.49679399 | 0.619334334 | 1 | 0.66984408 |
| ENSMUSG00000040374 | Pex2 | 19302 | protein_coding | 728.9120177 | 0.670146187 | 0.085682873 | -7.821238485 | 5.23E-15 | 1.90E-13 | 0.670146187 |
| ENSMUSG00000037725 | Ckap2 | 80986 | protein_coding | 181.2682928 | 0.670434946 | 0.165247834 | -4.057148151 | 4.97E-05 | 0.000464257 | 0.670434946 |
| ENSMUSG00000079165 | Sap25 | 751865 | protein_coding | 0.735624731 | -0.670437387 | 1.388589437 | 0.482819017 | 0.629224247 | 1 | 0.670437387 |
| ENSMUSG00000028840 | Zfp593 | 68040 | protein_coding | 11.38025386 | -0.67065384 | 0.326638696 | 2.053197763 | 0.040053401 | 0.125988927 | 0.67065384 |
| ENSMUSG00000050967 | Creg2 | 263764 | protein_coding | 2.556436647 | 0.670662515 | 1.547350195 | -0.433426458 | 0.664704982 | 1 | 0.670662515 |
| ENSMUSG00000025196 | Cpn1 | 93721 | protein_coding | 168.405754 | 0.670747331 | 0.297968508 | -2.251067856 | 0.02438124 | 0.085735413 | 0.670747331 |
| ENSMUSG00000093147 | Gm23369 | 115489667 | snRNA | 0.555228781 | -0.67088004 | 1.638467477 | 0.409455817 | 0.682205185 | 1 | 0.67088004 |
| ENSMUSG00000105504 | Gbp5 | 229898 | protein_coding | 5.248963721 | -0.671134055 | 0.524191609 | 1.280322011 | 0.20043191 | 0.401690844 | 0.671134055 |
| ENSMUSG00000043415 | Otud1 | 71198 | protein_coding | 18.99428439 | -0.671341156 | 0.304246275 | 2.206571488 | 0.027344011 | 0.093703879 | 0.671341156 |
| ENSMUSG00000028402 | Mpdz | 17475 | protein_coding | 48.18381883 | 0.671926585 | 0.225635767 | -2.97792586 | 0.002902062 | 0.015454578 | 0.671926585 |
| ENSMUSG00000040017 | Saa4 | 20211 | protein_coding | 2.317774396 | -0.671977931 | 1.479775911 | 0.454107899 | 0.649751168 | 1 | 0.671977931 |
| ENSMUSG00000022360 | Atad2 | 70472 | protein_coding | 429.3671829 | 0.67201081 | 0.195077002 | -3.444848974 | 0.000571379 | 0.003964599 | 0.67201081 |
| ENSMUSG00000024778 | Fas | 14102 | protein_coding | 88.53613991 | 0.672031196 | 0.158024859 | -4.252692893 | 2.11E-05 | 0.000216378 | 0.672031196 |
| ENSMUSG00000031163 | Glod5 | 69824 | protein_coding | 229.1093992 | -0.672657069 | 0.100888619 | 6.667323569 | 2.61E-11 | 6.98E-10 | 0.672657069 |
| ENSMUSG00000021013 | Ttc8 | 76260 | protein_coding | 13.33887964 | 0.672749058 | 0.283612012 | -2.372075335 | 0.017688484 | 0.066630388 | 0.672749058 |
| ENSMUSG00000026620 | Mark1 | 226778 | protein_coding | 81.08278774 | 0.672924741 | 0.205274407 | -3.278171638 | 0.001044818 | 0.006600663 | 0.672924741 |
| ENSMUSG00000042367 | Gjb3 | 14620 | protein_coding | 513.046649 | -0.673217077 | 0.135404073 | 4.971911554 | 6.63E-07 | 9.51E-06 | 0.673217077 |
| ENSMUSG00000021086 | Ccdc175 | 73936 | protein_coding | 2.016558434 | -0.673301145 | 2.093389082 | 0.321632109 | 0.747731416 | 1 | 0.673301145 |
| ENSMUSG00000037544 | Dlgap5 | 218977 | protein_coding | 259.382654 | 0.673400227 | 0.148036676 | -4.548874269 | 5.39E-06 | 6.42E-05 | 0.673400227 |
| ENSMUSG00000106951 | 5930430L01Rik | 319982 | lncRNA | 12.19395557 | -0.673478517 | 0.300049241 | 2.244559972 | 0.024796395 | 0.086811038 | 0.673478517 |
| ENSMUSG00000109336 | Samd4b | 233033 | protein_coding | 498.7022059 | 0.67355613 | 0.127016317 | -5.302910276 | 1.14E-07 | 1.86E-06 | 0.67355613 |
| ENSMUSG00000004562 | Arhgef40 | 268739 | protein_coding | 312.7839211 | 0.673755234 | 0.130849373 | -5.149090284 | 2.62E-07 | 4.03E-06 | 0.673755234 |
| ENSMUSG00000027379 | Bub1 | 12235 | protein_coding | 173.944463 | 0.673809074 | 0.210080834 | -3.207380046 | 0.001339499 | 0.008117753 | 0.673809074 |
| ENSMUSG00000026605 | Cenpf | 108000 | protein_coding | 651.819101 | 0.673809703 | 0.154118969 | -4.372010178 | 1.23E-05 | 0.000134753 | 0.673809703 |
| ENSMUSG00000009646 | Pla2g12b | 69836 | protein_coding | 40.08664343 | 0.673847463 | 0.335246957 | -2.010003219 | 0.044430848 | 0.136254601 | 0.673847463 |
| ENSMUSG00000057706 | Mex3b | 108797 | protein_coding | 19.78365968 | 0.673855731 | 0.288546114 | -2.335348489 | 0.019525223 | 0.07201649 | 0.673855731 |
| ENSMUSG00000030110 | Ret | 19713 | protein_coding | 37.35007588 | 0.673916258 | 0.221148893 | -3.047341765 | 0.00230875 | 0.012721623 | 0.673916258 |
| ENSMUSG00000077347 | Gm24671 | 115490298 | snRNA | 0.932372629 | 0.674119013 | 1.670948229 | -0.403435008 | 0.686628237 | 1 | 0.674119013 |
| ENSMUSG00000054545 | Ugt1a6a | 94284 | protein_coding | 68.29045528 | -0.674198423 | 0.203228037 | 3.317447899 | 0.000908439 | 0.005867014 | 0.674198423 |

| ensembl_gene_id | external_gene_name | entrezgene_id | gene_biotype | baseMean | log2FoldChange | lfcSE | stat | pvalue | padj | absLog2FC |
|---|---|---|---|---|---|---|---|---|---|---|
| ENSMUSG00000006435 | Neurl1a | 18011 | protein_coding | 35.51293804 | -0.674204907 | 0.213235421 | 3.161786656 | 0.001568044 | 0.009251404 | 0.674204907 |
| ENSMUSG00000051726 | Kcnf1 | 382571 | protein_coding | 192.5344743 | -0.674251787 | 0.22134925 | 3.046099264 | 0.002318313 | 0.012765795 | 0.674251787 |
| ENSMUSG00000063447 | Ube2d2b | 73318 | protein_coding | 2.758437576 | -0.674261056 | 0.724970537 | 0.930053046 | 0.35234362 | 1 | 0.674261056 |
| ENSMUSG00000074274 | D930028M14Rik | 434147 | lncRNA | 1.725418167 | -0.674264019 | 2.055187723 | 0.328079042 | 0.7428519 | 1 | 0.674264019 |
| ENSMUSG00000054293 | P2ry10b | 213438 | protein_coding | 1.343789643 | -0.674508828 | 1.364377323 | 0.494371181 | 0.621044063 | 1 | 0.674508828 |
| ENSMUSG00000033634 | Nat8f2 | 93673 | protein_coding | 3.314840522 | -0.67456905 | 0.73573416 | 0.916865204 | 0.35921328 | 1 | 0.67456905 |
| ENSMUSG00000040414 | Slc25a28 | 246696 | protein_coding | 282.292355 | 0.674816129 | 0.102947308 | -6.554966234 | 5.57E-11 | 1.45E-09 | 0.674816129 |
| ENSMUSG00000090125 | Pou3f1 | 18991 | protein_coding | 2.865635174 | 0.674972472 | 1.316171911 | -0.512830024 | 0.608070226 | 1 | 0.674972472 |
| ENSMUSG00000096696 | Zfp960 | 449000 | protein_coding | 25.23734932 | -0.675086537 | 0.233015965 | 2.897168595 | 0.003765473 | 0.019221956 | 0.675086537 |
| ENSMUSG00000055322 | Tns1 | 21961 | protein_coding | 12.62030252 | 0.675183528 | 1.074364987 | -0.628448932 | 0.529709891 | 0.727211667 | 0.675183528 |
| ENSMUSG00000081058 | H3c15 | 97114 | protein_coding | 16.72699827 | 0.675248334 | 0.395157221 | -1.708809298 | 0.08748628 | 0.224444373 | 0.675248334 |
| ENSMUSG00000026536 | Ifi211 | 381308 | protein_coding | 1.191964284 | 0.675399589 | 1.850570007 | -0.364968408 | 0.715134989 | 1 | 0.675399589 |
| ENSMUSG00000096233 | Gm13238 | 236069 | processed_pseudogene | 2.842766401 | -0.675491177 | 1.047218572 | 0.645033611 | 0.518905405 | 1 | 0.675491177 |
| ENSMUSG00000074264 | Amy1 | 11722 | protein_coding | 1.167192673 | -0.675516803 | 1.181361672 | 0.57181202 | 0.567449333 | 1 | 0.675516803 |
| ENSMUSG00000021087 | Rtn1 | 104001 | protein_coding | 8.019271532 | -0.675599867 | 0.542518044 | 1.245303957 | 0.213020042 | 0.417785903 | 0.675599867 |
| ENSMUSG00000038457 | Tmem255b | 272465 | protein_coding | 2.897960663 | -0.675734396 | 0.922409585 | 0.73257521 | 0.463817555 | 1 | 0.675734396 |
| ENSMUSG00000065611 | Mir23a | 387216 | miRNA | 24.74079527 | -0.676001431 | 0.277906154 | 2.432480968 | 0.01499578 | 0.058405732 | 0.676001431 |
| ENSMUSG00000032334 | Loxl1 | 16949 | protein_coding | 2.292342528 | 0.676035745 | 1.501063229 | -0.450371265 | 0.652442761 | 1 | 0.676035745 |
| ENSMUSG00000100281 | 4930422M22Rik | 73774 | lncRNA | 3.874454342 | 0.676126437 | 0.543162282 | -1.244796371 | 0.213206611 | 1 | 0.676126437 |
| ENSMUSG00000086943 | 4732414G09Rik | 100043309 | lncRNA | 4.267584755 | -0.676140942 | 0.685337776 | 0.986580582 | 0.323848293 | 1 | 0.676140942 |
| ENSMUSG00000091650 | Apol11a | 626615 | protein_coding | 3.990374108 | 0.676187961 | 0.642561246 | -1.05233231 | 0.292647113 | 1 | 0.676187961 |
| ENSMUSG00000063605 | Ccdc102a | 234582 | protein_coding | 7.563374887 | -0.676321316 | 0.534729143 | 1.264792323 | 0.205945784 | 0.408576247 | 0.676321316 |
| ENSMUSG00000023084 | Lrrc71 | 74485 | protein_coding | 13.3858846 | 0.676374413 | 0.342914209 | -1.972430407 | 0.048560493 | 0.145821162 | 0.676374413 |
| ENSMUSG00000022454 | Nell2 | 54003 | protein_coding | 1.483175664 | -0.676671672 | 1.500512997 | 0.450960221 | 0.652018219 | 1 | 0.676671672 |
| ENSMUSG00000047528 | Flacc1 | 108812 | protein_coding | 13.5179533 | -0.676813731 | 0.379863361 | 1.781729433 | 0.074793365 | 0.19999509 | 0.676813731 |
| ENSMUSG00000011267 | Zfp296 | 63872 | protein_coding | 9.346339981 | -0.677560724 | 0.474125151 | 1.429075682 | 0.152982482 | 0.333996921 | 0.677560724 |
| ENSMUSG00000070530 | Wfdc16 | 277345 | protein_coding | 0.846280467 | -0.677600376 | 1.512867412 | 0.447891448 | 0.654231541 | 1 | 0.677600376 |
| ENSMUSG00000041609 | Bicdl1 | 75665 | protein_coding | 257.3946061 | 0.677614384 | 0.11066902 | -6.122891322 | 9.19E-10 | 2.03E-08 | 0.677614384 |
| ENSMUSG00000038065 | Mturn | 68235 | protein_coding | 14.75344753 | 0.677700404 | 0.317120538 | -2.137043557 | 0.032594449 | 0.107588657 | 0.677700404 |
| ENSMUSG00000066512 | Klk1b5 | 16622 | protein_coding | 10.00698869 | 0.677859422 | 0.437941613 | -1.547830585 | 0.121663091 | 0.284460467 | 0.677859422 |
| ENSMUSG00000050747 | Trim15 | 69097 | protein_coding | 284.0155782 | 0.677927972 | 0.260936579 | -2.598056484 | 0.009375307 | 0.040476816 | 0.677927972 |
| ENSMUSG00000025192 | Entpd7 | 93685 | protein_coding | 306.4757184 | 0.677968586 | 0.117632907 | -5.763426276 | 8.24E-09 | 1.61E-07 | 0.677968586 |
| ENSMUSG00000038349 | Plcl1 | 227120 | protein_coding | 8.362428651 | -0.677985861 | 0.450560465 | 1.504761101 | 0.132385508 | 0.301713259 | 0.677985861 |
| ENSMUSG00000035296 | Sgcg | 24053 | protein_coding | 2.002224169 | -0.678024298 | 1.292599626 | 0.524543165 | 0.599900806 | 1 | 0.678024298 |
| ENSMUSG00000024236 | Svil | 225115 | protein_coding | 130.6999116 | 0.678116001 | 0.146216532 | -4.637751907 | 3.52E-06 | 4.35E-05 | 0.678116001 |
| ENSMUSG00000032852 | Rspo4 | 228770 | protein_coding | 1.994095376 | -0.678272211 | 2.080709858 | 0.325981159 | 0.74443861 | 1 | 0.678272211 |
| ENSMUSG00000033342 | Plppr5 | 75769 | protein_coding | 1.182405043 | -0.678334025 | 1.826239467 | 0.371437611 | 0.710311616 | 1 | 0.678334025 |
| ENSMUSG00000035458 | Tnni3 | 21954 | protein_coding | 3.010333834 | 0.678355474 | 0.75190418 | -0.902183406 | 0.366959448 | 1 | 0.678355474 |
| ENSMUSG00000091811 | Inafm1 | 66300 | protein_coding | 200.7490186 | -0.678508769 | 0.116027364 | 5.847834028 | 4.98E-09 | 1.01E-07 | 0.678508769 |

Supplementary Table 1

| ensembl_gene_id | external_gene_name | entrezgene_id | gene_biotype | baseMean | log2FoldChange | lfcSE | stat | pvalue | padj | absLog2FC |
|---|---|---|---|---|---|---|---|---|---|---|
| ENSMUSG00000028034 | Fubp1 | 51886 | protein_coding | 614.4888061 | -0.678694877 | 0.073174177 | 9.275059923 | 1.78E-20 | 9.22E-19 | 0.678694877 |
| ENSMUSG00000021219 | Rgs6 | 50779 | protein_coding | 5.117930432 | 0.679586254 | 1.466568924 | -0.463385145 | 0.643088324 | 0.805274537 | 0.679586254 |
| ENSMUSG00000028849 | Map7d1 | 245877 | protein_coding | 131.0317005 | 0.679739643 | 0.173075676 | -3.927412897 | 8.59E-05 | 0.000754237 | 0.679739643 |
| ENSMUSG00000078794 | Dact3 | 629378 | protein_coding | 5.837009658 | 0.679847497 | 0.604289653 | -1.125035807 | 0.260573861 | 0.474068167 | 0.679847497 |
| ENSMUSG00000033949 | Trim36 | 28105 | protein_coding | 252.949328 | -0.679962767 | 0.107348564 | 6.334158014 | 2.39E-10 | 5.70E-09 | 0.679962767 |
| ENSMUSG00000014030 | Pax5 | 18507 | protein_coding | 5.158380758 | 0.680120049 | 1.256080026 | -0.541462355 | 0.588188938 | 0.769374426 | 0.680120049 |
| ENSMUSG00000040740 | Slc25a34 | 384071 | protein_coding | 517.6660588 | -0.680270527 | 0.129963518 | 5.234319115 | 1.66E-07 | 2.63E-06 | 0.680270527 |
| ENSMUSG00000028785 | Hpca | 15444 | protein_coding | 30.00077495 | 0.680336069 | 0.324225226 | -2.098344038 | 0.035874765 | 0.115554504 | 0.680336069 |
| ENSMUSG00000021223 | Papln | 170721 | protein_coding | 28.1754834 | 0.680509893 | 0.31997322 | -2.126771399 | 0.033439082 | 0.109621469 | 0.680509893 |
| ENSMUSG00000057667 | Bloc1s3 | 232946 | protein_coding | 257.8843336 | 0.680519184 | 0.0816105 | -8.338622844 | 7.52E-17 | 3.18E-15 | 0.680519184 |
| ENSMUSG00000022057 | Adamdec1 | 58860 | protein_coding | 2.740548806 | -0.680777323 | 0.967808712 | 0.703421362 | 0.481793198 | 1 | 0.680777323 |
| ENSMUSG00000040084 | Bub1b | 12236 | protein_coding | 483.1748062 | 0.681261323 | 0.1656731 | -4.112081699 | 3.92E-05 | 0.000377711 | 0.681261323 |
| ENSMUSG00000020651 | Slc26a4 | 23985 | protein_coding | 3.315395007 | -0.68139464 | 0.706285978 | 0.964757423 | 0.334666326 | 1 | 0.68139464 |
| ENSMUSG00000020330 | Hmmr | 15366 | protein_coding | 352.4948247 | 0.681653547 | 0.1794545 | -3.798475645 | 0.000145589 | 0.001199022 | 0.681653547 |
| ENSMUSG00000090698 | Apold1 | 381823 | protein_coding | 5.969794208 | -0.681969887 | 0.563420305 | 1.210410559 | 0.226121392 | 0.433866874 | 0.681969887 |
| ENSMUSG00000032218 | Ccnb2 | 12442 | protein_coding | 449.6226443 | 0.68207059 | 0.123419742 | -5.526430201 | 3.27E-08 | 5.78E-07 | 0.68207059 |
| ENSMUSG00000029361 | Nos1 | 18125 | protein_coding | 7.136223485 | 0.682122187 | 1.247393736 | -0.546837913 | 0.584490094 | 0.766833571 | 0.682122187 |
| ENSMUSG00000073421 | H2-Ab1 | 14961 | protein_coding | 34.5850375 | -0.682182876 | 0.22862449 | 2.983857396 | 0.002846394 | 0.015226871 | 0.682182876 |
| ENSMUSG00000030677 | Kif22 | 110033 | protein_coding | 266.5618183 | 0.68239347 | 0.170337762 | -4.006119732 | 6.17E-05 | 0.000563781 | 0.68239347 |
| ENSMUSG00000065878 | Snord34 | 27210 | snoRNA | 5.071679028 | 0.682707826 | 0.640569437 | -1.065782704 | 0.286521881 | 0.505083513 | 0.682707826 |
| ENSMUSG00000033715 | Akr1c14 | 105387 | protein_coding | 1841.771821 | -0.68282765 | 0.182969688 | 3.731916778 | 0.000190028 | 0.00151381 | 0.68282765 |
| ENSMUSG00000025602 | Zfp202 | 80902 | protein_coding | 21.47535058 | -0.683067687 | 0.233362482 | 2.927067291 | 0.003421748 | 0.017747096 | 0.683067687 |
| ENSMUSG00000085207 | Gm11767 | 102640451 | lncRNA | 4.933233166 | -0.683087242 | 0.485406529 | 1.407247742 | 0.159353939 | 0.343481339 | 0.683087242 |
| ENSMUSG00000087303 | Lipo2 | 101055671 | protein_coding | 1.386425299 | 0.683327339 | 1.542640529 | -0.44295954 | 0.657795 | 1 | 0.683327339 |
| ENSMUSG00000026729 | 4930562F07Rik | 75255 | lncRNA | 0.924259535 | -0.683328084 | 1.465176127 | 0.466379483 | 0.640943895 | 1 | 0.683328084 |
| ENSMUSG00000005611 | Irag1 | 17540 | protein_coding | 6.192747272 | 0.683581231 | 0.679748615 | -1.005638284 | 0.314589601 | 0.535801636 | 0.683581231 |
| ENSMUSG00000027719 | Adad1 | 21744 | protein_coding | 1.038165905 | 0.683646242 | 2.233067964 | -0.306146635 | 0.759493002 | 1 | 0.683646242 |
| ENSMUSG00000027865 | Gdap2 | 14547 | protein_coding | 293.076671 | 0.683727153 | 0.085562278 | -7.990988215 | 1.34E-15 | 5.13E-14 | 0.683727153 |
| ENSMUSG00000035775 | Krt20 | 66809 | protein_coding | 9465.065044 | 0.683740509 | 0.166824343 | -4.098565566 | 4.16E-05 | 0.000397981 | 0.683740509 |
| ENSMUSG00000008206 | Cers4 | 67260 | protein_coding | 451.4739938 | -0.683809359 | 0.131594377 | 5.196341786 | 2.03E-07 | 3.18E-06 | 0.683809359 |
| ENSMUSG00000046280 | She | 214547 | protein_coding | 1.72963949 | 0.683899665 | 2.220273564 | -0.308024955 | 0.758063343 | 1 | 0.683899665 |
| ENSMUSG00000032515 | Csrnp1 | 215418 | protein_coding | 407.0648489 | -0.684087076 | 0.268499958 | 2.54781074 | 0.010840126 | 0.045342919 | 0.684087076 |
| ENSMUSG00000022978 | Mis18a | 66578 | protein_coding | 132.4271329 | 0.684100018 | 0.166427538 | -4.110497741 | 3.95E-05 | 0.000379723 | 0.684100018 |
| ENSMUSG00000084990 | Gm14549 | 69591 | lncRNA | 3.641884317 | 0.684145136 | 0.63832263 | -1.071785808 | 0.283816249 | 1 | 0.684145136 |
| ENSMUSG00000026826 | Nr4a2 | 18227 | protein_coding | 100.3007906 | 0.684145793 | 0.173310839 | -3.947507258 | 7.90E-05 | 0.000701882 | 0.684145793 |
| ENSMUSG00000020912 | Krt12 | 268482 | protein_coding | 2.823754465 | -0.684400181 | 0.898126633 | 0.762030827 | 0.446041605 | 1 | 0.684400181 |
| ENSMUSG00000044949 | Ubtd2 | 327900 | protein_coding | 30.59278233 | -0.684438756 | 0.217284425 | 3.149966945 | 0.001632889 | 0.00957243 | 0.684438756 |
| ENSMUSG00000078932 | CN725425 | 545126 | protein_coding | 2.555088192 | 0.684532733 | 0.996966367 | -0.686615673 | 0.492324949 | 1 | 0.684532733 |
| ENSMUSG00000023886 | Smoc2 | 64074 | protein_coding | 382.7234019 | 0.684906305 | 0.214493621 | -3.193131348 | 0.001407389 | 0.00846073 | 0.684906305 |

| ensembl_gene_id | external_gene_name | entrezgene_id | gene_biotype | baseMean | log2FoldChange | lfcSE | stat | pvalue | padj | absLog2FC |
|---|---|---|---|---|---|---|---|---|---|---|
| ENSMUSG00000010841 | Cfap97d1 | 75437 | protein_coding | 5.514899277 | -0.685184653 | 0.485153212 | 1.412305712 | 0.157859965 | 0.341395949 | 0.685184653 |
| ENSMUSG00000110266 | Gm32742 | 102635385 | protein_coding | 19.06453728 | -0.685222245 | 0.310962923 | 2.203549666 | 0.02755603 | 0.094313013 | 0.685222245 |
| ENSMUSG00000023083 | H2-M10.2 | 333715 | protein_coding | 1.164192601 | 0.685612198 | 1.31145538 | -0.522787286 | 0.601122288 | 1 | 0.685612198 |
| ENSMUSG00000024155 | Meiob | 75178 | protein_coding | 0.725018898 | -0.685618257 | 1.280453599 | 0.535449514 | 0.592339059 | 1 | 0.685618257 |
| ENSMUSG00000030365 | Clec2i | 93675 | protein_coding | 4.696856304 | 0.685734495 | 0.799455505 | -0.857751921 | 0.391029463 | 0.613316042 | 0.685734495 |
| ENSMUSG00000004319 | Clcn3 | 12725 | protein_coding | 1895.240943 | 0.685873399 | 0.055079019 | -12.45253478 | 1.35E-35 | 1.22E-33 | 0.685873399 |
| ENSMUSG00000019944 | Rhobtb1 | 69288 | protein_coding | 36.90927689 | -0.685886305 | 0.21558017 | 3.181583473 | 0.001464723 | 0.008704007 | 0.685886305 |
| ENSMUSG00000114133 | Btf3l4b | 70533 | protein_coding | 2.381445567 | 0.686085123 | 0.741828087 | -0.924857302 | 0.355040139 | 1 | 0.686085123 |
| ENSMUSG00000035638 | Muc20 | 224116 | protein_coding | 15.75762468 | 0.686103536 | 0.340505503 | -2.014955795 | 0.043909281 | 0.135006119 | 0.686103536 |
| ENSMUSG00000047250 | Ptgs1 | 19224 | protein_coding | 51.13845478 | -0.686245094 | 0.223444409 | 3.071211752 | 0.002131919 | 0.011942524 | 0.686245094 |
| ENSMUSG00000116895 | Gm3435 | 100041621 | protein_coding | 4.714525196 | -0.686471088 | 0.527041459 | 1.302499218 | 0.192745785 | 0.390530809 | 0.686471088 |
| ENSMUSG00000050288 | Fzd2 | 57265 | protein_coding | 31.47110478 | 0.686624366 | 0.274109787 | -2.504924665 | 0.012247748 | 0.049834789 | 0.686624366 |
| ENSMUSG00000072941 | Sod3 | 20657 | protein_coding | 11.02025193 | 0.686956963 | 0.511836062 | -1.342142563 | 0.179549776 | 0.372872861 | 0.686956963 |
| ENSMUSG00000118047 | 4930554I06Rik | 75354 | lncRNA | 9.261617242 | 0.68705105 | 0.54418387 | -1.262534757 | 0.206756428 | 0.40949576 | 0.68705105 |
| ENSMUSG00000001248 | Gramd1a | 52857 | protein_coding | 89.18754716 | 0.687257921 | 0.171421511 | -4.009169665 | 6.09E-05 | 0.000557783 | 0.687257921 |
| ENSMUSG00000009394 | Syn2 | 20965 | protein_coding | 33.12159644 | -0.687581563 | 0.243135136 | 2.827981075 | 0.004684258 | 0.023015718 | 0.687581563 |
| ENSMUSG00000019762 | Iyd | 70337 | protein_coding | 69.62237479 | -0.687705804 | 0.198845336 | 3.458496025 | 0.0005432 | 0.003805572 | 0.687705804 |
| ENSMUSG00000029727 | Cyp3a13 | 13113 | protein_coding | 1090.854129 | -0.687923692 | 0.151800279 | 4.531768293 | 5.85E-06 | 6.88E-05 | 0.687923692 |
| ENSMUSG00000030492 | Slc7a9 | 30962 | protein_coding | 103.4800641 | 0.688291243 | 0.245387248 | -2.804918549 | 0.00503293 | 0.024481237 | 0.688291243 |
| ENSMUSG00000114004 | Gm48552 | 117934532 | protein_coding | 1.483776741 | -0.68831078 | 0.841550584 | 0.817907792 | 0.413409841 | 1 | 0.68831078 |
| ENSMUSG00000022434 | Fam118a | 73225 | protein_coding | 104.5885983 | 0.688325738 | 0.149470043 | -4.605108326 | 4.12E-06 | 5.03E-05 | 0.688325738 |
| ENSMUSG00000050138 | Kcnk12 | 210741 | protein_coding | 1.269557569 | -0.688441877 | 1.634828101 | 0.421109642 | 0.673675021 | 1 | 0.688441877 |
| ENSMUSG00000034701 | Neurod1 | 18012 | protein_coding | 18.77043516 | -0.688459106 | 0.25064235 | 2.746778848 | 0.00601837 | 0.02823687 | 0.688459106 |
| ENSMUSG00000042993 | Ifnk | 387510 | protein_coding | 1.695738342 | -0.688539642 | 0.874590624 | 0.787270779 | 0.431123372 | 1 | 0.688539642 |
| ENSMUSG00000026934 | Lhx3 | 16871 | protein_coding | 1.845954372 | 0.688571746 | 1.490184517 | -0.462071467 | 0.644030071 | 1 | 0.688571746 |
| ENSMUSG00000059395 | Nkapl | 66707 | protein_coding | 5.77744354 | -0.688594206 | 0.394427466 | 1.745806939 | 0.080844505 | 0.211378428 | 0.688594206 |
| ENSMUSG00000006389 | Mpl | 17480 | protein_coding | 2.525270532 | -0.688658869 | 1.586613718 | 0.434043184 | 0.664257083 | 1 | 0.688658869 |
| ENSMUSG00000021069 | Pygl | 110095 | protein_coding | 103.5293441 | -0.688715445 | 0.143906076 | 4.785867715 | 1.70E-06 | 2.24E-05 | 0.688715445 |
| ENSMUSG00000097711 | Gm5523 | 433273 | processed_pseudogene | 1.107845151 | 0.688863983 | 1.338378703 | -0.514700347 | 0.606762431 | 1 | 0.688863983 |
| ENSMUSG00000076315 | Mir343 | 100124438 | miRNA | 1.009625905 | 0.689070346 | 1.034420343 | -0.666141526 | 0.505320644 | 1 | 0.689070346 |
| ENSMUSG00000053279 | Aldh1a1 | 11668 | protein_coding | 1832.860746 | -0.689081368 | 0.189749357 | 3.631534662 | 0.000281741 | 0.002154013 | 0.689081368 |
| ENSMUSG00000031543 | Ank1 | 11733 | protein_coding | 8.45764758 | 0.689200661 | 0.778847327 | -0.884898282 | 0.376211495 | 0.59846773 | 0.689200661 |
| ENSMUSG00000053687 | Dpep2 | 319446 | protein_coding | 1.506191763 | 0.689446734 | 1.401974161 | -0.491768503 | 0.622883005 | 1 | 0.689446734 |
| ENSMUSG00000067578 | Cbln4 | 228942 | protein_coding | 1.718497212 | 0.689447912 | 2.106639937 | -0.327273731 | 0.743460859 | 1 | 0.689447912 |
| ENSMUSG00000054958 | Nt5c1a | 230718 | protein_coding | 18.68926142 | -0.689556623 | 0.412131813 | 1.673145824 | 0.094298592 | 0.236536038 | 0.689556623 |
| ENSMUSG00000037106 | Fer1l6 | 631797 | protein_coding | 1184.647486 | -0.689638183 | 0.218020994 | 3.163173287 | 0.001560594 | 0.009213125 | 0.689638183 |
| ENSMUSG00000040231 | Syngr4 | 58867 | protein_coding | 4.643474836 | 0.689814553 | 0.509827153 | -1.35303612 | 0.176044091 | 0.368018849 | 0.689814553 |
| ENSMUSG00000047604 | Frat2 | 212398 | protein_coding | 401.2641992 | -0.689864681 | 0.110354704 | 6.251339145 | 4.07E-10 | 9.40E-09 | 0.689864681 |
| ENSMUSG00000008734 | Gprc5b | 64297 | protein_coding | 6.533451463 | 0.689910258 | 0.602621687 | -1.144848042 | 0.25227211 | 0.464617944 | 0.689910258 |

| ensembl_gene_id | external_gene_name | entrezgene_id | gene_biotype | baseMean | log2FoldChange | lfcSE | stat | pvalue | padj | absLog2FC |
|---|---|---|---|---|---|---|---|---|---|---|
| ENSMUSG00000045328 | Cenpe | 229841 | protein_coding | 531.9434516 | 0.690159773 | 0.152217397 | -4.534040036 | 5.79E-06 | 6.82E-05 | 0.690159773 |
| ENSMUSG00000045333 | Zfp423 | 94187 | protein_coding | 4.55567773 | 0.69066913 | 1.065402188 | -0.648270801 | 0.516809816 | 0.717554842 | 0.69066913 |
| ENSMUSG00000026304 | Rab17 | 19329 | protein_coding | 195.2623877 | -0.690678672 | 0.155649514 | 4.437396916 | 9.11E-06 | 0.000103273 | 0.690678672 |
| ENSMUSG00000075031 | H2bc3 | 319178 | protein_coding | 1438.114097 | 0.691172554 | 0.204344202 | -3.382393768 | 0.000718571 | 0.004789552 | 0.691172554 |
| ENSMUSG00000006469 | Slc34a3 | 142681 | protein_coding | 2.24971595 | 0.691375077 | 1.919464377 | -0.360191669 | 0.718703805 | 1 | 0.691375077 |
| ENSMUSG00000035407 | Kank4 | 242553 | protein_coding | 4.536010713 | -0.691700084 | 0.540987009 | 1.278589083 | 0.201041796 | 0.402386658 | 0.691700084 |
| ENSMUSG00000098719 | Mir7212 | 102465698 | miRNA | 2.317151312 | -0.691836813 | 0.805679589 | 0.85869969 | 0.390506222 | 1 | 0.691836813 |
| ENSMUSG00000026312 | Cdh7 | 241201 | protein_coding | 1.152454083 | -0.691919105 | 1.919868113 | 0.36039929 | 0.718548557 | 1 | 0.691919105 |
| ENSMUSG00000034810 | Scn7a | 20272 | protein_coding | 1.700606522 | 0.692091772 | 1.759104722 | -0.393434094 | 0.693998892 | 1 | 0.692091772 |
| ENSMUSG00000086765 | Gm11827 | 100503518 | lncRNA | 1.202628472 | -0.692179427 | 0.985700144 | 0.702221087 | 0.482541297 | 1 | 0.692179427 |
| ENSMUSG00000033486 | Catsper2 | 212670 | protein_coding | 28.94336997 | 0.692327015 | 0.282680618 | -2.449149221 | 0.014319411 | 0.056215521 | 0.692327015 |
| ENSMUSG00000074736 | Syndig1 | 433485 | protein_coding | 1.471183746 | -0.693014013 | 1.043814704 | 0.663924364 | 0.506738722 | 1 | 0.693014013 |
| ENSMUSG00000063511 | Snrnp70 | 20637 | protein_coding | 2217.416876 | 0.693055886 | 0.136710846 | -5.069501837 | 3.99E-07 | 5.99E-06 | 0.693055886 |
| ENSMUSG00000078864 | Gm14322 | 626802 | protein_coding | 4.021611183 | -0.693163296 | 0.539987461 | 1.283665541 | 0.199259007 | 1 | 0.693163296 |
| ENSMUSG00000044647 | Csrnp3 | 77771 | protein_coding | 2.941844716 | 0.693193961 | 2.409746695 | -0.287662584 | 0.773605034 | 1 | 0.693193961 |
| ENSMUSG00000023367 | Tmem176a | 66058 | protein_coding | 454.5118373 | 0.693464475 | 0.099862718 | -6.944177886 | 3.81E-12 | 1.10E-10 | 0.693464475 |
| ENSMUSG00000030877 | Mfsd13b | 74466 | protein_coding | 6.952660022 | 0.693531729 | 0.454541107 | -1.525784396 | 0.127063568 | 0.293636687 | 0.693531729 |
| ENSMUSG00000030345 | Dyrk4 | 101320 | protein_coding | 2.562463564 | -0.693741968 | 1.088607331 | 0.637274753 | 0.523945893 | 1 | 0.693741968 |
| ENSMUSG00000059301 | Gm5434 | 432649 | transcribed_processed_pseudogene | 0.970868571 | -0.69404751 | 1.550617654 | 0.447594227 | 0.65444607 | 1 | 0.69404751 |
| ENSMUSG00000018449 | Rpain | 69723 | protein_coding | 118.9757296 | 0.694130771 | 0.138991407 | -4.994055279 | 5.91E-07 | 8.58E-06 | 0.694130771 |
| ENSMUSG00000026226 | Spata3 | 70060 | protein_coding | 2.582682865 | -0.694495665 | 0.955383092 | 0.726928989 | 0.467269458 | 1 | 0.694495665 |
| ENSMUSG00000028176 | Lrrc7 | 242274 | protein_coding | 2.609519552 | 0.694548127 | 1.9630635 | -0.353808283 | 0.723482565 | 1 | 0.694548127 |
| ENSMUSG00000110101 | Gm6249 | 621677 | lncRNA | 1.677010997 | -0.694728932 | 1.812044359 | 0.383395102 | 0.701426835 | 1 | 0.694728932 |
| ENSMUSG00000085995 | Gm2788 | 108167326 | lncRNA | 9.298319009 | 0.694743551 | 0.521622701 | -1.331889026 | 0.182896658 | 0.377599236 | 0.694743551 |
| ENSMUSG00000010435 | Spaca7 | 78634 | protein_coding | 1.452120146 | 0.69474401 | 1.15755144 | -0.600184135 | 0.548383525 | 1 | 0.69474401 |
| ENSMUSG00000039497 | Dse | 212898 | protein_coding | 2.44121662 | 0.694907274 | 0.878963503 | -0.790598554 | 0.429178291 | 1 | 0.694907274 |
| ENSMUSG00000069267 | H3c2 | 319150 | protein_coding | 716.3759691 | 0.695142861 | 0.189954617 | -3.659520752 | 0.000252687 | 0.001949033 | 0.695142861 |
| ENSMUSG00000041737 | Tmem45b | 235135 | protein_coding | 3465.302885 | -0.695348159 | 0.102094778 | 6.810810237 | 9.71E-12 | 2.72E-10 | 0.695348159 |
| ENSMUSG00000024420 | Zfp521 | 225207 | protein_coding | 2.838786157 | -0.695476014 | 1.842797436 | 0.377402312 | 0.705874649 | 1 | 0.695476014 |
| ENSMUSG00000088148 | Mir1983 | 100316716 | miRNA | 1.036085414 | 0.695727226 | 1.240068574 | -0.561039317 | 0.574770735 | 1 | 0.695727226 |
| ENSMUSG00000065219 | Snord32a | 27209 | snoRNA | 46.78920833 | 0.695830019 | 0.370198318 | -1.879614208 | 0.060160676 | 0.171114949 | 0.695830019 |
| ENSMUSG00000002664 | Pspn | 19197 | protein_coding | 5.959940618 | -0.696026783 | 0.44668196 | 1.558215566 | 0.119182156 | 0.280659184 | 0.696026783 |
| ENSMUSG00000034551 | Hdx | 245596 | protein_coding | 1.377569994 | 0.696171344 | 2.37074032 | -0.293651455 | 0.769024253 | 1 | 0.696171344 |
| ENSMUSG00000044197 | Gpr146 | 80290 | protein_coding | 8.878868125 | -0.696318171 | 0.485212426 | 1.435079015 | 0.151264583 | 0.331256586 | 0.696318171 |
| ENSMUSG00000030579 | Tyrobp | 22177 | protein_coding | 15.05920529 | 0.69656227 | 0.515047812 | -1.35242254 | 0.176240183 | 0.368253661 | 0.69656227 |
| ENSMUSG00000054169 | Ceacam10 | 26366 | protein_coding | 260.1530742 | -0.696602865 | 0.248778171 | 2.800096417 | 0.005108734 | 0.024791549 | 0.696602865 |
| ENSMUSG00000023021 | Cers5 | 71949 | protein_coding | 339.1916331 | -0.696678289 | 0.110897792 | 6.28216555 | 3.34E-10 | 7.84E-09 | 0.696678289 |
| ENSMUSG00000002055 | Spag5 | 54141 | protein_coding | 302.4265706 | 0.69667876 | 0.136497517 | -5.103966536 | 3.33E-07 | 5.06E-06 | 0.69667876 |
| ENSMUSG00000032611 | 1700102P08Rik | 112418 | protein_coding | 2.858730142 | -0.696883922 | 0.74863337 | 0.930874778 | 0.351918343 | 1 | 0.696883922 |

Supplementary Table 1

| ensembl_gene_id | external_gene_name | entrezgene_id | gene_biotype | baseMean | log2FoldChange | lfcSE | stat | pvalue | padj | absLog2FC |
|---|---|---|---|---|---|---|---|---|---|---|
| ENSMUSG00000024620 | Pdgfrb | 18596 | protein_coding | 103.3903192 | -0.697187524 | 0.201220217 | 3.464798583 | 0.000530629 | 0.00373044 | 0.697187524 |
| ENSMUSG00000078810 | Gp6 | 243816 | protein_coding | 7.771605336 | 0.697207306 | 0.555728944 | -1.254581598 | 0.209630687 | 0.41315684 | 0.697207306 |
| ENSMUSG00000066797 | Zfp648 | 100503355 | protein_coding | 1.136391595 | 0.697276486 | 1.538012022 | -0.453362182 | 0.650287964 | 1 | 0.697276486 |
| ENSMUSG00000068396 | Rpl34-ps1 | 619547 | processed_pseudogene | 4.332464098 | 0.69729104 | 0.474952222 | -1.468128807 | 0.142069238 | 1 | 0.69729104 |
| ENSMUSG00000027400 | Pdyn | 18610 | protein_coding | 1.229788694 | -0.697317117 | 1.516637274 | 0.459778438 | 0.645675261 | 1 | 0.697317117 |
| ENSMUSG00000078878 | Gm14305 | 100043387 | protein_coding | 24.69198039 | -0.697588185 | 0.226641599 | 3.077935332 | 0.002084401 | 0.01173204 | 0.697588185 |
| ENSMUSG00000044080 | S100a1 | 20193 | protein_coding | 747.4106129 | -0.697742949 | 0.108969047 | 6.403129772 | 1.52E-10 | 3.68E-09 | 0.697742949 |
| ENSMUSG00000104763 | 9830132P13Rik | 329753 | lncRNA | 0.78867778 | -0.697927385 | 1.857666796 | 0.37570106 | 0.707139149 | 1 | 0.697927385 |
| ENSMUSG00000096255 | Dynlt1b | 21648 | protein_coding | 20.60136743 | -0.698202161 | 0.292251448 | 2.389046026 | 0.016892186 | 0.064246329 | 0.698202161 |
| ENSMUSG00000022206 | Npr3 | 18162 | protein_coding | 2.070341876 | 0.69827775 | 2.410145821 | -0.289724275 | 0.772027184 | 1 | 0.69827775 |
| ENSMUSG00000022144 | Gdnf | 14573 | protein_coding | 1.569304991 | 0.698601184 | 1.749821672 | -0.39924136 | 0.689715371 | 1 | 0.698601184 |
| ENSMUSG00000028707 | Dmbx1 | 140477 | protein_coding | 1.563368295 | 0.698631533 | 2.384220672 | -0.293023016 | 0.769504559 | 1 | 0.698631533 |
| ENSMUSG00000019122 | Ccl9 | 20308 | protein_coding | 200.8183241 | 0.698750125 | 0.149895753 | -4.661573875 | 3.14E-06 | 3.91E-05 | 0.698750125 |
| ENSMUSG00000061479 | Snrpa | 53607 | protein_coding | 477.8438418 | 0.698754122 | 0.090354832 | -7.733445012 | 1.05E-14 | 3.73E-13 | 0.698754122 |
| ENSMUSG00000051599 | Pcdhb2 | 93873 | protein_coding | 1.235197465 | -0.699082623 | 1.960260176 | 0.356627468 | 0.721370711 | 1 | 0.699082623 |
| ENSMUSG00000052384 | Nrros | 224109 | protein_coding | 16.75134277 | 0.699136484 | 0.290474444 | -2.40687778 | 0.016089553 | 0.061791369 | 0.699136484 |
| ENSMUSG00000030124 | Lag3 | 16768 | protein_coding | 6.751555444 | 0.699407977 | 0.621289776 | -1.125735533 | 0.260277479 | 0.473946662 | 0.699407977 |
| ENSMUSG00000000739 | Sult5a1 | 57429 | protein_coding | 1.503717611 | -0.699424803 | 1.651345368 | 0.423548469 | 0.671895143 | 1 | 0.699424803 |
| ENSMUSG00000079057 | Cyp4v3 | 102294 | protein_coding | 102.9281843 | -0.699599762 | 0.130605257 | 5.356597282 | 8.48E-08 | 1.42E-06 | 0.699599762 |
| ENSMUSG00000035969 | Rusc2 | 100213 | protein_coding | 69.5228638 | -0.699834816 | 0.135665883 | 5.158517375 | 2.49E-07 | 3.84E-06 | 0.699834816 |
| ENSMUSG00000003070 | Efna2 | 13637 | protein_coding | 6.387303279 | 0.699955963 | 0.55048124 | -1.271534637 | 0.203538513 | 0.405848226 | 0.699955963 |
| ENSMUSG00000021986 | Amer2 | 72125 | protein_coding | 5.102432429 | 0.7001257 | 0.73207117 | -0.956362891 | 0.338888926 | 0.560514185 | 0.7001257 |
| ENSMUSG00000001999 | Blvra | 109778 | protein_coding | 242.776673 | 0.700368889 | 0.085882804 | -8.154937379 | 3.49E-16 | 1.41E-14 | 0.700368889 |
| ENSMUSG00000001119 | Col6a1 | 12833 | protein_coding | 10.8219581 | 0.700468095 | 0.793539469 | -0.882713617 | 0.377391023 | 0.599541641 | 0.700468095 |
| ENSMUSG00000055200 | Sertad3 | 170742 | protein_coding | 238.2824106 | 0.700486639 | 0.130377335 | -5.372763902 | 7.75E-08 | 1.30E-06 | 0.700486639 |
| ENSMUSG00000052182 | Or7g30 | 258520 | protein_coding | 1.048491331 | -0.700878591 | 2.253437159 | 0.311026464 | 0.755780503 | 1 | 0.700878591 |
| ENSMUSG00000002227 | Mov10 | 17454 | protein_coding | 423.6765833 | 0.700981112 | 0.237608727 | -2.950148854 | 0.003176209 | 0.016683177 | 0.700981112 |
| ENSMUSG00000078427 | Sarnp | 66118 | protein_coding | 368.4773167 | 0.701059762 | 0.1281801 | -5.469333877 | 4.52E-08 | 7.80E-07 | 0.701059762 |
| ENSMUSG00000053411 | Cbx7 | 52609 | protein_coding | 149.1066235 | -0.701451449 | 0.111879872 | 6.26968405 | 3.62E-10 | 8.41E-09 | 0.701451449 |
| ENSMUSG00000056091 | St3gal5 | 20454 | protein_coding | 22.56561683 | 0.701585407 | 0.730586493 | -0.960304376 | 0.336902048 | 0.559233675 | 0.701585407 |
| ENSMUSG00000049685 | Cyp2g1 | 13108 | protein_coding | 1.242960117 | 0.701620711 | 2.320089484 | -0.302411056 | 0.762338728 | 1 | 0.701620711 |
| ENSMUSG00000103906 | Tigd5 | 105734 | protein_coding | 59.37208472 | 0.701811316 | 0.174100479 | -4.031070563 | 5.55E-05 | 0.000512242 | 0.701811316 |
| ENSMUSG00000030249 | Abcc9 | 20928 | protein_coding | 3.19976439 | 0.702004972 | 1.859629373 | -0.377497249 | 0.705804109 | 1 | 0.702004972 |
| ENSMUSG00000076438 | Oxct2b | 353371 | protein_coding | 2.077470882 | 0.702050896 | 1.1743177 | -0.597837276 | 0.549948515 | 1 | 0.702050896 |
| ENSMUSG00000050963 | Kcns2 | 16539 | protein_coding | 1.311591693 | 0.702092941 | 1.365790767 | -0.514056002 | 0.607212838 | 1 | 0.702092941 |
| ENSMUSG00000043286 | Pnpla1 | 433091 | protein_coding | 26.56506494 | 0.702543249 | 0.288801721 | -2.432614477 | 0.014990252 | 0.058397977 | 0.702543249 |
| ENSMUSG00000035181 | Heatr5a | 320487 | protein_coding | 1275.972393 | 0.702670673 | 0.086406531 | -8.13214774 | 4.22E-16 | 1.69E-14 | 0.702670673 |
| ENSMUSG00000110648 | Gm38574 | 102641752 | processed_pseudogene | 0.647974005 | -0.702678958 | 1.858706901 | 0.37804721 | 0.705395523 | 1 | 0.702678958 |
| ENSMUSG00000045034 | Ankrd34b | 218440 | protein_coding | 2.301334256 | 0.703121814 | 1.36903975 | -0.513587581 | 0.607540364 | 1 | 0.703121814 |

| ensembl_gene_id | external_gene_name | entrezgene_id | gene_biotype | baseMean | log2FoldChange | lfcSE | stat | pvalue | padj | absLog2FC |
|---|---|---|---|---|---|---|---|---|---|---|
| ENSMUSG00000029134 | Plb1 | 665270 | protein_coding | 13.22080584 | 0.703147441 | 0.863879671 | -0.813941414 | 0.415678514 | 0.635956548 | 0.703147441 |
| ENSMUSG00000065089 | Gm26493 | 115487044 | snoRNA | 13.70524452 | 0.703163892 | 0.558300251 | -1.259472643 | 0.207859665 | 0.410793982 | 0.703163892 |
| ENSMUSG00000037440 | Vnn1 | 22361 | protein_coding | 44.50027169 | -0.703707486 | 0.258920585 | 2.717850672 | 0.006570749 | 0.030225443 | 0.703707486 |
| ENSMUSG00000027360 | Hdc | 15186 | protein_coding | 3.95339027 | 0.703835459 | 0.857833966 | -0.820479821 | 0.411942628 | 1 | 0.703835459 |
| ENSMUSG00000073494 | Sh2d1b2 | 545378 | protein_coding | 1.637664584 | -0.703977447 | 2.04983086 | 0.343431968 | 0.731273513 | 1 | 0.703977447 |
| ENSMUSG00000036820 | Amdhd2 | 245847 | protein_coding | 355.2405695 | -0.704273072 | 0.10176849 | 6.920345081 | 4.51E-12 | 1.29E-10 | 0.704273072 |
| ENSMUSG00000020089 | Ppa1 | 67895 | protein_coding | 3529.416273 | 0.704278251 | 0.122648688 | -5.742240379 | 9.34E-09 | 1.81E-07 | 0.704278251 |
| ENSMUSG00000000320 | Alox12 | 11684 | protein_coding | 3.163026191 | -0.704508725 | 1.224848999 | 0.575180064 | 0.565169524 | 1 | 0.704508725 |
| ENSMUSG00000099583 | H3c4 | 319149 | protein_coding | 390.7107621 | 0.704687814 | 0.207746055 | -3.392063527 | 0.000693684 | 0.004647131 | 0.704687814 |
| ENSMUSG00000067231 | Cyp2c65 | 72303 | protein_coding | 1436.364532 | -0.704725297 | 0.182180596 | 3.868278578 | 0.000109606 | 0.000933973 | 0.704725297 |
| ENSMUSG00000026418 | Tnni1 | 21952 | protein_coding | 456.0632588 | -0.704847077 | 0.23125035 | 3.047982751 | 0.002303832 | 0.012707239 | 0.704847077 |
| ENSMUSG00000054422 | Fabp1 | 14080 | protein_coding | 14.01599686 | -0.705644428 | 0.637879945 | 1.106233915 | 0.268625287 | 0.483654382 | 0.705644428 |
| ENSMUSG00000079644 | Gm1110 | 382064 | protein_coding | 1.817636962 | 0.705828063 | 2.153183478 | -0.327806743 | 0.743057789 | 1 | 0.705828063 |
| ENSMUSG00000090077 | Lime1 | 72699 | protein_coding | 3.823533065 | 0.705840509 | 0.586160739 | -1.204175685 | 0.228521682 | 1 | 0.705840509 |
| ENSMUSG00000023919 | Cenpq | 83815 | protein_coding | 88.67687877 | 0.705969229 | 0.203558757 | -3.468134894 | 0.000524084 | 0.003689141 | 0.705969229 |
| ENSMUSG00000046223 | Plaur | 18793 | protein_coding | 102.5729411 | 0.706273731 | 0.318432915 | -2.217967109 | 0.026557075 | 0.091519937 | 0.706273731 |
| ENSMUSG00000091076 | Vmn2r115 | 638102 | protein_coding | 3.061286323 | 0.706310531 | 3.48494292 | -0.202674921 | 0.839389127 | 1 | 0.706310531 |
| ENSMUSG00000068762 | Gstm6 | 14867 | protein_coding | 163.4574193 | 0.706378274 | 0.129839178 | -5.440409329 | 5.32E-08 | 9.08E-07 | 0.706378274 |
| ENSMUSG00000028681 | Ptch2 | 19207 | protein_coding | 5.652803692 | 0.70652013 | 0.811866395 | -0.87024187 | 0.384168239 | 0.606515707 | 0.70652013 |
| ENSMUSG00000040154 | Wfdc5 | 209232 | protein_coding | 0.746647184 | -0.706835166 | 1.665979928 | 0.424275919 | 0.671364599 | 1 | 0.706835166 |
| ENSMUSG00000029659 | Medag | 70717 | protein_coding | 1.16279286 | -0.707063821 | 1.628790038 | 0.434103724 | 0.664213121 | 1 | 0.707063821 |
| ENSMUSG00000000253 | Gmpr | 66355 | protein_coding | 139.1062491 | -0.707126804 | 0.184730405 | 3.827885301 | 0.000129249 | 0.001083461 | 0.707126804 |
| ENSMUSG00000078657 | Crnn | 381457 | protein_coding | 3.316092295 | -0.707221292 | 0.616402957 | 1.147335982 | 0.251242788 | 1 | 0.707221292 |
| ENSMUSG00000021384 | Susd3 | 66329 | protein_coding | 3.442095675 | -0.707360942 | 0.819746141 | 0.862902436 | 0.388191111 | 1 | 0.707360942 |
| ENSMUSG00000031397 | Tktl1 | 83553 | protein_coding | 1.19442217 | -0.707389892 | 2.314956858 | 0.305573682 | 0.759929261 | 1 | 0.707389892 |
| ENSMUSG00000097789 | Ctxnd1 | 100039239 | protein_coding | 116.5042636 | -0.707410548 | 0.165292888 | 4.279739786 | 1.87E-05 | 0.000194686 | 0.707410548 |
| ENSMUSG00000007594 | Hapln4 | 330790 | protein_coding | 17.67739972 | 0.707464389 | 0.321848719 | -2.198127095 | 0.02794005 | 0.095316478 | 0.707464389 |
| ENSMUSG00000043342 | Hoxd9 | 15438 | protein_coding | 3.886867536 | -0.707503725 | 0.670043523 | 1.055907119 | 0.291010652 | 1 | 0.707503725 |
| ENSMUSG00000072663 | Spef2 | 320277 | protein_coding | 1.577480953 | -0.707580343 | 2.316838184 | 0.305407753 | 0.760055618 | 1 | 0.707580343 |
| ENSMUSG00000026209 | Dnpep | 13437 | protein_coding | 1526.068741 | 0.707757182 | 0.066657465 | -10.61782324 | 2.46E-26 | 1.68E-24 | 0.707757182 |
| ENSMUSG00000024935 | Slc1a1 | 20510 | protein_coding | 352.5273427 | -0.707788677 | 0.193895665 | 3.650358443 | 0.000261875 | 0.002014251 | 0.707788677 |
| ENSMUSG00000034317 | Trim59 | 66949 | protein_coding | 143.0528307 | 0.707844549 | 0.131447875 | -5.3849828 | 7.25E-08 | 1.22E-06 | 0.707844549 |
| ENSMUSG00000020930 | Ccdc103 | 73293 | protein_coding | 2.444682377 | -0.707948487 | 1.020648913 | 0.693625867 | 0.487916868 | 1 | 0.707948487 |
| ENSMUSG00000089011 | Gm24879 | 115488696 | snoRNA | 0.74238337 | 0.708201874 | 1.601109545 | -0.442319438 | 0.658258067 | 1 | 0.708201874 |
| ENSMUSG00000028687 | Mutyh | 70603 | protein_coding | 24.27760917 | 0.708388173 | 0.2800969 | -2.52908252 | 0.011436114 | 0.047391714 | 0.708388173 |
| ENSMUSG00000067399 | Trim43c | 666731 | protein_coding | 3.188868115 | -0.708585903 | 0.669237465 | 1.058795929 | 0.289692728 | 1 | 0.708585903 |
| ENSMUSG00000031586 | Rbpms | 19663 | protein_coding | 38.30443584 | -0.708620316 | 0.297486438 | 2.382025615 | 0.0172177 | 0.065243941 | 0.708620316 |
| ENSMUSG00000028207 | Asph | 65973 | protein_coding | 1427.49301 | 0.708677891 | 0.141383236 | -5.012460552 | 5.37E-07 | 7.88E-06 | 0.708677891 |
| ENSMUSG00000031216 | Stard8 | 236920 | protein_coding | 42.62866094 | -0.708807823 | 0.180564679 | 3.925506519 | 8.65E-05 | 0.000759025 | 0.708807823 |

Supplementary Table 1

| ensembl_gene_id | external_gene_name | entrezgene_id | gene_biotype | baseMean | log2FoldChange | lfcSE | stat | pvalue | padj | absLog2FC |
|---|---|---|---|---|---|---|---|---|---|---|
| ENSMUSG00000061171 | Slc38a11 | 320106 | protein_coding | 17.35433068 | -0.70890356 | 0.266210037 | 2.662948274 | 0.007745932 | 0.034686086 | 0.70890356 |
| ENSMUSG00000029455 | Aldh2 | 11669 | protein_coding | 5982.856659 | -0.709613984 | 0.133087342 | 5.331941973 | 9.72E-08 | 1.60E-06 | 0.709613984 |
| ENSMUSG00000043445 | Pgp | 67078 | protein_coding | 1456.921143 | 0.709649617 | 0.08427878 | -8.420264427 | 3.76E-17 | 1.61E-15 | 0.709649617 |
| ENSMUSG00000005864 | Cnga2 | 12789 | protein_coding | 1.295251898 | -0.709673445 | 2.33815514 | 0.303518545 | 0.761494717 | 1 | 0.709673445 |
| ENSMUSG00000065818 | Snord35a | 27211 | snoRNA | 5.094177049 | 0.709690897 | 0.522632048 | -1.357916911 | 0.174490051 | 0.365696631 | 0.709690897 |
| ENSMUSG00000040105 | Plpp6 | 74411 | protein_coding | 574.4336533 | 0.709911967 | 0.071986694 | -9.861710859 | 6.10E-23 | 3.62E-21 | 0.709911967 |
| ENSMUSG00000110218 | Tincr | 100504425 | protein_coding | 26.82754692 | -0.710006041 | 0.200493527 | 3.541291584 | 0.000398173 | 0.002918524 | 0.710006041 |
| ENSMUSG00000049608 | Gpr55 | 227326 | protein_coding | 19.6109337 | 0.710177886 | 0.321872469 | -2.20639525 | 0.027356337 | 0.093726671 | 0.710177886 |
| ENSMUSG00000041608 | Entpd3 | 215446 | protein_coding | 7.126294398 | -0.710390906 | 0.455384094 | 1.559981817 | 0.118764178 | 0.279846882 | 0.710390906 |
| ENSMUSG00000062866 | Phactr2 | 215789 | protein_coding | 273.9435609 | -0.710429049 | 0.115668037 | 6.141965128 | 8.15E-10 | 1.81E-08 | 0.710429049 |
| ENSMUSG00000021216 | Tubal3 | 238463 | protein_coding | 212.1758307 | -0.710688962 | 0.106551942 | 6.669882777 | 2.56E-11 | 6.87E-10 | 0.710688962 |
| ENSMUSG00000022445 | Cyp2d26 | 76279 | protein_coding | 302.3539409 | -0.71099169 | 0.148127845 | 4.799851726 | 1.59E-06 | 2.10E-05 | 0.71099169 |
| ENSMUSG00000050234 | Gja4 | 14612 | protein_coding | 2.842089547 | -0.711099432 | 0.779343425 | 0.912433992 | 0.361540302 | 1 | 0.711099432 |
| ENSMUSG00000008193 | Spib | 272382 | protein_coding | 98.43904131 | -0.711271608 | 0.172974906 | 4.111993028 | 3.92E-05 | 0.000377711 | 0.711271608 |
| ENSMUSG00000026580 | Selp | 20344 | protein_coding | 2.319285088 | 0.71136946 | 1.798030727 | -0.3956381 | 0.692372026 | 1 | 0.71136946 |
| ENSMUSG00000036768 | Kif15 | 209737 | protein_coding | 297.2208949 | 0.711403928 | 0.166426867 | -4.27457381 | 1.92E-05 | 0.000197903 | 0.711403928 |
| ENSMUSG00000114456 | H2bc9 | 319182 | protein_coding | 597.8077163 | 0.711734107 | 0.199670541 | -3.564542394 | 0.000364492 | 0.002708917 | 0.711734107 |
| ENSMUSG00000115958 | Gm28040 | 280287 | protein_coding | 2.388417087 | -0.712226352 | 0.749003791 | 0.950898194 | 0.341656059 | 1 | 0.712226352 |
| ENSMUSG00000035722 | Abca7 | 27403 | protein_coding | 1165.123436 | 0.712226479 | 0.121323073 | -5.870494902 | 4.34E-09 | 8.87E-08 | 0.712226479 |
| ENSMUSG00000035277 | Arx | 11878 | protein_coding | 5.754611271 | 0.712244639 | 0.640649166 | -1.111754571 | 0.266243689 | 0.481307968 | 0.712244639 |
| ENSMUSG00000021665 | Hexb | 15212 | protein_coding | 878.0686 | -0.712402786 | 0.274416988 | 2.596059347 | 0.009429978 | 0.0406597 | 0.712402786 |
| ENSMUSG00000025355 | Mmp19 | 58223 | protein_coding | 1.66531588 | -0.712763869 | 1.423843966 | 0.500591277 | 0.616658803 | 1 | 0.712763869 |
| ENSMUSG00000088273 | Gm23123 | 115488841 | snoRNA | 1.461333329 | -0.712879617 | 1.027594265 | 0.693736469 | 0.487847491 | 1 | 0.712879617 |
| ENSMUSG00000059479 | B3gnt8 | 232984 | protein_coding | 15.04385182 | -0.712960945 | 0.288589311 | 2.470503646 | 0.013492294 | 0.053754342 | 0.712960945 |
| ENSMUSG00000078870 | Gm14410 | 100043403 | protein_coding | 21.51785742 | -0.713054723 | 0.282147858 | 2.527237766 | 0.011496364 | 0.047593622 | 0.713054723 |
| ENSMUSG00000026110 | Mgat4a | 269181 | protein_coding | 4434.819478 | -0.713087409 | 0.150723773 | 4.731087832 | 2.23E-06 | 2.88E-05 | 0.713087409 |
| ENSMUSG00000069272 | H2ac8 | 319166 | protein_coding | 208.9253535 | 0.713094142 | 0.222149425 | -3.209975181 | 0.001327464 | 0.008065396 | 0.713094142 |
| ENSMUSG00000039154 | Shd | 20420 | protein_coding | 1.847551975 | -0.713116005 | 1.038529662 | 0.686659255 | 0.492297478 | 1 | 0.713116005 |
| ENSMUSG00000024989 | Cep55 | 74107 | protein_coding | 167.5618759 | 0.71328156 | 0.158692854 | -4.494730163 | 6.97E-06 | 8.11E-05 | 0.71328156 |
| ENSMUSG00000021537 | Cetn3 | 12626 | protein_coding | 423.6078351 | 0.713338286 | 0.110094925 | -6.479302186 | 9.21E-11 | 2.32E-09 | 0.713338286 |
| ENSMUSG00000026723 | Trdmt1 | 13434 | protein_coding | 44.98627528 | -0.713854819 | 0.219101633 | 3.258099042 | 0.001121613 | 0.007010715 | 0.713854819 |
| ENSMUSG00000040037 | Negr1 | 320840 | protein_coding | 5.331344972 | 0.713977144 | 1.119375436 | -0.637835279 | 0.523580912 | 0.722169631 | 0.713977144 |
| ENSMUSG00000016552 | Foxred2 | 239554 | protein_coding | 37.36586961 | -0.714373142 | 0.213992925 | 3.338302619 | 0.000842919 | 0.005507949 | 0.714373142 |
| ENSMUSG00000043535 | Setx | 269254 | protein_coding | 1294.166083 | 0.714456063 | 0.08819925 | -8.10047776 | 5.47E-16 | 2.18E-14 | 0.714456063 |
| ENSMUSG00000070448 | Vmn2r89 | 22301 | protein_coding | 1.75490326 | -0.714519352 | 2.391813594 | 0.298735384 | 0.765141957 | 1 | 0.714519352 |
| ENSMUSG00000091119 | Ccdc152 | 100039139 | protein_coding | 2.963592668 | -0.714612007 | 0.740727324 | 0.964743683 | 0.33467321 | 1 | 0.714612007 |
| ENSMUSG00000085253 | Gm16833 | 100579151 | lncRNA | 2.63384545 | 0.714626949 | 0.823309202 | -0.867993395 | 0.385397948 | 1 | 0.714626949 |
| ENSMUSG00000048895 | Cdk5r1 | 12569 | protein_coding | 15.95983843 | 0.715021065 | 0.37296045 | -1.917149833 | 0.055218899 | 0.160458367 | 0.715021065 |
| ENSMUSG00000053297 | AI854703 | 243373 | protein_coding | 363.1784811 | -0.715041722 | 0.328129515 | 2.179144787 | 0.029320914 | 0.099059146 | 0.715041722 |

Supplementary Table 1

| ensembl_gene_id | external_gene_name | entrezgene_id | gene_biotype | baseMean | log2FoldChange | lfcSE | stat | pvalue | padj | absLog2FC |
|---|---|---|---|---|---|---|---|---|---|---|
| ENSMUSG00000086247 | Gm15787 | 100504007 | lncRNA | 18.8369068 | -0.715375681 | 0.316709871 | 2.258772919 | 0.023897514 | 0.084361596 | 0.715375681 |
| ENSMUSG00000058441 | Panx2 | 406218 | protein_coding | 3.878337886 | 0.715493541 | 1.039504041 | -0.688302799 | 0.49126212 | 1 | 0.715493541 |
| ENSMUSG00000064715 | Gm24207 | 115488457 | snRNA | 1.47605639 | 0.715602241 | 0.930139389 | -0.769349465 | 0.44168588 | 1 | 0.715602241 |
| ENSMUSG00000074476 | Spc24 | 67629 | protein_coding | 193.9586513 | 0.715621535 | 0.159278476 | -4.492895426 | 7.03E-06 | 8.17E-05 | 0.715621535 |
| ENSMUSG00000077192 | Snora17 | 100306954 | snoRNA | 57.94828107 | 0.715808121 | 0.356117822 | -2.010031729 | 0.044427831 | 0.136254601 | 0.715808121 |
| ENSMUSG00000062275 | Fbxw24 | 382106 | protein_coding | 1.754639829 | -0.715863867 | 1.13232075 | 0.63220944 | 0.52725003 | 1 | 0.715863867 |
| ENSMUSG00000030142 | Clec4e | 56619 | protein_coding | 2.998063051 | 0.716084033 | 1.868845529 | -0.383169193 | 0.701594319 | 1 | 0.716084033 |
| ENSMUSG00000049985 | Ankrd55 | 77318 | protein_coding | 1.677526752 | 0.71616302 | 2.329016505 | -0.307495897 | 0.758465945 | 1 | 0.71616302 |
| ENSMUSG00000098528 | Zfp572 | 100416830 | transcribed_processed_pseudogene | 0.793265449 | -0.716412748 | 1.676979073 | 0.427204346 | 0.669230496 | 1 | 0.716412748 |
| ENSMUSG00000096630 | Vmn2r26 | 56552 | protein_coding | 1.182361604 | -0.716570326 | 1.783707636 | 0.401730817 | 0.687882142 | 1 | 0.716570326 |
| ENSMUSG00000098839 | Mir7054 | 102466218 | miRNA | 1.011930231 | -0.716588185 | 1.515114768 | 0.472959673 | 0.63624195 | 1 | 0.716588185 |
| ENSMUSG00000036295 | Lrrn3 | 16981 | protein_coding | 14.01747565 | -0.716843402 | 0.333966943 | 2.146450175 | 0.03183708 | 0.105701154 | 0.716843402 |
| ENSMUSG00000067653 | Ankrd23 | 78321 | protein_coding | 6.136854854 | 0.716991519 | 0.483679671 | -1.482368522 | 0.138242265 | 0.310941537 | 0.716991519 |
| ENSMUSG00000003352 | Cacnb3 | 12297 | protein_coding | 412.4363593 | -0.717183612 | 0.100737605 | 7.119323649 | 1.08E-12 | 3.26E-11 | 0.717183612 |
| ENSMUSG00000039315 | Clnk | 27278 | protein_coding | 2.698549657 | 0.717331722 | 1.189086851 | -0.603262681 | 0.546333955 | 1 | 0.717331722 |
| ENSMUSG00000026683 | Nuf2 | 66977 | protein_coding | 135.7842561 | 0.717918232 | 0.193745772 | -3.705465281 | 0.000211003 | 0.001655346 | 0.717918232 |
| ENSMUSG00000061126 | Cyp4f39 | 320997 | protein_coding | 6.299517384 | 0.718004828 | 0.449046547 | -1.598954123 | 0.109830797 | 0.264732086 | 0.718004828 |
| ENSMUSG00000112160 | BC024063 | 666584 | protein_coding | 19.91307506 | -0.718209455 | 0.276522514 | 2.59729107 | 0.009396226 | 0.04055653 | 0.718209455 |
| ENSMUSG00000020327 | Fgf22 | 67112 | protein_coding | 2.344949523 | 0.718487953 | 1.092229094 | -0.657817995 | 0.51065509 | 1 | 0.718487953 |
| ENSMUSG00000051951 | Xkr4 | 497097 | protein_coding | 15.25570992 | -0.718605246 | 0.331754194 | 2.166077353 | 0.030305276 | 0.101719784 | 0.718605246 |
| ENSMUSG00000024049 | Myom1 | 17929 | protein_coding | 27.78955333 | 0.718970636 | 0.30968393 | -2.3216272 | 0.020253018 | 0.07410998 | 0.718970636 |
| ENSMUSG00000047022 | Mipol1 | 73490 | protein_coding | 32.29576228 | -0.719164259 | 0.202261301 | 3.555619657 | 0.000377089 | 0.002788738 | 0.719164259 |
| ENSMUSG00000054362 | Lexm | 242602 | protein_coding | 2.144027163 | -0.719216256 | 1.19976593 | 0.59946381 | 0.548863636 | 1 | 0.719216256 |
| ENSMUSG00000031387 | Renbp | 19703 | protein_coding | 8.05201361 | -0.71923513 | 0.493322317 | 1.457941604 | 0.14485664 | 0.321916639 | 0.71923513 |
| ENSMUSG00000027331 | Knstrn | 51944 | protein_coding | 325.4157188 | 0.719249197 | 0.121964366 | -5.897207673 | 3.70E-09 | 7.60E-08 | 0.719249197 |
| ENSMUSG00000071633 | Gm4952 | 240549 | protein_coding | 7.227470519 | -0.719381502 | 0.45159989 | 1.592962085 | 0.111168697 | 0.26676599 | 0.719381502 |
| ENSMUSG00000022491 | Glycam1 | 14663 | protein_coding | 1.166851689 | -0.719425423 | 2.324739483 | 0.309464965 | 0.756967859 | 1 | 0.719425423 |
| ENSMUSG00000022659 | Gcsam | 14525 | protein_coding | 6.615030478 | 0.719682967 | 0.547491451 | -1.314509962 | 0.188674648 | 0.385300595 | 0.719682967 |
| ENSMUSG00000033389 | Arhgap44 | 216831 | protein_coding | 6.489462148 | -0.719747777 | 0.66988525 | 1.07443443 | 0.282628018 | 0.50078531 | 0.719747777 |
| ENSMUSG00000021728 | Emb | 13723 | protein_coding | 15.23099721 | 0.719954507 | 0.361009176 | -1.994283124 | 0.046121122 | 0.140291806 | 0.719954507 |
| ENSMUSG00000096959 | 4930509G22Rik | 105246071 | lncRNA | 5.380531304 | 0.720232678 | 0.631388722 | -1.140711979 | 0.2539898 | 0.466665282 | 0.720232678 |
| ENSMUSG00000048399 | Tprg | 71338 | protein_coding | 0.9376923 | 0.720976388 | 1.378887663 | -0.522868111 | 0.601066037 | 1 | 0.720976388 |
| ENSMUSG00000042096 | Dao | 13142 | protein_coding | 2.709141352 | 0.720995547 | 1.70247761 | -0.423497815 | 0.671932093 | 1 | 0.720995547 |
| ENSMUSG00000046532 | Ar | 11835 | protein_coding | 4.481725136 | 0.721440641 | 1.490099023 | -0.484156173 | 0.628275038 | 0.79681545 | 0.721440641 |
| ENSMUSG00000031149 | Praf2 | 54637 | protein_coding | 111.1176351 | 0.721498041 | 0.1334663 | -5.405844315 | 6.45E-08 | 1.09E-06 | 0.721498041 |
| ENSMUSG00000031750 | Il34 | 76527 | protein_coding | 36.3091249 | -0.721594153 | 0.22168431 | 3.255052891 | 0.001133713 | 0.00707029 | 0.721594153 |
| ENSMUSG00000039648 | Kyat1 | 70266 | protein_coding | 270.3906992 | -0.721862158 | 0.211946668 | 3.405866986 | 0.000659543 | 0.004469372 | 0.721862158 |
| ENSMUSG00000086199 | Bcas3os1 | 71489 | lncRNA | 4.952625343 | -0.721916753 | 0.595660967 | 1.211959138 | 0.225528023 | 0.433049501 | 0.721916753 |
| ENSMUSG00000020774 | Aspa | 11484 | protein_coding | 52.32122426 | -0.722136299 | 0.24726277 | 2.920521761 | 0.003494458 | 0.018083738 | 0.722136299 |

Supplementary Table 1

| ensembl_gene_id | external_gene_name | entrezgene_id | gene_biotype | baseMean | log2FoldChange | lfcSE | stat | pvalue | padj | absLog2FC |
|---|---|---|---|---|---|---|---|---|---|---|
| ENSMUSG00000062151 | Unc13c | 208898 | protein_coding | 3.157958279 | 0.722218921 | 1.436219802 | -0.502860997 | 0.615062005 | 1 | 0.722218921 |
| ENSMUSG00000044408 | Sptssa | 104725 | protein_coding | 692.0618823 | 0.722456142 | 0.081766978 | -8.835548977 | 9.96E-19 | 4.73E-17 | 0.722456142 |
| ENSMUSG00000049295 | Zfp219 | 69890 | protein_coding | 694.8309438 | 0.722558574 | 0.074886636 | -9.648698499 | 4.98E-22 | 2.81E-20 | 0.722558574 |
| ENSMUSG00000025081 | Tdrd1 | 83561 | protein_coding | 30.86681396 | -0.722718153 | 0.253915274 | 2.846296493 | 0.004423099 | 0.021981059 | 0.722718153 |
| ENSMUSG00000100372 | 1700003I22Rik | 69356 | lncRNA | 9.38019227 | 0.722723789 | 0.460434011 | -1.569657695 | 0.116494768 | 0.275852395 | 0.722723789 |
| ENSMUSG00000052508 | Or12j5 | 258513 | protein_coding | 2.101741393 | 0.722958352 | 2.168248867 | -0.3334296 | 0.738810023 | 1 | 0.722958352 |
| ENSMUSG00000052752 | Traf7 | 224619 | protein_coding | 1148.490105 | 0.723122986 | 0.063243122 | -11.43401787 | 2.83E-30 | 2.23E-28 | 0.723122986 |
| ENSMUSG00000087338 | Scpep1os | 73921 | lncRNA | 1.485941852 | -0.723731374 | 1.637140487 | 0.442070415 | 0.658438252 | 1 | 0.723731374 |
| ENSMUSG00000028718 | Stil | 20460 | protein_coding | 141.3899666 | 0.724403745 | 0.114986716 | -6.299890712 | 2.98E-10 | 7.04E-09 | 0.724403745 |
| ENSMUSG00000026482 | Rgl1 | 19731 | protein_coding | 17.23871253 | 0.724411577 | 0.467863846 | -1.548338439 | 0.121540834 | 0.284375933 | 0.724411577 |
| ENSMUSG00000022176 | Rem2 | 140743 | protein_coding | 12.26362075 | 0.724522061 | 0.45873177 | -1.579402404 | 0.114243786 | 0.271847534 | 0.724522061 |
| ENSMUSG00000028544 | Slc5a9 | 230612 | protein_coding | 312.6270619 | 0.72465246 | 0.137822407 | -5.257871179 | 1.46E-07 | 2.34E-06 | 0.72465246 |
| ENSMUSG00000033386 | Frrs1 | 20321 | protein_coding | 391.0381167 | -0.7246727 | 0.127164546 | 5.698700798 | 1.21E-08 | 2.30E-07 | 0.7246727 |
| ENSMUSG00000060985 | Tdrd5 | 214575 | protein_coding | 3.250673721 | -0.725452455 | 0.819113749 | 0.885655327 | 0.375803288 | 1 | 0.725452455 |
| ENSMUSG00000040189 | Odad1 | 211535 | protein_coding | 83.59739836 | 0.725680838 | 0.19266097 | -3.76662091 | 0.000165472 | 0.001338605 | 0.725680838 |
| ENSMUSG00000038916 | Soga3 | 67412 | protein_coding | 1.417184603 | 0.725883553 | 1.814011213 | -0.40015384 | 0.689043211 | 1 | 0.725883553 |
| ENSMUSG00000040327 | Cul9 | 78309 | protein_coding | 45.87880793 | -0.726153633 | 0.393741012 | 1.8442418 | 0.065147909 | 0.181269925 | 0.726153633 |
| ENSMUSG00000098531 | Mir7041 | 102466992 | miRNA | 0.776761256 | -0.726398949 | 1.312146844 | 0.553595775 | 0.579855518 | 1 | 0.726398949 |
| ENSMUSG00000031847 | 1700030J22Rik | 69528 | protein_coding | 15.71810491 | 0.726408419 | 0.324255062 | -2.240237711 | 0.025075495 | 0.087546875 | 0.726408419 |
| ENSMUSG00000089647 | Gm2245 | 102632231 | lncRNA | 1.352004336 | 0.726659171 | 1.261798032 | -0.575891825 | 0.564688302 | 1 | 0.726659171 |
| ENSMUSG00000030000 | Add2 | 11519 | protein_coding | 3.961899617 | -0.726759089 | 1.114240193 | 0.652246341 | 0.51424227 | 1 | 0.726759089 |
| ENSMUSG00000027939 | Nup210l | 77595 | protein_coding | 38.63952817 | 0.72693561 | 0.255584147 | -2.844212435 | 0.004452135 | 0.022092116 | 0.72693561 |
| ENSMUSG00000051043 | Gprc5c | 70355 | protein_coding | 14.86465604 | -0.727045487 | 0.315530328 | 2.304201604 | 0.021211329 | 0.076884085 | 0.727045487 |
| ENSMUSG00000020700 | Map3k3 | 26406 | protein_coding | 419.1637913 | 0.727066462 | 0.102045074 | -7.124954046 | 1.04E-12 | 3.14E-11 | 0.727066462 |
| ENSMUSG00000029120 | Ppp2r2c | 269643 | protein_coding | 3.857666317 | -0.727229716 | 0.784733883 | 0.926721443 | 0.354071181 | 1 | 0.727229716 |
| ENSMUSG00000064215 | Ifi27 | 52668 | protein_coding | 326.2044664 | -0.727313479 | 0.148620685 | 4.89375673 | 9.89E-07 | 1.37E-05 | 0.727313479 |
| ENSMUSG00000034157 | Cipc | 217732 | protein_coding | 1198.683423 | -0.727524199 | 0.138257091 | 5.262111278 | 1.42E-07 | 2.29E-06 | 0.727524199 |
| ENSMUSG00000033788 | Dysf | 26903 | protein_coding | 5.230792788 | 0.727573224 | 1.821631526 | -0.399407462 | 0.689592997 | 0.834370416 | 0.727573224 |
| ENSMUSG00000003402 | Prkcsh | 19089 | protein_coding | 2063.630554 | 0.7277426 | 0.054495663 | -13.35413788 | 1.12E-40 | 1.21E-38 | 0.7277426 |
| ENSMUSG00000040627 | Aicda | 11628 | protein_coding | 2.472378032 | 0.727753185 | 0.814961737 | -0.892990618 | 0.371862197 | 1 | 0.727753185 |
| ENSMUSG00000055159 | 4930583K01Rik | 67745 | lncRNA | 1.04923397 | 0.727923933 | 1.750276317 | -0.415890866 | 0.677489864 | 1 | 0.727923933 |
| ENSMUSG00000019831 | Wasf1 | 83767 | protein_coding | 2.002579354 | 0.728032456 | 1.650939757 | -0.440980631 | 0.659227019 | 1 | 0.728032456 |
| ENSMUSG00000021898 | Asb14 | 142687 | protein_coding | 3.653430498 | 0.72819116 | 0.753260186 | -0.966719301 | 0.333684376 | 1 | 0.72819116 |
| ENSMUSG00000030606 | Hapln3 | 67666 | protein_coding | 1.638031347 | -0.728246668 | 1.913012399 | 0.380680579 | 0.703440281 | 1 | 0.728246668 |
| ENSMUSG00000050132 | Sarm1 | 237868 | protein_coding | 3.377446834 | -0.728251423 | 1.599798854 | 0.455214367 | 0.648955025 | 1 | 0.728251423 |
| ENSMUSG00000038775 | Vill | 22351 | protein_coding | 85.52752566 | -0.728347765 | 0.151538757 | 4.806346454 | 1.54E-06 | 2.04E-05 | 0.728347765 |
| ENSMUSG00000030373 | Psg28 | 114871 | protein_coding | 4.079928972 | 0.7285754 | 0.831877455 | -0.875820586 | 0.381127577 | 1 | 0.7285754 |
| ENSMUSG00000071073 | Lrrc73 | 224813 | protein_coding | 4.616912156 | 0.7286128 | 0.808049603 | -0.90169316 | 0.367219887 | 0.589852079 | 0.7286128 |
| ENSMUSG00000020262 | Adarb1 | 110532 | protein_coding | 7.044377083 | 0.728747298 | 0.635408121 | -1.146896419 | 0.251424433 | 0.463839794 | 0.728747298 |

| ensembl_gene_id | external_gene_name | entrezgene_id | gene_biotype | baseMean | log2FoldChange | lfcSE | stat | pvalue | padj | absLog2FC |
|---|---|---|---|---|---|---|---|---|---|---|
| ENSMUSG00000020891 | Alox8 | 11688 | protein_coding | 4.190079984 | -0.729160547 | 1.056426883 | 0.690213926 | 0.490059667 | 1 | 0.729160547 |
| ENSMUSG00000021273 | Fdft1 | 14137 | protein_coding | 1345.198279 | 0.729386456 | 0.179167958 | -4.070964838 | 4.68E-05 | 0.000440521 | 0.729386456 |
| ENSMUSG00000037973 | Itprid1 | 232016 | protein_coding | 13.61429065 | -0.729826658 | 0.415952298 | 1.7545922 | 0.079329087 | 0.208654479 | 0.729826658 |
| ENSMUSG00000024534 | Sncaip | 67847 | protein_coding | 1.055030345 | -0.729945512 | 1.823195993 | 0.400365904 | 0.688887033 | 1 | 0.729945512 |
| ENSMUSG00000034563 | Ccpg1 | 72278 | protein_coding | 14.94869243 | 0.730085713 | 0.357294519 | -2.043372273 | 0.0410156 | 0.128452801 | 0.730085713 |
| ENSMUSG00000027797 | Dclk1 | 13175 | protein_coding | 440.3316582 | -0.730229081 | 0.153943248 | 4.74349535 | 2.10E-06 | 2.73E-05 | 0.730229081 |
| ENSMUSG00000024795 | Kif20b | 240641 | protein_coding | 346.102757 | 0.730400234 | 0.169839679 | -4.300527633 | 1.70E-05 | 0.000179112 | 0.730400234 |
| ENSMUSG00000064899 | Snord118 | 100216530 | snoRNA | 65.62870326 | 0.730752057 | 0.293831302 | -2.486978252 | 0.012883327 | 0.051803085 | 0.730752057 |
| ENSMUSG00000009941 | Nxf2 | 83454 | protein_coding | 1.930549897 | -0.730955916 | 2.609729171 | 0.280088802 | 0.779409376 | 1 | 0.730955916 |
| ENSMUSG00000051900 | Abca16 | 233810 | protein_coding | 2.690907905 | -0.731130138 | 2.612449029 | 0.279863886 | 0.779581935 | 1 | 0.731130138 |
| ENSMUSG00000025037 | Maoa | 17161 | protein_coding | 8022.229441 | -0.731364272 | 0.100802606 | 7.255410364 | 4.00E-13 | 1.26E-11 | 0.731364272 |
| ENSMUSG00000047237 | Fbxw21 | 320082 | protein_coding | 1.352801136 | -0.731469516 | 2.617956638 | 0.279404749 | 0.779934227 | 1 | 0.731469516 |
| ENSMUSG00000075104 | Or4c114 | 258901 | protein_coding | 1.516089767 | -0.731612278 | 2.620216832 | 0.279218219 | 0.780077363 | 1 | 0.731612278 |
| ENSMUSG00000110235 | Gm5086 | 328314 | lncRNA | 1.261930988 | -0.731865427 | 2.624331178 | 0.278876932 | 0.780339272 | 1 | 0.731865427 |
| ENSMUSG00000050164 | Mchr1 | 207911 | protein_coding | 1.368525737 | 0.731948563 | 1.081511518 | -0.676782957 | 0.498543673 | 1 | 0.731948563 |
| ENSMUSG00000027196 | Alkbh3os1 | 70852 | lncRNA | 2.864751267 | -0.731974064 | 0.633734773 | 1.155016412 | 0.248083706 | 1 | 0.731974064 |
| ENSMUSG00000046699 | Slitrk4 | 245446 | protein_coding | 1.796122981 | -0.732025908 | 2.626860803 | 0.27866947 | 0.780498494 | 1 | 0.732025908 |
| ENSMUSG00000033220 | Rac2 | 19354 | protein_coding | 58.77148461 | -0.73216623 | 0.208594798 | 3.509992759 | 0.000448119 | 0.003232956 | 0.73216623 |
| ENSMUSG00000032997 | Chpf | 74241 | protein_coding | 258.0815831 | 0.732468606 | 0.1132012 | -6.47050212 | 9.77E-11 | 2.43E-09 | 0.732468606 |
| ENSMUSG00000029309 | Sparcl1 | 13602 | protein_coding | 1.394765044 | 0.732545491 | 1.452876548 | -0.504203534 | 0.614118357 | 1 | 0.732545491 |
| ENSMUSG00000092627 | D130058E05Rik | 102637806 | lncRNA | 0.984081372 | -0.732796041 | 2.639572269 | 0.277619238 | 0.781304663 | 1 | 0.732796041 |
| ENSMUSG00000020061 | Mybpc1 | 109272 | protein_coding | 1.863709106 | -0.733071893 | 2.644016393 | 0.27725694 | 0.781582821 | 1 | 0.733071893 |
| ENSMUSG00000070337 | Gpr179 | 217143 | protein_coding | 6.956047734 | -0.733098874 | 0.96158482 | 0.762386073 | 0.445829616 | 0.662983366 | 0.733098874 |
| ENSMUSG00000028246 | Faxc | 76132 | protein_coding | 8.961409532 | 0.733172212 | 0.524861916 | -1.396885904 | 0.162447884 | 0.347924739 | 0.733172212 |
| ENSMUSG00000099033 | Mir7013 | 102465613 | miRNA | 0.764889643 | 0.733729113 | 1.330438968 | -0.551494003 | 0.581295076 | 1 | 0.733729113 |
| ENSMUSG00000027583 | Zbtb46 | 72147 | protein_coding | 9.741448853 | 0.733796889 | 0.58069294 | -1.263657327 | 0.206353048 | 0.408991146 | 0.733796889 |
| ENSMUSG00000033182 | Kbtbd12 | 74589 | protein_coding | 4.017975713 | 0.734030631 | 1.015258392 | -0.722998832 | 0.469680602 | 1 | 0.734030631 |
| ENSMUSG00000066026 | Dhrs3 | 20148 | protein_coding | 388.7246952 | -0.734067566 | 0.11030136 | 6.655108952 | 2.83E-11 | 7.56E-10 | 0.734067566 |
| ENSMUSG00000031651 | Triml1 | 244448 | protein_coding | 1.39049674 | -0.734087227 | 2.66113292 | 0.275855152 | 0.782659321 | 1 | 0.734087227 |
| ENSMUSG00000096918 | Gm16863 | 328646 | lncRNA | 7.172909391 | -0.734238546 | 0.394845456 | 1.859559312 | 0.0629479 | 0.176609179 | 0.734238546 |
| ENSMUSG00000048503 | Tlcd5 | 235300 | protein_coding | 2.188502089 | -0.734238609 | 2.663636166 | 0.27565274 | 0.782814797 | 1 | 0.734238609 |
| ENSMUSG00000050700 | Emilin3 | 280635 | protein_coding | 4.053930536 | 0.734449063 | 0.673223098 | -1.090944541 | 0.275297287 | 1 | 0.734449063 |
| ENSMUSG00000021356 | Irf4 | 16364 | protein_coding | 2.614477214 | 0.734570363 | 1.172614104 | -0.626438281 | 0.531027508 | 1 | 0.734570363 |
| ENSMUSG00000103613 | Gm34882 | 102638281 | lncRNA | 3.103780197 | -0.734658745 | 0.71820424 | 1.022910621 | 0.306350107 | 1 | 0.734658745 |
| ENSMUSG00000022101 | Fgf17 | 14171 | protein_coding | 1.267216914 | -0.734715794 | 1.422759299 | 0.516402033 | 0.605573647 | 1 | 0.734715794 |
| ENSMUSG00000046447 | Camk2n1 | 66259 | protein_coding | 834.7942457 | 0.735002729 | 0.108621689 | -6.766629531 | 1.32E-11 | 3.64E-10 | 0.735002729 |
| ENSMUSG00000038663 | Fsd2 | 244091 | protein_coding | 2.56538954 | 0.735248667 | 1.207818666 | -0.608740937 | 0.542696168 | 1 | 0.735248667 |
| ENSMUSG00000034416 | Pkd1l2 | 76645 | protein_coding | 2.387269694 | 0.735509992 | 1.957116799 | -0.375813029 | 0.707055901 | 1 | 0.735509992 |
| ENSMUSG00000007827 | Ankrd26 | 232339 | protein_coding | 115.9899591 | 0.73556234 | 0.138720042 | -5.302495084 | 1.14E-07 | 1.86E-06 | 0.73556234 |

| ensembl_gene_id | external_gene_name | entrezgene_id | gene_biotype | baseMean | log2FoldChange | lfcSE | stat | pvalue | padj | absLog2FC |
|---|---|---|---|---|---|---|---|---|---|---|
| ENSMUSG00000016664 | Pacsin2 | 23970 | protein_coding | 2407.086493 | 0.735922133 | 0.05661882 | -12.99783602 | 1.26E-38 | 1.22E-36 | 0.735922133 |
| ENSMUSG00000000266 | Mid2 | 23947 | protein_coding | 972.4627311 | 0.736201028 | 0.078660899 | -9.359173792 | 8.04E-21 | 4.31E-19 | 0.736201028 |
| ENSMUSG00000031727 | Pmfbp1 | 56523 | protein_coding | 1.991317566 | 0.736366729 | 1.974910493 | -0.372860812 | 0.70925204 | 1 | 0.736366729 |
| ENSMUSG00000093668 | Pou5f2 | 75507 | protein_coding | 2.76743137 | -0.736468543 | 0.805239658 | 0.914595469 | 0.360404038 | 1 | 0.736468543 |
| ENSMUSG00000021610 | Clptm1l | 218335 | protein_coding | 2091.230721 | 0.736727029 | 0.093606737 | -7.870448806 | 3.53E-15 | 1.31E-13 | 0.736727029 |
| ENSMUSG00000076361 | Mir182 | 387177 | miRNA | 0.579058261 | 0.736889213 | 1.334781536 | -0.552067281 | 0.580902258 | 1 | 0.736889213 |
| ENSMUSG00000110058 | Gm5907 | 546036 | lncRNA | 0.979412889 | -0.736955739 | 1.723007474 | 0.427714766 | 0.668858798 | 1 | 0.736955739 |
| ENSMUSG00000055978 | Fut2 | 14344 | protein_coding | 3248.129495 | 0.737016806 | 0.115052623 | -6.405910529 | 1.49E-10 | 3.62E-09 | 0.737016806 |
| ENSMUSG00000048174 | Tmem81 | 74626 | protein_coding | 22.79520356 | -0.737090088 | 0.223948715 | 3.291334308 | 0.000997133 | 0.006345534 | 0.737090088 |
| ENSMUSG00000027643 | Ghrh | 14601 | protein_coding | 0.854998285 | -0.73719167 | 1.241858849 | 0.593619533 | 0.552766613 | 1 | 0.73719167 |
| ENSMUSG00000023015 | Racgap1 | 26934 | protein_coding | 531.8861177 | 0.737301553 | 0.148739299 | -4.957005707 | 7.16E-07 | 1.02E-05 | 0.737301553 |
| ENSMUSG00000045004 | Spata21 | 329972 | protein_coding | 7.410684015 | 0.737819138 | 0.711075876 | -1.037609576 | 0.299451858 | 0.519448318 | 0.737819138 |
| ENSMUSG00000024301 | Kifc5b | 16580 | protein_coding | 70.6636411 | 0.738020856 | 0.204318169 | -3.612115657 | 0.000303709 | 0.002303836 | 0.738020856 |
| ENSMUSG00000052477 | C130026I21Rik | 620078 | protein_coding | 10.83158238 | 0.738071763 | 2.614393891 | -0.28231085 | 0.777705166 | 0.888933284 | 0.738071763 |
| ENSMUSG00000031371 | Haus7 | 73738 | protein_coding | 125.7349425 | 0.738310518 | 0.104945492 | -7.0351809 | 1.99E-12 | 5.88E-11 | 0.738310518 |
| ENSMUSG00000035505 | Cox18 | 231430 | protein_coding | 104.3028089 | 0.738350057 | 0.156380527 | -4.721496168 | 2.34E-06 | 3.00E-05 | 0.738350057 |
| ENSMUSG00000020377 | Ltc4s | 17001 | protein_coding | 17.35564522 | -0.738415656 | 0.240450428 | 3.070968364 | 0.002133658 | 0.011948193 | 0.738415656 |
| ENSMUSG00000051378 | Kif18b | 70218 | protein_coding | 154.6123751 | 0.739049054 | 0.186100145 | -3.971243838 | 7.15E-05 | 0.000642047 | 0.739049054 |
| ENSMUSG00000070345 | Hsf5 | 327992 | protein_coding | 5.185704809 | 0.739107375 | 0.518851885 | -1.424505522 | 0.154300188 | 0.336218865 | 0.739107375 |
| ENSMUSG00000040033 | Stat2 | 20847 | protein_coding | 362.4693765 | 0.739313993 | 0.170582823 | -4.334047118 | 1.46E-05 | 0.000156354 | 0.739313993 |
| ENSMUSG00000044303 | Cdkn2a | 12578 | protein_coding | 0.655284349 | 0.739565451 | 1.474746717 | -0.50148642 | 0.616028833 | 1 | 0.739565451 |
| ENSMUSG00000039913 | Pak5 | 241656 | protein_coding | 2.115974716 | 0.739587048 | 1.786103828 | -0.414078418 | 0.678816678 | 1 | 0.739587048 |
| ENSMUSG00000028766 | Alpl | 11647 | protein_coding | 2.907812959 | 0.739587178 | 1.5567985 | -0.475069302 | 0.634737578 | 1 | 0.739587178 |
| ENSMUSG00000042567 | Nek10 | 674895 | protein_coding | 6.616717388 | 0.739633735 | 0.648991922 | -1.139665549 | 0.254425665 | 0.467050404 | 0.739633735 |
| ENSMUSG00000036731 | Cysrt1 | 67859 | protein_coding | 1.570898927 | 0.739654525 | 1.082259094 | -0.683435722 | 0.494331554 | 1 | 0.739654525 |
| ENSMUSG00000068696 | Gpr88 | 64378 | protein_coding | 1.482903341 | 0.739710373 | 1.91420896 | -0.386431361 | 0.699177228 | 1 | 0.739710373 |
| ENSMUSG00000084918 | Gm12708 | 105244624 | lncRNA | 1.777333952 | -0.739802039 | 0.910209421 | 0.812782226 | 0.416342928 | 1 | 0.739802039 |
| ENSMUSG00000039989 | Cbx4 | 12418 | protein_coding | 291.9492419 | 0.739831594 | 0.126316134 | -5.856984154 | 4.71E-09 | 9.54E-08 | 0.739831594 |
| ENSMUSG00000028088 | Fmo5 | 14263 | protein_coding | 303.2130556 | -0.739886122 | 0.289295305 | 2.557546243 | 0.010541354 | 0.044453504 | 0.739886122 |
| ENSMUSG00000097336 | Fendrr | 68790 | lncRNA | 1.133544756 | -0.739928294 | 2.219817911 | 0.333328374 | 0.738886424 | 1 | 0.739928294 |
| ENSMUSG00000021136 | Smoc1 | 64075 | protein_coding | 6.699990323 | 0.739957025 | 0.676746707 | -1.093403215 | 0.274216803 | 0.490619315 | 0.739957025 |
| ENSMUSG00000027326 | Knl1 | 76464 | protein_coding | 327.2909352 | 0.739977036 | 0.175755156 | -4.210272149 | 2.55E-05 | 0.000256367 | 0.739977036 |
| ENSMUSG00000074170 | Plekhf1 | 72287 | protein_coding | 29.22201622 | 0.740234207 | 0.296193719 | -2.499155655 | 0.012448962 | 0.050506026 | 0.740234207 |
| ENSMUSG00000021484 | Lman2 | 66890 | protein_coding | 4296.146306 | 0.740249406 | 0.086656958 | -8.542296246 | 1.32E-17 | 5.83E-16 | 0.740249406 |
| ENSMUSG00000038180 | Spag4 | 245865 | protein_coding | 6.504183518 | 0.740488015 | 0.681992691 | -1.085771188 | 0.277580237 | 0.494519255 | 0.740488015 |
| ENSMUSG00000075297 | H60b | 667281 | protein_coding | 0.933758146 | -0.74055595 | 2.128969604 | 0.347847122 | 0.727954999 | 1 | 0.74055595 |
| ENSMUSG00000094796 | BC147527 | 625360 | protein_coding | 1.240410664 | -0.740644823 | 1.641041469 | 0.451326086 | 0.651754547 | 1 | 0.740644823 |
| ENSMUSG00000065651 | Gm25973 | 115488840 | snoRNA | 2.514717472 | 0.740675743 | 0.810546905 | -0.91379751 | 0.360823255 | 1 | 0.740675743 |
| ENSMUSG00000070527 | Mkrn3 | 22652 | protein_coding | 2.677596994 | -0.740938962 | 0.597592952 | 1.239872324 | 0.215022622 | 1 | 0.740938962 |

| ensembl_gene_id | external_gene_name | entrezgene_id | gene_biotype | baseMean | log2FoldChange | lfcSE | stat | pvalue | padj | absLog2FC |
|---|---|---|---|---|---|---|---|---|---|---|
| ENSMUSG00000075270 | Pde11a | 241489 | protein_coding | 50.6237745 | 0.740958076 | 0.359525875 | -2.060931154 | 0.039309609 | 0.124359938 | 0.740958076 |
| ENSMUSG00000078773 | Rad54b | 623474 | protein_coding | 53.64673628 | 0.740981422 | 0.228367684 | -3.244685981 | 0.001175802 | 0.007288737 | 0.740981422 |
| ENSMUSG00000048728 | Zfp454 | 237758 | protein_coding | 6.400888208 | -0.74106707 | 0.509787418 | 1.453678618 | 0.146035429 | 0.323442761 | 0.74106707 |
| ENSMUSG00000035984 | Nme5 | 75533 | protein_coding | 0.967152058 | 0.741144604 | 1.275273588 | -0.581165179 | 0.561129133 | 1 | 0.741144604 |
| ENSMUSG00000048058 | Ldlrad3 | 241576 | protein_coding | 231.7042446 | -0.741166255 | 0.180032021 | 4.116857948 | 3.84E-05 | 0.000370911 | 0.741166255 |
| ENSMUSG00000051054 | 1700014D04Rik | 102638268 | protein_coding | 1.252987466 | 0.742119339 | 1.816792144 | -0.408477844 | 0.682922893 | 1 | 0.742119339 |
| ENSMUSG00000080575 | Mir1190 | 100316805 | miRNA | 2.239724748 | 0.742350495 | 0.721265321 | -1.02923359 | 0.303369922 | 1 | 0.742350495 |
| ENSMUSG00000048482 | Bdnf | 12064 | protein_coding | 2.05735647 | -0.742673487 | 1.11032892 | 0.668877009 | 0.503573937 | 1 | 0.742673487 |
| ENSMUSG00000028152 | Tspan5 | 56224 | protein_coding | 118.7681467 | 0.743071238 | 0.188259164 | -3.947065436 | 7.91E-05 | 0.000702799 | 0.743071238 |
| ENSMUSG00000021032 | Ngb | 64242 | protein_coding | 1.264347725 | -0.743138469 | 1.218290733 | 0.609984505 | 0.541872072 | 1 | 0.743138469 |
| ENSMUSG00000021974 | Fgf9 | 14180 | protein_coding | 24.87874268 | -0.74336075 | 0.306696449 | 2.423767058 | 0.015360455 | 0.059503296 | 0.74336075 |
| ENSMUSG00000031209 | Heph | 15203 | protein_coding | 3221.233656 | 0.74345289 | 0.081993984 | -9.067163896 | 1.22E-19 | 6.08E-18 | 0.74345289 |
| ENSMUSG00000003354 | Ccdc65 | 105833 | protein_coding | 0.684996149 | 0.743502144 | 1.791741631 | -0.414960579 | 0.678170762 | 1 | 0.743502144 |
| ENSMUSG00000025473 | Adam8 | 11501 | protein_coding | 11.03977602 | 0.743601814 | 0.545752738 | -1.362525119 | 0.173032222 | 0.363688435 | 0.743601814 |
| ENSMUSG00000020326 | Ccng1 | 12450 | protein_coding | 1949.460338 | 0.743828896 | 0.094507433 | -7.870586198 | 3.53E-15 | 1.31E-13 | 0.743828896 |
| ENSMUSG00000037628 | Cdkn3 | 72391 | protein_coding | 107.8548362 | 0.744056177 | 0.170128676 | -4.373490679 | 1.22E-05 | 0.00013417 | 0.744056177 |
| ENSMUSG00000046317 | BC107364 | 329716 | lncRNA | 2.805873428 | 0.744191668 | 0.790231095 | -0.941739286 | 0.346326136 | 1 | 0.744191668 |
| ENSMUSG00000085793 | Lin52 | 217708 | protein_coding | 105.8696119 | 0.744236466 | 0.114986642 | -6.472373225 | 9.65E-11 | 2.41E-09 | 0.744236466 |
| ENSMUSG00000070324 | Fbxw22 | 382156 | protein_coding | 7.015422647 | -0.744284082 | 0.466743954 | 1.594630366 | 0.110794918 | 0.266249604 | 0.744284082 |
| ENSMUSG00000090124 | Ugt1a7c | 394432 | protein_coding | 1658.51012 | -0.744390399 | 0.09667421 | 7.699989458 | 1.36E-14 | 4.79E-13 | 0.744390399 |
| ENSMUSG00000042616 | Oscp1 | 230751 | protein_coding | 24.37547571 | 0.744950987 | 0.290163607 | -2.567348105 | 0.010247968 | 0.043415838 | 0.744950987 |
| ENSMUSG00000046997 | Spsb4 | 211949 | protein_coding | 37.36389492 | -0.744951603 | 0.262460102 | 2.83834227 | 0.004534853 | 0.022441881 | 0.744951603 |
| ENSMUSG00000029234 | Tmem165 | 21982 | protein_coding | 659.6300594 | 0.745192944 | 0.101711113 | -7.326563622 | 2.36E-13 | 7.57E-12 | 0.745192944 |
| ENSMUSG00000035158 | Mitf | 17342 | protein_coding | 28.3140433 | 0.745375964 | 0.238255399 | -3.128474601 | 0.001757162 | 0.010174535 | 0.745375964 |
| ENSMUSG00000097407 | 4933408J17Rik | 74454 | lncRNA | 1.218896472 | 0.745713988 | 1.643765033 | -0.453662156 | 0.650072009 | 1 | 0.745713988 |
| ENSMUSG00000031772 | Cntnap4 | 170571 | protein_coding | 2.619368162 | -0.745755914 | 1.766773937 | 0.42210036 | 0.672951765 | 1 | 0.745755914 |
| ENSMUSG00000033470 | Cysltr2 | 70086 | protein_coding | 2.136066913 | 0.745826543 | 0.978130227 | -0.762502295 | 0.445760274 | 1 | 0.745826543 |
| ENSMUSG00000044352 | Sowaha | 237761 | protein_coding | 60.85489567 | -0.74609658 | 0.184478968 | 4.044344932 | 5.25E-05 | 0.000487336 | 0.74609658 |
| ENSMUSG00000043501 | Lgals2 | 107753 | protein_coding | 41.65236951 | 0.746127372 | 0.242304239 | -3.079299711 | 0.002074878 | 0.011690393 | 0.746127372 |
| ENSMUSG00000026768 | Itga8 | 241226 | protein_coding | 7.270783937 | 0.746145337 | 0.434464764 | -1.717389761 | 0.085907981 | 0.221358153 | 0.746145337 |
| ENSMUSG00000025154 | Arhgap19 | 71085 | protein_coding | 176.9217266 | 0.746595061 | 0.149976606 | -4.97807679 | 6.42E-07 | 9.23E-06 | 0.746595061 |
| ENSMUSG00000070942 | Il1rl2 | 107527 | protein_coding | 20.89931386 | -0.746598523 | 0.309965601 | 2.408649602 | 0.016011662 | 0.061535163 | 0.746598523 |
| ENSMUSG00000068101 | Cenpm | 66570 | protein_coding | 52.53687813 | 0.746714064 | 0.189732726 | -3.935610273 | 8.30E-05 | 0.000732064 | 0.746714064 |
| ENSMUSG00000026644 | Acbd7 | 78245 | protein_coding | 0.641220612 | 0.747419608 | 1.397460061 | -0.534841481 | 0.592759477 | 1 | 0.747419608 |
| ENSMUSG00000031821 | Gins2 | 272551 | protein_coding | 105.9639338 | 0.747439482 | 0.170317755 | -4.388500072 | 1.14E-05 | 0.000125991 | 0.747439482 |
| ENSMUSG00000042042 | Csgalnact2 | 78752 | protein_coding | 147.716343 | 0.747535854 | 0.124606522 | -5.99917117 | 1.98E-09 | 4.22E-08 | 0.747535854 |
| ENSMUSG00000027371 | Fahd2a | 68126 | protein_coding | 103.1198052 | -0.747796711 | 0.110543375 | 6.764735665 | 1.34E-11 | 3.68E-10 | 0.747796711 |
| ENSMUSG00000100277 | 1810053B23Rik | 69857 | lncRNA | 2.134820563 | 0.748033393 | 2.153198675 | -0.347405653 | 0.728286587 | 1 | 0.748033393 |
| ENSMUSG00000098207 | Arl14 | 71619 | protein_coding | 174.8610679 | -0.748107572 | 0.223898436 | 3.341280913 | 0.000833928 | 0.005458378 | 0.748107572 |

| ensembl_gene_id | external_gene_name | entrezgene_id | gene_biotype | baseMean | log2FoldChange | lfcSE | stat | pvalue | padj | absLog2FC |
|---|---|---|---|---|---|---|---|---|---|---|
| ENSMUSG00000052760 | A630001G21Rik | 319997 | protein_coding | 20.71523597 | 0.74812005 | 0.259284306 | -2.885327165 | 0.003910071 | 0.01984355 | 0.74812005 |
| ENSMUSG00000078503 | Zfp990 | 101056073 | protein_coding | 1.390484899 | 0.748196038 | 1.085249394 | -0.689423134 | 0.49055703 | 1 | 0.748196038 |
| ENSMUSG00000028716 | Pdzk1ip1 | 67182 | protein_coding | 495.7795002 | -0.748271959 | 0.18324561 | 4.083437296 | 4.44E-05 | 0.000421391 | 0.748271959 |
| ENSMUSG00000056629 | Fkbp2 | 14227 | protein_coding | 639.4060372 | 0.748481867 | 0.051954389 | -14.40651853 | 4.71E-47 | 5.98E-45 | 0.748481867 |
| ENSMUSG00000026039 | Sgo2a | 68549 | protein_coding | 75.76017775 | 0.748554341 | 0.180353237 | -4.150490196 | 3.32E-05 | 0.000325728 | 0.748554341 |
| ENSMUSG00000113585 | Gm19605 | 102638940 | lncRNA | 53.06296798 | 0.748616688 | 0.167590866 | -4.466930125 | 7.94E-06 | 9.12E-05 | 0.748616688 |
| ENSMUSG00000026222 | Sp100 | 20684 | protein_coding | 39.38899044 | -0.748641141 | 0.350427952 | 2.136362518 | 0.032649877 | 0.107730217 | 0.748641141 |
| ENSMUSG00000052305 | Hbb-bs | 15129 | protein_coding | 270.8742913 | -0.748975479 | 0.404755454 | 1.850439501 | 0.06425023 | 0.179449017 | 0.748975479 |
| ENSMUSG00000029875 | Ccdc184 | 239650 | protein_coding | 1.947557843 | -0.749186619 | 0.816168341 | 0.917931487 | 0.358654736 | 1 | 0.749186619 |
| ENSMUSG00000035901 | Dennd5a | 19347 | protein_coding | 32.92020856 | 0.749302045 | 0.275407305 | -2.720705045 | 0.006514286 | 0.030010701 | 0.749302045 |
| ENSMUSG00000046634 | Pkd1l1 | 171395 | protein_coding | 6.017908548 | 0.749413234 | 1.076768609 | -0.695983545 | 0.486439135 | 0.695624496 | 0.749413234 |
| ENSMUSG00000026669 | Mcm10 | 70024 | protein_coding | 232.452994 | 0.749621096 | 0.170419437 | -4.398683098 | 1.09E-05 | 0.000120951 | 0.749621096 |
| ENSMUSG00000015880 | Ncapg | 54392 | protein_coding | 193.8870386 | 0.749739646 | 0.216417387 | -3.464322603 | 0.000531569 | 0.003735457 | 0.749739646 |
| ENSMUSG00000053141 | Ptprt | 19281 | protein_coding | 31.69981587 | -0.749785017 | 0.252001826 | 2.975315814 | 0.002926871 | 0.015546587 | 0.749785017 |
| ENSMUSG00000099110 | Mir6951 | 102465574 | miRNA | 1.223741064 | -0.749998798 | 1.048309817 | 0.715436206 | 0.474339547 | 1 | 0.749998798 |
| ENSMUSG00000026764 | Kif5c | 16574 | protein_coding | 34.25034619 | 0.750374352 | 0.290642807 | -2.581775068 | 0.009829363 | 0.042030578 | 0.750374352 |
| ENSMUSG00000024163 | Mapk8ip3 | 30957 | protein_coding | 484.698594 | 0.750426366 | 0.160301197 | -4.681352224 | 2.85E-06 | 3.57E-05 | 0.750426366 |
| ENSMUSG00000003992 | Ssbp2 | 66970 | protein_coding | 35.39019771 | -0.750447022 | 0.285772703 | 2.626027658 | 0.00863878 | 0.037851103 | 0.750447022 |
| ENSMUSG00000020424 | Castor1 | 71962 | protein_coding | 21.10635436 | 0.750939663 | 0.244719035 | -3.068578886 | 0.002150795 | 0.012023776 | 0.750939663 |
| ENSMUSG00000006411 | Nectin4 | 71740 | protein_coding | 50.71587179 | -0.751072462 | 0.186842336 | 4.019819481 | 5.82E-05 | 0.000535833 | 0.751072462 |
| ENSMUSG00000039683 | Sdk1 | 330222 | protein_coding | 8.306857674 | 0.751303373 | 0.782727746 | -0.959852742 | 0.337129334 | 0.559327232 | 0.751303373 |
| ENSMUSG00000038077 | Kcna6 | 16494 | protein_coding | 1.772962577 | -0.751719705 | 3.044902823 | 0.246878061 | 0.805002596 | 1 | 0.751719705 |
| ENSMUSG00000018381 | Abi3 | 66610 | protein_coding | 17.06137001 | -0.75173191 | 0.330834063 | 2.272232498 | 0.023072471 | 0.08204539 | 0.75173191 |
| ENSMUSG00000032946 | Rasgrp2 | 19395 | protein_coding | 9.749022798 | 0.751939502 | 0.863468569 | -0.870835985 | 0.383843714 | 0.606235185 | 0.751939502 |
| ENSMUSG00000035523 | Vmn1r172 | 81010 | protein_coding | 1.681968812 | -0.752018177 | 3.053333921 | 0.246294115 | 0.805454565 | 1 | 0.752018177 |
| ENSMUSG00000073747 | Zpld2 | 665186 | protein_coding | 1.729529425 | -0.752194537 | 3.05833442 | 0.24594908 | 0.80572165 | 1 | 0.752194537 |
| ENSMUSG00000029544 | Cabp1 | 29867 | protein_coding | 7.138514036 | 0.752225396 | 0.823504643 | -0.913444026 | 0.36100906 | 0.583394032 | 0.752225396 |
| ENSMUSG00000072875 | Gpr27 | 14761 | protein_coding | 15.51402996 | 0.75227351 | 0.49139802 | -1.530884293 | 0.125797992 | 0.291527931 | 0.75227351 |
| ENSMUSG00000070583 | Fv1 | 14349 | protein_coding | 47.87796091 | -0.752482083 | 0.14932629 | 5.039180203 | 4.68E-07 | 6.95E-06 | 0.752482083 |
| ENSMUSG00000025573 | 6030468B19Rik | 77727 | protein_coding | 1.803334231 | 0.752525261 | 1.101747141 | -0.683029011 | 0.494588511 | 1 | 0.752525261 |
| ENSMUSG00000055125 | M5C1000I18Rik | 102633151 | lncRNA | 7.352589234 | 0.752624423 | 0.514563526 | -1.462646272 | 0.143564195 | 0.319862266 | 0.752624423 |
| ENSMUSG00000052160 | Pld4 | 104759 | protein_coding | 2.90903091 | 0.752684462 | 1.496035714 | -0.503119314 | 0.614880388 | 1 | 0.752684462 |
| ENSMUSG00000041930 | Fam222a | 433940 | protein_coding | 68.24357454 | 0.752740648 | 0.186138516 | -4.043981141 | 5.26E-05 | 0.000487819 | 0.752740648 |
| ENSMUSG00000020044 | Timp3 | 21859 | protein_coding | 37.94463458 | -0.752776143 | 0.248540421 | 3.02878759 | 0.002455373 | 0.013413173 | 0.752776143 |
| ENSMUSG00000024347 | Psd2 | 74002 | protein_coding | 3.118210899 | 0.753055402 | 1.82902445 | -0.41172517 | 0.680540875 | 1 | 0.753055402 |
| ENSMUSG00000078598 | Skint5 | 242627 | protein_coding | 1.419998537 | 0.753063658 | 3.231069646 | -0.23306946 | 0.815707469 | 1 | 0.753063658 |
| ENSMUSG00000044086 | Lmod3 | 320502 | protein_coding | 1.29720156 | 0.75306431 | 3.220972646 | -0.233800281 | 0.815140031 | 1 | 0.75306431 |
| ENSMUSG00000018634 | Crhr1 | 12921 | protein_coding | 1.415871077 | 0.753064844 | 3.195054973 | -0.235696991 | 0.813667805 | 1 | 0.753064844 |
| ENSMUSG00000049561 | Or10c1 | 258506 | protein_coding | 1.506864841 | 0.753064876 | 3.175858456 | -0.237121675 | 0.812562398 | 1 | 0.753064876 |

| ensembl_gene_id | external_gene_name | entrezgene_id | gene_biotype | baseMean | log2FoldChange | lfcSE | stat | pvalue | padj | absLog2FC |
|---|---|---|---|---|---|---|---|---|---|---|
| ENSMUSG00000059203 | Il1rapl2 | 60367 | protein_coding | 1.186322936 | 0.753064925 | 3.246291617 | -0.23197698 | 0.816555893 | 1 | 0.753064925 |
| ENSMUSG00000023802 | Nox3 | 224480 | protein_coding | 1.368310465 | 0.753064977 | 3.200652102 | -0.235284858 | 0.813987647 | 1 | 0.753064977 |
| ENSMUSG00000066748 | Or8b8 | 258310 | protein_coding | 1.091201712 | 0.753065022 | 3.268642747 | -0.23039074 | 0.817788156 | 1 | 0.753065022 |
| ENSMUSG00000100969 | 1700030N03Rik | 70011 | lncRNA | 1.091201712 | 0.753065022 | 3.268642747 | -0.23039074 | 0.817788156 | 1 | 0.753065022 |
| ENSMUSG00000021404 | Serpinb9c | 20707 | protein_coding | 1.138762324 | 0.753065077 | 3.253923062 | -0.23143297 | 0.816978454 | 1 | 0.753065077 |
| ENSMUSG00000049691 | Nkx3-2 | 12020 | protein_coding | 1.41574746 | 0.753080655 | 2.671329605 | -0.281912293 | 0.778010763 | 1 | 0.753080655 |
| ENSMUSG00000036139 | Hoxc9 | 15427 | protein_coding | 1.292098567 | 0.753080948 | 2.66797747 | -0.282266607 | 0.777739089 | 1 | 0.753080948 |
| ENSMUSG00000030500 | Slc17a6 | 140919 | protein_coding | 1.458439748 | 0.753081018 | 2.650052567 | -0.284175879 | 0.776275599 | 1 | 0.753081018 |
| ENSMUSG00000070368 | Prok1 | 246691 | protein_coding | 1.063753426 | 0.753081191 | 2.70718169 | -0.278179035 | 0.780874927 | 1 | 0.753081191 |
| ENSMUSG00000021799 | Opn4 | 30044 | protein_coding | 1.01545195 | 0.753081543 | 2.704770258 | -0.278427175 | 0.780684461 | 1 | 0.753081543 |
| ENSMUSG00000102439 | Flg | 14246 | protein_coding | 1.229632471 | 0.753081607 | 2.670997658 | -0.281947685 | 0.777983625 | 1 | 0.753081607 |
| ENSMUSG00000026167 | Wnt10a | 22409 | protein_coding | 1.006492197 | 0.753082058 | 2.685890627 | -0.280384484 | 0.77918254 | 1 | 0.753082058 |
| ENSMUSG00000019102 | Aldh3a1 | 11670 | protein_coding | 1.090337231 | 0.753082129 | 2.682564499 | -0.280732161 | 0.778915839 | 1 | 0.753082129 |
| ENSMUSG00000111658 | B930092H01Rik | 622037 | lncRNA | 1.043517482 | 0.753082267 | 2.688145085 | -0.280149413 | 0.779362876 | 1 | 0.753082267 |
| ENSMUSG00000020636 | Allc | 94041 | protein_coding | 1.320502618 | 0.753082329 | 2.626145592 | -0.286763358 | 0.774293521 | 1 | 0.753082329 |
| ENSMUSG00000091318 | Semp2l1 | 408191 | protein_coding | 1.198056725 | 0.753082336 | 2.634413593 | -0.285863366 | 0.774982773 | 1 | 0.753082336 |
| ENSMUSG00000024738 | Pga5 | 58803 | protein_coding | 0.997998308 | 0.753082542 | 2.663956907 | -0.282693215 | 0.777412018 | 1 | 0.753082542 |
| ENSMUSG00000094891 | Or10h1b | 100038859 | protein_coding | 1.225381394 | 0.753083001 | 2.618075258 | -0.287647576 | 0.773616523 | 1 | 0.753083001 |
| ENSMUSG00000108379 | A730082K24Rik | 101786 | lncRNA | 0.853027799 | 0.753083675 | 2.686006119 | -0.28037303 | 0.779191326 | 1 | 0.753083675 |
| ENSMUSG00000024768 | Lipf | 67717 | protein_coding | 1.075973911 | 0.753092189 | 2.270863771 | -0.331632482 | 0.740166789 | 1 | 0.753092189 |
| ENSMUSG00000020407 | Upp1 | 22271 | protein_coding | 210.1279515 | -0.753128231 | 0.219553836 | 3.430266785 | 0.000602988 | 0.004145606 | 0.753128231 |
| ENSMUSG00000024034 | Tmprss3 | 140765 | protein_coding | 2.056431671 | -0.75327634 | 0.872125256 | 0.863724946 | 0.387739006 | 1 | 0.75327634 |
| ENSMUSG00000114206 | Gm30551 | 102632492 | lncRNA | 1.698443274 | -0.753303576 | 1.831652402 | 0.411269942 | 0.680874608 | 1 | 0.753303576 |
| ENSMUSG00000001039 | B9d1 | 27078 | protein_coding | 41.88111224 | 0.753368555 | 0.175936529 | -4.282047385 | 1.85E-05 | 0.000192981 | 0.753368555 |
| ENSMUSG00000039196 | Orm1 | 18405 | protein_coding | 1.036757542 | -0.753554427 | 3.098252442 | 0.243219183 | 0.807835607 | 1 | 0.753554427 |
| ENSMUSG00000025013 | Tll2 | 24087 | protein_coding | 1.874858862 | 0.753644777 | 1.735688524 | -0.434205082 | 0.664139523 | 1 | 0.753644777 |
| ENSMUSG00000026437 | Cdk18 | 18557 | protein_coding | 342.7601526 | 0.753736377 | 0.145926171 | -5.165189863 | 2.40E-07 | 3.71E-06 | 0.753736377 |
| ENSMUSG00000057406 | Nsd2 | 107823 | protein_coding | 1022.044019 | 0.753945756 | 0.085939988 | -8.772932997 | 1.74E-18 | 8.12E-17 | 0.753945756 |
| ENSMUSG00000062296 | Trank1 | 320429 | protein_coding | 5.233053163 | 0.754303953 | 0.776468116 | -0.971455154 | 0.331321675 | 0.553734049 | 0.754303953 |
| ENSMUSG00000049280 | Or10ab5 | 258369 | protein_coding | 0.847331937 | -0.754422907 | 2.231890207 | 0.338019722 | 0.735348326 | 1 | 0.754422907 |
| ENSMUSG00000015478 | Rnf5 | 54197 | protein_coding | 225.3125357 | 0.754494379 | 0.109064464 | -6.917875472 | 4.58E-12 | 1.31E-10 | 0.754494379 |
| ENSMUSG00000037568 | Vash2 | 226841 | protein_coding | 5.034158216 | 0.754765662 | 0.641007362 | -1.177468009 | 0.239008759 | 0.449185233 | 0.754765662 |
| ENSMUSG00000073000 | Gm10451 | 100041694 | lncRNA | 18.12713057 | -0.754872609 | 0.326364672 | 2.312972797 | 0.020724135 | 0.075366299 | 0.754872609 |
| ENSMUSG00000001663 | Gstt1 | 14871 | protein_coding | 197.0863371 | -0.755201517 | 0.122926508 | 6.143520489 | 8.07E-10 | 1.80E-08 | 0.755201517 |
| ENSMUSG00000040152 | Thbs1 | 21825 | protein_coding | 1516.517856 | -0.755254636 | 0.237809516 | 3.175880638 | 0.001493824 | 0.008870554 | 0.755254636 |
| ENSMUSG00000046460 | Sh2d7 | 244885 | protein_coding | 184.0292158 | -0.755955034 | 0.138087024 | 5.474482781 | 4.39E-08 | 7.60E-07 | 0.755955034 |
| ENSMUSG00000045091 | Aqp12 | 208760 | protein_coding | 9.985188665 | 0.755955577 | 0.384261643 | -1.967293877 | 0.049149341 | 0.147109787 | 0.755955577 |
| ENSMUSG00000034330 | Plcg2 | 234779 | protein_coding | 149.1557825 | -0.756420364 | 0.128997089 | 5.863856063 | 4.52E-09 | 9.17E-08 | 0.756420364 |
| ENSMUSG00000073700 | Klhl21 | 242785 | protein_coding | 350.2190592 | 0.756443518 | 0.104849404 | -7.214571449 | 5.41E-13 | 1.68E-11 | 0.756443518 |

| ensembl_gene_id | external_gene_name | entrezgene_id | gene_biotype | baseMean | log2FoldChange | lfcSE | stat | pvalue | padj | absLog2FC |
|---|---|---|---|---|---|---|---|---|---|---|
| ENSMUSG00000021490 | Slc34a1 | 20505 | protein_coding | 12.11044439 | 0.756463897 | 0.458552974 | -1.649676132 | 0.099009195 | 0.245833384 | 0.756463897 |
| ENSMUSG00000025265 | Fgd1 | 14163 | protein_coding | 45.16787277 | 0.757031003 | 0.216465998 | -3.497228244 | 0.000470119 | 0.003368135 | 0.757031003 |
| ENSMUSG00000020325 | Fstl3 | 83554 | protein_coding | 7.044908056 | -0.757088872 | 0.41922652 | 1.805918366 | 0.070931092 | 0.192815811 | 0.757088872 |
| ENSMUSG00000030098 | Grip2 | 243547 | protein_coding | 6.524718652 | -0.757109098 | 0.840240387 | 0.901062493 | 0.367555094 | 0.590046157 | 0.757109098 |
| ENSMUSG00000078143 | Gm17344 | 115486898 | lncRNA | 2.069501695 | -0.757220093 | 0.77208507 | 0.98074697 | 0.326717535 | 1 | 0.757220093 |
| ENSMUSG00000115021 | Vmn1r173 | 545934 | protein_coding | 1.26858609 | -0.75749223 | 2.312017436 | 0.327632577 | 0.743189488 | 1 | 0.75749223 |
| ENSMUSG00000104063 | Pcdhgb7 | 93704 | protein_coding | 9.126358381 | 0.757529337 | 0.349630733 | -2.166655461 | 0.030261135 | 0.101654269 | 0.757529337 |
| ENSMUSG00000015749 | Anp32e | 66471 | protein_coding | 719.6446732 | 0.757539136 | 0.10662917 | -7.104426855 | 1.21E-12 | 3.63E-11 | 0.757539136 |
| ENSMUSG00000071553 | Cpa2 | 232680 | protein_coding | 29.58209556 | 0.757549091 | 0.300133684 | -2.524038894 | 0.011601509 | 0.047864939 | 0.757549091 |
| ENSMUSG00000089704 | Galnt2 | 108148 | protein_coding | 13.69144186 | 0.757551198 | 0.299998105 | -2.525186609 | 0.011563687 | 0.047764563 | 0.757551198 |
| ENSMUSG00000085316 | D330050G23Rik | 320975 | lncRNA | 0.90845457 | -0.757668725 | 2.201030107 | 0.344233694 | 0.730670545 | 1 | 0.757668725 |
| ENSMUSG00000020335 | Zfp354b | 27274 | protein_coding | 3.64545709 | -0.757913318 | 0.747760201 | 1.013578039 | 0.310784141 | 1 | 0.757913318 |
| ENSMUSG00000020312 | Shc2 | 216148 | protein_coding | 6.335960864 | -0.757915509 | 0.585244817 | 1.295040103 | 0.195306396 | 0.39448261 | 0.757915509 |
| ENSMUSG00000047777 | Phf13 | 230936 | protein_coding | 225.8884621 | 0.758029762 | 0.11703417 | -6.476995218 | 9.36E-11 | 2.35E-09 | 0.758029762 |
| ENSMUSG00000073197 | 5730507C01Rik | 236366 | protein_coding | 31.49056813 | 0.758035392 | 0.349574081 | -2.168454223 | 0.030124144 | 0.101317742 | 0.758035392 |
| ENSMUSG00000030724 | Cd19 | 12478 | protein_coding | 2.506124742 | 0.758548989 | 1.491973938 | -0.508419732 | 0.611159018 | 1 | 0.758548989 |
| ENSMUSG00000032175 | Tyk2 | 54721 | protein_coding | 991.1469097 | 0.758586458 | 0.042127702 | -18.006832 | 1.72E-72 | 3.90E-70 | 0.758586458 |
| ENSMUSG00000022759 | Lrrc74b | 74685 | protein_coding | 2.83550102 | -0.758613426 | 0.995329308 | 0.762173302 | 0.445956578 | 1 | 0.758613426 |
| ENSMUSG00000028957 | Per3 | 18628 | protein_coding | 168.1158561 | -0.758708222 | 0.267285439 | 2.83856923 | 0.004531629 | 0.022432649 | 0.758708222 |
| ENSMUSG00000001870 | Ltbp1 | 268977 | protein_coding | 4.901571028 | 0.758745419 | 1.221930125 | -0.620940105 | 0.534639033 | 0.730517043 | 0.758745419 |
| ENSMUSG00000055114 | Anxa13 | 69787 | protein_coding | 106.1500459 | -0.759014254 | 0.252138946 | 3.010301528 | 0.002609885 | 0.014131027 | 0.759014254 |
| ENSMUSG00000028456 | Unc13b | 22249 | protein_coding | 990.8787079 | 0.7592246 | 0.100686814 | -7.540457083 | 4.68E-14 | 1.58E-12 | 0.7592246 |
| ENSMUSG00000002688 | Prkd1 | 18760 | protein_coding | 10.7909257 | 0.75932913 | 0.331224516 | -2.292490723 | 0.021877342 | 0.078813794 | 0.75932913 |
| ENSMUSG00000110803 | Gm20275 | 100504539 | lncRNA | 34.40315835 | -0.759355806 | 0.216693862 | 3.504279259 | 0.000457845 | 0.003290188 | 0.759355806 |
| ENSMUSG00000024391 | Apom | 55938 | protein_coding | 3.222019937 | 0.759570988 | 0.622629112 | -1.219941332 | 0.222487116 | 1 | 0.759570988 |
| ENSMUSG00000074219 | Gm10644 | 100126034 | lncRNA | 4.836806679 | 0.759749918 | 0.521460852 | -1.456964439 | 0.145126197 | 0.322298818 | 0.759749918 |
| ENSMUSG00000038094 | Atp13a4 | 224079 | protein_coding | 6.291538004 | 0.759910716 | 0.774190292 | -0.98155547 | 0.326318895 | 0.548314024 | 0.759910716 |
| ENSMUSG00000032066 | Bco2 | 170752 | protein_coding | 161.2332639 | -0.75998136 | 0.206939405 | 3.672482577 | 0.000240206 | 0.001860579 | 0.75998136 |
| ENSMUSG00000060509 | Xcr1 | 23832 | protein_coding | 2.714583009 | 0.760224937 | 0.984960564 | -0.771832868 | 0.440213422 | 1 | 0.760224937 |
| ENSMUSG00000105676 | Gm43403 | 108168904 | lncRNA | 2.81320065 | 0.76046668 | 1.080592941 | -0.703749443 | 0.481588824 | 1 | 0.76046668 |
| ENSMUSG00000074491 | Clec4g | 75863 | protein_coding | 1.645961332 | -0.76055164 | 1.853095275 | 0.410422308 | 0.681496185 | 1 | 0.76055164 |
| ENSMUSG00000032932 | Hspa13 | 110920 | protein_coding | 570.0108027 | 0.76101825 | 0.109965134 | -6.9205413 | 4.50E-12 | 1.29E-10 | 0.76101825 |
| ENSMUSG00000087476 | Rap1gapos | 69808 | lncRNA | 23.90551408 | 0.761083792 | 0.238703657 | -3.188404408 | 0.001430603 | 0.008565982 | 0.761083792 |
| ENSMUSG00000025935 | Tram1 | 72265 | protein_coding | 2565.951392 | 0.761235499 | 0.084053885 | -9.056517692 | 1.35E-19 | 6.66E-18 | 0.761235499 |
| ENSMUSG00000026782 | Abi2 | 329165 | protein_coding | 73.31485424 | 0.762509824 | 0.158016432 | -4.825509678 | 1.40E-06 | 1.87E-05 | 0.762509824 |
| ENSMUSG00000037370 | Enpp1 | 18605 | protein_coding | 315.6153885 | 0.762544857 | 0.137264078 | -5.555312555 | 2.77E-08 | 4.98E-07 | 0.762544857 |
| ENSMUSG00000025887 | Casp12 | 12364 | protein_coding | 10.92329521 | 0.762765804 | 0.432559239 | -1.763378827 | 0.077836618 | 0.205992615 | 0.762765804 |
| ENSMUSG00000020857 | Nme2 | 18103 | protein_coding | 161.3207077 | 0.762905256 | 0.17711678 | -4.307357303 | 1.65E-05 | 0.000174227 | 0.762905256 |
| ENSMUSG00000064916 | Gm22573 | 115490009 | snRNA | 2.278168866 | -0.763009021 | 0.770277998 | 0.990563178 | 0.321898925 | 1 | 0.763009021 |

Supplementary Table 1

| ensembl_gene_id | external_gene_name | entrezgene_id | gene_biotype | baseMean | log2FoldChange | lfcSE | stat | pvalue | padj | absLog2FC |
|---|---|---|---|---|---|---|---|---|---|---|
| ENSMUSG00000055116 | Arntl | 11865 | protein_coding | 750.4460266 | 0.763245459 | 0.187326705 | -4.07440819 | 4.61E-05 | 0.00043511 | 0.763245459 |
| ENSMUSG00000035983 | Gm7008 | 629820 | lncRNA | 31.17281901 | 0.763345877 | 0.366404254 | -2.083343379 | 0.037219935 | 0.119049005 | 0.763345877 |
| ENSMUSG00000039457 | Ppl | 19041 | protein_coding | 143.9378075 | -0.763349293 | 0.113811194 | 6.707154747 | 1.98E-11 | 5.39E-10 | 0.763349293 |
| ENSMUSG00000013584 | Aldh1a2 | 19378 | protein_coding | 1.01460204 | 0.76356262 | 1.172819206 | -0.651048871 | 0.51501494 | 1 | 0.76356262 |
| ENSMUSG00000038600 | Atp6v0a4 | 140494 | protein_coding | 5.136026768 | -0.763894332 | 0.79879974 | 0.956302679 | 0.338919337 | 0.560514185 | 0.763894332 |
| ENSMUSG00000003934 | Efnb3 | 13643 | protein_coding | 9.53507863 | 0.763982366 | 0.620677813 | -1.230883964 | 0.218366266 | 0.42452042 | 0.763982366 |
| ENSMUSG00000032118 | Fez1 | 235180 | protein_coding | 2.231747615 | -0.764174255 | 1.768301575 | 0.432151543 | 0.665631275 | 1 | 0.764174255 |
| ENSMUSG00000053080 | Zfta | 108899 | protein_coding | 77.84172197 | 0.764391558 | 0.185021992 | -4.131355146 | 3.61E-05 | 0.00035094 | 0.764391558 |
| ENSMUSG00000052133 | Sema5b | 20357 | protein_coding | 3.17455905 | 0.764906121 | 1.577282926 | -0.484951754 | 0.627710569 | 1 | 0.764906121 |
| ENSMUSG00000030930 | Chst15 | 77590 | protein_coding | 84.20484861 | -0.76491172 | 0.222429576 | 3.438893939 | 0.000584096 | 0.004039264 | 0.76491172 |
| ENSMUSG00000090145 | Ugt1a6b | 394435 | protein_coding | 13.28534899 | -0.765005367 | 0.365979469 | 2.090295855 | 0.036591231 | 0.117379098 | 0.765005367 |
| ENSMUSG00000025026 | Add3 | 27360 | protein_coding | 1158.670657 | -0.765035749 | 0.10227157 | 7.48043422 | 7.41E-14 | 2.46E-12 | 0.765035749 |
| ENSMUSG00000063919 | Srrm4 | 68955 | protein_coding | 3.841048211 | -0.76527639 | 1.999929404 | 0.382651702 | 0.70197803 | 1 | 0.76527639 |
| ENSMUSG00000056133 | Unc93a2 | 667055 | protein_coding | 2.385986467 | -0.765646819 | 1.196473524 | 0.639919567 | 0.522224892 | 1 | 0.765646819 |
| ENSMUSG00000044522 | A730020M07Rik | 100503044 | lncRNA | 1.273576458 | -0.766115511 | 1.913472621 | 0.400379657 | 0.688876906 | 1 | 0.766115511 |
| ENSMUSG00000026814 | Eng | 13805 | protein_coding | 5.317911667 | 0.766382352 | 1.173286435 | -0.653192885 | 0.513631936 | 0.715488638 | 0.766382352 |
| ENSMUSG00000014592 | Camta1 | 100072 | protein_coding | 108.8959647 | 0.76677968 | 0.167647135 | -4.573771455 | 4.79E-06 | 5.74E-05 | 0.76677968 |
| ENSMUSG00000052551 | Adarb2 | 94191 | protein_coding | 2.043814421 | -0.767194834 | 2.627209866 | 0.292018862 | 0.770272203 | 1 | 0.767194834 |
| ENSMUSG00000031492 | Chrnb3 | 108043 | protein_coding | 1.034628094 | -0.767200405 | 2.627891326 | 0.291945255 | 0.770328481 | 1 | 0.767200405 |
| ENSMUSG00000029868 | Trpv6 | 64177 | protein_coding | 285.9276155 | -0.767245018 | 0.35426707 | 2.165724913 | 0.030332214 | 0.101750358 | 0.767245018 |
| ENSMUSG00000081895 | Rpl17-ps10 | 100042880 | processed_pseudogene | 0.859311523 | -0.767486617 | 1.030074125 | 0.745079017 | 0.456223953 | 1 | 0.767486617 |
| ENSMUSG00000073928 | Or52h9 | 258809 | protein_coding | 1.004115007 | -0.767829364 | 1.230998298 | 0.623745268 | 0.532794889 | 1 | 0.767829364 |
| ENSMUSG00000116852 | 1700102H20Rik | 68230 | lncRNA | 2.775433742 | 0.767975854 | 0.715813031 | -1.072872133 | 0.283328491 | 1 | 0.767975854 |
| ENSMUSG00000027699 | Ect2 | 13605 | protein_coding | 490.2862361 | 0.768012157 | 0.173225965 | -4.433585669 | 9.27E-06 | 0.0001049 | 0.768012157 |
| ENSMUSG00000069910 | Spdl1 | 70385 | protein_coding | 112.3313432 | 0.768058449 | 0.188455699 | -4.075538457 | 4.59E-05 | 0.000434207 | 0.768058449 |
| ENSMUSG00000050241 | Klre1 | 243655 | protein_coding | 4.214391113 | 0.768127214 | 0.751471656 | -1.02216392 | 0.306703324 | 1 | 0.768127214 |
| ENSMUSG00000069998 | Or2z8 | 404315 | protein_coding | 1.179439379 | -0.768176934 | 3.692070821 | 0.208061267 | 0.83518113 | 1 | 0.768176934 |
| ENSMUSG00000089265 | Gm22888 | 115490072 | snoRNA | 0.66325591 | -0.768504889 | 1.844516941 | 0.416642901 | 0.676939625 | 1 | 0.768504889 |
| ENSMUSG00000022237 | Ankrd33b | 67434 | protein_coding | 5.796227934 | -0.768668314 | 0.623151285 | 1.23351798 | 0.217382573 | 0.423132462 | 0.768668314 |
| ENSMUSG00000086569 | Rab11fip4os2 | 72218 | lncRNA | 1.999645031 | -0.76954059 | 1.453064585 | 0.529598339 | 0.596390446 | 1 | 0.76954059 |
| ENSMUSG00000075588 | Hoxb2 | 103889 | protein_coding | 2.336705894 | 0.769742092 | 0.918200918 | -0.838315533 | 0.401853514 | 1 | 0.769742092 |
| ENSMUSG00000097387 | 4930563E18Rik | 75370 | lncRNA | 1.736407383 | 0.769756192 | 1.418694842 | -0.542580525 | 0.587418651 | 1 | 0.769756192 |
| ENSMUSG00000087107 | AI662270 | 100043636 | lncRNA | 4.889006123 | 0.770152443 | 0.790544488 | -0.974205063 | 0.329954731 | 0.552120015 | 0.770152443 |
| ENSMUSG00000038508 | Gdf15 | 23886 | protein_coding | 219.52447 | 0.770431739 | 0.200880296 | -3.835277796 | 0.000125422 | 0.001055133 | 0.770431739 |
| ENSMUSG00000064036 | Mro | 71263 | protein_coding | 5.191746966 | -0.770684672 | 0.722093936 | 1.067291433 | 0.285840255 | 0.504312604 | 0.770684672 |
| ENSMUSG00000066687 | Zbtb16 | 235320 | protein_coding | 17.50127262 | -0.770884114 | 0.425829337 | 1.810312364 | 0.070247361 | 0.191493053 | 0.770884114 |
| ENSMUSG00000029231 | Pdgfra | 18595 | protein_coding | 3.368832764 | 0.771108222 | 1.255596437 | -0.614136994 | 0.539124804 | 1 | 0.771108222 |
| ENSMUSG00000018654 | Ikzf1 | 22778 | protein_coding | 12.01023704 | 0.771194244 | 0.448901702 | -1.717957943 | 0.085804287 | 0.22115998 | 0.771194244 |
| ENSMUSG00000024579 | Pcyox1l | 240334 | protein_coding | 126.580784 | 0.771842139 | 0.152692148 | -5.054890829 | 4.31E-07 | 6.44E-06 | 0.771842139 |

Supplementary Table 1

| ensembl_gene_id | external_gene_name | entrezgene_id | gene_biotype | baseMean | log2FoldChange | lfcSE | stat | pvalue | padj | absLog2FC |
|---|---|---|---|---|---|---|---|---|---|---|
| ENSMUSG00000030878 | Cdr2 | 12585 | protein_coding | 107.6180362 | 0.772096875 | 0.166685022 | -4.63207111 | 3.62E-06 | 4.46E-05 | 0.772096875 |
| ENSMUSG00000026586 | Prrx1 | 18933 | protein_coding | 4.163786285 | 0.772199795 | 1.701344514 | -0.453876207 | 0.64991793 | 1 | 0.772199795 |
| ENSMUSG00000041695 | Kcnj2 | 16518 | protein_coding | 16.54287282 | -0.77304442 | 0.321582321 | 2.403877233 | 0.01622222 | 0.06221406 | 0.77304442 |
| ENSMUSG00000024774 | Ankrd22 | 52024 | protein_coding | 70.17844162 | 0.773094589 | 0.229544448 | -3.367951593 | 0.000757289 | 0.005018407 | 0.773094589 |
| ENSMUSG00000009246 | Trpm5 | 56843 | protein_coding | 308.3272368 | -0.773240639 | 0.165615101 | 4.66890177 | 3.03E-06 | 3.78E-05 | 0.773240639 |
| ENSMUSG00000040181 | Fmo1 | 14261 | protein_coding | 7.128585037 | -0.773329869 | 0.482080805 | 1.604149888 | 0.108681018 | 0.262814222 | 0.773329869 |
| ENSMUSG00000039607 | Rbms3 | 207181 | protein_coding | 33.29377222 | -0.773368007 | 0.253420571 | 3.051717554 | 0.002275361 | 0.012571198 | 0.773368007 |
| ENSMUSG00000051586 | Mical3 | 194401 | protein_coding | 425.1387203 | 0.773468916 | 0.175154811 | -4.415915897 | 1.01E-05 | 0.000112689 | 0.773468916 |
| ENSMUSG00000101583 | Gm20753 | 626049 | lncRNA | 2.123882543 | -0.773583356 | 0.722468552 | 1.070750213 | 0.284281759 | 1 | 0.773583356 |
| ENSMUSG00000073804 | Nps | 100043254 | protein_coding | 0.819994804 | -0.773638031 | 1.659644499 | 0.466146835 | 0.641110401 | 1 | 0.773638031 |
| ENSMUSG00000020255 | Nopchap1 | 28109 | protein_coding | 640.5576279 | 0.773755075 | 0.070977198 | -10.90145982 | 1.13E-27 | 8.11E-26 | 0.773755075 |
| ENSMUSG00000001440 | Kpnb1 | 16211 | protein_coding | 3575.146901 | 0.774223651 | 0.093501107 | -8.280368836 | 1.23E-16 | 5.15E-15 | 0.774223651 |
| ENSMUSG00000086993 | Rsf1os2 | 102641859 | lncRNA | 3.961004583 | 0.774227792 | 0.736750181 | -1.050868818 | 0.293318843 | 1 | 0.774227792 |
| ENSMUSG00000048572 | Tmem252 | 226040 | protein_coding | 322.2576806 | -0.774429328 | 0.226144282 | 3.42449219 | 0.00061595 | 0.004224166 | 0.774429328 |
| ENSMUSG00000028873 | Cdca8 | 52276 | protein_coding | 229.6369128 | 0.774494429 | 0.154876808 | -5.000712766 | 5.71E-07 | 8.31E-06 | 0.774494429 |
| ENSMUSG00000001082 | Mfsd10 | 68294 | protein_coding | 262.9569254 | 0.774678351 | 0.105333982 | -7.354496032 | 1.92E-13 | 6.21E-12 | 0.774678351 |
| ENSMUSG00000031883 | Car7 | 12354 | protein_coding | 6.066227449 | -0.774814816 | 0.565576503 | 1.369955809 | 0.170700697 | 0.360508479 | 0.774814816 |
| ENSMUSG00000065519 | Mir10a | 723893 | miRNA | 1.288041616 | -0.774890507 | 1.086997283 | 0.712872533 | 0.475924637 | 1 | 0.774890507 |
| ENSMUSG00000075610 | Tmem92 | 544806 | protein_coding | 0.949471929 | 0.775109979 | 1.811323992 | -0.427924536 | 0.668706063 | 1 | 0.775109979 |
| ENSMUSG00000027075 | Slc43a1 | 72401 | protein_coding | 75.47221184 | 0.775252327 | 0.186306675 | -4.161162381 | 3.17E-05 | 0.000311984 | 0.775252327 |
| ENSMUSG00000058260 | Serpina9 | 71907 | protein_coding | 22.33095573 | -0.775279575 | 0.388924047 | 1.993395832 | 0.04621812 | 0.140457496 | 0.775279575 |
| ENSMUSG00000049548 | Krt82 | 114566 | protein_coding | 0.993154313 | -0.775473998 | 1.619234344 | 0.478914001 | 0.63199981 | 1 | 0.775473998 |
| ENSMUSG00000053040 | Aph1c | 68318 | protein_coding | 6.957583161 | 0.775495552 | 0.591489862 | -1.311088493 | 0.189827868 | 0.387122324 | 0.775495552 |
| ENSMUSG00000033453 | Adamts15 | 235130 | protein_coding | 462.8922811 | 0.77554865 | 0.19777287 | -3.921410701 | 8.80E-05 | 0.000770817 | 0.77554865 |
| ENSMUSG00000020653 | Klf11 | 194655 | protein_coding | 242.8304928 | -0.775805632 | 0.201643672 | 3.847408775 | 0.000119374 | 0.001007326 | 0.775805632 |
| ENSMUSG00000041649 | Klf8 | 245671 | protein_coding | 10.34636672 | -0.775871472 | 0.416653551 | 1.862150151 | 0.062581929 | 0.175901555 | 0.775871472 |
| ENSMUSG00000021416 | Eci3 | 69123 | protein_coding | 105.7107236 | -0.775927052 | 0.117011163 | 6.631222441 | 3.33E-11 | 8.85E-10 | 0.775927052 |
| ENSMUSG00000018830 | Myh11 | 17880 | protein_coding | 11.02298904 | 0.776147317 | 0.812200033 | -0.955611039 | 0.339268782 | 0.560708667 | 0.776147317 |
| ENSMUSG00000033676 | Gabrb3 | 14402 | protein_coding | 3.352355448 | -0.776599867 | 0.856315182 | 0.906908908 | 0.36445497 | 1 | 0.776599867 |
| ENSMUSG00000039364 | Sectm1b | 58210 | protein_coding | 1409.422283 | -0.777139271 | 0.239487243 | 3.24501323 | 0.001174452 | 0.0072831 | 0.777139271 |
| ENSMUSG00000021953 | Tdh | 58865 | protein_coding | 1.033885078 | -0.777251667 | 1.964247332 | 0.39569949 | 0.692326732 | 1 | 0.777251667 |
| ENSMUSG00000069289 | Vmn1r203 | 171270 | protein_coding | 1.28616848 | -0.777640852 | 1.966053252 | 0.395533972 | 0.692448855 | 1 | 0.777640852 |
| ENSMUSG00000085845 | Gm13944 | 100503106 | lncRNA | 1.523016106 | -0.777688981 | 1.189106311 | 0.654011314 | 0.513104517 | 1 | 0.777688981 |
| ENSMUSG00000044349 | Snhg11 | 319317 | protein_coding | 2.92207971 | 0.777843895 | 1.465708117 | -0.530694949 | 0.595630187 | 1 | 0.777843895 |
| ENSMUSG00000046961 | Gpr156 | 239845 | protein_coding | 1.473209449 | -0.777991208 | 2.71245016 | 0.286822305 | 0.774248383 | 1 | 0.777991208 |
| ENSMUSG00000018599 | Mief2 | 237781 | protein_coding | 225.3873829 | 0.778155293 | 0.097475814 | -7.98306019 | 1.43E-15 | 5.42E-14 | 0.778155293 |
| ENSMUSG00000064109 | Hcst | 23900 | protein_coding | 11.26370107 | -0.778403859 | 0.418030122 | 1.862076005 | 0.062592378 | 0.175901555 | 0.778403859 |
| ENSMUSG00000029563 | Foxp2 | 114142 | protein_coding | 32.72702625 | -0.778780122 | 0.254935613 | 3.054811029 | 0.002252024 | 0.012461318 | 0.778780122 |
| ENSMUSG00000107875 | Gm54719 | 100503622 | lncRNA | 0.932683463 | -0.778992644 | 1.600153842 | 0.486823594 | 0.626383349 | 1 | 0.778992644 |

Supplementary Table 1

| ensembl_gene_id | external_gene_name | entrezgene_id | gene_biotype | baseMean | log2FoldChange | lfcSE | stat | pvalue | padj | absLog2FC |
|---|---|---|---|---|---|---|---|---|---|---|
| ENSMUSG00000094306 | Gm24924 | 115488991 | snRNA | 1.749886423 | 0.779017831 | 0.943040634 | -0.82607027 | 0.408764231 | 1 | 0.779017831 |
| ENSMUSG00000032221 | Mns1 | 17427 | protein_coding | 31.0927865 | 0.779031703 | 0.236321457 | -3.296491617 | 0.000979005 | 0.006255193 | 0.779031703 |
| ENSMUSG00000074916 | Chst14 | 72136 | protein_coding | 9.06137033 | 0.77921039 | 0.350801395 | -2.221229448 | 0.026335425 | 0.090888861 | 0.77921039 |
| ENSMUSG00000027870 | Hao2 | 56185 | protein_coding | 6630.787091 | -0.779232826 | 0.155973786 | 4.995921737 | 5.86E-07 | 8.50E-06 | 0.779232826 |
| ENSMUSG00000034883 | Lrr1 | 69706 | protein_coding | 16.95945544 | 0.779290882 | 0.35739753 | -2.180459621 | 0.029223409 | 0.098851981 | 0.779290882 |
| ENSMUSG00000021367 | Edn1 | 13614 | protein_coding | 184.7476636 | -0.779291268 | 0.379145774 | 2.055386925 | 0.039841647 | 0.125633944 | 0.779291268 |
| ENSMUSG00000022092 | Ppp3cc | 19057 | protein_coding | 53.17450084 | -0.779574594 | 0.169438831 | 4.600920514 | 4.21E-06 | 5.12E-05 | 0.779574594 |
| ENSMUSG00000047511 | Or2v2 | 258334 | protein_coding | 2.216747535 | 0.779642849 | 1.65148929 | -0.472084714 | 0.636866323 | 1 | 0.779642849 |
| ENSMUSG00000004558 | Ndrg2 | 29811 | protein_coding | 1375.628626 | -0.780119675 | 0.178698541 | 4.365562632 | 1.27E-05 | 0.000137975 | 0.780119675 |
| ENSMUSG00000027803 | Wwtr1 | 97064 | protein_coding | 10.19186426 | -0.780126584 | 0.388110843 | 2.010061297 | 0.044424702 | 0.136254601 | 0.780126584 |
| ENSMUSG00000027995 | Tlr2 | 24088 | protein_coding | 227.4663169 | -0.780169134 | 0.145244843 | 5.371406783 | 7.81E-08 | 1.31E-06 | 0.780169134 |
| ENSMUSG00000039438 | Ttc36 | 192653 | protein_coding | 4.484189511 | -0.780288156 | 0.539132345 | 1.44730355 | 0.147811925 | 0.32611438 | 0.780288156 |
| ENSMUSG00000032503 | Arpp21 | 74100 | protein_coding | 6.671532813 | -0.78068266 | 2.471020003 | 0.315935387 | 0.752051552 | 0.875590759 | 0.78068266 |
| ENSMUSG00000003813 | Rad23a | 19358 | protein_coding | 330.5583871 | -0.780733242 | 0.0813395 | 9.59845147 | 8.12E-22 | 4.54E-20 | 0.780733242 |
| ENSMUSG00000036912 | Piwil4 | 330890 | protein_coding | 86.54179918 | 0.780901194 | 0.184630692 | -4.229530787 | 2.34E-05 | 0.000237545 | 0.780901194 |
| ENSMUSG00000034023 | Fancd2 | 211651 | protein_coding | 128.1096091 | 0.781603972 | 0.179503411 | -4.354256943 | 1.34E-05 | 0.000144113 | 0.781603972 |
| ENSMUSG00000100094 | 1810008I18Rik | 100503969 | lncRNA | 20.35480377 | 0.781882129 | 0.326406489 | -2.395424585 | 0.016601129 | 0.063431523 | 0.781882129 |
| ENSMUSG00000066721 | Zfp575 | 101544 | protein_coding | 3.386435498 | -0.781952796 | 1.114721414 | 0.701478222 | 0.483004621 | 1 | 0.781952796 |
| ENSMUSG00000035836 | Ugt2b1 | 71773 | protein_coding | 3.072124748 | -0.782022428 | 0.900441867 | 0.868487413 | 0.38512756 | 1 | 0.782022428 |
| ENSMUSG00000024905 | Tesmin | 17771 | protein_coding | 5.353132072 | 0.782094265 | 0.569609976 | -1.373034703 | 0.169741559 | 0.359143126 | 0.782094265 |
| ENSMUSG00000032060 | Cryab | 12955 | protein_coding | 1.057537823 | -0.782106905 | 1.203227692 | 0.650007401 | 0.515687441 | 1 | 0.782106905 |
| ENSMUSG00000029672 | Fam3c | 27999 | protein_coding | 234.844579 | 0.782354572 | 0.15624817 | -5.007127916 | 5.52E-07 | 8.08E-06 | 0.782354572 |
| ENSMUSG00000020396 | Nefh | 380684 | protein_coding | 5.76925256 | -0.782462685 | 0.520979671 | 1.501906366 | 0.133121292 | 0.302752759 | 0.782462685 |
| ENSMUSG00000102785 | Gm2447 | 329639 | lncRNA | 5.262390408 | 0.782942564 | 0.593836464 | -1.31844811 | 0.187353685 | 0.383501006 | 0.782942564 |
| ENSMUSG00000067081 | Asb18 | 208372 | protein_coding | 1.314077999 | 0.782981531 | 2.34762086 | -0.333521287 | 0.738740823 | 1 | 0.782981531 |
| ENSMUSG00000085719 | 4933424L21Rik | 78762 | lncRNA | 9.922626978 | -0.782993526 | 0.352928247 | 2.218562929 | 0.026516474 | 0.091399092 | 0.782993526 |
| ENSMUSG00000089809 | Rasgef1b | 320292 | protein_coding | 501.914255 | -0.783195592 | 0.16095489 | 4.865932268 | 1.14E-06 | 1.55E-05 | 0.783195592 |
| ENSMUSG00000039096 | Rsad1 | 237926 | protein_coding | 57.40585639 | -0.783252417 | 0.231572149 | 3.382325645 | 0.000718749 | 0.004789552 | 0.783252417 |
| ENSMUSG00000025986 | Slc39a10 | 227059 | protein_coding | 164.593612 | 0.783428555 | 0.134590862 | -5.820815367 | 5.86E-09 | 1.17E-07 | 0.783428555 |
| ENSMUSG00000113434 | 3200001D21Rik | 71818 | lncRNA | 0.768482422 | -0.783450942 | 1.421030758 | 0.551325816 | 0.581410345 | 1 | 0.783450942 |
| ENSMUSG00000017716 | Birc5 | 11799 | protein_coding | 323.9370588 | 0.783701832 | 0.170165965 | -4.605514563 | 4.11E-06 | 5.03E-05 | 0.783701832 |
| ENSMUSG00000038147 | Cd84 | 12523 | protein_coding | 5.671423288 | 0.78385679 | 0.874540049 | -0.896307484 | 0.370088563 | 0.59267286 | 0.78385679 |
| ENSMUSG00000030306 | Tmtc1 | 387314 | protein_coding | 6.160166696 | 0.783941547 | 0.671098071 | -1.16814752 | 0.242747265 | 0.453835428 | 0.783941547 |
| ENSMUSG00000075122 | Cd80 | 12519 | protein_coding | 3.215071418 | -0.784101544 | 0.863796353 | 0.907738892 | 0.364016191 | 1 | 0.784101544 |
| ENSMUSG00000029346 | Srrd | 70118 | protein_coding | 90.28501651 | 0.78413154 | 0.131864665 | -5.946487175 | 2.74E-09 | 5.73E-08 | 0.78413154 |
| ENSMUSG00000047898 | Ccr4 | 12773 | protein_coding | 2.340950906 | 0.78448959 | 1.007865626 | -0.778367245 | 0.436352543 | 1 | 0.78448959 |
| ENSMUSG00000042641 | Rgsl1 | 240816 | protein_coding | 6.732943043 | -0.784618321 | 0.776698275 | 1.010197069 | 0.312400883 | 0.533456622 | 0.784618321 |
| ENSMUSG00000038179 | Slamf7 | 75345 | protein_coding | 5.782829618 | 0.785058983 | 0.695230342 | -1.12920702 | 0.258810513 | 0.472212663 | 0.785058983 |
| ENSMUSG00000038793 | Lefty1 | 13590 | protein_coding | 52.38452636 | -0.785383162 | 0.217313933 | 3.614048819 | 0.000301452 | 0.002287768 | 0.785383162 |

Supplementary Table 1

| ensembl_gene_id | external_gene_name | entrezgene_id | gene_biotype | baseMean | log2FoldChange | lfcSE | stat | pvalue | padj | absLog2FC |
|---|---|---|---|---|---|---|---|---|---|---|
| ENSMUSG00000041084 | Ostc | 66357 | protein_coding | 1224.91313 | 0.785823961 | 0.117206378 | -6.70461776 | 2.02E-11 | 5.48E-10 | 0.785823961 |
| ENSMUSG00000022322 | Shcbp1 | 20419 | protein_coding | 151.4677924 | 0.785869111 | 0.179598863 | -4.375690911 | 1.21E-05 | 0.000132912 | 0.785869111 |
| ENSMUSG00000031262 | Cenpi | 102920 | protein_coding | 84.9920606 | 0.785999121 | 0.181545676 | -4.329484119 | 1.49E-05 | 0.000159134 | 0.785999121 |
| ENSMUSG00000024883 | Rin1 | 225870 | protein_coding | 90.52885993 | -0.786167929 | 0.161377624 | 4.871604303 | 1.11E-06 | 1.51E-05 | 0.786167929 |
| ENSMUSG00000035211 | Xrra1 | 446101 | protein_coding | 4.196301237 | 0.786231082 | 0.834337989 | -0.942341224 | 0.346017969 | 1 | 0.786231082 |
| ENSMUSG00000049565 | Aknad1 | 329738 | protein_coding | 2.90642305 | 0.78643505 | 1.554855158 | -0.505793125 | 0.613001885 | 1 | 0.78643505 |
| ENSMUSG00000024785 | Rcl1 | 59028 | protein_coding | 189.9528577 | 0.786481933 | 0.143501243 | -5.480662881 | 4.24E-08 | 7.37E-07 | 0.786481933 |
| ENSMUSG00000000731 | Aire | 11634 | protein_coding | 3.399576755 | 0.786550303 | 1.049521089 | -0.749437349 | 0.453593647 | 1 | 0.786550303 |
| ENSMUSG00000000049 | Apoh | 11818 | protein_coding | 4.907260399 | -0.786712196 | 0.538477974 | 1.460992342 | 0.144017543 | 0.320612794 | 0.786712196 |
| ENSMUSG00000086429 | Gt(ROSA)26Sor | 14910 | lncRNA | 79.28454346 | 0.787152757 | 0.184204636 | -4.273251621 | 1.93E-05 | 0.000198956 | 0.787152757 |
| ENSMUSG00000027822 | Slc33a1 | 11416 | protein_coding | 602.2725848 | 0.787198327 | 0.117075557 | -6.723848678 | 1.77E-11 | 4.83E-10 | 0.787198327 |
| ENSMUSG00000098896 | Mir3473g | 102466258 | miRNA | 1.585258213 | -0.787378117 | 0.890509868 | 0.884187975 | 0.376594749 | 1 | 0.787378117 |
| ENSMUSG00000027134 | Lpcat4 | 99010 | protein_coding | 230.7852871 | 0.787475518 | 0.257070464 | -3.063267193 | 0.002189345 | 0.01218155 | 0.787475518 |
| ENSMUSG00000050556 | Kcnb1 | 16500 | protein_coding | 7.981042505 | -0.787476332 | 0.549762185 | 1.432394504 | 0.152030951 | 0.332667168 | 0.787476332 |
| ENSMUSG00000065474 | Mir141 | 387159 | miRNA | 0.777955732 | -0.787516947 | 1.531515361 | 0.514207671 | 0.607106805 | 1 | 0.787516947 |
| ENSMUSG00000090260 | 2810442N19Rik | 72753 | lncRNA | 1.622784427 | 0.787779122 | 0.903834802 | -0.871596358 | 0.383428618 | 1 | 0.787779122 |
| ENSMUSG00000055148 | Klf2 | 16598 | protein_coding | 435.5870511 | -0.788038073 | 0.179743639 | 4.384233451 | 1.16E-05 | 0.000128314 | 0.788038073 |
| ENSMUSG00000065505 | Mir148a | 387166 | miRNA | 0.698630019 | -0.788245168 | 1.369763493 | 0.57546078 | 0.564979708 | 1 | 0.788245168 |
| ENSMUSG00000077222 | Gm22270 | 115490266 | snoRNA | 17.05790229 | 0.788358311 | 0.370616655 | -2.127152949 | 0.033407377 | 0.109549131 | 0.788358311 |
| ENSMUSG00000069266 | H4c2 | 326620 | protein_coding | 432.5986085 | 0.788550487 | 0.193501889 | -4.075156535 | 4.60E-05 | 0.000434422 | 0.788550487 |
| ENSMUSG00000024936 | Kcnk7 | 16530 | protein_coding | 6.921433339 | -0.788808758 | 0.473255963 | 1.666769824 | 0.095560183 | 0.23910316 | 0.788808758 |
| ENSMUSG00000020486 | Septin4 | 18952 | protein_coding | 15.38208882 | 0.78933852 | 0.474916687 | -1.662056822 | 0.096501382 | 0.240801423 | 0.78933852 |
| ENSMUSG00000111259 | Or10ak7 | 258394 | protein_coding | 1.593641439 | -0.78967306 | 2.363897956 | 0.334055477 | 0.738337696 | 1 | 0.78967306 |
| ENSMUSG00000098790 | Mir6942 | 102465568 | miRNA | 0.564389452 | 0.789824306 | 1.299059789 | -0.607996886 | 0.543189539 | 1 | 0.789824306 |
| ENSMUSG00000024521 | Pmaip1 | 58801 | protein_coding | 254.1196987 | -0.790233511 | 0.187958274 | 4.204302866 | 2.62E-05 | 0.000261952 | 0.790233511 |
| ENSMUSG00000099050 | Mir7655 | 102466840 | miRNA | 0.557593203 | 0.790572641 | 1.637784004 | -0.482708733 | 0.629302562 | 1 | 0.790572641 |
| ENSMUSG00000099599 | Gm28548 | 102636046 | lncRNA | 0.833297795 | -0.790606981 | 1.447309421 | 0.546259818 | 0.584887353 | 1 | 0.790606981 |
| ENSMUSG00000115526 | 4930413F20Rik | 71637 | lncRNA | 6.131407444 | -0.790673755 | 0.490576837 | 1.61172256 | 0.107022326 | 0.260075004 | 0.790673755 |
| ENSMUSG00000020333 | Acsl6 | 216739 | protein_coding | 4.008057295 | 0.790782355 | 1.083226185 | -0.730025147 | 0.465374813 | 1 | 0.790782355 |
| ENSMUSG00000046351 | Zfp322a | 218100 | protein_coding | 287.6581332 | 0.790797446 | 0.096017953 | -8.235933177 | 1.78E-16 | 7.41E-15 | 0.790797446 |
| ENSMUSG00000061462 | Obscn | 380698 | protein_coding | 21.36229633 | -0.790919634 | 0.972748492 | 0.813077215 | 0.416173789 | 0.636419479 | 0.790919634 |
| ENSMUSG00000055775 | Myh8 | 17885 | protein_coding | 2.467284199 | -0.791007773 | 2.330930443 | 0.3393528 | 0.734343971 | 1 | 0.791007773 |
| ENSMUSG00000097360 | 9430065F17Rik | 77857 | lncRNA | 3.718283469 | -0.791010115 | 0.63473826 | 1.24619889 | 0.212691386 | 1 | 0.791010115 |
| ENSMUSG00000021684 | Pde8b | 218461 | protein_coding | 218.9961605 | -0.791094994 | 0.145426548 | 5.439825155 | 5.33E-08 | 9.10E-07 | 0.791094994 |
| ENSMUSG00000072623 | Zfp9 | 22750 | protein_coding | 88.42920703 | 0.791141113 | 0.180557195 | -4.381664832 | 1.18E-05 | 0.000129591 | 0.791141113 |
| ENSMUSG00000024986 | Hhex | 15242 | protein_coding | 9.485713269 | -0.791146785 | 0.436521558 | 1.812388804 | 0.069926142 | 0.190837928 | 0.791146785 |
| ENSMUSG00000042707 | Dnali1 | 75563 | protein_coding | 2.003719877 | -0.791343041 | 0.871433884 | 0.908093036 | 0.36382907 | 1 | 0.791343041 |
| ENSMUSG00000055629 | B4galnt4 | 330671 | protein_coding | 74.00112826 | 0.79146115 | 0.233075064 | -3.395734997 | 0.000684446 | 0.004600301 | 0.79146115 |
| ENSMUSG00000035164 | Zc3h12c | 244871 | protein_coding | 26.38892439 | 0.791537276 | 0.28528971 | -2.774503416 | 0.005528605 | 0.026425293 | 0.791537276 |

Supplementary Table 1

| ensembl_gene_id | external_gene_name | entrezgene_id | gene_biotype | baseMean | log2FoldChange | lfcSE | stat | pvalue | padj | absLog2FC |
|---|---|---|---|---|---|---|---|---|---|---|
| ENSMUSG00000038210 | Hoxa11 | 15396 | protein_coding | 33.32439088 | 0.791592521 | 0.286858222 | -2.759525297 | 0.005788541 | 0.027440166 | 0.791592521 |
| ENSMUSG00000037974 | Muc5ac | 17833 | protein_coding | 6.039470359 | 0.792015668 | 2.13264039 | -0.371377974 | 0.710356028 | 0.848979902 | 0.792015668 |
| ENSMUSG00000009292 | Trpm2 | 28240 | protein_coding | 8.280910736 | 0.792160215 | 1.035001341 | -0.765371197 | 0.444050539 | 0.661290522 | 0.792160215 |
| ENSMUSG00000024885 | Aldh3b1 | 67689 | protein_coding | 82.53551938 | -0.79242233 | 0.169503215 | 4.674969308 | 2.94E-06 | 3.67E-05 | 0.79242233 |
| ENSMUSG00000026043 | Col3a1 | 12825 | protein_coding | 35.37996377 | 0.792475145 | 0.477726852 | -1.658845721 | 0.097146884 | 0.241953298 | 0.792475145 |
| ENSMUSG00000036151 | Tm6sf2 | 107770 | protein_coding | 97.66243065 | -0.792491103 | 0.160709315 | 4.931208263 | 8.17E-07 | 1.15E-05 | 0.792491103 |
| ENSMUSG00000075256 | Cerkl | 228094 | protein_coding | 3.641984776 | -0.792504828 | 0.705883034 | 1.122714089 | 0.261558942 | 1 | 0.792504828 |
| ENSMUSG00000058935 | Tex48 | 75524 | protein_coding | 1.119114851 | 0.793401284 | 1.896605185 | -0.418327067 | 0.675708003 | 1 | 0.793401284 |
| ENSMUSG00000077649 | Gm24989 | 115487904 | snoRNA | 0.919737772 | 0.793613855 | 1.151814506 | -0.68901186 | 0.490815804 | 1 | 0.793613855 |
| ENSMUSG00000112116 | Gm46234 | 115487405 | processed_pseudogene | 2.807318897 | -0.793691152 | 0.739198969 | 1.073717882 | 0.282949144 | 1 | 0.793691152 |
| ENSMUSG00000063683 | Glyat | 107146 | protein_coding | 1.406861049 | -0.794286382 | 1.474173451 | 0.538801171 | 0.590024055 | 1 | 0.794286382 |
| ENSMUSG00000029108 | Pcdh7 | 54216 | protein_coding | 2.95821086 | -0.794458759 | 0.878291077 | 0.904550645 | 0.365703493 | 1 | 0.794458759 |
| ENSMUSG00000027978 | Prss12 | 19142 | protein_coding | 17.893595 | 0.794510139 | 0.578235179 | -1.374025945 | 0.169433628 | 0.358767397 | 0.794510139 |
| ENSMUSG00000033961 | Zfp446 | 269870 | protein_coding | 67.56370258 | 0.794564905 | 0.161688849 | -4.914160204 | 8.92E-07 | 1.24E-05 | 0.794564905 |
| ENSMUSG00000056411 | Gm12500 | 791415 | lncRNA | 11.80236121 | 0.794660878 | 0.40198355 | -1.976849247 | 0.048058673 | 0.14477215 | 0.794660878 |
| ENSMUSG00000028438 | Kif24 | 109242 | protein_coding | 99.85738525 | 0.79513207 | 0.140157801 | -5.673120318 | 1.40E-08 | 2.64E-07 | 0.79513207 |
| ENSMUSG00000030643 | Rab30 | 75985 | protein_coding | 7.383821387 | -0.795345059 | 0.444941166 | 1.787528601 | 0.073852094 | 0.19856618 | 0.795345059 |
| ENSMUSG00000000552 | Zfp385a | 29813 | protein_coding | 6.680385606 | 0.79534756 | 0.659804085 | -1.205429882 | 0.228037392 | 0.436620149 | 0.79534756 |
| ENSMUSG00000032437 | Stt3b | 68292 | protein_coding | 4888.16124 | 0.795394821 | 0.080038928 | -9.937599693 | 2.86E-23 | 1.71E-21 | 0.795394821 |
| ENSMUSG00000024379 | Tslp | 53603 | protein_coding | 1.119181256 | -0.795754367 | 1.51056475 | 0.526792623 | 0.598337606 | 1 | 0.795754367 |
| ENSMUSG00000023467 | Tulp2 | 56734 | protein_coding | 22.13331569 | 0.795796329 | 0.450383228 | -1.766931537 | 0.077239695 | 0.204938354 | 0.795796329 |
| ENSMUSG00000027762 | Sucnr1 | 84112 | protein_coding | 0.906918879 | -0.795847975 | 1.048287827 | 0.759188416 | 0.447739854 | 1 | 0.795847975 |
| ENSMUSG00000043903 | Zfp469 | 195209 | protein_coding | 5.849227309 | -0.796086782 | 0.78226202 | 1.017672802 | 0.308833477 | 0.529547921 | 0.796086782 |
| ENSMUSG00000042647 | Acad12 | 338350 | protein_coding | 119.8691015 | -0.796147124 | 0.141485356 | 5.627063781 | 1.83E-08 | 3.39E-07 | 0.796147124 |
| ENSMUSG00000011305 | Plin5 | 66968 | protein_coding | 6.169962176 | -0.796227926 | 0.588205822 | 1.353655295 | 0.175846377 | 0.367865096 | 0.796227926 |
| ENSMUSG00000020140 | Lgr5 | 14160 | protein_coding | 145.3544012 | 0.796231067 | 0.127304394 | -6.254545046 | 3.99E-10 | 9.23E-09 | 0.796231067 |
| ENSMUSG00000106519 | Gm38509 | 102638436 | lncRNA | 1.062861818 | -0.796645841 | 1.455851216 | 0.547202785 | 0.584239423 | 1 | 0.796645841 |
| ENSMUSG00000028019 | Pdgfc | 54635 | protein_coding | 47.77438322 | 0.79671124 | 0.262516838 | -3.034895759 | 0.002406188 | 0.013179368 | 0.79671124 |
| ENSMUSG00000040473 | Cfap69 | 207686 | protein_coding | 5.058347316 | -0.796738326 | 0.699013851 | 1.139803346 | 0.25436824 | 0.467048822 | 0.796738326 |
| ENSMUSG00000021569 | Trip13 | 69716 | protein_coding | 91.99914439 | 0.796745829 | 0.17771331 | -4.483321088 | 7.35E-06 | 8.50E-05 | 0.796745829 |
| ENSMUSG00000025170 | Rab40b | 217371 | protein_coding | 4.733202722 | -0.79681305 | 0.589684906 | 1.35125224 | 0.176614646 | 0.368771559 | 0.79681305 |
| ENSMUSG00000021252 | Erg28 | 58520 | protein_coding | 518.8931298 | 0.797165457 | 0.10644574 | -7.488937159 | 6.94E-14 | 2.32E-12 | 0.797165457 |
| ENSMUSG00000006398 | Cdc20 | 107995 | protein_coding | 593.4840608 | 0.797609987 | 0.139561321 | -5.715122085 | 1.10E-08 | 2.10E-07 | 0.797609987 |
| ENSMUSG00000097121 | D130020L05Rik | 319760 | lncRNA | 9.191067748 | -0.79781833 | 0.353569714 | 2.256466825 | 0.024041411 | 0.084706605 | 0.79781833 |
| ENSMUSG00000029177 | Cenpa | 12615 | protein_coding | 333.1267272 | 0.797910241 | 0.121332125 | -6.576248799 | 4.82E-11 | 1.27E-09 | 0.797910241 |
| ENSMUSG00000032783 | Troap | 78733 | protein_coding | 60.85643386 | 0.798095327 | 0.20355332 | -3.920817041 | 8.82E-05 | 0.00077231 | 0.798095327 |
| ENSMUSG00000101355 | H3c10 | 319152 | protein_coding | 297.9310675 | 0.798153293 | 0.212423375 | -3.757370366 | 0.000171708 | 0.001385242 | 0.798153293 |
| ENSMUSG00000053522 | Lgals7 | 16858 | protein_coding | 0.875718005 | 0.798506541 | 1.615212255 | -0.494366322 | 0.621047494 | 1 | 0.798506541 |
| ENSMUSG00000034156 | Tspoap1 | 207777 | protein_coding | 6.160133066 | 0.798509669 | 0.773817822 | -1.031909121 | 0.302114693 | 0.522421155 | 0.798509669 |

| ensembl_gene_id | external_gene_name | entrezgene_id | gene_biotype | baseMean | log2FoldChange | lfcSE | stat | pvalue | padj | absLog2FC |
|---|---|---|---|---|---|---|---|---|---|---|
| ENSMUSG00000027793 | Ccna1 | 12427 | protein_coding | 8.705915958 | 0.798676731 | 0.407287733 | -1.960964367 | 0.04988318 | 0.148614618 | 0.798676731 |
| ENSMUSG00000028362 | Tnfsf8 | 21949 | protein_coding | 2.833850481 | -0.798742015 | 0.812249703 | 0.98337003 | 0.325425355 | 1 | 0.798742015 |
| ENSMUSG00000026726 | Cubn | 65969 | protein_coding | 175.9985602 | -0.798777366 | 0.34454116 | 2.318380093 | 0.020428673 | 0.07460396 | 0.798777366 |
| ENSMUSG00000036908 | Unc93b1 | 54445 | protein_coding | 353.3453942 | -0.798877665 | 0.097283593 | 8.211843764 | 2.18E-16 | 8.97E-15 | 0.798877665 |
| ENSMUSG00000088464 | Mir1970 | 100316820 | miRNA | 0.906408079 | 0.799150847 | 1.755501494 | -0.455226526 | 0.648946279 | 1 | 0.799150847 |
| ENSMUSG00000025537 | Phkg1 | 18682 | protein_coding | 6.365441379 | 0.799155089 | 1.058596241 | -0.754919636 | 0.450297203 | 0.667103975 | 0.799155089 |
| ENSMUSG00000036777 | Anln | 68743 | protein_coding | 364.7767225 | 0.799567589 | 0.171616094 | -4.65904783 | 3.18E-06 | 3.95E-05 | 0.799567589 |
| ENSMUSG00000063583 | Or9s14 | 258484 | protein_coding | 2.378952621 | 0.79986321 | 2.734598104 | -0.292497537 | 0.769906243 | 1 | 0.79986321 |
| ENSMUSG00000092083 | Kcnb2 | 98741 | protein_coding | 10.56729762 | -0.800238989 | 0.600238463 | 1.333201783 | 0.182465594 | 0.377007156 | 0.800238989 |
| ENSMUSG00000041479 | Syt15 | 319508 | protein_coding | 1.878936266 | 0.800489971 | 1.44239257 | -0.55497372 | 0.578912636 | 1 | 0.800489971 |
| ENSMUSG00000020576 | Nbas | 71169 | protein_coding | 1021.431277 | 0.80055727 | 0.082098257 | -9.751209068 | 1.82E-22 | 1.04E-20 | 0.80055727 |
| ENSMUSG00000026500 | Cox20 | 66359 | protein_coding | 17.29135705 | 0.800782202 | 0.39153216 | -2.045252685 | 0.040829953 | 0.127992756 | 0.800782202 |
| ENSMUSG00000042607 | Asb4 | 65255 | protein_coding | 14.32072124 | -0.800936018 | 0.313007244 | 2.558841795 | 0.010502152 | 0.044310818 | 0.800936018 |
| ENSMUSG00000019989 | Enpp3 | 209558 | protein_coding | 25.25783311 | 0.801189222 | 0.305585965 | -2.621812887 | 0.008746344 | 0.038220989 | 0.801189222 |
| ENSMUSG00000034463 | Scara3 | 219151 | protein_coding | 3.157599495 | 0.801492164 | 0.925342241 | -0.866157545 | 0.386403771 | 1 | 0.801492164 |
| ENSMUSG00000026185 | Igfbp5 | 16011 | protein_coding | 3.785776911 | -0.801514977 | 1.114870794 | 0.718930823 | 0.472183544 | 1 | 0.801514977 |
| ENSMUSG00000062410 | Hsd3b3 | 15494 | protein_coding | 1743.86409 | -0.801792929 | 0.144832919 | 5.535985439 | 3.09E-08 | 5.49E-07 | 0.801792929 |
| ENSMUSG00000034792 | Gna15 | 14676 | protein_coding | 2.095354787 | 0.802036857 | 0.982200733 | -0.816571226 | 0.414173505 | 1 | 0.802036857 |
| ENSMUSG00000034829 | Nxnl1 | 234404 | protein_coding | 0.995384592 | 0.80254352 | 1.778880392 | -0.451150917 | 0.651880782 | 1 | 0.80254352 |
| ENSMUSG00000044860 | Gm1123 | 382097 | protein_coding | 1054.063647 | -0.802903998 | 0.34847525 | 2.304048844 | 0.021219902 | 0.07689828 | 0.802903998 |
| ENSMUSG00000037362 | Ccn3 | 18133 | protein_coding | 1922.651078 | -0.803678623 | 0.435729677 | 1.844443163 | 0.065118582 | 0.181269925 | 0.803678623 |
| ENSMUSG00000092049 | Vmn2r4 | 637053 | protein_coding | 2.703745603 | 0.803971406 | 3.098193565 | -0.259496829 | 0.795251928 | 1 | 0.803971406 |
| ENSMUSG00000056498 | Tmem154 | 320782 | protein_coding | 9.560591606 | 0.80414709 | 0.414299572 | -1.940979777 | 0.05226073 | 0.154444113 | 0.80414709 |
| ENSMUSG00000030556 | Lrrc28 | 67867 | protein_coding | 103.8905676 | 0.804404788 | 0.141318821 | -5.692127783 | 1.25E-08 | 2.38E-07 | 0.804404788 |
| ENSMUSG00000045031 | Cetn4 | 207175 | protein_coding | 3.797766521 | 0.804612862 | 0.747372479 | -1.076588829 | 0.281663997 | 1 | 0.804612862 |
| ENSMUSG00000052415 | Tchh | 99681 | protein_coding | 16.86703708 | 0.804877404 | 0.292736164 | -2.749497678 | 0.005968669 | 0.028073652 | 0.804877404 |
| ENSMUSG00000059659 | Gm10069 | 791299 | lncRNA | 9.724633139 | 0.804937656 | 0.463087601 | -1.738197382 | 0.08217604 | 0.214114094 | 0.804937656 |
| ENSMUSG00000025405 | Inhbc | 16325 | protein_coding | 0.810973008 | -0.804998899 | 1.302864875 | 0.617868295 | 0.53666216 | 1 | 0.804998899 |
| ENSMUSG00000073409 | H2-Q6 | 110557 | protein_coding | 30.54011091 | 0.805305401 | 0.322313881 | -2.498512935 | 0.012471559 | 0.05055359 | 0.805305401 |
| ENSMUSG00000062074 | Ccn6 | 327743 | protein_coding | 1.193280374 | 0.805433347 | 1.27728578 | -0.630581941 | 0.52831391 | 1 | 0.805433347 |
| ENSMUSG00000067341 | H2-Eb2 | 381091 | protein_coding | 2.637587753 | -0.805621535 | 1.026015078 | 0.785194635 | 0.43233946 | 1 | 0.805621535 |
| ENSMUSG00000028528 | Dnajc6 | 72685 | protein_coding | 6.159581735 | 0.805730111 | 0.736988187 | -1.09327412 | 0.274273462 | 0.490623547 | 0.805730111 |
| ENSMUSG00000074607 | Tox2 | 269389 | protein_coding | 2.021577112 | 0.805782095 | 1.366609368 | -0.589621375 | 0.555444518 | 1 | 0.805782095 |
| ENSMUSG00000030075 | Cntn3 | 18488 | protein_coding | 23.84261652 | 0.805965459 | 0.320356468 | -2.515839506 | 0.011874924 | 0.048696921 | 0.805965459 |
| ENSMUSG00000031661 | Nkd1 | 93960 | protein_coding | 12.3261097 | -0.806444547 | 0.456936463 | 1.764894273 | 0.077581536 | 0.205647109 | 0.806444547 |
| ENSMUSG00000045752 | Tssc4 | 56844 | protein_coding | 285.1346503 | 0.806488763 | 0.107639456 | -7.49250126 | 6.76E-14 | 2.26E-12 | 0.806488763 |
| ENSMUSG00000042788 | Fam166b | 329831 | protein_coding | 1.137139471 | -0.806588798 | 1.606848523 | 0.501969405 | 0.615689042 | 1 | 0.806588798 |
| ENSMUSG00000038811 | Gngt2 | 14710 | protein_coding | 30.83603385 | -0.806668866 | 0.27020366 | 2.985410578 | 0.002831979 | 0.015169415 | 0.806668866 |
| ENSMUSG00000001025 | S100a6 | 20200 | protein_coding | 7206.257691 | -0.806963128 | 0.144103066 | 5.599902578 | 2.14E-08 | 3.92E-07 | 0.806963128 |

| ensembl_gene_id | external_gene_name | entrezgene_id | gene_biotype | baseMean | log2FoldChange | lfcSE | stat | pvalue | padj | absLog2FC |
|---|---|---|---|---|---|---|---|---|---|---|
| ENSMUSG00000039563 | 2210406O10Rik | 76710 | lncRNA | 5.618896014 | 0.807557196 | 0.587248159 | -1.375154921 | 0.16908342 | 0.358301502 | 0.807557196 |
| ENSMUSG00000001155 | Ftcd | 14317 | protein_coding | 3.166969344 | 0.807758637 | 3.650113684 | -0.221296843 | 0.824861308 | 1 | 0.807758637 |
| ENSMUSG00000027469 | Tpx2 | 72119 | protein_coding | 479.4230331 | 0.80776252 | 0.157509607 | -5.128338097 | 2.92E-07 | 4.48E-06 | 0.80776252 |
| ENSMUSG00000043953 | Ccrl2 | 54199 | protein_coding | 12.95127113 | -0.807817187 | 0.300721855 | 2.686260325 | 0.007225677 | 0.032763544 | 0.807817187 |
| ENSMUSG00000105096 | Gbp10 | 626578 | protein_coding | 0.939074118 | 0.807870341 | 1.673735051 | -0.482675165 | 0.6293264 | 1 | 0.807870341 |
| ENSMUSG00000039313 | Minar1 | 209743 | protein_coding | 4.346009945 | -0.808148857 | 0.794830273 | 1.016756513 | 0.309269274 | 1 | 0.808148857 |
| ENSMUSG00000048327 | Ckap2l | 70466 | protein_coding | 203.8442515 | 0.808237041 | 0.168210752 | -4.804907134 | 1.55E-06 | 2.05E-05 | 0.808237041 |
| ENSMUSG00000052688 | Rab7b | 226421 | protein_coding | 2.158186525 | 0.80869267 | 1.784469713 | -0.453183747 | 0.650416435 | 1 | 0.80869267 |
| ENSMUSG00000042156 | Dzip1 | 66573 | protein_coding | 8.89916678 | 0.808872666 | 0.494323824 | -1.636321431 | 0.101772333 | 0.250788441 | 0.808872666 |
| ENSMUSG00000028989 | Angptl7 | 654812 | protein_coding | 4.141959174 | -0.808886853 | 0.564727815 | 1.432348171 | 0.152044203 | 1 | 0.808886853 |
| ENSMUSG00000065208 | Gm24616 | 115488691 | snoRNA | 12.83668375 | 0.809216592 | 0.439306608 | -1.842031458 | 0.065470548 | 0.181863858 | 0.809216592 |
| ENSMUSG00000093221 | Mir3105 | 100526562 | miRNA | 1.036170765 | -0.809404852 | 1.372281844 | 0.589824062 | 0.555308609 | 1 | 0.809404852 |
| ENSMUSG00000023033 | Scn8a | 20273 | protein_coding | 7.603286458 | 0.809520025 | 1.020917562 | -0.792933784 | 0.427816403 | 0.647057325 | 0.809520025 |
| ENSMUSG00000028626 | Col9a2 | 12840 | protein_coding | 4.533143605 | 0.81004282 | 1.410030938 | -0.574485849 | 0.565639074 | 0.752299084 | 0.81004282 |
| ENSMUSG00000067288 | Rps28 | 54127 | protein_coding | 2361.15818 | -0.810081391 | 0.195742171 | 4.138512335 | 3.50E-05 | 0.000341177 | 0.810081391 |
| ENSMUSG00000051678 | Pcdhb6 | 93877 | protein_coding | 1.010581475 | 0.810143946 | 2.016660172 | -0.401725565 | 0.687886008 | 1 | 0.810143946 |
| ENSMUSG00000047496 | Rnf152 | 320311 | protein_coding | 147.6260935 | -0.810218324 | 0.26082169 | 3.106407003 | 0.001893759 | 0.010862636 | 0.810218324 |
| ENSMUSG00000063021 | H2ac15 | 319169 | protein_coding | 458.5741112 | 0.810684394 | 0.181527998 | -4.465891774 | 7.97E-06 | 9.15E-05 | 0.810684394 |
| ENSMUSG00000042213 | Zfand4 | 67492 | protein_coding | 8.995264689 | 0.810873923 | 0.364122004 | -2.226929203 | 0.025952006 | 0.089941537 | 0.810873923 |
| ENSMUSG00000051048 | P4ha3 | 320452 | protein_coding | 1.869271865 | -0.811239627 | 1.659878194 | 0.488734433 | 0.625029723 | 1 | 0.811239627 |
| ENSMUSG00000060314 | Zfp941 | 407812 | protein_coding | 8.956359564 | -0.811410641 | 0.349437427 | 2.322048468 | 0.020230326 | 0.074043352 | 0.811410641 |
| ENSMUSG00000108387 | Gm31510 | 102633760 | lncRNA | 1.560394462 | -0.811424788 | 1.016739756 | 0.798065369 | 0.424832557 | 1 | 0.811424788 |
| ENSMUSG00000032332 | Col12a1 | 12816 | protein_coding | 4.718114654 | 0.81143091 | 1.159400302 | -0.699871225 | 0.484007729 | 0.694394109 | 0.81143091 |
| ENSMUSG00000055489 | Ano5 | 233246 | protein_coding | 1.39033625 | 0.811558457 | 2.280114588 | -0.355928803 | 0.721893885 | 1 | 0.811558457 |
| ENSMUSG00000038155 | Gstp2 | 14869 | protein_coding | 269.4735942 | -0.811597913 | 0.152949204 | 5.306323215 | 1.12E-07 | 1.83E-06 | 0.811597913 |
| ENSMUSG00000069268 | H2bc7 | 319180 | protein_coding | 236.067513 | 0.811753814 | 0.197383572 | -4.112570291 | 3.91E-05 | 0.000377208 | 0.811753814 |
| ENSMUSG00000115426 | Gm19510 | 100503021 | lncRNA | 4.115912671 | -0.812006855 | 0.943462953 | 0.860666391 | 0.389421809 | 1 | 0.812006855 |
| ENSMUSG00000027434 | Nkx2-2 | 18088 | protein_coding | 9.816591688 | -0.812151927 | 0.410308758 | 1.97936776 | 0.047774616 | 0.144047837 | 0.812151927 |
| ENSMUSG00000071229 | Timm8a2 | 223262 | protein_coding | 1.203574468 | -0.812283377 | 1.58189884 | 0.513486296 | 0.607611194 | 1 | 0.812283377 |
| ENSMUSG00000021876 | Rnase4 | 58809 | protein_coding | 1187.383618 | -0.812448532 | 0.111169244 | 7.308213155 | 2.71E-13 | 8.63E-12 | 0.812448532 |
| ENSMUSG00000022091 | Sorbs3 | 20410 | protein_coding | 26.93343357 | -0.813059834 | 0.26227274 | 3.100054671 | 0.001934849 | 0.011044625 | 0.813059834 |
| ENSMUSG00000022464 | Slc38a4 | 69354 | protein_coding | 2.873573907 | 0.813506162 | 0.712132458 | -1.142352314 | 0.253307604 | 1 | 0.813506162 |
| ENSMUSG00000041014 | Nrg3 | 18183 | protein_coding | 2.268478313 | 0.81376651 | 2.14454008 | -0.379459688 | 0.704346534 | 1 | 0.81376651 |
| ENSMUSG00000019990 | Pde7b | 29863 | protein_coding | 20.21932846 | 0.81384172 | 0.290630397 | -2.800263598 | 0.005106089 | 0.024790788 | 0.81384172 |
| ENSMUSG00000025176 | Hoga1 | 67432 | protein_coding | 38.09262461 | 0.814151831 | 0.182044507 | -4.472268048 | 7.74E-06 | 8.92E-05 | 0.814151831 |
| ENSMUSG00000077611 | Gm23946 | 115488402 | snoRNA | 22.75662737 | 0.814534465 | 0.397488817 | -2.049200958 | 0.040442468 | 0.127019194 | 0.814534465 |
| ENSMUSG00000042388 | Dlgap3 | 242667 | protein_coding | 5.406559448 | -0.814930409 | 0.691091831 | 1.179192652 | 0.238321472 | 0.448589299 | 0.814930409 |
| ENSMUSG00000028150 | Rorc | 19885 | protein_coding | 619.868161 | 0.814993003 | 0.192515737 | -4.233383816 | 2.30E-05 | 0.000233941 | 0.814993003 |
| ENSMUSG00000060090 | Rp2 | 19889 | protein_coding | 962.164712 | 0.81513182 | 0.082994883 | -9.821470746 | 9.10E-23 | 5.37E-21 | 0.81513182 |

| ensembl_gene_id | external_gene_name | entrezgene_id | gene_biotype | baseMean | log2FoldChange | lfcSE | stat | pvalue | padj | absLog2FC |
|---|---|---|---|---|---|---|---|---|---|---|
| ENSMUSG00000075289 | Carns1 | 107239 | protein_coding | 6.320387753 | -0.81518033 | 0.444732076 | 1.832969498 | 0.066807107 | 0.184552119 | 0.81518033 |
| ENSMUSG00000022376 | Adcy8 | 11514 | protein_coding | 11.0963666 | 0.815576031 | 0.475845418 | -1.713951632 | 0.086537609 | 0.222529131 | 0.815576031 |
| ENSMUSG00000044067 | Gpr22 | 73010 | protein_coding | 10.30356145 | -0.815791147 | 0.426541002 | 1.912573804 | 0.055802636 | 0.161727752 | 0.815791147 |
| ENSMUSG00000071347 | C1qtnf9 | 239126 | protein_coding | 2.909102921 | -0.81580376 | 0.881269054 | 0.925714748 | 0.354594242 | 1 | 0.81580376 |
| ENSMUSG00000004698 | Hdac9 | 79221 | protein_coding | 25.12745919 | -0.816184413 | 0.320376591 | 2.547578182 | 0.010847354 | 0.045361664 | 0.816184413 |
| ENSMUSG00000028262 | Clca3a2 | 80797 | protein_coding | 107.2189887 | 0.816244025 | 0.196868197 | -4.146144658 | 3.38E-05 | 0.000331378 | 0.816244025 |
| ENSMUSG00000037759 | Ptger2 | 19217 | protein_coding | 2.754444709 | 0.816401256 | 1.778162129 | -0.459126444 | 0.646143368 | 1 | 0.816401256 |
| ENSMUSG00000050150 | Slc9b1 | 74446 | protein_coding | 3.423086411 | -0.816758519 | 0.862597776 | 0.94685906 | 0.343710602 | 1 | 0.816758519 |
| ENSMUSG00000104870 | 4930448I18Rik | 73965 | lncRNA | 0.634329308 | -0.816877869 | 1.74121606 | 0.469142163 | 0.638968023 | 1 | 0.816877869 |
| ENSMUSG00000028635 | Edn2 | 13615 | protein_coding | 203.492268 | -0.817004158 | 0.167678538 | 4.872443229 | 1.10E-06 | 1.50E-05 | 0.817004158 |
| ENSMUSG00000040219 | Ttc12 | 235330 | protein_coding | 2.192753691 | 0.817411464 | 1.702206921 | -0.480206874 | 0.6310803 | 1 | 0.817411464 |
| ENSMUSG00000015709 | Arnt2 | 11864 | protein_coding | 42.85276453 | 0.817537702 | 0.308204994 | -2.652577731 | 0.007987973 | 0.035574288 | 0.817537702 |
| ENSMUSG00000034037 | Fgd5 | 232237 | protein_coding | 4.088386543 | 0.817557059 | 0.93404448 | -0.875287073 | 0.381417726 | 1 | 0.817557059 |
| ENSMUSG00000097756 | A730056A06Rik | 319783 | lncRNA | 1.23118541 | -0.818023029 | 0.974587406 | 0.83935317 | 0.401271155 | 1 | 0.818023029 |
| ENSMUSG00000116165 | Pdxp | 57028 | protein_coding | 5.427494701 | 0.818321475 | 0.49197496 | -1.663339686 | 0.096244459 | 0.240265168 | 0.818321475 |
| ENSMUSG00000058672 | Tubb2a | 22151 | protein_coding | 164.6785891 | -0.818704134 | 0.26079709 | 3.139237999 | 0.001693878 | 0.009870396 | 0.818704134 |
| ENSMUSG00000055745 | Rtl6 | 223732 | protein_coding | 19.38419741 | 0.819193917 | 0.287924414 | -2.845170043 | 0.004438772 | 0.022045679 | 0.819193917 |
| ENSMUSG00000074619 | Fundc2b | 73297 | protein_coding | 1.409904985 | -0.819223261 | 1.057677292 | 0.774549352 | 0.438605995 | 1 | 0.819223261 |
| ENSMUSG00000035692 | Isg15 | 100038882 | protein_coding | 133.168886 | 0.81941044 | 0.453730594 | -1.805940467 | 0.070927639 | 0.192815811 | 0.81941044 |
| ENSMUSG00000055994 | Nod2 | 257632 | protein_coding | 44.44103282 | -0.819480454 | 0.168102011 | 4.874899779 | 1.09E-06 | 1.49E-05 | 0.819480454 |
| ENSMUSG00000031825 | Crispld2 | 78892 | protein_coding | 2.673111387 | 0.819694251 | 2.152340553 | -0.380838548 | 0.703323054 | 1 | 0.819694251 |
| ENSMUSG00000045312 | Lhfpl2 | 218454 | protein_coding | 677.5260724 | -0.819858507 | 0.284918924 | 2.877515104 | 0.004008207 | 0.02019882 | 0.819858507 |
| ENSMUSG00000091387 | Gcnt4 | 218476 | protein_coding | 91.09705272 | -0.820072591 | 0.151246739 | 5.422084433 | 5.89E-08 | 9.96E-07 | 0.820072591 |
| ENSMUSG00000031671 | Setd6 | 66083 | protein_coding | 73.00580257 | -0.820344816 | 0.12591121 | 6.515264317 | 7.26E-11 | 1.85E-09 | 0.820344816 |
| ENSMUSG00000048416 | Mlf1 | 17349 | protein_coding | 2.540526191 | 0.820599083 | 0.761640285 | -1.077410293 | 0.281297008 | 1 | 0.820599083 |
| ENSMUSG00000095385 | D630033O11Rik | 235302 | lncRNA | 20.20825997 | -0.820620043 | 0.380977092 | 2.15398789 | 0.031241122 | 0.10409925 | 0.820620043 |
| ENSMUSG00000097693 | 4930471M09Rik | 75787 | transcribed_processed_pseudogene | 3.237348436 | 0.821056796 | 1.180781688 | -0.695350211 | 0.486835855 | 1 | 0.821056796 |
| ENSMUSG00000042500 | Ago4 | 76850 | protein_coding | 76.99962059 | -0.821601984 | 0.167597737 | 4.902226004 | 9.48E-07 | 1.32E-05 | 0.821601984 |
| ENSMUSG00000075549 | Gm6878 | 628416 | lncRNA | 0.802981568 | 0.821627741 | 1.269743958 | -0.647081434 | 0.51757924 | 1 | 0.821627741 |
| ENSMUSG00000031304 | Il2rg | 16186 | protein_coding | 7.160349965 | 0.821649756 | 0.590110678 | -1.392365511 | 0.163811743 | 0.350004803 | 0.821649756 |
| ENSMUSG00000035365 | Parpbp | 75317 | protein_coding | 92.84846355 | 0.821651173 | 0.165452149 | -4.966095505 | 6.83E-07 | 9.77E-06 | 0.821651173 |
| ENSMUSG00000015363 | Trabd | 67976 | protein_coding | 1534.05495 | 0.821830893 | 0.071403459 | -11.50967894 | 1.18E-30 | 9.45E-29 | 0.821830893 |
| ENSMUSG00000029757 | Dync1i1 | 13426 | protein_coding | 47.94950442 | 0.822266764 | 0.179805077 | -4.573100934 | 4.81E-06 | 5.75E-05 | 0.822266764 |
| ENSMUSG00000040086 | Tnni3k | 435766 | protein_coding | 1.233104172 | 0.822403492 | 1.854729145 | -0.443408944 | 0.657469968 | 1 | 0.822403492 |
| ENSMUSG00000027490 | E2f1 | 13555 | protein_coding | 75.26688246 | 0.822778533 | 0.205361014 | -4.006498205 | 6.16E-05 | 0.00056319 | 0.822778533 |
| ENSMUSG00000032238 | Rora | 19883 | protein_coding | 12.71383875 | -0.823184713 | 0.475257122 | 1.732082854 | 0.083258811 | 0.216212948 | 0.823184713 |
| ENSMUSG00000033530 | Ttc7b | 104718 | protein_coding | 53.40634262 | 0.823325488 | 0.179699244 | -4.581685867 | 4.61E-06 | 5.56E-05 | 0.823325488 |
| ENSMUSG00000069300 | H2bc11 | 319183 | protein_coding | 371.0803477 | 0.82338366 | 0.238316625 | -3.454998822 | 0.000550295 | 0.003837987 | 0.82338366 |
| ENSMUSG00000108436 | Gm44851 | 115486493 | lncRNA | 1.420523148 | -0.823506544 | 1.612376311 | 0.51074091 | 0.609532491 | 1 | 0.823506544 |

| ensembl_gene_id | external_gene_name | entrezgene_id | gene_biotype | baseMean | log2FoldChange | lfcSE | stat | pvalue | padj | absLog2FC |
|---|---|---|---|---|---|---|---|---|---|---|
| ENSMUSG00000062995 | Ica1 | 15893 | protein_coding | 589.5023067 | 0.823699402 | 0.096053391 | -8.575432854 | 9.87E-18 | 4.45E-16 | 0.823699402 |
| ENSMUSG00000013707 | Tnfaip8l2 | 69769 | protein_coding | 3.3468647 | 0.823804893 | 0.84447328 | -0.975525114 | 0.32929985 | 1 | 0.823804893 |
| ENSMUSG00000030474 | Siglece | 83382 | protein_coding | 3.630155314 | 0.823851366 | 1.220558653 | -0.674978924 | 0.499689156 | 1 | 0.823851366 |
| ENSMUSG00000062727 | H2bc12 | 319184 | protein_coding | 477.5647243 | 0.823893867 | 0.200085618 | -4.117706589 | 3.83E-05 | 0.000369764 | 0.823893867 |
| ENSMUSG00000039977 | Deup1 | 234964 | protein_coding | 1.683828483 | -0.823905113 | 1.416632715 | 0.581594018 | 0.560840173 | 1 | 0.823905113 |
| ENSMUSG00000050592 | Fam78a | 241303 | protein_coding | 44.67359107 | -0.823949326 | 0.287903319 | 2.861895894 | 0.004211152 | 0.02103538 | 0.823949326 |
| ENSMUSG00000027716 | Trpc3 | 22065 | protein_coding | 2.624456884 | 0.824005884 | 1.853007638 | -0.444685638 | 0.65654695 | 1 | 0.824005884 |
| ENSMUSG00000039704 | Lmbrd2 | 320506 | protein_coding | 1024.845087 | 0.82400734 | 0.092648053 | -8.893952001 | 5.90E-19 | 2.85E-17 | 0.82400734 |
| ENSMUSG00000062232 | Rapgef2 | 76089 | protein_coding | 606.1887291 | 0.824427974 | 0.10312879 | -7.994159304 | 1.30E-15 | 5.01E-14 | 0.824427974 |
| ENSMUSG00000063406 | Tmed5 | 73130 | protein_coding | 491.5779843 | 0.82449592 | 0.104925048 | -7.857951357 | 3.90E-15 | 1.44E-13 | 0.82449592 |
| ENSMUSG00000039187 | Fanci | 208836 | protein_coding | 75.03646047 | 0.824528163 | 0.22444383 | -3.673650382 | 0.00023911 | 0.001853831 | 0.824528163 |
| ENSMUSG00000087903 | Gm22039 | 115486273 | snoRNA | 0.567056251 | -0.82491739 | 1.466142426 | 0.562644785 | 0.573676789 | 1 | 0.82491739 |
| ENSMUSG00000033031 | Cip2a | 224171 | protein_coding | 337.4175032 | 0.824935044 | 0.186217975 | -4.429943164 | 9.43E-06 | 0.000106469 | 0.824935044 |
| ENSMUSG00000053117 | E330013P04Rik | 107376 | lncRNA | 1.552983027 | 0.825004905 | 1.564474355 | -0.527336803 | 0.597959721 | 1 | 0.825004905 |
| ENSMUSG00000049580 | Tsku | 244152 | protein_coding | 638.4472535 | -0.825561487 | 0.235109686 | 3.511388666 | 0.000445772 | 0.003220245 | 0.825561487 |
| ENSMUSG00000040978 | Gm11992 | 626870 | protein_coding | 18.16829737 | 0.825601471 | 0.314375027 | -2.626167477 | 0.008635232 | 0.037845599 | 0.825601471 |
| ENSMUSG00000053030 | Spink2 | 69982 | protein_coding | 0.906821977 | -0.825759144 | 2.19237159 | 0.376651088 | 0.706432917 | 1 | 0.825759144 |
| ENSMUSG00000044276 | 4933427E11Rik | 66769 | lncRNA | 1.591464973 | 0.825864539 | 1.152513591 | -0.716576833 | 0.473635244 | 1 | 0.825864539 |
| ENSMUSG00000005233 | Spc25 | 66442 | protein_coding | 160.6373034 | 0.825928893 | 0.186619247 | -4.42574337 | 9.61E-06 | 0.000108339 | 0.825928893 |
| ENSMUSG00000035594 | Chrna5 | 110835 | protein_coding | 1.585535967 | -0.826074686 | 0.957699535 | 0.862561436 | 0.388378641 | 1 | 0.826074686 |
| ENSMUSG00000052921 | Arhgef15 | 442801 | protein_coding | 5.186223843 | 0.826076915 | 1.362515831 | -0.606287939 | 0.544323566 | 0.736796542 | 0.826076915 |
| ENSMUSG00000039234 | Sec24d | 69608 | protein_coding | 1087.561682 | 0.82611861 | 0.119101921 | -6.936232433 | 4.03E-12 | 1.16E-10 | 0.82611861 |
| ENSMUSG00000000982 | Ccl3 | 20302 | protein_coding | 3.518700217 | 0.826193839 | 0.953550549 | -0.866439477 | 0.386249203 | 1 | 0.826193839 |
| ENSMUSG00000031422 | Morf4l2 | 56397 | protein_coding | 2154.517806 | 0.826444388 | 0.093658487 | -8.824020241 | 1.10E-18 | 5.20E-17 | 0.826444388 |
| ENSMUSG00000073399 | Trim40 | 195359 | protein_coding | 226.2988507 | 0.826453357 | 0.253369197 | -3.261854118 | 0.001106861 | 0.006936888 | 0.826453357 |
| ENSMUSG00000059136 | Or13a25 | 258963 | protein_coding | 2.271078468 | -0.826950243 | 1.924124957 | 0.429779906 | 0.667355751 | 1 | 0.826950243 |
| ENSMUSG00000027559 | Car3 | 12350 | protein_coding | 10.54240956 | 0.827418706 | 0.376532833 | -2.197467615 | 0.027987067 | 0.095392863 | 0.827418706 |
| ENSMUSG00000035337 | Uchl4 | 93841 | protein_coding | 12.45606218 | 0.827439991 | 0.396594133 | -2.08636468 | 0.036945602 | 0.118331976 | 0.827439991 |
| ENSMUSG00000030351 | Tspan11 | 68498 | protein_coding | 1.390949004 | 0.827598697 | 1.881865058 | -0.439775792 | 0.660099502 | 1 | 0.827598697 |
| ENSMUSG00000059187 | Tafa1 | 320265 | protein_coding | 4.452405898 | -0.827809935 | 0.572754097 | 1.445314733 | 0.148369501 | 0.326976379 | 0.827809935 |
| ENSMUSG00000080645 | Mir1198 | 100316807 | miRNA | 0.75582156 | 0.82796429 | 1.612418919 | -0.513492046 | 0.607607173 | 1 | 0.82796429 |
| ENSMUSG00000032105 | Pdzd3 | 170761 | protein_coding | 242.0550632 | -0.828028067 | 0.118975194 | 6.959669833 | 3.41E-12 | 9.92E-11 | 0.828028067 |
| ENSMUSG00000010505 | Myt1 | 17932 | protein_coding | 17.77281974 | -0.82811733 | 0.376826487 | 2.197609137 | 0.027976972 | 0.095392863 | 0.82811733 |
| ENSMUSG00000023046 | Igfbp6 | 16012 | protein_coding | 1.825661631 | 0.828263467 | 1.269537588 | -0.652413505 | 0.514134454 | 1 | 0.828263467 |
| ENSMUSG00000043897 | Vmn2r2 | 100125586 | protein_coding | 1.071849107 | -0.828300908 | 2.207004382 | 0.375305511 | 0.707433268 | 1 | 0.828300908 |
| ENSMUSG00000025464 | Paox | 212503 | protein_coding | 79.039433 | -0.828651482 | 0.181926152 | 4.554878312 | 5.24E-06 | 6.25E-05 | 0.828651482 |
| ENSMUSG00000056508 | 1700001K19Rik | 66323 | protein_coding | 4.544709364 | 0.828732565 | 0.665772313 | -1.244768742 | 0.213216769 | 0.417979787 | 0.828732565 |
| ENSMUSG00000041406 | BC055324 | 381306 | protein_coding | 60.78556636 | 0.828738254 | 0.220811866 | -3.753141837 | 0.000174632 | 0.001407454 | 0.828738254 |
| ENSMUSG00000065528 | Mir320 | 723838 | miRNA | 3.035914841 | 0.828807476 | 1.115529167 | -0.742972484 | 0.45749834 | 1 | 0.828807476 |

| ensembl_gene_id | external_gene_name | entrezgene_id | gene_biotype | baseMean | log2FoldChange | lfcSE | stat | pvalue | padj | absLog2FC |
|---|---|---|---|---|---|---|---|---|---|---|
| ENSMUSG00000032925 | Itgbl1 | 223272 | protein_coding | 1.015849592 | -0.828869955 | 1.639850158 | 0.505454691 | 0.613239514 | 1 | 0.828869955 |
| ENSMUSG00000037846 | Rtkn2 | 170799 | protein_coding | 7.352238622 | 0.829087997 | 0.487797678 | -1.69965548 | 0.089195748 | 0.227241374 | 0.829087997 |
| ENSMUSG00000037313 | Tacc3 | 21335 | protein_coding | 336.3982531 | 0.829187605 | 0.150751412 | -5.500363755 | 3.79E-08 | 6.64E-07 | 0.829187605 |
| ENSMUSG00000065952 | Rps23rg1 | 546049 | protein_coding | 18.99579822 | 0.829323784 | 0.290409587 | -2.85570388 | 0.004294154 | 0.021404668 | 0.829323784 |
| ENSMUSG00000079277 | Hoxd3 | 15434 | protein_coding | 2.96914499 | 0.830018519 | 0.819682698 | -1.012609539 | 0.311246702 | 1 | 0.830018519 |
| ENSMUSG00000045733 | Sprn | 212518 | protein_coding | 1.253320976 | -0.830262047 | 2.245511677 | 0.369742921 | 0.711574048 | 1 | 0.830262047 |
| ENSMUSG00000002732 | Fkbp7 | 14231 | protein_coding | 27.79535589 | 0.830365917 | 0.217225578 | -3.82259735 | 0.000132053 | 0.001102492 | 0.830365917 |
| ENSMUSG00000048126 | Col6a3 | 12835 | protein_coding | 7.820893227 | 0.830407275 | 0.931001985 | -0.891950058 | 0.372419702 | 0.594675977 | 0.830407275 |
| ENSMUSG00000046959 | Slc26a1 | 231583 | protein_coding | 23.89182027 | 0.830408345 | 0.511782584 | -1.62258031 | 0.104679152 | 0.255796319 | 0.830408345 |
| ENSMUSG00000066392 | Nrxn3 | 18191 | protein_coding | 3.953590791 | 0.830424244 | 2.261110554 | -0.367263884 | 0.713422191 | 1 | 0.830424244 |
| ENSMUSG00000068324 | Gm10029 | 100039532 | processed_pseudogene | 0.564267939 | -0.830492162 | 1.493419533 | 0.556101044 | 0.578141781 | 1 | 0.830492162 |
| ENSMUSG00000097515 | 1700040D17Rik | 76602 | lncRNA | 0.54206279 | 0.83076115 | 1.511186142 | -0.549741112 | 0.582496954 | 1 | 0.83076115 |
| ENSMUSG00000027227 | Sord | 20322 | protein_coding | 933.4683339 | -0.83089097 | 0.072995938 | 11.38270144 | 5.10E-30 | 3.99E-28 | 0.83089097 |
| ENSMUSG00000005045 | Chd5 | 269610 | protein_coding | 19.58670874 | 0.831028301 | 0.67238467 | -1.235941774 | 0.216480208 | 0.421873277 | 0.831028301 |
| ENSMUSG00000068086 | Cyp2d9 | 13105 | protein_coding | 76.83012812 | -0.831074683 | 0.362144756 | 2.294868749 | 0.021740647 | 0.078372637 | 0.831074683 |
| ENSMUSG00000028926 | Cdk14 | 18647 | protein_coding | 44.42700412 | 0.83154056 | 0.211699054 | -3.927937066 | 8.57E-05 | 0.000752996 | 0.83154056 |
| ENSMUSG00000036381 | P2ry14 | 140795 | protein_coding | 2.124715904 | 0.83183165 | 2.176942494 | -0.38211007 | 0.702379722 | 1 | 0.83183165 |
| ENSMUSG00000060843 | Ctnna3 | 216033 | protein_coding | 1.953341753 | 0.831900577 | 1.658313082 | -0.501654715 | 0.615910424 | 1 | 0.831900577 |
| ENSMUSG00000022204 | Ngdn | 68966 | protein_coding | 511.3402791 | 0.832229001 | 0.071420438 | -11.65253281 | 2.23E-31 | 1.81E-29 | 0.832229001 |
| ENSMUSG00000025489 | Ifitm5 | 73835 | protein_coding | 1.808478824 | 0.83240566 | 0.837018592 | -0.994488854 | 0.319984929 | 1 | 0.83240566 |
| ENSMUSG00000110234 | Gm45799 | 14356 | protein_coding | 1.922101391 | 0.832422557 | 0.796759233 | -1.044760479 | 0.296133685 | 1 | 0.832422557 |
| ENSMUSG00000046337 | Fam178b | 381337 | protein_coding | 3.009687603 | 0.8328237 | 1.030948048 | -0.80782315 | 0.419192396 | 1 | 0.8328237 |
| ENSMUSG00000028884 | Rpa2 | 19891 | protein_coding | 188.1033474 | 0.832862305 | 0.17488848 | -4.76224794 | 1.91E-06 | 2.50E-05 | 0.832862305 |
| ENSMUSG00000026837 | Col5a1 | 12831 | protein_coding | 14.61466657 | 0.832879521 | 0.909506423 | -0.915748916 | 0.359798602 | 0.582349713 | 0.832879521 |
| ENSMUSG00000027715 | Ccna2 | 12428 | protein_coding | 972.1351248 | 0.833112983 | 0.166982011 | -4.989237917 | 6.06E-07 | 8.78E-06 | 0.833112983 |
| ENSMUSG00000057816 | Cfap299 | 75784 | protein_coding | 3.04171589 | -0.833362312 | 0.676152832 | 1.232505838 | 0.217760187 | 1 | 0.833362312 |
| ENSMUSG00000005667 | Mthfd2 | 17768 | protein_coding | 126.7750024 | 0.833954647 | 0.174316766 | -4.784133325 | 1.72E-06 | 2.25E-05 | 0.833954647 |
| ENSMUSG00000038552 | Fndc4 | 64339 | protein_coding | 3.827422378 | 0.834881009 | 0.787362931 | -1.060350921 | 0.288984981 | 1 | 0.834881009 |
| ENSMUSG00000062309 | Rpp25 | 102614 | protein_coding | 9.747760252 | -0.835035541 | 0.351200773 | 2.377658611 | 0.017422948 | 0.065855474 | 0.835035541 |
| ENSMUSG00000052726 | Kcnt2 | 240776 | protein_coding | 1.46541367 | 0.835195026 | 1.789510507 | -0.466717028 | 0.640702345 | 1 | 0.835195026 |
| ENSMUSG00000031232 | Magt1 | 67075 | protein_coding | 2538.563773 | 0.835283714 | 0.111474373 | -7.493055946 | 6.73E-14 | 2.25E-12 | 0.835283714 |
| ENSMUSG00000043925 | Or55b4 | 257926 | protein_coding | 0.814797316 | -0.835362868 | 1.976574048 | 0.422631709 | 0.672563988 | 1 | 0.835362868 |
| ENSMUSG00000021714 | Cenpk | 60411 | protein_coding | 55.95381863 | 0.835550994 | 0.192554306 | -4.339300486 | 1.43E-05 | 0.000153277 | 0.835550994 |
| ENSMUSG00000000628 | Hk2 | 15277 | protein_coding | 3103.883855 | 0.835561713 | 0.133256928 | -6.27030599 | 3.60E-10 | 8.39E-09 | 0.835561713 |
| ENSMUSG00000059742 | Kcnh7 | 170738 | protein_coding | 20.14406236 | -0.835632275 | 0.395853243 | 2.110964833 | 0.034775335 | 0.112936062 | 0.835632275 |
| ENSMUSG00000074577 | Ripor3 | 69553 | protein_coding | 3.064814158 | -0.835678921 | 1.226965493 | 0.68109407 | 0.495811969 | 1 | 0.835678921 |
| ENSMUSG00000035932 | Or6s1 | 404319 | protein_coding | 1.83839487 | -0.835769643 | 1.446468445 | 0.577800121 | 0.563399073 | 1 | 0.835769643 |
| ENSMUSG00000047992 | Dipk1c | 240479 | protein_coding | 10.51887901 | 0.836215055 | 0.448216966 | -1.865647931 | 0.062090639 | 0.17492772 | 0.836215055 |
| ENSMUSG00000033427 | Upb1 | 103149 | protein_coding | 41.53578032 | -0.836362977 | 0.177678939 | 4.70715877 | 2.51E-06 | 3.20E-05 | 0.836362977 |

Supplementary Table 1

| ensembl_gene_id | external_gene_name | entrezgene_id | gene_biotype | baseMean | log2FoldChange | lfcSE | stat | pvalue | padj | absLog2FC |
|---|---|---|---|---|---|---|---|---|---|---|
| ENSMUSG00000022821 | Hgd | 15233 | protein_coding | 1.780698357 | 0.836459555 | 1.472680031 | -0.567984584 | 0.570045436 | 1 | 0.836459555 |
| ENSMUSG00000072624 | Gm5460 | 432838 | protein_coding | 1.398361739 | -0.836693937 | 2.072192226 | 0.403772356 | 0.686380126 | 1 | 0.836693937 |
| ENSMUSG00000032502 | Stac | 20840 | protein_coding | 1.040561826 | -0.836827619 | 1.476224067 | 0.566870327 | 0.570802286 | 1 | 0.836827619 |
| ENSMUSG00000030507 | Dbx1 | 13172 | protein_coding | 1.111564919 | -0.83683862 | 2.083495575 | 0.401651258 | 0.687940701 | 1 | 0.83683862 |
| ENSMUSG00000043789 | Vwce | 71768 | protein_coding | 6.552449894 | -0.836916392 | 0.932679382 | 0.897324856 | 0.369545597 | 0.592205336 | 0.836916392 |
| ENSMUSG00000053702 | Nebl | 74103 | protein_coding | 80.46216277 | -0.837211613 | 0.196812533 | 4.253853155 | 2.10E-05 | 0.000215526 | 0.837211613 |
| ENSMUSG00000029463 | Fam216a | 68948 | protein_coding | 15.51279345 | 0.838462575 | 0.303326089 | -2.76422835 | 0.005705759 | 0.027169813 | 0.838462575 |
| ENSMUSG00000044390 | Tigd3 | 332359 | protein_coding | 4.549708274 | -0.838814522 | 0.558495458 | 1.501918252 | 0.133118222 | 0.302752759 | 0.838814522 |
| ENSMUSG00000069920 | B3gnt9 | 97440 | protein_coding | 51.63010782 | -0.838919716 | 0.160030598 | 5.242245723 | 1.59E-07 | 2.53E-06 | 0.838919716 |
| ENSMUSG00000090436 | Vmn2r75 | 546981 | protein_coding | 1.02643232 | -0.839228664 | 2.297811978 | 0.365229476 | 0.714940117 | 1 | 0.839228664 |
| ENSMUSG00000017167 | Cntnap1 | 53321 | protein_coding | 91.68977028 | -0.83941222 | 0.233059333 | 3.601710391 | 0.00031613 | 0.002383826 | 0.83941222 |
| ENSMUSG00000067951 | Vmn1r227 | 171229 | protein_coding | 1.43385017 | -0.839498973 | 2.325992012 | 0.360920833 | 0.71815863 | 1 | 0.839498973 |
| ENSMUSG00000025758 | Plk4 | 20873 | protein_coding | 196.21913 | 0.839877049 | 0.160727898 | -5.225459037 | 1.74E-07 | 2.75E-06 | 0.839877049 |
| ENSMUSG00000032586 | Traip | 22036 | protein_coding | 67.34436992 | 0.839908926 | 0.25271271 | -3.323572149 | 0.000888724 | 0.005762227 | 0.839908926 |
| ENSMUSG00000091972 | Tmem207 | 100043057 | protein_coding | 0.987373863 | -0.83994021 | 2.374742975 | 0.353697313 | 0.723565736 | 1 | 0.83994021 |
| ENSMUSG00000039384 | Dusp10 | 63953 | protein_coding | 76.64763615 | 0.839982418 | 0.251655981 | -3.3378202 | 0.000844384 | 0.005513702 | 0.839982418 |
| ENSMUSG00000105078 | Vamp9 | 102639650 | protein_coding | 1.645411585 | -0.840073629 | 1.029059156 | 0.816351154 | 0.414299325 | 1 | 0.840073629 |
| ENSMUSG00000028278 | Rragd | 52187 | protein_coding | 6.281571403 | -0.840124674 | 0.611755841 | 1.373300615 | 0.169658911 | 0.359060267 | 0.840124674 |
| ENSMUSG00000028669 | Pithd1 | 66193 | protein_coding | 616.9311948 | 0.840196534 | 0.098457491 | -8.533596803 | 1.42E-17 | 6.25E-16 | 0.840196534 |
| ENSMUSG00000096373 | Vmn2r31 | 100042591 | protein_coding | 0.843951163 | -0.840316958 | 2.418690882 | 0.347426355 | 0.728271036 | 1 | 0.840316958 |
| ENSMUSG00000028777 | Gnat3 | 242851 | protein_coding | 44.86707627 | -0.840455148 | 0.235030546 | 3.575940074 | 0.000348972 | 0.002606475 | 0.840455148 |
| ENSMUSG00000092120 | Vmn2r90 | 626942 | protein_coding | 1.082811775 | -0.84047043 | 2.437040587 | 0.344873382 | 0.730189562 | 1 | 0.84047043 |
| ENSMUSG00000043122 | A530016L24Rik | 319942 | protein_coding | 1.221366151 | -0.84048971 | 2.43929584 | 0.344562433 | 0.730423352 | 1 | 0.84048971 |
| ENSMUSG00000086822 | 5330413P13Rik | 414081 | lncRNA | 1.092252686 | -0.84056344 | 2.448467774 | 0.343301819 | 0.731371413 | 1 | 0.84056344 |
| ENSMUSG00000038704 | Aspdh | 68352 | protein_coding | 1.33206591 | 0.840618654 | 1.870206284 | -0.449479109 | 0.653086075 | 1 | 0.840618654 |
| ENSMUSG00000098442 | Mir6959 | 102465981 | miRNA | 0.787381622 | 0.840944612 | 1.55211209 | -0.541806624 | 0.587951727 | 1 | 0.840944612 |
| ENSMUSG00000037010 | Apln | 30878 | protein_coding | 3.604274032 | -0.840980496 | 0.566801467 | 1.483730274 | 0.137880492 | 1 | 0.840980496 |
| ENSMUSG00000022871 | Fetub | 59083 | protein_coding | 1.879419247 | 0.841283981 | 1.196054154 | -0.703382852 | 0.48181719 | 1 | 0.841283981 |
| ENSMUSG00000027209 | Fam227b | 75823 | protein_coding | 0.720571216 | 0.841863078 | 1.358382198 | -0.619754204 | 0.535419623 | 1 | 0.841863078 |
| ENSMUSG00000064267 | Hvcn1 | 74096 | protein_coding | 6.312655177 | 0.842072143 | 0.548813734 | -1.534349619 | 0.124943665 | 0.289914061 | 0.842072143 |
| ENSMUSG00000020658 | Efr3b | 668212 | protein_coding | 15.88214032 | 0.842615219 | 0.395225522 | -2.131985844 | 0.03300801 | 0.108572726 | 0.842615219 |
| ENSMUSG00000063935 | Zar1 | 317755 | protein_coding | 0.93292608 | -0.842847936 | 2.789133167 | 0.302189923 | 0.762507286 | 1 | 0.842847936 |
| ENSMUSG00000090949 | Vmn2r77 | 546983 | protein_coding | 1.262047447 | -0.842855656 | 2.790016533 | 0.302097011 | 0.762578111 | 1 | 0.842855656 |
| ENSMUSG00000035557 | Krt17 | 16667 | protein_coding | 0.980486692 | -0.842889762 | 2.796621946 | 0.301395676 | 0.763112791 | 1 | 0.842889762 |
| ENSMUSG00000058200 | Or51b4 | 18367 | protein_coding | 1.166725298 | -0.842949384 | 2.80720886 | 0.300280252 | 0.763963395 | 1 | 0.842949384 |
| ENSMUSG00000061602 | Vmn1r78 | 171242 | protein_coding | 1.071604074 | -0.842962086 | 2.809682496 | 0.300020407 | 0.76416159 | 1 | 0.842962086 |
| ENSMUSG00000032064 | Dixdc1 | 330938 | protein_coding | 123.7202671 | -0.843020208 | 0.216831348 | 3.887907424 | 0.000101112 | 0.000869217 | 0.843020208 |
| ENSMUSG00000097074 | 4833428L15Rik | 74602 | lncRNA | 1.027430058 | -0.843025252 | 2.82146748 | 0.29878964 | 0.765100556 | 1 | 0.843025252 |
| ENSMUSG00000058252 | Tcp11x2 | 71841 | protein_coding | 0.98061031 | -0.843032531 | 2.822903638 | 0.298640209 | 0.765214583 | 1 | 0.843032531 |

| ensembl_gene_id | external_gene_name | entrezgene_id | gene_biotype | baseMean | log2FoldChange | lfcSE | stat | pvalue | padj | absLog2FC |
|---|---|---|---|---|---|---|---|---|---|---|
| ENSMUSG00000103585 | Pcdhgb4 | 93701 | protein_coding | 8.719215681 | -0.843404031 | 0.409226862 | 2.060969376 | 0.039305962 | 0.124359938 | 0.843404031 |
| ENSMUSG00000103965 | Gm30173 | 102631981 | lncRNA | 1.332126257 | 0.843957941 | 1.282862037 | -0.657871163 | 0.510620922 | 1 | 0.843957941 |
| ENSMUSG00000048520 | Fbxl13 | 320118 | protein_coding | 2.420966362 | -0.844673761 | 1.076530723 | 0.784625782 | 0.432673009 | 1 | 0.844673761 |
| ENSMUSG00000030223 | Ptpro | 19277 | protein_coding | 58.13810814 | 0.844764507 | 0.26915223 | -3.138612324 | 0.001697499 | 0.009881034 | 0.844764507 |
| ENSMUSG00000027794 | Sohlh2 | 74434 | protein_coding | 1.166848915 | -0.845309728 | 3.370509209 | 0.250795852 | 0.801971949 | 1 | 0.845309728 |
| ENSMUSG00000091956 | C2cd4b | 75697 | protein_coding | 81.53374362 | 0.845433632 | 0.19598731 | -4.313716192 | 1.61E-05 | 0.000169939 | 0.845433632 |
| ENSMUSG00000061259 | Tmprss11d | 231382 | protein_coding | 1.170764491 | 0.845470933 | 2.282820694 | -0.370362392 | 0.711112491 | 1 | 0.845470933 |
| ENSMUSG00000041498 | Kif14 | 381293 | protein_coding | 137.3750944 | 0.845498246 | 0.176671885 | -4.785697775 | 1.70E-06 | 2.24E-05 | 0.845498246 |
| ENSMUSG00000028539 | Artn | 11876 | protein_coding | 4.683478556 | -0.845795797 | 0.807527127 | 1.047389949 | 0.294919765 | 0.514070615 | 0.845795797 |
| ENSMUSG00000044043 | Pcdhb14 | 93885 | protein_coding | 2.816344365 | 0.845800061 | 0.742319774 | -1.139401227 | 0.254535845 | 1 | 0.845800061 |
| ENSMUSG00000024965 | Fermt3 | 108101 | protein_coding | 47.77513215 | -0.845895366 | 0.202767208 | 4.171756233 | 3.02E-05 | 0.000298838 | 0.845895366 |
| ENSMUSG00000022582 | Ly6g | 546644 | protein_coding | 193.1269671 | -0.845915893 | 0.309860784 | 2.729986936 | 0.006333684 | 0.029306374 | 0.845915893 |
| ENSMUSG00000001670 | Tat | 234724 | protein_coding | 278.3606293 | 0.845930388 | 0.290916865 | -2.907807999 | 0.003639717 | 0.018707218 | 0.845930388 |
| ENSMUSG00000029664 | Tfpi2 | 21789 | protein_coding | 147.5778619 | 0.845948693 | 0.118239676 | -7.154524795 | 8.40E-13 | 2.57E-11 | 0.845948693 |
| ENSMUSG00000028475 | Spaar | 71406 | protein_coding | 1.891674938 | 0.846003123 | 1.931715368 | -0.437954337 | 0.661419385 | 1 | 0.846003123 |
| ENSMUSG00000027496 | Aurka | 20878 | protein_coding | 262.7375393 | 0.846052167 | 0.173576978 | -4.874218793 | 1.09E-06 | 1.49E-05 | 0.846052167 |
| ENSMUSG00000039693 | Msantd3 | 66665 | protein_coding | 27.89403101 | 0.846224709 | 0.238045143 | -3.554891725 | 0.000378135 | 0.002795218 | 0.846224709 |
| ENSMUSG00000028439 | Fam219a | 71901 | protein_coding | 40.7588163 | 0.846253661 | 0.25700068 | -3.292807085 | 0.000991925 | 0.006324576 | 0.846253661 |
| ENSMUSG00000035683 | Melk | 17279 | protein_coding | 299.1245604 | 0.846576541 | 0.178926494 | -4.73142084 | 2.23E-06 | 2.88E-05 | 0.846576541 |
| ENSMUSG00000030255 | Sspn | 16651 | protein_coding | 1.13038802 | -0.846591811 | 1.591886867 | 0.531816569 | 0.594853046 | 1 | 0.846591811 |
| ENSMUSG00000042104 | Uggt2 | 66435 | protein_coding | 22.86580189 | 0.846760359 | 0.314990093 | -2.688212671 | 0.007183563 | 0.032644292 | 0.846760359 |
| ENSMUSG00000050961 | AY358078 | 278676 | protein_coding | 0.71286111 | -0.846770523 | 1.831593808 | 0.462313489 | 0.643856528 | 1 | 0.846770523 |
| ENSMUSG00000038060 | Dlec1 | 320256 | protein_coding | 10.58001559 | 0.846796256 | 0.572984568 | -1.47786922 | 0.139442786 | 0.312959795 | 0.846796256 |
| ENSMUSG00000079025 | Gsdmc | 83492 | protein_coding | 2.15302564 | -0.84690243 | 1.438192781 | 0.588865722 | 0.555951354 | 1 | 0.84690243 |
| ENSMUSG00000032015 | Pou2f3 | 18988 | protein_coding | 27.26030594 | -0.847087731 | 0.249519757 | 3.394872376 | 0.000686606 | 0.004612942 | 0.847087731 |
| ENSMUSG00000097523 | 1700022N22Rik | 69431 | lncRNA | 4.345936645 | -0.847121419 | 0.594280206 | 1.425457906 | 0.15402488 | 1 | 0.847121419 |
| ENSMUSG00000018648 | Dusp14 | 56405 | protein_coding | 25.55030887 | 0.847410589 | 0.221800379 | -3.82060028 | 0.000133127 | 0.001109775 | 0.847410589 |
| ENSMUSG00000029707 | Fscn3 | 56223 | protein_coding | 2.824585127 | 0.847417067 | 0.939705819 | -0.901789741 | 0.367168571 | 1 | 0.847417067 |
| ENSMUSG00000065599 | Mir23b | 387217 | miRNA | 1.369274349 | 0.848265384 | 0.988671833 | -0.857984778 | 0.390900869 | 1 | 0.848265384 |
| ENSMUSG00000029910 | Mad2l1 | 56150 | protein_coding | 337.8419011 | 0.848555841 | 0.158704597 | -5.346762832 | 8.95E-08 | 1.49E-06 | 0.848555841 |
| ENSMUSG00000020076 | Ddx50 | 94213 | protein_coding | 1060.681269 | 0.848648336 | 0.080933025 | -10.48581022 | 1.00E-25 | 6.55E-24 | 0.848648336 |
| ENSMUSG00000058420 | Syt17 | 110058 | protein_coding | 6.413009431 | 0.849161784 | 0.548423651 | -1.548368276 | 0.121533655 | 0.284375933 | 0.849161784 |
| ENSMUSG00000071398 | 2410004P03Rik | 73667 | protein_coding | 2.507335265 | -0.849415362 | 1.074160951 | 0.79077103 | 0.429077618 | 1 | 0.849415362 |
| ENSMUSG00000110618 | Gm39822 | 105244162 | lncRNA | 1.792067955 | 0.849780402 | 1.159589953 | -0.732828358 | 0.463663123 | 1 | 0.849780402 |
| ENSMUSG00000015452 | Ager | 11596 | protein_coding | 12.31828457 | 0.850429362 | 0.417784303 | -2.035570403 | 0.041793512 | 0.130148607 | 0.850429362 |
| ENSMUSG00000066513 | Klk1b4 | 18048 | protein_coding | 6.573114099 | -0.851031913 | 0.448305663 | 1.898329607 | 0.057652676 | 0.165809177 | 0.851031913 |
| ENSMUSG00000025723 | Nmb | 68039 | protein_coding | 5.341506322 | 0.851596353 | 0.523726654 | -1.626032105 | 0.103942826 | 0.254637565 | 0.851596353 |
| ENSMUSG00000026715 | Serpinc1 | 11905 | protein_coding | 8.740107616 | -0.851634686 | 0.586972755 | 1.450893044 | 0.146809645 | 0.324596932 | 0.851634686 |
| ENSMUSG00000049723 | Mmp12 | 17381 | protein_coding | 1.421653398 | -0.851981335 | 2.201318749 | 0.387032244 | 0.698732337 | 1 | 0.851981335 |

Supplementary Table 1

| ensembl_gene_id | external_gene_name | entrezgene_id | gene_biotype | baseMean | log2FoldChange | lfcSE | stat | pvalue | padj | absLog2FC |
|---|---|---|---|---|---|---|---|---|---|---|
| ENSMUSG00000101695 | 1700094J05Rik | 67523 | lncRNA | 0.876121951 | 0.85199325 | 1.345478476 | -0.633226964 | 0.52658544 | 1 | 0.85199325 |
| ENSMUSG00000041075 | Fzd7 | 14369 | protein_coding | 224.2961143 | 0.852001918 | 0.142481785 | -5.979725197 | 2.24E-09 | 4.74E-08 | 0.852001918 |
| ENSMUSG00000018451 | 6330403K07Rik | 103712 | protein_coding | 2.146832265 | 0.852242589 | 0.935348006 | -0.911150271 | 0.362216202 | 1 | 0.852242589 |
| ENSMUSG00000038384 | Setd1b | 208043 | protein_coding | 300.7859195 | 0.852446958 | 0.339917984 | -2.507801877 | 0.012148476 | 0.049584759 | 0.852446958 |
| ENSMUSG00000041064 | Pif1 | 208084 | protein_coding | 98.1646192 | 0.852496082 | 0.180473426 | -4.723665416 | 2.32E-06 | 2.97E-05 | 0.852496082 |
| ENSMUSG00000038765 | Lmx1b | 16917 | protein_coding | 3.880269747 | 0.852629933 | 1.929497485 | -0.441892223 | 0.658567198 | 1 | 0.852629933 |
| ENSMUSG00000026969 | Fam166a | 68222 | protein_coding | 25.9896537 | 0.852705991 | 0.47758011 | -1.78547216 | 0.074184763 | 0.199168782 | 0.852705991 |
| ENSMUSG00000116953 | 9030025P20Rik | 100041574 | protein_coding | 5.250331518 | -0.852711362 | 0.448751755 | 1.90018502 | 0.057408843 | 0.165344954 | 0.852711362 |
| ENSMUSG00000055041 | Commd5 | 66398 | protein_coding | 9.564109128 | 0.853262403 | 0.457449478 | -1.865260414 | 0.062144911 | 0.174980573 | 0.853262403 |
| ENSMUSG00000050158 | Or2m13 | 258458 | protein_coding | 45.08214283 | 0.853536366 | 0.256653498 | -3.325636986 | 0.000882167 | 0.005729834 | 0.853536366 |
| ENSMUSG00000053678 | Defb40 | 360217 | protein_coding | 3.909170304 | 0.853607398 | 0.606777279 | -1.406788665 | 0.159490065 | 1 | 0.853607398 |
| ENSMUSG00000046688 | Tifa | 211550 | protein_coding | 624.9896872 | 0.854114926 | 0.162424881 | -5.258522708 | 1.45E-07 | 2.33E-06 | 0.854114926 |
| ENSMUSG00000031298 | Adgrg2 | 237175 | protein_coding | 11.79124585 | -0.85450892 | 0.486903952 | 1.754984563 | 0.079261948 | 0.208627959 | 0.85450892 |
| ENSMUSG00000041633 | Kctd12b | 207474 | protein_coding | 2.681469528 | -0.855116095 | 0.860172145 | 0.994122048 | 0.320163452 | 1 | 0.855116095 |
| ENSMUSG00000024687 | Osbp | 76303 | protein_coding | 1875.761977 | 0.85541264 | 0.058474123 | -14.62890921 | 1.84E-48 | 2.37E-46 | 0.85541264 |
| ENSMUSG00000030109 | Slc6a12 | 14411 | protein_coding | 4.11555343 | -0.855559377 | 1.228589073 | 0.696375538 | 0.486193678 | 1 | 0.855559377 |
| ENSMUSG00000044201 | Cdc25c | 12532 | protein_coding | 90.48994671 | 0.855671356 | 0.177547383 | -4.81939718 | 1.44E-06 | 1.92E-05 | 0.855671356 |
| ENSMUSG00000026609 | Ush2a | 22283 | protein_coding | 5.340961539 | 0.855882495 | 1.66424971 | -0.514275286 | 0.607059538 | 0.781796866 | 0.855882495 |
| ENSMUSG00000029778 | Adcyap1r1 | 11517 | protein_coding | 3.672173207 | 0.856252958 | 1.024232304 | -0.835994876 | 0.403157786 | 1 | 0.856252958 |
| ENSMUSG00000102099 | 1700011B04Rik | 100503727 | lncRNA | 2.506237076 | 0.856308328 | 2.148188919 | -0.398618725 | 0.690174162 | 1 | 0.856308328 |
| ENSMUSG00000020901 | Pik3r5 | 320207 | protein_coding | 131.0857158 | -0.856503145 | 0.206860768 | 4.140481325 | 3.47E-05 | 0.000338705 | 0.856503145 |
| ENSMUSG00000081021 | Gm11964 | 665181 | processed_pseudogene | 1.805774536 | 0.857269134 | 0.917385401 | -0.934469998 | 0.350061505 | 1 | 0.857269134 |
| ENSMUSG00000022099 | Dmtn | 13829 | protein_coding | 9.160489153 | 0.85744874 | 0.584968042 | -1.465804417 | 0.142701577 | 0.318422618 | 0.85744874 |
| ENSMUSG00000020492 | Ska2 | 66140 | protein_coding | 69.49626496 | 0.857616272 | 0.17383743 | -4.933438519 | 8.08E-07 | 1.14E-05 | 0.857616272 |
| ENSMUSG00000063446 | Plppr1 | 272031 | protein_coding | 1.561956354 | 0.857958096 | 1.729457477 | -0.496085106 | 0.619834367 | 1 | 0.857958096 |
| ENSMUSG00000026856 | Dolpp1 | 57170 | protein_coding | 807.7070774 | 0.858164341 | 0.09198692 | -9.329199601 | 1.07E-20 | 5.65E-19 | 0.858164341 |
| ENSMUSG00000021589 | Rhobtb3 | 73296 | protein_coding | 135.2314156 | 0.858384416 | 0.123018778 | -6.977669814 | 3.00E-12 | 8.80E-11 | 0.858384416 |
| ENSMUSG00000015396 | Cd83 | 12522 | protein_coding | 42.02743464 | -0.858396212 | 0.248892617 | 3.448861699 | 0.000562955 | 0.003915665 | 0.858396212 |
| ENSMUSG00000029419 | Ajm1 | 381353 | protein_coding | 4.098499593 | 0.860207084 | 0.775078335 | -1.109832446 | 0.267071235 | 1 | 0.860207084 |
| ENSMUSG00000039860 | Srrm3 | 58212 | protein_coding | 5.042427596 | 0.860489351 | 1.049678597 | -0.819764596 | 0.412350318 | 0.632329199 | 0.860489351 |
| ENSMUSG00000039620 | Trmt9b | 319582 | protein_coding | 12.84758383 | -0.86052575 | 0.357918292 | 2.404251943 | 0.0162056 | 0.062193652 | 0.86052575 |
| ENSMUSG00000112398 | 9230102K24Rik | 109335 | lncRNA | 1.42269633 | 0.861089921 | 0.908151438 | -0.948178778 | 0.343038449 | 1 | 0.861089921 |
| ENSMUSG00000022383 | Ppara | 19013 | protein_coding | 297.9127248 | -0.861231484 | 0.16286715 | 5.287938555 | 1.24E-07 | 2.01E-06 | 0.861231484 |
| ENSMUSG00000031877 | Ces2g | 72361 | protein_coding | 143.9702376 | -0.861311136 | 0.187381204 | 4.596571679 | 4.29E-06 | 5.21E-05 | 0.861311136 |
| ENSMUSG00000068745 | Mybphl | 68753 | protein_coding | 0.954424056 | -0.861348063 | 1.676462019 | 0.51378919 | 0.607399386 | 1 | 0.861348063 |
| ENSMUSG00000022490 | Ppp1r1a | 58200 | protein_coding | 9.543870674 | 0.861377823 | 0.335954142 | -2.563974413 | 0.010348117 | 0.043750335 | 0.861377823 |
| ENSMUSG00000023993 | Treml1 | 71326 | protein_coding | 1.469089361 | -0.861609348 | 2.293800784 | 0.375625187 | 0.707195563 | 1 | 0.861609348 |
| ENSMUSG00000030270 | Cpne9 | 211232 | protein_coding | 5.721950469 | 0.862176842 | 1.250344988 | -0.689551164 | 0.490476488 | 0.69793377 | 0.862176842 |
| ENSMUSG00000024245 | Tmem178 | 68027 | protein_coding | 0.775802785 | 0.862232353 | 1.396695258 | -0.617337496 | 0.53701214 | 1 | 0.862232353 |

| ensembl_gene_id | external_gene_name | entrezgene_id | gene_biotype | baseMean | log2FoldChange | lfcSE | stat | pvalue | padj | absLog2FC |
|---|---|---|---|---|---|---|---|---|---|---|
| ENSMUSG00000073940 | Hbb-bt | 101488143 | protein_coding | 42.74359198 | -0.862253027 | 0.509341959 | 1.692876488 | 0.09047898 | 0.229589716 | 0.862253027 |
| ENSMUSG00000117148 | Vmn1r229 | 171224 | protein_coding | 2.313201204 | -0.862351878 | 2.140790935 | 0.402819287 | 0.687081171 | 1 | 0.862351878 |
| ENSMUSG00000031880 | Rrad | 56437 | protein_coding | 1.705298179 | -0.862494169 | 1.381290454 | 0.62441188 | 0.532357125 | 1 | 0.862494169 |
| ENSMUSG00000033585 | Ndn | 17984 | protein_coding | 0.970024152 | -0.862686505 | 1.573535297 | 0.548247317 | 0.583522096 | 1 | 0.862686505 |
| ENSMUSG00000001225 | Slc26a3 | 13487 | protein_coding | 6410.810467 | -0.862914477 | 0.165081383 | 5.227206479 | 1.72E-07 | 2.73E-06 | 0.862914477 |
| ENSMUSG00000032038 | St3gal4 | 20443 | protein_coding | 2783.252604 | -0.86388726 | 0.26291521 | 3.285801759 | 0.001016926 | 0.006446644 | 0.86388726 |
| ENSMUSG00000028794 | A3galt2 | 215493 | protein_coding | 3.052823951 | 0.864096823 | 1.036869573 | -0.833370798 | 0.404635639 | 1 | 0.864096823 |
| ENSMUSG00000067261 | Foxd3 | 15221 | protein_coding | 1.232439437 | -0.864261797 | 2.210650846 | 0.39095355 | 0.69583157 | 1 | 0.864261797 |
| ENSMUSG00000085111 | Ascl4 | 67341 | protein_coding | 1.683833416 | -0.864292022 | 1.065991537 | 0.810786945 | 0.417488033 | 1 | 0.864292022 |
| ENSMUSG00000023191 | P3h3 | 14789 | protein_coding | 6.563482881 | 0.864477728 | 0.646854875 | -1.33643227 | 0.181408025 | 0.375503154 | 0.864477728 |
| ENSMUSG00000050357 | Carmil2 | 234695 | protein_coding | 9.01163276 | 0.864751451 | 0.640949006 | -1.349173558 | 0.177281226 | 0.369555941 | 0.864751451 |
| ENSMUSG00000049176 | Frmpd4 | 333605 | protein_coding | 4.28076162 | 0.864934006 | 1.22213643 | -0.707722955 | 0.479117319 | 1 | 0.864934006 |
| ENSMUSG00000056671 | Prelid2 | 77619 | protein_coding | 265.9862469 | -0.865078089 | 0.114445605 | 7.558858092 | 4.07E-14 | 1.39E-12 | 0.865078089 |
| ENSMUSG00000023092 | Fhl1 | 14199 | protein_coding | 1250.548017 | -0.865110859 | 0.178776572 | 4.839061674 | 1.30E-06 | 1.76E-05 | 0.865110859 |
| ENSMUSG00000053510 | Nrd1 | 230598 | protein_coding | 1383.750769 | 0.865141669 | 0.07255369 | -11.92415812 | 8.86E-33 | 7.46E-31 | 0.865141669 |
| ENSMUSG00000031485 | Plpbp | 114863 | protein_coding | 711.8853031 | 0.86566634 | 0.069849116 | -12.39337571 | 2.84E-35 | 2.52E-33 | 0.86566634 |
| ENSMUSG00000002032 | Tmem25 | 71687 | protein_coding | 64.40587943 | -0.865702973 | 0.132891381 | 6.514365066 | 7.30E-11 | 1.86E-09 | 0.865702973 |
| ENSMUSG00000089678 | Agxt2 | 268782 | protein_coding | 0.984586596 | -0.865770964 | 1.569380515 | 0.551664147 | 0.581178478 | 1 | 0.865770964 |
| ENSMUSG00000032519 | Slc25a38 | 208638 | protein_coding | 336.3145906 | 0.866194331 | 0.08099274 | -10.69471574 | 1.08E-26 | 7.51E-25 | 0.866194331 |
| ENSMUSG00000055027 | Smyd1 | 12180 | protein_coding | 4.93864477 | -0.866298295 | 0.877923264 | 0.986758559 | 0.323761015 | 0.545570347 | 0.866298295 |
| ENSMUSG00000027435 | Cd93 | 17064 | protein_coding | 2.998357799 | 0.866845278 | 1.633202876 | -0.530763992 | 0.595582335 | 1 | 0.866845278 |
| ENSMUSG00000060613 | Cyp2c70 | 226105 | protein_coding | 1.650630731 | 0.866854574 | 1.052781258 | -0.823394763 | 0.41028354 | 1 | 0.866854574 |
| ENSMUSG00000063903 | Klk1 | 16612 | protein_coding | 14711.77968 | 0.867460145 | 0.143532031 | -6.043669421 | 1.51E-09 | 3.26E-08 | 0.867460145 |
| ENSMUSG00000020656 | Grhl1 | 195733 | protein_coding | 31.26846692 | 0.867555985 | 0.26752975 | -3.24283929 | 0.001183449 | 0.00731937 | 0.867555985 |
| ENSMUSG00000041763 | Tpp2 | 22019 | protein_coding | 1375.97512 | 0.868185121 | 0.058163042 | -14.92674888 | 2.21E-50 | 3.14E-48 | 0.868185121 |
| ENSMUSG00000033831 | Fgb | 110135 | protein_coding | 1.480723938 | 0.868392624 | 1.694718878 | -0.512411017 | 0.608363382 | 1 | 0.868392624 |
| ENSMUSG00000015854 | Cd5l | 11801 | protein_coding | 1.554716401 | 0.868511853 | 1.374983228 | -0.631652689 | 0.527613849 | 1 | 0.868511853 |
| ENSMUSG00000037965 | Zc3h7a | 106205 | protein_coding | 1310.124333 | 0.868546701 | 0.108226353 | -8.025279191 | 1.01E-15 | 3.94E-14 | 0.868546701 |
| ENSMUSG00000020937 | Plcd3 | 72469 | protein_coding | 105.8770761 | -0.868692095 | 0.235434814 | 3.689735097 | 0.000224488 | 0.001752808 | 0.868692095 |
| ENSMUSG00000038387 | Rras | 20130 | protein_coding | 51.31889774 | -0.869216014 | 0.177055535 | 4.909284625 | 9.14E-07 | 1.27E-05 | 0.869216014 |
| ENSMUSG00000001053 | N4bp3 | 212706 | protein_coding | 93.42746185 | -0.869541783 | 0.124889549 | 6.962486397 | 3.34E-12 | 9.74E-11 | 0.869541783 |
| ENSMUSG00000040415 | Dtx3 | 80904 | protein_coding | 32.9598307 | -0.869589677 | 0.215703977 | 4.031403082 | 5.54E-05 | 0.000511803 | 0.869589677 |
| ENSMUSG00000017724 | Etv4 | 18612 | protein_coding | 217.1597308 | 0.86964432 | 0.236643676 | -3.674910452 | 0.000237933 | 0.001846439 | 0.86964432 |
| ENSMUSG00000052631 | Sh2d6 | 71130 | protein_coding | 82.68713291 | -0.869713 | 0.209844735 | 4.144554784 | 3.40E-05 | 0.000333487 | 0.869713 |
| ENSMUSG00000058966 | Tlcd3b | 68952 | protein_coding | 3.796366675 | 0.869991788 | 1.318802691 | -0.659683055 | 0.509457243 | 1 | 0.869991788 |
| ENSMUSG00000079550 | Mpp4 | 227157 | protein_coding | 3.487038461 | 0.870450345 | 1.146579516 | -0.759171373 | 0.447750048 | 1 | 0.870450345 |
| ENSMUSG00000054675 | Tmem119 | 231633 | protein_coding | 2.075467373 | -0.870950036 | 0.977900353 | 0.890632704 | 0.373126248 | 1 | 0.870950036 |
| ENSMUSG00000066595 | Flvcr1 | 226844 | protein_coding | 206.2036846 | 0.871017881 | 0.083048334 | -10.48808368 | 9.80E-26 | 6.45E-24 | 0.871017881 |
| ENSMUSG00000025909 | Sntg1 | 71096 | protein_coding | 6.059685555 | 0.871409962 | 1.146746627 | -0.759897558 | 0.447315821 | 0.664595004 | 0.871409962 |

Supplementary Table 1

| ensembl_gene_id | external_gene_name | entrezgene_id | gene_biotype | baseMean | log2FoldChange | lfcSE | stat | pvalue | padj | absLog2FC |
|---|---|---|---|---|---|---|---|---|---|---|
| ENSMUSG00000032593 | Amigo3 | 320844 | protein_coding | 184.2765389 | 0.871866059 | 0.109026414 | -7.996833337 | 1.28E-15 | 4.91E-14 | 0.871866059 |
| ENSMUSG00000043385 | Or2k2 | 258922 | protein_coding | 2.078834221 | 0.872158961 | 1.254162073 | -0.695411685 | 0.486797339 | 1 | 0.872158961 |
| ENSMUSG00000026531 | Mptx1 | 66289 | protein_coding | 7708.20089 | -0.872531338 | 0.683822608 | 1.275961526 | 0.20196912 | 0.403987166 | 0.872531338 |
| ENSMUSG00000034974 | Dapk3 | 13144 | protein_coding | 517.7636994 | 0.872676292 | 0.084889427 | -10.28015289 | 8.66E-25 | 5.50E-23 | 0.872676292 |
| ENSMUSG00000040328 | Or2v1 | 18356 | protein_coding | 11.63006848 | 0.872817425 | 0.617935095 | -1.412474274 | 0.15781036 | 0.341333371 | 0.872817425 |
| ENSMUSG00000042097 | Zfp239 | 22685 | protein_coding | 170.868314 | 0.872887562 | 0.138696582 | -6.293504487 | 3.10E-10 | 7.31E-09 | 0.872887562 |
| ENSMUSG00000051079 | Rgs13 | 246709 | protein_coding | 100.2860238 | -0.873037886 | 0.195285462 | 4.470572857 | 7.80E-06 | 8.98E-05 | 0.873037886 |
| ENSMUSG00000039179 | Tekt5 | 70426 | protein_coding | 2.496910863 | -0.873051686 | 0.663895081 | 1.315044667 | 0.188494891 | 1 | 0.873051686 |
| ENSMUSG00000051118 | Or7d10 | 258336 | protein_coding | 3.039272196 | 0.873163834 | 1.291834693 | -0.675909881 | 0.499097864 | 1 | 0.873163834 |
| ENSMUSG00000099722 | Gm29154 | 102635534 | lncRNA | 2.623801618 | -0.873329638 | 2.169258902 | 0.402593548 | 0.687247257 | 1 | 0.873329638 |
| ENSMUSG00000039629 | Strip2 | 320609 | protein_coding | 127.78592 | -0.873347058 | 0.127603271 | 6.84423723 | 7.69E-12 | 2.16E-10 | 0.873347058 |
| ENSMUSG00000023940 | Sgo1 | 72415 | protein_coding | 141.2356348 | 0.873678062 | 0.197572643 | -4.422059903 | 9.78E-06 | 0.000110053 | 0.873678062 |
| ENSMUSG00000025240 | Sacm1l | 83493 | protein_coding | 948.6485837 | 0.873944915 | 0.07184883 | -12.16366247 | 4.85E-34 | 4.20E-32 | 0.873944915 |
| ENSMUSG00000043418 | Lrit2 | 239038 | protein_coding | 0.811784668 | 0.874209496 | 1.470026135 | -0.594689765 | 0.552050865 | 1 | 0.874209496 |
| ENSMUSG00000045954 | Cavin2 | 20324 | protein_coding | 1.22723889 | -0.874371668 | 1.248761886 | 0.700190867 | 0.483808114 | 1 | 0.874371668 |
| ENSMUSG00000094806 | Cyp2d10 | 13101 | protein_coding | 169.128154 | -0.874775705 | 0.194610763 | 4.495001659 | 6.96E-06 | 8.10E-05 | 0.874775705 |
| ENSMUSG00000028940 | Hes2 | 15206 | protein_coding | 70.59928249 | 0.875382448 | 0.13823025 | -6.332784979 | 2.41E-10 | 5.75E-09 | 0.875382448 |
| ENSMUSG00000021614 | Vcan | 13003 | protein_coding | 113.2966437 | 0.875473918 | 0.146869017 | -5.960916299 | 2.51E-09 | 5.27E-08 | 0.875473918 |
| ENSMUSG00000042800 | Spata46 | 76925 | protein_coding | 4.658535862 | -0.875483492 | 0.601316138 | 1.455945445 | 0.145407702 | 0.322533618 | 0.875483492 |
| ENSMUSG00000064982 | Gm25259 | 115486693 | snRNA | 1.073007794 | 0.875495501 | 1.339655611 | -0.653522811 | 0.513419288 | 1 | 0.875495501 |
| ENSMUSG00000075463 | 4930594M22Rik | 105245866 | lncRNA | 10.14005916 | 0.875631463 | 0.429192526 | -2.040183392 | 0.041332063 | 0.129149989 | 0.875631463 |
| ENSMUSG00000001517 | Foxm1 | 14235 | protein_coding | 191.2555756 | 0.876514164 | 0.177417509 | -4.940403973 | 7.80E-07 | 1.10E-05 | 0.876514164 |
| ENSMUSG00000006014 | Prg4 | 96875 | protein_coding | 7.357783812 | 0.876745562 | 0.810601045 | -1.081599348 | 0.279430595 | 0.496825672 | 0.876745562 |
| ENSMUSG00000094145 | Vmn2r20 | 667180 | protein_coding | 4.890982006 | 0.876866601 | 2.100692055 | -0.417417964 | 0.676372717 | 0.826766769 | 0.876866601 |
| ENSMUSG00000022768 | Ccdc116 | 76872 | protein_coding | 159.0838222 | 0.877013755 | 0.248132781 | -3.534453416 | 0.000408619 | 0.00298846 | 0.877013755 |
| ENSMUSG00000001525 | Tubb5 | 22154 | protein_coding | 3287.877962 | 0.877286824 | 0.152815197 | -5.740834938 | 9.42E-09 | 1.82E-07 | 0.877286824 |
| ENSMUSG00000035373 | Ccl7 | 20306 | protein_coding | 0.972433854 | 0.877295989 | 2.135322357 | -0.410849438 | 0.68118294 | 1 | 0.877295989 |
| ENSMUSG00000085720 | Gm7854 | 665934 | lncRNA | 7.457090887 | -0.877493508 | 0.413074743 | 2.124297174 | 0.033645302 | 0.110042513 | 0.877493508 |
| ENSMUSG00000102918 | Pcdhgc3 | 93706 | protein_coding | 65.14849887 | 0.87752363 | 0.204399342 | -4.293182264 | 1.76E-05 | 0.000184206 | 0.87752363 |
| ENSMUSG00000034295 | Fhod3 | 225288 | protein_coding | 4.163953113 | -0.877619708 | 1.048836899 | 0.836755179 | 0.402730196 | 1 | 0.877619708 |
| ENSMUSG00000024798 | Htr7 | 15566 | protein_coding | 1.274491575 | 0.877723763 | 2.121239488 | -0.413778722 | 0.679036169 | 1 | 0.877723763 |
| ENSMUSG00000072952 | Gm5878 | 545861 | lncRNA | 1.582645534 | 0.878218646 | 1.478190496 | -0.594117367 | 0.552433616 | 1 | 0.878218646 |
| ENSMUSG00000004098 | Col5a3 | 53867 | protein_coding | 5.370239543 | 0.878548374 | 1.217129519 | -0.721819954 | 0.47040518 | 0.683706314 | 0.878548374 |
| ENSMUSG00000090369 | 4933411K16Rik | 66765 | protein_coding | 3.809414086 | 0.878711015 | 0.686871716 | -1.279294219 | 0.200793468 | 1 | 0.878711015 |
| ENSMUSG00000034000 | Neu4 | 241159 | protein_coding | 2.017603113 | -0.879094113 | 1.575443856 | 0.557997741 | 0.576845927 | 1 | 0.879094113 |
| ENSMUSG00000052981 | Ube2ql1 | 76980 | protein_coding | 1.656060936 | 0.879376288 | 1.717235416 | -0.512088372 | 0.608589162 | 1 | 0.879376288 |
| ENSMUSG00000028996 | Rbp7 | 63954 | protein_coding | 7.820896658 | -0.879505238 | 0.354098685 | 2.48378567 | 0.012999402 | 0.052155522 | 0.879505238 |
| ENSMUSG00000028542 | Slc6a9 | 14664 | protein_coding | 165.4468447 | 0.879632617 | 0.126595902 | -6.948349875 | 3.70E-12 | 1.07E-10 | 0.879632617 |
| ENSMUSG00000027857 | Tshb | 22094 | protein_coding | 1.282364304 | -0.879648951 | 2.554433255 | 0.344361689 | 0.730574296 | 1 | 0.879648951 |

| ensembl_gene_id | external_gene_name | entrezgene_id | gene_biotype | baseMean | log2FoldChange | lfcSE | stat | pvalue | padj | absLog2FC |
|---|---|---|---|---|---|---|---|---|---|---|
| ENSMUSG00000091519 | Skor2 | 664805 | protein_coding | 1.187243079 | -0.879755962 | 2.555293154 | 0.344287684 | 0.730629946 | 1 | 0.879755962 |
| ENSMUSG00000027827 | Kcnab1 | 16497 | protein_coding | 12.83275938 | 0.87976944 | 0.410561761 | -2.142843107 | 0.032125699 | 0.106381349 | 0.87976944 |
| ENSMUSG00000107722 | 2900060B14Rik | 68204 | lncRNA | 13.95944525 | 0.880518929 | 0.544512849 | -1.617076493 | 0.10586177 | 0.258037088 | 0.880518929 |
| ENSMUSG00000051617 | Krt9 | 107656 | protein_coding | 2.61542308 | -0.880557222 | 2.131290544 | 0.413156819 | 0.679491723 | 1 | 0.880557222 |
| ENSMUSG00000044824 | Or55b10 | 258837 | protein_coding | 0.874700101 | 0.88104969 | 1.997307243 | -0.441118758 | 0.659127024 | 1 | 0.88104969 |
| ENSMUSG00000039405 | Prss23 | 76453 | protein_coding | 153.1168113 | -0.8811276 | 0.228994842 | 3.84780545 | 0.000119181 | 0.001006726 | 0.8811276 |
| ENSMUSG00000062524 | Ncr1 | 17086 | protein_coding | 2.122727119 | 0.882272511 | 0.90342708 | -0.976584088 | 0.3287751 | 1 | 0.882272511 |
| ENSMUSG00000004939 | Nmrk2 | 69564 | protein_coding | 1.463340597 | 0.882488534 | 1.894586706 | -0.465794746 | 0.641362426 | 1 | 0.882488534 |
| ENSMUSG00000044951 | Mylk4 | 238564 | protein_coding | 3.428080827 | 0.882503618 | 1.857929094 | -0.474993163 | 0.634791846 | 1 | 0.882503618 |
| ENSMUSG00000085828 | Gm15612 | 100038356 | lncRNA | 2.639865129 | -0.88259587 | 1.024866519 | 0.861181289 | 0.389138204 | 1 | 0.88259587 |
| ENSMUSG00000045275 | Lca5l | 385668 | protein_coding | 4.884784607 | -0.882604924 | 0.562003345 | 1.570462048 | 0.116307657 | 0.275527851 | 0.882604924 |
| ENSMUSG00000027068 | Dhrs9 | 241452 | protein_coding | 423.4059394 | 0.88284513 | 0.253806421 | -3.478419206 | 0.00050438 | 0.00357329 | 0.88284513 |
| ENSMUSG00000037784 | Dzip1l | 72507 | protein_coding | 24.11279294 | -0.883189611 | 0.283422761 | 3.116156259 | 0.001832252 | 0.010564875 | 0.883189611 |
| ENSMUSG00000031951 | Tmem231 | 234740 | protein_coding | 50.17073503 | 0.883196714 | 0.243984909 | -3.619882544 | 0.000294737 | 0.002244022 | 0.883196714 |
| ENSMUSG00000059852 | Kcng2 | 240444 | protein_coding | 2.361725731 | -0.883927013 | 1.016207949 | 0.869828871 | 0.384393931 | 1 | 0.883927013 |
| ENSMUSG00000055333 | Fat2 | 245827 | protein_coding | 32.51733075 | 0.884051914 | 0.240042651 | -3.682895146 | 0.0002306 | 0.00179544 | 0.884051914 |
| ENSMUSG00000018581 | Dnah11 | 13411 | protein_coding | 7.838305433 | 0.884201969 | 0.8657885 | -1.02126786 | 0.307127549 | 0.52766379 | 0.884201969 |
| ENSMUSG00000036760 | Kcnk9 | 223604 | protein_coding | 3.242229228 | 0.884268093 | 2.213383096 | -0.399509735 | 0.689517652 | 1 | 0.884268093 |
| ENSMUSG00000035275 | Raver2 | 242570 | protein_coding | 8.108418499 | 0.884457391 | 0.61055511 | -1.448611888 | 0.147445999 | 0.325550242 | 0.884457391 |
| ENSMUSG00000079056 | Kcnip3 | 56461 | protein_coding | 1.980060328 | -0.884593993 | 1.771436801 | 0.499365257 | 0.61752209 | 1 | 0.884593993 |
| ENSMUSG00000025153 | Fasn | 14104 | protein_coding | 5581.1222 | 0.884991564 | 0.076731413 | -11.53362792 | 8.93E-31 | 7.19E-29 | 0.884991564 |
| ENSMUSG00000004110 | Cacna1e | 12290 | protein_coding | 44.42068949 | 0.885026089 | 0.654109994 | -1.353023341 | 0.176048174 | 0.368018849 | 0.885026089 |
| ENSMUSG00000118314 | 1700001K23Rik | 69319 | lncRNA | 3.073035639 | -0.88525214 | 0.633976214 | 1.396349139 | 0.162609383 | 1 | 0.88525214 |
| ENSMUSG00000030217 | Art4 | 109978 | protein_coding | 0.854167799 | 0.885539678 | 1.545443165 | -0.573000481 | 0.566644368 | 1 | 0.885539678 |
| ENSMUSG00000077271 | Gm22781 | 115486264 | snoRNA | 0.634872311 | 0.885654256 | 1.695418473 | -0.522380917 | 0.60140514 | 1 | 0.885654256 |
| ENSMUSG00000041180 | Hectd2 | 226098 | protein_coding | 7.925248884 | 0.885775373 | 0.509961644 | -1.736945088 | 0.082396864 | 0.214486414 | 0.885775373 |
| ENSMUSG00000031760 | Mt3 | 17751 | protein_coding | 31.60021426 | -0.886009337 | 0.308921727 | 2.868070649 | 0.004129833 | 0.020729502 | 0.886009337 |
| ENSMUSG00000021846 | Peli2 | 93834 | protein_coding | 156.3276866 | -0.88642446 | 0.154539827 | 5.735896559 | 9.70E-09 | 1.87E-07 | 0.88642446 |
| ENSMUSG00000070953 | Rabepk | 227746 | protein_coding | 404.9203853 | 0.886602355 | 0.068553939 | -12.9329163 | 2.93E-38 | 2.80E-36 | 0.886602355 |
| ENSMUSG00000074067 | Gm10619 | 100579147 | lncRNA | 9.026875496 | 0.886641803 | 0.470078902 | -1.88615528 | 0.059274033 | 0.169380754 | 0.886641803 |
| ENSMUSG00000039377 | Hlx | 15284 | protein_coding | 1.34073304 | 0.886774714 | 1.851685213 | -0.47890144 | 0.632008747 | 1 | 0.886774714 |
| ENSMUSG00000001510 | Dlx3 | 13393 | protein_coding | 0.920157312 | 0.887261424 | 1.873693831 | -0.473535969 | 0.635830845 | 1 | 0.887261424 |
| ENSMUSG00000035632 | Cnot3 | 232791 | protein_coding | 418.0640472 | 0.88729008 | 0.140502274 | -6.315129688 | 2.70E-10 | 6.41E-09 | 0.88729008 |
| ENSMUSG00000035227 | Spcs2 | 66624 | protein_coding | 1342.238089 | 0.887327469 | 0.108352149 | -8.189292778 | 2.63E-16 | 1.07E-14 | 0.887327469 |
| ENSMUSG00000115317 | Gm32618 | 102635226 | lncRNA | 2.099169076 | -0.887351607 | 0.762823583 | 1.163246164 | 0.244729642 | 1 | 0.887351607 |
| ENSMUSG00000063434 | Sorcs3 | 66673 | protein_coding | 5.591286696 | 0.887401022 | 0.716448112 | -1.238611711 | 0.215489329 | 0.420598049 | 0.887401022 |
| ENSMUSG00000039349 | C130074G19Rik | 226777 | protein_coding | 354.5277655 | -0.887416913 | 0.159018694 | 5.580582307 | 2.40E-08 | 4.35E-07 | 0.887416913 |
| ENSMUSG00000070390 | Nlrp1b | 637515 | protein_coding | 197.3508606 | -0.88771264 | 0.160013366 | 5.547740556 | 2.89E-08 | 5.17E-07 | 0.88771264 |
| ENSMUSG00000096878 | Gm21083 | 105242399 | protein_coding | 3.252788797 | -0.887844969 | 0.678107816 | 1.309297648 | 0.190433546 | 1 | 0.887844969 |

Supplementary Table 1

| ensembl_gene_id | external_gene_name | entrezgene_id | gene_biotype | baseMean | log2FoldChange | lfcSE | stat | pvalue | padj | absLog2FC |
|---|---|---|---|---|---|---|---|---|---|---|
| ENSMUSG00000112026 | Gm6653 | 626115 | TEC | 2.637082024 | 0.887860094 | 0.839622606 | -1.057451393 | 0.290305627 | 1 | 0.887860094 |
| ENSMUSG00000068373 | D430041D05Rik | 241589 | protein_coding | 5.44791237 | 0.887869405 | 2.028073658 | -0.437789526 | 0.661538865 | 0.816921981 | 0.887869405 |
| ENSMUSG00000098144 | 1700029N11Rik | 69491 | lncRNA | 1.129807505 | 0.888022141 | 1.702452229 | -0.521613544 | 0.601939431 | 1 | 0.888022141 |
| ENSMUSG00000065643 | Gm25897 | 115488597 | snRNA | 2.557494962 | -0.888372739 | 0.743879064 | 1.194243503 | 0.23238273 | 1 | 0.888372739 |
| ENSMUSG00000041696 | Rasl12 | 70784 | protein_coding | 16.33849234 | 0.888436555 | 0.431468209 | -2.059100849 | 0.039484579 | 0.124778744 | 0.888436555 |
| ENSMUSG00000034449 | Dhrs11 | 192970 | protein_coding | 1767.260651 | 0.888715601 | 0.103730925 | -8.567508704 | 1.06E-17 | 4.76E-16 | 0.888715601 |
| ENSMUSG00000019933 | Mrln | 69563 | protein_coding | 0.772196186 | 0.88909289 | 1.456721925 | -0.610338099 | 0.541637864 | 1 | 0.88909289 |
| ENSMUSG00000055172 | C1ra | 50909 | protein_coding | 15.93808932 | -0.889107569 | 0.304453571 | 2.92033878 | 0.003496511 | 0.018083738 | 0.889107569 |
| ENSMUSG00000029390 | Tmed2 | 56334 | protein_coding | 1392.810284 | 0.889451881 | 0.10951159 | -8.121988529 | 4.59E-16 | 1.83E-14 | 0.889451881 |
| ENSMUSG00000085667 | Gm12992 | 545681 | lncRNA | 22.1545187 | -0.88962174 | 0.257550253 | 3.454167609 | 0.000551994 | 0.003846258 | 0.88962174 |
| ENSMUSG00000022498 | Txndc11 | 106200 | protein_coding | 867.8486049 | 0.889811171 | 0.080213362 | -11.09305411 | 1.36E-28 | 9.95E-27 | 0.889811171 |
| ENSMUSG00000074141 | Il4i1 | 14204 | protein_coding | 2.297388094 | -0.889929086 | 0.698431522 | 1.274182304 | 0.202598815 | 1 | 0.889929086 |
| ENSMUSG00000047014 | Catsperb | 271036 | protein_coding | 4.23567943 | -0.890147185 | 0.932517118 | 0.954563909 | 0.339798276 | 1 | 0.890147185 |
| ENSMUSG00000099032 | Tcf24 | 100039596 | protein_coding | 16.99108351 | 0.890246541 | 0.370180683 | -2.404897342 | 0.016177009 | 0.062098357 | 0.890246541 |
| ENSMUSG00000034460 | Six4 | 20474 | protein_coding | 6.507819987 | -0.890458781 | 0.807298989 | 1.103009904 | 0.270022864 | 0.485271256 | 0.890458781 |
| ENSMUSG00000017316 | Ppy | 19064 | protein_coding | 14.38665488 | -0.890628762 | 0.467435984 | 1.905349165 | 0.056734699 | 0.163972315 | 0.890628762 |
| ENSMUSG00000090588 | Muc21 | 672682 | protein_coding | 7.389502334 | 0.890640229 | 1.811713513 | -0.49160103 | 0.623001415 | 0.793281643 | 0.890640229 |
| ENSMUSG00000023132 | Gzma | 14938 | protein_coding | 67.47670878 | 0.890961746 | 0.384460745 | -2.317432292 | 0.020480195 | 0.074722558 | 0.890961746 |
| ENSMUSG00000048003 | Catsper4 | 329954 | protein_coding | 12.24183026 | 0.891191736 | 0.468943 | -1.900426566 | 0.057377163 | 0.165304689 | 0.891191736 |
| ENSMUSG00000048582 | Gja3 | 14611 | protein_coding | 1.191024674 | 0.891256148 | 2.051350275 | -0.434472922 | 0.663945054 | 1 | 0.891256148 |
| ENSMUSG00000073125 | Xlr3b | 574437 | protein_coding | 21.55213485 | 0.891345466 | 0.745095482 | -1.196283548 | 0.231585927 | 0.440448001 | 0.891345466 |
| ENSMUSG00000013338 | Fer1l4 | 74562 | protein_coding | 83.39029113 | 0.891451016 | 0.259706884 | -3.432527478 | 0.000597983 | 0.004119773 | 0.891451016 |
| ENSMUSG00000056162 | Cndp1 | 338403 | protein_coding | 3.002278768 | -0.891830348 | 1.3129161 | 0.679274439 | 0.496963989 | 1 | 0.891830348 |
| ENSMUSG00000019579 | Mydgf | 28106 | protein_coding | 1338.01011 | 0.891882137 | 0.072992629 | -12.21879736 | 2.47E-34 | 2.16E-32 | 0.891882137 |
| ENSMUSG00000058908 | Pla2g2a | 18780 | protein_coding | 31.06132765 | 0.892005369 | 0.382253968 | -2.333541164 | 0.01961976 | 0.072289318 | 0.892005369 |
| ENSMUSG00000101605 | Ace3 | 217246 | protein_coding | 6.214007142 | 0.89215915 | 0.525222163 | -1.698631955 | 0.089388552 | 0.227592128 | 0.89215915 |
| ENSMUSG00000039278 | Pcsk1n | 30052 | protein_coding | 28.08511742 | -0.893124549 | 0.345957912 | 2.581598856 | 0.009834382 | 0.0420321 | 0.893124549 |
| ENSMUSG00000025739 | Gng13 | 64337 | protein_coding | 3.213966485 | 0.893145496 | 0.611677662 | -1.460157123 | 0.144246896 | 1 | 0.893145496 |
| ENSMUSG00000066760 | Psg16 | 26436 | protein_coding | 2.220503981 | 0.893284555 | 1.492728499 | -0.598423997 | 0.549557058 | 1 | 0.893284555 |
| ENSMUSG00000025213 | Kazald1 | 107250 | protein_coding | 19.54428874 | -0.893457257 | 0.276858496 | 3.22712602 | 0.001250404 | 0.007657016 | 0.893457257 |
| ENSMUSG00000052861 | Dnah6 | 330355 | protein_coding | 5.863796241 | 0.893647282 | 1.575046081 | -0.567378499 | 0.570457054 | 0.756061621 | 0.893647282 |
| ENSMUSG00000032517 | Mobp | 17433 | protein_coding | 2.620676955 | 0.893784971 | 2.42426148 | -0.368683402 | 0.712363722 | 1 | 0.893784971 |
| ENSMUSG00000111474 | Gm32926 | 102635639 | lncRNA | 2.788717273 | -0.893874928 | 0.712967096 | 1.253739386 | 0.209936747 | 1 | 0.893874928 |
| ENSMUSG00000031548 | Sfrp1 | 20377 | protein_coding | 2.946759209 | 0.894065914 | 1.099027649 | -0.813506298 | 0.415927837 | 1 | 0.894065914 |
| ENSMUSG00000089914 | 4930486I03Rik | 75028 | lncRNA | 0.798898479 | -0.894128781 | 1.702009345 | 0.525337175 | 0.599348819 | 1 | 0.894128781 |
| ENSMUSG00000044337 | Ackr3 | 12778 | protein_coding | 14.22355352 | 0.894161161 | 0.317773766 | -2.813829384 | 0.00489552 | 0.023887859 | 0.894161161 |
| ENSMUSG00000046804 | Phgr1 | 53906 | protein_coding | 2861.453242 | -0.894531734 | 0.115274279 | 7.76002889 | 8.49E-15 | 3.06E-13 | 0.894531734 |
| ENSMUSG00000062252 | Lhfpl5 | 328789 | protein_coding | 4.352694892 | -0.89457328 | 0.765335583 | 1.16886409 | 0.242458394 | 1 | 0.89457328 |
| ENSMUSG00000025389 | Mip | 17339 | protein_coding | 1.226734597 | 0.894638867 | 1.462869309 | -0.611564452 | 0.540825964 | 1 | 0.894638867 |

Supplementary Table 1

| ensembl_gene_id | external_gene_name | entrezgene_id | gene_biotype | baseMean | log2FoldChange | lfcSE | stat | pvalue | padj | absLog2FC |
|---|---|---|---|---|---|---|---|---|---|---|
| ENSMUSG00000046591 | Ticrr | 77011 | protein_coding | 94.99401508 | 0.895032284 | 0.200341183 | -4.467540175 | 7.91E-06 | 9.10E-05 | 0.895032284 |
| ENSMUSG00000014867 | Surf4 | 20932 | protein_coding | 4301.477393 | 0.895151842 | 0.057673239 | -15.52109547 | 2.50E-54 | 3.72E-52 | 0.895151842 |
| ENSMUSG00000106563 | Mir6962 | 102466774 | miRNA | 0.566983279 | -0.895239167 | 2.017289695 | 0.443783146 | 0.657199375 | 1 | 0.895239167 |
| ENSMUSG00000035355 | Kcnh4 | 380728 | protein_coding | 4.026444399 | 0.89565282 | 2.22757046 | -0.40207609 | 0.687628029 | 1 | 0.89565282 |
| ENSMUSG00000037610 | Kcnmb2 | 72413 | protein_coding | 1.4833374 | -0.895929959 | 1.117840176 | 0.801483055 | 0.42285205 | 1 | 0.895929959 |
| ENSMUSG00000008461 | Fut1 | 14343 | protein_coding | 26.62113007 | 0.896002854 | 0.231909836 | -3.86358281 | 0.000111736 | 0.000950159 | 0.896002854 |
| ENSMUSG00000038296 | Galnt18 | 233733 | protein_coding | 12.43508696 | 0.896714807 | 0.346543282 | -2.587598297 | 0.009664762 | 0.041412525 | 0.896714807 |
| ENSMUSG00000020712 | Tcam1 | 75870 | protein_coding | 1.119484996 | 0.896858759 | 1.491295769 | -0.60139563 | 0.547576508 | 1 | 0.896858759 |
| ENSMUSG00000091694 | Apol11b | 328563 | protein_coding | 1.43221576 | -0.897561346 | 1.347092172 | 0.666295421 | 0.505222292 | 1 | 0.897561346 |
| ENSMUSG00000078851 | H2ac25 | 319162 | protein_coding | 71.44591117 | 0.897845862 | 0.206164372 | -4.355000103 | 1.33E-05 | 0.000143719 | 0.897845862 |
| ENSMUSG00000021280 | Exoc3l4 | 74190 | protein_coding | 33.69190549 | 0.898266752 | 0.27425398 | -3.275309818 | 0.001055462 | 0.006655171 | 0.898266752 |
| ENSMUSG00000089943 | Ugt1a5 | 394433 | protein_coding | 10.13874202 | -0.898306638 | 0.503257872 | 1.784982783 | 0.07426411 | 0.199317001 | 0.898306638 |
| ENSMUSG00000020469 | Myl7 | 17898 | protein_coding | 32.03313184 | -0.89851253 | 0.215865134 | 4.162379138 | 3.15E-05 | 0.000310511 | 0.89851253 |
| ENSMUSG00000048967 | Yjefn3 | 234365 | protein_coding | 7.290232773 | 0.89861066 | 0.452937972 | -1.983959651 | 0.047260338 | 0.142911457 | 0.89861066 |
| ENSMUSG00000057967 | Fgf18 | 14172 | protein_coding | 2.487191531 | -0.898909891 | 0.892426286 | 1.007265143 | 0.313807376 | 1 | 0.898909891 |
| ENSMUSG00000022041 | Chrna2 | 110902 | protein_coding | 3.291524608 | 0.899385714 | 1.120735623 | -0.802495875 | 0.422266174 | 1 | 0.899385714 |
| ENSMUSG00000028177 | 1810013D15Rik | 69042 | lncRNA | 1.662153128 | -0.899543876 | 0.942542839 | 0.954379831 | 0.339891412 | 1 | 0.899543876 |
| ENSMUSG00000047953 | Gp5 | 14729 | protein_coding | 2.515938636 | 0.899600929 | 0.893547624 | -1.006774463 | 0.314043169 | 1 | 0.899600929 |
| ENSMUSG00000024357 | Sil1 | 81500 | protein_coding | 844.4621551 | 0.899634941 | 0.100906693 | -8.915513062 | 4.86E-19 | 2.35E-17 | 0.899634941 |
| ENSMUSG00000030742 | Lat | 16797 | protein_coding | 4.390917037 | -0.89969107 | 0.663579469 | 1.35581511 | 0.175158002 | 0.366750254 | 0.89969107 |
| ENSMUSG00000020704 | Asic2 | 11418 | protein_coding | 1.891745513 | 0.899740166 | 1.362611926 | -0.660305512 | 0.509057794 | 1 | 0.899740166 |
| ENSMUSG00000030713 | Klk7 | 23993 | protein_coding | 1.493489569 | -0.899818639 | 2.597408963 | 0.346429327 | 0.729020088 | 1 | 0.899818639 |
| ENSMUSG00000051652 | Lrrc3 | 237387 | protein_coding | 4.470782871 | -0.900688253 | 0.54833168 | 1.642597511 | 0.100466236 | 0.24866598 | 0.900688253 |
| ENSMUSG00000029403 | Cdkl2 | 53886 | protein_coding | 169.3151874 | 0.901167182 | 0.125603909 | -7.174674638 | 7.25E-13 | 2.25E-11 | 0.901167182 |
| ENSMUSG00000055561 | Spink5 | 72432 | protein_coding | 2.865403289 | 0.901246433 | 0.84310673 | -1.068958889 | 0.285088194 | 1 | 0.901246433 |
| ENSMUSG00000027201 | Myef2 | 17876 | protein_coding | 129.8639349 | 0.902434657 | 0.140002627 | -6.445840864 | 1.15E-10 | 2.83E-09 | 0.902434657 |
| ENSMUSG00000036551 | Akap14 | 434756 | protein_coding | 1.277358606 | -0.902647614 | 1.274324347 | 0.708334276 | 0.478737697 | 1 | 0.902647614 |
| ENSMUSG00000055926 | Gm14137 | 623781 | protein_coding | 41.82508548 | 0.902850641 | 0.397825194 | -2.269465722 | 0.023240019 | 0.082552308 | 0.902850641 |
| ENSMUSG00000054417 | Cyp3a44 | 337924 | protein_coding | 13.76216655 | -0.903055216 | 0.532771911 | 1.695012814 | 0.09007299 | 0.228901189 | 0.903055216 |
| ENSMUSG00000048371 | Pdp2 | 382051 | protein_coding | 419.2272262 | -0.903063787 | 0.149924861 | 6.023442554 | 1.71E-09 | 3.66E-08 | 0.903063787 |
| ENSMUSG00000106779 | Gm8579 | 667335 | lncRNA | 3.073366769 | 0.903075423 | 0.607571952 | -1.486367861 | 0.137181846 | 1 | 0.903075423 |
| ENSMUSG00000060550 | H2-Q7 | 15018 | protein_coding | 43.91490727 | 0.903470608 | 0.288684684 | -3.129610468 | 0.001750382 | 0.010149514 | 0.903470608 |
| ENSMUSG00000039286 | Fndc3b | 72007 | protein_coding | 893.4618893 | 0.903510838 | 0.083963347 | -10.76077684 | 5.27E-27 | 3.69E-25 | 0.903510838 |
| ENSMUSG00000039814 | Xkr5 | 319581 | protein_coding | 22.83596202 | 0.903514758 | 0.292186911 | -3.092249253 | 0.00198646 | 0.011271717 | 0.903514758 |
| ENSMUSG00000046805 | Mpeg1 | 17476 | protein_coding | 3.633003675 | 0.903525304 | 0.690013778 | -1.309430817 | 0.190388458 | 1 | 0.903525304 |
| ENSMUSG00000027611 | Procr | 19124 | protein_coding | 1.718653031 | 0.903527141 | 1.742306287 | -0.518581117 | 0.60405288 | 1 | 0.903527141 |
| ENSMUSG00000030590 | Fam98c | 73833 | protein_coding | 450.9332754 | 0.90413415 | 0.090563698 | -9.983405767 | 1.80E-23 | 1.09E-21 | 0.90413415 |
| ENSMUSG00000064057 | Scgb3a1 | 68662 | protein_coding | 1.016014019 | 0.904192022 | 1.439271538 | -0.628228932 | 0.52985398 | 1 | 0.904192022 |
| ENSMUSG00000085424 | Fhad1os1 | 545691 | lncRNA | 4.345573064 | -0.904299492 | 0.64063196 | 1.411574115 | 0.158075399 | 1 | 0.904299492 |

| ensembl_gene_id | external_gene_name | entrezgene_id | gene_biotype | baseMean | log2FoldChange | lfcSE | stat | pvalue | padj | absLog2FC |
|---|---|---|---|---|---|---|---|---|---|---|
| ENSMUSG00000033578 | Tmem35a | 67564 | protein_coding | 34.76250575 | -0.904730809 | 0.269819171 | 3.353100541 | 0.000799117 | 0.005253429 | 0.904730809 |
| ENSMUSG00000021238 | Aldh6a1 | 104776 | protein_coding | 830.2804486 | 0.905107601 | 0.091819916 | -9.857421309 | 6.37E-23 | 3.77E-21 | 0.905107601 |
| ENSMUSG00000061653 | Vmn1r46 | 113856 | protein_coding | 3.176155807 | 0.905140064 | 3.635552382 | -0.248969061 | 0.803384713 | 1 | 0.905140064 |
| ENSMUSG00000106004 | Gm43391 | 115490167 | lncRNA | 3.13996604 | 0.905333829 | 1.709299878 | -0.529651842 | 0.596353343 | 1 | 0.905333829 |
| ENSMUSG00000062438 | Adam1b | 280667 | protein_coding | 3.159963387 | 0.905424426 | 1.100559731 | -0.822694489 | 0.41068175 | 1 | 0.905424426 |
| ENSMUSG00000045790 | Ccdc149 | 100503884 | protein_coding | 211.2559477 | 0.905499763 | 0.092295736 | -9.810851495 | 1.01E-22 | 5.90E-21 | 0.905499763 |
| ENSMUSG00000095664 | Vmn2r67 | 620672 | protein_coding | 2.847561738 | -0.905728555 | 2.685068123 | 0.337320513 | 0.735875298 | 1 | 0.905728555 |
| ENSMUSG00000053916 | Nanp | 67311 | protein_coding | 1.951593483 | -0.906082559 | 0.8370008 | 1.082534879 | 0.279014926 | 1 | 0.906082559 |
| ENSMUSG00000027808 | Serp1 | 28146 | protein_coding | 4420.331383 | 0.906184556 | 0.092918912 | -9.752423282 | 1.80E-22 | 1.03E-20 | 0.906184556 |
| ENSMUSG00000027187 | Cat | 12359 | protein_coding | 1806.295625 | 0.906511613 | 0.061012073 | -14.85790554 | 6.18E-50 | 8.65E-48 | 0.906511613 |
| ENSMUSG00000104401 | Gm20750 | 622124 | lncRNA | 1.796185904 | 0.906918411 | 2.365887613 | -0.383331146 | 0.701474248 | 1 | 0.906918411 |
| ENSMUSG00000023505 | Cdca3 | 14793 | protein_coding | 545.7531951 | 0.907724988 | 0.146142188 | -6.211245379 | 5.26E-10 | 1.19E-08 | 0.907724988 |
| ENSMUSG00000067615 | Krt81 | 64818 | protein_coding | 0.706425736 | -0.908235054 | 1.656924362 | 0.548145151 | 0.58359224 | 1 | 0.908235054 |
| ENSMUSG00000098544 | Mir7211 | 102466817 | miRNA | 6.181784015 | -0.908344037 | 0.477905488 | 1.900677143 | 0.057344314 | 0.165238877 | 0.908344037 |
| ENSMUSG00000020654 | Adcy3 | 104111 | protein_coding | 18.41608331 | 0.909493205 | 0.280446924 | -3.243013658 | 0.001182725 | 0.00731937 | 0.909493205 |
| ENSMUSG00000009545 | Kcnq1 | 16535 | protein_coding | 765.1325257 | -0.909804822 | 0.096005842 | 9.476556855 | 2.63E-21 | 1.44E-19 | 0.909804822 |
| ENSMUSG00000052955 | Cpvl | 71287 | protein_coding | 3.018423606 | -0.910128918 | 0.720133872 | 1.263832953 | 0.206289991 | 1 | 0.910128918 |
| ENSMUSG00000059303 | Or7g21 | 258557 | protein_coding | 1.040319469 | -0.910240167 | 1.991182256 | 0.457135536 | 0.64757363 | 1 | 0.910240167 |
| ENSMUSG00000049038 | Mterf2 | 74238 | protein_coding | 56.8828101 | -0.910419172 | 0.156048298 | 5.834214051 | 5.40E-09 | 1.09E-07 | 0.910419172 |
| ENSMUSG00000046718 | Bst2 | 69550 | protein_coding | 202.5090878 | 0.911582482 | 0.263720113 | -3.456628585 | 0.000546978 | 0.003827457 | 0.911582482 |
| ENSMUSG00000022951 | Rcan1 | 54720 | protein_coding | 445.7967919 | 0.911665296 | 0.068668343 | -13.27635499 | 3.17E-40 | 3.36E-38 | 0.911665296 |
| ENSMUSG00000025092 | Hspa12a | 73442 | protein_coding | 39.74384708 | 0.911709111 | 0.268356452 | -3.397380999 | 0.000680342 | 0.00458204 | 0.911709111 |
| ENSMUSG00000030786 | Itgam | 16409 | protein_coding | 13.83432117 | 0.911775563 | 0.354591504 | -2.57134069 | 0.010130561 | 0.043017765 | 0.911775563 |
| ENSMUSG00000050463 | Krt78 | 332131 | protein_coding | 10.83314129 | 0.911815445 | 0.470109936 | -1.939579181 | 0.052430853 | 0.154753013 | 0.911815445 |
| ENSMUSG00000030324 | Rho | 212541 | protein_coding | 1.821536625 | 0.911831868 | 1.922141847 | -0.474383235 | 0.635226646 | 1 | 0.911831868 |
| ENSMUSG00000051503 | Ccdc121rt3 | 625424 | protein_coding | 4.45303544 | 0.9125183 | 0.666098193 | -1.369945618 | 0.170703878 | 0.360508479 | 0.9125183 |
| ENSMUSG00000065934 | Gm25630 | 115487300 | snRNA | 0.391082598 | -0.912751106 | 1.977039804 | 0.461675635 | 0.644313946 | 1 | 0.912751106 |
| ENSMUSG00000039391 | Ccdc81 | 70884 | protein_coding | 1.827659788 | 0.913024815 | 1.718585057 | -0.531265422 | 0.595234863 | 1 | 0.913024815 |
| ENSMUSG00000023411 | Nfatc4 | 73181 | protein_coding | 2.609552167 | 0.913160423 | 2.146390149 | -0.425440092 | 0.670515885 | 1 | 0.913160423 |
| ENSMUSG00000047904 | Sstr2 | 20606 | protein_coding | 31.65515154 | -0.913204287 | 0.293072638 | 3.115965702 | 0.001833436 | 0.010565985 | 0.913204287 |
| ENSMUSG00000029026 | Trp73 | 22062 | protein_coding | 1.848421696 | -0.913422601 | 1.509905085 | 0.604953656 | 0.54520979 | 1 | 0.913422601 |
| ENSMUSG00000042216 | Sgsm1 | 52850 | protein_coding | 430.9706152 | 0.913595208 | 0.133593381 | -6.8386263 | 8.00E-12 | 2.25E-10 | 0.913595208 |
| ENSMUSG00000022456 | Septin3 | 24050 | protein_coding | 3.084908777 | 0.914127346 | 1.339910463 | -0.682230172 | 0.495093419 | 1 | 0.914127346 |
| ENSMUSG00000069273 | H3c6 | 319151 | protein_coding | 1601.102433 | 0.914352674 | 0.160679208 | -5.690547542 | 1.27E-08 | 2.40E-07 | 0.914352674 |
| ENSMUSG00000063428 | Ddo | 70503 | protein_coding | 3.450322174 | 0.91452303 | 1.828270984 | -0.50021197 | 0.616925831 | 1 | 0.91452303 |
| ENSMUSG00000037035 | Inhbb | 16324 | protein_coding | 2.108778922 | 0.914841623 | 1.295423079 | -0.706210688 | 0.480057125 | 1 | 0.914841623 |
| ENSMUSG00000047669 | Msl3l2 | 73390 | protein_coding | 20.52007085 | 0.915372446 | 0.25903924 | -3.533721164 | 0.000409753 | 0.002995424 | 0.915372446 |
| ENSMUSG00000060268 | Armh1 | 381544 | protein_coding | 1.138930417 | -0.915641134 | 1.603913245 | 0.570879464 | 0.568081352 | 1 | 0.915641134 |
| ENSMUSG00000031668 | Eif2ak3 | 13666 | protein_coding | 811.907448 | 0.915805282 | 0.08815928 | -10.38807583 | 2.81E-25 | 1.80E-23 | 0.915805282 |

Supplementary Table 1

| ensembl_gene_id | external_gene_name | entrezgene_id | gene_biotype | baseMean | log2FoldChange | lfcSE | stat | pvalue | padj | absLog2FC |
|---|---|---|---|---|---|---|---|---|---|---|
| ENSMUSG00000078249 | Hmga1b | 111241 | protein_coding | 8.746298292 | 0.915938278 | 0.447514117 | -2.046724883 | 0.040685105 | 0.127638694 | 0.915938278 |
| ENSMUSG00000027293 | Ehd4 | 98878 | protein_coding | 937.3700393 | 0.915940471 | 0.09803294 | -9.34319088 | 9.35E-21 | 5.00E-19 | 0.915940471 |
| ENSMUSG00000017720 | Trp53tg5 | 73603 | protein_coding | 2.292161434 | 0.91613651 | 0.797812418 | -1.148310668 | 0.250840337 | 1 | 0.91613651 |
| ENSMUSG00000048031 | Fcrl5 | 329693 | protein_coding | 1.331317144 | 0.916206348 | 1.723813513 | -0.53149969 | 0.595072556 | 1 | 0.916206348 |
| ENSMUSG00000000594 | Gm2a | 14667 | protein_coding | 149.926243 | -0.916925 | 0.156234572 | 5.868899498 | 4.39E-09 | 8.92E-08 | 0.916925 |
| ENSMUSG00000031552 | Adam18 | 13524 | protein_coding | 4.828176156 | -0.917033044 | 0.543688223 | 1.686689182 | 0.091663133 | 0.231775376 | 0.917033044 |
| ENSMUSG00000035910 | Dcdc2a | 195208 | protein_coding | 15.34025911 | 0.917097499 | 0.356415645 | -2.573112354 | 0.010078848 | 0.042864306 | 0.917097499 |
| ENSMUSG00000022306 | Zfpm2 | 22762 | protein_coding | 2.714652083 | 0.917159338 | 2.732873133 | -0.335602603 | 0.737170561 | 1 | 0.917159338 |
| ENSMUSG00000025104 | Hdgfl3 | 29877 | protein_coding | 2.365758986 | -0.917410416 | 1.468169575 | 0.624866795 | 0.532058487 | 1 | 0.917410416 |
| ENSMUSG00000037469 | Acp7 | 101744 | protein_coding | 1.894656639 | 0.917437094 | 2.419712406 | -0.379151296 | 0.704575516 | 1 | 0.917437094 |
| ENSMUSG00000033510 | Otud7a | 170711 | protein_coding | 3.969137226 | -0.917623281 | 0.718403477 | 1.277309075 | 0.201493151 | 1 | 0.917623281 |
| ENSMUSG00000034923 | Ly6g6f | 433099 | protein_coding | 20.00284082 | -0.917626737 | 0.271031099 | 3.385687993 | 0.000710001 | 0.004741185 | 0.917626737 |
| ENSMUSG00000052736 | Klrc2 | 16642 | protein_coding | 2.874017702 | 0.918261555 | 0.86277791 | -1.064308142 | 0.287189131 | 1 | 0.918261555 |
| ENSMUSG00000079259 | Trim71 | 636931 | protein_coding | 7.678685006 | 0.919146257 | 0.670186592 | -1.371478135 | 0.170225955 | 0.359891361 | 0.919146257 |
| ENSMUSG00000025202 | Scd3 | 30049 | protein_coding | 5.544345619 | 0.919558287 | 0.727554771 | -1.263902491 | 0.206265028 | 0.408887615 | 0.919558287 |
| ENSMUSG00000060992 | Copz1 | 56447 | protein_coding | 1860.73397 | 0.919571028 | 0.070311447 | -13.07853939 | 4.37E-39 | 4.32E-37 | 0.919571028 |
| ENSMUSG00000054115 | Skp2 | 27401 | protein_coding | 80.47192549 | 0.919778558 | 0.176412894 | -5.213783072 | 1.85E-07 | 2.92E-06 | 0.919778558 |
| ENSMUSG00000056427 | Slit3 | 20564 | protein_coding | 3.280919435 | -0.920284163 | 1.843337435 | 0.499248887 | 0.617604059 | 1 | 0.920284163 |
| ENSMUSG00000054855 | Rnd1 | 223881 | protein_coding | 7.370571184 | 0.920303649 | 0.694505411 | -1.325120918 | 0.185131049 | 0.380539475 | 0.920303649 |
| ENSMUSG00000006056 | Calcoco2 | 76815 | protein_coding | 3.305015249 | -0.920366153 | 0.632208045 | 1.45579633 | 0.145448931 | 1 | 0.920366153 |
| ENSMUSG00000032400 | Zwilch | 68014 | protein_coding | 107.7737516 | 0.92085516 | 0.156210727 | -5.894954703 | 3.75E-09 | 7.70E-08 | 0.92085516 |
| ENSMUSG00000051159 | Cited1 | 12705 | protein_coding | 2.633182679 | -0.920942237 | 0.950857391 | 0.968538758 | 0.332775372 | 1 | 0.920942237 |
| ENSMUSG00000049336 | Tenm2 | 23964 | protein_coding | 4.360040465 | -0.921013558 | 1.889080454 | 0.487545968 | 0.625871476 | 1 | 0.921013558 |
| ENSMUSG00000069265 | H3c1 | 360198 | protein_coding | 614.4360551 | 0.921117764 | 0.191440771 | -4.811502569 | 1.50E-06 | 1.99E-05 | 0.921117764 |
| ENSMUSG00000067996 | Bpifb9b | 433492 | protein_coding | 1.322171592 | -0.921339891 | 1.317456332 | 0.699332394 | 0.484344327 | 1 | 0.921339891 |
| ENSMUSG00000038292 | Kash5 | 384619 | protein_coding | 2.972438753 | -0.921523872 | 1.478441464 | 0.623307648 | 0.533082373 | 1 | 0.921523872 |
| ENSMUSG00000062510 | Nsl1 | 381318 | protein_coding | 81.17290267 | 0.921670427 | 0.236144733 | -3.902989556 | 9.50E-05 | 0.000823203 | 0.921670427 |
| ENSMUSG00000048922 | Cdca2 | 108912 | protein_coding | 217.1866461 | 0.921813687 | 0.167714338 | -5.496332019 | 3.88E-08 | 6.78E-07 | 0.921813687 |
| ENSMUSG00000036251 | Trpm8 | 171382 | protein_coding | 3.852696925 | -0.921916708 | 2.131105096 | 0.432600302 | 0.66530517 | 1 | 0.921916708 |
| ENSMUSG00000075705 | Msrb1 | 27361 | protein_coding | 738.9115506 | 0.922126158 | 0.066976351 | -13.76793667 | 3.97E-43 | 4.53E-41 | 0.922126158 |
| ENSMUSG00000046031 | Calhm6 | 215900 | protein_coding | 1.296655829 | -0.922164913 | 1.176210775 | 0.784013319 | 0.433032295 | 1 | 0.922164913 |
| ENSMUSG00000045777 | Ifitm10 | 320802 | protein_coding | 3.433056983 | 0.92268299 | 1.499911186 | -0.615158416 | 0.538450107 | 1 | 0.92268299 |
| ENSMUSG00000114627 | B230220B15Rik | 102634835 | lncRNA | 0.596480381 | 0.922977537 | 1.828718431 | -0.504712766 | 0.613760594 | 1 | 0.922977537 |
| ENSMUSG00000045689 | Pcdhb4 | 93875 | protein_coding | 3.525510215 | 0.923275498 | 0.680568776 | -1.356623357 | 0.174900916 | 1 | 0.923275498 |
| ENSMUSG00000110206 | Flt3l | 14256 | protein_coding | 10.86710417 | -0.923323817 | 0.37181273 | 2.4833034 | 0.013017017 | 0.052213508 | 0.923323817 |
| ENSMUSG00000061603 | Akap6 | 238161 | protein_coding | 4.425319714 | -0.923650455 | 1.506083746 | 0.613279612 | 0.539691472 | 0.734194205 | 0.923650455 |
| ENSMUSG00000004961 | Syt5 | 53420 | protein_coding | 9.090668649 | -0.923814517 | 0.620549398 | 1.488704236 | 0.136565269 | 0.308556417 | 0.923814517 |
| ENSMUSG00000048038 | Ccdc187 | 329366 | protein_coding | 3.935716715 | -0.923928782 | 0.998806234 | 0.925033056 | 0.354948713 | 1 | 0.923928782 |
| ENSMUSG00000056724 | Nbeal2 | 235627 | protein_coding | 1585.722251 | 0.923956219 | 0.099804089 | -9.257699057 | 2.09E-20 | 1.07E-18 | 0.923956219 |

Supplementary Table 1

| ensembl_gene_id | external_gene_name | entrezgene_id | gene_biotype | baseMean | log2FoldChange | lfcSE | stat | pvalue | padj | absLog2FC |
|---|---|---|---|---|---|---|---|---|---|---|
| ENSMUSG00000068617 | Efcab1 | 66793 | protein_coding | 4.921577667 | -0.924080464 | 0.519991528 | 1.777106767 | 0.075550675 | 0.201525415 | 0.924080464 |
| ENSMUSG00000043029 | Trpv3 | 246788 | protein_coding | 197.7952676 | -0.924358444 | 0.2655786 | 3.480545663 | 0.000500394 | 0.003550608 | 0.924358444 |
| ENSMUSG00000066158 | AY512931 | 434310 | lncRNA | 3.156913574 | 0.924378699 | 1.025738779 | -0.90118334 | 0.367490847 | 1 | 0.924378699 |
| ENSMUSG00000034311 | Kif4 | 16571 | protein_coding | 419.1810353 | 0.924811695 | 0.147207587 | -6.282364343 | 3.33E-10 | 7.84E-09 | 0.924811695 |
| ENSMUSG00000071516 | H2ac13 | 319191 | protein_coding | 196.425756 | 0.925024897 | 0.192046317 | -4.81667605 | 1.46E-06 | 1.95E-05 | 0.925024897 |
| ENSMUSG00000058743 | Kcnj14 | 211480 | protein_coding | 1.777182778 | 0.925178029 | 1.087726191 | -0.850561508 | 0.395012979 | 1 | 0.925178029 |
| ENSMUSG00000035493 | Tgfbi | 21810 | protein_coding | 44.19936347 | -0.925286928 | 0.383179952 | 2.414758193 | 0.015745659 | 0.060682336 | 0.925286928 |
| ENSMUSG00000015812 | Gnrh1 | 14714 | protein_coding | 4.843757724 | 0.925424042 | 0.470205792 | -1.968125566 | 0.049053592 | 0.146938331 | 0.925424042 |
| ENSMUSG00000029061 | Mmp23 | 26561 | protein_coding | 4.153076524 | 0.925558047 | 0.757074823 | -1.22254501 | 0.221501594 | 1 | 0.925558047 |
| ENSMUSG00000000290 | Itgb2 | 16414 | protein_coding | 18.72504132 | 0.925698425 | 0.390345565 | -2.371484417 | 0.017716794 | 0.066683451 | 0.925698425 |
| ENSMUSG00000036256 | Igfbp7 | 29817 | protein_coding | 3.103257863 | -0.925772055 | 0.84356988 | 1.097445602 | 0.272446648 | 1 | 0.925772055 |
| ENSMUSG00000111746 | 4933422A05Rik | 71133 | lncRNA | 0.974314677 | -0.926425276 | 1.407998134 | 0.657973369 | 0.510555244 | 1 | 0.926425276 |
| ENSMUSG00000010830 | Kdelr3 | 105785 | protein_coding | 290.2955037 | 0.926542015 | 0.148407241 | -6.243239925 | 4.29E-10 | 9.86E-09 | 0.926542015 |
| ENSMUSG00000037627 | Rgs22 | 626596 | protein_coding | 6.192424931 | -0.926915135 | 0.771063422 | 1.202125673 | 0.229314841 | 0.438111017 | 0.926915135 |
| ENSMUSG00000113005 | Gm34408 | 102637649 | lncRNA | 0.971578612 | -0.927042832 | 1.737543005 | 0.533536625 | 0.593662165 | 1 | 0.927042832 |
| ENSMUSG00000026399 | Cd55 | 13136 | protein_coding | 199.0815865 | -0.92717424 | 0.122506273 | 7.568381766 | 3.78E-14 | 1.29E-12 | 0.92717424 |
| ENSMUSG00000097184 | 4632428C04Rik | 100043102 | lncRNA | 29.18475993 | -0.92722336 | 0.224555533 | 4.129149468 | 3.64E-05 | 0.000353491 | 0.92722336 |
| ENSMUSG00000026832 | Cytip | 227929 | protein_coding | 57.45871774 | -0.927509643 | 0.20222466 | 4.586530861 | 4.51E-06 | 5.45E-05 | 0.927509643 |
| ENSMUSG00000069302 | H2ac12 | 319168 | protein_coding | 57.15601589 | 0.927697782 | 0.270706921 | -3.426945188 | 0.000610412 | 0.004191413 | 0.927697782 |
| ENSMUSG00000086999 | Bcas1os2 | 73303 | lncRNA | 10.25563317 | -0.92797127 | 0.459999244 | 2.017332162 | 0.043660861 | 0.134442563 | 0.92797127 |
| ENSMUSG00000010663 | Fads1 | 76267 | protein_coding | 188.2102292 | 0.927989838 | 0.20399162 | -4.549156671 | 5.39E-06 | 6.42E-05 | 0.927989838 |
| ENSMUSG00000027202 | Slc12a1 | 20495 | protein_coding | 1.400085374 | -0.928362295 | 1.816207735 | 0.511154246 | 0.609243054 | 1 | 0.928362295 |
| ENSMUSG00000064354 | mt-Co2 | 17709 | protein_coding | 11.69372133 | 0.928392591 | 0.600483335 | -1.546075532 | 0.122086327 | 0.285167413 | 0.928392591 |
| ENSMUSG00000061186 | Sfmbt2 | 353282 | protein_coding | 8.074001509 | 0.928450763 | 0.755860419 | -1.228336263 | 0.219320763 | 0.425536825 | 0.928450763 |
| ENSMUSG00000030584 | Dpf1 | 29861 | protein_coding | 3.920199872 | -0.928771927 | 1.570571675 | 0.591359148 | 0.554279807 | 1 | 0.928771927 |
| ENSMUSG00000005470 | Asf1b | 66929 | protein_coding | 146.115357 | 0.928813938 | 0.215458762 | -4.310866397 | 1.63E-05 | 0.000171923 | 0.928813938 |
| ENSMUSG00000000028 | Cdc45 | 12544 | protein_coding | 132.7624057 | 0.928910028 | 0.212587155 | -4.369549181 | 1.25E-05 | 0.000136072 | 0.928910028 |
| ENSMUSG00000030108 | Slc6a13 | 14412 | protein_coding | 2.102181007 | -0.928911132 | 1.197347603 | 0.775807401 | 0.437862713 | 1 | 0.928911132 |
| ENSMUSG00000042743 | Sgtb | 218544 | protein_coding | 25.45449063 | -0.929523953 | 0.225845994 | 4.115742484 | 3.86E-05 | 0.000372492 | 0.929523953 |
| ENSMUSG00000029552 | Tes | 21753 | protein_coding | 3057.395813 | 0.929949386 | 0.051343948 | -18.11215196 | 2.56E-73 | 5.86E-71 | 0.929949386 |
| ENSMUSG00000086969 | 4930443O20Rik | 74687 | lncRNA | 3.8147469 | -0.930107353 | 0.618012206 | 1.504998354 | 0.1323245 | 1 | 0.930107353 |
| ENSMUSG00000026223 | Itm2c | 64294 | protein_coding | 836.4428093 | 0.930299717 | 0.085212665 | -10.91738785 | 9.52E-28 | 6.86E-26 | 0.930299717 |
| ENSMUSG00000039601 | Rcan2 | 53901 | protein_coding | 3.388186972 | 0.930345112 | 1.066347619 | -0.872459501 | 0.382957753 | 1 | 0.930345112 |
| ENSMUSG00000031085 | Acte1 | 102636989 | protein_coding | 1.601192077 | -0.930937674 | 1.855793724 | 0.501638551 | 0.615921797 | 1 | 0.930937674 |
| ENSMUSG00000026011 | Ctla4 | 12477 | protein_coding | 2.235826088 | 0.931539923 | 1.093224174 | -0.852103297 | 0.394156762 | 1 | 0.931539923 |
| ENSMUSG00000030790 | Adm | 11535 | protein_coding | 65.33720203 | 0.932003634 | 0.406652587 | -2.291891562 | 0.021911901 | 0.07888094 | 0.932003634 |
| ENSMUSG00000007944 | Ttc9b | 73032 | protein_coding | 1.330203125 | -0.932056829 | 1.468083349 | 0.634880049 | 0.52550665 | 1 | 0.932056829 |
| ENSMUSG00000065429 | Mir345 | 723946 | miRNA | 1.196599494 | 0.932349776 | 1.008041987 | -0.924911648 | 0.355011867 | 1 | 0.932349776 |
| ENSMUSG00000006345 | Ggt1 | 14598 | protein_coding | 299.2761748 | 0.932430644 | 0.457345784 | -2.038787011 | 0.041471289 | 0.129452831 | 0.932430644 |

| ensembl_gene_id | external_gene_name | entrezgene_id | gene_biotype | baseMean | log2FoldChange | lfcSE | stat | pvalue | padj | absLog2FC |
|---|---|---|---|---|---|---|---|---|---|---|
| ENSMUSG00000025383 | Il23a | 83430 | protein_coding | 2.125475796 | 0.932442036 | 1.203701235 | -0.774645742 | 0.43854902 | 1 | 0.932442036 |
| ENSMUSG00000041794 | Myrip | 245049 | protein_coding | 60.78178667 | -0.93296989 | 0.172347863 | 5.413295373 | 6.19E-08 | 1.04E-06 | 0.93296989 |
| ENSMUSG00000074882 | Cyp2c68 | 433247 | protein_coding | 587.5409233 | -0.933341443 | 0.243379707 | 3.834918918 | 0.000125606 | 0.001056137 | 0.933341443 |
| ENSMUSG00000026450 | Chit1 | 71884 | protein_coding | 10.73420166 | -0.93353076 | 0.410514722 | 2.274049406 | 0.022963016 | 0.081727701 | 0.93353076 |
| ENSMUSG00000069917 | Hba-a2 | 110257 | protein_coding | 25.34066783 | -0.933618823 | 0.55573128 | 1.679982497 | 0.092960721 | 0.234124348 | 0.933618823 |
| ENSMUSG00000026981 | Il1rn | 16181 | protein_coding | 390.0520056 | -0.933977923 | 0.230150726 | 4.058114167 | 4.95E-05 | 0.000462865 | 0.933977923 |
| ENSMUSG00000003411 | Rab3b | 69908 | protein_coding | 17.63819048 | 0.934226819 | 0.349239092 | -2.675035072 | 0.007472143 | 0.033659295 | 0.934226819 |
| ENSMUSG00000085912 | Trp53cor1 | 100504267 | lncRNA | 0.98376516 | 0.934282916 | 1.564817857 | -0.597055377 | 0.550470409 | 1 | 0.934282916 |
| ENSMUSG00000115432 | D130009I18Rik | 320249 | lncRNA | 2.205453243 | 0.934309771 | 1.955382555 | -0.477814313 | 0.632782373 | 1 | 0.934309771 |
| ENSMUSG00000026471 | Mr1 | 15064 | protein_coding | 2.014602042 | 0.934627716 | 0.956433149 | -0.977201299 | 0.328469505 | 1 | 0.934627716 |
| ENSMUSG00000041889 | Shisa4 | 77552 | protein_coding | 2.120577396 | -0.934890531 | 1.345944584 | 0.694598086 | 0.487307212 | 1 | 0.934890531 |
| ENSMUSG00000029811 | Aoc1 | 76507 | protein_coding | 2480.256173 | -0.93507092 | 0.18394713 | 5.083367816 | 3.71E-07 | 5.60E-06 | 0.93507092 |
| ENSMUSG00000041736 | Tspo | 12257 | protein_coding | 643.7073545 | -0.935116904 | 0.104580018 | 8.941640334 | 3.83E-19 | 1.87E-17 | 0.935116904 |
| ENSMUSG00000022021 | Diaph3 | 56419 | protein_coding | 143.5065713 | 0.935228457 | 0.151605591 | -6.168825659 | 6.88E-10 | 1.54E-08 | 0.935228457 |
| ENSMUSG00000053552 | Ebf4 | 228598 | protein_coding | 1.915609598 | 0.93536173 | 1.661635812 | -0.562916208 | 0.573491943 | 1 | 0.93536173 |
| ENSMUSG00000110310 | 4930518J21Rik | 75123 | lncRNA | 1.038590796 | -0.935628363 | 1.498501155 | 0.624376138 | 0.532380592 | 1 | 0.935628363 |
| ENSMUSG00000006378 | Gcat | 26912 | protein_coding | 74.259972 | 0.936088891 | 0.145013277 | -6.455194371 | 1.08E-10 | 2.67E-09 | 0.936088891 |
| ENSMUSG00000035539 | Ccdc180 | 381522 | protein_coding | 3.600471411 | 0.936220928 | 1.355543798 | -0.690660773 | 0.489778746 | 1 | 0.936220928 |
| ENSMUSG00000024378 | Stard4 | 170459 | protein_coding | 3236.942073 | 0.936429016 | 0.099492056 | -9.41209831 | 4.86E-21 | 2.63E-19 | 0.936429016 |
| ENSMUSG00000049916 | 2610318N02Rik | 70458 | protein_coding | 38.41473532 | 0.936618224 | 0.247241402 | -3.788274193 | 0.000151697 | 0.001240778 | 0.936618224 |
| ENSMUSG00000038173 | Enpp6 | 320981 | protein_coding | 1.784015021 | 0.936693165 | 1.40981118 | -0.664410368 | 0.506427699 | 1 | 0.936693165 |
| ENSMUSG00000041538 | H2-Ob | 15002 | protein_coding | 3.943691014 | 0.936784017 | 1.099307267 | -0.852158487 | 0.394126134 | 1 | 0.936784017 |
| ENSMUSG00000020627 | Klhl29 | 208439 | protein_coding | 2.951502377 | 0.936787367 | 1.011033973 | -0.926563687 | 0.354153116 | 1 | 0.936787367 |
| ENSMUSG00000090386 | Mir99ahg | 77994 | lncRNA | 8.765535866 | 0.936906105 | 0.630505403 | -1.485960469 | 0.137289577 | 0.309726787 | 0.936906105 |
| ENSMUSG00000021747 | Cfap20dc | 74430 | protein_coding | 22.05423216 | 0.936929998 | 0.278661591 | -3.362250232 | 0.0007731 | 0.005117025 | 0.936929998 |
| ENSMUSG00000052430 | Bmpr1b | 12167 | protein_coding | 1.660760341 | -0.937098979 | 1.896874526 | 0.494022649 | 0.621290184 | 1 | 0.937098979 |
| ENSMUSG00000117199 | Gm4566 | 100043644 | lncRNA | 0.982872845 | 0.937127572 | 1.807962821 | -0.518333431 | 0.604225652 | 1 | 0.937127572 |
| ENSMUSG00000073176 | Zfp449 | 78619 | protein_coding | 3.671912275 | 0.937339356 | 0.603818608 | -1.55235255 | 0.12057789 | 1 | 0.937339356 |
| ENSMUSG00000030043 | Tacr1 | 21336 | protein_coding | 1.533014142 | 0.93761448 | 2.335796189 | -0.401411084 | 0.68811749 | 1 | 0.93761448 |
| ENSMUSG00000041544 | Disp3 | 242748 | protein_coding | 3.263716855 | 0.937641839 | 1.750554024 | -0.535625765 | 0.592217218 | 1 | 0.937641839 |
| ENSMUSG00000039742 | Garin1b | 330277 | protein_coding | 2.579561144 | 0.937657205 | 1.185569434 | -0.790891852 | 0.429007104 | 1 | 0.937657205 |
| ENSMUSG00000032420 | Nt5e | 23959 | protein_coding | 465.8474689 | -0.937884042 | 0.211600334 | 4.432337245 | 9.32E-06 | 0.000105365 | 0.937884042 |
| ENSMUSG00000029307 | Dmp1 | 13406 | protein_coding | 1.076306715 | -0.938138181 | 2.414705497 | 0.388510393 | 0.697638364 | 1 | 0.938138181 |
| ENSMUSG00000049709 | Nlrp10 | 244202 | protein_coding | 33.1620934 | -0.938338023 | 0.244008969 | 3.845506282 | 0.000120304 | 0.001014656 | 0.938338023 |
| ENSMUSG00000024225 | Clps | 109791 | protein_coding | 80.06163289 | -0.938701414 | 0.286671655 | 3.274482842 | 0.001058556 | 0.006667458 | 0.938701414 |
| ENSMUSG00000020319 | Wdpcp | 216560 | protein_coding | 47.47150097 | 0.9387164 | 0.16256534 | -5.774394471 | 7.72E-09 | 1.51E-07 | 0.9387164 |
| ENSMUSG00000036446 | Lum | 17022 | protein_coding | 0.734222075 | -0.939013694 | 1.624651805 | 0.577978426 | 0.563278684 | 1 | 0.939013694 |
| ENSMUSG00000051732 | Pabpc2 | 18459 | protein_coding | 0.947493842 | 0.939223255 | 1.954700054 | -0.480494822 | 0.630875584 | 1 | 0.939223255 |
| ENSMUSG00000040612 | Ildr2 | 100039795 | protein_coding | 4.462379886 | 0.939251552 | 0.936458273 | -1.002982813 | 0.315869154 | 0.536652594 | 0.939251552 |

| ensembl_gene_id | external_gene_name | entrezgene_id | gene_biotype | baseMean | log2FoldChange | lfcSE | stat | pvalue | padj | absLog2FC |
|---|---|---|---|---|---|---|---|---|---|---|
| ENSMUSG00000030008 | Pradc1 | 73327 | protein_coding | 115.1479085 | 0.939259026 | 0.158692838 | -5.918723466 | 3.24E-09 | 6.71E-08 | 0.939259026 |
| ENSMUSG00000037578 | Pkd2l1 | 329064 | protein_coding | 1.832040884 | 0.939420356 | 1.791204055 | -0.524463058 | 0.599956508 | 1 | 0.939420356 |
| ENSMUSG00000066677 | Ifi208 | 100033459 | protein_coding | 1.611742688 | 0.939569331 | 1.374040884 | -0.683800127 | 0.494101386 | 1 | 0.939569331 |
| ENSMUSG00000022574 | Naprt | 223646 | protein_coding | 633.6110978 | -0.939836336 | 0.082225472 | 11.42999017 | 2.96E-30 | 2.33E-28 | 0.939836336 |
| ENSMUSG00000024786 | Majin | 622554 | protein_coding | 3.125068539 | 0.940030791 | 0.681724321 | -1.37890165 | 0.167925079 | 1 | 0.940030791 |
| ENSMUSG00000022385 | Gtse1 | 29870 | protein_coding | 148.4137754 | 0.940092203 | 0.216914407 | -4.333931594 | 1.46E-05 | 0.000156354 | 0.940092203 |
| ENSMUSG00000033849 | B3galt2 | 26878 | protein_coding | 105.4182544 | -0.940432669 | 0.247937678 | 3.793020386 | 0.000148826 | 0.001219773 | 0.940432669 |
| ENSMUSG00000070690 | Albfm1 | 622307 | protein_coding | 1.600181766 | 0.940571418 | 1.289738548 | -0.729272936 | 0.465834724 | 1 | 0.940571418 |
| ENSMUSG00000023247 | Guca2a | 14915 | protein_coding | 21304.74436 | 0.940713103 | 0.101243731 | -9.291568914 | 1.52E-20 | 7.95E-19 | 0.940713103 |
| ENSMUSG00000049252 | Lrp1b | 94217 | protein_coding | 4.913694617 | 0.940811512 | 2.060142987 | -0.456672919 | 0.647906158 | 0.8085485 | 0.940811512 |
| ENSMUSG00000037509 | Arhgef4 | 226970 | protein_coding | 20.636965 | 0.940814601 | 0.393579989 | -2.390402527 | 0.016829916 | 0.06408329 | 0.940814601 |
| ENSMUSG00000034206 | Polq | 77782 | protein_coding | 116.2133589 | 0.941252196 | 0.194694858 | -4.834499517 | 1.33E-06 | 1.79E-05 | 0.941252196 |
| ENSMUSG00000046841 | Ckap4 | 216197 | protein_coding | 592.23081 | 0.941361207 | 0.116670957 | -8.068513658 | 7.12E-16 | 2.82E-14 | 0.941361207 |
| ENSMUSG00000046387 | Pcdhb17 | 93888 | protein_coding | 5.60911885 | 0.941706286 | 0.557460425 | -1.689279173 | 0.091165943 | 0.230830869 | 0.941706286 |
| ENSMUSG00000043631 | Ecm2 | 407800 | protein_coding | 5.454978834 | 0.942085417 | 0.490276852 | -1.921537624 | 0.054663964 | 0.159434952 | 0.942085417 |
| ENSMUSG00000018669 | Cdk5rap3 | 80280 | protein_coding | 641.5429663 | 0.942495702 | 0.076581273 | -12.30713027 | 8.29E-35 | 7.28E-33 | 0.942495702 |
| ENSMUSG00000037287 | Tbcel | 272589 | protein_coding | 162.0431613 | -0.94252204 | 0.092810719 | 10.15531446 | 3.14E-24 | 1.94E-22 | 0.94252204 |
| ENSMUSG00000070806 | Zmynd12 | 332934 | protein_coding | 7.591215054 | 0.943086626 | 0.552850986 | -1.705860439 | 0.088034067 | 0.22515016 | 0.943086626 |
| ENSMUSG00000006651 | Aplp1 | 11803 | protein_coding | 252.9501547 | 0.943827175 | 0.309394097 | -3.050566202 | 0.002284103 | 0.012608423 | 0.943827175 |
| ENSMUSG00000072082 | Ccnf | 12449 | protein_coding | 249.1684051 | 0.944200406 | 0.184720122 | -5.11151895 | 3.20E-07 | 4.87E-06 | 0.944200406 |
| ENSMUSG00000018774 | Cd68 | 12514 | protein_coding | 26.20783283 | 0.94424811 | 0.500945451 | -1.884931997 | 0.059439019 | 0.169705389 | 0.94424811 |
| ENSMUSG00000029375 | Cxcl15 | 20309 | protein_coding | 0.656534616 | -0.94435704 | 1.756242398 | 0.537714521 | 0.590774153 | 1 | 0.94435704 |
| ENSMUSG00000015484 | Fam163a | 329274 | protein_coding | 2.70250079 | 0.944359006 | 1.992284796 | -0.474008037 | 0.635494176 | 1 | 0.944359006 |
| ENSMUSG00000034059 | Ypel4 | 241525 | protein_coding | 3.057546915 | 0.944432389 | 1.284811505 | -0.735074666 | 0.462294022 | 1 | 0.944432389 |
| ENSMUSG00000049037 | Clec4a1 | 269799 | protein_coding | 6.059127643 | -0.94474507 | 0.512016904 | 1.845144296 | 0.065016551 | 0.181090121 | 0.94474507 |
| ENSMUSG00000031089 | Slc6a14 | 56774 | protein_coding | 10412.40315 | -0.944806572 | 0.143053497 | 6.604568169 | 3.99E-11 | 1.05E-09 | 0.944806572 |
| ENSMUSG00000037386 | Rims2 | 116838 | protein_coding | 61.74669544 | -0.944820178 | 0.488281813 | 1.934989495 | 0.052991585 | 0.155962044 | 0.944820178 |
| ENSMUSG00000021287 | Xrcc3 | 74335 | protein_coding | 127.8274425 | 0.944856866 | 0.146056852 | -6.469103335 | 9.86E-11 | 2.45E-09 | 0.944856866 |
| ENSMUSG00000022894 | Adamts5 | 23794 | protein_coding | 2.404696156 | 0.945221396 | 1.525392834 | -0.619657688 | 0.535483178 | 1 | 0.945221396 |
| ENSMUSG00000039013 | Siglecf | 233186 | protein_coding | 57.85647359 | -0.945298827 | 0.221517085 | 4.26738564 | 1.98E-05 | 0.000203749 | 0.945298827 |
| ENSMUSG00000024210 | Ip6k3 | 271424 | protein_coding | 1.043216031 | -0.945389086 | 2.01611017 | 0.468917374 | 0.639128697 | 1 | 0.945389086 |
| ENSMUSG00000050052 | Tdrp | 72148 | protein_coding | 5.81539658 | -0.9455264 | 0.680709467 | 1.389030778 | 0.164823391 | 0.351529573 | 0.9455264 |
| ENSMUSG00000021966 | Prss52 | 73382 | protein_coding | 0.760939114 | -0.946620721 | 1.667069473 | 0.567835196 | 0.570146879 | 1 | 0.946620721 |
| ENSMUSG00000030607 | Acan | 11595 | protein_coding | 2.520783879 | 0.94676266 | 1.339866055 | -0.70660993 | 0.479808916 | 1 | 0.94676266 |
| ENSMUSG00000072589 | Gm10371 | 100038654 | lncRNA | 1.125412863 | 0.946806439 | 2.343097914 | -0.404083173 | 0.686151559 | 1 | 0.946806439 |
| ENSMUSG00000017204 | Gsdma | 57911 | protein_coding | 6.611336722 | -0.946939128 | 0.468935426 | 2.019338005 | 0.0434521 | 0.133877139 | 0.946939128 |
| ENSMUSG00000040298 | Btbd16 | 330660 | protein_coding | 0.981940403 | -0.947102528 | 1.923207767 | 0.492459808 | 0.622394328 | 1 | 0.947102528 |
| ENSMUSG00000044071 | Tafa2 | 268354 | protein_coding | 1.65440969 | 0.947459468 | 1.147997002 | -0.825315281 | 0.409192618 | 1 | 0.947459468 |
| ENSMUSG00000030662 | Ipo5 | 70572 | protein_coding | 2081.544373 | 0.947801492 | 0.132289353 | -7.16460901 | 7.80E-13 | 2.41E-11 | 0.947801492 |

| ensembl_gene_id | external_gene_name | entrezgene_id | gene_biotype | baseMean | log2FoldChange | lfcSE | stat | pvalue | padj | absLog2FC |
|---|---|---|---|---|---|---|---|---|---|---|
| ENSMUSG00000072884 | Gm10433 | 100038378 | lncRNA | 18.71970895 | 0.948988533 | 0.261148519 | -3.633903566 | 0.000279165 | 0.002138281 | 0.948988533 |
| ENSMUSG00000030870 | Ubfd1 | 28018 | protein_coding | 961.1161676 | 0.949461642 | 0.090556287 | -10.48476776 | 1.01E-25 | 6.60E-24 | 0.949461642 |
| ENSMUSG00000038379 | Ttk | 22137 | protein_coding | 146.7198981 | 0.949546046 | 0.192481396 | -4.933183505 | 8.09E-07 | 1.14E-05 | 0.949546046 |
| ENSMUSG00000022436 | Sh3bp1 | 20401 | protein_coding | 9.320177425 | -0.949594192 | 0.459992075 | 2.064370766 | 0.038982576 | 0.123467254 | 0.949594192 |
| ENSMUSG00000029335 | Bmp3 | 110075 | protein_coding | 867.3475676 | -0.949790815 | 0.155935825 | 6.090908326 | 1.12E-09 | 2.46E-08 | 0.949790815 |
| ENSMUSG00000031837 | Necab2 | 117148 | protein_coding | 2.328946054 | 0.949846895 | 1.175341108 | -0.808145728 | 0.419006696 | 1 | 0.949846895 |
| ENSMUSG00000056749 | Nfil3 | 18030 | protein_coding | 338.0493873 | 0.95002452 | 0.285589857 | -3.326534526 | 0.000879331 | 0.005716696 | 0.95002452 |
| ENSMUSG00000102416 | 4933424G06Rik | 71166 | protein_coding | 1.08156994 | 0.950292739 | 2.529215438 | -0.375726292 | 0.707120389 | 1 | 0.950292739 |
| ENSMUSG00000016200 | Syt14 | 329324 | protein_coding | 5.564507452 | -0.951024434 | 0.696771611 | 1.364901237 | 0.172284093 | 0.36280925 | 0.951024434 |
| ENSMUSG00000059455 | Plac8l1 | 69401 | protein_coding | 2.379133967 | 0.951392175 | 1.316311669 | -0.722771208 | 0.469820459 | 1 | 0.951392175 |
| ENSMUSG00000028758 | Kif17 | 16559 | protein_coding | 6.786367263 | 0.951424786 | 0.827591508 | -1.149630919 | 0.250295918 | 0.462658024 | 0.951424786 |
| ENSMUSG00000054988 | Agtr1b | 11608 | protein_coding | 1.628622275 | -0.951521472 | 0.971029956 | 0.979909494 | 0.327130796 | 1 | 0.951521472 |
| ENSMUSG00000036357 | Gpr101 | 245424 | protein_coding | 1.783952033 | 0.951758258 | 2.388467316 | -0.398480754 | 0.690275842 | 1 | 0.951758258 |
| ENSMUSG00000094152 | Slc6a16 | 381884 | protein_coding | 2.511902147 | 0.952145941 | 2.407587307 | -0.395477222 | 0.692490728 | 1 | 0.952145941 |
| ENSMUSG00000070423 | Or51e1 | 259097 | protein_coding | 3.947811015 | -0.952791337 | 0.644044261 | 1.479387978 | 0.139036651 | 1 | 0.952791337 |
| ENSMUSG00000116597 | Gm536 | 224098 | lncRNA | 1.417716346 | -0.953342058 | 1.940001817 | 0.491412972 | 0.623134391 | 1 | 0.953342058 |
| ENSMUSG00000037143 | Cfap61 | 78774 | protein_coding | 3.376924266 | -0.953992007 | 1.237415849 | 0.770955058 | 0.440733571 | 1 | 0.953992007 |
| ENSMUSG00000046240 | Hepacam | 72927 | protein_coding | 8.786582047 | -0.95402362 | 0.676403944 | 1.41043474 | 0.158411354 | 0.342262172 | 0.95402362 |
| ENSMUSG00000049173 | Myoz3 | 170947 | protein_coding | 2.507813864 | 0.954263278 | 1.761304533 | -0.541793461 | 0.587960797 | 1 | 0.954263278 |
| ENSMUSG00000096822 | Or1j15 | 258621 | protein_coding | 1.837566876 | -0.954317649 | 2.545910882 | 0.374843305 | 0.707777006 | 1 | 0.954317649 |
| ENSMUSG00000029254 | Stap1 | 56792 | protein_coding | 3.57354415 | 0.955510688 | 0.898714122 | -1.06319759 | 0.287692355 | 1 | 0.955510688 |
| ENSMUSG00000065444 | Mir27a | 387220 | miRNA | 22.26108567 | -0.955578888 | 0.333306112 | 2.866970792 | 0.004144213 | 0.02077642 | 0.955578888 |
| ENSMUSG00000031079 | Zfp300 | 245368 | protein_coding | 1.109131055 | -0.955795056 | 1.190703099 | 0.802714847 | 0.422139569 | 1 | 0.955795056 |
| ENSMUSG00000032500 | Dclk3 | 245038 | protein_coding | 40.14956701 | -0.955859058 | 0.220622443 | 4.332555861 | 1.47E-05 | 0.000157233 | 0.955859058 |
| ENSMUSG00000038209 | Itln1 | 16429 | protein_coding | 14.49399076 | -0.955861504 | 0.299849074 | 3.187808757 | 0.001433553 | 0.008569101 | 0.955861504 |
| ENSMUSG00000013353 | 4931406B18Rik | 74054 | protein_coding | 3.123792507 | 0.956031026 | 2.645897475 | -0.361325802 | 0.717855908 | 1 | 0.956031026 |
| ENSMUSG00000028757 | Ddost | 13200 | protein_coding | 4763.674709 | 0.956099464 | 0.109646486 | -8.719836802 | 2.79E-18 | 1.29E-16 | 0.956099464 |
| ENSMUSG00000021922 | Itih4 | 16427 | protein_coding | 2.534273152 | 0.956234193 | 2.661097272 | -0.359338309 | 0.719342019 | 1 | 0.956234193 |
| ENSMUSG00000096465 | Or5p64 | 258727 | protein_coding | 2.253160556 | 0.956246733 | 2.661979374 | -0.359223945 | 0.719427565 | 1 | 0.956246733 |
| ENSMUSG00000095448 | Or7e168 | 258519 | protein_coding | 2.075300488 | 0.956437907 | 2.674051624 | -0.357673688 | 0.720587526 | 1 | 0.956437907 |
| ENSMUSG00000027889 | Ampd2 | 109674 | protein_coding | 551.5635042 | 0.95706967 | 0.087721959 | -10.91026324 | 1.03E-27 | 7.39E-26 | 0.95706967 |
| ENSMUSG00000022367 | Has2 | 15117 | protein_coding | 1.428202454 | 0.957222445 | 1.035379116 | -0.924513959 | 0.355218787 | 1 | 0.957222445 |
| ENSMUSG00000009614 | Sardh | 192166 | protein_coding | 113.1940832 | 0.95751099 | 0.183889971 | -5.206977751 | 1.92E-07 | 3.01E-06 | 0.95751099 |
| ENSMUSG00000087412 | Gm15501 | 100040298 | transcribed_processed_pseudogene | 9.064612415 | 0.957525634 | 0.544547528 | -1.758387623 | 0.078681578 | 0.207566563 | 0.957525634 |
| ENSMUSG00000075027 | 4631405J19Rik | 433456 | lncRNA | 3.787641474 | 0.957865558 | 0.757100533 | -1.265176177 | 0.20580818 | 1 | 0.957865558 |
| ENSMUSG00000053121 | Gm5129 | 332993 | lncRNA | 2.27489155 | 0.958048792 | 0.773578099 | -1.238464214 | 0.215543983 | 1 | 0.958048792 |
| ENSMUSG00000006360 | Crip1 | 12925 | protein_coding | 4187.172091 | -0.9582824 | 0.07705189 | 12.43684478 | 1.65E-35 | 1.47E-33 | 0.9582824 |
| ENSMUSG00000030510 | Cers3 | 545975 | protein_coding | 8.092370443 | 0.958378092 | 0.443303385 | -2.161901137 | 0.030625797 | 0.102483165 | 0.958378092 |
| ENSMUSG00000021485 | Mxd3 | 17121 | protein_coding | 60.73504001 | 0.958441155 | 0.216762622 | -4.421616359 | 9.80E-06 | 0.000110204 | 0.958441155 |

| ensembl_gene_id | external_gene_name | entrezgene_id | gene_biotype | baseMean | log2FoldChange | lfcSE | stat | pvalue | padj | absLog2FC |
|---|---|---|---|---|---|---|---|---|---|---|
| ENSMUSG00000078127 | Fam170b | 105511 | protein_coding | 1.015300078 | -0.958881852 | 2.142851723 | 0.447479329 | 0.654529009 | 1 | 0.958881852 |
| ENSMUSG00000053046 | Brsk2 | 75770 | protein_coding | 18.22537147 | 0.959062086 | 0.361279755 | -2.654624489 | 0.007939674 | 0.035388872 | 0.959062086 |
| ENSMUSG00000079516 | Reg3a | 19694 | protein_coding | 17.76257254 | 0.959509462 | 0.543146111 | -1.766577063 | 0.077299085 | 0.205062987 | 0.959509462 |
| ENSMUSG00000019027 | Dnah1 | 110084 | protein_coding | 10.09079515 | 0.959520353 | 0.99410085 | -0.965214296 | 0.334437488 | 0.556711723 | 0.959520353 |
| ENSMUSG00000070419 | Cyp3a57 | 622127 | protein_coding | 1.660511348 | -0.959803648 | 2.181458399 | 0.439982559 | 0.659949739 | 1 | 0.959803648 |
| ENSMUSG00000024107 | Lhcgr | 16867 | protein_coding | 2.629765228 | 0.960046248 | 3.054144032 | -0.314342165 | 0.753261177 | 1 | 0.960046248 |
| ENSMUSG00000035681 | Kcnc2 | 268345 | protein_coding | 2.02798711 | 0.960275758 | 3.085349717 | -0.311237249 | 0.755620268 | 1 | 0.960275758 |
| ENSMUSG00000022321 | Cdh10 | 320873 | protein_coding | 1.755005817 | 0.96033746 | 3.093939144 | -0.310393132 | 0.756262017 | 1 | 0.96033746 |
| ENSMUSG00000021415 | Tex56 | 66761 | protein_coding | 1.017824647 | -0.960358877 | 1.266155395 | 0.758484212 | 0.448161161 | 1 | 0.960358877 |
| ENSMUSG00000023902 | Zscan10 | 332221 | protein_coding | 1.848399832 | 0.960847928 | 1.575285764 | -0.609951508 | 0.54189393 | 1 | 0.960847928 |
| ENSMUSG00000057722 | Lepr | 16847 | protein_coding | 1.367608714 | -0.96087342 | 1.625727922 | 0.591041961 | 0.554492306 | 1 | 0.96087342 |
| ENSMUSG00000014782 | Plekhg4 | 102075 | protein_coding | 7.013817307 | 0.960990363 | 0.596327128 | -1.611515421 | 0.107067429 | 0.260075004 | 0.960990363 |
| ENSMUSG00000047307 | Pcdhb13 | 93884 | protein_coding | 2.060195448 | 0.961070074 | 1.754496732 | -0.547775357 | 0.583846161 | 1 | 0.961070074 |
| ENSMUSG00000023906 | Cldn6 | 54419 | protein_coding | 1.620593519 | 0.961390733 | 1.397074046 | -0.688145869 | 0.491360928 | 1 | 0.961390733 |
| ENSMUSG00000039376 | Synpo2l | 68760 | protein_coding | 3.648933496 | 0.961563283 | 1.21546857 | -0.791105016 | 0.428882712 | 1 | 0.961563283 |
| ENSMUSG00000009115 | Spatc1l | 76573 | protein_coding | 0.895676099 | 0.961753691 | 1.097817143 | -0.876060004 | 0.380997414 | 1 | 0.961753691 |
| ENSMUSG00000041205 | Map6d1 | 208158 | protein_coding | 3.612044941 | -0.962309642 | 0.751616022 | 1.28032082 | 0.200432328 | 1 | 0.962309642 |
| ENSMUSG00000052234 | Epx | 13861 | protein_coding | 2.005686318 | -0.962323132 | 1.082763162 | 0.888766043 | 0.374128829 | 1 | 0.962323132 |
| ENSMUSG00000034533 | Scn10a | 20264 | protein_coding | 3.971953148 | 0.962467514 | 3.490088431 | -0.27577167 | 0.782723445 | 1 | 0.962467514 |
| ENSMUSG00000035896 | Rnase1 | 19752 | protein_coding | 2.183185476 | -0.962718244 | 0.757644436 | 1.270672889 | 0.203845044 | 1 | 0.962718244 |
| ENSMUSG00000025993 | Slc40a1 | 53945 | protein_coding | 2723.548767 | -0.962772625 | 0.12480765 | 7.714051399 | 1.22E-14 | 4.31E-13 | 0.962772625 |
| ENSMUSG00000108530 | 4930429H19Rik | 100505025 | lncRNA | 1.169473235 | 0.96373696 | 1.937019507 | -0.497536012 | 0.618811115 | 1 | 0.96373696 |
| ENSMUSG00000040552 | C3ar1 | 12267 | protein_coding | 2.335045362 | 0.963772876 | 1.972532561 | -0.488596688 | 0.625127258 | 1 | 0.963772876 |
| ENSMUSG00000046650 | Or5an6 | 258679 | protein_coding | 1.723503826 | 0.963848087 | 2.676736819 | -0.360083248 | 0.71878488 | 1 | 0.963848087 |
| ENSMUSG00000033697 | Arhgap39 | 223666 | protein_coding | 655.3354561 | 0.963859618 | 0.091797799 | -10.49981184 | 8.66E-26 | 5.74E-24 | 0.963859618 |
| ENSMUSG00000113087 | 4930473H19Rik | 74926 | lncRNA | 1.840333636 | -0.963910746 | 0.890228714 | 1.082767531 | 0.278911621 | 1 | 0.963910746 |
| ENSMUSG00000086179 | Gm14317 | 670507 | lncRNA | 0.949110067 | 0.964046719 | 1.903931677 | -0.506345228 | 0.612614318 | 1 | 0.964046719 |
| ENSMUSG00000110333 | Gm45861 | 70952 | protein_coding | 2.143294415 | 0.964101211 | 2.703576105 | -0.356602209 | 0.721389624 | 1 | 0.964101211 |
| ENSMUSG00000113584 | Gm46355 | 108168006 | lncRNA | 0.892364482 | 0.964211859 | 1.886981896 | -0.51098098 | 0.609364375 | 1 | 0.964211859 |
| ENSMUSG00000073207 | Ccdc160 | 434778 | protein_coding | 1.879208188 | -0.964315377 | 0.846873101 | 1.138677537 | 0.254837675 | 1 | 0.964315377 |
| ENSMUSG00000021961 | 4930578I06Rik | 67750 | protein_coding | 0.979458253 | 0.96446495 | 1.861948253 | -0.517986979 | 0.604467355 | 1 | 0.96446495 |
| ENSMUSG00000004473 | Clec11a | 20256 | protein_coding | 22.86244042 | 0.964584379 | 0.284005284 | -3.396360685 | 0.000682883 | 0.004593536 | 0.964584379 |
| ENSMUSG00000092353 | Gm3363 | 100041479 | lncRNA | 2.081493996 | 0.965089634 | 1.043966151 | -0.924445331 | 0.355254502 | 1 | 0.965089634 |
| ENSMUSG00000048347 | Pcdhb18 | 93889 | protein_coding | 2.735370035 | 0.965208586 | 0.661226524 | -1.459724545 | 0.144365793 | 1 | 0.965208586 |
| ENSMUSG00000113640 | Adat3 | 100113398 | protein_coding | 0.877233394 | 0.965237276 | 1.641633593 | -0.587973638 | 0.556549988 | 1 | 0.965237276 |
| ENSMUSG00000029304 | Spp1 | 20750 | protein_coding | 0.548265828 | 0.965265726 | 1.790376044 | -0.539141333 | 0.589789338 | 1 | 0.965265726 |
| ENSMUSG00000072115 | Ang | 11727 | protein_coding | 184.3952563 | -0.965311403 | 0.129312838 | 7.464930912 | 8.33E-14 | 2.76E-12 | 0.965311403 |
| ENSMUSG00000020368 | Canx | 12330 | protein_coding | 12654.15881 | 0.965704219 | 0.103442619 | -9.335651311 | 1.00E-20 | 5.33E-19 | 0.965704219 |
| ENSMUSG00000048230 | Fbxo43 | 78803 | protein_coding | 2.173950662 | 0.96571948 | 1.267286809 | -0.762037033 | 0.446037902 | 1 | 0.96571948 |

| ensembl_gene_id | external_gene_name | entrezgene_id | gene_biotype | baseMean | log2FoldChange | lfcSE | stat | pvalue | padj | absLog2FC |
|---|---|---|---|---|---|---|---|---|---|---|
| ENSMUSG00000079180 | Mptx2 | 100504232 | protein_coding | 0.949429968 | -0.966091112 | 1.340866757 | 0.720497474 | 0.471218754 | 1 | 0.966091112 |
| ENSMUSG00000025731 | Mettl26 | 68347 | protein_coding | 533.4987083 | -0.966139697 | 0.071679051 | 13.47868988 | 2.09E-41 | 2.28E-39 | 0.966139697 |
| ENSMUSG00000100550 | 2310039L15Rik | 69629 | lncRNA | 1.537399791 | 0.966435978 | 1.864276539 | -0.518397329 | 0.604181078 | 1 | 0.966435978 |
| ENSMUSG00000041556 | Fbxo2 | 230904 | protein_coding | 0.759064909 | 0.966532944 | 1.95649302 | -0.494012978 | 0.621297014 | 1 | 0.966532944 |
| ENSMUSG00000000617 | Grm6 | 108072 | protein_coding | 2.381213811 | 0.966675221 | 1.931656267 | -0.500438529 | 0.61676633 | 1 | 0.966675221 |
| ENSMUSG00000061991 | H2ac10 | 319173 | protein_coding | 130.666431 | 0.966766535 | 0.208225103 | -4.642891383 | 3.44E-06 | 4.25E-05 | 0.966766535 |
| ENSMUSG00000032735 | Ablim3 | 319713 | protein_coding | 4.259875416 | 0.966780389 | 1.436547236 | -0.672988931 | 0.500954333 | 1 | 0.966780389 |
| ENSMUSG00000043880 | Or11l3 | 258373 | protein_coding | 0.933041095 | 0.966989826 | 1.670807767 | -0.578755884 | 0.562753902 | 1 | 0.966989826 |
| ENSMUSG00000052613 | Pcdh15 | 11994 | protein_coding | 6.182321586 | 0.967512286 | 2.809915186 | -0.344320815 | 0.730605032 | 0.861253255 | 0.967512286 |
| ENSMUSG00000004451 | Ralb | 64143 | protein_coding | 1702.902925 | 0.967559894 | 0.072970425 | -13.25961715 | 3.97E-40 | 4.17E-38 | 0.967559894 |
| ENSMUSG00000085582 | 3110099E03Rik | 100043232 | lncRNA | 0.754589194 | 0.967940753 | 1.872573799 | -0.516903928 | 0.605223225 | 1 | 0.967940753 |
| ENSMUSG00000044084 | Spem2 | 108803 | protein_coding | 1.224979978 | 0.968345084 | 1.593031459 | -0.607863127 | 0.543278257 | 1 | 0.968345084 |
| ENSMUSG00000087306 | A230004M16Rik | 102637814 | lncRNA | 1.371250162 | -0.968538837 | 2.303739534 | 0.420420287 | 0.67417845 | 1 | 0.968538837 |
| ENSMUSG00000055409 | Nell1 | 338352 | protein_coding | 2.000295891 | 0.968872108 | 2.675686914 | -0.362102196 | 0.717275662 | 1 | 0.968872108 |
| ENSMUSG00000048217 | Nags | 217214 | protein_coding | 101.5841799 | -0.968948311 | 0.245779919 | 3.942341239 | 8.07E-05 | 0.000714486 | 0.968948311 |
| ENSMUSG00000026572 | Tbx19 | 83993 | protein_coding | 1.398517773 | 0.969179695 | 2.704589563 | -0.358346312 | 0.720084164 | 1 | 0.969179695 |
| ENSMUSG00000023984 | Gm20517 | 112694759 | protein_coding | 9.475238661 | 0.969326668 | 0.373601613 | -2.59454626 | 0.009471587 | 0.040777919 | 0.969326668 |
| ENSMUSG00000025194 | Abcc2 | 12780 | protein_coding | 30.31560026 | -0.969937068 | 0.474033311 | 2.046136939 | 0.0407429 | 0.127792639 | 0.969937068 |
| ENSMUSG00000003585 | Sec14l2 | 67815 | protein_coding | 86.86997598 | 0.970173458 | 0.183164736 | -5.296726215 | 1.18E-07 | 1.92E-06 | 0.970173458 |
| ENSMUSG00000060181 | Slc35e3 | 215436 | protein_coding | 108.1242363 | -0.970216943 | 0.181811932 | 5.336376614 | 9.48E-08 | 1.57E-06 | 0.970216943 |
| ENSMUSG00000028970 | Abcb1b | 18669 | protein_coding | 7.266506589 | 0.970374285 | 0.815529562 | -1.189870153 | 0.234097431 | 0.443634979 | 0.970374285 |
| ENSMUSG00000027831 | Veph1 | 72789 | protein_coding | 1.347966141 | -0.970589916 | 2.341617423 | 0.414495514 | 0.678511252 | 1 | 0.970589916 |
| ENSMUSG00000057335 | Cep170 | 545389 | protein_coding | 20.78159206 | 0.970867246 | 0.354826426 | -2.73617514 | 0.006215793 | 0.02891482 | 0.970867246 |
| ENSMUSG00000064408 | Gm23924 | 115490511 | snRNA | 1.25727908 | 0.971201145 | 2.065634089 | -0.470170951 | 0.638232886 | 1 | 0.971201145 |
| ENSMUSG00000071661 | Zbtb3 | 75291 | protein_coding | 40.71348849 | 0.971423216 | 0.171511406 | -5.663898634 | 1.48E-08 | 2.77E-07 | 0.971423216 |
| ENSMUSG00000025203 | Scd2 | 20250 | protein_coding | 13269.83872 | 0.971433267 | 0.184467564 | -5.266146775 | 1.39E-07 | 2.24E-06 | 0.971433267 |
| ENSMUSG00000037843 | Vstm2l | 277432 | protein_coding | 11.69681895 | -0.972144085 | 0.364634395 | 2.666078952 | 0.007674167 | 0.034419117 | 0.972144085 |
| ENSMUSG00000017499 | Cdc6 | 23834 | protein_coding | 122.4242328 | 0.972588609 | 0.199914152 | -4.86503131 | 1.14E-06 | 1.56E-05 | 0.972588609 |
| ENSMUSG00000010362 | Rdm1 | 66599 | protein_coding | 125.0990552 | -0.972595756 | 0.10493121 | 9.268889178 | 1.88E-20 | 9.74E-19 | 0.972595756 |
| ENSMUSG00000036712 | Cyld | 74256 | protein_coding | 1036.180658 | 0.972709757 | 0.068797547 | -14.13872728 | 2.19E-45 | 2.66E-43 | 0.972709757 |
| ENSMUSG00000060594 | Layn | 244864 | protein_coding | 4.398900833 | -0.973655345 | 0.683579985 | 1.424347358 | 0.154345945 | 0.336218865 | 0.973655345 |
| ENSMUSG00000026070 | Il18r1 | 16182 | protein_coding | 9.550786184 | 0.973780101 | 0.487366375 | -1.998045314 | 0.045711748 | 0.139328867 | 0.973780101 |
| ENSMUSG00000004415 | Col26a1 | 140709 | protein_coding | 2.805519866 | 0.974052972 | 1.480591302 | -0.657881058 | 0.510614563 | 1 | 0.974052972 |
| ENSMUSG00000021919 | Chat | 12647 | protein_coding | 20.88413105 | -0.974403112 | 0.409171613 | 2.381404479 | 0.017246763 | 0.065324093 | 0.974403112 |
| ENSMUSG00000062184 | Hs6st2 | 50786 | protein_coding | 2.272444417 | 0.974471121 | 1.518149796 | -0.641880744 | 0.520950618 | 1 | 0.974471121 |
| ENSMUSG00000040136 | Abcc8 | 20927 | protein_coding | 39.34365287 | -0.97453043 | 0.269892656 | 3.610807505 | 0.000305245 | 0.002313363 | 0.97453043 |
| ENSMUSG00000096847 | Tmem151b | 210573 | protein_coding | 2.629673582 | 0.974568529 | 1.449669088 | -0.672269649 | 0.501412048 | 1 | 0.974568529 |
| ENSMUSG00000063971 | Krt88 | 66322 | protein_coding | 2.102955163 | 0.975155482 | 1.590382877 | -0.613157685 | 0.539772081 | 1 | 0.975155482 |
| ENSMUSG00000024737 | Slc15a3 | 65221 | protein_coding | 2.885534822 | 0.975199804 | 1.616788237 | -0.603171016 | 0.546394927 | 1 | 0.975199804 |

| ensembl_gene_id | external_gene_name | entrezgene_id | gene_biotype | baseMean | log2FoldChange | lfcSE | stat | pvalue | padj | absLog2FC |
|---|---|---|---|---|---|---|---|---|---|---|
| ENSMUSG00000019846 | Lama4 | 16775 | protein_coding | 12.26718949 | 0.975308897 | 0.527155807 | -1.850134027 | 0.064294235 | 0.179528541 | 0.975308897 |
| ENSMUSG00000027210 | Meis2 | 17536 | protein_coding | 3.498110347 | 0.975309738 | 1.254657006 | -0.777351686 | 0.436951309 | 1 | 0.975309738 |
| ENSMUSG00000025656 | Arhgef9 | 236915 | protein_coding | 15.61680783 | 0.975329241 | 0.324750785 | -3.003316037 | 0.002670549 | 0.014421666 | 0.975329241 |
| ENSMUSG00000078566 | Bnip3 | 12176 | protein_coding | 45.60563044 | -0.975338454 | 0.204188272 | 4.776662453 | 1.78E-06 | 2.33E-05 | 0.975338454 |
| ENSMUSG00000056004 | Elapor2 | 231014 | protein_coding | 15.10042806 | -0.975407897 | 0.511314694 | 1.907646912 | 0.056436867 | 0.163322332 | 0.975407897 |
| ENSMUSG00000025468 | Caly | 68566 | protein_coding | 1.950624214 | -0.975462 | 1.443249993 | 0.675878749 | 0.499117631 | 1 | 0.975462 |
| ENSMUSG00000113198 | Gm6988 | 629595 | processed_pseudogene | 3.159242804 | 0.975708232 | 0.788830415 | -1.236904934 | 0.216122378 | 1 | 0.975708232 |
| ENSMUSG00000006724 | Cyp27b1 | 13115 | protein_coding | 3.929718646 | 0.975864231 | 0.916335853 | -1.064963492 | 0.286892451 | 1 | 0.975864231 |
| ENSMUSG00000080712 | H2bc26-ps | 382522 | protein_coding | 25.5453595 | 0.97613251 | 0.238186397 | -4.09818748 | 4.16E-05 | 0.000398401 | 0.97613251 |
| ENSMUSG00000032121 | Tmem218 | 66279 | protein_coding | 114.216747 | 0.976386686 | 0.129221141 | -7.555936122 | 4.16E-14 | 1.42E-12 | 0.976386686 |
| ENSMUSG00000034266 | Batf | 53314 | protein_coding | 2.474182659 | 0.976447262 | 1.105449519 | -0.883303349 | 0.377072395 | 1 | 0.976447262 |
| ENSMUSG00000000392 | Fap | 14089 | protein_coding | 39.05958091 | 0.97657379 | 0.213765474 | -4.568435537 | 4.91E-06 | 5.88E-05 | 0.97657379 |
| ENSMUSG00000039556 | Ppp1r3f | 54646 | protein_coding | 8.031685437 | 0.976829071 | 0.524940789 | -1.860836673 | 0.062767245 | 0.176252046 | 0.976829071 |
| ENSMUSG00000046922 | Gpr6 | 140741 | protein_coding | 0.779852189 | -0.977102935 | 1.300830419 | 0.751137828 | 0.452569712 | 1 | 0.977102935 |
| ENSMUSG00000093930 | Hmgcs1 | 208715 | protein_coding | 5326.572769 | 0.977432719 | 0.152040036 | -6.428785087 | 1.29E-10 | 3.14E-09 | 0.977432719 |
| ENSMUSG00000101337 | Dnah7c | 100101919 | protein_coding | 3.549310999 | 0.977453185 | 1.483795406 | -0.658751996 | 0.510055038 | 1 | 0.977453185 |
| ENSMUSG00000086645 | Gm15743 | 100047123 | lncRNA | 9.340923178 | -0.978236137 | 0.41079966 | 2.381297338 | 0.017251781 | 0.065328114 | 0.978236137 |
| ENSMUSG00000032083 | Apoa1 | 11806 | protein_coding | 80.42771587 | -0.978576495 | 0.904465923 | 1.08193849 | 0.27927986 | 0.496688848 | 0.978576495 |
| ENSMUSG00000109816 | 4930483O08Rik | 74970 | lncRNA | 1.2397932 | -0.97887499 | 1.096479838 | 0.892743265 | 0.371994675 | 1 | 0.97887499 |
| ENSMUSG00000034774 | Dsg1c | 211924 | protein_coding | 3.042626449 | 0.978897477 | 1.045029952 | -0.936717149 | 0.348904072 | 1 | 0.978897477 |
| ENSMUSG00000096655 | 1700065D16Rik | 73410 | lncRNA | 1.654415432 | -0.978931096 | 1.728687314 | 0.56628581 | 0.571199505 | 1 | 0.978931096 |
| ENSMUSG00000054453 | Sytl5 | 236643 | protein_coding | 15.32614495 | 0.979379169 | 0.300819611 | -3.255702535 | 0.001131122 | 0.007064806 | 0.979379169 |
| ENSMUSG00000117655 | 6720468P15Rik | 78073 | lncRNA | 16.60521996 | 0.980118197 | 0.275962156 | -3.551639877 | 0.000382838 | 0.00281989 | 0.980118197 |
| ENSMUSG00000025190 | Got1 | 14718 | protein_coding | 897.606946 | 0.980651375 | 0.11845222 | -8.278877118 | 1.24E-16 | 5.20E-15 | 0.980651375 |
| ENSMUSG00000028343 | Erp44 | 76299 | protein_coding | 1300.849203 | 0.980774741 | 0.084137802 | -11.65676687 | 2.12E-31 | 1.73E-29 | 0.980774741 |
| ENSMUSG00000028641 | P3h1 | 56401 | protein_coding | 23.68035117 | 0.980897231 | 0.286492137 | -3.423819026 | 0.000617477 | 0.004231128 | 0.980897231 |
| ENSMUSG00000096257 | Ccer2 | 100504112 | protein_coding | 1.489337209 | -0.980973987 | 2.10961595 | 0.465001218 | 0.641930584 | 1 | 0.980973987 |
| ENSMUSG00000078771 | Evi2a | 14017 | protein_coding | 3.210513949 | -0.981349888 | 0.692021984 | 1.418090625 | 0.156164308 | 1 | 0.981349888 |
| ENSMUSG00000031111 | Igsf1 | 209268 | protein_coding | 1.98885688 | -0.982004774 | 2.66680766 | 0.368232321 | 0.712700012 | 1 | 0.982004774 |
| ENSMUSG00000042751 | Nmnat2 | 226518 | protein_coding | 5.84288799 | -0.982069418 | 0.694642877 | 1.413775986 | 0.157427687 | 0.340684159 | 0.982069418 |
| ENSMUSG00000058589 | Anks1b | 77531 | protein_coding | 7.384845685 | 0.982115054 | 1.317868298 | -0.745230047 | 0.456132662 | 0.671412316 | 0.982115054 |
| ENSMUSG00000052087 | Rgs14 | 51791 | protein_coding | 33.69130535 | 0.982329999 | 0.197802586 | -4.966214126 | 6.83E-07 | 9.77E-06 | 0.982329999 |
| ENSMUSG00000009378 | Slc16a12 | 240638 | protein_coding | 82.30781094 | 0.982372195 | 0.411695234 | -2.386163633 | 0.017025174 | 0.064634295 | 0.982372195 |
| ENSMUSG00000059741 | Myl3 | 17897 | protein_coding | 1.631178626 | 0.982393655 | 1.929747673 | -0.509078813 | 0.610696982 | 1 | 0.982393655 |
| ENSMUSG00000023267 | Gabrr2 | 14409 | protein_coding | 2.720234021 | -0.982730161 | 1.068229206 | 0.919961891 | 0.357592674 | 1 | 0.982730161 |
| ENSMUSG00000019737 | Syne4 | 233066 | protein_coding | 41.87995172 | 0.983151858 | 0.229565483 | -4.282664115 | 1.85E-05 | 0.000192892 | 0.983151858 |
| ENSMUSG00000061615 | H2ac4 | 319172 | protein_coding | 461.7683185 | 0.983314476 | 0.169776423 | -5.791819981 | 6.96E-09 | 1.37E-07 | 0.983314476 |
| ENSMUSG00000007670 | Khsrp | 16549 | protein_coding | 1485.620321 | 0.983493967 | 0.096623233 | -10.17864887 | 2.47E-24 | 1.55E-22 | 0.983493967 |
| ENSMUSG00000096228 | Or6z7 | 18349 | protein_coding | 1.304656606 | -0.983619775 | 2.663380532 | 0.369312519 | 0.711894795 | 1 | 0.983619775 |

| ensembl_gene_id | external_gene_name | entrezgene_id | gene_biotype | baseMean | log2FoldChange | lfcSE | stat | pvalue | padj | absLog2FC |
|---|---|---|---|---|---|---|---|---|---|---|
| ENSMUSG00000037411 | Serpine1 | 18787 | protein_coding | 6.785568064 | -0.983902733 | 0.659313599 | 1.492313726 | 0.13561692 | 0.306987369 | 0.983902733 |
| ENSMUSG00000055184 | Fam72a | 108900 | protein_coding | 18.270299 | 0.984316701 | 0.316739268 | -3.107656043 | 0.001885774 | 0.010831886 | 0.984316701 |
| ENSMUSG00000021927 | Hnrnpa1l2-ps2 | 545091 | processed_pseudogene | 0.696269369 | -0.984326775 | 1.707339439 | 0.576526701 | 0.564259227 | 1 | 0.984326775 |
| ENSMUSG00000046546 | Fam43a | 224093 | protein_coding | 173.1558597 | -0.984463149 | 0.375138886 | 2.624263135 | 0.008683667 | 0.038017517 | 0.984463149 |
| ENSMUSG00000032006 | Pdgfd | 71785 | protein_coding | 14.68256889 | -0.984806165 | 0.308854715 | 3.188574166 | 0.001429763 | 0.008564059 | 0.984806165 |
| ENSMUSG00000103577 | 9330162B11Rik | 320272 | lncRNA | 1.34936452 | -0.98507199 | 1.10687254 | 0.889959732 | 0.373487508 | 1 | 0.98507199 |
| ENSMUSG00000060257 | Scrt2 | 545474 | protein_coding | 1.848438153 | 0.985359467 | 1.886237633 | -0.52239413 | 0.601395942 | 1 | 0.985359467 |
| ENSMUSG00000032679 | Cd59a | 12509 | protein_coding | 12.54357314 | -0.985896279 | 0.382582871 | 2.576948298 | 0.009967686 | 0.042468102 | 0.985896279 |
| ENSMUSG00000031613 | Hpgd | 15446 | protein_coding | 1424.13286 | -0.985941018 | 0.11212485 | 8.793242714 | 1.45E-18 | 6.82E-17 | 0.985941018 |
| ENSMUSG00000115779 | 4930592A05Rik | 75880 | lncRNA | 0.740941633 | -0.985945577 | 1.382485129 | 0.713169029 | 0.475741169 | 1 | 0.985945577 |
| ENSMUSG00000042834 | Nrep | 27528 | protein_coding | 207.6623153 | -0.986072859 | 0.152232903 | 6.477396388 | 9.33E-11 | 2.35E-09 | 0.986072859 |
| ENSMUSG00000049577 | Zfpm1 | 22761 | protein_coding | 67.61942736 | 0.986308896 | 0.18773497 | -5.253730281 | 1.49E-07 | 2.39E-06 | 0.986308896 |
| ENSMUSG00000036862 | Dchs1 | 233651 | protein_coding | 4.169660462 | 0.986682179 | 1.634640504 | -0.603608057 | 0.546104255 | 1 | 0.986682179 |
| ENSMUSG00000005540 | Fcer2a | 14128 | protein_coding | 2.68224642 | -0.986683609 | 1.241564824 | 0.7947097 | 0.426782388 | 1 | 0.986683609 |
| ENSMUSG00000099153 | Mir7671 | 102465772 | miRNA | 0.572092067 | 0.986894554 | 1.62757808 | -0.606357733 | 0.544277229 | 1 | 0.986894554 |
| ENSMUSG00000026610 | Esrrg | 26381 | protein_coding | 56.6915269 | -0.987263703 | 0.247305246 | 3.992085571 | 6.55E-05 | 0.000594601 | 0.987263703 |
| ENSMUSG00000021573 | Tppp | 72948 | protein_coding | 1194.098799 | -0.987291325 | 0.116236924 | 8.493783999 | 2.00E-17 | 8.69E-16 | 0.987291325 |
| ENSMUSG00000015093 | Clic3 | 69454 | protein_coding | 7.538459001 | 0.987438365 | 0.561227935 | -1.759424832 | 0.078505377 | 0.207264236 | 0.987438365 |
| ENSMUSG00000113581 | Dio3os | 353504 | lncRNA | 8.252586447 | -0.987752801 | 0.525676764 | 1.879011722 | 0.060242893 | 0.171319292 | 0.987752801 |
| ENSMUSG00000050014 | Apol10b | 328561 | protein_coding | 1.846358182 | 0.988000448 | 1.119619051 | -0.882443405 | 0.377537072 | 1 | 0.988000448 |
| ENSMUSG00000001021 | S100a3 | 20197 | protein_coding | 2.361254018 | -0.988179204 | 1.467658271 | 0.673303332 | 0.500754334 | 1 | 0.988179204 |
| ENSMUSG00000031351 | Zfp185 | 22673 | protein_coding | 10.13083706 | 0.988693105 | 0.393938417 | -2.509765646 | 0.012081131 | 0.049395172 | 0.988693105 |
| ENSMUSG00000115656 | 4930556J02Rik | 75349 | lncRNA | 3.215219475 | 0.988758756 | 1.66523107 | -0.593766699 | 0.552668165 | 1 | 0.988758756 |
| ENSMUSG00000030116 | Mfap5 | 50530 | protein_coding | 1.879753104 | -0.98878853 | 1.721580495 | 0.574349287 | 0.565731463 | 1 | 0.98878853 |
| ENSMUSG00000020334 | Slc22a4 | 30805 | protein_coding | 8.25118927 | -0.988814526 | 0.629387015 | 1.571075512 | 0.11616511 | 0.275269138 | 0.988814526 |
| ENSMUSG00000024970 | Spindoc | 68229 | protein_coding | 258.7757514 | 0.988863189 | 0.113478245 | -8.714121248 | 2.93E-18 | 1.36E-16 | 0.988863189 |
| ENSMUSG00000021207 | Akr1c21 | 77337 | protein_coding | 2.778013954 | -0.989285106 | 0.718923818 | 1.376063891 | 0.168801854 | 1 | 0.989285106 |
| ENSMUSG00000027833 | Shox2 | 20429 | protein_coding | 1.502480663 | -0.989413416 | 1.860584995 | 0.531775446 | 0.59488153 | 1 | 0.989413416 |
| ENSMUSG00000041801 | Phlda3 | 27280 | protein_coding | 28.49183817 | 0.989443335 | 0.254538163 | -3.88721016 | 0.000101403 | 0.00087081 | 0.989443335 |
| ENSMUSG00000066975 | Cryba4 | 12959 | protein_coding | 0.697634057 | -0.989497186 | 1.838433011 | 0.538228579 | 0.590419252 | 1 | 0.989497186 |
| ENSMUSG00000035845 | Alg12 | 223774 | protein_coding | 326.9791188 | 0.989732587 | 0.086055725 | -11.50106621 | 1.30E-30 | 1.04E-28 | 0.989732587 |
| ENSMUSG00000022623 | Shank3 | 58234 | protein_coding | 6.453151712 | -0.989739293 | 0.7515996 | 1.316843826 | 0.187890979 | 0.384217774 | 0.989739293 |
| ENSMUSG00000100782 | Gm28231 | 102640900 | lncRNA | 1.058330007 | -0.989917373 | 1.773417023 | 0.558197739 | 0.576709364 | 1 | 0.989917373 |
| ENSMUSG00000097735 | D930032P07Rik | 320176 | lncRNA | 0.751066119 | -0.99010684 | 1.66046047 | 0.5962845 | 0.550985184 | 1 | 0.99010684 |
| ENSMUSG00000040380 | Cbln3 | 56410 | protein_coding | 51.64592137 | 0.990469348 | 0.177361598 | -5.584463385 | 2.34E-08 | 4.27E-07 | 0.990469348 |
| ENSMUSG00000056716 | Gm5420 | 432436 | lncRNA | 4.502971794 | 0.990660471 | 0.609338815 | -1.625795775 | 0.103993108 | 0.254685181 | 0.990660471 |
| ENSMUSG00000107768 | Gm36582 | 102640545 | lncRNA | 5.667175383 | 0.990838683 | 0.672582112 | -1.473186197 | 0.140700836 | 0.315055404 | 0.990838683 |
| ENSMUSG00000042524 | Sun2 | 223697 | protein_coding | 584.4713947 | -0.99105957 | 0.21484718 | 4.612858177 | 3.97E-06 | 4.87E-05 | 0.99105957 |
| ENSMUSG00000026676 | Ccdc3 | 74186 | protein_coding | 20.24451956 | 0.991462961 | 0.285699238 | -3.470303135 | 0.000519871 | 0.003668869 | 0.991462961 |

| ensembl_gene_id | external_gene_name | entrezgene_id | gene_biotype | baseMean | log2FoldChange | lfcSE | stat | pvalue | padj | absLog2FC |
|---|---|---|---|---|---|---|---|---|---|---|
| ENSMUSG00000024065 | Ehd3 | 57440 | protein_coding | 3.12178363 | 0.991488977 | 1.225931942 | -0.808763475 | 0.418651209 | 1 | 0.991488977 |
| ENSMUSG00000048782 | Insc | 233752 | protein_coding | 23.3846095 | 0.992289325 | 0.244306634 | -4.061655253 | 4.87E-05 | 0.000456675 | 0.992289325 |
| ENSMUSG00000023236 | Scg5 | 20394 | protein_coding | 11.38167759 | -0.992461146 | 0.368322653 | 2.69454278 | 0.007048528 | 0.032074783 | 0.992461146 |
| ENSMUSG00000079547 | H2-DMb1 | 14999 | protein_coding | 8.498323706 | -0.99268495 | 0.374258331 | 2.652405752 | 0.007992044 | 0.035574288 | 0.99268495 |
| ENSMUSG00000021965 | Ska3 | 219114 | protein_coding | 75.92955948 | 0.993051087 | 0.210564218 | -4.716143591 | 2.40E-06 | 3.07E-05 | 0.993051087 |
| ENSMUSG00000019992 | Mtfr2 | 71804 | protein_coding | 19.97044172 | 0.993166528 | 0.311633048 | -3.186974339 | 0.001437695 | 0.008577348 | 0.993166528 |
| ENSMUSG00000108929 | Cc2d2b | 668310 | protein_coding | 2.578495212 | 0.994195709 | 1.487982108 | -0.668150312 | 0.504037649 | 1 | 0.994195709 |
| ENSMUSG00000023973 | Cnpy3 | 72029 | protein_coding | 585.0795 | 0.994884343 | 0.071774976 | -13.86115893 | 1.09E-43 | 1.27E-41 | 0.994884343 |
| ENSMUSG00000044176 | Spink10 | 328971 | protein_coding | 3.209722827 | 0.995247095 | 1.286923688 | -0.773353622 | 0.43931313 | 1 | 0.995247095 |
| ENSMUSG00000037071 | Scd1 | 20249 | protein_coding | 601.5407759 | 0.995777204 | 0.127912934 | -7.78480468 | 6.98E-15 | 2.53E-13 | 0.995777204 |
| ENSMUSG00000102014 | 2900009J06Rik | 100502745 | lncRNA | 1.21559364 | 0.996028858 | 0.904746578 | -1.10089265 | 0.270943381 | 1 | 0.996028858 |
| ENSMUSG00000052316 | Lrrc15 | 74488 | protein_coding | 1.567483495 | -0.996420324 | 1.689115806 | 0.589906459 | 0.555253364 | 1 | 0.996420324 |
| ENSMUSG00000020279 | Il9r | 16199 | protein_coding | 3.933690668 | -0.99642705 | 0.691193001 | 1.441604657 | 0.149413934 | 1 | 0.99642705 |
| ENSMUSG00000040760 | Appl1 | 72993 | protein_coding | 1323.095092 | 0.996494263 | 0.119556068 | -8.334953467 | 7.75E-17 | 3.27E-15 | 0.996494263 |
| ENSMUSG00000050671 | Ism2 | 217738 | protein_coding | 1.413586723 | 0.996553454 | 1.369644083 | -0.727600306 | 0.466858293 | 1 | 0.996553454 |
| ENSMUSG00000092094 | Zfp804b | 207618 | protein_coding | 2.091184583 | 0.996597415 | 1.777567698 | -0.560652298 | 0.575034592 | 1 | 0.996597415 |
| ENSMUSG00000051236 | Msrb3 | 320183 | protein_coding | 6.985378875 | 0.9966448 | 0.548209294 | -1.818000553 | 0.069064048 | 0.189172946 | 0.9966448 |
| ENSMUSG00000026208 | Des | 13346 | protein_coding | 4.32587105 | -0.996714308 | 1.053542589 | 0.946059817 | 0.344118078 | 1 | 0.996714308 |
| ENSMUSG00000028108 | Ecm1 | 13601 | protein_coding | 11.4525023 | 0.996754086 | 0.404411066 | -2.464705268 | 0.013712599 | 0.054382773 | 0.996754086 |
| ENSMUSG00000046589 | Lrrc8e | 72267 | protein_coding | 235.2794838 | -0.997279514 | 0.125779304 | 7.928804526 | 2.21E-15 | 8.30E-14 | 0.997279514 |
| ENSMUSG00000053746 | Ptrh1 | 329384 | protein_coding | 175.0420117 | 0.99739694 | 0.221147467 | -4.510098862 | 6.48E-06 | 7.57E-05 | 0.99739694 |
| ENSMUSG00000118215 | Vmn1r55 | 384522 | protein_coding | 4.334099297 | 0.99794293 | 2.065824565 | -0.483072448 | 0.629044296 | 1 | 0.99794293 |
| ENSMUSG00000043157 | Arl11 | 219144 | protein_coding | 0.846748815 | 0.997964852 | 1.29067828 | -0.773209612 | 0.439398338 | 1 | 0.997964852 |
| ENSMUSG00000006442 | Srm | 20810 | protein_coding | 271.8742474 | 0.998393584 | 0.152327426 | -6.554260196 | 5.59E-11 | 1.46E-09 | 0.998393584 |
| ENSMUSG00000052852 | Reep1 | 52250 | protein_coding | 5.833800541 | 0.998569451 | 0.502273732 | -1.988098098 | 0.046800842 | 0.141758822 | 0.998569451 |
| ENSMUSG00000038045 | Sult6b1 | 73671 | protein_coding | 8.461225997 | 0.998647806 | 0.41968961 | -2.379491374 | 0.017336549 | 0.065598201 | 0.998647806 |
| ENSMUSG00000050468 | Astl | 215095 | protein_coding | 2.182897188 | -0.998828865 | 2.106767003 | 0.474105045 | 0.635425001 | 1 | 0.998828865 |
| ENSMUSG00000049571 | Cfap46 | 212124 | protein_coding | 5.769030504 | 0.999641983 | 1.514670376 | -0.659973285 | 0.509270973 | 0.712720411 | 0.999641983 |
| ENSMUSG00000030222 | Rerg | 232441 | protein_coding | 7.864340589 | -0.999650084 | 0.488661496 | 2.045690305 | 0.04078685 | 0.127906199 | 0.999650084 |
| ENSMUSG00000007872 | Id3 | 15903 | protein_coding | 2189.869434 | 0.999719376 | 0.130511094 | -7.660033709 | 1.86E-14 | 6.50E-13 | 0.999719376 |
| ENSMUSG00000048996 | Or1f12 | 258280 | protein_coding | 1.661769324 | 0.999963486 | 1.279058762 | -0.781796361 | 0.43433426 | 1 | 0.999963486 |
| ENSMUSG00000064246 | Chil1 | 12654 | protein_coding | 2.918566326 | -1.000239043 | 0.783751769 | 1.276219184 | 0.201878049 | 1 | 1.000239043 |
| ENSMUSG00000025586 | Cpeb1 | 12877 | protein_coding | 2.297014863 | -1.001113327 | 1.680038362 | 0.595887183 | 0.551250596 | 1 | 1.001113327 |
| ENSMUSG00000024063 | Lbh | 77889 | protein_coding | 228.1660139 | -1.001280825 | 0.153464138 | 6.524526429 | 6.82E-11 | 1.75E-09 | 1.001280825 |
| ENSMUSG00000118204 | 4930414N06Rik | 73894 | lncRNA | 7.497994076 | 1.002130212 | 0.457610632 | -2.189918984 | 0.028530113 | 0.097003559 | 1.002130212 |
| ENSMUSG00000042195 | Slc35f2 | 72022 | protein_coding | 260.4485413 | 1.003947697 | 0.116646728 | -8.606736926 | 7.52E-18 | 3.42E-16 | 1.003947697 |
| ENSMUSG00000097028 | Ptgs2os | 320019 | lncRNA | 1.470534059 | -1.003959845 | 1.880433994 | 0.533897945 | 0.593412144 | 1 | 1.003959845 |
| ENSMUSG00000113237 | Gm6109 | 619883 | processed_pseudogene | 1.150575352 | 1.004038693 | 1.148509481 | -0.874210191 | 0.382003797 | 1 | 1.004038693 |
| ENSMUSG00000061947 | Serpina10 | 217847 | protein_coding | 8.771655444 | 1.004039323 | 0.402733617 | -2.493060624 | 0.012664722 | 0.051148256 | 1.004039323 |

| ensembl_gene_id | external_gene_name | entrezgene_id | gene_biotype | baseMean | log2FoldChange | lfcSE | stat | pvalue | padj | absLog2FC |
|---|---|---|---|---|---|---|---|---|---|---|
| ENSMUSG00000079853 | Klra1 | 16627 | protein_coding | 2.968174022 | 1.004058926 | 0.857673006 | -1.170678008 | 0.241728228 | 1 | 1.004058926 |
| ENSMUSG00000073422 | H2-Ke6 | 14979 | protein_coding | 6.692357316 | 1.004192732 | 0.520421994 | -1.929573967 | 0.053659647 | 0.157297398 | 1.004192732 |
| ENSMUSG00000087478 | 4930506C21Rik | 75060 | lncRNA | 4.825503213 | -1.006204124 | 0.564489036 | 1.782504284 | 0.074667033 | 0.199909758 | 1.006204124 |
| ENSMUSG00000045378 | Pxt1 | 69307 | protein_coding | 0.927924133 | -1.006532126 | 1.91089598 | 0.526733081 | 0.59837896 | 1 | 1.006532126 |
| ENSMUSG00000019429 | Ffar3 | 233080 | protein_coding | 24.75084574 | -1.007173257 | 0.312414569 | 3.223835754 | 0.001264859 | 0.007727668 | 1.007173257 |
| ENSMUSG00000031173 | Otc | 18416 | protein_coding | 252.5555206 | -1.00734844 | 0.133061549 | 7.570544967 | 3.72E-14 | 1.28E-12 | 1.00734844 |
| ENSMUSG00000093536 | Smim17 | 100042450 | protein_coding | 1.401735342 | -1.007487242 | 1.398722918 | 0.720290795 | 0.471345971 | 1 | 1.007487242 |
| ENSMUSG00000034412 | Tbc1d10a | 103724 | protein_coding | 271.0056764 | 1.007618313 | 0.092580443 | -10.88370585 | 1.38E-27 | 9.77E-26 | 1.007618313 |
| ENSMUSG00000054404 | Slfn5 | 327978 | protein_coding | 9.852801262 | -1.007657471 | 0.663368849 | 1.519000286 | 0.128762424 | 0.29613919 | 1.007657471 |
| ENSMUSG00000049932 | H2ax | 15270 | protein_coding | 859.8472606 | 1.007692222 | 0.152262283 | -6.618134175 | 3.64E-11 | 9.64E-10 | 1.007692222 |
| ENSMUSG00000024011 | Pi16 | 74116 | protein_coding | 3.432557598 | 1.007885888 | 1.229539029 | -0.819726633 | 0.412371964 | 1 | 1.007885888 |
| ENSMUSG00000038067 | Csf3 | 12985 | protein_coding | 1.262812832 | 1.008190233 | 1.87230767 | -0.538474659 | 0.590249396 | 1 | 1.008190233 |
| ENSMUSG00000034837 | Gnat1 | 14685 | protein_coding | 3.219100382 | 1.008387162 | 0.957613499 | -1.053021039 | 0.29233135 | 1 | 1.008387162 |
| ENSMUSG00000020099 | Unc5b | 107449 | protein_coding | 1756.998296 | 1.008440363 | 0.15043846 | -6.703341436 | 2.04E-11 | 5.51E-10 | 1.008440363 |
| ENSMUSG00000110353 | Gm33543 | 102636491 | lncRNA | 1.521795702 | 1.009284215 | 1.174426781 | -0.859384537 | 0.390128398 | 1 | 1.009284215 |
| ENSMUSG00000026387 | Sctr | 319229 | protein_coding | 1.207189779 | -1.009464984 | 1.923937665 | 0.524686949 | 0.599800831 | 1 | 1.009464984 |
| ENSMUSG00000050395 | Tnfsf15 | 326623 | protein_coding | 21.25981356 | -1.009490378 | 0.259149965 | 3.895390754 | 9.80E-05 | 0.000845894 | 1.009490378 |
| ENSMUSG00000038264 | Sema7a | 20361 | protein_coding | 27.77534126 | -1.009881115 | 0.380740163 | 2.652415512 | 0.007991813 | 0.035574288 | 1.009881115 |
| ENSMUSG00000001497 | Pax9 | 18511 | protein_coding | 1.34097199 | -1.009967112 | 1.931162929 | 0.522983896 | 0.60098546 | 1 | 1.009967112 |
| ENSMUSG00000028100 | Nudt17 | 78373 | protein_coding | 18.06698675 | -1.010072993 | 0.28016049 | 3.605337041 | 0.000311748 | 0.002358318 | 1.010072993 |
| ENSMUSG00000088490 | Gm24400 | 115489076 | snRNA | 2.378864076 | -1.010788454 | 0.763167213 | 1.32446525 | 0.185348577 | 1 | 1.010788454 |
| ENSMUSG00000109305 | Smim38 | 100040049 | protein_coding | 27.76607016 | -1.01107479 | 0.312104537 | 3.239538909 | 0.001197231 | 0.007382777 | 1.01107479 |
| ENSMUSG00000091680 | Klhdc7b | 546648 | protein_coding | 2.294371842 | -1.011203112 | 1.43975033 | 0.702346158 | 0.482463314 | 1 | 1.011203112 |
| ENSMUSG00000061356 | Nuggc | 100503545 | protein_coding | 8.665569564 | 1.011525183 | 0.601093715 | -1.682807784 | 0.092412308 | 0.233151171 | 1.011525183 |
| ENSMUSG00000071715 | Ncf4 | 17972 | protein_coding | 3.384765413 | -1.011756188 | 0.765078224 | 1.322421886 | 0.186027705 | 1 | 1.011756188 |
| ENSMUSG00000032826 | Ank2 | 109676 | protein_coding | 13.2407126 | 1.011881213 | 0.80125522 | -1.262870042 | 0.206635888 | 0.409413929 | 1.011881213 |
| ENSMUSG00000031450 | Grk1 | 24013 | protein_coding | 3.160882627 | 1.012291144 | 1.909261767 | -0.530200291 | 0.595973069 | 1 | 1.012291144 |
| ENSMUSG00000038070 | Cntln | 338349 | protein_coding | 37.38236327 | 1.012444101 | 0.193800475 | -5.224156977 | 1.75E-07 | 2.77E-06 | 1.012444101 |
| ENSMUSG00000002068 | Ccne1 | 12447 | protein_coding | 87.16057659 | 1.013376708 | 0.196163337 | -5.165984242 | 2.39E-07 | 3.70E-06 | 1.013376708 |
| ENSMUSG00000059631 | 1500035N22Rik | 70258 | lncRNA | 1.566515793 | -1.013603681 | 1.278718655 | 0.79267138 | 0.427969309 | 1 | 1.013603681 |
| ENSMUSG00000064080 | Fbln2 | 14115 | protein_coding | 3.510008876 | 1.013754555 | 1.152464308 | -0.879640738 | 0.379053962 | 1 | 1.013754555 |
| ENSMUSG00000036027 | Pou2af2 | 69809 | protein_coding | 15.73937043 | -1.013917519 | 0.31001759 | 3.270516103 | 0.001073514 | 0.006748388 | 1.013917519 |
| ENSMUSG00000085415 | Selenok-ps1 | 100134990 | lncRNA | 0.988741977 | 1.01395509 | 1.98546391 | -0.510689258 | 0.609568664 | 1 | 1.01395509 |
| ENSMUSG00000056895 | H2bc27 | 78303 | protein_coding | 20.82596893 | 1.013956342 | 0.287268434 | -3.529647607 | 0.000416114 | 0.003035203 | 1.013956342 |
| ENSMUSG00000013611 | Snx31 | 66696 | protein_coding | 1.264935449 | -1.014241654 | 2.07628371 | 0.488488952 | 0.625203549 | 1 | 1.014241654 |
| ENSMUSG00000034612 | Chst11 | 58250 | protein_coding | 146.8054588 | 1.014261935 | 0.262958457 | -3.857118527 | 0.000114732 | 0.000972627 | 1.014261935 |
| ENSMUSG00000091006 | Vmn2r69 | 330581 | protein_coding | 3.033432157 | 1.014302054 | 2.188704282 | -0.46342581 | 0.643059182 | 1 | 1.014302054 |
| ENSMUSG00000039200 | Atf7ip2 | 75329 | protein_coding | 11.83117191 | 1.014448056 | 0.362123956 | -2.80138345 | 0.005088402 | 0.024729213 | 1.014448056 |
| ENSMUSG00000088113 | Mir1960 | 100316709 | miRNA | 0.923241583 | -1.014584629 | 1.122826178 | 0.903599016 | 0.366208063 | 1 | 1.014584629 |

| ensembl_gene_id | external_gene_name | entrezgene_id | gene_biotype | baseMean | log2FoldChange | lfcSE | stat | pvalue | padj | absLog2FC |
|---|---|---|---|---|---|---|---|---|---|---|
| ENSMUSG00000050762 | Prss27 | 213171 | protein_coding | 15.66893216 | 1.014806203 | 0.375816251 | -2.700272276 | 0.006928275 | 0.031640911 | 1.014806203 |
| ENSMUSG00000093979 | Gm2237 | 100039441 | protein_coding | 4.183305565 | -1.014885436 | 0.548402478 | 1.850621534 | 0.06422402 | 1 | 1.014885436 |
| ENSMUSG00000089324 | Mir1936 | 100316812 | miRNA | 2.926399572 | 1.015485646 | 0.786768293 | -1.290704843 | 0.196806045 | 1 | 1.015485646 |
| ENSMUSG00000043795 | Prr33 | 677289 | protein_coding | 2.541813794 | -1.015677768 | 2.129431078 | 0.476971421 | 0.633382472 | 1 | 1.015677768 |
| ENSMUSG00000086554 | 9530034E10Rik | 78615 | lncRNA | 6.188339404 | -1.016354722 | 0.458042331 | 2.218910028 | 0.026492846 | 0.091355786 | 1.016354722 |
| ENSMUSG00000068117 | Mei1 | 74369 | protein_coding | 2.070138029 | 1.016396951 | 1.804332557 | -0.563309102 | 0.573224422 | 1 | 1.016396951 |
| ENSMUSG00000074743 | Thbd | 21824 | protein_coding | 2.048047125 | -1.016596215 | 0.970568201 | 1.047423781 | 0.294904168 | 1 | 1.016596215 |
| ENSMUSG00000045027 | Prss22 | 70835 | protein_coding | 0.983303418 | 1.016690661 | 2.160802195 | -0.470515378 | 0.63798685 | 1 | 1.016690661 |
| ENSMUSG00000100593 | 1700119H24Rik | 76649 | lncRNA | 1.729793988 | 1.016862852 | 1.131688128 | -0.89853629 | 0.368899707 | 1 | 1.016862852 |
| ENSMUSG00000013415 | Igf2bp1 | 140486 | protein_coding | 22.12356443 | -1.017506402 | 0.484087815 | 2.101904596 | 0.035561634 | 0.114834733 | 1.017506402 |
| ENSMUSG00000084938 | BB557941 | 767815 | lncRNA | 0.639158009 | 1.017664986 | 1.539945378 | -0.660844859 | 0.508711811 | 1 | 1.017664986 |
| ENSMUSG00000108356 | 9130221F21Rik | 330350 | lncRNA | 5.351754279 | 1.017750544 | 0.968528492 | -1.050821481 | 0.293340588 | 0.512561987 | 1.017750544 |
| ENSMUSG00000030630 | Fah | 14085 | protein_coding | 50.13163681 | -1.018137022 | 0.175505323 | 5.801174585 | 6.59E-09 | 1.30E-07 | 1.018137022 |
| ENSMUSG00000101734 | 4933400C23Rik | 331003 | lncRNA | 1.200308645 | 1.018426558 | 1.948946556 | -0.522552327 | 0.601285823 | 1 | 1.018426558 |
| ENSMUSG00000040770 | Il25 | 140806 | protein_coding | 4.038908582 | -1.018520451 | 0.604912623 | 1.683748053 | 0.09223037 | 1 | 1.018520451 |
| ENSMUSG00000064272 | Gpbar1 | 227289 | protein_coding | 2.662213592 | 1.019410479 | 0.630147403 | -1.617733365 | 0.105720072 | 1 | 1.019410479 |
| ENSMUSG00000096549 | Prickle4 | 381104 | protein_coding | 0.907940844 | 1.019418525 | 1.508636694 | -0.675721682 | 0.499217367 | 1 | 1.019418525 |
| ENSMUSG00000006463 | Zdhhc24 | 70605 | protein_coding | 174.0705022 | 1.01971294 | 0.109207892 | -9.337355732 | 9.88E-21 | 5.26E-19 | 1.01971294 |
| ENSMUSG00000029752 | Asns | 27053 | protein_coding | 538.4258996 | 1.019765167 | 0.127713307 | -7.984799647 | 1.41E-15 | 5.36E-14 | 1.019765167 |
| ENSMUSG00000091983 | Or2r3 | 258989 | protein_coding | 0.962073513 | -1.019904873 | 2.145176756 | 0.475440949 | 0.634472713 | 1 | 1.019904873 |
| ENSMUSG00000040866 | Rsph6a | 83434 | protein_coding | 1.781342278 | 1.020074748 | 1.751209236 | -0.582497355 | 0.560231723 | 1 | 1.020074748 |
| ENSMUSG00000036278 | Macrod1 | 107227 | protein_coding | 93.03129762 | -1.020699659 | 0.140482897 | 7.265650718 | 3.71E-13 | 1.17E-11 | 1.020699659 |
| ENSMUSG00000090777 | Ccdc188 | 102638083 | protein_coding | 1.681076274 | 1.020863456 | 1.402398195 | -0.727941222 | 0.466649568 | 1 | 1.020863456 |
| ENSMUSG00000099140 | Mir7237 | 102466915 | miRNA | 0.637770026 | -1.020940057 | 1.572709628 | 0.64915992 | 0.516235015 | 1 | 1.020940057 |
| ENSMUSG00000053219 | Raet1e | 379043 | protein_coding | 52.56260457 | -1.021362618 | 0.246943817 | 4.136012118 | 3.53E-05 | 0.00034471 | 1.021362618 |
| ENSMUSG00000031893 | Tsnaxip1 | 72236 | protein_coding | 1.663524739 | 1.021507129 | 1.038605178 | -0.98353749 | 0.325342973 | 1 | 1.021507129 |
| ENSMUSG00000044816 | D630023F18Rik | 98303 | protein_coding | 1.204243049 | 1.022015606 | 1.279649232 | -0.798668558 | 0.424482623 | 1 | 1.022015606 |
| ENSMUSG00000039959 | Hip1 | 215114 | protein_coding | 6.756327492 | 1.022275715 | 0.92359234 | -1.106847329 | 0.268359941 | 0.483440174 | 1.022275715 |
| ENSMUSG00000057455 | Rit2 | 19762 | protein_coding | 1.546870365 | 1.022349239 | 2.339294164 | -0.437033211 | 0.662087263 | 1 | 1.022349239 |
| ENSMUSG00000025492 | Ifitm3 | 66141 | protein_coding | 113.9444478 | 1.022503434 | 0.237144671 | -4.311728495 | 1.62E-05 | 0.000171364 | 1.022503434 |
| ENSMUSG00000023571 | C1qtnf12 | 67389 | protein_coding | 76.9623644 | -1.022506899 | 0.168613287 | 6.064213071 | 1.33E-09 | 2.89E-08 | 1.022506899 |
| ENSMUSG00000096351 | Samd11 | 231004 | protein_coding | 3.066177427 | -1.022603565 | 0.747888997 | 1.367319976 | 0.171525033 | 1 | 1.022603565 |
| ENSMUSG00000103037 | Pcdhgb1 | 93699 | protein_coding | 3.735897389 | 1.022930332 | 0.730380627 | -1.400544174 | 0.161350425 | 1 | 1.022930332 |
| ENSMUSG00000015843 | Rxrg | 20183 | protein_coding | 1.196647782 | 1.023015222 | 1.848849892 | -0.553325192 | 0.580040754 | 1 | 1.023015222 |
| ENSMUSG00000039474 | Wfs1 | 22393 | protein_coding | 211.8816001 | 1.023103783 | 0.111435399 | -9.181138053 | 4.27E-20 | 2.16E-18 | 1.023103783 |
| ENSMUSG00000032473 | Cldn18 | 56492 | protein_coding | 1.370312051 | 1.023383311 | 2.345333911 | -0.436348661 | 0.662583782 | 1 | 1.023383311 |
| ENSMUSG00000027072 | Prg3 | 53856 | protein_coding | 1.124320065 | 1.023410674 | 2.345480895 | -0.436332982 | 0.662595155 | 1 | 1.023410674 |
| ENSMUSG00000040599 | Mis12 | 67139 | protein_coding | 332.4310078 | 1.023423967 | 0.116982788 | -8.748500423 | 2.16E-18 | 1.01E-16 | 1.023423967 |
| ENSMUSG00000031841 | Cdh13 | 12554 | protein_coding | 4.287904803 | 1.024353603 | 2.032569619 | -0.50396975 | 0.614282634 | 1 | 1.024353603 |

Supplementary Table 1

| ensembl_gene_id | external_gene_name | entrezgene_id | gene_biotype | baseMean | log2FoldChange | lfcSE | stat | pvalue | padj | absLog2FC |
|---|---|---|---|---|---|---|---|---|---|---|
| ENSMUSG00000052928 | Ctif | 269037 | protein_coding | 174.8560154 | 1.024441371 | 0.144440352 | -7.092487375 | 1.32E-12 | 3.93E-11 | 1.024441371 |
| ENSMUSG00000043496 | Tril | 66873 | protein_coding | 0.967800269 | 1.024883515 | 1.709124174 | -0.599654215 | 0.548736707 | 1 | 1.024883515 |
| ENSMUSG00000017652 | Cd40 | 21939 | protein_coding | 2.047414775 | 1.024912637 | 1.645440945 | -0.622880231 | 0.53336323 | 1 | 1.024912637 |
| ENSMUSG00000046709 | Mapk10 | 26414 | protein_coding | 4.533521575 | 1.025018132 | 1.789383095 | -0.572833249 | 0.566757604 | 0.753213832 | 1.025018132 |
| ENSMUSG00000056596 | Trnp1 | 69539 | protein_coding | 2.888899018 | 1.025049646 | 1.075908872 | -0.952729058 | 0.340727367 | 1 | 1.025049646 |
| ENSMUSG00000049103 | Ccr2 | 12772 | protein_coding | 5.502213209 | 1.025063209 | 0.902293119 | -1.136064531 | 0.255929558 | 0.468613008 | 1.025063209 |
| ENSMUSG00000001566 | Mnx1 | 15285 | protein_coding | 12.05929935 | -1.025079076 | 0.30519739 | 3.358741292 | 0.000782983 | 0.005170946 | 1.025079076 |
| ENSMUSG00000067276 | Capn6 | 12338 | protein_coding | 19.46148672 | 1.025249949 | 0.320002303 | -3.203883032 | 0.001355875 | 0.008201805 | 1.025249949 |
| ENSMUSG00000037446 | Tulp1 | 22157 | protein_coding | 4.329697642 | 1.025749071 | 1.522746941 | -0.673617555 | 0.50055449 | 1 | 1.025749071 |
| ENSMUSG00000038199 | Iqca1l | 231045 | protein_coding | 2.620025589 | 1.025800211 | 1.83001665 | -0.560541464 | 0.575110166 | 1 | 1.025800211 |
| ENSMUSG00000021904 | Sema3g | 218877 | protein_coding | 4.338332114 | -1.025899667 | 0.836920244 | 1.225803384 | 0.220272674 | 1 | 1.025899667 |
| ENSMUSG00000047720 | Clec2m | 381816 | protein_coding | 1.185814661 | 1.025959506 | 1.306141204 | -0.785488968 | 0.432166936 | 1 | 1.025959506 |
| ENSMUSG00000031933 | Izumo1r | 64931 | protein_coding | 1.607807739 | -1.026264772 | 1.462237342 | 0.70184555 | 0.482775489 | 1 | 1.026264772 |
| ENSMUSG00000071478 | H2ac7 | 319165 | protein_coding | 102.5493046 | 1.026333837 | 0.218594714 | -4.695144809 | 2.66E-06 | 3.36E-05 | 1.026333837 |
| ENSMUSG00000074224 | Wdr87-ps | 114675 | transcribed_unitary_pseudogene | 7.213990272 | 1.027017861 | 1.739439492 | -0.590430346 | 0.554902171 | 0.744703672 | 1.027017861 |
| ENSMUSG00000096950 | Gm9530 | 671336 | lncRNA | 9.219619077 | 1.027491299 | 0.663690118 | -1.548149161 | 0.121586388 | 0.284401928 | 1.027491299 |
| ENSMUSG00000079484 | Phyhd1 | 227696 | protein_coding | 2.814627113 | 1.027627162 | 0.776695216 | -1.323076467 | 0.18580995 | 1 | 1.027627162 |
| ENSMUSG00000086604 | Gm15510 | 102640192 | lncRNA | 1.070889616 | -1.027656121 | 1.211195056 | 0.848464594 | 0.396179283 | 1 | 1.027656121 |
| ENSMUSG00000006386 | Tek | 21687 | protein_coding | 7.95376847 | -1.02794468 | 0.516734165 | 1.989310461 | 0.046666946 | 0.141461059 | 1.02794468 |
| ENSMUSG00000039419 | Cntnap2 | 66797 | protein_coding | 3.617849012 | 1.028146151 | 1.391919071 | -0.738653685 | 0.460117316 | 1 | 1.028146151 |
| ENSMUSG00000056832 | Ttc26 | 264134 | protein_coding | 32.30517238 | 1.028678067 | 0.226852726 | -4.53456339 | 5.77E-06 | 6.80E-05 | 1.028678067 |
| ENSMUSG00000036896 | C1qc | 12262 | protein_coding | 3.688015561 | 1.029034737 | 0.845927667 | -1.216457124 | 0.223810835 | 1 | 1.029034737 |
| ENSMUSG00000024409 | Psors1c2 | 57390 | protein_coding | 0.757523077 | 1.029340736 | 1.481807616 | -0.694652076 | 0.487273368 | 1 | 1.029340736 |
| ENSMUSG00000050103 | Agmo | 319660 | protein_coding | 46.16293494 | -1.029451912 | 0.193652743 | 5.315968645 | 1.06E-07 | 1.74E-06 | 1.029451912 |
| ENSMUSG00000067916 | Zfp991 | 666532 | protein_coding | 29.52672378 | -1.029466165 | 0.274634393 | 3.748496877 | 0.000177898 | 0.001430282 | 1.029466165 |
| ENSMUSG00000108322 | 5430431A17Rik | 71368 | lncRNA | 2.840347924 | -1.029501664 | 0.79240167 | 1.299216929 | 0.193869493 | 1 | 1.029501664 |
| ENSMUSG00000022439 | Parvg | 64099 | protein_coding | 7.792162565 | 1.029974551 | 0.747229628 | -1.378390942 | 0.168082619 | 0.356592544 | 1.029974551 |
| ENSMUSG00000000901 | Mmp11 | 17385 | protein_coding | 7.621487165 | 1.030768831 | 0.55801676 | -1.847200485 | 0.064718089 | 0.180532859 | 1.030768831 |
| ENSMUSG00000019893 | Ros1 | 19886 | protein_coding | 10.72741042 | 1.031369965 | 0.502237916 | -2.053548589 | 0.040019402 | 0.125929956 | 1.031369965 |
| ENSMUSG00000031932 | Gpr83 | 14608 | protein_coding | 7.526606506 | -1.031394105 | 0.565020162 | 1.825411154 | 0.067939007 | 0.186897346 | 1.031394105 |
| ENSMUSG00000036192 | Rorb | 225998 | protein_coding | 9.942951382 | 1.03142929 | 0.44445732 | -2.320648672 | 0.020305813 | 0.074253808 | 1.03142929 |
| ENSMUSG00000079694 | Gm14862 | 102636575 | lncRNA | 1.476579108 | -1.031560513 | 1.104411523 | 0.934036355 | 0.35028514 | 1 | 1.031560513 |
| ENSMUSG00000078700 | D030028A08Rik | 319371 | lncRNA | 150.8927474 | 1.031585129 | 0.129969023 | -7.937161557 | 2.07E-15 | 7.78E-14 | 1.031585129 |
| ENSMUSG00000058571 | Gpc6 | 23888 | protein_coding | 15.91621578 | 1.031953406 | 0.344545515 | -2.995114899 | 0.002743415 | 0.014762057 | 1.031953406 |
| ENSMUSG00000089762 | Ier5l | 72500 | protein_coding | 29.7155191 | 1.032298831 | 0.499585496 | -2.066310649 | 0.038799157 | 0.123098803 | 1.032298831 |
| ENSMUSG00000077567 | Gm23200 | 115485981 | snoRNA | 0.640060562 | -1.032476832 | 1.427205023 | 0.723425727 | 0.46941837 | 1 | 1.032476832 |
| ENSMUSG00000028487 | Bnc2 | 242509 | protein_coding | 2.494232254 | 1.032553777 | 1.786656306 | -0.577925241 | 0.563314593 | 1 | 1.032553777 |
| ENSMUSG00000111825 | Gm10646 | 102632690 | lncRNA | 2.054933219 | -1.032727423 | 1.489972868 | 0.693118274 | 0.48823533 | 1 | 1.032727423 |
| ENSMUSG00000025189 | Cnnm1 | 83674 | protein_coding | 15.67078358 | -1.032811549 | 0.409893298 | 2.519708311 | 0.011745212 | 0.048307661 | 1.032811549 |

| ensembl_gene_id | external_gene_name | entrezgene_id | gene_biotype | baseMean | log2FoldChange | lfcSE | stat | pvalue | padj | absLog2FC |
|---|---|---|---|---|---|---|---|---|---|---|
| ENSMUSG00000056856 | Jakmip3 | 74004 | protein_coding | 2.360731223 | 1.033016072 | 2.409931906 | -0.428649486 | 0.668178327 | 1 | 1.033016072 |
| ENSMUSG00000029882 | 2210010C04Rik | 67373 | protein_coding | 1.592661647 | -1.033043676 | 2.089349936 | 0.494433057 | 0.621000373 | 1 | 1.033043676 |
| ENSMUSG00000022358 | Fbxo32 | 67731 | protein_coding | 505.7744424 | -1.033438972 | 0.209471076 | 4.933564059 | 8.07E-07 | 1.14E-05 | 1.033438972 |
| ENSMUSG00000046191 | Pcdhb20 | 93891 | protein_coding | 1.854801716 | 1.033570169 | 0.988360203 | -1.045742398 | 0.295679982 | 1 | 1.033570169 |
| ENSMUSG00000010064 | Slc38a3 | 76257 | protein_coding | 3.04480249 | -1.03389027 | 1.360121957 | 0.760145269 | 0.447167756 | 1 | 1.03389027 |
| ENSMUSG00000079442 | St6galnac4 | 20448 | protein_coding | 556.8388708 | 1.034643947 | 0.141121521 | -7.331581602 | 2.27E-13 | 7.32E-12 | 1.034643947 |
| ENSMUSG00000034110 | Kctd7 | 212919 | protein_coding | 84.96835073 | 1.034920894 | 0.153714098 | -6.732764953 | 1.66E-11 | 4.55E-10 | 1.034920894 |
| ENSMUSG00000024018 | Ccdc167 | 68597 | protein_coding | 77.92610278 | 1.034996974 | 0.163937896 | -6.313347906 | 2.73E-10 | 6.48E-09 | 1.034996974 |
| ENSMUSG00000045064 | Zc2hc1c | 72350 | protein_coding | 12.14374177 | -1.035058853 | 0.378868924 | 2.73197084 | 0.006295671 | 0.029171357 | 1.035058853 |
| ENSMUSG00000026797 | Stxbp1 | 20910 | protein_coding | 96.57447746 | -1.035159519 | 0.176347792 | 5.869988545 | 4.36E-09 | 8.87E-08 | 1.035159519 |
| ENSMUSG00000085260 | Med9os | 100504121 | lncRNA | 2.45585152 | 1.03519968 | 0.835846414 | -1.238504662 | 0.215528995 | 1 | 1.03519968 |
| ENSMUSG00000100911 | 1700027F09Rik | 69999 | lncRNA | 0.647689803 | 1.035366799 | 1.506673833 | -0.687187085 | 0.491964842 | 1 | 1.035366799 |
| ENSMUSG00000003865 | Gys1 | 14936 | protein_coding | 476.9571757 | 1.035904028 | 0.085046747 | -12.18040744 | 3.95E-34 | 3.44E-32 | 1.035904028 |
| ENSMUSG00000026496 | Parp1 | 11545 | protein_coding | 1069.117495 | 1.036221179 | 0.119943028 | -8.639278131 | 5.66E-18 | 2.60E-16 | 1.036221179 |
| ENSMUSG00000032135 | Mcam | 84004 | protein_coding | 6.506628612 | 1.03622197 | 0.588326197 | -1.761305166 | 0.078186766 | 0.206753604 | 1.03622197 |
| ENSMUSG00000075701 | Selenos | 109815 | protein_coding | 905.1635689 | 1.036688427 | 0.100494471 | -10.31587526 | 5.97E-25 | 3.81E-23 | 1.036688427 |
| ENSMUSG00000029608 | Rph3a | 19894 | protein_coding | 3.556773402 | 1.036920696 | 3.480098972 | -0.297957243 | 0.765735797 | 1 | 1.036920696 |
| ENSMUSG00000002111 | Spi1 | 20375 | protein_coding | 2.288731882 | 1.037104012 | 0.903415352 | -1.14798139 | 0.250976247 | 1 | 1.037104012 |
| ENSMUSG00000033740 | St18 | 240690 | protein_coding | 41.14720508 | -1.037112138 | 0.238051995 | 4.356662238 | 1.32E-05 | 0.000143007 | 1.037112138 |
| ENSMUSG00000026668 | Ucma | 68527 | protein_coding | 1.268470488 | 1.037143363 | 1.996362006 | -0.519516681 | 0.603400484 | 1 | 1.037143363 |
| ENSMUSG00000030671 | Pde3b | 18576 | protein_coding | 184.9802439 | 1.03724742 | 0.139875123 | -7.415524646 | 1.21E-13 | 3.98E-12 | 1.03724742 |
| ENSMUSG00000002204 | Napsa | 16541 | protein_coding | 3.439953133 | 1.037288404 | 0.861604471 | -1.203903228 | 0.228626984 | 1 | 1.037288404 |
| ENSMUSG00000009108 | Gnat2 | 14686 | protein_coding | 21.52868654 | 1.038008072 | 0.260248097 | -3.988532804 | 6.65E-05 | 0.000602545 | 1.038008072 |
| ENSMUSG00000059173 | Pde1a | 18573 | protein_coding | 2.500559513 | 1.038124009 | 2.171874528 | -0.477985259 | 0.632660697 | 1 | 1.038124009 |
| ENSMUSG00000046207 | Pik3r6 | 104709 | protein_coding | 2.817933509 | 1.038782641 | 1.551004199 | -0.669748439 | 0.503018167 | 1 | 1.038782641 |
| ENSMUSG00000098651 | Mir6999 | 102465604 | miRNA | 0.687314813 | 1.039126156 | 1.637194769 | -0.634699167 | 0.525624637 | 1 | 1.039126156 |
| ENSMUSG00000033770 | Clcnka | 12733 | protein_coding | 2.705041664 | 1.039181163 | 2.381004732 | -0.436446492 | 0.662512813 | 1 | 1.039181163 |
| ENSMUSG00000044365 | Cxxc4 | 319478 | protein_coding | 16.70290867 | 1.039667668 | 0.394437125 | -2.635826102 | 0.008393273 | 0.036961738 | 1.039667668 |
| ENSMUSG00000066515 | Klk1b3 | 18050 | protein_coding | 1.59349339 | -1.040056777 | 1.136713448 | 0.914968305 | 0.36020827 | 1 | 1.040056777 |
| ENSMUSG00000020728 | Cep112 | 76380 | protein_coding | 16.91182805 | 1.040148282 | 0.310160668 | -3.35357893 | 0.000797737 | 0.005248535 | 1.040148282 |
| ENSMUSG00000070822 | Zscan18 | 232875 | protein_coding | 7.950824682 | 1.040685346 | 0.489404607 | -2.126431447 | 0.033467352 | 0.109658699 | 1.040685346 |
| ENSMUSG00000038252 | Ncapd2 | 68298 | protein_coding | 874.3516631 | 1.040700797 | 0.137152093 | -7.587932281 | 3.25E-14 | 1.12E-12 | 1.040700797 |
| ENSMUSG00000030376 | Slc8a2 | 110891 | protein_coding | 3.685089387 | 1.040798406 | 1.800789978 | -0.57796768 | 0.56328594 | 1 | 1.040798406 |
| ENSMUSG00000006179 | Prss16 | 54373 | protein_coding | 24.62870081 | 1.041026345 | 0.306366066 | -3.397981896 | 0.000678849 | 0.00457572 | 1.041026345 |
| ENSMUSG00000099907 | Gm10421 | 100038689 | lncRNA | 4.154959 | -1.041185052 | 0.512224929 | 2.032671575 | 0.042085715 | 1 | 1.041185052 |
| ENSMUSG00000070284 | Gmppb | 331026 | protein_coding | 1042.655344 | 1.041276537 | 0.093878956 | -11.09169271 | 1.38E-28 | 1.01E-26 | 1.041276537 |
| ENSMUSG00000033420 | Antxr1 | 69538 | protein_coding | 2.616965095 | 1.041365176 | 1.998454795 | -0.521085179 | 0.602307435 | 1 | 1.041365176 |
| ENSMUSG00000058290 | Espl1 | 105988 | protein_coding | 248.9180209 | 1.041687552 | 0.150095786 | -6.940151887 | 3.92E-12 | 1.13E-10 | 1.041687552 |
| ENSMUSG00000117401 | 4833419F23Rik | 73915 | lncRNA | 1.047220863 | -1.04309595 | 1.503651718 | 0.693708482 | 0.487865046 | 1 | 1.04309595 |

| ensembl_gene_id | external_gene_name | entrezgene_id | gene_biotype | baseMean | log2FoldChange | lfcSE | stat | pvalue | padj | absLog2FC |
|---|---|---|---|---|---|---|---|---|---|---|
| ENSMUSG00000020808 | Pimreg | 109212 | protein_coding | 42.71480641 | 1.043536489 | 0.221339613 | -4.714639527 | 2.42E-06 | 3.09E-05 | 1.043536489 |
| ENSMUSG00000005268 | Prlr | 19116 | protein_coding | 172.6672719 | -1.043673282 | 0.157343669 | 6.633080884 | 3.29E-11 | 8.76E-10 | 1.043673282 |
| ENSMUSG00000072214 | Septin5 | 18951 | protein_coding | 450.3284197 | 1.043895836 | 0.139529881 | -7.48152172 | 7.35E-14 | 2.45E-12 | 1.043895836 |
| ENSMUSG00000098785 | Mir6395 | 102465211 | miRNA | 1.330650311 | 1.044246177 | 1.037752363 | -1.006257576 | 0.314291682 | 1 | 1.044246177 |
| ENSMUSG00000030897 | Cnga4 | 233649 | protein_coding | 1.856142183 | 1.044741107 | 1.478412681 | -0.70666406 | 0.479775269 | 1 | 1.044741107 |
| ENSMUSG00000063751 | Platr8 | 100038454 | lncRNA | 4.397248038 | -1.045495156 | 0.508535329 | 2.055894832 | 0.039792654 | 0.125530417 | 1.045495156 |
| ENSMUSG00000008129 | Brme1 | 74666 | protein_coding | 5.524226962 | -1.045973464 | 0.447704646 | 2.336302455 | 0.019475484 | 0.071918188 | 1.045973464 |
| ENSMUSG00000074818 | Pdzd7 | 100503041 | protein_coding | 108.7182815 | -1.046215738 | 0.239934246 | 4.36042688 | 1.30E-05 | 0.000141031 | 1.046215738 |
| ENSMUSG00000035262 | Amh | 11705 | protein_coding | 2.337177898 | 1.046311148 | 0.830705295 | -1.259545538 | 0.207833353 | 1 | 1.046311148 |
| ENSMUSG00000038628 | Polr3k | 67005 | protein_coding | 621.5584978 | 1.046362015 | 0.096287815 | -10.86702415 | 1.66E-27 | 1.17E-25 | 1.046362015 |
| ENSMUSG00000090489 | Gm17415 | 108167670 | processed_pseudogene | 23.67723775 | 1.046532765 | 0.266012262 | -3.93415235 | 8.35E-05 | 0.000735343 | 1.046532765 |
| ENSMUSG00000089759 | 3632454L22Rik | 320606 | lncRNA | 3.050229483 | 1.046795245 | 1.092275078 | -0.95836229 | 0.337880104 | 1 | 1.046795245 |
| ENSMUSG00000073906 | Or52w1 | 258352 | protein_coding | 0.830372288 | 1.047075729 | 1.523246305 | -0.687397518 | 0.491832262 | 1 | 1.047075729 |
| ENSMUSG00000079055 | Slc8a3 | 110893 | protein_coding | 2.600064984 | 1.047270662 | 1.611478562 | -0.649881845 | 0.515768546 | 1 | 1.047270662 |
| ENSMUSG00000032243 | Itga11 | 319480 | protein_coding | 5.530027971 | -1.047320393 | 0.861664884 | 1.215461385 | 0.224190168 | 0.431299678 | 1.047320393 |
| ENSMUSG00000097098 | 9330111N05Rik | 319983 | lncRNA | 2.297000103 | -1.047617789 | 1.18428723 | 0.884597724 | 0.376373635 | 1 | 1.047617789 |
| ENSMUSG00000030319 | Cand2 | 67088 | protein_coding | 48.18391624 | 1.047967066 | 0.167030851 | -6.274092828 | 3.52E-10 | 8.22E-09 | 1.047967066 |
| ENSMUSG00000039275 | Foxk2 | 68837 | protein_coding | 757.8738102 | 1.047988767 | 0.079542241 | -13.17524818 | 1.22E-39 | 1.24E-37 | 1.047988767 |
| ENSMUSG00000106350 | Kif19b | 101055939 | protein_coding | 1.466090362 | -1.048103914 | 1.226866772 | 0.854293178 | 0.392942566 | 1 | 1.048103914 |
| ENSMUSG00000018776 | Slc35g3 | 56293 | protein_coding | 4.702657727 | 1.049026466 | 0.79102461 | -1.326161605 | 0.184786174 | 0.380019786 | 1.049026466 |
| ENSMUSG00000037032 | Apbb1 | 11785 | protein_coding | 10.87718926 | 1.049191431 | 0.477968647 | -2.195105134 | 0.028156057 | 0.095909473 | 1.049191431 |
| ENSMUSG00000042514 | Klhl14 | 225266 | protein_coding | 3.028715646 | 1.049548395 | 0.851029453 | -1.233269179 | 0.217475353 | 1 | 1.049548395 |
| ENSMUSG00000003283 | Hck | 15162 | protein_coding | 216.8266574 | -1.049626166 | 0.184015326 | 5.704014916 | 1.17E-08 | 2.23E-07 | 1.049626166 |
| ENSMUSG00000019872 | Smpdl3a | 57319 | protein_coding | 1779.391504 | -1.049860095 | 0.15164054 | 6.923347108 | 4.41E-12 | 1.26E-10 | 1.049860095 |
| ENSMUSG00000067006 | Serpinb5 | 20724 | protein_coding | 734.3123053 | -1.050430025 | 0.126386386 | 8.311259302 | 9.47E-17 | 3.98E-15 | 1.050430025 |
| ENSMUSG00000038044 | Cct8l1 | 242891 | protein_coding | 2.315754239 | 1.050995534 | 1.076500864 | -0.976307191 | 0.328912258 | 1 | 1.050995534 |
| ENSMUSG00000097519 | 4930558J18Rik | 75323 | lncRNA | 6.448856873 | 1.051116878 | 0.500419715 | -2.100470557 | 0.035687468 | 0.115106025 | 1.051116878 |
| ENSMUSG00000060441 | Trim5 | 667823 | protein_coding | 15.48361495 | -1.051174323 | 0.382731658 | 2.746504766 | 0.006023401 | 0.028250624 | 1.051174323 |
| ENSMUSG00000089512 | Gm26287 | 115487683 | snoRNA | 1.499426341 | -1.051399812 | 0.861813027 | 1.219985983 | 0.222470189 | 1 | 1.051399812 |
| ENSMUSG00000051537 | Gm5124 | 331392 | transcribed_processed_pseudogene | 2.701661716 | 1.051752145 | 0.884990563 | -1.188433176 | 0.234662792 | 1 | 1.051752145 |
| ENSMUSG00000015405 | Ace2 | 70008 | protein_coding | 101.9547383 | -1.052320961 | 0.172165151 | 6.112276236 | 9.82E-10 | 2.16E-08 | 1.052320961 |
| ENSMUSG00000041710 | Trpc5 | 22067 | protein_coding | 1.784773037 | -1.052590348 | 2.121884976 | 0.496063811 | 0.619849391 | 1 | 1.052590348 |
| ENSMUSG00000074115 | Saa1 | 20208 | protein_coding | 1938.291749 | -1.052855875 | 0.209416019 | 5.027580423 | 4.97E-07 | 7.34E-06 | 1.052855875 |
| ENSMUSG00000019232 | Etnppl | 71760 | protein_coding | 55.82349107 | 1.05308989 | 0.218558578 | -4.818341616 | 1.45E-06 | 1.93E-05 | 1.05308989 |
| ENSMUSG00000027306 | Nusap1 | 108907 | protein_coding | 427.0270659 | 1.053367507 | 0.192935108 | -5.459698438 | 4.77E-08 | 8.20E-07 | 1.053367507 |
| ENSMUSG00000044092 | C130050O18Rik | 319772 | protein_coding | 2.417991228 | -1.053775367 | 0.789573008 | 1.334614223 | 0.18200264 | 1 | 1.053775367 |
| ENSMUSG00000023068 | Nus1 | 52014 | protein_coding | 2088.492465 | 1.054118295 | 0.066424481 | -15.86942466 | 1.03E-56 | 1.65E-54 | 1.054118295 |
| ENSMUSG00000045658 | Pid1 | 98496 | protein_coding | 3.072644188 | 1.055212606 | 2.007977076 | -0.525510285 | 0.599228505 | 1 | 1.055212606 |
| ENSMUSG00000063672 | Nkx6-3 | 74561 | protein_coding | 5.016029602 | -1.05575962 | 0.534458075 | 1.975383419 | 0.048224653 | 0.145192693 | 1.05575962 |

| ensembl_gene_id | external_gene_name | entrezgene_id | gene_biotype | baseMean | log2FoldChange | lfcSE | stat | pvalue | padj | absLog2FC |
|---|---|---|---|---|---|---|---|---|---|---|
| ENSMUSG00000030737 | Slco2b1 | 101488 | protein_coding | 108.1070481 | -1.056128001 | 0.130925923 | 8.066607236 | 7.23E-16 | 2.86E-14 | 1.056128001 |
| ENSMUSG00000024042 | Sik1 | 17691 | protein_coding | 301.8979605 | -1.056526046 | 0.166192478 | 6.357243487 | 2.05E-10 | 4.94E-09 | 1.056526046 |
| ENSMUSG00000048794 | Cfap100 | 243538 | protein_coding | 6.558066075 | -1.056600907 | 0.661267857 | 1.597841019 | 0.110078363 | 0.265144996 | 1.056600907 |
| ENSMUSG00000038572 | Bpifb5 | 228802 | protein_coding | 1.173157065 | 1.056793073 | 2.087410425 | -0.506269903 | 0.612667189 | 1 | 1.056793073 |
| ENSMUSG00000059994 | Fcrl1 | 229499 | protein_coding | 2.776623629 | 1.057159491 | 1.431785616 | -0.738350406 | 0.460301543 | 1 | 1.057159491 |
| ENSMUSG00000015312 | Gadd45b | 17873 | protein_coding | 218.6795878 | 1.057294086 | 0.266272337 | -3.970724471 | 7.17E-05 | 0.000643098 | 1.057294086 |
| ENSMUSG00000045969 | Ing1 | 26356 | protein_coding | 532.1579162 | 1.057661626 | 0.113771746 | -9.296346994 | 1.45E-20 | 7.62E-19 | 1.057661626 |
| ENSMUSG00000037855 | Zfp365 | 216049 | protein_coding | 1.385184169 | 1.058126211 | 1.888843365 | -0.560197966 | 0.575344414 | 1 | 1.058126211 |
| ENSMUSG00000013076 | Amotl1 | 75723 | protein_coding | 201.4995652 | 1.058620504 | 0.172215843 | -6.147056422 | 7.89E-10 | 1.77E-08 | 1.058620504 |
| ENSMUSG00000047619 | Ddi1 | 71829 | protein_coding | 4.194305521 | -1.059268557 | 0.861665365 | 1.229327068 | 0.218949202 | 1 | 1.059268557 |
| ENSMUSG00000027971 | Ndst4 | 64580 | protein_coding | 3.903615224 | 1.059986814 | 1.848867999 | -0.573316653 | 0.566430312 | 1 | 1.059986814 |
| ENSMUSG00000073489 | Ifi204 | 15951 | protein_coding | 7.395893328 | 1.06005395 | 0.770039076 | -1.37662358 | 0.168628657 | 0.357475435 | 1.06005395 |
| ENSMUSG00000009114 | 2610028H24Rik | 76964 | protein_coding | 0.911536257 | -1.060316066 | 1.691152316 | 0.626978455 | 0.530673358 | 1 | 1.060316066 |
| ENSMUSG00000001946 | Esam | 69524 | protein_coding | 7.517013388 | -1.060789067 | 0.416899431 | 2.544472334 | 0.010944298 | 0.045686082 | 1.060789067 |
| ENSMUSG00000019577 | Pdk4 | 27273 | protein_coding | 174.1233768 | 1.061065535 | 0.387627384 | -2.737333786 | 0.006193941 | 0.028870044 | 1.061065535 |
| ENSMUSG00000038503 | Mesd | 67943 | protein_coding | 495.4703791 | 1.061246267 | 0.100099792 | -10.60188283 | 2.92E-26 | 1.98E-24 | 1.061246267 |
| ENSMUSG00000021541 | Trpc7 | 26946 | protein_coding | 3.018433355 | 1.061401138 | 1.188138817 | -0.893330916 | 0.371679986 | 1 | 1.061401138 |
| ENSMUSG00000079478 | Znrd2 | 56390 | protein_coding | 1.554802687 | 1.061652254 | 0.942175918 | -1.126808946 | 0.259823267 | 1 | 1.061652254 |
| ENSMUSG00000020080 | Hkdc1 | 216019 | protein_coding | 403.7883211 | -1.061656752 | 0.302542603 | 3.509114887 | 0.000449601 | 0.003240814 | 1.061656752 |
| ENSMUSG00000026958 | Dpp7 | 83768 | protein_coding | 136.1466012 | -1.061779565 | 0.095261155 | 11.14598673 | 7.49E-29 | 5.55E-27 | 1.061779565 |
| ENSMUSG00000024806 | Mlana | 77836 | protein_coding | 0.963233465 | 1.061896477 | 2.135852503 | -0.497176877 | 0.619064326 | 1 | 1.061896477 |
| ENSMUSG00000028637 | Ccdc30 | 73332 | protein_coding | 8.089794126 | 1.062025068 | 0.812109409 | -1.307736441 | 0.190962719 | 0.388417088 | 1.062025068 |
| ENSMUSG00000028747 | Htr6 | 15565 | protein_coding | 1.336170526 | 1.062206945 | 2.146549616 | -0.494843882 | 0.620710326 | 1 | 1.062206945 |
| ENSMUSG00000032690 | Oas2 | 246728 | protein_coding | 49.47240418 | -1.062640973 | 0.57707093 | 1.841439098 | 0.065557238 | 0.182012815 | 1.062640973 |
| ENSMUSG00000114824 | 3110015C05Rik | 73129 | lncRNA | 1.274158908 | 1.062689261 | 1.453400747 | -0.731174292 | 0.464672701 | 1 | 1.062689261 |
| ENSMUSG00000042761 | Mrap2 | 244958 | protein_coding | 1.873223583 | 1.062779156 | 1.706354936 | -0.622835926 | 0.533392347 | 1 | 1.062779156 |
| ENSMUSG00000032649 | Colgalt2 | 269132 | protein_coding | 1.594784637 | 1.06357362 | 2.097531023 | -0.50705978 | 0.612112874 | 1 | 1.06357362 |
| ENSMUSG00000045559 | Or9k7 | 258297 | protein_coding | 3.255477821 | 1.063591577 | 2.240546016 | -0.474701956 | 0.634999423 | 1 | 1.063591577 |
| ENSMUSG00000040373 | Cacng5 | 140723 | protein_coding | 1.389222314 | 1.063592358 | 1.651835529 | -0.643885144 | 0.519649916 | 1 | 1.063592358 |
| ENSMUSG00000019935 | Slc17a8 | 216227 | protein_coding | 19.46158074 | 1.063607573 | 0.379953866 | -2.79930715 | 0.00512124 | 0.024837637 | 1.063607573 |
| ENSMUSG00000016498 | Pdcd1lg2 | 58205 | protein_coding | 1.205869745 | 1.063629578 | 1.883316371 | -0.564764155 | 0.572234191 | 1 | 1.063629578 |
| ENSMUSG00000093865 | Lrit3 | 242235 | protein_coding | 7.006902233 | -1.063750479 | 0.483877445 | 2.198388227 | 0.027921452 | 0.09528722 | 1.063750479 |
| ENSMUSG00000035187 | Nkx6-1 | 18096 | protein_coding | 1.286926248 | 1.063796885 | 2.203777385 | -0.48271522 | 0.629297955 | 1 | 1.063796885 |
| ENSMUSG00000035237 | Lcat | 16816 | protein_coding | 1.121814515 | 1.063799522 | 2.203851425 | -0.482700199 | 0.629308622 | 1 | 1.063799522 |
| ENSMUSG00000032987 | Or8s8 | 258277 | protein_coding | 0.991260596 | 1.064130695 | 1.893064739 | -0.562120604 | 0.574033851 | 1 | 1.064130695 |
| ENSMUSG00000099238 | Mir7001 | 102465605 | miRNA | 2.336993821 | 1.064197432 | 0.804566219 | -1.322697134 | 0.185936117 | 1 | 1.064197432 |
| ENSMUSG00000085900 | A930041C12Rik | 78443 | lncRNA | 1.580966804 | 1.064542774 | 0.963406113 | -1.104978223 | 0.269169026 | 1 | 1.064542774 |
| ENSMUSG00000117571 | 4933433H22Rik | 74473 | lncRNA | 1.014421 | 1.064556182 | 1.901492573 | -0.559852927 | 0.575579759 | 1 | 1.064556182 |
| ENSMUSG00000042604 | Kcna4 | 16492 | protein_coding | 1.349387215 | 1.064627075 | 2.674001572 | -0.398140033 | 0.690526966 | 1 | 1.064627075 |

Supplementary Table 1

| ensembl_gene_id | external_gene_name | entrezgene_id | gene_biotype | baseMean | log2FoldChange | lfcSE | stat | pvalue | padj | absLog2FC |
|---|---|---|---|---|---|---|---|---|---|---|
| ENSMUSG00000034872 | Gipc3 | 209047 | protein_coding | 7.255905914 | 1.064726957 | 0.694367803 | -1.533376047 | 0.125183227 | 0.290347713 | 1.064726957 |
| ENSMUSG00000032125 | Robo4 | 74144 | protein_coding | 3.260660695 | -1.064969045 | 1.832277891 | 0.581226816 | 0.561087597 | 1 | 1.064969045 |
| ENSMUSG00000030062 | Rpn1 | 103963 | protein_coding | 6294.839139 | 1.065153387 | 0.090734483 | -11.73923465 | 8.02E-32 | 6.66E-30 | 1.065153387 |
| ENSMUSG00000017765 | Slc12a4 | 20498 | protein_coding | 7.312649835 | -1.06520896 | 0.909568736 | 1.171114308 | 0.241552833 | 0.452165438 | 1.06520896 |
| ENSMUSG00000043924 | Ncmap | 230822 | protein_coding | 24.80342236 | 1.066016624 | 0.365371066 | -2.917627374 | 0.003527056 | 0.018218892 | 1.066016624 |
| ENSMUSG00000029700 | Slc13a1 | 55961 | protein_coding | 190.2455883 | -1.066229136 | 0.212174507 | 5.025246212 | 5.03E-07 | 7.42E-06 | 1.066229136 |
| ENSMUSG00000031381 | Piga | 18700 | protein_coding | 606.0125098 | 1.066284886 | 0.084260855 | -12.65456997 | 1.06E-36 | 9.68E-35 | 1.066284886 |
| ENSMUSG00000098879 | Mir7675 | 102465775 | miRNA | 0.580880825 | -1.066476856 | 1.457352667 | 0.731790513 | 0.46429644 | 1 | 1.066476856 |
| ENSMUSG00000085715 | Tsix | 22097 | lncRNA | 1.450534001 | -1.067387222 | 1.277628573 | 0.835444075 | 0.403467722 | 1 | 1.067387222 |
| ENSMUSG00000072294 | Klf12 | 16597 | protein_coding | 193.846867 | 1.067585736 | 0.126102138 | -8.466039906 | 2.54E-17 | 1.10E-15 | 1.067585736 |
| ENSMUSG00000079608 | Stard6 | 170461 | protein_coding | 0.763707544 | 1.068018184 | 1.690746505 | -0.631684396 | 0.527593127 | 1 | 1.068018184 |
| ENSMUSG00000043410 | Hfm1 | 330149 | protein_coding | 4.26068948 | 1.068213355 | 1.132692098 | -0.943074783 | 0.345642653 | 1 | 1.068213355 |
| ENSMUSG00000022754 | Tmem45a | 56277 | protein_coding | 11.02791037 | -1.068391002 | 0.321428409 | 3.323884801 | 0.000887729 | 0.005758032 | 1.068391002 |
| ENSMUSG00000073600 | Prob1 | 381148 | protein_coding | 14.30419953 | 1.068446447 | 0.350574006 | -3.047705841 | 0.002305955 | 0.012714707 | 1.068446447 |
| ENSMUSG00000046159 | Chrm3 | 12671 | protein_coding | 26.27882804 | -1.06863104 | 0.229409747 | 4.658176274 | 3.19E-06 | 3.96E-05 | 1.06863104 |
| ENSMUSG00000039691 | Tspan10 | 208634 | protein_coding | 1.308783227 | 1.068710871 | 2.110823425 | -0.50630046 | 0.61264574 | 1 | 1.068710871 |
| ENSMUSG00000064451 | Snora23 | 100379145 | snoRNA | 27.84922239 | 1.069993549 | 0.461843066 | -2.316790328 | 0.020515156 | 0.074820514 | 1.069993549 |
| ENSMUSG00000111901 | Gm6150 | 620417 | transcribed_processed_pseudogene | 1.754038488 | 1.070091284 | 2.392355395 | -0.447296119 | 0.654661269 | 1 | 1.070091284 |
| ENSMUSG00000065701 | Rny1 | 19872 | misc_RNA | 167.2176973 | 1.070092384 | 0.291839598 | -3.666714152 | 0.000245687 | 0.001900364 | 1.070092384 |
| ENSMUSG00000005716 | Pvalb | 19293 | protein_coding | 1.024180815 | 1.070357869 | 2.504560876 | -0.427363487 | 0.669114598 | 1 | 1.070357869 |
| ENSMUSG00000102408 | 4930568G15Rik | 75857 | lncRNA | 1.066949922 | 1.07138629 | 2.301408052 | -0.465535127 | 0.641548288 | 1 | 1.07138629 |
| ENSMUSG00000032191 | Bcl2l10 | 12049 | protein_coding | 0.720436533 | -1.071651321 | 1.258513628 | 0.851521427 | 0.394479766 | 1 | 1.071651321 |
| ENSMUSG00000027777 | Schip1 | 30953 | protein_coding | 5.23146396 | -1.071693818 | 0.997558245 | 1.074317037 | 0.282680612 | 0.500824763 | 1.071693818 |
| ENSMUSG00000001444 | Tbx21 | 57765 | protein_coding | 8.721004406 | 1.072024528 | 0.748473311 | -1.43228157 | 0.152063255 | 0.332693773 | 1.072024528 |
| ENSMUSG00000030035 | Wbp1 | 22377 | protein_coding | 1160.275275 | 1.073003377 | 0.062792611 | -17.08805166 | 1.82E-65 | 3.42E-63 | 1.073003377 |
| ENSMUSG00000025197 | Cyp2c23 | 226143 | protein_coding | 3.936912414 | -1.073332183 | 0.792085836 | 1.355070542 | 0.175395083 | 1 | 1.073332183 |
| ENSMUSG00000043263 | Ifi209 | 236312 | protein_coding | 2.528926754 | -1.074049797 | 0.791624012 | 1.356767581 | 0.174855071 | 1 | 1.074049797 |
| ENSMUSG00000022389 | Tef | 21685 | protein_coding | 305.5009118 | -1.075105086 | 0.488720849 | 2.199834705 | 0.027818625 | 0.095014843 | 1.075105086 |
| ENSMUSG00000085843 | Kank4os | 399570 | lncRNA | 1.164981074 | -1.075519595 | 2.167961432 | 0.496097199 | 0.619825835 | 1 | 1.075519595 |
| ENSMUSG00000041377 | Ninj2 | 29862 | protein_coding | 5.087118005 | -1.076285516 | 0.538753869 | 1.997731389 | 0.045745789 | 0.139398299 | 1.076285516 |
| ENSMUSG00000042682 | Selenok | 80795 | protein_coding | 943.3814367 | 1.076384718 | 0.083371642 | -12.91068159 | 3.92E-38 | 3.72E-36 | 1.076384718 |
| ENSMUSG00000037762 | Slc16a9 | 66859 | protein_coding | 183.7161167 | -1.076899986 | 0.159604861 | 6.747288122 | 1.51E-11 | 4.13E-10 | 1.076899986 |
| ENSMUSG00000069309 | H2ac22 | 319170 | protein_coding | 56.29840226 | 1.07703827 | 0.19524374 | -5.516377993 | 3.46E-08 | 6.11E-07 | 1.07703827 |
| ENSMUSG00000032740 | Ccdc88a | 108686 | protein_coding | 16.83647864 | 1.07759181 | 0.339492699 | -3.174123661 | 0.001502896 | 0.008911609 | 1.07759181 |
| ENSMUSG00000072789 | Gm10420 | 108169197 | processed_pseudogene | 1.829803266 | 1.077718361 | 0.735826986 | -1.464635548 | 0.143020378 | 1 | 1.077718361 |
| ENSMUSG00000020427 | Igfbp3 | 16009 | protein_coding | 5.043983344 | 1.078757978 | 0.59031044 | -1.827441808 | 0.067633368 | 0.186304828 | 1.078757978 |
| ENSMUSG00000094921 | Vmn2r112 | 628185 | protein_coding | 3.091818679 | -1.078810251 | 2.168597431 | 0.497469118 | 0.618858275 | 1 | 1.078810251 |
| ENSMUSG00000027702 | Lrrc34 | 71827 | protein_coding | 0.927153245 | 1.07886645 | 1.292887498 | -0.834462745 | 0.404020273 | 1 | 1.07886645 |
| ENSMUSG00000079605 | Zbtb9 | 474156 | protein_coding | 0.88446424 | 1.079541446 | 1.512520965 | -0.713736517 | 0.475390121 | 1 | 1.079541446 |

| ensembl_gene_id | external_gene_name | entrezgene_id | gene_biotype | baseMean | log2FoldChange | lfcSE | stat | pvalue | padj | absLog2FC |
|---|---|---|---|---|---|---|---|---|---|---|
| ENSMUSG00000024165 | Jpt2 | 52009 | protein_coding | 1386.424313 | 1.080039887 | 0.109349198 | -9.876980433 | 5.24E-23 | 3.12E-21 | 1.080039887 |
| ENSMUSG00000071001 | Hrct1 | 100039781 | protein_coding | 1.316522158 | -1.081968297 | 1.290478795 | 0.838423925 | 0.401792657 | 1 | 1.081968297 |
| ENSMUSG00000040667 | Nup88 | 19069 | protein_coding | 1070.10689 | 1.082361609 | 0.057126899 | -18.94661927 | 4.71E-80 | 1.23E-77 | 1.082361609 |
| ENSMUSG00000001333 | Sync | 68828 | protein_coding | 8.655730584 | -1.082370216 | 0.367880627 | 2.942177804 | 0.003259127 | 0.017053622 | 1.082370216 |
| ENSMUSG00000051497 | Kcnj16 | 16517 | protein_coding | 1.647151887 | -1.082939246 | 1.434201668 | 0.75508157 | 0.450200041 | 1 | 1.082939246 |
| ENSMUSG00000097740 | E030044B06Rik | 381223 | lncRNA | 1.571784178 | 1.083040003 | 1.916099849 | -0.565231506 | 0.57191631 | 1 | 1.083040003 |
| ENSMUSG00000037990 | Sh3rf3 | 237353 | protein_coding | 3.321923312 | 1.08309574 | 2.177186821 | -0.497474874 | 0.618854217 | 1 | 1.08309574 |
| ENSMUSG00000102080 | Gm5092 | 328766 | lncRNA | 1.03432649 | 1.083734455 | 1.467824265 | -0.738327115 | 0.460315693 | 1 | 1.083734455 |
| ENSMUSG00000078137 | Ankrd63 | 383787 | protein_coding | 1.787174797 | -1.083809335 | 1.167464374 | 0.928344675 | 0.353228807 | 1 | 1.083809335 |
| ENSMUSG00000075302 | Erich2 | 66748 | protein_coding | 2.265685584 | 1.083888953 | 1.066099304 | -1.016686671 | 0.309302509 | 1 | 1.083888953 |
| ENSMUSG00000031274 | Col4a5 | 12830 | protein_coding | 2.845161868 | 1.084335922 | 1.151583115 | -0.941604569 | 0.34639513 | 1 | 1.084335922 |
| ENSMUSG00000086008 | Gm8817 | 667794 | lncRNA | 1.076001287 | 1.084478118 | 1.660787712 | -0.652990211 | 0.51376259 | 1 | 1.084478118 |
| ENSMUSG00000085794 | Vax2os | 574519 | lncRNA | 1.350690637 | 1.084556347 | 2.516547058 | -0.430970024 | 0.666490166 | 1 | 1.084556347 |
| ENSMUSG00000068037 | Mas1 | 17171 | protein_coding | 3.083250005 | -1.084883191 | 0.841413386 | 1.289358131 | 0.19727361 | 1 | 1.084883191 |
| ENSMUSG00000044364 | Tmem74b | 108832 | protein_coding | 2.298938206 | -1.086741543 | 0.697979748 | 1.556981482 | 0.119474882 | 1 | 1.086741543 |
| ENSMUSG00000076460 | Mir744 | 791070 | miRNA | 0.86372534 | -1.086848862 | 1.261293316 | 0.861693984 | 0.388855937 | 1 | 1.086848862 |
| ENSMUSG00000021880 | Rnase6 | 78416 | protein_coding | 0.863394161 | 1.086885289 | 1.94055939 | -0.56008865 | 0.575418972 | 1 | 1.086885289 |
| ENSMUSG00000075119 | Or4a27 | 433449 | protein_coding | 0.983882487 | 1.087327096 | 1.89038779 | -0.575187325 | 0.565164614 | 1 | 1.087327096 |
| ENSMUSG00000085982 | 9530051G07Rik | 319781 | lncRNA | 1.496083801 | -1.087849259 | 1.602061389 | 0.679030945 | 0.497118255 | 1 | 1.087849259 |
| ENSMUSG00000030283 | St8sia1 | 20449 | protein_coding | 5.160526307 | 1.088038813 | 0.944681744 | -1.151751708 | 0.249423117 | 0.461664802 | 1.088038813 |
| ENSMUSG00000020717 | Pecam1 | 18613 | protein_coding | 24.33524162 | -1.088047264 | 0.390086803 | 2.789243973 | 0.005283125 | 0.025510388 | 1.088047264 |
| ENSMUSG00000076441 | Ass1 | 11898 | protein_coding | 138.3492925 | -1.088070062 | 0.239295919 | 4.546964554 | 5.44E-06 | 6.46E-05 | 1.088070062 |
| ENSMUSG00000022512 | Cldn1 | 12737 | protein_coding | 1.391492363 | 1.088268146 | 1.901652592 | -0.572274952 | 0.567135716 | 1 | 1.088268146 |
| ENSMUSG00000073405 | H2-T-ps | 667803 | unprocessed_pseudogene | 23.57573572 | 1.088520406 | 0.340867726 | -3.193380663 | 0.001406174 | 0.008459586 | 1.088520406 |
| ENSMUSG00000068855 | H2ac20 | 319176 | protein_coding | 348.8333784 | 1.088888715 | 0.140865171 | -7.730006694 | 1.08E-14 | 3.83E-13 | 1.088888715 |
| ENSMUSG00000086150 | Bach2os | 414068 | lncRNA | 1.143012776 | 1.089512293 | 1.096053147 | -0.994032356 | 0.320207115 | 1 | 1.089512293 |
| ENSMUSG00000022449 | Adamts20 | 223838 | protein_coding | 3.511654743 | 1.089896321 | 3.461736258 | -0.314840947 | 0.752882421 | 1 | 1.089896321 |
| ENSMUSG00000042254 | Cilp | 214425 | protein_coding | 1.765316426 | -1.090286275 | 2.038046129 | 0.534966436 | 0.592673067 | 1 | 1.090286275 |
| ENSMUSG00000032110 | Acrv1 | 11451 | protein_coding | 1.030702236 | 1.090631609 | 1.930617937 | -0.564913227 | 0.572132787 | 1 | 1.090631609 |
| ENSMUSG00000062200 | Vmn2r7 | 319217 | protein_coding | 2.508187437 | 1.090796954 | 2.361424178 | -0.461923344 | 0.644136293 | 1 | 1.090796954 |
| ENSMUSG00000040693 | Slco4c1 | 227394 | protein_coding | 2.005149577 | -1.091536828 | 1.041727667 | 1.047813995 | 0.294724313 | 1 | 1.091536828 |
| ENSMUSG00000004231 | Pax2 | 18504 | protein_coding | 3.27269671 | 1.091797027 | 2.45167396 | -0.44532717 | 0.656083336 | 1 | 1.091797027 |
| ENSMUSG00000032878 | Ccdc85a | 216613 | protein_coding | 1.497429698 | -1.091858734 | 1.773136073 | 0.61577831 | 0.538040844 | 1 | 1.091858734 |
| ENSMUSG00000097362 | Gm26544 | 102633872 | lncRNA | 1.766263381 | 1.091902825 | 2.704666706 | -0.403710676 | 0.686425488 | 1 | 1.091902825 |
| ENSMUSG00000058057 | Mettl7a3 | 668178 | protein_coding | 7.427656575 | -1.091920857 | 0.412453273 | 2.647380752 | 0.008111796 | 0.036011679 | 1.091920857 |
| ENSMUSG00000030214 | Plbd1 | 66857 | protein_coding | 554.7303668 | -1.092954827 | 0.205267828 | 5.324530588 | 1.01E-07 | 1.67E-06 | 1.092954827 |
| ENSMUSG00000037972 | Snn | 20621 | protein_coding | 22.11014123 | 1.093110381 | 0.225834366 | -4.840319037 | 1.30E-06 | 1.75E-05 | 1.093110381 |
| ENSMUSG00000042870 | Tom1 | 21968 | protein_coding | 105.6242584 | -1.09325751 | 0.138391108 | 7.899766995 | 2.79E-15 | 1.04E-13 | 1.09325751 |
| ENSMUSG00000020160 | Meis1 | 17268 | protein_coding | 2.373586282 | 1.093330825 | 1.357405671 | -0.805456209 | 0.420556467 | 1 | 1.093330825 |

| ensembl_gene_id | external_gene_name | entrezgene_id | gene_biotype | baseMean | log2FoldChange | lfcSE | stat | pvalue | padj | absLog2FC |
|---|---|---|---|---|---|---|---|---|---|---|
| ENSMUSG00000074682 | Zcchc3 | 67917 | protein_coding | 45.07086244 | 1.093491132 | 0.28339024 | -3.858605474 | 0.000114036 | 0.000967723 | 1.093491132 |
| ENSMUSG00000024083 | Pja2 | 224938 | protein_coding | 1134.847888 | 1.094724871 | 0.0868125 | -12.61022163 | 1.85E-36 | 1.69E-34 | 1.094724871 |
| ENSMUSG00000033219 | Gm9758 | 381714 | protein_coding | 1.578732301 | 1.094934712 | 1.986381341 | -0.5512208 | 0.581482323 | 1 | 1.094934712 |
| ENSMUSG00000025212 | Sfxn3 | 94280 | protein_coding | 135.73936 | -1.095297615 | 0.175862648 | 6.228142397 | 4.72E-10 | 1.08E-08 | 1.095297615 |
| ENSMUSG00000040726 | Hesx1 | 15209 | protein_coding | 5.647284974 | 1.095944839 | 0.811362224 | -1.35074669 | 0.17677659 | 0.368923116 | 1.095944839 |
| ENSMUSG00000032850 | Rnft2 | 269695 | protein_coding | 11.75205608 | 1.095945607 | 0.39436984 | -2.778979259 | 0.005453001 | 0.026177574 | 1.095945607 |
| ENSMUSG00000032648 | Pygm | 19309 | protein_coding | 13.19698902 | 1.09596419 | 0.387304508 | -2.829722265 | 0.004658843 | 0.022923163 | 1.09596419 |
| ENSMUSG00000062624 | Cyp2c67 | 545288 | protein_coding | 2.556855024 | 1.096178691 | 0.75962035 | -1.443061248 | 0.149003219 | 1 | 1.096178691 |
| ENSMUSG00000068289 | Cma2 | 545055 | protein_coding | 1.000822077 | -1.097806656 | 1.899305914 | 0.578004127 | 0.563261333 | 1 | 1.097806656 |
| ENSMUSG00000031302 | Nlgn3 | 245537 | protein_coding | 2.229201866 | 1.098322756 | 1.805124334 | -0.60844715 | 0.542890948 | 1 | 1.098322756 |
| ENSMUSG00000087549 | Gm13427 | 108168742 | lncRNA | 2.23138122 | 1.098950187 | 0.935536637 | -1.174673598 | 0.240125326 | 1 | 1.098950187 |
| ENSMUSG00000030934 | Oat | 18242 | protein_coding | 4309.075333 | 1.099014182 | 0.098589414 | -11.14738525 | 7.37E-29 | 5.49E-27 | 1.099014182 |
| ENSMUSG00000064156 | Prol1 | 17830 | protein_coding | 1.798257674 | 1.099242836 | 2.395810775 | -0.458818721 | 0.64636435 | 1 | 1.099242836 |
| ENSMUSG00000022229 | Atp12a | 192113 | protein_coding | 1789.486123 | -1.099257927 | 0.331459597 | 3.316416049 | 0.0009118 | 0.005884119 | 1.099257927 |
| ENSMUSG00000091239 | Vmn2r76 | 675969 | protein_coding | 4.652423004 | 1.099472178 | 1.803185891 | -0.609738676 | 0.54203493 | 0.735735661 | 1.099472178 |
| ENSMUSG00000104835 | Gm5547 | 433637 | lncRNA | 0.590415846 | 1.099586072 | 1.845313804 | -0.595880261 | 0.55125522 | 1 | 1.099586072 |
| ENSMUSG00000025425 | St8sia5 | 225742 | protein_coding | 42.71196 | -1.09961008 | 0.473760079 | 2.321027307 | 0.02028537 | 0.074195482 | 1.09961008 |
| ENSMUSG00000109819 | B930018H19Rik | 330734 | lncRNA | 0.757806583 | -1.099667559 | 1.329079986 | 0.827390053 | 0.408016017 | 1 | 1.099667559 |
| ENSMUSG00000020897 | Aurkb | 20877 | protein_coding | 325.5785265 | 1.100891597 | 0.218815931 | -5.031130927 | 4.88E-07 | 7.22E-06 | 1.100891597 |
| ENSMUSG00000070719 | Pla2g4d | 78390 | protein_coding | 3.273305171 | -1.101214934 | 0.957544548 | 1.15004042 | 0.250127224 | 1 | 1.101214934 |
| ENSMUSG00000042895 | Abra | 223513 | protein_coding | 1.462069542 | 1.101451601 | 1.053875984 | -1.045143468 | 0.295956667 | 1 | 1.101451601 |
| ENSMUSG00000068417 | Pnp2 | 667034 | protein_coding | 166.5160982 | -1.102290385 | 0.156526229 | 7.042208809 | 1.89E-12 | 5.60E-11 | 1.102290385 |
| ENSMUSG00000025038 | Efhc2 | 74405 | protein_coding | 1.227769599 | 1.102592126 | 2.287812084 | -0.481941735 | 0.629847337 | 1 | 1.102592126 |
| ENSMUSG00000028730 | Cfap57 | 68625 | protein_coding | 1.812722053 | 1.102986402 | 1.506901692 | -0.731956443 | 0.464195154 | 1 | 1.102986402 |
| ENSMUSG00000045498 | Pcdhb3 | 93874 | protein_coding | 1.234049734 | 1.103005211 | 1.318983616 | -0.836253914 | 0.403012074 | 1 | 1.103005211 |
| ENSMUSG00000114442 | F630042J09Rik | 100038627 | lncRNA | 6.005072814 | -1.103014102 | 0.524454048 | 2.103166344 | 0.035451231 | 0.114635135 | 1.103014102 |
| ENSMUSG00000038248 | Sobp | 109205 | protein_coding | 19.17780741 | -1.103276618 | 0.432760226 | 2.54939468 | 0.01079101 | 0.045254367 | 1.103276618 |
| ENSMUSG00000039954 | Stk32a | 269019 | protein_coding | 17.43231905 | -1.103318249 | 0.428534484 | 2.57463119 | 0.010034702 | 0.04269855 | 1.103318249 |
| ENSMUSG00000053303 | Slc22a26 | 236149 | protein_coding | 2.12859128 | 1.103702916 | 1.253853195 | -0.880248916 | 0.378724479 | 1 | 1.103702916 |
| ENSMUSG00000069733 | Ube2u | 381534 | protein_coding | 1.880984458 | 1.103988387 | 1.279173503 | -0.863048198 | 0.388110967 | 1 | 1.103988387 |
| ENSMUSG00000021730 | Hcn1 | 15165 | protein_coding | 4.924871364 | 1.10458982 | 1.422143119 | -0.776707917 | 0.437331115 | 0.655599676 | 1.10458982 |
| ENSMUSG00000026791 | Slc2a8 | 56017 | protein_coding | 209.3253654 | 1.104816081 | 0.097400398 | -11.34303456 | 8.03E-30 | 6.23E-28 | 1.104816081 |
| ENSMUSG00000084781 | D930015M05Rik | 320507 | lncRNA | 8.377485496 | -1.105131125 | 0.473173764 | 2.335571431 | 0.019513589 | 0.071994506 | 1.105131125 |
| ENSMUSG00000032487 | Ptgs2 | 19225 | protein_coding | 1.875862736 | 1.105462752 | 1.017026267 | -1.08695595 | 0.277056278 | 1 | 1.105462752 |
| ENSMUSG00000030747 | Dgat2 | 67800 | protein_coding | 664.0565304 | 1.105499176 | 0.17763257 | -6.223516179 | 4.86E-10 | 1.11E-08 | 1.105499176 |
| ENSMUSG00000047911 | Npm2 | 328440 | protein_coding | 1.223955447 | -1.105558784 | 1.158238624 | 0.954517283 | 0.339821865 | 1 | 1.105558784 |
| ENSMUSG00000047025 | Ccer1 | 66716 | protein_coding | 1.173374432 | 1.106208618 | 2.154131228 | -0.51352889 | 0.607581407 | 1 | 1.106208618 |
| ENSMUSG00000031444 | F10 | 14058 | protein_coding | 2.346386647 | 1.106337538 | 0.905388564 | -1.221947771 | 0.221727379 | 1 | 1.106337538 |
| ENSMUSG00000037196 | Pacrg | 69310 | protein_coding | 1.286609154 | -1.107267657 | 1.010760982 | 1.095479225 | 0.273306742 | 1 | 1.107267657 |

Supplementary Table 1

| ensembl_gene_id | external_gene_name | entrezgene_id | gene_biotype | baseMean | log2FoldChange | lfcSE | stat | pvalue | padj | absLog2FC |
|---|---|---|---|---|---|---|---|---|---|---|
| ENSMUSG00000016349 | Eef1a2 | 13628 | protein_coding | 1.221093564 | 1.107311508 | 2.172561353 | -0.509680201 | 0.610275526 | 1 | 1.107311508 |
| ENSMUSG00000020401 | Garin3 | 432552 | protein_coding | 1.282139927 | 1.108383589 | 2.190870782 | -0.505910069 | 0.612919783 | 1 | 1.108383589 |
| ENSMUSG00000057461 | Or52s1b | 258139 | protein_coding | 1.076326485 | 1.108501925 | 2.192897972 | -0.505496352 | 0.61321026 | 1 | 1.108501925 |
| ENSMUSG00000057195 | Ceacam13 | 69785 | protein_coding | 1.028645879 | 1.108765806 | 1.08155067 | -1.025163071 | 0.30528625 | 1 | 1.108765806 |
| ENSMUSG00000065642 | Snora69 | 104369 | snoRNA | 1.656353426 | 1.108873608 | 0.923233292 | -1.20107628 | 0.229721613 | 1 | 1.108873608 |
| ENSMUSG00000079679 | Vwde | 232585 | protein_coding | 1.387876632 | 1.108894451 | 2.607784818 | -0.425224675 | 0.670672898 | 1 | 1.108894451 |
| ENSMUSG00000043119 | Or2a5 | 258270 | protein_coding | 1.224208379 | 1.108956321 | 2.200807945 | -0.503886004 | 0.614341486 | 1 | 1.108956321 |
| ENSMUSG00000046993 | Gm5637 | 434782 | processed_pseudogene | 7.16589828 | 1.109095769 | 0.487073982 | -2.277058127 | 0.022782755 | 0.081330396 | 1.109095769 |
| ENSMUSG00000058831 | Opn1sw | 12057 | protein_coding | 5.692580221 | 1.109177683 | 0.505573073 | -2.193901817 | 0.028242469 | 0.096124511 | 1.109177683 |
| ENSMUSG00000045136 | Tubb2b | 73710 | protein_coding | 7.522600076 | -1.109314908 | 0.474935215 | 2.335718372 | 0.019505924 | 0.071982309 | 1.109314908 |
| ENSMUSG00000032372 | Plscr2 | 18828 | protein_coding | 18.38289137 | -1.109332403 | 0.364903649 | 3.040069362 | 0.002365237 | 0.012989531 | 1.109332403 |
| ENSMUSG00000025946 | Pth2r | 213527 | protein_coding | 1.174824666 | 1.109359464 | 2.20788559 | -0.502453329 | 0.615348674 | 1 | 1.109359464 |
| ENSMUSG00000067242 | Lgi1 | 56839 | protein_coding | 4.473153138 | 1.109477055 | 1.546780557 | -0.717281485 | 0.473200429 | 0.685335967 | 1.109477055 |
| ENSMUSG00000074852 | Hpse2 | 545291 | protein_coding | 1.357211614 | 1.109713154 | 2.214151593 | -0.501191137 | 0.616236611 | 1 | 1.109713154 |
| ENSMUSG00000004462 | Tbccd1 | 70573 | protein_coding | 606.0550707 | 1.109836739 | 0.108505846 | -10.22835891 | 1.48E-24 | 9.33E-23 | 1.109836739 |
| ENSMUSG00000053797 | Krt16 | 16666 | protein_coding | 1.033283113 | 1.110058955 | 2.220281289 | -0.499963208 | 0.617100984 | 1 | 1.110058955 |
| ENSMUSG00000079698 | Vmn2r93 | 627132 | protein_coding | 1.765711771 | 1.110242233 | 2.2236229 | -0.499294297 | 0.617572072 | 1 | 1.110242233 |
| ENSMUSG00000029636 | Wasf3 | 245880 | protein_coding | 6.001062622 | 1.110273449 | 1.333020795 | -0.832900322 | 0.404900948 | 0.626138614 | 1.110273449 |
| ENSMUSG00000048978 | Nrsn1 | 22360 | protein_coding | 1.978546535 | -1.110320857 | 1.160367651 | 0.956869881 | 0.338632935 | 1 | 1.110320857 |
| ENSMUSG00000042109 | Csdc2 | 105859 | protein_coding | 4.082365645 | 1.11045444 | 0.847371477 | -1.310469458 | 0.19003707 | 1 | 1.11045444 |
| ENSMUSG00000075113 | Or4c29 | 258453 | protein_coding | 1.46064369 | 1.110667627 | 2.231270869 | -0.497773552 | 0.618643659 | 1 | 1.110667627 |
| ENSMUSG00000062037 | Or8s16 | 257943 | protein_coding | 1.793123252 | 1.110748032 | 2.232754519 | -0.497478797 | 0.618851451 | 1 | 1.110748032 |
| ENSMUSG00000021362 | Gcm2 | 107889 | protein_coding | 1.769098367 | 1.110818634 | 2.234031643 | -0.497226007 | 0.619029683 | 1 | 1.110818634 |
| ENSMUSG00000000247 | Lhx2 | 16870 | protein_coding | 1.425867308 | 1.110950746 | 2.236410458 | -0.496756193 | 0.61936099 | 1 | 1.110950746 |
| ENSMUSG00000089871 | Speer4cos | 75858 | lncRNA | 1.594580859 | 1.110965784 | 2.236701123 | -0.496698362 | 0.619401778 | 1 | 1.110965784 |
| ENSMUSG00000098120 | Gm5914 | 546100 | lncRNA | 1.277599038 | -1.1112734 | 1.121508246 | 0.990874034 | 0.321747093 | 1 | 1.1112734 |
| ENSMUSG00000061937 | Csn1s2a | 12993 | protein_coding | 1.194521301 | 1.111548892 | 2.24735353 | -0.494603486 | 0.620880041 | 1 | 1.111548892 |
| ENSMUSG00000026692 | Fmo4 | 226564 | protein_coding | 26.36301374 | -1.111644376 | 0.250400609 | 4.439463556 | 9.02E-06 | 0.000102427 | 1.111644376 |
| ENSMUSG00000023176 | Cpn2 | 71756 | protein_coding | 55.73556121 | 1.111705476 | 0.410637209 | -2.70726922 | 0.006783922 | 0.031110715 | 1.111705476 |
| ENSMUSG00000087223 | 4930442L01Rik | 67583 | lncRNA | 1.290661621 | 1.111705803 | 2.250265679 | -0.494033133 | 0.62128278 | 1 | 1.111705803 |
| ENSMUSG00000087200 | Lrp8os3 | 105244644 | lncRNA | 1.353776399 | 1.112036257 | 1.299661412 | -0.855635357 | 0.392199506 | 1 | 1.112036257 |
| ENSMUSG00000050628 | Ubald2 | 319370 | protein_coding | 824.1410093 | 1.112699959 | 0.173674248 | -6.406821822 | 1.49E-10 | 3.61E-09 | 1.112699959 |
| ENSMUSG00000025044 | Msr1 | 20288 | protein_coding | 5.025618693 | -1.112999253 | 0.61233034 | 1.817645117 | 0.069118391 | 0.189251307 | 1.112999253 |
| ENSMUSG00000028631 | Kcnq4 | 60613 | protein_coding | 6.240579701 | -1.113046098 | 0.69005633 | 1.612978606 | 0.106749156 | 0.259548816 | 1.113046098 |
| ENSMUSG00000022212 | Cpne6 | 12891 | protein_coding | 2.71543377 | 1.113268318 | 2.279783307 | -0.488321989 | 0.625321789 | 1 | 1.113268318 |
| ENSMUSG00000032511 | Scn5a | 20271 | protein_coding | 3.789095886 | 1.113268785 | 1.658414297 | -0.671285086 | 0.502038933 | 1 | 1.113268785 |
| ENSMUSG00000028644 | Ermap | 27028 | protein_coding | 1.903807022 | 1.114474874 | 1.348456688 | -0.826481773 | 0.408530853 | 1 | 1.114474874 |
| ENSMUSG00000002847 | Pla1a | 85031 | protein_coding | 1.214363268 | 1.115108744 | 1.762414466 | -0.632716518 | 0.526918781 | 1 | 1.115108744 |
| ENSMUSG00000083282 | Ctsf | 56464 | protein_coding | 72.27810975 | -1.115800778 | 0.196364036 | 5.682307229 | 1.33E-08 | 2.51E-07 | 1.115800778 |

| ensembl_gene_id | external_gene_name | entrezgene_id | gene_biotype | baseMean | log2FoldChange | lfcSE | stat | pvalue | padj | absLog2FC |
|---|---|---|---|---|---|---|---|---|---|---|
| ENSMUSG00000000902 | Smarcb1 | 20587 | protein_coding | 948.1252589 | 1.115809849 | 0.070511104 | -15.82459774 | 2.11E-56 | 3.28E-54 | 1.115809849 |
| ENSMUSG00000088037 | Gm24992 | 115485640 | snoRNA | 0.648178139 | -1.115995594 | 1.461691043 | 0.763496226 | 0.445167512 | 1 | 1.115995594 |
| ENSMUSG00000069919 | Hba-a1 | 15122 | protein_coding | 56.68320283 | -1.11603886 | 0.516725044 | 2.159831177 | 0.030785741 | 0.102914115 | 1.11603886 |
| ENSMUSG00000046881 | Or1ab2 | 258335 | protein_coding | 3.380569712 | -1.116369007 | 1.628676674 | 0.685445445 | 0.493062876 | 1 | 1.116369007 |
| ENSMUSG00000044199 | S1pr4 | 13611 | protein_coding | 3.48921088 | -1.116589841 | 0.822508343 | 1.357542267 | 0.174608973 | 1 | 1.116589841 |
| ENSMUSG00000015001 | Oc90 | 18256 | protein_coding | 1.592416588 | 1.11660654 | 1.609447326 | -0.693782593 | 0.48781856 | 1 | 1.11660654 |
| ENSMUSG00000056973 | Ces1d | 104158 | protein_coding | 413.5835353 | -1.116794538 | 0.257078497 | 4.344177179 | 1.40E-05 | 0.000150204 | 1.116794538 |
| ENSMUSG00000037797 | Adh4 | 26876 | protein_coding | 1.372866811 | -1.117511579 | 1.445074751 | 0.77332441 | 0.439330413 | 1 | 1.117511579 |
| ENSMUSG00000043168 | 4930426D05Rik | 74644 | lncRNA | 2.021611427 | -1.117854557 | 1.70866331 | 0.654227518 | 0.512965235 | 1 | 1.117854557 |
| ENSMUSG00000053181 | A830005F24Rik | 442803 | lncRNA | 3.012495249 | 1.118184821 | 0.685664426 | -1.630804777 | 0.102931519 | 1 | 1.118184821 |
| ENSMUSG00000055748 | Gsdmc4 | 74548 | protein_coding | 8462.866959 | -1.118235936 | 0.388412116 | 2.878993447 | 0.003989466 | 0.020128948 | 1.118235936 |
| ENSMUSG00000030302 | Atp2b2 | 11941 | protein_coding | 5.766618605 | 1.11847907 | 2.690531445 | -0.415709347 | 0.677622701 | 0.827306109 | 1.11847907 |
| ENSMUSG00000103144 | Pcdhga1 | 93709 | protein_coding | 1.996710564 | -1.118800445 | 0.880492224 | 1.270653407 | 0.203851978 | 1 | 1.118800445 |
| ENSMUSG00000029371 | Cxcl5 | 20311 | protein_coding | 8.668298889 | 1.118980437 | 0.544954313 | -2.053347243 | 0.040038912 | 0.125967345 | 1.118980437 |
| ENSMUSG00000007682 | Dio2 | 13371 | protein_coding | 3.752759342 | 1.119132346 | 0.738773845 | -1.514851065 | 0.129810125 | 1 | 1.119132346 |
| ENSMUSG00000027397 | Slc20a1 | 20515 | protein_coding | 5439.840287 | 1.119804598 | 0.201770312 | -5.549897727 | 2.86E-08 | 5.11E-07 | 1.119804598 |
| ENSMUSG00000029712 | Actl6b | 83766 | protein_coding | 3.207834568 | 1.120034938 | 1.161365503 | -0.964412095 | 0.334839361 | 1 | 1.120034938 |
| ENSMUSG00000026829 | Gbgt1 | 227671 | protein_coding | 5.894197939 | -1.120366474 | 0.54658845 | 2.049744143 | 0.040389405 | 0.126876666 | 1.120366474 |
| ENSMUSG00000034910 | Pygo1 | 72135 | protein_coding | 2.18091613 | -1.120501233 | 1.720030007 | 0.65144284 | 0.514760664 | 1 | 1.120501233 |
| ENSMUSG00000056656 | Apol8 | 239552 | protein_coding | 3.550221738 | 1.121016114 | 0.949257065 | -1.180940501 | 0.237626361 | 1 | 1.121016114 |
| ENSMUSG00000089686 | Mnd1-ps | 100042415 | transcribed_processed_pseudogene | 0.925155222 | 1.121896485 | 1.214838174 | -0.923494593 | 0.355749517 | 1 | 1.121896485 |
| ENSMUSG00000111997 | Gm5176 | 382421 | transcribed_processed_pseudogene | 0.962253979 | 1.122930086 | 1.175720294 | -0.955099688 | 0.339527287 | 1 | 1.122930086 |
| ENSMUSG00000040569 | Slc26a7 | 208890 | protein_coding | 2.100192767 | 1.124207517 | 2.518957406 | -0.44629874 | 0.655381466 | 1 | 1.124207517 |
| ENSMUSG00000116560 | Gm2808 | 100040500 | unprocessed_pseudogene | 1.452123949 | -1.124255208 | 1.134936224 | 0.990588885 | 0.321886367 | 1 | 1.124255208 |
| ENSMUSG00000049797 | Or51a10 | 258326 | protein_coding | 1.35722418 | 1.124319204 | 2.521673309 | -0.445862356 | 0.655696675 | 1 | 1.124319204 |
| ENSMUSG00000074591 | Ankub1 | 242037 | protein_coding | 1.333816541 | 1.124473076 | 2.525526995 | -0.445242945 | 0.656144195 | 1 | 1.124473076 |
| ENSMUSG00000060600 | Eno3 | 13808 | protein_coding | 1001.010361 | -1.12447332 | 0.263288333 | 4.270881682 | 1.95E-05 | 0.000200831 | 1.12447332 |
| ENSMUSG00000015721 | Nlrp5 | 23968 | protein_coding | 1.472370917 | 1.124480316 | 2.525729776 | -0.445210064 | 0.656167954 | 1 | 1.124480316 |
| ENSMUSG00000026204 | Ptprn | 19275 | protein_coding | 21.85626068 | -1.124548324 | 0.418470907 | 2.687279581 | 0.007203663 | 0.032709605 | 1.124548324 |
| ENSMUSG00000043999 | Gpr75 | 237716 | protein_coding | 1.491024004 | -1.12463216 | 1.617161703 | 0.69543581 | 0.486782225 | 1 | 1.12463216 |
| ENSMUSG00000075394 | Hoxc4 | 15423 | protein_coding | 1.267433415 | 1.124943979 | 2.537463388 | -0.443334073 | 0.657524114 | 1 | 1.124943979 |
| ENSMUSG00000097960 | A330074K22Rik | 434353 | lncRNA | 1.594738873 | 1.124985255 | 2.538568284 | -0.443157374 | 0.657651908 | 1 | 1.124985255 |
| ENSMUSG00000025175 | Fn3k | 63828 | protein_coding | 4.828758837 | -1.12502843 | 0.560375258 | 2.007634017 | 0.044682197 | 0.136786828 | 1.12502843 |
| ENSMUSG00000072188 | Gm10354 | 100503549 | protein_coding | 1.264021018 | 1.125208798 | 2.544253567 | -0.442254975 | 0.658304708 | 1 | 1.125208798 |
| ENSMUSG00000029658 | Wdr95 | 381693 | protein_coding | 2.67192331 | 1.125535048 | 2.552802882 | -0.440901668 | 0.659284186 | 1 | 1.125535048 |
| ENSMUSG00000057050 | Or1e1f | 258343 | protein_coding | 1.636602079 | 1.125714211 | 2.557380887 | -0.44018246 | 0.659804962 | 1 | 1.125714211 |
| ENSMUSG00000091722 | Siah3 | 380918 | protein_coding | 1.658609523 | 1.125866932 | 2.561358623 | -0.439558491 | 0.66025691 | 1 | 1.125866932 |
| ENSMUSG00000075502 | Kbtbd6 | 432879 | protein_coding | 8.364918238 | 1.125895724 | 0.474170921 | -2.374451226 | 0.017575058 | 0.066324156 | 1.125895724 |
| ENSMUSG00000112071 | 5430401H09Rik | 100504461 | lncRNA | 1.289766142 | 1.125912001 | 2.562483493 | -0.439383124 | 0.660383953 | 1 | 1.125912001 |

Supplementary Table 1

| ensembl_gene_id | external_gene_name | entrezgene_id | gene_biotype | baseMean | log2FoldChange | lfcSE | stat | pvalue | padj | absLog2FC |
|---|---|---|---|---|---|---|---|---|---|---|
| ENSMUSG00000041986 | Elmod1 | 270162 | protein_coding | 1.290507006 | 1.125916043 | 2.562589518 | -0.439366521 | 0.660395981 | 1 | 1.125916043 |
| ENSMUSG00000024837 | Dmrt1 | 50796 | protein_coding | 1.429061382 | 1.125952056 | 2.563556721 | -0.439214801 | 0.660505901 | 1 | 1.125952056 |
| ENSMUSG00000056564 | Or2ab1 | 257892 | protein_coding | 1.471753671 | 1.125963198 | 2.563856481 | -0.439167795 | 0.660539958 | 1 | 1.125963198 |
| ENSMUSG00000044903 | Psg22 | 243862 | protein_coding | 1.198772378 | 1.125963653 | 2.563819501 | -0.439174307 | 0.66053524 | 1 | 1.125963653 |
| ENSMUSG00000027355 | Tmco5 | 67356 | protein_coding | 1.472494534 | 1.125966105 | 2.56393287 | -0.439155845 | 0.660548617 | 1 | 1.125966105 |
| ENSMUSG00000049491 | Slc36a3 | 215332 | protein_coding | 1.433389767 | 1.125970652 | 2.564045324 | -0.439138358 | 0.660561287 | 1 | 1.125970652 |
| ENSMUSG00000022061 | Nkx3-1 | 18095 | protein_coding | 1.151952629 | 1.126019775 | 2.565284293 | -0.438945413 | 0.66070109 | 1 | 1.126019775 |
| ENSMUSG00000051414 | Or7g17 | 259070 | protein_coding | 1.151952629 | 1.126019775 | 2.565284293 | -0.438945413 | 0.66070109 | 1 | 1.126019775 |
| ENSMUSG00000000263 | Glra1 | 14654 | protein_coding | 1.162076331 | 1.126067804 | 2.566561146 | -0.438745753 | 0.660845771 | 1 | 1.126067804 |
| ENSMUSG00000026582 | Sele | 20339 | protein_coding | 1.323827916 | 1.126089588 | 2.567153885 | -0.438652936 | 0.660913034 | 1 | 1.126089588 |
| ENSMUSG00000109561 | Ankrd31 | 625662 | protein_coding | 1.323827916 | 1.126089588 | 2.567153885 | -0.438652936 | 0.660913034 | 1 | 1.126089588 |
| ENSMUSG00000033044 | Dhrs7c | 68460 | protein_coding | 1.338067618 | 1.126107492 | 2.567627129 | -0.43857906 | 0.660966573 | 1 | 1.126107492 |
| ENSMUSG00000049041 | Or1r1 | 258705 | protein_coding | 1.338067618 | 1.126107492 | 2.567627129 | -0.43857906 | 0.660966573 | 1 | 1.126107492 |
| ENSMUSG00000109097 | Gm29683 | 624549 | lncRNA | 1.242946394 | 1.126149062 | 2.568707064 | -0.438410856 | 0.661088479 | 1 | 1.126149062 |
| ENSMUSG00000047085 | Lrrc4b | 272381 | protein_coding | 1.790004438 | 1.12616829 | 2.569298481 | -0.438317423 | 0.661156197 | 1 | 1.12616829 |
| ENSMUSG00000066197 | Gpr139 | 209776 | protein_coding | 2.260387641 | 1.126174692 | 2.56950845 | -0.438284097 | 0.661180352 | 1 | 1.126174692 |
| ENSMUSG00000096703 | Or4k40 | 277562 | protein_coding | 1.197907898 | 1.126183031 | 2.569591719 | -0.43827314 | 0.661188295 | 1 | 1.126183031 |
| ENSMUSG00000041912 | Tdrkh | 72634 | protein_coding | 7.797796152 | 1.126280298 | 0.50121648 | -2.247093506 | 0.024634052 | 0.086352522 | 1.126280298 |
| ENSMUSG00000029379 | Cxcl3 | 330122 | protein_coding | 1.108858236 | 1.126282296 | 2.595094062 | -0.434004421 | 0.664285231 | 1 | 1.126282296 |
| ENSMUSG00000054690 | Emcn | 59308 | protein_coding | 1.615176371 | 1.126343937 | 2.573918549 | -0.437598904 | 0.661677066 | 1 | 1.126343937 |
| ENSMUSG00000115701 | Vmn1r6 | 171193 | protein_coding | 1.753730747 | 1.126419763 | 2.575949007 | -0.437283409 | 0.661905826 | 1 | 1.126419763 |
| ENSMUSG00000059206 | Vmn1r71 | 252910 | protein_coding | 1.24633299 | 1.126460431 | 2.576952714 | -0.437128871 | 0.66201789 | 1 | 1.126460431 |
| ENSMUSG00000105923 | A830019L24Rik | 320072 | lncRNA | 1.24633299 | 1.126460431 | 2.576952714 | -0.437128871 | 0.66201789 | 1 | 1.126460431 |
| ENSMUSG00000041911 | Dlx1 | 13390 | protein_coding | 1.330908157 | 1.126575062 | 2.580031019 | -0.436651751 | 0.662363926 | 1 | 1.126575062 |
| ENSMUSG00000054134 | Umodl1 | 52020 | protein_coding | 1.706495419 | 1.126682841 | 2.582966568 | -0.436197222 | 0.662693643 | 1 | 1.126682841 |
| ENSMUSG00000073722 | 4931408C20Rik | 210940 | protein_coding | 1.939845736 | 1.127113663 | 2.594593054 | -0.434408649 | 0.663991718 | 1 | 1.127113663 |
| ENSMUSG00000022366 | Slc22a22 | 210463 | protein_coding | 1.006238791 | 1.127383413 | 2.601785078 | -0.433311507 | 0.664788479 | 1 | 1.127383413 |
| ENSMUSG00000017692 | Rhbdl3 | 246104 | protein_coding | 51.97220181 | -1.127755409 | 0.239897548 | 4.700987635 | 2.59E-06 | 3.28E-05 | 1.127755409 |
| ENSMUSG00000086509 | Nkx2-2os | 100313531 | lncRNA | 1.294634466 | 1.127803748 | 2.61338636 | -0.4315488 | 0.666069377 | 1 | 1.127803748 |
| ENSMUSG00000089964 | Gm16582 | 102634844 | lncRNA | 1.433188843 | 1.128017972 | 2.619351678 | -0.430647775 | 0.666724498 | 1 | 1.128017972 |
| ENSMUSG00000006675 | P4htm | 74443 | protein_coding | 5.318041176 | 1.129003225 | 0.64810678 | -1.742001875 | 0.081508116 | 0.21257502 | 1.129003225 |
| ENSMUSG00000071356 | Reg3b | 18489 | protein_coding | 23075.53129 | 1.129174827 | 0.359338567 | -3.142370261 | 0.00167586 | 0.009789581 | 1.129174827 |
| ENSMUSG00000064202 | Spata6l | 381218 | protein_coding | 17.16941398 | 1.129651021 | 0.304645065 | -3.708089021 | 0.000208829 | 0.001642193 | 1.129651021 |
| ENSMUSG00000028859 | Csf3r | 12986 | protein_coding | 1.843534831 | 1.129838539 | 1.879102184 | -0.601265088 | 0.547663439 | 1 | 1.129838539 |
| ENSMUSG00000032454 | Rbp2 | 19660 | protein_coding | 78.33389971 | -1.13032031 | 0.413309607 | 2.734802897 | 0.006241764 | 0.028986639 | 1.13032031 |
| ENSMUSG00000064225 | Paqr9 | 75552 | protein_coding | 1.730957532 | -1.130436016 | 1.679075361 | 0.673249124 | 0.500788814 | 1 | 1.130436016 |
| ENSMUSG00000026770 | Il2ra | 16184 | protein_coding | 3.278618378 | 1.130463219 | 1.976421268 | -0.57197483 | 0.567339027 | 1 | 1.130463219 |
| ENSMUSG00000020732 | Rab37 | 58222 | protein_coding | 36.96562988 | -1.130496265 | 0.291549672 | 3.877542565 | 0.000105517 | 0.000903321 | 1.130496265 |
| ENSMUSG00000022603 | Mroh4 | 69439 | protein_coding | 2.789480315 | 1.130555512 | 1.796384779 | -0.629350418 | 0.52911967 | 1 | 1.130555512 |

Supplementary Table 1

| ensembl_gene_id | external_gene_name | entrezgene_id | gene_biotype | baseMean | log2FoldChange | lfcSE | stat | pvalue | padj | absLog2FC |
|---|---|---|---|---|---|---|---|---|---|---|
| ENSMUSG00000027513 | Pck1 | 18534 | protein_coding | 2387.009888 | -1.130813902 | 0.16176533 | 6.990458978 | 2.74E-12 | 8.05E-11 | 1.130813902 |
| ENSMUSG00000074502 | Ubtfl1 | 546118 | protein_coding | 2.605852313 | 1.130925961 | 1.82595567 | -0.619361126 | 0.535678484 | 1 | 1.130925961 |
| ENSMUSG00000053825 | Ppfia2 | 327814 | protein_coding | 2.870836442 | 1.13104319 | 2.018919137 | -0.560222136 | 0.57532793 | 1 | 1.13104319 |
| ENSMUSG00000048528 | Nkx1-2 | 20231 | protein_coding | 1.84015924 | 1.131110074 | 1.635210812 | -0.691721255 | 0.489112397 | 1 | 1.131110074 |
| ENSMUSG00000086862 | Gm13546 | 100042926 | lncRNA | 0.60471289 | 1.131367261 | 1.870630167 | -0.604805419 | 0.545308293 | 1 | 1.131367261 |
| ENSMUSG00000032940 | Rbm11 | 224344 | protein_coding | 7.209591826 | 1.131888598 | 0.667796394 | -1.694960633 | 0.090082889 | 0.228901189 | 1.131888598 |
| ENSMUSG00000087633 | Gm14455 | 118568483 | lncRNA | 1.779607244 | -1.132605876 | 1.639424594 | 0.690855731 | 0.489656208 | 1 | 1.132605876 |
| ENSMUSG00000112833 | Gm36595 | 102640560 | lncRNA | 1.558037718 | 1.132689476 | 1.389542939 | -0.815152555 | 0.414984992 | 1 | 1.132689476 |
| ENSMUSG00000030589 | Rasgrp4 | 233046 | protein_coding | 12.86743386 | 1.133039009 | 0.718242678 | -1.577515572 | 0.114676942 | 0.272799628 | 1.133039009 |
| ENSMUSG00000056880 | Gadl1 | 73748 | protein_coding | 2.478266253 | 1.133173231 | 2.431484542 | -0.466041717 | 0.64118564 | 1 | 1.133173231 |
| ENSMUSG00000055789 | Ftdc2 | 224247 | protein_coding | 1.158393678 | 1.134223596 | 1.172039741 | -0.967734759 | 0.333176854 | 1 | 1.134223596 |
| ENSMUSG00000056258 | Kcnq3 | 110862 | protein_coding | 4.546959142 | 1.134495652 | 1.938164961 | -0.585345249 | 0.558315595 | 0.747162726 | 1.134495652 |
| ENSMUSG00000044220 | Nkx2-3 | 18089 | protein_coding | 1.197480529 | 1.135351955 | 1.555650926 | -0.729824369 | 0.465497547 | 1 | 1.135351955 |
| ENSMUSG00000040254 | Sema3d | 108151 | protein_coding | 5.289663635 | 1.135429577 | 0.566469142 | -2.004397932 | 0.045027451 | 0.137751634 | 1.135429577 |
| ENSMUSG00000028370 | Pappa | 18491 | protein_coding | 7.579438758 | 1.135636128 | 0.653735854 | -1.737148299 | 0.082360998 | 0.214426852 | 1.135636128 |
| ENSMUSG00000048218 | Amigo2 | 105827 | protein_coding | 15.85397742 | -1.13580945 | 0.36138999 | 3.142891283 | 0.00167288 | 0.009775631 | 1.13580945 |
| ENSMUSG00000021758 | Ddx4 | 13206 | protein_coding | 1.842554698 | 1.135881427 | 1.467608214 | -0.773967749 | 0.438949863 | 1 | 1.135881427 |
| ENSMUSG00000032131 | Abcg4 | 192663 | protein_coding | 3.254173442 | 1.135931174 | 0.845773655 | -1.343067577 | 0.179250093 | 1 | 1.135931174 |
| ENSMUSG00000040364 | Sec1 | 56546 | protein_coding | 11.13729007 | 1.13705854 | 0.412378473 | -2.757317885 | 0.005827767 | 0.027575856 | 1.13705854 |
| ENSMUSG00000055435 | Maf | 17132 | protein_coding | 88.64283171 | 1.137136474 | 0.341936043 | -3.325582365 | 0.00088234 | 0.005729834 | 1.137136474 |
| ENSMUSG00000112117 | Rmst | 110333 | lncRNA | 1.829230211 | 1.137525318 | 2.112036732 | -0.538591636 | 0.59016866 | 1 | 1.137525318 |
| ENSMUSG00000040297 | Suco | 226551 | protein_coding | 1031.14491 | 1.137598835 | 0.07672422 | -14.82711493 | 9.78E-50 | 1.34E-47 | 1.137598835 |
| ENSMUSG00000036676 | Tmtc3 | 237500 | protein_coding | 383.2074714 | 1.137726409 | 0.119424301 | -9.52675794 | 1.62E-21 | 8.99E-20 | 1.137726409 |
| ENSMUSG00000031443 | F7 | 14068 | protein_coding | 3.261901926 | 1.137943524 | 0.927050412 | -1.22748829 | 0.21963912 | 1 | 1.137943524 |
| ENSMUSG00000056605 | Krt72 | 105866 | protein_coding | 1.710421213 | 1.138368292 | 2.959384456 | -0.384663875 | 0.700486463 | 1 | 1.138368292 |
| ENSMUSG00000102260 | D330025C20Rik | 665054 | TEC | 1.524306224 | 1.138383346 | 2.959923665 | -0.384598887 | 0.700534619 | 1 | 1.138383346 |
| ENSMUSG00000021478 | Drd1 | 13488 | protein_coding | 1.571866836 | 1.138520494 | 2.965293259 | -0.383948701 | 0.70101647 | 1 | 1.138520494 |
| ENSMUSG00000027584 | Oprl1 | 18389 | protein_coding | 1.571866836 | 1.138520494 | 2.965293259 | -0.383948701 | 0.70101647 | 1 | 1.138520494 |
| ENSMUSG00000056910 | Vmn2r107 | 22312 | protein_coding | 1.571866836 | 1.138520494 | 2.965293259 | -0.383948701 | 0.70101647 | 1 | 1.138520494 |
| ENSMUSG00000067597 | Dgat2l6 | 668257 | protein_coding | 1.338191235 | 1.138642844 | 2.970025917 | -0.383378084 | 0.701439451 | 1 | 1.138642844 |
| ENSMUSG00000073956 | Or52j3 | 404317 | protein_coding | 1.477811759 | 1.138666993 | 2.9710332 | -0.383256233 | 0.701529787 | 1 | 1.138666993 |
| ENSMUSG00000022342 | Kcnv1 | 67498 | protein_coding | 1.43331246 | 1.138682638 | 2.971621624 | -0.383185608 | 0.701582148 | 1 | 1.138682638 |
| ENSMUSG00000036972 | Zic4 | 22774 | protein_coding | 1.43331246 | 1.138682638 | 2.971621624 | -0.383185608 | 0.701582148 | 1 | 1.138682638 |
| ENSMUSG00000057000 | Nxf3 | 245610 | protein_coding | 1.43331246 | 1.138682638 | 2.971621624 | -0.383185608 | 0.701582148 | 1 | 1.138682638 |
| ENSMUSG00000061469 | Msantd5 | 622699 | protein_coding | 1.43331246 | 1.138682638 | 2.971621624 | -0.383185608 | 0.701582148 | 1 | 1.138682638 |
| ENSMUSG00000072259 | Gm5592 | 434172 | protein_coding | 1.43331246 | 1.138682638 | 2.971621624 | -0.383185608 | 0.701582148 | 1 | 1.138682638 |
| ENSMUSG00000107770 | Gm44126 | 118567384 | processed_pseudogene | 1.364406687 | 1.138814671 | 0.869070494 | -1.31038239 | 0.190066508 | 1 | 1.138814671 |
| ENSMUSG00000058537 | AW822073 | 100504180 | protein_coding | 1.345271476 | 1.138850891 | 2.978255804 | -0.382388541 | 0.702173189 | 1 | 1.138850891 |
| ENSMUSG00000019785 | Clvs2 | 215890 | protein_coding | 1.392515401 | 1.138948657 | 2.982152761 | -0.381921634 | 0.702519493 | 1 | 1.138948657 |

| ensembl_gene_id | external_gene_name | entrezgene_id | gene_biotype | baseMean | log2FoldChange | lfcSE | stat | pvalue | padj | absLog2FC |
|---|---|---|---|---|---|---|---|---|---|---|
| ENSMUSG00000091345 | Col6a5 | 665033 | protein_coding | 8.000442835 | -1.139000414 | 0.896120966 | 1.271034221 | 0.203716474 | 0.405918699 | 1.139000414 |
| ENSMUSG00000054141 | Or1m1 | 18322 | protein_coding | 1.156203707 | 1.139010406 | 2.984508309 | -0.38164089 | 0.70272775 | 1 | 1.139010406 |
| ENSMUSG00000105218 | Gm38413 | 330133 | lncRNA | 1.156203707 | 1.139010406 | 2.984508309 | -0.38164089 | 0.70272775 | 1 | 1.139010406 |
| ENSMUSG00000050063 | Klk6 | 19144 | protein_coding | 1.199636859 | 1.13903185 | 2.98538511 | -0.381535986 | 0.702805574 | 1 | 1.13903185 |
| ENSMUSG00000072980 | Oip5 | 70645 | protein_coding | 46.78739142 | 1.139072548 | 0.250843635 | -4.540966518 | 5.60E-06 | 6.62E-05 | 1.139072548 |
| ENSMUSG00000036744 | Or2ag2b | 66786 | protein_coding | 1.108643095 | 1.139130168 | 2.98927341 | -0.381072592 | 0.703149384 | 1 | 1.139130168 |
| ENSMUSG00000033882 | Rbm46 | 633285 | protein_coding | 1.658943545 | 1.139196962 | 2.113790977 | -0.538935483 | 0.589931372 | 1 | 1.139196962 |
| ENSMUSG00000043635 | Adamts3 | 330119 | protein_coding | 28.39528672 | 1.139227627 | 0.339177109 | -3.358798678 | 0.000782821 | 0.005170946 | 1.139227627 |
| ENSMUSG00000091859 | Vmn2r100 | 627537 | protein_coding | 1.067846036 | 1.139495317 | 3.003997024 | -0.37932638 | 0.704445512 | 1 | 1.139495317 |
| ENSMUSG00000028373 | Astn2 | 56079 | protein_coding | 3.317228366 | -1.139684944 | 0.85932397 | 1.326257598 | 0.184754387 | 1 | 1.139684944 |
| ENSMUSG00000055937 | Krt28 | 70843 | protein_coding | 1.061082482 | 1.139836645 | 3.017922337 | -0.377689191 | 0.705661499 | 1 | 1.139836645 |
| ENSMUSG00000010651 | Acaa1b | 235674 | protein_coding | 101.4526814 | 1.14002688 | 0.291419566 | -3.911977827 | 9.15E-05 | 0.000797335 | 1.14002688 |
| ENSMUSG00000069301 | H2ac11 | 319167 | protein_coding | 65.76920573 | 1.140268207 | 0.292634878 | -3.896556058 | 9.76E-05 | 0.000842717 | 1.140268207 |
| ENSMUSG00000053128 | Rnf26 | 213211 | protein_coding | 44.37152282 | 1.140423057 | 0.195478272 | -5.834014429 | 5.41E-09 | 1.09E-07 | 1.140423057 |
| ENSMUSG00000100335 | 2310008N11Rik | 72316 | lncRNA | 1.056291633 | 1.141135797 | 1.323857297 | -0.861977948 | 0.388699653 | 1 | 1.141135797 |
| ENSMUSG00000056821 | 1700028I16Rik | 70003 | lncRNA | 0.701106157 | 1.141468221 | 1.680697559 | -0.679163372 | 0.497034353 | 1 | 1.141468221 |
| ENSMUSG00000073950 | Or52ab4 | 258726 | protein_coding | 1.122532989 | -1.141734515 | 2.432434589 | 0.469379329 | 0.638798521 | 1 | 1.141734515 |
| ENSMUSG00000024526 | Cidea | 12683 | protein_coding | 0.876036513 | -1.141977658 | 1.279318979 | 0.892644975 | 0.372047326 | 1 | 1.141977658 |
| ENSMUSG00000043773 | 1700048O20Rik | 69430 | lncRNA | 1.764472031 | -1.142351728 | 1.309278013 | 0.872505089 | 0.382932893 | 1 | 1.142351728 |
| ENSMUSG00000032525 | Nktr | 18087 | protein_coding | 2064.586928 | 1.143056506 | 0.083658514 | -13.66336134 | 1.68E-42 | 1.89E-40 | 1.143056506 |
| ENSMUSG00000037892 | Pcdh18 | 73173 | protein_coding | 46.51217088 | 1.14329604 | 0.183560737 | -6.228434574 | 4.71E-10 | 1.08E-08 | 1.14329604 |
| ENSMUSG00000094595 | Fsbp | 100503583 | protein_coding | 4.164047763 | 1.143429475 | 0.511060587 | -2.237365793 | 0.025262442 | 1 | 1.143429475 |
| ENSMUSG00000050558 | Prokr2 | 246313 | protein_coding | 1.911047607 | 1.143461477 | 2.096850755 | -0.545323254 | 0.585531213 | 1 | 1.143461477 |
| ENSMUSG00000087361 | 0610043K17Rik | 68400 | lncRNA | 4.346671463 | 1.143868581 | 0.755238734 | -1.514578809 | 0.129879102 | 1 | 1.143868581 |
| ENSMUSG00000035711 | Dok3 | 27261 | protein_coding | 9.638565797 | 1.144354927 | 0.449387249 | -2.54647841 | 0.010881594 | 0.045481814 | 1.144354927 |
| ENSMUSG00000026701 | Prdx6 | 11758 | protein_coding | 11486.11579 | -1.144400305 | 0.168582057 | 6.788387366 | 1.13E-11 | 3.16E-10 | 1.144400305 |
| ENSMUSG00000034762 | Glis1 | 230587 | protein_coding | 3.106514726 | -1.144989295 | 1.506663825 | 0.75995008 | 0.447284425 | 1 | 1.144989295 |
| ENSMUSG00000021379 | Id4 | 15904 | protein_coding | 64.14524043 | 1.145024577 | 0.203727552 | -5.620371739 | 1.91E-08 | 3.51E-07 | 1.145024577 |
| ENSMUSG00000038068 | Rnf144b | 218215 | protein_coding | 5.622971562 | -1.145133281 | 0.565533173 | 2.024873757 | 0.042880329 | 0.132607819 | 1.145133281 |
| ENSMUSG00000037820 | Tgm2 | 21817 | protein_coding | 407.9042 | 1.145795117 | 0.184571575 | -6.207863356 | 5.37E-10 | 1.22E-08 | 1.145795117 |
| ENSMUSG00000073968 | Trim68 | 101700 | protein_coding | 199.5714907 | 1.145814899 | 0.112822938 | -10.15586832 | 3.12E-24 | 1.94E-22 | 1.145814899 |
| ENSMUSG00000070526 | Peg12 | 27412 | protein_coding | 1.145599865 | 1.146495271 | 1.471748633 | -0.779002097 | 0.43597848 | 1 | 1.146495271 |
| ENSMUSG00000099398 | Ms4a14 | 383435 | protein_coding | 1.869561207 | 1.146881338 | 1.872207608 | -0.612582351 | 0.540152531 | 1 | 1.146881338 |
| ENSMUSG00000086784 | Isoc2a | 664994 | protein_coding | 438.8102715 | 1.146927963 | 0.129882246 | -8.830521457 | 1.04E-18 | 4.93E-17 | 1.146927963 |
| ENSMUSG00000047564 | Krtap3-1 | 69473 | protein_coding | 2.047059248 | 1.147415063 | 1.296181132 | -0.885227407 | 0.376033993 | 1 | 1.147415063 |
| ENSMUSG00000033857 | Engase | 217364 | protein_coding | 653.1490532 | 1.147639622 | 0.080934171 | -14.17991442 | 1.22E-45 | 1.49E-43 | 1.147639622 |
| ENSMUSG00000047757 | Fancb | 237211 | protein_coding | 41.87289519 | 1.147873799 | 0.214576943 | -5.349474101 | 8.82E-08 | 1.47E-06 | 1.147873799 |
| ENSMUSG00000051036 | Ttc24 | 214191 | transcribed_unitary_pseudogene | 1.599131474 | 1.147935066 | 2.37896062 | -0.482536388 | 0.629424955 | 1 | 1.147935066 |
| ENSMUSG00000035270 | Impg2 | 224224 | protein_coding | 5.859581265 | 1.149517962 | 0.629425523 | -1.826297028 | 0.067805533 | 0.186623428 | 1.149517962 |

| ensembl_gene_id | external_gene_name | entrezgene_id | gene_biotype | baseMean | log2FoldChange | lfcSE | stat | pvalue | padj | absLog2FC |
|---|---|---|---|---|---|---|---|---|---|---|
| ENSMUSG00000029909 | Prss37 | 67690 | protein_coding | 1.123168613 | 1.149806115 | 2.32742157 | -0.494025719 | 0.621288016 | 1 | 1.149806115 |
| ENSMUSG00000073007 | Tent5d | 213449 | protein_coding | 1.07396951 | 1.149881521 | 1.811383889 | -0.634808297 | 0.525553452 | 1 | 1.149881521 |
| ENSMUSG00000027642 | Rpn2 | 20014 | protein_coding | 7435.18169 | 1.15081477 | 0.08397508 | -13.70424146 | 9.58E-43 | 1.08E-40 | 1.15081477 |
| ENSMUSG00000074121 | Ntf5 | 78405 | protein_coding | 1.535739354 | -1.150908368 | 1.448810226 | 0.794381726 | 0.426973239 | 1 | 1.150908368 |
| ENSMUSG00000042793 | Lgr6 | 329252 | protein_coding | 2.555733301 | -1.15113013 | 1.682811379 | 0.684051786 | 0.493942465 | 1 | 1.15113013 |
| ENSMUSG00000029452 | Tmem116 | 77462 | protein_coding | 34.06035139 | 1.151470567 | 0.282675945 | -4.073464994 | 4.63E-05 | 0.000436592 | 1.151470567 |
| ENSMUSG00000019564 | Arid3a | 13496 | protein_coding | 132.5735754 | 1.152348473 | 0.219415435 | -5.251902498 | 1.51E-07 | 2.41E-06 | 1.152348473 |
| ENSMUSG00000037416 | Dmxl1 | 240283 | protein_coding | 1928.995809 | 1.15238916 | 0.066613385 | -17.29966377 | 4.73E-67 | 9.41E-65 | 1.15238916 |
| ENSMUSG00000111290 | Gm5122 | 330956 | lncRNA | 1.532825559 | -1.152563293 | 2.290484058 | 0.503196383 | 0.614826208 | 1 | 1.152563293 |
| ENSMUSG00000087626 | Gm15050 | 100504154 | lncRNA | 5.89763718 | 1.152847595 | 0.456752481 | -2.524009488 | 0.01160248 | 0.047864939 | 1.152847595 |
| ENSMUSG00000031937 | Vstm5 | 69137 | protein_coding | 115.1570515 | -1.153349091 | 0.154432275 | 7.468316389 | 8.12E-14 | 2.69E-12 | 1.153349091 |
| ENSMUSG00000029407 | Uso1 | 56041 | protein_coding | 1283.99674 | 1.153679853 | 0.090562491 | -12.73904729 | 3.59E-37 | 3.35E-35 | 1.153679853 |
| ENSMUSG00000060371 | Caln1 | 140904 | protein_coding | 4.412567298 | 1.153848912 | 3.084343749 | -0.374098676 | 0.708330904 | 0.847451753 | 1.153848912 |
| ENSMUSG00000031532 | Saraf | 67887 | protein_coding | 2457.699748 | 1.154226225 | 0.0645665 | -17.87654942 | 1.80E-71 | 4.01E-69 | 1.154226225 |
| ENSMUSG00000054074 | Skida1 | 72668 | protein_coding | 14.2785909 | -1.155102758 | 0.392502826 | 2.942915771 | 0.003251369 | 0.017018416 | 1.155102758 |
| ENSMUSG00000068085 | Cyp2d11 | 545123 | protein_coding | 2.561363036 | 1.155641404 | 1.052056169 | -1.098459795 | 0.272003765 | 1 | 1.155641404 |
| ENSMUSG00000038233 | Gask1a | 245050 | protein_coding | 1.690684584 | -1.156578451 | 1.053120607 | 1.098239311 | 0.272100005 | 1 | 1.156578451 |
| ENSMUSG00000078907 | Fam186b | 545136 | protein_coding | 2.445287512 | 1.156829676 | 1.258797593 | -0.91899578 | 0.358097779 | 1 | 1.156829676 |
| ENSMUSG00000112739 | Gm20597 | 100502868 | lncRNA | 1.727897744 | -1.156997188 | 2.28623998 | 0.506069878 | 0.612807596 | 1 | 1.156997188 |
| ENSMUSG00000095002 | Or6c66 | 258549 | protein_coding | 1.142523136 | -1.157309222 | 2.125653489 | 0.544448673 | 0.586132759 | 1 | 1.157309222 |
| ENSMUSG00000073922 | Or52n20 | 259052 | protein_coding | 2.005006497 | -1.157533771 | 2.294041467 | 0.504582758 | 0.613851922 | 1 | 1.157533771 |
| ENSMUSG00000065494 | Mir28a | 723830 | miRNA | 0.789145927 | -1.157564374 | 1.608709721 | 0.71956075 | 0.471795485 | 1 | 1.157564374 |
| ENSMUSG00000024186 | Rgs11 | 50782 | protein_coding | 27.10616722 | -1.157942608 | 0.251691524 | 4.600642042 | 4.21E-06 | 5.13E-05 | 1.157942608 |
| ENSMUSG00000114780 | AI197445 | 105120 | lncRNA | 1.281077512 | -1.158789892 | 2.134856642 | 0.542795178 | 0.587270833 | 1 | 1.158789892 |
| ENSMUSG00000030069 | Prok2 | 50501 | protein_coding | 1.071758937 | 1.158870617 | 2.095625757 | -0.552995025 | 0.580266817 | 1 | 1.158870617 |
| ENSMUSG00000037795 | N4bp2 | 333789 | protein_coding | 637.3054201 | 1.159345913 | 0.089126262 | -13.00790464 | 1.10E-38 | 1.07E-36 | 1.159345913 |
| ENSMUSG00000115792 | Vmn1r16 | 171202 | protein_coding | 1.26892508 | -1.159690208 | 2.326681574 | 0.498430993 | 0.618180296 | 1 | 1.159690208 |
| ENSMUSG00000021359 | Tfap2a | 21418 | protein_coding | 2.611212316 | -1.16027404 | 2.335762882 | 0.496743076 | 0.619370241 | 1 | 1.16027404 |
| ENSMUSG00000023257 | Cypt1 | 66742 | protein_coding | 0.627707281 | -1.160285344 | 1.657791911 | 0.699898061 | 0.483990969 | 1 | 1.160285344 |
| ENSMUSG00000072915 | Gm12258 | 108167848 | protein_coding | 31.2099288 | 1.160413347 | 0.272359916 | -4.260587849 | 2.04E-05 | 0.000209523 | 1.160413347 |
| ENSMUSG00000024901 | Peli3 | 240518 | protein_coding | 16.67314905 | -1.160630367 | 0.382535968 | 3.034042455 | 0.002413005 | 0.013212321 | 1.160630367 |
| ENSMUSG00000038527 | C1rl | 232371 | protein_coding | 12.12900054 | 1.161072494 | 0.317132595 | -3.661157864 | 0.000251078 | 0.001937524 | 1.161072494 |
| ENSMUSG00000027401 | Tgm3 | 21818 | protein_coding | 7317.414329 | -1.161717891 | 0.335115216 | 3.466622328 | 0.000527042 | 0.003708381 | 1.161717891 |
| ENSMUSG00000024030 | Abcg1 | 11307 | protein_coding | 52.96791243 | 1.161719463 | 0.180671333 | -6.430015466 | 1.28E-10 | 3.12E-09 | 1.161719463 |
| ENSMUSG00000048602 | Morc2b | 240069 | protein_coding | 2.229639652 | 1.16228452 | 1.701126284 | -0.683244114 | 0.494452602 | 1 | 1.16228452 |
| ENSMUSG00000043518 | Rai2 | 24004 | protein_coding | 9.409735152 | -1.162377293 | 0.534283474 | 2.175581595 | 0.029586561 | 0.099855399 | 1.162377293 |
| ENSMUSG00000065610 | Mir29a | 387222 | miRNA | 3.90238733 | 1.162624841 | 0.577481578 | -2.013267409 | 0.044086505 | 1 | 1.162624841 |
| ENSMUSG00000024670 | Cd6 | 12511 | protein_coding | 4.60419838 | -1.162925426 | 1.307444114 | 0.88946473 | 0.37375337 | 0.595844868 | 1.162925426 |
| ENSMUSG00000034088 | Hdlbp | 110611 | protein_coding | 9771.880304 | 1.163137222 | 0.067353687 | -17.26909511 | 8.04E-67 | 1.58E-64 | 1.163137222 |

| ensembl_gene_id | external_gene_name | entrezgene_id | gene_biotype | baseMean | log2FoldChange | lfcSE | stat | pvalue | padj | absLog2FC |
|---|---|---|---|---|---|---|---|---|---|---|
| ENSMUSG00000048489 | Depp1 | 213393 | protein_coding | 34.7543765 | -1.163145511 | 0.438000665 | 2.655579325 | 0.007917231 | 0.035343465 | 1.163145511 |
| ENSMUSG00000062017 | Abca14 | 67928 | protein_coding | 7.440051131 | 1.163201987 | 0.760636597 | -1.529247989 | 0.126202978 | 0.292138489 | 1.163201987 |
| ENSMUSG00000059864 | Or2y1g | 258463 | protein_coding | 1.345339101 | -1.163448717 | 1.230486704 | 0.94551913 | 0.344393909 | 1 | 1.163448717 |
| ENSMUSG00000041132 | N4bp2l1 | 100637 | protein_coding | 16.88668614 | -1.163475781 | 0.274688959 | 4.235611741 | 2.28E-05 | 0.000232062 | 1.163475781 |
| ENSMUSG00000040829 | Zmynd15 | 574428 | protein_coding | 4.395792338 | 1.163703358 | 1.066298432 | -1.091348654 | 0.275119496 | 0.491701663 | 1.163703358 |
| ENSMUSG00000025804 | Ccr1 | 12768 | protein_coding | 11.80676246 | 1.163831839 | 0.446393461 | -2.607188369 | 0.009128912 | 0.039588986 | 1.163831839 |
| ENSMUSG00000040649 | Rimklb | 108653 | protein_coding | 4.499548343 | 1.163856609 | 0.692271282 | -1.681214633 | 0.092721232 | 0.233671624 | 1.163856609 |
| ENSMUSG00000001435 | Col18a1 | 12822 | protein_coding | 14.35811765 | 1.164229736 | 0.546218654 | -2.131435328 | 0.033053295 | 0.108625296 | 1.164229736 |
| ENSMUSG00000044988 | Ucn3 | 83428 | protein_coding | 1.475330145 | 1.164759944 | 1.621776223 | -0.718200161 | 0.472633879 | 1 | 1.164759944 |
| ENSMUSG00000093599 | Mir5620 | 100885832 | miRNA | 1.42403302 | 1.164823929 | 1.114630346 | -1.045031596 | 0.296008367 | 1 | 1.164823929 |
| ENSMUSG00000092008 | Cyp2c69 | 100043108 | protein_coding | 61.01010921 | -1.165017536 | 0.282925428 | 4.117754788 | 3.83E-05 | 0.000369764 | 1.165017536 |
| ENSMUSG00000057176 | Ccdc189 | 233899 | protein_coding | 14.8431521 | -1.16506603 | 0.271049866 | 4.298345712 | 1.72E-05 | 0.000180539 | 1.16506603 |
| ENSMUSG00000061578 | Ksr2 | 333050 | protein_coding | 17.48390569 | 1.165772419 | 0.746841747 | -1.560936333 | 0.118538774 | 0.279490192 | 1.165772419 |
| ENSMUSG00000034664 | Itga2b | 16399 | protein_coding | 22.77104122 | 1.165930621 | 0.550389077 | -2.11837529 | 0.034143297 | 0.111321304 | 1.165930621 |
| ENSMUSG00000088217 | Gm22079 | 115488044 | snRNA | 2.013328377 | 1.167142163 | 0.837901353 | -1.39293505 | 0.163639432 | 1 | 1.167142163 |
| ENSMUSG00000056025 | Clca3a1 | 12722 | protein_coding | 33.72346554 | 1.168566652 | 0.264948451 | -4.41054344 | 1.03E-05 | 0.000114898 | 1.168566652 |
| ENSMUSG00000055022 | Cntn1 | 12805 | protein_coding | 6.678921354 | 1.168633666 | 1.106878493 | -1.055792188 | 0.291063168 | 0.510000436 | 1.168633666 |
| ENSMUSG00000087203 | Gm13986 | 329496 | lncRNA | 6.891501347 | -1.168677992 | 0.448649804 | 2.604877975 | 0.009190699 | 0.039794231 | 1.168677992 |
| ENSMUSG00000073514 | Dok6 | 623279 | protein_coding | 2.055798935 | -1.16894898 | 2.154948487 | 0.542448688 | 0.587509447 | 1 | 1.16894898 |
| ENSMUSG00000027776 | Il12a | 16159 | protein_coding | 1.709490469 | 1.170102447 | 1.188288679 | -0.984695443 | 0.324773692 | 1 | 1.170102447 |
| ENSMUSG00000053519 | Kcnip1 | 70357 | protein_coding | 3.176975618 | 1.170568849 | 1.366693949 | -0.856496694 | 0.391723101 | 1 | 1.170568849 |
| ENSMUSG00000048621 | Gm6377 | 622976 | protein_coding | 1.012911658 | 1.17149212 | 1.436200838 | -0.815688231 | 0.414678472 | 1 | 1.17149212 |
| ENSMUSG00000039519 | Cyp7b1 | 13123 | protein_coding | 1.741779011 | 1.171675566 | 1.367471359 | -0.85681909 | 0.391544874 | 1 | 1.171675566 |
| ENSMUSG00000047712 | Ust | 338362 | protein_coding | 2.284219334 | -1.172020684 | 1.419479651 | 0.825669239 | 0.408991746 | 1 | 1.172020684 |
| ENSMUSG00000026360 | Rgs2 | 19735 | protein_coding | 210.3181491 | 1.172183453 | 0.135873638 | -8.627011628 | 6.30E-18 | 2.88E-16 | 1.172183453 |
| ENSMUSG00000019433 | Gipc1 | 67903 | protein_coding | 1283.217229 | 1.173235741 | 0.081951463 | -14.316227 | 1.73E-46 | 2.17E-44 | 1.173235741 |
| ENSMUSG00000047361 | Gm973 | 381260 | protein_coding | 1.915779883 | -1.173282438 | 1.395136326 | 0.840980495 | 0.400358862 | 1 | 1.173282438 |
| ENSMUSG00000068323 | Slc4a5 | 232156 | protein_coding | 8.157313142 | 1.173971074 | 0.84401368 | -1.390938444 | 0.164244094 | 0.350538094 | 1.173971074 |
| ENSMUSG00000031595 | Pdgfrl | 68797 | protein_coding | 6.452215804 | 1.174078807 | 0.584082981 | -2.010123296 | 0.044418141 | 0.136254601 | 1.174078807 |
| ENSMUSG00000027858 | Tspan2 | 70747 | protein_coding | 2.615115794 | -1.174150591 | 0.917497033 | 1.279732303 | 0.2006393 | 1 | 1.174150591 |
| ENSMUSG00000059908 | Mug1 | 17836 | protein_coding | 2.34994861 | -1.174602832 | 2.622463951 | 0.447900469 | 0.65422503 | 1 | 1.174602832 |
| ENSMUSG00000048004 | Tmem196 | 217951 | protein_coding | 1.456366685 | -1.174892918 | 2.630009724 | 0.446725693 | 0.655073128 | 1 | 1.174892918 |
| ENSMUSG00000030708 | Dnajb13 | 69387 | protein_coding | 63.53329363 | -1.175555297 | 0.244412664 | 4.80971517 | 1.51E-06 | 2.01E-05 | 1.175555297 |
| ENSMUSG00000077148 | Gm22935 | 115489782 | snRNA | 3.788447406 | 1.1756056 | 0.614752978 | -1.912321928 | 0.055834915 | 1 | 1.1756056 |
| ENSMUSG00000050198 | Or6c38 | 258863 | protein_coding | 1.680385395 | -1.175617556 | 2.648945801 | 0.443805817 | 0.657182982 | 1 | 1.175617556 |
| ENSMUSG00000021294 | Kif26a | 668303 | protein_coding | 6.26295292 | -1.175628621 | 0.916943034 | 1.282117402 | 0.199801465 | 0.400865192 | 1.175628621 |
| ENSMUSG00000027674 | Pex5l | 58869 | protein_coding | 25.35398338 | -1.175691362 | 0.295878277 | 3.973564319 | 7.08E-05 | 0.000636513 | 1.175691362 |
| ENSMUSG00000020963 | Tshr | 22095 | protein_coding | 2.514206573 | 1.175977866 | 1.911698175 | -0.615148291 | 0.538456793 | 1 | 1.175977866 |
| ENSMUSG00000048304 | Slitrk3 | 386750 | protein_coding | 2.124651546 | -1.176043197 | 2.660286829 | 0.442073834 | 0.658435778 | 1 | 1.176043197 |

| ensembl_gene_id | external_gene_name | entrezgene_id | gene_biotype | baseMean | log2FoldChange | lfcSE | stat | pvalue | padj | absLog2FC |
|---|---|---|---|---|---|---|---|---|---|---|
| ENSMUSG00000017723 | Wfdc2 | 67701 | protein_coding | 1741.428165 | -1.176280719 | 0.204317382 | 5.757125047 | 8.56E-09 | 1.67E-07 | 1.176280719 |
| ENSMUSG00000031559 | 4930555F03Rik | 75359 | lncRNA | 1.617994653 | -1.176299481 | 2.66732159 | 0.441003996 | 0.659210104 | 1 | 1.176299481 |
| ENSMUSG00000063529 | Stmnd1 | 380842 | protein_coding | 1.194880588 | -1.177684599 | 1.192616037 | 0.987480096 | 0.323407336 | 1 | 1.177684599 |
| ENSMUSG00000064373 | Selenop | 20363 | protein_coding | 3119.956012 | -1.17778829 | 0.181299797 | 6.496357455 | 8.23E-11 | 2.09E-09 | 1.17778829 |
| ENSMUSG00000049241 | Hcar1 | 243270 | protein_coding | 2.123854713 | 1.178505623 | 0.90334514 | -1.30460172 | 0.192028502 | 1 | 1.178505623 |
| ENSMUSG00000049653 | Spatc1 | 74281 | protein_coding | 1.041622957 | 1.178829447 | 2.325107517 | -0.506999972 | 0.612154838 | 1 | 1.178829447 |
| ENSMUSG00000043833 | 2900005J15Rik | 67261 | lncRNA | 8.956474367 | -1.179084545 | 0.472162255 | 2.497202037 | 0.012517761 | 0.050703535 | 1.179084545 |
| ENSMUSG00000026622 | Nek2 | 18005 | protein_coding | 522.0315641 | 1.179542151 | 0.147200268 | -8.013179371 | 1.12E-15 | 4.32E-14 | 1.179542151 |
| ENSMUSG00000053914 | Kdm4d | 244694 | protein_coding | 2.531244883 | 1.18001425 | 0.840281948 | -1.40430751 | 0.160227299 | 1 | 1.18001425 |
| ENSMUSG00000030830 | Itgal | 16408 | protein_coding | 13.05881218 | 1.180513538 | 0.750897775 | -1.572136151 | 0.115918979 | 0.274894315 | 1.180513538 |
| ENSMUSG00000022053 | Ebf2 | 13592 | protein_coding | 1.502080244 | -1.181254602 | 2.056885025 | 0.574292966 | 0.565769568 | 1 | 1.181254602 |
| ENSMUSG00000087067 | Gm11532 | 100127433 | lncRNA | 0.934898644 | 1.181269444 | 1.751794975 | -0.674319462 | 0.500108234 | 1 | 1.181269444 |
| ENSMUSG00000048065 | Cyb5r2 | 320635 | protein_coding | 2.050234291 | 1.181372371 | 1.257252107 | -0.939646364 | 0.347398986 | 1 | 1.181372371 |
| ENSMUSG00000030359 | Pzp | 11287 | protein_coding | 3.128232823 | 1.181376703 | 1.530950631 | -0.771662181 | 0.440314536 | 1 | 1.181376703 |
| ENSMUSG00000033024 | Klra9 | 16640 | protein_coding | 0.73761659 | 1.181396493 | 1.870174211 | -0.631703981 | 0.527580326 | 1 | 1.181396493 |
| ENSMUSG00000112323 | Gm32828 | 102635512 | lncRNA | 0.828194259 | -1.181428387 | 1.660275486 | 0.711585756 | 0.47672133 | 1 | 1.181428387 |
| ENSMUSG00000034687 | Fras1 | 231470 | protein_coding | 11.11929574 | 1.18215911 | 0.814032417 | -1.452226086 | 0.146438752 | 0.32403719 | 1.18215911 |
| ENSMUSG00000030826 | Bcat2 | 12036 | protein_coding | 793.1458859 | 1.182250361 | 0.120538064 | -9.808108064 | 1.04E-22 | 6.04E-21 | 1.182250361 |
| ENSMUSG00000015829 | Tnr | 21960 | protein_coding | 6.201201677 | -1.182667262 | 0.911454375 | 1.297560574 | 0.194438377 | 0.393348786 | 1.182667262 |
| ENSMUSG00000089997 | 1810020O05Rik | 613264 | protein_coding | 1.07426788 | 1.183364729 | 2.03897544 | -0.580372233 | 0.561663626 | 1 | 1.183364729 |
| ENSMUSG00000030344 | Akap3 | 11642 | protein_coding | 1.364076959 | 1.183520711 | 2.0544673 | -0.576071817 | 0.56456664 | 1 | 1.183520711 |
| ENSMUSG00000088139 | Gm27343 | 115489597 | snRNA | 1.315873515 | 1.18378172 | 0.989953555 | -1.195795211 | 0.231776485 | 1 | 1.18378172 |
| ENSMUSG00000052469 | Tcp10c | 100041352 | protein_coding | 2.049598054 | 1.183851457 | 2.088374468 | -0.566877002 | 0.57079775 | 1 | 1.183851457 |
| ENSMUSG00000042846 | Lrrtm3 | 216028 | protein_coding | 0.710531819 | -1.184325255 | 1.791319625 | 0.66114681 | 0.508518169 | 1 | 1.184325255 |
| ENSMUSG00000104900 | 4930596I21Rik | 78229 | lncRNA | 6.868502406 | -1.184357647 | 0.555399685 | 2.132442059 | 0.032970523 | 0.108526787 | 1.184357647 |
| ENSMUSG00000020218 | Wif1 | 24117 | protein_coding | 9.280728156 | -1.184644382 | 0.404593538 | 2.927986414 | 0.00341165 | 0.017711405 | 1.184644382 |
| ENSMUSG00000043629 | 1700019D03Rik | 67080 | protein_coding | 225.3993205 | 1.185231406 | 0.159970218 | -7.409075384 | 1.27E-13 | 4.16E-12 | 1.185231406 |
| ENSMUSG00000031539 | Ap3m2 | 64933 | protein_coding | 17.15577005 | 1.185365564 | 0.303380346 | -3.907193006 | 9.34E-05 | 0.000811145 | 1.185365564 |
| ENSMUSG00000032484 | Ngp | 18054 | protein_coding | 1.220067842 | 1.185907612 | 2.342516522 | -0.506253681 | 0.612678575 | 1 | 1.185907612 |
| ENSMUSG00000032523 | Hhatl | 74770 | protein_coding | 1.236762575 | 1.186014542 | 2.358446299 | -0.502879605 | 0.615048921 | 1 | 1.186014542 |
| ENSMUSG00000015401 | Cltrn | 57394 | protein_coding | 0.983397733 | 1.186047621 | 2.363234877 | -0.501874626 | 0.615755715 | 1 | 1.186047621 |
| ENSMUSG00000084876 | Gm14965 | 105246848 | lncRNA | 0.933002146 | 1.186053269 | 2.364093976 | -0.501694637 | 0.615882338 | 1 | 1.186053269 |
| ENSMUSG00000061723 | Tnnt3 | 21957 | protein_coding | 1.837799829 | 1.186097182 | 2.371321527 | -0.50018404 | 0.616945495 | 1 | 1.186097182 |
| ENSMUSG00000030111 | A2m | 232345 | protein_coding | 2.722425821 | -1.186150004 | 3.007716491 | 0.394368953 | 0.693308662 | 1 | 1.186150004 |
| ENSMUSG00000023159 | Psg29 | 114872 | protein_coding | 0.98611401 | 1.186161563 | 2.380534268 | -0.498275357 | 0.618289974 | 1 | 1.186161563 |
| ENSMUSG00000114744 | Gm38409 | 328436 | lncRNA | 3.904286808 | 1.186512027 | 0.620435187 | -1.912386742 | 0.055826607 | 1 | 1.186512027 |
| ENSMUSG00000064201 | Krt2 | 16681 | protein_coding | 2.081342012 | -1.186921824 | 3.042661873 | 0.390093239 | 0.696467602 | 1 | 1.186921824 |
| ENSMUSG00000062609 | Kcnj15 | 16516 | protein_coding | 1.847666411 | -1.187286212 | 3.059727468 | 0.38803659 | 0.697988956 | 1 | 1.187286212 |
| ENSMUSG00000018263 | Tbx5 | 21388 | protein_coding | 1.946915095 | -1.18738031 | 3.064149174 | 0.387507345 | 0.698380649 | 1 | 1.18738031 |

| ensembl_gene_id | external_gene_name | entrezgene_id | gene_biotype | baseMean | log2FoldChange | lfcSE | stat | pvalue | padj | absLog2FC |
|---|---|---|---|---|---|---|---|---|---|---|
| ENSMUSG00000110027 | C030029H02Rik | 77383 | lncRNA | 1.392697589 | -1.1877723 | 3.083199965 | 0.385240112 | 0.700059527 | 1 | 1.1877723 |
| ENSMUSG00000023945 | Slc5a7 | 63993 | protein_coding | 3.132386009 | -1.187814625 | 1.158049288 | 1.025702997 | 0.305031601 | 1 | 1.187814625 |
| ENSMUSG00000086308 | G630016G05Rik | 108168785 | lncRNA | 1.751446922 | 1.188038012 | 2.736829046 | -0.434092884 | 0.664220993 | 1 | 1.188038012 |
| ENSMUSG00000022986 | Krt75 | 109052 | protein_coding | 1.508545406 | 1.188064011 | 2.742691848 | -0.433174442 | 0.664888045 | 1 | 1.188064011 |
| ENSMUSG00000112858 | Rnf212b | 102632837 | protein_coding | 1.679009509 | 1.188112109 | 2.754196226 | -0.43138252 | 0.666190258 | 1 | 1.188112109 |
| ENSMUSG00000040759 | Cmtm5 | 67272 | protein_coding | 1.541195996 | 1.188148867 | 2.762847565 | -0.430045031 | 0.667162885 | 1 | 1.188148867 |
| ENSMUSG00000045225 | Or5w19 | 258103 | protein_coding | 1.430053102 | 1.188183564 | 2.771179188 | -0.42876461 | 0.668094536 | 1 | 1.188183564 |
| ENSMUSG00000067338 | Tuba3b | 22147 | protein_coding | 1.244790028 | 1.188192682 | 2.773139093 | -0.42846487 | 0.668312704 | 1 | 1.188192682 |
| ENSMUSG00000095959 | Gm10845 | 100038734 | lncRNA | 1.355081008 | 1.18822375 | 2.782023833 | -0.427107682 | 0.669300898 | 1 | 1.18822375 |
| ENSMUSG00000072778 | Vmn2r27 | 232367 | protein_coding | 1.311647856 | 1.188228368 | 2.781869191 | -0.427133084 | 0.669282397 | 1 | 1.188228368 |
| ENSMUSG00000068048 | Rhox9 | 104384 | protein_coding | 1.12128179 | 1.18831899 | 2.804094019 | -0.423780009 | 0.671726258 | 1 | 1.18831899 |
| ENSMUSG00000060735 | Rxfp3 | 239336 | protein_coding | 1.0864281 | 1.188325957 | 2.805755112 | -0.423531602 | 0.671907447 | 1 | 1.188325957 |
| ENSMUSG00000005892 | Trh | 22044 | protein_coding | 1.129660327 | 1.188326988 | 2.806094619 | -0.423480727 | 0.671944557 | 1 | 1.188326988 |
| ENSMUSG00000023903 | Mmp25 | 240047 | protein_coding | 2.148233304 | -1.188458081 | 1.87001813 | 0.635532919 | 0.525080904 | 1 | 1.188458081 |
| ENSMUSG00000100830 | Gm28441 | 102633688 | lncRNA | 0.957196193 | 1.188533176 | 2.309274641 | -0.51467814 | 0.606777952 | 1 | 1.188533176 |
| ENSMUSG00000027500 | Stmn2 | 20257 | protein_coding | 1.10551655 | -1.18856102 | 1.776148778 | 0.66917875 | 0.50338146 | 1 | 1.18856102 |
| ENSMUSG00000056569 | Mpz | 17528 | protein_coding | 2.398946162 | -1.189239056 | 1.137571859 | 1.045418842 | 0.295829432 | 1 | 1.189239056 |
| ENSMUSG00000073067 | 9130019P16Rik | 100042056 | lncRNA | 6.66660187 | -1.189320866 | 0.67681223 | 1.75723903 | 0.078877075 | 0.207979566 | 1.189320866 |
| ENSMUSG00000061816 | Myl1 | 17901 | protein_coding | 1.064676334 | 1.189773279 | 1.727938628 | -0.688550658 | 0.491106081 | 1 | 1.189773279 |
| ENSMUSG00000079324 | 4932414N04Rik | 75721 | protein_coding | 1.636440838 | 1.189810608 | 3.279200328 | -0.362835597 | 0.716727699 | 1 | 1.189810608 |
| ENSMUSG00000032680 | 6820408C15Rik | 228778 | protein_coding | 4.402507368 | 1.189830978 | 0.756158313 | -1.57352099 | 0.115598232 | 0.274358896 | 1.189830978 |
| ENSMUSG00000059319 | Or14c41 | 258850 | protein_coding | 1.541319613 | 1.189849873 | 3.29492847 | -0.36111554 | 0.718013077 | 1 | 1.189849873 |
| ENSMUSG00000034402 | Kcnh5 | 238271 | protein_coding | 1.497886461 | 1.189865565 | 3.301241951 | -0.360429675 | 0.718525838 | 1 | 1.189865565 |
| ENSMUSG00000091897 | Gm17019 | 66773 | protein_coding | 1.450325849 | 1.189885147 | 3.309202954 | -0.359568501 | 0.719169842 | 1 | 1.189885147 |
| ENSMUSG00000024857 | Cabp2 | 29866 | protein_coding | 1.359332085 | 1.189927792 | 3.326745722 | -0.357685225 | 0.720578891 | 1 | 1.189927792 |
| ENSMUSG00000091541 | Vmn1r9 | 171203 | protein_coding | 1.359332085 | 1.189927792 | 3.326745722 | -0.357685225 | 0.720578891 | 1 | 1.189927792 |
| ENSMUSG00000047225 | Or56a4 | 244187 | protein_coding | 1.363459545 | 1.189960122 | 3.340475188 | -0.356224805 | 0.721672217 | 1 | 1.189960122 |
| ENSMUSG00000000094 | Tbx4 | 21387 | protein_coding | 1.411020157 | 1.189983526 | 3.350693961 | -0.355145394 | 0.722480671 | 1 | 1.189983526 |
| ENSMUSG00000039661 | Dusp26 | 66959 | protein_coding | 1.220777709 | 1.190007392 | 3.360312125 | -0.35413597 | 0.723236987 | 1 | 1.190007392 |
| ENSMUSG00000107008 | Gm2762 | 100040421 | lncRNA | 2.074642318 | -1.190240776 | 1.338430963 | 0.889280664 | 0.373852261 | 1 | 1.190240776 |
| ENSMUSG00000025407 | Gli1 | 14632 | protein_coding | 3.729663767 | -1.19063743 | 1.133593838 | 1.050321014 | 0.293570547 | 1 | 1.19063743 |
| ENSMUSG00000108348 | Pnma8c | 105247240 | protein_coding | 3.315992708 | 1.190920687 | 1.405362574 | -0.847411699 | 0.396765686 | 1 | 1.190920687 |
| ENSMUSG00000078627 | Marchf10 | 632687 | protein_coding | 2.066801307 | 1.191213477 | 2.197779978 | -0.542007612 | 0.587813262 | 1 | 1.191213477 |
| ENSMUSG00000030125 | Lrrc23 | 16977 | protein_coding | 0.918791717 | -1.191492795 | 2.162912128 | 0.550874342 | 0.581719818 | 1 | 1.191492795 |
| ENSMUSG00000069893 | 9930111J21Rik1 | 667214 | protein_coding | 1.784316176 | 1.19189141 | 1.151375562 | -1.035189081 | 0.300580617 | 1 | 1.19189141 |
| ENSMUSG00000096993 | Gm26787 | 102637672 | lncRNA | 1.441036915 | 1.192116789 | 1.182442437 | -1.008181668 | 0.313367257 | 1 | 1.192116789 |
| ENSMUSG00000099639 | 1700084F23Rik | 74280 | lncRNA | 1.095750569 | 1.192777164 | 2.327432073 | -0.512486348 | 0.608310672 | 1 | 1.192777164 |
| ENSMUSG00000064263 | Platr26 | 381371 | lncRNA | 2.65316534 | -1.193284126 | 0.649037901 | 1.838543055 | 0.065982424 | 1 | 1.193284126 |
| ENSMUSG00000026904 | Slc4a10 | 94229 | protein_coding | 2.567275284 | 1.193339323 | 1.133089196 | -1.053173332 | 0.292261559 | 1 | 1.193339323 |

| ensembl_gene_id | external_gene_name | entrezgene_id | gene_biotype | baseMean | log2FoldChange | lfcSE | stat | pvalue | padj | absLog2FC |
|---|---|---|---|---|---|---|---|---|---|---|
| ENSMUSG00000038028 | Tigar | 319801 | protein_coding | 322.2821316 | 1.193386506 | 0.11359748 | -10.50539596 | 8.16E-26 | 5.43E-24 | 1.193386506 |
| ENSMUSG00000109473 | B930025P03Rik | 320014 | lncRNA | 1.492416162 | -1.193927635 | 1.892213627 | 0.630968733 | 0.528060969 | 1 | 1.193927635 |
| ENSMUSG00000048852 | Gm12185 | 620913 | protein_coding | 5.01144173 | 1.194531311 | 0.720392292 | -1.658167812 | 0.0972836 | 0.242240857 | 1.194531311 |
| ENSMUSG00000028717 | Tal1 | 21349 | protein_coding | 1.284625268 | 1.195108172 | 2.33751077 | -0.511273867 | 0.609159301 | 1 | 1.195108172 |
| ENSMUSG00000028255 | Clca1 | 23844 | protein_coding | 27967.43667 | -1.195284189 | 0.375607322 | 3.182270738 | 0.001461252 | 0.008692083 | 1.195284189 |
| ENSMUSG00000031481 | Tpte | 234129 | protein_coding | 1.902296711 | -1.196166348 | 1.675154012 | 0.714063507 | 0.475187911 | 1 | 1.196166348 |
| ENSMUSG00000041309 | Nkx6-2 | 14912 | protein_coding | 1.586151292 | 1.19622767 | 1.183080669 | -1.011112514 | 0.311962582 | 1 | 1.19622767 |
| ENSMUSG00000005640 | Insrr | 23920 | protein_coding | 62.9208825 | -1.197216284 | 0.280791687 | 4.263716983 | 2.01E-05 | 0.000206866 | 1.197216284 |
| ENSMUSG00000030160 | Tmem52b | 330428 | protein_coding | 1.859072729 | 1.197273211 | 1.59183198 | -0.752135418 | 0.451969627 | 1 | 1.197273211 |
| ENSMUSG00000022790 | Igsf11 | 207683 | protein_coding | 12.71342862 | 1.197544608 | 0.36273451 | -3.301435551 | 0.000961914 | 0.006161774 | 1.197544608 |
| ENSMUSG00000056436 | Cyct | 13067 | protein_coding | 0.633386396 | -1.197981398 | 1.317456989 | 0.909313479 | 0.363184678 | 1 | 1.197981398 |
| ENSMUSG00000052563 | D930048N14Rik | 97775 | lncRNA | 19.75854223 | -1.198068452 | 0.314306308 | 3.81178621 | 0.000137966 | 0.001144911 | 1.198068452 |
| ENSMUSG00000062713 | Sim2 | 20465 | protein_coding | 1.132432299 | -1.198755949 | 2.204612594 | 0.543749025 | 0.586614191 | 1 | 1.198755949 |
| ENSMUSG00000036899 | Trpv5 | 194352 | protein_coding | 1.649258072 | 1.198801452 | 1.634393933 | -0.733483787 | 0.463263413 | 1 | 1.198801452 |
| ENSMUSG00000030543 | Mesp2 | 17293 | protein_coding | 4.134368213 | 1.199060956 | 0.555089787 | -2.160120729 | 0.030763325 | 1 | 1.199060956 |
| ENSMUSG00000110125 | Gm33148 | 102635939 | lncRNA | 9.379756762 | 1.199610873 | 2.956412702 | -0.405765701 | 0.684914762 | 0.832085503 | 1.199610873 |
| ENSMUSG00000018442 | Derl2 | 116891 | protein_coding | 884.9787376 | 1.199752212 | 0.072992141 | -16.43673138 | 1.04E-60 | 1.76E-58 | 1.199752212 |
| ENSMUSG00000021318 | Gli3 | 14634 | protein_coding | 5.314552774 | 1.200233499 | 0.892979917 | -1.344076698 | 0.178923586 | 0.371853021 | 1.200233499 |
| ENSMUSG00000027886 | Cfap276 | 75504 | protein_coding | 1.104808018 | 1.200593111 | 1.605844113 | -0.747639887 | 0.454677399 | 1 | 1.200593111 |
| ENSMUSG00000039167 | Adgrl4 | 170757 | protein_coding | 1.827828143 | 1.200819769 | 2.362505494 | -0.508282318 | 0.611255368 | 1 | 1.200819769 |
| ENSMUSG00000087289 | 4933424M12Rik | 67548 | lncRNA | 1.29419339 | -1.201097389 | 2.218681615 | 0.541356354 | 0.588261985 | 1 | 1.201097389 |
| ENSMUSG00000033063 | Cntnap3 | 238680 | protein_coding | 1.47629197 | -1.201421019 | 2.220647315 | 0.541022886 | 0.588491808 | 1 | 1.201421019 |
| ENSMUSG00000084764 | Grip1os2 | 75321 | lncRNA | 0.833396706 | -1.202361699 | 1.62418933 | 0.7402842 | 0.459127566 | 1 | 1.202361699 |
| ENSMUSG00000109394 | A230057D06Rik | 319893 | lncRNA | 2.470246202 | 1.202386851 | 2.195084245 | -0.547763419 | 0.58385436 | 1 | 1.202386851 |
| ENSMUSG00000040867 | Begain | 380785 | protein_coding | 1.818218047 | 1.202884266 | 1.504191028 | -0.7996885 | 0.423891298 | 1 | 1.202884266 |
| ENSMUSG00000018983 | E2f2 | 242705 | protein_coding | 573.9730946 | 1.203261947 | 0.163370774 | -7.365221556 | 1.77E-13 | 5.76E-12 | 1.203261947 |
| ENSMUSG00000100182 | 1810006J02Rik | 69030 | lncRNA | 4.775930747 | -1.203398319 | 0.569765375 | 2.112094506 | 0.034678344 | 0.112709737 | 1.203398319 |
| ENSMUSG00000069378 | Prdm6 | 225518 | protein_coding | 1.554182422 | -1.203400411 | 2.45254945 | 0.49067325 | 0.623657569 | 1 | 1.203400411 |
| ENSMUSG00000048096 | Lmod1 | 93689 | protein_coding | 1.612696374 | -1.203429859 | 1.37553372 | 0.874882121 | 0.381638048 | 1 | 1.203429859 |
| ENSMUSG00000092111 | Vmn2r113 | 434701 | protein_coding | 3.035457291 | -1.204086316 | 2.23683011 | 0.538300298 | 0.590369745 | 1 | 1.204086316 |
| ENSMUSG00000070866 | Zfp804a | 241514 | protein_coding | 1.525803625 | -1.204094225 | 2.3099901 | 0.521255145 | 0.602189043 | 1 | 1.204094225 |
| ENSMUSG00000089357 | Mir2137 | 100316779 | miRNA | 1.449322262 | 1.204326561 | 1.15357931 | -1.043991124 | 0.296489496 | 1 | 1.204326561 |
| ENSMUSG00000035566 | Pcdh17 | 219228 | protein_coding | 267.7558049 | 1.204923316 | 0.171693423 | -7.017876956 | 2.25E-12 | 6.63E-11 | 1.204923316 |
| ENSMUSG00000046886 | Zfp474 | 66758 | protein_coding | 1.118115289 | -1.205298933 | 2.244780076 | 0.536934083 | 0.591313147 | 1 | 1.205298933 |
| ENSMUSG00000033196 | Myh2 | 17882 | protein_coding | 2.279354832 | -1.205535809 | 2.246253601 | 0.536687313 | 0.591483621 | 1 | 1.205535809 |
| ENSMUSG00000051067 | Lingo3 | 237403 | protein_coding | 4.314685758 | 1.205660165 | 1.147726146 | -1.050477214 | 0.293498762 | 1 | 1.205660165 |
| ENSMUSG00000026567 | Adcy10 | 271639 | protein_coding | 4.906317918 | 1.205827181 | 1.040315708 | -1.159097351 | 0.246416505 | 0.458272537 | 1.205827181 |
| ENSMUSG00000044966 | Fbxo48 | 319701 | protein_coding | 8.525308757 | 1.205929978 | 0.50898018 | -2.369306361 | 0.017821484 | 0.066917689 | 1.205929978 |
| ENSMUSG00000030074 | Gxylt2 | 232313 | protein_coding | 4.366874963 | -1.206228992 | 0.961495608 | 1.254534063 | 0.209647953 | 1 | 1.206228992 |

| ensembl_gene_id | external_gene_name | entrezgene_id | gene_biotype | baseMean | log2FoldChange | lfcSE | stat | pvalue | padj | absLog2FC |
|---|---|---|---|---|---|---|---|---|---|---|
| ENSMUSG00000010796 | Asz1 | 74068 | protein_coding | 1.486623387 | 1.206608612 | 1.672440981 | -0.721465586 | 0.470623107 | 1 | 1.206608612 |
| ENSMUSG00000022537 | Tmem44 | 224090 | protein_coding | 67.53219862 | -1.206751334 | 0.184175399 | 6.552185258 | 5.67E-11 | 1.48E-09 | 1.206751334 |
| ENSMUSG00000085891 | Gm14634 | 552913 | lncRNA | 1.716162412 | 1.208210341 | 1.063377418 | -1.136200863 | 0.255872509 | 1 | 1.208210341 |
| ENSMUSG00000096330 | Gm21976 | 102637808 | transcribed_unprocessed_pseudogene | 1.711828582 | 1.20822665 | 1.552099988 | -0.778446401 | 0.436305893 | 1 | 1.20822665 |
| ENSMUSG00000112068 | Gm9182 | 668459 | processed_pseudogene | 0.534171647 | 1.208387936 | 1.648650725 | -0.732955694 | 0.463585453 | 1 | 1.208387936 |
| ENSMUSG00000042066 | Tmcc2 | 68875 | protein_coding | 4.433574502 | -1.209653231 | 0.915062374 | 1.321935275 | 0.186189705 | 0.38181259 | 1.209653231 |
| ENSMUSG00000074061 | Fbxw19 | 235612 | protein_coding | 1.514745747 | -1.209762248 | 2.273718785 | 0.532063269 | 0.594682176 | 1 | 1.209762248 |
| ENSMUSG00000074505 | Fat3 | 270120 | protein_coding | 6.805762269 | 1.210281989 | 2.761277366 | -0.438305114 | 0.66116512 | 0.816884691 | 1.210281989 |
| ENSMUSG00000112188 | Gm47708 | 102641383 | lncRNA | 15.32279974 | -1.210424796 | 0.367747421 | 3.291456926 | 0.000996699 | 0.006345534 | 1.210424796 |
| ENSMUSG00000108094 | Gm31108 | 102633311 | lncRNA | 2.214570217 | -1.210524524 | 0.839743629 | 1.441540586 | 0.14943202 | 1 | 1.210524524 |
| ENSMUSG00000112762 | 4930459C07Rik | 74881 | lncRNA | 1.470904163 | 1.210937295 | 1.158708541 | -1.045074971 | 0.295988321 | 1 | 1.210937295 |
| ENSMUSG00000097321 | 1700028E10Rik | 625175 | lncRNA | 9.161399138 | 1.211132988 | 0.583600698 | -2.075276796 | 0.037960889 | 0.120950437 | 1.211132988 |
| ENSMUSG00000069515 | Lyz1 | 17110 | protein_coding | 2.8487006 | 1.211750338 | 1.231876136 | -0.983662482 | 0.325281492 | 1 | 1.211750338 |
| ENSMUSG00000027350 | Chgb | 12653 | protein_coding | 1087.661617 | -1.211921318 | 0.185315124 | 6.539786366 | 6.16E-11 | 1.59E-09 | 1.211921318 |
| ENSMUSG00000023918 | Adgrf4 | 78249 | protein_coding | 1.048269821 | -1.212652595 | 1.611642818 | 0.752432599 | 0.451790949 | 1 | 1.212652595 |
| ENSMUSG00000029408 | Abcb9 | 56325 | protein_coding | 246.0695413 | 1.213290249 | 0.094089725 | -12.89503449 | 4.80E-38 | 4.53E-36 | 1.213290249 |
| ENSMUSG00000033361 | Prrg3 | 208748 | protein_coding | 3.094172138 | 1.213488638 | 2.388341984 | -0.508088308 | 0.611391414 | 1 | 1.213488638 |
| ENSMUSG00000054072 | Iigp1 | 60440 | protein_coding | 24.98528187 | 1.213593757 | 0.74456915 | -1.629927531 | 0.103116814 | 0.253177382 | 1.213593757 |
| ENSMUSG00000042662 | Dusp15 | 252864 | protein_coding | 2.937767333 | 1.21380942 | 2.255752203 | -0.538095195 | 0.59051133 | 1 | 1.21380942 |
| ENSMUSG00000003617 | Cp | 12870 | protein_coding | 5.063433856 | 1.214256651 | 0.567300382 | -2.140412189 | 0.03232147 | 0.10685821 | 1.214256651 |
| ENSMUSG00000025724 | Sec11a | 56529 | protein_coding | 1057.144432 | 1.214274262 | 0.074923107 | -16.20693942 | 4.50E-59 | 7.37E-57 | 1.214274262 |
| ENSMUSG00000016283 | H2-M2 | 14990 | protein_coding | 1.575688809 | 1.21442421 | 1.225699294 | -0.990801101 | 0.321782712 | 1 | 1.21442421 |
| ENSMUSG00000027801 | Tm4sf4 | 229302 | protein_coding | 10.90364825 | -1.214895959 | 0.436896837 | 2.780738738 | 0.005423537 | 0.026043699 | 1.214895959 |
| ENSMUSG00000103770 | Pcdha9 | 192161 | protein_coding | 0.537960488 | -1.215008066 | 2.032428658 | 0.59781093 | 0.549966097 | 1 | 1.215008066 |
| ENSMUSG00000093619 | 4930535L15Rik | 78027 | lncRNA | 0.871011464 | 1.215218266 | 1.356266853 | -0.896002334 | 0.370251517 | 1 | 1.215218266 |
| ENSMUSG00000028348 | Cavin4 | 68016 | protein_coding | 3.240434741 | 1.215392238 | 0.759259575 | -1.600759843 | 0.109430121 | 1 | 1.215392238 |
| ENSMUSG00000051980 | Casr | 12374 | protein_coding | 5.167958659 | 1.215662138 | 0.804118978 | -1.511793866 | 0.13058631 | 0.298680373 | 1.215662138 |
| ENSMUSG00000071547 | Nt5dc2 | 70021 | protein_coding | 48.13390787 | 1.216262177 | 0.239186745 | -5.084989869 | 3.68E-07 | 5.56E-06 | 1.216262177 |
| ENSMUSG00000086266 | Igf2os | 111975 | lncRNA | 2.352740016 | 1.216269991 | 1.551591942 | -0.78388522 | 0.433107463 | 1 | 1.216269991 |
| ENSMUSG00000001642 | Akr1b3 | 11677 | protein_coding | 798.9666704 | 1.2164239 | 0.153468064 | -7.926234755 | 2.26E-15 | 8.46E-14 | 1.2164239 |
| ENSMUSG00000073786 | Gm7579 | 105243090 | protein_coding | 0.723080373 | 1.216541821 | 1.681780855 | -0.723365245 | 0.469455518 | 1 | 1.216541821 |
| ENSMUSG00000051811 | Cox6b2 | 333182 | protein_coding | 14.39543612 | -1.217674865 | 0.404222448 | 3.012388029 | 0.00259201 | 0.014043458 | 1.217674865 |
| ENSMUSG00000039781 | Cep131 | 12009 | protein_coding | 295.5806933 | 1.218231392 | 0.127735184 | -9.537163955 | 1.47E-21 | 8.16E-20 | 1.218231392 |
| ENSMUSG00000029428 | Stx2 | 13852 | protein_coding | 11.03347127 | 1.218298713 | 0.388844746 | -3.133123757 | 0.001729565 | 0.01004291 | 1.218298713 |
| ENSMUSG00000044534 | Ackr2 | 59289 | protein_coding | 6.775507274 | 1.218547651 | 0.589323021 | -2.067707534 | 0.038667533 | 0.122751949 | 1.218547651 |
| ENSMUSG00000028036 | Ptgfr | 19220 | protein_coding | 5.358244265 | -1.218943506 | 0.683950496 | 1.782210134 | 0.074714971 | 0.199909758 | 1.218943506 |
| ENSMUSG00000046130 | Vmn1r184 | 100312477 | protein_coding | 1.267043637 | 1.219610627 | 2.37920306 | -0.512613088 | 0.608221996 | 1 | 1.219610627 |
| ENSMUSG00000041044 | Lrit1 | 239037 | protein_coding | 0.79425627 | -1.219807861 | 2.128351519 | 0.573123307 | 0.566561207 | 1 | 1.219807861 |
| ENSMUSG00000028602 | Tnfrsf8 | 21941 | protein_coding | 18.00678159 | 1.220317078 | 0.393429802 | -3.101740318 | 0.001923866 | 0.01100288 | 1.220317078 |

| ensembl_gene_id | external_gene_name | entrezgene_id | gene_biotype | baseMean | log2FoldChange | lfcSE | stat | pvalue | padj | absLog2FC |
|---|---|---|---|---|---|---|---|---|---|---|
| ENSMUSG00000068859 | Sp9 | 381373 | protein_coding | 23.26318018 | 1.221029954 | 0.578883588 | -2.109284112 | 0.034920065 | 0.113250188 | 1.221029954 |
| ENSMUSG00000039787 | Cercam | 99151 | protein_coding | 2.366456539 | 1.221558226 | 1.026539068 | -1.189977337 | 0.2340553 | 1 | 1.221558226 |
| ENSMUSG00000033526 | Ppip5k1 | 327655 | protein_coding | 2217.254359 | 1.221655722 | 0.052384692 | -23.32085327 | 2.72E-120 | 1.18E-117 | 1.221655722 |
| ENSMUSG00000034987 | Hrh2 | 15466 | protein_coding | 4.722039259 | 1.221696325 | 1.16684896 | -1.047004682 | 0.295097419 | 0.514264394 | 1.221696325 |
| ENSMUSG00000055838 | Or1q1 | 258616 | protein_coding | 1.03759238 | -1.222111822 | 2.210046627 | 0.552980108 | 0.580277032 | 1 | 1.222111822 |
| ENSMUSG00000044041 | Krt13 | 16663 | protein_coding | 3.217450758 | 1.223667455 | 0.703561844 | -1.739246473 | 0.081991418 | 1 | 1.223667455 |
| ENSMUSG00000038048 | Cntnap5c | 620292 | protein_coding | 2.153394808 | 1.223894397 | 1.508386654 | -0.811393017 | 0.417140008 | 1 | 1.223894397 |
| ENSMUSG00000048402 | Gli2 | 14633 | protein_coding | 3.5801337 | 1.224118591 | 1.826941704 | -0.670037029 | 0.502834185 | 1 | 1.224118591 |
| ENSMUSG00000090061 | Nwd2 | 319807 | protein_coding | 2.528404405 | 1.225175003 | 1.194749241 | -1.025466233 | 0.30514325 | 1 | 1.225175003 |
| ENSMUSG00000053334 | Ficd | 231630 | protein_coding | 205.1061131 | 1.225347353 | 0.109914181 | -11.14821896 | 7.31E-29 | 5.46E-27 | 1.225347353 |
| ENSMUSG00000053044 | Cd8b1 | 12526 | protein_coding | 1.728799973 | -1.226081399 | 1.157541804 | 1.059211334 | 0.289503544 | 1 | 1.226081399 |
| ENSMUSG00000033965 | Slc16a2 | 20502 | protein_coding | 20.62565035 | -1.226098715 | 0.299720222 | 4.090810784 | 4.30E-05 | 0.000409864 | 1.226098715 |
| ENSMUSG00000025007 | Aldh18a1 | 56454 | protein_coding | 1100.503522 | 1.226820179 | 0.122039862 | -10.05261857 | 8.95E-24 | 5.51E-22 | 1.226820179 |
| ENSMUSG00000091685 | Gm17359 | 100233207 | protein_coding | 1.418095149 | -1.226871752 | 1.550269503 | 0.791392561 | 0.42871495 | 1 | 1.226871752 |
| ENSMUSG00000009654 | Oit3 | 18302 | protein_coding | 1.969659998 | 1.22717128 | 0.966711746 | -1.269428333 | 0.204288336 | 1 | 1.22717128 |
| ENSMUSG00000028356 | Ambp | 11699 | protein_coding | 20.03339464 | -1.228134005 | 0.333905927 | 3.678083872 | 0.000234993 | 0.001826197 | 1.228134005 |
| ENSMUSG00000035486 | Plk5 | 216166 | protein_coding | 3.017239998 | -1.228137737 | 0.881245635 | 1.393638377 | 0.163426833 | 1 | 1.228137737 |
| ENSMUSG00000017400 | Stac2 | 217154 | protein_coding | 2.710132417 | 1.228324303 | 2.205990386 | -0.556813081 | 0.577655146 | 1 | 1.228324303 |
| ENSMUSG00000026347 | Tmem163 | 72160 | protein_coding | 5.181298039 | -1.228395303 | 0.605045303 | 2.030253431 | 0.042330785 | 0.131303585 | 1.228395303 |
| ENSMUSG00000111611 | Or13c7c | 100040268 | protein_coding | 1.234839611 | -1.22994011 | 2.072229935 | 0.593534573 | 0.552823452 | 1 | 1.22994011 |
| ENSMUSG00000101009 | 1700108F19Rik | 73272 | lncRNA | 7.277219143 | 1.230152226 | 0.429476402 | -2.864306911 | 0.004179229 | 0.020901206 | 1.230152226 |
| ENSMUSG00000096718 | Zfp781 | 331188 | protein_coding | 1.066010125 | -1.230604802 | 1.413694413 | 0.870488552 | 0.384033473 | 1 | 1.230604802 |
| ENSMUSG00000021176 | Efcab11 | 78767 | protein_coding | 35.44715669 | 1.230798197 | 0.213051323 | -5.777003299 | 7.60E-09 | 1.49E-07 | 1.230798197 |
| ENSMUSG00000032818 | Loxhd1 | 240411 | protein_coding | 3.898593083 | 1.230875395 | 1.837364867 | -0.669913427 | 0.50291298 | 1 | 1.230875395 |
| ENSMUSG00000039577 | Nphp4 | 260305 | protein_coding | 6.583370241 | 1.230992545 | 0.604427126 | -2.036626903 | 0.041687445 | 0.129891785 | 1.230992545 |
| ENSMUSG00000030017 | Reg3g | 19695 | protein_coding | 2678.6097 | 1.231647202 | 0.517558154 | -2.379727172 | 0.01732546 | 0.065583696 | 1.231647202 |
| ENSMUSG00000017195 | Zpbp2 | 69376 | protein_coding | 1.450859149 | -1.23186727 | 1.152904387 | 1.068490401 | 0.285299358 | 1 | 1.23186727 |
| ENSMUSG00000036634 | Mag | 17136 | protein_coding | 4.267552005 | -1.232184668 | 0.806503293 | 1.527811081 | 0.12655945 | 1 | 1.232184668 |
| ENSMUSG00000070799 | Psg26 | 574429 | protein_coding | 3.023911641 | 1.23256742 | 0.928662736 | -1.327249788 | 0.184426068 | 1 | 1.23256742 |
| ENSMUSG00000040828 | Catsperd | 106757 | protein_coding | 5.903762155 | -1.233214455 | 0.680921421 | 1.811096577 | 0.070125903 | 0.191288267 | 1.233214455 |
| ENSMUSG00000036211 | H1f6 | 107970 | protein_coding | 15.14887436 | 1.233225083 | 0.532151879 | -2.317430664 | 0.020480284 | 0.074722558 | 1.233225083 |
| ENSMUSG00000006948 | Klk4 | 56640 | protein_coding | 1.616021246 | 1.233947612 | 2.112994189 | -0.583980599 | 0.559233365 | 1 | 1.233947612 |
| ENSMUSG00000073154 | 9330158H04Rik | 319472 | lncRNA | 4.421931364 | 1.234040885 | 3.08346926 | -0.400211833 | 0.6890005 | 0.833930977 | 1.234040885 |
| ENSMUSG00000027022 | Xirp2 | 241431 | protein_coding | 2.46397427 | 1.234386702 | 2.003832318 | -0.616012973 | 0.537885958 | 1 | 1.234386702 |
| ENSMUSG00000031519 | Asb5 | 76294 | protein_coding | 4.57211953 | 1.234702054 | 0.68687805 | -1.797556428 | 0.072247328 | 0.195556857 | 1.234702054 |
| ENSMUSG00000060429 | Sntb1 | 20649 | protein_coding | 12.72165708 | 1.234797897 | 0.428726536 | -2.880152717 | 0.003974826 | 0.020091911 | 1.234797897 |
| ENSMUSG00000014846 | Tppp3 | 67971 | protein_coding | 538.3517422 | -1.23506993 | 0.177064884 | 6.975239265 | 3.05E-12 | 8.94E-11 | 1.23506993 |
| ENSMUSG00000044141 | E130201H02Rik | 78552 | processed_pseudogene | 0.967876224 | 1.235167738 | 2.226932285 | -0.554649886 | 0.57913416 | 1 | 1.235167738 |
| ENSMUSG00000022441 | Efcab6 | 77627 | protein_coding | 4.145569297 | 1.235209262 | 3.556413944 | -0.347318755 | 0.728351862 | 1 | 1.235209262 |

| ensembl_gene_id | external_gene_name | entrezgene_id | gene_biotype | baseMean | log2FoldChange | lfcSE | stat | pvalue | padj | absLog2FC |
|---|---|---|---|---|---|---|---|---|---|---|
| ENSMUSG00000078938 | Syce3 | 75459 | protein_coding | 5.405622151 | -1.235332601 | 0.544654992 | 2.268101128 | 0.023323044 | 0.082758217 | 1.235332601 |
| ENSMUSG00000023120 | Ldc1 | 332942 | protein_coding | 1.543450454 | -1.235433671 | 2.256631987 | 0.547467943 | 0.584057289 | 1 | 1.235433671 |
| ENSMUSG00000072662 | Mansc4 | 545893 | protein_coding | 2.334223581 | -1.236534074 | 0.812546049 | 1.521801842 | 0.12805874 | 1 | 1.236534074 |
| ENSMUSG00000020363 | Gfpt2 | 14584 | protein_coding | 5.261656329 | 1.236551779 | 0.86014884 | -1.437602101 | 0.150546985 | 0.330514877 | 1.236551779 |
| ENSMUSG00000028003 | Lrat | 79235 | protein_coding | 107.6049396 | -1.236684633 | 0.156130462 | 7.920841434 | 2.36E-15 | 8.81E-14 | 1.236684633 |
| ENSMUSG00000087695 | Gm16291 | 100503072 | lncRNA | 1.506243438 | 1.237182731 | 3.127107382 | -0.395631675 | 0.692376767 | 1 | 1.237182731 |
| ENSMUSG00000073386 | 9830107B12Rik | 328829 | protein_coding | 1.725754572 | -1.237245317 | 2.487048956 | 0.497475256 | 0.618853948 | 1 | 1.237245317 |
| ENSMUSG00000094520 | Or51q1 | 259122 | protein_coding | 2.520673683 | 1.238197624 | 2.710750819 | -0.456772941 | 0.647834257 | 1 | 1.238197624 |
| ENSMUSG00000111602 | Gm32014 | 102634431 | lncRNA | 1.452896897 | -1.239686079 | 2.509619709 | 0.493973678 | 0.621324769 | 1 | 1.239686079 |
| ENSMUSG00000048814 | Lonrf2 | 381338 | protein_coding | 1.543890661 | -1.239691759 | 2.509668164 | 0.493966405 | 0.621329906 | 1 | 1.239691759 |
| ENSMUSG00000078670 | Fam174b | 100038347 | protein_coding | 1603.348307 | 1.239832475 | 0.123562102 | -10.03408372 | 1.08E-23 | 6.63E-22 | 1.239832475 |
| ENSMUSG00000070601 | Vmn2r84 | 625068 | protein_coding | 1.630967395 | -1.239944091 | 2.512030622 | 0.493602299 | 0.621587077 | 1 | 1.239944091 |
| ENSMUSG00000046934 | Csl | 71832 | protein_coding | 1.879308628 | 1.240892662 | 0.970863101 | -1.278133509 | 0.201202356 | 1 | 1.240892662 |
| ENSMUSG00000053137 | Mapk11 | 19094 | protein_coding | 10.61569818 | -1.24135249 | 0.523132361 | 2.372922383 | 0.017647972 | 0.066568892 | 1.24135249 |
| ENSMUSG00000073102 | Drc1 | 381738 | protein_coding | 10.62338251 | -1.241354517 | 0.378884747 | 3.276338063 | 0.001051626 | 0.006636053 | 1.241354517 |
| ENSMUSG00000026955 | Sapcd2 | 72080 | protein_coding | 131.2306225 | 1.241717499 | 0.191467215 | -6.485274766 | 8.86E-11 | 2.24E-09 | 1.241717499 |
| ENSMUSG00000111035 | Gm33054 | 102635808 | lncRNA | 1.346050731 | -1.241731032 | 1.527413663 | 0.812963156 | 0.416239183 | 1 | 1.241731032 |
| ENSMUSG00000053049 | Gm15413 | 791381 | lncRNA | 1.238418841 | 1.241807211 | 1.471716158 | -0.843781733 | 0.398791385 | 1 | 1.241807211 |
| ENSMUSG00000014529 | Tmbim7 | 75010 | protein_coding | 1.668118468 | 1.241949519 | 2.116562282 | -0.586776742 | 0.557353659 | 1 | 1.241949519 |
| ENSMUSG00000050321 | Neto1 | 246317 | protein_coding | 1.85861407 | 1.242454041 | 2.11741653 | -0.586778285 | 0.557352622 | 1 | 1.242454041 |
| ENSMUSG00000041653 | Pnpla3 | 116939 | protein_coding | 2.903852172 | -1.242609594 | 0.89508616 | 1.388256963 | 0.165058812 | 1 | 1.242609594 |
| ENSMUSG00000115705 | Vmn1r28 | 171198 | protein_coding | 2.054057768 | -1.242695851 | 2.538412394 | 0.489556328 | 0.624447886 | 1 | 1.242695851 |
| ENSMUSG00000046750 | Selenov | 280621 | protein_coding | 1.825250491 | -1.243153468 | 2.542904197 | 0.488871531 | 0.624932652 | 1 | 1.243153468 |
| ENSMUSG00000045648 | Vwc2l | 320460 | protein_coding | 1.409587362 | -1.24353889 | 2.546722354 | 0.488289934 | 0.625344491 | 1 | 1.24353889 |
| ENSMUSG00000026983 | Il36rn | 54450 | protein_coding | 1.22685897 | -1.243701273 | 2.548341217 | 0.488043463 | 0.625519056 | 1 | 1.243701273 |
| ENSMUSG00000036356 | Csgalnact1 | 234356 | protein_coding | 3.403085889 | 1.243753377 | 1.115198551 | -1.115275281 | 0.264732475 | 1 | 1.243753377 |
| ENSMUSG00000097682 | 4930554H23Rik | 77631 | lncRNA | 1.295520006 | -1.244078224 | 2.552072232 | 0.487477669 | 0.625919864 | 1 | 1.244078224 |
| ENSMUSG00000041380 | Htr2c | 15560 | protein_coding | 1.227476216 | -1.244300687 | 2.55429903 | 0.487139788 | 0.626159272 | 1 | 1.244300687 |
| ENSMUSG00000098414 | Mir6909 | 102465548 | miRNA | 0.905462659 | -1.244729877 | 1.321663936 | 0.941789999 | 0.346300166 | 1 | 1.244729877 |
| ENSMUSG00000026807 | Ak8 | 68870 | protein_coding | 1.907461003 | 1.244833698 | 1.487203847 | -0.837029638 | 0.402575909 | 1 | 1.244833698 |
| ENSMUSG00000045780 | Or51v8 | 258189 | protein_coding | 1.275160445 | -1.244885241 | 2.561436695 | 0.486010544 | 0.62695969 | 1 | 1.244885241 |
| ENSMUSG00000057003 | Myh4 | 17884 | protein_coding | 1.275160445 | -1.244885241 | 2.561436695 | 0.486010544 | 0.62695969 | 1 | 1.244885241 |
| ENSMUSG00000066058 | Cldn19 | 242653 | protein_coding | 2.74171 | 1.246162585 | 2.393992397 | -0.520537403 | 0.602689066 | 1 | 1.246162585 |
| ENSMUSG00000065232 | Gm22973 | 115488085 | snRNA | 128.4120056 | 1.246793204 | 0.256210551 | -4.866283604 | 1.14E-06 | 1.55E-05 | 1.246793204 |
| ENSMUSG00000025151 | Maged1 | 94275 | protein_coding | 1644.000355 | 1.247008094 | 0.106709678 | -11.68598871 | 1.50E-31 | 1.24E-29 | 1.247008094 |
| ENSMUSG00000068874 | Selenbp1 | 20341 | protein_coding | 10787.17891 | -1.247250538 | 0.122602491 | 10.17312559 | 2.61E-24 | 1.64E-22 | 1.247250538 |
| ENSMUSG00000032087 | Dscaml1 | 114873 | protein_coding | 6.729054019 | -1.247281642 | 0.955370367 | 1.305547759 | 0.191706396 | 0.389463531 | 1.247281642 |
| ENSMUSG00000020321 | Mdh1 | 17449 | protein_coding | 8006.177835 | 1.247367001 | 0.074124194 | -16.82806829 | 1.52E-63 | 2.73E-61 | 1.247367001 |
| ENSMUSG00000039488 | Cntn5 | 244682 | protein_coding | 1.377344507 | -1.247960141 | 2.141456233 | 0.582762385 | 0.56005327 | 1 | 1.247960141 |

| ensembl_gene_id | external_gene_name | entrezgene_id | gene_biotype | baseMean | log2FoldChange | lfcSE | stat | pvalue | padj | absLog2FC |
|---|---|---|---|---|---|---|---|---|---|---|
| ENSMUSG00000034706 | Dnai2 | 432611 | protein_coding | 2.734410656 | 1.248816367 | 1.183901476 | -1.054831329 | 0.291502474 | 1 | 1.248816367 |
| ENSMUSG00000095524 | Gm15682 | 108169053 | processed_pseudogene | 0.991781519 | -1.24901795 | 1.619129129 | 0.771413427 | 0.440461919 | 1 | 1.24901795 |
| ENSMUSG00000025478 | Dpysl4 | 26757 | protein_coding | 4.029433614 | 1.24919737 | 2.300978213 | -0.542898391 | 0.587199764 | 1 | 1.24919737 |
| ENSMUSG00000067855 | Speer3 | 71026 | protein_coding | 1.421518522 | -1.249233464 | 2.14810884 | 0.581550358 | 0.560869589 | 1 | 1.249233464 |
| ENSMUSG00000075224 | Lrrc55 | 241528 | protein_coding | 2.138400625 | 1.249482307 | 2.3741942 | -0.526276371 | 0.598696198 | 1 | 1.249482307 |
| ENSMUSG00000002603 | Tgfb1 | 21803 | protein_coding | 25.90605244 | -1.249659055 | 0.445905546 | 2.802519648 | 0.005070513 | 0.024649528 | 1.249659055 |
| ENSMUSG00000019039 | Dalrd3 | 67789 | protein_coding | 638.3880739 | 1.249801693 | 0.094259136 | -13.25921018 | 3.99E-40 | 4.17E-38 | 1.249801693 |
| ENSMUSG00000033987 | Dnah17 | 69926 | protein_coding | 31.74854177 | 1.25007412 | 0.455863771 | -2.742209842 | 0.006102734 | 0.028549731 | 1.25007412 |
| ENSMUSG00000029299 | Abcg3 | 27405 | protein_coding | 2.725246323 | 1.250368351 | 1.037320778 | -1.205382537 | 0.22805566 | 1 | 1.250368351 |
| ENSMUSG00000029212 | Gabrb1 | 14400 | protein_coding | 6.446166089 | 1.250902715 | 1.049471066 | -1.191936354 | 0.233286202 | 0.442718029 | 1.250902715 |
| ENSMUSG00000074802 | Gas2l3 | 237436 | protein_coding | 212.9989843 | 1.250982448 | 0.135691917 | -9.219284932 | 2.99E-20 | 1.52E-18 | 1.250982448 |
| ENSMUSG00000073430 | Gm10505 | 100038429 | lncRNA | 3.90665951 | 1.251399223 | 1.404524023 | -0.890977443 | 0.372941272 | 1 | 1.251399223 |
| ENSMUSG00000078302 | Foxd1 | 15229 | protein_coding | 2.335361797 | 1.251565036 | 1.824324361 | -0.686043043 | 0.492685966 | 1 | 1.251565036 |
| ENSMUSG00000030410 | Dmwd | 13401 | protein_coding | 23.0299163 | 1.252016689 | 0.263756796 | -4.746860394 | 2.07E-06 | 2.68E-05 | 1.252016689 |
| ENSMUSG00000090173 | Fbxw10 | 213980 | protein_coding | 14.87623505 | 1.252068254 | 0.548082497 | -2.284452178 | 0.02234497 | 0.080148748 | 1.252068254 |
| ENSMUSG00000079492 | Gm11127 | 100529082 | protein_coding | 0.656671815 | -1.252173552 | 1.868686197 | 0.670082304 | 0.502805325 | 1 | 1.252173552 |
| ENSMUSG00000029767 | Calu | 12321 | protein_coding | 3095.978611 | 1.25245281 | 0.06116152 | -20.47779086 | 3.40E-93 | 1.06E-90 | 1.25245281 |
| ENSMUSG00000100863 | Gm12669 | 620016 | processed_pseudogene | 6.621959398 | 1.252864203 | 0.494868475 | -2.531711486 | 0.011350734 | 0.047144373 | 1.252864203 |
| ENSMUSG00000040706 | Agmat | 75986 | protein_coding | 12.25459479 | 1.252877927 | 0.3680603 | -3.40400181 | 0.000664063 | 0.004494401 | 1.252877927 |
| ENSMUSG00000040310 | Alx4 | 11695 | protein_coding | 3.617152238 | 1.254356079 | 1.451295433 | -0.864300989 | 0.387422567 | 1 | 1.254356079 |
| ENSMUSG00000073555 | Gm4951 | 240327 | protein_coding | 0.897012077 | 1.25478849 | 1.432176007 | -0.876141259 | 0.380953245 | 1 | 1.25478849 |
| ENSMUSG00000028845 | Tekt2 | 24084 | protein_coding | 2.777894932 | 1.255356072 | 1.145978175 | -1.095445009 | 0.273321725 | 1 | 1.255356072 |
| ENSMUSG00000023031 | Cela1 | 109901 | protein_coding | 1240.7771 | -1.255656282 | 0.200065304 | 6.276232106 | 3.47E-10 | 8.13E-09 | 1.255656282 |
| ENSMUSG00000086918 | 4930429F24Rik | 74633 | lncRNA | 2.431995993 | 1.256206269 | 0.971780072 | -1.292685767 | 0.196119762 | 1 | 1.256206269 |
| ENSMUSG00000005583 | Mef2c | 17260 | protein_coding | 4.513671626 | 1.256342981 | 1.596186672 | -0.787090259 | 0.431229032 | 0.649597228 | 1.256342981 |
| ENSMUSG00000017002 | Slpi | 20568 | protein_coding | 30.54990675 | 1.256774603 | 0.46833807 | -2.683477352 | 0.007286091 | 0.032955362 | 1.256774603 |
| ENSMUSG00000066537 | Vmn2r57 | 269902 | protein_coding | 1.217187066 | -1.257027768 | 0.955293013 | 1.315855712 | 0.188222476 | 1 | 1.257027768 |
| ENSMUSG00000028222 | Calb1 | 12307 | protein_coding | 0.952296021 | -1.257227664 | 2.195887583 | 0.572537353 | 0.566957988 | 1 | 1.257227664 |
| ENSMUSG00000026058 | Khdrbs2 | 170771 | protein_coding | 2.440254699 | 1.257611701 | 2.165377866 | -0.580781637 | 0.561387633 | 1 | 1.257611701 |
| ENSMUSG00000037159 | Wee2 | 381759 | protein_coding | 1.846762458 | -1.257624009 | 1.007721741 | 1.247987373 | 0.212035681 | 1 | 1.257624009 |
| ENSMUSG00000096986 | 4930509E16Rik | 75115 | lncRNA | 20.61589951 | 1.258872688 | 0.260898934 | -4.825135416 | 1.40E-06 | 1.87E-05 | 1.258872688 |
| ENSMUSG00000062542 | Syt9 | 60510 | protein_coding | 2.483758161 | 1.259744827 | 1.285679609 | -0.979827959 | 0.327171048 | 1 | 1.259744827 |
| ENSMUSG00000021364 | Elovl2 | 54326 | protein_coding | 2.457132217 | 1.259802323 | 1.301400582 | -0.968035777 | 0.333026502 | 1 | 1.259802323 |
| ENSMUSG00000048502 | Duxbl1 | 278672 | protein_coding | 0.741456417 | -1.259965774 | 1.283401075 | 0.98173969 | 0.326228107 | 1 | 1.259965774 |
| ENSMUSG00000087434 | Rab11fip4os1 | 790912 | lncRNA | 1.052441034 | 1.260024323 | 2.651635557 | -0.475187595 | 0.634653268 | 1 | 1.260024323 |
| ENSMUSG00000117662 | Gm4013 | 100042757 | lncRNA | 3.048474415 | -1.260162072 | 0.647881487 | 1.945050287 | 0.051768926 | 1 | 1.260162072 |
| ENSMUSG00000051242 | Pcdhb9 | 93880 | protein_coding | 1.106149877 | -1.260373131 | 2.209974482 | 0.570311169 | 0.568466667 | 1 | 1.260373131 |
| ENSMUSG00000038151 | Prdm1 | 12142 | protein_coding | 51.08002697 | 1.262922 | 0.367358088 | -3.43784999 | 0.000586352 | 0.004051748 | 1.262922 |
| ENSMUSG00000045672 | Col27a1 | 373864 | protein_coding | 19.56430984 | 1.263023566 | 0.567103347 | -2.227148848 | 0.025937327 | 0.089909535 | 1.263023566 |

Supplementary Table 1

| ensembl_gene_id | external_gene_name | entrezgene_id | gene_biotype | baseMean | log2FoldChange | lfcSE | stat | pvalue | padj | absLog2FC |
|---|---|---|---|---|---|---|---|---|---|---|
| ENSMUSG00000063063 | Ctnna2 | 12386 | protein_coding | 5.422728015 | -1.263074066 | 0.767568674 | 1.64555187 | 0.099856055 | 0.247378172 | 1.263074066 |
| ENSMUSG00000102692 | Dchs2 | 100534287 | protein_coding | 3.704670798 | 1.263166277 | 1.592297616 | -0.793297851 | 0.427604308 | 1 | 1.263166277 |
| ENSMUSG00000085831 | Lncbate3 | 72229 | lncRNA | 0.595141039 | -1.263541308 | 1.424039188 | 0.887293916 | 0.37492068 | 1 | 1.263541308 |
| ENSMUSG00000024940 | Ltbp3 | 16998 | protein_coding | 9.181812642 | 1.263996846 | 0.784600802 | -1.611006315 | 0.107178346 | 0.26018046 | 1.263996846 |
| ENSMUSG00000078875 | Gm14419 | 628147 | protein_coding | 0.475615093 | -1.264025665 | 1.730807135 | 0.730309946 | 0.46520075 | 1 | 1.264025665 |
| ENSMUSG00000021719 | Rgs7bp | 52882 | protein_coding | 6.301719493 | 1.264429286 | 0.600766439 | -2.104693609 | 0.035317987 | 0.114316197 | 1.264429286 |
| ENSMUSG00000025875 | Tspan17 | 74257 | protein_coding | 71.78974895 | -1.264995287 | 0.13906667 | 9.096322577 | 9.34E-20 | 4.68E-18 | 1.264995287 |
| ENSMUSG00000052336 | Cx3cr1 | 13051 | protein_coding | 6.695689402 | 1.265325119 | 0.672308656 | -1.882059837 | 0.059827892 | 0.170406218 | 1.265325119 |
| ENSMUSG00000021732 | Fgf10 | 14165 | protein_coding | 1.816483526 | 1.26652749 | 1.365218718 | -0.927710317 | 0.353557855 | 1 | 1.26652749 |
| ENSMUSG00000032554 | Trf | 22041 | protein_coding | 375.7941244 | 1.266750057 | 0.491021531 | -2.579825889 | 0.009885014 | 0.042192071 | 1.266750057 |
| ENSMUSG00000010592 | Dazl | 13164 | protein_coding | 3.708868644 | 1.266857763 | 0.615782676 | -2.057313095 | 0.039656119 | 1 | 1.266857763 |
| ENSMUSG00000116138 | C030006K11Rik | 223665 | protein_coding | 66.762265 | -1.266885626 | 0.147222945 | 8.605218591 | 7.62E-18 | 3.46E-16 | 1.266885626 |
| ENSMUSG00000077380 | Gm22661 | 115485922 | snoRNA | 1.8084343 | 1.266921841 | 0.89711969 | -1.412210495 | 0.157887991 | 1 | 1.266921841 |
| ENSMUSG00000074782 | 4833422C13Rik | 373852 | lncRNA | 1.658387158 | -1.267038999 | 1.481655548 | 0.855150848 | 0.392467643 | 1 | 1.267038999 |
| ENSMUSG00000076288 | Mir301b | 791069 | miRNA | 0.967675365 | 1.267997826 | 1.374607869 | -0.922443305 | 0.35629739 | 1 | 1.267997826 |
| ENSMUSG00000109392 | Gm5737 | 436008 | protein_coding | 0.956681534 | -1.268120971 | 1.957668673 | 0.647770988 | 0.517133082 | 1 | 1.268120971 |
| ENSMUSG00000025433 | Crisp3 | 11572 | protein_coding | 21.36333756 | -1.26819492 | 0.423376769 | 2.995428691 | 0.002740594 | 0.014751683 | 1.26819492 |
| ENSMUSG00000075528 | Aarsd1 | 69684 | protein_coding | 6.161400735 | 1.268423171 | 0.506288474 | -2.505336853 | 0.012233482 | 0.049808611 | 1.268423171 |
| ENSMUSG00000028864 | Hgf | 15234 | protein_coding | 6.532978304 | 1.269059236 | 0.634682908 | -1.999516956 | 0.045552449 | 0.139022943 | 1.269059236 |
| ENSMUSG00000049436 | Upk1b | 22268 | protein_coding | 33.04489601 | 1.269106945 | 0.247401812 | -5.129739885 | 2.90E-07 | 4.45E-06 | 1.269106945 |
| ENSMUSG00000053024 | Cntn2 | 21367 | protein_coding | 5.572201462 | 1.26930858 | 1.517849034 | -0.836254826 | 0.403011561 | 0.624327666 | 1.26930858 |
| ENSMUSG00000008318 | Relt | 320100 | protein_coding | 7.807805873 | 1.269546485 | 0.588301158 | -2.157987397 | 0.030928812 | 0.103308737 | 1.269546485 |
| ENSMUSG00000075102 | Or4c116 | 258904 | protein_coding | 1.201437499 | 1.269555286 | 2.326978761 | -0.545580951 | 0.585354021 | 1 | 1.269555286 |
| ENSMUSG00000069816 | Or1e17 | 18321 | protein_coding | 1.190229798 | 1.269716052 | 2.327598028 | -0.545504867 | 0.585406334 | 1 | 1.269716052 |
| ENSMUSG00000099974 | Bcl2a1d | 12047 | protein_coding | 0.715647515 | 1.269723467 | 1.421543371 | -0.893200653 | 0.371749728 | 1 | 1.269723467 |
| ENSMUSG00000036223 | Ska1 | 66468 | protein_coding | 49.0740378 | 1.269787229 | 0.248577254 | -5.108219716 | 3.25E-07 | 4.95E-06 | 1.269787229 |
| ENSMUSG00000066456 | Hmgn3 | 94353 | protein_coding | 4.905763004 | -1.269921347 | 0.554373629 | 2.290731881 | 0.021978926 | 0.079007396 | 1.269921347 |
| ENSMUSG00000094230 | Gm21847 | 74062 | lncRNA | 1.402458198 | -1.270105843 | 2.002659978 | 0.63420943 | 0.525944154 | 1 | 1.270105843 |
| ENSMUSG00000074489 | Bglap3 | 12095 | protein_coding | 18.76705753 | -1.27119305 | 0.371846261 | 3.418598448 | 0.000629445 | 0.004304208 | 1.27119305 |
| ENSMUSG00000009628 | Tex15 | 104271 | protein_coding | 4.695713567 | 1.271517038 | 0.692858135 | -1.835176603 | 0.066479527 | 0.184012672 | 1.271517038 |
| ENSMUSG00000053025 | Sv2b | 64176 | protein_coding | 2.436886744 | 1.272617246 | 1.599761687 | -0.795504266 | 0.426320231 | 1 | 1.272617246 |
| ENSMUSG00000030717 | Nupr1 | 56312 | protein_coding | 2990.504117 | 1.272679167 | 0.126897254 | -10.02920967 | 1.13E-23 | 6.91E-22 | 1.272679167 |
| ENSMUSG00000074203 | G430095P16Rik | 619292 | lncRNA | 1.814556294 | -1.27278996 | 0.9393059 | 1.355032434 | 0.175407224 | 1 | 1.27278996 |
| ENSMUSG00000070605 | Zfp992 | 433791 | protein_coding | 23.06652346 | -1.272961389 | 0.297559969 | 4.277999473 | 1.89E-05 | 0.000195572 | 1.272961389 |
| ENSMUSG00000055368 | Slc6a2 | 20538 | protein_coding | 2.102482861 | -1.273007098 | 2.911175485 | 0.437282845 | 0.661906235 | 1 | 1.273007098 |
| ENSMUSG00000059956 | Serpinb12 | 71869 | protein_coding | 0.679472352 | -1.273034978 | 1.57214774 | 0.809742587 | 0.418088136 | 1 | 1.273034978 |
| ENSMUSG00000026786 | Apbb1ip | 54519 | protein_coding | 21.52594642 | 1.273555414 | 0.382618937 | -3.328521646 | 0.000873082 | 0.005685363 | 1.273555414 |
| ENSMUSG00000055691 | Gja6 | 414089 | protein_coding | 1.299556463 | 1.273668807 | 1.058000416 | -1.20384528 | 0.228649385 | 1 | 1.273668807 |
| ENSMUSG00000065423 | Mir181a-2 | 387176 | miRNA | 0.775110588 | -1.275426418 | 1.411501077 | 0.903595781 | 0.366209779 | 1 | 1.275426418 |

Supplementary Table 1

| ensembl_gene_id | external_gene_name | entrezgene_id | gene_biotype | baseMean | log2FoldChange | lfcSE | stat | pvalue | padj | absLog2FC |
|---|---|---|---|---|---|---|---|---|---|---|
| ENSMUSG00000022309 | Angpt1 | 11600 | protein_coding | 1.322844675 | -1.276635485 | 2.969592939 | 0.429902519 | 0.667266553 | 1 | 1.276635485 |
| ENSMUSG00000051180 | Or6ae1 | 258954 | protein_coding | 1.322844675 | -1.276635485 | 2.969592939 | 0.429902519 | 0.667266553 | 1 | 1.276635485 |
| ENSMUSG00000032691 | Nlrp3 | 216799 | protein_coding | 2.109322129 | 1.276667741 | 1.600844332 | -0.797496493 | 0.425162739 | 1 | 1.276667741 |
| ENSMUSG00000031489 | Adrb3 | 11556 | protein_coding | 1.167092946 | 1.276710067 | 1.387686311 | -0.92002786 | 0.3575582 | 1 | 1.276710067 |
| ENSMUSG00000023011 | Faim2 | 72393 | protein_coding | 10.73124388 | 1.276768397 | 1.078371696 | -1.183978031 | 0.236421759 | 0.446602599 | 1.276768397 |
| ENSMUSG00000087095 | Emx2os | 329078 | lncRNA | 1.184290298 | -1.277066262 | 2.976810765 | 0.429004852 | 0.667919694 | 1 | 1.277066262 |
| ENSMUSG00000000183 | Fgf6 | 14177 | protein_coding | 1.370405287 | -1.277286216 | 2.980475376 | 0.428551172 | 0.668249886 | 1 | 1.277286216 |
| ENSMUSG00000108367 | Erfl | 232974 | protein_coding | 1.264432301 | 1.277489581 | 1.447124791 | -0.88277776 | 0.377356359 | 1 | 1.277489581 |
| ENSMUSG00000091059 | Vmn2r14 | 231591 | protein_coding | 1.093296534 | -1.277819923 | 2.989557878 | 0.427427725 | 0.669067817 | 1 | 1.277819923 |
| ENSMUSG00000020673 | Tpo | 22018 | protein_coding | 1.653577157 | 1.278355093 | 2.361323726 | -0.541372231 | 0.588251043 | 1 | 1.278355093 |
| ENSMUSG00000091043 | Glyatl3 | 435528 | protein_coding | 0.734993937 | -1.278598168 | 1.818733712 | 0.703015598 | 0.482046029 | 1 | 1.278598168 |
| ENSMUSG00000060568 | Fam78b | 226610 | protein_coding | 2.081315922 | -1.27901746 | 1.740819297 | 0.734721554 | 0.46250909 | 1 | 1.27901746 |
| ENSMUSG00000016763 | Scube1 | 64706 | protein_coding | 3.76272964 | 1.279159745 | 2.234302692 | -0.572509602 | 0.566976783 | 1 | 1.279159745 |
| ENSMUSG00000062329 | Cytl1 | 231162 | protein_coding | 1.140857146 | -1.27955412 | 3.019536911 | 0.423758397 | 0.671742022 | 1 | 1.27955412 |
| ENSMUSG00000031486 | Adgra2 | 78560 | protein_coding | 16.84748743 | -1.280550721 | 0.516376888 | 2.479876134 | 0.013142803 | 0.052654112 | 1.280550721 |
| ENSMUSG00000068606 | Gm4841 | 225594 | protein_coding | 1.667666566 | 1.281040714 | 1.425664847 | -0.89855671 | 0.368888826 | 1 | 1.281040714 |
| ENSMUSG00000086727 | 4931428L18Rik | 70988 | protein_coding | 4.003611084 | 1.281104565 | 1.085489815 | -1.180208739 | 0.237917204 | 1 | 1.281104565 |
| ENSMUSG00000095040 | 1700001J03Rik | 69282 | protein_coding | 0.703495518 | -1.281488592 | 1.316779393 | 0.973199154 | 0.330454331 | 1 | 1.281488592 |
| ENSMUSG00000025515 | Muc2 | 17831 | protein_coding | 35461.6037 | -1.282370647 | 0.170162308 | 7.536161594 | 4.84E-14 | 1.63E-12 | 1.282370647 |
| ENSMUSG00000043017 | Ptgir | 19222 | protein_coding | 1.60040382 | 1.283809451 | 1.724919843 | -0.744271948 | 0.456711969 | 1 | 1.283809451 |
| ENSMUSG00000024112 | Cacna1h | 58226 | protein_coding | 182.2006993 | -1.284594407 | 0.235122097 | 5.463520541 | 4.67E-08 | 8.05E-07 | 1.284594407 |
| ENSMUSG00000112920 | Spata22 | 380709 | protein_coding | 0.493882762 | -1.284935711 | 1.333182822 | 0.963810582 | 0.335140901 | 1 | 1.284935711 |
| ENSMUSG00000065574 | Mir203 | 387199 | miRNA | 2.538507363 | -1.28526275 | 0.857085626 | 1.499573334 | 0.133724959 | 1 | 1.28526275 |
| ENSMUSG00000107153 | Gm38404 | 232386 | lncRNA | 0.809912943 | 1.285355556 | 2.118446765 | -0.606744327 | 0.5440206 | 1 | 1.285355556 |
| ENSMUSG00000117666 | Gm550 | 225852 | protein_coding | 2.498124762 | 1.285725628 | 0.904162222 | -1.422007684 | 0.15502402 | 1 | 1.285725628 |
| ENSMUSG00000028591 | Pramel13 | 77632 | protein_coding | 1.577806364 | -1.285831172 | 1.826411776 | 0.704020413 | 0.481420062 | 1 | 1.285831172 |
| ENSMUSG00000025150 | Cbr2 | 12409 | protein_coding | 442.8742127 | 1.286626969 | 0.248614785 | -5.17518285 | 2.28E-07 | 3.53E-06 | 1.286626969 |
| ENSMUSG00000061911 | Myt1l | 17933 | protein_coding | 4.608234262 | 1.286638922 | 2.064638484 | -0.623178795 | 0.533167035 | 0.729592345 | 1.286638922 |
| ENSMUSG00000036545 | Adamts2 | 216725 | protein_coding | 2.922393836 | 1.286669117 | 1.656773697 | -0.776611265 | 0.437388153 | 1 | 1.286669117 |
| ENSMUSG00000039452 | Snx22 | 382083 | protein_coding | 10.07826281 | 1.28669393 | 0.487883014 | -2.637300118 | 0.008356886 | 0.036850631 | 1.28669393 |
| ENSMUSG00000007209 | Ceacam9 | 26368 | protein_coding | 2.180513666 | 1.288357957 | 0.843326539 | -1.527709491 | 0.126584683 | 1 | 1.288357957 |
| ENSMUSG00000029082 | Bst1 | 12182 | protein_coding | 4.70120023 | 1.290982422 | 1.054167331 | -1.224646585 | 0.220708408 | 0.42723932 | 1.290982422 |
| ENSMUSG00000034780 | B3galt1 | 26877 | protein_coding | 1.355365934 | 1.291168274 | 1.882310163 | -0.685948735 | 0.492745437 | 1 | 1.291168274 |
| ENSMUSG00000106139 | Gm30648 | 102632621 | lncRNA | 2.074863045 | 1.291642925 | 1.062265707 | -1.215932056 | 0.224010805 | 1 | 1.291642925 |
| ENSMUSG00000028044 | Cks1b | 54124 | protein_coding | 132.4972807 | 1.292757517 | 0.18050126 | -7.162041518 | 7.95E-13 | 2.45E-11 | 1.292757517 |
| ENSMUSG00000084890 | A830036E02Rik | 353123 | lncRNA | 7.387549701 | 1.29324271 | 0.727428096 | -1.777828925 | 0.075431956 | 0.201273764 | 1.29324271 |
| ENSMUSG00000032268 | Tmprss5 | 80893 | protein_coding | 3.750828305 | 1.293268614 | 1.872000771 | -0.690848334 | 0.489660857 | 1 | 1.293268614 |
| ENSMUSG00000099210 | Mir7051 | 102465636 | miRNA | 0.83141899 | 1.293319458 | 1.386823239 | -0.932577001 | 0.351038413 | 1 | 1.293319458 |
| ENSMUSG00000026272 | Agxt | 11611 | protein_coding | 3.375930418 | 1.29344034 | 0.639782761 | -2.021686765 | 0.043208722 | 1 | 1.29344034 |

| ensembl_gene_id | external_gene_name | entrezgene_id | gene_biotype | baseMean | log2FoldChange | lfcSE | stat | pvalue | padj | absLog2FC |
|---|---|---|---|---|---|---|---|---|---|---|
| ENSMUSG00000020038 | Cry1 | 12952 | protein_coding | 338.0161705 | 1.293845492 | 0.256103127 | -5.052048782 | 4.37E-07 | 6.52E-06 | 1.293845492 |
| ENSMUSG00000100816 | Gm28321 | 102633780 | lncRNA | 1.300368808 | 1.294214421 | 1.875165726 | -0.690186688 | 0.490076794 | 1 | 1.294214421 |
| ENSMUSG00000020701 | Tmem132e | 270893 | protein_coding | 1.888676653 | -1.294783933 | 2.451389254 | 0.528183736 | 0.597371817 | 1 | 1.294783933 |
| ENSMUSG00000029184 | Or12d17 | 258832 | protein_coding | 1.6416415 | 1.294830433 | 2.142877125 | -0.604248568 | 0.545678395 | 1 | 1.294830433 |
| ENSMUSG00000030443 | Zfp583 | 213011 | protein_coding | 2.243630584 | -1.295036748 | 1.065827387 | 1.215052985 | 0.224345883 | 1 | 1.295036748 |
| ENSMUSG00000017309 | Cd300lg | 52685 | protein_coding | 1.377892299 | -1.29520975 | 2.454401746 | 0.527708942 | 0.597701366 | 1 | 1.29520975 |
| ENSMUSG00000017733 | Eppin | 75526 | protein_coding | 0.96635663 | -1.295271625 | 2.454871975 | 0.527633065 | 0.597754039 | 1 | 1.295271625 |
| ENSMUSG00000098904 | Mir6998 | 102466214 | miRNA | 0.855144504 | 1.295541747 | 1.628080523 | -0.795747955 | 0.426178547 | 1 | 1.295541747 |
| ENSMUSG00000078439 | Smim24 | 72273 | protein_coding | 1309.80598 | -1.295617344 | 0.165411502 | 7.832691953 | 4.78E-15 | 1.74E-13 | 1.295617344 |
| ENSMUSG00000043747 | 1520401A03Rik | 320309 | protein_coding | 6.545721704 | 1.296001179 | 0.717609338 | -1.805998208 | 0.070918619 | 0.192815811 | 1.296001179 |
| ENSMUSG00000039375 | Wdr17 | 244484 | protein_coding | 1.981981873 | 1.296274498 | 1.883153379 | -0.688353117 | 0.49123044 | 1 | 1.296274498 |
| ENSMUSG00000062939 | Stat4 | 20849 | protein_coding | 5.782274757 | 1.297457492 | 0.62729082 | -2.068350837 | 0.038607045 | 0.122583491 | 1.297457492 |
| ENSMUSG00000075402 | Krt76 | 77055 | protein_coding | 1.161176042 | 1.298234557 | 2.332224479 | -0.556650772 | 0.577766057 | 1 | 1.298234557 |
| ENSMUSG00000025650 | Col7a1 | 12836 | protein_coding | 57.54535633 | 1.299171714 | 0.440543587 | -2.949019692 | 0.003187837 | 0.01673893 | 1.299171714 |
| ENSMUSG00000028351 | Brinp1 | 56710 | protein_coding | 1.936093755 | 1.299248433 | 2.418666934 | -0.537175423 | 0.591146446 | 1 | 1.299248433 |
| ENSMUSG00000032394 | Igdcc3 | 19289 | protein_coding | 2.567633234 | 1.29941813 | 1.492433428 | -0.870670748 | 0.383933955 | 1 | 1.29941813 |
| ENSMUSG00000113949 | Scamp4 | 56214 | protein_coding | 13.94017375 | 1.299766174 | 0.450858454 | -2.882869693 | 0.003940704 | 0.01996653 | 1.299766174 |
| ENSMUSG00000053310 | Nrgn | 64011 | protein_coding | 9.246224713 | -1.300353337 | 0.471399907 | 2.758492985 | 0.005806855 | 0.027492662 | 1.300353337 |
| ENSMUSG00000027070 | Lrp2 | 14725 | protein_coding | 5.824612325 | 1.300846099 | 3.140928128 | -0.414159779 | 0.678757096 | 0.827904179 | 1.300846099 |
| ENSMUSG00000092243 | Gm7030 | 630294 | protein_coding | 1.070524021 | -1.300991763 | 1.399707623 | 0.929473943 | 0.352643523 | 1 | 1.300991763 |
| ENSMUSG00000056486 | Chn1 | 108699 | protein_coding | 4.945451873 | 1.301111944 | 0.721319716 | -1.803793679 | 0.071263657 | 0.193498767 | 1.301111944 |
| ENSMUSG00000028452 | Vcp | 269523 | protein_coding | 4257.9378 | 1.301855175 | 0.11668905 | -11.15661819 | 6.65E-29 | 4.99E-27 | 1.301855175 |
| ENSMUSG00000027199 | Gatm | 67092 | protein_coding | 260.1047481 | 1.302213334 | 0.37917077 | -3.434371625 | 0.000593929 | 0.004096971 | 1.302213334 |
| ENSMUSG00000067064 | Or6b2 | 259040 | protein_coding | 1.525151099 | 1.303200074 | 2.350876105 | -0.554346557 | 0.579341694 | 1 | 1.303200074 |
| ENSMUSG00000035383 | Pmch | 110312 | protein_coding | 1.182713764 | 1.303899154 | 1.139569553 | -1.144203221 | 0.252539368 | 1 | 1.303899154 |
| ENSMUSG00000076121 | Mir687 | 751541 | miRNA | 0.493020421 | 1.304343362 | 1.455754493 | -0.895991301 | 0.370257409 | 1 | 1.304343362 |
| ENSMUSG00000091898 | Tnnc1 | 21924 | protein_coding | 1.886381164 | -1.304404379 | 0.798882723 | 1.632785816 | 0.102514051 | 1 | 1.304404379 |
| ENSMUSG00000056947 | Mab21l1 | 17116 | protein_coding | 0.719101275 | -1.304444185 | 1.402999227 | 0.92975403 | 0.352498453 | 1 | 1.304444185 |
| ENSMUSG00000043811 | Rtn4r | 65079 | protein_coding | 40.09472912 | -1.304953068 | 0.20187218 | 6.46425409 | 1.02E-10 | 2.52E-09 | 1.304953068 |
| ENSMUSG00000039059 | Hrh3 | 99296 | protein_coding | 0.885163616 | -1.305387711 | 2.000780726 | 0.652439167 | 0.514117904 | 1 | 1.305387711 |
| ENSMUSG00000090174 | Gm10612 | 102638502 | lncRNA | 4.17904695 | -1.308492143 | 0.587860155 | 2.225856152 | 0.026023818 | 1 | 1.308492143 |
| ENSMUSG00000023274 | Cd4 | 12504 | protein_coding | 2.670283595 | -1.30866602 | 1.868570264 | 0.700356869 | 0.483704465 | 1 | 1.30866602 |
| ENSMUSG00000040026 | Saa3 | 20210 | protein_coding | 51.57212172 | 1.308699365 | 0.313199947 | -4.178478885 | 2.93E-05 | 0.000291068 | 1.308699365 |
| ENSMUSG00000064936 | Gm23722 | 115485772 | snoRNA | 1.612400982 | -1.309404659 | 0.830881987 | 1.575921345 | 0.115043931 | 1 | 1.309404659 |
| ENSMUSG00000048458 | Inka2 | 109050 | protein_coding | 8.823187016 | 1.309648323 | 0.575416123 | -2.276002133 | 0.022845882 | 0.081458356 | 1.309648323 |
| ENSMUSG00000022076 | Klhl1 | 93688 | protein_coding | 1.391576098 | 1.310104526 | 2.347370915 | -0.558115685 | 0.576765391 | 1 | 1.310104526 |
| ENSMUSG00000020622 | Nt5c1b | 70881 | protein_coding | 1.509630368 | 1.310147648 | 1.75535734 | -0.746370906 | 0.45544339 | 1 | 1.310147648 |
| ENSMUSG00000111731 | Gm48727 | 118567821 | lncRNA | 4.702865355 | 1.310352523 | 0.517765876 | -2.53078193 | 0.011380858 | 0.047245724 | 1.310352523 |
| ENSMUSG00000003227 | Edar | 13608 | protein_coding | 1.95276358 | -1.310724405 | 1.487846493 | 0.880954057 | 0.378342687 | 1 | 1.310724405 |

| ensembl_gene_id | external_gene_name | entrezgene_id | gene_biotype | baseMean | log2FoldChange | lfcSE | stat | pvalue | padj | absLog2FC |
|---|---|---|---|---|---|---|---|---|---|---|
| ENSMUSG00000025482 | Odf3 | 69287 | protein_coding | 1.199900588 | -1.31171762 | 1.305952872 | 1.004414208 | 0.315179004 | 1 | 1.31171762 |
| ENSMUSG00000065692 | Gm25410 | 115487519 | snoRNA | 0.650040771 | -1.312277136 | 1.772828919 | 0.740216454 | 0.459168665 | 1 | 1.312277136 |
| ENSMUSG00000094112 | 9330182O14Rik | 328531 | protein_coding | 3.739038721 | -1.312916722 | 0.693008814 | 1.894516629 | 0.058156469 | 1 | 1.312916722 |
| ENSMUSG00000087236 | Kif28 | 383592 | protein_coding | 1.812247188 | -1.313023083 | 2.085332142 | 0.629646979 | 0.52892558 | 1 | 1.313023083 |
| ENSMUSG00000059434 | Gckr | 231103 | protein_coding | 2.319597694 | 1.313548046 | 1.309599074 | -1.003015405 | 0.315853429 | 1 | 1.313548046 |
| ENSMUSG00000068231 | Vmn1r43 | 113847 | protein_coding | 3.751051414 | -1.313915695 | 1.729510519 | 0.759703789 | 0.447431663 | 1 | 1.313915695 |
| ENSMUSG00000088054 | Mir1968 | 100316713 | miRNA | 4.621668315 | -1.313951572 | 0.490318325 | 2.679792913 | 0.007366772 | 0.03324812 | 1.313951572 |
| ENSMUSG00000065845 | Gm25189 | 115489620 | snRNA | 1.961048524 | 1.313997563 | 1.403400665 | -0.936295383 | 0.349121125 | 1 | 1.313997563 |
| ENSMUSG00000013083 | 2200002J24Rik | 69147 | protein_coding | 5.475432379 | -1.314634992 | 0.429618836 | 3.060003156 | 0.002213347 | 0.012282071 | 1.314634992 |
| ENSMUSG00000023047 | Amhr2 | 110542 | protein_coding | 4.921331539 | -1.315120014 | 0.611067325 | 2.152168771 | 0.031384064 | 0.104512288 | 1.315120014 |
| ENSMUSG00000056921 | Or1e34 | 259009 | protein_coding | 1.207004072 | 1.315583716 | 2.3454089 | -0.560918702 | 0.57485296 | 1 | 1.315583716 |
| ENSMUSG00000067220 | Cnga1 | 12788 | protein_coding | 2.081763483 | 1.315867317 | 2.404544663 | -0.547241787 | 0.584212632 | 1 | 1.315867317 |
| ENSMUSG00000024500 | Ppp2r2b | 72930 | protein_coding | 2.391533627 | 1.315927179 | 1.222523925 | -1.076401984 | 0.281747515 | 1 | 1.315927179 |
| ENSMUSG00000035232 | Pdk3 | 236900 | protein_coding | 564.8219954 | 1.316803663 | 0.091585493 | -14.37786292 | 7.13E-47 | 8.98E-45 | 1.316803663 |
| ENSMUSG00000076144 | Mir709 | 735271 | miRNA | 1.112672845 | -1.316915071 | 1.175001243 | 1.120777598 | 0.262382542 | 1 | 1.316915071 |
| ENSMUSG00000092837 | Rpph1 | 85029 | ribozyme | 15421.6678 | 1.317071875 | 0.771479395 | -1.707202919 | 0.087784343 | 0.224929501 | 1.317071875 |
| ENSMUSG00000035941 | Ibtk | 108837 | protein_coding | 2286.021099 | 1.317246199 | 0.085787801 | -15.354703 | 3.29E-53 | 4.77E-51 | 1.317246199 |
| ENSMUSG00000050075 | Gpr171 | 229323 | protein_coding | 7.443320086 | 1.317424213 | 0.669298022 | -1.968367108 | 0.049025814 | 0.146881764 | 1.317424213 |
| ENSMUSG00000031765 | Mt1 | 17748 | protein_coding | 1293.91968 | -1.317617924 | 0.177820218 | 7.409831918 | 1.26E-13 | 4.14E-12 | 1.317617924 |
| ENSMUSG00000032625 | Thsd7a | 330267 | protein_coding | 5.362482199 | -1.318107703 | 1.176952443 | 1.11993285 | 0.262742378 | 0.476857637 | 1.318107703 |
| ENSMUSG00000045417 | Vmn1r230 | 171231 | protein_coding | 2.047707677 | 1.318322364 | 2.731834536 | -0.482577677 | 0.629395632 | 1 | 1.318322364 |
| ENSMUSG00000116270 | Gm1600 | 381033 | lncRNA | 3.820175216 | 1.318854328 | 0.990962174 | -1.330882615 | 0.183227639 | 1 | 1.318854328 |
| ENSMUSG00000045776 | Lrtm1 | 319476 | protein_coding | 2.406980641 | 1.318958263 | 2.699295075 | -0.488630634 | 0.625103221 | 1 | 1.318958263 |
| ENSMUSG00000049098 | Or8b3 | 258869 | protein_coding | 1.623214994 | 1.318994972 | 2.697078209 | -0.489045875 | 0.624809219 | 1 | 1.318994972 |
| ENSMUSG00000067215 | Usp51 | 635253 | protein_coding | 1.501259302 | 1.319605335 | 1.434499095 | -0.919906704 | 0.357621515 | 1 | 1.319605335 |
| ENSMUSG00000078962 | 4932414J04Rik | 237694 | lncRNA | 1.218074183 | -1.319943525 | 1.299128666 | 1.016022169 | 0.30961883 | 1 | 1.319943525 |
| ENSMUSG00000053063 | Clec12a | 232413 | protein_coding | 1.810883211 | 1.321871466 | 1.795149246 | -0.736357419 | 0.461513211 | 1 | 1.321871466 |
| ENSMUSG00000100512 | Ovol3 | 381867 | protein_coding | 0.877042772 | 1.322417362 | 1.770783641 | -0.746797819 | 0.455185612 | 1 | 1.322417362 |
| ENSMUSG00000034891 | Sncb | 104069 | protein_coding | 1.753211964 | -1.324200569 | 1.892076188 | 0.69986641 | 0.484010736 | 1 | 1.324200569 |
| ENSMUSG00000108841 | Frmpd2 | 268729 | protein_coding | 2.838802323 | 1.325247131 | 2.22242766 | -0.596306082 | 0.550970769 | 1 | 1.325247131 |
| ENSMUSG00000028072 | Ntrk1 | 18211 | protein_coding | 1.408227333 | 1.326691038 | 1.608466067 | -0.824817549 | 0.409475182 | 1 | 1.326691038 |
| ENSMUSG00000049092 | Gpr137c | 70713 | protein_coding | 11.36982868 | 1.326743355 | 0.511849723 | -2.59205641 | 0.009540413 | 0.040996715 | 1.326743355 |
| ENSMUSG00000027489 | Necab3 | 56846 | protein_coding | 4.82304847 | 1.327227824 | 0.922703252 | -1.438412427 | 0.150317069 | 0.330166076 | 1.327227824 |
| ENSMUSG00000055150 | Zfp78 | 330463 | protein_coding | 8.328791311 | -1.327313105 | 0.380094592 | 3.492059963 | 0.000479311 | 0.003417675 | 1.327313105 |
| ENSMUSG00000006344 | Ggt5 | 23887 | protein_coding | 3.094254498 | 1.327704489 | 1.94899498 | -0.681225197 | 0.495729007 | 1 | 1.327704489 |
| ENSMUSG00000059412 | Fxyd2 | 11936 | protein_coding | 2.584936753 | 1.327827551 | 2.35010978 | -0.565006606 | 0.572069271 | 1 | 1.327827551 |
| ENSMUSG00000031965 | Tbx20 | 57246 | protein_coding | 4.316289689 | 1.328530785 | 2.05250435 | -0.647273067 | 0.517455229 | 1 | 1.328530785 |
| ENSMUSG00000052374 | Actn2 | 11472 | protein_coding | 2.805641165 | 1.329739681 | 2.010597093 | -0.661365565 | 0.508377904 | 1 | 1.329739681 |
| ENSMUSG00000061702 | Tmem91 | 320208 | protein_coding | 2.135265814 | -1.330371853 | 0.817100488 | 1.628161863 | 0.103490569 | 1 | 1.330371853 |

Supplementary Table 1

| ensembl_gene_id | external_gene_name | entrezgene_id | gene_biotype | baseMean | log2FoldChange | lfcSE | stat | pvalue | padj | absLog2FC |
|---|---|---|---|---|---|---|---|---|---|---|
| ENSMUSG00000021266 | Wars1 | 22375 | protein_coding | 2693.535193 | 1.330417026 | 0.097405775 | -13.65850248 | 1.80E-42 | 2.00E-40 | 1.330417026 |
| ENSMUSG00000046908 | Ltb4r1 | 16995 | protein_coding | 3.441244688 | 1.332118186 | 0.791655469 | -1.682699405 | 0.092433297 | 1 | 1.332118186 |
| ENSMUSG00000090273 | Prr22 | 100504446 | protein_coding | 5.427799555 | 1.332598275 | 0.520321479 | -2.56110564 | 0.010433962 | 0.044068147 | 1.332598275 |
| ENSMUSG00000031098 | Syt8 | 55925 | protein_coding | 30.00663858 | 1.333122407 | 0.282161967 | -4.724670799 | 2.30E-06 | 2.96E-05 | 1.333122407 |
| ENSMUSG00000043719 | Col6a6 | 245026 | protein_coding | 2.910626222 | 1.333209728 | 1.382179891 | -0.96457034 | 0.334760061 | 1 | 1.333209728 |
| ENSMUSG00000114412 | 4930405A10Rik | 74896 | lncRNA | 1.200568465 | 1.333243514 | 1.849029638 | -0.721050375 | 0.470878522 | 1 | 1.333243514 |
| ENSMUSG00000090122 | Kcne1l | 66240 | protein_coding | 3.170870024 | -1.333296389 | 0.695284521 | 1.917627028 | 0.055158321 | 1 | 1.333296389 |
| ENSMUSG00000046678 | Or10g6 | 258283 | protein_coding | 2.100067754 | 1.334053501 | 1.940278027 | -0.687557908 | 0.491731222 | 1 | 1.334053501 |
| ENSMUSG00000097804 | Gm16685 | 102634900 | lncRNA | 5.574497802 | 1.334652559 | 0.549825232 | -2.427412353 | 0.015206961 | 0.059061089 | 1.334652559 |
| ENSMUSG00000037913 | Tmem156 | 243025 | protein_coding | 0.938023564 | -1.335395988 | 1.531718143 | 0.871828798 | 0.383301782 | 1 | 1.335395988 |
| ENSMUSG00000002459 | Rgs20 | 58175 | protein_coding | 2.375310497 | 1.335433942 | 1.642023087 | -0.813285728 | 0.416054258 | 1 | 1.335433942 |
| ENSMUSG00000080115 | Eef1akmt3 | 100504608 | protein_coding | 20.24250162 | 1.335518002 | 0.300331362 | -4.44681498 | 8.72E-06 | 9.92E-05 | 1.335518002 |
| ENSMUSG00000027225 | Duoxa2 | 66811 | protein_coding | 150.7933131 | 1.336060422 | 0.325971791 | -4.098699517 | 4.15E-05 | 0.000397981 | 1.336060422 |
| ENSMUSG00000040466 | Blvrb | 233016 | protein_coding | 741.4191753 | 1.337292591 | 0.090099629 | -14.8423762 | 7.79E-50 | 1.08E-47 | 1.337292591 |
| ENSMUSG00000031824 | 6430548M08Rik | 234797 | protein_coding | 1672.247659 | 1.339283153 | 0.078730508 | -17.01098065 | 6.81E-65 | 1.25E-62 | 1.339283153 |
| ENSMUSG00000005364 | Il5ra | 16192 | protein_coding | 1.109050155 | 1.339631674 | 1.988336663 | -0.673744894 | 0.500473515 | 1 | 1.339631674 |
| ENSMUSG00000028560 | Usp1 | 230484 | protein_coding | 855.9865594 | 1.34005618 | 0.105404267 | -12.71349081 | 4.98E-37 | 4.62E-35 | 1.34005618 |
| ENSMUSG00000095917 | Or11g7 | 258661 | protein_coding | 1.746158171 | -1.340102802 | 2.117685139 | 0.632814944 | 0.526854497 | 1 | 1.340102802 |
| ENSMUSG00000094732 | 1500015L24Rik | 68994 | lncRNA | 0.856282743 | -1.340397925 | 1.7934379 | 0.747390208 | 0.454828054 | 1 | 1.340397925 |
| ENSMUSG00000057715 | A830018L16Rik | 320492 | protein_coding | 1.379155131 | -1.340466696 | 2.122914108 | 0.631427664 | 0.527760933 | 1 | 1.340466696 |
| ENSMUSG00000090685 | Atp6v1fnb | 668210 | protein_coding | 0.922069716 | -1.340707109 | 1.683151567 | 0.79654568 | 0.425714936 | 1 | 1.340707109 |
| ENSMUSG00000025321 | Itgb8 | 320910 | protein_coding | 9.07461707 | -1.340739274 | 0.490300353 | 2.734526429 | 0.006247008 | 0.02899794 | 1.340739274 |
| ENSMUSG00000059483 | Gm5528 | 433319 | processed_pseudogene | 0.714474069 | 1.341060391 | 1.571613483 | -0.853301658 | 0.393492041 | 1 | 1.341060391 |
| ENSMUSG00000097178 | 2310002F09Rik | 100504720 | lncRNA | 1.511502101 | 1.341503604 | 1.938500199 | -0.692031708 | 0.488917417 | 1 | 1.341503604 |
| ENSMUSG00000039911 | Spsb1 | 74646 | protein_coding | 31.4637716 | -1.343072901 | 0.193873712 | 6.927565807 | 4.28E-12 | 1.23E-10 | 1.343072901 |
| ENSMUSG00000065541 | Mir24-2 | 723960 | miRNA | 0.891803635 | 1.343537881 | 1.294103402 | -1.038199791 | 0.29917705 | 1 | 1.343537881 |
| ENSMUSG00000050035 | Fhl4 | 14202 | protein_coding | 1.906564582 | 1.343979684 | 1.129536372 | -1.189850737 | 0.234105064 | 1 | 1.343979684 |
| ENSMUSG00000090219 | Or51a42 | 259081 | protein_coding | 2.069363903 | 1.345514257 | 2.658438747 | -0.506129494 | 0.612765747 | 1 | 1.345514257 |
| ENSMUSG00000022658 | Tagln3 | 56370 | protein_coding | 1.339698058 | 1.346798899 | 1.072093342 | -1.256232873 | 0.209031552 | 1 | 1.346798899 |
| ENSMUSG00000061376 | Vmn1r219 | 171272 | protein_coding | 1.56683447 | 1.348536948 | 2.673527208 | -0.504403674 | 0.613977737 | 1 | 1.348536948 |
| ENSMUSG00000096002 | Vmn2r53 | 637908 | protein_coding | 4.779415338 | 1.348644659 | 2.113294082 | -0.638171786 | 0.523361861 | 0.722098569 | 1.348644659 |
| ENSMUSG00000041134 | Cyyr1 | 224405 | protein_coding | 2.216381283 | 1.34891835 | 2.718159333 | -0.496261692 | 0.61970979 | 1 | 1.34891835 |
| ENSMUSG00000030641 | Ddias | 74041 | protein_coding | 65.38817405 | 1.349093189 | 0.214040914 | -6.302968723 | 2.92E-10 | 6.91E-09 | 1.349093189 |
| ENSMUSG00000095276 | Gfy | 100039953 | protein_coding | 2.106499397 | -1.34954337 | 2.313799805 | 0.583258485 | 0.559719305 | 1 | 1.34954337 |
| ENSMUSG00000022534 | Mefv | 54483 | protein_coding | 1.044044985 | -1.350050069 | 1.372347248 | 0.983752524 | 0.325237208 | 1 | 1.350050069 |
| ENSMUSG00000084336 | Or4x6 | 258339 | protein_coding | 1.153499454 | -1.350414722 | 2.33671623 | 0.577911303 | 0.563324003 | 1 | 1.350414722 |
| ENSMUSG00000028979 | Masp2 | 17175 | protein_coding | 3.080674155 | 1.350561953 | 0.828295417 | -1.630531723 | 0.102989166 | 1 | 1.350561953 |
| ENSMUSG00000078808 | Vmn1r58 | 81014 | protein_coding | 2.210122776 | -1.351214375 | 2.358329349 | 0.57295406 | 0.5666758 | 1 | 1.351214375 |
| ENSMUSG00000035831 | Krt25 | 70810 | protein_coding | 0.733329392 | -1.351789474 | 2.160411983 | 0.625709117 | 0.531505752 | 1 | 1.351789474 |

| ensembl_gene_id | external_gene_name | entrezgene_id | gene_biotype | baseMean | log2FoldChange | lfcSE | stat | pvalue | padj | absLog2FC |
|---|---|---|---|---|---|---|---|---|---|---|
| ENSMUSG00000038754 | Elovl3 | 12686 | protein_coding | 0.854622111 | -1.352096778 | 2.444186712 | 0.553188826 | 0.580134117 | 1 | 1.352096778 |
| ENSMUSG00000037406 | Htra4 | 330723 | protein_coding | 1.703607945 | -1.352391009 | 0.940764467 | 1.437544738 | 0.15056327 | 1 | 1.352391009 |
| ENSMUSG00000093007 | Mir15a | 387174 | miRNA | 0.50139606 | -1.353013989 | 1.8337761 | 0.737829438 | 0.460618102 | 1 | 1.353013989 |
| ENSMUSG00000001334 | Fndc5 | 384061 | protein_coding | 181.3808573 | -1.354117352 | 0.237782827 | 5.694765132 | 1.24E-08 | 2.35E-07 | 1.354117352 |
| ENSMUSG00000090411 | Vmn1r174 | 404291 | protein_coding | 1.803642615 | 1.354336613 | 2.715038727 | -0.498827733 | 0.617900749 | 1 | 1.354336613 |
| ENSMUSG00000024757 | Slc22a19 | 207151 | protein_coding | 1.529796841 | 1.354439053 | 2.705292071 | -0.500662781 | 0.61660847 | 1 | 1.354439053 |
| ENSMUSG00000038517 | Tbkbp1 | 73174 | protein_coding | 5.639165413 | -1.355016289 | 0.814197876 | 1.664234615 | 0.096065553 | 0.239895137 | 1.355016289 |
| ENSMUSG00000054966 | Lmntd1 | 74071 | protein_coding | 4.169359959 | 1.356304939 | 3.048881921 | -0.44485322 | 0.656425831 | 1 | 1.356304939 |
| ENSMUSG00000036854 | Hspb6 | 243912 | protein_coding | 27.87116913 | 1.356332125 | 0.267081688 | -5.078341887 | 3.81E-07 | 5.73E-06 | 1.356332125 |
| ENSMUSG00000090307 | 1700071M16Rik | 73504 | lncRNA | 1.176712509 | -1.356464193 | 2.160064764 | 0.627973853 | 0.530021069 | 1 | 1.356464193 |
| ENSMUSG00000034818 | Celf5 | 319586 | protein_coding | 42.74545032 | -1.356624573 | 0.277636315 | 4.886336907 | 1.03E-06 | 1.41E-05 | 1.356624573 |
| ENSMUSG00000039347 | Atp6v0e2 | 76252 | protein_coding | 179.014244 | -1.35714368 | 0.147065407 | 9.228163909 | 2.75E-20 | 1.41E-18 | 1.35714368 |
| ENSMUSG00000047497 | Adamts12 | 239337 | protein_coding | 3.088452668 | 1.357397254 | 1.523791439 | -0.89080252 | 0.373035123 | 1 | 1.357397254 |
| ENSMUSG00000026532 | Spta1 | 20739 | protein_coding | 15.91432325 | -1.357739883 | 0.678953046 | 1.999755197 | 0.045526705 | 0.138995748 | 1.357739883 |
| ENSMUSG00000021612 | Slc6a18 | 22598 | protein_coding | 2.232102891 | 1.358238592 | 2.467571054 | -0.550435454 | 0.582020737 | 1 | 1.358238592 |
| ENSMUSG00000016758 | Bik | 12124 | protein_coding | 38.00543226 | -1.3583055 | 0.191970357 | 7.075600217 | 1.49E-12 | 4.43E-11 | 1.3583055 |
| ENSMUSG00000047959 | Kcna3 | 16491 | protein_coding | 2.117055531 | 1.358778294 | 0.995549952 | -1.36485195 | 0.172299587 | 1 | 1.358778294 |
| ENSMUSG00000075044 | Slc22a29 | 236293 | protein_coding | 1.495327564 | 1.358885987 | 1.112028507 | -1.221988445 | 0.221711997 | 1 | 1.358885987 |
| ENSMUSG00000068758 | Il3ra | 16188 | protein_coding | 24.05957096 | 1.359127637 | 0.398777233 | -3.408237797 | 0.000653839 | 0.00443976 | 1.359127637 |
| ENSMUSG00000028023 | Pitx2 | 18741 | protein_coding | 36.81005491 | 1.35931468 | 0.895961889 | -1.517156809 | 0.129227098 | 0.296850229 | 1.35931468 |
| ENSMUSG00000094185 | Gm5445 | 432725 | processed_pseudogene | 0.7192246 | 1.359978428 | 1.656729512 | -0.820881392 | 0.411713831 | 1 | 1.359978428 |
| ENSMUSG00000088308 | Gm24507 | 115488238 | snoRNA | 1.53655147 | 1.360243336 | 0.832546881 | -1.633833923 | 0.102293727 | 1 | 1.360243336 |
| ENSMUSG00000026523 | Wdr64 | 75820 | protein_coding | 2.349241075 | -1.360991672 | 2.666318928 | 0.510438439 | 0.609744333 | 1 | 1.360991672 |
| ENSMUSG00000001986 | Gria3 | 53623 | protein_coding | 3.40732184 | 1.361010784 | 1.586047109 | -0.85811498 | 0.390828976 | 1 | 1.361010784 |
| ENSMUSG00000037625 | Cldn11 | 18417 | protein_coding | 1.096405706 | 1.361671607 | 1.430342198 | -0.951990096 | 0.341102005 | 1 | 1.361671607 |
| ENSMUSG00000099170 | 5730403I07Rik | 70487 | lncRNA | 1.060928392 | 1.361727134 | 1.615917343 | -0.842696033 | 0.399398467 | 1 | 1.361727134 |
| ENSMUSG00000074102 | Rbm15b | 109095 | protein_coding | 1876.903628 | 1.364934648 | 0.087066536 | -15.67691456 | 2.18E-55 | 3.36E-53 | 1.364934648 |
| ENSMUSG00000041247 | Lamp3 | 239739 | protein_coding | 0.941961287 | 1.365030713 | 1.934282218 | -0.705704007 | 0.480372228 | 1 | 1.365030713 |
| ENSMUSG00000096965 | 3300005D01Rik | 78512 | lncRNA | 1.048048394 | 1.366490021 | 1.277730319 | -1.069466695 | 0.284859428 | 1 | 1.366490021 |
| ENSMUSG00000031380 | Vegfd | 14205 | protein_coding | 1.236594314 | -1.367551276 | 1.224464524 | 1.116856593 | 0.264055647 | 1 | 1.367551276 |
| ENSMUSG00000021423 | Ly86 | 17084 | protein_coding | 1.350186813 | -1.367667454 | 1.67667823 | 0.815700609 | 0.41467139 | 1 | 1.367667454 |
| ENSMUSG00000050148 | Ubqln2 | 54609 | protein_coding | 386.2619821 | 1.367800581 | 0.104909688 | -13.0378863 | 7.45E-39 | 7.28E-37 | 1.367800581 |
| ENSMUSG00000020914 | Top2a | 21973 | protein_coding | 3209.300281 | 1.36788918 | 0.193796391 | -7.058383151 | 1.68E-12 | 5.00E-11 | 1.36788918 |
| ENSMUSG00000032807 | Alox12b | 11686 | protein_coding | 2.043157412 | 1.368025689 | 1.547574177 | -0.88398069 | 0.376706637 | 1 | 1.368025689 |
| ENSMUSG00000031709 | Tbc1d9 | 71310 | protein_coding | 104.0620815 | 1.368201876 | 0.248528595 | -5.505209072 | 3.69E-08 | 6.48E-07 | 1.368201876 |
| ENSMUSG00000099063 | Mir7031 | 102465624 | miRNA | 0.678560098 | 1.368541916 | 1.83962469 | -0.74392452 | 0.45692214 | 1 | 1.368541916 |
| ENSMUSG00000036432 | Siah2 | 20439 | protein_coding | 378.4284064 | 1.369804084 | 0.10263217 | -13.34673212 | 1.24E-40 | 1.33E-38 | 1.369804084 |
| ENSMUSG00000046334 | Gm6195 | 108167613 | processed_pseudogene | 0.56538195 | 1.370201114 | 2.05553156 | -0.666592107 | 0.505032712 | 1 | 1.370201114 |
| ENSMUSG00000021950 | Anxa8 | 11752 | protein_coding | 227.1840179 | -1.370937501 | 0.313866739 | 4.367896717 | 1.25E-05 | 0.000136924 | 1.370937501 |

Supplementary Table 1

| ensembl_gene_id | external_gene_name | entrezgene_id | gene_biotype | baseMean | log2FoldChange | lfcSE | stat | pvalue | padj | absLog2FC |
|---|---|---|---|---|---|---|---|---|---|---|
| ENSMUSG00000037185 | Krt80 | 74127 | protein_coding | 6.269389088 | 1.371451277 | 0.76401475 | -1.795058639 | 0.072644358 | 0.196184284 | 1.371451277 |
| ENSMUSG00000079017 | Ifi27l2a | 76933 | protein_coding | 11.20778046 | -1.371512322 | 0.686553154 | 1.997678277 | 0.045751551 | 0.139398299 | 1.371512322 |
| ENSMUSG00000058099 | Nfam1 | 74039 | protein_coding | 11.76340877 | 1.372024828 | 0.808642662 | -1.696701018 | 0.0897532 | 0.228322063 | 1.372024828 |
| ENSMUSG00000063130 | Calml3 | 70405 | protein_coding | 0.908659759 | -1.372412961 | 1.547135768 | 0.88706692 | 0.375042872 | 1 | 1.372412961 |
| ENSMUSG00000040046 | Tph1 | 21990 | protein_coding | 192.2960384 | -1.372544829 | 0.292266222 | 4.696214366 | 2.65E-06 | 3.34E-05 | 1.372544829 |
| ENSMUSG00000004035 | Gstm7 | 68312 | protein_coding | 214.6190818 | 1.372928085 | 0.130917456 | -10.48697499 | 9.92E-26 | 6.50E-24 | 1.372928085 |
| ENSMUSG00000044927 | H1f10 | 243529 | protein_coding | 14.42064635 | 1.374035982 | 0.290020951 | -4.73771284 | 2.16E-06 | 2.80E-05 | 1.374035982 |
| ENSMUSG00000030256 | Bhlhe41 | 79362 | protein_coding | 1.945207451 | 1.374163123 | 1.271623982 | -1.080636369 | 0.279858898 | 1 | 1.374163123 |
| ENSMUSG00000031372 | Trex2 | 24102 | protein_coding | 1.322354558 | -1.37509257 | 1.286587972 | 1.068790164 | 0.285164232 | 1 | 1.37509257 |
| ENSMUSG00000027869 | Hsd3b6 | 15497 | protein_coding | 1.123723208 | -1.375449579 | 1.67490145 | 0.821212244 | 0.411525383 | 1 | 1.375449579 |
| ENSMUSG00000040372 | Gpr63 | 81006 | protein_coding | 1.489400742 | 1.375531859 | 1.371013749 | -1.003295452 | 0.31571833 | 1 | 1.375531859 |
| ENSMUSG00000037716 | Ccdc33 | 382077 | protein_coding | 2.588575124 | 1.375786748 | 1.843768604 | -0.746181893 | 0.455557544 | 1 | 1.375786748 |
| ENSMUSG00000098453 | Mir6997 | 102465603 | miRNA | 0.576033571 | -1.375935707 | 1.710118433 | 0.804585039 | 0.421059178 | 1 | 1.375935707 |
| ENSMUSG00000066268 | Or51a5 | 259115 | protein_coding | 1.162832834 | -1.377079016 | 2.430117373 | 0.566671812 | 0.570937175 | 1 | 1.377079016 |
| ENSMUSG00000027368 | Dusp2 | 13537 | protein_coding | 6.741259317 | 1.378170869 | 0.556709354 | -2.475566215 | 0.013302509 | 0.053152102 | 1.378170869 |
| ENSMUSG00000023439 | Gnb3 | 14695 | protein_coding | 2.912181327 | 1.37850255 | 1.155456777 | -1.193036881 | 0.232854929 | 1 | 1.37850255 |
| ENSMUSG00000025916 | Ppp1r42 | 69312 | protein_coding | 1.167585683 | 1.379349928 | 1.731294124 | -0.796716115 | 0.425615923 | 1 | 1.379349928 |
| ENSMUSG00000065298 | Gm26121 | 115487853 | snRNA | 0.484840685 | -1.379535719 | 1.387111787 | 0.994538242 | 0.319960897 | 1 | 1.379535719 |
| ENSMUSG00000034310 | Tmem132d | 243274 | protein_coding | 2.80699348 | 1.379590756 | 1.931301732 | -0.714332066 | 0.475021869 | 1 | 1.379590756 |
| ENSMUSG00000032978 | Guca2b | 14916 | protein_coding | 129.0855957 | -1.380488377 | 0.162418684 | 8.499566339 | 1.90E-17 | 8.29E-16 | 1.380488377 |
| ENSMUSG00000035041 | Creb3l3 | 208677 | protein_coding | 52.74436221 | 1.380649509 | 0.482927946 | -2.858914091 | 0.004250939 | 0.02121487 | 1.380649509 |
| ENSMUSG00000022064 | Pibf1 | 52023 | protein_coding | 407.7244951 | 1.380945398 | 0.077343429 | -17.85472165 | 2.66E-71 | 5.85E-69 | 1.380945398 |
| ENSMUSG00000049556 | Lingo1 | 235402 | protein_coding | 1.687008891 | 1.380998427 | 1.594104458 | -0.866316144 | 0.386316816 | 1 | 1.380998427 |
| ENSMUSG00000030399 | Ckm | 12715 | protein_coding | 0.97462739 | 1.38112741 | 1.878348542 | -0.735288142 | 0.462164028 | 1 | 1.38112741 |
| ENSMUSG00000025889 | Snca | 20617 | protein_coding | 8.58898549 | 1.381159212 | 0.448282262 | -3.081003489 | 0.002063042 | 0.011643568 | 1.381159212 |
| ENSMUSG00000066240 | Or10a49 | 258136 | protein_coding | 1.110662649 | 1.38120202 | 2.686711337 | -0.514086497 | 0.607191518 | 1 | 1.38120202 |
| ENSMUSG00000016024 | Lbp | 16803 | protein_coding | 39.78019434 | 1.381355931 | 0.4244141 | -3.254736188 | 0.001134977 | 0.007072837 | 1.381355931 |
| ENSMUSG00000054408 | Spcs3 | 76687 | protein_coding | 2180.067051 | 1.38179639 | 0.121510778 | -11.37180099 | 5.78E-30 | 4.50E-28 | 1.38179639 |
| ENSMUSG00000086264 | Gm15850 | 329253 | lncRNA | 2.484565507 | -1.382216818 | 0.946412507 | 1.460480295 | 0.144158119 | 1 | 1.382216818 |
| ENSMUSG00000036814 | Slc6a20a | 102680 | protein_coding | 21.3240507 | -1.383476719 | 0.339198026 | 4.078669723 | 4.53E-05 | 0.000429384 | 1.383476719 |
| ENSMUSG00000097934 | 6720483E21Rik | 77741 | lncRNA | 0.835988522 | -1.385363293 | 1.851740792 | 0.748141046 | 0.454375086 | 1 | 1.385363293 |
| ENSMUSG00000094053 | Scgb2b7 | 100043836 | protein_coding | 1.161706416 | -1.386150638 | 1.000487227 | 1.385475597 | 0.165907089 | 1 | 1.386150638 |
| ENSMUSG00000044339 | Alkbh2 | 231642 | protein_coding | 55.86575836 | 1.386266255 | 0.194492163 | -7.127620115 | 1.02E-12 | 3.09E-11 | 1.386266255 |
| ENSMUSG00000093539 | Mir5621 | 100885833 | miRNA | 0.672346343 | -1.386967519 | 1.5517895 | 0.893785864 | 0.371436474 | 1 | 1.386967519 |
| ENSMUSG00000028773 | Fabp3 | 14077 | protein_coding | 1.808511328 | -1.387040579 | 0.980598759 | 1.414483311 | 0.157220044 | 1 | 1.387040579 |
| ENSMUSG00000092867 | Mir3058 | 100526463 | miRNA | 2.436790921 | -1.387245404 | 0.763488069 | 1.816983737 | 0.069219604 | 1 | 1.387245404 |
| ENSMUSG00000022094 | Slc39a14 | 213053 | protein_coding | 983.5419524 | 1.387311401 | 0.126131894 | -10.99889458 | 3.87E-28 | 2.80E-26 | 1.387311401 |
| ENSMUSG00000004842 | Pou1f1 | 18736 | protein_coding | 3.034851355 | 1.387638956 | 1.409209904 | -0.984692878 | 0.324774952 | 1 | 1.387638956 |
| ENSMUSG00000018169 | Mfng | 17305 | protein_coding | 7.17005072 | 1.388906932 | 0.714216068 | -1.944659319 | 0.051815995 | 0.1532944 | 1.388906932 |

Supplementary Table 1

| ensembl_gene_id | external_gene_name | entrezgene_id | gene_biotype | baseMean | log2FoldChange | lfcSE | stat | pvalue | padj | absLog2FC |
|---|---|---|---|---|---|---|---|---|---|---|
| ENSMUSG00000029377 | Ereg | 13874 | protein_coding | 53.57728224 | 1.389298054 | 0.346692752 | -4.007289007 | 6.14E-05 | 0.000561931 | 1.389298054 |
| ENSMUSG00000118505 | Etv3l | 546801 | protein_coding | 1.976489853 | -1.389501579 | 1.324966897 | 1.048706637 | 0.294313159 | 1 | 1.389501579 |
| ENSMUSG00000038408 | 1700018A04Rik | 71307 | lncRNA | 2.679266158 | 1.389553854 | 1.471389351 | -0.944382161 | 0.344974394 | 1 | 1.389553854 |
| ENSMUSG00000064105 | Cnnm2 | 94219 | protein_coding | 142.4711345 | 1.389694696 | 0.130187379 | -10.67457314 | 1.34E-26 | 9.25E-25 | 1.389694696 |
| ENSMUSG00000023828 | Slc22a3 | 20519 | protein_coding | 7.233927993 | -1.389787566 | 0.481047991 | 2.889082985 | 0.003863671 | 0.019656396 | 1.389787566 |
| ENSMUSG00000036097 | Slf2 | 226151 | protein_coding | 882.297754 | 1.390677724 | 0.097314268 | -14.29058401 | 2.50E-46 | 3.11E-44 | 1.390677724 |
| ENSMUSG00000026527 | Rgs7 | 24012 | protein_coding | 3.877653452 | -1.391578431 | 0.64208258 | 2.167288872 | 0.030212834 | 1 | 1.391578431 |
| ENSMUSG00000024731 | Ms4a10 | 69826 | protein_coding | 25.92276679 | 1.391867146 | 0.330428203 | -4.212313391 | 2.53E-05 | 0.000254828 | 1.391867146 |
| ENSMUSG00000024538 | Ppic | 19038 | protein_coding | 26.3647279 | -1.395054202 | 0.348838877 | 3.999136254 | 6.36E-05 | 0.000580034 | 1.395054202 |
| ENSMUSG00000038421 | Fcrla | 98752 | protein_coding | 1.271922671 | 1.395232939 | 1.429570523 | -0.975980489 | 0.329074133 | 1 | 1.395232939 |
| ENSMUSG00000022757 | Tfg | 21787 | protein_coding | 1336.417049 | 1.395751208 | 0.067897134 | -20.55685023 | 6.68E-94 | 2.12E-91 | 1.395751208 |
| ENSMUSG00000072919 | Noxred1 | 71275 | protein_coding | 5.062051655 | 1.395870908 | 0.684529851 | -2.039167329 | 0.04143333 | 0.129368504 | 1.395870908 |
| ENSMUSG00000067571 | Ms4a15 | 545279 | protein_coding | 3.807789308 | 1.396012 | 0.749088702 | -1.863613743 | 0.062375966 | 1 | 1.396012 |
| ENSMUSG00000100417 | Gm28578 | 102639746 | lncRNA | 2.518686723 | 1.396256738 | 2.011594195 | -0.694104577 | 0.487616628 | 1 | 1.396256738 |
| ENSMUSG00000086564 | Cd101 | 630146 | protein_coding | 6.926529101 | 1.397339725 | 0.492162444 | -2.839183978 | 0.004522907 | 0.022396188 | 1.397339725 |
| ENSMUSG00000028197 | Col24a1 | 71355 | protein_coding | 4.766832061 | 1.399130236 | 2.264796798 | -0.617772966 | 0.536725006 | 0.732579283 | 1.399130236 |
| ENSMUSG00000077155 | Gm24582 | 115487856 | snRNA | 1.288019414 | -1.399136093 | 1.112866244 | 1.257236528 | 0.208668 | 1 | 1.399136093 |
| ENSMUSG00000027849 | Syt6 | 54524 | protein_coding | 10.69454274 | -1.400524197 | 0.375271769 | 3.73202653 | 0.000189945 | 0.00151381 | 1.400524197 |
| ENSMUSG00000039057 | Myo16 | 244281 | protein_coding | 3.805085077 | -1.400991757 | 1.977385323 | 0.70850721 | 0.478630337 | 1 | 1.400991757 |
| ENSMUSG00000010461 | Eya4 | 14051 | protein_coding | 2.26956525 | 1.401733296 | 2.410745025 | -0.581452323 | 0.560935642 | 1 | 1.401733296 |
| ENSMUSG00000070979 | Actl7a | 11470 | protein_coding | 0.72778767 | 1.401832785 | 1.8738287 | -0.748111492 | 0.454392911 | 1 | 1.401832785 |
| ENSMUSG00000055102 | Zfp819 | 74400 | protein_coding | 0.711663201 | 1.402109172 | 1.677824574 | -0.8356709 | 0.40334007 | 1 | 1.402109172 |
| ENSMUSG00000043873 | Chil5 | 229687 | protein_coding | 1.180878237 | 1.4026869 | 1.131219831 | -1.23997729 | 0.214983794 | 1 | 1.4026869 |
| ENSMUSG00000043165 | Lor | 16939 | protein_coding | 3.38469467 | 1.402764016 | 0.71027309 | -1.974964328 | 0.048272196 | 1 | 1.402764016 |
| ENSMUSG00000054871 | Tmem158 | 72309 | protein_coding | 8.922913228 | 1.403701197 | 0.416181814 | -3.372807628 | 0.000744059 | 0.004936679 | 1.403701197 |
| ENSMUSG00000048483 | Zdhhc22 | 238331 | protein_coding | 2.825565216 | -1.404058128 | 0.929237794 | 1.510978284 | 0.130793984 | 1 | 1.404058128 |
| ENSMUSG00000044854 | 1700056E22Rik | 73363 | protein_coding | 16.15835165 | 1.405396514 | 0.41029548 | -3.425327801 | 0.000614058 | 0.004212943 | 1.405396514 |
| ENSMUSG00000112666 | Gm48485 | 382395 | lncRNA | 1.362269868 | -1.40578781 | 1.661026745 | 0.84633665 | 0.397364968 | 1 | 1.40578781 |
| ENSMUSG00000051727 | Kctd14 | 233529 | protein_coding | 187.8613581 | 1.405995648 | 0.154831196 | -9.08082921 | 1.08E-19 | 5.38E-18 | 1.405995648 |
| ENSMUSG00000020067 | Mypn | 68802 | protein_coding | 1.876153238 | 1.408791457 | 2.367224129 | -0.595123816 | 0.55176071 | 1 | 1.408791457 |
| ENSMUSG00000005696 | Sh2d1a | 20400 | protein_coding | 0.924569147 | 1.408920385 | 1.885558338 | -0.747216544 | 0.454932859 | 1 | 1.408920385 |
| ENSMUSG00000041399 | 1700013G24Rik | 69380 | protein_coding | 1.726562693 | -1.409123325 | 1.295774707 | 1.08747556 | 0.276826694 | 1 | 1.409123325 |
| ENSMUSG00000052271 | Bhlha15 | 17341 | protein_coding | 67.61474922 | 1.40918012 | 0.224605389 | -6.274026322 | 3.52E-10 | 8.22E-09 | 1.40918012 |
| ENSMUSG00000085723 | Gm15915 | 100504069 | lncRNA | 1.044800893 | -1.409322879 | 1.253135793 | 1.124637 | 0.260742888 | 1 | 1.409322879 |
| ENSMUSG00000066510 | Ankdd1a | 330963 | protein_coding | 1.625457394 | 1.41183606 | 1.910796115 | -0.738873211 | 0.459983991 | 1 | 1.41183606 |
| ENSMUSG00000073119 | Speer4a | 75657 | protein_coding | 1.981369028 | 1.411869924 | 2.395449855 | -0.589396568 | 0.555595278 | 1 | 1.411869924 |
| ENSMUSG00000031131 | Vgll1 | 170828 | protein_coding | 0.635092792 | -1.412665438 | 1.561048324 | 0.904946642 | 0.365493656 | 1 | 1.412665438 |
| ENSMUSG00000024330 | Col11a2 | 12815 | protein_coding | 55.52793372 | 1.412690863 | 0.416635954 | -3.390707997 | 0.000697123 | 0.004664625 | 1.412690863 |
| ENSMUSG00000034224 | Slc38a8 | 234788 | protein_coding | 1.18216771 | 1.413872086 | 1.815349234 | -0.778843023 | 0.436072191 | 1 | 1.413872086 |

| ensembl_gene_id | external_gene_name | entrezgene_id | gene_biotype | baseMean | log2FoldChange | lfcSE | stat | pvalue | padj | absLog2FC |
|---|---|---|---|---|---|---|---|---|---|---|
| ENSMUSG00000006216 | Clcnkb | 56365 | protein_coding | 3.013964932 | -1.413948679 | 1.447561575 | 0.976779643 | 0.328678256 | 1 | 1.413948679 |
| ENSMUSG00000026520 | Pycr2 | 69051 | protein_coding | 593.5837822 | 1.415453693 | 0.085859368 | -16.48572227 | 4.65E-61 | 8.08E-59 | 1.415453693 |
| ENSMUSG00000043932 | Klri2 | 320407 | protein_coding | 1.450010585 | 1.416073998 | 1.104439037 | -1.282165833 | 0.199784479 | 1 | 1.416073998 |
| ENSMUSG00000039760 | Il22ra2 | 237310 | protein_coding | 3.20345551 | -1.417435554 | 1.302635827 | 1.088128796 | 0.276538253 | 1 | 1.417435554 |
| ENSMUSG00000097845 | A230020J21Rik | 100125339 | lncRNA | 1.420325988 | 1.417747328 | 1.267896099 | -1.118188887 | 0.26348633 | 1 | 1.417747328 |
| ENSMUSG00000037138 | Aff3 | 16764 | protein_coding | 3.14464763 | 1.418782707 | 1.283668566 | -1.105256251 | 0.269048569 | 1 | 1.418782707 |
| ENSMUSG00000041670 | Rims1 | 116837 | protein_coding | 3.779882932 | 1.420724864 | 1.376488566 | -1.032137062 | 0.302007914 | 1 | 1.420724864 |
| ENSMUSG00000037548 | H2-DMb2 | 15000 | protein_coding | 1.489243693 | -1.422224579 | 0.880457175 | 1.615325105 | 0.10624031 | 1 | 1.422224579 |
| ENSMUSG00000031362 | Xlr4c | 72891 | protein_coding | 1.160002016 | -1.42268944 | 1.408668129 | 1.009953594 | 0.312517524 | 1 | 1.42268944 |
| ENSMUSG00000031448 | Adprhl1 | 234072 | protein_coding | 4.529116466 | 1.422988507 | 0.599736743 | -2.372688555 | 0.017659147 | 0.066595834 | 1.422988507 |
| ENSMUSG00000079177 | Fam228a | 74855 | protein_coding | 2.117430516 | 1.423202442 | 1.355117602 | -1.050242754 | 0.293606518 | 1 | 1.423202442 |
| ENSMUSG00000020873 | Slc35b1 | 110172 | protein_coding | 1347.80781 | 1.423523792 | 0.080543403 | -17.6739962 | 6.65E-70 | 1.43E-67 | 1.423523792 |
| ENSMUSG00000000686 | Abhd15 | 67477 | protein_coding | 5.984832882 | 1.424560034 | 0.517384947 | -2.753385157 | 0.005898246 | 0.02782961 | 1.424560034 |
| ENSMUSG00000038860 | Garnl3 | 99326 | protein_coding | 10.45907661 | -1.424585141 | 0.61255682 | 2.325637549 | 0.020037895 | 0.073420411 | 1.424585141 |
| ENSMUSG00000050505 | Pcdh20 | 219257 | protein_coding | 11.08969581 | -1.427886537 | 0.439747976 | 3.247056529 | 0.001166052 | 0.007239168 | 1.427886537 |
| ENSMUSG00000021613 | Hapln1 | 12950 | protein_coding | 15.81057969 | 1.428737819 | 0.332555268 | -4.296241724 | 1.74E-05 | 0.000182029 | 1.428737819 |
| ENSMUSG00000075410 | Prcd | 100038570 | protein_coding | 1.918687223 | -1.429746274 | 1.074354405 | 1.330795747 | 0.183256228 | 1 | 1.429746274 |
| ENSMUSG00000079076 | Gm3086 | 100040999 | transcribed_processed_pseudogene | 0.785873882 | -1.429940603 | 1.391445115 | 1.027665833 | 0.304107045 | 1 | 1.429940603 |
| ENSMUSG00000028906 | Epb41 | 269587 | protein_coding | 2266.175467 | 1.430082918 | 0.058170386 | -24.58438086 | 1.86E-133 | 8.51E-131 | 1.430082918 |
| ENSMUSG00000032126 | Hmbs | 15288 | protein_coding | 776.9031489 | 1.43232264 | 0.08539061 | -16.77377222 | 3.80E-63 | 6.67E-61 | 1.43232264 |
| ENSMUSG00000054901 | Arhgef33 | 381112 | protein_coding | 4.922610245 | 1.432715792 | 0.836521732 | -1.712706002 | 0.08676664 | 0.223013899 | 1.432715792 |
| ENSMUSG00000019178 | Styxl1 | 76571 | protein_coding | 2.402072907 | -1.433052406 | 0.781330113 | 1.834119001 | 0.066636332 | 1 | 1.433052406 |
| ENSMUSG00000029663 | Gngt1 | 14699 | protein_coding | 1.227679324 | 1.433507511 | 1.27034098 | -1.128443097 | 0.259132835 | 1 | 1.433507511 |
| ENSMUSG00000031963 | Bmper | 73230 | protein_coding | 13.81115173 | 1.43390898 | 0.435334385 | -3.293810525 | 0.000988391 | 0.006304476 | 1.43390898 |
| ENSMUSG00000021848 | Otx2 | 18424 | protein_coding | 2.226124182 | 1.434093305 | 1.794532554 | -0.799145885 | 0.424205826 | 1 | 1.434093305 |
| ENSMUSG00000054320 | Lrrc36 | 270091 | protein_coding | 2.089028927 | -1.434309532 | 0.897359348 | 1.598366959 | 0.109961333 | 1 | 1.434309532 |
| ENSMUSG00000055827 | Gsdmc3 | 270328 | protein_coding | 997.6114602 | -1.434940877 | 0.27960675 | 5.131996554 | 2.87E-07 | 4.40E-06 | 1.434940877 |
| ENSMUSG00000091906 | 1700099I09Rik | 76624 | lncRNA | 0.795227036 | 1.435015476 | 1.527267338 | -0.939596782 | 0.347424428 | 1 | 1.435015476 |
| ENSMUSG00000087252 | Gm14379 | 100270707 | lncRNA | 1.267069922 | 1.437009527 | 1.041380809 | -1.379907824 | 0.167615027 | 1 | 1.437009527 |
| ENSMUSG00000017491 | Rarb | 218772 | protein_coding | 2.095730838 | 1.437374105 | 1.222436816 | -1.175826911 | 0.23966405 | 1 | 1.437374105 |
| ENSMUSG00000020490 | Btnl10 | 192194 | protein_coding | 2.317928247 | 1.437913241 | 1.63992996 | -0.876813813 | 0.380587775 | 1 | 1.437913241 |
| ENSMUSG00000027828 | Ssr3 | 67437 | protein_coding | 4947.580592 | 1.438327976 | 0.096752022 | -14.86612839 | 5.47E-50 | 7.72E-48 | 1.438327976 |
| ENSMUSG00000092586 | Ly6g6c | 68468 | protein_coding | 5.057591496 | -1.438722695 | 0.778138496 | 1.848928826 | 0.064468091 | 0.179926661 | 1.438722695 |
| ENSMUSG00000030732 | Chrdl2 | 69121 | protein_coding | 2.388850947 | -1.438763147 | 2.001529405 | 0.718831881 | 0.472244512 | 1 | 1.438763147 |
| ENSMUSG00000042678 | Myo15a | 17910 | protein_coding | 8.583695686 | 1.439644038 | 1.232640795 | -1.167934766 | 0.242833079 | 0.453944473 | 1.439644038 |
| ENSMUSG00000028417 | Tal2 | 21350 | protein_coding | 2.110814455 | -1.439919656 | 0.947505866 | 1.519694715 | 0.128587721 | 1 | 1.439919656 |
| ENSMUSG00000036053 | Fmnl2 | 71409 | protein_coding | 47.23679329 | 1.439968829 | 0.223168368 | -6.452387695 | 1.10E-10 | 2.71E-09 | 1.439968829 |
| ENSMUSG00000027474 | Ccm2l | 228788 | protein_coding | 2.396508759 | -1.440489346 | 1.595241563 | 0.902991359 | 0.366530481 | 1 | 1.440489346 |
| ENSMUSG00000038540 | Tmc3 | 233424 | protein_coding | 3.191572806 | 1.441333575 | 0.74141566 | -1.944029041 | 0.051891949 | 1 | 1.441333575 |

Supplementary Table 1

| ensembl_gene_id | external_gene_name | entrezgene_id | gene_biotype | baseMean | log2FoldChange | lfcSE | stat | pvalue | padj | absLog2FC |
|---|---|---|---|---|---|---|---|---|---|---|
| ENSMUSG00000094626 | Tmem121b | 94047 | protein_coding | 1.39922162 | -1.442346854 | 1.092443928 | 1.320293717 | 0.186736973 | 1 | 1.442346854 |
| ENSMUSG00000051373 | Plpp7 | 227721 | protein_coding | 3.422705866 | -1.442717587 | 0.642805501 | 2.244407655 | 0.024806184 | 1 | 1.442717587 |
| ENSMUSG00000074060 | Fbxw15 | 382105 | protein_coding | 1.194524895 | 1.442983445 | 1.73301989 | -0.832641018 | 0.405047218 | 1 | 1.442983445 |
| ENSMUSG00000001943 | Vsig2 | 57276 | protein_coding | 89.50811043 | -1.443213287 | 0.147087763 | 9.811919493 | 1.00E-22 | 5.86E-21 | 1.443213287 |
| ENSMUSG00000029287 | Tgfbr3 | 21814 | protein_coding | 2.827024144 | 1.443911404 | 1.183528544 | -1.22000556 | 0.222462767 | 1 | 1.443911404 |
| ENSMUSG00000064867 | Gm25039 | 115489772 | snRNA | 0.706690239 | -1.444944969 | 1.270490844 | 1.137312383 | 0.25540772 | 1 | 1.444944969 |
| ENSMUSG00000022211 | Carmil3 | 268747 | protein_coding | 11.82666795 | -1.445423401 | 0.610780797 | 2.366517429 | 0.017956329 | 0.067342119 | 1.445423401 |
| ENSMUSG00000023041 | Krt6b | 16688 | protein_coding | 1.164485416 | -1.445619089 | 1.198704527 | 1.205984508 | 0.227823464 | 1 | 1.445619089 |
| ENSMUSG00000029206 | Nsun7 | 70918 | protein_coding | 1.938649695 | 1.446011681 | 1.367864672 | -1.057130658 | 0.290451961 | 1 | 1.446011681 |
| ENSMUSG00000051674 | Dcun1d4 | 100737 | protein_coding | 228.4345946 | 1.447035708 | 0.10476689 | -13.81195628 | 2.16E-43 | 2.49E-41 | 1.447035708 |
| ENSMUSG00000045474 | Or2ad1 | 258527 | protein_coding | 0.740054721 | -1.44729684 | 1.625969492 | 0.890113158 | 0.373405128 | 1 | 1.44729684 |
| ENSMUSG00000058287 | Gm12253 | 624860 | protein_coding | 3.504813299 | 1.447521761 | 0.724018358 | -1.999288754 | 0.04557712 | 1 | 1.447521761 |
| ENSMUSG00000033227 | Wnt6 | 22420 | protein_coding | 5.359641403 | -1.448265448 | 0.602610826 | 2.403318004 | 0.016247052 | 0.062280366 | 1.448265448 |
| ENSMUSG00000097365 | C030034L19Rik | 320088 | lncRNA | 2.708077629 | 1.448493493 | 1.176272739 | -1.23142656 | 0.218163368 | 1 | 1.448493493 |
| ENSMUSG00000047182 | Irs3 | 16369 | protein_coding | 1.416035437 | -1.449030346 | 1.215703141 | 1.19192778 | 0.233289564 | 1 | 1.449030346 |
| ENSMUSG00000056863 | Or13n4 | 258590 | protein_coding | 0.909430882 | -1.449131639 | 1.87469998 | 0.772993895 | 0.439525994 | 1 | 1.449131639 |
| ENSMUSG00000042845 | Wfdc12 | 192200 | protein_coding | 0.772048828 | 1.449344582 | 1.276919904 | -1.13503171 | 0.256362032 | 1 | 1.449344582 |
| ENSMUSG00000046275 | Trarg1 | 237858 | protein_coding | 1.134324471 | -1.450210411 | 1.439554199 | 1.007402439 | 0.31374142 | 1 | 1.450210411 |
| ENSMUSG00000100937 | Nscme3l | 75555 | protein_coding | 0.753743667 | 1.450356195 | 1.517577883 | -0.955704621 | 0.339221487 | 1 | 1.450356195 |
| ENSMUSG00000049100 | Pcdh10 | 18526 | protein_coding | 2.872992009 | 1.45087418 | 1.845784127 | -0.786047598 | 0.431839602 | 1 | 1.45087418 |
| ENSMUSG00000030219 | Erp27 | 69187 | protein_coding | 0.816285465 | 1.4516082 | 1.447340067 | -1.00294895 | 0.315885493 | 1 | 1.4516082 |
| ENSMUSG00000097486 | Gm17733 | 100503783 | lncRNA | 1.332890145 | 1.452666802 | 0.993398443 | -1.462320393 | 0.143653433 | 1 | 1.452666802 |
| ENSMUSG00000050382 | Kif7 | 16576 | protein_coding | 4.083267246 | 1.454037969 | 0.805020109 | -1.806213227 | 0.07088504 | 1 | 1.454037969 |
| ENSMUSG00000024352 | Spata24 | 71242 | protein_coding | 12.41998409 | 1.454309395 | 0.359053568 | -4.050396726 | 5.11E-05 | 0.000475973 | 1.454309395 |
| ENSMUSG00000092515 | C87198 | 97836 | lncRNA | 1.496234869 | 1.457311324 | 2.383530159 | -0.611408804 | 0.540928976 | 1 | 1.457311324 |
| ENSMUSG00000035745 | Grin3b | 170483 | protein_coding | 4.292617236 | 1.457565652 | 0.748494487 | -1.947329843 | 0.051495203 | 1 | 1.457565652 |
| ENSMUSG00000029591 | Ung | 22256 | protein_coding | 322.0791014 | 1.458232303 | 0.113816133 | -12.81217582 | 1.40E-37 | 1.32E-35 | 1.458232303 |
| ENSMUSG00000064140 | Trim38 | 214158 | protein_coding | 20.05467493 | -1.458358196 | 0.315372949 | 4.624233631 | 3.76E-06 | 4.63E-05 | 1.458358196 |
| ENSMUSG00000040640 | Erc2 | 238988 | protein_coding | 6.195600758 | 1.459473638 | 0.959657896 | -1.520827 | 0.128303257 | 0.2954831 | 1.459473638 |
| ENSMUSG00000043605 | Or2a7 | 18310 | protein_coding | 1.160465358 | -1.459618934 | 1.803816889 | 0.809183539 | 0.418409582 | 1 | 1.459618934 |
| ENSMUSG00000021804 | Rgr | 57811 | protein_coding | 1.353429415 | 1.459943037 | 2.117943761 | -0.689320965 | 0.490621308 | 1 | 1.459943037 |
| ENSMUSG00000043687 | 1190005I06Rik | 68918 | protein_coding | 2.790128414 | 1.459993733 | 0.777661443 | -1.877415612 | 0.060461153 | 1 | 1.459993733 |
| ENSMUSG00000099950 | 9130227L01Rik | 329159 | lncRNA | 27.40535469 | 1.460080735 | 0.295949128 | -4.933553095 | 8.07E-07 | 1.14E-05 | 1.460080735 |
| ENSMUSG00000018543 | 1700001P01Rik | 72215 | protein_coding | 1.319689198 | -1.460328718 | 1.172901119 | 1.245056974 | 0.213110808 | 1 | 1.460328718 |
| ENSMUSG00000036961 | Wnt8b | 22423 | protein_coding | 10.46295172 | -1.46161032 | 0.592658888 | 2.466191514 | 0.01365583 | 0.05423578 | 1.46161032 |
| ENSMUSG00000048387 | Osr1 | 23967 | protein_coding | 0.861549015 | 1.461714268 | 1.950901254 | -0.749250771 | 0.453706073 | 1 | 1.461714268 |
| ENSMUSG00000001403 | Ube2c | 68612 | protein_coding | 600.5691237 | 1.462138743 | 0.159933247 | -9.142181318 | 6.12E-20 | 3.09E-18 | 1.462138743 |
| ENSMUSG00000098731 | Mir7027 | 102465621 | miRNA | 0.747789477 | -1.462499108 | 1.35738511 | 1.077438596 | 0.281284369 | 1 | 1.462499108 |
| ENSMUSG00000006403 | Adamts4 | 240913 | protein_coding | 4.033668262 | 1.462598416 | 0.994750542 | -1.470316782 | 0.141475978 | 1 | 1.462598416 |

Supplementary Table 1

| ensembl_gene_id | external_gene_name | entrezgene_id | gene_biotype | baseMean | log2FoldChange | lfcSE | stat | pvalue | padj | absLog2FC |
|---|---|---|---|---|---|---|---|---|---|---|
| ENSMUSG00000087591 | Gm14635 | 100043946 | lncRNA | 0.750415115 | 1.462847708 | 1.891994981 | -0.773177373 | 0.439417416 | 1 | 1.462847708 |
| ENSMUSG00000046182 | Gsg1l | 269994 | protein_coding | 3.720392912 | -1.463495699 | 1.959771509 | 0.746768535 | 0.455203292 | 1 | 1.463495699 |
| ENSMUSG00000024175 | Tekt4 | 71840 | protein_coding | 3.371651136 | 1.463829371 | 1.065378097 | -1.373999874 | 0.169441722 | 1 | 1.463829371 |
| ENSMUSG00000046807 | Lrrc75b | 192734 | protein_coding | 16.02166863 | 1.464763071 | 0.535217008 | -2.736764805 | 0.006204664 | 0.028888694 | 1.464763071 |
| ENSMUSG00000067702 | Tuba3a | 22144 | protein_coding | 1.93284952 | 1.464830046 | 1.280352329 | -1.144083556 | 0.252588986 | 1 | 1.464830046 |
| ENSMUSG00000021903 | Galnt15 | 78754 | protein_coding | 2.356348179 | 1.465996439 | 2.044145201 | -0.717168447 | 0.473270166 | 1 | 1.465996439 |
| ENSMUSG00000028008 | Asic5 | 58170 | protein_coding | 1.465550213 | -1.466059043 | 1.988869797 | 0.737131734 | 0.461042242 | 1 | 1.466059043 |
| ENSMUSG00000085824 | Platr9 | 73696 | lncRNA | 1.629678579 | -1.467125609 | 2.202642992 | 0.666075081 | 0.505363111 | 1 | 1.467125609 |
| ENSMUSG00000070687 | Htr1d | 15552 | protein_coding | 2.206250495 | -1.467256229 | 1.331801934 | 1.101707537 | 0.270588839 | 1 | 1.467256229 |
| ENSMUSG00000004031 | Brinp2 | 240843 | protein_coding | 3.108091818 | 1.467778209 | 1.696785295 | -0.86503473 | 0.387019727 | 1 | 1.467778209 |
| ENSMUSG00000079262 | Slco1a6 | 28254 | protein_coding | 1.668431151 | -1.468932821 | 1.383853941 | 1.061479667 | 0.288471968 | 1 | 1.468932821 |
| ENSMUSG00000063230 | Or13a22 | 258956 | protein_coding | 1.879730895 | 1.469317749 | 3.121915194 | -0.470646273 | 0.637893358 | 1 | 1.469317749 |
| ENSMUSG00000087125 | A230108P19Rik | 320132 | lncRNA | 1.044039301 | 1.470027507 | 1.52062678 | -0.966724726 | 0.333681663 | 1 | 1.470027507 |
| ENSMUSG00000025370 | Cdh9 | 12565 | protein_coding | 1.199092929 | -1.470249984 | 2.231304606 | 0.658919441 | 0.509947501 | 1 | 1.470249984 |
| ENSMUSG00000054568 | Usp17la | 13531 | protein_coding | 1.313194681 | -1.470286713 | 2.248250786 | 0.653969176 | 0.513131665 | 1 | 1.470286713 |
| ENSMUSG00000046585 | Cfap58 | 381229 | protein_coding | 1.549914938 | 1.470520149 | 2.020127108 | -0.727934467 | 0.466653703 | 1 | 1.470520149 |
| ENSMUSG00000067971 | Or5o1 | 236785 | protein_coding | 1.862543863 | -1.471149208 | 2.256580396 | 0.651937423 | 0.514441542 | 1 | 1.471149208 |
| ENSMUSG00000058246 | Gm10037 | 102637366 | protein_coding | 2.830242565 | -1.471512165 | 0.998556035 | 1.473640049 | 0.140578532 | 1 | 1.471512165 |
| ENSMUSG00000094822 | Or52a20 | 436002 | protein_coding | 1.404505133 | -1.471584607 | 2.243935704 | 0.655805157 | 0.511949498 | 1 | 1.471584607 |
| ENSMUSG00000005220 | Corin | 53419 | protein_coding | 1.586245427 | -1.472269972 | 2.250515379 | 0.654192363 | 0.512987882 | 1 | 1.472269972 |
| ENSMUSG00000068196 | Col8a1 | 12837 | protein_coding | 374.0047587 | -1.472900312 | 0.299587579 | 4.916426494 | 8.81E-07 | 1.23E-05 | 1.472900312 |
| ENSMUSG00000035576 | L3mbtl1 | 241764 | protein_coding | 5.853526412 | 1.472901145 | 1.004058245 | -1.466947911 | 0.142390226 | 0.317889981 | 1.472901145 |
| ENSMUSG00000117800 | 1700122C19Rik | 78640 | lncRNA | 1.863601414 | -1.472916094 | 2.256777376 | 0.652663444 | 0.513973275 | 1 | 1.472916094 |
| ENSMUSG00000057933 | Gsta2 | 14858 | protein_coding | 0.826317336 | 1.475057786 | 1.874009434 | -0.787113319 | 0.431215533 | 1 | 1.475057786 |
| ENSMUSG00000016346 | Kcnq2 | 16536 | protein_coding | 6.284709395 | 1.47574415 | 1.54801961 | -0.953311018 | 0.340432511 | 0.561998621 | 1.47574415 |
| ENSMUSG00000093839 | Or4d2b | 258406 | protein_coding | 1.285503888 | -1.476051154 | 1.628807753 | 0.906215698 | 0.364821696 | 1 | 1.476051154 |
| ENSMUSG00000048376 | F2r | 14062 | protein_coding | 6.51169626 | 1.476879918 | 0.573044934 | -2.57724976 | 0.009958996 | 0.042442029 | 1.476879918 |
| ENSMUSG00000022103 | Gfra2 | 14586 | protein_coding | 2.511571686 | 1.477486661 | 2.433342096 | -0.607184113 | 0.543728735 | 1 | 1.477486661 |
| ENSMUSG00000030731 | Syt3 | 20981 | protein_coding | 8.027651423 | -1.477556395 | 0.616381002 | 2.397147852 | 0.016523255 | 0.063163803 | 1.477556395 |
| ENSMUSG00000024518 | Rax | 19434 | protein_coding | 2.147473012 | 1.477811694 | 2.412305722 | -0.612613766 | 0.540131753 | 1 | 1.477811694 |
| ENSMUSG00000033854 | Kcnk10 | 72258 | protein_coding | 7.008110588 | -1.477889475 | 0.502785701 | 2.939402357 | 0.003288459 | 0.017169019 | 1.477889475 |
| ENSMUSG00000033061 | Resp18 | 19711 | protein_coding | 19.2440005 | -1.478248205 | 0.329229753 | 4.490020084 | 7.12E-06 | 8.26E-05 | 1.478248205 |
| ENSMUSG00000118454 | Wdr88 | 384605 | protein_coding | 1.57190938 | 1.47853495 | 1.934901187 | -0.764139771 | 0.444783954 | 1 | 1.47853495 |
| ENSMUSG00000085957 | Syna | 214292 | protein_coding | 10.83686801 | 1.478676801 | 0.420492891 | -3.51653222 | 0.000437224 | 0.003173763 | 1.478676801 |
| ENSMUSG00000006649 | Nphs1 | 54631 | protein_coding | 2.546233201 | -1.47870466 | 2.098109008 | 0.704779711 | 0.480947337 | 1 | 1.47870466 |
| ENSMUSG00000068099 | Smim45 | 76505 | protein_coding | 1.959199709 | -1.479193358 | 1.713263977 | 0.863377377 | 0.387930014 | 1 | 1.479193358 |
| ENSMUSG00000021751 | Acox2 | 93732 | protein_coding | 73.27784375 | 1.479468001 | 0.226332577 | -6.536699329 | 6.29E-11 | 1.63E-09 | 1.479468001 |
| ENSMUSG00000111296 | 2010001M07Rik | 115487174 | lncRNA | 8.936056747 | -1.480191538 | 0.409293541 | 3.616454673 | 0.000298666 | 0.002269749 | 1.480191538 |
| ENSMUSG00000029245 | Epha5 | 13839 | protein_coding | 5.090829212 | 1.480855931 | 1.20771782 | -1.226160537 | 0.220138269 | 0.426485613 | 1.480855931 |

| ensembl_gene_id | external_gene_name | entrezgene_id | gene_biotype | baseMean | log2FoldChange | lfcSE | stat | pvalue | padj | absLog2FC |
|---|---|---|---|---|---|---|---|---|---|---|
| ENSMUSG00000024059 | Clip4 | 78785 | protein_coding | 2.721624859 | 1.480995889 | 2.001791389 | -0.739835278 | 0.45939995 | 1 | 1.480995889 |
| ENSMUSG00000029616 | Erp29 | 67397 | protein_coding | 1751.466239 | 1.481045072 | 0.058820813 | -25.17892878 | 6.82E-140 | 3.52E-137 | 1.481045072 |
| ENSMUSG00000036052 | Dnajb5 | 56323 | protein_coding | 80.9883854 | 1.481651878 | 0.174270917 | -8.502003098 | 1.86E-17 | 8.14E-16 | 1.481651878 |
| ENSMUSG00000033368 | Trim69 | 70928 | protein_coding | 1.096058246 | 1.482180746 | 1.365525799 | -1.085428593 | 0.277731874 | 1 | 1.482180746 |
| ENSMUSG00000002769 | Gnmt | 14711 | protein_coding | 4.511727037 | 1.483097264 | 0.644201825 | -2.302224562 | 0.021322512 | 0.077202358 | 1.483097264 |
| ENSMUSG00000059776 | Rpl13-ps6 | 100040416 | processed_pseudogene | 1.330390968 | 1.483493716 | 1.211670297 | -1.224337775 | 0.220824832 | 1 | 1.483493716 |
| ENSMUSG00000000805 | Car4 | 12351 | protein_coding | 1755.379997 | -1.484209494 | 0.397379027 | 3.734997049 | 0.000187717 | 0.001499741 | 1.484209494 |
| ENSMUSG00000010307 | Tmem86a | 67893 | protein_coding | 21.31619087 | -1.48442056 | 0.257734941 | 5.759485138 | 8.44E-09 | 1.64E-07 | 1.48442056 |
| ENSMUSG00000006369 | Fbln1 | 14114 | protein_coding | 62.41598309 | 1.484863979 | 0.237647251 | -6.248184962 | 4.15E-10 | 9.56E-09 | 1.484863979 |
| ENSMUSG00000108228 | 6430584L05Rik | 330324 | lncRNA | 1.554186567 | -1.485086758 | 2.026239526 | 0.732927543 | 0.463602623 | 1 | 1.485086758 |
| ENSMUSG00000040554 | Aipl1 | 114230 | protein_coding | 1.766193046 | -1.485127502 | 1.323625326 | 1.122015024 | 0.261856052 | 1 | 1.485127502 |
| ENSMUSG00000021306 | Gpr137b | 83924 | protein_coding | 81.03492382 | -1.485338598 | 0.260256555 | 5.707209183 | 1.15E-08 | 2.20E-07 | 1.485338598 |
| ENSMUSG00000071185 | Or1i2 | 257883 | protein_coding | 1.09173548 | -1.485558275 | 2.032032841 | 0.731070013 | 0.46473639 | 1 | 1.485558275 |
| ENSMUSG00000032968 | Inha | 16322 | protein_coding | 5.133330042 | 1.486002057 | 0.550952629 | -2.697150318 | 0.006993569 | 0.031886196 | 1.486002057 |
| ENSMUSG00000027015 | Cybrd1 | 73649 | protein_coding | 2.124995997 | 1.486270984 | 1.030298368 | -1.44256366 | 0.149143427 | 1 | 1.486270984 |
| ENSMUSG00000032174 | Icam5 | 15898 | protein_coding | 1.033507203 | 1.487315035 | 1.434245904 | -1.037001417 | 0.299735197 | 1 | 1.487315035 |
| ENSMUSG00000056061 | Gata5os | 100036520 | lncRNA | 1.956617672 | -1.488677876 | 1.770371566 | 0.840884425 | 0.400412685 | 1 | 1.488677876 |
| ENSMUSG00000089555 | Mir1933 | 100316753 | miRNA | 1.485615234 | 1.488759848 | 0.979750839 | -1.519529036 | 0.128629385 | 1 | 1.488759848 |
| ENSMUSG00000090417 | Vmn2r94 | 665227 | protein_coding | 1.576440034 | -1.488972246 | 2.076159224 | 0.717176327 | 0.473265304 | 1 | 1.488972246 |
| ENSMUSG00000049036 | Tmem121 | 69195 | protein_coding | 2.154146926 | -1.490423079 | 0.97678906 | 1.525839243 | 0.127049905 | 1 | 1.490423079 |
| ENSMUSG00000039153 | Runx2 | 12393 | protein_coding | 5.549929042 | 1.490647522 | 1.261288747 | -1.18184478 | 0.237267296 | 0.44767168 | 1.490647522 |
| ENSMUSG00000034730 | Adgrb1 | 107831 | protein_coding | 10.56038952 | -1.491196479 | 0.572303246 | 2.60560549 | 0.009171203 | 0.039741076 | 1.491196479 |
| ENSMUSG00000050192 | Eif5a2 | 208691 | protein_coding | 8.905403062 | 1.492319253 | 0.393439607 | -3.793007176 | 0.000148834 | 0.001219773 | 1.492319253 |
| ENSMUSG00000021760 | Gpx8 | 69590 | protein_coding | 7.983836285 | 1.492645568 | 0.415315485 | -3.59400413 | 0.000325635 | 0.00244767 | 1.492645568 |
| ENSMUSG00000044576 | Garem2 | 242915 | protein_coding | 3.971275409 | 1.493682623 | 1.013805745 | -1.473342039 | 0.140658831 | 1 | 1.493682623 |
| ENSMUSG00000029516 | Cit | 12704 | protein_coding | 336.8555674 | 1.495129073 | 0.177681762 | -8.414645711 | 3.94E-17 | 1.68E-15 | 1.495129073 |
| ENSMUSG00000033597 | Caskin1 | 268932 | protein_coding | 11.79892737 | 1.496391769 | 0.590449129 | -2.534328016 | 0.011266322 | 0.046817321 | 1.496391769 |
| ENSMUSG00000056978 | Hamp2 | 66438 | protein_coding | 1.25414867 | -1.497278567 | 1.289837813 | 1.160827006 | 0.245712258 | 1 | 1.497278567 |
| ENSMUSG00000030617 | Ccdc83 | 75338 | protein_coding | 1.313634986 | -1.49781879 | 2.556908103 | 0.585792969 | 0.558014649 | 1 | 1.49781879 |
| ENSMUSG00000074336 | Apoc4 | 11425 | protein_coding | 2.429506656 | -1.498522387 | 2.238531275 | 0.669422136 | 0.503226235 | 1 | 1.498522387 |
| ENSMUSG00000028332 | Hemgn | 93966 | protein_coding | 1.361316184 | -1.498533719 | 2.238673449 | 0.669384684 | 0.503250119 | 1 | 1.498533719 |
| ENSMUSG00000075837 | Gm22005 | 115489653 | snRNA | 0.926332705 | -1.49996275 | 1.2190344 | 1.230451536 | 0.218528065 | 1 | 1.49996275 |
| ENSMUSG00000047394 | Odf3b | 70113 | protein_coding | 36.33959206 | -1.500682038 | 0.280444357 | 5.351086583 | 8.74E-08 | 1.46E-06 | 1.500682038 |
| ENSMUSG00000028033 | Kcnq5 | 226922 | protein_coding | 2.732777938 | -1.500964582 | 2.292928801 | 0.654605839 | 0.512721564 | 1 | 1.500964582 |
| ENSMUSG00000097694 | G730013B05Rik | 619312 | lncRNA | 1.672676035 | -1.501040426 | 1.769625782 | 0.848224772 | 0.396312804 | 1 | 1.501040426 |
| ENSMUSG00000024076 | Vit | 74199 | protein_coding | 1.163894017 | 1.501892911 | 2.229536626 | -0.673634554 | 0.500543679 | 1 | 1.501892911 |
| ENSMUSG00000003477 | Inmt | 21743 | protein_coding | 1.233689018 | -1.502168681 | 2.302499336 | 0.652407867 | 0.51413809 | 1 | 1.502168681 |
| ENSMUSG00000111994 | A330049N07Rik | 327768 | lncRNA | 1.253838149 | -1.503002112 | 2.321036467 | 0.64755644 | 0.517271879 | 1 | 1.503002112 |
| ENSMUSG00000002279 | Lmf1 | 76483 | protein_coding | 772.2195881 | 1.503087321 | 0.0786466 | -19.11191737 | 2.01E-81 | 5.35E-79 | 1.503087321 |

| ensembl_gene_id | external_gene_name | entrezgene_id | gene_biotype | baseMean | log2FoldChange | lfcSE | stat | pvalue | padj | absLog2FC |
|---|---|---|---|---|---|---|---|---|---|---|
| ENSMUSG00000026875 | Traf1 | 22029 | protein_coding | 178.166264 | -1.50315082 | 0.182663339 | 8.229077764 | 1.89E-16 | 7.83E-15 | 1.50315082 |
| ENSMUSG00000032368 | Zic1 | 22771 | protein_coding | 2.223568625 | -1.504027329 | 2.344681671 | 0.641463337 | 0.521221694 | 1 | 1.504027329 |
| ENSMUSG00000017493 | Igfbp4 | 16010 | protein_coding | 31.72924206 | 1.504387411 | 0.278814183 | -5.395663143 | 6.83E-08 | 1.15E-06 | 1.504387411 |
| ENSMUSG00000039109 | F13a1 | 74145 | protein_coding | 4.684665715 | 1.504490327 | 1.157669745 | -1.299585079 | 0.193743217 | 0.392188708 | 1.504490327 |
| ENSMUSG00000054910 | 4931415C17Rik | 70966 | lncRNA | 1.08518329 | -1.505069897 | 2.252095068 | 0.668297675 | 0.503943597 | 1 | 1.505069897 |
| ENSMUSG00000030605 | Mfge8 | 17304 | protein_coding | 336.1873617 | 1.505564026 | 0.085722691 | -17.5631914 | 4.71E-69 | 9.73E-67 | 1.505564026 |
| ENSMUSG00000045102 | Poln | 272158 | protein_coding | 2.764073629 | 1.50646146 | 1.023116968 | -1.472423493 | 0.140906553 | 1 | 1.50646146 |
| ENSMUSG00000015852 | Fcrls | 80891 | protein_coding | 0.949679416 | 1.507009139 | 2.037783604 | -0.73953345 | 0.459583136 | 1 | 1.507009139 |
| ENSMUSG00000056487 | Mettl7a2 | 393082 | protein_coding | 14.34553942 | -1.507073865 | 0.368881825 | 4.085519434 | 4.40E-05 | 0.000418111 | 1.507073865 |
| ENSMUSG00000106581 | Mir7042 | 102465631 | miRNA | 0.69313603 | -1.507267075 | 1.370101899 | 1.10011312 | 0.271282838 | 1 | 1.507267075 |
| ENSMUSG00000000693 | Loxl3 | 16950 | protein_coding | 12.03641953 | 1.507388832 | 0.392019342 | -3.845189945 | 0.000120459 | 0.001015447 | 1.507388832 |
| ENSMUSG00000078491 | Gm13090 | 105704528 | lncRNA | 0.568901581 | 1.507819475 | 1.609147181 | -0.937030182 | 0.348743033 | 1 | 1.507819475 |
| ENSMUSG00000071392 | Ect2l | 100045792 | protein_coding | 3.193328013 | 1.508894171 | 1.530913637 | -0.985616781 | 0.3243212 | 1 | 1.508894171 |
| ENSMUSG00000074004 | B3gnt6 | 272411 | protein_coding | 201.67482 | -1.509727715 | 0.267668414 | 5.640290875 | 1.70E-08 | 3.16E-07 | 1.509727715 |
| ENSMUSG00000020642 | Rnf144a | 108089 | protein_coding | 7.277914865 | 1.511757856 | 0.935383184 | -1.616190971 | 0.10605303 | 0.258312646 | 1.511757856 |
| ENSMUSG00000021725 | Parp8 | 52552 | protein_coding | 5.408940678 | 1.511803912 | 0.71313397 | -2.119943764 | 0.034010788 | 0.110941817 | 1.511803912 |
| ENSMUSG00000028460 | Sit1 | 54390 | protein_coding | 1.390308435 | -1.512099421 | 1.928311681 | 0.784157165 | 0.432947896 | 1 | 1.512099421 |
| ENSMUSG00000025963 | Mdh1b | 76668 | protein_coding | 1.117959979 | 1.512890776 | 1.146905437 | -1.319106813 | 0.187133406 | 1 | 1.512890776 |
| ENSMUSG00000048834 | Vstm2a | 211739 | protein_coding | 1.653392795 | -1.514616261 | 2.681648059 | 0.564807994 | 0.572204369 | 1 | 1.514616261 |
| ENSMUSG00000031870 | Pgr | 18667 | protein_coding | 1.65772118 | -1.514627453 | 2.682131781 | 0.564710304 | 0.572270825 | 1 | 1.514627453 |
| ENSMUSG00000037206 | Islr | 26968 | protein_coding | 2.391972035 | 1.515931075 | 1.661203066 | -0.91255013 | 0.361479193 | 1 | 1.515931075 |
| ENSMUSG00000092171 | 4833427F10Rik | 74601 | lncRNA | 0.816273238 | 1.516596284 | 1.915249907 | -0.791852947 | 0.428446425 | 1 | 1.516596284 |
| ENSMUSG00000031239 | Itm2a | 16431 | protein_coding | 2.384756999 | 1.517055771 | 0.773665223 | -1.960868507 | 0.049894364 | 1 | 1.517055771 |
| ENSMUSG00000020052 | Ascl1 | 17172 | protein_coding | 2.779823532 | 1.517245158 | 1.260606011 | -1.203583947 | 0.228750427 | 1 | 1.517245158 |
| ENSMUSG00000040836 | Gpr161 | 240888 | protein_coding | 10.51745527 | 1.518304923 | 0.53546934 | -2.835465653 | 0.004575894 | 0.022584072 | 1.518304923 |
| ENSMUSG00000028476 | Reck | 53614 | protein_coding | 1.749086321 | -1.5185078 | 1.448876077 | 1.048059129 | 0.294611365 | 1 | 1.5185078 |
| ENSMUSG00000021070 | Bdkrb2 | 12062 | protein_coding | 1.111804548 | 1.518598082 | 2.039575092 | -0.744565909 | 0.456534184 | 1 | 1.518598082 |
| ENSMUSG00000022816 | Fstl1 | 14314 | protein_coding | 2.664327143 | 1.519231558 | 0.992959969 | -1.530002825 | 0.12601603 | 1 | 1.519231558 |
| ENSMUSG00000050097 | Ces2b | 234669 | protein_coding | 99.06141893 | -1.519518927 | 0.282762117 | 5.373841945 | 7.71E-08 | 1.29E-06 | 1.519518927 |
| ENSMUSG00000017969 | Ptgis | 19223 | protein_coding | 2.391293286 | 1.520044328 | 1.430997979 | -1.062226747 | 0.28813276 | 1 | 1.520044328 |
| ENSMUSG00000092210 | A930009A15Rik | 77798 | protein_coding | 0.944554913 | 1.520258197 | 1.242547195 | -1.223501372 | 0.221140386 | 1 | 1.520258197 |
| ENSMUSG00000026401 | Cd55b | 13137 | protein_coding | 0.706331492 | 1.520645824 | 1.743737001 | -0.872061454 | 0.383174853 | 1 | 1.520645824 |
| ENSMUSG00000094747 | Or4f14b | 257956 | protein_coding | 2.433154598 | -1.520902013 | 3.037481919 | 0.500711462 | 0.616574204 | 1 | 1.520902013 |
| ENSMUSG00000020473 | Aebp1 | 11568 | protein_coding | 6.717578233 | 1.521022646 | 0.871422209 | -1.745448567 | 0.080906818 | 0.211474389 | 1.521022646 |
| ENSMUSG00000042849 | Or6b3 | 259041 | protein_coding | 2.25116707 | -1.521030556 | 3.047254844 | 0.499147801 | 0.617675264 | 1 | 1.521030556 |
| ENSMUSG00000040899 | Ccr6 | 12458 | protein_coding | 1.041834475 | 1.521196223 | 1.358287735 | -1.119936656 | 0.262740757 | 1 | 1.521196223 |
| ENSMUSG00000075068 | Or4c3d | 57272 | protein_coding | 2.017491469 | -1.521262145 | 3.065189772 | 0.49630276 | 0.619680819 | 1 | 1.521262145 |
| ENSMUSG00000093942 | Or13a18 | 18345 | protein_coding | 2.278120161 | -1.52153055 | 2.014736703 | 0.755200691 | 0.450128574 | 1 | 1.52153055 |
| ENSMUSG00000079343 | C1s2 | 317677 | protein_coding | 1.241980743 | -1.52158501 | 3.091491691 | 0.492184732 | 0.622588757 | 1 | 1.52158501 |

| ensembl_gene_id | external_gene_name | entrezgene_id | gene_biotype | baseMean | log2FoldChange | lfcSE | stat | pvalue | padj | absLog2FC |
|---|---|---|---|---|---|---|---|---|---|---|
| ENSMUSG00000024856 | Cdk2ap2 | 52004 | protein_coding | 1918.135023 | 1.523140514 | 0.088917951 | -17.12973023 | 8.91E-66 | 1.71E-63 | 1.523140514 |
| ENSMUSG00000006356 | Crip2 | 68337 | protein_coding | 15.39349312 | -1.523506153 | 0.351086033 | 4.339409747 | 1.43E-05 | 0.000153277 | 1.523506153 |
| ENSMUSG00000050272 | Dscam | 13508 | protein_coding | 4.52067108 | 1.523701076 | 0.962000289 | -1.583888378 | 0.113219123 | 0.270127107 | 1.523701076 |
| ENSMUSG00000036110 | Slc17a2 | 218103 | protein_coding | 15.48565496 | -1.523801525 | 0.465356006 | 3.274485567 | 0.001058545 | 0.006667458 | 1.523801525 |
| ENSMUSG00000035606 | Ky | 16716 | protein_coding | 3.013351237 | -1.524005603 | 1.396695962 | 1.091150576 | 0.275206631 | 1 | 1.524005603 |
| ENSMUSG00000044985 | Or2b4 | 259064 | protein_coding | 0.998205607 | -1.524561413 | 2.547495947 | 0.598454893 | 0.549536448 | 1 | 1.524561413 |
| ENSMUSG00000015568 | Lpl | 16956 | protein_coding | 255.3866161 | -1.525640468 | 0.182405308 | 8.364013548 | 6.06E-17 | 2.57E-15 | 1.525640468 |
| ENSMUSG00000042988 | Notum | 77583 | protein_coding | 3.01378135 | 1.526587289 | 0.646812422 | -2.360170022 | 0.018266561 | 1 | 1.526587289 |
| ENSMUSG00000084902 | Gm281 | 238939 | protein_coding | 1.413868737 | -1.526649531 | 2.57912181 | 0.591926106 | 0.553900071 | 1 | 1.526649531 |
| ENSMUSG00000026574 | Dpt | 56429 | protein_coding | 0.869910225 | -1.526889254 | 2.058109791 | 0.741889116 | 0.458154521 | 1 | 1.526889254 |
| ENSMUSG00000029467 | Atp2a2 | 11938 | protein_coding | 8557.699925 | 1.529326602 | 0.053943607 | -28.35046974 | 8.26E-177 | 4.70E-174 | 1.529326602 |
| ENSMUSG00000023045 | Soat2 | 223920 | protein_coding | 30.27267725 | 1.530451805 | 0.327555764 | -4.67233972 | 2.98E-06 | 3.72E-05 | 1.530451805 |
| ENSMUSG00000043388 | Tmem130 | 243339 | protein_coding | 3.05141181 | 1.531816577 | 1.013231787 | -1.511812595 | 0.130581544 | 1 | 1.531816577 |
| ENSMUSG00000097182 | A830009L08Rik | 320297 | lncRNA | 1.115381204 | -1.532900623 | 1.283198152 | 1.194593852 | 0.232245753 | 1 | 1.532900623 |
| ENSMUSG00000020399 | Havcr2 | 171285 | protein_coding | 1.94173486 | 1.533316849 | 1.392451965 | -1.101163191 | 0.270825639 | 1 | 1.533316849 |
| ENSMUSG00000057454 | Lypd3 | 72434 | protein_coding | 1.424492852 | 1.533659827 | 1.500850748 | -1.021860321 | 0.306847015 | 1 | 1.533659827 |
| ENSMUSG00000108976 | Gm29797 | 101056205 | protein_coding | 0.74720138 | -1.534429571 | 1.298782533 | 1.18143687 | 0.237429218 | 1 | 1.534429571 |
| ENSMUSG00000032383 | Ppib | 19035 | protein_coding | 2565.998304 | 1.535563757 | 0.086581901 | -17.7353897 | 2.24E-70 | 4.86E-68 | 1.535563757 |
| ENSMUSG00000017817 | Jph2 | 59091 | protein_coding | 2.98533298 | 1.536784356 | 1.702111123 | -0.902869581 | 0.366595117 | 1 | 1.536784356 |
| ENSMUSG00000020439 | Smtn | 29856 | protein_coding | 254.1411983 | 1.537415836 | 0.111615633 | -13.77419806 | 3.64E-43 | 4.18E-41 | 1.537415836 |
| ENSMUSG00000107902 | Gm32592 | 102635190 | lncRNA | 0.984131415 | 1.537619392 | 2.454053203 | -0.626563185 | 0.530945607 | 1 | 1.537619392 |
| ENSMUSG00000100890 | 1700085C21Rik | 74283 | lncRNA | 1.089913642 | 1.537751903 | 2.422709912 | -0.634723908 | 0.525608498 | 1 | 1.537751903 |
| ENSMUSG00000072423 | Psmb11 | 73902 | protein_coding | 1.501526619 | 1.539283735 | 2.134947796 | -0.720993618 | 0.470913441 | 1 | 1.539283735 |
| ENSMUSG00000089996 | Tmsb15b2 | 100034363 | protein_coding | 1.057519194 | 1.539293299 | 1.232011882 | -1.249414329 | 0.21151357 | 1 | 1.539293299 |
| ENSMUSG00000052949 | Rnf157 | 217340 | protein_coding | 28.46411907 | 1.53950685 | 0.304008964 | -5.064017941 | 4.11E-07 | 6.16E-06 | 1.53950685 |
| ENSMUSG00000032864 | Rag2 | 19374 | protein_coding | 1.052142684 | 1.539687032 | 2.076369453 | -0.741528455 | 0.458373085 | 1 | 1.539687032 |
| ENSMUSG00000050761 | Gp1bb | 14724 | protein_coding | 2.705159908 | 1.539807729 | 0.691594557 | -2.22646016 | 0.025983375 | 1 | 1.539807729 |
| ENSMUSG00000102224 | 4930447F24Rik | 76873 | lncRNA | 11.71898204 | -1.539977251 | 0.309394522 | 4.977390165 | 6.44E-07 | 9.25E-06 | 1.539977251 |
| ENSMUSG00000027233 | Patl2 | 67578 | protein_coding | 1.440517498 | 1.54008357 | 2.158840245 | -0.713384686 | 0.475607747 | 1 | 1.54008357 |
| ENSMUSG00000079563 | Pglyrp2 | 57757 | protein_coding | 4.50768986 | 1.54295114 | 0.792979024 | -1.945765391 | 0.051682928 | 0.152928125 | 1.54295114 |
| ENSMUSG00000056370 | Sftpb | 20388 | protein_coding | 2.264522191 | 1.542953256 | 1.648981425 | -0.93570081 | 0.349427253 | 1 | 1.542953256 |
| ENSMUSG00000085589 | A430078I02Rik | 320486 | lncRNA | 46.8815522 | 1.543569914 | 0.201651765 | -7.654631301 | 1.94E-14 | 6.75E-13 | 1.543569914 |
| ENSMUSG00000009876 | Cox4i2 | 84682 | protein_coding | 0.888447236 | 1.54376765 | 1.689545425 | -0.913717753 | 0.360865173 | 1 | 1.54376765 |
| ENSMUSG00000036856 | Wnt4 | 22417 | protein_coding | 2.267780592 | 1.543925521 | 1.325142193 | -1.165101774 | 0.243977799 | 1 | 1.543925521 |
| ENSMUSG00000079598 | Clec2l | 665180 | protein_coding | 0.876886607 | 1.545017955 | 1.783765465 | -0.866155324 | 0.386404989 | 1 | 1.545017955 |
| ENSMUSG00000016529 | Il10 | 16153 | protein_coding | 1.164046517 | 1.545476946 | 2.218925793 | -0.696497806 | 0.486117131 | 1 | 1.545476946 |
| ENSMUSG00000078926 | Cdc20b | 238896 | protein_coding | 1.370094339 | -1.546126028 | 1.855546508 | 0.833245635 | 0.40470621 | 1 | 1.546126028 |
| ENSMUSG00000050359 | Sprr1a | 20753 | protein_coding | 91.83138017 | 1.547902425 | 0.504003881 | -3.071211319 | 0.002131922 | 0.011942524 | 1.547902425 |
| ENSMUSG00000031189 | Aff2 | 14266 | protein_coding | 1.475336757 | 1.54864986 | 1.965727354 | -0.78782536 | 0.430798865 | 1 | 1.54864986 |

| ensembl_gene_id | external_gene_name | entrezgene_id | gene_biotype | baseMean | log2FoldChange | lfcSE | stat | pvalue | padj | absLog2FC |
|---|---|---|---|---|---|---|---|---|---|---|
| ENSMUSG00000028736 | Pax7 | 18509 | protein_coding | 3.372566864 | 1.549802494 | 2.219897395 | -0.698141498 | 0.485088708 | 1 | 1.549802494 |
| ENSMUSG00000055385 | Rnf212 | 671564 | protein_coding | 18.75186574 | -1.550448166 | 0.316892902 | 4.892656659 | 9.95E-07 | 1.37E-05 | 1.550448166 |
| ENSMUSG00000001270 | Ckb | 12709 | protein_coding | 290.1346698 | 1.553830894 | 0.225667287 | -6.885494621 | 5.76E-12 | 1.63E-10 | 1.553830894 |
| ENSMUSG00000070031 | Sp140 | 434484 | protein_coding | 16.7963753 | -1.55503074 | 0.416085386 | 3.737287567 | 0.000186016 | 0.001489031 | 1.55503074 |
| ENSMUSG00000025854 | Fam20c | 80752 | protein_coding | 4.921503952 | -1.55512212 | 0.814837466 | 1.90850591 | 0.05632586 | 0.16307245 | 1.55512212 |
| ENSMUSG00000021194 | Chga | 12652 | protein_coding | 525.1160137 | -1.555183787 | 0.228427605 | 6.808212996 | 9.88E-12 | 2.77E-10 | 1.555183787 |
| ENSMUSG00000093565 | Rab26os | 75614 | lncRNA | 50.61403494 | 1.55530935 | 0.32488406 | -4.787275037 | 1.69E-06 | 2.22E-05 | 1.55530935 |
| ENSMUSG00000030711 | Sult1a1 | 20887 | protein_coding | 2454.819297 | -1.555326645 | 0.164939543 | 9.429677189 | 4.11E-21 | 2.24E-19 | 1.555326645 |
| ENSMUSG00000052922 | Bpi | 329547 | protein_coding | 2.147286072 | 1.556172221 | 2.14823728 | -0.724394943 | 0.468823304 | 1 | 1.556172221 |
| ENSMUSG00000118339 | 4930592I03Rik | 75853 | lncRNA | 10.46911776 | 1.557897497 | 0.382111367 | -4.077077082 | 4.56E-05 | 0.000431592 | 1.557897497 |
| ENSMUSG00000096209 | Or5p81 | 258308 | protein_coding | 1.168938964 | 1.558262138 | 2.002161398 | -0.778289972 | 0.436398086 | 1 | 1.558262138 |
| ENSMUSG00000025243 | Slc6a20b | 22599 | protein_coding | 10.83303496 | 1.558737821 | 0.509321738 | -3.060418797 | 0.002210277 | 0.012277333 | 1.558737821 |
| ENSMUSG00000025469 | Msx3 | 17703 | protein_coding | 0.532807464 | -1.558754245 | 1.363545891 | 1.143162291 | 0.252971215 | 1 | 1.558754245 |
| ENSMUSG00000097650 | 4921507G05Rik | 66709 | lncRNA | 1.176898447 | -1.560512382 | 1.111154851 | 1.404405858 | 0.160198028 | 1 | 1.560512382 |
| ENSMUSG00000026544 | Dusp23 | 68440 | protein_coding | 176.2129485 | 1.560770569 | 0.112525559 | -13.87036495 | 9.58E-44 | 1.12E-41 | 1.560770569 |
| ENSMUSG00000024600 | Slc27a6 | 225579 | protein_coding | 1.171699383 | 1.561182698 | 2.102070052 | -0.742688236 | 0.457670454 | 1 | 1.561182698 |
| ENSMUSG00000114384 | 4930455J16Rik | 67585 | lncRNA | 4.498433408 | 1.561255414 | 2.271913041 | -0.687198579 | 0.4919576 | 0.698788598 | 1.561255414 |
| ENSMUSG00000020542 | Myocd | 214384 | protein_coding | 1.514612092 | -1.562769459 | 2.084620261 | 0.749666253 | 0.453455738 | 1 | 1.562769459 |
| ENSMUSG00000100625 | 1700016G22Rik | 71867 | lncRNA | 2.320333188 | 1.563008269 | 1.144835188 | -1.365269241 | 0.172168443 | 1 | 1.563008269 |
| ENSMUSG00000065513 | Mir26a-1 | 387218 | miRNA | 0.925367559 | 1.5634573 | 1.442029741 | -1.084206002 | 0.278273471 | 1 | 1.5634573 |
| ENSMUSG00000062170 | Fmr1nb | 207854 | protein_coding | 2.005371012 | 1.565300093 | 1.715030864 | -0.912694999 | 0.361402975 | 1 | 1.565300093 |
| ENSMUSG00000036040 | Adamtsl2 | 77794 | protein_coding | 3.177897747 | 1.566025908 | 1.327980501 | -1.179253691 | 0.238297172 | 1 | 1.566025908 |
| ENSMUSG00000043986 | Spata31d1d | 238663 | protein_coding | 2.48663289 | 1.566407838 | 1.982924183 | -0.789948426 | 0.429557888 | 1 | 1.566407838 |
| ENSMUSG00000067158 | Col4a4 | 12829 | protein_coding | 8.091920229 | 1.567069878 | 0.802822556 | -1.951950485 | 0.050944084 | 0.15122966 | 1.567069878 |
| ENSMUSG00000079419 | Ms4a6c | 73656 | protein_coding | 0.951629368 | 1.567563559 | 1.45415807 | -1.077987044 | 0.281039538 | 1 | 1.567563559 |
| ENSMUSG00000048040 | Arxes2 | 76976 | protein_coding | 5.370210505 | 1.567775663 | 0.500988851 | -3.129362381 | 0.001751861 | 0.010154522 | 1.567775663 |
| ENSMUSG00000110001 | C86187 | 97402 | lncRNA | 1.101541491 | 1.569448922 | 1.84403727 | -0.851093927 | 0.394717179 | 1 | 1.569448922 |
| ENSMUSG00000063506 | Arhgap22 | 239027 | protein_coding | 1.831021899 | 1.57032748 | 1.301201812 | -1.206828538 | 0.227498183 | 1 | 1.57032748 |
| ENSMUSG00000015981 | Stk32c | 57740 | protein_coding | 2.944920689 | 1.570988283 | 1.031603021 | -1.522861266 | 0.127793418 | 1 | 1.570988283 |
| ENSMUSG00000056043 | Rgs9bp | 243923 | protein_coding | 7.952739008 | -1.571111221 | 0.548951341 | 2.862022742 | 0.004209467 | 0.021033325 | 1.571111221 |
| ENSMUSG00000031722 | Hp | 15439 | protein_coding | 0.89285198 | 1.571229055 | 2.203256809 | -0.713139316 | 0.475759553 | 1 | 1.571229055 |
| ENSMUSG00000041895 | Wipi1 | 52639 | protein_coding | 439.6169679 | 1.571726892 | 0.084949074 | -18.50198977 | 1.99E-76 | 4.83E-74 | 1.571726892 |
| ENSMUSG00000100510 | Hand2os1 | 102636514 | lncRNA | 4.93251411 | 1.57308725 | 1.918478182 | -0.819966192 | 0.41223538 | 0.632329199 | 1.57308725 |
| ENSMUSG00000065462 | Mir200c | 723944 | miRNA | 1.109437848 | 1.573195802 | 1.08133035 | -1.454870662 | 0.145705071 | 1 | 1.573195802 |
| ENSMUSG00000022510 | Trp63 | 22061 | protein_coding | 3.823168405 | 1.573767554 | 3.494094757 | -0.450407806 | 0.652416417 | 1 | 1.573767554 |
| ENSMUSG00000060780 | Lrrtm1 | 74342 | protein_coding | 5.360158616 | 1.573849661 | 0.782104808 | -2.012325772 | 0.044185607 | 0.135678899 | 1.573849661 |
| ENSMUSG00000036790 | Slitrk2 | 245450 | protein_coding | 1.775178802 | 1.57425834 | 1.938921222 | -0.811924859 | 0.416834749 | 1 | 1.57425834 |
| ENSMUSG00000043230 | Fam124b | 241128 | protein_coding | 1.256898066 | 1.574723239 | 2.118810727 | -0.743210905 | 0.457354002 | 1 | 1.574723239 |
| ENSMUSG00000044083 | Efcab8 | 100504221 | protein_coding | 6.132004675 | 1.574916656 | 1.414500747 | -1.113408147 | 0.265533177 | 0.480535393 | 1.574916656 |

| ensembl_gene_id | external_gene_name | entrezgene_id | gene_biotype | baseMean | log2FoldChange | lfcSE | stat | pvalue | padj | absLog2FC |
|---|---|---|---|---|---|---|---|---|---|---|
| ENSMUSG00000000223 | Drp2 | 13497 | protein_coding | 3.71775744 | 1.578727141 | 1.293372086 | -1.220628741 | 0.222226619 | 1 | 1.578727141 |
| ENSMUSG00000014905 | Dnajb9 | 27362 | protein_coding | 1088.57999 | 1.57992868 | 0.086868407 | -18.18760974 | 6.47E-74 | 1.53E-71 | 1.57992868 |
| ENSMUSG00000050702 | 4930563M21Rik | 75258 | lncRNA | 1.246568595 | -1.580045301 | 1.728640332 | 0.914039359 | 0.360696164 | 1 | 1.580045301 |
| ENSMUSG00000065417 | Mir340 | 723845 | miRNA | 0.719584947 | -1.58077567 | 1.470596362 | 1.074921515 | 0.282409868 | 1 | 1.58077567 |
| ENSMUSG00000045896 | Paip2b | 232164 | protein_coding | 711.2078065 | 1.583098427 | 0.086492926 | -18.3032127 | 7.80E-75 | 1.87E-72 | 1.583098427 |
| ENSMUSG00000028011 | Tdo2 | 56720 | protein_coding | 4.874504031 | -1.583198028 | 0.49712858 | 3.184685191 | 0.001449116 | 0.008633007 | 1.583198028 |
| ENSMUSG00000023964 | Calcr | 12311 | protein_coding | 1.104986587 | 1.58359596 | 1.806102976 | -0.876802696 | 0.380593814 | 1 | 1.58359596 |
| ENSMUSG00000006457 | Actn3 | 11474 | protein_coding | 2.331919116 | 1.584770026 | 0.963444127 | -1.64490081 | 0.099990268 | 1 | 1.584770026 |
| ENSMUSG00000024843 | Chka | 12660 | protein_coding | 982.6937598 | 1.587314036 | 0.156149727 | -10.16533339 | 2.83E-24 | 1.76E-22 | 1.587314036 |
| ENSMUSG00000047986 | Palm3 | 74337 | protein_coding | 3.546188805 | 1.587685761 | 1.707436567 | -0.929865151 | 0.352440909 | 1 | 1.587685761 |
| ENSMUSG00000023913 | Pla2g7 | 27226 | protein_coding | 4.759585918 | -1.58846117 | 0.633842694 | 2.506081061 | 0.012207763 | 0.049740685 | 1.58846117 |
| ENSMUSG00000067750 | Khdc1a | 368204 | protein_coding | 2.464113338 | -1.588670808 | 0.866979114 | 1.832421085 | 0.066888709 | 1 | 1.588670808 |
| ENSMUSG00000028883 | Sema3a | 20346 | protein_coding | 2.758412931 | -1.588980932 | 1.399982927 | 1.135000221 | 0.256375226 | 1 | 1.588980932 |
| ENSMUSG00000034739 | Mfrp | 259172 | protein_coding | 6.255097946 | 1.589140364 | 0.553896729 | -2.869019225 | 0.004117468 | 0.020673719 | 1.589140364 |
| ENSMUSG00000031253 | Srpx2 | 68792 | protein_coding | 12.37077782 | -1.589164766 | 0.438504048 | 3.624059511 | 0.000290015 | 0.00221215 | 1.589164766 |
| ENSMUSG00000091382 | Vmn1r18 | 171199 | protein_coding | 5.325501715 | 1.58922688 | 3.62499689 | -0.438407791 | 0.6610907 | 0.816884691 | 1.58922688 |
| ENSMUSG00000035186 | Ubd | 24108 | protein_coding | 2.316445506 | -1.589533426 | 0.827098423 | 1.921818955 | 0.054628542 | 1 | 1.589533426 |
| ENSMUSG00000046402 | Rbp1 | 19659 | protein_coding | 9.054772806 | 1.589564337 | 0.607229349 | -2.617733053 | 0.008851602 | 0.038619645 | 1.589564337 |
| ENSMUSG00000077639 | Gm22772 | 115485835 | snoRNA | 0.957745487 | 1.592772848 | 1.137176046 | -1.400638761 | 0.161322124 | 1 | 1.592772848 |
| ENSMUSG00000056822 | Or2l13 | 259071 | protein_coding | 3.200206606 | 1.592853683 | 0.714196012 | -2.23027524 | 0.025729176 | 1 | 1.592853683 |
| ENSMUSG00000048022 | Tmem229a | 319832 | protein_coding | 7.12343124 | -1.594094465 | 0.574221772 | 2.776095478 | 0.005501605 | 0.026334349 | 1.594094465 |
| ENSMUSG00000048626 | Klf17 | 75753 | protein_coding | 0.848925364 | -1.594409391 | 1.308618378 | 1.21839141 | 0.223075268 | 1 | 1.594409391 |
| ENSMUSG00000038550 | Ciart | 229599 | protein_coding | 24.6468893 | -1.595751519 | 0.616904685 | 2.586706761 | 0.009689802 | 0.041509049 | 1.595751519 |
| ENSMUSG00000044056 | Efcab9 | 69306 | protein_coding | 1.643128523 | -1.595917759 | 0.955309575 | 1.670576535 | 0.094805351 | 1 | 1.595917759 |
| ENSMUSG00000032079 | Apoa5 | 66113 | protein_coding | 1.121948254 | -1.597352446 | 1.422754371 | 1.122718354 | 0.26155713 | 1 | 1.597352446 |
| ENSMUSG00000087782 | Gm23169 | 115487625 | snRNA | 0.681615295 | 1.600986716 | 1.603356114 | -0.998522226 | 0.318026192 | 1 | 1.600986716 |
| ENSMUSG00000079669 | Gm17396 | 102635502 | lncRNA | 1.827915405 | -1.601129965 | 1.359386767 | 1.177832537 | 0.238863374 | 1 | 1.601129965 |
| ENSMUSG00000105816 | D030025E07Rik | 402774 | lncRNA | 0.811085833 | -1.601539286 | 2.072966029 | 0.772583469 | 0.439768931 | 1 | 1.601539286 |
| ENSMUSG00000036106 | Prr5 | 109270 | protein_coding | 376.9379439 | 1.601752306 | 0.151880554 | -10.54613155 | 5.29E-26 | 3.54E-24 | 1.601752306 |
| ENSMUSG00000047220 | Iho1 | 434438 | protein_coding | 1.262328665 | 1.602152381 | 2.221318796 | -0.721261794 | 0.470748459 | 1 | 1.602152381 |
| ENSMUSG00000063804 | Lin28b | 380669 | protein_coding | 2.713004432 | 1.602866662 | 2.399927585 | -0.667881261 | 0.50420939 | 1 | 1.602866662 |
| ENSMUSG00000041202 | Pla2g2d | 18782 | protein_coding | 1.081022436 | 1.60320806 | 1.748658199 | -0.916821859 | 0.359235996 | 1 | 1.60320806 |
| ENSMUSG00000054594 | Oscar | 232790 | protein_coding | 2.264428269 | 1.604558453 | 1.803737461 | -0.88957428 | 0.373694522 | 1 | 1.604558453 |
| ENSMUSG00000113211 | 4921525O09Rik | 74050 | lncRNA | 1.006981735 | 1.604988559 | 1.732413832 | -0.926446401 | 0.35421404 | 1 | 1.604988559 |
| ENSMUSG00000049133 | Flg2 | 229574 | protein_coding | 1.35350317 | 1.605992036 | 2.664608868 | -0.602712111 | 0.546700223 | 1 | 1.605992036 |
| ENSMUSG00000004631 | Sgce | 20392 | protein_coding | 6.827481344 | 1.606217148 | 0.764813161 | -2.100143185 | 0.035716247 | 0.115153868 | 1.606217148 |
| ENSMUSG00000079227 | Ccr5 | 12774 | protein_coding | 4.525417897 | 1.607539957 | 0.837140326 | -1.92027538 | 0.054823125 | 0.159701725 | 1.607539957 |
| ENSMUSG00000024150 | Mcfd2 | 193813 | protein_coding | 1934.0395 | 1.607591366 | 0.056847921 | -28.27880652 | 6.30E-176 | 3.47E-173 | 1.607591366 |
| ENSMUSG00000014453 | Blk | 12143 | protein_coding | 1.495648026 | 1.60759819 | 1.798761914 | -0.893724832 | 0.371469136 | 1 | 1.60759819 |

Supplementary Table 1

| ensembl_gene_id | external_gene_name | entrezgene_id | gene_biotype | baseMean | log2FoldChange | lfcSE | stat | pvalue | padj | absLog2FC |
|---|---|---|---|---|---|---|---|---|---|---|
| ENSMUSG00000060487 | Samd5 | 320825 | protein_coding | 21.40565657 | -1.608218769 | 0.338562379 | 4.750140212 | 2.03E-06 | 2.65E-05 | 1.608218769 |
| ENSMUSG00000032323 | Cyp11a1 | 13070 | protein_coding | 3.341467068 | -1.608325729 | 1.426896849 | 1.127149261 | 0.259679379 | 1 | 1.608325729 |
| ENSMUSG00000087213 | 2810408I11Rik | 69941 | lncRNA | 16.88457376 | 1.608529862 | 0.394476466 | -4.077632006 | 4.55E-05 | 0.00043081 | 1.608529862 |
| ENSMUSG00000054580 | Pla2r1 | 18779 | protein_coding | 4.328538578 | 1.609042273 | 1.119290508 | -1.437555542 | 0.150560203 | 1 | 1.609042273 |
| ENSMUSG00000037463 | Fbxo27 | 233040 | protein_coding | 1.81436392 | -1.609386994 | 1.555638011 | 1.034551086 | 0.300878607 | 1 | 1.609386994 |
| ENSMUSG00000115008 | 4933429O19Rik | 66770 | lncRNA | 0.999144203 | 1.612286333 | 2.268370056 | -0.710768655 | 0.477227608 | 1 | 1.612286333 |
| ENSMUSG00000025479 | Cyp2e1 | 13106 | protein_coding | 1.331730435 | -1.612844944 | 2.11031259 | 0.764268266 | 0.444707392 | 1 | 1.612844944 |
| ENSMUSG00000021852 | Slc35f4 | 75288 | protein_coding | 1.599706277 | 1.613061579 | 2.27250929 | -0.709815175 | 0.477818757 | 1 | 1.613061579 |
| ENSMUSG00000042349 | Ikbke | 56489 | protein_coding | 1169.969864 | 1.613529755 | 0.082981232 | -19.44451425 | 3.24E-84 | 8.78E-82 | 1.613529755 |
| ENSMUSG00000042812 | Foxf1 | 15227 | protein_coding | 0.794667908 | -1.61587138 | 1.615994007 | 0.999924117 | 0.317347232 | 1 | 1.61587138 |
| ENSMUSG00000050147 | F2rl3 | 14065 | protein_coding | 0.897805272 | 1.616760303 | 1.913338942 | -0.844994197 | 0.398114079 | 1 | 1.616760303 |
| ENSMUSG00000090774 | Vmn2r74 | 546980 | protein_coding | 1.881574924 | -1.618722037 | 1.839935711 | 0.879770976 | 0.378983391 | 1 | 1.618722037 |
| ENSMUSG00000022225 | Cma1 | 17228 | protein_coding | 5.304280926 | 1.620911491 | 0.68021469 | -2.382941027 | 0.017174945 | 0.065126762 | 1.620911491 |
| ENSMUSG00000112846 | 4930432O09Rik | 74647 | lncRNA | 1.222190825 | 1.623123746 | 2.228904347 | -0.728215972 | 0.46648139 | 1 | 1.623123746 |
| ENSMUSG00000065590 | Mir212 | 387208 | miRNA | 0.702287118 | -1.624004428 | 1.227210404 | 1.323330068 | 0.185725637 | 1 | 1.624004428 |
| ENSMUSG00000110018 | 5430437J10Rik | 71432 | transcribed_processed_pseudogene | 0.852991338 | 1.624788215 | 2.146863234 | -0.756819619 | 0.449157937 | 1 | 1.624788215 |
| ENSMUSG00000072955 | Tmsb15l | 399591 | protein_coding | 1.849181547 | 1.625101251 | 0.7727844 | -2.102916741 | 0.035473048 | 1 | 1.625101251 |
| ENSMUSG00000085967 | Gm12530 | 329851 | lncRNA | 0.842801087 | -1.626224128 | 1.747276308 | 0.930719498 | 0.351998681 | 1 | 1.626224128 |
| ENSMUSG00000075073 | Or4a47 | 258985 | protein_coding | 2.397534451 | -1.626290069 | 2.129229825 | 0.76379264 | 0.444990824 | 1 | 1.626290069 |
| ENSMUSG00000042282 | Gucy2f | 245650 | protein_coding | 2.617130037 | -1.627217453 | 2.184752811 | 0.744806206 | 0.456388883 | 1 | 1.627217453 |
| ENSMUSG00000031097 | Tnni2 | 21953 | protein_coding | 3.546624319 | 1.62776654 | 1.34451763 | -1.210669539 | 0.226022081 | 1 | 1.62776654 |
| ENSMUSG00000052520 | Cyp2j5 | 13109 | protein_coding | 1.117519737 | 1.628366068 | 1.850312319 | -0.880049303 | 0.378832601 | 1 | 1.628366068 |
| ENSMUSG00000097691 | 9030616G12Rik | 77699 | lncRNA | 14.62413308 | -1.629224121 | 0.354011403 | 4.602179777 | 4.18E-06 | 5.10E-05 | 1.629224121 |
| ENSMUSG00000117864 | Gm19500 | 100503007 | lncRNA | 1.823040052 | -1.630155916 | 1.343013077 | 1.213804947 | 0.224822217 | 1 | 1.630155916 |
| ENSMUSG00000023153 | Tmem52 | 69671 | protein_coding | 3.636642104 | -1.631235298 | 0.571932052 | 2.852148768 | 0.004342477 | 1 | 1.631235298 |
| ENSMUSG00000035394 | Cfap53 | 74453 | protein_coding | 3.097028199 | -1.632188502 | 0.753017538 | 2.167530528 | 0.030194425 | 1 | 1.632188502 |
| ENSMUSG00000027581 | Stmn3 | 20262 | protein_coding | 1.184027352 | -1.632379905 | 1.185641956 | 1.376789929 | 0.168577205 | 1 | 1.632379905 |
| ENSMUSG00000021281 | Tnfaip2 | 21928 | protein_coding | 15.51361534 | 1.634960263 | 0.597409272 | -2.736750734 | 0.006204929 | 0.028888694 | 1.634960263 |
| ENSMUSG00000032478 | Nme6 | 54369 | protein_coding | 247.9663488 | 1.635022815 | 0.146702296 | -11.14517537 | 7.56E-29 | 5.57E-27 | 1.635022815 |
| ENSMUSG00000068600 | Gml2 | 15202 | protein_coding | 1.075338652 | 1.635441028 | 2.477898384 | -0.660011338 | 0.509246554 | 1 | 1.635441028 |
| ENSMUSG00000102697 | Pcdhac2 | 353237 | protein_coding | 1.698360118 | -1.636900501 | 1.793689491 | 0.912588555 | 0.361458976 | 1 | 1.636900501 |
| ENSMUSG00000043857 | Mgat5b | 268510 | protein_coding | 3.055257769 | -1.637941138 | 1.899603754 | 0.862254107 | 0.388547701 | 1 | 1.637941138 |
| ENSMUSG00000066682 | Pilrb2 | 545812 | protein_coding | 1.091249373 | 1.638416066 | 1.975756396 | -0.82926016 | 0.406957209 | 1 | 1.638416066 |
| ENSMUSG00000021130 | Galnt16 | 108760 | protein_coding | 2.937144961 | 1.638631185 | 2.047695529 | -0.800231852 | 0.423576478 | 1 | 1.638631185 |
| ENSMUSG00000020229 | Slc5a4a | 64452 | protein_coding | 4.635481713 | 1.640813392 | 0.75127936 | -2.184025649 | 0.028960363 | 0.09822375 | 1.640813392 |
| ENSMUSG00000022678 | Nde1 | 67203 | protein_coding | 536.4673697 | 1.640820249 | 0.105945627 | -15.48738067 | 4.22E-54 | 6.17E-52 | 1.640820249 |
| ENSMUSG00000040462 | Os9 | 216440 | protein_coding | 4563.620233 | 1.641010024 | 0.052697839 | -31.13998722 | 6.93E-213 | 5.20E-210 | 1.641010024 |
| ENSMUSG00000044165 | Bcl2l15 | 229672 | protein_coding | 1.332152851 | 1.641477059 | 1.316670027 | -1.246688256 | 0.212511827 | 1 | 1.641477059 |
| ENSMUSG00000050424 | Pnma5 | 385377 | protein_coding | 1.119572458 | 1.642304202 | 1.665985024 | -0.985785693 | 0.324238288 | 1 | 1.642304202 |

| ensembl_gene_id | external_gene_name | entrezgene_id | gene_biotype | baseMean | log2FoldChange | lfcSE | stat | pvalue | padj | absLog2FC |
|---|---|---|---|---|---|---|---|---|---|---|
| ENSMUSG00000026241 | Nppc | 18159 | protein_coding | 0.791732761 | 1.643149699 | 1.854136588 | -0.886207472 | 0.375505738 | 1 | 1.643149699 |
| ENSMUSG00000054976 | Nyap2 | 241134 | protein_coding | 2.2299075 | -1.643388806 | 1.81942921 | 0.903244159 | 0.366396326 | 1 | 1.643388806 |
| ENSMUSG00000074899 | Sptbn5 | 640524 | protein_coding | 44.67714294 | 1.644642133 | 0.441909957 | -3.721667971 | 0.000197911 | 0.00156603 | 1.644642133 |
| ENSMUSG00000065586 | Mir96 | 723886 | miRNA | 0.747392125 | -1.644944728 | 1.346013904 | 1.222085985 | 0.221675112 | 1 | 1.644944728 |
| ENSMUSG00000032172 | Olfm2 | 244723 | protein_coding | 3.961481435 | 1.644969003 | 0.920555251 | -1.786931313 | 0.073948591 | 1 | 1.644969003 |
| ENSMUSG00000063129 | Aldoart2 | 79459 | protein_coding | 1.40420477 | 1.644978029 | 1.241891934 | -1.324574211 | 0.185312414 | 1 | 1.644978029 |
| ENSMUSG00000020893 | Per1 | 18626 | protein_coding | 707.770219 | 1.645047361 | 0.203430215 | -8.086543892 | 6.14E-16 | 2.44E-14 | 1.645047361 |
| ENSMUSG00000036960 | Clca2 | 229933 | protein_coding | 8.742140021 | -1.645417404 | 0.508222986 | 3.237589502 | 0.001205441 | 0.007425086 | 1.645417404 |
| ENSMUSG00000031762 | Mt2 | 17750 | protein_coding | 126.8137351 | -1.645693236 | 0.366323744 | 4.492455817 | 7.04E-06 | 8.18E-05 | 1.645693236 |
| ENSMUSG00000025105 | Bnc1 | 12173 | protein_coding | 1.184808448 | -1.645889739 | 2.142406377 | 0.768243484 | 0.442342544 | 1 | 1.645889739 |
| ENSMUSG00000098553 | Mir3569 | 102465774 | miRNA | 1.529273112 | 1.646146288 | 1.384848632 | -1.188683189 | 0.234564358 | 1 | 1.646146288 |
| ENSMUSG00000099284 | Mir7026 | 102466216 | miRNA | 0.701201205 | -1.646319427 | 1.331945 | 1.236026584 | 0.216448682 | 1 | 1.646319427 |
| ENSMUSG00000029007 | Agtrap | 11610 | protein_coding | 793.2425748 | 1.646386517 | 0.154371358 | -10.66510357 | 1.48E-26 | 1.02E-24 | 1.646386517 |
| ENSMUSG00000009900 | Wnt3a | 22416 | protein_coding | 0.988659055 | 1.646428577 | 2.450290032 | -0.671932121 | 0.501626911 | 1 | 1.646428577 |
| ENSMUSG00000116376 | Tmem249 | 666504 | protein_coding | 1.208094954 | 1.646538625 | 2.456164227 | -0.670369923 | 0.502622003 | 1 | 1.646538625 |
| ENSMUSG00000047104 | Pbp2 | 76400 | protein_coding | 2.705330516 | 1.646822378 | 0.967350748 | -1.702404614 | 0.088679546 | 1 | 1.646822378 |
| ENSMUSG00000097622 | A330033J07Rik | 320614 | lncRNA | 1.095971695 | -1.646876128 | 1.840213607 | 0.89493748 | 0.37082051 | 1 | 1.646876128 |
| ENSMUSG00000100291 | 2310069B03Rik | 69652 | lncRNA | 11.5266965 | 1.648332961 | 0.551922092 | -2.986531947 | 0.002821614 | 0.015118795 | 1.648332961 |
| ENSMUSG00000049420 | Tmem200a | 77220 | protein_coding | 2.422448423 | 1.648698593 | 1.457038677 | -1.13154072 | 0.257827575 | 1 | 1.648698593 |
| ENSMUSG00000049907 | Rasl11b | 68939 | protein_coding | 33.47025534 | 1.649003205 | 0.333204928 | -4.94891601 | 7.46E-07 | 1.06E-05 | 1.649003205 |
| ENSMUSG00000075570 | Krt26 | 320864 | protein_coding | 2.253135895 | 1.649237956 | 2.431389589 | -0.678310857 | 0.497574616 | 1 | 1.649237956 |
| ENSMUSG00000012520 | Phox2b | 18935 | protein_coding | 1.826713544 | 1.649931959 | 2.041950146 | -0.808017748 | 0.419080365 | 1 | 1.649931959 |
| ENSMUSG00000031142 | Cacna1f | 54652 | protein_coding | 4.670372964 | 1.650002328 | 1.725879499 | -0.95603565 | 0.339054224 | 0.560532006 | 1.650002328 |
| ENSMUSG00000057802 | Gm10030 | 791282 | lncRNA | 2.243205438 | -1.650015921 | 1.229622273 | 1.341888445 | 0.179632169 | 1 | 1.650015921 |
| ENSMUSG00000028441 | 1110017D15Rik | 73721 | protein_coding | 1.659356735 | 1.650452932 | 1.436037342 | -1.149310595 | 0.250427931 | 1 | 1.650452932 |
| ENSMUSG00000059743 | Fdps | 110196 | protein_coding | 546.6658641 | 1.650641319 | 0.11235026 | -14.69192253 | 7.26E-49 | 9.59E-47 | 1.650641319 |
| ENSMUSG00000020799 | Tekt1 | 21689 | protein_coding | 2.329578106 | -1.650667186 | 0.996904352 | 1.655792937 | 0.097763759 | 1 | 1.650667186 |
| ENSMUSG00000043301 | Kcnj6 | 16522 | protein_coding | 3.137313558 | 1.651940298 | 2.223637598 | -0.742899967 | 0.457542246 | 1 | 1.651940298 |
| ENSMUSG00000040212 | Emp3 | 13732 | protein_coding | 7.728487166 | 1.652262873 | 0.662735674 | -2.493094816 | 0.012663502 | 0.051148256 | 1.652262873 |
| ENSMUSG00000054537 | Tmprss11e | 243084 | protein_coding | 1.947199286 | 1.652429091 | 1.162169765 | -1.421848288 | 0.155070298 | 1 | 1.652429091 |
| ENSMUSG00000037470 | Uggt1 | 320011 | protein_coding | 4572.442825 | 1.65250563 | 0.11197383 | -14.75796285 | 2.73E-49 | 3.67E-47 | 1.65250563 |
| ENSMUSG00000057280 | Musk | 18198 | protein_coding | 1.864394443 | 1.652618261 | 2.171667081 | -0.760990612 | 0.446662675 | 1 | 1.652618261 |
| ENSMUSG00000086687 | 4930547E08Rik | 78031 | lncRNA | 1.004695204 | 1.654933233 | 2.183535162 | -0.757914625 | 0.448502094 | 1 | 1.654933233 |
| ENSMUSG00000095928 | Or5ac22 | 258994 | protein_coding | 2.466686909 | -1.655030558 | 2.359542158 | 0.701420211 | 0.483040813 | 1 | 1.655030558 |
| ENSMUSG00000026751 | Nr5a1 | 26423 | protein_coding | 2.289270758 | -1.655457959 | 2.370131575 | 0.698466691 | 0.484885381 | 1 | 1.655457959 |
| ENSMUSG00000032171 | Pin1 | 23988 | protein_coding | 569.3852515 | 1.655655389 | 0.100442365 | -16.48363599 | 4.81E-61 | 8.27E-59 | 1.655655389 |
| ENSMUSG00000064259 | Vmn1r13 | 113862 | protein_coding | 1.733951663 | -1.656147324 | 2.205127223 | 0.751043888 | 0.452626243 | 1 | 1.656147324 |
| ENSMUSG00000028148 | Them5 | 66198 | protein_coding | 1.562047736 | 1.656179979 | 1.330397727 | -1.244875833 | 0.213177395 | 1 | 1.656179979 |
| ENSMUSG00000097146 | Gm4211 | 100043074 | lncRNA | 0.807808091 | 1.65750748 | 1.951403755 | -0.84939238 | 0.395662992 | 1 | 1.65750748 |

Supplementary Table 1

| ensembl_gene_id | external_gene_name | entrezgene_id | gene_biotype | baseMean | log2FoldChange | lfcSE | stat | pvalue | padj | absLog2FC |
|---|---|---|---|---|---|---|---|---|---|---|
| ENSMUSG00000028445 | Enho | 69638 | protein_coding | 4.451091078 | 1.65802145 | 0.760875967 | -2.17909557 | 0.029324569 | 0.099059146 | 1.65802145 |
| ENSMUSG00000001128 | Cfp | 18636 | protein_coding | 31.41813094 | 1.658699354 | 0.272525484 | -6.086400918 | 1.15E-09 | 2.53E-08 | 1.658699354 |
| ENSMUSG00000107736 | Gm44148 | 115490371 | lncRNA | 1.464336435 | 1.659992933 | 0.998893018 | -1.661832552 | 0.096546353 | 1 | 1.659992933 |
| ENSMUSG00000030468 | Siglecg | 243958 | protein_coding | 2.828013711 | 1.660256434 | 2.002395772 | -0.829135008 | 0.407028016 | 1 | 1.660256434 |
| ENSMUSG00000080461 | Gm22488 | 115488567 | snoRNA | 0.952585193 | 1.660440712 | 1.405751904 | -1.181176215 | 0.237532728 | 1 | 1.660440712 |
| ENSMUSG00000117853 | Vmn1r88 | 100312474 | protein_coding | 2.839608038 | -1.660833345 | 2.668474024 | 0.622390674 | 0.533685012 | 1 | 1.660833345 |
| ENSMUSG00000039555 | Cylc2 | 74914 | protein_coding | 2.14853541 | -1.661403185 | 2.248961214 | 0.73874248 | 0.460063385 | 1 | 1.661403185 |
| ENSMUSG00000000560 | Gabra2 | 14395 | protein_coding | 1.48090822 | -1.662054173 | 2.41240487 | 0.688961539 | 0.490847471 | 1 | 1.662054173 |
| ENSMUSG00000032591 | Mst1 | 15235 | protein_coding | 16.36394741 | 1.662335211 | 0.338740455 | -4.907400897 | 9.23E-07 | 1.29E-05 | 1.662335211 |
| ENSMUSG00000095730 | Vmn2r29 | 76229 | protein_coding | 1.027932652 | -1.662582196 | 1.288597805 | 1.290225848 | 0.196972254 | 1 | 1.662582196 |
| ENSMUSG00000074930 | Gm13981 | 108168832 | processed_pseudogene | 0.485898037 | 1.6646438 | 1.954872662 | -0.851535669 | 0.394471858 | 1 | 1.6646438 |
| ENSMUSG00000060034 | Ctf2 | 244218 | protein_coding | 1.41623436 | -1.665536768 | 1.111602009 | 1.498321121 | 0.134049839 | 1 | 1.665536768 |
| ENSMUSG00000047094 | Ofcc1 | 218165 | protein_coding | 1.322721141 | 1.665550793 | 2.57774238 | -0.646127715 | 0.518196648 | 1 | 1.665550793 |
| ENSMUSG00000020581 | Agr2 | 23795 | protein_coding | 31390.40055 | 1.666179595 | 0.194612511 | -8.561523531 | 1.11E-17 | 4.98E-16 | 1.666179595 |
| ENSMUSG00000065034 | Gm22363 | 115490517 | snRNA | 0.856727798 | 1.666874867 | 1.523907975 | -1.093815962 | 0.274035702 | 1 | 1.666874867 |
| ENSMUSG00000094898 | Vmn1r201 | 171255 | protein_coding | 0.958728596 | -1.666912953 | 2.451371204 | 0.679992059 | 0.496509489 | 1 | 1.666912953 |
| ENSMUSG00000031927 | 1700012B09Rik | 69325 | protein_coding | 2.086642866 | 1.667487262 | 0.824561525 | -2.022271489 | 0.043148312 | 1 | 1.667487262 |
| ENSMUSG00000039539 | Sgcz | 244431 | protein_coding | 1.323444517 | -1.668166758 | 2.461756313 | 0.677632774 | 0.498004561 | 1 | 1.668166758 |
| ENSMUSG00000054641 | Mmrn1 | 70945 | protein_coding | 1.614957062 | -1.668772642 | 1.809105307 | 0.922429798 | 0.356304433 | 1 | 1.668772642 |
| ENSMUSG00000044309 | Apol7c | 108956 | protein_coding | 2.459340203 | 1.668776024 | 0.962383674 | -1.73400284 | 0.082917579 | 1 | 1.668776024 |
| ENSMUSG00000062987 | Olfr715b | 384732 | protein_coding | 1.284138825 | -1.669249363 | 2.47086928 | 0.675571701 | 0.499312614 | 1 | 1.669249363 |
| ENSMUSG00000046768 | Rhoj | 80837 | protein_coding | 4.303915075 | 1.66964858 | 0.806533323 | -2.07015449 | 0.03843788 | 1 | 1.66964858 |
| ENSMUSG00000031554 | Adam5 | 11499 | protein_coding | 2.496808195 | 1.670046009 | 1.856777205 | -0.899432633 | 0.368422264 | 1 | 1.670046009 |
| ENSMUSG00000059343 | Aldoart1 | 353204 | protein_coding | 1.548039415 | 1.671136987 | 1.37186426 | -1.218150393 | 0.223166827 | 1 | 1.671136987 |
| ENSMUSG00000009216 | Fam163b | 109349 | protein_coding | 2.084115323 | -1.672344119 | 2.346696033 | 0.712637724 | 0.476069962 | 1 | 1.672344119 |
| ENSMUSG00000026170 | Cyp27a1 | 104086 | protein_coding | 97.45629417 | -1.67242376 | 0.203256694 | 8.2281362 | 1.90E-16 | 7.87E-15 | 1.67242376 |
| ENSMUSG00000006095 | Tbcb | 66411 | protein_coding | 1360.913785 | 1.673441515 | 0.079346685 | -21.09025124 | 9.77E-99 | 3.43E-96 | 1.673441515 |
| ENSMUSG00000057060 | Slc35f3 | 210027 | protein_coding | 2.171607717 | 1.673745869 | 1.527623994 | -1.095653037 | 0.273230642 | 1 | 1.673745869 |
| ENSMUSG00000031168 | Ebp | 13595 | protein_coding | 1934.119922 | 1.674542875 | 0.097775703 | -17.12637002 | 9.44E-66 | 1.79E-63 | 1.674542875 |
| ENSMUSG00000031924 | Cyb5b | 66427 | protein_coding | 8929.821707 | 1.674686139 | 0.076331699 | -21.9395896 | 1.09E-106 | 4.00E-104 | 1.674686139 |
| ENSMUSG00000022596 | Slurp1 | 57277 | protein_coding | 1.768607031 | -1.674784377 | 1.860930877 | 0.899971298 | 0.368135525 | 1 | 1.674784377 |
| ENSMUSG00000086254 | 3100003L05Rik | 73076 | lncRNA | 5.134287854 | -1.677080597 | 0.636727707 | 2.633905481 | 0.008440898 | 0.037106194 | 1.677080597 |
| ENSMUSG00000022468 | Endou | 19011 | protein_coding | 1.416982213 | -1.677713795 | 1.471589551 | 1.140069114 | 0.254257508 | 1 | 1.677713795 |
| ENSMUSG00000074156 | Ces1h | 75704 | protein_coding | 1.11941248 | 1.678194257 | 1.592499087 | -1.053811755 | 0.291969111 | 1 | 1.678194257 |
| ENSMUSG00000098944 | Mir8120 | 102465905 | miRNA | 1.826996774 | -1.678379569 | 1.315332859 | 1.276011283 | 0.201951531 | 1 | 1.678379569 |
| ENSMUSG00000037994 | Slc9b2 | 97086 | protein_coding | 1.913103347 | -1.678385259 | 1.292822201 | 1.29823363 | 0.194207065 | 1 | 1.678385259 |
| ENSMUSG00000097545 | Mir124a-1hg | 268755 | lncRNA | 2.112642991 | -1.678386512 | 2.683687988 | 0.625402998 | 0.531706595 | 1 | 1.678386512 |
| ENSMUSG00000095616 | Gm26244 | 115489000 | snRNA | 3.328595567 | 1.678864229 | 1.001751672 | -1.675928551 | 0.093752189 | 1 | 1.678864229 |
| ENSMUSG00000051728 | Fam243 | 75328 | protein_coding | 1.020577594 | -1.678951224 | 1.839515253 | 0.912713945 | 0.361393008 | 1 | 1.678951224 |

| ensembl_gene_id | external_gene_name | entrezgene_id | gene_biotype | baseMean | log2FoldChange | lfcSE | stat | pvalue | padj | absLog2FC |
|---|---|---|---|---|---|---|---|---|---|---|
| ENSMUSG00000116504 | I730030J21Rik | 619313 | lncRNA | 1.722520867 | -1.679347449 | 1.229274551 | 1.366128866 | 0.17189852 | 1 | 1.679347449 |
| ENSMUSG00000035916 | Ptprq | 237523 | protein_coding | 3.221817005 | -1.681115405 | 1.574608116 | 1.067640506 | 0.285682705 | 1 | 1.681115405 |
| ENSMUSG00000021098 | 4930447C04Rik | 75801 | protein_coding | 1.775683144 | 1.681489086 | 2.125086244 | -0.791256869 | 0.428794112 | 1 | 1.681489086 |
| ENSMUSG00000000157 | Itgb2l | 16415 | protein_coding | 208.5838436 | 1.681635834 | 0.171921473 | -9.781418229 | 1.35E-22 | 7.84E-21 | 1.681635834 |
| ENSMUSG00000048856 | Slc25a47 | 104910 | protein_coding | 144.3245194 | 1.682153755 | 0.136000016 | -12.36877617 | 3.86E-35 | 3.41E-33 | 1.682153755 |
| ENSMUSG00000042386 | Tex13b | 83555 | protein_coding | 2.091685664 | 1.682932149 | 1.988874137 | -0.846173279 | 0.397456086 | 1 | 1.682932149 |
| ENSMUSG00000047935 | Sox1ot | 434280 | lncRNA | 12.81236859 | 1.683586674 | 0.436361759 | -3.85823606 | 0.000114208 | 0.000968688 | 1.683586674 |
| ENSMUSG00000096316 | Faiml | 623459 | protein_coding | 0.915868232 | -1.684239532 | 2.150240706 | 0.783279531 | 0.433462981 | 1 | 1.684239532 |
| ENSMUSG00000015354 | Pcolce2 | 76477 | protein_coding | 1.785602918 | 1.684339247 | 1.474486887 | -1.142322297 | 0.253320076 | 1 | 1.684339247 |
| ENSMUSG00000087159 | Gm15246 | 101055907 | lncRNA | 1.317208781 | 1.684357323 | 1.408847288 | -1.195557061 | 0.231869456 | 1 | 1.684357323 |
| ENSMUSG00000018238 | Gdf9 | 14566 | protein_coding | 1.442918153 | -1.684854258 | 1.392431143 | 1.210009031 | 0.226275428 | 1 | 1.684854258 |
| ENSMUSG00000024440 | Pcdh12 | 53601 | protein_coding | 2.019395633 | 1.684924886 | 2.181810535 | -0.772259946 | 0.439960483 | 1 | 1.684924886 |
| ENSMUSG00000031132 | Cd40lg | 21947 | protein_coding | 1.046661126 | 1.686112304 | 1.98911073 | -0.847671414 | 0.396620991 | 1 | 1.686112304 |
| ENSMUSG00000027887 | Sypl2 | 17306 | protein_coding | 3.93571929 | -1.686480295 | 1.130715318 | 1.491516271 | 0.135826003 | 1 | 1.686480295 |
| ENSMUSG00000030368 | Ceacam11 | 66996 | protein_coding | 0.952414023 | 1.686550943 | 1.990575985 | -0.847267804 | 0.396845868 | 1 | 1.686550943 |
| ENSMUSG00000074895 | Eif4e1b | 218268 | protein_coding | 0.890644381 | -1.687093936 | 2.520136534 | 0.66944545 | 0.503211367 | 1 | 1.687093936 |
| ENSMUSG00000087382 | Ctcflos | 74161 | lncRNA | 1.918184437 | -1.688395339 | 2.217542015 | 0.761381443 | 0.446429268 | 1 | 1.688395339 |
| ENSMUSG00000085008 | Dbhos | 329371 | lncRNA | 1.442856055 | -1.688416624 | 2.217851441 | 0.761284815 | 0.446486968 | 1 | 1.688416624 |
| ENSMUSG00000086029 | Pax6os1 | 402728 | lncRNA | 1.243207204 | -1.688626267 | 1.732013875 | 0.974949619 | 0.329585252 | 1 | 1.688626267 |
| ENSMUSG00000030815 | Phkg2 | 68961 | protein_coding | 280.7459859 | 1.689228497 | 0.09574097 | -17.64373699 | 1.14E-69 | 2.41E-67 | 1.689228497 |
| ENSMUSG00000001657 | Hoxc8 | 15426 | protein_coding | 0.908058845 | 1.690130105 | 2.178728139 | -0.775741624 | 0.437901558 | 1 | 1.690130105 |
| ENSMUSG00000028492 | Saxo1 | 75811 | protein_coding | 2.851222891 | 1.690229777 | 1.223362378 | -1.381626415 | 0.167086439 | 1 | 1.690229777 |
| ENSMUSG00000026840 | Lamc3 | 23928 | protein_coding | 2.324198594 | 1.690276694 | 2.189965212 | -0.771828102 | 0.440216245 | 1 | 1.690276694 |
| ENSMUSG00000061165 | Or8a1b | 80706 | protein_coding | 1.305690264 | -1.690285874 | 2.562539052 | 0.659613703 | 0.509501758 | 1 | 1.690285874 |
| ENSMUSG00000095093 | Vmn2r111 | 210876 | protein_coding | 1.258870515 | -1.690361931 | 2.563586087 | 0.659373968 | 0.509655654 | 1 | 1.690361931 |
| ENSMUSG00000096852 | Cyp2d12 | 380997 | protein_coding | 95.21325453 | -1.690796304 | 0.358702466 | 4.713645611 | 2.43E-06 | 3.10E-05 | 1.690796304 |
| ENSMUSG00000071686 | Tex13a | 67944 | protein_coding | 1.120316139 | -1.691445467 | 2.578615932 | 0.65595091 | 0.51185571 | 1 | 1.691445467 |
| ENSMUSG00000023999 | Kif6 | 319991 | protein_coding | 2.140156151 | 1.691944921 | 2.405405983 | -0.703392663 | 0.481811077 | 1 | 1.691944921 |
| ENSMUSG00000046844 | Vat1l | 270097 | protein_coding | 1.792597677 | -1.691999242 | 2.056976132 | 0.822566298 | 0.410754671 | 1 | 1.691999242 |
| ENSMUSG00000031491 | Chrna6 | 11440 | protein_coding | 1.131881682 | 1.692075025 | 2.54620739 | -0.664547213 | 0.506340141 | 1 | 1.692075025 |
| ENSMUSG00000079105 | C7 | 109828 | protein_coding | 1.342631701 | -1.692868931 | 2.211809329 | 0.765377426 | 0.444046831 | 1 | 1.692868931 |
| ENSMUSG00000114767 | Gm46392 | 108168052 | lncRNA | 1.840480499 | 1.694049482 | 1.739845469 | -0.97367813 | 0.33021638 | 1 | 1.694049482 |
| ENSMUSG00000040367 | Lrrd1 | 242838 | protein_coding | 3.684387875 | 1.694227463 | 0.904337332 | -1.873446339 | 0.061006776 | 1 | 1.694227463 |
| ENSMUSG00000079357 | Gm11100 | 102633458 | protein_coding | 2.07547091 | 1.694253631 | 1.051906809 | -1.610649934 | 0.107256044 | 1 | 1.694253631 |
| ENSMUSG00000101969 | Gm20125 | 100504235 | lncRNA | 4.53349698 | 1.694363094 | 1.17940811 | -1.436621539 | 0.15082556 | 0.33090651 | 1.694363094 |
| ENSMUSG00000013629 | Cad | 69719 | protein_coding | 1246.413542 | 1.69495055 | 0.120335264 | -14.08523567 | 4.68E-45 | 5.60E-43 | 1.69495055 |
| ENSMUSG00000077460 | Gm26419 | 115486011 | snoRNA | 0.594830023 | -1.695196852 | 1.380669818 | 1.227807569 | 0.219519214 | 1 | 1.695196852 |
| ENSMUSG00000027931 | Npr1 | 18160 | protein_coding | 8.778388908 | 1.696611654 | 0.410701134 | -4.131012829 | 3.61E-05 | 0.000351256 | 1.696611654 |
| ENSMUSG00000005800 | Mmp8 | 17394 | protein_coding | 0.927679158 | 1.696703949 | 2.411011449 | -0.703731187 | 0.481600195 | 1 | 1.696703949 |

Supplementary Table 1

| ensembl_gene_id | external_gene_name | entrezgene_id | gene_biotype | baseMean | log2FoldChange | lfcSE | stat | pvalue | padj | absLog2FC |
|---|---|---|---|---|---|---|---|---|---|---|
| ENSMUSG00000018906 | P4ha2 | 18452 | protein_coding | 94.93944511 | 1.696743554 | 0.178824324 | -9.488326385 | 2.35E-21 | 1.29E-19 | 1.696743554 |
| ENSMUSG00000075065 | Or4b1 | 258987 | protein_coding | 1.500641871 | 1.696776738 | 2.412693238 | -0.703270814 | 0.481886995 | 1 | 1.696776738 |
| ENSMUSG00000022577 | Ly6h | 23934 | protein_coding | 1.616295203 | -1.697476823 | 2.386806124 | 0.711191749 | 0.476965421 | 1 | 1.697476823 |
| ENSMUSG00000063409 | Lrrc43 | 381741 | protein_coding | 1.001036898 | 1.69759759 | 2.431103541 | -0.698282719 | 0.485000404 | 1 | 1.69759759 |
| ENSMUSG00000058295 | Prp2 | 83380 | protein_coding | 1.136666814 | 1.697698262 | 2.433416909 | -0.697660255 | 0.485389689 | 1 | 1.697698262 |
| ENSMUSG00000088544 | Mir1945 | 100316833 | miRNA | 1.083253035 | -1.698283058 | 1.438745516 | 1.180391556 | 0.237844519 | 1 | 1.698283058 |
| ENSMUSG00000064682 | Gm25813 | 115486988 | snRNA | 1.554988413 | 1.698522938 | 1.125563316 | -1.509042551 | 0.131287909 | 1 | 1.698522938 |
| ENSMUSG00000105662 | Gm6639 | 625963 | lncRNA | 1.370666956 | 1.698703331 | 2.456652596 | -0.691470717 | 0.489269777 | 1 | 1.698703331 |
| ENSMUSG00000089645 | Gm5766 | 436332 | transcribed_processed_pseudogene | 0.800393058 | -1.699022636 | 1.819914519 | 0.933572769 | 0.350524317 | 1 | 1.699022636 |
| ENSMUSG00000086449 | 9030204H09Rik | 77665 | lncRNA | 2.349136011 | -1.699102955 | 0.809559331 | 2.098799791 | 0.035834553 | 1 | 1.699102955 |
| ENSMUSG00000042401 | Crtac1 | 72832 | protein_coding | 1.073585362 | 1.699497858 | 2.475389926 | -0.686557637 | 0.492361532 | 1 | 1.699497858 |
| ENSMUSG00000055540 | Epha6 | 13840 | protein_coding | 0.977012013 | 1.69958101 | 2.47733625 | -0.686051807 | 0.49268044 | 1 | 1.69958101 |
| ENSMUSG00000021081 | Serpina1f | 68348 | protein_coding | 1.187151887 | 1.699617885 | 2.478266355 | -0.685809208 | 0.492833429 | 1 | 1.699617885 |
| ENSMUSG00000030945 | Acsm2 | 233799 | protein_coding | 1.254244383 | 1.700816114 | 2.507410485 | -0.678315786 | 0.497571492 | 1 | 1.700816114 |
| ENSMUSG00000027505 | Fam209 | 76426 | protein_coding | 2.115032382 | 1.701305958 | 0.84697731 | -2.008679499 | 0.044571134 | 1 | 1.701305958 |
| ENSMUSG00000032965 | Ift57 | 73916 | protein_coding | 4.124261059 | 1.702393937 | 0.58725677 | -2.898891972 | 0.003744839 | 1 | 1.702393937 |
| ENSMUSG00000071341 | Egr4 | 13656 | protein_coding | 1.877420068 | 1.703080131 | 1.190387377 | -1.430694045 | 0.152517921 | 1 | 1.703080131 |
| ENSMUSG00000005360 | Slc1a3 | 20512 | protein_coding | 13.01389393 | -1.703176661 | 0.534144604 | 3.188605947 | 0.001429606 | 0.008564059 | 1.703176661 |
| ENSMUSG00000047636 | Cdcp2 | 242603 | protein_coding | 1.648237499 | -1.703511717 | 2.837036804 | 0.600454571 | 0.548203328 | 1 | 1.703511717 |
| ENSMUSG00000050541 | Adra1b | 11548 | protein_coding | 1.473310619 | 1.703945403 | 2.050071303 | -0.83116397 | 0.405881006 | 1 | 1.703945403 |
| ENSMUSG00000029155 | Spata18 | 73472 | protein_coding | 2.135824644 | -1.704813955 | 2.854756173 | 0.597183735 | 0.550384717 | 1 | 1.704813955 |
| ENSMUSG00000060565 | Gm5591 | 434171 | protein_coding | 1.375256206 | -1.705001228 | 2.857400203 | 0.596696685 | 0.550709906 | 1 | 1.705001228 |
| ENSMUSG00000041809 | Efhc1 | 71877 | protein_coding | 9.171101619 | 1.705478032 | 0.420232294 | -4.058417352 | 4.94E-05 | 0.000462789 | 1.705478032 |
| ENSMUSG00000028280 | Gabrr1 | 14408 | protein_coding | 1.416325104 | -1.705626555 | 2.866105098 | 0.595102586 | 0.5517749 | 1 | 1.705626555 |
| ENSMUSG00000032262 | Elovl4 | 83603 | protein_coding | 0.914198284 | 1.706124205 | 2.000537662 | -0.852832834 | 0.393752013 | 1 | 1.706124205 |
| ENSMUSG00000098590 | Gm1113 | 115487164 | protein_coding | 1.626039014 | 1.706285365 | 1.960654647 | -0.870263087 | 0.384156647 | 1 | 1.706285365 |
| ENSMUSG00000028386 | Slc46a2 | 30936 | protein_coding | 1.545880134 | 1.706333729 | 1.393206313 | -1.224753085 | 0.220668266 | 1 | 1.706333729 |
| ENSMUSG00000043274 | Or10ag2 | 258347 | protein_coding | 1.054714302 | -1.707211135 | 2.889551194 | 0.590822249 | 0.554639526 | 1 | 1.707211135 |
| ENSMUSG00000001076 | C1ql4 | 239659 | protein_coding | 1.024039591 | 1.707575636 | 1.462879277 | -1.16727037 | 0.2431012 | 1 | 1.707575636 |
| ENSMUSG00000020020 | Usp44 | 327799 | protein_coding | 2.258511807 | 1.707728716 | 1.251492693 | -1.364553486 | 0.172393433 | 1 | 1.707728716 |
| ENSMUSG00000046623 | Gjb4 | 14621 | protein_coding | 4.764586647 | -1.70857351 | 0.704738104 | 2.424409154 | 0.015333319 | 0.059481896 | 1.70857351 |
| ENSMUSG00000062483 | Vmn1r171 | 81012 | protein_coding | 0.959593077 | -1.709825175 | 2.927859825 | 0.583984643 | 0.559230644 | 1 | 1.709825175 |
| ENSMUSG00000066122 | Or13a17 | 18344 | protein_coding | 0.842069657 | -1.710424779 | 2.43815069 | 0.701525458 | 0.482975153 | 1 | 1.710424779 |
| ENSMUSG00000027204 | Fbn1 | 14118 | protein_coding | 29.8110868 | -1.710901044 | 0.344927044 | 4.960182378 | 7.04E-07 | 1.01E-05 | 1.710901044 |
| ENSMUSG00000085682 | Gm14267 | 383773 | transcribed_processed_pseudogene | 1.487668278 | 1.711609317 | 2.820079158 | -0.606936622 | 0.543892973 | 1 | 1.711609317 |
| ENSMUSG00000100658 | F730311O21Rik | 622304 | lncRNA | 2.095832242 | -1.71170909 | 1.26421727 | 1.353967495 | 0.175746748 | 1 | 1.71170909 |
| ENSMUSG00000048173 | Or1aa2 | 258289 | protein_coding | 1.900196517 | 1.712137037 | 2.838268889 | -0.603232852 | 0.546353796 | 1 | 1.712137037 |
| ENSMUSG00000015619 | Gata3 | 14462 | protein_coding | 2.83432994 | 1.712323393 | 0.957662038 | -1.788024715 | 0.07377202 | 1 | 1.712323393 |
| ENSMUSG00000085034 | Gm11240 | 667768 | lncRNA | 11.3444304 | 1.712387185 | 0.354957099 | -4.824208868 | 1.41E-06 | 1.88E-05 | 1.712387185 |

| ensembl_gene_id | external_gene_name | entrezgene_id | gene_biotype | baseMean | log2FoldChange | lfcSE | stat | pvalue | padj | absLog2FC |
|---|---|---|---|---|---|---|---|---|---|---|
| ENSMUSG00000096099 | Vmn1r220 | 171271 | protein_coding | 1.555501638 | 1.712407551 | 2.84764644 | -0.601341349 | 0.547612655 | 1 | 1.712407551 |
| ENSMUSG00000040127 | Sdr9c7 | 70061 | protein_coding | 5.67817302 | 1.713045767 | 0.557399558 | -3.073281533 | 0.002117186 | 0.011888207 | 1.713045767 |
| ENSMUSG00000081945 | Or52d13 | 546989 | protein_coding | 1.80932637 | 1.71324786 | 2.877523372 | -0.595389729 | 0.55158299 | 1 | 1.71324786 |
| ENSMUSG00000090206 | Tepp | 73407 | protein_coding | 1.159740405 | 1.713294283 | 2.87904212 | -0.595091774 | 0.551782127 | 1 | 1.713294283 |
| ENSMUSG00000046450 | Or13j1 | 56015 | protein_coding | 1.143965969 | 1.713576648 | 2.88929467 | -0.593077842 | 0.553129059 | 1 | 1.713576648 |
| ENSMUSG00000058398 | Prss43 | 272643 | protein_coding | 1.623211381 | 1.713591987 | 2.889961641 | -0.592946274 | 0.553217108 | 1 | 1.713591987 |
| ENSMUSG00000010803 | Gabra1 | 14394 | protein_coding | 1.483916141 | 1.713967395 | 2.903710231 | -0.590268057 | 0.555010951 | 1 | 1.713967395 |
| ENSMUSG00000098627 | 4930524O08Rik | 74700 | lncRNA | 1.726345334 | -1.714020408 | 2.981691294 | 0.574848379 | 0.565393844 | 1 | 1.714020408 |
| ENSMUSG00000075387 | Or1j13 | 258952 | protein_coding | 1.397049837 | 1.714324617 | 2.916968316 | -0.587707658 | 0.556728535 | 1 | 1.714324617 |
| ENSMUSG00000021508 | Cxcl14 | 57266 | protein_coding | 1.292557234 | 1.714569563 | 2.926144032 | -0.585948451 | 0.557910157 | 1 | 1.714569563 |
| ENSMUSG00000025902 | Sox17 | 20671 | protein_coding | 1.119941084 | 1.714807483 | 2.935103622 | -0.584240866 | 0.55905827 | 1 | 1.714807483 |
| ENSMUSG00000048456 | Or5b12 | 258696 | protein_coding | 1.119941084 | 1.714807483 | 2.935103622 | -0.584240866 | 0.55905827 | 1 | 1.714807483 |
| ENSMUSG00000073400 | Trim10 | 19824 | protein_coding | 3.584357818 | 1.714977473 | 0.855230377 | -2.005281289 | 0.044932984 | 1 | 1.714977473 |
| ENSMUSG00000063188 | Or1o1 | 258504 | protein_coding | 1.16824256 | 1.715033981 | 2.943778667 | -0.582596103 | 0.560165229 | 1 | 1.715033981 |
| ENSMUSG00000025652 | Tmem89 | 69384 | protein_coding | 1.248629959 | -1.715545412 | 2.085435746 | 0.822631632 | 0.410717505 | 1 | 1.715545412 |
| ENSMUSG00000091679 | Vmn2r96 | 433070 | protein_coding | 1.207548252 | 1.715561361 | 2.964237384 | -0.578753028 | 0.56275583 | 1 | 1.715561361 |
| ENSMUSG00000094347 | Or6c208 | 258724 | protein_coding | 0.898498368 | 1.716841879 | 1.701864795 | -1.008800396 | 0.313070371 | 1 | 1.716841879 |
| ENSMUSG00000041468 | Gpr12 | 14738 | protein_coding | 1.550232034 | 1.717066658 | 1.588062131 | -1.08123393 | 0.279593069 | 1 | 1.717066658 |
| ENSMUSG00000050737 | Ptges | 64292 | protein_coding | 42.44999067 | 1.718113539 | 0.195730921 | -8.777936218 | 1.67E-18 | 7.79E-17 | 1.718113539 |
| ENSMUSG00000070225 | n-R5s33 | 115488675 | rRNA | 1.465861387 | 1.718356607 | 1.179621526 | -1.456701637 | 0.145198758 | 1 | 1.718356607 |
| ENSMUSG00000054003 | Tdrd9 | 74691 | protein_coding | 3.921535678 | 1.719050937 | 1.571765866 | -1.09370675 | 0.274083613 | 1 | 1.719050937 |
| ENSMUSG00000042918 | Mamstr | 74490 | protein_coding | 21.79031266 | 1.719682403 | 0.403393592 | -4.263038471 | 2.02E-05 | 0.000207367 | 1.719682403 |
| ENSMUSG00000041193 | Pla2g5 | 18784 | protein_coding | 303.0682084 | 1.719711858 | 0.321717164 | -5.345415322 | 9.02E-08 | 1.50E-06 | 1.719711858 |
| ENSMUSG00000085298 | F730035M05Rik | 414124 | lncRNA | 0.857886436 | 1.720338855 | 1.569447272 | -1.096143136 | 0.27301614 | 1 | 1.720338855 |
| ENSMUSG00000054746 | Abca15 | 320631 | protein_coding | 3.595481892 | 1.722440106 | 1.400795373 | -1.229615788 | 0.218841015 | 1 | 1.722440106 |
| ENSMUSG00000002341 | Ncan | 13004 | protein_coding | 3.02609161 | 1.72445485 | 2.244600621 | -0.768268009 | 0.442327977 | 1 | 1.72445485 |
| ENSMUSG00000098474 | Mir6919 | 102466765 | miRNA | 0.799119027 | -1.725202543 | 1.226946968 | 1.406093815 | 0.159696269 | 1 | 1.725202543 |
| ENSMUSG00000069823 | Or1e16 | 258923 | protein_coding | 1.584029306 | 1.725211797 | 3.426430099 | -0.503501238 | 0.61461191 | 1 | 1.725211797 |
| ENSMUSG00000069171 | Nr2f1 | 13865 | protein_coding | 1.350353705 | 1.725835269 | 3.463590869 | -0.498279195 | 0.61828727 | 1 | 1.725835269 |
| ENSMUSG00000040808 | S100g | 12309 | protein_coding | 1030.286379 | -1.725854045 | 0.236275697 | 7.3044078 | 2.78E-13 | 8.84E-12 | 1.725854045 |
| ENSMUSG00000022315 | Slc30a8 | 239436 | protein_coding | 1.306920553 | 1.72597809 | 3.472234747 | -0.497079897 | 0.61913271 | 1 | 1.72597809 |
| ENSMUSG00000070378 | Or1a1 | 404316 | protein_coding | 1.306920553 | 1.72597809 | 3.472234747 | -0.497079897 | 0.61913271 | 1 | 1.72597809 |
| ENSMUSG00000075062 | Or4b12 | 258789 | protein_coding | 1.306920553 | 1.72597809 | 3.472234747 | -0.497079897 | 0.61913271 | 1 | 1.72597809 |
| ENSMUSG00000107070 | Gm35191 | 102638690 | lncRNA | 1.259359941 | 1.726125194 | 3.48123375 | -0.4958372 | 0.620009277 | 1 | 1.726125194 |
| ENSMUSG00000094992 | Fbxw25 | 546160 | protein_coding | 1.302793093 | 1.726152933 | 3.482977837 | -0.495596875 | 0.620178858 | 1 | 1.726152933 |
| ENSMUSG00000079168 | Cd209g | 70192 | protein_coding | 1.215926789 | 1.726335262 | 3.494211184 | -0.494055789 | 0.62126678 | 1 | 1.726335262 |
| ENSMUSG00000027261 | Hao1 | 15112 | protein_coding | 1.168366177 | 1.726431943 | 3.500209988 | -0.493236677 | 0.621845365 | 1 | 1.726431943 |
| ENSMUSG00000058670 | Dmtf1l | 237029 | protein_coding | 1.263487401 | 1.72643412 | 3.500418808 | -0.493207875 | 0.621865714 | 1 | 1.72643412 |
| ENSMUSG00000037171 | Nodal | 18119 | protein_coding | 1.26342737 | 1.726719411 | 2.123631911 | -0.813097318 | 0.416162264 | 1 | 1.726719411 |

Supplementary Table 1

| ensembl_gene_id | external_gene_name | entrezgene_id | gene_biotype | baseMean | log2FoldChange | lfcSE | stat | pvalue | padj | absLog2FC |
|---|---|---|---|---|---|---|---|---|---|---|
| ENSMUSG00000030310 | Slc6a1 | 232333 | protein_coding | 6.445095494 | 1.727679261 | 0.917593807 | -1.882836663 | 0.059722507 | 0.170249866 | 1.727679261 |
| ENSMUSG00000086594 | Nudt12os | 75039 | lncRNA | 0.595039682 | -1.728314046 | 1.760850214 | 0.981522467 | 0.326335161 | 1 | 1.728314046 |
| ENSMUSG00000072437 | Nanos1 | 332397 | protein_coding | 30.73834153 | 1.72893917 | 0.266900841 | -6.477833353 | 9.30E-11 | 2.34E-09 | 1.72893917 |
| ENSMUSG00000040875 | Osbpl10 | 74486 | protein_coding | 590.4721796 | 1.729881543 | 0.131536394 | -13.15135295 | 1.67E-39 | 1.69E-37 | 1.729881543 |
| ENSMUSG00000039476 | Prrx2 | 20204 | protein_coding | 1.424749118 | 1.730054115 | 2.069630795 | -0.83592403 | 0.403197643 | 1 | 1.730054115 |
| ENSMUSG00000047773 | Ankfn1 | 382543 | protein_coding | 2.921823894 | 1.730458213 | 1.684208314 | -1.027460914 | 0.304203481 | 1 | 1.730458213 |
| ENSMUSG00000010342 | Tex14 | 83560 | protein_coding | 13.2168582 | 1.730554086 | 0.713969505 | -2.423848741 | 0.015357 | 0.059503296 | 1.730554086 |
| ENSMUSG00000020241 | Col6a2 | 12834 | protein_coding | 15.17225406 | 1.731689136 | 0.845773661 | -2.047461651 | 0.040612778 | 0.127457131 | 1.731689136 |
| ENSMUSG00000013483 | Card14 | 170720 | protein_coding | 504.6635359 | 1.734860479 | 0.130294904 | -13.31487592 | 1.90E-40 | 2.02E-38 | 1.734860479 |
| ENSMUSG00000097662 | Gm2093 | 328489 | lncRNA | 1.613337266 | -1.734872852 | 1.123671414 | 1.543932533 | 0.122604678 | 1 | 1.734872852 |
| ENSMUSG00000033191 | Tie1 | 21846 | protein_coding | 3.849708213 | 1.735062795 | 2.44542175 | -0.709514747 | 0.478005103 | 1 | 1.735062795 |
| ENSMUSG00000104109 | Gm30292 | 102632139 | lncRNA | 1.657643086 | 1.735683327 | 2.141658038 | -0.81043906 | 0.417687876 | 1 | 1.735683327 |
| ENSMUSG00000029656 | C8b | 110382 | protein_coding | 1.777951398 | -1.736324486 | 2.231920874 | 0.777950736 | 0.436598056 | 1 | 1.736324486 |
| ENSMUSG00000118075 | Gm38576 | 102641800 | lncRNA | 0.718946968 | -1.736379047 | 1.527711849 | 1.13658806 | 0.255710534 | 1 | 1.736379047 |
| ENSMUSG00000015890 | Amdhd1 | 71761 | protein_coding | 0.87544079 | 1.736470849 | 1.529943137 | -1.134990449 | 0.25637932 | 1 | 1.736470849 |
| ENSMUSG00000031202 | Rab39b | 67790 | protein_coding | 1.31375042 | 1.739826812 | 1.282269109 | -1.356834381 | 0.174833841 | 1 | 1.739826812 |
| ENSMUSG00000056293 | Gsdmc2 | 331063 | protein_coding | 2425.080819 | -1.744107979 | 0.390826836 | 4.462610597 | 8.10E-06 | 9.28E-05 | 1.744107979 |
| ENSMUSG00000046694 | Tent5b | 100342 | protein_coding | 25.08560603 | 1.744553054 | 0.240747113 | -7.246413181 | 4.28E-13 | 1.34E-11 | 1.744553054 |
| ENSMUSG00000033210 | Slc9c1 | 208169 | protein_coding | 2.338392076 | 1.744890679 | 1.760378147 | -0.991202193 | 0.32158686 | 1 | 1.744890679 |
| ENSMUSG00000001027 | Scn4a | 110880 | protein_coding | 2.847666275 | -1.745866859 | 3.722672941 | 0.468982069 | 0.639082453 | 1 | 1.745866859 |
| ENSMUSG00000103114 | Gm32200 | 102634668 | lncRNA | 1.992093908 | 1.74690046 | 2.175311843 | -0.803057486 | 0.421941509 | 1 | 1.74690046 |
| ENSMUSG00000116660 | 9030404E10Rik | 71528 | lncRNA | 11.05824844 | -1.7471369 | 0.381582877 | 4.57865645 | 4.68E-06 | 5.62E-05 | 1.7471369 |
| ENSMUSG00000038295 | Atg9b | 213948 | protein_coding | 12.58316035 | 1.748086466 | 0.384695861 | -4.544074016 | 5.52E-06 | 6.54E-05 | 1.748086466 |
| ENSMUSG00000055809 | Dnaaf3 | 436022 | protein_coding | 6.903880774 | 1.749390357 | 0.541017741 | -3.233517544 | 0.001222758 | 0.007512139 | 1.749390357 |
| ENSMUSG00000018893 | Mb | 17189 | protein_coding | 0.833646693 | 1.750024173 | 1.687092424 | -1.037301897 | 0.299595182 | 1 | 1.750024173 |
| ENSMUSG00000044405 | Adig | 246747 | protein_coding | 5.956431003 | 1.751419294 | 0.517178507 | -3.386488943 | 0.000707931 | 0.004731193 | 1.751419294 |
| ENSMUSG00000052562 | Slc22a30 | 319800 | protein_coding | 1.015078818 | 1.75393082 | 2.236263925 | -0.784312979 | 0.432856486 | 1 | 1.75393082 |
| ENSMUSG00000027790 | Sis | 69983 | protein_coding | 142.5817754 | -1.755067977 | 0.413323465 | 4.246233576 | 2.17E-05 | 0.000222179 | 1.755067977 |
| ENSMUSG00000035472 | Slc25a21 | 217593 | protein_coding | 1.909982451 | 1.755596301 | 1.623182934 | -1.081576367 | 0.27944081 | 1 | 1.755596301 |
| ENSMUSG00000070369 | Itgad | 381924 | protein_coding | 5.846513244 | 1.755928615 | 1.033281806 | -1.699370495 | 0.089249398 | 0.227342982 | 1.755928615 |
| ENSMUSG00000045514 | Or9a4 | 258381 | protein_coding | 2.095266658 | 1.756343426 | 2.336516672 | -0.751693085 | 0.45223565 | 1 | 1.756343426 |
| ENSMUSG00000020019 | Ntn4 | 57764 | protein_coding | 72.20600955 | -1.758079437 | 0.232459418 | 7.562952083 | 3.94E-14 | 1.35E-12 | 1.758079437 |
| ENSMUSG00000074473 | A230072I06Rik | 619290 | protein_coding | 1.939787474 | 1.758684461 | 1.062002945 | -1.65600714 | 0.097720373 | 1 | 1.758684461 |
| ENSMUSG00000044629 | Cnrip1 | 380686 | protein_coding | 2.434863205 | -1.75885346 | 1.899870487 | 0.925775452 | 0.354562687 | 1 | 1.75885346 |
| ENSMUSG00000048655 | Ccdc169 | 320604 | protein_coding | 1.560562524 | 1.759312202 | 1.484225864 | -1.185339944 | 0.235883067 | 1 | 1.759312202 |
| ENSMUSG00000023259 | Slc26a6 | 171429 | protein_coding | 66.45492768 | 1.760706816 | 0.179301498 | -9.81981094 | 9.25E-23 | 5.44E-21 | 1.760706816 |
| ENSMUSG00000023943 | Sult1c1 | 20888 | protein_coding | 0.766045571 | 1.761381212 | 1.45378066 | -1.211586631 | 0.225670655 | 1 | 1.761381212 |
| ENSMUSG00000028451 | 1700022I11Rik | 67317 | protein_coding | 7.241526235 | 1.76341462 | 0.610755648 | -2.887266988 | 0.003886043 | 0.019745882 | 1.76341462 |
| ENSMUSG00000052180 | Serpinb6c | 97848 | protein_coding | 1.06022881 | 1.764262344 | 2.204166783 | -0.800421437 | 0.423466665 | 1 | 1.764262344 |

| ensembl_gene_id | external_gene_name | entrezgene_id | gene_biotype | baseMean | log2FoldChange | lfcSE | stat | pvalue | padj | absLog2FC |
|---|---|---|---|---|---|---|---|---|---|---|
| ENSMUSG00000099353 | 1700031M16Rik | 73302 | transcribed_unitary_pseudogene | 1.430001084 | 1.766556168 | 2.287465129 | -0.772276764 | 0.439950524 | 1 | 1.766556168 |
| ENSMUSG00000087143 | A830082K12Rik | 320174 | lncRNA | 3.493732586 | 1.7677283 | 2.482729034 | -0.712010161 | 0.476458484 | 1 | 1.7677283 |
| ENSMUSG00000038168 | P3h2 | 210530 | protein_coding | 1.748796732 | 1.767823792 | 1.544446746 | -1.144632404 | 0.252361463 | 1 | 1.767823792 |
| ENSMUSG00000089261 | Mir669n | 100316705 | miRNA | 1.260013236 | -1.768401455 | 0.915071868 | 1.932527396 | 0.053294444 | 1 | 1.768401455 |
| ENSMUSG00000027525 | Phactr3 | 74189 | protein_coding | 1.86303038 | -1.76958553 | 1.880570128 | 0.940983537 | 0.346713296 | 1 | 1.76958553 |
| ENSMUSG00000068078 | 2310034C09Rik | 117172 | protein_coding | 1.755080449 | 1.771357605 | 1.023594863 | -1.73052608 | 0.083536326 | 1 | 1.771357605 |
| ENSMUSG00000027895 | Kcnc4 | 99738 | protein_coding | 1.54942769 | 1.771908437 | 2.35587692 | -0.752122669 | 0.451977293 | 1 | 1.771908437 |
| ENSMUSG00000060988 | Galnt13 | 271786 | protein_coding | 2.412489234 | 1.772255301 | 1.846133981 | -0.959981951 | 0.337064299 | 1 | 1.772255301 |
| ENSMUSG00000071714 | Csf2rb2 | 12984 | protein_coding | 3.317638884 | 1.772790059 | 1.628528882 | -1.088583739 | 0.276337491 | 1 | 1.772790059 |
| ENSMUSG00000033007 | Asic4 | 241118 | protein_coding | 0.621939364 | -1.772895472 | 1.503970697 | 1.17880985 | 0.238473901 | 1 | 1.772895472 |
| ENSMUSG00000037003 | Tns2 | 209039 | protein_coding | 481.5427784 | 1.773785721 | 0.114412564 | -15.50341726 | 3.29E-54 | 4.85E-52 | 1.773785721 |
| ENSMUSG00000089679 | Gm16299 | 102641509 | lncRNA | 2.225774973 | -1.774042802 | 0.812807379 | 2.182611585 | 0.029064426 | 1 | 1.774042802 |
| ENSMUSG00000042985 | Upk3b | 100647 | protein_coding | 1.982125444 | -1.774337348 | 1.418233607 | 1.251089623 | 0.210901781 | 1 | 1.774337348 |
| ENSMUSG00000063820 | Arl9 | 384185 | protein_coding | 1.145129567 | 1.774396938 | 1.265912987 | -1.401673697 | 0.161012708 | 1 | 1.774396938 |
| ENSMUSG00000049112 | Oxtr | 18430 | protein_coding | 4.939084958 | -1.774407639 | 0.825723272 | 2.148913201 | 0.031641281 | 0.105177962 | 1.774407639 |
| ENSMUSG00000042817 | Flt3 | 14255 | protein_coding | 2.952763407 | 1.774485732 | 1.263454012 | -1.40447196 | 0.160178356 | 1 | 1.774485732 |
| ENSMUSG00000057716 | Tmem178b | 434008 | protein_coding | 4.504256061 | 1.775235684 | 1.34163592 | -1.323187355 | 0.18577308 | 0.381242282 | 1.775235684 |
| ENSMUSG00000032623 | Oas1d | 100535 | protein_coding | 0.944432125 | -1.775494602 | 1.027693039 | 1.727650704 | 0.084050867 | 1 | 1.775494602 |
| ENSMUSG00000019124 | Scrn1 | 69938 | protein_coding | 4.429790437 | -1.775852493 | 0.795780301 | 2.231586395 | 0.025642312 | 0.089020995 | 1.775852493 |
| ENSMUSG00000070331 | Qrich2 | 217341 | protein_coding | 19.63959544 | 1.776460859 | 0.371334989 | -4.783984575 | 1.72E-06 | 2.25E-05 | 1.776460859 |
| ENSMUSG00000085918 | Gm13032 | 100049161 | lncRNA | 1.21577693 | -1.776593998 | 1.7083567 | 1.039943238 | 0.298366273 | 1 | 1.776593998 |
| ENSMUSG00000028236 | Sdr16c5 | 242285 | protein_coding | 2.698036104 | 1.776692413 | 0.862288823 | -2.060437716 | 0.039356714 | 1 | 1.776692413 |
| ENSMUSG00000049988 | Lrrc25 | 211228 | protein_coding | 2.452242588 | 1.777352927 | 1.498440805 | -1.186134895 | 0.235569032 | 1 | 1.777352927 |
| ENSMUSG00000026424 | Gpr37l1 | 171469 | protein_coding | 1.488717797 | 1.779000667 | 1.41289888 | -1.259113934 | 0.207989182 | 1 | 1.779000667 |
| ENSMUSG00000025400 | Tac2 | 21334 | protein_coding | 1.644232227 | 1.77978055 | 2.36572426 | -0.752319524 | 0.451858929 | 1 | 1.77978055 |
| ENSMUSG00000045991 | Onecut2 | 225631 | protein_coding | 5.167880145 | 1.780050087 | 1.657020021 | -1.074247785 | 0.282711641 | 0.500826007 | 1.780050087 |
| ENSMUSG00000063689 | H2ac21 | 621893 | protein_coding | 37.3246904 | 1.780404437 | 0.24953301 | -7.134945532 | 9.68E-13 | 2.94E-11 | 1.780404437 |
| ENSMUSG00000037451 | Slc22a20 | 381203 | protein_coding | 1.601222434 | 1.78169292 | 2.363259593 | -0.753913334 | 0.450901263 | 1 | 1.78169292 |
| ENSMUSG00000038594 | Cep85l | 100038725 | protein_coding | 2.535226668 | 1.782352283 | 1.131935374 | -1.574606046 | 0.115347406 | 1 | 1.782352283 |
| ENSMUSG00000038453 | Srcin1 | 56013 | protein_coding | 15.39055459 | 1.783261259 | 0.788717768 | -2.260962453 | 0.023761582 | 0.084025676 | 1.783261259 |
| ENSMUSG00000040125 | Gpr26 | 233919 | protein_coding | 1.927997582 | -1.784331711 | 2.108250206 | 0.846356711 | 0.39735378 | 1 | 1.784331711 |
| ENSMUSG00000075081 | Or4a74 | 258968 | protein_coding | 1.374267554 | 1.78446677 | 2.037662209 | -0.875742192 | 0.381170203 | 1 | 1.78446677 |
| ENSMUSG00000029122 | Evc | 59056 | protein_coding | 2.893943827 | -1.78517203 | 2.102075953 | 0.849242401 | 0.395746424 | 1 | 1.78517203 |
| ENSMUSG00000026211 | Obsl1 | 98733 | protein_coding | 25.18796974 | 1.786067647 | 0.287399026 | -6.214591867 | 5.15E-10 | 1.17E-08 | 1.786067647 |
| ENSMUSG00000041700 | Lhfpl1 | 237091 | protein_coding | 1.046904357 | 1.787295405 | 1.413226844 | -1.264691095 | 0.205982083 | 1 | 1.787295405 |
| ENSMUSG00000064964 | Gm25147 | 115489572 | snRNA | 0.812799278 | -1.787372406 | 1.187273685 | 1.505442619 | 0.132210319 | 1 | 1.787372406 |
| ENSMUSG00000095864 | Vmn1r77 | 628946 | protein_coding | 2.689475109 | -1.787413755 | 2.629283017 | 0.67981033 | 0.496624565 | 1 | 1.787413755 |
| ENSMUSG00000013367 | Iglon5 | 210094 | protein_coding | 1.936884795 | -1.787632997 | 1.627339489 | 1.098500349 | 0.271986065 | 1 | 1.787632997 |
| ENSMUSG00000048332 | Lhfp | 108927 | protein_coding | 2.21052086 | 1.788228128 | 1.770408363 | -1.010065342 | 0.312463986 | 1 | 1.788228128 |

| ensembl_gene_id | external_gene_name | entrezgene_id | gene_biotype | baseMean | log2FoldChange | lfcSE | stat | pvalue | padj | absLog2FC |
|---|---|---|---|---|---|---|---|---|---|---|
| ENSMUSG00000111205 | Gm39307 | 105243360 | lncRNA | 1.095184969 | 1.788397209 | 1.552955436 | -1.15160884 | 0.249481847 | 1 | 1.788397209 |
| ENSMUSG00000011486 | Slc25a41 | 103775 | protein_coding | 1.470966689 | 1.788816301 | 1.463615125 | -1.222190364 | 0.221635647 | 1 | 1.788816301 |
| ENSMUSG00000030847 | Bag3 | 29810 | protein_coding | 979.8572491 | 1.789015341 | 0.112753526 | -15.86660216 | 1.08E-56 | 1.71E-54 | 1.789015341 |
| ENSMUSG00000041828 | Abca8a | 217258 | protein_coding | 3.79430756 | 1.789090661 | 1.633066322 | -1.095540724 | 0.273279814 | 1 | 1.789090661 |
| ENSMUSG00000047554 | Tmem41b | 233724 | protein_coding | 941.7348586 | 1.789340151 | 0.081078267 | -22.06929438 | 6.24E-108 | 2.34E-105 | 1.789340151 |
| ENSMUSG00000026841 | Fibcd1 | 98970 | protein_coding | 2.180060934 | 1.789429852 | 1.512440744 | -1.18314047 | 0.236753481 | 1 | 1.789429852 |
| ENSMUSG00000040732 | Erg | 13876 | protein_coding | 2.503108683 | 1.789852734 | 2.378382291 | -0.75255048 | 0.451720085 | 1 | 1.789852734 |
| ENSMUSG00000045466 | Zfp956 | 101197 | protein_coding | 202.3735133 | 1.790174789 | 0.106130805 | -16.86762662 | 7.79E-64 | 1.41E-61 | 1.790174789 |
| ENSMUSG00000045322 | Tlr9 | 81897 | protein_coding | 3.469959679 | 1.790593195 | 0.94027681 | -1.904325594 | 0.056867793 | 1 | 1.790593195 |
| ENSMUSG00000006221 | Hspb7 | 29818 | protein_coding | 5.120149543 | -1.79071562 | 0.703334103 | 2.546038379 | 0.010895321 | 0.045516147 | 1.79071562 |
| ENSMUSG00000115744 | Vmn1r80 | 171238 | protein_coding | 2.46442528 | -1.792124438 | 3.049597347 | 0.587659364 | 0.556760956 | 1 | 1.792124438 |
| ENSMUSG00000039838 | Slc45a1 | 242773 | protein_coding | 2.947341486 | 1.792506424 | 1.574727739 | -1.138296088 | 0.254996867 | 1 | 1.792506424 |
| ENSMUSG00000029564 | 4930519G04Rik | 67593 | protein_coding | 1.909338251 | 1.792641517 | 2.381893788 | -0.752611861 | 0.451683188 | 1 | 1.792641517 |
| ENSMUSG00000020607 | Lratd1 | 105005 | protein_coding | 146.5985149 | 1.792746634 | 0.14775324 | -12.13338288 | 7.03E-34 | 6.01E-32 | 1.792746634 |
| ENSMUSG00000020212 | Mdm1 | 17245 | protein_coding | 115.1161457 | 1.793302828 | 0.136607816 | -13.12738087 | 2.29E-39 | 2.30E-37 | 1.793302828 |
| ENSMUSG00000097825 | 9630001P10Rik | 319251 | lncRNA | 0.806898538 | -1.794985629 | 1.342196977 | 1.337348884 | 0.181108781 | 1 | 1.794985629 |
| ENSMUSG00000028661 | Epha8 | 13842 | protein_coding | 2.172585038 | 1.795448077 | 2.195600899 | -0.817747924 | 0.413501139 | 1 | 1.795448077 |
| ENSMUSG00000057606 | Colq | 382864 | protein_coding | 1.221843556 | -1.796561033 | 1.368526039 | 1.312770807 | 0.18926019 | 1 | 1.796561033 |
| ENSMUSG00000020892 | Aloxe3 | 23801 | protein_coding | 6.821553453 | 1.798020349 | 1.001021043 | -1.796186367 | 0.072464882 | 0.195920934 | 1.798020349 |
| ENSMUSG00000025716 | Myo3a | 667663 | protein_coding | 103.1884015 | -1.798091251 | 0.180073781 | 9.985302904 | 1.77E-23 | 1.07E-21 | 1.798091251 |
| ENSMUSG00000029477 | Morn3 | 74890 | protein_coding | 1.017912165 | 1.799007657 | 1.904079462 | -0.944817531 | 0.34475204 | 1 | 1.799007657 |
| ENSMUSG00000025461 | Scart1 | 244233 | protein_coding | 2.249551205 | 1.800876169 | 1.334379013 | -1.349598691 | 0.177144744 | 1 | 1.800876169 |
| ENSMUSG00000074179 | Gsta5 | 100042314 | protein_coding | 1.574548748 | 1.801877266 | 1.085121139 | -1.660530979 | 0.096807681 | 1 | 1.801877266 |
| ENSMUSG00000075033 | Nxpe3 | 385658 | protein_coding | 1.727059567 | 1.802162014 | 1.088600114 | -1.655485784 | 0.097825998 | 1 | 1.802162014 |
| ENSMUSG00000036832 | Lpar3 | 65086 | protein_coding | 10.06346832 | 1.802357487 | 0.582596094 | -3.093665586 | 0.001977002 | 0.011230894 | 1.802357487 |
| ENSMUSG00000097683 | Gm26644 | 102639186 | lncRNA | 1.958953109 | 1.802502561 | 2.338189412 | -0.770896725 | 0.440768148 | 1 | 1.802502561 |
| ENSMUSG00000086851 | A430108G06Rik | 319650 | lncRNA | 0.874557467 | 1.802917078 | 1.661021801 | -1.085426499 | 0.277732801 | 1 | 1.802917078 |
| ENSMUSG00000050211 | Pla2g4e | 329502 | protein_coding | 2.037779409 | -1.804091334 | 1.840132271 | 0.980413942 | 0.32688183 | 1 | 1.804091334 |
| ENSMUSG00000075023 | Accsl | 381411 | protein_coding | 1.465098338 | 1.805063169 | 1.977951603 | -0.912592182 | 0.361457068 | 1 | 1.805063169 |
| ENSMUSG00000092216 | Gm19345 | 100502736 | protein_coding | 1.69735501 | 1.806140241 | 0.935917895 | -1.929806291 | 0.053630844 | 1 | 1.806140241 |
| ENSMUSG00000025557 | Slc15a1 | 56643 | protein_coding | 562.7978312 | -1.811072607 | 0.305256907 | 5.932945551 | 2.98E-09 | 6.20E-08 | 1.811072607 |
| ENSMUSG00000090362 | Vmn2r79 | 621430 | protein_coding | 2.659282482 | -1.811200482 | 1.873617561 | 0.966686329 | 0.333700863 | 1 | 1.811200482 |
| ENSMUSG00000036599 | Chst12 | 59031 | protein_coding | 16.08188541 | 1.811759974 | 0.354015684 | -5.117739281 | 3.09E-07 | 4.73E-06 | 1.811759974 |
| ENSMUSG00000112129 | Pbld1 | 68371 | protein_coding | 61.51373259 | -1.812103519 | 0.226930627 | 7.985275257 | 1.40E-15 | 5.35E-14 | 1.812103519 |
| ENSMUSG00000071552 | Tigit | 100043314 | protein_coding | 1.684418387 | -1.812117387 | 1.130578045 | 1.60282379 | 0.108973563 | 1 | 1.812117387 |
| ENSMUSG00000006313 | Upk1a | 109637 | protein_coding | 214.3405732 | 1.812515097 | 0.212033026 | -8.548267822 | 1.25E-17 | 5.56E-16 | 1.812515097 |
| ENSMUSG00000067010 | Vmn2r10 | 22307 | protein_coding | 1.408245817 | 1.813472748 | 2.351614621 | -0.771160688 | 0.440611693 | 1 | 1.813472748 |
| ENSMUSG00000048142 | Nat8l | 269642 | protein_coding | 3.743541736 | 1.815597316 | 1.306364932 | -1.389808675 | 0.164586983 | 1 | 1.815597316 |
| ENSMUSG00000020323 | Prss57 | 73106 | protein_coding | 1.282880418 | -1.815899003 | 1.604504997 | 1.131750295 | 0.25773943 | 1 | 1.815899003 |

| ensembl_gene_id | external_gene_name | entrezgene_id | gene_biotype | baseMean | log2FoldChange | lfcSE | stat | pvalue | padj | absLog2FC |
|---|---|---|---|---|---|---|---|---|---|---|
| ENSMUSG00000090509 | Sfta2 | 433102 | protein_coding | 1.933473359 | 1.815927125 | 1.403821656 | -1.293559704 | 0.195817549 | 1 | 1.815927125 |
| ENSMUSG00000066060 | Gm12866 | 433751 | lncRNA | 1.158410958 | 1.816136735 | 1.417401089 | -1.281314618 | 0.20008318 | 1 | 1.816136735 |
| ENSMUSG00000084967 | Gm12296 | 100093714 | lncRNA | 1.107698731 | 1.816185032 | 2.041041781 | -0.889832364 | 0.373555906 | 1 | 1.816185032 |
| ENSMUSG00000059043 | Or2c1 | 18312 | protein_coding | 1.577657758 | 1.81995627 | 2.051650339 | -0.887069417 | 0.375041528 | 1 | 1.81995627 |
| ENSMUSG00000075249 | Fsip2 | 241516 | protein_coding | 2.344747739 | 1.822595495 | 1.801516835 | -1.011700506 | 0.311681273 | 1 | 1.822595495 |
| ENSMUSG00000059708 | Akap17b | 338351 | protein_coding | 63.83536063 | 1.822624667 | 0.189435199 | -9.621362202 | 6.50E-22 | 3.65E-20 | 1.822624667 |
| ENSMUSG00000040164 | Kcns1 | 16538 | protein_coding | 1.341891355 | 1.823164588 | 2.709281383 | -0.672932904 | 0.500989978 | 1 | 1.823164588 |
| ENSMUSG00000088984 | Gm25604 | 115486291 | snoRNA | 1.158564567 | -1.823492891 | 1.19336473 | 1.528026466 | 0.126505967 | 1 | 1.823492891 |
| ENSMUSG00000022558 | Mroh1 | 223658 | protein_coding | 1898.531982 | 1.824102527 | 0.078305509 | -23.29468974 | 5.02E-120 | 2.12E-117 | 1.824102527 |
| ENSMUSG00000092090 | Gm3294 | 118567373 | lncRNA | 1.710381201 | -1.824151793 | 1.968553103 | 0.926645966 | 0.354110381 | 1 | 1.824151793 |
| ENSMUSG00000037347 | Chst7 | 60322 | protein_coding | 2.156117117 | 1.825584606 | 0.916177479 | -1.992610218 | 0.046304145 | 1 | 1.825584606 |
| ENSMUSG00000025268 | Maged2 | 80884 | protein_coding | 433.9503508 | 1.826406771 | 0.162478349 | -11.24092401 | 2.57E-29 | 1.94E-27 | 1.826406771 |
| ENSMUSG00000027867 | Spag17 | 74362 | protein_coding | 43.0121278 | 1.828327058 | 0.322566728 | -5.66805842 | 1.44E-08 | 2.71E-07 | 1.828327058 |
| ENSMUSG00000075266 | Cenpw | 66311 | protein_coding | 118.3599257 | 1.828719621 | 0.214255924 | -8.53521146 | 1.40E-17 | 6.18E-16 | 1.828719621 |
| ENSMUSG00000078632 | Lrrc37a | 237954 | protein_coding | 3.670422294 | 1.829987391 | 2.225019406 | -0.82245907 | 0.410815672 | 1 | 1.829987391 |
| ENSMUSG00000022388 | Ttll8 | 239591 | protein_coding | 2.229759468 | -1.83029565 | 2.257105942 | 0.81090374 | 0.417420952 | 1 | 1.83029565 |
| ENSMUSG00000041731 | Pgm5 | 226041 | protein_coding | 4.95938998 | 1.830298862 | 0.75094003 | -2.437343581 | 0.014795614 | 0.057707786 | 1.830298862 |
| ENSMUSG00000046856 | Gpr1 | 241070 | protein_coding | 3.197883767 | 1.831010212 | 1.129990971 | -1.620375968 | 0.105151539 | 1 | 1.831010212 |
| ENSMUSG00000074987 | Wt1os | 329480 | lncRNA | 1.33574434 | 1.831123527 | 2.583338627 | -0.708820558 | 0.478435839 | 1 | 1.831123527 |
| ENSMUSG00000038648 | Creb3l2 | 208647 | protein_coding | 244.0800547 | 1.831449675 | 0.122647627 | -14.93261403 | 2.02E-50 | 2.90E-48 | 1.831449675 |
| ENSMUSG00000031698 | Mylk3 | 213435 | protein_coding | 0.986593266 | -1.832049159 | 1.969634162 | 0.930146925 | 0.352295018 | 1 | 1.832049159 |
| ENSMUSG00000047565 | Acot10 | 64833 | protein_coding | 0.624020166 | 1.835977672 | 1.46409852 | -1.253998721 | 0.20984247 | 1 | 1.835977672 |
| ENSMUSG00000048915 | Efna5 | 13640 | protein_coding | 1.377707406 | -1.837166519 | 1.732368607 | 1.060494004 | 0.288919917 | 1 | 1.837166519 |
| ENSMUSG00000043673 | Kcns3 | 238076 | protein_coding | 2.534782994 | -1.838433926 | 0.913928113 | 2.011573885 | 0.044264875 | 1 | 1.838433926 |
| ENSMUSG00000064294 | Aox3 | 71724 | protein_coding | 3.073610297 | 1.841671413 | 2.419615732 | -0.761142106 | 0.446572194 | 1 | 1.841671413 |
| ENSMUSG00000109713 | Pvrig | 102640920 | protein_coding | 0.913316106 | 1.84312758 | 1.339893647 | -1.37557752 | 0.168952471 | 1 | 1.84312758 |
| ENSMUSG00000028359 | Orm3 | 18407 | protein_coding | 1.300426822 | 1.845715913 | 1.570445446 | -1.175281776 | 0.239882003 | 1 | 1.845715913 |
| ENSMUSG00000034634 | Ly6d | 17068 | protein_coding | 1.735673877 | 1.847478298 | 1.098635389 | -1.681611859 | 0.09264413 | 1 | 1.847478298 |
| ENSMUSG00000086111 | Gm15326 | 102636620 | lncRNA | 1.529739134 | -1.848257659 | 1.116756929 | 1.655022334 | 0.097919968 | 1 | 1.848257659 |
| ENSMUSG00000038216 | Pnmt | 18948 | protein_coding | 1.028984202 | 1.848857723 | 2.243311076 | -0.824164665 | 0.409846001 | 1 | 1.848857723 |
| ENSMUSG00000042474 | Fcmr | 69169 | protein_coding | 2.668622608 | -1.848994113 | 1.241007088 | 1.489914225 | 0.136246791 | 1 | 1.848994113 |
| ENSMUSG00000034854 | Mfsd12 | 73822 | protein_coding | 451.0646259 | 1.849561526 | 0.126120997 | -14.66497709 | 1.08E-48 | 1.41E-46 | 1.849561526 |
| ENSMUSG00000070291 | Ddx43 | 100048658 | protein_coding | 30.02497646 | 1.849815186 | 0.393064011 | -4.706142347 | 2.52E-06 | 3.21E-05 | 1.849815186 |
| ENSMUSG00000040963 | Asgr2 | 11890 | protein_coding | 0.90896979 | -1.85002358 | 2.183304002 | 0.847350428 | 0.396799827 | 1 | 1.85002358 |
| ENSMUSG00000005917 | Otx1 | 18423 | protein_coding | 2.490066582 | 1.850603353 | 1.991993435 | -0.92902081 | 0.352878302 | 1 | 1.850603353 |
| ENSMUSG00000070645 | Ren1 | 19701 | protein_coding | 1.568401795 | 1.851075657 | 1.718756394 | -1.076985466 | 0.281486759 | 1 | 1.851075657 |
| ENSMUSG00000028541 | B4galt2 | 53418 | protein_coding | 10.01798148 | 1.851139228 | 0.522926925 | -3.539957766 | 0.000400191 | 0.002930712 | 1.851139228 |
| ENSMUSG00000021775 | Nr1d2 | 353187 | protein_coding | 375.1461095 | -1.852928612 | 0.269672214 | 6.871040148 | 6.37E-12 | 1.81E-10 | 1.852928612 |
| ENSMUSG00000108709 | 4933431G14Rik | 71265 | lncRNA | 1.112326512 | -1.854498387 | 1.975583771 | 0.938709061 | 0.34788014 | 1 | 1.854498387 |

| ensembl_gene_id | external_gene_name | entrezgene_id | gene_biotype | baseMean | log2FoldChange | lfcSE | stat | pvalue | padj | absLog2FC |
|---|---|---|---|---|---|---|---|---|---|---|
| ENSMUSG00000041358 | Nutm1 | 213765 | protein_coding | 2.92105929 | 1.854810498 | 1.106157032 | -1.676805774 | 0.093580468 | 1 | 1.854810498 |
| ENSMUSG00000065495 | Mir150 | 387168 | miRNA | 0.547578905 | -1.85534298 | 1.943048813 | 0.954861745 | 0.339647618 | 1 | 1.85534298 |
| ENSMUSG00000032274 | Cyp19a1 | 13075 | protein_coding | 2.171857232 | -1.855903884 | 2.248874387 | 0.825259025 | 0.409224549 | 1 | 1.855903884 |
| ENSMUSG00000025900 | Rp1 | 19888 | protein_coding | 2.460416395 | 1.856703098 | 2.628874851 | -0.706272913 | 0.480018435 | 1 | 1.856703098 |
| ENSMUSG00000062132 | Arhgap33os | 100504207 | protein_coding | 1.378518632 | 1.857519393 | 2.273731837 | -0.816947435 | 0.413958468 | 1 | 1.857519393 |
| ENSMUSG00000042436 | Mfap4 | 76293 | protein_coding | 1.155069473 | -1.857721957 | 2.071103081 | 0.896972234 | 0.369733734 | 1 | 1.857721957 |
| ENSMUSG00000074625 | Arhgap40 | 545481 | protein_coding | 0.988762075 | 1.858120793 | 2.196589742 | -0.845911623 | 0.397602048 | 1 | 1.858120793 |
| ENSMUSG00000057203 | Vmn1r234 | 171232 | protein_coding | 1.637345056 | 1.860728975 | 2.660131784 | -0.699487516 | 0.484247412 | 1 | 1.860728975 |
| ENSMUSG00000045989 | 4930451I11Rik | 78118 | protein_coding | 1.281682541 | 1.862060497 | 2.670389518 | -0.697299208 | 0.485615563 | 1 | 1.862060497 |
| ENSMUSG00000046480 | Scn4b | 399548 | protein_coding | 2.144125567 | 1.862211987 | 2.671638159 | -0.697030015 | 0.48578401 | 1 | 1.862211987 |
| ENSMUSG00000068947 | Or1af1 | 236509 | protein_coding | 1.273694541 | 1.862437419 | 2.673379342 | -0.696660362 | 0.486015371 | 1 | 1.862437419 |
| ENSMUSG00000096735 | Vmn1r79 | 100042437 | protein_coding | 1.589908061 | 1.862605678 | 2.674749107 | -0.696366501 | 0.486199336 | 1 | 1.862605678 |
| ENSMUSG00000036647 | Or6b9 | 233670 | protein_coding | 1.439578083 | 1.862621 | 2.29199412 | -0.812663952 | 0.416410754 | 1 | 1.862621 |
| ENSMUSG00000021721 | Htr1a | 15550 | protein_coding | 1.684288422 | 1.863475044 | 2.681735061 | -0.694876638 | 0.487132615 | 1 | 1.863475044 |
| ENSMUSG00000050224 | Krtap13 | 16699 | protein_coding | 1.859208947 | 1.863795433 | 1.075226782 | -1.733397516 | 0.083025039 | 1 | 1.863795433 |
| ENSMUSG00000073924 | Or52p1 | 259078 | protein_coding | 1.648799992 | 1.863801358 | 2.684371975 | -0.694315607 | 0.487484305 | 1 | 1.863801358 |
| ENSMUSG00000054555 | Adam12 | 11489 | protein_coding | 4.432815474 | 1.866050382 | 1.397363493 | -1.335407996 | 0.181742849 | 0.375961595 | 1.866050382 |
| ENSMUSG00000044681 | Cnpy1 | 269637 | protein_coding | 2.485031096 | -1.868147361 | 1.135324976 | 1.645473676 | 0.099872167 | 1 | 1.868147361 |
| ENSMUSG00000084904 | Gm14827 | 100503393 | lncRNA | 1.937509671 | 1.86889453 | 1.485592054 | -1.258013278 | 0.208386954 | 1 | 1.86889453 |
| ENSMUSG00000107660 | Gm6559 | 625167 | lncRNA | 0.639590743 | 1.86910452 | 1.489244702 | -1.255068772 | 0.209453796 | 1 | 1.86910452 |
| ENSMUSG00000031523 | Dlc1 | 50768 | protein_coding | 12.3235363 | -1.87019999 | 0.594220391 | 3.147317085 | 0.001647762 | 0.009652762 | 1.87019999 |
| ENSMUSG00000067229 | Cyp2c66 | 69888 | protein_coding | 20.23187044 | -1.871767168 | 0.493087253 | 3.796016136 | 0.00014704 | 0.00120867 | 1.871767168 |
| ENSMUSG00000060890 | Arr3 | 170735 | protein_coding | 3.591829684 | 1.871818715 | 1.262298153 | -1.482865764 | 0.138110079 | 1 | 1.871818715 |
| ENSMUSG00000079450 | Cldn34c1 | 73061 | protein_coding | 12.99366535 | 1.871961866 | 0.84235799 | -2.222287777 | 0.026263864 | 0.09075569 | 1.871961866 |
| ENSMUSG00000040118 | Cacna2d1 | 12293 | protein_coding | 77.60562488 | 1.873495576 | 0.186773022 | -10.03086826 | 1.12E-23 | 6.82E-22 | 1.873495576 |
| ENSMUSG00000068615 | Gjd2 | 14617 | protein_coding | 1.041075763 | -1.874870124 | 1.499977615 | 1.249932069 | 0.211324363 | 1 | 1.874870124 |
| ENSMUSG00000028005 | Gucy1b1 | 54195 | protein_coding | 1.70654028 | 1.874933106 | 1.951027259 | -0.960997904 | 0.336553222 | 1 | 1.874933106 |
| ENSMUSG00000116331 | B230214G05Rik | 328579 | lncRNA | 1.027580278 | 1.875157026 | 2.243086369 | -0.835971834 | 0.403170748 | 1 | 1.875157026 |
| ENSMUSG00000044441 | Or5b94 | 258692 | protein_coding | 0.98093061 | -1.876205865 | 2.544710291 | 0.737296451 | 0.46094209 | 1 | 1.876205865 |
| ENSMUSG00000095139 | Pou3f2 | 18992 | protein_coding | 1.201783526 | -1.876940692 | 2.583200825 | 0.726594957 | 0.467474118 | 1 | 1.876940692 |
| ENSMUSG00000055944 | E130018O15Rik | 442795 | lncRNA | 1.808129409 | -1.877823634 | 2.57443673 | 0.729411452 | 0.465750015 | 1 | 1.877823634 |
| ENSMUSG00000049741 | Rxfp4 | 242093 | protein_coding | 1.697568257 | -1.879954002 | 0.949626921 | 1.979676398 | 0.047739902 | 1 | 1.879954002 |
| ENSMUSG00000026307 | Scly | 50880 | protein_coding | 377.344321 | 1.880656428 | 0.100518461 | -18.70956246 | 4.14E-78 | 1.02E-75 | 1.880656428 |
| ENSMUSG00000020672 | Sntg2 | 268534 | protein_coding | 1.826491309 | 1.881174622 | 1.542671024 | -1.21942695 | 0.222682186 | 1 | 1.881174622 |
| ENSMUSG00000027322 | Siglec1 | 20612 | protein_coding | 2.699973324 | 1.882178768 | 2.164228162 | -0.869676682 | 0.384477119 | 1 | 1.882178768 |
| ENSMUSG00000095105 | Edaradd | 171211 | protein_coding | 20.48558172 | 1.882179929 | 0.376266873 | -5.002247241 | 5.67E-07 | 8.27E-06 | 1.882179929 |
| ENSMUSG00000112805 | Gm34574 | 102637874 | lncRNA | 1.506275045 | -1.882236476 | 1.467754498 | 1.282391897 | 0.199705205 | 1 | 1.882236476 |
| ENSMUSG00000043670 | Diras1 | 208666 | protein_coding | 1.772263321 | -1.882348289 | 1.663243524 | 1.131733424 | 0.257746525 | 1 | 1.882348289 |
| ENSMUSG00000074771 | Ankef1 | 319196 | protein_coding | 1.765473859 | -1.883164419 | 1.988210851 | 0.947165346 | 0.343554531 | 1 | 1.883164419 |

| ensembl_gene_id | external_gene_name | entrezgene_id | gene_biotype | baseMean | log2FoldChange | lfcSE | stat | pvalue | padj | absLog2FC |
|---|---|---|---|---|---|---|---|---|---|---|
| ENSMUSG00000097502 | 4930528D03Rik | 75223 | lncRNA | 1.721383691 | -1.883261748 | 2.322936098 | 0.810724733 | 0.417523766 | 1 | 1.883261748 |
| ENSMUSG00000004929 | Thop1 | 50492 | protein_coding | 1186.463533 | 1.884272595 | 0.092801877 | -20.30425083 | 1.18E-91 | 3.48E-89 | 1.884272595 |
| ENSMUSG00000023949 | Tcte1 | 21645 | protein_coding | 1.065115972 | -1.884324805 | 2.182807305 | 0.863257513 | 0.387995899 | 1 | 1.884324805 |
| ENSMUSG00000071178 | Serpina1b | 20701 | protein_coding | 5.03749461 | 1.885338241 | 0.676883337 | -2.785322283 | 0.005347454 | 0.025753241 | 1.885338241 |
| ENSMUSG00000016327 | Atp1b4 | 67821 | protein_coding | 1.577670323 | 1.885784707 | 2.279126721 | -0.82741547 | 0.408001615 | 1 | 1.885784707 |
| ENSMUSG00000031509 | Spaca7b | 76413 | protein_coding | 0.867514331 | -1.885829728 | 2.09778527 | 0.898962232 | 0.368672778 | 1 | 1.885829728 |
| ENSMUSG00000021750 | Fam107a | 268709 | protein_coding | 1.625869325 | 1.886446816 | 1.847419175 | -1.021125493 | 0.307194986 | 1 | 1.886446816 |
| ENSMUSG00000072753 | 9230020A06Rik | 619299 | lncRNA | 1.429003705 | 1.886601444 | 2.281937141 | -0.826754344 | 0.408376312 | 1 | 1.886601444 |
| ENSMUSG00000026725 | Tnn | 329278 | protein_coding | 2.573041967 | 1.887874966 | 1.821495729 | -1.036442159 | 0.299995911 | 1 | 1.887874966 |
| ENSMUSG00000070166 | Gm25821 | 115485864 | snoRNA | 0.678075579 | 1.888197479 | 2.107378072 | -0.895993701 | 0.370256127 | 1 | 1.888197479 |
| ENSMUSG00000073988 | Ttpa | 50500 | protein_coding | 44.95072213 | 1.888397246 | 0.219823044 | -8.590533604 | 8.66E-18 | 3.92E-16 | 1.888397246 |
| ENSMUSG00000030792 | Dkkl1 | 50722 | protein_coding | 5.588067453 | -1.888776311 | 0.511006067 | 3.69619156 | 0.000218858 | 0.001712088 | 1.888776311 |
| ENSMUSG00000105102 | Gm35507 | 102639119 | lncRNA | 1.136058071 | -1.889119112 | 2.134605812 | 0.884996706 | 0.376158408 | 1 | 1.889119112 |
| ENSMUSG00000030562 | Nox4 | 50490 | protein_coding | 1.905292469 | 1.890026489 | 2.293811789 | -0.823967554 | 0.409957994 | 1 | 1.890026489 |
| ENSMUSG00000046607 | Hrk | 12123 | protein_coding | 2.038007121 | -1.890499214 | 1.866014002 | 1.013121666 | 0.311002051 | 1 | 1.890499214 |
| ENSMUSG00000032572 | Col6a4 | 68553 | protein_coding | 3.871615153 | 1.890961287 | 1.844699739 | -1.025078091 | 0.305326342 | 1 | 1.890961287 |
| ENSMUSG00000033214 | Slitrk5 | 75409 | protein_coding | 1.631570313 | 1.891770092 | 2.299893633 | -0.822546776 | 0.410765776 | 1 | 1.891770092 |
| ENSMUSG00000106406 | Mir7682 | 102466847 | miRNA | 0.605938876 | -1.891808405 | 1.63701515 | 1.15564502 | 0.247826386 | 1 | 1.891808405 |
| ENSMUSG00000106795 | Gm43050 | 433945 | lncRNA | 1.179807911 | -1.892045847 | 2.143808098 | 0.882563066 | 0.377472392 | 1 | 1.892045847 |
| ENSMUSG00000036699 | Zcchc12 | 72693 | protein_coding | 1.029421666 | 1.892164847 | 1.958429501 | -0.966164391 | 0.333961928 | 1 | 1.892164847 |
| ENSMUSG00000061489 | Or13a19 | 258958 | protein_coding | 1.318362288 | -1.892296424 | 2.145722691 | 0.881892349 | 0.377835025 | 1 | 1.892296424 |
| ENSMUSG00000044624 | Gm4922 | 237300 | protein_coding | 0.758146106 | -1.89231043 | 1.633160744 | 1.15867984 | 0.246586711 | 1 | 1.89231043 |
| ENSMUSG00000047990 | C2cd4a | 244911 | protein_coding | 162.3551021 | 1.895176934 | 0.210119777 | -9.01950762 | 1.89E-19 | 9.31E-18 | 1.895176934 |
| ENSMUSG00000022842 | Ece2 | 107522 | protein_coding | 1.344770381 | 1.896474723 | 1.909132182 | -0.993370046 | 0.320529653 | 1 | 1.896474723 |
| ENSMUSG00000074139 | 1700057G04Rik | 78459 | protein_coding | 2.193419412 | -1.896645588 | 0.950605186 | 1.995198024 | 0.046021287 | 1 | 1.896645588 |
| ENSMUSG00000044708 | Kcnj10 | 16513 | protein_coding | 3.586444014 | 1.8966475 | 1.616571762 | -1.173252896 | 0.240694406 | 1 | 1.8966475 |
| ENSMUSG00000020185 | E2f7 | 52679 | protein_coding | 71.79391763 | 1.897761946 | 0.214542803 | -8.845609935 | 9.10E-19 | 4.37E-17 | 1.897761946 |
| ENSMUSG00000048620 | Or6z3 | 258917 | protein_coding | 2.460663629 | 1.898019955 | 3.026298114 | -0.627175474 | 0.530544218 | 1 | 1.898019955 |
| ENSMUSG00000057337 | Chst3 | 53374 | protein_coding | 2.051846763 | -1.898568919 | 1.384442221 | 1.37136017 | 0.170262707 | 1 | 1.898568919 |
| ENSMUSG00000105076 | A930003O13Rik | 330052 | lncRNA | 2.612253663 | -1.898806956 | 1.134333216 | 1.673941068 | 0.094142182 | 1 | 1.898806956 |
| ENSMUSG00000034755 | Pcdh11x | 245578 | protein_coding | 2.191809796 | 1.899158723 | 3.040451546 | -0.624630485 | 0.532213607 | 1 | 1.899158723 |
| ENSMUSG00000024211 | Grm8 | 14823 | protein_coding | 2.144249184 | 1.899189798 | 3.040837721 | -0.624561378 | 0.532258975 | 1 | 1.899189798 |
| ENSMUSG00000031997 | Trpc6 | 22068 | protein_coding | 1.492700031 | -1.899902658 | 2.589310714 | 0.733748425 | 0.463102079 | 1 | 1.899902658 |
| ENSMUSG00000098752 | Mir6897 | 102465540 | miRNA | 0.809225168 | 1.900073427 | 1.291698492 | -1.470988345 | 0.141294269 | 1 | 1.900073427 |
| ENSMUSG00000093460 | Six3os1 | 100043902 | lncRNA | 2.679277322 | -1.900201875 | 2.115544794 | 0.898209237 | 0.36907401 | 1 | 1.900201875 |
| ENSMUSG00000073761 | 4933427I04Rik | 115489902 | lncRNA | 0.704563645 | 1.90063952 | 1.766582808 | -1.07588476 | 0.281978799 | 1 | 1.90063952 |
| ENSMUSG00000096497 | Or6c5c | 258069 | protein_coding | 1.13493924 | 1.901119173 | 3.065240641 | -0.620218572 | 0.535113896 | 1 | 1.901119173 |
| ENSMUSG00000038498 | Catsper1 | 225865 | protein_coding | 1.918828504 | 1.901344307 | 3.068283173 | -0.619676933 | 0.535470505 | 1 | 1.901344307 |
| ENSMUSG00000060284 | Sp7 | 170574 | protein_coding | 1.594159139 | 1.90180325 | 3.074205675 | -0.618632405 | 0.536158549 | 1 | 1.90180325 |

Supplementary Table 1

| ensembl_gene_id | external_gene_name | entrezgene_id | gene_biotype | baseMean | log2FoldChange | lfcSE | stat | pvalue | padj | absLog2FC |
|---|---|---|---|---|---|---|---|---|---|---|
| ENSMUSG00000111008 | Gm31562 | 102633831 | lncRNA | 1.594159139 | 1.90180325 | 3.074205675 | -0.618632405 | 0.536158549 | 1 | 1.90180325 |
| ENSMUSG00000024471 | Myot | 58916 | protein_coding | 4.27956282 | 1.904010835 | 0.650154562 | -2.928551064 | 0.003405459 | 1 | 1.904010835 |
| ENSMUSG00000002900 | Lamb1 | 16777 | protein_coding | 20.38195747 | -1.904299167 | 0.419715329 | 4.537120847 | 5.70E-06 | 6.73E-05 | 1.904299167 |
| ENSMUSG00000052303 | Mrgpra6 | 381886 | protein_coding | 1.412047993 | 1.905376038 | 3.121989923 | -0.610308196 | 0.541657669 | 1 | 1.905376038 |
| ENSMUSG00000075115 | Or4a67 | 257887 | protein_coding | 1.321177846 | 1.905578109 | 3.124751905 | -0.60983341 | 0.541972167 | 1 | 1.905578109 |
| ENSMUSG00000100253 | 1700020N18Rik | 67086 | lncRNA | 0.737112081 | -1.907470546 | 1.749138104 | 1.090520264 | 0.275484032 | 1 | 1.907470546 |
| ENSMUSG00000062712 | Or2j6 | 258955 | protein_coding | 1.307034583 | -1.908412185 | 2.662847043 | 0.716681114 | 0.473570882 | 1 | 1.908412185 |
| ENSMUSG00000020932 | Gfap | 14580 | protein_coding | 2.357800631 | -1.909475485 | 0.842268215 | 2.267063448 | 0.02338635 | 1 | 1.909475485 |
| ENSMUSG00000040413 | Timd2 | 171284 | protein_coding | 1.865819854 | -1.9095795 | 2.695839534 | 0.708343162 | 0.478732179 | 1 | 1.9095795 |
| ENSMUSG00000107252 | Litafd | 436336 | protein_coding | 1.689845478 | 1.910191893 | 0.935278664 | -2.042377278 | 0.041114122 | 1 | 1.910191893 |
| ENSMUSG00000018620 | Mmp20 | 30800 | protein_coding | 2.565965331 | 1.910997214 | 1.503099657 | -1.271370933 | 0.203596718 | 1 | 1.910997214 |
| ENSMUSG00000094845 | Tmem95 | 100503361 | protein_coding | 2.710423574 | -1.912138793 | 0.812484803 | 2.353445609 | 0.018600329 | 1 | 1.912138793 |
| ENSMUSG00000025519 | Tktl2 | 74419 | protein_coding | 1.03214268 | -1.913432422 | 2.182012101 | 0.876911921 | 0.380534481 | 1 | 1.913432422 |
| ENSMUSG00000040564 | Apoc1 | 11812 | protein_coding | 7.587308294 | -1.913938463 | 0.416237774 | 4.598185415 | 4.26E-06 | 5.18E-05 | 1.913938463 |
| ENSMUSG00000090254 | Gm1965 | 434065 | lncRNA | 21.59687128 | -1.914866615 | 0.261390775 | 7.325685515 | 2.38E-13 | 7.61E-12 | 1.914866615 |
| ENSMUSG00000067878 | Map7d3 | 320923 | protein_coding | 0.993107537 | 1.916388381 | 2.273371986 | -0.842971758 | 0.399244239 | 1 | 1.916388381 |
| ENSMUSG00000032103 | Pus3 | 67049 | protein_coding | 0.668902929 | 1.916473964 | 1.49208236 | -1.284429074 | 0.198991865 | 1 | 1.916473964 |
| ENSMUSG00000030519 | Apba2 | 11784 | protein_coding | 1.351231163 | -1.916925739 | 1.249889526 | 1.533676136 | 0.125109347 | 1 | 1.916925739 |
| ENSMUSG00000029866 | Kel | 23925 | protein_coding | 2.30543132 | 1.917557068 | 2.183048822 | -0.878384876 | 0.379734889 | 1 | 1.917557068 |
| ENSMUSG00000114136 | 4921511I17Rik | 70858 | lncRNA | 1.429040361 | -1.919793723 | 0.951680424 | 2.01726722 | 0.043667635 | 1 | 1.919793723 |
| ENSMUSG00000024256 | Adcyap1 | 11516 | protein_coding | 1.469101025 | 1.920768299 | 1.950229962 | -0.984893237 | 0.324676516 | 1 | 1.920768299 |
| ENSMUSG00000021991 | Cacna2d3 | 12294 | protein_coding | 2.682889411 | 1.921670804 | 1.900319151 | -1.011235825 | 0.311903573 | 1 | 1.921670804 |
| ENSMUSG00000094985 | Topaz1 | 671232 | protein_coding | 2.230660307 | -1.923698605 | 2.12563933 | 0.904997653 | 0.365466631 | 1 | 1.923698605 |
| ENSMUSG00000092958 | Mir5128 | 100628597 | miRNA | 0.713739827 | -1.92398528 | 2.039810163 | 0.943217813 | 0.345569504 | 1 | 1.92398528 |
| ENSMUSG00000038651 | Sycp2l | 637277 | protein_coding | 0.997627312 | -1.925459713 | 2.489918873 | 0.773302188 | 0.439343561 | 1 | 1.925459713 |
| ENSMUSG00000031725 | Ces1f | 234564 | protein_coding | 939.6115003 | -1.926624979 | 0.160280951 | 12.02029913 | 2.78E-33 | 2.37E-31 | 1.926624979 |
| ENSMUSG00000047592 | Nxpe5 | 381680 | protein_coding | 1.333837311 | -1.927312311 | 1.550356944 | 1.243141019 | 0.213815873 | 1 | 1.927312311 |
| ENSMUSG00000046196 | Ttc39d | 67737 | protein_coding | 1.373307662 | 1.927578355 | 1.519355134 | -1.268681898 | 0.204554541 | 1 | 1.927578355 |
| ENSMUSG00000028226 | Mmp16 | 17389 | protein_coding | 1.964909524 | 1.929269821 | 1.986536045 | -0.971172824 | 0.331462224 | 1 | 1.929269821 |
| ENSMUSG00000078554 | Fam229a | 68233 | protein_coding | 0.691824586 | 1.929741742 | 1.430109125 | -1.349366777 | 0.177219186 | 1 | 1.929741742 |
| ENSMUSG00000060816 | Vmn1r52 | 113849 | protein_coding | 0.997675982 | 1.931241477 | 2.15907162 | -0.894477728 | 0.371066341 | 1 | 1.931241477 |
| ENSMUSG00000046550 | Spin2c | 278240 | protein_coding | 23.92087193 | 1.931255725 | 0.302148749 | -6.391738282 | 1.64E-10 | 3.95E-09 | 1.931255725 |
| ENSMUSG00000107797 | Gm10445 | 100038556 | lncRNA | 1.029011844 | -1.932391573 | 2.084361929 | 0.927090227 | 0.35387969 | 1 | 1.932391573 |
| ENSMUSG00000044499 | Hs3st5 | 319415 | protein_coding | 1.105897553 | -1.93326248 | 1.656924332 | 1.166777772 | 0.243300126 | 1 | 1.93326248 |
| ENSMUSG00000075316 | Scn9a | 20274 | protein_coding | 2.707773312 | -1.934109711 | 1.967739882 | 0.982909239 | 0.32565211 | 1 | 1.934109711 |
| ENSMUSG00000030187 | Klra2 | 16633 | protein_coding | 1.709707953 | 1.934622735 | 1.441175723 | -1.342391982 | 0.179468933 | 1 | 1.934622735 |
| ENSMUSG00000032595 | Cdhr4 | 69398 | protein_coding | 1.434356749 | 1.935370195 | 1.545106584 | -1.252580382 | 0.210358458 | 1 | 1.935370195 |
| ENSMUSG00000080152 | H3f4 | 382523 | protein_coding | 0.907914008 | 1.935772068 | 1.426536721 | -1.356973178 | 0.174789733 | 1 | 1.935772068 |
| ENSMUSG00000093757 | Mir5623 | 100885835 | miRNA | 2.219527052 | -1.936346 | 0.798708831 | 2.4243453 | 0.015336016 | 1 | 1.936346 |

Supplementary Table 1

| ensembl_gene_id | external_gene_name | entrezgene_id | gene_biotype | baseMean | log2FoldChange | lfcSE | stat | pvalue | padj | absLog2FC |
|---|---|---|---|---|---|---|---|---|---|---|
| ENSMUSG00000001493 | Meox1 | 17285 | protein_coding | 12.81190615 | 1.936826516 | 0.351496271 | -5.510233468 | 3.58E-08 | 6.31E-07 | 1.936826516 |
| ENSMUSG00000056600 | Or2h2 | 258469 | protein_coding | 1.213530516 | 1.936964785 | 2.59308075 | -0.746974341 | 0.455079049 | 1 | 1.936964785 |
| ENSMUSG00000015053 | Gata2 | 14461 | protein_coding | 3.282991381 | -1.937533679 | 1.033779784 | 1.874222836 | 0.060899718 | 1 | 1.937533679 |
| ENSMUSG00000049999 | Ppp1r3d | 228966 | protein_coding | 100.0635891 | 1.937579872 | 0.185770255 | -10.42997908 | 1.81E-25 | 1.17E-23 | 1.937579872 |
| ENSMUSG00000055653 | Gpc3 | 14734 | protein_coding | 25.4969663 | 1.937841215 | 0.325106472 | -5.960635605 | 2.51E-09 | 5.27E-08 | 1.937841215 |
| ENSMUSG00000026247 | Ecel1 | 13599 | protein_coding | 2.953874207 | 1.937850432 | 2.207475275 | -0.877858273 | 0.380020636 | 1 | 1.937850432 |
| ENSMUSG00000064084 | Or4c105 | 258454 | protein_coding | 1.397009411 | 1.938225234 | 2.598855016 | -0.745799678 | 0.455788435 | 1 | 1.938225234 |
| ENSMUSG00000026249 | Serpine2 | 20720 | protein_coding | 17.65839528 | 1.938263017 | 0.44702457 | -4.335920542 | 1.45E-05 | 0.000155248 | 1.938263017 |
| ENSMUSG00000114832 | Gm21388 | 100861998 | lncRNA | 2.054296953 | 1.939937029 | 1.922892605 | -1.00886395 | 0.313039886 | 1 | 1.939937029 |
| ENSMUSG00000091034 | Gm17660 | 100271704 | protein_coding | 1.187654386 | 1.941557954 | 1.193306263 | -1.62704078 | 0.103728437 | 1 | 1.941557954 |
| ENSMUSG00000048699 | Krt90 | 239673 | protein_coding | 2.689786949 | 1.943619113 | 1.092107286 | -1.779696132 | 0.075125704 | 1 | 1.943619113 |
| ENSMUSG00000113492 | Gm47792 | 115488144 | lncRNA | 1.07692062 | 1.943642764 | 1.98884243 | -0.97727338 | 0.328433828 | 1 | 1.943642764 |
| ENSMUSG00000042078 | Svop | 68666 | protein_coding | 2.897932798 | 1.947115718 | 1.44000241 | -1.35216143 | 0.176323679 | 1 | 1.947115718 |
| ENSMUSG00000092021 | Gbp11 | 634650 | protein_coding | 0.85210347 | 1.947530208 | 1.59512653 | -1.220925219 | 0.222114335 | 1 | 1.947530208 |
| ENSMUSG00000030553 | Pgpep1l | 78444 | protein_coding | 0.765633035 | -1.948582462 | 2.413066391 | 0.807512992 | 0.419370992 | 1 | 1.948582462 |
| ENSMUSG00000025666 | Tmem47 | 192216 | protein_coding | 4.550306146 | 1.949627087 | 0.634504195 | -3.072678009 | 0.002121473 | 0.011900366 | 1.949627087 |
| ENSMUSG00000064567 | Gm24143 | 115487889 | snoRNA | 0.906609459 | -1.949725517 | 1.401676214 | 1.390995651 | 0.164226746 | 1 | 1.949725517 |
| ENSMUSG00000042655 | Shisal2b | 77803 | protein_coding | 1.132666357 | 1.952191268 | 1.932367486 | -1.010258805 | 0.312371312 | 1 | 1.952191268 |
| ENSMUSG00000027398 | Il1b | 16176 | protein_coding | 5.801640884 | 1.952304339 | 0.890619033 | -2.192075698 | 0.028374041 | 0.096552424 | 1.952304339 |
| ENSMUSG00000029154 | Cwh43 | 231293 | protein_coding | 188.9365669 | -1.955044962 | 0.186326939 | 10.49255129 | 9.35E-26 | 6.17E-24 | 1.955044962 |
| ENSMUSG00000023978 | Prph2 | 19133 | protein_coding | 1.284169123 | -1.956777461 | 2.518253483 | 0.777037528 | 0.43713663 | 1 | 1.956777461 |
| ENSMUSG00000027574 | Nkain4 | 58237 | protein_coding | 1.473337701 | 1.957364045 | 1.924933354 | -1.016847696 | 0.309225888 | 1 | 1.957364045 |
| ENSMUSG00000030104 | Edem1 | 192193 | protein_coding | 4827.665362 | 1.957591901 | 0.065144946 | -30.04978922 | 2.20E-198 | 1.45E-195 | 1.957591901 |
| ENSMUSG00000044937 | Ttc41 | 103220 | protein_coding | 53.45757419 | 1.957895381 | 0.221385551 | -8.843826399 | 9.25E-19 | 4.41E-17 | 1.957895381 |
| ENSMUSG00000070692 | Rps12-ps10 | 670832 | processed_pseudogene | 1.849907745 | 1.958815428 | 0.875570383 | -2.237187857 | 0.025274065 | 1 | 1.958815428 |
| ENSMUSG00000049372 | Or4c31 | 258522 | protein_coding | 3.677677944 | 1.963010059 | 2.155668509 | -0.910627052 | 0.362491912 | 1 | 1.963010059 |
| ENSMUSG00000047586 | Nccrp1 | 233038 | protein_coding | 7.803414341 | -1.963401359 | 0.560542287 | 3.502681966 | 0.000460599 | 0.003307101 | 1.963401359 |
| ENSMUSG00000097993 | Ptprv | 13924 | protein_coding | 3.46093666 | 1.963763043 | 1.909850071 | -1.028228903 | 0.303842166 | 1 | 1.963763043 |
| ENSMUSG00000052013 | Btla | 208154 | protein_coding | 0.92462114 | -1.964049897 | 1.255486734 | 1.564373278 | 0.117729932 | 1 | 1.964049897 |
| ENSMUSG00000079845 | Xlr4a | 434794 | protein_coding | 2.156822309 | -1.964222429 | 1.323821227 | 1.483752027 | 0.137874718 | 1 | 1.964222429 |
| ENSMUSG00000016255 | Tubb1 | 545486 | protein_coding | 1.41118484 | 1.965823582 | 1.833449467 | -1.072199489 | 0.28363044 | 1 | 1.965823582 |
| ENSMUSG00000020826 | Nos2 | 18126 | protein_coding | 40.50911209 | 1.966028867 | 0.660488536 | -2.976628298 | 0.002914371 | 0.015492034 | 1.966028867 |
| ENSMUSG00000085247 | 4930545L23Rik | 78805 | lncRNA | 2.555180713 | -1.966032462 | 1.082661762 | 1.815924909 | 0.069381892 | 1 | 1.966032462 |
| ENSMUSG00000116557 | 4933432I09Rik | 66779 | lncRNA | 1.581356791 | 1.966476678 | 2.252882227 | -0.872871495 | 0.382733125 | 1 | 1.966476678 |
| ENSMUSG00000039706 | Ldb2 | 16826 | protein_coding | 1.803214621 | 1.968033359 | 2.600377367 | -0.756826061 | 0.449154077 | 1 | 1.968033359 |
| ENSMUSG00000001508 | Sgca | 20391 | protein_coding | 1.613302564 | 1.969962105 | 1.743085758 | -1.13015788 | 0.258409704 | 1 | 1.969962105 |
| ENSMUSG00000006204 | Cdcp3 | 71395 | protein_coding | 4.665347757 | 1.971597298 | 2.054111039 | -0.959829951 | 0.337140806 | 0.559327232 | 1.971597298 |
| ENSMUSG00000116628 | 4930478L05Rik | 75818 | lncRNA | 1.196048723 | 1.971979442 | 1.403247328 | -1.405297129 | 0.159932942 | 1 | 1.971979442 |
| ENSMUSG00000029343 | Crybb1 | 12960 | protein_coding | 4.50255753 | 1.972646556 | 0.715116272 | -2.758497651 | 0.005806772 | 0.027492662 | 1.972646556 |

| ensembl_gene_id | external_gene_name | entrezgene_id | gene_biotype | baseMean | log2FoldChange | lfcSE | stat | pvalue | padj | absLog2FC |
|---|---|---|---|---|---|---|---|---|---|---|
| ENSMUSG00000033006 | Sox10 | 20665 | protein_coding | 3.068402095 | 1.97282262 | 1.529163595 | -1.290131825 | 0.197004892 | 1 | 1.97282262 |
| ENSMUSG00000096356 | Or8b40 | 258475 | protein_coding | 0.764152489 | 1.973544628 | 1.411868124 | -1.397825048 | 0.162165611 | 1 | 1.973544628 |
| ENSMUSG00000064581 | Gm25133 | 115485933 | snoRNA | 1.280229282 | 1.974405589 | 2.087538867 | -0.945805427 | 0.344247837 | 1 | 1.974405589 |
| ENSMUSG00000075091 | Or4c125 | 258974 | protein_coding | 1.832773746 | 1.974427311 | 2.170186626 | -0.909796092 | 0.362930056 | 1 | 1.974427311 |
| ENSMUSG00000026774 | Potegl | 70981 | protein_coding | 0.657443273 | 1.97504166 | 1.622358643 | -1.217389058 | 0.223456224 | 1 | 1.97504166 |
| ENSMUSG00000039070 | Cpa4 | 71791 | protein_coding | 1.418665399 | -1.976174131 | 2.229161245 | 0.886510177 | 0.375342672 | 1 | 1.976174131 |
| ENSMUSG00000044556 | Tex38 | 75173 | protein_coding | 2.301008153 | 1.977302665 | 0.940430005 | -2.102551656 | 0.03550498 | 1 | 1.977302665 |
| ENSMUSG00000015222 | Map2 | 17756 | protein_coding | 98.29582906 | 1.977926904 | 0.157931614 | -12.52394537 | 5.52E-36 | 4.98E-34 | 1.977926904 |
| ENSMUSG00000028264 | Spaca1 | 67652 | protein_coding | 1.634956951 | 1.979825483 | 1.626376268 | -1.217323151 | 0.223481289 | 1 | 1.979825483 |
| ENSMUSG00000064356 | mt-Atp8 | 17706 | protein_coding | 4.467185373 | 1.980570156 | 1.05741014 | -1.87303874 | 0.061063036 | 0.173046478 | 1.980570156 |
| ENSMUSG00000044313 | Mab21l3 | 242125 | protein_coding | 4.961882878 | 1.980674387 | 0.785680739 | -2.520965944 | 0.011703318 | 0.048196656 | 1.980674387 |
| ENSMUSG00000074281 | Gm12522 | 100038453 | lncRNA | 2.213051958 | 1.980889094 | 1.491031071 | -1.328536429 | 0.184000957 | 1 | 1.980889094 |
| ENSMUSG00000043843 | Tmem145 | 330485 | protein_coding | 2.330901189 | 1.981393479 | 2.148254762 | -0.922327051 | 0.356358008 | 1 | 1.981393479 |
| ENSMUSG00000054986 | Sec14l3 | 380683 | protein_coding | 1.255042638 | 1.982754913 | 2.628570672 | -0.754309151 | 0.450663609 | 1 | 1.982754913 |
| ENSMUSG00000042286 | Stab1 | 192187 | protein_coding | 8.825527858 | 1.982919276 | 1.174524099 | -1.688274662 | 0.091358517 | 0.231252996 | 1.982919276 |
| ENSMUSG00000095257 | Scgb1b3 | 384585 | protein_coding | 1.289066569 | -1.982963126 | 0.955883894 | 2.074481156 | 0.038034647 | 1 | 1.982963126 |
| ENSMUSG00000029913 | Prdm5 | 70779 | protein_coding | 25.08320851 | 1.987029568 | 0.256239493 | -7.754579689 | 8.86E-15 | 3.18E-13 | 1.987029568 |
| ENSMUSG00000045440 | Insm2 | 56856 | protein_coding | 1.249283102 | -1.987203215 | 1.387514038 | 1.432204044 | 0.152085435 | 1 | 1.987203215 |
| ENSMUSG00000034732 | Pabpc5 | 93728 | protein_coding | 1.133298407 | 1.987224228 | 1.333479217 | -1.490255118 | 0.136157169 | 1 | 1.987224228 |
| ENSMUSG00000118233 | Gm30381 | 102632253 | lncRNA | 0.758894835 | -1.987988258 | 1.39019836 | 1.430003311 | 0.152716069 | 1 | 1.987988258 |
| ENSMUSG00000002190 | Clgn | 12745 | protein_coding | 1.726076114 | 1.991358969 | 2.159406377 | -0.922178887 | 0.356435274 | 1 | 1.991358969 |
| ENSMUSG00000089768 | Tmsb15b1 | 666244 | protein_coding | 4.686064062 | 1.993279538 | 0.584333073 | -3.411204379 | 0.000646766 | 0.004397156 | 1.993279538 |
| ENSMUSG00000053216 | Btn2a2 | 238555 | protein_coding | 2.186935584 | 1.994908742 | 1.420669461 | -1.404203298 | 0.16025832 | 1 | 1.994908742 |
| ENSMUSG00000062391 | 4932435O22Rik | 319819 | lncRNA | 1.093767868 | 1.995642052 | 2.332722211 | -0.855499228 | 0.392274831 | 1 | 1.995642052 |
| ENSMUSG00000022025 | Cnmd | 16840 | protein_coding | 0.791998831 | -1.995936297 | 1.459140614 | 1.36788482 | 0.171348132 | 1 | 1.995936297 |
| ENSMUSG00000026421 | Csrp1 | 13007 | protein_coding | 910.7115381 | 1.997140601 | 0.128563523 | -15.53427091 | 2.03E-54 | 3.08E-52 | 1.997140601 |
| ENSMUSG00000051596 | Otop1 | 21906 | protein_coding | 1.706295518 | 1.997219044 | 1.742076657 | -1.14645876 | 0.251605382 | 1 | 1.997219044 |
| ENSMUSG00000074813 | Morrbid | 100043424 | lncRNA | 2.202520868 | -1.999139567 | 1.076423466 | 1.857205487 | 0.063281925 | 1 | 1.999139567 |
| ENSMUSG00000118298 | Vmn1r60 | 636697 | protein_coding | 1.86989584 | 1.999557035 | 2.357964837 | -0.84800121 | 0.396437297 | 1 | 1.999557035 |
| ENSMUSG00000025408 | Ddit3 | 13198 | protein_coding | 765.5283613 | 2.000643843 | 0.174376579 | -11.47312247 | 1.80E-30 | 1.43E-28 | 2.000643843 |
| ENSMUSG00000095957 | Or7g19 | 258075 | protein_coding | 1.415050636 | 2.001442369 | 2.370416885 | -0.844341931 | 0.398478362 | 1 | 2.001442369 |
| ENSMUSG00000047940 | Stpg2 | 381476 | protein_coding | 1.477427379 | -2.003403038 | 1.627083232 | 1.231284914 | 0.218216322 | 1 | 2.003403038 |
| ENSMUSG00000043698 | Or13p10 | 18363 | protein_coding | 1.514500245 | 2.003864557 | 2.386806115 | -0.839559001 | 0.401155695 | 1 | 2.003864557 |
| ENSMUSG00000029660 | Tex26 | 75860 | protein_coding | 0.513340874 | -2.00420288 | 1.66863251 | 1.201105017 | 0.229710467 | 1 | 2.00420288 |
| ENSMUSG00000038917 | 3930402G23Rik | 665306 | lncRNA | 30.09622536 | 2.004322372 | 0.313331878 | -6.396803243 | 1.59E-10 | 3.83E-09 | 2.004322372 |
| ENSMUSG00000112967 | 4930428E07Rik | 73867 | lncRNA | 1.601042008 | 2.004454413 | 2.390863592 | -0.838380918 | 0.401816803 | 1 | 2.004454413 |
| ENSMUSG00000066372 | Vmn2r65 | 100009609 | protein_coding | 2.365673846 | -2.004490308 | 2.216892805 | 0.904189099 | 0.365895139 | 1 | 2.004490308 |
| ENSMUSG00000039252 | Lgi2 | 246316 | protein_coding | 5.918062341 | 2.005416571 | 0.894128324 | -2.242873329 | 0.024904984 | 0.087084073 | 2.005416571 |
| ENSMUSG00000036915 | Kirrel2 | 243911 | protein_coding | 5.912960758 | 2.006208597 | 1.080082613 | -1.85745847 | 0.063245955 | 0.177264714 | 2.006208597 |

Supplementary Table 1

| ensembl_gene_id | external_gene_name | entrezgene_id | gene_biotype | baseMean | log2FoldChange | lfcSE | stat | pvalue | padj | absLog2FC |
|---|---|---|---|---|---|---|---|---|---|---|
| ENSMUSG00000097584 | 1700125G22Rik | 76657 | lncRNA | 0.64565357 | 2.007137762 | 1.851110539 | -1.084288442 | 0.278236928 | 1 | 2.007137762 |
| ENSMUSG00000032017 | Grik4 | 110637 | protein_coding | 2.529816125 | -2.008843408 | 2.059290255 | 0.9755028 | 0.329310914 | 1 | 2.008843408 |
| ENSMUSG00000036872 | Abcc12 | 244562 | protein_coding | 3.84591453 | 2.009529999 | 2.232904995 | -0.899962158 | 0.368140389 | 1 | 2.009529999 |
| ENSMUSG00000019906 | Lin7a | 108030 | protein_coding | 2.91834616 | -2.010177424 | 1.028642947 | 1.954203283 | 0.050677181 | 1 | 2.010177424 |
| ENSMUSG00000024781 | Lipa | 16889 | protein_coding | 1004.41932 | 2.010288107 | 0.088608149 | -22.68739546 | 5.97E-114 | 2.35E-111 | 2.010288107 |
| ENSMUSG00000045518 | Onecut3 | 246086 | protein_coding | 1.582101785 | 2.010440725 | 1.613252405 | -1.246203457 | 0.21268971 | 1 | 2.010440725 |
| ENSMUSG00000046790 | Or13p3 | 258852 | protein_coding | 1.180176574 | 2.010475249 | 2.66771017 | -0.753633311 | 0.451069436 | 1 | 2.010475249 |
| ENSMUSG00000029695 | Aass | 30956 | protein_coding | 1.840842085 | 2.010819434 | 2.378462688 | -0.845428202 | 0.397871804 | 1 | 2.010819434 |
| ENSMUSG00000059481 | Plg | 18815 | protein_coding | 2.063387292 | 2.012945105 | 2.393074408 | -0.841154416 | 0.400261434 | 1 | 2.012945105 |
| ENSMUSG00000020703 | 5530401A14Rik | 71388 | lncRNA | 1.088083224 | 2.014045302 | 1.63559693 | -1.231382418 | 0.21817987 | 1 | 2.014045302 |
| ENSMUSG00000079657 | Rab26 | 328778 | protein_coding | 18.38948082 | 2.014125402 | 0.617650786 | -3.260945259 | 0.001110415 | 0.006956521 | 2.014125402 |
| ENSMUSG00000040134 | Rdh7 | 54150 | protein_coding | 2.924586971 | -2.014574096 | 1.111620736 | 1.812285459 | 0.0699421 | 1 | 2.014574096 |
| ENSMUSG00000066475 | Gm10268 | 118568354 | processed_pseudogene | 1.751293317 | 2.016146467 | 0.941337155 | -2.14178996 | 0.032210388 | 1 | 2.016146467 |
| ENSMUSG00000026678 | Rgs5 | 19737 | protein_coding | 1.199382429 | -2.018579441 | 2.589950484 | 0.779389202 | 0.435750482 | 1 | 2.018579441 |
| ENSMUSG00000028427 | Aqp7 | 11832 | protein_coding | 2.938632879 | 2.019265119 | 1.289875023 | -1.565473462 | 0.117471935 | 1 | 2.019265119 |
| ENSMUSG00000056679 | Gpr173 | 70771 | protein_coding | 2.198892052 | 2.019432514 | 2.437574809 | -0.828459708 | 0.407410204 | 1 | 2.019432514 |
| ENSMUSG00000000159 | Igsf5 | 72058 | protein_coding | 621.900734 | 2.020057239 | 0.069854224 | -28.91818311 | 7.05E-184 | 4.31E-181 | 2.020057239 |
| ENSMUSG00000029813 | Aoc1l2 | 69761 | protein_coding | 1.636183675 | -2.020317047 | 2.224011515 | 0.908411235 | 0.363660992 | 1 | 2.020317047 |
| ENSMUSG00000046323 | Dppa3 | 73708 | protein_coding | 0.943876819 | -2.020539754 | 1.510424322 | 1.337729884 | 0.180984506 | 1 | 2.020539754 |
| ENSMUSG00000054106 | Try4 | 22074 | protein_coding | 1.097619225 | -2.021452192 | 2.484523413 | 0.813617687 | 0.415864002 | 1 | 2.021452192 |
| ENSMUSG00000032315 | Cyp1a1 | 13076 | protein_coding | 1.168527418 | 2.021657461 | 3.051780841 | -0.662451718 | 0.507681768 | 1 | 2.021657461 |
| ENSMUSG00000090958 | Lrrc32 | 434215 | protein_coding | 2.209698396 | -2.021999258 | 2.249584909 | 0.898832158 | 0.368742068 | 1 | 2.021999258 |
| ENSMUSG00000046378 | Asphd1 | 233879 | protein_coding | 2.150588055 | -2.023317429 | 1.023893378 | 1.976101685 | 0.048143262 | 1 | 2.023317429 |
| ENSMUSG00000098505 | Gm14496 | 672125 | protein_coding | 1.980783631 | -2.024152856 | 2.245747152 | 0.901327139 | 0.367414408 | 1 | 2.024152856 |
| ENSMUSG00000022964 | Tmem50b | 77975 | protein_coding | 1732.943302 | 2.028956625 | 0.081506276 | -24.89325626 | 8.80E-137 | 4.41E-134 | 2.028956625 |
| ENSMUSG00000035780 | Ugt2a3 | 72094 | protein_coding | 1.943497268 | 2.029206963 | 1.271529868 | -1.595878331 | 0.11051596 | 1 | 2.029206963 |
| ENSMUSG00000038805 | Six3 | 20473 | protein_coding | 2.576436277 | -2.030156388 | 2.561136522 | 0.792677927 | 0.427965494 | 1 | 2.030156388 |
| ENSMUSG00000010311 | Optc | 269120 | protein_coding | 1.638094584 | -2.030235629 | 2.562572266 | 0.792264732 | 0.428206331 | 1 | 2.030235629 |
| ENSMUSG00000097378 | B230208H11Rik | 320273 | lncRNA | 2.673238331 | 2.031497015 | 1.007488377 | -2.01639747 | 0.04375843 | 1 | 2.031497015 |
| ENSMUSG00000041096 | Tspyl2 | 52808 | protein_coding | 158.4810633 | 2.032960203 | 0.153570709 | -13.23794243 | 5.30E-40 | 5.47E-38 | 2.032960203 |
| ENSMUSG00000117752 | Gm35554 | 102639185 | lncRNA | 1.968509832 | 2.03695203 | 2.739820815 | -0.743461769 | 0.457202159 | 1 | 2.03695203 |
| ENSMUSG00000060024 | Vmn1r213 | 171249 | protein_coding | 1.663859428 | -2.038265033 | 2.565994276 | 0.794337327 | 0.426999079 | 1 | 2.038265033 |
| ENSMUSG00000044976 | Wdr72 | 546144 | protein_coding | 1.925454658 | 2.040252137 | 2.000615532 | -1.019812205 | 0.307817534 | 1 | 2.040252137 |
| ENSMUSG00000056900 | Usp13 | 72607 | protein_coding | 3.797437613 | 2.040913255 | 1.885823678 | -1.082239702 | 0.279146031 | 1 | 2.040913255 |
| ENSMUSG00000109857 | Gm53058 | 118027462 | lncRNA | 0.728296352 | -2.042177828 | 2.016832657 | 1.012566819 | 0.311267115 | 1 | 2.042177828 |
| ENSMUSG00000097462 | 9530026P05Rik | 330385 | lncRNA | 3.307202525 | -2.042783673 | 0.894429741 | 2.283895068 | 0.022377699 | 1 | 2.042783673 |
| ENSMUSG00000060924 | Csmd1 | 94109 | protein_coding | 6.638093961 | 2.043782723 | 1.912108884 | -1.068863149 | 0.285131339 | 0.503546031 | 2.043782723 |
| ENSMUSG00000032374 | Plod2 | 26432 | protein_coding | 16.00044787 | 2.044241093 | 0.39282749 | -5.203915575 | 1.95E-07 | 3.06E-06 | 2.044241093 |
| ENSMUSG00000094878 | Or2g7 | 258480 | protein_coding | 2.149949568 | 2.045237447 | 2.455067924 | -0.833067561 | 0.404806627 | 1 | 2.045237447 |

Supplementary Table 1

| ensembl_gene_id | external_gene_name | entrezgene_id | gene_biotype | baseMean | log2FoldChange | lfcSE | stat | pvalue | padj | absLog2FC |
|---|---|---|---|---|---|---|---|---|---|---|
| ENSMUSG00000050654 | Or6d15 | 258438 | protein_coding | 1.881322456 | 2.047445973 | 2.687619029 | -0.761806622 | 0.446175428 | 1 | 2.047445973 |
| ENSMUSG00000031271 | Serpina7 | 331535 | protein_coding | 0.838584848 | 2.049734378 | 1.760871844 | -1.164045178 | 0.244405704 | 1 | 2.049734378 |
| ENSMUSG00000006542 | Prkag3 | 241113 | protein_coding | 2.177976946 | -2.051285163 | 1.333818714 | 1.53790402 | 0.124072084 | 1 | 2.051285163 |
| ENSMUSG00000018479 | Chct1 | 73634 | protein_coding | 1.002697249 | 2.051427047 | 2.21596897 | -0.92574719 | 0.354577378 | 1 | 2.051427047 |
| ENSMUSG00000035179 | Ppp1r32 | 67752 | protein_coding | 4.017200302 | 2.053977901 | 1.131303875 | -1.815584606 | 0.069434117 | 1 | 2.053977901 |
| ENSMUSG00000020889 | Nr1d1 | 217166 | protein_coding | 506.307353 | -2.054181308 | 0.536192664 | 3.831050747 | 0.000127597 | 0.001071245 | 2.054181308 |
| ENSMUSG00000027933 | Ints3 | 229543 | protein_coding | 2575.201486 | 2.054884213 | 0.08277408 | -24.82521355 | 4.79E-136 | 2.33E-133 | 2.054884213 |
| ENSMUSG00000002104 | Rapsn | 19400 | protein_coding | 1.120229785 | 2.055925908 | 1.884840057 | -1.090769426 | 0.275374353 | 1 | 2.055925908 |
| ENSMUSG00000022996 | Wnt10b | 22410 | protein_coding | 2.622384157 | -2.057002847 | 0.99805109 | 2.061019588 | 0.039301171 | 1 | 2.057002847 |
| ENSMUSG00000035431 | Sstr1 | 20605 | protein_coding | 199.9470164 | -2.058046773 | 0.233073843 | 8.83002032 | 1.05E-18 | 4.94E-17 | 2.058046773 |
| ENSMUSG00000009097 | Tbx1 | 21380 | protein_coding | 1.109379945 | -2.059808424 | 1.749901086 | 1.177099918 | 0.239155628 | 1 | 2.059808424 |
| ENSMUSG00000089728 | Clec2f | 435921 | protein_coding | 0.631808872 | 2.059892812 | 1.779336592 | -1.157674619 | 0.246996845 | 1 | 2.059892812 |
| ENSMUSG00000063060 | Sox7 | 20680 | protein_coding | 1.354826577 | -2.060849111 | 1.928615225 | 1.068564161 | 0.285266105 | 1 | 2.060849111 |
| ENSMUSG00000045331 | 2310079G19Rik | 69699 | protein_coding | 15.92157396 | -2.063323298 | 0.543101081 | 3.799151521 | 0.000145192 | 0.00119646 | 2.063323298 |
| ENSMUSG00000040663 | Clcf1 | 56708 | protein_coding | 6.206698911 | 2.063995502 | 0.667401117 | -3.092586227 | 0.001984206 | 0.011264068 | 2.063995502 |
| ENSMUSG00000034683 | Ppp1r1c | 75276 | protein_coding | 1.576279664 | 2.067822763 | 1.907345819 | -1.08413626 | 0.278304388 | 1 | 2.067822763 |
| ENSMUSG00000087198 | Tmem274 | 102636051 | protein_coding | 1.98395137 | 2.069685424 | 1.299905303 | -1.592181691 | 0.111343886 | 1 | 2.069685424 |
| ENSMUSG00000040694 | Apobec2 | 11811 | protein_coding | 57.2694951 | 2.070852417 | 0.41538877 | -4.985335583 | 6.19E-07 | 8.94E-06 | 2.070852417 |
| ENSMUSG00000032259 | Drd2 | 13489 | protein_coding | 1.45587309 | -2.071360181 | 2.390238165 | 0.866591544 | 0.386165849 | 1 | 2.071360181 |
| ENSMUSG00000094248 | H2ac23 | 665433 | protein_coding | 0.759102023 | -2.072760337 | 2.046725206 | 1.012720385 | 0.311193738 | 1 | 2.072760337 |
| ENSMUSG00000093755 | Vmn1r39 | 546903 | protein_coding | 1.42667964 | -2.073017988 | 2.402866932 | 0.86272692 | 0.388287627 | 1 | 2.073017988 |
| ENSMUSG00000074472 | Zfp872 | 619310 | protein_coding | 4.044127317 | 2.073949137 | 0.688132491 | -3.01388056 | 0.002579293 | 1 | 2.073949137 |
| ENSMUSG00000054406 | Or3a1d | 258704 | protein_coding | 0.937298041 | -2.074528272 | 1.683849061 | 1.232015576 | 0.217943266 | 1 | 2.074528272 |
| ENSMUSG00000069855 | Slc47a2 | 380701 | protein_coding | 1.252686932 | 2.074607811 | 2.177547235 | -0.952726892 | 0.340728464 | 1 | 2.074607811 |
| ENSMUSG00000096937 | Gm10825 | 100038752 | lncRNA | 2.014084384 | 2.075940687 | 1.930442965 | -1.075370122 | 0.282209053 | 1 | 2.075940687 |
| ENSMUSG00000067279 | Ppp1r3c | 53412 | protein_coding | 2.342006724 | 2.077360725 | 1.116247588 | -1.861021468 | 0.062741145 | 1 | 2.077360725 |
| ENSMUSG00000032318 | Isl2 | 104360 | protein_coding | 0.916663748 | 2.080642764 | 2.61473829 | -0.795736526 | 0.426185192 | 1 | 2.080642764 |
| ENSMUSG00000043943 | Naalad2 | 72560 | protein_coding | 1.557646707 | -2.080708858 | 1.2857668 | 1.618263015 | 0.105605928 | 1 | 2.080708858 |
| ENSMUSG00000025726 | Slc28a1 | 434203 | protein_coding | 0.930621151 | -2.080974366 | 1.875006352 | 1.109849235 | 0.267063999 | 1 | 2.080974366 |
| ENSMUSG00000037636 | Slc25a43 | 194744 | protein_coding | 2.476021895 | 2.082121554 | 0.923096389 | -2.255584119 | 0.024096689 | 1 | 2.082121554 |
| ENSMUSG00000021499 | Catsper3 | 76856 | protein_coding | 2.228763712 | 2.08287833 | 1.484801833 | -1.402798868 | 0.160676825 | 1 | 2.08287833 |
| ENSMUSG00000042425 | Frmpd3 | 245643 | protein_coding | 3.48523498 | 2.084547673 | 3.383058125 | -0.616172586 | 0.53778062 | 1 | 2.084547673 |
| ENSMUSG00000013419 | Zfp651 | 270210 | protein_coding | 13.07934384 | 2.084936086 | 0.614222092 | -3.394433565 | 0.000687707 | 0.004614708 | 2.084936086 |
| ENSMUSG00000079462 | Gm15737 | 115490424 | lncRNA | 1.251163056 | -2.086205873 | 1.911160708 | 1.091591023 | 0.275012904 | 1 | 2.086205873 |
| ENSMUSG00000030600 | Lrfn1 | 80749 | protein_coding | 9.1078257 | 2.086664679 | 0.481795195 | -4.331020115 | 1.48E-05 | 0.00015813 | 2.086664679 |
| ENSMUSG00000056529 | Ptafr | 19204 | protein_coding | 2.507861368 | 2.086781472 | 1.477197421 | -1.412662548 | 0.157754969 | 1 | 2.086781472 |
| ENSMUSG00000032179 | Bmp5 | 12160 | protein_coding | 1.11260868 | -2.087453219 | 1.920165483 | 1.08712152 | 0.276983108 | 1 | 2.087453219 |
| ENSMUSG00000029683 | Lmod2 | 93677 | protein_coding | 1.118144104 | 2.088936213 | 1.505179148 | -1.387832282 | 0.165188122 | 1 | 2.088936213 |
| ENSMUSG00000017588 | Krt27 | 16675 | protein_coding | 0.874335508 | -2.09216905 | 1.565040772 | 1.336814406 | 0.181283226 | 1 | 2.09216905 |

| ensembl_gene_id | external_gene_name | entrezgene_id | gene_biotype | baseMean | log2FoldChange | lfcSE | stat | pvalue | padj | absLog2FC |
|---|---|---|---|---|---|---|---|---|---|---|
| ENSMUSG00000062393 | Dgkk | 331374 | protein_coding | 3.991247926 | 2.094390115 | 2.055274699 | -1.019031722 | 0.308187907 | 1 | 2.094390115 |
| ENSMUSG00000022126 | Acod1 | 16365 | protein_coding | 0.656399127 | 2.094562872 | 2.441092063 | -0.858043375 | 0.390868513 | 1 | 2.094562872 |
| ENSMUSG00000043067 | Dpy19l1 | 244745 | protein_coding | 179.6476948 | 2.094741287 | 0.128817572 | -16.2613008 | 1.86E-59 | 3.07E-57 | 2.094741287 |
| ENSMUSG00000079051 | Vinac1 | 668894 | protein_coding | 2.14474796 | 2.09604109 | 1.015202733 | -2.064652727 | 0.03895587 | 1 | 2.09604109 |
| ENSMUSG00000078896 | Zfp965 | 100503949 | protein_coding | 0.576830807 | -2.09702636 | 2.240874747 | 0.935807038 | 0.349372547 | 1 | 2.09702636 |
| ENSMUSG00000034641 | Cd300ld | 217305 | protein_coding | 1.31647079 | 2.099740825 | 1.624983848 | -1.292161044 | 0.196301379 | 1 | 2.099740825 |
| ENSMUSG00000070424 | Art5 | 11875 | protein_coding | 9.628297338 | 2.100060943 | 0.434231403 | -4.836271464 | 1.32E-06 | 1.78E-05 | 2.100060943 |
| ENSMUSG00000036083 | Slc17a3 | 105355 | protein_coding | 4.764414265 | 2.100075948 | 1.152503322 | -1.82218646 | 0.068426698 | 0.187894662 | 2.100075948 |
| ENSMUSG00000026621 | Mtarc1 | 66112 | protein_coding | 2.239215393 | 2.100892514 | 1.219809137 | -1.722312491 | 0.085012926 | 1 | 2.100892514 |
| ENSMUSG00000075304 | Sp5 | 64406 | protein_coding | 9.528753322 | 2.102335219 | 0.430713928 | -4.881047684 | 1.06E-06 | 1.45E-05 | 2.102335219 |
| ENSMUSG00000020566 | Atp6v1c2 | 68775 | protein_coding | 5.309616044 | 2.103292525 | 0.582335622 | -3.61182185 | 0.000304053 | 0.002305389 | 2.103292525 |
| ENSMUSG00000058818 | Pirb | 18733 | protein_coding | 4.256042392 | 2.104256686 | 1.038452614 | -2.026338668 | 0.042730091 | 1 | 2.104256686 |
| ENSMUSG00000033475 | Tomm6 | 66119 | protein_coding | 2.148574934 | 2.105559045 | 0.853364665 | -2.467361412 | 0.01361129 | 1 | 2.105559045 |
| ENSMUSG00000040489 | Sox30 | 214105 | protein_coding | 1.215987905 | 2.106010415 | 1.638124725 | -1.285622751 | 0.198574753 | 1 | 2.106010415 |
| ENSMUSG00000069299 | Vmn1r188 | 252912 | protein_coding | 2.411338521 | -2.111927361 | 2.414034218 | 0.874853946 | 0.38165338 | 1 | 2.111927361 |
| ENSMUSG00000053368 | Rxfp2 | 140498 | protein_coding | 2.367905369 | -2.112133138 | 2.416189751 | 0.874158636 | 0.382031869 | 1 | 2.112133138 |
| ENSMUSG00000006310 | Zbtb32 | 58206 | protein_coding | 142.188018 | 2.11216938 | 0.363017389 | -5.818369706 | 5.94E-09 | 1.19E-07 | 2.11216938 |
| ENSMUSG00000085562 | 2610028E06Rik | 72395 | lncRNA | 1.040105695 | -2.112189026 | 2.136357146 | 0.988687229 | 0.322816193 | 1 | 2.112189026 |
| ENSMUSG00000029123 | Stk32b | 64293 | protein_coding | 2.692574734 | -2.11233539 | 2.418307145 | 0.873476884 | 0.3824032 | 1 | 2.11233539 |
| ENSMUSG00000025130 | P4hb | 18453 | protein_coding | 28433.61093 | 2.113431818 | 0.05672226 | -37.25930203 | 7.49E-304 | 1.12E-300 | 2.113431818 |
| ENSMUSG00000045928 | 4933440M02Rik | 71208 | lncRNA | 1.19799598 | -2.114423789 | 2.819928052 | 0.7498148 | 0.453366254 | 1 | 2.114423789 |
| ENSMUSG00000054034 | Tceal5 | 331532 | protein_coding | 1.293117205 | -2.115599068 | 2.835603441 | 0.746084251 | 0.455616523 | 1 | 2.115599068 |
| ENSMUSG00000094970 | Or8g26 | 258323 | protein_coding | 1.059441604 | -2.115707245 | 2.837085271 | 0.745732695 | 0.455828905 | 1 | 2.115707245 |
| ENSMUSG00000006517 | Mvd | 192156 | protein_coding | 1543.999878 | 2.1157787 | 0.136281731 | -15.52503542 | 2.35E-54 | 3.53E-52 | 2.1157787 |
| ENSMUSG00000091375 | Vmn2r15 | 211223 | protein_coding | 1.037559891 | -2.116757596 | 2.466300782 | 0.85827228 | 0.390742132 | 1 | 2.116757596 |
| ENSMUSG00000078591 | Hs3st4 | 628779 | protein_coding | 1.176855131 | -2.116946309 | 2.468417619 | 0.857612704 | 0.391106358 | 1 | 2.116946309 |
| ENSMUSG00000014351 | Gip | 14607 | protein_coding | 0.812890937 | -2.1170191 | 2.413264575 | 0.877242853 | 0.380354745 | 1 | 2.1170191 |
| ENSMUSG00000087341 | 0610040F04Rik | 75394 | lncRNA | 19.2546907 | 2.117110607 | 0.349099931 | -6.064483032 | 1.32E-09 | 2.89E-08 | 2.117110607 |
| ENSMUSG00000004951 | Hspb1 | 15507 | protein_coding | 6.546042992 | -2.117482207 | 0.489924103 | 4.322061711 | 1.55E-05 | 0.000164055 | 2.117482207 |
| ENSMUSG00000061561 | Or2z9 | 258532 | protein_coding | 0.9806279 | -2.117521358 | 2.474964465 | 0.855576469 | 0.39223209 | 1 | 2.117521358 |
| ENSMUSG00000065518 | Mir423 | 751519 | miRNA | 0.949150108 | -2.118775262 | 1.323844535 | 1.600471359 | 0.109494056 | 1 | 2.118775262 |
| ENSMUSG00000098181 | Rps12-ps24 | 671641 | processed_pseudogene | 0.644031632 | 2.118912747 | 1.309851767 | -1.617673694 | 0.105732938 | 1 | 2.118912747 |
| ENSMUSG00000020374 | Rasgef1c | 74563 | protein_coding | 1.169146913 | -2.119102208 | 2.493195579 | 0.849954262 | 0.395350516 | 1 | 2.119102208 |
| ENSMUSG00000032537 | Ephb1 | 270190 | protein_coding | 3.317372227 | -2.11929441 | 2.407219856 | 0.880390881 | 0.378647595 | 1 | 2.11929441 |
| ENSMUSG00000047031 | Vmn1r68 | 628580 | protein_coding | 2.295233568 | 2.120520709 | 2.041438211 | -1.038738619 | 0.298926315 | 1 | 2.120520709 |
| ENSMUSG00000058806 | Col13a1 | 12817 | protein_coding | 3.552852709 | -2.120531382 | 1.865617826 | 1.136637607 | 0.255689812 | 1 | 2.120531382 |
| ENSMUSG00000096014 | Sox1 | 20664 | protein_coding | 3.1589844 | 2.120602075 | 1.000234491 | -2.12010493 | 0.033997197 | 1 | 2.120602075 |
| ENSMUSG00000078897 | Gm4724 | 100043915 | protein_coding | 0.991493397 | -2.123962033 | 1.109962567 | 1.91354384 | 0.055678467 | 1 | 2.123962033 |
| ENSMUSG00000037477 | Tbx10 | 109575 | protein_coding | 2.564404702 | 2.126269847 | 1.715160931 | -1.239691162 | 0.215089647 | 1 | 2.126269847 |

| ensembl_gene_id | external_gene_name | entrezgene_id | gene_biotype | baseMean | log2FoldChange | lfcSE | stat | pvalue | padj | absLog2FC |
|---|---|---|---|---|---|---|---|---|---|---|
| ENSMUSG00000099630 | 1700096J18Rik | 67927 | lncRNA | 0.674377641 | -2.127527413 | 1.738747378 | 1.223597769 | 0.221104002 | 1 | 2.127527413 |
| ENSMUSG00000062421 | Arf2 | 11841 | protein_coding | 438.5736976 | 2.128595638 | 0.101332125 | -21.0061285 | 5.76E-98 | 1.94E-95 | 2.128595638 |
| ENSMUSG00000022435 | Upk3a | 22270 | protein_coding | 1.862724539 | 2.13019944 | 1.768020749 | -1.204849796 | 0.228261292 | 1 | 2.13019944 |
| ENSMUSG00000026246 | Alppl2 | 11650 | protein_coding | 2.87851433 | -2.130648232 | 0.9681039 | 2.200846658 | 0.027746881 | 1 | 2.130648232 |
| ENSMUSG00000045515 | Pou3f3 | 18993 | protein_coding | 3.137119342 | -2.131243689 | 1.97090629 | 1.081352117 | 0.279540513 | 1 | 2.131243689 |
| ENSMUSG00000044581 | 4932415D10Rik | 102635990 | protein_coding | 2.001077426 | 2.132221063 | 1.629050547 | -1.308873482 | 0.19057721 | 1 | 2.132221063 |
| ENSMUSG00000091510 | Mtag2 | 50994 | lncRNA | 0.960741581 | 2.132252114 | 1.674514586 | -1.273355355 | 0.202891972 | 1 | 2.132252114 |
| ENSMUSG00000045036 | Tmem232 | 381107 | protein_coding | 0.853674019 | -2.132963237 | 2.426501826 | 0.879028078 | 0.379386052 | 1 | 2.132963237 |
| ENSMUSG00000091151 | Vmn1r224 | 665525 | protein_coding | 0.964334103 | -2.138834453 | 2.474716816 | 0.864274425 | 0.387437156 | 1 | 2.138834453 |
| ENSMUSG00000008482 | Rnf151 | 67504 | protein_coding | 4.484393227 | 2.138904507 | 1.429633311 | -1.496121062 | 0.13462211 | 0.305377683 | 2.138904507 |
| ENSMUSG00000118170 | Gm30094 | 102631870 | lncRNA | 1.004403102 | 2.140424283 | 1.504200593 | -1.422964658 | 0.154746401 | 1 | 2.140424283 |
| ENSMUSG00000037437 | Adam32 | 353188 | protein_coding | 1.853698868 | 2.140478075 | 1.699012324 | -1.259836697 | 0.207728278 | 1 | 2.140478075 |
| ENSMUSG00000051323 | Pcdh19 | 279653 | protein_coding | 3.141386727 | 2.140747598 | 1.391393084 | -1.538564208 | 0.123910722 | 1 | 2.140747598 |
| ENSMUSG00000051969 | Tlr11 | 239081 | protein_coding | 1.797335178 | 2.141013465 | 2.16425564 | -0.989260892 | 0.322535513 | 1 | 2.141013465 |
| ENSMUSG00000086155 | 9430041J12Rik | 77323 | lncRNA | 1.241055284 | -2.142364928 | 1.459999686 | 1.467373554 | 0.142274464 | 1 | 2.142364928 |
| ENSMUSG00000049123 | Catsperg2 | 76718 | protein_coding | 2.651333232 | 2.142666942 | 2.469251101 | -0.867739592 | 0.385536907 | 1 | 2.142666942 |
| ENSMUSG00000032224 | Fam81a | 76886 | protein_coding | 1.48911921 | 2.14292827 | 1.201008685 | -1.784273749 | 0.074379194 | 1 | 2.14292827 |
| ENSMUSG00000054204 | Alkal2 | 100294583 | protein_coding | 1.325333189 | 2.143833607 | 1.845827901 | -1.161448261 | 0.245459652 | 1 | 2.143833607 |
| ENSMUSG00000053287 | Or9g19 | 258757 | protein_coding | 1.778756866 | -2.143880682 | 2.857766269 | 0.75019455 | 0.45313754 | 1 | 2.143880682 |
| ENSMUSG00000047531 | Rtp2 | 224055 | protein_coding | 1.384234586 | -2.144749463 | 3.352457119 | 0.639754481 | 0.52233223 | 1 | 2.144749463 |
| ENSMUSG00000080968 | Atf1-ps | 100040260 | processed_pseudogene | 1.070902097 | 2.145544971 | 1.246778622 | -1.720870838 | 0.085274264 | 1 | 2.145544971 |
| ENSMUSG00000025905 | Oprk1 | 18387 | protein_coding | 1.133545596 | -2.146036644 | 2.898618836 | 0.740365245 | 0.459078401 | 1 | 2.146036644 |
| ENSMUSG00000080673 | Gm23336 | 115489848 | snRNA | 1.22737115 | -2.147190622 | 1.196495619 | 1.794566222 | 0.072722839 | 1 | 2.147190622 |
| ENSMUSG00000094133 | Or5an9 | 258409 | protein_coding | 1.107125833 | -2.147498509 | 3.418999613 | 0.628107269 | 0.529933672 | 1 | 2.147498509 |
| ENSMUSG00000056184 | Or8s2 | 259038 | protein_coding | 1.111253293 | -2.148288159 | 3.438904421 | 0.624701328 | 0.532167101 | 1 | 2.148288159 |
| ENSMUSG00000020598 | Nrcam | 319504 | protein_coding | 5.359418216 | 2.148555058 | 1.356199943 | -1.584246534 | 0.113137628 | 0.270127107 | 2.148555058 |
| ENSMUSG00000026251 | Chrnd | 11447 | protein_coding | 1.016132069 | -2.148612902 | 3.447229196 | 0.62328693 | 0.533095985 | 1 | 2.148612902 |
| ENSMUSG00000057654 | Or2t47 | 258495 | protein_coding | 0.968571457 | -2.149421667 | 3.468192496 | 0.619752701 | 0.535420613 | 1 | 2.149421667 |
| ENSMUSG00000050876 | Spata31d1a | 72219 | protein_coding | 2.446861868 | 2.150362598 | 2.021525014 | -1.063732866 | 0.287449731 | 1 | 2.150362598 |
| ENSMUSG00000019767 | Ccdc170 | 100504234 | protein_coding | 1.348133778 | -2.151079925 | 1.81917152 | 1.182450308 | 0.237027071 | 1 | 2.151079925 |
| ENSMUSG00000068428 | Gmnc | 239789 | protein_coding | 1.822031596 | 2.15297232 | 2.400539406 | -0.896870226 | 0.369788169 | 1 | 2.15297232 |
| ENSMUSG00000085514 | Bcas3os2 | 78420 | lncRNA | 0.757506933 | -2.153939561 | 1.261698543 | 1.707174486 | 0.087789626 | 1 | 2.153939561 |
| ENSMUSG00000015947 | Fcgr1 | 14129 | protein_coding | 1.160598722 | 2.155812569 | 2.56089209 | -0.841820933 | 0.399888193 | 1 | 2.155812569 |
| ENSMUSG00000042268 | Slc26a9 | 320718 | protein_coding | 3.21869917 | 2.155889178 | 2.418539987 | -0.891401089 | 0.372714034 | 1 | 2.155889178 |
| ENSMUSG00000052854 | Nrk | 27206 | protein_coding | 4.386396352 | 2.156498531 | 0.99645274 | -2.164175423 | 0.030450889 | 0.102046874 | 2.156498531 |
| ENSMUSG00000070471 | Erich6 | 545527 | protein_coding | 2.244469628 | 2.157317454 | 1.578455232 | -1.366727045 | 0.171710878 | 1 | 2.157317454 |
| ENSMUSG00000084739 | Gm22067 | 115488600 | snRNA | 1.344928801 | 2.157382834 | 1.338014743 | -1.612375981 | 0.106880148 | 1 | 2.157382834 |
| ENSMUSG00000038539 | Atf5 | 107503 | protein_coding | 205.8861594 | 2.157902914 | 0.191063935 | -11.29414042 | 1.40E-29 | 1.07E-27 | 2.157902914 |
| ENSMUSG00000072845 | Tmprss11a | 194597 | protein_coding | 2.146387386 | 2.161332341 | 1.343217535 | -1.609070969 | 0.107600824 | 1 | 2.161332341 |

Supplementary Table 1

| ensembl_gene_id | external_gene_name | entrezgene_id | gene_biotype | baseMean | log2FoldChange | lfcSE | stat | pvalue | padj | absLog2FC |
|---|---|---|---|---|---|---|---|---|---|---|
| ENSMUSG00000020836 | Coro6 | 216961 | protein_coding | 1.835076816 | 2.162038031 | 1.850130625 | -1.168586694 | 0.242570192 | 1 | 2.162038031 |
| ENSMUSG00000097441 | Gm26633 | 102633600 | lncRNA | 1.628399687 | 2.163836013 | 2.284899503 | -0.947015836 | 0.34363071 | 1 | 2.163836013 |
| ENSMUSG00000028655 | Mfsd2a | 76574 | protein_coding | 110.9739732 | 2.168458122 | 0.185999421 | -11.65841328 | 2.08E-31 | 1.71E-29 | 2.168458122 |
| ENSMUSG00000111544 | 4930534H03Rik | 102633753 | lncRNA | 1.485801046 | -2.169559466 | 1.766971185 | 1.227840887 | 0.219506704 | 1 | 2.169559466 |
| ENSMUSG00000025353 | Ormdl2 | 66844 | protein_coding | 1517.4769 | 2.174429942 | 0.078354843 | -27.75106004 | 1.69E-169 | 9.01E-167 | 2.174429942 |
| ENSMUSG00000064565 | Gm24145 | 115488697 | snRNA | 1.528018518 | 2.17764609 | 1.144614205 | -1.902515345 | 0.057103815 | 1 | 2.17764609 |
| ENSMUSG00000022425 | Enpp2 | 18606 | protein_coding | 2.515366962 | -2.178395692 | 1.799083038 | 1.210836658 | 0.225958013 | 1 | 2.178395692 |
| ENSMUSG00000029867 | Llcfc1 | 76606 | protein_coding | 1.334096438 | 2.17907174 | 2.214611471 | -0.98395216 | 0.325139036 | 1 | 2.17907174 |
| ENSMUSG00000059910 | Or4x11 | 258340 | protein_coding | 1.677946135 | 2.179641232 | 2.381749781 | -0.915142829 | 0.360116653 | 1 | 2.179641232 |
| ENSMUSG00000028031 | Dkk2 | 56811 | protein_coding | 1.631126387 | 2.18049326 | 2.387422221 | -0.91332536 | 0.361071448 | 1 | 2.18049326 |
| ENSMUSG00000068407 | Rnase12 | 497106 | protein_coding | 2.072990218 | 2.18143211 | 1.011208019 | -2.157253571 | 0.030985912 | 1 | 2.18143211 |
| ENSMUSG00000057400 | Ces1c | 13884 | protein_coding | 1.789376869 | -2.181689658 | 1.001026044 | 2.179453443 | 0.029298 | 1 | 2.181689658 |
| ENSMUSG00000040204 | Pclaf | 68026 | protein_coding | 782.6050096 | 2.184214986 | 0.210177712 | -10.39222934 | 2.69E-25 | 1.73E-23 | 2.184214986 |
| ENSMUSG00000048721 | Fndc9 | 320116 | protein_coding | 2.253871979 | 2.185315 | 1.59627512 | -1.369008996 | 0.170996463 | 1 | 2.185315 |
| ENSMUSG00000032204 | Aqp9 | 64008 | protein_coding | 1.603985615 | 2.191951725 | 1.462466884 | -1.498804348 | 0.133924396 | 1 | 2.191951725 |
| ENSMUSG00000059668 | Krt4 | 16682 | protein_coding | 3.275363154 | 2.193154889 | 0.983896266 | -2.229050931 | 0.025810516 | 1 | 2.193154889 |
| ENSMUSG00000028001 | Fga | 14161 | protein_coding | 1.839029647 | 2.196240183 | 1.562909805 | -1.405225161 | 0.159954334 | 1 | 2.196240183 |
| ENSMUSG00000071158 | Klrh1 | 232415 | protein_coding | 3.13573094 | -2.197977345 | 0.824588341 | 2.665545021 | 0.007686364 | 1 | 2.197977345 |
| ENSMUSG00000030996 | Art1 | 11870 | protein_coding | 2.183349891 | 2.199066123 | 1.059518977 | -2.075532549 | 0.037937205 | 1 | 2.199066123 |
| ENSMUSG00000023000 | Dhh | 13363 | protein_coding | 2.4690369 | 2.201512004 | 1.624065443 | -1.355556215 | 0.175240411 | 1 | 2.201512004 |
| ENSMUSG00000097738 | 4930445N18Rik | 73985 | lncRNA | 1.052513023 | 2.20293302 | 1.738340504 | -1.26726209 | 0.205061591 | 1 | 2.20293302 |
| ENSMUSG00000025386 | Pde6g | 18588 | protein_coding | 3.292241309 | -2.204093648 | 1.241873734 | 1.774813 | 0.075928769 | 1 | 2.204093648 |
| ENSMUSG00000030862 | Cpxm2 | 55987 | protein_coding | 1.772154349 | -2.204652933 | 1.174259265 | 1.877483959 | 0.060451794 | 1 | 2.204652933 |
| ENSMUSG00000060332 | Tmc2 | 192140 | protein_coding | 2.354600762 | 2.205978462 | 2.67061977 | -0.826017424 | 0.408794208 | 1 | 2.205978462 |
| ENSMUSG00000020908 | Myh3 | 17883 | protein_coding | 3.4792745 | 2.207408066 | 1.219745768 | -1.809727998 | 0.070337979 | 1 | 2.207408066 |
| ENSMUSG00000032558 | Nphp3 | 74025 | protein_coding | 239.9656916 | 2.207919757 | 0.126748296 | -17.41971947 | 5.85E-68 | 1.19E-65 | 2.207919757 |
| ENSMUSG00000113998 | Gm40909 | 105245459 | lncRNA | 1.914009535 | 2.209802617 | 2.176359948 | -1.015366332 | 0.309931236 | 1 | 2.209802617 |
| ENSMUSG00000115422 | 4930452G13Rik | 73989 | lncRNA | 1.085287418 | 2.211969154 | 2.640372086 | -0.83774903 | 0.40217167 | 1 | 2.211969154 |
| ENSMUSG00000075070 | 4932412D23Rik | 75722 | lncRNA | 1.624115599 | 2.213380584 | 2.654254287 | -0.833899222 | 0.404337776 | 1 | 2.213380584 |
| ENSMUSG00000074206 | Adh6b | 100417514 | protein_coding | 1.082728633 | 2.213571397 | 2.656043922 | -0.833409184 | 0.404613997 | 1 | 2.213571397 |
| ENSMUSG00000044362 | Ccdc89 | 70054 | protein_coding | 26.71076845 | 2.214460412 | 0.262679399 | -8.430278206 | 3.45E-17 | 1.48E-15 | 2.214460412 |
| ENSMUSG00000051313 | Or4c127 | 258976 | protein_coding | 1.726687916 | 2.216602049 | 2.68677242 | -0.825005509 | 0.409368463 | 1 | 2.216602049 |
| ENSMUSG00000034913 | Cby2 | 67926 | protein_coding | 1.500950548 | 2.217268286 | 2.693652945 | -0.823145495 | 0.41042526 | 1 | 2.217268286 |
| ENSMUSG00000043308 | Vmn1r75 | 171241 | protein_coding | 1.216954625 | 2.217322124 | 2.694161437 | -0.823010118 | 0.410502239 | 1 | 2.217322124 |
| ENSMUSG00000073913 | Or52e5 | 258753 | protein_coding | 1.260387777 | 2.217399472 | 2.694981766 | -0.822788302 | 0.41062839 | 1 | 2.217399472 |
| ENSMUSG00000004500 | Zfp324 | 243834 | protein_coding | 436.1867514 | 2.218343206 | 0.113930334 | -19.47104982 | 1.93E-84 | 5.32E-82 | 2.218343206 |
| ENSMUSG00000053399 | Adamts18 | 208936 | protein_coding | 20.35323875 | 2.218359916 | 0.527633866 | -4.204354683 | 2.62E-05 | 0.000261952 | 2.218359916 |
| ENSMUSG00000105602 | Gm10636 | 100038607 | lncRNA | 1.279441879 | -2.21994762 | 1.634810479 | 1.357923532 | 0.17448795 | 1 | 2.21994762 |
| ENSMUSG00000049511 | Htr1b | 15551 | protein_coding | 1.454198598 | -2.22265961 | 1.4207572 | 1.564419036 | 0.117719193 | 1 | 2.22265961 |

| ensembl_gene_id | external_gene_name | entrezgene_id | gene_biotype | baseMean | log2FoldChange | lfcSE | stat | pvalue | padj | absLog2FC |
|---|---|---|---|---|---|---|---|---|---|---|
| ENSMUSG00000020940 | Efcab15 | 69441 | protein_coding | 1.033187403 | 2.223069118 | 1.738120578 | -1.279007421 | 0.200894443 | 1 | 2.223069118 |
| ENSMUSG00000084601 | Gm23111 | 115488885 | snRNA | 0.761352641 | 2.224028278 | 1.67056773 | -1.331300873 | 0.183090031 | 1 | 2.224028278 |
| ENSMUSG00000067616 | Klk11 | 56538 | protein_coding | 1.363775975 | 2.226265413 | 2.384513824 | -0.933634936 | 0.350492237 | 1 | 2.226265413 |
| ENSMUSG00000031297 | Slc7a3 | 11989 | protein_coding | 4.109787946 | 2.228272041 | 1.195717909 | -1.863543253 | 0.062385873 | 1 | 2.228272041 |
| ENSMUSG00000024459 | H2-M5 | 240095 | protein_coding | 1.657307949 | 2.23071889 | 2.081971484 | -1.071445458 | 0.283969182 | 1 | 2.23071889 |
| ENSMUSG00000050751 | Pgbd5 | 209966 | protein_coding | 3.654208061 | 2.230887137 | 0.852783583 | -2.616006195 | 0.008896494 | 1 | 2.230887137 |
| ENSMUSG00000053398 | Phgdh | 236539 | protein_coding | 358.9443354 | 2.231199673 | 0.192117396 | -11.61373059 | 3.51E-31 | 2.84E-29 | 2.231199673 |
| ENSMUSG00000084504 | Gm25323 | 115490016 | snRNA | 0.584916339 | -2.231512989 | 2.010993813 | 1.109656815 | 0.267146939 | 1 | 2.231512989 |
| ENSMUSG00000031737 | Irx5 | 54352 | protein_coding | 1.152772466 | 2.23283134 | 2.516413311 | -0.887307077 | 0.374913596 | 1 | 2.23283134 |
| ENSMUSG00000111767 | 4930546K05Rik | 78194 | lncRNA | 1.346741435 | 2.232943354 | 2.039422321 | -1.094890122 | 0.273564777 | 1 | 2.232943354 |
| ENSMUSG00000027419 | Pcsk2 | 18549 | protein_coding | 2.372377142 | 2.233604541 | 2.137961121 | -1.044735809 | 0.29614509 | 1 | 2.233604541 |
| ENSMUSG00000006529 | Itih1 | 16424 | protein_coding | 3.076036894 | 2.233801704 | 1.86422764 | -1.198245137 | 0.230821598 | 1 | 2.233801704 |
| ENSMUSG00000096167 | Or8b39 | 258415 | protein_coding | 1.443177793 | 2.235844678 | 1.464338377 | -1.526863403 | 0.126794982 | 1 | 2.235844678 |
| ENSMUSG00000009406 | Elk1 | 13712 | protein_coding | 156.04832 | 2.237651202 | 0.17672448 | -12.66180669 | 9.63E-37 | 8.88E-35 | 2.237651202 |
| ENSMUSG00000085811 | Cep112it | 629945 | lncRNA | 2.942791185 | 2.239808779 | 1.291278218 | -1.734567149 | 0.082817502 | 1 | 2.239808779 |
| ENSMUSG00000031382 | Asb11 | 68854 | protein_coding | 12.71839321 | -2.240258943 | 0.435499445 | 5.144114348 | 2.69E-07 | 4.14E-06 | 2.240258943 |
| ENSMUSG00000034488 | Edil3 | 13612 | protein_coding | 2.033474163 | -2.240747123 | 1.351429885 | 1.658056514 | 0.09730606 | 1 | 2.240747123 |
| ENSMUSG00000047026 | Acsm4 | 233801 | protein_coding | 1.440547255 | -2.242821818 | 1.377428778 | 1.628266996 | 0.103468284 | 1 | 2.242821818 |
| ENSMUSG00000095401 | Or6c6c | 258068 | protein_coding | 2.776354614 | 2.244169188 | 3.039421248 | -0.738354116 | 0.46029929 | 1 | 2.244169188 |
| ENSMUSG00000046178 | Nxph1 | 18231 | protein_coding | 1.683941677 | 2.244214743 | 1.292893922 | -1.735807327 | 0.082597908 | 1 | 2.244214743 |
| ENSMUSG00000033080 | Vsx1 | 114889 | protein_coding | 2.273825181 | 2.245633944 | 3.063010702 | -0.733145967 | 0.463469407 | 1 | 2.245633944 |
| ENSMUSG00000075176 | Or8k38 | 258583 | protein_coding | 1.8147289 | 2.246973209 | 3.085122726 | -0.728325389 | 0.466414424 | 1 | 2.246973209 |
| ENSMUSG00000087333 | Gm13652 | 102634853 | lncRNA | 1.862289512 | 2.247026709 | 3.08602763 | -0.728129161 | 0.466534525 | 1 | 2.247026709 |
| ENSMUSG00000072944 | Nup62cl | 279706 | protein_coding | 1.893555825 | 2.247351896 | 1.0996789 | -2.043643736 | 0.040988755 | 1 | 2.247351896 |
| ENSMUSG00000063827 | Or2y12 | 257971 | protein_coding | 1.355632618 | 2.247911264 | 3.100879636 | -0.724926965 | 0.468496837 | 1 | 2.247911264 |
| ENSMUSG00000061311 | Rag1 | 19373 | protein_coding | 1.537620147 | 2.247946433 | 3.101531845 | -0.724785862 | 0.46858341 | 1 | 2.247946433 |
| ENSMUSG00000116455 | Gm35974 | 102639732 | lncRNA | 1.161507731 | -2.248304343 | 2.113723753 | 1.063669905 | 0.287478263 | 1 | 2.248304343 |
| ENSMUSG00000031636 | Pdlim3 | 53318 | protein_coding | 2.157500147 | 2.252568822 | 1.980088595 | -1.137610119 | 0.255283319 | 1 | 2.252568822 |
| ENSMUSG00000027470 | Mylk2 | 228785 | protein_coding | 1.398138984 | 2.253635255 | 1.86034647 | -1.211406204 | 0.225739763 | 1 | 2.253635255 |
| ENSMUSG00000027248 | Pdia3 | 14827 | protein_coding | 19064.86349 | 2.25766793 | 0.110343787 | -20.46030861 | 4.86E-93 | 1.49E-90 | 2.25766793 |
| ENSMUSG00000078949 | R3hdml | 100043899 | protein_coding | 2.058042485 | 2.257677207 | 1.045850263 | -2.158700234 | 0.030873431 | 1 | 2.257677207 |
| ENSMUSG00000075139 | Or5d14 | 258105 | protein_coding | 1.500969268 | 2.26025389 | 2.220648141 | -1.01783522 | 0.308756271 | 1 | 2.26025389 |
| ENSMUSG00000009734 | Pou6f2 | 218030 | protein_coding | 3.555745566 | 2.261513575 | 3.368164889 | -0.671437905 | 0.501941605 | 1 | 2.261513575 |
| ENSMUSG00000060180 | Myh13 | 544791 | protein_coding | 2.655657182 | -2.263295364 | 2.038440624 | 1.110307231 | 0.266866658 | 1 | 2.263295364 |
| ENSMUSG00000092695 | Mir3092 | 100526525 | miRNA | 0.9658349 | 2.266345986 | 1.249658752 | -1.813571891 | 0.069743661 | 1 | 2.266345986 |
| ENSMUSG00000021047 | Nova1 | 664883 | protein_coding | 2.566761501 | 2.269888519 | 1.793637017 | -1.265522789 | 0.205683985 | 1 | 2.269888519 |
| ENSMUSG00000079699 | Gm6592 | 625480 | protein_coding | 0.775993164 | 2.272307913 | 2.110809727 | -1.076510063 | 0.281699203 | 1 | 2.272307913 |
| ENSMUSG00000032122 | Slc37a2 | 56857 | protein_coding | 1389.494996 | -2.273101445 | 0.504273742 | 4.507673621 | 6.55E-06 | 7.65E-05 | 2.273101445 |
| ENSMUSG00000027408 | Cpxm1 | 56264 | protein_coding | 2.225637867 | 2.274936513 | 1.395013751 | -1.630762787 | 0.102940382 | 1 | 2.274936513 |

| ensembl_gene_id | external_gene_name | entrezgene_id | gene_biotype | baseMean | log2FoldChange | lfcSE | stat | pvalue | padj | absLog2FC |
|---|---|---|---|---|---|---|---|---|---|---|
| ENSMUSG00000048424 | Ranbp3l | 223332 | protein_coding | 1.818577795 | -2.28055445 | 1.726415935 | 1.320976251 | 0.186509283 | 1 | 2.28055445 |
| ENSMUSG00000076376 | Mir674 | 732489 | miRNA | 0.863131832 | 2.284359581 | 1.702568727 | -1.34171358 | 0.179688882 | 1 | 2.284359581 |
| ENSMUSG00000068105 | Tnfrsf13c | 72049 | protein_coding | 1.379964495 | 2.285709147 | 1.664470416 | -1.37323507 | 0.169679281 | 1 | 2.285709147 |
| ENSMUSG00000049409 | Prokr1 | 58182 | protein_coding | 2.503919813 | 2.287565719 | 1.487678445 | -1.537674843 | 0.124128138 | 1 | 2.287565719 |
| ENSMUSG00000060044 | Tmem26 | 327766 | protein_coding | 2.319482096 | 2.287621514 | 0.897658269 | -2.548432509 | 0.010820822 | 1 | 2.287621514 |
| ENSMUSG00000042064 | Myo3b | 329421 | protein_coding | 3.806122935 | -2.290872175 | 2.171188055 | 1.055123793 | 0.291368712 | 1 | 2.290872175 |
| ENSMUSG00000049630 | C1ql3 | 227580 | protein_coding | 0.841764876 | -2.290888723 | 1.365332328 | 1.677898249 | 0.093366964 | 1 | 2.290888723 |
| ENSMUSG00000025551 | Fgf14 | 14169 | protein_coding | 3.088081523 | 2.291718408 | 0.883675795 | -2.59339276 | 0.009503417 | 1 | 2.291718408 |
| ENSMUSG00000003505 | Psg18 | 26438 | protein_coding | 0.847224122 | -2.295132616 | 2.103019069 | 1.09135131 | 0.275118328 | 1 | 2.295132616 |
| ENSMUSG00000024650 | Slc22a6 | 18399 | protein_coding | 2.648069704 | 2.29795567 | 2.203786261 | -1.042730736 | 0.297073019 | 1 | 2.29795567 |
| ENSMUSG00000030483 | Cyp2b10 | 13088 | protein_coding | 2.495119218 | 2.299619235 | 1.416383018 | -1.623585716 | 0.104464256 | 1 | 2.299619235 |
| ENSMUSG00000048292 | Or10v5 | 258938 | protein_coding | 1.33079128 | 2.300764664 | 1.275442107 | -1.803895803 | 0.071247643 | 1 | 2.300764664 |
| ENSMUSG00000036602 | Alx1 | 216285 | protein_coding | 1.136233738 | -2.302676554 | 2.131964026 | 1.080072894 | 0.280109721 | 1 | 2.302676554 |
| ENSMUSG00000003355 | Fkbp11 | 66120 | protein_coding | 258.7692062 | 2.304107505 | 0.165811442 | -13.89594996 | 6.70E-44 | 7.91E-42 | 2.304107505 |
| ENSMUSG00000042684 | Npl | 74091 | protein_coding | 1.831442703 | -2.304508471 | 1.180018564 | 1.952942557 | 0.050826402 | 1 | 2.304508471 |
| ENSMUSG00000004460 | Dnajb11 | 67838 | protein_coding | 6658.618525 | 2.304742478 | 0.069686767 | -33.07288568 | 7.29E-240 | 6.02E-237 | 2.304742478 |
| ENSMUSG00000024807 | Syvn1 | 74126 | protein_coding | 1412.965039 | 2.305526002 | 0.144773477 | -15.92505784 | 4.25E-57 | 6.87E-55 | 2.305526002 |
| ENSMUSG00000118318 | Vmn1r183 | 209824 | protein_coding | 2.707682716 | -2.305957133 | 2.697220323 | 0.854938365 | 0.39258527 | 1 | 2.305957133 |
| ENSMUSG00000115253 | Vmn1r20 | 434017 | protein_coding | 1.186984963 | -2.306086571 | 2.580556387 | 0.893639287 | 0.371514919 | 1 | 2.306086571 |
| ENSMUSG00000037922 | Bank1 | 242248 | protein_coding | 1.952636252 | -2.306229062 | 1.49639806 | 1.541186883 | 0.123271309 | 1 | 2.306229062 |
| ENSMUSG00000062036 | 4932415M13Rik | 211496 | transcribed_processed_pseudogene | 1.120378224 | 2.308104942 | 2.251408412 | -1.025182694 | 0.305276992 | 1 | 2.308104942 |
| ENSMUSG00000038135 | Crygn | 214301 | protein_coding | 0.919232971 | -2.308656895 | 1.577674826 | 1.463328727 | 0.143377451 | 1 | 2.308656895 |
| ENSMUSG00000030769 | Slc5a11 | 233836 | protein_coding | 4.045484028 | 2.309067048 | 1.29718221 | -1.780063764 | 0.075065526 | 1 | 2.309067048 |
| ENSMUSG00000078648 | Gm17546 | 102634389 | lncRNA | 1.509505042 | 2.312299902 | 1.054548449 | -2.192691957 | 0.028329581 | 1 | 2.312299902 |
| ENSMUSG00000022997 | Wnt1 | 22408 | protein_coding | 1.361035129 | 2.312847993 | 2.287360502 | -1.011142752 | 0.311948111 | 1 | 2.312847993 |
| ENSMUSG00000050742 | Or2m12 | 258443 | protein_coding | 0.648316627 | -2.312964932 | 2.006330834 | 1.152833268 | 0.248978827 | 1 | 2.312964932 |
| ENSMUSG00000066279 | Chrna10 | 504186 | protein_coding | 8.29671613 | 2.315703964 | 0.633824166 | -3.653543188 | 0.000258646 | 0.001991275 | 2.315703964 |
| ENSMUSG00000020990 | Cdkl1 | 71091 | protein_coding | 11.26615912 | -2.319142869 | 0.513261517 | 4.518442922 | 6.23E-06 | 7.30E-05 | 2.319142869 |
| ENSMUSG00000030284 | Creld1 | 171508 | protein_coding | 171.3548715 | 2.319294108 | 0.110090963 | -21.06707075 | 1.59E-98 | 5.49E-96 | 2.319294108 |
| ENSMUSG00000021255 | Esrrb | 26380 | protein_coding | 2.33888031 | -2.31996059 | 2.251186722 | 1.03055005 | 0.30275187 | 1 | 2.31996059 |
| ENSMUSG00000000120 | Ngfr | 18053 | protein_coding | 1.219304145 | 2.321363915 | 2.275430292 | -1.020186786 | 0.307639884 | 1 | 2.321363915 |
| ENSMUSG00000107108 | Gm9936 | 791292 | lncRNA | 1.419218522 | 2.32214194 | 2.294317642 | -1.012127483 | 0.311477103 | 1 | 2.32214194 |
| ENSMUSG00000042477 | Tfap2e | 332937 | protein_coding | 1.277585963 | 2.322938793 | 1.692873219 | -1.3721871 | 0.1700052 | 1 | 2.322938793 |
| ENSMUSG00000053153 | Spag16 | 66722 | protein_coding | 3.626241794 | 2.323477309 | 1.976682957 | -1.175442577 | 0.239817698 | 1 | 2.323477309 |
| ENSMUSG00000070683 | Lactbl1 | 242707 | protein_coding | 2.089174855 | 2.323796427 | 2.32340817 | -1.000167107 | 0.317229645 | 1 | 2.323796427 |
| ENSMUSG00000117994 | Gm5820 | 545253 | lncRNA | 1.512952352 | 2.326315868 | 1.980359905 | -1.17469348 | 0.240117369 | 1 | 2.326315868 |
| ENSMUSG00000029885 | Moxd2 | 194357 | protein_coding | 2.42679538 | 2.326914189 | 2.01324808 | -1.155801022 | 0.247762555 | 1 | 2.326914189 |
| ENSMUSG00000055088 | Or1n2 | 258941 | protein_coding | 2.236657652 | 2.328247069 | 2.423812828 | -0.960572138 | 0.336767344 | 1 | 2.328247069 |
| ENSMUSG00000064325 | Hhip | 15245 | protein_coding | 2.146404751 | 2.328780537 | 2.428463306 | -0.958952326 | 0.337582762 | 1 | 2.328780537 |

Supplementary Table 1

| ensembl_gene_id | external_gene_name | entrezgene_id | gene_biotype | baseMean | log2FoldChange | lfcSE | stat | pvalue | padj | absLog2FC |
|---|---|---|---|---|---|---|---|---|---|---|
| ENSMUSG00000031957 | Ctrb1 | 66473 | protein_coding | 1.776070224 | -2.328967293 | 2.177965045 | 1.069331805 | 0.284920184 | 1 | 2.328967293 |
| ENSMUSG00000048080 | Or4k6 | 258360 | protein_coding | 1.416200576 | 2.331234072 | 2.393190154 | -0.974111509 | 0.330001176 | 1 | 2.331234072 |
| ENSMUSG00000115267 | Vmn1r76 | 171239 | protein_coding | 1.751671191 | 2.337000462 | 2.838179084 | -0.823415434 | 0.410271789 | 1 | 2.337000462 |
| ENSMUSG00000004540 | Psg17 | 26437 | protein_coding | 2.822348249 | 2.338145053 | 0.840586419 | -2.781564156 | 0.005409764 | 1 | 2.338145053 |
| ENSMUSG00000097882 | 0610038B21Rik | 70345 | lncRNA | 1.225505123 | 2.338476216 | 1.49459295 | -1.564624145 | 0.117671064 | 1 | 2.338476216 |
| ENSMUSG00000063531 | Sema3e | 20349 | protein_coding | 6.736655096 | 2.340503899 | 0.625993244 | -3.738864473 | 0.000184853 | 0.001482597 | 2.340503899 |
| ENSMUSG00000060615 | Ang4 | 219033 | protein_coding | 483.2323146 | -2.341384895 | 0.626658515 | 3.736301092 | 0.000186747 | 0.00149302 | 2.341384895 |
| ENSMUSG00000022995 | Enah | 13800 | protein_coding | 11.06829015 | 2.342642835 | 0.857717801 | -2.731251273 | 0.006309435 | 0.029218733 | 2.342642835 |
| ENSMUSG00000012017 | Scarf2 | 224024 | protein_coding | 58.69664758 | 2.342736508 | 0.265087409 | -8.83760008 | 9.78E-19 | 4.65E-17 | 2.342736508 |
| ENSMUSG00000053963 | Stum | 381310 | protein_coding | 2.672275416 | 2.342773121 | 1.987704993 | -1.178632206 | 0.238544661 | 1 | 2.342773121 |
| ENSMUSG00000069189 | Zdhhc11 | 71164 | protein_coding | 0.988278029 | 2.3523225 | 2.576000905 | -0.913168352 | 0.361154006 | 1 | 2.3523225 |
| ENSMUSG00000022610 | Mapk12 | 29857 | protein_coding | 6.442245898 | -2.353015938 | 0.532761864 | 4.416637337 | 1.00E-05 | 0.00011239 | 2.353015938 |
| ENSMUSG00000014813 | Stc1 | 20855 | protein_coding | 2.051407144 | 2.354935918 | 2.706446927 | -0.870120856 | 0.384234361 | 1 | 2.354935918 |
| ENSMUSG00000058881 | Zfp516 | 329003 | protein_coding | 783.2719658 | 2.356007356 | 0.158868373 | -14.82993317 | 9.38E-50 | 1.29E-47 | 2.356007356 |
| ENSMUSG00000006143 | Upk3bl | 69665 | protein_coding | 1.159790034 | 2.356717939 | 1.680382397 | -1.402489066 | 0.160769253 | 1 | 2.356717939 |
| ENSMUSG00000076433 | Cep295nl | 58251 | protein_coding | 1.418008659 | -2.358989387 | 1.008730106 | 2.338573393 | 0.019357523 | 1 | 2.358989387 |
| ENSMUSG00000089730 | 1700007P06Rik | 69308 | lncRNA | 1.356231813 | 2.35989102 | 1.629289053 | -1.448417649 | 0.147500282 | 1 | 2.35989102 |
| ENSMUSG00000046415 | B430212C06Rik | 338360 | lncRNA | 1.269789906 | 2.361786765 | 2.276280952 | -1.037563822 | 0.299473168 | 1 | 2.361786765 |
| ENSMUSG00000109487 | Or4k48 | 258888 | protein_coding | 1.102043604 | -2.362074013 | 2.851138799 | 0.828466861 | 0.407406155 | 1 | 2.362074013 |
| ENSMUSG00000115027 | Vmn1r81 | 171244 | protein_coding | 1.404015898 | 2.367254853 | 2.293373125 | -1.032215311 | 0.301971264 | 1 | 2.367254853 |
| ENSMUSG00000000318 | Clec10a | 17312 | protein_coding | 1.391029638 | 2.367967802 | 2.534545722 | -0.934277011 | 0.35016102 | 1 | 2.367967802 |
| ENSMUSG00000104093 | A330015K06Rik | 102637797 | lncRNA | 1.894612944 | 2.373037377 | 1.301686032 | -1.823048968 | 0.068295975 | 1 | 2.373037377 |
| ENSMUSG00000084984 | Far1os | 69295 | lncRNA | 77.84502988 | 2.37354346 | 0.20030717 | -11.84951822 | 2.16E-32 | 1.81E-30 | 2.37354346 |
| ENSMUSG00000038973 | Cldnd2 | 74276 | protein_coding | 2.776073242 | 2.373808506 | 1.564089356 | -1.517693664 | 0.129091642 | 1 | 2.373808506 |
| ENSMUSG00000052295 | 8030423F21Rik | 211001 | lncRNA | 1.971166009 | 2.374134929 | 1.839587909 | -1.290579764 | 0.196849437 | 1 | 2.374134929 |
| ENSMUSG00000023930 | Crisp2 | 22024 | protein_coding | 1.670913336 | 2.375057318 | 1.663989634 | -1.427326991 | 0.153485664 | 1 | 2.375057318 |
| ENSMUSG00000035107 | Dcbld2 | 73379 | protein_coding | 275.2317904 | 2.375864873 | 0.103027281 | -23.06054143 | 1.15E-117 | 4.64E-115 | 2.375864873 |
| ENSMUSG00000092519 | Actl9 | 69481 | protein_coding | 1.065087047 | 2.376594709 | 1.794218709 | -1.324584732 | 0.185308923 | 1 | 2.376594709 |
| ENSMUSG00000028020 | Glrb | 14658 | protein_coding | 2.001576762 | 2.37693728 | 1.949316881 | -1.219369361 | 0.222704033 | 1 | 2.37693728 |
| ENSMUSG00000059089 | Fcgr4 | 246256 | protein_coding | 1.133214581 | 2.377806092 | 3.075152883 | -0.77323183 | 0.439385192 | 1 | 2.377806092 |
| ENSMUSG00000028613 | Lrp8 | 16975 | protein_coding | 141.3040671 | 2.380478509 | 0.181065663 | -13.14704548 | 1.77E-39 | 1.78E-37 | 2.380478509 |
| ENSMUSG00000043458 | Pcdhb12 | 93883 | protein_coding | 1.016042382 | 2.383213333 | 1.443444028 | -1.65106044 | 0.098726234 | 1 | 2.383213333 |
| ENSMUSG00000084782 | Tex53 | 670833 | protein_coding | 1.209100941 | 2.389306901 | 1.302594953 | -1.834266973 | 0.066614374 | 1 | 2.389306901 |
| ENSMUSG00000034898 | Filip1 | 70598 | protein_coding | 3.019608489 | -2.390012578 | 2.39270999 | 0.998872654 | 0.317856385 | 1 | 2.390012578 |
| ENSMUSG00000038011 | Dnah10 | 56087 | protein_coding | 29.43307011 | 2.390287275 | 0.567476913 | -4.212131315 | 2.53E-05 | 0.000254828 | 2.390287275 |
| ENSMUSG00000018459 | Slc13a3 | 114644 | protein_coding | 2.052631512 | 2.390989553 | 2.475944092 | -0.965688022 | 0.334200315 | 1 | 2.390989553 |
| ENSMUSG00000021933 | Gucy1b2 | 239134 | protein_coding | 1.91372087 | 2.391332646 | 2.309555947 | -1.035407975 | 0.300478423 | 1 | 2.391332646 |
| ENSMUSG00000081675 | Gm12712 | 108167910 | processed_pseudogene | 0.81118014 | 2.391500119 | 1.334226088 | -1.792424943 | 0.073064922 | 1 | 2.391500119 |
| ENSMUSG00000115529 | 9630013A20Rik | 319903 | lncRNA | 2.926466202 | 2.396717721 | 0.913278147 | -2.624302057 | 0.008682675 | 1 | 2.396717721 |

| ensembl_gene_id | external_gene_name | entrezgene_id | gene_biotype | baseMean | log2FoldChange | lfcSE | stat | pvalue | padj | absLog2FC |
|---|---|---|---|---|---|---|---|---|---|---|
| ENSMUSG00000031343 | Gabra3 | 14396 | protein_coding | 2.05094863 | 2.398653068 | 2.274442111 | -1.054611615 | 0.291602989 | 1 | 2.398653068 |
| ENSMUSG00000068114 | Ccdc134 | 76457 | protein_coding | 442.9924051 | 2.400035664 | 0.127175365 | -18.87185989 | 1.94E-79 | 4.94E-77 | 2.400035664 |
| ENSMUSG00000043807 | Ly6g5b | 266614 | protein_coding | 3.674937721 | 2.40468178 | 0.836032571 | -2.876301551 | 0.004023651 | 1 | 2.40468178 |
| ENSMUSG00000021743 | Fezf2 | 54713 | protein_coding | 1.151428981 | 2.406512542 | 1.397895602 | -1.721525226 | 0.085155559 | 1 | 2.406512542 |
| ENSMUSG00000112148 | Lilrb4a | 14728 | protein_coding | 1.471910516 | 2.413618952 | 1.556420119 | -1.550750291 | 0.120961537 | 1 | 2.413618952 |
| ENSMUSG00000070476 | Fam217b | 71532 | protein_coding | 75.9920063 | 2.414020268 | 0.215472384 | -11.2033859 | 3.92E-29 | 2.96E-27 | 2.414020268 |
| ENSMUSG00000039728 | Slc6a5 | 104245 | protein_coding | 3.061404324 | -2.414635842 | 2.302301625 | 1.048792137 | 0.294273797 | 1 | 2.414635842 |
| ENSMUSG00000031329 | Tsx | 22127 | protein_coding | 0.893155954 | 2.416128035 | 1.720542588 | -1.40428261 | 0.160234711 | 1 | 2.416128035 |
| ENSMUSG00000071324 | Armc2 | 213402 | protein_coding | 2.391436592 | 2.42060778 | 1.8842145 | -1.284677397 | 0.19890504 | 1 | 2.42060778 |
| ENSMUSG00000032081 | Apoc3 | 11814 | protein_coding | 2.899894246 | -2.422425335 | 1.196552771 | 2.024503553 | 0.042918367 | 1 | 2.422425335 |
| ENSMUSG00000026468 | Lhx4 | 16872 | protein_coding | 1.892947254 | 2.423887142 | 2.154448563 | -1.125061505 | 0.260562972 | 1 | 2.423887142 |
| ENSMUSG00000059824 | Dbp | 13170 | protein_coding | 103.2492282 | -2.424805752 | 0.681594648 | 3.557548112 | 0.000374333 | 0.002774564 | 2.424805752 |
| ENSMUSG00000051906 | Cd209f | 69142 | protein_coding | 1.124202969 | 2.425418171 | 2.565110234 | -0.945541497 | 0.344382496 | 1 | 2.425418171 |
| ENSMUSG00000030087 | Klf15 | 66277 | protein_coding | 6.574130605 | -2.428406813 | 0.55871905 | 4.346382703 | 1.38E-05 | 0.00014909 | 2.428406813 |
| ENSMUSG00000048752 | Prss50 | 235631 | protein_coding | 1.218586414 | 2.429177992 | 1.838583795 | -1.321222345 | 0.186427238 | 1 | 2.429177992 |
| ENSMUSG00000110404 | Rnf223 | 100045778 | protein_coding | 13.24846164 | 2.432406653 | 0.383234171 | -6.347050551 | 2.19E-10 | 5.26E-09 | 2.432406653 |
| ENSMUSG00000062563 | Cys1 | 12879 | protein_coding | 3.339574449 | 2.433837704 | 1.835975267 | -1.325637522 | 0.184959791 | 1 | 2.433837704 |
| ENSMUSG00000031283 | Chrdl1 | 83453 | protein_coding | 0.99564604 | 2.438136991 | 2.196895723 | -1.109810067 | 0.26708088 | 1 | 2.438136991 |
| ENSMUSG00000078964 | Ces1b | 382044 | protein_coding | 2.73582639 | -2.440151183 | 0.984458944 | 2.478672368 | 0.013187238 | 1 | 2.440151183 |
| ENSMUSG00000011751 | Sptbn4 | 80297 | protein_coding | 100.9281533 | 2.443241539 | 0.262331932 | -9.313549904 | 1.24E-20 | 6.50E-19 | 2.443241539 |
| ENSMUSG00000020571 | Pdia6 | 71853 | protein_coding | 10911.70418 | 2.446372013 | 0.121134763 | -20.19545799 | 1.07E-90 | 3.11E-88 | 2.446372013 |
| ENSMUSG00000034391 | Fbxo15 | 50764 | protein_coding | 2.508116419 | 2.447794607 | 1.264804741 | -1.935314225 | 0.052951748 | 1 | 2.447794607 |
| ENSMUSG00000024552 | Slc14a2 | 27411 | protein_coding | 4.273637793 | 2.451011219 | 1.829883227 | -1.339435863 | 0.180428821 | 1 | 2.451011219 |
| ENSMUSG00000068647 | Or4f54 | 258389 | protein_coding | 2.016128316 | 2.452014823 | 2.558989046 | -0.958196686 | 0.337963588 | 1 | 2.452014823 |
| ENSMUSG00000097881 | Celrr | 98452 | lncRNA | 1.415831925 | 2.45218749 | 2.560087406 | -0.957853034 | 0.338136872 | 1 | 2.45218749 |
| ENSMUSG00000090762 | Vmn2r56 | 629079 | protein_coding | 1.23839037 | 2.45271743 | 2.333118715 | -1.051261307 | 0.293138593 | 1 | 2.45271743 |
| ENSMUSG00000091631 | Vmn2r95 | 328759 | protein_coding | 3.455675756 | -2.453315439 | 2.397943297 | 1.023091514 | 0.306264579 | 1 | 2.453315439 |
| ENSMUSG00000022155 | Mroh2b | 223825 | protein_coding | 3.165474364 | 2.453442628 | 2.568319258 | -0.955271671 | 0.33944033 | 1 | 2.453442628 |
| ENSMUSG00000053461 | Hhipl2 | 78772 | protein_coding | 2.886168343 | 2.4540206 | 1.136014084 | -2.160202619 | 0.030756987 | 1 | 2.4540206 |
| ENSMUSG00000045812 | Or4d5 | 258601 | protein_coding | 2.662944931 | 2.455065641 | 2.579709238 | -0.951683083 | 0.341257731 | 1 | 2.455065641 |
| ENSMUSG00000085263 | 1700125G02Rik | 73628 | lncRNA | 1.098800234 | 2.455495525 | 2.582459328 | -0.950836088 | 0.34168759 | 1 | 2.455495525 |
| ENSMUSG00000075132 | Or5d43 | 404329 | protein_coding | 1.189793999 | 2.455920285 | 2.585249365 | -0.949974234 | 0.342125345 | 1 | 2.455920285 |
| ENSMUSG00000056380 | Gpr50 | 14765 | protein_coding | 1.710531629 | -2.456642222 | 2.441963779 | 1.006010918 | 0.314410318 | 1 | 2.456642222 |
| ENSMUSG00000111174 | Or4a68 | 258804 | protein_coding | 1.149930874 | 2.457185998 | 2.593606656 | -0.947401177 | 0.343434392 | 1 | 2.457185998 |
| ENSMUSG00000068999 | Vmn2r5 | 667060 | protein_coding | 1.467643615 | 2.457219744 | 2.593846502 | -0.947326583 | 0.343472389 | 1 | 2.457219744 |
| ENSMUSG00000041624 | Gucy1a2 | 234889 | protein_coding | 1.675308175 | 2.457228199 | 1.88492043 | -1.303624365 | 0.192361689 | 1 | 2.457228199 |
| ENSMUSG00000079451 | Tmprss11g | 320454 | protein_coding | 1.371781527 | 2.457270557 | 2.594179881 | -0.947224429 | 0.34352443 | 1 | 2.457270557 |
| ENSMUSG00000030114 | Klrg1 | 50928 | protein_coding | 1.239990704 | 2.458022535 | 2.599190183 | -0.945687834 | 0.344307829 | 1 | 2.458022535 |
| ENSMUSG00000061387 | Or10w1 | 258098 | protein_coding | 1.703531133 | 2.459059124 | 2.606280375 | -0.943512888 | 0.345418626 | 1 | 2.459059124 |

Supplementary Table 1

| ensembl_gene_id | external_gene_name | entrezgene_id | gene_biotype | baseMean | log2FoldChange | lfcSE | stat | pvalue | padj | absLog2FC |
|---|---|---|---|---|---|---|---|---|---|---|
| ENSMUSG00000054236 | Or5p4 | 258927 | protein_coding | 1.14710171 | 2.460072759 | 2.613295671 | -0.941367939 | 0.346516338 | 1 | 2.460072759 |
| ENSMUSG00000048391 | Or7g25 | 258560 | protein_coding | 1.362788672 | -2.460143839 | 2.491398365 | 0.987455027 | 0.32341962 | 1 | 2.460143839 |
| ENSMUSG00000004892 | Bcan | 12032 | protein_coding | 6.60520812 | -2.462469808 | 1.069920838 | 2.301543928 | 0.021360905 | 0.077273602 | 2.462469808 |
| ENSMUSG00000020388 | Pdlim4 | 30794 | protein_coding | 6.616996864 | -2.46353675 | 0.563907745 | 4.368687558 | 1.25E-05 | 0.000136519 | 2.46353675 |
| ENSMUSG00000110412 | Gm33175 | 102635976 | lncRNA | 0.658866448 | 2.465236438 | 1.455265313 | -1.69401168 | 0.090263064 | 1 | 2.465236438 |
| ENSMUSG00000000244 | Tspan32 | 27027 | protein_coding | 5.360928028 | 2.46870981 | 1.348681604 | -1.830461544 | 0.067180952 | 0.185243988 | 2.46870981 |
| ENSMUSG00000074037 | Mc1r | 17199 | protein_coding | 0.984086368 | 2.469570157 | 2.938075256 | -0.84054013 | 0.400605611 | 1 | 2.469570157 |
| ENSMUSG00000066672 | Or10x1 | 258238 | protein_coding | 1.12248613 | 2.471018784 | 2.947638436 | -0.838304574 | 0.401859668 | 1 | 2.471018784 |
| ENSMUSG00000091358 | Speer5-ps1 | 70365 | transcribed_processed_pseudogene | 3.378551934 | 2.477627628 | 2.056381823 | -1.204848049 | 0.228261967 | 1 | 2.477627628 |
| ENSMUSG00000072492 | Gm10364 | 100151693 | transcribed_processed_pseudogene | 1.118725492 | 2.4776382 | 1.275400483 | -1.942635457 | 0.05206022 | 1 | 2.4776382 |
| ENSMUSG00000057836 | Xlr3a | 22445 | protein_coding | 116.6037453 | 2.479948509 | 0.289591494 | -8.563609638 | 1.09E-17 | 4.91E-16 | 2.479948509 |
| ENSMUSG00000047861 | Foxi1 | 14233 | protein_coding | 1.210458269 | 2.480647131 | 2.018441906 | -1.228991096 | 0.219075144 | 1 | 2.480647131 |
| ENSMUSG00000068234 | Vmn1r44 | 113854 | protein_coding | 0.982828749 | -2.482423969 | 2.915032114 | 0.851594038 | 0.39443945 | 1 | 2.482423969 |
| ENSMUSG00000052382 | Rnase9 | 328401 | protein_coding | 0.956279958 | 2.485086843 | 2.467249776 | -1.007229534 | 0.313824483 | 1 | 2.485086843 |
| ENSMUSG00000041423 | Paqr6 | 68957 | protein_coding | 1.394394725 | 2.486459092 | 2.017636899 | -1.232362024 | 0.21781388 | 1 | 2.486459092 |
| ENSMUSG00000065905 | Gm26110 | 115487642 | snRNA | 1.642160596 | 2.48896694 | 1.228705978 | -2.025681476 | 0.042797436 | 1 | 2.48896694 |
| ENSMUSG00000087149 | Itih6 | 634882 | transcribed_unprocessed_pseudogene | 4.170828151 | 2.494147815 | 1.31593563 | -1.895341807 | 0.058047133 | 1 | 2.494147815 |
| ENSMUSG00000062905 | Vmn1r32 | 171188 | protein_coding | 1.321893713 | 2.494522379 | 1.956727517 | -1.274844023 | 0.202364455 | 1 | 2.494522379 |
| ENSMUSG00000013158 | 4933405L10Rik | 71046 | protein_coding | 1.202939684 | 2.494906496 | 1.582478036 | -1.576582069 | 0.114891721 | 1 | 2.494906496 |
| ENSMUSG00000056158 | Car10 | 72605 | protein_coding | 2.651198109 | 2.497700069 | 2.431089707 | -1.027399385 | 0.304232441 | 1 | 2.497700069 |
| ENSMUSG00000045421 | Or2y17 | 259068 | protein_coding | 1.299687337 | 2.499963404 | 2.530384657 | -0.987977617 | 0.32316361 | 1 | 2.499963404 |
| ENSMUSG00000039904 | Gpr37 | 14763 | protein_coding | 1.848936438 | 2.50012791 | 2.6890158 | -0.929755753 | 0.35249756 | 1 | 2.50012791 |
| ENSMUSG00000020155 | Kcnmb1 | 16533 | protein_coding | 1.451740553 | 2.500739607 | 2.53376935 | -0.986964187 | 0.323660196 | 1 | 2.500739607 |
| ENSMUSG00000033910 | Gucy1a1 | 60596 | protein_coding | 1.667072526 | 2.501936273 | 2.712112162 | -0.922504721 | 0.356265369 | 1 | 2.501936273 |
| ENSMUSG00000066820 | Vmn2r28 | 665255 | protein_coding | 1.653882221 | 2.502049708 | 2.983535398 | -0.838619079 | 0.401683101 | 1 | 2.502049708 |
| ENSMUSG00000073907 | Or56b35 | 258745 | protein_coding | 1.653882221 | 2.502049708 | 2.983535398 | -0.838619079 | 0.401683101 | 1 | 2.502049708 |
| ENSMUSG00000091407 | Vmn2r117 | 619788 | protein_coding | 1.576078762 | 2.502079956 | 2.713964823 | -0.921927924 | 0.356566172 | 1 | 2.502079956 |
| ENSMUSG00000075204 | Or5al5 | 257950 | protein_coding | 1.515327844 | 2.502591161 | 2.989498218 | -0.837127497 | 0.402520906 | 1 | 2.502591161 |
| ENSMUSG00000087148 | 1700030C14Rik | 75662 | lncRNA | 1.42433408 | 2.503383793 | 2.99832299 | -0.834927992 | 0.403758253 | 1 | 2.503383793 |
| ENSMUSG00000064194 | Zfp936 | 668620 | protein_coding | 1.749003445 | 2.503728193 | 3.002264291 | -0.833946632 | 0.404311058 | 1 | 2.503728193 |
| ENSMUSG00000114189 | 9330199G10Rik | 319466 | lncRNA | 1.238219091 | 2.504132081 | 3.006820487 | -0.832817287 | 0.404947783 | 1 | 2.504132081 |
| ENSMUSG00000044120 | Or56a5 | 259047 | protein_coding | 1.471894692 | 2.504393294 | 3.009842397 | -0.832067917 | 0.40537061 | 1 | 2.504393294 |
| ENSMUSG00000021872 | Rnase10 | 75019 | protein_coding | 1.333340316 | 2.504628202 | 3.012527557 | -0.831404246 | 0.405745301 | 1 | 2.504628202 |
| ENSMUSG00000026835 | Fcnb | 14134 | protein_coding | 1.194785939 | 2.504712705 | 3.013481711 | -0.831169041 | 0.405878141 | 1 | 2.504712705 |
| ENSMUSG00000079157 | Nalf1 | 270028 | protein_coding | 1.675129858 | -2.505024354 | 1.919822439 | 1.304820854 | 0.191953856 | 1 | 2.505024354 |
| ENSMUSG00000019139 | Isyna1 | 71780 | protein_coding | 2345.385463 | 2.505085881 | 0.120376411 | -20.81043843 | 3.48E-96 | 1.15E-93 | 2.505085881 |
| ENSMUSG00000024640 | Psat1 | 107272 | protein_coding | 251.7123618 | 2.505342338 | 0.169618399 | -14.7704633 | 2.27E-49 | 3.07E-47 | 2.505342338 |
| ENSMUSG00000066273 | Or51a39 | 18332 | protein_coding | 1.242346552 | 2.507241405 | 3.043137371 | -0.823900173 | 0.409996282 | 1 | 2.507241405 |
| ENSMUSG00000092048 | Vmn2r85 | 623734 | protein_coding | 1.242346552 | 2.507241405 | 3.043137371 | -0.823900173 | 0.409996282 | 1 | 2.507241405 |

| ensembl_gene_id | external_gene_name | entrezgene_id | gene_biotype | baseMean | log2FoldChange | lfcSE | stat | pvalue | padj | absLog2FC |
|---|---|---|---|---|---|---|---|---|---|---|
| ENSMUSG00000022136 | Dnajc3 | 100037258 | protein_coding | 7203.861503 | 2.507501229 | 0.107908658 | -23.237257 | 1.91E-119 | 7.90E-117 | 2.507501229 |
| ENSMUSG00000071361 | Mcpt9 | 17232 | protein_coding | 1.477753106 | -2.511280455 | 1.574139116 | 1.59533578 | 0.110637168 | 1 | 2.511280455 |
| ENSMUSG00000034118 | Tpst1 | 22021 | protein_coding | 67.59800108 | 2.511594419 | 0.160405709 | -15.65776205 | 2.94E-55 | 4.50E-53 | 2.511594419 |
| ENSMUSG00000054666 | Or10h28 | 258939 | protein_coding | 1.762093202 | 2.511981446 | 2.147456453 | -1.169747327 | 0.242102666 | 1 | 2.511981446 |
| ENSMUSG00000030046 | Bmp10 | 12154 | protein_coding | 1.74409962 | 2.512713418 | 2.041928118 | -1.230559193 | 0.218487775 | 1 | 2.512713418 |
| ENSMUSG00000018623 | Mmp7 | 17393 | protein_coding | 39.38128061 | 2.514588385 | 0.307063147 | -8.189157219 | 2.63E-16 | 1.07E-14 | 2.514588385 |
| ENSMUSG00000066607 | Insyn1 | 319477 | protein_coding | 0.888927597 | 2.516609316 | 2.181402328 | -1.153665825 | 0.2486372 | 1 | 2.516609316 |
| ENSMUSG00000040340 | Tex45 | 76406 | protein_coding | 2.038782304 | 2.517929105 | 1.465753964 | -1.717838851 | 0.085826013 | 1 | 2.517929105 |
| ENSMUSG00000035131 | Brinp3 | 215378 | protein_coding | 1.175064768 | 2.52104946 | 2.283681435 | -1.103940953 | 0.269618752 | 1 | 2.52104946 |
| ENSMUSG00000059049 | Frem1 | 329872 | protein_coding | 4.065767285 | 2.525741788 | 1.266739847 | -1.99389148 | 0.046163915 | 1 | 2.525741788 |
| ENSMUSG00000068392 | Rnase13 | 497071 | protein_coding | 2.144050299 | 2.526564878 | 2.520416398 | -1.00243947 | 0.316131387 | 1 | 2.526564878 |
| ENSMUSG00000022494 | Shisa9 | 72555 | protein_coding | 3.052274989 | 2.527057037 | 1.98598595 | -1.272444569 | 0.203215207 | 1 | 2.527057037 |
| ENSMUSG00000029205 | Chrna9 | 231252 | protein_coding | 1.556400848 | 2.527773417 | 2.013218819 | -1.255588013 | 0.20926538 | 1 | 2.527773417 |
| ENSMUSG00000006519 | Cyba | 13057 | protein_coding | 378.4742548 | -2.5291 | 0.147106902 | 17.19225921 | 3.03E-66 | 5.90E-64 | 2.5291 |
| ENSMUSG00000059901 | Adamts14 | 237360 | protein_coding | 2.804147831 | 2.530860724 | 1.625170618 | -1.557289244 | 0.119401828 | 1 | 2.530860724 |
| ENSMUSG00000041120 | Nbl1 | 17965 | protein_coding | 54.51293891 | -2.534240257 | 0.297347401 | 8.522826327 | 1.56E-17 | 6.82E-16 | 2.534240257 |
| ENSMUSG00000108676 | 1700123J17Rik | 76728 | lncRNA | 1.637504458 | 2.536036035 | 2.525507289 | -1.004168963 | 0.315297179 | 1 | 2.536036035 |
| ENSMUSG00000046932 | Vmn1r193 | 171259 | protein_coding | 2.20169665 | 2.537236116 | 2.199104934 | -1.153758549 | 0.248599173 | 1 | 2.537236116 |
| ENSMUSG00000111715 | Or4a73 | 258788 | protein_coding | 1.679783874 | 2.537407025 | 2.341942604 | -1.083462515 | 0.27860318 | 1 | 2.537407025 |
| ENSMUSG00000002266 | Zim1 | 22776 | protein_coding | 3.629382414 | 2.537534922 | 3.387829635 | -0.749014914 | 0.453848217 | 1 | 2.537534922 |
| ENSMUSG00000072791 | Abcb5 | 77706 | protein_coding | 1.469781778 | 2.539944654 | 2.056521524 | -1.235068354 | 0.216805066 | 1 | 2.539944654 |
| ENSMUSG00000111657 | Gm31592 | 102633867 | lncRNA | 1.289907164 | 2.542699857 | 3.602993856 | -0.705718622 | 0.480363138 | 1 | 2.542699857 |
| ENSMUSG00000030162 | Olr1 | 108078 | protein_coding | 1.531607777 | -2.545697056 | 1.66766521 | 1.526503666 | 0.126884478 | 1 | 2.545697056 |
| ENSMUSG00000068950 | Or1j10 | 258949 | protein_coding | 1.506251798 | -2.55196577 | 2.784672676 | 0.916432941 | 0.359439865 | 1 | 2.55196577 |
| ENSMUSG00000031617 | Tmem184c | 234463 | protein_coding | 178.5112509 | 2.554492321 | 0.147238804 | -17.34931452 | 2.00E-67 | 4.02E-65 | 2.554492321 |
| ENSMUSG00000085058 | 8030453O22Rik | 77209 | lncRNA | 0.908724757 | -2.55726856 | 2.880073365 | 0.887917854 | 0.37458494 | 1 | 2.55726856 |
| ENSMUSG00000045083 | Lingo2 | 242384 | protein_coding | 5.95735192 | 2.557594767 | 0.900423367 | -2.840435801 | 0.004505194 | 0.022335267 | 2.557594767 |
| ENSMUSG00000117172 | A230051N06Rik | 320593 | lncRNA | 0.906541609 | -2.559196265 | 1.163351636 | 2.199847566 | 0.027817712 | 1 | 2.559196265 |
| ENSMUSG00000036570 | Fxyd1 | 56188 | protein_coding | 2.274365764 | 2.559203218 | 2.165767547 | -1.18166108 | 0.237340207 | 1 | 2.559203218 |
| ENSMUSG00000071491 | Vmn1r209 | 432736 | protein_coding | 0.900537944 | 2.560800022 | 2.531061018 | -1.011749619 | 0.311657784 | 1 | 2.560800022 |
| ENSMUSG00000043310 | Or51s1 | 259089 | protein_coding | 0.950285717 | 2.560830032 | 2.173355896 | -1.178283795 | 0.238683486 | 1 | 2.560830032 |
| ENSMUSG00000025892 | Gria4 | 14802 | protein_coding | 1.578808371 | 2.5633511 | 1.937488058 | -1.323028077 | 0.185826041 | 1 | 2.5633511 |
| ENSMUSG00000022296 | Baalc | 118452 | protein_coding | 1.647419361 | 2.564408721 | 2.265691008 | -1.131843977 | 0.257700035 | 1 | 2.564408721 |
| ENSMUSG00000047295 | Smgc | 223809 | protein_coding | 1.088198464 | 2.568849921 | 2.594096963 | -0.990267503 | 0.322043386 | 1 | 2.568849921 |
| ENSMUSG00000100857 | 1700041M19Rik | 76601 | lncRNA | 1.467037177 | 2.569049561 | 2.290566636 | -1.121578181 | 0.262041833 | 1 | 2.569049561 |
| ENSMUSG00000072683 | Gm7457 | 665037 | lncRNA | 2.483052308 | 2.570592013 | 2.71898761 | -0.945422481 | 0.344443229 | 1 | 2.570592013 |
| ENSMUSG00000055312 | Them7 | 74088 | protein_coding | 1.409401352 | 2.572121689 | 1.3209055 | -1.947241259 | 0.051505817 | 1 | 2.572121689 |
| ENSMUSG00000061100 | Retnla | 57262 | protein_coding | 0.985336487 | 2.572701581 | 2.140303987 | -1.202026253 | 0.229353357 | 1 | 2.572701581 |
| ENSMUSG00000014158 | Trpv4 | 63873 | protein_coding | 4.612178587 | 2.573554024 | 1.009312917 | -2.549807875 | 0.01077823 | 0.045227455 | 2.573554024 |

| ensembl_gene_id | external_gene_name | entrezgene_id | gene_biotype | baseMean | log2FoldChange | lfcSE | stat | pvalue | padj | absLog2FC |
|---|---|---|---|---|---|---|---|---|---|---|
| ENSMUSG00000100586 | Vmn1r90 | 627280 | protein_coding | 1.736047173 | -2.575331535 | 3.312665454 | 0.777419746 | 0.436911166 | 1 | 2.575331535 |
| ENSMUSG00000064580 | Gm25134 | 115486604 | snRNA | 0.921904654 | 2.577702347 | 2.132108346 | -1.208992194 | 0.226665844 | 1 | 2.577702347 |
| ENSMUSG00000068180 | AY512915 | 414067 | lncRNA | 1.452605033 | 2.578156424 | 2.924945102 | -0.881437543 | 0.378081045 | 1 | 2.578156424 |
| ENSMUSG00000015665 | Awat1 | 245533 | protein_coding | 1.003969599 | -2.580250757 | 3.473052999 | 0.742934461 | 0.457521361 | 1 | 2.580250757 |
| ENSMUSG00000046287 | Pnma3 | 245468 | protein_coding | 0.956408987 | -2.58068551 | 3.488501165 | 0.739769141 | 0.459440087 | 1 | 2.58068551 |
| ENSMUSG00000118384 | 9030625G05Rik | 71545 | lncRNA | 22.21720899 | 2.582208515 | 0.342192193 | -7.546076647 | 4.49E-14 | 1.52E-12 | 2.582208515 |
| ENSMUSG00000092998 | Mir5099 | 100628578 | miRNA | 1.180760878 | 2.582476486 | 1.281040364 | -2.015921245 | 0.043808212 | 1 | 2.582476486 |
| ENSMUSG00000022857 | Tmprss15 | 19146 | protein_coding | 1.871652568 | 2.584731255 | 2.000806857 | -1.29184446 | 0.196411015 | 1 | 2.584731255 |
| ENSMUSG00000087090 | Nctc1 | 330677 | lncRNA | 1.740854099 | 2.587280141 | 2.12636808 | -1.216760242 | 0.223695451 | 1 | 2.587280141 |
| ENSMUSG00000022044 | Stmn4 | 56471 | protein_coding | 1.939248618 | -2.589492777 | 2.201967021 | 1.175990718 | 0.239598585 | 1 | 2.589492777 |
| ENSMUSG00000029423 | Piwil1 | 57749 | protein_coding | 2.821819716 | 2.589536514 | 2.4132521 | -1.073048487 | 0.283249362 | 1 | 2.589536514 |
| ENSMUSG00000027238 | Frmd5 | 228564 | protein_coding | 22.74504677 | 2.591404496 | 0.353157699 | -7.337811143 | 2.17E-13 | 7.02E-12 | 2.591404496 |
| ENSMUSG00000070858 | Gm1673 | 381633 | protein_coding | 9.042779233 | 2.592594912 | 0.471289181 | -5.501070294 | 3.77E-08 | 6.62E-07 | 2.592594912 |
| ENSMUSG00000009588 | St6galnac1 | 20445 | protein_coding | 1.206120811 | 2.593412692 | 2.301002031 | -1.127079706 | 0.259708783 | 1 | 2.593412692 |
| ENSMUSG00000026237 | Nmur1 | 14767 | protein_coding | 1.019453813 | 2.594022889 | 2.246131871 | -1.154884503 | 0.248137726 | 1 | 2.594022889 |
| ENSMUSG00000037482 | Erv3 | 71995 | protein_coding | 1.980103996 | 2.597638802 | 2.278196234 | -1.140217319 | 0.254195774 | 1 | 2.597638802 |
| ENSMUSG00000020913 | Krt24 | 75706 | protein_coding | 1.523960495 | 2.597766629 | 2.279346983 | -1.139697751 | 0.254412245 | 1 | 2.597766629 |
| ENSMUSG00000058132 | Vmn1r82 | 171268 | protein_coding | 1.569420242 | 2.598371764 | 2.269703043 | -1.144806926 | 0.252289146 | 1 | 2.598371764 |
| ENSMUSG00000017211 | Gsdma2 | 76758 | protein_coding | 2.03322546 | -2.600893784 | 1.941164206 | 1.339862839 | 0.180289942 | 1 | 2.600893784 |
| ENSMUSG00000089309 | Gm24838 | 115485663 | snoRNA | 1.002290665 | -2.602411195 | 1.406869334 | 1.849788842 | 0.06434399 | 1 | 2.602411195 |
| ENSMUSG00000099207 | Gm10637 | 100038706 | lncRNA | 1.530439092 | 2.604287209 | 2.302448083 | -1.131094867 | 0.258015164 | 1 | 2.604287209 |
| ENSMUSG00000047658 | Gal3st3 | 545276 | protein_coding | 1.122157984 | 2.606721513 | 1.368228868 | -1.905179444 | 0.056756749 | 1 | 2.606721513 |
| ENSMUSG00000037034 | Pax1 | 18503 | protein_coding | 2.126348673 | -2.606920469 | 2.200336741 | 1.184782502 | 0.236103453 | 1 | 2.606920469 |
| ENSMUSG00000054446 | Cpa1 | 109697 | protein_coding | 1.401163015 | -2.616585831 | 1.720369179 | 1.520944379 | 0.128273796 | 1 | 2.616585831 |
| ENSMUSG00000034968 | Lbx2 | 16815 | protein_coding | 0.790296951 | 2.619933692 | 1.867730763 | -1.402736274 | 0.160695496 | 1 | 2.619933692 |
| ENSMUSG00000064006 | Or5h22 | 258392 | protein_coding | 1.479024796 | 2.636725021 | 2.605805505 | -1.011865627 | 0.311602306 | 1 | 2.636725021 |
| ENSMUSG00000033633 | Clec18a | 353287 | protein_coding | 1.913030253 | 2.639638734 | 1.64011443 | -1.609423517 | 0.107523766 | 1 | 2.639638734 |
| ENSMUSG00000020176 | Grb10 | 14783 | protein_coding | 130.5387459 | 2.643037735 | 0.182052623 | -14.51798763 | 9.32E-48 | 1.19E-45 | 2.643037735 |
| ENSMUSG00000022026 | Olfm4 | 380924 | protein_coding | 6.078221276 | 2.643416656 | 1.222506774 | -2.162292032 | 0.030595674 | 0.102444638 | 2.643416656 |
| ENSMUSG00000062142 | Or52r1 | 259092 | protein_coding | 1.386828032 | 2.644748562 | 2.60556674 | -1.015037735 | 0.31008784 | 1 | 2.644748562 |
| ENSMUSG00000049530 | Clrn2 | 624224 | protein_coding | 1.147044032 | 2.645497698 | 2.612208418 | -1.01274373 | 0.311182584 | 1 | 2.645497698 |
| ENSMUSG00000086644 | Gm13470 | 102633065 | lncRNA | 0.969723728 | 2.649169289 | 1.680688973 | -1.57624006 | 0.114970489 | 1 | 2.649169289 |
| ENSMUSG00000100486 | Gm4131 | 100042960 | protein_coding | 1.185239067 | -2.650755502 | 1.383834401 | 1.915514963 | 0.055426863 | 1 | 2.650755502 |
| ENSMUSG00000045875 | Adra1a | 11549 | protein_coding | 3.156887139 | 2.655655369 | 2.074916602 | -1.279885354 | 0.200585459 | 1 | 2.655655369 |
| ENSMUSG00000036510 | Cdh8 | 12564 | protein_coding | 1.255101457 | 2.655855407 | 1.769539392 | -1.500873854 | 0.133388192 | 1 | 2.655855407 |
| ENSMUSG00000070816 | Vmn1r86 | 100312473 | protein_coding | 2.592470414 | 2.657012505 | 2.200543846 | -1.207434476 | 0.227264865 | 1 | 2.657012505 |
| ENSMUSG00000039097 | Rln1 | 19773 | protein_coding | 2.029057241 | 2.659173826 | 1.679883796 | -1.582951054 | 0.113432621 | 1 | 2.659173826 |
| ENSMUSG00000027596 | a | 50518 | protein_coding | 2.052040171 | 2.660518464 | 1.366767399 | -1.946577351 | 0.051585426 | 1 | 2.660518464 |
| ENSMUSG00000028642 | Tmem269 | 75180 | protein_coding | 1.133936134 | 2.662417007 | 2.03059183 | -1.311153215 | 0.189806005 | 1 | 2.662417007 |

| ensembl_gene_id | external_gene_name | entrezgene_id | gene_biotype | baseMean | log2FoldChange | lfcSE | stat | pvalue | padj | absLog2FC |
|---|---|---|---|---|---|---|---|---|---|---|
| ENSMUSG00000031770 | Herpud1 | 64209 | protein_coding | 9449.185792 | 2.667464595 | 0.078381712 | -34.0317215 | 7.57E-254 | 6.94E-251 | 2.667464595 |
| ENSMUSG00000037542 | Aldh8a1 | 237320 | protein_coding | 2.022008372 | 2.667769673 | 1.739308794 | -1.53381026 | 0.125076338 | 1 | 2.667769673 |
| ENSMUSG00000034532 | Fbxo16 | 50759 | protein_coding | 8.54648418 | 2.669648191 | 0.572083192 | -4.666538415 | 3.06E-06 | 3.82E-05 | 2.669648191 |
| ENSMUSG00000043740 | B430306N03Rik | 320148 | protein_coding | 1.608536593 | 2.673643059 | 2.124039374 | -1.258754 | 0.2081192 | 1 | 2.673643059 |
| ENSMUSG00000100241 | Slc18a3 | 20508 | protein_coding | 1.457378626 | -2.674159457 | 1.2826737 | 2.08483222 | 0.037084533 | 1 | 2.674159457 |
| ENSMUSG00000036770 | Stpg3 | 74472 | protein_coding | 3.168420655 | 2.674368333 | 1.373060745 | -1.947742183 | 0.05144582 | 1 | 2.674368333 |
| ENSMUSG00000089947 | Gm15509 | 105244050 | lncRNA | 0.96126654 | 2.679467287 | 1.501663047 | -1.784333238 | 0.074369532 | 1 | 2.679467287 |
| ENSMUSG00000031616 | Ednra | 13617 | protein_coding | 2.464623679 | 2.679482922 | 2.188187875 | -1.224521419 | 0.220755591 | 1 | 2.679482922 |
| ENSMUSG00000102424 | Paupar | 103164605 | lncRNA | 1.130360578 | -2.684916714 | 1.856274317 | 1.446400831 | 0.148064809 | 1 | 2.684916714 |
| ENSMUSG00000038311 | 2410017I17Rik | 675325 | lncRNA | 2.837668903 | 2.690516879 | 1.070904255 | -2.512378549 | 0.011992037 | 1 | 2.690516879 |
| ENSMUSG00000087668 | Gm11186 | 790909 | lncRNA | 1.004719691 | 2.695091457 | 2.222924554 | -1.212407975 | 0.22535625 | 1 | 2.695091457 |
| ENSMUSG00000090619 | Vmn2r60 | 637898 | protein_coding | 2.528235778 | 2.695336073 | 1.990892689 | -1.353832925 | 0.175789687 | 1 | 2.695336073 |
| ENSMUSG00000030759 | Far1 | 67420 | protein_coding | 10211.71228 | 2.696355798 | 0.084078707 | -32.06942527 | 1.18E-225 | 9.26E-223 | 2.696355798 |
| ENSMUSG00000050830 | Vwc2 | 319922 | protein_coding | 1.433686909 | 2.696470522 | 2.06851755 | -1.303576333 | 0.192378075 | 1 | 2.696470522 |
| ENSMUSG00000112778 | Gm33677 | 102636674 | lncRNA | 8.580437133 | -2.697642509 | 0.451157579 | 5.979379788 | 2.24E-09 | 4.74E-08 | 2.697642509 |
| ENSMUSG00000004542 | Psg19 | 26439 | protein_coding | 1.835784679 | 2.702518162 | 2.342637342 | -1.153622079 | 0.248655142 | 1 | 2.702518162 |
| ENSMUSG00000021999 | Cpb2 | 56373 | protein_coding | 3.780998952 | -2.70406252 | 0.770931332 | 3.507527074 | 0.000452292 | 1 | 2.70406252 |
| ENSMUSG00000030022 | Adamts9 | 101401 | protein_coding | 12.81110704 | -2.704969699 | 0.856622338 | 3.157715576 | 0.001590106 | 0.009348171 | 2.704969699 |
| ENSMUSG00000030994 | D7Ertd443e | 71007 | protein_coding | 3.147675411 | 2.70830429 | 2.240516282 | -1.20878581 | 0.226745144 | 1 | 2.70830429 |
| ENSMUSG00000029522 | Pla2g1b | 18778 | protein_coding | 0.9748227 | 2.708893965 | 1.999730975 | -1.354629197 | 0.175535726 | 1 | 2.708893965 |
| ENSMUSG00000080411 | Mir1193 | 100316673 | miRNA | 0.597903929 | -2.7119888 | 1.533130393 | 1.76892247 | 0.076906814 | 1 | 2.7119888 |
| ENSMUSG00000098380 | Mir6939 | 102465566 | miRNA | 1.03187834 | -2.71723194 | 1.104195805 | 2.460824365 | 0.013861821 | 1 | 2.71723194 |
| ENSMUSG00000023809 | Rps6ka2 | 20112 | protein_coding | 98.11929368 | 2.721769871 | 0.205940385 | -13.2162998 | 7.07E-40 | 7.25E-38 | 2.721769871 |
| ENSMUSG00000065341 | Gm25579 | 115486570 | snRNA | 0.588176796 | 2.726379442 | 1.531483897 | -1.78022077 | 0.075039838 | 1 | 2.726379442 |
| ENSMUSG00000029373 | Pf4 | 56744 | protein_coding | 10.6125054 | 2.726948516 | 0.411718527 | -6.623332051 | 3.51E-11 | 9.32E-10 | 2.726948516 |
| ENSMUSG00000029641 | Rasl11a | 68895 | protein_coding | 20.87936103 | 2.728717738 | 0.347300201 | -7.856942588 | 3.94E-15 | 1.44E-13 | 2.728717738 |
| ENSMUSG00000027270 | Lamp5 | 76161 | protein_coding | 2.039045789 | 2.731035183 | 2.101602763 | -1.299501139 | 0.193772003 | 1 | 2.731035183 |
| ENSMUSG00000026663 | Atf6 | 226641 | protein_coding | 5122.74007 | 2.735749772 | 0.06347466 | -43.0998729 | 0 | 0 | 2.735749772 |
| ENSMUSG00000009070 | Rsph14 | 71236 | protein_coding | 1.610885608 | -2.736533952 | 2.500807395 | 1.094260181 | 0.273840884 | 1 | 2.736533952 |
| ENSMUSG00000064329 | Scn1a | 20265 | protein_coding | 3.228701838 | 2.738480306 | 2.711985564 | -1.0097695 | 0.312605736 | 1 | 2.738480306 |
| ENSMUSG00000074444 | Defa30 | 100038927 | protein_coding | 1.025541095 | 2.738502405 | 2.478887147 | -1.104730568 | 0.269276354 | 1 | 2.738502405 |
| ENSMUSG00000039759 | Thap3 | 69876 | protein_coding | 545.423784 | 2.738998392 | 0.136140953 | -20.11884247 | 5.05E-90 | 1.44E-87 | 2.738998392 |
| ENSMUSG00000073458 | Smok2a | 27263 | protein_coding | 3.125635892 | 2.739184389 | 2.722860529 | -1.005995114 | 0.31441792 | 1 | 2.739184389 |
| ENSMUSG00000091879 | Vmn2r17 | 384221 | protein_coding | 3.0863302 | 2.739951092 | 2.73487817 | -1.001854899 | 0.316413678 | 1 | 2.739951092 |
| ENSMUSG00000090942 | F830016B08Rik | 240328 | protein_coding | 0.89775056 | 2.7462775 | 2.134533685 | -1.286593657 | 0.198235956 | 1 | 2.7462775 |
| ENSMUSG00000028185 | Dnase2b | 56629 | protein_coding | 1.806855131 | 2.746451838 | 1.558253168 | -1.762519656 | 0.077981538 | 1 | 2.746451838 |
| ENSMUSG00000041681 | Iapp | 15874 | protein_coding | 9.709227391 | -2.747862814 | 0.503923889 | 5.452932226 | 4.95E-08 | 8.50E-07 | 2.747862814 |
| ENSMUSG00000022146 | Osmr | 18414 | protein_coding | 22.02018873 | 2.754197203 | 0.645835478 | -4.264549249 | 2.00E-05 | 0.000206225 | 2.754197203 |
| ENSMUSG00000073975 | Or51r1 | 259108 | protein_coding | 1.338768796 | -2.754611189 | 2.917250731 | 0.944249035 | 0.345042403 | 1 | 2.754611189 |

Supplementary Table 1

| ensembl_gene_id | external_gene_name | entrezgene_id | gene_biotype | baseMean | log2FoldChange | lfcSE | stat | pvalue | padj | absLog2FC |
|---|---|---|---|---|---|---|---|---|---|---|
| ENSMUSG00000074001 | Klhl40 | 72330 | protein_coding | 1.715579423 | 2.755708408 | 2.290851769 | -1.202918689 | 0.229007787 | 1 | 2.755708408 |
| ENSMUSG00000021322 | Aoah | 27052 | protein_coding | 4.532656972 | 2.75909519 | 1.023388202 | -2.696039669 | 0.007016931 | 0.031966234 | 2.75909519 |
| ENSMUSG00000044485 | Klk1b11 | 16613 | protein_coding | 1.133748276 | 2.760133999 | 1.340694787 | -2.058734042 | 0.039519723 | 1 | 2.760133999 |
| ENSMUSG00000097003 | D930007P13Rik | 432989 | lncRNA | 1.622385447 | 2.764143018 | 2.244712019 | -1.231402066 | 0.218172525 | 1 | 2.764143018 |
| ENSMUSG00000087658 | Hotairm1 | 105242658 | lncRNA | 26.70848337 | 2.764419192 | 0.378143477 | -7.310503446 | 2.66E-13 | 8.50E-12 | 2.764419192 |
| ENSMUSG00000047261 | Gap43 | 14432 | protein_coding | 1.258204603 | 2.767139052 | 2.269318898 | -1.219369853 | 0.222703847 | 1 | 2.767139052 |
| ENSMUSG00000030268 | Bcat1 | 12035 | protein_coding | 2.54318371 | 2.770558915 | 2.305780062 | -1.2015712 | 0.229529705 | 1 | 2.770558915 |
| ENSMUSG00000043939 | A530064D06Rik | 328830 | protein_coding | 1.447254696 | 2.772972669 | 2.319610761 | -1.195447407 | 0.231912273 | 1 | 2.772972669 |
| ENSMUSG00000031962 | Cdh15 | 12555 | protein_coding | 1.99951263 | 2.777955963 | 2.330634245 | -1.191931325 | 0.233288174 | 1 | 2.777955963 |
| ENSMUSG00000000903 | Vpreb3 | 22364 | protein_coding | 0.919996442 | 2.7830779 | 1.887259982 | -1.474665879 | 0.140302394 | 1 | 2.7830779 |
| ENSMUSG00000029348 | Asphd2 | 72898 | protein_coding | 2.318213574 | 2.786634646 | 1.672024847 | -1.666622748 | 0.095589443 | 1 | 2.786634646 |
| ENSMUSG00000026452 | Syt2 | 20980 | protein_coding | 29.8285408 | -2.793741512 | 0.323736635 | 8.629673665 | 6.15E-18 | 2.82E-16 | 2.793741512 |
| ENSMUSG00000000248 | Clec2g | 70809 | protein_coding | 1.108922244 | 2.798263689 | 1.435901353 | -1.948785467 | 0.05132105 | 1 | 2.798263689 |
| ENSMUSG00000049515 | Espnl | 227357 | protein_coding | 6.52033437 | 2.80141191 | 0.778100491 | -3.600321478 | 0.000317824 | 0.002393317 | 2.80141191 |
| ENSMUSG00000078796 | Zfp541 | 666528 | protein_coding | 2.541123602 | 2.801478497 | 1.975112764 | -1.418389141 | 0.156077184 | 1 | 2.801478497 |
| ENSMUSG00000027932 | Slc27a3 | 26568 | protein_coding | 82.75319936 | 2.805375146 | 0.312293621 | -8.983133043 | 2.63E-19 | 1.29E-17 | 2.805375146 |
| ENSMUSG00000053852 | Adgrg4 | 236798 | protein_coding | 4.377480532 | 2.809868612 | 2.12343702 | -1.323264399 | 0.185747466 | 1 | 2.809868612 |
| ENSMUSG00000056995 | Or4p7 | 258203 | protein_coding | 1.133872253 | 2.810080953 | 2.573397747 | -1.091973037 | 0.274844953 | 1 | 2.810080953 |
| ENSMUSG00000045132 | Or51v14 | 258808 | protein_coding | 1.592968534 | 2.811015351 | 2.581388386 | -1.08895483 | 0.276173805 | 1 | 2.811015351 |
| ENSMUSG00000050366 | Or6b13 | 258055 | protein_coding | 1.406853546 | 2.813734259 | 2.604564909 | -1.080308749 | 0.280004715 | 1 | 2.813734259 |
| ENSMUSG00000074365 | Crxos | 546024 | lncRNA | 1.458665235 | 2.814719353 | 2.613156437 | -1.077133888 | 0.281420456 | 1 | 2.814719353 |
| ENSMUSG00000070280 | Slc22a14 | 382113 | protein_coding | 1.692541761 | 2.815111987 | 2.616623496 | -1.075856726 | 0.281991338 | 1 | 2.815111987 |
| ENSMUSG00000022485 | Hoxc5 | 15424 | protein_coding | 1.146188638 | 2.815461532 | 2.619669149 | -1.074739355 | 0.282491439 | 1 | 2.815461532 |
| ENSMUSG00000117570 | Gm38410 | 328919 | lncRNA | 1.448552993 | 2.81927659 | 2.654255744 | -1.062172172 | 0.28815753 | 1 | 2.81927659 |
| ENSMUSG00000051817 | Sox12 | 20667 | protein_coding | 51.88746045 | 2.823591774 | 0.289496069 | -9.753471914 | 1.78E-22 | 1.03E-20 | 2.823591774 |
| ENSMUSG00000037095 | Lrg1 | 76905 | protein_coding | 27.38105458 | 2.82359387 | 0.691541365 | -4.083044073 | 4.44E-05 | 0.000421772 | 2.82359387 |
| ENSMUSG00000022864 | D16Ertd472e | 67102 | protein_coding | 46.38016138 | 2.825325856 | 0.216167244 | -13.07009244 | 4.88E-39 | 4.80E-37 | 2.825325856 |
| ENSMUSG00000027048 | Abcb11 | 27413 | protein_coding | 3.575152327 | 2.828112913 | 1.181407796 | -2.393849882 | 0.016672571 | 1 | 2.828112913 |
| ENSMUSG00000053469 | Tg | 21819 | protein_coding | 3.499896575 | -2.830101356 | 2.035572382 | 1.390322143 | 0.164431077 | 1 | 2.830101356 |
| ENSMUSG00000048636 | A730049H05Rik | 74516 | lncRNA | 0.929405761 | -2.831541923 | 1.80118163 | 1.572046859 | 0.115939684 | 1 | 2.831541923 |
| ENSMUSG00000018930 | Ccl4 | 20303 | protein_coding | 2.443623543 | 2.831592737 | 1.177916033 | -2.403900327 | 0.016221195 | 1 | 2.831592737 |
| ENSMUSG00000042357 | Gjb5 | 14622 | protein_coding | 14.32411666 | -2.840614381 | 0.504506745 | 5.630478496 | 1.80E-08 | 3.32E-07 | 2.840614381 |
| ENSMUSG00000019368 | Sec14l4 | 103655 | protein_coding | 7.10685565 | -2.84276611 | 0.523933992 | 5.425809648 | 5.77E-08 | 9.77E-07 | 2.84276611 |
| ENSMUSG00000101429 | BC055402 | 407809 | lncRNA | 2.05231205 | 2.849042648 | 2.994802458 | -0.951329074 | 0.341437353 | 1 | 2.849042648 |
| ENSMUSG00000056300 | Zfp981 | 100041433 | protein_coding | 1.042261243 | 2.84992038 | 3.007206784 | -0.947696845 | 0.343283808 | 1 | 2.84992038 |
| ENSMUSG00000040576 | Tmem270 | 76629 | protein_coding | 1.506349465 | 2.850318779 | 3.012235436 | -0.946247011 | 0.344022613 | 1 | 2.850318779 |
| ENSMUSG00000091962 | Vmn2r78 | 637896 | protein_coding | 1.4153557 | 2.850605432 | 3.016352303 | -0.94505056 | 0.344633064 | 1 | 2.850605432 |
| ENSMUSG00000022431 | Ribc2 | 67747 | protein_coding | 1.18580756 | 2.850701429 | 3.018488571 | -0.944413524 | 0.344958373 | 1 | 2.850701429 |
| ENSMUSG00000054763 | Defb42 | 619548 | protein_coding | 1.18580756 | 2.850701429 | 3.018488571 | -0.944413524 | 0.344958373 | 1 | 2.850701429 |

Supplementary Table 1

| ensembl_gene_id | external_gene_name | entrezgene_id | gene_biotype | baseMean | log2FoldChange | lfcSE | stat | pvalue | padj | absLog2FC |
|---|---|---|---|---|---|---|---|---|---|---|
| ENSMUSG00000073006 | Wmp | 213450 | protein_coding | 1.18580756 | 2.850701429 | 3.018488571 | -0.944413524 | 0.344958373 | 1 | 2.850701429 |
| ENSMUSG00000007279 | Scube2 | 56788 | protein_coding | 1.276801324 | 2.85073109 | 3.018143248 | -0.944531407 | 0.34489816 | 1 | 2.85073109 |
| ENSMUSG00000066263 | Or51k1 | 259088 | protein_coding | 1.55654617 | 2.850733646 | 3.018235103 | -0.944503509 | 0.34491241 | 1 | 2.850733646 |
| ENSMUSG00000004872 | Pax3 | 18505 | protein_coding | 2.556308184 | 2.851252177 | 1.95264037 | -1.460203436 | 0.144234171 | 1 | 2.851252177 |
| ENSMUSG00000040901 | Kcnk18 | 332396 | protein_coding | 1.379987856 | 2.851253078 | 3.025760002 | -0.942326251 | 0.346025632 | 1 | 2.851253078 |
| ENSMUSG00000081724 | Or10al7 | 258324 | protein_coding | 1.462916313 | 2.851312973 | 3.02664268 | -0.942071224 | 0.346156176 | 1 | 2.851312973 |
| ENSMUSG00000095698 | Rhox2d | 434760 | protein_coding | 1.283564877 | 2.851937291 | 3.035791827 | -0.9394377 | 0.347506065 | 1 | 2.851937291 |
| ENSMUSG00000072295 | C2cd6 | 73463 | protein_coding | 1.554976224 | 2.852027029 | 3.037165303 | -0.939042411 | 0.34770897 | 1 | 2.852027029 |
| ENSMUSG00000117744 | Vmn1r59 | 404284 | protein_coding | 1.497970927 | 2.852361715 | 3.042118572 | -0.937623451 | 0.348437955 | 1 | 2.852361715 |
| ENSMUSG00000107385 | C330024D21Rik | 320479 | lncRNA | 1.41122824 | 2.852576153 | 3.045299188 | -0.936714581 | 0.348905394 | 1 | 2.852576153 |
| ENSMUSG00000084174 | Sycn | 68416 | protein_coding | 1507.143309 | -2.854419298 | 0.349662913 | 8.163345871 | 3.26E-16 | 1.32E-14 | 2.854419298 |
| ENSMUSG00000058013 | Septin11 | 52398 | protein_coding | 2283.520203 | 2.855106132 | 0.070553526 | -40.4672351 | 0 | 0 | 2.855106132 |
| ENSMUSG00000037161 | Mgarp | 67749 | protein_coding | 0.895182777 | 2.855639196 | 1.437800801 | -1.986116014 | 0.047020443 | 1 | 2.855639196 |
| ENSMUSG00000050445 | Cyp8b1 | 13124 | protein_coding | 4.453082542 | 2.855657613 | 0.834525525 | -3.421893672 | 0.000621866 | 0.00425767 | 2.855657613 |
| ENSMUSG00000057069 | Ero1b | 67475 | protein_coding | 446.6238052 | 2.855756525 | 0.11556066 | -24.7121859 | 7.91E-135 | 3.73E-132 | 2.855756525 |
| ENSMUSG00000038026 | Kcnj9 | 16524 | protein_coding | 3.226009129 | 2.856955964 | 1.617137856 | -1.766674346 | 0.077282782 | 1 | 2.856955964 |
| ENSMUSG00000058755 | Osm | 18413 | protein_coding | 0.988059283 | 2.857925381 | 1.73638071 | -1.645909428 | 0.099782408 | 1 | 2.857925381 |
| ENSMUSG00000117221 | Gm17830 | 100415784 | lncRNA | 1.490755753 | 2.862075234 | 2.307663647 | -1.240248005 | 0.214883678 | 1 | 2.862075234 |
| ENSMUSG00000060534 | Dcc | 13176 | protein_coding | 3.834657279 | 2.862261079 | 3.515825119 | -0.814107921 | 0.415583128 | 1 | 2.862261079 |
| ENSMUSG00000025576 | Rbfox3 | 52897 | protein_coding | 5.470736693 | 2.862996577 | 1.424808438 | -2.009390526 | 0.044495734 | 0.13639879 | 2.862996577 |
| ENSMUSG00000079654 | Prrt4 | 101359 | protein_coding | 2.257794755 | 2.863236197 | 1.451702291 | -1.972330149 | 0.04857193 | 1 | 2.863236197 |
| ENSMUSG00000096960 | A230028O05Rik | 319487 | lncRNA | 2.248775099 | 2.865786344 | 1.68750595 | -1.698237772 | 0.089462894 | 1 | 2.865786344 |
| ENSMUSG00000022548 | Apod | 11815 | protein_coding | 8.646028955 | 2.874570049 | 0.583621587 | -4.925400486 | 8.42E-07 | 1.18E-05 | 2.874570049 |
| ENSMUSG00000062546 | V1ra8 | 113850 | protein_coding | 0.795913552 | 2.877682788 | 1.589082588 | -1.810908262 | 0.070155054 | 1 | 2.877682788 |
| ENSMUSG00000040405 | Havcr1 | 171283 | protein_coding | 1.090686335 | 2.880499286 | 3.599258361 | -0.800303562 | 0.42353494 | 1 | 2.880499286 |
| ENSMUSG00000109761 | 5430403N17Rik | 330737 | lncRNA | 1.280928784 | 2.880854377 | 3.608637447 | -0.798321921 | 0.4246837 | 1 | 2.880854377 |
| ENSMUSG00000009350 | Mpo | 17523 | protein_coding | 2.296447025 | -2.88398533 | 1.632378396 | 1.766738237 | 0.077272076 | 1 | 2.88398533 |
| ENSMUSG00000022901 | Cd86 | 12524 | protein_coding | 1.068685954 | 2.884195661 | 1.916605368 | -1.504845864 | 0.132363709 | 1 | 2.884195661 |
| ENSMUSG00000113218 | 4933409F18Rik | 74668 | lncRNA | 1.135676435 | 2.887985439 | 2.265801333 | -1.27459782 | 0.20245163 | 1 | 2.887985439 |
| ENSMUSG00000077314 | Gm24171 | 115485789 | snoRNA | 0.868678166 | 2.888230242 | 1.862726808 | -1.550538828 | 0.121012241 | 1 | 2.888230242 |
| ENSMUSG00000020175 | Rab36 | 76877 | protein_coding | 12.14025347 | 2.892530458 | 0.475827737 | -6.078944609 | 1.21E-09 | 2.64E-08 | 2.892530458 |
| ENSMUSG00000071342 | Lsmem1 | 380755 | protein_coding | 1.537363816 | 2.89443152 | 1.390810317 | -2.081111625 | 0.03742369 | 1 | 2.89443152 |
| ENSMUSG00000040289 | Hey1 | 15213 | protein_coding | 4.51076458 | 2.895731197 | 0.764599524 | -3.787252155 | 0.000152322 | 0.001244042 | 2.895731197 |
| ENSMUSG00000015243 | Abca1 | 11303 | protein_coding | 431.1207672 | 2.896717358 | 0.136536373 | -21.21571922 | 6.84E-100 | 2.45E-97 | 2.896717358 |
| ENSMUSG00000063632 | Sox11 | 20666 | protein_coding | 2.024035584 | 2.906239593 | 1.926420265 | -1.508621793 | 0.131395462 | 1 | 2.906239593 |
| ENSMUSG00000059430 | Actg2 | 11468 | protein_coding | 3.138972869 | 2.909093797 | 1.093222836 | -2.661025457 | 0.007790307 | 1 | 2.909093797 |
| ENSMUSG00000071176 | Arhgef10 | 234094 | protein_coding | 187.9321794 | 2.916024923 | 0.17703162 | -16.47177451 | 5.85E-61 | 9.96E-59 | 2.916024923 |
| ENSMUSG00000109748 | Gm35392 | 102638957 | lncRNA | 1.604349715 | 2.91851994 | 2.318964384 | -1.25854453 | 0.208194894 | 1 | 2.91851994 |
| ENSMUSG00000099413 | Gm17767 | 100045066 | lncRNA | 11.09812632 | 2.923418399 | 0.486018376 | -6.015036762 | 1.80E-09 | 3.84E-08 | 2.923418399 |

| ensembl_gene_id | external_gene_name | entrezgene_id | gene_biotype | baseMean | log2FoldChange | lfcSE | stat | pvalue | padj | absLog2FC |
|---|---|---|---|---|---|---|---|---|---|---|
| ENSMUSG00000027485 | Bpifb1 | 228801 | protein_coding | 1.975253076 | 2.924583418 | 2.314797231 | -1.263429633 | 0.20643482 | 1 | 2.924583418 |
| ENSMUSG00000051046 | Or6d14 | 258754 | protein_coding | 1.220303711 | 2.925654998 | 2.545096274 | -1.149526259 | 0.250339045 | 1 | 2.925654998 |
| ENSMUSG00000038143 | Stox2 | 71069 | protein_coding | 479.3872002 | 2.930234456 | 0.088206578 | -33.22013528 | 5.51E-242 | 4.79E-239 | 2.930234456 |
| ENSMUSG00000058981 | Or10j7 | 258758 | protein_coding | 1.003439699 | 2.931478928 | 2.198878855 | -1.33316982 | 0.182476081 | 1 | 2.931478928 |
| ENSMUSG00000045349 | Sh2d5 | 230863 | protein_coding | 15.98718625 | 2.936378027 | 0.529019326 | -5.550606343 | 2.85E-08 | 5.10E-07 | 2.936378027 |
| ENSMUSG00000046259 | Sprr2h | 20762 | protein_coding | 1.819321232 | 2.93974439 | 1.730856437 | -1.698433404 | 0.089425992 | 1 | 2.93974439 |
| ENSMUSG00000091620 | Vmn2r23 | 435916 | protein_coding | 3.441461911 | 2.951668584 | 2.04157828 | -1.445777814 | 0.148239531 | 1 | 2.951668584 |
| ENSMUSG00000003469 | Phyhip | 105653 | protein_coding | 1.468056436 | 2.952805668 | 1.771933194 | -1.666431713 | 0.095627459 | 1 | 2.952805668 |
| ENSMUSG00000114491 | A230103O09Rik | 77753 | lncRNA | 1.470046416 | 2.953373818 | 2.617333221 | -1.128390453 | 0.259155058 | 1 | 2.953373818 |
| ENSMUSG00000089992 | G6pd2 | 14380 | protein_coding | 0.811677511 | 2.957963311 | 1.834689252 | -1.612242132 | 0.10690926 | 1 | 2.957963311 |
| ENSMUSG00000001655 | Hoxc13 | 15422 | protein_coding | 1.088023538 | 2.959253997 | 2.53780121 | -1.166070055 | 0.243586122 | 1 | 2.959253997 |
| ENSMUSG00000030137 | Tuba8 | 53857 | protein_coding | 1.213589323 | 2.959832988 | 2.036284842 | -1.453545657 | 0.146072313 | 1 | 2.959832988 |
| ENSMUSG00000057981 | Vmn1r12 | 626397 | protein_coding | 2.211812005 | 2.961051418 | 2.630094801 | -1.125834482 | 0.260235586 | 1 | 2.961051418 |
| ENSMUSG00000093789 | Methig1 | 554292 | protein_coding | 1.173004538 | -2.962297115 | 1.098367142 | 2.697000851 | 0.006996709 | 1 | 2.962297115 |
| ENSMUSG00000116831 | Gm30505 | 102632430 | lncRNA | 6.183064907 | 2.97379987 | 0.715521477 | -4.156129433 | 3.24E-05 | 0.000318174 | 2.97379987 |
| ENSMUSG00000109093 | Gm39079 | 105243043 | lncRNA | 1.653473661 | 2.974169408 | 1.561546807 | -1.904630329 | 0.056828141 | 1 | 2.974169408 |
| ENSMUSG00000091572 | Vmn2r3 | 637004 | protein_coding | 3.347070133 | 2.974902894 | 3.712296898 | -0.801364486 | 0.422920669 | 1 | 2.974902894 |
| ENSMUSG00000019876 | Pkib | 18768 | protein_coding | 64.13752951 | 2.977799279 | 0.237507283 | -12.53771778 | 4.64E-36 | 4.21E-34 | 2.977799279 |
| ENSMUSG00000056966 | Gjc3 | 118446 | protein_coding | 1.250022869 | -2.989140361 | 2.078747853 | 1.43795235 | 0.150447575 | 1 | 2.989140361 |
| ENSMUSG00000040724 | Kcna2 | 16490 | protein_coding | 4.247071685 | 2.991693874 | 2.203524314 | -1.357685892 | 0.174563375 | 1 | 2.991693874 |
| ENSMUSG00000005232 | G6pc2 | 14378 | protein_coding | 0.586036909 | -2.998009211 | 1.366234723 | 2.194358818 | 0.028209624 | 1 | 2.998009211 |
| ENSMUSG00000087075 | Lbhd2 | 380787 | protein_coding | 1.114878833 | 3.000480696 | 1.322154861 | -2.26938673 | 0.023244818 | 1 | 3.000480696 |
| ENSMUSG00000064923 | Gm22042 | 115490033 | snRNA | 1.356069965 | 3.002056631 | 1.732985456 | -1.732303419 | 0.083219554 | 1 | 3.002056631 |
| ENSMUSG00000073768 | Or10ak16 | 258331 | protein_coding | 1.679047418 | 3.007526732 | 2.516899959 | -1.194932966 | 0.232113222 | 1 | 3.007526732 |
| ENSMUSG00000042686 | Jph1 | 57339 | protein_coding | 11.4129647 | 3.009706267 | 0.528108635 | -5.699028703 | 1.20E-08 | 2.30E-07 | 3.009706267 |
| ENSMUSG00000117763 | 9830166K06Rik | 402757 | lncRNA | 1.31890718 | -3.014151465 | 1.934060226 | 1.558457914 | 0.119124737 | 1 | 3.014151465 |
| ENSMUSG00000056054 | S100a8 | 20201 | protein_coding | 1.097998078 | 3.014794401 | 1.369958782 | -2.200646064 | 0.02776109 | 1 | 3.014794401 |
| ENSMUSG00000023484 | Prph | 19132 | protein_coding | 2.081738818 | 3.019039839 | 2.496621981 | -1.209249883 | 0.226566859 | 1 | 3.019039839 |
| ENSMUSG00000003814 | Calr | 12317 | protein_coding | 43330.08781 | 3.022189297 | 0.080641702 | -37.47675463 | 2.20E-307 | 3.64E-304 | 3.022189297 |
| ENSMUSG00000064115 | Cadm2 | 239857 | protein_coding | 1.30536543 | -3.022991786 | 2.085698928 | 1.449390295 | 0.147228617 | 1 | 3.022991786 |
| ENSMUSG00000077980 | Mir654 | 100124453 | miRNA | 0.707828139 | -3.026472096 | 1.587952391 | 1.905895991 | 0.056663701 | 1 | 3.026472096 |
| ENSMUSG00000117146 | Fut4-ps1 | 14346 | transcribed_processed_pseudogene | 1.115908335 | 3.030827604 | 2.570335761 | -1.179156299 | 0.238335944 | 1 | 3.030827604 |
| ENSMUSG00000046500 | Tafa4 | 320701 | protein_coding | 1.367028758 | 3.03441491 | 2.575075324 | -1.178379087 | 0.238645511 | 1 | 3.03441491 |
| ENSMUSG00000055523 | Gucy2g | 73707 | protein_coding | 2.123023022 | -3.039404474 | 1.49612713 | 2.031514845 | 0.042202795 | 1 | 3.039404474 |
| ENSMUSG00000089437 | Gm23199 | 115489195 | snRNA | 1.311443321 | 3.041684341 | 1.251657398 | -2.430125325 | 0.015093603 | 1 | 3.041684341 |
| ENSMUSG00000064223 | Or2ag1b | 258444 | protein_coding | 1.507295749 | 3.043560767 | 2.901138154 | -1.049091979 | 0.294135787 | 1 | 3.043560767 |
| ENSMUSG00000034456 | Uroc1 | 243537 | protein_coding | 2.659577961 | -3.043725342 | 2.018461626 | 1.50794313 | 0.131569083 | 1 | 3.043725342 |
| ENSMUSG00000054626 | Xlr | 22441 | protein_coding | 0.891782102 | 3.046641756 | 2.965285215 | -1.027436329 | 0.304215052 | 1 | 3.046641756 |
| ENSMUSG00000070330 | Tmem235 | 546519 | protein_coding | 0.965831543 | 3.047925878 | 2.499933547 | -1.219202759 | 0.222767245 | 1 | 3.047925878 |

| ensembl_gene_id | external_gene_name | entrezgene_id | gene_biotype | baseMean | log2FoldChange | lfcSE | stat | pvalue | padj | absLog2FC |
|---|---|---|---|---|---|---|---|---|---|---|
| ENSMUSG00000054477 | Kcnn2 | 140492 | protein_coding | 2.14156442 | -3.048209015 | 1.995604023 | 1.527461851 | 0.126646206 | 1 | 3.048209015 |
| ENSMUSG00000031155 | Pim2 | 18715 | protein_coding | 149.7827691 | 3.051110609 | 0.161766431 | -18.86120986 | 2.38E-79 | 5.95E-77 | 3.051110609 |
| ENSMUSG00000028435 | Aqp3 | 11828 | protein_coding | 3.378587511 | 3.053359222 | 1.054800336 | -2.894727197 | 0.003794882 | 1 | 3.053359222 |
| ENSMUSG00000038576 | Susd4 | 96935 | protein_coding | 40.69916672 | 3.056436622 | 0.312702807 | -9.774253886 | 1.45E-22 | 8.39E-21 | 3.056436622 |
| ENSMUSG00000058579 | Cela2a | 13706 | protein_coding | 2.254777082 | -3.058893584 | 1.483256467 | 2.062282318 | 0.039180865 | 1 | 3.058893584 |
| ENSMUSG00000033777 | Tlr13 | 279572 | protein_coding | 1.474016976 | 3.072997645 | 1.534865548 | -2.002128232 | 0.045270942 | 1 | 3.072997645 |
| ENSMUSG00000066319 | Rtp3 | 235636 | protein_coding | 1.618360461 | 3.074243075 | 2.930527751 | -1.049040765 | 0.294159357 | 1 | 3.074243075 |
| ENSMUSG00000084973 | Gm13848 | 434003 | lncRNA | 1.388812321 | 3.075914829 | 2.949014425 | -1.043031463 | 0.296933721 | 1 | 3.075914829 |
| ENSMUSG00000110629 | 4930567H12Rik | 75930 | lncRNA | 1.387879953 | -3.077310183 | 1.859687552 | 1.65474581 | 0.09797607 | 1 | 3.077310183 |
| ENSMUSG00000028354 | Fmn2 | 54418 | protein_coding | 18.35850277 | 3.082736865 | 0.555954522 | -5.544944316 | 2.94E-08 | 5.24E-07 | 3.082736865 |
| ENSMUSG00000007877 | Tcap | 21393 | protein_coding | 1.378018697 | 3.090445101 | 1.929829954 | -1.601407987 | 0.109286585 | 1 | 3.090445101 |
| ENSMUSG00000032564 | Cpne4 | 74020 | protein_coding | 1.871975278 | 3.092917108 | 2.125362376 | -1.455242241 | 0.14560221 | 1 | 3.092917108 |
| ENSMUSG00000035860 | Cdhr3 | 68764 | protein_coding | 2.648157204 | 3.094690106 | 2.242704772 | -1.379891881 | 0.167619937 | 1 | 3.094690106 |
| ENSMUSG00000020964 | Sel1l | 20338 | protein_coding | 7712.920593 | 3.095023395 | 0.076121755 | -40.65885471 | 0 | 0 | 3.095023395 |
| ENSMUSG00000063234 | Gpr84 | 80910 | protein_coding | 1.401138146 | 3.09912862 | 2.609034557 | -1.187844987 | 0.234894487 | 1 | 3.09912862 |
| ENSMUSG00000045341 | Or2l5 | 258937 | protein_coding | 2.860382672 | 3.100410928 | 1.880612841 | -1.648617334 | 0.099226055 | 1 | 3.100410928 |
| ENSMUSG00000021090 | Lrrc9 | 78257 | protein_coding | 1.774557145 | 3.101538723 | 2.634815285 | -1.177137062 | 0.239140804 | 1 | 3.101538723 |
| ENSMUSG00000050134 | Or6n2 | 258713 | protein_coding | 1.865349985 | 3.102277734 | 2.642900363 | -1.173815622 | 0.240468885 | 1 | 3.102277734 |
| ENSMUSG00000043144 | Aqp6 | 11831 | protein_coding | 2.003904362 | 3.102667482 | 2.647206164 | -1.172053587 | 0.241175542 | 1 | 3.102667482 |
| ENSMUSG00000091013 | Vmn1r1 | 625823 | protein_coding | 1.406253704 | 3.102670041 | 2.647181488 | -1.17206548 | 0.241170768 | 1 | 3.102670041 |
| ENSMUSG00000063823 | Or13a21 | 259028 | protein_coding | 1.54480808 | 3.104176405 | 2.664041971 | -1.165213025 | 0.243932774 | 1 | 3.104176405 |
| ENSMUSG00000030559 | Rab38 | 72433 | protein_coding | 120.3891284 | 3.109626238 | 0.164386373 | -18.91656947 | 8.33E-80 | 2.15E-77 | 3.109626238 |
| ENSMUSG00000029248 | Thegl | 71868 | protein_coding | 1.554850919 | 3.112721026 | 1.344013507 | -2.315989392 | 0.020558848 | 1 | 3.112721026 |
| ENSMUSG00000026220 | Slc16a14 | 71781 | protein_coding | 2.423115718 | 3.113163616 | 1.86208003 | -1.67187423 | 0.094549125 | 1 | 3.113163616 |
| ENSMUSG00000046167 | Gldn | 235379 | protein_coding | 1.396159138 | 3.114168152 | 3.01466225 | -1.033007314 | 0.301600476 | 1 | 3.114168152 |
| ENSMUSG00000033207 | Mamdc2 | 71738 | protein_coding | 1.297942174 | 3.115193563 | 3.510891757 | -0.887294106 | 0.374920578 | 1 | 3.115193563 |
| ENSMUSG00000096757 | Or8c11 | 258874 | protein_coding | 1.297942174 | 3.115193563 | 3.510891757 | -0.887294106 | 0.374920578 | 1 | 3.115193563 |
| ENSMUSG00000043468 | Adam30 | 71078 | protein_coding | 1.250381562 | 3.115917285 | 3.524957908 | -0.883958721 | 0.376718496 | 1 | 3.115917285 |
| ENSMUSG00000094200 | Or2a14 | 258648 | protein_coding | 1.202820949 | 3.117693405 | 3.560305466 | -0.875681436 | 0.38120324 | 1 | 3.117693405 |
| ENSMUSG00000039632 | Odad3 | 77609 | protein_coding | 1.359064216 | 3.128090538 | 1.41700514 | -2.207536479 | 0.027276601 | 1 | 3.128090538 |
| ENSMUSG00000030145 | Zfp248 | 72720 | protein_coding | 8.737570432 | 3.129633977 | 0.561103745 | -5.577638727 | 2.44E-08 | 4.41E-07 | 3.129633977 |
| ENSMUSG00000079606 | Fsip2l | 209005 | protein_coding | 1.640052922 | 3.132041262 | 3.065671365 | -1.021649384 | 0.306946875 | 1 | 3.132041262 |
| ENSMUSG00000037335 | Hand1 | 15110 | protein_coding | 1.530679452 | -3.141029182 | 1.977561255 | 1.588334709 | 0.112210673 | 1 | 3.141029182 |
| ENSMUSG00000036587 | Fut7 | 14347 | protein_coding | 1.46712333 | 3.141080564 | 2.293292888 | -1.369681378 | 0.170786384 | 1 | 3.141080564 |
| ENSMUSG00000023992 | Trem2 | 83433 | protein_coding | 2.166294336 | 3.142978889 | 2.360282138 | -1.331611522 | 0.182987877 | 1 | 3.142978889 |
| ENSMUSG00000048067 | Or10am5 | 269862 | protein_coding | 1.963473853 | 3.186890288 | 2.351221812 | -1.355418817 | 0.175284158 | 1 | 3.186890288 |
| ENSMUSG00000091805 | Vmn2r108 | 627805 | protein_coding | 2.568204752 | 3.18949291 | 2.373584345 | -1.343745343 | 0.179030749 | 1 | 3.18949291 |
| ENSMUSG00000084487 | Gm23787 | 115488058 | snoRNA | 0.46560142 | -3.190571029 | 1.99116454 | 1.602364328 | 0.109075068 | 1 | 3.190571029 |
| ENSMUSG00000097910 | Hdnr | 75996 | lncRNA | 2.941941109 | 3.194290096 | 2.761132072 | -1.156876967 | 0.24732263 | 1 | 3.194290096 |

| ensembl_gene_id | external_gene_name | entrezgene_id | gene_biotype | baseMean | log2FoldChange | lfcSE | stat | pvalue | padj | absLog2FC |
|---|---|---|---|---|---|---|---|---|---|---|
| ENSMUSG00000061815 | Rufy4 | 435626 | protein_coding | 1.533898483 | 3.211935434 | 2.504667209 | -1.28238012 | 0.199709335 | 1 | 3.211935434 |
| ENSMUSG00000041534 | Rbp3 | 19661 | protein_coding | 1.599622413 | 3.214069076 | 2.631452819 | -1.221404789 | 0.221932795 | 1 | 3.214069076 |
| ENSMUSG00000029211 | Gabra4 | 14397 | protein_coding | 1.872603706 | 3.214903647 | 2.642134202 | -1.216782874 | 0.223686838 | 1 | 3.214903647 |
| ENSMUSG00000019146 | Cacng2 | 12300 | protein_coding | 2.244833683 | 3.215651127 | 2.652063644 | -1.212509034 | 0.225317587 | 1 | 3.215651127 |
| ENSMUSG00000115424 | D130079A08Rik | 320671 | lncRNA | 1.897273397 | 3.217437259 | 2.932079479 | -1.097322662 | 0.272500368 | 1 | 3.217437259 |
| ENSMUSG00000070868 | Skint3 | 195564 | protein_coding | 1.481610267 | 3.218740681 | 2.949144426 | -1.091415074 | 0.275090283 | 1 | 3.218740681 |
| ENSMUSG00000047696 | Ccdc144b | 241943 | protein_coding | 1.759223602 | 3.219147383 | 1.850354165 | -1.739746608 | 0.081903521 | 1 | 3.219147383 |
| ENSMUSG00000057116 | Semp2l2a | 231201 | protein_coding | 1.493123629 | 3.221296485 | 2.600975379 | -1.23849557 | 0.215532364 | 1 | 3.221296485 |
| ENSMUSG00000085041 | BB031773 | 100473 | lncRNA | 1.115348207 | 3.225384238 | 2.968713197 | -1.086458686 | 0.27727611 | 1 | 3.225384238 |
| ENSMUSG00000117450 | 4933424G05Rik | 74456 | lncRNA | 1.208455557 | 3.23222123 | 1.462304828 | -2.210360773 | 0.027080134 | 1 | 3.23222123 |
| ENSMUSG00000022886 | Prl2a1 | 56635 | protein_coding | 0.841972588 | 3.232987752 | 2.447956523 | -1.320688387 | 0.186605288 | 1 | 3.232987752 |
| ENSMUSG00000045326 | Fndc7 | 320181 | protein_coding | 2.935659946 | 3.23877555 | 2.0051632 | -1.615217928 | 0.106263509 | 1 | 3.23877555 |
| ENSMUSG00000064213 | Defa24 | 503491 | protein_coding | 2.147764605 | 3.239752492 | 2.27226248 | -1.425782682 | 0.153931081 | 1 | 3.239752492 |
| ENSMUSG00000020000 | Moxd1 | 59012 | protein_coding | 5.667723667 | 3.240806601 | 0.87873926 | -3.688018448 | 0.000226007 | 0.001763006 | 3.240806601 |
| ENSMUSG00000076439 | Mog | 17441 | protein_coding | 2.150472807 | 3.244261967 | 1.487288388 | -2.181326764 | 0.029159256 | 1 | 3.244261967 |
| ENSMUSG00000012705 | Retn | 57264 | protein_coding | 1.746472428 | 3.249344393 | 2.094745825 | -1.551187907 | 0.120856659 | 1 | 3.249344393 |
| ENSMUSG00000094553 | Vmn1r50 | 113852 | protein_coding | 1.218205973 | 3.249360954 | 3.00613685 | -1.080909192 | 0.279737509 | 1 | 3.249360954 |
| ENSMUSG00000086236 | 5830418P13Rik | 100529079 | lncRNA | 1.949080548 | 3.251495258 | 2.92662256 | -1.111006012 | 0.266565762 | 1 | 3.251495258 |
| ENSMUSG00000109449 | Or4k36 | 258910 | protein_coding | 1.541672339 | 3.253726912 | 2.956686431 | -1.100463978 | 0.271130016 | 1 | 3.253726912 |
| ENSMUSG00000062785 | Kcnc3 | 16504 | protein_coding | 48.50791653 | 3.261193207 | 0.398783098 | -8.177862158 | 2.89E-16 | 1.18E-14 | 3.261193207 |
| ENSMUSG00000105406 | Gm43247 | 108169063 | processed_pseudogene | 0.911015902 | 3.271859191 | 1.258223414 | -2.60038015 | 0.009312054 | 1 | 3.271859191 |
| ENSMUSG00000054036 | Or11m3 | 258502 | protein_coding | 4.447745271 | 3.273841532 | 2.078201296 | -1.575324555 | 0.115181549 | 0.273724003 | 3.273841532 |
| ENSMUSG00000053184 | Spaca3 | 75622 | protein_coding | 1.534727658 | 3.277256133 | 1.52144125 | -2.154047113 | 0.031236477 | 1 | 3.277256133 |
| ENSMUSG00000022803 | Popdc2 | 64082 | protein_coding | 1.338266551 | 3.295246756 | 1.705584815 | -1.932033358 | 0.053355389 | 1 | 3.295246756 |
| ENSMUSG00000004891 | Nes | 18008 | protein_coding | 32.68802317 | 3.296137289 | 0.424808332 | -7.759116386 | 8.55E-15 | 3.08E-13 | 3.296137289 |
| ENSMUSG00000044938 | Klhl31 | 244923 | protein_coding | 10.28790493 | 3.305540366 | 0.686509134 | -4.814998375 | 1.47E-06 | 1.96E-05 | 3.305540366 |
| ENSMUSG00000058427 | Cxcl2 | 20310 | protein_coding | 3.11855461 | 3.307404296 | 1.071710454 | -3.086098753 | 0.002028015 | 1 | 3.307404296 |
| ENSMUSG00000057074 | Ces1g | 12623 | protein_coding | 342.3975464 | -3.33361884 | 0.311998218 | 10.6847368 | 1.20E-26 | 8.33E-25 | 3.33361884 |
| ENSMUSG00000101500 | Btbd35f12 | 100040606 | protein_coding | 0.949028367 | 3.349923482 | 1.572643922 | -2.130122042 | 0.033161539 | 1 | 3.349923482 |
| ENSMUSG00000079186 | Gzmc | 14940 | protein_coding | 1.503630383 | 3.354197328 | 1.462722902 | -2.293118761 | 0.021841169 | 1 | 3.354197328 |
| ENSMUSG00000030247 | Kcnj8 | 16523 | protein_coding | 3.156900283 | -3.385334928 | 1.049471833 | 3.225751106 | 0.001256426 | 1 | 3.385334928 |
| ENSMUSG00000046186 | Cd109 | 235505 | protein_coding | 2.025122519 | 3.385951792 | 2.563731111 | -1.320712526 | 0.186597236 | 1 | 3.385951792 |
| ENSMUSG00000027313 | Chac1 | 69065 | protein_coding | 67.10614678 | 3.388295162 | 0.409914596 | -8.265856329 | 1.39E-16 | 5.78E-15 | 3.388295162 |
| ENSMUSG00000011350 | Gm5893 | 545936 | lncRNA | 2.019601046 | 3.394628528 | 1.258140282 | -2.698131977 | 0.006972979 | 1 | 3.394628528 |
| ENSMUSG00000037577 | Ephx3 | 71932 | protein_coding | 2.808474926 | 3.397864952 | 1.507322381 | -2.254239036 | 0.024181134 | 1 | 3.397864952 |
| ENSMUSG00000000791 | Il12rb1 | 16161 | protein_coding | 20.43910081 | 3.409782077 | 0.520641052 | -6.549199423 | 5.78E-11 | 1.50E-09 | 3.409782077 |
| ENSMUSG00000031665 | Sall1 | 58198 | protein_coding | 2.418139042 | 3.410062904 | 2.470465076 | -1.380332366 | 0.167484334 | 1 | 3.410062904 |
| ENSMUSG00000072900 | Mir540 | 723880 | miRNA | 0.63160829 | -3.415096596 | 1.36053899 | 2.510105643 | 0.012069505 | 1 | 3.415096596 |
| ENSMUSG00000034239 | Gm884 | 380730 | protein_coding | 2.713566016 | 3.417462665 | 2.880394115 | -1.18645662 | 0.235442023 | 1 | 3.417462665 |

Supplementary Table 1

| ensembl_gene_id | external_gene_name | entrezgene_id | gene_biotype | baseMean | log2FoldChange | lfcSE | stat | pvalue | padj | absLog2FC |
|---|---|---|---|---|---|---|---|---|---|---|
| ENSMUSG00000029816 | Gpnmb | 93695 | protein_coding | 2.020176888 | 3.421621748 | 2.933483606 | -1.166402206 | 0.243451867 | 1 | 3.421621748 |
| ENSMUSG00000086219 | Srrm4os | 76798 | lncRNA | 1.609258465 | 3.422263582 | 2.941578494 | -1.163410594 | 0.244662953 | 1 | 3.422263582 |
| ENSMUSG00000024184 | Pdia2 | 69191 | protein_coding | 17.94633075 | 3.422777481 | 0.403633709 | -8.4799099 | 2.25E-17 | 9.77E-16 | 3.422777481 |
| ENSMUSG00000046974 | BC053393 | 407814 | protein_coding | 1.609382082 | 3.423462447 | 2.957608593 | -1.157510313 | 0.247063928 | 1 | 3.423462447 |
| ENSMUSG00000115855 | Gm34643 | 102637961 | lncRNA | 1.336400789 | 3.427245298 | 3.010212594 | -1.138539286 | 0.254895364 | 1 | 3.427245298 |
| ENSMUSG00000046486 | Or6k6 | 404222 | protein_coding | 1.691614593 | 3.43693629 | 2.011254449 | -1.708852051 | 0.087478358 | 1 | 3.43693629 |
| ENSMUSG00000090223 | Pcp4 | 18546 | protein_coding | 1.049712151 | 3.437466542 | 1.474506257 | -2.331266161 | 0.019739329 | 1 | 3.437466542 |
| ENSMUSG00000038738 | Shank1 | 243961 | protein_coding | 50.71131737 | 3.441199298 | 0.467232404 | -7.36506986 | 1.77E-13 | 5.76E-12 | 3.441199298 |
| ENSMUSG00000019834 | Slc22a16 | 70840 | protein_coding | 0.986841564 | 3.448979755 | 2.045673734 | -1.685987212 | 0.091798262 | 1 | 3.448979755 |
| ENSMUSG00000022715 | Tmem114 | 74720 | protein_coding | 2.211283817 | 3.450120327 | 3.419704948 | -1.008894153 | 0.3130254 | 1 | 3.450120327 |
| ENSMUSG00000042256 | Ptchd4 | 627626 | protein_coding | 1.475078783 | 3.45400293 | 3.509571082 | -0.984166683 | 0.325033564 | 1 | 3.45400293 |
| ENSMUSG00000045613 | Chrm2 | 243764 | protein_coding | 1.470951323 | 3.454143579 | 3.512982227 | -0.983251083 | 0.325483879 | 1 | 3.454143579 |
| ENSMUSG00000027718 | Il21 | 60505 | protein_coding | 1.336524406 | 3.454724777 | 3.527143366 | -0.979468204 | 0.327348692 | 1 | 3.454724777 |
| ENSMUSG00000033082 | Clec1a | 243653 | protein_coding | 1.288963794 | 3.455051354 | 3.535193752 | -0.977330126 | 0.328405743 | 1 | 3.455051354 |
| ENSMUSG00000098128 | Gm3693 | 100042143 | lncRNA | 1.570200007 | 3.455364677 | 3.543091099 | -0.975240145 | 0.329441153 | 1 | 3.455364677 |
| ENSMUSG00000019803 | Nr2e1 | 21907 | protein_coding | 1.245530642 | 3.455526097 | 3.547011307 | -0.974207804 | 0.32995337 | 1 | 3.455526097 |
| ENSMUSG00000070407 | Hs3st3b1 | 54710 | protein_coding | 878.6500474 | 3.460242948 | 0.114919403 | -30.11017151 | 3.57E-199 | 2.45E-196 | 3.460242948 |
| ENSMUSG00000020357 | Flt4 | 14257 | protein_coding | 3.555858054 | 3.465477832 | 1.738286092 | -1.993617649 | 0.046193855 | 1 | 3.465477832 |
| ENSMUSG00000057244 | Gm6139 | 108169072 | processed_pseudogene | 0.796821494 | 3.484855576 | 1.869335196 | -1.864221881 | 0.062290552 | 1 | 3.484855576 |
| ENSMUSG00000032115 | Hyou1 | 12282 | protein_coding | 17034.26386 | 3.4882642 | 0.091778356 | -38.00748182 | 0 | 0 | 3.4882642 |
| ENSMUSG00000032575 | Manf | 74840 | protein_coding | 8107.677243 | 3.49242431 | 0.096514809 | -36.18537259 | 1.03E-286 | 1.22E-283 | 3.49242431 |
| ENSMUSG00000017754 | Pltp | 18830 | protein_coding | 24.38449358 | 3.502336863 | 0.47022949 | -7.448143809 | 9.47E-14 | 3.13E-12 | 3.502336863 |
| ENSMUSG00000072592 | Gm10373 | 100038728 | lncRNA | 0.839192953 | 3.502981163 | 2.030358033 | -1.725302191 | 0.084473027 | 1 | 3.502981163 |
| ENSMUSG00000003341 | Atp8b3 | 67331 | protein_coding | 3.506164913 | 3.505344774 | 2.504869077 | -1.399412371 | 0.161689359 | 1 | 3.505344774 |
| ENSMUSG00000075604 | Cyp11b1 | 110115 | protein_coding | 1.139066784 | 3.505722 | 2.973108656 | -1.179143585 | 0.238341006 | 1 | 3.505722 |
| ENSMUSG00000006764 | Tph2 | 216343 | protein_coding | 1.067096086 | 3.508829846 | 2.15904516 | -1.62517668 | 0.104124918 | 1 | 3.508829846 |
| ENSMUSG00000023968 | Crip3 | 114570 | protein_coding | 1.539855139 | 3.513912835 | 1.994323205 | -1.761957554 | 0.078076469 | 1 | 3.513912835 |
| ENSMUSG00000111448 | Or8b48 | 258806 | protein_coding | 1.570067528 | 3.515786957 | 2.070793005 | -1.697797389 | 0.089546009 | 1 | 3.515786957 |
| ENSMUSG00000080406 | Mir1188 | 100316669 | miRNA | 0.865863334 | -3.520060915 | 1.29616674 | 2.715746983 | 0.006612643 | 1 | 3.520060915 |
| ENSMUSG00000064452 | Gm24564 | 115488063 | snoRNA | 0.805957084 | -3.521273579 | 1.46478141 | 2.403958403 | 0.016218619 | 1 | 3.521273579 |
| ENSMUSG00000090581 | Vmn2r6 | 667069 | protein_coding | 2.313060444 | 3.535143065 | 2.578886305 | -1.370802217 | 0.17043662 | 1 | 3.535143065 |
| ENSMUSG00000078817 | Nlrp12 | 378425 | protein_coding | 2.547245349 | 3.547553688 | 2.701255715 | -1.313297985 | 0.189082557 | 1 | 3.547553688 |
| ENSMUSG00000105265 | Sox2ot | 320478 | lncRNA | 2.219623203 | 3.548386848 | 2.714470316 | -1.307211513 | 0.191140887 | 1 | 3.548386848 |
| ENSMUSG00000095483 | Or6c206 | 404321 | protein_coding | 1.209227594 | 3.559037022 | 3.004118808 | -1.18471913 | 0.236128516 | 1 | 3.559037022 |
| ENSMUSG00000053626 | Tll1 | 21892 | protein_coding | 1.895077455 | 3.564592338 | 3.032014561 | -1.175651457 | 0.239734184 | 1 | 3.564592338 |
| ENSMUSG00000074665 | Bpifb4 | 381399 | protein_coding | 3.451145275 | 3.564681205 | 2.003505841 | -1.779221769 | 0.075203411 | 1 | 3.564681205 |
| ENSMUSG00000057465 | Saa2 | 20209 | protein_coding | 9.166922642 | 3.567110288 | 0.973299495 | -3.66496675 | 0.000247371 | 0.001911596 | 3.567110288 |
| ENSMUSG00000093910 | Zfp853 | 330230 | protein_coding | 1.793099944 | 3.580591507 | 1.881505765 | -1.903045727 | 0.057034579 | 1 | 3.580591507 |
| ENSMUSG00000026864 | Hspa5 | 14828 | protein_coding | 77842.70454 | 3.582512118 | 0.095426568 | -37.54208273 | 0 | 0 | 3.582512118 |

| ensembl_gene_id | external_gene_name | entrezgene_id | gene_biotype | baseMean | log2FoldChange | lfcSE | stat | pvalue | padj | absLog2FC |
|---|---|---|---|---|---|---|---|---|---|---|
| ENSMUSG00000000031 | H19 | 14955 | lncRNA | 3.551274218 | 3.583524102 | 1.251637401 | -2.863068889 | 0.004195593 | 1 | 3.583524102 |
| ENSMUSG00000078349 | AW011738 | 100382 | lncRNA | 90.65410283 | 3.604268243 | 0.296588436 | -12.15242338 | 5.57E-34 | 4.79E-32 | 3.604268243 |
| ENSMUSG00000025017 | Pik3ap1 | 83490 | protein_coding | 99.57623848 | 3.609080189 | 0.278593673 | -12.95463802 | 2.21E-38 | 2.12E-36 | 3.609080189 |
| ENSMUSG00000056071 | S100a9 | 20202 | protein_coding | 1.728403674 | 3.622646086 | 1.613588421 | -2.245086813 | 0.02476256 | 1 | 3.622646086 |
| ENSMUSG00000095201 | Vmn1r91 | 667067 | protein_coding | 2.101688949 | 3.624601194 | 1.225544944 | -2.957542448 | 0.00310102 | 1 | 3.624601194 |
| ENSMUSG00000070101 | Mir341 | 723846 | miRNA | 0.637875245 | -3.625453265 | 1.790230005 | 2.025132667 | 0.042853743 | 1 | 3.625453265 |
| ENSMUSG00000074968 | Ano3 | 228432 | protein_coding | 2.320047104 | 3.627038299 | 1.959099378 | -1.851380456 | 0.064114839 | 1 | 3.627038299 |
| ENSMUSG00000046008 | Pnlip | 69060 | protein_coding | 2.301583049 | -3.632750527 | 2.46689869 | 1.472598183 | 0.140859415 | 1 | 3.632750527 |
| ENSMUSG00000074195 | Clca4b | 99709 | protein_coding | 368.2115338 | 3.640561962 | 0.463827746 | -7.848952535 | 4.20E-15 | 1.54E-13 | 3.640561962 |
| ENSMUSG00000073898 | Or10a5 | 259036 | protein_coding | 1.331581821 | 3.642885794 | 2.030381896 | -1.794187488 | 0.072783249 | 1 | 3.642885794 |
| ENSMUSG00000011463 | Cpb1 | 76703 | protein_coding | 2.334780951 | -3.666506859 | 2.148511488 | 1.706533514 | 0.087908792 | 1 | 3.666506859 |
| ENSMUSG00000063594 | Gng8 | 14709 | protein_coding | 2.136130428 | 3.673976391 | 2.586018745 | -1.420707564 | 0.155401793 | 1 | 3.673976391 |
| ENSMUSG00000046152 | Fut10 | 171167 | protein_coding | 1.904647431 | 3.688512711 | 1.359581142 | -2.712977252 | 0.006668168 | 1 | 3.688512711 |
| ENSMUSG00000062380 | Tubb3 | 22152 | protein_coding | 5.22139331 | 3.689688608 | 0.890333125 | -4.144166383 | 3.41E-05 | 0.000333855 | 3.689688608 |
| ENSMUSG00000112201 | 4930532I03Rik | 75833 | lncRNA | 1.509409938 | 3.704126399 | 2.978958186 | -1.243430142 | 0.213709369 | 1 | 3.704126399 |
| ENSMUSG00000091556 | Gm14569 | 101055983 | protein_coding | 1.428268196 | 3.724276707 | 1.491186532 | -2.497525713 | 0.012506339 | 1 | 3.724276707 |
| ENSMUSG00000077925 | Mir453 | 100124484 | miRNA | 0.81287917 | -3.726601777 | 1.259314212 | 2.959231097 | 0.003084077 | 1 | 3.726601777 |
| ENSMUSG00000059887 | Or4e5 | 57269 | protein_coding | 1.236552262 | 3.729508932 | 3.545236343 | -1.051977519 | 0.292809864 | 1 | 3.729508932 |
| ENSMUSG00000053550 | Shisa7 | 232813 | protein_coding | 134.0532363 | 3.730227742 | 0.186997965 | -19.94795899 | 1.56E-88 | 4.37E-86 | 3.730227742 |
| ENSMUSG00000016458 | Wt1 | 22431 | protein_coding | 3.387359626 | 3.734779569 | 2.242600935 | -1.665378584 | 0.095837249 | 1 | 3.734779569 |
| ENSMUSG00000022769 | Sdf2l1 | 64136 | protein_coding | 4723.134471 | 3.763644763 | 0.103595037 | -36.33035792 | 5.37E-289 | 6.82E-286 | 3.763644763 |
| ENSMUSG00000037624 | Kcnk2 | 16526 | protein_coding | 2.917036586 | 3.766619529 | 2.205644451 | -1.707718362 | 0.087688613 | 1 | 3.766619529 |
| ENSMUSG00000025475 | Adgra1 | 52389 | protein_coding | 2.481179221 | 3.76713279 | 2.399414764 | -1.57002151 | 0.116410107 | 1 | 3.76713279 |
| ENSMUSG00000028713 | Cyp4b1 | 13120 | protein_coding | 54.77585792 | -3.768033417 | 0.469438219 | 8.0266865 | 1.00E-15 | 3.91E-14 | 3.768033417 |
| ENSMUSG00000045761 | Togaram2 | 320159 | protein_coding | 2.664280745 | 3.787723339 | 2.589704576 | -1.462608274 | 0.143574598 | 1 | 3.787723339 |
| ENSMUSG00000090961 | Vmn2r8 | 627479 | protein_coding | 1.931000172 | 3.793663065 | 2.595704541 | -1.461515748 | 0.143873957 | 1 | 3.793663065 |
| ENSMUSG00000004360 | 9330159F19Rik | 212448 | protein_coding | 1.645641036 | 3.795683596 | 2.983081721 | -1.272403491 | 0.203229794 | 1 | 3.795683596 |
| ENSMUSG00000018845 | Unc45b | 217012 | protein_coding | 1.881157094 | 3.798580525 | 2.972296626 | -1.2779951 | 0.201251154 | 1 | 3.798580525 |
| ENSMUSG00000054999 | Naaladl1 | 381204 | protein_coding | 3521.256213 | 3.800819753 | 0.183883707 | -20.66969285 | 6.49E-95 | 2.10E-92 | 3.800819753 |
| ENSMUSG00000020695 | Mrc2 | 17534 | protein_coding | 3.742270726 | 3.800970518 | 1.870008457 | -2.032595363 | 0.04209342 | 1 | 3.800970518 |
| ENSMUSG00000094672 | Vmn2r25 | 545874 | protein_coding | 1.797899872 | 3.813378746 | 2.981034312 | -1.279213302 | 0.200821953 | 1 | 3.813378746 |
| ENSMUSG00000056367 | Actr3b | 242894 | protein_coding | 9.737522965 | 3.817230303 | 0.839498142 | -4.547038417 | 5.44E-06 | 6.46E-05 | 3.817230303 |
| ENSMUSG00000076145 | Mir679 | 751539 | miRNA | 0.779082315 | -3.825435563 | 1.399596062 | 2.733242589 | 0.006271412 | 1 | 3.825435563 |
| ENSMUSG00000026866 | Kynu | 70789 | protein_coding | 2.25158955 | -3.833638328 | 1.137856077 | 3.369176827 | 0.00075393 | 1 | 3.833638328 |
| ENSMUSG00000036800 | Fam135b | 70363 | protein_coding | 5.563073501 | 3.836297985 | 2.016352869 | -1.902592569 | 0.05709373 | 0.164631718 | 3.836297985 |
| ENSMUSG00000086514 | Gm11747 | 100126224 | lncRNA | 7.349085309 | 3.836795888 | 0.791061576 | -4.850186137 | 1.23E-06 | 1.67E-05 | 3.836795888 |
| ENSMUSG00000050663 | Trhde | 237553 | protein_coding | 3.067800083 | 3.882054496 | 2.653850858 | -1.462800551 | 0.143521962 | 1 | 3.882054496 |
| ENSMUSG00000030659 | Nucb2 | 53322 | protein_coding | 833.795018 | 3.907121229 | 0.114552338 | -34.10773873 | 5.66E-255 | 5.50E-252 | 3.907121229 |
| ENSMUSG00000058152 | Chsy3 | 78923 | protein_coding | 3.439127211 | 3.921729767 | 1.249030311 | -3.139819533 | 0.001690519 | 1 | 3.921729767 |

Supplementary Table 1

| ensembl_gene_id | external_gene_name | entrezgene_id | gene_biotype | baseMean | log2FoldChange | lfcSE | stat | pvalue | padj | absLog2FC |
|---|---|---|---|---|---|---|---|---|---|---|
| ENSMUSG00000004988 | Fxyd4 | 108017 | protein_coding | 83.2017336 | -3.931769573 | 0.431989376 | 9.101542279 | 8.91E-20 | 4.47E-18 | 3.931769573 |
| ENSMUSG00000049583 | Grm5 | 108071 | protein_coding | 2.093831139 | 3.935968825 | 2.938545872 | -1.339427389 | 0.180431578 | 1 | 3.935968825 |
| ENSMUSG00000040452 | Cdh12 | 215654 | protein_coding | 1.958786976 | 3.93793523 | 2.975526721 | -1.323441393 | 0.185688634 | 1 | 3.93793523 |
| ENSMUSG00000087035 | Tmem74bos | 228767 | protein_coding | 1.191270834 | 4.027070767 | 2.901113475 | -1.388112117 | 0.165102907 | 1 | 4.027070767 |
| ENSMUSG00000060671 | Atp8b2 | 54667 | protein_coding | 496.6384482 | 4.04944735 | 0.104603706 | -38.71227408 | 0 | 0 | 4.04944735 |
| ENSMUSG00000020048 | Hsp90b1 | 22027 | protein_coding | 59760.54841 | 4.054929711 | 0.110367273 | -36.74032707 | 1.66E-295 | 2.28E-292 | 4.054929711 |
| ENSMUSG00000032098 | Treh | 58866 | protein_coding | 79.90698834 | 4.05579021 | 0.285839403 | -14.18905218 | 1.07E-45 | 1.32E-43 | 4.05579021 |
| ENSMUSG00000059891 | Tsks | 22116 | protein_coding | 3.274513272 | 4.064381306 | 1.389442344 | -2.925188889 | 0.003442472 | 1 | 4.064381306 |
| ENSMUSG00000000215 | Ins2 | 16334 | protein_coding | 1.772390915 | 4.066791811 | 2.869914931 | -1.417042633 | 0.156470463 | 1 | 4.066791811 |
| ENSMUSG00000047591 | Mafa | 378435 | protein_coding | 2.112846556 | 4.067167623 | 1.838838179 | -2.211813779 | 0.026979535 | 1 | 4.067167623 |
| ENSMUSG00000047842 | Diras2 | 68203 | protein_coding | 4.314317771 | 4.087707454 | 1.164884594 | -3.509109377 | 0.00044961 | 1 | 4.087707454 |
| ENSMUSG00000085925 | Rtl1 | 353326 | protein_coding | 24.43262334 | -4.134738625 | 0.395626333 | 10.45112085 | 1.45E-25 | 9.38E-24 | 4.134738625 |
| ENSMUSG00000094379 | Vmn1r202 | 171258 | protein_coding | 2.227534596 | 4.138463609 | 2.959307281 | -1.398456874 | 0.161975915 | 1 | 4.138463609 |
| ENSMUSG00000044749 | Abca6 | 76184 | protein_coding | 2.686754494 | 4.13881597 | 2.96711889 | -1.394893876 | 0.163047844 | 1 | 4.13881597 |
| ENSMUSG00000048108 | Tmem72 | 319776 | protein_coding | 10.90615682 | 4.15025903 | 0.944282158 | -4.395147145 | 1.11E-05 | 0.000122689 | 4.15025903 |
| ENSMUSG00000094430 | Smbd1 | 381043 | protein_coding | 83.15669502 | 4.16818348 | 0.236994669 | -17.58766769 | 3.06E-69 | 6.40E-67 | 4.16818348 |
| ENSMUSG00000021268 | Meg3 | 17263 | lncRNA | 287.9468587 | -4.175766366 | 0.256300792 | 16.29244426 | 1.12E-59 | 1.86E-57 | 4.175766366 |
| ENSMUSG00000027737 | Slc7a11 | 26570 | protein_coding | 273.2204548 | 4.180723167 | 0.205228229 | -20.37109216 | 3.02E-92 | 9.06E-90 | 4.180723167 |
| ENSMUSG00000097725 | 4930593A02Rik | 71301 | lncRNA | 2.245280061 | 4.188850243 | 1.264654926 | -3.3122476 | 0.000925496 | 1 | 4.188850243 |
| ENSMUSG00000055193 | Klk15 | 317652 | protein_coding | 3.645113934 | -4.277574446 | 0.921931075 | 4.639798532 | 3.49E-06 | 1 | 4.277574446 |
| ENSMUSG00000032875 | Arhgef17 | 207212 | protein_coding | 251.2761479 | 4.282848538 | 0.174675533 | -24.51888058 | 9.29E-133 | 4.15E-130 | 4.282848538 |
| ENSMUSG00000068341 | Reg3d | 30053 | protein_coding | 7.973044311 | 4.285612835 | 0.789505442 | -5.428224562 | 5.69E-08 | 9.65E-07 | 4.285612835 |
| ENSMUSG00000023272 | Creld2 | 76737 | protein_coding | 5943.83244 | 4.29069154 | 0.12206831 | -35.14992168 | 1.17E-270 | 1.28E-267 | 4.29069154 |
| ENSMUSG00000072720 | Myo18b | 74376 | protein_coding | 49.10286963 | 4.299491807 | 0.389218795 | -11.04646504 | 2.28E-28 | 1.66E-26 | 4.299491807 |
| ENSMUSG00000007783 | Cpt1c | 78070 | protein_coding | 21.34409874 | 4.30389438 | 0.527330097 | -8.161670281 | 3.30E-16 | 1.34E-14 | 4.30389438 |
| ENSMUSG00000029814 | Igf2bp3 | 140488 | protein_coding | 140.8419519 | 4.3207522 | 0.294557043 | -14.66864331 | 1.02E-48 | 1.34E-46 | 4.3207522 |
| ENSMUSG00000026833 | Olfm1 | 56177 | protein_coding | 47.68135729 | 4.321240481 | 0.330192372 | -13.08703911 | 3.91E-39 | 3.89E-37 | 4.321240481 |
| ENSMUSG00000033768 | Nrxn2 | 18190 | protein_coding | 45.06359578 | 4.321630962 | 0.51846812 | -8.335384172 | 7.72E-17 | 3.26E-15 | 4.321630962 |
| ENSMUSG00000020081 | Tacr2 | 21337 | protein_coding | 5.301864875 | 4.323780638 | 0.919348144 | -4.703093888 | 2.56E-06 | 3.25E-05 | 4.323780638 |
| ENSMUSG00000097393 | D030068K23Rik | 628686 | lncRNA | 2.040943382 | 4.331839848 | 3.502723129 | -1.236706325 | 0.21619613 | 1 | 4.331839848 |
| ENSMUSG00000097391 | Mirg | 100040724 | lncRNA | 36.35077887 | -4.369007668 | 0.405739676 | 10.76800698 | 4.87E-27 | 3.43E-25 | 4.369007668 |
| ENSMUSG00000021622 | Ckmt2 | 76722 | protein_coding | 745.7908349 | 4.372820079 | 0.150672951 | -29.02193162 | 3.48E-185 | 2.21E-182 | 4.372820079 |
| ENSMUSG00000101126 | Gm10538 | 100038653 | lncRNA | 2.801567302 | 4.380230648 | 1.063960731 | -4.116910069 | 3.84E-05 | 1 | 4.380230648 |
| ENSMUSG00000054136 | Adm2 | 223780 | protein_coding | 1.729834424 | 4.448707334 | 1.458523237 | -3.050144983 | 0.002287309 | 1 | 4.448707334 |
| ENSMUSG00000025823 | Pdia4 | 12304 | protein_coding | 34193.82101 | 4.504158879 | 0.099246691 | -45.38346634 | 0 | 0 | 4.504158879 |
| ENSMUSG00000053395 | Cacng8 | 81905 | protein_coding | 2.953047103 | 4.543590723 | 1.642707044 | -2.765916625 | 0.005676305 | 1 | 4.543590723 |
| ENSMUSG00000069303 | H2bc24 | 665622 | protein_coding | 2.446340669 | -4.555701871 | 2.344055322 | 1.943512949 | 0.051954212 | 1 | 4.555701871 |
| ENSMUSG00000040856 | Dlk1 | 13386 | protein_coding | 31.60494253 | -4.57398896 | 0.578180771 | 7.911001521 | 2.55E-15 | 9.52E-14 | 4.57398896 |
| ENSMUSG00000029576 | Radil | 231858 | protein_coding | 3.757863904 | 4.591896027 | 1.253789373 | -3.662414218 | 0.000249849 | 1 | 4.591896027 |

Supplementary Table 1

| ensembl_gene_id | external_gene_name | entrezgene_id | gene_biotype | baseMean | log2FoldChange | lfcSE | stat | pvalue | padj | absLog2FC |
|---|---|---|---|---|---|---|---|---|---|---|
| ENSMUSG00000042821 | Snai1 | 20613 | protein_coding | 23.52100224 | 4.616014404 | 0.58982338 | -7.826096018 | 5.03E-15 | 1.83E-13 | 4.616014404 |
| ENSMUSG00000042448 | Hoxd1 | 15429 | protein_coding | 7.645814686 | 4.628251173 | 0.847583752 | -5.460523706 | 4.75E-08 | 8.17E-07 | 4.628251173 |
| ENSMUSG00000076060 | Mir704 | 735289 | miRNA | 2.370790226 | 4.66228559 | 1.107485032 | -4.209795579 | 2.56E-05 | 1 | 4.66228559 |
| ENSMUSG00000098343 | Mir6240 | 102466620 | miRNA | 84.15836697 | 4.708234746 | 1.073463964 | -4.386020306 | 1.15E-05 | 0.00012735 | 4.708234746 |
| ENSMUSG00000027221 | Chst1 | 76969 | protein_coding | 3.37736889 | 4.771634465 | 1.261474241 | -3.782585731 | 0.000155208 | 1 | 4.771634465 |
| ENSMUSG00000031734 | Irx3 | 16373 | protein_coding | 10.81906627 | 4.824045787 | 0.844808289 | -5.710225443 | 1.13E-08 | 2.16E-07 | 4.824045787 |
| ENSMUSG00000070436 | Serpinh1 | 12406 | protein_coding | 59.50824199 | 4.872520578 | 0.411915941 | -11.82891967 | 2.77E-32 | 2.31E-30 | 4.872520578 |
| ENSMUSG00000112035 | Gm30539 | 102632477 | lncRNA | 5.64650565 | 4.876007816 | 1.126888429 | -4.32696591 | 1.51E-05 | 0.00016086 | 4.876007816 |
| ENSMUSG00000041329 | Atp1b2 | 11932 | protein_coding | 3.845489952 | 4.884642499 | 2.128267387 | -2.295126321 | 0.021725886 | 1 | 4.884642499 |
| ENSMUSG00000029697 | Fezf1 | 73191 | protein_coding | 9.49620915 | 5.005238748 | 1.069326539 | -4.680739294 | 2.86E-06 | 3.57E-05 | 5.005238748 |
| ENSMUSG00000029844 | Hoxa1 | 15394 | protein_coding | 58.85934614 | 5.076982057 | 0.372718868 | -13.62147854 | 2.98E-42 | 3.31E-40 | 5.076982057 |
| ENSMUSG00000028972 | Car6 | 12353 | protein_coding | 6.389151187 | 5.11003164 | 1.034846658 | -4.937960227 | 7.89E-07 | 1.12E-05 | 5.11003164 |
| ENSMUSG00000018340 | Anxa6 | 11749 | protein_coding | 1417.445842 | 5.119469989 | 0.111455702 | -45.93277779 | 0 | 0 | 5.119469989 |
| ENSMUSG00000036067 | Slc2a6 | 227659 | protein_coding | 565.2035175 | 5.548418663 | 0.192945942 | -28.7563377 | 7.55E-182 | 4.45E-179 | 5.548418663 |
| ENSMUSG00000036036 | Zfp57 | 22715 | protein_coding | 62.33106599 | 5.830791664 | 0.414430534 | -14.06940655 | 5.86E-45 | 6.96E-43 | 5.830791664 |
| ENSMUSG00000032715 | Trib3 | 228775 | protein_coding | 73.55303011 | 5.837363353 | 0.433752163 | -13.4578311 | 2.77E-41 | 3.01E-39 | 5.837363353 |
| ENSMUSG00000026442 | Nfasc | 269116 | protein_coding | 376.7559273 | 6.144157452 | 0.338221834 | -18.16605798 | 9.58E-74 | 2.23E-71 | 6.144157452 |
| ENSMUSG00000060445 | Sycp2 | 320558 | protein_coding | 27.25336193 | 6.203540691 | 0.756543159 | -8.199850356 | 2.41E-16 | 9.86E-15 | 6.203540691 |
| ENSMUSG00000019789 | Hey2 | 15214 | protein_coding | 25.83898677 | 6.282968506 | 0.736942868 | -8.525719947 | 1.52E-17 | 6.67E-16 | 6.282968506 |
| ENSMUSG00000001120 | Pcbp3 | 59093 | protein_coding | 49.68484743 | 6.462378314 | 0.570805968 | -11.32149745 | 1.03E-29 | 7.93E-28 | 6.462378314 |
| ENSMUSG00000052584 | Serp2 | 72661 | protein_coding | 321.3873242 | 7.399481288 | 0.334588282 | -22.11518361 | 2.26E-108 | 8.67E-106 | 7.399481288 |
| ENSMUSG00000029151 | Slc30a3 | 22784 | protein_coding | 114.4813816 | 7.414473821 | 0.503768938 | -14.71800514 | 4.94E-49 | 6.58E-47 | 7.414473821 |
| ENSMUSG00000017446 | C1qtnf1 | 56745 | protein_coding | 1030.062915 | 7.710628117 | 0.222320727 | -34.68245276 | 1.45E-263 | 1.50E-260 | 7.710628117 |
| ENSMUSG00000098682 | Otx2os1 | 606497 | lncRNA | 72.0894564 | 8.586684591 | 0.930039432 | -9.232602719 | 2.64E-20 | 1.35E-18 | 8.586684591 |
| ENSMUSG00000009092 | Derl3 | 70377 | protein_coding | 6340.735785 | 9.247893935 | 0.177753681 | -52.02645535 | 0 | 0 | 9.247893935 |
| ENSMUSG00000028558 | Calr4 | 108802 | protein_coding | 350.804723 | 10.47456851 | 0.74195555 | -14.11751488 | 2.96E-45 | 3.57E-43 | 10.47456851 |

| ensembl_gene_id | external_gene_name | entrezgene_id | gene_biotype | baseMean | log2FoldChange | lfcSE | stat | pvalue | padj | absLog2FC |
|---|---|---|---|---|---|---|---|---|---|---|
| ENSMUSG00000026380 | Tfcp2l1 | 81879 | protein_coding | 11754.64326 | -1.21E-05 | 0.071476548 | -0.000168855 | 0.999865273 | 0.999895545 | 1.21E-05 |
| ENSMUSG00000046157 | Tmem229b | 268567 | protein_coding | 63.02585261 | 2.20E-05 | 0.168239215 | 0.000130915 | 0.999895545 | 0.999895545 | 2.20E-05 |
| ENSMUSG00000022546 | Gpt | 76282 | protein_coding | 1171.811069 | -5.90E-05 | 0.106183976 | -0.000555958 | 0.99955641 | 0.999680386 | 5.90E-05 |
| ENSMUSG00000011306 | Sugp1 | 70616 | protein_coding | 441.657895 | -5.99E-05 | 0.064579181 | -0.000926982 | 0.999260375 | 0.999454275 | 5.99E-05 |
| ENSMUSG00000041765 | Ubac2 | 68889 | protein_coding | 791.370507 | 7.12E-05 | 0.077648992 | 0.000916984 | 0.999268353 | 0.999454275 | 7.12E-05 |
| ENSMUSG00000021772 | Nkiras1 | 69721 | protein_coding | 163.2385985 | -9.60E-05 | 0.096800315 | -0.000991319 | 0.999209042 | 0.999454275 | 9.60E-05 |
| ENSMUSG00000067851 | Arfgef1 | 211673 | protein_coding | 4780.279692 | -0.000115976 | 0.052708267 | -0.002200331 | 0.998244391 | 0.999077457 | 0.000115976 |
| ENSMUSG00000027875 | Hmgcs2 | 15360 | protein_coding | 28847.52922 | 0.000128146 | 0.076153208 | 0.001682737 | 0.998657371 | 0.99920261 | 0.000128146 |
| ENSMUSG00000064302 | Clasp1 | 76707 | protein_coding | 942.4859238 | -0.000133487 | 0.053593797 | -0.002490711 | 0.998012702 | 0.999077457 | 0.000133487 |
| ENSMUSG00000005774 | Rfx5 | 53970 | protein_coding | 249.5306544 | -0.000152778 | 0.094272347 | -0.001620607 | 0.998706943 | 0.99920261 | 0.000152778 |
| ENSMUSG00000038023 | Atp6v0a2 | 21871 | protein_coding | 3841.516773 | 0.000154578 | 0.049399464 | 0.003129151 | 0.997503303 | 0.998865928 | 0.000154578 |
| ENSMUSG00000032366 | Tpm1 | 22003 | protein_coding | 5572.894198 | 0.000174761 | 0.042656573 | 0.004096938 | 0.996731126 | 0.998588735 | 0.000174761 |
| ENSMUSG00000021892 | Sh3bp5 | 24056 | protein_coding | 160.9328122 | 0.000180989 | 0.13488544 | 0.001341797 | 0.998929401 | 0.99936318 | 0.000180989 |
| ENSMUSG00000033628 | Pik3c3 | 225326 | protein_coding | 1248.986673 | 0.00020322 | 0.052191194 | 0.00389376 | 0.996893237 | 0.998627072 | 0.00020322 |
| ENSMUSG00000041974 | Spidr | 224008 | protein_coding | 271.8335082 | 0.000211538 | 0.082368652 | 0.002568191 | 0.997950883 | 0.999077457 | 0.000211538 |
| ENSMUSG00000047423 | AI837181 | 107242 | protein_coding | 695.7399487 | -0.000211904 | 0.091632161 | -0.00231255 | 0.998154853 | 0.999077457 | 0.000211904 |
| ENSMUSG00000027522 | Stx16 | 228960 | protein_coding | 994.3717352 | -0.000216831 | 0.052188536 | -0.004154758 | 0.996684992 | 0.998588735 | 0.000216831 |
| ENSMUSG00000040565 | Btaf1 | 107182 | protein_coding | 1324.300267 | 0.000223565 | 0.048410454 | 0.004618111 | 0.996315294 | 0.998544326 | 0.000223565 |
| ENSMUSG00000024066 | Xdh | 22436 | protein_coding | 1788.766446 | -0.000265221 | 0.123785858 | -0.002142582 | 0.998290468 | 0.999077457 | 0.000265221 |
| ENSMUSG00000020741 | Cluh | 74148 | protein_coding | 6022.584014 | -0.000280286 | 0.059701236 | -0.00469481 | 0.996254097 | 0.998544326 | 0.000280286 |
| ENSMUSG00000041482 | Piezo2 | 667742 | protein_coding | 54.58723582 | -0.000295643 | 0.165622117 | -0.001785048 | 0.998575738 | 0.999195317 | 0.000295643 |
| ENSMUSG00000022905 | Kpna1 | 16646 | protein_coding | 1318.736395 | -0.000302912 | 0.049431576 | -0.006127899 | 0.995110675 | 0.997828443 | 0.000302912 |
| ENSMUSG00000002221 | Paxip1 | 55982 | protein_coding | 623.8967958 | -0.000330023 | 0.059160089 | -0.005578474 | 0.995549045 | 0.998061632 | 0.000330023 |
| ENSMUSG00000053436 | Mapk14 | 26416 | protein_coding | 1306.104845 | 0.000347677 | 0.042284742 | 0.008222273 | 0.99343965 | 0.996976705 | 0.000347677 |
| ENSMUSG00000031453 | Rasa3 | 19414 | protein_coding | 732.860376 | 0.000358156 | 0.072554541 | 0.004936362 | 0.996061369 | 0.998475988 | 0.000358156 |
| ENSMUSG00000025366 | Esyt1 | 23943 | protein_coding | 870.8850817 | 0.000387152 | 0.063926794 | 0.00605618 | 0.995167897 | 0.997828443 | 0.000387152 |
| ENSMUSG00000046691 | Chtf8 | 214987 | protein_coding | 93.94865187 | -0.000388059 | 0.135874317 | -0.002856016 | 0.997721232 | 0.998960099 | 0.000388059 |
| ENSMUSG00000062647 | Rpl7a | 27176 | protein_coding | 17599.97174 | -0.000394989 | 0.038495549 | -0.010260633 | 0.991813343 | 0.996742599 | 0.000394989 |
| ENSMUSG00000020707 | Rnf135 | 71956 | protein_coding | 167.6783903 | -0.000400349 | 0.09507123 | -0.004211044 | 0.996640083 | 0.998588735 | 0.000400349 |
| ENSMUSG00000041840 | Haus1 | 225745 | protein_coding | 138.4234126 | -0.000418266 | 0.1324879 | -0.003157014 | 0.997481072 | 0.998865928 | 0.000418266 |
| ENSMUSG00000042520 | Ubap2l | 74383 | protein_coding | 1377.64275 | 0.00042534 | 0.049225536 | 0.008640634 | 0.993105857 | 0.996814475 | 0.00042534 |
| ENSMUSG00000003863 | Ppfia3 | 76787 | protein_coding | 845.9599737 | -0.000427279 | 0.145646837 | -0.002933667 | 0.997659276 | 0.998960086 | 0.000427279 |
| ENSMUSG00000028865 | Cd164l2 | 69655 | protein_coding | 9.658006999 | 0.000429984 | 0.390694659 | 0.001100563 | 0.999121878 | 0.999454275 | 0.000429984 |
| ENSMUSG00000093565 | Rab26os | 75614 | lncRNA | 47.30626966 | 0.000430628 | 0.179965303 | 0.002392838 | 0.998090794 | 0.999077457 | 0.000430628 |
| ENSMUSG00000002413 | Braf | 109880 | protein_coding | 723.4896987 | -0.000460328 | 0.060683214 | -0.007585747 | 0.993947508 | 0.996976705 | 0.000460328 |
| ENSMUSG00000071708 | Sms | 20603 | protein_coding | 226.8657504 | 0.000489199 | 0.100933792 | 0.00484673 | 0.996132884 | 0.998485612 | 0.000489199 |
| ENSMUSG00000034453 | Polr3b | 70428 | protein_coding | 790.7529613 | 0.000558377 | 0.064573768 | 0.008647125 | 0.993100678 | 0.996814475 | 0.000558377 |
| ENSMUSG00000022312 | Eif3h | 68135 | protein_coding | 3600.103474 | 0.000566799 | 0.033719716 | 0.016809133 | 0.986588884 | 0.995437687 | 0.000566799 |
| ENSMUSG00000022538 | Lsg1 | 224092 | protein_coding | 464.0898922 | -0.00058928 | 0.068029915 | -0.008662071 | 0.993088754 | 0.996814475 | 0.00058928 |

Supplementary Table 1

| ensembl_gene_id | external_gene_name | entrezgene_id | gene_biotype | baseMean | log2FoldChange | lfcSE | stat | pvalue | padj | absLog2FC |
|---|---|---|---|---|---|---|---|---|---|---|
| ENSMUSG00000057738 | Sptan1 | 20740 | protein_coding | 11347.39028 | 0.000600789 | 0.04161762 | 0.014435918 | 0.988482204 | 0.995768162 | 0.000600789 |
| ENSMUSG00000028639 | Ybx1 | 22608 | protein_coding | 10303.14643 | -0.000613886 | 0.043386659 | -0.014149179 | 0.988710965 | 0.995814509 | 0.000613886 |
| ENSMUSG00000027427 | Polr3f | 70408 | protein_coding | 381.2727915 | 0.000615642 | 0.069031888 | 0.008918226 | 0.99288438 | 0.996814475 | 0.000615642 |
| ENSMUSG00000032369 | Plscr1 | 22038 | protein_coding | 3284.047765 | 0.000630915 | 0.031967621 | 0.019736066 | 0.98425392 | 0.994488063 | 0.000630915 |
| ENSMUSG00000030557 | Mef2a | 17258 | protein_coding | 669.9157798 | 0.000634756 | 0.051864269 | 0.012238794 | 0.990235099 | 0.996133239 | 0.000634756 |
| ENSMUSG00000015846 | Rxra | 20181 | protein_coding | 3161.578839 | 0.000640474 | 0.056517784 | 0.011332265 | 0.990958354 | 0.996376861 | 0.000640474 |
| ENSMUSG00000004285 | Atp6v1f | 66144 | protein_coding | 1276.113512 | -0.000671817 | 0.060398782 | -0.011123025 | 0.991125293 | 0.996376861 | 0.000671817 |
| ENSMUSG00000026988 | Wdsub1 | 72137 | protein_coding | 383.4631974 | 0.000674914 | 0.077126721 | 0.008750721 | 0.993018024 | 0.996814475 | 0.000674914 |
| ENSMUSG00000075000 | Nrbf2 | 641340 | protein_coding | 211.3849624 | 0.000675345 | 0.088773357 | 0.007607514 | 0.99393014 | 0.996976705 | 0.000675345 |
| ENSMUSG00000056167 | Cnot10 | 78893 | protein_coding | 856.6398813 | -0.000677669 | 0.050887916 | -0.013316891 | 0.989374973 | 0.995858831 | 0.000677669 |
| ENSMUSG00000041143 | Tmco4 | 77056 | protein_coding | 1834.900975 | 0.000679374 | 0.054475906 | 0.012471096 | 0.990049763 | 0.996040707 | 0.000679374 |
| ENSMUSG00000024790 | Sac3d1 | 66406 | protein_coding | 302.7697338 | -0.000689468 | 0.099091323 | -0.00695791 | 0.994448436 | 0.997293074 | 0.000689468 |
| ENSMUSG00000035067 | Xkr6 | 219149 | protein_coding | 56.02946711 | -0.00069728 | 0.194658038 | -0.003582076 | 0.997141923 | 0.998690088 | 0.00069728 |
| ENSMUSG00000075706 | Gpx4 | 625249 | protein_coding | 1875.350095 | 0.000699556 | 0.062411619 | 0.011208746 | 0.991056902 | 0.996376861 | 0.000699556 |
| ENSMUSG00000019982 | Myb | 17863 | protein_coding | 2255.890316 | -0.000705347 | 0.05446349 | -0.012950814 | 0.989667034 | 0.995924128 | 0.000705347 |
| ENSMUSG00000035305 | Ror1 | 26563 | protein_coding | 265.4081853 | -0.000708577 | 0.128087082 | -0.005531997 | 0.995586127 | 0.998061632 | 0.000708577 |
| ENSMUSG00000021548 | Ccnh | 66671 | protein_coding | 289.6817721 | 0.000721356 | 0.087324245 | 0.008260664 | 0.993409019 | 0.996976705 | 0.000721356 |
| ENSMUSG00000041815 | Poldip3 | 73826 | protein_coding | 2107.209698 | 0.000762807 | 0.049218619 | 0.015498339 | 0.987634609 | 0.995704119 | 0.000762807 |
| ENSMUSG00000038416 | Cdc16 | 69957 | protein_coding | 836.5785316 | 0.000799524 | 0.058715832 | 0.013616839 | 0.98913567 | 0.995858831 | 0.000799524 |
| ENSMUSG00000063457 | Rps15 | 20054 | protein_coding | 11076.38609 | -0.000801574 | 0.056926368 | -0.014080891 | 0.988765445 | 0.995814509 | 0.000801574 |
| ENSMUSG00000026761 | Orc4 | 26428 | protein_coding | 550.0737892 | -0.000813545 | 0.063238354 | -0.012864733 | 0.989735712 | 0.995924128 | 0.000813545 |
| ENSMUSG00000058239 | Usf2 | 22282 | protein_coding | 469.9276711 | 0.000823907 | 0.085719684 | 0.009611643 | 0.992331137 | 0.996791885 | 0.000823907 |
| ENSMUSG00000057219 | Armc7 | 276905 | protein_coding | 135.0818283 | 0.000824526 | 0.127158281 | 0.006484252 | 0.994826352 | 0.997610035 | 0.000824526 |
| ENSMUSG00000030208 | Emp1 | 13730 | protein_coding | 15700.59051 | -0.000855183 | 0.109375844 | -0.007818759 | 0.993761597 | 0.996976705 | 0.000855183 |
| ENSMUSG00000032185 | Carm1 | 59035 | protein_coding | 524.409919 | 0.000858502 | 0.066817037 | 0.012848552 | 0.989748621 | 0.995924128 | 0.000858502 |
| ENSMUSG00000003345 | Csnk1g2 | 103236 | protein_coding | 2537.695543 | -0.000862321 | 0.055902082 | -0.015425562 | 0.98769267 | 0.995704119 | 0.000862321 |
| ENSMUSG00000028028 | Alpk1 | 71481 | protein_coding | 528.9714794 | -0.000864327 | 0.070840797 | -0.012200971 | 0.990265275 | 0.996133239 | 0.000864327 |
| ENSMUSG00000024366 | Gfra3 | 14587 | protein_coding | 43.24504721 | 0.000870388 | 0.202541851 | 0.004297323 | 0.996571243 | 0.998588735 | 0.000870388 |
| ENSMUSG00000059263 | Usp47 | 74996 | protein_coding | 3849.494114 | 0.000889072 | 0.048122083 | 0.018475348 | 0.985259644 | 0.994696524 | 0.000889072 |
| ENSMUSG00000006019 | Dhx34 | 71723 | protein_coding | 378.6029968 | -0.00089372 | 0.078583452 | -0.011372878 | 0.990925952 | 0.996376861 | 0.00089372 |
| ENSMUSG00000005142 | Man2b1 | 17159 | protein_coding | 3675.859186 | 0.000908646 | 0.061435569 | 0.014790232 | 0.988199532 | 0.995768162 | 0.000908646 |
| ENSMUSG00000024359 | Hspa9 | 15526 | protein_coding | 7800.321488 | 0.000929149 | 0.063325849 | 0.014672509 | 0.988293452 | 0.995768162 | 0.000929149 |
| ENSMUSG00000018160 | Med1 | 19014 | protein_coding | 1441.925936 | 0.000934104 | 0.05566311 | 0.01678138 | 0.986611024 | 0.995437687 | 0.000934104 |
| ENSMUSG00000021033 | Gstz1 | 14874 | protein_coding | 671.0824384 | -0.000943207 | 0.065096837 | -0.014489296 | 0.988439619 | 0.995768162 | 0.000943207 |
| ENSMUSG00000026849 | Tor1a | 30931 | protein_coding | 658.3548079 | 0.000950382 | 0.054552519 | 0.017421409 | 0.986100429 | 0.995233846 | 0.000950382 |
| ENSMUSG00000029759 | Pon3 | 269823 | protein_coding | 1422.037604 | 0.000950405 | 0.058426884 | 0.016266578 | 0.987021721 | 0.995550783 | 0.000950405 |
| ENSMUSG00000032643 | Fhl3 | 14201 | protein_coding | 7.071041189 | -0.000964884 | 0.462118099 | -0.002087961 | 0.99833405 | 0.999077457 | 0.000964884 |
| ENSMUSG00000079523 | Tmsb10 | 19240 | protein_coding | 2864.562861 | 0.000970429 | 0.097917402 | 0.009910685 | 0.992092547 | 0.996791885 | 0.000970429 |
| ENSMUSG00000032397 | Tipin | 66131 | protein_coding | 182.5781554 | 0.000979167 | 0.12624045 | 0.007756366 | 0.993811377 | 0.996976705 | 0.000979167 |

| ensembl_gene_id | external_gene_name | entrezgene_id | gene_biotype | baseMean | log2FoldChange | lfcSE | stat | pvalue | padj | absLog2FC |
|---|---|---|---|---|---|---|---|---|---|---|
| ENSMUSG00000005813 | Metap1 | 75624 | protein_coding | 1398.602137 | 0.000984097 | 0.04593116 | 0.021425483 | 0.982906246 | 0.993782244 | 0.000984097 |
| ENSMUSG00000026589 | Sec16b | 89867 | protein_coding | 992.6831237 | 0.000987343 | 0.049119691 | 0.020100759 | 0.983962994 | 0.994383457 | 0.000987343 |
| ENSMUSG00000024457 | Trim26 | 22670 | protein_coding | 1454.533359 | -0.001008389 | 0.051509758 | -0.019576664 | 0.984381079 | 0.994488063 | 0.001008389 |
| ENSMUSG00000064030 | Pym1 | 78428 | protein_coding | 346.9619857 | 0.001026696 | 0.088216004 | 0.011638428 | 0.990714088 | 0.996376861 | 0.001026696 |
| ENSMUSG00000031862 | Atp13a1 | 170759 | protein_coding | 1930.433393 | 0.001041603 | 0.058362994 | 0.017846985 | 0.985760922 | 0.995078017 | 0.001041603 |
| ENSMUSG00000046185 | Zfp84 | 74352 | protein_coding | 623.8029242 | -0.001070028 | 0.057684457 | -0.018549672 | 0.985200352 | 0.994696524 | 0.001070028 |
| ENSMUSG00000035601 | Trmt10b | 69934 | protein_coding | 219.9716141 | -0.001073061 | 0.090716275 | -0.011828758 | 0.990562237 | 0.996369812 | 0.001073061 |
| ENSMUSG00000038903 | Ccdc68 | 381175 | protein_coding | 607.5004811 | -0.001098253 | 0.078284247 | -0.014029043 | 0.98880681 | 0.995814509 | 0.001098253 |
| ENSMUSG00000035649 | Zcchc7 | 319885 | protein_coding | 659.7974899 | 0.001100138 | 0.106620385 | 0.010318269 | 0.991767358 | 0.996742599 | 0.001100138 |
| ENSMUSG00000037946 | Fgd3 | 30938 | protein_coding | 342.6110418 | 0.001128342 | 0.080907755 | 0.013946033 | 0.988873036 | 0.995814509 | 0.001128342 |
| ENSMUSG00000047804 | Akap10 | 56697 | protein_coding | 813.6157288 | -0.001149055 | 0.045924633 | -0.025020459 | 0.980038644 | 0.992512654 | 0.001149055 |
| ENSMUSG00000005370 | Msh6 | 17688 | protein_coding | 851.9500647 | -0.001160685 | 0.079125655 | -0.014668889 | 0.988296339 | 0.995768162 | 0.001160685 |
| ENSMUSG00000049550 | Clip1 | 56430 | protein_coding | 1319.809676 | -0.001170575 | 0.059239288 | -0.019760104 | 0.984234744 | 0.994488063 | 0.001170575 |
| ENSMUSG00000031487 | Brf2 | 66653 | protein_coding | 146.6319461 | 0.001176739 | 0.105424806 | 0.011161876 | 0.991094296 | 0.996376861 | 0.001176739 |
| ENSMUSG00000024219 | Anks1 | 224650 | protein_coding | 1152.202554 | -0.001199218 | 0.050746437 | -0.023631578 | 0.981146483 | 0.993012764 | 0.001199218 |
| ENSMUSG00000027222 | Pex16 | 18633 | protein_coding | 771.1033814 | -0.001213648 | 0.052160255 | -0.023267684 | 0.981436749 | 0.993012764 | 0.001213648 |
| ENSMUSG00000004667 | Polr2e | 66420 | protein_coding | 995.3631852 | -0.001264243 | 0.055838016 | -0.022641261 | 0.98193643 | 0.993218186 | 0.001264243 |
| ENSMUSG00000054733 | Msra | 110265 | protein_coding | 1060.339061 | -0.001269945 | 0.061079311 | -0.020791739 | 0.983411788 | 0.993888695 | 0.001269945 |
| ENSMUSG00000038331 | Satb2 | 212712 | protein_coding | 3711.925154 | 0.001281624 | 0.075801078 | 0.016907728 | 0.986510228 | 0.995437687 | 0.001281624 |
| ENSMUSG00000018882 | Mrpl45 | 67036 | protein_coding | 875.7065123 | -0.001287105 | 0.056174632 | -0.022912565 | 0.981720017 | 0.993174752 | 0.001287105 |
| ENSMUSG00000078656 | Vps25 | 28084 | protein_coding | 14.1198262 | 0.001379515 | 0.315910634 | 0.00436679 | 0.996515817 | 0.998588735 | 0.001379515 |
| ENSMUSG00000003581 | Rnf215 | 71673 | protein_coding | 407.4063625 | 0.001445959 | 0.069512375 | 0.020801456 | 0.983404037 | 0.993888695 | 0.001445959 |
| ENSMUSG00000059995 | Atxn7l3 | 217218 | protein_coding | 1289.67391 | -0.001453197 | 0.068591612 | -0.021186225 | 0.983097103 | 0.993782244 | 0.001453197 |
| ENSMUSG00000035439 | Haus8 | 76478 | protein_coding | 310.7206883 | -0.001469142 | 0.100290247 | -0.014648904 | 0.988312283 | 0.995768162 | 0.001469142 |
| ENSMUSG00000039737 | Prkrip1 | 66801 | protein_coding | 324.5302456 | 0.001478032 | 0.088949026 | 0.016616615 | 0.986742469 | 0.995445753 | 0.001478032 |
| ENSMUSG00000032743 | Katnip | 233865 | protein_coding | 1109.104742 | -0.001481274 | 0.043885931 | -0.033752814 | 0.973074264 | 0.989206294 | 0.001481274 |
| ENSMUSG00000022792 | Yars2 | 70120 | protein_coding | 314.9582418 | 0.001485373 | 0.093320055 | 0.015916981 | 0.987300623 | 0.995635139 | 0.001485373 |
| ENSMUSG00000020386 | Sar1b | 66397 | protein_coding | 1188.098951 | -0.001500016 | 0.040168812 | -0.037342813 | 0.970211669 | 0.987977748 | 0.001500016 |
| ENSMUSG00000012296 | Tjap1 | 74094 | protein_coding | 306.0110781 | -0.001521391 | 0.071914579 | -0.021155525 | 0.983121592 | 0.993782244 | 0.001521391 |
| ENSMUSG00000069833 | Ahnak | 66395 | protein_coding | 28042.74733 | 0.00153445 | 0.059886911 | 0.025622453 | 0.979558477 | 0.992419874 | 0.00153445 |
| ENSMUSG00000037349 | Nudt22 | 68323 | protein_coding | 422.6651034 | 0.001536847 | 0.073418044 | 0.02093283 | 0.983299238 | 0.993888695 | 0.001536847 |
| ENSMUSG00000042404 | Dennd4b | 229541 | protein_coding | 58.57574249 | -0.001587861 | 0.165965789 | -0.0095674 | 0.992366436 | 0.996791885 | 0.001587861 |
| ENSMUSG00000033124 | Atg9a | 245860 | protein_coding | 986.4001755 | 0.001590501 | 0.063468551 | 0.025059668 | 0.98000737 | 0.992512654 | 0.001590501 |
| ENSMUSG00000041939 | Mvk | 17855 | protein_coding | 883.7542433 | -0.001602989 | 0.117002555 | -0.013700458 | 0.989068958 | 0.995858831 | 0.001602989 |
| ENSMUSG00000037740 | Mrps26 | 99045 | protein_coding | 706.3473377 | 0.001606974 | 0.066907951 | 0.024017681 | 0.980838505 | 0.992886774 | 0.001606974 |
| ENSMUSG00000025226 | Fbxl15 | 68431 | protein_coding | 129.1990097 | 0.00161151 | 0.119681508 | 0.013464988 | 0.989256819 | 0.995858831 | 0.00161151 |
| ENSMUSG00000026608 | Kctd3 | 226823 | protein_coding | 1232.434299 | 0.001617417 | 0.045017586 | 0.035928558 | 0.971339324 | 0.988536579 | 0.001617417 |
| ENSMUSG00000020122 | Egfr | 13649 | protein_coding | 1273.731996 | 0.001623348 | 0.054483809 | 0.029795062 | 0.976230497 | 0.990541665 | 0.001623348 |
| ENSMUSG00000001473 | Tubb6 | 67951 | protein_coding | 104.5760407 | 0.001623426 | 0.123108663 | 0.013186939 | 0.98947865 | 0.995901029 | 0.001623426 |

Supplementary Table 1

| ensembl_gene_id | external_gene_name | entrezgene_id | gene_biotype | baseMean | log2FoldChange | lfcSE | stat | pvalue | padj | absLog2FC |
|---|---|---|---|---|---|---|---|---|---|---|
| ENSMUSG00000003429 | Rps11 | 27207 | protein_coding | 8695.968871 | -0.001639166 | 0.039646266 | -0.041344784 | 0.967021031 | 0.986846159 | 0.001639166 |
| ENSMUSG00000003812 | Dnase2a | 13423 | protein_coding | 169.2910587 | -0.001668937 | 0.114059849 | -0.014632116 | 0.988325677 | 0.995768162 | 0.001668937 |
| ENSMUSG00000039480 | Nt5dc1 | 319638 | protein_coding | 522.215936 | 0.00168836 | 0.072526584 | 0.023279194 | 0.981427568 | 0.993012764 | 0.00168836 |
| ENSMUSG00000032261 | Sh3bgrl2 | 212531 | protein_coding | 4721.044623 | -0.001695585 | 0.037770229 | -0.044892102 | 0.964193312 | 0.985482308 | 0.001695585 |
| ENSMUSG00000020740 | Gga3 | 260302 | protein_coding | 256.2717694 | -0.001697461 | 0.104963186 | -0.016171968 | 0.987097199 | 0.995554504 | 0.001697461 |
| ENSMUSG00000012114 | Med15 | 94112 | protein_coding | 910.2966171 | -0.001709473 | 0.047344733 | -0.036106937 | 0.971197091 | 0.988509005 | 0.001709473 |
| ENSMUSG00000065219 | Snord32a | 27209 | snoRNA | 42.06216601 | -0.001727441 | 0.462191581 | -0.0037375 | 0.997017913 | 0.998662378 | 0.001727441 |
| ENSMUSG00000026205 | Slc23a3 | 22626 | protein_coding | 90.09911208 | -0.001781408 | 0.128031128 | -0.013913863 | 0.988898702 | 0.995814509 | 0.001781408 |
| ENSMUSG00000040774 | Cept1 | 99712 | protein_coding | 2307.868852 | 0.001790117 | 0.062371983 | 0.028700659 | 0.977103331 | 0.990867082 | 0.001790117 |
| ENSMUSG00000028923 | Necap2 | 66147 | protein_coding | 1578.426413 | 0.001808751 | 0.062211769 | 0.029074096 | 0.976805495 | 0.990867082 | 0.001808751 |
| ENSMUSG00000053134 | Supt7l | 72195 | protein_coding | 394.3845298 | 0.001818282 | 0.07381524 | 0.024632881 | 0.980347792 | 0.992621208 | 0.001818282 |
| ENSMUSG00000036061 | Smug1 | 71726 | protein_coding | 450.2211036 | 0.001856402 | 0.063022092 | 0.02945637 | 0.976500616 | 0.990691082 | 0.001856402 |
| ENSMUSG00000030536 | Iqgap1 | 29875 | protein_coding | 10856.57056 | 0.001857129 | 0.061177729 | 0.030356293 | 0.975782902 | 0.990489989 | 0.001857129 |
| ENSMUSG00000021660 | Btf3 | 218490 | protein_coding | 3763.168586 | 0.001866035 | 0.049113965 | 0.037993981 | 0.969692481 | 0.987827196 | 0.001866035 |
| ENSMUSG00000042659 | Arrdc4 | 66412 | protein_coding | 1667.333264 | 0.001899018 | 0.109813292 | 0.017293152 | 0.986202749 | 0.995274827 | 0.001899018 |
| ENSMUSG00000039005 | Tlr4 | 21898 | protein_coding | 757.5086041 | -0.001905813 | 0.06765819 | -0.028168256 | 0.977527955 | 0.991109854 | 0.001905813 |
| ENSMUSG00000029490 | Mfsd7a | 243197 | protein_coding | 343.8921466 | -0.001935392 | 0.145111221 | -0.013337302 | 0.989358688 | 0.995858831 | 0.001935392 |
| ENSMUSG00000025287 | Acot9 | 56360 | protein_coding | 385.2525184 | 0.001935648 | 0.078927178 | 0.024524484 | 0.980434254 | 0.992621208 | 0.001935648 |
| ENSMUSG00000101892 | 9130401M01Rik | 75758 | protein_coding | 116.1769866 | 0.001952592 | 0.128118554 | 0.015240507 | 0.987840306 | 0.995743522 | 0.001952592 |
| ENSMUSG00000036450 | Hif1an | 319594 | protein_coding | 1670.439019 | -0.001954198 | 0.041533257 | -0.047051392 | 0.962472268 | 0.984572804 | 0.001954198 |
| ENSMUSG00000040390 | Map3k10 | 269881 | protein_coding | 270.2060729 | 0.001987673 | 0.099582237 | 0.019960117 | 0.984075188 | 0.994434523 | 0.001987673 |
| ENSMUSG00000095325 | Zfp870 | 240066 | protein_coding | 163.5301616 | -0.001990273 | 0.122315653 | -0.016271613 | 0.987017704 | 0.995550783 | 0.001990273 |
| ENSMUSG00000039087 | Rreb1 | 68750 | protein_coding | 1766.660802 | 0.001997574 | 0.080006842 | 0.024967535 | 0.980080859 | 0.992512654 | 0.001997574 |
| ENSMUSG00000078546 | Zfp995 | 70081 | protein_coding | 244.455019 | 0.002009456 | 0.105879121 | 0.018978778 | 0.984858035 | 0.994664675 | 0.002009456 |
| ENSMUSG00000059291 | Rpl11 | 67025 | protein_coding | 6888.704768 | -0.002012401 | 0.072671315 | -0.027691823 | 0.977907945 | 0.991283179 | 0.002012401 |
| ENSMUSG00000041390 | Mdfic | 16543 | protein_coding | 83.24915771 | 0.002052883 | 0.128625899 | 0.015960104 | 0.98726622 | 0.995635139 | 0.002052883 |
| ENSMUSG00000022812 | Gsk3b | 56637 | protein_coding | 1881.561988 | -0.002073558 | 0.048356534 | -0.042880623 | 0.965796695 | 0.98615951 | 0.002073558 |
| ENSMUSG00000040354 | Mars1 | 216443 | protein_coding | 679.5794216 | -0.00207404 | 0.084457637 | -0.024557167 | 0.980408185 | 0.992621208 | 0.00207404 |
| ENSMUSG00000048787 | Dcun1d3 | 233805 | protein_coding | 221.8555 | 0.002125296 | 0.091296041 | 0.023279167 | 0.98142759 | 0.993012764 | 0.002125296 |
| ENSMUSG00000038212 | Mfsd14b | 66631 | protein_coding | 2258.979896 | 0.002130437 | 0.043308906 | 0.04919167 | 0.960766549 | 0.984076351 | 0.002130437 |
| ENSMUSG00000024925 | Rnaseh2c | 68209 | protein_coding | 382.6051731 | 0.002167329 | 0.11045194 | 0.019622375 | 0.984344614 | 0.994488063 | 0.002167329 |
| ENSMUSG00000027351 | Spred1 | 114715 | protein_coding | 944.3910606 | -0.002187157 | 0.056842108 | -0.038477766 | 0.969306758 | 0.987804745 | 0.002187157 |
| ENSMUSG00000003604 | Aven | 74268 | protein_coding | 556.3996901 | 0.002189158 | 0.066662789 | 0.032839276 | 0.973802757 | 0.989317668 | 0.002189158 |
| ENSMUSG00000027215 | Cd82 | 12521 | protein_coding | 2972.096362 | 0.002235224 | 0.054585077 | 0.040949359 | 0.967336268 | 0.986873905 | 0.002235224 |
| ENSMUSG00000041351 | Rap1gap | 110351 | protein_coding | 3203.868555 | -0.002236174 | 0.074655633 | -0.029953179 | 0.976104394 | 0.990526145 | 0.002236174 |
| ENSMUSG00000023088 | Abcc1 | 17250 | protein_coding | 1121.375765 | -0.002246529 | 0.053071357 | -0.042330351 | 0.96623535 | 0.986357608 | 0.002246529 |
| ENSMUSG00000063320 | 1190007I07Rik | 544717 | protein_coding | 185.367215 | 0.002251578 | 0.122388145 | 0.018397028 | 0.985322124 | 0.994697332 | 0.002251578 |
| ENSMUSG00000029475 | Kdm2b | 30841 | protein_coding | 367.843293 | 0.002259943 | 0.072660494 | 0.031102775 | 0.975187576 | 0.990174853 | 0.002259943 |
| ENSMUSG00000020079 | Supv3l1 | 338359 | protein_coding | 561.2131534 | -0.002261204 | 0.06560613 | -0.034466355 | 0.972505271 | 0.989126672 | 0.002261204 |

| ensembl_gene_id | external_gene_name | entrezgene_id | gene_biotype | baseMean | log2FoldChange | lfcSE | stat | pvalue | padj | absLog2FC |
|---|---|---|---|---|---|---|---|---|---|---|
| ENSMUSG00000039615 | Stub1 | 56424 | protein_coding | 693.1991944 | 0.002320696 | 0.07020065 | 0.033058041 | 0.973628303 | 0.989317668 | 0.002320696 |
| ENSMUSG00000039496 | Cdnf | 227526 | protein_coding | 31.73883782 | -0.002363128 | 0.258218258 | -0.00915167 | 0.992698126 | 0.996814475 | 0.002363128 |
| ENSMUSG00000038286 | Bphl | 68021 | protein_coding | 674.6780808 | -0.00237313 | 0.057982625 | -0.040928303 | 0.967353054 | 0.986873905 | 0.00237313 |
| ENSMUSG00000038418 | Egr1 | 13653 | protein_coding | 6445.216921 | 0.002376746 | 0.101642864 | 0.023383302 | 0.981344525 | 0.993012764 | 0.002376746 |
| ENSMUSG00000025507 | Pidd1 | 57913 | protein_coding | 74.69894114 | 0.002386633 | 0.146833796 | 0.016253973 | 0.987031777 | 0.995550783 | 0.002386633 |
| ENSMUSG00000049090 | Zadh2 | 225791 | protein_coding | 817.0312584 | 0.002411735 | 0.075611158 | 0.031896544 | 0.974554555 | 0.989650608 | 0.002411735 |
| ENSMUSG00000029575 | Mmab | 77697 | protein_coding | 436.1521879 | 0.002412084 | 0.113554532 | 0.021241634 | 0.983052903 | 0.993782244 | 0.002412084 |
| ENSMUSG00000041303 | Gtf3c3 | 98488 | protein_coding | 597.3344818 | 0.002434008 | 0.061894142 | 0.03932534 | 0.968631004 | 0.987552927 | 0.002434008 |
| ENSMUSG00000018398 | Septin8 | 20362 | protein_coding | 957.1035994 | 0.002442205 | 0.049077792 | 0.049761914 | 0.960312117 | 0.983923349 | 0.002442205 |
| ENSMUSG00000020064 | Herc4 | 67345 | protein_coding | 1226.071132 | -0.00246071 | 0.062983402 | -0.039069182 | 0.968835231 | 0.987573843 | 0.00246071 |
| ENSMUSG00000025857 | Dnaaf5 | 433956 | protein_coding | 267.6934646 | 0.00246135 | 0.084802629 | 0.029024449 | 0.976845091 | 0.990867082 | 0.00246135 |
| ENSMUSG00000021221 | Dpf3 | 70127 | protein_coding | 88.57188053 | -0.002471397 | 0.156699773 | -0.015771542 | 0.987416652 | 0.995640723 | 0.002471397 |
| ENSMUSG00000025939 | Ube2w | 66799 | protein_coding | 484.1278789 | 0.002474827 | 0.077361837 | 0.031990277 | 0.974479805 | 0.989637016 | 0.002474827 |
| ENSMUSG00000027739 | Rab33b | 19338 | protein_coding | 392.8861893 | 0.002505682 | 0.068707585 | 0.036468783 | 0.97090857 | 0.988437757 | 0.002505682 |
| ENSMUSG00000034656 | Cacna1a | 12286 | protein_coding | 177.1023236 | 0.002510558 | 0.118431664 | 0.021198371 | 0.983087414 | 0.993782244 | 0.002510558 |
| ENSMUSG00000024608 | Rps14 | 20044 | protein_coding | 13378.14538 | -0.002534027 | 0.060821959 | -0.041663022 | 0.966767333 | 0.986650853 | 0.002534027 |
| ENSMUSG00000001761 | Smo | 319757 | protein_coding | 611.794794 | -0.002538367 | 0.07749971 | -0.032753241 | 0.973871366 | 0.989317668 | 0.002538367 |
| ENSMUSG00000031349 | Nsdhl | 18194 | protein_coding | 910.7261921 | 0.00254693 | 0.08501411 | 0.029958904 | 0.976099828 | 0.990526145 | 0.00254693 |
| ENSMUSG00000006392 | Med8 | 80509 | protein_coding | 749.3392881 | 0.002554166 | 0.058764238 | 0.043464633 | 0.965331157 | 0.986105006 | 0.002554166 |
| ENSMUSG00000034607 | Pof1b | 69693 | protein_coding | 4272.153654 | 0.002564847 | 0.071100538 | 0.036073521 | 0.971223736 | 0.988509005 | 0.002564847 |
| ENSMUSG00000021870 | Slmap | 83997 | protein_coding | 2185.494995 | 0.00258687 | 0.053423864 | 0.048421628 | 0.961380223 | 0.984186478 | 0.00258687 |
| ENSMUSG00000028991 | Mtor | 56717 | protein_coding | 1709.35894 | -0.002591451 | 0.044244655 | -0.058570934 | 0.953293862 | 0.981022852 | 0.002591451 |
| ENSMUSG00000024294 | Mib1 | 225164 | protein_coding | 2923.710874 | -0.00259509 | 0.036638249 | -0.070830077 | 0.943532994 | 0.975407474 | 0.00259509 |
| ENSMUSG00000050350 | Gpr18 | 110168 | protein_coding | 16.04349452 | 0.002609207 | 0.339119585 | 0.007694062 | 0.993861087 | 0.996976705 | 0.002609207 |
| ENSMUSG00000050705 | 2310061I04Rik | 69662 | protein_coding | 404.0587812 | 0.002615354 | 0.076352004 | 0.034253906 | 0.972674681 | 0.989174207 | 0.002615354 |
| ENSMUSG00000030761 | Myo7a | 17921 | protein_coding | 330.6452076 | -0.002624858 | 0.07874501 | -0.033333648 | 0.973408521 | 0.989317668 | 0.002624858 |
| ENSMUSG00000014550 | Rbsn | 78287 | protein_coding | 793.9864116 | -0.002634575 | 0.051244786 | -0.051411567 | 0.958997568 | 0.983542327 | 0.002634575 |
| ENSMUSG00000014956 | Ppp1cb | 19046 | protein_coding | 5990.905221 | -0.0026398 | 0.045243396 | -0.058346644 | 0.953472514 | 0.981022852 | 0.0026398 |
| ENSMUSG00000051444 | Bbs12 | 241950 | protein_coding | 18.61158718 | 0.002640996 | 0.34042832 | 0.007757861 | 0.993810184 | 0.996976705 | 0.002640996 |
| ENSMUSG00000040848 | Sft2d2 | 108735 | protein_coding | 5224.248219 | -0.002641072 | 0.040258305 | -0.065603152 | 0.947693779 | 0.978148441 | 0.002641072 |
| ENSMUSG00000039782 | Cpeb2 | 231207 | protein_coding | 477.9494294 | 0.002686943 | 0.086868476 | 0.030931162 | 0.975324438 | 0.990245355 | 0.002686943 |
| ENSMUSG00000057346 | Apol9a | 223672 | protein_coding | 42.75666991 | 0.002700789 | 0.239883195 | 0.011258768 | 0.991016992 | 0.996376861 | 0.002700789 |
| ENSMUSG00000022641 | Bbx | 70508 | protein_coding | 879.1924625 | 0.002700817 | 0.052990278 | 0.050968165 | 0.959350888 | 0.983542327 | 0.002700817 |
| ENSMUSG00000004044 | Cavin1 | 19285 | protein_coding | 63.87889341 | -0.002701836 | 0.195418946 | -0.013825866 | 0.988968906 | 0.995823024 | 0.002701836 |
| ENSMUSG00000028331 | Trmo | 74753 | protein_coding | 127.5748246 | 0.002705286 | 0.112889187 | 0.02396408 | 0.98088126 | 0.992886774 | 0.002705286 |
| ENSMUSG00000046541 | Zfp526 | 210172 | protein_coding | 287.5893566 | -0.002726304 | 0.070067501 | -0.038909674 | 0.968962404 | 0.98764105 | 0.002726304 |
| ENSMUSG00000051343 | Rab11fip5 | 52055 | protein_coding | 101.5800159 | 0.002748451 | 0.147952083 | 0.018576629 | 0.985178847 | 0.994696524 | 0.002748451 |
| ENSMUSG00000020850 | Prpf8 | 192159 | protein_coding | 5516.10043 | 0.002756997 | 0.064075587 | 0.043027261 | 0.965679802 | 0.986120526 | 0.002756997 |
| ENSMUSG00000021314 | Amph | 218038 | protein_coding | 55.80932304 | -0.002762149 | 0.179746702 | -0.015366896 | 0.987739474 | 0.995704119 | 0.002762149 |

| ensembl_gene_id | external_gene_name | entrezgene_id | gene_biotype | baseMean | log2FoldChange | lfcSE | stat | pvalue | padj | absLog2FC |
|---|---|---|---|---|---|---|---|---|---|---|
| ENSMUSG00000032462 | Pik3cb | 74769 | protein_coding | 367.4809589 | -0.002771492 | 0.0754987 | -0.036709133 | 0.970716926 | 0.988305042 | 0.002771492 |
| ENSMUSG00000027018 | Hat1 | 107435 | protein_coding | 405.6664317 | -0.002776559 | 0.109736115 | -0.025302147 | 0.979813962 | 0.992429328 | 0.002776559 |
| ENSMUSG00000039899 | Fgl2 | 14190 | protein_coding | 1387.616199 | 0.002779094 | 0.099705952 | 0.027872899 | 0.977763524 | 0.991278195 | 0.002779094 |
| ENSMUSG00000015971 | Actr8 | 56249 | protein_coding | 680.0409524 | -0.002784633 | 0.05723966 | -0.048648663 | 0.961199289 | 0.984186478 | 0.002784633 |
| ENSMUSG00000054199 | Gon4l | 76022 | protein_coding | 556.1595944 | -0.002792139 | 0.05425298 | -0.051465166 | 0.958954858 | 0.983542327 | 0.002792139 |
| ENSMUSG00000003721 | Insig2 | 72999 | protein_coding | 507.8037242 | -0.002800177 | 0.058798788 | -0.047623046 | 0.962016665 | 0.984481423 | 0.002800177 |
| ENSMUSG00000032313 | Tmem266 | 244886 | protein_coding | 11.7751275 | -0.002807403 | 0.384200544 | -0.007307129 | 0.994169806 | 0.997075651 | 0.002807403 |
| ENSMUSG00000073910 | Mob3b | 214944 | protein_coding | 579.0317446 | 0.002818683 | 0.083092358 | 0.033922295 | 0.972939115 | 0.989206294 | 0.002818683 |
| ENSMUSG00000045545 | Krt14 | 16664 | protein_coding | 20.59932586 | -0.002839186 | 0.264923071 | -0.010717019 | 0.991449219 | 0.996516146 | 0.002839186 |
| ENSMUSG00000026280 | Atg4b | 66615 | protein_coding | 1118.025193 | 0.002850874 | 0.048767876 | 0.058458022 | 0.953383799 | 0.981022852 | 0.002850874 |
| ENSMUSG00000047810 | Ccdc88b | 78317 | protein_coding | 34.59672349 | -0.002873605 | 0.214087106 | -0.013422599 | 0.989290637 | 0.995858831 | 0.002873605 |
| ENSMUSG00000031163 | Glod5 | 69824 | protein_coding | 669.9779766 | 0.002882312 | 0.063755442 | 0.045208879 | 0.963940817 | 0.985325065 | 0.002882312 |
| ENSMUSG00000020905 | Usp43 | 216835 | protein_coding | 723.59526 | 0.002905215 | 0.075805989 | 0.038324349 | 0.969429077 | 0.987807993 | 0.002905215 |
| ENSMUSG00000018102 | H2bc4 | 68024 | protein_coding | 3855.905773 | 0.002913564 | 0.064944228 | 0.044862557 | 0.964216862 | 0.985482308 | 0.002913564 |
| ENSMUSG00000046679 | C87436 | 232196 | protein_coding | 297.1354016 | 0.002928692 | 0.081611451 | 0.035885796 | 0.971373422 | 0.988536579 | 0.002928692 |
| ENSMUSG00000001847 | Rac1 | 19353 | protein_coding | 6348.716552 | 0.002947968 | 0.047342867 | 0.062268473 | 0.950349035 | 0.979743296 | 0.002947968 |
| ENSMUSG00000006333 | Rps9 | 76846 | protein_coding | 15223.70804 | -0.002960038 | 0.038186136 | -0.077516048 | 0.938213025 | 0.97297734 | 0.002960038 |
| ENSMUSG00000021115 | Vrk1 | 22367 | protein_coding | 349.0779726 | 0.002960707 | 0.088814987 | 0.033335672 | 0.973406907 | 0.989317668 | 0.002960707 |
| ENSMUSG00000026531 | Mptx1 | 66289 | protein_coding | 37542.21937 | -0.00296231 | 0.136290665 | -0.021735239 | 0.982659154 | 0.993688498 | 0.00296231 |
| ENSMUSG00000021643 | Serf1 | 20365 | protein_coding | 148.7954083 | 0.003011611 | 0.133249122 | 0.022601358 | 0.981968261 | 0.993218186 | 0.003011611 |
| ENSMUSG00000026470 | Stx6 | 58244 | protein_coding | 688.2917063 | 0.00301465 | 0.055911486 | 0.053918252 | 0.957000295 | 0.982772999 | 0.00301465 |
| ENSMUSG00000021748 | Pdhb | 68263 | protein_coding | 2054.385807 | -0.003021887 | 0.055124038 | -0.054819759 | 0.956282058 | 0.982572197 | 0.003021887 |
| ENSMUSG00000027900 | Dram2 | 67171 | protein_coding | 3469.474183 | 0.003023299 | 0.06261751 | 0.048282012 | 0.96149149 | 0.984193404 | 0.003023299 |
| ENSMUSG00000057315 | Arhgap24 | 231532 | protein_coding | 398.7307083 | -0.003027956 | 0.0973769 | -0.031095222 | 0.9751936 | 0.990174853 | 0.003027956 |
| ENSMUSG00000046675 | Tmem251 | 320351 | protein_coding | 210.5057612 | 0.003054112 | 0.091820511 | 0.033261766 | 0.973465843 | 0.989317668 | 0.003054112 |
| ENSMUSG00000024960 | Plcb3 | 18797 | protein_coding | 10399.46812 | -0.003069004 | 0.037187384 | -0.082528101 | 0.934226773 | 0.971391049 | 0.003069004 |
| ENSMUSG00000078695 | Cisd3 | 217149 | protein_coding | 2178.657008 | 0.00310569 | 0.04970645 | 0.062480614 | 0.950180099 | 0.979641635 | 0.00310569 |
| ENSMUSG00000029862 | Clcn1 | 12723 | protein_coding | 9.473347663 | -0.003107555 | 0.392600064 | -0.007915321 | 0.993684554 | 0.996976705 | 0.003107555 |
| ENSMUSG00000020737 | Jpt1 | 15374 | protein_coding | 4035.856041 | -0.003117462 | 0.067569614 | -0.046137041 | 0.963201023 | 0.984779784 | 0.003117462 |
| ENSMUSG00000050122 | Vwa3b | 70853 | protein_coding | 33.54286614 | 0.003131455 | 0.215264727 | 0.014546997 | 0.988393585 | 0.995768162 | 0.003131455 |
| ENSMUSG00000036819 | Jmjd4 | 194952 | protein_coding | 313.8025765 | 0.003168609 | 0.0820265 | 0.038629087 | 0.969186111 | 0.987804745 | 0.003168609 |
| ENSMUSG00000020698 | Cct6b | 12467 | protein_coding | 10.60114099 | 0.003205206 | 0.35813232 | 0.008949783 | 0.992859202 | 0.996814475 | 0.003205206 |
| ENSMUSG00000021340 | Gpld1 | 14756 | protein_coding | 265.2403547 | 0.003208076 | 0.090141542 | 0.035589316 | 0.971609828 | 0.988590012 | 0.003208076 |
| ENSMUSG00000026740 | Dnajc1 | 13418 | protein_coding | 665.3816317 | 0.003211441 | 0.052004694 | 0.061752914 | 0.950759601 | 0.979948241 | 0.003211441 |
| ENSMUSG00000022023 | Wbp4 | 22380 | protein_coding | 623.6202941 | 0.00322161 | 0.064319683 | 0.050087472 | 0.960052683 | 0.983921291 | 0.00322161 |
| ENSMUSG00000068523 | Gng5 | 14707 | protein_coding | 1548.697458 | -0.003241255 | 0.06253056 | -0.05183474 | 0.958660374 | 0.983480652 | 0.003241255 |
| ENSMUSG00000020846 | Rflnb | 76566 | protein_coding | 2805.346479 | 0.003253746 | 0.060861667 | 0.053461333 | 0.957364338 | 0.982963945 | 0.003253746 |
| ENSMUSG00000031755 | Bbs2 | 67378 | protein_coding | 247.5762591 | 0.003257395 | 0.097784777 | 0.033311884 | 0.973425877 | 0.989317668 | 0.003257395 |
| ENSMUSG00000024726 | Carnmt1 | 67383 | protein_coding | 217.8037944 | 0.003292959 | 0.10740732 | 0.030658605 | 0.975541804 | 0.990403694 | 0.003292959 |

| ensembl_gene_id | external_gene_name | entrezgene_id | gene_biotype | baseMean | log2FoldChange | lfcSE | stat | pvalue | padj | absLog2FC |
|---|---|---|---|---|---|---|---|---|---|---|
| ENSMUSG00000027164 | Traf6 | 22034 | protein_coding | 759.9804682 | -0.003366071 | 0.046553281 | -0.072305765 | 0.942358577 | 0.974755742 | 0.003366071 |
| ENSMUSG00000021595 | Nsun2 | 28114 | protein_coding | 1730.713585 | -0.003373871 | 0.073173235 | -0.04610799 | 0.963224178 | 0.984779784 | 0.003373871 |
| ENSMUSG00000019088 | Dnase1l1 | 69537 | protein_coding | 299.0227996 | -0.003380789 | 0.078613444 | -0.043005225 | 0.965697369 | 0.986120526 | 0.003380789 |
| ENSMUSG00000028874 | Fgr | 14191 | protein_coding | 29.59075345 | -0.003389926 | 0.232311995 | -0.014592127 | 0.98835758 | 0.995768162 | 0.003389926 |
| ENSMUSG00000051212 | Gpr183 | 321019 | protein_coding | 17.04502735 | -0.003406663 | 0.333913495 | -0.010202231 | 0.991859939 | 0.996742599 | 0.003406663 |
| ENSMUSG00000033557 | Fam20b | 215015 | protein_coding | 4879.827914 | 0.003407306 | 0.0322832 | 0.105544247 | 0.915943962 | 0.962496141 | 0.003407306 |
| ENSMUSG00000020949 | Fkbp3 | 30795 | protein_coding | 314.2183046 | -0.003411916 | 0.085791799 | -0.03976972 | 0.968276717 | 0.987316554 | 0.003411916 |
| ENSMUSG00000004789 | Dlst | 78920 | protein_coding | 4053.479686 | 0.003440301 | 0.045845946 | 0.075040463 | 0.940182518 | 0.974004206 | 0.003440301 |
| ENSMUSG00000031708 | Tecr | 106529 | protein_coding | 2248.525034 | 0.0034539 | 0.062589074 | 0.055183755 | 0.955992071 | 0.982430646 | 0.0034539 |
| ENSMUSG00000037070 | Rbmxl1 | 19656 | protein_coding | 230.4123019 | -0.003466884 | 0.088699794 | -0.039085597 | 0.968822144 | 0.987573843 | 0.003466884 |
| ENSMUSG00000018919 | Tm4sf5 | 75604 | protein_coding | 73.35241992 | -0.00349677 | 0.222003026 | -0.015751004 | 0.987433037 | 0.995640723 | 0.00349677 |
| ENSMUSG00000042558 | Adprs | 100206 | protein_coding | 483.3787787 | -0.00350442 | 0.074426538 | -0.047085622 | 0.962444987 | 0.984572804 | 0.00350442 |
| ENSMUSG00000026356 | Dars | 226414 | protein_coding | 1453.48605 | 0.003515696 | 0.060019628 | 0.058575777 | 0.953290004 | 0.981022852 | 0.003515696 |
| ENSMUSG00000026480 | Ncf2 | 17970 | protein_coding | 711.5264363 | -0.00352661 | 0.066991567 | -0.052642594 | 0.958016679 | 0.983146794 | 0.00352661 |
| ENSMUSG00000017404 | Rpl19 | 19921 | protein_coding | 9931.430153 | -0.003529862 | 0.054353104 | -0.064943157 | 0.948219259 | 0.978324393 | 0.003529862 |
| ENSMUSG00000038844 | Kif16b | 16558 | protein_coding | 2008.994504 | -0.003532403 | 0.038250693 | -0.09234874 | 0.926420965 | 0.967733155 | 0.003532403 |
| ENSMUSG00000029723 | Spacdr | 78829 | protein_coding | 1782.077413 | -0.003566595 | 0.108293753 | -0.032934446 | 0.973726864 | 0.989317668 | 0.003566595 |
| ENSMUSG00000044881 | Coa4 | 68185 | protein_coding | 399.8563174 | -0.003571867 | 0.073848393 | -0.048367559 | 0.961423313 | 0.984186478 | 0.003571867 |
| ENSMUSG00000068749 | Psma5 | 26442 | protein_coding | 1242.792565 | 0.003589303 | 0.070320326 | 0.051042188 | 0.959291904 | 0.983542327 | 0.003589303 |
| ENSMUSG00000023932 | Cdc5l | 71702 | protein_coding | 1542.51961 | 0.003607547 | 0.053548644 | 0.067369539 | 0.946287519 | 0.977252742 | 0.003607547 |
| ENSMUSG00000058908 | Pla2g2a | 18780 | protein_coding | 18.1672606 | -0.00361564 | 0.448912617 | -0.008054217 | 0.993573734 | 0.996976705 | 0.00361564 |
| ENSMUSG00000033400 | Agl | 77559 | protein_coding | 742.8279805 | -0.003629915 | 0.065308174 | -0.055581331 | 0.955675338 | 0.982360082 | 0.003629915 |
| ENSMUSG00000024889 | Rce1 | 19671 | protein_coding | 647.2703219 | 0.003649718 | 0.070675305 | 0.051640642 | 0.958815035 | 0.983514188 | 0.003649718 |
| ENSMUSG00000072966 | Gprasp2 | 245607 | protein_coding | 9.334423081 | -0.003652389 | 0.381842783 | -0.009565165 | 0.992368219 | 0.996791885 | 0.003652389 |
| ENSMUSG00000036023 | Parp2 | 11546 | protein_coding | 504.17717 | -0.003656843 | 0.07797871 | -0.046895401 | 0.962596594 | 0.984637528 | 0.003656843 |
| ENSMUSG00000031536 | Polb | 18970 | protein_coding | 351.498832 | 0.003682856 | 0.073290436 | 0.050250163 | 0.959923038 | 0.98389976 | 0.003682856 |
| ENSMUSG00000046338 | Gpat2 | 215456 | protein_coding | 11.58583641 | 0.003718316 | 0.406036316 | 0.009157594 | 0.992693399 | 0.996814475 | 0.003718316 |
| ENSMUSG00000019809 | Pex3 | 56535 | protein_coding | 362.6984886 | 0.003723074 | 0.074994865 | 0.049644387 | 0.960405775 | 0.983956796 | 0.003723074 |
| ENSMUSG00000028796 | Phc2 | 54383 | protein_coding | 1163.864316 | -0.003732978 | 0.064641695 | -0.057748757 | 0.953948756 | 0.981205596 | 0.003732978 |
| ENSMUSG00000036975 | Tmem177 | 66343 | protein_coding | 215.4892902 | 0.003759263 | 0.087313515 | 0.043054769 | 0.965657875 | 0.986120526 | 0.003759263 |
| ENSMUSG00000051285 | Pcmtd1 | 319263 | protein_coding | 1144.423482 | 0.003768615 | 0.065344436 | 0.057673079 | 0.954009038 | 0.981205596 | 0.003768615 |
| ENSMUSG00000017670 | Elmo2 | 140579 | protein_coding | 327.0622186 | 0.003787102 | 0.089773618 | 0.042185021 | 0.966351203 | 0.986375338 | 0.003787102 |
| ENSMUSG00000028089 | Chd1l | 68058 | protein_coding | 478.7569057 | -0.003793524 | 0.071073386 | -0.05337474 | 0.957433331 | 0.982972201 | 0.003793524 |
| ENSMUSG00000025025 | Mxi1 | 17859 | protein_coding | 1094.725814 | 0.003795184 | 0.067758565 | 0.056010399 | 0.955333523 | 0.982192001 | 0.003795184 |
| ENSMUSG00000024795 | Kif20b | 240641 | protein_coding | 457.2162781 | 0.003796644 | 0.110594073 | 0.03432954 | 0.972614369 | 0.989174207 | 0.003796644 |
| ENSMUSG00000037563 | Rps16 | 20055 | protein_coding | 2687.487713 | 0.00380724 | 0.085010228 | 0.044785669 | 0.964278148 | 0.98548249 | 0.00380724 |
| ENSMUSG00000048922 | Cdca2 | 108912 | protein_coding | 286.2037547 | -0.00384345 | 0.113696841 | -0.033804366 | 0.973033154 | 0.989206294 | 0.00384345 |
| ENSMUSG00000028497 | Hacd4 | 66775 | protein_coding | 46.69915389 | -0.003880678 | 0.219827056 | -0.017653324 | 0.985915417 | 0.99517168 | 0.003880678 |
| ENSMUSG00000027523 | Gnas | 14683 | protein_coding | 378.3664994 | 0.003921972 | 0.201202992 | 0.019492614 | 0.984448129 | 0.994488063 | 0.003921972 |

| ensembl_gene_id | external_gene_name | entrezgene_id | gene_biotype | baseMean | log2FoldChange | lfcSE | stat | pvalue | padj | absLog2FC |
|---|---|---|---|---|---|---|---|---|---|---|
| ENSMUSG00000028291 | Akirin2 | 433693 | protein_coding | 493.6331375 | -0.003941959 | 0.069250335 | -0.056923325 | 0.954606274 | 0.981694642 | 0.003941959 |
| ENSMUSG00000030850 | Ate1 | 11907 | protein_coding | 874.6384533 | 0.00395873 | 0.054226107 | 0.073004141 | 0.941802823 | 0.974493399 | 0.00395873 |
| ENSMUSG00000026439 | Rbbp5 | 213464 | protein_coding | 689.5946119 | -0.004011686 | 0.048866026 | -0.08209561 | 0.934570684 | 0.97153938 | 0.004011686 |
| ENSMUSG00000038058 | Nod1 | 107607 | protein_coding | 410.6481218 | 0.004021853 | 0.067186213 | 0.05986129 | 0.952266111 | 0.98066383 | 0.004021853 |
| ENSMUSG00000029394 | Cdk2ap1 | 13445 | protein_coding | 311.7977171 | -0.004023139 | 0.073832013 | -0.05449044 | 0.956544425 | 0.982572197 | 0.004023139 |
| ENSMUSG00000034685 | Fam171a2 | 217219 | protein_coding | 7.327059926 | -0.004029965 | 0.423271106 | -0.009521003 | 0.992403454 | 0.996791885 | 0.004029965 |
| ENSMUSG00000071669 | Snx29 | 74478 | protein_coding | 143.5494163 | 0.004035044 | 0.119280193 | 0.033828279 | 0.973014086 | 0.989206294 | 0.004035044 |
| ENSMUSG00000031833 | Mast3 | 546071 | protein_coding | 373.5509927 | -0.004051121 | 0.072025571 | -0.056245589 | 0.955146164 | 0.982061982 | 0.004051121 |
| ENSMUSG00000071072 | Ptges3 | 56351 | protein_coding | 1812.707729 | -0.004068695 | 0.052739285 | -0.07714733 | 0.938506341 | 0.97297734 | 0.004068695 |
| ENSMUSG00000024613 | Tcof1 | 21453 | protein_coding | 640.29202 | 0.004127824 | 0.07553751 | 0.054646016 | 0.956420478 | 0.982572197 | 0.004127824 |
| ENSMUSG00000033542 | Arhgef5 | 54324 | protein_coding | 3199.701774 | -0.00413404 | 0.035370126 | -0.116879432 | 0.906955597 | 0.95873684 | 0.00413404 |
| ENSMUSG00000025525 | Apool | 68117 | protein_coding | 698.0183244 | 0.004146278 | 0.053140736 | 0.078024473 | 0.937808586 | 0.97291985 | 0.004146278 |
| ENSMUSG00000033961 | Zfp446 | 269870 | protein_coding | 85.96656364 | 0.004168697 | 0.158545291 | 0.026293415 | 0.979023307 | 0.992064648 | 0.004168697 |
| ENSMUSG00000046791 | Riox1 | 71952 | protein_coding | 292.25153 | -0.004198858 | 0.084282094 | -0.049819103 | 0.960266544 | 0.983923349 | 0.004198858 |
| ENSMUSG00000020652 | Cenpo | 52504 | protein_coding | 190.291192 | -0.00420699 | 0.094632609 | -0.044456027 | 0.964540902 | 0.985688558 | 0.00420699 |
| ENSMUSG00000026491 | Ahctf1 | 226747 | protein_coding | 1606.69868 | -0.004241082 | 0.051081824 | -0.083025271 | 0.933831445 | 0.971167841 | 0.004241082 |
| ENSMUSG00000030834 | Abcc6 | 27421 | protein_coding | 311.5703091 | -0.004265959 | 0.101196712 | -0.042155114 | 0.966375044 | 0.986375338 | 0.004265959 |
| ENSMUSG00000027893 | Ahcyl1 | 229709 | protein_coding | 1934.22033 | -0.004273065 | 0.053179766 | -0.080351325 | 0.935957839 | 0.972062858 | 0.004273065 |
| ENSMUSG00000031781 | Ciapin1 | 109006 | protein_coding | 456.4442863 | -0.00428067 | 0.082238235 | -0.052052063 | 0.95848721 | 0.983365559 | 0.00428067 |
| ENSMUSG00000007411 | Mark3 | 17169 | protein_coding | 1463.420507 | -0.004300511 | 0.041282853 | -0.104171849 | 0.917032974 | 0.963263907 | 0.004300511 |
| ENSMUSG00000029169 | Dhx15 | 13204 | protein_coding | 2975.047443 | -0.004303618 | 0.039240255 | -0.10967354 | 0.912668286 | 0.961085859 | 0.004303618 |
| ENSMUSG00000079465 | Col4a3 | 12828 | protein_coding | 6.921312202 | 0.004304196 | 0.528155306 | 0.008149489 | 0.993497721 | 0.996976705 | 0.004304196 |
| ENSMUSG00000023908 | Pkmyt1 | 268930 | protein_coding | 244.2681478 | 0.004311928 | 0.121984111 | 0.035348277 | 0.971802028 | 0.988660819 | 0.004311928 |
| ENSMUSG00000063334 | Krr1 | 52705 | protein_coding | 573.6675673 | 0.0043348 | 0.062055642 | 0.069853433 | 0.944310318 | 0.975710693 | 0.0043348 |
| ENSMUSG00000102106 | 2310043O21Rik | 69679 | lncRNA | 8.032737612 | -0.004344803 | 0.451762811 | -0.009617443 | 0.992326509 | 0.996791885 | 0.004344803 |
| ENSMUSG00000028132 | Tlcd4 | 99887 | protein_coding | 818.3312909 | -0.004357619 | 0.050744099 | -0.085874407 | 0.931566256 | 0.970125857 | 0.004357619 |
| ENSMUSG00000019718 | L3hypdh | 67217 | protein_coding | 18.1344457 | 0.004388626 | 0.308442583 | 0.014228341 | 0.98864781 | 0.995814509 | 0.004388626 |
| ENSMUSG00000023078 | Cxcl13 | 55985 | protein_coding | 9.186716749 | 0.004392253 | 0.404344572 | 0.010862648 | 0.991333031 | 0.996471562 | 0.004392253 |
| ENSMUSG00000023072 | Cep89 | 72140 | protein_coding | 437.3232289 | -0.004392567 | 0.071763931 | -0.061208566 | 0.951193108 | 0.980184745 | 0.004392567 |
| ENSMUSG00000031600 | Vps37a | 52348 | protein_coding | 933.2468252 | 0.004397294 | 0.053942379 | 0.081518357 | 0.935029727 | 0.971724733 | 0.004397294 |
| ENSMUSG00000024251 | Thada | 240174 | protein_coding | 408.7432581 | -0.004401462 | 0.09348725 | -0.047080882 | 0.962448764 | 0.984572804 | 0.004401462 |
| ENSMUSG00000038534 | Osbpl7 | 71240 | protein_coding | 1029.582329 | -0.00442811 | 0.070627423 | -0.062696753 | 0.950007983 | 0.979588542 | 0.00442811 |
| ENSMUSG00000037119 | Fam91a1 | 210998 | protein_coding | 2636.576853 | -0.004449109 | 0.04960547 | -0.089689895 | 0.928533646 | 0.968593927 | 0.004449109 |
| ENSMUSG00000025779 | Ly96 | 17087 | protein_coding | 74.86793191 | 0.004483941 | 0.149959787 | 0.029900953 | 0.976146045 | 0.990526145 | 0.004483941 |
| ENSMUSG00000044502 | Bod1 | 69556 | protein_coding | 768.1474389 | 0.004512988 | 0.049912256 | 0.090418426 | 0.927954715 | 0.968383271 | 0.004512988 |
| ENSMUSG00000018501 | Ncor1 | 20185 | protein_coding | 5163.05737 | 0.004526002 | 0.030639772 | 0.147716557 | 0.882566464 | 0.947613141 | 0.004526002 |
| ENSMUSG00000009291 | Pttg1ip | 108705 | protein_coding | 5506.426288 | -0.004533418 | 0.049678019 | -0.091256008 | 0.927289173 | 0.968095473 | 0.004533418 |
| ENSMUSG00000022386 | Trmu | 72026 | protein_coding | 231.8850884 | 0.004587323 | 0.087029972 | 0.052709695 | 0.957963214 | 0.983146794 | 0.004587323 |
| ENSMUSG00000028179 | Cth | 107869 | protein_coding | 2135.827816 | -0.00460922 | 0.061847576 | -0.074525468 | 0.940592277 | 0.974178386 | 0.00460922 |

| ensembl_gene_id | external_gene_name | entrezgene_id | gene_biotype | baseMean | log2FoldChange | lfcSE | stat | pvalue | padj | absLog2FC |
|---|---|---|---|---|---|---|---|---|---|---|
| ENSMUSG00000041849 | Card6 | 239319 | protein_coding | 88.30516104 | -0.004618891 | 0.160269471 | -0.028819532 | 0.977008523 | 0.990867082 | 0.004618891 |
| ENSMUSG00000078762 | Haus5 | 71909 | protein_coding | 20.7205249 | 0.004673898 | 0.274256616 | 0.017042062 | 0.98640306 | 0.995414689 | 0.004673898 |
| ENSMUSG00000035284 | Vps13c | 320528 | protein_coding | 1736.929624 | -0.004683516 | 0.094689271 | -0.049461948 | 0.960551162 | 0.984043233 | 0.004683516 |
| ENSMUSG00000020803 | Txndc17 | 52700 | protein_coding | 2267.304276 | -0.004691368 | 0.045185569 | -0.103824466 | 0.917308651 | 0.963427979 | 0.004691368 |
| ENSMUSG00000028797 | Tmem234 | 76799 | protein_coding | 2211.182036 | 0.004712252 | 0.0392851 | 0.119950118 | 0.904522663 | 0.95823668 | 0.004712252 |
| ENSMUSG00000045968 | Teddm2 | 240817 | protein_coding | 18.13921942 | 0.004724072 | 0.301320771 | 0.015677882 | 0.987491372 | 0.995640723 | 0.004724072 |
| ENSMUSG00000057230 | Aak1 | 269774 | protein_coding | 909.5781963 | 0.004726204 | 0.074702851 | 0.06326672 | 0.949554117 | 0.979309282 | 0.004726204 |
| ENSMUSG00000054693 | Adam10 | 11487 | protein_coding | 2501.420079 | -0.004735461 | 0.051727058 | -0.091547079 | 0.927057899 | 0.968095473 | 0.004735461 |
| ENSMUSG00000042073 | Abhd14b | 76491 | protein_coding | 731.2999692 | 0.004743612 | 0.063862487 | 0.074278529 | 0.940788761 | 0.974240941 | 0.004743612 |
| ENSMUSG00000055963 | Triqk | 208820 | protein_coding | 175.0862862 | -0.004748623 | 0.102553748 | -0.046303753 | 0.963068148 | 0.98474512 | 0.004748623 |
| ENSMUSG00000029416 | Slc15a4 | 100561 | protein_coding | 420.3381091 | 0.004775708 | 0.100140152 | 0.047690243 | 0.96196311 | 0.984481423 | 0.004775708 |
| ENSMUSG00000028468 | Rgp1 | 242406 | protein_coding | 1574.24844 | -0.004813723 | 0.046916693 | -0.102601488 | 0.918279262 | 0.963947444 | 0.004813723 |
| ENSMUSG00000028857 | Tmem222 | 52174 | protein_coding | 651.884505 | 0.00482125 | 0.054725272 | 0.088099155 | 0.929797868 | 0.96903517 | 0.00482125 |
| ENSMUSG00000025006 | Sorbs1 | 20411 | protein_coding | 295.6495054 | -0.00482459 | 0.104020909 | -0.046380963 | 0.96300661 | 0.984744632 | 0.00482459 |
| ENSMUSG00000005442 | Cic | 71722 | protein_coding | 1073.365929 | -0.004847122 | 0.080509791 | -0.060205378 | 0.951992062 | 0.980585772 | 0.004847122 |
| ENSMUSG00000006273 | Atp6v1b2 | 11966 | protein_coding | 1555.023669 | -0.004877092 | 0.04872172 | -0.100100971 | 0.920264164 | 0.964898263 | 0.004877092 |
| ENSMUSG00000021983 | Atp8a2 | 50769 | protein_coding | 1069.54646 | -0.004879828 | 0.061267193 | -0.079648299 | 0.93651698 | 0.972438509 | 0.004879828 |
| ENSMUSG00000051721 | Wdcp | 238037 | protein_coding | 54.13120297 | -0.004888179 | 0.170717293 | -0.028633178 | 0.97715715 | 0.990867082 | 0.004888179 |
| ENSMUSG00000031590 | Frg1 | 14300 | protein_coding | 1045.103476 | 0.004916617 | 0.066229772 | 0.074235748 | 0.940822802 | 0.974240941 | 0.004916617 |
| ENSMUSG00000067722 | BC003965 | 214489 | protein_coding | 552.0255043 | 0.004921455 | 0.071470163 | 0.068860267 | 0.945100845 | 0.976374805 | 0.004921455 |
| ENSMUSG00000026527 | Rgs7 | 24012 | protein_coding | 10.14005902 | 0.004952926 | 0.383332234 | 0.012920715 | 0.989691048 | 0.995924128 | 0.004952926 |
| ENSMUSG00000023977 | Ubr2 | 224826 | protein_coding | 1650.180557 | 0.004954338 | 0.062597381 | 0.079146087 | 0.936916427 | 0.972619969 | 0.004954338 |
| ENSMUSG00000034445 | Cyb561a3 | 225912 | protein_coding | 543.5201059 | -0.004969864 | 0.070332815 | -0.070662091 | 0.943666692 | 0.975483158 | 0.004969864 |
| ENSMUSG00000029119 | Man2b2 | 17160 | protein_coding | 1230.295988 | -0.004971366 | 0.070831644 | -0.070185661 | 0.944045887 | 0.975687517 | 0.004971366 |
| ENSMUSG00000057367 | Birc2 | 11797 | protein_coding | 846.3148465 | 0.00497476 | 0.067366636 | 0.073846044 | 0.941132889 | 0.974315 | 0.00497476 |
| ENSMUSG00000028729 | Ebna1bp2 | 69072 | protein_coding | 860.4382981 | 0.005004029 | 0.062834788 | 0.079637878 | 0.936525269 | 0.972438509 | 0.005004029 |
| ENSMUSG00000026893 | Gca | 227960 | protein_coding | 300.2758911 | 0.005006621 | 0.08826098 | 0.056725196 | 0.954764103 | 0.981794343 | 0.005006621 |
| ENSMUSG00000056666 | Retsat | 67442 | protein_coding | 2139.748381 | -0.005007752 | 0.042529607 | -0.117747434 | 0.906267781 | 0.958651438 | 0.005007752 |
| ENSMUSG00000000881 | Dlg3 | 53310 | protein_coding | 1418.426983 | -0.005017136 | 0.047595998 | -0.105410877 | 0.916049786 | 0.962544624 | 0.005017136 |
| ENSMUSG00000068747 | Sort1 | 20661 | protein_coding | 12952.96722 | -0.005026595 | 0.041720565 | -0.120482439 | 0.90410099 | 0.958041702 | 0.005026595 |
| ENSMUSG00000040010 | Slc7a5 | 20539 | protein_coding | 208.2534624 | -0.005037079 | 0.099149686 | -0.050802774 | 0.95948268 | 0.983542327 | 0.005037079 |
| ENSMUSG00000027086 | Fastkd1 | 320720 | protein_coding | 646.0247306 | 0.005046252 | 0.054611755 | 0.09240231 | 0.926378404 | 0.967733155 | 0.005046252 |
| ENSMUSG00000025227 | Mfsd13a | 75146 | protein_coding | 283.8542199 | -0.005056182 | 0.077788315 | -0.064999254 | 0.948174594 | 0.978324393 | 0.005056182 |
| ENSMUSG00000029076 | Sdf4 | 20318 | protein_coding | 4177.616894 | 0.005075558 | 0.04282845 | 0.118509019 | 0.905664349 | 0.958526272 | 0.005075558 |
| ENSMUSG00000020402 | Vdac1 | 22333 | protein_coding | 6796.538933 | 0.005085033 | 0.045775629 | 0.111086033 | 0.911548124 | 0.960412417 | 0.005085033 |
| ENSMUSG00000035824 | Tk2 | 57813 | protein_coding | 339.4255843 | -0.005115085 | 0.079598271 | -0.064261258 | 0.948762201 | 0.978805528 | 0.005115085 |
| ENSMUSG00000035078 | Mtmr9 | 210376 | protein_coding | 693.2998145 | 0.005122729 | 0.07653998 | 0.066928799 | 0.946638387 | 0.9774899 | 0.005122729 |
| ENSMUSG00000030835 | Nomo1 | 211548 | protein_coding | 2484.722659 | 0.005149472 | 0.043554681 | 0.11823005 | 0.905885381 | 0.958628969 | 0.005149472 |
| ENSMUSG00000026181 | Ppm1f | 68606 | protein_coding | 166.8409865 | 0.005158475 | 0.11073805 | 0.046582678 | 0.962845838 | 0.984687419 | 0.005158475 |

| ensembl_gene_id | external_gene_name | entrezgene_id | gene_biotype | baseMean | log2FoldChange | lfcSE | stat | pvalue | padj | absLog2FC |
|---|---|---|---|---|---|---|---|---|---|---|
| ENSMUSG00000039244 | Ints15 | 231868 | protein_coding | 402.4850662 | 0.005158947 | 0.098173643 | 0.052549204 | 0.95809109 | 0.983146794 | 0.005158947 |
| ENSMUSG00000034473 | Sec22a | 317717 | protein_coding | 626.9845342 | -0.005175908 | 0.066823484 | -0.07745642 | 0.938260459 | 0.97297734 | 0.005175908 |
| ENSMUSG00000034042 | Gpbp1l1 | 77110 | protein_coding | 3984.374111 | -0.005195138 | 0.030009031 | -0.173119154 | 0.86255777 | 0.937787345 | 0.005195138 |
| ENSMUSG00000018012 | Rac3 | 170758 | protein_coding | 89.76455595 | 0.005201577 | 0.126561253 | 0.041099283 | 0.967216746 | 0.986873905 | 0.005201577 |
| ENSMUSG00000030534 | Vps33b | 233405 | protein_coding | 459.8272831 | -0.00526021 | 0.062429537 | -0.084258355 | 0.932851023 | 0.970732425 | 0.00526021 |
| ENSMUSG00000033793 | Atp6v1h | 108664 | protein_coding | 1385.082926 | 0.005263944 | 0.05116428 | 0.102883181 | 0.918055687 | 0.963937797 | 0.005263944 |
| ENSMUSG00000051890 | Klhdc1 | 271005 | protein_coding | 77.77252059 | -0.005265537 | 0.140998033 | -0.037344758 | 0.970210119 | 0.987977748 | 0.005265537 |
| ENSMUSG00000071074 | Yipf3 | 28064 | protein_coding | 961.1332651 | -0.005272246 | 0.074240667 | -0.071015601 | 0.943385339 | 0.97537988 | 0.005272246 |
| ENSMUSG00000070426 | Rnf121 | 75212 | protein_coding | 744.8418932 | 0.00532126 | 0.05586781 | 0.095247344 | 0.924118366 | 0.966739549 | 0.00532126 |
| ENSMUSG00000046836 | Brox | 71678 | protein_coding | 1269.708098 | -0.005321807 | 0.048538901 | -0.109640039 | 0.912694855 | 0.961085859 | 0.005321807 |
| ENSMUSG00000030271 | Ogg1 | 18294 | protein_coding | 172.1453066 | 0.005329973 | 0.097841594 | 0.054475532 | 0.956556302 | 0.982572197 | 0.005329973 |
| ENSMUSG00000024560 | Cxxc1 | 74322 | protein_coding | 1017.016512 | 0.00533706 | 0.063633111 | 0.083872381 | 0.933157899 | 0.970843006 | 0.00533706 |
| ENSMUSG00000048992 | Prss32 | 69814 | protein_coding | 12108.1062 | -0.005357824 | 0.067654373 | -0.079194056 | 0.936878273 | 0.972619969 | 0.005357824 |
| ENSMUSG00000049939 | Lrrc4 | 192198 | protein_coding | 7.902610027 | 0.005381834 | 0.479486529 | 0.011224161 | 0.991044604 | 0.996376861 | 0.005381834 |
| ENSMUSG00000020541 | Tom1l1 | 71943 | protein_coding | 1652.856315 | 0.005398141 | 0.040285957 | 0.133995609 | 0.893406047 | 0.952355017 | 0.005398141 |
| ENSMUSG00000031828 | Klhl36 | 234796 | protein_coding | 130.2941204 | 0.005403564 | 0.113201702 | 0.047733944 | 0.961928282 | 0.984481423 | 0.005403564 |
| ENSMUSG00000029836 | Cbx3 | 12417 | protein_coding | 485.5526053 | -0.005461956 | 0.092070053 | -0.059323917 | 0.952694112 | 0.980775949 | 0.005461956 |
| ENSMUSG00000028670 | Lypla2 | 26394 | protein_coding | 1660.798664 | 0.005490099 | 0.082392321 | 0.066633624 | 0.946873378 | 0.977654576 | 0.005490099 |
| ENSMUSG00000050994 | Adgb | 215772 | protein_coding | 7.088935564 | -0.005493614 | 0.554796611 | -0.009902033 | 0.99209945 | 0.996791885 | 0.005493614 |
| ENSMUSG00000040006 | Ginm1 | 215751 | protein_coding | 1412.412525 | -0.005502526 | 0.043370125 | -0.126873655 | 0.899040398 | 0.955322545 | 0.005502526 |
| ENSMUSG00000032047 | Acat1 | 110446 | protein_coding | 4547.879751 | -0.005511861 | 0.051100269 | -0.107863648 | 0.914103853 | 0.961718903 | 0.005511861 |
| ENSMUSG00000024105 | Themis3 | 74556 | protein_coding | 1306.072794 | -0.005556861 | 0.07486832 | -0.074221791 | 0.940833907 | 0.974240941 | 0.005556861 |
| ENSMUSG00000020109 | Dnajb12 | 56709 | protein_coding | 1373.455602 | -0.005573064 | 0.061764416 | -0.090230986 | 0.928103661 | 0.968383271 | 0.005573064 |
| ENSMUSG00000047141 | Zfp654 | 72020 | protein_coding | 630.1082841 | 0.00557657 | 0.064829264 | 0.086019325 | 0.931451055 | 0.970068528 | 0.00557657 |
| ENSMUSG00000020941 | Map3k14 | 53859 | protein_coding | 178.7296387 | -0.005593645 | 0.102049961 | -0.054812805 | 0.956287598 | 0.982572197 | 0.005593645 |
| ENSMUSG00000021557 | Agtpbp1 | 67269 | protein_coding | 115.6541182 | 0.005607412 | 0.164235798 | 0.034142448 | 0.97276356 | 0.989202215 | 0.005607412 |
| ENSMUSG00000024505 | Dtwd2 | 68857 | protein_coding | 78.69131636 | 0.005609584 | 0.155139241 | 0.036158381 | 0.971156071 | 0.988509005 | 0.005609584 |
| ENSMUSG00000040681 | Hmgn1 | 15312 | protein_coding | 1334.651235 | -0.005610552 | 0.065768493 | -0.085307592 | 0.932016856 | 0.970344469 | 0.005610552 |
| ENSMUSG00000019710 | Mrpl24 | 67707 | protein_coding | 1118.303472 | 0.005625076 | 0.045780426 | 0.122870766 | 0.902209436 | 0.956874924 | 0.005625076 |
| ENSMUSG00000039313 | Minar1 | 209743 | protein_coding | 12.16256732 | 0.00562987 | 0.518879098 | 0.010850062 | 0.991343073 | 0.996471562 | 0.00562987 |
| ENSMUSG00000079610 | Ankrd39 | 109346 | protein_coding | 77.28136093 | 0.005647455 | 0.1603188 | 0.035226408 | 0.971899205 | 0.988697312 | 0.005647455 |
| ENSMUSG00000036242 | Armh4 | 67419 | protein_coding | 194.5977947 | -0.005678561 | 0.093997367 | -0.060411921 | 0.951827565 | 0.980585772 | 0.005678561 |
| ENSMUSG00000051934 | Spats2 | 72572 | protein_coding | 272.4168841 | 0.005708931 | 0.080500698 | 0.070917781 | 0.943463191 | 0.975397839 | 0.005708931 |
| ENSMUSG00000040738 | Ints8 | 72656 | protein_coding | 681.5915757 | 0.005716057 | 0.061354731 | 0.093164074 | 0.925773215 | 0.967733155 | 0.005716057 |
| ENSMUSG00000061462 | Obscn | 380698 | protein_coding | 10.72755727 | -0.005724735 | 0.649871105 | -0.008809031 | 0.992971501 | 0.996814475 | 0.005724735 |
| ENSMUSG00000009630 | Ppp2cb | 19053 | protein_coding | 2093.851863 | -0.005739511 | 0.056779293 | -0.101084585 | 0.919483315 | 0.964769513 | 0.005739511 |
| ENSMUSG00000037989 | Wnk2 | 75607 | protein_coding | 350.3441063 | 0.005757186 | 0.114981179 | 0.050070679 | 0.960066065 | 0.983921291 | 0.005757186 |
| ENSMUSG00000006345 | Ggt1 | 14598 | protein_coding | 167.4475109 | 0.005757236 | 0.249597665 | 0.023066066 | 0.981597574 | 0.993113179 | 0.005757236 |
| ENSMUSG00000025538 | Sumf2 | 67902 | protein_coding | 114.8073756 | 0.00576434 | 0.10887896 | 0.052942646 | 0.957777606 | 0.983146794 | 0.00576434 |

| ensembl_gene_id | external_gene_name | entrezgene_id | gene_biotype | baseMean | log2FoldChange | lfcSE | stat | pvalue | padj | absLog2FC |
|---|---|---|---|---|---|---|---|---|---|---|
| ENSMUSG00000042510 | AA986860 | 212439 | protein_coding | 835.7260222 | 0.005764742 | 0.081682809 | 0.070574728 | 0.943736224 | 0.975492506 | 0.005764742 |
| ENSMUSG00000038784 | Cnot4 | 53621 | protein_coding | 350.0175766 | 0.005766732 | 0.068308241 | 0.084422203 | 0.932720755 | 0.970701318 | 0.005766732 |
| ENSMUSG00000057541 | Pus7 | 78697 | protein_coding | 426.0753863 | -0.005771113 | 0.093286577 | -0.061864345 | 0.950670861 | 0.979948241 | 0.005771113 |
| ENSMUSG00000026803 | Ttf1 | 22130 | protein_coding | 586.4944769 | 0.005794814 | 0.063442909 | 0.09133904 | 0.927223199 | 0.968095473 | 0.005794814 |
| ENSMUSG00000025208 | Mrpl43 | 94067 | protein_coding | 576.9221585 | 0.005816787 | 0.06453502 | 0.090133806 | 0.928180885 | 0.968383271 | 0.005816787 |
| ENSMUSG00000022914 | Brwd1 | 93871 | protein_coding | 1710.29353 | -0.005822807 | 0.06542222 | -0.089003509 | 0.929079121 | 0.96881133 | 0.005822807 |
| ENSMUSG00000059119 | Nap1l4 | 17955 | protein_coding | 1590.957255 | -0.005833465 | 0.042502402 | -0.137250233 | 0.890833007 | 0.95112373 | 0.005833465 |
| ENSMUSG00000030706 | Mrpl48 | 52443 | protein_coding | 756.7610318 | -0.005888035 | 0.058673922 | -0.100351814 | 0.920065023 | 0.96489645 | 0.005888035 |
| ENSMUSG00000079557 | Marchf2 | 224703 | protein_coding | 1123.890608 | -0.005888084 | 0.051594947 | -0.114121331 | 0.909141613 | 0.959446759 | 0.005888084 |
| ENSMUSG00000024095 | Hnrnpll | 72692 | protein_coding | 1299.707466 | -0.005896485 | 0.038881245 | -0.151653719 | 0.879460062 | 0.945762765 | 0.005896485 |
| ENSMUSG00000069208 | Zfp825 | 235956 | protein_coding | 97.38151331 | 0.005896617 | 0.147640589 | 0.039938998 | 0.96814176 | 0.987251745 | 0.005896617 |
| ENSMUSG00000035877 | Zhx3 | 320799 | protein_coding | 424.8978733 | -0.005897987 | 0.066619674 | -0.088532211 | 0.929453684 | 0.96881133 | 0.005897987 |
| ENSMUSG00000039529 | Atp8b1 | 54670 | protein_coding | 14225.0637 | 0.005914658 | 0.043401625 | 0.136277347 | 0.891602032 | 0.951425975 | 0.005914658 |
| ENSMUSG00000031731 | Ap1g1 | 11765 | protein_coding | 2139.57543 | 0.005923171 | 0.052685222 | 0.112425659 | 0.910485912 | 0.959960814 | 0.005923171 |
| ENSMUSG00000063801 | Ap3s2 | 11778 | protein_coding | 609.0597297 | 0.005930702 | 0.066998014 | 0.08852056 | 0.929462944 | 0.96881133 | 0.005930702 |
| ENSMUSG00000038400 | Pmepa1 | 65112 | protein_coding | 121.5939774 | 0.005942254 | 0.137503674 | 0.04321524 | 0.965529957 | 0.986120526 | 0.005942254 |
| ENSMUSG00000019837 | Gtf3c6 | 67371 | protein_coding | 670.3070914 | -0.00594293 | 0.073793394 | -0.080534712 | 0.93581199 | 0.971973978 | 0.00594293 |
| ENSMUSG00000004233 | Wars2 | 70560 | protein_coding | 437.3227505 | 0.005944568 | 0.068284328 | 0.087056106 | 0.930626915 | 0.969585908 | 0.005944568 |
| ENSMUSG00000054226 | Tprkb | 69786 | protein_coding | 396.3740475 | -0.005950904 | 0.076973579 | -0.077310998 | 0.938376142 | 0.97297734 | 0.005950904 |
| ENSMUSG00000016028 | Celsr1 | 12614 | protein_coding | 2536.167618 | 0.005956883 | 0.083048281 | 0.071727948 | 0.942818414 | 0.975106302 | 0.005956883 |
| ENSMUSG00000071655 | Ubxn1 | 225896 | protein_coding | 1826.989547 | 0.005957206 | 0.041472819 | 0.143641215 | 0.885783793 | 0.949475732 | 0.005957206 |
| ENSMUSG00000033953 | Ppp3r1 | 19058 | protein_coding | 1296.254714 | 0.005958748 | 0.043026643 | 0.138489732 | 0.889853384 | 0.950815976 | 0.005958748 |
| ENSMUSG00000041133 | Smc1a | 24061 | protein_coding | 2286.241446 | -0.005970248 | 0.082305932 | -0.072537279 | 0.94217434 | 0.974690203 | 0.005970248 |
| ENSMUSG00000024766 | Lipo3 | 381236 | protein_coding | 585.1564816 | 0.005982834 | 0.076349322 | 0.078361325 | 0.937540637 | 0.972883156 | 0.005982834 |
| ENSMUSG00000067148 | Polr1c | 20016 | protein_coding | 993.3938912 | 0.00598634 | 0.056264021 | 0.106397306 | 0.915267131 | 0.962265867 | 0.00598634 |
| ENSMUSG00000022721 | Trmt2a | 15547 | protein_coding | 567.1316987 | 0.005989835 | 0.064777957 | 0.092467173 | 0.926326871 | 0.967733155 | 0.005989835 |
| ENSMUSG00000035868 | Zfp983 | 73229 | protein_coding | 142.9816919 | 0.006000228 | 0.109766208 | 0.054663705 | 0.956406385 | 0.982572197 | 0.006000228 |
| ENSMUSG00000026049 | Tex30 | 75623 | protein_coding | 118.8023846 | -0.006030106 | 0.108889457 | -0.055378232 | 0.955837137 | 0.982430646 | 0.006030106 |
| ENSMUSG00000032015 | Pou2f3 | 18988 | protein_coding | 47.4628815 | -0.006096009 | 0.186466208 | -0.032692297 | 0.973919967 | 0.989317668 | 0.006096009 |
| ENSMUSG00000031077 | Fadd | 14082 | protein_coding | 349.8808156 | -0.006130736 | 0.075990343 | -0.08067783 | 0.935698168 | 0.971973978 | 0.006130736 |
| ENSMUSG00000036112 | Metap2 | 56307 | protein_coding | 2591.999724 | 0.006163073 | 0.055840546 | 0.110369137 | 0.912116628 | 0.960727899 | 0.006163073 |
| ENSMUSG00000036887 | C1qa | 12259 | protein_coding | 36.75796352 | -0.006175594 | 0.257395748 | -0.023992602 | 0.98085851 | 0.992886774 | 0.006175594 |
| ENSMUSG00000029541 | Cyp2w1 | 545817 | protein_coding | 127.6453635 | -0.00617727 | 0.173500501 | -0.035603761 | 0.971598309 | 0.988590012 | 0.00617727 |
| ENSMUSG00000049657 | Zbtb5 | 230119 | protein_coding | 241.7381037 | 0.006187539 | 0.081175799 | 0.076223938 | 0.939240938 | 0.973466491 | 0.006187539 |
| ENSMUSG00000000282 | Mnt | 17428 | protein_coding | 338.6954663 | -0.006208252 | 0.096950486 | -0.064035281 | 0.948942134 | 0.97886592 | 0.006208252 |
| ENSMUSG00000056260 | Lrif1 | 321000 | protein_coding | 309.3418561 | -0.006236693 | 0.097306923 | -0.064093008 | 0.948896169 | 0.97886592 | 0.006236693 |
| ENSMUSG00000039202 | Abhd2 | 54608 | protein_coding | 13390.89582 | -0.006242627 | 0.050974721 | -0.122465146 | 0.902530647 | 0.957069421 | 0.006242627 |
| ENSMUSG00000026826 | Nr4a2 | 18227 | protein_coding | 129.9230514 | -0.006251482 | 0.190648262 | -0.032790659 | 0.973841528 | 0.989317668 | 0.006251482 |
| ENSMUSG00000032393 | Dpp8 | 74388 | protein_coding | 3152.267276 | -0.006278571 | 0.045159367 | -0.139031424 | 0.889425317 | 0.950733597 | 0.006278571 |

| ensembl_gene_id | external_gene_name | entrezgene_id | gene_biotype | baseMean | log2FoldChange | lfcSE | stat | pvalue | padj | absLog2FC |
|---|---|---|---|---|---|---|---|---|---|---|
| ENSMUSG00000026879 | Gsn | 227753 | protein_coding | 5093.651536 | -0.006309044 | 0.106512645 | -0.059232819 | 0.95276667 | 0.980775949 | 0.006309044 |
| ENSMUSG00000038009 | Dnajc22 | 72778 | protein_coding | 1551.51495 | -0.00631637 | 0.051008057 | -0.123830834 | 0.901449218 | 0.956441676 | 0.00631637 |
| ENSMUSG00000042167 | Tent2 | 100715 | protein_coding | 510.6320592 | -0.006317061 | 0.064574133 | -0.097826495 | 0.922070069 | 0.96563322 | 0.006317061 |
| ENSMUSG00000033983 | Coil | 12812 | protein_coding | 175.6018221 | 0.006367972 | 0.118258893 | 0.053847725 | 0.957056486 | 0.982772999 | 0.006367972 |
| ENSMUSG00000026832 | Cytip | 227929 | protein_coding | 256.155809 | -0.006394898 | 0.106712694 | -0.05992631 | 0.952214325 | 0.98066383 | 0.006394898 |
| ENSMUSG00000113204 | Gm46430 | 108168101 | lncRNA | 168.1787354 | -0.006395 | 0.158836624 | -0.040261494 | 0.967884652 | 0.987251745 | 0.006395 |
| ENSMUSG00000053119 | Chmp3 | 66700 | protein_coding | 3483.000181 | 0.006401249 | 0.044883998 | 0.14261762 | 0.886592181 | 0.949785579 | 0.006401249 |
| ENSMUSG00000041915 | Ammecr1l | 225339 | protein_coding | 618.0489396 | 0.006412054 | 0.05543173 | 0.115674804 | 0.907910276 | 0.959122007 | 0.006412054 |
| ENSMUSG00000027397 | Slc20a1 | 20515 | protein_coding | 8494.564857 | 0.00641526 | 0.093499332 | 0.068612894 | 0.945297755 | 0.976480713 | 0.00641526 |
| ENSMUSG00000023032 | Slc4a8 | 59033 | protein_coding | 35.52119206 | 0.006417865 | 0.227535264 | 0.028206023 | 0.977497834 | 0.991109854 | 0.006417865 |
| ENSMUSG00000025036 | Sfxn2 | 94279 | protein_coding | 328.8566605 | -0.006425742 | 0.084719086 | -0.075847632 | 0.93954032 | 0.973589073 | 0.006425742 |
| ENSMUSG00000046034 | Otulin | 432940 | protein_coding | 429.0113638 | -0.00643081 | 0.067916363 | -0.094687198 | 0.924563286 | 0.966993877 | 0.00643081 |
| ENSMUSG00000038495 | Otud7b | 229603 | protein_coding | 1028.728514 | 0.006437615 | 0.053309451 | 0.120759356 | 0.903881643 | 0.957904159 | 0.006437615 |
| ENSMUSG00000035047 | Kri1 | 215194 | protein_coding | 222.7337585 | -0.006466419 | 0.109248315 | -0.059190102 | 0.952800694 | 0.980775949 | 0.006466419 |
| ENSMUSG00000018372 | Cep95 | 320162 | protein_coding | 237.335236 | 0.006475667 | 0.127255807 | 0.050887006 | 0.95941556 | 0.983542327 | 0.006475667 |
| ENSMUSG00000044352 | Sowaha | 237761 | protein_coding | 201.8737439 | 0.006486606 | 0.099912035 | 0.064923173 | 0.94823517 | 0.978324393 | 0.006486606 |
| ENSMUSG00000021981 | Cab39l | 69008 | protein_coding | 942.9451527 | 0.006518155 | 0.059745597 | 0.109098496 | 0.913124368 | 0.961349829 | 0.006518155 |
| ENSMUSG00000047215 | Rpl9 | 20005 | protein_coding | 13449.72019 | 0.006521279 | 0.054738548 | 0.11913503 | 0.905168378 | 0.958435249 | 0.006521279 |
| ENSMUSG00000081534 | Slc48a1 | 67739 | protein_coding | 599.3157332 | -0.006522163 | 0.061142322 | -0.106671816 | 0.915049342 | 0.962265867 | 0.006522163 |
| ENSMUSG00000020841 | Cpd | 12874 | protein_coding | 10240.4945 | 0.006533607 | 0.047692691 | 0.136993874 | 0.891035637 | 0.95113395 | 0.006533607 |
| ENSMUSG00000032411 | Tfdp2 | 211586 | protein_coding | 1518.500227 | -0.006550036 | 0.053137168 | -0.123266566 | 0.901896016 | 0.956772045 | 0.006550036 |
| ENSMUSG00000069539 | Scyl2 | 213326 | protein_coding | 1402.042051 | -0.006557174 | 0.044102599 | -0.148679998 | 0.881806145 | 0.947331698 | 0.006557174 |
| ENSMUSG00000028514 | Usp24 | 329908 | protein_coding | 2188.99998 | 0.006558273 | 0.063831745 | 0.102743132 | 0.91816684 | 0.963947444 | 0.006558273 |
| ENSMUSG00000024588 | Fech | 14151 | protein_coding | 1160.148849 | -0.006585873 | 0.045472773 | -0.144831124 | 0.884844208 | 0.948995857 | 0.006585873 |
| ENSMUSG00000042121 | Ssh1 | 231637 | protein_coding | 239.7356745 | 0.006597424 | 0.097458472 | 0.067694724 | 0.946028649 | 0.977047968 | 0.006597424 |
| ENSMUSG00000061607 | Mdc1 | 240087 | protein_coding | 405.4240501 | -0.00660703 | 0.086479818 | -0.076399674 | 0.939101129 | 0.973407543 | 0.00660703 |
| ENSMUSG00000024247 | Pkdcc | 106522 | protein_coding | 2083.454792 | -0.006607031 | 0.071874685 | -0.091924312 | 0.926758175 | 0.96786874 | 0.006607031 |
| ENSMUSG00000029826 | Zc3hav1 | 78781 | protein_coding | 2233.636711 | 0.006612933 | 0.040078182 | 0.165000828 | 0.868943331 | 0.940691929 | 0.006612933 |
| ENSMUSG00000117545 | Gm30794 | 102632818 | lncRNA | 6.760273536 | -0.006614464 | 0.440413711 | -0.015018751 | 0.988017221 | 0.995768162 | 0.006614464 |
| ENSMUSG00000021598 | Med10 | 28077 | protein_coding | 430.3409868 | 0.006625028 | 0.074555024 | 0.088860917 | 0.929192445 | 0.96881133 | 0.006625028 |
| ENSMUSG00000035585 | Tsen34 | 66078 | protein_coding | 792.6314925 | 0.006665259 | 0.058445642 | 0.114042023 | 0.909204481 | 0.959446759 | 0.006665259 |
| ENSMUSG00000019087 | Atp6ap1 | 54411 | protein_coding | 2449.723934 | 0.006695717 | 0.055159795 | 0.121387642 | 0.903384005 | 0.957533608 | 0.006695717 |
| ENSMUSG00000011179 | Odc1 | 18263 | protein_coding | 4262.508307 | 0.006730577 | 0.071196409 | 0.094535344 | 0.924683908 | 0.967017533 | 0.006730577 |
| ENSMUSG00000020522 | Mfap3 | 216760 | protein_coding | 521.6332616 | 0.006740403 | 0.055834416 | 0.120721292 | 0.903911793 | 0.957904159 | 0.006740403 |
| ENSMUSG00000068921 | Dap3 | 65111 | protein_coding | 1376.653754 | 0.006752024 | 0.05215425 | 0.129462581 | 0.896991632 | 0.954400543 | 0.006752024 |
| ENSMUSG00000015850 | Adamtsl4 | 229595 | protein_coding | 74.10238594 | -0.006777271 | 0.171339752 | -0.039554573 | 0.968448245 | 0.987429022 | 0.006777271 |
| ENSMUSG00000032470 | Mras | 17532 | protein_coding | 30.93562915 | -0.006780486 | 0.22362354 | -0.030320987 | 0.975811059 | 0.990489989 | 0.006780486 |
| ENSMUSG00000029684 | Wasl | 73178 | protein_coding | 1416.503896 | 0.006781836 | 0.0510299 | 0.132899252 | 0.894273058 | 0.952773623 | 0.006781836 |
| ENSMUSG00000045795 | Whamm | 434204 | protein_coding | 423.2190114 | -0.006782649 | 0.067585287 | -0.100356884 | 0.920060998 | 0.96489645 | 0.006782649 |

| ensembl_gene_id | external_gene_name | entrezgene_id | gene_biotype | baseMean | log2FoldChange | lfcSE | stat | pvalue | padj | absLog2FC |
|---|---|---|---|---|---|---|---|---|---|---|
| ENSMUSG00000019945 | Cabcoco1 | 73287 | protein_coding | 78.86171766 | -0.006811754 | 0.141305353 | -0.048205917 | 0.961552134 | 0.984193404 | 0.006811754 |
| ENSMUSG00000047260 | Emc6 | 66048 | protein_coding | 991.8483294 | -0.006824462 | 0.083831924 | -0.081406484 | 0.935118692 | 0.971726183 | 0.006824462 |
| ENSMUSG00000038910 | Plcl2 | 224860 | protein_coding | 2437.021813 | -0.006826999 | 0.058579144 | -0.116543169 | 0.907222076 | 0.958828915 | 0.006826999 |
| ENSMUSG00000033596 | Rfwd3 | 234736 | protein_coding | 1099.937196 | 0.006845477 | 0.066720854 | 0.102598759 | 0.918281428 | 0.963947444 | 0.006845477 |
| ENSMUSG00000047824 | Pygo2 | 68911 | protein_coding | 1032.316463 | 0.006858589 | 0.07186596 | 0.095435855 | 0.923968638 | 0.966739549 | 0.006858589 |
| ENSMUSG00000037408 | Cnnm4 | 94220 | protein_coding | 11187.28379 | -0.006866383 | 0.077567962 | -0.088520864 | 0.929462702 | 0.96881133 | 0.006866383 |
| ENSMUSG00000039680 | Mrps6 | 121022 | protein_coding | 506.7011222 | 0.006871836 | 0.089105141 | 0.077120537 | 0.938527655 | 0.97297734 | 0.006871836 |
| ENSMUSG00000069495 | Epc2 | 227867 | protein_coding | 802.7090769 | 0.006888882 | 0.053562806 | 0.128613168 | 0.897663746 | 0.954463189 | 0.006888882 |
| ENSMUSG00000028098 | Rnf115 | 67845 | protein_coding | 1075.2273 | -0.006895923 | 0.061290574 | -0.112511964 | 0.910417484 | 0.959960814 | 0.006895923 |
| ENSMUSG00000043290 | Zfp784 | 654801 | protein_coding | 12.51028878 | 0.006897438 | 0.368172953 | 0.018734234 | 0.985053118 | 0.994674825 | 0.006897438 |
| ENSMUSG00000034758 | Tle6 | 114606 | protein_coding | 56.56745644 | -0.006907385 | 0.162826835 | -0.04242166 | 0.966162562 | 0.986345738 | 0.006907385 |
| ENSMUSG00000042272 | Sestd1 | 228071 | protein_coding | 1064.17975 | 0.006917749 | 0.0596306 | 0.11601006 | 0.907644569 | 0.95908688 | 0.006917749 |
| ENSMUSG00000019731 | Slc35e1 | 270066 | protein_coding | 2853.98038 | 0.006973308 | 0.043442346 | 0.160518672 | 0.872472514 | 0.942710088 | 0.006973308 |
| ENSMUSG00000034602 | Mon2 | 67074 | protein_coding | 2925.037003 | 0.007005166 | 0.029067709 | 0.240994793 | 0.809559152 | 0.91448208 | 0.007005166 |
| ENSMUSG00000015112 | Slc25a13 | 50799 | protein_coding | 1199.889435 | 0.007019326 | 0.058935784 | 0.119101255 | 0.905195137 | 0.958435249 | 0.007019326 |
| ENSMUSG00000031641 | Cbr4 | 234309 | protein_coding | 478.0456561 | 0.007031667 | 0.078810769 | 0.089222163 | 0.928905352 | 0.968789653 | 0.007031667 |
| ENSMUSG00000021065 | Fut8 | 53618 | protein_coding | 2663.95983 | -0.007044428 | 0.05931566 | -0.118761682 | 0.905464167 | 0.958468846 | 0.007044428 |
| ENSMUSG00000041219 | Arhgap11a | 228482 | protein_coding | 730.5134964 | -0.007060627 | 0.072789034 | -0.097001247 | 0.922725404 | 0.966069983 | 0.007060627 |
| ENSMUSG00000027651 | Rprd1b | 70470 | protein_coding | 714.8083131 | 0.007105867 | 0.059361803 | 0.119704358 | 0.904717348 | 0.95837997 | 0.007105867 |
| ENSMUSG00000021577 | Sdha | 66945 | protein_coding | 11435.91125 | 0.007112932 | 0.035796124 | 0.198706774 | 0.842492125 | 0.929981434 | 0.007112932 |
| ENSMUSG00000032577 | Mapkapk3 | 102626 | protein_coding | 2071.085847 | -0.007118919 | 0.051092685 | -0.139333433 | 0.889186672 | 0.950733597 | 0.007118919 |
| ENSMUSG00000027706 | Sec62 | 69276 | protein_coding | 1759.640372 | 0.007129617 | 0.040437233 | 0.176313177 | 0.860047911 | 0.93678189 | 0.007129617 |
| ENSMUSG00000039828 | Wdr70 | 545085 | protein_coding | 226.3564461 | 0.007162453 | 0.097015131 | 0.073828201 | 0.941147087 | 0.974315 | 0.007162453 |
| ENSMUSG00000027540 | Ptpn1 | 19246 | protein_coding | 672.8117071 | -0.007174458 | 0.07136001 | -0.100538917 | 0.919916488 | 0.96489645 | 0.007174458 |
| ENSMUSG00000021809 | Nudt13 | 67725 | protein_coding | 304.7382874 | -0.007196025 | 0.072955549 | -0.09863574 | 0.921427492 | 0.965302141 | 0.007196025 |
| ENSMUSG00000000826 | Dnajc5 | 13002 | protein_coding | 3311.45584 | -0.007196878 | 0.039870384 | -0.180506858 | 0.856754671 | 0.935731633 | 0.007196878 |
| ENSMUSG00000020349 | Ppp2ca | 19052 | protein_coding | 5582.565026 | -0.007199844 | 0.038299248 | -0.187989167 | 0.850885143 | 0.933103815 | 0.007199844 |
| ENSMUSG00000090862 | Rps13 | 68052 | protein_coding | 5340.033952 | 0.00721387 | 0.05756684 | 0.125312938 | 0.90027581 | 0.955895949 | 0.00721387 |
| ENSMUSG00000002006 | Pdzd4 | 245469 | protein_coding | 32.50950678 | 0.007253531 | 0.218549156 | 0.03318947 | 0.973523496 | 0.989317668 | 0.007253531 |
| ENSMUSG00000025337 | Sbds | 66711 | protein_coding | 1752.285325 | 0.007263176 | 0.042614556 | 0.170438845 | 0.864665025 | 0.938514797 | 0.007263176 |
| ENSMUSG00000037525 | Bcdin3d | 75284 | protein_coding | 127.2202309 | -0.007289107 | 0.1211107 | -0.06018549 | 0.952007902 | 0.980585772 | 0.007289107 |
| ENSMUSG00000022111 | Uchl3 | 50933 | protein_coding | 389.3536078 | 0.007300042 | 0.080057792 | 0.091184649 | 0.927345872 | 0.968095473 | 0.007300042 |
| ENSMUSG00000029387 | Gtf2h3 | 209357 | protein_coding | 666.6355144 | 0.007307476 | 0.0744569 | 0.098143711 | 0.921818178 | 0.965541569 | 0.007307476 |
| ENSMUSG00000039753 | Fbxl5 | 242960 | protein_coding | 1849.338044 | 0.007313617 | 0.066618056 | 0.109784299 | 0.912580444 | 0.961085859 | 0.007313617 |
| ENSMUSG00000026915 | Strbp | 20744 | protein_coding | 3184.169121 | 0.007322063 | 0.040077369 | 0.182698196 | 0.855034832 | 0.935174414 | 0.007322063 |
| ENSMUSG00000027109 | Sp3 | 20687 | protein_coding | 3347.482353 | 0.007326108 | 0.040477807 | 0.180990729 | 0.856374853 | 0.935610703 | 0.007326108 |
| ENSMUSG00000032773 | Chrm1 | 12669 | protein_coding | 157.4542286 | -0.007342247 | 0.132112324 | -0.055575792 | 0.95567975 | 0.982360082 | 0.007342247 |
| ENSMUSG00000031938 | 4931406C07Rik | 70984 | protein_coding | 1456.68651 | -0.007346798 | 0.050407532 | -0.145748023 | 0.884120309 | 0.948596624 | 0.007346798 |
| ENSMUSG00000013495 | Tmem175 | 72392 | protein_coding | 358.0651037 | -0.007396884 | 0.066532672 | -0.111176712 | 0.911476219 | 0.960412417 | 0.007396884 |

| ensembl_gene_id | external_gene_name | entrezgene_id | gene_biotype | baseMean | log2FoldChange | lfcSE | stat | pvalue | padj | absLog2FC |
|---|---|---|---|---|---|---|---|---|---|---|
| ENSMUSG00000063730 | Hsd3b2 | 15493 | protein_coding | 674.5124179 | -0.007398809 | 0.08344998 | -0.088661603 | 0.929350849 | 0.96881133 | 0.007398809 |
| ENSMUSG00000024727 | Trpm6 | 225997 | protein_coding | 2780.023476 | -0.007436124 | 0.088988614 | -0.08356264 | 0.933404172 | 0.970931898 | 0.007436124 |
| ENSMUSG00000049482 | Ctu2 | 66965 | protein_coding | 291.0271638 | -0.007449946 | 0.07551608 | -0.098653777 | 0.92141317 | 0.965302141 | 0.007449946 |
| ENSMUSG00000024590 | Lmnb1 | 16906 | protein_coding | 986.2121552 | 0.007453836 | 0.084753136 | 0.087947616 | 0.929918311 | 0.969098068 | 0.007453836 |
| ENSMUSG00000003970 | Rpl8 | 26961 | protein_coding | 17578.45413 | 0.007463784 | 0.048759435 | 0.153073632 | 0.878340209 | 0.945542552 | 0.007463784 |
| ENSMUSG00000054814 | Usp46 | 69727 | protein_coding | 610.9770046 | 0.007468484 | 0.053636537 | 0.139242474 | 0.889258546 | 0.950733597 | 0.007468484 |
| ENSMUSG00000014856 | Tmem208 | 66320 | protein_coding | 851.9496604 | -0.007472408 | 0.06676305 | -0.111924301 | 0.910883429 | 0.960244284 | 0.007472408 |
| ENSMUSG00000033379 | Atp6v0b | 114143 | protein_coding | 1999.602987 | -0.007479081 | 0.064067182 | -0.116738102 | 0.907067595 | 0.95873684 | 0.007479081 |
| ENSMUSG00000029141 | Slc4a1ap | 20534 | protein_coding | 426.4727698 | -0.007491604 | 0.067310082 | -0.111299872 | 0.911378557 | 0.960412417 | 0.007491604 |
| ENSMUSG00000075232 | Amd1 | 11702 | protein_coding | 748.0632616 | 0.007495812 | 0.064509944 | 0.116196231 | 0.907497024 | 0.958993808 | 0.007495812 |
| ENSMUSG00000037742 | Eef1a1 | 13627 | protein_coding | 144081.2469 | 0.007509912 | 0.060115085 | 0.124925575 | 0.900582471 | 0.955984485 | 0.007509912 |
| ENSMUSG00000000743 | Chmp1a | 234852 | protein_coding | 5184.923432 | -0.007516446 | 0.051065236 | -0.147193008 | 0.882979679 | 0.947930588 | 0.007516446 |
| ENSMUSG00000037613 | Tnfrsf23 | 79201 | protein_coding | 32.04355773 | 0.007517315 | 0.27191676 | 0.027645647 | 0.977944775 | 0.991283179 | 0.007517315 |
| ENSMUSG00000028898 | Trnau1ap | 71787 | protein_coding | 475.4821161 | -0.007540297 | 0.059584933 | -0.126547037 | 0.899298918 | 0.955338492 | 0.007540297 |
| ENSMUSG00000027883 | Gpsm2 | 76123 | protein_coding | 1134.963387 | -0.007542348 | 0.065171588 | -0.115730621 | 0.907866037 | 0.959122007 | 0.007542348 |
| ENSMUSG00000033020 | Polr2f | 69833 | protein_coding | 871.7245282 | 0.007556276 | 0.099296559 | 0.076098062 | 0.939341082 | 0.973498396 | 0.007556276 |
| ENSMUSG00000029624 | Ptcd1 | 71799 | protein_coding | 570.1224166 | 0.007556724 | 0.056215324 | 0.134424634 | 0.893066804 | 0.952330613 | 0.007556724 |
| ENSMUSG00000025508 | Rplp2 | 67186 | protein_coding | 8088.820208 | -0.007567894 | 0.083897827 | -0.090203693 | 0.92812535 | 0.968383271 | 0.007567894 |
| ENSMUSG00000015536 | Mocs2 | 17434 | protein_coding | 808.9360285 | 0.007579278 | 0.052663023 | 0.143920304 | 0.885563402 | 0.949314078 | 0.007579278 |
| ENSMUSG00000015994 | Fnta | 14272 | protein_coding | 1142.034353 | 0.007590019 | 0.046727662 | 0.162430966 | 0.870966488 | 0.941678503 | 0.007590019 |
| ENSMUSG00000049760 | Micos13 | 224904 | protein_coding | 2031.535876 | 0.007600401 | 0.056029194 | 0.135650732 | 0.892097397 | 0.951700386 | 0.007600401 |
| ENSMUSG00000026455 | Klhl12 | 240756 | protein_coding | 479.9754685 | -0.007601445 | 0.057257785 | -0.132758278 | 0.894384552 | 0.952805789 | 0.007601445 |
| ENSMUSG00000098176 | Ccdc166 | 223648 | protein_coding | 164.5070889 | 0.007612584 | 0.131899482 | 0.057715042 | 0.953975612 | 0.981205596 | 0.007612584 |
| ENSMUSG00000038020 | Rapgefl1 | 268480 | protein_coding | 1874.766448 | 0.007638355 | 0.076132934 | 0.100329183 | 0.92008299 | 0.96489645 | 0.007638355 |
| ENSMUSG00000033862 | Cdk10 | 234854 | protein_coding | 435.6324802 | 0.007651667 | 0.067042376 | 0.114131793 | 0.90913332 | 0.959446759 | 0.007651667 |
| ENSMUSG00000053390 | Zfp952 | 240067 | protein_coding | 222.9210544 | 0.007656885 | 0.088966701 | 0.086064612 | 0.931415054 | 0.970068528 | 0.007656885 |
| ENSMUSG00000069206 | Zfp874a | 238692 | protein_coding | 269.40755 | -0.007692009 | 0.109332366 | -0.070354364 | 0.943911614 | 0.975611267 | 0.007692009 |
| ENSMUSG00000036661 | Dennd3 | 105841 | protein_coding | 385.4863485 | 0.007721142 | 0.06086091 | 0.126865369 | 0.899046957 | 0.955322545 | 0.007721142 |
| ENSMUSG00000023707 | Ogfod2 | 66627 | protein_coding | 517.5390707 | 0.00775277 | 0.081919136 | 0.094639309 | 0.924601326 | 0.966993877 | 0.00775277 |
| ENSMUSG00000028657 | Ppt1 | 19063 | protein_coding | 838.547964 | 0.00775434 | 0.055704172 | 0.139205732 | 0.889287579 | 0.950733597 | 0.00775434 |
| ENSMUSG00000012405 | Rpl15 | 66480 | protein_coding | 5229.264587 | -0.007768434 | 0.063794293 | -0.121773182 | 0.903078653 | 0.957335794 | 0.007768434 |
| ENSMUSG00000024312 | Wdr46 | 57315 | protein_coding | 478.2557671 | -0.007811509 | 0.100716793 | -0.077559149 | 0.938178739 | 0.97297734 | 0.007811509 |
| ENSMUSG00000025732 | Mcrip2 | 68241 | protein_coding | 591.7554871 | 0.0078126 | 0.099571181 | 0.078462464 | 0.937460188 | 0.972883156 | 0.0078126 |
| ENSMUSG00000042271 | Nxt2 | 237082 | protein_coding | 505.4183546 | 0.007830408 | 0.060673502 | 0.129058116 | 0.897311663 | 0.954463189 | 0.007830408 |
| ENSMUSG00000031029 | Eif3f | 66085 | protein_coding | 4310.793601 | 0.007832665 | 0.069286192 | 0.113047987 | 0.909992511 | 0.959845275 | 0.007832665 |
| ENSMUSG00000028914 | Casp9 | 12371 | protein_coding | 558.5411452 | -0.00783596 | 0.070313509 | -0.111443173 | 0.911264926 | 0.960412417 | 0.00783596 |
| ENSMUSG00000031309 | Rps6ka3 | 110651 | protein_coding | 2909.804155 | -0.007861748 | 0.078765022 | -0.099812686 | 0.920493036 | 0.965012754 | 0.007861748 |
| ENSMUSG00000031984 | 2810004N23Rik | 66523 | protein_coding | 492.4978964 | -0.007867208 | 0.084774367 | -0.092801736 | 0.926061072 | 0.967733155 | 0.007867208 |
| ENSMUSG00000056486 | Chn1 | 108699 | protein_coding | 8.728587949 | 0.007871509 | 0.448556019 | 0.017548553 | 0.985998999 | 0.995193757 | 0.007871509 |

| ensembl_gene_id | external_gene_name | entrezgene_id | gene_biotype | baseMean | log2FoldChange | lfcSE | stat | pvalue | padj | absLog2FC |
|---|---|---|---|---|---|---|---|---|---|---|
| ENSMUSG00000070002 | Ell | 13716 | protein_coding | 518.4702949 | 0.00787292 | 0.101534084 | 0.077539673 | 0.938194232 | 0.97297734 | 0.00787292 |
| ENSMUSG00000007646 | Rad51c | 114714 | protein_coding | 30.03587976 | 0.007909402 | 0.267968978 | 0.029516111 | 0.97645297 | 0.990691082 | 0.007909402 |
| ENSMUSG00000032328 | Tmem30a | 69981 | protein_coding | 3053.325539 | 0.007928218 | 0.04321254 | 0.183470314 | 0.854429009 | 0.934988261 | 0.007928218 |
| ENSMUSG00000028572 | Hook1 | 77963 | protein_coding | 3971.043145 | -0.007938464 | 0.059074421 | -0.134380731 | 0.893101518 | 0.952330613 | 0.007938464 |
| ENSMUSG00000021716 | Srek1ip1 | 67288 | protein_coding | 558.8685276 | 0.007938777 | 0.057304485 | 0.138536753 | 0.889816225 | 0.950815976 | 0.007938777 |
| ENSMUSG00000033352 | Map2k4 | 26398 | protein_coding | 1114.641972 | 0.007955567 | 0.04706176 | 0.169045251 | 0.865761049 | 0.939297866 | 0.007955567 |
| ENSMUSG00000097290 | 1300002E11Rik | 100043489 | lncRNA | 208.9580578 | -0.007963036 | 0.103967441 | -0.07659163 | 0.938948418 | 0.973350976 | 0.007963036 |
| ENSMUSG00000035151 | Elmod2 | 244548 | protein_coding | 670.9467457 | 0.00797479 | 0.052821935 | 0.150974961 | 0.879995468 | 0.946112461 | 0.00797479 |
| ENSMUSG00000004099 | Dnmt1 | 13433 | protein_coding | 1722.984998 | 0.00801515 | 0.109569538 | 0.073151263 | 0.941685749 | 0.974493399 | 0.00801515 |
| ENSMUSG00000022307 | Oxr1 | 170719 | protein_coding | 1082.495242 | 0.008018403 | 0.048074508 | 0.166791168 | 0.867534368 | 0.939984329 | 0.008018403 |
| ENSMUSG00000016495 | Plgrkt | 67759 | protein_coding | 1804.589363 | -0.008023239 | 0.074195983 | -0.108135759 | 0.913888002 | 0.961718903 | 0.008023239 |
| ENSMUSG00000020024 | Cep83 | 77048 | protein_coding | 448.7322954 | -0.008031807 | 0.095758193 | -0.083875925 | 0.933155081 | 0.970843006 | 0.008031807 |
| ENSMUSG00000047388 | Atmin | 234776 | protein_coding | 803.0228848 | -0.00805549 | 0.059370426 | -0.135681865 | 0.892072785 | 0.951700386 | 0.00805549 |
| ENSMUSG00000021681 | Aggf1 | 66549 | protein_coding | 1161.152579 | -0.008060927 | 0.060435169 | -0.133381386 | 0.893891765 | 0.95260639 | 0.008060927 |
| ENSMUSG00000035365 | Parpbp | 75317 | protein_coding | 129.9731238 | -0.008087062 | 0.13648115 | -0.059254056 | 0.952749755 | 0.980775949 | 0.008087062 |
| ENSMUSG00000034518 | Hmgxb4 | 70823 | protein_coding | 444.7781059 | 0.008100152 | 0.067387883 | 0.120201913 | 0.904323202 | 0.958088312 | 0.008100152 |
| ENSMUSG00000042724 | Map3k9 | 338372 | protein_coding | 539.4111104 | -0.008154142 | 0.069595679 | -0.117164484 | 0.906729711 | 0.95873684 | 0.008154142 |
| ENSMUSG00000029145 | Eif2b4 | 13667 | protein_coding | 726.3278958 | 0.008185848 | 0.069396923 | 0.117956925 | 0.906101788 | 0.958651438 | 0.008185848 |
| ENSMUSG00000026377 | Nifk | 67949 | protein_coding | 655.6930215 | 0.008188754 | 0.075994637 | 0.107754374 | 0.914190536 | 0.961718903 | 0.008188754 |
| ENSMUSG00000038365 | Fbxo25 | 66822 | protein_coding | 2354.958789 | -0.008206931 | 0.053697491 | -0.152836403 | 0.87852729 | 0.945542552 | 0.008206931 |
| ENSMUSG00000032373 | Car12 | 76459 | protein_coding | 1719.349796 | -0.008249467 | 0.079811691 | -0.103361632 | 0.917675962 | 0.963750993 | 0.008249467 |
| ENSMUSG00000058317 | Ube2e2 | 218793 | protein_coding | 164.1993688 | 0.008253982 | 0.100602295 | 0.082045664 | 0.934610401 | 0.97153938 | 0.008253982 |
| ENSMUSG00000039201 | Tbc1d25 | 209815 | protein_coding | 175.5333433 | -0.008255495 | 0.090408061 | -0.091313703 | 0.92724333 | 0.968095473 | 0.008255495 |
| ENSMUSG00000051550 | Zfp579 | 68490 | protein_coding | 101.0317093 | -0.008292404 | 0.129687485 | -0.063941439 | 0.949016856 | 0.978880386 | 0.008292404 |
| ENSMUSG00000021615 | Xrcc4 | 108138 | protein_coding | 182.6197827 | 0.008292962 | 0.089311335 | 0.092854524 | 0.926019134 | 0.967733155 | 0.008292962 |
| ENSMUSG00000021118 | Plek2 | 27260 | protein_coding | 678.6255267 | -0.008330771 | 0.088943986 | -0.093663118 | 0.925376769 | 0.967491161 | 0.008330771 |
| ENSMUSG00000039611 | Pgap4 | 67063 | protein_coding | 1706.338027 | -0.008333452 | 0.059919834 | -0.139076686 | 0.889389551 | 0.950733597 | 0.008333452 |
| ENSMUSG00000096199 | Ptrhd1 | 69709 | protein_coding | 123.5625129 | 0.00837101 | 0.11302723 | 0.074061885 | 0.940961143 | 0.974310135 | 0.00837101 |
| ENSMUSG00000027652 | Ralgapb | 228850 | protein_coding | 2990.282704 | -0.008385306 | 0.046200242 | -0.181499176 | 0.85597578 | 0.935604164 | 0.008385306 |
| ENSMUSG00000024976 | Shoc2 | 56392 | protein_coding | 1946.533952 | -0.008387047 | 0.036682701 | -0.228637662 | 0.81915055 | 0.918795445 | 0.008387047 |
| ENSMUSG00000022860 | Chodl | 246048 | protein_coding | 37.15775076 | 0.008401291 | 0.290617041 | 0.028908459 | 0.976937599 | 0.990867082 | 0.008401291 |
| ENSMUSG00000004849 | Ap1s1 | 11769 | protein_coding | 1370.938289 | 0.008416966 | 0.073890074 | 0.113911996 | 0.909307556 | 0.959446759 | 0.008416966 |
| ENSMUSG00000018707 | Dync1h1 | 13424 | protein_coding | 11361.27253 | -0.008457339 | 0.057553497 | -0.14694744 | 0.883173507 | 0.948012456 | 0.008457339 |
| ENSMUSG00000031066 | Usp11 | 236733 | protein_coding | 61.84948675 | 0.008460865 | 0.1483019 | 0.057051626 | 0.954504071 | 0.981652136 | 0.008460865 |
| ENSMUSG00000000631 | Myo18a | 360013 | protein_coding | 4967.056212 | -0.008487628 | 0.066100076 | -0.128405721 | 0.897827904 | 0.954467075 | 0.008487628 |
| ENSMUSG00000033009 | Ogfod1 | 270086 | protein_coding | 570.4757215 | 0.008509135 | 0.05891448 | 0.144431979 | 0.885159366 | 0.949133318 | 0.008509135 |
| ENSMUSG00000034290 | Nek9 | 217718 | protein_coding | 1369.404442 | -0.008510698 | 0.040682035 | -0.209200407 | 0.834291795 | 0.926479877 | 0.008510698 |
| ENSMUSG00000037239 | Spred3 | 101809 | protein_coding | 8.542695315 | -0.008526121 | 0.511251743 | -0.016676952 | 0.986694334 | 0.995445753 | 0.008526121 |
| ENSMUSG00000044080 | S100a1 | 20193 | protein_coding | 2147.259551 | 0.008568052 | 0.060777946 | 0.140973048 | 0.887891234 | 0.950419616 | 0.008568052 |

| ensembl_gene_id | external_gene_name | entrezgene_id | gene_biotype | baseMean | log2FoldChange | lfcSE | stat | pvalue | padj | absLog2FC |
|---|---|---|---|---|---|---|---|---|---|---|
| ENSMUSG00000063236 | 1110038F14Rik | 117171 | protein_coding | 296.7984991 | 0.008570106 | 0.086797056 | 0.098737286 | 0.921346863 | 0.965302141 | 0.008570106 |
| ENSMUSG00000092118 | Fancf | 100040608 | protein_coding | 54.86084835 | -0.008582728 | 0.164582136 | -0.052148603 | 0.958410286 | 0.983349197 | 0.008582728 |
| ENSMUSG00000029166 | Mapre3 | 100732 | protein_coding | 112.8103491 | -0.008593024 | 0.144381844 | -0.059515962 | 0.952541152 | 0.980775949 | 0.008593024 |
| ENSMUSG00000058835 | Abi1 | 11308 | protein_coding | 2821.708663 | -0.008599637 | 0.031497541 | -0.273025675 | 0.784833479 | 0.899716112 | 0.008599637 |
| ENSMUSG00000024182 | Axin1 | 12005 | protein_coding | 1358.591946 | -0.00860959 | 0.051586806 | -0.166895203 | 0.867452508 | 0.939958785 | 0.00860959 |
| ENSMUSG00000032396 | Dis3l | 213550 | protein_coding | 565.5587709 | -0.008620749 | 0.056400724 | -0.152848197 | 0.878517988 | 0.945542552 | 0.008620749 |
| ENSMUSG00000032036 | Kirrel3 | 67703 | protein_coding | 17.04791071 | 0.008623825 | 0.332850568 | 0.025909 | 0.979329922 | 0.992312996 | 0.008623825 |
| ENSMUSG00000020648 | Dus4l | 71916 | protein_coding | 65.10068618 | 0.00862889 | 0.176205529 | 0.048970595 | 0.96094273 | 0.984093538 | 0.00862889 |
| ENSMUSG00000038217 | Tlcd2 | 380712 | protein_coding | 91.3672586 | -0.008631241 | 0.131544726 | -0.065614499 | 0.947684746 | 0.978148441 | 0.008631241 |
| ENSMUSG00000057193 | Slc44a2 | 68682 | protein_coding | 835.9343377 | 0.008645031 | 0.049600659 | 0.174292662 | 0.861635466 | 0.937561241 | 0.008645031 |
| ENSMUSG00000047517 | Dmbt1 | 12945 | protein_coding | 120136.3745 | -0.008646519 | 0.062985005 | -0.137279006 | 0.890810264 | 0.95112373 | 0.008646519 |
| ENSMUSG00000040940 | Arhgef1 | 16801 | protein_coding | 886.7255431 | 0.008653726 | 0.064670065 | 0.133813466 | 0.893550079 | 0.952419871 | 0.008653726 |
| ENSMUSG00000019370 | Calm3 | 12315 | protein_coding | 11699.26123 | -0.008653794 | 0.075217279 | -0.115050613 | 0.908405005 | 0.959234074 | 0.008653794 |
| ENSMUSG00000047728 | Ly6g2 | 223631 | protein_coding | 236.805451 | -0.008679383 | 0.155182701 | -0.055930093 | 0.955397497 | 0.982195158 | 0.008679383 |
| ENSMUSG00000044030 | Irf2bp1 | 272359 | protein_coding | 974.1064454 | -0.008702203 | 0.088161425 | -0.098707608 | 0.921370428 | 0.965302141 | 0.008702203 |
| ENSMUSG00000001166 | Oas1c | 114643 | protein_coding | 66.27179873 | -0.008705713 | 0.15746332 | -0.055287245 | 0.955909624 | 0.982430646 | 0.008705713 |
| ENSMUSG00000043059 | Zfp513 | 101023 | protein_coding | 358.0764777 | -0.008716559 | 0.077016843 | -0.113177309 | 0.909889985 | 0.959845275 | 0.008716559 |
| ENSMUSG00000036352 | Ubac1 | 98766 | protein_coding | 886.9556062 | 0.008754317 | 0.065972449 | 0.132696561 | 0.894433363 | 0.952805789 | 0.008754317 |
| ENSMUSG00000031928 | Mre11a | 17535 | protein_coding | 475.849883 | 0.008777201 | 0.078490107 | 0.11182557 | 0.910961713 | 0.96026404 | 0.008777201 |
| ENSMUSG00000027326 | Knl1 | 76464 | protein_coding | 448.9620453 | -0.008793207 | 0.09759912 | -0.090095144 | 0.928211608 | 0.968383271 | 0.008793207 |
| ENSMUSG00000032561 | Acpp | 56318 | protein_coding | 1327.621265 | 0.00880552 | 0.094709461 | 0.092974025 | 0.925924197 | 0.967733155 | 0.00880552 |
| ENSMUSG00000042043 | Tbca | 21371 | protein_coding | 1126.412517 | -0.008809951 | 0.051965432 | -0.169534832 | 0.865375977 | 0.939033668 | 0.008809951 |
| ENSMUSG00000024277 | Mapre2 | 212307 | protein_coding | 1663.470862 | -0.008854871 | 0.044753028 | -0.197860826 | 0.843153955 | 0.930064557 | 0.008854871 |
| ENSMUSG00000036529 | Sbf1 | 77980 | protein_coding | 2511.90701 | 0.008857593 | 0.058031035 | 0.152635456 | 0.878685763 | 0.94557102 | 0.008857593 |
| ENSMUSG00000019432 | Ddx39b | 53817 | protein_coding | 2705.202988 | 0.008868806 | 0.040764047 | 0.217564424 | 0.827768506 | 0.923349179 | 0.008868806 |
| ENSMUSG00000053226 | Dand5 | 23863 | protein_coding | 63.04847891 | 0.008880744 | 0.153928053 | 0.057694126 | 0.953992273 | 0.981205596 | 0.008880744 |
| ENSMUSG00000063659 | Zbtb18 | 30928 | protein_coding | 875.6504996 | -0.008894069 | 0.045601565 | -0.195038674 | 0.845362672 | 0.930668864 | 0.008894069 |
| ENSMUSG00000036275 | 9530068E07Rik | 213673 | protein_coding | 2130.972465 | 0.00893473 | 0.042353608 | 0.210955587 | 0.832921928 | 0.92587823 | 0.00893473 |
| ENSMUSG00000023944 | Hsp90ab1 | 15516 | protein_coding | 19488.02028 | 0.008947963 | 0.078257362 | 0.114340205 | 0.908968112 | 0.959446759 | 0.008947963 |
| ENSMUSG00000024764 | Naa40 | 70999 | protein_coding | 787.7756936 | -0.0089611 | 0.066828575 | -0.134090837 | 0.893330746 | 0.952355017 | 0.0089611 |
| ENSMUSG00000039671 | Zmynd8 | 228880 | protein_coding | 1578.427575 | 0.008980416 | 0.045788228 | 0.196129352 | 0.844508923 | 0.930210801 | 0.008980416 |
| ENSMUSG00000024136 | Dnase1l2 | 66705 | protein_coding | 149.8717343 | -0.009007812 | 0.229811927 | -0.039196451 | 0.968733763 | 0.987573843 | 0.009007812 |
| ENSMUSG00000025521 | Tmem192 | 73067 | protein_coding | 256.4669966 | -0.009010916 | 0.097611266 | -0.092314299 | 0.926448328 | 0.967733155 | 0.009010916 |
| ENSMUSG00000029363 | Rfc5 | 72151 | protein_coding | 480.1851887 | 0.009019422 | 0.102116385 | 0.088324927 | 0.929618428 | 0.968910773 | 0.009019422 |
| ENSMUSG00000041966 | Dcaf17 | 75763 | protein_coding | 488.976871 | -0.00903054 | 0.064727526 | -0.139516229 | 0.889042233 | 0.950733597 | 0.00903054 |
| ENSMUSG00000034308 | Sdr42e1 | 74032 | protein_coding | 910.4211945 | 0.009042693 | 0.072669481 | 0.124435911 | 0.900970141 | 0.956248672 | 0.009042693 |
| ENSMUSG00000024833 | Pola2 | 18969 | protein_coding | 422.5284741 | -0.009115638 | 0.099151301 | -0.091936641 | 0.92674838 | 0.96786874 | 0.009115638 |
| ENSMUSG00000031667 | Aktip | 14339 | protein_coding | 985.71029 | -0.009135782 | 0.048068677 | -0.190056863 | 0.849264572 | 0.931896969 | 0.009135782 |
| ENSMUSG00000041515 | Irf8 | 15900 | protein_coding | 1806.76639 | 0.009146406 | 0.076030709 | 0.120298837 | 0.904246426 | 0.95806991 | 0.009146406 |

| ensembl_gene_id | external_gene_name | entrezgene_id | gene_biotype | baseMean | log2FoldChange | lfcSE | stat | pvalue | padj | absLog2FC |
|---|---|---|---|---|---|---|---|---|---|---|
| ENSMUSG00000067567 | Hdac8 | 70315 | protein_coding | 165.9538032 | -0.009148822 | 0.091932006 | -0.09951727 | 0.920727577 | 0.965070185 | 0.009148822 |
| ENSMUSG00000051329 | Nup160 | 59015 | protein_coding | 558.6437427 | -0.009167239 | 0.064977451 | -0.141083385 | 0.887804069 | 0.950389394 | 0.009167239 |
| ENSMUSG00000044477 | Zfand3 | 21769 | protein_coding | 1407.546405 | 0.009187713 | 0.055270536 | 0.166231654 | 0.867974649 | 0.940208703 | 0.009187713 |
| ENSMUSG00000045467 | Ttll13 | 269954 | protein_coding | 10.83892303 | 0.009209768 | 0.408417056 | 0.02254991 | 0.9820093 | 0.993218186 | 0.009209768 |
| ENSMUSG00000047434 | Xxylt1 | 268880 | protein_coding | 281.6160468 | 0.009218478 | 0.075614458 | 0.121914222 | 0.902966952 | 0.957280308 | 0.009218478 |
| ENSMUSG00000003199 | Mpnd | 68047 | protein_coding | 1707.264168 | -0.009235861 | 0.057716218 | -0.160021936 | 0.872863794 | 0.942844904 | 0.009235861 |
| ENSMUSG00000021868 | Ppif | 105675 | protein_coding | 2198.671173 | -0.009243221 | 0.054721145 | -0.168914982 | 0.865863514 | 0.939297866 | 0.009243221 |
| ENSMUSG00000034035 | Ccdc17 | 622665 | protein_coding | 81.20564577 | -0.00924559 | 0.136252482 | -0.067856306 | 0.945900021 | 0.97697769 | 0.00924559 |
| ENSMUSG00000018196 | Glrx2 | 69367 | protein_coding | 376.7285253 | -0.009262089 | 0.073820445 | -0.125467795 | 0.900153219 | 0.955868241 | 0.009262089 |
| ENSMUSG00000055670 | Zzef1 | 195018 | protein_coding | 4640.831356 | -0.009275707 | 0.04034434 | -0.229913469 | 0.81815901 | 0.91832199 | 0.009275707 |
| ENSMUSG00000025157 | Zdhhc16 | 74168 | protein_coding | 293.3301368 | 0.00927652 | 0.071968728 | 0.128896537 | 0.897439517 | 0.954463189 | 0.00927652 |
| ENSMUSG00000034022 | Cpsf1 | 94230 | protein_coding | 1704.82501 | -0.009285849 | 0.054235751 | -0.171212686 | 0.864056533 | 0.938233215 | 0.009285849 |
| ENSMUSG00000038622 | Med30 | 69790 | protein_coding | 299.9388882 | 0.009287471 | 0.075300436 | 0.12333887 | 0.901838762 | 0.956772045 | 0.009287471 |
| ENSMUSG00000028969 | Cdk5 | 12568 | protein_coding | 605.8935129 | 0.009287536 | 0.060889492 | 0.152531012 | 0.878768133 | 0.94557102 | 0.009287536 |
| ENSMUSG00000058407 | Txndc9 | 98258 | protein_coding | 1326.088204 | 0.00932256 | 0.064939668 | 0.143557253 | 0.885850098 | 0.949475732 | 0.00932256 |
| ENSMUSG00000074238 | Ap1ar | 211556 | protein_coding | 2199.509251 | 0.009347994 | 0.035211912 | 0.265478171 | 0.790641124 | 0.902382831 | 0.009347994 |
| ENSMUSG00000041645 | Ddx24 | 27225 | protein_coding | 1436.608923 | 0.009356728 | 0.04858861 | 0.192570389 | 0.847295437 | 0.930926537 | 0.009356728 |
| ENSMUSG00000024969 | Mark2 | 13728 | protein_coding | 2016.506481 | 0.009385599 | 0.060161382 | 0.156007044 | 0.876027469 | 0.944301517 | 0.009385599 |
| ENSMUSG00000029617 | Ccz1 | 231874 | protein_coding | 1605.779324 | -0.009396705 | 0.051617032 | -0.182046598 | 0.855546157 | 0.935543319 | 0.009396705 |
| ENSMUSG00000016946 | Kctd5 | 69259 | protein_coding | 1496.081759 | -0.009399514 | 0.056767941 | -0.165577848 | 0.868489182 | 0.940513365 | 0.009399514 |
| ENSMUSG00000025781 | Atp5c1 | 11949 | protein_coding | 10135.04681 | 0.00947542 | 0.039625816 | 0.239122402 | 0.811010671 | 0.914838553 | 0.00947542 |
| ENSMUSG00000033200 | Tpsg1 | 26945 | protein_coding | 1050.929815 | -0.009496049 | 0.089293172 | -0.106346865 | 0.915307149 | 0.962265867 | 0.009496049 |
| ENSMUSG00000075600 | Zc3h3 | 223642 | protein_coding | 122.4101091 | 0.009502294 | 0.133207917 | 0.071334301 | 0.943131696 | 0.975305217 | 0.009502294 |
| ENSMUSG00000028992 | Nmnat1 | 66454 | protein_coding | 294.5351832 | 0.009509649 | 0.085382312 | 0.111377277 | 0.911317178 | 0.960412417 | 0.009509649 |
| ENSMUSG00000022811 | Zfp148 | 22661 | protein_coding | 1813.825335 | -0.009514759 | 0.046611284 | -0.204129949 | 0.838251951 | 0.928098115 | 0.009514759 |
| ENSMUSG00000043153 | Crppa | 75847 | protein_coding | 76.89278917 | 0.009523983 | 0.133691597 | 0.071238453 | 0.943207978 | 0.975321561 | 0.009523983 |
| ENSMUSG00000047866 | Lonp2 | 66887 | protein_coding | 2191.567305 | 0.009524548 | 0.036268003 | 0.262615717 | 0.792846785 | 0.903748946 | 0.009524548 |
| ENSMUSG00000029577 | Ube3b | 117146 | protein_coding | 3330.735631 | 0.009525579 | 0.04381816 | 0.217388847 | 0.827905322 | 0.923349179 | 0.009525579 |
| ENSMUSG00000020827 | Mink1 | 50932 | protein_coding | 1409.552016 | -0.00952658 | 0.052168371 | -0.182612179 | 0.855102327 | 0.935184812 | 0.00952658 |
| ENSMUSG00000015672 | Mrpl32 | 75398 | protein_coding | 521.5952726 | -0.009550435 | 0.063663497 | -0.150014301 | 0.880753332 | 0.946674819 | 0.009550435 |
| ENSMUSG00000002550 | Uck1 | 22245 | protein_coding | 737.7276223 | -0.00958019 | 0.058343848 | -0.164202231 | 0.869571945 | 0.940691929 | 0.00958019 |
| ENSMUSG00000059447 | Hadhb | 231086 | protein_coding | 3089.472787 | -0.009588851 | 0.037600733 | -0.255017661 | 0.798709459 | 0.907948215 | 0.009588851 |
| ENSMUSG00000024483 | Ankhd1 | 108857 | protein_coding | 1619.351303 | 0.009611177 | 0.05429976 | 0.177002203 | 0.859506659 | 0.936612161 | 0.009611177 |
| ENSMUSG00000038646 | Ramac | 67148 | protein_coding | 680.9866336 | 0.009620983 | 0.085362463 | 0.112707422 | 0.910262516 | 0.959960814 | 0.009620983 |
| ENSMUSG00000054309 | Cpsf3 | 54451 | protein_coding | 1099.2927 | -0.00962105 | 0.048485987 | -0.198429496 | 0.842709042 | 0.929981914 | 0.00962105 |
| ENSMUSG00000022401 | Xpnpep3 | 321003 | protein_coding | 323.0424059 | -0.009626095 | 0.070471014 | -0.136596518 | 0.891349729 | 0.951406253 | 0.009626095 |
| ENSMUSG00000024661 | Fth1 | 14319 | protein_coding | 18754.76393 | -0.009641676 | 0.070276284 | -0.137196724 | 0.8908753 | 0.95112373 | 0.009641676 |
| ENSMUSG00000020303 | Stc2 | 20856 | protein_coding | 347.8830588 | 0.009655512 | 0.073701081 | 0.131009098 | 0.89576811 | 0.953534806 | 0.009655512 |
| ENSMUSG00000001521 | Tulp3 | 22158 | protein_coding | 42.37604983 | 0.009657428 | 0.189214597 | 0.05103955 | 0.959294005 | 0.983542327 | 0.009657428 |

| ensembl_gene_id | external_gene_name | entrezgene_id | gene_biotype | baseMean | log2FoldChange | lfcSE | stat | pvalue | padj | absLog2FC |
|---|---|---|---|---|---|---|---|---|---|---|
| ENSMUSG00000031703 | Itfg1 | 71927 | protein_coding | 2342.784319 | -0.009744068 | 0.045434432 | -0.214464392 | 0.830184932 | 0.924804338 | 0.009744068 |
| ENSMUSG00000030224 | Strap | 20901 | protein_coding | 1880.122403 | -0.009760071 | 0.048081054 | -0.202992028 | 0.839141263 | 0.928687288 | 0.009760071 |
| ENSMUSG00000048012 | Zfp473 | 243963 | protein_coding | 35.47947497 | -0.009766561 | 0.205812602 | -0.047453659 | 0.962151664 | 0.984557099 | 0.009766561 |
| ENSMUSG00000041741 | Pde3a | 54611 | protein_coding | 2259.401854 | -0.00977798 | 0.080789788 | -0.121029899 | 0.903667353 | 0.957770991 | 0.00977798 |
| ENSMUSG00000040771 | Oard1 | 106821 | protein_coding | 553.8262742 | 0.009779875 | 0.082735017 | 0.118207201 | 0.905903484 | 0.958628969 | 0.009779875 |
| ENSMUSG00000062328 | Rpl17 | 319195 | protein_coding | 11883.24483 | -0.009801983 | 0.045163663 | -0.217032511 | 0.828183009 | 0.923583729 | 0.009801983 |
| ENSMUSG00000040365 | Trim41 | 211007 | protein_coding | 946.648708 | 0.009827476 | 0.055113963 | 0.178311907 | 0.858478028 | 0.93615636 | 0.009827476 |
| ENSMUSG00000020875 | Hoxb9 | 15417 | protein_coding | 971.1611994 | -0.009857788 | 0.065511389 | -0.150474421 | 0.88039033 | 0.946410802 | 0.009857788 |
| ENSMUSG00000021807 | Rtraf | 68045 | protein_coding | 2163.173447 | 0.009860917 | 0.049700674 | 0.1984061 | 0.842727345 | 0.929981914 | 0.009860917 |
| ENSMUSG00000053886 | Sh2d4a | 72281 | protein_coding | 2449.126498 | -0.009868743 | 0.049496755 | -0.199381617 | 0.841964239 | 0.929706079 | 0.009868743 |
| ENSMUSG00000009566 | Fpgs | 14287 | protein_coding | 375.6649581 | 0.009878254 | 0.103870918 | 0.095101246 | 0.924234409 | 0.96679823 | 0.009878254 |
| ENSMUSG00000024747 | Aldh1a7 | 26358 | protein_coding | 576.6682284 | 0.009884788 | 0.051324335 | 0.192594558 | 0.847276507 | 0.930926537 | 0.009884788 |
| ENSMUSG00000079355 | Ackr4 | 252837 | protein_coding | 72.13706401 | -0.00991172 | 0.17584087 | -0.056367557 | 0.955049002 | 0.982024691 | 0.00991172 |
| ENSMUSG00000038122 | Tbc1d32 | 544696 | protein_coding | 682.5553451 | -0.009939585 | 0.054727406 | -0.181619881 | 0.855881046 | 0.935592329 | 0.009939585 |
| ENSMUSG00000058454 | Dhcr7 | 13360 | protein_coding | 1510.630774 | -0.009941021 | 0.086209218 | -0.115312738 | 0.908197243 | 0.959153588 | 0.009941021 |
| ENSMUSG00000002748 | Baz1b | 22385 | protein_coding | 2422.248118 | 0.009960309 | 0.042058331 | 0.236821317 | 0.812795413 | 0.915984616 | 0.009960309 |
| ENSMUSG00000028647 | Mycbp | 56309 | protein_coding | 337.4425824 | -0.009989717 | 0.071630165 | -0.139462435 | 0.889084738 | 0.950733597 | 0.009989717 |
| ENSMUSG00000044201 | Cdc25c | 12532 | protein_coding | 120.1887593 | 0.009989952 | 0.124727034 | 0.080094519 | 0.936162082 | 0.97221237 | 0.009989952 |
| ENSMUSG00000037787 | Coa8 | 68020 | protein_coding | 406.3436651 | -0.010014294 | 0.068483383 | -0.146229543 | 0.883740184 | 0.948431353 | 0.010014294 |
| ENSMUSG00000024360 | Etf1 | 225363 | protein_coding | 2815.100579 | -0.010034306 | 0.044096689 | -0.227552362 | 0.819994258 | 0.919348861 | 0.010034306 |
| ENSMUSG00000051359 | Ncald | 52589 | protein_coding | 560.5876927 | 0.010040899 | 0.065386263 | 0.153562827 | 0.877954449 | 0.945241431 | 0.010040899 |
| ENSMUSG00000035199 | Arl6ip5 | 65106 | protein_coding | 775.3665879 | 0.010063047 | 0.053838402 | 0.186912064 | 0.85172958 | 0.933520082 | 0.010063047 |
| ENSMUSG00000024048 | Myl12a | 67268 | protein_coding | 1092.715017 | 0.010063324 | 0.054097264 | 0.186022783 | 0.852426895 | 0.933715176 | 0.010063324 |
| ENSMUSG00000020275 | Rel | 19696 | protein_coding | 151.6339734 | -0.01007701 | 0.116184959 | -0.086732481 | 0.930884158 | 0.969665987 | 0.01007701 |
| ENSMUSG00000037966 | Ninj1 | 18081 | protein_coding | 412.2355012 | 0.010080464 | 0.070290934 | 0.143410586 | 0.885965923 | 0.949475732 | 0.010080464 |
| ENSMUSG00000031660 | Brd7 | 26992 | protein_coding | 805.6757991 | 0.010095861 | 0.045777229 | 0.2205433 | 0.825448054 | 0.922363751 | 0.010095861 |
| ENSMUSG00000024787 | Snx15 | 69024 | protein_coding | 831.2866502 | 0.010098896 | 0.078116003 | 0.129280758 | 0.897135496 | 0.954412098 | 0.010098896 |
| ENSMUSG00000037935 | Smarce1 | 57376 | protein_coding | 1294.943681 | 0.010107409 | 0.047673467 | 0.212013299 | 0.832096659 | 0.925655158 | 0.010107409 |
| ENSMUSG00000040009 | Gnaz | 14687 | protein_coding | 21.62080494 | -0.010123719 | 0.252731852 | -0.040057154 | 0.96804756 | 0.987251745 | 0.010123719 |
| ENSMUSG00000010663 | Fads1 | 76267 | protein_coding | 257.6010743 | -0.01012782 | 0.101021732 | -0.100253878 | 0.920142773 | 0.96489645 | 0.01012782 |
| ENSMUSG00000013622 | Atraid | 381629 | protein_coding | 1200.045094 | 0.010172936 | 0.045302039 | 0.224558006 | 0.822323137 | 0.920659938 | 0.010172936 |
| ENSMUSG00000022109 | Med4 | 67381 | protein_coding | 267.5845369 | -0.010179859 | 0.075471945 | -0.134882694 | 0.892704623 | 0.952159223 | 0.010179859 |
| ENSMUSG00000034127 | Tspan8 | 216350 | protein_coding | 13797.91255 | 0.010200172 | 0.037607127 | 0.271229753 | 0.786214331 | 0.900404409 | 0.010200172 |
| ENSMUSG00000031885 | Cbfb | 12400 | protein_coding | 933.9949922 | -0.010249337 | 0.062411161 | -0.164222817 | 0.86955574 | 0.940691929 | 0.010249337 |
| ENSMUSG00000028532 | Cachd1 | 320508 | protein_coding | 1663.792892 | 0.010276012 | 0.051801479 | 0.198372954 | 0.842753276 | 0.929981914 | 0.010276012 |
| ENSMUSG00000017760 | Ctsa | 19025 | protein_coding | 2871.954277 | -0.01027971 | 0.051441152 | -0.19983436 | 0.841610128 | 0.929633324 | 0.01027971 |
| ENSMUSG00000025364 | Pa2g4 | 18813 | protein_coding | 1414.33975 | 0.010279857 | 0.063714145 | 0.16134341 | 0.871822935 | 0.94222547 | 0.010279857 |
| ENSMUSG00000041236 | Vps41 | 218035 | protein_coding | 1724.174962 | -0.010283225 | 0.046873177 | -0.21938399 | 0.826350941 | 0.922892079 | 0.010283225 |
| ENSMUSG00000005899 | Smpd4 | 77626 | protein_coding | 520.5589316 | 0.010303143 | 0.064442056 | 0.159882283 | 0.872973805 | 0.94290058 | 0.010303143 |

| ensembl_gene_id | external_gene_name | entrezgene_id | gene_biotype | baseMean | log2FoldChange | lfcSE | stat | pvalue | padj | absLog2FC |
|---|---|---|---|---|---|---|---|---|---|---|
| ENSMUSG00000029298 | Gbp9 | 236573 | protein_coding | 14.69172207 | 0.010314168 | 0.407116164 | 0.025334705 | 0.979787992 | 0.992429328 | 0.010314168 |
| ENSMUSG00000062762 | Ei24 | 13663 | protein_coding | 2152.176276 | -0.010319696 | 0.038186769 | -0.270242702 | 0.786973543 | 0.900576045 | 0.010319696 |
| ENSMUSG00000039176 | Polg | 18975 | protein_coding | 1638.381292 | -0.010338634 | 0.049006597 | -0.210964118 | 0.832915271 | 0.92587823 | 0.010338634 |
| ENSMUSG00000040659 | Efhd2 | 27984 | protein_coding | 2373.948734 | -0.010340418 | 0.061578715 | -0.16792195 | 0.866644682 | 0.939714838 | 0.010340418 |
| ENSMUSG00000038708 | Golga4 | 54214 | protein_coding | 6255.044664 | 0.010354162 | 0.031463074 | 0.329089323 | 0.742088174 | 0.876754335 | 0.010354162 |
| ENSMUSG00000013858 | Tmem259 | 216157 | protein_coding | 1946.87524 | 0.01036586 | 0.096034468 | 0.107938958 | 0.914044114 | 0.961718903 | 0.01036586 |
| ENSMUSG00000041817 | Fam169a | 320557 | protein_coding | 8.859353093 | 0.010390626 | 0.441335306 | 0.023543609 | 0.981216653 | 0.993012764 | 0.010390626 |
| ENSMUSG00000038342 | Mlxip | 208104 | protein_coding | 1342.46124 | 0.010440718 | 0.075185404 | 0.138866286 | 0.889555813 | 0.950733597 | 0.010440718 |
| ENSMUSG00000031299 | Pdha1 | 18597 | protein_coding | 6357.747772 | -0.010442997 | 0.039311325 | -0.265648575 | 0.790509872 | 0.902360752 | 0.010442997 |
| ENSMUSG00000052031 | Tagap1 | 380608 | protein_coding | 179.2757917 | 0.010449543 | 0.110888848 | 0.094234386 | 0.92492297 | 0.967142108 | 0.010449543 |
| ENSMUSG00000031969 | Acad8 | 66948 | protein_coding | 1052.147496 | 0.010485824 | 0.051959888 | 0.201806131 | 0.840068288 | 0.929183351 | 0.010485824 |
| ENSMUSG00000028030 | Tbck | 271981 | protein_coding | 966.3067492 | -0.010532376 | 0.047468026 | -0.221883596 | 0.824404501 | 0.921867382 | 0.010532376 |
| ENSMUSG00000013033 | Adgrl1 | 330814 | protein_coding | 50.80916427 | 0.010537046 | 0.170775258 | 0.061701245 | 0.950800748 | 0.979948241 | 0.010537046 |
| ENSMUSG00000038181 | Chpf2 | 100910 | protein_coding | 537.7992237 | 0.010538846 | 0.089346918 | 0.117954214 | 0.906103936 | 0.958651438 | 0.010538846 |
| ENSMUSG00000034041 | Lyl1 | 17095 | protein_coding | 15.14606701 | 0.010541453 | 0.368291901 | 0.02862255 | 0.977165627 | 0.990867082 | 0.010541453 |
| ENSMUSG00000029089 | 5730480H06Rik | 66768 | protein_coding | 138.4564664 | -0.010559856 | 0.113981951 | -0.092644985 | 0.926185604 | 0.967733155 | 0.010559856 |
| ENSMUSG00000029202 | Pds5a | 71521 | protein_coding | 2695.584808 | -0.010564782 | 0.061923499 | -0.170610228 | 0.864530255 | 0.938494846 | 0.010564782 |
| ENSMUSG00000022338 | Eny2 | 223527 | protein_coding | 887.7227957 | -0.010566382 | 0.056692474 | -0.186380674 | 0.852146247 | 0.933591927 | 0.010566382 |
| ENSMUSG00000027955 | Gask1b | 68659 | protein_coding | 10.11425271 | 0.010583854 | 0.430960577 | 0.024558752 | 0.980406921 | 0.992621208 | 0.010583854 |
| ENSMUSG00000064140 | Trim38 | 214158 | protein_coding | 35.51391051 | -0.010592717 | 0.216138339 | -0.049008968 | 0.960912149 | 0.984093538 | 0.010592717 |
| ENSMUSG00000071266 | Zfp946 | 74149 | protein_coding | 452.8718363 | -0.010609234 | 0.088954631 | -0.11926567 | 0.90506488 | 0.958435249 | 0.010609234 |
| ENSMUSG00000033732 | Sf3b3 | 101943 | protein_coding | 2308.225315 | -0.010610936 | 0.058540812 | -0.18125707 | 0.856165801 | 0.935604164 | 0.010610936 |
| ENSMUSG00000005312 | Ubqln1 | 56085 | protein_coding | 3038.807738 | 0.010613703 | 0.047170239 | 0.225008454 | 0.821972698 | 0.920488417 | 0.010613703 |
| ENSMUSG00000002010 | Idh3g | 15929 | protein_coding | 3375.05443 | 0.010626492 | 0.053601094 | 0.198251406 | 0.842848369 | 0.929981914 | 0.010626492 |
| ENSMUSG00000021497 | Txndc15 | 69672 | protein_coding | 836.4972397 | 0.010662658 | 0.06174404 | 0.172691285 | 0.862894095 | 0.937787345 | 0.010662658 |
| ENSMUSG00000006191 | Cdkal1 | 68916 | protein_coding | 394.3643463 | -0.010663236 | 0.068662574 | -0.155299105 | 0.876585521 | 0.944776763 | 0.010663236 |
| ENSMUSG00000040782 | Cop1 | 26374 | protein_coding | 1948.5793 | 0.01068701 | 0.047342646 | 0.225737479 | 0.821405606 | 0.920176481 | 0.01068701 |
| ENSMUSG00000003363 | Pld3 | 18807 | protein_coding | 619.1650488 | -0.010696368 | 0.076754018 | -0.139359057 | 0.889166424 | 0.950733597 | 0.010696368 |
| ENSMUSG00000057858 | Fam204a | 76539 | protein_coding | 301.7730492 | 0.010700625 | 0.087105578 | 0.122846607 | 0.902228567 | 0.956874924 | 0.010700625 |
| ENSMUSG00000037104 | Socs5 | 56468 | protein_coding | 293.8084727 | 0.010709545 | 0.085525443 | 0.125220571 | 0.900348932 | 0.955895949 | 0.010709545 |
| ENSMUSG00000015837 | Sqstm1 | 18412 | protein_coding | 4454.718468 | -0.010714872 | 0.046204998 | -0.231898539 | 0.816616819 | 0.917421069 | 0.010714872 |
| ENSMUSG00000038039 | Gcc2 | 70297 | protein_coding | 3118.523691 | 0.010725164 | 0.06146693 | 0.174486733 | 0.861482956 | 0.937521805 | 0.010725164 |
| ENSMUSG00000027091 | Zc3h15 | 69082 | protein_coding | 1488.357582 | 0.010734086 | 0.045403791 | 0.23641388 | 0.813111527 | 0.915984616 | 0.010734086 |
| ENSMUSG00000019726 | Lyst | 17101 | protein_coding | 543.6970812 | -0.010758233 | 0.06970255 | -0.154344896 | 0.8773378 | 0.945176551 | 0.010758233 |
| ENSMUSG00000048307 | Ankrd46 | 68839 | protein_coding | 510.866131 | 0.010767743 | 0.055889434 | 0.192661511 | 0.847224068 | 0.930926537 | 0.010767743 |
| ENSMUSG00000037570 | Mcrs1 | 51812 | protein_coding | 1075.483908 | 0.010780024 | 0.047070849 | 0.22901698 | 0.818855719 | 0.918528635 | 0.010780024 |
| ENSMUSG00000028646 | Rragc | 54170 | protein_coding | 1108.805111 | 0.010807099 | 0.054189417 | 0.199431918 | 0.841924895 | 0.929706079 | 0.010807099 |
| ENSMUSG00000032702 | Kank1 | 107351 | protein_coding | 970.1750046 | 0.010813502 | 0.056312186 | 0.192027752 | 0.847720467 | 0.931084542 | 0.010813502 |
| ENSMUSG00000057054 | Inca1 | 103844 | protein_coding | 25.75476167 | -0.01083707 | 0.246057372 | -0.044042857 | 0.964870242 | 0.985900171 | 0.01083707 |

Supplementary Table 1

| ensembl_gene_id | external_gene_name | entrezgene_id | gene_biotype | baseMean | log2FoldChange | lfcSE | stat | pvalue | padj | absLog2FC |
|---|---|---|---|---|---|---|---|---|---|---|
| ENSMUSG00000039048 | Foxred1 | 235169 | protein_coding | 642.7471054 | 0.010842957 | 0.05228811 | 0.207369457 | 0.835721334 | 0.927001716 | 0.010842957 |
| ENSMUSG00000022671 | Mzt2 | 72083 | protein_coding | 307.2836092 | 0.010888529 | 0.081270795 | 0.133978378 | 0.893419672 | 0.952355017 | 0.010888529 |
| ENSMUSG00000053929 | Cyhr1 | 54151 | protein_coding | 1920.16303 | 0.010889099 | 0.043533819 | 0.250129665 | 0.802487076 | 0.910121497 | 0.010889099 |
| ENSMUSG00000061374 | Fiz1 | 23877 | protein_coding | 577.1740572 | -0.010905135 | 0.079509445 | -0.137155216 | 0.890908109 | 0.95112373 | 0.010905135 |
| ENSMUSG00000028954 | Nub1 | 53312 | protein_coding | 1007.77178 | -0.010907752 | 0.058457447 | -0.18659302 | 0.851979739 | 0.933591927 | 0.010907752 |
| ENSMUSG00000115987 | Vps28 | 66914 | protein_coding | 2506.400493 | -0.010931251 | 0.043021081 | -0.25409058 | 0.799425581 | 0.908422797 | 0.010931251 |
| ENSMUSG00000020682 | Mmp28 | 118453 | protein_coding | 269.3240672 | -0.010942685 | 0.08682784 | -0.126027373 | 0.899710257 | 0.955523695 | 0.010942685 |
| ENSMUSG00000020954 | Strn3 | 94186 | protein_coding | 1143.572384 | 0.010970032 | 0.047394927 | 0.231460055 | 0.816957414 | 0.917675853 | 0.010970032 |
| ENSMUSG00000039298 | Cdk5rap2 | 214444 | protein_coding | 457.8810722 | 0.010988013 | 0.073564414 | 0.149365879 | 0.881264934 | 0.947098467 | 0.010988013 |
| ENSMUSG00000004642 | Slbp | 20492 | protein_coding | 805.2571639 | -0.010996369 | 0.066793383 | -0.164632609 | 0.869233163 | 0.940691929 | 0.010996369 |
| ENSMUSG00000030744 | Rps3 | 27050 | protein_coding | 18014.1617 | -0.01104179 | 0.038464728 | -0.287062726 | 0.774064292 | 0.893779608 | 0.01104179 |
| ENSMUSG00000023965 | Fbxl17 | 50758 | protein_coding | 502.4785369 | -0.011068408 | 0.063756115 | -0.173605434 | 0.862175561 | 0.937787345 | 0.011068408 |
| ENSMUSG00000029179 | Zcchc4 | 78796 | protein_coding | 233.6676225 | -0.011069765 | 0.081225252 | -0.136284768 | 0.891596165 | 0.951425975 | 0.011069765 |
| ENSMUSG00000028530 | Jak1 | 16451 | protein_coding | 2868.893616 | 0.011114197 | 0.040278159 | 0.275936072 | 0.782597167 | 0.898825528 | 0.011114197 |
| ENSMUSG00000022419 | Deptor | 97998 | protein_coding | 2966.076843 | -0.011124762 | 0.062068964 | -0.179232275 | 0.857755322 | 0.935951904 | 0.011124762 |
| ENSMUSG00000038533 | Cbfa2t2 | 12396 | protein_coding | 679.719309 | -0.011130241 | 0.053956202 | -0.206282885 | 0.836569946 | 0.927560229 | 0.011130241 |
| ENSMUSG00000060798 | Intu | 380614 | protein_coding | 186.8730928 | -0.011134488 | 0.133583023 | -0.083352569 | 0.933571202 | 0.971022429 | 0.011134488 |
| ENSMUSG00000003166 | Dgcr2 | 13356 | protein_coding | 1722.824032 | 0.011136397 | 0.034885603 | 0.319226164 | 0.749555018 | 0.880475742 | 0.011136397 |
| ENSMUSG00000061718 | Ppp1r1b | 19049 | protein_coding | 5854.236936 | -0.011149977 | 0.069808862 | -0.159721508 | 0.873100458 | 0.942905245 | 0.011149977 |
| ENSMUSG00000079509 | Zfx | 22764 | protein_coding | 979.641838 | -0.011171933 | 0.053384416 | -0.209273306 | 0.834234889 | 0.926479877 | 0.011171933 |
| ENSMUSG00000022387 | Brd1 | 223770 | protein_coding | 2598.813314 | 0.011175455 | 0.042971816 | 0.260064755 | 0.794813824 | 0.904781532 | 0.011175455 |
| ENSMUSG00000021485 | Mxd3 | 17121 | protein_coding | 99.39513902 | -0.011186101 | 0.15406986 | -0.072604082 | 0.942121178 | 0.974690203 | 0.011186101 |
| ENSMUSG00000034377 | Tulp4 | 68842 | protein_coding | 1741.650797 | -0.011219019 | 0.050040892 | -0.224197018 | 0.822604004 | 0.920659938 | 0.011219019 |
| ENSMUSG00000032279 | Idh3a | 67834 | protein_coding | 3519.719908 | -0.011230754 | 0.057036894 | -0.196903328 | 0.843903189 | 0.930177636 | 0.011230754 |
| ENSMUSG00000005656 | Snx6 | 72183 | protein_coding | 1189.584051 | 0.011230832 | 0.055765527 | 0.201393804 | 0.840390659 | 0.929368454 | 0.011230832 |
| ENSMUSG00000032763 | Ilvbl | 216136 | protein_coding | 1734.027086 | -0.01124185 | 0.0375744 | -0.299189083 | 0.764795778 | 0.889439786 | 0.01124185 |
| ENSMUSG00000003161 | Sri | 109552 | protein_coding | 7752.268231 | 0.011263711 | 0.046743058 | 0.240970775 | 0.809577767 | 0.91448208 | 0.011263711 |
| ENSMUSG00000033623 | Pcgf3 | 69587 | protein_coding | 1718.923153 | 0.011297191 | 0.054941978 | 0.205620388 | 0.837087451 | 0.927812059 | 0.011297191 |
| ENSMUSG00000059070 | Rpl18 | 19899 | protein_coding | 10348.25043 | 0.011315896 | 0.0394326 | 0.286968042 | 0.774136791 | 0.893793243 | 0.011315896 |
| ENSMUSG00000027243 | Harbi1 | 241547 | protein_coding | 319.54709 | -0.01132589 | 0.085739483 | -0.13209655 | 0.894907925 | 0.953178075 | 0.01132589 |
| ENSMUSG00000009246 | Trpm5 | 56843 | protein_coding | 625.1741404 | 0.011326344 | 0.087869089 | 0.128900212 | 0.897436609 | 0.954463189 | 0.011326344 |
| ENSMUSG00000025555 | Farp1 | 223254 | protein_coding | 953.0074658 | -0.01133691 | 0.054204871 | -0.209149293 | 0.834331695 | 0.926479877 | 0.01133691 |
| ENSMUSG00000024962 | Vegfb | 22340 | protein_coding | 176.2104328 | -0.011342893 | 0.139407412 | -0.081365065 | 0.935151631 | 0.971726183 | 0.011342893 |
| ENSMUSG00000042041 | 2010003K11Rik | 69861 | protein_coding | 300.5214271 | -0.011348222 | 0.150595359 | -0.075355724 | 0.939931686 | 0.973932042 | 0.011348222 |
| ENSMUSG00000004356 | Utp20 | 70683 | protein_coding | 1993.381638 | 0.011363866 | 0.068472968 | 0.165961341 | 0.868187374 | 0.940312812 | 0.011363866 |
| ENSMUSG00000090935 | Synj2bp | 24071 | protein_coding | 1418.960019 | 0.011367841 | 0.056848152 | 0.19996853 | 0.841505194 | 0.929613548 | 0.011367841 |
| ENSMUSG00000034282 | Evpl | 14027 | protein_coding | 1707.640566 | 0.011369439 | 0.103951647 | 0.109372381 | 0.912907139 | 0.961246633 | 0.011369439 |
| ENSMUSG00000074406 | Zfp628 | 232816 | protein_coding | 198.9805559 | 0.011370127 | 0.12299224 | 0.09244589 | 0.926343781 | 0.967733155 | 0.011370127 |
| ENSMUSG00000024818 | Slc25a45 | 107375 | protein_coding | 62.14265635 | 0.011371911 | 0.22755858 | 0.049973553 | 0.960143464 | 0.983923349 | 0.011371911 |

| ensembl_gene_id | external_gene_name | entrezgene_id | gene_biotype | baseMean | log2FoldChange | lfcSE | stat | pvalue | padj | absLog2FC |
|---|---|---|---|---|---|---|---|---|---|---|
| ENSMUSG00000040850 | Psme4 | 103554 | protein_coding | 3767.134059 | 0.011372384 | 0.048401599 | 0.234958842 | 0.814240678 | 0.916408517 | 0.011372384 |
| ENSMUSG00000028696 | Ipp | 16351 | protein_coding | 260.7919574 | 0.011387126 | 0.088732236 | 0.128331328 | 0.897886774 | 0.954467075 | 0.011387126 |
| ENSMUSG00000031154 | Otud5 | 54644 | protein_coding | 1581.336664 | -0.011401235 | 0.058014733 | -0.196523104 | 0.844200751 | 0.930210801 | 0.011401235 |
| ENSMUSG00000092819 | Gm23639 | 115489002 | snoRNA | 84.72523307 | -0.011401854 | 0.307140468 | -0.037122604 | 0.970387249 | 0.988094151 | 0.011401854 |
| ENSMUSG00000021816 | Ppp3cb | 19056 | protein_coding | 914.8690978 | -0.011409798 | 0.0567961 | -0.200890515 | 0.840784185 | 0.929613548 | 0.011409798 |
| ENSMUSG00000022683 | Pla2g10 | 26565 | protein_coding | 1307.432302 | 0.011442735 | 0.090109648 | 0.126986792 | 0.898950853 | 0.955322545 | 0.011442735 |
| ENSMUSG00000024142 | Mlst8 | 56716 | protein_coding | 530.5823296 | -0.011454022 | 0.057997204 | -0.197492656 | 0.843442028 | 0.930151569 | 0.011454022 |
| ENSMUSG00000032320 | Rcn2 | 26611 | protein_coding | 488.7727376 | 0.01145587 | 0.063353708 | 0.180823985 | 0.856505736 | 0.935610703 | 0.01145587 |
| ENSMUSG00000029614 | Rpl6 | 19988 | protein_coding | 8528.063613 | -0.011499601 | 0.048937727 | -0.234984376 | 0.81422086 | 0.916408517 | 0.011499601 |
| ENSMUSG00000010358 | Ifi35 | 70110 | protein_coding | 460.5320864 | -0.01151072 | 0.079911067 | -0.144044131 | 0.885465621 | 0.949313881 | 0.01151072 |
| ENSMUSG00000025979 | Mob4 | 19070 | protein_coding | 1390.879673 | -0.011515134 | 0.047472101 | -0.242566355 | 0.808341348 | 0.913597373 | 0.011515134 |
| ENSMUSG00000032812 | Arap1 | 69710 | protein_coding | 1767.803903 | -0.011535088 | 0.064225356 | -0.179603334 | 0.857463987 | 0.935863679 | 0.011535088 |
| ENSMUSG00000028556 | Dock7 | 67299 | protein_coding | 1426.48365 | -0.011535625 | 0.045508139 | -0.253484868 | 0.799893553 | 0.908570455 | 0.011535625 |
| ENSMUSG00000040616 | Tmem51 | 214359 | protein_coding | 627.6445851 | 0.01155948 | 0.06298038 | 0.183540967 | 0.854373578 | 0.934988261 | 0.01155948 |
| ENSMUSG00000028860 | Sytl1 | 269589 | protein_coding | 1586.397848 | -0.011611926 | 0.084179333 | -0.137942717 | 0.89028569 | 0.951039003 | 0.011611926 |
| ENSMUSG00000022706 | Mrpl40 | 18100 | protein_coding | 682.1692079 | -0.011629426 | 0.058907945 | -0.197416934 | 0.843501279 | 0.930151569 | 0.011629426 |
| ENSMUSG00000048442 | Smim5 | 66528 | protein_coding | 168.2881222 | -0.01163331 | 0.101419412 | -0.114704966 | 0.908678977 | 0.959237145 | 0.01163331 |
| ENSMUSG00000032899 | Styk1 | 243659 | protein_coding | 2983.998261 | 0.011645626 | 0.042021763 | 0.277133197 | 0.781677832 | 0.898575693 | 0.011645626 |
| ENSMUSG00000029518 | Rab35 | 77407 | protein_coding | 1383.056227 | 0.011665415 | 0.0487301 | 0.239388282 | 0.810804515 | 0.914838553 | 0.011665415 |
| ENSMUSG00000009470 | Tnpo1 | 238799 | protein_coding | 3349.540777 | -0.011667316 | 0.038426639 | -0.303625723 | 0.761413052 | 0.887492981 | 0.011667316 |
| ENSMUSG00000020205 | Phlda1 | 21664 | protein_coding | 159.6244832 | -0.01167935 | 0.110277757 | -0.105908483 | 0.915654963 | 0.962443303 | 0.01167935 |
| ENSMUSG00000042492 | Tbc1d10b | 68449 | protein_coding | 1216.620327 | 0.011681286 | 0.08391547 | 0.139203013 | 0.889289728 | 0.950733597 | 0.011681286 |
| ENSMUSG00000004508 | Gab2 | 14389 | protein_coding | 398.4034902 | -0.011701776 | 0.089708327 | -0.130442475 | 0.896216364 | 0.953823091 | 0.011701776 |
| ENSMUSG00000018841 | Rad51d | 19364 | protein_coding | 555.9245629 | -0.011702285 | 0.055317042 | -0.211549371 | 0.832458611 | 0.92580236 | 0.011702285 |
| ENSMUSG00000091512 | Lamtor3 | 56692 | protein_coding | 615.7603768 | 0.011705786 | 0.071954938 | 0.162682184 | 0.870768675 | 0.941634557 | 0.011705786 |
| ENSMUSG00000041625 | Ggact | 223267 | protein_coding | 1171.923515 | 0.011717842 | 0.078870985 | 0.148569747 | 0.881893147 | 0.947331698 | 0.011717842 |
| ENSMUSG00000025270 | Alas2 | 11656 | protein_coding | 11.02592473 | -0.011765011 | 0.386552597 | -0.030435733 | 0.975719548 | 0.990489989 | 0.011765011 |
| ENSMUSG00000000740 | Rpl13 | 270106 | protein_coding | 9809.773084 | -0.011773789 | 0.059007161 | -0.199531522 | 0.841846988 | 0.929706079 | 0.011773789 |
| ENSMUSG00000027646 | Src | 20779 | protein_coding | 4070.598431 | -0.011777311 | 0.060845045 | -0.193562371 | 0.846518564 | 0.930889346 | 0.011777311 |
| ENSMUSG00000030554 | Synm | 233335 | protein_coding | 7.287821166 | -0.011777855 | 0.432684198 | -0.027220441 | 0.978283912 | 0.991564619 | 0.011777855 |
| ENSMUSG00000025283 | Sat1 | 20229 | protein_coding | 4205.940966 | 0.011777997 | 0.069726272 | 0.168917641 | 0.865861423 | 0.939297866 | 0.011777997 |
| ENSMUSG00000028988 | Ctnnbip1 | 67087 | protein_coding | 474.0006961 | -0.011814996 | 0.068599697 | -0.172231026 | 0.863255907 | 0.937932225 | 0.011814996 |
| ENSMUSG00000040383 | Aqr | 11834 | protein_coding | 1252.414799 | 0.011828648 | 0.059038523 | 0.200354737 | 0.841203156 | 0.929613548 | 0.011828648 |
| ENSMUSG00000066191 | Anks6 | 75691 | protein_coding | 234.0509958 | 0.011836234 | 0.102683585 | 0.115268996 | 0.908231912 | 0.959153588 | 0.011836234 |
| ENSMUSG00000027661 | Slc2a10 | 170441 | protein_coding | 245.3526745 | -0.011846457 | 0.088601086 | -0.133705554 | 0.893635413 | 0.952419871 | 0.011846457 |
| ENSMUSG00000051295 | 9630028B13Rik | 319562 | lncRNA | 19.29205771 | 0.011857861 | 0.312137819 | 0.037989183 | 0.969696307 | 0.987827196 | 0.011857861 |
| ENSMUSG00000020647 | Ncoa1 | 17977 | protein_coding | 1444.53198 | -0.011864467 | 0.065464003 | -0.181236499 | 0.856181947 | 0.935604164 | 0.011864467 |
| ENSMUSG00000029480 | Dhx37 | 208144 | protein_coding | 314.5740099 | 0.011866022 | 0.077067979 | 0.15396825 | 0.87763477 | 0.945241431 | 0.011866022 |
| ENSMUSG00000079614 | Seh1l | 72124 | protein_coding | 992.1312654 | -0.011896375 | 0.055764142 | -0.213333781 | 0.831066626 | 0.925277452 | 0.011896375 |

| ensembl_gene_id | external_gene_name | entrezgene_id | gene_biotype | baseMean | log2FoldChange | lfcSE | stat | pvalue | padj | absLog2FC |
|---|---|---|---|---|---|---|---|---|---|---|
| ENSMUSG00000039804 | Ncoa5 | 228869 | protein_coding | 572.6236835 | -0.011900864 | 0.061147629 | -0.194625115 | 0.84568644 | 0.93082665 | 0.011900864 |
| ENSMUSG00000063065 | Mapk3 | 26417 | protein_coding | 6651.125902 | 0.011913119 | 0.061754892 | 0.192909719 | 0.847029674 | 0.930926537 | 0.011913119 |
| ENSMUSG00000032939 | Nup93 | 71805 | protein_coding | 491.1770719 | 0.011914215 | 0.105433453 | 0.11300223 | 0.910028787 | 0.959845275 | 0.011914215 |
| ENSMUSG00000043895 | S1pr2 | 14739 | protein_coding | 42.88185754 | 0.011920691 | 0.195882278 | 0.060856403 | 0.95147357 | 0.98032045 | 0.011920691 |
| ENSMUSG00000029050 | Ski | 20481 | protein_coding | 1501.001504 | -0.011935623 | 0.072460523 | -0.164718971 | 0.869165184 | 0.940691929 | 0.011935623 |
| ENSMUSG00000042772 | Smg7 | 226517 | protein_coding | 1465.86608 | 0.011954445 | 0.042941808 | 0.278387093 | 0.780715226 | 0.898301545 | 0.011954445 |
| ENSMUSG00000040549 | Ckap5 | 75786 | protein_coding | 1862.536055 | -0.011959338 | 0.06615212 | -0.180785403 | 0.856536021 | 0.935610703 | 0.011959338 |
| ENSMUSG00000028036 | Ptgfr | 19220 | protein_coding | 10.39641451 | -0.011960119 | 0.367757656 | -0.032521741 | 0.974055979 | 0.989393511 | 0.011960119 |
| ENSMUSG00000067369 | Trmt2b | 215201 | protein_coding | 1081.463072 | 0.011967114 | 0.058490867 | 0.20459799 | 0.837886226 | 0.928042853 | 0.011967114 |
| ENSMUSG00000068267 | Cenpb | 12616 | protein_coding | 5368.168038 | -0.011980699 | 0.055557786 | -0.215643927 | 0.829265313 | 0.924226793 | 0.011980699 |
| ENSMUSG00000025236 | Adpgk | 72141 | protein_coding | 367.8735136 | 0.011993654 | 0.083278046 | 0.144019401 | 0.88548515 | 0.949313881 | 0.011993654 |
| ENSMUSG00000020516 | Rps6kb1 | 72508 | protein_coding | 863.4655372 | 0.011999363 | 0.054356027 | 0.220754969 | 0.825283228 | 0.922274452 | 0.011999363 |
| ENSMUSG00000001082 | Mfsd10 | 68294 | protein_coding | 412.0089489 | 0.012005185 | 0.108340679 | 0.110809577 | 0.911767351 | 0.960548214 | 0.012005185 |
| ENSMUSG00000025893 | Kbtbd3 | 69149 | protein_coding | 104.2225246 | 0.012015116 | 0.165979938 | 0.072388965 | 0.942292366 | 0.974749775 | 0.012015116 |
| ENSMUSG00000029499 | Pxmp2 | 19301 | protein_coding | 90.8503633 | -0.012058627 | 0.149295379 | -0.080770265 | 0.935624656 | 0.971967201 | 0.012058627 |
| ENSMUSG00000066152 | Slc31a2 | 20530 | protein_coding | 297.0700486 | -0.012070714 | 0.086947047 | -0.13882834 | 0.889585799 | 0.950733597 | 0.012070714 |
| ENSMUSG00000034949 | Zfr2 | 103406 | protein_coding | 82.60606958 | 0.012098022 | 0.174791285 | 0.069214102 | 0.944819198 | 0.976173951 | 0.012098022 |
| ENSMUSG00000084957 | Bbip1 | 100503572 | protein_coding | 424.9475326 | 0.012103677 | 0.071248038 | 0.169880841 | 0.865103849 | 0.938898215 | 0.012103677 |
| ENSMUSG00000010601 | Apol7a | 75761 | protein_coding | 171.2347016 | -0.012103721 | 0.201955708 | -0.05993255 | 0.952209355 | 0.98066383 | 0.012103721 |
| ENSMUSG00000049643 | 2310022A10Rik | 66367 | protein_coding | 141.2781274 | 0.012106397 | 0.112306127 | 0.107798189 | 0.91415578 | 0.961718903 | 0.012106397 |
| ENSMUSG00000022614 | Lmf2 | 105847 | protein_coding | 415.2571004 | -0.012119355 | 0.085449069 | -0.141831334 | 0.887213233 | 0.95019842 | 0.012119355 |
| ENSMUSG00000070394 | Tmem256 | 69186 | protein_coding | 637.6479997 | 0.01214586 | 0.087172902 | 0.139330686 | 0.889188842 | 0.950733597 | 0.01214586 |
| ENSMUSG00000022898 | Vps26c | 13185 | protein_coding | 934.7764884 | 0.012150757 | 0.049324856 | 0.246341462 | 0.805417917 | 0.911698936 | 0.012150757 |
| ENSMUSG00000037519 | Ppfia1 | 233977 | protein_coding | 1539.399692 | 0.012159637 | 0.044547029 | 0.272961788 | 0.784882589 | 0.899716112 | 0.012159637 |
| ENSMUSG00000053347 | Zfp943 | 74670 | protein_coding | 442.2036345 | -0.012179161 | 0.070870015 | -0.171852102 | 0.863553803 | 0.93800311 | 0.012179161 |
| ENSMUSG00000047832 | Cdca4 | 71963 | protein_coding | 505.1237653 | 0.012255075 | 0.068537381 | 0.178808625 | 0.858087972 | 0.936097283 | 0.012255075 |
| ENSMUSG00000060121 | Gemin2 | 66603 | protein_coding | 136.7651441 | -0.012282577 | 0.113961307 | -0.107778489 | 0.914171407 | 0.961718903 | 0.012282577 |
| ENSMUSG00000050471 | Fam118b | 109229 | protein_coding | 555.7898894 | 0.012283833 | 0.06651801 | 0.184669284 | 0.85348844 | 0.934372006 | 0.012283833 |
| ENSMUSG00000029670 | Ing3 | 71777 | protein_coding | 332.2267791 | -0.012309321 | 0.07325672 | -0.168029927 | 0.866559735 | 0.93968591 | 0.012309321 |
| ENSMUSG00000058799 | Nap1l1 | 53605 | protein_coding | 887.308128 | -0.012310756 | 0.095603923 | -0.128768318 | 0.897540975 | 0.954463189 | 0.012310756 |
| ENSMUSG00000021039 | Snw1 | 66354 | protein_coding | 1034.705337 | 0.012315243 | 0.052301927 | 0.23546442 | 0.813848292 | 0.916289542 | 0.012315243 |
| ENSMUSG00000034285 | Nipsnap1 | 18082 | protein_coding | 1179.984991 | 0.012323111 | 0.051657992 | 0.238551884 | 0.811453079 | 0.915190139 | 0.012323111 |
| ENSMUSG00000030727 | Rabep2 | 70314 | protein_coding | 673.4129974 | 0.012347323 | 0.080398333 | 0.153576854 | 0.877943388 | 0.945241431 | 0.012347323 |
| ENSMUSG00000002804 | Nudt14 | 66174 | protein_coding | 613.002454 | -0.012350454 | 0.080969725 | -0.152531754 | 0.878767547 | 0.94557102 | 0.012350454 |
| ENSMUSG00000042502 | Cd2bp2 | 70233 | protein_coding | 1012.180928 | 0.012351829 | 0.056622868 | 0.218142057 | 0.82731843 | 0.923333171 | 0.012351829 |
| ENSMUSG00000006169 | Clint1 | 216705 | protein_coding | 5832.715313 | -0.01235669 | 0.031022269 | -0.398316757 | 0.69039671 | 0.84457466 | 0.01235669 |
| ENSMUSG00000022255 | Mtdh | 67154 | protein_coding | 2552.051989 | 0.012360083 | 0.049035722 | 0.252062839 | 0.800992493 | 0.90924234 | 0.012360083 |
| ENSMUSG00000052085 | Dock8 | 76088 | protein_coding | 738.8119635 | -0.01237756 | 0.059462067 | -0.208158918 | 0.835104885 | 0.926699249 | 0.01237756 |
| ENSMUSG00000049687 | Pheta2 | 338368 | protein_coding | 12.66499769 | 0.012385508 | 0.345887546 | 0.035807905 | 0.97143553 | 0.988537404 | 0.012385508 |

| ensembl_gene_id | external_gene_name | entrezgene_id | gene_biotype | baseMean | log2FoldChange | lfcSE | stat | pvalue | padj | absLog2FC |
|---|---|---|---|---|---|---|---|---|---|---|
| ENSMUSG00000024853 | Sf3b2 | 319322 | protein_coding | 4147.220471 | 0.012386413 | 0.037962342 | 0.326281584 | 0.74421132 | 0.87823035 | 0.012386413 |
| ENSMUSG00000018965 | Ywhah | 22629 | protein_coding | 3169.557421 | 0.012407369 | 0.051761726 | 0.239701613 | 0.810561585 | 0.914760439 | 0.012407369 |
| ENSMUSG00000068101 | Cenpm | 66570 | protein_coding | 88.29798397 | -0.012408949 | 0.157289699 | -0.078892317 | 0.937118275 | 0.972704281 | 0.012408949 |
| ENSMUSG00000004626 | Stxbp2 | 20911 | protein_coding | 2190.843106 | -0.01241311 | 0.039776942 | -0.31206797 | 0.754988868 | 0.883718994 | 0.01241311 |
| ENSMUSG00000024908 | Ppp6r3 | 52036 | protein_coding | 4321.623844 | -0.012434281 | 0.033801827 | -0.367858259 | 0.712978925 | 0.858612846 | 0.012434281 |
| ENSMUSG00000020460 | Rps27a | 78294 | protein_coding | 7043.723448 | 0.012450772 | 0.042009038 | 0.296383163 | 0.766937482 | 0.89053739 | 0.012450772 |
| ENSMUSG00000074480 | Mex3a | 72640 | protein_coding | 45.34969672 | 0.012486778 | 0.181447038 | 0.068817754 | 0.945134686 | 0.976374805 | 0.012486778 |
| ENSMUSG00000004561 | Mettl17 | 52535 | protein_coding | 355.3852878 | 0.012541132 | 0.073317671 | 0.171051969 | 0.864182902 | 0.938307255 | 0.012541132 |
| ENSMUSG00000041268 | Dmxl2 | 235380 | protein_coding | 44.99554193 | 0.012573969 | 0.191915476 | 0.065518264 | 0.947761365 | 0.978148441 | 0.012573969 |
| ENSMUSG00000004221 | Ikbkg | 16151 | protein_coding | 1312.375591 | 0.012607956 | 0.044757489 | 0.28169489 | 0.778177473 | 0.896853291 | 0.012607956 |
| ENSMUSG00000036805 | Noxa1 | 241275 | protein_coding | 1304.09463 | 0.01264149 | 0.066616701 | 0.189764572 | 0.849493619 | 0.931958067 | 0.01264149 |
| ENSMUSG00000030613 | Ccdc90b | 66365 | protein_coding | 374.3158606 | -0.012747305 | 0.070174143 | -0.181652457 | 0.855855479 | 0.935592329 | 0.012747305 |
| ENSMUSG00000057375 | Yipf1 | 230584 | protein_coding | 2752.10921 | 0.012756556 | 0.036079387 | 0.353569095 | 0.723661839 | 0.865314338 | 0.012756556 |
| ENSMUSG00000020844 | Nxn | 18230 | protein_coding | 37.53723495 | -0.012780739 | 0.219306468 | -0.058277986 | 0.953527203 | 0.981022852 | 0.012780739 |
| ENSMUSG00000024621 | Csf1r | 12978 | protein_coding | 13.58463931 | -0.012790863 | 0.335100816 | -0.038170194 | 0.969551985 | 0.987807993 | 0.012790863 |
| ENSMUSG00000029507 | Pus1 | 56361 | protein_coding | 564.1534333 | 0.012808024 | 0.079074542 | 0.161974052 | 0.871326287 | 0.941878077 | 0.012808024 |
| ENSMUSG00000034432 | Cops8 | 108679 | protein_coding | 579.7784296 | 0.012815825 | 0.058943562 | 0.217425353 | 0.827876875 | 0.923349179 | 0.012815825 |
| ENSMUSG00000045817 | Zfp36l2 | 12193 | protein_coding | 4263.999185 | 0.012827492 | 0.097540302 | 0.13150966 | 0.895372146 | 0.953302079 | 0.012827492 |
| ENSMUSG00000001016 | Ilf2 | 67781 | protein_coding | 1039.137031 | 0.012840184 | 0.059048462 | 0.21745163 | 0.827856399 | 0.923349179 | 0.012840184 |
| ENSMUSG00000025602 | Zfp202 | 80902 | protein_coding | 49.92087229 | 0.012842682 | 0.191263717 | 0.067146462 | 0.946465106 | 0.977373552 | 0.012842682 |
| ENSMUSG00000044125 | 9530080O11Rik | 319247 | lncRNA | 15.0393513 | 0.012861131 | 0.295225424 | 0.043563766 | 0.965252135 | 0.986105006 | 0.012861131 |
| ENSMUSG00000020272 | Stk10 | 20868 | protein_coding | 856.8141746 | -0.012877627 | 0.11302881 | -0.11393225 | 0.909291501 | 0.959446759 | 0.012877627 |
| ENSMUSG00000018428 | Akap1 | 11640 | protein_coding | 2625.890662 | 0.012879271 | 0.041327871 | 0.311636453 | 0.755316828 | 0.88389772 | 0.012879271 |
| ENSMUSG00000020485 | Supt4a | 20922 | protein_coding | 538.7481255 | 0.012880701 | 0.061389456 | 0.20981943 | 0.833808607 | 0.926340413 | 0.012880701 |
| ENSMUSG00000028195 | Ccn1 | 16007 | protein_coding | 1299.536267 | 0.012916057 | 0.136927972 | 0.094327379 | 0.924849102 | 0.967127575 | 0.012916057 |
| ENSMUSG00000024758 | Rtn3 | 20168 | protein_coding | 3148.540067 | -0.012917455 | 0.042310141 | -0.305304005 | 0.760134626 | 0.886576344 | 0.012917455 |
| ENSMUSG00000022246 | Rai14 | 75646 | protein_coding | 790.1374309 | -0.012963959 | 0.057563707 | -0.225210637 | 0.821815415 | 0.920376194 | 0.012963959 |
| ENSMUSG00000028312 | Smc2 | 14211 | protein_coding | 698.2104771 | 0.012968537 | 0.133182453 | 0.097374215 | 0.922429221 | 0.965912346 | 0.012968537 |
| ENSMUSG00000004864 | Mapk13 | 26415 | protein_coding | 2608.353684 | 0.012989749 | 0.047926586 | 0.271034307 | 0.786364646 | 0.900451559 | 0.012989749 |
| ENSMUSG00000003135 | Cnot11 | 52846 | protein_coding | 594.3011609 | 0.013052918 | 0.065835075 | 0.198266927 | 0.842836226 | 0.929981914 | 0.013052918 |
| ENSMUSG00000020570 | Sypl | 19027 | protein_coding | 3372.28474 | -0.013064076 | 0.047493343 | -0.275071729 | 0.783261131 | 0.898858056 | 0.013064076 |
| ENSMUSG00000025154 | Arhgap19 | 71085 | protein_coding | 245.537507 | -0.013069522 | 0.094496699 | -0.138306647 | 0.889998073 | 0.950907574 | 0.013069522 |
| ENSMUSG00000018474 | Chd3 | 216848 | protein_coding | 1551.090043 | 0.01315917 | 0.059063343 | 0.222797574 | 0.823693056 | 0.921535618 | 0.01315917 |
| ENSMUSG00000061171 | Slc38a11 | 320106 | protein_coding | 44.33006898 | -0.013198536 | 0.202396443 | -0.065211304 | 0.948005761 | 0.978275461 | 0.013198536 |
| ENSMUSG00000055538 | Zcchc24 | 71918 | protein_coding | 10.4413369 | 0.013204664 | 0.441264889 | 0.029924573 | 0.976127208 | 0.990526145 | 0.013204664 |
| ENSMUSG00000106918 | Mrpl33 | 66845 | protein_coding | 518.6401027 | -0.013241138 | 0.056055975 | -0.236212791 | 0.813267555 | 0.915984616 | 0.013241138 |
| ENSMUSG00000030652 | Coq7 | 12850 | protein_coding | 572.0021228 | 0.013244741 | 0.067407636 | 0.196487251 | 0.844228811 | 0.930210801 | 0.013244741 |
| ENSMUSG00000032380 | Dapk2 | 13143 | protein_coding | 738.1852063 | -0.013246587 | 0.054078897 | -0.244949283 | 0.806495701 | 0.912178372 | 0.013246587 |
| ENSMUSG00000058174 | Gm5148 | 381438 | protein_coding | 37.22268029 | 0.013250384 | 0.208102845 | 0.063672287 | 0.949231172 | 0.979038828 | 0.013250384 |

| ensembl_gene_id | external_gene_name | entrezgene_id | gene_biotype | baseMean | log2FoldChange | lfcSE | stat | pvalue | padj | absLog2FC |
|---|---|---|---|---|---|---|---|---|---|---|
| ENSMUSG00000031595 | Pdgfrl | 68797 | protein_coding | 5.917490556 | 0.013258106 | 0.522991077 | 0.02535054 | 0.979775361 | 0.992429328 | 0.013258106 |
| ENSMUSG00000029352 | Crybb3 | 12962 | protein_coding | 9.739613936 | 0.01326666 | 0.410691788 | 0.032303203 | 0.974230255 | 0.989464121 | 0.01326666 |
| ENSMUSG00000040631 | Dok4 | 114255 | protein_coding | 2233.860263 | -0.013273997 | 0.064289395 | -0.206472575 | 0.836421785 | 0.927459717 | 0.013273997 |
| ENSMUSG00000051469 | Zfp24 | 59057 | protein_coding | 919.5289387 | 0.013277853 | 0.051819733 | 0.256231598 | 0.797772011 | 0.907119032 | 0.013277853 |
| ENSMUSG00000057363 | Uxs1 | 67883 | protein_coding | 445.0665956 | 0.013290698 | 0.065754464 | 0.20212617 | 0.83981809 | 0.928990077 | 0.013290698 |
| ENSMUSG00000066037 | Hnrnpr | 74326 | protein_coding | 1075.582507 | 0.013296779 | 0.057107245 | 0.232838744 | 0.815886626 | 0.917020505 | 0.013296779 |
| ENSMUSG00000038605 | Samd10 | 229011 | protein_coding | 330.1238188 | -0.013304481 | 0.073588259 | -0.18079625 | 0.856527507 | 0.935610703 | 0.013304481 |
| ENSMUSG00000062906 | Hdac10 | 170787 | protein_coding | 462.1370744 | -0.013343565 | 0.056376661 | -0.236685984 | 0.812900409 | 0.915984616 | 0.013343565 |
| ENSMUSG00000028792 | Ak2 | 11637 | protein_coding | 4768.728558 | 0.013352237 | 0.052937134 | 0.252228177 | 0.8008647 | 0.909225274 | 0.013352237 |
| ENSMUSG00000037890 | Wdr19 | 213081 | protein_coding | 198.2515355 | -0.01336592 | 0.109055488 | -0.122560725 | 0.902454957 | 0.957052087 | 0.01336592 |
| ENSMUSG00000066278 | Vps37b | 330192 | protein_coding | 440.1243185 | 0.013380319 | 0.081494647 | 0.164186477 | 0.869584346 | 0.940691929 | 0.013380319 |
| ENSMUSG00000071724 | Smpd5 | 100503915 | protein_coding | 8.327074563 | -0.013382082 | 0.414553695 | -0.032280697 | 0.974248203 | 0.989464121 | 0.013382082 |
| ENSMUSG00000036046 | 5031439G07Rik | 223739 | protein_coding | 1129.256739 | -0.013386917 | 0.061126203 | -0.219004562 | 0.826646494 | 0.923060252 | 0.013386917 |
| ENSMUSG00000018932 | Map2k3 | 26397 | protein_coding | 998.0935301 | 0.013390225 | 0.051482432 | 0.260093095 | 0.794791964 | 0.904781532 | 0.013390225 |
| ENSMUSG00000022412 | Mief1 | 239555 | protein_coding | 1192.973928 | -0.013392306 | 0.044303949 | -0.302282455 | 0.762436752 | 0.887976999 | 0.013392306 |
| ENSMUSG00000028494 | Plin2 | 11520 | protein_coding | 2360.643122 | -0.013404193 | 0.067466479 | -0.198679304 | 0.842513614 | 0.929981434 | 0.013404193 |
| ENSMUSG00000018189 | Uchl5 | 56207 | protein_coding | 355.2896221 | -0.01342008 | 0.070416446 | -0.190581611 | 0.848853398 | 0.931678839 | 0.01342008 |
| ENSMUSG00000035011 | Zbtb7a | 16969 | protein_coding | 3213.727122 | 0.013451285 | 0.079877935 | 0.168398 | 0.866270181 | 0.939561451 | 0.013451285 |
| ENSMUSG00000058440 | Nrf1 | 18181 | protein_coding | 644.7284194 | 0.013470554 | 0.068308675 | 0.197201223 | 0.843670074 | 0.930175172 | 0.013470554 |
| ENSMUSG00000024944 | Arl2 | 56327 | protein_coding | 326.5891396 | -0.013475607 | 0.068534919 | -0.196623956 | 0.844121822 | 0.930210801 | 0.013475607 |
| ENSMUSG00000028920 | Fbxo42 | 213499 | protein_coding | 759.515908 | 0.013480433 | 0.077973071 | 0.172885749 | 0.862741234 | 0.937787345 | 0.013480433 |
| ENSMUSG00000070371 | Prss36 | 77613 | protein_coding | 55.06141587 | 0.013483318 | 0.162752072 | 0.082845755 | 0.933974187 | 0.971253657 | 0.013483318 |
| ENSMUSG00000029135 | Fosl2 | 14284 | protein_coding | 5181.804156 | 0.013512824 | 0.063087813 | 0.214190725 | 0.830398329 | 0.924850404 | 0.013512824 |
| ENSMUSG00000071172 | Srsf3 | 20383 | protein_coding | 2921.495149 | -0.013523837 | 0.088824012 | -0.152254288 | 0.878986377 | 0.945595293 | 0.013523837 |
| ENSMUSG00000022471 | Xrcc6 | 14375 | protein_coding | 538.2495155 | 0.013532448 | 0.079260615 | 0.170733569 | 0.864433266 | 0.938452729 | 0.013532448 |
| ENSMUSG00000024892 | Pcx | 18563 | protein_coding | 2060.909106 | -0.013537566 | 0.124974394 | -0.108322722 | 0.913739699 | 0.961718903 | 0.013537566 |
| ENSMUSG00000013275 | Slc41a1 | 98396 | protein_coding | 377.3457772 | -0.013555249 | 0.065969803 | -0.20547658 | 0.837199795 | 0.927812059 | 0.013555249 |
| ENSMUSG00000033430 | Terf2ip | 57321 | protein_coding | 495.8531295 | -0.013580488 | 0.076952835 | -0.176478074 | 0.859918372 | 0.936765956 | 0.013580488 |
| ENSMUSG00000025437 | Usp33 | 170822 | protein_coding | 1837.124609 | -0.01358332 | 0.046834561 | -0.290027701 | 0.771795046 | 0.892559247 | 0.01358332 |
| ENSMUSG00000059187 | Tafa1 | 320265 | protein_coding | 6.35708538 | -0.013589642 | 0.501883438 | -0.027077287 | 0.978398091 | 0.991618025 | 0.013589642 |
| ENSMUSG00000034825 | Nrip3 | 78593 | protein_coding | 19.03583329 | 0.01359788 | 0.353118229 | 0.038508008 | 0.969282647 | 0.987804745 | 0.01359788 |
| ENSMUSG00000032518 | Rpsa | 16785 | protein_coding | 18706.98596 | 0.013614023 | 0.045871061 | 0.29678892 | 0.766627665 | 0.890414429 | 0.013614023 |
| ENSMUSG00000024750 | Zfand5 | 22682 | protein_coding | 2839.045946 | 0.013634615 | 0.097346957 | 0.140062056 | 0.88861096 | 0.950733597 | 0.013634615 |
| ENSMUSG00000054400 | Cklf | 75458 | protein_coding | 99.50669374 | 0.013687914 | 0.121454549 | 0.11269989 | 0.910268488 | 0.959960814 | 0.013687914 |
| ENSMUSG00000021023 | Prorp | 66132 | protein_coding | 381.0046622 | 0.013776189 | 0.075203455 | 0.183185584 | 0.854652405 | 0.935123077 | 0.013776189 |
| ENSMUSG00000035215 | Lsm7 | 66094 | protein_coding | 173.2655916 | 0.013785652 | 0.095736964 | 0.143995082 | 0.885504353 | 0.949313881 | 0.013785652 |
| ENSMUSG00000024258 | Polr2d | 69241 | protein_coding | 262.5820876 | -0.013810054 | 0.090748458 | -0.152179494 | 0.879045367 | 0.945595293 | 0.013810054 |
| ENSMUSG00000038525 | Armc10 | 67211 | protein_coding | 820.3632108 | -0.01384908 | 0.071555727 | -0.19354258 | 0.846534062 | 0.930889346 | 0.01384908 |
| ENSMUSG00000031246 | Sh3bgrl | 56726 | protein_coding | 3189.545552 | 0.013854821 | 0.051657899 | 0.268203337 | 0.788542806 | 0.901391398 | 0.013854821 |

Supplementary Table 1

| ensembl_gene_id | external_gene_name | entrezgene_id | gene_biotype | baseMean | log2FoldChange | lfcSE | stat | pvalue | padj | absLog2FC |
|---|---|---|---|---|---|---|---|---|---|---|
| ENSMUSG00000039813 | Tbc1d2 | 381605 | protein_coding | 1297.63663 | -0.01385606 | 0.078947978 | -0.175508736 | 0.860679907 | 0.93708084 | 0.01385606 |
| ENSMUSG00000036977 | Anapc10 | 68999 | protein_coding | 149.3084819 | 0.013875199 | 0.111796281 | 0.124111458 | 0.901227026 | 0.956378776 | 0.013875199 |
| ENSMUSG00000024556 | Me2 | 107029 | protein_coding | 4519.761871 | 0.013890999 | 0.070502179 | 0.197029355 | 0.843804566 | 0.930175172 | 0.013890999 |
| ENSMUSG00000043257 | Pigv | 230801 | protein_coding | 595.8852998 | 0.01390561 | 0.091226641 | 0.152429267 | 0.878848375 | 0.945572603 | 0.01390561 |
| ENSMUSG00000026174 | Cnot9 | 58184 | protein_coding | 817.0169777 | 0.013911104 | 0.064262063 | 0.2164746 | 0.828617822 | 0.923812345 | 0.013911104 |
| ENSMUSG00000037062 | Sh3glb1 | 54673 | protein_coding | 2724.751675 | -0.013912193 | 0.037736717 | -0.368664642 | 0.712377706 | 0.858248564 | 0.013912193 |
| ENSMUSG00000060938 | Rpl26 | 19941 | protein_coding | 9238.86842 | -0.013930301 | 0.031100243 | -0.447916142 | 0.654213718 | 0.822000293 | 0.013930301 |
| ENSMUSG00000028040 | Efna4 | 13639 | protein_coding | 142.9283339 | 0.013939896 | 0.114802789 | 0.121424715 | 0.903354642 | 0.957533608 | 0.013939896 |
| ENSMUSG00000005102 | Eif2ak4 | 27103 | protein_coding | 671.3975905 | 0.013945055 | 0.061647281 | 0.226207135 | 0.821040322 | 0.920019265 | 0.013945055 |
| ENSMUSG00000029708 | Gcc1 | 74375 | protein_coding | 636.970465 | -0.013951329 | 0.061217107 | -0.227899194 | 0.819724609 | 0.919197481 | 0.013951329 |
| ENSMUSG00000055560 | Zfp459 | 328274 | protein_coding | 65.53060045 | 0.01397118 | 0.142500261 | 0.098043187 | 0.921898 | 0.965544165 | 0.01397118 |
| ENSMUSG00000032898 | Fbxo21 | 231670 | protein_coding | 597.430174 | -0.013992447 | 0.073980868 | -0.189136021 | 0.84998621 | 0.932292793 | 0.013992447 |
| ENSMUSG00000072258 | Taf1a | 21339 | protein_coding | 160.6035268 | -0.014082114 | 0.129516916 | -0.108727993 | 0.913418238 | 0.961596444 | 0.014082114 |
| ENSMUSG00000070730 | Rmdn3 | 67809 | protein_coding | 1327.422578 | -0.014082821 | 0.065695523 | -0.21436501 | 0.830262425 | 0.924826781 | 0.014082821 |
| ENSMUSG00000026563 | Tada1 | 27878 | protein_coding | 218.8674738 | 0.014083193 | 0.085124087 | 0.1654431 | 0.868595232 | 0.940565051 | 0.014083193 |
| ENSMUSG00000032307 | Ube2q2 | 109161 | protein_coding | 613.1326643 | -0.014086107 | 0.056513552 | -0.249251852 | 0.803165973 | 0.910363905 | 0.014086107 |
| ENSMUSG00000027650 | Tti1 | 75425 | protein_coding | 398.840748 | -0.014094007 | 0.069616319 | -0.202452631 | 0.839562886 | 0.928898921 | 0.014094007 |
| ENSMUSG00000036435 | Exoc1 | 69940 | protein_coding | 781.2050696 | -0.014122408 | 0.06349513 | -0.222417187 | 0.823989133 | 0.921722463 | 0.014122408 |
| ENSMUSG00000030016 | Zfp638 | 18139 | protein_coding | 1667.005102 | 0.014141789 | 0.074194542 | 0.190604163 | 0.848835728 | 0.931678839 | 0.014141789 |
| ENSMUSG00000049739 | Zfp646 | 233905 | protein_coding | 1495.967033 | -0.014147511 | 0.045683728 | -0.309683805 | 0.75680142 | 0.884543883 | 0.014147511 |
| ENSMUSG00000050708 | Ftl1 | 14325 | protein_coding | 8913.151782 | -0.014167095 | 0.049670928 | -0.285219056 | 0.775476322 | 0.894974876 | 0.014167095 |
| ENSMUSG00000015214 | Mtmr1 | 53332 | protein_coding | 769.8733244 | 0.014172453 | 0.049066258 | 0.288843162 | 0.772701401 | 0.89315908 | 0.014172453 |
| ENSMUSG00000006673 | Qrich1 | 69232 | protein_coding | 2061.965849 | -0.014182418 | 0.038277979 | -0.37051115 | 0.711001671 | 0.857064391 | 0.014182418 |
| ENSMUSG00000022800 | Fyttd1 | 69823 | protein_coding | 916.2935619 | 0.014208881 | 0.064310079 | 0.220943298 | 0.825136584 | 0.922174476 | 0.014208881 |
| ENSMUSG00000030432 | Rpl28 | 19943 | protein_coding | 7852.409729 | -0.014260171 | 0.049014819 | -0.290935913 | 0.771100334 | 0.892331737 | 0.014260171 |
| ENSMUSG00000022620 | Arsa | 11883 | protein_coding | 508.657059 | -0.014273848 | 0.068051164 | -0.209751715 | 0.833861461 | 0.926340413 | 0.014273848 |
| ENSMUSG00000040321 | Zfp770 | 228491 | protein_coding | 134.0862642 | -0.014277979 | 0.121046771 | -0.117954235 | 0.90610392 | 0.958651438 | 0.014277979 |
| ENSMUSG00000031983 | 2310022B05Rik | 69551 | protein_coding | 458.6061429 | 0.014289442 | 0.082792407 | 0.172593631 | 0.862970859 | 0.937787345 | 0.014289442 |
| ENSMUSG00000022526 | Zfp251 | 71591 | protein_coding | 316.6942839 | -0.014338162 | 0.083887337 | -0.170921648 | 0.864285375 | 0.938355341 | 0.014338162 |
| ENSMUSG00000026155 | Smap1 | 98366 | protein_coding | 526.6963224 | -0.014381826 | 0.0597693 | -0.240622299 | 0.809847866 | 0.914534473 | 0.014381826 |
| ENSMUSG00000033047 | Eif3l | 223691 | protein_coding | 3691.626369 | 0.014393068 | 0.045789464 | 0.314331431 | 0.753269329 | 0.882654542 | 0.014393068 |
| ENSMUSG00000024604 | Rbm22 | 66810 | protein_coding | 835.894079 | -0.01441141 | 0.049596494 | -0.290573159 | 0.771377791 | 0.892332662 | 0.01441141 |
| ENSMUSG00000020246 | Hcfc2 | 67933 | protein_coding | 364.3084031 | -0.014421328 | 0.071479308 | -0.201755279 | 0.840108045 | 0.929183351 | 0.014421328 |
| ENSMUSG00000028042 | Zbtb7b | 22724 | protein_coding | 5616.685248 | 0.014441548 | 0.088742069 | 0.162736212 | 0.870726134 | 0.941634557 | 0.014441548 |
| ENSMUSG00000027185 | Nat10 | 98956 | protein_coding | 558.7958718 | -0.014467272 | 0.076038387 | -0.190262736 | 0.849103252 | 0.931783353 | 0.014467272 |
| ENSMUSG00000039318 | Rab3gap2 | 98732 | protein_coding | 1225.711659 | 0.014516371 | 0.04532309 | 0.320286437 | 0.748751204 | 0.880475742 | 0.014516371 |
| ENSMUSG00000021518 | Ptdss1 | 19210 | protein_coding | 2601.623433 | 0.014570814 | 0.04022939 | 0.362193251 | 0.717207622 | 0.861383838 | 0.014570814 |
| ENSMUSG00000032220 | Myo1e | 71602 | protein_coding | 2524.858336 | 0.01457361 | 0.061021687 | 0.238826727 | 0.811239944 | 0.915013749 | 0.01457361 |
| ENSMUSG00000061838 | Suclg2 | 20917 | protein_coding | 6152.38283 | -0.014574949 | 0.032018567 | -0.455203036 | 0.648963177 | 0.818730278 | 0.014574949 |

Supplementary Table 1

| ensembl_gene_id | external_gene_name | entrezgene_id | gene_biotype | baseMean | log2FoldChange | lfcSE | stat | pvalue | padj | absLog2FC |
|---|---|---|---|---|---|---|---|---|---|---|
| ENSMUSG00000024777 | Ppp2r5b | 225849 | protein_coding | 827.8140676 | 0.014596104 | 0.072774693 | 0.20056565 | 0.841038219 | 0.929613548 | 0.014596104 |
| ENSMUSG00000000001 | Gnai3 | 14679 | protein_coding | 4128.888081 | 0.014614911 | 0.047351651 | 0.308646284 | 0.757590609 | 0.885131169 | 0.014614911 |
| ENSMUSG00000044847 | Lsm11 | 72290 | protein_coding | 302.6693142 | -0.014641469 | 0.099036993 | -0.147838381 | 0.882470319 | 0.947572996 | 0.014641469 |
| ENSMUSG00000074282 | Zfp94 | 22756 | protein_coding | 52.41304029 | -0.014651552 | 0.164322687 | -0.089163292 | 0.928952138 | 0.968789653 | 0.014651552 |
| ENSMUSG00000028960 | Ube4b | 63958 | protein_coding | 1188.49136 | -0.014675284 | 0.051939352 | -0.282546542 | 0.777524464 | 0.89645106 | 0.014675284 |
| ENSMUSG00000003849 | Nqo1 | 18104 | protein_coding | 1209.302728 | -0.01468356 | 0.078590635 | -0.186836002 | 0.851789218 | 0.933520082 | 0.01468356 |
| ENSMUSG00000026568 | Mpc2 | 70456 | protein_coding | 1818.163765 | 0.014694575 | 0.073606759 | 0.199636225 | 0.841765094 | 0.929706079 | 0.014694575 |
| ENSMUSG00000021211 | Akr1c12 | 622402 | protein_coding | 3053.700821 | -0.014707207 | 0.048169747 | -0.305320419 | 0.760122126 | 0.886576344 | 0.014707207 |
| ENSMUSG00000028163 | Nfkb1 | 18033 | protein_coding | 2231.241845 | -0.014722639 | 0.035944053 | -0.409598755 | 0.68210031 | 0.839327919 | 0.014722639 |
| ENSMUSG00000015357 | Clpx | 270166 | protein_coding | 1266.693408 | 0.014743219 | 0.052424553 | 0.28122736 | 0.778536021 | 0.896941755 | 0.014743219 |
| ENSMUSG00000022952 | Runx1 | 12394 | protein_coding | 283.3262247 | -0.014808497 | 0.086074062 | -0.172043659 | 0.863403206 | 0.937932724 | 0.014808497 |
| ENSMUSG00000062031 | Pgghg | 212974 | protein_coding | 760.2734635 | -0.014821275 | 0.052349252 | -0.283122948 | 0.777082591 | 0.896141595 | 0.014821275 |
| ENSMUSG00000018322 | Tomm34 | 67145 | protein_coding | 511.9833101 | -0.014830103 | 0.070067436 | -0.211654718 | 0.832376417 | 0.92580236 | 0.014830103 |
| ENSMUSG00000029569 | Tmem168 | 101118 | protein_coding | 1902.703561 | 0.014838009 | 0.059342741 | 0.250039158 | 0.802557066 | 0.910121497 | 0.014838009 |
| ENSMUSG00000032456 | Nmnat3 | 74080 | protein_coding | 339.8426994 | -0.01485157 | 0.096110979 | -0.154525221 | 0.877195626 | 0.945176551 | 0.01485157 |
| ENSMUSG00000038446 | Cdc40 | 71713 | protein_coding | 523.7953046 | -0.01485905 | 0.064472409 | -0.230471457 | 0.817725441 | 0.918128862 | 0.01485905 |
| ENSMUSG00000026753 | Ppp6c | 67857 | protein_coding | 1466.011534 | 0.01486557 | 0.042403515 | 0.350574002 | 0.725907963 | 0.866137742 | 0.01486557 |
| ENSMUSG00000021213 | Akr1c13 | 27384 | protein_coding | 4571.086293 | 0.014871661 | 0.055656604 | 0.267203888 | 0.789312183 | 0.901682062 | 0.014871661 |
| ENSMUSG00000024425 | Ndfip1 | 65113 | protein_coding | 2343.778224 | 0.014877998 | 0.036964014 | 0.402499539 | 0.687316428 | 0.842728192 | 0.014877998 |
| ENSMUSG00000001942 | Siae | 22619 | protein_coding | 778.5246189 | -0.014887975 | 0.06708394 | -0.221930548 | 0.82436795 | 0.921867382 | 0.014887975 |
| ENSMUSG00000020715 | Ern1 | 78943 | protein_coding | 1023.222766 | -0.014897153 | 0.063709033 | -0.233831089 | 0.815116113 | 0.916810722 | 0.014897153 |
| ENSMUSG00000017550 | Atad5 | 237877 | protein_coding | 192.3187664 | -0.014898585 | 0.128888389 | -0.115592917 | 0.907975177 | 0.959122007 | 0.014898585 |
| ENSMUSG00000061286 | Exosc5 | 27998 | protein_coding | 435.0282441 | 0.01492352 | 0.088589935 | 0.168456162 | 0.866224428 | 0.939561451 | 0.01492352 |
| ENSMUSG00000038705 | Gmeb2 | 229004 | protein_coding | 676.7762233 | -0.014926059 | 0.052827546 | -0.282543113 | 0.777527093 | 0.89645106 | 0.014926059 |
| ENSMUSG00000018143 | Mafk | 17135 | protein_coding | 2511.649872 | -0.014939256 | 0.082589199 | -0.180886312 | 0.856456813 | 0.935610703 | 0.014939256 |
| ENSMUSG00000050271 | Prag1 | 244418 | protein_coding | 822.7523647 | 0.014983676 | 0.077399558 | 0.193588654 | 0.846497983 | 0.930889346 | 0.014983676 |
| ENSMUSG00000037280 | Galnt6 | 207839 | protein_coding | 3064.078109 | 0.015009955 | 0.056886091 | 0.263859847 | 0.791887916 | 0.903021982 | 0.015009955 |
| ENSMUSG00000031505 | Naxd | 69225 | protein_coding | 1125.227734 | 0.015045986 | 0.047766524 | 0.314990174 | 0.752769114 | 0.882492633 | 0.015045986 |
| ENSMUSG00000060923 | Acyp2 | 75572 | protein_coding | 86.83409667 | 0.015055746 | 0.173583892 | 0.086734696 | 0.930882397 | 0.969665987 | 0.015055746 |
| ENSMUSG00000009090 | Ap1b1 | 11764 | protein_coding | 3304.594307 | -0.015058311 | 0.054620543 | -0.275689509 | 0.782786554 | 0.898825528 | 0.015058311 |
| ENSMUSG00000080538 | Gm25541 | 115489638 | snRNA | 17.84605841 | -0.015061691 | 0.367222184 | -0.0410152 | 0.967283778 | 0.986873905 | 0.015061691 |
| ENSMUSG00000022855 | Senp2 | 75826 | protein_coding | 886.9897912 | -0.01507399 | 0.058431654 | -0.257976442 | 0.796425089 | 0.906034665 | 0.01507399 |
| ENSMUSG00000024561 | Mbd1 | 17190 | protein_coding | 1502.933686 | 0.015095492 | 0.046767225 | 0.322779294 | 0.746862395 | 0.879915444 | 0.015095492 |
| ENSMUSG00000002771 | Grin2d | 14814 | protein_coding | 60.08558339 | 0.015106749 | 0.172760911 | 0.087443097 | 0.930319314 | 0.969328051 | 0.015106749 |
| ENSMUSG00000068391 | Chrac1 | 93696 | protein_coding | 396.9064383 | -0.015110806 | 0.075527724 | -0.200069653 | 0.841426107 | 0.929613548 | 0.015110806 |
| ENSMUSG00000017837 | Nkiras2 | 71966 | protein_coding | 1913.325161 | 0.015163443 | 0.066774317 | 0.227084953 | 0.820357684 | 0.919509896 | 0.015163443 |
| ENSMUSG00000025357 | Dgka | 13139 | protein_coding | 2592.841556 | 0.015178228 | 0.061527957 | 0.246688309 | 0.805149455 | 0.911459024 | 0.015178228 |
| ENSMUSG00000062352 | Itgb1bp1 | 16413 | protein_coding | 130.4730006 | 0.015185729 | 0.12951044 | 0.11725486 | 0.906658094 | 0.95873684 | 0.015185729 |
| ENSMUSG00000034278 | Dnajc17 | 69408 | protein_coding | 159.3160762 | 0.015187535 | 0.101175217 | 0.150111221 | 0.880676867 | 0.946655725 | 0.015187535 |

| ensembl_gene_id | external_gene_name | entrezgene_id | gene_biotype | baseMean | log2FoldChange | lfcSE | stat | pvalue | padj | absLog2FC |
|---|---|---|---|---|---|---|---|---|---|---|
| ENSMUSG00000001247 | Lsr | 54135 | protein_coding | 1341.801982 | 0.015193491 | 0.074423433 | 0.204149294 | 0.838236834 | 0.928098115 | 0.015193491 |
| ENSMUSG00000041000 | Trim62 | 67525 | protein_coding | 114.2405753 | 0.0152605 | 0.133587371 | 0.1142361 | 0.909050636 | 0.959446759 | 0.0152605 |
| ENSMUSG00000061894 | Zscan20 | 269585 | protein_coding | 182.5730863 | 0.015267903 | 0.097151549 | 0.157155531 | 0.875122274 | 0.943767347 | 0.015267903 |
| ENSMUSG00000109901 | Chmp1b | 67064 | protein_coding | 1354.756192 | 0.015271892 | 0.051796114 | 0.294846287 | 0.768111305 | 0.891239011 | 0.015271892 |
| ENSMUSG00000078941 | Ak6 | 108143 | protein_coding | 199.2624337 | 0.015282838 | 0.097122594 | 0.157356153 | 0.874964167 | 0.943745489 | 0.015282838 |
| ENSMUSG00000034853 | Acot11 | 329910 | protein_coding | 1492.35631 | 0.015284428 | 0.063001794 | 0.242603062 | 0.808312909 | 0.913597373 | 0.015284428 |
| ENSMUSG00000025264 | Tsr2 | 69499 | protein_coding | 229.2444912 | 0.015291428 | 0.096832879 | 0.157915657 | 0.874523261 | 0.943745489 | 0.015291428 |
| ENSMUSG00000062949 | Atp11c | 320940 | protein_coding | 478.8987401 | -0.015303234 | 0.071281268 | -0.214688011 | 0.83001057 | 0.924673975 | 0.015303234 |
| ENSMUSG00000004994 | Ccdc130 | 67736 | protein_coding | 169.3090073 | 0.015327849 | 0.093757188 | 0.16348452 | 0.870136961 | 0.941223339 | 0.015327849 |
| ENSMUSG00000022897 | Dyrk1a | 13548 | protein_coding | 1651.447504 | -0.01536236 | 0.044377359 | -0.346175622 | 0.729210734 | 0.868664612 | 0.01536236 |
| ENSMUSG00000037337 | Map4k1 | 26411 | protein_coding | 23.76666343 | -0.015369928 | 0.254903212 | -0.060297115 | 0.951919 | 0.980585772 | 0.015369928 |
| ENSMUSG00000024479 | Mal2 | 105853 | protein_coding | 6615.065442 | 0.015401855 | 0.052719743 | 0.292145857 | 0.770175106 | 0.892032325 | 0.015401855 |
| ENSMUSG00000026883 | Dab2ip | 69601 | protein_coding | 3462.164233 | -0.015406487 | 0.05827149 | -0.2643915 | 0.791478261 | 0.902954861 | 0.015406487 |
| ENSMUSG00000059839 | Zfp874b | 408067 | protein_coding | 152.9945792 | -0.015407441 | 0.160480915 | -0.096007933 | 0.923514272 | 0.966483755 | 0.015407441 |
| ENSMUSG00000032705 | Exd2 | 97827 | protein_coding | 422.7790658 | -0.015432286 | 0.061521707 | -0.250842935 | 0.801935545 | 0.909865152 | 0.015432286 |
| ENSMUSG00000067629 | Syngap1 | 240057 | protein_coding | 11.48514055 | -0.015435288 | 0.454089422 | -0.033991738 | 0.972883739 | 0.989206294 | 0.015435288 |
| ENSMUSG00000026641 | Usf1 | 22278 | protein_coding | 875.5380273 | -0.015451141 | 0.058289142 | -0.265077507 | 0.790949754 | 0.90265349 | 0.015451141 |
| ENSMUSG00000020132 | Rab21 | 216344 | protein_coding | 947.647805 | 0.015452307 | 0.072550232 | 0.212987698 | 0.831336558 | 0.925325507 | 0.015452307 |
| ENSMUSG00000054469 | Lclat1 | 225010 | protein_coding | 1933.217533 | -0.015455889 | 0.043933014 | -0.351805792 | 0.724983915 | 0.865987203 | 0.015455889 |
| ENSMUSG00000051864 | Tbc1d22a | 223754 | protein_coding | 1608.96035 | -0.015456956 | 0.064215469 | -0.240704558 | 0.809784106 | 0.914534473 | 0.015456956 |
| ENSMUSG00000062006 | Rpl34 | 68436 | protein_coding | 5487.52856 | 0.015460329 | 0.034412934 | 0.449259255 | 0.653244647 | 0.821648745 | 0.015460329 |
| ENSMUSG00000031370 | Zrsr2 | 22184 | protein_coding | 437.5160725 | 0.015485158 | 0.063959843 | 0.242107512 | 0.808696858 | 0.913844956 | 0.015485158 |
| ENSMUSG00000002395 | Use1 | 67023 | protein_coding | 1671.963447 | -0.015499624 | 0.045691098 | -0.339226335 | 0.734439232 | 0.87231562 | 0.015499624 |
| ENSMUSG00000030929 | Eri2 | 71151 | protein_coding | 126.8820748 | 0.015527718 | 0.139829861 | 0.111047227 | 0.911578896 | 0.960412417 | 0.015527718 |
| ENSMUSG00000071649 | B3gat3 | 72727 | protein_coding | 732.4156407 | 0.015553289 | 0.083276583 | 0.186766662 | 0.851843587 | 0.933520082 | 0.015553289 |
| ENSMUSG00000052392 | Acot4 | 171282 | protein_coding | 141.1159512 | 0.015556872 | 0.108575255 | 0.143281925 | 0.88606753 | 0.949475732 | 0.015556872 |
| ENSMUSG00000078816 | Prkcg | 18752 | protein_coding | 7.124979475 | -0.015563684 | 0.470302305 | -0.033092937 | 0.973600475 | 0.989317668 | 0.015563684 |
| ENSMUSG00000017176 | Nt5c3b | 68106 | protein_coding | 168.6394125 | 0.015570356 | 0.095296101 | 0.163389227 | 0.870211985 | 0.94124136 | 0.015570356 |
| ENSMUSG00000026917 | Wdr5 | 140858 | protein_coding | 851.3673613 | -0.015582219 | 0.057453954 | -0.271212308 | 0.786227748 | 0.900404409 | 0.015582219 |
| ENSMUSG00000103793 | Pcdhga6 | 93714 | protein_coding | 9.319372705 | -0.015619082 | 0.391198532 | -0.039926228 | 0.968151941 | 0.987251745 | 0.015619082 |
| ENSMUSG00000029833 | Trim24 | 21848 | protein_coding | 1328.351072 | 0.015641113 | 0.06195875 | 0.25244397 | 0.800697917 | 0.909183504 | 0.015641113 |
| ENSMUSG00000033808 | Tmem87a | 211499 | protein_coding | 1679.328871 | -0.015656682 | 0.053944847 | -0.290234989 | 0.77163647 | 0.892439856 | 0.015656682 |
| ENSMUSG00000032601 | Prkar2a | 19087 | protein_coding | 3444.568676 | 0.01565785 | 0.054831499 | 0.285563047 | 0.77521281 | 0.894779343 | 0.01565785 |
| ENSMUSG00000024571 | Naa12 | 27366 | protein_coding | 1018.864887 | 0.01566572 | 0.056697881 | 0.276301685 | 0.782316362 | 0.898825528 | 0.01566572 |
| ENSMUSG00000022761 | Lztr1 | 66863 | protein_coding | 1457.782033 | 0.015684146 | 0.039345994 | 0.398621164 | 0.690172365 | 0.844556471 | 0.015684146 |
| ENSMUSG00000027663 | Zmat3 | 22401 | protein_coding | 359.8715644 | 0.015699372 | 0.128479392 | 0.122193695 | 0.90274562 | 0.957171506 | 0.015699372 |
| ENSMUSG00000038838 | Vars2 | 68915 | protein_coding | 759.2477359 | -0.015701888 | 0.066579452 | -0.235836852 | 0.813559271 | 0.916202738 | 0.015701888 |
| ENSMUSG00000058812 | 0610039K10Rik | 68386 | lncRNA | 25.25733136 | -0.015709102 | 0.258312951 | -0.060814224 | 0.951507162 | 0.98032045 | 0.015709102 |
| ENSMUSG00000044715 | Gskip | 66787 | protein_coding | 334.5258158 | 0.015717193 | 0.082859651 | 0.189684523 | 0.849556349 | 0.931963489 | 0.015717193 |

| ensembl_gene_id | external_gene_name | entrezgene_id | gene_biotype | baseMean | log2FoldChange | lfcSE | stat | pvalue | padj | absLog2FC |
|---|---|---|---|---|---|---|---|---|---|---|
| ENSMUSG00000027714 | Exosc9 | 50911 | protein_coding | 562.0209375 | 0.015718061 | 0.067375124 | 0.233291759 | 0.815534857 | 0.917017949 | 0.015718061 |
| ENSMUSG00000009995 | Tafazzin | 66826 | protein_coding | 1132.019473 | 0.015745521 | 0.047389533 | 0.332257364 | 0.739694932 | 0.875013582 | 0.015745521 |
| ENSMUSG00000044350 | Lacc1 | 210808 | protein_coding | 163.9844454 | 0.015747841 | 0.124928634 | 0.126054699 | 0.899688626 | 0.955523695 | 0.015747841 |
| ENSMUSG00000026792 | Lrsam1 | 227738 | protein_coding | 448.8868017 | 0.01575778 | 0.072663385 | 0.216859974 | 0.828317471 | 0.92361722 | 0.01575778 |
| ENSMUSG00000030086 | Chchd6 | 66098 | protein_coding | 81.2429276 | -0.015766157 | 0.147898544 | -0.106601164 | 0.915105395 | 0.962265867 | 0.015766157 |
| ENSMUSG00000049755 | Zfp672 | 319475 | protein_coding | 875.6108525 | 0.015833056 | 0.056765989 | 0.278917999 | 0.780307755 | 0.898056783 | 0.015833056 |
| ENSMUSG00000025138 | Sirt7 | 209011 | protein_coding | 1445.995555 | 0.01583515 | 0.067415468 | 0.234888968 | 0.814294913 | 0.916408517 | 0.01583515 |
| ENSMUSG00000069844 | Sco1 | 52892 | protein_coding | 1331.881583 | 0.015835198 | 0.053088201 | 0.298280926 | 0.765488761 | 0.889849511 | 0.015835198 |
| ENSMUSG00000051483 | Cbr1 | 12408 | protein_coding | 3699.770748 | -0.015836032 | 0.073062502 | -0.216746363 | 0.828406014 | 0.923652088 | 0.015836032 |
| ENSMUSG00000037012 | Hk1 | 15275 | protein_coding | 3764.315623 | 0.015852844 | 0.089016791 | 0.17808824 | 0.858653677 | 0.93621174 | 0.015852844 |
| ENSMUSG00000032012 | Nectin1 | 58235 | protein_coding | 247.545044 | 0.015879165 | 0.128895742 | 0.123193868 | 0.901953581 | 0.956772045 | 0.015879165 |
| ENSMUSG00000009356 | Lpo | 76113 | protein_coding | 350.6066817 | 0.015933684 | 0.205976879 | 0.077356661 | 0.938339817 | 0.97297734 | 0.015933684 |
| ENSMUSG00000004961 | Syt5 | 53420 | protein_coding | 17.89187114 | 0.015943047 | 0.381856973 | 0.041751357 | 0.966696913 | 0.986641422 | 0.015943047 |
| ENSMUSG00000074865 | Zfp934 | 77117 | protein_coding | 58.29997218 | -0.015958505 | 0.191033445 | -0.083537757 | 0.933423957 | 0.970931898 | 0.015958505 |
| ENSMUSG00000008690 | Ncaph2 | 52683 | protein_coding | 2136.740841 | 0.015964547 | 0.052442675 | 0.304419015 | 0.760808683 | 0.886968072 | 0.015964547 |
| ENSMUSG00000008976 | Gabpa | 14390 | protein_coding | 1121.159797 | -0.016018854 | 0.066642488 | -0.240369982 | 0.810043447 | 0.914623726 | 0.016018854 |
| ENSMUSG00000020098 | Pcbd1 | 13180 | protein_coding | 604.1338167 | 0.01602925 | 0.060046403 | 0.266947708 | 0.789509423 | 0.901729353 | 0.01602925 |
| ENSMUSG00000027387 | Zc3h8 | 57432 | protein_coding | 63.27149865 | -0.016041114 | 0.160230927 | -0.10011247 | 0.920255035 | 0.964898263 | 0.016041114 |
| ENSMUSG00000028840 | Zfp593 | 68040 | protein_coding | 24.93890524 | -0.016102496 | 0.298936649 | -0.053865916 | 0.957041992 | 0.982772999 | 0.016102496 |
| ENSMUSG00000037103 | Dcaf15 | 212123 | protein_coding | 998.4885949 | 0.01610444 | 0.057278628 | 0.281159671 | 0.778587935 | 0.896941755 | 0.01610444 |
| ENSMUSG00000029524 | Sirt4 | 75387 | protein_coding | 126.9188562 | -0.016123314 | 0.108969779 | -0.147961332 | 0.882373285 | 0.947541514 | 0.016123314 |
| ENSMUSG00000022325 | Pop1 | 67724 | protein_coding | 147.8791818 | -0.016140876 | 0.104513905 | -0.154437592 | 0.877264715 | 0.945176551 | 0.016140876 |
| ENSMUSG00000092607 | Scnm1 | 69269 | protein_coding | 339.7780199 | 0.016152056 | 0.065818876 | 0.245401583 | 0.806145503 | 0.912178372 | 0.016152056 |
| ENSMUSG00000030059 | Tmf1 | 232286 | protein_coding | 3071.706119 | -0.016153174 | 0.042853358 | -0.376940688 | 0.706217683 | 0.853889082 | 0.016153174 |
| ENSMUSG00000001674 | Ddx18 | 66942 | protein_coding | 322.1455382 | -0.016172962 | 0.100810139 | -0.160429919 | 0.872542423 | 0.942710088 | 0.016172962 |
| ENSMUSG00000039356 | Exosc2 | 227715 | protein_coding | 367.6193442 | 0.016191839 | 0.076326799 | 0.21213832 | 0.831999125 | 0.925610506 | 0.016191839 |
| ENSMUSG00000021932 | Rnaseh2b | 67153 | protein_coding | 290.112462 | 0.016192445 | 0.093722425 | 0.172770233 | 0.862832035 | 0.937787345 | 0.016192445 |
| ENSMUSG00000033382 | Trappc8 | 75964 | protein_coding | 1989.916138 | 0.016220829 | 0.043684171 | 0.371320508 | 0.710398825 | 0.856611363 | 0.016220829 |
| ENSMUSG00000005968 | Tuft1 | 22156 | protein_coding | 926.7984762 | -0.016231234 | 0.05089346 | -0.318925742 | 0.749782823 | 0.880488429 | 0.016231234 |
| ENSMUSG00000003344 | Btbd2 | 208198 | protein_coding | 842.8005535 | -0.016238602 | 0.076936432 | -0.211065183 | 0.832836408 | 0.92587823 | 0.016238602 |
| ENSMUSG00000015355 | Cd48 | 12506 | protein_coding | 10.25032244 | 0.016253496 | 0.380328591 | 0.042735403 | 0.965912457 | 0.986201549 | 0.016253496 |
| ENSMUSG00000036197 | Gxylt1 | 223827 | protein_coding | 781.5541187 | -0.016276305 | 0.048194906 | -0.337718359 | 0.735575439 | 0.872889311 | 0.016276305 |
| ENSMUSG00000022779 | Top3b | 21976 | protein_coding | 894.4704303 | 0.016283179 | 0.051654191 | 0.315234421 | 0.752583673 | 0.882492321 | 0.016283179 |
| ENSMUSG00000028975 | Pex14 | 56273 | protein_coding | 577.5958807 | -0.016296541 | 0.070680103 | -0.230567598 | 0.817650742 | 0.918128862 | 0.016296541 |
| ENSMUSG00000026638 | Irf6 | 54139 | protein_coding | 4674.851431 | -0.016297241 | 0.032244356 | -0.505429261 | 0.613257371 | 0.793911553 | 0.016297241 |
| ENSMUSG00000060636 | Rpl35a | 57808 | protein_coding | 4332.024529 | -0.016299107 | 0.048741301 | -0.334400316 | 0.738077501 | 0.874596535 | 0.016299107 |
| ENSMUSG00000031622 | Sin3b | 20467 | protein_coding | 1534.46523 | -0.016386154 | 0.047945722 | -0.341764674 | 0.732527997 | 0.871007815 | 0.016386154 |
| ENSMUSG00000040007 | Bahd1 | 228536 | protein_coding | 846.9515907 | -0.016388895 | 0.103218914 | -0.158778019 | 0.873843767 | 0.943461099 | 0.016388895 |
| ENSMUSG00000053565 | Eif3k | 73830 | protein_coding | 2515.140962 | 0.016416956 | 0.045611539 | 0.359929894 | 0.718899561 | 0.862409121 | 0.016416956 |

| ensembl_gene_id | external_gene_name | entrezgene_id | gene_biotype | baseMean | log2FoldChange | lfcSE | stat | pvalue | padj | absLog2FC |
|---|---|---|---|---|---|---|---|---|---|---|
| ENSMUSG00000042650 | Alkbh5 | 268420 | protein_coding | 3908.600341 | 0.016425196 | 0.052168653 | 0.314847993 | 0.752877071 | 0.882492633 | 0.016425196 |
| ENSMUSG00000013150 | Gfod2 | 70575 | protein_coding | 391.3261806 | 0.016451325 | 0.101630694 | 0.161873589 | 0.871405401 | 0.941900463 | 0.016451325 |
| ENSMUSG00000034807 | Colgalt1 | 234407 | protein_coding | 1166.167993 | -0.016480052 | 0.053438183 | -0.308394689 | 0.757782023 | 0.885240904 | 0.016480052 |
| ENSMUSG00000038024 | Dennd4c | 329877 | protein_coding | 2793.186327 | -0.016486977 | 0.037807841 | -0.436072956 | 0.662783798 | 0.827627866 | 0.016486977 |
| ENSMUSG00000026854 | Usp20 | 74270 | protein_coding | 365.8738969 | -0.016488462 | 0.080274382 | -0.205401298 | 0.837258606 | 0.927813478 | 0.016488462 |
| ENSMUSG00000039982 | Dtx4 | 207521 | protein_coding | 539.409458 | 0.01649212 | 0.091196441 | 0.180841703 | 0.856491829 | 0.935610703 | 0.01649212 |
| ENSMUSG00000084910 | C630043F03Rik | 68285 | lncRNA | 51.58932497 | 0.01650121 | 0.181391233 | 0.090970274 | 0.927516211 | 0.968095473 | 0.01650121 |
| ENSMUSG00000058503 | Fam133b | 68152 | protein_coding | 340.3001361 | -0.016543945 | 0.082543758 | -0.200426364 | 0.841147142 | 0.929613548 | 0.016543945 |
| ENSMUSG00000043716 | Rpl7 | 19989 | protein_coding | 18132.3568 | -0.016557714 | 0.051736331 | -0.320040351 | 0.748937742 | 0.880475742 | 0.016557714 |
| ENSMUSG00000043639 | Rbm20 | 73713 | protein_coding | 110.7157833 | -0.016562981 | 0.128877208 | -0.128517533 | 0.897739424 | 0.954463189 | 0.016562981 |
| ENSMUSG00000040268 | Plekha1 | 101476 | protein_coding | 1367.355255 | 0.016611594 | 0.07144151 | 0.232520193 | 0.816134006 | 0.917183108 | 0.016611594 |
| ENSMUSG00000049489 | Ccnq | 69109 | protein_coding | 269.6305694 | -0.016632906 | 0.114774876 | -0.144917654 | 0.884775887 | 0.948995857 | 0.016632906 |
| ENSMUSG00000113679 | Gm10432 | 100038713 | lncRNA | 8.157993875 | -0.016658518 | 0.470001479 | -0.035443543 | 0.971726064 | 0.988645905 | 0.016658518 |
| ENSMUSG00000000355 | Mcts1 | 68995 | protein_coding | 344.7453677 | -0.016659979 | 0.066899371 | -0.249030423 | 0.803337249 | 0.910371624 | 0.016659979 |
| ENSMUSG00000025486 | Sirt3 | 64384 | protein_coding | 362.0969324 | -0.016664431 | 0.072570148 | -0.229632033 | 0.818377713 | 0.918354815 | 0.016664431 |
| ENSMUSG00000032512 | Wdr48 | 67561 | protein_coding | 1466.688343 | -0.016727213 | 0.050547918 | -0.330917945 | 0.740706467 | 0.875888927 | 0.016727213 |
| ENSMUSG00000057278 | Snrpg | 68011 | protein_coding | 435.5138358 | 0.016749522 | 0.097112965 | 0.172474621 | 0.863064412 | 0.937787345 | 0.016749522 |
| ENSMUSG00000001441 | Npepps | 19155 | protein_coding | 1519.606116 | 0.01675255 | 0.048337042 | 0.346577895 | 0.728908455 | 0.868368667 | 0.01675255 |
| ENSMUSG00000037242 | Clic4 | 29876 | protein_coding | 1048.033903 | 0.016757388 | 0.088648366 | 0.189032109 | 0.850067651 | 0.932292793 | 0.016757388 |
| ENSMUSG00000025261 | Huwe1 | 59026 | protein_coding | 5702.752632 | 0.016760033 | 0.068280035 | 0.245460224 | 0.806100103 | 0.912178372 | 0.016760033 |
| ENSMUSG00000028889 | Yrdc | 230734 | protein_coding | 431.9452723 | 0.016761065 | 0.068912689 | 0.243221758 | 0.807833612 | 0.913151515 | 0.016761065 |
| ENSMUSG00000036748 | Cuedc2 | 67116 | protein_coding | 447.1596836 | 0.016773001 | 0.070630539 | 0.237475194 | 0.812288161 | 0.915791177 | 0.016773001 |
| ENSMUSG00000023156 | Rpp14 | 67053 | protein_coding | 430.3339744 | -0.016775335 | 0.062997719 | -0.266284806 | 0.790019874 | 0.902120505 | 0.016775335 |
| ENSMUSG00000008305 | Tle1 | 21885 | protein_coding | 1043.229056 | 0.016780845 | 0.050457053 | 0.332576794 | 0.739453763 | 0.874869968 | 0.016780845 |
| ENSMUSG00000040407 | Akap9 | 100986 | protein_coding | 5873.545347 | -0.016814941 | 0.050391629 | -0.333685196 | 0.738617122 | 0.874765244 | 0.016814941 |
| ENSMUSG00000040139 | 9430038I01Rik | 77252 | protein_coding | 255.7866196 | -0.01686264 | 0.102451663 | -0.164591182 | 0.869265772 | 0.940691929 | 0.01686264 |
| ENSMUSG00000024392 | Bag6 | 224727 | protein_coding | 2793.256188 | -0.016870197 | 0.057792932 | -0.291907622 | 0.770357256 | 0.892047592 | 0.016870197 |
| ENSMUSG00000056917 | Sipa1 | 20469 | protein_coding | 104.2250989 | -0.016870866 | 0.149260029 | -0.113030036 | 0.910006743 | 0.959845275 | 0.016870866 |
| ENSMUSG00000037461 | Ints7 | 77065 | protein_coding | 835.7947928 | -0.016873425 | 0.077292741 | -0.21830543 | 0.827191144 | 0.923255006 | 0.016873425 |
| ENSMUSG00000028407 | Smim27 | 100504309 | protein_coding | 117.75252 | 0.016879716 | 0.111507465 | 0.151377456 | 0.879677972 | 0.94589723 | 0.016879716 |
| ENSMUSG00000055725 | Paqr3 | 231474 | protein_coding | 10.8889883 | 0.016881214 | 0.361382826 | 0.046712829 | 0.962742106 | 0.984661462 | 0.016881214 |
| ENSMUSG00000005078 | Jkamp | 104771 | protein_coding | 291.3093697 | -0.016883173 | 0.078930518 | -0.213899173 | 0.830625685 | 0.925039736 | 0.016883173 |
| ENSMUSG00000045482 | Trrap | 100683 | protein_coding | 1654.757035 | 0.016900355 | 0.068199086 | 0.247809119 | 0.804282099 | 0.910796813 | 0.016900355 |
| ENSMUSG00000021218 | Gdi2 | 14569 | protein_coding | 8778.523657 | 0.016909712 | 0.035462901 | 0.476828215 | 0.633484453 | 0.808283189 | 0.016909712 |
| ENSMUSG00000022770 | Dlg1 | 13383 | protein_coding | 1852.385007 | -0.016924756 | 0.040701074 | -0.415830705 | 0.677533889 | 0.836508911 | 0.016924756 |
| ENSMUSG00000045005 | Fzd5 | 14367 | protein_coding | 4456.312501 | -0.016941434 | 0.064944733 | -0.260859241 | 0.794201058 | 0.904717134 | 0.016941434 |
| ENSMUSG00000046324 | Ermp1 | 226090 | protein_coding | 2631.405711 | -0.01697581 | 0.056911057 | -0.298286677 | 0.765484372 | 0.889849511 | 0.01697581 |
| ENSMUSG00000041720 | Pi4ka | 224020 | protein_coding | 1946.516069 | -0.016985043 | 0.057913199 | -0.293284486 | 0.769304711 | 0.891725514 | 0.016985043 |
| ENSMUSG00000020432 | Tcn2 | 21452 | protein_coding | 2266.262772 | 0.016998026 | 0.053426401 | 0.318157786 | 0.750365252 | 0.880695457 | 0.016998026 |

| ensembl_gene_id | external_gene_name | entrezgene_id | gene_biotype | baseMean | log2FoldChange | lfcSE | stat | pvalue | padj | absLog2FC |
|---|---|---|---|---|---|---|---|---|---|---|
| ENSMUSG00000021254 | Gpatch2l | 70373 | protein_coding | 459.9931161 | 0.0170005 | 0.060575898 | 0.280647928 | 0.778980452 | 0.897137595 | 0.0170005 |
| ENSMUSG00000029676 | Pot1a | 101185 | protein_coding | 304.6655585 | -0.017001392 | 0.099090501 | -0.171574391 | 0.863772142 | 0.938113902 | 0.017001392 |
| ENSMUSG00000005501 | Usp40 | 227334 | protein_coding | 705.0469364 | -0.017009712 | 0.05307023 | -0.320513255 | 0.748579284 | 0.880475742 | 0.017009712 |
| ENSMUSG00000027508 | Pag1 | 94212 | protein_coding | 709.4756751 | -0.017014802 | 0.062464909 | -0.272389759 | 0.785322346 | 0.900006643 | 0.017014802 |
| ENSMUSG00000024981 | Acsl5 | 433256 | protein_coding | 2865.246558 | 0.017014877 | 0.070723421 | 0.240583337 | 0.809878066 | 0.914534473 | 0.017014877 |
| ENSMUSG00000029322 | Plac8 | 231507 | protein_coding | 9492.014172 | -0.017017528 | 0.096839755 | -0.175728742 | 0.860507054 | 0.936965584 | 0.017017528 |
| ENSMUSG00000040506 | Ambra1 | 228361 | protein_coding | 1044.483497 | 0.017029913 | 0.086148211 | 0.197681564 | 0.843294215 | 0.930150544 | 0.017029913 |
| ENSMUSG00000022214 | Dcaf11 | 28199 | protein_coding | 2541.482336 | -0.0170441 | 0.035052455 | -0.486245536 | 0.626793089 | 0.803190139 | 0.0170441 |
| ENSMUSG00000029776 | Hibadh | 58875 | protein_coding | 1531.790423 | -0.01706337 | 0.049629091 | -0.343817893 | 0.730983243 | 0.870004926 | 0.01706337 |
| ENSMUSG00000024835 | Coro1b | 23789 | protein_coding | 6300.467157 | 0.017100411 | 0.055623228 | 0.307432922 | 0.758513872 | 0.885839179 | 0.017100411 |
| ENSMUSG00000060166 | Zdhhc8 | 27801 | protein_coding | 150.3820566 | -0.017109113 | 0.137004158 | -0.124880244 | 0.900618359 | 0.955984485 | 0.017109113 |
| ENSMUSG00000034329 | Brip1 | 237911 | protein_coding | 205.2095175 | 0.017112888 | 0.112423126 | 0.152218573 | 0.879014545 | 0.945595293 | 0.017112888 |
| ENSMUSG00000033033 | Calhm2 | 72691 | protein_coding | 46.57906902 | 0.017118107 | 0.190800766 | 0.089717182 | 0.928511962 | 0.968593927 | 0.017118107 |
| ENSMUSG00000000148 | Brat1 | 231841 | protein_coding | 450.8698807 | 0.017152271 | 0.075381533 | 0.227539426 | 0.820004315 | 0.919348861 | 0.017152271 |
| ENSMUSG00000028413 | B4galt1 | 14595 | protein_coding | 2386.84527 | -0.017156276 | 0.116711746 | -0.146996997 | 0.88313439 | 0.948012456 | 0.017156276 |
| ENSMUSG00000024603 | Dctn4 | 67665 | protein_coding | 1873.379204 | -0.017172059 | 0.039672747 | -0.432842707 | 0.665129045 | 0.82901007 | 0.017172059 |
| ENSMUSG00000061477 | Rps7 | 20115 | protein_coding | 9386.281285 | -0.017174898 | 0.047655612 | -0.360396129 | 0.718550921 | 0.862201627 | 0.017174898 |
| ENSMUSG00000038279 | Nop2 | 110109 | protein_coding | 735.426304 | -0.017177562 | 0.056946751 | -0.301642533 | 0.762924581 | 0.888341059 | 0.017177562 |
| ENSMUSG00000044709 | Gemin7 | 69731 | protein_coding | 494.0887042 | 0.017202157 | 0.06857445 | 0.250853742 | 0.80192719 | 0.909865152 | 0.017202157 |
| ENSMUSG00000022440 | C1qtnf6 | 72709 | protein_coding | 306.5959668 | 0.017205746 | 0.096011685 | 0.179204709 | 0.857776967 | 0.935951904 | 0.017205746 |
| ENSMUSG00000041058 | Wwp1 | 107568 | protein_coding | 1890.958689 | -0.01720818 | 0.062450514 | -0.275549063 | 0.782894437 | 0.898825528 | 0.01720818 |
| ENSMUSG00000025510 | Cd151 | 12476 | protein_coding | 2895.373785 | -0.017213804 | 0.053124472 | -0.324027756 | 0.745917021 | 0.879490473 | 0.017213804 |
| ENSMUSG00000028955 | Vamp3 | 22319 | protein_coding | 2326.806392 | -0.017230884 | 0.040301088 | -0.427553819 | 0.668975994 | 0.831002845 | 0.017230884 |
| ENSMUSG00000000168 | Dlat | 235339 | protein_coding | 2444.430189 | 0.017233716 | 0.06737357 | 0.255793416 | 0.798110358 | 0.907439773 | 0.017233716 |
| ENSMUSG00000038563 | Efl1 | 101592 | protein_coding | 625.2357787 | -0.017305975 | 0.06779037 | -0.255286624 | 0.798501731 | 0.907820755 | 0.017305975 |
| ENSMUSG00000022394 | L3mbtl2 | 214669 | protein_coding | 376.7268478 | 0.017356247 | 0.088391799 | 0.196355853 | 0.844331648 | 0.930210801 | 0.017356247 |
| ENSMUSG00000036292 | Gramd1c | 207798 | protein_coding | 301.1261953 | 0.017357502 | 0.090276979 | 0.192269415 | 0.847531175 | 0.931008464 | 0.017357502 |
| ENSMUSG00000030678 | Maz | 17188 | protein_coding | 1207.368976 | -0.017417391 | 0.081863674 | -0.212760929 | 0.83151344 | 0.925325507 | 0.017417391 |
| ENSMUSG00000032547 | Ryk | 20187 | protein_coding | 1329.422572 | 0.017430138 | 0.0406902 | 0.428362055 | 0.668387546 | 0.830822102 | 0.017430138 |
| ENSMUSG00000002265 | Peg3 | 18616 | protein_coding | 80.73624347 | 0.017464573 | 0.175625385 | 0.099442191 | 0.920787185 | 0.965070185 | 0.017464573 |
| ENSMUSG00000039960 | Rhou | 69581 | protein_coding | 977.7199676 | -0.017471374 | 0.080158658 | -0.217959915 | 0.827460344 | 0.923349179 | 0.017471374 |
| ENSMUSG00000002767 | Mrpl2 | 27398 | protein_coding | 1136.149174 | -0.017473015 | 0.072227029 | -0.241917959 | 0.808843734 | 0.913844956 | 0.017473015 |
| ENSMUSG00000054342 | Kcnn4 | 16534 | protein_coding | 969.1655238 | -0.017483453 | 0.059121531 | -0.295720575 | 0.767443483 | 0.890729692 | 0.017483453 |
| ENSMUSG00000022701 | Ccdc191 | 212153 | protein_coding | 171.4369846 | 0.017483707 | 0.100273329 | 0.17436049 | 0.861582163 | 0.937561241 | 0.017483707 |
| ENSMUSG00000005148 | Klf5 | 12224 | protein_coding | 7731.219532 | -0.017488871 | 0.064307863 | -0.271955408 | 0.785656306 | 0.90019746 | 0.017488871 |
| ENSMUSG00000030752 | Kdm8 | 77035 | protein_coding | 139.5756299 | -0.017532569 | 0.127798823 | -0.137188816 | 0.890881551 | 0.95112373 | 0.017532569 |
| ENSMUSG00000030060 | Hmces | 232210 | protein_coding | 239.6409077 | -0.01755366 | 0.085006428 | -0.20649803 | 0.836401903 | 0.927459717 | 0.01755366 |
| ENSMUSG00000024118 | Tedc2 | 72016 | protein_coding | 98.45065568 | 0.017554744 | 0.130974725 | 0.134031542 | 0.893377633 | 0.952355017 | 0.017554744 |
| ENSMUSG00000040040 | Ift88 | 21821 | protein_coding | 244.895424 | 0.017590001 | 0.086537438 | 0.203264641 | 0.838928191 | 0.928642661 | 0.017590001 |

| ensembl_gene_id | external_gene_name | entrezgene_id | gene_biotype | baseMean | log2FoldChange | lfcSE | stat | pvalue | padj | absLog2FC |
|---|---|---|---|---|---|---|---|---|---|---|
| ENSMUSG00000072872 | Rybp | 56353 | protein_coding | 482.1078409 | -0.017623747 | 0.05654481 | -0.311677541 | 0.755285598 | 0.88389772 | 0.017623747 |
| ENSMUSG00000030583 | Sipa1l3 | 74206 | protein_coding | 2014.767383 | -0.017642566 | 0.095537067 | -0.184667233 | 0.853490049 | 0.934372006 | 0.017642566 |
| ENSMUSG00000014668 | Chfr | 231600 | protein_coding | 1126.075288 | 0.017650742 | 0.043785475 | 0.403118653 | 0.686860939 | 0.842421961 | 0.017650742 |
| ENSMUSG00000001729 | Akt1 | 11651 | protein_coding | 3736.420702 | 0.017671615 | 0.041033548 | 0.430662611 | 0.666713709 | 0.829882036 | 0.017671615 |
| ENSMUSG00000041075 | Fzd7 | 14369 | protein_coding | 425.7079098 | 0.017675444 | 0.085073854 | 0.207765875 | 0.835411779 | 0.926722091 | 0.017675444 |
| ENSMUSG00000030609 | Aen | 68048 | protein_coding | 297.2641914 | 0.017675605 | 0.095909393 | 0.184294826 | 0.853782173 | 0.934564929 | 0.017675605 |
| ENSMUSG00000017478 | Zc3h18 | 76014 | protein_coding | 2080.322917 | 0.017679819 | 0.048604261 | 0.363750398 | 0.716044405 | 0.860552159 | 0.017679819 |
| ENSMUSG00000035770 | Dync1li2 | 234663 | protein_coding | 2270.793297 | 0.017704321 | 0.046004775 | 0.384836594 | 0.700358485 | 0.850619214 | 0.017704321 |
| ENSMUSG00000064289 | Tank | 21353 | protein_coding | 560.365913 | -0.017710367 | 0.083937368 | -0.210995024 | 0.832891154 | 0.92587823 | 0.017710367 |
| ENSMUSG00000027678 | Ncoa3 | 17979 | protein_coding | 1556.318634 | -0.017714116 | 0.050527084 | -0.350586549 | 0.725898549 | 0.866137742 | 0.017714116 |
| ENSMUSG00000003131 | Pafah1b2 | 18475 | protein_coding | 1324.012335 | -0.017722249 | 0.045005068 | -0.393783407 | 0.693740956 | 0.84654428 | 0.017722249 |
| ENSMUSG00000019838 | Slc16a10 | 72472 | protein_coding | 284.1930301 | -0.01776213 | 0.09615536 | -0.184723241 | 0.853446116 | 0.934372006 | 0.01776213 |
| ENSMUSG00000027993 | Trim2 | 80890 | protein_coding | 9044.758791 | -0.017780001 | 0.03207136 | -0.554388751 | 0.579312823 | 0.771609416 | 0.017780001 |
| ENSMUSG00000073838 | Tufm | 233870 | protein_coding | 2542.737431 | 0.017808811 | 0.085449479 | 0.208413331 | 0.834906248 | 0.926650089 | 0.017808811 |
| ENSMUSG00000005803 | Sqor | 59010 | protein_coding | 13752.9883 | -0.017815275 | 0.042806814 | -0.416178471 | 0.677279412 | 0.83633117 | 0.017815275 |
| ENSMUSG00000029797 | Sspo | 243369 | protein_coding | 122.4725184 | -0.01782939 | 0.12781033 | -0.139498817 | 0.889055991 | 0.950733597 | 0.01782939 |
| ENSMUSG00000037761 | Actr5 | 109275 | protein_coding | 239.6679649 | -0.017833327 | 0.099391064 | -0.179425855 | 0.857603332 | 0.935889087 | 0.017833327 |
| ENSMUSG00000005566 | Trim28 | 21849 | protein_coding | 3800.136546 | -0.017835447 | 0.047671604 | -0.374131465 | 0.70830651 | 0.855350218 | 0.017835447 |
| ENSMUSG00000072949 | Acot1 | 26897 | protein_coding | 114.9617178 | -0.017849342 | 0.160429691 | -0.111259592 | 0.911410497 | 0.960412417 | 0.017849342 |
| ENSMUSG00000027994 | Mcub | 66815 | protein_coding | 64.66016323 | -0.017877658 | 0.166952648 | -0.107082208 | 0.914723762 | 0.962154325 | 0.017877658 |
| ENSMUSG00000021012 | Zc3h14 | 75553 | protein_coding | 1434.733855 | 0.017892701 | 0.049436347 | 0.361934116 | 0.717401265 | 0.861469114 | 0.017892701 |
| ENSMUSG00000018765 | Fxr2 | 23879 | protein_coding | 1197.017763 | 0.017915151 | 0.056928017 | 0.314698312 | 0.752990726 | 0.882492633 | 0.017915151 |
| ENSMUSG00000035024 | Ncapd3 | 78658 | protein_coding | 668.9171511 | -0.01792702 | 0.083875049 | -0.21373484 | 0.830753841 | 0.925118574 | 0.01792702 |
| ENSMUSG00000039953 | Clstn1 | 65945 | protein_coding | 6362.750123 | -0.017932438 | 0.051932844 | -0.345300519 | 0.729868457 | 0.869255491 | 0.017932438 |
| ENSMUSG00000001156 | Mxd1 | 17119 | protein_coding | 7855.858527 | -0.017939278 | 0.086035873 | -0.208509275 | 0.834831342 | 0.926650089 | 0.017939278 |
| ENSMUSG00000052926 | Rnaseh2a | 69724 | protein_coding | 503.2697927 | 0.017954117 | 0.057059237 | 0.314657507 | 0.753021711 | 0.882492633 | 0.017954117 |
| ENSMUSG00000073643 | Wdfy1 | 69368 | protein_coding | 1701.461153 | -0.017988567 | 0.043463709 | -0.413875563 | 0.678965241 | 0.837322968 | 0.017988567 |
| ENSMUSG00000055991 | Zkscan5 | 22757 | protein_coding | 343.1848433 | 0.018051662 | 0.075193723 | 0.24006874 | 0.810276968 | 0.914739662 | 0.018051662 |
| ENSMUSG00000003779 | Kif20a | 19348 | protein_coding | 494.0032121 | 0.018077877 | 0.101389793 | 0.178300754 | 0.858486787 | 0.93615636 | 0.018077877 |
| ENSMUSG00000007415 | Gatad1 | 67210 | protein_coding | 1512.719861 | 0.018096274 | 0.045456891 | 0.398097495 | 0.690558321 | 0.844620107 | 0.018096274 |
| ENSMUSG00000025262 | Fam120c | 207375 | protein_coding | 207.3724657 | 0.018104296 | 0.108302264 | 0.167164523 | 0.867240598 | 0.939855462 | 0.018104296 |
| ENSMUSG00000021587 | Pcsk1 | 18548 | protein_coding | 488.5969926 | 0.01811608 | 0.073805225 | 0.245457956 | 0.806101859 | 0.912178372 | 0.01811608 |
| ENSMUSG00000022635 | Zcrb1 | 67197 | protein_coding | 483.4241406 | 0.018123561 | 0.059170343 | 0.306294673 | 0.759380295 | 0.886076667 | 0.018123561 |
| ENSMUSG00000024218 | Taf11 | 68776 | protein_coding | 285.5025789 | 0.018128783 | 0.077754256 | 0.233154867 | 0.815641151 | 0.917020505 | 0.018128783 |
| ENSMUSG00000000085 | Scmh1 | 29871 | protein_coding | 463.1941055 | -0.018133897 | 0.06432804 | -0.281897249 | 0.778022299 | 0.896738538 | 0.018133897 |
| ENSMUSG00000061315 | Naca | 17938 | protein_coding | 5790.01091 | -0.018136429 | 0.048268901 | -0.375737348 | 0.707112169 | 0.854394093 | 0.018136429 |
| ENSMUSG00000057234 | Mettl15 | 76894 | protein_coding | 99.33786146 | -0.018147875 | 0.12833823 | -0.141406616 | 0.887548728 | 0.95031765 | 0.018147875 |
| ENSMUSG00000017781 | Pitpna | 18738 | protein_coding | 2872.804791 | 0.018161467 | 0.041263865 | 0.44013006 | 0.659842912 | 0.826041967 | 0.018161467 |
| ENSMUSG00000041415 | Dicer1 | 192119 | protein_coding | 1433.122462 | -0.018162876 | 0.050900795 | -0.356828924 | 0.721219881 | 0.863755116 | 0.018162876 |

| ensembl_gene_id | external_gene_name | entrezgene_id | gene_biotype | baseMean | log2FoldChange | lfcSE | stat | pvalue | padj | absLog2FC |
|---|---|---|---|---|---|---|---|---|---|---|
| ENSMUSG00000026342 | Slc35f5 | 74150 | protein_coding | 3098.222867 | -0.01818859 | 0.064145506 | -0.283552057 | 0.776753682 | 0.895916653 | 0.01818859 |
| ENSMUSG00000004934 | Pias4 | 59004 | protein_coding | 694.2660831 | -0.018232283 | 0.087502022 | -0.208364139 | 0.834944655 | 0.926650089 | 0.018232283 |
| ENSMUSG00000042978 | Sbk1 | 104175 | protein_coding | 482.1706549 | 0.018246962 | 0.079265177 | 0.23020149 | 0.817935204 | 0.918198597 | 0.018246962 |
| ENSMUSG00000032485 | Scap | 235623 | protein_coding | 1614.989801 | 0.018280907 | 0.065811031 | 0.27777876 | 0.781182197 | 0.898390166 | 0.018280907 |
| ENSMUSG00000026116 | Tmem131 | 56030 | protein_coding | 2473.762134 | 0.01833023 | 0.070315848 | 0.260684195 | 0.794336055 | 0.904717934 | 0.01833023 |
| ENSMUSG00000041264 | Uspl1 | 231915 | protein_coding | 520.4754551 | -0.018331973 | 0.079041566 | -0.231928258 | 0.816593736 | 0.917421069 | 0.018331973 |
| ENSMUSG00000037487 | Ubr5 | 70790 | protein_coding | 4946.623168 | -0.018336342 | 0.051425927 | -0.356558311 | 0.721422492 | 0.863807468 | 0.018336342 |
| ENSMUSG00000039630 | Hnrnpu | 51810 | protein_coding | 7356.036937 | 0.018340487 | 0.044766244 | 0.409694577 | 0.682030008 | 0.839305452 | 0.018340487 |
| ENSMUSG00000046668 | Cxxc5 | 67393 | protein_coding | 292.7938882 | -0.018377094 | 0.094839782 | -0.193769893 | 0.846356062 | 0.930889346 | 0.018377094 |
| ENSMUSG00000049672 | Zbtb14 | 22666 | protein_coding | 345.1939002 | 0.018382044 | 0.071315593 | 0.257756303 | 0.796594991 | 0.906052077 | 0.018382044 |
| ENSMUSG00000029445 | Hpd | 15445 | protein_coding | 1348.972331 | -0.018411668 | 0.056843881 | -0.323898852 | 0.746014613 | 0.879490473 | 0.018411668 |
| ENSMUSG00000027162 | Lin7c | 22343 | protein_coding | 5777.154759 | -0.018422235 | 0.036341975 | -0.506913424 | 0.612215566 | 0.793255831 | 0.018422235 |
| ENSMUSG00000058301 | Upf1 | 19704 | protein_coding | 2481.530334 | 0.01842561 | 0.057194758 | 0.322155574 | 0.747334839 | 0.880070575 | 0.01842561 |
| ENSMUSG00000040147 | Maob | 109731 | protein_coding | 753.3853037 | 0.018433759 | 0.081466238 | 0.226274833 | 0.820987671 | 0.920019265 | 0.018433759 |
| ENSMUSG00000041870 | Ankrd13a | 68420 | protein_coding | 5636.599502 | 0.01846979 | 0.04881762 | 0.378342686 | 0.705176039 | 0.853461462 | 0.01846979 |
| ENSMUSG00000030610 | Det1 | 76375 | protein_coding | 285.9728029 | -0.018493738 | 0.091448978 | -0.202230118 | 0.839736829 | 0.928990077 | 0.018493738 |
| ENSMUSG00000079477 | Rab7 | 19349 | protein_coding | 2172.784607 | -0.018536061 | 0.035850837 | -0.517032864 | 0.605133217 | 0.789158883 | 0.018536061 |
| ENSMUSG00000067847 | Romo1 | 67067 | protein_coding | 691.8491886 | 0.018550332 | 0.107546792 | 0.172486152 | 0.863055348 | 0.937787345 | 0.018550332 |
| ENSMUSG00000099843 | Gm7160 | 635504 | lncRNA | 32.81640378 | 0.018569117 | 0.2401019 | 0.077338484 | 0.938354277 | 0.97297734 | 0.018569117 |
| ENSMUSG00000066613 | Zfp932 | 69504 | protein_coding | 135.2621996 | -0.018569994 | 0.157665559 | -0.117780916 | 0.906241251 | 0.958651438 | 0.018569994 |
| ENSMUSG00000002565 | Scin | 20259 | protein_coding | 6062.716884 | 0.018612252 | 0.048658978 | 0.38250397 | 0.702087585 | 0.851790685 | 0.018612252 |
| ENSMUSG00000040605 | Bace2 | 56175 | protein_coding | 1422.657837 | -0.018617247 | 0.073382341 | -0.253702006 | 0.799725784 | 0.908570455 | 0.018617247 |
| ENSMUSG00000075576 | Gm12359 | 100037262 | lncRNA | 13.56905214 | -0.018618007 | 0.346914544 | -0.053667415 | 0.957200145 | 0.982857935 | 0.018618007 |
| ENSMUSG00000028070 | Naxe | 246703 | protein_coding | 1296.786959 | -0.018624144 | 0.059467495 | -0.313181915 | 0.754142457 | 0.883228425 | 0.018624144 |
| ENSMUSG00000079560 | Hoxa3 | 15400 | protein_coding | 104.8741012 | -0.018628334 | 0.120230214 | -0.154938871 | 0.876869509 | 0.945019685 | 0.018628334 |
| ENSMUSG00000020925 | Ccdc43 | 52715 | protein_coding | 454.6544184 | -0.018634085 | 0.075177008 | -0.247869471 | 0.804235401 | 0.910796813 | 0.018634085 |
| ENSMUSG00000051022 | Hs3st1 | 15476 | protein_coding | 264.3759893 | -0.018656086 | 0.096147758 | -0.194035575 | 0.846148026 | 0.930889346 | 0.018656086 |
| ENSMUSG00000037703 | Lzts3 | 241638 | protein_coding | 2421.607101 | 0.018700218 | 0.06170253 | 0.303070518 | 0.761836121 | 0.887606361 | 0.018700218 |
| ENSMUSG00000053600 | Zfp472 | 224691 | protein_coding | 234.4442717 | 0.018704146 | 0.095499378 | 0.195856209 | 0.844722714 | 0.930261042 | 0.018704146 |
| ENSMUSG00000029265 | Dr1 | 13486 | protein_coding | 798.3699822 | -0.018707098 | 0.050493837 | -0.370482804 | 0.711022787 | 0.857064391 | 0.018707098 |
| ENSMUSG00000022369 | Mtbp | 105837 | protein_coding | 133.1824062 | -0.018735653 | 0.160506765 | -0.116728119 | 0.907075507 | 0.95873684 | 0.018735653 |
| ENSMUSG00000025010 | Ccnj | 240665 | protein_coding | 112.5470067 | 0.018739274 | 0.133461694 | 0.140409383 | 0.888336545 | 0.950656978 | 0.018739274 |
| ENSMUSG00000024527 | Afg3l2 | 69597 | protein_coding | 3294.797013 | -0.018740631 | 0.046036449 | -0.407082455 | 0.683947429 | 0.840783841 | 0.018740631 |
| ENSMUSG00000026707 | Nsun6 | 74455 | protein_coding | 240.8215942 | -0.018747025 | 0.095627431 | -0.196042341 | 0.844577026 | 0.930210801 | 0.018747025 |
| ENSMUSG00000041757 | Plekha6 | 240753 | protein_coding | 7690.839273 | 0.018752299 | 0.062365353 | 0.300684567 | 0.763655038 | 0.888754045 | 0.018752299 |
| ENSMUSG00000059890 | Ube4a | 140630 | protein_coding | 1035.445714 | 0.018772074 | 0.047456844 | 0.395560933 | 0.692428962 | 0.845713562 | 0.018772074 |
| ENSMUSG00000041429 | Nthl1 | 18207 | protein_coding | 50.05162075 | 0.018785644 | 0.166795486 | 0.112626814 | 0.910326426 | 0.959960814 | 0.018785644 |
| ENSMUSG00000041975 | Mettl8 | 228019 | protein_coding | 194.5183279 | 0.01882377 | 0.086560318 | 0.217464199 | 0.827846604 | 0.923349179 | 0.01882377 |
| ENSMUSG00000014418 | Hps5 | 246694 | protein_coding | 1368.658637 | 0.018837946 | 0.045168664 | 0.417057858 | 0.676636088 | 0.835792754 | 0.018837946 |

| ensembl_gene_id | external_gene_name | entrezgene_id | gene_biotype | baseMean | log2FoldChange | lfcSE | stat | pvalue | padj | absLog2FC |
|---|---|---|---|---|---|---|---|---|---|---|
| ENSMUSG00000006784 | Odad4 | 74407 | protein_coding | 28.6491394 | -0.018848342 | 0.232481503 | -0.081074588 | 0.935382635 | 0.971834101 | 0.018848342 |
| ENSMUSG00000031075 | Ano1 | 101772 | protein_coding | 3743.780343 | -0.018851375 | 0.123252496 | -0.152949236 | 0.878438308 | 0.945542552 | 0.018851375 |
| ENSMUSG00000026781 | Acbd5 | 74159 | protein_coding | 1886.726283 | -0.018855958 | 0.052946341 | -0.356133358 | 0.721740697 | 0.86404218 | 0.018855958 |
| ENSMUSG00000009035 | Tmem184b | 223693 | protein_coding | 483.9654337 | -0.018865936 | 0.077006396 | -0.244991799 | 0.806462781 | 0.912178372 | 0.018865936 |
| ENSMUSG00000028577 | Plaa | 18786 | protein_coding | 1220.357088 | 0.018873693 | 0.063238109 | 0.298454418 | 0.765356361 | 0.889849511 | 0.018873693 |
| ENSMUSG00000027346 | Gpcpd1 | 74182 | protein_coding | 807.457897 | -0.01889834 | 0.093769488 | -0.201540395 | 0.840276047 | 0.929305432 | 0.01889834 |
| ENSMUSG00000029735 | Tpk1 | 29807 | protein_coding | 862.4801584 | 0.018917117 | 0.068335295 | 0.276827913 | 0.781912247 | 0.898695579 | 0.018917117 |
| ENSMUSG00000030225 | Dera | 232449 | protein_coding | 1135.867635 | 0.018924551 | 0.060338011 | 0.313642275 | 0.753792748 | 0.883011233 | 0.018924551 |
| ENSMUSG00000058587 | Tmod3 | 50875 | protein_coding | 2541.660994 | 0.018933754 | 0.043487576 | 0.435383069 | 0.663284398 | 0.827892753 | 0.018933754 |
| ENSMUSG00000028634 | Hivep3 | 16656 | protein_coding | 127.9161205 | 0.018985701 | 0.123567178 | 0.153646795 | 0.877888238 | 0.945241431 | 0.018985701 |
| ENSMUSG00000041199 | Rpusd1 | 106707 | protein_coding | 103.6485057 | 0.018993444 | 0.130235178 | 0.145839584 | 0.884048026 | 0.948596624 | 0.018993444 |
| ENSMUSG00000022557 | Bop1 | 12181 | protein_coding | 966.804691 | 0.018996457 | 0.102258368 | 0.185769214 | 0.852625748 | 0.933806142 | 0.018996457 |
| ENSMUSG00000020153 | Ndufs7 | 75406 | protein_coding | 2113.575731 | 0.01902745 | 0.071193682 | 0.267263185 | 0.78926653 | 0.901682062 | 0.01902745 |
| ENSMUSG00000021832 | Psmc6 | 67089 | protein_coding | 1845.97422 | 0.019030556 | 0.056639731 | 0.335993049 | 0.736876108 | 0.873729946 | 0.019030556 |
| ENSMUSG00000105814 | Mir703 | 735265 | miRNA | 35.90538863 | -0.019042575 | 0.208824977 | -0.091189165 | 0.927342284 | 0.968095473 | 0.019042575 |
| ENSMUSG00000034613 | Ppm1h | 319468 | protein_coding | 1403.78231 | 0.019045096 | 0.067910942 | 0.280442233 | 0.779138239 | 0.897245014 | 0.019045096 |
| ENSMUSG00000021866 | Anxa11 | 11744 | protein_coding | 2409.661767 | 0.019047801 | 0.086762875 | 0.219538606 | 0.82623051 | 0.922846345 | 0.019047801 |
| ENSMUSG00000036206 | Sh3bp4 | 98402 | protein_coding | 352.1620808 | -0.019053287 | 0.085829158 | -0.221990841 | 0.824321013 | 0.921867382 | 0.019053287 |
| ENSMUSG00000041238 | Rbbp8 | 225182 | protein_coding | 764.0953031 | 0.019055387 | 0.081444698 | 0.233967185 | 0.815010454 | 0.91676947 | 0.019055387 |
| ENSMUSG00000022219 | Cideb | 12684 | protein_coding | 168.0478687 | -0.019081668 | 0.120660609 | -0.158143309 | 0.874343875 | 0.94368516 | 0.019081668 |
| ENSMUSG00000029221 | Slc30a9 | 109108 | protein_coding | 1399.042819 | -0.019104943 | 0.044519525 | -0.429136279 | 0.667824052 | 0.830562128 | 0.019104943 |
| ENSMUSG00000039285 | Azi2 | 27215 | protein_coding | 2720.803967 | 0.019124432 | 0.041947883 | 0.455909342 | 0.648455171 | 0.818405307 | 0.019124432 |
| ENSMUSG00000003421 | Nosip | 66394 | protein_coding | 767.7152873 | -0.019133609 | 0.066873364 | -0.286117037 | 0.774788483 | 0.89435358 | 0.019133609 |
| ENSMUSG00000026981 | Il1rn | 16181 | protein_coding | 924.2066487 | -0.019151679 | 0.126640346 | -0.151228891 | 0.879795161 | 0.945960168 | 0.019151679 |
| ENSMUSG00000027968 | Larp7 | 28036 | protein_coding | 431.378581 | 0.019168588 | 0.106262836 | 0.180388445 | 0.856847625 | 0.935731633 | 0.019168588 |
| ENSMUSG00000037692 | Ahdc1 | 230793 | protein_coding | 499.1700985 | 0.01917396 | 0.124342983 | 0.154202186 | 0.877450318 | 0.945203479 | 0.01917396 |
| ENSMUSG00000045410 | Akr1e1 | 56043 | protein_coding | 966.6367516 | -0.019204264 | 0.053634972 | -0.35805488 | 0.720302244 | 0.863217232 | 0.019204264 |
| ENSMUSG00000001783 | Rtcb | 28088 | protein_coding | 3558.993721 | -0.019222821 | 0.046103425 | -0.416949952 | 0.676715016 | 0.835804884 | 0.019222821 |
| ENSMUSG00000046722 | Cdc42se1 | 57912 | protein_coding | 1973.83167 | 0.019243044 | 0.039635448 | 0.485500864 | 0.6273211 | 0.803431336 | 0.019243044 |
| ENSMUSG00000021559 | Dapk1 | 69635 | protein_coding | 406.5488107 | 0.019316191 | 0.08256082 | 0.233963167 | 0.815013573 | 0.91676947 | 0.019316191 |
| ENSMUSG00000019969 | Psen1 | 19164 | protein_coding | 4266.122917 | 0.019370486 | 0.033687806 | 0.574999927 | 0.565291346 | 0.761818669 | 0.019370486 |
| ENSMUSG00000063160 | Numbl | 18223 | protein_coding | 8.939643113 | 0.01939756 | 0.397105213 | 0.048847406 | 0.961040903 | 0.984107349 | 0.01939756 |
| ENSMUSG00000097099 | Gm9917 | 100038755 | lncRNA | 186.758568 | -0.019411275 | 0.109804795 | -0.17677985 | 0.859681317 | 0.936698912 | 0.019411275 |
| ENSMUSG00000056537 | Rlim | 19820 | protein_coding | 2442.967857 | -0.019432014 | 0.039095182 | -0.497043695 | 0.619158238 | 0.798277593 | 0.019432014 |
| ENSMUSG00000107121 | 1110006O24Rik | 66123 | lncRNA | 10.42970638 | 0.019435629 | 0.371405848 | 0.052329895 | 0.958265833 | 0.983263542 | 0.019435629 |
| ENSMUSG00000027963 | Extl2 | 58193 | protein_coding | 237.7976423 | -0.019467602 | 0.096987441 | -0.200722917 | 0.840915239 | 0.929613548 | 0.019467602 |
| ENSMUSG00000029646 | Cdx2 | 12591 | protein_coding | 5451.34571 | 0.019495427 | 0.066536057 | 0.293005447 | 0.769517988 | 0.891758816 | 0.019495427 |
| ENSMUSG00000033111 | 3830406C13Rik | 218734 | protein_coding | 395.9719891 | -0.019500741 | 0.066373512 | -0.293803066 | 0.768908392 | 0.891610224 | 0.019500741 |
| ENSMUSG00000056698 | Elmod3 | 232089 | protein_coding | 259.1486843 | 0.019552016 | 0.090908636 | 0.215073251 | 0.829710208 | 0.924467081 | 0.019552016 |

| ensembl_gene_id | external_gene_name | entrezgene_id | gene_biotype | baseMean | log2FoldChange | lfcSE | stat | pvalue | padj | absLog2FC |
|---|---|---|---|---|---|---|---|---|---|---|
| ENSMUSG00000056342 | Usp34 | 17847 | protein_coding | 2451.276087 | -0.019581878 | 0.082474595 | -0.237429214 | 0.812323828 | 0.915791177 | 0.019581878 |
| ENSMUSG00000060206 | Zfp462 | 242466 | protein_coding | 393.673904 | 0.019586453 | 0.075946536 | 0.257897907 | 0.7964857 | 0.906039704 | 0.019586453 |
| ENSMUSG00000024566 | Atp9b | 50771 | protein_coding | 1234.803031 | -0.019591471 | 0.045075495 | -0.434636852 | 0.663826042 | 0.828295599 | 0.019591471 |
| ENSMUSG00000019841 | Rev3l | 19714 | protein_coding | 1140.393074 | 0.019593826 | 0.054570447 | 0.359055621 | 0.71955348 | 0.862768697 | 0.019593826 |
| ENSMUSG00000026083 | Eif5b | 226982 | protein_coding | 859.4383483 | -0.019619441 | 0.065419284 | -0.29990302 | 0.764251131 | 0.889251381 | 0.019619441 |
| ENSMUSG00000014402 | Tsg101 | 22088 | protein_coding | 1153.565744 | -0.01962499 | 0.044689664 | -0.43913936 | 0.660560561 | 0.826315558 | 0.01962499 |
| ENSMUSG00000021275 | Tecpr2 | 104859 | protein_coding | 491.0500359 | 0.019634524 | 0.060887009 | 0.322474762 | 0.747093054 | 0.880021159 | 0.019634524 |
| ENSMUSG00000036499 | Eea1 | 216238 | protein_coding | 2655.743089 | -0.019676013 | 0.051697604 | -0.38059816 | 0.703501447 | 0.852778701 | 0.019676013 |
| ENSMUSG00000066842 | Hmcn1 | 545370 | protein_coding | 810.2886748 | 0.019678527 | 0.110165035 | 0.178627706 | 0.858230038 | 0.936097283 | 0.019678527 |
| ENSMUSG00000032044 | Rpusd4 | 71989 | protein_coding | 327.3686114 | 0.019678999 | 0.0667136 | 0.294977316 | 0.768011208 | 0.891239011 | 0.019678999 |
| ENSMUSG00000027805 | Pfn2 | 18645 | protein_coding | 22.57842297 | 0.019695808 | 0.254954094 | 0.077252369 | 0.938422781 | 0.97297734 | 0.019695808 |
| ENSMUSG00000028274 | Rngtt | 24018 | protein_coding | 431.8288026 | 0.019708485 | 0.065350038 | 0.301583381 | 0.762969679 | 0.888341059 | 0.019708485 |
| ENSMUSG00000043909 | Trp53bp1 | 27223 | protein_coding | 639.3168341 | -0.019735828 | 0.059269452 | -0.332984823 | 0.73914574 | 0.87479925 | 0.019735828 |
| ENSMUSG00000036158 | Prickle1 | 106042 | protein_coding | 40.88612323 | 0.019747335 | 0.198164241 | 0.099651357 | 0.92062112 | 0.965021567 | 0.019747335 |
| ENSMUSG00000013997 | Nit1 | 27045 | protein_coding | 951.8520112 | 0.019760545 | 0.047315891 | 0.417630195 | 0.676217517 | 0.8354677 | 0.019760545 |
| ENSMUSG00000024963 | Dnajc4 | 57431 | protein_coding | 489.7703986 | -0.019777686 | 0.072694077 | -0.27206736 | 0.785570226 | 0.900162785 | 0.019777686 |
| ENSMUSG00000025409 | Mbd6 | 110962 | protein_coding | 233.693398 | -0.019802505 | 0.110329073 | -0.179485829 | 0.857556243 | 0.935889087 | 0.019802505 |
| ENSMUSG00000042590 | Ipo11 | 76582 | protein_coding | 537.8487382 | 0.019804962 | 0.075840798 | 0.261138626 | 0.793985605 | 0.904599495 | 0.019804962 |
| ENSMUSG00000026730 | Pter | 19212 | protein_coding | 297.4158081 | -0.019825437 | 0.093171712 | -0.212783866 | 0.831495549 | 0.925325507 | 0.019825437 |
| ENSMUSG00000023232 | Serinc2 | 230779 | protein_coding | 2399.606104 | 0.01988194 | 0.080136822 | 0.248099936 | 0.804057086 | 0.910778265 | 0.01988194 |
| ENSMUSG00000020982 | Nemf | 66244 | protein_coding | 1010.618528 | 0.019884161 | 0.068391916 | 0.290738474 | 0.771251344 | 0.892332662 | 0.019884161 |
| ENSMUSG00000044345 | Marveld1 | 277010 | protein_coding | 65.2307214 | -0.019886159 | 0.14721244 | -0.135084774 | 0.892544848 | 0.952051774 | 0.019886159 |
| ENSMUSG00000063480 | Snu13 | 20826 | protein_coding | 678.5960443 | 0.019887591 | 0.076655728 | 0.259440377 | 0.795295479 | 0.905108834 | 0.019887591 |
| ENSMUSG00000066900 | Suds3 | 71954 | protein_coding | 2313.879831 | 0.019891867 | 0.048239064 | 0.412360128 | 0.680075483 | 0.838147754 | 0.019891867 |
| ENSMUSG00000023940 | Sgo1 | 72415 | protein_coding | 183.2629102 | 0.019894355 | 0.144238146 | 0.137927143 | 0.890297999 | 0.951039003 | 0.019894355 |
| ENSMUSG00000038046 | Mrm3 | 67390 | protein_coding | 165.4003492 | -0.019896112 | 0.100795205 | -0.197391451 | 0.843521219 | 0.930151569 | 0.019896112 |
| ENSMUSG00000040929 | Rfx3 | 19726 | protein_coding | 193.7300401 | 0.019896965 | 0.126453375 | 0.157346252 | 0.87497197 | 0.943745489 | 0.019896965 |
| ENSMUSG00000020358 | Hnrnpab | 15384 | protein_coding | 3847.062723 | 0.019904093 | 0.055783253 | 0.356811252 | 0.721233112 | 0.863755116 | 0.019904093 |
| ENSMUSG00000045538 | Ddx28 | 71986 | protein_coding | 219.7750078 | 0.019921523 | 0.085272084 | 0.233623035 | 0.815277644 | 0.916810722 | 0.019921523 |
| ENSMUSG00000041733 | Coq5 | 52064 | protein_coding | 965.7329679 | -0.019922962 | 0.056998088 | -0.349537382 | 0.726685912 | 0.866745337 | 0.019922962 |
| ENSMUSG00000040701 | Ap1g2 | 11766 | protein_coding | 949.8501637 | 0.019927447 | 0.061289298 | 0.325137463 | 0.745077037 | 0.878866104 | 0.019927447 |
| ENSMUSG00000066232 | Ipo7 | 233726 | protein_coding | 3795.269798 | -0.019951062 | 0.045130765 | -0.44207231 | 0.658436881 | 0.825261698 | 0.019951062 |
| ENSMUSG00000071253 | Slc25a16 | 73132 | protein_coding | 839.981187 | -0.019951308 | 0.057909026 | -0.344528467 | 0.730448891 | 0.869497289 | 0.019951308 |
| ENSMUSG00000021975 | Ints9 | 210925 | protein_coding | 323.5842747 | 0.01995759 | 0.074726789 | 0.267074104 | 0.789412106 | 0.901682062 | 0.01995759 |
| ENSMUSG00000024989 | Cep55 | 74107 | protein_coding | 221.4997282 | -0.020008204 | 0.104662371 | -0.191169034 | 0.848393164 | 0.93163806 | 0.020008204 |
| ENSMUSG00000005262 | Ufd1 | 22230 | protein_coding | 680.783637 | -0.020043497 | 0.050855988 | -0.394122647 | 0.693490491 | 0.846497211 | 0.020043497 |
| ENSMUSG00000042507 | Mideas | 238317 | protein_coding | 1012.375355 | -0.020061701 | 0.074029195 | -0.270997151 | 0.786393223 | 0.900451559 | 0.020061701 |
| ENSMUSG00000022556 | Hsf1 | 15499 | protein_coding | 489.9276321 | 0.020080721 | 0.068635591 | 0.292570087 | 0.76985078 | 0.891927276 | 0.020080721 |
| ENSMUSG00000034674 | Tdg | 21665 | protein_coding | 529.4982458 | 0.0200968 | 0.072287231 | 0.278013143 | 0.78100227 | 0.898311362 | 0.0200968 |

| ensembl_gene_id | external_gene_name | entrezgene_id | gene_biotype | baseMean | log2FoldChange | lfcSE | stat | pvalue | padj | absLog2FC |
|---|---|---|---|---|---|---|---|---|---|---|
| ENSMUSG00000037306 | Man1c1 | 230815 | protein_coding | 31.9521849 | -0.020106121 | 0.248201828 | -0.081007143 | 0.935436272 | 0.971834101 | 0.020106121 |
| ENSMUSG00000029148 | Nrbp1 | 192292 | protein_coding | 1802.885462 | 0.020119655 | 0.043840638 | 0.458927057 | 0.646286548 | 0.816759123 | 0.020119655 |
| ENSMUSG00000030888 | Rrp8 | 101867 | protein_coding | 238.3513966 | -0.020125694 | 0.079902281 | -0.251878843 | 0.801134713 | 0.909275777 | 0.020125694 |
| ENSMUSG00000088108 | Snora47 | 100217450 | snoRNA | 8.737767831 | 0.020139518 | 0.403595952 | 0.049900199 | 0.960201919 | 0.983923349 | 0.020139518 |
| ENSMUSG00000005802 | Slc30a4 | 22785 | protein_coding | 3480.136441 | -0.020151714 | 0.047201075 | -0.426933376 | 0.669427855 | 0.831282284 | 0.020151714 |
| ENSMUSG00000017861 | Mybl2 | 17865 | protein_coding | 112.9737853 | -0.020153508 | 0.225504303 | -0.089370836 | 0.9287872 | 0.968789653 | 0.020153508 |
| ENSMUSG00000057816 | Cfap299 | 75784 | protein_coding | 7.902337804 | -0.020157668 | 0.534419665 | -0.037718799 | 0.969911887 | 0.987915089 | 0.020157668 |
| ENSMUSG00000006526 | Stimate | 69179 | protein_coding | 303.312117 | -0.020207626 | 0.08235341 | -0.245376917 | 0.8061646 | 0.912178372 | 0.020207626 |
| ENSMUSG00000002546 | Golga2 | 99412 | protein_coding | 2136.176421 | 0.020237861 | 0.038110288 | 0.531034062 | 0.595395176 | 0.782426499 | 0.020237861 |
| ENSMUSG00000062190 | Lancl2 | 71835 | protein_coding | 397.0985709 | -0.020254694 | 0.068089006 | -0.297473784 | 0.76610482 | 0.890191832 | 0.020254694 |
| ENSMUSG00000028868 | Wasf2 | 242687 | protein_coding | 2324.991726 | 0.020264575 | 0.068792549 | 0.294575139 | 0.768318456 | 0.891415233 | 0.020264575 |
| ENSMUSG00000046330 | Rpl37a | 19981 | protein_coding | 8766.022871 | 0.020267931 | 0.066380125 | 0.305331323 | 0.760113822 | 0.886576344 | 0.020267931 |
| ENSMUSG00000055976 | Cldn23 | 71908 | protein_coding | 2009.420861 | -0.020289917 | 0.084346244 | -0.24055507 | 0.809899977 | 0.914534473 | 0.020289917 |
| ENSMUSG00000030922 | Lyrm1 | 73919 | protein_coding | 36.85111044 | 0.020326733 | 0.197129977 | 0.103113353 | 0.917873007 | 0.963895161 | 0.020326733 |
| ENSMUSG00000034601 | 2700049A03Rik | 76967 | protein_coding | 265.7119357 | 0.020328932 | 0.079746531 | 0.25491933 | 0.798785406 | 0.907951244 | 0.020328932 |
| ENSMUSG00000003184 | Irf3 | 54131 | protein_coding | 1308.342732 | 0.020331693 | 0.063491554 | 0.320226741 | 0.748796454 | 0.880475742 | 0.020331693 |
| ENSMUSG00000015804 | Med28 | 66999 | protein_coding | 1025.008332 | 0.020338611 | 0.053571732 | 0.379651922 | 0.704203813 | 0.853157579 | 0.020338611 |
| ENSMUSG00000020549 | Elac2 | 68626 | protein_coding | 630.0038263 | 0.020345328 | 0.058667468 | 0.346790617 | 0.728748626 | 0.868306546 | 0.020345328 |
| ENSMUSG00000078622 | Ccdc47 | 67163 | protein_coding | 1943.798022 | 0.020357747 | 0.045553858 | 0.446894029 | 0.654951576 | 0.822489025 | 0.020357747 |
| ENSMUSG00000036591 | Arhgap21 | 71435 | protein_coding | 2435.822616 | 0.020374059 | 0.043465606 | 0.468739782 | 0.639255648 | 0.812170498 | 0.020374059 |
| ENSMUSG00000032298 | Neil1 | 72774 | protein_coding | 211.7050143 | -0.020381617 | 0.087534986 | -0.232839664 | 0.815885912 | 0.917020505 | 0.020381617 |
| ENSMUSG00000020493 | Prr11 | 270906 | protein_coding | 94.32568877 | -0.020384002 | 0.146847067 | -0.138811093 | 0.889599428 | 0.950733597 | 0.020384002 |
| ENSMUSG00000096687 | Mfsd4b4 | 213332 | protein_coding | 53.65463045 | -0.020391977 | 0.176190051 | -0.115738525 | 0.907859773 | 0.959122007 | 0.020391977 |
| ENSMUSG00000079037 | Prnp | 19122 | protein_coding | 2101.754467 | 0.020401131 | 0.073708925 | 0.27677966 | 0.781949301 | 0.898695579 | 0.020401131 |
| ENSMUSG00000033335 | Dnm2 | 13430 | protein_coding | 4594.267834 | -0.020429682 | 0.060070413 | -0.340095588 | 0.733784545 | 0.871859084 | 0.020429682 |
| ENSMUSG00000033306 | Lpp | 210126 | protein_coding | 3091.070053 | 0.020469817 | 0.087763184 | 0.233239229 | 0.815575646 | 0.917017949 | 0.020469817 |
| ENSMUSG00000015189 | Casd1 | 213819 | protein_coding | 1994.005184 | 0.020510328 | 0.078841766 | 0.260145471 | 0.794751564 | 0.904781532 | 0.020510328 |
| ENSMUSG00000032571 | Pik3r4 | 75669 | protein_coding | 810.694426 | 0.02051843 | 0.052141303 | 0.393515864 | 0.69393851 | 0.846659581 | 0.02051843 |
| ENSMUSG00000025207 | Sema4g | 26456 | protein_coding | 5667.824645 | 0.02059131 | 0.076020042 | 0.270866867 | 0.786493429 | 0.900451559 | 0.02059131 |
| ENSMUSG00000014075 | Dynlt2b | 66061 | protein_coding | 46.3833888 | -0.020593735 | 0.178571234 | -0.115325043 | 0.908187489 | 0.959153588 | 0.020593735 |
| ENSMUSG00000074916 | Chst14 | 72136 | protein_coding | 17.88579582 | 0.020599605 | 0.3109878 | 0.066239269 | 0.947187333 | 0.977863038 | 0.020599605 |
| ENSMUSG00000026095 | Asnsd1 | 70396 | protein_coding | 198.5989839 | 0.020602663 | 0.118773086 | 0.173462386 | 0.862287992 | 0.937787345 | 0.020602663 |
| ENSMUSG00000019210 | Atp6v1e1 | 11973 | protein_coding | 2380.417325 | 0.020604649 | 0.035998954 | 0.572367994 | 0.567072695 | 0.762827428 | 0.020604649 |
| ENSMUSG00000025995 | Wdr75 | 73674 | protein_coding | 517.7289492 | 0.020606873 | 0.070878471 | 0.290735287 | 0.771253781 | 0.892332662 | 0.020606873 |
| ENSMUSG00000042532 | Golga7b | 71146 | protein_coding | 6.534589551 | -0.02063322 | 0.475334103 | -0.043407826 | 0.965376439 | 0.986105006 | 0.02063322 |
| ENSMUSG00000024456 | Diaph1 | 13367 | protein_coding | 2473.383577 | -0.020633948 | 0.045218383 | -0.456317691 | 0.648161544 | 0.81829507 | 0.020633948 |
| ENSMUSG00000078578 | Ube2d3 | 66105 | protein_coding | 5922.156182 | -0.02063439 | 0.041734192 | -0.494424094 | 0.621006701 | 0.799982033 | 0.02063439 |
| ENSMUSG00000020868 | Xylt2 | 217119 | protein_coding | 393.1031172 | -0.020636008 | 0.082447116 | -0.250293878 | 0.80236009 | 0.910121497 | 0.020636008 |
| ENSMUSG00000003731 | Kpna6 | 16650 | protein_coding | 1462.90845 | 0.020646375 | 0.044817202 | 0.460679683 | 0.645028434 | 0.81580845 | 0.020646375 |

Supplementary Table 1

| ensembl_gene_id | external_gene_name | entrezgene_id | gene_biotype | baseMean | log2FoldChange | lfcSE | stat | pvalue | padj | absLog2FC |
|---|---|---|---|---|---|---|---|---|---|---|
| ENSMUSG00000037738 | Nek5 | 330721 | protein_coding | 23.41449985 | -0.020648177 | 0.248070472 | -0.083235125 | 0.933664584 | 0.971056929 | 0.020648177 |
| ENSMUSG00000066640 | Fbxl18 | 231863 | protein_coding | 604.6433498 | -0.020654199 | 0.066338817 | -0.3113441 | 0.755539045 | 0.883965335 | 0.020654199 |
| ENSMUSG00000086429 | Gt(ROSA)26Sor | 14910 | lncRNA | 118.6158612 | -0.020656026 | 0.186753349 | -0.110605921 | 0.911928852 | 0.960655601 | 0.020656026 |
| ENSMUSG00000041258 | Zfp236 | 329002 | protein_coding | 1042.703485 | 0.02066949 | 0.058873915 | 0.35108061 | 0.725527874 | 0.865987203 | 0.02066949 |
| ENSMUSG00000021546 | Hnrnpk | 15387 | protein_coding | 7838.238445 | 0.020671983 | 0.039072092 | 0.529072851 | 0.596754914 | 0.783256002 | 0.020671983 |
| ENSMUSG00000004085 | Map3k20 | 65964 | protein_coding | 592.3353293 | -0.02069078 | 0.071763563 | -0.288318743 | 0.773102761 | 0.893430901 | 0.02069078 |
| ENSMUSG00000031365 | Zfp275 | 27081 | protein_coding | 339.3631481 | 0.020755122 | 0.078496413 | 0.264408539 | 0.791465132 | 0.902954861 | 0.020755122 |
| ENSMUSG00000021669 | Cert1 | 68018 | protein_coding | 1555.974983 | -0.020778248 | 0.045411176 | -0.457558018 | 0.647270011 | 0.817489503 | 0.020778248 |
| ENSMUSG00000032097 | Ddx6 | 13209 | protein_coding | 3465.863932 | -0.02078931 | 0.062347632 | -0.333441858 | 0.738800771 | 0.87479925 | 0.02078931 |
| ENSMUSG00000021376 | Tpmt | 22017 | protein_coding | 328.6019597 | -0.020850137 | 0.078797757 | -0.264603182 | 0.791315169 | 0.902896542 | 0.020850137 |
| ENSMUSG00000025958 | Creb1 | 12912 | protein_coding | 1170.420312 | 0.020865993 | 0.060516056 | 0.344800937 | 0.730244028 | 0.869327992 | 0.020865993 |
| ENSMUSG00000033427 | Upb1 | 103149 | protein_coding | 86.18266386 | 0.020875187 | 0.179258917 | 0.116452714 | 0.90729376 | 0.958841839 | 0.020875187 |
| ENSMUSG00000049401 | Ogfr | 72075 | protein_coding | 1325.453634 | -0.020890993 | 0.065049203 | -0.321156781 | 0.748091582 | 0.880353749 | 0.020890993 |
| ENSMUSG00000034226 | Rhov | 228543 | protein_coding | 19.3139053 | 0.020895376 | 0.298885554 | 0.069910959 | 0.944264531 | 0.975710693 | 0.020895376 |
| ENSMUSG00000034021 | Pds5b | 100710 | protein_coding | 426.1611843 | 0.020919285 | 0.088117592 | 0.237401911 | 0.812345007 | 0.915791177 | 0.020919285 |
| ENSMUSG00000025155 | Dus1l | 68730 | protein_coding | 978.881969 | 0.020937621 | 0.06904675 | 0.303238333 | 0.761708238 | 0.887577222 | 0.020937621 |
| ENSMUSG00000030861 | Acadsb | 66885 | protein_coding | 470.1140762 | 0.020946855 | 0.060866864 | 0.344142181 | 0.730739362 | 0.869778854 | 0.020946855 |
| ENSMUSG00000028759 | Hp1bp3 | 15441 | protein_coding | 3192.544745 | 0.020947797 | 0.038758833 | 0.54046512 | 0.588876309 | 0.778426904 | 0.020947797 |
| ENSMUSG00000046070 | Igfals | 16005 | protein_coding | 44.02596396 | -0.02095236 | 0.193038602 | -0.108539743 | 0.913567556 | 0.96169086 | 0.02095236 |
| ENSMUSG00000042499 | Hoxd11 | 15431 | protein_coding | 22.04194119 | 0.020973607 | 0.44534568 | 0.047095118 | 0.962437418 | 0.984572804 | 0.020973607 |
| ENSMUSG00000065822 | Snord15a | 449630 | snoRNA | 182.7877098 | -0.020987295 | 0.179754576 | -0.116755274 | 0.907053987 | 0.95873684 | 0.020987295 |
| ENSMUSG00000028478 | Clta | 12757 | protein_coding | 3419.500874 | -0.020989682 | 0.050662897 | -0.414300857 | 0.678653786 | 0.837002876 | 0.020989682 |
| ENSMUSG00000020611 | Gna13 | 14674 | protein_coding | 4328.773329 | -0.02100295 | 0.045567177 | -0.460922775 | 0.644854011 | 0.81580845 | 0.02100295 |
| ENSMUSG00000090841 | Myl6 | 17904 | protein_coding | 12103.08424 | 0.021004684 | 0.038280181 | 0.548709127 | 0.583205081 | 0.774740391 | 0.021004684 |
| ENSMUSG00000021279 | Cdc42bpb | 217866 | protein_coding | 2676.4071 | 0.021010353 | 0.048966907 | 0.429072506 | 0.66787046 | 0.830562128 | 0.021010353 |
| ENSMUSG00000000049 | Apoh | 11818 | protein_coding | 13.06437118 | -0.021038035 | 0.355073118 | -0.059249867 | 0.952753091 | 0.980775949 | 0.021038035 |
| ENSMUSG00000057143 | Trim12c | 319236 | protein_coding | 509.0055595 | -0.02106087 | 0.080833694 | -0.260545678 | 0.794442885 | 0.904736983 | 0.02106087 |
| ENSMUSG00000040174 | Alkbh3 | 69113 | protein_coding | 271.5429139 | -0.021077101 | 0.078862267 | -0.267264705 | 0.78926536 | 0.901682062 | 0.021077101 |
| ENSMUSG00000000127 | Fer | 14158 | protein_coding | 350.1200063 | 0.021092624 | 0.076602699 | 0.275350917 | 0.783046649 | 0.898856746 | 0.021092624 |
| ENSMUSG00000001901 | Kcnh6 | 192775 | protein_coding | 99.77927617 | 0.021149409 | 0.167635183 | 0.126163304 | 0.899602658 | 0.955523695 | 0.021149409 |
| ENSMUSG00000046079 | Lrrc8d | 231549 | protein_coding | 675.3185309 | 0.021154268 | 0.073370032 | 0.288323007 | 0.773099498 | 0.893430901 | 0.021154268 |
| ENSMUSG00000039901 | Armh3 | 71617 | protein_coding | 445.9051558 | -0.021174524 | 0.059732814 | -0.354487293 | 0.722973725 | 0.864747998 | 0.021174524 |
| ENSMUSG00000028256 | Odf2l | 52184 | protein_coding | 222.8641198 | -0.021199939 | 0.111247471 | -0.190565577 | 0.848865961 | 0.931678839 | 0.021199939 |
| ENSMUSG00000027881 | Prpf38b | 66921 | protein_coding | 1324.436363 | 0.021208108 | 0.08507073 | 0.249299707 | 0.803128958 | 0.910363905 | 0.021208108 |
| ENSMUSG00000003062 | Stard3nl | 76205 | protein_coding | 796.5094104 | -0.021211216 | 0.067376384 | -0.314816784 | 0.752900767 | 0.882492633 | 0.021211216 |
| ENSMUSG00000022469 | Rapgef3 | 223864 | protein_coding | 416.8491053 | -0.021221089 | 0.078501661 | -0.270326629 | 0.786908981 | 0.900576045 | 0.021221089 |
| ENSMUSG00000021928 | Ebpl | 68177 | protein_coding | 352.9814578 | 0.021223067 | 0.102089052 | 0.207887788 | 0.835316584 | 0.926721051 | 0.021223067 |
| ENSMUSG00000028437 | Ubap1 | 67123 | protein_coding | 1253.133601 | 0.021225803 | 0.040671599 | 0.521882671 | 0.601752025 | 0.786805165 | 0.021225803 |
| ENSMUSG00000033554 | Dph5 | 69740 | protein_coding | 189.5398919 | -0.021268377 | 0.101460299 | -0.20962265 | 0.833962201 | 0.926388511 | 0.021268377 |

| ensembl_gene_id | external_gene_name | entrezgene_id | gene_biotype | baseMean | log2FoldChange | lfcSE | stat | pvalue | padj | absLog2FC |
|---|---|---|---|---|---|---|---|---|---|---|
| ENSMUSG00000019873 | Reep3 | 28193 | protein_coding | 1859.017921 | -0.021270848 | 0.044218577 | -0.481038729 | 0.630488974 | 0.805900922 | 0.021270848 |
| ENSMUSG00000021830 | Txndc16 | 70561 | protein_coding | 749.2120556 | 0.021279513 | 0.067814479 | 0.313790111 | 0.753680455 | 0.882943826 | 0.021279513 |
| ENSMUSG00000029063 | Nadk | 192185 | protein_coding | 4438.145693 | 0.021293977 | 0.056159462 | 0.379169886 | 0.704561713 | 0.853157579 | 0.021293977 |
| ENSMUSG00000036980 | Taf6 | 21343 | protein_coding | 367.6787548 | 0.021332921 | 0.087101209 | 0.244921072 | 0.806517545 | 0.912178372 | 0.021332921 |
| ENSMUSG00000033450 | Tagap | 72536 | protein_coding | 82.99867646 | 0.021379662 | 0.195714986 | 0.109238757 | 0.913013121 | 0.961295463 | 0.021379662 |
| ENSMUSG00000047767 | Atg16l2 | 73683 | protein_coding | 305.564446 | -0.02138076 | 0.144460058 | -0.148004649 | 0.8823391 | 0.947541514 | 0.02138076 |
| ENSMUSG00000010660 | Plcd1 | 18799 | protein_coding | 1214.94364 | -0.02140424 | 0.08304643 | -0.257738227 | 0.796608942 | 0.906052077 | 0.02140424 |
| ENSMUSG00000022314 | Rad21 | 19357 | protein_coding | 3106.301159 | 0.02140859 | 0.058816469 | 0.363989725 | 0.715865682 | 0.860522201 | 0.02140859 |
| ENSMUSG00000041638 | Gcn1 | 231659 | protein_coding | 2234.472926 | 0.021452074 | 0.064411294 | 0.333048326 | 0.739097805 | 0.87479925 | 0.021452074 |
| ENSMUSG00000063354 | Slc39a4 | 72027 | protein_coding | 2966.433117 | -0.021456631 | 0.108876001 | -0.197074018 | 0.843769615 | 0.930175172 | 0.021456631 |
| ENSMUSG00000022544 | Eef2kmt | 70511 | protein_coding | 337.5629431 | -0.021472175 | 0.082509138 | -0.260239959 | 0.794678683 | 0.904781532 | 0.021472175 |
| ENSMUSG00000042225 | Ammecr1 | 56068 | protein_coding | 499.2417365 | 0.021503841 | 0.086365642 | 0.248986056 | 0.803371568 | 0.910371624 | 0.021503841 |
| ENSMUSG00000047044 | D030056L22Rik | 225995 | protein_coding | 345.8820223 | 0.021509111 | 0.089669238 | 0.239871685 | 0.810429732 | 0.914739662 | 0.021509111 |
| ENSMUSG00000053477 | Tcf4 | 21413 | protein_coding | 498.2166716 | -0.021517507 | 0.071853491 | -0.299463628 | 0.76458632 | 0.889388602 | 0.021517507 |
| ENSMUSG00000031783 | Polr2c | 20021 | protein_coding | 779.2104926 | 0.021519975 | 0.056786078 | 0.378965686 | 0.704713345 | 0.853157579 | 0.021519975 |
| ENSMUSG00000015776 | Med22 | 20933 | protein_coding | 949.3411694 | -0.021548485 | 0.045088576 | -0.477914518 | 0.632711048 | 0.807820141 | 0.021548485 |
| ENSMUSG00000059552 | Trp53 | 22059 | protein_coding | 536.8038927 | -0.02155678 | 0.082915222 | -0.259985798 | 0.794874729 | 0.904781532 | 0.02155678 |
| ENSMUSG00000023806 | Rsph3b | 100037282 | protein_coding | 359.1943197 | 0.021558412 | 0.074926126 | 0.287728902 | 0.773554265 | 0.893440495 | 0.021558412 |
| ENSMUSG00000033039 | Micall1 | 27008 | protein_coding | 237.0167679 | -0.021612305 | 0.080918812 | -0.267086288 | 0.789402725 | 0.901682062 | 0.021612305 |
| ENSMUSG00000060038 | Dhps | 330817 | protein_coding | 365.5487343 | -0.021631727 | 0.069803621 | -0.309894056 | 0.756641523 | 0.884485201 | 0.021631727 |
| ENSMUSG00000001348 | Acp5 | 11433 | protein_coding | 3792.082989 | -0.021650261 | 0.067464373 | -0.320913993 | 0.74827557 | 0.880449447 | 0.021650261 |
| ENSMUSG00000033632 | AW554918 | 225289 | protein_coding | 163.9565084 | 0.021661254 | 0.120274758 | 0.180098091 | 0.857075561 | 0.935731633 | 0.021661254 |
| ENSMUSG00000072946 | Ptgr2 | 77219 | protein_coding | 1089.520503 | 0.021664151 | 0.05049646 | 0.429023156 | 0.667906373 | 0.830562128 | 0.021664151 |
| ENSMUSG00000004070 | Hmox2 | 15369 | protein_coding | 579.8344938 | 0.021665266 | 0.065055143 | 0.333029259 | 0.739112197 | 0.87479925 | 0.021665266 |
| ENSMUSG00000005506 | Celf1 | 13046 | protein_coding | 2630.293472 | 0.02167544 | 0.038012342 | 0.570221111 | 0.568527739 | 0.763734926 | 0.02167544 |
| ENSMUSG00000052738 | Suclg1 | 56451 | protein_coding | 5305.61958 | 0.021676683 | 0.057760345 | 0.375286591 | 0.707447338 | 0.854735032 | 0.021676683 |
| ENSMUSG00000022235 | Cmbl | 69574 | protein_coding | 999.6086442 | 0.02171213 | 0.062278894 | 0.348627411 | 0.727369046 | 0.867367688 | 0.02171213 |
| ENSMUSG00000064179 | Tnnt1 | 21955 | protein_coding | 39.1519516 | -0.021723113 | 0.210890059 | -0.103006815 | 0.917957562 | 0.963921188 | 0.021723113 |
| ENSMUSG00000020166 | Cnot2 | 72068 | protein_coding | 1172.381274 | -0.021728805 | 0.046382505 | -0.468469833 | 0.639448639 | 0.812190927 | 0.021728805 |
| ENSMUSG00000022272 | Myo10 | 17909 | protein_coding | 2886.730402 | -0.02172996 | 0.060869064 | -0.356995134 | 0.721095448 | 0.863755116 | 0.02172996 |
| ENSMUSG00000063172 | Hspb11 | 72938 | protein_coding | 246.2816567 | 0.021734234 | 0.087366708 | 0.248770207 | 0.803538538 | 0.910466241 | 0.021734234 |
| ENSMUSG00000003382 | Etv3 | 27049 | protein_coding | 1505.920615 | 0.021745113 | 0.041945034 | 0.518419245 | 0.60416579 | 0.788682346 | 0.021745113 |
| ENSMUSG00000049600 | Zbtb45 | 232879 | protein_coding | 232.4126724 | 0.021761523 | 0.129988203 | 0.167411521 | 0.867046261 | 0.939855462 | 0.021761523 |
| ENSMUSG00000026878 | Rab14 | 68365 | protein_coding | 2624.380076 | 0.021765536 | 0.034783339 | 0.625746016 | 0.531481546 | 0.740026398 | 0.021765536 |
| ENSMUSG00000062944 | 9130023H24Rik | 100043133 | protein_coding | 181.5543454 | 0.021786368 | 0.102159846 | 0.213257642 | 0.831126011 | 0.925277452 | 0.021786368 |
| ENSMUSG00000019715 | Gle1 | 74412 | protein_coding | 532.7417729 | 0.021794858 | 0.060697458 | 0.359073643 | 0.719539998 | 0.862768697 | 0.021794858 |
| ENSMUSG00000048373 | Fgfbp1 | 14181 | protein_coding | 2553.042684 | 0.021800543 | 0.050718348 | 0.429835435 | 0.667315355 | 0.830281488 | 0.021800543 |
| ENSMUSG00000057322 | Rpl38 | 67671 | protein_coding | 5343.425818 | 0.02182384 | 0.02950864 | 0.739574574 | 0.459558174 | 0.68653709 | 0.02182384 |
| ENSMUSG00000023967 | Mrps18a | 68565 | protein_coding | 741.1003389 | 0.021857218 | 0.051501552 | 0.424399206 | 0.6712747 | 0.832361742 | 0.021857218 |

| ensembl_gene_id | external_gene_name | entrezgene_id | gene_biotype | baseMean | log2FoldChange | lfcSE | stat | pvalue | padj | absLog2FC |
|---|---|---|---|---|---|---|---|---|---|---|
| ENSMUSG00000073433 | Arhgdig | 14570 | protein_coding | 14.81096011 | 0.021857998 | 0.334325775 | 0.065379339 | 0.947871974 | 0.978199995 | 0.021857998 |
| ENSMUSG00000028559 | Osbpl9 | 100273 | protein_coding | 1255.429961 | 0.021876838 | 0.060962892 | 0.358854981 | 0.71970358 | 0.862820371 | 0.021876838 |
| ENSMUSG00000021198 | Unc79 | 217843 | protein_coding | 45.97269724 | 0.021919019 | 0.185002291 | 0.118479719 | 0.905687564 | 0.958526272 | 0.021919019 |
| ENSMUSG00000017950 | Hnf4a | 15378 | protein_coding | 17341.66028 | 0.021932605 | 0.048036632 | 0.456580816 | 0.647972371 | 0.818120287 | 0.021932605 |
| ENSMUSG00000026554 | Dcaf8 | 98193 | protein_coding | 2631.85143 | -0.021944884 | 0.035536788 | -0.617525802 | 0.536887968 | 0.74340107 | 0.021944884 |
| ENSMUSG00000038611 | Phrf1 | 101471 | protein_coding | 1324.853349 | 0.021965877 | 0.058709358 | 0.374146098 | 0.708295624 | 0.855350218 | 0.021965877 |
| ENSMUSG00000093661 | Eif4e3 | 66892 | protein_coding | 94.46971647 | 0.021972343 | 0.171010251 | 0.128485529 | 0.897764749 | 0.954463189 | 0.021972343 |
| ENSMUSG00000043683 | Fem1a | 14154 | protein_coding | 1463.892559 | -0.021974261 | 0.07481183 | -0.293727086 | 0.768966455 | 0.891610224 | 0.021974261 |
| ENSMUSG00000020536 | Llgl1 | 16897 | protein_coding | 876.1712974 | -0.022058399 | 0.050908485 | -0.433295137 | 0.664800371 | 0.828854703 | 0.022058399 |
| ENSMUSG00000015733 | Capza2 | 12343 | protein_coding | 3071.57164 | 0.022076868 | 0.039037082 | 0.565535823 | 0.571709365 | 0.766604883 | 0.022076868 |
| ENSMUSG00000042903 | Foxo4 | 54601 | protein_coding | 822.567946 | -0.022090753 | 0.059781908 | -0.369522375 | 0.711738398 | 0.857670564 | 0.022090753 |
| ENSMUSG00000060012 | Kif13b | 16554 | protein_coding | 5770.800539 | 0.022107778 | 0.058529926 | 0.377717515 | 0.705640455 | 0.853575129 | 0.022107778 |
| ENSMUSG00000014496 | Ankrd28 | 105522 | protein_coding | 1101.466422 | 0.022125094 | 0.056488966 | 0.391671072 | 0.695301267 | 0.84768023 | 0.022125094 |
| ENSMUSG00000014355 | Anapc1 | 17222 | protein_coding | 2422.565396 | 0.022130974 | 0.044017368 | 0.502778215 | 0.615120212 | 0.795448934 | 0.022130974 |
| ENSMUSG00000037818 | Abhd18 | 269423 | protein_coding | 199.8434441 | 0.022141297 | 0.107755702 | 0.205476798 | 0.837199624 | 0.927812059 | 0.022141297 |
| ENSMUSG00000024975 | Pdcd4 | 18569 | protein_coding | 4404.176007 | 0.022143541 | 0.047156819 | 0.469572403 | 0.638660546 | 0.811701365 | 0.022143541 |
| ENSMUSG00000061288 | Taok3 | 330177 | protein_coding | 1284.120745 | 0.022148477 | 0.048048325 | 0.460962518 | 0.644825497 | 0.81580845 | 0.022148477 |
| ENSMUSG00000033498 | Strc | 140476 | protein_coding | 31.56398526 | 0.022152126 | 0.255936812 | 0.086553107 | 0.931026741 | 0.969751874 | 0.022152126 |
| ENSMUSG00000110605 | Gm32856 | 102635552 | lncRNA | 41.81719484 | -0.022166274 | 0.209668063 | -0.105720795 | 0.915803881 | 0.962474368 | 0.022166274 |
| ENSMUSG00000041231 | Ublcp1 | 79560 | protein_coding | 528.8961832 | 0.022186405 | 0.056340169 | 0.393793726 | 0.693733337 | 0.84654428 | 0.022186405 |
| ENSMUSG00000050866 | Clrn3 | 212070 | protein_coding | 2439.065672 | 0.022196248 | 0.052522813 | 0.422602033 | 0.672585643 | 0.833278687 | 0.022196248 |
| ENSMUSG00000021271 | Zfp839 | 72805 | protein_coding | 276.1169839 | -0.022210432 | 0.079741958 | -0.278528804 | 0.780606457 | 0.898240481 | 0.022210432 |
| ENSMUSG00000063052 | Lrrc40 | 67144 | protein_coding | 610.710975 | 0.022216081 | 0.069473237 | 0.319778976 | 0.749135886 | 0.880475742 | 0.022216081 |
| ENSMUSG00000020600 | Slc7a15 | 328059 | protein_coding | 13.17057822 | 0.022257867 | 0.551502234 | 0.040358617 | 0.967807222 | 0.987251745 | 0.022257867 |
| ENSMUSG00000074136 | 4930513N10Rik | 319960 | lncRNA | 33.93903853 | -0.022264259 | 0.199562633 | -0.111565268 | 0.911168112 | 0.960412417 | 0.022264259 |
| ENSMUSG00000048200 | Cracr2b | 213573 | protein_coding | 463.4622112 | 0.022270577 | 0.080840913 | 0.275486456 | 0.78294253 | 0.898825528 | 0.022270577 |
| ENSMUSG00000025234 | Arih1 | 23806 | protein_coding | 1910.427039 | -0.022301879 | 0.05133804 | -0.43441237 | 0.663989017 | 0.828333496 | 0.022301879 |
| ENSMUSG00000026655 | Fam107b | 66540 | protein_coding | 3488.101578 | -0.022333903 | 0.065724303 | -0.339811947 | 0.73399815 | 0.871991313 | 0.022333903 |
| ENSMUSG00000030935 | Acsm3 | 20216 | protein_coding | 3659.81364 | -0.022338417 | 0.082761243 | -0.26991399 | 0.787226424 | 0.900662858 | 0.022338417 |
| ENSMUSG00000002205 | Vrk3 | 101568 | protein_coding | 888.5802738 | -0.022342719 | 0.049307586 | -0.453129448 | 0.650455531 | 0.819843404 | 0.022342719 |
| ENSMUSG00000054280 | Prr14l | 215476 | protein_coding | 1953.089815 | -0.02235493 | 0.063697084 | -0.350956874 | 0.725620703 | 0.865987203 | 0.02235493 |
| ENSMUSG00000027698 | Nceh1 | 320024 | protein_coding | 2025.587611 | -0.022371463 | 0.110296696 | -0.202829853 | 0.839268023 | 0.928763838 | 0.022371463 |
| ENSMUSG00000024122 | Pdpk1 | 18607 | protein_coding | 2435.024147 | 0.022388146 | 0.035824314 | 0.624942752 | 0.532008632 | 0.740523322 | 0.022388146 |
| ENSMUSG00000041153 | Osgin2 | 209212 | protein_coding | 158.7158467 | 0.022407563 | 0.096253685 | 0.232796934 | 0.815919094 | 0.917020505 | 0.022407563 |
| ENSMUSG00000048758 | Rpl29 | 19944 | protein_coding | 5644.701177 | 0.022410562 | 0.076713237 | 0.292134221 | 0.770184003 | 0.892032325 | 0.022410562 |
| ENSMUSG00000024130 | Abca3 | 27410 | protein_coding | 2231.960297 | -0.02243156 | 0.055562784 | -0.403715552 | 0.686421902 | 0.842267824 | 0.02243156 |
| ENSMUSG00000021532 | Fastkd3 | 69577 | protein_coding | 463.2387998 | 0.022437459 | 0.067981772 | 0.330051111 | 0.741361343 | 0.876277805 | 0.022437459 |
| ENSMUSG00000027575 | Arfgap1 | 228998 | protein_coding | 1294.266641 | -0.022461393 | 0.041060209 | -0.547035527 | 0.584354325 | 0.775436698 | 0.022461393 |
| ENSMUSG00000025887 | Casp12 | 12364 | protein_coding | 12.95597718 | -0.022497636 | 0.426795294 | -0.052712943 | 0.957960626 | 0.983146794 | 0.022497636 |

| ensembl_gene_id | external_gene_name | entrezgene_id | gene_biotype | baseMean | log2FoldChange | lfcSE | stat | pvalue | padj | absLog2FC |
|---|---|---|---|---|---|---|---|---|---|---|
| ENSMUSG00000035834 | Polr3g | 67486 | protein_coding | 298.6977998 | -0.022503414 | 0.076669911 | -0.293510374 | 0.769132071 | 0.891717679 | 0.022503414 |
| ENSMUSG00000054079 | Utp18 | 217109 | protein_coding | 418.5230315 | -0.022514497 | 0.073340904 | -0.306984171 | 0.75885542 | 0.886038321 | 0.022514497 |
| ENSMUSG00000021215 | Net1 | 56349 | protein_coding | 4662.710406 | -0.022518526 | 0.078446499 | -0.287055846 | 0.77406956 | 0.893779608 | 0.022518526 |
| ENSMUSG00000035246 | Pcyt1b | 236899 | protein_coding | 8.065422084 | 0.022543321 | 0.481835123 | 0.04678638 | 0.962683485 | 0.984661462 | 0.022543321 |
| ENSMUSG00000051978 | Erich1 | 234086 | protein_coding | 176.8151259 | -0.022568929 | 0.114905692 | -0.196412633 | 0.844287209 | 0.930210801 | 0.022568929 |
| ENSMUSG00000020519 | Sap30l | 50724 | protein_coding | 644.2787455 | 0.022578327 | 0.086972309 | 0.259603633 | 0.795169533 | 0.905053219 | 0.022578327 |
| ENSMUSG00000063787 | Chchd1 | 66121 | protein_coding | 481.9832615 | 0.02258287 | 0.11406616 | 0.197980456 | 0.843060355 | 0.930024923 | 0.02258287 |
| ENSMUSG00000070047 | Fat1 | 14107 | protein_coding | 9998.343467 | -0.022591546 | 0.078180054 | -0.288968151 | 0.77260575 | 0.893129098 | 0.022591546 |
| ENSMUSG00000064120 | Mocs1 | 56738 | protein_coding | 609.4652753 | 0.022593475 | 0.077348005 | 0.29210158 | 0.770208959 | 0.892032325 | 0.022593475 |
| ENSMUSG00000015305 | Sash1 | 70097 | protein_coding | 2312.353634 | -0.022606644 | 0.050622268 | -0.446575102 | 0.655181876 | 0.822565162 | 0.022606644 |
| ENSMUSG00000111375 | Btbd8 | 100503185 | protein_coding | 206.8506513 | -0.022611 | 0.100160595 | -0.225747463 | 0.821397841 | 0.920176481 | 0.022611 |
| ENSMUSG00000030447 | Cyfip1 | 20430 | protein_coding | 3146.19939 | 0.022613271 | 0.040114366 | 0.563720026 | 0.572944686 | 0.767564688 | 0.022613271 |
| ENSMUSG00000057236 | Rbbp4 | 19646 | protein_coding | 2181.571772 | 0.02263105 | 0.046558856 | 0.486074027 | 0.626914681 | 0.803226458 | 0.02263105 |
| ENSMUSG00000087165 | 2010001A14Rik | 76516 | lncRNA | 156.0906246 | -0.022648818 | 0.118795092 | -0.190654493 | 0.848796294 | 0.931678839 | 0.022648818 |
| ENSMUSG00000016503 | Gtf3a | 66596 | protein_coding | 313.338373 | 0.022650499 | 0.070904483 | 0.319450875 | 0.749384638 | 0.880475742 | 0.022650499 |
| ENSMUSG00000051220 | Ercc6l | 236930 | protein_coding | 199.2001376 | -0.022651382 | 0.179434861 | -0.126237355 | 0.899544043 | 0.955523695 | 0.022651382 |
| ENSMUSG00000041958 | Pigs | 276846 | protein_coding | 1197.489025 | -0.022662051 | 0.075547243 | -0.299971914 | 0.764198579 | 0.889251381 | 0.022662051 |
| ENSMUSG00000025059 | Gk | 14933 | protein_coding | 1651.861149 | 0.022663772 | 0.117173151 | 0.193421202 | 0.846629112 | 0.930889346 | 0.022663772 |
| ENSMUSG00000031913 | Vps4a | 116733 | protein_coding | 1202.709291 | -0.022694006 | 0.053244394 | -0.426223394 | 0.669945072 | 0.831606417 | 0.022694006 |
| ENSMUSG00000040389 | Wdr47 | 99512 | protein_coding | 147.6621428 | 0.022705334 | 0.1064465 | 0.213302778 | 0.831090807 | 0.925277452 | 0.022705334 |
| ENSMUSG00000058761 | Rnf169 | 108937 | protein_coding | 650.7872555 | -0.022717433 | 0.058515805 | -0.388227297 | 0.697847834 | 0.849362552 | 0.022717433 |
| ENSMUSG00000058921 | Slc10a5 | 241877 | protein_coding | 1600.334757 | -0.022726188 | 0.066473694 | -0.341882432 | 0.732439371 | 0.871007815 | 0.022726188 |
| ENSMUSG00000025034 | Trim8 | 93679 | protein_coding | 910.3691087 | 0.022753312 | 0.091166961 | 0.249578488 | 0.802913336 | 0.910278712 | 0.022753312 |
| ENSMUSG00000017386 | Traf4 | 22032 | protein_coding | 1612.678944 | -0.022754971 | 0.06374282 | -0.356980927 | 0.721106084 | 0.863755116 | 0.022754971 |
| ENSMUSG00000040018 | Cox15 | 226139 | protein_coding | 1453.384696 | 0.022771869 | 0.056217408 | 0.405067931 | 0.685427578 | 0.841671981 | 0.022771869 |
| ENSMUSG00000038766 | Gabpb2 | 213054 | protein_coding | 1709.99401 | -0.022784506 | 0.047648016 | -0.478183741 | 0.632519434 | 0.80777961 | 0.022784506 |
| ENSMUSG00000029916 | Agk | 69923 | protein_coding | 177.5721315 | -0.022794539 | 0.099066234 | -0.230093925 | 0.818018785 | 0.918228506 | 0.022794539 |
| ENSMUSG00000029455 | Aldh2 | 11669 | protein_coding | 11156.56227 | -0.022803901 | 0.095356081 | -0.239144698 | 0.810993383 | 0.914838553 | 0.022803901 |
| ENSMUSG00000003541 | Ier3 | 15937 | protein_coding | 226.8157178 | -0.022814011 | 0.115201108 | -0.198036385 | 0.843016597 | 0.930024923 | 0.022814011 |
| ENSMUSG00000078945 | Naip2 | 17948 | protein_coding | 3633.883462 | 0.022821499 | 0.043242719 | 0.527753554 | 0.597670398 | 0.783947174 | 0.022821499 |
| ENSMUSG00000068923 | Syt11 | 229521 | protein_coding | 63.58561325 | 0.022834117 | 0.157087897 | 0.145358854 | 0.884427549 | 0.948750818 | 0.022834117 |
| ENSMUSG00000034659 | Tmem109 | 68539 | protein_coding | 702.89058 | 0.02283683 | 0.087832891 | 0.260003169 | 0.794861329 | 0.904781532 | 0.02283683 |
| ENSMUSG00000039410 | Prdm16 | 70673 | protein_coding | 181.6609531 | 0.022838748 | 0.166162638 | 0.137448155 | 0.89067657 | 0.95112373 | 0.022838748 |
| ENSMUSG00000040356 | Skiv2l | 108077 | protein_coding | 1972.027642 | 0.022875592 | 0.053903455 | 0.424380806 | 0.671288116 | 0.832361742 | 0.022875592 |
| ENSMUSG00000024191 | Bnip1 | 224630 | protein_coding | 468.5902547 | -0.02287707 | 0.06242413 | -0.366477991 | 0.714008433 | 0.85937861 | 0.02287707 |
| ENSMUSG00000073207 | Ccdc160 | 434778 | protein_coding | 9.361293812 | 0.022894655 | 0.395568156 | 0.057877902 | 0.953845886 | 0.981205596 | 0.022894655 |
| ENSMUSG00000030231 | Plekha5 | 109135 | protein_coding | 2211.709014 | 0.022920392 | 0.051752345 | 0.44288605 | 0.657848158 | 0.824906092 | 0.022920392 |
| ENSMUSG00000043822 | Adamtsl5 | 66548 | protein_coding | 139.7850293 | -0.02292235 | 0.150881145 | -0.151923226 | 0.879247489 | 0.945686559 | 0.02292235 |
| ENSMUSG00000035443 | Thyn1 | 77862 | protein_coding | 403.5141762 | 0.022939111 | 0.085758933 | 0.267483633 | 0.789096814 | 0.90164123 | 0.022939111 |

Supplementary Table 1

| ensembl_gene_id | external_gene_name | entrezgene_id | gene_biotype | baseMean | log2FoldChange | lfcSE | stat | pvalue | padj | absLog2FC |
|---|---|---|---|---|---|---|---|---|---|---|
| ENSMUSG00000018672 | Copz2 | 56358 | protein_coding | 127.9565464 | 0.022960363 | 0.110315222 | 0.208134129 | 0.835124239 | 0.926699249 | 0.022960363 |
| ENSMUSG00000062859 | Tcp11 | 21463 | protein_coding | 15.18008923 | -0.022975723 | 0.293894496 | -0.078176771 | 0.937687439 | 0.972883156 | 0.022975723 |
| ENSMUSG00000044465 | Fhip1b | 74349 | protein_coding | 1372.571146 | -0.023003667 | 0.051988099 | -0.442479475 | 0.65814228 | 0.825115883 | 0.023003667 |
| ENSMUSG00000054027 | Nt5dc3 | 103466 | protein_coding | 148.0004628 | 0.023010438 | 0.109081665 | 0.210946891 | 0.832928713 | 0.92587823 | 0.023010438 |
| ENSMUSG00000059475 | Zfp426 | 235028 | protein_coding | 410.5170213 | 0.023017565 | 0.069323161 | 0.332032822 | 0.739864475 | 0.875021589 | 0.023017565 |
| ENSMUSG00000022828 | Gtf2e1 | 74197 | protein_coding | 246.4283785 | -0.023045137 | 0.07418841 | -0.310629887 | 0.756082004 | 0.884312567 | 0.023045137 |
| ENSMUSG00000029003 | Mad2l2 | 71890 | protein_coding | 119.2909732 | -0.023052721 | 0.118073193 | -0.195240942 | 0.845204329 | 0.930607647 | 0.023052721 |
| ENSMUSG00000073468 | Sft2d1 | 106489 | protein_coding | 452.3875768 | 0.023065127 | 0.072689077 | 0.317312149 | 0.75100676 | 0.881219879 | 0.023065127 |
| ENSMUSG00000036572 | Upf3b | 68134 | protein_coding | 392.6185977 | -0.023086578 | 0.104711474 | -0.22047802 | 0.825498889 | 0.922363751 | 0.023086578 |
| ENSMUSG00000050174 | Nudt6 | 229228 | protein_coding | 47.1375717 | 0.023092125 | 0.167513781 | 0.137852089 | 0.890357316 | 0.951039372 | 0.023092125 |
| ENSMUSG00000054277 | Arfgap3 | 66251 | protein_coding | 3908.043441 | -0.023113053 | 0.041854119 | -0.552228883 | 0.580791548 | 0.772645663 | 0.023113053 |
| ENSMUSG00000007603 | Dus3l | 224907 | protein_coding | 670.4095629 | 0.023137162 | 0.060465172 | 0.382652701 | 0.701977289 | 0.851790685 | 0.023137162 |
| ENSMUSG00000049957 | Ccdc137 | 67291 | protein_coding | 348.1519212 | -0.023144972 | 0.075343731 | -0.307191743 | 0.75869743 | 0.885925238 | 0.023144972 |
| ENSMUSG00000028800 | Hdac1 | 433759 | protein_coding | 1350.989575 | -0.023149894 | 0.048252392 | -0.479766762 | 0.63139325 | 0.806596874 | 0.023149894 |
| ENSMUSG00000026240 | Cops7b | 26895 | protein_coding | 266.3850173 | -0.023157079 | 0.088755489 | -0.260908698 | 0.794162917 | 0.904717134 | 0.023157079 |
| ENSMUSG00000038722 | Bud31 | 231889 | protein_coding | 592.1039225 | 0.023190529 | 0.075146196 | 0.308605497 | 0.757621639 | 0.885131169 | 0.023190529 |
| ENSMUSG00000027601 | Mtfr1 | 67472 | protein_coding | 1086.346554 | 0.023192514 | 0.045892967 | 0.505360956 | 0.613305337 | 0.793911553 | 0.023192514 |
| ENSMUSG00000006599 | Gtf2h1 | 14884 | protein_coding | 1126.212114 | -0.023216763 | 0.052314279 | -0.443794 | 0.657191527 | 0.824377651 | 0.023216763 |
| ENSMUSG00000113800 | 2210039B01Rik | 73644 | lncRNA | 25.98920288 | -0.023228499 | 0.237545926 | -0.097785296 | 0.922102784 | 0.96563322 | 0.023228499 |
| ENSMUSG00000040435 | Ppp1r15a | 17872 | protein_coding | 1013.803922 | -0.023236505 | 0.131619955 | -0.176542416 | 0.859867829 | 0.936765956 | 0.023236505 |
| ENSMUSG00000056602 | Fry | 320365 | protein_coding | 329.487435 | 0.023247966 | 0.072298417 | 0.321555673 | 0.74778933 | 0.880198417 | 0.023247966 |
| ENSMUSG00000023175 | Bsg | 12215 | protein_coding | 13939.09043 | -0.023285095 | 0.071682914 | -0.324834661 | 0.74530621 | 0.879007843 | 0.023285095 |
| ENSMUSG00000025217 | Btrc | 12234 | protein_coding | 401.618769 | -0.023291725 | 0.07456115 | -0.312384197 | 0.754748559 | 0.883681575 | 0.023291725 |
| ENSMUSG00000017999 | Ddx27 | 228889 | protein_coding | 806.3786624 | -0.023307944 | 0.060961744 | -0.382337231 | 0.702211242 | 0.851790951 | 0.023307944 |
| ENSMUSG00000035919 | Bbs9 | 319845 | protein_coding | 107.3787562 | 0.023323887 | 0.138129651 | 0.168855033 | 0.86591067 | 0.939297866 | 0.023323887 |
| ENSMUSG00000024054 | Smchd1 | 74355 | protein_coding | 1419.510274 | -0.023332004 | 0.055504621 | -0.420361475 | 0.674221407 | 0.834301331 | 0.023332004 |
| ENSMUSG00000029249 | Rest | 19712 | protein_coding | 1326.632703 | -0.023332428 | 0.039016525 | -0.598013992 | 0.549830597 | 0.751487589 | 0.023332428 |
| ENSMUSG00000027479 | Mapre1 | 13589 | protein_coding | 2029.564786 | 0.023340318 | 0.055910221 | 0.417460657 | 0.676341496 | 0.835556864 | 0.023340318 |
| ENSMUSG00000020409 | Slu7 | 193116 | protein_coding | 1413.501165 | 0.023407089 | 0.046013194 | 0.508703848 | 0.610959825 | 0.792706869 | 0.023407089 |
| ENSMUSG00000029432 | Nipsnap2 | 14467 | protein_coding | 2257.35148 | 0.023418947 | 0.053595399 | 0.436958159 | 0.662141693 | 0.827333871 | 0.023418947 |
| ENSMUSG00000020677 | Ddx52 | 78394 | protein_coding | 640.3717847 | -0.023430788 | 0.086543408 | -0.270740293 | 0.786590784 | 0.900451559 | 0.023430788 |
| ENSMUSG00000040325 | Dcaf1 | 321006 | protein_coding | 1239.049562 | 0.023463013 | 0.043049926 | 0.545018657 | 0.585740686 | 0.776292513 | 0.023463013 |
| ENSMUSG00000024811 | Tnks2 | 74493 | protein_coding | 3303.136836 | -0.023464546 | 0.032332264 | -0.725731616 | 0.468003316 | 0.693047027 | 0.023464546 |
| ENSMUSG00000027905 | Ddx20 | 53975 | protein_coding | 337.7890851 | 0.023474827 | 0.069298668 | 0.338748607 | 0.73479912 | 0.872486041 | 0.023474827 |
| ENSMUSG00000029571 | Tmem106b | 71900 | protein_coding | 1856.745763 | 0.023499807 | 0.047932913 | 0.490264521 | 0.623946729 | 0.801464668 | 0.023499807 |
| ENSMUSG00000028669 | Pithd1 | 66193 | protein_coding | 1026.827278 | 0.023500805 | 0.053226595 | 0.441523738 | 0.65883388 | 0.825416419 | 0.023500805 |
| ENSMUSG00000032842 | Abcc10 | 224814 | protein_coding | 527.8463525 | 0.023528692 | 0.092422909 | 0.254576409 | 0.799050282 | 0.908143613 | 0.023528692 |
| ENSMUSG00000021575 | Ahrr | 11624 | protein_coding | 6.982422438 | -0.023554922 | 0.46143693 | -0.051046893 | 0.959288154 | 0.983542327 | 0.023554922 |
| ENSMUSG00000024006 | Stk38 | 106504 | protein_coding | 1667.481187 | -0.023566665 | 0.038863202 | -0.606400504 | 0.544248834 | 0.748889318 | 0.023566665 |

| ensembl_gene_id | external_gene_name | entrezgene_id | gene_biotype | baseMean | log2FoldChange | lfcSE | stat | pvalue | padj | absLog2FC |
|---|---|---|---|---|---|---|---|---|---|---|
| ENSMUSG00000057092 | Fxyd3 | 17178 | protein_coding | 586.917976 | 0.023577083 | 0.087788695 | 0.268566281 | 0.788263463 | 0.901263727 | 0.023577083 |
| ENSMUSG00000019984 | Med23 | 70208 | protein_coding | 821.5645234 | -0.023584464 | 0.048756971 | -0.483714711 | 0.628588351 | 0.804287872 | 0.023584464 |
| ENSMUSG00000038872 | Zfhx3 | 11906 | protein_coding | 410.511644 | 0.023592605 | 0.141026317 | 0.167292214 | 0.86714013 | 0.939855462 | 0.023592605 |
| ENSMUSG00000033417 | Cacul1 | 78832 | protein_coding | 1286.356452 | -0.023650711 | 0.041556528 | -0.569121438 | 0.569273731 | 0.764610432 | 0.023650711 |
| ENSMUSG00000017778 | Cox7c | 12867 | protein_coding | 4072.833726 | 0.023696739 | 0.105044961 | 0.225586635 | 0.821522936 | 0.920176481 | 0.023696739 |
| ENSMUSG00000002496 | Tsc2 | 22084 | protein_coding | 1542.571659 | -0.023712553 | 0.042242581 | -0.561342422 | 0.574564127 | 0.768227276 | 0.023712553 |
| ENSMUSG00000006335 | Tfpt | 69714 | protein_coding | 232.6194749 | -0.023736474 | 0.111880507 | -0.212159158 | 0.831982869 | 0.925610506 | 0.023736474 |
| ENSMUSG00000038415 | Foxq1 | 15220 | protein_coding | 95.15406128 | 0.023752625 | 0.183600988 | 0.129370898 | 0.897064174 | 0.954410472 | 0.023752625 |
| ENSMUSG00000072694 | 1500011B03Rik | 66236 | protein_coding | 116.048609 | 0.023792841 | 0.13408867 | 0.177441095 | 0.85916193 | 0.936398271 | 0.023792841 |
| ENSMUSG00000057229 | Dmac2 | 66349 | protein_coding | 1109.327263 | 0.023823418 | 0.049246079 | 0.483762733 | 0.628554266 | 0.804287872 | 0.023823418 |
| ENSMUSG00000002881 | Nab1 | 17936 | protein_coding | 1450.439984 | -0.02383056 | 0.048855791 | -0.4877735 | 0.625710284 | 0.8024647 | 0.02383056 |
| ENSMUSG00000054008 | Ndst1 | 15531 | protein_coding | 1310.929006 | 0.023844764 | 0.063659473 | 0.374567405 | 0.707982218 | 0.855125017 | 0.023844764 |
| ENSMUSG00000039347 | Atp6v0e2 | 76252 | protein_coding | 619.6323272 | 0.023848709 | 0.072241128 | 0.33012647 | 0.741304403 | 0.876274728 | 0.023848709 |
| ENSMUSG00000097316 | Gm10516 | 100038712 | lncRNA | 261.5888007 | -0.023883412 | 0.074777309 | -0.319393847 | 0.749427876 | 0.880475742 | 0.023883412 |
| ENSMUSG00000012429 | Mplkip | 66308 | protein_coding | 270.8579814 | -0.023893358 | 0.10065746 | -0.237372949 | 0.812367474 | 0.915791177 | 0.023893358 |
| ENSMUSG00000018677 | Slc25a39 | 68066 | protein_coding | 3783.454535 | 0.023898731 | 0.061167699 | 0.390708349 | 0.696012827 | 0.847971508 | 0.023898731 |
| ENSMUSG00000057691 | Zfp746 | 69228 | protein_coding | 783.4012096 | -0.023910968 | 0.063410973 | -0.377079347 | 0.706114639 | 0.853850569 | 0.023910968 |
| ENSMUSG00000027770 | Dhx36 | 72162 | protein_coding | 1069.950053 | -0.023912705 | 0.056331993 | -0.424495988 | 0.67120413 | 0.832361742 | 0.023912705 |
| ENSMUSG00000043587 | Pxylp1 | 235534 | protein_coding | 1271.454833 | -0.023922539 | 0.057572786 | -0.415518177 | 0.677762611 | 0.836585484 | 0.023922539 |
| ENSMUSG00000025873 | Faf2 | 76577 | protein_coding | 1573.066436 | 0.023931613 | 0.034175225 | 0.700262048 | 0.483763669 | 0.705835346 | 0.023931613 |
| ENSMUSG00000038250 | Usp38 | 74841 | protein_coding | 920.7595825 | -0.023933846 | 0.053649275 | -0.446116868 | 0.655512829 | 0.82274538 | 0.023933846 |
| ENSMUSG00000037465 | Klf10 | 21847 | protein_coding | 2446.073866 | 0.023942226 | 0.135301103 | 0.176955146 | 0.859543622 | 0.936612161 | 0.023942226 |
| ENSMUSG00000029780 | Nt5c3 | 107569 | protein_coding | 1366.79893 | -0.023968006 | 0.074516987 | -0.321644864 | 0.747721752 | 0.880183116 | 0.023968006 |
| ENSMUSG00000039850 | Endov | 338371 | protein_coding | 425.7585262 | -0.023975177 | 0.085939814 | -0.278976366 | 0.780262963 | 0.898056783 | 0.023975177 |
| ENSMUSG00000033114 | Slc35d2 | 70484 | protein_coding | 118.0339017 | -0.02398491 | 0.134292979 | -0.178601366 | 0.858250722 | 0.936097283 | 0.02398491 |
| ENSMUSG00000028544 | Slc5a9 | 230612 | protein_coding | 314.011766 | -0.023993281 | 0.124632631 | -0.192512032 | 0.847341144 | 0.930926537 | 0.023993281 |
| ENSMUSG00000049871 | Nlrc3 | 268857 | protein_coding | 21.37177158 | -0.024009033 | 0.247628586 | -0.096955821 | 0.922761479 | 0.966069983 | 0.024009033 |
| ENSMUSG00000047264 | Zfp358 | 140482 | protein_coding | 312.1030422 | -0.024014069 | 0.090955636 | -0.264019581 | 0.791764831 | 0.903021982 | 0.024014069 |
| ENSMUSG00000070691 | Runx3 | 12399 | protein_coding | 8.855762698 | -0.024014911 | 0.380270583 | -0.063152165 | 0.949645335 | 0.979340729 | 0.024014911 |
| ENSMUSG00000026492 | Tfb2m | 15278 | protein_coding | 307.3707714 | -0.024018505 | 0.081182083 | -0.295859675 | 0.767337248 | 0.890725386 | 0.024018505 |
| ENSMUSG00000028292 | Rars2 | 109093 | protein_coding | 858.0489348 | 0.024030385 | 0.060088691 | 0.399915278 | 0.689218919 | 0.844094282 | 0.024030385 |
| ENSMUSG00000025064 | Col17a1 | 12821 | protein_coding | 14.45708904 | -0.024041161 | 0.327813956 | -0.073337819 | 0.941537297 | 0.974493399 | 0.024041161 |
| ENSMUSG00000055900 | Tmem69 | 230657 | protein_coding | 210.3731876 | 0.024050822 | 0.098129792 | 0.245091954 | 0.806385233 | 0.912178372 | 0.024050822 |
| ENSMUSG00000030045 | Mrpl19 | 56284 | protein_coding | 545.0271255 | 0.024066371 | 0.078587807 | 0.306235433 | 0.759425396 | 0.886076667 | 0.024066371 |
| ENSMUSG00000047368 | Abhd17b | 226016 | protein_coding | 1278.414313 | 0.02406725 | 0.052345372 | 0.459777998 | 0.645675577 | 0.816434847 | 0.02406725 |
| ENSMUSG00000008892 | Vdac3 | 22335 | protein_coding | 4481.661301 | -0.024081128 | 0.045424878 | -0.530130822 | 0.596021231 | 0.782866459 | 0.024081128 |
| ENSMUSG00000064138 | Fam172a | 68675 | protein_coding | 366.1229135 | -0.024103108 | 0.068922243 | -0.349714507 | 0.726552965 | 0.866650863 | 0.024103108 |
| ENSMUSG00000020346 | Mgat1 | 17308 | protein_coding | 2170.849918 | -0.0241046 | 0.04080966 | -0.590659177 | 0.554748806 | 0.754782249 | 0.0241046 |
| ENSMUSG00000053333 | Dis3l2 | 208718 | protein_coding | 475.8075859 | -0.024129396 | 0.0658797 | -0.366264502 | 0.714167715 | 0.859442037 | 0.024129396 |

| ensembl_gene_id | external_gene_name | entrezgene_id | gene_biotype | baseMean | log2FoldChange | lfcSE | stat | pvalue | padj | absLog2FC |
|---|---|---|---|---|---|---|---|---|---|---|
| ENSMUSG00000078350 | Smim1 | 68859 | protein_coding | 16.34252865 | -0.024159526 | 0.288636033 | -0.08370239 | 0.933293057 | 0.970920986 | 0.024159526 |
| ENSMUSG00000047284 | Neurl4 | 216860 | protein_coding | 1133.951833 | -0.024165533 | 0.071900258 | -0.336098002 | 0.736796965 | 0.873700342 | 0.024165533 |
| ENSMUSG00000045912 | C2cd4c | 237397 | protein_coding | 56.67267002 | -0.024167028 | 0.206398215 | -0.117089328 | 0.906789266 | 0.95873684 | 0.024167028 |
| ENSMUSG00000051346 | Spryd4 | 66701 | protein_coding | 369.486449 | 0.024167038 | 0.087437243 | 0.276392949 | 0.782246271 | 0.898825528 | 0.024167038 |
| ENSMUSG00000023861 | Mpc1 | 55951 | protein_coding | 1071.391295 | -0.024178774 | 0.074141404 | -0.326117027 | 0.744335815 | 0.878312994 | 0.024178774 |
| ENSMUSG00000020430 | Pes1 | 64934 | protein_coding | 1094.914889 | 0.02421089 | 0.049186509 | 0.492226243 | 0.622559415 | 0.800830795 | 0.02421089 |
| ENSMUSG00000024190 | Dusp1 | 19252 | protein_coding | 7898.143906 | 0.024221244 | 0.116234015 | 0.208383441 | 0.834929584 | 0.926650089 | 0.024221244 |
| ENSMUSG00000002679 | Med6 | 69792 | protein_coding | 307.1649783 | 0.024225235 | 0.072565301 | 0.333840484 | 0.738499932 | 0.874754933 | 0.024225235 |
| ENSMUSG00000041483 | Zfp281 | 226442 | protein_coding | 201.3959857 | 0.024225868 | 0.121054434 | 0.200123755 | 0.841383795 | 0.929613548 | 0.024225868 |
| ENSMUSG00000074811 | Hps6 | 20170 | protein_coding | 230.3611707 | 0.024236662 | 0.100559914 | 0.241017136 | 0.809541835 | 0.91448208 | 0.024236662 |
| ENSMUSG00000026857 | Ntmt1 | 66617 | protein_coding | 354.6772645 | 0.024270736 | 0.065219261 | 0.37214061 | 0.709788162 | 0.856292985 | 0.024270736 |
| ENSMUSG00000098132 | Rassf10 | 78748 | protein_coding | 121.737374 | -0.024279104 | 0.163262329 | -0.148712226 | 0.881780713 | 0.947331698 | 0.024279104 |
| ENSMUSG00000004865 | Srpk1 | 20815 | protein_coding | 1920.676047 | -0.024288968 | 0.052080005 | -0.466377989 | 0.640944964 | 0.813324566 | 0.024288968 |
| ENSMUSG00000005043 | Sgsh | 27029 | protein_coding | 380.9630696 | 0.02428903 | 0.088638127 | 0.274024635 | 0.78406569 | 0.89946133 | 0.02428903 |
| ENSMUSG00000031441 | Atp11a | 50770 | protein_coding | 1157.922202 | -0.024302255 | 0.079110824 | -0.30719254 | 0.758696824 | 0.885925238 | 0.024302255 |
| ENSMUSG00000037754 | Ppp1r16b | 228852 | protein_coding | 30.53647749 | 0.024308258 | 0.254939082 | 0.09534928 | 0.924037401 | 0.966739549 | 0.024308258 |
| ENSMUSG00000051319 | Mtln | 67885 | protein_coding | 367.9044541 | -0.024341342 | 0.07840694 | -0.310448825 | 0.75621967 | 0.884312567 | 0.024341342 |
| ENSMUSG00000002949 | Timm44 | 21856 | protein_coding | 1393.452483 | -0.024372033 | 0.047644608 | -0.511538115 | 0.608974306 | 0.791308423 | 0.024372033 |
| ENSMUSG00000026516 | Nvl | 67459 | protein_coding | 730.800254 | -0.024375457 | 0.066989556 | -0.363869519 | 0.715955447 | 0.860552159 | 0.024375457 |
| ENSMUSG00000037366 | Pafah2 | 100163 | protein_coding | 1821.460262 | 0.024387803 | 0.076556084 | 0.318561272 | 0.750059225 | 0.880597382 | 0.024387803 |
| ENSMUSG00000032560 | Dnajc13 | 235567 | protein_coding | 2732.506108 | 0.024397433 | 0.034892016 | 0.699226819 | 0.484410293 | 0.706263486 | 0.024397433 |
| ENSMUSG00000037949 | Ano10 | 102566 | protein_coding | 1894.581731 | -0.02443645 | 0.042636717 | -0.573131592 | 0.566555598 | 0.762547332 | 0.02443645 |
| ENSMUSG00000039086 | Ss18l1 | 269397 | protein_coding | 186.9815677 | 0.02444352 | 0.091853179 | 0.26611512 | 0.790150551 | 0.902120505 | 0.02444352 |
| ENSMUSG00000037971 | 1110032A03Rik | 68721 | protein_coding | 497.8308197 | 0.024447154 | 0.063929381 | 0.382408746 | 0.702158204 | 0.851790685 | 0.024447154 |
| ENSMUSG00000059920 | 4930453N24Rik | 67609 | protein_coding | 679.3782536 | -0.024466984 | 0.065401754 | -0.374102871 | 0.708327783 | 0.855350218 | 0.024466984 |
| ENSMUSG00000043987 | Cep164 | 214552 | protein_coding | 870.6757887 | 0.024488091 | 0.058183851 | 0.420874363 | 0.673846827 | 0.834200336 | 0.024488091 |
| ENSMUSG00000053749 | Gm9920 | 100503794 | lncRNA | 61.41776284 | 0.024520168 | 0.185726178 | 0.13202322 | 0.894965925 | 0.953178075 | 0.024520168 |
| ENSMUSG00000039041 | Adrm1 | 56436 | protein_coding | 897.5297334 | 0.024553532 | 0.057083477 | 0.430133787 | 0.667098323 | 0.830161713 | 0.024553532 |
| ENSMUSG00000029275 | Gfi1 | 14581 | protein_coding | 253.6994806 | 0.0245731 | 0.111849304 | 0.219698285 | 0.82610614 | 0.922846345 | 0.0245731 |
| ENSMUSG00000017264 | Exosc10 | 50912 | protein_coding | 810.0520522 | 0.024575567 | 0.069361877 | 0.354309429 | 0.723107002 | 0.864843268 | 0.024575567 |
| ENSMUSG00000023022 | Lima1 | 65970 | protein_coding | 8448.746121 | -0.024610022 | 0.063858477 | -0.385383797 | 0.699953085 | 0.850330953 | 0.024610022 |
| ENSMUSG00000001052 | Sec24b | 99683 | protein_coding | 564.5568141 | 0.024636896 | 0.097087491 | 0.253759739 | 0.799681179 | 0.908570455 | 0.024636896 |
| ENSMUSG00000052914 | Cyp2j6 | 13110 | protein_coding | 1924.817341 | -0.024650198 | 0.051205697 | -0.481395612 | 0.630235355 | 0.805755951 | 0.024650198 |
| ENSMUSG00000029458 | Brap | 72399 | protein_coding | 1352.918328 | -0.024679534 | 0.040599021 | -0.607884962 | 0.543263774 | 0.748182313 | 0.024679534 |
| ENSMUSG00000029196 | Tada2b | 231151 | protein_coding | 407.4696463 | -0.024681566 | 0.062688926 | -0.393714934 | 0.693791514 | 0.84654428 | 0.024681566 |
| ENSMUSG00000028289 | Epha7 | 13841 | protein_coding | 35.53305524 | -0.024693979 | 0.211021577 | -0.11702111 | 0.906843325 | 0.95873684 | 0.024693979 |
| ENSMUSG00000035842 | Ddx11 | 320209 | protein_coding | 197.5535233 | -0.024707316 | 0.110858658 | -0.222872231 | 0.82363495 | 0.921535618 | 0.024707316 |
| ENSMUSG00000043262 | Uevld | 54122 | protein_coding | 514.5483491 | 0.0247268 | 0.056428979 | 0.438193291 | 0.661246171 | 0.826855248 | 0.0247268 |
| ENSMUSG00000011008 | Mcoln2 | 68279 | protein_coding | 719.2993864 | -0.024741246 | 0.059590994 | -0.415184322 | 0.678006975 | 0.836653033 | 0.024741246 |

Supplementary Table 1

| ensembl_gene_id | external_gene_name | entrezgene_id | gene_biotype | baseMean | log2FoldChange | lfcSE | stat | pvalue | padj | absLog2FC |
|---|---|---|---|---|---|---|---|---|---|---|
| ENSMUSG00000062376 | Borcs7 | 66439 | protein_coding | 134.8127994 | -0.024771539 | 0.123608878 | -0.200402585 | 0.841165738 | 0.929613548 | 0.024771539 |
| ENSMUSG00000004897 | Hdgf | 15191 | protein_coding | 3101.318031 | -0.024788074 | 0.05260116 | -0.471245768 | 0.637465238 | 0.810693313 | 0.024788074 |
| ENSMUSG00000036368 | Rmdn2 | 381110 | protein_coding | 129.012376 | -0.02482815 | 0.135353172 | -0.183432349 | 0.854458796 | 0.934988261 | 0.02482815 |
| ENSMUSG00000022809 | Nr1i2 | 18171 | protein_coding | 1935.656451 | -0.024862982 | 0.036478064 | -0.681587207 | 0.495500006 | 0.714177192 | 0.024862982 |
| ENSMUSG00000000628 | Hk2 | 15277 | protein_coding | 3225.279098 | 0.024897186 | 0.105917634 | 0.235061764 | 0.814160795 | 0.916408517 | 0.024897186 |
| ENSMUSG00000007036 | Abhd16a | 193742 | protein_coding | 1688.883664 | -0.024923576 | 0.047588931 | -0.523726324 | 0.600468904 | 0.786019643 | 0.024923576 |
| ENSMUSG00000033253 | Szt2 | 230676 | protein_coding | 947.6875435 | 0.024949745 | 0.083463454 | 0.298930172 | 0.764993325 | 0.889541232 | 0.024949745 |
| ENSMUSG00000048490 | Nrip1 | 268903 | protein_coding | 3123.497934 | 0.02495644 | 0.055586393 | 0.448966705 | 0.653455675 | 0.821663954 | 0.02495644 |
| ENSMUSG00000000131 | Xpo6 | 74204 | protein_coding | 1155.340375 | -0.024963471 | 0.047463096 | -0.525955391 | 0.598919202 | 0.784827956 | 0.024963471 |
| ENSMUSG00000024387 | Csnk2b | 13001 | protein_coding | 1613.844335 | 0.024966996 | 0.063196838 | 0.395067162 | 0.692793323 | 0.845902325 | 0.024966996 |
| ENSMUSG00000042215 | Bag2 | 213539 | protein_coding | 46.16617871 | 0.024975271 | 0.19605531 | 0.127388902 | 0.8986326 | 0.955133984 | 0.024975271 |
| ENSMUSG00000004393 | Ddx56 | 52513 | protein_coding | 649.6631877 | -0.024979307 | 0.070928396 | -0.352176398 | 0.724705976 | 0.865841503 | 0.024979307 |
| ENSMUSG00000023923 | Tbc1d5 | 72238 | protein_coding | 558.7528234 | -0.025022498 | 0.055709554 | -0.449159911 | 0.653316305 | 0.821648745 | 0.025022498 |
| ENSMUSG00000025809 | Itgb1 | 16412 | protein_coding | 5150.013078 | 0.025047464 | 0.032546904 | 0.769580559 | 0.441548741 | 0.672888818 | 0.025047464 |
| ENSMUSG00000002985 | Apoe | 11816 | protein_coding | 50.30866242 | -0.02505071 | 0.187396767 | -0.133677389 | 0.893657686 | 0.952419871 | 0.02505071 |
| ENSMUSG00000118423 | Lrrc70 | 113523645 | transcribed_unitary_pseudogene | 5.868177594 | 0.025071829 | 0.667267695 | 0.037573869 | 0.970027443 | 0.987915089 | 0.025071829 |
| ENSMUSG00000031585 | Gtf2e2 | 68153 | protein_coding | 396.3493929 | 0.025084793 | 0.06881235 | 0.364539114 | 0.715455471 | 0.860253025 | 0.025084793 |
| ENSMUSG00000031352 | Hccs | 15159 | protein_coding | 630.4734913 | -0.02511108 | 0.07334838 | -0.342353573 | 0.732084823 | 0.870787945 | 0.02511108 |
| ENSMUSG00000024286 | Ccny | 67974 | protein_coding | 2371.309348 | 0.025112548 | 0.031865441 | 0.788080989 | 0.430649334 | 0.665528957 | 0.025112548 |
| ENSMUSG00000050158 | Or2m13 | 258458 | protein_coding | 106.7384643 | -0.02511315 | 0.158260808 | -0.158682053 | 0.873919379 | 0.943461975 | 0.02511315 |
| ENSMUSG00000007589 | Tinf2 | 28113 | protein_coding | 417.3772067 | 0.025121474 | 0.080991869 | 0.310172796 | 0.756429557 | 0.884429744 | 0.025121474 |
| ENSMUSG00000030722 | Nfatc2ip | 18020 | protein_coding | 100.2990169 | -0.025137909 | 0.141379669 | -0.177804272 | 0.858876692 | 0.936328291 | 0.025137909 |
| ENSMUSG00000027367 | Stard7 | 99138 | protein_coding | 3923.571922 | 0.025142544 | 0.046979989 | 0.535175613 | 0.592528428 | 0.780262005 | 0.025142544 |
| ENSMUSG00000032278 | Paqr5 | 74090 | protein_coding | 1429.150044 | 0.025144295 | 0.057168437 | 0.439828281 | 0.660061483 | 0.826041967 | 0.025144295 |
| ENSMUSG00000026972 | Arrdc1 | 215705 | protein_coding | 1479.991208 | 0.025180171 | 0.056534584 | 0.445394114 | 0.656034965 | 0.823272576 | 0.025180171 |
| ENSMUSG00000051034 | Zfp11 | 22648 | protein_coding | 53.43006874 | -0.025206412 | 0.162201322 | -0.155402015 | 0.876504395 | 0.944752465 | 0.025206412 |
| ENSMUSG00000020679 | Hnf1b | 21410 | protein_coding | 418.0191574 | -0.02522667 | 0.092115947 | -0.27385779 | 0.784193911 | 0.899522993 | 0.02522667 |
| ENSMUSG00000025075 | Habp2 | 226243 | protein_coding | 9.438393705 | 0.025233479 | 0.456856359 | 0.055232851 | 0.955952958 | 0.982430646 | 0.025233479 |
| ENSMUSG00000018923 | Med11 | 66172 | protein_coding | 309.9226023 | 0.025271182 | 0.080296805 | 0.314722131 | 0.752972639 | 0.882492633 | 0.025271182 |
| ENSMUSG00000019054 | Fis1 | 66437 | protein_coding | 1893.46314 | 0.025308394 | 0.054799321 | 0.461837741 | 0.644197684 | 0.815589264 | 0.025308394 |
| ENSMUSG00000019143 | Hars2 | 70791 | protein_coding | 561.1044729 | -0.02531406 | 0.056566426 | -0.447510326 | 0.654506634 | 0.822141364 | 0.02531406 |
| ENSMUSG00000020287 | Mpg | 268395 | protein_coding | 136.8050639 | -0.025317842 | 0.119258147 | -0.212294444 | 0.83187733 | 0.925602711 | 0.025317842 |
| ENSMUSG00000049562 | Ap5b1 | 381201 | protein_coding | 394.873441 | -0.025335305 | 0.097670831 | -0.259394797 | 0.795330644 | 0.905108834 | 0.025335305 |
| ENSMUSG00000033434 | Gtpbp6 | 107999 | protein_coding | 164.3338671 | 0.025340562 | 0.13600556 | 0.186320041 | 0.852193792 | 0.933591927 | 0.025340562 |
| ENSMUSG00000021277 | Traf3 | 22031 | protein_coding | 628.3136608 | -0.025348087 | 0.052019586 | -0.487279675 | 0.626060149 | 0.802772682 | 0.025348087 |
| ENSMUSG00000031169 | Porcn | 53627 | protein_coding | 62.18146637 | 0.025390952 | 0.1562458 | 0.162506462 | 0.87090704 | 0.941677361 | 0.025390952 |
| ENSMUSG00000028102 | Pex11b | 18632 | protein_coding | 223.3519898 | 0.025394051 | 0.086307457 | 0.294227777 | 0.768583857 | 0.891466618 | 0.025394051 |
| ENSMUSG00000054630 | Ugt2b5 | 22238 | protein_coding | 1096.11934 | -0.025425709 | 0.090306345 | -0.281549531 | 0.778288944 | 0.896895273 | 0.025425709 |
| ENSMUSG00000022960 | Donson | 60364 | protein_coding | 154.333857 | 0.025448633 | 0.109776407 | 0.231822429 | 0.816675936 | 0.917423574 | 0.025448633 |

| ensembl_gene_id | external_gene_name | entrezgene_id | gene_biotype | baseMean | log2FoldChange | lfcSE | stat | pvalue | padj | absLog2FC |
|---|---|---|---|---|---|---|---|---|---|---|
| ENSMUSG00000078580 | E430018J23Rik | 101604 | protein_coding | 263.1448256 | 0.025471651 | 0.079254832 | 0.321389248 | 0.74791543 | 0.880274689 | 0.025471651 |
| ENSMUSG00000059288 | Cdyl | 12593 | protein_coding | 302.0807361 | -0.025479499 | 0.080094678 | -0.31811725 | 0.750395999 | 0.880695457 | 0.025479499 |
| ENSMUSG00000033105 | Lss | 16987 | protein_coding | 1844.916826 | -0.0254795 | 0.135822733 | -0.187593778 | 0.851195104 | 0.933253344 | 0.0254795 |
| ENSMUSG00000053646 | Plxnb1 | 235611 | protein_coding | 242.3951786 | -0.025556903 | 0.084399232 | -0.30280966 | 0.762034921 | 0.887701328 | 0.025556903 |
| ENSMUSG00000038949 | Cnst | 226744 | protein_coding | 1248.838335 | -0.025562082 | 0.078117045 | -0.327227964 | 0.743495472 | 0.877686912 | 0.025562082 |
| ENSMUSG00000022772 | Senp5 | 320213 | protein_coding | 786.4633528 | -0.025570676 | 0.060047416 | -0.425841406 | 0.670223413 | 0.831693828 | 0.025570676 |
| ENSMUSG00000009907 | Vps4b | 20479 | protein_coding | 3040.588126 | 0.025575736 | 0.035143867 | 0.727743922 | 0.466770358 | 0.692404394 | 0.025575736 |
| ENSMUSG00000097121 | D130020L05Rik | 319760 | lncRNA | 23.8372966 | -0.025580858 | 0.240122918 | -0.106532345 | 0.915159994 | 0.962265867 | 0.025580858 |
| ENSMUSG00000020224 | Llph | 66225 | protein_coding | 262.3954932 | -0.025602199 | 0.08556427 | -0.299216004 | 0.764775239 | 0.889439786 | 0.025602199 |
| ENSMUSG00000031433 | Rbm41 | 237073 | protein_coding | 141.572584 | 0.02560472 | 0.118867365 | 0.2154058 | 0.829450949 | 0.924270223 | 0.02560472 |
| ENSMUSG00000074622 | Mafb | 16658 | protein_coding | 204.9657792 | -0.025630895 | 0.168804826 | -0.151837455 | 0.87931514 | 0.94569625 | 0.025630895 |
| ENSMUSG00000022160 | Mettl3 | 56335 | protein_coding | 392.9328883 | -0.025631631 | 0.077782666 | -0.329528831 | 0.741756006 | 0.876551558 | 0.025631631 |
| ENSMUSG00000026659 | Dusp12 | 80915 | protein_coding | 112.6147166 | -0.025638146 | 0.121400173 | -0.211187066 | 0.832741303 | 0.92587823 | 0.025638146 |
| ENSMUSG00000027454 | Gins1 | 69270 | protein_coding | 57.80099185 | -0.025656256 | 0.215423838 | -0.119096643 | 0.90519879 | 0.958435249 | 0.025656256 |
| ENSMUSG00000022378 | Cyrib | 223601 | protein_coding | 734.8968066 | -0.025676005 | 0.067132385 | -0.382468238 | 0.702114084 | 0.851790685 | 0.025676005 |
| ENSMUSG00000032508 | Myd88 | 17874 | protein_coding | 581.7805982 | -0.025696049 | 0.053021997 | -0.484629965 | 0.627938854 | 0.803865895 | 0.025696049 |
| ENSMUSG00000030753 | Thap12 | 72981 | protein_coding | 1202.343065 | -0.025697468 | 0.044943184 | -0.571776762 | 0.567473222 | 0.762827428 | 0.025697468 |
| ENSMUSG00000039449 | Prpf18 | 67229 | protein_coding | 953.9978541 | 0.025697499 | 0.055340106 | 0.464355791 | 0.642392858 | 0.814098293 | 0.025697499 |
| ENSMUSG00000034218 | Atm | 11920 | protein_coding | 1256.847584 | -0.025702976 | 0.071189618 | -0.361049501 | 0.718062443 | 0.861878015 | 0.025702976 |
| ENSMUSG00000001785 | Pwp1 | 103136 | protein_coding | 553.0608686 | -0.025720456 | 0.065864608 | -0.390504957 | 0.696163191 | 0.848090631 | 0.025720456 |
| ENSMUSG00000047454 | Gphn | 268566 | protein_coding | 505.9439786 | -0.025728672 | 0.07915947 | -0.325023298 | 0.745163439 | 0.878903736 | 0.025728672 |
| ENSMUSG00000029263 | Pigg | 433931 | protein_coding | 632.85612 | -0.025741606 | 0.051285273 | -0.501929792 | 0.615716908 | 0.795599668 | 0.025741606 |
| ENSMUSG00000061517 | Sox21 | 223227 | protein_coding | 109.3483412 | 0.025755337 | 0.168510609 | 0.152841043 | 0.87852363 | 0.945542552 | 0.025755337 |
| ENSMUSG00000030917 | Tmem159 | 233806 | protein_coding | 520.8255854 | 0.025761491 | 0.068251628 | 0.377448744 | 0.705840148 | 0.853688621 | 0.025761491 |
| ENSMUSG00000026615 | Eprs | 107508 | protein_coding | 3760.875993 | 0.025774122 | 0.070419027 | 0.366010769 | 0.71435704 | 0.859482312 | 0.025774122 |
| ENSMUSG00000059363 | Fxn | 14297 | protein_coding | 153.9351339 | -0.025780897 | 0.101663139 | -0.253591389 | 0.79981125 | 0.908570455 | 0.025780897 |
| ENSMUSG00000010097 | Nxf1 | 53319 | protein_coding | 1261.589932 | -0.025785915 | 0.065911971 | -0.391217472 | 0.695636495 | 0.847823651 | 0.025785915 |
| ENSMUSG00000026896 | Ifih1 | 71586 | protein_coding | 1854.829164 | -0.025804075 | 0.073915758 | -0.349101132 | 0.727013387 | 0.867071801 | 0.025804075 |
| ENSMUSG00000065990 | Aurkaip1 | 66077 | protein_coding | 1459.533668 | -0.025807339 | 0.064173515 | -0.402149379 | 0.687574094 | 0.842976085 | 0.025807339 |
| ENSMUSG00000033752 | Mnd1 | 76915 | protein_coding | 32.78455821 | -0.025857832 | 0.204063677 | -0.126714524 | 0.89916635 | 0.955338492 | 0.025857832 |
| ENSMUSG00000042595 | Fam199x | 245622 | protein_coding | 792.4954528 | 0.025886193 | 0.056495614 | 0.458198278 | 0.646809997 | 0.817036486 | 0.025886193 |
| ENSMUSG00000028187 | Rpf1 | 70285 | protein_coding | 648.7463758 | -0.025911169 | 0.074679198 | -0.346966359 | 0.728616592 | 0.868277527 | 0.025911169 |
| ENSMUSG00000039000 | Ube3c | 100763 | protein_coding | 2280.797176 | 0.025916857 | 0.043190312 | 0.600061827 | 0.548465031 | 0.750864769 | 0.025916857 |
| ENSMUSG00000031916 | Cog8 | 97484 | protein_coding | 798.1636126 | 0.025930483 | 0.066927404 | 0.38744193 | 0.698429067 | 0.849567474 | 0.025930483 |
| ENSMUSG00000029613 | Eif2ak1 | 15467 | protein_coding | 1171.16748 | 0.025939372 | 0.051494502 | 0.5037309 | 0.614450491 | 0.794774067 | 0.025939372 |
| ENSMUSG00000021038 | Vipas39 | 104799 | protein_coding | 1386.074329 | 0.025964955 | 0.055759051 | 0.465663498 | 0.641456384 | 0.813708274 | 0.025964955 |
| ENSMUSG00000028538 | St3gal3 | 20441 | protein_coding | 1365.384708 | -0.02599885 | 0.083315307 | -0.312053706 | 0.754999708 | 0.883718994 | 0.02599885 |
| ENSMUSG00000014074 | Rnf168 | 70238 | protein_coding | 628.793404 | -0.026038099 | 0.058027242 | -0.448721985 | 0.653632223 | 0.821663954 | 0.026038099 |
| ENSMUSG00000042608 | Stk40 | 74178 | protein_coding | 908.2833889 | 0.026065799 | 0.071936626 | 0.362343918 | 0.717095043 | 0.861383838 | 0.026065799 |

| ensembl_gene_id | external_gene_name | entrezgene_id | gene_biotype | baseMean | log2FoldChange | lfcSE | stat | pvalue | padj | absLog2FC |
|---|---|---|---|---|---|---|---|---|---|---|
| ENSMUSG00000005672 | Kit | 16590 | protein_coding | 1857.603508 | -0.026094604 | 0.083358426 | -0.313040985 | 0.754249524 | 0.883289673 | 0.026094604 |
| ENSMUSG00000017747 | Ghdc | 80860 | protein_coding | 319.71196 | -0.026114598 | 0.078146141 | -0.334176427 | 0.738246431 | 0.874711645 | 0.026114598 |
| ENSMUSG00000004788 | Eif2b2 | 217715 | protein_coding | 596.0921085 | 0.026122724 | 0.082401256 | 0.31701852 | 0.75122955 | 0.881353045 | 0.026122724 |
| ENSMUSG00000034931 | Dhx8 | 217207 | protein_coding | 1108.226748 | 0.026127793 | 0.048736482 | 0.536103376 | 0.591887107 | 0.780262005 | 0.026127793 |
| ENSMUSG00000022262 | Dnah5 | 110082 | protein_coding | 301.3294012 | 0.026130532 | 0.110618412 | 0.236222262 | 0.813260206 | 0.915984616 | 0.026130532 |
| ENSMUSG00000020392 | Cdkn2aipnl | 52626 | protein_coding | 728.3058755 | 0.026134247 | 0.060545131 | 0.431649034 | 0.665996515 | 0.829512341 | 0.026134247 |
| ENSMUSG00000002728 | Naa20 | 67877 | protein_coding | 873.8470551 | 0.026145362 | 0.064533994 | 0.405140927 | 0.685373924 | 0.841671981 | 0.026145362 |
| ENSMUSG00000113902 | Ndufb1 | 102631912 | protein_coding | 1412.631476 | 0.026159666 | 0.077970529 | 0.335507102 | 0.737242588 | 0.873998696 | 0.026159666 |
| ENSMUSG00000026927 | Entr1 | 68112 | protein_coding | 716.1977914 | 0.026167652 | 0.053269867 | 0.491228039 | 0.62326517 | 0.801099657 | 0.026167652 |
| ENSMUSG00000003269 | Cyth2 | 19158 | protein_coding | 1251.979583 | 0.026218965 | 0.05103221 | 0.513772877 | 0.607410793 | 0.790170193 | 0.026218965 |
| ENSMUSG00000021798 | Ldb3 | 24131 | protein_coding | 233.3272027 | -0.026219487 | 0.126064538 | -0.207984637 | 0.835240963 | 0.926721051 | 0.026219487 |
| ENSMUSG00000085451 | Gm40614 | 105245120 | lncRNA | 24.33186679 | -0.026240674 | 0.278921275 | -0.094079142 | 0.92504629 | 0.967208345 | 0.026240674 |
| ENSMUSG00000037029 | Zfp146 | 26465 | protein_coding | 625.9813757 | -0.026245377 | 0.082269688 | -0.31901637 | 0.749714099 | 0.880475742 | 0.026245377 |
| ENSMUSG00000060002 | Chpt1 | 212862 | protein_coding | 2595.642199 | -0.026248151 | 0.049488986 | -0.530383684 | 0.595845936 | 0.78269996 | 0.026248151 |
| ENSMUSG00000038121 | Fam210a | 108654 | protein_coding | 1096.967151 | -0.026253526 | 0.056946553 | -0.461020464 | 0.644783923 | 0.81580845 | 0.026253526 |
| ENSMUSG00000097180 | 2700038G22Rik | 67194 | lncRNA | 12.81134882 | -0.026261739 | 0.357269352 | -0.073506833 | 0.941402807 | 0.974454626 | 0.026261739 |
| ENSMUSG00000040128 | Pnrc1 | 108767 | protein_coding | 2272.412 | -0.026269657 | 0.087364662 | -0.300689733 | 0.763651098 | 0.888754045 | 0.026269657 |
| ENSMUSG00000026111 | Unc50 | 67387 | protein_coding | 948.6608288 | 0.02629243 | 0.047017791 | 0.559201727 | 0.576024055 | 0.768874353 | 0.02629243 |
| ENSMUSG00000021338 | Carmil1 | 68732 | protein_coding | 961.6419512 | 0.026334077 | 0.047121756 | 0.558851772 | 0.576262886 | 0.769065836 | 0.026334077 |
| ENSMUSG00000025745 | Hadha | 97212 | protein_coding | 6603.189608 | 0.026362601 | 0.036476083 | 0.722736614 | 0.469841717 | 0.694009261 | 0.026362601 |
| ENSMUSG00000020171 | Yeats4 | 64050 | protein_coding | 668.6562468 | 0.026384859 | 0.064984938 | 0.40601499 | 0.684731586 | 0.841281905 | 0.026384859 |
| ENSMUSG00000026102 | Inpp1 | 16329 | protein_coding | 467.4132394 | -0.026390792 | 0.0743047 | -0.355169888 | 0.722462322 | 0.864328625 | 0.026390792 |
| ENSMUSG00000050447 | Lypd6 | 320343 | protein_coding | 10.39429931 | -0.026391662 | 0.356818138 | -0.073963903 | 0.941039108 | 0.974315 | 0.026391662 |
| ENSMUSG00000020397 | Med7 | 66213 | protein_coding | 286.3837386 | -0.026394478 | 0.073055383 | -0.361294085 | 0.717879615 | 0.861722706 | 0.026394478 |
| ENSMUSG00000031446 | Cul4a | 99375 | protein_coding | 2062.427583 | -0.026401693 | 0.037885841 | -0.696874948 | 0.485881057 | 0.707074879 | 0.026401693 |
| ENSMUSG00000042447 | Mios | 252875 | protein_coding | 379.5493055 | -0.026413235 | 0.065204166 | -0.405085083 | 0.685414971 | 0.841671981 | 0.026413235 |
| ENSMUSG00000022698 | Naa50 | 72117 | protein_coding | 1549.069905 | 0.026432968 | 0.063913593 | 0.413573488 | 0.679186493 | 0.837472006 | 0.026432968 |
| ENSMUSG00000028807 | Zbtb8a | 73680 | protein_coding | 425.0726704 | -0.026446838 | 0.072582829 | -0.364367693 | 0.715583457 | 0.860311198 | 0.026446838 |
| ENSMUSG00000040818 | Dennd6a | 211922 | protein_coding | 1262.335085 | -0.026457915 | 0.053582087 | -0.49378283 | 0.62145956 | 0.800250831 | 0.026457915 |
| ENSMUSG00000024163 | Mapk8ip3 | 30957 | protein_coding | 592.9219741 | -0.026485005 | 0.102618428 | -0.258092091 | 0.796335835 | 0.906028181 | 0.026485005 |
| ENSMUSG00000042111 | Ccdc115 | 69668 | protein_coding | 357.5582307 | 0.026524308 | 0.078446736 | 0.33811869 | 0.735273747 | 0.87279256 | 0.026524308 |
| ENSMUSG00000008373 | Prpf31 | 68988 | protein_coding | 650.5866765 | 0.02657101 | 0.081113421 | 0.327578463 | 0.743230408 | 0.877650787 | 0.02657101 |
| ENSMUSG00000044857 | Lemd2 | 224640 | protein_coding | 1017.0115 | -0.026580708 | 0.069497464 | -0.382470191 | 0.702112635 | 0.851790685 | 0.026580708 |
| ENSMUSG00000097779 | 4833407H14Rik | 73899 | lncRNA | 14.95738931 | -0.02658418 | 0.312213589 | -0.085147416 | 0.932144195 | 0.970414397 | 0.02658418 |
| ENSMUSG00000054737 | Zfp182 | 319535 | protein_coding | 236.1457725 | 0.026610404 | 0.137963159 | 0.192880505 | 0.847052553 | 0.930926537 | 0.026610404 |
| ENSMUSG00000045598 | Zfp553 | 233887 | protein_coding | 444.2146131 | -0.026627138 | 0.066304071 | -0.401591299 | 0.687984835 | 0.843351431 | 0.026627138 |
| ENSMUSG00000037197 | Rbm17 | 76938 | protein_coding | 1067.080139 | 0.02662752 | 0.053110822 | 0.501357721 | 0.616119389 | 0.795599668 | 0.02662752 |
| ENSMUSG00000090176 | Cd200r2 | 271375 | protein_coding | 162.916783 | 0.026644458 | 0.137839463 | 0.193300649 | 0.846723518 | 0.930889346 | 0.026644458 |
| ENSMUSG00000020271 | Fbxw11 | 103583 | protein_coding | 1816.5951 | -0.026652858 | 0.061620526 | -0.43253214 | 0.665354698 | 0.82916121 | 0.026652858 |

Supplementary Table 1

| ensembl_gene_id | external_gene_name | entrezgene_id | gene_biotype | baseMean | log2FoldChange | lfcSE | stat | pvalue | padj | absLog2FC |
|---|---|---|---|---|---|---|---|---|---|---|
| ENSMUSG00000049811 | Fam161a | 73873 | protein_coding | 202.5628436 | 0.026661675 | 0.088069312 | 0.30273513 | 0.762091723 | 0.887703374 | 0.026661675 |
| ENSMUSG00000038178 | Slc43a2 | 215113 | protein_coding | 429.3549354 | 0.026683498 | 0.130155872 | 0.205011865 | 0.837562855 | 0.927834878 | 0.026683498 |
| ENSMUSG00000032666 | 1700025G04Rik | 69399 | protein_coding | 393.0707463 | 0.026699007 | 0.098231265 | 0.271797448 | 0.785777767 | 0.900219089 | 0.026699007 |
| ENSMUSG00000019948 | Actr6 | 67019 | protein_coding | 76.76747269 | -0.026727529 | 0.146971317 | -0.181855408 | 0.8556962 | 0.935592329 | 0.026727529 |
| ENSMUSG00000032309 | Fbxo22 | 71999 | protein_coding | 642.6701402 | 0.02672797 | 0.050845469 | 0.525670641 | 0.599117066 | 0.784887158 | 0.02672797 |
| ENSMUSG00000018900 | Slc22a5 | 20520 | protein_coding | 816.6189885 | 0.026739105 | 0.063551952 | 0.420744038 | 0.673942 | 0.834254117 | 0.026739105 |
| ENSMUSG00000035933 | Cog5 | 238123 | protein_coding | 1095.426178 | 0.02675109 | 0.046650088 | 0.573441377 | 0.566345881 | 0.762477743 | 0.02675109 |
| ENSMUSG00000022820 | Ndufb4 | 68194 | protein_coding | 651.6794776 | -0.026779342 | 0.063698994 | -0.420404472 | 0.674190002 | 0.834301331 | 0.026779342 |
| ENSMUSG00000028833 | Ncdn | 26562 | protein_coding | 665.5534149 | -0.026781565 | 0.078186687 | -0.342533569 | 0.731949386 | 0.870769235 | 0.026781565 |
| ENSMUSG00000026087 | Mrpl30 | 107734 | protein_coding | 1338.740576 | 0.026794904 | 0.048448494 | 0.553059583 | 0.580222611 | 0.772159116 | 0.026794904 |
| ENSMUSG00000114744 | Gm38409 | 328436 | lncRNA | 8.088757166 | -0.026874237 | 0.428503702 | -0.062716464 | 0.949992287 | 0.979588542 | 0.026874237 |
| ENSMUSG00000079427 | Mthfsl | 100039707 | protein_coding | 143.3765699 | -0.026874759 | 0.100154434 | -0.268333196 | 0.788442857 | 0.901341033 | 0.026874759 |
| ENSMUSG00000051839 | Gypa | 14934 | protein_coding | 13.57534153 | 0.02688851 | 0.357275909 | 0.07525979 | 0.940008014 | 0.973941151 | 0.02688851 |
| ENSMUSG00000022969 | Il10rb | 16155 | protein_coding | 997.1621315 | -0.026891985 | 0.058545051 | -0.459338306 | 0.645991244 | 0.816513895 | 0.026891985 |
| ENSMUSG00000035596 | Mboat7 | 77582 | protein_coding | 923.1980337 | 0.026918441 | 0.067524301 | 0.398648203 | 0.690152439 | 0.844556471 | 0.026918441 |
| ENSMUSG00000073639 | Rab18 | 19330 | protein_coding | 1364.430411 | 0.026918959 | 0.046850746 | 0.574568427 | 0.56558321 | 0.76190043 | 0.026918959 |
| ENSMUSG00000028284 | Map3k7 | 26409 | protein_coding | 1136.702492 | 0.026920009 | 0.053218563 | 0.505838695 | 0.612969892 | 0.793750237 | 0.026920009 |
| ENSMUSG00000031320 | Rps4x | 20102 | protein_coding | 15845.47727 | -0.026933377 | 0.041856734 | -0.643465792 | 0.519921905 | 0.731656244 | 0.026933377 |
| ENSMUSG00000022426 | Josd1 | 74158 | protein_coding | 1208.536396 | -0.026949815 | 0.05463299 | -0.493288294 | 0.621808898 | 0.800376096 | 0.026949815 |
| ENSMUSG00000061207 | Stk19 | 54402 | protein_coding | 298.7215081 | -0.026952585 | 0.10011765 | -0.269209126 | 0.787768758 | 0.900953603 | 0.026952585 |
| ENSMUSG00000042115 | Klhdc8a | 213417 | protein_coding | 212.5342555 | -0.026957058 | 0.110414152 | -0.244144957 | 0.807118555 | 0.912727084 | 0.026957058 |
| ENSMUSG00000056820 | Tsnax | 53424 | protein_coding | 1138.445208 | -0.026966157 | 0.056978325 | -0.473270441 | 0.636020248 | 0.809697277 | 0.026966157 |
| ENSMUSG00000085772 | D630024D03Rik | 414116 | lncRNA | 14.6634157 | -0.026985581 | 0.341944891 | -0.078917923 | 0.937097907 | 0.972704281 | 0.026985581 |
| ENSMUSG00000046010 | Zfp830 | 66983 | protein_coding | 289.7427217 | -0.027009123 | 0.080774449 | -0.334377059 | 0.738095048 | 0.874596535 | 0.027009123 |
| ENSMUSG00000002963 | Pnkp | 59047 | protein_coding | 696.647657 | -0.027037874 | 0.068630983 | -0.393960177 | 0.693610441 | 0.84654428 | 0.027037874 |
| ENSMUSG00000041408 | Wapl | 218914 | protein_coding | 3598.955053 | -0.027083769 | 0.049703482 | -0.544906878 | 0.585817566 | 0.776292513 | 0.027083769 |
| ENSMUSG00000072582 | Ptrh2 | 217057 | protein_coding | 300.4366064 | 0.02710693 | 0.078618821 | 0.344789319 | 0.730252763 | 0.869327992 | 0.02710693 |
| ENSMUSG00000030768 | Disp1 | 68897 | protein_coding | 209.3004749 | -0.027120512 | 0.085965721 | -0.315480542 | 0.752396822 | 0.882337373 | 0.027120512 |
| ENSMUSG00000008730 | Hipk1 | 15257 | protein_coding | 3562.676528 | 0.027127874 | 0.059853878 | 0.453235028 | 0.650379512 | 0.819811661 | 0.027127874 |
| ENSMUSG00000031257 | Nox1 | 237038 | protein_coding | 576.6914214 | 0.027147427 | 0.102811193 | 0.264051277 | 0.791740407 | 0.903021982 | 0.027147427 |
| ENSMUSG00000042638 | Gucy2c | 14917 | protein_coding | 9828.4039 | -0.027164983 | 0.025666886 | -1.058366928 | 0.289888191 | 0.532281323 | 0.027164983 |
| ENSMUSG00000097047 | 1110020A21Rik | 68531 | lncRNA | 40.60191902 | -0.027172428 | 0.20165296 | -0.134748469 | 0.89281075 | 0.952209442 | 0.027172428 |
| ENSMUSG00000037845 | Fdxacb1 | 382137 | protein_coding | 89.22319614 | 0.027175256 | 0.140384335 | 0.193577551 | 0.846506678 | 0.930889346 | 0.027175256 |
| ENSMUSG00000034107 | Ano7 | 404545 | protein_coding | 778.4874258 | 0.027178105 | 0.099576395 | 0.272937228 | 0.784901469 | 0.899716112 | 0.027178105 |
| ENSMUSG00000073062 | Zxdb | 668166 | protein_coding | 100.2353003 | -0.027206937 | 0.124437924 | -0.218638629 | 0.826931559 | 0.923060252 | 0.027206937 |
| ENSMUSG00000048039 | Isg20l2 | 229504 | protein_coding | 622.5802833 | -0.027221368 | 0.062956278 | -0.432385276 | 0.665461418 | 0.829230125 | 0.027221368 |
| ENSMUSG00000034321 | Exosc1 | 66583 | protein_coding | 178.9877114 | -0.02722551 | 0.117838958 | -0.231039977 | 0.817283745 | 0.917914545 | 0.02722551 |
| ENSMUSG00000001419 | Mef2d | 17261 | protein_coding | 1096.902832 | -0.027228341 | 0.066291863 | -0.410734291 | 0.68126738 | 0.838859992 | 0.027228341 |
| ENSMUSG00000028049 | Scamp3 | 24045 | protein_coding | 258.6934462 | -0.027235111 | 0.095823137 | -0.284222698 | 0.776239722 | 0.895541399 | 0.027235111 |

| ensembl_gene_id | external_gene_name | entrezgene_id | gene_biotype | baseMean | log2FoldChange | lfcSE | stat | pvalue | padj | absLog2FC |
|---|---|---|---|---|---|---|---|---|---|---|
| ENSMUSG00000061787 | Rps17 | 20068 | protein_coding | 7449.168491 | -0.027262186 | 0.067732952 | -0.402495167 | 0.687319644 | 0.842728192 | 0.027262186 |
| ENSMUSG00000040725 | Hnrnpul1 | 232989 | protein_coding | 3768.967224 | -0.027291744 | 0.053499396 | -0.510131806 | 0.609959124 | 0.79214131 | 0.027291744 |
| ENSMUSG00000036850 | Mrpl41 | 107733 | protein_coding | 467.695435 | 0.027309532 | 0.066596235 | 0.410076221 | 0.681750035 | 0.839028452 | 0.027309532 |
| ENSMUSG00000039428 | Tmem135 | 72759 | protein_coding | 1099.12072 | 0.027324982 | 0.049944196 | 0.547110254 | 0.584302988 | 0.77543238 | 0.027324982 |
| ENSMUSG00000027884 | Clcc1 | 229725 | protein_coding | 712.2150391 | 0.027333246 | 0.066091641 | 0.413565857 | 0.679192083 | 0.837472006 | 0.027333246 |
| ENSMUSG00000073434 | Wdr90 | 106618 | protein_coding | 269.3946834 | 0.027334711 | 0.107806503 | 0.253553448 | 0.799840565 | 0.908570455 | 0.027334711 |
| ENSMUSG00000006763 | Saal1 | 78935 | protein_coding | 258.0429509 | 0.027349006 | 0.079057852 | 0.345936624 | 0.729390344 | 0.868814394 | 0.027349006 |
| ENSMUSG00000084990 | Gm14549 | 69591 | lncRNA | 6.163610018 | -0.027365165 | 0.529540763 | -0.051677164 | 0.958785934 | 0.983514188 | 0.027365165 |
| ENSMUSG00000098905 | Zfp953 | 629016 | protein_coding | 197.7766165 | 0.027403161 | 0.095756613 | 0.286175131 | 0.77474399 | 0.89435358 | 0.027403161 |
| ENSMUSG00000042694 | Stn1 | 108689 | protein_coding | 208.3077358 | 0.027447513 | 0.099853047 | 0.274879076 | 0.783409143 | 0.898963943 | 0.027447513 |
| ENSMUSG00000028300 | C9orf72 | 73205 | protein_coding | 267.9608266 | -0.027479312 | 0.081631083 | -0.336628049 | 0.736397309 | 0.873419092 | 0.027479312 |
| ENSMUSG00000035329 | Fbxo33 | 70611 | protein_coding | 643.2274718 | -0.027480163 | 0.069380639 | -0.396078265 | 0.692047292 | 0.845589334 | 0.027480163 |
| ENSMUSG00000021290 | Atp5mpl | 70257 | protein_coding | 3420.929521 | 0.027480404 | 0.045718739 | 0.601075282 | 0.547789846 | 0.750761183 | 0.027480404 |
| ENSMUSG00000052151 | Plpp2 | 50784 | protein_coding | 5083.433231 | 0.027494519 | 0.053068307 | 0.518096783 | 0.604390745 | 0.788784457 | 0.027494519 |
| ENSMUSG00000019470 | Xab2 | 67439 | protein_coding | 984.4576911 | -0.027498088 | 0.047843558 | -0.574750078 | 0.565460334 | 0.761899808 | 0.027498088 |
| ENSMUSG00000024866 | Acy3 | 71670 | protein_coding | 438.5774404 | 0.027498788 | 0.088219811 | 0.311707632 | 0.755262727 | 0.88389772 | 0.027498788 |
| ENSMUSG00000004264 | Phb2 | 12034 | protein_coding | 3810.976743 | 0.027499434 | 0.052262825 | 0.526175812 | 0.598766058 | 0.784827956 | 0.027499434 |
| ENSMUSG00000021241 | Isca2 | 74316 | protein_coding | 392.0088388 | -0.027500433 | 0.084393688 | -0.325858883 | 0.744531126 | 0.878396765 | 0.027500433 |
| ENSMUSG00000006276 | Eps15l1 | 13859 | protein_coding | 1038.471572 | -0.027504225 | 0.057712329 | -0.476574502 | 0.633665144 | 0.808283189 | 0.027504225 |
| ENSMUSG00000025103 | Btbd1 | 83962 | protein_coding | 1384.779183 | 0.027519191 | 0.043374182 | 0.634460171 | 0.525780552 | 0.736366735 | 0.027519191 |
| ENSMUSG00000028003 | Lrat | 79235 | protein_coding | 409.8131884 | -0.027522613 | 0.109366085 | -0.251655834 | 0.801307098 | 0.909407429 | 0.027522613 |
| ENSMUSG00000059355 | Wdr83os | 414077 | protein_coding | 333.8068478 | 0.027543611 | 0.100696927 | 0.273529805 | 0.784445985 | 0.899577643 | 0.027543611 |
| ENSMUSG00000023015 | Racgap1 | 26934 | protein_coding | 797.2257622 | 0.027552732 | 0.094418633 | 0.291814562 | 0.770428411 | 0.892047592 | 0.027552732 |
| ENSMUSG00000031701 | Dnaja2 | 56445 | protein_coding | 3709.967553 | -0.027630756 | 0.043682107 | -0.632541731 | 0.527032948 | 0.737165628 | 0.027630756 |
| ENSMUSG00000030619 | Eed | 13626 | protein_coding | 665.4080625 | 0.027658654 | 0.056840608 | 0.48660025 | 0.626541646 | 0.803150559 | 0.027658654 |
| ENSMUSG00000024581 | Napg | 108123 | protein_coding | 853.265854 | -0.027689633 | 0.047976018 | -0.577155707 | 0.563834276 | 0.76110784 | 0.027689633 |
| ENSMUSG00000015575 | Atp6v0e | 11974 | protein_coding | 961.0883463 | -0.027702415 | 0.057636433 | -0.4806407 | 0.630771884 | 0.805994625 | 0.027702415 |
| ENSMUSG00000029273 | Sult1d1 | 53315 | protein_coding | 9897.488937 | -0.027707539 | 0.051119316 | -0.542017018 | 0.587806783 | 0.777778141 | 0.027707539 |
| ENSMUSG00000025786 | Zdhhc3 | 69035 | protein_coding | 2691.459323 | -0.027714915 | 0.035186708 | -0.787652978 | 0.430899717 | 0.665624496 | 0.027714915 |
| ENSMUSG00000043411 | Usp48 | 170707 | protein_coding | 1336.610149 | 0.027723415 | 0.048517623 | 0.571409174 | 0.567722312 | 0.763029223 | 0.027723415 |
| ENSMUSG00000032064 | Dixdc1 | 330938 | protein_coding | 208.3168456 | -0.027723978 | 0.142157851 | -0.195022486 | 0.845375345 | 0.930668864 | 0.027723978 |
| ENSMUSG00000029032 | Arhgef16 | 230972 | protein_coding | 3059.730839 | 0.027725921 | 0.058531302 | 0.473693897 | 0.635718206 | 0.80962603 | 0.027725921 |
| ENSMUSG00000027939 | Nup210l | 77595 | protein_coding | 40.56360465 | -0.027737386 | 0.19981344 | -0.13881642 | 0.889595219 | 0.950733597 | 0.027737386 |
| ENSMUSG00000006732 | Mettl1 | 17299 | protein_coding | 253.4030794 | -0.027759237 | 0.123093603 | -0.225513235 | 0.821580031 | 0.920176481 | 0.027759237 |
| ENSMUSG00000061028 | Clasrp | 53609 | protein_coding | 582.6658551 | -0.027778897 | 0.107646518 | -0.258056622 | 0.796363209 | 0.906028181 | 0.027778897 |
| ENSMUSG00000028081 | Rps3a1 | 20091 | protein_coding | 10847.70502 | -0.027797102 | 0.04235409 | -0.65630266 | 0.511629406 | 0.725473655 | 0.027797102 |
| ENSMUSG00000037234 | Hook3 | 320191 | protein_coding | 578.8577203 | -0.027816529 | 0.088755796 | -0.313405209 | 0.753972827 | 0.883093891 | 0.027816529 |
| ENSMUSG00000023953 | Polh | 80905 | protein_coding | 316.5232181 | 0.027826749 | 0.07952032 | 0.349932557 | 0.726389313 | 0.866519746 | 0.027826749 |
| ENSMUSG00000035293 | G2e3 | 217558 | protein_coding | 587.2886495 | -0.027827946 | 0.080707635 | -0.344799427 | 0.730245164 | 0.869327992 | 0.027827946 |

Supplementary Table 1

| ensembl_gene_id | external_gene_name | entrezgene_id | gene_biotype | baseMean | log2FoldChange | lfcSE | stat | pvalue | padj | absLog2FC |
|---|---|---|---|---|---|---|---|---|---|---|
| ENSMUSG00000058006 | Mdn1 | 100019 | protein_coding | 674.6067082 | 0.02783681 | 0.11154 | 0.249567961 | 0.802921478 | 0.910278712 | 0.02783681 |
| ENSMUSG00000029326 | Enoph1 | 67870 | protein_coding | 633.6403834 | 0.027843209 | 0.070826361 | 0.393119297 | 0.694231372 | 0.846888755 | 0.027843209 |
| ENSMUSG00000027079 | Clp1 | 98985 | protein_coding | 315.9327799 | -0.027849732 | 0.086490278 | -0.321998408 | 0.7474539 | 0.880070575 | 0.027849732 |
| ENSMUSG00000022075 | Rhobtb2 | 246710 | protein_coding | 406.0712934 | -0.027872929 | 0.070676351 | -0.394374193 | 0.693304794 | 0.846398669 | 0.027872929 |
| ENSMUSG00000030301 | Ccdc91 | 67015 | protein_coding | 1049.941904 | 0.027978482 | 0.047015525 | 0.595090277 | 0.551783128 | 0.752522003 | 0.027978482 |
| ENSMUSG00000008683 | Rps15a | 267019 | protein_coding | 6014.38729 | -0.028019068 | 0.046214842 | -0.606278561 | 0.544329793 | 0.748889318 | 0.028019068 |
| ENSMUSG00000109511 | Nup62 | 18226 | protein_coding | 678.4337932 | 0.028037203 | 0.075382433 | 0.371932851 | 0.709942846 | 0.856338689 | 0.028037203 |
| ENSMUSG00000056598 | Drc3 | 74665 | protein_coding | 37.31363584 | 0.02804929 | 0.199378862 | 0.14068337 | 0.888120083 | 0.950538398 | 0.02804929 |
| ENSMUSG00000029097 | Trmt44 | 78890 | protein_coding | 187.8352622 | -0.028065558 | 0.119872091 | -0.234129212 | 0.814884668 | 0.916752358 | 0.028065558 |
| ENSMUSG00000010048 | Ifrd2 | 15983 | protein_coding | 641.3759005 | -0.028115141 | 0.078025802 | -0.360331331 | 0.718599372 | 0.862201627 | 0.028115141 |
| ENSMUSG00000058318 | Phf21a | 192285 | protein_coding | 347.132187 | 0.02812946 | 0.077772757 | 0.36168783 | 0.717585322 | 0.861553924 | 0.02812946 |
| ENSMUSG00000040331 | Nsmce4a | 67872 | protein_coding | 1092.79671 | -0.028140175 | 0.064930579 | -0.433388642 | 0.664732451 | 0.828854703 | 0.028140175 |
| ENSMUSG00000034636 | Zyg11b | 414872 | protein_coding | 1883.760765 | 0.028161196 | 0.044927663 | 0.626811948 | 0.53078251 | 0.73971984 | 0.028161196 |
| ENSMUSG00000036270 | Edc4 | 234699 | protein_coding | 964.8285251 | -0.028163552 | 0.04325531 | -0.651100449 | 0.514981647 | 0.72762476 | 0.028163552 |
| ENSMUSG00000050623 | Catsperz | 67077 | protein_coding | 13.17867611 | -0.028207369 | 0.335146938 | -0.084164187 | 0.932925892 | 0.970732425 | 0.028207369 |
| ENSMUSG00000034729 | Mrps10 | 64657 | protein_coding | 361.263575 | 0.028207914 | 0.077549239 | 0.363741988 | 0.716050686 | 0.860552159 | 0.028207914 |
| ENSMUSG00000046456 | Tmem150b | 330460 | protein_coding | 420.2982438 | -0.028212076 | 0.08000101 | -0.352646503 | 0.72435347 | 0.865627893 | 0.028212076 |
| ENSMUSG00000054920 | Klhl5 | 71778 | protein_coding | 410.7559509 | -0.028223626 | 0.097095722 | -0.290678369 | 0.771297316 | 0.892332662 | 0.028223626 |
| ENSMUSG00000021096 | Ppm1a | 19042 | protein_coding | 2738.107797 | -0.028243871 | 0.039051995 | -0.723237594 | 0.469533926 | 0.69386728 | 0.028243871 |
| ENSMUSG00000031955 | Bcar1 | 12927 | protein_coding | 1552.521278 | 0.028330674 | 0.075340868 | 0.376033284 | 0.706892152 | 0.854251145 | 0.028330674 |
| ENSMUSG00000035206 | Sppl2b | 73218 | protein_coding | 920.8887838 | -0.028342467 | 0.072959111 | -0.38847056 | 0.697667837 | 0.849281085 | 0.028342467 |
| ENSMUSG00000049721 | Gal3st1 | 53897 | protein_coding | 1430.423181 | -0.028374718 | 0.076269853 | -0.372030592 | 0.709870072 | 0.856314957 | 0.028374718 |
| ENSMUSG00000058290 | Espl1 | 105988 | protein_coding | 327.20995 | 0.028398243 | 0.104583575 | 0.271536362 | 0.785978537 | 0.900310808 | 0.028398243 |
| ENSMUSG00000049305 | Ccdc71 | 72454 | protein_coding | 750.1162696 | 0.028407375 | 0.052539904 | 0.540681897 | 0.588726858 | 0.778426904 | 0.028407375 |
| ENSMUSG00000042790 | Rnf214 | 235315 | protein_coding | 449.7588158 | -0.028408597 | 0.065105748 | -0.436345455 | 0.662586107 | 0.827608087 | 0.028408597 |
| ENSMUSG00000024056 | Ndc80 | 67052 | protein_coding | 189.4404655 | -0.028441135 | 0.164621825 | -0.172766493 | 0.862834976 | 0.937787345 | 0.028441135 |
| ENSMUSG00000036632 | Alg5 | 66248 | protein_coding | 721.6591496 | -0.028442865 | 0.060896456 | -0.467069304 | 0.640450295 | 0.812950957 | 0.028442865 |
| ENSMUSG00000027673 | Ndufb5 | 66046 | protein_coding | 2610.100169 | 0.028448271 | 0.047173262 | 0.603059232 | 0.546469286 | 0.749779664 | 0.028448271 |
| ENSMUSG00000035773 | Kiss1r | 114229 | protein_coding | 38.96234885 | -0.028448351 | 0.20263238 | -0.140393906 | 0.888348772 | 0.950656978 | 0.028448351 |
| ENSMUSG00000019773 | Fbxo5 | 67141 | protein_coding | 147.1090578 | 0.028497533 | 0.171063895 | 0.166589995 | 0.867692666 | 0.940092686 | 0.028497533 |
| ENSMUSG00000085492 | Trmt61b | 68789 | lncRNA | 101.6939186 | 0.028531102 | 0.118987848 | 0.239781646 | 0.810499537 | 0.914754428 | 0.028531102 |
| ENSMUSG00000020864 | Ankrd40 | 71452 | protein_coding | 1297.871602 | -0.028546281 | 0.06211808 | -0.459548668 | 0.645840211 | 0.816513895 | 0.028546281 |
| ENSMUSG00000008435 | Rdh13 | 108841 | protein_coding | 923.2714512 | 0.028564296 | 0.055548211 | 0.514225301 | 0.607094481 | 0.790011514 | 0.028564296 |
| ENSMUSG00000034892 | Rps29 | 20090 | protein_coding | 8313.104543 | 0.028587536 | 0.107651898 | 0.265555333 | 0.79058169 | 0.90237886 | 0.028587536 |
| ENSMUSG00000038876 | Rnf146 | 68031 | protein_coding | 300.4933778 | -0.028601506 | 0.076763215 | -0.372593905 | 0.70945071 | 0.856166251 | 0.028601506 |
| ENSMUSG00000026078 | Pdcl3 | 68833 | protein_coding | 898.0175088 | 0.028620534 | 0.053378781 | 0.536178119 | 0.591835454 | 0.780262005 | 0.028620534 |
| ENSMUSG00000110185 | Igip | 109169 | protein_coding | 269.1562317 | -0.028623631 | 0.093377164 | -0.30653781 | 0.759195196 | 0.886076667 | 0.028623631 |
| ENSMUSG00000059273 | Zc3h4 | 330474 | protein_coding | 1452.131835 | -0.028625121 | 0.068422475 | -0.418358457 | 0.675685055 | 0.835024575 | 0.028625121 |
| ENSMUSG00000033022 | Cdo1 | 12583 | protein_coding | 745.2753708 | -0.02865719 | 0.075549273 | -0.379317874 | 0.704451828 | 0.853157579 | 0.02865719 |

| ensembl_gene_id | external_gene_name | entrezgene_id | gene_biotype | baseMean | log2FoldChange | lfcSE | stat | pvalue | padj | absLog2FC |
|---|---|---|---|---|---|---|---|---|---|---|
| ENSMUSG00000090137 | Uba52 | 22186 | protein_coding | 163.1870326 | -0.028666324 | 0.156798901 | -0.182822226 | 0.854937509 | 0.93513139 | 0.028666324 |
| ENSMUSG00000002015 | Bcap31 | 27061 | protein_coding | 2873.99124 | -0.028719561 | 0.051379027 | -0.558974394 | 0.576179196 | 0.769017785 | 0.028719561 |
| ENSMUSG00000030057 | Cnbp | 12785 | protein_coding | 4365.685372 | 0.028726754 | 0.036135128 | 0.794981382 | 0.426624332 | 0.662000443 | 0.028726754 |
| ENSMUSG00000037339 | Fam53a | 74504 | protein_coding | 621.344418 | 0.028752831 | 0.055328041 | 0.51967917 | 0.603287207 | 0.788045747 | 0.028752831 |
| ENSMUSG00000052681 | Rap1b | 215449 | protein_coding | 1831.782141 | 0.028796971 | 0.040361821 | 0.713470541 | 0.475554636 | 0.698858175 | 0.028796971 |
| ENSMUSG00000029413 | Naaa | 67111 | protein_coding | 524.9167103 | 0.028802217 | 0.080363245 | 0.358400381 | 0.720043707 | 0.863035664 | 0.028802217 |
| ENSMUSG00000021067 | Sav1 | 64010 | protein_coding | 498.0107348 | 0.028806657 | 0.058498271 | 0.492436044 | 0.622411124 | 0.800805946 | 0.028806657 |
| ENSMUSG00000027242 | Wdr76 | 241627 | protein_coding | 239.0216171 | -0.028837161 | 0.124765389 | -0.231131094 | 0.81721296 | 0.917898969 | 0.028837161 |
| ENSMUSG00000003031 | Cdkn1b | 12576 | protein_coding | 1600.150973 | -0.028892607 | 0.043785307 | -0.659869921 | 0.509337308 | 0.72429229 | 0.028892607 |
| ENSMUSG00000028282 | Casp8ap2 | 26885 | protein_coding | 587.6864336 | 0.028944724 | 0.088014321 | 0.3288638 | 0.742258637 | 0.876765146 | 0.028944724 |
| ENSMUSG00000022197 | Pdzd2 | 68070 | protein_coding | 70.36713722 | 0.028948052 | 0.175939567 | 0.164534065 | 0.869310732 | 0.940691929 | 0.028948052 |
| ENSMUSG00000035620 | Ric8b | 237422 | protein_coding | 446.2603154 | -0.029001349 | 0.063463736 | -0.456975123 | 0.647688926 | 0.817954527 | 0.029001349 |
| ENSMUSG00000024542 | Cep192 | 70799 | protein_coding | 661.1532996 | 0.029005338 | 0.078745693 | 0.368341895 | 0.712618318 | 0.858410189 | 0.029005338 |
| ENSMUSG00000052139 | Babam2 | 107976 | protein_coding | 647.2294747 | 0.029005972 | 0.050264608 | 0.577065508 | 0.563895204 | 0.761121151 | 0.029005972 |
| ENSMUSG00000017843 | Ppp2r5c | 26931 | protein_coding | 3048.441549 | -0.029010862 | 0.041901558 | -0.692357595 | 0.488712789 | 0.708970987 | 0.029010862 |
| ENSMUSG00000002968 | Med25 | 75613 | protein_coding | 1415.099331 | 0.029013023 | 0.061177469 | 0.474243594 | 0.63532621 | 0.809423077 | 0.029013023 |
| ENSMUSG00000030983 | Bccip | 66165 | protein_coding | 779.6343108 | -0.029026379 | 0.078337066 | -0.37053186 | 0.710986242 | 0.857064391 | 0.029026379 |
| ENSMUSG00000041608 | Entpd3 | 215446 | protein_coding | 9.691520428 | 0.029065099 | 0.386540642 | 0.075192866 | 0.940061261 | 0.973941151 | 0.029065099 |
| ENSMUSG00000041841 | Rpl37 | 67281 | protein_coding | 6936.603043 | -0.029066293 | 0.055646074 | -0.522342197 | 0.601432094 | 0.786583124 | 0.029066293 |
| ENSMUSG00000000194 | Gpr107 | 277463 | protein_coding | 1957.647329 | 0.029081133 | 0.04208501 | 0.69100928 | 0.489559709 | 0.709610771 | 0.029081133 |
| ENSMUSG00000014791 | Elmo3 | 234683 | protein_coding | 1886.977251 | -0.029106239 | 0.033478301 | -0.869406115 | 0.384625039 | 0.624972092 | 0.029106239 |
| ENSMUSG00000033565 | Rbfox2 | 93686 | protein_coding | 638.7849821 | -0.029153559 | 0.061611032 | -0.47318732 | 0.636079544 | 0.809697277 | 0.029153559 |
| ENSMUSG00000034653 | Ythdc2 | 240255 | protein_coding | 802.381325 | 0.029169691 | 0.055929387 | 0.521544973 | 0.601987185 | 0.786991399 | 0.029169691 |
| ENSMUSG00000042046 | Dstyk | 213452 | protein_coding | 408.9436905 | -0.029180871 | 0.06706376 | -0.435121317 | 0.663474371 | 0.828033678 | 0.029180871 |
| ENSMUSG00000028164 | Manba | 110173 | protein_coding | 810.8374384 | 0.029185794 | 0.053684153 | 0.543657527 | 0.586677165 | 0.776984696 | 0.029185794 |
| ENSMUSG00000021715 | Cwc27 | 67285 | protein_coding | 242.536991 | -0.029192653 | 0.080728778 | -0.361613958 | 0.717640532 | 0.861553924 | 0.029192653 |
| ENSMUSG00000027597 | Ahcy | 269378 | protein_coding | 2421.241997 | 0.029195641 | 0.047576442 | 0.613657512 | 0.539441671 | 0.745464938 | 0.029195641 |
| ENSMUSG00000034868 | Myl12b | 67938 | protein_coding | 815.0875665 | 0.02920562 | 0.08780859 | 0.332605496 | 0.739432094 | 0.874869968 | 0.02920562 |
| ENSMUSG00000025609 | Mkln1 | 27418 | protein_coding | 1702.244675 | -0.029209741 | 0.055251534 | -0.528668427 | 0.597035483 | 0.783432973 | 0.029209741 |
| ENSMUSG00000062312 | Erbb2 | 13866 | protein_coding | 6032.060125 | -0.029216252 | 0.065146191 | -0.448472152 | 0.653812479 | 0.821762439 | 0.029216252 |
| ENSMUSG00000040843 | Tiprl | 226591 | protein_coding | 584.3789205 | 0.029225685 | 0.06014461 | 0.4859236 | 0.627021335 | 0.803226458 | 0.029225685 |
| ENSMUSG00000029060 | Mib2 | 76580 | protein_coding | 836.4918205 | 0.029232572 | 0.068628507 | 0.425953781 | 0.670141525 | 0.831656204 | 0.029232572 |
| ENSMUSG00000005951 | Shpk | 74637 | protein_coding | 606.4789026 | -0.029233927 | 0.057117471 | -0.511821104 | 0.608776219 | 0.791306043 | 0.029233927 |
| ENSMUSG00000026479 | Lamc2 | 16782 | protein_coding | 1678.222926 | -0.029240769 | 0.062893641 | -0.464924093 | 0.641985816 | 0.813747171 | 0.029240769 |
| ENSMUSG00000032246 | Calml4 | 75600 | protein_coding | 2866.157059 | -0.02924818 | 0.051693817 | -0.56579648 | 0.57153214 | 0.766430968 | 0.02924818 |
| ENSMUSG00000044320 | 1700001O22Rik | 73598 | protein_coding | 39.37118882 | -0.02924916 | 0.206042333 | -0.141957041 | 0.887113939 | 0.95019842 | 0.02924916 |
| ENSMUSG00000021326 | Trim27 | 19720 | protein_coding | 920.0004723 | 0.029252778 | 0.053788322 | 0.543849972 | 0.586544718 | 0.776873083 | 0.029252778 |
| ENSMUSG00000022568 | Scrib | 105782 | protein_coding | 2151.828075 | -0.029266398 | 0.060578814 | -0.483112771 | 0.629015667 | 0.804579288 | 0.029266398 |
| ENSMUSG00000006307 | Kmt2b | 75410 | protein_coding | 924.1076227 | -0.029277554 | 0.073008672 | -0.40101475 | 0.688409264 | 0.843368224 | 0.029277554 |

| ensembl_gene_id | external_gene_name | entrezgene_id | gene_biotype | baseMean | log2FoldChange | lfcSE | stat | pvalue | padj | absLog2FC |
|---|---|---|---|---|---|---|---|---|---|---|
| ENSMUSG00000017686 | Rhot1 | 59040 | protein_coding | 805.4859617 | 0.029278086 | 0.061595695 | 0.475326828 | 0.63455404 | 0.80894922 | 0.029278086 |
| ENSMUSG00000039382 | Wdr45 | 54636 | protein_coding | 280.2296888 | 0.02927996 | 0.081954296 | 0.357271814 | 0.720888328 | 0.863727048 | 0.02927996 |
| ENSMUSG00000043760 | Pkhd1 | 241035 | protein_coding | 293.8979831 | -0.029286546 | 0.106852014 | -0.274085111 | 0.784019215 | 0.89946133 | 0.029286546 |
| ENSMUSG00000036918 | Ttc7 | 225049 | protein_coding | 2505.717723 | 0.029330086 | 0.046553271 | 0.630032762 | 0.52867315 | 0.738046389 | 0.029330086 |
| ENSMUSG00000029312 | Klhl8 | 246293 | protein_coding | 98.40437373 | -0.029332831 | 0.117208004 | -0.250263037 | 0.802383939 | 0.910121497 | 0.029332831 |
| ENSMUSG00000030494 | Rhpn2 | 52428 | protein_coding | 3259.266916 | -0.029353071 | 0.050173384 | -0.585032719 | 0.558525716 | 0.756983295 | 0.029353071 |
| ENSMUSG00000022035 | Ccdc25 | 67179 | protein_coding | 1037.979569 | 0.029355347 | 0.079891001 | 0.367442474 | 0.713288995 | 0.858813602 | 0.029355347 |
| ENSMUSG00000046971 | Pla2g4f | 271844 | protein_coding | 222.6292913 | 0.029365408 | 0.19722387 | 0.14889378 | 0.881637449 | 0.947331698 | 0.029365408 |
| ENSMUSG00000028041 | Adam15 | 11490 | protein_coding | 797.846257 | 0.029365927 | 0.096718675 | 0.303622094 | 0.761415817 | 0.887492981 | 0.029365927 |
| ENSMUSG00000058260 | Serpina9 | 71907 | protein_coding | 98.04316709 | 0.029367223 | 0.192565037 | 0.152505477 | 0.878788271 | 0.94557102 | 0.029367223 |
| ENSMUSG00000040021 | Lats1 | 16798 | protein_coding | 2069.818179 | -0.02939956 | 0.043928884 | -0.669253521 | 0.50333377 | 0.719943844 | 0.02939956 |
| ENSMUSG00000003435 | Supt5 | 20924 | protein_coding | 1897.299762 | 0.029441006 | 0.034019314 | 0.865420326 | 0.386808129 | 0.626877167 | 0.029441006 |
| ENSMUSG00000061904 | Slc25a3 | 18674 | protein_coding | 13844.42013 | -0.029442042 | 0.044395527 | -0.663175859 | 0.507217931 | 0.72271052 | 0.029442042 |
| ENSMUSG00000112489 | 9230116L04Rik | 321017 | lncRNA | 119.7450514 | 0.029446066 | 0.135320464 | 0.217602464 | 0.827738865 | 0.923349179 | 0.029446066 |
| ENSMUSG00000001962 | Fam50a | 108160 | protein_coding | 357.5095652 | 0.029465248 | 0.089978206 | 0.327470946 | 0.743311714 | 0.877651795 | 0.029465248 |
| ENSMUSG00000022149 | C9 | 12279 | protein_coding | 14.10181921 | -0.029493164 | 0.387896995 | -0.076033495 | 0.93939245 | 0.973498396 | 0.029493164 |
| ENSMUSG00000036104 | Rab3gap1 | 226407 | protein_coding | 1955.382624 | 0.029523714 | 0.047507786 | 0.621449993 | 0.534303587 | 0.741606925 | 0.029523714 |
| ENSMUSG00000045394 | Epcam | 17075 | protein_coding | 36243.08521 | -0.029549763 | 0.053496439 | -0.552368784 | 0.580695713 | 0.772616101 | 0.029549763 |
| ENSMUSG00000032637 | Atxn2l | 233871 | protein_coding | 926.7346314 | -0.02955792 | 0.07148264 | -0.413497871 | 0.679241882 | 0.837472006 | 0.02955792 |
| ENSMUSG00000031819 | Emc8 | 18117 | protein_coding | 396.4461033 | -0.029602231 | 0.068652628 | -0.431188611 | 0.666331235 | 0.829602333 | 0.029602231 |
| ENSMUSG00000033578 | Tmem35a | 67564 | protein_coding | 61.64890463 | 0.029614605 | 0.152718353 | 0.193916478 | 0.846241281 | 0.930889346 | 0.029614605 |
| ENSMUSG00000030340 | Scnn1a | 20276 | protein_coding | 1424.428655 | 0.029626244 | 0.10830828 | 0.273536276 | 0.784441011 | 0.899577643 | 0.029626244 |
| ENSMUSG00000015533 | Itga2 | 16398 | protein_coding | 1784.048336 | -0.029636086 | 0.101014225 | -0.29338527 | 0.769227683 | 0.891725514 | 0.029636086 |
| ENSMUSG00000028830 | AU040320 | 100317 | protein_coding | 2965.964862 | -0.029647803 | 0.033308938 | -0.890085504 | 0.373419976 | 0.615257862 | 0.029647803 |
| ENSMUSG00000021375 | Kif13a | 16553 | protein_coding | 2603.778283 | -0.029680686 | 0.051255345 | -0.579074944 | 0.562538606 | 0.76019675 | 0.029680686 |
| ENSMUSG00000058444 | Map2k5 | 23938 | protein_coding | 280.1712271 | 0.029708791 | 0.07870814 | 0.377455134 | 0.7058354 | 0.853688621 | 0.029708791 |
| ENSMUSG00000000346 | Dazap2 | 23994 | protein_coding | 4296.866058 | 0.029746781 | 0.046952442 | 0.633551316 | 0.526373681 | 0.736749554 | 0.029746781 |
| ENSMUSG00000067194 | Eif1ax | 66235 | protein_coding | 1599.616341 | -0.029759124 | 0.054808293 | -0.542967534 | 0.587152156 | 0.777161429 | 0.029759124 |
| ENSMUSG00000028458 | Tesk1 | 21754 | protein_coding | 631.7314248 | -0.029763345 | 0.089832014 | -0.331322253 | 0.740401084 | 0.875592013 | 0.029763345 |
| ENSMUSG00000029725 | Ppp1r35 | 69871 | protein_coding | 197.8983023 | -0.029772577 | 0.119172321 | -0.249827953 | 0.802720402 | 0.910242717 | 0.029772577 |
| ENSMUSG00000031967 | Afg3l1 | 114896 | protein_coding | 844.2084811 | 0.02978646 | 0.078116775 | 0.381306824 | 0.702975591 | 0.85233346 | 0.02978646 |
| ENSMUSG00000058638 | Zfp110 | 65020 | protein_coding | 993.2023378 | -0.029789935 | 0.084113012 | -0.3541656 | 0.723214782 | 0.864908031 | 0.029789935 |
| ENSMUSG00000010122 | Slc47a1 | 67473 | protein_coding | 36.66977193 | -0.029797053 | 0.21737701 | -0.137075455 | 0.890971153 | 0.951128072 | 0.029797053 |
| ENSMUSG00000050944 | Efcab5 | 319634 | protein_coding | 80.24587247 | 0.029803591 | 0.14946231 | 0.199405397 | 0.841945639 | 0.929706079 | 0.029803591 |
| ENSMUSG00000024576 | Csnk1a1 | 93687 | protein_coding | 9076.423508 | -0.02984851 | 0.029979703 | -0.995623956 | 0.31943289 | 0.562943309 | 0.02984851 |
| ENSMUSG00000019261 | Map1s | 270058 | protein_coding | 220.7760525 | 0.029860329 | 0.116210494 | 0.256950369 | 0.797217087 | 0.906615892 | 0.029860329 |
| ENSMUSG00000017667 | Zfp334 | 228876 | protein_coding | 73.09845227 | 0.029863403 | 0.152352024 | 0.196015796 | 0.844597803 | 0.930210801 | 0.029863403 |
| ENSMUSG00000049493 | Pls1 | 102502 | protein_coding | 7024.107374 | 0.029876049 | 0.073259615 | 0.407810626 | 0.683412712 | 0.840526771 | 0.029876049 |
| ENSMUSG00000019978 | Epb41l2 | 13822 | protein_coding | 758.4060339 | 0.029878522 | 0.059410057 | 0.502920273 | 0.615020327 | 0.795383546 | 0.029878522 |

| ensembl_gene_id | external_gene_name | entrezgene_id | gene_biotype | baseMean | log2FoldChange | lfcSE | stat | pvalue | padj | absLog2FC |
|---|---|---|---|---|---|---|---|---|---|---|
| ENSMUSG00000024990 | Rbp4 | 19662 | protein_coding | 102.2192687 | 0.029910832 | 0.128668568 | 0.232464168 | 0.816177515 | 0.917183108 | 0.029910832 |
| ENSMUSG00000018566 | Slc2a4 | 20528 | protein_coding | 20.45343126 | -0.029928441 | 0.294884096 | -0.101492218 | 0.919159735 | 0.964681052 | 0.029928441 |
| ENSMUSG00000000567 | Sox9 | 20682 | protein_coding | 858.2004304 | -0.029944657 | 0.104248575 | -0.28724284 | 0.773926386 | 0.893742272 | 0.029944657 |
| ENSMUSG00000010080 | Epn3 | 71889 | protein_coding | 873.7805924 | 0.029951414 | 0.078183226 | 0.383092578 | 0.701651122 | 0.851559877 | 0.029951414 |
| ENSMUSG00000040666 | Sh3bgr | 50795 | protein_coding | 95.26237782 | -0.029958591 | 0.171097633 | -0.175096466 | 0.861003835 | 0.937253381 | 0.029958591 |
| ENSMUSG00000022120 | Obi1 | 72486 | protein_coding | 257.1475362 | -0.02999084 | 0.086511024 | -0.346670734 | 0.728838699 | 0.868349712 | 0.02999084 |
| ENSMUSG00000031584 | Gsr | 14782 | protein_coding | 2927.466852 | 0.029996727 | 0.065466108 | 0.45820238 | 0.64680705 | 0.817036486 | 0.029996727 |
| ENSMUSG00000032217 | Rnf111 | 93836 | protein_coding | 790.2189465 | -0.030002161 | 0.056173844 | -0.534094852 | 0.593275911 | 0.780786732 | 0.030002161 |
| ENSMUSG00000078515 | Ddi2 | 68817 | protein_coding | 2362.147762 | 0.030006891 | 0.053000141 | 0.566166244 | 0.571280775 | 0.766221312 | 0.030006891 |
| ENSMUSG00000085112 | A530072M11Rik | 100415915 | lncRNA | 31.89279567 | -0.030008375 | 0.225751525 | -0.132926564 | 0.894251458 | 0.952773623 | 0.030008375 |
| ENSMUSG00000023047 | Amhr2 | 110542 | protein_coding | 8.126746293 | 0.030015746 | 0.453613966 | 0.066170242 | 0.947242288 | 0.977863038 | 0.030015746 |
| ENSMUSG00000038074 | Fkbp14 | 231997 | protein_coding | 68.01850738 | -0.03002431 | 0.172867241 | -0.173684209 | 0.862113649 | 0.937787345 | 0.03002431 |
| ENSMUSG00000042729 | Wdr74 | 107071 | protein_coding | 453.3123046 | 0.030028172 | 0.069877488 | 0.429725987 | 0.667394978 | 0.830281488 | 0.030028172 |
| ENSMUSG00000055485 | Soga1 | 320706 | protein_coding | 98.21506342 | -0.030039548 | 0.142536099 | -0.210750455 | 0.833082001 | 0.925920981 | 0.030039548 |
| ENSMUSG00000034863 | Ano8 | 382014 | protein_coding | 389.0665042 | -0.030121936 | 0.077320618 | -0.389571848 | 0.696853173 | 0.848535732 | 0.030121936 |
| ENSMUSG00000026784 | Pdss1 | 56075 | protein_coding | 592.1258792 | -0.030211995 | 0.098494102 | -0.306739131 | 0.759041942 | 0.886076667 | 0.030211995 |
| ENSMUSG00000024513 | Mbd2 | 17191 | protein_coding | 2441.34664 | -0.03022258 | 0.050021671 | -0.604189728 | 0.545717509 | 0.749130599 | 0.03022258 |
| ENSMUSG00000020888 | Dvl2 | 13543 | protein_coding | 256.4348904 | -0.030229011 | 0.085896573 | -0.351923361 | 0.724895739 | 0.86595508 | 0.030229011 |
| ENSMUSG00000068290 | Ddrgk1 | 77006 | protein_coding | 1864.764104 | -0.030254201 | 0.040224647 | -0.752130935 | 0.451972322 | 0.680967534 | 0.030254201 |
| ENSMUSG00000097119 | B230354K17Rik | 105246278 | lncRNA | 253.5165439 | 0.030263435 | 0.082832858 | 0.365355444 | 0.714846096 | 0.859709385 | 0.030263435 |
| ENSMUSG00000022817 | Itgb5 | 16419 | protein_coding | 791.1114498 | 0.030270647 | 0.068567106 | 0.44147477 | 0.658869323 | 0.825416419 | 0.030270647 |
| ENSMUSG00000049792 | Bag5 | 70369 | protein_coding | 369.7450126 | -0.030280454 | 0.07966046 | -0.380119 | 0.703857083 | 0.852953349 | 0.030280454 |
| ENSMUSG00000000876 | Pxmp4 | 59038 | protein_coding | 721.7849186 | 0.030283912 | 0.058366872 | 0.518854463 | 0.603862235 | 0.78841372 | 0.030283912 |
| ENSMUSG00000015755 | Tab2 | 68652 | protein_coding | 1667.760446 | -0.03029978 | 0.04506465 | -0.672362488 | 0.501352957 | 0.718418124 | 0.03029978 |
| ENSMUSG00000051355 | Commd1 | 17846 | protein_coding | 591.2162582 | 0.030316014 | 0.070556682 | 0.429668932 | 0.667436486 | 0.830281488 | 0.030316014 |
| ENSMUSG00000060261 | Gtf2i | 14886 | protein_coding | 1428.460071 | -0.030330156 | 0.060858821 | -0.498369098 | 0.618223913 | 0.797543961 | 0.030330156 |
| ENSMUSG00000030780 | Rusf1 | 233913 | protein_coding | 598.9962351 | -0.030352346 | 0.069561045 | -0.43634114 | 0.662589237 | 0.827608087 | 0.030352346 |
| ENSMUSG00000047635 | Mtrfr | 72650 | protein_coding | 143.705539 | -0.030360874 | 0.115433712 | -0.263015661 | 0.792538508 | 0.90358915 | 0.030360874 |
| ENSMUSG00000013155 | Enkd1 | 102124 | protein_coding | 34.5135105 | -0.030364043 | 0.214864131 | -0.141317411 | 0.887619195 | 0.95031765 | 0.030364043 |
| ENSMUSG00000001056 | Nhp2 | 52530 | protein_coding | 579.8378146 | -0.030365401 | 0.089582518 | -0.338965707 | 0.734635565 | 0.872420306 | 0.030365401 |
| ENSMUSG00000053110 | Yap1 | 22601 | protein_coding | 1266.006134 | -0.03036947 | 0.042483402 | -0.714854946 | 0.474698679 | 0.698573273 | 0.03036947 |
| ENSMUSG00000034120 | Srsf2 | 20382 | protein_coding | 2616.920625 | 0.030375871 | 0.054028238 | 0.562222127 | 0.573964688 | 0.76815045 | 0.030375871 |
| ENSMUSG00000099881 | 2810013P06Rik | 100503178 | lncRNA | 409.8478985 | -0.030388896 | 0.082553343 | -0.368112238 | 0.712789546 | 0.85855232 | 0.030388896 |
| ENSMUSG00000032468 | Armc8 | 74125 | protein_coding | 1000.942996 | 0.030392345 | 0.0597488 | 0.508668708 | 0.610984459 | 0.792706869 | 0.030392345 |
| ENSMUSG00000031012 | Cask | 12361 | protein_coding | 1836.495362 | 0.030398506 | 0.046022073 | 0.660520128 | 0.508920106 | 0.724071862 | 0.030398506 |
| ENSMUSG00000028573 | Fggy | 75578 | protein_coding | 196.6674974 | -0.030450891 | 0.103492079 | -0.29423403 | 0.768579079 | 0.891466618 | 0.030450891 |
| ENSMUSG00000045427 | Hnrnph2 | 56258 | protein_coding | 1179.465488 | 0.030466365 | 0.065391903 | 0.465904245 | 0.641284042 | 0.813625029 | 0.030466365 |
| ENSMUSG00000038564 | Ift172 | 67661 | protein_coding | 639.1041269 | 0.030473508 | 0.069115969 | 0.440904006 | 0.659282493 | 0.825741594 | 0.030473508 |
| ENSMUSG00000063704 | Mapk15 | 332110 | protein_coding | 57.03785252 | 0.030475057 | 0.170794856 | 0.178430766 | 0.858384689 | 0.93615636 | 0.030475057 |

| ensembl_gene_id | external_gene_name | entrezgene_id | gene_biotype | baseMean | log2FoldChange | lfcSE | stat | pvalue | padj | absLog2FC |
|---|---|---|---|---|---|---|---|---|---|---|
| ENSMUSG00000092183 | 4930515G01Rik | 67642 | lncRNA | 25.4682919 | -0.030527139 | 0.261505813 | -0.116735985 | 0.907069274 | 0.95873684 | 0.030527139 |
| ENSMUSG00000030452 | Nipa2 | 93790 | protein_coding | 956.3134467 | -0.030543438 | 0.051935106 | -0.588107755 | 0.556459968 | 0.755437307 | 0.030543438 |
| ENSMUSG00000020523 | Fam114a2 | 67726 | protein_coding | 1722.126465 | 0.030546342 | 0.050739192 | 0.602026583 | 0.547156443 | 0.750223246 | 0.030546342 |
| ENSMUSG00000034795 | Ccdc122 | 108811 | protein_coding | 188.2871002 | -0.030561567 | 0.103650305 | -0.294852651 | 0.768106444 | 0.891239011 | 0.030561567 |
| ENSMUSG00000025733 | Rhot2 | 214952 | protein_coding | 1054.32239 | -0.030562419 | 0.052085663 | -0.586772198 | 0.557356711 | 0.756034291 | 0.030562419 |
| ENSMUSG00000030522 | Mtmr10 | 233315 | protein_coding | 422.8857374 | -0.030572998 | 0.062883091 | -0.486187902 | 0.626833948 | 0.803190139 | 0.030572998 |
| ENSMUSG00000036211 | H1f6 | 107970 | protein_coding | 6.322153476 | 0.030594349 | 0.597807926 | 0.051177557 | 0.959184036 | 0.983542327 | 0.030594349 |
| ENSMUSG00000061111 | Mcrip1 | 192173 | protein_coding | 207.9628015 | -0.030599162 | 0.100375389 | -0.304847257 | 0.760482488 | 0.886731192 | 0.030599162 |
| ENSMUSG00000035000 | Dpp4 | 13482 | protein_coding | 1670.832643 | 0.030637048 | 0.073676876 | 0.415829898 | 0.677534479 | 0.836508911 | 0.030637048 |
| ENSMUSG00000045095 | Magi1 | 14924 | protein_coding | 1727.578416 | 0.030672877 | 0.045728628 | 0.670758729 | 0.502374243 | 0.718956777 | 0.030672877 |
| ENSMUSG00000020225 | Tmbim4 | 68212 | protein_coding | 1426.822493 | -0.030692353 | 0.067994721 | -0.451393174 | 0.651706202 | 0.820714191 | 0.030692353 |
| ENSMUSG00000050530 | Fam171a1 | 269233 | protein_coding | 592.0767379 | -0.030742607 | 0.076076693 | -0.404100205 | 0.686139034 | 0.841984797 | 0.030742607 |
| ENSMUSG00000026279 | Thap4 | 67026 | protein_coding | 1780.195735 | 0.030749534 | 0.055143899 | 0.557623509 | 0.5771015 | 0.769727784 | 0.030749534 |
| ENSMUSG00000041161 | Otud3 | 73162 | protein_coding | 405.4758746 | -0.030789604 | 0.086410454 | -0.356318038 | 0.721602402 | 0.863940753 | 0.030789604 |
| ENSMUSG00000005204 | Senp3 | 80886 | protein_coding | 1093.38676 | -0.030795438 | 0.047256959 | -0.651659323 | 0.514620969 | 0.727380907 | 0.030795438 |
| ENSMUSG00000045767 | B230219D22Rik | 78521 | protein_coding | 1114.166651 | 0.030813338 | 0.041015556 | 0.751259794 | 0.452496321 | 0.68117317 | 0.030813338 |
| ENSMUSG00000033871 | Ppargc1b | 170826 | protein_coding | 996.3010776 | -0.030823074 | 0.049729402 | -0.619815897 | 0.535379001 | 0.74228379 | 0.030823074 |
| ENSMUSG00000074221 | Zfp568 | 243905 | protein_coding | 526.9070645 | -0.030882332 | 0.080003724 | -0.386011182 | 0.699488387 | 0.850145091 | 0.030882332 |
| ENSMUSG00000026627 | Pacc1 | 66950 | protein_coding | 131.7580072 | -0.030890641 | 0.121141823 | -0.254995673 | 0.798726442 | 0.907948215 | 0.030890641 |
| ENSMUSG00000006676 | Usp19 | 71472 | protein_coding | 2299.39109 | 0.030901045 | 0.051783342 | 0.596737162 | 0.550682877 | 0.752041897 | 0.030901045 |
| ENSMUSG00000027378 | Nphp1 | 53885 | protein_coding | 94.82701487 | 0.030904824 | 0.128669659 | 0.240187344 | 0.810185024 | 0.914719539 | 0.030904824 |
| ENSMUSG00000056952 | Tatdn2 | 381801 | protein_coding | 1316.67756 | -0.030917973 | 0.044273269 | -0.698344025 | 0.484962073 | 0.706263486 | 0.030917973 |
| ENSMUSG00000020183 | Cpm | 70574 | protein_coding | 1774.534885 | -0.030922791 | 0.067040653 | -0.461254314 | 0.644616158 | 0.81580845 | 0.030922791 |
| ENSMUSG00000056144 | Trim34a | 94094 | protein_coding | 134.7625026 | 0.030923916 | 0.106289886 | 0.290939402 | 0.771097665 | 0.892331737 | 0.030923916 |
| ENSMUSG00000019917 | Septin10 | 103080 | protein_coding | 414.6623885 | -0.030936813 | 0.07485777 | -0.413274569 | 0.67940546 | 0.837609652 | 0.030936813 |
| ENSMUSG00000034168 | Irf2bpl | 238330 | protein_coding | 1158.102247 | 0.03093705 | 0.115149595 | 0.268668332 | 0.788184924 | 0.901237824 | 0.03093705 |
| ENSMUSG00000096950 | Gm9530 | 671336 | lncRNA | 7.549685837 | 0.030954021 | 0.415058031 | 0.074577574 | 0.940550817 | 0.974178386 | 0.030954021 |
| ENSMUSG00000046591 | Ticrr | 77011 | protein_coding | 113.7757192 | 0.030963246 | 0.147608565 | 0.209765918 | 0.833850375 | 0.926340413 | 0.030963246 |
| ENSMUSG00000021111 | Papola | 18789 | protein_coding | 2845.474856 | 0.030979496 | 0.047120222 | 0.657456508 | 0.510887427 | 0.725207424 | 0.030979496 |
| ENSMUSG00000028430 | Nol6 | 230082 | protein_coding | 689.3739776 | -0.030987897 | 0.078360103 | -0.395455029 | 0.692507105 | 0.845744951 | 0.030987897 |
| ENSMUSG00000025451 | Paip1 | 218693 | protein_coding | 1105.927031 | 0.031017359 | 0.043483764 | 0.713308972 | 0.475654587 | 0.698941369 | 0.031017359 |
| ENSMUSG00000027312 | Atrn | 11990 | protein_coding | 1537.686603 | -0.031036519 | 0.049536599 | -0.626537129 | 0.530962691 | 0.73971984 | 0.031036519 |
| ENSMUSG00000007029 | Vars | 22321 | protein_coding | 2144.964443 | -0.031050522 | 0.086819126 | -0.357646103 | 0.720608172 | 0.86351969 | 0.031050522 |
| ENSMUSG00000079334 | Naa80 | 56441 | protein_coding | 229.7507526 | 0.031059731 | 0.086741486 | 0.358072382 | 0.720289146 | 0.863217232 | 0.031059731 |
| ENSMUSG00000001098 | Kctd10 | 330171 | protein_coding | 1843.424542 | 0.031072311 | 0.071293705 | 0.435835267 | 0.662956254 | 0.827707324 | 0.031072311 |
| ENSMUSG00000009927 | Rps25 | 75617 | protein_coding | 7408.073978 | -0.031076721 | 0.033386137 | -0.930827097 | 0.351943011 | 0.594351535 | 0.031076721 |
| ENSMUSG00000036452 | Arhgap26 | 71302 | protein_coding | 790.6098366 | 0.031082905 | 0.071935996 | 0.432091116 | 0.665675191 | 0.829315377 | 0.031082905 |
| ENSMUSG00000008859 | Rala | 56044 | protein_coding | 1446.928223 | 0.031122797 | 0.06102742 | 0.50998055 | 0.610065088 | 0.792215128 | 0.031122797 |
| ENSMUSG00000034883 | Lrr1 | 69706 | protein_coding | 22.03022863 | 0.031123183 | 0.260828339 | 0.119324391 | 0.90501836 | 0.958435249 | 0.031123183 |

Supplementary Table 1

| ensembl_gene_id | external_gene_name | entrezgene_id | gene_biotype | baseMean | log2FoldChange | lfcSE | stat | pvalue | padj | absLog2FC |
|---|---|---|---|---|---|---|---|---|---|---|
| ENSMUSG00000001815 | Evx2 | 14029 | protein_coding | 16.32823215 | -0.031151105 | 0.444900272 | -0.070018175 | 0.944179193 | 0.975710693 | 0.031151105 |
| ENSMUSG00000021958 | Pinx1 | 72400 | protein_coding | 100.578978 | -0.031153389 | 0.132199159 | -0.23565497 | 0.813700415 | 0.916250984 | 0.031153389 |
| ENSMUSG00000042202 | Slc35e2 | 320541 | protein_coding | 663.9759504 | 0.031187961 | 0.062206061 | 0.50136531 | 0.616114049 | 0.795599668 | 0.031187961 |
| ENSMUSG00000099759 | 1700030C10Rik | 69513 | lncRNA | 32.86140683 | -0.031205394 | 0.262568729 | -0.118846575 | 0.905396908 | 0.958468846 | 0.031205394 |
| ENSMUSG00000061950 | Ppp4r1 | 70351 | protein_coding | 1570.861122 | -0.031227375 | 0.048790737 | -0.640026718 | 0.522155229 | 0.733199598 | 0.031227375 |
| ENSMUSG00000060733 | Ipmk | 69718 | protein_coding | 5371.765814 | 0.031227868 | 0.051576523 | 0.60546671 | 0.544868937 | 0.749012844 | 0.031227868 |
| ENSMUSG00000025484 | Bet1l | 54399 | protein_coding | 467.8241878 | 0.031230252 | 0.062656906 | 0.498432722 | 0.618179078 | 0.797543961 | 0.031230252 |
| ENSMUSG00000078652 | Psme3 | 19192 | protein_coding | 1226.103748 | 0.031251142 | 0.063044687 | 0.495698264 | 0.620107312 | 0.79914261 | 0.031251142 |
| ENSMUSG00000005505 | Kbtbd4 | 67136 | protein_coding | 131.213463 | 0.031267549 | 0.134212774 | 0.232969992 | 0.815784707 | 0.917020505 | 0.031267549 |
| ENSMUSG00000050989 | Selenon | 74777 | protein_coding | 61.49417382 | 0.031282514 | 0.197709741 | 0.158224443 | 0.874279944 | 0.943679315 | 0.031282514 |
| ENSMUSG00000028223 | Decr1 | 67460 | protein_coding | 1816.710025 | 0.031306602 | 0.061367061 | 0.510153187 | 0.609944146 | 0.79214131 | 0.031306602 |
| ENSMUSG00000057134 | Ado | 211488 | protein_coding | 756.5616023 | 0.031329216 | 0.060719413 | 0.515967037 | 0.605877431 | 0.789638382 | 0.031329216 |
| ENSMUSG00000045094 | Arhgef37 | 328967 | protein_coding | 236.0423585 | -0.031353896 | 0.083376687 | -0.376051114 | 0.706878897 | 0.854251145 | 0.031353896 |
| ENSMUSG00000022808 | Snx4 | 69150 | protein_coding | 1476.436312 | -0.031363088 | 0.043384254 | -0.722914081 | 0.469732672 | 0.694009261 | 0.031363088 |
| ENSMUSG00000025369 | Smarcc2 | 68094 | protein_coding | 1934.947015 | -0.031364099 | 0.040882854 | -0.767169992 | 0.442980452 | 0.674020733 | 0.031364099 |
| ENSMUSG00000039208 | Metrnl | 210029 | protein_coding | 1276.188735 | 0.031378602 | 0.071380319 | 0.439597392 | 0.66022873 | 0.826041967 | 0.031378602 |
| ENSMUSG00000029634 | Rnf6 | 74132 | protein_coding | 1575.931135 | 0.031428426 | 0.043729692 | 0.718697634 | 0.472327241 | 0.696728273 | 0.031428426 |
| ENSMUSG00000029270 | Dipk1a | 67266 | protein_coding | 180.8978617 | 0.031438312 | 0.106810576 | 0.294337075 | 0.768500345 | 0.891466618 | 0.031438312 |
| ENSMUSG00000071456 | 1110002L01Rik | 100043040 | lncRNA | 33.01951049 | 0.031448648 | 0.31344425 | 0.100332507 | 0.920080351 | 0.96489645 | 0.031448648 |
| ENSMUSG00000001150 | Mcm3ap | 54387 | protein_coding | 1051.289293 | 0.031522069 | 0.049151207 | 0.641328488 | 0.521309284 | 0.732548832 | 0.031522069 |
| ENSMUSG00000096188 | Cmtm4 | 97487 | protein_coding | 7874.380901 | -0.031550727 | 0.04317275 | -0.73080188 | 0.464900175 | 0.690945086 | 0.031550727 |
| ENSMUSG00000021621 | Zcchc9 | 69085 | protein_coding | 509.4175795 | 0.031565537 | 0.071449702 | 0.441786824 | 0.658643474 | 0.825261698 | 0.031565537 |
| ENSMUSG00000071645 | Tut1 | 70044 | protein_coding | 463.8840882 | -0.031577829 | 0.07466766 | -0.42291172 | 0.672359671 | 0.833136568 | 0.031577829 |
| ENSMUSG00000021460 | Auh | 11992 | protein_coding | 608.1253208 | -0.031588953 | 0.069168933 | -0.456692789 | 0.647891875 | 0.818082701 | 0.031588953 |
| ENSMUSG00000031633 | Slc25a4 | 11739 | protein_coding | 573.2163923 | 0.031609781 | 0.083432404 | 0.378866959 | 0.704786661 | 0.853182291 | 0.031609781 |
| ENSMUSG00000026273 | Mterf4 | 69821 | protein_coding | 287.1486797 | 0.031627382 | 0.08949332 | 0.353404949 | 0.723784876 | 0.865397294 | 0.031627382 |
| ENSMUSG00000069808 | Tlcd3a | 116972 | protein_coding | 755.8428514 | -0.031654977 | 0.06437171 | -0.491752931 | 0.622894014 | 0.800888833 | 0.031654977 |
| ENSMUSG00000027498 | Cstf1 | 67337 | protein_coding | 566.2674765 | 0.031668241 | 0.056256523 | 0.562925676 | 0.573485496 | 0.767901078 | 0.031668241 |
| ENSMUSG00000042548 | Asxl1 | 228790 | protein_coding | 1230.021212 | -0.031678484 | 0.055926614 | -0.5664295 | 0.571101846 | 0.766045036 | 0.031678484 |
| ENSMUSG00000040250 | Ints13 | 71177 | protein_coding | 573.0208136 | -0.031679953 | 0.059457072 | -0.532820598 | 0.594157774 | 0.781437157 | 0.031679953 |
| ENSMUSG00000055114 | Anxa13 | 69787 | protein_coding | 464.611152 | -0.031754465 | 0.088936648 | -0.357045893 | 0.721057449 | 0.863755116 | 0.031754465 |
| ENSMUSG00000031753 | Cog4 | 102339 | protein_coding | 1621.365221 | 0.031775365 | 0.054081743 | 0.587543298 | 0.55683888 | 0.755776857 | 0.031775365 |
| ENSMUSG00000016833 | Mrps18c | 68735 | protein_coding | 421.3699323 | 0.031787123 | 0.07285078 | 0.436331956 | 0.662595899 | 0.827608087 | 0.031787123 |
| ENSMUSG00000020899 | Pfas | 237823 | protein_coding | 792.5213882 | -0.03179887 | 0.073484678 | -0.432727902 | 0.665212457 | 0.829048013 | 0.03179887 |
| ENSMUSG00000046159 | Chrm3 | 12671 | protein_coding | 69.75531085 | -0.031809737 | 0.174975141 | -0.181795751 | 0.85574302 | 0.935592329 | 0.031809737 |
| ENSMUSG00000102047 | 2010010A06Rik | 100503117 | transcribed_processed_pseudogene | 17.85057214 | 0.03184769 | 0.32142972 | 0.099081348 | 0.921073681 | 0.965195323 | 0.03184769 |
| ENSMUSG00000020664 | Dld | 13382 | protein_coding | 3165.785129 | 0.031860566 | 0.051834478 | 0.614659723 | 0.538779464 | 0.74493282 | 0.031860566 |
| ENSMUSG00000000884 | Gnb1l | 13972 | protein_coding | 62.96194735 | -0.031870696 | 0.174321584 | -0.182827022 | 0.854933745 | 0.93513139 | 0.031870696 |
| ENSMUSG00000052751 | Repin1 | 58887 | protein_coding | 504.4688766 | -0.031886667 | 0.100171538 | -0.318320625 | 0.750241741 | 0.880642591 | 0.031886667 |

| ensembl_gene_id | external_gene_name | entrezgene_id | gene_biotype | baseMean | log2FoldChange | lfcSE | stat | pvalue | padj | absLog2FC |
|---|---|---|---|---|---|---|---|---|---|---|
| ENSMUSG00000020475 | Pgam2 | 56012 | protein_coding | 26.56148584 | -0.031891519 | 0.250943296 | -0.127086555 | 0.898871893 | 0.955322545 | 0.031891519 |
| ENSMUSG00000026074 | Map4k4 | 26921 | protein_coding | 504.8092287 | 0.031917094 | 0.070494678 | 0.452758913 | 0.650722353 | 0.81998745 | 0.031917094 |
| ENSMUSG00000033773 | Rpap2 | 231571 | protein_coding | 290.1861041 | 0.03192769 | 0.091831033 | 0.347678658 | 0.728081526 | 0.867832282 | 0.03192769 |
| ENSMUSG00000021094 | Dhrs7 | 66375 | protein_coding | 585.1028637 | -0.031939164 | 0.075270663 | -0.424324195 | 0.671329396 | 0.832361742 | 0.031939164 |
| ENSMUSG00000052533 | Nup188 | 227699 | protein_coding | 548.6520262 | 0.031947623 | 0.087041094 | 0.367040689 | 0.713588668 | 0.859001601 | 0.031947623 |
| ENSMUSG00000028678 | Kif2c | 73804 | protein_coding | 281.9012498 | 0.031950852 | 0.120044898 | 0.266157518 | 0.790117899 | 0.902120505 | 0.031950852 |
| ENSMUSG00000052446 | Zfp961 | 234413 | protein_coding | 298.676573 | -0.031951816 | 0.089534429 | -0.356866246 | 0.72119194 | 0.863755116 | 0.031951816 |
| ENSMUSG00000023991 | Foxp4 | 74123 | protein_coding | 827.4454318 | -0.03196049 | 0.093507386 | -0.341796419 | 0.732504105 | 0.871007815 | 0.03196049 |
| ENSMUSG00000036009 | Mettl25 | 216292 | protein_coding | 114.7235631 | 0.031971106 | 0.121119516 | 0.263963292 | 0.791808205 | 0.903021982 | 0.031971106 |
| ENSMUSG00000018569 | Cldn7 | 53624 | protein_coding | 13194.96427 | 0.031978583 | 0.074264513 | 0.430603815 | 0.666756467 | 0.829882036 | 0.031978583 |
| ENSMUSG00000018167 | Stard3 | 59045 | protein_coding | 1182.456222 | 0.031988189 | 0.048818617 | 0.655245707 | 0.51230957 | 0.725948198 | 0.031988189 |
| ENSMUSG00000020196 | Cabin1 | 104248 | protein_coding | 1534.199358 | 0.032008529 | 0.048268742 | 0.663131627 | 0.507246257 | 0.72271052 | 0.032008529 |
| ENSMUSG00000030341 | Tnfrsf1a | 21937 | protein_coding | 687.5634932 | 0.032008689 | 0.061460406 | 0.520801777 | 0.602504865 | 0.7874413 | 0.032008689 |
| ENSMUSG00000037279 | Ovol2 | 107586 | protein_coding | 399.0935415 | -0.032027816 | 0.079169574 | -0.404547027 | 0.685810503 | 0.841836579 | 0.032027816 |
| ENSMUSG00000038406 | Scaf1 | 233208 | protein_coding | 1238.653395 | 0.032043321 | 0.096004945 | 0.333767399 | 0.738555086 | 0.874756013 | 0.032043321 |
| ENSMUSG00000015880 | Ncapg | 54392 | protein_coding | 263.4153455 | 0.032048153 | 0.128131478 | 0.25011928 | 0.802495106 | 0.910121497 | 0.032048153 |
| ENSMUSG00000021125 | Arg2 | 11847 | protein_coding | 183.7630464 | 0.032066411 | 0.180538674 | 0.177615189 | 0.859025195 | 0.936390154 | 0.032066411 |
| ENSMUSG00000024283 | Wac | 225131 | protein_coding | 1563.439635 | 0.032098476 | 0.042602294 | 0.753444776 | 0.451182685 | 0.680343771 | 0.032098476 |
| ENSMUSG00000034006 | Slc66a2 | 66943 | protein_coding | 1085.346494 | 0.032104755 | 0.051251863 | 0.626411473 | 0.531045086 | 0.73971984 | 0.032104755 |
| ENSMUSG00000053286 | Trmt1l | 98685 | protein_coding | 705.5149855 | -0.032115563 | 0.077848025 | -0.412541785 | 0.679942361 | 0.838143427 | 0.032115563 |
| ENSMUSG00000022913 | Psmg1 | 56088 | protein_coding | 280.7933793 | 0.032121836 | 0.086250546 | 0.372424732 | 0.709576643 | 0.856166251 | 0.032121836 |
| ENSMUSG00000072964 | Bhlhb9 | 70237 | protein_coding | 39.12667142 | 0.032150894 | 0.185551549 | 0.173272032 | 0.862437607 | 0.937787345 | 0.032150894 |
| ENSMUSG00000026721 | Rabgap1l | 29809 | protein_coding | 2010.27851 | -0.032177866 | 0.060522113 | -0.531671222 | 0.594953727 | 0.782189365 | 0.032177866 |
| ENSMUSG00000038812 | Trmt112 | 67674 | protein_coding | 190.5110758 | -0.032178092 | 0.089370172 | -0.360054046 | 0.718806718 | 0.862386247 | 0.032178092 |
| ENSMUSG00000050555 | Hyls1 | 76832 | protein_coding | 134.7961318 | -0.032200159 | 0.106265671 | -0.303015629 | 0.761877951 | 0.887606361 | 0.032200159 |
| ENSMUSG00000019849 | Prep | 19072 | protein_coding | 1016.69387 | 0.032207345 | 0.063778915 | 0.504984203 | 0.613569931 | 0.794017195 | 0.032207345 |
| ENSMUSG00000070056 | Mfhas1 | 52065 | protein_coding | 1049.06245 | 0.032211281 | 0.073972045 | 0.435452081 | 0.663234314 | 0.827892753 | 0.032211281 |
| ENSMUSG00000025794 | Rpl14 | 67115 | protein_coding | 9148.604759 | -0.03223023 | 0.034875544 | -0.924149888 | 0.355408282 | 0.598108042 | 0.03223023 |
| ENSMUSG00000000326 | Comt | 12846 | protein_coding | 1400.479668 | -0.032234386 | 0.063093458 | -0.51089902 | 0.609421767 | 0.791668558 | 0.032234386 |
| ENSMUSG00000024293 | Esco1 | 77805 | protein_coding | 983.2510309 | 0.032241039 | 0.059946515 | 0.537830076 | 0.590694366 | 0.779660849 | 0.032241039 |
| ENSMUSG00000045140 | Pigw | 70325 | protein_coding | 146.2842894 | -0.032256007 | 0.128720438 | -0.25058963 | 0.8021314 | 0.910022729 | 0.032256007 |
| ENSMUSG00000039157 | Fam102a | 98952 | protein_coding | 2326.996263 | -0.03229166 | 0.085600743 | -0.377235741 | 0.705998421 | 0.853816014 | 0.03229166 |
| ENSMUSG00000087679 | Tmem250-ps | 109299 | protein_coding | 1433.828745 | 0.032306804 | 0.065184123 | 0.495623814 | 0.620159848 | 0.799146453 | 0.032306804 |
| ENSMUSG00000027589 | Pcmtd2 | 245867 | protein_coding | 1340.93978 | -0.032324288 | 0.049829712 | -0.648695052 | 0.516535503 | 0.729245212 | 0.032324288 |
| ENSMUSG00000022561 | Gpaa1 | 14731 | protein_coding | 1100.63087 | -0.032332473 | 0.084297682 | -0.383551151 | 0.701311152 | 0.851297936 | 0.032332473 |
| ENSMUSG00000034112 | Atp2c2 | 69047 | protein_coding | 1565.806238 | 0.032352086 | 0.087781918 | 0.368550682 | 0.712462661 | 0.858286796 | 0.032352086 |
| ENSMUSG00000020100 | Slc29a3 | 71279 | protein_coding | 179.8589067 | -0.032356183 | 0.110600414 | -0.292550284 | 0.769865919 | 0.891927276 | 0.032356183 |
| ENSMUSG00000077450 | Rab11b | 19326 | protein_coding | 1368.522575 | 0.03237332 | 0.042429214 | 0.762995997 | 0.445465783 | 0.676462024 | 0.03237332 |
| ENSMUSG00000005465 | Il27ra | 50931 | protein_coding | 13.00316677 | -0.03238044 | 0.377657448 | -0.085740239 | 0.931672914 | 0.970174281 | 0.03238044 |

| ensembl_gene_id | external_gene_name | entrezgene_id | gene_biotype | baseMean | log2FoldChange | lfcSE | stat | pvalue | padj | absLog2FC |
|---|---|---|---|---|---|---|---|---|---|---|
| ENSMUSG00000037343 | Taf2 | 319944 | protein_coding | 723.9394314 | -0.032453381 | 0.0627202 | -0.517431079 | 0.604855269 | 0.789053031 | 0.032453381 |
| ENSMUSG00000034639 | Setmar | 74729 | protein_coding | 95.08905651 | -0.032496598 | 0.141509216 | -0.229642979 | 0.818369206 | 0.918354815 | 0.032496598 |
| ENSMUSG00000062040 | Zfp27 | 22689 | protein_coding | 219.947347 | 0.032502773 | 0.083919716 | 0.387307946 | 0.698528244 | 0.849624028 | 0.032502773 |
| ENSMUSG00000000544 | Gpa33 | 59290 | protein_coding | 12096.52343 | -0.032509133 | 0.064320167 | -0.505426747 | 0.613259136 | 0.793911553 | 0.032509133 |
| ENSMUSG00000025504 | Eps8l2 | 98845 | protein_coding | 3175.745158 | -0.032511231 | 0.072368451 | -0.449245912 | 0.653254271 | 0.821648745 | 0.032511231 |
| ENSMUSG00000074923 | Pak6 | 214230 | protein_coding | 48.29214832 | -0.032516762 | 0.179226785 | -0.181428025 | 0.856031623 | 0.935604164 | 0.032516762 |
| ENSMUSG00000043991 | Pura | 19290 | protein_coding | 1469.209915 | -0.032528413 | 0.06205723 | -0.52416798 | 0.60016171 | 0.78574508 | 0.032528413 |
| ENSMUSG00000020280 | Pus10 | 74467 | protein_coding | 607.7788078 | -0.032558069 | 0.077031668 | -0.422658232 | 0.672544634 | 0.833278687 | 0.032558069 |
| ENSMUSG00000005981 | Trap1 | 68015 | protein_coding | 2447.158982 | -0.032558206 | 0.053821573 | -0.604928539 | 0.54522648 | 0.749029274 | 0.032558206 |
| ENSMUSG00000002661 | Alkbh7 | 66400 | protein_coding | 411.0756575 | -0.032565094 | 0.111404933 | -0.292312851 | 0.770047434 | 0.892032325 | 0.032565094 |
| ENSMUSG00000020124 | Usp15 | 14479 | protein_coding | 1860.231521 | 0.032570829 | 0.056591805 | 0.575539676 | 0.564926365 | 0.761555988 | 0.032570829 |
| ENSMUSG00000032602 | Slc25a20 | 57279 | protein_coding | 3438.772782 | -0.032581235 | 0.083319431 | -0.391040061 | 0.695767624 | 0.847823651 | 0.032581235 |
| ENSMUSG00000020766 | Galk1 | 14635 | protein_coding | 491.8889333 | -0.032606936 | 0.074022635 | -0.440499535 | 0.659575349 | 0.82591596 | 0.032606936 |
| ENSMUSG00000001415 | Smg5 | 229512 | protein_coding | 1589.893379 | 0.032624411 | 0.061847012 | 0.527501811 | 0.597845159 | 0.784048864 | 0.032624411 |
| ENSMUSG00000007892 | Rplp1 | 56040 | protein_coding | 18547.18147 | -0.032635291 | 0.075697198 | -0.431129443 | 0.666374253 | 0.829602333 | 0.032635291 |
| ENSMUSG00000025139 | Tollip | 54473 | protein_coding | 1520.36128 | -0.03263821 | 0.036846004 | -0.885800534 | 0.375725022 | 0.617288396 | 0.03263821 |
| ENSMUSG00000111080 | Gm20300 | 100504586 | lncRNA | 381.5290214 | 0.032640803 | 0.081720597 | 0.399419541 | 0.689584098 | 0.844292391 | 0.032640803 |
| ENSMUSG00000010751 | Tnfrsf22 | 79202 | protein_coding | 8.783716553 | -0.032656638 | 0.396767345 | -0.082306768 | 0.934402773 | 0.971511412 | 0.032656638 |
| ENSMUSG00000029650 | Slc46a3 | 71706 | protein_coding | 144.7742647 | -0.032674397 | 0.190392476 | -0.171616007 | 0.863739422 | 0.938113902 | 0.032674397 |
| ENSMUSG00000001065 | Zfp276 | 57247 | protein_coding | 520.8390278 | -0.032695753 | 0.070757289 | -0.462083184 | 0.644021669 | 0.815430435 | 0.032695753 |
| ENSMUSG00000006740 | Kif5b | 16573 | protein_coding | 6105.510191 | 0.032697984 | 0.045624203 | 0.716680651 | 0.473571167 | 0.697472342 | 0.032697984 |
| ENSMUSG00000028337 | Coro2a | 107684 | protein_coding | 6815.493113 | 0.032707664 | 0.054044278 | 0.60520123 | 0.545045299 | 0.749012844 | 0.032707664 |
| ENSMUSG00000021635 | Rad17 | 19356 | protein_coding | 454.5764683 | -0.032731103 | 0.059592229 | -0.549251197 | 0.582833074 | 0.774433683 | 0.032731103 |
| ENSMUSG00000018509 | Cenpv | 73139 | protein_coding | 698.1629178 | -0.032749546 | 0.066605876 | -0.491691534 | 0.622937424 | 0.800888833 | 0.032749546 |
| ENSMUSG00000066861 | Oas1g | 23960 | protein_coding | 286.7379323 | -0.032752489 | 0.101929133 | -0.321326079 | 0.747963295 | 0.880274689 | 0.032752489 |
| ENSMUSG00000034573 | Ptpn13 | 19249 | protein_coding | 708.0459808 | -0.032766183 | 0.072701936 | -0.450692022 | 0.652211533 | 0.820965922 | 0.032766183 |
| ENSMUSG00000003660 | Snrnp200 | 320632 | protein_coding | 3249.260086 | 0.032786663 | 0.048375788 | 0.677749439 | 0.497930574 | 0.715687493 | 0.032786663 |
| ENSMUSG00000032612 | Usp4 | 22258 | protein_coding | 2607.301184 | 0.0327883 | 0.040309317 | 0.813417413 | 0.415978779 | 0.653976386 | 0.0327883 |
| ENSMUSG00000019790 | Stxbp5 | 78808 | protein_coding | 901.4778849 | 0.032805698 | 0.048766915 | 0.672703979 | 0.501135636 | 0.718355879 | 0.032805698 |
| ENSMUSG00000061601 | Pclo | 26875 | protein_coding | 36.10964783 | -0.032834386 | 0.247689647 | -0.132562612 | 0.894539303 | 0.952855703 | 0.032834386 |
| ENSMUSG00000038451 | Spsb2 | 14794 | protein_coding | 370.3444943 | -0.032856701 | 0.078033714 | -0.421057766 | 0.6737129 | 0.834153989 | 0.032856701 |
| ENSMUSG00000003437 | Paf1 | 54624 | protein_coding | 737.114219 | 0.032870827 | 0.058256094 | 0.564247019 | 0.572586033 | 0.7673338 | 0.032870827 |
| ENSMUSG00000030697 | Ppp4c | 56420 | protein_coding | 2201.552871 | 0.032876592 | 0.037189274 | 0.884034242 | 0.376677728 | 0.618288216 | 0.032876592 |
| ENSMUSG00000030729 | Pgm2l1 | 70974 | protein_coding | 548.7572811 | -0.032882959 | 0.073641503 | -0.446527537 | 0.655216226 | 0.822565162 | 0.032882959 |
| ENSMUSG00000032563 | Mrpl3 | 94062 | protein_coding | 697.3093997 | 0.032899995 | 0.058068388 | 0.566573246 | 0.571004156 | 0.765987453 | 0.032899995 |
| ENSMUSG00000071256 | Zfp213 | 449521 | protein_coding | 280.8449593 | -0.032942664 | 0.080473675 | -0.409359509 | 0.68227585 | 0.839415825 | 0.032942664 |
| ENSMUSG00000031575 | Ash2l | 23808 | protein_coding | 1178.327719 | -0.032952938 | 0.042934194 | -0.767521998 | 0.44277122 | 0.673829523 | 0.032952938 |
| ENSMUSG00000062054 | Iah1 | 67732 | protein_coding | 742.9682862 | 0.032955928 | 0.053484921 | 0.616172325 | 0.537780792 | 0.744190049 | 0.032955928 |
| ENSMUSG00000027167 | Elp4 | 77766 | protein_coding | 292.6042972 | -0.032956461 | 0.086953204 | -0.379013763 | 0.704677644 | 0.853157579 | 0.032956461 |

| ensembl_gene_id | external_gene_name | entrezgene_id | gene_biotype | baseMean | log2FoldChange | lfcSE | stat | pvalue | padj | absLog2FC |
|---|---|---|---|---|---|---|---|---|---|---|
| ENSMUSG00000056209 | Npm3 | 18150 | protein_coding | 533.943988 | 0.032980884 | 0.100028925 | 0.329713469 | 0.741616475 | 0.876450894 | 0.032980884 |
| ENSMUSG00000008384 | Sertad1 | 55942 | protein_coding | 1303.60274 | 0.032983 | 0.070385289 | 0.468606446 | 0.639350969 | 0.812170498 | 0.032983 |
| ENSMUSG00000047248 | C2cd3 | 277939 | protein_coding | 671.7088057 | 0.032989153 | 0.069992115 | 0.471326701 | 0.637407451 | 0.810693313 | 0.032989153 |
| ENSMUSG00000026970 | Rbms1 | 56878 | protein_coding | 372.5361435 | 0.033022401 | 0.076428478 | 0.432069324 | 0.665691029 | 0.829315377 | 0.033022401 |
| ENSMUSG00000045327 | 6330549D23Rik | 229613 | transcribed_processed_pseudogene | 19.37355832 | 0.033023247 | 0.266602688 | 0.123866895 | 0.901420665 | 0.956441676 | 0.033023247 |
| ENSMUSG00000030279 | C2cd5 | 74741 | protein_coding | 633.8214575 | 0.033024057 | 0.050573001 | 0.652997766 | 0.513757719 | 0.727042009 | 0.033024057 |
| ENSMUSG00000029017 | Pmpcb | 73078 | protein_coding | 1822.086804 | 0.033025446 | 0.049424942 | 0.668193912 | 0.504009821 | 0.720199042 | 0.033025446 |
| ENSMUSG00000000827 | Tpd52l2 | 66314 | protein_coding | 1062.250297 | 0.033031005 | 0.065092309 | 0.507448662 | 0.611840049 | 0.793051316 | 0.033031005 |
| ENSMUSG00000034757 | Tmub2 | 72053 | protein_coding | 1049.302717 | 0.033067129 | 0.053052904 | 0.623285943 | 0.533096634 | 0.741142191 | 0.033067129 |
| ENSMUSG00000029869 | Ephb6 | 13848 | protein_coding | 239.5502097 | 0.033099113 | 0.310012163 | 0.106767142 | 0.914973716 | 0.962265867 | 0.033099113 |
| ENSMUSG00000030374 | Strn4 | 97387 | protein_coding | 1988.123836 | 0.033113354 | 0.043300492 | 0.764733912 | 0.444430008 | 0.675397921 | 0.033113354 |
| ENSMUSG00000024525 | Impa2 | 114663 | protein_coding | 535.9590256 | -0.033133777 | 0.073229306 | -0.452466081 | 0.650933251 | 0.820125043 | 0.033133777 |
| ENSMUSG00000026313 | Hdac4 | 208727 | protein_coding | 1129.251108 | -0.033140189 | 0.047422761 | -0.698824541 | 0.48466169 | 0.706263486 | 0.033140189 |
| ENSMUSG00000026626 | Ppp2r5a | 226849 | protein_coding | 2961.008038 | -0.033156065 | 0.045048769 | -0.73600381 | 0.461728379 | 0.68813359 | 0.033156065 |
| ENSMUSG00000024259 | Slc25a46 | 67453 | protein_coding | 1284.537706 | 0.033160981 | 0.045613945 | 0.726992162 | 0.467230757 | 0.692786125 | 0.033160981 |
| ENSMUSG00000047557 | Lxn | 17035 | protein_coding | 281.33328 | -0.033197897 | 0.077456367 | -0.428601263 | 0.668213427 | 0.830797774 | 0.033197897 |
| ENSMUSG00000020250 | Txnrd1 | 50493 | protein_coding | 5613.858694 | 0.033205148 | 0.04055631 | 0.818741852 | 0.412933713 | 0.650647971 | 0.033205148 |
| ENSMUSG00000042506 | Usp22 | 216825 | protein_coding | 1985.592904 | -0.033236301 | 0.055335939 | -0.600627766 | 0.54808794 | 0.750796946 | 0.033236301 |
| ENSMUSG00000072825 | Cep170b | 217882 | protein_coding | 4076.158104 | -0.033238658 | 0.058643144 | -0.566795288 | 0.570853273 | 0.765902723 | 0.033238658 |
| ENSMUSG00000038126 | Mphosph9 | 269702 | protein_coding | 238.1242875 | 0.033255569 | 0.088198238 | 0.377054794 | 0.706132885 | 0.853850569 | 0.033255569 |
| ENSMUSG00000076431 | Sox4 | 20677 | protein_coding | 1219.680067 | -0.033256984 | 0.067656245 | -0.491558233 | 0.623031676 | 0.800888833 | 0.033256984 |
| ENSMUSG00000018446 | C1qbp | 12261 | protein_coding | 1209.494532 | -0.033264997 | 0.076754877 | -0.433392616 | 0.664729564 | 0.828854703 | 0.033264997 |
| ENSMUSG00000026632 | Tatdn3 | 68972 | protein_coding | 104.2147283 | -0.033283615 | 0.141837335 | -0.234660464 | 0.814472276 | 0.916470201 | 0.033283615 |
| ENSMUSG00000053819 | Camk2d | 108058 | protein_coding | 2256.711103 | 0.03328644 | 0.0341428 | 0.974918294 | 0.329600791 | 0.57340582 | 0.03328644 |
| ENSMUSG00000022553 | Maf1 | 68877 | protein_coding | 1630.492654 | 0.033354131 | 0.042888032 | 0.77770252 | 0.436744406 | 0.669904607 | 0.033354131 |
| ENSMUSG00000030188 | Magohb | 66441 | protein_coding | 174.005029 | -0.03335796 | 0.101493069 | -0.328672295 | 0.742403397 | 0.876866822 | 0.03335796 |
| ENSMUSG00000028228 | Cpne3 | 70568 | protein_coding | 3337.131398 | -0.033364447 | 0.041725514 | -0.799617418 | 0.423932493 | 0.659840391 | 0.033364447 |
| ENSMUSG00000018362 | Kpna2 | 16647 | protein_coding | 2203.822756 | 0.033401994 | 0.057267526 | 0.58326239 | 0.559716677 | 0.758094025 | 0.033401994 |
| ENSMUSG00000053730 | Tmem39b | 230770 | protein_coding | 119.616781 | -0.033427375 | 0.163508735 | -0.204437853 | 0.838011352 | 0.92807369 | 0.033427375 |
| ENSMUSG00000058600 | Rpl30 | 19946 | protein_coding | 3708.147471 | -0.033430158 | 0.044195286 | -0.756419101 | 0.449397959 | 0.679246742 | 0.033430158 |
| ENSMUSG00000003458 | Ncstn | 59287 | protein_coding | 3129.971349 | -0.033434093 | 0.03552596 | -0.941117211 | 0.346644796 | 0.588828801 | 0.033434093 |
| ENSMUSG00000023004 | Tuba1b | 22143 | protein_coding | 891.3776986 | 0.033442759 | 0.10415276 | 0.32109335 | 0.748139649 | 0.880353749 | 0.033442759 |
| ENSMUSG00000027774 | Gfm1 | 28030 | protein_coding | 1394.456509 | 0.033485968 | 0.081189414 | 0.41244254 | 0.680015088 | 0.838147754 | 0.033485968 |
| ENSMUSG00000026150 | Mff | 75734 | protein_coding | 1216.420794 | 0.033487908 | 0.053974088 | 0.620444175 | 0.534965397 | 0.741885541 | 0.033487908 |
| ENSMUSG00000032382 | Snx1 | 56440 | protein_coding | 2804.487591 | 0.033510062 | 0.030949589 | 1.082730418 | 0.278928098 | 0.52178462 | 0.033510062 |
| ENSMUSG00000030189 | Ybx3 | 56449 | protein_coding | 3865.706083 | -0.033518503 | 0.056920146 | -0.58886889 | 0.555949228 | 0.755288004 | 0.033518503 |
| ENSMUSG00000085704 | 4921531C22Rik | 66730 | lncRNA | 111.2646913 | 0.033520251 | 0.152704872 | 0.219510031 | 0.826252767 | 0.922846345 | 0.033520251 |
| ENSMUSG00000028540 | Dph2 | 67728 | protein_coding | 150.8488786 | -0.033546047 | 0.099314886 | -0.337774617 | 0.73553304 | 0.872889311 | 0.033546047 |
| ENSMUSG00000007891 | Ctsd | 13033 | protein_coding | 3631.420744 | -0.033546272 | 0.049567245 | -0.676783078 | 0.498543596 | 0.716014372 | 0.033546272 |

| ensembl_gene_id | external_gene_name | entrezgene_id | gene_biotype | baseMean | log2FoldChange | lfcSE | stat | pvalue | padj | absLog2FC |
|---|---|---|---|---|---|---|---|---|---|---|
| ENSMUSG00000074754 | Smim26 | 228715 | protein_coding | 154.4465976 | -0.033561887 | 0.096612444 | -0.347386789 | 0.728300757 | 0.868029437 | 0.033561887 |
| ENSMUSG00000037286 | Stag1 | 20842 | protein_coding | 1220.898842 | 0.033623616 | 0.062531034 | 0.537710856 | 0.590776684 | 0.779660849 | 0.033623616 |
| ENSMUSG00000034024 | Cct2 | 12461 | protein_coding | 4259.724947 | -0.033648567 | 0.04402955 | -0.764226913 | 0.444732031 | 0.675687484 | 0.033648567 |
| ENSMUSG00000002763 | Pex6 | 224824 | protein_coding | 1497.42991 | 0.033650251 | 0.090634023 | 0.371276145 | 0.710431864 | 0.856611363 | 0.033650251 |
| ENSMUSG00000035735 | Dagla | 269060 | protein_coding | 307.4205265 | -0.03366864 | 0.095842416 | -0.351291647 | 0.725369561 | 0.865987203 | 0.03366864 |
| ENSMUSG00000118537 | Shld3 | 113002583 | protein_coding | 54.16900461 | -0.033671352 | 0.173012316 | -0.194618236 | 0.845691826 | 0.93082665 | 0.033671352 |
| ENSMUSG00000028010 | Gar1 | 68147 | protein_coding | 151.7030258 | -0.033714383 | 0.107566395 | -0.313428581 | 0.753955073 | 0.883093891 | 0.033714383 |
| ENSMUSG00000001517 | Foxm1 | 14235 | protein_coding | 241.1956073 | -0.033716795 | 0.121276095 | -0.278016829 | 0.78099944 | 0.898311362 | 0.033716795 |
| ENSMUSG00000029276 | Glmn | 170823 | protein_coding | 92.99304605 | -0.033723886 | 0.145316075 | -0.232072649 | 0.816481588 | 0.917421069 | 0.033723886 |
| ENSMUSG00000028156 | Eif4e | 13684 | protein_coding | 955.2405837 | 0.033739781 | 0.064385268 | 0.524029514 | 0.600258012 | 0.785807367 | 0.033739781 |
| ENSMUSG00000075467 | Dnlz | 52838 | protein_coding | 545.1276186 | -0.033794993 | 0.0747448 | -0.45213839 | 0.651169289 | 0.820358341 | 0.033794993 |
| ENSMUSG00000030801 | Kat8 | 67773 | protein_coding | 342.2406773 | 0.033839713 | 0.069723543 | 0.485341274 | 0.627434282 | 0.80351248 | 0.033839713 |
| ENSMUSG00000023243 | Kcnk5 | 16529 | protein_coding | 1936.287645 | -0.033851234 | 0.100091004 | -0.338204561 | 0.735209039 | 0.87277999 | 0.033851234 |
| ENSMUSG00000022749 | Tbc1d23 | 67581 | protein_coding | 774.180313 | -0.033879551 | 0.075796788 | -0.446978712 | 0.65489043 | 0.822476285 | 0.033879551 |
| ENSMUSG00000005698 | Ctcf | 13018 | protein_coding | 1621.184086 | -0.033880545 | 0.040010009 | -0.84680174 | 0.397105638 | 0.636592707 | 0.033880545 |
| ENSMUSG00000034442 | Trmt5 | 76357 | protein_coding | 165.3056458 | -0.033884836 | 0.100476241 | -0.337242272 | 0.735934273 | 0.87299831 | 0.033884836 |
| ENSMUSG00000022656 | Nectin3 | 58998 | protein_coding | 2984.09497 | 0.033886341 | 0.048050576 | 0.70522238 | 0.480671856 | 0.702983134 | 0.033886341 |
| ENSMUSG00000021540 | Smad5 | 17129 | protein_coding | 1033.788126 | -0.033898943 | 0.05684179 | -0.5963736 | 0.550925673 | 0.752092411 | 0.033898943 |
| ENSMUSG00000028085 | Gatb | 229487 | protein_coding | 394.6361142 | 0.033903019 | 0.069507704 | 0.487759157 | 0.625720444 | 0.8024647 | 0.033903019 |
| ENSMUSG00000012428 | Steap4 | 117167 | protein_coding | 30.0583112 | -0.033918623 | 0.212520465 | -0.159601679 | 0.873194857 | 0.942905245 | 0.033918623 |
| ENSMUSG00000047747 | Rnf150 | 330812 | protein_coding | 300.8535454 | 0.033952305 | 0.113774089 | 0.298418604 | 0.765383692 | 0.889849511 | 0.033952305 |
| ENSMUSG00000059479 | B3gnt8 | 232984 | protein_coding | 27.41918809 | 0.033956001 | 0.282025672 | 0.120400391 | 0.904165981 | 0.95804762 | 0.033956001 |
| ENSMUSG00000021488 | Nsd1 | 18193 | protein_coding | 4882.28284 | -0.033965273 | 0.04827118 | -0.703634617 | 0.481660347 | 0.7037268 | 0.033965273 |
| ENSMUSG00000078765 | U2af1l4 | 233073 | protein_coding | 6.672626584 | 0.033980415 | 0.464245174 | 0.073194977 | 0.941650964 | 0.974493399 | 0.033980415 |
| ENSMUSG00000028318 | Polr1e | 64424 | protein_coding | 174.4327591 | 0.0339843 | 0.102054247 | 0.333002308 | 0.739132541 | 0.87479925 | 0.0339843 |
| ENSMUSG00000038861 | Pi4kb | 107650 | protein_coding | 1051.387276 | 0.034005003 | 0.051945478 | 0.654628739 | 0.512706816 | 0.726301064 | 0.034005003 |
| ENSMUSG00000025991 | Cps1 | 227231 | protein_coding | 1391.834556 | 0.034030406 | 0.129889316 | 0.261995422 | 0.793324971 | 0.904038427 | 0.034030406 |
| ENSMUSG00000021982 | Cdadc1 | 71891 | protein_coding | 453.338674 | -0.0340309 | 0.074916648 | -0.454250175 | 0.649648773 | 0.819274771 | 0.0340309 |
| ENSMUSG00000054428 | Atpif1 | 11983 | protein_coding | 3044.944896 | 0.034056279 | 0.038755374 | 0.878749848 | 0.379536925 | 0.62053716 | 0.034056279 |
| ENSMUSG00000024491 | Rbm27 | 225432 | protein_coding | 1003.609304 | -0.034058713 | 0.050400793 | -0.67575747 | 0.499194641 | 0.716301449 | 0.034058713 |
| ENSMUSG00000005699 | Pard6a | 56513 | protein_coding | 94.77358874 | 0.034063765 | 0.14529142 | 0.234451316 | 0.814634624 | 0.916534992 | 0.034063765 |
| ENSMUSG00000026213 | Stk11ip | 71728 | protein_coding | 467.0426554 | 0.034071226 | 0.061542635 | 0.553619874 | 0.579839022 | 0.771934807 | 0.034071226 |
| ENSMUSG00000021483 | Cdk20 | 105278 | protein_coding | 51.241847 | 0.034082984 | 0.178650337 | 0.19078041 | 0.848697637 | 0.931678839 | 0.034082984 |
| ENSMUSG00000066151 | Fkbp15 | 338355 | protein_coding | 1279.368466 | 0.034102147 | 0.043379574 | 0.786133747 | 0.431789135 | 0.666296372 | 0.034102147 |
| ENSMUSG00000089847 | Timm10b | 14356 | protein_coding | 170.3618827 | -0.034121145 | 0.10214206 | -0.334055777 | 0.73833747 | 0.874754933 | 0.034121145 |
| ENSMUSG00000029447 | Cct6a | 12466 | protein_coding | 2883.975525 | 0.034121246 | 0.055648299 | 0.613158823 | 0.539771329 | 0.745625688 | 0.034121246 |
| ENSMUSG00000047539 | Fbxo28 | 67948 | protein_coding | 738.7194024 | 0.034134234 | 0.061318461 | 0.556671407 | 0.577751956 | 0.770224502 | 0.034134234 |
| ENSMUSG00000062202 | Btbd9 | 224671 | protein_coding | 847.0603454 | -0.034147701 | 0.051720273 | -0.660238216 | 0.509100971 | 0.724201408 | 0.034147701 |
| ENSMUSG00000024844 | Banf1 | 23825 | protein_coding | 1048.459829 | -0.03414904 | 0.079914935 | -0.427317371 | 0.669148182 | 0.831111145 | 0.03414904 |

| ensembl_gene_id | external_gene_name | entrezgene_id | gene_biotype | baseMean | log2FoldChange | lfcSE | stat | pvalue | padj | absLog2FC |
|---|---|---|---|---|---|---|---|---|---|---|
| ENSMUSG00000041782 | Lad1 | 16763 | protein_coding | 8295.327296 | 0.034173644 | 0.046959348 | 0.727728257 | 0.466779949 | 0.692404394 | 0.034173644 |
| ENSMUSG00000043881 | Kbtbd7 | 211255 | protein_coding | 353.5000494 | -0.034191505 | 0.070969928 | -0.481774552 | 0.629966109 | 0.805574037 | 0.034191505 |
| ENSMUSG00000048410 | Zfp407 | 240476 | protein_coding | 599.3789822 | 0.034192955 | 0.071113225 | 0.480824144 | 0.630641489 | 0.805987676 | 0.034192955 |
| ENSMUSG00000019797 | Mtres1 | 67851 | protein_coding | 533.1165315 | 0.03420632 | 0.075835274 | 0.451060807 | 0.651945725 | 0.820951722 | 0.03420632 |
| ENSMUSG00000020770 | Unk | 217331 | protein_coding | 357.360786 | 0.03422936 | 0.076154189 | 0.449474426 | 0.653089452 | 0.821622092 | 0.03422936 |
| ENSMUSG00000004633 | Chn2 | 69993 | protein_coding | 544.9647269 | -0.03428856 | 0.106965401 | -0.320557488 | 0.748545758 | 0.880475742 | 0.03428856 |
| ENSMUSG00000022752 | Tomm70a | 28185 | protein_coding | 2605.065081 | 0.034299949 | 0.043706776 | 0.784774168 | 0.432585988 | 0.666799776 | 0.034299949 |
| ENSMUSG00000002107 | Celf2 | 14007 | protein_coding | 118.2872183 | 0.034306606 | 0.111321486 | 0.308175966 | 0.75794844 | 0.885371178 | 0.034306606 |
| ENSMUSG00000056201 | Cfl1 | 12631 | protein_coding | 6112.452355 | 0.034313199 | 0.053768581 | 0.638164484 | 0.523366614 | 0.734516873 | 0.034313199 |
| ENSMUSG00000031148 | Gpkow | 209416 | protein_coding | 474.2947446 | -0.034335865 | 0.071614739 | -0.479452486 | 0.631616762 | 0.806746273 | 0.034335865 |
| ENSMUSG00000040997 | Abhd4 | 105501 | protein_coding | 1206.927206 | -0.034408674 | 0.057952055 | -0.59374381 | 0.552683476 | 0.753179518 | 0.034408674 |
| ENSMUSG00000046314 | Stxbp6 | 217517 | protein_coding | 494.0099391 | 0.034420753 | 0.06771331 | 0.50833068 | 0.611221458 | 0.792823007 | 0.034420753 |
| ENSMUSG00000031314 | Taf1 | 270627 | protein_coding | 1607.141629 | -0.034431082 | 0.050352949 | -0.683794747 | 0.494104784 | 0.713490595 | 0.034431082 |
| ENSMUSG00000035642 | Aamdc | 66273 | protein_coding | 716.0217684 | 0.034431211 | 0.061510315 | 0.559763206 | 0.575640964 | 0.76862707 | 0.034431211 |
| ENSMUSG00000045165 | AI467606 | 101602 | protein_coding | 51.28644042 | 0.034460916 | 0.177345807 | 0.194314803 | 0.845929395 | 0.93086312 | 0.034460916 |
| ENSMUSG00000035392 | Dennd1a | 227801 | protein_coding | 265.286984 | 0.034463947 | 0.074728544 | 0.461188527 | 0.644663352 | 0.81580845 | 0.034463947 |
| ENSMUSG00000005982 | Naa60 | 74763 | protein_coding | 655.9005934 | 0.03451585 | 0.060364509 | 0.571790457 | 0.567463943 | 0.762827428 | 0.03451585 |
| ENSMUSG00000028463 | Car9 | 230099 | protein_coding | 840.2259086 | 0.034534329 | 0.057585833 | 0.599701817 | 0.548704977 | 0.751121736 | 0.034534329 |
| ENSMUSG00000045867 | Cradd | 12905 | protein_coding | 151.9337758 | -0.03453577 | 0.10089405 | -0.342297389 | 0.7321271 | 0.870787945 | 0.03453577 |
| ENSMUSG00000027801 | Tm4sf4 | 229302 | protein_coding | 28.34123321 | 0.034553259 | 0.341824683 | 0.101084741 | 0.919483191 | 0.964769513 | 0.034553259 |
| ENSMUSG00000024797 | Vps51 | 68505 | protein_coding | 919.4214516 | -0.034585515 | 0.069973996 | -0.494262396 | 0.621120879 | 0.800065209 | 0.034585515 |
| ENSMUSG00000036644 | Tbc1d9b | 76795 | protein_coding | 2074.584847 | 0.034589051 | 0.047846538 | 0.722916478 | 0.469731199 | 0.694009261 | 0.034589051 |
| ENSMUSG00000033285 | Wdr3 | 269470 | protein_coding | 530.2752719 | -0.03459532 | 0.097303524 | -0.355540261 | 0.722184888 | 0.864247815 | 0.03459532 |
| ENSMUSG00000064068 | Mtx1 | 17827 | protein_coding | 655.372335 | 0.03459677 | 0.068216006 | 0.507164998 | 0.612039052 | 0.793245503 | 0.03459677 |
| ENSMUSG00000026889 | Rbm18 | 67889 | protein_coding | 427.1124889 | 0.034597691 | 0.082712026 | 0.418290937 | 0.675734415 | 0.835024575 | 0.034597691 |
| ENSMUSG00000025421 | Hdhd2 | 76987 | protein_coding | 252.4123846 | -0.034603153 | 0.085397294 | -0.405201987 | 0.685329045 | 0.841671981 | 0.034603153 |
| ENSMUSG00000029176 | Anapc4 | 52206 | protein_coding | 759.9966304 | 0.034620216 | 0.078707353 | 0.439859993 | 0.660038513 | 0.826041967 | 0.034620216 |
| ENSMUSG00000096370 | Gm21992 | 102902673 | protein_coding | 84.4004912 | 0.034626451 | 0.147054491 | 0.235466804 | 0.813846441 | 0.916289542 | 0.034626451 |
| ENSMUSG00000022899 | Slc15a2 | 57738 | protein_coding | 149.6970645 | -0.034651535 | 0.708028148 | -0.048940901 | 0.960966394 | 0.984093538 | 0.034651535 |
| ENSMUSG00000007613 | Tgfbr1 | 21812 | protein_coding | 1201.2978 | 0.034673774 | 0.04681863 | 0.740597804 | 0.45893734 | 0.685946212 | 0.034673774 |
| ENSMUSG00000033396 | Spg11 | 214585 | protein_coding | 1362.165188 | -0.034689225 | 0.038182318 | -0.908515425 | 0.363605968 | 0.606076959 | 0.034689225 |
| ENSMUSG00000025173 | Wdr45b | 66840 | protein_coding | 896.7430825 | 0.034698828 | 0.053055313 | 0.65401231 | 0.513103875 | 0.726547391 | 0.034698828 |
| ENSMUSG00000049551 | Fzd9 | 14371 | protein_coding | 25.84064787 | 0.034706009 | 0.274150637 | 0.126594668 | 0.899261217 | 0.955338492 | 0.034706009 |
| ENSMUSG00000060475 | Wtap | 60532 | protein_coding | 1344.92652 | -0.034713955 | 0.049406193 | -0.70262356 | 0.482290376 | 0.704519646 | 0.034713955 |
| ENSMUSG00000032398 | Snapc5 | 330959 | protein_coding | 816.5739986 | -0.034722927 | 0.052561752 | -0.660612051 | 0.508861138 | 0.724051842 | 0.034722927 |
| ENSMUSG00000002820 | Atg4d | 235040 | protein_coding | 2118.531685 | 0.03478408 | 0.065818006 | 0.528488806 | 0.597160114 | 0.783469018 | 0.03478408 |
| ENSMUSG00000037857 | Nufip2 | 68564 | protein_coding | 2016.247857 | -0.034802909 | 0.057279375 | -0.607599324 | 0.543453249 | 0.74831545 | 0.034802909 |
| ENSMUSG00000027134 | Lpcat4 | 99010 | protein_coding | 150.9581943 | 0.034809425 | 0.176943382 | 0.196726347 | 0.844041691 | 0.930210801 | 0.034809425 |
| ENSMUSG00000090330 | 9130221H12Rik | 77124 | lncRNA | 56.49872484 | 0.03481705 | 0.181750221 | 0.191565382 | 0.848082662 | 0.93136051 | 0.03481705 |

| ensembl_gene_id | external_gene_name | entrezgene_id | gene_biotype | baseMean | log2FoldChange | lfcSE | stat | pvalue | padj | absLog2FC |
|---|---|---|---|---|---|---|---|---|---|---|
| ENSMUSG00000014907 | Naf1 | 234344 | protein_coding | 157.0251814 | 0.03481986 | 0.114285923 | 0.304673219 | 0.760615049 | 0.886815999 | 0.03481986 |
| ENSMUSG00000027997 | Casp6 | 12368 | protein_coding | 584.5516644 | 0.034830033 | 0.054171363 | 0.642960257 | 0.520249889 | 0.731862348 | 0.034830033 |
| ENSMUSG00000016018 | Mtrex | 72198 | protein_coding | 979.0412492 | 0.034845388 | 0.065067347 | 0.535528023 | 0.592284785 | 0.780262005 | 0.034845388 |
| ENSMUSG00000042419 | Nfkbil1 | 18038 | protein_coding | 318.5673814 | 0.034865424 | 0.119373548 | 0.29206993 | 0.770233158 | 0.892032325 | 0.034865424 |
| ENSMUSG00000020114 | Cand1 | 71902 | protein_coding | 2442.001353 | -0.034867378 | 0.044344066 | -0.786291861 | 0.43169652 | 0.666254386 | 0.034867378 |
| ENSMUSG00000009681 | Bcr | 110279 | protein_coding | 913.0732894 | -0.034870618 | 0.069945051 | -0.498543028 | 0.618101349 | 0.797513438 | 0.034870618 |
| ENSMUSG00000069020 | Urm1 | 68205 | protein_coding | 574.1554875 | 0.034871375 | 0.084117939 | 0.414553364 | 0.678468894 | 0.836920218 | 0.034871375 |
| ENSMUSG00000020629 | Adi1 | 104923 | protein_coding | 388.5054058 | -0.034881861 | 0.0775504 | -0.449796018 | 0.652857529 | 0.82145077 | 0.034881861 |
| ENSMUSG00000048170 | Mcmbp | 210711 | protein_coding | 1697.620784 | -0.034919177 | 0.048614502 | -0.71828726 | 0.472580184 | 0.696772776 | 0.034919177 |
| ENSMUSG00000021339 | Mrs2 | 380836 | protein_coding | 1582.366579 | 0.034941305 | 0.055571805 | 0.628759583 | 0.529506464 | 0.738762068 | 0.034941305 |
| ENSMUSG00000028619 | Tceanc2 | 66526 | protein_coding | 433.8040275 | -0.034944652 | 0.067483506 | -0.51782509 | 0.604580311 | 0.788968011 | 0.034944652 |
| ENSMUSG00000048310 | Pskh1 | 244631 | protein_coding | 774.9072313 | 0.034952185 | 0.06305501 | 0.554312581 | 0.579364942 | 0.771609416 | 0.034952185 |
| ENSMUSG00000022295 | Atp6v1c1 | 66335 | protein_coding | 857.2139171 | 0.034982169 | 0.048799929 | 0.716848775 | 0.473467412 | 0.697472342 | 0.034982169 |
| ENSMUSG00000044533 | Rps2 | 16898 | protein_coding | 17828.37307 | 0.034990821 | 0.051005721 | 0.686017578 | 0.492702024 | 0.712161455 | 0.034990821 |
| ENSMUSG00000061959 | Ces1e | 13897 | protein_coding | 590.8899155 | -0.035022184 | 0.093064184 | -0.376322904 | 0.706676854 | 0.854251145 | 0.035022184 |
| ENSMUSG00000027635 | Dsn1 | 66934 | protein_coding | 151.1415341 | 0.03506237 | 0.108563293 | 0.322967083 | 0.746720171 | 0.879838986 | 0.03506237 |
| ENSMUSG00000026171 | Rnf25 | 57751 | protein_coding | 403.3705823 | 0.035066012 | 0.064772962 | 0.541368045 | 0.588253928 | 0.778114427 | 0.035066012 |
| ENSMUSG00000015337 | Endog | 13804 | protein_coding | 308.7362877 | -0.035140828 | 0.105256294 | -0.333859631 | 0.738485483 | 0.874754933 | 0.035140828 |
| ENSMUSG00000024997 | Prdx3 | 11757 | protein_coding | 2769.810415 | 0.035159369 | 0.042147882 | 0.834190651 | 0.404173558 | 0.643509426 | 0.035159369 |
| ENSMUSG00000054514 | Atad3aos | 70448 | lncRNA | 11.19753798 | -0.035176586 | 0.381971618 | -0.092092146 | 0.926624829 | 0.967854832 | 0.035176586 |
| ENSMUSG00000041362 | Shtn1 | 71653 | protein_coding | 1117.573472 | 0.035176756 | 0.05666323 | 0.620803928 | 0.534728638 | 0.741748559 | 0.035176756 |
| ENSMUSG00000079283 | 2310009B15Rik | 69549 | protein_coding | 140.8827145 | -0.035179593 | 0.12339916 | -0.285087786 | 0.775576887 | 0.895007403 | 0.035179593 |
| ENSMUSG00000028990 | Lzic | 69151 | protein_coding | 544.67338 | -0.035183496 | 0.079854253 | -0.440596397 | 0.659505212 | 0.82591596 | 0.035183496 |
| ENSMUSG00000061613 | U2af1 | 108121 | protein_coding | 805.1214888 | 0.035196856 | 0.065735342 | 0.535432759 | 0.592350643 | 0.780262005 | 0.035196856 |
| ENSMUSG00000023755 | Rhebl1 | 69159 | protein_coding | 36.11159317 | -0.035214727 | 0.227781591 | -0.154598651 | 0.877137734 | 0.945176551 | 0.035214727 |
| ENSMUSG00000074639 | Rdh16f2 | 216454 | protein_coding | 6.838450491 | -0.035216912 | 0.60285625 | -0.058416765 | 0.953416661 | 0.981022852 | 0.035216912 |
| ENSMUSG00000017831 | Rab5a | 271457 | protein_coding | 1614.439472 | 0.03522377 | 0.046524471 | 0.757101989 | 0.448988762 | 0.679126034 | 0.03522377 |
| ENSMUSG00000007833 | Aldh16a1 | 69748 | protein_coding | 2202.119707 | 0.035252042 | 0.084802001 | 0.415698237 | 0.677630831 | 0.836508911 | 0.035252042 |
| ENSMUSG00000044968 | Napepld | 242864 | protein_coding | 316.9207128 | 0.035254108 | 0.091599394 | 0.384872725 | 0.700331714 | 0.850619214 | 0.035254108 |
| ENSMUSG00000033107 | Rnf125 | 67664 | protein_coding | 163.9680788 | -0.035269747 | 0.14372738 | -0.245393376 | 0.806151857 | 0.912178372 | 0.035269747 |
| ENSMUSG00000031792 | Usb1 | 101985 | protein_coding | 337.3823601 | 0.035274943 | 0.070339458 | 0.501495796 | 0.616022236 | 0.795599668 | 0.035274943 |
| ENSMUSG00000028126 | Pip5k1a | 18720 | protein_coding | 455.9606543 | -0.035300327 | 0.072135105 | -0.489364041 | 0.624583989 | 0.801920996 | 0.035300327 |
| ENSMUSG00000030107 | Usp18 | 24110 | protein_coding | 47.49630432 | -0.035317872 | 0.314177958 | -0.11241359 | 0.910495481 | 0.959960814 | 0.035317872 |
| ENSMUSG00000037953 | A4gnt | 333424 | protein_coding | 394.5299214 | 0.035335478 | 0.112223528 | 0.314866936 | 0.752862687 | 0.882492633 | 0.035335478 |
| ENSMUSG00000022949 | Clic6 | 209195 | protein_coding | 1017.277705 | 0.035337185 | 0.082884673 | 0.426341608 | 0.669858944 | 0.831606417 | 0.035337185 |
| ENSMUSG00000024073 | Birc6 | 12211 | protein_coding | 5017.43343 | 0.035370156 | 0.048768714 | 0.725263242 | 0.468290551 | 0.693047027 | 0.035370156 |
| ENSMUSG00000053253 | Ndfip2 | 76273 | protein_coding | 3513.652354 | -0.035377422 | 0.061382258 | -0.576346051 | 0.564381302 | 0.76133645 | 0.035377422 |
| ENSMUSG00000064127 | Med14 | 26896 | protein_coding | 976.7164646 | 0.03541227 | 0.058945998 | 0.600757837 | 0.54800129 | 0.750796946 | 0.03541227 |
| ENSMUSG00000048661 | Lemd3 | 380664 | protein_coding | 571.364804 | -0.035442154 | 0.067316557 | -0.526499802 | 0.59854099 | 0.784685171 | 0.035442154 |

Supplementary Table 1

| ensembl_gene_id | external_gene_name | entrezgene_id | gene_biotype | baseMean | log2FoldChange | lfcSE | stat | pvalue | padj | absLog2FC |
|---|---|---|---|---|---|---|---|---|---|---|
| ENSMUSG00000022000 | Zc3h13 | 67302 | protein_coding | 481.5171336 | 0.035470408 | 0.066249142 | 0.535409319 | 0.592366848 | 0.780262005 | 0.035470408 |
| ENSMUSG00000067336 | Bmpr2 | 12168 | protein_coding | 1225.749269 | -0.035483842 | 0.080281215 | -0.441994335 | 0.658493306 | 0.825261698 | 0.035483842 |
| ENSMUSG00000021639 | Gtf2h2 | 23894 | protein_coding | 303.8567959 | -0.03549478 | 0.073719714 | -0.481482881 | 0.630173344 | 0.805740546 | 0.03549478 |
| ENSMUSG00000022682 | Rrn3 | 106298 | protein_coding | 879.569597 | 0.035496475 | 0.051595645 | 0.687974248 | 0.491468998 | 0.711036201 | 0.035496475 |
| ENSMUSG00000037031 | Tspan15 | 70423 | protein_coding | 3363.560987 | -0.035504325 | 0.047536307 | -0.746888584 | 0.455130818 | 0.683035055 | 0.035504325 |
| ENSMUSG00000030029 | Lrig1 | 16206 | protein_coding | 3861.244413 | -0.035539391 | 0.0507356 | -0.700482323 | 0.483626141 | 0.705703834 | 0.035539391 |
| ENSMUSG00000055373 | Fut9 | 14348 | protein_coding | 346.5881305 | 0.03554098 | 0.169262439 | 0.209975588 | 0.833686727 | 0.926340413 | 0.03554098 |
| ENSMUSG00000030884 | Uqcrc2 | 67003 | protein_coding | 8167.318044 | 0.035564401 | 0.053298213 | 0.667271918 | 0.504598461 | 0.720720873 | 0.035564401 |
| ENSMUSG00000029851 | Tcaf2 | 232748 | protein_coding | 765.3230303 | 0.035579419 | 0.058822585 | 0.604859835 | 0.545272132 | 0.749029274 | 0.035579419 |
| ENSMUSG00000028064 | Sema4a | 20351 | protein_coding | 3864.545863 | -0.035587764 | 0.052633288 | -0.676145558 | 0.498948234 | 0.716075302 | 0.035587764 |
| ENSMUSG00000029404 | Arl6ip4 | 65105 | protein_coding | 567.4205872 | -0.035590913 | 0.054636591 | -0.651411663 | 0.514780784 | 0.727432033 | 0.035590913 |
| ENSMUSG00000022234 | Cct5 | 12465 | protein_coding | 5079.840118 | 0.035602458 | 0.051071708 | 0.697107251 | 0.485735677 | 0.706927105 | 0.035602458 |
| ENSMUSG00000020743 | Mif4gd | 69674 | protein_coding | 915.2798807 | -0.035611644 | 0.053231367 | -0.66899736 | 0.503497162 | 0.719946838 | 0.035611644 |
| ENSMUSG00000050107 | Haspin | 14841 | protein_coding | 171.7238119 | 0.035635709 | 0.156826335 | 0.227230391 | 0.820244597 | 0.919448845 | 0.035635709 |
| ENSMUSG00000033054 | Npat | 244879 | protein_coding | 581.2925811 | -0.035663684 | 0.070953277 | -0.50263618 | 0.615220087 | 0.795450521 | 0.035663684 |
| ENSMUSG00000024079 | Eif2ak2 | 19106 | protein_coding | 982.1250039 | 0.03566993 | 0.073319895 | 0.486497287 | 0.626614628 | 0.80316437 | 0.03566993 |
| ENSMUSG00000027679 | Dnajc19 | 67713 | protein_coding | 504.4536459 | 0.03567288 | 0.063979645 | 0.557566089 | 0.577140718 | 0.769727784 | 0.03567288 |
| ENSMUSG00000030816 | Rnf40 | 233900 | protein_coding | 1470.842172 | 0.035702232 | 0.045810183 | 0.779351434 | 0.435772724 | 0.66917794 | 0.035702232 |
| ENSMUSG00000079414 | Gm11110 | 100169874 | lncRNA | 6.317098937 | -0.035730409 | 0.476918602 | -0.074919302 | 0.940278919 | 0.974024613 | 0.035730409 |
| ENSMUSG00000066798 | Zbtb6 | 241322 | protein_coding | 705.5477975 | -0.035748151 | 0.062050727 | -0.576111727 | 0.564539666 | 0.761422697 | 0.035748151 |
| ENSMUSG00000079508 | Apoo | 68316 | protein_coding | 297.1198592 | 0.035802527 | 0.082094616 | 0.436112968 | 0.662754769 | 0.827627866 | 0.035802527 |
| ENSMUSG00000045822 | Zswim3 | 67538 | protein_coding | 118.9946016 | 0.03583183 | 0.117729381 | 0.304357584 | 0.760855479 | 0.886968072 | 0.03583183 |
| ENSMUSG00000036120 | Rfxank | 19727 | protein_coding | 373.9757733 | 0.035833601 | 0.06833446 | 0.524385513 | 0.60001043 | 0.785648606 | 0.035833601 |
| ENSMUSG00000019842 | Traf3ip2 | 103213 | protein_coding | 576.9816695 | -0.035836933 | 0.065216281 | -0.549508994 | 0.582656194 | 0.774266351 | 0.035836933 |
| ENSMUSG00000020590 | Snx13 | 217463 | protein_coding | 3148.918735 | 0.035852045 | 0.033922635 | 1.056876784 | 0.290567825 | 0.53310436 | 0.035852045 |
| ENSMUSG00000028890 | Mtf1 | 17764 | protein_coding | 597.9529878 | -0.03585673 | 0.061685664 | -0.581281403 | 0.561050812 | 0.759206347 | 0.03585673 |
| ENSMUSG00000086401 | Gm15559 | 100504049 | lncRNA | 7.093594047 | 0.035874628 | 0.469739659 | 0.076371299 | 0.939123703 | 0.973407543 | 0.035874628 |
| ENSMUSG00000033658 | Ddx19b | 234733 | protein_coding | 901.5505531 | 0.035899729 | 0.056011593 | 0.640933901 | 0.521565629 | 0.732626853 | 0.035899729 |
| ENSMUSG00000002635 | Pdcd2l | 68079 | protein_coding | 260.8838544 | 0.035902263 | 0.083976758 | 0.427526181 | 0.66899612 | 0.831002845 | 0.035902263 |
| ENSMUSG00000037157 | Il22ra1 | 230828 | protein_coding | 850.2423717 | -0.035912029 | 0.060916739 | -0.58952646 | 0.555508168 | 0.75524772 | 0.035912029 |
| ENSMUSG00000001472 | Tcf25 | 66855 | protein_coding | 4709.99587 | -0.035937236 | 0.034846456 | -1.031302478 | 0.302398997 | 0.546174109 | 0.035937236 |
| ENSMUSG00000033706 | Smyd5 | 232187 | protein_coding | 220.647682 | 0.03594867 | 0.09343449 | 0.38474732 | 0.700424633 | 0.850619214 | 0.03594867 |
| ENSMUSG00000030654 | Arl6ip1 | 54208 | protein_coding | 3576.186528 | 0.035957772 | 0.038706468 | 0.928986141 | 0.352896269 | 0.595378328 | 0.035957772 |
| ENSMUSG00000001138 | Cnnm3 | 94218 | protein_coding | 511.5287701 | -0.035961643 | 0.070792328 | -0.507987859 | 0.611461858 | 0.792879744 | 0.035961643 |
| ENSMUSG00000043085 | Tmem82 | 213989 | protein_coding | 311.3140878 | -0.03597663 | 0.09091172 | -0.395731486 | 0.692303125 | 0.84565495 | 0.03597663 |
| ENSMUSG00000042208 | Sanbr | 71675 | protein_coding | 558.648778 | -0.035986473 | 0.061594013 | -0.584252779 | 0.559050257 | 0.757536946 | 0.035986473 |
| ENSMUSG00000026333 | Gin1 | 252876 | protein_coding | 117.1373791 | 0.036007489 | 0.116672541 | 0.308620079 | 0.757610545 | 0.885131169 | 0.036007489 |
| ENSMUSG00000108084 | Gm7932 | 666105 | lncRNA | 27.03613338 | 0.036026413 | 0.248921499 | 0.144730017 | 0.884924039 | 0.94900718 | 0.036026413 |
| ENSMUSG00000043241 | Upf2 | 326622 | protein_coding | 607.1692923 | 0.036036457 | 0.060430129 | 0.596332619 | 0.550953044 | 0.752092411 | 0.036036457 |

| ensembl_gene_id | external_gene_name | entrezgene_id | gene_biotype | baseMean | log2FoldChange | lfcSE | stat | pvalue | padj | absLog2FC |
|---|---|---|---|---|---|---|---|---|---|---|
| ENSMUSG00000041702 | Btbd7 | 238386 | protein_coding | 1812.083747 | -0.036044747 | 0.039793657 | -0.905791265 | 0.365046345 | 0.60723078 | 0.036044747 |
| ENSMUSG00000028436 | Dcaf12 | 68970 | protein_coding | 1231.379911 | 0.036052418 | 0.059963665 | 0.601237737 | 0.547681653 | 0.750740503 | 0.036052418 |
| ENSMUSG00000000056 | Narf | 67608 | protein_coding | 1543.444662 | -0.036101036 | 0.043513768 | -0.829646282 | 0.406738801 | 0.645607651 | 0.036101036 |
| ENSMUSG00000042826 | Fgf11 | 14166 | protein_coding | 33.19729354 | 0.036108974 | 0.217690426 | 0.165873044 | 0.868256862 | 0.940324922 | 0.036108974 |
| ENSMUSG00000042997 | Nhlrc3 | 212114 | protein_coding | 1518.806219 | 0.036130667 | 0.040628674 | 0.889289845 | 0.373847328 | 0.615647488 | 0.036130667 |
| ENSMUSG00000061013 | Mkx | 210719 | protein_coding | 331.0359491 | 0.036139275 | 0.090497429 | 0.399340347 | 0.689642442 | 0.844292391 | 0.036139275 |
| ENSMUSG00000033728 | Lrrc14 | 223664 | protein_coding | 372.8654221 | 0.036147691 | 0.106699454 | 0.338780465 | 0.734775119 | 0.872486041 | 0.036147691 |
| ENSMUSG00000001418 | Glmp | 56700 | protein_coding | 1421.218642 | -0.036148703 | 0.066881628 | -0.540487781 | 0.588860685 | 0.778426904 | 0.036148703 |
| ENSMUSG00000039063 | Echdc3 | 67856 | protein_coding | 261.9657748 | 0.036158695 | 0.082920531 | 0.436064445 | 0.662789973 | 0.827627866 | 0.036158695 |
| ENSMUSG00000066148 | Prpf4 | 70052 | protein_coding | 458.0092634 | 0.036165311 | 0.07030463 | 0.514408678 | 0.606966294 | 0.789908442 | 0.036165311 |
| ENSMUSG00000050608 | Micos10 | 433771 | protein_coding | 1092.669839 | 0.03616618 | 0.064207858 | 0.563267201 | 0.573252949 | 0.767653434 | 0.03616618 |
| ENSMUSG00000050796 | B3galt6 | 117592 | protein_coding | 175.6764167 | -0.036208536 | 0.108719871 | -0.33304432 | 0.739100829 | 0.87479925 | 0.036208536 |
| ENSMUSG00000074886 | Grk6 | 26385 | protein_coding | 1956.580394 | 0.036221758 | 0.053836967 | 0.672804592 | 0.501071616 | 0.718355583 | 0.036221758 |
| ENSMUSG00000036940 | Kdm1a | 99982 | protein_coding | 1496.759571 | -0.036233151 | 0.041142418 | -0.880676279 | 0.378493059 | 0.619709567 | 0.036233151 |
| ENSMUSG00000026754 | Golga1 | 76899 | protein_coding | 838.657347 | 0.036249645 | 0.059716624 | 0.607027705 | 0.543832526 | 0.748688924 | 0.036249645 |
| ENSMUSG00000033964 | Zbtb41 | 226470 | protein_coding | 827.5110879 | 0.036296521 | 0.054871529 | 0.661481865 | 0.508303341 | 0.723769025 | 0.036296521 |
| ENSMUSG00000031129 | Slc9a9 | 331004 | protein_coding | 47.16944869 | 0.036341647 | 0.185614989 | 0.195790476 | 0.844774165 | 0.930261042 | 0.036341647 |
| ENSMUSG00000028847 | Trappc3 | 27096 | protein_coding | 946.3364603 | 0.036351274 | 0.057333627 | 0.634030602 | 0.526060851 | 0.736439527 | 0.036351274 |
| ENSMUSG00000022142 | Nup155 | 170762 | protein_coding | 483.3999382 | 0.036357938 | 0.08217549 | 0.442442606 | 0.658168954 | 0.825115883 | 0.036357938 |
| ENSMUSG00000061859 | Patj | 12695 | protein_coding | 2948.677287 | -0.036380672 | 0.037745137 | -0.963850567 | 0.33512085 | 0.578701569 | 0.036380672 |
| ENSMUSG00000062691 | Cebpzos | 68554 | protein_coding | 257.7995173 | -0.036397516 | 0.09020635 | -0.403491734 | 0.686586515 | 0.84234163 | 0.036397516 |
| ENSMUSG00000027566 | Psma7 | 26444 | protein_coding | 2377.171063 | 0.036399636 | 0.048507695 | 0.750388908 | 0.453020509 | 0.681834975 | 0.036399636 |
| ENSMUSG00000041215 | Yeats2 | 208146 | protein_coding | 738.1378949 | 0.036402167 | 0.058847329 | 0.618586561 | 0.536188758 | 0.742878353 | 0.036402167 |
| ENSMUSG00000033618 | Map3k13 | 71751 | protein_coding | 544.5973281 | -0.036403122 | 0.063397977 | -0.574200051 | 0.565832435 | 0.762042164 | 0.036403122 |
| ENSMUSG00000026643 | Nmt2 | 18108 | protein_coding | 864.3167082 | -0.036426916 | 0.050982837 | -0.71449371 | 0.474921944 | 0.698573273 | 0.036426916 |
| ENSMUSG00000025076 | Casp7 | 12369 | protein_coding | 3948.604367 | -0.036500005 | 0.088581473 | -0.412050108 | 0.680302696 | 0.838321829 | 0.036500005 |
| ENSMUSG00000028096 | Gpr89 | 67549 | protein_coding | 1223.498597 | 0.036501448 | 0.08016396 | 0.455334889 | 0.64886833 | 0.818686277 | 0.036501448 |
| ENSMUSG00000037992 | Rara | 19401 | protein_coding | 550.7823504 | -0.036503326 | 0.094306908 | -0.387069479 | 0.698704772 | 0.849656295 | 0.036503326 |
| ENSMUSG00000031776 | Arl2bp | 107566 | protein_coding | 1901.587232 | -0.036532664 | 0.045681906 | -0.799718474 | 0.423873927 | 0.659827685 | 0.036532664 |
| ENSMUSG00000078153 | Psme2b | 621823 | protein_coding | 52.15445398 | 0.036538995 | 0.218031914 | 0.167585535 | 0.866909352 | 0.939855462 | 0.036538995 |
| ENSMUSG00000022095 | Fhip2b | 239170 | protein_coding | 462.2672901 | -0.036550562 | 0.072823268 | -0.50190774 | 0.615732421 | 0.795599668 | 0.036550562 |
| ENSMUSG00000041498 | Kif14 | 381293 | protein_coding | 194.8392448 | -0.036555199 | 0.128292383 | -0.28493663 | 0.775692692 | 0.895076992 | 0.036555199 |
| ENSMUSG00000035311 | Gnptab | 432486 | protein_coding | 1150.583714 | -0.036572247 | 0.076604145 | -0.47741864 | 0.633064043 | 0.808091169 | 0.036572247 |
| ENSMUSG00000031357 | Syap1 | 67043 | protein_coding | 1585.643043 | -0.03658505 | 0.053391027 | -0.685228446 | 0.493199776 | 0.712517495 | 0.03658505 |
| ENSMUSG00000020257 | Wdr82 | 77305 | protein_coding | 1819.891623 | -0.036606341 | 0.049020474 | -0.746756157 | 0.455210765 | 0.683091468 | 0.036606341 |
| ENSMUSG00000043872 | Zmym1 | 68310 | protein_coding | 92.86367388 | -0.036610866 | 0.123735261 | -0.295880623 | 0.76732125 | 0.890725386 | 0.036610866 |
| ENSMUSG00000021694 | Ercc8 | 71991 | protein_coding | 160.853464 | -0.036615398 | 0.105573263 | -0.346824535 | 0.728723144 | 0.868306546 | 0.036615398 |
| ENSMUSG00000007476 | Lrrc8a | 241296 | protein_coding | 314.4697236 | 0.036629818 | 0.08550397 | 0.42839903 | 0.66836063 | 0.830822102 | 0.036629818 |
| ENSMUSG00000098207 | Arl14 | 71619 | protein_coding | 349.8798096 | -0.036724802 | 0.133049131 | -0.276024364 | 0.782529353 | 0.898825528 | 0.036724802 |

| ensembl_gene_id | external_gene_name | entrezgene_id | gene_biotype | baseMean | log2FoldChange | lfcSE | stat | pvalue | padj | absLog2FC |
|---|---|---|---|---|---|---|---|---|---|---|
| ENSMUSG00000029461 | Fam168a | 319604 | protein_coding | 340.5796122 | -0.036749059 | 0.121184387 | -0.30324912 | 0.761700018 | 0.887577222 | 0.036749059 |
| ENSMUSG00000049950 | Rpp38 | 227522 | protein_coding | 81.75156051 | 0.036818089 | 0.15559233 | 0.236631776 | 0.812942467 | 0.915984616 | 0.036818089 |
| ENSMUSG00000031995 | St14 | 19143 | protein_coding | 7901.428213 | 0.0368259 | 0.069579975 | 0.529260035 | 0.596625075 | 0.783232176 | 0.0368259 |
| ENSMUSG00000021713 | Ppwd1 | 238831 | protein_coding | 284.7126839 | -0.036841331 | 0.098909396 | -0.372475543 | 0.709538818 | 0.856166251 | 0.036841331 |
| ENSMUSG00000037531 | Mrpl47 | 74600 | protein_coding | 411.4819625 | 0.036849917 | 0.06837353 | 0.538950049 | 0.589921322 | 0.77918632 | 0.036849917 |
| ENSMUSG00000045690 | Wdr89 | 72338 | protein_coding | 39.03055304 | 0.036850193 | 0.194057078 | 0.189893579 | 0.849392524 | 0.931939078 | 0.036850193 |
| ENSMUSG00000028068 | Iqgap3 | 404710 | protein_coding | 555.1810029 | -0.036853683 | 0.098256107 | -0.375077785 | 0.707602618 | 0.854813943 | 0.036853683 |
| ENSMUSG00000073608 | Gal3st2c | 619597 | protein_coding | 2340.228074 | 0.036910939 | 0.123255905 | 0.299465891 | 0.764584594 | 0.889388602 | 0.036910939 |
| ENSMUSG00000000605 | Clcn4 | 12727 | protein_coding | 2534.084587 | 0.03692713 | 0.037964066 | 0.972686376 | 0.330709198 | 0.574280957 | 0.03692713 |
| ENSMUSG00000042207 | Kdm5b | 75605 | protein_coding | 1872.87503 | -0.036929266 | 0.044957476 | -0.821426585 | 0.411403326 | 0.649442192 | 0.036929266 |
| ENSMUSG00000087030 | Gm16143 | 102632251 | lncRNA | 16.18190678 | -0.036967413 | 0.345526536 | -0.106988638 | 0.914797994 | 0.96216965 | 0.036967413 |
| ENSMUSG00000020903 | Stx8 | 55943 | protein_coding | 534.3895012 | 0.03698872 | 0.06793139 | 0.544501146 | 0.58609666 | 0.776407166 | 0.03698872 |
| ENSMUSG00000022148 | Fyb | 23880 | protein_coding | 116.6767731 | 0.037000217 | 0.162138567 | 0.228201211 | 0.81948982 | 0.919112062 | 0.037000217 |
| ENSMUSG00000043421 | Hilpda | 69573 | protein_coding | 144.7324054 | -0.037053705 | 0.0966343 | -0.383442574 | 0.701391642 | 0.851309024 | 0.037053705 |
| ENSMUSG00000074892 | B3galt5 | 93961 | protein_coding | 6176.595667 | 0.037057767 | 0.097712653 | 0.379252494 | 0.704500374 | 0.853157579 | 0.037057767 |
| ENSMUSG00000035150 | Eif2s3x | 26905 | protein_coding | 1605.459159 | 0.037075708 | 0.057795266 | 0.641500768 | 0.521197382 | 0.732548832 | 0.037075708 |
| ENSMUSG00000042628 | Zfyve1 | 217695 | protein_coding | 1589.390797 | -0.037078982 | 0.056806896 | -0.65271972 | 0.513936987 | 0.727104289 | 0.037078982 |
| ENSMUSG00000050296 | Abca12 | 74591 | protein_coding | 890.6797164 | 0.037079256 | 0.135719288 | 0.273205502 | 0.784695251 | 0.899716112 | 0.037079256 |
| ENSMUSG00000022500 | Litaf | 56722 | protein_coding | 2184.817579 | 0.037128145 | 0.070100965 | 0.529638149 | 0.596362839 | 0.78312381 | 0.037128145 |
| ENSMUSG00000023087 | Noct | 12457 | protein_coding | 627.8739148 | -0.037148162 | 0.09354779 | -0.397103573 | 0.691291085 | 0.8450944 | 0.037148162 |
| ENSMUSG00000089911 | Mfsd14a | 15247 | protein_coding | 926.6039843 | -0.037158656 | 0.059797827 | -0.621404776 | 0.53433333 | 0.741606925 | 0.037158656 |
| ENSMUSG00000031353 | Rbbp7 | 245688 | protein_coding | 2593.083733 | 0.037164099 | 0.064319481 | 0.577804714 | 0.563395973 | 0.760707204 | 0.037164099 |
| ENSMUSG00000003039 | Fam32a | 67922 | protein_coding | 2466.6327 | -0.037202712 | 0.056533272 | -0.658067555 | 0.510494723 | 0.724852046 | 0.037202712 |
| ENSMUSG00000075595 | Zfp652 | 268469 | protein_coding | 1629.856418 | -0.037217088 | 0.049524865 | -0.75148288 | 0.4523621 | 0.68117317 | 0.037217088 |
| ENSMUSG00000004364 | Cul3 | 26554 | protein_coding | 2208.721128 | -0.037231126 | 0.034509741 | -1.078858462 | 0.280650828 | 0.523508209 | 0.037231126 |
| ENSMUSG00000022663 | Atg3 | 67841 | protein_coding | 756.2612321 | 0.037252492 | 0.059865196 | 0.622272954 | 0.533762402 | 0.741516545 | 0.037252492 |
| ENSMUSG00000020821 | Kif1c | 16562 | protein_coding | 8654.842115 | -0.03726898 | 0.050704927 | -0.735016926 | 0.462329186 | 0.688774391 | 0.03726898 |
| ENSMUSG00000036873 | 2410004B18Rik | 66421 | protein_coding | 302.9525823 | -0.037294249 | 0.081179432 | -0.459405152 | 0.645943249 | 0.816513895 | 0.037294249 |
| ENSMUSG00000024776 | Stambpl1 | 76630 | protein_coding | 506.2848479 | -0.037311037 | 0.075845156 | -0.491936974 | 0.622763899 | 0.800888833 | 0.037311037 |
| ENSMUSG00000048047 | Zbtb33 | 56805 | protein_coding | 446.0531236 | 0.037311718 | 0.088890028 | 0.419751443 | 0.67466704 | 0.834405417 | 0.037311718 |
| ENSMUSG00000059277 | R74862 | 97423 | lncRNA | 9.211226506 | -0.037325118 | 0.429468069 | -0.086910111 | 0.930742962 | 0.96964417 | 0.037325118 |
| ENSMUSG00000073423 | Zfp414 | 328801 | protein_coding | 140.7496422 | -0.03735005 | 0.116052373 | -0.321837882 | 0.747575514 | 0.880075211 | 0.03735005 |
| ENSMUSG00000096995 | 2810029C07Rik | 100504687 | lncRNA | 8.454324579 | -0.037350413 | 0.512242393 | -0.072915506 | 0.941873355 | 0.974503855 | 0.037350413 |
| ENSMUSG00000041879 | Ipo9 | 226432 | protein_coding | 761.4894185 | 0.037352307 | 0.077528934 | 0.481785374 | 0.629958421 | 0.805574037 | 0.037352307 |
| ENSMUSG00000029559 | 2210016L21Rik | 72357 | protein_coding | 570.3675881 | 0.037382526 | 0.069717079 | 0.536203269 | 0.591818075 | 0.780262005 | 0.037382526 |
| ENSMUSG00000024899 | Papss2 | 23972 | protein_coding | 15614.98001 | 0.037382941 | 0.056771322 | 0.658482828 | 0.510227928 | 0.724823423 | 0.037382941 |
| ENSMUSG00000026667 | Uhmk1 | 16589 | protein_coding | 3129.961618 | -0.037399376 | 0.034036374 | -1.098806124 | 0.27185264 | 0.51439253 | 0.037399376 |
| ENSMUSG00000039221 | Rpl22l1 | 68028 | protein_coding | 1134.452875 | 0.037411119 | 0.076494653 | 0.489068422 | 0.624793257 | 0.801977146 | 0.037411119 |
| ENSMUSG00000027998 | Plrg1 | 53317 | protein_coding | 810.5736832 | 0.037412753 | 0.058918292 | 0.634993847 | 0.525432428 | 0.735943093 | 0.037412753 |

| ensembl_gene_id | external_gene_name | entrezgene_id | gene_biotype | baseMean | log2FoldChange | lfcSE | stat | pvalue | padj | absLog2FC |
|---|---|---|---|---|---|---|---|---|---|---|
| ENSMUSG00000027488 | Snta1 | 20648 | protein_coding | 138.8122496 | 0.037443825 | 0.133259667 | 0.280983929 | 0.778722726 | 0.896968888 | 0.037443825 |
| ENSMUSG00000039831 | Arhgap29 | 214137 | protein_coding | 269.2316347 | 0.037444646 | 0.094104157 | 0.397906396 | 0.690699185 | 0.844688387 | 0.037444646 |
| ENSMUSG00000003068 | Stk11 | 20869 | protein_coding | 1523.984528 | 0.037453105 | 0.049993249 | 0.74916326 | 0.45375881 | 0.682563971 | 0.037453105 |
| ENSMUSG00000020691 | Mettl2 | 52686 | protein_coding | 284.3361363 | 0.037454979 | 0.084308833 | 0.444259254 | 0.656855157 | 0.824173587 | 0.037454979 |
| ENSMUSG00000036371 | Serbp1 | 66870 | protein_coding | 5852.381365 | 0.037464868 | 0.050710974 | 0.738792112 | 0.460033242 | 0.686940379 | 0.037464868 |
| ENSMUSG00000039841 | Zfp800 | 627049 | protein_coding | 565.8528718 | -0.037466082 | 0.064322622 | -0.582471318 | 0.560249256 | 0.758554256 | 0.037466082 |
| ENSMUSG00000071604 | Entrep1 | 381217 | protein_coding | 1165.295035 | 0.037466244 | 0.065110313 | 0.575427186 | 0.565002423 | 0.761555988 | 0.037466244 |
| ENSMUSG00000062116 | Zfp954 | 232853 | protein_coding | 417.0596065 | 0.037507653 | 0.066720276 | 0.562162731 | 0.574005151 | 0.76815045 | 0.037507653 |
| ENSMUSG00000033157 | Abhd10 | 213012 | protein_coding | 238.7837686 | 0.037532099 | 0.098737034 | 0.380121796 | 0.703855007 | 0.852953349 | 0.037532099 |
| ENSMUSG00000034334 | Fam151b | 73942 | protein_coding | 69.50251651 | 0.037532805 | 0.176284624 | 0.21291026 | 0.83139696 | 0.925325507 | 0.037532805 |
| ENSMUSG00000020134 | Peli1 | 67245 | protein_coding | 1362.117274 | -0.037537371 | 0.050806069 | -0.73883635 | 0.460006376 | 0.686940379 | 0.037537371 |
| ENSMUSG00000006215 | Zbtb17 | 22642 | protein_coding | 570.6636509 | -0.037549841 | 0.070233146 | -0.534645577 | 0.592894961 | 0.780422622 | 0.037549841 |
| ENSMUSG00000044674 | Fzd1 | 14362 | protein_coding | 730.0170567 | 0.037567966 | 0.090112066 | 0.416902723 | 0.676749561 | 0.835804884 | 0.037567966 |
| ENSMUSG00000006390 | Elovl1 | 54325 | protein_coding | 1440.587847 | 0.037600491 | 0.069518901 | 0.540867173 | 0.588599139 | 0.778379525 | 0.037600491 |
| ENSMUSG00000039349 | C130074G19Rik | 226777 | protein_coding | 825.4418802 | -0.03761876 | 0.083245885 | -0.451899337 | 0.651341502 | 0.8204259 | 0.03761876 |
| ENSMUSG00000074384 | AI429214 | 621080 | protein_coding | 11.63382388 | 0.037619946 | 0.362227326 | 0.103857285 | 0.917282606 | 0.963427979 | 0.037619946 |
| ENSMUSG00000025066 | Sfr1 | 67788 | protein_coding | 790.1959927 | -0.037631113 | 0.055627186 | -0.676487811 | 0.498730982 | 0.716014372 | 0.037631113 |
| ENSMUSG00000032872 | Cyb5r4 | 266690 | protein_coding | 1554.490009 | 0.037656244 | 0.045609938 | 0.825614892 | 0.409022585 | 0.647331425 | 0.037656244 |
| ENSMUSG00000038128 | Camk4 | 12326 | protein_coding | 6.451739508 | 0.037675573 | 0.460402098 | 0.081831888 | 0.934780398 | 0.971653463 | 0.037675573 |
| ENSMUSG00000021432 | Slc35b3 | 108652 | protein_coding | 1065.191361 | -0.037689504 | 0.049919686 | -0.755002826 | 0.450247286 | 0.679535938 | 0.037689504 |
| ENSMUSG00000022231 | Sema5a | 20356 | protein_coding | 2707.282104 | -0.037690131 | 0.072097302 | -0.522767557 | 0.601136019 | 0.786473529 | 0.037690131 |
| ENSMUSG00000025403 | Shmt2 | 108037 | protein_coding | 336.2876629 | 0.037694832 | 0.08814465 | 0.427647421 | 0.668907835 | 0.831002845 | 0.037694832 |
| ENSMUSG00000020463 | Ppp4r3b | 104570 | protein_coding | 2358.098733 | -0.03771254 | 0.042312729 | -0.891281207 | 0.372778328 | 0.614514576 | 0.03771254 |
| ENSMUSG00000021549 | Rasa1 | 218397 | protein_coding | 1084.331622 | -0.037715882 | 0.063485772 | -0.59408401 | 0.552455925 | 0.752996679 | 0.037715882 |
| ENSMUSG00000057554 | Lgals8 | 56048 | protein_coding | 1267.316103 | 0.037740791 | 0.05144009 | 0.733684387 | 0.463141116 | 0.689357631 | 0.037740791 |
| ENSMUSG00000043929 | Klhl15 | 236904 | protein_coding | 262.6649182 | -0.037759081 | 0.077543721 | -0.486939251 | 0.626301382 | 0.803018158 | 0.037759081 |
| ENSMUSG00000060397 | Zfp128 | 243833 | protein_coding | 31.90224583 | 0.037763347 | 0.201999781 | 0.186947466 | 0.851701823 | 0.933520082 | 0.037763347 |
| ENSMUSG00000028706 | Nsun4 | 72181 | protein_coding | 303.2363324 | -0.037771013 | 0.075936548 | -0.497402296 | 0.618905387 | 0.798167707 | 0.037771013 |
| ENSMUSG00000021978 | Extl3 | 54616 | protein_coding | 1023.317064 | -0.037776932 | 0.057591456 | -0.655946817 | 0.511858344 | 0.725570423 | 0.037776932 |
| ENSMUSG00000035367 | Rmi1 | 74386 | protein_coding | 219.83431 | -0.037779386 | 0.089926168 | -0.420115595 | 0.674401011 | 0.834309996 | 0.037779386 |
| ENSMUSG00000029152 | Ociad1 | 68095 | protein_coding | 2019.501914 | 0.037786817 | 0.04384063 | 0.861913179 | 0.388735296 | 0.628610661 | 0.037786817 |
| ENSMUSG00000075229 | Ccdc58 | 381045 | protein_coding | 416.4169544 | -0.037792764 | 0.065049159 | -0.580987738 | 0.561248718 | 0.759206347 | 0.037792764 |
| ENSMUSG00000026234 | Ncl | 17975 | protein_coding | 3745.181122 | 0.037831047 | 0.079949047 | 0.473189464 | 0.636078014 | 0.809697277 | 0.037831047 |
| ENSMUSG00000054256 | Msi1 | 17690 | protein_coding | 22.16275974 | 0.037833241 | 0.254564143 | 0.148619678 | 0.881853745 | 0.947331698 | 0.037833241 |
| ENSMUSG00000025766 | D3Ertd751e | 73852 | protein_coding | 264.5519131 | 0.037843248 | 0.098119955 | 0.385683505 | 0.699731081 | 0.850189342 | 0.037843248 |
| ENSMUSG00000046223 | Plaur | 18793 | protein_coding | 113.7722688 | 0.03784533 | 0.175699655 | 0.215397862 | 0.829457137 | 0.924270223 | 0.03784533 |
| ENSMUSG00000030421 | Uri1 | 19777 | protein_coding | 823.6005673 | 0.037862479 | 0.065453802 | 0.578461111 | 0.562952845 | 0.760490915 | 0.037862479 |
| ENSMUSG00000024619 | Cdx1 | 12590 | protein_coding | 7030.688933 | 0.037864239 | 0.080237606 | 0.471901408 | 0.636997163 | 0.810530941 | 0.037864239 |
| ENSMUSG00000065687 | Gm23734 | 115487646 | snoRNA | 22.98776328 | 0.03788655 | 0.363066233 | 0.104351621 | 0.916890314 | 0.963217713 | 0.03788655 |

| ensembl_gene_id | external_gene_name | entrezgene_id | gene_biotype | baseMean | log2FoldChange | lfcSE | stat | pvalue | padj | absLog2FC |
|---|---|---|---|---|---|---|---|---|---|---|
| ENSMUSG00000031918 | Mtmr2 | 77116 | protein_coding | 1282.784608 | 0.037901784 | 0.056629654 | 0.669292171 | 0.50330912 | 0.719943844 | 0.037901784 |
| ENSMUSG00000032570 | Atp2c1 | 235574 | protein_coding | 1985.573376 | 0.037905433 | 0.037313159 | 1.015873052 | 0.309689843 | 0.553189591 | 0.037905433 |
| ENSMUSG00000052302 | Tbc1d30 | 74694 | protein_coding | 532.0028265 | -0.037911384 | 0.062030781 | -0.611170519 | 0.541086698 | 0.746586685 | 0.037911384 |
| ENSMUSG00000051403 | Ppp1r37 | 232947 | protein_coding | 2668.405913 | -0.037919534 | 0.063770448 | -0.594625485 | 0.552093841 | 0.752630378 | 0.037919534 |
| ENSMUSG00000030681 | Mvp | 78388 | protein_coding | 9080.792531 | 0.037919828 | 0.059951558 | 0.632507806 | 0.527055109 | 0.737165628 | 0.037919828 |
| ENSMUSG00000020288 | Ahsa2 | 268390 | protein_coding | 270.1515299 | 0.037927049 | 0.109082775 | 0.347690542 | 0.7280726 | 0.867832282 | 0.037927049 |
| ENSMUSG00000052299 | Ltn1 | 78913 | protein_coding | 1493.939912 | -0.037941596 | 0.039915763 | -0.950541683 | 0.341837085 | 0.584451398 | 0.037941596 |
| ENSMUSG00000071553 | Cpa2 | 232680 | protein_coding | 36.58526143 | 0.03794639 | 0.24686707 | 0.153711833 | 0.877836954 | 0.945241431 | 0.03794639 |
| ENSMUSG00000049191 | Rtl5 | 331474 | protein_coding | 8.366534851 | -0.037958971 | 0.425780853 | -0.089151427 | 0.928961567 | 0.968789653 | 0.037958971 |
| ENSMUSG00000042055 | Wdr11 | 207425 | protein_coding | 1335.584199 | -0.037977194 | 0.045819317 | -0.828846788 | 0.407191109 | 0.645827204 | 0.037977194 |
| ENSMUSG00000029192 | Tbc1d14 | 100855 | protein_coding | 727.0874547 | 0.0379774 | 0.050095144 | 0.758105414 | 0.448387879 | 0.678988684 | 0.0379774 |
| ENSMUSG00000026867 | Gapvd1 | 66691 | protein_coding | 1971.251018 | 0.037985522 | 0.053574546 | 0.709021827 | 0.478310932 | 0.700798986 | 0.037985522 |
| ENSMUSG00000021377 | Dek | 110052 | protein_coding | 1284.838988 | -0.037986254 | 0.084463847 | -0.449733884 | 0.652902336 | 0.82145077 | 0.037986254 |
| ENSMUSG00000046756 | Mrps7 | 50529 | protein_coding | 851.5280491 | 0.038005707 | 0.071817203 | 0.529200601 | 0.5966663 | 0.783232176 | 0.038005707 |
| ENSMUSG00000051671 | Coa6 | 67892 | protein_coding | 317.5462031 | 0.038043437 | 0.089649402 | 0.424357952 | 0.671304781 | 0.832361742 | 0.038043437 |
| ENSMUSG00000069565 | Dazap1 | 70248 | protein_coding | 1636.573242 | 0.038054967 | 0.094535143 | 0.402548361 | 0.687280504 | 0.842728192 | 0.038054967 |
| ENSMUSG00000026605 | Cenpf | 108000 | protein_coding | 880.4784462 | -0.03806544 | 0.089474136 | -0.425435119 | 0.67051951 | 0.831904926 | 0.03806544 |
| ENSMUSG00000040479 | Dgkz | 104418 | protein_coding | 3652.643693 | 0.038070893 | 0.042510304 | 0.895568597 | 0.370483214 | 0.61248414 | 0.038070893 |
| ENSMUSG00000040795 | Iqcc | 230767 | protein_coding | 186.2181197 | -0.038071023 | 0.097230964 | -0.391552459 | 0.695388921 | 0.84768023 | 0.038071023 |
| ENSMUSG00000008822 | Acyp1 | 66204 | protein_coding | 168.3781209 | 0.038087012 | 0.094237139 | 0.404161375 | 0.686094055 | 0.841984797 | 0.038087012 |
| ENSMUSG00000081984 | Dnajb3 | 15504 | protein_coding | 14.81795471 | -0.038107772 | 0.33158804 | -0.114925048 | 0.908504531 | 0.959234074 | 0.038107772 |
| ENSMUSG00000032216 | Nedd4 | 17999 | protein_coding | 7482.217092 | -0.038111169 | 0.050108459 | -0.760573567 | 0.446911813 | 0.677828158 | 0.038111169 |
| ENSMUSG00000030199 | Etv6 | 14011 | protein_coding | 835.9112364 | -0.03812555 | 0.055729202 | -0.684121588 | 0.493898391 | 0.713271187 | 0.03812555 |
| ENSMUSG00000026176 | Ctdsp1 | 227292 | protein_coding | 1546.988928 | -0.038133287 | 0.056058011 | -0.680246875 | 0.496348156 | 0.714747243 | 0.038133287 |
| ENSMUSG00000040084 | Bub1b | 12236 | protein_coding | 702.3342428 | 0.038161129 | 0.097474017 | 0.391500525 | 0.6954273 | 0.84768023 | 0.038161129 |
| ENSMUSG00000039069 | Mtg2 | 52856 | protein_coding | 301.7666399 | 0.038173652 | 0.087172875 | 0.437907457 | 0.66145337 | 0.826919264 | 0.038173652 |
| ENSMUSG00000065701 | Rny1 | 19872 | misc_RNA | 115.6277207 | 0.038181234 | 0.25684954 | 0.148652142 | 0.881828127 | 0.947331698 | 0.038181234 |
| ENSMUSG00000049728 | Zfp668 | 244219 | protein_coding | 166.6860287 | 0.038188504 | 0.102799818 | 0.371484164 | 0.710276949 | 0.856611363 | 0.038188504 |
| ENSMUSG00000073982 | Rhog | 56212 | protein_coding | 430.7531288 | 0.038193604 | 0.092078449 | 0.414794172 | 0.678292587 | 0.836813383 | 0.038193604 |
| ENSMUSG00000028677 | Rnf220 | 66743 | protein_coding | 938.1872585 | -0.038194738 | 0.065090763 | -0.586791981 | 0.557343423 | 0.756034291 | 0.038194738 |
| ENSMUSG00000031511 | Arhgef7 | 54126 | protein_coding | 856.4170728 | -0.038210501 | 0.058393134 | -0.654366331 | 0.512875821 | 0.726387506 | 0.038210501 |
| ENSMUSG00000037361 | Sf3b6 | 66055 | protein_coding | 844.3635248 | 0.038231281 | 0.05272321 | 0.725131882 | 0.468371127 | 0.693047027 | 0.038231281 |
| ENSMUSG00000009772 | Nuak2 | 74137 | protein_coding | 887.1200974 | -0.038241614 | 0.080923266 | -0.472566362 | 0.636522586 | 0.810069424 | 0.038241614 |
| ENSMUSG00000022517 | Mgrn1 | 17237 | protein_coding | 2480.04756 | 0.038254211 | 0.049031601 | 0.780195019 | 0.435276094 | 0.668797405 | 0.038254211 |
| ENSMUSG00000039660 | Spout1 | 227695 | protein_coding | 359.3337943 | 0.038267041 | 0.068140649 | 0.561589033 | 0.574396054 | 0.768186995 | 0.038267041 |
| ENSMUSG00000063316 | Rpl27 | 19942 | protein_coding | 2644.283861 | 0.03833573 | 0.056098461 | 0.683365097 | 0.494376169 | 0.713491312 | 0.03833573 |
| ENSMUSG00000036427 | Gpi1 | 14751 | protein_coding | 11649.02313 | 0.038359187 | 0.042078925 | 0.911600917 | 0.361978839 | 0.604873354 | 0.038359187 |
| ENSMUSG00000028433 | Ubap2 | 68926 | protein_coding | 541.6370963 | -0.038361316 | 0.070196782 | -0.546482537 | 0.584734289 | 0.775689464 | 0.038361316 |
| ENSMUSG00000021552 | Gkap1 | 56278 | protein_coding | 68.48523658 | -0.038369386 | 0.152618009 | -0.251407983 | 0.801498696 | 0.909560866 | 0.038369386 |

| ensembl_gene_id | external_gene_name | entrezgene_id | gene_biotype | baseMean | log2FoldChange | lfcSE | stat | pvalue | padj | absLog2FC |
|---|---|---|---|---|---|---|---|---|---|---|
| ENSMUSG00000063358 | Mapk1 | 26413 | protein_coding | 3144.332631 | 0.038372451 | 0.030078743 | 1.275733218 | 0.202049842 | 0.435870434 | 0.038372451 |
| ENSMUSG00000022533 | Atp13a3 | 224088 | protein_coding | 4554.918868 | 0.038375578 | 0.057186597 | 0.671058949 | 0.502182977 | 0.718950524 | 0.038375578 |
| ENSMUSG00000024824 | Rad9a | 19367 | protein_coding | 162.1018008 | -0.038390984 | 0.10917421 | -0.35164884 | 0.725101633 | 0.865987203 | 0.038390984 |
| ENSMUSG00000042744 | Hectd4 | 269700 | protein_coding | 2001.676902 | -0.038396959 | 0.06439868 | -0.596238288 | 0.551016051 | 0.75210185 | 0.038396959 |
| ENSMUSG00000028405 | Aco1 | 11428 | protein_coding | 3383.486473 | 0.038401257 | 0.06416363 | 0.598489466 | 0.549513386 | 0.751462933 | 0.038401257 |
| ENSMUSG00000034800 | Zfp661 | 72180 | protein_coding | 102.5668593 | 0.038411472 | 0.135691159 | 0.283080135 | 0.777115409 | 0.896141595 | 0.038411472 |
| ENSMUSG00000035086 | Becn1 | 56208 | protein_coding | 1966.629603 | 0.038447794 | 0.046675427 | 0.823726663 | 0.410094886 | 0.64854129 | 0.038447794 |
| ENSMUSG00000053897 | Slc39a8 | 67547 | protein_coding | 1053.03996 | -0.038454649 | 0.059784216 | -0.643224116 | 0.520078687 | 0.73174917 | 0.038454649 |
| ENSMUSG00000103906 | Tigd5 | 105734 | protein_coding | 105.2750644 | 0.038455314 | 0.133874218 | 0.287249585 | 0.773921222 | 0.893742272 | 0.038455314 |
| ENSMUSG00000020738 | Sumo2 | 170930 | protein_coding | 320.1647124 | -0.038462935 | 0.118603498 | -0.324298487 | 0.745712065 | 0.879357888 | 0.038462935 |
| ENSMUSG00000029250 | Polr2b | 231329 | protein_coding | 1603.370856 | 0.038464323 | 0.06256608 | 0.614779181 | 0.53870056 | 0.74493282 | 0.038464323 |
| ENSMUSG00000032119 | Hinfp | 102423 | protein_coding | 542.8959849 | 0.038465951 | 0.064677478 | 0.594734864 | 0.552020713 | 0.752630378 | 0.038465951 |
| ENSMUSG00000033294 | Noc4l | 100608 | protein_coding | 294.0812124 | -0.038469432 | 0.081223573 | -0.473623978 | 0.635768074 | 0.80962603 | 0.038469432 |
| ENSMUSG00000063727 | Tnfrsf11b | 18383 | protein_coding | 17.16488544 | 0.03847715 | 0.299086494 | 0.128648906 | 0.897635466 | 0.954463189 | 0.03847715 |
| ENSMUSG00000019966 | Kitl | 17311 | protein_coding | 3163.918242 | -0.038491684 | 0.052352175 | -0.735245171 | 0.462190194 | 0.688630936 | 0.038491684 |
| ENSMUSG00000071037 | Camkmt | 73582 | protein_coding | 164.1301293 | -0.038493199 | 0.094917637 | -0.405543167 | 0.685078295 | 0.84151555 | 0.038493199 |
| ENSMUSG00000021731 | Mrps30 | 59054 | protein_coding | 477.2865345 | -0.038509113 | 0.072721576 | -0.529541778 | 0.596429671 | 0.783147802 | 0.038509113 |
| ENSMUSG00000042496 | Prdm10 | 382066 | protein_coding | 387.9794078 | 0.038511564 | 0.088782697 | 0.433773312 | 0.664453063 | 0.828806137 | 0.038511564 |
| ENSMUSG00000049354 | Dcaf7 | 71833 | protein_coding | 2373.19358 | -0.038515857 | 0.051263599 | -0.751329569 | 0.452454338 | 0.68117317 | 0.038515857 |
| ENSMUSG00000025894 | Aasdhppt | 67618 | protein_coding | 415.7689424 | -0.038527182 | 0.075559985 | -0.509888688 | 0.610129447 | 0.792234911 | 0.038527182 |
| ENSMUSG00000071893 | Vmn1r4 | 171194 | protein_coding | 6.645570397 | 0.038540896 | 0.466575565 | 0.08260376 | 0.934166611 | 0.971391049 | 0.038540896 |
| ENSMUSG00000002055 | Spag5 | 54141 | protein_coding | 431.5489821 | 0.038544153 | 0.103578127 | 0.372126373 | 0.709798761 | 0.856292985 | 0.038544153 |
| ENSMUSG00000037643 | Prkci | 18759 | protein_coding | 2370.482641 | 0.038555824 | 0.053271306 | 0.723763432 | 0.469210983 | 0.693580708 | 0.038555824 |
| ENSMUSG00000062075 | Lmnb2 | 16907 | protein_coding | 1725.364427 | -0.038593427 | 0.055403651 | -0.696586343 | 0.486061705 | 0.707210133 | 0.038593427 |
| ENSMUSG00000027305 | Ndufaf1 | 69702 | protein_coding | 328.9150074 | -0.038597893 | 0.074110858 | -0.520812934 | 0.602497092 | 0.7874413 | 0.038597893 |
| ENSMUSG00000061762 | Tac1 | 21333 | protein_coding | 78.82276646 | 0.038642983 | 0.161053131 | 0.239939347 | 0.810377277 | 0.914739662 | 0.038642983 |
| ENSMUSG00000032733 | Snx33 | 235406 | protein_coding | 721.6669141 | -0.038684929 | 0.058166819 | -0.665068677 | 0.506006568 | 0.721828109 | 0.038684929 |
| ENSMUSG00000035451 | Foxa1 | 15375 | protein_coding | 948.8604737 | -0.03871866 | 0.075750775 | -0.511132197 | 0.609258492 | 0.791550125 | 0.03871866 |
| ENSMUSG00000052566 | Hook2 | 170833 | protein_coding | 2240.555354 | -0.038733394 | 0.061241848 | -0.632466125 | 0.527082337 | 0.737165628 | 0.038733394 |
| ENSMUSG00000051951 | Xkr4 | 497097 | protein_coding | 40.79179399 | -0.038781479 | 0.19397436 | -0.199930955 | 0.841534581 | 0.929613548 | 0.038781479 |
| ENSMUSG00000038546 | Ranbp9 | 56705 | protein_coding | 853.2581332 | -0.038784478 | 0.049051516 | -0.790688663 | 0.429125693 | 0.664262236 | 0.038784478 |
| ENSMUSG00000030281 | Il17rc | 171095 | protein_coding | 3117.02493 | 0.038796117 | 0.084565219 | 0.458771557 | 0.646398222 | 0.816823161 | 0.038796117 |
| ENSMUSG00000079434 | Neu2 | 23956 | protein_coding | 44.19452019 | -0.038799384 | 0.187471412 | -0.206961604 | 0.836039845 | 0.927227465 | 0.038799384 |
| ENSMUSG00000063550 | Nup98 | 269966 | protein_coding | 2154.817461 | 0.038803471 | 0.048097063 | 0.806774225 | 0.419796571 | 0.657014281 | 0.038803471 |
| ENSMUSG00000020736 | Nt5c | 50773 | protein_coding | 1948.081933 | 0.038850355 | 0.067779979 | 0.573183339 | 0.566520564 | 0.762547332 | 0.038850355 |
| ENSMUSG00000028967 | Errfi1 | 74155 | protein_coding | 2980.266333 | -0.038868333 | 0.071265692 | -0.545400336 | 0.585478209 | 0.776097903 | 0.038868333 |
| ENSMUSG00000031578 | Mak16 | 67920 | protein_coding | 255.7577206 | 0.038931157 | 0.111827207 | 0.348136725 | 0.727737505 | 0.867614603 | 0.038931157 |
| ENSMUSG00000030882 | Dnhd1 | 77505 | protein_coding | 12.41613221 | -0.038934366 | 0.458912579 | -0.084840485 | 0.932388207 | 0.970417852 | 0.038934366 |
| ENSMUSG00000019923 | Zwint | 52696 | protein_coding | 673.3679052 | 0.038937189 | 0.085515704 | 0.455322093 | 0.648877534 | 0.818686277 | 0.038937189 |

| ensembl_gene_id | external_gene_name | entrezgene_id | gene_biotype | baseMean | log2FoldChange | lfcSE | stat | pvalue | padj | absLog2FC |
|---|---|---|---|---|---|---|---|---|---|---|
| ENSMUSG00000023387 | Kcnk16 | 74571 | protein_coding | 6.446964703 | 0.038944953 | 0.520210149 | 0.074863885 | 0.940323011 | 0.974024613 | 0.038944953 |
| ENSMUSG00000050565 | Tor1aip2 | 240832 | protein_coding | 4785.069276 | 0.038945606 | 0.044428519 | 0.876590253 | 0.380709235 | 0.621615656 | 0.038945606 |
| ENSMUSG00000029994 | Anxa4 | 11746 | protein_coding | 6829.929305 | -0.03895433 | 0.0438914 | -0.887516232 | 0.374801031 | 0.616422248 | 0.03895433 |
| ENSMUSG00000041313 | Slc7a1 | 11987 | protein_coding | 712.5366101 | -0.038999742 | 0.069444817 | -0.561593263 | 0.574393172 | 0.768186995 | 0.038999742 |
| ENSMUSG00000018736 | Ndel1 | 83431 | protein_coding | 1331.560441 | 0.039008143 | 0.053339055 | 0.731324229 | 0.464581134 | 0.690725542 | 0.039008143 |
| ENSMUSG00000022684 | Bfar | 67118 | protein_coding | 1455.092384 | -0.039009791 | 0.044409263 | -0.878415642 | 0.379718199 | 0.62056297 | 0.039009791 |
| ENSMUSG00000024921 | Smarca2 | 67155 | protein_coding | 1002.498804 | -0.039034943 | 0.044181297 | -0.883517364 | 0.376956805 | 0.618431576 | 0.039034943 |
| ENSMUSG00000023353 | Agap3 | 213990 | protein_coding | 813.5905075 | -0.039061897 | 0.058859257 | -0.663649172 | 0.506914878 | 0.72262143 | 0.039061897 |
| ENSMUSG00000020961 | Ston2 | 108800 | protein_coding | 701.9688585 | 0.039068712 | 0.082315347 | 0.474622447 | 0.635056104 | 0.809382719 | 0.039068712 |
| ENSMUSG00000029106 | Add1 | 11518 | protein_coding | 6594.611938 | -0.03907109 | 0.031169392 | -1.25350825 | 0.210020798 | 0.445219276 | 0.03907109 |
| ENSMUSG00000025188 | Hps1 | 192236 | protein_coding | 439.6814604 | 0.039080607 | 0.067554708 | 0.578503084 | 0.562924516 | 0.760490915 | 0.039080607 |
| ENSMUSG00000018995 | Nars2 | 244141 | protein_coding | 341.7349554 | 0.039093028 | 0.075343022 | 0.518867275 | 0.6038533 | 0.78841372 | 0.039093028 |
| ENSMUSG00000053559 | Smagp | 207818 | protein_coding | 2365.17353 | 0.039095369 | 0.060258538 | 0.648793846 | 0.516471635 | 0.729218881 | 0.039095369 |
| ENSMUSG00000002280 | Ciao3 | 67563 | protein_coding | 642.6628959 | 0.039109789 | 0.055488044 | 0.704832711 | 0.480914349 | 0.703210238 | 0.039109789 |
| ENSMUSG00000032562 | Gnai2 | 14678 | protein_coding | 2679.379262 | 0.039116055 | 0.064719629 | 0.604392444 | 0.545582758 | 0.749073143 | 0.039116055 |
| ENSMUSG00000050910 | Cdr2l | 237988 | protein_coding | 214.9833241 | 0.039146199 | 0.133154947 | 0.293989819 | 0.768765683 | 0.891549271 | 0.039146199 |
| ENSMUSG00000051007 | Gatd1 | 213350 | protein_coding | 761.5164547 | 0.039161434 | 0.052409396 | 0.747221637 | 0.454929786 | 0.682988492 | 0.039161434 |
| ENSMUSG00000030275 | Etnk1 | 75320 | protein_coding | 1698.875794 | 0.039166613 | 0.044377866 | 0.882570907 | 0.377468153 | 0.61867166 | 0.039166613 |
| ENSMUSG00000043333 | Rhbdl2 | 230726 | protein_coding | 951.5861202 | 0.039208336 | 0.078124942 | 0.501867067 | 0.615761033 | 0.795599668 | 0.039208336 |
| ENSMUSG00000021395 | Spin1 | 20729 | protein_coding | 1368.873282 | -0.039214282 | 0.049831224 | -0.786941976 | 0.431315833 | 0.666012107 | 0.039214282 |
| ENSMUSG00000022768 | Ccdc116 | 76872 | protein_coding | 248.7422475 | -0.03922473 | 0.131791311 | -0.297627585 | 0.765987418 | 0.890119548 | 0.03922473 |
| ENSMUSG00000031304 | Il2rg | 16186 | protein_coding | 11.55673787 | 0.039241375 | 0.392848467 | 0.099889341 | 0.920432178 | 0.965011685 | 0.039241375 |
| ENSMUSG00000008958 | Vps72 | 21427 | protein_coding | 727.0433704 | -0.039257332 | 0.063555042 | -0.617690283 | 0.536779519 | 0.74336929 | 0.039257332 |
| ENSMUSG00000037503 | Fam168b | 214469 | protein_coding | 2778.78184 | -0.039298373 | 0.034447724 | -1.140811911 | 0.253948203 | 0.495031103 | 0.039298373 |
| ENSMUSG00000028743 | Akr7a5 | 110198 | protein_coding | 1135.526266 | 0.039301719 | 0.085365311 | 0.460394489 | 0.64523309 | 0.816003296 | 0.039301719 |
| ENSMUSG00000033712 | Ccar2 | 219158 | protein_coding | 1504.082908 | -0.039310425 | 0.048550536 | -0.80968055 | 0.418123799 | 0.655860109 | 0.039310425 |
| ENSMUSG00000030505 | Prmt3 | 71974 | protein_coding | 322.6522622 | -0.039313102 | 0.091180862 | -0.431155189 | 0.666355534 | 0.829602333 | 0.039313102 |
| ENSMUSG00000020150 | Gamt | 14431 | protein_coding | 27.18167341 | -0.03932705 | 0.22646511 | -0.173656112 | 0.862135731 | 0.937787345 | 0.03932705 |
| ENSMUSG00000023104 | Rfc2 | 19718 | protein_coding | 905.5446528 | 0.039357257 | 0.070093879 | 0.561493498 | 0.574461162 | 0.768186995 | 0.039357257 |
| ENSMUSG00000068039 | Tcp1 | 21454 | protein_coding | 3526.555189 | 0.039359158 | 0.054943117 | 0.716361951 | 0.473767883 | 0.697669633 | 0.039359158 |
| ENSMUSG00000000958 | Slc7a7 | 20540 | protein_coding | 870.0284404 | 0.039368698 | 0.059975962 | 0.656407934 | 0.511561687 | 0.725473655 | 0.039368698 |
| ENSMUSG00000057788 | Ddx49 | 234374 | protein_coding | 587.4592984 | 0.039375673 | 0.071203019 | 0.553005669 | 0.580259528 | 0.772159116 | 0.039375673 |
| ENSMUSG00000031592 | Pcm1 | 18536 | protein_coding | 1563.300163 | -0.039397711 | 0.047434104 | -0.830577744 | 0.406212212 | 0.645353595 | 0.039397711 |
| ENSMUSG00000024896 | Minpp1 | 17330 | protein_coding | 780.3538516 | -0.039429361 | 0.051982953 | -0.758505601 | 0.448148361 | 0.678921982 | 0.039429361 |
| ENSMUSG00000030822 | Prr14 | 233895 | protein_coding | 666.1240213 | -0.039449632 | 0.057105868 | -0.690815731 | 0.489681348 | 0.709700079 | 0.039449632 |
| ENSMUSG00000024067 | Dpy30 | 66310 | protein_coding | 805.0896453 | -0.039457144 | 0.052367895 | -0.753460576 | 0.451173193 | 0.680343771 | 0.039457144 |
| ENSMUSG00000021785 | Ngly1 | 59007 | protein_coding | 1706.287363 | 0.039474684 | 0.045957382 | 0.858941098 | 0.390373014 | 0.630176941 | 0.039474684 |
| ENSMUSG00000028673 | Fuca1 | 71665 | protein_coding | 5623.153821 | -0.039481174 | 0.036158719 | -1.091885319 | 0.274883511 | 0.517285207 | 0.039481174 |
| ENSMUSG00000010517 | Faf1 | 14084 | protein_coding | 1619.447687 | 0.039492311 | 0.06298588 | 0.62700261 | 0.530657524 | 0.73971984 | 0.039492311 |

| ensembl_gene_id | external_gene_name | entrezgene_id | gene_biotype | baseMean | log2FoldChange | lfcSE | stat | pvalue | padj | absLog2FC |
|---|---|---|---|---|---|---|---|---|---|---|
| ENSMUSG00000021456 | Fbp2 | 14120 | protein_coding | 3503.052528 | 0.039533903 | 0.039836169 | 0.992412263 | 0.32099646 | 0.564526707 | 0.039533903 |
| ENSMUSG00000019214 | Chtf18 | 214901 | protein_coding | 155.9691526 | 0.039544844 | 0.146827667 | 0.269328287 | 0.787677066 | 0.90092814 | 0.039544844 |
| ENSMUSG00000023939 | Mrpl14 | 68463 | protein_coding | 622.3101942 | 0.039558899 | 0.093700463 | 0.422184665 | 0.672890233 | 0.833463963 | 0.039558899 |
| ENSMUSG00000001672 | Marveld3 | 73608 | protein_coding | 1346.080542 | 0.039578778 | 0.069149036 | 0.572369188 | 0.567071886 | 0.762827428 | 0.039578778 |
| ENSMUSG00000097207 | 6030443J06Rik | 320719 | lncRNA | 28.37909258 | 0.039584795 | 0.252616573 | 0.15669912 | 0.87548198 | 0.943965895 | 0.039584795 |
| ENSMUSG00000048251 | Bcl11b | 58208 | protein_coding | 576.303651 | -0.03958566 | 0.104434344 | -0.379048291 | 0.704652004 | 0.853157579 | 0.03958566 |
| ENSMUSG00000031516 | Dctn6 | 22428 | protein_coding | 871.1569736 | -0.039595038 | 0.056482101 | -0.701019206 | 0.483291029 | 0.705342483 | 0.039595038 |
| ENSMUSG00000035561 | Aldh1b1 | 72535 | protein_coding | 9507.387734 | -0.039600019 | 0.050577774 | -0.782952969 | 0.43365473 | 0.667785655 | 0.039600019 |
| ENSMUSG00000048796 | Cyb561d1 | 72023 | protein_coding | 700.7706666 | -0.039611483 | 0.062843 | -0.630324516 | 0.528482286 | 0.737950908 | 0.039611483 |
| ENSMUSG00000054717 | Hmgb2 | 97165 | protein_coding | 1795.817526 | 0.039626409 | 0.091989052 | 0.430773096 | 0.666633364 | 0.829856909 | 0.039626409 |
| ENSMUSG00000028862 | Map3k6 | 53608 | protein_coding | 221.074691 | 0.039687867 | 0.096363887 | 0.411854158 | 0.680446323 | 0.838364494 | 0.039687867 |
| ENSMUSG00000044224 | Dnajc21 | 78244 | protein_coding | 583.9715274 | 0.039691214 | 0.061331777 | 0.647155782 | 0.517531126 | 0.729947916 | 0.039691214 |
| ENSMUSG00000028587 | Orc1 | 18392 | protein_coding | 204.483967 | 0.039693573 | 0.125248508 | 0.316918527 | 0.751305423 | 0.881377942 | 0.039693573 |
| ENSMUSG00000021235 | Coq6 | 217707 | protein_coding | 534.5398962 | -0.039738703 | 0.070106395 | -0.56683421 | 0.570826826 | 0.765902723 | 0.039738703 |
| ENSMUSG00000004552 | Ctse | 13034 | protein_coding | 1059.108182 | 0.039746411 | 0.151443087 | 0.262451139 | 0.79297365 | 0.903818244 | 0.039746411 |
| ENSMUSG00000027610 | Gss | 14854 | protein_coding | 3486.620016 | -0.039769051 | 0.042070776 | -0.945289231 | 0.344511235 | 0.58688999 | 0.039769051 |
| ENSMUSG00000028798 | Eif3i | 54709 | protein_coding | 3018.613874 | 0.039807169 | 0.040646739 | 0.979344721 | 0.327409681 | 0.571134226 | 0.039807169 |
| ENSMUSG00000026519 | Tmem63a | 208795 | protein_coding | 2693.781655 | -0.039808638 | 0.059644379 | -0.667433193 | 0.50449547 | 0.72070143 | 0.039808638 |
| ENSMUSG00000049164 | Zfp518a | 72672 | protein_coding | 922.6001572 | -0.039830795 | 0.058559475 | -0.680176782 | 0.496392532 | 0.714747243 | 0.039830795 |
| ENSMUSG00000031751 | Amfr | 23802 | protein_coding | 4573.508954 | -0.039848863 | 0.035468308 | -1.123506183 | 0.261222576 | 0.503084198 | 0.039848863 |
| ENSMUSG00000024614 | Tmx3 | 67988 | protein_coding | 1003.574183 | -0.039865274 | 0.049253297 | -0.809392997 | 0.418289129 | 0.655863747 | 0.039865274 |
| ENSMUSG00000026112 | Coa5 | 76178 | protein_coding | 2637.55292 | 0.03989132 | 0.046874477 | 0.851024314 | 0.394755846 | 0.634212745 | 0.03989132 |
| ENSMUSG00000022370 | Mrpl13 | 68537 | protein_coding | 600.7331984 | 0.039891737 | 0.077410807 | 0.51532517 | 0.606325813 | 0.789753 | 0.039891737 |
| ENSMUSG00000005625 | Psmd4 | 19185 | protein_coding | 1899.128741 | 0.039900826 | 0.039821122 | 1.002001547 | 0.316342846 | 0.560212157 | 0.039900826 |
| ENSMUSG00000027170 | Eif3m | 98221 | protein_coding | 1806.335323 | -0.039908411 | 0.069983931 | -0.570251057 | 0.568507432 | 0.763734926 | 0.039908411 |
| ENSMUSG00000073802 | Cdkn2b | 12579 | protein_coding | 1305.088861 | -0.039909521 | 0.078595379 | -0.507784566 | 0.611604435 | 0.792937111 | 0.039909521 |
| ENSMUSG00000028643 | Svbp | 69216 | protein_coding | 347.9548655 | 0.039943165 | 0.101029547 | 0.395361219 | 0.692576325 | 0.84576544 | 0.039943165 |
| ENSMUSG00000012126 | Ubxn11 | 67586 | protein_coding | 29.73676064 | -0.039997078 | 0.228796514 | -0.174815068 | 0.861224948 | 0.937367533 | 0.039997078 |
| ENSMUSG00000038274 | Fau | 14109 | protein_coding | 8783.830539 | -0.040005702 | 0.05149891 | -0.776826191 | 0.437261322 | 0.670378681 | 0.040005702 |
| ENSMUSG00000002504 | Slc9a3r2 | 65962 | protein_coding | 215.6759746 | -0.04003166 | 0.105746009 | -0.37856426 | 0.705011466 | 0.853326322 | 0.04003166 |
| ENSMUSG00000020808 | Pimreg | 109212 | protein_coding | 69.27250595 | 0.040044245 | 0.147903857 | 0.270745106 | 0.786587083 | 0.900451559 | 0.040044245 |
| ENSMUSG00000052397 | Ezr | 22350 | protein_coding | 13741.01855 | 0.040062963 | 0.078252193 | 0.511972398 | 0.608670327 | 0.791295959 | 0.040062963 |
| ENSMUSG00000046410 | Kcnk6 | 52150 | protein_coding | 919.5873441 | -0.040081954 | 0.091719965 | -0.437003585 | 0.662108748 | 0.827333871 | 0.040081954 |
| ENSMUSG00000033540 | Idua | 15932 | protein_coding | 239.7730039 | -0.040098514 | 0.118513071 | -0.338346767 | 0.735101885 | 0.872717027 | 0.040098514 |
| ENSMUSG00000063077 | Kif1b | 16561 | protein_coding | 2149.552239 | -0.040112595 | 0.03787201 | -1.059162022 | 0.289525998 | 0.531747525 | 0.040112595 |
| ENSMUSG00000068184 | Ndufaf2 | 75597 | protein_coding | 255.2218449 | 0.040115859 | 0.088991425 | 0.450783417 | 0.652145654 | 0.820965922 | 0.040115859 |
| ENSMUSG00000027206 | Cops2 | 12848 | protein_coding | 1481.180954 | -0.04012126 | 0.069111336 | -0.580530805 | 0.561556719 | 0.759431884 | 0.04012126 |
| ENSMUSG00000027509 | Rae1 | 66679 | protein_coding | 744.8407091 | 0.040125018 | 0.051334832 | 0.781633378 | 0.434430065 | 0.668221346 | 0.040125018 |
| ENSMUSG00000026843 | Fubp3 | 320267 | protein_coding | 997.8825802 | -0.040133616 | 0.054451353 | -0.737054529 | 0.46108919 | 0.687717915 | 0.040133616 |

| ensembl_gene_id | external_gene_name | entrezgene_id | gene_biotype | baseMean | log2FoldChange | lfcSE | stat | pvalue | padj | absLog2FC |
|---|---|---|---|---|---|---|---|---|---|---|
| ENSMUSG00000055720 | Ubl7 | 69459 | protein_coding | 535.947959 | 0.040148962 | 0.063197351 | 0.635295013 | 0.525236026 | 0.735923665 | 0.040148962 |
| ENSMUSG00000060579 | Fhit | 14198 | protein_coding | 154.8410574 | 0.040159627 | 0.093992512 | 0.427264115 | 0.669186967 | 0.831111145 | 0.040159627 |
| ENSMUSG00000049287 | Iba57 | 216792 | protein_coding | 486.3748887 | -0.040163629 | 0.087171846 | -0.460740836 | 0.644984554 | 0.81580845 | 0.040163629 |
| ENSMUSG00000004952 | Rasa4 | 54153 | protein_coding | 908.3196916 | 0.040187709 | 0.08011908 | 0.501599728 | 0.615949111 | 0.795599668 | 0.040187709 |
| ENSMUSG00000037958 | Nsrp1 | 237859 | protein_coding | 410.7633294 | 0.040189771 | 0.085498256 | 0.470065392 | 0.638308299 | 0.811381567 | 0.040189771 |
| ENSMUSG00000044700 | Tmem201 | 230917 | protein_coding | 358.6301948 | -0.040265939 | 0.104947263 | -0.383677842 | 0.701217237 | 0.851297936 | 0.040265939 |
| ENSMUSG00000032412 | Atp1b3 | 11933 | protein_coding | 944.8700944 | -0.040309713 | 0.06805372 | -0.592321962 | 0.553635012 | 0.75391672 | 0.040309713 |
| ENSMUSG00000038717 | Atp5l | 27425 | protein_coding | 546.0965077 | -0.040331212 | 0.086742339 | -0.464954168 | 0.641964278 | 0.813747171 | 0.040331212 |
| ENSMUSG00000042680 | Garem1 | 381126 | protein_coding | 656.0746363 | 0.040340678 | 0.058088115 | 0.69447388 | 0.487385076 | 0.708115319 | 0.040340678 |
| ENSMUSG00000028691 | Prdx1 | 18477 | protein_coding | 7680.352198 | 0.040343385 | 0.04484496 | 0.899619141 | 0.368322968 | 0.610412891 | 0.040343385 |
| ENSMUSG00000060657 | Marf1 | 223989 | protein_coding | 1498.14342 | 0.040372299 | 0.050421077 | 0.80070283 | 0.423303704 | 0.659294819 | 0.040372299 |
| ENSMUSG00000028672 | Hmgcl | 15356 | protein_coding | 1813.43248 | 0.040373853 | 0.046038397 | 0.876960444 | 0.380508124 | 0.621413115 | 0.040373853 |
| ENSMUSG00000031109 | Enox2 | 209224 | protein_coding | 630.4852472 | -0.040377769 | 0.051157039 | -0.789290566 | 0.4299422 | 0.664948752 | 0.040377769 |
| ENSMUSG00000048429 | Timm29 | 69773 | protein_coding | 592.268464 | -0.040395218 | 0.067447294 | -0.598915321 | 0.549229354 | 0.751378251 | 0.040395218 |
| ENSMUSG00000026187 | Xrcc5 | 22596 | protein_coding | 647.2918501 | -0.040400755 | 0.068134619 | -0.592954885 | 0.553211345 | 0.753561 | 0.040400755 |
| ENSMUSG00000018547 | Pip4k2b | 108083 | protein_coding | 232.7403444 | 0.040421614 | 0.088039245 | 0.459131767 | 0.646139546 | 0.81663734 | 0.040421614 |
| ENSMUSG00000028273 | Pdlim5 | 56376 | protein_coding | 2619.229116 | 0.040429642 | 0.04160607 | 0.971724619 | 0.331187566 | 0.574740328 | 0.040429642 |
| ENSMUSG00000044155 | Lsm8 | 76522 | protein_coding | 487.0163468 | 0.040444408 | 0.066155856 | 0.611350382 | 0.540967644 | 0.746550877 | 0.040444408 |
| ENSMUSG00000044719 | E230025N22Rik | 240216 | protein_coding | 84.18820843 | 0.040466291 | 0.14639311 | 0.2764221 | 0.782223884 | 0.898825528 | 0.040466291 |
| ENSMUSG00000031851 | Ntpcr | 66566 | protein_coding | 339.779986 | 0.040472179 | 0.072130911 | 0.561093414 | 0.574733858 | 0.768233148 | 0.040472179 |
| ENSMUSG00000021027 | Ralgapa1 | 56784 | protein_coding | 1671.234157 | -0.040498309 | 0.06878272 | -0.588786089 | 0.556004779 | 0.755288004 | 0.040498309 |
| ENSMUSG00000020253 | Ppm1m | 67905 | protein_coding | 63.26643608 | 0.040593067 | 0.145555687 | 0.278883418 | 0.780334295 | 0.898056783 | 0.040593067 |
| ENSMUSG00000040148 | Hmx3 | 15373 | protein_coding | 20.69108631 | 0.040603948 | 0.251411309 | 0.161504065 | 0.87169641 | 0.942151867 | 0.040603948 |
| ENSMUSG00000030397 | Mark4 | 232944 | protein_coding | 444.6524757 | -0.040605495 | 0.099950519 | -0.406255966 | 0.684554536 | 0.841256649 | 0.040605495 |
| ENSMUSG00000035776 | Cd99l2 | 171486 | protein_coding | 575.4278973 | -0.040622659 | 0.092235424 | -0.440423616 | 0.659630324 | 0.82592067 | 0.040622659 |
| ENSMUSG00000026499 | Acbd3 | 170760 | protein_coding | 1855.826086 | 0.040647375 | 0.039480731 | 1.029549707 | 0.303221434 | 0.547169304 | 0.040647375 |
| ENSMUSG00000051351 | Zfp46 | 22704 | protein_coding | 452.4897348 | 0.040653079 | 0.086451513 | 0.47024138 | 0.638182573 | 0.811381567 | 0.040653079 |
| ENSMUSG00000060301 | 2610008E11Rik | 72128 | protein_coding | 326.9461947 | -0.040668509 | 0.110557163 | -0.367850509 | 0.712984704 | 0.858612846 | 0.040668509 |
| ENSMUSG00000069270 | H2ac6 | 319164 | protein_coding | 398.0509542 | -0.04070623 | 0.099213228 | -0.410290354 | 0.681592967 | 0.839021966 | 0.04070623 |
| ENSMUSG00000054702 | Ap1s3 | 252903 | protein_coding | 231.5740612 | 0.040758907 | 0.155993902 | 0.261285257 | 0.793872535 | 0.904534575 | 0.040758907 |
| ENSMUSG00000031327 | Chic1 | 12212 | protein_coding | 424.2087982 | 0.040760959 | 0.087934185 | 0.463539391 | 0.642977787 | 0.814430263 | 0.040760959 |
| ENSMUSG00000001054 | Rmnd5b | 66089 | protein_coding | 428.5776461 | -0.040765576 | 0.070437903 | -0.578744886 | 0.562761324 | 0.760423283 | 0.040765576 |
| ENSMUSG00000030094 | Xpc | 22591 | protein_coding | 555.1647035 | -0.040780085 | 0.053580467 | -0.761099835 | 0.446597439 | 0.677580465 | 0.040780085 |
| ENSMUSG00000054099 | Slc25a40 | 319653 | protein_coding | 497.3669328 | -0.04081537 | 0.092365505 | -0.441889753 | 0.658568986 | 0.825261698 | 0.04081537 |
| ENSMUSG00000026829 | Gbgt1 | 227671 | protein_coding | 8.301771464 | 0.040837875 | 0.478430993 | 0.08535792 | 0.931976846 | 0.970344469 | 0.040837875 |
| ENSMUSG00000024769 | Cdc42bpg | 240505 | protein_coding | 3115.557309 | -0.040856294 | 0.054693092 | -0.747010139 | 0.455057441 | 0.682988492 | 0.040856294 |
| ENSMUSG00000031639 | Tlr3 | 142980 | protein_coding | 573.5978939 | 0.040867211 | 0.072934405 | 0.560328301 | 0.575255527 | 0.76842093 | 0.040867211 |
| ENSMUSG00000092837 | Rpph1 | 85029 | ribozyme | 484.8215962 | -0.040883423 | 0.307662165 | -0.132884142 | 0.894285008 | 0.952773623 | 0.040883423 |
| ENSMUSG00000053297 | AI854703 | 243373 | protein_coding | 920.7848088 | -0.040907212 | 0.16484965 | -0.248148611 | 0.804019425 | 0.910778265 | 0.040907212 |

Supplementary Table 1

| ensembl_gene_id | external_gene_name | entrezgene_id | gene_biotype | baseMean | log2FoldChange | lfcSE | stat | pvalue | padj | absLog2FC |
|---|---|---|---|---|---|---|---|---|---|---|
| ENSMUSG00000020486 | Septin4 | 18952 | protein_coding | 16.49797349 | 0.04092998 | 0.329484197 | 0.12422441 | 0.901137596 | 0.956346802 | 0.04092998 |
| ENSMUSG00000022008 | Gpalpp1 | 67467 | protein_coding | 1003.581465 | -0.040945317 | 0.062574552 | -0.654344558 | 0.512889845 | 0.726387506 | 0.040945317 |
| ENSMUSG00000001739 | Cldn15 | 60363 | protein_coding | 4001.978258 | 0.040972792 | 0.052363054 | 0.782475228 | 0.433935338 | 0.668045287 | 0.040972792 |
| ENSMUSG00000001767 | Crnkl1 | 66877 | protein_coding | 677.4505514 | 0.040989995 | 0.067832328 | 0.604284065 | 0.545654799 | 0.749108278 | 0.040989995 |
| ENSMUSG00000026618 | Iars2 | 381314 | protein_coding | 3069.82433 | -0.041033883 | 0.047026305 | -0.872572981 | 0.382895873 | 0.623481598 | 0.041033883 |
| ENSMUSG00000030509 | Asb7 | 117589 | protein_coding | 831.4359383 | 0.041056929 | 0.075373277 | 0.544714661 | 0.58594978 | 0.77637824 | 0.041056929 |
| ENSMUSG00000025577 | Cbx2 | 12416 | protein_coding | 169.8439873 | 0.041084752 | 0.1004197 | 0.409130396 | 0.682443971 | 0.839494578 | 0.041084752 |
| ENSMUSG00000024654 | Asrgl1 | 66514 | protein_coding | 107.0901896 | 0.041108346 | 0.147285588 | 0.279106368 | 0.780163196 | 0.898056783 | 0.041108346 |
| ENSMUSG00000000275 | Trim25 | 217069 | protein_coding | 1902.763695 | 0.04114853 | 0.03828666 | 1.074748512 | 0.282487338 | 0.525210203 | 0.04114853 |
| ENSMUSG00000031386 | Hcfc1 | 15161 | protein_coding | 2266.949253 | 0.041148605 | 0.068745818 | 0.598561577 | 0.549465285 | 0.751460876 | 0.041148605 |
| ENSMUSG00000002365 | Snx9 | 66616 | protein_coding | 2183.096351 | -0.041161091 | 0.037587503 | -1.095073838 | 0.273484289 | 0.51596644 | 0.041161091 |
| ENSMUSG00000034485 | Uaca | 72565 | protein_coding | 1982.248882 | 0.04118369 | 0.051169667 | 0.804845778 | 0.420908681 | 0.657926757 | 0.04118369 |
| ENSMUSG00000049553 | Polr1a | 20019 | protein_coding | 673.7645794 | -0.041189048 | 0.063214753 | -0.651573345 | 0.514676448 | 0.727384723 | 0.041189048 |
| ENSMUSG00000037773 | Pced1a | 319513 | protein_coding | 176.9160356 | 0.041214715 | 0.102741867 | 0.4011482 | 0.688311015 | 0.843368224 | 0.041214715 |
| ENSMUSG00000068115 | Ninl | 78177 | protein_coding | 59.79514354 | 0.041216152 | 0.152748667 | 0.269829865 | 0.787291146 | 0.900662858 | 0.041216152 |
| ENSMUSG00000026779 | Mastl | 67121 | protein_coding | 338.687962 | -0.041216428 | 0.116741471 | -0.353057294 | 0.724045488 | 0.865580106 | 0.041216428 |
| ENSMUSG00000045725 | Prr15 | 78004 | protein_coding | 1284.219727 | -0.041252005 | 0.103292222 | -0.399371843 | 0.689619238 | 0.844292391 | 0.041252005 |
| ENSMUSG00000033060 | Lmo7 | 380928 | protein_coding | 5742.868139 | -0.04128025 | 0.048789527 | -0.846088348 | 0.397503461 | 0.637040476 | 0.04128025 |
| ENSMUSG00000036241 | Ube2r2 | 67615 | protein_coding | 2105.28539 | 0.041293145 | 0.058037675 | 0.711488615 | 0.476781504 | 0.699625498 | 0.041293145 |
| ENSMUSG00000007836 | Hnrnpa0 | 77134 | protein_coding | 2537.202804 | 0.041305289 | 0.09196608 | 0.44913613 | 0.653333459 | 0.821648745 | 0.041305289 |
| ENSMUSG00000015474 | Ppt2 | 54397 | protein_coding | 296.9139288 | -0.041319306 | 0.098146444 | -0.420996462 | 0.673757665 | 0.834153989 | 0.041319306 |
| ENSMUSG00000028854 | Slc9a1 | 20544 | protein_coding | 830.3509819 | -0.041326918 | 0.08004479 | -0.516297415 | 0.605646701 | 0.789401467 | 0.041326918 |
| ENSMUSG00000030232 | Aebp2 | 11569 | protein_coding | 935.9808129 | -0.041342152 | 0.053544169 | -0.772113058 | 0.440047469 | 0.671885771 | 0.041342152 |
| ENSMUSG00000040188 | Scamp2 | 24044 | protein_coding | 4544.489727 | 0.041356009 | 0.050247728 | 0.823042379 | 0.410483894 | 0.648632239 | 0.041356009 |
| ENSMUSG00000047731 | Wbp1l | 226178 | protein_coding | 1665.521099 | 0.041367036 | 0.04868917 | 0.849614741 | 0.395539314 | 0.634775849 | 0.041367036 |
| ENSMUSG00000038346 | Zfp384 | 269800 | protein_coding | 603.5700065 | -0.041378184 | 0.063051948 | -0.656255437 | 0.511659785 | 0.725473655 | 0.041378184 |
| ENSMUSG00000009076 | Zmat5 | 67178 | protein_coding | 240.1403864 | 0.041397726 | 0.086860346 | 0.476600977 | 0.633646288 | 0.808283189 | 0.041397726 |
| ENSMUSG00000006134 | Crkl | 12929 | protein_coding | 1012.692274 | -0.041424788 | 0.057102159 | -0.725450477 | 0.468175716 | 0.693047027 | 0.041424788 |
| ENSMUSG00000021963 | Sap18 | 20220 | protein_coding | 1084.685829 | 0.041444611 | 0.05658278 | 0.732459801 | 0.46388797 | 0.690013032 | 0.041444611 |
| ENSMUSG00000020962 | Gtf2a1 | 83602 | protein_coding | 911.9191156 | -0.041454019 | 0.046884797 | -0.884167602 | 0.376605745 | 0.618232985 | 0.041454019 |
| ENSMUSG00000056752 | Dnah9 | 237806 | protein_coding | 24.95498355 | -0.041462362 | 0.281304088 | -0.147393386 | 0.882821526 | 0.947823896 | 0.041462362 |
| ENSMUSG00000010609 | Psen2 | 19165 | protein_coding | 392.9078289 | 0.041469878 | 0.082185435 | 0.504589139 | 0.613847439 | 0.794312578 | 0.041469878 |
| ENSMUSG00000038764 | Ptpn3 | 545622 | protein_coding | 2345.066908 | 0.041493495 | 0.062070028 | 0.668494865 | 0.503817759 | 0.720116015 | 0.041493495 |
| ENSMUSG00000007908 | Hmgcll1 | 208982 | protein_coding | 8.667200272 | 0.041493648 | 0.397820142 | 0.104302531 | 0.91692927 | 0.963217713 | 0.041493648 |
| ENSMUSG00000006262 | Mob1b | 68473 | protein_coding | 2054.719637 | 0.04149861 | 0.052152485 | 0.795716825 | 0.426196645 | 0.66169488 | 0.04149861 |
| ENSMUSG00000002455 | Prpf6 | 68879 | protein_coding | 1333.30771 | 0.041513865 | 0.05544507 | 0.748738602 | 0.454014772 | 0.682675952 | 0.041513865 |
| ENSMUSG00000014602 | Kif1a | 16560 | protein_coding | 39.10437881 | -0.041560283 | 0.210970661 | -0.196995557 | 0.843831014 | 0.930175172 | 0.041560283 |
| ENSMUSG00000024442 | Dele1 | 66839 | protein_coding | 1054.144172 | 0.041571542 | 0.054129636 | 0.767999645 | 0.442487396 | 0.673641376 | 0.041571542 |
| ENSMUSG00000039738 | Slx4 | 52864 | protein_coding | 218.0980467 | -0.041602384 | 0.108140282 | -0.384707561 | 0.700454093 | 0.850619214 | 0.041602384 |

| ensembl_gene_id | external_gene_name | entrezgene_id | gene_biotype | baseMean | log2FoldChange | lfcSE | stat | pvalue | padj | absLog2FC |
|---|---|---|---|---|---|---|---|---|---|---|
| ENSMUSG00000068154 | Insm1 | 53626 | protein_coding | 49.8958271 | -0.04160265 | 0.197566098 | -0.210575855 | 0.833218255 | 0.9260086 | 0.04160265 |
| ENSMUSG00000039911 | Spsb1 | 74646 | protein_coding | 55.37931034 | 0.041606737 | 0.195079699 | 0.213280713 | 0.831108016 | 0.925277452 | 0.041606737 |
| ENSMUSG00000034903 | Cobll1 | 319876 | protein_coding | 528.8294649 | -0.041625606 | 0.059978646 | -0.694007099 | 0.487677757 | 0.708241492 | 0.041625606 |
| ENSMUSG00000022074 | Tnfrsf10b | 21933 | protein_coding | 199.0953333 | 0.041634776 | 0.110740295 | 0.375967721 | 0.706940893 | 0.854251145 | 0.041634776 |
| ENSMUSG00000042211 | Fbxo38 | 107035 | protein_coding | 1555.525363 | -0.041668509 | 0.043263312 | -0.963137268 | 0.335478641 | 0.579071395 | 0.041668509 |
| ENSMUSG00000035382 | Pcsk7 | 18554 | protein_coding | 3566.62748 | 0.041712171 | 0.040988625 | 1.017652381 | 0.308843185 | 0.552491851 | 0.041712171 |
| ENSMUSG00000002372 | Ranbp3 | 71810 | protein_coding | 1192.871377 | 0.041725741 | 0.042303984 | 0.986331252 | 0.323970588 | 0.567961048 | 0.041725741 |
| ENSMUSG00000034211 | Mrps17 | 66258 | protein_coding | 889.5296042 | 0.041731732 | 0.049208387 | 0.848061377 | 0.39640379 | 0.635720358 | 0.041731732 |
| ENSMUSG00000050730 | Arhgap42 | 71544 | protein_coding | 692.254974 | -0.041742922 | 0.059234848 | -0.704702102 | 0.480995643 | 0.703236518 | 0.041742922 |
| ENSMUSG00000035273 | Hpse | 15442 | protein_coding | 341.8800546 | -0.041768304 | 0.082671303 | -0.505233407 | 0.613394909 | 0.793911553 | 0.041768304 |
| ENSMUSG00000026098 | Pms1 | 227099 | protein_coding | 111.740534 | 0.041834889 | 0.128825509 | 0.324740722 | 0.745377312 | 0.879027415 | 0.041834889 |
| ENSMUSG00000020610 | Amz2 | 13929 | protein_coding | 736.6109017 | 0.041840953 | 0.055669118 | 0.75160078 | 0.452291174 | 0.68117317 | 0.041840953 |
| ENSMUSG00000021905 | Dph3 | 105638 | protein_coding | 647.4011457 | 0.041858882 | 0.066687268 | 0.627689257 | 0.530207524 | 0.739484281 | 0.041858882 |
| ENSMUSG00000047719 | Ubiad1 | 71707 | protein_coding | 431.3963752 | 0.041877951 | 0.077114504 | 0.543061932 | 0.587087162 | 0.777161429 | 0.041877951 |
| ENSMUSG00000033454 | Zbtb1 | 268564 | protein_coding | 617.8109867 | -0.041890503 | 0.062852318 | -0.666490979 | 0.505097328 | 0.721241774 | 0.041890503 |
| ENSMUSG00000031467 | Agpat5 | 52123 | protein_coding | 1231.132905 | -0.041905316 | 0.064091323 | -0.653837587 | 0.513216448 | 0.726594825 | 0.041905316 |
| ENSMUSG00000001445 | Mrpl10 | 107732 | protein_coding | 1252.670822 | 0.041922016 | 0.05538432 | 0.756929337 | 0.449092198 | 0.679215578 | 0.041922016 |
| ENSMUSG00000107476 | Zfp862-ps | 58894 | transcribed_unitary_pseudogene | 164.6675811 | -0.041948365 | 0.142633009 | -0.294099981 | 0.768681506 | 0.89151576 | 0.041948365 |
| ENSMUSG00000032737 | Inppl1 | 16332 | protein_coding | 2290.16191 | -0.041965715 | 0.059934088 | -0.700197774 | 0.483803802 | 0.705835346 | 0.041965715 |
| ENSMUSG00000010067 | Rassf1 | 56289 | protein_coding | 377.7283052 | -0.04196739 | 0.090417804 | -0.464149625 | 0.64254055 | 0.814130378 | 0.04196739 |
| ENSMUSG00000028549 | Itgb3bp | 67733 | protein_coding | 101.7718842 | -0.041989042 | 0.133286707 | -0.31502798 | 0.75274041 | 0.882492633 | 0.041989042 |
| ENSMUSG00000057604 | Lmcd1 | 30937 | protein_coding | 15.79808483 | -0.042005835 | 0.363268779 | -0.115632935 | 0.907943459 | 0.959122007 | 0.042005835 |
| ENSMUSG00000021113 | Snapc1 | 75627 | protein_coding | 239.0603058 | 0.042036014 | 0.103382756 | 0.406605661 | 0.684297639 | 0.841005032 | 0.042036014 |
| ENSMUSG00000031921 | Terf2 | 21750 | protein_coding | 351.6389812 | -0.042053524 | 0.070006509 | -0.600708777 | 0.548033972 | 0.750796946 | 0.042053524 |
| ENSMUSG00000018347 | Zkscan6 | 52712 | protein_coding | 429.9570381 | 0.042056284 | 0.067961473 | 0.618825378 | 0.536031402 | 0.742852833 | 0.042056284 |
| ENSMUSG00000028066 | Pmf1 | 67037 | protein_coding | 383.5567821 | 0.042069629 | 0.092211155 | 0.456231453 | 0.64822355 | 0.818309291 | 0.042069629 |
| ENSMUSG00000021371 | Mcur1 | 76137 | protein_coding | 1381.792804 | -0.042079815 | 0.062335242 | -0.675056583 | 0.499639816 | 0.716748916 | 0.042079815 |
| ENSMUSG00000022019 | Tdrd3 | 219249 | protein_coding | 825.7930018 | -0.042083302 | 0.062620934 | -0.672032496 | 0.501563009 | 0.718549049 | 0.042083302 |
| ENSMUSG00000020776 | Fbf1 | 217335 | protein_coding | 304.4078956 | 0.042101333 | 0.086906159 | 0.484445907 | 0.628069445 | 0.803942848 | 0.042101333 |
| ENSMUSG00000025869 | Nop16 | 28126 | protein_coding | 417.1724032 | 0.042106416 | 0.067749649 | 0.621500138 | 0.534270604 | 0.741606925 | 0.042106416 |
| ENSMUSG00000068394 | Cep152 | 99100 | protein_coding | 267.6105607 | -0.042113449 | 0.094033299 | -0.447856765 | 0.654256573 | 0.822000293 | 0.042113449 |
| ENSMUSG00000021420 | Fars2 | 69955 | protein_coding | 948.1326165 | 0.04211495 | 0.058068763 | 0.725259986 | 0.468292549 | 0.693047027 | 0.04211495 |
| ENSMUSG00000033931 | Rbm34 | 52202 | protein_coding | 324.4340699 | 0.04213053 | 0.088478049 | 0.476169297 | 0.633953771 | 0.808458996 | 0.04213053 |
| ENSMUSG00000029290 | Zfp326 | 54367 | protein_coding | 397.0080688 | -0.042132149 | 0.078653439 | -0.535668231 | 0.592187864 | 0.780262005 | 0.042132149 |
| ENSMUSG00000031198 | Fundc2 | 67391 | protein_coding | 394.3551709 | 0.042134126 | 0.084760467 | 0.497096433 | 0.61912105 | 0.798277593 | 0.042134126 |
| ENSMUSG00000006418 | Rnf114 | 81018 | protein_coding | 1973.228461 | 0.042147565 | 0.053897901 | 0.781988978 | 0.434221051 | 0.668067439 | 0.042147565 |
| ENSMUSG00000025068 | Gsto1 | 14873 | protein_coding | 9619.509042 | 0.042178558 | 0.051722324 | 0.815480734 | 0.414797188 | 0.65259636 | 0.042178558 |
| ENSMUSG00000040957 | Cables1 | 63955 | protein_coding | 848.5973669 | 0.042242083 | 0.115029921 | 0.367226915 | 0.713449764 | 0.858898503 | 0.042242083 |
| ENSMUSG00000037376 | Trmt6 | 66926 | protein_coding | 374.7864373 | 0.04225769 | 0.089889028 | 0.47010954 | 0.638276758 | 0.811381567 | 0.04225769 |

| ensembl_gene_id | external_gene_name | entrezgene_id | gene_biotype | baseMean | log2FoldChange | lfcSE | stat | pvalue | padj | absLog2FC |
|---|---|---|---|---|---|---|---|---|---|---|
| ENSMUSG00000015013 | Trappc2l | 59005 | protein_coding | 562.7051424 | -0.042270872 | 0.068663435 | -0.615624202 | 0.538142574 | 0.744499039 | 0.042270872 |
| ENSMUSG00000042396 | Rbm7 | 67010 | protein_coding | 1189.222736 | -0.042287667 | 0.052759399 | -0.801519114 | 0.422831183 | 0.658779673 | 0.042287667 |
| ENSMUSG00000037896 | Rcor1 | 217864 | protein_coding | 1675.333339 | -0.042296389 | 0.068669923 | -0.615937612 | 0.537935696 | 0.744340538 | 0.042296389 |
| ENSMUSG00000027593 | Raly | 19383 | protein_coding | 2649.260525 | 0.042297008 | 0.054756763 | 0.772452684 | 0.43984636 | 0.671740242 | 0.042297008 |
| ENSMUSG00000028495 | Rps6 | 20104 | protein_coding | 5095.549808 | -0.04232644 | 0.054832135 | -0.771927628 | 0.440157293 | 0.671885771 | 0.04232644 |
| ENSMUSG00000033355 | Rtp4 | 67775 | protein_coding | 123.1173825 | 0.042352721 | 0.231302655 | 0.18310521 | 0.854715468 | 0.935123077 | 0.042352721 |
| ENSMUSG00000033589 | Reep4 | 72549 | protein_coding | 775.6907187 | 0.042360383 | 0.071023145 | 0.596430688 | 0.550887545 | 0.752092411 | 0.042360383 |
| ENSMUSG00000027751 | Supt20 | 56790 | protein_coding | 866.2611281 | -0.042370906 | 0.081909 | -0.517292433 | 0.604952036 | 0.789053031 | 0.042370906 |
| ENSMUSG00000043311 | D17H6S53E | 114585 | protein_coding | 848.2488211 | -0.042378651 | 0.058359435 | -0.726166222 | 0.467736876 | 0.692926016 | 0.042378651 |
| ENSMUSG00000013593 | Ndufs2 | 226646 | protein_coding | 7485.975685 | 0.042399742 | 0.066150676 | 0.640957051 | 0.521550588 | 0.732626853 | 0.042399742 |
| ENSMUSG00000053046 | Brsk2 | 75770 | protein_coding | 12.66631512 | -0.042436208 | 0.356772634 | -0.118944685 | 0.90531918 | 0.95845089 | 0.042436208 |
| ENSMUSG00000075284 | Wipf1 | 215280 | protein_coding | 277.6544795 | -0.042445868 | 0.097210989 | -0.436636525 | 0.662374969 | 0.827558191 | 0.042445868 |
| ENSMUSG00000073448 | Gm10509 | 100310809 | protein_coding | 47.17184708 | -0.042485037 | 0.21353152 | -0.198963775 | 0.842291081 | 0.929889314 | 0.042485037 |
| ENSMUSG00000036894 | Rap2b | 74012 | protein_coding | 57.7886551 | 0.042517381 | 0.202043824 | 0.21043643 | 0.833327062 | 0.926065706 | 0.042517381 |
| ENSMUSG00000032480 | Dhx30 | 72831 | protein_coding | 1020.119941 | 0.042530556 | 0.054065593 | 0.786647355 | 0.43148833 | 0.666166426 | 0.042530556 |
| ENSMUSG00000014426 | Map3k4 | 26407 | protein_coding | 1431.746641 | 0.042535893 | 0.039013316 | 1.090291666 | 0.275584685 | 0.518231602 | 0.042535893 |
| ENSMUSG00000021911 | Parg | 26430 | protein_coding | 1125.621832 | 0.042544844 | 0.062454501 | 0.681213413 | 0.495736462 | 0.714326476 | 0.042544844 |
| ENSMUSG00000029535 | Triap1 | 69076 | protein_coding | 480.9105483 | 0.042561491 | 0.079263574 | 0.53696154 | 0.59129418 | 0.780129105 | 0.042561491 |
| ENSMUSG00000033545 | Znrf1 | 170737 | protein_coding | 401.77805 | 0.042567935 | 0.072743161 | 0.585181263 | 0.558425842 | 0.756975167 | 0.042567935 |
| ENSMUSG00000031402 | Mpp1 | 17524 | protein_coding | 783.3484831 | -0.042641623 | 0.085509395 | -0.498677643 | 0.618006497 | 0.797454855 | 0.042641623 |
| ENSMUSG00000015176 | Nolc1 | 70769 | protein_coding | 790.4195416 | 0.042648985 | 0.088831679 | 0.480110082 | 0.631149119 | 0.806348875 | 0.042648985 |
| ENSMUSG00000040396 | Abhd13 | 68904 | protein_coding | 1952.692828 | 0.04264926 | 0.036631019 | 1.16429356 | 0.244305065 | 0.485573395 | 0.04264926 |
| ENSMUSG00000026932 | Nacc2 | 67991 | protein_coding | 845.3330657 | 0.042660779 | 0.061141661 | 0.697736674 | 0.485341888 | 0.706609067 | 0.042660779 |
| ENSMUSG00000048000 | Gigyf2 | 227331 | protein_coding | 1993.506873 | -0.042677629 | 0.040208396 | -1.061410877 | 0.288503216 | 0.530912668 | 0.042677629 |
| ENSMUSG00000055128 | Cgrrf1 | 68755 | protein_coding | 377.1374161 | -0.042678216 | 0.081842459 | -0.521467908 | 0.602040856 | 0.786991399 | 0.042678216 |
| ENSMUSG00000025759 | Mfsd8 | 72175 | protein_coding | 333.3039651 | 0.042679389 | 0.078533921 | 0.543451652 | 0.58681887 | 0.777011446 | 0.042679389 |
| ENSMUSG00000097360 | 9430065F17Rik | 77857 | lncRNA | 8.14836902 | 0.042699878 | 0.424430773 | 0.100605047 | 0.91986399 | 0.96489645 | 0.042699878 |
| ENSMUSG00000038267 | Slc22a23 | 73102 | protein_coding | 3031.160286 | -0.042705595 | 0.058700833 | -0.727512583 | 0.46691201 | 0.692467352 | 0.042705595 |
| ENSMUSG00000041431 | Ccnb1 | 268697 | protein_coding | 534.8218152 | 0.042717589 | 0.082884614 | 0.515386232 | 0.606283151 | 0.789753 | 0.042717589 |
| ENSMUSG00000067424 | Zfp563 | 240068 | protein_coding | 198.5357355 | 0.042734021 | 0.093202443 | 0.458507525 | 0.646587857 | 0.816941518 | 0.042734021 |
| ENSMUSG00000029098 | Acox3 | 80911 | protein_coding | 977.1495714 | -0.042741208 | 0.067122673 | -0.636762607 | 0.524279486 | 0.735121083 | 0.042741208 |
| ENSMUSG00000087305 | A430035B10Rik | 320312 | lncRNA | 54.79422919 | 0.042742088 | 0.172765044 | 0.247400093 | 0.804598602 | 0.911091255 | 0.042742088 |
| ENSMUSG00000090258 | Churc1 | 211151 | protein_coding | 346.6529118 | -0.042765125 | 0.116543945 | -0.366944205 | 0.713660637 | 0.859024115 | 0.042765125 |
| ENSMUSG00000027677 | Ttc14 | 67120 | protein_coding | 638.499456 | -0.042767893 | 0.090167388 | -0.474316646 | 0.635274123 | 0.809423077 | 0.042767893 |
| ENSMUSG00000056019 | Zfp709 | 236193 | protein_coding | 146.4413787 | -0.042771741 | 0.13236278 | -0.323140244 | 0.746589034 | 0.879811357 | 0.042771741 |
| ENSMUSG00000000339 | Rtca | 66368 | protein_coding | 814.2373449 | -0.042788682 | 0.059951864 | -0.713717282 | 0.475402018 | 0.698824933 | 0.042788682 |
| ENSMUSG00000051390 | Zbtb22 | 81630 | protein_coding | 1021.651493 | 0.04279762 | 0.081394363 | 0.525805705 | 0.59902321 | 0.784827956 | 0.04279762 |
| ENSMUSG00000026946 | Nmi | 64685 | protein_coding | 612.7743186 | 0.042801732 | 0.079254467 | 0.540054498 | 0.589159449 | 0.77864167 | 0.042801732 |
| ENSMUSG00000097133 | Gm26628 | 102632638 | lncRNA | 7.880971753 | -0.04280511 | 0.457778367 | -0.093506188 | 0.925501434 | 0.967558773 | 0.04280511 |

| ensembl_gene_id | external_gene_name | entrezgene_id | gene_biotype | baseMean | log2FoldChange | lfcSE | stat | pvalue | padj | absLog2FC |
|---|---|---|---|---|---|---|---|---|---|---|
| ENSMUSG00000028063 | Lmna | 16905 | protein_coding | 10178.87829 | 0.042819691 | 0.083194242 | 0.514695363 | 0.606765914 | 0.789775133 | 0.042819691 |
| ENSMUSG00000018572 | Phf23 | 78246 | protein_coding | 728.5366938 | -0.042820112 | 0.073826132 | -0.580012938 | 0.561905893 | 0.759659123 | 0.042820112 |
| ENSMUSG00000040749 | Siah1b | 20438 | protein_coding | 98.87508803 | 0.042827573 | 0.120120556 | 0.356538248 | 0.721437514 | 0.863807468 | 0.042827573 |
| ENSMUSG00000052539 | Magi3 | 99470 | protein_coding | 2795.893288 | 0.042870749 | 0.046006393 | 0.931843283 | 0.351417522 | 0.593808716 | 0.042870749 |
| ENSMUSG00000025243 | Slc6a20b | 22599 | protein_coding | 9.114246783 | -0.042875098 | 0.425494223 | -0.100765405 | 0.91973669 | 0.96489645 | 0.042875098 |
| ENSMUSG00000042251 | Pm20d1 | 212933 | protein_coding | 655.4499458 | -0.042885839 | 0.064463574 | -0.665272433 | 0.505876259 | 0.721715006 | 0.042885839 |
| ENSMUSG00000031950 | Gabarapl2 | 93739 | protein_coding | 997.4302354 | 0.042889652 | 0.061892353 | 0.692971753 | 0.488327279 | 0.708843746 | 0.042889652 |
| ENSMUSG00000037058 | Paip2 | 67869 | protein_coding | 1666.032918 | 0.042925276 | 0.038800933 | 1.106294934 | 0.268598884 | 0.511015761 | 0.042925276 |
| ENSMUSG00000022972 | Cfap298 | 68001 | protein_coding | 136.9837704 | 0.042930367 | 0.130874905 | 0.328025964 | 0.742892031 | 0.87731545 | 0.042930367 |
| ENSMUSG00000002332 | Dhrs1 | 52585 | protein_coding | 717.779302 | 0.042931473 | 0.075011535 | 0.572331607 | 0.56709734 | 0.762827428 | 0.042931473 |
| ENSMUSG00000029592 | Usp30 | 100756 | protein_coding | 453.8278939 | 0.042942563 | 0.0816616 | 0.525859924 | 0.598985536 | 0.784827956 | 0.042942563 |
| ENSMUSG00000020585 | Laptm4a | 17775 | protein_coding | 4173.159363 | 0.042955925 | 0.040877903 | 1.050834853 | 0.293334445 | 0.536288924 | 0.042955925 |
| ENSMUSG00000025439 | Clns1a | 12729 | protein_coding | 945.2556938 | -0.042963552 | 0.058576341 | -0.733462537 | 0.463276369 | 0.689357631 | 0.042963552 |
| ENSMUSG00000017119 | Nbr1 | 17966 | protein_coding | 5942.23308 | 0.042963801 | 0.034548484 | 1.243579932 | 0.213654206 | 0.449303803 | 0.042963801 |
| ENSMUSG00000028948 | Nol9 | 74035 | protein_coding | 615.2934254 | -0.043076397 | 0.058936853 | -0.730890698 | 0.464845918 | 0.690944422 | 0.043076397 |
| ENSMUSG00000054263 | Lifr | 16880 | protein_coding | 256.8867928 | 0.04307868 | 0.155370496 | 0.277264223 | 0.781577229 | 0.898525697 | 0.04307868 |
| ENSMUSG00000015291 | Gdi1 | 14567 | protein_coding | 1333.368723 | -0.043089034 | 0.055719432 | -0.773321481 | 0.439332146 | 0.671700891 | 0.043089034 |
| ENSMUSG00000020630 | Rnaseh1 | 19819 | protein_coding | 232.6261133 | -0.043099951 | 0.088939067 | -0.484600892 | 0.627959481 | 0.803865895 | 0.043099951 |
| ENSMUSG00000050390 | C77080 | 97130 | protein_coding | 4726.701024 | -0.043102055 | 0.074857261 | -0.575789899 | 0.564757202 | 0.761525033 | 0.043102055 |
| ENSMUSG00000019794 | Katna1 | 23924 | protein_coding | 777.734732 | 0.043104397 | 0.071771393 | 0.600579077 | 0.548120377 | 0.750796946 | 0.043104397 |
| ENSMUSG00000048486 | Fitm2 | 228859 | protein_coding | 465.2113898 | -0.043124741 | 0.074549866 | -0.578468388 | 0.562947934 | 0.760490915 | 0.043124741 |
| ENSMUSG00000031604 | Msmo1 | 66234 | protein_coding | 3529.185536 | -0.043147713 | 0.126456773 | -0.341205238 | 0.73294908 | 0.871251553 | 0.043147713 |
| ENSMUSG00000031644 | Nek1 | 18004 | protein_coding | 496.106739 | -0.043166869 | 0.077737558 | -0.555289748 | 0.57869649 | 0.771101239 | 0.043166869 |
| ENSMUSG00000073563 | Csnk1g3 | 70425 | protein_coding | 1214.565355 | -0.043185754 | 0.053644124 | -0.805041652 | 0.420795644 | 0.65789349 | 0.043185754 |
| ENSMUSG00000018171 | Vmp1 | 75909 | protein_coding | 2262.308183 | -0.043206484 | 0.085394885 | -0.505961027 | 0.612884009 | 0.793748541 | 0.043206484 |
| ENSMUSG00000000581 | C1d | 57316 | protein_coding | 744.2283463 | 0.043231868 | 0.068179579 | 0.63408822 | 0.52602325 | 0.736439527 | 0.043231868 |
| ENSMUSG00000030172 | Erc1 | 111173 | protein_coding | 631.6127576 | 0.043258457 | 0.057293836 | 0.75502812 | 0.45023211 | 0.679535938 | 0.043258457 |
| ENSMUSG00000074748 | Atxn7l3b | 382423 | protein_coding | 2607.662675 | 0.043263967 | 0.04162146 | 1.039462967 | 0.298589473 | 0.542138138 | 0.043263967 |
| ENSMUSG00000044433 | Camsap3 | 69697 | protein_coding | 1439.163603 | 0.043266978 | 0.089486533 | 0.483502677 | 0.62873886 | 0.804352815 | 0.043266978 |
| ENSMUSG00000025133 | Ints4 | 101861 | protein_coding | 877.5222639 | 0.04328654 | 0.051358047 | 0.842838501 | 0.399318772 | 0.638789048 | 0.04328654 |
| ENSMUSG00000021276 | Cinp | 67236 | protein_coding | 282.9427598 | 0.043297287 | 0.083769434 | 0.516862591 | 0.605252084 | 0.789158883 | 0.043297287 |
| ENSMUSG00000045659 | Plekha7 | 233765 | protein_coding | 1709.287454 | -0.043324005 | 0.065628306 | -0.660142056 | 0.509162673 | 0.724225297 | 0.043324005 |
| ENSMUSG00000033377 | Palmd | 114301 | protein_coding | 32.00110075 | 0.04332589 | 0.220101223 | 0.196845293 | 0.843948606 | 0.930177636 | 0.04332589 |
| ENSMUSG00000048003 | Catsper4 | 329954 | protein_coding | 15.09241451 | 0.043326805 | 0.313667032 | 0.138129929 | 0.890137733 | 0.950993787 | 0.043326805 |
| ENSMUSG00000040648 | Ppip5k2 | 227399 | protein_coding | 2589.884656 | 0.04332747 | 0.044548572 | 0.972589416 | 0.330757405 | 0.574302828 | 0.04332747 |
| ENSMUSG00000027551 | Zfp64 | 22722 | protein_coding | 515.9656806 | 0.043344377 | 0.072456314 | 0.598213935 | 0.549697195 | 0.751487589 | 0.043344377 |
| ENSMUSG00000039616 | Mocos | 68591 | protein_coding | 234.5276684 | 0.043392099 | 0.137141812 | 0.316403131 | 0.751696541 | 0.881708496 | 0.043392099 |
| ENSMUSG00000073000 | Gm10451 | 100041694 | lncRNA | 53.34063065 | 0.043408878 | 0.256244232 | 0.169404311 | 0.865478633 | 0.939081875 | 0.043408878 |
| ENSMUSG00000029440 | Psmd9 | 67151 | protein_coding | 432.9224352 | 0.043412091 | 0.06659496 | 0.651882532 | 0.514476955 | 0.727325341 | 0.043412091 |

| ensembl_gene_id | external_gene_name | entrezgene_id | gene_biotype | baseMean | log2FoldChange | lfcSE | stat | pvalue | padj | absLog2FC |
|---|---|---|---|---|---|---|---|---|---|---|
| ENSMUSG00000039108 | Lsm14b | 241846 | protein_coding | 744.8215934 | -0.043448589 | 0.063939079 | -0.679531038 | 0.496801448 | 0.715005734 | 0.043448589 |
| ENSMUSG00000019782 | Rwdd1 | 66521 | protein_coding | 329.314269 | 0.043471344 | 0.102919456 | 0.422382181 | 0.672746082 | 0.833349418 | 0.043471344 |
| ENSMUSG00000062997 | Rpl35 | 66489 | protein_coding | 2485.6013 | -0.043477861 | 0.042503679 | -1.022919956 | 0.306345693 | 0.550527858 | 0.043477861 |
| ENSMUSG00000010066 | Cacna2d2 | 56808 | protein_coding | 226.2881725 | 0.043497671 | 0.088804273 | 0.489815075 | 0.624264762 | 0.801681623 | 0.043497671 |
| ENSMUSG00000020745 | Pafah1b1 | 18472 | protein_coding | 3790.609009 | -0.043499469 | 0.036046317 | -1.206765989 | 0.227522278 | 0.4661143 | 0.043499469 |
| ENSMUSG00000039018 | Mtg1 | 212508 | protein_coding | 219.1600126 | 0.043527113 | 0.093241266 | 0.466822417 | 0.640626936 | 0.813111176 | 0.043527113 |
| ENSMUSG00000021820 | Camk2g | 12325 | protein_coding | 290.3638877 | 0.043533876 | 0.0780972 | 0.557431967 | 0.57723233 | 0.769786305 | 0.043533876 |
| ENSMUSG00000025607 | Copg2 | 54160 | protein_coding | 1454.337946 | 0.043557045 | 0.051218522 | 0.850415882 | 0.395093909 | 0.634439856 | 0.043557045 |
| ENSMUSG00000015869 | Prpsap1 | 67763 | protein_coding | 2297.106181 | 0.043559988 | 0.079482136 | 0.548047528 | 0.583659268 | 0.775075241 | 0.043559988 |
| ENSMUSG00000101942 | Gm19582 | 100503175 | lncRNA | 5.949597013 | 0.043587576 | 0.479276193 | 0.090944588 | 0.927536621 | 0.968095473 | 0.043587576 |
| ENSMUSG00000048874 | Phf3 | 213109 | protein_coding | 1922.051219 | -0.043596527 | 0.045344217 | -0.961457275 | 0.336322298 | 0.57990695 | 0.043596527 |
| ENSMUSG00000024248 | Cox7a2l | 20463 | protein_coding | 4558.363263 | 0.04370629 | 0.050302995 | 0.868860587 | 0.384923389 | 0.625286182 | 0.04370629 |
| ENSMUSG00000036893 | Ehmt1 | 77683 | protein_coding | 1277.938736 | -0.04372091 | 0.038538538 | -1.134472458 | 0.25659642 | 0.498509875 | 0.04372091 |
| ENSMUSG00000068328 | Aup1 | 11993 | protein_coding | 1813.673692 | 0.043730231 | 0.041551335 | 1.052438636 | 0.29259835 | 0.535307271 | 0.043730231 |
| ENSMUSG00000024180 | Pgap6 | 60455 | protein_coding | 2785.598745 | -0.043737078 | 0.054144908 | -0.807778228 | 0.419218261 | 0.656573069 | 0.043737078 |
| ENSMUSG00000043140 | Tmem186 | 66690 | protein_coding | 550.3949899 | -0.043761382 | 0.060394121 | -0.724596727 | 0.468699468 | 0.693332995 | 0.043761382 |
| ENSMUSG00000006728 | Cdk4 | 12567 | protein_coding | 1644.727038 | -0.043786077 | 0.07385859 | -0.592836625 | 0.553290494 | 0.753561 | 0.043786077 |
| ENSMUSG00000021782 | Dlg5 | 71228 | protein_coding | 571.1558387 | -0.043792544 | 0.076762721 | -0.570492336 | 0.568343819 | 0.763615826 | 0.043792544 |
| ENSMUSG00000036550 | Cnot1 | 234594 | protein_coding | 6210.085133 | 0.043793603 | 0.043785443 | 1.000186352 | 0.317220333 | 0.561067373 | 0.043793603 |
| ENSMUSG00000039105 | Atp6v1g1 | 66290 | protein_coding | 1574.967351 | 0.043800896 | 0.045742134 | 0.957561267 | 0.338284039 | 0.582029301 | 0.043800896 |
| ENSMUSG00000067288 | Rps28 | 54127 | protein_coding | 6866.492219 | -0.043802027 | 0.050386715 | -0.869316988 | 0.384673773 | 0.624988308 | 0.043802027 |
| ENSMUSG00000036606 | Plxnb2 | 140570 | protein_coding | 11288.63635 | -0.043815313 | 0.0512602 | -0.854762813 | 0.392682469 | 0.632267311 | 0.043815313 |
| ENSMUSG00000002210 | Smg9 | 71997 | protein_coding | 581.5774253 | 0.043817544 | 0.08686167 | 0.504452011 | 0.613943777 | 0.794327773 | 0.043817544 |
| ENSMUSG00000024137 | E4f1 | 13560 | protein_coding | 460.0685753 | 0.04381931 | 0.081483891 | 0.537766537 | 0.590738237 | 0.779660849 | 0.04381931 |
| ENSMUSG00000037221 | Mospd3 | 68929 | protein_coding | 82.74803662 | -0.043830235 | 0.177274487 | -0.247245027 | 0.8047186 | 0.911163158 | 0.043830235 |
| ENSMUSG00000040124 | Gorab | 98376 | protein_coding | 347.0632346 | -0.04383345 | 0.086188437 | -0.508576918 | 0.611048812 | 0.792726586 | 0.04383345 |
| ENSMUSG00000036721 | Zscan12 | 22758 | protein_coding | 128.6066407 | -0.043850654 | 0.117616728 | -0.372826681 | 0.709277444 | 0.856166251 | 0.043850654 |
| ENSMUSG00000037331 | Larp1 | 73158 | protein_coding | 3427.57479 | -0.043870954 | 0.049114175 | -0.893244236 | 0.371726393 | 0.613533062 | 0.043870954 |
| ENSMUSG00000060126 | Tpt1 | 22070 | protein_coding | 18661.19369 | -0.043903256 | 0.042364377 | -1.036324832 | 0.300050625 | 0.543683373 | 0.043903256 |
| ENSMUSG00000024528 | Srfbp1 | 67222 | protein_coding | 228.5609991 | -0.043915661 | 0.1046948 | -0.419463631 | 0.674877329 | 0.834451176 | 0.043915661 |
| ENSMUSG00000025785 | Exosc7 | 66446 | protein_coding | 489.503031 | 0.043926573 | 0.075586689 | 0.581141647 | 0.561144992 | 0.759206347 | 0.043926573 |
| ENSMUSG00000021175 | Cdca7l | 217946 | protein_coding | 218.4071309 | -0.043958602 | 0.139039071 | -0.316160064 | 0.751881018 | 0.88186074 | 0.043958602 |
| ENSMUSG00000037410 | Tbc1d2b | 67016 | protein_coding | 638.2330396 | 0.04396021 | 0.085077475 | 0.516707977 | 0.605360027 | 0.789158883 | 0.04396021 |
| ENSMUSG00000038797 | Zscan2 | 22691 | protein_coding | 71.51568947 | -0.043960322 | 0.148052526 | -0.296923821 | 0.76652467 | 0.890414429 | 0.043960322 |
| ENSMUSG00000059851 | Kmt5c | 232811 | protein_coding | 923.0082367 | 0.043962067 | 0.058622993 | 0.749911672 | 0.453307904 | 0.682136666 | 0.043962067 |
| ENSMUSG00000073016 | Uprt | 331487 | protein_coding | 462.0945202 | -0.043976962 | 0.085174174 | -0.516318035 | 0.605632302 | 0.789401467 | 0.043976962 |
| ENSMUSG00000024335 | Brd2 | 14312 | protein_coding | 4518.702436 | -0.043983697 | 0.051993395 | -0.845947785 | 0.397581873 | 0.637102829 | 0.043983697 |
| ENSMUSG00000054423 | Cadps | 27062 | protein_coding | 37.91678274 | -0.044025046 | 0.221078373 | -0.199137732 | 0.842155007 | 0.92985306 | 0.044025046 |
| ENSMUSG00000021686 | Ap3b1 | 11774 | protein_coding | 3217.072545 | -0.044088579 | 0.030235427 | -1.458176195 | 0.144791984 | 0.35731604 | 0.044088579 |

| ensembl_gene_id | external_gene_name | entrezgene_id | gene_biotype | baseMean | log2FoldChange | lfcSE | stat | pvalue | padj | absLog2FC |
|---|---|---|---|---|---|---|---|---|---|---|
| ENSMUSG00000062961 | Ccdc177 | 380768 | protein_coding | 17.93331775 | -0.044092463 | 0.282146057 | -0.156275313 | 0.875816014 | 0.944199804 | 0.044092463 |
| ENSMUSG00000002608 | Ccdc97 | 52132 | protein_coding | 392.4912336 | -0.044105912 | 0.093398345 | -0.472234403 | 0.636759486 | 0.81030697 | 0.044105912 |
| ENSMUSG00000033769 | Exoc6b | 75914 | protein_coding | 1066.468994 | -0.044110703 | 0.054919234 | -0.803192247 | 0.421863626 | 0.65828686 | 0.044110703 |
| ENSMUSG00000022889 | Mrpl39 | 27393 | protein_coding | 778.7315465 | -0.044126334 | 0.059927951 | -0.736323086 | 0.4615341 | 0.687971202 | 0.044126334 |
| ENSMUSG00000035726 | Supt16 | 114741 | protein_coding | 1836.370675 | -0.044128965 | 0.067507477 | -0.653690043 | 0.51331152 | 0.726665632 | 0.044128965 |
| ENSMUSG00000043668 | Tox3 | 244579 | protein_coding | 319.3986277 | -0.044150541 | 0.086219027 | -0.512074218 | 0.608599068 | 0.791267105 | 0.044150541 |
| ENSMUSG00000025269 | Apex2 | 77622 | protein_coding | 115.4674338 | -0.04420282 | 0.128799744 | -0.343190274 | 0.731455321 | 0.870438309 | 0.04420282 |
| ENSMUSG00000053684 | Kics2 | 270802 | protein_coding | 68.34026917 | -0.044203723 | 0.146291671 | -0.302161584 | 0.762528888 | 0.888020174 | 0.044203723 |
| ENSMUSG00000031922 | Cep57 | 74360 | protein_coding | 453.9121468 | 0.044256208 | 0.081263266 | 0.544602866 | 0.586026683 | 0.77637824 | 0.044256208 |
| ENSMUSG00000022300 | Dcaf13 | 223499 | protein_coding | 877.9867663 | 0.044277559 | 0.063746753 | 0.694585329 | 0.487315209 | 0.708115319 | 0.044277559 |
| ENSMUSG00000046460 | Sh2d7 | 244885 | protein_coding | 306.3022913 | 0.044290542 | 0.102147125 | 0.433595576 | 0.664582148 | 0.828854703 | 0.044290542 |
| ENSMUSG00000063281 | Zfp35 | 22694 | protein_coding | 255.2502166 | 0.044297988 | 0.092701225 | 0.477857636 | 0.632751536 | 0.807820141 | 0.044297988 |
| ENSMUSG00000005882 | Uqcc1 | 56046 | protein_coding | 1217.321358 | 0.044331967 | 0.04656759 | 0.951991865 | 0.341101108 | 0.584025646 | 0.044331967 |
| ENSMUSG00000026648 | Dclre1c | 227525 | protein_coding | 191.4488169 | -0.044346909 | 0.098002813 | -0.45250649 | 0.650904147 | 0.820125043 | 0.044346909 |
| ENSMUSG00000001755 | Coasy | 71743 | protein_coding | 862.602837 | 0.044374985 | 0.069372083 | 0.639666319 | 0.522389557 | 0.733400921 | 0.044374985 |
| ENSMUSG00000018809 | Smyd4 | 319822 | protein_coding | 155.1645806 | 0.044393428 | 0.107964405 | 0.411185779 | 0.680936315 | 0.838663506 | 0.044393428 |
| ENSMUSG00000021901 | Bap1 | 104416 | protein_coding | 948.1737082 | -0.044397478 | 0.050717069 | -0.875395198 | 0.381358912 | 0.622235448 | 0.044397478 |
| ENSMUSG00000029055 | Plch2 | 269615 | protein_coding | 1552.25367 | 0.044415576 | 0.063620577 | 0.698132241 | 0.485094497 | 0.706376429 | 0.044415576 |
| ENSMUSG00000031790 | Mmp15 | 17388 | protein_coding | 3377.667469 | -0.044418687 | 0.089055884 | -0.498773182 | 0.617939182 | 0.797435496 | 0.044418687 |
| ENSMUSG00000040151 | Hs2st1 | 23908 | protein_coding | 475.7889405 | -0.04444109 | 0.065577268 | -0.677690479 | 0.497967964 | 0.715687493 | 0.04444109 |
| ENSMUSG00000044447 | Dock5 | 68813 | protein_coding | 1987.172011 | -0.04445215 | 0.036412709 | -1.220786672 | 0.222166801 | 0.459226353 | 0.04445215 |
| ENSMUSG00000021311 | Mtr | 238505 | protein_coding | 219.6150562 | 0.044465113 | 0.09977087 | 0.445672297 | 0.655833978 | 0.823084402 | 0.044465113 |
| ENSMUSG00000009079 | Ewsr1 | 14030 | protein_coding | 3647.300289 | -0.044476196 | 0.053648306 | -0.82903262 | 0.407085949 | 0.645753068 | 0.044476196 |
| ENSMUSG00000042462 | Dctpp1 | 66422 | protein_coding | 451.543416 | -0.044505787 | 0.067324276 | -0.661065959 | 0.508570015 | 0.723987816 | 0.044505787 |
| ENSMUSG00000049881 | 2810025M15Rik | 69953 | transcribed_processed_pseudogene | 118.3164437 | -0.044541777 | 0.123429371 | -0.360868541 | 0.718197722 | 0.861944627 | 0.044541777 |
| ENSMUSG00000028788 | Ptp4a2 | 19244 | protein_coding | 9612.990944 | 0.04456096 | 0.03790745 | 1.175519852 | 0.239786799 | 0.480291336 | 0.04456096 |
| ENSMUSG00000031479 | Vps36 | 70160 | protein_coding | 998.9908588 | -0.04457709 | 0.046855649 | -0.951370655 | 0.341416251 | 0.584146725 | 0.04457709 |
| ENSMUSG00000049598 | Vsig8 | 240916 | protein_coding | 51.28237945 | 0.04458569 | 0.196411023 | 0.227001973 | 0.820422208 | 0.919518309 | 0.04458569 |
| ENSMUSG00000030726 | Pold3 | 67967 | protein_coding | 644.2717475 | -0.044595631 | 0.070948941 | -0.628559502 | 0.52963748 | 0.738880938 | 0.044595631 |
| ENSMUSG00000035378 | Shq1 | 72171 | protein_coding | 176.4813324 | 0.044599904 | 0.106694114 | 0.418016535 | 0.675935028 | 0.835182669 | 0.044599904 |
| ENSMUSG00000032498 | Mlh1 | 17350 | protein_coding | 378.8094182 | 0.04461973 | 0.077361332 | 0.576770456 | 0.56409453 | 0.761174286 | 0.04461973 |
| ENSMUSG00000021823 | Vcl | 22330 | protein_coding | 1469.247674 | 0.04466052 | 0.059267348 | 0.753543427 | 0.451123425 | 0.680343771 | 0.04466052 |
| ENSMUSG00000021285 | Ppp1r13b | 21981 | protein_coding | 1036.31534 | -0.044665638 | 0.049164756 | -0.90848897 | 0.363619939 | 0.606076959 | 0.044665638 |
| ENSMUSG00000019810 | Fuca2 | 66848 | protein_coding | 1982.560987 | 0.044665865 | 0.039634806 | 1.126935372 | 0.259769807 | 0.50123327 | 0.044665865 |
| ENSMUSG00000011960 | Ccnt1 | 12455 | protein_coding | 988.5690918 | -0.044673091 | 0.075789818 | -0.589433938 | 0.555570216 | 0.75524772 | 0.044673091 |
| ENSMUSG00000034487 | Poglut3 | 68304 | protein_coding | 328.4443197 | 0.044677268 | 0.078654599 | 0.568018509 | 0.5700224 | 0.765233601 | 0.044677268 |
| ENSMUSG00000038296 | Galnt18 | 233733 | protein_coding | 18.34576946 | -0.044682515 | 0.312909982 | -0.142796707 | 0.886450738 | 0.949760235 | 0.044682515 |
| ENSMUSG00000034563 | Ccpg1 | 72278 | protein_coding | 14.945109 | -0.044686481 | 0.317340675 | -0.140815483 | 0.888015711 | 0.950489771 | 0.044686481 |
| ENSMUSG00000033653 | Vps8 | 209018 | protein_coding | 541.3514739 | -0.044688108 | 0.070963698 | -0.62973195 | 0.528869975 | 0.73819336 | 0.044688108 |

| ensembl_gene_id | external_gene_name | entrezgene_id | gene_biotype | baseMean | log2FoldChange | lfcSE | stat | pvalue | padj | absLog2FC |
|---|---|---|---|---|---|---|---|---|---|---|
| ENSMUSG00000027782 | Kpna4 | 16649 | protein_coding | 1500.351733 | 0.044699931 | 0.039580561 | 1.129340519 | 0.258754214 | 0.500569859 | 0.044699931 |
| ENSMUSG00000042292 | Mrtfa | 223701 | protein_coding | 592.1688526 | -0.04470101 | 0.096013484 | -0.465570126 | 0.641523231 | 0.813708274 | 0.04470101 |
| ENSMUSG00000021636 | Marveld2 | 218518 | protein_coding | 1077.158293 | -0.044710534 | 0.046698533 | -0.957429091 | 0.338350721 | 0.582029301 | 0.044710534 |
| ENSMUSG00000001988 | Npas1 | 18142 | protein_coding | 9.070617259 | 0.044711622 | 0.405066503 | 0.110380942 | 0.912107267 | 0.960727899 | 0.044711622 |
| ENSMUSG00000037373 | Ctbp1 | 13016 | protein_coding | 4192.838128 | 0.044720765 | 0.050744008 | 0.881301385 | 0.378154717 | 0.619451611 | 0.044720765 |
| ENSMUSG00000022247 | Brix1 | 67832 | protein_coding | 475.2004645 | -0.044740496 | 0.067308145 | -0.664711472 | 0.506235055 | 0.722035264 | 0.044740496 |
| ENSMUSG00000024212 | Mllt1 | 64144 | protein_coding | 1158.681237 | 0.044744178 | 0.062988035 | 0.710359966 | 0.477480942 | 0.700094114 | 0.044744178 |
| ENSMUSG00000038014 | Fam120a | 218236 | protein_coding | 11840.7778 | 0.044748805 | 0.048295898 | 0.926554981 | 0.354157638 | 0.596379092 | 0.044748805 |
| ENSMUSG00000020705 | Ddx42 | 72047 | protein_coding | 1564.617796 | -0.044761841 | 0.049747509 | -0.899780551 | 0.368237046 | 0.610334928 | 0.044761841 |
| ENSMUSG00000063229 | Ldha | 16828 | protein_coding | 19574.97522 | 0.044766108 | 0.068309526 | 0.655342102 | 0.512247519 | 0.725924055 | 0.044766108 |
| ENSMUSG00000057342 | Sphk2 | 56632 | protein_coding | 2151.125635 | 0.044782755 | 0.075928044 | 0.589805197 | 0.555321258 | 0.755247648 | 0.044782755 |
| ENSMUSG00000024169 | Ift140 | 106633 | protein_coding | 519.1945417 | 0.044834254 | 0.068653731 | 0.653049062 | 0.51372465 | 0.727042009 | 0.044834254 |
| ENSMUSG00000040220 | Gas8 | 104346 | protein_coding | 151.9939691 | 0.044854489 | 0.112596988 | 0.398363129 | 0.690362532 | 0.84457466 | 0.044854489 |
| ENSMUSG00000036273 | Lrrk2 | 66725 | protein_coding | 457.1370513 | 0.044855001 | 0.120445177 | 0.372410106 | 0.709587531 | 0.856166251 | 0.044855001 |
| ENSMUSG00000028029 | Aimp1 | 13722 | protein_coding | 976.9580043 | -0.044862245 | 0.05992592 | -0.7486284 | 0.45408121 | 0.682675952 | 0.044862245 |
| ENSMUSG00000066324 | Bpnt2 | 242291 | protein_coding | 4629.680185 | -0.044873146 | 0.037639605 | -1.192178967 | 0.233191079 | 0.472624422 | 0.044873146 |
| ENSMUSG00000044150 | Bclaf3 | 382252 | protein_coding | 172.6374937 | -0.044885843 | 0.105675522 | -0.424751563 | 0.671017791 | 0.83235935 | 0.044885843 |
| ENSMUSG00000022179 | 4931414P19Rik | 74359 | protein_coding | 174.8002084 | 0.044887035 | 0.102033581 | 0.439924136 | 0.659992055 | 0.826041967 | 0.044887035 |
| ENSMUSG00000029924 | Slc37a3 | 72144 | protein_coding | 903.0413189 | 0.044920239 | 0.051447358 | 0.873130125 | 0.382592156 | 0.62329844 | 0.044920239 |
| ENSMUSG00000031200 | Mtcp1 | 17763 | protein_coding | 62.39332108 | -0.044928295 | 0.231250822 | -0.194283828 | 0.845953648 | 0.93086312 | 0.044928295 |
| ENSMUSG00000109293 | Dcst2 | 329702 | protein_coding | 32.22692205 | -0.044956355 | 0.262415087 | -0.171317721 | 0.863973947 | 0.93820671 | 0.044956355 |
| ENSMUSG00000024991 | Eif3a | 13669 | protein_coding | 4818.007579 | -0.044956911 | 0.058692905 | -0.765968411 | 0.443695098 | 0.674737597 | 0.044956911 |
| ENSMUSG00000030213 | Atf7ip | 54343 | protein_coding | 1240.745819 | -0.044964597 | 0.057988116 | -0.775410544 | 0.438097106 | 0.671081619 | 0.044964597 |
| ENSMUSG00000038697 | Taf5l | 102162 | protein_coding | 939.9678148 | -0.044970759 | 0.047282946 | -0.951098929 | 0.341554158 | 0.584243096 | 0.044970759 |
| ENSMUSG00000020325 | Fstl3 | 83554 | protein_coding | 10.11941022 | -0.044977375 | 0.359214907 | -0.125210213 | 0.900357132 | 0.955895949 | 0.044977375 |
| ENSMUSG00000024521 | Pmaip1 | 58801 | protein_coding | 582.490537 | 0.044978516 | 0.144725414 | 0.310785194 | 0.755963926 | 0.884269981 | 0.044978516 |
| ENSMUSG00000029501 | Ankle2 | 71782 | protein_coding | 1032.644759 | 0.044981294 | 0.044628278 | 1.007910139 | 0.313497604 | 0.55760602 | 0.044981294 |
| ENSMUSG00000091625 | Lsm5 | 66373 | protein_coding | 144.658522 | -0.044987767 | 0.136194324 | -0.330320428 | 0.741157859 | 0.876229953 | 0.044987767 |
| ENSMUSG00000096054 | Syne1 | 64009 | protein_coding | 295.0464183 | 0.045006905 | 0.077364146 | 0.581754053 | 0.560732357 | 0.759080896 | 0.045006905 |
| ENSMUSG00000026484 | Rnf2 | 19821 | protein_coding | 589.8218569 | -0.045007404 | 0.064550522 | -0.697243061 | 0.485650695 | 0.706867216 | 0.045007404 |
| ENSMUSG00000036167 | Pphln1 | 223828 | protein_coding | 395.3058025 | 0.045039411 | 0.072842341 | 0.618313611 | 0.536368632 | 0.742937123 | 0.045039411 |
| ENSMUSG00000047603 | Zfp235 | 56525 | protein_coding | 236.2118637 | -0.045042175 | 0.096397756 | -0.467253359 | 0.640318622 | 0.812911785 | 0.045042175 |
| ENSMUSG00000012535 | Tnpo3 | 320938 | protein_coding | 1638.611975 | 0.045047091 | 0.046938354 | 0.959707501 | 0.337202447 | 0.580796825 | 0.045047091 |
| ENSMUSG00000055717 | Slain1 | 105439 | protein_coding | 138.9114792 | 0.045048325 | 0.119610581 | 0.376624916 | 0.706452368 | 0.854108805 | 0.045048325 |
| ENSMUSG00000028952 | Zbtb48 | 100090 | protein_coding | 190.4076557 | -0.045051664 | 0.113050892 | -0.398507812 | 0.690255901 | 0.84457466 | 0.045051664 |
| ENSMUSG00000001467 | Cyp51 | 13121 | protein_coding | 4063.669965 | -0.045056294 | 0.114827453 | -0.392382595 | 0.694775544 | 0.847296219 | 0.045056294 |
| ENSMUSG00000072884 | Gm10433 | 100038378 | lncRNA | 39.48224906 | -0.04507411 | 0.191781671 | -0.235028246 | 0.81418681 | 0.916408517 | 0.04507411 |
| ENSMUSG00000022125 | Cln5 | 211286 | protein_coding | 639.0940121 | 0.045093422 | 0.068368051 | 0.659568632 | 0.509530689 | 0.72436245 | 0.045093422 |
| ENSMUSG00000056310 | Tyw1 | 100929 | protein_coding | 536.9057007 | -0.045108219 | 0.063471952 | -0.710679563 | 0.477282827 | 0.699994557 | 0.045108219 |

| ensembl_gene_id | external_gene_name | entrezgene_id | gene_biotype | baseMean | log2FoldChange | lfcSE | stat | pvalue | padj | absLog2FC |
|---|---|---|---|---|---|---|---|---|---|---|
| ENSMUSG00000074576 | Mocs3 | 69372 | protein_coding | 232.7715816 | 0.045166936 | 0.11263452 | 0.401004383 | 0.688416897 | 0.843368224 | 0.045166936 |
| ENSMUSG00000048379 | Socs4 | 67296 | protein_coding | 851.031211 | -0.045168797 | 0.044098441 | -1.02427198 | 0.305706829 | 0.54999264 | 0.045168797 |
| ENSMUSG00000030051 | Aplf | 72103 | protein_coding | 247.2303443 | -0.045179839 | 0.083218887 | -0.542903666 | 0.587196132 | 0.777161429 | 0.045179839 |
| ENSMUSG00000006373 | Pgrmc1 | 53328 | protein_coding | 2881.78645 | -0.045199207 | 0.038262737 | -1.181285243 | 0.237489427 | 0.47743605 | 0.045199207 |
| ENSMUSG00000021806 | Nid2 | 18074 | protein_coding | 216.5552445 | -0.045224505 | 0.097258099 | -0.464994744 | 0.641935221 | 0.813747171 | 0.045224505 |
| ENSMUSG00000034101 | Ctnnd1 | 12388 | protein_coding | 1347.097713 | 0.045231344 | 0.044168641 | 1.024060119 | 0.30580688 | 0.550111272 | 0.045231344 |
| ENSMUSG00000054128 | H2-T3 | 15043 | protein_coding | 1386.77966 | -0.045265277 | 0.109615189 | -0.412947127 | 0.679645352 | 0.837841354 | 0.045265277 |
| ENSMUSG00000021770 | Samd8 | 67630 | protein_coding | 816.9898504 | 0.045282343 | 0.064012458 | 0.707398907 | 0.479318616 | 0.701704005 | 0.045282343 |
| ENSMUSG00000044149 | Nkrf | 77286 | protein_coding | 112.9290779 | 0.045319687 | 0.128173677 | 0.353580296 | 0.723653443 | 0.865314338 | 0.045319687 |
| ENSMUSG00000024695 | Zfp91 | 109910 | protein_coding | 2031.814719 | -0.045333424 | 0.047425354 | -0.955890051 | 0.339127786 | 0.58240762 | 0.045333424 |
| ENSMUSG00000025858 | Get4 | 67604 | protein_coding | 540.2709505 | -0.045376001 | 0.053751005 | -0.844188886 | 0.398563864 | 0.638169126 | 0.045376001 |
| ENSMUSG00000002396 | Ocel1 | 77090 | protein_coding | 689.1855999 | -0.045399068 | 0.071110166 | -0.638432875 | 0.523191936 | 0.734335627 | 0.045399068 |
| ENSMUSG00000041605 | Inava | 67313 | protein_coding | 2571.129454 | 0.045402256 | 0.063773742 | 0.711927106 | 0.476509916 | 0.699497125 | 0.045402256 |
| ENSMUSG00000015757 | Ppil4 | 67418 | protein_coding | 647.2393918 | -0.045412747 | 0.065944551 | -0.688650483 | 0.491043244 | 0.710842238 | 0.045412747 |
| ENSMUSG00000025533 | Asl | 109900 | protein_coding | 821.9833761 | -0.045429463 | 0.06253475 | -0.726467486 | 0.467552233 | 0.692926016 | 0.045429463 |
| ENSMUSG00000025245 | Lztfl1 | 93730 | protein_coding | 405.2542056 | 0.045434874 | 0.075563001 | 0.601284665 | 0.547650401 | 0.750740503 | 0.045434874 |
| ENSMUSG00000021929 | Kpna3 | 16648 | protein_coding | 785.5813082 | -0.045436281 | 0.053335529 | -0.851895191 | 0.394272266 | 0.633814677 | 0.045436281 |
| ENSMUSG00000057469 | E2f6 | 50496 | protein_coding | 289.6896783 | 0.045443554 | 0.080987785 | 0.561116143 | 0.574718364 | 0.768233148 | 0.045443554 |
| ENSMUSG00000033940 | Brk1 | 101314 | protein_coding | 1645.680923 | 0.045463047 | 0.047212265 | 0.962949924 | 0.335572654 | 0.579171681 | 0.045463047 |
| ENSMUSG00000079215 | Zfp664 | 269704 | protein_coding | 1376.579283 | 0.045471665 | 0.04259455 | 1.067546564 | 0.285725099 | 0.528548826 | 0.045471665 |
| ENSMUSG00000096959 | 4930509G22Rik | 105246071 | lncRNA | 9.334327379 | -0.045518389 | 0.380716381 | -0.119559839 | 0.904831835 | 0.958435249 | 0.045518389 |
| ENSMUSG00000044860 | Gm1123 | 382097 | protein_coding | 4029.860657 | -0.045568575 | 0.054180694 | -0.841048194 | 0.400320936 | 0.63958547 | 0.045568575 |
| ENSMUSG00000033805 | Ephx4 | 384214 | protein_coding | 58.86653372 | 0.045592091 | 0.164397819 | 0.277327835 | 0.781528388 | 0.898525697 | 0.045592091 |
| ENSMUSG00000054161 | Fam83e | 73813 | protein_coding | 9218.78209 | -0.045680492 | 0.033802693 | -1.351386195 | 0.176571754 | 0.403338905 | 0.045680492 |
| ENSMUSG00000026227 | 2810459M11Rik | 72792 | protein_coding | 970.4914402 | -0.045692404 | 0.078513401 | -0.58196949 | 0.560587232 | 0.758948144 | 0.045692404 |
| ENSMUSG00000027570 | Col9a3 | 12841 | protein_coding | 7.733474144 | 0.045708823 | 0.639345063 | 0.071493197 | 0.943005239 | 0.975236981 | 0.045708823 |
| ENSMUSG00000052997 | Uba2 | 50995 | protein_coding | 1619.835755 | 0.045754966 | 0.05321247 | 0.859854208 | 0.389869413 | 0.629623876 | 0.045754966 |
| ENSMUSG00000010392 | Gosr1 | 53334 | protein_coding | 1384.816367 | 0.045805609 | 0.045039635 | 1.017006672 | 0.309150256 | 0.552606516 | 0.045805609 |
| ENSMUSG00000037885 | Stk35 | 67333 | protein_coding | 701.8278992 | -0.045812649 | 0.066407689 | -0.689869656 | 0.490276159 | 0.710266225 | 0.045812649 |
| ENSMUSG00000024968 | Rcor2 | 104383 | protein_coding | 15.82528327 | -0.045829141 | 0.293641342 | -0.156071828 | 0.875976404 | 0.944301517 | 0.045829141 |
| ENSMUSG00000033739 | Fkbpl | 56299 | protein_coding | 114.6191589 | 0.045850793 | 0.109484209 | 0.418789103 | 0.67537027 | 0.834932644 | 0.045850793 |
| ENSMUSG00000039043 | Arpin | 70420 | protein_coding | 1523.584586 | 0.045871472 | 0.045338767 | 1.011749422 | 0.311657878 | 0.555505998 | 0.045871472 |
| ENSMUSG00000018583 | G3bp1 | 27041 | protein_coding | 3433.707756 | 0.045873423 | 0.063297605 | 0.724726041 | 0.468620116 | 0.693332995 | 0.045873423 |
| ENSMUSG00000025816 | Sec61a2 | 57743 | protein_coding | 156.7431889 | -0.045932912 | 0.142322902 | -0.322737322 | 0.746894184 | 0.879915444 | 0.045932912 |
| ENSMUSG00000039013 | Siglecf | 233186 | protein_coding | 80.02513234 | 0.045948583 | 0.156445208 | 0.293703994 | 0.768984102 | 0.891610224 | 0.045948583 |
| ENSMUSG00000039851 | 4932438H23Rik | 74387 | protein_coding | 24.22898044 | 0.04597711 | 0.286006945 | 0.160755223 | 0.872286193 | 0.942599802 | 0.04597711 |
| ENSMUSG00000029191 | Rfc1 | 19687 | protein_coding | 1104.238758 | 0.045983837 | 0.076717606 | 0.599390934 | 0.548912221 | 0.751266959 | 0.045983837 |
| ENSMUSG00000038780 | Smurf1 | 75788 | protein_coding | 815.0425299 | 0.045986178 | 0.06681073 | 0.688305284 | 0.491260555 | 0.710862177 | 0.045986178 |
| ENSMUSG00000047466 | 8030462N17Rik | 212163 | protein_coding | 596.9065813 | -0.045987141 | 0.052771344 | -0.871441542 | 0.383513112 | 0.624309975 | 0.045987141 |

Supplementary Table 1

| ensembl_gene_id | external_gene_name | entrezgene_id | gene_biotype | baseMean | log2FoldChange | lfcSE | stat | pvalue | padj | absLog2FC |
|---|---|---|---|---|---|---|---|---|---|---|
| ENSMUSG00000035772 | Mrps2 | 118451 | protein_coding | 604.8586088 | 0.046022034 | 0.076688818 | 0.600114002 | 0.548430262 | 0.750864769 | 0.046022034 |
| ENSMUSG00000039159 | Ube2h | 22214 | protein_coding | 1409.853485 | 0.046044293 | 0.044325805 | 1.038769478 | 0.29891196 | 0.542488541 | 0.046044293 |
| ENSMUSG00000021619 | Atg10 | 66795 | protein_coding | 122.8995113 | -0.046076383 | 0.109581037 | -0.420477705 | 0.674136513 | 0.834301331 | 0.046076383 |
| ENSMUSG00000045252 | Zfp574 | 232976 | protein_coding | 449.2778333 | 0.046105014 | 0.088769064 | 0.519381544 | 0.603494699 | 0.788189101 | 0.046105014 |
| ENSMUSG00000028885 | Smpdl3b | 100340 | protein_coding | 66.41340294 | 0.046123399 | 0.220173012 | 0.20948707 | 0.834068028 | 0.926442254 | 0.046123399 |
| ENSMUSG00000033192 | Lpcat2 | 270084 | protein_coding | 353.6826434 | 0.046123674 | 0.086014435 | 0.536231781 | 0.591798372 | 0.780262005 | 0.046123674 |
| ENSMUSG00000027695 | Pld1 | 18805 | protein_coding | 3205.56042 | -0.046139602 | 0.048527216 | -0.950798441 | 0.341706704 | 0.584442043 | 0.046139602 |
| ENSMUSG00000035476 | Tab3 | 66724 | protein_coding | 266.0231336 | -0.046162947 | 0.084255254 | -0.547893989 | 0.583764697 | 0.775075241 | 0.046162947 |
| ENSMUSG00000024867 | Pip5k1b | 18719 | protein_coding | 1576.696316 | -0.046175052 | 0.037636385 | -1.226872664 | 0.219870454 | 0.456057983 | 0.046175052 |
| ENSMUSG00000027556 | Car1 | 12346 | protein_coding | 151271.3177 | 0.046202497 | 0.086899133 | 0.531679609 | 0.594947917 | 0.782189365 | 0.046202497 |
| ENSMUSG00000024037 | Wdr4 | 57773 | protein_coding | 339.2994507 | -0.046207746 | 0.07635657 | -0.605157437 | 0.545074393 | 0.749012844 | 0.046207746 |
| ENSMUSG00000032475 | Nck1 | 17973 | protein_coding | 734.2023528 | -0.046213236 | 0.064302341 | -0.718686689 | 0.472333987 | 0.696728273 | 0.046213236 |
| ENSMUSG00000015889 | Lta4h | 16993 | protein_coding | 2356.882691 | 0.046218769 | 0.050613496 | 0.913170854 | 0.36115269 | 0.604118809 | 0.046218769 |
| ENSMUSG00000006398 | Cdc20 | 107995 | protein_coding | 858.1396461 | 0.046228924 | 0.100037905 | 0.46211408 | 0.643999514 | 0.815430435 | 0.046228924 |
| ENSMUSG00000043284 | Tmem11 | 216821 | protein_coding | 750.4467361 | 0.046235497 | 0.060948543 | 0.75859889 | 0.448092537 | 0.678921982 | 0.046235497 |
| ENSMUSG00000027951 | Adar | 56417 | protein_coding | 1310.384491 | -0.046246612 | 0.059910694 | -0.771925826 | 0.44015836 | 0.671885771 | 0.046246612 |
| ENSMUSG00000021774 | Ube2e1 | 22194 | protein_coding | 359.5685267 | 0.046254914 | 0.074035069 | 0.624770322 | 0.532121812 | 0.740616938 | 0.046254914 |
| ENSMUSG00000026604 | Ptpn14 | 19250 | protein_coding | 89.76059304 | -0.04625591 | 0.18030636 | -0.256540644 | 0.7975334 | 0.906911659 | 0.04625591 |
| ENSMUSG00000000416 | Cttnbp2 | 30785 | protein_coding | 296.9640892 | -0.046359111 | 0.10822205 | -0.428370294 | 0.668381549 | 0.830822102 | 0.046359111 |
| ENSMUSG00000029815 | Malsu1 | 75593 | protein_coding | 308.1406713 | 0.046366761 | 0.073484585 | 0.630972611 | 0.528058433 | 0.737831675 | 0.046366761 |
| ENSMUSG00000025373 | Rnf41 | 67588 | protein_coding | 652.922441 | -0.046379262 | 0.067904535 | -0.683006838 | 0.494602522 | 0.713491312 | 0.046379262 |
| ENSMUSG00000058298 | Mcm9 | 71567 | protein_coding | 345.0994147 | -0.046394695 | 0.062874433 | -0.737894436 | 0.460578601 | 0.687305492 | 0.046394695 |
| ENSMUSG00000017858 | Ift52 | 245866 | protein_coding | 693.207133 | 0.04641137 | 0.052553463 | 0.883126766 | 0.377167784 | 0.61858892 | 0.04641137 |
| ENSMUSG00000022414 | Tab1 | 66513 | protein_coding | 724.9388648 | -0.046440832 | 0.053900578 | -0.861601746 | 0.38890671 | 0.628768653 | 0.046440832 |
| ENSMUSG00000018574 | Acadvl | 11370 | protein_coding | 2886.134019 | 0.046453909 | 0.039451888 | 1.177482513 | 0.239002973 | 0.479283878 | 0.046453909 |
| ENSMUSG00000045854 | Lyrm2 | 108755 | protein_coding | 249.4607824 | 0.046457103 | 0.095222567 | 0.487879126 | 0.625635461 | 0.8024647 | 0.046457103 |
| ENSMUSG00000037296 | Lsm1 | 67207 | protein_coding | 378.0254451 | 0.046457326 | 0.075577078 | 0.614701271 | 0.53875202 | 0.74493282 | 0.046457326 |
| ENSMUSG00000027562 | Car2 | 12349 | protein_coding | 21238.55832 | -0.046503666 | 0.068580537 | -0.678088393 | 0.497715649 | 0.715643748 | 0.046503666 |
| ENSMUSG00000056608 | Chd9 | 109151 | protein_coding | 1560.614956 | 0.046507442 | 0.040175147 | 1.157617244 | 0.247020269 | 0.487958829 | 0.046507442 |
| ENSMUSG00000003134 | Tbc1d8 | 54610 | protein_coding | 434.2902497 | -0.046554288 | 0.074415622 | -0.625598311 | 0.531578447 | 0.740026398 | 0.046554288 |
| ENSMUSG00000021597 | Slf1 | 105377 | protein_coding | 350.3923888 | 0.04657385 | 0.071920285 | 0.647575995 | 0.517259228 | 0.729755889 | 0.04657385 |
| ENSMUSG00000034566 | Atp5h | 71679 | protein_coding | 5673.053286 | 0.046578339 | 0.04455004 | 1.045528554 | 0.295778751 | 0.538805075 | 0.046578339 |
| ENSMUSG00000018861 | Fdxr | 14149 | protein_coding | 358.9352568 | 0.046579814 | 0.102829819 | 0.452979641 | 0.650563402 | 0.819915284 | 0.046579814 |
| ENSMUSG00000091337 | Eid1 | 58521 | protein_coding | 440.4233924 | -0.046638934 | 0.085130546 | -0.547851937 | 0.583793573 | 0.775075241 | 0.046638934 |
| ENSMUSG00000100153 | Ppp1ccb | 434233 | transcribed_processed_pseudogene | 16.61455337 | -0.046720153 | 0.3308022 | -0.141232898 | 0.887685958 | 0.950326038 | 0.046720153 |
| ENSMUSG00000035095 | Fam167a | 219148 | protein_coding | 23.7638578 | -0.046724801 | 0.235246599 | -0.198620516 | 0.842559604 | 0.929981434 | 0.046724801 |
| ENSMUSG00000015957 | Wnt11 | 22411 | protein_coding | 12.65077317 | -0.046732532 | 0.488080039 | -0.095747682 | 0.92372097 | 0.966637342 | 0.046732532 |
| ENSMUSG00000031425 | Plp1 | 18823 | protein_coding | 13.42159781 | 0.046743947 | 0.377540059 | 0.123811887 | 0.90146422 | 0.956441676 | 0.046743947 |
| ENSMUSG00000051705 | Senp8 | 71599 | protein_coding | 157.8777461 | 0.046745931 | 0.128245337 | 0.364503945 | 0.715481728 | 0.860253025 | 0.046745931 |

| ensembl_gene_id | external_gene_name | entrezgene_id | gene_biotype | baseMean | log2FoldChange | lfcSE | stat | pvalue | padj | absLog2FC |
|---|---|---|---|---|---|---|---|---|---|---|
| ENSMUSG00000029267 | Mtf2 | 17765 | protein_coding | 737.3101086 | 0.046771727 | 0.075417094 | 0.620174082 | 0.535143183 | 0.742068282 | 0.046771727 |
| ENSMUSG00000010290 | AI597479 | 98404 | protein_coding | 377.3356804 | -0.046781023 | 0.077249196 | -0.605585898 | 0.544789769 | 0.749012844 | 0.046781023 |
| ENSMUSG00000048924 | Ccdc125 | 76041 | protein_coding | 168.1343968 | 0.04678464 | 0.118704538 | 0.394126806 | 0.693487421 | 0.846497211 | 0.04678464 |
| ENSMUSG00000022848 | Slc49a4 | 224132 | protein_coding | 959.651483 | -0.04680286 | 0.066420769 | -0.704641941 | 0.481033091 | 0.703236518 | 0.04680286 |
| ENSMUSG00000028886 | Eya3 | 14050 | protein_coding | 847.2999321 | 0.046809023 | 0.047860323 | 0.978033996 | 0.32805751 | 0.571645972 | 0.046809023 |
| ENSMUSG00000038515 | Grtp1 | 66790 | protein_coding | 1983.434321 | -0.046826169 | 0.048830021 | -0.958962706 | 0.337577533 | 0.581263386 | 0.046826169 |
| ENSMUSG00000096935 | 1700113A16Rik | 76642 | lncRNA | 23.23731978 | -0.046860491 | 0.260839184 | -0.179652806 | 0.857425146 | 0.935863679 | 0.046860491 |
| ENSMUSG00000072647 | Adam1a | 280668 | protein_coding | 9.218727764 | -0.046866296 | 0.417300131 | -0.112308367 | 0.910578908 | 0.959986013 | 0.046866296 |
| ENSMUSG00000038683 | Pak1ip1 | 68083 | protein_coding | 887.7325182 | 0.046870809 | 0.05846663 | 0.801667705 | 0.422745202 | 0.658779673 | 0.046870809 |
| ENSMUSG00000029030 | Tprgl | 67808 | protein_coding | 2008.97226 | -0.046909907 | 0.048823547 | -0.960804977 | 0.336650236 | 0.580252873 | 0.046909907 |
| ENSMUSG00000026198 | Abcb6 | 74104 | protein_coding | 104.579877 | -0.046912656 | 0.132961526 | -0.352828803 | 0.724216789 | 0.865592838 | 0.046912656 |
| ENSMUSG00000059981 | Taok2 | 381921 | protein_coding | 1423.810148 | -0.046980671 | 0.072212627 | -0.650588032 | 0.515312458 | 0.727964612 | 0.046980671 |
| ENSMUSG00000002660 | Clpp | 53895 | protein_coding | 673.5877864 | 0.046985686 | 0.088029895 | 0.533746922 | 0.593516641 | 0.780976082 | 0.046985686 |
| ENSMUSG00000024942 | Capn1 | 12333 | protein_coding | 3650.929263 | 0.047001423 | 0.056672713 | 0.829348395 | 0.406907294 | 0.645631043 | 0.047001423 |
| ENSMUSG00000032186 | Tmod2 | 50876 | protein_coding | 14.04107979 | -0.04702038 | 0.35998672 | -0.130616986 | 0.896078305 | 0.953802048 | 0.04702038 |
| ENSMUSG00000033799 | Tasor2 | 105203 | protein_coding | 829.7059828 | -0.047023493 | 0.074369056 | -0.632299182 | 0.527191398 | 0.737190295 | 0.047023493 |
| ENSMUSG00000022853 | Ehhadh | 74147 | protein_coding | 1595.708123 | 0.047026789 | 0.052236333 | 0.900269716 | 0.367976733 | 0.610154305 | 0.047026789 |
| ENSMUSG00000028333 | Anp32b | 67628 | protein_coding | 653.8095973 | -0.047068436 | 0.090138136 | -0.522181159 | 0.601544203 | 0.786660993 | 0.047068436 |
| ENSMUSG00000028391 | Wdr31 | 71354 | protein_coding | 58.1240448 | -0.047087591 | 0.212532178 | -0.221555117 | 0.824660226 | 0.92202835 | 0.047087591 |
| ENSMUSG00000020526 | Znhit3 | 448850 | protein_coding | 100.2568881 | 0.047103173 | 0.130092593 | 0.362074209 | 0.717296576 | 0.861407542 | 0.047103173 |
| ENSMUSG00000026433 | Rab29 | 226422 | protein_coding | 262.118719 | -0.047113939 | 0.087594124 | -0.537866434 | 0.590669263 | 0.779660849 | 0.047113939 |
| ENSMUSG00000096971 | 4930556M19Rik | 75259 | lncRNA | 33.61986627 | -0.047114914 | 0.235180622 | -0.200335017 | 0.841218578 | 0.929613548 | 0.047114914 |
| ENSMUSG00000037997 | Parp11 | 101187 | protein_coding | 201.5974371 | -0.047129953 | 0.092712381 | -0.508345835 | 0.611210831 | 0.792823007 | 0.047129953 |
| ENSMUSG00000024160 | Spsb3 | 79043 | protein_coding | 173.2543707 | -0.047206389 | 0.128337292 | -0.367830643 | 0.712999518 | 0.858612846 | 0.047206389 |
| ENSMUSG00000028790 | Khdrbs1 | 20218 | protein_coding | 1551.948038 | -0.0472111 | 0.037276839 | -1.266499563 | 0.205334286 | 0.439838414 | 0.0472111 |
| ENSMUSG00000020362 | Cnot6 | 104625 | protein_coding | 1107.106546 | 0.047219139 | 0.055445581 | 0.851630342 | 0.394419293 | 0.633886008 | 0.047219139 |
| ENSMUSG00000053768 | Chchd3 | 66075 | protein_coding | 3843.906498 | 0.047220721 | 0.044037434 | 1.072285928 | 0.283591625 | 0.526203742 | 0.047220721 |
| ENSMUSG00000026245 | Farsb | 23874 | protein_coding | 1184.264281 | 0.047223078 | 0.067035187 | 0.704452101 | 0.481151269 | 0.703237858 | 0.047223078 |
| ENSMUSG00000050071 | Bex1 | 19716 | protein_coding | 38.9325028 | -0.047227834 | 0.210525521 | -0.22433306 | 0.822498153 | 0.920659938 | 0.047227834 |
| ENSMUSG00000034616 | Ssh3 | 245857 | protein_coding | 2167.220246 | -0.047272045 | 0.04037766 | -1.170747507 | 0.241700283 | 0.482707243 | 0.047272045 |
| ENSMUSG00000026039 | Sgo2a | 68549 | protein_coding | 111.9833691 | 0.047295149 | 0.171800648 | 0.275290866 | 0.783092781 | 0.898856746 | 0.047295149 |
| ENSMUSG00000022971 | Ifnar2 | 15976 | protein_coding | 43.00018823 | 0.047311288 | 0.185913224 | 0.254480487 | 0.799124377 | 0.908144516 | 0.047311288 |
| ENSMUSG00000026259 | Ngef | 53972 | protein_coding | 1435.62439 | -0.047329378 | 0.0587055 | -0.806217098 | 0.420117682 | 0.657152071 | 0.047329378 |
| ENSMUSG00000028879 | Stx12 | 100226 | protein_coding | 1675.788973 | -0.047340118 | 0.04435484 | -1.067304438 | 0.285834385 | 0.5285691 | 0.047340118 |
| ENSMUSG00000028057 | Rit1 | 19769 | protein_coding | 855.7998272 | 0.047340151 | 0.047531082 | 0.995983016 | 0.319258397 | 0.562758791 | 0.047340151 |
| ENSMUSG00000027957 | Slc35a3 | 229782 | protein_coding | 7506.637129 | 0.047364205 | 0.050429123 | 0.939223261 | 0.347616129 | 0.589851392 | 0.047364205 |
| ENSMUSG00000052763 | Zfp212 | 232784 | protein_coding | 339.1680898 | -0.04738031 | 0.069317603 | -0.68352493 | 0.494275203 | 0.713491312 | 0.04738031 |
| ENSMUSG00000027309 | Dnaaf9 | 228602 | protein_coding | 431.783011 | -0.047389369 | 0.073621677 | -0.643687718 | 0.519777956 | 0.731645204 | 0.047389369 |
| ENSMUSG00000036330 | Slc18a1 | 110877 | protein_coding | 119.0493777 | -0.047389811 | 0.126023707 | -0.376038859 | 0.706888007 | 0.854251145 | 0.047389811 |

| ensembl_gene_id | external_gene_name | entrezgene_id | gene_biotype | baseMean | log2FoldChange | lfcSE | stat | pvalue | padj | absLog2FC |
|---|---|---|---|---|---|---|---|---|---|---|
| ENSMUSG00000044501 | Zfp758 | 224598 | protein_coding | 342.6893329 | -0.047390544 | 0.105713106 | -0.448293934 | 0.653941078 | 0.821860019 | 0.047390544 |
| ENSMUSG00000062300 | Nectin2 | 19294 | protein_coding | 2839.214503 | 0.047417412 | 0.064056334 | 0.740245485 | 0.459151053 | 0.686131304 | 0.047417412 |
| ENSMUSG00000037253 | Mex3c | 240396 | protein_coding | 1304.013384 | -0.047456019 | 0.04110145 | -1.154606929 | 0.248251428 | 0.489252203 | 0.047456019 |
| ENSMUSG00000039686 | Zer1 | 227693 | protein_coding | 507.3880223 | -0.04746548 | 0.082129835 | -0.577932248 | 0.563309862 | 0.760667465 | 0.04746548 |
| ENSMUSG00000033577 | Myo6 | 17920 | protein_coding | 7587.199177 | -0.047473091 | 0.042450721 | -1.118310592 | 0.263434365 | 0.504916665 | 0.047473091 |
| ENSMUSG00000021999 | Cpb2 | 56373 | protein_coding | 10.3149247 | 0.047496207 | 0.463737529 | 0.102420451 | 0.918422951 | 0.964033255 | 0.047496207 |
| ENSMUSG00000038383 | Pigu | 228812 | protein_coding | 606.9898368 | 0.047535282 | 0.059422148 | 0.799959004 | 0.42373455 | 0.659792516 | 0.047535282 |
| ENSMUSG00000052656 | Rnf103 | 22644 | protein_coding | 2506.572553 | 0.047536516 | 0.052697637 | 0.902061629 | 0.367024131 | 0.609389288 | 0.047536516 |
| ENSMUSG00000038324 | Trpc4ap | 56407 | protein_coding | 2244.249248 | 0.04757323 | 0.043079405 | 1.104314908 | 0.269456558 | 0.511930778 | 0.04757323 |
| ENSMUSG00000027852 | Nras | 18176 | protein_coding | 1263.112679 | -0.047618054 | 0.05441473 | -0.87509493 | 0.381522256 | 0.622373434 | 0.047618054 |
| ENSMUSG00000061778 | Mospd2 | 76763 | protein_coding | 512.7968242 | -0.047626401 | 0.079000107 | -0.602865016 | 0.546598491 | 0.749829352 | 0.047626401 |
| ENSMUSG00000030876 | Mettl9 | 59052 | protein_coding | 1537.402321 | 0.047717505 | 0.071691103 | 0.665598701 | 0.505667637 | 0.721608883 | 0.047717505 |
| ENSMUSG00000032352 | Lrrc1 | 214345 | protein_coding | 1457.294569 | 0.047729355 | 0.046724962 | 1.02149586 | 0.30701957 | 0.550940759 | 0.047729355 |
| ENSMUSG00000024208 | Uqcc2 | 67267 | protein_coding | 702.7759192 | 0.047737309 | 0.079800184 | 0.598210515 | 0.549699476 | 0.751487589 | 0.047737309 |
| ENSMUSG00000001034 | Mapk7 | 23939 | protein_coding | 304.099102 | 0.047738869 | 0.082104475 | 0.58144052 | 0.560943595 | 0.759175676 | 0.047738869 |
| ENSMUSG00000044475 | Ascc1 | 69090 | protein_coding | 509.241608 | 0.04778616 | 0.061274447 | 0.779870926 | 0.435466853 | 0.668899318 | 0.04778616 |
| ENSMUSG00000027331 | Knstrn | 51944 | protein_coding | 474.6064173 | -0.047793213 | 0.103717865 | -0.460800201 | 0.644941958 | 0.81580845 | 0.047793213 |
| ENSMUSG00000014980 | Tsen15 | 66637 | protein_coding | 114.4244227 | 0.047798854 | 0.120687674 | 0.396054146 | 0.692065084 | 0.845589334 | 0.047798854 |
| ENSMUSG00000029772 | Ahcyl2 | 74340 | protein_coding | 16470.82021 | 0.047813183 | 0.040537362 | 1.179484321 | 0.238205376 | 0.478577066 | 0.047813183 |
| ENSMUSG00000036639 | Nudt1 | 17766 | protein_coding | 350.6565492 | 0.047832304 | 0.086132439 | 0.555334373 | 0.578665972 | 0.771101239 | 0.047832304 |
| ENSMUSG00000028411 | Aptx | 66408 | protein_coding | 244.6416575 | -0.047866339 | 0.083814618 | -0.571097744 | 0.567933388 | 0.763127957 | 0.047866339 |
| ENSMUSG00000063200 | Nol7 | 70078 | protein_coding | 938.3348491 | 0.0478743 | 0.057301135 | 0.835486065 | 0.40344409 | 0.642601761 | 0.0478743 |
| ENSMUSG00000024403 | Atp6v1g2 | 66237 | protein_coding | 25.28024982 | -0.047876259 | 0.234059786 | -0.204547135 | 0.837925962 | 0.928042853 | 0.047876259 |
| ENSMUSG00000036298 | Slc2a13 | 239606 | protein_coding | 1260.995527 | -0.047887967 | 0.060119061 | -0.796552139 | 0.425711183 | 0.661307016 | 0.047887967 |
| ENSMUSG00000062198 | 2700097O09Rik | 72658 | protein_coding | 111.3245247 | -0.047901429 | 0.140486754 | -0.340967585 | 0.733127984 | 0.871318547 | 0.047901429 |
| ENSMUSG00000022098 | Bmp1 | 12153 | protein_coding | 225.9266385 | 0.047938675 | 0.168967979 | 0.283714552 | 0.776629142 | 0.895837077 | 0.047938675 |
| ENSMUSG00000027582 | Zgpat | 229007 | protein_coding | 646.173453 | -0.047957688 | 0.059244451 | -0.809488265 | 0.41823435 | 0.655860109 | 0.047957688 |
| ENSMUSG00000027349 | Fam98b | 68215 | protein_coding | 180.7442124 | 0.04799915 | 0.09573979 | 0.501350066 | 0.616124776 | 0.795599668 | 0.04799915 |
| ENSMUSG00000032322 | Pstpip1 | 19200 | protein_coding | 27.32814813 | 0.048006436 | 0.222166138 | 0.216083499 | 0.828922662 | 0.923972613 | 0.048006436 |
| ENSMUSG00000069308 | H2bc22 | 319188 | protein_coding | 65.6374283 | 0.048008487 | 0.197196886 | 0.243454588 | 0.807653261 | 0.91309405 | 0.048008487 |
| ENSMUSG00000022685 | Parn | 74108 | protein_coding | 477.1747282 | 0.048016999 | 0.067819854 | 0.708007994 | 0.478940293 | 0.701468541 | 0.048016999 |
| ENSMUSG00000029313 | Aff1 | 17355 | protein_coding | 2085.450133 | -0.048032998 | 0.060627213 | -0.792267949 | 0.428204456 | 0.663421426 | 0.048032998 |
| ENSMUSG00000027882 | Stxbp3 | 20912 | protein_coding | 1455.137622 | 0.048037641 | 0.041551262 | 1.156105465 | 0.247638021 | 0.488580666 | 0.048037641 |
| ENSMUSG00000005886 | Ncoa2 | 17978 | protein_coding | 1510.820631 | -0.048070093 | 0.056096278 | -0.85692126 | 0.391488403 | 0.631323671 | 0.048070093 |
| ENSMUSG00000028121 | Bcar3 | 29815 | protein_coding | 1739.526747 | 0.048071953 | 0.057260946 | 0.839524245 | 0.40117519 | 0.640442713 | 0.048071953 |
| ENSMUSG00000030019 | Fbxl14 | 101358 | protein_coding | 1057.848989 | -0.048090012 | 0.067101408 | -0.71667664 | 0.473573643 | 0.697472342 | 0.048090012 |
| ENSMUSG00000027583 | Zbtb46 | 72147 | protein_coding | 8.95018578 | -0.048098302 | 0.504688617 | -0.095302926 | 0.924074219 | 0.966739549 | 0.048098302 |
| ENSMUSG00000039656 | Rxrb | 20182 | protein_coding | 1116.254672 | -0.048106917 | 0.070814109 | -0.679340846 | 0.496921922 | 0.715076276 | 0.048106917 |
| ENSMUSG00000036461 | Elf1 | 13709 | protein_coding | 2714.267743 | -0.048107511 | 0.036861409 | -1.305091489 | 0.191861697 | 0.423218928 | 0.048107511 |

| ensembl_gene_id | external_gene_name | entrezgene_id | gene_biotype | baseMean | log2FoldChange | lfcSE | stat | pvalue | padj | absLog2FC |
|---|---|---|---|---|---|---|---|---|---|---|
| ENSMUSG00000046862 | Pramel12 | 242736 | protein_coding | 452.8001043 | -0.048119565 | 0.064618727 | -0.744669037 | 0.456471822 | 0.684487275 | 0.048119565 |
| ENSMUSG00000028945 | Rheb | 19744 | protein_coding | 1046.843166 | -0.048148817 | 0.045245433 | -1.064169643 | 0.287251857 | 0.529611374 | 0.048148817 |
| ENSMUSG00000060288 | Ppih | 66101 | protein_coding | 308.5743271 | 0.048164945 | 0.099558909 | 0.48378338 | 0.628539611 | 0.804287872 | 0.048164945 |
| ENSMUSG00000015748 | Prpf3 | 70767 | protein_coding | 308.2176989 | 0.048169164 | 0.099474286 | 0.484237343 | 0.628217437 | 0.804068461 | 0.048169164 |
| ENSMUSG00000024941 | Scyl1 | 78891 | protein_coding | 1479.442308 | 0.048182838 | 0.056947746 | 0.846088602 | 0.397503319 | 0.637040476 | 0.048182838 |
| ENSMUSG00000025872 | Thoc3 | 73666 | protein_coding | 538.8035671 | 0.048183157 | 0.077215241 | 0.624010964 | 0.532620385 | 0.740799392 | 0.048183157 |
| ENSMUSG00000037475 | Thoc2 | 331401 | protein_coding | 1369.410041 | 0.048210728 | 0.055913041 | 0.86224479 | 0.388552827 | 0.628430063 | 0.048210728 |
| ENSMUSG00000029420 | Rimbp2 | 231760 | protein_coding | 68.38845539 | -0.048216328 | 0.150798935 | -0.319739181 | 0.749166055 | 0.880475742 | 0.048216328 |
| ENSMUSG00000013465 | Nelfb | 58202 | protein_coding | 1547.3158 | 0.048231709 | 0.060939749 | 0.791465496 | 0.428672403 | 0.663709663 | 0.048231709 |
| ENSMUSG00000031834 | Pik3r2 | 18709 | protein_coding | 1329.141709 | 0.04824657 | 0.069546315 | 0.693732949 | 0.487849699 | 0.708341774 | 0.04824657 |
| ENSMUSG00000034109 | Golim4 | 73124 | protein_coding | 2047.713173 | 0.048261796 | 0.051746854 | 0.932651779 | 0.35099979 | 0.593363209 | 0.048261796 |
| ENSMUSG00000026276 | Septin2 | 18000 | protein_coding | 59.16968352 | -0.048270976 | 0.211823235 | -0.22788329 | 0.819736973 | 0.919197481 | 0.048270976 |
| ENSMUSG00000039230 | Tbcd | 108903 | protein_coding | 1675.337485 | 0.048286527 | 0.051280119 | 0.941622766 | 0.34638581 | 0.588629676 | 0.048286527 |
| ENSMUSG00000037933 | Bicd2 | 76895 | protein_coding | 455.5368775 | -0.048293974 | 0.094188176 | -0.512739242 | 0.608133736 | 0.790834872 | 0.048293974 |
| ENSMUSG00000025362 | Rps26 | 27370 | protein_coding | 7875.559901 | -0.048309994 | 0.054443071 | -0.8873488 | 0.37489114 | 0.616422248 | 0.048309994 |
| ENSMUSG00000029110 | Rnf4 | 19822 | protein_coding | 2794.497394 | -0.048310454 | 0.049751025 | -0.971044399 | 0.33152617 | 0.574895496 | 0.048310454 |
| ENSMUSG00000039703 | Nploc4 | 217365 | protein_coding | 1602.603498 | 0.048318995 | 0.047427921 | 1.018787961 | 0.308303642 | 0.552028135 | 0.048318995 |
| ENSMUSG00000031333 | Abcb7 | 11306 | protein_coding | 1908.4992 | 0.048332514 | 0.049508754 | 0.976241774 | 0.328944666 | 0.5725114 | 0.048332514 |
| ENSMUSG00000028165 | Cisd2 | 67006 | protein_coding | 1234.578808 | 0.048345957 | 0.042054659 | 1.149598114 | 0.250309435 | 0.491446343 | 0.048345957 |
| ENSMUSG00000002409 | Dyrk1b | 13549 | protein_coding | 297.9291866 | -0.048353837 | 0.124588507 | -0.388108326 | 0.697935871 | 0.849362552 | 0.048353837 |
| ENSMUSG00000021969 | Zdhhc20 | 75965 | protein_coding | 1775.823225 | -0.048391763 | 0.050317799 | -0.961722571 | 0.336188981 | 0.579801059 | 0.048391763 |
| ENSMUSG00000022634 | Yaf2 | 67057 | protein_coding | 202.9601066 | 0.048392957 | 0.098388233 | 0.49185716 | 0.622820325 | 0.800888833 | 0.048392957 |
| ENSMUSG00000039849 | Pcif1 | 228866 | protein_coding | 1130.247894 | -0.048405745 | 0.061660606 | -0.785035188 | 0.432432938 | 0.666779423 | 0.048405745 |
| ENSMUSG00000027012 | Dync1i2 | 13427 | protein_coding | 2063.511262 | 0.048407928 | 0.045993483 | 1.052495374 | 0.292572332 | 0.535307271 | 0.048407928 |
| ENSMUSG00000078963 | Hsbp1l1 | 66255 | protein_coding | 77.16587175 | 0.048456738 | 0.190925038 | 0.253799807 | 0.799650222 | 0.908570455 | 0.048456738 |
| ENSMUSG00000034889 | Cactin | 70312 | protein_coding | 578.1396548 | 0.048469813 | 0.082810392 | 0.585310757 | 0.558338783 | 0.756920776 | 0.048469813 |
| ENSMUSG00000016128 | Stard13 | 243362 | protein_coding | 618.3918531 | -0.048469823 | 0.059035946 | -0.821022214 | 0.411633615 | 0.649614964 | 0.048469823 |
| ENSMUSG00000001844 | Zdhhc4 | 72881 | protein_coding | 432.2182126 | -0.048492444 | 0.066054869 | -0.734123689 | 0.462873357 | 0.689266725 | 0.048492444 |
| ENSMUSG00000047417 | Rexo1 | 66932 | protein_coding | 807.4231883 | -0.048500081 | 0.0734864 | -0.659987165 | 0.509262066 | 0.72429229 | 0.048500081 |
| ENSMUSG00000062044 | Lmtk3 | 381983 | protein_coding | 7.520984015 | 0.04850014 | 0.55361026 | 0.08760701 | 0.93018903 | 0.969254926 | 0.04850014 |
| ENSMUSG00000056220 | Pla2g4a | 18783 | protein_coding | 1386.72854 | 0.048509747 | 0.054960426 | 0.882630466 | 0.377435963 | 0.61867166 | 0.048509747 |
| ENSMUSG00000011096 | Akt1s1 | 67605 | protein_coding | 1064.905509 | 0.048510063 | 0.101899636 | 0.476057272 | 0.634033576 | 0.808496836 | 0.048510063 |
| ENSMUSG00000028668 | Eloa | 27224 | protein_coding | 1810.980407 | -0.048512587 | 0.039539626 | -1.226935916 | 0.219846678 | 0.456057983 | 0.048512587 |
| ENSMUSG00000051375 | Pcdh1 | 75599 | protein_coding | 2702.649747 | -0.04852536 | 0.062714342 | -0.773752197 | 0.439077346 | 0.6716305 | 0.04852536 |
| ENSMUSG00000038745 | Nlrp6 | 101613 | protein_coding | 5498.201563 | -0.048527791 | 0.061887547 | -0.784128528 | 0.432964698 | 0.667195279 | 0.048527791 |
| ENSMUSG00000051147 | Nat2 | 17961 | protein_coding | 524.9156296 | -0.048569187 | 0.087593606 | -0.554483243 | 0.579248171 | 0.771609416 | 0.048569187 |
| ENSMUSG00000023938 | Aars2 | 224805 | protein_coding | 505.4678218 | -0.048592374 | 0.091410971 | -0.531581418 | 0.595015938 | 0.782189365 | 0.048592374 |
| ENSMUSG00000032299 | Commd4 | 66199 | protein_coding | 503.5130016 | -0.048594956 | 0.063211141 | -0.768772003 | 0.442028672 | 0.673207706 | 0.048594956 |
| ENSMUSG00000107283 | Mpv17 | 17527 | protein_coding | 285.439136 | 0.048596811 | 0.084456456 | 0.575406705 | 0.565016271 | 0.761555988 | 0.048596811 |

| ensembl_gene_id | external_gene_name | entrezgene_id | gene_biotype | baseMean | log2FoldChange | lfcSE | stat | pvalue | padj | absLog2FC |
|---|---|---|---|---|---|---|---|---|---|---|
| ENSMUSG00000034994 | Eef2 | 13629 | protein_coding | 66323.27302 | -0.048622895 | 0.044806551 | -1.085173789 | 0.277844691 | 0.520660159 | 0.048622895 |
| ENSMUSG00000051306 | Usp42 | 76800 | protein_coding | 422.0970022 | 0.04865099 | 0.077279991 | 0.629541863 | 0.528994371 | 0.738303091 | 0.04865099 |
| ENSMUSG00000051768 | Xrcc1 | 22594 | protein_coding | 493.0351046 | 0.04869957 | 0.056255337 | 0.86568801 | 0.386661277 | 0.626702152 | 0.04869957 |
| ENSMUSG00000021973 | Micu2 | 68514 | protein_coding | 1680.656611 | -0.048719087 | 0.058997556 | -0.82578144 | 0.408928084 | 0.64731555 | 0.048719087 |
| ENSMUSG00000029456 | Acad10 | 71985 | protein_coding | 510.6881452 | -0.04872944 | 0.059567938 | -0.81804814 | 0.413329699 | 0.651017488 | 0.04872944 |
| ENSMUSG00000087497 | 2810001G20Rik | 66456 | lncRNA | 112.88297 | 0.048741097 | 0.151956914 | 0.320756034 | 0.74839528 | 0.880475742 | 0.048741097 |
| ENSMUSG00000034723 | Tmx4 | 52837 | protein_coding | 361.0676284 | 0.048759085 | 0.071400064 | 0.682899733 | 0.494670203 | 0.713491312 | 0.048759085 |
| ENSMUSG00000059878 | Zfp422 | 67255 | protein_coding | 696.6907822 | -0.048761246 | 0.061281283 | -0.795695582 | 0.426208995 | 0.66169488 | 0.048761246 |
| ENSMUSG00000031021 | Tmem9b | 56786 | protein_coding | 1299.045096 | 0.048783132 | 0.044114329 | 1.105834166 | 0.268798304 | 0.511131971 | 0.048783132 |
| ENSMUSG00000060591 | Ifitm2 | 80876 | protein_coding | 467.8071791 | -0.048786773 | 0.103169763 | -0.4728786 | 0.636299794 | 0.809849797 | 0.048786773 |
| ENSMUSG00000030555 | Ttc23 | 67009 | protein_coding | 263.0194978 | 0.04881697 | 0.083361707 | 0.585604251 | 0.558141491 | 0.756755317 | 0.04881697 |
| ENSMUSG00000030846 | Tial1 | 21843 | protein_coding | 735.3097918 | -0.048840436 | 0.058110433 | -0.84047619 | 0.400641446 | 0.639843989 | 0.048840436 |
| ENSMUSG00000018931 | Natd1 | 24083 | protein_coding | 298.7830796 | -0.048897332 | 0.087197137 | -0.560767639 | 0.574955951 | 0.768361739 | 0.048897332 |
| ENSMUSG00000001062 | Vps9d1 | 72325 | protein_coding | 441.4522006 | -0.048904619 | 0.06354992 | -0.769546512 | 0.441568944 | 0.672888818 | 0.048904619 |
| ENSMUSG00000001227 | Sema6b | 20359 | protein_coding | 86.16001909 | 0.048921428 | 0.123587291 | 0.395845137 | 0.692219276 | 0.84565495 | 0.048921428 |
| ENSMUSG00000035105 | Egln3 | 112407 | protein_coding | 410.2181816 | 0.048927514 | 0.157851854 | 0.309958439 | 0.756592562 | 0.884485201 | 0.048927514 |
| ENSMUSG00000050552 | Lamtor4 | 66096 | protein_coding | 969.8812904 | 0.048957575 | 0.079159162 | 0.618470097 | 0.536265504 | 0.742878353 | 0.048957575 |
| ENSMUSG00000004328 | Hif3a | 53417 | protein_coding | 95.05614212 | -0.049002361 | 0.153825176 | -0.318558783 | 0.750061113 | 0.880597382 | 0.049002361 |
| ENSMUSG00000029625 | Cpsf4 | 54188 | protein_coding | 345.6166633 | -0.049007417 | 0.071685601 | -0.683643796 | 0.494200122 | 0.713491312 | 0.049007417 |
| ENSMUSG00000024085 | Man2a1 | 17158 | protein_coding | 3383.945225 | 0.049009358 | 0.042935719 | 1.141458886 | 0.253679012 | 0.494901086 | 0.049009358 |
| ENSMUSG00000002342 | Tmem161a | 234371 | protein_coding | 656.3931997 | 0.049012021 | 0.069198389 | 0.708282693 | 0.478769722 | 0.701282408 | 0.049012021 |
| ENSMUSG00000050244 | Heatr1 | 217995 | protein_coding | 736.3768311 | 0.049077105 | 0.092795581 | 0.528873301 | 0.596893344 | 0.783337403 | 0.049077105 |
| ENSMUSG00000047414 | Flrt2 | 399558 | protein_coding | 45.17578426 | -0.04911742 | 0.177923866 | -0.276058635 | 0.782503031 | 0.898825528 | 0.04911742 |
| ENSMUSG00000025466 | Fuom | 69064 | protein_coding | 774.2379053 | -0.049145912 | 0.070738321 | -0.694756551 | 0.487207881 | 0.708115319 | 0.049145912 |
| ENSMUSG00000031545 | Gpat4 | 102247 | protein_coding | 1975.554388 | 0.049164082 | 0.049857962 | 0.986082867 | 0.32409245 | 0.567992068 | 0.049164082 |
| ENSMUSG00000045975 | C2cd2 | 207781 | protein_coding | 488.4816609 | -0.049180417 | 0.058785451 | -0.836608646 | 0.402812584 | 0.642103246 | 0.049180417 |
| ENSMUSG00000034621 | Gpatch8 | 237943 | protein_coding | 1081.396148 | -0.049191968 | 0.072013216 | -0.683096381 | 0.494545942 | 0.713491312 | 0.049191968 |
| ENSMUSG00000038374 | Rbm8a | 60365 | protein_coding | 879.2084314 | 0.04919398 | 0.054102726 | 0.909269895 | 0.363207677 | 0.605769124 | 0.04919398 |
| ENSMUSG00000028998 | Tomm7 | 66169 | protein_coding | 2203.537886 | 0.04924947 | 0.057790246 | 0.852210779 | 0.394097115 | 0.633722622 | 0.04924947 |
| ENSMUSG00000047721 | Bola2 | 66162 | protein_coding | 559.6961969 | -0.049257527 | 0.0707102 | -0.696611334 | 0.486046061 | 0.707210133 | 0.049257527 |
| ENSMUSG00000038777 | Sema6c | 20360 | protein_coding | 9.943775037 | 0.049280537 | 0.429334015 | 0.114783677 | 0.908616588 | 0.959234074 | 0.049280537 |
| ENSMUSG00000079555 | Haus3 | 231123 | protein_coding | 138.2948841 | 0.049300457 | 0.12693201 | 0.388400508 | 0.697719668 | 0.849281085 | 0.049300457 |
| ENSMUSG00000032018 | Sc5d | 235293 | protein_coding | 1823.595659 | -0.04930298 | 0.062919699 | -0.783585767 | 0.43328321 | 0.667452319 | 0.04930298 |
| ENSMUSG00000037149 | Ddx1 | 104721 | protein_coding | 2680.295314 | 0.049318333 | 0.054100278 | 0.911609606 | 0.361974264 | 0.604873354 | 0.049318333 |
| ENSMUSG00000033256 | Shf | 435684 | protein_coding | 150.7176251 | -0.049337773 | 0.109972892 | -0.44863577 | 0.653694425 | 0.821678098 | 0.049337773 |
| ENSMUSG00000031971 | Ccsap | 73420 | protein_coding | 53.55203112 | -0.049338858 | 0.173562765 | -0.284270982 | 0.776202722 | 0.895541399 | 0.049338858 |
| ENSMUSG00000019278 | Dpep1 | 13479 | protein_coding | 6664.989719 | -0.04935544 | 0.117476199 | -0.420131398 | 0.674389467 | 0.834309996 | 0.04935544 |
| ENSMUSG00000029705 | Cux1 | 13047 | protein_coding | 2742.980103 | -0.049358922 | 0.039089909 | -1.262702377 | 0.206696159 | 0.441566958 | 0.049358922 |
| ENSMUSG00000028266 | Lmo4 | 16911 | protein_coding | 729.5465403 | -0.049360149 | 0.08544692 | -0.577670306 | 0.56348673 | 0.760766053 | 0.049360149 |

| ensembl_gene_id | external_gene_name | entrezgene_id | gene_biotype | baseMean | log2FoldChange | lfcSE | stat | pvalue | padj | absLog2FC |
|---|---|---|---|---|---|---|---|---|---|---|
| ENSMUSG00000030660 | Pik3c2a | 18704 | protein_coding | 2454.740967 | 0.049376505 | 0.046150018 | 1.069913012 | 0.284658465 | 0.527543906 | 0.049376505 |
| ENSMUSG00000044768 | Macir | 52392 | protein_coding | 1336.511373 | -0.049379818 | 0.045497201 | -1.085337487 | 0.277772209 | 0.520660159 | 0.049379818 |
| ENSMUSG00000015522 | Arnt | 11863 | protein_coding | 853.4470701 | -0.049432705 | 0.049441609 | -0.999819908 | 0.31739767 | 0.561196384 | 0.049432705 |
| ENSMUSG00000024242 | Map4k3 | 225028 | protein_coding | 2316.389713 | -0.049437735 | 0.05400256 | -0.915470206 | 0.359944836 | 0.603099387 | 0.049437735 |
| ENSMUSG00000029472 | Anapc5 | 59008 | protein_coding | 3550.473011 | -0.049442318 | 0.037811365 | -1.307604666 | 0.191007433 | 0.422142919 | 0.049442318 |
| ENSMUSG00000006219 | Fblim1 | 74202 | protein_coding | 1289.582546 | 0.04945616 | 0.07308199 | 0.676721581 | 0.498582621 | 0.716014372 | 0.04945616 |
| ENSMUSG00000067928 | Zfp760 | 240034 | protein_coding | 140.5278106 | 0.049476279 | 0.140919018 | 0.351097249 | 0.725515392 | 0.865987203 | 0.049476279 |
| ENSMUSG00000040560 | Wdr7 | 104082 | protein_coding | 723.1442309 | 0.04950601 | 0.05332418 | 0.928397018 | 0.353201665 | 0.595699984 | 0.04950601 |
| ENSMUSG00000029802 | Abcg2 | 26357 | protein_coding | 2501.437211 | 0.049506368 | 0.061367219 | 0.806723343 | 0.419825892 | 0.657014281 | 0.049506368 |
| ENSMUSG00000052419 | Smim30 | 66520 | protein_coding | 495.391579 | 0.049508067 | 0.066176019 | 0.748126996 | 0.45438356 | 0.682675952 | 0.049508067 |
| ENSMUSG00000039067 | Psmd7 | 17463 | protein_coding | 1076.773486 | 0.049514565 | 0.064681254 | 0.765516471 | 0.443964062 | 0.674900387 | 0.049514565 |
| ENSMUSG00000030560 | Ctsc | 13032 | protein_coding | 1839.122181 | -0.049557434 | 0.049713469 | -0.996861314 | 0.318831832 | 0.562506362 | 0.049557434 |
| ENSMUSG00000010406 | Mrpl52 | 68836 | protein_coding | 940.1608575 | 0.049584263 | 0.084973448 | 0.583526551 | 0.559538888 | 0.75798157 | 0.049584263 |
| ENSMUSG00000030852 | Tacc2 | 57752 | protein_coding | 1273.938984 | -0.049584554 | 0.07141771 | -0.694289334 | 0.487500778 | 0.708115319 | 0.049584554 |
| ENSMUSG00000024949 | Sf1 | 22668 | protein_coding | 2068.386327 | -0.049624557 | 0.068500105 | -0.72444497 | 0.4687926 | 0.693362933 | 0.049624557 |
| ENSMUSG00000003464 | Pex19 | 19298 | protein_coding | 1909.022158 | -0.049651165 | 0.040870466 | -1.21484214 | 0.224426305 | 0.462439318 | 0.049651165 |
| ENSMUSG00000061032 | Rrp1 | 18114 | protein_coding | 2121.205603 | 0.049665562 | 0.039883872 | 1.24525429 | 0.213038292 | 0.448715514 | 0.049665562 |
| ENSMUSG00000028737 | Aldh4a1 | 212647 | protein_coding | 451.5642594 | 0.049668995 | 0.074072373 | 0.670546833 | 0.502509262 | 0.719074257 | 0.049668995 |
| ENSMUSG00000054065 | Pkp3 | 56460 | protein_coding | 2160.173023 | 0.049683659 | 0.072980972 | 0.680775525 | 0.496013539 | 0.714476238 | 0.049683659 |
| ENSMUSG00000039263 | Npepl1 | 228961 | protein_coding | 1475.985083 | -0.049694634 | 0.039961541 | -1.243561495 | 0.213660995 | 0.449303803 | 0.049694634 |
| ENSMUSG00000030400 | Ercc2 | 13871 | protein_coding | 420.1011259 | -0.049741102 | 0.069439833 | -0.716319445 | 0.473794123 | 0.697669633 | 0.049741102 |
| ENSMUSG00000024732 | Ccdc86 | 108673 | protein_coding | 227.4379996 | -0.049744085 | 0.111115443 | -0.447679315 | 0.654384652 | 0.822097164 | 0.049744085 |
| ENSMUSG00000027425 | Kat14 | 228714 | protein_coding | 684.1609382 | -0.049776441 | 0.059027992 | -0.843268425 | 0.399078338 | 0.638678506 | 0.049776441 |
| ENSMUSG00000024436 | Mrps18b | 66973 | protein_coding | 339.3609313 | -0.049779088 | 0.076508531 | -0.650634469 | 0.515282474 | 0.727964612 | 0.049779088 |
| ENSMUSG00000039810 | Zc3h10 | 103284 | protein_coding | 412.6209557 | -0.049793903 | 0.072825691 | -0.683740884 | 0.494138802 | 0.713490595 | 0.049793903 |
| ENSMUSG00000017716 | Birc5 | 11799 | protein_coding | 428.0634328 | 0.049813034 | 0.116378529 | 0.428025981 | 0.668632205 | 0.830998117 | 0.049813034 |
| ENSMUSG00000001855 | Nup214 | 227720 | protein_coding | 540.7781479 | -0.049822586 | 0.093456835 | -0.533107993 | 0.593958827 | 0.781302936 | 0.049822586 |
| ENSMUSG00000009647 | Mcu | 215999 | protein_coding | 3234.900363 | 0.049830001 | 0.060477081 | 0.823948509 | 0.409968816 | 0.64854129 | 0.049830001 |
| ENSMUSG00000020372 | Rack1 | 14694 | protein_coding | 23979.32173 | -0.049835692 | 0.044743698 | -1.113803613 | 0.265363447 | 0.507231991 | 0.049835692 |
| ENSMUSG00000022672 | Prkdc | 19090 | protein_coding | 654.9422203 | 0.049847057 | 0.05212664 | 0.956268356 | 0.338936673 | 0.58240762 | 0.049847057 |
| ENSMUSG00000096472 | Cdkn2d | 12581 | protein_coding | 138.8133715 | 0.049852824 | 0.122063573 | 0.408416886 | 0.682967638 | 0.840074678 | 0.049852824 |
| ENSMUSG00000047989 | Ino80c | 225280 | protein_coding | 583.4440365 | 0.0498758 | 0.056205768 | 0.887378666 | 0.374875065 | 0.616422248 | 0.0498758 |
| ENSMUSG00000022827 | Rabl3 | 67657 | protein_coding | 238.9626167 | -0.049881975 | 0.10160554 | -0.490937549 | 0.623470618 | 0.801299862 | 0.049881975 |
| ENSMUSG00000030717 | Nupr1 | 56312 | protein_coding | 3236.218412 | -0.049882077 | 0.115685611 | -0.431186523 | 0.666332753 | 0.829602333 | 0.049882077 |
| ENSMUSG00000031065 | Cdk16 | 18555 | protein_coding | 2809.227593 | 0.049891034 | 0.051040446 | 0.97748037 | 0.328331391 | 0.571937821 | 0.049891034 |
| ENSMUSG00000030166 | Rad52 | 19365 | protein_coding | 212.517667 | -0.049897873 | 0.091134563 | -0.54751865 | 0.584022461 | 0.775222412 | 0.049897873 |
| ENSMUSG00000006538 | Ihh | 16147 | protein_coding | 2310.363777 | 0.049913921 | 0.087263547 | 0.57199052 | 0.567328397 | 0.762827428 | 0.049913921 |
| ENSMUSG00000115219 | Eef1akmt4 | 110599566 | protein_coding | 88.1650512 | -0.049919365 | 0.155083253 | -0.32188753 | 0.747537901 | 0.880075211 | 0.049919365 |
| ENSMUSG00000007867 | Ift43 | 76411 | protein_coding | 366.6390066 | -0.0499243 | 0.079633602 | -0.62692505 | 0.530708366 | 0.73971984 | 0.0499243 |

| ensembl_gene_id | external_gene_name | entrezgene_id | gene_biotype | baseMean | log2FoldChange | lfcSE | stat | pvalue | padj | absLog2FC |
|---|---|---|---|---|---|---|---|---|---|---|
| ENSMUSG00000067150 | Xpo5 | 72322 | protein_coding | 874.4123295 | 0.049927349 | 0.057086485 | 0.874591393 | 0.381796272 | 0.622494568 | 0.049927349 |
| ENSMUSG00000029550 | Sppl3 | 74585 | protein_coding | 1097.406527 | -0.049938767 | 0.057424004 | -0.869649677 | 0.384491881 | 0.624972092 | 0.049938767 |
| ENSMUSG00000030866 | Ern2 | 26918 | protein_coding | 2602.852658 | 0.049944931 | 0.069024286 | 0.723584897 | 0.469320616 | 0.693679184 | 0.049944931 |
| ENSMUSG00000030342 | Cd9 | 12527 | protein_coding | 4061.599057 | -0.049971522 | 0.06622258 | -0.754599444 | 0.450489357 | 0.679801803 | 0.049971522 |
| ENSMUSG00000042472 | Zfp410 | 52708 | protein_coding | 615.3730442 | 0.050008056 | 0.061636082 | 0.811343844 | 0.417168238 | 0.655085793 | 0.050008056 |
| ENSMUSG00000036810 | Cnep1r1 | 382030 | protein_coding | 359.4816159 | 0.050030183 | 0.075301492 | 0.664398298 | 0.506435422 | 0.722065604 | 0.050030183 |
| ENSMUSG00000020457 | Drg1 | 13494 | protein_coding | 989.9854793 | 0.050054872 | 0.064774789 | 0.772752372 | 0.439668945 | 0.671740242 | 0.050054872 |
| ENSMUSG00000008167 | Fbxw9 | 68628 | protein_coding | 446.1315136 | -0.050056616 | 0.085681724 | -0.584215794 | 0.559075137 | 0.757536946 | 0.050056616 |
| ENSMUSG00000041444 | Arhgap32 | 330914 | protein_coding | 3101.58187 | -0.050072462 | 0.047476391 | -1.054681321 | 0.291571098 | 0.534094399 | 0.050072462 |
| ENSMUSG00000042535 | Gtpbp1 | 14904 | protein_coding | 973.7259009 | 0.050077499 | 0.05700677 | 0.878448282 | 0.379700492 | 0.62056297 | 0.050077499 |
| ENSMUSG00000027809 | Etfdh | 66841 | protein_coding | 2587.197293 | 0.050111533 | 0.042742381 | 1.172408545 | 0.24103307 | 0.481643597 | 0.050111533 |
| ENSMUSG00000029173 | Sepsecs | 211006 | protein_coding | 1192.468518 | -0.050117136 | 0.058569215 | -0.85569077 | 0.392168847 | 0.632008293 | 0.050117136 |
| ENSMUSG00000041712 | Ubr7 | 66622 | protein_coding | 575.6265077 | -0.05012047 | 0.082783815 | -0.605438031 | 0.544887988 | 0.749012844 | 0.05012047 |
| ENSMUSG00000102869 | Norad | 347740 | lncRNA | 2340.696494 | 0.05013851 | 0.054210671 | 0.924882653 | 0.355026951 | 0.597653407 | 0.05013851 |
| ENSMUSG00000017776 | Crk | 12928 | protein_coding | 3785.440917 | 0.050149731 | 0.048142847 | 1.041686014 | 0.297557275 | 0.540774324 | 0.050149731 |
| ENSMUSG00000006288 | Ttc5 | 219022 | protein_coding | 660.0601572 | -0.050195058 | 0.056828159 | -0.88327792 | 0.37708613 | 0.618517903 | 0.050195058 |
| ENSMUSG00000098274 | Rpl24 | 68193 | protein_coding | 270.9978384 | 0.050239361 | 0.115935863 | 0.433337534 | 0.664769574 | 0.828854703 | 0.050239361 |
| ENSMUSG00000074925 | Ptar1 | 72351 | protein_coding | 867.3459681 | 0.050267641 | 0.060031316 | 0.837356961 | 0.402391951 | 0.641658935 | 0.050267641 |
| ENSMUSG00000034235 | Usp54 | 78787 | protein_coding | 187.4287097 | 0.050287888 | 0.097021581 | 0.518316515 | 0.604237452 | 0.788707774 | 0.050287888 |
| ENSMUSG00000042406 | Atf4 | 11911 | protein_coding | 5586.477198 | 0.050290044 | 0.054443176 | 0.923716207 | 0.355634092 | 0.598408466 | 0.050290044 |
| ENSMUSG00000039770 | Ypel5 | 383295 | protein_coding | 904.7887759 | 0.050321253 | 0.057540528 | 0.874535826 | 0.381826519 | 0.622494568 | 0.050321253 |
| ENSMUSG00000021242 | Npc2 | 67963 | protein_coding | 4876.935422 | -0.050421368 | 0.06898789 | -0.730872746 | 0.464856884 | 0.690944422 | 0.050421368 |
| ENSMUSG00000030741 | Spns1 | 73658 | protein_coding | 859.4613015 | 0.050431023 | 0.087209599 | 0.578273768 | 0.563079302 | 0.760578285 | 0.050431023 |
| ENSMUSG00000010760 | Phlda2 | 22113 | protein_coding | 24.35050193 | 0.050452633 | 0.287024396 | 0.175778204 | 0.860468193 | 0.936965584 | 0.050452633 |
| ENSMUSG00000027087 | Itgav | 16410 | protein_coding | 818.1378438 | 0.050469839 | 0.061808545 | 0.816551152 | 0.41418498 | 0.651982545 | 0.050469839 |
| ENSMUSG00000034789 | Rab24 | 19336 | protein_coding | 227.0525023 | 0.050538422 | 0.100321552 | 0.50376436 | 0.614426975 | 0.794774067 | 0.050538422 |
| ENSMUSG00000022529 | Zfp263 | 74120 | protein_coding | 671.8461837 | -0.05057112 | 0.059338457 | -0.852248655 | 0.394076097 | 0.633722622 | 0.05057112 |
| ENSMUSG00000024446 | Rpp21 | 67676 | protein_coding | 464.7145561 | 0.050624377 | 0.0721102 | 0.70204184 | 0.482653071 | 0.70472411 | 0.050624377 |
| ENSMUSG00000028629 | Exo5 | 73172 | protein_coding | 273.9662108 | -0.050646935 | 0.089136124 | -0.568197634 | 0.569900778 | 0.765233601 | 0.050646935 |
| ENSMUSG00000040112 | Mrps35 | 232536 | protein_coding | 1131.875009 | -0.050651821 | 0.044455503 | -1.139382486 | 0.254543658 | 0.495656311 | 0.050651821 |
| ENSMUSG00000038827 | Abitram | 230234 | protein_coding | 166.8530207 | -0.050660915 | 0.090451707 | -0.56008799 | 0.575419422 | 0.768546546 | 0.050660915 |
| ENSMUSG00000020956 | Dtd2 | 328092 | protein_coding | 227.924918 | 0.05066607 | 0.118026065 | 0.429278654 | 0.667720449 | 0.830562128 | 0.05066607 |
| ENSMUSG00000031388 | Naa10 | 56292 | protein_coding | 541.2199307 | 0.050669338 | 0.073980862 | 0.684897918 | 0.49340834 | 0.712754953 | 0.050669338 |
| ENSMUSG00000041488 | Stx3 | 20908 | protein_coding | 1550.632183 | -0.050718473 | 0.04447536 | -1.140372401 | 0.254131186 | 0.495032448 | 0.050718473 |
| ENSMUSG00000032469 | Dbr1 | 83703 | protein_coding | 312.7480603 | 0.050739526 | 0.068131118 | 0.744733503 | 0.456432841 | 0.684487275 | 0.050739526 |
| ENSMUSG00000058594 | Fbh1 | 50755 | protein_coding | 2598.678653 | 0.050752531 | 0.039215137 | 1.29420768 | 0.195593695 | 0.428130377 | 0.050752531 |
| ENSMUSG00000034210 | Efcab14 | 230648 | protein_coding | 1551.972298 | -0.050767869 | 0.055924192 | -0.907797989 | 0.363984961 | 0.606215581 | 0.050767869 |
| ENSMUSG00000026709 | Dars2 | 226539 | protein_coding | 596.7051648 | 0.050776514 | 0.08153265 | 0.622775212 | 0.53343225 | 0.741375537 | 0.050776514 |
| ENSMUSG00000006021 | Kptn | 70394 | protein_coding | 358.6664758 | -0.050778632 | 0.081639529 | -0.62198585 | 0.533951172 | 0.741606925 | 0.050778632 |

| ensembl_gene_id | external_gene_name | entrezgene_id | gene_biotype | baseMean | log2FoldChange | lfcSE | stat | pvalue | padj | absLog2FC |
|---|---|---|---|---|---|---|---|---|---|---|
| ENSMUSG00000049517 | Rps23 | 66475 | protein_coding | 7359.417134 | -0.050799038 | 0.032330175 | -1.571257762 | 0.116122788 | 0.314900319 | 0.050799038 |
| ENSMUSG00000073411 | H2-D1 | 14964 | protein_coding | 11623.21074 | -0.050835664 | 0.115595288 | -0.439772807 | 0.660101665 | 0.826041967 | 0.050835664 |
| ENSMUSG00000115338 | Pnp | 18950 | protein_coding | 540.4321319 | 0.050836764 | 0.081182799 | 0.626201177 | 0.531182995 | 0.73982107 | 0.050836764 |
| ENSMUSG00000017421 | Zfp207 | 22680 | protein_coding | 2400.135728 | -0.05084641 | 0.048138854 | -1.056244697 | 0.290856435 | 0.5332246 | 0.05084641 |
| ENSMUSG00000025134 | Alyref | 21681 | protein_coding | 1135.677507 | 0.050857607 | 0.070109245 | 0.72540515 | 0.468203514 | 0.693047027 | 0.050857607 |
| ENSMUSG00000021678 | F2rl1 | 14063 | protein_coding | 2142.09343 | 0.050858159 | 0.047548473 | 1.069606568 | 0.284796437 | 0.527617708 | 0.050858159 |
| ENSMUSG00000101225 | 1700008J07Rik | 629159 | lncRNA | 47.30020159 | 0.050859765 | 0.17107628 | 0.297292911 | 0.766242893 | 0.890223985 | 0.050859765 |
| ENSMUSG00000035248 | Tut7 | 214290 | protein_coding | 2267.647116 | 0.050886141 | 0.040082168 | 1.269545623 | 0.20424653 | 0.438338886 | 0.050886141 |
| ENSMUSG00000023027 | Atf1 | 11908 | protein_coding | 455.8125989 | 0.050897792 | 0.064265188 | 0.791996311 | 0.428362827 | 0.663421426 | 0.050897792 |
| ENSMUSG00000022771 | Ppil2 | 66053 | protein_coding | 1542.781286 | 0.050926306 | 0.044652975 | 1.14049078 | 0.254081892 | 0.495031103 | 0.050926306 |
| ENSMUSG00000004768 | Rab23 | 19335 | protein_coding | 120.3777999 | 0.050932387 | 0.117952526 | 0.43180412 | 0.665883785 | 0.829435992 | 0.050932387 |
| ENSMUSG00000047213 | Ythdf3 | 229096 | protein_coding | 2759.976849 | -0.050952351 | 0.044560549 | -1.143440837 | 0.252855605 | 0.493979691 | 0.050952351 |
| ENSMUSG00000040624 | Plekhg1 | 213783 | protein_coding | 502.5969816 | -0.050959743 | 0.067057579 | -0.759940106 | 0.447290387 | 0.678210988 | 0.050959743 |
| ENSMUSG00000041977 | Arhgef11 | 213498 | protein_coding | 2117.170304 | -0.050959747 | 0.035335817 | -1.442155635 | 0.149258473 | 0.36399386 | 0.050959747 |
| ENSMUSG00000030905 | Crym | 12971 | protein_coding | 68.02999031 | -0.050964119 | 0.159701268 | -0.319121568 | 0.74963433 | 0.880475742 | 0.050964119 |
| ENSMUSG00000030313 | Dennd5b | 320560 | protein_coding | 536.5643939 | -0.05097585 | 0.075763299 | -0.672830397 | 0.501055197 | 0.718355583 | 0.05097585 |
| ENSMUSG00000085741 | 5430405H02Rik | 74487 | lncRNA | 34.76271921 | -0.050983281 | 0.20470257 | -0.249060286 | 0.803314148 | 0.910371624 | 0.050983281 |
| ENSMUSG00000028016 | Ints12 | 71793 | protein_coding | 414.3552563 | -0.050984612 | 0.071054805 | -0.717539249 | 0.473041427 | 0.697134159 | 0.050984612 |
| ENSMUSG00000000303 | Cdh1 | 12550 | protein_coding | 22312.45681 | 0.051015315 | 0.0626519 | 0.814266046 | 0.415492556 | 0.653466789 | 0.051015315 |
| ENSMUSG00000055065 | Ddx17 | 67040 | protein_coding | 4839.981075 | -0.051028951 | 0.048576965 | -1.050476299 | 0.293499182 | 0.536379174 | 0.051028951 |
| ENSMUSG00000043279 | Trim56 | 384309 | protein_coding | 1374.413071 | 0.051033881 | 0.071759188 | 0.711182529 | 0.476971133 | 0.699728324 | 0.051033881 |
| ENSMUSG00000040697 | Dnajc16 | 214063 | protein_coding | 668.8194647 | 0.051046723 | 0.063370987 | 0.805521986 | 0.420518524 | 0.657587729 | 0.051046723 |
| ENSMUSG00000040721 | Zfhx2 | 239102 | protein_coding | 76.61657075 | -0.051059615 | 0.182414554 | -0.279909765 | 0.779546736 | 0.897539087 | 0.051059615 |
| ENSMUSG00000015937 | Macroh2a1 | 26914 | protein_coding | 2169.218201 | 0.051067068 | 0.045456813 | 1.123419461 | 0.261259388 | 0.503084198 | 0.051067068 |
| ENSMUSG00000028107 | Tars2 | 71807 | protein_coding | 677.1332563 | 0.051082065 | 0.07633198 | 0.669209223 | 0.503362024 | 0.719943844 | 0.051082065 |
| ENSMUSG00000047515 | BC049715 | 320135 | protein_coding | 68.80653866 | 0.051164752 | 0.15629863 | 0.327352528 | 0.743401267 | 0.877659754 | 0.051164752 |
| ENSMUSG00000034158 | Lrrc58 | 320184 | protein_coding | 1681.626317 | 0.05117212 | 0.053966871 | 0.948213581 | 0.343020735 | 0.585633643 | 0.05117212 |
| ENSMUSG00000026853 | Crat | 12908 | protein_coding | 1562.436276 | -0.051184262 | 0.051909642 | -0.986026108 | 0.324120301 | 0.567992068 | 0.051184262 |
| ENSMUSG00000059316 | Slc27a4 | 26569 | protein_coding | 3505.501961 | 0.051192058 | 0.067665418 | 0.756546825 | 0.449321409 | 0.679246742 | 0.051192058 |
| ENSMUSG00000027598 | Itch | 16396 | protein_coding | 2849.789746 | -0.051198392 | 0.051270906 | -0.99858568 | 0.31799544 | 0.561627805 | 0.051198392 |
| ENSMUSG00000069237 | Fam8a1 | 97863 | protein_coding | 1768.329251 | -0.051198419 | 0.066161764 | -0.773836973 | 0.439027205 | 0.671617505 | 0.051198419 |
| ENSMUSG00000032297 | Celf6 | 76183 | protein_coding | 26.93076524 | -0.051212022 | 0.248555107 | -0.206038904 | 0.836760522 | 0.927643978 | 0.051212022 |
| ENSMUSG00000030811 | Fbxl19 | 233902 | protein_coding | 1038.633133 | 0.051327807 | 0.099142351 | 0.517718281 | 0.604654842 | 0.789001427 | 0.051327807 |
| ENSMUSG00000002297 | Dbf4 | 27214 | protein_coding | 402.949182 | 0.051333766 | 0.084726214 | 0.605878197 | 0.544595638 | 0.749012844 | 0.051333766 |
| ENSMUSG00000037523 | Mavs | 228607 | protein_coding | 1128.045096 | 0.051352512 | 0.059438794 | 0.863956159 | 0.387611975 | 0.627536092 | 0.051352512 |
| ENSMUSG00000040820 | Hlcs | 110948 | protein_coding | 350.0446519 | -0.051358489 | 0.067064496 | -0.765807429 | 0.443790893 | 0.674808197 | 0.051358489 |
| ENSMUSG00000052144 | Ppp4r2 | 232314 | protein_coding | 1008.303979 | -0.051381866 | 0.061364902 | -0.837316851 | 0.40241449 | 0.641658935 | 0.051381866 |
| ENSMUSG00000089682 | Bcl2l2 | 12050 | protein_coding | 1208.552718 | 0.051389118 | 0.045464877 | 1.130303688 | 0.258348281 | 0.50034619 | 0.051389118 |
| ENSMUSG00000022619 | Mapk8ip2 | 60597 | protein_coding | 12.82110536 | -0.051409185 | 0.362784536 | -0.141707211 | 0.887311279 | 0.950240321 | 0.051409185 |

| ensembl_gene_id | external_gene_name | entrezgene_id | gene_biotype | baseMean | log2FoldChange | lfcSE | stat | pvalue | padj | absLog2FC |
|---|---|---|---|---|---|---|---|---|---|---|
| ENSMUSG00000038880 | Mrps34 | 79044 | protein_coding | 297.5085833 | 0.05141641 | 0.107253528 | 0.479391314 | 0.631660271 | 0.806746273 | 0.05141641 |
| ENSMUSG00000039458 | Mtmr12 | 268783 | protein_coding | 984.0349524 | -0.051439901 | 0.045799784 | -1.123147254 | 0.26137496 | 0.50318658 | 0.051439901 |
| ENSMUSG00000041203 | Trir | 68544 | protein_coding | 1528.053965 | 0.05144631 | 0.078904682 | 0.65200579 | 0.514397437 | 0.727325341 | 0.05144631 |
| ENSMUSG00000061136 | Prpf40a | 56194 | protein_coding | 2199.447507 | 0.051446553 | 0.055288732 | 0.93050701 | 0.352108636 | 0.594540465 | 0.051446553 |
| ENSMUSG00000022191 | Drosha | 14000 | protein_coding | 950.0647298 | 0.051457273 | 0.047749001 | 1.077661769 | 0.281184726 | 0.523633495 | 0.051457273 |
| ENSMUSG00000008140 | Emc10 | 69683 | protein_coding | 1392.69591 | 0.051476691 | 0.095135361 | 0.541088931 | 0.588446287 | 0.778305034 | 0.051476691 |
| ENSMUSG00000073676 | Hspe1 | 15528 | protein_coding | 1524.124444 | 0.051477939 | 0.08959855 | 0.574539873 | 0.565602527 | 0.76190043 | 0.051477939 |
| ENSMUSG00000027465 | Tbc1d20 | 67231 | protein_coding | 1918.239383 | -0.051491203 | 0.044148534 | -1.166317388 | 0.243486145 | 0.484584527 | 0.051491203 |
| ENSMUSG00000000385 | Tmprss2 | 50528 | protein_coding | 10376.90045 | 0.05153712 | 0.072750096 | 0.708413082 | 0.478688771 | 0.701227523 | 0.05153712 |
| ENSMUSG00000019373 | Cops3 | 26572 | protein_coding | 1008.288879 | 0.051551779 | 0.064386425 | 0.800662244 | 0.423327206 | 0.659294819 | 0.051551779 |
| ENSMUSG00000016253 | Nelfcd | 57314 | protein_coding | 978.5049416 | 0.051561729 | 0.073670218 | 0.699899236 | 0.483990234 | 0.706043466 | 0.051561729 |
| ENSMUSG00000047126 | Cltc | 67300 | protein_coding | 12153.9342 | 0.05158223 | 0.045722078 | 1.128168974 | 0.259248564 | 0.500967327 | 0.05158223 |
| ENSMUSG00000033106 | Slc7a6os | 66432 | protein_coding | 385.3830647 | -0.051594226 | 0.091354822 | -0.564767408 | 0.572231978 | 0.767059504 | 0.051594226 |
| ENSMUSG00000021496 | Pcbd2 | 72562 | protein_coding | 253.8663453 | -0.051618187 | 0.105656197 | -0.488548595 | 0.625161314 | 0.802110238 | 0.051618187 |
| ENSMUSG00000079144 | A130010J15Rik | 319266 | protein_coding | 293.6435822 | 0.05167485 | 0.085233823 | 0.606271648 | 0.544334382 | 0.748889318 | 0.05167485 |
| ENSMUSG00000054426 | A930005H10Rik | 68161 | lncRNA | 31.20445033 | -0.051707834 | 0.223948223 | -0.230891915 | 0.817398772 | 0.917979805 | 0.051707834 |
| ENSMUSG00000047888 | Tnrc6b | 213988 | protein_coding | 1995.45199 | 0.051725802 | 0.070740162 | 0.731208418 | 0.464651859 | 0.690767011 | 0.051725802 |
| ENSMUSG00000024400 | Wdr33 | 74320 | protein_coding | 1287.82072 | -0.051744099 | 0.049991591 | -1.035056036 | 0.300642743 | 0.54415999 | 0.051744099 |
| ENSMUSG00000037674 | Rfx7 | 319758 | protein_coding | 732.100262 | 0.051764346 | 0.070289683 | 0.736443013 | 0.461461136 | 0.687971202 | 0.051764346 |
| ENSMUSG00000041420 | Meis3 | 17537 | protein_coding | 114.655548 | 0.051772254 | 0.130375625 | 0.397100715 | 0.691293193 | 0.8450944 | 0.051772254 |
| ENSMUSG00000078619 | Smarcd2 | 83796 | protein_coding | 2093.497011 | -0.051773297 | 0.059412049 | -0.871427572 | 0.383520737 | 0.624309975 | 0.051773297 |
| ENSMUSG00000041272 | Tox | 252838 | protein_coding | 364.2796022 | -0.051807382 | 0.085401923 | -0.606630155 | 0.544096383 | 0.748817407 | 0.051807382 |
| ENSMUSG00000060227 | Golm2 | 319996 | protein_coding | 406.0269443 | -0.05183845 | 0.084448802 | -0.613844712 | 0.539317948 | 0.745357832 | 0.05183845 |
| ENSMUSG00000057101 | Zfp180 | 210135 | protein_coding | 695.5563252 | -0.051900024 | 0.066131492 | -0.784800432 | 0.432570587 | 0.666799776 | 0.051900024 |
| ENSMUSG00000054517 | Trim65 | 338364 | protein_coding | 301.1906598 | 0.051901708 | 0.097055155 | 0.534765084 | 0.59281231 | 0.780422622 | 0.051901708 |
| ENSMUSG00000023345 | Poc1a | 70235 | protein_coding | 325.2245249 | 0.051918778 | 0.092750966 | 0.559765357 | 0.575639496 | 0.76862707 | 0.051918778 |
| ENSMUSG00000074093 | Svip | 75744 | protein_coding | 556.6924898 | 0.051932186 | 0.06151077 | 0.844277935 | 0.398514113 | 0.638169126 | 0.051932186 |
| ENSMUSG00000030878 | Cdr2 | 12585 | protein_coding | 167.7797904 | 0.051970463 | 0.11439033 | 0.454325664 | 0.649594447 | 0.819270326 | 0.051970463 |
| ENSMUSG00000011257 | Pabpc4 | 230721 | protein_coding | 1685.065092 | -0.051998837 | 0.065633366 | -0.792262236 | 0.428207786 | 0.663421426 | 0.051998837 |
| ENSMUSG00000002103 | Acp2 | 11432 | protein_coding | 665.0897554 | 0.052012085 | 0.055178691 | 0.942611792 | 0.345879506 | 0.588148333 | 0.052012085 |
| ENSMUSG00000029687 | Ezh2 | 14056 | protein_coding | 699.8101137 | -0.052039713 | 0.078176761 | -0.665667294 | 0.505623783 | 0.721608883 | 0.052039713 |
| ENSMUSG00000031820 | Babam1 | 68251 | protein_coding | 1207.922644 | 0.05204648 | 0.065286512 | 0.79720112 | 0.425334235 | 0.661079917 | 0.05204648 |
| ENSMUSG00000028136 | Snx27 | 76742 | protein_coding | 899.2270579 | -0.052061659 | 0.06105833 | -0.852654494 | 0.393850933 | 0.633492143 | 0.052061659 |
| ENSMUSG00000055239 | Kcmf1 | 74287 | protein_coding | 1176.184575 | -0.052062153 | 0.049539272 | -1.050926893 | 0.293292168 | 0.536272425 | 0.052062153 |
| ENSMUSG00000033916 | Chmp2a | 68953 | protein_coding | 2939.933514 | 0.052062804 | 0.050854949 | 1.023750998 | 0.305952901 | 0.55012849 | 0.052062804 |
| ENSMUSG00000028902 | Sf3a3 | 75062 | protein_coding | 723.1248856 | 0.052065322 | 0.074229601 | 0.701409154 | 0.483047711 | 0.705097682 | 0.052065322 |
| ENSMUSG00000020532 | Acaca | 107476 | protein_coding | 1891.731976 | 0.052071554 | 0.058078458 | 0.896572587 | 0.369947031 | 0.612036907 | 0.052071554 |
| ENSMUSG00000033826 | Dnah8 | 13417 | protein_coding | 730.4913877 | -0.05209125 | 0.130056877 | -0.400526685 | 0.688768633 | 0.843735036 | 0.05209125 |
| ENSMUSG00000059714 | Flot1 | 14251 | protein_coding | 679.0433638 | -0.052108598 | 0.072608763 | -0.71766265 | 0.472965318 | 0.697134159 | 0.052108598 |

| ensembl_gene_id | external_gene_name | entrezgene_id | gene_biotype | baseMean | log2FoldChange | lfcSE | stat | pvalue | padj | absLog2FC |
|---|---|---|---|---|---|---|---|---|---|---|
| ENSMUSG00000040687 | Madd | 228355 | protein_coding | 902.1077389 | -0.052112601 | 0.065750934 | -0.792575823 | 0.428024999 | 0.663407897 | 0.052112601 |
| ENSMUSG00000005609 | Ctr9 | 22083 | protein_coding | 1578.890832 | 0.05211946 | 0.049301295 | 1.057162087 | 0.29043762 | 0.533047399 | 0.05211946 |
| ENSMUSG00000063800 | Prpf38a | 230596 | protein_coding | 753.9679489 | 0.052155298 | 0.055179164 | 0.94519912 | 0.344557228 | 0.58688999 | 0.052155298 |
| ENSMUSG00000078681 | Tm2d3 | 68634 | protein_coding | 587.4853605 | 0.052166 | 0.054673773 | 0.954132079 | 0.34001679 | 0.583162735 | 0.052166 |
| ENSMUSG00000009563 | Tor2a | 30933 | protein_coding | 877.5105028 | 0.052183816 | 0.05165711 | 1.010196207 | 0.312401296 | 0.556203985 | 0.052183816 |
| ENSMUSG00000033066 | Gas7 | 14457 | protein_coding | 256.7229631 | -0.052216775 | 0.092592929 | -0.563939124 | 0.572795563 | 0.767550813 | 0.052216775 |
| ENSMUSG00000071660 | Ttc9c | 70387 | protein_coding | 877.175446 | 0.052296811 | 0.051596991 | 1.013563196 | 0.310791227 | 0.554623228 | 0.052296811 |
| ENSMUSG00000052798 | Nup107 | 103468 | protein_coding | 334.3737263 | 0.052311359 | 0.083429698 | 0.627011249 | 0.530651861 | 0.73971984 | 0.052311359 |
| ENSMUSG00000056770 | Setd3 | 52690 | protein_coding | 2932.740243 | 0.05232164 | 0.038609061 | 1.355164784 | 0.175365062 | 0.401707581 | 0.05232164 |
| ENSMUSG00000030877 | Mfsd13b | 74466 | protein_coding | 9.664631094 | 0.052344413 | 0.365012754 | 0.143404341 | 0.885970855 | 0.949475732 | 0.052344413 |
| ENSMUSG00000078995 | Zfp456 | 408065 | protein_coding | 75.31277972 | -0.052348735 | 0.160071134 | -0.327034201 | 0.743642018 | 0.877751213 | 0.052348735 |
| ENSMUSG00000025199 | Chuk | 12675 | protein_coding | 2087.866656 | -0.052349486 | 0.051931634 | -1.008046183 | 0.313432292 | 0.55760602 | 0.052349486 |
| ENSMUSG00000017765 | Slc12a4 | 20498 | protein_coding | 9.37770529 | -0.052375285 | 0.446603305 | -0.117274737 | 0.906642343 | 0.95873684 | 0.052375285 |
| ENSMUSG00000025225 | Nfkb2 | 18034 | protein_coding | 1036.319255 | -0.052377446 | 0.057749218 | -0.906981035 | 0.364416826 | 0.606684232 | 0.052377446 |
| ENSMUSG00000020074 | Ccar1 | 67500 | protein_coding | 1382.489702 | 0.052403531 | 0.065154029 | 0.804302234 | 0.421222448 | 0.657986673 | 0.052403531 |
| ENSMUSG00000020116 | Pno1 | 66249 | protein_coding | 493.467445 | 0.052425028 | 0.086158731 | 0.60847029 | 0.542875605 | 0.747967026 | 0.052425028 |
| ENSMUSG00000038034 | Igsf8 | 140559 | protein_coding | 701.7192551 | -0.052432356 | 0.078123004 | -0.671151308 | 0.502124144 | 0.718950524 | 0.052432356 |
| ENSMUSG00000062127 | Cttnbp2nl | 80281 | protein_coding | 1594.969548 | -0.052432759 | 0.065185939 | -0.804356889 | 0.421190891 | 0.657986673 | 0.052432759 |
| ENSMUSG00000072772 | Grcc10 | 14790 | protein_coding | 390.8893221 | -0.052443653 | 0.087613602 | -0.598578891 | 0.549453736 | 0.751460876 | 0.052443653 |
| ENSMUSG00000039183 | Nubp2 | 26426 | protein_coding | 498.1691293 | 0.052459813 | 0.091035485 | 0.576256749 | 0.564441652 | 0.761354176 | 0.052459813 |
| ENSMUSG00000028580 | Pum1 | 80912 | protein_coding | 1924.4453 | -0.052465278 | 0.04994863 | -1.050384723 | 0.293541267 | 0.536379174 | 0.052465278 |
| ENSMUSG00000062234 | Gak | 231580 | protein_coding | 4665.634062 | 0.052486201 | 0.042522348 | 1.234320395 | 0.217083539 | 0.45330399 | 0.052486201 |
| ENSMUSG00000025645 | Ccdc51 | 66658 | protein_coding | 363.1201782 | 0.052496355 | 0.070827839 | 0.741182504 | 0.45858279 | 0.685788934 | 0.052496355 |
| ENSMUSG00000051378 | Kif18b | 70218 | protein_coding | 218.5112553 | 0.052499901 | 0.135368658 | 0.387829071 | 0.69814253 | 0.849391002 | 0.052499901 |
| ENSMUSG00000019795 | Pcmt1 | 18537 | protein_coding | 916.8140497 | -0.052506048 | 0.044667793 | -1.175478892 | 0.239803177 | 0.480291336 | 0.052506048 |
| ENSMUSG00000039166 | Akap7 | 432442 | protein_coding | 1169.486919 | 0.052542626 | 0.041007412 | 1.281295833 | 0.200089776 | 0.433483049 | 0.052542626 |
| ENSMUSG00000040034 | Nup43 | 69912 | protein_coding | 121.4314769 | 0.052557692 | 0.136147069 | 0.386036161 | 0.699469888 | 0.850145091 | 0.052557692 |
| ENSMUSG00000032540 | Abhd5 | 67469 | protein_coding | 635.0953119 | 0.052574476 | 0.053111641 | 0.989886108 | 0.32222979 | 0.566263466 | 0.052574476 |
| ENSMUSG00000024570 | Rbfa | 68731 | protein_coding | 245.9470588 | 0.052583571 | 0.089323094 | 0.588689536 | 0.556069558 | 0.75530479 | 0.052583571 |
| ENSMUSG00000037772 | Mrpl23 | 19935 | protein_coding | 224.9154069 | -0.052593239 | 0.119729429 | -0.439267433 | 0.660467769 | 0.826263572 | 0.052593239 |
| ENSMUSG00000030750 | Nsmce1 | 67711 | protein_coding | 356.3675695 | 0.052625714 | 0.077376898 | 0.680121788 | 0.49642735 | 0.714747243 | 0.052625714 |
| ENSMUSG00000028651 | Ppie | 56031 | protein_coding | 363.1008572 | 0.052634498 | 0.086397706 | 0.60921175 | 0.542384093 | 0.747864433 | 0.052634498 |
| ENSMUSG00000033295 | Ptprf | 19268 | protein_coding | 9802.722659 | -0.052636025 | 0.068188295 | -0.771921713 | 0.440160797 | 0.671885771 | 0.052636025 |
| ENSMUSG00000045180 | Shroom2 | 110380 | protein_coding | 1086.877073 | 0.052689295 | 0.061158303 | 0.861523171 | 0.388949965 | 0.628768653 | 0.052689295 |
| ENSMUSG00000008604 | Ubqln4 | 94232 | protein_coding | 1096.146397 | -0.052692273 | 0.065088321 | -0.809550346 | 0.418198656 | 0.655860109 | 0.052692273 |
| ENSMUSG00000026021 | Sumo1 | 22218 | protein_coding | 1124.437461 | 0.052766999 | 0.059252201 | 0.89054918 | 0.373171073 | 0.614910585 | 0.052766999 |
| ENSMUSG00000019295 | Tmem129 | 68366 | protein_coding | 577.6540189 | 0.05277593 | 0.066624964 | 0.792134472 | 0.428282272 | 0.663421426 | 0.05277593 |
| ENSMUSG00000001289 | Pfdn5 | 56612 | protein_coding | 2011.95132 | -0.05277769 | 0.045203475 | -1.167558228 | 0.242985008 | 0.484078966 | 0.05277769 |
| ENSMUSG00000067714 | Lpar5 | 381810 | protein_coding | 161.0443606 | 0.052795213 | 0.125784732 | 0.419726722 | 0.674685102 | 0.834405417 | 0.052795213 |

| ensembl_gene_id | external_gene_name | entrezgene_id | gene_biotype | baseMean | log2FoldChange | lfcSE | stat | pvalue | padj | absLog2FC |
|---|---|---|---|---|---|---|---|---|---|---|
| ENSMUSG00000032199 | Polr2m | 28015 | protein_coding | 2121.642354 | 0.052800091 | 0.040934651 | 1.289863001 | 0.197098229 | 0.429540965 | 0.052800091 |
| ENSMUSG00000033016 | Nfatc1 | 18018 | protein_coding | 242.7046207 | 0.052801145 | 0.10396617 | 0.507868517 | 0.611545555 | 0.792924519 | 0.052801145 |
| ENSMUSG00000072640 | Lyrm9 | 66274 | protein_coding | 212.3628194 | -0.052842624 | 0.088535533 | -0.596852157 | 0.550606092 | 0.752000715 | 0.052842624 |
| ENSMUSG00000023092 | Fhl1 | 14199 | protein_coding | 3691.881003 | -0.052855569 | 0.109718844 | -0.481736471 | 0.629993164 | 0.805574037 | 0.052855569 |
| ENSMUSG00000049307 | Fut4 | 14345 | protein_coding | 2775.705849 | -0.052873552 | 0.100265245 | -0.527336786 | 0.597959733 | 0.784071601 | 0.052873552 |
| ENSMUSG00000022710 | Usp7 | 252870 | protein_coding | 3735.217627 | 0.052880262 | 0.039665038 | 1.333170595 | 0.182475827 | 0.410430635 | 0.052880262 |
| ENSMUSG00000032477 | Cdc25a | 12530 | protein_coding | 1110.83324 | 0.052886786 | 0.073561096 | 0.71895049 | 0.472171426 | 0.696651125 | 0.052886786 |
| ENSMUSG00000041623 | D11Wsu47e | 276852 | protein_coding | 199.9368914 | -0.052904802 | 0.105084087 | -0.503452079 | 0.614646464 | 0.794963793 | 0.052904802 |
| ENSMUSG00000032077 | Bud13 | 215051 | protein_coding | 203.9840156 | -0.052924182 | 0.085128603 | -0.621696827 | 0.534141238 | 0.741606925 | 0.052924182 |
| ENSMUSG00000021245 | Mlh3 | 217716 | protein_coding | 316.9895491 | 0.052949271 | 0.076655133 | 0.690746576 | 0.489724814 | 0.709700079 | 0.052949271 |
| ENSMUSG00000054422 | Fabp1 | 14080 | protein_coding | 74.40702574 | -0.052956757 | 0.858999357 | -0.061649356 | 0.950842071 | 0.979948241 | 0.052956757 |
| ENSMUSG00000043702 | Pde12 | 211948 | protein_coding | 698.3319461 | -0.052975704 | 0.060850528 | -0.870587426 | 0.383979465 | 0.624863562 | 0.052975704 |
| ENSMUSG00000002076 | Hsf2bp | 74377 | protein_coding | 9.726976815 | 0.052999112 | 0.412146271 | 0.128592968 | 0.897679731 | 0.954463189 | 0.052999112 |
| ENSMUSG00000040312 | Cchcr1 | 240084 | protein_coding | 268.1869173 | 0.053008476 | 0.091303358 | 0.580575313 | 0.561526715 | 0.759431884 | 0.053008476 |
| ENSMUSG00000041126 | H2az2 | 77605 | protein_coding | 1094.229428 | -0.053036828 | 0.052864131 | -1.0032668 | 0.31573215 | 0.559908993 | 0.053036828 |
| ENSMUSG00000030600 | Lrfn1 | 80749 | protein_coding | 7.56643879 | 0.053036862 | 0.461268554 | 0.114980442 | 0.908460624 | 0.959234074 | 0.053036862 |
| ENSMUSG00000027273 | Snap25 | 20614 | protein_coding | 25.39511213 | -0.053085115 | 0.283254764 | -0.187411199 | 0.851338243 | 0.933346828 | 0.053085115 |
| ENSMUSG00000008601 | Rab25 | 53868 | protein_coding | 2825.792454 | 0.053087042 | 0.064485438 | 0.82324078 | 0.410371083 | 0.648598559 | 0.053087042 |
| ENSMUSG00000029314 | Gpat3 | 231510 | protein_coding | 235.1507429 | 0.053105431 | 0.129434674 | 0.410287521 | 0.681595045 | 0.839021966 | 0.053105431 |
| ENSMUSG00000052837 | Junb | 16477 | protein_coding | 14786.23949 | -0.053110371 | 0.112803974 | -0.470820037 | 0.637769254 | 0.810952043 | 0.053110371 |
| ENSMUSG00000039652 | Cpeb3 | 208922 | protein_coding | 149.8614049 | -0.053125153 | 0.130538117 | -0.406970425 | 0.68402971 | 0.840803897 | 0.053125153 |
| ENSMUSG00000007080 | Pole | 18973 | protein_coding | 364.897946 | 0.053136234 | 0.170168877 | 0.312255889 | 0.75484606 | 0.883718994 | 0.053136234 |
| ENSMUSG00000054452 | Tle5 | 14797 | protein_coding | 8889.892468 | 0.053159356 | 0.066026045 | 0.805127064 | 0.42074636 | 0.657880216 | 0.053159356 |
| ENSMUSG00000010057 | Nprl2 | 56032 | protein_coding | 365.2575792 | 0.053166564 | 0.071472364 | 0.743875826 | 0.456951602 | 0.685107692 | 0.053166564 |
| ENSMUSG00000023971 | Rrp36 | 224823 | protein_coding | 606.4483594 | 0.053174821 | 0.063246564 | 0.840754306 | 0.400485591 | 0.639735319 | 0.053174821 |
| ENSMUSG00000026305 | Lrrfip1 | 16978 | protein_coding | 2172.078832 | 0.05319863 | 0.046957232 | 1.132916652 | 0.257249256 | 0.499176845 | 0.05319863 |
| ENSMUSG00000079197 | Psme2 | 19188 | protein_coding | 2218.157977 | -0.053205952 | 0.101309013 | -0.525184783 | 0.599454742 | 0.785201968 | 0.053205952 |
| ENSMUSG00000045215 | Asxl3 | 211961 | protein_coding | 56.6781081 | 0.053222746 | 0.156981657 | 0.339037992 | 0.73458111 | 0.87241988 | 0.053222746 |
| ENSMUSG00000056116 | H2-T22 | 15039 | protein_coding | 844.8585317 | 0.053223533 | 0.086646081 | 0.614263587 | 0.539041161 | 0.745103008 | 0.053223533 |
| ENSMUSG00000057894 | Zfp329 | 67230 | protein_coding | 232.6874684 | -0.053236613 | 0.10883524 | -0.489148671 | 0.624736446 | 0.801977146 | 0.053236613 |
| ENSMUSG00000029587 | Zfp12 | 231866 | protein_coding | 437.562556 | -0.053245186 | 0.066620208 | -0.799234764 | 0.424154298 | 0.660009298 | 0.053245186 |
| ENSMUSG00000032609 | Klhdc8b | 78267 | protein_coding | 64.79670537 | -0.053245894 | 0.166629712 | -0.319546219 | 0.749312349 | 0.880475742 | 0.053245894 |
| ENSMUSG00000031988 | Vps26b | 69091 | protein_coding | 1233.703098 | 0.053263889 | 0.049283026 | 1.080775548 | 0.279796968 | 0.522981653 | 0.053263889 |
| ENSMUSG00000027878 | Notch2 | 18129 | protein_coding | 357.4500861 | -0.053291589 | 0.080197975 | -0.664500429 | 0.506370075 | 0.722065604 | 0.053291589 |
| ENSMUSG00000027630 | Tbl1xr1 | 81004 | protein_coding | 2450.517361 | 0.053294731 | 0.045618996 | 1.168257426 | 0.242702943 | 0.483875678 | 0.053294731 |
| ENSMUSG00000038510 | Rpf2 | 67239 | protein_coding | 206.9796022 | -0.053304906 | 0.094993608 | -0.56114203 | 0.574700718 | 0.768233148 | 0.053304906 |
| ENSMUSG00000045210 | Vcpip1 | 70675 | protein_coding | 1081.336962 | -0.053326408 | 0.049004793 | -1.088187601 | 0.276512297 | 0.519430846 | 0.053326408 |
| ENSMUSG00000020412 | Ascc2 | 75452 | protein_coding | 1379.330252 | 0.053351731 | 0.047041797 | 1.134134628 | 0.25673808 | 0.498604892 | 0.053351731 |
| ENSMUSG00000030084 | Plxna1 | 18844 | protein_coding | 867.8943064 | -0.053371331 | 0.058980893 | -0.904891871 | 0.365522674 | 0.60759958 | 0.053371331 |

| ensembl_gene_id | external_gene_name | entrezgene_id | gene_biotype | baseMean | log2FoldChange | lfcSE | stat | pvalue | padj | absLog2FC |
|---|---|---|---|---|---|---|---|---|---|---|
| ENSMUSG00000033720 | Sfxn5 | 94282 | protein_coding | 653.9762778 | 0.053373207 | 0.090914518 | 0.587070227 | 0.557156543 | 0.755953522 | 0.053373207 |
| ENSMUSG00000038506 | Dcun1d2 | 102323 | protein_coding | 524.6008499 | 0.053375227 | 0.06359023 | 0.839362072 | 0.401266161 | 0.640499152 | 0.053375227 |
| ENSMUSG00000025047 | Pdcd11 | 18572 | protein_coding | 734.9005843 | -0.053376351 | 0.058619028 | -0.910563577 | 0.362525369 | 0.605347548 | 0.053376351 |
| ENSMUSG00000048495 | Tyw5 | 68736 | protein_coding | 82.26580202 | 0.053379518 | 0.128246638 | 0.41622548 | 0.677245016 | 0.83633117 | 0.053379518 |
| ENSMUSG00000014599 | Csf1 | 12977 | protein_coding | 13.54171463 | 0.05338784 | 0.341494088 | 0.156336058 | 0.875768136 | 0.944199804 | 0.05338784 |
| ENSMUSG00000029153 | Ociad2 | 433904 | protein_coding | 4839.268858 | -0.053388273 | 0.041533581 | -1.285424265 | 0.198644067 | 0.431859377 | 0.053388273 |
| ENSMUSG00000044949 | Ubtd2 | 327900 | protein_coding | 66.40633591 | -0.053399702 | 0.179305731 | -0.297813695 | 0.765845362 | 0.890018603 | 0.053399702 |
| ENSMUSG00000022565 | Plec | 18810 | protein_coding | 21719.75114 | 0.053419854 | 0.110123522 | 0.485090316 | 0.627612281 | 0.803676612 | 0.053419854 |
| ENSMUSG00000041734 | Kirrel | 170643 | protein_coding | 7.502872767 | 0.053438532 | 0.493579665 | 0.108267289 | 0.913783669 | 0.961718903 | 0.053438532 |
| ENSMUSG00000027959 | Sass6 | 72776 | protein_coding | 98.687071 | -0.053462738 | 0.146029903 | -0.366108151 | 0.714284375 | 0.859482312 | 0.053462738 |
| ENSMUSG00000038305 | Spats2l | 67198 | protein_coding | 284.2581883 | 0.05348531 | 0.177647114 | 0.301076156 | 0.763356423 | 0.888663035 | 0.05348531 |
| ENSMUSG00000038527 | C1rl | 232371 | protein_coding | 13.40473704 | -0.053495082 | 0.364849103 | -0.146622484 | 0.883430006 | 0.948224673 | 0.053495082 |
| ENSMUSG00000001507 | Itga3 | 16400 | protein_coding | 3183.191338 | -0.053525573 | 0.063970609 | -0.836721337 | 0.402749223 | 0.642103246 | 0.053525573 |
| ENSMUSG00000021413 | Prpf4b | 19134 | protein_coding | 2102.139258 | -0.053525745 | 0.061200049 | -0.87460298 | 0.381789966 | 0.622494568 | 0.053525745 |
| ENSMUSG00000018565 | Elp5 | 54351 | protein_coding | 743.4281133 | 0.053549372 | 0.059702531 | 0.896936369 | 0.369752872 | 0.611904008 | 0.053549372 |
| ENSMUSG00000039450 | Dcxr | 67880 | protein_coding | 1212.632516 | 0.053554456 | 0.078997992 | 0.677921739 | 0.497821316 | 0.715668065 | 0.053554456 |
| ENSMUSG00000039001 | Rps21 | 66481 | protein_coding | 5094.177052 | -0.053580187 | 0.088650581 | -0.604397473 | 0.545579415 | 0.749073143 | 0.053580187 |
| ENSMUSG00000022938 | Fam3b | 52793 | protein_coding | 1587.825296 | 0.053582563 | 0.069954899 | 0.765958695 | 0.44370088 | 0.674737597 | 0.053582563 |
| ENSMUSG00000063450 | Syne2 | 319565 | protein_coding | 5365.333986 | -0.053600051 | 0.038732698 | -1.383844992 | 0.166405923 | 0.389893694 | 0.053600051 |
| ENSMUSG00000049091 | Sephs2 | 20768 | protein_coding | 1837.729442 | -0.053616289 | 0.048895166 | -1.096556028 | 0.272835519 | 0.515284976 | 0.053616289 |
| ENSMUSG00000005447 | Pafah1b3 | 18476 | protein_coding | 1557.832301 | -0.053631625 | 0.083478429 | -0.642460885 | 0.520573979 | 0.732190524 | 0.053631625 |
| ENSMUSG00000074646 | 6430550D23Rik | 320095 | protein_coding | 20.37103517 | 0.053658052 | 0.27618782 | 0.194281023 | 0.845955844 | 0.93086312 | 0.053658052 |
| ENSMUSG00000045775 | Slc16a5 | 217316 | protein_coding | 2677.74362 | 0.053690945 | 0.073996693 | 0.725585734 | 0.468092769 | 0.693047027 | 0.053690945 |
| ENSMUSG00000023980 | Taf8 | 63856 | protein_coding | 458.821568 | 0.053698178 | 0.061754352 | 0.86954484 | 0.384549193 | 0.624972092 | 0.053698178 |
| ENSMUSG00000037089 | Slc35b2 | 73836 | protein_coding | 1225.571072 | 0.05370043 | 0.044282497 | 1.212678472 | 0.225252774 | 0.463289311 | 0.05370043 |
| ENSMUSG00000046519 | Golph3l | 229593 | protein_coding | 8485.250843 | 0.053708236 | 0.046314303 | 1.159646853 | 0.246192617 | 0.487131402 | 0.053708236 |
| ENSMUSG00000024393 | Prrc2a | 53761 | protein_coding | 3113.514653 | 0.05370955 | 0.104330054 | 0.5148042 | 0.60668985 | 0.789765867 | 0.05370955 |
| ENSMUSG00000039771 | Polr2j | 20022 | protein_coding | 472.1608043 | 0.05372676 | 0.058999323 | 0.910633498 | 0.362488514 | 0.605347548 | 0.05372676 |
| ENSMUSG00000075273 | Ttc30b | 72421 | protein_coding | 295.6607405 | 0.053729414 | 0.071072069 | 0.755984938 | 0.449658227 | 0.679246742 | 0.053729414 |
| ENSMUSG00000033809 | Alg3 | 208624 | protein_coding | 143.0376155 | -0.053735549 | 0.126770103 | -0.423881876 | 0.671651962 | 0.832569654 | 0.053735549 |
| ENSMUSG00000057522 | Spop | 20747 | protein_coding | 1278.944296 | 0.053759532 | 0.050786459 | 1.058540662 | 0.289809023 | 0.532196551 | 0.053759532 |
| ENSMUSG00000066036 | Ubr4 | 69116 | protein_coding | 4856.114158 | 0.053774122 | 0.067896226 | 0.792004576 | 0.428358008 | 0.663421426 | 0.053774122 |
| ENSMUSG00000032384 | Csnk1g1 | 214897 | protein_coding | 683.8874807 | -0.053784801 | 0.054717638 | -0.982951806 | 0.325631159 | 0.56923128 | 0.053784801 |
| ENSMUSG00000025920 | Stau2 | 29819 | protein_coding | 352.2834319 | -0.053795987 | 0.085969445 | -0.625757057 | 0.531474303 | 0.740026398 | 0.053795987 |
| ENSMUSG00000046449 | Nexmif | 245555 | protein_coding | 90.61841437 | 0.05380788 | 0.123328137 | 0.436298488 | 0.662620179 | 0.827608087 | 0.05380788 |
| ENSMUSG00000020628 | Trappc12 | 217449 | protein_coding | 944.321451 | 0.053838987 | 0.050793337 | 1.059961604 | 0.289162067 | 0.531418201 | 0.053838987 |
| ENSMUSG00000001983 | Taco1 | 70207 | protein_coding | 293.4040308 | 0.053865899 | 0.09148766 | 0.58877776 | 0.556010367 | 0.755288004 | 0.053865899 |
| ENSMUSG00000037415 | Ranbp10 | 74334 | protein_coding | 663.9310439 | -0.0539148 | 0.04952257 | -1.08869148 | 0.27628996 | 0.519134124 | 0.0539148 |
| ENSMUSG00000056553 | Ptprn2 | 19276 | protein_coding | 1208.920178 | 0.053948857 | 0.079460011 | 0.678943479 | 0.497173675 | 0.71518329 | 0.053948857 |

| ensembl_gene_id | external_gene_name | entrezgene_id | gene_biotype | baseMean | log2FoldChange | lfcSE | stat | pvalue | padj | absLog2FC |
|---|---|---|---|---|---|---|---|---|---|---|
| ENSMUSG00000057841 | Rpl32 | 19951 | protein_coding | 11264.87513 | -0.053975317 | 0.037158436 | -1.452572358 | 0.146342526 | 0.35932794 | 0.053975317 |
| ENSMUSG00000044452 | Zfp507 | 668501 | protein_coding | 786.512592 | -0.053975395 | 0.055682816 | -0.969336661 | 0.332377243 | 0.575751644 | 0.053975395 |
| ENSMUSG00000027366 | Sppl2a | 66552 | protein_coding | 5899.143408 | 0.05399622 | 0.062467973 | 0.864382452 | 0.38737783 | 0.627358857 | 0.05399622 |
| ENSMUSG00000078517 | Emc1 | 230866 | protein_coding | 1568.823811 | 0.053998958 | 0.048474996 | 1.113954853 | 0.265298556 | 0.507219974 | 0.053998958 |
| ENSMUSG00000023723 | Mrps23 | 64656 | protein_coding | 1208.798397 | 0.054015823 | 0.04547258 | 1.187876808 | 0.234881948 | 0.474717773 | 0.054015823 |
| ENSMUSG00000020305 | Asb3 | 65257 | protein_coding | 228.0813133 | 0.054028851 | 0.111037333 | 0.48658275 | 0.626554051 | 0.803150559 | 0.054028851 |
| ENSMUSG00000005640 | Insrr | 23920 | protein_coding | 187.0559268 | -0.054045773 | 0.122324194 | -0.441824068 | 0.65861652 | 0.825261698 | 0.054045773 |
| ENSMUSG00000030306 | Tmtc1 | 387314 | protein_coding | 6.50254276 | -0.054048904 | 0.508593348 | -0.106271355 | 0.915367058 | 0.96226612 | 0.054048904 |
| ENSMUSG00000022185 | Acin1 | 56215 | protein_coding | 2556.580248 | 0.054053822 | 0.041499487 | 1.30251784 | 0.192739424 | 0.42444695 | 0.054053822 |
| ENSMUSG00000027628 | Aar2 | 68295 | protein_coding | 721.049062 | 0.054056359 | 0.07409106 | 0.729593536 | 0.465638674 | 0.691532821 | 0.054056359 |
| ENSMUSG00000071646 | Mta2 | 23942 | protein_coding | 2525.723157 | 0.054131935 | 0.039836618 | 1.358848656 | 0.174194552 | 0.400176002 | 0.054131935 |
| ENSMUSG00000029538 | Srsf9 | 108014 | protein_coding | 565.3958622 | 0.054143712 | 0.085541902 | 0.632949584 | 0.526766567 | 0.737129071 | 0.054143712 |
| ENSMUSG00000026584 | Scyl3 | 240880 | protein_coding | 630.3095026 | -0.054166976 | 0.060153892 | -0.900473336 | 0.367868409 | 0.610100147 | 0.054166976 |
| ENSMUSG00000049969 | Plekhf2 | 71801 | protein_coding | 1727.125302 | -0.054199257 | 0.042357247 | -1.279574577 | 0.200694796 | 0.434385314 | 0.054199257 |
| ENSMUSG00000031902 | Nfatc3 | 18021 | protein_coding | 1477.010698 | -0.054207736 | 0.061427598 | -0.882465493 | 0.377525132 | 0.61867166 | 0.054207736 |
| ENSMUSG00000038806 | Sde2 | 208768 | protein_coding | 544.5953528 | -0.05421556 | 0.057319342 | -0.945851058 | 0.344224559 | 0.586633147 | 0.05421556 |
| ENSMUSG00000022350 | Washc5 | 223593 | protein_coding | 1167.963105 | -0.054223704 | 0.059338437 | -0.91380404 | 0.360819823 | 0.604050305 | 0.054223704 |
| ENSMUSG00000036339 | Tmem260 | 218989 | protein_coding | 765.8509953 | 0.054226612 | 0.046626048 | 1.163011114 | 0.244824993 | 0.486063359 | 0.054226612 |
| ENSMUSG00000057969 | Sema3b | 20347 | protein_coding | 2097.40291 | -0.054290559 | 0.050861804 | -1.067413161 | 0.285785308 | 0.5285691 | 0.054290559 |
| ENSMUSG00000027713 | 1810062G17Rik | 72282 | protein_coding | 21.62872564 | 0.054302293 | 0.271467606 | 0.200032314 | 0.841455309 | 0.929613548 | 0.054302293 |
| ENSMUSG00000020793 | Galr2 | 14428 | protein_coding | 8.910233679 | 0.054310968 | 0.395704097 | 0.137251468 | 0.89083203 | 0.95112373 | 0.054310968 |
| ENSMUSG00000015217 | Hmgb3 | 15354 | protein_coding | 471.3457038 | -0.054335884 | 0.074491724 | -0.729421762 | 0.46574371 | 0.691625121 | 0.054335884 |
| ENSMUSG00000030980 | Knop1 | 66356 | protein_coding | 672.9360138 | -0.054339129 | 0.065503419 | -0.829561716 | 0.40678663 | 0.645607651 | 0.054339129 |
| ENSMUSG00000050912 | Tmem123 | 71929 | protein_coding | 585.250068 | 0.054387286 | 0.068335189 | 0.795889884 | 0.426096042 | 0.661690021 | 0.054387286 |
| ENSMUSG00000052632 | Asap2 | 211914 | protein_coding | 810.6844874 | 0.054387731 | 0.055016891 | 0.988564228 | 0.322876395 | 0.566904861 | 0.054387731 |
| ENSMUSG00000001670 | Tat | 234724 | protein_coding | 334.0960948 | 0.054397568 | 0.132142384 | 0.411658744 | 0.680589568 | 0.838364494 | 0.054397568 |
| ENSMUSG00000031782 | Coq9 | 67914 | protein_coding | 2323.334978 | 0.054398449 | 0.047859488 | 1.136628308 | 0.255693701 | 0.497115408 | 0.054398449 |
| ENSMUSG00000021143 | Pacs2 | 217893 | protein_coding | 1116.142058 | -0.054410553 | 0.043704318 | -1.244969749 | 0.21314287 | 0.448799461 | 0.054410553 |
| ENSMUSG00000050556 | Kcnb1 | 16500 | protein_coding | 13.82814119 | 0.054413892 | 0.343027236 | 0.158628488 | 0.873961583 | 0.943461975 | 0.054413892 |
| ENSMUSG00000020069 | Hnrnph3 | 432467 | protein_coding | 284.2300922 | -0.054424041 | 0.092113763 | -0.590835063 | 0.554630939 | 0.754685552 | 0.054424041 |
| ENSMUSG00000052395 | Rft1 | 328370 | protein_coding | 393.6520358 | 0.054440578 | 0.0771671 | 0.705489489 | 0.480505671 | 0.702867569 | 0.054440578 |
| ENSMUSG00000012443 | Kif11 | 16551 | protein_coding | 1019.844333 | 0.054493777 | 0.110700367 | 0.492263748 | 0.622532904 | 0.800830795 | 0.054493777 |
| ENSMUSG00000034957 | Cebpa | 12606 | protein_coding | 1400.302819 | -0.054499887 | 0.089020504 | -0.612217229 | 0.540394044 | 0.746077797 | 0.054499887 |
| ENSMUSG00000020220 | Vps13d | 230895 | protein_coding | 5071.894005 | -0.054505518 | 0.039643991 | -1.374874656 | 0.169170308 | 0.393510681 | 0.054505518 |
| ENSMUSG00000026966 | Ssna1 | 68475 | protein_coding | 625.9940147 | -0.054533965 | 0.066189067 | -0.823911977 | 0.409989574 | 0.64854129 | 0.054533965 |
| ENSMUSG00000002250 | Ppard | 19015 | protein_coding | 2033.978939 | -0.054551391 | 0.085001005 | -0.64177348 | 0.521020271 | 0.732526772 | 0.054551391 |
| ENSMUSG00000006527 | Sfmbt1 | 54650 | protein_coding | 276.1760398 | -0.054561526 | 0.082436465 | -0.661861546 | 0.508059958 | 0.723614143 | 0.054561526 |
| ENSMUSG00000006642 | Tcf23 | 69852 | protein_coding | 107.7520881 | 0.054567551 | 0.137588773 | 0.396598862 | 0.691663292 | 0.845418668 | 0.054567551 |
| ENSMUSG00000041164 | Zmiz2 | 52915 | protein_coding | 777.4122677 | -0.054584338 | 0.090799082 | -0.601155177 | 0.547736636 | 0.750752059 | 0.054584338 |

| ensembl_gene_id | external_gene_name | entrezgene_id | gene_biotype | baseMean | log2FoldChange | lfcSE | stat | pvalue | padj | absLog2FC |
|---|---|---|---|---|---|---|---|---|---|---|
| ENSMUSG00000029203 | Ube2k | 53323 | protein_coding | 979.6001612 | 0.05459042 | 0.055277479 | 0.987570713 | 0.323362935 | 0.567389191 | 0.05459042 |
| ENSMUSG00000063047 | Zfp780b | 338354 | protein_coding | 242.2516718 | -0.054637882 | 0.102210032 | -0.534564768 | 0.592950852 | 0.780422622 | 0.054637882 |
| ENSMUSG00000001305 | Rrp15 | 67223 | protein_coding | 218.9795098 | -0.054640736 | 0.128446864 | -0.425395643 | 0.670548281 | 0.831904926 | 0.054640736 |
| ENSMUSG00000052384 | Nrros | 224109 | protein_coding | 27.77380015 | -0.05464207 | 0.283584467 | -0.192683579 | 0.847206784 | 0.930926537 | 0.05464207 |
| ENSMUSG00000025899 | Alkbh8 | 67667 | protein_coding | 377.3725801 | 0.054647555 | 0.084926634 | 0.643467815 | 0.519920592 | 0.731656244 | 0.054647555 |
| ENSMUSG00000027536 | Chmp4c | 66371 | protein_coding | 4528.670519 | -0.054681632 | 0.039069888 | -1.399585066 | 0.161637609 | 0.382218433 | 0.054681632 |
| ENSMUSG00000029528 | Pxn | 19303 | protein_coding | 3342.665433 | 0.054684918 | 0.062547477 | 0.87429456 | 0.381957861 | 0.622645752 | 0.054684918 |
| ENSMUSG00000021520 | Uqcrb | 67530 | protein_coding | 5442.090595 | 0.054710579 | 0.042285663 | 1.293832815 | 0.195723175 | 0.428222446 | 0.054710579 |
| ENSMUSG00000014547 | Wdfy2 | 268752 | protein_coding | 390.3074252 | 0.054741741 | 0.088355493 | 0.619562394 | 0.535545932 | 0.742435249 | 0.054741741 |
| ENSMUSG00000039509 | Nup133 | 234865 | protein_coding | 455.4402739 | 0.054756003 | 0.098907912 | 0.553605893 | 0.579848592 | 0.771934807 | 0.054756003 |
| ENSMUSG00000005686 | Ampd3 | 11717 | protein_coding | 1183.500494 | 0.054781071 | 0.044097129 | 1.242282048 | 0.214132516 | 0.449826116 | 0.054781071 |
| ENSMUSG00000030786 | Itgam | 16409 | protein_coding | 20.43307064 | 0.054788228 | 0.283036601 | 0.193572944 | 0.846510285 | 0.930889346 | 0.054788228 |
| ENSMUSG00000045671 | Spred2 | 114716 | protein_coding | 552.2014748 | 0.054797079 | 0.099920916 | 0.548404495 | 0.583414191 | 0.774890517 | 0.054797079 |
| ENSMUSG00000022110 | Sucla2 | 20916 | protein_coding | 2512.027372 | -0.054801844 | 0.044535487 | -1.230520815 | 0.218502137 | 0.454666616 | 0.054801844 |
| ENSMUSG00000110231 | Gm6329 | 622523 | lncRNA | 11.24690102 | -0.054818595 | 0.360497643 | -0.15206367 | 0.879136718 | 0.945630484 | 0.054818595 |
| ENSMUSG00000040690 | Col16a1 | 107581 | protein_coding | 561.6021552 | 0.054883512 | 0.077085513 | 0.711982186 | 0.476475807 | 0.699497125 | 0.054883512 |
| ENSMUSG00000008475 | Arpc5 | 67771 | protein_coding | 5361.206663 | 0.054896847 | 0.037683746 | 1.456777855 | 0.145177711 | 0.357829885 | 0.054896847 |
| ENSMUSG00000019979 | Apaf1 | 11783 | protein_coding | 791.6974848 | 0.054897219 | 0.067749028 | 0.810302692 | 0.417766228 | 0.655527867 | 0.054897219 |
| ENSMUSG00000071657 | Bscl2 | 14705 | protein_coding | 1043.532266 | -0.054911779 | 0.091213442 | -0.602014112 | 0.547164744 | 0.750223246 | 0.054911779 |
| ENSMUSG00000035245 | Eogt | 101351 | protein_coding | 339.337902 | 0.054915675 | 0.08120929 | 0.676224047 | 0.498898407 | 0.716067516 | 0.054915675 |
| ENSMUSG00000024800 | Rpp30 | 54364 | protein_coding | 111.5256016 | 0.054919576 | 0.150173701 | 0.365707016 | 0.71458371 | 0.859633975 | 0.054919576 |
| ENSMUSG00000053289 | Ddx10 | 77591 | protein_coding | 646.2760667 | 0.054924213 | 0.084922502 | 0.646756888 | 0.517789299 | 0.730184332 | 0.054924213 |
| ENSMUSG00000033319 | Fem1c | 240263 | protein_coding | 925.5527227 | -0.054933242 | 0.066448231 | -0.826707361 | 0.408402948 | 0.646860933 | 0.054933242 |
| ENSMUSG00000037822 | Smim14 | 68552 | protein_coding | 2771.719554 | 0.054938841 | 0.03891516 | 1.411759358 | 0.158020829 | 0.377781372 | 0.054938841 |
| ENSMUSG00000044864 | Ankrd50 | 99696 | protein_coding | 1252.511427 | -0.054956341 | 0.049819354 | -1.103112272 | 0.269978412 | 0.512409304 | 0.054956341 |
| ENSMUSG00000074746 | Pdzd8 | 107368 | protein_coding | 2168.264626 | -0.054960979 | 0.040232726 | -1.366076428 | 0.171914976 | 0.396668438 | 0.054960979 |
| ENSMUSG00000034893 | Cog3 | 338337 | protein_coding | 1926.921166 | -0.054964707 | 0.049905228 | -1.101381736 | 0.270730551 | 0.513373374 | 0.054964707 |
| ENSMUSG00000001630 | Stk38l | 232533 | protein_coding | 1766.523088 | -0.054982217 | 0.06460533 | -0.851047699 | 0.394742857 | 0.634212745 | 0.054982217 |
| ENSMUSG00000030533 | Unc45a | 101869 | protein_coding | 1048.208837 | -0.054995609 | 0.051387751 | -1.07020851 | 0.284525465 | 0.527473335 | 0.054995609 |
| ENSMUSG00000034083 | Ccdc174 | 232236 | protein_coding | 405.6204545 | -0.055070009 | 0.077335941 | -0.712088171 | 0.476410179 | 0.699497125 | 0.055070009 |
| ENSMUSG00000026355 | Mcm6 | 17219 | protein_coding | 1221.683172 | 0.055108134 | 0.154844015 | 0.355894507 | 0.72191957 | 0.864080715 | 0.055108134 |
| ENSMUSG00000050428 | Fbxo46 | 243867 | protein_coding | 460.2674906 | 0.055117021 | 0.096120232 | 0.573417481 | 0.566362057 | 0.762477743 | 0.055117021 |
| ENSMUSG00000048807 | Slc35e4 | 103710 | protein_coding | 323.2401839 | 0.055142803 | 0.088865624 | 0.620518949 | 0.534916183 | 0.741885541 | 0.055142803 |
| ENSMUSG00000028247 | Coq3 | 230027 | protein_coding | 224.3113861 | -0.055156309 | 0.102343778 | -0.538931733 | 0.58993396 | 0.77918632 | 0.055156309 |
| ENSMUSG00000021133 | Susd6 | 217684 | protein_coding | 2285.469294 | -0.05516206 | 0.05133807 | -1.074486439 | 0.282604719 | 0.52530732 | 0.05516206 |
| ENSMUSG00000002052 | Supt6 | 20926 | protein_coding | 2061.014621 | 0.05517267 | 0.031047147 | 1.777060869 | 0.075558225 | 0.241780511 | 0.05517267 |
| ENSMUSG00000028358 | Zfp618 | 72701 | protein_coding | 847.4936896 | -0.055182037 | 0.079722441 | -0.692176968 | 0.4888262 | 0.708993627 | 0.055182037 |
| ENSMUSG00000021697 | Depdc1b | 218581 | protein_coding | 74.61853744 | 0.055183693 | 0.206107119 | 0.267742782 | 0.788897316 | 0.901477241 | 0.055183693 |
| ENSMUSG00000029036 | Atad3a | 108888 | protein_coding | 1397.707414 | -0.055228587 | 0.061034516 | -0.904874671 | 0.365531787 | 0.60759958 | 0.055228587 |

Supplementary Table 1

| ensembl_gene_id | external_gene_name | entrezgene_id | gene_biotype | baseMean | log2FoldChange | lfcSE | stat | pvalue | padj | absLog2FC |
|---|---|---|---|---|---|---|---|---|---|---|
| ENSMUSG00000026192 | Atic | 108147 | protein_coding | 1673.451067 | 0.055256514 | 0.083929646 | 0.658367059 | 0.510302297 | 0.724823423 | 0.055256514 |
| ENSMUSG00000089872 | Rps6kc1 | 320119 | protein_coding | 648.3693113 | -0.055260951 | 0.063941923 | -0.864236612 | 0.387457924 | 0.627362845 | 0.055260951 |
| ENSMUSG00000024063 | Lbh | 77889 | protein_coding | 578.8980798 | -0.055305162 | 0.111064373 | -0.497955918 | 0.618515113 | 0.797791988 | 0.055305162 |
| ENSMUSG00000026271 | Gpr35 | 64095 | protein_coding | 691.5356184 | 0.055307426 | 0.05552404 | 0.996098737 | 0.319202173 | 0.562721191 | 0.055307426 |
| ENSMUSG00000026153 | Fam135a | 68187 | protein_coding | 1716.642824 | -0.055368329 | 0.041271115 | -1.341575814 | 0.179733573 | 0.406831399 | 0.055368329 |
| ENSMUSG00000029207 | Apbb2 | 11787 | protein_coding | 505.7499349 | -0.055368712 | 0.056658748 | -0.977231461 | 0.328454576 | 0.572028827 | 0.055368712 |
| ENSMUSG00000028821 | Syf2 | 68592 | protein_coding | 1122.591926 | -0.055378675 | 0.049506991 | -1.118603124 | 0.26330949 | 0.504916665 | 0.055378675 |
| ENSMUSG00000027552 | E2f5 | 13559 | protein_coding | 640.4621413 | -0.055392972 | 0.081787718 | -0.677277388 | 0.498229975 | 0.715879721 | 0.055392972 |
| ENSMUSG00000047675 | Rps8 | 20116 | protein_coding | 14898.33237 | -0.055399533 | 0.046219997 | -1.198605273 | 0.230681467 | 0.469603637 | 0.055399533 |
| ENSMUSG00000042275 | Pelo | 105083 | protein_coding | 60.97317834 | 0.055428431 | 0.152626584 | 0.363163672 | 0.716482623 | 0.860942945 | 0.055428431 |
| ENSMUSG00000041836 | Ptpre | 19267 | protein_coding | 318.2101298 | 0.055462839 | 0.089545479 | 0.619381794 | 0.535664872 | 0.742536306 | 0.055462839 |
| ENSMUSG00000032507 | Fbxl2 | 72179 | protein_coding | 11.88599669 | -0.055470061 | 0.34240961 | -0.161999136 | 0.871306534 | 0.941878077 | 0.055470061 |
| ENSMUSG00000032239 | Rp9 | 55934 | protein_coding | 937.4792874 | -0.055479034 | 0.059407187 | -0.933877472 | 0.350367101 | 0.592592578 | 0.055479034 |
| ENSMUSG00000028403 | Zdhhc21 | 68268 | protein_coding | 1486.453375 | 0.055487635 | 0.070246368 | 0.789900422 | 0.429585925 | 0.664608067 | 0.055487635 |
| ENSMUSG00000022516 | Nudt16l1 | 66911 | protein_coding | 291.7245617 | 0.055492452 | 0.07339162 | 0.756114288 | 0.449580676 | 0.679246742 | 0.055492452 |
| ENSMUSG00000042712 | Tceal9 | 22381 | protein_coding | 1535.466412 | 0.055495501 | 0.070581588 | 0.786260315 | 0.431714997 | 0.666254386 | 0.055495501 |
| ENSMUSG00000032534 | Cep63 | 28135 | protein_coding | 614.5789901 | -0.055554043 | 0.061269645 | -0.906713965 | 0.364558077 | 0.606794107 | 0.055554043 |
| ENSMUSG00000032002 | Dcun1d5 | 76863 | protein_coding | 672.0092771 | 0.055569767 | 0.059699525 | 0.93082427 | 0.351944473 | 0.594351535 | 0.055569767 |
| ENSMUSG00000024826 | Dpf2 | 19708 | protein_coding | 836.2521474 | -0.055619614 | 0.049281757 | -1.128604519 | 0.259064704 | 0.500771482 | 0.055619614 |
| ENSMUSG00000019814 | Ltv1 | 353258 | protein_coding | 500.299451 | -0.055631286 | 0.090072114 | -0.617630519 | 0.536818922 | 0.74336929 | 0.055631286 |
| ENSMUSG00000015149 | Sirt2 | 64383 | protein_coding | 802.5304125 | 0.055664435 | 0.064621299 | 0.861394557 | 0.389020773 | 0.628820087 | 0.055664435 |
| ENSMUSG00000018415 | Gid4 | 66771 | protein_coding | 889.8235405 | -0.055665302 | 0.074404671 | -0.748142569 | 0.454374167 | 0.682675952 | 0.055665302 |
| ENSMUSG00000018846 | Pank3 | 211347 | protein_coding | 8343.986935 | -0.055694927 | 0.044374263 | -1.255117791 | 0.209436003 | 0.444801234 | 0.055694927 |
| ENSMUSG00000036180 | Gatad2a | 234366 | protein_coding | 1891.078395 | -0.05570851 | 0.06208196 | -0.897338124 | 0.369538519 | 0.611674813 | 0.05570851 |
| ENSMUSG00000031004 | Mki67 | 17345 | protein_coding | 4387.058722 | 0.055757457 | 0.099459811 | 0.560602888 | 0.575068283 | 0.768361739 | 0.055757457 |
| ENSMUSG00000060703 | Cd302 | 66205 | protein_coding | 69.17163269 | 0.05576789 | 0.163404166 | 0.341288053 | 0.732886741 | 0.871241669 | 0.05576789 |
| ENSMUSG00000024044 | Epb41l3 | 13823 | protein_coding | 2650.074328 | -0.055796132 | 0.0382913 | -1.457149057 | 0.145075239 | 0.357631975 | 0.055796132 |
| ENSMUSG00000038206 | Fbxo8 | 50753 | protein_coding | 827.3885377 | 0.055801805 | 0.052288198 | 1.067196937 | 0.285882915 | 0.528598232 | 0.055801805 |
| ENSMUSG00000001018 | Snapin | 20615 | protein_coding | 443.4478738 | -0.055808139 | 0.072780386 | -0.766801908 | 0.443199302 | 0.674226502 | 0.055808139 |
| ENSMUSG00000036817 | Sun1 | 77053 | protein_coding | 2154.881326 | -0.055819724 | 0.044481334 | -1.254902206 | 0.209514263 | 0.444801234 | 0.055819724 |
| ENSMUSG00000020986 | Sec23a | 20334 | protein_coding | 2663.964067 | -0.055826465 | 0.050699781 | -1.101118473 | 0.270845098 | 0.513373374 | 0.055826465 |
| ENSMUSG00000067916 | Zfp991 | 666532 | protein_coding | 66.00067954 | -0.055828102 | 0.190352556 | -0.293287903 | 0.769302099 | 0.891725514 | 0.055828102 |
| ENSMUSG00000041439 | Mfsd6 | 98682 | protein_coding | 4826.519124 | 0.055839726 | 0.048437015 | 1.152831709 | 0.248979467 | 0.489898763 | 0.055839726 |
| ENSMUSG00000033952 | Aspm | 12316 | protein_coding | 590.2736744 | -0.055840132 | 0.092486349 | -0.603766203 | 0.545999092 | 0.749262029 | 0.055840132 |
| ENSMUSG00000032218 | Ccnb2 | 12442 | protein_coding | 650.9684352 | -0.055861675 | 0.09205443 | -0.606833097 | 0.543961681 | 0.748759819 | 0.055861675 |
| ENSMUSG00000036155 | Mgat5 | 107895 | protein_coding | 3347.958078 | -0.055869883 | 0.042827562 | -1.304531028 | 0.192052587 | 0.423524146 | 0.055869883 |
| ENSMUSG00000022843 | Clcn2 | 12724 | protein_coding | 5974.813572 | -0.055894302 | 0.080502814 | -0.69431489 | 0.487484755 | 0.708115319 | 0.055894302 |
| ENSMUSG00000071064 | Zfp827 | 622675 | protein_coding | 558.7007252 | -0.055922549 | 0.070929748 | -0.788421648 | 0.430450112 | 0.66537611 | 0.055922549 |
| ENSMUSG00000061533 | Cep128 | 75216 | protein_coding | 148.3666298 | 0.055944943 | 0.11760884 | 0.475686549 | 0.634297705 | 0.808705755 | 0.055944943 |

Supplementary Table 1

| ensembl_gene_id | external_gene_name | entrezgene_id | gene_biotype | baseMean | log2FoldChange | lfcSE | stat | pvalue | padj | absLog2FC |
|---|---|---|---|---|---|---|---|---|---|---|
| ENSMUSG00000026975 | Dph7 | 67228 | protein_coding | 156.7287949 | 0.055949955 | 0.125287493 | 0.446572549 | 0.65518372 | 0.822565162 | 0.055949955 |
| ENSMUSG00000021068 | Nin | 18080 | protein_coding | 183.7051947 | -0.055962907 | 0.088634507 | -0.631389615 | 0.527785804 | 0.737765595 | 0.055962907 |
| ENSMUSG00000038215 | Cep44 | 382010 | protein_coding | 170.9013972 | -0.055996471 | 0.124226921 | -0.450759547 | 0.65216286 | 0.820965922 | 0.055996471 |
| ENSMUSG00000029246 | Ppat | 231327 | protein_coding | 377.7583341 | -0.056007285 | 0.118272283 | -0.473545313 | 0.635824181 | 0.80962603 | 0.056007285 |
| ENSMUSG00000006930 | Hap1 | 15114 | protein_coding | 64.13436864 | -0.056031112 | 0.168093183 | -0.333333635 | 0.738882453 | 0.87479925 | 0.056031112 |
| ENSMUSG00000070923 | Klhl9 | 242521 | protein_coding | 2504.2355 | 0.05605799 | 0.043579171 | 1.286348228 | 0.198321558 | 0.431310511 | 0.05605799 |
| ENSMUSG00000031886 | Ces2e | 234673 | protein_coding | 6047.70709 | -0.056059885 | 0.072384909 | -0.774469234 | 0.438653355 | 0.671427662 | 0.056059885 |
| ENSMUSG00000060679 | Mrps9 | 69527 | protein_coding | 870.560546 | -0.05606034 | 0.069202103 | -0.810095906 | 0.417885057 | 0.655631123 | 0.05606034 |
| ENSMUSG00000021501 | Caml | 12328 | protein_coding | 473.3559053 | 0.056077531 | 0.059788804 | 0.937926962 | 0.348281946 | 0.590676511 | 0.056077531 |
| ENSMUSG00000034906 | Ncaph | 215387 | protein_coding | 327.8990201 | 0.056092225 | 0.137675548 | 0.407423295 | 0.683697121 | 0.840651377 | 0.056092225 |
| ENSMUSG00000036718 | Micall2 | 231830 | protein_coding | 604.7159948 | 0.056100646 | 0.075435545 | 0.743689808 | 0.457064157 | 0.685107692 | 0.056100646 |
| ENSMUSG00000062580 | Timm17a | 21854 | protein_coding | 1281.253597 | 0.056128299 | 0.053229224 | 1.054463975 | 0.291670546 | 0.534215888 | 0.056128299 |
| ENSMUSG00000028973 | Abcb8 | 74610 | protein_coding | 1769.945772 | 0.056137976 | 0.060124606 | 0.933693883 | 0.350461821 | 0.592690624 | 0.056137976 |
| ENSMUSG00000036565 | Ttyh3 | 78339 | protein_coding | 792.3966863 | -0.056201887 | 0.107115562 | -0.524684608 | 0.599802459 | 0.785533185 | 0.056201887 |
| ENSMUSG00000025762 | Larp1b | 214048 | protein_coding | 1436.866015 | 0.056223688 | 0.041890015 | 1.342173987 | 0.179539589 | 0.406691669 | 0.056223688 |
| ENSMUSG00000040811 | Eml2 | 72205 | protein_coding | 379.80007 | -0.056227539 | 0.08426129 | -0.667299768 | 0.504580675 | 0.720720873 | 0.056227539 |
| ENSMUSG00000050747 | Trim15 | 69097 | protein_coding | 270.197577 | -0.056250141 | 0.105559265 | -0.532877346 | 0.594118488 | 0.781437157 | 0.056250141 |
| ENSMUSG00000024498 | Tcerg1 | 56070 | protein_coding | 742.4922429 | -0.056319211 | 0.066824883 | -0.842788028 | 0.399347005 | 0.638789048 | 0.056319211 |
| ENSMUSG00000046027 | Stard5 | 170460 | protein_coding | 1106.475301 | -0.056337417 | 0.051313779 | -1.097900358 | 0.272248001 | 0.514726356 | 0.056337417 |
| ENSMUSG00000060450 | Rnf14 | 56736 | protein_coding | 1599.069213 | 0.05635692 | 0.038441707 | 1.466035831 | 0.142638526 | 0.354359599 | 0.05635692 |
| ENSMUSG00000063455 | D630045J12Rik | 330286 | protein_coding | 22.56277908 | -0.056364592 | 0.24567093 | -0.229431265 | 0.818533738 | 0.918422987 | 0.056364592 |
| ENSMUSG00000042599 | Kdm7a | 338523 | protein_coding | 2082.036676 | -0.056367844 | 0.057345313 | -0.982954698 | 0.325629735 | 0.56923128 | 0.056367844 |
| ENSMUSG00000029279 | Brdt | 114642 | protein_coding | 22.39982154 | -0.05637002 | 0.248926673 | -0.22645231 | 0.820849647 | 0.919933444 | 0.05637002 |
| ENSMUSG00000024082 | Ndufaf7 | 73694 | protein_coding | 445.8602891 | 0.056390263 | 0.057324187 | 0.983708038 | 0.325259087 | 0.56903126 | 0.056390263 |
| ENSMUSG00000026502 | Desi2 | 78825 | protein_coding | 726.9520717 | -0.056408816 | 0.059452178 | -0.948809913 | 0.342717299 | 0.585270749 | 0.056408816 |
| ENSMUSG00000030770 | Parva | 57342 | protein_coding | 2368.518033 | 0.056430569 | 0.056638093 | 0.996335964 | 0.319086935 | 0.562649386 | 0.056430569 |
| ENSMUSG00000039623 | Ap5z1 | 231855 | protein_coding | 396.4794378 | -0.056444604 | 0.078557016 | -0.718517672 | 0.472438155 | 0.696772776 | 0.056444604 |
| ENSMUSG00000040181 | Fmo1 | 14261 | protein_coding | 18.21410448 | 0.056473119 | 0.314139094 | 0.179771063 | 0.857332302 | 0.935846624 | 0.056473119 |
| ENSMUSG00000046792 | Zfp787 | 67109 | protein_coding | 663.0576826 | -0.056485535 | 0.090039128 | -0.627344321 | 0.530433557 | 0.739671592 | 0.056485535 |
| ENSMUSG00000015970 | Chdh | 218865 | protein_coding | 3200.604105 | 0.05648888 | 0.065695651 | 0.859857225 | 0.38986775 | 0.629623876 | 0.05648888 |
| ENSMUSG00000028917 | Plekhm2 | 69582 | protein_coding | 414.3745988 | -0.056506605 | 0.068685814 | -0.822682328 | 0.410688667 | 0.64875856 | 0.056506605 |
| ENSMUSG00000042105 | Inpp5f | 101490 | protein_coding | 294.7098768 | -0.056533862 | 0.087558028 | -0.645673082 | 0.518491097 | 0.730599032 | 0.056533862 |
| ENSMUSG00000000134 | Tfe3 | 209446 | protein_coding | 465.0957125 | -0.056549234 | 0.073336938 | -0.771088009 | 0.440654767 | 0.672385224 | 0.056549234 |
| ENSMUSG00000038319 | Kcnh2 | 16511 | protein_coding | 65.86804207 | 0.056550166 | 0.157380854 | 0.35932049 | 0.719355347 | 0.862768697 | 0.056550166 |
| ENSMUSG00000024858 | Grk2 | 110355 | protein_coding | 2222.723001 | 0.056585386 | 0.047736875 | 1.185360077 | 0.23587511 | 0.476068282 | 0.056585386 |
| ENSMUSG00000062014 | Gmfb | 63985 | protein_coding | 1847.796782 | -0.056609791 | 0.045555155 | -1.242664888 | 0.213991348 | 0.449718678 | 0.056609791 |
| ENSMUSG00000006301 | Tmbim1 | 69660 | protein_coding | 3989.149518 | -0.056617746 | 0.070598512 | -0.801967971 | 0.422571486 | 0.658754022 | 0.056617746 |
| ENSMUSG00000096740 | Lbhd1 | 102308570 | protein_coding | 94.38755867 | 0.056713499 | 0.120422559 | 0.470954109 | 0.637673506 | 0.810894231 | 0.056713499 |
| ENSMUSG00000040177 | 2310057M21Rik | 68277 | protein_coding | 121.3040017 | -0.056750608 | 0.11023357 | -0.514821464 | 0.606677785 | 0.789765867 | 0.056750608 |

| ensembl_gene_id | external_gene_name | entrezgene_id | gene_biotype | baseMean | log2FoldChange | lfcSE | stat | pvalue | padj | absLog2FC |
|---|---|---|---|---|---|---|---|---|---|---|
| ENSMUSG00000090659 | Zfp493 | 72958 | protein_coding | 57.67323528 | -0.056764095 | 0.172490163 | -0.329085985 | 0.742090697 | 0.876754335 | 0.056764095 |
| ENSMUSG00000027901 | Dennd2d | 72121 | protein_coding | 1490.0236 | 0.056769031 | 0.069267219 | 0.819565614 | 0.412463783 | 0.650098067 | 0.056769031 |
| ENSMUSG00000031583 | Wrn | 22427 | protein_coding | 713.7227059 | -0.056793086 | 0.058042428 | -0.978475374 | 0.327839265 | 0.571636266 | 0.056793086 |
| ENSMUSG00000034867 | Ankrd27 | 245886 | protein_coding | 828.5293948 | -0.056804745 | 0.046813135 | -1.213436044 | 0.224963153 | 0.46304796 | 0.056804745 |
| ENSMUSG00000048109 | Rbm15 | 229700 | protein_coding | 1074.657072 | -0.056833885 | 0.052914985 | -1.074060305 | 0.282795654 | 0.525455543 | 0.056833885 |
| ENSMUSG00000032594 | Ip6k1 | 27399 | protein_coding | 1828.321435 | -0.056873558 | 0.065761267 | -0.864848886 | 0.387121735 | 0.627196326 | 0.056873558 |
| ENSMUSG00000027014 | Cwc22 | 80744 | protein_coding | 561.9859471 | 0.056873939 | 0.070520262 | 0.80649075 | 0.419959939 | 0.657096531 | 0.056873939 |
| ENSMUSG00000026219 | Trip12 | 14897 | protein_coding | 4887.489545 | -0.056884653 | 0.033668443 | -1.689554022 | 0.091113308 | 0.272007465 | 0.056884653 |
| ENSMUSG00000037957 | Wdr20 | 69641 | protein_coding | 630.010093 | 0.056894698 | 0.051514056 | 1.104449968 | 0.269397995 | 0.511910154 | 0.056894698 |
| ENSMUSG00000025795 | Rassf3 | 192678 | protein_coding | 1447.195657 | 0.056908126 | 0.054538411 | 1.043450399 | 0.296739742 | 0.539639357 | 0.056908126 |
| ENSMUSG00000028073 | Pear1 | 73182 | protein_coding | 41.27318874 | 0.056922348 | 0.206833852 | 0.275208081 | 0.783156379 | 0.898858056 | 0.056922348 |
| ENSMUSG00000048701 | Ccdc6 | 76551 | protein_coding | 1956.272439 | 0.056939524 | 0.059250925 | 0.960989621 | 0.336557386 | 0.580250264 | 0.056939524 |
| ENSMUSG00000003452 | Bicd1 | 12121 | protein_coding | 382.4305591 | 0.056955681 | 0.076756832 | 0.742027503 | 0.458070672 | 0.685606255 | 0.056955681 |
| ENSMUSG00000002326 | Gmpr2 | 105446 | protein_coding | 77.50923644 | -0.056957884 | 0.1485111 | -0.383526105 | 0.701329719 | 0.851297936 | 0.056957884 |
| ENSMUSG00000056216 | Cebpg | 12611 | protein_coding | 2091.172597 | 0.056988073 | 0.056541102 | 1.00790524 | 0.313499956 | 0.55760602 | 0.056988073 |
| ENSMUSG00000039725 | Trp53rka | 381406 | protein_coding | 218.3847644 | -0.056996691 | 0.108042522 | -0.527539435 | 0.597819039 | 0.784048864 | 0.056996691 |
| ENSMUSG00000029502 | Golga3 | 269682 | protein_coding | 2482.602904 | -0.057003105 | 0.047792979 | -1.192708767 | 0.232983451 | 0.472500517 | 0.057003105 |
| ENSMUSG00000048271 | Rbm33 | 381626 | protein_coding | 1043.699058 | -0.057012424 | 0.064179029 | -0.888334158 | 0.374361031 | 0.616178847 | 0.057012424 |
| ENSMUSG00000022024 | Sugt1 | 67955 | protein_coding | 1197.991512 | -0.057023615 | 0.046831562 | -1.217632122 | 0.223363802 | 0.460873597 | 0.057023615 |
| ENSMUSG00000071019 | Sdr16c6 | 242286 | protein_coding | 24.71083685 | 0.057035765 | 0.277057885 | 0.205862268 | 0.8368985 | 0.927733167 | 0.057035765 |
| ENSMUSG00000037458 | Azin1 | 54375 | protein_coding | 3803.221268 | -0.057069698 | 0.07706401 | -0.740549296 | 0.458966761 | 0.685946212 | 0.057069698 |
| ENSMUSG00000032855 | Pkd1 | 18763 | protein_coding | 1340.604436 | -0.05714344 | 0.072007947 | -0.793571307 | 0.427445042 | 0.66295501 | 0.05714344 |
| ENSMUSG00000003527 | Ess2 | 27886 | protein_coding | 341.6934144 | -0.057188279 | 0.078603704 | -0.727551956 | 0.4668879 | 0.692467352 | 0.057188279 |
| ENSMUSG00000074405 | Zfp865 | 319748 | protein_coding | 247.5547412 | -0.057193085 | 0.120793818 | -0.473476918 | 0.635872965 | 0.80962603 | 0.057193085 |
| ENSMUSG00000036875 | Dna2 | 327762 | protein_coding | 113.5704411 | -0.057257611 | 0.146763094 | -0.390136305 | 0.696435759 | 0.848230457 | 0.057257611 |
| ENSMUSG00000008226 | Scrn3 | 74616 | protein_coding | 294.8492392 | 0.05728596 | 0.080634359 | 0.710441067 | 0.477430664 | 0.700084044 | 0.05728596 |
| ENSMUSG00000045128 | Rpl18a | 76808 | protein_coding | 10460.48521 | -0.057295544 | 0.053351773 | -1.073920143 | 0.282858474 | 0.525476168 | 0.057295544 |
| ENSMUSG00000020694 | Tlk2 | 24086 | protein_coding | 838.8449411 | -0.057316558 | 0.051202099 | -1.119418135 | 0.262961798 | 0.504855347 | 0.057316558 |
| ENSMUSG00000035495 | Tstd2 | 272027 | protein_coding | 633.4004486 | -0.057319432 | 0.059424678 | -0.964572881 | 0.334758788 | 0.578382994 | 0.057319432 |
| ENSMUSG00000042197 | Zfp451 | 98403 | protein_coding | 503.1043272 | -0.057319495 | 0.063081306 | -0.908660565 | 0.363529326 | 0.606076959 | 0.057319495 |
| ENSMUSG00000025423 | Pias2 | 17344 | protein_coding | 780.5303393 | -0.05733236 | 0.048793046 | -1.175010866 | 0.239990369 | 0.480533622 | 0.05733236 |
| ENSMUSG00000038344 | Txlng | 353170 | protein_coding | 477.9667877 | -0.057351407 | 0.086800425 | -0.660727256 | 0.508787241 | 0.724049267 | 0.057351407 |
| ENSMUSG00000048668 | Rhno1 | 72440 | protein_coding | 207.9778824 | -0.057358212 | 0.090623881 | -0.632926025 | 0.526781952 | 0.737129071 | 0.057358212 |
| ENSMUSG00000040093 | Bmf | 171543 | protein_coding | 370.8207087 | -0.057387485 | 0.099282604 | -0.578021556 | 0.563249566 | 0.760667465 | 0.057387485 |
| ENSMUSG00000029925 | Tbxas1 | 21391 | protein_coding | 11.96844829 | -0.057399373 | 0.400812684 | -0.143207477 | 0.886126325 | 0.949475732 | 0.057399373 |
| ENSMUSG00000020974 | Pole2 | 18974 | protein_coding | 86.23903292 | 0.057434367 | 0.170186882 | 0.337478224 | 0.735756425 | 0.872915755 | 0.057434367 |
| ENSMUSG00000039740 | Alg2 | 56737 | protein_coding | 354.6126748 | 0.05743982 | 0.083483214 | 0.688040356 | 0.491427368 | 0.711036201 | 0.05743982 |
| ENSMUSG00000034354 | Mtmr3 | 74302 | protein_coding | 2480.462988 | -0.057477966 | 0.042854762 | -1.34122706 | 0.179846743 | 0.406900734 | 0.057477966 |
| ENSMUSG00000001518 | Itfg2 | 101142 | protein_coding | 275.5431388 | 0.057496451 | 0.089212704 | 0.644487262 | 0.519259515 | 0.731043055 | 0.057496451 |

Supplementary Table 1

| ensembl_gene_id | external_gene_name | entrezgene_id | gene_biotype | baseMean | log2FoldChange | lfcSE | stat | pvalue | padj | absLog2FC |
|---|---|---|---|---|---|---|---|---|---|---|
| ENSMUSG00000000374 | Trappc10 | 216131 | protein_coding | 3771.745847 | 0.057500295 | 0.037341424 | 1.539852752 | 0.12359625 | 0.326278724 | 0.057500295 |
| ENSMUSG00000021103 | Mnat1 | 17420 | protein_coding | 267.1109824 | -0.057502296 | 0.084926861 | -0.677080199 | 0.498355071 | 0.715904735 | 0.057502296 |
| ENSMUSG00000056531 | Ccdc18 | 73254 | protein_coding | 123.1284377 | -0.057506541 | 0.117273206 | -0.490363849 | 0.623876454 | 0.801438232 | 0.057506541 |
| ENSMUSG00000064655 | Gm25788 | 115490049 | snoRNA | 308.6164222 | 0.057511858 | 0.38846829 | 0.148047754 | 0.882305081 | 0.947541514 | 0.057511858 |
| ENSMUSG00000059939 | 9430015G10Rik | 230996 | protein_coding | 173.8171338 | -0.0575129 | 0.125197311 | -0.459378076 | 0.645962689 | 0.816513895 | 0.0575129 |
| ENSMUSG00000024841 | Eif1ad | 69860 | protein_coding | 574.1223637 | 0.057541171 | 0.067589057 | 0.85133856 | 0.394581311 | 0.634058669 | 0.057541171 |
| ENSMUSG00000042751 | Nmnat2 | 226518 | protein_coding | 29.4259967 | 0.057590662 | 0.239417299 | 0.240545116 | 0.809907692 | 0.914534473 | 0.057590662 |
| ENSMUSG00000020496 | Rnf187 | 108660 | protein_coding | 1648.056749 | 0.05760207 | 0.069752257 | 0.825809414 | 0.408912213 | 0.64731555 | 0.05760207 |
| ENSMUSG00000004394 | Tmed4 | 103694 | protein_coding | 1378.7146 | 0.057640197 | 0.041174017 | 1.399916764 | 0.161538245 | 0.382095524 | 0.057640197 |
| ENSMUSG00000041237 | Pklr | 18770 | protein_coding | 1378.167856 | 0.057686017 | 0.064067445 | 0.900395157 | 0.367909997 | 0.610106378 | 0.057686017 |
| ENSMUSG00000038542 | Pcid2 | 234069 | protein_coding | 346.1512303 | -0.057712939 | 0.076149705 | -0.757887896 | 0.448518097 | 0.678988684 | 0.057712939 |
| ENSMUSG00000009112 | Bcl2l13 | 94044 | protein_coding | 1273.659815 | 0.057731801 | 0.052581711 | 1.097944503 | 0.272228724 | 0.514726356 | 0.057731801 |
| ENSMUSG00000005045 | Chd5 | 269610 | protein_coding | 17.58736631 | -0.05774311 | 0.366295532 | -0.15764077 | 0.874739875 | 0.943745489 | 0.05774311 |
| ENSMUSG00000038366 | Lasp1 | 16796 | protein_coding | 8964.184206 | 0.05775725 | 0.052953622 | 1.090713874 | 0.275398804 | 0.518002859 | 0.05775725 |
| ENSMUSG00000004127 | Trmt10a | 108943 | protein_coding | 79.89613019 | 0.05775902 | 0.146646878 | 0.39386464 | 0.693680977 | 0.84654428 | 0.05775902 |
| ENSMUSG00000023947 | Nfkbie | 18037 | protein_coding | 65.19471502 | 0.057762297 | 0.149345026 | 0.386770809 | 0.69892589 | 0.849723168 | 0.057762297 |
| ENSMUSG00000027406 | Idh3b | 170718 | protein_coding | 5980.841317 | -0.057790397 | 0.035007421 | -1.650804183 | 0.098778566 | 0.285896663 | 0.057790397 |
| ENSMUSG00000030417 | Pdcd5 | 56330 | protein_coding | 611.2797238 | 0.057821227 | 0.066029391 | 0.875689247 | 0.381198993 | 0.6220843 | 0.057821227 |
| ENSMUSG00000006705 | Pknox1 | 18771 | protein_coding | 502.7006986 | 0.057823426 | 0.079542372 | 0.726951243 | 0.467255824 | 0.692786125 | 0.057823426 |
| ENSMUSG00000027288 | Zfp106 | 20402 | protein_coding | 3648.704745 | 0.057834768 | 0.052872646 | 1.093850454 | 0.274020572 | 0.516555203 | 0.057834768 |
| ENSMUSG00000025485 | Ric8a | 101489 | protein_coding | 1421.405548 | -0.057841591 | 0.038548419 | -1.500491907 | 0.133487028 | 0.340947717 | 0.057841591 |
| ENSMUSG00000073155 | 1810058I24Rik | 67705 | protein_coding | 263.7896051 | -0.05784669 | 0.079059102 | -0.731689185 | 0.464358299 | 0.690585235 | 0.05784669 |
| ENSMUSG00000026037 | Orc2 | 18393 | protein_coding | 289.7730289 | 0.057868068 | 0.089191233 | 0.648808923 | 0.51646189 | 0.729218881 | 0.057868068 |
| ENSMUSG00000054894 | Dmac2l | 68055 | protein_coding | 229.2125797 | -0.057873607 | 0.099833436 | -0.57970164 | 0.562115837 | 0.759869414 | 0.057873607 |
| ENSMUSG00000019737 | Syne4 | 233066 | protein_coding | 26.91644472 | 0.057897246 | 0.252732059 | 0.229085483 | 0.818802477 | 0.918528635 | 0.057897246 |
| ENSMUSG00000071652 | Ints5 | 109077 | protein_coding | 467.5463296 | 0.057919582 | 0.100874471 | 0.574174828 | 0.565849502 | 0.762042164 | 0.057919582 |
| ENSMUSG00000059842 | Zfp341 | 228807 | protein_coding | 74.79722819 | 0.05792708 | 0.135256402 | 0.428276066 | 0.668450142 | 0.830835872 | 0.05792708 |
| ENSMUSG00000036902 | Neto2 | 74513 | protein_coding | 50.76354352 | 0.05793062 | 0.181885115 | 0.318501158 | 0.750104817 | 0.880597382 | 0.05793062 |
| ENSMUSG00000025410 | Dctn2 | 69654 | protein_coding | 3054.433987 | 0.057978942 | 0.046903768 | 1.236125467 | 0.21641193 | 0.45231664 | 0.057978942 |
| ENSMUSG00000031729 | Ist1 | 71955 | protein_coding | 2858.429476 | -0.05801543 | 0.041367215 | -1.402449507 | 0.160781059 | 0.381255129 | 0.05801543 |
| ENSMUSG00000020782 | Llgl2 | 217325 | protein_coding | 5447.990786 | 0.05803336 | 0.067985398 | 0.853615066 | 0.393318308 | 0.632975187 | 0.05803336 |
| ENSMUSG00000028232 | Tmem68 | 72098 | protein_coding | 311.9742558 | -0.058100778 | 0.072055434 | -0.80633444 | 0.420050038 | 0.657152071 | 0.058100778 |
| ENSMUSG00000044916 | 1700029I15Rik | 75641 | protein_coding | 19.79388739 | 0.05810105 | 0.272933227 | 0.212876427 | 0.83142335 | 0.925325507 | 0.05810105 |
| ENSMUSG00000044296 | Zfp879 | 214779 | protein_coding | 15.6910385 | 0.058126807 | 0.300949267 | 0.19314487 | 0.846845513 | 0.930889346 | 0.058126807 |
| ENSMUSG00000059970 | Hspa2 | 15512 | protein_coding | 46.60033542 | -0.058133228 | 0.204554766 | -0.284193954 | 0.776261748 | 0.895541399 | 0.058133228 |
| ENSMUSG00000024477 | Pggt1b | 225467 | protein_coding | 1100.631665 | 0.058155367 | 0.06235575 | 0.932638404 | 0.351006698 | 0.593363209 | 0.058155367 |
| ENSMUSG00000041881 | Ndufa7 | 66416 | protein_coding | 3148.524995 | 0.05816463 | 0.05616988 | 1.035512805 | 0.30042949 | 0.544018232 | 0.05816463 |
| ENSMUSG00000033955 | Tnks1bp1 | 228140 | protein_coding | 3557.258668 | -0.058179162 | 0.067350298 | -0.863829322 | 0.387681657 | 0.627536092 | 0.058179162 |
| ENSMUSG00000059323 | Tonsl | 72749 | protein_coding | 437.4230429 | 0.058186663 | 0.064603803 | 0.900669318 | 0.367764167 | 0.609989995 | 0.058186663 |

Supplementary Table 1

| ensembl_gene_id | external_gene_name | entrezgene_id | gene_biotype | baseMean | log2FoldChange | lfcSE | stat | pvalue | padj | absLog2FC |
|---|---|---|---|---|---|---|---|---|---|---|
| ENSMUSG00000020640 | Itsn2 | 20403 | protein_coding | 2244.352923 | -0.058206402 | 0.043388981 | -1.341501952 | 0.179757536 | 0.406831399 | 0.058206402 |
| ENSMUSG00000070372 | Capza1 | 12340 | protein_coding | 1608.066857 | 0.058243782 | 0.05607392 | 1.038696454 | 0.298945931 | 0.542489144 | 0.058243782 |
| ENSMUSG00000051435 | Fhad1 | 329977 | protein_coding | 63.75980524 | 0.058271551 | 0.165233002 | 0.352662908 | 0.72434117 | 0.865627893 | 0.058271551 |
| ENSMUSG00000029777 | Gars | 353172 | protein_coding | 3042.155796 | 0.0582762 | 0.047789868 | 1.219425847 | 0.222682605 | 0.460024268 | 0.0582762 |
| ENSMUSG00000030528 | Blm | 12144 | protein_coding | 109.7482815 | 0.058279288 | 0.144028016 | 0.404638553 | 0.685743216 | 0.841819353 | 0.058279288 |
| ENSMUSG00000029370 | Rassf6 | 73246 | protein_coding | 1362.268453 | 0.058294913 | 0.068541663 | 0.850503334 | 0.395045308 | 0.634427961 | 0.058294913 |
| ENSMUSG00000090150 | Acad11 | 102632 | protein_coding | 3299.832236 | 0.058327576 | 0.049730059 | 1.17288371 | 0.240842443 | 0.481535591 | 0.058327576 |
| ENSMUSG00000049775 | Tmsb4x | 19241 | protein_coding | 6987.089141 | 0.058341 | 0.0624382 | 0.934379914 | 0.350107955 | 0.592368513 | 0.058341 |
| ENSMUSG00000022756 | Slc7a4 | 224022 | protein_coding | 580.9429866 | -0.058343889 | 0.069178276 | -0.843384546 | 0.399013412 | 0.638678506 | 0.058343889 |
| ENSMUSG00000035299 | Mid1 | 17318 | protein_coding | 113.0924273 | -0.058351247 | 0.138448255 | -0.421466104 | 0.673414758 | 0.833985547 | 0.058351247 |
| ENSMUSG00000025001 | Hells | 15201 | protein_coding | 360.0486979 | 0.058360566 | 0.165367388 | 0.352914601 | 0.724152464 | 0.865580106 | 0.058360566 |
| ENSMUSG00000028149 | Rap1gds1 | 229877 | protein_coding | 1108.017426 | -0.058393746 | 0.084371626 | -0.692101695 | 0.488873467 | 0.708998418 | 0.058393746 |
| ENSMUSG00000075703 | Selenoi | 28042 | protein_coding | 1791.93295 | 0.058398733 | 0.03558107 | 1.64128661 | 0.100737934 | 0.289486009 | 0.058398733 |
| ENSMUSG00000022946 | Dop1b | 70028 | protein_coding | 2133.356668 | 0.058421479 | 0.056606698 | 1.032059488 | 0.302044251 | 0.545716741 | 0.058421479 |
| ENSMUSG00000019297 | Nop9 | 67842 | protein_coding | 429.5642958 | 0.058425638 | 0.070795205 | 0.825276774 | 0.409214475 | 0.647571567 | 0.058425638 |
| ENSMUSG00000038417 | Fig4 | 103199 | protein_coding | 650.1997699 | 0.058429448 | 0.057183823 | 1.021782827 | 0.306883699 | 0.550870027 | 0.058429448 |
| ENSMUSG00000051154 | Commd3 | 12238 | protein_coding | 664.0963659 | 0.058434629 | 0.069455117 | 0.84132936 | 0.400163447 | 0.639482053 | 0.058434629 |
| ENSMUSG00000025145 | Lrrc45 | 217366 | protein_coding | 677.4818066 | 0.058439525 | 0.065235775 | 0.895820191 | 0.370348804 | 0.612387488 | 0.058439525 |
| ENSMUSG00000024220 | Zfp523 | 224656 | protein_coding | 386.3499719 | 0.058454989 | 0.064290632 | 0.909230273 | 0.363228587 | 0.605769124 | 0.058454989 |
| ENSMUSG00000032290 | Ptpn9 | 56294 | protein_coding | 1327.409544 | 0.058460432 | 0.050488995 | 1.157884638 | 0.246911118 | 0.487862729 | 0.058460432 |
| ENSMUSG00000020755 | Sap30bp | 57230 | protein_coding | 743.5208058 | 0.058474977 | 0.059888341 | 0.976400011 | 0.328866276 | 0.572498536 | 0.058474977 |
| ENSMUSG00000030127 | Cops7a | 26894 | protein_coding | 1662.870671 | 0.05849606 | 0.060625311 | 0.964878525 | 0.334605659 | 0.578183377 | 0.05849606 |
| ENSMUSG00000022833 | Ccdc14 | 239839 | protein_coding | 139.6362925 | -0.058497246 | 0.126568184 | -0.462179702 | 0.643952458 | 0.815430435 | 0.058497246 |
| ENSMUSG00000020454 | Eif4enif1 | 74203 | protein_coding | 756.0355327 | -0.058508868 | 0.052841688 | -1.107248271 | 0.268186603 | 0.510669852 | 0.058508868 |
| ENSMUSG00000044279 | Crb3 | 224912 | protein_coding | 1614.334562 | 0.058516155 | 0.074275127 | 0.787829748 | 0.430796298 | 0.665561246 | 0.058516155 |
| ENSMUSG00000022788 | Fgd4 | 224014 | protein_coding | 2207.092523 | -0.058550204 | 0.035494328 | -1.649565073 | 0.099031923 | 0.286216457 | 0.058550204 |
| ENSMUSG00000052760 | A630001G21Rik | 319997 | protein_coding | 30.62110626 | -0.058566439 | 0.209238102 | -0.279903316 | 0.779551683 | 0.897539087 | 0.058566439 |
| ENSMUSG00000021711 | Trappc13 | 66975 | protein_coding | 860.0627619 | -0.058592987 | 0.060519025 | -0.968174667 | 0.332957145 | 0.576322837 | 0.058592987 |
| ENSMUSG00000036188 | Ankmy2 | 217473 | protein_coding | 851.5358701 | 0.058596348 | 0.052166991 | 1.123245686 | 0.261333164 | 0.503166182 | 0.058596348 |
| ENSMUSG00000031818 | Cox4i1 | 12857 | protein_coding | 18792.15285 | 0.05859943 | 0.049224821 | 1.190444757 | 0.23387163 | 0.473349371 | 0.05859943 |
| ENSMUSG00000027342 | Pcna | 18538 | protein_coding | 1528.680775 | 0.058615899 | 0.087650323 | 0.668747102 | 0.503656816 | 0.720013604 | 0.058615899 |
| ENSMUSG00000027304 | Rtf1 | 76246 | protein_coding | 646.5211803 | 0.058675513 | 0.052159779 | 1.124918754 | 0.260623464 | 0.502482139 | 0.058675513 |
| ENSMUSG00000009376 | Met | 17295 | protein_coding | 1735.387668 | -0.058679495 | 0.054857792 | -1.069665633 | 0.284769841 | 0.527617708 | 0.058679495 |
| ENSMUSG00000052423 | B4galt3 | 57370 | protein_coding | 422.725206 | -0.05869437 | 0.088144298 | -0.665889583 | 0.505481679 | 0.721608883 | 0.05869437 |
| ENSMUSG00000039878 | Slc39a5 | 72002 | protein_coding | 2792.38861 | -0.058694761 | 0.07770478 | -0.755355858 | 0.450035492 | 0.679535938 | 0.058694761 |
| ENSMUSG00000045071 | E130308A19Rik | 230259 | protein_coding | 197.8040664 | -0.058694932 | 0.089309633 | -0.657207175 | 0.511047713 | 0.725307266 | 0.058694932 |
| ENSMUSG00000029405 | G3bp2 | 23881 | protein_coding | 2908.010397 | 0.05869899 | 0.047983863 | 1.223306898 | 0.221213803 | 0.457725814 | 0.05869899 |
| ENSMUSG00000034848 | Ttc21b | 73668 | protein_coding | 367.7274966 | 0.058706589 | 0.072671929 | 0.80783033 | 0.419188263 | 0.656573069 | 0.058706589 |
| ENSMUSG00000052812 | Atad2b | 320817 | protein_coding | 963.9245052 | -0.058727989 | 0.042495086 | -1.381994823 | 0.166973291 | 0.390767415 | 0.058727989 |

| ensembl_gene_id | external_gene_name | entrezgene_id | gene_biotype | baseMean | log2FoldChange | lfcSE | stat | pvalue | padj | absLog2FC |
|---|---|---|---|---|---|---|---|---|---|---|
| ENSMUSG00000024712 | Rfk | 54391 | protein_coding | 11821.06646 | -0.058734468 | 0.073698507 | -0.796956013 | 0.425476578 | 0.661237426 | 0.058734468 |
| ENSMUSG00000035765 | Dym | 69190 | protein_coding | 1185.594126 | -0.0587499 | 0.061921916 | -0.948773945 | 0.342735596 | 0.585270749 | 0.0587499 |
| ENSMUSG00000003038 | Hmgn2 | 15331 | protein_coding | 1105.809108 | -0.058751419 | 0.068907672 | -0.852610707 | 0.393875223 | 0.633492143 | 0.058751419 |
| ENSMUSG00000034751 | Mast4 | 328329 | protein_coding | 477.399801 | -0.058774954 | 0.063774305 | -0.921608694 | 0.356732722 | 0.599586098 | 0.058774954 |
| ENSMUSG00000022668 | Gtpbp8 | 66067 | protein_coding | 560.5927909 | -0.058776709 | 0.075206576 | -0.781536827 | 0.434486825 | 0.668221346 | 0.058776709 |
| ENSMUSG00000057719 | Sh3rf2 | 269016 | protein_coding | 716.9794244 | -0.058796141 | 0.083455519 | -0.704520703 | 0.481108562 | 0.703237858 | 0.058796141 |
| ENSMUSG00000062203 | Gspt1 | 14852 | protein_coding | 1543.243498 | 0.05881637 | 0.05157305 | 1.140447772 | 0.2540998 | 0.495031103 | 0.05881637 |
| ENSMUSG00000028047 | Thbs3 | 21827 | protein_coding | 17.88146134 | -0.058856153 | 0.374270861 | -0.157255504 | 0.875043486 | 0.943745489 | 0.058856153 |
| ENSMUSG00000038299 | Wdr36 | 225348 | protein_coding | 1070.768867 | -0.058887975 | 0.050883057 | -1.157319908 | 0.247141683 | 0.488015969 | 0.058887975 |
| ENSMUSG00000039231 | Suv39h1 | 20937 | protein_coding | 254.2745481 | -0.058902138 | 0.102974196 | -0.572008717 | 0.567316069 | 0.762827428 | 0.058902138 |
| ENSMUSG00000079316 | Rab9 | 56382 | protein_coding | 637.1989138 | -0.058914344 | 0.061869179 | -0.952240598 | 0.340974976 | 0.583871675 | 0.058914344 |
| ENSMUSG00000048833 | Slc39a9 | 328133 | protein_coding | 3014.724141 | 0.058930601 | 0.044950514 | 1.311010613 | 0.189854178 | 0.420477604 | 0.058930601 |
| ENSMUSG00000032459 | Mrps22 | 64655 | protein_coding | 461.9044049 | 0.058956591 | 0.076506534 | 0.77060857 | 0.44093898 | 0.672622244 | 0.058956591 |
| ENSMUSG00000042505 | Sdhaf3 | 71238 | protein_coding | 113.4423678 | 0.05897817 | 0.112635286 | 0.523620724 | 0.600542365 | 0.786052002 | 0.05897817 |
| ENSMUSG00000037816 | Fbxw17 | 109082 | protein_coding | 148.8171714 | -0.058989339 | 0.105434745 | -0.559486714 | 0.575829596 | 0.768742044 | 0.058989339 |
| ENSMUSG00000022718 | Dgcr8 | 94223 | protein_coding | 430.0380304 | -0.059007238 | 0.087776252 | -0.672246044 | 0.501427073 | 0.718418124 | 0.059007238 |
| ENSMUSG00000019818 | Cd164 | 53599 | protein_coding | 5867.744973 | -0.059024269 | 0.027826944 | -2.12111934 | 0.033911761 | 0.141755849 | 0.059024269 |
| ENSMUSG00000018339 | Gpx3 | 14778 | protein_coding | 20.60328407 | -0.059031759 | 0.286486532 | -0.206054221 | 0.836748558 | 0.927643978 | 0.059031759 |
| ENSMUSG00000047879 | Usp14 | 59025 | protein_coding | 1011.705889 | 0.059045722 | 0.053964483 | 1.094158943 | 0.273885275 | 0.516481271 | 0.059045722 |
| ENSMUSG00000014551 | Mrps25 | 64658 | protein_coding | 529.4785724 | 0.059050907 | 0.085724145 | 0.688848007 | 0.490918921 | 0.710750466 | 0.059050907 |
| ENSMUSG00000054874 | Pcnx3 | 104401 | protein_coding | 1853.947203 | -0.059063909 | 0.076443997 | -0.772642869 | 0.439733766 | 0.671740242 | 0.059063909 |
| ENSMUSG00000026235 | Epha4 | 13838 | protein_coding | 495.8531177 | 0.05907579 | 0.085125688 | 0.693983114 | 0.487692799 | 0.708241492 | 0.05907579 |
| ENSMUSG00000022515 | Anks3 | 72615 | protein_coding | 441.0474417 | 0.059101625 | 0.077251106 | 0.765058622 | 0.444236638 | 0.67516768 | 0.059101625 |
| ENSMUSG00000055013 | Agap1 | 347722 | protein_coding | 1668.819993 | -0.059106151 | 0.053022156 | -1.114744394 | 0.264959971 | 0.506761084 | 0.059106151 |
| ENSMUSG00000048772 | Tmem53 | 68777 | protein_coding | 116.7278512 | -0.059127959 | 0.148143155 | -0.399127177 | 0.689799498 | 0.844356471 | 0.059127959 |
| ENSMUSG00000020783 | Ncbp3 | 66874 | protein_coding | 829.780252 | -0.059165513 | 0.054769328 | -1.080267272 | 0.280023179 | 0.523118689 | 0.059165513 |
| ENSMUSG00000040731 | Eif4h | 22384 | protein_coding | 6503.973136 | -0.059195588 | 0.039261352 | -1.50773175 | 0.131623196 | 0.337846137 | 0.059195588 |
| ENSMUSG00000039536 | Stau1 | 20853 | protein_coding | 876.5378487 | -0.059199578 | 0.047696532 | -1.241171532 | 0.214542389 | 0.450335169 | 0.059199578 |
| ENSMUSG00000005615 | Pcyt1a | 13026 | protein_coding | 1470.873504 | 0.059224945 | 0.046070591 | 1.285526055 | 0.198608518 | 0.431840309 | 0.059224945 |
| ENSMUSG00000042331 | Specc1 | 432572 | protein_coding | 1368.917736 | -0.0592308 | 0.055348493 | -1.070142946 | 0.284554971 | 0.527473335 | 0.0592308 |
| ENSMUSG00000033904 | Ccp110 | 101565 | protein_coding | 71.95325323 | 0.059231465 | 0.196895505 | 0.300826904 | 0.763546492 | 0.888754045 | 0.059231465 |
| ENSMUSG00000037720 | Tmem33 | 67878 | protein_coding | 1359.035581 | 0.059246937 | 0.050486277 | 1.173525557 | 0.240585114 | 0.481155234 | 0.059246937 |
| ENSMUSG00000055125 | M5C1000I18Rik | 102633151 | lncRNA | 7.245659263 | -0.059249173 | 0.471340452 | -0.125703562 | 0.899966582 | 0.955732983 | 0.059249173 |
| ENSMUSG00000028099 | Polr3c | 74414 | protein_coding | 892.6566118 | -0.059269515 | 0.052592079 | -1.126966565 | 0.259756618 | 0.50123327 | 0.059269515 |
| ENSMUSG00000057614 | Gnai1 | 14677 | protein_coding | 293.8848457 | 0.059290704 | 0.098940615 | 0.599255462 | 0.549002543 | 0.751273993 | 0.059290704 |
| ENSMUSG00000074604 | Mgst2 | 211666 | protein_coding | 1710.057055 | -0.059326714 | 0.061049698 | -0.971777352 | 0.331161325 | 0.574740328 | 0.059326714 |
| ENSMUSG00000061983 | Rps12 | 20042 | protein_coding | 11664.89276 | -0.059332646 | 0.074569155 | -0.795672764 | 0.426222261 | 0.66169488 | 0.059332646 |
| ENSMUSG00000023007 | Prpf40b | 54614 | protein_coding | 189.2212544 | -0.059335675 | 0.112515203 | -0.527356959 | 0.597945727 | 0.784071601 | 0.059335675 |
| ENSMUSG00000078202 | Nrarp | 67122 | protein_coding | 125.9844038 | 0.059345576 | 0.154978592 | 0.382927572 | 0.701773467 | 0.85164427 | 0.059345576 |

| ensembl_gene_id | external_gene_name | entrezgene_id | gene_biotype | baseMean | log2FoldChange | lfcSE | stat | pvalue | padj | absLog2FC |
|---|---|---|---|---|---|---|---|---|---|---|
| ENSMUSG00000039621 | Prex1 | 277360 | protein_coding | 153.1616668 | -0.059349183 | 0.10240305 | -0.579564599 | 0.562208272 | 0.759930668 | 0.059349183 |
| ENSMUSG00000037826 | Ppm1k | 243382 | protein_coding | 435.4296436 | -0.05936382 | 0.060306275 | -0.984372197 | 0.324932543 | 0.568614628 | 0.05936382 |
| ENSMUSG00000056919 | Cep162 | 382090 | protein_coding | 333.9664286 | 0.059391766 | 0.077176703 | 0.769555628 | 0.441563535 | 0.672888818 | 0.059391766 |
| ENSMUSG00000037536 | Fbxo34 | 78938 | protein_coding | 791.9684963 | 0.059416712 | 0.04839866 | 1.227652008 | 0.219577629 | 0.455743685 | 0.059416712 |
| ENSMUSG00000030469 | Zfp719 | 210105 | protein_coding | 232.0254333 | -0.059436358 | 0.110982167 | -0.535548721 | 0.592270477 | 0.780262005 | 0.059436358 |
| ENSMUSG00000067713 | Prkag1 | 19082 | protein_coding | 603.8429729 | 0.059478358 | 0.062097313 | 0.957824991 | 0.338151015 | 0.582002286 | 0.059478358 |
| ENSMUSG00000021009 | Ptpn21 | 24000 | protein_coding | 259.6871679 | -0.059483958 | 0.092072632 | -0.646054714 | 0.518243922 | 0.730504162 | 0.059483958 |
| ENSMUSG00000021188 | Trip11 | 109181 | protein_coding | 2005.42713 | 0.05949174 | 0.046418883 | 1.281627991 | 0.199973176 | 0.433436829 | 0.05949174 |
| ENSMUSG00000018412 | Kansl1 | 76719 | protein_coding | 1329.76492 | -0.05949535 | 0.055485964 | -1.072259454 | 0.283603513 | 0.526203742 | 0.05949535 |
| ENSMUSG00000042857 | Gm9776 | 328309 | lncRNA | 50.43292434 | 0.059504957 | 0.17733811 | 0.335545228 | 0.737213833 | 0.873998696 | 0.059504957 |
| ENSMUSG00000060402 | Chst8 | 68947 | protein_coding | 47.84645818 | 0.059569065 | 0.211611253 | 0.281502347 | 0.778325128 | 0.896895273 | 0.059569065 |
| ENSMUSG00000026657 | Frmd4a | 209630 | protein_coding | 251.0135111 | 0.059591119 | 0.080116295 | 0.743807727 | 0.456992805 | 0.685107692 | 0.059591119 |
| ENSMUSG00000019577 | Pdk4 | 27273 | protein_coding | 235.4183172 | -0.059596969 | 0.398632589 | -0.149503506 | 0.881156343 | 0.947044875 | 0.059596969 |
| ENSMUSG00000052942 | Glis3 | 226075 | protein_coding | 258.8832188 | -0.059601924 | 0.085262986 | -0.699036318 | 0.484529334 | 0.706263486 | 0.059601924 |
| ENSMUSG00000027463 | Slc52a3 | 69698 | protein_coding | 1128.986763 | 0.059629446 | 0.079522325 | 0.749845352 | 0.453347851 | 0.682136666 | 0.059629446 |
| ENSMUSG00000048755 | Mcat | 223722 | protein_coding | 422.8393103 | 0.059634031 | 0.068229195 | 0.874025131 | 0.382104568 | 0.622821949 | 0.059634031 |
| ENSMUSG00000029681 | Bcl7b | 12054 | protein_coding | 466.2121096 | -0.0596417 | 0.064299026 | -0.927567705 | 0.353631856 | 0.595956629 | 0.0596417 |
| ENSMUSG00000030339 | Ltbr | 17000 | protein_coding | 2878.82204 | 0.059646763 | 0.078133983 | 0.763390786 | 0.445230374 | 0.676259511 | 0.059646763 |
| ENSMUSG00000056962 | Jmjd6 | 107817 | protein_coding | 667.8368474 | -0.059682711 | 0.062235031 | -0.958989015 | 0.337564279 | 0.581263386 | 0.059682711 |
| ENSMUSG00000005846 | Rsl1d1 | 66409 | protein_coding | 1090.802512 | -0.059694578 | 0.065465947 | -0.911841666 | 0.361852073 | 0.604844153 | 0.059694578 |
| ENSMUSG00000067367 | Lyar | 17089 | protein_coding | 287.3492655 | 0.059699093 | 0.098637186 | 0.605239213 | 0.545020065 | 0.749012844 | 0.059699093 |
| ENSMUSG00000042284 | Itga1 | 109700 | protein_coding | 421.6923048 | 0.059700077 | 0.066520416 | 0.897469982 | 0.369468184 | 0.611621167 | 0.059700077 |
| ENSMUSG00000035621 | Midn | 59090 | protein_coding | 3340.769782 | 0.059722041 | 0.106303821 | 0.561805214 | 0.574248739 | 0.768186995 | 0.059722041 |
| ENSMUSG00000070738 | Dgkd | 227333 | protein_coding | 3467.832048 | -0.059757102 | 0.040630975 | -1.470727722 | 0.141364766 | 0.352354239 | 0.059757102 |
| ENSMUSG00000039164 | Naif1 | 71254 | protein_coding | 216.2923737 | -0.059762452 | 0.105395011 | -0.567033029 | 0.570691742 | 0.76587715 | 0.059762452 |
| ENSMUSG00000039047 | Pigk | 329777 | protein_coding | 1043.388308 | 0.059770579 | 0.048196446 | 1.24014494 | 0.214921789 | 0.45050036 | 0.059770579 |
| ENSMUSG00000021234 | Fam161b | 217705 | protein_coding | 44.94923907 | 0.059793817 | 0.197598321 | 0.30260286 | 0.762192535 | 0.887756682 | 0.059793817 |
| ENSMUSG00000038291 | Snx25 | 102141 | protein_coding | 759.0103534 | -0.059806413 | 0.072430875 | -0.825703308 | 0.408972415 | 0.64731555 | 0.059806413 |
| ENSMUSG00000059669 | Taf1b | 21340 | protein_coding | 113.8449186 | 0.059811714 | 0.135774001 | 0.440524058 | 0.659557592 | 0.82591596 | 0.059811714 |
| ENSMUSG00000001285 | Myg1 | 60315 | protein_coding | 430.5460412 | -0.059811773 | 0.072837433 | -0.821168054 | 0.411550551 | 0.649560515 | 0.059811773 |
| ENSMUSG00000006221 | Hspb7 | 29818 | protein_coding | 16.3681925 | 0.059827491 | 0.322241627 | 0.185660342 | 0.85271113 | 0.933836235 | 0.059827491 |
| ENSMUSG00000035473 | Galm | 319625 | protein_coding | 1583.455717 | -0.059835859 | 0.039191782 | -1.526745038 | 0.126824423 | 0.331045348 | 0.059835859 |
| ENSMUSG00000027937 | Jtb | 23922 | protein_coding | 881.9206922 | 0.059848798 | 0.06029348 | 0.992624712 | 0.320892877 | 0.564467652 | 0.059848798 |
| ENSMUSG00000043940 | Wdfy3 | 72145 | protein_coding | 1154.766084 | -0.059869928 | 0.069845852 | -0.857172278 | 0.391349681 | 0.631257944 | 0.059869928 |
| ENSMUSG00000028249 | Sdcbp | 53378 | protein_coding | 5330.501658 | 0.059897077 | 0.035841746 | 1.671153993 | 0.094691265 | 0.278512863 | 0.059897077 |
| ENSMUSG00000032741 | Tpcn1 | 252972 | protein_coding | 1947.74874 | 0.059906584 | 0.095967394 | 0.624238937 | 0.532470679 | 0.740799392 | 0.059906584 |
| ENSMUSG00000073702 | Rpl31 | 114641 | protein_coding | 7718.934749 | -0.059917433 | 0.043494032 | -1.377601254 | 0.168326434 | 0.392283294 | 0.059917433 |
| ENSMUSG00000030595 | Nfkbib | 18036 | protein_coding | 407.554474 | 0.059937923 | 0.106903043 | 0.56067556 | 0.575018731 | 0.768361739 | 0.059937923 |
| ENSMUSG00000056211 | R3hdm1 | 226412 | protein_coding | 461.9568853 | -0.059945728 | 0.084959219 | -0.705582378 | 0.480447886 | 0.702846794 | 0.059945728 |

| ensembl_gene_id | external_gene_name | entrezgene_id | gene_biotype | baseMean | log2FoldChange | lfcSE | stat | pvalue | padj | absLog2FC |
|---|---|---|---|---|---|---|---|---|---|---|
| ENSMUSG00000022503 | Nubp1 | 26425 | protein_coding | 472.8346193 | 0.059964037 | 0.074166324 | 0.808507605 | 0.41879843 | 0.656297957 | 0.059964037 |
| ENSMUSG00000074129 | Rpl13a | 22121 | protein_coding | 519.6940849 | -0.060003242 | 0.066615126 | -0.900745004 | 0.367723914 | 0.609985966 | 0.060003242 |
| ENSMUSG00000030204 | Ddx47 | 67755 | protein_coding | 894.2073472 | -0.060009706 | 0.05719637 | -1.049187325 | 0.294091911 | 0.537064913 | 0.060009706 |
| ENSMUSG00000027132 | Katnbl1 | 72425 | protein_coding | 238.6627351 | 0.060025729 | 0.092737581 | 0.647264334 | 0.517460881 | 0.729947916 | 0.060025729 |
| ENSMUSG00000037533 | Rapgef6 | 192786 | protein_coding | 1813.792858 | 0.060025758 | 0.034407372 | 1.744560936 | 0.081061326 | 0.252369884 | 0.060025758 |
| ENSMUSG00000029730 | Mcm7 | 17220 | protein_coding | 893.5019832 | 0.060047087 | 0.140882467 | 0.426221149 | 0.669946708 | 0.831606417 | 0.060047087 |
| ENSMUSG00000035873 | Pawr | 114774 | protein_coding | 1043.414206 | 0.060058276 | 0.067497701 | 0.889782547 | 0.37358266 | 0.615400159 | 0.060058276 |
| ENSMUSG00000020400 | Tnip1 | 57783 | protein_coding | 1814.24532 | -0.060091477 | 0.068234944 | -0.880655473 | 0.378504324 | 0.619709567 | 0.060091477 |
| ENSMUSG00000025868 | Higd2a | 67044 | protein_coding | 3178.332253 | -0.06010211 | 0.077907224 | -0.771457467 | 0.440435824 | 0.672114736 | 0.06010211 |
| ENSMUSG00000047371 | Zfp768 | 233890 | protein_coding | 893.5781284 | 0.060133871 | 0.056616105 | 1.062133655 | 0.288175014 | 0.530646089 | 0.060133871 |
| ENSMUSG00000041775 | Mapk1ip1 | 69546 | protein_coding | 139.7663446 | -0.060173257 | 0.108603211 | -0.554065171 | 0.579534245 | 0.771707437 | 0.060173257 |
| ENSMUSG00000029530 | Ccr9 | 12769 | protein_coding | 14.05516907 | 0.060216686 | 0.337175559 | 0.178591493 | 0.858258475 | 0.936097283 | 0.060216686 |
| ENSMUSG00000071632 | 2510002D24Rik | 72307 | protein_coding | 370.487674 | 0.060226093 | 0.076866927 | 0.783511127 | 0.433327022 | 0.667456054 | 0.060226093 |
| ENSMUSG00000028294 | Cfap206 | 69329 | protein_coding | 11.38482484 | 0.060245344 | 0.392173297 | 0.153619191 | 0.877910005 | 0.945241431 | 0.060245344 |
| ENSMUSG00000031799 | Tpm4 | 326618 | protein_coding | 2426.311649 | 0.060259719 | 0.073493491 | 0.81993274 | 0.412254451 | 0.649970343 | 0.060259719 |
| ENSMUSG00000110444 | Gm10033 | 378466 | protein_coding | 125.9453581 | -0.060277532 | 0.148922444 | -0.404757878 | 0.685655493 | 0.841775742 | 0.060277532 |
| ENSMUSG00000028243 | Ubxn2b | 68053 | protein_coding | 179.6749346 | 0.060278313 | 0.09725474 | 0.619798203 | 0.535390652 | 0.74228379 | 0.060278313 |
| ENSMUSG00000032182 | Yipf2 | 74766 | protein_coding | 396.8847266 | 0.060309457 | 0.070218802 | 0.858879037 | 0.390407256 | 0.630176941 | 0.060309457 |
| ENSMUSG00000004056 | Akt2 | 11652 | protein_coding | 838.1490948 | -0.060310637 | 0.058995009 | -1.022300672 | 0.306638615 | 0.550747405 | 0.060310637 |
| ENSMUSG00000075254 | Heg1 | 77446 | protein_coding | 159.0952948 | -0.060332753 | 0.105337886 | -0.572754547 | 0.566810899 | 0.762762443 | 0.060332753 |
| ENSMUSG00000023942 | Slc29a1 | 63959 | protein_coding | 381.5687671 | 0.060337523 | 0.10686646 | 0.564606738 | 0.572341281 | 0.767069546 | 0.060337523 |
| ENSMUSG00000030867 | Plk1 | 18817 | protein_coding | 1137.552458 | 0.060347036 | 0.060198025 | 1.002475349 | 0.316114067 | 0.560212157 | 0.060347036 |
| ENSMUSG00000028393 | Alad | 17025 | protein_coding | 784.6483227 | -0.060352274 | 0.060847898 | -0.991854703 | 0.321268408 | 0.564943367 | 0.060352274 |
| ENSMUSG00000029086 | Prom1 | 19126 | protein_coding | 8264.55482 | 0.06036985 | 0.088235207 | 0.684192304 | 0.493853741 | 0.713270579 | 0.06036985 |
| ENSMUSG00000021124 | Vti1b | 53612 | protein_coding | 1312.163098 | 0.060375026 | 0.04083257 | 1.478599717 | 0.139247328 | 0.34961375 | 0.060375026 |
| ENSMUSG00000022824 | Muc13 | 17063 | protein_coding | 24954.14071 | 0.060396153 | 0.042273765 | 1.428691117 | 0.153093032 | 0.3701874 | 0.060396153 |
| ENSMUSG00000038072 | Galnt11 | 231050 | protein_coding | 435.3819668 | 0.060453344 | 0.061161507 | 0.988421426 | 0.322946299 | 0.566904861 | 0.060453344 |
| ENSMUSG00000046605 | B3gntl1 | 210004 | protein_coding | 164.830416 | -0.060458011 | 0.097638676 | -0.619201463 | 0.535783648 | 0.742637121 | 0.060458011 |
| ENSMUSG00000090000 | Ier3ip1 | 66191 | protein_coding | 683.8735454 | 0.060468597 | 0.05407695 | 1.118195404 | 0.263483547 | 0.504916665 | 0.060468597 |
| ENSMUSG00000001687 | Ubl3 | 24109 | protein_coding | 2912.006782 | -0.060475888 | 0.038472624 | -1.571920047 | 0.115969095 | 0.314662099 | 0.060475888 |
| ENSMUSG00000025933 | Tmem14a | 75712 | protein_coding | 66.50063751 | 0.060489897 | 0.171786893 | 0.352121728 | 0.724746974 | 0.865841503 | 0.060489897 |
| ENSMUSG00000004054 | Map3k11 | 26403 | protein_coding | 1916.463885 | 0.060500569 | 0.079878082 | 0.757411388 | 0.448803436 | 0.678988684 | 0.060500569 |
| ENSMUSG00000025232 | Hexa | 15211 | protein_coding | 3283.52816 | -0.060517539 | 0.045578341 | -1.327769694 | 0.184254202 | 0.412933228 | 0.060517539 |
| ENSMUSG00000025580 | Eif4a3 | 192170 | protein_coding | 1750.132921 | 0.060526604 | 0.050800948 | 1.191446351 | 0.233478406 | 0.472837299 | 0.060526604 |
| ENSMUSG00000062661 | Ncs1 | 14299 | protein_coding | 7.037560729 | -0.060552383 | 0.437189384 | -0.138503781 | 0.889842281 | 0.950815976 | 0.060552383 |
| ENSMUSG00000032388 | Spg21 | 27965 | protein_coding | 869.7031205 | 0.06055846 | 0.046585646 | 1.2999382 | 0.193622151 | 0.425472739 | 0.06055846 |
| ENSMUSG00000020876 | Snx11 | 74479 | protein_coding | 316.0740513 | 0.060559187 | 0.085412318 | 0.709021701 | 0.47831101 | 0.700798986 | 0.060559187 |
| ENSMUSG00000037112 | Sik2 | 235344 | protein_coding | 352.5457435 | 0.060583136 | 0.072650743 | 0.833895623 | 0.404339804 | 0.643710564 | 0.060583136 |
| ENSMUSG00000029227 | Fip1l1 | 66899 | protein_coding | 975.4283124 | -0.06059264 | 0.045907963 | -1.319872089 | 0.186877728 | 0.416448019 | 0.06059264 |

| ensembl_gene_id | external_gene_name | entrezgene_id | gene_biotype | baseMean | log2FoldChange | lfcSE | stat | pvalue | padj | absLog2FC |
|---|---|---|---|---|---|---|---|---|---|---|
| ENSMUSG00000031848 | Lsm4 | 50783 | protein_coding | 1017.716281 | 0.060605607 | 0.093282854 | 0.649697176 | 0.515887848 | 0.728649792 | 0.060605607 |
| ENSMUSG00000024039 | Cbs | 12411 | protein_coding | 1684.166676 | 0.060671206 | 0.069467662 | 0.873373366 | 0.382459603 | 0.623211682 | 0.060671206 |
| ENSMUSG00000020802 | Ube2o | 217342 | protein_coding | 405.254483 | -0.060671728 | 0.092292514 | -0.657385147 | 0.5109333 | 0.725208707 | 0.060671728 |
| ENSMUSG00000029775 | Klhdc10 | 76788 | protein_coding | 1267.148256 | 0.060702343 | 0.040174229 | 1.510977157 | 0.130794271 | 0.336624075 | 0.060702343 |
| ENSMUSG00000021702 | Thbs4 | 21828 | protein_coding | 5058.649593 | 0.060703891 | 0.128080003 | 0.473952915 | 0.635533484 | 0.809577291 | 0.060703891 |
| ENSMUSG00000004565 | Pnpla6 | 50767 | protein_coding | 1340.111751 | 0.060846299 | 0.08097435 | 0.751426827 | 0.452395823 | 0.68117317 | 0.060846299 |
| ENSMUSG00000031513 | Leprotl1 | 68192 | protein_coding | 991.5325303 | 0.060855552 | 0.064558719 | 0.942638783 | 0.345865695 | 0.588148333 | 0.060855552 |
| ENSMUSG00000040260 | Daam2 | 76441 | protein_coding | 11.84619726 | -0.060862968 | 0.395862466 | -0.153747762 | 0.877808624 | 0.945241431 | 0.060862968 |
| ENSMUSG00000034928 | Rnf44 | 105239 | protein_coding | 1898.261534 | 0.060884118 | 0.092023231 | 0.661616828 | 0.50821682 | 0.723769025 | 0.060884118 |
| ENSMUSG00000030942 | Thumpd1 | 233802 | protein_coding | 658.742652 | -0.060891914 | 0.054580733 | -1.115630185 | 0.264580466 | 0.5061553 | 0.060891914 |
| ENSMUSG00000033790 | Tubgcp5 | 233276 | protein_coding | 282.3057634 | 0.060904593 | 0.086034255 | 0.707910969 | 0.479000548 | 0.701493083 | 0.060904593 |
| ENSMUSG00000027953 | Slc50a1 | 19729 | protein_coding | 785.4812927 | -0.060911595 | 0.073222945 | -0.831864866 | 0.405485225 | 0.644516088 | 0.060911595 |
| ENSMUSG00000033187 | BC016579 | 212998 | protein_coding | 688.4754207 | 0.060916398 | 0.066853465 | 0.911192827 | 0.362193783 | 0.605044452 | 0.060916398 |
| ENSMUSG00000028496 | Mllt3 | 70122 | protein_coding | 1004.595756 | -0.060923257 | 0.059826543 | -1.018331561 | 0.308520415 | 0.552220724 | 0.060923257 |
| ENSMUSG00000063364 | 3300002I08Rik | 69277 | protein_coding | 23.74206964 | 0.060924157 | 0.269600711 | 0.225979214 | 0.821217587 | 0.920153966 | 0.060924157 |
| ENSMUSG00000025086 | Trub1 | 72133 | protein_coding | 203.2242625 | -0.060949642 | 0.086350387 | -0.705840985 | 0.480287031 | 0.70267522 | 0.060949642 |
| ENSMUSG00000032076 | Cadm1 | 54725 | protein_coding | 64.39159579 | -0.060963401 | 0.20442342 | -0.298221218 | 0.765534328 | 0.889849511 | 0.060963401 |
| ENSMUSG00000022404 | Slc25a17 | 20524 | protein_coding | 1103.960961 | -0.060963477 | 0.052874291 | -1.152989017 | 0.248914893 | 0.489898763 | 0.060963477 |
| ENSMUSG00000047021 | Cfap65 | 241116 | protein_coding | 7.306249133 | -0.060976536 | 0.554396463 | -0.109987238 | 0.912419496 | 0.960984144 | 0.060976536 |
| ENSMUSG00000050017 | Pitpnb | 56305 | protein_coding | 532.7876199 | 0.060978381 | 0.078122614 | 0.780547114 | 0.435068907 | 0.66862102 | 0.060978381 |
| ENSMUSG00000022443 | Myh9 | 17886 | protein_coding | 21838.68706 | 0.060982838 | 0.061936088 | 0.984609134 | 0.324816101 | 0.568516308 | 0.060982838 |
| ENSMUSG00000016831 | Tox4 | 268741 | protein_coding | 1409.4623 | 0.060995726 | 0.044869965 | 1.359388764 | 0.17402343 | 0.399914072 | 0.060995726 |
| ENSMUSG00000057835 | Zfp119a | 104349 | protein_coding | 119.1471963 | -0.06101545 | 0.122477754 | -0.498175774 | 0.618360156 | 0.797655914 | 0.06101545 |
| ENSMUSG00000022100 | Xpo7 | 65246 | protein_coding | 1358.415344 | -0.061016912 | 0.057473051 | -1.061661262 | 0.288389491 | 0.5307986 | 0.061016912 |
| ENSMUSG00000029610 | Aimp2 | 231872 | protein_coding | 248.9455887 | -0.061020376 | 0.081931006 | -0.744777567 | 0.456406198 | 0.684487275 | 0.061020376 |
| ENSMUSG00000004880 | Lbr | 98386 | protein_coding | 1975.253136 | -0.061029657 | 0.064798149 | -0.941842592 | 0.346273235 | 0.588629676 | 0.061029657 |
| ENSMUSG00000037098 | Rab11fip3 | 215445 | protein_coding | 441.7468359 | 0.061039901 | 0.066012409 | 0.924673125 | 0.355135963 | 0.59777452 | 0.061039901 |
| ENSMUSG00000030647 | Ndufc2 | 68197 | protein_coding | 2202.180989 | 0.061041736 | 0.068609074 | 0.889703535 | 0.373625095 | 0.615407202 | 0.061041736 |
| ENSMUSG00000041360 | Pum3 | 52874 | protein_coding | 765.0937424 | -0.061053924 | 0.066617544 | -0.916484149 | 0.359413017 | 0.602834501 | 0.061053924 |
| ENSMUSG00000006717 | Acot13 | 66834 | protein_coding | 716.2574296 | 0.061062679 | 0.061990006 | 0.985040709 | 0.324604075 | 0.568485113 | 0.061062679 |
| ENSMUSG00000007987 | Ift22 | 67286 | protein_coding | 154.6846125 | 0.061076062 | 0.118050596 | 0.517371908 | 0.604896566 | 0.789053031 | 0.061076062 |
| ENSMUSG00000022400 | Rbx1 | 56438 | protein_coding | 1053.531879 | 0.061109795 | 0.060442031 | 1.011048016 | 0.311993449 | 0.555723255 | 0.061109795 |
| ENSMUSG00000085347 | Hk1os | 70162 | lncRNA | 10.20576546 | 0.061180086 | 0.360269404 | 0.1698176 | 0.865153585 | 0.938898215 | 0.061180086 |
| ENSMUSG00000037600 | Kdf1 | 69073 | protein_coding | 680.3297656 | 0.061186235 | 0.092237449 | 0.663355673 | 0.507102789 | 0.722697656 | 0.061186235 |
| ENSMUSG00000004455 | Ppp1cc | 19047 | protein_coding | 1342.290306 | -0.061191705 | 0.052344628 | -1.169015939 | 0.24239721 | 0.483565042 | 0.061191705 |
| ENSMUSG00000040029 | Ipo8 | 320727 | protein_coding | 1707.409237 | 0.061199586 | 0.040600867 | 1.507346797 | 0.131721788 | 0.337937843 | 0.061199586 |
| ENSMUSG00000029686 | Cul1 | 26965 | protein_coding | 2582.584636 | 0.061233941 | 0.056747898 | 1.079052135 | 0.280564487 | 0.523452524 | 0.061233941 |
| ENSMUSG00000021550 | 2210016F16Rik | 70153 | protein_coding | 1599.19812 | 0.061237244 | 0.064227101 | 0.953448671 | 0.340362791 | 0.583213766 | 0.061237244 |
| ENSMUSG00000029513 | Prkab1 | 19079 | protein_coding | 2690.051053 | 0.061317892 | 0.045171719 | 1.357439854 | 0.174641492 | 0.40092091 | 0.061317892 |

| ensembl_gene_id | external_gene_name | entrezgene_id | gene_biotype | baseMean | log2FoldChange | lfcSE | stat | pvalue | padj | absLog2FC |
|---|---|---|---|---|---|---|---|---|---|---|
| ENSMUSG00000016940 | Kctd2 | 70382 | protein_coding | 519.0442773 | -0.061334118 | 0.071651114 | -0.856010665 | 0.39199188 | 0.631786234 | 0.061334118 |
| ENSMUSG00000005103 | Wdr1 | 22388 | protein_coding | 8781.18912 | 0.061382985 | 0.027551592 | 2.22792879 | 0.025885264 | 0.117532596 | 0.061382985 |
| ENSMUSG00000078861 | Zfp931 | 353208 | protein_coding | 102.1709872 | 0.061389104 | 0.140142151 | 0.438048821 | 0.661350893 | 0.826855248 | 0.061389104 |
| ENSMUSG00000041697 | Cox6a1 | 12861 | protein_coding | 11353.01035 | 0.061391353 | 0.084988677 | 0.722347441 | 0.470080892 | 0.694251203 | 0.061391353 |
| ENSMUSG00000045519 | Zfp560 | 434377 | protein_coding | 105.2032602 | 0.061411935 | 0.165524672 | 0.371013784 | 0.710627265 | 0.856779748 | 0.061411935 |
| ENSMUSG00000032434 | Cmtm6 | 67213 | protein_coding | 3343.868393 | 0.061416749 | 0.042340651 | 1.450538613 | 0.14690838 | 0.360168963 | 0.061416749 |
| ENSMUSG00000032024 | Clmp | 71566 | protein_coding | 122.6601355 | 0.061474225 | 0.213625433 | 0.287766418 | 0.773525545 | 0.893440495 | 0.061474225 |
| ENSMUSG00000027709 | Mccc1 | 72039 | protein_coding | 559.3617272 | -0.06151777 | 0.064250574 | -0.957466467 | 0.338331864 | 0.582029301 | 0.06151777 |
| ENSMUSG00000021576 | Pdcd6 | 18570 | protein_coding | 1560.115006 | 0.061527724 | 0.037391552 | 1.645498025 | 0.09986715 | 0.287678519 | 0.061527724 |
| ENSMUSG00000024104 | Washc2 | 28006 | protein_coding | 1221.980652 | -0.061544858 | 0.040840937 | -1.5069404 | 0.131825935 | 0.33815124 | 0.061544858 |
| ENSMUSG00000040472 | Rabggta | 56187 | protein_coding | 478.6232362 | 0.061552993 | 0.077713109 | 0.792054179 | 0.428329086 | 0.663421426 | 0.061552993 |
| ENSMUSG00000042293 | Gm5617 | 434402 | protein_coding | 169.9649127 | 0.061557346 | 0.106602201 | 0.577449111 | 0.563636106 | 0.760904026 | 0.061557346 |
| ENSMUSG00000033885 | Pxk | 218699 | protein_coding | 1202.479991 | -0.06156009 | 0.045255693 | -1.360272832 | 0.173743602 | 0.399424529 | 0.06156009 |
| ENSMUSG00000034681 | Rnps1 | 19826 | protein_coding | 876.0279774 | -0.061572381 | 0.064319257 | -0.957293087 | 0.338419344 | 0.582029301 | 0.061572381 |
| ENSMUSG00000007656 | Arpp19 | 59046 | protein_coding | 1563.135271 | 0.061579484 | 0.051699047 | 1.191114488 | 0.233608643 | 0.472995177 | 0.061579484 |
| ENSMUSG00000055531 | Cpsf6 | 432508 | protein_coding | 1423.002125 | 0.061582926 | 0.051672635 | 1.191789927 | 0.233343627 | 0.47275536 | 0.061582926 |
| ENSMUSG00000021076 | Actr10 | 56444 | protein_coding | 1146.824749 | 0.061597581 | 0.056812287 | 1.084229916 | 0.278262871 | 0.521098971 | 0.061597581 |
| ENSMUSG00000098766 | Mir6386 | 102465203 | miRNA | 12.17270658 | -0.061639671 | 0.391480536 | -0.15745271 | 0.874888075 | 0.943745489 | 0.061639671 |
| ENSMUSG00000024121 | Atp6v0c | 11984 | protein_coding | 1237.013173 | 0.061655289 | 0.069378474 | 0.888680389 | 0.374174874 | 0.616119646 | 0.061655289 |
| ENSMUSG00000039068 | Zzz3 | 108946 | protein_coding | 1490.920055 | -0.061679852 | 0.049903333 | -1.235986636 | 0.216463531 | 0.452365863 | 0.061679852 |
| ENSMUSG00000071644 | Eef1g | 67160 | protein_coding | 8648.127777 | 0.061705019 | 0.052485629 | 1.175655525 | 0.239732558 | 0.480269188 | 0.061705019 |
| ENSMUSG00000029427 | Zcchc8 | 70650 | protein_coding | 834.5869524 | -0.061707296 | 0.057937775 | -1.065061554 | 0.286848076 | 0.529236838 | 0.061707296 |
| ENSMUSG00000040434 | Large2 | 228366 | protein_coding | 162.0728891 | 0.061729899 | 0.094001676 | 0.656689351 | 0.511380683 | 0.725473655 | 0.061729899 |
| ENSMUSG00000026432 | Avpr1b | 26361 | protein_coding | 14.59638045 | -0.061731411 | 0.366802377 | -0.168296103 | 0.86635034 | 0.9395852 | 0.061731411 |
| ENSMUSG00000041957 | Pkp2 | 67451 | protein_coding | 1354.513385 | 0.061766744 | 0.05573376 | 1.108246493 | 0.267755377 | 0.510292007 | 0.061766744 |
| ENSMUSG00000057863 | Rpl36 | 54217 | protein_coding | 5268.680207 | -0.061768588 | 0.09555454 | -0.646422327 | 0.518005885 | 0.730398705 | 0.061768588 |
| ENSMUSG00000026434 | Nucks1 | 98415 | protein_coding | 3002.144151 | -0.061800462 | 0.067298105 | -0.918309086 | 0.358457072 | 0.601570652 | 0.061800462 |
| ENSMUSG00000028062 | Lamtor2 | 83409 | protein_coding | 938.2107506 | 0.061802295 | 0.057874172 | 1.067873507 | 0.285577575 | 0.528457779 | 0.061802295 |
| ENSMUSG00000030733 | Sh2b1 | 20399 | protein_coding | 840.8437469 | -0.061824408 | 0.057658729 | -1.072247154 | 0.283609036 | 0.526203742 | 0.061824408 |
| ENSMUSG00000019189 | Rnf145 | 74315 | protein_coding | 1607.082212 | 0.061838575 | 0.069533102 | 0.889340089 | 0.373820333 | 0.615647488 | 0.061838575 |
| ENSMUSG00000027942 | 4933434E20Rik | 99650 | protein_coding | 197.3099817 | -0.06187387 | 0.104263392 | -0.593438106 | 0.552887993 | 0.753394564 | 0.06187387 |
| ENSMUSG00000021891 | Mettl6 | 67011 | protein_coding | 284.3738343 | 0.061874015 | 0.078353739 | 0.789675329 | 0.429717403 | 0.664689483 | 0.061874015 |
| ENSMUSG00000023572 | Ccndbp1 | 17151 | protein_coding | 669.1381918 | -0.061896535 | 0.049615049 | -1.247535489 | 0.212201216 | 0.447576381 | 0.061896535 |
| ENSMUSG00000062444 | Ap3b2 | 11775 | protein_coding | 10.140235 | 0.061909272 | 0.519483391 | 0.11917469 | 0.905136958 | 0.958435249 | 0.061909272 |
| ENSMUSG00000041740 | Rnf10 | 50849 | protein_coding | 2643.414388 | 0.061917049 | 0.078871138 | 0.785040641 | 0.432429741 | 0.666779423 | 0.061917049 |
| ENSMUSG00000022329 | Stk3 | 56274 | protein_coding | 543.3423107 | 0.061939427 | 0.058012033 | 1.067699646 | 0.285656018 | 0.528542291 | 0.061939427 |
| ENSMUSG00000022514 | Il1rap | 16180 | protein_coding | 262.6009899 | 0.0619793 | 0.095627658 | 0.648131524 | 0.516899886 | 0.7296319 | 0.0619793 |
| ENSMUSG00000022248 | Rad1 | 19355 | protein_coding | 136.189199 | 0.061987174 | 0.099603427 | 0.622339767 | 0.533718478 | 0.741516545 | 0.061987174 |
| ENSMUSG00000014859 | E2f4 | 104394 | protein_coding | 763.1198753 | 0.061996447 | 0.06787502 | 0.913391212 | 0.361036826 | 0.604050305 | 0.061996447 |

| ensembl_gene_id | external_gene_name | entrezgene_id | gene_biotype | baseMean | log2FoldChange | lfcSE | stat | pvalue | padj | absLog2FC |
|---|---|---|---|---|---|---|---|---|---|---|
| ENSMUSG00000002812 | Flii | 14248 | protein_coding | 5200.292094 | 0.062001156 | 0.042100132 | 1.47270693 | 0.140830077 | 0.351955161 | 0.062001156 |
| ENSMUSG00000034525 | Ice1 | 218333 | protein_coding | 1019.344573 | -0.062023252 | 0.049981388 | -1.24092695 | 0.214632736 | 0.450343118 | 0.062023252 |
| ENSMUSG00000041064 | Pif1 | 208084 | protein_coding | 122.5459387 | 0.062044851 | 0.149248002 | 0.41571646 | 0.677617495 | 0.836508911 | 0.062044851 |
| ENSMUSG00000046364 | Rpl27a | 26451 | protein_coding | 5748.500865 | -0.062054795 | 0.080837885 | -0.767644959 | 0.442698145 | 0.673781897 | 0.062054795 |
| ENSMUSG00000103711 | Tstd1 | 226654 | protein_coding | 2542.79014 | 0.062069494 | 0.049012434 | 1.266402997 | 0.205368839 | 0.439838414 | 0.062069494 |
| ENSMUSG00000035781 | R3hdm4 | 109284 | protein_coding | 2208.414254 | -0.062071583 | 0.076126505 | -0.815374143 | 0.414858181 | 0.65259636 | 0.062071583 |
| ENSMUSG00000036707 | Cab39 | 12283 | protein_coding | 7020.978477 | 0.062074117 | 0.047424092 | 1.308915236 | 0.190563065 | 0.421276292 | 0.062074117 |
| ENSMUSG00000031760 | Mt3 | 17751 | protein_coding | 47.07017003 | -0.062075935 | 0.206448216 | -0.300685255 | 0.763654513 | 0.888754045 | 0.062075935 |
| ENSMUSG00000028224 | Nbn | 27354 | protein_coding | 625.4384645 | 0.062094568 | 0.061056419 | 1.017003115 | 0.309151947 | 0.552606516 | 0.062094568 |
| ENSMUSG00000079418 | Atg4a | 666468 | protein_coding | 389.2501809 | 0.06209608 | 0.091117531 | 0.681494325 | 0.495558756 | 0.71419804 | 0.06209608 |
| ENSMUSG00000090069 | E430024P14Rik | 100040822 | lncRNA | 23.24972135 | -0.062114083 | 0.284141204 | -0.21860287 | 0.826959417 | 0.923060252 | 0.062114083 |
| ENSMUSG00000039046 | Usp6nl | 98910 | protein_coding | 1154.620689 | -0.0621388 | 0.050074785 | -1.240919969 | 0.214635315 | 0.450343118 | 0.0621388 |
| ENSMUSG00000021930 | Spryd7 | 66674 | protein_coding | 604.7841461 | -0.06214845 | 0.086166097 | -0.721263383 | 0.470747481 | 0.695023769 | 0.06214845 |
| ENSMUSG00000033624 | Pdpr | 319518 | protein_coding | 1302.772979 | -0.062161632 | 0.047466545 | -1.309588312 | 0.190335145 | 0.421061026 | 0.062161632 |
| ENSMUSG00000021257 | Angel1 | 68737 | protein_coding | 143.8238603 | 0.062163046 | 0.110655163 | 0.561772671 | 0.574270914 | 0.768186995 | 0.062163046 |
| ENSMUSG00000025793 | Hgs | 15239 | protein_coding | 1645.097568 | 0.062177632 | 0.080229776 | 0.774994458 | 0.438342934 | 0.671207415 | 0.062177632 |
| ENSMUSG00000015023 | Ddx19a | 13680 | protein_coding | 486.5489612 | 0.062182506 | 0.068909177 | 0.90238353 | 0.366853167 | 0.609230875 | 0.062182506 |
| ENSMUSG00000028893 | Sesn2 | 230784 | protein_coding | 439.9364278 | 0.062190118 | 0.078439727 | 0.792839549 | 0.427871311 | 0.663355008 | 0.062190118 |
| ENSMUSG00000046441 | Cmtr2 | 234728 | protein_coding | 246.2704034 | -0.062217957 | 0.083120517 | -0.748527078 | 0.454142299 | 0.682675952 | 0.062217957 |
| ENSMUSG00000027236 | Eif3j1 | 78655 | protein_coding | 461.8274619 | -0.062254701 | 0.076952808 | -0.808998427 | 0.418516051 | 0.655993003 | 0.062254701 |
| ENSMUSG00000024972 | Lgals12 | 56072 | protein_coding | 454.6117431 | 0.062275988 | 0.086005889 | 0.724089812 | 0.469010599 | 0.693475193 | 0.062275988 |
| ENSMUSG00000071866 | Ppia | 268373 | protein_coding | 9042.055939 | 0.062289844 | 0.067041558 | 0.929122846 | 0.352825426 | 0.595376755 | 0.062289844 |
| ENSMUSG00000024325 | Ring1 | 19763 | protein_coding | 505.438204 | -0.062289888 | 0.072474383 | -0.859474559 | 0.39007875 | 0.629835803 | 0.062289888 |
| ENSMUSG00000056592 | Zfp658 | 210104 | protein_coding | 68.08135601 | -0.062305607 | 0.143723095 | -0.43351145 | 0.66464325 | 0.828854703 | 0.062305607 |
| ENSMUSG00000022881 | Rfc4 | 106344 | protein_coding | 184.2207013 | 0.062312903 | 0.166142835 | 0.375056214 | 0.70761866 | 0.854813943 | 0.062312903 |
| ENSMUSG00000055313 | Pgbd1 | 319207 | protein_coding | 23.34195979 | 0.062320845 | 0.263847038 | 0.236200661 | 0.813276967 | 0.915984616 | 0.062320845 |
| ENSMUSG00000032777 | Gtf3c1 | 233863 | protein_coding | 2642.722689 | 0.06233291 | 0.047318562 | 1.317303564 | 0.187736891 | 0.417546937 | 0.06233291 |
| ENSMUSG00000025743 | Sdc3 | 20970 | protein_coding | 7.04674264 | -0.062341915 | 0.481275175 | -0.129534865 | 0.896934439 | 0.954400543 | 0.062341915 |
| ENSMUSG00000032555 | Topbp1 | 235559 | protein_coding | 1104.652532 | 0.062346294 | 0.109006862 | 0.571948342 | 0.567356972 | 0.762827428 | 0.062346294 |
| ENSMUSG00000047146 | Tet1 | 52463 | protein_coding | 62.34891242 | -0.062348208 | 0.19828209 | -0.314441953 | 0.753185397 | 0.882620324 | 0.062348208 |
| ENSMUSG00000022106 | Rcbtb2 | 105670 | protein_coding | 389.8669135 | 0.062387728 | 0.070412965 | 0.886026151 | 0.375603435 | 0.617215876 | 0.062387728 |
| ENSMUSG00000024501 | Dpysl3 | 22240 | protein_coding | 267.3312313 | 0.062404523 | 0.081002693 | 0.770400599 | 0.441062298 | 0.67268883 | 0.062404523 |
| ENSMUSG00000062101 | Zfp119b | 240120 | protein_coding | 82.34435645 | -0.062410023 | 0.145857519 | -0.427883479 | 0.668735956 | 0.831002845 | 0.062410023 |
| ENSMUSG00000026254 | Eif4e2 | 26987 | protein_coding | 1574.460824 | -0.062410737 | 0.043522949 | -1.433973061 | 0.151579951 | 0.367724228 | 0.062410737 |
| ENSMUSG00000020893 | Per1 | 18626 | protein_coding | 809.1827955 | 0.062444023 | 0.282110312 | 0.22134612 | 0.824822942 | 0.92202835 | 0.062444023 |
| ENSMUSG00000074358 | Ccdc61 | 232933 | protein_coding | 369.1920541 | 0.062470442 | 0.079881834 | 0.782035652 | 0.434193621 | 0.668067439 | 0.062470442 |
| ENSMUSG00000024782 | Ak3 | 56248 | protein_coding | 5121.753956 | 0.062495205 | 0.05863136 | 1.065900657 | 0.286468552 | 0.529013894 | 0.062495205 |
| ENSMUSG00000087651 | 1500009L16Rik | 69784 | protein_coding | 79.95310396 | -0.062496624 | 0.140859431 | -0.443680795 | 0.657273382 | 0.824377651 | 0.062496624 |
| ENSMUSG00000027870 | Hao2 | 56185 | protein_coding | 12498.10664 | -0.062517881 | 0.070475451 | -0.887087344 | 0.375031877 | 0.61652794 | 0.062517881 |

| ensembl_gene_id | external_gene_name | entrezgene_id | gene_biotype | baseMean | log2FoldChange | lfcSE | stat | pvalue | padj | absLog2FC |
|---|---|---|---|---|---|---|---|---|---|---|
| ENSMUSG00000067825 | Pex26 | 74043 | protein_coding | 683.7613275 | 0.062529652 | 0.066290907 | 0.943261377 | 0.345547226 | 0.587831236 | 0.062529652 |
| ENSMUSG00000036989 | Trim3 | 55992 | protein_coding | 150.0834952 | -0.06263372 | 0.105010449 | -0.596452264 | 0.550873135 | 0.752092411 | 0.06263372 |
| ENSMUSG00000024234 | Mtpap | 67440 | protein_coding | 517.4789505 | -0.062637456 | 0.073844586 | -0.848233559 | 0.396307911 | 0.635629805 | 0.062637456 |
| ENSMUSG00000033446 | Lpar6 | 67168 | protein_coding | 315.2092553 | -0.062644056 | 0.116992246 | -0.535454766 | 0.592335428 | 0.780262005 | 0.062644056 |
| ENSMUSG00000066441 | Rdh11 | 17252 | protein_coding | 875.6107739 | -0.062653103 | 0.102891458 | -0.608924239 | 0.542574657 | 0.747864433 | 0.062653103 |
| ENSMUSG00000026127 | Imp4 | 27993 | protein_coding | 951.6699077 | 0.062654481 | 0.048692483 | 1.286738265 | 0.198185531 | 0.431211287 | 0.062654481 |
| ENSMUSG00000001542 | Ell2 | 192657 | protein_coding | 1283.859603 | 0.062656726 | 0.057171689 | 1.095939747 | 0.273105144 | 0.515514161 | 0.062656726 |
| ENSMUSG00000079020 | Slc45a4 | 106068 | protein_coding | 542.9217249 | 0.062661787 | 0.063048261 | 0.993870194 | 0.320286067 | 0.563769199 | 0.062661787 |
| ENSMUSG00000020669 | Sh3yl1 | 24057 | protein_coding | 761.4195722 | 0.062669368 | 0.053928533 | 1.162081816 | 0.245202233 | 0.486452996 | 0.062669368 |
| ENSMUSG00000028614 | Ndc1 | 72787 | protein_coding | 407.2751477 | 0.062685065 | 0.10807116 | 0.580035089 | 0.561890955 | 0.759659123 | 0.062685065 |
| ENSMUSG00000059495 | Arhgef12 | 69632 | protein_coding | 2117.726522 | -0.062707939 | 0.037029408 | -1.69346317 | 0.09036734 | 0.270581895 | 0.062707939 |
| ENSMUSG00000061244 | Exoc5 | 105504 | protein_coding | 1451.905251 | 0.062717664 | 0.045387633 | 1.381822761 | 0.167026129 | 0.390777655 | 0.062717664 |
| ENSMUSG00000030921 | Trim30a | 20128 | protein_coding | 106.2347037 | -0.062722623 | 0.211932191 | -0.29595609 | 0.767263615 | 0.890725386 | 0.062722623 |
| ENSMUSG00000024231 | Cul2 | 71745 | protein_coding | 765.8067495 | 0.062723972 | 0.057411205 | 1.092538839 | 0.274596329 | 0.517029701 | 0.062723972 |
| ENSMUSG00000026035 | Ppil3 | 70225 | protein_coding | 219.2717572 | -0.062776471 | 0.080577189 | -0.779084901 | 0.435929704 | 0.669302737 | 0.062776471 |
| ENSMUSG00000036087 | Slain2 | 75991 | protein_coding | 1847.800273 | 0.062789283 | 0.036237541 | 1.732713665 | 0.083146575 | 0.256583392 | 0.062789283 |
| ENSMUSG00000008763 | Man1a2 | 17156 | protein_coding | 3561.149513 | 0.062790674 | 0.053026246 | 1.184143297 | 0.236356343 | 0.476551574 | 0.062790674 |
| ENSMUSG00000026134 | Prim2 | 19076 | protein_coding | 154.5869009 | 0.06279289 | 0.160000134 | 0.392455235 | 0.694721881 | 0.847294848 | 0.06279289 |
| ENSMUSG00000043866 | Taf10 | 24075 | protein_coding | 1497.357257 | 0.062819379 | 0.082567617 | 0.760823449 | 0.446762527 | 0.677665469 | 0.062819379 |
| ENSMUSG00000077167 | Gm24119 | 115487534 | snoRNA | 41.08343861 | -0.06284744 | 0.217365782 | -0.289132169 | 0.772480237 | 0.893129098 | 0.06284744 |
| ENSMUSG00000024240 | Epc1 | 13831 | protein_coding | 786.3245112 | -0.062848192 | 0.054723915 | -1.148459348 | 0.250778986 | 0.491745579 | 0.062848192 |
| ENSMUSG00000049521 | Cdc42ep1 | 104445 | protein_coding | 500.3876673 | -0.062866229 | 0.084698707 | -0.742233635 | 0.457945793 | 0.685606255 | 0.062866229 |
| ENSMUSG00000020376 | Rnf130 | 59044 | protein_coding | 1539.248013 | -0.062871397 | 0.051829661 | -1.213038953 | 0.225114928 | 0.463123925 | 0.062871397 |
| ENSMUSG00000001173 | Ocrl | 320634 | protein_coding | 532.6929775 | -0.062902602 | 0.066212875 | -0.950005598 | 0.342109408 | 0.584809039 | 0.062902602 |
| ENSMUSG00000075289 | Carns1 | 107239 | protein_coding | 16.65861277 | 0.06290916 | 0.355438019 | 0.176990521 | 0.859515835 | 0.936612161 | 0.06290916 |
| ENSMUSG00000024012 | Mtch1 | 56462 | protein_coding | 3606.124365 | 0.0629558 | 0.059736037 | 1.053899846 | 0.291928773 | 0.534455336 | 0.0629558 |
| ENSMUSG00000054889 | Dsp | 109620 | protein_coding | 18783.24729 | -0.062975099 | 0.042814014 | -1.470899203 | 0.141318378 | 0.352354239 | 0.062975099 |
| ENSMUSG00000043924 | Ncmap | 230822 | protein_coding | 16.03191965 | -0.062987022 | 0.308456522 | -0.204200648 | 0.838196705 | 0.928098115 | 0.062987022 |
| ENSMUSG00000085826 | Gm15638 | 100503378 | lncRNA | 31.28340878 | -0.063004777 | 0.23200099 | -0.271571154 | 0.785951782 | 0.900310808 | 0.063004777 |
| ENSMUSG00000027424 | Mgme1 | 74528 | protein_coding | 237.113921 | 0.063072997 | 0.136470235 | 0.462174018 | 0.643956534 | 0.815430435 | 0.063072997 |
| ENSMUSG00000022358 | Fbxo32 | 67731 | protein_coding | 1332.308396 | 0.063073876 | 0.087563668 | 0.720320163 | 0.471327893 | 0.695753311 | 0.063073876 |
| ENSMUSG00000017688 | Hnf4g | 30942 | protein_coding | 858.505688 | -0.063075835 | 0.048689298 | -1.295476359 | 0.195155952 | 0.427445339 | 0.063075835 |
| ENSMUSG00000026790 | Odf2 | 18286 | protein_coding | 691.6466781 | -0.06310009 | 0.057697521 | -1.093636057 | 0.274114628 | 0.516672115 | 0.06310009 |
| ENSMUSG00000038984 | Tspyl5 | 239364 | protein_coding | 12.0709235 | -0.063111718 | 0.353142885 | -0.178714397 | 0.858161964 | 0.936097283 | 0.063111718 |
| ENSMUSG00000031808 | Slc27a1 | 26457 | protein_coding | 648.4059483 | -0.063124376 | 0.058520534 | -1.078670543 | 0.280734622 | 0.523508209 | 0.063124376 |
| ENSMUSG00000048779 | P2ry6 | 233571 | protein_coding | 347.625561 | -0.06313575 | 0.086477446 | -0.730083433 | 0.465339187 | 0.691455871 | 0.06313575 |
| ENSMUSG00000053329 | Gatd3a | 28295 | protein_coding | 1227.080731 | 0.063164096 | 0.066864925 | 0.944652156 | 0.34483649 | 0.58705564 | 0.063164096 |
| ENSMUSG00000042626 | Shc1 | 20416 | protein_coding | 1700.11327 | 0.063168123 | 0.03720574 | 1.697805849 | 0.089544412 | 0.268966796 | 0.063168123 |
| ENSMUSG00000020459 | Mtif2 | 76784 | protein_coding | 744.8900736 | 0.063189636 | 0.061447135 | 1.028357729 | 0.303781585 | 0.547921576 | 0.063189636 |

| ensembl_gene_id | external_gene_name | entrezgene_id | gene_biotype | baseMean | log2FoldChange | lfcSE | stat | pvalue | padj | absLog2FC |
|---|---|---|---|---|---|---|---|---|---|---|
| ENSMUSG00000020605 | Hs1bp3 | 58240 | protein_coding | 428.0508134 | 0.063198468 | 0.073262327 | 0.862632552 | 0.388339527 | 0.628274245 | 0.063198468 |
| ENSMUSG00000028703 | Lrrc41 | 230654 | protein_coding | 707.5896859 | 0.063225827 | 0.061458892 | 1.028749872 | 0.303597228 | 0.547724856 | 0.063225827 |
| ENSMUSG00000047067 | Dusp28 | 67446 | protein_coding | 169.6435058 | 0.06323959 | 0.12679131 | 0.498769116 | 0.617942047 | 0.797435496 | 0.06323959 |
| ENSMUSG00000097145 | 9230114K14Rik | 414108 | lncRNA | 26.26494894 | -0.063248474 | 0.254599478 | -0.248423426 | 0.80380681 | 0.910706227 | 0.063248474 |
| ENSMUSG00000036403 | Cep135 | 381644 | protein_coding | 203.6173337 | -0.063275651 | 0.107413727 | -0.589083463 | 0.555805286 | 0.755288004 | 0.063275651 |
| ENSMUSG00000021640 | Naip1 | 17940 | protein_coding | 4826.272231 | 0.063294038 | 0.0469451 | 1.348256545 | 0.177575883 | 0.404658227 | 0.063294038 |
| ENSMUSG00000042444 | Mindy2 | 235461 | protein_coding | 699.3595904 | -0.063300739 | 0.060259836 | -1.050463177 | 0.293505212 | 0.536379174 | 0.063300739 |
| ENSMUSG00000046909 | Tefm | 68550 | protein_coding | 164.504113 | 0.063355383 | 0.13648802 | 0.464182739 | 0.642516827 | 0.814130378 | 0.063355383 |
| ENSMUSG00000056692 | Ilrun | 224647 | protein_coding | 3065.185921 | 0.063373245 | 0.055255243 | 1.146918219 | 0.251415422 | 0.492478624 | 0.063373245 |
| ENSMUSG00000035133 | Arhgap5 | 11855 | protein_coding | 6546.504154 | 0.06338212 | 0.046705507 | 1.357058838 | 0.174762516 | 0.400966723 | 0.06338212 |
| ENSMUSG00000026275 | Ppp1r7 | 66385 | protein_coding | 834.7565716 | -0.063410891 | 0.058494883 | -1.084041678 | 0.27834632 | 0.521115754 | 0.063410891 |
| ENSMUSG00000020451 | Limk2 | 16886 | protein_coding | 4403.632039 | -0.063419671 | 0.055453677 | -1.143651338 | 0.252768262 | 0.493868883 | 0.063419671 |
| ENSMUSG00000027803 | Wwtr1 | 97064 | protein_coding | 18.04973198 | -0.063421741 | 0.309103468 | -0.20517965 | 0.837431768 | 0.927834878 | 0.063421741 |
| ENSMUSG00000095677 | Dynlt1f | 100040531 | protein_coding | 124.9823569 | -0.063424574 | 0.114628525 | -0.553305336 | 0.580054348 | 0.772081254 | 0.063424574 |
| ENSMUSG00000029147 | Ppm1g | 14208 | protein_coding | 1538.170676 | 0.063431298 | 0.07092683 | 0.894320223 | 0.371150583 | 0.613223192 | 0.063431298 |
| ENSMUSG00000051495 | Irf2bp2 | 270110 | protein_coding | 4893.013835 | -0.063434638 | 0.075244106 | -0.843051248 | 0.399199783 | 0.638743416 | 0.063434638 |
| ENSMUSG00000051413 | Plagl2 | 54711 | protein_coding | 627.3668377 | -0.06347308 | 0.081626773 | -0.777601245 | 0.436804127 | 0.669932492 | 0.06347308 |
| ENSMUSG00000052117 | D630039A03Rik | 242484 | protein_coding | 1161.470423 | -0.063540445 | 0.108891905 | -0.583518541 | 0.559544279 | 0.75798157 | 0.063540445 |
| ENSMUSG00000024078 | Ttc27 | 74196 | protein_coding | 299.0689246 | -0.063547489 | 0.082629602 | -0.769064443 | 0.441855055 | 0.673006845 | 0.063547489 |
| ENSMUSG00000097357 | Gm16793 | 100504714 | lncRNA | 12.35259504 | 0.06355902 | 0.353327734 | 0.179886871 | 0.857241382 | 0.93581072 | 0.06355902 |
| ENSMUSG00000089824 | Rbm12 | 75710 | protein_coding | 217.7592657 | -0.063566492 | 0.103147543 | -0.61626763 | 0.537717899 | 0.744166872 | 0.063566492 |
| ENSMUSG00000032333 | Stoml1 | 69106 | protein_coding | 281.3910126 | 0.063577506 | 0.086205849 | 0.737508035 | 0.460813459 | 0.687468885 | 0.063577506 |
| ENSMUSG00000027207 | Galk2 | 69976 | protein_coding | 1220.150706 | 0.063581872 | 0.056446293 | 1.126413599 | 0.259990493 | 0.501479092 | 0.063581872 |
| ENSMUSG00000032908 | Sgpp2 | 433323 | protein_coding | 2882.15734 | 0.06358506 | 0.058485254 | 1.087198142 | 0.276949252 | 0.519767321 | 0.06358506 |
| ENSMUSG00000034379 | Wdr5b | 69544 | protein_coding | 103.6328611 | 0.0636153 | 0.138875602 | 0.458073979 | 0.646899293 | 0.817085283 | 0.0636153 |
| ENSMUSG00000040712 | Camta2 | 216874 | protein_coding | 939.9594517 | -0.063615957 | 0.065702526 | -0.968242182 | 0.332923433 | 0.576322837 | 0.063615957 |
| ENSMUSG00000028850 | Gpatch3 | 242691 | protein_coding | 134.862866 | 0.063641224 | 0.106268304 | 0.598873057 | 0.549257539 | 0.751378251 | 0.063641224 |
| ENSMUSG00000085282 | Gm15663 | 327812 | lncRNA | 39.4850946 | 0.063650533 | 0.218295603 | 0.291579546 | 0.770608116 | 0.892082197 | 0.063650533 |
| ENSMUSG00000051224 | Tceanc | 245695 | protein_coding | 148.072243 | -0.063653016 | 0.113502947 | -0.56080496 | 0.574930505 | 0.768361739 | 0.063653016 |
| ENSMUSG00000032497 | Lrrfip2 | 71268 | protein_coding | 2216.143133 | -0.063657739 | 0.042248542 | -1.506744042 | 0.131876278 | 0.33822658 | 0.063657739 |
| ENSMUSG00000038042 | Ptpdc1 | 218232 | protein_coding | 19.92906671 | 0.063661532 | 0.271150677 | 0.234782863 | 0.81437727 | 0.91643725 | 0.063661532 |
| ENSMUSG00000029703 | Lrwd1 | 71735 | protein_coding | 337.3548787 | 0.063686489 | 0.070697045 | 0.900836659 | 0.367675173 | 0.609985966 | 0.063686489 |
| ENSMUSG00000039210 | Gpatch2 | 67769 | protein_coding | 1063.27503 | -0.063705146 | 0.06226736 | -1.023090529 | 0.306265045 | 0.550505615 | 0.063705146 |
| ENSMUSG00000031483 | Erlin2 | 244373 | protein_coding | 3491.732457 | 0.06371924 | 0.030924763 | 2.060460078 | 0.039354579 | 0.15670896 | 0.06371924 |
| ENSMUSG00000020993 | Trappc6b | 78232 | protein_coding | 569.787852 | 0.063743324 | 0.056684376 | 1.124530755 | 0.260787931 | 0.502716586 | 0.063743324 |
| ENSMUSG00000003534 | Ddr1 | 12305 | protein_coding | 2845.444057 | 0.063747477 | 0.08573233 | 0.743564024 | 0.457140275 | 0.685158105 | 0.063747477 |
| ENSMUSG00000029177 | Cenpa | 12615 | protein_coding | 466.8434655 | 0.063755451 | 0.085808619 | 0.742995886 | 0.457484171 | 0.68535506 | 0.063755451 |
| ENSMUSG00000015087 | Rabl6 | 227624 | protein_coding | 1303.383892 | 0.063784096 | 0.052129814 | 1.223562715 | 0.221117232 | 0.457721775 | 0.063784096 |
| ENSMUSG00000019845 | Tube1 | 71924 | protein_coding | 76.84780841 | 0.063797489 | 0.151917076 | 0.419949425 | 0.6745224 | 0.834396161 | 0.063797489 |

| ensembl_gene_id | external_gene_name | entrezgene_id | gene_biotype | baseMean | log2FoldChange | lfcSE | stat | pvalue | padj | absLog2FC |
|---|---|---|---|---|---|---|---|---|---|---|
| ENSMUSG00000007610 | Gtpbp3 | 70359 | protein_coding | 442.7707757 | 0.063798982 | 0.075573714 | 0.84419541 | 0.398560219 | 0.638169126 | 0.063798982 |
| ENSMUSG00000025290 | Rps24 | 20088 | protein_coding | 12648.95154 | -0.063817472 | 0.045471809 | -1.403451376 | 0.160482282 | 0.380826627 | 0.063817472 |
| ENSMUSG00000020218 | Wif1 | 24117 | protein_coding | 24.43853295 | 0.063826571 | 0.277247779 | 0.230214906 | 0.817924779 | 0.918198597 | 0.063826571 |
| ENSMUSG00000025204 | Ndufb8 | 67264 | protein_coding | 4150.622357 | 0.063873269 | 0.072935279 | 0.875752712 | 0.381164482 | 0.6220843 | 0.063873269 |
| ENSMUSG00000056121 | Fez2 | 225020 | protein_coding | 581.8452029 | -0.063892356 | 0.069284562 | -0.922173042 | 0.356438322 | 0.599216181 | 0.063892356 |
| ENSMUSG00000102758 | Naaladl2 | 635702 | protein_coding | 319.3579656 | -0.063905599 | 0.099842877 | -0.640061675 | 0.522132504 | 0.733199598 | 0.063905599 |
| ENSMUSG00000023286 | Ube2j2 | 140499 | protein_coding | 1111.056052 | 0.063937894 | 0.061031736 | 1.047617158 | 0.294815029 | 0.537837327 | 0.063937894 |
| ENSMUSG00000060675 | Plaat3 | 225845 | protein_coding | 933.4307808 | -0.063956566 | 0.079922492 | -0.800232379 | 0.423576173 | 0.659618863 | 0.063956566 |
| ENSMUSG00000021670 | Hmgcr | 15357 | protein_coding | 5760.375956 | 0.063973964 | 0.118355756 | 0.540522631 | 0.588836658 | 0.778426904 | 0.063973964 |
| ENSMUSG00000038902 | Pogz | 229584 | protein_coding | 318.137582 | -0.063987155 | 0.077454718 | -0.826123405 | 0.408734092 | 0.647240398 | 0.063987155 |
| ENSMUSG00000032348 | Gsta4 | 14860 | protein_coding | 631.6131484 | 0.063988554 | 0.094917427 | 0.674149688 | 0.500216152 | 0.71738425 | 0.063988554 |
| ENSMUSG00000041168 | Lonp1 | 74142 | protein_coding | 1963.140078 | -0.064025561 | 0.051579445 | -1.241299927 | 0.214494972 | 0.450294247 | 0.064025561 |
| ENSMUSG00000020361 | Hspa4 | 15525 | protein_coding | 3972.962406 | 0.064028805 | 0.057650822 | 1.110631251 | 0.266727105 | 0.508753167 | 0.064028805 |
| ENSMUSG00000065546 | Mir196a-1 | 387191 | miRNA | 75.48013189 | -0.064093227 | 0.151556159 | -0.422900841 | 0.672367609 | 0.833136568 | 0.064093227 |
| ENSMUSG00000041115 | Iqsec2 | 245666 | protein_coding | 402.618051 | -0.064131887 | 0.107780409 | -0.595023604 | 0.551827693 | 0.752522003 | 0.064131887 |
| ENSMUSG00000027439 | Gzf1 | 74533 | protein_coding | 698.5628459 | -0.064155812 | 0.053000571 | -1.210473984 | 0.226097068 | 0.464315218 | 0.064155812 |
| ENSMUSG00000032905 | Atg12 | 67526 | protein_coding | 1045.422264 | 0.064177434 | 0.048365114 | 1.326936463 | 0.184529701 | 0.413206122 | 0.064177434 |
| ENSMUSG00000024101 | Washc1 | 68767 | protein_coding | 598.1209256 | 0.064202442 | 0.061048284 | 1.051666612 | 0.292952534 | 0.535833676 | 0.064202442 |
| ENSMUSG00000027176 | Cstf3 | 228410 | protein_coding | 441.6528361 | 0.064214451 | 0.073771557 | 0.870449979 | 0.384054544 | 0.624863562 | 0.064214451 |
| ENSMUSG00000112627 | 4933412E12Rik | 71086 | lncRNA | 9.345067986 | -0.06426783 | 0.390163099 | -0.164720422 | 0.869164042 | 0.940691929 | 0.06426783 |
| ENSMUSG00000045294 | Insig1 | 231070 | protein_coding | 4180.642676 | 0.064269303 | 0.119067262 | 0.539773084 | 0.589353532 | 0.778738583 | 0.064269303 |
| ENSMUSG00000029012 | Orc5 | 26429 | protein_coding | 313.3230449 | -0.064281296 | 0.076624499 | -0.838913107 | 0.401518072 | 0.640672993 | 0.064281296 |
| ENSMUSG00000030965 | Abraxas2 | 109359 | protein_coding | 696.2138931 | 0.064332218 | 0.049007926 | 1.312690074 | 0.189287404 | 0.419779698 | 0.064332218 |
| ENSMUSG00000038227 | Hoxa9 | 15405 | protein_coding | 457.9818794 | 0.064333391 | 0.079310031 | 0.811163357 | 0.417271865 | 0.655115204 | 0.064333391 |
| ENSMUSG00000032184 | Lysmd2 | 70082 | protein_coding | 30.04875376 | 0.064337046 | 0.222076352 | 0.289706875 | 0.772040497 | 0.892779083 | 0.064337046 |
| ENSMUSG00000029922 | Mkrn1 | 54484 | protein_coding | 5320.43201 | -0.064346278 | 0.051102945 | -1.259150092 | 0.207976124 | 0.443185909 | 0.064346278 |
| ENSMUSG00000032966 | Fkbp1a | 14225 | protein_coding | 846.3119359 | -0.064347356 | 0.075560287 | -0.851602851 | 0.394434557 | 0.633886008 | 0.064347356 |
| ENSMUSG00000060743 | H3f3a | 15078 | protein_coding | 3852.026632 | -0.064380106 | 0.051354317 | -1.253645463 | 0.209970899 | 0.445201247 | 0.064380106 |
| ENSMUSG00000018040 | Rrp7a | 74778 | protein_coding | 1141.924954 | 0.064392018 | 0.060740092 | 1.060123824 | 0.289088271 | 0.531357026 | 0.064392018 |
| ENSMUSG00000022427 | Tomm22 | 223696 | protein_coding | 2229.554477 | 0.064404177 | 0.052211714 | 1.233519694 | 0.217381934 | 0.453693043 | 0.064404177 |
| ENSMUSG00000038116 | Phf20 | 228829 | protein_coding | 372.1032646 | 0.064405418 | 0.079226836 | 0.812924268 | 0.41626148 | 0.654207378 | 0.064405418 |
| ENSMUSG00000028115 | Bnipl | 171388 | protein_coding | 200.408843 | 0.064406072 | 0.135792186 | 0.474298808 | 0.635286842 | 0.809423077 | 0.064406072 |
| ENSMUSG00000038213 | Tapbpl | 213233 | protein_coding | 876.720879 | -0.064412735 | 0.095862921 | -0.671925431 | 0.501631169 | 0.718582863 | 0.064412735 |
| ENSMUSG00000020448 | Rnf185 | 193670 | protein_coding | 698.3952703 | 0.064416528 | 0.052752593 | 1.221106395 | 0.222045739 | 0.459085065 | 0.064416528 |
| ENSMUSG00000023017 | Asic1 | 11419 | protein_coding | 116.0437531 | -0.064423037 | 0.114644525 | -0.561937319 | 0.574158726 | 0.768186995 | 0.064423037 |
| ENSMUSG00000021771 | Vdac2 | 22334 | protein_coding | 9725.99548 | 0.064429386 | 0.045565007 | 1.414010235 | 0.157358897 | 0.376573221 | 0.064429386 |
| ENSMUSG00000055567 | Unc80 | 329178 | protein_coding | 19.38666035 | 0.064467672 | 0.275783574 | 0.233761826 | 0.815169887 | 0.916810722 | 0.064467672 |
| ENSMUSG00000034163 | Zfc3h1 | 216345 | protein_coding | 1138.320595 | -0.064497735 | 0.063427762 | -1.016869162 | 0.309215675 | 0.552606516 | 0.064497735 |
| ENSMUSG00000051278 | Zgrf1 | 71643 | protein_coding | 86.82887421 | 0.064530906 | 0.191500023 | 0.336975972 | 0.736135014 | 0.87317221 | 0.064530906 |

| ensembl_gene_id | external_gene_name | entrezgene_id | gene_biotype | baseMean | log2FoldChange | lfcSE | stat | pvalue | padj | absLog2FC |
|---|---|---|---|---|---|---|---|---|---|---|
| ENSMUSG00000029599 | Ddx54 | 71990 | protein_coding | 1877.922539 | 0.064543985 | 0.055465347 | 1.163681268 | 0.244553204 | 0.485822803 | 0.064543985 |
| ENSMUSG00000021481 | Zfp346 | 26919 | protein_coding | 163.683366 | 0.064555969 | 0.096197077 | 0.671080356 | 0.50216934 | 0.718950524 | 0.064555969 |
| ENSMUSG00000022680 | Pdxdc1 | 94184 | protein_coding | 14368.00701 | 0.064556478 | 0.042657373 | 1.513372087 | 0.130185171 | 0.336231563 | 0.064556478 |
| ENSMUSG00000015143 | Actn1 | 109711 | protein_coding | 2097.555229 | 0.064592504 | 0.051291488 | 1.259322081 | 0.207914021 | 0.443170686 | 0.064592504 |
| ENSMUSG00000074715 | Ccl28 | 56838 | protein_coding | 3318.517178 | 0.0645963 | 0.077920734 | 0.829000146 | 0.407104325 | 0.645753068 | 0.0645963 |
| ENSMUSG00000001833 | Septin7 | 235072 | protein_coding | 1750.533816 | 0.064611083 | 0.051514612 | 1.254228275 | 0.209759046 | 0.44497561 | 0.064611083 |
| ENSMUSG00000036737 | Oxsr1 | 108737 | protein_coding | 1826.418985 | -0.064633599 | 0.040088148 | -1.612286988 | 0.106899503 | 0.299347968 | 0.064633599 |
| ENSMUSG00000038544 | Inip | 66209 | protein_coding | 353.6698372 | 0.064636858 | 0.066087077 | 0.978055933 | 0.328046661 | 0.571645972 | 0.064636858 |
| ENSMUSG00000054648 | Zfp869 | 66869 | protein_coding | 753.2489668 | -0.06464077 | 0.093693407 | -0.689918028 | 0.490245738 | 0.710266225 | 0.06464077 |
| ENSMUSG00000039477 | Tnrc18 | 231861 | protein_coding | 2418.856606 | -0.064697063 | 0.088257562 | -0.733048383 | 0.463528921 | 0.689542561 | 0.064697063 |
| ENSMUSG00000023147 | Get1 | 71446 | protein_coding | 174.6505929 | 0.064711177 | 0.094707902 | 0.68327115 | 0.494435521 | 0.713491312 | 0.064711177 |
| ENSMUSG00000016554 | Eif3d | 55944 | protein_coding | 1601.958839 | 0.064782901 | 0.050437931 | 1.284408372 | 0.198999105 | 0.432388349 | 0.064782901 |
| ENSMUSG00000035152 | Ap2b1 | 71770 | protein_coding | 4023.890484 | 0.064792468 | 0.048469706 | 1.336762132 | 0.181300294 | 0.408299098 | 0.064792468 |
| ENSMUSG00000019851 | Perp | 64058 | protein_coding | 5288.329306 | 0.064794212 | 0.054104943 | 1.197565484 | 0.23108622 | 0.469953022 | 0.064794212 |
| ENSMUSG00000042015 | Wdr41 | 218460 | protein_coding | 707.0272287 | -0.064794654 | 0.050787082 | -1.275809742 | 0.202022783 | 0.435870434 | 0.064794654 |
| ENSMUSG00000025246 | Tbl1x | 21372 | protein_coding | 1787.976596 | -0.064807642 | 0.051791249 | -1.251324172 | 0.21081623 | 0.445849236 | 0.064807642 |
| ENSMUSG00000042350 | Arel1 | 68497 | protein_coding | 1335.993979 | -0.064821284 | 0.044323746 | -1.462450476 | 0.143617806 | 0.355746355 | 0.064821284 |
| ENSMUSG00000042133 | Ppig | 228005 | protein_coding | 1042.418885 | -0.064851124 | 0.052050605 | -1.245924487 | 0.212792119 | 0.448412192 | 0.064851124 |
| ENSMUSG00000033004 | Mycbp2 | 105689 | protein_coding | 1707.796924 | 0.064865024 | 0.062905004 | 1.031158416 | 0.302466538 | 0.546234922 | 0.064865024 |
| ENSMUSG00000021952 | Xpo4 | 57258 | protein_coding | 586.3717704 | -0.064877292 | 0.064229961 | -1.010078349 | 0.312457754 | 0.556243095 | 0.064877292 |
| ENSMUSG00000015968 | Cacna1d | 12289 | protein_coding | 31.35281265 | 0.064883721 | 0.236234261 | 0.274658387 | 0.783578704 | 0.89909454 | 0.064883721 |
| ENSMUSG00000003033 | Ap1m1 | 11767 | protein_coding | 246.0704197 | -0.064904365 | 0.083331545 | -0.778869097 | 0.43605683 | 0.669302737 | 0.064904365 |
| ENSMUSG00000023882 | Zfp54 | 22712 | protein_coding | 272.0585252 | 0.064954701 | 0.110199498 | 0.589428279 | 0.555574011 | 0.75524772 | 0.064954701 |
| ENSMUSG00000008333 | Snrpb2 | 20639 | protein_coding | 438.4844746 | 0.064960855 | 0.078288874 | 0.829758451 | 0.406675366 | 0.645607651 | 0.064960855 |
| ENSMUSG00000034189 | Hsdl1 | 72552 | protein_coding | 696.2602214 | -0.064972748 | 0.062170595 | -1.045072002 | 0.295989693 | 0.538897588 | 0.064972748 |
| ENSMUSG00000047844 | Bex4 | 406217 | protein_coding | 37.28369475 | -0.064987512 | 0.216744169 | -0.299835111 | 0.764302932 | 0.889251381 | 0.064987512 |
| ENSMUSG00000028622 | Mrpl37 | 56280 | protein_coding | 817.1179889 | 0.065002025 | 0.074945508 | 0.867323838 | 0.385764599 | 0.625831733 | 0.065002025 |
| ENSMUSG00000031750 | Il34 | 76527 | protein_coding | 82.53125553 | 0.065032865 | 0.203329477 | 0.31983983 | 0.749089753 | 0.880475742 | 0.065032865 |
| ENSMUSG00000049823 | Zbtb12 | 193736 | protein_coding | 72.51746741 | 0.065034219 | 0.150181477 | 0.433037551 | 0.66498749 | 0.828976491 | 0.065034219 |
| ENSMUSG00000001280 | Sp1 | 20683 | protein_coding | 2800.777475 | -0.065079014 | 0.051888858 | -1.25420016 | 0.209769262 | 0.44497561 | 0.065079014 |
| ENSMUSG00000019889 | Ptprk | 19272 | protein_coding | 4162.379501 | -0.065089827 | 0.041668643 | -1.562081779 | 0.118268725 | 0.317992618 | 0.065089827 |
| ENSMUSG00000028385 | Snx30 | 209131 | protein_coding | 412.5144334 | -0.065107285 | 0.111546442 | -0.583678726 | 0.559436483 | 0.75796288 | 0.065107285 |
| ENSMUSG00000031696 | Vps35 | 65114 | protein_coding | 2872.789066 | 0.065124942 | 0.051500113 | 1.264559214 | 0.206029382 | 0.440784802 | 0.065124942 |
| ENSMUSG00000042670 | Immp1l | 66541 | protein_coding | 420.192417 | 0.065136794 | 0.066394983 | 0.98104994 | 0.326568115 | 0.57003615 | 0.065136794 |
| ENSMUSG00000020929 | Eftud2 | 20624 | protein_coding | 1120.044374 | 0.065148518 | 0.085773904 | 0.759537754 | 0.447530938 | 0.678448152 | 0.065148518 |
| ENSMUSG00000031631 | Cfap97 | 66756 | protein_coding | 430.5282139 | -0.065174735 | 0.067163659 | -0.970386904 | 0.331853674 | 0.575141012 | 0.065174735 |
| ENSMUSG00000019494 | Cops6 | 26893 | protein_coding | 847.187876 | 0.065192417 | 0.07144487 | 0.912485626 | 0.361513133 | 0.604649053 | 0.065192417 |
| ENSMUSG00000040896 | Kcnd3 | 56543 | protein_coding | 10.39493296 | 0.065195752 | 0.369246763 | 0.176564179 | 0.859850733 | 0.936765956 | 0.065195752 |
| ENSMUSG00000026510 | Trp53bp2 | 209456 | protein_coding | 1121.849477 | 0.065203456 | 0.058289148 | 1.118620839 | 0.263301929 | 0.504916665 | 0.065203456 |

| ensembl_gene_id | external_gene_name | entrezgene_id | gene_biotype | baseMean | log2FoldChange | lfcSE | stat | pvalue | padj | absLog2FC |
|---|---|---|---|---|---|---|---|---|---|---|
| ENSMUSG00000022199 | Slc22a17 | 59049 | protein_coding | 15.78205901 | 0.065221644 | 0.312492792 | 0.208714074 | 0.834671453 | 0.926650089 | 0.065221644 |
| ENSMUSG00000029598 | Plbd2 | 71772 | protein_coding | 515.5759352 | -0.065255464 | 0.07391218 | -0.882878367 | 0.377301993 | 0.61867166 | 0.065255464 |
| ENSMUSG00000061755 | Bod1l | 665775 | protein_coding | 1347.394422 | -0.065274845 | 0.058708994 | -1.111837243 | 0.266208136 | 0.508185669 | 0.065274845 |
| ENSMUSG00000028467 | Gba2 | 230101 | protein_coding | 373.0284813 | -0.065295986 | 0.08427019 | -0.774840855 | 0.438433704 | 0.671251818 | 0.065295986 |
| ENSMUSG00000083649 | Rasl2-9 | 19428 | protein_coding | 10.34049317 | 0.065326904 | 0.375999312 | 0.173742084 | 0.862068162 | 0.937787345 | 0.065326904 |
| ENSMUSG00000014601 | Strip1 | 229707 | protein_coding | 857.9422366 | 0.065343876 | 0.050527462 | 1.293234891 | 0.195929832 | 0.428500189 | 0.065343876 |
| ENSMUSG00000032281 | Acsbg1 | 94180 | protein_coding | 8.043578474 | 0.06536155 | 0.460704003 | 0.141873197 | 0.887180166 | 0.95019842 | 0.06536155 |
| ENSMUSG00000074529 | Zfp972 | 78703 | protein_coding | 12.66736771 | -0.065390621 | 0.3685498 | -0.177426824 | 0.859173139 | 0.936398271 | 0.065390621 |
| ENSMUSG00000024414 | Mrpl27 | 94064 | protein_coding | 626.3995107 | 0.065412555 | 0.069463338 | 0.94168458 | 0.346354152 | 0.588629676 | 0.065412555 |
| ENSMUSG00000034108 | Ccs | 12460 | protein_coding | 960.7919621 | -0.065430163 | 0.0563481 | -1.161177799 | 0.245569601 | 0.486881861 | 0.065430163 |
| ENSMUSG00000007216 | Zfp775 | 243372 | protein_coding | 187.8478532 | -0.065450895 | 0.136247027 | -0.480384019 | 0.630954355 | 0.806163912 | 0.065450895 |
| ENSMUSG00000060216 | Arrb2 | 216869 | protein_coding | 65.67551295 | 0.065461979 | 0.155454214 | 0.421101345 | 0.67368108 | 0.834153989 | 0.065461979 |
| ENSMUSG00000035517 | Tdrd7 | 100121 | protein_coding | 1499.490553 | 0.065464452 | 0.054412847 | 1.203106531 | 0.228935098 | 0.468116689 | 0.065464452 |
| ENSMUSG00000018209 | Stk4 | 58231 | protein_coding | 1444.8088 | -0.065472087 | 0.043972945 | -1.488917493 | 0.136509097 | 0.345547355 | 0.065472087 |
| ENSMUSG00000055254 | Ntrk2 | 18212 | protein_coding | 168.3765779 | 0.065472394 | 0.136551464 | 0.479470464 | 0.631603975 | 0.806746273 | 0.065472394 |
| ENSMUSG00000046111 | Cep295 | 319675 | protein_coding | 311.9980765 | -0.06547475 | 0.109323475 | -0.598908417 | 0.549233958 | 0.751378251 | 0.06547475 |
| ENSMUSG00000058291 | Zfp68 | 24135 | protein_coding | 727.862817 | -0.065482856 | 0.063712124 | -1.0277927 | 0.304047351 | 0.548046454 | 0.065482856 |
| ENSMUSG00000038848 | Ythdf1 | 228994 | protein_coding | 789.7711906 | -0.065508591 | 0.047318342 | -1.384422767 | 0.166229042 | 0.389715871 | 0.065508591 |
| ENSMUSG00000066829 | Zfp810 | 235050 | protein_coding | 295.5486683 | -0.065512783 | 0.11714554 | -0.559242657 | 0.575996124 | 0.768874353 | 0.065512783 |
| ENSMUSG00000036985 | Zdhhc9 | 208884 | protein_coding | 618.0682612 | 0.065515618 | 0.081451865 | 0.804347673 | 0.421196212 | 0.657986673 | 0.065515618 |
| ENSMUSG00000028741 | Mrto4 | 69902 | protein_coding | 277.7191806 | -0.065527087 | 0.089195167 | -0.734648401 | 0.462553653 | 0.688854258 | 0.065527087 |
| ENSMUSG00000031527 | Eri1 | 67276 | protein_coding | 659.3776838 | 0.065530259 | 0.078628912 | 0.833411746 | 0.404612553 | 0.644017631 | 0.065530259 |
| ENSMUSG00000069895 | Atxn1l | 52335 | protein_coding | 907.3617514 | -0.065533374 | 0.052467263 | -1.249033601 | 0.211652784 | 0.44700425 | 0.065533374 |
| ENSMUSG00000050945 | Zfp438 | 240186 | protein_coding | 79.61078781 | 0.06555289 | 0.126188083 | 0.519485581 | 0.603422165 | 0.788158197 | 0.06555289 |
| ENSMUSG00000057649 | Brd9 | 105246 | protein_coding | 758.1672499 | -0.06560713 | 0.084935816 | -0.772431855 | 0.439858693 | 0.671740242 | 0.06560713 |
| ENSMUSG00000025702 | Marchf8 | 71779 | protein_coding | 3214.32669 | 0.065619257 | 0.039140939 | 1.676486516 | 0.093642935 | 0.276589673 | 0.065619257 |
| ENSMUSG00000051502 | Ufsp1 | 70240 | protein_coding | 95.20019607 | 0.06562736 | 0.128941976 | 0.50896816 | 0.610774542 | 0.792689605 | 0.06562736 |
| ENSMUSG00000020921 | Tmem101 | 76547 | protein_coding | 209.1935388 | 0.065640372 | 0.101230572 | 0.648424387 | 0.516710502 | 0.729428419 | 0.065640372 |
| ENSMUSG00000024096 | Ralbp1 | 19765 | protein_coding | 2346.362678 | 0.065662549 | 0.055623142 | 1.180489756 | 0.237805482 | 0.477892712 | 0.065662549 |
| ENSMUSG00000000308 | Ckmt1 | 12716 | protein_coding | 45544.62588 | 0.065674125 | 0.096330632 | 0.681757436 | 0.495392342 | 0.714113831 | 0.065674125 |
| ENSMUSG00000021585 | Cast | 12380 | protein_coding | 4203.303572 | 0.06572192 | 0.042126028 | 1.560126198 | 0.118730062 | 0.318807809 | 0.06572192 |
| ENSMUSG00000002031 | Ift46 | 76568 | protein_coding | 656.6187124 | -0.065726371 | 0.051442576 | -1.277664837 | 0.201367629 | 0.43512903 | 0.065726371 |
| ENSMUSG00000002257 | Def6 | 23853 | protein_coding | 27.04788124 | 0.065726463 | 0.28628663 | 0.229582721 | 0.818416035 | 0.918354815 | 0.065726463 |
| ENSMUSG00000020456 | Ogdh | 18293 | protein_coding | 8189.91885 | 0.065737314 | 0.050901793 | 1.291453801 | 0.196546365 | 0.429092085 | 0.065737314 |
| ENSMUSG00000057572 | Zbtb8os | 67106 | protein_coding | 352.9866963 | -0.065746536 | 0.071734504 | -0.916525972 | 0.359391092 | 0.602834501 | 0.065746536 |
| ENSMUSG00000046949 | Nqo2 | 18105 | protein_coding | 767.4612356 | 0.065760028 | 0.065572837 | 1.002854708 | 0.315930968 | 0.560076808 | 0.065760028 |
| ENSMUSG00000029474 | Rnf34 | 80751 | protein_coding | 550.098211 | -0.065798873 | 0.060803637 | -1.082153568 | 0.279184297 | 0.522078519 | 0.065798873 |
| ENSMUSG00000014850 | Msh3 | 17686 | protein_coding | 649.7553618 | -0.065806181 | 0.066828673 | -0.984699788 | 0.324771557 | 0.568500043 | 0.065806181 |
| ENSMUSG00000006304 | Arpc2 | 76709 | protein_coding | 4874.363566 | 0.065821321 | 0.030950569 | 2.126659451 | 0.033448389 | 0.140437951 | 0.065821321 |

| ensembl_gene_id | external_gene_name | entrezgene_id | gene_biotype | baseMean | log2FoldChange | lfcSE | stat | pvalue | padj | absLog2FC |
|---|---|---|---|---|---|---|---|---|---|---|
| ENSMUSG00000029178 | Klf3 | 16599 | protein_coding | 4881.398087 | -0.065828869 | 0.056148546 | -1.172405599 | 0.241034252 | 0.481643597 | 0.065828869 |
| ENSMUSG00000042572 | Ube2q1 | 70093 | protein_coding | 1661.38685 | -0.065865824 | 0.045915218 | -1.434509651 | 0.151426877 | 0.3675902 | 0.065865824 |
| ENSMUSG00000020085 | Aifm2 | 71361 | protein_coding | 272.1373992 | -0.065879914 | 0.089789186 | -0.733717685 | 0.463120818 | 0.689357631 | 0.065879914 |
| ENSMUSG00000037938 | Chchd5 | 66170 | protein_coding | 280.9923018 | 0.065897003 | 0.090096245 | 0.731406763 | 0.464530735 | 0.690725542 | 0.065897003 |
| ENSMUSG00000042709 | Atpaf2 | 246782 | protein_coding | 642.3990789 | 0.065912146 | 0.06213548 | 1.060781146 | 0.288789373 | 0.531170873 | 0.065912146 |
| ENSMUSG00000025980 | Hspd1 | 15510 | protein_coding | 5450.164036 | 0.065966786 | 0.097683337 | 0.675312571 | 0.499477199 | 0.716579378 | 0.065966786 |
| ENSMUSG00000031537 | Ikbkb | 16150 | protein_coding | 962.3159913 | -0.065968838 | 0.05242791 | -1.258277092 | 0.208291563 | 0.443565039 | 0.065968838 |
| ENSMUSG00000039457 | Ppl | 19041 | protein_coding | 403.181207 | -0.065981621 | 0.063178421 | -1.044369591 | 0.296314427 | 0.539082898 | 0.065981621 |
| ENSMUSG00000071415 | Rpl23 | 65019 | protein_coding | 10233.36841 | -0.065981742 | 0.045214458 | -1.459306267 | 0.144480831 | 0.357032789 | 0.065981742 |
| ENSMUSG00000026787 | Gad2 | 14417 | protein_coding | 49.03848615 | 0.065986314 | 0.187702856 | 0.351546673 | 0.725178264 | 0.865987203 | 0.065986314 |
| ENSMUSG00000063362 | Alg11 | 207958 | protein_coding | 700.438172 | 0.066008625 | 0.051918788 | 1.271382254 | 0.203592693 | 0.437895353 | 0.066008625 |
| ENSMUSG00000034850 | Tmem127 | 69470 | protein_coding | 2838.129532 | -0.066062044 | 0.040382122 | -1.635923033 | 0.101855697 | 0.291161534 | 0.066062044 |
| ENSMUSG00000039735 | Fnbp1l | 214459 | protein_coding | 1427.065927 | -0.0660674 | 0.067893673 | -0.973100993 | 0.330503111 | 0.574170383 | 0.0660674 |
| ENSMUSG00000038900 | Rpl12 | 269261 | protein_coding | 11072.38035 | -0.066073871 | 0.043614361 | -1.51495674 | 0.12978336 | 0.335814568 | 0.066073871 |
| ENSMUSG00000059183 | Mtfmt | 69606 | protein_coding | 307.9134649 | -0.066077451 | 0.07215417 | -0.915781457 | 0.359781531 | 0.603018132 | 0.066077451 |
| ENSMUSG00000053617 | Sh3pxd2a | 14218 | protein_coding | 563.1854121 | 0.066094932 | 0.077280386 | 0.855261411 | 0.392406445 | 0.632120905 | 0.066094932 |
| ENSMUSG00000042814 | Mcts2 | 66405 | protein_coding | 127.1101068 | 0.06609701 | 0.135073533 | 0.489340946 | 0.624600337 | 0.801920996 | 0.06609701 |
| ENSMUSG00000006058 | Snf8 | 27681 | protein_coding | 981.80885 | 0.066099071 | 0.068656579 | 0.962749268 | 0.335673366 | 0.579221524 | 0.066099071 |
| ENSMUSG00000027752 | Exosc8 | 69639 | protein_coding | 180.1950793 | 0.066132144 | 0.105577985 | 0.626381951 | 0.531064445 | 0.73971984 | 0.066132144 |
| ENSMUSG00000031835 | Mbtps1 | 56453 | protein_coding | 3238.03473 | 0.06613986 | 0.034872889 | 1.89659825 | 0.057880982 | 0.203825108 | 0.06613986 |
| ENSMUSG00000079003 | Samd1 | 666704 | protein_coding | 407.747908 | -0.06617706 | 0.086029155 | -0.769239921 | 0.441750896 | 0.672975316 | 0.06617706 |
| ENSMUSG00000049265 | Kcnk3 | 16527 | protein_coding | 17.11612835 | 0.066187515 | 0.305940045 | 0.216341456 | 0.828721597 | 0.923812345 | 0.066187515 |
| ENSMUSG00000073139 | Tmem185a | 236848 | protein_coding | 261.3595698 | 0.066199296 | 0.084752175 | 0.781092594 | 0.434748038 | 0.668304413 | 0.066199296 |
| ENSMUSG00000028295 | Smim8 | 66291 | protein_coding | 260.1431107 | 0.066208692 | 0.080078837 | 0.826793869 | 0.408353906 | 0.646860933 | 0.066208692 |
| ENSMUSG00000028653 | Trit1 | 66966 | protein_coding | 252.9279367 | -0.066214958 | 0.082290247 | -0.804651343 | 0.421020905 | 0.657926757 | 0.066214958 |
| ENSMUSG00000068547 | Clca4a | 99663 | protein_coding | 1108.165054 | -0.066249501 | 0.169237616 | -0.391458485 | 0.695458369 | 0.84768023 | 0.066249501 |
| ENSMUSG00000034781 | Gna11 | 14672 | protein_coding | 8179.563994 | 0.066284839 | 0.056246861 | 1.178462904 | 0.238612113 | 0.478916931 | 0.066284839 |
| ENSMUSG00000028841 | Cnksr1 | 194231 | protein_coding | 898.3284052 | 0.066285152 | 0.053988787 | 1.227757745 | 0.219537922 | 0.455719922 | 0.066285152 |
| ENSMUSG00000027546 | Atp9a | 11981 | protein_coding | 7863.933739 | -0.066293593 | 0.051431805 | -1.288961029 | 0.197411635 | 0.429875431 | 0.066293593 |
| ENSMUSG00000016757 | Ttll12 | 223723 | protein_coding | 574.5943106 | 0.066294999 | 0.077436227 | 0.856123831 | 0.391929288 | 0.631748489 | 0.066294999 |
| ENSMUSG00000001376 | Vps50 | 73288 | protein_coding | 1002.060515 | 0.066349335 | 0.06604624 | 1.004589133 | 0.315094732 | 0.559370353 | 0.066349335 |
| ENSMUSG00000002981 | Clptm1 | 56457 | protein_coding | 4688.823747 | -0.066371263 | 0.074778757 | -0.88756841 | 0.374772953 | 0.616422248 | 0.066371263 |
| ENSMUSG00000059674 | Cdh24 | 239096 | protein_coding | 42.9478366 | 0.066388703 | 0.228634922 | 0.290369914 | 0.771533258 | 0.892439856 | 0.066388703 |
| ENSMUSG00000034201 | Gas2l1 | 78926 | protein_coding | 1048.434102 | -0.066403082 | 0.101789661 | -0.652355858 | 0.514171633 | 0.727308651 | 0.066403082 |
| ENSMUSG00000034544 | Rsrc1 | 66880 | protein_coding | 399.3219345 | -0.0664263 | 0.068756686 | -0.966106775 | 0.333990755 | 0.577368303 | 0.0664263 |
| ENSMUSG00000047037 | Nipa1 | 233280 | protein_coding | 87.87019073 | -0.066433457 | 0.156633443 | -0.424133291 | 0.671468608 | 0.832445548 | 0.066433457 |
| ENSMUSG00000023074 | Mospd1 | 70380 | protein_coding | 470.5711509 | -0.066438556 | 0.066805638 | -0.994505236 | 0.319976958 | 0.563409586 | 0.066438556 |
| ENSMUSG00000026688 | Mgst3 | 66447 | protein_coding | 11790.48778 | 0.066445421 | 0.078276106 | 0.848859569 | 0.395959439 | 0.635262971 | 0.066445421 |
| ENSMUSG00000003032 | Klf4 | 16600 | protein_coding | 13173.6835 | 0.066453915 | 0.072429904 | 0.917492802 | 0.358884463 | 0.602198495 | 0.066453915 |

| ensembl_gene_id | external_gene_name | entrezgene_id | gene_biotype | baseMean | log2FoldChange | lfcSE | stat | pvalue | padj | absLog2FC |
|---|---|---|---|---|---|---|---|---|---|---|
| ENSMUSG00000092260 | Zfp963 | 620419 | protein_coding | 52.85618206 | -0.066466565 | 0.158126603 | -0.420337653 | 0.674238807 | 0.834301331 | 0.066466565 |
| ENSMUSG00000021464 | Ror2 | 26564 | protein_coding | 136.8178137 | 0.066474987 | 0.117940733 | 0.563630439 | 0.573005667 | 0.767564688 | 0.066474987 |
| ENSMUSG00000043155 | Hpdl | 242642 | protein_coding | 164.0287863 | 0.06649976 | 0.124564353 | 0.533858671 | 0.593439318 | 0.780938056 | 0.06649976 |
| ENSMUSG00000024217 | Snrpc | 20630 | protein_coding | 493.4819398 | 0.066534671 | 0.096085893 | 0.692449937 | 0.488654815 | 0.708970987 | 0.066534671 |
| ENSMUSG00000051786 | Tubgcp6 | 328580 | protein_coding | 467.737269 | -0.066544628 | 0.069649395 | -0.955422911 | 0.339363872 | 0.582473517 | 0.066544628 |
| ENSMUSG00000036114 | Rpp25l | 69961 | protein_coding | 410.5508269 | 0.066550515 | 0.097496103 | 0.68259667 | 0.494861739 | 0.713639923 | 0.066550515 |
| ENSMUSG00000045328 | Cenpe | 229841 | protein_coding | 745.2533332 | -0.066552637 | 0.120107957 | -0.554106811 | 0.579505749 | 0.771707437 | 0.066552637 |
| ENSMUSG00000039670 | Oxld1 | 66431 | protein_coding | 229.1961652 | 0.066555763 | 0.103151425 | 0.645223887 | 0.518782108 | 0.730713836 | 0.066555763 |
| ENSMUSG00000035629 | Rubcn | 100502698 | protein_coding | 831.5292042 | 0.066604101 | 0.062342911 | 1.068350834 | 0.285362286 | 0.528301869 | 0.066604101 |
| ENSMUSG00000057409 | Zfp53 | 24132 | protein_coding | 321.7369042 | 0.066622024 | 0.103161679 | 0.645802057 | 0.518407556 | 0.730599032 | 0.066622024 |
| ENSMUSG00000020289 | Nprl3 | 17168 | protein_coding | 254.187853 | 0.066649317 | 0.092003234 | 0.724423635 | 0.468805694 | 0.693362933 | 0.066649317 |
| ENSMUSG00000029310 | Nudt9 | 74167 | protein_coding | 760.5708645 | 0.066651858 | 0.052815767 | 1.261968959 | 0.206959956 | 0.442013404 | 0.066651858 |
| ENSMUSG00000035578 | Iqcg | 69707 | protein_coding | 37.25943763 | 0.066692854 | 0.199381812 | 0.334498181 | 0.738003663 | 0.874596535 | 0.066692854 |
| ENSMUSG00000021240 | Abcd4 | 19300 | protein_coding | 271.0986349 | -0.066738956 | 0.081408167 | -0.819806635 | 0.412326348 | 0.649970343 | 0.066738956 |
| ENSMUSG00000022962 | Gart | 14450 | protein_coding | 923.0363905 | -0.066775408 | 0.077412494 | -0.862592122 | 0.388361763 | 0.628274245 | 0.066775408 |
| ENSMUSG00000040688 | Tbl3 | 213773 | protein_coding | 715.7322926 | -0.066879979 | 0.073526996 | -0.909597596 | 0.363034767 | 0.605769124 | 0.066879979 |
| ENSMUSG00000035509 | Fbxl21 | 213311 | protein_coding | 11.86758875 | -0.06688675 | 0.351033913 | -0.190542133 | 0.848884331 | 0.931678839 | 0.06688675 |
| ENSMUSG00000032193 | Ldlr | 16835 | protein_coding | 8697.44174 | -0.06691988 | 0.099750297 | -0.670873994 | 0.502300805 | 0.718956777 | 0.06691988 |
| ENSMUSG00000032187 | Smarca4 | 20586 | protein_coding | 3144.754154 | 0.066920064 | 0.06878033 | 0.972953516 | 0.330576405 | 0.57417401 | 0.066920064 |
| ENSMUSG00000019254 | Ppp1r12c | 232807 | protein_coding | 665.003818 | -0.066926713 | 0.053906271 | -1.241538547 | 0.214406867 | 0.450167889 | 0.066926713 |
| ENSMUSG00000034274 | Thoc5 | 107829 | protein_coding | 689.8523629 | 0.06692675 | 0.065586037 | 1.020442049 | 0.30751886 | 0.551334926 | 0.06692675 |
| ENSMUSG00000026319 | Relch | 227446 | protein_coding | 2457.479036 | -0.066938564 | 0.040968487 | -1.633903741 | 0.102279064 | 0.291835539 | 0.066938564 |
| ENSMUSG00000020186 | Csrp2 | 13008 | protein_coding | 4514.424571 | -0.066946959 | 0.058190482 | -1.150479557 | 0.249946409 | 0.490972685 | 0.066946959 |
| ENSMUSG00000024983 | Vti1a | 53611 | protein_coding | 661.1393793 | 0.066964333 | 0.056090871 | 1.193854401 | 0.232534927 | 0.47182823 | 0.066964333 |
| ENSMUSG00000058997 | Vwa8 | 219189 | protein_coding | 4296.606379 | 0.066968971 | 0.042334493 | 1.581900858 | 0.113672205 | 0.310710449 | 0.066968971 |
| ENSMUSG00000043501 | Lgals2 | 107753 | protein_coding | 179.3752102 | 0.066977321 | 0.376383937 | 0.177949466 | 0.858762663 | 0.93626727 | 0.066977321 |
| ENSMUSG00000031378 | Abcd1 | 11666 | protein_coding | 510.0160125 | 0.066992129 | 0.079606007 | 0.841546152 | 0.400042041 | 0.639482053 | 0.066992129 |
| ENSMUSG00000048826 | Dact2 | 240025 | protein_coding | 800.1593724 | 0.067006694 | 0.066374712 | 1.009521433 | 0.312724628 | 0.556656741 | 0.067006694 |
| ENSMUSG00000023883 | Phf10 | 72057 | protein_coding | 610.4991305 | 0.067011707 | 0.063798466 | 1.050365485 | 0.293550108 | 0.536379174 | 0.067011707 |
| ENSMUSG00000022945 | Chaf1b | 110749 | protein_coding | 205.0598509 | -0.067041923 | 0.176037728 | -0.380838376 | 0.703323181 | 0.852626696 | 0.067041923 |
| ENSMUSG00000025354 | Dnajc14 | 74330 | protein_coding | 2155.868063 | -0.067049994 | 0.041593524 | -1.612029648 | 0.106955488 | 0.299347968 | 0.067049994 |
| ENSMUSG00000047881 | Rell1 | 100532 | protein_coding | 1184.515549 | -0.067068947 | 0.060539374 | -1.107856636 | 0.267923736 | 0.510492213 | 0.067068947 |
| ENSMUSG00000020267 | Hint1 | 15254 | protein_coding | 7116.418938 | 0.067119826 | 0.070611335 | 0.950553132 | 0.34183127 | 0.584451398 | 0.067119826 |
| ENSMUSG00000034327 | Kctd9 | 105440 | protein_coding | 377.8408301 | 0.067163797 | 0.068435663 | 0.981415147 | 0.32638806 | 0.569880807 | 0.067163797 |
| ENSMUSG00000073413 | Ly6g6d | 114654 | protein_coding | 63.72810175 | 0.067167311 | 0.154448243 | 0.434885561 | 0.663645495 | 0.828183154 | 0.067167311 |
| ENSMUSG00000032238 | Rora | 19883 | protein_coding | 42.48214454 | -0.067191648 | 0.197276647 | -0.340596056 | 0.733407699 | 0.871475535 | 0.067191648 |
| ENSMUSG00000028617 | Lrrc42 | 77809 | protein_coding | 1324.542807 | 0.067195131 | 0.064939687 | 1.034731372 | 0.30079438 | 0.544312273 | 0.067195131 |
| ENSMUSG00000045284 | Dcaf12l1 | 245404 | protein_coding | 45.24644554 | -0.067199252 | 0.17414771 | -0.385875024 | 0.699589229 | 0.850145091 | 0.067199252 |
| ENSMUSG00000040043 | Rbms2 | 56516 | protein_coding | 241.3062505 | -0.067230239 | 0.103121195 | -0.651953639 | 0.51443108 | 0.727325341 | 0.067230239 |

| ensembl_gene_id | external_gene_name | entrezgene_id | gene_biotype | baseMean | log2FoldChange | lfcSE | stat | pvalue | padj | absLog2FC |
|---|---|---|---|---|---|---|---|---|---|---|
| ENSMUSG00000038987 | Cfap157 | 227736 | protein_coding | 92.24251885 | 0.067257365 | 0.197011019 | 0.341388848 | 0.732810869 | 0.871241669 | 0.067257365 |
| ENSMUSG00000041453 | Rpl21 | 19933 | protein_coding | 9570.242765 | -0.067269984 | 0.04335711 | -1.551532932 | 0.120774022 | 0.321618667 | 0.067269984 |
| ENSMUSG00000020917 | Acly | 104112 | protein_coding | 11875.36971 | -0.067272685 | 0.112078555 | -0.600227983 | 0.548354307 | 0.750864769 | 0.067272685 |
| ENSMUSG00000030314 | Atg7 | 74244 | protein_coding | 886.612739 | -0.067289815 | 0.067045946 | -1.003637351 | 0.315553444 | 0.559838328 | 0.067289815 |
| ENSMUSG00000015437 | Gzmb | 14939 | protein_coding | 16.65674048 | -0.067290805 | 0.475229564 | -0.14159642 | 0.887398794 | 0.95027094 | 0.067290805 |
| ENSMUSG00000066415 | Msl2 | 77853 | protein_coding | 1676.821914 | -0.067359191 | 0.055702891 | -1.209258444 | 0.226563571 | 0.465036363 | 0.067359191 |
| ENSMUSG00000027377 | Mall | 228576 | protein_coding | 2205.153007 | 0.06736362 | 0.099463515 | 0.677269648 | 0.498234885 | 0.715879721 | 0.06736362 |
| ENSMUSG00000066306 | Numa1 | 101706 | protein_coding | 5061.713595 | -0.067392855 | 0.048277229 | -1.395955334 | 0.162727946 | 0.383952244 | 0.067392855 |
| ENSMUSG00000043131 | Mob1a | 232157 | protein_coding | 3353.137625 | -0.067408519 | 0.035485521 | -1.899606292 | 0.057484806 | 0.202990467 | 0.067408519 |
| ENSMUSG00000014313 | Cox6c | 12864 | protein_coding | 7482.751568 | 0.067433411 | 0.043430502 | 1.552674 | 0.120501037 | 0.321263055 | 0.067433411 |
| ENSMUSG00000027519 | Rab22a | 19334 | protein_coding | 1457.890299 | -0.067444875 | 0.047156828 | -1.430225015 | 0.152652448 | 0.369310836 | 0.067444875 |
| ENSMUSG00000069089 | Cdk7 | 12572 | protein_coding | 523.6591878 | 0.067449185 | 0.065090616 | 1.036235165 | 0.300092445 | 0.543683373 | 0.067449185 |
| ENSMUSG00000107002 | 0610012G03Rik | 106264 | protein_coding | 905.3325972 | 0.067449961 | 0.062424678 | 1.08050155 | 0.279918897 | 0.523118689 | 0.067449961 |
| ENSMUSG00000030272 | Camk1 | 52163 | protein_coding | 408.5216485 | -0.067481551 | 0.085827012 | -0.786250726 | 0.431720613 | 0.666254386 | 0.067481551 |
| ENSMUSG00000057388 | Mrpl18 | 67681 | protein_coding | 931.6133874 | 0.067522396 | 0.068856715 | 0.980621803 | 0.326779278 | 0.570343011 | 0.067522396 |
| ENSMUSG00000021417 | Eci2 | 23986 | protein_coding | 1001.66927 | 0.067528164 | 0.055477998 | 1.217206215 | 0.223525766 | 0.461089796 | 0.067528164 |
| ENSMUSG00000058192 | Zfp846 | 244721 | protein_coding | 184.6933917 | 0.067533946 | 0.092954686 | 0.726525454 | 0.467516709 | 0.692926016 | 0.067533946 |
| ENSMUSG00000061759 | Armt1 | 73419 | protein_coding | 298.4441289 | 0.06757031 | 0.076553851 | 0.88265071 | 0.377425021 | 0.61867166 | 0.06757031 |
| ENSMUSG00000038025 | Phf2 | 18676 | protein_coding | 616.6892996 | -0.067574252 | 0.071422214 | -0.946123741 | 0.344085476 | 0.586511041 | 0.067574252 |
| ENSMUSG00000047502 | Mroh7 | 381538 | protein_coding | 8.65178968 | 0.067595378 | 0.429713609 | 0.157303321 | 0.875005803 | 0.943745489 | 0.067595378 |
| ENSMUSG00000043962 | Thrap3 | 230753 | protein_coding | 2608.140387 | 0.067602351 | 0.03997233 | 1.691228701 | 0.090793131 | 0.271210133 | 0.067602351 |
| ENSMUSG00000037805 | Rpl10a | 19896 | protein_coding | 7623.640764 | -0.067611243 | 0.069887833 | -0.967425085 | 0.333331575 | 0.576562634 | 0.067611243 |
| ENSMUSG00000032435 | Dync1li1 | 235661 | protein_coding | 1284.46324 | 0.067626533 | 0.054729125 | 1.235658955 | 0.21658536 | 0.452561817 | 0.067626533 |
| ENSMUSG00000037606 | Osbpl5 | 79196 | protein_coding | 908.697353 | -0.067635201 | 0.077970856 | -0.867442065 | 0.385699842 | 0.625831733 | 0.067635201 |
| ENSMUSG00000019590 | Cyb561 | 13056 | protein_coding | 2385.594906 | 0.067669742 | 0.056222687 | 1.203602062 | 0.228743422 | 0.467843395 | 0.067669742 |
| ENSMUSG00000051235 | Gen1 | 209334 | protein_coding | 137.094883 | 0.067684122 | 0.125513874 | 0.5392561 | 0.589710156 | 0.779018323 | 0.067684122 |
| ENSMUSG00000004591 | Pkn2 | 109333 | protein_coding | 2962.904411 | -0.067698469 | 0.042433313 | -1.595408544 | 0.110620907 | 0.305372024 | 0.067698469 |
| ENSMUSG00000038253 | Hoxa5 | 15402 | protein_coding | 116.2703775 | -0.067712549 | 0.122141353 | -0.554378572 | 0.579319788 | 0.771609416 | 0.067712549 |
| ENSMUSG00000059811 | Atl2 | 56298 | protein_coding | 1759.269137 | 0.067722106 | 0.041041141 | 1.650102882 | 0.098921895 | 0.286052251 | 0.067722106 |
| ENSMUSG00000026960 | Arl6ip6 | 65103 | protein_coding | 346.0280322 | -0.067748849 | 0.077206347 | -0.877503612 | 0.380213158 | 0.621060566 | 0.067748849 |
| ENSMUSG00000114797 | Gm49336 | 71844 | protein_coding | 1134.567235 | -0.067770397 | 0.043987936 | -1.540658719 | 0.123399869 | 0.326064971 | 0.067770397 |
| ENSMUSG00000029486 | Mrpl1 | 94061 | protein_coding | 522.4679018 | 0.067778175 | 0.082507613 | 0.821477836 | 0.411374144 | 0.649442192 | 0.067778175 |
| ENSMUSG00000029561 | Oasl2 | 23962 | protein_coding | 345.4446251 | -0.067782672 | 0.322995415 | -0.209856451 | 0.833779713 | 0.926340413 | 0.067782672 |
| ENSMUSG00000003360 | Ddx23 | 74351 | protein_coding | 1807.51617 | 0.067786327 | 0.040482316 | 1.674467604 | 0.094038736 | 0.27725095 | 0.067786327 |
| ENSMUSG00000058325 | Dock1 | 330662 | protein_coding | 3112.718577 | -0.067795732 | 0.036191895 | -1.873229666 | 0.061036677 | 0.210508661 | 0.067795732 |
| ENSMUSG00000049680 | Urgcp | 72046 | protein_coding | 1859.89463 | 0.067799942 | 0.070913172 | 0.956097998 | 0.339022726 | 0.58240762 | 0.067799942 |
| ENSMUSG00000031858 | Mau2 | 74549 | protein_coding | 2361.646436 | -0.067826452 | 0.033956015 | -1.997479763 | 0.045773091 | 0.174387582 | 0.067826452 |
| ENSMUSG00000031629 | Cenpu | 71876 | protein_coding | 70.50407413 | 0.067833188 | 0.190860349 | 0.355407438 | 0.722284377 | 0.864247815 | 0.067833188 |
| ENSMUSG00000028799 | Zfp362 | 230761 | protein_coding | 124.8308591 | -0.067835063 | 0.150103297 | -0.451922542 | 0.651324785 | 0.8204259 | 0.067835063 |

| ensembl_gene_id | external_gene_name | entrezgene_id | gene_biotype | baseMean | log2FoldChange | lfcSE | stat | pvalue | padj | absLog2FC |
|---|---|---|---|---|---|---|---|---|---|---|
| ENSMUSG00000058022 | Adtrp | 109254 | protein_coding | 947.2351031 | -0.067843682 | 0.074048067 | -0.916211392 | 0.359556031 | 0.603010752 | 0.067843682 |
| ENSMUSG00000005355 | Casp14 | 12365 | protein_coding | 326.6508932 | 0.067866395 | 0.207889191 | 0.32645466 | 0.744080388 | 0.878140096 | 0.067866395 |
| ENSMUSG00000038520 | Tbc1d17 | 233204 | protein_coding | 903.2590546 | -0.067874251 | 0.059496385 | -1.14081302 | 0.253947741 | 0.495031103 | 0.067874251 |
| ENSMUSG00000004568 | Arhgef18 | 102098 | protein_coding | 1341.548359 | -0.067920018 | 0.060275057 | -1.126834565 | 0.259812434 | 0.501255547 | 0.067920018 |
| ENSMUSG00000026003 | Acadl | 11363 | protein_coding | 1964.043545 | -0.067923973 | 0.065523246 | -1.03663931 | 0.299903986 | 0.543615578 | 0.067923973 |
| ENSMUSG00000041598 | Cdc42ep4 | 56699 | protein_coding | 2144.111774 | 0.067925248 | 0.061467162 | 1.105065628 | 0.269131153 | 0.511644242 | 0.067925248 |
| ENSMUSG00000097787 | 2700046G09Rik | 67188 | lncRNA | 34.94365899 | -0.067939078 | 0.23073195 | -0.294450241 | 0.768413881 | 0.891461812 | 0.067939078 |
| ENSMUSG00000004270 | Lpcat3 | 14792 | protein_coding | 1648.559085 | 0.06794259 | 0.054118573 | 1.25543942 | 0.209319287 | 0.444790913 | 0.06794259 |
| ENSMUSG00000039242 | B3galnt2 | 97884 | protein_coding | 615.5045713 | 0.067961198 | 0.065574038 | 1.036404034 | 0.300013689 | 0.543683373 | 0.067961198 |
| ENSMUSG00000015697 | Setdb1 | 84505 | protein_coding | 853.3192538 | -0.067974911 | 0.056461645 | -1.203913056 | 0.228623185 | 0.467716111 | 0.067974911 |
| ENSMUSG00000016308 | Ube2a | 22209 | protein_coding | 724.8905244 | 0.067980419 | 0.075730303 | 0.897664685 | 0.369364342 | 0.611593275 | 0.067980419 |
| ENSMUSG00000052296 | Ppp6r1 | 243819 | protein_coding | 3485.663608 | 0.067992014 | 0.055847293 | 1.217463027 | 0.223428096 | 0.460947281 | 0.067992014 |
| ENSMUSG00000054604 | Cggbp1 | 106143 | protein_coding | 2377.683335 | 0.068005981 | 0.042765939 | 1.59019031 | 0.111791914 | 0.3074195 | 0.068005981 |
| ENSMUSG00000005054 | Cstb | 13014 | protein_coding | 1004.474777 | 0.068010895 | 0.081145535 | 0.838134774 | 0.401955015 | 0.641247695 | 0.068010895 |
| ENSMUSG00000054302 | Eapp | 66266 | protein_coding | 677.3814749 | -0.06801909 | 0.053314166 | -1.275816443 | 0.202020414 | 0.435870434 | 0.06801909 |
| ENSMUSG00000020829 | Slc46a1 | 52466 | protein_coding | 479.5193646 | 0.068019212 | 0.16682105 | 0.407737583 | 0.683466342 | 0.840526771 | 0.068019212 |
| ENSMUSG00000039065 | Atpsckmt | 68073 | protein_coding | 275.5965525 | 0.068080172 | 0.089563624 | 0.760131946 | 0.447175719 | 0.678100876 | 0.068080172 |
| ENSMUSG00000007034 | Slc44a4 | 70129 | protein_coding | 9826.938109 | -0.068115908 | 0.063153713 | -1.078573286 | 0.280777995 | 0.523508209 | 0.068115908 |
| ENSMUSG00000024382 | Ercc3 | 13872 | protein_coding | 819.4025411 | 0.068138166 | 0.047736721 | 1.427374258 | 0.153472047 | 0.370570998 | 0.068138166 |
| ENSMUSG00000056429 | Tgoln1 | 22134 | protein_coding | 20571.51448 | 0.068138369 | 0.064225225 | 1.060928462 | 0.288722414 | 0.531168876 | 0.068138369 |
| ENSMUSG00000046761 | Fam83h | 105732 | protein_coding | 3446.431762 | -0.068159032 | 0.07000979 | -0.973564296 | 0.330272921 | 0.573894128 | 0.068159032 |
| ENSMUSG00000031979 | Cog2 | 76332 | protein_coding | 1289.530416 | 0.068160814 | 0.05397728 | 1.262768589 | 0.206672356 | 0.441566958 | 0.068160814 |
| ENSMUSG00000035575 | Utp6 | 216987 | protein_coding | 860.7083533 | 0.06816802 | 0.06920971 | 0.984948781 | 0.324649231 | 0.568485113 | 0.06816802 |
| ENSMUSG00000024902 | Mrpl11 | 66419 | protein_coding | 659.207711 | 0.068180961 | 0.06325177 | 1.077929691 | 0.281065134 | 0.523522715 | 0.068180961 |
| ENSMUSG00000028414 | Fktn | 246179 | protein_coding | 474.836242 | 0.068195946 | 0.057603545 | 1.183884536 | 0.236458772 | 0.476619666 | 0.068195946 |
| ENSMUSG00000021114 | Atp6v1d | 73834 | protein_coding | 785.2877586 | 0.068233955 | 0.054634501 | 1.248916962 | 0.211695447 | 0.447035809 | 0.068233955 |
| ENSMUSG00000031849 | Comp | 12845 | protein_coding | 8.637899773 | 0.068275357 | 0.388025999 | 0.175955624 | 0.860328805 | 0.936965584 | 0.068275357 |
| ENSMUSG00000015002 | Efr3a | 76740 | protein_coding | 2091.279097 | -0.068313822 | 0.041512767 | -1.645609958 | 0.099844088 | 0.287678519 | 0.068313822 |
| ENSMUSG00000001157 | Gmcl1 | 23885 | protein_coding | 913.2544379 | 0.068355951 | 0.05248235 | 1.302455986 | 0.192760555 | 0.42444695 | 0.068355951 |
| ENSMUSG00000024548 | Setbp1 | 240427 | protein_coding | 806.0882433 | 0.068374282 | 0.071419395 | 0.957362945 | 0.338384095 | 0.582029301 | 0.068374282 |
| ENSMUSG00000032803 | Cdv3 | 321022 | protein_coding | 3188.98483 | 0.068382765 | 0.047255152 | 1.447096501 | 0.147869898 | 0.361756348 | 0.068382765 |
| ENSMUSG00000040532 | Abhd11 | 68758 | protein_coding | 941.7592217 | -0.068446349 | 0.055057843 | -1.243171638 | 0.213804592 | 0.449547151 | 0.068446349 |
| ENSMUSG00000071359 | Tbpl1 | 237336 | protein_coding | 395.1671979 | 0.068451501 | 0.070028197 | 0.977484829 | 0.328329185 | 0.571937821 | 0.068451501 |
| ENSMUSG00000028484 | Psip1 | 101739 | protein_coding | 346.1568134 | 0.068461392 | 0.078926929 | 0.867402202 | 0.385721676 | 0.625831733 | 0.068461392 |
| ENSMUSG00000004980 | Hnrnpa2b1 | 53379 | protein_coding | 6789.60166 | -0.068469144 | 0.074450664 | -0.919657931 | 0.357751543 | 0.600860147 | 0.068469144 |
| ENSMUSG00000038141 | Tmem181a | 77106 | protein_coding | 884.9784344 | -0.068481011 | 0.045232854 | -1.513966176 | 0.130034418 | 0.33596044 | 0.068481011 |
| ENSMUSG00000020513 | Tubd1 | 56427 | protein_coding | 94.34804529 | 0.068537546 | 0.138501716 | 0.494849794 | 0.620706153 | 0.799662139 | 0.068537546 |
| ENSMUSG00000026938 | Fcna | 14133 | protein_coding | 94.11533723 | 0.068546516 | 0.150501958 | 0.455452651 | 0.648783625 | 0.818686277 | 0.068546516 |
| ENSMUSG00000037843 | Vstm2l | 277432 | protein_coding | 13.03514947 | -0.068547931 | 0.318905571 | -0.214947425 | 0.829808309 | 0.924512511 | 0.068547931 |

| ensembl_gene_id | external_gene_name | entrezgene_id | gene_biotype | baseMean | log2FoldChange | lfcSE | stat | pvalue | padj | absLog2FC |
|---|---|---|---|---|---|---|---|---|---|---|
| ENSMUSG00000033249 | Hsf4 | 26386 | protein_coding | 11.8740466 | -0.068554879 | 0.394974748 | -0.173567753 | 0.862205177 | 0.937787345 | 0.068554879 |
| ENSMUSG00000051232 | Tmem199 | 195040 | protein_coding | 220.0869582 | 0.068557159 | 0.08532312 | 0.803500373 | 0.421685581 | 0.65828686 | 0.068557159 |
| ENSMUSG00000021668 | Polk | 27015 | protein_coding | 270.2662767 | 0.06855992 | 0.099525458 | 0.688868171 | 0.490906231 | 0.710750466 | 0.06855992 |
| ENSMUSG00000027412 | Lpin3 | 64899 | protein_coding | 1254.853886 | 0.06856619 | 0.054734863 | 1.252696842 | 0.210316056 | 0.445347562 | 0.06856619 |
| ENSMUSG00000061689 | Dlgap4 | 228836 | protein_coding | 505.7442983 | 0.068575337 | 0.084876776 | 0.807939936 | 0.41912516 | 0.65655478 | 0.068575337 |
| ENSMUSG00000038712 | Mindy1 | 75007 | protein_coding | 2499.240177 | -0.068585418 | 0.046522039 | -1.474256482 | 0.140412548 | 0.351472935 | 0.068585418 |
| ENSMUSG00000022895 | Ets2 | 23872 | protein_coding | 5917.526309 | -0.068600402 | 0.037331823 | -1.837585103 | 0.066123566 | 0.221791752 | 0.068600402 |
| ENSMUSG00000070939 | Tgfbrap1 | 73122 | protein_coding | 688.3388875 | -0.068616572 | 0.051230869 | -1.339359899 | 0.180453538 | 0.40741624 | 0.068616572 |
| ENSMUSG00000062098 | Btbd3 | 228662 | protein_coding | 674.9230253 | -0.068634335 | 0.058211139 | -1.179058448 | 0.238374903 | 0.478679126 | 0.068634335 |
| ENSMUSG00000027361 | Gabpb1 | 14391 | protein_coding | 316.7552915 | -0.06863622 | 0.080023965 | -0.857695818 | 0.39106045 | 0.630978677 | 0.06863622 |
| ENSMUSG00000034902 | Pip5k1c | 18717 | protein_coding | 947.6476951 | -0.068649223 | 0.050707241 | -1.353834692 | 0.175789123 | 0.402176854 | 0.068649223 |
| ENSMUSG00000046230 | Vps13a | 271564 | protein_coding | 1855.315735 | -0.068667688 | 0.060015652 | -1.144162989 | 0.252556049 | 0.493633669 | 0.068667688 |
| ENSMUSG00000039130 | Zc3hc1 | 232679 | protein_coding | 236.7370388 | -0.068713704 | 0.093102202 | -0.738045959 | 0.460486521 | 0.687265524 | 0.068713704 |
| ENSMUSG00000039182 | AW209491 | 105351 | protein_coding | 237.9650585 | 0.068763296 | 0.08612343 | 0.798427278 | 0.424622579 | 0.660610489 | 0.068763296 |
| ENSMUSG00000038080 | Kdm1b | 218214 | protein_coding | 866.4063068 | -0.068791784 | 0.062324847 | -1.103761787 | 0.269696485 | 0.512235923 | 0.068791784 |
| ENSMUSG00000006931 | P3h4 | 66180 | protein_coding | 43.73416874 | 0.068794209 | 0.222526544 | 0.309150576 | 0.757206988 | 0.884953772 | 0.068794209 |
| ENSMUSG00000074628 | Tldc2 | 383766 | protein_coding | 187.1323178 | 0.068798987 | 0.102461288 | 0.671463225 | 0.501925479 | 0.71887675 | 0.068798987 |
| ENSMUSG00000039954 | Stk32a | 269019 | protein_coding | 16.39934198 | -0.068822945 | 0.309345776 | -0.222479021 | 0.823941003 | 0.921722463 | 0.068822945 |
| ENSMUSG00000043733 | Ptpn11 | 19247 | protein_coding | 1988.517537 | 0.068832257 | 0.048619873 | 1.415722689 | 0.156856713 | 0.375594388 | 0.068832257 |
| ENSMUSG00000027605 | Acss2 | 60525 | protein_coding | 3731.393429 | -0.068872246 | 0.076418233 | -0.901254104 | 0.36745323 | 0.609913364 | 0.068872246 |
| ENSMUSG00000020515 | Cnot8 | 69125 | protein_coding | 832.1647045 | 0.068903556 | 0.049264521 | 1.398644595 | 0.161919586 | 0.382604713 | 0.068903556 |
| ENSMUSG00000032215 | Rsl24d1 | 225215 | protein_coding | 353.0182754 | -0.068922028 | 0.072916648 | -0.945216634 | 0.344548289 | 0.58688999 | 0.068922028 |
| ENSMUSG00000072568 | Lratd2 | 399603 | protein_coding | 1004.621476 | 0.068928657 | 0.055675212 | 1.238049309 | 0.215697778 | 0.451467627 | 0.068928657 |
| ENSMUSG00000040720 | Virma | 66185 | protein_coding | 1191.97507 | 0.06893088 | 0.051907186 | 1.327964109 | 0.184189964 | 0.412891303 | 0.06893088 |
| ENSMUSG00000025165 | Sectm1a | 209588 | protein_coding | 83.25853037 | -0.068939876 | 0.146289991 | -0.471254908 | 0.637458712 | 0.810693313 | 0.068939876 |
| ENSMUSG00000035840 | Lysmd3 | 80289 | protein_coding | 856.1396787 | 0.068943679 | 0.070058598 | 0.984085917 | 0.325073271 | 0.568781234 | 0.068943679 |
| ENSMUSG00000025888 | Casp1 | 12362 | protein_coding | 3594.585228 | -0.068946394 | 0.071931618 | -0.95849914 | 0.337811125 | 0.581541419 | 0.068946394 |
| ENSMUSG00000030364 | Clec2h | 94071 | protein_coding | 7146.742001 | 0.068974844 | 0.078010579 | 0.884172955 | 0.376602856 | 0.618232985 | 0.068974844 |
| ENSMUSG00000031652 | N4bp1 | 80750 | protein_coding | 926.0278943 | -0.06899533 | 0.048805354 | -1.413683629 | 0.157454814 | 0.376690964 | 0.06899533 |
| ENSMUSG00000018849 | Wwc1 | 211652 | protein_coding | 1295.692422 | -0.068997562 | 0.044108708 | -1.564261674 | 0.117756129 | 0.317250963 | 0.068997562 |
| ENSMUSG00000026277 | Stk25 | 59041 | protein_coding | 4079.540216 | 0.069005854 | 0.051800241 | 1.332153159 | 0.182809865 | 0.410843172 | 0.069005854 |
| ENSMUSG00000030446 | Zfp273 | 212569 | protein_coding | 101.0586142 | -0.06909897 | 0.127317191 | -0.542730874 | 0.587315114 | 0.777255116 | 0.06909897 |
| ENSMUSG00000028137 | Celf3 | 78784 | protein_coding | 40.0128466 | -0.069100689 | 0.196800242 | -0.351120955 | 0.725497608 | 0.865987203 | 0.069100689 |
| ENSMUSG00000001786 | Fbxo7 | 69754 | protein_coding | 364.6211951 | -0.069142032 | 0.068997303 | -1.002097607 | 0.316296454 | 0.560212157 | 0.069142032 |
| ENSMUSG00000070520 | Nsmce3 | 66647 | protein_coding | 391.0953528 | 0.069162216 | 0.088520212 | 0.781315521 | 0.434616943 | 0.668304413 | 0.069162216 |
| ENSMUSG00000020777 | Acox1 | 11430 | protein_coding | 7565.454514 | -0.069167977 | 0.055304898 | -1.250666391 | 0.211056217 | 0.4461694 | 0.069167977 |
| ENSMUSG00000028832 | Stmn1 | 16765 | protein_coding | 982.3978169 | -0.069190441 | 0.103940002 | -0.665676732 | 0.505617749 | 0.721608883 | 0.069190441 |
| ENSMUSG00000117924 | Tmem223 | 66836 | protein_coding | 359.4507478 | -0.069223005 | 0.113663583 | -0.609016562 | 0.542513461 | 0.747864433 | 0.069223005 |
| ENSMUSG00000063870 | Chd4 | 107932 | protein_coding | 5819.999648 | -0.069271488 | 0.045357633 | -1.527228902 | 0.126704102 | 0.330800882 | 0.069271488 |

| ensembl_gene_id | external_gene_name | entrezgene_id | gene_biotype | baseMean | log2FoldChange | lfcSE | stat | pvalue | padj | absLog2FC |
|---|---|---|---|---|---|---|---|---|---|---|
| ENSMUSG00000050029 | Rap2c | 72065 | protein_coding | 860.2625721 | 0.069299196 | 0.066072553 | 1.048834846 | 0.294254137 | 0.537241256 | 0.069299196 |
| ENSMUSG00000035212 | Leprot | 230514 | protein_coding | 1618.206367 | 0.069299347 | 0.061001741 | 1.136022433 | 0.255947176 | 0.497488262 | 0.069299347 |
| ENSMUSG00000033186 | Mzt1 | 76789 | protein_coding | 987.8005453 | 0.069371682 | 0.063377209 | 1.094584038 | 0.273698912 | 0.516310954 | 0.069371682 |
| ENSMUSG00000019338 | Zfp687 | 78266 | protein_coding | 785.2486529 | 0.069418867 | 0.058357012 | 1.189554856 | 0.234221398 | 0.473819429 | 0.069418867 |
| ENSMUSG00000053536 | Cstf2t | 83410 | protein_coding | 1259.340214 | 0.069423618 | 0.044007355 | 1.577545789 | 0.114669995 | 0.312220666 | 0.069423618 |
| ENSMUSG00000058258 | Idi1 | 319554 | protein_coding | 584.8318058 | 0.069448247 | 0.122205999 | 0.568288365 | 0.569839178 | 0.765233601 | 0.069448247 |
| ENSMUSG00000025925 | Terf1 | 21749 | protein_coding | 197.9625614 | 0.069477045 | 0.096826583 | 0.717541018 | 0.473040337 | 0.697134159 | 0.069477045 |
| ENSMUSG00000041040 | Fam117b | 72750 | protein_coding | 1283.649466 | -0.069493858 | 0.050101127 | -1.387071764 | 0.165419881 | 0.388484989 | 0.069493858 |
| ENSMUSG00000087687 | Pet100 | 100503890 | protein_coding | 696.0417367 | 0.069502909 | 0.078650724 | 0.883690642 | 0.376863233 | 0.61835018 | 0.069502909 |
| ENSMUSG00000027394 | Ttl | 69737 | protein_coding | 188.2783224 | -0.069507689 | 0.100633515 | -0.690701186 | 0.489753344 | 0.709700079 | 0.069507689 |
| ENSMUSG00000086199 | Bcas3os1 | 71489 | lncRNA | 7.238525441 | 0.069511015 | 0.500678364 | 0.13883367 | 0.889581587 | 0.950733597 | 0.069511015 |
| ENSMUSG00000074272 | Ceacam1 | 26365 | protein_coding | 32617.697 | -0.069517857 | 0.039666168 | -1.752573077 | 0.079675316 | 0.249693708 | 0.069517857 |
| ENSMUSG00000024099 | Ndufv2 | 72900 | protein_coding | 3675.809478 | 0.069528296 | 0.064229799 | 1.082492826 | 0.279033601 | 0.521917755 | 0.069528296 |
| ENSMUSG00000032244 | Fem1b | 14155 | protein_coding | 1416.408129 | -0.069533812 | 0.050886614 | -1.366446053 | 0.171799002 | 0.396593546 | 0.069533812 |
| ENSMUSG00000031754 | Nudt21 | 68219 | protein_coding | 566.4438602 | -0.069541745 | 0.068758202 | -1.011395636 | 0.311827109 | 0.555549689 | 0.069541745 |
| ENSMUSG00000033326 | Kdm4a | 230674 | protein_coding | 1903.28713 | 0.069584601 | 0.049340707 | 1.410287867 | 0.1584547 | 0.378465483 | 0.069584601 |
| ENSMUSG00000020923 | Ubtf | 21429 | protein_coding | 2874.644163 | 0.069600166 | 0.036213485 | 1.921940577 | 0.054613235 | 0.196288755 | 0.069600166 |
| ENSMUSG00000037815 | Ctnna1 | 12385 | protein_coding | 12193.49002 | 0.069616413 | 0.040967042 | 1.699327295 | 0.089257532 | 0.268368093 | 0.069616413 |
| ENSMUSG00000021710 | Nln | 75805 | protein_coding | 1246.893044 | -0.069624712 | 0.043524493 | -1.599667389 | 0.10967239 | 0.303689326 | 0.069624712 |
| ENSMUSG00000022982 | Sod1 | 20655 | protein_coding | 5505.137904 | -0.069643084 | 0.052772058 | -1.319696181 | 0.186936476 | 0.416513477 | 0.069643084 |
| ENSMUSG00000057551 | Zfp317 | 244713 | protein_coding | 638.4790922 | -0.069678509 | 0.070501046 | -0.988332993 | 0.322989592 | 0.56691915 | 0.069678509 |
| ENSMUSG00000039853 | Trim14 | 74735 | protein_coding | 404.1393579 | 0.069682355 | 0.074519598 | 0.935087633 | 0.34974314 | 0.592096117 | 0.069682355 |
| ENSMUSG00000034940 | Synrg | 217030 | protein_coding | 1170.032487 | -0.069696659 | 0.052056752 | -1.33885915 | 0.18061653 | 0.407535035 | 0.069696659 |
| ENSMUSG00000029338 | Antxr2 | 71914 | protein_coding | 1240.135398 | -0.069700638 | 0.057700996 | -1.207962484 | 0.227061693 | 0.465615713 | 0.069700638 |
| ENSMUSG00000038152 | 5033430I15Rik | 76000 | lncRNA | 48.79116022 | 0.069700684 | 0.192237291 | 0.362576288 | 0.716921426 | 0.861277699 | 0.069700684 |
| ENSMUSG00000021139 | Gm20498 | 105940408 | protein_coding | 8.783688609 | 0.069717568 | 0.383645081 | 0.181724129 | 0.855799229 | 0.935592329 | 0.069717568 |
| ENSMUSG00000034875 | Nudt19 | 110959 | protein_coding | 2035.611663 | 0.06972894 | 0.058485323 | 1.192246816 | 0.233164482 | 0.472624422 | 0.06972894 |
| ENSMUSG00000015711 | Prune1 | 229589 | protein_coding | 422.9259967 | -0.069739525 | 0.059982494 | -1.162664648 | 0.24496559 | 0.486120986 | 0.069739525 |
| ENSMUSG00000054000 | Tusc1 | 69136 | protein_coding | 112.4164407 | -0.069746631 | 0.141326528 | -0.493514078 | 0.621649395 | 0.800298539 | 0.069746631 |
| ENSMUSG00000021890 | Eaf1 | 74427 | protein_coding | 862.3747692 | 0.069758597 | 0.053107684 | 1.313531148 | 0.189004032 | 0.419381952 | 0.069758597 |
| ENSMUSG00000058558 | Rpl5 | 100503670 | protein_coding | 9162.413494 | -0.069761572 | 0.041924814 | -1.663968541 | 0.096118716 | 0.281223973 | 0.069761572 |
| ENSMUSG00000005553 | Atp4a | 11944 | protein_coding | 8.006315452 | 0.069784141 | 0.420111261 | 0.166108713 | 0.868071398 | 0.940250348 | 0.069784141 |
| ENSMUSG00000057897 | Camk2b | 12323 | protein_coding | 581.2042371 | 0.069786659 | 0.130022375 | 0.536728075 | 0.59145546 | 0.7802374 | 0.069786659 |
| ENSMUSG00000032839 | Trpc1 | 22063 | protein_coding | 21.51218448 | -0.069792734 | 0.276821329 | -0.252121951 | 0.800946804 | 0.90924234 | 0.069792734 |
| ENSMUSG00000045282 | Tmem86b | 68255 | protein_coding | 50.07949328 | 0.06981397 | 0.202094631 | 0.345451878 | 0.729754682 | 0.869184177 | 0.06981397 |
| ENSMUSG00000028700 | Pomgnt1 | 68273 | protein_coding | 894.2777539 | -0.069825088 | 0.064471416 | -1.083039471 | 0.278790904 | 0.521705837 | 0.069825088 |
| ENSMUSG00000063019 | Manbal | 69161 | protein_coding | 308.4789883 | 0.069852527 | 0.106537668 | 0.655660367 | 0.512042675 | 0.725697532 | 0.069852527 |
| ENSMUSG00000063273 | Naa15 | 74838 | protein_coding | 839.861232 | -0.069872477 | 0.079633209 | -0.87742887 | 0.380253738 | 0.621060566 | 0.069872477 |
| ENSMUSG00000078653 | Cntd1 | 68107 | protein_coding | 20.79323523 | -0.069879345 | 0.251521973 | -0.277826005 | 0.781145928 | 0.898390166 | 0.069879345 |

| ensembl_gene_id | external_gene_name | entrezgene_id | gene_biotype | baseMean | log2FoldChange | lfcSE | stat | pvalue | padj | absLog2FC |
|---|---|---|---|---|---|---|---|---|---|---|
| ENSMUSG00000041997 | Tlk1 | 228012 | protein_coding | 2200.118106 | -0.06988164 | 0.052195209 | -1.338851612 | 0.180618985 | 0.407535035 | 0.06988164 |
| ENSMUSG00000039031 | Arhgap18 | 73910 | protein_coding | 1464.127892 | -0.069901207 | 0.042603265 | -1.640747644 | 0.10084981 | 0.289568133 | 0.069901207 |
| ENSMUSG00000035725 | Prkx | 19108 | protein_coding | 733.4614637 | -0.069928885 | 0.075424722 | -0.927134799 | 0.35385655 | 0.596118728 | 0.069928885 |
| ENSMUSG00000005980 | Dnase1 | 13419 | protein_coding | 27.27541661 | -0.06998811 | 0.299514585 | -0.233671793 | 0.815239787 | 0.916810722 | 0.06998811 |
| ENSMUSG00000020832 | Eral1 | 57837 | protein_coding | 367.1977176 | 0.069992884 | 0.074330692 | 0.941641756 | 0.346376084 | 0.588629676 | 0.069992884 |
| ENSMUSG00000025646 | Atrip | 235610 | protein_coding | 260.227753 | -0.069995317 | 0.084115056 | -0.832137798 | 0.405331169 | 0.644334723 | 0.069995317 |
| ENSMUSG00000032624 | Eml4 | 78798 | protein_coding | 2145.03016 | -0.069997285 | 0.043570066 | -1.606545319 | 0.108154146 | 0.301243853 | 0.069997285 |
| ENSMUSG00000096299 | Gm21814 | 436100 | lncRNA | 10.068879 | -0.069998852 | 0.356768133 | -0.196202646 | 0.844451557 | 0.930210801 | 0.069998852 |
| ENSMUSG00000025277 | Abhd6 | 66082 | protein_coding | 693.4713083 | -0.070007649 | 0.062983161 | -1.111529627 | 0.266340445 | 0.508211544 | 0.070007649 |
| ENSMUSG00000024817 | Uhrf2 | 109113 | protein_coding | 2170.252106 | 0.070047233 | 0.039096357 | 1.791656279 | 0.073188042 | 0.236781989 | 0.070047233 |
| ENSMUSG00000028194 | Ddah1 | 69219 | protein_coding | 1151.556159 | 0.070063807 | 0.075105043 | 0.932877522 | 0.350883209 | 0.593278833 | 0.070063807 |
| ENSMUSG00000028958 | Tmub1 | 64295 | protein_coding | 548.8253981 | 0.07006898 | 0.090512763 | 0.774133694 | 0.438851734 | 0.671540176 | 0.07006898 |
| ENSMUSG00000091803 | Cox16 | 66272 | protein_coding | 217.7202337 | 0.070077855 | 0.126676851 | 0.553201747 | 0.580125271 | 0.772111929 | 0.070077855 |
| ENSMUSG00000046598 | Bdh1 | 71911 | protein_coding | 3282.827759 | -0.070085212 | 0.050837883 | -1.378602076 | 0.168017476 | 0.391953183 | 0.070085212 |
| ENSMUSG00000047501 | Cldn4 | 12740 | protein_coding | 903.0662761 | 0.070098567 | 0.104141725 | 0.673107409 | 0.50087896 | 0.718207077 | 0.070098567 |
| ENSMUSG00000038351 | Sgsm2 | 97761 | protein_coding | 886.719475 | -0.070101252 | 0.061525193 | -1.139391014 | 0.254540102 | 0.495656311 | 0.070101252 |
| ENSMUSG00000020521 | Rnft1 | 76892 | protein_coding | 1045.275972 | 0.070127899 | 0.058204089 | 1.204862073 | 0.228256552 | 0.467203125 | 0.070127899 |
| ENSMUSG00000017286 | Glod4 | 67201 | protein_coding | 1255.644555 | 0.070159854 | 0.056098912 | 1.25064554 | 0.211063828 | 0.4461694 | 0.070159854 |
| ENSMUSG00000039294 | Cybc1 | 217370 | protein_coding | 460.0227802 | -0.070171815 | 0.074646894 | -0.940050021 | 0.347191903 | 0.589385665 | 0.070171815 |
| ENSMUSG00000028455 | Stoml2 | 66592 | protein_coding | 927.4103576 | 0.070177885 | 0.054397748 | 1.290088069 | 0.197020082 | 0.429486734 | 0.070177885 |
| ENSMUSG00000053644 | Aldh7a1 | 110695 | protein_coding | 459.6613404 | 0.070196987 | 0.069028429 | 1.016928646 | 0.309187375 | 0.552606516 | 0.070196987 |
| ENSMUSG00000029048 | Rer1 | 67830 | protein_coding | 3308.600757 | 0.070207712 | 0.048161737 | 1.457748775 | 0.144909803 | 0.357552079 | 0.070207712 |
| ENSMUSG00000034312 | Iqsec1 | 232227 | protein_coding | 388.6270955 | -0.070208071 | 0.120956692 | -0.58043974 | 0.561618112 | 0.759451224 | 0.070208071 |
| ENSMUSG00000022054 | Nefm | 18040 | protein_coding | 10.00760544 | 0.070210356 | 0.422239877 | 0.166280731 | 0.867936029 | 0.940208703 | 0.070210356 |
| ENSMUSG00000040498 | Igsf23 | 70080 | protein_coding | 99.54634826 | 0.070214379 | 0.158564752 | 0.442812029 | 0.657901702 | 0.824909093 | 0.070214379 |
| ENSMUSG00000036528 | Ppfibp2 | 19024 | protein_coding | 817.188188 | 0.070219244 | 0.071760185 | 0.978526527 | 0.327813978 | 0.571636266 | 0.070219244 |
| ENSMUSG00000009863 | Sdhb | 67680 | protein_coding | 5065.313854 | 0.070223022 | 0.052433851 | 1.339268819 | 0.180483176 | 0.407426116 | 0.070223022 |
| ENSMUSG00000025395 | Prim1 | 19075 | protein_coding | 241.1216864 | 0.070240465 | 0.152392326 | 0.460918649 | 0.644856971 | 0.81580845 | 0.070240465 |
| ENSMUSG00000057411 | Antkmt | 214917 | protein_coding | 381.4944221 | -0.070302696 | 0.08568679 | -0.820461314 | 0.411953174 | 0.649970343 | 0.070302696 |
| ENSMUSG00000059423 | Zfp933 | 242747 | protein_coding | 188.4703151 | -0.070306274 | 0.092349085 | -0.761309917 | 0.446471979 | 0.677479639 | 0.070306274 |
| ENSMUSG00000014763 | Fam120b | 67544 | protein_coding | 1144.144129 | 0.070329115 | 0.051691626 | 1.360551417 | 0.173655494 | 0.399424529 | 0.070329115 |
| ENSMUSG00000056459 | Zbtb25 | 109929 | protein_coding | 50.63729727 | 0.070335677 | 0.160897181 | 0.437146732 | 0.662004938 | 0.827333871 | 0.070335677 |
| ENSMUSG00000028420 | Tmem38b | 52076 | protein_coding | 250.0232274 | 0.070348654 | 0.113635955 | 0.619070385 | 0.535869992 | 0.742692967 | 0.070348654 |
| ENSMUSG00000013787 | Ehmt2 | 110147 | protein_coding | 4025.651046 | -0.070348695 | 0.054356634 | -1.294206232 | 0.195594195 | 0.428130377 | 0.070348695 |
| ENSMUSG00000064558 | Gm24620 | 115488217 | snoRNA | 18.31330268 | 0.070349972 | 0.533132881 | 0.131955792 | 0.895019259 | 0.953178075 | 0.070349972 |
| ENSMUSG00000001383 | Zmat2 | 66492 | protein_coding | 961.3995351 | 0.070362816 | 0.049602871 | 1.418523039 | 0.156038117 | 0.374523994 | 0.070362816 |
| ENSMUSG00000014077 | Chp1 | 56398 | protein_coding | 6948.533843 | -0.070379803 | 0.048735307 | -1.444123532 | 0.14870423 | 0.363026509 | 0.070379803 |
| ENSMUSG00000036086 | Zranb3 | 226409 | protein_coding | 103.6845357 | 0.070398327 | 0.117461115 | 0.599333042 | 0.548950818 | 0.751266959 | 0.070398327 |
| ENSMUSG00000028108 | Ecm1 | 13601 | protein_coding | 14.86118267 | 0.070401379 | 0.342339082 | 0.205648092 | 0.837065808 | 0.927812059 | 0.070401379 |

| ensembl_gene_id | external_gene_name | entrezgene_id | gene_biotype | baseMean | log2FoldChange | lfcSE | stat | pvalue | padj | absLog2FC |
|---|---|---|---|---|---|---|---|---|---|---|
| ENSMUSG00000013878 | Rnf170 | 77733 | protein_coding | 391.3097034 | 0.070425328 | 0.06667047 | 1.056319657 | 0.290822198 | 0.5332246 | 0.070425328 |
| ENSMUSG00000040195 | Nemp1 | 72243 | protein_coding | 222.059791 | -0.070431606 | 0.084568395 | -0.832836032 | 0.40493721 | 0.644152928 | 0.070431606 |
| ENSMUSG00000040242 | Fgfr1op2 | 67529 | protein_coding | 1686.404142 | 0.070442462 | 0.045465353 | 1.549365778 | 0.121293815 | 0.322629946 | 0.070442462 |
| ENSMUSG00000071014 | Ndufb6 | 230075 | protein_coding | 1008.360822 | 0.070451655 | 0.102527535 | 0.687148626 | 0.491989075 | 0.711469495 | 0.070451655 |
| ENSMUSG00000029669 | Tspan12 | 269831 | protein_coding | 1311.000146 | 0.070457867 | 0.064529365 | 1.091872929 | 0.274888958 | 0.517285207 | 0.070457867 |
| ENSMUSG00000024319 | Vps52 | 224705 | protein_coding | 932.9336406 | 0.070465749 | 0.091006875 | 0.77429039 | 0.438759085 | 0.671462115 | 0.070465749 |
| ENSMUSG00000020634 | Ubxn2a | 217379 | protein_coding | 1818.952105 | 0.070466092 | 0.048780261 | 1.444561607 | 0.148581063 | 0.362780743 | 0.070466092 |
| ENSMUSG00000060032 | H2aj | 232440 | protein_coding | 1569.212783 | 0.070478289 | 0.082440215 | 0.85490181 | 0.392605508 | 0.632267311 | 0.070478289 |
| ENSMUSG00000020137 | Thap2 | 66816 | protein_coding | 411.2028308 | 0.070488299 | 0.078360561 | 0.899537947 | 0.368366193 | 0.610412891 | 0.070488299 |
| ENSMUSG00000110218 | Tincr | 100504425 | protein_coding | 57.9753527 | 0.070498424 | 0.185811638 | 0.379408013 | 0.704384901 | 0.853157579 | 0.070498424 |
| ENSMUSG00000018651 | Tada2a | 217031 | protein_coding | 187.5804061 | -0.070505092 | 0.099287056 | -0.710113632 | 0.477633674 | 0.700254387 | 0.070505092 |
| ENSMUSG00000004931 | Apba3 | 57267 | protein_coding | 830.6904177 | 0.070536709 | 0.082693659 | 0.852988119 | 0.393665893 | 0.633344957 | 0.070536709 |
| ENSMUSG00000022554 | Hgh1 | 59053 | protein_coding | 162.19985 | -0.070576488 | 0.094283303 | -0.748557649 | 0.454123866 | 0.682675952 | 0.070576488 |
| ENSMUSG00000116903 | Gm19522 | 100503040 | lncRNA | 15.43472871 | -0.070585967 | 0.366958752 | -0.192353953 | 0.847464959 | 0.930999141 | 0.070585967 |
| ENSMUSG00000021840 | Mapk1ip1l | 218975 | protein_coding | 1657.019915 | 0.070594003 | 0.059109817 | 1.194285595 | 0.232366271 | 0.471545344 | 0.070594003 |
| ENSMUSG00000010277 | Bltp2 | 72503 | protein_coding | 9268.436772 | 0.070606581 | 0.043508292 | 1.622830436 | 0.104625657 | 0.29617307 | 0.070606581 |
| ENSMUSG00000031985 | Gnpat | 14712 | protein_coding | 824.8307923 | 0.070614624 | 0.059741148 | 1.18200983 | 0.237201801 | 0.47721475 | 0.070614624 |
| ENSMUSG00000022151 | Ttc33 | 67515 | protein_coding | 483.1587033 | -0.070662715 | 0.067502211 | -1.046820745 | 0.295182261 | 0.538150682 | 0.070662715 |
| ENSMUSG00000019763 | Rmnd1 | 66084 | protein_coding | 436.5447812 | -0.070677869 | 0.070731494 | -0.999241851 | 0.317677547 | 0.561449273 | 0.070677869 |
| ENSMUSG00000021987 | Mtmr6 | 219135 | protein_coding | 3075.849476 | -0.070687962 | 0.042788969 | -1.652013666 | 0.098531765 | 0.285589823 | 0.070687962 |
| ENSMUSG00000040430 | Pitpnc1 | 71795 | protein_coding | 61.98934417 | -0.070693995 | 0.174778576 | -0.404477461 | 0.685861649 | 0.841836579 | 0.070693995 |
| ENSMUSG00000020170 | Frs2 | 327826 | protein_coding | 671.8587811 | -0.070739669 | 0.082693794 | -0.855441084 | 0.392307008 | 0.632104617 | 0.070739669 |
| ENSMUSG00000015605 | Srf | 20807 | protein_coding | 1111.653875 | -0.070755832 | 0.082777748 | -0.854768758 | 0.392679177 | 0.632267311 | 0.070755832 |
| ENSMUSG00000023051 | Tarbp2 | 21357 | protein_coding | 471.8119818 | 0.070758706 | 0.089661418 | 0.789176741 | 0.430008714 | 0.664948752 | 0.070758706 |
| ENSMUSG00000036442 | Thap11 | 59016 | protein_coding | 573.3753513 | -0.070761316 | 0.073935598 | -0.95706693 | 0.338533474 | 0.5820991 | 0.070761316 |
| ENSMUSG00000022636 | Alcam | 11658 | protein_coding | 1216.241668 | 0.070786373 | 0.061107313 | 1.158394474 | 0.246703093 | 0.487630933 | 0.070786373 |
| ENSMUSG00000025503 | Taldo1 | 21351 | protein_coding | 7129.386245 | 0.070805752 | 0.06705077 | 1.056002066 | 0.290967271 | 0.53329119 | 0.070805752 |
| ENSMUSG00000031068 | Glrx3 | 30926 | protein_coding | 1284.879014 | 0.070811727 | 0.042616717 | 1.661595075 | 0.096593991 | 0.282102734 | 0.070811727 |
| ENSMUSG00000020375 | Rufy1 | 216724 | protein_coding | 611.2996891 | -0.070816548 | 0.061365967 | -1.154003612 | 0.248498688 | 0.4893628 | 0.070816548 |
| ENSMUSG00000002781 | Tmem143 | 70209 | protein_coding | 603.8726302 | 0.070819056 | 0.069707462 | 1.015946566 | 0.309654833 | 0.553189591 | 0.070819056 |
| ENSMUSG00000038007 | Acer2 | 230379 | protein_coding | 172.799268 | -0.070852783 | 0.099166289 | -0.714484564 | 0.474927597 | 0.698573273 | 0.070852783 |
| ENSMUSG00000054237 | Fra10ac1 | 70567 | protein_coding | 186.3892708 | -0.070852908 | 0.10026909 | -0.706627615 | 0.479797923 | 0.702150736 | 0.070852908 |
| ENSMUSG00000020491 | 2810021J22Rik | 69944 | protein_coding | 146.5877454 | -0.070875923 | 0.105878253 | -0.669409635 | 0.503234207 | 0.719943844 | 0.070875923 |
| ENSMUSG00000037628 | Cdkn3 | 72391 | protein_coding | 141.6121008 | 0.070916221 | 0.135486447 | 0.523419295 | 0.6006825 | 0.786125192 | 0.070916221 |
| ENSMUSG00000033102 | Cdc14b | 218294 | protein_coding | 499.3165025 | -0.070934644 | 0.08106567 | -0.875026922 | 0.381559257 | 0.622373434 | 0.070934644 |
| ENSMUSG00000091586 | Cyp4f17 | 208285 | protein_coding | 74.89926826 | -0.070938262 | 0.1632756 | -0.434469459 | 0.663947568 | 0.828333496 | 0.070938262 |
| ENSMUSG00000028709 | Mob3c | 100465 | protein_coding | 427.3118137 | 0.070943615 | 0.07363424 | 0.963459594 | 0.335316932 | 0.578888054 | 0.070943615 |
| ENSMUSG00000000088 | Cox5a | 12858 | protein_coding | 5225.418266 | 0.070953636 | 0.075378702 | 0.941295539 | 0.346553428 | 0.588735609 | 0.070953636 |
| ENSMUSG00000005299 | Letm1 | 56384 | protein_coding | 3434.915564 | 0.070983213 | 0.071037423 | 0.99923688 | 0.317679954 | 0.561449273 | 0.070983213 |

Supplementary Table 1

| ensembl_gene_id | external_gene_name | entrezgene_id | gene_biotype | baseMean | log2FoldChange | lfcSE | stat | pvalue | padj | absLog2FC |
|---|---|---|---|---|---|---|---|---|---|---|
| ENSMUSG00000026458 | Ppfia4 | 68507 | protein_coding | 27.9877043 | -0.071010891 | 0.263500516 | -0.269490518 | 0.787552238 | 0.900897641 | 0.071010891 |
| ENSMUSG00000054792 | Klhl18 | 270201 | protein_coding | 489.8896627 | 0.071012502 | 0.064989302 | 1.092679881 | 0.274534376 | 0.516979903 | 0.071012502 |
| ENSMUSG00000030766 | Arhgap17 | 70497 | protein_coding | 588.2043078 | -0.07102633 | 0.059444619 | -1.194831959 | 0.232152691 | 0.471289835 | 0.07102633 |
| ENSMUSG00000072437 | Nanos1 | 332397 | protein_coding | 38.45942071 | -0.071032692 | 0.222494235 | -0.319256328 | 0.749532147 | 0.880475742 | 0.071032692 |
| ENSMUSG00000040158 | Tax1bp3 | 76281 | protein_coding | 1258.383676 | 0.071041876 | 0.081067608 | 0.876328759 | 0.380851335 | 0.621784722 | 0.071041876 |
| ENSMUSG00000057497 | Fam136a | 66488 | protein_coding | 890.9092843 | 0.071075078 | 0.055521566 | 1.280134611 | 0.200497798 | 0.434133726 | 0.071075078 |
| ENSMUSG00000047821 | Trim16 | 94092 | protein_coding | 3214.197678 | -0.071092463 | 0.059455175 | -1.19573214 | 0.231801105 | 0.470931774 | 0.071092463 |
| ENSMUSG00000040229 | Gpr34 | 23890 | protein_coding | 9.477416838 | 0.071103813 | 0.400395015 | 0.177584161 | 0.859049564 | 0.936390154 | 0.071103813 |
| ENSMUSG00000028397 | Kdm4c | 76804 | protein_coding | 768.7029009 | -0.071109199 | 0.045969551 | -1.546876084 | 0.121893129 | 0.323423905 | 0.071109199 |
| ENSMUSG00000025737 | Wdr24 | 268933 | protein_coding | 755.8034993 | 0.071110264 | 0.061288337 | 1.160257682 | 0.245943909 | 0.487025714 | 0.071110264 |
| ENSMUSG00000031392 | Irak1 | 16179 | protein_coding | 1087.725127 | 0.071117705 | 0.063051911 | 1.127923085 | 0.259352404 | 0.500967327 | 0.071117705 |
| ENSMUSG00000040822 | 1700123O20Rik | 58248 | protein_coding | 587.5751261 | 0.071140616 | 0.059516204 | 1.195315084 | 0.231963949 | 0.471166492 | 0.071140616 |
| ENSMUSG00000045160 | Bola3 | 78653 | protein_coding | 599.6715267 | 0.071149594 | 0.078245831 | 0.909308437 | 0.363187338 | 0.605769124 | 0.071149594 |
| ENSMUSG00000040661 | Rad54l2 | 81000 | protein_coding | 791.0158215 | -0.071173501 | 0.049234505 | -1.445602053 | 0.14828885 | 0.362451393 | 0.071173501 |
| ENSMUSG00000026698 | Pigc | 67292 | protein_coding | 523.5109902 | 0.071185865 | 0.069598022 | 1.022814479 | 0.306395571 | 0.550544997 | 0.071185865 |
| ENSMUSG00000016534 | Lamp2 | 16784 | protein_coding | 2806.040394 | -0.071185985 | 0.044848175 | -1.587266036 | 0.112452404 | 0.30863821 | 0.071185985 |
| ENSMUSG00000059866 | Tnip2 | 231130 | protein_coding | 247.3101628 | -0.071191597 | 0.102028549 | -0.697761533 | 0.485326339 | 0.706609067 | 0.071191597 |
| ENSMUSG00000050079 | Rspry1 | 67610 | protein_coding | 797.2591999 | 0.071224872 | 0.050613786 | 1.407222765 | 0.159361343 | 0.379450817 | 0.071224872 |
| ENSMUSG00000096010 | H4c16 | 320332 | protein_coding | 1323.460439 | 0.07123465 | 0.115171397 | 0.618509902 | 0.536239272 | 0.742878353 | 0.07123465 |
| ENSMUSG00000029545 | Acads | 11409 | protein_coding | 6645.015954 | -0.071243848 | 0.059790035 | -1.19156725 | 0.233430973 | 0.472813526 | 0.071243848 |
| ENSMUSG00000041297 | Cdk13 | 69562 | protein_coding | 1961.374899 | -0.071251389 | 0.038526996 | -1.849388647 | 0.064401714 | 0.218011427 | 0.071251389 |
| ENSMUSG00000022407 | Adsl | 11564 | protein_coding | 1061.781756 | 0.071256259 | 0.052635899 | 1.353757818 | 0.175813655 | 0.402176854 | 0.071256259 |
| ENSMUSG00000014301 | Pam16 | 66449 | protein_coding | 199.8772453 | 0.071316714 | 0.088386943 | 0.806869343 | 0.419741763 | 0.657014281 | 0.071316714 |
| ENSMUSG00000020818 | Mfsd11 | 69900 | protein_coding | 454.7260345 | 0.071323162 | 0.076086294 | 0.937398294 | 0.348553718 | 0.591013123 | 0.071323162 |
| ENSMUSG00000055415 | Atp10b | 319767 | protein_coding | 4995.132391 | 0.071334594 | 0.048255787 | 1.47825989 | 0.139338229 | 0.349744377 | 0.071334594 |
| ENSMUSG00000021222 | Dcaf4 | 73828 | protein_coding | 339.5230915 | -0.071352726 | 0.083304766 | -0.856526337 | 0.391706711 | 0.631515959 | 0.071352726 |
| ENSMUSG00000042705 | Commd10 | 69456 | protein_coding | 443.2333597 | -0.071356234 | 0.065365441 | -1.091650769 | 0.274986632 | 0.517408635 | 0.071356234 |
| ENSMUSG00000023915 | Tnfrsf21 | 94185 | protein_coding | 1854.636561 | 0.071364503 | 0.058770162 | 1.214298223 | 0.224633864 | 0.462665431 | 0.071364503 |
| ENSMUSG00000027333 | Smox | 228608 | protein_coding | 440.4315907 | 0.071403652 | 0.082098168 | 0.869735027 | 0.384445226 | 0.624972092 | 0.071403652 |
| ENSMUSG00000028234 | Rps20 | 67427 | protein_coding | 9398.83035 | -0.07145416 | 0.07118644 | -1.003760827 | 0.31549391 | 0.559794288 | 0.07145416 |
| ENSMUSG00000046380 | Jrk | 16469 | protein_coding | 106.49231 | 0.071462298 | 0.128777477 | 0.554928546 | 0.578943535 | 0.771366688 | 0.071462298 |
| ENSMUSG00000024953 | Prdx5 | 54683 | protein_coding | 3697.874623 | -0.071472287 | 0.059520806 | -1.200795001 | 0.229830731 | 0.469064473 | 0.071472287 |
| ENSMUSG00000002325 | Irf9 | 16391 | protein_coding | 623.4022474 | -0.071473493 | 0.122980444 | -0.581177712 | 0.561120687 | 0.759206347 | 0.071473493 |
| ENSMUSG00000048118 | Arid4a | 238247 | protein_coding | 1096.509086 | -0.071506956 | 0.062501822 | -1.144077952 | 0.25259131 | 0.493642761 | 0.071506956 |
| ENSMUSG00000060098 | Prmt7 | 214572 | protein_coding | 506.492333 | -0.071525957 | 0.078740973 | -0.908370247 | 0.36368264 | 0.606076959 | 0.071525957 |
| ENSMUSG00000023805 | Synj2 | 20975 | protein_coding | 951.5134327 | -0.07154079 | 0.063814284 | -1.121078006 | 0.26225466 | 0.504097842 | 0.07154079 |
| ENSMUSG00000027654 | Fam83d | 71878 | protein_coding | 244.472248 | 0.07155131 | 0.115213645 | 0.621031568 | 0.534578853 | 0.741732183 | 0.07155131 |
| ENSMUSG00000006731 | B4galnt1 | 14421 | protein_coding | 2273.580744 | 0.071596362 | 0.094918437 | 0.754293517 | 0.450672994 | 0.679951459 | 0.071596362 |
| ENSMUSG00000048899 | Rimkla | 194237 | protein_coding | 29.24759613 | 0.071599835 | 0.256805705 | 0.278809361 | 0.78039113 | 0.898056783 | 0.071599835 |

| ensembl_gene_id | external_gene_name | entrezgene_id | gene_biotype | baseMean | log2FoldChange | lfcSE | stat | pvalue | padj | absLog2FC |
|---|---|---|---|---|---|---|---|---|---|---|
| ENSMUSG00000032902 | Slc16a1 | 20501 | protein_coding | 9067.029237 | -0.071603917 | 0.08325521 | -0.860053295 | 0.389759665 | 0.629572728 | 0.071603917 |
| ENSMUSG00000018661 | Cog1 | 16834 | protein_coding | 1685.608419 | 0.071612649 | 0.051332812 | 1.395065785 | 0.162996003 | 0.384472215 | 0.071612649 |
| ENSMUSG00000028322 | Exosc3 | 66362 | protein_coding | 384.5329422 | 0.07161386 | 0.067382541 | 1.062795474 | 0.287874713 | 0.530274756 | 0.07161386 |
| ENSMUSG00000061887 | Ssbp3 | 72475 | protein_coding | 1682.303176 | 0.071636046 | 0.061106354 | 1.17231748 | 0.241069616 | 0.481643597 | 0.071636046 |
| ENSMUSG00000003072 | Atp5d | 66043 | protein_coding | 8280.533133 | 0.071644825 | 0.097658129 | 0.733628893 | 0.463174947 | 0.689357631 | 0.071644825 |
| ENSMUSG00000071654 | Uqcc3 | 107197 | protein_coding | 517.4169361 | -0.071645023 | 0.082575635 | -0.867629095 | 0.385597413 | 0.625831733 | 0.071645023 |
| ENSMUSG00000034333 | Zbed4 | 223773 | protein_coding | 331.8912156 | 0.071718832 | 0.073922707 | 0.970186771 | 0.331953404 | 0.575141012 | 0.071718832 |
| ENSMUSG00000025470 | Zfp511 | 69752 | protein_coding | 205.0495987 | -0.071724825 | 0.087834468 | -0.816590881 | 0.414162268 | 0.651982545 | 0.071724825 |
| ENSMUSG00000026413 | Pkp1 | 18772 | protein_coding | 946.5471928 | 0.071758131 | 0.098858577 | 0.725866518 | 0.467920604 | 0.693047027 | 0.071758131 |
| ENSMUSG00000030315 | Vgll4 | 232334 | protein_coding | 1246.008166 | -0.071787944 | 0.066790526 | -1.074822247 | 0.282454318 | 0.525210203 | 0.071787944 |
| ENSMUSG00000021936 | Mapk8 | 26419 | protein_coding | 569.356394 | 0.071830183 | 0.064044848 | 1.121560676 | 0.262049279 | 0.503943326 | 0.071830183 |
| ENSMUSG00000051113 | Garin5a | 75538 | protein_coding | 15.21538105 | -0.071848235 | 0.367663547 | -0.195418434 | 0.845065387 | 0.930518195 | 0.071848235 |
| ENSMUSG00000065208 | Gm24616 | 115488691 | snoRNA | 11.6936305 | -0.071860048 | 0.35399422 | -0.202997799 | 0.839136752 | 0.928687288 | 0.071860048 |
| ENSMUSG00000032582 | Rbm6 | 19654 | protein_coding | 1635.682956 | -0.071862693 | 0.063920459 | -1.124251842 | 0.260906202 | 0.502824369 | 0.071862693 |
| ENSMUSG00000032010 | Usp2 | 53376 | protein_coding | 241.9360373 | -0.07186737 | 0.108913271 | -0.659858704 | 0.509344507 | 0.72429229 | 0.07186737 |
| ENSMUSG00000018168 | Ikzf3 | 22780 | protein_coding | 29.73657611 | 0.071872248 | 0.240359107 | 0.299020283 | 0.764924569 | 0.88952542 | 0.071872248 |
| ENSMUSG00000067219 | Nipal1 | 70701 | protein_coding | 1386.889923 | -0.071884416 | 0.067001095 | -1.07288419 | 0.283323081 | 0.525975748 | 0.071884416 |
| ENSMUSG00000058297 | Spock2 | 94214 | protein_coding | 19.88770225 | -0.071890262 | 0.299564149 | -0.239982863 | 0.810343542 | 0.914739662 | 0.071890262 |
| ENSMUSG00000026274 | Pask | 269224 | protein_coding | 152.5241035 | 0.071962844 | 0.120344533 | 0.59797352 | 0.549857602 | 0.751487589 | 0.071962844 |
| ENSMUSG00000041028 | Ghitm | 66092 | protein_coding | 6884.118098 | 0.072004634 | 0.039960771 | 1.801883016 | 0.071563813 | 0.233254754 | 0.072004634 |
| ENSMUSG00000038167 | Plekhg6 | 213522 | protein_coding | 764.0886129 | 0.07202724 | 0.119936082 | 0.600546881 | 0.548141827 | 0.750796946 | 0.07202724 |
| ENSMUSG00000032783 | Troap | 78733 | protein_coding | 77.92843424 | 0.072029071 | 0.168800542 | 0.426711135 | 0.669589739 | 0.831366151 | 0.072029071 |
| ENSMUSG00000017009 | Sdc4 | 20971 | protein_coding | 10370.9468 | -0.072036033 | 0.04416133 | -1.631201611 | 0.102847785 | 0.292733187 | 0.072036033 |
| ENSMUSG00000027122 | Arl14ep | 212772 | protein_coding | 305.57393 | 0.072038106 | 0.094369732 | 0.763360287 | 0.445248557 | 0.676259511 | 0.072038106 |
| ENSMUSG00000001380 | Hars | 15115 | protein_coding | 925.1461395 | 0.072059353 | 0.054825983 | 1.314328529 | 0.188735671 | 0.419197611 | 0.072059353 |
| ENSMUSG00000025142 | Aspscr1 | 68938 | protein_coding | 915.5116987 | 0.072080997 | 0.045710317 | 1.576908715 | 0.114816531 | 0.312566879 | 0.072080997 |
| ENSMUSG00000027180 | Fbxo3 | 57443 | protein_coding | 1367.962251 | -0.072092829 | 0.044575905 | -1.617304896 | 0.105812483 | 0.297926072 | 0.072092829 |
| ENSMUSG00000031442 | Mcf2l | 17207 | protein_coding | 2396.196093 | 0.072141464 | 0.062155134 | 1.160667817 | 0.245777015 | 0.486955333 | 0.072141464 |
| ENSMUSG00000005804 | Bloc1s6 | 18457 | protein_coding | 661.1455277 | 0.072165291 | 0.057091233 | 1.264034549 | 0.206217627 | 0.440953549 | 0.072165291 |
| ENSMUSG00000068246 | Apol9b | 71898 | protein_coding | 21.75464977 | -0.072175675 | 0.336814206 | -0.214289283 | 0.830321474 | 0.924828677 | 0.072175675 |
| ENSMUSG00000039745 | Htatip2 | 53415 | protein_coding | 509.175463 | 0.072198056 | 0.08055416 | 0.896267247 | 0.370110048 | 0.612180999 | 0.072198056 |
| ENSMUSG00000037017 | Zscan21 | 22697 | protein_coding | 414.9582595 | -0.072214029 | 0.075043349 | -0.962297525 | 0.335900172 | 0.579426898 | 0.072214029 |
| ENSMUSG00000056394 | Lig1 | 16881 | protein_coding | 574.2814962 | 0.072218887 | 0.128618064 | 0.561498789 | 0.574457555 | 0.768186995 | 0.072218887 |
| ENSMUSG00000055827 | Gsdmc3 | 270328 | protein_coding | 3381.904728 | 0.07222239 | 0.103286576 | 0.699242753 | 0.484400336 | 0.706263486 | 0.07222239 |
| ENSMUSG00000073926 | Or52d3 | 57250 | protein_coding | 6.290045628 | -0.072222799 | 0.606718767 | -0.119038347 | 0.905244975 | 0.958435249 | 0.072222799 |
| ENSMUSG00000021794 | Glud1 | 14661 | protein_coding | 5687.144328 | -0.072227506 | 0.040296228 | -1.792413577 | 0.073066741 | 0.236530829 | 0.072227506 |
| ENSMUSG00000020380 | Rad50 | 19360 | protein_coding | 789.0084981 | -0.072228188 | 0.061478263 | -1.174857323 | 0.240051802 | 0.480533622 | 0.072228188 |
| ENSMUSG00000039512 | Uhrf1bp1 | 224648 | protein_coding | 418.1490799 | -0.07225639 | 0.087891566 | -0.822108339 | 0.41101524 | 0.649146117 | 0.07225639 |
| ENSMUSG00000021051 | Ppp2r5e | 26932 | protein_coding | 715.0802635 | -0.072287311 | 0.072636155 | -0.995197379 | 0.319640276 | 0.563143068 | 0.072287311 |

| ensembl_gene_id | external_gene_name | entrezgene_id | gene_biotype | baseMean | log2FoldChange | lfcSE | stat | pvalue | padj | absLog2FC |
|---|---|---|---|---|---|---|---|---|---|---|
| ENSMUSG00000006575 | Rundc3a | 51799 | protein_coding | 144.3957086 | -0.072308745 | 0.097448555 | -0.742019668 | 0.458075419 | 0.685606255 | 0.072308745 |
| ENSMUSG00000022037 | Clu | 12759 | protein_coding | 50.48251049 | 0.072310952 | 0.187683496 | 0.385281359 | 0.700028971 | 0.850359085 | 0.072310952 |
| ENSMUSG00000049300 | Prmt6 | 99890 | protein_coding | 187.9594902 | 0.072351803 | 0.090203973 | 0.802091088 | 0.42250027 | 0.65874317 | 0.072351803 |
| ENSMUSG00000073609 | D2hgdh | 98314 | protein_coding | 480.9967172 | -0.072362271 | 0.064977218 | -1.113656041 | 0.265426775 | 0.507292913 | 0.072362271 |
| ENSMUSG00000044792 | Isca1 | 69046 | protein_coding | 1104.461335 | 0.07236851 | 0.066540082 | 1.087592738 | 0.276774938 | 0.519742597 | 0.07236851 |
| ENSMUSG00000086003 | B230206L02Rik | 100039440 | lncRNA | 10.43505499 | -0.072393183 | 0.461692989 | -0.156799398 | 0.875402947 | 0.94394379 | 0.072393183 |
| ENSMUSG00000048351 | Coa7 | 69893 | protein_coding | 411.0547759 | -0.072424773 | 0.070983053 | -1.020310773 | 0.307581095 | 0.551334926 | 0.072424773 |
| ENSMUSG00000005683 | Cs | 12974 | protein_coding | 13556.93333 | 0.072430036 | 0.078751558 | 0.919728296 | 0.357714762 | 0.600860147 | 0.072430036 |
| ENSMUSG00000027341 | Tmem230 | 70612 | protein_coding | 286.6197525 | 0.072455149 | 0.078696734 | 0.920688134 | 0.357213273 | 0.600206132 | 0.072455149 |
| ENSMUSG00000021458 | Aopep | 72061 | protein_coding | 1341.307013 | -0.072464524 | 0.067596439 | -1.072016874 | 0.283712453 | 0.526335066 | 0.072464524 |
| ENSMUSG00000042428 | Mgat3 | 17309 | protein_coding | 3873.099402 | 0.07247461 | 0.048535757 | 1.493220985 | 0.135379351 | 0.343658553 | 0.07247461 |
| ENSMUSG00000045078 | Rnf216 | 108086 | protein_coding | 949.4962526 | -0.072497797 | 0.05203984 | -1.393121052 | 0.163583188 | 0.385406293 | 0.072497797 |
| ENSMUSG00000049323 | Smcr8 | 237782 | protein_coding | 955.7643231 | -0.072557944 | 0.047173189 | -1.538118275 | 0.124019698 | 0.326892409 | 0.072557944 |
| ENSMUSG00000024913 | Lrp5 | 16973 | protein_coding | 1528.185535 | -0.072566604 | 0.078915436 | -0.919548926 | 0.357808527 | 0.600893275 | 0.072566604 |
| ENSMUSG00000027030 | Stk39 | 53416 | protein_coding | 937.2649351 | 0.0725777 | 0.073343371 | 0.989560464 | 0.322389002 | 0.566481525 | 0.0725777 |
| ENSMUSG00000044982 | Sft2d3 | 67158 | protein_coding | 147.6193523 | 0.072587411 | 0.115012079 | 0.631128592 | 0.527956448 | 0.737831675 | 0.072587411 |
| ENSMUSG00000046201 | Scaf8 | 106583 | protein_coding | 1261.229579 | -0.072594358 | 0.042255568 | -1.717983245 | 0.085799671 | 0.261666282 | 0.072594358 |
| ENSMUSG00000030423 | Pop4 | 66161 | protein_coding | 298.0433583 | -0.072601166 | 0.095962017 | -0.75656149 | 0.44931262 | 0.679246742 | 0.072601166 |
| ENSMUSG00000004096 | Cwc15 | 66070 | protein_coding | 1249.07324 | 0.072620532 | 0.04911653 | 1.478535477 | 0.139264509 | 0.34961375 | 0.072620532 |
| ENSMUSG00000031311 | Nono | 53610 | protein_coding | 3542.252465 | -0.072621574 | 0.038627384 | -1.880054175 | 0.060100695 | 0.208317411 | 0.072621574 |
| ENSMUSG00000090553 | Snrpe | 20643 | protein_coding | 732.6715904 | -0.072664216 | 0.049686647 | -1.462449584 | 0.14361805 | 0.355746355 | 0.072664216 |
| ENSMUSG00000031072 | LTO1 | 72284 | protein_coding | 320.4131202 | 0.072721639 | 0.078340459 | 0.928276907 | 0.35326395 | 0.595742729 | 0.072721639 |
| ENSMUSG00000022787 | Wdr53 | 68980 | protein_coding | 193.299878 | -0.07273243 | 0.087588467 | -0.830388203 | 0.406319333 | 0.645460194 | 0.07273243 |
| ENSMUSG00000030990 | Pgap2 | 233575 | protein_coding | 1082.866198 | 0.072743277 | 0.055941419 | 1.300347361 | 0.193481943 | 0.425338507 | 0.072743277 |
| ENSMUSG00000034930 | Rtkn | 20166 | protein_coding | 330.929451 | -0.072751035 | 0.074520374 | -0.976256979 | 0.328937133 | 0.5725114 | 0.072751035 |
| ENSMUSG00000033933 | Vhl | 22346 | protein_coding | 466.6812907 | -0.072778228 | 0.067046659 | -1.085486265 | 0.277706344 | 0.520659264 | 0.072778228 |
| ENSMUSG00000055240 | Zfp101 | 22643 | protein_coding | 192.6068253 | -0.072781261 | 0.13268283 | -0.54853564 | 0.583324165 | 0.774834759 | 0.072781261 |
| ENSMUSG00000027076 | Timm10 | 30059 | protein_coding | 329.5906178 | -0.072784465 | 0.085360802 | -0.852668475 | 0.393843178 | 0.633492143 | 0.072784465 |
| ENSMUSG00000029397 | Rchy1 | 68098 | protein_coding | 778.4475501 | 0.072801737 | 0.051348636 | 1.417792984 | 0.156251213 | 0.374715872 | 0.072801737 |
| ENSMUSG00000005510 | Ndufs3 | 68349 | protein_coding | 2151.886316 | 0.07287984 | 0.061669387 | 1.181783107 | 0.237291772 | 0.477271565 | 0.07287984 |
| ENSMUSG00000045205 | Dpy19l4 | 381510 | protein_coding | 675.4080739 | -0.072891452 | 0.05880719 | -1.239498964 | 0.215160772 | 0.450636075 | 0.072891452 |
| ENSMUSG00000024290 | Rock1 | 19877 | protein_coding | 2602.894341 | -0.072893633 | 0.04331596 | -1.682835438 | 0.092406953 | 0.274093604 | 0.072893633 |
| ENSMUSG00000012848 | Rps5 | 20103 | protein_coding | 13687.54028 | -0.072926864 | 0.047173502 | -1.545928555 | 0.122121823 | 0.323870851 | 0.072926864 |
| ENSMUSG00000064363 | mt-Nd4 | 17719 | protein_coding | 15778.44669 | 0.072927317 | 0.129449377 | 0.563365533 | 0.573186002 | 0.767653434 | 0.072927317 |
| ENSMUSG00000028329 | Xpa | 22590 | protein_coding | 234.0005773 | -0.072971322 | 0.100889353 | -0.723280702 | 0.469507447 | 0.69386728 | 0.072971322 |
| ENSMUSG00000021796 | Bmpr1a | 12166 | protein_coding | 2269.826181 | -0.072975821 | 0.043916034 | -1.661712438 | 0.096570446 | 0.282085054 | 0.072975821 |
| ENSMUSG00000019820 | Utrn | 22288 | protein_coding | 3604.273103 | 0.072985437 | 0.060992924 | 1.196621378 | 0.231454164 | 0.470523296 | 0.072985437 |
| ENSMUSG00000023505 | Cdca3 | 14793 | protein_coding | 716.6600448 | 0.072986606 | 0.098450916 | 0.741350196 | 0.458481134 | 0.685788934 | 0.072986606 |
| ENSMUSG00000001017 | Chtop | 66511 | protein_coding | 2125.670565 | 0.073005357 | 0.033459478 | 2.181903674 | 0.029116643 | 0.127460395 | 0.073005357 |

| ensembl_gene_id | external_gene_name | entrezgene_id | gene_biotype | baseMean | log2FoldChange | lfcSE | stat | pvalue | padj | absLog2FC |
|---|---|---|---|---|---|---|---|---|---|---|
| ENSMUSG00000050312 | Nsun3 | 106338 | protein_coding | 389.251106 | 0.073024305 | 0.078386554 | 0.931592242 | 0.351547294 | 0.593903542 | 0.073024305 |
| ENSMUSG00000042672 | Dcst1 | 77772 | protein_coding | 32.83900435 | 0.073044596 | 0.21171203 | 0.345018636 | 0.73008036 | 0.869327992 | 0.073044596 |
| ENSMUSG00000051065 | Mb21d2 | 239796 | protein_coding | 378.9046386 | -0.073082442 | 0.088283604 | -0.827814436 | 0.407775598 | 0.646388536 | 0.073082442 |
| ENSMUSG00000032481 | Smarcc1 | 20588 | protein_coding | 1197.039314 | -0.073089825 | 0.064238244 | -1.137793002 | 0.255206928 | 0.496528185 | 0.073089825 |
| ENSMUSG00000020863 | Luc7l3 | 67684 | protein_coding | 1180.919717 | -0.073166205 | 0.087557122 | -0.835639681 | 0.403357638 | 0.642544198 | 0.073166205 |
| ENSMUSG00000036552 | Ermard | 381062 | protein_coding | 119.3238535 | 0.073188708 | 0.115807981 | 0.631983281 | 0.527397802 | 0.737351049 | 0.073188708 |
| ENSMUSG00000022957 | Itsn1 | 16443 | protein_coding | 830.2811027 | -0.073226785 | 0.056461316 | -1.296937274 | 0.19465277 | 0.426574973 | 0.073226785 |
| ENSMUSG00000054150 | Syne3 | 212073 | protein_coding | 205.5619568 | -0.07323533 | 0.122557183 | -0.597560482 | 0.550133239 | 0.751671802 | 0.07323533 |
| ENSMUSG00000021250 | Fos | 14281 | protein_coding | 43639.58709 | 0.073249089 | 0.058864591 | 1.244365872 | 0.213364937 | 0.449133255 | 0.073249089 |
| ENSMUSG00000026806 | Ddx31 | 227674 | protein_coding | 179.8470098 | -0.073254301 | 0.104734769 | -0.699426776 | 0.484285359 | 0.706263486 | 0.073254301 |
| ENSMUSG00000029594 | Rbm19 | 74111 | protein_coding | 354.872076 | 0.073279958 | 0.092537367 | 0.791895865 | 0.428421398 | 0.663448424 | 0.073279958 |
| ENSMUSG00000069631 | Strada | 72149 | protein_coding | 147.730039 | -0.073289848 | 0.1171594 | -0.625556706 | 0.531605743 | 0.740026398 | 0.073289848 |
| ENSMUSG00000059796 | Eif4a1 | 13681 | protein_coding | 7872.416921 | 0.073301574 | 0.048754085 | 1.503496051 | 0.132711175 | 0.339503983 | 0.073301574 |
| ENSMUSG00000010755 | Cars | 27267 | protein_coding | 513.3403579 | 0.073328969 | 0.071618378 | 1.02388481 | 0.305889686 | 0.55012849 | 0.073328969 |
| ENSMUSG00000046671 | Mtfr1l | 76824 | protein_coding | 918.7271201 | -0.073380812 | 0.044937284 | -1.632960542 | 0.102477296 | 0.292221615 | 0.073380812 |
| ENSMUSG00000061455 | Stx17 | 67727 | protein_coding | 1070.265352 | -0.073394324 | 0.064106656 | -1.144878374 | 0.252259543 | 0.493470298 | 0.073394324 |
| ENSMUSG00000040521 | Tsfm | 66399 | protein_coding | 450.2280951 | 0.073418353 | 0.10158462 | 0.722730989 | 0.469845173 | 0.694009261 | 0.073418353 |
| ENSMUSG00000032254 | Kif23 | 71819 | protein_coding | 668.11012 | 0.073433885 | 0.09021894 | 0.813951985 | 0.415672458 | 0.653685981 | 0.073433885 |
| ENSMUSG00000049339 | Retreg2 | 227298 | protein_coding | 2000.210739 | -0.073445076 | 0.04846462 | -1.515436959 | 0.129661784 | 0.335750736 | 0.073445076 |
| ENSMUSG00000022681 | Ntan1 | 18203 | protein_coding | 518.5829104 | -0.073471712 | 0.069111577 | -1.063088345 | 0.28774189 | 0.530090639 | 0.073471712 |
| ENSMUSG00000034216 | Vps18 | 228545 | protein_coding | 603.6169108 | 0.073478888 | 0.074441898 | 0.987063609 | 0.323611456 | 0.567701725 | 0.073478888 |
| ENSMUSG00000058230 | Arhgap35 | 232906 | protein_coding | 1889.823107 | -0.073487169 | 0.063221257 | -1.16238069 | 0.245080863 | 0.486272033 | 0.073487169 |
| ENSMUSG00000078813 | Leng1 | 69757 | protein_coding | 350.0395529 | 0.073497775 | 0.084486577 | 0.869934348 | 0.384336283 | 0.624972092 | 0.073497775 |
| ENSMUSG00000039308 | Ndst2 | 17423 | protein_coding | 475.1592626 | -0.073509151 | 0.07338551 | -1.001684814 | 0.316495843 | 0.560339056 | 0.073509151 |
| ENSMUSG00000025722 | Wdr73 | 71968 | protein_coding | 262.3969313 | -0.073557268 | 0.082421244 | -0.892455209 | 0.372148991 | 0.61393518 | 0.073557268 |
| ENSMUSG00000042320 | Prox2 | 73422 | protein_coding | 15.31322311 | -0.073565774 | 0.321195147 | -0.229037627 | 0.818839671 | 0.918528635 | 0.073565774 |
| ENSMUSG00000002416 | Ndufb2 | 68198 | protein_coding | 758.7271518 | 0.073602989 | 0.080762045 | 0.91135618 | 0.362107734 | 0.605026049 | 0.073602989 |
| ENSMUSG00000051177 | Plcb1 | 18795 | protein_coding | 218.1473041 | 0.073643825 | 0.106485061 | 0.691588322 | 0.489195897 | 0.709266447 | 0.073643825 |
| ENSMUSG00000027363 | Usp8 | 84092 | protein_coding | 1355.505355 | 0.073647868 | 0.044063239 | 1.671412942 | 0.094640141 | 0.278464068 | 0.073647868 |
| ENSMUSG00000026409 | Pfkfb2 | 18640 | protein_coding | 599.4444976 | 0.073648312 | 0.06946241 | 1.060261388 | 0.2890257 | 0.531357026 | 0.073648312 |
| ENSMUSG00000026317 | Cln8 | 26889 | protein_coding | 424.7029999 | -0.073656686 | 0.069725634 | -1.056378867 | 0.290795157 | 0.5332246 | 0.073656686 |
| ENSMUSG00000029642 | Polr1d | 20018 | protein_coding | 1820.519019 | 0.073692563 | 0.072386222 | 1.018046819 | 0.308655707 | 0.552278995 | 0.073692563 |
| ENSMUSG00000022200 | Golph3 | 66629 | protein_coding | 3880.636828 | 0.073693809 | 0.02772924 | 2.6576209 | 0.007869436 | 0.0501028 | 0.073693809 |
| ENSMUSG00000031816 | Mthfsd | 234814 | protein_coding | 411.3365424 | 0.073703179 | 0.094989192 | 0.775911209 | 0.437801414 | 0.670824076 | 0.073703179 |
| ENSMUSG00000030650 | Tmc5 | 74424 | protein_coding | 2638.893693 | -0.073741011 | 0.078317928 | -0.941559775 | 0.346418072 | 0.588629676 | 0.073741011 |
| ENSMUSG00000036636 | Clcn7 | 26373 | protein_coding | 720.0028555 | -0.073781572 | 0.06381486 | -1.156181675 | 0.247606854 | 0.488580666 | 0.073781572 |
| ENSMUSG00000021572 | Cep72 | 74470 | protein_coding | 130.450248 | -0.073783091 | 0.123982357 | -0.595109603 | 0.55177021 | 0.752522003 | 0.073783091 |
| ENSMUSG00000020167 | Tcf3 | 21423 | protein_coding | 922.1197606 | 0.073795555 | 0.071796009 | 1.02785038 | 0.304020214 | 0.548046454 | 0.073795555 |
| ENSMUSG00000034570 | Inpp5j | 170835 | protein_coding | 1304.098683 | -0.073810113 | 0.071639508 | -1.030299005 | 0.302869666 | 0.546718055 | 0.073810113 |

| ensembl_gene_id | external_gene_name | entrezgene_id | gene_biotype | baseMean | log2FoldChange | lfcSE | stat | pvalue | padj | absLog2FC |
|---|---|---|---|---|---|---|---|---|---|---|
| ENSMUSG00000005893 | Nr2c2 | 22026 | protein_coding | 1094.214911 | -0.073826137 | 0.046634232 | -1.58308895 | 0.113401192 | 0.310127356 | 0.073826137 |
| ENSMUSG00000040669 | Phc1 | 13619 | protein_coding | 252.4513375 | -0.073842598 | 0.085788831 | -0.860748393 | 0.389376634 | 0.62920611 | 0.073842598 |
| ENSMUSG00000034923 | Ly6g6f | 433099 | protein_coding | 38.52260051 | -0.073851324 | 0.204192554 | -0.361674914 | 0.717594976 | 0.861553924 | 0.073851324 |
| ENSMUSG00000021451 | Sema4d | 20354 | protein_coding | 175.4750873 | 0.073869949 | 0.118632137 | 0.622680754 | 0.533494333 | 0.741375537 | 0.073869949 |
| ENSMUSG00000023025 | Larp4 | 207214 | protein_coding | 3180.795274 | -0.073909771 | 0.057175395 | -1.292684925 | 0.196120054 | 0.428683669 | 0.073909771 |
| ENSMUSG00000070733 | Fryl | 72313 | protein_coding | 4298.088923 | 0.073920834 | 0.064077916 | 1.153608589 | 0.248660676 | 0.489520351 | 0.073920834 |
| ENSMUSG00000024773 | Atg2a | 329015 | protein_coding | 1355.369237 | -0.073923708 | 0.065262109 | -1.132720174 | 0.257331783 | 0.499216262 | 0.073923708 |
| ENSMUSG00000026172 | Bcs1l | 66821 | protein_coding | 269.8400287 | 0.073941201 | 0.077886146 | 0.949349859 | 0.342442703 | 0.585130336 | 0.073941201 |
| ENSMUSG00000038524 | Fchsd1 | 319262 | protein_coding | 171.5707954 | 0.073947217 | 0.097702302 | 0.75686258 | 0.449132196 | 0.679215578 | 0.073947217 |
| ENSMUSG00000033544 | Angptl1 | 72713 | protein_coding | 9.016518769 | -0.073953243 | 0.414950656 | -0.178221777 | 0.858548808 | 0.936160692 | 0.073953243 |
| ENSMUSG00000032551 | 1110059G10Rik | 66202 | protein_coding | 265.7798406 | -0.073959329 | 0.094755144 | -0.780531019 | 0.435078377 | 0.66862102 | 0.073959329 |
| ENSMUSG00000061118 | Dnajc30 | 66114 | protein_coding | 274.1704207 | -0.07399976 | 0.07451272 | -0.993115811 | 0.320653519 | 0.564169682 | 0.07399976 |
| ENSMUSG00000044573 | Acp1 | 11431 | protein_coding | 1003.463387 | 0.07407245 | 0.052038881 | 1.423405881 | 0.154618529 | 0.372364958 | 0.07407245 |
| ENSMUSG00000030216 | Wbp11 | 60321 | protein_coding | 883.0821427 | 0.07409043 | 0.058813382 | 1.25975462 | 0.207757895 | 0.443013562 | 0.07409043 |
| ENSMUSG00000035572 | Dcaf10 | 242418 | protein_coding | 734.1954079 | 0.074115905 | 0.053702037 | 1.3801321 | 0.167545976 | 0.391491529 | 0.074115905 |
| ENSMUSG00000075012 | Fjx1 | 14221 | protein_coding | 10.14508869 | 0.074142588 | 0.374451321 | 0.198003276 | 0.843042501 | 0.930024923 | 0.074142588 |
| ENSMUSG00000029863 | Casp2 | 12366 | protein_coding | 1054.377855 | -0.074154421 | 0.054308522 | -1.365428811 | 0.172118314 | 0.396989709 | 0.074154421 |
| ENSMUSG00000022351 | Sqle | 20775 | protein_coding | 3135.131492 | -0.074209507 | 0.124489032 | -0.59611281 | 0.551099867 | 0.75210185 | 0.074209507 |
| ENSMUSG00000031389 | Arhgap4 | 171207 | protein_coding | 58.1666985 | 0.074213259 | 0.159927255 | 0.464043851 | 0.642616328 | 0.814162427 | 0.074213259 |
| ENSMUSG00000071573 | Rnls | 67795 | protein_coding | 92.87648231 | 0.074220635 | 0.134877756 | 0.55028077 | 0.582126812 | 0.773817927 | 0.074220635 |
| ENSMUSG00000020530 | Ggnbp2 | 217039 | protein_coding | 2211.121154 | -0.074248197 | 0.059543074 | -1.246966127 | 0.212409918 | 0.447899418 | 0.074248197 |
| ENSMUSG00000001100 | Poldip2 | 67811 | protein_coding | 1312.200325 | 0.074256598 | 0.049505658 | 1.499961831 | 0.13362429 | 0.34107542 | 0.074256598 |
| ENSMUSG00000068882 | Ssb | 20823 | protein_coding | 1550.202882 | 0.074277082 | 0.074470017 | 0.997409229 | 0.318565913 | 0.562331745 | 0.074277082 |
| ENSMUSG00000031697 | Orc6 | 56452 | protein_coding | 195.0304577 | 0.074285679 | 0.12093369 | 0.614267856 | 0.53903834 | 0.745103008 | 0.074285679 |
| ENSMUSG00000037188 | Grhl3 | 230824 | protein_coding | 10.56802967 | 0.074291077 | 0.39242565 | 0.18931249 | 0.849847907 | 0.932219915 | 0.074291077 |
| ENSMUSG00000032085 | Tagln | 21345 | protein_coding | 27.5638147 | -0.07431353 | 0.229872153 | -0.323282003 | 0.746481682 | 0.879750811 | 0.07431353 |
| ENSMUSG00000049184 | Purg | 75029 | protein_coding | 26.06278218 | -0.074318614 | 0.237722903 | -0.312627069 | 0.75456401 | 0.883529645 | 0.074318614 |
| ENSMUSG00000029389 | Ddx55 | 67848 | protein_coding | 283.5615636 | -0.074335296 | 0.086121288 | -0.863146597 | 0.388056871 | 0.628080406 | 0.074335296 |
| ENSMUSG00000040936 | Ulk4 | 209012 | protein_coding | 28.80092384 | -0.07438774 | 0.235540302 | -0.315817461 | 0.752141064 | 0.882101588 | 0.07438774 |
| ENSMUSG00000020290 | Xpo1 | 103573 | protein_coding | 1833.576877 | 0.074404103 | 0.047467506 | 1.567474457 | 0.117003832 | 0.31618145 | 0.074404103 |
| ENSMUSG00000032212 | Sltm | 66660 | protein_coding | 1262.555168 | -0.074424845 | 0.068673135 | -1.083754869 | 0.2784735 | 0.521232839 | 0.074424845 |
| ENSMUSG00000057637 | Prdm2 | 110593 | protein_coding | 812.8017737 | -0.074467349 | 0.059304728 | -1.255673069 | 0.209234527 | 0.444692306 | 0.074467349 |
| ENSMUSG00000012350 | Ehf | 13661 | protein_coding | 10922.52958 | -0.074477483 | 0.048754841 | -1.527591537 | 0.126613984 | 0.330619127 | 0.074477483 |
| ENSMUSG00000060601 | Nr1h2 | 22260 | protein_coding | 1147.610467 | -0.074499611 | 0.058068554 | -1.282959624 | 0.199506222 | 0.433033222 | 0.074499611 |
| ENSMUSG00000035949 | Fbxw2 | 30050 | protein_coding | 847.9005557 | -0.074559898 | 0.050920095 | -1.464252935 | 0.143124852 | 0.355049143 | 0.074559898 |
| ENSMUSG00000028907 | Utp11 | 67205 | protein_coding | 570.7745029 | 0.074577497 | 0.07538414 | 0.989299565 | 0.322516597 | 0.5665985 | 0.074577497 |
| ENSMUSG00000052488 | Cherp | 27967 | protein_coding | 262.3175321 | -0.074605481 | 0.103024077 | -0.72415578 | 0.468970103 | 0.693475193 | 0.074605481 |
| ENSMUSG00000055817 | Mta3 | 116871 | protein_coding | 699.4249057 | 0.074605516 | 0.056316118 | 1.324763125 | 0.185249729 | 0.414300704 | 0.074605516 |
| ENSMUSG00000037916 | Ndufv1 | 17995 | protein_coding | 5582.836797 | 0.074622474 | 0.063032413 | 1.183874617 | 0.236462699 | 0.476619666 | 0.074622474 |

| ensembl_gene_id | external_gene_name | entrezgene_id | gene_biotype | baseMean | log2FoldChange | lfcSE | stat | pvalue | padj | absLog2FC |
|---|---|---|---|---|---|---|---|---|---|---|
| ENSMUSG00000021044 | Adck1 | 72113 | protein_coding | 328.4048362 | 0.074626708 | 0.072172288 | 1.034007797 | 0.301132518 | 0.544679691 | 0.074626708 |
| ENSMUSG00000024222 | Fkbp5 | 14229 | protein_coding | 4767.851704 | 0.074710269 | 0.119714955 | 0.624067974 | 0.532582945 | 0.740799392 | 0.074710269 |
| ENSMUSG00000040209 | Zfp704 | 170753 | protein_coding | 1453.998372 | 0.074762115 | 0.064881522 | 1.152286702 | 0.24920328 | 0.489990404 | 0.074762115 |
| ENSMUSG00000015165 | Hnrnpl | 15388 | protein_coding | 4900.739916 | 0.074796949 | 0.050375868 | 1.484777392 | 0.137602803 | 0.347443307 | 0.074796949 |
| ENSMUSG00000020477 | Mrps24 | 64660 | protein_coding | 1124.975783 | 0.0748136 | 0.069009504 | 1.084105752 | 0.278317913 | 0.521115754 | 0.0748136 |
| ENSMUSG00000037646 | Vps13b | 666173 | protein_coding | 3133.105796 | -0.074816921 | 0.041159221 | -1.81774384 | 0.069103294 | 0.228065548 | 0.074816921 |
| ENSMUSG00000029782 | Tmem209 | 72649 | protein_coding | 504.1454351 | -0.074824542 | 0.090893181 | -0.823214033 | 0.41038629 | 0.648598559 | 0.074824542 |
| ENSMUSG00000016756 | Cmah | 12763 | protein_coding | 647.5596994 | -0.07482571 | 0.180147947 | -0.415356993 | 0.677880585 | 0.836585484 | 0.07482571 |
| ENSMUSG00000118100 | Gm50340 | 115489003 | lncRNA | 20.37140467 | 0.07491098 | 0.277595154 | 0.269856943 | 0.787270313 | 0.900662858 | 0.07491098 |
| ENSMUSG00000035051 | Dhx57 | 106794 | protein_coding | 445.6540018 | 0.074930181 | 0.064505195 | 1.16161467 | 0.245392021 | 0.486649916 | 0.074930181 |
| ENSMUSG00000017830 | Dhx58 | 80861 | protein_coding | 154.6706682 | 0.075025012 | 0.154264965 | 0.486338629 | 0.626727095 | 0.803180854 | 0.075025012 |
| ENSMUSG00000017718 | Afmid | 71562 | protein_coding | 118.3002021 | 0.075026371 | 0.125281626 | 0.598861727 | 0.549265096 | 0.751378251 | 0.075026371 |
| ENSMUSG00000043633 | Fam221b | 242408 | protein_coding | 21.95604174 | 0.075062442 | 0.277648318 | 0.27035079 | 0.786890395 | 0.900576045 | 0.075062442 |
| ENSMUSG00000027580 | Helz2 | 229003 | protein_coding | 2041.345373 | -0.075076356 | 0.068270801 | -1.099684704 | 0.271469522 | 0.513977697 | 0.075076356 |
| ENSMUSG00000024862 | Klc2 | 16594 | protein_coding | 324.7457046 | -0.075142561 | 0.075248338 | -0.99859429 | 0.317991267 | 0.561627805 | 0.075142561 |
| ENSMUSG00000024146 | Cript | 56724 | protein_coding | 909.1040009 | 0.075149209 | 0.045602408 | 1.647921967 | 0.099368685 | 0.286840423 | 0.075149209 |
| ENSMUSG00000031949 | Adat1 | 30947 | protein_coding | 55.71685584 | 0.075178088 | 0.170059988 | 0.442068052 | 0.658439962 | 0.825261698 | 0.075178088 |
| ENSMUSG00000025648 | Pfkfb4 | 270198 | protein_coding | 656.8089347 | -0.075185709 | 0.075461273 | -0.996348272 | 0.319080957 | 0.562649386 | 0.075185709 |
| ENSMUSG00000026991 | Pkp4 | 227937 | protein_coding | 1796.336273 | 0.075188676 | 0.043645129 | 1.722727783 | 0.084937764 | 0.260026017 | 0.075188676 |
| ENSMUSG00000024151 | Msh2 | 17685 | protein_coding | 755.8760574 | 0.075201615 | 0.059764546 | 1.258298107 | 0.208283966 | 0.443565039 | 0.075201615 |
| ENSMUSG00000021192 | Golga5 | 27277 | protein_coding | 1329.768012 | 0.075267229 | 0.059051477 | 1.274603667 | 0.202449559 | 0.436583513 | 0.075267229 |
| ENSMUSG00000031591 | Asah1 | 11886 | protein_coding | 2932.492355 | 0.075351744 | 0.060589386 | 1.243645941 | 0.2136299 | 0.449303803 | 0.075351744 |
| ENSMUSG00000022748 | Cmss1 | 66497 | protein_coding | 115.32092 | 0.075386174 | 0.128083185 | 0.588571984 | 0.556148432 | 0.755348304 | 0.075386174 |
| ENSMUSG00000024027 | Glp1r | 14652 | protein_coding | 25.6071525 | 0.075425097 | 0.278010067 | 0.271303475 | 0.786157634 | 0.900404409 | 0.075425097 |
| ENSMUSG00000039934 | Gsap | 212167 | protein_coding | 55.28903548 | 0.075428498 | 0.187928314 | 0.40136846 | 0.688148867 | 0.843368224 | 0.075428498 |
| ENSMUSG00000030788 | Rnf141 | 67150 | protein_coding | 1909.943203 | -0.07543804 | 0.049854811 | -1.513154672 | 0.130240375 | 0.336231563 | 0.07543804 |
| ENSMUSG00000022263 | Trio | 223435 | protein_coding | 2045.778622 | -0.075438624 | 0.05356707 | -1.408302239 | 0.159041593 | 0.37913729 | 0.075438624 |
| ENSMUSG00000030126 | Tmcc1 | 330401 | protein_coding | 804.9597656 | -0.075439585 | 0.046444872 | -1.624282325 | 0.104315568 | 0.295551199 | 0.075439585 |
| ENSMUSG00000041263 | Rusc1 | 72296 | protein_coding | 161.0980112 | -0.075446633 | 0.099785912 | -0.756085015 | 0.449598226 | 0.679246742 | 0.075446633 |
| ENSMUSG00000018381 | Abi3 | 66610 | protein_coding | 46.62282128 | 0.075452918 | 0.183707577 | 0.410722949 | 0.681275698 | 0.838859992 | 0.075452918 |
| ENSMUSG00000032422 | Snx14 | 244962 | protein_coding | 1169.824563 | -0.075453992 | 0.051841695 | -1.455469232 | 0.145539402 | 0.358228623 | 0.075453992 |
| ENSMUSG00000110424 | Zbed5-ps | 75479 | lncRNA | 14.81388293 | 0.075469438 | 0.338343725 | 0.223055528 | 0.82349229 | 0.921486272 | 0.075469438 |
| ENSMUSG00000062760 | Shisal1 | 72301 | protein_coding | 11.11722243 | -0.075498143 | 0.371564598 | -0.203189817 | 0.838986672 | 0.928643655 | 0.075498143 |
| ENSMUSG00000025917 | Cops5 | 26754 | protein_coding | 1124.731568 | 0.07551957 | 0.050913909 | 1.483279749 | 0.1380001 | 0.347998271 | 0.07551957 |
| ENSMUSG00000049516 | Spty2d1 | 101685 | protein_coding | 1110.451053 | 0.075534749 | 0.046130233 | 1.637423963 | 0.101541915 | 0.290503186 | 0.075534749 |
| ENSMUSG00000022594 | Lynx1 | 23936 | protein_coding | 238.3665241 | -0.075547531 | 0.097714815 | -0.773143063 | 0.439437718 | 0.671700891 | 0.075547531 |
| ENSMUSG00000062825 | Actg1 | 11465 | protein_coding | 27525.30669 | 0.075553108 | 0.049165188 | 1.536719597 | 0.12436199 | 0.32728228 | 0.075553108 |
| ENSMUSG00000024845 | Tmem134 | 66990 | protein_coding | 1466.212126 | 0.075561583 | 0.073942305 | 1.021899205 | 0.306828609 | 0.550870027 | 0.075561583 |
| ENSMUSG00000050144 | Slc25a44 | 229517 | protein_coding | 1115.007919 | 0.075588169 | 0.04713146 | 1.603773106 | 0.108764075 | 0.302003828 | 0.075588169 |

| ensembl_gene_id | external_gene_name | entrezgene_id | gene_biotype | baseMean | log2FoldChange | lfcSE | stat | pvalue | padj | absLog2FC |
|---|---|---|---|---|---|---|---|---|---|---|
| ENSMUSG00000093606 | B130034C11Rik | 414095 | lncRNA | 19.48169682 | 0.075595927 | 0.284201121 | 0.265994472 | 0.790243466 | 0.902120505 | 0.075595927 |
| ENSMUSG00000091931 | Gon7 | 100233175 | protein_coding | 69.22645195 | 0.075626655 | 0.164963788 | 0.458443979 | 0.646633501 | 0.816941518 | 0.075626655 |
| ENSMUSG00000036561 | Ppp6r2 | 71474 | protein_coding | 935.6724115 | -0.075659028 | 0.058218596 | -1.299568062 | 0.193749052 | 0.425519673 | 0.075659028 |
| ENSMUSG00000024213 | Nudt3 | 56409 | protein_coding | 1243.8527 | -0.07569025 | 0.074820103 | -1.011629861 | 0.311715063 | 0.555505998 | 0.07569025 |
| ENSMUSG00000034341 | Wbp2 | 22378 | protein_coding | 1391.929029 | 0.075708743 | 0.064878656 | 1.166928361 | 0.243239301 | 0.484226665 | 0.075708743 |
| ENSMUSG00000020594 | Pum2 | 80913 | protein_coding | 3177.052766 | -0.075741244 | 0.049530539 | -1.529182717 | 0.126219154 | 0.329962116 | 0.075741244 |
| ENSMUSG00000025235 | Bbs4 | 102774 | protein_coding | 110.4224522 | -0.075758752 | 0.122816402 | -0.616845555 | 0.537336601 | 0.743740778 | 0.075758752 |
| ENSMUSG00000070780 | Rbm47 | 245945 | protein_coding | 6341.065376 | 0.075766647 | 0.080309633 | 0.943431613 | 0.34546018 | 0.587779496 | 0.075766647 |
| ENSMUSG00000023066 | Rttn | 246102 | protein_coding | 251.2125024 | 0.075770916 | 0.077802652 | 0.973886025 | 0.330113133 | 0.573740112 | 0.075770916 |
| ENSMUSG00000008668 | Rps18 | 20084 | protein_coding | 12833.17735 | -0.07580844 | 0.045688899 | -1.659231066 | 0.09706924 | 0.282795761 | 0.07580844 |
| ENSMUSG00000037251 | Pomk | 74653 | protein_coding | 304.1848811 | 0.075838658 | 0.093820898 | 0.80833439 | 0.41889811 | 0.656326613 | 0.075838658 |
| ENSMUSG00000005371 | Fbxo11 | 225055 | protein_coding | 706.3296926 | -0.075869079 | 0.071122668 | -1.066735557 | 0.286091266 | 0.528797792 | 0.075869079 |
| ENSMUSG00000038781 | Stap2 | 106766 | protein_coding | 1937.373695 | -0.075896965 | 0.055961599 | -1.356232953 | 0.17502506 | 0.401397774 | 0.075896965 |
| ENSMUSG00000066150 | Slc31a1 | 20529 | protein_coding | 2252.843429 | -0.075916572 | 0.049844653 | -1.523063518 | 0.127742815 | 0.332543725 | 0.075916572 |
| ENSMUSG00000030265 | Kras | 16653 | protein_coding | 2247.784232 | -0.07591955 | 0.053978437 | -1.406479207 | 0.159581875 | 0.379596103 | 0.07591955 |
| ENSMUSG00000079104 | Prps1l3 | 328099 | protein_coding | 522.7082409 | 0.075929136 | 0.065880776 | 1.152523402 | 0.249106059 | 0.489920593 | 0.075929136 |
| ENSMUSG00000048756 | Foxo3 | 56484 | protein_coding | 3764.104447 | -0.07593379 | 0.056562632 | -1.342472713 | 0.179442772 | 0.406671386 | 0.07593379 |
| ENSMUSG00000031948 | Kars | 85305 | protein_coding | 2781.912232 | -0.075949448 | 0.059304025 | -1.280679488 | 0.200306268 | 0.433794437 | 0.075949448 |
| ENSMUSG00000039958 | Etfbkmt | 320204 | protein_coding | 326.707607 | -0.075970381 | 0.088113582 | -0.862186957 | 0.388584646 | 0.628430063 | 0.075970381 |
| ENSMUSG00000039715 | Dync2i2 | 71820 | protein_coding | 178.1727756 | 0.075992324 | 0.107589848 | 0.706315 | 0.479992267 | 0.702307684 | 0.075992324 |
| ENSMUSG00000034738 | Nostrin | 329416 | protein_coding | 1259.972754 | 0.076014343 | 0.074466789 | 1.020781794 | 0.307357832 | 0.551334926 | 0.076014343 |
| ENSMUSG00000074649 | BC029722 | 613262 | protein_coding | 159.7362336 | -0.076016341 | 0.120684731 | -0.629875378 | 0.528776123 | 0.738126248 | 0.076016341 |
| ENSMUSG00000019988 | Nedd1 | 17997 | protein_coding | 485.9185446 | 0.076025621 | 0.07269367 | 1.045835499 | 0.295636988 | 0.538727425 | 0.076025621 |
| ENSMUSG00000049134 | Nrap | 18175 | protein_coding | 547.2566681 | 0.076042618 | 0.114298673 | 0.665297469 | 0.505860249 | 0.721715006 | 0.076042618 |
| ENSMUSG00000026096 | Osgepl1 | 72085 | protein_coding | 157.3204222 | -0.07606155 | 0.101487917 | -0.749464095 | 0.453577532 | 0.682418589 | 0.07606155 |
| ENSMUSG00000018541 | Cwc25 | 67480 | protein_coding | 398.5675773 | -0.076070272 | 0.065604636 | -1.159525847 | 0.246241907 | 0.487137296 | 0.076070272 |
| ENSMUSG00000048644 | Ctxn1 | 330695 | protein_coding | 10.01237802 | -0.076086838 | 0.393685305 | -0.193268169 | 0.846748954 | 0.930889346 | 0.076086838 |
| ENSMUSG00000008429 | Herpud2 | 80517 | protein_coding | 631.948656 | 0.076107447 | 0.056514918 | 1.346678899 | 0.178083671 | 0.405350432 | 0.076107447 |
| ENSMUSG00000097804 | Gm16685 | 102634900 | lncRNA | 8.979898482 | -0.076124137 | 0.454885301 | -0.167347983 | 0.867096252 | 0.939855462 | 0.076124137 |
| ENSMUSG00000062270 | Morf4l1 | 21761 | protein_coding | 1405.136851 | 0.076125396 | 0.040642902 | 1.873030504 | 0.061064173 | 0.210558459 | 0.076125396 |
| ENSMUSG00000023307 | Marchf5 | 69104 | protein_coding | 840.3596493 | -0.076169506 | 0.04764705 | -1.598619533 | 0.109905167 | 0.304125022 | 0.076169506 |
| ENSMUSG00000041426 | Hibch | 227095 | protein_coding | 659.8932899 | 0.076196073 | 0.069351368 | 1.098696022 | 0.271900678 | 0.514423069 | 0.076196073 |
| ENSMUSG00000020898 | Ctc1 | 68964 | protein_coding | 448.3936552 | -0.076206528 | 0.087014311 | -0.875793044 | 0.381142552 | 0.6220843 | 0.076206528 |
| ENSMUSG00000031161 | Hdac6 | 15185 | protein_coding | 395.9907235 | 0.076207283 | 0.06171746 | 1.234776715 | 0.216913616 | 0.453012934 | 0.076207283 |
| ENSMUSG00000038507 | Parp12 | 243771 | protein_coding | 1628.143952 | -0.076215675 | 0.112566234 | -0.67707404 | 0.498358979 | 0.715904735 | 0.076215675 |
| ENSMUSG00000034192 | Lsm3 | 67678 | protein_coding | 292.0623427 | 0.076221168 | 0.085037866 | 0.896320333 | 0.370081702 | 0.612180999 | 0.076221168 |
| ENSMUSG00000030738 | Eif3c | 56347 | protein_coding | 6792.968996 | -0.076269668 | 0.033999787 | -2.243239551 | 0.024881371 | 0.114352201 | 0.076269668 |
| ENSMUSG00000030643 | Rab30 | 75985 | protein_coding | 26.17696207 | -0.076285445 | 0.242810762 | -0.314176541 | 0.753386959 | 0.88272824 | 0.076285445 |
| ENSMUSG00000035754 | Wdr18 | 216156 | protein_coding | 757.6932289 | 0.076287009 | 0.073395129 | 1.039401522 | 0.298618037 | 0.542138138 | 0.076287009 |

| ensembl_gene_id | external_gene_name | entrezgene_id | gene_biotype | baseMean | log2FoldChange | lfcSE | stat | pvalue | padj | absLog2FC |
|---|---|---|---|---|---|---|---|---|---|---|
| ENSMUSG00000025266 | Gnl3l | 237107 | protein_coding | 1304.509168 | 0.076291132 | 0.050846837 | 1.500410576 | 0.133508082 | 0.340947717 | 0.076291132 |
| ENSMUSG00000036160 | Surf6 | 20935 | protein_coding | 693.0310133 | 0.076341392 | 0.069250954 | 1.10238758 | 0.270293209 | 0.512797437 | 0.076341392 |
| ENSMUSG00000026812 | Tsc1 | 64930 | protein_coding | 856.6478439 | -0.076360686 | 0.055156967 | -1.384425047 | 0.166228344 | 0.389715871 | 0.076360686 |
| ENSMUSG00000031826 | Usp10 | 22224 | protein_coding | 760.4985866 | 0.076362792 | 0.081921689 | 0.932143775 | 0.351262228 | 0.593670714 | 0.076362792 |
| ENSMUSG00000040423 | Rc3h1 | 381305 | protein_coding | 1764.7895 | -0.076370168 | 0.056412346 | -1.35378465 | 0.175805092 | 0.402176854 | 0.076370168 |
| ENSMUSG00000045664 | Cdc42ep2 | 104252 | protein_coding | 249.9953519 | 0.076376365 | 0.111224377 | 0.686687281 | 0.492279814 | 0.711826106 | 0.076376365 |
| ENSMUSG00000024778 | Fas | 14102 | protein_coding | 166.0767377 | -0.076384209 | 0.116075569 | -0.658055865 | 0.510502234 | 0.724852046 | 0.076384209 |
| ENSMUSG00000060036 | Rpl3 | 27367 | protein_coding | 15285.94483 | -0.076403747 | 0.043168507 | -1.76989554 | 0.076744544 | 0.244162411 | 0.076403747 |
| ENSMUSG00000033222 | Ttf2 | 74044 | protein_coding | 353.1284661 | 0.076415743 | 0.08237439 | 0.927663839 | 0.353581971 | 0.595956629 | 0.076415743 |
| ENSMUSG00000029175 | Slc35f6 | 74919 | protein_coding | 942.4406833 | 0.076417332 | 0.06065549 | 1.259858456 | 0.207720428 | 0.442992242 | 0.076417332 |
| ENSMUSG00000002274 | Metrn | 70083 | protein_coding | 30.08803533 | -0.076445594 | 0.276128275 | -0.276848122 | 0.781896729 | 0.898695579 | 0.076445594 |
| ENSMUSG00000028688 | Toe1 | 68276 | protein_coding | 270.6779852 | 0.076450699 | 0.088805955 | 0.860873569 | 0.38930768 | 0.629157727 | 0.076450699 |
| ENSMUSG00000040363 | Bcor | 71458 | protein_coding | 1003.480255 | -0.076452568 | 0.066963519 | -1.141704747 | 0.253576767 | 0.494893811 | 0.076452568 |
| ENSMUSG00000051950 | B3glct | 381694 | protein_coding | 426.4114222 | -0.076456827 | 0.097883707 | -0.781098609 | 0.4347445 | 0.668304413 | 0.076456827 |
| ENSMUSG00000023952 | Gtpbp2 | 56055 | protein_coding | 2443.780522 | -0.076477405 | 0.064710067 | -1.181847106 | 0.237266373 | 0.477271565 | 0.076477405 |
| ENSMUSG00000025008 | Tctn3 | 67590 | protein_coding | 100.4185003 | -0.076488061 | 0.138329718 | -0.552940193 | 0.580304364 | 0.772159116 | 0.076488061 |
| ENSMUSG00000026255 | Efhd1 | 98363 | protein_coding | 6.173887242 | 0.0765026 | 0.477268727 | 0.160292506 | 0.872650662 | 0.942740972 | 0.0765026 |
| ENSMUSG00000028434 | Epb41l4b | 54357 | protein_coding | 5658.731026 | 0.076504315 | 0.041514245 | 1.842844907 | 0.065351658 | 0.220072289 | 0.076504315 |
| ENSMUSG00000054978 | Kbtbd13 | 74492 | protein_coding | 53.47366998 | 0.076539808 | 0.170054029 | 0.450091116 | 0.652644743 | 0.821331915 | 0.076539808 |
| ENSMUSG00000033610 | Pank1 | 75735 | protein_coding | 1019.982578 | -0.076546218 | 0.080255505 | -0.953781527 | 0.340194242 | 0.583162735 | 0.076546218 |
| ENSMUSG00000025260 | Hsd17b10 | 15108 | protein_coding | 1448.023398 | -0.076569135 | 0.052621899 | -1.455081187 | 0.145646787 | 0.358438231 | 0.076569135 |
| ENSMUSG00000024982 | Zdhhc6 | 66980 | protein_coding | 800.7976229 | 0.076582918 | 0.066080931 | 1.158926142 | 0.246486292 | 0.487381613 | 0.076582918 |
| ENSMUSG00000028901 | Gmeb1 | 56809 | protein_coding | 349.7980312 | -0.076583756 | 0.074184502 | -1.03234172 | 0.301912064 | 0.545539031 | 0.076583756 |
| ENSMUSG00000020097 | Sgpl1 | 20397 | protein_coding | 3025.056932 | -0.076607192 | 0.051526573 | -1.486751152 | 0.137080547 | 0.346504386 | 0.076607192 |
| ENSMUSG00000036526 | Card11 | 108723 | protein_coding | 72.99793487 | -0.07660744 | 0.18063961 | -0.424089935 | 0.671500225 | 0.832445548 | 0.07660744 |
| ENSMUSG00000035900 | Gramd4 | 223752 | protein_coding | 976.7798736 | -0.076609352 | 0.059006791 | -1.298314152 | 0.194179405 | 0.426123143 | 0.076609352 |
| ENSMUSG00000047230 | Cldn2 | 12738 | protein_coding | 3432.887114 | -0.076611861 | 0.093784833 | -0.816889666 | 0.413991484 | 0.65186884 | 0.076611861 |
| ENSMUSG00000002393 | Nr2f6 | 13864 | protein_coding | 1387.744244 | 0.076615962 | 0.106425369 | 0.719903183 | 0.471584608 | 0.696004848 | 0.076615962 |
| ENSMUSG00000033499 | Larp4b | 217980 | protein_coding | 3903.447447 | -0.076640113 | 0.032988926 | -2.323207269 | 0.02016802 | 0.098830041 | 0.076640113 |
| ENSMUSG00000020553 | Pctp | 18559 | protein_coding | 549.7720363 | -0.076695362 | 0.05352124 | -1.432989236 | 0.151860913 | 0.36822447 | 0.076695362 |
| ENSMUSG00000042410 | Agps | 228061 | protein_coding | 1647.895551 | -0.076722645 | 0.073882976 | -1.03843469 | 0.299067726 | 0.542649102 | 0.076722645 |
| ENSMUSG00000009292 | Trpm2 | 28240 | protein_coding | 13.90204053 | -0.076723433 | 0.374379836 | -0.204934735 | 0.837623116 | 0.927834878 | 0.076723433 |
| ENSMUSG00000035139 | Secisbp2 | 75420 | protein_coding | 547.4308604 | -0.076754552 | 0.058152721 | -1.319878956 | 0.186875435 | 0.416448019 | 0.076754552 |
| ENSMUSG00000034543 | Morc2a | 74522 | protein_coding | 741.4895414 | -0.07677572 | 0.05005112 | -1.533946076 | 0.125042919 | 0.328198932 | 0.07677572 |
| ENSMUSG00000027680 | Fxr1 | 14359 | protein_coding | 866.0200368 | -0.076783083 | 0.077153488 | -0.995199114 | 0.319639432 | 0.563143068 | 0.076783083 |
| ENSMUSG00000030814 | Bcl7c | 12055 | protein_coding | 207.5113194 | 0.076798913 | 0.105297734 | 0.729350104 | 0.465787531 | 0.691625579 | 0.076798913 |
| ENSMUSG00000029376 | Mthfd2l | 665563 | protein_coding | 193.1347792 | -0.076808392 | 0.08802346 | -0.872590016 | 0.382886584 | 0.623481598 | 0.076808392 |
| ENSMUSG00000027115 | Kif18a | 228421 | protein_coding | 167.4846915 | 0.076811704 | 0.149288023 | 0.514520202 | 0.60688834 | 0.789870733 | 0.076811704 |
| ENSMUSG00000025616 | Usp16 | 74112 | protein_coding | 1012.411584 | -0.076853351 | 0.052895207 | -1.452936006 | 0.146241523 | 0.359247178 | 0.076853351 |

Supplementary Table 1

| ensembl_gene_id | external_gene_name | entrezgene_id | gene_biotype | baseMean | log2FoldChange | lfcSE | stat | pvalue | padj | absLog2FC |
|---|---|---|---|---|---|---|---|---|---|---|
| ENSMUSG00000058135 | Gstm1 | 14862 | protein_coding | 3230.276673 | 0.076871883 | 0.084038115 | 0.914726401 | 0.360335281 | 0.603559666 | 0.076871883 |
| ENSMUSG00000039515 | Ptpa | 110854 | protein_coding | 2085.946992 | 0.076879718 | 0.05998004 | 1.281755043 | 0.199928589 | 0.433436829 | 0.076879718 |
| ENSMUSG00000027573 | Gid8 | 76425 | protein_coding | 1362.683035 | 0.076889498 | 0.045777289 | 1.679642884 | 0.093026819 | 0.275392886 | 0.076889498 |
| ENSMUSG00000070583 | Fv1 | 14349 | protein_coding | 106.5318632 | -0.076890402 | 0.145136539 | -0.52977977 | 0.596264633 | 0.783092209 | 0.076890402 |
| ENSMUSG00000049313 | Sorl1 | 20660 | protein_coding | 4499.477109 | -0.076893832 | 0.072179647 | -1.065311831 | 0.286734841 | 0.529168268 | 0.076893832 |
| ENSMUSG00000026526 | Fh1 | 14194 | protein_coding | 3247.080047 | 0.076903082 | 0.052802384 | 1.45643201 | 0.145273232 | 0.358010607 | 0.076903082 |
| ENSMUSG00000021573 | Tppp | 72948 | protein_coding | 2890.691781 | -0.076937967 | 0.130773069 | -0.588331895 | 0.556309541 | 0.755428309 | 0.076937967 |
| ENSMUSG00000043065 | Spice1 | 212514 | protein_coding | 286.5843726 | 0.076976745 | 0.080531725 | 0.955856162 | 0.339144909 | 0.58240762 | 0.076976745 |
| ENSMUSG00000000555 | Itga5 | 16402 | protein_coding | 105.2804875 | -0.077006798 | 0.124718924 | -0.617442776 | 0.536942714 | 0.743413046 | 0.077006798 |
| ENSMUSG00000026918 | Brd3 | 67382 | protein_coding | 797.5760511 | -0.077011969 | 0.065524819 | -1.175309908 | 0.239870752 | 0.480367021 | 0.077011969 |
| ENSMUSG00000049800 | Sertad2 | 58172 | protein_coding | 1325.997427 | -0.077016165 | 0.060097995 | -1.281509718 | 0.200014689 | 0.433436829 | 0.077016165 |
| ENSMUSG00000024854 | Pold4 | 69745 | protein_coding | 291.4788572 | 0.077043884 | 0.077045425 | 0.999979995 | 0.317320189 | 0.561120909 | 0.077043884 |
| ENSMUSG00000030124 | Lag3 | 16768 | protein_coding | 7.649502125 | -0.077047615 | 0.410416455 | -0.187730325 | 0.851088057 | 0.933199421 | 0.077047615 |
| ENSMUSG00000019303 | Psmc3ip | 19183 | protein_coding | 30.82410044 | -0.077084294 | 0.241878252 | -0.31869047 | 0.749961242 | 0.880597382 | 0.077084294 |
| ENSMUSG00000047045 | Tmem164 | 209497 | protein_coding | 3398.625804 | -0.077103314 | 0.045265316 | -1.703364079 | 0.088499956 | 0.266988405 | 0.077103314 |
| ENSMUSG00000035914 | Cd276 | 102657 | protein_coding | 83.32795702 | -0.077120008 | 0.149655649 | -0.51531638 | 0.606331955 | 0.789753 | 0.077120008 |
| ENSMUSG00000049536 | Tceal1 | 237052 | protein_coding | 25.47653097 | 0.077150013 | 0.251808918 | 0.306383163 | 0.759312926 | 0.886076667 | 0.077150013 |
| ENSMUSG00000024422 | Dhx16 | 69192 | protein_coding | 1230.844151 | 0.077152724 | 0.069507429 | 1.109992491 | 0.267002262 | 0.509217772 | 0.077152724 |
| ENSMUSG00000037608 | Bclaf1 | 72567 | protein_coding | 2650.740182 | -0.077248736 | 0.071965859 | -1.073408097 | 0.283088053 | 0.5257021 | 0.077248736 |
| ENSMUSG00000025968 | Ndufs1 | 227197 | protein_coding | 4296.406878 | 0.077276452 | 0.04400823 | 1.755954555 | 0.079096169 | 0.248419031 | 0.077276452 |
| ENSMUSG00000036257 | Pnpla8 | 67452 | protein_coding | 1653.174067 | 0.077306578 | 0.052088072 | 1.484151283 | 0.137768791 | 0.347590314 | 0.077306578 |
| ENSMUSG00000060152 | Pop5 | 117109 | protein_coding | 444.581546 | -0.077317758 | 0.060751582 | -1.272687156 | 0.203129077 | 0.437185296 | 0.077317758 |
| ENSMUSG00000028882 | Ppp1r8 | 100336 | protein_coding | 623.2244955 | 0.077344334 | 0.065640506 | 1.178301932 | 0.238676258 | 0.478917898 | 0.077344334 |
| ENSMUSG00000030518 | Entrep2 | 70638 | protein_coding | 80.52500474 | -0.077381988 | 0.15481601 | -0.499831948 | 0.617193413 | 0.796788198 | 0.077381988 |
| ENSMUSG00000022992 | Kansl2 | 69612 | protein_coding | 459.7158235 | -0.077391313 | 0.079865905 | -0.969015665 | 0.332537373 | 0.575843468 | 0.077391313 |
| ENSMUSG00000022437 | Samm50 | 68653 | protein_coding | 2969.471381 | 0.077402846 | 0.061683054 | 1.254847818 | 0.20953401 | 0.444801234 | 0.077402846 |
| ENSMUSG00000028645 | Slc2a1 | 20525 | protein_coding | 2328.864375 | 0.077414947 | 0.069220745 | 1.118377838 | 0.263405655 | 0.504916665 | 0.077414947 |
| ENSMUSG00000027177 | Hipk3 | 15259 | protein_coding | 2379.359459 | -0.077418124 | 0.040595282 | -1.907071978 | 0.056511267 | 0.200916492 | 0.077418124 |
| ENSMUSG00000027613 | Eif6 | 16418 | protein_coding | 4924.355052 | 0.077443991 | 0.062941605 | 1.230410165 | 0.218543549 | 0.454666616 | 0.077443991 |
| ENSMUSG00000024120 | Lrpprc | 72416 | protein_coding | 2054.75608 | -0.077451353 | 0.057883446 | -1.338057047 | 0.180877841 | 0.407746288 | 0.077451353 |
| ENSMUSG00000020955 | Ap4s1 | 11782 | protein_coding | 286.0928618 | -0.077472723 | 0.072221567 | -1.072708965 | 0.283401717 | 0.526061175 | 0.077472723 |
| ENSMUSG00000029482 | Aacs | 78894 | protein_coding | 3388.331425 | -0.077484205 | 0.145272269 | -0.533372308 | 0.593775883 | 0.781164996 | 0.077484205 |
| ENSMUSG00000035764 | Fbxo45 | 268882 | protein_coding | 468.4031039 | -0.077485214 | 0.063156078 | -1.226884517 | 0.219865999 | 0.456057983 | 0.077485214 |
| ENSMUSG00000020455 | Trim11 | 94091 | protein_coding | 539.0003464 | -0.077493812 | 0.059742173 | -1.29713748 | 0.194583888 | 0.426539942 | 0.077493812 |
| ENSMUSG00000020840 | Blmh | 104184 | protein_coding | 1247.841807 | 0.077508915 | 0.069387939 | 1.117037287 | 0.263978383 | 0.505363174 | 0.077508915 |
| ENSMUSG00000028776 | Tinagl1 | 94242 | protein_coding | 1257.492893 | -0.077555542 | 0.093437344 | -0.830027254 | 0.406523375 | 0.645466424 | 0.077555542 |
| ENSMUSG00000073791 | Efcab7 | 230500 | protein_coding | 63.51312088 | 0.077621902 | 0.149566271 | 0.518979993 | 0.603774693 | 0.78841372 | 0.077621902 |
| ENSMUSG00000020592 | Sdc1 | 20969 | protein_coding | 6244.246209 | -0.077622603 | 0.04628661 | -1.676999086 | 0.093542661 | 0.276344108 | 0.077622603 |
| ENSMUSG00000024317 | Rnf138 | 56515 | protein_coding | 418.8565374 | -0.077659672 | 0.068199651 | -1.138710694 | 0.254823841 | 0.496022219 | 0.077659672 |

| ensembl_gene_id | external_gene_name | entrezgene_id | gene_biotype | baseMean | log2FoldChange | lfcSE | stat | pvalue | padj | absLog2FC |
|---|---|---|---|---|---|---|---|---|---|---|
| ENSMUSG00000094724 | Rnaset2b | 68195 | protein_coding | 13.29538287 | 0.07767442 | 0.311463496 | 0.24938531 | 0.803062747 | 0.910363905 | 0.07767442 |
| ENSMUSG00000066838 | Zfp772 | 232855 | protein_coding | 117.850269 | -0.077694965 | 0.130135316 | -0.597032133 | 0.550485927 | 0.751963963 | 0.077694965 |
| ENSMUSG00000055371 | Stam2 | 56324 | protein_coding | 1796.540256 | 0.077707224 | 0.041999899 | 1.850176458 | 0.064288121 | 0.217764025 | 0.077707224 |
| ENSMUSG00000021392 | Nol8 | 70930 | protein_coding | 366.0819392 | -0.077720008 | 0.081871522 | -0.949292324 | 0.342471957 | 0.585130336 | 0.077720008 |
| ENSMUSG00000036111 | Lmo1 | 109594 | protein_coding | 7.927176085 | 0.077725052 | 0.416076031 | 0.186804927 | 0.851813584 | 0.933520082 | 0.077725052 |
| ENSMUSG00000050473 | Slc35d3 | 76157 | protein_coding | 9.315391888 | 0.077734134 | 0.422161829 | 0.184133496 | 0.853908729 | 0.934631853 | 0.077734134 |
| ENSMUSG00000025381 | Cnpy2 | 56530 | protein_coding | 1562.000456 | 0.077738258 | 0.049563721 | 1.568450795 | 0.116775965 | 0.316003891 | 0.077738258 |
| ENSMUSG00000070777 | Ceacam20 | 71601 | protein_coding | 2126.762759 | -0.07774078 | 0.102811354 | -0.756149753 | 0.449559415 | 0.679246742 | 0.07774078 |
| ENSMUSG00000027864 | Ptgfrn | 19221 | protein_coding | 4127.076701 | 0.077750338 | 0.06506685 | 1.194930108 | 0.232114338 | 0.4712713 | 0.077750338 |
| ENSMUSG00000038084 | Opa1 | 74143 | protein_coding | 2806.739128 | 0.077766031 | 0.042772887 | 1.818115075 | 0.069046546 | 0.22803884 | 0.077766031 |
| ENSMUSG00000079480 | Pin4 | 69713 | protein_coding | 233.9592271 | 0.077782084 | 0.097517423 | 0.797622424 | 0.425089634 | 0.66096665 | 0.077782084 |
| ENSMUSG00000031278 | Acsl4 | 50790 | protein_coding | 425.4465285 | 0.077787081 | 0.079104624 | 0.983344299 | 0.325438015 | 0.569172418 | 0.077787081 |
| ENSMUSG00000021706 | Zfyve16 | 218441 | protein_coding | 824.4693428 | 0.077839478 | 0.055333489 | 1.406733608 | 0.159506396 | 0.379596103 | 0.077839478 |
| ENSMUSG00000036231 | Agr3 | 403205 | protein_coding | 31.98981051 | -0.077845397 | 0.261353938 | -0.297854308 | 0.765814363 | 0.890018603 | 0.077845397 |
| ENSMUSG00000020022 | Ndufa12 | 66414 | protein_coding | 1963.904724 | 0.077846423 | 0.058533163 | 1.329954156 | 0.183533377 | 0.411947497 | 0.077846423 |
| ENSMUSG00000034501 | Pcnx4 | 67708 | protein_coding | 112.1237204 | -0.077890959 | 0.110070284 | -0.707647474 | 0.479164204 | 0.701661251 | 0.077890959 |
| ENSMUSG00000014786 | Slc9a5 | 277973 | protein_coding | 77.01675935 | 0.077901791 | 0.15712075 | 0.495808425 | 0.62002958 | 0.799106292 | 0.077901791 |
| ENSMUSG00000019942 | Cdk1 | 12534 | protein_coding | 615.9011158 | 0.077923474 | 0.127116576 | 0.613007965 | 0.539871073 | 0.745625688 | 0.077923474 |
| ENSMUSG00000095026 | Gm3336 | 100502950 | protein_coding | 1100.192915 | 0.077938323 | 0.084972918 | 0.917213679 | 0.35903068 | 0.602381167 | 0.077938323 |
| ENSMUSG00000065954 | Tacc1 | 320165 | protein_coding | 242.1739964 | 0.078002675 | 0.103646617 | 0.752582933 | 0.451700577 | 0.680801421 | 0.078002675 |
| ENSMUSG00000040620 | Dhx33 | 216877 | protein_coding | 558.846741 | -0.078006162 | 0.071780144 | -1.086737328 | 0.277152912 | 0.51990697 | 0.078006162 |
| ENSMUSG00000066440 | Zfyve26 | 211978 | protein_coding | 583.4170965 | -0.078011842 | 0.06179247 | -1.262481374 | 0.206775624 | 0.441678211 | 0.078011842 |
| ENSMUSG00000031787 | Katnb1 | 74187 | protein_coding | 436.2457936 | 0.078012329 | 0.097930005 | 0.796613142 | 0.425675743 | 0.661307016 | 0.078012329 |
| ENSMUSG00000117648 | Gm32282 | 102634780 | lncRNA | 5.893364455 | -0.078014331 | 0.496765008 | -0.15704474 | 0.875209588 | 0.943798397 | 0.078014331 |
| ENSMUSG00000031103 | Elf4 | 56501 | protein_coding | 1086.508689 | -0.078017594 | 0.06416659 | -1.215860058 | 0.224038236 | 0.461733499 | 0.078017594 |
| ENSMUSG00000022114 | Spry2 | 24064 | protein_coding | 942.2411461 | -0.07801814 | 0.067603449 | -1.154055628 | 0.248477363 | 0.4893628 | 0.07801814 |
| ENSMUSG00000086837 | Gm16618 | 100504291 | lncRNA | 7.434403094 | -0.078037007 | 0.489015192 | -0.159579924 | 0.873211994 | 0.942905245 | 0.078037007 |
| ENSMUSG00000028389 | Zfp37 | 22696 | protein_coding | 172.1924072 | 0.078049546 | 0.173794163 | 0.449091875 | 0.653365382 | 0.821648745 | 0.078049546 |
| ENSMUSG00000032536 | Trak1 | 67095 | protein_coding | 6651.598047 | -0.078051279 | 0.063287019 | -1.233290487 | 0.217467406 | 0.453693043 | 0.078051279 |
| ENSMUSG00000046865 | Fbl | 14113 | protein_coding | 563.2335507 | -0.07806102 | 0.080958192 | -0.964213976 | 0.334938659 | 0.578448892 | 0.07806102 |
| ENSMUSG00000039865 | Slc44a3 | 213603 | protein_coding | 2268.725471 | -0.078061315 | 0.051052777 | -1.529031704 | 0.126256584 | 0.330006473 | 0.078061315 |
| ENSMUSG00000016528 | Mapkapk2 | 17164 | protein_coding | 2269.314674 | 0.078072177 | 0.056510668 | 1.381547584 | 0.167110658 | 0.390918709 | 0.078072177 |
| ENSMUSG00000074283 | Zfp109 | 56869 | protein_coding | 104.547153 | -0.078090865 | 0.125765738 | -0.620923205 | 0.534650152 | 0.741735868 | 0.078090865 |
| ENSMUSG00000021243 | Fcf1 | 73736 | protein_coding | 511.7068921 | 0.078093141 | 0.05671988 | 1.376821342 | 0.168567491 | 0.392565467 | 0.078093141 |
| ENSMUSG00000017428 | Psmd11 | 69077 | protein_coding | 1002.889736 | 0.078138592 | 0.061100163 | 1.278860608 | 0.200946146 | 0.434666577 | 0.078138592 |
| ENSMUSG00000016409 | Nkap | 67050 | protein_coding | 359.1092201 | -0.0781592 | 0.068521915 | -1.1406453 | 0.254017558 | 0.495031103 | 0.0781592 |
| ENSMUSG00000039523 | Cep104 | 230967 | protein_coding | 410.6575893 | 0.078197199 | 0.090255002 | 0.866402935 | 0.386269235 | 0.626292308 | 0.078197199 |
| ENSMUSG00000003233 | Dvl3 | 13544 | protein_coding | 384.5125161 | -0.07820639 | 0.117927751 | -0.663172062 | 0.507220362 | 0.72271052 | 0.07820639 |
| ENSMUSG00000071862 | Lrrtm2 | 107065 | protein_coding | 49.07890712 | -0.078229649 | 0.267179273 | -0.292798344 | 0.769676294 | 0.891835724 | 0.078229649 |

| ensembl_gene_id | external_gene_name | entrezgene_id | gene_biotype | baseMean | log2FoldChange | lfcSE | stat | pvalue | padj | absLog2FC |
|---|---|---|---|---|---|---|---|---|---|---|
| ENSMUSG00000024127 | Prepl | 213760 | protein_coding | 863.1376562 | 0.078244259 | 0.050288365 | 1.555911767 | 0.119729075 | 0.320093959 | 0.078244259 |
| ENSMUSG00000000959 | Oxa1l | 69089 | protein_coding | 1447.030039 | -0.078256328 | 0.054209909 | -1.443579758 | 0.148857222 | 0.363290016 | 0.078256328 |
| ENSMUSG00000071550 | Cfap44 | 212517 | protein_coding | 9.249351336 | 0.078291079 | 0.452375952 | 0.173066404 | 0.862599232 | 0.937787345 | 0.078291079 |
| ENSMUSG00000006423 | Steep1 | 77644 | protein_coding | 722.5787869 | 0.078324674 | 0.055256451 | 1.417475645 | 0.15634391 | 0.37481169 | 0.078324674 |
| ENSMUSG00000100164 | 2610306M01Rik | 67170 | lncRNA | 82.88366105 | 0.078333066 | 0.173714837 | 0.450929048 | 0.652040687 | 0.820965922 | 0.078333066 |
| ENSMUSG00000025220 | Oga | 76055 | protein_coding | 2870.842952 | -0.078347286 | 0.050069804 | -1.564761186 | 0.117638916 | 0.317145235 | 0.078347286 |
| ENSMUSG00000019132 | BC005537 | 79555 | protein_coding | 2627.424485 | -0.078348412 | 0.05665921 | -1.382800998 | 0.166725893 | 0.39047175 | 0.078348412 |
| ENSMUSG00000019927 | Ube2d1 | 216080 | protein_coding | 214.7581334 | -0.078357696 | 0.092897607 | -0.843484545 | 0.398957506 | 0.638672592 | 0.078357696 |
| ENSMUSG00000058546 | Rpl23a | 268449 | protein_coding | 4028.283848 | -0.078357852 | 0.072007514 | -1.08818993 | 0.276511269 | 0.519430846 | 0.078357852 |
| ENSMUSG00000003746 | Man1a | 17155 | protein_coding | 2465.111352 | -0.078374306 | 0.048865332 | -1.603883609 | 0.10873971 | 0.302001032 | 0.078374306 |
| ENSMUSG00000052658 | 5830454E08Rik | 76100 | lncRNA | 13.32125629 | 0.078417619 | 0.316163534 | 0.248028664 | 0.804112229 | 0.910778265 | 0.078417619 |
| ENSMUSG00000022089 | Bin3 | 57784 | protein_coding | 578.5103797 | 0.07842396 | 0.054207134 | 1.44674609 | 0.147968051 | 0.361886677 | 0.07842396 |
| ENSMUSG00000040102 | Klhl42 | 232539 | protein_coding | 571.246961 | -0.078450845 | 0.077752537 | -1.008981158 | 0.31298367 | 0.556994886 | 0.078450845 |
| ENSMUSG00000019951 | Uhrf1bp1l | 75089 | protein_coding | 1043.14993 | -0.078466064 | 0.065919483 | -1.190331911 | 0.233915963 | 0.473379688 | 0.078466064 |
| ENSMUSG00000036879 | Phkb | 102093 | protein_coding | 1262.070409 | 0.078489936 | 0.049553358 | 1.58394787 | 0.113205583 | 0.309749988 | 0.078489936 |
| ENSMUSG00000024132 | Eci1 | 13177 | protein_coding | 1425.541617 | 0.07850359 | 0.045582188 | 1.722242681 | 0.085025566 | 0.260141777 | 0.07850359 |
| ENSMUSG00000009741 | Ubp1 | 22221 | protein_coding | 1079.769852 | -0.078511063 | 0.047265327 | -1.661070978 | 0.096699192 | 0.282265491 | 0.078511063 |
| ENSMUSG00000058388 | Phtf1 | 18685 | protein_coding | 343.3375714 | 0.078592047 | 0.072238798 | 1.087947882 | 0.276618116 | 0.519508601 | 0.078592047 |
| ENSMUSG00000027487 | Cdk5rap1 | 66971 | protein_coding | 422.6568641 | -0.078600736 | 0.167185664 | -0.470140405 | 0.638254709 | 0.811381567 | 0.078600736 |
| ENSMUSG00000050373 | Snx21 | 101113 | protein_coding | 346.4802711 | 0.078601505 | 0.070827898 | 1.109753464 | 0.267105278 | 0.509293782 | 0.078601505 |
| ENSMUSG00000073436 | Eme2 | 193838 | protein_coding | 176.5017767 | -0.078610407 | 0.129984365 | -0.604768171 | 0.545333045 | 0.749049145 | 0.078610407 |
| ENSMUSG00000063564 | Col23a1 | 237759 | protein_coding | 9.83794335 | -0.078614808 | 0.409307946 | -0.192067632 | 0.847689229 | 0.931084542 | 0.078614808 |
| ENSMUSG00000027947 | Il6ra | 16194 | protein_coding | 85.68045764 | -0.078616003 | 0.190911663 | -0.411792563 | 0.680491473 | 0.838364494 | 0.078616003 |
| ENSMUSG00000033088 | Triobp | 110253 | protein_coding | 1425.90652 | 0.078620071 | 0.071727327 | 1.09609648 | 0.273036555 | 0.515483555 | 0.078620071 |
| ENSMUSG00000056383 | AI987944 | 233168 | protein_coding | 292.1102492 | -0.078623509 | 0.105245385 | -0.747049464 | 0.455033704 | 0.682988492 | 0.078623509 |
| ENSMUSG00000004951 | Hspb1 | 15507 | protein_coding | 24.38892048 | -0.078632522 | 0.265134579 | -0.296575883 | 0.766790326 | 0.89053739 | 0.078632522 |
| ENSMUSG00000020156 | Pwwp3a | 68114 | protein_coding | 592.3895035 | -0.07864046 | 0.053782753 | -1.462187344 | 0.143689879 | 0.355746355 | 0.07864046 |
| ENSMUSG00000068250 | Amn1 | 232566 | protein_coding | 256.7969022 | -0.078652825 | 0.101085279 | -0.778083861 | 0.436519577 | 0.669750853 | 0.078652825 |
| ENSMUSG00000036057 | Ptpn23 | 104831 | protein_coding | 380.564618 | -0.078681794 | 0.095175202 | -0.826704771 | 0.408404416 | 0.646860933 | 0.078681794 |
| ENSMUSG00000085667 | Gm12992 | 545681 | lncRNA | 31.40578596 | 0.078724715 | 0.211327583 | 0.37252456 | 0.70950233 | 0.856166251 | 0.078724715 |
| ENSMUSG00000035885 | Cox8a | 12868 | protein_coding | 12364.7691 | 0.078738147 | 0.075184888 | 1.047260276 | 0.294979552 | 0.538015747 | 0.078738147 |
| ENSMUSG00000021156 | Zmynd11 | 66505 | protein_coding | 1158.18267 | -0.078770593 | 0.046204957 | -1.704808279 | 0.088230187 | 0.26640858 | 0.078770593 |
| ENSMUSG00000024843 | Chka | 12660 | protein_coding | 909.6901128 | 0.078786145 | 0.095676821 | 0.823461155 | 0.410245798 | 0.648598559 | 0.078786145 |
| ENSMUSG00000074826 | Gm10767 | 100038538 | protein_coding | 57.03817601 | 0.078797314 | 0.188047988 | 0.419027693 | 0.675195894 | 0.834781062 | 0.078797314 |
| ENSMUSG00000028465 | Tln1 | 21894 | protein_coding | 3504.892571 | 0.078811928 | 0.067518171 | 1.167269887 | 0.243101395 | 0.484226665 | 0.078811928 |
| ENSMUSG00000023904 | Hcfc1r1 | 353502 | protein_coding | 895.8606156 | 0.078878535 | 0.067471412 | 1.169066027 | 0.24237703 | 0.483565042 | 0.078878535 |
| ENSMUSG00000027088 | Phospho2 | 73373 | protein_coding | 523.6417915 | 0.078904396 | 0.066482032 | 1.186852952 | 0.235285628 | 0.475376011 | 0.078904396 |
| ENSMUSG00000000751 | Rpa1 | 68275 | protein_coding | 930.0360745 | 0.078936245 | 0.101147589 | 0.780406588 | 0.435151591 | 0.668669815 | 0.078936245 |
| ENSMUSG00000017631 | Abr | 109934 | protein_coding | 5158.268617 | 0.078938958 | 0.049177656 | 1.605179354 | 0.10845434 | 0.301715221 | 0.078938958 |

| ensembl_gene_id | external_gene_name | entrezgene_id | gene_biotype | baseMean | log2FoldChange | lfcSE | stat | pvalue | padj | absLog2FC |
|---|---|---|---|---|---|---|---|---|---|---|
| ENSMUSG00000022466 | Rpap3 | 71919 | protein_coding | 251.6981776 | -0.078979368 | 0.076811978 | -1.028216815 | 0.303847851 | 0.547921576 | 0.078979368 |
| ENSMUSG00000039193 | Nlrc4 | 268973 | protein_coding | 1378.273269 | -0.078979881 | 0.050590248 | -1.5611681 | 0.118484094 | 0.318412428 | 0.078979881 |
| ENSMUSG00000022421 | Nptxr | 73340 | protein_coding | 12.94580315 | -0.07900707 | 0.420485254 | -0.187894984 | 0.850958975 | 0.933121329 | 0.07900707 |
| ENSMUSG00000022283 | Pabpc1 | 18458 | protein_coding | 6736.50605 | -0.079036635 | 0.09789501 | -0.807361223 | 0.419458401 | 0.656821597 | 0.079036635 |
| ENSMUSG00000003235 | Eif2b5 | 224045 | protein_coding | 1413.205498 | 0.079058888 | 0.061247282 | 1.29081463 | 0.196767963 | 0.429227235 | 0.079058888 |
| ENSMUSG00000015126 | Tsr3 | 68327 | protein_coding | 156.0048332 | -0.079064406 | 0.103689084 | -0.762514261 | 0.445753135 | 0.67671134 | 0.079064406 |
| ENSMUSG00000034194 | R3hcc1 | 71843 | protein_coding | 314.6573211 | -0.079066956 | 0.075299074 | -1.050038887 | 0.293700234 | 0.536541439 | 0.079066956 |
| ENSMUSG00000041769 | Ppp2r2d | 52432 | protein_coding | 683.3298605 | -0.079103868 | 0.050540472 | -1.565158876 | 0.117545661 | 0.317074504 | 0.079103868 |
| ENSMUSG00000017715 | Pgs1 | 74451 | protein_coding | 771.1826785 | 0.079120533 | 0.060667213 | 1.304172862 | 0.19217465 | 0.423671625 | 0.079120533 |
| ENSMUSG00000027306 | Nusap1 | 108907 | protein_coding | 571.5373197 | 0.079155985 | 0.133142624 | 0.594520243 | 0.552164207 | 0.75266268 | 0.079155985 |
| ENSMUSG00000032641 | Gpr19 | 14760 | protein_coding | 82.18598613 | 0.079176214 | 0.154082035 | 0.513857531 | 0.607351602 | 0.790170193 | 0.079176214 |
| ENSMUSG00000074909 | Ranbp6 | 240614 | protein_coding | 427.1090647 | -0.079182979 | 0.094200251 | -0.840581401 | 0.400582482 | 0.639813181 | 0.079182979 |
| ENSMUSG00000047945 | Marcksl1 | 17357 | protein_coding | 352.8486996 | 0.079206026 | 0.076331025 | 1.037664913 | 0.299426085 | 0.543015856 | 0.079206026 |
| ENSMUSG00000070697 | Utp3 | 65961 | protein_coding | 506.1940909 | -0.079229342 | 0.065345064 | -1.212476306 | 0.225330107 | 0.463330185 | 0.079229342 |
| ENSMUSG00000022789 | Dnm1l | 74006 | protein_coding | 1985.41689 | 0.079250085 | 0.052509055 | 1.509265113 | 0.131231046 | 0.337215278 | 0.079250085 |
| ENSMUSG00000022956 | Atp5o | 28080 | protein_coding | 5760.161945 | 0.079265548 | 0.048636922 | 1.629740225 | 0.103156412 | 0.293497536 | 0.079265548 |
| ENSMUSG00000021408 | Ripk1 | 19766 | protein_coding | 1291.821647 | -0.079266697 | 0.044442251 | -1.783588725 | 0.074490519 | 0.239447599 | 0.079266697 |
| ENSMUSG00000031775 | Pllp | 67801 | protein_coding | 477.7626793 | 0.079273414 | 0.124263772 | 0.637944697 | 0.523509681 | 0.734525893 | 0.079273414 |
| ENSMUSG00000021536 | Adcy2 | 210044 | protein_coding | 27.96037064 | -0.079307358 | 0.224574432 | -0.353145092 | 0.72397967 | 0.865566026 | 0.079307358 |
| ENSMUSG00000018293 | Pfn1 | 18643 | protein_coding | 11077.01888 | 0.079339244 | 0.092212872 | 0.860392289 | 0.389572835 | 0.629397025 | 0.079339244 |
| ENSMUSG00000074912 | Gm14207 | 100043609 | lncRNA | 112.923311 | -0.07935841 | 0.133759808 | -0.593290402 | 0.552986821 | 0.753401915 | 0.07935841 |
| ENSMUSG00000018858 | Mrpl58 | 68572 | protein_coding | 696.8354203 | 0.079371441 | 0.073474557 | 1.080257492 | 0.280027533 | 0.523118689 | 0.079371441 |
| ENSMUSG00000097333 | Zfp87 | 170763 | protein_coding | 216.3477938 | 0.079378331 | 0.08888226 | 0.893072821 | 0.371818177 | 0.61362175 | 0.079378331 |
| ENSMUSG00000079658 | Eloc | 67923 | protein_coding | 651.5490921 | 0.079395489 | 0.064812864 | 1.224995842 | 0.220576787 | 0.45711152 | 0.079395489 |
| ENSMUSG00000000732 | Icosl | 50723 | protein_coding | 151.522842 | -0.079406465 | 0.115538045 | -0.687275479 | 0.491909148 | 0.711469495 | 0.079406465 |
| ENSMUSG00000073557 | Ppp1r12b | 329251 | protein_coding | 214.3339965 | -0.079443244 | 0.101236325 | -0.784730622 | 0.432611525 | 0.666799776 | 0.079443244 |
| ENSMUSG00000028309 | Rnf20 | 109331 | protein_coding | 1329.872543 | 0.079448575 | 0.049136971 | 1.616879793 | 0.105904231 | 0.297926072 | 0.079448575 |
| ENSMUSG00000005907 | Pex1 | 71382 | protein_coding | 333.591544 | -0.079452561 | 0.074928546 | -1.060377734 | 0.288972788 | 0.531357026 | 0.079452561 |
| ENSMUSG00000022422 | Dscc1 | 72107 | protein_coding | 39.13574937 | 0.079468911 | 0.268987853 | 0.295436802 | 0.767660223 | 0.890907917 | 0.079468911 |
| ENSMUSG00000034247 | Plekhm1 | 353047 | protein_coding | 898.4863993 | -0.079482774 | 0.068721059 | -1.156599949 | 0.247435843 | 0.488420788 | 0.079482774 |
| ENSMUSG00000039678 | Tbc1d13 | 70296 | protein_coding | 2536.222812 | 0.079492394 | 0.062556111 | 1.270737459 | 0.203822065 | 0.437979805 | 0.079492394 |
| ENSMUSG00000070000 | Fcho1 | 74015 | protein_coding | 90.44077545 | 0.079508191 | 0.162019396 | 0.490732549 | 0.623615622 | 0.801324379 | 0.079508191 |
| ENSMUSG00000068823 | Csde1 | 229663 | protein_coding | 8236.470114 | -0.079517237 | 0.043870868 | -1.812529393 | 0.069904437 | 0.229602617 | 0.079517237 |
| ENSMUSG00000004677 | Myo9b | 17925 | protein_coding | 979.1998747 | -0.079521708 | 0.053394682 | -1.489318885 | 0.136403417 | 0.345432975 | 0.079521708 |
| ENSMUSG00000039414 | Heatr5b | 320473 | protein_coding | 761.212645 | 0.079548328 | 0.063124527 | 1.260180973 | 0.207604085 | 0.442919841 | 0.079548328 |
| ENSMUSG00000038241 | Cep250 | 16328 | protein_coding | 404.6225795 | -0.079551031 | 0.077855267 | -1.021780987 | 0.30688457 | 0.550870027 | 0.079551031 |
| ENSMUSG00000071054 | Safb | 224903 | protein_coding | 1125.786528 | -0.079569421 | 0.081653641 | -0.974474873 | 0.329820809 | 0.573541103 | 0.079569421 |
| ENSMUSG00000043384 | Gprasp1 | 67298 | protein_coding | 199.7403395 | -0.079570401 | 0.103021888 | -0.772364039 | 0.439898846 | 0.671740242 | 0.079570401 |
| ENSMUSG00000091537 | Tma7 | 66167 | protein_coding | 1447.308379 | 0.079571347 | 0.052169347 | 1.525250961 | 0.127196514 | 0.331496151 | 0.079571347 |

| ensembl_gene_id | external_gene_name | entrezgene_id | gene_biotype | baseMean | log2FoldChange | lfcSE | stat | pvalue | padj | absLog2FC |
|---|---|---|---|---|---|---|---|---|---|---|
| ENSMUSG00000036687 | Tmem184a | 231832 | protein_coding | 855.6545342 | -0.079601173 | 0.053338156 | -1.492387035 | 0.135597712 | 0.344158687 | 0.079601173 |
| ENSMUSG00000027108 | Ola1 | 67059 | protein_coding | 1701.159005 | 0.079602956 | 0.059344676 | 1.341366426 | 0.179801512 | 0.406855478 | 0.079602956 |
| ENSMUSG00000020790 | Ankfy1 | 11736 | protein_coding | 2861.636035 | 0.079610179 | 0.041070782 | 1.938365303 | 0.05257867 | 0.191494178 | 0.079610179 |
| ENSMUSG00000029366 | Dck | 13178 | protein_coding | 255.4938968 | 0.079635969 | 0.101709892 | 0.78297172 | 0.433643719 | 0.667785655 | 0.079635969 |
| ENSMUSG00000060681 | Slc9a6 | 236794 | protein_coding | 489.1512564 | 0.079668699 | 0.057618002 | 1.382704981 | 0.166755344 | 0.390484018 | 0.079668699 |
| ENSMUSG00000020801 | Med31 | 67279 | protein_coding | 102.0350682 | 0.079691935 | 0.153180633 | 0.520248111 | 0.602890656 | 0.787655348 | 0.079691935 |
| ENSMUSG00000067786 | Nnat | 18111 | protein_coding | 48.61372497 | -0.079701565 | 0.231112697 | -0.344860174 | 0.730199493 | 0.869327992 | 0.079701565 |
| ENSMUSG00000000399 | Ndufa9 | 66108 | protein_coding | 3703.29624 | 0.079708306 | 0.056051669 | 1.422050519 | 0.155011586 | 0.372948208 | 0.079708306 |
| ENSMUSG00000035173 | Ccdc186 | 213993 | protein_coding | 1668.226008 | -0.079721446 | 0.042466322 | -1.877286325 | 0.060478861 | 0.209031847 | 0.079721446 |
| ENSMUSG00000031337 | Mtm1 | 17772 | protein_coding | 1617.337038 | -0.079741196 | 0.0538959 | -1.47954103 | 0.138995774 | 0.349210912 | 0.079741196 |
| ENSMUSG00000031488 | Rab11fip1 | 75767 | protein_coding | 4272.956402 | -0.079767768 | 0.041244139 | -1.934038881 | 0.053108348 | 0.192640198 | 0.079767768 |
| ENSMUSG00000056941 | Commd7 | 99311 | protein_coding | 766.7591796 | 0.079770213 | 0.058649142 | 1.36012582 | 0.173790111 | 0.399474505 | 0.079770213 |
| ENSMUSG00000026596 | Abl2 | 11352 | protein_coding | 641.8075908 | -0.079792769 | 0.05243691 | -1.521690884 | 0.128086553 | 0.333247419 | 0.079792769 |
| ENSMUSG00000045435 | Tmem60 | 212090 | protein_coding | 240.5995015 | 0.079800421 | 0.090664294 | 0.880174731 | 0.37876466 | 0.619821175 | 0.079800421 |
| ENSMUSG00000067787 | Blcap | 53619 | protein_coding | 724.2464334 | -0.079802807 | 0.055994878 | -1.425180461 | 0.154105043 | 0.371487597 | 0.079802807 |
| ENSMUSG00000054716 | Zfp771 | 244216 | protein_coding | 253.1911667 | -0.079882488 | 0.15763432 | -0.506758222 | 0.612324473 | 0.793255831 | 0.079882488 |
| ENSMUSG00000028082 | Sh3d19 | 27059 | protein_coding | 2707.312705 | -0.079903039 | 0.046856199 | -1.705282133 | 0.088141818 | 0.26624145 | 0.079903039 |
| ENSMUSG00000048376 | F2r | 14062 | protein_coding | 6.869758375 | -0.079930466 | 0.496547008 | -0.160972606 | 0.872114977 | 0.942477935 | 0.079930466 |
| ENSMUSG00000022022 | Mtrf1 | 211253 | protein_coding | 125.8268423 | 0.07993111 | 0.119167543 | 0.670745639 | 0.502382583 | 0.718956777 | 0.07993111 |
| ENSMUSG00000031756 | Cenpn | 72155 | protein_coding | 64.7349245 | 0.079955444 | 0.204481904 | 0.391014765 | 0.695786322 | 0.847823651 | 0.079955444 |
| ENSMUSG00000020520 | Galnt10 | 171212 | protein_coding | 4212.895858 | 0.07995861 | 0.055818691 | 1.43247017 | 0.152009309 | 0.368528883 | 0.07995861 |
| ENSMUSG00000071379 | Hpcal1 | 53602 | protein_coding | 1613.348445 | 0.079971504 | 0.056401882 | 1.417887158 | 0.156223712 | 0.374715872 | 0.079971504 |
| ENSMUSG00000020877 | Scrn2 | 217140 | protein_coding | 450.8929946 | 0.079982366 | 0.095993363 | 0.833207247 | 0.404727856 | 0.644137584 | 0.079982366 |
| ENSMUSG00000070814 | Zswim9 | 321008 | protein_coding | 87.28893798 | 0.080046226 | 0.143462885 | 0.557957731 | 0.576873248 | 0.769561988 | 0.080046226 |
| ENSMUSG00000022840 | Adcy5 | 224129 | protein_coding | 111.9446353 | 0.08005878 | 0.144017307 | 0.555896942 | 0.578281309 | 0.770802766 | 0.08005878 |
| ENSMUSG00000045973 | Slc25a51 | 230125 | protein_coding | 1754.21045 | -0.080088495 | 0.042498595 | -1.884497492 | 0.059497714 | 0.207110275 | 0.080088495 |
| ENSMUSG00000079083 | Jrkl | 77532 | protein_coding | 248.6902665 | -0.080089178 | 0.098803849 | -0.810587631 | 0.417602522 | 0.655442335 | 0.080089178 |
| ENSMUSG00000067365 | Tmem128 | 66309 | protein_coding | 638.4017564 | 0.080106954 | 0.061708469 | 1.298151706 | 0.19423521 | 0.426123143 | 0.080106954 |
| ENSMUSG00000004798 | Ulk2 | 29869 | protein_coding | 575.7831521 | 0.080113202 | 0.056909705 | 1.407724775 | 0.159212583 | 0.379320626 | 0.080113202 |
| ENSMUSG00000024620 | Pdgfrb | 18596 | protein_coding | 312.2939867 | -0.080117742 | 0.13528886 | -0.59219763 | 0.553718256 | 0.75391672 | 0.080117742 |
| ENSMUSG00000022836 | Mylk | 107589 | protein_coding | 1596.004332 | -0.080171218 | 0.081931769 | -0.978511978 | 0.32782117 | 0.571636266 | 0.080171218 |
| ENSMUSG00000074227 | Spint2 | 20733 | protein_coding | 14689.20282 | -0.080189531 | 0.074363103 | -1.078351054 | 0.280877121 | 0.523508209 | 0.080189531 |
| ENSMUSG00000026525 | Opn3 | 13603 | protein_coding | 33.98931926 | 0.080197003 | 0.218914694 | 0.366339058 | 0.714112088 | 0.859439227 | 0.080197003 |
| ENSMUSG00000026576 | Atp1b1 | 11931 | protein_coding | 18745.16933 | -0.080197965 | 0.064665613 | -1.240194927 | 0.214903304 | 0.45050036 | 0.080197965 |
| ENSMUSG00000066235 | Pomgnt2 | 215494 | protein_coding | 226.4898717 | 0.080209932 | 0.112894873 | 0.710483393 | 0.477404426 | 0.700084044 | 0.080209932 |
| ENSMUSG00000036282 | Naa30 | 70646 | protein_coding | 861.181332 | -0.080216464 | 0.067719322 | -1.184543218 | 0.236198099 | 0.47644362 | 0.080216464 |
| ENSMUSG00000026269 | Rnpepl1 | 108657 | protein_coding | 2137.810474 | 0.080235095 | 0.074415438 | 1.078204962 | 0.280942298 | 0.523508209 | 0.080235095 |
| ENSMUSG00000018363 | Smurf2 | 66313 | protein_coding | 1576.896616 | -0.080238855 | 0.038686981 | -2.074053134 | 0.038074376 | 0.153429653 | 0.080238855 |
| ENSMUSG00000026939 | Tmem141 | 51875 | protein_coding | 187.5278199 | 0.080268541 | 0.097651114 | 0.821993085 | 0.411080833 | 0.649146117 | 0.080268541 |

Supplementary Table 1

| ensembl_gene_id | external_gene_name | entrezgene_id | gene_biotype | baseMean | log2FoldChange | lfcSE | stat | pvalue | padj | absLog2FC |
|---|---|---|---|---|---|---|---|---|---|---|
| ENSMUSG00000029622 | Arpc1b | 11867 | protein_coding | 7368.809638 | 0.080274428 | 0.071952729 | 1.115655079 | 0.264569806 | 0.5061553 | 0.080274428 |
| ENSMUSG00000027067 | Ssrp1 | 20833 | protein_coding | 2156.525474 | 0.080289575 | 0.064486994 | 1.245050662 | 0.213113128 | 0.448795432 | 0.080289575 |
| ENSMUSG00000005069 | Pex5 | 19305 | protein_coding | 1116.681199 | 0.080296477 | 0.054178956 | 1.482060229 | 0.13832427 | 0.348392238 | 0.080296477 |
| ENSMUSG00000062510 | Nsl1 | 381318 | protein_coding | 124.0504699 | 0.080316223 | 0.176192579 | 0.455843391 | 0.648502599 | 0.818405307 | 0.080316223 |
| ENSMUSG00000029388 | Eif2b1 | 209354 | protein_coding | 1584.056213 | 0.080353989 | 0.052959045 | 1.517285457 | 0.129194628 | 0.334810892 | 0.080353989 |
| ENSMUSG00000029998 | Pcyox1 | 66881 | protein_coding | 2061.517412 | -0.080378684 | 0.047322856 | -1.698517177 | 0.089410194 | 0.268663721 | 0.080378684 |
| ENSMUSG00000032578 | Cish | 12700 | protein_coding | 235.9549585 | -0.080389371 | 0.121670654 | -0.660712904 | 0.508796447 | 0.724049267 | 0.080389371 |
| ENSMUSG00000036323 | Srp72 | 66661 | protein_coding | 2023.883563 | 0.080395383 | 0.059875281 | 1.342714071 | 0.179364577 | 0.406551304 | 0.080395383 |
| ENSMUSG00000047641 | Krt87 | 406219 | protein_coding | 64.52615967 | 0.080413056 | 0.168731547 | 0.476573929 | 0.633665552 | 0.808283189 | 0.080413056 |
| ENSMUSG00000036199 | Ndufa13 | 67184 | protein_coding | 3848.666388 | 0.080418122 | 0.080185443 | 1.002901752 | 0.315908267 | 0.560076808 | 0.080418122 |
| ENSMUSG00000021504 | B4galt7 | 218271 | protein_coding | 420.3617592 | 0.080423019 | 0.080396121 | 1.000334566 | 0.317148625 | 0.56100207 | 0.080423019 |
| ENSMUSG00000062028 | Irgc1 | 210145 | protein_coding | 9.017988631 | 0.080476545 | 0.416634648 | 0.19315855 | 0.846834799 | 0.930889346 | 0.080476545 |
| ENSMUSG00000031078 | Cttn | 13043 | protein_coding | 6350.237463 | 0.080486236 | 0.051345903 | 1.567529847 | 0.116990895 | 0.31618145 | 0.080486236 |
| ENSMUSG00000036376 | Abt1 | 30946 | protein_coding | 308.3446969 | -0.080490319 | 0.072937758 | -1.103548022 | 0.269789249 | 0.512256092 | 0.080490319 |
| ENSMUSG00000062604 | Srpk2 | 20817 | protein_coding | 449.6683176 | -0.080499032 | 0.078551673 | -1.024790796 | 0.30546191 | 0.549797347 | 0.080499032 |
| ENSMUSG00000016256 | Ctsz | 64138 | protein_coding | 1504.898137 | -0.080501289 | 0.058473431 | -1.376715673 | 0.168600171 | 0.392580849 | 0.080501289 |
| ENSMUSG00000000759 | Tubgcp3 | 259279 | protein_coding | 776.8779788 | -0.080518461 | 0.069354276 | -1.160973266 | 0.245652772 | 0.486927513 | 0.080518461 |
| ENSMUSG00000016541 | Atxn10 | 54138 | protein_coding | 456.8062738 | 0.080557336 | 0.085648338 | 0.940559237 | 0.34693078 | 0.589066402 | 0.080557336 |
| ENSMUSG00000031497 | Tnfsf13b | 24099 | protein_coding | 65.77571629 | 0.080560791 | 0.165237957 | 0.487544103 | 0.625872797 | 0.802596262 | 0.080560791 |
| ENSMUSG00000033751 | Gadd45gip1 | 102060 | protein_coding | 350.2562655 | 0.080585471 | 0.092957483 | 0.866906764 | 0.385993097 | 0.625996649 | 0.080585471 |
| ENSMUSG00000039200 | Atf7ip2 | 75329 | protein_coding | 16.68760039 | -0.080592515 | 0.371442009 | -0.216971999 | 0.828230166 | 0.923583729 | 0.080592515 |
| ENSMUSG00000074211 | Sdhaf1 | 68332 | protein_coding | 135.4172161 | 0.080598996 | 0.115363042 | 0.69865526 | 0.484767501 | 0.706263486 | 0.080598996 |
| ENSMUSG00000116632 | Magef1 | 76222 | transcribed_processed_pseudogene | 61.23448723 | 0.080609017 | 0.177729841 | 0.453548015 | 0.650154177 | 0.819655754 | 0.080609017 |
| ENSMUSG00000037638 | Zbtb42 | 382639 | protein_coding | 726.8928028 | -0.080633003 | 0.077257191 | -1.043695753 | 0.296626175 | 0.539493665 | 0.080633003 |
| ENSMUSG00000039191 | Rbpj | 19664 | protein_coding | 470.4742274 | -0.080645484 | 0.058453713 | -1.379646905 | 0.167695388 | 0.391604912 | 0.080645484 |
| ENSMUSG00000024401 | Tnf | 21926 | protein_coding | 20.6084377 | -0.080691879 | 0.327020319 | -0.246748824 | 0.805102619 | 0.911459024 | 0.080691879 |
| ENSMUSG00000024830 | Rps6kb2 | 58988 | protein_coding | 314.7560558 | -0.080704942 | 0.089569161 | -0.90103492 | 0.367569753 | 0.609972551 | 0.080704942 |
| ENSMUSG00000024927 | Rela | 19697 | protein_coding | 771.0559787 | -0.080728099 | 0.084480678 | -0.955580629 | 0.339284152 | 0.582473517 | 0.080728099 |
| ENSMUSG00000019864 | Rtn4ip1 | 170728 | protein_coding | 450.0007793 | 0.080743656 | 0.076338226 | 1.057709357 | 0.290187968 | 0.532649825 | 0.080743656 |
| ENSMUSG00000039834 | Zfp335 | 329559 | protein_coding | 559.4970809 | 0.080744781 | 0.086881884 | 0.929362689 | 0.352701157 | 0.595229357 | 0.080744781 |
| ENSMUSG00000033740 | St18 | 240690 | protein_coding | 74.03360225 | -0.080751287 | 0.165894076 | -0.486764138 | 0.626425487 | 0.803113429 | 0.080751287 |
| ENSMUSG00000117284 | Gm7072 | 631624 | protein_coding | 608.5882381 | -0.080753274 | 0.119703737 | -0.674609467 | 0.499923917 | 0.717028905 | 0.080753274 |
| ENSMUSG00000002731 | Prkra | 23992 | protein_coding | 794.2208793 | -0.080802614 | 0.066453095 | -1.215934544 | 0.224009858 | 0.461733499 | 0.080802614 |
| ENSMUSG00000000915 | Hip1r | 29816 | protein_coding | 2567.304855 | -0.080805352 | 0.046973779 | -1.720222504 | 0.085392004 | 0.260866991 | 0.080805352 |
| ENSMUSG00000037216 | Lipt1 | 623661 | protein_coding | 67.16687189 | -0.080830033 | 0.157906466 | -0.511885521 | 0.608731132 | 0.791306043 | 0.080830033 |
| ENSMUSG00000040410 | Fbxl4 | 269514 | protein_coding | 381.2490805 | 0.080839257 | 0.062611826 | 1.291118034 | 0.196662752 | 0.429095149 | 0.080839257 |
| ENSMUSG00000015750 | Aph1a | 226548 | protein_coding | 1656.82943 | 0.080857669 | 0.047557513 | 1.700208102 | 0.089091789 | 0.268056582 | 0.080857669 |
| ENSMUSG00000062309 | Rpp25 | 102614 | protein_coding | 16.55181603 | -0.080873541 | 0.296165631 | -0.273068624 | 0.784800465 | 0.899716112 | 0.080873541 |
| ENSMUSG00000020787 | P2rx1 | 18436 | protein_coding | 13.38304514 | -0.080897888 | 0.339258041 | -0.23845533 | 0.811527957 | 0.915210585 | 0.080897888 |

| ensembl_gene_id | external_gene_name | entrezgene_id | gene_biotype | baseMean | log2FoldChange | lfcSE | stat | pvalue | padj | absLog2FC |
|---|---|---|---|---|---|---|---|---|---|---|
| ENSMUSG00000020231 | Dip2a | 64451 | protein_coding | 1937.468052 | -0.080913689 | 0.06068622 | -1.3333124 | 0.182429306 | 0.410383236 | 0.080913689 |
| ENSMUSG00000028191 | Bcl10 | 12042 | protein_coding | 1462.673111 | 0.080915947 | 0.053841398 | 1.502857454 | 0.132875806 | 0.339763458 | 0.080915947 |
| ENSMUSG00000026827 | Gpd2 | 14571 | protein_coding | 1385.831618 | 0.080920908 | 0.064622688 | 1.252205862 | 0.21049486 | 0.445550678 | 0.080920908 |
| ENSMUSG00000034382 | AI661453 | 224833 | protein_coding | 616.0950241 | 0.080932002 | 0.068119317 | 1.188091805 | 0.234797242 | 0.474627116 | 0.080932002 |
| ENSMUSG00000021373 | Cap2 | 67252 | protein_coding | 15.45244207 | -0.080938864 | 0.338394479 | -0.239184943 | 0.810962178 | 0.914838553 | 0.080938864 |
| ENSMUSG00000030880 | Polr3e | 26939 | protein_coding | 609.9839369 | -0.080952416 | 0.071704551 | -1.128971804 | 0.258909728 | 0.500729794 | 0.080952416 |
| ENSMUSG00000083307 | AA414768 | 245350 | processed_pseudogene | 16.59800822 | -0.080972827 | 0.295928611 | -0.273622841 | 0.784374479 | 0.899577643 | 0.080972827 |
| ENSMUSG00000025613 | Cct8 | 12469 | protein_coding | 2726.438624 | 0.080977882 | 0.059622334 | 1.358180355 | 0.174406463 | 0.400571839 | 0.080977882 |
| ENSMUSG00000031980 | Agt | 11606 | protein_coding | 230.4795673 | 0.080979908 | 0.091885439 | 0.881313827 | 0.378147984 | 0.619451611 | 0.080979908 |
| ENSMUSG00000050812 | Ecpas | 230249 | protein_coding | 3532.407523 | -0.080999664 | 0.046566138 | -1.739454201 | 0.081954901 | 0.254268314 | 0.080999664 |
| ENSMUSG00000026457 | Adipor1 | 72674 | protein_coding | 5562.459663 | 0.081018573 | 0.062228819 | 1.301946175 | 0.192934786 | 0.424714619 | 0.081018573 |
| ENSMUSG00000025782 | Taf3 | 209361 | protein_coding | 133.2785533 | -0.081067825 | 0.111175189 | -0.729189897 | 0.46588551 | 0.691634539 | 0.081067825 |
| ENSMUSG00000026032 | Ndufb3 | 66495 | protein_coding | 1063.183266 | 0.081079465 | 0.063483031 | 1.277183264 | 0.201537554 | 0.435332994 | 0.081079465 |
| ENSMUSG00000033883 | Zfp267 | 241944 | protein_coding | 317.8690552 | -0.081079864 | 0.098829486 | -0.820401561 | 0.411987226 | 0.649970343 | 0.081079864 |
| ENSMUSG00000017291 | Taok1 | 216965 | protein_coding | 2827.099879 | -0.081116227 | 0.034021426 | -2.384268879 | 0.017113094 | 0.088190332 | 0.081116227 |
| ENSMUSG00000021494 | Ddx41 | 72935 | protein_coding | 755.3584462 | 0.081126868 | 0.050546305 | 1.605000962 | 0.108493593 | 0.301772366 | 0.081126868 |
| ENSMUSG00000027340 | Slc23a2 | 54338 | protein_coding | 1724.567805 | -0.081149071 | 0.046314988 | -1.752112544 | 0.079754458 | 0.249745186 | 0.081149071 |
| ENSMUSG00000032869 | Psmf1 | 228769 | protein_coding | 613.4230905 | 0.081164553 | 0.050318779 | 1.613007198 | 0.106742944 | 0.299280839 | 0.081164553 |
| ENSMUSG00000016344 | Ppdpf | 66496 | protein_coding | 1255.92703 | 0.08119838 | 0.074673161 | 1.08738373 | 0.276867259 | 0.519767321 | 0.08119838 |
| ENSMUSG00000002105 | Slc39a13 | 68427 | protein_coding | 123.7773071 | -0.081214601 | 0.122043326 | -0.665457124 | 0.505758158 | 0.721674201 | 0.081214601 |
| ENSMUSG00000028576 | Ift74 | 67694 | protein_coding | 192.6287405 | -0.081218754 | 0.13354225 | -0.608187702 | 0.54306299 | 0.748033553 | 0.081218754 |
| ENSMUSG00000034620 | Rxylt1 | 216395 | protein_coding | 516.2829905 | -0.081246086 | 0.07134483 | -1.138780292 | 0.254794804 | 0.496022219 | 0.081246086 |
| ENSMUSG00000060950 | Trmt61a | 328162 | protein_coding | 139.0313937 | -0.081253918 | 0.131383613 | -0.61844789 | 0.536280138 | 0.742878353 | 0.081253918 |
| ENSMUSG00000018481 | Appbp2 | 66884 | protein_coding | 1410.316751 | 0.081257614 | 0.052479231 | 1.548376603 | 0.121531651 | 0.323030784 | 0.081257614 |
| ENSMUSG00000047714 | Ppp1r2 | 66849 | protein_coding | 1302.426247 | 0.081276648 | 0.042547014 | 1.910278515 | 0.056097363 | 0.19992976 | 0.081276648 |
| ENSMUSG00000019822 | Smpd2 | 20598 | protein_coding | 523.3196589 | -0.081288914 | 0.064820399 | -1.254063764 | 0.20981883 | 0.44497561 | 0.081288914 |
| ENSMUSG00000024974 | Smc3 | 13006 | protein_coding | 1641.548256 | -0.081290574 | 0.066485722 | -1.222677168 | 0.221451654 | 0.458159182 | 0.081290574 |
| ENSMUSG00000015289 | Lage3 | 66192 | protein_coding | 315.8922774 | -0.081305237 | 0.081816848 | -0.993746873 | 0.320346116 | 0.56381336 | 0.081305237 |
| ENSMUSG00000029311 | Hsd17b11 | 114664 | protein_coding | 1554.093612 | -0.081323198 | 0.071369337 | -1.139469712 | 0.254507294 | 0.495656311 | 0.081323198 |
| ENSMUSG00000027411 | Vps16 | 80743 | protein_coding | 789.9439341 | -0.081341455 | 0.053554205 | -1.518862138 | 0.128797201 | 0.334230211 | 0.081341455 |
| ENSMUSG00000038371 | Sbf2 | 319934 | protein_coding | 1647.010677 | -0.081354917 | 0.057766472 | -1.408341462 | 0.159029984 | 0.37913729 | 0.081354917 |
| ENSMUSG00000011837 | Snapc2 | 102209 | protein_coding | 277.2278456 | 0.081358415 | 0.081311863 | 1.000572511 | 0.317033526 | 0.56085999 | 0.081358415 |
| ENSMUSG00000029199 | Lias | 79464 | protein_coding | 702.754717 | 0.081374221 | 0.062572025 | 1.300488853 | 0.193433474 | 0.425320463 | 0.081374221 |
| ENSMUSG00000019961 | Tmpo | 21917 | protein_coding | 2717.018818 | 0.081375419 | 0.069991501 | 1.162647148 | 0.244972693 | 0.486120986 | 0.081375419 |
| ENSMUSG00000022223 | Sdr39u1 | 654795 | protein_coding | 134.0635536 | -0.081388751 | 0.117006924 | -0.695589183 | 0.486686142 | 0.707799388 | 0.081388751 |
| ENSMUSG00000040760 | Appl1 | 72993 | protein_coding | 1453.849545 | 0.081411319 | 0.058650038 | 1.388086382 | 0.165110743 | 0.387871952 | 0.081411319 |
| ENSMUSG00000031832 | Taf1c | 21341 | protein_coding | 383.8104687 | 0.081430787 | 0.093723571 | 0.868839989 | 0.384934657 | 0.625286182 | 0.081430787 |
| ENSMUSG00000049421 | Zfp260 | 26466 | protein_coding | 801.1753105 | 0.081439006 | 0.052568297 | 1.549203809 | 0.121332734 | 0.322680245 | 0.081439006 |
| ENSMUSG00000002718 | Cse1l | 110750 | protein_coding | 1062.282136 | 0.08144951 | 0.073198427 | 1.112722126 | 0.26582779 | 0.507758471 | 0.08144951 |

| ensembl_gene_id | external_gene_name | entrezgene_id | gene_biotype | baseMean | log2FoldChange | lfcSE | stat | pvalue | padj | absLog2FC |
|---|---|---|---|---|---|---|---|---|---|---|
| ENSMUSG00000044026 | Slc35g1 | 240660 | protein_coding | 3406.300601 | -0.081451855 | 0.034569934 | -2.356147263 | 0.018465596 | 0.093148158 | 0.081451855 |
| ENSMUSG00000058254 | Tspan7 | 21912 | protein_coding | 1331.166862 | -0.081455316 | 0.055569203 | -1.465835615 | 0.142693076 | 0.354359599 | 0.081455316 |
| ENSMUSG00000110277 | Gm45871 | 108168395 | protein_coding | 478.2705456 | -0.081549327 | 0.101676035 | -0.802050619 | 0.422523678 | 0.65874317 | 0.081549327 |
| ENSMUSG00000029505 | Ep400 | 75560 | protein_coding | 1795.600175 | -0.081558828 | 0.073914173 | -1.103426116 | 0.269842161 | 0.51227128 | 0.081558828 |
| ENSMUSG00000015745 | Plekho1 | 67220 | protein_coding | 22.19228266 | 0.081572099 | 0.29611824 | 0.275471374 | 0.782954115 | 0.898825528 | 0.081572099 |
| ENSMUSG00000026821 | Ralgds | 19730 | protein_coding | 2064.160567 | 0.081587777 | 0.059568134 | 1.369654733 | 0.170794705 | 0.395236937 | 0.081587777 |
| ENSMUSG00000001323 | Srr | 27364 | protein_coding | 375.5259594 | -0.081599087 | 0.065311355 | -1.24938592 | 0.211523956 | 0.446836226 | 0.081599087 |
| ENSMUSG00000032607 | Amt | 434437 | protein_coding | 410.4334304 | 0.081601897 | 0.076802042 | 1.062496455 | 0.288010366 | 0.530464045 | 0.081601897 |
| ENSMUSG00000026708 | Cenpl | 70454 | protein_coding | 155.5861664 | -0.081639684 | 0.117512512 | -0.694731839 | 0.487223371 | 0.708115319 | 0.081639684 |
| ENSMUSG00000027938 | Creb3l4 | 78284 | protein_coding | 1142.456967 | 0.081662925 | 0.068679174 | 1.18904932 | 0.23442026 | 0.473983893 | 0.081662925 |
| ENSMUSG00000020235 | Fzr1 | 56371 | protein_coding | 1062.345714 | 0.08166583 | 0.06358561 | 1.284344517 | 0.199021437 | 0.432388349 | 0.08166583 |
| ENSMUSG00000045996 | Polr2k | 17749 | protein_coding | 165.4308593 | 0.08168923 | 0.101587495 | 0.804126832 | 0.42132373 | 0.658017412 | 0.08168923 |
| ENSMUSG00000036309 | Skp1 | 21402 | protein_coding | 3951.385384 | 0.081729396 | 0.041769768 | 1.956663876 | 0.050387 | 0.186070341 | 0.081729396 |
| ENSMUSG00000066043 | Phactr4 | 100169 | protein_coding | 824.1897284 | -0.081757786 | 0.056587612 | -1.444800084 | 0.148514047 | 0.362726949 | 0.081757786 |
| ENSMUSG00000039148 | Sart1 | 20227 | protein_coding | 1199.727599 | 0.081764342 | 0.047889012 | 1.707371661 | 0.087752994 | 0.265315436 | 0.081764342 |
| ENSMUSG00000030304 | Ergic2 | 67456 | protein_coding | 556.2131322 | 0.08178287 | 0.058041085 | 1.40905136 | 0.158819982 | 0.379001161 | 0.08178287 |
| ENSMUSG00000033126 | Ybey | 216119 | protein_coding | 165.8813412 | -0.081815042 | 0.099801412 | -0.819778407 | 0.412342443 | 0.649970343 | 0.081815042 |
| ENSMUSG00000005628 | Tmod4 | 50874 | protein_coding | 6.583328635 | 0.081838691 | 0.465452049 | 0.175826256 | 0.860430442 | 0.936965584 | 0.081838691 |
| ENSMUSG00000029761 | Cald1 | 109624 | protein_coding | 22.358419 | -0.081943214 | 0.282974299 | -0.289578289 | 0.772138878 | 0.89282883 | 0.081943214 |
| ENSMUSG00000003662 | Ciao1 | 26371 | protein_coding | 861.3665193 | 0.081963815 | 0.058823798 | 1.393378501 | 0.163505363 | 0.385391844 | 0.081963815 |
| ENSMUSG00000035354 | Uvrag | 78610 | protein_coding | 447.7332801 | -0.081963972 | 0.075987457 | -1.07865134 | 0.280743185 | 0.523508209 | 0.081963972 |
| ENSMUSG00000018666 | Cbx1 | 12412 | protein_coding | 1076.023337 | 0.081965743 | 0.059798299 | 1.370703597 | 0.170467373 | 0.394876088 | 0.081965743 |
| ENSMUSG00000097610 | A930012L18Rik | 626275 | lncRNA | 6.440484825 | -0.082051261 | 0.497321369 | -0.164986397 | 0.868954689 | 0.940691929 | 0.082051261 |
| ENSMUSG00000029790 | Cep41 | 83922 | protein_coding | 83.62492551 | 0.08205331 | 0.145665357 | 0.563300097 | 0.573230552 | 0.767653434 | 0.08205331 |
| ENSMUSG00000031897 | Psmb10 | 19171 | protein_coding | 2702.249972 | -0.082094062 | 0.101778756 | -0.806593299 | 0.419900836 | 0.657067803 | 0.082094062 |
| ENSMUSG00000056665 | Them6 | 223626 | protein_coding | 33.31079489 | 0.082099408 | 0.207924918 | 0.394851222 | 0.69295269 | 0.846032859 | 0.082099408 |
| ENSMUSG00000026556 | Vangl2 | 93840 | protein_coding | 40.39981294 | -0.082102474 | 0.205283326 | -0.399947117 | 0.689195468 | 0.844094282 | 0.082102474 |
| ENSMUSG00000066877 | Nck2 | 17974 | protein_coding | 1743.739128 | -0.082112224 | 0.053558795 | -1.533123078 | 0.125245533 | 0.328321636 | 0.082112224 |
| ENSMUSG00000030327 | Necap1 | 67602 | protein_coding | 804.8785025 | 0.082121153 | 0.053385386 | 1.538270284 | 0.123982542 | 0.326892409 | 0.082121153 |
| ENSMUSG00000039016 | Timm8b | 30057 | protein_coding | 832.7413201 | 0.082125227 | 0.06209595 | 1.322553661 | 0.185983853 | 0.415160239 | 0.082125227 |
| ENSMUSG00000030930 | Chst15 | 77590 | protein_coding | 207.7738506 | -0.082153031 | 0.12466037 | -0.65901482 | 0.509886252 | 0.724638828 | 0.082153031 |
| ENSMUSG00000026353 | Ubxn4 | 67812 | protein_coding | 1674.255567 | 0.082173355 | 0.037204074 | 2.208719252 | 0.027194174 | 0.121104897 | 0.082173355 |
| ENSMUSG00000045624 | Esf1 | 66580 | protein_coding | 441.0361087 | -0.082187722 | 0.082960958 | -0.99067952 | 0.321842094 | 0.565705445 | 0.082187722 |
| ENSMUSG00000028718 | Stil | 20460 | protein_coding | 189.2403036 | 0.082189368 | 0.124278865 | 0.661330211 | 0.508400571 | 0.723843561 | 0.082189368 |
| ENSMUSG00000011427 | Zfp790 | 233056 | protein_coding | 431.4958953 | 0.0821949 | 0.079961056 | 1.027936644 | 0.303979631 | 0.548046454 | 0.0821949 |
| ENSMUSG00000035498 | Cdcp1 | 109332 | protein_coding | 2845.909634 | 0.082207527 | 0.038944146 | 2.11090843 | 0.034780183 | 0.143968177 | 0.082207527 |
| ENSMUSG00000006005 | Tpr | 108989 | protein_coding | 3990.15253 | -0.082234486 | 0.064708723 | -1.270840815 | 0.203785285 | 0.437979805 | 0.082234486 |
| ENSMUSG00000045679 | Pqlc3 | 217430 | protein_coding | 563.2116077 | 0.082237134 | 0.069548611 | 1.182441059 | 0.237030739 | 0.477153008 | 0.082237134 |
| ENSMUSG00000047547 | Cltb | 74325 | protein_coding | 3073.761225 | 0.082246788 | 0.049380199 | 1.665582329 | 0.095796633 | 0.280581712 | 0.082246788 |

Supplementary Table 1

| ensembl_gene_id | external_gene_name | entrezgene_id | gene_biotype | baseMean | log2FoldChange | lfcSE | stat | pvalue | padj | absLog2FC |
|---|---|---|---|---|---|---|---|---|---|---|
| ENSMUSG00000038736 | Nudcd1 | 67429 | protein_coding | 753.7416692 | -0.082259723 | 0.069569146 | -1.182416734 | 0.237040386 | 0.477153008 | 0.082259723 |
| ENSMUSG00000043964 | Orai3 | 269999 | protein_coding | 416.3985912 | -0.082263472 | 0.071601888 | -1.148900882 | 0.250596854 | 0.491645551 | 0.082263472 |
| ENSMUSG00000044991 | Shld1 | 73747 | protein_coding | 118.9101781 | 0.082288814 | 0.137585887 | 0.598090516 | 0.549779538 | 0.751487589 | 0.082288814 |
| ENSMUSG00000027797 | Dclk1 | 13175 | protein_coding | 764.289101 | 0.082326204 | 0.081805002 | 1.006371273 | 0.314237007 | 0.558485807 | 0.082326204 |
| ENSMUSG00000055835 | Zfp1 | 22640 | protein_coding | 127.6838174 | -0.082335063 | 0.119337479 | -0.689934661 | 0.490235278 | 0.710266225 | 0.082335063 |
| ENSMUSG00000039840 | Epg5 | 100502841 | protein_coding | 1017.979371 | -0.082335421 | 0.051914594 | -1.585978347 | 0.112744221 | 0.309011907 | 0.082335421 |
| ENSMUSG00000096001 | 2610528A11Rik | 70045 | protein_coding | 7187.632697 | -0.082362696 | 0.078442251 | -1.049978737 | 0.293727889 | 0.536541439 | 0.082362696 |
| ENSMUSG00000030168 | Adipor2 | 68465 | protein_coding | 4744.085296 | 0.082371032 | 0.046469413 | 1.772586002 | 0.076297333 | 0.243298134 | 0.082371032 |
| ENSMUSG00000024069 | Slc30a6 | 210148 | protein_coding | 468.5272812 | 0.08238613 | 0.064623943 | 1.274854578 | 0.202360719 | 0.43646801 | 0.08238613 |
| ENSMUSG00000079487 | Med12 | 59024 | protein_coding | 1037.280769 | -0.082417207 | 0.070623392 | -1.166995873 | 0.243212036 | 0.484226665 | 0.082417207 |
| ENSMUSG00000035007 | Rundc1 | 217201 | protein_coding | 476.075365 | 0.082500066 | 0.08239028 | 1.001332512 | 0.31666608 | 0.560517382 | 0.082500066 |
| ENSMUSG00000019082 | Slc25a22 | 68267 | protein_coding | 325.8763636 | -0.082514496 | 0.101997276 | -0.808987254 | 0.418522478 | 0.655993003 | 0.082514496 |
| ENSMUSG00000095115 | Itpripl2 | 319622 | protein_coding | 561.9074749 | -0.082520355 | 0.133750483 | -0.616972388 | 0.537252938 | 0.743714861 | 0.082520355 |
| ENSMUSG00000038619 | Ensa | 56205 | protein_coding | 2297.462619 | 0.082521455 | 0.031997462 | 2.579000047 | 0.009908677 | 0.058857178 | 0.082521455 |
| ENSMUSG00000028080 | Lrba | 80877 | protein_coding | 5545.724375 | -0.082577189 | 0.054932065 | -1.503260239 | 0.132771949 | 0.339551732 | 0.082577189 |
| ENSMUSG00000028470 | Hint2 | 68917 | protein_coding | 747.0492091 | 0.082593331 | 0.076948534 | 1.073358088 | 0.283110482 | 0.5257021 | 0.082593331 |
| ENSMUSG00000073889 | Il11ra1 | 16157 | protein_coding | 207.4866621 | -0.082608507 | 0.091389222 | -0.903919574 | 0.366038049 | 0.608127704 | 0.082608507 |
| ENSMUSG00000016200 | Syt14 | 329324 | protein_coding | 21.93862656 | -0.082621454 | 0.297659926 | -0.277569962 | 0.781342493 | 0.898510439 | 0.082621454 |
| ENSMUSG00000046694 | Tent5b | 100342 | protein_coding | 17.5018302 | 0.082636928 | 0.30333433 | 0.272428539 | 0.785292531 | 0.900006643 | 0.082636928 |
| ENSMUSG00000047446 | Arl4a | 11861 | protein_coding | 2453.731603 | -0.082642534 | 0.086016804 | -0.96077196 | 0.33666684 | 0.580252873 | 0.082642534 |
| ENSMUSG00000074466 | Gm15417 | 545539 | lncRNA | 141.4471912 | 0.082645733 | 0.125510539 | 0.658476439 | 0.510232032 | 0.724823423 | 0.082645733 |
| ENSMUSG00000048967 | Yjefn3 | 234365 | protein_coding | 7.810180122 | 0.082647032 | 0.433960549 | 0.190448262 | 0.848957882 | 0.931687225 | 0.082647032 |
| ENSMUSG00000034152 | Exoc3 | 211446 | protein_coding | 1372.251699 | 0.082677008 | 0.048746257 | 1.696068864 | 0.089872839 | 0.269652976 | 0.082677008 |
| ENSMUSG00000034438 | Gbp8 | 76074 | protein_coding | 14.37991872 | -0.082700513 | 0.491647722 | -0.168210915 | 0.866417355 | 0.939594693 | 0.082700513 |
| ENSMUSG00000025949 | Pikfyve | 18711 | protein_coding | 982.7229069 | -0.082730682 | 0.050221444 | -1.647317865 | 0.099492728 | 0.286931191 | 0.082730682 |
| ENSMUSG00000041781 | Cpsf2 | 51786 | protein_coding | 1173.48622 | 0.082747753 | 0.077339463 | 1.069929242 | 0.284651159 | 0.527543906 | 0.082747753 |
| ENSMUSG00000061143 | Maml3 | 433586 | protein_coding | 573.3134804 | -0.08277399 | 0.077782256 | -1.064175751 | 0.28724909 | 0.529611374 | 0.08277399 |
| ENSMUSG00000031715 | Smarca5 | 93762 | protein_coding | 2716.673198 | 0.082775759 | 0.054705128 | 1.513126134 | 0.130247622 | 0.336231563 | 0.082775759 |
| ENSMUSG00000097431 | Gm26782 | 102640338 | lncRNA | 55.48010996 | -0.082805667 | 0.159149584 | -0.52030087 | 0.602853889 | 0.787655348 | 0.082805667 |
| ENSMUSG00000070699 | Sars2 | 71984 | protein_coding | 271.2656046 | 0.082813981 | 0.102125991 | 0.810900151 | 0.417423014 | 0.655224931 | 0.082813981 |
| ENSMUSG00000006315 | Tmem147 | 69804 | protein_coding | 962.6487525 | -0.082832164 | 0.070422381 | -1.176219301 | 0.239507254 | 0.480032716 | 0.082832164 |
| ENSMUSG00000024639 | Gnaq | 14682 | protein_coding | 1001.010414 | 0.082877791 | 0.057800493 | 1.4338596 | 0.151612333 | 0.367724228 | 0.082877791 |
| ENSMUSG00000045248 | Med26 | 70625 | protein_coding | 254.1028468 | 0.082882229 | 0.097542223 | 0.849706174 | 0.395488465 | 0.634757412 | 0.082882229 |
| ENSMUSG00000036073 | Galt | 14430 | protein_coding | 539.5135617 | -0.082893705 | 0.066405061 | -1.248304011 | 0.211919746 | 0.447440166 | 0.082893705 |
| ENSMUSG00000034254 | Agpat1 | 55979 | protein_coding | 2496.171575 | 0.082906753 | 0.103215572 | 0.803238803 | 0.421836722 | 0.65828686 | 0.082906753 |
| ENSMUSG00000033364 | Usp37 | 319651 | protein_coding | 833.7891055 | -0.082917753 | 0.052431812 | -1.581439779 | 0.113777518 | 0.310840257 | 0.082917753 |
| ENSMUSG00000003420 | Fcgrt | 14132 | protein_coding | 1383.773066 | 0.082926018 | 0.101110552 | 0.820151974 | 0.412129477 | 0.649970343 | 0.082926018 |
| ENSMUSG00000021476 | Habp4 | 56541 | protein_coding | 305.5150973 | 0.082946817 | 0.076940341 | 1.078066667 | 0.281004006 | 0.523508209 | 0.082946817 |
| ENSMUSG00000022744 | Cldnd1 | 224250 | protein_coding | 492.0369519 | 0.08295654 | 0.072481045 | 1.144527369 | 0.252404994 | 0.493560011 | 0.08295654 |

| ensembl_gene_id | external_gene_name | entrezgene_id | gene_biotype | baseMean | log2FoldChange | lfcSE | stat | pvalue | padj | absLog2FC |
|---|---|---|---|---|---|---|---|---|---|---|
| ENSMUSG00000002910 | Arrdc2 | 70807 | protein_coding | 430.2613629 | 0.082969615 | 0.085826142 | 0.966717286 | 0.333685383 | 0.577025968 | 0.082969615 |
| ENSMUSG00000027131 | Emc4 | 68032 | protein_coding | 562.779769 | 0.082996114 | 0.072055361 | 1.151838153 | 0.249387586 | 0.490195121 | 0.082996114 |
| ENSMUSG00000002308 | Cd320 | 54219 | protein_coding | 575.2425979 | -0.083011808 | 0.08305142 | -0.99952305 | 0.317541379 | 0.561388929 | 0.083011808 |
| ENSMUSG00000037860 | Aim2 | 383619 | protein_coding | 878.4251201 | 0.083018641 | 0.067266449 | 1.234176062 | 0.217137305 | 0.45330399 | 0.083018641 |
| ENSMUSG00000024050 | Wiz | 22404 | protein_coding | 469.6025278 | -0.083059728 | 0.08779515 | -0.946062828 | 0.344116542 | 0.586511041 | 0.083059728 |
| ENSMUSG00000008301 | Phax | 56698 | protein_coding | 583.1608674 | -0.083062282 | 0.059960903 | -1.385274029 | 0.165968692 | 0.389376941 | 0.083062282 |
| ENSMUSG00000028563 | Tm2d1 | 94043 | protein_coding | 343.8920738 | 0.083074808 | 0.066311292 | 1.252800309 | 0.210278389 | 0.445347562 | 0.083074808 |
| ENSMUSG00000046807 | Lrrc75b | 192734 | protein_coding | 8.337432527 | -0.083082996 | 0.480732267 | -0.172825919 | 0.862788263 | 0.937787345 | 0.083082996 |
| ENSMUSG00000040167 | Ikzf5 | 67143 | protein_coding | 75.03737237 | -0.083133322 | 0.17860747 | -0.465452658 | 0.641607332 | 0.813714227 | 0.083133322 |
| ENSMUSG00000025950 | Idh1 | 15926 | protein_coding | 5411.309787 | -0.083146037 | 0.056556407 | -1.470143558 | 0.141522877 | 0.352392829 | 0.083146037 |
| ENSMUSG00000030614 | Tmem126b | 68472 | protein_coding | 418.9231656 | 0.083147821 | 0.0733318 | 1.133857625 | 0.256854274 | 0.498650401 | 0.083147821 |
| ENSMUSG00000023845 | Lnpep | 240028 | protein_coding | 4725.998499 | -0.083216856 | 0.046333585 | -1.796037498 | 0.072488554 | 0.235215876 | 0.083216856 |
| ENSMUSG00000037440 | Vnn1 | 22361 | protein_coding | 121.5435144 | -0.083227209 | 0.209897014 | -0.396514497 | 0.691725516 | 0.845430647 | 0.083227209 |
| ENSMUSG00000042757 | Tmem108 | 81907 | protein_coding | 7.592258569 | -0.083239402 | 0.455153489 | -0.182882048 | 0.854890569 | 0.93513139 | 0.083239402 |
| ENSMUSG00000019857 | Asf1a | 66403 | protein_coding | 206.7589656 | -0.083249872 | 0.097883448 | -0.850499997 | 0.395047162 | 0.634427961 | 0.083249872 |
| ENSMUSG00000056204 | Pgpep1 | 66522 | protein_coding | 1321.060323 | 0.083257765 | 0.048680854 | 1.710277407 | 0.087214588 | 0.264480943 | 0.083257765 |
| ENSMUSG00000034371 | Tkfc | 225913 | protein_coding | 384.0694689 | 0.083267526 | 0.081609006 | 1.020322757 | 0.307575414 | 0.551334926 | 0.083267526 |
| ENSMUSG00000064125 | Prr36 | 73072 | protein_coding | 11.34804877 | 0.083306605 | 0.418734877 | 0.198948331 | 0.842303163 | 0.929889314 | 0.083306605 |
| ENSMUSG00000020009 | Ifngr1 | 15979 | protein_coding | 5076.184879 | -0.083310883 | 0.059941515 | -1.389869503 | 0.164568508 | 0.386825021 | 0.083310883 |
| ENSMUSG00000038485 | Socs7 | 192157 | protein_coding | 630.4672633 | -0.083335364 | 0.058854138 | -1.415964385 | 0.156785932 | 0.375480655 | 0.083335364 |
| ENSMUSG00000062397 | Zfp706 | 68036 | protein_coding | 2748.982793 | 0.083358886 | 0.045926475 | 1.815050811 | 0.069516101 | 0.228944671 | 0.083358886 |
| ENSMUSG00000039599 | Fam149b | 105428 | protein_coding | 745.283733 | 0.083362777 | 0.050522527 | 1.650012021 | 0.098940477 | 0.286054693 | 0.083362777 |
| ENSMUSG00000037262 | Kin | 16588 | protein_coding | 271.2689346 | 0.083375041 | 0.080758264 | 1.032402592 | 0.301883558 | 0.545539031 | 0.083375041 |
| ENSMUSG00000046032 | Snx12 | 55988 | protein_coding | 581.3944253 | 0.083391225 | 0.072724495 | 1.146673144 | 0.251516734 | 0.492557422 | 0.083391225 |
| ENSMUSG00000016559 | H3f3b | 15081 | protein_coding | 16667.82968 | 0.083404565 | 0.042870953 | 1.945479586 | 0.051717284 | 0.189254514 | 0.083404565 |
| ENSMUSG00000032399 | Rpl4 | 67891 | protein_coding | 23523.03356 | -0.08342418 | 0.040587007 | -2.055440575 | 0.03983647 | 0.158042883 | 0.08342418 |
| ENSMUSG00000025077 | Dclre1a | 55947 | protein_coding | 414.3810065 | -0.083454952 | 0.067772079 | -1.231406112 | 0.218171012 | 0.454403192 | 0.083454952 |
| ENSMUSG00000061991 | H2ac10 | 319173 | protein_coding | 153.9502687 | 0.083458572 | 0.164633655 | 0.506935061 | 0.612200384 | 0.793255831 | 0.083458572 |
| ENSMUSG00000030530 | Furin | 18550 | protein_coding | 1415.933637 | -0.083543802 | 0.071449063 | -1.169277785 | 0.242291731 | 0.483474231 | 0.083543802 |
| ENSMUSG00000066406 | Akap13 | 75547 | protein_coding | 4199.58043 | -0.083630324 | 0.038570706 | -2.168234224 | 0.03014087 | 0.130601934 | 0.083630324 |
| ENSMUSG00000000600 | Krit1 | 79264 | protein_coding | 527.340102 | -0.083639042 | 0.090173581 | -0.927533772 | 0.353649466 | 0.595956629 | 0.083639042 |
| ENSMUSG00000005718 | Tfap4 | 83383 | protein_coding | 438.5264792 | 0.083660421 | 0.062838621 | 1.331353534 | 0.183072711 | 0.411180542 | 0.083660421 |
| ENSMUSG00000062683 | Atp5g2 | 67942 | protein_coding | 1339.386866 | 0.083694918 | 0.108749152 | 0.769614445 | 0.441528634 | 0.672888818 | 0.083694918 |
| ENSMUSG00000029260 | Ugt2b34 | 100727 | protein_coding | 14639.97807 | -0.08370317 | 0.05002225 | -1.673318775 | 0.094264558 | 0.277610848 | 0.08370317 |
| ENSMUSG00000032314 | Etfa | 110842 | protein_coding | 2173.800123 | -0.083715794 | 0.042370606 | -1.975798837 | 0.048177565 | 0.180310883 | 0.083715794 |
| ENSMUSG00000031557 | Plekha2 | 83436 | protein_coding | 1514.739505 | 0.083715984 | 0.06679546 | 1.253318468 | 0.21008983 | 0.445219276 | 0.083715984 |
| ENSMUSG00000031681 | Smad1 | 17125 | protein_coding | 1333.155743 | -0.083753104 | 0.045059766 | -1.858711472 | 0.063068046 | 0.21475895 | 0.083753104 |
| ENSMUSG00000019977 | Hbs1l | 56422 | protein_coding | 1021.161292 | 0.083763501 | 0.062029455 | 1.350382673 | 0.176893266 | 0.40378736 | 0.083763501 |
| ENSMUSG00000022490 | Ppp1r1a | 58200 | protein_coding | 15.66321474 | 0.083804921 | 0.292764592 | 0.286253611 | 0.774683885 | 0.89435358 | 0.083804921 |

| ensembl_gene_id | external_gene_name | entrezgene_id | gene_biotype | baseMean | log2FoldChange | lfcSE | stat | pvalue | padj | absLog2FC |
|---|---|---|---|---|---|---|---|---|---|---|
| ENSMUSG00000025340 | Rabgef1 | 56715 | protein_coding | 687.3466847 | 0.083830788 | 0.057859041 | 1.448879666 | 0.147371191 | 0.360974361 | 0.083830788 |
| ENSMUSG00000036617 | Etl4 | 208618 | protein_coding | 2835.553725 | -0.083839798 | 0.07149408 | -1.172681685 | 0.240923478 | 0.481593365 | 0.083839798 |
| ENSMUSG00000025147 | Mob2 | 101513 | protein_coding | 1194.754095 | 0.083842755 | 0.038797081 | 2.161058304 | 0.030690836 | 0.132269137 | 0.083842755 |
| ENSMUSG00000003500 | Impdh1 | 23917 | protein_coding | 375.7183992 | -0.083844343 | 0.066606847 | -1.258794649 | 0.208104514 | 0.44340091 | 0.083844343 |
| ENSMUSG00000028367 | Txn1 | 22166 | protein_coding | 12599.91845 | 0.083912223 | 0.071846978 | 1.167929759 | 0.242835099 | 0.483959669 | 0.083912223 |
| ENSMUSG00000039361 | Picalm | 233489 | protein_coding | 4852.572984 | -0.083926527 | 0.029067139 | -2.887333656 | 0.00388522 | 0.02955516 | 0.083926527 |
| ENSMUSG00000028233 | Tgs1 | 116940 | protein_coding | 252.9251366 | -0.083938997 | 0.098209632 | -0.854692099 | 0.392721626 | 0.632267311 | 0.083938997 |
| ENSMUSG00000099950 | 9130227L01Rik | 329159 | lncRNA | 22.43097876 | 0.083943842 | 0.25585271 | 0.328094401 | 0.742840287 | 0.87731545 | 0.083943842 |
| ENSMUSG00000032226 | Gcnt3 | 72077 | protein_coding | 8588.32536 | -0.083945407 | 0.087204801 | -0.962623693 | 0.335736403 | 0.579268318 | 0.083945407 |
| ENSMUSG00000018821 | Avpi1 | 69534 | protein_coding | 229.1325904 | 0.08396414 | 0.08442282 | 0.994566872 | 0.319946967 | 0.563409586 | 0.08396414 |
| ENSMUSG00000030165 | Klrd1 | 16643 | protein_coding | 28.42197345 | 0.083986225 | 0.252367232 | 0.332793699 | 0.739290015 | 0.87485545 | 0.083986225 |
| ENSMUSG00000021134 | Srsf5 | 20384 | protein_coding | 6387.042743 | -0.083998336 | 0.093925763 | -0.894305601 | 0.371158404 | 0.613223192 | 0.083998336 |
| ENSMUSG00000057177 | Gsk3a | 606496 | protein_coding | 767.614457 | 0.084003617 | 0.054624507 | 1.537837539 | 0.124088343 | 0.326989003 | 0.084003617 |
| ENSMUSG00000065610 | Mir29a | 387222 | miRNA | 8.025010484 | -0.084015978 | 0.523780556 | -0.16040301 | 0.872563619 | 0.942710088 | 0.084015978 |
| ENSMUSG00000035376 | Hacd2 | 70757 | protein_coding | 410.8212686 | 0.084029586 | 0.083629813 | 1.004780271 | 0.315002666 | 0.559370353 | 0.084029586 |
| ENSMUSG00000036295 | Lrrn3 | 16981 | protein_coding | 23.65954017 | 0.084051837 | 0.304923062 | 0.275649328 | 0.782817418 | 0.898825528 | 0.084051837 |
| ENSMUSG00000039518 | Cdsn | 386463 | protein_coding | 12.76845461 | -0.084063503 | 0.310748293 | -0.270519596 | 0.786760543 | 0.900566916 | 0.084063503 |
| ENSMUSG00000068130 | Zfp442 | 668923 | protein_coding | 64.98250253 | 0.08414459 | 0.195836688 | 0.429667143 | 0.667437788 | 0.830281488 | 0.08414459 |
| ENSMUSG00000019907 | Ppp1r12a | 17931 | protein_coding | 2100.338474 | -0.084155613 | 0.039978205 | -2.105037305 | 0.035288061 | 0.145361573 | 0.084155613 |
| ENSMUSG00000036678 | Aaas | 223921 | protein_coding | 228.5538266 | 0.084180238 | 0.114091884 | 0.737828456 | 0.460618699 | 0.687305492 | 0.084180238 |
| ENSMUSG00000021692 | Dimt1 | 66254 | protein_coding | 93.15408834 | -0.084189781 | 0.138299023 | -0.608751813 | 0.542688958 | 0.747901626 | 0.084189781 |
| ENSMUSG00000054057 | A930004D18Rik | 77940 | lncRNA | 22.78430486 | -0.084227814 | 0.284169761 | -0.296399638 | 0.766924902 | 0.89053739 | 0.084227814 |
| ENSMUSG00000022055 | Nefl | 18039 | protein_coding | 8.629481818 | -0.084234628 | 0.4333661 | -0.194372905 | 0.845883904 | 0.93086312 | 0.084234628 |
| ENSMUSG00000029319 | Coq2 | 71883 | protein_coding | 1068.444719 | 0.084239014 | 0.050671674 | 1.662447809 | 0.09642302 | 0.281807546 | 0.084239014 |
| ENSMUSG00000041852 | Tcf20 | 21411 | protein_coding | 5648.96735 | -0.084254903 | 0.036329886 | -2.31916237 | 0.020386234 | 0.099626908 | 0.084254903 |
| ENSMUSG00000059923 | Grb2 | 14784 | protein_coding | 1180.514155 | -0.084258518 | 0.046453773 | -1.813814315 | 0.069706318 | 0.229325538 | 0.084258518 |
| ENSMUSG00000026027 | Stradb | 227154 | protein_coding | 438.143473 | -0.084311796 | 0.07385213 | -1.14162985 | 0.253607911 | 0.494893811 | 0.084311796 |
| ENSMUSG00000035759 | Bbs10 | 71769 | protein_coding | 41.11098454 | 0.084329789 | 0.197250958 | 0.427525371 | 0.66899671 | 0.831002845 | 0.084329789 |
| ENSMUSG00000034252 | Senp6 | 215351 | protein_coding | 2060.744664 | -0.084347651 | 0.044761196 | -1.88439225 | 0.059511937 | 0.207110275 | 0.084347651 |
| ENSMUSG00000048087 | Ahcyl | 11615 | protein_coding | 18.33759786 | -0.084348696 | 0.27184231 | -0.310285386 | 0.756343943 | 0.884393762 | 0.084348696 |
| ENSMUSG00000041361 | Myzap | 102371 | protein_coding | 298.6056389 | 0.084358164 | 0.111416086 | 0.757145283 | 0.448962827 | 0.679126034 | 0.084358164 |
| ENSMUSG00000078440 | Dohh | 102115 | protein_coding | 196.4142748 | 0.084390343 | 0.147545077 | 0.571963126 | 0.567346956 | 0.762827428 | 0.084390343 |
| ENSMUSG00000060708 | Bloc1s4 | 117197 | protein_coding | 184.3409859 | -0.084397941 | 0.123628325 | -0.68267479 | 0.494812363 | 0.713632533 | 0.084397941 |
| ENSMUSG00000036777 | Anln | 68743 | protein_coding | 505.1368881 | 0.084415565 | 0.111559257 | 0.756688119 | 0.449236735 | 0.679246742 | 0.084415565 |
| ENSMUSG00000000384 | Tbrg4 | 21379 | protein_coding | 1804.794517 | 0.084416864 | 0.060126323 | 1.40399179 | 0.160321294 | 0.380724711 | 0.084416864 |
| ENSMUSG00000051427 | Ccdc157 | 216516 | protein_coding | 139.5644851 | 0.084478494 | 0.120449876 | 0.701358084 | 0.483079574 | 0.705097682 | 0.084478494 |
| ENSMUSG00000024410 | Rmc1 | 76482 | protein_coding | 808.3089448 | 0.084487694 | 0.054952606 | 1.537464731 | 0.124179545 | 0.327122432 | 0.084487694 |
| ENSMUSG00000003531 | Dgcr6 | 13353 | protein_coding | 475.9240064 | -0.084518937 | 0.072591688 | -1.164305985 | 0.244300032 | 0.485573395 | 0.084518937 |
| ENSMUSG00000004945 | Tmem242 | 70544 | protein_coding | 912.3111903 | 0.084532188 | 0.050084079 | 1.687805582 | 0.091448556 | 0.272408741 | 0.084532188 |

| ensembl_gene_id | external_gene_name | entrezgene_id | gene_biotype | baseMean | log2FoldChange | lfcSE | stat | pvalue | padj | absLog2FC |
|---|---|---|---|---|---|---|---|---|---|---|
| ENSMUSG00000021116 | Eif2s1 | 13665 | protein_coding | 1195.053538 | 0.084557673 | 0.063250703 | 1.336865338 | 0.181266597 | 0.408280224 | 0.084557673 |
| ENSMUSG00000052911 | Lamb2 | 16779 | protein_coding | 538.6165599 | 0.084563872 | 0.059319467 | 1.42556696 | 0.153993379 | 0.371384959 | 0.084563872 |
| ENSMUSG00000026504 | Sdccag8 | 76816 | protein_coding | 203.5301532 | 0.08456559 | 0.088652296 | 0.953901852 | 0.340133326 | 0.583162735 | 0.08456559 |
| ENSMUSG00000033014 | Trim33 | 94093 | protein_coding | 1150.676257 | -0.08456679 | 0.055708665 | -1.518018605 | 0.129009709 | 0.334558164 | 0.08456679 |
| ENSMUSG00000070319 | Eif3g | 53356 | protein_coding | 1685.596604 | 0.084586283 | 0.067698429 | 1.24945709 | 0.211497939 | 0.446836226 | 0.084586283 |
| ENSMUSG00000014349 | Ube2z | 268470 | protein_coding | 2336.60057 | 0.084600231 | 0.043700014 | 1.935931466 | 0.052876095 | 0.192230115 | 0.084600231 |
| ENSMUSG00000035390 | Brsk1 | 381979 | protein_coding | 25.92824922 | 0.084695971 | 0.256499666 | 0.330199146 | 0.741249492 | 0.876274048 | 0.084695971 |
| ENSMUSG00000036813 | Entpd8 | 72090 | protein_coding | 2508.245211 | 0.084700627 | 0.066780407 | 1.268345475 | 0.204674604 | 0.43881778 | 0.084700627 |
| ENSMUSG00000029401 | Rilpl2 | 80291 | protein_coding | 194.4707719 | 0.0847055 | 0.091891587 | 0.921798204 | 0.356633844 | 0.599482386 | 0.0847055 |
| ENSMUSG00000021947 | Cryl1 | 68631 | protein_coding | 2637.355342 | -0.084724572 | 0.053372799 | -1.587411086 | 0.11241957 | 0.30863821 | 0.084724572 |
| ENSMUSG00000040728 | Esrp1 | 207920 | protein_coding | 5573.930204 | 0.08482125 | 0.03610151 | 2.349520872 | 0.01879759 | 0.094291987 | 0.08482125 |
| ENSMUSG00000026887 | Mrrf | 67871 | protein_coding | 361.6871646 | 0.084824321 | 0.081941127 | 1.035186168 | 0.300581977 | 0.544111073 | 0.084824321 |
| ENSMUSG00000029128 | Rab28 | 100972 | protein_coding | 586.3797001 | 0.084825041 | 0.055996464 | 1.514828521 | 0.129815836 | 0.335814568 | 0.084825041 |
| ENSMUSG00000031642 | Sh3rf1 | 59009 | protein_coding | 969.9254426 | 0.084833373 | 0.075713863 | 1.120447039 | 0.262523309 | 0.504493972 | 0.084833373 |
| ENSMUSG00000024301 | Kifc5b | 16580 | protein_coding | 93.17728915 | 0.084846283 | 0.172584392 | 0.491621992 | 0.622986593 | 0.800888833 | 0.084846283 |
| ENSMUSG00000057982 | Zfp809 | 235047 | protein_coding | 400.8730588 | -0.084859166 | 0.071984783 | -1.178848668 | 0.238458441 | 0.47878726 | 0.084859166 |
| ENSMUSG00000049233 | Apoo-ps | 621156 | processed_pseudogene | 10.28708345 | 0.084882224 | 0.401453424 | 0.211437289 | 0.832546062 | 0.92583577 | 0.084882224 |
| ENSMUSG00000036912 | Piwil4 | 330890 | protein_coding | 121.7542731 | 0.084897729 | 0.172376742 | 0.492512668 | 0.622356969 | 0.800805946 | 0.084897729 |
| ENSMUSG00000024143 | Rhoq | 104215 | protein_coding | 886.5154544 | -0.08491413 | 0.07900709 | -1.07476595 | 0.282479528 | 0.525210203 | 0.08491413 |
| ENSMUSG00000027671 | Actl6a | 56456 | protein_coding | 738.4814395 | 0.084922321 | 0.079977091 | 1.061833084 | 0.288311468 | 0.530775478 | 0.084922321 |
| ENSMUSG00000072501 | Phf20l1 | 239510 | protein_coding | 1051.401755 | -0.084927614 | 0.049498812 | -1.715750559 | 0.086207707 | 0.262563114 | 0.084927614 |
| ENSMUSG00000026646 | Suv39h2 | 64707 | protein_coding | 63.04405999 | 0.084946741 | 0.160116558 | 0.530530645 | 0.595744067 | 0.78269996 | 0.084946741 |
| ENSMUSG00000037846 | Rtkn2 | 170799 | protein_coding | 10.1210222 | 0.084952092 | 0.383512876 | 0.221510402 | 0.824695038 | 0.92202835 | 0.084952092 |
| ENSMUSG00000024883 | Rin1 | 225870 | protein_coding | 217.9981836 | -0.085001607 | 0.097960147 | -0.867716204 | 0.385549713 | 0.625831733 | 0.085001607 |
| ENSMUSG00000029213 | Commd8 | 27784 | protein_coding | 597.7496108 | 0.085041492 | 0.069150469 | 1.229803534 | 0.218770684 | 0.454982371 | 0.085041492 |
| ENSMUSG00000028809 | Srrm1 | 51796 | protein_coding | 1010.557215 | -0.085079469 | 0.057598995 | -1.477099885 | 0.139648864 | 0.350415857 | 0.085079469 |
| ENSMUSG00000052915 | Msl1 | 74026 | protein_coding | 1305.916581 | -0.085125627 | 0.060768515 | -1.400817952 | 0.161268519 | 0.381891659 | 0.085125627 |
| ENSMUSG00000028104 | Polr3gl | 69870 | protein_coding | 190.8411464 | 0.085126747 | 0.093809552 | 0.90744221 | 0.364172997 | 0.606340896 | 0.085126747 |
| ENSMUSG00000029557 | Mrm2 | 68017 | protein_coding | 164.9703748 | 0.085129971 | 0.112370842 | 0.757580605 | 0.448702095 | 0.678988684 | 0.085129971 |
| ENSMUSG00000003873 | Bax | 12028 | protein_coding | 410.0898433 | 0.085139168 | 0.078873915 | 1.079433771 | 0.280394403 | 0.523431015 | 0.085139168 |
| ENSMUSG00000029578 | Wipi2 | 74781 | protein_coding | 2148.141255 | -0.085143672 | 0.042020746 | -2.026229412 | 0.042741281 | 0.166294002 | 0.085143672 |
| ENSMUSG00000024736 | Tmem132a | 98170 | protein_coding | 95.76712029 | -0.085151135 | 0.147049815 | -0.579063188 | 0.562546538 | 0.76019675 | 0.085151135 |
| ENSMUSG00000041607 | Mbp | 17196 | protein_coding | 1906.97969 | -0.085155373 | 0.044127237 | -1.929768988 | 0.053635468 | 0.193941522 | 0.085155373 |
| ENSMUSG00000043183 | Simc1 | 319719 | protein_coding | 76.09151034 | -0.085198221 | 0.144190429 | -0.590872931 | 0.554605564 | 0.754685552 | 0.085198221 |
| ENSMUSG00000044361 | BC024139 | 271278 | protein_coding | 37.85811589 | -0.085198274 | 0.220117251 | -0.387058597 | 0.698712828 | 0.849656295 | 0.085198274 |
| ENSMUSG00000006010 | Odr4 | 226499 | protein_coding | 876.162754 | 0.085222569 | 0.056575494 | 1.506351308 | 0.131977013 | 0.338431117 | 0.085222569 |
| ENSMUSG00000001700 | Gramd3 | 107022 | protein_coding | 1665.267728 | -0.085248241 | 0.072758626 | -1.1716582 | 0.241334311 | 0.482040926 | 0.085248241 |
| ENSMUSG00000020029 | Nudt4 | 71207 | protein_coding | 9940.939529 | -0.085268457 | 0.063057989 | -1.352222915 | 0.176304015 | 0.402927855 | 0.085268457 |
| ENSMUSG00000033543 | Gtf2a2 | 235459 | protein_coding | 550.2389229 | 0.085292912 | 0.077483171 | 1.100792729 | 0.270986877 | 0.513537646 | 0.085292912 |

| ensembl_gene_id | external_gene_name | entrezgene_id | gene_biotype | baseMean | log2FoldChange | lfcSE | stat | pvalue | padj | absLog2FC |
|---|---|---|---|---|---|---|---|---|---|---|
| ENSMUSG00000023452 | Pisd | 320951 | protein_coding | 1169.239844 | 0.085310455 | 0.059216186 | 1.440661095 | 0.149680451 | 0.364802272 | 0.085310455 |
| ENSMUSG00000019813 | Cep57l1 | 103268 | protein_coding | 71.41485786 | 0.085327904 | 0.139136997 | 0.613265379 | 0.539700881 | 0.745625688 | 0.085327904 |
| ENSMUSG00000060519 | Tor3a | 30935 | protein_coding | 746.0961137 | 0.085348294 | 0.057672443 | 1.479879993 | 0.138905276 | 0.349146725 | 0.085348294 |
| ENSMUSG00000050714 | Zbtb26 | 320633 | protein_coding | 174.2360363 | 0.085359804 | 0.120818235 | 0.706514241 | 0.4798684 | 0.702190153 | 0.085359804 |
| ENSMUSG00000022141 | Nipbl | 71175 | protein_coding | 2089.699998 | -0.085364442 | 0.051663007 | -1.652332052 | 0.098466879 | 0.285546433 | 0.085364442 |
| ENSMUSG00000027193 | Api5 | 11800 | protein_coding | 2605.590811 | 0.085367157 | 0.047678701 | 1.790467326 | 0.073378815 | 0.237154121 | 0.085367157 |
| ENSMUSG00000021918 | Nek4 | 23955 | protein_coding | 382.8543132 | 0.085373233 | 0.065400981 | 1.305381537 | 0.191762962 | 0.423163427 | 0.085373233 |
| ENSMUSG00000001930 | Vwf | 22371 | protein_coding | 80.11171565 | -0.085392131 | 0.158482297 | -0.538811796 | 0.590016723 | 0.779231815 | 0.085392131 |
| ENSMUSG00000024309 | Pfdn6 | 14976 | protein_coding | 606.1815433 | 0.085393164 | 0.083660844 | 1.02070646 | 0.307393533 | 0.551334926 | 0.085393164 |
| ENSMUSG00000038010 | Ccdc138 | 76138 | protein_coding | 95.26672209 | 0.085405686 | 0.122982796 | 0.694452307 | 0.4873986 | 0.708115319 | 0.085405686 |
| ENSMUSG00000021500 | Ddx46 | 212880 | protein_coding | 1619.762394 | -0.085434804 | 0.056637728 | -1.508443357 | 0.131441093 | 0.337593648 | 0.085434804 |
| ENSMUSG00000049940 | Pgrmc2 | 70804 | protein_coding | 780.5000075 | 0.085438517 | 0.07923431 | 1.078302019 | 0.280898996 | 0.523508209 | 0.085438517 |
| ENSMUSG00000045098 | Kmt5b | 225888 | protein_coding | 1648.07626 | -0.085444159 | 0.049167686 | -1.737811256 | 0.082244077 | 0.254626651 | 0.085444159 |
| ENSMUSG00000026260 | Ndufa10 | 67273 | protein_coding | 4568.683964 | 0.085456137 | 0.05459875 | 1.565166551 | 0.117543862 | 0.317074504 | 0.085456137 |
| ENSMUSG00000044702 | Palb2 | 233826 | protein_coding | 83.85141908 | -0.085488435 | 0.159319013 | -0.536586522 | 0.591553256 | 0.780262005 | 0.085488435 |
| ENSMUSG00000009535 | Rnmt | 67897 | protein_coding | 472.4887606 | -0.085529381 | 0.066881346 | -1.27882267 | 0.200959509 | 0.434666577 | 0.085529381 |
| ENSMUSG00000027428 | Rbbp9 | 26450 | protein_coding | 252.4209485 | 0.085543888 | 0.07586455 | 1.127587095 | 0.259494339 | 0.501083584 | 0.085543888 |
| ENSMUSG00000032754 | Slc8b1 | 170756 | protein_coding | 1197.143725 | -0.085553322 | 0.082905382 | -1.031939306 | 0.302100551 | 0.545733091 | 0.085553322 |
| ENSMUSG00000052459 | Atp6v1a | 11964 | protein_coding | 2065.216008 | 0.085569364 | 0.044967051 | 1.902934737 | 0.057049062 | 0.201938151 | 0.085569364 |
| ENSMUSG00000034981 | Parm1 | 231440 | protein_coding | 3198.726119 | -0.085569379 | 0.073742026 | -1.160388239 | 0.245890773 | 0.487025714 | 0.085569379 |
| ENSMUSG00000031959 | Wdr59 | 319481 | protein_coding | 417.7778072 | 0.085582107 | 0.089426495 | 0.957010637 | 0.338561886 | 0.5820991 | 0.085582107 |
| ENSMUSG00000029066 | Mrpl20 | 66448 | protein_coding | 789.5708591 | 0.085649631 | 0.068247058 | 1.254993739 | 0.209481033 | 0.444801234 | 0.085649631 |
| ENSMUSG00000022615 | Tymp | 72962 | protein_coding | 105.7457485 | -0.085678956 | 0.137617263 | -0.622588723 | 0.533554823 | 0.741395712 | 0.085678956 |
| ENSMUSG00000010607 | Pigyl | 66268 | protein_coding | 231.9797028 | 0.085685169 | 0.097177778 | 0.881736245 | 0.377919456 | 0.619192021 | 0.085685169 |
| ENSMUSG00000003346 | Abhd17a | 216169 | protein_coding | 1840.328431 | 0.085695051 | 0.093940334 | 0.912228511 | 0.361648438 | 0.604649053 | 0.085695051 |
| ENSMUSG00000015714 | Cers2 | 76893 | protein_coding | 1505.548803 | -0.085698662 | 0.04895264 | -1.750644332 | 0.080007194 | 0.250247482 | 0.085698662 |
| ENSMUSG00000045730 | Adrb2 | 11555 | protein_coding | 32.96468252 | -0.085701998 | 0.277720973 | -0.308590297 | 0.757633203 | 0.885131169 | 0.085701998 |
| ENSMUSG00000038888 | Ctu1 | 233189 | protein_coding | 331.8518359 | 0.085721425 | 0.088787295 | 0.965469501 | 0.334309705 | 0.577733884 | 0.085721425 |
| ENSMUSG00000089774 | Slc5a3 | 53881 | protein_coding | 1771.110882 | -0.085730863 | 0.092843326 | -0.923392846 | 0.355802519 | 0.598521667 | 0.085730863 |
| ENSMUSG00000038909 | Kat7 | 217127 | protein_coding | 1599.785251 | -0.085793414 | 0.046165727 | -1.858378899 | 0.063115227 | 0.214828885 | 0.085793414 |
| ENSMUSG00000022194 | Pabpn1 | 12050 | protein_coding | 338.2430014 | -0.085803339 | 0.096724148 | -0.887093254 | 0.375028696 | 0.61652794 | 0.085803339 |
| ENSMUSG00000063663 | Brwd3 | 382236 | protein_coding | 644.0308735 | 0.085857851 | 0.05667786 | 1.514839331 | 0.129813097 | 0.335814568 | 0.085857851 |
| ENSMUSG00000026611 | Spata17 | 74717 | protein_coding | 18.62801955 | -0.085861224 | 0.281333795 | -0.305193422 | 0.760218842 | 0.886610448 | 0.085861224 |
| ENSMUSG00000075054 | Yae1d1 | 67008 | protein_coding | 275.47079 | -0.085890167 | 0.078223 | -1.098016778 | 0.272197163 | 0.514726356 | 0.085890167 |
| ENSMUSG00000000184 | Ccnd2 | 12444 | protein_coding | 3339.203519 | -0.085901987 | 0.114218877 | -0.752082229 | 0.45200161 | 0.680967534 | 0.085901987 |
| ENSMUSG00000029064 | Gnb1 | 14688 | protein_coding | 5972.384697 | 0.08590422 | 0.032507494 | 2.64259745 | 0.008227279 | 0.051687309 | 0.08590422 |
| ENSMUSG00000039801 | Cplane1 | 73692 | protein_coding | 143.616064 | 0.085908959 | 0.113261796 | 0.758499003 | 0.44815231 | 0.678921982 | 0.085908959 |
| ENSMUSG00000021974 | Fgf9 | 14180 | protein_coding | 44.65432313 | -0.085930842 | 0.175077878 | -0.49081496 | 0.623557328 | 0.801324379 | 0.085930842 |
| ENSMUSG00000043003 | Rasef | 242505 | protein_coding | 3309.895878 | 0.085935729 | 0.056475208 | 1.521654051 | 0.128095786 | 0.333247419 | 0.085935729 |

| ensembl_gene_id | external_gene_name | entrezgene_id | gene_biotype | baseMean | log2FoldChange | lfcSE | stat | pvalue | padj | absLog2FC |
|---|---|---|---|---|---|---|---|---|---|---|
| ENSMUSG00000026888 | Grb14 | 50915 | protein_coding | 42.30796864 | 0.0859915 | 0.19445092 | 0.442227273 | 0.658324753 | 0.825247049 | 0.0859915 |
| ENSMUSG00000004895 | Prcc | 94315 | protein_coding | 964.2509344 | 0.086003825 | 0.0553801 | 1.552973447 | 0.120429478 | 0.321178468 | 0.086003825 |
| ENSMUSG00000025728 | Pigq | 14755 | protein_coding | 1505.213528 | 0.086035987 | 0.058033129 | 1.482532283 | 0.13819872 | 0.348184778 | 0.086035987 |
| ENSMUSG00000022536 | Glyr1 | 74022 | protein_coding | 3458.389261 | 0.086076185 | 0.031549759 | 2.728267582 | 0.006366794 | 0.042790188 | 0.086076185 |
| ENSMUSG00000040746 | Rnf167 | 70510 | protein_coding | 1397.811677 | -0.086093327 | 0.067046726 | -1.284079512 | 0.199114136 | 0.432432462 | 0.086093327 |
| ENSMUSG00000039637 | Coro7 | 78885 | protein_coding | 467.5431097 | 0.086096341 | 0.07975559 | 1.079502281 | 0.280363877 | 0.523431015 | 0.086096341 |
| ENSMUSG00000078713 | Tomm5 | 68512 | protein_coding | 864.2605923 | 0.086117497 | 0.061001136 | 1.411735942 | 0.158027726 | 0.377781372 | 0.086117497 |
| ENSMUSG00000041506 | Rrp9 | 27966 | protein_coding | 367.0405062 | 0.08614918 | 0.074095807 | 1.162672815 | 0.244962275 | 0.486120986 | 0.08614918 |
| ENSMUSG00000026662 | Sephs1 | 109079 | protein_coding | 744.1512052 | 0.086175605 | 0.066297704 | 1.299827893 | 0.193659963 | 0.425497851 | 0.086175605 |
| ENSMUSG00000090925 | 1810064F22Rik | 69862 | lncRNA | 11.02420066 | -0.086196049 | 0.363047942 | -0.237423325 | 0.812328396 | 0.915791177 | 0.086196049 |
| ENSMUSG00000040025 | Ythdf2 | 213541 | protein_coding | 1601.078051 | -0.086206114 | 0.034768209 | -2.479452233 | 0.013158435 | 0.072948122 | 0.086206114 |
| ENSMUSG00000031358 | Msl3 | 17692 | protein_coding | 1145.270097 | 0.086214849 | 0.04583408 | 1.881020594 | 0.059969118 | 0.207983217 | 0.086214849 |
| ENSMUSG00000042579 | 4632404H12Rik | 74034 | lncRNA | 116.8842074 | -0.086223153 | 0.113683143 | -0.758451525 | 0.448180722 | 0.678921982 | 0.086223153 |
| ENSMUSG00000023026 | Dip2b | 239667 | protein_coding | 1397.269446 | -0.086227911 | 0.054002556 | -1.596737601 | 0.11032421 | 0.304905289 | 0.086227911 |
| ENSMUSG00000027160 | Ccdc34 | 68201 | protein_coding | 360.0240436 | 0.086229607 | 0.091039248 | 0.947169586 | 0.343552371 | 0.586128211 | 0.086229607 |
| ENSMUSG00000016382 | Pls3 | 102866 | protein_coding | 345.3583197 | 0.086263057 | 0.122637836 | 0.703396765 | 0.481808522 | 0.703879521 | 0.086263057 |
| ENSMUSG00000054520 | Sh3bp2 | 24055 | protein_coding | 138.0642733 | -0.08629368 | 0.116750276 | -0.739130416 | 0.459827809 | 0.686760796 | 0.08629368 |
| ENSMUSG00000027475 | Kif3b | 16569 | protein_coding | 1308.036238 | 0.08634493 | 0.064095956 | 1.34711977 | 0.177941662 | 0.405210667 | 0.08634493 |
| ENSMUSG00000026466 | Tor1aip1 | 208263 | protein_coding | 531.3336651 | -0.086345288 | 0.061374811 | -1.406852219 | 0.159471214 | 0.379596103 | 0.086345288 |
| ENSMUSG00000002320 | Tm9sf1 | 74140 | protein_coding | 1865.654562 | 0.086358753 | 0.062080637 | 1.391073901 | 0.164203018 | 0.38635024 | 0.086358753 |
| ENSMUSG00000029563 | Foxp2 | 114142 | protein_coding | 93.17452335 | -0.086369196 | 0.206394373 | -0.418466818 | 0.675605842 | 0.835024575 | 0.086369196 |
| ENSMUSG00000003923 | Tfam | 21780 | protein_coding | 884.254757 | -0.086376478 | 0.058147151 | -1.485480814 | 0.137416502 | 0.347081587 | 0.086376478 |
| ENSMUSG00000048038 | Ccdc187 | 329366 | protein_coding | 8.034804628 | -0.086385478 | 0.431758578 | -0.200078198 | 0.841419424 | 0.929613548 | 0.086385478 |
| ENSMUSG00000032782 | Cntrob | 216846 | protein_coding | 299.6242497 | -0.08639919 | 0.09503635 | -0.909117305 | 0.363288209 | 0.605805909 | 0.08639919 |
| ENSMUSG00000016481 | Cr1l | 12946 | protein_coding | 1861.582562 | -0.086400594 | 0.03559261 | -2.427486896 | 0.015203836 | 0.080948255 | 0.086400594 |
| ENSMUSG00000030510 | Cers3 | 545975 | protein_coding | 9.034044924 | 0.086422272 | 0.431361275 | 0.200347776 | 0.8412086 | 0.929613548 | 0.086422272 |
| ENSMUSG00000043207 | Zmpste24 | 230709 | protein_coding | 1044.735578 | 0.086428171 | 0.051185465 | 1.688529581 | 0.091309616 | 0.272341441 | 0.086428171 |
| ENSMUSG00000041886 | Macc1 | 238455 | protein_coding | 144.2633505 | 0.086458258 | 0.135722201 | 0.637023694 | 0.52410941 | 0.734983692 | 0.086458258 |
| ENSMUSG00000007739 | Cct4 | 12464 | protein_coding | 3612.577628 | -0.086500533 | 0.043382267 | -1.99391455 | 0.046161394 | 0.175287213 | 0.086500533 |
| ENSMUSG00000002222 | Rmnd5a | 68477 | protein_coding | 4570.504383 | 0.086510105 | 0.03630471 | 2.382889331 | 0.017177357 | 0.088419803 | 0.086510105 |
| ENSMUSG00000021007 | Spata7 | 104871 | protein_coding | 96.03447975 | -0.086546231 | 0.140681779 | -0.61519147 | 0.53842828 | 0.744805917 | 0.086546231 |
| ENSMUSG00000034362 | Csta1 | 209294 | protein_coding | 25.93375796 | -0.086566333 | 0.267267757 | -0.323893665 | 0.74601854 | 0.879490473 | 0.086566333 |
| ENSMUSG00000035027 | Map2k2 | 26396 | protein_coding | 2741.976355 | 0.086569153 | 0.095894927 | 0.902750075 | 0.366658554 | 0.609033113 | 0.086569153 |
| ENSMUSG00000021112 | Pals1 | 56217 | protein_coding | 1704.908859 | -0.086591685 | 0.057012078 | -1.518830539 | 0.128805157 | 0.334230211 | 0.086591685 |
| ENSMUSG00000005836 | Gata6 | 14465 | protein_coding | 1321.853667 | -0.086631413 | 0.079002313 | -1.096568056 | 0.272830258 | 0.515284976 | 0.086631413 |
| ENSMUSG00000067931 | Zfp948 | 381066 | protein_coding | 344.4809098 | -0.08664093 | 0.093226179 | -0.929362668 | 0.352701168 | 0.595229357 | 0.08664093 |
| ENSMUSG00000032407 | U2surp | 67958 | protein_coding | 2124.199117 | -0.08665238 | 0.064098274 | -1.351867604 | 0.176417673 | 0.40304403 | 0.08665238 |
| ENSMUSG00000050106 | Tmc8 | 217356 | protein_coding | 214.460994 | 0.086658887 | 0.123449479 | 0.701978559 | 0.482692535 | 0.70472411 | 0.086658887 |
| ENSMUSG00000002814 | Top3a | 21975 | protein_coding | 236.1575646 | -0.086661361 | 0.077075066 | -1.124376082 | 0.260853515 | 0.502782913 | 0.086661361 |

| ensembl_gene_id | external_gene_name | entrezgene_id | gene_biotype | baseMean | log2FoldChange | lfcSE | stat | pvalue | padj | absLog2FC |
|---|---|---|---|---|---|---|---|---|---|---|
| ENSMUSG00000024816 | Frmd8 | 67457 | protein_coding | 1031.491957 | 0.086714467 | 0.090889055 | 0.954069404 | 0.340048512 | 0.583162735 | 0.086714467 |
| ENSMUSG00000052428 | Tmco1 | 68944 | protein_coding | 1052.246896 | 0.086716492 | 0.044496353 | 1.948844937 | 0.051313945 | 0.188419854 | 0.086716492 |
| ENSMUSG00000030982 | Vps35l | 71517 | protein_coding | 1209.225843 | 0.086721151 | 0.046894855 | 1.8492679 | 0.064419138 | 0.218015518 | 0.086721151 |
| ENSMUSG00000015656 | Hspa8 | 15481 | protein_coding | 24157.29732 | 0.086731449 | 0.072265929 | 1.200170675 | 0.230073062 | 0.46920123 | 0.086731449 |
| ENSMUSG00000028140 | Mrpl9 | 78523 | protein_coding | 1551.031363 | -0.086731484 | 0.068496928 | -1.266209827 | 0.20543797 | 0.439869642 | 0.086731484 |
| ENSMUSG00000020894 | Vamp2 | 22318 | protein_coding | 311.9836693 | -0.086750076 | 0.110397527 | -0.785797273 | 0.431986264 | 0.666473021 | 0.086750076 |
| ENSMUSG00000036402 | Gng12 | 14701 | protein_coding | 5368.329225 | -0.086774087 | 0.040489582 | -2.143121347 | 0.032103356 | 0.136424458 | 0.086774087 |
| ENSMUSG00000043190 | Rfesd | 218341 | protein_coding | 195.8086443 | 0.086777457 | 0.100061762 | 0.86723895 | 0.385811099 | 0.625831733 | 0.086777457 |
| ENSMUSG00000057440 | Mpp7 | 75739 | protein_coding | 482.7189728 | 0.08680784 | 0.082677662 | 1.049955176 | 0.293738722 | 0.536541439 | 0.08680784 |
| ENSMUSG00000020329 | Polrmt | 216151 | protein_coding | 663.0297393 | -0.086815349 | 0.080744358 | -1.075187806 | 0.282290654 | 0.525086663 | 0.086815349 |
| ENSMUSG00000029500 | Pgam5 | 72542 | protein_coding | 677.7011759 | 0.08685203 | 0.07312292 | 1.187753855 | 0.234930399 | 0.474717773 | 0.08685203 |
| ENSMUSG00000029186 | Pi4k2b | 67073 | protein_coding | 3591.161315 | 0.08685914 | 0.034565656 | 2.512874063 | 0.011975208 | 0.067977533 | 0.08685914 |
| ENSMUSG00000022802 | Lmln | 239833 | protein_coding | 148.2620908 | -0.086870358 | 0.126775473 | -0.685230004 | 0.493198794 | 0.712517495 | 0.086870358 |
| ENSMUSG00000090626 | Tex9 | 21778 | protein_coding | 159.3368482 | -0.086900581 | 0.160659622 | -0.540898705 | 0.588577404 | 0.778379525 | 0.086900581 |
| ENSMUSG00000024254 | Abcg8 | 67470 | protein_coding | 48.34614226 | 0.08692652 | 0.298512827 | 0.291198608 | 0.770899427 | 0.892291327 | 0.08692652 |
| ENSMUSG00000022177 | Haus4 | 219072 | protein_coding | 323.747546 | 0.086972221 | 0.095947582 | 0.906455576 | 0.364694768 | 0.606833732 | 0.086972221 |
| ENSMUSG00000027935 | Rab13 | 68328 | protein_coding | 212.7746171 | 0.086992442 | 0.091425365 | 0.951513203 | 0.341343919 | 0.584131301 | 0.086992442 |
| ENSMUSG00000033161 | Atp1a1 | 11928 | protein_coding | 72551.99321 | -0.0869929 | 0.056875326 | -1.529536738 | 0.126131439 | 0.32978627 | 0.0869929 |
| ENSMUSG00000021180 | Rps6ka5 | 73086 | protein_coding | 90.22176257 | 0.087004378 | 0.122334813 | 0.711198847 | 0.476961023 | 0.699728324 | 0.087004378 |
| ENSMUSG00000024135 | Srbd1 | 78586 | protein_coding | 405.5671744 | 0.087007969 | 0.09522316 | 0.91372696 | 0.360860334 | 0.604050305 | 0.087007969 |
| ENSMUSG00000015542 | Nat9 | 66176 | protein_coding | 442.6850923 | 0.087026755 | 0.057881893 | 1.503522976 | 0.132704237 | 0.339503983 | 0.087026755 |
| ENSMUSG00000038976 | Ppp1r9b | 217124 | protein_coding | 1306.629486 | -0.087077841 | 0.073538688 | -1.184109255 | 0.236369817 | 0.476551574 | 0.087077841 |
| ENSMUSG00000020070 | Rufy2 | 70432 | protein_coding | 174.4861261 | -0.08708217 | 0.133592722 | -0.651848158 | 0.514499131 | 0.727325341 | 0.08708217 |
| ENSMUSG00000042660 | Wdr55 | 67936 | protein_coding | 396.0827238 | -0.087084017 | 0.073263521 | -1.188640884 | 0.234581012 | 0.474249466 | 0.087084017 |
| ENSMUSG00000070737 | Tmem35b | 100039968 | protein_coding | 251.2668853 | -0.087131348 | 0.111740932 | -0.779762138 | 0.435530896 | 0.668933977 | 0.087131348 |
| ENSMUSG00000020248 | Nfyb | 18045 | protein_coding | 524.6423744 | 0.087134665 | 0.064682536 | 1.347112691 | 0.177943941 | 0.405210667 | 0.087134665 |
| ENSMUSG00000029468 | P2rx7 | 18439 | protein_coding | 88.61571253 | -0.087134982 | 0.162760823 | -0.535355989 | 0.592403717 | 0.780262005 | 0.087134982 |
| ENSMUSG00000047409 | Ctdspl | 69274 | protein_coding | 2433.107579 | 0.087139275 | 0.046480988 | 1.874729394 | 0.060829961 | 0.209930404 | 0.087139275 |
| ENSMUSG00000030403 | Vasp | 22323 | protein_coding | 9133.837631 | 0.087160716 | 0.086216871 | 1.010947338 | 0.312041635 | 0.555731141 | 0.087160716 |
| ENSMUSG00000091896 | Ube2d2a | 56550 | protein_coding | 833.289169 | 0.087168824 | 0.051771869 | 1.683710204 | 0.092237688 | 0.273701805 | 0.087168824 |
| ENSMUSG00000089844 | A530032D15Rik | 381287 | protein_coding | 7.247121267 | 0.087194178 | 0.48088889 | 0.181318762 | 0.85611738 | 0.935604164 | 0.087194178 |
| ENSMUSG00000030499 | Kctd15 | 233107 | protein_coding | 31.57305888 | 0.087213379 | 0.224720842 | 0.388096531 | 0.697944599 | 0.849362552 | 0.087213379 |
| ENSMUSG00000023019 | Gpd1 | 14555 | protein_coding | 592.747653 | -0.087227761 | 0.055985577 | -1.558039873 | 0.119223796 | 0.319397925 | 0.087227761 |
| ENSMUSG00000018999 | Slc35b4 | 58246 | protein_coding | 411.6615208 | 0.087252451 | 0.075146102 | 1.161104144 | 0.24559955 | 0.486881861 | 0.087252451 |
| ENSMUSG00000025531 | Chm | 12662 | protein_coding | 571.053248 | 0.087255868 | 0.073805774 | 1.18223634 | 0.237111937 | 0.477153008 | 0.087255868 |
| ENSMUSG00000026029 | Casp8 | 12370 | protein_coding | 1818.028213 | 0.087262213 | 0.05998066 | 1.454839146 | 0.145713798 | 0.358494177 | 0.087262213 |
| ENSMUSG00000026014 | Raph1 | 77300 | protein_coding | 939.1249145 | -0.087269472 | 0.096460035 | -0.904721546 | 0.365612924 | 0.607609194 | 0.087269472 |
| ENSMUSG00000002129 | Sf3a1 | 67465 | protein_coding | 1407.122878 | 0.087274527 | 0.061416225 | 1.421033732 | 0.155306953 | 0.373324674 | 0.087274527 |
| ENSMUSG00000034793 | G6pc3 | 68401 | protein_coding | 737.6368292 | -0.087286785 | 0.069427239 | -1.257241194 | 0.208666311 | 0.444245755 | 0.087286785 |

| ensembl_gene_id | external_gene_name | entrezgene_id | gene_biotype | baseMean | log2FoldChange | lfcSE | stat | pvalue | padj | absLog2FC |
|---|---|---|---|---|---|---|---|---|---|---|
| ENSMUSG00000059834 | Sclt1 | 67161 | protein_coding | 261.9522652 | 0.087303859 | 0.089729471 | 0.972967492 | 0.330569459 | 0.57417401 | 0.087303859 |
| ENSMUSG00000024667 | Tmem216 | 68642 | protein_coding | 94.60318143 | 0.087315621 | 0.141139463 | 0.618647817 | 0.536148394 | 0.742878353 | 0.087315621 |
| ENSMUSG00000037148 | Arhgap10 | 78514 | protein_coding | 50.60846996 | 0.087323682 | 0.165871813 | 0.526452808 | 0.598573633 | 0.784685171 | 0.087323682 |
| ENSMUSG00000032403 | 2300009A05Rik | 69478 | protein_coding | 96.58946551 | 0.087350495 | 0.146618495 | 0.595767236 | 0.551330734 | 0.752162317 | 0.087350495 |
| ENSMUSG00000059734 | Ndufs8 | 225887 | protein_coding | 2573.962104 | 0.087350797 | 0.068472085 | 1.275714005 | 0.202056636 | 0.435870434 | 0.087350797 |
| ENSMUSG00000066090 | Insl5 | 23919 | protein_coding | 798.7445656 | -0.087353316 | 0.147232196 | -0.593303087 | 0.552978333 | 0.753401915 | 0.087353316 |
| ENSMUSG00000088054 | Mir1968 | 100316713 | miRNA | 9.577740046 | 0.087361401 | 0.422567905 | 0.20673932 | 0.836213447 | 0.927356228 | 0.087361401 |
| ENSMUSG00000032657 | Entrep3 | 68521 | protein_coding | 39.63784697 | -0.0873982 | 0.196828455 | -0.444032344 | 0.6570192 | 0.824251158 | 0.0873982 |
| ENSMUSG00000030264 | Thumpd3 | 14911 | protein_coding | 1093.257982 | -0.087434703 | 0.07399182 | -1.181680663 | 0.237332434 | 0.477271565 | 0.087434703 |
| ENSMUSG00000057895 | Zfp105 | 22646 | protein_coding | 19.84327054 | -0.087443232 | 0.294040037 | -0.297385462 | 0.766172242 | 0.890206033 | 0.087443232 |
| ENSMUSG00000004530 | Coro1c | 23790 | protein_coding | 3260.48773 | 0.087460592 | 0.050640903 | 1.727074108 | 0.084154355 | 0.258604665 | 0.087460592 |
| ENSMUSG00000044018 | Mrpl50 | 28028 | protein_coding | 793.7386171 | 0.087506202 | 0.057751297 | 1.515224896 | 0.12971546 | 0.335781899 | 0.087506202 |
| ENSMUSG00000042380 | Smim12 | 80284 | protein_coding | 389.7860794 | 0.087520556 | 0.096548918 | 0.90648925 | 0.364676952 | 0.606833732 | 0.087520556 |
| ENSMUSG00000027655 | Dhx35 | 71715 | protein_coding | 531.9312635 | -0.08752314 | 0.057964287 | -1.509949401 | 0.131056336 | 0.336981111 | 0.08752314 |
| ENSMUSG00000097523 | 1700022N22Rik | 69431 | lncRNA | 7.884840221 | 0.087527891 | 0.419175588 | 0.208809611 | 0.834596869 | 0.926646726 | 0.087527891 |
| ENSMUSG00000004567 | Mcoln1 | 94178 | protein_coding | 675.9391852 | -0.08754103 | 0.071952448 | -1.216651164 | 0.223736967 | 0.461466437 | 0.08754103 |
| ENSMUSG00000030067 | Foxp1 | 108655 | protein_coding | 2487.98024 | -0.087549243 | 0.060334381 | -1.451067231 | 0.14676114 | 0.359972154 | 0.087549243 |
| ENSMUSG00000034064 | Poglut1 | 224143 | protein_coding | 336.8003271 | 0.087559231 | 0.070651783 | 1.239306738 | 0.215231924 | 0.450726561 | 0.087559231 |
| ENSMUSG00000053898 | Ech1 | 51798 | protein_coding | 5228.573245 | 0.087564943 | 0.038388928 | 2.28099474 | 0.022548758 | 0.106976995 | 0.087564943 |
| ENSMUSG00000034317 | Trim59 | 66949 | protein_coding | 190.4164998 | 0.087572995 | 0.106756926 | 0.820302698 | 0.412043569 | 0.649970343 | 0.087572995 |
| ENSMUSG00000003868 | Ruvbl2 | 20174 | protein_coding | 1024.273663 | 0.087600223 | 0.088961158 | 0.98470191 | 0.324770514 | 0.568500043 | 0.087600223 |
| ENSMUSG00000060621 | Nkpd1 | 69547 | protein_coding | 10.25720303 | -0.087605073 | 0.402456873 | -0.217675678 | 0.827681816 | 0.923349179 | 0.087605073 |
| ENSMUSG00000074217 | Misp3 | 70134 | protein_coding | 97.72572846 | 0.087613199 | 0.138999875 | 0.630311349 | 0.528490899 | 0.737950908 | 0.087613199 |
| ENSMUSG00000042210 | Abhd14a | 68644 | protein_coding | 239.9684106 | 0.08761883 | 0.084310597 | 1.039238637 | 0.298693765 | 0.542214583 | 0.08761883 |
| ENSMUSG00000024926 | Kat5 | 81601 | protein_coding | 617.7374186 | -0.087631631 | 0.055251102 | -1.586061244 | 0.112725417 | 0.309011907 | 0.087631631 |
| ENSMUSG00000042323 | Pbrm1 | 66923 | protein_coding | 2454.189234 | -0.087636948 | 0.047232141 | -1.855451516 | 0.063531777 | 0.215882209 | 0.087636948 |
| ENSMUSG00000041688 | Amot | 27494 | protein_coding | 961.5370493 | 0.087660751 | 0.050832578 | 1.724499424 | 0.084617723 | 0.259435364 | 0.087660751 |
| ENSMUSG00000038570 | Saxo2 | 330577 | protein_coding | 11.78900272 | 0.087667888 | 0.355230598 | 0.246791487 | 0.805069599 | 0.911459024 | 0.087667888 |
| ENSMUSG00000021910 | Nisch | 64652 | protein_coding | 2608.209435 | -0.087670627 | 0.054888825 | -1.597240008 | 0.110212218 | 0.304713258 | 0.087670627 |
| ENSMUSG00000061666 | Gdpd1 | 66569 | protein_coding | 2347.359049 | -0.087672697 | 0.050506501 | -1.735869547 | 0.082586903 | 0.255198121 | 0.087672697 |
| ENSMUSG00000019471 | Cdc37 | 12539 | protein_coding | 2288.168005 | 0.087709231 | 0.05958788 | 1.471930716 | 0.141039588 | 0.352096817 | 0.087709231 |
| ENSMUSG00000031601 | Cnot7 | 18983 | protein_coding | 846.8601408 | -0.087726446 | 0.067910472 | -1.291795553 | 0.196427956 | 0.428914823 | 0.087726446 |
| ENSMUSG00000048930 | Tada3 | 101206 | protein_coding | 559.4347541 | 0.087726565 | 0.066998777 | 1.309375615 | 0.190407148 | 0.421109924 | 0.087726565 |
| ENSMUSG00000032127 | Vps11 | 71732 | protein_coding | 823.9256203 | 0.087740087 | 0.06081482 | 1.442741858 | 0.149093203 | 0.363700815 | 0.087740087 |
| ENSMUSG00000046020 | Pofut1 | 140484 | protein_coding | 1361.084323 | 0.087751856 | 0.054217766 | 1.618507404 | 0.105553293 | 0.297761537 | 0.087751856 |
| ENSMUSG00000041718 | Alg13 | 67574 | protein_coding | 84.69087915 | 0.087761888 | 0.146202689 | 0.60027547 | 0.548322664 | 0.750864769 | 0.087761888 |
| ENSMUSG00000031217 | Efnb1 | 13641 | protein_coding | 3974.796207 | 0.087763651 | 0.095920784 | 0.914959691 | 0.360212792 | 0.603422841 | 0.087763651 |
| ENSMUSG00000020781 | Tsen54 | 76265 | protein_coding | 451.2838206 | -0.087771978 | 0.076355161 | -1.149522529 | 0.250340582 | 0.491447666 | 0.087771978 |
| ENSMUSG00000047514 | Tspyl1 | 22110 | protein_coding | 907.405768 | 0.087780366 | 0.049022886 | 1.790599717 | 0.073357552 | 0.237154121 | 0.087780366 |

| ensembl_gene_id | external_gene_name | entrezgene_id | gene_biotype | baseMean | log2FoldChange | lfcSE | stat | pvalue | padj | absLog2FC |
|---|---|---|---|---|---|---|---|---|---|---|
| ENSMUSG00000034394 | Lif | 16878 | protein_coding | 79.90048885 | -0.087796263 | 0.152998892 | -0.573835937 | 0.566078828 | 0.7622237 | 0.087796263 |
| ENSMUSG00000057286 | St6galnac2 | 20446 | protein_coding | 1152.355534 | -0.087820164 | 0.059200853 | -1.483427351 | 0.137960905 | 0.347997676 | 0.087820164 |
| ENSMUSG00000038497 | Tmco3 | 234076 | protein_coding | 1896.139938 | 0.087821138 | 0.06592454 | 1.332146382 | 0.182812092 | 0.410843172 | 0.087821138 |
| ENSMUSG00000059820 | Nkapd1 | 270156 | protein_coding | 389.4297711 | -0.087876196 | 0.074158319 | -1.184980952 | 0.236024978 | 0.47621354 | 0.087876196 |
| ENSMUSG00000045667 | Smtnl2 | 276829 | protein_coding | 21.57839107 | -0.087886909 | 0.258863214 | -0.339511 | 0.734224812 | 0.872125178 | 0.087886909 |
| ENSMUSG00000021177 | Tdp1 | 104884 | protein_coding | 92.04461934 | -0.087902621 | 0.129660484 | -0.677944571 | 0.497806839 | 0.715668065 | 0.087902621 |
| ENSMUSG00000030682 | Cdipt | 52858 | protein_coding | 1695.146627 | 0.0880111 | 0.058167626 | 1.513059872 | 0.130264451 | 0.336231563 | 0.0880111 |
| ENSMUSG00000031935 | Med17 | 234959 | protein_coding | 673.9460665 | -0.088016168 | 0.052225878 | -1.685298015 | 0.091931089 | 0.273384904 | 0.088016168 |
| ENSMUSG00000017132 | Cyth1 | 19157 | protein_coding | 1168.440636 | 0.08803147 | 0.060476917 | 1.455620997 | 0.14549742 | 0.358198217 | 0.08803147 |
| ENSMUSG00000075592 | Nynrin | 277154 | protein_coding | 60.54756738 | -0.088041063 | 0.168414999 | -0.522762604 | 0.601139466 | 0.786473529 | 0.088041063 |
| ENSMUSG00000021996 | Esd | 13885 | protein_coding | 3733.463229 | 0.088046486 | 0.042907315 | 2.052015738 | 0.040168133 | 0.159045294 | 0.088046486 |
| ENSMUSG00000038756 | Ttll6 | 237930 | protein_coding | 656.7588645 | -0.08806575 | 0.089700249 | -0.981778218 | 0.326209121 | 0.569717836 | 0.08806575 |
| ENSMUSG00000005161 | Prdx2 | 21672 | protein_coding | 3350.539143 | 0.088066523 | 0.050449602 | 1.74563367 | 0.080874628 | 0.252031907 | 0.088066523 |
| ENSMUSG00000042541 | Sem1 | 20422 | protein_coding | 2039.422931 | 0.088078388 | 0.085002222 | 1.036189237 | 0.300113867 | 0.543683373 | 0.088078388 |
| ENSMUSG00000047407 | Tgif1 | 21815 | protein_coding | 1017.802198 | -0.088104525 | 0.0881659 | -0.999303869 | 0.317647512 | 0.561449273 | 0.088104525 |
| ENSMUSG00000022540 | Rogdi | 66049 | protein_coding | 169.3463686 | 0.088114338 | 0.121082392 | 0.727722148 | 0.46678369 | 0.692404394 | 0.088114338 |
| ENSMUSG00000020152 | Actr2 | 66713 | protein_coding | 4780.013989 | 0.08811549 | 0.040927938 | 2.152942309 | 0.031323213 | 0.133956365 | 0.08811549 |
| ENSMUSG00000024493 | Lars | 107045 | protein_coding | 1283.447177 | -0.088141763 | 0.07352073 | -1.198869535 | 0.230578679 | 0.469603637 | 0.088141763 |
| ENSMUSG00000002803 | Btbd6 | 399566 | protein_coding | 585.2193426 | -0.088149745 | 0.057097755 | -1.543839064 | 0.122627325 | 0.32462424 | 0.088149745 |
| ENSMUSG00000078651 | Aoc2 | 237940 | protein_coding | 24.21737195 | -0.088158142 | 0.283017759 | -0.311493323 | 0.755425618 | 0.883960887 | 0.088158142 |
| ENSMUSG00000024507 | Hsd17b4 | 15488 | protein_coding | 2208.532314 | 0.088214679 | 0.036976382 | 2.385703377 | 0.017046494 | 0.088027156 | 0.088214679 |
| ENSMUSG00000025903 | Lypla1 | 18777 | protein_coding | 2098.679775 | 0.08821677 | 0.055984705 | 1.575729822 | 0.115088082 | 0.313094719 | 0.08821677 |
| ENSMUSG00000017485 | Top2b | 21974 | protein_coding | 3365.140103 | -0.088219329 | 0.042049243 | -2.098000413 | 0.035905109 | 0.146964896 | 0.088219329 |
| ENSMUSG00000062210 | Tnfaip8 | 106869 | protein_coding | 827.5201403 | 0.088220478 | 0.070219656 | 1.256350193 | 0.208989031 | 0.44452204 | 0.088220478 |
| ENSMUSG00000028755 | Cda | 72269 | protein_coding | 383.2199939 | -0.088266065 | 0.109688357 | -0.804698577 | 0.420993641 | 0.657926757 | 0.088266065 |
| ENSMUSG00000019302 | Atp6v0a1 | 11975 | protein_coding | 1211.272788 | -0.088272778 | 0.07162201 | -1.23248116 | 0.2177694 | 0.453918459 | 0.088272778 |
| ENSMUSG00000068580 | Zfyve19 | 72008 | protein_coding | 592.5819675 | 0.0883055 | 0.070038897 | 1.260806549 | 0.207378554 | 0.442731525 | 0.0883055 |
| ENSMUSG00000000441 | Raf1 | 110157 | protein_coding | 1392.496532 | 0.088313426 | 0.040725118 | 2.168524733 | 0.030118785 | 0.13057141 | 0.088313426 |
| ENSMUSG00000078676 | Casc3 | 192160 | protein_coding | 949.9811816 | -0.088315628 | 0.043842589 | -2.014379844 | 0.043969669 | 0.169803364 | 0.088315628 |
| ENSMUSG00000100084 | 2310068J16Rik | 70281 | lncRNA | 16.18720694 | -0.088345007 | 0.311752975 | -0.28338144 | 0.776884454 | 0.896003404 | 0.088345007 |
| ENSMUSG00000042364 | Snx18 | 170625 | protein_coding | 440.523073 | -0.088367593 | 0.059720085 | -1.479696383 | 0.138954291 | 0.349163297 | 0.088367593 |
| ENSMUSG00000036955 | Kifbp | 72320 | protein_coding | 439.7618242 | 0.08837406 | 0.076319973 | 1.157941451 | 0.246887931 | 0.487862729 | 0.08837406 |
| ENSMUSG00000030243 | Recql | 19691 | protein_coding | 268.8810432 | 0.088386552 | 0.092981776 | 0.950579303 | 0.341817979 | 0.584451398 | 0.088386552 |
| ENSMUSG00000004846 | Plod3 | 26433 | protein_coding | 519.2679531 | 0.08839554 | 0.070537577 | 1.253169505 | 0.210144025 | 0.445254566 | 0.08839554 |
| ENSMUSG00000079685 | Ulbp1 | 77777 | protein_coding | 75.94272772 | 0.088399295 | 0.284740565 | 0.310455572 | 0.75621454 | 0.884312567 | 0.088399295 |
| ENSMUSG00000021263 | Degs2 | 70059 | protein_coding | 4851.732133 | 0.088419681 | 0.060659681 | 1.457635107 | 0.144941148 | 0.357574711 | 0.088419681 |
| ENSMUSG00000064090 | Vrk2 | 69922 | protein_coding | 155.0092302 | 0.088436137 | 0.106611731 | 0.829516003 | 0.406812485 | 0.645607651 | 0.088436137 |
| ENSMUSG00000024563 | Smad2 | 17126 | protein_coding | 1214.787824 | -0.088464535 | 0.042049468 | -2.103820535 | 0.035394105 | 0.145575294 | 0.088464535 |
| ENSMUSG00000037075 | Rnf139 | 75841 | protein_coding | 627.6497009 | -0.088467997 | 0.058422833 | -1.514270917 | 0.129957142 | 0.335868401 | 0.088467997 |

| ensembl_gene_id | external_gene_name | entrezgene_id | gene_biotype | baseMean | log2FoldChange | lfcSE | stat | pvalue | padj | absLog2FC |
|---|---|---|---|---|---|---|---|---|---|---|
| ENSMUSG00000002996 | Hbp1 | 73389 | protein_coding | 1072.332669 | -0.08847068 | 0.078944278 | -1.120672489 | 0.262427297 | 0.504369564 | 0.08847068 |
| ENSMUSG00000028618 | Tmem59 | 56374 | protein_coding | 8277.366202 | -0.088482321 | 0.033903891 | -2.609798372 | 0.00905956 | 0.05546831 | 0.088482321 |
| ENSMUSG00000018820 | Zfyve27 | 319740 | protein_coding | 779.2007931 | -0.088482829 | 0.063355529 | -1.396607841 | 0.162531531 | 0.383544922 | 0.088482829 |
| ENSMUSG00000048720 | Tbc1d12 | 209478 | protein_coding | 275.1592702 | -0.088486779 | 0.086604764 | -1.021731078 | 0.306908198 | 0.550870027 | 0.088486779 |
| ENSMUSG00000032267 | Usp28 | 235323 | protein_coding | 475.6855916 | -0.088517136 | 0.064251161 | -1.377673716 | 0.16830405 | 0.392283294 | 0.088517136 |
| ENSMUSG00000086382 | Chrna1os | 102633437 | lncRNA | 14.41782977 | 0.088533722 | 0.335522911 | 0.263867888 | 0.791881721 | 0.903021982 | 0.088533722 |
| ENSMUSG00000062519 | Zfp398 | 272347 | protein_coding | 430.1169694 | -0.0885506 | 0.071800815 | -1.233281266 | 0.217470845 | 0.453693043 | 0.0885506 |
| ENSMUSG00000040302 | Rbm48 | 269623 | protein_coding | 114.3613694 | -0.088619337 | 0.130977488 | -0.676599763 | 0.498659929 | 0.716014372 | 0.088619337 |
| ENSMUSG00000030805 | Stx4a | 20909 | protein_coding | 526.258828 | -0.088648523 | 0.05687039 | -1.558781714 | 0.119048054 | 0.319180794 | 0.088648523 |
| ENSMUSG00000053754 | Chd8 | 67772 | protein_coding | 1320.455177 | -0.088670121 | 0.053394306 | -1.660666216 | 0.096780502 | 0.282442845 | 0.088670121 |
| ENSMUSG00000087060 | Eldr | 72735 | lncRNA | 70.32622968 | -0.088676171 | 0.157346021 | -0.563574284 | 0.573043892 | 0.767564688 | 0.088676171 |
| ENSMUSG00000022587 | Ly6e | 17069 | protein_coding | 2620.702974 | -0.088685042 | 0.12216075 | -0.725970021 | 0.467857149 | 0.693040529 | 0.088685042 |
| ENSMUSG00000032120 | C2cd2l | 71764 | protein_coding | 1254.477343 | 0.088694175 | 0.085565265 | 1.036567514 | 0.29993746 | 0.543615578 | 0.088694175 |
| ENSMUSG00000042246 | Tmc7 | 209760 | protein_coding | 147.9858628 | -0.088721409 | 0.119490509 | -0.742497541 | 0.457785941 | 0.685488753 | 0.088721409 |
| ENSMUSG00000020817 | Rabep1 | 54189 | protein_coding | 872.5426563 | -0.088730479 | 0.059764901 | -1.484658671 | 0.137634265 | 0.347468345 | 0.088730479 |
| ENSMUSG00000020680 | Taf15 | 70439 | protein_coding | 510.7125085 | 0.088730997 | 0.055391225 | 1.601896273 | 0.109178549 | 0.302685656 | 0.088730997 |
| ENSMUSG00000028995 | Hycc1 | 84652 | protein_coding | 549.5514313 | -0.088735923 | 0.114647466 | -0.7739894 | 0.438937059 | 0.671607017 | 0.088735923 |
| ENSMUSG00000063873 | Slc24a3 | 94249 | protein_coding | 135.1775093 | 0.088761033 | 0.179769436 | 0.4937493 | 0.621483243 | 0.800250831 | 0.088761033 |
| ENSMUSG00000033159 | Cnppd1 | 69171 | protein_coding | 1685.610652 | -0.088762068 | 0.047616613 | -1.864098728 | 0.062307841 | 0.213115282 | 0.088762068 |
| ENSMUSG00000023055 | Calcoco1 | 67488 | protein_coding | 426.3746051 | -0.088763544 | 0.077443593 | -1.146170277 | 0.251724704 | 0.492727559 | 0.088763544 |
| ENSMUSG00000087370 | Tmem170b | 621976 | protein_coding | 551.7790662 | -0.088771832 | 0.0733922 | -1.209554044 | 0.226450062 | 0.464862544 | 0.088771832 |
| ENSMUSG00000025934 | Gsta3 | 14859 | protein_coding | 153.3847995 | -0.088777704 | 0.230053239 | -0.385900691 | 0.699570219 | 0.850145091 | 0.088777704 |
| ENSMUSG00000024097 | Srsf7 | 225027 | protein_coding | 1699.329153 | 0.088802645 | 0.076196671 | 1.165439962 | 0.243840949 | 0.485005302 | 0.088802645 |
| ENSMUSG00000031119 | Gpc4 | 14735 | protein_coding | 526.2281066 | 0.088817162 | 0.119723182 | 0.741854342 | 0.458175592 | 0.685606255 | 0.088817162 |
| ENSMUSG00000042854 | Trp53rkb | 76367 | protein_coding | 60.78552996 | -0.088840622 | 0.152318531 | -0.583255507 | 0.55972131 | 0.758094025 | 0.088840622 |
| ENSMUSG00000027618 | Nfs1 | 18041 | protein_coding | 942.6215954 | -0.088873092 | 0.052737035 | -1.685212148 | 0.091947648 | 0.273384904 | 0.088873092 |
| ENSMUSG00000037275 | Gemin5 | 216766 | protein_coding | 579.0609938 | -0.088887919 | 0.063097303 | -1.40874355 | 0.158911012 | 0.379061279 | 0.088887919 |
| ENSMUSG00000014769 | Psmb1 | 19170 | protein_coding | 3123.870888 | 0.088926982 | 0.03923769 | 2.266366378 | 0.02342896 | 0.109582031 | 0.088926982 |
| ENSMUSG00000025228 | Actr1a | 54130 | protein_coding | 3814.814326 | 0.088969965 | 0.049071488 | 1.813068431 | 0.069821267 | 0.229476957 | 0.088969965 |
| ENSMUSG00000030079 | Ruvbl1 | 56505 | protein_coding | 293.2476004 | 0.088977078 | 0.082517879 | 1.078276357 | 0.280910445 | 0.523508209 | 0.088977078 |
| ENSMUSG00000025898 | Cwf19l2 | 244672 | protein_coding | 519.5896211 | -0.088984105 | 0.080391618 | -1.106882879 | 0.268344569 | 0.510774618 | 0.088984105 |
| ENSMUSG00000034424 | Gcsh | 68133 | protein_coding | 983.4866492 | 0.088989524 | 0.057182891 | 1.556226395 | 0.119654267 | 0.320010674 | 0.088989524 |
| ENSMUSG00000069114 | Zbtb10 | 229055 | protein_coding | 865.6129452 | -0.089011334 | 0.056182998 | -1.584310866 | 0.113122994 | 0.309576536 | 0.089011334 |
| ENSMUSG00000020495 | Smg8 | 74133 | protein_coding | 364.1008329 | 0.089014734 | 0.075099798 | 1.185285935 | 0.235904412 | 0.476068282 | 0.089014734 |
| ENSMUSG00000047084 | Ngrn | 83485 | protein_coding | 465.9302104 | 0.089015822 | 0.070113954 | 1.269587811 | 0.204231493 | 0.438338886 | 0.089015822 |
| ENSMUSG00000031146 | Plp2 | 18824 | protein_coding | 1907.620691 | -0.089028295 | 0.082716263 | -1.076309441 | 0.281788888 | 0.524516319 | 0.089028295 |
| ENSMUSG00000039952 | Dag1 | 13138 | protein_coding | 4301.698289 | -0.089059446 | 0.058897806 | -1.512101253 | 0.130508106 | 0.336485813 | 0.089059446 |
| ENSMUSG00000054611 | Kdm2a | 225876 | protein_coding | 2362.837737 | -0.089078511 | 0.040348 | -2.207755317 | 0.027261334 | 0.121280976 | 0.089078511 |
| ENSMUSG00000061411 | Nol4l | 329540 | protein_coding | 1148.141344 | 0.089094614 | 0.05505658 | 1.618237341 | 0.105611459 | 0.297761537 | 0.089094614 |

| ensembl_gene_id | external_gene_name | entrezgene_id | gene_biotype | baseMean | log2FoldChange | lfcSE | stat | pvalue | padj | absLog2FC |
|---|---|---|---|---|---|---|---|---|---|---|
| ENSMUSG00000037573 | Tob1 | 22057 | protein_coding | 4278.822106 | 0.08910464 | 0.057852198 | 1.540211826 | 0.123508728 | 0.326177557 | 0.08910464 |
| ENSMUSG00000037010 | Apln | 30878 | protein_coding | 8.601614382 | 0.089106737 | 0.421131593 | 0.21158882 | 0.832427832 | 0.92580236 | 0.089106737 |
| ENSMUSG00000063849 | Ppcdc | 66812 | protein_coding | 329.9747324 | -0.089125353 | 0.087716959 | -1.016056124 | 0.309602661 | 0.553175506 | 0.089125353 |
| ENSMUSG00000078908 | Mon1b | 270096 | protein_coding | 854.7609549 | 0.089126724 | 0.087237914 | 1.021651244 | 0.306945995 | 0.550870027 | 0.089126724 |
| ENSMUSG00000032567 | Aste1 | 66595 | protein_coding | 314.0664722 | -0.089191063 | 0.076789912 | -1.161494533 | 0.245440845 | 0.486686894 | 0.089191063 |
| ENSMUSG00000028517 | Plpp3 | 67916 | protein_coding | 129.2476231 | 0.089198541 | 0.115305629 | 0.773583577 | 0.439177087 | 0.671631718 | 0.089198541 |
| ENSMUSG00000003200 | Sh3gl1 | 20405 | protein_coding | 769.9976403 | 0.089214493 | 0.05909094 | 1.509782944 | 0.131098819 | 0.336982889 | 0.089214493 |
| ENSMUSG00000024072 | Yipf4 | 67864 | protein_coding | 1272.295975 | 0.089224857 | 0.059262035 | 1.505598947 | 0.132170159 | 0.338818654 | 0.089224857 |
| ENSMUSG00000032579 | Hemk1 | 69536 | protein_coding | 208.2884048 | -0.089229113 | 0.104944487 | -0.850250596 | 0.395185777 | 0.634524196 | 0.089229113 |
| ENSMUSG00000041684 | Bivm | 246229 | protein_coding | 406.9979432 | 0.089236706 | 0.074511169 | 1.197628588 | 0.231061642 | 0.469953022 | 0.089236706 |
| ENSMUSG00000028878 | Fam76a | 230789 | protein_coding | 610.6786002 | -0.08924207 | 0.052969245 | -1.684790301 | 0.092029037 | 0.273475636 | 0.08924207 |
| ENSMUSG00000030521 | Mphosph10 | 67973 | protein_coding | 352.3022308 | -0.089300355 | 0.082385828 | -1.0839286 | 0.278396458 | 0.521149121 | 0.089300355 |
| ENSMUSG00000027952 | Pmvk | 68603 | protein_coding | 712.7496651 | -0.089323358 | 0.10179697 | -0.877465788 | 0.380233693 | 0.621060566 | 0.089323358 |
| ENSMUSG00000051185 | Fam174a | 67698 | protein_coding | 325.0715368 | 0.089328323 | 0.070566814 | 1.265868717 | 0.205560088 | 0.440072685 | 0.089328323 |
| ENSMUSG00000000171 | Sdhd | 66925 | protein_coding | 7124.902787 | 0.08935273 | 0.050353811 | 1.774497866 | 0.075980835 | 0.242642163 | 0.08935273 |
| ENSMUSG00000033323 | Ctdp1 | 67655 | protein_coding | 668.2646083 | -0.089370042 | 0.065802845 | -1.358148606 | 0.174416535 | 0.400571839 | 0.089370042 |
| ENSMUSG00000028693 | Nasp | 50927 | protein_coding | 788.4319239 | -0.089408152 | 0.096243504 | -0.928978566 | 0.352900195 | 0.595378328 | 0.089408152 |
| ENSMUSG00000055660 | Mettl4 | 76781 | protein_coding | 544.8847511 | 0.08943668 | 0.066249237 | 1.350003165 | 0.177014968 | 0.403950811 | 0.08943668 |
| ENSMUSG00000031478 | Nek3 | 23954 | protein_coding | 248.1172305 | -0.0894494 | 0.091788762 | -0.974513642 | 0.329801569 | 0.573541103 | 0.0894494 |
| ENSMUSG00000031823 | Zdhhc7 | 102193 | protein_coding | 1747.235267 | 0.089452762 | 0.061123831 | 1.463467856 | 0.143339403 | 0.355303993 | 0.089452762 |
| ENSMUSG00000073664 | Nbeal1 | 269198 | protein_coding | 4265.321712 | 0.089478497 | 0.032649251 | 2.740598747 | 0.006132735 | 0.041645508 | 0.089478497 |
| ENSMUSG00000036840 | Siah1a | 20437 | protein_coding | 390.5532017 | -0.0894902 | 0.073967961 | -1.209850849 | 0.22633613 | 0.464746979 | 0.0894902 |
| ENSMUSG00000049225 | Pdp1 | 381511 | protein_coding | 437.1118194 | 0.089525128 | 0.098003817 | 0.913486134 | 0.360986923 | 0.604050305 | 0.089525128 |
| ENSMUSG00000036932 | Aifm1 | 26926 | protein_coding | 1720.064411 | 0.089578525 | 0.066622405 | 1.344570557 | 0.178763957 | 0.405874466 | 0.089578525 |
| ENSMUSG00000020092 | Pald1 | 27355 | protein_coding | 26.73361797 | 0.089592302 | 0.274985617 | 0.325807228 | 0.74457021 | 0.878396765 | 0.089592302 |
| ENSMUSG00000014554 | Dguok | 27369 | protein_coding | 367.4847477 | 0.089610769 | 0.072470312 | 1.236516948 | 0.21626647 | 0.45218843 | 0.089610769 |
| ENSMUSG00000026020 | Nop58 | 55989 | protein_coding | 1231.96921 | -0.089641762 | 0.098289663 | -0.912016163 | 0.361760209 | 0.60475867 | 0.089641762 |
| ENSMUSG00000024384 | Iws1 | 73473 | protein_coding | 1359.439707 | -0.089699038 | 0.057188697 | -1.56847494 | 0.116770334 | 0.316003891 | 0.089699038 |
| ENSMUSG00000035021 | Baz1a | 217578 | protein_coding | 688.4585606 | -0.089710107 | 0.066246273 | -1.354191 | 0.17567545 | 0.402088841 | 0.089710107 |
| ENSMUSG00000014932 | Yes1 | 22612 | protein_coding | 1626.801672 | -0.089729681 | 0.047727837 | -1.880028236 | 0.06010423 | 0.208317411 | 0.089729681 |
| ENSMUSG00000049658 | Bdp1 | 544971 | protein_coding | 1322.98729 | -0.089732755 | 0.043381741 | -2.068445227 | 0.038598176 | 0.154536441 | 0.089732755 |
| ENSMUSG00000008575 | Nfib | 18028 | protein_coding | 1043.300943 | -0.089741899 | 0.067786648 | -1.323887543 | 0.185540396 | 0.414720716 | 0.089741899 |
| ENSMUSG00000041354 | Rgl2 | 19732 | protein_coding | 799.3045047 | -0.089742643 | 0.080104594 | -1.120318313 | 0.262578141 | 0.50453922 | 0.089742643 |
| ENSMUSG00000041629 | Fam104a | 28081 | protein_coding | 346.8892059 | -0.089743824 | 0.113258156 | -0.792382881 | 0.428137458 | 0.663421426 | 0.089743824 |
| ENSMUSG00000005873 | Reep5 | 13476 | protein_coding | 881.9603636 | 0.089754572 | 0.056996689 | 1.574733094 | 0.115318066 | 0.3135617 | 0.089754572 |
| ENSMUSG00000032030 | Cul5 | 75717 | protein_coding | 848.1018552 | -0.089764904 | 0.061042714 | -1.470526109 | 0.14141932 | 0.352354239 | 0.089764904 |
| ENSMUSG00000031295 | Phka2 | 110094 | protein_coding | 386.8475156 | -0.08978872 | 0.067084778 | -1.338436573 | 0.180754163 | 0.407666398 | 0.08978872 |
| ENSMUSG00000034932 | Mrpl54 | 66047 | protein_coding | 990.7216025 | 0.089789639 | 0.094721211 | 0.947935928 | 0.343162073 | 0.585688936 | 0.089789639 |
| ENSMUSG00000022911 | Arl13b | 68146 | protein_coding | 335.8643091 | 0.089799453 | 0.074354995 | 1.207712457 | 0.227157885 | 0.465663557 | 0.089799453 |

Supplementary Table 1

| ensembl_gene_id | external_gene_name | entrezgene_id | gene_biotype | baseMean | log2FoldChange | lfcSE | stat | pvalue | padj | absLog2FC |
|---|---|---|---|---|---|---|---|---|---|---|
| ENSMUSG00000096956 | Snhg18 | 100616095 | lncRNA | 770.3328677 | 0.08984265 | 0.06333207 | 1.418596446 | 0.156016703 | 0.374523994 | 0.08984265 |
| ENSMUSG00000068264 | Ap5s1 | 69596 | protein_coding | 386.9073822 | 0.089853039 | 0.090295167 | 0.995103524 | 0.319685917 | 0.563143068 | 0.089853039 |
| ENSMUSG00000031445 | Proz | 66901 | protein_coding | 28.24132967 | -0.089883437 | 0.268277522 | -0.335039017 | 0.737595654 | 0.87419748 | 0.089883437 |
| ENSMUSG00000034269 | Setd5 | 72895 | protein_coding | 2229.632829 | -0.089891213 | 0.054031716 | -1.663674949 | 0.096177405 | 0.281344642 | 0.089891213 |
| ENSMUSG00000029641 | Rasl11a | 68895 | protein_coding | 15.01213577 | -0.089892271 | 0.296685082 | -0.302988848 | 0.761898361 | 0.887606361 | 0.089892271 |
| ENSMUSG00000053004 | Hrh1 | 15465 | protein_coding | 195.3920197 | 0.08990705 | 0.094290303 | 0.953513218 | 0.340330102 | 0.583213766 | 0.08990705 |
| ENSMUSG00000038975 | Rabggtb | 19352 | protein_coding | 819.6036247 | -0.0899075 | 0.051260126 | -1.753946133 | 0.079439738 | 0.249100651 | 0.0899075 |
| ENSMUSG00000033147 | Slc22a15 | 242126 | protein_coding | 360.647623 | -0.089908152 | 0.078101269 | -1.151174018 | 0.249660652 | 0.490471109 | 0.089908152 |
| ENSMUSG00000034297 | Med13 | 327987 | protein_coding | 2105.592239 | -0.089914875 | 0.046063337 | -1.951983532 | 0.05094016 | 0.18743143 | 0.089914875 |
| ENSMUSG00000022978 | Mis18a | 66578 | protein_coding | 209.1412249 | 0.089925834 | 0.137927449 | 0.651979243 | 0.514414563 | 0.727325341 | 0.089925834 |
| ENSMUSG00000033373 | Fntb | 110606 | protein_coding | 360.9401654 | 0.089988272 | 0.088318048 | 1.01891146 | 0.308245003 | 0.552028135 | 0.089988272 |
| ENSMUSG00000034435 | Tmem30b | 238257 | protein_coding | 5996.78663 | 0.089994696 | 0.074899712 | 1.201535941 | 0.229543373 | 0.468848355 | 0.089994696 |
| ENSMUSG00000031221 | Igbp1 | 18518 | protein_coding | 1811.560949 | -0.090027177 | 0.047519729 | -1.894522092 | 0.058155745 | 0.204158306 | 0.090027177 |
| ENSMUSG00000062908 | Acadm | 11364 | protein_coding | 3732.920055 | -0.090048028 | 0.042160055 | -2.135861243 | 0.032690725 | 0.138156009 | 0.090048028 |
| ENSMUSG00000020229 | Slc5a4a | 64452 | protein_coding | 19.76763906 | -0.090062796 | 0.538074016 | -0.167379939 | 0.867071109 | 0.939855462 | 0.090062796 |
| ENSMUSG00000026925 | Inpp5e | 64436 | protein_coding | 424.1773238 | -0.090077253 | 0.069953249 | -1.287677894 | 0.197858112 | 0.430673204 | 0.090077253 |
| ENSMUSG00000052102 | Gnpda1 | 26384 | protein_coding | 365.5166565 | 0.090092549 | 0.088874987 | 1.013699716 | 0.31072606 | 0.554568301 | 0.090092549 |
| ENSMUSG00000027327 | 1700037H04Rik | 67326 | protein_coding | 350.0848146 | 0.090094361 | 0.072351604 | 1.245229628 | 0.213047355 | 0.448715514 | 0.090094361 |
| ENSMUSG00000062867 | Impdh2 | 23918 | protein_coding | 2126.043063 | -0.090122 | 0.056471772 | -1.595876967 | 0.110516265 | 0.305187638 | 0.090122 |
| ENSMUSG00000019312 | Grb7 | 14786 | protein_coding | 1968.775322 | -0.09012894 | 0.064184421 | -1.40421832 | 0.160253848 | 0.380620591 | 0.09012894 |
| ENSMUSG00000074582 | Arfgef2 | 99371 | protein_coding | 2540.286862 | -0.090129764 | 0.052953329 | -1.702060381 | 0.08874405 | 0.267559412 | 0.090129764 |
| ENSMUSG00000027602 | Map1lc3a | 66734 | protein_coding | 720.9421254 | -0.090136802 | 0.103481763 | -0.871040462 | 0.383732061 | 0.624590931 | 0.090136802 |
| ENSMUSG00000040424 | Hipk4 | 233020 | protein_coding | 9.579298495 | -0.09017885 | 0.379936702 | -0.2373523 | 0.812383492 | 0.915791177 | 0.09017885 |
| ENSMUSG00000024535 | Snx24 | 69226 | protein_coding | 380.4061884 | 0.090209707 | 0.074001186 | 1.219030551 | 0.222832597 | 0.460189691 | 0.090209707 |
| ENSMUSG00000038990 | Cables2 | 252966 | protein_coding | 485.3918982 | 0.090248567 | 0.054267011 | 1.663046574 | 0.096303113 | 0.281610209 | 0.090248567 |
| ENSMUSG00000027223 | Mapk8ip1 | 19099 | protein_coding | 67.91136489 | -0.090256902 | 0.140740734 | -0.641299075 | 0.52132839 | 0.732548832 | 0.090256902 |
| ENSMUSG00000041837 | Pdcd7 | 50996 | protein_coding | 327.275543 | -0.09031673 | 0.104167389 | -0.867034595 | 0.385923055 | 0.625946003 | 0.09031673 |
| ENSMUSG00000002768 | Mea1 | 17256 | protein_coding | 1364.092994 | 0.090327254 | 0.066853263 | 1.351127071 | 0.176654731 | 0.403453687 | 0.090327254 |
| ENSMUSG00000019877 | Serinc1 | 56442 | protein_coding | 2351.457821 | -0.090327685 | 0.04281104 | -2.109915677 | 0.03486562 | 0.144247781 | 0.090327685 |
| ENSMUSG00000039328 | Rnf122 | 68867 | protein_coding | 70.6225058 | -0.090336738 | 0.167719566 | -0.538617765 | 0.590150627 | 0.779344838 | 0.090336738 |
| ENSMUSG00000029062 | Cdk11b | 12537 | protein_coding | 892.5389435 | 0.090350365 | 0.060341217 | 1.497324198 | 0.134308921 | 0.342017997 | 0.090350365 |
| ENSMUSG00000046994 | Mars2 | 212679 | protein_coding | 528.7492811 | -0.090377221 | 0.074535492 | -1.212539408 | 0.225305967 | 0.463330185 | 0.090377221 |
| ENSMUSG00000046818 | Ddit4l | 73284 | protein_coding | 7.438214372 | 0.090419093 | 0.621832179 | 0.145407549 | 0.884389105 | 0.948750818 | 0.090419093 |
| ENSMUSG00000061665 | Cd2ap | 12488 | protein_coding | 8607.737249 | 0.090419148 | 0.046191732 | 1.957474713 | 0.050291682 | 0.185896496 | 0.090419148 |
| ENSMUSG00000059554 | Ccdc28a | 215814 | protein_coding | 360.7379257 | -0.090422405 | 0.075582079 | -1.196347154 | 0.231561115 | 0.470652152 | 0.090422405 |
| ENSMUSG00000022884 | Eif4a2 | 13682 | protein_coding | 7450.276255 | 0.090422901 | 0.051029196 | 1.771983641 | 0.076397273 | 0.243489885 | 0.090422901 |
| ENSMUSG00000018761 | Mpdu1 | 24070 | protein_coding | 1180.422815 | -0.090471322 | 0.044597333 | -2.028626311 | 0.042496367 | 0.165460865 | 0.090471322 |
| ENSMUSG00000039983 | Ccdc32 | 269336 | protein_coding | 969.6613918 | 0.090480817 | 0.075613493 | 1.19662264 | 0.231453672 | 0.470523296 | 0.090480817 |
| ENSMUSG00000007950 | Abhd8 | 64296 | protein_coding | 173.3457497 | 0.090485515 | 0.149460408 | 0.605414615 | 0.544903543 | 0.749012844 | 0.090485515 |

| ensembl_gene_id | external_gene_name | entrezgene_id | gene_biotype | baseMean | log2FoldChange | lfcSE | stat | pvalue | padj | absLog2FC |
|---|---|---|---|---|---|---|---|---|---|---|
| ENSMUSG00000020464 | Pnpt1 | 71701 | protein_coding | 781.0756543 | 0.090496093 | 0.071205545 | 1.270913564 | 0.2037594 | 0.437979805 | 0.090496093 |
| ENSMUSG00000028394 | Pole3 | 59001 | protein_coding | 578.57777 | 0.090538568 | 0.070529158 | 1.283704086 | 0.199245514 | 0.432642037 | 0.090538568 |
| ENSMUSG00000038174 | Hycc2 | 213056 | protein_coding | 818.1878252 | -0.090539854 | 0.065759367 | -1.376835849 | 0.168563004 | 0.392565467 | 0.090539854 |
| ENSMUSG00000062646 | Ganc | 76051 | protein_coding | 464.6730888 | -0.090556492 | 0.080914681 | -1.119160225 | 0.263071791 | 0.504886203 | 0.090556492 |
| ENSMUSG00000041440 | Gk5 | 235533 | protein_coding | 553.9079756 | 0.090588178 | 0.061637853 | 1.469684184 | 0.141647308 | 0.35263139 | 0.090588178 |
| ENSMUSG00000022551 | Cyc1 | 66445 | protein_coding | 6885.534556 | 0.090614571 | 0.073959929 | 1.225184672 | 0.220505648 | 0.457022824 | 0.090614571 |
| ENSMUSG00000027189 | Trim44 | 80985 | protein_coding | 1181.330406 | -0.090615189 | 0.061712649 | -1.468340626 | 0.142011721 | 0.353156981 | 0.090615189 |
| ENSMUSG00000042305 | Tmem183a | 57439 | protein_coding | 3575.940882 | 0.090625444 | 0.038596698 | 2.348010266 | 0.018874001 | 0.094455763 | 0.090625444 |
| ENSMUSG00000043705 | Capn13 | 381122 | protein_coding | 1107.854556 | 0.090634685 | 0.117233698 | 0.773111206 | 0.43945657 | 0.671700891 | 0.090634685 |
| ENSMUSG00000028648 | Ndufs5 | 595136 | protein_coding | 628.5230027 | 0.090677601 | 0.133509205 | 0.679186138 | 0.497019929 | 0.71512751 | 0.090677601 |
| ENSMUSG00000037514 | Pank2 | 74450 | protein_coding | 939.2974168 | 0.090742612 | 0.063612323 | 1.426494229 | 0.153725731 | 0.370961375 | 0.090742612 |
| ENSMUSG00000047617 | Paxx | 227622 | protein_coding | 268.4298184 | -0.09077895 | 0.075484116 | -1.20262322 | 0.229122158 | 0.468321044 | 0.09077895 |
| ENSMUSG00000025651 | Uqcrc1 | 22273 | protein_coding | 10987.88426 | 0.090827433 | 0.071278338 | 1.274264184 | 0.202569805 | 0.436627004 | 0.090827433 |
| ENSMUSG00000036766 | Dner | 227325 | protein_coding | 11.15834354 | -0.090834809 | 0.405173167 | -0.224187622 | 0.822611314 | 0.920659938 | 0.090834809 |
| ENSMUSG00000042229 | Rabif | 98710 | protein_coding | 669.7842752 | 0.090857559 | 0.047482245 | 1.913505966 | 0.055683311 | 0.198761565 | 0.090857559 |
| ENSMUSG00000047036 | Zfp445 | 235682 | protein_coding | 1720.826361 | -0.090924274 | 0.06792887 | -1.338521804 | 0.180726397 | 0.407666398 | 0.090924274 |
| ENSMUSG00000039756 | Dnttip2 | 99480 | protein_coding | 850.5114784 | -0.09092489 | 0.072604229 | -1.252335995 | 0.210447458 | 0.445508816 | 0.09092489 |
| ENSMUSG00000030030 | 1700003E16Rik | 71837 | protein_coding | 80.8342114 | 0.090991975 | 0.182826355 | 0.497696159 | 0.618698216 | 0.797964342 | 0.090991975 |
| ENSMUSG00000021018 | Polr2h | 245841 | protein_coding | 441.0959651 | -0.091009316 | 0.078913149 | -1.153284549 | 0.24879361 | 0.489722268 | 0.091009316 |
| ENSMUSG00000020780 | Srp68 | 217337 | protein_coding | 1110.806672 | 0.091044872 | 0.050857142 | 1.790208182 | 0.07342045 | 0.237154121 | 0.091044872 |
| ENSMUSG00000022552 | Sharpin | 106025 | protein_coding | 744.2661273 | -0.091058072 | 0.052344258 | -1.739600019 | 0.081929276 | 0.25423772 | 0.091058072 |
| ENSMUSG00000032228 | Tcf12 | 21406 | protein_coding | 1574.648476 | -0.091068378 | 0.053797267 | -1.692806769 | 0.090492254 | 0.27082801 | 0.091068378 |
| ENSMUSG00000003778 | Brd8 | 78656 | protein_coding | 1265.511365 | -0.091080444 | 0.065405409 | -1.392552173 | 0.163755254 | 0.385586361 | 0.091080444 |
| ENSMUSG00000038914 | Dido1 | 23856 | protein_coding | 1982.673881 | -0.091083333 | 0.043003426 | -2.11804831 | 0.034170977 | 0.142359943 | 0.091083333 |
| ENSMUSG00000021474 | Sfxn1 | 14057 | protein_coding | 1310.660077 | -0.091095446 | 0.056086737 | -1.624188735 | 0.104335534 | 0.295551199 | 0.091095446 |
| ENSMUSG00000063808 | Gpatch1 | 67471 | protein_coding | 429.6530022 | -0.091125476 | 0.062191346 | -1.465243664 | 0.142854451 | 0.35463345 | 0.091125476 |
| ENSMUSG00000054612 | Mgmt | 17314 | protein_coding | 64.71597167 | 0.091130699 | 0.15386053 | 0.592294195 | 0.553653602 | 0.75391672 | 0.091130699 |
| ENSMUSG00000052934 | Fbxo31 | 76454 | protein_coding | 559.1322548 | -0.091131306 | 0.085509393 | -1.065746151 | 0.286538409 | 0.529021743 | 0.091131306 |
| ENSMUSG00000020561 | Polr1f | 28071 | protein_coding | 382.5988666 | 0.091134672 | 0.083390405 | 1.092867603 | 0.274451934 | 0.5169296 | 0.091134672 |
| ENSMUSG00000024074 | Crim1 | 50766 | protein_coding | 1729.923744 | -0.09115024 | 0.082176797 | -1.109196801 | 0.267345292 | 0.509691161 | 0.09115024 |
| ENSMUSG00000087819 | Gm25117 | 115487522 | snoRNA | 450.5973317 | 0.091154999 | 0.298561685 | 0.305313788 | 0.760127176 | 0.886576344 | 0.091154999 |
| ENSMUSG00000020572 | Nampt | 59027 | protein_coding | 1704.309069 | -0.091186842 | 0.078047255 | -1.168354252 | 0.2426639 | 0.483875678 | 0.091186842 |
| ENSMUSG00000036019 | Tmtc2 | 278279 | protein_coding | 1032.864722 | 0.091189382 | 0.048720011 | 1.871702793 | 0.061247733 | 0.210875374 | 0.091189382 |
| ENSMUSG00000066551 | Hmgb1 | 15289 | protein_coding | 2911.958128 | 0.091208525 | 0.05501279 | 1.657951278 | 0.097327301 | 0.283295883 | 0.091208525 |
| ENSMUSG00000052056 | Zfp217 | 228913 | protein_coding | 1337.083432 | -0.091222817 | 0.080086691 | -1.139050886 | 0.254681931 | 0.495865688 | 0.091222817 |
| ENSMUSG00000028211 | Trp53inp1 | 60599 | protein_coding | 1949.577206 | -0.091238493 | 0.096543498 | -0.945050623 | 0.344633032 | 0.586900547 | 0.091238493 |
| ENSMUSG00000014039 | Prdm15 | 114604 | protein_coding | 412.6186567 | -0.091240763 | 0.070632638 | -1.291764901 | 0.196438574 | 0.428914823 | 0.091240763 |
| ENSMUSG00000027291 | Vps39 | 269338 | protein_coding | 753.6748327 | -0.091259269 | 0.052485089 | -1.738765632 | 0.082075996 | 0.254497133 | 0.091259269 |
| ENSMUSG00000027531 | Impa1 | 55980 | protein_coding | 1747.423727 | 0.091261492 | 0.048918818 | 1.865570247 | 0.062101516 | 0.21271653 | 0.091261492 |

| ensembl_gene_id | external_gene_name | entrezgene_id | gene_biotype | baseMean | log2FoldChange | lfcSE | stat | pvalue | padj | absLog2FC |
|---|---|---|---|---|---|---|---|---|---|---|
| ENSMUSG00000040613 | Apobec1 | 11810 | protein_coding | 519.9618853 | 0.091265848 | 0.125646364 | 0.726370785 | 0.467611496 | 0.692926016 | 0.091265848 |
| ENSMUSG00000006850 | Tmco6 | 71983 | protein_coding | 415.4451753 | 0.091266267 | 0.074049412 | 1.232504958 | 0.217760516 | 0.453918459 | 0.091266267 |
| ENSMUSG00000027699 | Ect2 | 13605 | protein_coding | 709.0598969 | 0.091333147 | 0.109226529 | 0.836180986 | 0.403053093 | 0.642359644 | 0.091333147 |
| ENSMUSG00000042564 | Fam227a | 75729 | protein_coding | 19.8797062 | 0.091344287 | 0.294744328 | 0.309910245 | 0.756629212 | 0.884485201 | 0.091344287 |
| ENSMUSG00000038323 | 1700066M21Rik | 73467 | protein_coding | 172.3859742 | 0.091347847 | 0.094584474 | 0.96578056 | 0.334153998 | 0.577547724 | 0.091347847 |
| ENSMUSG00000022476 | Polr3h | 78929 | protein_coding | 187.5731793 | -0.091375418 | 0.11506785 | -0.794100338 | 0.427137021 | 0.662540996 | 0.091375418 |
| ENSMUSG00000021893 | Capn7 | 12339 | protein_coding | 1584.115181 | 0.091386527 | 0.045301139 | 2.017311933 | 0.043662971 | 0.168942595 | 0.091386527 |
| ENSMUSG00000007815 | Rhoa | 11848 | protein_coding | 5159.047789 | 0.091398515 | 0.053352124 | 1.713118595 | 0.086690724 | 0.263635924 | 0.091398515 |
| ENSMUSG00000025780 | Itih5 | 209378 | protein_coding | 6179.285831 | -0.091442958 | 0.112617803 | -0.811976048 | 0.416805376 | 0.654765273 | 0.091442958 |
| ENSMUSG00000019558 | Slc6a8 | 102857 | protein_coding | 6574.189475 | 0.091449399 | 0.111549379 | 0.819810923 | 0.412323903 | 0.649970343 | 0.091449399 |
| ENSMUSG00000075376 | Rc3h2 | 319817 | protein_coding | 1389.969293 | -0.091454532 | 0.056719681 | -1.61239504 | 0.106876003 | 0.299347968 | 0.091454532 |
| ENSMUSG00000030655 | Smg1 | 233789 | protein_coding | 3789.613032 | -0.091473808 | 0.053500511 | -1.709774473 | 0.087307585 | 0.26456259 | 0.091473808 |
| ENSMUSG00000006154 | Eps8l1 | 67425 | protein_coding | 120.0986656 | 0.09149838 | 0.260341317 | 0.351455472 | 0.725246673 | 0.865987203 | 0.09149838 |
| ENSMUSG00000040455 | Usp45 | 77593 | protein_coding | 681.8809375 | -0.091499984 | 0.059750527 | -1.531366972 | 0.125678722 | 0.328975938 | 0.091499984 |
| ENSMUSG00000064853 | Gm23442 | 115488838 | snoRNA | 390.7981028 | 0.091508489 | 0.172502174 | 0.530477307 | 0.595781038 | 0.78269996 | 0.091508489 |
| ENSMUSG00000079435 | Rpl36a | 19982 | protein_coding | 4179.141073 | -0.091523223 | 0.060832744 | -1.504505896 | 0.132451156 | 0.339377153 | 0.091523223 |
| ENSMUSG00000092417 | Gpank1 | 81845 | protein_coding | 255.160605 | -0.091552537 | 0.086635918 | -1.056750351 | 0.290625539 | 0.533149593 | 0.091552537 |
| ENSMUSG00000074579 | Lekr1 | 624866 | protein_coding | 53.09646675 | -0.091573762 | 0.187329387 | -0.488838208 | 0.624956245 | 0.802084296 | 0.091573762 |
| ENSMUSG00000020814 | Mxra7 | 67622 | protein_coding | 22.17668631 | -0.091613362 | 0.278150865 | -0.329365727 | 0.74187927 | 0.876632986 | 0.091613362 |
| ENSMUSG00000071252 | 2210408I21Rik | 72371 | protein_coding | 82.57218647 | 0.091629161 | 0.139303761 | 0.657765163 | 0.510689043 | 0.725053455 | 0.091629161 |
| ENSMUSG00000023994 | Nfya | 18044 | protein_coding | 525.7040315 | 0.091650274 | 0.074109129 | 1.236693454 | 0.21620091 | 0.452109968 | 0.091650274 |
| ENSMUSG00000029090 | Adgra3 | 70693 | protein_coding | 2856.484563 | 0.091658459 | 0.054965326 | 1.667568731 | 0.09540137 | 0.279964463 | 0.091658459 |
| ENSMUSG00000033287 | Kctd17 | 72844 | protein_coding | 410.5990617 | -0.09171887 | 0.090430881 | -1.014242797 | 0.310466912 | 0.554289814 | 0.09171887 |
| ENSMUSG00000046731 | Kctd11 | 216858 | protein_coding | 194.6830791 | -0.091725622 | 0.106712704 | -0.859556726 | 0.390033438 | 0.629825698 | 0.091725622 |
| ENSMUSG00000037640 | Zfp60 | 22718 | protein_coding | 237.71336 | -0.091738007 | 0.129325851 | -0.709355525 | 0.47810388 | 0.700752637 | 0.091738007 |
| ENSMUSG00000016984 | Etaa1 | 68145 | protein_coding | 240.7595466 | -0.091755595 | 0.090070486 | -1.018708788 | 0.308341239 | 0.552028135 | 0.091755595 |
| ENSMUSG00000038260 | Trpm4 | 68667 | protein_coding | 6880.710251 | -0.091790749 | 0.090430644 | -1.015040314 | 0.310086611 | 0.553733449 | 0.091790749 |
| ENSMUSG00000030303 | Far2 | 330450 | protein_coding | 340.9585754 | -0.091790939 | 0.123837424 | -0.741221321 | 0.458559258 | 0.685788934 | 0.091790939 |
| ENSMUSG00000024847 | Aip | 11632 | protein_coding | 513.975205 | 0.091806027 | 0.074308937 | 1.235464138 | 0.216657815 | 0.452654565 | 0.091806027 |
| ENSMUSG00000038235 | F11r | 16456 | protein_coding | 4620.029018 | 0.091807021 | 0.043267842 | 2.121830367 | 0.033851986 | 0.141616335 | 0.091807021 |
| ENSMUSG00000047187 | Rab2a | 59021 | protein_coding | 3661.39881 | 0.091814207 | 0.047323086 | 1.940156811 | 0.052360636 | 0.190915661 | 0.091814207 |
| ENSMUSG00000096255 | Dynlt1b | 21648 | protein_coding | 66.24227508 | -0.091844905 | 0.143249027 | -0.641155523 | 0.521421643 | 0.732616034 | 0.091844905 |
| ENSMUSG00000027502 | Rtf2 | 66404 | protein_coding | 1659.047221 | 0.091876037 | 0.044841847 | 2.048890542 | 0.040472819 | 0.159898373 | 0.091876037 |
| ENSMUSG00000032344 | Cgas | 214763 | protein_coding | 69.02251774 | -0.091923075 | 0.205419592 | -0.447489325 | 0.654521794 | 0.822141364 | 0.091923075 |
| ENSMUSG00000022286 | Grhl2 | 252973 | protein_coding | 1580.161424 | -0.091935993 | 0.049844719 | -1.844448016 | 0.065117875 | 0.21965195 | 0.091935993 |
| ENSMUSG00000039648 | Kyat1 | 70266 | protein_coding | 606.4013812 | -0.091945837 | 0.080300369 | -1.145023849 | 0.252199278 | 0.493414747 | 0.091945837 |
| ENSMUSG00000024909 | Efemp2 | 58859 | protein_coding | 18.80363057 | 0.091972557 | 0.276575903 | 0.332540022 | 0.739481524 | 0.874869968 | 0.091972557 |
| ENSMUSG00000046352 | Gjb2 | 14619 | protein_coding | 218.2053065 | 0.091978231 | 0.099685031 | 0.922688487 | 0.356169567 | 0.598889231 | 0.091978231 |
| ENSMUSG00000073684 | Faap20 | 67513 | protein_coding | 286.2306524 | -0.091993956 | 0.083015898 | -1.108148662 | 0.267797618 | 0.510312204 | 0.091993956 |

| ensembl_gene_id | external_gene_name | entrezgene_id | gene_biotype | baseMean | log2FoldChange | lfcSE | stat | pvalue | padj | absLog2FC |
|---|---|---|---|---|---|---|---|---|---|---|
| ENSMUSG00000005621 | Zfp592 | 233410 | protein_coding | 978.5946884 | -0.092025066 | 0.054380841 | -1.692233238 | 0.090601511 | 0.270930943 | 0.092025066 |
| ENSMUSG00000023030 | Slc11a2 | 18174 | protein_coding | 864.0218342 | -0.09204488 | 0.093964485 | -0.979570948 | 0.327297951 | 0.571042203 | 0.09204488 |
| ENSMUSG00000028252 | Ccnc | 51813 | protein_coding | 448.7372023 | 0.092044916 | 0.072851484 | 1.263459716 | 0.206424015 | 0.44133635 | 0.092044916 |
| ENSMUSG00000112980 | D430020J02Rik | 319545 | lncRNA | 51.16705218 | 0.092049359 | 0.208560023 | 0.441356679 | 0.6589548 | 0.82545938 | 0.092049359 |
| ENSMUSG00000026640 | Plxna2 | 18845 | protein_coding | 1307.032618 | -0.092060185 | 0.053325595 | -1.726378963 | 0.084279258 | 0.258850238 | 0.092060185 |
| ENSMUSG00000046947 | Adck2 | 57869 | protein_coding | 440.1329092 | 0.092090553 | 0.078400698 | 1.174613943 | 0.240149202 | 0.480566595 | 0.092090553 |
| ENSMUSG00000022261 | Sdc2 | 15529 | protein_coding | 942.4529997 | 0.09212769 | 0.044027779 | 2.092490069 | 0.036394699 | 0.148403872 | 0.09212769 |
| ENSMUSG00000001998 | Ap4e1 | 108011 | protein_coding | 276.72402 | -0.092164935 | 0.09514633 | -0.968665157 | 0.332712282 | 0.576084494 | 0.092164935 |
| ENSMUSG00000029028 | Lrrc47 | 72946 | protein_coding | 602.9134061 | 0.092165035 | 0.064909003 | 1.419911419 | 0.155633471 | 0.373998061 | 0.092165035 |
| ENSMUSG00000025730 | Rab40c | 224624 | protein_coding | 632.8194517 | 0.092172162 | 0.063636871 | 1.44840815 | 0.147502937 | 0.361113508 | 0.092172162 |
| ENSMUSG00000014470 | Rnf166 | 68718 | protein_coding | 461.3288165 | 0.092232569 | 0.08649214 | 1.066369371 | 0.286256702 | 0.528881227 | 0.092232569 |
| ENSMUSG00000047443 | Erfe | 227358 | protein_coding | 164.9382155 | -0.092242458 | 0.099177746 | -0.930072136 | 0.352333737 | 0.594858267 | 0.092242458 |
| ENSMUSG00000006818 | Sod2 | 20656 | protein_coding | 2588.737438 | 0.092258713 | 0.055065868 | 1.675424662 | 0.093850941 | 0.276898911 | 0.092258713 |
| ENSMUSG00000023984 | Gm20517 | 112694759 | protein_coding | 8.821529013 | -0.092258915 | 0.495179951 | -0.186313915 | 0.852198596 | 0.933591927 | 0.092258915 |
| ENSMUSG00000039988 | Ankrd13c | 433667 | protein_coding | 1962.917783 | 0.092276126 | 0.042497868 | 2.171311893 | 0.029907604 | 0.129900331 | 0.092276126 |
| ENSMUSG00000022755 | Adgrg7 | 239853 | protein_coding | 7974.158338 | -0.092277791 | 0.052013587 | -1.77410937 | 0.076045061 | 0.242751129 | 0.092277791 |
| ENSMUSG00000040412 | Elapor1 | 229722 | protein_coding | 7435.577599 | 0.092318053 | 0.066234252 | 1.393811355 | 0.163374578 | 0.385139866 | 0.092318053 |
| ENSMUSG00000021759 | Plpp1 | 19012 | protein_coding | 397.2706485 | 0.092337994 | 0.085645604 | 1.078140496 | 0.280971062 | 0.523508209 | 0.092337994 |
| ENSMUSG00000061119 | Prcp | 72461 | protein_coding | 1065.614211 | -0.09233818 | 0.053015917 | -1.741706758 | 0.081559769 | 0.253334823 | 0.09233818 |
| ENSMUSG00000020612 | Prkar1a | 19084 | protein_coding | 1483.972339 | 0.092395911 | 0.04596258 | 2.010242034 | 0.044405579 | 0.170791501 | 0.092395911 |
| ENSMUSG00000038195 | Rilp | 280408 | protein_coding | 334.3046177 | -0.09240423 | 0.086686683 | -1.065956467 | 0.286443321 | 0.529013894 | 0.09240423 |
| ENSMUSG00000036304 | Zdhhc23 | 332175 | protein_coding | 544.3541013 | -0.092422672 | 0.061654721 | -1.499036414 | 0.133864186 | 0.341208744 | 0.092422672 |
| ENSMUSG00000054659 | Pm20d2 | 242377 | protein_coding | 55.07272079 | -0.092437548 | 0.179503124 | -0.514963448 | 0.606578562 | 0.789765867 | 0.092437548 |
| ENSMUSG00000019897 | Ccdc59 | 52713 | protein_coding | 367.3389625 | -0.092442734 | 0.076271596 | -1.212020447 | 0.225504554 | 0.463452522 | 0.092442734 |
| ENSMUSG00000033938 | Ndufb7 | 66916 | protein_coding | 2170.049918 | 0.092444737 | 0.089400268 | 1.034054351 | 0.301110755 | 0.544679691 | 0.092444737 |
| ENSMUSG00000018326 | Ywhab | 54401 | protein_coding | 3871.108269 | 0.092455492 | 0.066626289 | 1.387672847 | 0.165236688 | 0.388111283 | 0.092455492 |
| ENSMUSG00000003380 | Rabac1 | 14470 | protein_coding | 908.0679345 | 0.092469701 | 0.094094812 | 0.982729004 | 0.325740832 | 0.56923128 | 0.092469701 |
| ENSMUSG00000028656 | Cap1 | 12331 | protein_coding | 5362.598461 | 0.092472603 | 0.048231129 | 1.917280525 | 0.055202303 | 0.19765709 | 0.092472603 |
| ENSMUSG00000101304 | Plet1os | 69631 | lncRNA | 28.40265222 | -0.092489789 | 0.243767905 | -0.379417418 | 0.704377918 | 0.853157579 | 0.092489789 |
| ENSMUSG00000060716 | Plekhh1 | 211945 | protein_coding | 2857.225998 | -0.092501693 | 0.061599264 | -1.501668814 | 0.133182662 | 0.340305926 | 0.092501693 |
| ENSMUSG00000094112 | 9330182O14Rik | 328531 | protein_coding | 14.03027731 | 0.092504803 | 0.32124313 | 0.287958851 | 0.773378237 | 0.893440495 | 0.092504803 |
| ENSMUSG00000103472 | Pcdhga7 | 93715 | protein_coding | 21.36669956 | 0.09252037 | 0.315617796 | 0.293140536 | 0.769414734 | 0.891758816 | 0.09252037 |
| ENSMUSG00000048329 | Mfsd6l | 215723 | protein_coding | 321.6820525 | 0.092525014 | 0.073672899 | 1.255889409 | 0.209156069 | 0.444642753 | 0.092525014 |
| ENSMUSG00000044952 | Kctd21 | 622320 | protein_coding | 192.4172276 | -0.092569024 | 0.112472755 | -0.823035088 | 0.41048804 | 0.648632239 | 0.092569024 |
| ENSMUSG00000026926 | Pmpca | 66865 | protein_coding | 1250.552948 | 0.092608533 | 0.046001736 | 2.013153032 | 0.044098533 | 0.170138047 | 0.092608533 |
| ENSMUSG00000024965 | Fermt3 | 108101 | protein_coding | 98.60745974 | -0.092611251 | 0.138440587 | -0.668960261 | 0.503520828 | 0.719946838 | 0.092611251 |
| ENSMUSG00000057147 | Dph6 | 66632 | protein_coding | 105.8721288 | -0.092627134 | 0.136658239 | -0.677801313 | 0.497897679 | 0.715687493 | 0.092627134 |
| ENSMUSG00000036327 | Qsox2 | 227638 | protein_coding | 725.2447731 | 0.092635482 | 0.065398503 | 1.416477108 | 0.156635862 | 0.375325502 | 0.092635482 |
| ENSMUSG00000058385 | H2bc8 | 319181 | protein_coding | 1386.690579 | 0.092638205 | 0.130475328 | 0.710005532 | 0.477700706 | 0.700289 | 0.092638205 |

| ensembl_gene_id | external_gene_name | entrezgene_id | gene_biotype | baseMean | log2FoldChange | lfcSE | stat | pvalue | padj | absLog2FC |
|---|---|---|---|---|---|---|---|---|---|---|
| ENSMUSG00000037656 | Slc20a2 | 20516 | protein_coding | 506.5359681 | -0.092658075 | 0.072191693 | -1.283500517 | 0.199316779 | 0.432738515 | 0.092658075 |
| ENSMUSG00000073077 | Cfap47 | 636104 | protein_coding | 62.96104897 | -0.092662574 | 0.184443185 | -0.502390882 | 0.615392592 | 0.795545998 | 0.092662574 |
| ENSMUSG00000022708 | Zbtb20 | 56490 | protein_coding | 471.6384552 | -0.092692546 | 0.107649125 | -0.861061773 | 0.389204022 | 0.629053243 | 0.092692546 |
| ENSMUSG00000021998 | Lcp1 | 18826 | protein_coding | 403.3208827 | 0.092696994 | 0.083996399 | 1.103582943 | 0.269774094 | 0.512256092 | 0.092696994 |
| ENSMUSG00000049624 | Slc17a5 | 235504 | protein_coding | 732.8610878 | -0.092758649 | 0.071029495 | -1.305917336 | 0.19158067 | 0.423061957 | 0.092758649 |
| ENSMUSG00000050212 | Eva1b | 230752 | protein_coding | 30.8329032 | 0.092770442 | 0.248715983 | 0.372997507 | 0.709150299 | 0.856166251 | 0.092770442 |
| ENSMUSG00000020297 | Nsg2 | 18197 | protein_coding | 14.5328332 | -0.092798576 | 0.357940067 | -0.259257302 | 0.795436722 | 0.90516568 | 0.092798576 |
| ENSMUSG00000087259 | 2610035D17Rik | 72386 | lncRNA | 452.1426421 | -0.092816354 | 0.084930317 | -1.092853036 | 0.274458331 | 0.5169296 | 0.092816354 |
| ENSMUSG00000039989 | Cbx4 | 12418 | protein_coding | 424.192825 | 0.092827094 | 0.102075676 | 0.909394853 | 0.363141738 | 0.605769124 | 0.092827094 |
| ENSMUSG00000026341 | Actr3 | 74117 | protein_coding | 7128.256176 | 0.092853447 | 0.051558861 | 1.800921231 | 0.071715296 | 0.233599794 | 0.092853447 |
| ENSMUSG00000025742 | Prps2 | 110639 | protein_coding | 984.6314022 | -0.092873193 | 0.055430291 | -1.675495329 | 0.093837087 | 0.276898911 | 0.092873193 |
| ENSMUSG00000032479 | Map4 | 17758 | protein_coding | 1821.886644 | 0.092901523 | 0.043008894 | 2.160053731 | 0.03076851 | 0.132451326 | 0.092901523 |
| ENSMUSG00000030748 | Il4ra | 16190 | protein_coding | 803.1976697 | 0.092925509 | 0.059724004 | 1.555915609 | 0.119728161 | 0.320093959 | 0.092925509 |
| ENSMUSG00000020978 | Klhdc2 | 69554 | protein_coding | 699.1983407 | -0.092955996 | 0.058747623 | -1.582293735 | 0.11358253 | 0.310570612 | 0.092955996 |
| ENSMUSG00000021196 | Pfkp | 56421 | protein_coding | 4337.99084 | -0.093020613 | 0.079711608 | -1.166964457 | 0.243224724 | 0.484226665 | 0.093020613 |
| ENSMUSG00000060181 | Slc35e3 | 215436 | protein_coding | 264.9905313 | -0.093037025 | 0.113107958 | -0.822550657 | 0.410763568 | 0.648813327 | 0.093037025 |
| ENSMUSG00000039206 | Daglb | 231871 | protein_coding | 1009.710856 | 0.09304166 | 0.079949342 | 1.163757672 | 0.24452223 | 0.485821117 | 0.09304166 |
| ENSMUSG00000103421 | Golt1a | 68338 | protein_coding | 80.51241213 | -0.093049585 | 0.157329283 | -0.591432079 | 0.554230952 | 0.754395894 | 0.093049585 |
| ENSMUSG00000025959 | Klf7 | 93691 | protein_coding | 460.2133241 | 0.093103615 | 0.074244473 | 1.254014076 | 0.209836889 | 0.44497561 | 0.093103615 |
| ENSMUSG00000024592 | C330018D20Rik | 77422 | protein_coding | 322.161861 | 0.093108966 | 0.09558761 | 0.974069396 | 0.330022084 | 0.573705524 | 0.093108966 |
| ENSMUSG00000035049 | Rrp12 | 107094 | protein_coding | 294.6896016 | 0.093117299 | 0.090463366 | 1.029337102 | 0.303321294 | 0.547288265 | 0.093117299 |
| ENSMUSG00000042178 | Armc5 | 233912 | protein_coding | 641.6155004 | 0.093131826 | 0.117101889 | 0.795305927 | 0.426435567 | 0.661834894 | 0.093131826 |
| ENSMUSG00000026289 | Atg16l1 | 77040 | protein_coding | 691.749872 | -0.093159017 | 0.069835977 | -1.333968844 | 0.182214067 | 0.410013432 | 0.093159017 |
| ENSMUSG00000047583 | Tyw3 | 209584 | protein_coding | 67.77009538 | 0.093179995 | 0.164065552 | 0.567943688 | 0.570073205 | 0.765238104 | 0.093179995 |
| ENSMUSG00000040596 | Pogk | 71592 | protein_coding | 608.2806157 | -0.093180387 | 0.058798516 | -1.584740469 | 0.113025313 | 0.309466761 | 0.093180387 |
| ENSMUSG00000025436 | Atp23 | 68876 | protein_coding | 90.22324257 | 0.093230965 | 0.137207623 | 0.679488228 | 0.496828564 | 0.715005734 | 0.093230965 |
| ENSMUSG00000027099 | Mtx2 | 53375 | protein_coding | 1175.536554 | 0.093235624 | 0.050545218 | 1.844598326 | 0.065095991 | 0.219624067 | 0.093235624 |
| ENSMUSG00000020687 | Cdc27 | 217232 | protein_coding | 1280.165585 | 0.093242737 | 0.06112821 | 1.525363449 | 0.12716847 | 0.331476631 | 0.093242737 |
| ENSMUSG00000052406 | Rexo4 | 227656 | protein_coding | 692.6236238 | 0.093261768 | 0.059581704 | 1.565275271 | 0.117518379 | 0.317074504 | 0.093261768 |
| ENSMUSG00000045438 | Cox19 | 68033 | protein_coding | 513.6911895 | 0.093269892 | 0.060213482 | 1.548986849 | 0.121384882 | 0.322765702 | 0.093269892 |
| ENSMUSG00000004771 | Rab11a | 53869 | protein_coding | 2287.455602 | 0.093278055 | 0.041659437 | 2.239061831 | 0.025151893 | 0.115148818 | 0.093278055 |
| ENSMUSG00000055733 | Nap1l3 | 54561 | protein_coding | 22.55764252 | -0.093318087 | 0.323855314 | -0.288147464 | 0.773233862 | 0.893440495 | 0.093318087 |
| ENSMUSG00000115902 | D730005E14Rik | 109361 | lncRNA | 41.12810359 | -0.093345033 | 0.18335898 | -0.509083509 | 0.61069369 | 0.792648462 | 0.093345033 |
| ENSMUSG00000001774 | Chordc1 | 66917 | protein_coding | 866.8152275 | 0.093353034 | 0.080560726 | 1.158790871 | 0.24654144 | 0.48740394 | 0.093353034 |
| ENSMUSG00000019362 | D8Ertd738e | 101966 | protein_coding | 1499.924647 | 0.093357719 | 0.050475449 | 1.849566875 | 0.064376001 | 0.217970138 | 0.093357719 |
| ENSMUSG00000030061 | Uba3 | 22200 | protein_coding | 1063.016798 | -0.093358319 | 0.046532011 | -2.006324616 | 0.044821625 | 0.171818006 | 0.093358319 |
| ENSMUSG00000030612 | Mrpl46 | 67308 | protein_coding | 636.600651 | 0.093373929 | 0.07283054 | 1.282071086 | 0.199817711 | 0.433436829 | 0.093373929 |
| ENSMUSG00000022346 | Myc | 17869 | protein_coding | 469.4790481 | -0.093394222 | 0.08653485 | -1.079267161 | 0.280468648 | 0.523452524 | 0.093394222 |
| ENSMUSG00000045691 | Thtpa | 105663 | protein_coding | 186.0464166 | 0.093398292 | 0.103644375 | 0.901141932 | 0.36751286 | 0.609949562 | 0.093398292 |

| ensembl_gene_id | external_gene_name | entrezgene_id | gene_biotype | baseMean | log2FoldChange | lfcSE | stat | pvalue | padj | absLog2FC |
|---|---|---|---|---|---|---|---|---|---|---|
| ENSMUSG00000042750 | Bex2 | 12069 | protein_coding | 59.10710766 | 0.093403861 | 0.159054163 | 0.587245627 | 0.557038753 | 0.755953522 | 0.093403861 |
| ENSMUSG00000034349 | Smc4 | 70099 | protein_coding | 1488.890016 | 0.093420988 | 0.089403222 | 1.044939836 | 0.296050777 | 0.538897588 | 0.093420988 |
| ENSMUSG00000051223 | Bzw1 | 66882 | protein_coding | 5127.73916 | 0.093427661 | 0.058923619 | 1.585572333 | 0.112836357 | 0.309131951 | 0.093427661 |
| ENSMUSG00000045251 | Zfp688 | 69234 | protein_coding | 186.5053197 | -0.093438125 | 0.114247046 | -0.817860321 | 0.41343695 | 0.651059241 | 0.093438125 |
| ENSMUSG00000022353 | Mtss1 | 211401 | protein_coding | 1224.76573 | 0.093445867 | 0.059765841 | 1.563533048 | 0.117927269 | 0.317445011 | 0.093445867 |
| ENSMUSG00000047694 | Yipf6 | 77929 | protein_coding | 3011.393806 | 0.093457161 | 0.046863264 | 1.994252041 | 0.046124518 | 0.175229704 | 0.093457161 |
| ENSMUSG00000024911 | Fibp | 58249 | protein_coding | 929.2968389 | -0.093516047 | 0.05541982 | -1.687411602 | 0.091524234 | 0.272527941 | 0.093516047 |
| ENSMUSG00000069729 | Arid1b | 239985 | protein_coding | 1784.772773 | -0.093588609 | 0.058968325 | -1.587099672 | 0.112490072 | 0.30863821 | 0.093588609 |
| ENSMUSG00000078773 | Rad54b | 623474 | protein_coding | 63.97944752 | 0.093598149 | 0.171741788 | 0.54499345 | 0.585758023 | 0.776292513 | 0.093598149 |
| ENSMUSG00000005610 | Eif4g2 | 13690 | protein_coding | 12640.40256 | 0.093606258 | 0.0496024 | 1.887131616 | 0.059142625 | 0.206435622 | 0.093606258 |
| ENSMUSG00000028217 | Cdh17 | 12557 | protein_coding | 19175.63553 | 0.093606626 | 0.057133177 | 1.638393485 | 0.101339637 | 0.290200492 | 0.093606626 |
| ENSMUSG00000009073 | Nf2 | 18016 | protein_coding | 2693.236017 | 0.093632989 | 0.052069796 | 1.798220769 | 0.07214203 | 0.234551147 | 0.093632989 |
| ENSMUSG00000047907 | Tshz2 | 228911 | protein_coding | 266.2698423 | -0.093649313 | 0.104072194 | -0.899849514 | 0.36820034 | 0.610334928 | 0.093649313 |
| ENSMUSG00000020843 | Timm22 | 56322 | protein_coding | 446.670494 | 0.093652965 | 0.085955282 | 1.089554505 | 0.275909433 | 0.518721313 | 0.093652965 |
| ENSMUSG00000043162 | Pyurf | 66459 | protein_coding | 422.9147735 | 0.093671601 | 0.07307163 | 1.281914758 | 0.19987255 | 0.433436829 | 0.093671601 |
| ENSMUSG00000041241 | Mul1 | 68350 | protein_coding | 1580.833606 | 0.093681376 | 0.049397251 | 1.896489644 | 0.057895328 | 0.203825108 | 0.093681376 |
| ENSMUSG00000038214 | Bend3 | 331623 | protein_coding | 259.7477368 | -0.093686818 | 0.104006403 | -0.90077933 | 0.367705659 | 0.609985966 | 0.093686818 |
| ENSMUSG00000025347 | Mettl7b | 71664 | protein_coding | 6535.84077 | -0.093690049 | 0.088865178 | -1.054294287 | 0.291748204 | 0.534297443 | 0.093690049 |
| ENSMUSG00000039942 | Ptger4 | 19219 | protein_coding | 1009.98899 | -0.093699357 | 0.101248159 | -0.925442573 | 0.354735742 | 0.597225526 | 0.093699357 |
| ENSMUSG00000005687 | Bcas2 | 68183 | protein_coding | 840.6314416 | -0.093711751 | 0.059815222 | -1.566687336 | 0.117187792 | 0.316511058 | 0.093711751 |
| ENSMUSG00000048546 | Tob2 | 57259 | protein_coding | 881.6967576 | -0.093720405 | 0.064236563 | -1.458988463 | 0.144568283 | 0.357091852 | 0.093720405 |
| ENSMUSG00000013418 | B4galnt2 | 14422 | protein_coding | 9793.31216 | 0.093749172 | 0.062041294 | 1.511076982 | 0.130768838 | 0.336624075 | 0.093749172 |
| ENSMUSG00000001524 | Gtf2h4 | 14885 | protein_coding | 291.3949104 | 0.093772343 | 0.077774624 | 1.205693299 | 0.22793577 | 0.46690209 | 0.093772343 |
| ENSMUSG00000068587 | Mgam | 232714 | protein_coding | 12724.39917 | -0.093826197 | 0.049454318 | -1.897229627 | 0.057797638 | 0.203782797 | 0.093826197 |
| ENSMUSG00000050240 | Hic2 | 58180 | protein_coding | 259.2076505 | -0.093855912 | 0.106765775 | -0.879082383 | 0.379356609 | 0.620491412 | 0.093855912 |
| ENSMUSG00000028488 | Sh3gl2 | 20404 | protein_coding | 310.3601731 | -0.09386989 | 0.092490679 | -1.014911895 | 0.310147828 | 0.553781446 | 0.09386989 |
| ENSMUSG00000031917 | Nip7 | 66164 | protein_coding | 352.7701145 | -0.093873251 | 0.092660047 | -1.013093062 | 0.311015712 | 0.554839644 | 0.093873251 |
| ENSMUSG00000020537 | Drg2 | 13495 | protein_coding | 804.2049774 | 0.093875485 | 0.063242347 | 1.484377 | 0.137708933 | 0.347583794 | 0.093875485 |
| ENSMUSG00000046962 | Zbtb21 | 114565 | protein_coding | 459.2201784 | -0.093879585 | 0.062550682 | -1.500856299 | 0.133392733 | 0.340761066 | 0.093879585 |
| ENSMUSG00000029102 | Hgfac | 54426 | protein_coding | 4588.941344 | 0.093883508 | 0.076031777 | 1.234793022 | 0.216907545 | 0.453012934 | 0.093883508 |
| ENSMUSG00000037750 | Fam222b | 216971 | protein_coding | 159.0756446 | 0.093887532 | 0.114251028 | 0.821765317 | 0.411210476 | 0.649264867 | 0.093887532 |
| ENSMUSG00000043460 | Elfn2 | 207393 | protein_coding | 39.3498843 | -0.093900865 | 0.202166405 | -0.464473141 | 0.642308798 | 0.81409258 | 0.093900865 |
| ENSMUSG00000026473 | Glul | 14645 | protein_coding | 11647.78785 | 0.093902379 | 0.060026257 | 1.564355063 | 0.117734208 | 0.317250963 | 0.093902379 |
| ENSMUSG00000026456 | Cyb5r1 | 72017 | protein_coding | 253.241843 | -0.093910889 | 0.120255967 | -0.780924982 | 0.434846618 | 0.668392242 | 0.093910889 |
| ENSMUSG00000030838 | Ush1c | 72088 | protein_coding | 4847.724636 | 0.093952392 | 0.042674432 | 2.201608489 | 0.027692976 | 0.122794781 | 0.093952392 |
| ENSMUSG00000028552 | Eps15 | 13858 | protein_coding | 1250.725814 | -0.093963604 | 0.062989952 | -1.491723701 | 0.135771593 | 0.34449158 | 0.093963604 |
| ENSMUSG00000025085 | Ablim1 | 226251 | protein_coding | 2466.455773 | -0.093986194 | 0.082682943 | -1.136705963 | 0.255661226 | 0.497115408 | 0.093986194 |
| ENSMUSG00000050213 | Snip1 | 76793 | protein_coding | 506.7798382 | -0.093993296 | 0.05746726 | -1.635597318 | 0.101923892 | 0.291285948 | 0.093993296 |
| ENSMUSG00000030064 | Frmd4b | 232288 | protein_coding | 1216.480599 | -0.094008558 | 0.063108119 | -1.489642852 | 0.136318168 | 0.345280838 | 0.094008558 |

| ensembl_gene_id | external_gene_name | entrezgene_id | gene_biotype | baseMean | log2FoldChange | lfcSE | stat | pvalue | padj | absLog2FC |
|---|---|---|---|---|---|---|---|---|---|---|
| ENSMUSG00000049686 | Orai1 | 109305 | protein_coding | 533.2629879 | 0.094009616 | 0.124773982 | 0.753439256 | 0.451186001 | 0.680343771 | 0.094009616 |
| ENSMUSG00000033502 | Cdc14a | 229776 | protein_coding | 1039.979223 | -0.094049274 | 0.059553011 | -1.57925304 | 0.114278028 | 0.311837862 | 0.094049274 |
| ENSMUSG00000045374 | Wdr81 | 192652 | protein_coding | 854.3361796 | -0.094052525 | 0.067707383 | -1.389102936 | 0.164801451 | 0.387258196 | 0.094052525 |
| ENSMUSG00000061410 | Zcchc14 | 142682 | protein_coding | 999.8314756 | -0.094058652 | 0.068339059 | -1.376352754 | 0.168712448 | 0.392724841 | 0.094058652 |
| ENSMUSG00000095567 | Noc2l | 57741 | protein_coding | 881.7256778 | -0.094134797 | 0.070913937 | -1.327451286 | 0.184359444 | 0.413054287 | 0.094134797 |
| ENSMUSG00000027835 | Pdcd10 | 56426 | protein_coding | 965.4768986 | 0.094145054 | 0.06886075 | 1.367180199 | 0.17156883 | 0.396321966 | 0.094145054 |
| ENSMUSG00000007850 | Hnrnph1 | 59013 | protein_coding | 2907.599007 | -0.094151145 | 0.090252543 | -1.043196588 | 0.296857253 | 0.539792189 | 0.094151145 |
| ENSMUSG00000101856 | 1700096K18Rik | 73571 | lncRNA | 87.27346549 | 0.094157937 | 0.13660699 | 0.689261484 | 0.490658731 | 0.710501334 | 0.094157937 |
| ENSMUSG00000025967 | Eef1b2 | 55949 | protein_coding | 8310.399602 | -0.094183988 | 0.033692168 | -2.795426727 | 0.005183125 | 0.036667193 | 0.094183988 |
| ENSMUSG00000002957 | Ap2a2 | 11772 | protein_coding | 2720.911833 | 0.094203861 | 0.048576034 | 1.939307352 | 0.052463924 | 0.191162608 | 0.094203861 |
| ENSMUSG00000027469 | Tpx2 | 72119 | protein_coding | 649.7266448 | 0.094204772 | 0.13467544 | 0.699494814 | 0.484242853 | 0.706263486 | 0.094204772 |
| ENSMUSG00000059439 | Bcas3 | 192197 | protein_coding | 484.8418332 | -0.094207862 | 0.096539682 | -0.975845991 | 0.32914079 | 0.572749056 | 0.094207862 |
| ENSMUSG00000014195 | Dnajc7 | 56354 | protein_coding | 1607.528815 | -0.094283937 | 0.058619667 | -1.60840111 | 0.107747359 | 0.300630044 | 0.094283937 |
| ENSMUSG00000024740 | Ddb1 | 13194 | protein_coding | 8906.32028 | 0.094295268 | 0.047095716 | 2.002204785 | 0.045262711 | 0.172892408 | 0.094295268 |
| ENSMUSG00000001420 | Tmem79 | 71913 | protein_coding | 583.7669836 | 0.094297861 | 0.078570631 | 1.200166784 | 0.230074573 | 0.46920123 | 0.094297861 |
| ENSMUSG00000035683 | Melk | 17279 | protein_coding | 418.8656985 | 0.094331771 | 0.136815691 | 0.689480647 | 0.490520848 | 0.710394783 | 0.094331771 |
| ENSMUSG00000041147 | Brca2 | 12190 | protein_coding | 140.8456694 | -0.094368698 | 0.159817601 | -0.590477503 | 0.554870564 | 0.754884223 | 0.094368698 |
| ENSMUSG00000027102 | Hoxd8 | 15437 | protein_coding | 38.26769881 | -0.094373331 | 0.216776949 | -0.43534763 | 0.663310117 | 0.827892753 | 0.094373331 |
| ENSMUSG00000009640 | Dmap1 | 66233 | protein_coding | 388.5997175 | 0.094378048 | 0.067005153 | 1.408519259 | 0.158977368 | 0.37913729 | 0.094378048 |
| ENSMUSG00000000439 | Mkrn2 | 67027 | protein_coding | 449.4457673 | -0.094395165 | 0.064777456 | -1.45722249 | 0.145054975 | 0.357631975 | 0.094395165 |
| ENSMUSG00000020580 | Rock2 | 19878 | protein_coding | 2866.116863 | -0.09445508 | 0.059312992 | -1.592485517 | 0.111275655 | 0.306582205 | 0.09445508 |
| ENSMUSG00000036093 | Arl5a | 75423 | protein_coding | 1192.360681 | 0.09446128 | 0.040430118 | 2.336408694 | 0.019469951 | 0.096286999 | 0.09446128 |
| ENSMUSG00000036986 | Pml | 18854 | protein_coding | 674.4423781 | -0.094466946 | 0.063392744 | -1.490185478 | 0.136175474 | 0.345087178 | 0.094466946 |
| ENSMUSG00000107709 | 9430018G01Rik | 319812 | lncRNA | 6.804790323 | -0.094496208 | 0.497714143 | -0.189860403 | 0.849418522 | 0.931939078 | 0.094496208 |
| ENSMUSG00000045930 | Clec14a | 66864 | protein_coding | 497.7759464 | -0.094529378 | 0.080552885 | -1.173507042 | 0.240592535 | 0.481155234 | 0.094529378 |
| ENSMUSG00000042918 | Mamstr | 74490 | protein_coding | 16.77213961 | 0.094548793 | 0.296380066 | 0.319011986 | 0.749717424 | 0.880475742 | 0.094548793 |
| ENSMUSG00000089984 | Fbxo24 | 71176 | protein_coding | 27.20717331 | 0.094564321 | 0.236167491 | 0.400412101 | 0.688853013 | 0.843774308 | 0.094564321 |
| ENSMUSG00000032387 | Rbpms2 | 71973 | protein_coding | 8.37879915 | 0.094587276 | 0.419162527 | 0.225657759 | 0.821467614 | 0.920176481 | 0.094587276 |
| ENSMUSG00000074785 | Plxnc1 | 54712 | protein_coding | 8.123016586 | 0.094602802 | 0.513952556 | 0.184069134 | 0.85395922 | 0.934631853 | 0.094602802 |
| ENSMUSG00000037022 | Mmaa | 109136 | protein_coding | 452.0639041 | 0.094672232 | 0.075422524 | 1.255224921 | 0.209397121 | 0.444801234 | 0.094672232 |
| ENSMUSG00000044359 | P2ry4 | 57385 | protein_coding | 187.7314485 | 0.094677352 | 0.122518696 | 0.772758408 | 0.439665372 | 0.671740242 | 0.094677352 |
| ENSMUSG00000040464 | Gtpbp10 | 207704 | protein_coding | 381.9418431 | -0.094705034 | 0.07708275 | -1.228615151 | 0.219216132 | 0.455544189 | 0.094705034 |
| ENSMUSG00000042050 | Dync2i1 | 217935 | protein_coding | 186.4426411 | 0.094708326 | 0.104644066 | 0.905052045 | 0.365437816 | 0.60759958 | 0.094708326 |
| ENSMUSG00000027430 | Dtd1 | 66044 | protein_coding | 217.8371079 | -0.094765362 | 0.10344775 | -0.916069825 | 0.359630273 | 0.603010752 | 0.094765362 |
| ENSMUSG00000006941 | Eif1b | 68969 | protein_coding | 255.1798281 | -0.094778663 | 0.083811233 | -1.130858714 | 0.258114563 | 0.500132153 | 0.094778663 |
| ENSMUSG00000033096 | Apmap | 71881 | protein_coding | 807.9956262 | 0.094813779 | 0.067794308 | 1.398550733 | 0.161947749 | 0.382615199 | 0.094813779 |
| ENSMUSG00000029470 | P2rx4 | 18438 | protein_coding | 2976.163163 | 0.09482482 | 0.050010821 | 1.896086064 | 0.057948665 | 0.203825108 | 0.09482482 |
| ENSMUSG00000027405 | Nop56 | 67134 | protein_coding | 743.8161817 | -0.094829852 | 0.087464657 | -1.084207671 | 0.278272731 | 0.521098971 | 0.094829852 |
| ENSMUSG00000038776 | Ephx1 | 13849 | protein_coding | 284.7541262 | 0.094898139 | 0.081238994 | 1.168135342 | 0.242752177 | 0.483886312 | 0.094898139 |

| ensembl_gene_id | external_gene_name | entrezgene_id | gene_biotype | baseMean | log2FoldChange | lfcSE | stat | pvalue | padj | absLog2FC |
|---|---|---|---|---|---|---|---|---|---|---|
| ENSMUSG00000053799 | Exoc6 | 107371 | protein_coding | 972.9241298 | -0.094916023 | 0.052201771 | -1.818252925 | 0.069025484 | 0.228015974 | 0.094916023 |
| ENSMUSG00000027298 | Tyro3 | 22174 | protein_coding | 323.404432 | 0.094987818 | 0.096292525 | 0.986450589 | 0.323912051 | 0.567946164 | 0.094987818 |
| ENSMUSG00000027624 | Epb41l1 | 13821 | protein_coding | 1969.390938 | -0.094988442 | 0.073613464 | -1.290367778 | 0.196922994 | 0.429391175 | 0.094988442 |
| ENSMUSG00000063882 | Uqcrh | 66576 | protein_coding | 8263.218242 | 0.095022083 | 0.052107273 | 1.823585807 | 0.068214714 | 0.226218114 | 0.095022083 |
| ENSMUSG00000033478 | Fhip2a | 226252 | protein_coding | 578.5233442 | -0.095035905 | 0.058859275 | -1.614629207 | 0.106391016 | 0.298608218 | 0.095035905 |
| ENSMUSG00000020849 | Ywhae | 22627 | protein_coding | 2973.639222 | 0.09506141 | 0.053963442 | 1.761589079 | 0.07813875 | 0.246748312 | 0.09506141 |
| ENSMUSG00000042079 | Hnrnpf | 98758 | protein_coding | 2117.788111 | 0.095079944 | 0.070270785 | 1.353050843 | 0.176039388 | 0.402521936 | 0.095079944 |
| ENSMUSG00000055493 | Epm2a | 13853 | protein_coding | 38.94863772 | -0.095097673 | 0.209056019 | -0.454890864 | 0.649187756 | 0.818885486 | 0.095097673 |
| ENSMUSG00000049122 | Frmd3 | 242506 | protein_coding | 107.9307662 | 0.095117103 | 0.139382606 | 0.682417311 | 0.494975112 | 0.713739596 | 0.095117103 |
| ENSMUSG00000010830 | Kdelr3 | 105785 | protein_coding | 341.0734309 | 0.095123861 | 0.100135897 | 0.949947651 | 0.342138853 | 0.584809039 | 0.095123861 |
| ENSMUSG00000024174 | Pot1b | 72836 | protein_coding | 103.2186029 | -0.09513617 | 0.122144375 | -0.778882937 | 0.436048676 | 0.669302737 | 0.09513617 |
| ENSMUSG00000026797 | Stxbp1 | 20910 | protein_coding | 222.4390942 | 0.095136887 | 0.159073831 | 0.598067492 | 0.5497949 | 0.751487589 | 0.095136887 |
| ENSMUSG00000033166 | Dis3 | 72662 | protein_coding | 359.0079114 | -0.095145134 | 0.080327841 | -1.184460254 | 0.236230921 | 0.476450233 | 0.095145134 |
| ENSMUSG00000014778 | Fhod1 | 234686 | protein_coding | 435.6156024 | 0.09515011 | 0.113553157 | 0.837934518 | 0.402067481 | 0.641349295 | 0.09515011 |
| ENSMUSG00000038119 | Cdon | 57810 | protein_coding | 565.0693725 | 0.09515671 | 0.079315569 | 1.199722982 | 0.230246944 | 0.469307431 | 0.09515671 |
| ENSMUSG00000046982 | Tshz1 | 110796 | protein_coding | 1888.852217 | -0.095199533 | 0.071874663 | -1.324521456 | 0.185329922 | 0.414422581 | 0.095199533 |
| ENSMUSG00000040618 | Pck2 | 74551 | protein_coding | 822.0221984 | -0.095219555 | 0.061057161 | -1.559514941 | 0.118874551 | 0.319001795 | 0.095219555 |
| ENSMUSG00000029149 | Krtcap3 | 69815 | protein_coding | 1296.603736 | -0.095221307 | 0.063643126 | -1.496175843 | 0.134607838 | 0.342321275 | 0.095221307 |
| ENSMUSG00000031877 | Ces2g | 72361 | protein_coding | 293.9943641 | -0.095229512 | 0.136271733 | -0.698820731 | 0.484664071 | 0.706263486 | 0.095229512 |
| ENSMUSG00000019779 | Frk | 14302 | protein_coding | 3053.897394 | -0.095235058 | 0.06671855 | -1.427414982 | 0.153460315 | 0.370570998 | 0.095235058 |
| ENSMUSG00000031107 | Rbmx2 | 209003 | protein_coding | 70.45930447 | -0.095235721 | 0.158376188 | -0.601326012 | 0.547622868 | 0.750740503 | 0.095235721 |
| ENSMUSG00000025860 | Xiap | 11798 | protein_coding | 2793.85521 | -0.095239324 | 0.049337275 | -1.930372599 | 0.053560687 | 0.19375801 | 0.095239324 |
| ENSMUSG00000038489 | Polr2l | 66491 | protein_coding | 183.3086247 | 0.095246432 | 0.149456864 | 0.637283759 | 0.523940028 | 0.734827388 | 0.095246432 |
| ENSMUSG00000042148 | Cox10 | 70383 | protein_coding | 1006.683064 | 0.095269278 | 0.064622996 | 1.474231838 | 0.14041918 | 0.351472935 | 0.095269278 |
| ENSMUSG00000028101 | Pias3 | 229615 | protein_coding | 328.6345129 | -0.095279649 | 0.086643516 | -1.099674312 | 0.271474051 | 0.513977697 | 0.095279649 |
| ENSMUSG00000074182 | Znhit6 | 229937 | protein_coding | 104.864322 | 0.095299113 | 0.11855413 | 0.803844739 | 0.421486648 | 0.658144381 | 0.095299113 |
| ENSMUSG00000049606 | Zfp644 | 52397 | protein_coding | 2112.824777 | -0.095325873 | 0.060265221 | -1.581772545 | 0.113701505 | 0.310737869 | 0.095325873 |
| ENSMUSG00000024404 | Riok3 | 66878 | protein_coding | 2838.727971 | 0.095332886 | 0.033703566 | 2.828569731 | 0.004675651 | 0.033935297 | 0.095332886 |
| ENSMUSG00000063605 | Ccdc102a | 234582 | protein_coding | 7.550347702 | 0.095371823 | 0.458869883 | 0.207840668 | 0.835353377 | 0.926721051 | 0.095371823 |
| ENSMUSG00000041827 | Oasl1 | 231655 | protein_coding | 321.1758243 | 0.095375344 | 0.09795394 | 0.973675419 | 0.330217726 | 0.57386005 | 0.095375344 |
| ENSMUSG00000016181 | Utp25 | 215193 | protein_coding | 156.7392619 | -0.09539084 | 0.106068 | -0.899336651 | 0.368473371 | 0.610431648 | 0.09539084 |
| ENSMUSG00000050954 | Zfp169 | 67911 | protein_coding | 176.6038267 | -0.095405306 | 0.122388986 | -0.779525264 | 0.435670361 | 0.66908446 | 0.095405306 |
| ENSMUSG00000029999 | Tgfa | 21802 | protein_coding | 987.7754914 | -0.095453742 | 0.061965087 | -1.540443922 | 0.123452182 | 0.326111931 | 0.095453742 |
| ENSMUSG00000036391 | Sec24a | 77371 | protein_coding | 972.8543038 | 0.095459491 | 0.073883343 | 1.292029936 | 0.196346778 | 0.428914823 | 0.095459491 |
| ENSMUSG00000004366 | Sst | 20604 | protein_coding | 128.3738041 | 0.095484759 | 0.127771247 | 0.747310222 | 0.454876324 | 0.682988492 | 0.095484759 |
| ENSMUSG00000060260 | Pwwp2b | 101631 | protein_coding | 413.4800063 | 0.095490906 | 0.077017108 | 1.239866165 | 0.2150249 | 0.45050036 | 0.095490906 |
| ENSMUSG00000041225 | Arhgap12 | 75415 | protein_coding | 2340.666029 | -0.095533354 | 0.035492409 | -2.691655943 | 0.007109825 | 0.046364799 | 0.095533354 |
| ENSMUSG00000039253 | Fn3krp | 238024 | protein_coding | 327.043538 | 0.095610389 | 0.069121115 | 1.383229846 | 0.166594401 | 0.390220465 | 0.095610389 |
| ENSMUSG00000025171 | Ubtd1 | 226122 | protein_coding | 126.4537099 | 0.095616242 | 0.127153851 | 0.751972839 | 0.452067393 | 0.680972431 | 0.095616242 |

| ensembl_gene_id | external_gene_name | entrezgene_id | gene_biotype | baseMean | log2FoldChange | lfcSE | stat | pvalue | padj | absLog2FC |
|---|---|---|---|---|---|---|---|---|---|---|
| ENSMUSG00000002249 | Tead3 | 21678 | protein_coding | 367.2212118 | -0.095635054 | 0.10701371 | -0.893671051 | 0.371497919 | 0.613407078 | 0.095635054 |
| ENSMUSG00000022801 | Lrch3 | 70144 | protein_coding | 538.6387634 | -0.095638987 | 0.057405582 | -1.66602243 | 0.095708947 | 0.280483044 | 0.095638987 |
| ENSMUSG00000078786 | BC024978 | 414069 | protein_coding | 123.9121305 | 0.095645169 | 0.138896549 | 0.68860724 | 0.491070464 | 0.710842238 | 0.095645169 |
| ENSMUSG00000055737 | Ghr | 14600 | protein_coding | 538.4068906 | 0.095666966 | 0.104923695 | 0.911776556 | 0.361886354 | 0.604844153 | 0.095666966 |
| ENSMUSG00000031807 | Pgls | 66171 | protein_coding | 1520.195909 | 0.095673649 | 0.071531918 | 1.337495922 | 0.181060813 | 0.407987666 | 0.095673649 |
| ENSMUSG00000021803 | Cdhr1 | 170677 | protein_coding | 118.4515462 | -0.095678063 | 0.109527805 | -0.873550452 | 0.382363119 | 0.623117424 | 0.095678063 |
| ENSMUSG00000028069 | Gpatch4 | 66614 | protein_coding | 181.8161235 | 0.095684937 | 0.093064551 | 1.028156655 | 0.303876144 | 0.547921576 | 0.095684937 |
| ENSMUSG00000021264 | Yy1 | 22632 | protein_coding | 2513.561031 | -0.095714058 | 0.049814127 | -1.921423975 | 0.054678278 | 0.196434974 | 0.095714058 |
| ENSMUSG00000068876 | Cgn | 70737 | protein_coding | 4100.879679 | 0.09572526 | 0.048878667 | 1.958426156 | 0.050180028 | 0.185651138 | 0.09572526 |
| ENSMUSG00000039473 | Ubn1 | 170644 | protein_coding | 1532.963617 | -0.095727862 | 0.044646238 | -2.144141754 | 0.03202153 | 0.13614849 | 0.095727862 |
| ENSMUSG00000029817 | Tra2a | 101214 | protein_coding | 1155.886375 | 0.095741971 | 0.088134422 | 1.086317571 | 0.277338516 | 0.520134695 | 0.095741971 |
| ENSMUSG00000024947 | Men1 | 17283 | protein_coding | 739.7970902 | 0.095751833 | 0.076575277 | 1.250427519 | 0.211143417 | 0.446279147 | 0.095751833 |
| ENSMUSG00000062470 | Fbxl12os | 66662 | lncRNA | 16.23184872 | -0.095758216 | 0.319504545 | -0.299708461 | 0.764399544 | 0.889299628 | 0.095758216 |
| ENSMUSG00000027195 | Hsd17b12 | 56348 | protein_coding | 1982.845617 | 0.095796001 | 0.065206566 | 1.469115882 | 0.141801361 | 0.35279706 | 0.095796001 |
| ENSMUSG00000005150 | Wdr83 | 67836 | protein_coding | 282.9702744 | 0.095826456 | 0.078789484 | 1.216234078 | 0.223895766 | 0.461675874 | 0.095826456 |
| ENSMUSG00000049539 | H1f1 | 80838 | protein_coding | 1586.57083 | 0.095842155 | 0.123951079 | 0.773225661 | 0.439388842 | 0.671700891 | 0.095842155 |
| ENSMUSG00000018076 | Med13l | 76199 | protein_coding | 1127.970167 | -0.095847203 | 0.058849489 | -1.628683694 | 0.103379996 | 0.293833134 | 0.095847203 |
| ENSMUSG00000079563 | Pglyrp2 | 57757 | protein_coding | 9.006336026 | 0.09586742 | 0.40796263 | 0.234990689 | 0.81421596 | 0.916408517 | 0.09586742 |
| ENSMUSG00000045576 | St7l | 229681 | protein_coding | 600.2273777 | -0.095881465 | 0.06633747 | -1.44535908 | 0.148357051 | 0.362508206 | 0.095881465 |
| ENSMUSG00000095687 | Rnaset2a | 100037283 | protein_coding | 6.878532627 | -0.095889101 | 0.499526792 | -0.191959876 | 0.847773635 | 0.931084542 | 0.095889101 |
| ENSMUSG00000049299 | Trappc1 | 245828 | protein_coding | 636.9635468 | 0.095895319 | 0.06281268 | 1.526687277 | 0.126838793 | 0.331045348 | 0.095895319 |
| ENSMUSG00000095193 | Gm20939 | 100044193 | lncRNA | 63.95273423 | 0.095907729 | 0.16051713 | 0.597492179 | 0.550178827 | 0.751671802 | 0.095907729 |
| ENSMUSG00000024140 | Epas1 | 13819 | protein_coding | 2135.54812 | -0.095909412 | 0.06101227 | -1.571969227 | 0.115957688 | 0.314662099 | 0.095909412 |
| ENSMUSG00000060376 | Bckdha | 12039 | protein_coding | 422.4976045 | -0.095947316 | 0.078336656 | -1.224807388 | 0.220647801 | 0.457199933 | 0.095947316 |
| ENSMUSG00000027303 | Ptpra | 19262 | protein_coding | 1096.84251 | -0.095955577 | 0.047105723 | -2.037025879 | 0.041647449 | 0.163100633 | 0.095955577 |
| ENSMUSG00000046873 | Mbtps2 | 270669 | protein_coding | 295.6377963 | -0.095986365 | 0.085832386 | -1.118300088 | 0.263438849 | 0.504916665 | 0.095986365 |
| ENSMUSG00000031634 | Ufsp2 | 192169 | protein_coding | 860.4063048 | 0.095993436 | 0.05993324 | 1.601672733 | 0.109227998 | 0.302750973 | 0.095993436 |
| ENSMUSG00000017299 | Dnttip1 | 76233 | protein_coding | 446.164987 | 0.09601431 | 0.062610716 | 1.533512418 | 0.125149649 | 0.328283734 | 0.09601431 |
| ENSMUSG00000079429 | Mroh2a | 100040766 | protein_coding | 172.9788646 | -0.096041348 | 0.123721185 | -0.776272459 | 0.437588131 | 0.67075219 | 0.096041348 |
| ENSMUSG00000022742 | Cpox | 12892 | protein_coding | 521.6393339 | 0.096053924 | 0.066714927 | 1.439766608 | 0.149933441 | 0.365277364 | 0.096053924 |
| ENSMUSG00000011034 | Slc5a1 | 20537 | protein_coding | 6022.663108 | -0.096053979 | 0.042117022 | -2.280645087 | 0.022569457 | 0.106976995 | 0.096053979 |
| ENSMUSG00000003234 | Abcf3 | 27406 | protein_coding | 1099.350772 | 0.096058088 | 0.049735419 | 1.931381876 | 0.053435845 | 0.193436559 | 0.096058088 |
| ENSMUSG00000037270 | Bltp1 | 229227 | protein_coding | 2348.178834 | 0.096072874 | 0.07125259 | 1.348342196 | 0.177548347 | 0.404658227 | 0.096072874 |
| ENSMUSG00000014303 | Glis2 | 83396 | protein_coding | 11.91947138 | 0.096081595 | 0.345421416 | 0.278157608 | 0.780891375 | 0.898311362 | 0.096081595 |
| ENSMUSG00000022323 | Rida | 15473 | protein_coding | 372.3896178 | 0.096109752 | 0.066264964 | 1.450385635 | 0.146951011 | 0.360218718 | 0.096109752 |
| ENSMUSG00000044617 | Zbtb39 | 320080 | protein_coding | 429.289149 | -0.096140387 | 0.091473289 | -1.051021435 | 0.293248745 | 0.536272425 | 0.096140387 |
| ENSMUSG00000039968 | Rsbn1l | 242860 | protein_coding | 511.6922774 | -0.096237712 | 0.073164338 | -1.315363673 | 0.188387708 | 0.418734297 | 0.096237712 |
| ENSMUSG00000039100 | Marchf6 | 223455 | protein_coding | 2877.6699 | -0.096259302 | 0.040821712 | -2.358041747 | 0.018371627 | 0.092761185 | 0.096259302 |
| ENSMUSG00000028565 | Nfia | 18027 | protein_coding | 641.2648539 | -0.096298891 | 0.077166766 | -1.247932187 | 0.212055892 | 0.447440166 | 0.096298891 |

| ensembl_gene_id | external_gene_name | entrezgene_id | gene_biotype | baseMean | log2FoldChange | lfcSE | stat | pvalue | padj | absLog2FC |
|---|---|---|---|---|---|---|---|---|---|---|
| ENSMUSG00000007338 | Mrpl49 | 18120 | protein_coding | 788.6021502 | -0.096301653 | 0.061391915 | -1.568637383 | 0.116732457 | 0.316003891 | 0.096301653 |
| ENSMUSG00000047921 | Trappc9 | 76510 | protein_coding | 1027.396552 | 0.096302333 | 0.065949077 | 1.460252927 | 0.144220574 | 0.356724723 | 0.096302333 |
| ENSMUSG00000031255 | Sytl4 | 27359 | protein_coding | 595.167488 | 0.096309692 | 0.071524248 | 1.346532048 | 0.178130992 | 0.405350432 | 0.096309692 |
| ENSMUSG00000028861 | Mrps15 | 66407 | protein_coding | 659.2143993 | 0.096347462 | 0.073946781 | 1.302929773 | 0.192598738 | 0.424322384 | 0.096347462 |
| ENSMUSG00000089669 | Tnfsf13 | 69583 | protein_coding | 203.7982983 | 0.096399956 | 0.118079531 | 0.816398537 | 0.414272233 | 0.652056247 | 0.096399956 |
| ENSMUSG00000023170 | Gps2 | 56310 | protein_coding | 492.9776875 | -0.096403034 | 0.0605382 | -1.592433116 | 0.11128742 | 0.306582205 | 0.096403034 |
| ENSMUSG00000042737 | Dpm3 | 68563 | protein_coding | 442.4892339 | 0.096410354 | 0.106142424 | 0.908311209 | 0.363713822 | 0.606076959 | 0.096410354 |
| ENSMUSG00000026558 | Uck2 | 80914 | protein_coding | 1549.973761 | -0.096412496 | 0.06850563 | -1.407366036 | 0.159318877 | 0.379408267 | 0.096412496 |
| ENSMUSG00000056153 | Socs6 | 54607 | protein_coding | 1168.582916 | 0.096421165 | 0.038068656 | 2.532822944 | 0.011314809 | 0.064914238 | 0.096421165 |
| ENSMUSG00000074794 | Arrdc3 | 105171 | protein_coding | 1696.893083 | 0.096484195 | 0.123964367 | 0.778322007 | 0.436379205 | 0.669599185 | 0.096484195 |
| ENSMUSG00000040945 | Rcc2 | 108911 | protein_coding | 1849.938429 | -0.09651153 | 0.081427993 | -1.185237734 | 0.235923464 | 0.476068282 | 0.09651153 |
| ENSMUSG00000091144 | Phf11c | 628705 | protein_coding | 23.81103061 | -0.096538371 | 0.28598634 | -0.337562875 | 0.735692623 | 0.872904276 | 0.096538371 |
| ENSMUSG00000058331 | Zfp85 | 22746 | protein_coding | 68.28784641 | -0.096552715 | 0.176337053 | -0.547546383 | 0.584003413 | 0.775222412 | 0.096552715 |
| ENSMUSG00000027708 | Dcun1d1 | 114893 | protein_coding | 850.8229543 | -0.096554595 | 0.052452423 | -1.840803342 | 0.065650383 | 0.220801611 | 0.096554595 |
| ENSMUSG00000001482 | Def8 | 23854 | protein_coding | 427.4572347 | 0.096564578 | 0.062979345 | 1.53327377 | 0.125208415 | 0.328321636 | 0.096564578 |
| ENSMUSG00000060678 | H4c3 | 319155 | protein_coding | 5516.316755 | 0.096591692 | 0.078652301 | 1.228084757 | 0.219415153 | 0.455582357 | 0.096591692 |
| ENSMUSG00000047656 | Trpt1 | 107328 | protein_coding | 182.5330176 | -0.096635145 | 0.116015802 | -0.832948125 | 0.404873986 | 0.644152928 | 0.096635145 |
| ENSMUSG00000026922 | Agpat2 | 67512 | protein_coding | 2419.550101 | 0.096688219 | 0.0596272 | 1.621545529 | 0.104900694 | 0.296691247 | 0.096688219 |
| ENSMUSG00000027198 | Ext2 | 14043 | protein_coding | 1206.050667 | 0.096707689 | 0.054619308 | 1.770576958 | 0.076631076 | 0.244015888 | 0.096707689 |
| ENSMUSG00000018433 | Nol11 | 68979 | protein_coding | 476.3521739 | 0.096719252 | 0.066323112 | 1.458303899 | 0.144756796 | 0.357283876 | 0.096719252 |
| ENSMUSG00000034820 | Cpsf7 | 269061 | protein_coding | 804.0339985 | -0.096730054 | 0.066453999 | -1.45559418 | 0.145504838 | 0.358198217 | 0.096730054 |
| ENSMUSG00000040234 | Tm7sf3 | 67623 | protein_coding | 1302.811035 | -0.096733271 | 0.055065346 | -1.756699602 | 0.078969025 | 0.248300541 | 0.096733271 |
| ENSMUSG00000030431 | Tmem238 | 664968 | protein_coding | 671.6306076 | -0.096733603 | 0.122473024 | -0.789835992 | 0.429623556 | 0.664608067 | 0.096733603 |
| ENSMUSG00000044921 | Rassf9 | 237504 | protein_coding | 173.2639525 | 0.096809198 | 0.121697719 | 0.795489012 | 0.4263291 | 0.661733339 | 0.096809198 |
| ENSMUSG00000039990 | Edrf1 | 214764 | protein_coding | 641.118783 | 0.096809213 | 0.083733928 | 1.156152777 | 0.247618672 | 0.488580666 | 0.096809213 |
| ENSMUSG00000024906 | Mus81 | 71711 | protein_coding | 316.3864575 | 0.096813102 | 0.073852335 | 1.310901037 | 0.1898912 | 0.420477604 | 0.096813102 |
| ENSMUSG00000020315 | Sptbn1 | 20742 | protein_coding | 17802.06153 | -0.096828237 | 0.035130209 | -2.756267046 | 0.005846524 | 0.040237686 | 0.096828237 |
| ENSMUSG00000037852 | Cpe | 12876 | protein_coding | 159.9748394 | 0.096887012 | 0.116502678 | 0.831629054 | 0.405618357 | 0.644664161 | 0.096887012 |
| ENSMUSG00000038332 | Sesn1 | 140742 | protein_coding | 426.4251518 | -0.096913204 | 0.103999204 | -0.931864864 | 0.351406368 | 0.593808716 | 0.096913204 |
| ENSMUSG00000025089 | Gfra1 | 14585 | protein_coding | 6.399042355 | -0.096928741 | 0.491773392 | -0.197100418 | 0.843748956 | 0.930175172 | 0.096928741 |
| ENSMUSG00000038528 | Mfsd4b5 | 215928 | protein_coding | 17.30092655 | -0.096933419 | 0.277729134 | -0.349021427 | 0.727073223 | 0.867079041 | 0.096933419 |
| ENSMUSG00000060512 | 0610040J01Rik | 76261 | protein_coding | 1696.42688 | 0.096941667 | 0.050452349 | 1.921450023 | 0.054674997 | 0.196434974 | 0.096941667 |
| ENSMUSG00000020671 | Rab10 | 19325 | protein_coding | 3463.923946 | 0.096946861 | 0.045857453 | 2.114091723 | 0.034507434 | 0.143269254 | 0.096946861 |
| ENSMUSG00000032841 | Prr5l | 72446 | protein_coding | 97.92329789 | 0.096954262 | 0.138782881 | 0.6986039 | 0.484799606 | 0.706263486 | 0.096954262 |
| ENSMUSG00000034640 | Tiparp | 99929 | protein_coding | 574.2644605 | -0.097013687 | 0.078742317 | -1.232040033 | 0.21793413 | 0.454144426 | 0.097013687 |
| ENSMUSG00000018377 | Vezf1 | 22344 | protein_coding | 1016.54907 | -0.097025831 | 0.055695326 | -1.742082126 | 0.081494075 | 0.253277114 | 0.097025831 |
| ENSMUSG00000006589 | Aprt | 11821 | protein_coding | 1675.718489 | -0.097048216 | 0.07022246 | -1.382011047 | 0.166968309 | 0.390767415 | 0.097048216 |
| ENSMUSG00000068663 | Clec16a | 74374 | protein_coding | 675.3573828 | 0.097074309 | 0.050012589 | 1.940997488 | 0.052258582 | 0.190759202 | 0.097074309 |
| ENSMUSG00000029233 | Srd5a3 | 57357 | protein_coding | 372.7602803 | -0.097107732 | 0.066416487 | -1.462102809 | 0.143713039 | 0.355746355 | 0.097107732 |

| ensembl_gene_id | external_gene_name | entrezgene_id | gene_biotype | baseMean | log2FoldChange | lfcSE | stat | pvalue | padj | absLog2FC |
|---|---|---|---|---|---|---|---|---|---|---|
| ENSMUSG00000074698 | Csnk2a1 | 12995 | protein_coding | 2821.917674 | -0.097123262 | 0.039068947 | -2.485945212 | 0.012920786 | 0.071852934 | 0.097123262 |
| ENSMUSG00000029434 | Vps33a | 77573 | protein_coding | 796.6280574 | 0.097147704 | 0.057745636 | 1.682338464 | 0.092503227 | 0.274255759 | 0.097147704 |
| ENSMUSG00000031171 | Ftsj1 | 54632 | protein_coding | 289.6587367 | 0.097184449 | 0.103154551 | 0.942124693 | 0.346128804 | 0.588510197 | 0.097184449 |
| ENSMUSG00000041372 | B4galnt3 | 330406 | protein_coding | 1950.426862 | -0.097202088 | 0.083198794 | -1.16831126 | 0.242681235 | 0.483875678 | 0.097202088 |
| ENSMUSG00000042647 | Acad12 | 338350 | protein_coding | 398.7329717 | -0.097207857 | 0.093884214 | -1.035401504 | 0.300481444 | 0.544051223 | 0.097207857 |
| ENSMUSG00000039431 | Mtmr7 | 54384 | protein_coding | 369.3437202 | 0.09726939 | 0.070912333 | 1.3716851 | 0.170161489 | 0.394462093 | 0.09726939 |
| ENSMUSG00000025862 | Stag2 | 20843 | protein_coding | 3437.464298 | -0.097272791 | 0.058990944 | -1.64894449 | 0.099159007 | 0.286429753 | 0.097272791 |
| ENSMUSG00000026743 | Mllt10 | 17354 | protein_coding | 1004.81128 | -0.097278676 | 0.053354338 | -1.823257091 | 0.068264462 | 0.226290026 | 0.097278676 |
| ENSMUSG00000020919 | Stat5b | 20851 | protein_coding | 667.2244643 | -0.097287398 | 0.058825026 | -1.653843694 | 0.098159275 | 0.284919841 | 0.097287398 |
| ENSMUSG00000019943 | Atp2b1 | 67972 | protein_coding | 6798.450046 | 0.097296621 | 0.064794134 | 1.501627007 | 0.133193465 | 0.340305926 | 0.097296621 |
| ENSMUSG00000020121 | Srgap1 | 117600 | protein_coding | 337.0804432 | -0.097316198 | 0.101755934 | -0.956368781 | 0.338885952 | 0.58240762 | 0.097316198 |
| ENSMUSG00000015341 | Golga7 | 57437 | protein_coding | 785.4601452 | -0.097340489 | 0.059973433 | -1.623060152 | 0.104576547 | 0.29613801 | 0.097340489 |
| ENSMUSG00000035004 | Igsf6 | 80719 | protein_coding | 8.381052382 | 0.097347746 | 0.412869393 | 0.235783391 | 0.813600757 | 0.916202738 | 0.097347746 |
| ENSMUSG00000021097 | Clmn | 94040 | protein_coding | 4230.784349 | -0.097399713 | 0.063928999 | -1.523560741 | 0.127618476 | 0.332327332 | 0.097399713 |
| ENSMUSG00000040471 | Ggt6 | 71522 | protein_coding | 1408.068129 | 0.097418738 | 0.068856715 | 1.414803743 | 0.157126046 | 0.376183454 | 0.097418738 |
| ENSMUSG00000037348 | Paqr7 | 71904 | protein_coding | 88.55162801 | -0.097510604 | 0.143132486 | -0.681261163 | 0.495706253 | 0.714326476 | 0.097510604 |
| ENSMUSG00000036775 | Decr2 | 26378 | protein_coding | 1543.195639 | 0.097558086 | 0.04730532 | 2.062306874 | 0.039178529 | 0.156316709 | 0.097558086 |
| ENSMUSG00000022814 | Umps | 22247 | protein_coding | 642.4384813 | -0.09760476 | 0.058018989 | -1.682289899 | 0.092512639 | 0.274255759 | 0.09760476 |
| ENSMUSG00000048439 | Nupl2 | 231042 | protein_coding | 190.190052 | 0.097606748 | 0.110199064 | 0.885731189 | 0.375762397 | 0.617288396 | 0.097606748 |
| ENSMUSG00000031993 | Snx19 | 102607 | protein_coding | 1082.984765 | 0.097611204 | 0.050207945 | 1.944138605 | 0.051878739 | 0.189587225 | 0.097611204 |
| ENSMUSG00000064145 | Arih2 | 23807 | protein_coding | 1462.42876 | -0.097611889 | 0.052475893 | -1.86012822 | 0.062867387 | 0.214392548 | 0.097611889 |
| ENSMUSG00000032897 | Nfyc | 18046 | protein_coding | 701.9718518 | 0.097638113 | 0.053111434 | 1.838363327 | 0.066008885 | 0.221624129 | 0.097638113 |
| ENSMUSG00000031605 | Klhl2 | 77113 | protein_coding | 890.3273794 | 0.097644546 | 0.058218213 | 1.677216477 | 0.093500158 | 0.276314934 | 0.097644546 |
| ENSMUSG00000029171 | Pgm2 | 66681 | protein_coding | 1859.470334 | -0.097650068 | 0.045069104 | -2.166674272 | 0.0302597 | 0.130971062 | 0.097650068 |
| ENSMUSG00000043252 | Tmem64 | 100201 | protein_coding | 311.3284074 | 0.097676592 | 0.125885308 | 0.775917327 | 0.437797801 | 0.670824076 | 0.097676592 |
| ENSMUSG00000003226 | Ranbp2 | 19386 | protein_coding | 4207.505876 | -0.09770662 | 0.063484177 | -1.539070429 | 0.123787103 | 0.326515312 | 0.09770662 |
| ENSMUSG00000025813 | Homer2 | 26557 | protein_coding | 603.3230146 | -0.097709384 | 0.112746061 | -0.866632353 | 0.386143481 | 0.626177569 | 0.097709384 |
| ENSMUSG00000030967 | Zranb1 | 360216 | protein_coding | 1348.252433 | -0.097740711 | 0.058368717 | -1.674539314 | 0.094024655 | 0.27725095 | 0.097740711 |
| ENSMUSG00000036099 | Vezt | 215008 | protein_coding | 419.1149419 | -0.097760923 | 0.077854573 | -1.255686329 | 0.209229718 | 0.444692306 | 0.097760923 |
| ENSMUSG00000029125 | Stx18 | 71116 | protein_coding | 420.3940803 | 0.097767344 | 0.070855748 | 1.379808237 | 0.167645696 | 0.391604912 | 0.097767344 |
| ENSMUSG00000031533 | Mrps31 | 57312 | protein_coding | 892.2459822 | -0.09776784 | 0.052394534 | -1.865993116 | 0.062042329 | 0.2125678 | 0.09776784 |
| ENSMUSG00000044017 | Adgrd1 | 243277 | protein_coding | 10.5137751 | 0.097773614 | 0.36748188 | 0.266063769 | 0.790190098 | 0.902120505 | 0.097773614 |
| ENSMUSG00000037106 | Fer1l6 | 631797 | protein_coding | 3285.269541 | 0.097803715 | 0.065437964 | 1.494602058 | 0.135018329 | 0.342958039 | 0.097803715 |
| ENSMUSG00000066892 | Fbxl12 | 30843 | protein_coding | 154.6336024 | 0.097803952 | 0.100227156 | 0.975822879 | 0.329152245 | 0.572749056 | 0.097803952 |
| ENSMUSG00000072692 | Rpl37rt | 100502825 | transcribed_processed_pseudogene | 151.5190578 | -0.097807174 | 0.189715264 | -0.515547203 | 0.606170694 | 0.789753 | 0.097807174 |
| ENSMUSG00000030695 | Aldoa | 11674 | protein_coding | 24536.7546 | 0.097854916 | 0.071344624 | 1.371580798 | 0.170193975 | 0.394469421 | 0.097854916 |
| ENSMUSG00000037730 | Mynn | 80732 | protein_coding | 448.3766122 | -0.097876477 | 0.0686577 | -1.425571751 | 0.153991995 | 0.371384959 | 0.097876477 |
| ENSMUSG00000047153 | Khnyn | 219094 | protein_coding | 2218.118684 | -0.097886701 | 0.056301171 | -1.738626361 | 0.082100507 | 0.254517969 | 0.097886701 |
| ENSMUSG00000025016 | Tm9sf3 | 107358 | protein_coding | 12862.13672 | 0.097907814 | 0.032691663 | 2.994886284 | 0.002745472 | 0.022452446 | 0.097907814 |

| ensembl_gene_id | external_gene_name | entrezgene_id | gene_biotype | baseMean | log2FoldChange | lfcSE | stat | pvalue | padj | absLog2FC |
|---|---|---|---|---|---|---|---|---|---|---|
| ENSMUSG00000025280 | Polr3a | 218832 | protein_coding | 544.2718488 | -0.097921881 | 0.06087965 | -1.608450132 | 0.10773663 | 0.300630044 | 0.097921881 |
| ENSMUSG00000022408 | Fam83f | 213956 | protein_coding | 862.8986591 | 0.097935246 | 0.06204614 | 1.578426088 | 0.114467756 | 0.31208616 | 0.097935246 |
| ENSMUSG00000073725 | Lmbrd1 | 68421 | protein_coding | 1467.713556 | -0.097965806 | 0.049826728 | -1.966129644 | 0.049283637 | 0.1835442 | 0.097965806 |
| ENSMUSG00000040998 | Npnt | 114249 | protein_coding | 7.915015324 | 0.098000114 | 0.448074967 | 0.218713656 | 0.826873111 | 0.923060252 | 0.098000114 |
| ENSMUSG00000052776 | Oas1a | 246730 | protein_coding | 714.8763145 | 0.098013029 | 0.09853831 | 0.994669276 | 0.319897142 | 0.563409586 | 0.098013029 |
| ENSMUSG00000038859 | Baiap2l1 | 66898 | protein_coding | 2675.81696 | 0.098017865 | 0.057194334 | 1.713768802 | 0.086571195 | 0.263421446 | 0.098017865 |
| ENSMUSG00000028582 | Cc2d1b | 319965 | protein_coding | 968.9979436 | 0.098030709 | 0.044620775 | 2.196974607 | 0.02802226 | 0.12411837 | 0.098030709 |
| ENSMUSG00000037204 | Atg101 | 68118 | protein_coding | 878.2310809 | 0.098070236 | 0.079699715 | 1.23049669 | 0.218511166 | 0.454666616 | 0.098070236 |
| ENSMUSG00000063383 | Zfp947 | 210853 | protein_coding | 113.438088 | 0.098090141 | 0.154462245 | 0.635042831 | 0.525400481 | 0.735943093 | 0.098090141 |
| ENSMUSG00000021831 | Ero1a | 50527 | protein_coding | 917.4120309 | 0.098092995 | 0.065224817 | 1.503921359 | 0.132601617 | 0.339503983 | 0.098092995 |
| ENSMUSG00000042719 | Naa25 | 231713 | protein_coding | 600.8952387 | -0.09811257 | 0.073555294 | -1.333861437 | 0.182249272 | 0.410035436 | 0.09811257 |
| ENSMUSG00000055305 | Zfp93 | 22755 | protein_coding | 143.497562 | 0.098129424 | 0.122023217 | 0.804186502 | 0.421289273 | 0.658017412 | 0.098129424 |
| ENSMUSG00000038893 | Fam117a | 215512 | protein_coding | 224.5824927 | 0.098217024 | 0.092642428 | 1.060173251 | 0.289065788 | 0.531357026 | 0.098217024 |
| ENSMUSG00000035891 | Cerk | 223753 | protein_coding | 603.5821051 | 0.098225312 | 0.060791235 | 1.615780836 | 0.106141706 | 0.298245186 | 0.098225312 |
| ENSMUSG00000026197 | Zfand2b | 68818 | protein_coding | 634.8684052 | -0.098238048 | 0.057358876 | -1.712691318 | 0.086769343 | 0.263676126 | 0.098238048 |
| ENSMUSG00000042032 | Mat2b | 108645 | protein_coding | 2220.227247 | 0.098248902 | 0.039939114 | 2.459967012 | 0.013894979 | 0.075935046 | 0.098248902 |
| ENSMUSG00000009731 | Kcnd1 | 16506 | protein_coding | 8.961459514 | -0.098257683 | 0.43108216 | -0.227932613 | 0.819698628 | 0.919197481 | 0.098257683 |
| ENSMUSG00000050914 | Ankrd37 | 654824 | protein_coding | 150.0483515 | 0.098273398 | 0.139980888 | 0.702048682 | 0.482648804 | 0.70472411 | 0.098273398 |
| ENSMUSG00000040734 | Ppp1r13l | 333654 | protein_coding | 86.06012706 | 0.098276256 | 0.160746433 | 0.611374413 | 0.540951738 | 0.746550877 | 0.098276256 |
| ENSMUSG00000032667 | Pon2 | 330260 | protein_coding | 3831.130422 | 0.098278491 | 0.033535177 | 2.930609008 | 0.003382983 | 0.026613348 | 0.098278491 |
| ENSMUSG00000045039 | Megf8 | 269878 | protein_coding | 1008.464248 | -0.098287749 | 0.05609945 | -1.752026956 | 0.079769173 | 0.249745186 | 0.098287749 |
| ENSMUSG00000025885 | Myo5b | 17919 | protein_coding | 6976.466234 | 0.098316266 | 0.042654531 | 2.304943086 | 0.021169761 | 0.102159742 | 0.098316266 |
| ENSMUSG00000034617 | Mtrr | 210009 | protein_coding | 246.4418405 | 0.098333007 | 0.083493607 | 1.177730967 | 0.238903878 | 0.479144737 | 0.098333007 |
| ENSMUSG00000015787 | Abo | 80908 | protein_coding | 289.6000575 | 0.098346782 | 0.106869771 | 0.920248829 | 0.357442743 | 0.60052913 | 0.098346782 |
| ENSMUSG00000070972 | Dnajc25 | 72429 | protein_coding | 166.3469249 | 0.098351962 | 0.099556083 | 0.987905101 | 0.323199127 | 0.567163473 | 0.098351962 |
| ENSMUSG00000021189 | Atxn3 | 110616 | protein_coding | 1030.222231 | -0.09837654 | 0.055802027 | -1.762956375 | 0.077907848 | 0.246356835 | 0.09837654 |
| ENSMUSG00000037375 | Hhat | 226861 | protein_coding | 82.31476197 | -0.09841639 | 0.147286444 | -0.668197202 | 0.504007721 | 0.720199042 | 0.09841639 |
| ENSMUSG00000004558 | Ndrg2 | 29811 | protein_coding | 3548.296934 | -0.098432663 | 0.08072274 | -1.219391997 | 0.222695446 | 0.460024268 | 0.098432663 |
| ENSMUSG00000022205 | Sub1 | 20024 | protein_coding | 1350.745183 | 0.09844884 | 0.058173687 | 1.692325933 | 0.090583846 | 0.270928354 | 0.09844884 |
| ENSMUSG00000021216 | Tubal3 | 238463 | protein_coding | 516.1802396 | -0.098455266 | 0.069439176 | -1.417863393 | 0.156230652 | 0.374715872 | 0.098455266 |
| ENSMUSG00000036214 | Polr1has | 76416 | protein_coding | 107.5993309 | 0.098476931 | 0.131077306 | 0.751288949 | 0.452478779 | 0.68117317 | 0.098476931 |
| ENSMUSG00000089712 | Gm15889 | 115486954 | lncRNA | 34.05886317 | 0.09847704 | 0.244112993 | 0.403407615 | 0.686648386 | 0.842345237 | 0.09847704 |
| ENSMUSG00000052062 | Pard3b | 72823 | protein_coding | 939.5678554 | -0.098515182 | 0.063976697 | -1.539860393 | 0.123594387 | 0.326278724 | 0.098515182 |
| ENSMUSG00000022545 | Ercc4 | 50505 | protein_coding | 317.6782984 | -0.098524652 | 0.07283832 | -1.352648596 | 0.17616792 | 0.402701636 | 0.098524652 |
| ENSMUSG00000044312 | Neurog3 | 11925 | protein_coding | 19.02818804 | 0.098526626 | 0.259671825 | 0.379427478 | 0.704370449 | 0.853157579 | 0.098526626 |
| ENSMUSG00000086390 | 1810019D21Rik | 69771 | lncRNA | 176.3330519 | 0.098526861 | 0.144743635 | 0.680699092 | 0.496061911 | 0.714476238 | 0.098526861 |
| ENSMUSG00000050064 | Zfp697 | 242109 | protein_coding | 89.0981503 | -0.09854274 | 0.145196653 | -0.678684647 | 0.497337696 | 0.715355425 | 0.09854274 |
| ENSMUSG00000037134 | Prmt9 | 102182 | protein_coding | 392.4649677 | 0.098551517 | 0.076851273 | 1.282366752 | 0.199714022 | 0.4333093 | 0.098551517 |
| ENSMUSG00000031298 | Adgrg2 | 237175 | protein_coding | 22.94808192 | 0.098571485 | 0.247573843 | 0.398149834 | 0.690519742 | 0.844620107 | 0.098571485 |

Supplementary Table 1

| ensembl_gene_id | external_gene_name | entrezgene_id | gene_biotype | baseMean | log2FoldChange | lfcSE | stat | pvalue | padj | absLog2FC |
|---|---|---|---|---|---|---|---|---|---|---|
| ENSMUSG00000027314 | Dll4 | 54485 | protein_coding | 300.07577 | 0.098591005 | 0.12613773 | 0.781613915 | 0.434441507 | 0.668221346 | 0.098591005 |
| ENSMUSG00000053560 | Ier2 | 15936 | protein_coding | 10475.61915 | 0.098596901 | 0.094372278 | 1.04476551 | 0.296131359 | 0.538897588 | 0.098596901 |
| ENSMUSG00000058729 | Lin9 | 72568 | protein_coding | 189.5149962 | -0.098626763 | 0.121372028 | -0.812598794 | 0.416448123 | 0.654395292 | 0.098626763 |
| ENSMUSG00000029726 | Mepce | 231803 | protein_coding | 928.3313985 | -0.098666334 | 0.069653518 | -1.416530511 | 0.156620237 | 0.375325502 | 0.098666334 |
| ENSMUSG00000007041 | Clic1 | 114584 | protein_coding | 3880.716699 | 0.098675052 | 0.054502894 | 1.810455272 | 0.070225215 | 0.230421574 | 0.098675052 |
| ENSMUSG00000037833 | Sh2d4b | 328381 | protein_coding | 8.959617497 | 0.098677451 | 0.396372778 | 0.248951131 | 0.803398583 | 0.910371624 | 0.098677451 |
| ENSMUSG00000041945 | Mfsd9 | 211798 | protein_coding | 315.7909904 | -0.09868565 | 0.087456807 | -1.128393019 | 0.259153975 | 0.500884007 | 0.09868565 |
| ENSMUSG00000038604 | Ripor1 | 75687 | protein_coding | 109.4765574 | -0.098735291 | 0.136936594 | -0.72102926 | 0.470891513 | 0.695172778 | 0.098735291 |
| ENSMUSG00000026976 | Pax8 | 18510 | protein_coding | 75.23807368 | -0.09874561 | 0.165720579 | -0.595856051 | 0.551271395 | 0.752144991 | 0.09874561 |
| ENSMUSG00000038398 | Upf3a | 67031 | protein_coding | 754.503671 | -0.09879831 | 0.053308252 | -1.853339891 | 0.063833657 | 0.21663413 | 0.09879831 |
| ENSMUSG00000019791 | Hint3 | 66847 | protein_coding | 644.9765697 | 0.09881966 | 0.060281263 | 1.639309711 | 0.101148775 | 0.290022548 | 0.09881966 |
| ENSMUSG00000007989 | Fzd3 | 14365 | protein_coding | 269.9281018 | 0.098822359 | 0.077819395 | 1.269893697 | 0.204122498 | 0.438306743 | 0.098822359 |
| ENSMUSG00000037126 | Psd | 73728 | protein_coding | 116.1955169 | 0.098853348 | 0.138815434 | 0.712120727 | 0.47639002 | 0.699497125 | 0.098853348 |
| ENSMUSG00000035142 | Nubpl | 76826 | protein_coding | 174.6991446 | -0.098877764 | 0.105505865 | -0.937177883 | 0.348667065 | 0.591143162 | 0.098877764 |
| ENSMUSG00000057133 | Chd6 | 71389 | protein_coding | 1178.337717 | -0.098883232 | 0.067981573 | -1.454559341 | 0.145791294 | 0.358583208 | 0.098883232 |
| ENSMUSG00000066233 | Tmem42 | 66079 | protein_coding | 143.6468358 | 0.098925675 | 0.113761618 | 0.869587451 | 0.384525898 | 0.624972092 | 0.098925675 |
| ENSMUSG00000025505 | Tmem80 | 71448 | protein_coding | 231.3330003 | 0.098928132 | 0.089546973 | 1.104762432 | 0.269262543 | 0.511744874 | 0.098928132 |
| ENSMUSG00000035342 | Lzts2 | 226154 | protein_coding | 1713.589253 | -0.098928921 | 0.072114805 | -1.371825402 | 0.170117798 | 0.394462093 | 0.098928921 |
| ENSMUSG00000050890 | Pdik1l | 230809 | protein_coding | 701.2301604 | -0.098946942 | 0.059595168 | -1.660318205 | 0.096850455 | 0.282560439 | 0.098946942 |
| ENSMUSG00000023044 | Csad | 246277 | protein_coding | 735.1475238 | -0.09896481 | 0.065361604 | -1.514112323 | 0.129997354 | 0.335918494 | 0.09896481 |
| ENSMUSG00000020935 | Dcakd | 68087 | protein_coding | 253.9037955 | 0.098983014 | 0.096563489 | 1.025056316 | 0.305336616 | 0.549639002 | 0.098983014 |
| ENSMUSG00000057530 | Ece1 | 230857 | protein_coding | 3468.660843 | -0.098988518 | 0.055295053 | -1.790187593 | 0.073423759 | 0.237154121 | 0.098988518 |
| ENSMUSG00000066568 | Lsm14a | 67070 | protein_coding | 994.5396484 | 0.099036069 | 0.063788395 | 1.552571887 | 0.120525446 | 0.32127502 | 0.099036069 |
| ENSMUSG00000037152 | Ndufc1 | 66377 | protein_coding | 1474.19918 | 0.099059838 | 0.097390831 | 1.017137204 | 0.309088164 | 0.552606516 | 0.099059838 |
| ENSMUSG00000022340 | Sybu | 319613 | protein_coding | 253.7803739 | 0.099066976 | 0.097432047 | 1.016780197 | 0.309258005 | 0.552620924 | 0.099066976 |
| ENSMUSG00000020273 | Papolg | 216578 | protein_coding | 458.9739514 | -0.099133824 | 0.069344785 | -1.429578657 | 0.152837985 | 0.369704242 | 0.099133824 |
| ENSMUSG00000015501 | Hivep2 | 15273 | protein_coding | 669.2393066 | -0.099138798 | 0.054933246 | -1.804714016 | 0.071119445 | 0.232409989 | 0.099138798 |
| ENSMUSG00000040964 | Arhgef10l | 72754 | protein_coding | 1185.515778 | 0.099156557 | 0.070954221 | 1.397472274 | 0.162271598 | 0.383043628 | 0.099156557 |
| ENSMUSG00000000276 | Dgke | 56077 | protein_coding | 225.8213826 | -0.099218223 | 0.078116469 | -1.270131945 | 0.204037634 | 0.438306743 | 0.099218223 |
| ENSMUSG00000022865 | Cxadr | 13052 | protein_coding | 5800.296354 | -0.09922747 | 0.053004974 | -1.872040725 | 0.061200969 | 0.210759777 | 0.09922747 |
| ENSMUSG00000032305 | Fam219b | 78323 | protein_coding | 284.1195752 | -0.099249386 | 0.077870279 | -1.274547706 | 0.202469377 | 0.436583513 | 0.099249386 |
| ENSMUSG00000035666 | Gtf3c4 | 269252 | protein_coding | 803.0607786 | -0.099250531 | 0.051008871 | -1.945750399 | 0.051684729 | 0.189222442 | 0.099250531 |
| ENSMUSG00000028603 | Scp2 | 20280 | protein_coding | 3922.224466 | 0.099263455 | 0.042995415 | 2.308698613 | 0.020960311 | 0.101551787 | 0.099263455 |
| ENSMUSG00000018659 | Pnpo | 103711 | protein_coding | 1481.743116 | 0.099291546 | 0.069500553 | 1.428643972 | 0.153106589 | 0.3701874 | 0.099291546 |
| ENSMUSG00000038538 | Ubn2 | 320538 | protein_coding | 942.8239315 | -0.099303855 | 0.072542906 | -1.368898223 | 0.171031091 | 0.395624278 | 0.099303855 |
| ENSMUSG00000040548 | Tex2 | 21763 | protein_coding | 1449.826986 | 0.099305252 | 0.045770745 | 2.169622785 | 0.030035434 | 0.130315158 | 0.099305252 |
| ENSMUSG00000037552 | Plekhg2 | 101497 | protein_coding | 243.9131619 | -0.0993448 | 0.108649087 | -0.914363872 | 0.360525679 | 0.60375884 | 0.0993448 |
| ENSMUSG00000039155 | Cdh26 | 381409 | protein_coding | 42.94432692 | 0.099348453 | 0.17842172 | 0.556818156 | 0.577651678 | 0.770206405 | 0.099348453 |
| ENSMUSG00000034917 | Tjp3 | 27375 | protein_coding | 7733.621218 | 0.099354724 | 0.061379696 | 1.618690381 | 0.105513897 | 0.297761537 | 0.099354724 |

| ensembl_gene_id | external_gene_name | entrezgene_id | gene_biotype | baseMean | log2FoldChange | lfcSE | stat | pvalue | padj | absLog2FC |
|---|---|---|---|---|---|---|---|---|---|---|
| ENSMUSG00000002778 | Kdelr1 | 68137 | protein_coding | 2429.291124 | 0.099356678 | 0.066894022 | 1.485284865 | 0.137468379 | 0.347158246 | 0.099356678 |
| ENSMUSG00000031303 | Map3k15 | 270672 | protein_coding | 72.82941853 | 0.099399781 | 0.199278001 | 0.49879957 | 0.617920591 | 0.797435496 | 0.099399781 |
| ENSMUSG00000032504 | Pdcd6ip | 18571 | protein_coding | 4984.59472 | 0.099405599 | 0.059493341 | 1.67086934 | 0.094747489 | 0.278626509 | 0.099405599 |
| ENSMUSG00000000776 | Polr3d | 67065 | protein_coding | 385.1699863 | -0.099416826 | 0.096343871 | -1.031895702 | 0.30212098 | 0.545733091 | 0.099416826 |
| ENSMUSG00000001366 | Fbxo9 | 71538 | protein_coding | 1046.983892 | 0.099521951 | 0.051896385 | 1.917704874 | 0.055148443 | 0.197639766 | 0.099521951 |
| ENSMUSG00000011589 | Fsd1 | 240121 | protein_coding | 7.610516347 | -0.099562057 | 0.431662112 | -0.230648125 | 0.817588177 | 0.91812858 | 0.099562057 |
| ENSMUSG00000040857 | Erf | 13875 | protein_coding | 764.5730392 | -0.099589927 | 0.086895099 | -1.146093718 | 0.251756377 | 0.492727559 | 0.099589927 |
| ENSMUSG00000002058 | Unc119 | 22248 | protein_coding | 504.243212 | -0.099598179 | 0.129376699 | -0.769830887 | 0.441400215 | 0.672888818 | 0.099598179 |
| ENSMUSG00000024957 | Kcnk4 | 16528 | protein_coding | 12.32069173 | 0.099607311 | 0.329979145 | 0.301859412 | 0.762759238 | 0.888224293 | 0.099607311 |
| ENSMUSG00000021226 | Acot2 | 171210 | protein_coding | 174.1523262 | 0.099613176 | 0.136996749 | 0.727120731 | 0.467152 | 0.692759569 | 0.099613176 |
| ENSMUSG00000000563 | Atp5pb | 11950 | protein_coding | 8778.419265 | 0.099615879 | 0.050317141 | 1.979760327 | 0.047730466 | 0.179428725 | 0.099615879 |
| ENSMUSG00000020413 | Hus1 | 15574 | protein_coding | 184.4827047 | -0.099646275 | 0.088126908 | -1.130713397 | 0.258175741 | 0.500132153 | 0.099646275 |
| ENSMUSG00000041685 | Fcho2 | 218503 | protein_coding | 1209.196585 | -0.099661377 | 0.072385414 | -1.37681574 | 0.168569223 | 0.392565467 | 0.099661377 |
| ENSMUSG00000020075 | Ddx21 | 56200 | protein_coding | 1843.440722 | -0.099670647 | 0.067537269 | -1.475787334 | 0.140000992 | 0.350881819 | 0.099670647 |
| ENSMUSG00000046804 | Phgr1 | 53906 | protein_coding | 8218.843449 | -0.099696897 | 0.110168967 | -0.904945376 | 0.365494327 | 0.60759958 | 0.099696897 |
| ENSMUSG00000028214 | Gem | 14579 | protein_coding | 19.16559768 | 0.099757166 | 0.326076451 | 0.305931831 | 0.759656549 | 0.886275134 | 0.099757166 |
| ENSMUSG00000017978 | Cadps2 | 320405 | protein_coding | 2186.684236 | -0.099763139 | 0.052295905 | -1.907666358 | 0.056434353 | 0.20073154 | 0.099763139 |
| ENSMUSG00000034748 | Sirt6 | 50721 | protein_coding | 266.738495 | 0.09980029 | 0.086186734 | 1.157954185 | 0.246882734 | 0.487862729 | 0.09980029 |
| ENSMUSG00000020973 | Dnaaf2 | 109065 | protein_coding | 35.95797389 | -0.099807439 | 0.189459149 | -0.526801895 | 0.598331167 | 0.784494856 | 0.099807439 |
| ENSMUSG00000036138 | Acaa1a | 113868 | protein_coding | 2200.669494 | -0.099826229 | 0.056474041 | -1.767648059 | 0.077119758 | 0.244881547 | 0.099826229 |
| ENSMUSG00000032330 | Cox7a2 | 12866 | protein_coding | 6215.693765 | 0.099837342 | 0.061357781 | 1.627134162 | 0.103708607 | 0.294300318 | 0.099837342 |
| ENSMUSG00000064367 | mt-Nd5 | 17721 | protein_coding | 18290.69205 | 0.09983848 | 0.116740547 | 0.855216825 | 0.392431123 | 0.632120905 | 0.09983848 |
| ENSMUSG00000025481 | Urah | 76974 | protein_coding | 218.4085165 | -0.099842259 | 0.115071568 | -0.867653591 | 0.385583999 | 0.625831733 | 0.099842259 |
| ENSMUSG00000021178 | Psmc1 | 19179 | protein_coding | 1982.585704 | 0.099863642 | 0.056099429 | 1.780118682 | 0.07505654 | 0.240742704 | 0.099863642 |
| ENSMUSG00000044927 | H1f10 | 243529 | protein_coding | 15.39811949 | 0.099876715 | 0.305029096 | 0.327433405 | 0.743340104 | 0.877651795 | 0.099876715 |
| ENSMUSG00000018559 | Ctdnep1 | 67181 | protein_coding | 551.6295199 | 0.09989042 | 0.06385922 | 1.564228632 | 0.117763885 | 0.317250963 | 0.09989042 |
| ENSMUSG00000022051 | Bnip3l | 12177 | protein_coding | 1458.910389 | 0.099925689 | 0.048555119 | 2.05798464 | 0.039591608 | 0.157342006 | 0.099925689 |
| ENSMUSG00000007888 | Crlf1 | 12931 | protein_coding | 7.570413759 | 0.09995769 | 0.433791463 | 0.230427979 | 0.817759222 | 0.918128862 | 0.09995769 |
| ENSMUSG00000041278 | Ttc1 | 66827 | protein_coding | 563.6472412 | 0.099975837 | 0.073734398 | 1.355891401 | 0.175133724 | 0.401524931 | 0.099975837 |
| ENSMUSG00000074030 | Exoc8 | 102058 | protein_coding | 932.2790468 | -0.099984591 | 0.061626323 | -1.622433175 | 0.104710631 | 0.296257604 | 0.099984591 |
| ENSMUSG00000035064 | Eef2k | 13631 | protein_coding | 2053.983254 | 0.099985236 | 0.06198973 | 1.612932273 | 0.106759223 | 0.299280839 | 0.099985236 |
| ENSMUSG00000010936 | Vac14 | 234729 | protein_coding | 560.7851693 | 0.099995117 | 0.063685995 | 1.570127262 | 0.116385507 | 0.315453626 | 0.099995117 |
| ENSMUSG00000031657 | Heatr3 | 234549 | protein_coding | 412.8020134 | -0.100034534 | 0.09463603 | -1.057044914 | 0.29049109 | 0.5330518 | 0.100034534 |
| ENSMUSG00000028869 | Gnl2 | 230737 | protein_coding | 685.2477294 | -0.100052177 | 0.067128783 | -1.490451225 | 0.136105632 | 0.345013444 | 0.100052177 |
| ENSMUSG00000044813 | Shb | 230126 | protein_coding | 544.0059044 | 0.100085333 | 0.109270007 | 0.915945149 | 0.359695665 | 0.603018132 | 0.100085333 |
| ENSMUSG00000043336 | Filip1l | 78749 | protein_coding | 753.5564112 | -0.100101559 | 0.107945053 | -0.927338086 | 0.353751026 | 0.596003217 | 0.100101559 |
| ENSMUSG00000028578 | Caap1 | 67770 | protein_coding | 225.424921 | 0.100131022 | 0.089387551 | 1.12018979 | 0.262632894 | 0.504584307 | 0.100131022 |
| ENSMUSG00000044795 | Cyb5d1 | 327951 | protein_coding | 153.9984792 | 0.100158663 | 0.105547477 | 0.948944167 | 0.342649009 | 0.585270749 | 0.100158663 |
| ENSMUSG00000003154 | Foxj2 | 60611 | protein_coding | 440.0900233 | -0.10016592 | 0.103873683 | -0.964305078 | 0.334892996 | 0.578448892 | 0.10016592 |

| ensembl_gene_id | external_gene_name | entrezgene_id | gene_biotype | baseMean | log2FoldChange | lfcSE | stat | pvalue | padj | absLog2FC |
|---|---|---|---|---|---|---|---|---|---|---|
| ENSMUSG00000035401 | Emsy | 233545 | protein_coding | 447.7571035 | -0.100168006 | 0.069645238 | -1.438260651 | 0.150360112 | 0.365794955 | 0.100168006 |
| ENSMUSG00000035569 | Ankrd11 | 77087 | protein_coding | 4161.305427 | -0.100168149 | 0.037799513 | -2.649985209 | 0.008049529 | 0.050867853 | 0.100168149 |
| ENSMUSG00000045319 | Proser2 | 227545 | protein_coding | 294.0444823 | -0.100192986 | 0.089256278 | -1.122531521 | 0.261636513 | 0.503509791 | 0.100192986 |
| ENSMUSG00000066438 | Plekhd1 | 217682 | protein_coding | 109.4744354 | -0.100225759 | 0.127703275 | -0.784833115 | 0.432551422 | 0.666799776 | 0.100225759 |
| ENSMUSG00000047648 | Fbxo30 | 71865 | protein_coding | 337.0242075 | -0.100228435 | 0.080833206 | -1.239941346 | 0.21499709 | 0.45050036 | 0.100228435 |
| ENSMUSG00000017548 | Suz12 | 52615 | protein_coding | 1189.074197 | -0.100240953 | 0.059389736 | -1.68784977 | 0.091440071 | 0.272408741 | 0.100240953 |
| ENSMUSG00000002733 | Plekha3 | 83435 | protein_coding | 960.9163794 | -0.100252437 | 0.061330481 | -1.634626611 | 0.102127346 | 0.291609013 | 0.100252437 |
| ENSMUSG00000044636 | Csrnp2 | 207785 | protein_coding | 140.5722934 | -0.100257474 | 0.14346317 | -0.698837712 | 0.484653457 | 0.706263486 | 0.100257474 |
| ENSMUSG00000057181 | 5730455P16Rik | 70591 | protein_coding | 279.8985515 | -0.100275845 | 0.081740967 | -1.226751388 | 0.219916047 | 0.456081872 | 0.100275845 |
| ENSMUSG00000035580 | Kcnh8 | 211468 | protein_coding | 19.44779366 | 0.100303006 | 0.314101471 | 0.319333129 | 0.749473913 | 0.880475742 | 0.100303006 |
| ENSMUSG00000054474 | Thnsl2 | 232078 | protein_coding | 625.6849067 | 0.10031645 | 0.095948288 | 1.045526216 | 0.29577983 | 0.538805075 | 0.10031645 |
| ENSMUSG00000016995 | Matn4 | 17183 | protein_coding | 10.15802312 | 0.100340371 | 0.40432066 | 0.248170278 | 0.804002662 | 0.910778265 | 0.100340371 |
| ENSMUSG00000053113 | Socs3 | 12702 | protein_coding | 417.4413466 | -0.100345624 | 0.177645227 | -0.564865297 | 0.572165389 | 0.767059504 | 0.100345624 |
| ENSMUSG00000022175 | Lrp10 | 65107 | protein_coding | 6293.530499 | 0.100365697 | 0.078649737 | 1.276109768 | 0.201916719 | 0.435860116 | 0.100365697 |
| ENSMUSG00000030486 | Zfp108 | 54678 | protein_coding | 61.28232191 | -0.100367651 | 0.185738363 | -0.54037114 | 0.588941106 | 0.778448752 | 0.100367651 |
| ENSMUSG00000021948 | Prkcd | 18753 | protein_coding | 4648.861259 | 0.100379648 | 0.055706503 | 1.8019377 | 0.071555208 | 0.233254754 | 0.100379648 |
| ENSMUSG00000091754 | Gm3636 | 100041151 | protein_coding | 5.943887358 | -0.100380017 | 0.51861593 | -0.193553672 | 0.846525377 | 0.930889346 | 0.100380017 |
| ENSMUSG00000029729 | Zkscan1 | 74570 | protein_coding | 2902.083401 | -0.100383908 | 0.055831829 | -1.797969191 | 0.07218189 | 0.234551147 | 0.100383908 |
| ENSMUSG00000010175 | Prox1 | 19130 | protein_coding | 42.25801733 | 0.100394825 | 0.18355517 | 0.546946325 | 0.584415609 | 0.775454214 | 0.100394825 |
| ENSMUSG00000022890 | Atp5j | 11957 | protein_coding | 2633.374197 | 0.100399656 | 0.04449041 | 2.256658385 | 0.024029429 | 0.111677984 | 0.100399656 |
| ENSMUSG00000028699 | Tspan1 | 66805 | protein_coding | 24397.95993 | 0.100422856 | 0.054820439 | 1.831850646 | 0.066973675 | 0.223527413 | 0.100422856 |
| ENSMUSG00000010047 | Hyal2 | 15587 | protein_coding | 875.8033843 | 0.100424302 | 0.063632908 | 1.578181864 | 0.114523835 | 0.31208616 | 0.100424302 |
| ENSMUSG00000016510 | Mtif3 | 76366 | protein_coding | 530.2569136 | 0.100439354 | 0.080632234 | 1.245647663 | 0.212893776 | 0.448543447 | 0.100439354 |
| ENSMUSG00000022489 | Pde1b | 18574 | protein_coding | 145.0274544 | -0.100519018 | 0.117824046 | -0.853128207 | 0.393588211 | 0.633283156 | 0.100519018 |
| ENSMUSG00000023186 | Vwa5a | 67776 | protein_coding | 2092.57454 | -0.100535499 | 0.067326306 | -1.493257322 | 0.135369842 | 0.343658553 | 0.100535499 |
| ENSMUSG00000036822 | Topors | 106021 | protein_coding | 985.0865133 | -0.100580427 | 0.060390378 | -1.665504189 | 0.095812209 | 0.280581712 | 0.100580427 |
| ENSMUSG00000028986 | Klhl7 | 52323 | protein_coding | 479.0244898 | 0.100583068 | 0.07340793 | 1.370193492 | 0.170626509 | 0.395074475 | 0.100583068 |
| ENSMUSG00000095440 | Fignl2 | 668225 | protein_coding | 63.36369628 | -0.100586835 | 0.172925788 | -0.581676315 | 0.560784728 | 0.759088073 | 0.100586835 |
| ENSMUSG00000031996 | Aplp2 | 11804 | protein_coding | 4608.702902 | -0.100599568 | 0.047543163 | -2.11596289 | 0.034347964 | 0.142802169 | 0.100599568 |
| ENSMUSG00000031482 | Slc25a15 | 18408 | protein_coding | 880.3070023 | -0.100635312 | 0.064206185 | -1.567377227 | 0.117026544 | 0.31618145 | 0.100635312 |
| ENSMUSG00000043467 | Zbtb37 | 240869 | protein_coding | 247.8518744 | -0.100655671 | 0.094050271 | -1.070232655 | 0.284514599 | 0.527473335 | 0.100655671 |
| ENSMUSG00000022361 | Zhx1 | 22770 | protein_coding | 964.4948513 | -0.100676968 | 0.057926656 | -1.738007583 | 0.082209478 | 0.254568403 | 0.100676968 |
| ENSMUSG00000033597 | Caskin1 | 268932 | protein_coding | 8.000712012 | -0.100682798 | 0.411088401 | -0.244917632 | 0.806520209 | 0.912178372 | 0.100682798 |
| ENSMUSG00000064254 | Ethe1 | 66071 | protein_coding | 11033.95335 | -0.100686877 | 0.082765382 | -1.216533701 | 0.223781681 | 0.461499638 | 0.100686877 |
| ENSMUSG00000021786 | Oxsm | 71147 | protein_coding | 631.1596716 | 0.100687322 | 0.067910479 | 1.482647789 | 0.138168013 | 0.348161805 | 0.100687322 |
| ENSMUSG00000000420 | Galnt1 | 14423 | protein_coding | 2491.823271 | 0.100758705 | 0.052942205 | 1.903183014 | 0.057016669 | 0.201934601 | 0.100758705 |
| ENSMUSG00000019528 | Gyg | 27357 | protein_coding | 521.0371463 | -0.1008605 | 0.065880457 | -1.530962356 | 0.125778696 | 0.329184199 | 0.1008605 |
| ENSMUSG00000045106 | Ccdc73 | 211936 | protein_coding | 22.91000491 | -0.100881423 | 0.303411889 | -0.332490014 | 0.739519279 | 0.874869968 | 0.100881423 |
| ENSMUSG00000048782 | Insc | 233752 | protein_coding | 44.8554101 | 0.100893891 | 0.184287629 | 0.547480544 | 0.584048633 | 0.775222412 | 0.100893891 |

| ensembl_gene_id | external_gene_name | entrezgene_id | gene_biotype | baseMean | log2FoldChange | lfcSE | stat | pvalue | padj | absLog2FC |
|---|---|---|---|---|---|---|---|---|---|---|
| ENSMUSG00000022607 | Ptk2 | 14083 | protein_coding | 1552.534188 | 0.100899101 | 0.046847795 | 2.153764124 | 0.031258675 | 0.133784989 | 0.100899101 |
| ENSMUSG00000006641 | Slc5a6 | 330064 | protein_coding | 991.9592202 | -0.10090552 | 0.080591397 | -1.252063172 | 0.210546846 | 0.445602229 | 0.10090552 |
| ENSMUSG00000025571 | Tnrc6c | 217351 | protein_coding | 1106.500941 | -0.100914084 | 0.059923234 | -1.68405604 | 0.092170839 | 0.273594537 | 0.100914084 |
| ENSMUSG00000033909 | Usp36 | 72344 | protein_coding | 518.3703154 | -0.100939942 | 0.101433208 | -0.995137042 | 0.319669617 | 0.563143068 | 0.100939942 |
| ENSMUSG00000060279 | Ap2a1 | 11771 | protein_coding | 1286.214258 | 0.100948008 | 0.069073778 | 1.461451965 | 0.143891449 | 0.356019852 | 0.100948008 |
| ENSMUSG00000018733 | Pex12 | 103737 | protein_coding | 158.613777 | 0.100978691 | 0.109084228 | 0.92569469 | 0.354604668 | 0.597067183 | 0.100978691 |
| ENSMUSG00000021428 | Riok1 | 71340 | protein_coding | 369.0993706 | -0.101050907 | 0.093269991 | -1.083423572 | 0.278620458 | 0.521447386 | 0.101050907 |
| ENSMUSG00000024900 | Cpt1a | 12894 | protein_coding | 8303.878728 | -0.101053588 | 0.059950159 | -1.685626703 | 0.091867723 | 0.27324802 | 0.101053588 |
| ENSMUSG00000034947 | Tmem106a | 217203 | protein_coding | 387.3027348 | -0.101058893 | 0.078132174 | -1.293435056 | 0.195860632 | 0.428406946 | 0.101058893 |
| ENSMUSG00000001333 | Sync | 68828 | protein_coding | 17.34568487 | -0.101081428 | 0.278628566 | -0.362781998 | 0.716767741 | 0.861157219 | 0.101081428 |
| ENSMUSG00000048897 | Zfp710 | 209225 | protein_coding | 992.2038307 | 0.101092097 | 0.089324266 | 1.131742825 | 0.257742571 | 0.499713206 | 0.101092097 |
| ENSMUSG00000046688 | Tifa | 211550 | protein_coding | 665.5205666 | -0.101206939 | 0.085859233 | -1.17875429 | 0.238496031 | 0.478803123 | 0.101206939 |
| ENSMUSG00000026104 | Stat1 | 20846 | protein_coding | 412.9392895 | -0.101221413 | 0.156480691 | -0.646862003 | 0.51772126 | 0.73015223 | 0.101221413 |
| ENSMUSG00000018287 | Spag7 | 216873 | protein_coding | 826.393763 | 0.101261159 | 0.053252936 | 1.901513173 | 0.057234829 | 0.20232926 | 0.101261159 |
| ENSMUSG00000043391 | 2510009E07Rik | 72190 | protein_coding | 747.8166156 | 0.101267705 | 0.058046871 | 1.744585061 | 0.081057123 | 0.252369884 | 0.101267705 |
| ENSMUSG00000005936 | Kctd20 | 66989 | protein_coding | 908.2690999 | -0.101274469 | 0.04727962 | -2.142032218 | 0.03219089 | 0.136724381 | 0.101274469 |
| ENSMUSG00000031438 | Rnf128 | 66889 | protein_coding | 5088.272396 | -0.101316709 | 0.051396302 | -1.971284016 | 0.048691399 | 0.181685836 | 0.101316709 |
| ENSMUSG00000032745 | Gpbp1 | 73274 | protein_coding | 1408.525062 | -0.101357128 | 0.064851208 | -1.562918131 | 0.118071852 | 0.31772177 | 0.101357128 |
| ENSMUSG00000007617 | Homer1 | 26556 | protein_coding | 722.950989 | 0.10140058 | 0.067777039 | 1.496090435 | 0.13463009 | 0.342321275 | 0.10140058 |
| ENSMUSG00000033392 | Clasp2 | 76499 | protein_coding | 536.9749834 | -0.101424943 | 0.06067617 | -1.671577872 | 0.094607591 | 0.278419092 | 0.101424943 |
| ENSMUSG00000024287 | Thoc1 | 225160 | protein_coding | 379.2668862 | -0.101434738 | 0.08930017 | -1.135885158 | 0.256004631 | 0.497539971 | 0.101434738 |
| ENSMUSG00000037993 | Dhx38 | 64340 | protein_coding | 1103.9477 | 0.101498371 | 0.059664901 | 1.701140348 | 0.088916636 | 0.267808218 | 0.101498371 |
| ENSMUSG00000021468 | Sptlc1 | 268656 | protein_coding | 1517.191384 | 0.101499111 | 0.045497871 | 2.23085409 | 0.025690796 | 0.116930286 | 0.101499111 |
| ENSMUSG00000060739 | Nsa2 | 59050 | protein_coding | 1377.692 | -0.101509938 | 0.057735288 | -1.75819575 | 0.078714208 | 0.247934382 | 0.101509938 |
| ENSMUSG00000041037 | Irgq | 210146 | protein_coding | 826.9700815 | 0.101510162 | 0.068307889 | 1.486067919 | 0.137261157 | 0.346747943 | 0.101510162 |
| ENSMUSG00000010538 | Tsacc | 76927 | protein_coding | 35.95251204 | -0.101515388 | 0.222638538 | -0.45596503 | 0.648415125 | 0.818405307 | 0.101515388 |
| ENSMUSG00000020794 | Ube2g1 | 67128 | protein_coding | 1094.185212 | -0.101520683 | 0.043911445 | -2.311941276 | 0.02078092 | 0.10100479 | 0.101520683 |
| ENSMUSG00000022602 | Arc | 11838 | protein_coding | 545.5203689 | 0.101533266 | 0.187388285 | 0.541833585 | 0.587933153 | 0.777817716 | 0.101533266 |
| ENSMUSG00000032028 | Nxpe2 | 78252 | protein_coding | 1737.017119 | 0.101567931 | 0.15754017 | 0.64471132 | 0.519114279 | 0.730966208 | 0.101567931 |
| ENSMUSG00000018567 | Gabarap | 56486 | protein_coding | 4809.918297 | 0.101580864 | 0.041712515 | 2.435261057 | 0.014881049 | 0.079756433 | 0.101580864 |
| ENSMUSG00000037214 | Thap1 | 73754 | protein_coding | 159.3893599 | 0.10160865 | 0.094161175 | 1.079092837 | 0.280546344 | 0.523452524 | 0.10160865 |
| ENSMUSG00000021900 | Btd | 26363 | protein_coding | 314.3406705 | 0.101608866 | 0.087667933 | 1.159019756 | 0.246448132 | 0.487365914 | 0.101608866 |
| ENSMUSG00000038690 | Atp5j2 | 57423 | protein_coding | 6365.309529 | 0.101610396 | 0.080210581 | 1.266795413 | 0.205228453 | 0.439737247 | 0.101610396 |
| ENSMUSG00000058704 | Memo1 | 76890 | protein_coding | 635.3067195 | 0.101637071 | 0.062888016 | 1.616159607 | 0.106059809 | 0.298192616 | 0.101637071 |
| ENSMUSG00000029131 | Dnajb6 | 23950 | protein_coding | 789.8660609 | -0.101641935 | 0.061399839 | -1.655410456 | 0.097841266 | 0.284213967 | 0.101641935 |
| ENSMUSG00000032253 | Phip | 83946 | protein_coding | 1436.602601 | -0.101663901 | 0.057704975 | -1.761787465 | 0.078105213 | 0.246739036 | 0.101663901 |
| ENSMUSG00000045594 | Glb1 | 12091 | protein_coding | 2111.311296 | -0.101701662 | 0.033754635 | -3.012968758 | 0.002587055 | 0.021395612 | 0.101701662 |
| ENSMUSG00000020608 | Smc6 | 67241 | protein_coding | 1330.378462 | -0.101703616 | 0.05772202 | -1.761955242 | 0.07807686 | 0.24669779 | 0.101703616 |
| ENSMUSG00000047604 | Frat2 | 212398 | protein_coding | 1013.010481 | -0.101744635 | 0.085783261 | -1.18606631 | 0.235596114 | 0.475764905 | 0.101744635 |

| ensembl_gene_id | external_gene_name | entrezgene_id | gene_biotype | baseMean | log2FoldChange | lfcSE | stat | pvalue | padj | absLog2FC |
|---|---|---|---|---|---|---|---|---|---|---|
| ENSMUSG00000029840 | Mtpn | 14489 | protein_coding | 3728.317596 | 0.101754226 | 0.040979507 | 2.483051501 | 0.013026225 | 0.07233951 | 0.101754226 |
| ENSMUSG00000034190 | Chmp7 | 105513 | protein_coding | 995.8156025 | 0.10177912 | 0.049187798 | 2.069194486 | 0.03852784 | 0.15440817 | 0.10177912 |
| ENSMUSG00000048027 | Rgmb | 68799 | protein_coding | 1679.348233 | -0.10179101 | 0.072112071 | -1.411566865 | 0.158077535 | 0.377840372 | 0.10179101 |
| ENSMUSG00000024878 | Cbwd1 | 226043 | protein_coding | 254.7565215 | 0.101811079 | 0.082571658 | 1.233002721 | 0.217574749 | 0.453747313 | 0.101811079 |
| ENSMUSG00000097404 | Gm10814 | 100502985 | lncRNA | 13.37892569 | -0.101831358 | 0.324286574 | -0.314016573 | 0.753508452 | 0.882806451 | 0.101831358 |
| ENSMUSG00000005881 | Ergic3 | 66366 | protein_coding | 2786.886992 | -0.101845848 | 0.05349205 | -1.903943627 | 0.056917526 | 0.20173823 | 0.101845848 |
| ENSMUSG00000035530 | Eif1 | 20918 | protein_coding | 3445.426758 | -0.101892817 | 0.073853111 | -1.379668574 | 0.167688713 | 0.391604912 | 0.101892817 |
| ENSMUSG00000015839 | Nfe2l2 | 18024 | protein_coding | 5758.776475 | -0.101898412 | 0.056202844 | -1.813047242 | 0.069824535 | 0.229476957 | 0.101898412 |
| ENSMUSG00000015016 | Acsf3 | 257633 | protein_coding | 636.0060546 | 0.101925238 | 0.060909987 | 1.6733748 | 0.094253536 | 0.277610848 | 0.101925238 |
| ENSMUSG00000038268 | Ovca2 | 246257 | protein_coding | 170.639542 | 0.101979648 | 0.103915194 | 0.98137379 | 0.326408446 | 0.569880807 | 0.101979648 |
| ENSMUSG00000028822 | Tmem50a | 71817 | protein_coding | 1345.204297 | -0.101981496 | 0.053485903 | -1.906698583 | 0.056559631 | 0.200957912 | 0.101981496 |
| ENSMUSG00000029911 | Ssbp1 | 381760 | protein_coding | 397.5279738 | 0.101996567 | 0.079726108 | 1.279337097 | 0.200778374 | 0.434462179 | 0.101996567 |
| ENSMUSG00000047797 | Gjb1 | 14618 | protein_coding | 1115.217246 | 0.102000767 | 0.077125965 | 1.322521772 | 0.185994464 | 0.415160239 | 0.102000767 |
| ENSMUSG00000029655 | N4bp2l2 | 381695 | protein_coding | 1231.626414 | -0.10200446 | 0.08267374 | -1.233819346 | 0.217270229 | 0.453522778 | 0.10200446 |
| ENSMUSG00000036503 | Rnf13 | 24017 | protein_coding | 1325.670749 | -0.102022082 | 0.058592287 | -1.741220343 | 0.081644963 | 0.253501793 | 0.102022082 |
| ENSMUSG00000014748 | Tex261 | 21766 | protein_coding | 1872.412481 | -0.102054304 | 0.048806314 | -2.091006158 | 0.036527512 | 0.148719814 | 0.102054304 |
| ENSMUSG00000026196 | Bard1 | 12021 | protein_coding | 104.2910811 | 0.102057612 | 0.137565413 | 0.741884236 | 0.458157478 | 0.685606255 | 0.102057612 |
| ENSMUSG00000022450 | Ndufa6 | 67130 | protein_coding | 4250.187849 | 0.102064987 | 0.059223074 | 1.723399025 | 0.084816391 | 0.259748184 | 0.102064987 |
| ENSMUSG00000074212 | Dnajb14 | 70604 | protein_coding | 674.2004069 | 0.102119158 | 0.078370855 | 1.303024675 | 0.192566338 | 0.424308967 | 0.102119158 |
| ENSMUSG00000005871 | Apc | 11789 | protein_coding | 3084.850722 | -0.102119607 | 0.039176441 | -2.606658585 | 0.009143047 | 0.055884001 | 0.102119607 |
| ENSMUSG00000019578 | Ubxn6 | 66530 | protein_coding | 506.5219661 | 0.102152293 | 0.075887232 | 1.346106453 | 0.178268188 | 0.405548182 | 0.102152293 |
| ENSMUSG00000067071 | Hes6 | 55927 | protein_coding | 1290.55205 | -0.102191047 | 0.090342218 | -1.131154956 | 0.257989877 | 0.499952265 | 0.102191047 |
| ENSMUSG00000019774 | Mtrf1l | 108853 | protein_coding | 195.1758978 | -0.102196511 | 0.093252364 | -1.095913346 | 0.273116698 | 0.515514161 | 0.102196511 |
| ENSMUSG00000048222 | Mfap1b | 100034361 | protein_coding | 236.1310923 | -0.102199498 | 0.085232673 | -1.199064803 | 0.230502749 | 0.469603637 | 0.102199498 |
| ENSMUSG00000019863 | Qrsl1 | 76563 | protein_coding | 382.7857063 | 0.102207582 | 0.073541826 | 1.389788461 | 0.164593123 | 0.386825021 | 0.102207582 |
| ENSMUSG00000053838 | Nudcd3 | 209586 | protein_coding | 1138.353947 | -0.102222922 | 0.05119219 | -1.996846047 | 0.04584191 | 0.174452968 | 0.102222922 |
| ENSMUSG00000033128 | Gga1 | 106039 | protein_coding | 1164.810221 | 0.102224899 | 0.086649448 | 1.179752451 | 0.238098686 | 0.478422316 | 0.102224899 |
| ENSMUSG00000043323 | Fbrsl1 | 381668 | protein_coding | 585.0885491 | -0.102238677 | 0.087334553 | -1.170655516 | 0.241737273 | 0.482707243 | 0.102238677 |
| ENSMUSG00000020260 | Pofut2 | 80294 | protein_coding | 761.0848643 | 0.102240786 | 0.061580024 | 1.660291433 | 0.096855838 | 0.282560439 | 0.102240786 |
| ENSMUSG00000056973 | Ces1d | 104158 | protein_coding | 1236.788458 | -0.102260822 | 0.076047371 | -1.344698967 | 0.178722468 | 0.405874466 | 0.102260822 |
| ENSMUSG00000029551 | Psmg3 | 66506 | protein_coding | 166.5300931 | 0.102271623 | 0.104436363 | 0.979272157 | 0.327445524 | 0.571134973 | 0.102271623 |
| ENSMUSG00000035898 | Uba6 | 231380 | protein_coding | 591.2586242 | 0.102296374 | 0.062648988 | 1.632849582 | 0.102500636 | 0.292221615 | 0.102296374 |
| ENSMUSG00000044221 | Grsf1 | 231413 | protein_coding | 1487.62347 | -0.10231611 | 0.052013299 | -1.967114424 | 0.049170021 | 0.183344491 | 0.10231611 |
| ENSMUSG00000009905 | Kdsr | 70750 | protein_coding | 905.0432749 | -0.10234948 | 0.061438254 | -1.665891734 | 0.09573498 | 0.280508363 | 0.10234948 |
| ENSMUSG00000031403 | Dkc1 | 245474 | protein_coding | 577.7967008 | -0.102355044 | 0.090623117 | -1.129458434 | 0.258704494 | 0.500565425 | 0.102355044 |
| ENSMUSG00000037697 | Ddhd1 | 114874 | protein_coding | 349.7055413 | -0.102363654 | 0.080887595 | -1.265504979 | 0.205690365 | 0.440293139 | 0.102363654 |
| ENSMUSG00000015961 | Adss | 11566 | protein_coding | 1662.678524 | -0.102429683 | 0.040200693 | -2.547958133 | 0.010835547 | 0.06269999 | 0.102429683 |
| ENSMUSG00000042605 | Atxn2 | 20239 | protein_coding | 653.2483785 | -0.102430172 | 0.081526499 | -1.256403424 | 0.208969741 | 0.44452204 | 0.102430172 |
| ENSMUSG00000036948 | Trappc14 | 231807 | protein_coding | 445.6975772 | -0.102449211 | 0.079304818 | -1.291840945 | 0.196412233 | 0.428914823 | 0.102449211 |

| ensembl_gene_id | external_gene_name | entrezgene_id | gene_biotype | baseMean | log2FoldChange | lfcSE | stat | pvalue | padj | absLog2FC |
|---|---|---|---|---|---|---|---|---|---|---|
| ENSMUSG00000051391 | Ywhag | 22628 | protein_coding | 1226.994568 | -0.102469942 | 0.078525219 | -1.304930362 | 0.191916561 | 0.423282055 | 0.102469942 |
| ENSMUSG00000022020 | Naa16 | 66897 | protein_coding | 204.6079971 | -0.102497903 | 0.090809853 | -1.128709057 | 0.259020587 | 0.500746225 | 0.102497903 |
| ENSMUSG00000034528 | Hsd17b13 | 243168 | protein_coding | 380.5871195 | 0.102535446 | 0.158840804 | 0.645523335 | 0.518588101 | 0.730671877 | 0.102535446 |
| ENSMUSG00000030691 | Fchsd2 | 207278 | protein_coding | 1295.363503 | -0.102627111 | 0.078822273 | -1.302006491 | 0.192914167 | 0.424714619 | 0.102627111 |
| ENSMUSG00000034203 | Chchd4 | 72170 | protein_coding | 588.7637757 | 0.102629368 | 0.075129709 | 1.366029097 | 0.171929831 | 0.396668438 | 0.102629368 |
| ENSMUSG00000042642 | Flad1 | 319945 | protein_coding | 817.4934417 | 0.10265741 | 0.06585831 | 1.558761667 | 0.1190528 | 0.319180794 | 0.10265741 |
| ENSMUSG00000056091 | St3gal5 | 20454 | protein_coding | 316.0579147 | 0.102663203 | 0.169786282 | 0.604661354 | 0.545404031 | 0.749069007 | 0.102663203 |
| ENSMUSG00000031458 | Coprs | 66423 | protein_coding | 265.6489921 | -0.102665484 | 0.075422725 | -1.361200933 | 0.173450199 | 0.399148312 | 0.102665484 |
| ENSMUSG00000031758 | Cdyl2 | 75796 | protein_coding | 53.87327235 | -0.102694144 | 0.17837802 | -0.575710753 | 0.564810706 | 0.761533505 | 0.102694144 |
| ENSMUSG00000045316 | Fahd1 | 68636 | protein_coding | 1658.551601 | 0.102713305 | 0.068052077 | 1.509333865 | 0.131213485 | 0.337215278 | 0.102713305 |
| ENSMUSG00000030034 | Ino80b | 70020 | protein_coding | 337.6287535 | 0.10272661 | 0.091836654 | 1.11857962 | 0.263319522 | 0.504916665 | 0.10272661 |
| ENSMUSG00000039981 | Zc3h12d | 237256 | protein_coding | 441.1424853 | 0.102738398 | 0.08513491 | 1.206771674 | 0.227520088 | 0.4661143 | 0.102738398 |
| ENSMUSG00000002064 | Sdf2 | 20316 | protein_coding | 1220.424088 | 0.102748223 | 0.063675202 | 1.61363012 | 0.10660768 | 0.299002096 | 0.102748223 |
| ENSMUSG00000028599 | Tnfrsf1b | 21938 | protein_coding | 392.0120334 | -0.102752022 | 0.080535253 | -1.2758639 | 0.202003634 | 0.435870434 | 0.102752022 |
| ENSMUSG00000019854 | Reps1 | 19707 | protein_coding | 365.6237359 | 0.102760957 | 0.080183585 | 1.281571005 | 0.199993177 | 0.433436829 | 0.102760957 |
| ENSMUSG00000069769 | Msi2 | 76626 | protein_coding | 2262.63657 | 0.10276885 | 0.05074834 | 2.025068213 | 0.04286036 | 0.1666367 | 0.10276885 |
| ENSMUSG00000032531 | Amotl2 | 56332 | protein_coding | 266.3662324 | 0.102797238 | 0.100527321 | 1.022580096 | 0.306506424 | 0.550571297 | 0.102797238 |
| ENSMUSG00000034032 | Rpap1 | 68925 | protein_coding | 648.2610779 | 0.102798607 | 0.077686489 | 1.323249499 | 0.18575242 | 0.415061056 | 0.102798607 |
| ENSMUSG00000024241 | Sos1 | 20662 | protein_coding | 729.9030464 | -0.102800081 | 0.068741014 | -1.495469375 | 0.134791987 | 0.342598955 | 0.102800081 |
| ENSMUSG00000031812 | Map1lc3b | 67443 | protein_coding | 2353.254584 | -0.102849253 | 0.050362046 | -2.04219769 | 0.041131926 | 0.161710035 | 0.102849253 |
| ENSMUSG00000026142 | Rhbdd1 | 76867 | protein_coding | 595.6308222 | 0.102868665 | 0.062101821 | 1.656451664 | 0.097630386 | 0.283793301 | 0.102868665 |
| ENSMUSG00000039050 | Osbpl2 | 228983 | protein_coding | 2055.751955 | 0.102915908 | 0.04689512 | 2.194597398 | 0.02819249 | 0.124462166 | 0.102915908 |
| ENSMUSG00000032584 | Mst1r | 19882 | protein_coding | 1312.760846 | -0.102931031 | 0.067842431 | -1.5172073 | 0.129214354 | 0.334810892 | 0.102931031 |
| ENSMUSG00000042082 | Arsb | 11881 | protein_coding | 430.1141447 | -0.102938014 | 0.069998167 | -1.47058157 | 0.141404311 | 0.352354239 | 0.102938014 |
| ENSMUSG00000022208 | Jph4 | 319984 | protein_coding | 6.952863266 | -0.10298732 | 0.469035928 | -0.219572349 | 0.826204228 | 0.922846345 | 0.10298732 |
| ENSMUSG00000044950 | Pwwp2a | 70802 | protein_coding | 464.3495002 | -0.102988218 | 0.081028469 | -1.271012755 | 0.203724111 | 0.437979805 | 0.102988218 |
| ENSMUSG00000033228 | Scaf11 | 72193 | protein_coding | 2553.247336 | -0.103061642 | 0.039613495 | -2.601680094 | 0.009276834 | 0.056413087 | 0.103061642 |
| ENSMUSG00000021703 | Serinc5 | 218442 | protein_coding | 2784.106585 | -0.103076177 | 0.057903643 | -1.780132858 | 0.07505422 | 0.240742704 | 0.103076177 |
| ENSMUSG00000026429 | Ube2t | 67196 | protein_coding | 80.46475523 | 0.103130058 | 0.145269944 | 0.709920135 | 0.477753663 | 0.700302975 | 0.103130058 |
| ENSMUSG00000034538 | Zfp418 | 232854 | protein_coding | 73.94693718 | -0.103149099 | 0.151201338 | -0.682197002 | 0.49511439 | 0.713812781 | 0.103149099 |
| ENSMUSG00000024683 | Mrpl16 | 94063 | protein_coding | 471.4688689 | 0.10324499 | 0.075122847 | 1.374348742 | 0.169333441 | 0.393611906 | 0.10324499 |
| ENSMUSG00000039640 | Mrpl12 | 56282 | protein_coding | 1745.813242 | 0.103269775 | 0.058274578 | 1.772123951 | 0.076373984 | 0.243463775 | 0.103269775 |
| ENSMUSG00000021040 | Slirp | 380773 | protein_coding | 176.5088869 | 0.103285026 | 0.14176059 | 0.728587725 | 0.466253889 | 0.692064102 | 0.103285026 |
| ENSMUSG00000068551 | Zfp467 | 68910 | protein_coding | 708.6986187 | -0.103305492 | 0.076366722 | -1.352755363 | 0.176133797 | 0.402680713 | 0.103305492 |
| ENSMUSG00000064177 | Ghrl | 58991 | protein_coding | 14.35635407 | -0.103309432 | 0.394855878 | -0.261638329 | 0.793600291 | 0.90428827 | 0.103309432 |
| ENSMUSG00000033055 | Ankrd54 | 223690 | protein_coding | 348.3009102 | 0.103338831 | 0.098527464 | 1.048832751 | 0.294255101 | 0.537241256 | 0.103338831 |
| ENSMUSG00000063904 | Dpp3 | 75221 | protein_coding | 3451.024044 | 0.103348754 | 0.06309296 | 1.6380394 | 0.101413475 | 0.290238707 | 0.103348754 |
| ENSMUSG00000030649 | Anapc15 | 75430 | protein_coding | 101.1740919 | -0.103356871 | 0.120653877 | -0.856639446 | 0.391644179 | 0.631478271 | 0.103356871 |
| ENSMUSG00000032463 | Faim | 23873 | protein_coding | 197.2194177 | -0.103381731 | 0.084666785 | -1.221042361 | 0.222069981 | 0.459085065 | 0.103381731 |

| ensembl_gene_id | external_gene_name | entrezgene_id | gene_biotype | baseMean | log2FoldChange | lfcSE | stat | pvalue | padj | absLog2FC |
|---|---|---|---|---|---|---|---|---|---|---|
| ENSMUSG00000022750 | Klhl22 | 224023 | protein_coding | 866.9193873 | 0.103392827 | 0.05052004 | 2.046570592 | 0.040700265 | 0.160521685 | 0.103392827 |
| ENSMUSG00000052962 | Mrpl35 | 66223 | protein_coding | 547.6067161 | 0.103398039 | 0.062835132 | 1.645545027 | 0.099857465 | 0.287678519 | 0.103398039 |
| ENSMUSG00000002324 | Rec8 | 56739 | protein_coding | 65.54027566 | -0.103436119 | 0.163984024 | -0.630769489 | 0.528191256 | 0.737884649 | 0.103436119 |
| ENSMUSG00000038696 | Mapkap1 | 227743 | protein_coding | 1709.367379 | 0.103437994 | 0.046657637 | 2.216957405 | 0.026626002 | 0.119243971 | 0.103437994 |
| ENSMUSG00000026343 | Gpr39 | 71111 | protein_coding | 193.7194979 | 0.103487009 | 0.089867556 | 1.151550283 | 0.249505921 | 0.490346331 | 0.103487009 |
| ENSMUSG00000026072 | Il1r1 | 16177 | protein_coding | 766.1927093 | 0.103507765 | 0.074005031 | 1.398658493 | 0.161915417 | 0.382604713 | 0.103507765 |
| ENSMUSG00000035673 | Sbno2 | 216161 | protein_coding | 1641.837518 | -0.103519215 | 0.05939283 | -1.742958106 | 0.081340933 | 0.252899642 | 0.103519215 |
| ENSMUSG00000019998 | Stx7 | 53331 | protein_coding | 2056.999284 | 0.103524953 | 0.044697428 | 2.316127727 | 0.020551296 | 0.100099894 | 0.103524953 |
| ENSMUSG00000014243 | Zswim7 | 69747 | protein_coding | 137.4601393 | -0.103554197 | 0.103310985 | -1.002354169 | 0.316172569 | 0.560212157 | 0.103554197 |
| ENSMUSG00000024955 | Esrra | 26379 | protein_coding | 4300.205703 | 0.103594251 | 0.076009692 | 1.362908448 | 0.172911366 | 0.398306185 | 0.103594251 |
| ENSMUSG00000051000 | Fhip1a | 229488 | protein_coding | 647.0327026 | 0.103633584 | 0.071043849 | 1.458727045 | 0.144640249 | 0.357160205 | 0.103633584 |
| ENSMUSG00000047789 | Slc38a9 | 268706 | protein_coding | 198.9157373 | -0.103643661 | 0.128426972 | -0.807024093 | 0.419652603 | 0.657004431 | 0.103643661 |
| ENSMUSG00000026126 | Ptpn18 | 19253 | protein_coding | 984.4999502 | 0.103717146 | 0.050882203 | 2.038377645 | 0.04151218 | 0.162728957 | 0.103717146 |
| ENSMUSG00000031428 | Zcchc18 | 66995 | protein_coding | 5.914014497 | -0.103748189 | 0.509568764 | -0.203599979 | 0.83866611 | 0.928416279 | 0.103748189 |
| ENSMUSG00000024260 | Sap130 | 269003 | protein_coding | 632.7740805 | 0.103767858 | 0.064162761 | 1.617259857 | 0.105822201 | 0.297926072 | 0.103767858 |
| ENSMUSG00000023150 | Ivns1abp | 117198 | protein_coding | 8623.957061 | 0.103782616 | 0.059756633 | 1.736754756 | 0.082430468 | 0.254910098 | 0.103782616 |
| ENSMUSG00000055943 | Emc7 | 73024 | protein_coding | 936.698344 | 0.103791456 | 0.05334271 | 1.945747703 | 0.051685053 | 0.189222442 | 0.103791456 |
| ENSMUSG00000067873 | Htatsf1 | 72459 | protein_coding | 723.8599848 | 0.103795449 | 0.081018939 | 1.281125755 | 0.200149499 | 0.433554193 | 0.103795449 |
| ENSMUSG00000027722 | Spata5 | 57815 | protein_coding | 346.8631293 | -0.103799143 | 0.088720741 | -1.169953526 | 0.242019672 | 0.482991121 | 0.103799143 |
| ENSMUSG00000069135 | Cep43 | 75296 | protein_coding | 327.0405555 | -0.103844947 | 0.091999733 | -1.128752702 | 0.25900217 | 0.500746225 | 0.103844947 |
| ENSMUSG00000026603 | Smyd2 | 226830 | protein_coding | 414.1998081 | -0.103855593 | 0.074142396 | -1.400758522 | 0.161286296 | 0.381891659 | 0.103855593 |
| ENSMUSG00000040524 | Zfp609 | 214812 | protein_coding | 494.1882802 | -0.103862646 | 0.093764658 | -1.107695037 | 0.267993543 | 0.510504592 | 0.103862646 |
| ENSMUSG00000057789 | Bak1 | 12018 | protein_coding | 1739.768703 | 0.103863267 | 0.099165648 | 1.04737143 | 0.294928303 | 0.537983118 | 0.103863267 |
| ENSMUSG00000030725 | Lipt2 | 67164 | protein_coding | 254.5625104 | 0.103874956 | 0.08598197 | 1.208101607 | 0.227008182 | 0.465615713 | 0.103874956 |
| ENSMUSG00000056999 | Ide | 15925 | protein_coding | 1360.632494 | 0.103919667 | 0.046866723 | 2.217344427 | 0.026599564 | 0.119191767 | 0.103919667 |
| ENSMUSG00000020283 | Pex13 | 72129 | protein_coding | 836.5183737 | -0.10394578 | 0.05204902 | -1.997074665 | 0.045817073 | 0.174452968 | 0.10394578 |
| ENSMUSG00000066880 | Zfp617 | 170938 | protein_coding | 295.5113773 | -0.103952394 | 0.111131006 | -0.935404056 | 0.349580107 | 0.592006307 | 0.103952394 |
| ENSMUSG00000102805 | Gm37240 | 99889 | protein_coding | 11.49552514 | -0.103952404 | 0.356313796 | -0.291743978 | 0.770482382 | 0.892047592 | 0.103952404 |
| ENSMUSG00000084128 | Esrp2 | 77411 | protein_coding | 1618.572433 | 0.103980914 | 0.055835628 | 1.862268204 | 0.062565295 | 0.213633392 | 0.103980914 |
| ENSMUSG00000036686 | Cc2d1a | 212139 | protein_coding | 1372.929763 | 0.104017345 | 0.056416915 | 1.843726216 | 0.06522305 | 0.219914726 | 0.104017345 |
| ENSMUSG00000062410 | Hsd3b3 | 15494 | protein_coding | 3420.133741 | -0.104067237 | 0.064191936 | -1.621188643 | 0.104977188 | 0.296855533 | 0.104067237 |
| ENSMUSG00000041375 | Ccdc9 | 243846 | protein_coding | 1329.579324 | -0.104069246 | 0.055093367 | -1.88896145 | 0.058896994 | 0.20590328 | 0.104069246 |
| ENSMUSG00000024910 | Ctsw | 13041 | protein_coding | 74.77416359 | 0.104072472 | 0.146205619 | 0.711822659 | 0.476574599 | 0.699528403 | 0.104072472 |
| ENSMUSG00000031266 | Gla | 11605 | protein_coding | 453.9661174 | 0.104123188 | 0.068858167 | 1.512139976 | 0.130498257 | 0.336485813 | 0.104123188 |
| ENSMUSG00000037617 | Spag1 | 26942 | protein_coding | 38.39168398 | -0.104148081 | 0.249001796 | -0.418262368 | 0.6757553 | 0.835024575 | 0.104148081 |
| ENSMUSG00000030532 | Hddc3 | 68695 | protein_coding | 51.0570364 | 0.104155896 | 0.161136713 | 0.646382152 | 0.518031896 | 0.730398705 | 0.104155896 |
| ENSMUSG00000048078 | Tenm4 | 23966 | protein_coding | 286.6376351 | 0.104175135 | 0.093056468 | 1.119483011 | 0.262934135 | 0.504855347 | 0.104175135 |
| ENSMUSG00000047109 | Cldn14 | 56173 | protein_coding | 41.91575625 | -0.104176736 | 0.253802915 | -0.410463119 | 0.681466253 | 0.838995821 | 0.104176736 |
| ENSMUSG00000026623 | Lpgat1 | 226856 | protein_coding | 1300.164391 | 0.104245896 | 0.038400839 | 2.714677547 | 0.006634033 | 0.044045718 | 0.104245896 |

| ensembl_gene_id | external_gene_name | entrezgene_id | gene_biotype | baseMean | log2FoldChange | lfcSE | stat | pvalue | padj | absLog2FC |
|---|---|---|---|---|---|---|---|---|---|---|
| ENSMUSG00000078249 | Hmga1b | 111241 | protein_coding | 10.73268 | 0.104249096 | 0.351763893 | 0.296360991 | 0.766954413 | 0.89053739 | 0.104249096 |
| ENSMUSG00000027495 | Fam210b | 67017 | protein_coding | 741.2482962 | -0.104255096 | 0.064084248 | -1.626844341 | 0.103770162 | 0.294423187 | 0.104255096 |
| ENSMUSG00000038732 | Mboat1 | 218121 | protein_coding | 5735.268256 | 0.104255909 | 0.086301482 | 1.20804308 | 0.227030692 | 0.465615713 | 0.104255909 |
| ENSMUSG00000023169 | Slc38a1 | 105727 | protein_coding | 1126.776357 | 0.104256037 | 0.051641413 | 2.018845562 | 0.043503274 | 0.168526855 | 0.104256037 |
| ENSMUSG00000023249 | Parp3 | 235587 | protein_coding | 130.7378395 | -0.104264731 | 0.196981696 | -0.529311776 | 0.596589188 | 0.783232176 | 0.104264731 |
| ENSMUSG00000021140 | Pcnx | 54604 | protein_coding | 1211.058102 | -0.104267826 | 0.053382643 | -1.953215892 | 0.050794019 | 0.187149906 | 0.104267826 |
| ENSMUSG00000024145 | Pigf | 18701 | protein_coding | 304.0034627 | -0.10431209 | 0.074989666 | -1.391019526 | 0.164219506 | 0.38635024 | 0.10431209 |
| ENSMUSG00000034255 | Arhgap27 | 544817 | protein_coding | 2073.80969 | -0.10432422 | 0.082826089 | -1.259557477 | 0.207829043 | 0.443094402 | 0.10432422 |
| ENSMUSG00000028004 | Npy2r | 18167 | protein_coding | 22.78093374 | -0.104359682 | 0.280062703 | -0.372629704 | 0.709424062 | 0.856166251 | 0.104359682 |
| ENSMUSG00000117048 | Gm41584 | 105246280 | lncRNA | 6.480461603 | 0.104370031 | 0.50029192 | 0.208618263 | 0.834746252 | 0.926650089 | 0.104370031 |
| ENSMUSG00000038633 | Degs1 | 13244 | protein_coding | 370.9612044 | 0.104370341 | 0.093085479 | 1.121231177 | 0.262189472 | 0.504092706 | 0.104370341 |
| ENSMUSG00000020482 | Ccdc117 | 104479 | protein_coding | 646.1848402 | -0.104408871 | 0.056611581 | -1.844302349 | 0.065139089 | 0.21967756 | 0.104408871 |
| ENSMUSG00000053641 | Dennd4a | 102442 | protein_coding | 293.2233494 | -0.104422532 | 0.086876592 | -1.201963947 | 0.229377497 | 0.468724137 | 0.104422532 |
| ENSMUSG00000038256 | Bcl9 | 77578 | protein_coding | 272.2584076 | -0.104423431 | 0.09560245 | -1.092267313 | 0.274715623 | 0.517140055 | 0.104423431 |
| ENSMUSG00000021127 | Zfp36l1 | 12192 | protein_coding | 663.4354608 | -0.104448598 | 0.07360182 | -1.419103473 | 0.155868851 | 0.374284837 | 0.104448598 |
| ENSMUSG00000070327 | Rnf213 | 672511 | protein_coding | 3234.992395 | -0.104452486 | 0.078125966 | -1.336975293 | 0.181230702 | 0.408256395 | 0.104452486 |
| ENSMUSG00000027860 | Vangl1 | 229658 | protein_coding | 296.8400888 | -0.104493661 | 0.08152316 | -1.281766568 | 0.199924545 | 0.433436829 | 0.104493661 |
| ENSMUSG00000055923 | Aasdh | 231326 | protein_coding | 204.3100079 | -0.104531653 | 0.100122059 | -1.044042177 | 0.296465876 | 0.539262936 | 0.104531653 |
| ENSMUSG00000003208 | Yju2 | 72886 | protein_coding | 193.065644 | 0.104555535 | 0.085238721 | 1.226620176 | 0.219965382 | 0.456081872 | 0.104555535 |
| ENSMUSG00000029253 | Cenpc1 | 12617 | protein_coding | 478.351256 | 0.104577181 | 0.072271573 | 1.447002977 | 0.14789609 | 0.361765545 | 0.104577181 |
| ENSMUSG00000036580 | Spg20 | 229285 | protein_coding | 307.1639141 | 0.104581662 | 0.078295917 | 1.33572306 | 0.181639809 | 0.408949491 | 0.104581662 |
| ENSMUSG00000057058 | Skap1 | 78473 | protein_coding | 24.27139328 | -0.104597421 | 0.297038745 | -0.352133931 | 0.724737823 | 0.865841503 | 0.104597421 |
| ENSMUSG00000038013 | Wipf2 | 68524 | protein_coding | 947.0360195 | -0.104647266 | 0.067577969 | -1.548541148 | 0.121492063 | 0.322997444 | 0.104647266 |
| ENSMUSG00000032329 | Hmg20a | 66867 | protein_coding | 486.5931799 | -0.10465718 | 0.076989753 | -1.359365053 | 0.174030939 | 0.399914072 | 0.10465718 |
| ENSMUSG00000024539 | Ptpn2 | 19255 | protein_coding | 716.9112158 | 0.104723667 | 0.056349714 | 1.858459598 | 0.063103776 | 0.214828885 | 0.104723667 |
| ENSMUSG00000014245 | Pigl | 327942 | protein_coding | 147.597803 | 0.104736302 | 0.112087672 | 0.934414107 | 0.350090323 | 0.592368513 | 0.104736302 |
| ENSMUSG00000001910 | Nacc1 | 66830 | protein_coding | 2216.690742 | 0.104780846 | 0.06916042 | 1.515040628 | 0.129762116 | 0.335814568 | 0.104780846 |
| ENSMUSG00000025156 | Gps1 | 209318 | protein_coding | 1225.645565 | 0.104821999 | 0.069695924 | 1.503990376 | 0.132583845 | 0.339503983 | 0.104821999 |
| ENSMUSG00000035236 | Scai | 320271 | protein_coding | 151.8381249 | -0.104824934 | 0.141442373 | -0.74111408 | 0.458624273 | 0.685788934 | 0.104824934 |
| ENSMUSG00000024013 | Fgd2 | 26382 | protein_coding | 42.34845238 | 0.104846558 | 0.192389552 | 0.544970122 | 0.585774067 | 0.776292513 | 0.104846558 |
| ENSMUSG00000020638 | Cmpk2 | 22169 | protein_coding | 280.2830065 | -0.104860333 | 0.136131867 | -0.770284986 | 0.441130861 | 0.672729777 | 0.104860333 |
| ENSMUSG00000047496 | Rnf152 | 320311 | protein_coding | 527.6974406 | -0.1048647 | 0.108435751 | -0.967067586 | 0.333510248 | 0.576784962 | 0.1048647 |
| ENSMUSG00000037316 | Bag4 | 67384 | protein_coding | 616.5495583 | -0.104920703 | 0.059422766 | -1.76566507 | 0.077452056 | 0.245557111 | 0.104920703 |
| ENSMUSG00000025140 | Pycr1 | 209027 | protein_coding | 97.6900751 | 0.104959768 | 0.154692981 | 0.678503753 | 0.497452345 | 0.71539272 | 0.104959768 |
| ENSMUSG00000045103 | Dmd | 13405 | protein_coding | 483.3514333 | 0.104964119 | 0.073173038 | 1.434464414 | 0.151439777 | 0.3675902 | 0.104964119 |
| ENSMUSG00000062797 | Hikeshi | 67669 | protein_coding | 346.3945462 | 0.104965723 | 0.077960717 | 1.34639248 | 0.178175975 | 0.405395591 | 0.104965723 |
| ENSMUSG00000017774 | Myo1c | 17913 | protein_coding | 2109.212417 | -0.104970532 | 0.049038564 | -2.140571087 | 0.032308643 | 0.137116178 | 0.104970532 |
| ENSMUSG00000001569 | Nom1 | 433864 | protein_coding | 780.7151397 | -0.105001432 | 0.066525647 | -1.578360177 | 0.114482888 | 0.31208616 | 0.105001432 |
| ENSMUSG00000045757 | Zfp764 | 233893 | protein_coding | 211.3844571 | -0.10504913 | 0.089321357 | -1.176080766 | 0.239562603 | 0.480047975 | 0.10504913 |

| ensembl_gene_id | external_gene_name | entrezgene_id | gene_biotype | baseMean | log2FoldChange | lfcSE | stat | pvalue | padj | absLog2FC |
|---|---|---|---|---|---|---|---|---|---|---|
| ENSMUSG00000034330 | Plcg2 | 234779 | protein_coding | 317.4351317 | -0.105066525 | 0.08825279 | -1.190517886 | 0.233842904 | 0.473349371 | 0.105066525 |
| ENSMUSG00000020775 | Mrpl38 | 60441 | protein_coding | 773.247234 | 0.105083241 | 0.095900923 | 1.09574796 | 0.273189088 | 0.515530122 | 0.105083241 |
| ENSMUSG00000054823 | Nsd3 | 234135 | protein_coding | 1017.125827 | -0.105133277 | 0.048734899 | -2.157248287 | 0.030986324 | 0.133044847 | 0.105133277 |
| ENSMUSG00000021519 | Mterf3 | 66410 | protein_coding | 378.7798122 | -0.105153996 | 0.084620929 | -1.242647619 | 0.213997714 | 0.449718678 | 0.105153996 |
| ENSMUSG00000039099 | Wdr93 | 626359 | protein_coding | 10.0917364 | -0.105183506 | 0.443440204 | -0.237198849 | 0.81250253 | 0.915797337 | 0.105183506 |
| ENSMUSG00000019647 | Sema6a | 20358 | protein_coding | 488.0893107 | -0.105188983 | 0.114877071 | -0.91566561 | 0.359842307 | 0.603052779 | 0.105188983 |
| ENSMUSG00000012483 | Rpa3 | 68240 | protein_coding | 290.9046843 | 0.105218935 | 0.083061218 | 1.266763686 | 0.2052398 | 0.439737247 | 0.105218935 |
| ENSMUSG00000030101 | Sumf1 | 58911 | protein_coding | 829.5279285 | 0.105237718 | 0.054593285 | 1.927667788 | 0.053896461 | 0.194492779 | 0.105237718 |
| ENSMUSG00000023829 | Slc22a1 | 20517 | protein_coding | 894.6073677 | -0.105253847 | 0.06101979 | -1.724913309 | 0.084543097 | 0.259255851 | 0.105253847 |
| ENSMUSG00000024891 | Slc29a2 | 13340 | protein_coding | 35.24216155 | -0.105300918 | 0.2132715 | -0.493741161 | 0.621488992 | 0.800250831 | 0.105300918 |
| ENSMUSG00000040813 | Tex264 | 21767 | protein_coding | 1381.796015 | -0.105312247 | 0.091763274 | -1.147651372 | 0.251112513 | 0.492145102 | 0.105312247 |
| ENSMUSG00000026088 | Mitd1 | 69028 | protein_coding | 330.3005394 | -0.10532328 | 0.087942255 | -1.197641341 | 0.231056674 | 0.469953022 | 0.10532328 |
| ENSMUSG00000013646 | Sh3bp5l | 79566 | protein_coding | 863.519958 | 0.105347163 | 0.060760667 | 1.733805264 | 0.082952641 | 0.256181012 | 0.105347163 |
| ENSMUSG00000026819 | Slc25a25 | 227731 | protein_coding | 1459.79084 | -0.105365287 | 0.093234487 | -1.130110646 | 0.258429604 | 0.500383507 | 0.105365287 |
| ENSMUSG00000028896 | Rcc1 | 100088 | protein_coding | 745.0541364 | 0.105414127 | 0.077115141 | 1.36697055 | 0.171634537 | 0.396321966 | 0.105414127 |
| ENSMUSG00000052087 | Rgs14 | 51791 | protein_coding | 52.76356936 | 0.105423131 | 0.162758794 | 0.647726174 | 0.517162072 | 0.729722934 | 0.105423131 |
| ENSMUSG00000087175 | Gm15133 | 622070 | lncRNA | 26.77509044 | 0.105490065 | 0.278577449 | 0.378674101 | 0.704929888 | 0.853291624 | 0.105490065 |
| ENSMUSG00000030663 | 1110004F10Rik | 56372 | protein_coding | 877.0737439 | 0.105536565 | 0.048054493 | 2.19618516 | 0.028078693 | 0.12424678 | 0.105536565 |
| ENSMUSG00000042743 | Sgtb | 218544 | protein_coding | 61.99734239 | -0.105539617 | 0.160010991 | -0.659577301 | 0.509525125 | 0.72436245 | 0.105539617 |
| ENSMUSG00000065304 | Gm23245 | 115490273 | snoRNA | 7.137053341 | -0.105550254 | 0.514955006 | -0.204969856 | 0.837595676 | 0.927834878 | 0.105550254 |
| ENSMUSG00000007659 | Bcl2l1 | 12048 | protein_coding | 921.6549234 | 0.105568158 | 0.049877734 | 2.116538783 | 0.03429901 | 0.142635415 | 0.105568158 |
| ENSMUSG00000000861 | Bcl11a | 14025 | protein_coding | 100.9104012 | -0.105581954 | 0.127098728 | -0.830708186 | 0.406138501 | 0.645300059 | 0.105581954 |
| ENSMUSG00000029364 | Wsb2 | 59043 | protein_coding | 1790.289616 | -0.105637528 | 0.04569502 | -2.311795198 | 0.020788972 | 0.101013484 | 0.105637528 |
| ENSMUSG00000027710 | Acad9 | 229211 | protein_coding | 1028.674927 | -0.10564063 | 0.058780071 | -1.797218476 | 0.072300942 | 0.234890673 | 0.10564063 |
| ENSMUSG00000031827 | Cotl1 | 72042 | protein_coding | 1229.280046 | -0.105643466 | 0.09440849 | -1.119003872 | 0.263138487 | 0.504894037 | 0.105643466 |
| ENSMUSG00000032342 | Mto1 | 68291 | protein_coding | 332.7649061 | 0.10565396 | 0.081748676 | 1.292424106 | 0.196210313 | 0.428764731 | 0.10565396 |
| ENSMUSG00000025630 | Hprt | 15452 | protein_coding | 1564.128804 | 0.10565983 | 0.040239062 | 2.625802523 | 0.008644495 | 0.05353678 | 0.10565983 |
| ENSMUSG00000018425 | Dhx40 | 67487 | protein_coding | 803.80028 | 0.10566586 | 0.067215594 | 1.572043842 | 0.115940384 | 0.314662099 | 0.10566586 |
| ENSMUSG00000044600 | Smim7 | 66818 | protein_coding | 2434.154867 | 0.105694009 | 0.055967704 | 1.888482148 | 0.058961252 | 0.205949343 | 0.105694009 |
| ENSMUSG00000056493 | Foxk1 | 17425 | protein_coding | 805.7163654 | -0.105723292 | 0.097505071 | -1.084285065 | 0.278238425 | 0.521098971 | 0.105723292 |
| ENSMUSG00000063296 | Tmem117 | 320709 | protein_coding | 961.9962883 | -0.105730417 | 0.069895376 | -1.512695445 | 0.130357037 | 0.336416697 | 0.105730417 |
| ENSMUSG00000026347 | Tmem163 | 72160 | protein_coding | 11.44317793 | -0.105740633 | 0.363332081 | -0.291030269 | 0.77102817 | 0.892331737 | 0.105740633 |
| ENSMUSG00000034560 | Washc4 | 319277 | protein_coding | 761.3809536 | 0.105778783 | 0.060339285 | 1.75306657 | 0.079590581 | 0.249525136 | 0.105778783 |
| ENSMUSG00000038005 | Hpf1 | 72612 | protein_coding | 246.8500424 | -0.105791443 | 0.094363164 | -1.121109533 | 0.262241241 | 0.504097842 | 0.105791443 |
| ENSMUSG00000023905 | Tnfrsf12a | 27279 | protein_coding | 123.1093847 | 0.105878368 | 0.159354903 | 0.664418645 | 0.506422403 | 0.722065604 | 0.105878368 |
| ENSMUSG00000002603 | Tgfb1 | 21803 | protein_coding | 23.56474418 | -0.105915855 | 0.250114047 | -0.423470238 | 0.671952208 | 0.832813808 | 0.105915855 |
| ENSMUSG00000059810 | Rgs3 | 50780 | protein_coding | 469.1045246 | 0.105962107 | 0.064010394 | 1.655389092 | 0.097845597 | 0.284213967 | 0.105962107 |
| ENSMUSG00000023067 | Cdkn1a | 12575 | protein_coding | 2669.327162 | 0.106005576 | 0.068807985 | 1.540599924 | 0.123414186 | 0.326064971 | 0.106005576 |
| ENSMUSG00000109865 | Hspa14 | 50497 | protein_coding | 578.4958397 | 0.106101572 | 0.064494125 | 1.645135462 | 0.099941879 | 0.287712011 | 0.106101572 |

| ensembl_gene_id | external_gene_name | entrezgene_id | gene_biotype | baseMean | log2FoldChange | lfcSE | stat | pvalue | padj | absLog2FC |
|---|---|---|---|---|---|---|---|---|---|---|
| ENSMUSG00000071528 | Atp5md | 66477 | protein_coding | 2782.792029 | 0.106102912 | 0.054987439 | 1.929584526 | 0.053658338 | 0.193980724 | 0.106102912 |
| ENSMUSG00000093674 | Rpl41 | 67945 | protein_coding | 14762.98973 | 0.106106885 | 0.06371561 | 1.665320072 | 0.095848916 | 0.280638248 | 0.106106885 |
| ENSMUSG00000089857 | Zfp882 | 382019 | protein_coding | 44.64462065 | 0.106114018 | 0.193312127 | 0.548925823 | 0.583056356 | 0.774606627 | 0.106114018 |
| ENSMUSG00000051977 | Prdm9 | 213389 | protein_coding | 144.1681813 | 0.106123632 | 0.100697184 | 1.053888779 | 0.291933841 | 0.534455336 | 0.106123632 |
| ENSMUSG00000074918 | Inafm2 | 100043272 | protein_coding | 561.4170589 | -0.106166494 | 0.071885617 | -1.476880883 | 0.13970757 | 0.350507776 | 0.106166494 |
| ENSMUSG00000040711 | Sh3pxd2b | 268396 | protein_coding | 184.5425391 | -0.106172889 | 0.112480836 | -0.943919811 | 0.345210627 | 0.587571366 | 0.106172889 |
| ENSMUSG00000027544 | Nfatc2 | 18019 | protein_coding | 93.02699283 | 0.106181578 | 0.188319124 | 0.563838529 | 0.572864028 | 0.767564688 | 0.106181578 |
| ENSMUSG00000087406 | E130215H24Rik | 109344 | lncRNA | 14.01549803 | -0.106205667 | 0.325183422 | -0.32660234 | 0.743968674 | 0.878072512 | 0.106205667 |
| ENSMUSG00000037400 | Atp11b | 76295 | protein_coding | 3498.200857 | -0.106232947 | 0.048728181 | -2.180113137 | 0.029249076 | 0.127901261 | 0.106232947 |
| ENSMUSG00000038671 | Arfrp1 | 76688 | protein_coding | 494.1163864 | -0.106255632 | 0.068655525 | -1.547663249 | 0.121703395 | 0.323186341 | 0.106255632 |
| ENSMUSG00000068479 | Mfap1a | 67532 | protein_coding | 390.014679 | -0.106265587 | 0.069855128 | -1.521228148 | 0.128202594 | 0.333363953 | 0.106265587 |
| ENSMUSG00000078532 | Nkain1 | 67149 | protein_coding | 138.8070255 | 0.106284886 | 0.119617022 | 0.888543153 | 0.374248654 | 0.616119646 | 0.106284886 |
| ENSMUSG00000029127 | Zbtb49 | 75079 | protein_coding | 141.960656 | -0.106286052 | 0.10550259 | -1.007425995 | 0.313730104 | 0.557830804 | 0.106286052 |
| ENSMUSG00000034673 | Pbx2 | 18515 | protein_coding | 780.8330925 | -0.106291191 | 0.052445448 | -2.026700019 | 0.0426931 | 0.166146629 | 0.106291191 |
| ENSMUSG00000062981 | Mrpl42 | 67270 | protein_coding | 1311.268081 | 0.106328449 | 0.060402693 | 1.760326314 | 0.078352495 | 0.247278021 | 0.106328449 |
| ENSMUSG00000022305 | Lrp12 | 239393 | protein_coding | 322.6872752 | -0.106332339 | 0.083455064 | -1.274126858 | 0.202618461 | 0.436673517 | 0.106332339 |
| ENSMUSG00000031586 | Rbpms | 19663 | protein_coding | 24.86599817 | 0.106349595 | 0.269114459 | 0.395183505 | 0.692707465 | 0.845861535 | 0.106349595 |
| ENSMUSG00000036667 | Tcaf1 | 77574 | protein_coding | 479.9053649 | 0.10635629 | 0.07849285 | 1.354980604 | 0.175423737 | 0.401740784 | 0.10635629 |
| ENSMUSG00000039501 | Znfx1 | 98999 | protein_coding | 1477.633342 | -0.106357965 | 0.058486614 | -1.818500963 | 0.0689876 | 0.227937517 | 0.106357965 |
| ENSMUSG00000020658 | Efr3b | 668212 | protein_coding | 20.64700209 | 0.106362379 | 0.316253934 | 0.336319543 | 0.736629914 | 0.873566485 | 0.106362379 |
| ENSMUSG00000025404 | R3hdm2 | 71750 | protein_coding | 560.7982742 | -0.106364693 | 0.090014436 | -1.181640393 | 0.237348419 | 0.477271565 | 0.106364693 |
| ENSMUSG00000025162 | Csnk1d | 104318 | protein_coding | 2966.251262 | 0.106372203 | 0.042633603 | 2.495031987 | 0.012594578 | 0.07062335 | 0.106372203 |
| ENSMUSG00000040046 | Tph1 | 21990 | protein_coding | 285.5669748 | -0.106381746 | 0.119008033 | -0.893903912 | 0.371373305 | 0.613347321 | 0.106381746 |
| ENSMUSG00000030120 | Mlf2 | 30853 | protein_coding | 3699.053848 | 0.106391309 | 0.077088454 | 1.380119896 | 0.167549732 | 0.391491529 | 0.106391309 |
| ENSMUSG00000049555 | Tmie | 20776 | protein_coding | 27.71727401 | -0.106396738 | 0.246658534 | -0.431352349 | 0.666212192 | 0.829602333 | 0.106396738 |
| ENSMUSG00000038384 | Setd1b | 208043 | protein_coding | 285.8355766 | -0.106418005 | 0.143605586 | -0.741043628 | 0.458666988 | 0.685789219 | 0.106418005 |
| ENSMUSG00000024077 | Strn | 268980 | protein_coding | 1639.597957 | -0.106465082 | 0.040270069 | -2.643776974 | 0.008198666 | 0.051608073 | 0.106465082 |
| ENSMUSG00000020668 | Kif3c | 16570 | protein_coding | 43.69597207 | -0.106500614 | 0.24998545 | -0.42602725 | 0.670087989 | 0.831653764 | 0.106500614 |
| ENSMUSG00000020716 | Nf1 | 18015 | protein_coding | 1181.421577 | -0.10655586 | 0.087841477 | -1.213047222 | 0.225111767 | 0.463123925 | 0.10655586 |
| ENSMUSG00000064181 | Rab3ip | 216363 | protein_coding | 926.2783138 | -0.106567565 | 0.0599991 | -1.776152722 | 0.075707748 | 0.242105663 | 0.106567565 |
| ENSMUSG00000068011 | Mkrn2os | 70291 | protein_coding | 743.0267154 | 0.10657744 | 0.063844521 | 1.669327899 | 0.095052414 | 0.27921863 | 0.10657744 |
| ENSMUSG00000030200 | Bcl2l14 | 66813 | protein_coding | 1916.519686 | 0.106604999 | 0.053464623 | 1.993935291 | 0.046159127 | 0.175287213 | 0.106604999 |
| ENSMUSG00000024759 | Atl3 | 109168 | protein_coding | 1767.383034 | -0.106665917 | 0.046780659 | -2.280128567 | 0.022600064 | 0.107008585 | 0.106665917 |
| ENSMUSG00000087174 | 5530601H04Rik | 71445 | lncRNA | 182.776096 | -0.106682922 | 0.101050217 | -1.055741644 | 0.291086266 | 0.53344866 | 0.106682922 |
| ENSMUSG00000025870 | Arl10 | 56795 | protein_coding | 275.3976306 | -0.106701458 | 0.078656804 | -1.356544541 | 0.174925974 | 0.401227589 | 0.106701458 |
| ENSMUSG00000052794 | 1700030K09Rik | 72254 | protein_coding | 103.0447763 | 0.106730885 | 0.117201377 | 0.910662379 | 0.362473292 | 0.605347548 | 0.106730885 |
| ENSMUSG00000021036 | Sptlc2 | 20773 | protein_coding | 5561.487687 | 0.106744545 | 0.041708634 | 2.559291337 | 0.01048858 | 0.061486489 | 0.106744545 |
| ENSMUSG00000034973 | Dop1a | 320615 | protein_coding | 1301.710839 | -0.106753892 | 0.047071239 | -2.267921847 | 0.023333971 | 0.109264501 | 0.106753892 |
| ENSMUSG00000047227 | Gm527 | 217648 | protein_coding | 42.13288076 | -0.10675656 | 0.195403755 | -0.546338325 | 0.584833397 | 0.775689464 | 0.10675656 |

| ensembl_gene_id | external_gene_name | entrezgene_id | gene_biotype | baseMean | log2FoldChange | lfcSE | stat | pvalue | padj | absLog2FC |
|---|---|---|---|---|---|---|---|---|---|---|
| ENSMUSG00000053291 | Rab4b | 19342 | protein_coding | 71.18323885 | 0.106824111 | 0.16218958 | 0.658637323 | 0.51012869 | 0.724823423 | 0.106824111 |
| ENSMUSG00000070934 | Rraga | 68441 | protein_coding | 567.5019925 | 0.106834895 | 0.063859562 | 1.672966303 | 0.09433393 | 0.277715094 | 0.106834895 |
| ENSMUSG00000020476 | Dbnl | 13169 | protein_coding | 2358.559524 | 0.106863436 | 0.060692424 | 1.760737639 | 0.078282819 | 0.247106483 | 0.106863436 |
| ENSMUSG00000019518 | Ap4m1 | 11781 | protein_coding | 521.9771563 | -0.106908506 | 0.075637972 | -1.413423737 | 0.15753117 | 0.376817737 | 0.106908506 |
| ENSMUSG00000037725 | Ckap2 | 80986 | protein_coding | 284.046654 | -0.106911282 | 0.108829629 | -0.982372938 | 0.325916153 | 0.569329485 | 0.106911282 |
| ENSMUSG00000010154 | Spire2 | 234857 | protein_coding | 1184.590728 | 0.106928943 | 0.059110335 | 1.808972025 | 0.07045535 | 0.230847912 | 0.106928943 |
| ENSMUSG00000031791 | Tmem38a | 74166 | protein_coding | 91.91451247 | -0.106930002 | 0.138695399 | -0.770970075 | 0.44072467 | 0.672428264 | 0.106930002 |
| ENSMUSG00000020189 | Osbpl8 | 237542 | protein_coding | 478.1395558 | -0.106931995 | 0.060079523 | -1.779840956 | 0.075101993 | 0.240748848 | 0.106931995 |
| ENSMUSG00000043782 | Bicdl2 | 212733 | protein_coding | 495.6563511 | 0.106963053 | 0.076167954 | 1.404305189 | 0.16022799 | 0.380615229 | 0.106963053 |
| ENSMUSG00000029104 | Htt | 15194 | protein_coding | 1804.378202 | 0.106970132 | 0.054701906 | 1.955510132 | 0.050522889 | 0.186363821 | 0.106970132 |
| ENSMUSG00000003316 | Glg1 | 20340 | protein_coding | 3663.756754 | -0.10701953 | 0.041481847 | -2.579912345 | 0.00988254 | 0.058770253 | 0.10701953 |
| ENSMUSG00000035878 | Hykk | 235386 | protein_coding | 237.0475234 | 0.107025401 | 0.107025156 | 1.000002293 | 0.317309398 | 0.561120909 | 0.107025401 |
| ENSMUSG00000009207 | Lnpk | 69605 | protein_coding | 364.1757886 | 0.107031355 | 0.088857039 | 1.204534339 | 0.228383118 | 0.467402861 | 0.107031355 |
| ENSMUSG00000037466 | Tedc1 | 104732 | protein_coding | 303.6994996 | 0.107031468 | 0.118705172 | 0.901657999 | 0.367238571 | 0.609619808 | 0.107031468 |
| ENSMUSG00000037166 | Ppp1r14a | 68458 | protein_coding | 28.83386631 | -0.107054617 | 0.274348911 | -0.390213383 | 0.696378766 | 0.848225103 | 0.107054617 |
| ENSMUSG00000028786 | Tmem54 | 66260 | protein_coding | 2648.958338 | 0.107074412 | 0.069285891 | 1.545399949 | 0.122249553 | 0.324102998 | 0.107074412 |
| ENSMUSG00000034116 | Vav1 | 22324 | protein_coding | 223.1672273 | -0.107083263 | 0.093085391 | -1.150376681 | 0.249988761 | 0.490996071 | 0.107083263 |
| ENSMUSG00000030102 | Itpr1 | 16438 | protein_coding | 862.6796355 | 0.107086242 | 0.056024717 | 1.911410677 | 0.055951826 | 0.199502233 | 0.107086242 |
| ENSMUSG00000025764 | Jade1 | 269424 | protein_coding | 1157.576151 | -0.107099261 | 0.055578523 | -1.926990055 | 0.053980869 | 0.194579678 | 0.107099261 |
| ENSMUSG00000029623 | Pdap1 | 231887 | protein_coding | 1174.461052 | 0.107103071 | 0.07071342 | 1.514607426 | 0.129871851 | 0.335820819 | 0.107103071 |
| ENSMUSG00000046798 | Cldn12 | 64945 | protein_coding | 2085.028588 | 0.107134975 | 0.041525658 | 2.579970555 | 0.009880874 | 0.058770253 | 0.107134975 |
| ENSMUSG00000079641 | Rpl39 | 67248 | protein_coding | 4584.868703 | -0.107137013 | 0.048244654 | -2.22070229 | 0.026371132 | 0.118795321 | 0.107137013 |
| ENSMUSG00000050229 | Pigm | 67556 | protein_coding | 607.9731433 | -0.107139413 | 0.059382664 | -1.804220372 | 0.071196767 | 0.232615531 | 0.107139413 |
| ENSMUSG00000032423 | Syncrip | 56403 | protein_coding | 1626.389706 | -0.107211958 | 0.063809695 | -1.680182895 | 0.092921736 | 0.275316708 | 0.107211958 |
| ENSMUSG00000055652 | Klhl25 | 207952 | protein_coding | 388.7229556 | 0.107219315 | 0.078358885 | 1.368310889 | 0.171214784 | 0.395821216 | 0.107219315 |
| ENSMUSG00000019792 | Trmt11 | 73681 | protein_coding | 282.6428812 | 0.107234283 | 0.084400045 | 1.270547693 | 0.203889605 | 0.438066569 | 0.107234283 |
| ENSMUSG00000022420 | Dnal4 | 54152 | protein_coding | 342.9594803 | 0.10724542 | 0.074681855 | 1.436030474 | 0.15099367 | 0.36685181 | 0.10724542 |
| ENSMUSG00000051517 | Arhgef39 | 230098 | protein_coding | 382.6926835 | -0.107247298 | 0.080764359 | -1.327903792 | 0.184209892 | 0.412891303 | 0.107247298 |
| ENSMUSG00000064109 | Hcst | 23900 | protein_coding | 39.86355254 | 0.10726657 | 0.183185636 | 0.585562123 | 0.558169808 | 0.756755317 | 0.10726657 |
| ENSMUSG00000017615 | Tnfaip1 | 21927 | protein_coding | 2519.831743 | 0.107277251 | 0.043017214 | 2.49382148 | 0.012637609 | 0.070741657 | 0.107277251 |
| ENSMUSG00000022434 | Fam118a | 73225 | protein_coding | 265.8139296 | 0.107279729 | 0.092006098 | 1.166006729 | 0.243611725 | 0.484728722 | 0.107279729 |
| ENSMUSG00000021638 | Ocln | 18260 | protein_coding | 3253.075347 | -0.107300104 | 0.036335815 | -2.953012188 | 0.003146895 | 0.025111322 | 0.107300104 |
| ENSMUSG00000031785 | Adgrg1 | 14766 | protein_coding | 1152.617439 | 0.10734413 | 0.073326279 | 1.463924409 | 0.143214603 | 0.355127279 | 0.10734413 |
| ENSMUSG00000036961 | Wnt8b | 22423 | protein_coding | 25.5357131 | 0.107363302 | 0.279922535 | 0.383546477 | 0.701314616 | 0.851297936 | 0.107363302 |
| ENSMUSG00000018796 | Acsl1 | 14081 | protein_coding | 993.857819 | 0.107364363 | 0.060376801 | 1.778238677 | 0.075364663 | 0.241391445 | 0.107364363 |
| ENSMUSG00000045665 | Mfsd5 | 106073 | protein_coding | 486.2126715 | 0.107404689 | 0.087226911 | 1.231325139 | 0.218201283 | 0.454407553 | 0.107404689 |
| ENSMUSG00000014226 | Cacybp | 12301 | protein_coding | 735.4686051 | 0.107457359 | 0.078656963 | 1.366151898 | 0.171891292 | 0.396668438 | 0.107457359 |
| ENSMUSG00000038347 | Tcte2 | 21646 | protein_coding | 39.09612211 | -0.107526625 | 0.215168232 | -0.499732809 | 0.617263227 | 0.796814542 | 0.107526625 |
| ENSMUSG00000031201 | Brcc3 | 210766 | protein_coding | 652.8652981 | 0.107546216 | 0.070014985 | 1.536045688 | 0.124527175 | 0.327433832 | 0.107546216 |

| ensembl_gene_id | external_gene_name | entrezgene_id | gene_biotype | baseMean | log2FoldChange | lfcSE | stat | pvalue | padj | absLog2FC |
|---|---|---|---|---|---|---|---|---|---|---|
| ENSMUSG00000029536 | Gatc | 384281 | protein_coding | 241.0485528 | 0.107555508 | 0.079959162 | 1.345130505 | 0.178583094 | 0.405715197 | 0.107555508 |
| ENSMUSG00000037999 | Arap2 | 212285 | protein_coding | 2322.971413 | -0.107560737 | 0.060453003 | -1.779245551 | 0.075199513 | 0.241001487 | 0.107560737 |
| ENSMUSG00000059895 | Ptp4a3 | 19245 | protein_coding | 145.3425634 | -0.10758598 | 0.107345182 | -1.002243211 | 0.316226143 | 0.560212157 | 0.10758598 |
| ENSMUSG00000021938 | Pspc1 | 66645 | protein_coding | 432.5725027 | 0.107589086 | 0.078176141 | 1.376239414 | 0.168747523 | 0.392724841 | 0.107589086 |
| ENSMUSG00000054021 | Sirt5 | 68346 | protein_coding | 248.8786031 | -0.107603766 | 0.093992883 | -1.144807591 | 0.25228887 | 0.493470298 | 0.107603766 |
| ENSMUSG00000045302 | Preb | 50907 | protein_coding | 1447.102616 | 0.107606104 | 0.052367675 | 2.054819195 | 0.039896472 | 0.158163814 | 0.107606104 |
| ENSMUSG00000021102 | Glrx5 | 73046 | protein_coding | 1065.243411 | 0.107657832 | 0.072148207 | 1.492176128 | 0.135652979 | 0.344244782 | 0.107657832 |
| ENSMUSG00000028383 | Hsdl2 | 72479 | protein_coding | 674.3376941 | 0.107710646 | 0.062644537 | 1.719394079 | 0.08554264 | 0.261228205 | 0.107710646 |
| ENSMUSG00000024036 | Slc37a1 | 224674 | protein_coding | 4140.341285 | -0.107713002 | 0.044504713 | -2.420260573 | 0.015509389 | 0.082141188 | 0.107713002 |
| ENSMUSG00000018378 | Cuedc1 | 70393 | protein_coding | 148.6399394 | -0.107713945 | 0.134069325 | -0.803419757 | 0.421732159 | 0.65828686 | 0.107713945 |
| ENSMUSG00000033237 | Arid2 | 77044 | protein_coding | 1135.192872 | -0.10772927 | 0.057954007 | -1.858875266 | 0.063044821 | 0.21475471 | 0.10772927 |
| ENSMUSG00000022863 | Btg3 | 12228 | protein_coding | 120.816504 | -0.107739216 | 0.145086337 | -0.742586918 | 0.457731811 | 0.685471346 | 0.107739216 |
| ENSMUSG00000032590 | Apeh | 235606 | protein_coding | 1501.656441 | 0.107746414 | 0.06848957 | 1.573179904 | 0.115677167 | 0.314273193 | 0.107746414 |
| ENSMUSG00000054200 | Ffar4 | 107221 | protein_coding | 453.2294546 | 0.107759449 | 0.072027761 | 1.496082168 | 0.134632244 | 0.342321275 | 0.107759449 |
| ENSMUSG00000038047 | Haus6 | 230376 | protein_coding | 226.8841526 | -0.107762146 | 0.116734513 | -0.923138697 | 0.355934931 | 0.598619525 | 0.107762146 |
| ENSMUSG00000027276 | Jag1 | 16449 | protein_coding | 615.7851225 | -0.107788852 | 0.079409596 | -1.357378167 | 0.174661082 | 0.40092091 | 0.107788852 |
| ENSMUSG00000100599 | 1700120C14Rik | 73600 | lncRNA | 11.37892411 | -0.10779957 | 0.344624894 | -0.312802622 | 0.754430623 | 0.883437602 | 0.10779957 |
| ENSMUSG00000023089 | Ndufa5 | 68202 | protein_coding | 1678.517188 | 0.107862619 | 0.062944188 | 1.713623181 | 0.086597953 | 0.263453159 | 0.107862619 |
| ENSMUSG00000024805 | Pcgf5 | 76073 | protein_coding | 721.4306933 | -0.107894918 | 0.066213399 | -1.629502787 | 0.103206625 | 0.293509716 | 0.107894918 |
| ENSMUSG00000046785 | Epm2aip1 | 77781 | protein_coding | 550.5111236 | -0.107924639 | 0.075568467 | -1.428170286 | 0.153242851 | 0.370189171 | 0.107924639 |
| ENSMUSG00000021944 | Gata4 | 14463 | protein_coding | 16.11900155 | -0.107926278 | 0.407230572 | -0.26502499 | 0.790990211 | 0.90265349 | 0.107926278 |
| ENSMUSG00000025825 | Iscu | 66383 | protein_coding | 747.8977413 | 0.107948005 | 0.059006183 | 1.829435473 | 0.067334398 | 0.224318083 | 0.107948005 |
| ENSMUSG00000017288 | Vps53 | 68299 | protein_coding | 1274.169592 | 0.107980235 | 0.04903216 | 2.202232889 | 0.027648862 | 0.122734158 | 0.107980235 |
| ENSMUSG00000018848 | Rars | 104458 | protein_coding | 1600.44984 | 0.108000953 | 0.064437598 | 1.676054915 | 0.093727437 | 0.276737893 | 0.108000953 |
| ENSMUSG00000038059 | Smim3 | 106878 | protein_coding | 166.2978623 | 0.108034355 | 0.120129544 | 0.899315452 | 0.368484659 | 0.610431648 | 0.108034355 |
| ENSMUSG00000038845 | Phb | 18673 | protein_coding | 1557.637506 | 0.10804158 | 0.070332711 | 1.536149793 | 0.124501646 | 0.327433832 | 0.10804158 |
| ENSMUSG00000020994 | Pnn | 18949 | protein_coding | 857.4680981 | 0.108113737 | 0.083085237 | 1.301238834 | 0.193176717 | 0.425015131 | 0.108113737 |
| ENSMUSG00000042198 | Chchd7 | 66433 | protein_coding | 744.4175744 | 0.108122114 | 0.068429056 | 1.580061445 | 0.114092796 | 0.311490522 | 0.108122114 |
| ENSMUSG00000048154 | Kmt2d | 381022 | protein_coding | 1593.833804 | -0.108156138 | 0.101810701 | -1.062325833 | 0.288087791 | 0.530546055 | 0.108156138 |
| ENSMUSG00000036054 | Sugp2 | 234373 | protein_coding | 442.0139589 | -0.108160207 | 0.07558762 | -1.430924893 | 0.152451742 | 0.369268435 | 0.108160207 |
| ENSMUSG00000019080 | Mfsd3 | 69572 | protein_coding | 109.1483535 | -0.10818402 | 0.132671832 | -0.815425691 | 0.414828684 | 0.65259636 | 0.10818402 |
| ENSMUSG00000027009 | Itga4 | 16401 | protein_coding | 28.69069504 | 0.108272825 | 0.227288718 | 0.476366913 | 0.633813003 | 0.8083434 | 0.108272825 |
| ENSMUSG00000025316 | Banp | 53325 | protein_coding | 211.8121927 | 0.108289039 | 0.11487246 | 0.942689303 | 0.345839846 | 0.588148333 | 0.108289039 |
| ENSMUSG00000037169 | Mycn | 18109 | protein_coding | 53.45912979 | 0.108293887 | 0.213313334 | 0.507675186 | 0.611681153 | 0.79296591 | 0.108293887 |
| ENSMUSG00000050855 | Zfp940 | 233057 | protein_coding | 25.78838871 | 0.108401671 | 0.298729299 | 0.362875926 | 0.716697571 | 0.861137068 | 0.108401671 |
| ENSMUSG00000036860 | Mrpl55 | 67212 | protein_coding | 519.8826096 | 0.108456664 | 0.086585472 | 1.252596556 | 0.210352569 | 0.445366401 | 0.108456664 |
| ENSMUSG00000044783 | Hjurp | 212427 | protein_coding | 3051.598201 | -0.108479102 | 0.066479092 | -1.631777728 | 0.102726319 | 0.29253061 | 0.108479102 |
| ENSMUSG00000026417 | Pigr | 18703 | protein_coding | 92541.70362 | 0.108490865 | 0.056536987 | 1.918936104 | 0.054992421 | 0.197299838 | 0.108490865 |
| ENSMUSG00000051234 | Rnf7 | 19823 | protein_coding | 876.5359601 | 0.108492009 | 0.061663014 | 1.759434087 | 0.078503806 | 0.247610187 | 0.108492009 |

| ensembl_gene_id | external_gene_name | entrezgene_id | gene_biotype | baseMean | log2FoldChange | lfcSE | stat | pvalue | padj | absLog2FC |
|---|---|---|---|---|---|---|---|---|---|---|
| ENSMUSG00000078572 | Ndufaf8 | 208501 | protein_coding | 549.0604842 | 0.108503904 | 0.085803729 | 1.264559295 | 0.206029352 | 0.440784802 | 0.108503904 |
| ENSMUSG00000020300 | Cpeb4 | 67579 | protein_coding | 1764.54932 | -0.108522247 | 0.052912257 | -2.050985026 | 0.040268405 | 0.159364066 | 0.108522247 |
| ENSMUSG00000028964 | Park7 | 57320 | protein_coding | 1441.131131 | 0.108531377 | 0.057918496 | 1.87386386 | 0.060949192 | 0.21029688 | 0.108531377 |
| ENSMUSG00000054455 | Vapb | 56491 | protein_coding | 1918.655275 | 0.108534447 | 0.057219223 | 1.896817912 | 0.057851974 | 0.203825108 | 0.108534447 |
| ENSMUSG00000029111 | Nelfa | 24116 | protein_coding | 489.7274244 | 0.108563353 | 0.068198142 | 1.591881387 | 0.111411359 | 0.306818815 | 0.108563353 |
| ENSMUSG00000025982 | Sf3b1 | 81898 | protein_coding | 6209.946708 | -0.108619237 | 0.053254803 | -2.039613905 | 0.041388796 | 0.162442714 | 0.108619237 |
| ENSMUSG00000040370 | Etfrf1 | 67636 | protein_coding | 536.1290986 | 0.108632624 | 0.062081506 | 1.749838738 | 0.080146144 | 0.250536319 | 0.108632624 |
| ENSMUSG00000026384 | Ptpn4 | 19258 | protein_coding | 691.3231883 | -0.10863398 | 0.052700912 | -2.061330182 | 0.039271551 | 0.156571643 | 0.10863398 |
| ENSMUSG00000038239 | Hrc | 15464 | protein_coding | 31.58028161 | -0.108638693 | 0.2645308 | -0.410684476 | 0.681303911 | 0.838859992 | 0.108638693 |
| ENSMUSG00000024370 | Cdc23 | 52563 | protein_coding | 548.2267758 | -0.108654647 | 0.061131184 | -1.777401309 | 0.075502236 | 0.241736064 | 0.108654647 |
| ENSMUSG00000021596 | Mctp1 | 78771 | protein_coding | 89.23213804 | -0.108693732 | 0.131819602 | -0.824564256 | 0.409619022 | 0.648148152 | 0.108693732 |
| ENSMUSG00000035506 | Slc12a8 | 171286 | protein_coding | 2462.37142 | 0.108712426 | 0.080629297 | 1.348299317 | 0.177562132 | 0.404658227 | 0.108712426 |
| ENSMUSG00000025417 | Pip4k2c | 117150 | protein_coding | 2000.176685 | 0.10872883 | 0.056519418 | 1.923742926 | 0.054386815 | 0.195649377 | 0.10872883 |
| ENSMUSG00000044134 | Pheta1 | 231717 | protein_coding | 1686.39436 | 0.108731556 | 0.066777358 | 1.628269814 | 0.103467687 | 0.293978751 | 0.108731556 |
| ENSMUSG00000041846 | Ppp4r3a | 68734 | protein_coding | 1841.455542 | -0.108732783 | 0.040082297 | -2.712738282 | 0.006672979 | 0.044230856 | 0.108732783 |
| ENSMUSG00000020544 | Cox11 | 69802 | protein_coding | 151.3529502 | -0.108737726 | 0.106807098 | -1.018075842 | 0.308641915 | 0.552278995 | 0.108737726 |
| ENSMUSG00000035863 | Palm | 18483 | protein_coding | 44.94193595 | -0.10876999 | 0.220344173 | -0.493636788 | 0.621562715 | 0.800250831 | 0.10876999 |
| ENSMUSG00000043372 | Hexim2 | 71059 | protein_coding | 74.65166703 | -0.108809234 | 0.173390656 | -0.627538047 | 0.530306605 | 0.73955851 | 0.108809234 |
| ENSMUSG00000049504 | Proser1 | 212127 | protein_coding | 112.7094649 | -0.108815065 | 0.143483201 | -0.758381915 | 0.448222381 | 0.678921982 | 0.108815065 |
| ENSMUSG00000037426 | Depdc5 | 277854 | protein_coding | 608.3293189 | -0.108828763 | 0.053778586 | -2.023644933 | 0.043006698 | 0.167125066 | 0.108828763 |
| ENSMUSG00000019826 | Zbtb24 | 268294 | protein_coding | 402.3233492 | 0.108848088 | 0.072649621 | 1.49826091 | 0.134065476 | 0.341505912 | 0.108848088 |
| ENSMUSG00000038425 | Poli | 26447 | protein_coding | 133.798292 | -0.108853186 | 0.125053169 | -0.870455238 | 0.384051671 | 0.624863562 | 0.108853186 |
| ENSMUSG00000005354 | Txn2 | 56551 | protein_coding | 2127.818453 | 0.108856828 | 0.067789394 | 1.605809134 | 0.108315853 | 0.301538022 | 0.108856828 |
| ENSMUSG00000071648 | Rom1 | 19881 | protein_coding | 57.54852112 | 0.108863362 | 0.168081489 | 0.647682043 | 0.517190621 | 0.729722934 | 0.108863362 |
| ENSMUSG00000051579 | Tceal8 | 66684 | protein_coding | 375.4507865 | 0.108897316 | 0.120218754 | 0.905826361 | 0.365027766 | 0.60723078 | 0.108897316 |
| ENSMUSG00000058655 | Eif4b | 75705 | protein_coding | 9540.579921 | -0.10894416 | 0.042205352 | -2.581287773 | 0.009843249 | 0.058646044 | 0.10894416 |
| ENSMUSG00000049882 | Vcpkmt | 207965 | protein_coding | 89.10306344 | -0.108959292 | 0.15302698 | -0.712026677 | 0.476448257 | 0.699497125 | 0.108959292 |
| ENSMUSG00000027787 | Nmd3 | 97112 | protein_coding | 408.2178343 | -0.109001087 | 0.087716279 | -1.242655167 | 0.213994931 | 0.449718678 | 0.109001087 |
| ENSMUSG00000052906 | Ubxn8 | 108159 | protein_coding | 592.3049649 | -0.109051438 | 0.061216191 | -1.781414955 | 0.074844687 | 0.240298679 | 0.109051438 |
| ENSMUSG00000030291 | Med21 | 108098 | protein_coding | 322.6533962 | 0.109110177 | 0.084493004 | 1.291351617 | 0.19658178 | 0.429095149 | 0.109110177 |
| ENSMUSG00000014498 | Ankrd52 | 237615 | protein_coding | 919.2215653 | -0.109112569 | 0.084393509 | -1.292902383 | 0.196044823 | 0.428635421 | 0.109112569 |
| ENSMUSG00000024792 | Zfpl1 | 81909 | protein_coding | 490.9091944 | 0.10915509 | 0.066192658 | 1.649051333 | 0.099137119 | 0.286417827 | 0.10915509 |
| ENSMUSG00000014232 | Cluap1 | 76779 | protein_coding | 132.4363551 | 0.109163401 | 0.125496369 | 0.869853064 | 0.384380708 | 0.624972092 | 0.109163401 |
| ENSMUSG00000044763 | Trmt10c | 52575 | protein_coding | 476.8964114 | -0.109165936 | 0.073976867 | -1.475676648 | 0.140030718 | 0.350881819 | 0.109165936 |
| ENSMUSG00000049439 | Cyp20a1 | 77951 | protein_coding | 492.650581 | 0.109190613 | 0.070298016 | 1.55325313 | 0.120362673 | 0.321053395 | 0.109190613 |
| ENSMUSG00000033083 | Tbc1d4 | 210789 | protein_coding | 755.3323445 | -0.109197363 | 0.071260302 | -1.532373004 | 0.125430415 | 0.328615354 | 0.109197363 |
| ENSMUSG00000028676 | Srsf10 | 14105 | protein_coding | 1180.822798 | 0.109211393 | 0.050526477 | 2.161468603 | 0.03065916 | 0.132167941 | 0.109211393 |
| ENSMUSG00000006699 | Cdc42 | 12540 | protein_coding | 8064.875603 | 0.109214932 | 0.051902836 | 2.104218958 | 0.035359352 | 0.145506574 | 0.109214932 |
| ENSMUSG00000014767 | Tbp | 21374 | protein_coding | 271.5123968 | -0.109240167 | 0.085514309 | -1.277448986 | 0.20144378 | 0.435188726 | 0.109240167 |

| ensembl_gene_id | external_gene_name | entrezgene_id | gene_biotype | baseMean | log2FoldChange | lfcSE | stat | pvalue | padj | absLog2FC |
|---|---|---|---|---|---|---|---|---|---|---|
| ENSMUSG00000034075 | Zdhhc5 | 228136 | protein_coding | 4071.462265 | 0.109262504 | 0.040576932 | 2.692724652 | 0.007087077 | 0.046272589 | 0.109262504 |
| ENSMUSG00000000532 | Acvr1b | 11479 | protein_coding | 2876.251557 | -0.109273679 | 0.05008784 | -2.181640854 | 0.029136049 | 0.127510737 | 0.109273679 |
| ENSMUSG00000031166 | Wdr13 | 73447 | protein_coding | 1985.903486 | -0.109278252 | 0.041206773 | -2.651948765 | 0.008002868 | 0.05073202 | 0.109278252 |
| ENSMUSG00000006920 | Ezh1 | 14055 | protein_coding | 1279.550981 | -0.109297676 | 0.045603928 | -2.396672431 | 0.016544707 | 0.086181037 | 0.109297676 |
| ENSMUSG00000020778 | Ten1 | 69535 | protein_coding | 445.0077143 | 0.109341087 | 0.083128547 | 1.315325373 | 0.188400574 | 0.418734297 | 0.109341087 |
| ENSMUSG00000034462 | Pkd2 | 18764 | protein_coding | 292.7604805 | -0.109350796 | 0.081705107 | -1.338359381 | 0.180779313 | 0.407666398 | 0.109350796 |
| ENSMUSG00000078784 | Rbis | 69126 | protein_coding | 452.7503972 | 0.109354051 | 0.095408739 | 1.146163888 | 0.251727347 | 0.492727559 | 0.109354051 |
| ENSMUSG00000034390 | Cmip | 74440 | protein_coding | 1265.214469 | -0.109357704 | 0.079335448 | -1.378421708 | 0.168073125 | 0.391964304 | 0.109357704 |
| ENSMUSG00000038970 | Lmtk2 | 231876 | protein_coding | 2715.504837 | 0.109376453 | 0.054525913 | 2.005953652 | 0.044861193 | 0.171848571 | 0.109376453 |
| ENSMUSG00000028426 | Rad23b | 19359 | protein_coding | 3003.430427 | 0.109382435 | 0.04279859 | 2.555748564 | 0.010595965 | 0.061835229 | 0.109382435 |
| ENSMUSG00000019878 | Hsf2 | 15500 | protein_coding | 172.9580981 | -0.109391193 | 0.129723321 | -0.843265436 | 0.399080009 | 0.638678506 | 0.109391193 |
| ENSMUSG00000108659 | Gm34121 | 102637256 | lncRNA | 14.09074831 | 0.109406728 | 0.303601465 | 0.360362978 | 0.718575708 | 0.862201627 | 0.109406728 |
| ENSMUSG00000004018 | Fancl | 67030 | protein_coding | 207.9073043 | 0.109455798 | 0.087095319 | 1.256735715 | 0.208849351 | 0.44445942 | 0.109455798 |
| ENSMUSG00000061650 | Med9 | 192191 | protein_coding | 442.1214608 | 0.109495058 | 0.085588444 | 1.279320578 | 0.200784189 | 0.434462179 | 0.109495058 |
| ENSMUSG00000035936 | Aldh5a1 | 214579 | protein_coding | 433.037571 | -0.109510643 | 0.076222801 | -1.436717642 | 0.15079824 | 0.366529497 | 0.109510643 |
| ENSMUSG00000024791 | Cdca5 | 67849 | protein_coding | 166.5752368 | 0.109547324 | 0.128383302 | 0.853283269 | 0.393502236 | 0.633207999 | 0.109547324 |
| ENSMUSG00000021000 | Mia2 | 338320 | protein_coding | 2617.718601 | -0.109597825 | 0.044588231 | -2.457998921 | 0.013971359 | 0.076221398 | 0.109597825 |
| ENSMUSG00000034343 | Ube2f | 67921 | protein_coding | 699.3168577 | 0.109598514 | 0.071414215 | 1.534687645 | 0.124860572 | 0.32795219 | 0.109598514 |
| ENSMUSG00000000916 | Nsun5 | 100609 | protein_coding | 167.7706249 | 0.109603976 | 0.102612875 | 1.068130835 | 0.285461498 | 0.528364236 | 0.109603976 |
| ENSMUSG00000006281 | Tep1 | 21745 | protein_coding | 4624.035102 | -0.109614101 | 0.05844783 | -1.875417809 | 0.060735267 | 0.20964847 | 0.109614101 |
| ENSMUSG00000022453 | Naga | 17939 | protein_coding | 2949.629593 | 0.109631541 | 0.058878079 | 1.862009481 | 0.062601754 | 0.213667404 | 0.109631541 |
| ENSMUSG00000074064 | Mlycd | 56690 | protein_coding | 887.0791769 | 0.109647787 | 0.057561042 | 1.904895814 | 0.056793615 | 0.201440865 | 0.109647787 |
| ENSMUSG00000025453 | Nnt | 18115 | protein_coding | 8.760388906 | 0.109658681 | 0.397391493 | 0.275946222 | 0.782589372 | 0.898825528 | 0.109658681 |
| ENSMUSG00000031490 | Eif4ebp1 | 13685 | protein_coding | 308.3156411 | 0.109671625 | 0.114713311 | 0.95604969 | 0.339047131 | 0.58240762 | 0.109671625 |
| ENSMUSG00000039763 | Dnajc28 | 246738 | protein_coding | 171.6169413 | -0.109685674 | 0.095620073 | -1.147098828 | 0.251340778 | 0.492421447 | 0.109685674 |
| ENSMUSG00000023143 | Nagpa | 27426 | protein_coding | 410.2612896 | 0.10969354 | 0.104696606 | 1.047727754 | 0.294764057 | 0.537823306 | 0.10969354 |
| ENSMUSG00000044328 | Trp53i13 | 216964 | protein_coding | 97.52818889 | -0.10972103 | 0.141058371 | -0.777841324 | 0.436662563 | 0.669842781 | 0.10972103 |
| ENSMUSG00000020785 | Camkk1 | 55984 | protein_coding | 50.73806835 | -0.109769116 | 0.166394959 | -0.659690157 | 0.509452685 | 0.72436245 | 0.109769116 |
| ENSMUSG00000039633 | Lonrf1 | 244421 | protein_coding | 118.1736643 | 0.10979104 | 0.118324133 | 0.92788375 | 0.353467874 | 0.595956629 | 0.10979104 |
| ENSMUSG00000036781 | Rps27l | 67941 | protein_coding | 2094.351835 | 0.109800184 | 0.047596038 | 2.306918552 | 0.021059361 | 0.101895312 | 0.109800184 |
| ENSMUSG00000090733 | Rps27 | 57294 | protein_coding | 10692.00633 | -0.109807393 | 0.048887262 | -2.246135067 | 0.024695358 | 0.113854212 | 0.109807393 |
| ENSMUSG00000027133 | Nop10 | 66181 | protein_coding | 709.3664169 | 0.109823 | 0.069705444 | 1.575529739 | 0.11513422 | 0.313114597 | 0.109823 |
| ENSMUSG00000036957 | Lrfn3 | 233067 | protein_coding | 215.6186769 | 0.10984194 | 0.099819316 | 1.10040766 | 0.271154542 | 0.513727015 | 0.10984194 |
| ENSMUSG00000028779 | Pef1 | 67898 | protein_coding | 1785.618212 | 0.109853562 | 0.061090169 | 1.798219968 | 0.072142156 | 0.234551147 | 0.109853562 |
| ENSMUSG00000062590 | Armc9 | 78795 | protein_coding | 122.160688 | 0.109887158 | 0.116242594 | 0.945326098 | 0.344492419 | 0.58688999 | 0.109887158 |
| ENSMUSG00000056124 | B4galt6 | 56386 | protein_coding | 550.9992617 | -0.109897281 | 0.066557745 | -1.651156912 | 0.098706539 | 0.285839532 | 0.109897281 |
| ENSMUSG00000049878 | Rlf | 109263 | protein_coding | 840.1403011 | -0.109903419 | 0.073886414 | -1.487464513 | 0.136892169 | 0.34629961 | 0.109903419 |
| ENSMUSG00000035161 | Ints6 | 18130 | protein_coding | 854.9390955 | 0.109920317 | 0.057763502 | 1.90293721 | 0.057048739 | 0.201938151 | 0.109920317 |
| ENSMUSG00000099988 | 1700054A03Rik | 78457 | lncRNA | 13.60239699 | -0.1099287 | 0.416141368 | -0.264161913 | 0.791655158 | 0.903021982 | 0.1099287 |

| ensembl_gene_id | external_gene_name | entrezgene_id | gene_biotype | baseMean | log2FoldChange | lfcSE | stat | pvalue | padj | absLog2FC |
|---|---|---|---|---|---|---|---|---|---|---|
| ENSMUSG00000049327 | Kmt5a | 67956 | protein_coding | 3329.170929 | 0.109954475 | 0.055439098 | 1.983338094 | 0.047329677 | 0.178465808 | 0.109954475 |
| ENSMUSG00000021707 | Dhfr | 13361 | protein_coding | 402.6121951 | 0.109971232 | 0.120374156 | 0.913578424 | 0.360938408 | 0.604050305 | 0.109971232 |
| ENSMUSG00000042303 | Sgsm3 | 105835 | protein_coding | 2736.985589 | 0.110010037 | 0.061402032 | 1.791635134 | 0.073191431 | 0.236781989 | 0.110010037 |
| ENSMUSG00000030512 | Snrpa1 | 68981 | protein_coding | 356.0915763 | 0.110012673 | 0.102735069 | 1.070838554 | 0.284242028 | 0.52707842 | 0.110012673 |
| ENSMUSG00000032092 | Mpzl2 | 14012 | protein_coding | 522.0334065 | -0.110016655 | 0.069635343 | -1.579896777 | 0.114130507 | 0.31154074 | 0.110016655 |
| ENSMUSG00000033444 | Specc1l | 74392 | protein_coding | 3378.837172 | -0.110022932 | 0.066954737 | -1.643243416 | 0.10033258 | 0.288630668 | 0.110022932 |
| ENSMUSG00000028409 | Smu1 | 74255 | protein_coding | 1780.693025 | 0.110042629 | 0.05608281 | 1.962145433 | 0.049745555 | 0.184509329 | 0.110042629 |
| ENSMUSG00000040359 | Ufl1 | 67490 | protein_coding | 690.6288679 | 0.110060888 | 0.067547163 | 1.629393198 | 0.103229808 | 0.293509716 | 0.110060888 |
| ENSMUSG00000097184 | 4632428C04Rik | 100043102 | lncRNA | 56.64806905 | -0.110084579 | 0.168523445 | -0.653230053 | 0.513607978 | 0.727021493 | 0.110084579 |
| ENSMUSG00000029992 | Gfpt1 | 14583 | protein_coding | 20690.20886 | -0.110103199 | 0.050585245 | -2.176587257 | 0.029511377 | 0.128599602 | 0.110103199 |
| ENSMUSG00000075470 | Alg10b | 380959 | protein_coding | 1408.761527 | 0.110138128 | 0.047520706 | 2.3176871 | 0.020466333 | 0.099836826 | 0.110138128 |
| ENSMUSG00000036534 | Slc38a7 | 234595 | protein_coding | 296.9109583 | -0.11017277 | 0.08023077 | -1.37319846 | 0.169690658 | 0.393925615 | 0.11017277 |
| ENSMUSG00000052369 | Tmem106c | 380967 | protein_coding | 358.3641191 | 0.110178296 | 0.065222505 | 1.689268071 | 0.091168069 | 0.272039484 | 0.110178296 |
| ENSMUSG00000055370 | Gm9968 | 791329 | lncRNA | 8.846297418 | -0.110209159 | 0.426784032 | -0.258231684 | 0.796228107 | 0.906002305 | 0.110209159 |
| ENSMUSG00000038658 | Ric1 | 226089 | protein_coding | 661.7404202 | -0.110240102 | 0.057388263 | -1.920952072 | 0.05473775 | 0.196492651 | 0.110240102 |
| ENSMUSG00000066357 | Wdr6 | 83669 | protein_coding | 556.6624649 | 0.110250387 | 0.069535267 | 1.58553194 | 0.112845526 | 0.309131951 | 0.110250387 |
| ENSMUSG00000039879 | Heca | 380629 | protein_coding | 1743.518011 | -0.110306094 | 0.052985413 | -2.081820014 | 0.037358913 | 0.150979148 | 0.110306094 |
| ENSMUSG00000028793 | Rnf19b | 75234 | protein_coding | 1261.657166 | -0.110317842 | 0.067777379 | -1.62764988 | 0.103599146 | 0.294196765 | 0.110317842 |
| ENSMUSG00000035112 | Wnk4 | 69847 | protein_coding | 382.7571247 | -0.110324088 | 0.100839807 | -1.094052942 | 0.27393176 | 0.516506461 | 0.110324088 |
| ENSMUSG00000050002 | Idnk | 75731 | protein_coding | 615.4205705 | -0.110325589 | 0.06884536 | -1.602513066 | 0.109042201 | 0.30241162 | 0.110325589 |
| ENSMUSG00000037851 | Iars | 105148 | protein_coding | 1110.660745 | 0.110346119 | 0.090511025 | 1.219145616 | 0.222788929 | 0.460158435 | 0.110346119 |
| ENSMUSG00000035234 | Abraxas1 | 70681 | protein_coding | 151.9793151 | -0.11035199 | 0.109588719 | -1.006964874 | 0.313951654 | 0.558101668 | 0.11035199 |
| ENSMUSG00000036941 | Elac1 | 114615 | protein_coding | 245.6195951 | -0.110370173 | 0.100305174 | -1.100343766 | 0.27118237 | 0.513727015 | 0.110370173 |
| ENSMUSG00000057329 | Bcl2 | 12043 | protein_coding | 401.0428329 | 0.110403863 | 0.097474185 | 1.132647208 | 0.257362436 | 0.499216262 | 0.110403863 |
| ENSMUSG00000020021 | Fgd6 | 13998 | protein_coding | 601.3875492 | -0.110405835 | 0.080392937 | -1.373327548 | 0.169650542 | 0.39391111 | 0.110405835 |
| ENSMUSG00000063317 | Usp31 | 76179 | protein_coding | 354.2550667 | -0.110425222 | 0.074402085 | -1.484168374 | 0.137764258 | 0.347590314 | 0.110425222 |
| ENSMUSG00000067942 | Zfp160 | 224585 | protein_coding | 416.0043342 | -0.11044099 | 0.110896802 | -0.995889763 | 0.31930371 | 0.562777151 | 0.11044099 |
| ENSMUSG00000042439 | Zfp532 | 328977 | protein_coding | 237.6521737 | -0.110504186 | 0.111532637 | -0.990778929 | 0.32179354 | 0.565681756 | 0.110504186 |
| ENSMUSG00000044496 | 2510039O18Rik | 77034 | protein_coding | 2665.299513 | 0.110507581 | 0.059941964 | 1.843576253 | 0.065244919 | 0.219942478 | 0.110507581 |
| ENSMUSG00000000706 | Btn1a1 | 12231 | protein_coding | 65.0222335 | 0.110610058 | 0.188485754 | 0.586835109 | 0.557314455 | 0.756034291 | 0.110610058 |
| ENSMUSG00000056234 | Ncoa4 | 27057 | protein_coding | 1301.601327 | 0.110631647 | 0.051303951 | 2.156396225 | 0.031052739 | 0.133152759 | 0.110631647 |
| ENSMUSG00000021687 | Scamp1 | 107767 | protein_coding | 3319.547082 | -0.110718194 | 0.049910047 | -2.218354819 | 0.026530649 | 0.119015238 | 0.110718194 |
| ENSMUSG00000019986 | Ahi1 | 52906 | protein_coding | 35.91479945 | -0.110729258 | 0.196473622 | -0.563583329 | 0.573037735 | 0.767564688 | 0.110729258 |
| ENSMUSG00000041353 | Tmem29 | 382245 | protein_coding | 167.7466898 | 0.110777316 | 0.099024507 | 1.118685852 | 0.263274183 | 0.504916665 | 0.110777316 |
| ENSMUSG00000089917 | Uckl1 | 68556 | protein_coding | 774.7635807 | -0.11079293 | 0.052888501 | -2.094839682 | 0.036185246 | 0.147736574 | 0.11079293 |
| ENSMUSG00000032263 | Bckdhb | 12040 | protein_coding | 175.3076527 | 0.110817775 | 0.110912739 | 0.999143794 | 0.317725039 | 0.561462863 | 0.110817775 |
| ENSMUSG00000044156 | Hepacam2 | 101202 | protein_coding | 4515.915794 | 0.11085751 | 0.067800869 | 1.635045553 | 0.102039499 | 0.29151302 | 0.11085751 |
| ENSMUSG00000046840 | Hnf4aos | 68314 | lncRNA | 7.456078038 | -0.110881833 | 0.483798322 | -0.229190197 | 0.818721091 | 0.918528635 | 0.110881833 |
| ENSMUSG00000010045 | Tmem115 | 56395 | protein_coding | 1148.206479 | 0.110904027 | 0.076831612 | 1.443468698 | 0.148888484 | 0.36331133 | 0.110904027 |

| ensembl_gene_id | external_gene_name | entrezgene_id | gene_biotype | baseMean | log2FoldChange | lfcSE | stat | pvalue | padj | absLog2FC |
|---|---|---|---|---|---|---|---|---|---|---|
| ENSMUSG00000034724 | Cnot6l | 231464 | protein_coding | 1002.611688 | -0.110928207 | 0.098385789 | -1.127482004 | 0.259538745 | 0.501087195 | 0.110928207 |
| ENSMUSG00000009549 | Srp14 | 20813 | protein_coding | 657.6224077 | 0.110992018 | 0.061221379 | 1.812961758 | 0.06983772 | 0.229476957 | 0.110992018 |
| ENSMUSG00000032515 | Csrnp1 | 215418 | protein_coding | 975.8237354 | -0.111042268 | 0.104038331 | -1.067320731 | 0.28582703 | 0.5285691 | 0.111042268 |
| ENSMUSG00000028175 | Depdc1a | 76131 | protein_coding | 86.41429475 | 0.111077161 | 0.160400173 | 0.692500257 | 0.488623224 | 0.708970987 | 0.111077161 |
| ENSMUSG00000026933 | Camsap1 | 227634 | protein_coding | 493.7875721 | -0.111100595 | 0.083613564 | -1.328738906 | 0.183934124 | 0.412502521 | 0.111100595 |
| ENSMUSG00000022961 | Son | 20658 | protein_coding | 6750.467307 | -0.11115393 | 0.029639907 | -3.750144379 | 0.000176733 | 0.002365286 | 0.11115393 |
| ENSMUSG00000092558 | Med20 | 56771 | protein_coding | 409.0838753 | -0.111157448 | 0.077435596 | -1.435482567 | 0.151149633 | 0.367102434 | 0.111157448 |
| ENSMUSG00000032911 | Cspg4 | 121021 | protein_coding | 50.79171106 | -0.111169173 | 0.215675125 | -0.515447357 | 0.606240447 | 0.789753 | 0.111169173 |
| ENSMUSG00000053977 | Cd8a | 12525 | protein_coding | 28.33101151 | -0.111184616 | 0.256871992 | -0.432840557 | 0.665130607 | 0.82901007 | 0.111184616 |
| ENSMUSG00000024068 | Spast | 50850 | protein_coding | 911.0826847 | -0.111217239 | 0.045993458 | -2.418109962 | 0.015601361 | 0.082474622 | 0.111217239 |
| ENSMUSG00000032288 | Imp3 | 102462 | protein_coding | 787.1156939 | -0.111223192 | 0.071271453 | -1.560557397 | 0.118628217 | 0.318746628 | 0.111223192 |
| ENSMUSG00000026858 | Miga2 | 108958 | protein_coding | 1285.710277 | 0.111235488 | 0.05930032 | 1.875799109 | 0.06068287 | 0.209557313 | 0.111235488 |
| ENSMUSG00000038482 | Tfdp1 | 21781 | protein_coding | 1096.247784 | 0.111244575 | 0.07828337 | 1.421049902 | 0.155302253 | 0.373324674 | 0.111244575 |
| ENSMUSG00000048175 | Asb8 | 78541 | protein_coding | 701.975224 | -0.111264801 | 0.077718579 | -1.431637103 | 0.152247705 | 0.368991352 | 0.111264801 |
| ENSMUSG00000020190 | Mknk2 | 17347 | protein_coding | 5357.613951 | -0.111271935 | 0.061811401 | -1.80018465 | 0.071831487 | 0.233883785 | 0.111271935 |
| ENSMUSG00000027184 | Caprin1 | 53872 | protein_coding | 7682.143896 | 0.111289825 | 0.043571015 | 2.55421696 | 0.010642692 | 0.061984359 | 0.111289825 |
| ENSMUSG00000027329 | Spef1 | 70997 | protein_coding | 146.114788 | 0.111316265 | 0.152958807 | 0.727753224 | 0.466764663 | 0.692404394 | 0.111316265 |
| ENSMUSG00000062963 | Ufc1 | 66155 | protein_coding | 1854.796629 | 0.111326712 | 0.06033479 | 1.845149571 | 0.065015784 | 0.219399361 | 0.111326712 |
| ENSMUSG00000026048 | Ercc5 | 22592 | protein_coding | 395.1554916 | -0.111341864 | 0.077637367 | -1.434127248 | 0.151535953 | 0.367724228 | 0.111341864 |
| ENSMUSG00000020697 | Lig3 | 16882 | protein_coding | 776.2874905 | 0.111351778 | 0.060558846 | 1.838736776 | 0.065953911 | 0.221545247 | 0.111351778 |
| ENSMUSG00000063785 | Utp14a | 72554 | protein_coding | 561.7677634 | -0.111415759 | 0.068889312 | -1.617315599 | 0.105810174 | 0.297926072 | 0.111415759 |
| ENSMUSG00000058076 | Sdhc | 66052 | protein_coding | 4574.262122 | 0.111422055 | 0.069044688 | 1.613767231 | 0.106577925 | 0.298970638 | 0.111422055 |
| ENSMUSG00000039117 | Taf4 | 228980 | protein_coding | 716.1376213 | -0.111435743 | 0.056131666 | -1.985256297 | 0.047115964 | 0.177906613 | 0.111435743 |
| ENSMUSG00000020307 | Cdc34 | 216150 | protein_coding | 973.557464 | 0.111482117 | 0.085646854 | 1.301648698 | 0.193036504 | 0.424880538 | 0.111482117 |
| ENSMUSG00000006715 | Gmnn | 57441 | protein_coding | 346.1465971 | 0.111496499 | 0.094024953 | 1.185818184 | 0.235694109 | 0.475903204 | 0.111496499 |
| ENSMUSG00000050732 | Vamp8 | 22320 | protein_coding | 2575.059243 | 0.111578339 | 0.052247455 | 2.135574616 | 0.032714102 | 0.138216735 | 0.111578339 |
| ENSMUSG00000057778 | Cyb5d2 | 192986 | protein_coding | 173.6995428 | 0.111628886 | 0.090976003 | 1.227014619 | 0.219817096 | 0.456057983 | 0.111628886 |
| ENSMUSG00000054381 | Zfp747 | 269997 | protein_coding | 338.2089091 | -0.111632177 | 0.070138171 | -1.591603768 | 0.111473763 | 0.306938257 | 0.111632177 |
| ENSMUSG00000026594 | Ralgps2 | 78255 | protein_coding | 1282.733592 | 0.11164818 | 0.078381354 | 1.424422703 | 0.154324146 | 0.371849022 | 0.11164818 |
| ENSMUSG00000028557 | Rnf11 | 29864 | protein_coding | 1614.313703 | 0.111665366 | 0.052177834 | 2.140092016 | 0.032347331 | 0.137199451 | 0.111665366 |
| ENSMUSG00000079317 | Trappc2 | 66226 | protein_coding | 55.41421045 | -0.111710274 | 0.178517467 | -0.625766632 | 0.531468021 | 0.740026398 | 0.111710274 |
| ENSMUSG00000032604 | Qars | 97541 | protein_coding | 3731.118454 | 0.111721109 | 0.039646752 | 2.817913258 | 0.004833687 | 0.034738353 | 0.111721109 |
| ENSMUSG00000024215 | Spdef | 30051 | protein_coding | 1994.533442 | 0.111757748 | 0.094857578 | 1.178163625 | 0.238731381 | 0.478917898 | 0.111757748 |
| ENSMUSG00000118559 | A930007A09Rik | 432999 | protein_coding | 13.85682779 | 0.111829908 | 0.343398396 | 0.325656467 | 0.744684285 | 0.878467081 | 0.111829908 |
| ENSMUSG00000061751 | Kalrn | 545156 | protein_coding | 819.8174809 | -0.111839386 | 0.08975832 | -1.246005786 | 0.212762271 | 0.448407886 | 0.111839386 |
| ENSMUSG00000029556 | Hnf1a | 21405 | protein_coding | 556.2304897 | 0.111854017 | 0.066501194 | 1.681985105 | 0.092571728 | 0.274380493 | 0.111854017 |
| ENSMUSG00000001995 | Sipa1l2 | 244668 | protein_coding | 654.5889706 | 0.11187899 | 0.051852093 | 2.15765619 | 0.030954572 | 0.13300792 | 0.11187899 |
| ENSMUSG00000058900 | Rsl1 | 380855 | protein_coding | 127.0037133 | -0.111880504 | 0.111232235 | -1.005828073 | 0.314498281 | 0.558770915 | 0.111880504 |
| ENSMUSG00000043366 | Or51e2 | 170639 | protein_coding | 17.89484671 | 0.111883458 | 0.273236807 | 0.409474328 | 0.682191603 | 0.83937621 | 0.111883458 |

Supplementary Table 1

| ensembl_gene_id | external_gene_name | entrezgene_id | gene_biotype | baseMean | log2FoldChange | lfcSE | stat | pvalue | padj | absLog2FC |
|---|---|---|---|---|---|---|---|---|---|---|
| ENSMUSG00000000804 | Usp32 | 237898 | protein_coding | 924.9673956 | -0.111897782 | 0.048036003 | -2.329456571 | 0.019834891 | 0.097583066 | 0.111897782 |
| ENSMUSG00000050199 | Lgr4 | 107515 | protein_coding | 6687.714829 | -0.111937966 | 0.071917192 | -1.5564841 | 0.119593021 | 0.319946717 | 0.111937966 |
| ENSMUSG00000001134 | Uxt | 22294 | protein_coding | 255.6413095 | -0.111949527 | 0.074773509 | -1.497181684 | 0.13434599 | 0.34205838 | 0.111949527 |
| ENSMUSG00000051853 | Arf3 | 11842 | protein_coding | 2865.829144 | -0.111963836 | 0.064549036 | -1.734554733 | 0.082819703 | 0.255819451 | 0.111963836 |
| ENSMUSG00000068798 | Rap1a | 109905 | protein_coding | 1060.833022 | 0.112016843 | 0.043239631 | 2.590605903 | 0.009580714 | 0.057587839 | 0.112016843 |
| ENSMUSG00000000486 | Septin1 | 54204 | protein_coding | 28.62377459 | 0.112025037 | 0.252254553 | 0.444095204 | 0.656973754 | 0.824251158 | 0.112025037 |
| ENSMUSG00000032860 | P2ry2 | 18442 | protein_coding | 477.6477863 | -0.112156193 | 0.066426213 | -1.688432733 | 0.091328191 | 0.272346476 | 0.112156193 |
| ENSMUSG00000074476 | Spc24 | 67629 | protein_coding | 283.9831449 | 0.112174329 | 0.114803813 | 0.977095845 | 0.328521705 | 0.572083958 | 0.112174329 |
| ENSMUSG00000058690 | Ccser2 | 72972 | protein_coding | 1332.627815 | 0.112174486 | 0.050434971 | 2.224140988 | 0.02613896 | 0.118377705 | 0.112174486 |
| ENSMUSG00000020184 | Mdm2 | 17246 | protein_coding | 848.6335143 | -0.112187792 | 0.060295682 | -1.860627291 | 0.062796828 | 0.214197218 | 0.112187792 |
| ENSMUSG00000087022 | 9130024F11Rik | 329160 | lncRNA | 276.8832859 | 0.112191757 | 0.090713479 | 1.236770525 | 0.216172288 | 0.452108738 | 0.112191757 |
| ENSMUSG00000026121 | Sema4c | 20353 | protein_coding | 87.74887623 | -0.11219304 | 0.147040484 | -0.763007823 | 0.445458731 | 0.676462024 | 0.11219304 |
| ENSMUSG00000026718 | Stam | 20844 | protein_coding | 1079.0415 | -0.112283111 | 0.046082269 | -2.436579493 | 0.01482691 | 0.079571907 | 0.112283111 |
| ENSMUSG00000040675 | Mthfd1l | 270685 | protein_coding | 379.1262715 | 0.112301874 | 0.103345434 | 1.086665078 | 0.277184853 | 0.51990697 | 0.112301874 |
| ENSMUSG00000015120 | Ube2i | 22196 | protein_coding | 1189.277998 | -0.112302596 | 0.069396288 | -1.618279592 | 0.105602357 | 0.297761537 | 0.112302596 |
| ENSMUSG00000050973 | Gdpgp1 | 269952 | protein_coding | 292.5225474 | -0.112322505 | 0.073903757 | -1.519848378 | 0.128549087 | 0.333871046 | 0.112322505 |
| ENSMUSG00000061518 | Cox5b | 12859 | protein_coding | 9645.226243 | 0.112356132 | 0.074945384 | 1.499173468 | 0.133828636 | 0.341184624 | 0.112356132 |
| ENSMUSG00000063884 | Ptcd3 | 69956 | protein_coding | 1020.229693 | -0.11237452 | 0.070766601 | -1.587959835 | 0.112295422 | 0.308484011 | 0.11237452 |
| ENSMUSG00000036537 | Rnf113a1 | 69942 | protein_coding | 43.46780054 | 0.112375217 | 0.20052498 | 0.560405075 | 0.575203171 | 0.76841464 | 0.112375217 |
| ENSMUSG00000033717 | Adra2a | 11551 | protein_coding | 919.4630524 | 0.11239924 | 0.146824294 | 0.765535706 | 0.443952613 | 0.674900387 | 0.11239924 |
| ENSMUSG00000003271 | Sult2b1 | 54200 | protein_coding | 453.5992742 | -0.112402154 | 0.112059635 | -1.003056573 | 0.315833566 | 0.560027259 | 0.112402154 |
| ENSMUSG00000019952 | Poc1b | 382406 | protein_coding | 486.5234626 | 0.112419814 | 0.071518931 | 1.571888912 | 0.115976316 | 0.314662099 | 0.112419814 |
| ENSMUSG00000073530 | Pappa2 | 23850 | protein_coding | 130.0568069 | 0.1124268 | 0.184650485 | 0.60886274 | 0.542615423 | 0.747864433 | 0.1124268 |
| ENSMUSG00000006585 | Cdt1 | 67177 | protein_coding | 382.4252058 | -0.112436622 | 0.135442397 | -0.830143475 | 0.406457669 | 0.645466424 | 0.112436622 |
| ENSMUSG00000036555 | Iqce | 74239 | protein_coding | 236.2279074 | 0.112443856 | 0.098027644 | 1.147062723 | 0.251355699 | 0.492421447 | 0.112443856 |
| ENSMUSG00000022696 | Sidt1 | 320007 | protein_coding | 720.9815981 | 0.112468369 | 0.073157258 | 1.537350791 | 0.12420743 | 0.327122701 | 0.112468369 |
| ENSMUSG00000003283 | Hck | 15162 | protein_coding | 384.8086378 | 0.112469414 | 0.104130004 | 1.080086521 | 0.280103653 | 0.523118689 | 0.112469414 |
| ENSMUSG00000039395 | Mreg | 381269 | protein_coding | 21.85997989 | -0.112471139 | 0.266203989 | -0.422499826 | 0.672660228 | 0.833307074 | 0.112471139 |
| ENSMUSG00000000530 | Acvrl1 | 11482 | protein_coding | 93.34048019 | 0.112533633 | 0.161619953 | 0.696285519 | 0.48625004 | 0.707337899 | 0.112533633 |
| ENSMUSG00000040717 | Il17rd | 171463 | protein_coding | 120.9545949 | -0.112562696 | 0.158357324 | -0.710814585 | 0.477199141 | 0.699935475 | 0.112562696 |
| ENSMUSG00000030527 | Crtc3 | 70461 | protein_coding | 292.8542321 | -0.112588221 | 0.0897126 | -1.254987828 | 0.209483179 | 0.444801234 | 0.112588221 |
| ENSMUSG00000042369 | Rbm45 | 241490 | protein_coding | 910.2816205 | 0.112590887 | 0.07883618 | 1.428162648 | 0.153245049 | 0.370189171 | 0.112590887 |
| ENSMUSG00000026672 | Optn | 71648 | protein_coding | 847.7412402 | -0.112601113 | 0.072406649 | -1.555121188 | 0.11991721 | 0.320286898 | 0.112601113 |
| ENSMUSG00000017801 | Mlx | 21428 | protein_coding | 2379.232202 | 0.112627273 | 0.056911386 | 1.978993657 | 0.04781672 | 0.179562869 | 0.112627273 |
| ENSMUSG00000030787 | Lyve1 | 114332 | protein_coding | 56.95669001 | 0.112642322 | 0.190323922 | 0.59184532 | 0.553954172 | 0.7540828 | 0.112642322 |
| ENSMUSG00000085436 | Zfp335os | 629777 | lncRNA | 27.6743042 | 0.112649778 | 0.256265296 | 0.43958265 | 0.660239409 | 0.826041967 | 0.112649778 |
| ENSMUSG00000071180 | Smim15 | 75616 | protein_coding | 1640.984198 | 0.112681967 | 0.050068539 | 2.250554311 | 0.02441378 | 0.112975905 | 0.112681967 |
| ENSMUSG00000057842 | Zfp595 | 218314 | protein_coding | 134.4592813 | -0.112685344 | 0.142523833 | -0.790642114 | 0.429152864 | 0.664262236 | 0.112685344 |
| ENSMUSG00000046567 | 4930430F08Rik | 68281 | protein_coding | 63.47376747 | 0.112739513 | 0.175405232 | 0.642737454 | 0.520394473 | 0.73200189 | 0.112739513 |

| ensembl_gene_id | external_gene_name | entrezgene_id | gene_biotype | baseMean | log2FoldChange | lfcSE | stat | pvalue | padj | absLog2FC |
|---|---|---|---|---|---|---|---|---|---|---|
| ENSMUSG00000070420 | Zscan25 | 666311 | protein_coding | 262.7870236 | -0.112771364 | 0.076142924 | -1.481048507 | 0.138593648 | 0.34868951 | 0.112771364 |
| ENSMUSG00000030616 | Sytl2 | 83671 | protein_coding | 2262.642344 | 0.112784757 | 0.06578003 | 1.714574415 | 0.086423283 | 0.263070646 | 0.112784757 |
| ENSMUSG00000049659 | Aftph | 216549 | protein_coding | 3147.482223 | 0.112815333 | 0.045977016 | 2.453733226 | 0.014138182 | 0.076795712 | 0.112815333 |
| ENSMUSG00000042557 | Sin3a | 20466 | protein_coding | 1191.079576 | -0.112825009 | 0.044089579 | -2.558994902 | 0.010497528 | 0.061495716 | 0.112825009 |
| ENSMUSG00000041144 | Dnah7b | 227058 | protein_coding | 29.21770566 | 0.112852098 | 0.266568256 | 0.423351599 | 0.672038753 | 0.832857063 | 0.112852098 |
| ENSMUSG00000030254 | Rad18 | 58186 | protein_coding | 142.1569796 | 0.112919721 | 0.15340968 | 0.736066466 | 0.461690249 | 0.68813359 | 0.112919721 |
| ENSMUSG00000059248 | Septin9 | 53860 | protein_coding | 955.6940439 | 0.112936104 | 0.086910182 | 1.29945769 | 0.193786905 | 0.425544854 | 0.112936104 |
| ENSMUSG00000041716 | Gm20604 | 100859931 | protein_coding | 87.81067902 | 0.113032501 | 0.12878551 | 0.877680271 | 0.380117254 | 0.621026335 | 0.113032501 |
| ENSMUSG00000058793 | Cds2 | 110911 | protein_coding | 1951.50855 | -0.11306649 | 0.047187183 | -2.396127122 | 0.016569343 | 0.086281495 | 0.11306649 |
| ENSMUSG00000018750 | Zbtb4 | 75580 | protein_coding | 553.5644986 | -0.11307386 | 0.070379852 | -1.606622592 | 0.108137184 | 0.301243853 | 0.11307386 |
| ENSMUSG00000028974 | Dffa | 13347 | protein_coding | 462.4978242 | -0.113073997 | 0.080387682 | -1.406608503 | 0.15954351 | 0.379596103 | 0.113073997 |
| ENSMUSG00000063856 | Gpx1 | 14775 | protein_coding | 4637.417581 | 0.113119445 | 0.071028896 | 1.592583453 | 0.111253668 | 0.306582205 | 0.113119445 |
| ENSMUSG00000055884 | Fancm | 104806 | protein_coding | 198.276768 | 0.113177562 | 0.091275286 | 1.239958453 | 0.214990762 | 0.45050036 | 0.113177562 |
| ENSMUSG00000039795 | Zfand1 | 66361 | protein_coding | 466.4388861 | -0.113200883 | 0.093201641 | -1.214580356 | 0.224526185 | 0.462553939 | 0.113200883 |
| ENSMUSG00000021470 | Ercc6l2 | 76251 | protein_coding | 594.100615 | -0.113204903 | 0.070569027 | -1.604172641 | 0.108676003 | 0.302001032 | 0.113204903 |
| ENSMUSG00000035614 | Togaram1 | 328108 | protein_coding | 978.3939816 | -0.113217308 | 0.048779542 | -2.320999822 | 0.020286853 | 0.099261555 | 0.113217308 |
| ENSMUSG00000020623 | Map2k6 | 26399 | protein_coding | 477.8712835 | 0.113238186 | 0.061631423 | 1.83734498 | 0.066158984 | 0.221864406 | 0.113238186 |
| ENSMUSG00000059456 | Ptk2b | 19229 | protein_coding | 1482.446186 | -0.113333654 | 0.04630405 | -2.447597005 | 0.014381241 | 0.077775409 | 0.113333654 |
| ENSMUSG00000048921 | Zfp689 | 71131 | protein_coding | 69.19522062 | -0.113337563 | 0.151081265 | -0.750176155 | 0.453148618 | 0.681964144 | 0.113337563 |
| ENSMUSG00000022337 | Emc2 | 66736 | protein_coding | 1027.088695 | 0.113341657 | 0.063841153 | 1.775369845 | 0.07583684 | 0.242470403 | 0.113341657 |
| ENSMUSG00000032652 | Crebl2 | 232430 | protein_coding | 507.8315535 | -0.113365345 | 0.078124779 | -1.451080521 | 0.14675744 | 0.359972154 | 0.113365345 |
| ENSMUSG00000024386 | Proc | 19123 | protein_coding | 61.23046621 | -0.113393731 | 0.211945131 | -0.535014562 | 0.592639788 | 0.780331635 | 0.113393731 |
| ENSMUSG00000004266 | Ptpn6 | 15170 | protein_coding | 665.7858621 | 0.113430341 | 0.062352265 | 1.819185578 | 0.068883122 | 0.227778986 | 0.113430341 |
| ENSMUSG00000098134 | Rnf113a2 | 66381 | protein_coding | 259.3378623 | 0.113444056 | 0.094634986 | 1.198753865 | 0.230623666 | 0.469603637 | 0.113444056 |
| ENSMUSG00000029462 | Vps29 | 56433 | protein_coding | 691.3126442 | 0.113481096 | 0.057344142 | 1.978948348 | 0.047821822 | 0.179562869 | 0.113481096 |
| ENSMUSG00000009828 | Cilk1 | 56542 | protein_coding | 2572.710339 | 0.113498529 | 0.055126032 | 2.058891691 | 0.039504615 | 0.157189965 | 0.113498529 |
| ENSMUSG00000030970 | Ctbp2 | 13017 | protein_coding | 1578.677971 | 0.113532005 | 0.039561185 | 2.869782743 | 0.004107539 | 0.030824702 | 0.113532005 |
| ENSMUSG00000020739 | Nup85 | 445007 | protein_coding | 560.9220618 | 0.113538859 | 0.093362683 | 1.216105359 | 0.22394479 | 0.461717926 | 0.113538859 |
| ENSMUSG00000031879 | Ciao2b | 68523 | protein_coding | 553.6725936 | 0.113554818 | 0.096658304 | 1.174806647 | 0.24007208 | 0.480533622 | 0.113554818 |
| ENSMUSG00000032370 | Lactb | 80907 | protein_coding | 382.6026075 | 0.113564176 | 0.088788612 | 1.279039882 | 0.200883012 | 0.4346177 | 0.113564176 |
| ENSMUSG00000001583 | Tnk1 | 83813 | protein_coding | 664.2239209 | -0.113571456 | 0.072423276 | -1.568162376 | 0.116843243 | 0.316056856 | 0.113571456 |
| ENSMUSG00000025937 | Lactb2 | 212442 | protein_coding | 595.5928639 | 0.113574721 | 0.058606789 | 1.937910651 | 0.052634124 | 0.191597869 | 0.113574721 |
| ENSMUSG00000092203 | 1110038B12Rik | 68763 | lncRNA | 1160.438621 | -0.113577749 | 0.077607788 | -1.463483907 | 0.143335014 | 0.355303993 | 0.113577749 |
| ENSMUSG00000020441 | 2310033P09Rik | 67862 | protein_coding | 186.2365677 | -0.113580045 | 0.107556182 | -1.056006669 | 0.290965168 | 0.53329119 | 0.113580045 |
| ENSMUSG00000018001 | Cyth3 | 19159 | protein_coding | 420.1947678 | -0.113582193 | 0.077226051 | -1.470775631 | 0.141351805 | 0.352354239 | 0.113582193 |
| ENSMUSG00000045211 | Nudt18 | 213484 | protein_coding | 150.4509288 | 0.113594408 | 0.116637335 | 0.973911209 | 0.330100628 | 0.573740112 | 0.113594408 |
| ENSMUSG00000028382 | Ptbp3 | 230257 | protein_coding | 7005.783861 | -0.113665584 | 0.059346838 | -1.915276165 | 0.055457294 | 0.198349916 | 0.113665584 |
| ENSMUSG00000035069 | Oma1 | 67013 | protein_coding | 955.8144026 | 0.113675378 | 0.074019966 | 1.535739398 | 0.124602308 | 0.327433832 | 0.113675378 |
| ENSMUSG00000022867 | Usp25 | 30940 | protein_coding | 1714.475882 | -0.113727563 | 0.058145552 | -1.955911661 | 0.050475561 | 0.186231841 | 0.113727563 |

| ensembl_gene_id | external_gene_name | entrezgene_id | gene_biotype | baseMean | log2FoldChange | lfcSE | stat | pvalue | padj | absLog2FC |
|---|---|---|---|---|---|---|---|---|---|---|
| ENSMUSG00000015597 | Zfp318 | 57908 | protein_coding | 453.9629346 | -0.113772221 | 0.061020835 | -1.864481556 | 0.062254109 | 0.21306899 | 0.113772221 |
| ENSMUSG00000021282 | Eif5 | 217869 | protein_coding | 5288.825179 | -0.113773438 | 0.066427568 | -1.712744304 | 0.086759591 | 0.263676126 | 0.113773438 |
| ENSMUSG00000057594 | Arl16 | 70317 | protein_coding | 147.0977952 | 0.113807848 | 0.109154228 | 1.042633443 | 0.297118094 | 0.540135201 | 0.113807848 |
| ENSMUSG00000021400 | Wrnip1 | 78903 | protein_coding | 696.4461873 | 0.113814978 | 0.057356494 | 1.984343354 | 0.047217576 | 0.178248561 | 0.113814978 |
| ENSMUSG00000015093 | Clic3 | 69454 | protein_coding | 9.169216026 | -0.113835536 | 0.383509712 | -0.296825692 | 0.766599589 | 0.890414429 | 0.113835536 |
| ENSMUSG00000037926 | Ssh2 | 237860 | protein_coding | 966.3408464 | 0.113849709 | 0.066367002 | 1.715456545 | 0.086261556 | 0.262627924 | 0.113849709 |
| ENSMUSG00000025428 | Atp5a1 | 11946 | protein_coding | 43047.00363 | 0.113881146 | 0.057598584 | 1.977151822 | 0.048024471 | 0.179863133 | 0.113881146 |
| ENSMUSG00000031328 | Flna | 192176 | protein_coding | 562.9492584 | 0.113943594 | 0.089503932 | 1.273056863 | 0.202997865 | 0.437065467 | 0.113943594 |
| ENSMUSG00000035890 | Rnf126 | 70294 | protein_coding | 572.6903305 | 0.113978293 | 0.076637491 | 1.487239365 | 0.136951603 | 0.346341304 | 0.113978293 |
| ENSMUSG00000030956 | Fam53b | 77938 | protein_coding | 691.6214351 | -0.113990926 | 0.07643086 | -1.491425395 | 0.135849845 | 0.344625981 | 0.113990926 |
| ENSMUSG00000050234 | Gja4 | 14612 | protein_coding | 8.655436655 | -0.11399545 | 0.413427475 | -0.275732642 | 0.782753423 | 0.898825528 | 0.11399545 |
| ENSMUSG00000028229 | Rmdn1 | 66302 | protein_coding | 241.6393808 | -0.114013167 | 0.091870763 | -1.241016873 | 0.214599516 | 0.450343118 | 0.114013167 |
| ENSMUSG00000054843 | Atrnl1 | 226255 | protein_coding | 3485.700512 | -0.114029987 | 0.041843102 | -2.725180068 | 0.006426643 | 0.043040891 | 0.114029987 |
| ENSMUSG00000024537 | Psmg2 | 107047 | protein_coding | 251.1046263 | 0.114053968 | 0.080962392 | 1.408727742 | 0.158915689 | 0.379061279 | 0.114053968 |
| ENSMUSG00000040918 | Slc19a2 | 116914 | protein_coding | 229.4757077 | -0.114070375 | 0.102152395 | -1.116668634 | 0.264136034 | 0.505545025 | 0.114070375 |
| ENSMUSG00000038112 | AW551984 | 244810 | protein_coding | 35.84562513 | 0.114107816 | 0.203212969 | 0.561518373 | 0.574444209 | 0.768186995 | 0.114107816 |
| ENSMUSG00000063275 | Hacd1 | 30963 | protein_coding | 498.0892173 | -0.114123457 | 0.073523819 | -1.552197073 | 0.120615076 | 0.321407689 | 0.114123457 |
| ENSMUSG00000005779 | Psmb4 | 19172 | protein_coding | 3072.579994 | 0.114131661 | 0.084854906 | 1.345021364 | 0.178618335 | 0.405715197 | 0.114131661 |
| ENSMUSG00000029146 | Snx17 | 266781 | protein_coding | 1272.297683 | 0.114144573 | 0.046906306 | 2.433459015 | 0.014955328 | 0.079994886 | 0.114144573 |
| ENSMUSG00000022601 | Zbtb11 | 271377 | protein_coding | 568.3770553 | -0.114149778 | 0.062835826 | -1.816635276 | 0.069272978 | 0.228412455 | 0.114149778 |
| ENSMUSG00000031400 | G6pdx | 14381 | protein_coding | 1262.303456 | 0.114159188 | 0.055785954 | 2.046378684 | 0.040719128 | 0.160556814 | 0.114159188 |
| ENSMUSG00000047507 | Baiap3 | 545192 | protein_coding | 78.55941026 | -0.114184226 | 0.15124475 | -0.754963241 | 0.450271039 | 0.679535938 | 0.114184226 |
| ENSMUSG00000074890 | Lcmt2 | 329504 | protein_coding | 234.5382526 | -0.114184642 | 0.095171626 | -1.199776092 | 0.230226312 | 0.469307431 | 0.114184642 |
| ENSMUSG00000036845 | Lin37 | 75660 | protein_coding | 229.4187805 | 0.114211785 | 0.104070192 | 1.097449549 | 0.272444924 | 0.514863213 | 0.114211785 |
| ENSMUSG00000059689 | Zfp637 | 232337 | protein_coding | 267.1246512 | 0.114218264 | 0.082039502 | 1.39223497 | 0.163851256 | 0.38575609 | 0.114218264 |
| ENSMUSG00000027810 | Eif2a | 229317 | protein_coding | 1291.177619 | -0.114219833 | 0.068903365 | -1.6576815 | 0.09738177 | 0.283377084 | 0.114219833 |
| ENSMUSG00000048280 | Zfp738 | 408068 | protein_coding | 270.8070885 | -0.11424159 | 0.090909826 | -1.256647326 | 0.208881369 | 0.444468907 | 0.11424159 |
| ENSMUSG00000075419 | Dolk | 227697 | protein_coding | 415.9288298 | 0.114283334 | 0.080013612 | 1.428298648 | 0.153205917 | 0.370189171 | 0.114283334 |
| ENSMUSG00000033880 | Lgals3bp | 19039 | protein_coding | 2230.069646 | 0.114328966 | 0.077181172 | 1.481306418 | 0.138524939 | 0.348679833 | 0.114328966 |
| ENSMUSG00000022847 | Thpo | 21832 | protein_coding | 11.13436587 | 0.114336476 | 0.370078861 | 0.308951653 | 0.757358304 | 0.885066476 | 0.114336476 |
| ENSMUSG00000071757 | Zhx2 | 387609 | protein_coding | 317.2680636 | 0.114339437 | 0.070676812 | 1.617778628 | 0.105710314 | 0.29784969 | 0.114339437 |
| ENSMUSG00000067995 | Gtf2f2 | 68705 | protein_coding | 410.9675287 | 0.114395845 | 0.067783864 | 1.687656007 | 0.091477281 | 0.272438433 | 0.114395845 |
| ENSMUSG00000046058 | Eid2 | 386655 | protein_coding | 14.26673189 | 0.114436519 | 0.302950795 | 0.377739623 | 0.70562403 | 0.853575129 | 0.114436519 |
| ENSMUSG00000030701 | Plekhb1 | 27276 | protein_coding | 21.71534653 | -0.11456694 | 0.4325759 | -0.26484818 | 0.791126419 | 0.902745048 | 0.11456694 |
| ENSMUSG00000002477 | Snrpd1 | 20641 | protein_coding | 476.0425174 | 0.114573003 | 0.097033688 | 1.18075491 | 0.237700101 | 0.47774047 | 0.114573003 |
| ENSMUSG00000016477 | E2f3 | 13557 | protein_coding | 365.5250215 | -0.114607574 | 0.087200896 | -1.314293542 | 0.18874744 | 0.419197611 | 0.114607574 |
| ENSMUSG00000021302 | Ggps1 | 14593 | protein_coding | 672.2626921 | -0.114645419 | 0.071251984 | -1.609013713 | 0.107613343 | 0.300613293 | 0.114645419 |
| ENSMUSG00000070167 | Snora57 | 118027464 | snoRNA | 315.1879448 | 0.114648365 | 0.428203548 | 0.267742678 | 0.788897396 | 0.901477241 | 0.114648365 |
| ENSMUSG00000037443 | Cep85 | 70012 | protein_coding | 686.3649981 | 0.114655366 | 0.076042804 | 1.507774046 | 0.131612367 | 0.337846137 | 0.114655366 |

| ensembl_gene_id | external_gene_name | entrezgene_id | gene_biotype | baseMean | log2FoldChange | lfcSE | stat | pvalue | padj | absLog2FC |
|---|---|---|---|---|---|---|---|---|---|---|
| ENSMUSG00000058240 | Cryzl1 | 66609 | protein_coding | 626.0804936 | -0.114678151 | 0.065853224 | -1.741420457 | 0.081609905 | 0.253441737 | 0.114678151 |
| ENSMUSG00000049106 | Dcaf5 | 320808 | protein_coding | 1037.803513 | 0.114680234 | 0.045665779 | 2.511294789 | 0.01202892 | 0.068138529 | 0.114680234 |
| ENSMUSG00000030798 | Cd37 | 12493 | protein_coding | 19.76387008 | 0.114688096 | 0.301407809 | 0.380508044 | 0.703568327 | 0.852795671 | 0.114688096 |
| ENSMUSG00000036027 | Pou2af2 | 69809 | protein_coding | 30.14322496 | -0.114742372 | 0.245947507 | -0.46653196 | 0.640834776 | 0.813310965 | 0.114742372 |
| ENSMUSG00000034799 | Unc13a | 382018 | protein_coding | 45.02006885 | -0.114765097 | 0.198583649 | -0.577918161 | 0.563319374 | 0.760667465 | 0.114765097 |
| ENSMUSG00000046707 | Csnk2a2 | 13000 | protein_coding | 554.0329747 | -0.114775532 | 0.074290525 | -1.544955189 | 0.122357103 | 0.324281511 | 0.114775532 |
| ENSMUSG00000026577 | Blzf1 | 66352 | protein_coding | 435.0848138 | 0.114795598 | 0.066288453 | 1.731758584 | 0.083316555 | 0.256862183 | 0.114795598 |
| ENSMUSG00000035372 | 1810055G02Rik | 72056 | protein_coding | 274.484897 | -0.114806817 | 0.086526438 | -1.326840892 | 0.18456132 | 0.413219549 | 0.114806817 |
| ENSMUSG00000035085 | 1700020L24Rik | 66330 | protein_coding | 39.36863281 | -0.114835026 | 0.2134607 | -0.537967997 | 0.590599143 | 0.779660849 | 0.114835026 |
| ENSMUSG00000013662 | Atad1 | 67979 | protein_coding | 2453.805426 | -0.114857453 | 0.046095297 | -2.49173908 | 0.012711938 | 0.071034452 | 0.114857453 |
| ENSMUSG00000023830 | Igf2r | 16004 | protein_coding | 5435.86328 | -0.114862674 | 0.037840688 | -3.035427744 | 0.002401948 | 0.02017511 | 0.114862674 |
| ENSMUSG00000041130 | Zfp598 | 213753 | protein_coding | 1534.549073 | -0.114869813 | 0.059457732 | -1.931957525 | 0.053364749 | 0.193265956 | 0.114869813 |
| ENSMUSG00000113450 | Zfp935 | 71508 | protein_coding | 71.9015106 | -0.114885458 | 0.159942152 | -0.718293811 | 0.472576145 | 0.696772776 | 0.114885458 |
| ENSMUSG00000043510 | Hscb | 100900 | protein_coding | 287.4229646 | -0.114890753 | 0.095439138 | -1.203811716 | 0.22866236 | 0.467736921 | 0.114890753 |
| ENSMUSG00000037020 | Wdr62 | 233064 | protein_coding | 115.7701478 | -0.114904602 | 0.126100411 | -0.911215126 | 0.362182036 | 0.605044452 | 0.114904602 |
| ENSMUSG00000028811 | Yars | 107271 | protein_coding | 1213.362023 | 0.114940847 | 0.072605164 | 1.583094667 | 0.113399889 | 0.310127356 | 0.114940847 |
| ENSMUSG00000038866 | Zcchc2 | 227449 | protein_coding | 621.8279493 | -0.114948923 | 0.054954669 | -2.091704402 | 0.036464967 | 0.148577695 | 0.114948923 |
| ENSMUSG00000035469 | Rcbtb1 | 71330 | protein_coding | 1288.057252 | 0.114958155 | 0.058638842 | 1.960443827 | 0.049943937 | 0.185048514 | 0.114958155 |
| ENSMUSG00000022014 | Epsti1 | 108670 | protein_coding | 878.2361956 | 0.114959519 | 0.068680221 | 1.673837352 | 0.094162569 | 0.277413181 | 0.114959519 |
| ENSMUSG00000021850 | ccdc198 | 67082 | protein_coding | 202.8420307 | -0.115008369 | 0.151130411 | -0.760987605 | 0.446664471 | 0.677580465 | 0.115008369 |
| ENSMUSG00000048327 | Ckap2l | 70466 | protein_coding | 278.1998949 | 0.115013527 | 0.117994582 | 0.974735659 | 0.3296914 | 0.573501586 | 0.115013527 |
| ENSMUSG00000015461 | Atf6b | 12915 | protein_coding | 788.8261548 | 0.115032203 | 0.079288448 | 1.450806589 | 0.146833724 | 0.360082332 | 0.115032203 |
| ENSMUSG00000025372 | Baiap2 | 108100 | protein_coding | 374.0147536 | 0.115050406 | 0.082027953 | 1.402575608 | 0.16074343 | 0.381255129 | 0.115050406 |
| ENSMUSG00000026199 | Ankzf1 | 52231 | protein_coding | 598.2659637 | -0.115054605 | 0.065503721 | -1.756459067 | 0.079010055 | 0.248330613 | 0.115054605 |
| ENSMUSG00000051518 | Rps19bp1 | 66538 | protein_coding | 271.0637763 | 0.115067522 | 0.076487124 | 1.504403836 | 0.132477417 | 0.339390517 | 0.115067522 |
| ENSMUSG00000021022 | Ppp2r3c | 59032 | protein_coding | 235.9779694 | -0.115093408 | 0.093518147 | -1.230706674 | 0.218432591 | 0.454666616 | 0.115093408 |
| ENSMUSG00000040761 | Spen | 56381 | protein_coding | 1097.681896 | -0.115105035 | 0.099740896 | -1.154040514 | 0.248483559 | 0.4893628 | 0.115105035 |
| ENSMUSG00000022464 | Slc38a4 | 69354 | protein_coding | 7.244294526 | 0.115106107 | 0.456138099 | 0.25234925 | 0.800771124 | 0.909183504 | 0.115106107 |
| ENSMUSG00000016637 | Ift27 | 67042 | protein_coding | 156.9098742 | 0.11510644 | 0.099186058 | 1.160510278 | 0.245841111 | 0.487001548 | 0.11510644 |
| ENSMUSG00000032294 | Pkm | 18746 | protein_coding | 17772.82479 | -0.115110082 | 0.081897897 | -1.405531603 | 0.159863259 | 0.380084737 | 0.115110082 |
| ENSMUSG00000013833 | Med16 | 216154 | protein_coding | 599.5480673 | -0.115143754 | 0.065328719 | -1.762528896 | 0.077979979 | 0.246536584 | 0.115143754 |
| ENSMUSG00000031845 | Bco1 | 63857 | protein_coding | 47.1785241 | -0.115151926 | 0.189186045 | -0.608670299 | 0.542742998 | 0.747912187 | 0.115151926 |
| ENSMUSG00000070565 | Rasal2 | 226525 | protein_coding | 254.1194698 | 0.115190572 | 0.085642117 | 1.345022469 | 0.178617979 | 0.405715197 | 0.115190572 |
| ENSMUSG00000022508 | Bcl6 | 12053 | protein_coding | 128.346317 | -0.115197262 | 0.18920079 | -0.608862478 | 0.542615597 | 0.747864433 | 0.115197262 |
| ENSMUSG00000043090 | Zfp866 | 330788 | protein_coding | 349.9903761 | -0.115211822 | 0.077013204 | -1.496000898 | 0.134653421 | 0.342321275 | 0.115211822 |
| ENSMUSG00000071078 | Nr2c2ap | 75692 | protein_coding | 46.32034747 | -0.115228462 | 0.190800551 | -0.603921014 | 0.545896157 | 0.749242602 | 0.115228462 |
| ENSMUSG00000028684 | Urod | 22275 | protein_coding | 651.3919561 | -0.11524026 | 0.069656758 | -1.654401709 | 0.09804592 | 0.284673902 | 0.11524026 |
| ENSMUSG00000022139 | Mbnl2 | 105559 | protein_coding | 4173.673694 | 0.115265557 | 0.047557708 | 2.423698725 | 0.015363345 | 0.081578678 | 0.115265557 |
| ENSMUSG00000019734 | Tmc4 | 353499 | protein_coding | 5058.002941 | -0.115322981 | 0.046833845 | -2.462385491 | 0.013801624 | 0.07559346 | 0.115322981 |

| ensembl_gene_id | external_gene_name | entrezgene_id | gene_biotype | baseMean | log2FoldChange | lfcSE | stat | pvalue | padj | absLog2FC |
|---|---|---|---|---|---|---|---|---|---|---|
| ENSMUSG00000001707 | Eef1e1 | 66143 | protein_coding | 252.9527416 | -0.115384526 | 0.078021158 | -1.478887638 | 0.139170348 | 0.349495708 | 0.115384526 |
| ENSMUSG00000048537 | Phldb1 | 102693 | protein_coding | 2531.374995 | -0.115397769 | 0.070496008 | -1.636940476 | 0.101642908 | 0.290740541 | 0.115397769 |
| ENSMUSG00000034522 | Zfp395 | 380912 | protein_coding | 232.9234263 | -0.115409369 | 0.114547235 | -1.007526448 | 0.313681854 | 0.557806513 | 0.115409369 |
| ENSMUSG00000024228 | Nudt12 | 67993 | protein_coding | 396.5349688 | 0.11541136 | 0.067165787 | 1.718305782 | 0.085740855 | 0.261585844 | 0.11541136 |
| ENSMUSG00000005732 | Ranbp1 | 19385 | protein_coding | 657.9035135 | 0.115414597 | 0.072080711 | 1.6011856 | 0.109335817 | 0.302965417 | 0.115414597 |
| ENSMUSG00000021072 | Tmx1 | 72736 | protein_coding | 1614.196528 | 0.115511001 | 0.048726151 | 2.37061619 | 0.017758461 | 0.090515394 | 0.115511001 |
| ENSMUSG00000073424 | Cyp4f15 | 106648 | protein_coding | 13.13898226 | -0.11551869 | 0.424428715 | -0.272174541 | 0.785487817 | 0.900132312 | 0.11551869 |
| ENSMUSG00000020180 | Snrpd3 | 67332 | protein_coding | 330.4741984 | 0.115579113 | 0.078449766 | 1.473288189 | 0.140673344 | 0.35172698 | 0.115579113 |
| ENSMUSG00000000568 | Hnrnpd | 11991 | protein_coding | 971.4367483 | -0.115631517 | 0.061579497 | -1.877760016 | 0.060414002 | 0.208897216 | 0.115631517 |
| ENSMUSG00000028007 | Snx7 | 76561 | protein_coding | 1294.20864 | 0.115633387 | 0.06934804 | 1.667435538 | 0.095427832 | 0.279964463 | 0.115633387 |
| ENSMUSG00000005823 | Gpr108 | 78308 | protein_coding | 2335.599919 | 0.11564627 | 0.052418995 | 2.20619012 | 0.027370691 | 0.121700339 | 0.11564627 |
| ENSMUSG00000024544 | Ldlrad4 | 52662 | protein_coding | 48.83612017 | 0.115671071 | 0.189612893 | 0.610038006 | 0.541836632 | 0.747365665 | 0.115671071 |
| ENSMUSG00000037434 | Slc30a1 | 22782 | protein_coding | 1477.377809 | -0.115672604 | 0.06334014 | -1.826213259 | 0.067818145 | 0.225459333 | 0.115672604 |
| ENSMUSG00000035769 | Xylb | 102448 | protein_coding | 113.7003527 | -0.115694421 | 0.141258672 | -0.819025263 | 0.412772 | 0.650456717 | 0.115694421 |
| ENSMUSG00000038095 | Sbno1 | 243272 | protein_coding | 2813.699416 | -0.115699154 | 0.033251909 | -3.479474074 | 0.000502399 | 0.005665866 | 0.115699154 |
| ENSMUSG00000021962 | Dcp1a | 75901 | protein_coding | 587.499181 | -0.115718053 | 0.064427553 | -1.796095723 | 0.072479295 | 0.235215876 | 0.115718053 |
| ENSMUSG00000031974 | Abcb10 | 56199 | protein_coding | 891.8371506 | -0.115726193 | 0.062996963 | -1.837012248 | 0.066208087 | 0.221890653 | 0.115726193 |
| ENSMUSG00000020901 | Pik3r5 | 320207 | protein_coding | 260.5384239 | -0.115731335 | 0.137835422 | -0.839634201 | 0.401113517 | 0.640407651 | 0.115731335 |
| ENSMUSG00000021913 | Ogdhl | 239017 | protein_coding | 29.72243456 | 0.115733661 | 0.275692533 | 0.419792512 | 0.674637035 | 0.834405417 | 0.115733661 |
| ENSMUSG00000031659 | Adcy7 | 11513 | protein_coding | 41.3380174 | 0.115734501 | 0.202589483 | 0.571275962 | 0.567812594 | 0.763029223 | 0.115734501 |
| ENSMUSG00000031256 | Cstf2 | 108062 | protein_coding | 1544.037696 | 0.115760328 | 0.051558141 | 2.245238609 | 0.024752819 | 0.114021339 | 0.115760328 |
| ENSMUSG00000035048 | Anapc13 | 69010 | protein_coding | 852.324121 | 0.115764813 | 0.048324662 | 2.395563863 | 0.016594823 | 0.086330552 | 0.115764813 |
| ENSMUSG00000028454 | Pigo | 56703 | protein_coding | 766.7917109 | 0.115777294 | 0.080552903 | 1.437282704 | 0.150637681 | 0.366304868 | 0.115777294 |
| ENSMUSG00000032078 | Zpr1 | 22687 | protein_coding | 627.336632 | -0.11578679 | 0.061737658 | -1.87546455 | 0.060728842 | 0.20964847 | 0.11578679 |
| ENSMUSG00000032000 | Birc3 | 11796 | protein_coding | 961.0959153 | -0.115838367 | 0.087132988 | -1.329443304 | 0.183701759 | 0.412210695 | 0.115838367 |
| ENSMUSG00000049604 | Hoxb13 | 15408 | protein_coding | 5840.33259 | 0.115844428 | 0.173980841 | 0.665845889 | 0.50550961 | 0.721608883 | 0.115844428 |
| ENSMUSG00000001300 | Efnb2 | 13642 | protein_coding | 1996.332697 | 0.115846689 | 0.07242712 | 1.599493241 | 0.109711049 | 0.303702082 | 0.115846689 |
| ENSMUSG00000041560 | Nop53 | 68077 | protein_coding | 3580.060043 | -0.115858217 | 0.046811475 | -2.474996077 | 0.013323764 | 0.073636854 | 0.115858217 |
| ENSMUSG00000049232 | Tigd2 | 68140 | protein_coding | 422.0222628 | -0.115869536 | 0.091313766 | -1.268916413 | 0.204470878 | 0.438439283 | 0.115869536 |
| ENSMUSG00000021767 | Kat6b | 54169 | protein_coding | 1050.005851 | 0.115898952 | 0.072920969 | 1.589377555 | 0.111975179 | 0.307740918 | 0.115898952 |
| ENSMUSG00000034744 | Nagk | 56174 | protein_coding | 964.4033226 | -0.11591289 | 0.050553057 | -2.292895751 | 0.021854008 | 0.104705757 | 0.11591289 |
| ENSMUSG00000031976 | Urb2 | 382038 | protein_coding | 552.7195253 | -0.115922192 | 0.05752272 | -2.015241854 | 0.043879314 | 0.169495017 | 0.115922192 |
| ENSMUSG00000022018 | Rgcc | 66214 | protein_coding | 168.6330801 | -0.115953095 | 0.123886693 | -0.935960854 | 0.349293343 | 0.591707327 | 0.115953095 |
| ENSMUSG00000046613 | Vwa5b2 | 328643 | protein_coding | 41.10479535 | -0.115957128 | 0.223062945 | -0.51984039 | 0.603174826 | 0.787962771 | 0.115957128 |
| ENSMUSG00000027374 | Mrps5 | 77721 | protein_coding | 849.248024 | 0.116011849 | 0.05997188 | 1.934437425 | 0.053059369 | 0.192513581 | 0.116011849 |
| ENSMUSG00000024429 | Gnl1 | 14670 | protein_coding | 780.4519865 | -0.116020201 | 0.058293665 | -1.990271169 | 0.046561072 | 0.176237021 | 0.116020201 |
| ENSMUSG00000060073 | Psma3 | 19167 | protein_coding | 1720.157886 | 0.116031094 | 0.049200354 | 2.358338623 | 0.018356939 | 0.092745101 | 0.116031094 |
| ENSMUSG00000017405 | Nek8 | 140859 | protein_coding | 325.0589074 | -0.116079828 | 0.084462177 | -1.374340949 | 0.169335859 | 0.393611906 | 0.116079828 |
| ENSMUSG00000002307 | Daxx | 13163 | protein_coding | 628.3407043 | -0.11608289 | 0.065626861 | -1.768831968 | 0.07692192 | 0.244581981 | 0.11608289 |

| ensembl_gene_id | external_gene_name | entrezgene_id | gene_biotype | baseMean | log2FoldChange | lfcSE | stat | pvalue | padj | absLog2FC |
|---|---|---|---|---|---|---|---|---|---|---|
| ENSMUSG00000025584 | Pde8a | 18584 | protein_coding | 176.6705852 | 0.116083408 | 0.098846546 | 1.174380012 | 0.240242848 | 0.480694343 | 0.116083408 |
| ENSMUSG00000031577 | Tti2 | 234138 | protein_coding | 269.85062 | -0.116221234 | 0.085724352 | -1.355755175 | 0.175177078 | 0.401524931 | 0.116221234 |
| ENSMUSG00000030591 | Psmd8 | 57296 | protein_coding | 2925.538578 | 0.116231461 | 0.057601393 | 2.017858522 | 0.043605998 | 0.168799213 | 0.116231461 |
| ENSMUSG00000028927 | Padi2 | 18600 | protein_coding | 150.2848708 | -0.116254268 | 0.141198899 | -0.823336929 | 0.410316419 | 0.648598559 | 0.116254268 |
| ENSMUSG00000037025 | Foxa2 | 15376 | protein_coding | 724.5729342 | 0.116274795 | 0.120307436 | 0.966480528 | 0.333803786 | 0.577106953 | 0.116274795 |
| ENSMUSG00000028820 | Sfpq | 71514 | protein_coding | 2839.574361 | 0.116275157 | 0.065670278 | 1.770590309 | 0.076628855 | 0.244015888 | 0.116275157 |
| ENSMUSG00000002524 | Puf60 | 67959 | protein_coding | 2383.461925 | 0.116299153 | 0.066713414 | 1.743264901 | 0.081287353 | 0.252829537 | 0.116299153 |
| ENSMUSG00000034111 | Tmed8 | 382620 | protein_coding | 131.5678074 | 0.116318941 | 0.103109974 | 1.12810562 | 0.259275316 | 0.500967327 | 0.116318941 |
| ENSMUSG00000043541 | Casc1 | 320662 | protein_coding | 9.009998985 | -0.116348267 | 0.432740293 | -0.268863957 | 0.788034374 | 0.901142999 | 0.116348267 |
| ENSMUSG00000000301 | Pemt | 18618 | protein_coding | 84.79501627 | 0.116351122 | 0.147433474 | 0.789177106 | 0.430008501 | 0.664948752 | 0.116351122 |
| ENSMUSG00000062169 | Cnih4 | 98417 | protein_coding | 1097.470897 | -0.116388232 | 0.05475853 | -2.125481295 | 0.033546473 | 0.14077647 | 0.116388232 |
| ENSMUSG00000031931 | Ankrd49 | 56503 | protein_coding | 419.1609655 | -0.116434127 | 0.092387948 | -1.26027398 | 0.207570543 | 0.442906874 | 0.116434127 |
| ENSMUSG00000020062 | Slc5a8 | 216225 | protein_coding | 8131.301159 | -0.116446218 | 0.100609545 | -1.157407261 | 0.247106009 | 0.488015969 | 0.116446218 |
| ENSMUSG00000030754 | Copb1 | 70349 | protein_coding | 4258.017376 | 0.11648617 | 0.046360489 | 2.512617371 | 0.011983923 | 0.067990196 | 0.11648617 |
| ENSMUSG00000045917 | Tmem268 | 230279 | protein_coding | 625.2459785 | 0.116487533 | 0.056284321 | 2.069626674 | 0.038487318 | 0.154396187 | 0.116487533 |
| ENSMUSG00000044791 | Setd2 | 235626 | protein_coding | 2039.712774 | -0.116488172 | 0.044378379 | -2.624885728 | 0.008667805 | 0.053657175 | 0.116488172 |
| ENSMUSG00000052291 | 5330438D12Rik | 102633868 | lncRNA | 33.13542352 | -0.116489364 | 0.207103713 | -0.562468736 | 0.573796698 | 0.76815045 | 0.116489364 |
| ENSMUSG00000028124 | Gclm | 14630 | protein_coding | 1647.271258 | -0.116526068 | 0.066092688 | -1.763070494 | 0.077888601 | 0.246344277 | 0.116526068 |
| ENSMUSG00000037627 | Rgs22 | 626596 | protein_coding | 8.82250396 | 0.116544366 | 0.444162214 | 0.262391448 | 0.793019666 | 0.903818244 | 0.116544366 |
| ENSMUSG00000028567 | Txndc12 | 66073 | protein_coding | 682.0436836 | 0.116565895 | 0.063885587 | 1.824603965 | 0.068060815 | 0.225933166 | 0.116565895 |
| ENSMUSG00000036291 | Ap5m1 | 74385 | protein_coding | 1082.070208 | -0.116605852 | 0.064026492 | -1.821212561 | 0.068574552 | 0.227084558 | 0.116605852 |
| ENSMUSG00000003680 | Taf6l | 225895 | protein_coding | 168.6001809 | 0.116614441 | 0.100562792 | 1.159618168 | 0.246204301 | 0.487131402 | 0.116614441 |
| ENSMUSG00000029821 | Gsdme | 54722 | protein_coding | 736.7766791 | -0.116618731 | 0.075038149 | -1.554125916 | 0.120154384 | 0.320656918 | 0.116618731 |
| ENSMUSG00000026175 | Vil1 | 22349 | protein_coding | 17508.8442 | 0.116644852 | 0.071060606 | 1.641484048 | 0.100696976 | 0.289420804 | 0.116644852 |
| ENSMUSG00000022217 | Emc9 | 85308 | protein_coding | 96.10175402 | 0.116690268 | 0.149989761 | 0.77798823 | 0.436575952 | 0.669773628 | 0.116690268 |
| ENSMUSG00000060477 | Irak2 | 108960 | protein_coding | 870.1068319 | -0.116693558 | 0.059930506 | -1.94714789 | 0.051517007 | 0.188907406 | 0.116693558 |
| ENSMUSG00000069184 | Zfp72 | 238722 | protein_coding | 77.50114978 | -0.116704814 | 0.161243721 | -0.72377897 | 0.469201442 | 0.693580708 | 0.116704814 |
| ENSMUSG00000000787 | Ddx3x | 13205 | protein_coding | 8563.423901 | -0.116769978 | 0.061916843 | -1.885916211 | 0.059306247 | 0.20675137 | 0.116769978 |
| ENSMUSG00000021764 | Ndufs4 | 17993 | protein_coding | 1929.937589 | 0.116881826 | 0.054897248 | 2.129101727 | 0.033245845 | 0.13980593 | 0.116881826 |
| ENSMUSG00000026229 | Psmd1 | 70247 | protein_coding | 2103.598645 | 0.116916348 | 0.055407335 | 2.110123991 | 0.034847677 | 0.144210545 | 0.116916348 |
| ENSMUSG00000025432 | Avil | 11567 | protein_coding | 294.4494936 | 0.116978292 | 0.103162519 | 1.133922406 | 0.256827098 | 0.498650401 | 0.116978292 |
| ENSMUSG00000030201 | Lrp6 | 16974 | protein_coding | 4103.289446 | -0.11698398 | 0.037848988 | -3.090808646 | 0.001996122 | 0.017495361 | 0.11698398 |
| ENSMUSG00000044308 | Ubr3 | 68795 | protein_coding | 2302.971987 | -0.11699455 | 0.038551808 | -3.034735791 | 0.002407465 | 0.020200409 | 0.11699455 |
| ENSMUSG00000042606 | Hirip3 | 233876 | protein_coding | 179.3533539 | 0.117098904 | 0.174562837 | 0.670812331 | 0.502340091 | 0.718956777 | 0.117098904 |
| ENSMUSG00000026131 | Dst | 13518 | protein_coding | 7488.165792 | -0.11711386 | 0.036943298 | -3.170097595 | 0.001523877 | 0.014180942 | 0.11711386 |
| ENSMUSG00000024780 | Cdc37l1 | 67072 | protein_coding | 787.5926906 | -0.117116577 | 0.066052856 | -1.773073632 | 0.076216507 | 0.243202137 | 0.117116577 |
| ENSMUSG00000030909 | Anks4b | 72074 | protein_coding | 3084.013414 | -0.117117332 | 0.083437831 | -1.403647856 | 0.160423737 | 0.380799766 | 0.117117332 |
| ENSMUSG00000020056 | Washc3 | 67282 | protein_coding | 532.0488683 | 0.11712269 | 0.059356198 | 1.973217516 | 0.048470786 | 0.181262767 | 0.11712269 |
| ENSMUSG00000027438 | Napb | 17957 | protein_coding | 41.76763845 | -0.117157634 | 0.21793342 | -0.537584526 | 0.590863916 | 0.779712165 | 0.117157634 |

| ensembl_gene_id | external_gene_name | entrezgene_id | gene_biotype | baseMean | log2FoldChange | lfcSE | stat | pvalue | padj | absLog2FC |
|---|---|---|---|---|---|---|---|---|---|---|
| ENSMUSG00000038668 | Lpar1 | 14745 | protein_coding | 1066.315572 | -0.117192841 | 0.053721857 | -2.181474102 | 0.029148368 | 0.127530041 | 0.117192841 |
| ENSMUSG00000021592 | Arsk | 77041 | protein_coding | 217.1233821 | 0.117205037 | 0.083346302 | 1.406241603 | 0.159652394 | 0.379695349 | 0.117205037 |
| ENSMUSG00000033685 | Ucp2 | 22228 | protein_coding | 5594.164277 | -0.117266359 | 0.054533095 | -2.150370485 | 0.031525921 | 0.134644738 | 0.117266359 |
| ENSMUSG00000028439 | Fam219a | 71901 | protein_coding | 49.82040665 | 0.11728104 | 0.187134897 | 0.626719239 | 0.53084329 | 0.73971984 | 0.11728104 |
| ENSMUSG00000010021 | Kif19a | 286942 | protein_coding | 17.61616529 | 0.117315802 | 0.28279834 | 0.414839076 | 0.678259712 | 0.836813383 | 0.117315802 |
| ENSMUSG00000026090 | Cracdl | 72097 | protein_coding | 1624.128694 | 0.117342573 | 0.056604974 | 2.073008151 | 0.038171521 | 0.153657835 | 0.117342573 |
| ENSMUSG00000047843 | Bri3 | 55950 | protein_coding | 482.6041292 | -0.11738326 | 0.128231827 | -0.915398798 | 0.359982309 | 0.603099387 | 0.11738326 |
| ENSMUSG00000045038 | Prkce | 18754 | protein_coding | 522.0282781 | -0.117383314 | 0.066531343 | -1.764331033 | 0.077676262 | 0.245920034 | 0.117383314 |
| ENSMUSG00000030056 | Isy1 | 57905 | protein_coding | 285.208334 | 0.11739028 | 0.075423727 | 1.556410484 | 0.119610514 | 0.319946717 | 0.11739028 |
| ENSMUSG00000060510 | Zfp266 | 77519 | protein_coding | 1293.845177 | -0.117416582 | 0.052424062 | -2.239745976 | 0.025107419 | 0.115030496 | 0.117416582 |
| ENSMUSG00000045348 | Nyap1 | 243300 | protein_coding | 144.1885086 | -0.117433389 | 0.14221584 | -0.825740572 | 0.408951272 | 0.64731555 | 0.117433389 |
| ENSMUSG00000021288 | Klc1 | 16593 | protein_coding | 1242.777099 | -0.117443155 | 0.070414343 | -1.667886818 | 0.095338197 | 0.279854223 | 0.117443155 |
| ENSMUSG00000032392 | Parp16 | 214424 | protein_coding | 341.6325559 | 0.117445826 | 0.068737209 | 1.7086208 | 0.087521213 | 0.264813247 | 0.117445826 |
| ENSMUSG00000028431 | Elp1 | 230233 | protein_coding | 491.0227315 | -0.117451506 | 0.077217885 | -1.52104018 | 0.128249755 | 0.333379077 | 0.117451506 |
| ENSMUSG00000048232 | Fbxo10 | 269529 | protein_coding | 101.1860273 | 0.117471747 | 0.120808623 | 0.972378832 | 0.330862119 | 0.574373898 | 0.117471747 |
| ENSMUSG00000024381 | Bin1 | 30948 | protein_coding | 261.5608855 | 0.117474447 | 0.090387796 | 1.299671552 | 0.193713565 | 0.425519673 | 0.117474447 |
| ENSMUSG00000053091 | Lins1 | 72635 | protein_coding | 176.4943532 | 0.117489932 | 0.110821431 | 1.060173384 | 0.289065727 | 0.531357026 | 0.117489932 |
| ENSMUSG00000008450 | Nutf2 | 68051 | protein_coding | 229.1948532 | 0.117560728 | 0.101364766 | 1.159779019 | 0.246138789 | 0.487131402 | 0.117560728 |
| ENSMUSG00000085628 | Appbp2os | 654810 | lncRNA | 24.41656241 | 0.117614338 | 0.230258904 | 0.510791703 | 0.60949692 | 0.791668558 | 0.117614338 |
| ENSMUSG00000024181 | Mrpl28 | 68611 | protein_coding | 1173.659948 | 0.117615945 | 0.062176443 | 1.891648025 | 0.058537893 | 0.20509246 | 0.117615945 |
| ENSMUSG00000021365 | Nedd9 | 18003 | protein_coding | 3181.846933 | -0.117623546 | 0.057487929 | -2.046056406 | 0.040750821 | 0.160603249 | 0.117623546 |
| ENSMUSG00000022390 | Zc3h7b | 20286 | protein_coding | 994.2363247 | -0.117631567 | 0.05965227 | -1.971954581 | 0.048614792 | 0.181568028 | 0.117631567 |
| ENSMUSG00000021224 | Numb | 18222 | protein_coding | 1220.05721 | 0.117671644 | 0.047128465 | 2.496827452 | 0.012530991 | 0.070364658 | 0.117671644 |
| ENSMUSG00000052310 | Slc39a1 | 30791 | protein_coding | 2847.365723 | 0.117693691 | 0.030556093 | 3.851725739 | 0.000117288 | 0.001697944 | 0.117693691 |
| ENSMUSG00000022195 | 6030458C11Rik | 77877 | protein_coding | 684.2555308 | 0.117725792 | 0.050432453 | 2.334326092 | 0.019578654 | 0.09661718 | 0.117725792 |
| ENSMUSG00000055553 | Kxd1 | 75620 | protein_coding | 1169.407083 | -0.117729317 | 0.057993573 | -2.030040776 | 0.042352395 | 0.165140974 | 0.117729317 |
| ENSMUSG00000040433 | Zbtb38 | 245007 | protein_coding | 1754.0452 | -0.11774366 | 0.050789628 | -2.318261905 | 0.020435091 | 0.099774968 | 0.11774366 |
| ENSMUSG00000028675 | Pnrc2 | 52830 | protein_coding | 2154.167467 | 0.117787357 | 0.056801047 | 2.073682858 | 0.038108774 | 0.153453234 | 0.117787357 |
| ENSMUSG00000030774 | Pak1 | 18479 | protein_coding | 1990.796141 | 0.117797484 | 0.047577842 | 2.475889613 | 0.013290466 | 0.073503206 | 0.117797484 |
| ENSMUSG00000086877 | A230072C01Rik | 320742 | lncRNA | 11.48874725 | -0.117818177 | 0.388367992 | -0.303367371 | 0.761609909 | 0.887577222 | 0.117818177 |
| ENSMUSG00000048905 | Bnip5 | 207819 | protein_coding | 1659.107952 | 0.117819065 | 0.076019408 | 1.549855064 | 0.121176307 | 0.322370555 | 0.117819065 |
| ENSMUSG00000075040 | Zfp408 | 381410 | protein_coding | 336.4199959 | -0.11785615 | 0.075659538 | -1.557717021 | 0.119300344 | 0.31943494 | 0.11785615 |
| ENSMUSG00000116165 | Pdxp | 57028 | protein_coding | 9.164189679 | -0.117872298 | 0.371805189 | -0.317027039 | 0.751223085 | 0.881353045 | 0.117872298 |
| ENSMUSG00000062175 | Tgif2 | 228839 | protein_coding | 7.301602307 | -0.117892021 | 0.43775475 | -0.26931066 | 0.787690629 | 0.90092814 | 0.117892021 |
| ENSMUSG00000024935 | Slc1a1 | 20510 | protein_coding | 793.7857887 | -0.117899445 | 0.107817085 | -1.093513567 | 0.274168375 | 0.516713029 | 0.117899445 |
| ENSMUSG00000016921 | Srsf6 | 67996 | protein_coding | 2658.534864 | -0.117906907 | 0.072711321 | -1.621575642 | 0.104894242 | 0.296691247 | 0.117906907 |
| ENSMUSG00000035351 | Nup37 | 69736 | protein_coding | 228.5176955 | 0.117913974 | 0.093642979 | 1.259186491 | 0.20796298 | 0.443185909 | 0.117913974 |
| ENSMUSG00000063810 | Alms1 | 236266 | protein_coding | 348.3581235 | -0.117920364 | 0.085305115 | -1.382336388 | 0.166868438 | 0.390635405 | 0.117920364 |
| ENSMUSG00000041187 | Prkd2 | 101540 | protein_coding | 391.8374372 | -0.117964581 | 0.095072667 | -1.240783334 | 0.214685799 | 0.450343118 | 0.117964581 |

| ensembl_gene_id | external_gene_name | entrezgene_id | gene_biotype | baseMean | log2FoldChange | lfcSE | stat | pvalue | padj | absLog2FC |
|---|---|---|---|---|---|---|---|---|---|---|
| ENSMUSG00000028114 | Mettl14 | 210529 | protein_coding | 322.0109495 | -0.117999308 | 0.08626348 | -1.367894132 | 0.171345217 | 0.39594273 | 0.117999308 |
| ENSMUSG00000003438 | Timm50 | 66525 | protein_coding | 671.6779351 | 0.11800201 | 0.103851182 | 1.136260628 | 0.255847503 | 0.49735447 | 0.11800201 |
| ENSMUSG00000005251 | Ripk4 | 72388 | protein_coding | 1541.896677 | 0.118039022 | 0.053944742 | 2.188146939 | 0.028658901 | 0.125951884 | 0.118039022 |
| ENSMUSG00000047030 | Spata2 | 263876 | protein_coding | 1377.91099 | -0.118040225 | 0.064681637 | -1.824941837 | 0.068009807 | 0.225910228 | 0.118040225 |
| ENSMUSG00000027254 | Map1a | 17754 | protein_coding | 30.40496678 | -0.118041624 | 0.248750558 | -0.474538126 | 0.635116217 | 0.809382719 | 0.118041624 |
| ENSMUSG00000039205 | Ciz1 | 68379 | protein_coding | 864.6895456 | -0.118063526 | 0.053462401 | -2.208346881 | 0.027220102 | 0.121164388 | 0.118063526 |
| ENSMUSG00000012819 | Cdh23 | 22295 | protein_coding | 8.85428525 | 0.118138914 | 0.71558727 | 0.165093649 | 0.868870272 | 0.940691929 | 0.118138914 |
| ENSMUSG00000035696 | Rnf38 | 73469 | protein_coding | 784.4083729 | -0.118147277 | 0.071475787 | -1.652969224 | 0.098337127 | 0.285282038 | 0.118147277 |
| ENSMUSG00000022507 | 1810013L24Rik | 69053 | protein_coding | 1354.760228 | -0.118147502 | 0.053236932 | -2.219277067 | 0.026467881 | 0.118917792 | 0.118147502 |
| ENSMUSG00000022201 | Zfr | 22763 | protein_coding | 1612.058684 | 0.118148699 | 0.044270901 | 2.668766497 | 0.007613036 | 0.048836685 | 0.118148699 |
| ENSMUSG00000023075 | Akirin1 | 68050 | protein_coding | 1163.591315 | 0.118155111 | 0.06623333 | 1.783922255 | 0.074436299 | 0.2394347 | 0.118155111 |
| ENSMUSG00000034707 | Gns | 75612 | protein_coding | 1517.6336 | -0.118169188 | 0.043649048 | -2.707256931 | 0.006784173 | 0.044729501 | 0.118169188 |
| ENSMUSG00000053470 | Kdm3a | 104263 | protein_coding | 1507.791014 | -0.118176142 | 0.062130745 | -1.902055756 | 0.057163866 | 0.202153934 | 0.118176142 |
| ENSMUSG00000038886 | Man2a2 | 140481 | protein_coding | 132.5345808 | 0.118181528 | 0.110545007 | 1.069080648 | 0.285033331 | 0.527935285 | 0.118181528 |
| ENSMUSG00000055204 | Ankrd17 | 81702 | protein_coding | 2968.741416 | -0.118243588 | 0.053268485 | -2.219766305 | 0.026434635 | 0.118917792 | 0.118243588 |
| ENSMUSG00000052748 | Swt1 | 66875 | protein_coding | 501.8842505 | -0.118246065 | 0.078317626 | -1.509826978 | 0.131087579 | 0.336982889 | 0.118246065 |
| ENSMUSG00000024346 | Pfdn1 | 67199 | protein_coding | 896.3483466 | 0.118257811 | 0.059021631 | 2.003635104 | 0.045109163 | 0.17246929 | 0.118257811 |
| ENSMUSG00000022841 | Ap2m1 | 11773 | protein_coding | 2886.444011 | 0.118287759 | 0.038676801 | 3.058364602 | 0.002225486 | 0.019080498 | 0.118287759 |
| ENSMUSG00000056076 | Eif3b | 27979 | protein_coding | 2505.737788 | 0.118304063 | 0.078275574 | 1.511379063 | 0.130691901 | 0.33658069 | 0.118304063 |
| ENSMUSG00000020219 | Timm13 | 30055 | protein_coding | 1477.744512 | 0.11831868 | 0.102056699 | 1.159342608 | 0.246316561 | 0.487165462 | 0.11831868 |
| ENSMUSG00000039462 | Col10a1 | 12813 | protein_coding | 8.752940167 | -0.118358102 | 0.406407183 | -0.291230339 | 0.770875161 | 0.892291327 | 0.118358102 |
| ENSMUSG00000073591 | Pcdhb22 | 93893 | protein_coding | 7.012279449 | 0.118364794 | 0.486230359 | 0.243433575 | 0.807669537 | 0.91309405 | 0.118364794 |
| ENSMUSG00000027799 | Nbea | 26422 | protein_coding | 485.2106287 | -0.118400138 | 0.113320887 | -1.044821844 | 0.296105317 | 0.538897588 | 0.118400138 |
| ENSMUSG00000020018 | Snrpf | 69878 | protein_coding | 199.2439052 | 0.118538268 | 0.118332708 | 1.001737142 | 0.316470563 | 0.560339056 | 0.118538268 |
| ENSMUSG00000040483 | Xaf1 | 327959 | protein_coding | 105.9158129 | -0.118553419 | 0.189891835 | -0.624320781 | 0.532416939 | 0.740799392 | 0.118553419 |
| ENSMUSG00000025371 | Chmp6 | 208092 | protein_coding | 1939.072047 | 0.118629045 | 0.065582299 | 1.80885767 | 0.070473118 | 0.230859227 | 0.118629045 |
| ENSMUSG00000062627 | Mysm1 | 320713 | protein_coding | 750.7978578 | -0.118631668 | 0.067119555 | -1.767468041 | 0.077149877 | 0.24490649 | 0.118631668 |
| ENSMUSG00000028837 | Psmb2 | 26445 | protein_coding | 1648.317141 | 0.118654475 | 0.056864563 | 2.086615428 | 0.036922912 | 0.149687228 | 0.118654475 |
| ENSMUSG00000028013 | Ppa2 | 74776 | protein_coding | 1695.704834 | -0.118692462 | 0.048730981 | -2.4356674 | 0.014864345 | 0.079719751 | 0.118692462 |
| ENSMUSG00000031534 | Smim19 | 102032 | protein_coding | 227.9467592 | -0.118704422 | 0.114967846 | -1.03250105 | 0.301837456 | 0.545539031 | 0.118704422 |
| ENSMUSG00000030083 | Abtb1 | 80283 | protein_coding | 708.2700477 | -0.118736868 | 0.08266103 | -1.436431028 | 0.150879729 | 0.366672302 | 0.118736868 |
| ENSMUSG00000054766 | Set | 56086 | protein_coding | 3191.433816 | 0.118743883 | 0.042155951 | 2.816776287 | 0.00485083 | 0.034815014 | 0.118743883 |
| ENSMUSG00000027615 | Hps3 | 12807 | protein_coding | 296.2577655 | -0.118755165 | 0.105570358 | -1.124891176 | 0.260635152 | 0.502482139 | 0.118755165 |
| ENSMUSG00000061559 | Wdr61 | 66317 | protein_coding | 956.7038998 | 0.118764734 | 0.05948004 | 1.996715771 | 0.045856068 | 0.174452968 | 0.118764734 |
| ENSMUSG00000040061 | Plcb2 | 18796 | protein_coding | 97.16957412 | -0.118771434 | 0.153542791 | -0.773539635 | 0.439203082 | 0.671631718 | 0.118771434 |
| ENSMUSG00000055210 | Foxd2 | 17301 | protein_coding | 1287.962927 | -0.11877692 | 0.117208827 | -1.013378621 | 0.310879347 | 0.554719101 | 0.11877692 |
| ENSMUSG00000034401 | Spata6 | 67946 | protein_coding | 101.0943074 | 0.118813303 | 0.155917688 | 0.762025813 | 0.446044598 | 0.676953924 | 0.118813303 |
| ENSMUSG00000058402 | Zfp420 | 233058 | protein_coding | 53.67005226 | -0.118853631 | 0.178016045 | -0.667656842 | 0.504352665 | 0.720625093 | 0.118853631 |
| ENSMUSG00000050505 | Pcdh20 | 219257 | protein_coding | 24.81648582 | -0.118870547 | 0.297777499 | -0.399192509 | 0.689751362 | 0.844356471 | 0.118870547 |

| ensembl_gene_id | external_gene_name | entrezgene_id | gene_biotype | baseMean | log2FoldChange | lfcSE | stat | pvalue | padj | absLog2FC |
|---|---|---|---|---|---|---|---|---|---|---|
| ENSMUSG00000022442 | Ttll1 | 319953 | protein_coding | 83.38819091 | -0.11889345 | 0.148890143 | -0.798531371 | 0.424562196 | 0.660580273 | 0.11889345 |
| ENSMUSG00000032198 | Dock6 | 319899 | protein_coding | 391.1601383 | -0.118909239 | 0.076034522 | -1.563884887 | 0.117844604 | 0.317275448 | 0.118909239 |
| ENSMUSG00000051627 | H1f4 | 50709 | protein_coding | 8084.648782 | 0.118913209 | 0.076251912 | 1.559478405 | 0.118883192 | 0.319001795 | 0.118913209 |
| ENSMUSG00000020003 | Pex7 | 18634 | protein_coding | 964.2093292 | 0.118961836 | 0.059159454 | 2.010867709 | 0.044339432 | 0.170740692 | 0.118961836 |
| ENSMUSG00000000278 | Scpep1 | 74617 | protein_coding | 623.8823903 | -0.118974722 | 0.091330794 | -1.302679153 | 0.192684322 | 0.42444695 | 0.118974722 |
| ENSMUSG00000042203 | Tbc1d22b | 381085 | protein_coding | 456.3474418 | 0.119010938 | 0.068664166 | 1.733232118 | 0.08305442 | 0.256348065 | 0.119010938 |
| ENSMUSG00000047735 | Samd9l | 209086 | protein_coding | 2966.124977 | 0.119019165 | 0.071022763 | 1.675789015 | 0.093779527 | 0.276801468 | 0.119019165 |
| ENSMUSG00000023079 | Gtf2ird1 | 57080 | protein_coding | 1370.848309 | 0.11903242 | 0.044379497 | 2.682148934 | 0.007315089 | 0.047263795 | 0.11903242 |
| ENSMUSG00000053012 | Krcc1 | 57896 | protein_coding | 1904.916064 | 0.119048563 | 0.082141593 | 1.449309158 | 0.147251265 | 0.360790207 | 0.119048563 |
| ENSMUSG00000003444 | Med29 | 67224 | protein_coding | 157.5643233 | 0.119053355 | 0.11526435 | 1.032872311 | 0.301663658 | 0.545395719 | 0.119053355 |
| ENSMUSG00000027395 | Polr1b | 20017 | protein_coding | 539.3344149 | -0.119077797 | 0.065966446 | -1.805126768 | 0.071054847 | 0.232315871 | 0.119077797 |
| ENSMUSG00000039163 | Cmc1 | 67899 | protein_coding | 452.4849921 | -0.119093119 | 0.069530147 | -1.712827093 | 0.086744354 | 0.263676126 | 0.119093119 |
| ENSMUSG00000002870 | Mcm2 | 17216 | protein_coding | 751.2016846 | 0.119114224 | 0.142620561 | 0.835182694 | 0.403614852 | 0.642746788 | 0.119114224 |
| ENSMUSG00000032952 | Ap4b1 | 67489 | protein_coding | 85.51665301 | 0.119125396 | 0.127183327 | 0.936643181 | 0.348942132 | 0.591360842 | 0.119125396 |
| ENSMUSG00000039754 | Alkbh4 | 72041 | protein_coding | 227.4014116 | -0.119178446 | 0.089794695 | -1.327232601 | 0.184431751 | 0.413101507 | 0.119178446 |
| ENSMUSG00000060860 | Ube2s | 77891 | protein_coding | 650.6102298 | 0.119236473 | 0.065905926 | 1.809191981 | 0.070421183 | 0.230782853 | 0.119236473 |
| ENSMUSG00000022237 | Ankrd33b | 67434 | protein_coding | 8.104506522 | -0.119238547 | 0.424208768 | -0.281084589 | 0.778645521 | 0.896944023 | 0.119238547 |
| ENSMUSG00000015671 | Psma2 | 19166 | protein_coding | 2272.479863 | 0.119284181 | 0.054313631 | 2.196210741 | 0.028076863 | 0.12424678 | 0.119284181 |
| ENSMUSG00000001036 | Epn2 | 13855 | protein_coding | 413.4343541 | -0.119319566 | 0.071091873 | -1.678385447 | 0.093271877 | 0.275847342 | 0.119319566 |
| ENSMUSG00000029229 | Chic2 | 74277 | protein_coding | 451.0657284 | -0.119331934 | 0.087470888 | -1.364247428 | 0.172489706 | 0.397562024 | 0.119331934 |
| ENSMUSG00000021583 | Erap1 | 80898 | protein_coding | 2781.874991 | -0.119386654 | 0.056115668 | -2.127510152 | 0.033377719 | 0.140323897 | 0.119386654 |
| ENSMUSG00000055932 | Fto | 26383 | protein_coding | 731.9134771 | 0.119426646 | 0.056311613 | 2.120817354 | 0.033937176 | 0.141755849 | 0.119426646 |
| ENSMUSG00000040652 | Oaz2 | 18247 | protein_coding | 503.1246349 | 0.119436185 | 0.066240116 | 1.803079354 | 0.071375753 | 0.233011492 | 0.119436185 |
| ENSMUSG00000085399 | Foxd2os | 100040736 | lncRNA | 599.6626863 | -0.119470931 | 0.068493932 | -1.744255702 | 0.081114512 | 0.252438005 | 0.119470931 |
| ENSMUSG00000034785 | Dio1 | 13370 | protein_coding | 1259.235549 | 0.119484545 | 0.06652464 | 1.79609456 | 0.072479479 | 0.235215876 | 0.119484545 |
| ENSMUSG00000020133 | 2310011J03Rik | 66374 | protein_coding | 828.7147474 | 0.1195003 | 0.073818024 | 1.618849879 | 0.105479567 | 0.297761537 | 0.1195003 |
| ENSMUSG00000023118 | Sympk | 68188 | protein_coding | 2075.850962 | 0.119519909 | 0.047212536 | 2.531529081 | 0.01135664 | 0.065084766 | 0.119519909 |
| ENSMUSG00000115529 | 9630013A20Rik | 319903 | lncRNA | 6.726888586 | -0.119523782 | 0.533228384 | -0.2241512 | 0.822639654 | 0.920659938 | 0.119523782 |
| ENSMUSG00000033272 | Slc35a4 | 67843 | protein_coding | 1243.642087 | 0.119527126 | 0.067761157 | 1.763947538 | 0.077740812 | 0.246021601 | 0.119527126 |
| ENSMUSG00000102440 | Pcdhga9 | 93717 | protein_coding | 14.1312792 | 0.119570029 | 0.336503492 | 0.355330723 | 0.722341842 | 0.8642486 | 0.119570029 |
| ENSMUSG00000031007 | Atp6ap2 | 70495 | protein_coding | 3706.078917 | 0.119583691 | 0.048416743 | 2.469883022 | 0.013515724 | 0.074453021 | 0.119583691 |
| ENSMUSG00000034926 | Dhcr24 | 74754 | protein_coding | 5603.842912 | 0.119597601 | 0.097264706 | 1.229609439 | 0.218843394 | 0.455009895 | 0.119597601 |
| ENSMUSG00000026585 | Kifap3 | 16579 | protein_coding | 393.1208871 | 0.119625983 | 0.069298023 | 1.726253919 | 0.084301742 | 0.258860279 | 0.119625983 |
| ENSMUSG00000013653 | 1810065E05Rik | 69864 | protein_coding | 15937.42802 | -0.119629502 | 0.125293245 | -0.954796098 | 0.339680821 | 0.582893445 | 0.119629502 |
| ENSMUSG00000087480 | Gm15910 | 100504616 | lncRNA | 8.004184345 | 0.119649074 | 0.413878143 | 0.289092516 | 0.772510581 | 0.893129098 | 0.119649074 |
| ENSMUSG00000036109 | Mbnl3 | 171170 | protein_coding | 627.4354211 | 0.11971456 | 0.077587755 | 1.542956871 | 0.122841241 | 0.325030466 | 0.11971456 |
| ENSMUSG00000036867 | Smad6 | 17130 | protein_coding | 499.7247918 | -0.119761213 | 0.073394664 | -1.631742774 | 0.102733685 | 0.29253061 | 0.119761213 |
| ENSMUSG00000049470 | Aff4 | 93736 | protein_coding | 3273.099727 | -0.1197671 | 0.042401684 | -2.824583562 | 0.00473421 | 0.034222527 | 0.1197671 |
| ENSMUSG00000040272 | Accs | 329470 | protein_coding | 115.4249283 | -0.119795866 | 0.132136556 | -0.906606542 | 0.364614901 | 0.606826059 | 0.119795866 |

| ensembl_gene_id | external_gene_name | entrezgene_id | gene_biotype | baseMean | log2FoldChange | lfcSE | stat | pvalue | padj | absLog2FC |
|---|---|---|---|---|---|---|---|---|---|---|
| ENSMUSG00000096916 | Zfp850 | 100043772 | protein_coding | 96.88373209 | -0.119799101 | 0.149133625 | -0.803300401 | 0.421801126 | 0.65828686 | 0.119799101 |
| ENSMUSG00000020953 | Coch | 12810 | protein_coding | 19.0718768 | 0.119817521 | 0.267484594 | 0.447941766 | 0.654195225 | 0.822000293 | 0.119817521 |
| ENSMUSG00000039699 | Batf2 | 74481 | protein_coding | 324.9974593 | -0.119844385 | 0.144233597 | -0.830904781 | 0.406027422 | 0.645187136 | 0.119844385 |
| ENSMUSG00000030630 | Fah | 14085 | protein_coding | 87.98822073 | -0.119865329 | 0.155816847 | -0.769270661 | 0.441732651 | 0.672975316 | 0.119865329 |
| ENSMUSG00000032400 | Zwilch | 68014 | protein_coding | 127.6281628 | 0.119877471 | 0.125781346 | 0.953062396 | 0.340558457 | 0.583344264 | 0.119877471 |
| ENSMUSG00000027811 | 4930579G24Rik | 75939 | protein_coding | 36.26710966 | 0.119915536 | 0.205684781 | 0.583006364 | 0.559889016 | 0.758250106 | 0.119915536 |
| ENSMUSG00000058833 | Rex1bd | 66462 | protein_coding | 592.2082565 | 0.11995543 | 0.087016763 | 1.378532438 | 0.16803896 | 0.391953183 | 0.11995543 |
| ENSMUSG00000078994 | Zfp429 | 72807 | protein_coding | 153.8953923 | -0.120003066 | 0.104867746 | -1.144327702 | 0.252487759 | 0.493560011 | 0.120003066 |
| ENSMUSG00000072969 | Armcx5 | 494468 | protein_coding | 97.16552286 | -0.120008522 | 0.138531461 | -0.866290742 | 0.386330742 | 0.626292308 | 0.120008522 |
| ENSMUSG00000078812 | Eif5a | 276770 | protein_coding | 5892.411861 | 0.120024772 | 0.072682034 | 1.651367815 | 0.098663493 | 0.28576619 | 0.120024772 |
| ENSMUSG00000048277 | Syngr2 | 20973 | protein_coding | 3675.356998 | 0.120073669 | 0.054688318 | 2.195599955 | 0.028120589 | 0.12424678 | 0.120073669 |
| ENSMUSG00000004151 | Etv1 | 14009 | protein_coding | 20.11361143 | 0.120138296 | 0.334495029 | 0.359163172 | 0.719473026 | 0.862768697 | 0.120138296 |
| ENSMUSG00000029673 | Auts2 | 319974 | protein_coding | 572.6654491 | -0.120144463 | 0.096138333 | -1.249704048 | 0.211407678 | 0.446720601 | 0.120144463 |
| ENSMUSG00000029071 | Dvl1 | 13542 | protein_coding | 1290.790067 | -0.120153824 | 0.057998454 | -2.071672892 | 0.038295957 | 0.153937911 | 0.120153824 |
| ENSMUSG00000044147 | Arf6 | 11845 | protein_coding | 4977.523786 | 0.12016459 | 0.068494022 | 1.754380693 | 0.079365297 | 0.24901248 | 0.12016459 |
| ENSMUSG00000068744 | Psrc1 | 56742 | protein_coding | 69.41519374 | -0.120164642 | 0.153578818 | -0.782429789 | 0.433962033 | 0.668045287 | 0.120164642 |
| ENSMUSG00000033760 | Rbm4b | 66704 | protein_coding | 166.1092543 | -0.120234022 | 0.131340252 | -0.915439254 | 0.359961079 | 0.603099387 | 0.120234022 |
| ENSMUSG00000025192 | Entpd7 | 93685 | protein_coding | 420.1853896 | 0.120351328 | 0.078458993 | 1.533939234 | 0.125044603 | 0.328198932 | 0.120351328 |
| ENSMUSG00000073728 | Tmem51os1 | 100038693 | lncRNA | 32.89101239 | -0.120396487 | 0.23360438 | -0.515386257 | 0.606283134 | 0.789753 | 0.120396487 |
| ENSMUSG00000074781 | Ube2n | 93765 | protein_coding | 901.9201519 | 0.120421151 | 0.058247519 | 2.067403932 | 0.038696108 | 0.1546981 | 0.120421151 |
| ENSMUSG00000034243 | Golgb1 | 224139 | protein_coding | 5209.74546 | 0.120449049 | 0.037800066 | 3.186477239 | 0.001440168 | 0.013511107 | 0.120449049 |
| ENSMUSG00000033356 | Pus7l | 78895 | protein_coding | 86.70154579 | 0.120449714 | 0.143190057 | 0.8411877 | 0.40024279 | 0.639523975 | 0.120449714 |
| ENSMUSG00000038481 | Cdk19 | 78334 | protein_coding | 758.5333779 | -0.120507977 | 0.056448083 | -2.134846244 | 0.032773572 | 0.138361096 | 0.120507977 |
| ENSMUSG00000056716 | Gm5420 | 432436 | lncRNA | 11.08191125 | 0.120526937 | 0.393586359 | 0.306227424 | 0.759431494 | 0.886076667 | 0.120526937 |
| ENSMUSG00000027954 | Efna1 | 13636 | protein_coding | 2532.608186 | 0.120533584 | 0.076850602 | 1.568414314 | 0.116784473 | 0.316003891 | 0.120533584 |
| ENSMUSG00000005611 | Irag1 | 17540 | protein_coding | 6.715870118 | -0.120545101 | 0.497270448 | -0.242413563 | 0.808459726 | 0.913667134 | 0.120545101 |
| ENSMUSG00000037085 | Trmt12 | 68260 | protein_coding | 149.0502799 | 0.120548429 | 0.114221099 | 1.055395455 | 0.291244501 | 0.533617367 | 0.120548429 |
| ENSMUSG00000014606 | Slc25a11 | 67863 | protein_coding | 2758.392436 | 0.120562006 | 0.052873566 | 2.280194338 | 0.022596165 | 0.107008585 | 0.120562006 |
| ENSMUSG00000056267 | Cep70 | 68121 | protein_coding | 212.9555191 | -0.12059684 | 0.107418673 | -1.122680414 | 0.261573249 | 0.503448119 | 0.12059684 |
| ENSMUSG00000028708 | Mknk1 | 17346 | protein_coding | 236.9271973 | -0.120615244 | 0.093220057 | -1.293876536 | 0.19570807 | 0.428222446 | 0.120615244 |
| ENSMUSG00000059534 | Uqcr10 | 66152 | protein_coding | 5164.142001 | 0.120645336 | 0.077138679 | 1.564005728 | 0.117816223 | 0.317252001 | 0.120645336 |
| ENSMUSG00000024045 | Akap8 | 56399 | protein_coding | 1528.919509 | -0.120657712 | 0.057988315 | -2.080724592 | 0.037459122 | 0.151214294 | 0.120657712 |
| ENSMUSG00000026036 | Nif3l1 | 65102 | protein_coding | 403.096819 | 0.120691861 | 0.063819574 | 1.891141764 | 0.058605423 | 0.20528446 | 0.120691861 |
| ENSMUSG00000039158 | Akna | 100182 | protein_coding | 152.9030715 | 0.120752725 | 0.119347408 | 1.011775015 | 0.311645639 | 0.555505998 | 0.120752725 |
| ENSMUSG00000030278 | Cidec | 14311 | protein_coding | 65.07112449 | 0.120783318 | 0.169267709 | 0.713563847 | 0.47549692 | 0.698858175 | 0.120783318 |
| ENSMUSG00000029185 | Fam114a1 | 68303 | protein_coding | 1709.398794 | 0.12086407 | 0.039611363 | 3.051247461 | 0.002278927 | 0.019455929 | 0.12086407 |
| ENSMUSG00000001288 | Rarg | 19411 | protein_coding | 256.0055252 | -0.120870123 | 0.111150498 | -1.087445625 | 0.276839917 | 0.519767321 | 0.120870123 |
| ENSMUSG00000028480 | Glipr2 | 384009 | protein_coding | 217.0285798 | -0.120920529 | 0.100032481 | -1.208812657 | 0.226734828 | 0.465277093 | 0.120920529 |
| ENSMUSG00000052593 | Adam17 | 11491 | protein_coding | 881.912893 | 0.120931127 | 0.049305294 | 2.452700654 | 0.014178828 | 0.076990557 | 0.120931127 |

| ensembl_gene_id | external_gene_name | entrezgene_id | gene_biotype | baseMean | log2FoldChange | lfcSE | stat | pvalue | padj | absLog2FC |
|---|---|---|---|---|---|---|---|---|---|---|
| ENSMUSG00000021182 | Ccdc88c | 68339 | protein_coding | 1520.011176 | 0.120954939 | 0.046883715 | 2.579892391 | 0.009883111 | 0.058770253 | 0.120954939 |
| ENSMUSG00000021336 | Slc17a4 | 319848 | protein_coding | 1872.890108 | -0.121037936 | 0.111603663 | -1.084533721 | 0.278128224 | 0.521009858 | 0.121037936 |
| ENSMUSG00000063488 | Zkscan7 | 382118 | protein_coding | 391.2935747 | -0.121055685 | 0.091184788 | -1.327586408 | 0.184314777 | 0.41301159 | 0.121055685 |
| ENSMUSG00000015083 | C8g | 69379 | protein_coding | 50.66032756 | -0.121069043 | 0.184283298 | -0.656972414 | 0.511198654 | 0.725329993 | 0.121069043 |
| ENSMUSG00000040225 | Prrc2c | 226562 | protein_coding | 2703.253119 | -0.121185754 | 0.063080801 | -1.921119446 | 0.054716651 | 0.196492651 | 0.121185754 |
| ENSMUSG00000055148 | Klf2 | 16598 | protein_coding | 1001.278737 | 0.12119691 | 0.136469955 | 0.888084932 | 0.374495067 | 0.61633656 | 0.12119691 |
| ENSMUSG00000040351 | Ankib1 | 70797 | protein_coding | 2470.308747 | -0.121260878 | 0.062938857 | -1.926645693 | 0.0540238 | 0.194647414 | 0.121260878 |
| ENSMUSG00000026810 | Dpm2 | 13481 | protein_coding | 472.1705992 | 0.121284502 | 0.078525059 | 1.544532457 | 0.122459395 | 0.324392683 | 0.121284502 |
| ENSMUSG00000026979 | Psd4 | 215632 | protein_coding | 2188.529956 | -0.121317418 | 0.039779393 | -3.049755367 | 0.002290278 | 0.019496529 | 0.121317418 |
| ENSMUSG00000073402 | Gm8909 | 667977 | protein_coding | 214.2620352 | -0.121442972 | 0.17663153 | -0.687549791 | 0.491736336 | 0.711295353 | 0.121442972 |
| ENSMUSG00000025978 | Rftn2 | 74013 | protein_coding | 142.9934696 | -0.121460091 | 0.129389488 | -0.938716841 | 0.347876144 | 0.590112399 | 0.121460091 |
| ENSMUSG00000021939 | Ctsb | 13030 | protein_coding | 7731.970497 | -0.121462723 | 0.03968692 | -3.060522773 | 0.00220951 | 0.018994009 | 0.121462723 |
| ENSMUSG00000003575 | Crtc1 | 382056 | protein_coding | 230.2602564 | -0.121496596 | 0.1037896 | -1.170604722 | 0.241757698 | 0.482707243 | 0.121496596 |
| ENSMUSG00000038759 | Nup205 | 70699 | protein_coding | 1114.592612 | 0.121518208 | 0.084922079 | 1.430937741 | 0.152448059 | 0.369268435 | 0.121518208 |
| ENSMUSG00000089809 | Rasgef1b | 320292 | protein_coding | 1043.811036 | -0.121536304 | 0.081632091 | -1.488829978 | 0.136532146 | 0.345551463 | 0.121536304 |
| ENSMUSG00000023764 | Sfi1 | 78887 | protein_coding | 622.6969289 | 0.12159198 | 0.088632547 | 1.371866025 | 0.170105149 | 0.394462093 | 0.12159198 |
| ENSMUSG00000027285 | Haus2 | 66296 | protein_coding | 319.9918244 | -0.121605848 | 0.097725544 | -1.244360915 | 0.213366761 | 0.449133255 | 0.121605848 |
| ENSMUSG00000030209 | Grin2b | 14812 | protein_coding | 20.99883967 | -0.121668724 | 0.355059398 | -0.342671465 | 0.731845632 | 0.870710034 | 0.121668724 |
| ENSMUSG00000038459 | Abhd17c | 70178 | protein_coding | 2745.495863 | 0.121676437 | 0.069418601 | 1.752793 | 0.079637545 | 0.249623847 | 0.121676437 |
| ENSMUSG00000020078 | Vps26a | 30930 | protein_coding | 1409.987101 | -0.121701146 | 0.040105836 | -3.0344997 | 0.00240935 | 0.020205713 | 0.121701146 |
| ENSMUSG00000046229 | Scand1 | 19018 | protein_coding | 549.6096786 | -0.121706478 | 0.120117758 | -1.013226351 | 0.310952057 | 0.554787456 | 0.121706478 |
| ENSMUSG00000036285 | Noa1 | 56412 | protein_coding | 383.4572324 | -0.121769876 | 0.082639059 | -1.473514807 | 0.140612274 | 0.351653742 | 0.121769876 |
| ENSMUSG00000102976 | Zc3h11a | 70579 | protein_coding | 200.1803874 | -0.12179385 | 0.110713794 | -1.100078373 | 0.271297976 | 0.513806497 | 0.12179385 |
| ENSMUSG00000019158 | Tmem160 | 69094 | protein_coding | 372.0256246 | -0.121840693 | 0.098929044 | -1.231596784 | 0.218099743 | 0.45437389 | 0.121840693 |
| ENSMUSG00000028910 | Mecr | 26922 | protein_coding | 1368.447388 | 0.121846607 | 0.067608642 | 1.802234204 | 0.071508565 | 0.233227308 | 0.121846607 |
| ENSMUSG00000020564 | Atxn7l1 | 380753 | protein_coding | 191.6918585 | -0.121892371 | 0.14583943 | -0.835798465 | 0.40326829 | 0.642512124 | 0.121892371 |
| ENSMUSG00000020732 | Rab37 | 58222 | protein_coding | 121.4676299 | 0.121895117 | 0.137659399 | 0.885483426 | 0.375895955 | 0.617324668 | 0.121895117 |
| ENSMUSG00000025792 | Slc25a10 | 27376 | protein_coding | 3437.356511 | 0.121914994 | 0.081747591 | 1.491358862 | 0.135867303 | 0.344625981 | 0.121914994 |
| ENSMUSG00000054321 | Taf4b | 72504 | protein_coding | 210.6573285 | -0.121926446 | 0.105475828 | -1.155965768 | 0.247695159 | 0.488633619 | 0.121926446 |
| ENSMUSG00000024168 | Tmem204 | 407831 | protein_coding | 8.894809791 | 0.121941583 | 0.400017127 | 0.304840904 | 0.760487327 | 0.886731192 | 0.121941583 |
| ENSMUSG00000040385 | Ppp1ca | 19045 | protein_coding | 4104.096674 | 0.121950396 | 0.052865584 | 2.306801274 | 0.021065901 | 0.101895312 | 0.121950396 |
| ENSMUSG00000034993 | Vat1 | 26949 | protein_coding | 1082.931815 | -0.121959224 | 0.057111054 | -2.135474933 | 0.032722236 | 0.138216735 | 0.121959224 |
| ENSMUSG00000098678 | Mroh6 | 223645 | protein_coding | 35.91473266 | 0.121963672 | 0.237728375 | 0.513037925 | 0.607924792 | 0.790772957 | 0.121963672 |
| ENSMUSG00000020708 | Psmc5 | 19184 | protein_coding | 1416.917433 | 0.121970362 | 0.060660906 | 2.010691404 | 0.044358063 | 0.170760157 | 0.121970362 |
| ENSMUSG00000024533 | Spire1 | 68166 | protein_coding | 100.9221865 | 0.12197994 | 0.130957482 | 0.931446893 | 0.351622444 | 0.593968278 | 0.12197994 |
| ENSMUSG00000031778 | Cx3cl1 | 20312 | protein_coding | 1922.938833 | 0.122038606 | 0.0820013 | 1.488252079 | 0.136684428 | 0.345828328 | 0.122038606 |
| ENSMUSG00000029465 | Arpc3 | 56378 | protein_coding | 3566.819913 | 0.122057323 | 0.064246242 | 1.899836006 | 0.057454644 | 0.202928394 | 0.122057323 |
| ENSMUSG00000043885 | Slc36a4 | 234967 | protein_coding | 338.397404 | 0.122061791 | 0.092964408 | 1.312994871 | 0.189184677 | 0.419609583 | 0.122061791 |
| ENSMUSG00000052631 | Sh2d6 | 71130 | protein_coding | 135.8145454 | -0.122072364 | 0.132196109 | -0.923418731 | 0.355789034 | 0.598521667 | 0.122072364 |

| ensembl_gene_id | external_gene_name | entrezgene_id | gene_biotype | baseMean | log2FoldChange | lfcSE | stat | pvalue | padj | absLog2FC |
|---|---|---|---|---|---|---|---|---|---|---|
| ENSMUSG00000024413 | Npc1 | 18145 | protein_coding | 787.3049975 | -0.122096319 | 0.06736229 | -1.812532182 | 0.069904006 | 0.229602617 | 0.122096319 |
| ENSMUSG00000089876 | Tmem102 | 380705 | protein_coding | 736.7126141 | 0.122120903 | 0.080490886 | 1.517201623 | 0.129215787 | 0.334810892 | 0.122120903 |
| ENSMUSG00000022906 | Parp9 | 80285 | protein_coding | 1102.475303 | -0.122147681 | 0.135878793 | -0.898945879 | 0.368681489 | 0.610694985 | 0.122147681 |
| ENSMUSG00000020390 | Ube2b | 22210 | protein_coding | 1269.99819 | -0.122165073 | 0.050093416 | -2.438745077 | 0.014738361 | 0.079254935 | 0.122165073 |
| ENSMUSG00000008136 | Fhl2 | 14200 | protein_coding | 320.849674 | 0.12217434 | 0.076770052 | 1.591432292 | 0.111512322 | 0.306958481 | 0.12217434 |
| ENSMUSG00000060427 | Zfp868 | 234362 | protein_coding | 309.9796225 | -0.122179521 | 0.073803347 | -1.655473982 | 0.09782839 | 0.284213967 | 0.122179521 |
| ENSMUSG00000049764 | Zfp280b | 64453 | protein_coding | 390.11449 | 0.122186318 | 0.076969373 | 1.587466723 | 0.112406978 | 0.30863821 | 0.122186318 |
| ENSMUSG00000024812 | Tjp2 | 21873 | protein_coding | 3890.483802 | 0.122211415 | 0.052042422 | 2.348303778 | 0.018859133 | 0.094449731 | 0.122211415 |
| ENSMUSG00000032712 | Resf1 | 67246 | protein_coding | 1013.72082 | -0.122234532 | 0.073661127 | -1.659417092 | 0.097031774 | 0.282766792 | 0.122234532 |
| ENSMUSG00000069456 | Rdh16 | 19683 | protein_coding | 484.7276353 | -0.122253608 | 0.139751268 | -0.874794264 | 0.381685859 | 0.622454024 | 0.122253608 |
| ENSMUSG00000055762 | Eef1d | 66656 | protein_coding | 2516.767667 | 0.122315878 | 0.061194411 | 1.998808005 | 0.045629131 | 0.174003547 | 0.122315878 |
| ENSMUSG00000042508 | Dmtf1 | 23857 | protein_coding | 501.4055387 | -0.122333206 | 0.093651164 | -1.306264656 | 0.191462572 | 0.423020709 | 0.122333206 |
| ENSMUSG00000020091 | Eif4ebp2 | 13688 | protein_coding | 2246.440632 | -0.122351161 | 0.087524168 | -1.397912867 | 0.162139234 | 0.382899316 | 0.122351161 |
| ENSMUSG00000068284 | Usf3 | 207806 | protein_coding | 4156.850407 | 0.122357077 | 0.078702744 | 1.55467358 | 0.12002383 | 0.3204676 | 0.122357077 |
| ENSMUSG00000047221 | Fam185a | 330050 | protein_coding | 278.588107 | 0.12237055 | 0.091464593 | 1.337900772 | 0.180928785 | 0.407747138 | 0.12237055 |
| ENSMUSG00000030683 | Sez6l2 | 233878 | protein_coding | 53.31520653 | -0.122371522 | 0.168517033 | -0.726167082 | 0.467736349 | 0.692926016 | 0.122371522 |
| ENSMUSG00000005897 | Nr2c1 | 22025 | protein_coding | 293.3948206 | -0.122384941 | 0.079632319 | -1.536875263 | 0.124323858 | 0.327235329 | 0.122384941 |
| ENSMUSG00000027011 | Ube2e3 | 22193 | protein_coding | 591.0888737 | 0.12241416 | 0.05920385 | 2.067672271 | 0.038670851 | 0.154660479 | 0.12241416 |
| ENSMUSG00000025135 | Anapc11 | 66156 | protein_coding | 494.308329 | 0.122418903 | 0.119505485 | 1.024378947 | 0.305656322 | 0.54999264 | 0.122418903 |
| ENSMUSG00000090100 | Ttbk2 | 140810 | protein_coding | 170.0859936 | -0.122443216 | 0.103580116 | -1.182111213 | 0.237161576 | 0.477193354 | 0.122443216 |
| ENSMUSG00000035203 | Epn1 | 13854 | protein_coding | 4317.53055 | 0.12246359 | 0.096255072 | 1.272281941 | 0.203272963 | 0.437382664 | 0.12246359 |
| ENSMUSG00000038056 | Kmt2c | 231051 | protein_coding | 2567.697905 | -0.122476838 | 0.060040426 | -2.03990621 | 0.041359668 | 0.162367908 | 0.122476838 |
| ENSMUSG00000061130 | Ppm1b | 19043 | protein_coding | 1970.489772 | 0.122496733 | 0.041933747 | 2.921196903 | 0.003486894 | 0.027205194 | 0.122496733 |
| ENSMUSG00000039156 | Stim2 | 116873 | protein_coding | 1038.137775 | -0.122505665 | 0.044698677 | -2.740700026 | 0.006130845 | 0.041645508 | 0.122505665 |
| ENSMUSG00000029381 | Shroom3 | 27428 | protein_coding | 6130.905129 | -0.122516164 | 0.074965424 | -1.634302296 | 0.102195392 | 0.291648396 | 0.122516164 |
| ENSMUSG00000029033 | Acap3 | 140500 | protein_coding | 258.2957808 | -0.122555449 | 0.103943166 | -1.179062107 | 0.238373446 | 0.478679126 | 0.122555449 |
| ENSMUSG00000073755 | 5730409E04Rik | 230757 | protein_coding | 15.73903125 | -0.122580165 | 0.28671028 | -0.427540182 | 0.668985925 | 0.831002845 | 0.122580165 |
| ENSMUSG00000022477 | Aco2 | 11429 | protein_coding | 13475.8801 | 0.122580597 | 0.057853598 | 2.11880679 | 0.034106799 | 0.142262102 | 0.122580597 |
| ENSMUSG00000027284 | Cdan1 | 68968 | protein_coding | 191.7866245 | -0.122582475 | 0.11358092 | -1.079252351 | 0.280475248 | 0.523452524 | 0.122582475 |
| ENSMUSG00000097115 | 1810019N24Rik | 69086 | lncRNA | 9.651079824 | -0.122586963 | 0.379140202 | -0.323328843 | 0.746446213 | 0.879750811 | 0.122586963 |
| ENSMUSG00000025200 | Cwf19l1 | 72502 | protein_coding | 244.0243633 | -0.122602088 | 0.089068108 | -1.376498175 | 0.168667452 | 0.392680814 | 0.122602088 |
| ENSMUSG00000059482 | Cfap418 | 67157 | protein_coding | 87.19395452 | 0.122615155 | 0.141833113 | 0.864503024 | 0.38731162 | 0.627314903 | 0.122615155 |
| ENSMUSG00000022849 | Hspbap1 | 66667 | protein_coding | 131.1602023 | -0.122643215 | 0.120252532 | -1.019880525 | 0.307785127 | 0.551639336 | 0.122643215 |
| ENSMUSG00000026047 | Poglut2 | 72050 | protein_coding | 156.9440884 | -0.122657361 | 0.105774911 | -1.159607314 | 0.246208722 | 0.487131402 | 0.122657361 |
| ENSMUSG00000032357 | Tinag | 26944 | protein_coding | 760.5882309 | -0.12266046 | 0.097619006 | -1.256522324 | 0.208926657 | 0.444506623 | 0.12266046 |
| ENSMUSG00000090247 | Bloc1s1 | 14533 | protein_coding | 208.2302548 | -0.122665264 | 0.121268386 | -1.011518901 | 0.31176814 | 0.555505998 | 0.122665264 |
| ENSMUSG00000031134 | Rbmx | 19655 | protein_coding | 713.592035 | -0.122717567 | 0.078730606 | -1.558702182 | 0.119066885 | 0.319180794 | 0.122717567 |
| ENSMUSG00000054939 | Zfp174 | 385674 | protein_coding | 45.80361105 | -0.122725925 | 0.207693075 | -0.590900418 | 0.554587146 | 0.754685552 | 0.122725925 |
| ENSMUSG00000027247 | Arhgap1 | 228359 | protein_coding | 2849.884603 | 0.122737061 | 0.060850614 | 2.017022575 | 0.043693157 | 0.168978309 | 0.122737061 |

| ensembl_gene_id | external_gene_name | entrezgene_id | gene_biotype | baseMean | log2FoldChange | lfcSE | stat | pvalue | padj | absLog2FC |
|---|---|---|---|---|---|---|---|---|---|---|
| ENSMUSG00000025856 | Pdgfa | 18590 | protein_coding | 876.2824889 | -0.12275375 | 0.088323311 | -1.389822784 | 0.164582698 | 0.386825021 | 0.12275375 |
| ENSMUSG00000024936 | Kcnk7 | 16530 | protein_coding | 9.516240565 | -0.122773412 | 0.39534454 | -0.310547888 | 0.756144349 | 0.884312567 | 0.122773412 |
| ENSMUSG00000029014 | Dnajc2 | 22791 | protein_coding | 748.6057222 | -0.122831885 | 0.060847824 | -2.018673422 | 0.043521174 | 0.16855571 | 0.122831885 |
| ENSMUSG00000026942 | Traf2 | 22030 | protein_coding | 562.5433511 | 0.122843284 | 0.079171253 | 1.551614751 | 0.120754431 | 0.321618667 | 0.122843284 |
| ENSMUSG00000014773 | Dll1 | 13388 | protein_coding | 504.5983501 | -0.122855935 | 0.113980526 | -1.077867759 | 0.281092775 | 0.523522715 | 0.122855935 |
| ENSMUSG00000011254 | Thg1l | 66628 | protein_coding | 185.1343491 | 0.122877852 | 0.103089844 | 1.191949143 | 0.233281187 | 0.472718587 | 0.122877852 |
| ENSMUSG00000032802 | Srxn1 | 76650 | protein_coding | 2765.780163 | 0.122884184 | 0.062025087 | 1.981201321 | 0.047568701 | 0.178987504 | 0.122884184 |
| ENSMUSG00000029832 | Nfe2l3 | 18025 | protein_coding | 61.87808569 | 0.122902652 | 0.169687687 | 0.724287389 | 0.468889318 | 0.693423019 | 0.122902652 |
| ENSMUSG00000041438 | Utp4 | 21771 | protein_coding | 376.72596 | -0.122908739 | 0.06671812 | -1.842209269 | 0.065444545 | 0.220339077 | 0.122908739 |
| ENSMUSG00000032977 | Fam207a | 108707 | protein_coding | 622.9514622 | 0.123038114 | 0.061281879 | 2.007740554 | 0.044670869 | 0.17144386 | 0.123038114 |
| ENSMUSG00000004069 | Dnaja3 | 83945 | protein_coding | 1815.920048 | 0.123070256 | 0.064697758 | 1.902233719 | 0.057140606 | 0.202129099 | 0.123070256 |
| ENSMUSG00000104011 | Gm32391 | 102634922 | lncRNA | 19.45869247 | 0.123110074 | 0.273584696 | 0.449988891 | 0.652718451 | 0.82134757 | 0.123110074 |
| ENSMUSG00000036478 | Btg1 | 12226 | protein_coding | 5381.48241 | -0.123131097 | 0.060669207 | -2.029548487 | 0.042402456 | 0.165184357 | 0.123131097 |
| ENSMUSG00000026113 | Inpp4a | 269180 | protein_coding | 303.5117901 | -0.123135472 | 0.098118299 | -1.254969491 | 0.209489835 | 0.444801234 | 0.123135472 |
| ENSMUSG00000029918 | Mrps33 | 14548 | protein_coding | 168.9041494 | 0.123142966 | 0.115177137 | 1.069161547 | 0.284996882 | 0.527928408 | 0.123142966 |
| ENSMUSG00000075031 | H2bc3 | 319178 | protein_coding | 2117.799012 | 0.123144154 | 0.130305153 | 0.945044396 | 0.344636211 | 0.586900547 | 0.123144154 |
| ENSMUSG00000073633 | Fbxo36 | 66153 | protein_coding | 100.5383728 | 0.123162077 | 0.120780001 | 1.019722441 | 0.307860116 | 0.551651121 | 0.123162077 |
| ENSMUSG00000046562 | Unc119b | 106840 | protein_coding | 1135.86388 | 0.123163204 | 0.062271537 | 1.977841087 | 0.047946637 | 0.179737215 | 0.123163204 |
| ENSMUSG00000032376 | Usp3 | 235441 | protein_coding | 2147.829903 | -0.123196959 | 0.056984437 | -2.161940459 | 0.030622766 | 0.132046349 | 0.123196959 |
| ENSMUSG00000033671 | Cep350 | 74081 | protein_coding | 2706.742497 | -0.123206796 | 0.047679897 | -2.584040706 | 0.009765027 | 0.05836938 | 0.123206796 |
| ENSMUSG00000048997 | Atxn7l2 | 72522 | protein_coding | 90.00244007 | -0.123224016 | 0.128667298 | -0.95769491 | 0.338216624 | 0.582029301 | 0.123224016 |
| ENSMUSG00000031839 | Hsbp1 | 68196 | protein_coding | 1472.634276 | 0.123243366 | 0.040412045 | 3.049669148 | 0.002290936 | 0.019496529 | 0.123243366 |
| ENSMUSG00000046822 | Slc39a3 | 106947 | protein_coding | 816.8395447 | 0.123306052 | 0.081412164 | 1.514590024 | 0.12987626 | 0.335820819 | 0.123306052 |
| ENSMUSG00000011877 | Git1 | 216963 | protein_coding | 1225.022146 | -0.123308262 | 0.059740709 | -2.064057561 | 0.039012259 | 0.155730372 | 0.123308262 |
| ENSMUSG00000031596 | Slc7a2 | 11988 | protein_coding | 32.11896759 | -0.123339125 | 0.242433071 | -0.508755364 | 0.61092371 | 0.792706869 | 0.123339125 |
| ENSMUSG00000034126 | Pomt2 | 217734 | protein_coding | 396.0584735 | 0.12334466 | 0.071316293 | 1.729543898 | 0.083711797 | 0.257588275 | 0.12334466 |
| ENSMUSG00000047534 | Mis18bp1 | 217653 | protein_coding | 209.1186494 | -0.1233667 | 0.131331072 | -0.93935653 | 0.347547723 | 0.589851392 | 0.1233667 |
| ENSMUSG00000046897 | Zfp740 | 68744 | protein_coding | 1884.672619 | -0.123374715 | 0.046542271 | -2.650809938 | 0.008029901 | 0.050786438 | 0.123374715 |
| ENSMUSG00000033918 | Parl | 381038 | protein_coding | 806.2016076 | -0.123413242 | 0.05216002 | -2.36605049 | 0.017978992 | 0.091350728 | 0.123413242 |
| ENSMUSG00000039262 | Prrc2b | 227723 | protein_coding | 1657.739321 | -0.123478904 | 0.07097262 | -1.739810418 | 0.081892312 | 0.254171925 | 0.123478904 |
| ENSMUSG00000028826 | Maco1 | 66146 | protein_coding | 1128.774766 | -0.123490329 | 0.046586215 | -2.650791168 | 0.008030348 | 0.050786438 | 0.123490329 |
| ENSMUSG00000048920 | Fkrp | 243853 | protein_coding | 475.3581784 | 0.123561366 | 0.075576305 | 1.634922027 | 0.102065394 | 0.291535355 | 0.123561366 |
| ENSMUSG00000042569 | Dhrs7b | 216820 | protein_coding | 1152.41712 | 0.123567591 | 0.071434938 | 1.729792086 | 0.083667429 | 0.257550033 | 0.123567591 |
| ENSMUSG00000020918 | Kat2a | 14534 | protein_coding | 484.6653708 | -0.123568916 | 0.084019211 | -1.470722165 | 0.141366269 | 0.352354239 | 0.123568916 |
| ENSMUSG00000033416 | Gucd1 | 68778 | protein_coding | 4121.759856 | 0.123576681 | 0.04657594 | 2.653230022 | 0.007972552 | 0.050599507 | 0.123576681 |
| ENSMUSG00000060093 | H4c1 | 326619 | protein_coding | 1044.067866 | 0.123599908 | 0.133247432 | 0.927596919 | 0.353616697 | 0.595956629 | 0.123599908 |
| ENSMUSG00000027353 | Mcm8 | 66634 | protein_coding | 104.227206 | 0.1236002 | 0.166194637 | 0.743707513 | 0.457053443 | 0.685107692 | 0.1236002 |
| ENSMUSG00000038664 | Herc1 | 235439 | protein_coding | 2500.688909 | -0.123636363 | 0.043058493 | -2.87135836 | 0.004087119 | 0.030728656 | 0.123636363 |
| ENSMUSG00000028710 | Atpaf1 | 230649 | protein_coding | 248.9649459 | -0.123657657 | 0.098679748 | -1.253120927 | 0.210161701 | 0.445254566 | 0.123657657 |

| ensembl_gene_id | external_gene_name | entrezgene_id | gene_biotype | baseMean | log2FoldChange | lfcSE | stat | pvalue | padj | absLog2FC |
|---|---|---|---|---|---|---|---|---|---|---|
| ENSMUSG00000021661 | Ankra2 | 68558 | protein_coding | 407.6606514 | 0.123657678 | 0.062000765 | 1.994454063 | 0.046102455 | 0.175187158 | 0.123657678 |
| ENSMUSG00000031555 | Adam9 | 11502 | protein_coding | 2117.881319 | 0.123665519 | 0.050096231 | 2.468559338 | 0.013565816 | 0.074591174 | 0.123665519 |
| ENSMUSG00000039131 | Gipc2 | 54120 | protein_coding | 3407.750517 | -0.123674391 | 0.053632716 | -2.305950524 | 0.021113397 | 0.102036486 | 0.123674391 |
| ENSMUSG00000020389 | Cdkl3 | 213084 | protein_coding | 51.92651627 | 0.123682711 | 0.195061159 | 0.634071444 | 0.526034198 | 0.736439527 | 0.123682711 |
| ENSMUSG00000021957 | Tkt | 21881 | protein_coding | 11364.20483 | 0.123715215 | 0.065356573 | 1.892926885 | 0.058367593 | 0.204673663 | 0.123715215 |
| ENSMUSG00000030868 | Dctn5 | 59288 | protein_coding | 877.256741 | 0.123717723 | 0.051886786 | 2.38437821 | 0.01710801 | 0.088190332 | 0.123717723 |
| ENSMUSG00000022474 | Pmm1 | 29858 | protein_coding | 563.9256784 | -0.123725811 | 0.070011572 | -1.767219434 | 0.077191486 | 0.244956139 | 0.123725811 |
| ENSMUSG00000089832 | Shkbp1 | 192192 | protein_coding | 476.9547881 | 0.12373849 | 0.102847864 | 1.203121636 | 0.228929254 | 0.468116689 | 0.12373849 |
| ENSMUSG00000031865 | Dctn1 | 13191 | protein_coding | 3157.999984 | 0.123746938 | 0.064727387 | 1.911817283 | 0.055899634 | 0.199401327 | 0.123746938 |
| ENSMUSG00000048799 | Cep120 | 225523 | protein_coding | 413.3352702 | -0.123762408 | 0.072363134 | -1.710296395 | 0.087211078 | 0.264480943 | 0.123762408 |
| ENSMUSG00000006241 | Ccdc159 | 67119 | protein_coding | 22.83722377 | -0.123791702 | 0.243067689 | -0.509289009 | 0.610549661 | 0.792525305 | 0.123791702 |
| ENSMUSG00000035623 | Rsf1 | 233532 | protein_coding | 938.3471752 | -0.123791991 | 0.05323562 | -2.325360172 | 0.020052709 | 0.098324732 | 0.123791991 |
| ENSMUSG00000038248 | Sobp | 109205 | protein_coding | 26.15262957 | -0.123801562 | 0.315943141 | -0.391847603 | 0.69517082 | 0.847585984 | 0.123801562 |
| ENSMUSG00000030769 | Slc5a11 | 233836 | protein_coding | 12.78487961 | -0.12383809 | 0.714129566 | -0.173411235 | 0.862328194 | 0.937787345 | 0.12383809 |
| ENSMUSG00000025332 | Kdm5c | 20591 | protein_coding | 4152.228972 | 0.123891695 | 0.070357781 | 1.760881226 | 0.078258508 | 0.247078105 | 0.123891695 |
| ENSMUSG00000064128 | Cenpj | 219103 | protein_coding | 106.6475796 | 0.123922483 | 0.111457597 | 1.111835235 | 0.266208999 | 0.508185669 | 0.123922483 |
| ENSMUSG00000037526 | Atg14 | 100504663 | protein_coding | 360.1641559 | -0.124002552 | 0.076685764 | -1.617021786 | 0.105873578 | 0.297926072 | 0.124002552 |
| ENSMUSG00000039220 | Ppp1r10 | 52040 | protein_coding | 1619.405436 | 0.124003975 | 0.045667629 | 2.715358319 | 0.00662041 | 0.044027772 | 0.124003975 |
| ENSMUSG00000040505 | Abcg5 | 27409 | protein_coding | 68.89778879 | 0.124011128 | 0.188966547 | 0.656259692 | 0.511657048 | 0.725473655 | 0.124011128 |
| ENSMUSG00000041112 | Elmo1 | 140580 | protein_coding | 297.9780637 | -0.124048784 | 0.082906387 | -1.496251238 | 0.134588197 | 0.342321275 | 0.124048784 |
| ENSMUSG00000035958 | Tdp2 | 56196 | protein_coding | 540.5155144 | -0.124049812 | 0.0589026 | -2.106015893 | 0.035202972 | 0.145122272 | 0.124049812 |
| ENSMUSG00000037049 | Smpd1 | 20597 | protein_coding | 826.6097534 | 0.124064183 | 0.081395892 | 1.524206939 | 0.127457025 | 0.332056189 | 0.124064183 |
| ENSMUSG00000090266 | Mettl23 | 74319 | protein_coding | 315.3942924 | -0.124091324 | 0.073255173 | -1.69395989 | 0.090272905 | 0.270399544 | 0.124091324 |
| ENSMUSG00000051146 | Camk2n2 | 73047 | protein_coding | 19.9879945 | -0.124124189 | 0.264655123 | -0.469003539 | 0.639067107 | 0.812026097 | 0.124124189 |
| ENSMUSG00000033365 | Ipo13 | 230673 | protein_coding | 316.2607296 | 0.124150206 | 0.095529049 | 1.299606848 | 0.193735752 | 0.425519673 | 0.124150206 |
| ENSMUSG00000026842 | Abl1 | 11350 | protein_coding | 2294.229351 | -0.124167307 | 0.057638507 | -2.154242262 | 0.031221178 | 0.133697276 | 0.124167307 |
| ENSMUSG00000030615 | Tmem126a | 66271 | protein_coding | 517.6311054 | 0.124180551 | 0.056207848 | 2.209309837 | 0.027153097 | 0.121033168 | 0.124180551 |
| ENSMUSG00000032349 | Elovl5 | 68801 | protein_coding | 1145.527155 | -0.124184993 | 0.066508908 | -1.867193392 | 0.061874586 | 0.212283335 | 0.124184993 |
| ENSMUSG00000032806 | Slc10a3 | 214601 | protein_coding | 479.5820045 | 0.12420706 | 0.079479256 | 1.56276072 | 0.118108887 | 0.31772177 | 0.12420706 |
| ENSMUSG00000052833 | Sae1 | 56459 | protein_coding | 1397.727878 | 0.124241495 | 0.077224002 | 1.608845581 | 0.107650111 | 0.300613293 | 0.124241495 |
| ENSMUSG00000065582 | Mir194-2 | 723957 | miRNA | 15.99487013 | 0.124261827 | 0.325811597 | 0.381391665 | 0.702912645 | 0.85232122 | 0.124261827 |
| ENSMUSG00000020437 | Myo1g | 246177 | protein_coding | 20.42594778 | 0.124325713 | 0.329133159 | 0.377736821 | 0.705626111 | 0.853575129 | 0.124325713 |
| ENSMUSG00000027300 | Ubox5 | 140629 | protein_coding | 259.1692687 | -0.124329803 | 0.097666636 | -1.273001793 | 0.203017406 | 0.437065467 | 0.124329803 |
| ENSMUSG00000029722 | Agfg2 | 231801 | protein_coding | 890.9337204 | -0.124331936 | 0.078683755 | -1.580147465 | 0.1140731 | 0.311489481 | 0.124331936 |
| ENSMUSG00000028959 | Fastk | 66587 | protein_coding | 1624.646423 | -0.124339813 | 0.053832756 | -2.309742666 | 0.020902405 | 0.101348119 | 0.124339813 |
| ENSMUSG00000026870 | Cutal | 77996 | protein_coding | 2298.636519 | 0.124362517 | 0.059209526 | 2.100380229 | 0.035695407 | 0.146403821 | 0.124362517 |
| ENSMUSG00000038677 | Scube3 | 268935 | protein_coding | 6.22302171 | -0.124419645 | 0.622065788 | -0.200010429 | 0.841472425 | 0.929613548 | 0.124419645 |
| ENSMUSG00000001248 | Gramd1a | 52857 | protein_coding | 140.4022645 | 0.124423755 | 0.135496728 | 0.918278666 | 0.358472993 | 0.601570652 | 0.124423755 |
| ENSMUSG00000027272 | Ubr1 | 22222 | protein_coding | 753.8941445 | -0.124446304 | 0.05607488 | -2.219287919 | 0.026467143 | 0.118917792 | 0.124446304 |

| ensembl_gene_id | external_gene_name | entrezgene_id | gene_biotype | baseMean | log2FoldChange | lfcSE | stat | pvalue | padj | absLog2FC |
|---|---|---|---|---|---|---|---|---|---|---|
| ENSMUSG00000011382 | Dhdh | 71755 | protein_coding | 37.34684785 | -0.124461869 | 0.20650789 | -0.602697889 | 0.546709686 | 0.749854321 | 0.124461869 |
| ENSMUSG00000027510 | Rbm38 | 56190 | protein_coding | 203.5580034 | -0.12446535 | 0.104141868 | -1.195151886 | 0.232027693 | 0.471169298 | 0.12446535 |
| ENSMUSG00000019996 | Map7 | 17761 | protein_coding | 3697.479472 | 0.124489249 | 0.05425788 | 2.294399437 | 0.021767565 | 0.104540062 | 0.124489249 |
| ENSMUSG00000003752 | Itpkc | 233011 | protein_coding | 784.5797981 | -0.124527782 | 0.073943008 | -1.684104901 | 0.092161397 | 0.273594537 | 0.124527782 |
| ENSMUSG00000027550 | Lrrcc1 | 71710 | protein_coding | 422.8067725 | -0.124543871 | 0.103704471 | -1.200949876 | 0.229770645 | 0.469064473 | 0.124543871 |
| ENSMUSG00000024767 | Otub1 | 107260 | protein_coding | 1574.926254 | 0.124550807 | 0.06083253 | 2.047437563 | 0.040615141 | 0.160285476 | 0.124550807 |
| ENSMUSG00000036568 | Bicral | 210982 | protein_coding | 652.5563369 | -0.124554382 | 0.091289801 | -1.364384425 | 0.172446607 | 0.397562024 | 0.124554382 |
| ENSMUSG00000024831 | Ighmbp2 | 20589 | protein_coding | 286.9408621 | -0.124590427 | 0.090746872 | -1.372944588 | 0.169769574 | 0.394052089 | 0.124590427 |
| ENSMUSG00000026820 | Ptges2 | 96979 | protein_coding | 1028.677339 | -0.124597103 | 0.065707941 | -1.896225931 | 0.057930176 | 0.203825108 | 0.124597103 |
| ENSMUSG00000027472 | Pdrg1 | 68559 | protein_coding | 389.496066 | 0.124599995 | 0.082429201 | 1.51160018 | 0.130635606 | 0.336509736 | 0.124599995 |
| ENSMUSG00000033257 | Ttll4 | 67534 | protein_coding | 602.5673072 | -0.124602161 | 0.091439555 | -1.362672427 | 0.172985771 | 0.398420671 | 0.124602161 |
| ENSMUSG00000035674 | Ndufa3 | 66091 | protein_coding | 2527.274151 | 0.124603872 | 0.050607601 | 2.46215727 | 0.013810409 | 0.07559346 | 0.124603872 |
| ENSMUSG00000003099 | Ppp5c | 19060 | protein_coding | 2092.930108 | 0.124617806 | 0.05759987 | 2.163508453 | 0.030502094 | 0.131772643 | 0.124617806 |
| ENSMUSG00000028161 | Ppp3ca | 19055 | protein_coding | 1915.147978 | -0.124624658 | 0.064610407 | -1.928863539 | 0.053747805 | 0.194173579 | 0.124624658 |
| ENSMUSG00000097519 | 4930558J18Rik | 75323 | lncRNA | 6.794829271 | -0.124637274 | 0.443243297 | -0.281193816 | 0.778561748 | 0.896941755 | 0.124637274 |
| ENSMUSG00000068036 | Afdn | 17356 | protein_coding | 4222.904122 | -0.124640936 | 0.035303321 | -3.53057258 | 0.000414661 | 0.004817897 | 0.124640936 |
| ENSMUSG00000061079 | Zfp143 | 20841 | protein_coding | 370.2705385 | 0.124649745 | 0.078376771 | 1.590391447 | 0.111746597 | 0.307374616 | 0.124649745 |
| ENSMUSG00000020733 | Slc9a3r1 | 26941 | protein_coding | 4649.825916 | -0.12466826 | 0.090778615 | -1.373321905 | 0.169652296 | 0.39391111 | 0.12466826 |
| ENSMUSG00000045980 | Tmem104 | 320534 | protein_coding | 159.4340228 | 0.124684324 | 0.126868253 | 0.982785855 | 0.325712845 | 0.56923128 | 0.124684324 |
| ENSMUSG00000025326 | Ube3a | 22215 | protein_coding | 1497.276403 | -0.124732046 | 0.046312771 | -2.693253776 | 0.007075839 | 0.046217923 | 0.124732046 |
| ENSMUSG00000038618 | Rassf7 | 66985 | protein_coding | 2064.715931 | 0.124740003 | 0.0473072 | 2.636808006 | 0.008369018 | 0.052253619 | 0.124740003 |
| ENSMUSG00000031568 | Rwdd4a | 192174 | protein_coding | 390.249511 | 0.124744505 | 0.063816675 | 1.954732139 | 0.050614695 | 0.186591593 | 0.124744505 |
| ENSMUSG00000006498 | Ptbp1 | 19205 | protein_coding | 4239.515519 | 0.124767292 | 0.048890232 | 2.551988144 | 0.010711016 | 0.062224986 | 0.124767292 |
| ENSMUSG00000028158 | Mttp | 17777 | protein_coding | 220.5905035 | 0.124771224 | 0.495354519 | 0.251882681 | 0.801131746 | 0.909275777 | 0.124771224 |
| ENSMUSG00000027255 | Arfgap2 | 77038 | protein_coding | 785.174657 | 0.124779795 | 0.060891832 | 2.049204174 | 0.040442154 | 0.159861993 | 0.124779795 |
| ENSMUSG00000004038 | Gstm3 | 14864 | protein_coding | 700.0052424 | 0.124784574 | 0.113197409 | 1.102362453 | 0.270304128 | 0.512797437 | 0.124784574 |
| ENSMUSG00000032046 | Abhd12 | 76192 | protein_coding | 967.7368168 | 0.124838885 | 0.079810534 | 1.564190574 | 0.11777282 | 0.317250963 | 0.124838885 |
| ENSMUSG00000042700 | Sipa1l1 | 217692 | protein_coding | 2229.117634 | -0.124855675 | 0.040725817 | -3.065762316 | 0.002171158 | 0.018714201 | 0.124855675 |
| ENSMUSG00000049502 | Dtx3l | 209200 | protein_coding | 1327.057784 | -0.124866264 | 0.121241997 | -1.029892833 | 0.303060315 | 0.546939761 | 0.124866264 |
| ENSMUSG00000022013 | Dnajc15 | 66148 | protein_coding | 637.6841826 | 0.12490322 | 0.069188415 | 1.80526206 | 0.071033684 | 0.232315871 | 0.12490322 |
| ENSMUSG00000038429 | Usp5 | 22225 | protein_coding | 2241.303325 | 0.124913802 | 0.063159692 | 1.977745581 | 0.047957415 | 0.179737215 | 0.124913802 |
| ENSMUSG00000079108 | Srp54c | 100101806 | protein_coding | 164.5464032 | 0.124925466 | 0.110326018 | 1.132330052 | 0.2574957 | 0.499365134 | 0.124925466 |
| ENSMUSG00000070003 | Ssbp4 | 76900 | protein_coding | 681.9806357 | 0.124926979 | 0.077598704 | 1.60991063 | 0.107417368 | 0.300176728 | 0.124926979 |
| ENSMUSG00000006050 | Sra1 | 24068 | protein_coding | 1526.577172 | 0.124927627 | 0.066424735 | 1.880739548 | 0.060007357 | 0.208071092 | 0.124927627 |
| ENSMUSG00000032245 | Cln6 | 76524 | protein_coding | 586.2327912 | -0.124975658 | 0.080127973 | -1.559700742 | 0.118830617 | 0.319001795 | 0.124975658 |
| ENSMUSG00000054715 | Zscan22 | 232878 | protein_coding | 118.7483404 | 0.125022599 | 0.129202571 | 0.967647923 | 0.333220235 | 0.576562634 | 0.125022599 |
| ENSMUSG00000000154 | Slc22a18 | 18400 | protein_coding | 1745.100204 | 0.125025249 | 0.086279012 | 1.449080668 | 0.147315056 | 0.360891677 | 0.125025249 |
| ENSMUSG00000026828 | Galnt5 | 241391 | protein_coding | 1263.907163 | 0.125112096 | 0.066971267 | 1.868145869 | 0.061741741 | 0.211988304 | 0.125112096 |
| ENSMUSG00000035285 | Nat14 | 269854 | protein_coding | 16.17551944 | -0.125121944 | 0.308424937 | -0.405680374 | 0.684977464 | 0.841455786 | 0.125121944 |

| ensembl_gene_id | external_gene_name | entrezgene_id | gene_biotype | baseMean | log2FoldChange | lfcSE | stat | pvalue | padj | absLog2FC |
|---|---|---|---|---|---|---|---|---|---|---|
| ENSMUSG00000020859 | Spag9 | 70834 | protein_coding | 2180.050569 | -0.125136874 | 0.044355048 | -2.821254421 | 0.004783625 | 0.03448615 | 0.125136874 |
| ENSMUSG00000090215 | Trim34b | 434218 | protein_coding | 14.90413496 | 0.125173222 | 0.347860518 | 0.359837394 | 0.718968738 | 0.862409121 | 0.125173222 |
| ENSMUSG00000025289 | Prdx4 | 53381 | protein_coding | 368.2466566 | 0.125233813 | 0.094052804 | 1.331526632 | 0.183015788 | 0.411129072 | 0.125233813 |
| ENSMUSG00000036264 | Fstl4 | 320027 | protein_coding | 47.0357717 | -0.125239023 | 0.23427958 | -0.534570801 | 0.592946679 | 0.780422622 | 0.125239023 |
| ENSMUSG00000061132 | Blnk | 17060 | protein_coding | 1868.720252 | -0.125248018 | 0.052660084 | -2.37842418 | 0.017386812 | 0.088986709 | 0.125248018 |
| ENSMUSG00000030800 | Prss8 | 76560 | protein_coding | 1414.788842 | 0.125262198 | 0.049281314 | 2.541778749 | 0.011028996 | 0.063591213 | 0.125262198 |
| ENSMUSG00000033762 | Recql4 | 79456 | protein_coding | 89.21406678 | 0.125276383 | 0.148387003 | 0.844254416 | 0.398527252 | 0.638169126 | 0.125276383 |
| ENSMUSG00000030055 | Rab43 | 69834 | protein_coding | 193.5654278 | -0.12528127 | 0.141759104 | -0.883761721 | 0.376824854 | 0.61835018 | 0.12528127 |
| ENSMUSG00000028602 | Tnfrsf8 | 21941 | protein_coding | 12.2854896 | 0.125291759 | 0.407342904 | 0.307583017 | 0.758399645 | 0.885834075 | 0.125291759 |
| ENSMUSG00000039254 | Pomt1 | 99011 | protein_coding | 588.5023644 | 0.125299488 | 0.060629794 | 2.066632275 | 0.038768817 | 0.154950363 | 0.125299488 |
| ENSMUSG00000039427 | Alg1 | 208211 | protein_coding | 275.742102 | -0.125324179 | 0.104452823 | -1.199816105 | 0.230210768 | 0.469307431 | 0.125324179 |
| ENSMUSG00000031813 | Mvb12a | 73711 | protein_coding | 1391.144555 | 0.125342573 | 0.076037327 | 1.648434758 | 0.099263488 | 0.286680207 | 0.125342573 |
| ENSMUSG00000046139 | Patl1 | 225929 | protein_coding | 305.1150603 | -0.125369244 | 0.093827737 | -1.336164 | 0.181495675 | 0.408682037 | 0.125369244 |
| ENSMUSG00000054191 | Klf1 | 16596 | protein_coding | 9.507048347 | -0.125552594 | 0.389930454 | -0.321987145 | 0.747462433 | 0.880070575 | 0.125552594 |
| ENSMUSG00000049916 | 2610318N02Rik | 70458 | protein_coding | 39.91146647 | 0.125554746 | 0.194008288 | 0.647161766 | 0.517527254 | 0.729947916 | 0.125554746 |
| ENSMUSG00000033429 | Mcee | 73724 | protein_coding | 734.6426627 | 0.125618958 | 0.052854823 | 2.37667918 | 0.017469275 | 0.089352043 | 0.125618958 |
| ENSMUSG00000042185 | Nfrkb | 235134 | protein_coding | 542.0170777 | -0.125642563 | 0.060208828 | -2.086779756 | 0.036908048 | 0.149664594 | 0.125642563 |
| ENSMUSG00000020709 | Adap2 | 216991 | protein_coding | 57.31976715 | -0.12566617 | 0.207423128 | -0.60584454 | 0.54461799 | 0.749012844 | 0.12566617 |
| ENSMUSG00000020120 | Plek | 56193 | protein_coding | 47.22900477 | 0.125666816 | 0.187908413 | 0.668766311 | 0.50364456 | 0.720013604 | 0.125666816 |
| ENSMUSG00000074634 | Tmem267 | 633640 | protein_coding | 72.67995584 | -0.125678643 | 0.565781558 | -0.222132803 | 0.824210503 | 0.921854467 | 0.125678643 |
| ENSMUSG00000048706 | Lurap1l | 52829 | protein_coding | 2999.327383 | -0.125680138 | 0.065559928 | -1.917026789 | 0.055234528 | 0.197684696 | 0.125680138 |
| ENSMUSG00000055197 | Fev | 260298 | protein_coding | 7.915239623 | -0.12569851 | 0.48208547 | -0.260739055 | 0.794293746 | 0.904717934 | 0.12569851 |
| ENSMUSG00000022711 | Pmm2 | 54128 | protein_coding | 4186.545791 | 0.125716332 | 0.052360128 | 2.400993612 | 0.016350622 | 0.085418361 | 0.125716332 |
| ENSMUSG00000030842 | Lamtor1 | 66508 | protein_coding | 1626.620524 | 0.125728649 | 0.077025692 | 1.632294956 | 0.102617365 | 0.292438637 | 0.125728649 |
| ENSMUSG00000037190 | Cyb561d2 | 56368 | protein_coding | 273.8317396 | 0.125739856 | 0.121028245 | 1.038929853 | 0.298837361 | 0.542414195 | 0.125739856 |
| ENSMUSG00000030672 | Mylpf | 17907 | protein_coding | 22.43112896 | 0.125792485 | 0.24894566 | 0.505300978 | 0.613347456 | 0.793911553 | 0.125792485 |
| ENSMUSG00000069305 | H4c18 | 319161 | protein_coding | 1351.211546 | 0.125808239 | 0.128009184 | 0.982806353 | 0.325702755 | 0.56923128 | 0.125808239 |
| ENSMUSG00000022570 | Gfus | 22122 | protein_coding | 2477.334247 | 0.125905045 | 0.043090304 | 2.921888067 | 0.003479166 | 0.027158036 | 0.125905045 |
| ENSMUSG00000025572 | Tmc6 | 217353 | protein_coding | 1603.280746 | 0.125925541 | 0.085385754 | 1.474783962 | 0.140270634 | 0.351340845 | 0.125925541 |
| ENSMUSG00000026478 | Lamc1 | 226519 | protein_coding | 274.455209 | -0.125936005 | 0.076654149 | -1.64291179 | 0.100401185 | 0.288725021 | 0.125936005 |
| ENSMUSG00000021054 | Sgpp1 | 81535 | protein_coding | 873.2974807 | -0.125976185 | 0.059590838 | -2.114019358 | 0.034513614 | 0.143269254 | 0.125976185 |
| ENSMUSG00000030704 | Rab6a | 19346 | protein_coding | 2176.926474 | 0.126005308 | 0.053822372 | 2.341132576 | 0.019225339 | 0.09547586 | 0.126005308 |
| ENSMUSG00000054855 | Rnd1 | 223881 | protein_coding | 6.818517589 | -0.12601168 | 0.466331933 | -0.270218853 | 0.786991889 | 0.900576045 | 0.12601168 |
| ENSMUSG00000039741 | Bahcc1 | 268515 | protein_coding | 396.0829402 | -0.126090457 | 0.105733945 | -1.192525804 | 0.233055139 | 0.472527059 | 0.126090457 |
| ENSMUSG00000036078 | Sigmar1 | 18391 | protein_coding | 967.8866989 | 0.126095709 | 0.071776634 | 1.756779346 | 0.078955427 | 0.248300541 | 0.126095709 |
| ENSMUSG00000032381 | Ciao2a | 68250 | protein_coding | 1561.495051 | 0.126140884 | 0.068719853 | 1.835581402 | 0.06641959 | 0.22236843 | 0.126140884 |
| ENSMUSG00000038943 | Prc1 | 233406 | protein_coding | 544.7218536 | 0.126143345 | 0.104467952 | 1.207483668 | 0.227245931 | 0.465784841 | 0.126143345 |
| ENSMUSG00000037710 | Cisd1 | 52637 | protein_coding | 1797.49998 | 0.126175726 | 0.065393571 | 1.929482131 | 0.053671036 | 0.193983147 | 0.126175726 |
| ENSMUSG00000021972 | Hmbox1 | 219150 | protein_coding | 439.9967213 | -0.126240368 | 0.083339962 | -1.514763936 | 0.129832196 | 0.335814568 | 0.126240368 |

| ensembl_gene_id | external_gene_name | entrezgene_id | gene_biotype | baseMean | log2FoldChange | lfcSE | stat | pvalue | padj | absLog2FC |
|---|---|---|---|---|---|---|---|---|---|---|
| ENSMUSG00000024668 | Sdhaf2 | 66072 | protein_coding | 794.852482 | 0.126251914 | 0.046617388 | 2.708257986 | 0.006763742 | 0.044625766 | 0.126251914 |
| ENSMUSG00000002428 | Hltf | 20585 | protein_coding | 960.6182773 | 0.126318335 | 0.061898617 | 2.040729516 | 0.04127772 | 0.162164626 | 0.126318335 |
| ENSMUSG00000020547 | Bzw2 | 66912 | protein_coding | 1107.787477 | -0.126325987 | 0.067478528 | -1.87209161 | 0.061193931 | 0.210759777 | 0.126325987 |
| ENSMUSG00000001520 | Nrip2 | 60345 | protein_coding | 11.04501256 | -0.126362587 | 0.41242399 | -0.306390002 | 0.759307719 | 0.886076667 | 0.126362587 |
| ENSMUSG00000022003 | Slc25a30 | 67554 | protein_coding | 114.1365709 | -0.126416655 | 0.117704453 | -1.074017608 | 0.28281479 | 0.525455543 | 0.126416655 |
| ENSMUSG00000002345 | Borcs8 | 72368 | protein_coding | 383.1991483 | 0.126456826 | 0.108364683 | 1.16695609 | 0.243228103 | 0.484226665 | 0.126456826 |
| ENSMUSG00000026620 | Mark1 | 226778 | protein_coding | 100.2854502 | 0.126457826 | 0.133614813 | 0.946435676 | 0.343926416 | 0.586381512 | 0.126457826 |
| ENSMUSG00000020635 | Fkbp1b | 14226 | protein_coding | 12.14794748 | 0.126504049 | 0.334848643 | 0.37779472 | 0.705583096 | 0.853575129 | 0.126504049 |
| ENSMUSG00000021171 | Esyt2 | 52635 | protein_coding | 3940.128579 | -0.126506323 | 0.043208379 | -2.927819208 | 0.003413485 | 0.026787965 | 0.126506323 |
| ENSMUSG00000022021 | Diaph3 | 56419 | protein_coding | 198.6480764 | 0.126525477 | 0.13868411 | 0.91232858 | 0.361595773 | 0.604649053 | 0.126525477 |
| ENSMUSG00000090213 | Peds1 | 407243 | protein_coding | 719.8789528 | 0.126535425 | 0.062755039 | 2.01633887 | 0.043764553 | 0.169132745 | 0.126535425 |
| ENSMUSG00000020393 | Kremen1 | 84035 | protein_coding | 721.034803 | -0.126536441 | 0.07129608 | -1.774802213 | 0.075930551 | 0.242581587 | 0.126536441 |
| ENSMUSG00000078789 | Dph1 | 116905 | protein_coding | 268.9947521 | -0.126542121 | 0.085995713 | -1.471493357 | 0.141157743 | 0.352173719 | 0.126542121 |
| ENSMUSG00000056851 | Pcbp2 | 18521 | protein_coding | 1172.848553 | -0.126612009 | 0.128852605 | -0.982611168 | 0.325798846 | 0.569247886 | 0.126612009 |
| ENSMUSG00000020483 | Dynll2 | 68097 | protein_coding | 1531.746304 | 0.126642896 | 0.07910712 | 1.600903891 | 0.109398208 | 0.303086221 | 0.126642896 |
| ENSMUSG00000003546 | Klc4 | 74764 | protein_coding | 4096.113936 | 0.126646867 | 0.059550238 | 2.126723115 | 0.033443096 | 0.140437951 | 0.126646867 |
| ENSMUSG00000056612 | Ppp1r14b | 18938 | protein_coding | 723.2365402 | 0.126693535 | 0.06218004 | 2.037527395 | 0.04159722 | 0.162956942 | 0.126693535 |
| ENSMUSG00000030413 | Pglyrp1 | 21946 | protein_coding | 1734.343361 | 0.126695621 | 0.08701948 | 1.455945503 | 0.145407685 | 0.358198217 | 0.126695621 |
| ENSMUSG00000003198 | Zfp959 | 224893 | protein_coding | 125.559672 | -0.126719661 | 0.106515137 | -1.189686882 | 0.234169483 | 0.473794416 | 0.126719661 |
| ENSMUSG00000045962 | Wnk1 | 232341 | protein_coding | 1767.529506 | -0.126726757 | 0.088861715 | -1.42611198 | 0.153836021 | 0.37117198 | 0.126726757 |
| ENSMUSG00000061360 | Phf5a | 68479 | protein_coding | 811.3100262 | 0.126764357 | 0.071669807 | 1.768727475 | 0.076939365 | 0.244589225 | 0.126764357 |
| ENSMUSG00000022858 | Tra2b | 20462 | protein_coding | 1843.66384 | -0.12678785 | 0.071461049 | -1.774223174 | 0.076026242 | 0.242739104 | 0.12678785 |
| ENSMUSG00000044340 | Phlpp1 | 98432 | protein_coding | 1009.784915 | -0.126799515 | 0.069607919 | -1.82162486 | 0.068511926 | 0.226923769 | 0.126799515 |
| ENSMUSG00000001665 | Gstt3 | 103140 | protein_coding | 1062.718684 | 0.126805394 | 0.059545441 | 2.12955671 | 0.033208228 | 0.139684167 | 0.126805394 |
| ENSMUSG00000023191 | P3h3 | 14789 | protein_coding | 8.573329392 | 0.126811895 | 0.398950598 | 0.317863655 | 0.750588363 | 0.880845945 | 0.126811895 |
| ENSMUSG00000019806 | Aig1 | 66253 | protein_coding | 1288.886616 | -0.126852306 | 0.04390457 | -2.889273414 | 0.003861332 | 0.02945681 | 0.126852306 |
| ENSMUSG00000058173 | Smco4 | 170748 | protein_coding | 318.7872574 | 0.126886454 | 0.073662742 | 1.722532328 | 0.084973132 | 0.260078507 | 0.126886454 |
| ENSMUSG00000042340 | Ctf1 | 13019 | protein_coding | 53.73088442 | 0.126896108 | 0.17113859 | 0.741481556 | 0.458401511 | 0.685788934 | 0.126896108 |
| ENSMUSG00000088008 | Gm25492 | 115490291 | scaRNA | 71.39076867 | -0.126896873 | 0.211173434 | -0.600913053 | 0.547897899 | 0.750796946 | 0.126896873 |
| ENSMUSG00000053070 | Cfap300 | 234912 | protein_coding | 28.80823827 | 0.12689744 | 0.238880849 | 0.531216466 | 0.595268784 | 0.782387912 | 0.12689744 |
| ENSMUSG00000040451 | Sgms1 | 208449 | protein_coding | 915.4012477 | -0.126912386 | 0.055529608 | -2.285490386 | 0.02228409 | 0.106135709 | 0.126912386 |
| ENSMUSG00000020128 | Vps54 | 245944 | protein_coding | 2556.25243 | -0.126988685 | 0.044821933 | -2.833181834 | 0.004608716 | 0.033555199 | 0.126988685 |
| ENSMUSG00000020284 | Cfap410 | 67884 | protein_coding | 204.9696702 | 0.12711332 | 0.110117377 | 1.154343875 | 0.248359215 | 0.4893628 | 0.12711332 |
| ENSMUSG00000036693 | Nop14 | 75416 | protein_coding | 321.4597055 | -0.127128483 | 0.099832012 | -1.273424033 | 0.202867614 | 0.436976894 | 0.127128483 |
| ENSMUSG00000022031 | Elp3 | 74195 | protein_coding | 1115.244592 | 0.127139327 | 0.047367744 | 2.684090807 | 0.007272735 | 0.047118814 | 0.127139327 |
| ENSMUSG00000053158 | Fes | 14159 | protein_coding | 185.9791271 | 0.127165905 | 0.099816516 | 1.273996624 | 0.202664613 | 0.436714619 | 0.127165905 |
| ENSMUSG00000060639 | H4c9 | 319158 | protein_coding | 200.692792 | 0.127220726 | 0.119053818 | 1.068598453 | 0.285250646 | 0.528173927 | 0.127220726 |
| ENSMUSG00000024187 | Fam234a | 106581 | protein_coding | 6499.988304 | -0.12722406 | 0.047132922 | -2.699261044 | 0.006949364 | 0.045592153 | 0.12722406 |
| ENSMUSG00000055447 | Cd47 | 16423 | protein_coding | 595.2606559 | -0.127290553 | 0.084208828 | -1.5116058 | 0.130634175 | 0.336509736 | 0.127290553 |

| ensembl_gene_id | external_gene_name | entrezgene_id | gene_biotype | baseMean | log2FoldChange | lfcSE | stat | pvalue | padj | absLog2FC |
|---|---|---|---|---|---|---|---|---|---|---|
| ENSMUSG00000041926 | Rnpep | 215615 | protein_coding | 1919.173187 | 0.127311218 | 0.074618763 | 1.706155566 | 0.087979119 | 0.265849589 | 0.127311218 |
| ENSMUSG00000034663 | Bmp2k | 140780 | protein_coding | 748.4745559 | -0.127334379 | 0.068478419 | -1.859481882 | 0.062958865 | 0.214568389 | 0.127334379 |
| ENSMUSG00000025758 | Plk4 | 20873 | protein_coding | 242.2981784 | 0.12733784 | 0.111271401 | 1.144389656 | 0.252462076 | 0.493560011 | 0.12733784 |
| ENSMUSG00000020130 | Tbc1d15 | 66687 | protein_coding | 1086.368974 | -0.12736551 | 0.083159317 | -1.531584371 | 0.125625032 | 0.328925409 | 0.12736551 |
| ENSMUSG00000030751 | Psma1 | 26440 | protein_coding | 2087.255071 | 0.127417394 | 0.079770755 | 1.597294574 | 0.11020006 | 0.304713258 | 0.127417394 |
| ENSMUSG00000037876 | Jmjd1c | 108829 | protein_coding | 2211.16076 | 0.127421502 | 0.055607729 | 2.291435095 | 0.021938262 | 0.104860209 | 0.127421502 |
| ENSMUSG00000052331 | Ankrd44 | 329154 | protein_coding | 424.553952 | -0.127423518 | 0.081578218 | -1.561979668 | 0.118292779 | 0.318004275 | 0.127423518 |
| ENSMUSG00000046516 | Cox17 | 12856 | protein_coding | 198.8241856 | -0.127440265 | 0.100720844 | -1.265281941 | 0.205770278 | 0.440405743 | 0.127440265 |
| ENSMUSG00000036768 | Kif15 | 209737 | protein_coding | 424.6432085 | -0.127440581 | 0.136115701 | -0.936266572 | 0.349135955 | 0.591564986 | 0.127440581 |
| ENSMUSG00000031153 | Gripap1 | 54645 | protein_coding | 504.2404178 | 0.127453031 | 0.084318355 | 1.511569234 | 0.130643483 | 0.336509736 | 0.127453031 |
| ENSMUSG00000025907 | Rb1cc1 | 12421 | protein_coding | 2151.364959 | -0.127528979 | 0.067750596 | -1.882329994 | 0.059791225 | 0.207678888 | 0.127528979 |
| ENSMUSG00000024855 | Pacs1 | 107975 | protein_coding | 216.5349649 | -0.127578134 | 0.09558287 | -1.334738473 | 0.181961957 | 0.409503276 | 0.127578134 |
| ENSMUSG00000039662 | Icmt | 57295 | protein_coding | 1392.211161 | 0.127605172 | 0.06959811 | 1.833457431 | 0.066734574 | 0.222950058 | 0.127605172 |
| ENSMUSG00000036904 | Fzd8 | 14370 | protein_coding | 593.1771458 | 0.127609455 | 0.08289464 | 1.539417452 | 0.123702416 | 0.326398702 | 0.127609455 |
| ENSMUSG00000002846 | Timmdc1 | 76916 | protein_coding | 532.0675755 | 0.127626576 | 0.081842388 | 1.559419007 | 0.118897241 | 0.319001795 | 0.127626576 |
| ENSMUSG00000027263 | Tubgcp4 | 51885 | protein_coding | 624.955427 | -0.127639252 | 0.073041874 | -1.747480533 | 0.080554019 | 0.251372806 | 0.127639252 |
| ENSMUSG00000040860 | Crocc | 230872 | protein_coding | 230.1307544 | -0.127667517 | 0.104800509 | -1.21819559 | 0.223149655 | 0.460551754 | 0.127667517 |
| ENSMUSG00000057948 | Unc13d | 70450 | protein_coding | 552.6464096 | -0.127695342 | 0.07345248 | -1.738475557 | 0.082127053 | 0.254517969 | 0.127695342 |
| ENSMUSG00000024515 | Smad4 | 17128 | protein_coding | 2281.909177 | -0.127704391 | 0.070371688 | -1.814712625 | 0.069568084 | 0.229057674 | 0.127704391 |
| ENSMUSG00000060244 | Alyref2 | 56009 | protein_coding | 47.04949035 | 0.127714643 | 0.227796117 | 0.560653291 | 0.575033915 | 0.768361739 | 0.127714643 |
| ENSMUSG00000029823 | Luc7l2 | 192196 | protein_coding | 1850.482448 | -0.127725419 | 0.0789963 | -1.616853188 | 0.105909975 | 0.297926072 | 0.127725419 |
| ENSMUSG00000039357 | Fut11 | 73068 | protein_coding | 266.3400948 | -0.127731653 | 0.080203619 | -1.592592125 | 0.111251721 | 0.306582205 | 0.127731653 |
| ENSMUSG00000025358 | Cdk2 | 12566 | protein_coding | 382.9271192 | -0.127738895 | 0.08680514 | -1.47155912 | 0.141139972 | 0.352173719 | 0.127738895 |
| ENSMUSG00000047139 | Cd24a | 12484 | protein_coding | 7261.723287 | 0.12779864 | 0.037992452 | 3.363790218 | 0.0007688 | 0.00809297 | 0.12779864 |
| ENSMUSG00000074259 | Gramd2 | 546134 | protein_coding | 83.71257406 | 0.127805169 | 0.198067201 | 0.64526165 | 0.518757641 | 0.730713836 | 0.127805169 |
| ENSMUSG00000079426 | Arpc4 | 68089 | protein_coding | 2073.420796 | 0.127812533 | 0.056658708 | 2.255832106 | 0.024081148 | 0.111811535 | 0.127812533 |
| ENSMUSG00000003810 | Mast2 | 17776 | protein_coding | 2417.276856 | -0.127812689 | 0.062765289 | -2.036359438 | 0.041714275 | 0.16319418 | 0.127812689 |
| ENSMUSG00000079043 | Fastkd5 | 380601 | protein_coding | 107.0962806 | -0.12784123 | 0.130711963 | -0.978037715 | 0.328055671 | 0.571645972 | 0.12784123 |
| ENSMUSG00000021431 | Snrnp48 | 67797 | protein_coding | 439.417541 | -0.127853551 | 0.124310485 | -1.028501747 | 0.30371387 | 0.547874002 | 0.127853551 |
| ENSMUSG00000038467 | Chmp4b | 75608 | protein_coding | 5279.7211 | 0.127940579 | 0.050123885 | 2.552487282 | 0.010695681 | 0.062180697 | 0.127940579 |
| ENSMUSG00000026944 | Abca2 | 11305 | protein_coding | 193.5523374 | 0.128016857 | 0.105076382 | 1.218321903 | 0.22310167 | 0.460551754 | 0.128016857 |
| ENSMUSG00000028716 | Pdzk1ip1 | 67182 | protein_coding | 1492.127879 | -0.128018139 | 0.1106298 | -1.157175906 | 0.247200499 | 0.488015969 | 0.128018139 |
| ENSMUSG00000056014 | A430033K04Rik | 243308 | protein_coding | 257.9151317 | -0.128026492 | 0.09079376 | -1.410080298 | 0.158515974 | 0.37855577 | 0.128026492 |
| ENSMUSG00000021482 | Prxl2c | 66129 | protein_coding | 1135.421479 | -0.128031059 | 0.066728699 | -1.918680584 | 0.05502477 | 0.197371991 | 0.128031059 |
| ENSMUSG00000070315 | 4930581F22Rik | 78934 | lncRNA | 52.85630851 | 0.12803697 | 0.200032446 | 0.640081012 | 0.522119933 | 0.733199598 | 0.12803697 |
| ENSMUSG00000017664 | Slc35c2 | 228875 | protein_coding | 438.92315 | 0.128063485 | 0.072488008 | 1.766685114 | 0.077280978 | 0.24519188 | 0.128063485 |
| ENSMUSG00000023921 | Mmut | 17850 | protein_coding | 2157.548723 | 0.128078751 | 0.047888139 | 2.67454013 | 0.007483182 | 0.048137725 | 0.128078751 |
| ENSMUSG00000041992 | Rapgef5 | 217944 | protein_coding | 925.683047 | -0.128092154 | 0.060376577 | -2.121553765 | 0.033875228 | 0.141652252 | 0.128092154 |
| ENSMUSG00000025035 | Arl3 | 56350 | protein_coding | 93.6306653 | 0.128202178 | 0.152361823 | 0.84143242 | 0.40010573 | 0.639482053 | 0.128202178 |

| ensembl_gene_id | external_gene_name | entrezgene_id | gene_biotype | baseMean | log2FoldChange | lfcSE | stat | pvalue | padj | absLog2FC |
|---|---|---|---|---|---|---|---|---|---|---|
| ENSMUSG00000027890 | Gstm4 | 14865 | protein_coding | 198.5397487 | 0.128271002 | 0.11162563 | 1.149117841 | 0.250507393 | 0.491535798 | 0.128271002 |
| ENSMUSG00000026851 | BC005624 | 227707 | protein_coding | 651.020001 | 0.128296483 | 0.055869397 | 2.296364212 | 0.021655065 | 0.104092769 | 0.128296483 |
| ENSMUSG00000041540 | Sox5 | 20678 | protein_coding | 10.59519129 | 0.128350811 | 0.354391767 | 0.362172103 | 0.717223425 | 0.861383838 | 0.128350811 |
| ENSMUSG00000052407 | Ccdc171 | 320226 | protein_coding | 118.9720312 | -0.128357499 | 0.143057965 | -0.897241196 | 0.369590227 | 0.611697618 | 0.128357499 |
| ENSMUSG00000021131 | Erh | 13877 | protein_coding | 444.3513287 | 0.12837483 | 0.095573354 | 1.343207333 | 0.179204848 | 0.406457009 | 0.12837483 |
| ENSMUSG00000050549 | Fam241a | 70617 | protein_coding | 310.3196655 | -0.128403424 | 0.081019681 | -1.584842384 | 0.11300215 | 0.309461501 | 0.128403424 |
| ENSMUSG00000021650 | Ptcd2 | 68927 | protein_coding | 1145.645102 | 0.128405612 | 0.04488326 | 2.860879822 | 0.004224672 | 0.031483955 | 0.128405612 |
| ENSMUSG00000000253 | Gmpr | 66355 | protein_coding | 317.2478096 | -0.128449026 | 0.0912534 | -1.407608114 | 0.159247143 | 0.379346925 | 0.128449026 |
| ENSMUSG00000086999 | Bcas1os2 | 73303 | lncRNA | 15.5724356 | -0.128467481 | 0.402069287 | -0.319515779 | 0.749335428 | 0.880475742 | 0.128467481 |
| ENSMUSG00000089715 | Cbx6 | 494448 | protein_coding | 141.7967123 | -0.128515031 | 0.125128056 | -1.027068074 | 0.304388411 | 0.548477307 | 0.128515031 |
| ENSMUSG00000023960 | Enpp5 | 83965 | protein_coding | 1061.473619 | 0.128525377 | 0.067782945 | 1.89613151 | 0.057942657 | 0.203825108 | 0.128525377 |
| ENSMUSG00000070871 | Ccnyl1 | 227210 | protein_coding | 223.5978617 | 0.128575198 | 0.087583001 | 1.468038287 | 0.142093823 | 0.353306674 | 0.128575198 |
| ENSMUSG00000038822 | Hace1 | 209462 | protein_coding | 449.1970491 | 0.128598653 | 0.071694885 | 1.79369356 | 0.072862094 | 0.236142883 | 0.128598653 |
| ENSMUSG00000037791 | Phf12 | 268448 | protein_coding | 585.706604 | -0.128645709 | 0.064647865 | -1.989945201 | 0.046596972 | 0.176277123 | 0.128645709 |
| ENSMUSG00000019359 | Gdpd2 | 71584 | protein_coding | 100.4610567 | -0.128673497 | 0.184452071 | -0.69759855 | 0.485428288 | 0.70667106 | 0.128673497 |
| ENSMUSG00000030435 | U2af2 | 22185 | protein_coding | 2492.415797 | 0.128716984 | 0.056875002 | 2.263155683 | 0.023626094 | 0.110184505 | 0.128716984 |
| ENSMUSG00000024993 | Dennd10 | 67894 | protein_coding | 488.9738219 | 0.128757652 | 0.091806654 | 1.402487147 | 0.160769826 | 0.381255129 | 0.128757652 |
| ENSMUSG00000024273 | 2700062C07Rik | 68046 | protein_coding | 180.7563315 | 0.128784488 | 0.10742466 | 1.198835432 | 0.230591942 | 0.469603637 | 0.128784488 |
| ENSMUSG00000037210 | Fam193a | 231128 | protein_coding | 726.138306 | -0.128818815 | 0.058975966 | -2.184259505 | 0.028943185 | 0.126769891 | 0.128818815 |
| ENSMUSG00000025239 | Limd1 | 29806 | protein_coding | 2256.272969 | -0.128822286 | 0.035403777 | -3.638659389 | 0.000274061 | 0.003428845 | 0.128822286 |
| ENSMUSG00000043487 | Acot6 | 217700 | protein_coding | 47.73719597 | 0.128837341 | 0.168982047 | 0.762432123 | 0.44580214 | 0.676717913 | 0.128837341 |
| ENSMUSG00000032249 | Anp32a | 11737 | protein_coding | 1804.504249 | 0.128879858 | 0.064354022 | 2.00266983 | 0.045212739 | 0.172742441 | 0.128879858 |
| ENSMUSG00000075010 | AW112010 | 107350 | lncRNA | 765.9710879 | -0.128883327 | 0.232048608 | -0.555415212 | 0.57861069 | 0.771101239 | 0.128883327 |
| ENSMUSG00000006299 | Aamp | 227290 | protein_coding | 2796.118943 | 0.128918155 | 0.042626092 | 3.024395375 | 0.002491306 | 0.020731319 | 0.128918155 |
| ENSMUSG00000022299 | Slc25a32 | 69906 | protein_coding | 543.5506929 | -0.128929931 | 0.057502756 | -2.242152204 | 0.024951537 | 0.114590169 | 0.128929931 |
| ENSMUSG00000043602 | Zfp3 | 193043 | protein_coding | 81.22099126 | 0.128949775 | 0.140263591 | 0.919338894 | 0.35791834 | 0.601015105 | 0.128949775 |
| ENSMUSG00000006435 | Neurl1a | 18011 | protein_coding | 67.26512898 | -0.128955386 | 0.185852838 | -0.693857501 | 0.487771578 | 0.708292116 | 0.128955386 |
| ENSMUSG00000038982 | Bloc1s5 | 17828 | protein_coding | 310.9296963 | 0.128957399 | 0.070622481 | 1.826010588 | 0.067848667 | 0.225514314 | 0.128957399 |
| ENSMUSG00000022354 | Ndufb9 | 66218 | protein_coding | 4487.408291 | 0.128958727 | 0.060761346 | 2.122380987 | 0.033805757 | 0.141484232 | 0.128958727 |
| ENSMUSG00000032850 | Rnft2 | 269695 | protein_coding | 12.76791507 | 0.128959659 | 0.356681916 | 0.36155368 | 0.717685584 | 0.861553924 | 0.128959659 |
| ENSMUSG00000018166 | Erbb3 | 13867 | protein_coding | 12730.86314 | -0.128965694 | 0.06723699 | -1.918076557 | 0.055101305 | 0.197514723 | 0.128965694 |
| ENSMUSG00000026321 | Tnfrsf11a | 21934 | protein_coding | 3390.040844 | 0.128980071 | 0.06648255 | 1.940059023 | 0.052372517 | 0.190915819 | 0.128980071 |
| ENSMUSG00000039814 | Xkr5 | 319581 | protein_coding | 31.70199538 | -0.12902729 | 0.231596304 | -0.557121542 | 0.57744439 | 0.770005431 | 0.12902729 |
| ENSMUSG00000044551 | 9930012K11Rik | 268759 | protein_coding | 37.06636325 | -0.129047999 | 0.222495726 | -0.580002148 | 0.561913169 | 0.759659123 | 0.129047999 |
| ENSMUSG00000021461 | Fancc | 14088 | protein_coding | 221.3597655 | -0.129089777 | 0.083711937 | -1.542071325 | 0.123056262 | 0.325332515 | 0.129089777 |
| ENSMUSG00000020922 | Lsm12 | 268490 | protein_coding | 900.9923238 | 0.129166427 | 0.064362204 | 2.006867674 | 0.044763755 | 0.171677781 | 0.129166427 |
| ENSMUSG00000020107 | Anapc16 | 52717 | protein_coding | 875.7089497 | 0.12918892 | 0.060241962 | 2.14450054 | 0.031992802 | 0.136098106 | 0.12918892 |
| ENSMUSG00000003423 | Pih1d1 | 68845 | protein_coding | 733.4412519 | 0.129221819 | 0.082509634 | 1.566142181 | 0.117315335 | 0.31680248 | 0.129221819 |
| ENSMUSG00000052712 | BC004004 | 80748 | protein_coding | 1167.256409 | -0.129233689 | 0.050664291 | -2.550784529 | 0.010748075 | 0.062318117 | 0.129233689 |

| ensembl_gene_id | external_gene_name | entrezgene_id | gene_biotype | baseMean | log2FoldChange | lfcSE | stat | pvalue | padj | absLog2FC |
|---|---|---|---|---|---|---|---|---|---|---|
| ENSMUSG00000039470 | Zdhhc2 | 70546 | protein_coding | 336.7632239 | 0.129235291 | 0.127003732 | 1.017570815 | 0.308881963 | 0.552499935 | 0.129235291 |
| ENSMUSG00000033313 | Fbxl8 | 50788 | protein_coding | 342.6660433 | 0.129281331 | 0.105022914 | 1.230982139 | 0.218329545 | 0.454615954 | 0.129281331 |
| ENSMUSG00000042302 | Ehbp1 | 216565 | protein_coding | 242.5651666 | -0.12931987 | 0.095505793 | -1.354052631 | 0.175719587 | 0.402132792 | 0.12931987 |
| ENSMUSG00000010914 | Pdhx | 27402 | protein_coding | 645.1570038 | 0.129328258 | 0.079425793 | 1.628290424 | 0.103463319 | 0.293978751 | 0.129328258 |
| ENSMUSG00000034858 | Atosa | 235493 | protein_coding | 366.4880615 | -0.129379737 | 0.095467335 | -1.355225187 | 0.175345822 | 0.401707581 | 0.129379737 |
| ENSMUSG00000020721 | Helz | 78455 | protein_coding | 646.0040743 | -0.129443226 | 0.080093185 | -1.6161578 | 0.1060602 | 0.298192616 | 0.129443226 |
| ENSMUSG00000063015 | Ccni | 12453 | protein_coding | 4734.467649 | -0.129458025 | 0.058222364 | -2.223510257 | 0.026181413 | 0.118482312 | 0.129458025 |
| ENSMUSG00000029547 | Ints1 | 68510 | protein_coding | 1636.110165 | 0.129466685 | 0.108040914 | 1.198311644 | 0.230795715 | 0.46971763 | 0.129466685 |
| ENSMUSG00000026961 | Lrrc26 | 227618 | protein_coding | 935.0352209 | 0.129514333 | 0.096229663 | 1.345887838 | 0.178338692 | 0.405619009 | 0.129514333 |
| ENSMUSG00000035829 | Ppp1r26 | 241289 | protein_coding | 163.7428465 | -0.129520784 | 0.10205594 | -1.269115593 | 0.20439984 | 0.438439283 | 0.129520784 |
| ENSMUSG00000023456 | Tpi1 | 21991 | protein_coding | 11705.22402 | 0.129524741 | 0.05747264 | 2.253676547 | 0.024216524 | 0.112320932 | 0.129524741 |
| ENSMUSG00000025821 | Zfp282 | 101095 | protein_coding | 707.8260456 | -0.129591454 | 0.066032837 | -1.962530455 | 0.049700759 | 0.184385584 | 0.129591454 |
| ENSMUSG00000024754 | Cemip2 | 83921 | protein_coding | 1242.838366 | -0.129599595 | 0.042833688 | -3.025646426 | 0.002481022 | 0.020709859 | 0.129599595 |
| ENSMUSG00000038879 | Nipal2 | 223473 | protein_coding | 772.4247896 | 0.129641159 | 0.05952175 | 2.17804682 | 0.029402551 | 0.128328807 | 0.129641159 |
| ENSMUSG00000033735 | Spr | 20751 | protein_coding | 1551.120645 | 0.129662849 | 0.067284721 | 1.927077162 | 0.053970014 | 0.194579678 | 0.129662849 |
| ENSMUSG00000028932 | Psmc2 | 19181 | protein_coding | 1293.544906 | 0.129697763 | 0.068190625 | 1.901988199 | 0.057172697 | 0.202153934 | 0.129697763 |
| ENSMUSG00000015396 | Cd83 | 12522 | protein_coding | 162.128706 | -0.129701799 | 0.129124293 | -1.004472482 | 0.315150929 | 0.559370353 | 0.129701799 |
| ENSMUSG00000032615 | Nt5m | 103850 | protein_coding | 324.5683149 | 0.129733122 | 0.072465705 | 1.790269233 | 0.07341064 | 0.237154121 | 0.129733122 |
| ENSMUSG00000078348 | Sf3b5 | 66125 | protein_coding | 628.6163021 | 0.1297563 | 0.102186634 | 1.26979718 | 0.204156886 | 0.438306743 | 0.1297563 |
| ENSMUSG00000026750 | Psmb7 | 19177 | protein_coding | 1255.843156 | 0.129775766 | 0.068128187 | 1.904876273 | 0.056796156 | 0.201440865 | 0.129775766 |
| ENSMUSG00000051451 | Crebzf | 233490 | protein_coding | 846.9178384 | -0.129856703 | 0.070008966 | -1.85485817 | 0.063616483 | 0.216033484 | 0.129856703 |
| ENSMUSG00000029649 | Pomp | 66537 | protein_coding | 1243.222614 | 0.12990278 | 0.082510481 | 1.574379148 | 0.115399822 | 0.313731106 | 0.12990278 |
| ENSMUSG00000022210 | Dhrs4 | 28200 | protein_coding | 952.6100448 | 0.129914699 | 0.065169487 | 1.993489673 | 0.046207853 | 0.175422328 | 0.129914699 |
| ENSMUSG00000019428 | Fkbp8 | 14232 | protein_coding | 6501.483681 | 0.129915137 | 0.060449837 | 2.149139593 | 0.031623336 | 0.13495357 | 0.129915137 |
| ENSMUSG00000027981 | Rnpc3 | 67225 | protein_coding | 214.5813551 | -0.129927739 | 0.106020355 | -1.225498052 | 0.220387623 | 0.456836916 | 0.129927739 |
| ENSMUSG00000079562 | Maea | 59003 | protein_coding | 2201.590108 | 0.129931683 | 0.055201234 | 2.353782217 | 0.018583495 | 0.093567289 | 0.129931683 |
| ENSMUSG00000050043 | Tmx2 | 66958 | protein_coding | 811.3876484 | 0.129990124 | 0.080348269 | 1.617833541 | 0.105698476 | 0.29784969 | 0.129990124 |
| ENSMUSG00000031540 | Kat6a | 244349 | protein_coding | 1594.752703 | -0.130026056 | 0.061831415 | -2.102912513 | 0.035473418 | 0.145789961 | 0.130026056 |
| ENSMUSG00000028630 | Dyrk2 | 69181 | protein_coding | 2583.587282 | -0.130047673 | 0.047216775 | -2.754268421 | 0.00588235 | 0.040385127 | 0.130047673 |
| ENSMUSG00000097852 | 4933405D12Rik | 74057 | lncRNA | 9.472888487 | 0.130069561 | 0.391602411 | 0.332146987 | 0.739778271 | 0.875021589 | 0.130069561 |
| ENSMUSG00000032998 | Foxj3 | 230700 | protein_coding | 750.7457832 | -0.130071194 | 0.052951193 | -2.456435571 | 0.014032296 | 0.076460123 | 0.130071194 |
| ENSMUSG00000004535 | Tax1bp1 | 52440 | protein_coding | 10462.33052 | -0.130188279 | 0.068399347 | -1.903355568 | 0.056994165 | 0.201924436 | 0.130188279 |
| ENSMUSG00000007050 | Lsm2 | 27756 | protein_coding | 180.0215583 | 0.130200849 | 0.124366931 | 1.046908919 | 0.295141588 | 0.538150682 | 0.130200849 |
| ENSMUSG00000003955 | Fam162a | 70186 | protein_coding | 2185.663168 | 0.13023726 | 0.081286489 | 1.602200588 | 0.10911126 | 0.302551116 | 0.13023726 |
| ENSMUSG00000051730 | Mettl5 | 75422 | protein_coding | 161.3963281 | -0.13024096 | 0.094360038 | -1.380255487 | 0.167507995 | 0.391491529 | 0.13024096 |
| ENSMUSG00000027637 | Rab5if | 67388 | protein_coding | 663.9622861 | 0.130253469 | 0.094316268 | 1.381028665 | 0.167270147 | 0.391235049 | 0.130253469 |
| ENSMUSG00000027365 | Trpm7 | 58800 | protein_coding | 2227.726553 | -0.130283993 | 0.057159061 | -2.279323532 | 0.02264784 | 0.107171866 | 0.130283993 |
| ENSMUSG00000062373 | Tmem65 | 74868 | protein_coding | 851.857614 | -0.130325321 | 0.079777849 | -1.633602836 | 0.102342272 | 0.291964235 | 0.130325321 |
| ENSMUSG00000041570 | Camsap2 | 67886 | protein_coding | 317.539647 | -0.130363215 | 0.123331824 | -1.05701197 | 0.290506124 | 0.5330518 | 0.130363215 |

| ensembl_gene_id | external_gene_name | entrezgene_id | gene_biotype | baseMean | log2FoldChange | lfcSE | stat | pvalue | padj | absLog2FC |
|---|---|---|---|---|---|---|---|---|---|---|
| ENSMUSG00000027668 | Mfn1 | 67414 | protein_coding | 1612.160017 | -0.130366199 | 0.047536339 | -2.742453477 | 0.006098209 | 0.041496124 | 0.130366199 |
| ENSMUSG00000097101 | 1810034E14Rik | 66503 | lncRNA | 193.5111078 | 0.130381737 | 0.091124082 | 1.430815361 | 0.152483139 | 0.369289021 | 0.130381737 |
| ENSMUSG00000035268 | Pkig | 18769 | protein_coding | 520.8218241 | 0.130388577 | 0.06904094 | 1.888568982 | 0.058949607 | 0.205949343 | 0.130388577 |
| ENSMUSG00000029868 | Trpv6 | 64177 | protein_coding | 249.2281666 | 0.130390552 | 0.39640156 | 0.328935518 | 0.742204427 | 0.876765146 | 0.130390552 |
| ENSMUSG00000024999 | Noc3l | 57753 | protein_coding | 293.1056599 | -0.130397523 | 0.086668976 | -1.504546712 | 0.132440655 | 0.339377153 | 0.130397523 |
| ENSMUSG00000034071 | Zfp551 | 619331 | protein_coding | 69.17147651 | -0.130414642 | 0.167772969 | -0.777328093 | 0.436965225 | 0.67011584 | 0.130414642 |
| ENSMUSG00000023827 | Agpat4 | 68262 | protein_coding | 1509.881289 | -0.130415332 | 0.087334465 | -1.493285986 | 0.135362342 | 0.343658553 | 0.130415332 |
| ENSMUSG00000055302 | Mrfap1 | 67568 | protein_coding | 2932.912582 | 0.130423601 | 0.067638547 | 1.928243692 | 0.053824822 | 0.194372111 | 0.130423601 |
| ENSMUSG00000006024 | Napa | 108124 | protein_coding | 1908.806574 | 0.130446842 | 0.044937726 | 2.902835867 | 0.003698004 | 0.028480281 | 0.130446842 |
| ENSMUSG00000057880 | Abat | 268860 | protein_coding | 7778.235662 | -0.130447697 | 0.079576862 | -1.639266659 | 0.101157736 | 0.290022548 | 0.130447697 |
| ENSMUSG00000022707 | Gbe1 | 74185 | protein_coding | 830.5170488 | -0.130450508 | 0.060403583 | -2.159648505 | 0.03079989 | 0.132491287 | 0.130450508 |
| ENSMUSG00000011658 | Fuz | 70300 | protein_coding | 123.1987709 | -0.130461669 | 0.127795848 | -1.020859996 | 0.307320775 | 0.551334926 | 0.130461669 |
| ENSMUSG00000058897 | Col25a1 | 77018 | protein_coding | 17.13973064 | 0.130491408 | 0.33647668 | 0.387817094 | 0.698151395 | 0.849391002 | 0.130491408 |
| ENSMUSG00000023330 | Dtwd1 | 69185 | protein_coding | 86.02303697 | 0.130497243 | 0.143344367 | 0.910375805 | 0.362624353 | 0.605450144 | 0.130497243 |
| ENSMUSG00000031145 | Prickle3 | 54630 | protein_coding | 193.9573705 | 0.130523722 | 0.098478185 | 1.325407469 | 0.185036041 | 0.414052481 | 0.130523722 |
| ENSMUSG00000027984 | Hadh | 15107 | protein_coding | 7466.255057 | -0.130525989 | 0.035735348 | -3.65257359 | 0.000259625 | 0.003276707 | 0.130525989 |
| ENSMUSG00000035847 | Ids | 15931 | protein_coding | 698.5186624 | 0.130529205 | 0.079142794 | 1.649287298 | 0.099088791 | 0.286329498 | 0.130529205 |
| ENSMUSG00000025024 | Smndc1 | 76479 | protein_coding | 488.0409953 | -0.130550982 | 0.065820766 | -1.98343151 | 0.04731925 | 0.178465808 | 0.130550982 |
| ENSMUSG00000040883 | Tmem205 | 235043 | protein_coding | 636.0595112 | -0.130558615 | 0.085734704 | -1.522821086 | 0.127803473 | 0.332647936 | 0.130558615 |
| ENSMUSG00000027203 | Dut | 110074 | protein_coding | 405.4655298 | 0.130566064 | 0.102453165 | 1.274397561 | 0.202522556 | 0.436583513 | 0.130566064 |
| ENSMUSG00000027684 | Mecom | 14013 | protein_coding | 3354.415181 | -0.130589943 | 0.055499386 | -2.352997998 | 0.018622734 | 0.093677117 | 0.130589943 |
| ENSMUSG00000036779 | Tent4b | 214627 | protein_coding | 777.2206961 | -0.13061483 | 0.068118625 | -1.917461336 | 0.055179348 | 0.19765709 | 0.13061483 |
| ENSMUSG00000008393 | Carhsp1 | 52502 | protein_coding | 2351.470672 | 0.130706306 | 0.055846091 | 2.340473666 | 0.019259297 | 0.095479458 | 0.130706306 |
| ENSMUSG00000062115 | Rai1 | 19377 | protein_coding | 377.5048645 | -0.130717789 | 0.086510296 | -1.511008447 | 0.130786299 | 0.336624075 | 0.130717789 |
| ENSMUSG00000050088 | 1600012H06Rik | 67912 | protein_coding | 772.1829746 | 0.130759811 | 0.070163263 | 1.863650669 | 0.062370777 | 0.213194897 | 0.130759811 |
| ENSMUSG00000028771 | Ptpn12 | 19248 | protein_coding | 522.4202545 | -0.130769145 | 0.059708463 | -2.190127484 | 0.028514992 | 0.125530222 | 0.130769145 |
| ENSMUSG00000031774 | Psme3ip1 | 102122 | protein_coding | 935.465168 | 0.130785143 | 0.045568375 | 2.870085751 | 0.004103605 | 0.030809515 | 0.130785143 |
| ENSMUSG00000071337 | Tia1 | 21841 | protein_coding | 482.9364109 | -0.130787967 | 0.099910402 | -1.309052555 | 0.190516549 | 0.421276292 | 0.130787967 |
| ENSMUSG00000029038 | Ssu72 | 68991 | protein_coding | 1236.706687 | 0.130816986 | 0.052922122 | 2.471877163 | 0.013440569 | 0.074161163 | 0.130816986 |
| ENSMUSG00000010453 | Kansl3 | 226976 | protein_coding | 1641.894593 | -0.130822449 | 0.053485909 | -2.445923629 | 0.01444816 | 0.078032645 | 0.130822449 |
| ENSMUSG00000073043 | Atoh1 | 11921 | protein_coding | 2131.931361 | 0.130837583 | 0.120170513 | 1.088766121 | 0.276257035 | 0.519132744 | 0.130837583 |
| ENSMUSG00000019920 | Lims1 | 110829 | protein_coding | 2293.027644 | 0.130886678 | 0.055922527 | 2.340500071 | 0.019257935 | 0.095479458 | 0.130886678 |
| ENSMUSG00000063063 | Ctnna2 | 12386 | protein_coding | 9.778251622 | 0.131032797 | 0.454963052 | 0.288007556 | 0.773340955 | 0.893440495 | 0.131032797 |
| ENSMUSG00000040482 | Dxo | 112403 | protein_coding | 452.7912365 | 0.131071236 | 0.075746602 | 1.730390962 | 0.083560448 | 0.25741348 | 0.131071236 |
| ENSMUSG00000057421 | Las1l | 76130 | protein_coding | 576.1158947 | 0.131087774 | 0.076987815 | 1.702708079 | 0.088622712 | 0.267243545 | 0.131087774 |
| ENSMUSG00000022129 | Dct | 13190 | protein_coding | 24.19578214 | -0.13111564 | 0.266262137 | -0.49243066 | 0.622414929 | 0.800805946 | 0.13111564 |
| ENSMUSG00000029430 | Ran | 19384 | protein_coding | 2576.913965 | 0.131123659 | 0.086031114 | 1.524142278 | 0.127473173 | 0.332056189 | 0.131123659 |
| ENSMUSG00000068566 | Myadm | 50918 | protein_coding | 1190.382868 | 0.131130278 | 0.0589755 | 2.223470378 | 0.026184099 | 0.118482312 | 0.131130278 |
| ENSMUSG00000038079 | Tmem237 | 381259 | protein_coding | 31.05312264 | 0.131137312 | 0.20727925 | 0.632660108 | 0.526955626 | 0.737165628 | 0.131137312 |

| ensembl_gene_id | external_gene_name | entrezgene_id | gene_biotype | baseMean | log2FoldChange | lfcSE | stat | pvalue | padj | absLog2FC |
|---|---|---|---|---|---|---|---|---|---|---|
| ENSMUSG00000030881 | Arfip2 | 76932 | protein_coding | 1165.565139 | 0.131203863 | 0.04485609 | 2.924995539 | 0.003444612 | 0.026953543 | 0.131203863 |
| ENSMUSG00000039483 | Asb6 | 72323 | protein_coding | 346.0452837 | 0.131238284 | 0.078215583 | 1.677904568 | 0.093365731 | 0.276023673 | 0.131238284 |
| ENSMUSG00000033032 | Afap1l1 | 106877 | protein_coding | 286.3501917 | -0.131241888 | 0.106336186 | -1.234216621 | 0.217122195 | 0.45330399 | 0.131241888 |
| ENSMUSG00000031521 | Aga | 11593 | protein_coding | 538.2062317 | 0.131242612 | 0.065487537 | 2.004085319 | 0.045060922 | 0.17240747 | 0.131242612 |
| ENSMUSG00000030091 | Nup210 | 54563 | protein_coding | 687.3030592 | -0.131267089 | 0.082515762 | -1.590812305 | 0.111651823 | 0.307265793 | 0.131267089 |
| ENSMUSG00000071533 | Pcnp | 76302 | protein_coding | 914.7367635 | 0.131326085 | 0.051465979 | 2.551706711 | 0.010719671 | 0.062252841 | 0.131326085 |
| ENSMUSG00000025878 | Uimc1 | 20184 | protein_coding | 430.3519265 | -0.131356845 | 0.076130391 | -1.725419299 | 0.084451936 | 0.259025555 | 0.131356845 |
| ENSMUSG00000042625 | Safb2 | 224902 | protein_coding | 980.70685 | -0.13141273 | 0.089387731 | -1.470142801 | 0.141523082 | 0.352392829 | 0.13141273 |
| ENSMUSG00000030516 | Tjp1 | 21872 | protein_coding | 2668.334302 | -0.131466988 | 0.03686734 | -3.56594722 | 0.000362544 | 0.004334138 | 0.131466988 |
| ENSMUSG00000001127 | Araf | 11836 | protein_coding | 1686.28399 | 0.131554583 | 0.045217543 | 2.909370441 | 0.003621575 | 0.028052417 | 0.131554583 |
| ENSMUSG00000035297 | Cops4 | 26891 | protein_coding | 1526.03095 | 0.131555717 | 0.049127137 | 2.677862497 | 0.007409363 | 0.047796318 | 0.131555717 |
| ENSMUSG00000026426 | Arl8a | 68724 | protein_coding | 261.417188 | -0.131557175 | 0.082214936 | -1.600161507 | 0.109562759 | 0.303437853 | 0.131557175 |
| ENSMUSG00000102252 | Snrpn | 20646 | protein_coding | 129.9990936 | 0.131631572 | 0.105023797 | 1.253349966 | 0.210078372 | 0.445219276 | 0.131631572 |
| ENSMUSG00000045636 | Mtus1 | 102103 | protein_coding | 5006.587527 | -0.131661922 | 0.043415386 | -3.032609763 | 0.002424489 | 0.020319562 | 0.131661922 |
| ENSMUSG00000018800 | Abca5 | 217265 | protein_coding | 59.54153987 | -0.131672822 | 0.158172633 | -0.832462729 | 0.405147807 | 0.644307787 | 0.131672822 |
| ENSMUSG00000038156 | Spon1 | 233744 | protein_coding | 16.18260672 | -0.131695439 | 0.332818859 | -0.395697045 | 0.692328536 | 0.84565495 | 0.131695439 |
| ENSMUSG00000025534 | Gusb | 110006 | protein_coding | 792.7229559 | 0.131699797 | 0.07285529 | 1.807690231 | 0.070654726 | 0.23126624 | 0.131699797 |
| ENSMUSG00000039578 | Ccser1 | 232035 | protein_coding | 638.0726866 | -0.13174636 | 0.073206764 | -1.799647359 | 0.071916338 | 0.234068382 | 0.13174636 |
| ENSMUSG00000025237 | Parp6 | 67287 | protein_coding | 531.6375644 | -0.131753347 | 0.068680765 | -1.918344211 | 0.05506738 | 0.197440646 | 0.131753347 |
| ENSMUSG00000036202 | Rif1 | 51869 | protein_coding | 1097.728195 | -0.131756877 | 0.071624357 | -1.839554047 | 0.065833736 | 0.221242374 | 0.131756877 |
| ENSMUSG00000057506 | Bloc1s2 | 73689 | protein_coding | 349.7613712 | 0.131786239 | 0.075348719 | 1.749017656 | 0.080287967 | 0.25082096 | 0.131786239 |
| ENSMUSG00000022781 | Pak2 | 224105 | protein_coding | 3526.06607 | -0.131801057 | 0.049601617 | -2.657192731 | 0.007879438 | 0.050113226 | 0.131801057 |
| ENSMUSG00000035242 | Oaz1 | 18245 | protein_coding | 3886.88215 | 0.131848898 | 0.055394413 | 2.380184044 | 0.017303993 | 0.088788259 | 0.131848898 |
| ENSMUSG00000032870 | Smap2 | 69780 | protein_coding | 387.6641104 | 0.131866153 | 0.080926147 | 1.629462877 | 0.103215067 | 0.293509716 | 0.131866153 |
| ENSMUSG00000023832 | Acat2 | 110460 | protein_coding | 758.5090975 | 0.131872501 | 0.112985541 | 1.167162627 | 0.2431447 | 0.484226665 | 0.131872501 |
| ENSMUSG00000070643 | Sox13 | 20668 | protein_coding | 920.8909489 | -0.131965615 | 0.073386499 | -1.79822741 | 0.072140978 | 0.234551147 | 0.131965615 |
| ENSMUSG00000037325 | Bbs7 | 71492 | protein_coding | 127.7653701 | 0.131972978 | 0.124935044 | 1.056332746 | 0.29081622 | 0.5332246 | 0.131972978 |
| ENSMUSG00000031231 | Cox7b | 66142 | protein_coding | 8634.171665 | 0.132039635 | 0.034675834 | 3.807828632 | 0.000140192 | 0.001965985 | 0.132039635 |
| ENSMUSG00000031878 | Nae1 | 234664 | protein_coding | 617.0049755 | -0.132070467 | 0.061962841 | -2.131446275 | 0.033052394 | 0.139211338 | 0.132070467 |
| ENSMUSG00000039456 | Morc3 | 338467 | protein_coding | 968.5688013 | -0.132073467 | 0.066635933 | -1.982015716 | 0.047477482 | 0.178786609 | 0.132073467 |
| ENSMUSG00000041995 | Zbed3 | 72114 | protein_coding | 188.7490943 | 0.132082473 | 0.108780553 | 1.214210353 | 0.224667408 | 0.46267543 | 0.132082473 |
| ENSMUSG00000043061 | Tmem18 | 211986 | protein_coding | 561.0716873 | 0.132116319 | 0.068730915 | 1.922225538 | 0.054577385 | 0.196218366 | 0.132116319 |
| ENSMUSG00000031245 | Hmgn5 | 50887 | protein_coding | 318.6730773 | 0.132127814 | 0.100812276 | 1.310632195 | 0.189982057 | 0.420568377 | 0.132127814 |
| ENSMUSG00000022313 | Utp23 | 78581 | protein_coding | 150.8250635 | 0.132132814 | 0.101624394 | 1.300207641 | 0.193529812 | 0.425385755 | 0.132132814 |
| ENSMUSG00000036315 | Polr1h | 66136 | protein_coding | 358.7030741 | 0.132154812 | 0.072028444 | 1.834758678 | 0.066541454 | 0.222414567 | 0.132154812 |
| ENSMUSG00000042742 | Bmt2 | 101148 | protein_coding | 571.98299 | -0.132156546 | 0.084947066 | -1.555751743 | 0.119767137 | 0.320093959 | 0.132156546 |
| ENSMUSG00000026000 | Lancl1 | 14768 | protein_coding | 505.2891482 | 0.132158897 | 0.065449272 | 2.019256954 | 0.043460519 | 0.168401681 | 0.132158897 |
| ENSMUSG00000035678 | Tnfsf9 | 21950 | protein_coding | 6.354123241 | -0.132159343 | 0.48167619 | -0.274373834 | 0.783797349 | 0.899281435 | 0.132159343 |
| ENSMUSG00000091705 | H2-Q2 | 15013 | protein_coding | 2384.734621 | 0.132176802 | 0.104368241 | 1.266446585 | 0.205353242 | 0.439838414 | 0.132176802 |

Supplementary Table 1

| ensembl_gene_id | external_gene_name | entrezgene_id | gene_biotype | baseMean | log2FoldChange | lfcSE | stat | pvalue | padj | absLog2FC |
|---|---|---|---|---|---|---|---|---|---|---|
| ENSMUSG00000038057 | Dbil5 | 13168 | protein_coding | 5.912537825 | 0.132208102 | 0.470647397 | 0.280906902 | 0.778781806 | 0.896972875 | 0.132208102 |
| ENSMUSG00000003228 | Grk5 | 14773 | protein_coding | 198.8226747 | 0.132214338 | 0.114558574 | 1.154119966 | 0.248450989 | 0.4893628 | 0.132214338 |
| ENSMUSG00000042029 | Ncapg2 | 76044 | protein_coding | 503.2576799 | 0.132219913 | 0.148938147 | 0.887750491 | 0.374674979 | 0.616422248 | 0.132219913 |
| ENSMUSG00000089417 | Gm22009 | 115489854 | scaRNA | 2873.605295 | -0.132223941 | 0.104839766 | -1.261200265 | 0.207236703 | 0.442487265 | 0.132223941 |
| ENSMUSG00000058355 | Abce1 | 24015 | protein_coding | 2093.05627 | 0.13224774 | 0.077192795 | 1.713213527 | 0.086673264 | 0.26363254 | 0.13224774 |
| ENSMUSG00000059897 | Zfp930 | 234358 | protein_coding | 191.6372032 | -0.13227372 | 0.164684803 | -0.803193241 | 0.421863052 | 0.65828686 | 0.13227372 |
| ENSMUSG00000046985 | Tapt1 | 231225 | protein_coding | 913.6570819 | -0.132310433 | 0.055404463 | -2.388082589 | 0.016936535 | 0.087561384 | 0.132310433 |
| ENSMUSG00000038685 | Rtel1 | 269400 | protein_coding | 468.5374585 | -0.132337392 | 0.071028254 | -1.863165493 | 0.062438986 | 0.213379724 | 0.132337392 |
| ENSMUSG00000071281 | Zfp65 | 235907 | protein_coding | 368.3123806 | -0.132417317 | 0.09841166 | -1.345544991 | 0.178449302 | 0.405619009 | 0.132417317 |
| ENSMUSG00000100094 | 1810008I18Rik | 100503969 | lncRNA | 28.04513104 | 0.132466572 | 0.220510418 | 0.600727045 | 0.548021803 | 0.750796946 | 0.132466572 |
| ENSMUSG00000054808 | Actn4 | 60595 | protein_coding | 7579.813363 | 0.132502188 | 0.079616557 | 1.664254191 | 0.096061643 | 0.281107986 | 0.132502188 |
| ENSMUSG00000019173 | Rab5c | 19345 | protein_coding | 3308.640386 | 0.132544837 | 0.079793819 | 1.66109153 | 0.096695065 | 0.282265491 | 0.132544837 |
| ENSMUSG00000021366 | Hivep1 | 110521 | protein_coding | 1359.786874 | -0.132547624 | 0.051972894 | -2.550322157 | 0.010762342 | 0.062357381 | 0.132547624 |
| ENSMUSG00000044641 | Pard6b | 58220 | protein_coding | 859.3025818 | -0.132549268 | 0.088587425 | -1.496253761 | 0.13458754 | 0.342321275 | 0.132549268 |
| ENSMUSG00000024645 | Timm21 | 67105 | protein_coding | 386.297907 | 0.132556455 | 0.06680786 | 1.984144592 | 0.047239723 | 0.178290432 | 0.132556455 |
| ENSMUSG00000026749 | Nek6 | 59126 | protein_coding | 1207.617892 | 0.132596966 | 0.065696077 | 2.018339159 | 0.043555951 | 0.168649897 | 0.132596966 |
| ENSMUSG00000037033 | Clca3b | 229927 | protein_coding | 6686.447948 | 0.13260466 | 0.095654096 | 1.386293585 | 0.165657275 | 0.388872616 | 0.13260466 |
| ENSMUSG00000031875 | Cmtm3 | 68119 | protein_coding | 16.02211818 | 0.132644296 | 0.29437194 | 0.450601017 | 0.652277134 | 0.820984417 | 0.132644296 |
| ENSMUSG00000050668 | Gpatch11 | 53951 | protein_coding | 317.5787387 | 0.132661286 | 0.086453464 | 1.534482009 | 0.124911116 | 0.327978113 | 0.132661286 |
| ENSMUSG00000024948 | Map4k2 | 26412 | protein_coding | 233.0291823 | -0.132712661 | 0.104928549 | -1.264790777 | 0.205946338 | 0.440724071 | 0.132712661 |
| ENSMUSG00000041530 | Ago1 | 236511 | protein_coding | 970.3989038 | -0.132738176 | 0.065930007 | -2.013319592 | 0.044081019 | 0.170111172 | 0.132738176 |
| ENSMUSG00000037108 | Zcwpw1 | 381678 | protein_coding | 21.25821296 | 0.132773943 | 0.288007011 | 0.461009411 | 0.644791853 | 0.81580845 | 0.132773943 |
| ENSMUSG00000050856 | Atp5k | 11958 | protein_coding | 4177.821316 | 0.132778129 | 0.093741779 | 1.416424252 | 0.156651327 | 0.375325502 | 0.132778129 |
| ENSMUSG00000033460 | Armcx1 | 78248 | protein_coding | 73.99092188 | 0.132808402 | 0.15873638 | 0.836660139 | 0.402783631 | 0.642103246 | 0.132808402 |
| ENSMUSG00000047181 | Samd14 | 217125 | protein_coding | 49.27887619 | -0.132835135 | 0.175267759 | -0.757898287 | 0.448511876 | 0.678988684 | 0.132835135 |
| ENSMUSG00000000409 | Lck | 16818 | protein_coding | 92.78218297 | 0.132848696 | 0.140796015 | 0.943554373 | 0.345397417 | 0.587762388 | 0.132848696 |
| ENSMUSG00000068922 | Msto1 | 229524 | protein_coding | 255.467002 | -0.132934813 | 0.094890891 | -1.400922807 | 0.161237158 | 0.381887449 | 0.132934813 |
| ENSMUSG00000037661 | Gpr160 | 71862 | protein_coding | 839.8438383 | 0.132974838 | 0.046173509 | 2.87989455 | 0.003978082 | 0.030176166 | 0.132974838 |
| ENSMUSG00000000823 | Zfp512b | 269401 | protein_coding | 492.2178144 | -0.132999682 | 0.086596579 | -1.535853773 | 0.124574248 | 0.327433832 | 0.132999682 |
| ENSMUSG00000039917 | Rhbdd2 | 215160 | protein_coding | 285.1751711 | -0.133034153 | 0.087014646 | -1.52887082 | 0.126296471 | 0.330057234 | 0.133034153 |
| ENSMUSG00000020105 | Lrig3 | 320398 | protein_coding | 2828.960895 | -0.133071984 | 0.063399364 | -2.098948245 | 0.035821463 | 0.146771529 | 0.133071984 |
| ENSMUSG00000031767 | Nudt7 | 67528 | protein_coding | 528.8006994 | -0.133101828 | 0.063046612 | -2.111165446 | 0.034758094 | 0.143913678 | 0.133101828 |
| ENSMUSG00000031105 | Slc25a14 | 20523 | protein_coding | 87.83579198 | 0.133126902 | 0.139653807 | 0.953263681 | 0.340456488 | 0.583293508 | 0.133126902 |
| ENSMUSG00000018921 | Pelp1 | 75273 | protein_coding | 379.9270729 | 0.133193085 | 0.106296114 | 1.253038147 | 0.210191824 | 0.445259891 | 0.133193085 |
| ENSMUSG00000068739 | Sars | 20226 | protein_coding | 2432.996049 | 0.133201229 | 0.049908498 | 2.668908814 | 0.007609811 | 0.048835423 | 0.133201229 |
| ENSMUSG00000029710 | Ephb4 | 13846 | protein_coding | 2828.491868 | -0.133203883 | 0.065617314 | -2.03001122 | 0.042355399 | 0.165140974 | 0.133203883 |
| ENSMUSG00000040842 | Szrd1 | 213491 | protein_coding | 2785.420522 | 0.133232631 | 0.060216269 | 2.21256869 | 0.026927396 | 0.12038719 | 0.133232631 |
| ENSMUSG00000062248 | Cks2 | 66197 | protein_coding | 570.9503371 | 0.133249272 | 0.070434541 | 1.891817134 | 0.05851535 | 0.205058028 | 0.133249272 |
| ENSMUSG00000029001 | Fbxo44 | 230903 | protein_coding | 56.37347359 | -0.133252717 | 0.176441535 | -0.755223069 | 0.450115149 | 0.679535938 | 0.133252717 |

| ensembl_gene_id | external_gene_name | entrezgene_id | gene_biotype | baseMean | log2FoldChange | lfcSE | stat | pvalue | padj | absLog2FC |
|---|---|---|---|---|---|---|---|---|---|---|
| ENSMUSG00000014164 | Klhl3 | 100503085 | protein_coding | 45.93291612 | 0.133323979 | 0.192609159 | 0.692199581 | 0.488812002 | 0.708993627 | 0.133323979 |
| ENSMUSG00000049303 | Syt12 | 171180 | protein_coding | 19.78605455 | -0.133377107 | 0.28027939 | -0.475871977 | 0.634165588 | 0.808601236 | 0.133377107 |
| ENSMUSG00000028466 | Creb3 | 12913 | protein_coding | 1048.823927 | 0.133381996 | 0.066506678 | 2.005542905 | 0.044905039 | 0.171933419 | 0.133381996 |
| ENSMUSG00000019808 | Adat2 | 66757 | protein_coding | 92.49338635 | 0.133385209 | 0.13330802 | 1.000579029 | 0.317030373 | 0.56085999 | 0.133385209 |
| ENSMUSG00000052675 | Zfp112 | 57745 | protein_coding | 42.37743302 | -0.133388834 | 0.210926112 | -0.632396023 | 0.527128132 | 0.737165747 | 0.133388834 |
| ENSMUSG00000050621 | Rps27rt | 100043813 | protein_coding | 23.4664327 | -0.133401062 | 0.239601953 | -0.556761161 | 0.577690623 | 0.770206405 | 0.133401062 |
| ENSMUSG00000031023 | Akip1 | 57373 | protein_coding | 221.2196189 | 0.13345751 | 0.087086544 | 1.532469941 | 0.12540651 | 0.328615354 | 0.13345751 |
| ENSMUSG00000025413 | Ttc4 | 72354 | protein_coding | 988.2768929 | 0.133458808 | 0.051502856 | 2.591289469 | 0.009561703 | 0.057537902 | 0.133458808 |
| ENSMUSG00000024665 | Fads2 | 56473 | protein_coding | 7086.346934 | -0.133526471 | 0.095223233 | -1.402246767 | 0.160841571 | 0.381342548 | 0.133526471 |
| ENSMUSG00000059518 | Znhit1 | 70103 | protein_coding | 492.7889635 | 0.133572277 | 0.097118274 | 1.375356788 | 0.169020859 | 0.393219762 | 0.133572277 |
| ENSMUSG00000040054 | Baz2a | 116848 | protein_coding | 1983.489472 | -0.133576608 | 0.055351148 | -2.413258129 | 0.015810618 | 0.083135909 | 0.133576608 |
| ENSMUSG00000029198 | Grpel1 | 17713 | protein_coding | 1748.504404 | 0.133577082 | 0.054066936 | 2.470587227 | 0.013489142 | 0.074372441 | 0.133577082 |
| ENSMUSG00000085683 | Tmem238l | 71576 | protein_coding | 1488.20701 | 0.133610826 | 0.057803521 | 2.311465166 | 0.020807175 | 0.10107148 | 0.133610826 |
| ENSMUSG00000025393 | Atp5b | 11947 | protein_coding | 52688.52356 | 0.133616379 | 0.055210783 | 2.420113821 | 0.01551565 | 0.082147368 | 0.133616379 |
| ENSMUSG00000031393 | Mecp2 | 17257 | protein_coding | 625.8325983 | -0.13363515 | 0.061153895 | -2.185227132 | 0.028872197 | 0.126596499 | 0.13363515 |
| ENSMUSG00000034430 | Zxdc | 80292 | protein_coding | 584.0646787 | -0.133708218 | 0.058015544 | -2.304696468 | 0.021183579 | 0.102159742 | 0.133708218 |
| ENSMUSG00000037572 | Wdhd1 | 218973 | protein_coding | 263.9324326 | 0.133719 | 0.126169833 | 1.059833376 | 0.28922041 | 0.531418201 | 0.133719 |
| ENSMUSG00000035529 | Prdm4 | 72843 | protein_coding | 702.6696071 | -0.133810893 | 0.05539478 | -2.415586673 | 0.015709883 | 0.082768143 | 0.133810893 |
| ENSMUSG00000076437 | Selenoh | 72657 | protein_coding | 266.749077 | -0.13390343 | 0.098435912 | -1.360310759 | 0.173731605 | 0.399424529 | 0.13390343 |
| ENSMUSG00000030802 | Bckdk | 12041 | protein_coding | 1343.672213 | 0.133956734 | 0.068105719 | 1.966894043 | 0.049195427 | 0.183396824 | 0.133956734 |
| ENSMUSG00000027455 | Nsfl1c | 386649 | protein_coding | 944.643655 | 0.133961137 | 0.058626533 | 2.284991608 | 0.02231332 | 0.106212193 | 0.133961137 |
| ENSMUSG00000030689 | Ino80e | 233875 | protein_coding | 315.5593759 | 0.133965526 | 0.071987106 | 1.860965571 | 0.062749039 | 0.214124788 | 0.133965526 |
| ENSMUSG00000001436 | Slc19a1 | 20509 | protein_coding | 232.326863 | -0.134016809 | 0.081211293 | -1.650223798 | 0.098897171 | 0.286052251 | 0.134016809 |
| ENSMUSG00000096986 | 4930509E16Rik | 75115 | lncRNA | 19.36663933 | 0.134056048 | 0.271736378 | 0.493331251 | 0.62177855 | 0.800376096 | 0.134056048 |
| ENSMUSG00000058773 | H1f5 | 56702 | protein_coding | 1986.303741 | 0.134084118 | 0.146221669 | 0.916992122 | 0.359146768 | 0.602513256 | 0.134084118 |
| ENSMUSG00000020111 | Micu1 | 216001 | protein_coding | 2726.808557 | -0.134095358 | 0.047284945 | -2.835899645 | 0.00456968 | 0.033346259 | 0.134095358 |
| ENSMUSG00000038593 | Tctn1 | 654470 | protein_coding | 104.8858292 | -0.134111533 | 0.137287195 | -0.976868481 | 0.328634268 | 0.572218186 | 0.134111533 |
| ENSMUSG00000019777 | Hdac2 | 15182 | protein_coding | 951.8837839 | 0.134145409 | 0.055193147 | 2.430472201 | 0.015079163 | 0.080364064 | 0.134145409 |
| ENSMUSG00000039987 | Phtf2 | 68770 | protein_coding | 89.32362116 | -0.134189408 | 0.142640081 | -0.94075527 | 0.346830288 | 0.589019803 | 0.134189408 |
| ENSMUSG00000020692 | Nle1 | 217011 | protein_coding | 145.6031539 | -0.134203303 | 0.101206263 | -1.326037524 | 0.184827268 | 0.413700119 | 0.134203303 |
| ENSMUSG00000030409 | Dmpk | 13400 | protein_coding | 132.319587 | -0.134302219 | 0.126762157 | -1.059481957 | 0.289380342 | 0.53165149 | 0.134302219 |
| ENSMUSG00000022297 | Fzd6 | 14368 | protein_coding | 290.4029036 | -0.134330875 | 0.078086365 | -1.720285924 | 0.085380481 | 0.260866991 | 0.134330875 |
| ENSMUSG00000026159 | Agfg1 | 15463 | protein_coding | 1033.214649 | -0.134338564 | 0.055665201 | -2.413331169 | 0.015807449 | 0.083135909 | 0.134338564 |
| ENSMUSG00000029713 | Gnb2 | 14693 | protein_coding | 5648.897138 | 0.134348176 | 0.064005415 | 2.09901264 | 0.035815786 | 0.146771529 | 0.134348176 |
| ENSMUSG00000021773 | Comtd1 | 69156 | protein_coding | 173.7818657 | 0.134385771 | 0.151360724 | 0.887851008 | 0.374620901 | 0.616422248 | 0.134385771 |
| ENSMUSG00000025583 | Rptor | 74370 | protein_coding | 1097.139412 | 0.134397631 | 0.069577377 | 1.931628304 | 0.0534054 | 0.193369753 | 0.134397631 |
| ENSMUSG00000020720 | Psmd12 | 66997 | protein_coding | 959.6429627 | 0.134399393 | 0.070647679 | 1.902389358 | 0.057120271 | 0.202129099 | 0.134399393 |
| ENSMUSG00000057396 | Zfp759 | 268670 | protein_coding | 152.6232966 | -0.134458913 | 0.122470286 | -1.097890093 | 0.272252484 | 0.514726356 | 0.134458913 |
| ENSMUSG00000022751 | Nit2 | 52633 | protein_coding | 394.0559193 | -0.134490495 | 0.079067133 | -1.700965875 | 0.088949395 | 0.267828024 | 0.134490495 |

| ensembl_gene_id | external_gene_name | entrezgene_id | gene_biotype | baseMean | log2FoldChange | lfcSE | stat | pvalue | padj | absLog2FC |
|---|---|---|---|---|---|---|---|---|---|---|
| ENSMUSG00000090035 | Galnt4 | 14426 | protein_coding | 6371.968825 | 0.134583936 | 0.037191252 | 3.618698671 | 0.000296088 | 0.00363672 | 0.134583936 |
| ENSMUSG00000024735 | Prpf19 | 28000 | protein_coding | 2040.540867 | 0.134725601 | 0.060933513 | 2.21102633 | 0.027034014 | 0.120635733 | 0.134725601 |
| ENSMUSG00000037313 | Tacc3 | 21335 | protein_coding | 470.1418165 | 0.134730051 | 0.101118821 | 1.33239341 | 0.182730946 | 0.410775296 | 0.134730051 |
| ENSMUSG00000057182 | Scn3a | 20269 | protein_coding | 54.34938226 | 0.134739978 | 0.16792403 | 0.802386523 | 0.422329407 | 0.658647373 | 0.134739978 |
| ENSMUSG00000067274 | Rplp0 | 11837 | protein_coding | 35001.84645 | 0.134776491 | 0.046305235 | 2.910610248 | 0.003607237 | 0.027981678 | 0.134776491 |
| ENSMUSG00000051314 | Ffar2 | 233079 | protein_coding | 475.2223315 | 0.134791443 | 0.112438972 | 1.198796474 | 0.230607094 | 0.469603637 | 0.134791443 |
| ENSMUSG00000019775 | Rgs17 | 56533 | protein_coding | 341.6348648 | -0.134813023 | 0.109142165 | -1.235205687 | 0.216753963 | 0.452796783 | 0.134813023 |
| ENSMUSG00000016427 | Ndufa1 | 54405 | protein_coding | 2404.879691 | 0.134838142 | 0.089923101 | 1.499482785 | 0.133748431 | 0.34107542 | 0.134838142 |
| ENSMUSG00000011831 | Evi5 | 14020 | protein_coding | 797.7157955 | -0.134893641 | 0.051359939 | -2.626436925 | 0.008628398 | 0.053478162 | 0.134893641 |
| ENSMUSG00000073616 | Cops9 | 66915 | protein_coding | 1128.410905 | 0.134951879 | 0.086833792 | 1.554140115 | 0.120150998 | 0.320656918 | 0.134951879 |
| ENSMUSG00000042359 | Osbpl6 | 99031 | protein_coding | 408.2485517 | -0.134963207 | 0.091140082 | -1.480832628 | 0.138651179 | 0.348725448 | 0.134963207 |
| ENSMUSG00000036430 | Tbcc | 72726 | protein_coding | 293.7252034 | 0.134969917 | 0.076442679 | 1.765635626 | 0.077456999 | 0.245557111 | 0.134969917 |
| ENSMUSG00000038002 | Cramp1 | 57354 | protein_coding | 831.1960318 | -0.135008322 | 0.079457442 | -1.699127468 | 0.089295169 | 0.268368093 | 0.135008322 |
| ENSMUSG00000000901 | Mmp11 | 17385 | protein_coding | 10.29507899 | 0.135010235 | 0.398690936 | 0.33863382 | 0.734885602 | 0.872524487 | 0.135010235 |
| ENSMUSG00000035559 | Mpv17l2 | 234384 | protein_coding | 656.6886245 | 0.135098276 | 0.090095791 | 1.499495976 | 0.133745012 | 0.34107542 | 0.135098276 |
| ENSMUSG00000038644 | Pold1 | 18971 | protein_coding | 560.0144285 | 0.135108847 | 0.139158279 | 0.970900535 | 0.331597812 | 0.574895496 | 0.135108847 |
| ENSMUSG00000049076 | Acap2 | 78618 | protein_coding | 2355.871891 | -0.135109074 | 0.068282402 | -1.978680749 | 0.047851962 | 0.179634215 | 0.135109074 |
| ENSMUSG00000078584 | AU022252 | 230696 | protein_coding | 394.2113416 | 0.135127074 | 0.064742774 | 2.087137537 | 0.036875704 | 0.149582964 | 0.135127074 |
| ENSMUSG00000075327 | Zbtb2 | 381990 | protein_coding | 377.427149 | -0.135262629 | 0.072961341 | -1.853894504 | 0.063754254 | 0.21645576 | 0.135262629 |
| ENSMUSG00000023106 | Denr | 68184 | protein_coding | 669.9715771 | 0.135309279 | 0.062861435 | 2.152500643 | 0.031357945 | 0.134033812 | 0.135309279 |
| ENSMUSG00000014353 | Tmem87b | 72477 | protein_coding | 2384.719166 | -0.135358864 | 0.0334547 | -4.04603437 | 5.21E-05 | 0.000847727 | 0.135358864 |
| ENSMUSG00000063235 | Ptpmt1 | 66461 | protein_coding | 647.2331054 | -0.135383309 | 0.065692879 | -2.060852119 | 0.03931715 | 0.156629283 | 0.135383309 |
| ENSMUSG00000037278 | Tmem97 | 69071 | protein_coding | 1158.442451 | 0.135388393 | 0.076508206 | 1.769593107 | 0.076794948 | 0.24422641 | 0.135388393 |
| ENSMUSG00000030032 | Wdr54 | 75659 | protein_coding | 25.93107467 | 0.135391088 | 0.270023799 | 0.501404277 | 0.61608663 | 0.795599668 | 0.135391088 |
| ENSMUSG00000025041 | Nt5c2 | 76952 | protein_coding | 424.9369649 | -0.13542341 | 0.066403174 | -2.039411701 | 0.041408956 | 0.162469023 | 0.13542341 |
| ENSMUSG00000053870 | Fpgt | 75540 | protein_coding | 1115.098432 | -0.135448211 | 0.065099812 | -2.080623678 | 0.037468365 | 0.151214294 | 0.135448211 |
| ENSMUSG00000023873 | 1700010I14Rik | 66931 | protein_coding | 73.48444349 | 0.13550312 | 0.142052371 | 0.953895519 | 0.340136532 | 0.583162735 | 0.13550312 |
| ENSMUSG00000056268 | Dennd1b | 329260 | protein_coding | 2053.453525 | -0.135603142 | 0.050487804 | -2.685859377 | 0.007234353 | 0.046962624 | 0.135603142 |
| ENSMUSG00000033423 | Eri3 | 140546 | protein_coding | 672.1543728 | 0.135605914 | 0.068411401 | 1.982212215 | 0.047455494 | 0.178786609 | 0.135605914 |
| ENSMUSG00000035960 | Apex1 | 11792 | protein_coding | 245.8976549 | 0.135627635 | 0.084186944 | 1.611029324 | 0.107173331 | 0.299598598 | 0.135627635 |
| ENSMUSG00000070705 | Eid2b | 434156 | protein_coding | 54.82947964 | 0.135705628 | 0.161176785 | 0.84196758 | 0.399806101 | 0.63926958 | 0.135705628 |
| ENSMUSG00000030286 | Emc3 | 66087 | protein_coding | 1523.50969 | 0.135723554 | 0.044063059 | 3.080211834 | 0.002068534 | 0.018022285 | 0.135723554 |
| ENSMUSG00000038187 | Btbd10 | 68815 | protein_coding | 329.134894 | -0.135763039 | 0.104019926 | -1.305163771 | 0.191837088 | 0.423218928 | 0.135763039 |
| ENSMUSG00000026895 | Ndufa8 | 68375 | protein_coding | 2377.155329 | 0.135764925 | 0.067136509 | 2.02222201 | 0.043153421 | 0.167533757 | 0.135764925 |
| ENSMUSG00000025509 | Pnpla2 | 66853 | protein_coding | 1651.354126 | -0.135825299 | 0.072935065 | -1.862277066 | 0.062564046 | 0.213633392 | 0.135825299 |
| ENSMUSG00000023020 | Cox14 | 66379 | protein_coding | 1573.462124 | 0.135827818 | 0.057819599 | 2.349165709 | 0.018815531 | 0.094323304 | 0.135827818 |
| ENSMUSG00000052033 | Pfdn4 | 109054 | protein_coding | 158.8511662 | 0.13583288 | 0.115329405 | 1.177781852 | 0.238883585 | 0.479144737 | 0.13583288 |
| ENSMUSG00000031156 | Slc35a2 | 22232 | protein_coding | 2288.019553 | 0.135858217 | 0.082030546 | 1.656190574 | 0.097683231 | 0.283895742 | 0.135858217 |
| ENSMUSG00000034300 | Fam53c | 66306 | protein_coding | 418.4930285 | 0.135911365 | 0.074945503 | 1.813469253 | 0.069759477 | 0.229360057 | 0.135911365 |

Supplementary Table 1

| ensembl_gene_id | external_gene_name | entrezgene_id | gene_biotype | baseMean | log2FoldChange | lfcSE | stat | pvalue | padj | absLog2FC |
|---|---|---|---|---|---|---|---|---|---|---|
| ENSMUSG00000037225 | Fgf2 | 14173 | protein_coding | 8.442280739 | -0.135941553 | 0.492097068 | -0.276249469 | 0.782356464 | 0.898825528 | 0.135941553 |
| ENSMUSG00000016619 | Nup50 | 18141 | protein_coding | 1443.796059 | 0.135956281 | 0.051467435 | 2.641598153 | 0.008251589 | 0.051819853 | 0.135956281 |
| ENSMUSG00000029647 | Pan3 | 72587 | protein_coding | 1763.591311 | -0.13597467 | 0.038637438 | -3.519246536 | 0.000432774 | 0.00500312 | 0.13597467 |
| ENSMUSG00000026860 | Sh3glb2 | 227700 | protein_coding | 1423.870331 | -0.136031542 | 0.058014026 | -2.344804356 | 0.019037066 | 0.094932209 | 0.136031542 |
| ENSMUSG00000029056 | Pank4 | 269614 | protein_coding | 451.1873456 | -0.136064688 | 0.078738098 | -1.728066737 | 0.08397626 | 0.258254223 | 0.136064688 |
| ENSMUSG00000024350 | Dnajc18 | 76594 | protein_coding | 144.6979786 | 0.136103753 | 0.115106063 | 1.182420369 | 0.237038945 | 0.477153008 | 0.136103753 |
| ENSMUSG00000045237 | Eola1 | 68618 | protein_coding | 335.5444103 | 0.136107526 | 0.085166756 | 1.59812975 | 0.110014104 | 0.304322033 | 0.136107526 |
| ENSMUSG00000017466 | Timp2 | 21858 | protein_coding | 119.9414448 | -0.136119638 | 0.152361361 | -0.893399985 | 0.371643011 | 0.613533062 | 0.136119638 |
| ENSMUSG00000029715 | Pop7 | 74097 | protein_coding | 338.5277811 | 0.136124441 | 0.082766244 | 1.644685484 | 0.100034688 | 0.287876412 | 0.136124441 |
| ENSMUSG00000056313 | Tcim | 69068 | protein_coding | 3331.531544 | 0.136161642 | 0.056365387 | 2.415696048 | 0.015705165 | 0.082768143 | 0.136161642 |
| ENSMUSG00000004947 | Dtx2 | 74198 | protein_coding | 189.9769822 | 0.136235038 | 0.107460377 | 1.267769969 | 0.20488011 | 0.43919999 | 0.136235038 |
| ENSMUSG00000055401 | Fbxo6 | 50762 | protein_coding | 512.7549602 | 0.136280815 | 0.088483928 | 1.540175927 | 0.123517476 | 0.326177557 | 0.136280815 |
| ENSMUSG00000047459 | Dynlrb1 | 67068 | protein_coding | 1154.430907 | 0.136295578 | 0.072936081 | 1.868698944 | 0.061664711 | 0.211904282 | 0.136295578 |
| ENSMUSG00000026755 | Arpc5l | 74192 | protein_coding | 775.0606387 | 0.136362599 | 0.061260752 | 2.225937404 | 0.026018375 | 0.117964106 | 0.136362599 |
| ENSMUSG00000069792 | Wfdc17 | 100034251 | protein_coding | 40.20989013 | 0.136388098 | 0.222488287 | 0.613012485 | 0.539868084 | 0.745625688 | 0.136388098 |
| ENSMUSG00000028551 | Cdkn2c | 12580 | protein_coding | 260.6591638 | 0.1363937 | 0.154016399 | 0.885579077 | 0.37584439 | 0.617324668 | 0.1363937 |
| ENSMUSG00000038302 | Afg1l | 215951 | protein_coding | 702.4319803 | 0.136402108 | 0.064150384 | 2.126286685 | 0.033479396 | 0.140531551 | 0.136402108 |
| ENSMUSG00000025487 | Psmd13 | 23997 | protein_coding | 1482.895693 | 0.136402357 | 0.07166442 | 1.903348368 | 0.056995103 | 0.201924436 | 0.136402357 |
| ENSMUSG00000036835 | Psenen | 66340 | protein_coding | 540.3615073 | 0.136421939 | 0.068973163 | 1.977898847 | 0.047940119 | 0.179737215 | 0.136421939 |
| ENSMUSG00000052698 | Tln2 | 70549 | protein_coding | 1438.323348 | 0.136491225 | 0.073667272 | 1.852806832 | 0.063910051 | 0.216756548 | 0.136491225 |
| ENSMUSG00000090110 | Cmc4 | 105886298 | protein_coding | 37.760376 | -0.136496686 | 0.219514445 | -0.621811864 | 0.534065584 | 0.741606925 | 0.136496686 |
| ENSMUSG00000089281 | Scarna6 | 100217415 | scaRNA | 831.7474847 | 0.136504661 | 0.118685026 | 1.150142238 | 0.250085292 | 0.491125853 | 0.136504661 |
| ENSMUSG00000021737 | Psmd6 | 66413 | protein_coding | 1168.072065 | 0.136546127 | 0.053123335 | 2.570360613 | 0.01015927 | 0.059943581 | 0.136546127 |
| ENSMUSG00000109336 | Samd4b | 233033 | protein_coding | 631.2284221 | 0.13654857 | 0.081006173 | 1.685656336 | 0.091862012 | 0.27324802 | 0.13654857 |
| ENSMUSG00000005378 | Bud23 | 66138 | protein_coding | 622.3666388 | 0.136559977 | 0.108381586 | 1.259992418 | 0.207672097 | 0.442947747 | 0.136559977 |
| ENSMUSG00000037351 | Actr1b | 226977 | protein_coding | 3764.3705 | 0.136587522 | 0.045204599 | 3.021540391 | 0.002514921 | 0.020906252 | 0.136587522 |
| ENSMUSG00000004637 | Wwox | 80707 | protein_coding | 225.0149923 | 0.136709534 | 0.097201049 | 1.406461511 | 0.159587126 | 0.379596103 | 0.136709534 |
| ENSMUSG00000050965 | Prkca | 18750 | protein_coding | 2968.346327 | -0.13672804 | 0.045132943 | -3.029451029 | 0.002449986 | 0.02049322 | 0.13672804 |
| ENSMUSG00000058709 | Egln2 | 112406 | protein_coding | 1789.299825 | 0.13676978 | 0.076362574 | 1.791057749 | 0.073284028 | 0.23698647 | 0.13676978 |
| ENSMUSG00000029627 | Zkscan14 | 67235 | protein_coding | 259.3398744 | -0.13684261 | 0.120851837 | -1.132317171 | 0.257501114 | 0.499365134 | 0.13684261 |
| ENSMUSG00000028180 | Zranb2 | 53861 | protein_coding | 1082.267197 | 0.13689921 | 0.074036737 | 1.84907136 | 0.064447509 | 0.218015518 | 0.13689921 |
| ENSMUSG00000030742 | Lat | 16797 | protein_coding | 10.70233218 | 0.136953732 | 0.354301525 | 0.386545701 | 0.699092563 | 0.849861734 | 0.136953732 |
| ENSMUSG00000022637 | Cblb | 208650 | protein_coding | 482.4962163 | -0.137004716 | 0.080952935 | -1.692399624 | 0.090569804 | 0.270928354 | 0.137004716 |
| ENSMUSG00000084883 | Ccdc85c | 668158 | protein_coding | 2295.103824 | 0.137013414 | 0.096908344 | 1.41384538 | 0.157407306 | 0.376633179 | 0.137013414 |
| ENSMUSG00000056899 | Immp2l | 93757 | protein_coding | 138.5176867 | -0.1370272 | 0.113699847 | -1.205166088 | 0.228139191 | 0.46714214 | 0.1370272 |
| ENSMUSG00000031987 | Egln1 | 112405 | protein_coding | 996.63819 | -0.137030049 | 0.050643218 | -2.705792684 | 0.006814157 | 0.044853842 | 0.137030049 |
| ENSMUSG00000034292 | Traf3ip1 | 74019 | protein_coding | 166.3273183 | 0.137055641 | 0.102816609 | 1.333010705 | 0.182528291 | 0.410491389 | 0.137055641 |
| ENSMUSG00000063268 | Parp10 | 671535 | protein_coding | 853.4606611 | -0.137065055 | 0.077541619 | -1.767632107 | 0.077122427 | 0.244881547 | 0.137065055 |
| ENSMUSG00000031963 | Bmper | 73230 | protein_coding | 47.52855414 | 0.13708823 | 0.321276106 | 0.426699115 | 0.669598495 | 0.831366151 | 0.13708823 |

| ensembl_gene_id | external_gene_name | entrezgene_id | gene_biotype | baseMean | log2FoldChange | lfcSE | stat | pvalue | padj | absLog2FC |
|---|---|---|---|---|---|---|---|---|---|---|
| ENSMUSG00000001025 | S100a6 | 20200 | protein_coding | 16578.6964 | 0.137185227 | 0.099573394 | 1.377729753 | 0.168286742 | 0.392283294 | 0.137185227 |
| ENSMUSG00000022403 | St13 | 70356 | protein_coding | 2702.060803 | 0.137224931 | 0.075211322 | 1.824524931 | 0.068072751 | 0.225933166 | 0.137224931 |
| ENSMUSG00000032766 | Gng11 | 66066 | protein_coding | 408.9458272 | 0.13723271 | 0.068732353 | 1.996624628 | 0.045865975 | 0.174452968 | 0.13723271 |
| ENSMUSG00000030138 | Bms1 | 213895 | protein_coding | 866.874899 | 0.137258389 | 0.049800549 | 2.756162169 | 0.005848399 | 0.040237686 | 0.137258389 |
| ENSMUSG00000026094 | Stk17b | 98267 | protein_coding | 1455.500826 | -0.137263869 | 0.12005556 | -1.143336211 | 0.252899026 | 0.494004675 | 0.137263869 |
| ENSMUSG00000054091 | 1810037I17Rik | 67704 | protein_coding | 1676.490562 | 0.137279326 | 0.054530856 | 2.517461437 | 0.011820391 | 0.067240721 | 0.137279326 |
| ENSMUSG00000072945 | Ripply1 | 622473 | protein_coding | 13.72381342 | -0.137309818 | 0.430217121 | -0.319164002 | 0.749602153 | 0.880475742 | 0.137309818 |
| ENSMUSG00000030869 | Ndufab1 | 70316 | protein_coding | 1583.34086 | 0.137344508 | 0.049301299 | 2.78581927 | 0.005339263 | 0.037453804 | 0.137344508 |
| ENSMUSG00000033075 | Senp1 | 223870 | protein_coding | 603.0704089 | -0.137345046 | 0.054267802 | -2.530875431 | 0.011377825 | 0.065183013 | 0.137345046 |
| ENSMUSG00000040364 | Sec1 | 56546 | protein_coding | 6.41466398 | 0.137347363 | 0.543082037 | 0.252903527 | 0.800342764 | 0.909016674 | 0.137347363 |
| ENSMUSG00000085208 | Brip1os | 74038 | lncRNA | 1455.287417 | 0.13737204 | 0.06247701 | 2.198761419 | 0.027894891 | 0.123622124 | 0.13737204 |
| ENSMUSG00000021967 | Mrpl57 | 67840 | protein_coding | 795.279037 | 0.137379168 | 0.08108068 | 1.694351457 | 0.090198518 | 0.270226918 | 0.137379168 |
| ENSMUSG00000023018 | Smarcd1 | 83797 | protein_coding | 269.9179863 | -0.137400369 | 0.086281855 | -1.592459603 | 0.111281473 | 0.306582205 | 0.137400369 |
| ENSMUSG00000029635 | Cdk8 | 264064 | protein_coding | 851.4461588 | -0.137413839 | 0.061919533 | -2.219232464 | 0.026470914 | 0.118917792 | 0.137413839 |
| ENSMUSG00000035093 | Secisbp2l | 70354 | protein_coding | 2272.158458 | 0.137469377 | 0.061873177 | 2.221792762 | 0.026297314 | 0.118778578 | 0.137469377 |
| ENSMUSG00000033985 | Tesk2 | 230661 | protein_coding | 639.2282251 | -0.137510805 | 0.061917628 | -2.220866802 | 0.026359984 | 0.118795321 | 0.137510805 |
| ENSMUSG00000058881 | Zfp516 | 329003 | protein_coding | 418.9866801 | 0.137511401 | 0.084961845 | 1.618507705 | 0.105553228 | 0.297761537 | 0.137511401 |
| ENSMUSG00000036151 | Tm6sf2 | 107770 | protein_coding | 244.5802478 | -0.137511864 | 0.142146356 | -0.967396335 | 0.333345941 | 0.576562634 | 0.137511864 |
| ENSMUSG00000004843 | Chmp2b | 68942 | protein_coding | 3268.56702 | 0.137514916 | 0.05569136 | 2.469232503 | 0.013540321 | 0.074501794 | 0.137514916 |
| ENSMUSG00000018678 | Sp2 | 78912 | protein_coding | 127.8007683 | 0.137577236 | 0.129083918 | 1.065796869 | 0.286515477 | 0.529021743 | 0.137577236 |
| ENSMUSG00000052595 | A1cf | 69865 | protein_coding | 576.7110155 | 0.137602643 | 0.075258591 | 1.828397815 | 0.067489869 | 0.224692221 | 0.137602643 |
| ENSMUSG00000036863 | Syde2 | 214804 | protein_coding | 447.3883065 | -0.137656631 | 0.065999801 | -2.085712811 | 0.037004645 | 0.149943194 | 0.137656631 |
| ENSMUSG00000061882 | Ccdc62 | 208908 | protein_coding | 34.5022473 | -0.137657539 | 0.293458036 | -0.469087646 | 0.639006989 | 0.812013688 | 0.137657539 |
| ENSMUSG00000118047 | 4930554I06Rik | 75354 | lncRNA | 16.42842386 | -0.137686739 | 0.320945713 | -0.429003205 | 0.667920893 | 0.830562128 | 0.137686739 |
| ENSMUSG00000033701 | Acbd6 | 72482 | protein_coding | 449.4503472 | -0.137718198 | 0.102452338 | -1.344217228 | 0.178878152 | 0.405982819 | 0.137718198 |
| ENSMUSG00000118346 | Tmem179b | 67706 | protein_coding | 399.1065962 | -0.137735198 | 0.070463861 | -1.954692739 | 0.050619348 | 0.186591593 | 0.137735198 |
| ENSMUSG00000041747 | Utp15 | 105372 | protein_coding | 315.5841883 | -0.137740017 | 0.08419875 | -1.635891469 | 0.101862304 | 0.291161534 | 0.137740017 |
| ENSMUSG00000025348 | Itga7 | 16404 | protein_coding | 548.7765317 | -0.137741072 | 0.087813206 | -1.568568989 | 0.116748403 | 0.316003891 | 0.137741072 |
| ENSMUSG00000014503 | Pkd2l2 | 53871 | protein_coding | 159.9761448 | -0.137758682 | 0.11987318 | -1.149203529 | 0.250472066 | 0.491535798 | 0.137758682 |
| ENSMUSG00000061024 | Rrs1 | 59014 | protein_coding | 345.1671781 | -0.137853377 | 0.095231711 | -1.44755749 | 0.147740847 | 0.361605197 | 0.137853377 |
| ENSMUSG00000098544 | Mir7211 | 102466817 | miRNA | 7.26265988 | -0.137857978 | 0.48836456 | -0.282284975 | 0.777725006 | 0.89645995 | 0.137857978 |
| ENSMUSG00000087598 | Zfp111 | 56707 | protein_coding | 258.0933699 | -0.137870393 | 0.082124788 | -1.678791458 | 0.093192694 | 0.275714286 | 0.137870393 |
| ENSMUSG00000030960 | Eef1akmt2 | 72096 | protein_coding | 227.3007669 | 0.137879525 | 0.102232294 | 1.348688549 | 0.177437025 | 0.404570466 | 0.137879525 |
| ENSMUSG00000003190 | Bcl2l12 | 75736 | protein_coding | 142.8056583 | 0.137888709 | 0.101706913 | 1.355745691 | 0.175180096 | 0.401524931 | 0.137888709 |
| ENSMUSG00000031310 | Zmym3 | 56364 | protein_coding | 518.3210291 | -0.137900301 | 0.060309521 | -2.28654281 | 0.022222523 | 0.105936335 | 0.137900301 |
| ENSMUSG00000040022 | Rab11fip2 | 74998 | protein_coding | 175.803639 | -0.137911079 | 0.114781747 | -1.201507049 | 0.229554574 | 0.468848355 | 0.137911079 |
| ENSMUSG00000025495 | Ptdss2 | 27388 | protein_coding | 598.6058565 | -0.137934569 | 0.058676929 | -2.350746201 | 0.018735808 | 0.094069857 | 0.137934569 |
| ENSMUSG00000020256 | Aldh1l2 | 216188 | protein_coding | 17.3200994 | -0.137945037 | 0.353992577 | -0.389683417 | 0.696770661 | 0.84851019 | 0.137945037 |
| ENSMUSG00000025195 | Dnmbp | 71972 | protein_coding | 4191.50958 | -0.137982156 | 0.058746875 | -2.34875739 | 0.018836175 | 0.094397452 | 0.137982156 |

| ensembl_gene_id | external_gene_name | entrezgene_id | gene_biotype | baseMean | log2FoldChange | lfcSE | stat | pvalue | padj | absLog2FC |
|---|---|---|---|---|---|---|---|---|---|---|
| ENSMUSG00000037706 | Cd81 | 12520 | protein_coding | 1580.495436 | -0.138012796 | 0.072749544 | -1.897094994 | 0.057815402 | 0.203800871 | 0.138012796 |
| ENSMUSG00000025209 | Twnk | 226153 | protein_coding | 410.8543516 | -0.138049407 | 0.083073843 | -1.661767425 | 0.096559416 | 0.282085054 | 0.138049407 |
| ENSMUSG00000040236 | Trappc5 | 66682 | protein_coding | 562.2081707 | 0.138068012 | 0.05543922 | 2.490439289 | 0.012758529 | 0.071196124 | 0.138068012 |
| ENSMUSG00000046556 | Zfp319 | 79233 | protein_coding | 392.2343613 | 0.138164549 | 0.096715951 | 1.428560109 | 0.153130707 | 0.370189171 | 0.138164549 |
| ENSMUSG00000031253 | Srpx2 | 68792 | protein_coding | 21.77828095 | 0.138180386 | 0.265593276 | 0.52027065 | 0.602874949 | 0.787655348 | 0.138180386 |
| ENSMUSG00000029521 | Chek2 | 50883 | protein_coding | 336.2782432 | 0.138206944 | 0.100638775 | 1.373297163 | 0.169659984 | 0.39391111 | 0.138206944 |
| ENSMUSG00000032091 | Tmprss4 | 214523 | protein_coding | 3886.87849 | 0.138218593 | 0.059529116 | 2.321865363 | 0.020240186 | 0.099093347 | 0.138218593 |
| ENSMUSG00000048007 | Timm8a1 | 30058 | protein_coding | 95.48195854 | 0.138237678 | 0.137151594 | 1.00791886 | 0.313493417 | 0.55760602 | 0.138237678 |
| ENSMUSG00000040123 | Zmym5 | 219105 | protein_coding | 682.3068614 | -0.138249484 | 0.094951299 | -1.456004133 | 0.145391477 | 0.358198217 | 0.138249484 |
| ENSMUSG00000033213 | AA467197 | 433470 | protein_coding | 3008.841912 | -0.138310364 | 0.077474122 | -1.785245965 | 0.074221429 | 0.238963663 | 0.138310364 |
| ENSMUSG00000032846 | Zswim6 | 67263 | protein_coding | 556.9697921 | -0.138390133 | 0.071041426 | -1.9480202 | 0.051412546 | 0.188695979 | 0.138390133 |
| ENSMUSG00000018974 | Sart3 | 53890 | protein_coding | 811.6565708 | 0.138403157 | 0.061715923 | 2.242584253 | 0.024923637 | 0.114513814 | 0.138403157 |
| ENSMUSG00000030089 | Slc41a3 | 71699 | protein_coding | 64.16795089 | -0.138443895 | 0.168816778 | -0.820083739 | 0.412168371 | 0.649970343 | 0.138443895 |
| ENSMUSG00000007458 | M6pr | 17113 | protein_coding | 4999.96995 | 0.13844694 | 0.038514935 | 3.594629941 | 0.000324853 | 0.003936066 | 0.13844694 |
| ENSMUSG00000049892 | Rasd1 | 19416 | protein_coding | 1024.450681 | 0.138448433 | 0.087442701 | 1.583304618 | 0.113352051 | 0.310098138 | 0.138448433 |
| ENSMUSG00000035967 | Ints6l | 236790 | protein_coding | 457.3416631 | -0.138499313 | 0.107418031 | -1.28934884 | 0.197276839 | 0.429755988 | 0.138499313 |
| ENSMUSG00000022674 | Ube2v2 | 70620 | protein_coding | 198.5481141 | -0.138520089 | 0.090760866 | -1.52620942 | 0.126957718 | 0.331141375 | 0.138520089 |
| ENSMUSG00000051076 | Vtcn1 | 242122 | protein_coding | 26.5781182 | -0.13852542 | 0.263332964 | -0.526046639 | 0.598855803 | 0.784827956 | 0.13852542 |
| ENSMUSG00000032336 | Nptn | 20320 | protein_coding | 1944.576422 | -0.138533877 | 0.040906644 | -3.386586256 | 0.00070768 | 0.007608506 | 0.138533877 |
| ENSMUSG00000024193 | Phf1 | 21652 | protein_coding | 279.0943389 | -0.138534165 | 0.116756736 | -1.186519686 | 0.235417132 | 0.475463004 | 0.138534165 |
| ENSMUSG00000037013 | Ss18 | 268996 | protein_coding | 1364.06111 | 0.138548366 | 0.044250301 | 3.131015214 | 0.001742031 | 0.015721175 | 0.138548366 |
| ENSMUSG00000032816 | Igdcc4 | 56741 | protein_coding | 1198.186125 | -0.138550683 | 0.062929275 | -2.201688854 | 0.027687295 | 0.122794781 | 0.138550683 |
| ENSMUSG00000025083 | Afap1l2 | 226250 | protein_coding | 78.27387799 | -0.138565677 | 0.14640945 | -0.946425775 | 0.343931464 | 0.586381512 | 0.138565677 |
| ENSMUSG00000032402 | Smad3 | 17127 | protein_coding | 1271.070222 | 0.138581402 | 0.058667025 | 2.36216857 | 0.018168379 | 0.092023067 | 0.138581402 |
| ENSMUSG00000006494 | Pdk1 | 228026 | protein_coding | 1849.08796 | -0.13858595 | 0.061436579 | -2.255756297 | 0.024085898 | 0.111811535 | 0.13858595 |
| ENSMUSG00000024914 | Drap1 | 66556 | protein_coding | 1553.333453 | 0.138592182 | 0.075194516 | 1.843115546 | 0.065312142 | 0.220031108 | 0.138592182 |
| ENSMUSG00000037679 | Inf2 | 70435 | protein_coding | 190.0725202 | -0.138601493 | 0.12423429 | -1.115646038 | 0.264573677 | 0.5061553 | 0.138601493 |
| ENSMUSG00000021792 | Prxl2a | 70564 | protein_coding | 210.9749432 | 0.138607338 | 0.117181062 | 1.182847597 | 0.236869553 | 0.477141554 | 0.138607338 |
| ENSMUSG00000054770 | Kctd18 | 51960 | protein_coding | 168.1523911 | -0.138612501 | 0.095770428 | -1.447341357 | 0.147801341 | 0.361643489 | 0.138612501 |
| ENSMUSG00000030515 | Tarsl2 | 272396 | protein_coding | 301.6366211 | -0.138685392 | 0.088799273 | -1.561785219 | 0.118338595 | 0.31807442 | 0.138685392 |
| ENSMUSG00000062619 | 2310039H08Rik | 67101 | protein_coding | 467.5891951 | 0.138693938 | 0.079156105 | 1.75215718 | 0.079746784 | 0.249745186 | 0.138693938 |
| ENSMUSG00000032096 | Arcn1 | 213827 | protein_coding | 5406.630497 | 0.138700807 | 0.048399597 | 2.865742977 | 0.004160319 | 0.031090574 | 0.138700807 |
| ENSMUSG00000040751 | Lat2 | 56743 | protein_coding | 9.024409133 | -0.138778854 | 0.395519588 | -0.350877322 | 0.725680385 | 0.865994345 | 0.138778854 |
| ENSMUSG00000026575 | Nme7 | 171567 | protein_coding | 198.6664514 | 0.138791304 | 0.103705548 | 1.338320921 | 0.180791845 | 0.407666398 | 0.138791304 |
| ENSMUSG00000038751 | Ptk6 | 20459 | protein_coding | 408.1966871 | 0.138837545 | 0.126312807 | 1.099156519 | 0.271699801 | 0.514223998 | 0.138837545 |
| ENSMUSG00000033902 | Mapkbp1 | 26390 | protein_coding | 248.7271009 | -0.138891685 | 0.0956373 | -1.452275257 | 0.146425085 | 0.359475923 | 0.138891685 |
| ENSMUSG00000039634 | Zfp189 | 230162 | protein_coding | 43.0352853 | 0.138962715 | 0.230729894 | 0.602274429 | 0.546991479 | 0.75017702 | 0.138962715 |
| ENSMUSG00000005580 | Adcy9 | 11515 | protein_coding | 507.4721186 | -0.138970757 | 0.066282429 | -2.096645504 | 0.036024968 | 0.147231287 | 0.138970757 |
| ENSMUSG00000054484 | Tmem62 | 96957 | protein_coding | 492.0472393 | 0.138970926 | 0.066453025 | 2.091265625 | 0.036504259 | 0.148662675 | 0.138970926 |

| ensembl_gene_id | external_gene_name | entrezgene_id | gene_biotype | baseMean | log2FoldChange | lfcSE | stat | pvalue | padj | absLog2FC |
|---|---|---|---|---|---|---|---|---|---|---|
| ENSMUSG00000035270 | Impg2 | 224224 | protein_coding | 7.490673847 | -0.139030646 | 0.452207941 | -0.307448484 | 0.758502029 | 0.885839179 | 0.139030646 |
| ENSMUSG00000021810 | Ecd | 70601 | protein_coding | 925.4304191 | 0.139086648 | 0.061113961 | 2.275857187 | 0.022854559 | 0.107689014 | 0.139086648 |
| ENSMUSG00000074519 | Zfp971 | 626848 | protein_coding | 140.843723 | -0.139091053 | 0.135683108 | -1.025116945 | 0.305308011 | 0.549639002 | 0.139091053 |
| ENSMUSG00000045257 | Morn2 | 378462 | protein_coding | 67.61465705 | 0.139109556 | 0.158974252 | 0.875044568 | 0.381549656 | 0.622373434 | 0.139109556 |
| ENSMUSG00000042570 | Mier2 | 70427 | protein_coding | 560.5337945 | 0.139125176 | 0.058992448 | 2.358355705 | 0.018356095 | 0.092745101 | 0.139125176 |
| ENSMUSG00000020492 | Ska2 | 66140 | protein_coding | 104.4505306 | 0.139126988 | 0.14537854 | 0.956998104 | 0.338568212 | 0.5820991 | 0.139126988 |
| ENSMUSG00000029269 | Sult1b1 | 56362 | protein_coding | 3926.132077 | 0.139146422 | 0.062396071 | 2.23005103 | 0.025744055 | 0.117115481 | 0.139146422 |
| ENSMUSG00000037486 | Asxl2 | 75302 | protein_coding | 1181.519915 | -0.139159343 | 0.037707355 | -3.690509288 | 0.000223806 | 0.002889761 | 0.139159343 |
| ENSMUSG00000015095 | Fbxw5 | 30839 | protein_coding | 994.7446767 | 0.139182677 | 0.059594848 | 2.335481704 | 0.019518271 | 0.09643724 | 0.139182677 |
| ENSMUSG00000116220 | A430088P11Rik | 106059 | lncRNA | 30.86201627 | -0.13918293 | 0.266462066 | -0.522336752 | 0.601435884 | 0.786583124 | 0.13918293 |
| ENSMUSG00000027315 | Spint1 | 20732 | protein_coding | 7138.280625 | -0.139203787 | 0.056191768 | -2.47729855 | 0.013238112 | 0.073314226 | 0.139203787 |
| ENSMUSG00000028953 | Abcf2 | 27407 | protein_coding | 871.823894 | 0.139206238 | 0.061334015 | 2.269641681 | 0.023229332 | 0.108932668 | 0.139206238 |
| ENSMUSG00000029763 | Exoc4 | 20336 | protein_coding | 1187.9556 | 0.139219012 | 0.065156069 | 2.136700621 | 0.032622349 | 0.13790318 | 0.139219012 |
| ENSMUSG00000022257 | Laptm4b | 114128 | protein_coding | 1753.738648 | 0.139232542 | 0.045782092 | 3.041200921 | 0.002356365 | 0.019885455 | 0.139232542 |
| ENSMUSG00000020458 | Rtn4 | 68585 | protein_coding | 2238.263537 | -0.13924008 | 0.062151482 | -2.240334048 | 0.025069244 | 0.114920894 | 0.13924008 |
| ENSMUSG00000040938 | Slc16a11 | 216867 | protein_coding | 71.76293035 | -0.139262729 | 0.175822111 | -0.792066078 | 0.428322148 | 0.663421426 | 0.139262729 |
| ENSMUSG00000026017 | Carf | 241066 | protein_coding | 146.0705375 | -0.139279036 | 0.1220538 | -1.14112822 | 0.253816569 | 0.495031103 | 0.139279036 |
| ENSMUSG00000026398 | Nr5a2 | 26424 | protein_coding | 2178.152486 | -0.139305013 | 0.033141093 | -4.203392246 | 2.63E-05 | 0.000459924 | 0.139305013 |
| ENSMUSG00000035278 | Plekhj1 | 78670 | protein_coding | 417.3274197 | 0.139311928 | 0.078504905 | 1.774563362 | 0.075970011 | 0.242642163 | 0.139311928 |
| ENSMUSG00000037493 | Cib2 | 56506 | protein_coding | 47.6363826 | 0.139322918 | 0.195833463 | 0.711435709 | 0.476814277 | 0.699625498 | 0.139322918 |
| ENSMUSG00000034471 | Caskin2 | 140721 | protein_coding | 1855.292105 | -0.139336939 | 0.095911154 | -1.452770959 | 0.146287358 | 0.359247178 | 0.139336939 |
| ENSMUSG00000038816 | Ctnnal1 | 54366 | protein_coding | 319.0375314 | -0.139371055 | 0.073432176 | -1.897956207 | 0.057701851 | 0.203588399 | 0.139371055 |
| ENSMUSG00000025956 | Mettl21a | 67099 | protein_coding | 176.9574265 | 0.139390432 | 0.096769957 | 1.440430852 | 0.14974554 | 0.364905761 | 0.139390432 |
| ENSMUSG00000035125 | Gcfc2 | 330361 | protein_coding | 109.9135487 | -0.139418815 | 0.140588766 | -0.991678207 | 0.321354525 | 0.564971593 | 0.139418815 |
| ENSMUSG00000024091 | Vapa | 30960 | protein_coding | 3113.904935 | 0.139471324 | 0.034607661 | 4.030070776 | 5.58E-05 | 0.000901045 | 0.139471324 |
| ENSMUSG00000018697 | Aatf | 56321 | protein_coding | 503.1932617 | -0.13947227 | 0.068503321 | -2.035992832 | 0.041751075 | 0.163268571 | 0.13947227 |
| ENSMUSG00000021555 | Naa35 | 78689 | protein_coding | 1088.297998 | 0.139511684 | 0.053727083 | 2.596673319 | 0.00941314 | 0.056941378 | 0.139511684 |
| ENSMUSG00000038102 | Trappc11 | 320714 | protein_coding | 1646.448588 | 0.139622133 | 0.050074742 | 2.788274634 | 0.00529896 | 0.037252106 | 0.139622133 |
| ENSMUSG00000020277 | Pfkl | 18641 | protein_coding | 6317.295433 | 0.139644106 | 0.068386213 | 2.041992091 | 0.041152316 | 0.161750767 | 0.139644106 |
| ENSMUSG00000028212 | Ccne2 | 12448 | protein_coding | 143.7572029 | 0.139710296 | 0.148761072 | 0.939158977 | 0.347649128 | 0.589851392 | 0.139710296 |
| ENSMUSG00000032454 | Rbp2 | 19660 | protein_coding | 239.4313845 | -0.139747806 | 0.20097594 | -0.695345954 | 0.486838522 | 0.707879718 | 0.139747806 |
| ENSMUSG00000032252 | Glce | 93683 | protein_coding | 1130.378164 | 0.139776222 | 0.052873265 | 2.643608694 | 0.008202742 | 0.051613588 | 0.139776222 |
| ENSMUSG00000005417 | Mprip | 26936 | protein_coding | 3028.28769 | -0.139788937 | 0.054229639 | -2.577722065 | 0.009945396 | 0.059031798 | 0.139788937 |
| ENSMUSG00000063714 | Sp3os | 100503849 | lncRNA | 54.45960512 | -0.139794521 | 0.175791634 | -0.795228518 | 0.426480586 | 0.661841071 | 0.139794521 |
| ENSMUSG00000034701 | Neurod1 | 18012 | protein_coding | 34.54050895 | -0.139912521 | 0.194425401 | -0.719620588 | 0.471758632 | 0.696197974 | 0.139912521 |
| ENSMUSG00000000708 | Kat2b | 18519 | protein_coding | 1304.285822 | -0.13994434 | 0.082134568 | -1.703842164 | 0.088410578 | 0.266803405 | 0.13994434 |
| ENSMUSG00000069601 | Ank3 | 11735 | protein_coding | 2081.684021 | -0.13994593 | 0.066435543 | -2.106491845 | 0.035161651 | 0.145026072 | 0.13994593 |
| ENSMUSG00000001123 | Lgals9 | 16859 | protein_coding | 1057.741648 | 0.139949287 | 0.09009846 | 1.553292777 | 0.120353205 | 0.321053395 | 0.139949287 |
| ENSMUSG00000030677 | Kif22 | 110033 | protein_coding | 374.8238092 | 0.139951825 | 0.099192082 | 1.410917315 | 0.158268996 | 0.378133941 | 0.139951825 |

Supplementary Table 1

| ensembl_gene_id | external_gene_name | entrezgene_id | gene_biotype | baseMean | log2FoldChange | lfcSE | stat | pvalue | padj | absLog2FC |
|---|---|---|---|---|---|---|---|---|---|---|
| ENSMUSG00000018126 | Baiap2l2 | 207495 | protein_coding | 1255.085309 | -0.139955413 | 0.098485771 | -1.421072426 | 0.155295705 | 0.373324674 | 0.139955413 |
| ENSMUSG00000021335 | Slc17a1 | 20504 | protein_coding | 42.6878332 | -0.140062268 | 0.198902694 | -0.704174816 | 0.481323912 | 0.703426437 | 0.140062268 |
| ENSMUSG00000002346 | Slc25a42 | 73095 | protein_coding | 153.7248768 | -0.140089903 | 0.156921261 | -0.892740108 | 0.371996366 | 0.613790199 | 0.140089903 |
| ENSMUSG00000038462 | Uqcrfs1 | 66694 | protein_coding | 6701.654449 | 0.140097064 | 0.052527422 | 2.667122403 | 0.00765038 | 0.048978834 | 0.140097064 |
| ENSMUSG00000000838 | Fmr1 | 14265 | protein_coding | 1308.400317 | -0.140102672 | 0.054663108 | -2.56302061 | 0.010376589 | 0.060985149 | 0.140102672 |
| ENSMUSG00000020265 | Sumo3 | 20610 | protein_coding | 1728.593782 | 0.140108143 | 0.048569843 | 2.884673611 | 0.003918197 | 0.029749885 | 0.140108143 |
| ENSMUSG00000036615 | Rfxap | 170767 | protein_coding | 234.99697 | 0.140126074 | 0.078119178 | 1.793747417 | 0.072853493 | 0.236142883 | 0.140126074 |
| ENSMUSG00000029291 | Rufy3 | 52822 | protein_coding | 618.1244566 | -0.140129674 | 0.052229218 | -2.682974781 | 0.007297049 | 0.047191414 | 0.140129674 |
| ENSMUSG00000040767 | Snrnp25 | 78372 | protein_coding | 310.8386063 | -0.140188681 | 0.10836416 | -1.293681244 | 0.195775547 | 0.428278926 | 0.140188681 |
| ENSMUSG00000021193 | Pitrm1 | 69617 | protein_coding | 995.8590964 | 0.14018935 | 0.062803539 | 2.232188679 | 0.025602495 | 0.116636001 | 0.14018935 |
| ENSMUSG00000042797 | Aqp11 | 66333 | protein_coding | 589.7658088 | -0.140199082 | 0.085672236 | -1.636458758 | 0.10174361 | 0.290939875 | 0.140199082 |
| ENSMUSG00000021010 | Npas3 | 27386 | protein_coding | 13.85576116 | -0.140253409 | 0.36163353 | -0.387832979 | 0.698139638 | 0.849391002 | 0.140253409 |
| ENSMUSG00000025591 | Tma16 | 66282 | protein_coding | 184.815133 | -0.140312849 | 0.104940984 | -1.337064358 | 0.18120163 | 0.408247931 | 0.140312849 |
| ENSMUSG00000028894 | Inpp5b | 16330 | protein_coding | 655.2561434 | 0.140324324 | 0.05535484 | 2.534996482 | 0.011244846 | 0.064641 | 0.140324324 |
| ENSMUSG00000042377 | Fam83g | 69640 | protein_coding | 1110.638393 | -0.140364886 | 0.054343183 | -2.582934562 | 0.00979639 | 0.058470165 | 0.140364886 |
| ENSMUSG00000027401 | Tgm3 | 21818 | protein_coding | 16941.04576 | -0.140385852 | 0.209208755 | -0.671032396 | 0.502199892 | 0.718950524 | 0.140385852 |
| ENSMUSG00000053175 | Bcl3 | 12051 | protein_coding | 343.6348678 | 0.140404896 | 0.112154363 | 1.251889736 | 0.210610045 | 0.445605776 | 0.140404896 |
| ENSMUSG00000010554 | Mettl16 | 67493 | protein_coding | 438.2572621 | 0.140435704 | 0.079220845 | 1.772711512 | 0.076276523 | 0.243297365 | 0.140435704 |
| ENSMUSG00000021055 | Esr2 | 13983 | protein_coding | 42.08122706 | 0.140447401 | 0.187655859 | 0.748430675 | 0.454200426 | 0.682675952 | 0.140447401 |
| ENSMUSG00000028271 | Gtf2b | 229906 | protein_coding | 735.4517653 | 0.140482945 | 0.056078315 | 2.505120664 | 0.012240963 | 0.069048619 | 0.140482945 |
| ENSMUSG00000020258 | Glyctk | 235582 | protein_coding | 29.05160712 | -0.140489279 | 0.218886488 | -0.641836234 | 0.520979521 | 0.732526772 | 0.140489279 |
| ENSMUSG00000034509 | Mad2l1bp | 66591 | protein_coding | 404.100652 | 0.140503457 | 0.070861269 | 1.982796236 | 0.047390195 | 0.178649292 | 0.140503457 |
| ENSMUSG00000017412 | Cacnb4 | 12298 | protein_coding | 8.078439415 | 0.140526826 | 0.41599783 | 0.337806632 | 0.735508912 | 0.872889311 | 0.140526826 |
| ENSMUSG00000027220 | Syt13 | 80976 | protein_coding | 44.97797102 | 0.140560881 | 0.214289375 | 0.655939571 | 0.511863006 | 0.725570423 | 0.140560881 |
| ENSMUSG00000034484 | Snx2 | 67804 | protein_coding | 1653.403486 | -0.140651115 | 0.050550159 | -2.78240696 | 0.005395733 | 0.037767792 | 0.140651115 |
| ENSMUSG00000024081 | Cebpz | 12607 | protein_coding | 741.14501 | -0.14065284 | 0.090896781 | -1.547390765 | 0.121769047 | 0.323307444 | 0.14065284 |
| ENSMUSG00000005763 | Cd247 | 12503 | protein_coding | 12.77964695 | 0.140662911 | 0.3919642 | 0.358866729 | 0.71969479 | 0.862820371 | 0.140662911 |
| ENSMUSG00000024070 | Prkd3 | 75292 | protein_coding | 332.6998382 | -0.140685168 | 0.097813989 | -1.438292919 | 0.15035096 | 0.365794955 | 0.140685168 |
| ENSMUSG00000024617 | Camk2a | 12322 | protein_coding | 16.83879506 | -0.140712549 | 0.313521032 | -0.448813748 | 0.65356602 | 0.821663954 | 0.140712549 |
| ENSMUSG00000032312 | Csk | 12988 | protein_coding | 1597.859819 | 0.140747908 | 0.103698642 | 1.357278212 | 0.174692828 | 0.40092091 | 0.140747908 |
| ENSMUSG00000040859 | Bsdc1 | 100383 | protein_coding | 2098.985936 | 0.140760077 | 0.044676803 | 3.150630014 | 0.001629187 | 0.014953844 | 0.140760077 |
| ENSMUSG00000026374 | Tsn | 22099 | protein_coding | 1110.114174 | 0.140792869 | 0.060729636 | 2.318355239 | 0.020430023 | 0.099774968 | 0.140792869 |
| ENSMUSG00000020747 | Tmem94 | 71947 | protein_coding | 3804.54981 | -0.140832669 | 0.065316193 | -2.156167758 | 0.031070568 | 0.1331711 | 0.140832669 |
| ENSMUSG00000053964 | Lgals4 | 16855 | protein_coding | 87252.07634 | 0.140842099 | 0.0600978 | 2.343548343 | 0.019101287 | 0.095193588 | 0.140842099 |
| ENSMUSG00000018334 | Ksr1 | 16706 | protein_coding | 519.6977718 | 0.140848002 | 0.058722432 | 2.398538305 | 0.016460655 | 0.085937513 | 0.140848002 |
| ENSMUSG00000049038 | Mterf2 | 74238 | protein_coding | 125.3847954 | -0.140852008 | 0.129608962 | -1.086745896 | 0.277149124 | 0.51990697 | 0.140852008 |
| ENSMUSG00000010376 | Nedd8 | 18002 | protein_coding | 1488.774258 | 0.140862016 | 0.069202795 | 2.035496063 | 0.041800984 | 0.163424115 | 0.140862016 |
| ENSMUSG00000037509 | Arhgef4 | 226970 | protein_coding | 14.54478829 | -0.140900752 | 0.330721524 | -0.426040467 | 0.670078359 | 0.831653764 | 0.140900752 |
| ENSMUSG00000041650 | Pcca | 110821 | protein_coding | 884.2010956 | -0.140910813 | 0.075495928 | -1.86646906 | 0.061975769 | 0.212475175 | 0.140910813 |

| ensembl_gene_id | external_gene_name | entrezgene_id | gene_biotype | baseMean | log2FoldChange | lfcSE | stat | pvalue | padj | absLog2FC |
|---|---|---|---|---|---|---|---|---|---|---|
| ENSMUSG00000036751 | Cox6b1 | 110323 | protein_coding | 9436.710609 | 0.140925728 | 0.052037382 | 2.708163296 | 0.006765672 | 0.044625766 | 0.140925728 |
| ENSMUSG00000060429 | Sntb1 | 20649 | protein_coding | 24.5714668 | 0.140952353 | 0.231341121 | 0.609283609 | 0.542336469 | 0.74786321 | 0.140952353 |
| ENSMUSG00000065036 | Gm22362 | 115486329 | snoRNA | 19.338799 | -0.141021789 | 0.36388553 | -0.387544372 | 0.698353242 | 0.849539318 | 0.141021789 |
| ENSMUSG00000025757 | Hspa4l | 18415 | protein_coding | 1586.704336 | 0.141037246 | 0.08931526 | 1.579094625 | 0.114314354 | 0.311884215 | 0.141037246 |
| ENSMUSG00000029923 | Rab19 | 19331 | protein_coding | 284.873147 | -0.14105066 | 0.084710716 | -1.665086395 | 0.095895521 | 0.280723736 | 0.14105066 |
| ENSMUSG00000071477 | Zfp777 | 72306 | protein_coding | 522.9631301 | -0.141071675 | 0.077678115 | -1.816105787 | 0.069354146 | 0.228633343 | 0.141071675 |
| ENSMUSG00000038145 | Snrk | 20623 | protein_coding | 1372.907514 | -0.141075564 | 0.056629068 | -2.491221704 | 0.012730465 | 0.071088718 | 0.141075564 |
| ENSMUSG00000021265 | Slc25a29 | 214663 | protein_coding | 114.4823144 | 0.141075714 | 0.14903909 | 0.946568538 | 0.343858682 | 0.586381407 | 0.141075714 |
| ENSMUSG00000024235 | Map3k8 | 26410 | protein_coding | 23.5923862 | -0.141107653 | 0.280713555 | -0.502674883 | 0.615192872 | 0.795450521 | 0.141107653 |
| ENSMUSG00000030008 | Pradc1 | 73327 | protein_coding | 196.2408755 | 0.14110836 | 0.102713431 | 1.373806306 | 0.169501823 | 0.393827388 | 0.14110836 |
| ENSMUSG00000021327 | Zkscan3 | 72739 | protein_coding | 1004.047403 | 0.14114423 | 0.065278951 | 2.162170616 | 0.030605028 | 0.132040471 | 0.14114423 |
| ENSMUSG00000014852 | Adamts13 | 279028 | protein_coding | 12.82633619 | 0.141149133 | 0.365914654 | 0.385743319 | 0.699686777 | 0.850189342 | 0.141149133 |
| ENSMUSG00000025558 | Dock9 | 105445 | protein_coding | 986.2930844 | -0.141207955 | 0.0592973 | -2.381355558 | 0.017249054 | 0.08870392 | 0.141207955 |
| ENSMUSG00000024887 | Asah2 | 54447 | protein_coding | 676.5934626 | 0.141220735 | 0.095303364 | 1.481802205 | 0.138392932 | 0.348456334 | 0.141220735 |
| ENSMUSG00000050697 | Prkaa1 | 105787 | protein_coding | 2019.552655 | -0.141253555 | 0.049312276 | -2.864470396 | 0.004177072 | 0.031201315 | 0.141253555 |
| ENSMUSG00000068742 | Cry2 | 12953 | protein_coding | 550.9762759 | -0.141311334 | 0.122474673 | -1.153800454 | 0.248581988 | 0.489425188 | 0.141311334 |
| ENSMUSG00000026869 | Psmd5 | 66998 | protein_coding | 1104.295688 | 0.14132311 | 0.052565408 | 2.688519217 | 0.00717697 | 0.04672709 | 0.14132311 |
| ENSMUSG00000028479 | Gne | 50798 | protein_coding | 11926.88181 | 0.141352691 | 0.063833755 | 2.214387855 | 0.02680211 | 0.119899479 | 0.141352691 |
| ENSMUSG00000016257 | Prelid3b | 66390 | protein_coding | 2903.678702 | 0.141374363 | 0.066081801 | 2.139384222 | 0.032404564 | 0.137270396 | 0.141374363 |
| ENSMUSG00000022664 | Slc35a5 | 74102 | protein_coding | 577.1784885 | 0.141385206 | 0.055067703 | 2.567479661 | 0.01024408 | 0.060338304 | 0.141385206 |
| ENSMUSG00000030688 | Stard10 | 56018 | protein_coding | 12368.6365 | 0.141389485 | 0.06375732 | 2.217619642 | 0.026580777 | 0.119173811 | 0.141389485 |
| ENSMUSG00000025474 | Tubgcp2 | 74237 | protein_coding | 392.3048681 | -0.141389715 | 0.091956206 | -1.53757664 | 0.124152163 | 0.327103729 | 0.141389715 |
| ENSMUSG00000090124 | Ugt1a7c | 394432 | protein_coding | 4137.303285 | -0.141392016 | 0.043671547 | -3.237623224 | 0.001205299 | 0.011764114 | 0.141392016 |
| ENSMUSG00000020706 | Ftsj3 | 56095 | protein_coding | 769.9289762 | -0.141409866 | 0.082408419 | -1.715963814 | 0.086168666 | 0.26249378 | 0.141409866 |
| ENSMUSG00000026834 | Acvr1c | 269275 | protein_coding | 597.7157018 | -0.141428692 | 0.10785599 | -1.311273412 | 0.189765408 | 0.42043505 | 0.141428692 |
| ENSMUSG00000001105 | Ift20 | 55978 | protein_coding | 722.9128204 | 0.141443449 | 0.06533383 | 2.1649343 | 0.030392717 | 0.131440962 | 0.141443449 |
| ENSMUSG00000090641 | Zfp712 | 78251 | protein_coding | 86.92576587 | -0.141490437 | 0.147944192 | -0.956377096 | 0.338881752 | 0.58240762 | 0.141490437 |
| ENSMUSG00000039768 | Dnajc11 | 230935 | protein_coding | 1759.174595 | 0.141498322 | 0.05371753 | 2.634118168 | 0.008435612 | 0.05258644 | 0.141498322 |
| ENSMUSG00000039354 | Smarcal1 | 54380 | protein_coding | 421.4076534 | 0.141505274 | 0.062211677 | 2.274577399 | 0.022931293 | 0.107974588 | 0.141505274 |
| ENSMUSG00000047412 | Zbtb44 | 235132 | protein_coding | 1117.350836 | -0.141509593 | 0.070817298 | -1.998234856 | 0.045691204 | 0.174157894 | 0.141509593 |
| ENSMUSG00000041594 | Tmtc4 | 70551 | protein_coding | 660.7512688 | 0.141514721 | 0.049929499 | 2.834290843 | 0.004592751 | 0.0334988 | 0.141514721 |
| ENSMUSG00000043068 | Fam89a | 69627 | protein_coding | 6.01217681 | -0.141521853 | 0.484986751 | -0.291805606 | 0.770435259 | 0.892047592 | 0.141521853 |
| ENSMUSG00000000938 | Hoxa10 | 15395 | protein_coding | 617.5526012 | 0.141526472 | 0.09549918 | 1.48196531 | 0.138349526 | 0.348401437 | 0.141526472 |
| ENSMUSG00000026999 | Nup35 | 69482 | protein_coding | 112.8129646 | 0.141527803 | 0.128780921 | 1.09898114 | 0.271776293 | 0.51430841 | 0.141527803 |
| ENSMUSG00000033579 | Fa2h | 338521 | protein_coding | 1629.743236 | 0.141528859 | 0.097047974 | 1.458339141 | 0.144747087 | 0.357283876 | 0.141528859 |
| ENSMUSG00000038836 | Agbl3 | 76223 | protein_coding | 32.60644634 | 0.14153424 | 0.20521103 | 0.689700937 | 0.490382277 | 0.710356146 | 0.14153424 |
| ENSMUSG00000074300 | Smim31 | 407795 | protein_coding | 2209.394031 | 0.141604997 | 0.071553497 | 1.979008745 | 0.047815022 | 0.179562869 | 0.141604997 |
| ENSMUSG00000068877 | Selenbp2 | 20342 | protein_coding | 173.0056924 | -0.141626712 | 0.126139113 | -1.122781892 | 0.261530138 | 0.503425225 | 0.141626712 |
| ENSMUSG00000046861 | Hectd3 | 76608 | protein_coding | 3314.690868 | -0.141640407 | 0.05475144 | -2.586971348 | 0.009682364 | 0.058047394 | 0.141640407 |

| ensembl_gene_id | external_gene_name | entrezgene_id | gene_biotype | baseMean | log2FoldChange | lfcSE | stat | pvalue | padj | absLog2FC |
|---|---|---|---|---|---|---|---|---|---|---|
| ENSMUSG00000008348 | Ubc | 22190 | protein_coding | 3111.556484 | 0.141642991 | 0.106951913 | 1.324361459 | 0.185383028 | 0.414426406 | 0.141642991 |
| ENSMUSG00000056050 | Mia3 | 338366 | protein_coding | 5625.842612 | 0.141740908 | 0.056612761 | 2.503691853 | 0.012290503 | 0.0692796 | 0.141740908 |
| ENSMUSG00000003559 | As3mt | 57344 | protein_coding | 1044.111142 | 0.141783584 | 0.062309093 | 2.275487846 | 0.022876681 | 0.1077489 | 0.141783584 |
| ENSMUSG00000031621 | Isx | 71597 | protein_coding | 2183.415965 | -0.141820597 | 0.212848347 | -0.666298793 | 0.505220137 | 0.721353267 | 0.141820597 |
| ENSMUSG00000099481 | Xndc1 | 102443350 | protein_coding | 55.08611999 | -0.141907563 | 0.167774281 | -0.845824298 | 0.397650768 | 0.637110813 | 0.141907563 |
| ENSMUSG00000046432 | Bex3 | 12070 | protein_coding | 312.2541068 | 0.141959517 | 0.076517313 | 1.855260108 | 0.063559092 | 0.215929529 | 0.141959517 |
| ENSMUSG00000027845 | Dclre1b | 140917 | protein_coding | 208.6947429 | 0.141971485 | 0.090451777 | 1.569582042 | 0.116512379 | 0.31570327 | 0.141971485 |
| ENSMUSG00000022560 | Slc52a2 | 52710 | protein_coding | 663.4333305 | -0.141989788 | 0.061558398 | -2.306586784 | 0.021077867 | 0.101895312 | 0.141989788 |
| ENSMUSG00000026848 | Tor1b | 30934 | protein_coding | 849.9419994 | -0.142008829 | 0.072026539 | -1.971618119 | 0.048653217 | 0.181585382 | 0.142008829 |
| ENSMUSG00000021945 | Zmym2 | 76007 | protein_coding | 1511.200437 | -0.142052977 | 0.058149017 | -2.442912763 | 0.014569259 | 0.078555143 | 0.142052977 |
| ENSMUSG00000026304 | Rab17 | 19329 | protein_coding | 618.0441364 | -0.142054884 | 0.104829483 | -1.355104299 | 0.175384329 | 0.401707581 | 0.142054884 |
| ENSMUSG00000070495 | Ctcfl | 664799 | protein_coding | 41.65435059 | -0.142090273 | 0.217867314 | -0.652187197 | 0.514280418 | 0.727325341 | 0.142090273 |
| ENSMUSG00000026546 | Cfap45 | 71870 | protein_coding | 47.36426677 | 0.142148077 | 0.183733365 | 0.773665018 | 0.439128912 | 0.671631718 | 0.142148077 |
| ENSMUSG00000053453 | Thoc7 | 66231 | protein_coding | 732.1599426 | -0.142187213 | 0.058417691 | -2.433975246 | 0.014934016 | 0.079926167 | 0.142187213 |
| ENSMUSG00000073481 | Mtarc2 | 67247 | protein_coding | 2289.79134 | -0.142204266 | 0.035984798 | -3.951787289 | 7.76E-05 | 0.001191396 | 0.142204266 |
| ENSMUSG00000057672 | Pkn1 | 320795 | protein_coding | 1944.432328 | -0.142274012 | 0.062640382 | -2.271282639 | 0.023129874 | 0.108600285 | 0.142274012 |
| ENSMUSG00000041729 | Coro2b | 235431 | protein_coding | 12.84256924 | -0.142284397 | 0.365288144 | -0.389512769 | 0.696896867 | 0.848535732 | 0.142284397 |
| ENSMUSG00000066839 | Ecsit | 26940 | protein_coding | 635.535588 | 0.14233255 | 0.055811335 | 2.550244509 | 0.010764739 | 0.062357381 | 0.14233255 |
| ENSMUSG00000025582 | Nptx1 | 18164 | protein_coding | 9.959601459 | 0.142419904 | 0.397314948 | 0.358455943 | 0.720002133 | 0.863035664 | 0.142419904 |
| ENSMUSG00000044786 | Zfp36 | 22695 | protein_coding | 9504.138378 | 0.14251203 | 0.082380908 | 1.729915743 | 0.08364533 | 0.257531164 | 0.14251203 |
| ENSMUSG00000037894 | H2az1 | 51788 | protein_coding | 2129.187349 | 0.142540942 | 0.070386882 | 2.02510664 | 0.042856415 | 0.1666367 | 0.142540942 |
| ENSMUSG00000028745 | Capzb | 12345 | protein_coding | 3874.065331 | 0.142552017 | 0.034452259 | 4.137668199 | 3.51E-05 | 0.000596283 | 0.142552017 |
| ENSMUSG00000031714 | Gab1 | 14388 | protein_coding | 737.0249218 | -0.142565183 | 0.074642064 | -1.909984457 | 0.056135215 | 0.200020461 | 0.142565183 |
| ENSMUSG00000106617 | Gm36266 | 108169076 | lncRNA | 54.05412963 | -0.142671668 | 0.169524075 | -0.841601219 | 0.400011207 | 0.639482053 | 0.142671668 |
| ENSMUSG00000022034 | Esco2 | 71988 | protein_coding | 212.1260635 | 0.142706674 | 0.144302881 | 0.988938495 | 0.322693235 | 0.566769091 | 0.142706674 |
| ENSMUSG00000037007 | Zfp113 | 56314 | protein_coding | 362.16028 | -0.142711633 | 0.091047408 | -1.567443117 | 0.117011152 | 0.31618145 | 0.142711633 |
| ENSMUSG00000041736 | Tspo | 12257 | protein_coding | 1557.428507 | -0.142729086 | 0.083899024 | -1.701200784 | 0.088905291 | 0.267808218 | 0.142729086 |
| ENSMUSG00000030763 | Lcmt1 | 30949 | protein_coding | 379.7553652 | 0.14274943 | 0.06563511 | 2.174894354 | 0.029638034 | 0.129042272 | 0.14274943 |
| ENSMUSG00000039842 | Mcph1 | 244329 | protein_coding | 282.5993146 | 0.142793609 | 0.08128273 | 1.756752128 | 0.078960068 | 0.248300541 | 0.142793609 |
| ENSMUSG00000070034 | Sp110 | 109032 | protein_coding | 62.60527608 | -0.142803512 | 0.29701911 | -0.480788971 | 0.63066649 | 0.805987676 | 0.142803512 |
| ENSMUSG00000023034 | Nr4a1 | 15370 | protein_coding | 3289.563969 | -0.142819307 | 0.181716153 | -0.785947228 | 0.431898404 | 0.666401221 | 0.142819307 |
| ENSMUSG00000096795 | Zfp433 | 73610 | protein_coding | 16.34875558 | 0.142861421 | 0.31621796 | 0.45178149 | 0.651426406 | 0.8204259 | 0.142861421 |
| ENSMUSG00000049288 | Lix1l | 280411 | protein_coding | 9.667775516 | 0.142869488 | 0.441317309 | 0.323734159 | 0.746139307 | 0.879490473 | 0.142869488 |
| ENSMUSG00000024033 | Rsph1 | 22092 | protein_coding | 30.0578783 | 0.142877077 | 0.219183967 | 0.65185916 | 0.514492033 | 0.727325341 | 0.142877077 |
| ENSMUSG00000022571 | Pycrl | 66194 | protein_coding | 1475.277711 | 0.142881892 | 0.072103573 | 1.981620139 | 0.047521771 | 0.178864648 | 0.142881892 |
| ENSMUSG00000021709 | Erbin | 59079 | protein_coding | 13712.05905 | -0.142892728 | 0.056358974 | -2.535403264 | 0.011231795 | 0.064598841 | 0.142892728 |
| ENSMUSG00000020696 | Rffl | 67338 | protein_coding | 596.4091256 | -0.142927334 | 0.067844034 | -2.106704527 | 0.0351432 | 0.14502415 | 0.142927334 |
| ENSMUSG00000029454 | Mapkapk5 | 17165 | protein_coding | 60.33964218 | -0.142958162 | 0.242718055 | -0.58898858 | 0.555868935 | 0.755288004 | 0.142958162 |
| ENSMUSG00000078201 | Tmem203 | 227615 | protein_coding | 87.16439786 | -0.142962409 | 0.159147254 | -0.898302705 | 0.369024191 | 0.611199869 | 0.142962409 |

| ensembl_gene_id | external_gene_name | entrezgene_id | gene_biotype | baseMean | log2FoldChange | lfcSE | stat | pvalue | padj | absLog2FC |
|---|---|---|---|---|---|---|---|---|---|---|
| ENSMUSG00000027434 | Nkx2-2 | 18088 | protein_coding | 14.52068377 | 0.142969706 | 0.358351283 | 0.398965243 | 0.689918815 | 0.844374344 | 0.142969706 |
| ENSMUSG00000040543 | Pitpnm3 | 327958 | protein_coding | 281.4610331 | -0.142998539 | 0.084888664 | -1.684542222 | 0.092076927 | 0.273529895 | 0.142998539 |
| ENSMUSG00000031144 | Syp | 20977 | protein_coding | 30.0788036 | -0.143014027 | 0.228151107 | -0.626839068 | 0.530764732 | 0.73971984 | 0.143014027 |
| ENSMUSG00000031843 | Mphosph6 | 68533 | protein_coding | 269.2348562 | 0.143022215 | 0.087602872 | 1.632620164 | 0.102548908 | 0.292295199 | 0.143022215 |
| ENSMUSG00000019320 | Noxo1 | 71893 | protein_coding | 77.21736224 | -0.14302457 | 0.150642918 | -0.949427775 | 0.342403089 | 0.585130336 | 0.14302457 |
| ENSMUSG00000028760 | Eif4g3 | 230861 | protein_coding | 1103.274505 | 0.143044502 | 0.059208873 | 2.415930155 | 0.015695072 | 0.082768143 | 0.143044502 |
| ENSMUSG00000034403 | Pja1 | 18744 | protein_coding | 1469.827873 | 0.143065569 | 0.042508338 | 3.365588392 | 0.000763806 | 0.008056179 | 0.143065569 |
| ENSMUSG00000034709 | Ppp1r21 | 73825 | protein_coding | 595.6619129 | 0.143097024 | 0.054715385 | 2.615297746 | 0.00891497 | 0.054811943 | 0.143097024 |
| ENSMUSG00000074277 | Phldb3 | 232970 | protein_coding | 160.4279757 | -0.143144196 | 0.108330637 | -1.321363921 | 0.18638005 | 0.415618234 | 0.143144196 |
| ENSMUSG00000035455 | Fignl1 | 60530 | protein_coding | 155.2474806 | 0.143148816 | 0.204920929 | 0.698556351 | 0.48482933 | 0.706263486 | 0.143148816 |
| ENSMUSG00000028689 | Ccdc163 | 68394 | protein_coding | 124.7672211 | -0.143182029 | 0.125541661 | -1.14051406 | 0.254072198 | 0.495031103 | 0.143182029 |
| ENSMUSG00000089862 | Umad1 | 100036521 | protein_coding | 496.7918888 | 0.143186969 | 0.074809992 | 1.914008614 | 0.055619055 | 0.198733926 | 0.143186969 |
| ENSMUSG00000025477 | Inpp5a | 212111 | protein_coding | 810.780977 | -0.143206427 | 0.052850631 | -2.709644609 | 0.006735534 | 0.044499775 | 0.143206427 |
| ENSMUSG00000021611 | Tert | 21752 | protein_coding | 51.14497107 | -0.143248921 | 0.168501187 | -0.850135976 | 0.395249492 | 0.634563321 | 0.143248921 |
| ENSMUSG00000041119 | Pde9a | 18585 | protein_coding | 2317.402494 | 0.143332635 | 0.075643653 | 1.894840204 | 0.058113575 | 0.20409356 | 0.143332635 |
| ENSMUSG00000049532 | Sall2 | 50524 | protein_coding | 18.43400278 | 0.143334413 | 0.280148257 | 0.511637711 | 0.608904588 | 0.791308423 | 0.143334413 |
| ENSMUSG00000044042 | Fmn1 | 14260 | protein_coding | 1027.052599 | 0.143334576 | 0.060086614 | 2.385466042 | 0.017057497 | 0.088049668 | 0.143334576 |
| ENSMUSG00000053716 | Dusp7 | 235584 | protein_coding | 147.7876195 | 0.143435449 | 0.14599862 | 0.982443869 | 0.325881223 | 0.569329485 | 0.143435449 |
| ENSMUSG00000036181 | H1f2 | 50708 | protein_coding | 13299.95221 | 0.14344203 | 0.060518061 | 2.370235058 | 0.017776779 | 0.090551523 | 0.14344203 |
| ENSMUSG00000039844 | Rapgef1 | 107746 | protein_coding | 1132.608971 | -0.143446639 | 0.045305714 | -3.166193086 | 0.001544482 | 0.014331339 | 0.143446639 |
| ENSMUSG00000026322 | Htr4 | 15562 | protein_coding | 717.2464724 | -0.14352396 | 0.096580107 | -1.486061309 | 0.137262905 | 0.346747943 | 0.14352396 |
| ENSMUSG00000023914 | Mep1a | 17287 | protein_coding | 13739.52837 | -0.143527755 | 0.082537504 | -1.738939866 | 0.082045341 | 0.254451003 | 0.143527755 |
| ENSMUSG00000035172 | Plekhh3 | 217198 | protein_coding | 588.4978781 | -0.143562268 | 0.100236103 | -1.432241112 | 0.15207483 | 0.368632313 | 0.143562268 |
| ENSMUSG00000027282 | Mtch2 | 56428 | protein_coding | 4274.608732 | 0.14358775 | 0.051823082 | 2.770729659 | 0.005593084 | 0.038894487 | 0.14358775 |
| ENSMUSG00000038298 | Pdzk1 | 59020 | protein_coding | 16.47171559 | 0.143610314 | 0.551969596 | 0.260177942 | 0.794726518 | 0.904781532 | 0.143610314 |
| ENSMUSG00000036438 | Calm2 | 12314 | protein_coding | 8382.215073 | -0.143641656 | 0.051511486 | -2.788536473 | 0.005294679 | 0.037252106 | 0.143641656 |
| ENSMUSG00000015224 | Cyp2j9 | 74519 | protein_coding | 59.76469778 | -0.143659404 | 0.219468339 | -0.654579173 | 0.512738737 | 0.726301064 | 0.143659404 |
| ENSMUSG00000020700 | Map3k3 | 26406 | protein_coding | 661.5361727 | 0.143677844 | 0.067404393 | 2.131579807 | 0.033041405 | 0.139211338 | 0.143677844 |
| ENSMUSG00000043518 | Rai2 | 24004 | protein_coding | 43.91693917 | -0.14369861 | 0.180147366 | -0.797672553 | 0.425060535 | 0.66096665 | 0.14369861 |
| ENSMUSG00000027001 | Dusp19 | 68082 | protein_coding | 188.6273685 | 0.143723081 | 0.113672448 | 1.264361623 | 0.206100261 | 0.440819483 | 0.143723081 |
| ENSMUSG00000103088 | Pcdhgb6 | 93703 | protein_coding | 38.08165121 | 0.143731982 | 0.190509921 | 0.7544593 | 0.450573475 | 0.679865123 | 0.143731982 |
| ENSMUSG00000023045 | Soat2 | 223920 | protein_coding | 42.56180463 | 0.143774938 | 0.268572661 | 0.535329761 | 0.592421851 | 0.780262005 | 0.143774938 |
| ENSMUSG00000031150 | Ccdc120 | 54648 | protein_coding | 300.1856542 | -0.143789703 | 0.10787637 | -1.332911958 | 0.182560698 | 0.410507024 | 0.143789703 |
| ENSMUSG00000036825 | Ssx2ip | 99167 | protein_coding | 1473.321163 | 0.143825468 | 0.053292025 | 2.698817873 | 0.006958625 | 0.04560043 | 0.143825468 |
| ENSMUSG00000042390 | Gatad2b | 229542 | protein_coding | 550.5270114 | -0.143835869 | 0.107435603 | -1.338810087 | 0.180632506 | 0.407535035 | 0.143835869 |
| ENSMUSG00000073542 | Cep76 | 225659 | protein_coding | 157.0984243 | -0.143868353 | 0.108771946 | -1.322660465 | 0.185948317 | 0.415160239 | 0.143868353 |
| ENSMUSG00000053604 | Rpia | 19895 | protein_coding | 607.5567551 | -0.143871602 | 0.061083699 | -2.355319088 | 0.018506806 | 0.093297675 | 0.143871602 |
| ENSMUSG00000001416 | Cct3 | 12462 | protein_coding | 3225.235356 | 0.143880042 | 0.080020092 | 1.798048953 | 0.072169251 | 0.234551147 | 0.143880042 |
| ENSMUSG00000078607 | 1810010H24Rik | 69066 | protein_coding | 39.47367682 | 0.143887211 | 0.269070356 | 0.534756831 | 0.592818018 | 0.780422622 | 0.143887211 |

| ensembl_gene_id | external_gene_name | entrezgene_id | gene_biotype | baseMean | log2FoldChange | lfcSE | stat | pvalue | padj | absLog2FC |
|---|---|---|---|---|---|---|---|---|---|---|
| ENSMUSG00000020758 | Itgb4 | 192897 | protein_coding | 4675.334187 | -0.143900311 | 0.078568152 | -1.831534892 | 0.067020745 | 0.223591963 | 0.143900311 |
| ENSMUSG00000019868 | Vta1 | 66201 | protein_coding | 655.9857836 | 0.14390091 | 0.06270995 | 2.294706208 | 0.021749967 | 0.104486659 | 0.14390091 |
| ENSMUSG00000025218 | Poll | 56626 | protein_coding | 257.475867 | 0.14396333 | 0.083711233 | 1.719761208 | 0.085475857 | 0.261073701 | 0.14396333 |
| ENSMUSG00000042102 | Dmgdh | 74129 | protein_coding | 12.42569325 | 0.144028574 | 0.446811654 | 0.322347398 | 0.74718953 | 0.880070519 | 0.144028574 |
| ENSMUSG00000024827 | Gldc | 104174 | protein_coding | 7.650448963 | 0.144032823 | 0.512281071 | 0.281159759 | 0.778587867 | 0.896941755 | 0.144032823 |
| ENSMUSG00000048732 | Klhl11 | 217194 | protein_coding | 247.4890993 | -0.14405373 | 0.084952689 | -1.695693592 | 0.089943923 | 0.269714697 | 0.14405373 |
| ENSMUSG00000097354 | 2310001H17Rik | 76432 | lncRNA | 165.4138853 | -0.144129208 | 0.09561315 | -1.507420342 | 0.131702948 | 0.337937843 | 0.144129208 |
| ENSMUSG00000074194 | Zfp791 | 244556 | protein_coding | 92.97598286 | 0.144150071 | 0.175545826 | 0.821153507 | 0.411558836 | 0.649560515 | 0.144150071 |
| ENSMUSG00000006932 | Ctnnb1 | 12387 | protein_coding | 18911.84763 | 0.144167153 | 0.03666487 | 3.932024149 | 8.42E-05 | 0.001273135 | 0.144167153 |
| ENSMUSG00000039278 | Pcsk1n | 30052 | protein_coding | 45.98324924 | -0.14418016 | 0.187804908 | -0.767712417 | 0.442658058 | 0.673781897 | 0.14418016 |
| ENSMUSG00000078453 | Abracl | 73112 | protein_coding | 743.237267 | 0.144183666 | 0.078715626 | 1.831703225 | 0.066995648 | 0.223554482 | 0.144183666 |
| ENSMUSG00000059975 | Zfp74 | 72723 | protein_coding | 90.77336185 | 0.144214757 | 0.131381402 | 1.097680148 | 0.272344181 | 0.514779021 | 0.144214757 |
| ENSMUSG00000022336 | Eif3e | 16341 | protein_coding | 2778.20861 | -0.144260595 | 0.079240465 | -1.820542009 | 0.068676505 | 0.227328817 | 0.144260595 |
| ENSMUSG00000050619 | Zscan29 | 99334 | protein_coding | 723.6371644 | -0.144314767 | 0.051797878 | -2.78611349 | 0.005334419 | 0.037436109 | 0.144314767 |
| ENSMUSG00000097494 | 4933406C10Rik | 74076 | lncRNA | 41.98430348 | 0.144387044 | 0.221182999 | 0.652794493 | 0.513888774 | 0.727099865 | 0.144387044 |
| ENSMUSG00000012609 | Ttll5 | 320244 | protein_coding | 428.4040942 | 0.144394003 | 0.05983724 | 2.413112672 | 0.015816929 | 0.083141987 | 0.144394003 |
| ENSMUSG00000078671 | Chd2 | 244059 | protein_coding | 1689.240914 | -0.144435834 | 0.068645804 | -2.104073727 | 0.035372016 | 0.145521558 | 0.144435834 |
| ENSMUSG00000027620 | Rbm39 | 170791 | protein_coding | 4349.599547 | -0.14443718 | 0.080550387 | -1.793128307 | 0.07295241 | 0.236340603 | 0.14443718 |
| ENSMUSG00000021895 | Arhgef3 | 71704 | protein_coding | 994.9712844 | 0.144448025 | 0.055588475 | 2.598524704 | 0.009362531 | 0.056752643 | 0.144448025 |
| ENSMUSG00000023919 | Cenpq | 83815 | protein_coding | 121.2462978 | 0.144448179 | 0.135510181 | 1.065958132 | 0.286442569 | 0.529013894 | 0.144448179 |
| ENSMUSG00000027649 | Ctnnbl1 | 66642 | protein_coding | 564.565615 | 0.144495387 | 0.067004827 | 2.156492195 | 0.031045253 | 0.133152759 | 0.144495387 |
| ENSMUSG00000086714 | 0610009E02Rik | 100125929 | lncRNA | 12.43099416 | 0.144578297 | 0.412512593 | 0.350482141 | 0.725976891 | 0.866155901 | 0.144578297 |
| ENSMUSG00000029426 | Scarb2 | 12492 | protein_coding | 3705.379669 | -0.144590406 | 0.052888595 | -2.733867384 | 0.006259525 | 0.042325937 | 0.144590406 |
| ENSMUSG00000032324 | Tspan3 | 56434 | protein_coding | 5736.756889 | 0.144597784 | 0.058079791 | 2.489640226 | 0.012787246 | 0.071277641 | 0.144597784 |
| ENSMUSG00000038240 | Pdss2 | 71365 | protein_coding | 675.7818515 | 0.144638206 | 0.071020204 | 2.036578311 | 0.041692318 | 0.16319418 | 0.144638206 |
| ENSMUSG00000028416 | Bag1 | 12017 | protein_coding | 4503.169911 | 0.144662724 | 0.037566861 | 3.850806814 | 0.000117729 | 0.001702799 | 0.144662724 |
| ENSMUSG00000037416 | Dmxl1 | 240283 | protein_coding | 2574.124701 | 0.144675175 | 0.039974608 | 3.619176791 | 0.000295542 | 0.003635136 | 0.144675175 |
| ENSMUSG00000028621 | Cyb5rl | 230582 | protein_coding | 116.0463671 | 0.14470006 | 0.127732152 | 1.132839754 | 0.257281554 | 0.499179454 | 0.14470006 |
| ENSMUSG00000040824 | Snrpd2 | 107686 | protein_coding | 833.902304 | 0.144719107 | 0.082964933 | 1.74434066 | 0.081099706 | 0.252438005 | 0.144719107 |
| ENSMUSG00000037553 | Zdhhc18 | 503610 | protein_coding | 405.5686539 | 0.144736461 | 0.145258823 | 0.996403926 | 0.319053926 | 0.562649386 | 0.144736461 |
| ENSMUSG00000019975 | Ikbip | 67454 | protein_coding | 220.0578268 | 0.144738822 | 0.089796417 | 1.611855207 | 0.106993452 | 0.299347968 | 0.144738822 |
| ENSMUSG00000033991 | Ttc37 | 218343 | protein_coding | 838.3368773 | 0.144795184 | 0.062718413 | 2.308655114 | 0.020962727 | 0.101551787 | 0.144795184 |
| ENSMUSG00000028199 | Cryz | 12972 | protein_coding | 415.7279573 | -0.144885667 | 0.066155426 | -2.190079863 | 0.028518445 | 0.125530222 | 0.144885667 |
| ENSMUSG00000005986 | Ankrd13d | 68423 | protein_coding | 9.338915113 | -0.144912412 | 0.369209589 | -0.392493632 | 0.694693516 | 0.847294848 | 0.144912412 |
| ENSMUSG00000035168 | Tanc1 | 66860 | protein_coding | 919.161833 | -0.144929167 | 0.046555456 | -3.113043618 | 0.001851686 | 0.016534966 | 0.144929167 |
| ENSMUSG00000033819 | Ppp1r16a | 73062 | protein_coding | 1281.792766 | 0.144977429 | 0.061982635 | 2.339000741 | 0.019335395 | 0.095738998 | 0.144977429 |
| ENSMUSG00000036641 | Ccdc148 | 227933 | protein_coding | 17.42031139 | -0.144980364 | 0.273977414 | -0.529169035 | 0.596688195 | 0.783232176 | 0.144980364 |
| ENSMUSG00000022724 | Riox2 | 67014 | protein_coding | 425.19772 | 0.144981542 | 0.084001128 | 1.725947571 | 0.084356845 | 0.25893088 | 0.144981542 |
| ENSMUSG00000025094 | Slc18a2 | 214084 | protein_coding | 60.16882192 | 0.144982722 | 0.210982147 | 0.68718005 | 0.491969275 | 0.711469495 | 0.144982722 |

| ensembl_gene_id | external_gene_name | entrezgene_id | gene_biotype | baseMean | log2FoldChange | lfcSE | stat | pvalue | padj | absLog2FC |
|---|---|---|---|---|---|---|---|---|---|---|
| ENSMUSG00000028262 | Clca3a2 | 80797 | protein_coding | 129.7237006 | 0.144995522 | 0.146607335 | 0.989005921 | 0.322660245 | 0.566769091 | 0.144995522 |
| ENSMUSG00000020609 | Apob | 238055 | protein_coding | 7356.826355 | -0.145044648 | 0.103473731 | -1.40175334 | 0.160988916 | 0.381467566 | 0.145044648 |
| ENSMUSG00000031529 | Tnks | 21951 | protein_coding | 1230.616412 | -0.145073592 | 0.074409357 | -1.949668656 | 0.051215624 | 0.188241916 | 0.145073592 |
| ENSMUSG00000015127 | Unkl | 74154 | protein_coding | 127.2188263 | -0.145075739 | 0.125857756 | -1.152696053 | 0.249035162 | 0.489898763 | 0.145075739 |
| ENSMUSG00000054931 | Zkscan4 | 544922 | protein_coding | 16.82274518 | 0.145085352 | 0.331625014 | 0.437498217 | 0.661750069 | 0.827097835 | 0.145085352 |
| ENSMUSG00000053414 | Hunk | 26559 | protein_coding | 777.3021538 | -0.145132865 | 0.062377119 | -2.326700362 | 0.019981219 | 0.098063637 | 0.145132865 |
| ENSMUSG00000031954 | Cfdp1 | 23837 | protein_coding | 1031.881379 | 0.145134781 | 0.049876975 | 2.909855334 | 0.003615961 | 0.028022394 | 0.145134781 |
| ENSMUSG00000039007 | Cpq | 54381 | protein_coding | 1541.437757 | -0.145204931 | 0.066700864 | -2.176957273 | 0.029483756 | 0.12855088 | 0.145204931 |
| ENSMUSG00000020415 | Pttg1 | 30939 | protein_coding | 532.9067519 | 0.145242455 | 0.069537564 | 2.0886906 | 0.036735584 | 0.149190319 | 0.145242455 |
| ENSMUSG00000022359 | Wdyhv1 | 76773 | protein_coding | 231.3032222 | 0.145245865 | 0.081134902 | 1.790177367 | 0.073425402 | 0.237154121 | 0.145245865 |
| ENSMUSG00000026495 | Efcab2 | 68226 | protein_coding | 95.9977522 | -0.145258856 | 0.129280417 | -1.123595196 | 0.261184795 | 0.503084198 | 0.145258856 |
| ENSMUSG00000093044 | Snord8 | 100217445 | snoRNA | 6.971609101 | 0.145274747 | 0.45891347 | 0.316562395 | 0.751575674 | 0.881630847 | 0.145274747 |
| ENSMUSG00000037960 | Card19 | 68480 | protein_coding | 778.9118645 | 0.145279912 | 0.090484661 | 1.605575033 | 0.108367315 | 0.301615868 | 0.145279912 |
| ENSMUSG00000032431 | Crtap | 56693 | protein_coding | 148.0048228 | -0.145282267 | 0.119831524 | -1.212387716 | 0.225364001 | 0.463340801 | 0.145282267 |
| ENSMUSG00000078954 | Arhgap8 | 73167 | protein_coding | 95.6682174 | -0.145423429 | 0.146207838 | -0.994634976 | 0.31991383 | 0.563409586 | 0.145423429 |
| ENSMUSG00000022377 | Asap1 | 13196 | protein_coding | 299.0688014 | -0.145426938 | 0.104028184 | -1.397957095 | 0.162125952 | 0.382899316 | 0.145426938 |
| ENSMUSG00000025626 | Phf6 | 70998 | protein_coding | 482.2020459 | 0.145456067 | 0.065855824 | 2.208704667 | 0.02719519 | 0.121104897 | 0.145456067 |
| ENSMUSG00000024985 | Tcf7l2 | 21416 | protein_coding | 752.4096369 | -0.145489161 | 0.082424062 | -1.765129726 | 0.077541966 | 0.245659762 | 0.145489161 |
| ENSMUSG00000056069 | Otulinl | 223433 | protein_coding | 33.10384506 | 0.145508765 | 0.284546572 | 0.511370647 | 0.609091545 | 0.791396982 | 0.145508765 |
| ENSMUSG00000022774 | Ncbp2 | 68092 | protein_coding | 304.8155914 | 0.145528352 | 0.08750545 | 1.663077587 | 0.096296906 | 0.281610209 | 0.145528352 |
| ENSMUSG00000046658 | Zfp316 | 54201 | protein_coding | 245.8508436 | -0.145619519 | 0.112684452 | -1.29227694 | 0.196261255 | 0.428817945 | 0.145619519 |
| ENSMUSG00000069814 | Ccdc92b | 432582 | protein_coding | 36.82085639 | -0.145637907 | 0.201186579 | -0.723894743 | 0.469130358 | 0.693580708 | 0.145637907 |
| ENSMUSG00000021024 | Psma6 | 26443 | protein_coding | 1769.483439 | 0.145669773 | 0.059773314 | 2.437036922 | 0.014808167 | 0.079524247 | 0.145669773 |
| ENSMUSG00000026031 | Cflar | 12633 | protein_coding | 1640.53401 | 0.145709794 | 0.043712678 | 3.333353161 | 0.00085806 | 0.008864236 | 0.145709794 |
| ENSMUSG00000031986 | Sprtn | 244666 | protein_coding | 482.1903025 | -0.14571561 | 0.068804691 | -2.117815055 | 0.034190734 | 0.142368697 | 0.14571561 |
| ENSMUSG00000026427 | Eif2d | 16865 | protein_coding | 883.145536 | 0.145727668 | 0.043308926 | 3.364841433 | 0.000765877 | 0.008068773 | 0.145727668 |
| ENSMUSG00000042992 | Borcs5 | 67774 | protein_coding | 199.7978307 | 0.14573533 | 0.09597867 | 1.518413723 | 0.128910134 | 0.3344488 | 0.14573533 |
| ENSMUSG00000055839 | Elob | 67673 | protein_coding | 2553.32258 | 0.145741942 | 0.058017447 | 2.512036467 | 0.012003668 | 0.06804329 | 0.145741942 |
| ENSMUSG00000059278 | Naa38 | 78304 | protein_coding | 442.1700529 | 0.145791269 | 0.103811585 | 1.40438342 | 0.160204705 | 0.380615229 | 0.145791269 |
| ENSMUSG00000110266 | Gm32742 | 102635385 | protein_coding | 23.47826126 | -0.14584228 | 0.279427395 | -0.521932649 | 0.601717225 | 0.786805165 | 0.14584228 |
| ENSMUSG00000057963 | Itpk1 | 217837 | protein_coding | 3818.864524 | 0.145857853 | 0.054125099 | 2.694828382 | 0.007042489 | 0.04601873 | 0.145857853 |
| ENSMUSG00000053965 | Pde5a | 242202 | protein_coding | 2977.04965 | -0.145911009 | 0.049863249 | -2.926223445 | 0.003431044 | 0.026860412 | 0.145911009 |
| ENSMUSG00000036022 | Fam122b | 78755 | protein_coding | 70.75009485 | -0.145927459 | 0.165464457 | -0.881926316 | 0.377816655 | 0.619086486 | 0.145927459 |
| ENSMUSG00000023852 | Chd1 | 12648 | protein_coding | 1620.125349 | -0.14600877 | 0.042071599 | -3.470483012 | 0.000519523 | 0.005846721 | 0.14600877 |
| ENSMUSG00000018585 | Atox1 | 11927 | protein_coding | 1298.721831 | 0.146050982 | 0.077477489 | 1.885076335 | 0.059419532 | 0.206976523 | 0.146050982 |
| ENSMUSG00000021486 | Prelid1 | 66494 | protein_coding | 4662.608457 | 0.146106012 | 0.050803653 | 2.875895762 | 0.004028827 | 0.030432271 | 0.146106012 |
| ENSMUSG00000025971 | Maip1 | 68115 | protein_coding | 221.017852 | -0.146128857 | 0.089786468 | -1.627515377 | 0.103627685 | 0.294226 | 0.146128857 |
| ENSMUSG00000064442 | Gm26225 | 115488410 | snoRNA | 8.13451578 | 0.146143422 | 0.506235718 | 0.288686508 | 0.772821288 | 0.893233635 | 0.146143422 |
| ENSMUSG00000026566 | Mpzl1 | 68481 | protein_coding | 341.3986677 | 0.146145181 | 0.096062354 | 1.521357492 | 0.12817015 | 0.333333335 | 0.146145181 |

| ensembl_gene_id | external_gene_name | entrezgene_id | gene_biotype | baseMean | log2FoldChange | lfcSE | stat | pvalue | padj | absLog2FC |
|---|---|---|---|---|---|---|---|---|---|---|
| ENSMUSG00000058672 | Tubb2a | 22151 | protein_coding | 298.1257541 | -0.14617618 | 0.100659513 | -1.452184447 | 0.146450326 | 0.359483167 | 0.14617618 |
| ENSMUSG00000020311 | Erlec1 | 66753 | protein_coding | 541.9178987 | -0.146215425 | 0.079587444 | -1.837166989 | 0.066185247 | 0.221867939 | 0.146215425 |
| ENSMUSG00000044712 | Slc38a6 | 625098 | protein_coding | 92.60947005 | -0.146244163 | 0.143005625 | -1.022646227 | 0.306475144 | 0.550571297 | 0.146244163 |
| ENSMUSG00000006906 | Stambp | 70527 | protein_coding | 231.3347549 | -0.146253609 | 0.099829455 | -1.465034639 | 0.142911467 | 0.35463345 | 0.146253609 |
| ENSMUSG00000027466 | Rbck1 | 24105 | protein_coding | 1953.790674 | 0.146270006 | 0.053697772 | 2.723949269 | 0.006450641 | 0.043147862 | 0.146270006 |
| ENSMUSG00000009569 | Mrtfb | 239719 | protein_coding | 1347.478727 | -0.146288998 | 0.086338257 | -1.694370521 | 0.090194898 | 0.270226918 | 0.146288998 |
| ENSMUSG00000032301 | Psma4 | 26441 | protein_coding | 2119.084811 | 0.146291432 | 0.06325173 | 2.312844744 | 0.020731177 | 0.10082379 | 0.146291432 |
| ENSMUSG00000031796 | Cfap20 | 14894 | protein_coding | 468.0414841 | -0.146297205 | 0.063138009 | -2.317101983 | 0.020498177 | 0.099931711 | 0.146297205 |
| ENSMUSG00000002329 | Mdp1 | 67881 | protein_coding | 1681.568668 | 0.146305181 | 0.061663518 | 2.372637586 | 0.017661584 | 0.090113624 | 0.146305181 |
| ENSMUSG00000079109 | Pms2 | 18861 | protein_coding | 194.8151046 | -0.146308923 | 0.10228793 | -1.430363517 | 0.152612714 | 0.369310836 | 0.146308923 |
| ENSMUSG00000024014 | Pim1 | 18712 | protein_coding | 421.4203403 | -0.146336835 | 0.09704345 | -1.507951697 | 0.13156689 | 0.337846137 | 0.146336835 |
| ENSMUSG00000009248 | Ascl2 | 17173 | protein_coding | 177.8006128 | -0.146346751 | 0.146198791 | -1.001012047 | 0.316820984 | 0.560658943 | 0.146346751 |
| ENSMUSG00000028863 | Meaf6 | 70088 | protein_coding | 332.8206621 | -0.146417812 | 0.075442593 | -1.940784453 | 0.052284428 | 0.190808975 | 0.146417812 |
| ENSMUSG00000033233 | Trim45 | 229644 | protein_coding | 60.7640279 | 0.14642419 | 0.155658704 | 0.940674605 | 0.346871637 | 0.589027996 | 0.14642419 |
| ENSMUSG00000026019 | Wdr12 | 57750 | protein_coding | 217.9690198 | -0.146508908 | 0.09040319 | -1.620616566 | 0.105099897 | 0.297150428 | 0.146508908 |
| ENSMUSG00000048481 | Mypop | 232934 | protein_coding | 30.73569404 | -0.14652809 | 0.216761869 | -0.675986467 | 0.499049237 | 0.716156527 | 0.14652809 |
| ENSMUSG00000032413 | Rasa2 | 114713 | protein_coding | 1375.799112 | -0.146548216 | 0.063153193 | -2.320519499 | 0.020312791 | 0.099328195 | 0.146548216 |
| ENSMUSG00000005034 | Prkacb | 18749 | protein_coding | 2209.094333 | -0.146560894 | 0.088817598 | -1.650133507 | 0.098915633 | 0.286052251 | 0.146560894 |
| ENSMUSG00000035235 | Trim13 | 66597 | protein_coding | 81.56941928 | 0.146593744 | 0.127720281 | 1.147771858 | 0.251062758 | 0.492145102 | 0.146593744 |
| ENSMUSG00000026436 | Elk4 | 13714 | protein_coding | 982.9660062 | -0.146634556 | 0.045761469 | -3.204323613 | 0.001353802 | 0.012865508 | 0.146634556 |
| ENSMUSG00000029101 | Rgs12 | 71729 | protein_coding | 130.6790736 | -0.146638083 | 0.126920527 | -1.155353558 | 0.247945672 | 0.488954885 | 0.146638083 |
| ENSMUSG00000085078 | C030013C21Rik | 77417 | lncRNA | 9.253589878 | -0.14663865 | 0.40639006 | -0.360832275 | 0.718224834 | 0.861944627 | 0.14663865 |
| ENSMUSG00000020381 | Mrnip | 68067 | protein_coding | 36.69408203 | -0.14671157 | 0.210784688 | -0.696025745 | 0.486412707 | 0.707465523 | 0.14671157 |
| ENSMUSG00000038582 | Pptc7 | 320717 | protein_coding | 1246.149289 | -0.146714531 | 0.056531994 | -2.595247775 | 0.009452275 | 0.057092449 | 0.146714531 |
| ENSMUSG00000055833 | 1700034H15Rik | 98736 | lncRNA | 7.753487425 | -0.146748859 | 0.436045607 | -0.336544749 | 0.736460113 | 0.873429345 | 0.146748859 |
| ENSMUSG00000066442 | Mthfs | 107885 | protein_coding | 53.59352145 | 0.146791826 | 0.228739321 | 0.641742859 | 0.521040156 | 0.732526772 | 0.146791826 |
| ENSMUSG00000024383 | Map3k2 | 26405 | protein_coding | 1617.276767 | -0.14683688 | 0.040787492 | -3.600046836 | 0.00031816 | 0.003863678 | 0.14683688 |
| ENSMUSG00000026309 | Ilkap | 67444 | protein_coding | 817.6380466 | -0.146848056 | 0.048541919 | -3.025180259 | 0.00248485 | 0.020731078 | 0.146848056 |
| ENSMUSG00000033943 | Mga | 29808 | protein_coding | 1425.905604 | -0.146861401 | 0.044602769 | -3.292652136 | 0.000992472 | 0.009984775 | 0.146861401 |
| ENSMUSG00000040170 | Fmo2 | 55990 | protein_coding | 22.07761492 | -0.1469008 | 0.266667741 | -0.550875781 | 0.581718831 | 0.773403098 | 0.1469008 |
| ENSMUSG00000022462 | Slc38a2 | 67760 | protein_coding | 4183.419202 | -0.146937723 | 0.052091269 | -2.820774471 | 0.004790787 | 0.034522352 | 0.146937723 |
| ENSMUSG00000045275 | Lca5l | 385668 | protein_coding | 6.626111287 | 0.146945657 | 0.461449843 | 0.3184434 | 0.750148622 | 0.880597382 | 0.146945657 |
| ENSMUSG00000034105 | Meak7 | 74347 | protein_coding | 478.623213 | 0.14695586 | 0.074844835 | 1.96347363 | 0.049591166 | 0.184148452 | 0.14695586 |
| ENSMUSG00000002147 | Stat6 | 20852 | protein_coding | 2203.74069 | -0.147000493 | 0.032822512 | -4.478648431 | 7.51E-06 | 0.000152571 | 0.147000493 |
| ENSMUSG00000012640 | Zfp715 | 69930 | protein_coding | 457.1675849 | -0.147006521 | 0.084607354 | -1.737514684 | 0.082296364 | 0.254690744 | 0.147006521 |
| ENSMUSG00000024524 | Gnal | 14680 | protein_coding | 49.11476471 | -0.14705251 | 0.211680787 | -0.694689925 | 0.487249643 | 0.708115319 | 0.14705251 |
| ENSMUSG00000028412 | Slc44a1 | 100434 | protein_coding | 10167.83415 | -0.147060075 | 0.054205218 | -2.713024323 | 0.006667221 | 0.044211463 | 0.147060075 |
| ENSMUSG00000034164 | Emid1 | 140703 | protein_coding | 10.40151991 | 0.14706716 | 0.456045217 | 0.322483724 | 0.747086266 | 0.880021159 | 0.14706716 |
| ENSMUSG00000028961 | Pgd | 110208 | protein_coding | 3676.075113 | 0.147092235 | 0.071562423 | 2.055439565 | 0.039836567 | 0.158042883 | 0.147092235 |

| ensembl_gene_id | external_gene_name | entrezgene_id | gene_biotype | baseMean | log2FoldChange | lfcSE | stat | pvalue | padj | absLog2FC |
|---|---|---|---|---|---|---|---|---|---|---|
| ENSMUSG00000038172 | Ttc39b | 69863 | protein_coding | 1490.246182 | -0.147124865 | 0.053848891 | -2.732180061 | 0.006291675 | 0.042447899 | 0.147124865 |
| ENSMUSG00000078185 | Chml | 12663 | protein_coding | 519.3532885 | 0.14716232 | 0.074594186 | 1.972839001 | 0.048513908 | 0.181275208 | 0.14716232 |
| ENSMUSG00000024589 | Nedd4l | 83814 | protein_coding | 1817.913973 | -0.14717753 | 0.053632729 | -2.744173828 | 0.006066341 | 0.04133159 | 0.14717753 |
| ENSMUSG00000009733 | Tfcp2 | 21422 | protein_coding | 260.8063456 | 0.147191037 | 0.078831832 | 1.867152302 | 0.061880322 | 0.212283335 | 0.147191037 |
| ENSMUSG00000024206 | Rfx2 | 19725 | protein_coding | 50.51285486 | 0.147222833 | 0.253477019 | 0.580813338 | 0.561366265 | 0.759301665 | 0.147222833 |
| ENSMUSG00000047098 | Rnf31 | 268749 | protein_coding | 731.9676748 | -0.147293351 | 0.07036581 | -2.093251676 | 0.036326694 | 0.148201516 | 0.147293351 |
| ENSMUSG00000019990 | Pde7b | 29863 | protein_coding | 25.03890092 | 0.147344448 | 0.311150405 | 0.473547343 | 0.635822733 | 0.80962603 | 0.147344448 |
| ENSMUSG00000042099 | Kank3 | 80880 | protein_coding | 46.47023108 | -0.147344487 | 0.190899255 | -0.771844225 | 0.440206695 | 0.671892236 | 0.147344487 |
| ENSMUSG00000030122 | Ptms | 69202 | protein_coding | 2936.552726 | 0.147375554 | 0.099624697 | 1.479307418 | 0.139058171 | 0.34931326 | 0.147375554 |
| ENSMUSG00000026469 | Xpr1 | 19775 | protein_coding | 2772.49084 | -0.147408611 | 0.039360564 | -3.745083812 | 0.000180334 | 0.002403504 | 0.147408611 |
| ENSMUSG00000068206 | Pick1 | 18693 | protein_coding | 313.2005569 | 0.147422677 | 0.087851713 | 1.678085399 | 0.093330429 | 0.275969898 | 0.147422677 |
| ENSMUSG00000020525 | Ppm1d | 53892 | protein_coding | 325.7705577 | -0.147432643 | 0.083836154 | -1.758580721 | 0.07864875 | 0.247826999 | 0.147432643 |
| ENSMUSG00000000538 | Tom1l2 | 216810 | protein_coding | 1748.738152 | -0.14744914 | 0.05970004 | -2.46983319 | 0.013517607 | 0.074453021 | 0.14744914 |
| ENSMUSG00000046623 | Gjb4 | 14621 | protein_coding | 15.36278575 | -0.147468061 | 0.370113701 | -0.398439887 | 0.690305961 | 0.84457466 | 0.147468061 |
| ENSMUSG00000059325 | Hopx | 74318 | protein_coding | 500.7817813 | -0.147557951 | 0.076195001 | -1.936583099 | 0.052796326 | 0.192070007 | 0.147557951 |
| ENSMUSG00000039774 | Galnt12 | 230145 | protein_coding | 2640.788684 | 0.147588331 | 0.058767503 | 2.511393613 | 0.012025553 | 0.068138529 | 0.147588331 |
| ENSMUSG00000031889 | Phaf1 | 234678 | protein_coding | 374.4099796 | -0.147597681 | 0.072615287 | -2.032597923 | 0.042093162 | 0.164406979 | 0.147597681 |
| ENSMUSG00000030123 | Plxnd1 | 67784 | protein_coding | 34.52240201 | 0.147617874 | 0.201125749 | 0.733958106 | 0.462974272 | 0.689353345 | 0.147617874 |
| ENSMUSG00000048581 | E130311K13Rik | 329659 | protein_coding | 76.25951373 | 0.147625074 | 0.141596949 | 1.042572417 | 0.29714637 | 0.540135201 | 0.147625074 |
| ENSMUSG00000040714 | Klc3 | 232943 | protein_coding | 31.04707405 | 0.147699894 | 0.212424361 | 0.695305819 | 0.486863668 | 0.707879718 | 0.147699894 |
| ENSMUSG00000080465 | Snord94 | 115490493 | snoRNA | 248.1457719 | 0.147847834 | 0.243579155 | 0.606980651 | 0.543863753 | 0.748688924 | 0.147847834 |
| ENSMUSG00000064307 | Lrrc51 | 69358 | protein_coding | 95.4712391 | -0.147945143 | 0.121304407 | -1.219618863 | 0.222609392 | 0.459964339 | 0.147945143 |
| ENSMUSG00000026669 | Mcm10 | 70024 | protein_coding | 314.1240435 | 0.14794643 | 0.12328043 | 1.200080418 | 0.23010811 | 0.469206409 | 0.14794643 |
| ENSMUSG00000044627 | Swi5 | 72931 | protein_coding | 1537.224079 | 0.147967152 | 0.068492032 | 2.160355689 | 0.030745145 | 0.132451326 | 0.147967152 |
| ENSMUSG00000044827 | Tlr1 | 21897 | protein_coding | 1407.191883 | 0.148008891 | 0.049163382 | 3.010551474 | 0.002607737 | 0.021533529 | 0.148008891 |
| ENSMUSG00000031563 | Wwc2 | 52357 | protein_coding | 494.4773002 | 0.148065684 | 0.141416495 | 1.047018484 | 0.295091053 | 0.538150682 | 0.148065684 |
| ENSMUSG00000074063 | Osgin1 | 71839 | protein_coding | 257.1326328 | -0.148077673 | 0.153527152 | -0.964504788 | 0.334792909 | 0.578382994 | 0.148077673 |
| ENSMUSG00000034265 | Zdhhc14 | 224454 | protein_coding | 169.9931369 | -0.148116557 | 0.127754636 | -1.159383032 | 0.24630009 | 0.487165462 | 0.148116557 |
| ENSMUSG00000021120 | Pigh | 110417 | protein_coding | 247.371129 | 0.148168817 | 0.09798131 | 1.512215111 | 0.130479148 | 0.336485813 | 0.148168817 |
| ENSMUSG00000032526 | Ss18l2 | 26901 | protein_coding | 332.2533235 | 0.148192375 | 0.087143522 | 1.70055527 | 0.089026529 | 0.267960217 | 0.148192375 |
| ENSMUSG00000037544 | Dlgap5 | 218977 | protein_coding | 341.0310534 | 0.148218648 | 0.083716778 | 1.770477206 | 0.076647678 | 0.244015888 | 0.148218648 |
| ENSMUSG00000063049 | Ing2 | 69260 | protein_coding | 217.1646885 | -0.148221281 | 0.082804184 | -1.790021639 | 0.073450433 | 0.237154121 | 0.148221281 |
| ENSMUSG00000071647 | Eml3 | 225898 | protein_coding | 581.9336782 | -0.148223611 | 0.06989873 | -2.120547976 | 0.03395986 | 0.141755849 | 0.148223611 |
| ENSMUSG00000023963 | Cyp39a1 | 56050 | protein_coding | 357.0692864 | -0.148259537 | 0.089319569 | -1.659877428 | 0.096939113 | 0.282652548 | 0.148259537 |
| ENSMUSG00000018417 | Myo1b | 17912 | protein_coding | 1562.398694 | -0.148260858 | 0.065950145 | -2.248074781 | 0.024571422 | 0.113433539 | 0.148260858 |
| ENSMUSG00000020462 | Cfap36 | 216618 | protein_coding | 252.1618793 | 0.148294545 | 0.090462947 | 1.639284933 | 0.101153932 | 0.290022548 | 0.148294545 |
| ENSMUSG00000042298 | Ttc19 | 72795 | protein_coding | 518.1087475 | -0.148350331 | 0.060661517 | -2.445542735 | 0.014463431 | 0.078088968 | 0.148350331 |
| ENSMUSG00000037972 | Snn | 20621 | protein_coding | 33.98464279 | 0.148355191 | 0.214008726 | 0.693220288 | 0.488171318 | 0.708681146 | 0.148355191 |
| ENSMUSG00000020719 | Ddx5 | 13207 | protein_coding | 16694.07306 | -0.148371295 | 0.050072162 | -2.96314936 | 0.003045088 | 0.024383381 | 0.148371295 |

Supplementary Table 1

| ensembl_gene_id | external_gene_name | entrezgene_id | gene_biotype | baseMean | log2FoldChange | lfcSE | stat | pvalue | padj | absLog2FC |
|---|---|---|---|---|---|---|---|---|---|---|
| ENSMUSG00000036990 | Otud4 | 73945 | protein_coding | 2013.057855 | -0.148397611 | 0.03883294 | -3.821436408 | 0.000132677 | 0.001878557 | 0.148397611 |
| ENSMUSG00000020916 | Krt36 | 16673 | protein_coding | 22.0566377 | -0.148562081 | 0.34785 | -0.42708662 | 0.669316238 | 0.831207683 | 0.148562081 |
| ENSMUSG00000028086 | Fbxw7 | 50754 | protein_coding | 428.8876476 | -0.148578069 | 0.078354507 | -1.896228745 | 0.057929804 | 0.203825108 | 0.148578069 |
| ENSMUSG00000027404 | Snrpb | 20638 | protein_coding | 2494.600354 | 0.1486154 | 0.074464952 | 1.995776477 | 0.045958258 | 0.174680375 | 0.1486154 |
| ENSMUSG00000026798 | Coq4 | 227683 | protein_coding | 300.5807557 | 0.148646582 | 0.072396464 | 2.053229851 | 0.040050291 | 0.158656604 | 0.148646582 |
| ENSMUSG00000041949 | Tango6 | 272538 | protein_coding | 129.1971305 | -0.148668586 | 0.110788876 | -1.341908964 | 0.179625515 | 0.406744622 | 0.148668586 |
| ENSMUSG00000013663 | Pten | 19211 | protein_coding | 2980.729802 | -0.148681897 | 0.039600136 | -3.754580457 | 0.000173632 | 0.00233347 | 0.148681897 |
| ENSMUSG00000036246 | Gmip | 78816 | protein_coding | 398.5402719 | -0.148744813 | 0.0877012 | -1.696040806 | 0.089878153 | 0.269652976 | 0.148744813 |
| ENSMUSG00000017679 | Ttpal | 76080 | protein_coding | 518.872018 | 0.148750869 | 0.068265397 | 2.179008321 | 0.02933105 | 0.128155471 | 0.148750869 |
| ENSMUSG00000040481 | Bptf | 207165 | protein_coding | 1692.867085 | -0.148798862 | 0.038956438 | -3.819621844 | 0.000133656 | 0.001890769 | 0.148798862 |
| ENSMUSG00000019823 | Mical1 | 171580 | protein_coding | 283.6859838 | -0.148801496 | 0.093708047 | -1.587926559 | 0.112302947 | 0.308484011 | 0.148801496 |
| ENSMUSG00000025812 | Pard3 | 93742 | protein_coding | 1589.531835 | -0.148843872 | 0.053040259 | -2.806243304 | 0.005012283 | 0.035735229 | 0.148843872 |
| ENSMUSG00000012076 | Brms1l | 52592 | protein_coding | 147.4039173 | 0.148912094 | 0.111006076 | 1.341476963 | 0.179765644 | 0.406831399 | 0.148912094 |
| ENSMUSG00000032112 | Trappc4 | 60409 | protein_coding | 645.072783 | 0.148932338 | 0.065758095 | 2.264851761 | 0.023521778 | 0.10995238 | 0.148932338 |
| ENSMUSG00000053483 | Usp21 | 30941 | protein_coding | 371.8710129 | 0.148979013 | 0.083835325 | 1.777043429 | 0.075561095 | 0.241780511 | 0.148979013 |
| ENSMUSG00000032648 | Pygm | 19309 | protein_coding | 7.909257825 | 0.148999856 | 0.419153592 | 0.355477941 | 0.722231568 | 0.864247815 | 0.148999856 |
| ENSMUSG00000061313 | Ddhd2 | 72108 | protein_coding | 410.6734525 | -0.149074523 | 0.080015186 | -1.863077871 | 0.062451311 | 0.213379724 | 0.149074523 |
| ENSMUSG00000030156 | Cd69 | 12515 | protein_coding | 32.20170623 | 0.149142116 | 0.290752095 | 0.512952853 | 0.607984301 | 0.790773109 | 0.149142116 |
| ENSMUSG00000014361 | Mertk | 17289 | protein_coding | 1298.160501 | -0.149150346 | 0.077101208 | -1.934474824 | 0.053054775 | 0.192513581 | 0.149150346 |
| ENSMUSG00000028330 | Ncbp1 | 433702 | protein_coding | 1031.21282 | 0.149162124 | 0.057258134 | 2.605081808 | 0.009185233 | 0.056046253 | 0.149162124 |
| ENSMUSG00000024098 | Twsg1 | 65960 | protein_coding | 961.6976978 | 0.149179715 | 0.062716765 | 2.37862578 | 0.017377307 | 0.088972317 | 0.149179715 |
| ENSMUSG00000025359 | Pmel | 20431 | protein_coding | 53.16187246 | -0.149214418 | 0.187404645 | -0.79621515 | 0.425906993 | 0.661460139 | 0.149214418 |
| ENSMUSG00000005483 | Dnajb1 | 81489 | protein_coding | 3005.014679 | 0.149265797 | 0.101269355 | 1.473948329 | 0.140495504 | 0.351500309 | 0.149265797 |
| ENSMUSG00000025092 | Hspa12a | 73442 | protein_coding | 36.09304756 | 0.149268888 | 0.328889049 | 0.453857884 | 0.649931118 | 0.819502669 | 0.149268888 |
| ENSMUSG00000020228 | Helb | 117599 | protein_coding | 693.4139595 | 0.14931088 | 0.071144957 | 2.09868537 | 0.035844645 | 0.146791923 | 0.14931088 |
| ENSMUSG00000022096 | Hr | 15460 | protein_coding | 788.5649692 | 0.149312192 | 0.090768154 | 1.644984339 | 0.099973041 | 0.287750353 | 0.149312192 |
| ENSMUSG00000052752 | Traf7 | 224619 | protein_coding | 1599.228335 | 0.149317469 | 0.046846571 | 3.187372417 | 0.001435718 | 0.013485044 | 0.149317469 |
| ENSMUSG00000026489 | Coq8a | 67426 | protein_coding | 1469.002652 | 0.149364005 | 0.063581695 | 2.349166753 | 0.018815478 | 0.094323304 | 0.149364005 |
| ENSMUSG00000027350 | Chgb | 12653 | protein_coding | 2400.558976 | -0.14937825 | 0.095212062 | -1.568900476 | 0.116671131 | 0.315962272 | 0.14937825 |
| ENSMUSG00000035910 | Dcdc2a | 195208 | protein_coding | 11.82495312 | 0.149391749 | 0.400618378 | 0.372902884 | 0.709220726 | 0.856166251 | 0.149391749 |
| ENSMUSG00000024851 | Pitpnm1 | 18739 | protein_coding | 468.5870764 | -0.149473915 | 0.087419259 | -1.709851092 | 0.087293412 | 0.26456259 | 0.149473915 |
| ENSMUSG00000020823 | Sec14l1 | 74136 | protein_coding | 776.1008351 | -0.149494967 | 0.086084182 | -1.736613672 | 0.082455385 | 0.254924035 | 0.149494967 |
| ENSMUSG00000037940 | Inpp4b | 234515 | protein_coding | 594.2516695 | 0.149540582 | 0.073039868 | 2.04738298 | 0.040620496 | 0.160285476 | 0.149540582 |
| ENSMUSG00000029238 | Clock | 12753 | protein_coding | 3161.413597 | -0.149581555 | 0.061891785 | -2.416824048 | 0.015656583 | 0.082730576 | 0.149581555 |
| ENSMUSG00000029047 | Pex10 | 668173 | protein_coding | 187.4831493 | -0.149586942 | 0.111553911 | -1.340938571 | 0.179940398 | 0.406927512 | 0.149586942 |
| ENSMUSG00000079038 | D130040H23Rik | 211135 | protein_coding | 35.00149604 | -0.149599722 | 0.266904643 | -0.560498763 | 0.575139283 | 0.768392944 | 0.149599722 |
| ENSMUSG00000027075 | Slc43a1 | 72401 | protein_coding | 96.97544182 | 0.149611326 | 0.125522235 | 1.19191095 | 0.233296164 | 0.472718587 | 0.149611326 |
| ENSMUSG00000024454 | Hdac3 | 15183 | protein_coding | 1273.360753 | 0.149613839 | 0.041735257 | 3.584830927 | 0.000337297 | 0.004065458 | 0.149613839 |
| ENSMUSG00000027080 | Med19 | 381379 | protein_coding | 433.4709491 | 0.149620662 | 0.070464515 | 2.123347624 | 0.033724733 | 0.141296696 | 0.149620662 |

| ensembl_gene_id | external_gene_name | entrezgene_id | gene_biotype | baseMean | log2FoldChange | lfcSE | stat | pvalue | padj | absLog2FC |
|---|---|---|---|---|---|---|---|---|---|---|
| ENSMUSG00000061979 | Rcc1l | 94254 | protein_coding | 367.3496964 | -0.149628108 | 0.093047339 | -1.608085829 | 0.107816382 | 0.300770593 | 0.149628108 |
| ENSMUSG00000024725 | Ostf1 | 20409 | protein_coding | 1844.413101 | 0.149664577 | 0.064143501 | 2.333277336 | 0.019633594 | 0.096777726 | 0.149664577 |
| ENSMUSG00000046879 | Irgm1 | 15944 | protein_coding | 1010.605842 | -0.149730198 | 0.215349019 | -0.69529083 | 0.48687306 | 0.707879718 | 0.149730198 |
| ENSMUSG00000026796 | Niban2 | 227737 | protein_coding | 2202.827988 | 0.149745263 | 0.066071703 | 2.266405384 | 0.023426574 | 0.109582031 | 0.149745263 |
| ENSMUSG00000066979 | Bub3 | 12237 | protein_coding | 1106.880298 | 0.149746883 | 0.056962393 | 2.62887276 | 0.008566841 | 0.05321343 | 0.149746883 |
| ENSMUSG00000031706 | Rfx1 | 19724 | protein_coding | 518.3256856 | 0.149770498 | 0.122951914 | 1.21812254 | 0.22317741 | 0.460551754 | 0.149770498 |
| ENSMUSG00000023110 | Prmt5 | 27374 | protein_coding | 806.3141549 | 0.14983586 | 0.077062988 | 1.944329751 | 0.051855699 | 0.18954598 | 0.14983586 |
| ENSMUSG00000027478 | Dnmt3b | 13436 | protein_coding | 66.05442829 | 0.149889586 | 0.180123423 | 0.832149332 | 0.405324659 | 0.644334723 | 0.149889586 |
| ENSMUSG00000026483 | Niban1 | 63913 | protein_coding | 2527.798227 | -0.149922715 | 0.069323527 | -2.162652734 | 0.030567899 | 0.131915576 | 0.149922715 |
| ENSMUSG00000066643 | Wdr35 | 74682 | protein_coding | 49.40298352 | 0.149973528 | 0.170387554 | 0.880190627 | 0.37875605 | 0.619821175 | 0.149973528 |
| ENSMUSG00000032359 | Ctsh | 13036 | protein_coding | 867.9629009 | 0.149973632 | 0.054623881 | 2.745568979 | 0.006040607 | 0.041260849 | 0.149973632 |
| ENSMUSG00000032105 | Pdzd3 | 170761 | protein_coding | 737.3111537 | -0.150001848 | 0.071765804 | -2.090157684 | 0.036603637 | 0.148841871 | 0.150001848 |
| ENSMUSG00000041079 | Rwdd2b | 53858 | protein_coding | 392.095398 | -0.150011181 | 0.082977686 | -1.807849657 | 0.070629903 | 0.231231921 | 0.150011181 |
| ENSMUSG00000096433 | Zfp994 | 240038 | protein_coding | 345.1138109 | -0.150138239 | 0.107927955 | -1.391096855 | 0.164196058 | 0.38635024 | 0.150138239 |
| ENSMUSG00000045414 | Dipk2a | 68861 | protein_coding | 641.081283 | 0.150173219 | 0.049960982 | 3.005809961 | 0.002648745 | 0.02180516 | 0.150173219 |
| ENSMUSG00000030587 | 2200002D01Rik | 72275 | protein_coding | 7758.343469 | -0.150190294 | 0.093375604 | -1.608453258 | 0.107735945 | 0.300630044 | 0.150190294 |
| ENSMUSG00000097254 | C430042M11Rik | 320021 | lncRNA | 50.51111572 | 0.150195909 | 0.168194427 | 0.892989803 | 0.371862633 | 0.613632322 | 0.150195909 |
| ENSMUSG00000061414 | Cracr2a | 381812 | protein_coding | 1532.048805 | -0.150219948 | 0.090510828 | -1.65969035 | 0.096976762 | 0.282657552 | 0.150219948 |
| ENSMUSG00000023892 | Zfp51 | 22709 | protein_coding | 311.5968847 | -0.150220491 | 0.092308971 | -1.62736611 | 0.103659365 | 0.294235769 | 0.150220491 |
| ENSMUSG00000032285 | Dnaja4 | 58233 | protein_coding | 62.16962055 | 0.150263935 | 0.196159195 | 0.766030544 | 0.443658128 | 0.674737597 | 0.150263935 |
| ENSMUSG00000043572 | Pars2 | 230577 | protein_coding | 186.4848747 | -0.150302963 | 0.106676712 | -1.408957594 | 0.158847708 | 0.379011242 | 0.150302963 |
| ENSMUSG00000058656 | Samd12 | 320679 | protein_coding | 362.3377298 | 0.150341835 | 0.126416328 | 1.189259624 | 0.234337518 | 0.473944903 | 0.150341835 |
| ENSMUSG00000009575 | Cbx5 | 12419 | protein_coding | 2486.737005 | 0.15034402 | 0.07653726 | 1.964324575 | 0.049492464 | 0.183993767 | 0.15034402 |
| ENSMUSG00000031626 | Sorbs2 | 234214 | protein_coding | 1121.487539 | -0.150355807 | 0.070760937 | -2.124841942 | 0.033599804 | 0.140900901 | 0.150355807 |
| ENSMUSG00000036898 | Zfp157 | 72154 | protein_coding | 227.9279129 | -0.150384873 | 0.103087899 | -1.45880239 | 0.144619505 | 0.357160205 | 0.150384873 |
| ENSMUSG00000104445 | Rhbg | 58176 | protein_coding | 51.82219952 | -0.150477556 | 0.204793173 | -0.734778183 | 0.462474596 | 0.688854258 | 0.150477556 |
| ENSMUSG00000005533 | Igf1r | 16001 | protein_coding | 527.3909509 | -0.150501391 | 0.084799321 | -1.774794758 | 0.075931782 | 0.242581587 | 0.150501391 |
| ENSMUSG00000031953 | Tmem170 | 66817 | protein_coding | 787.9728095 | 0.150522453 | 0.056046786 | 2.685657184 | 0.007238732 | 0.046962624 | 0.150522453 |
| ENSMUSG00000048234 | Rnf149 | 67702 | protein_coding | 1416.836112 | -0.150535951 | 0.06597104 | -2.281848995 | 0.022498257 | 0.10680877 | 0.150535951 |
| ENSMUSG00000028059 | Arhgef2 | 16800 | protein_coding | 1506.667211 | 0.150580774 | 0.086021788 | 1.750495736 | 0.080032809 | 0.25027906 | 0.150580774 |
| ENSMUSG00000023036 | Pcdhgc4 | 93707 | protein_coding | 178.8343881 | 0.150581183 | 0.117269903 | 1.284056508 | 0.199122184 | 0.432432462 | 0.150581183 |
| ENSMUSG00000032026 | Rexo2 | 104444 | protein_coding | 361.2068881 | 0.150596062 | 0.09819783 | 1.533598684 | 0.125128412 | 0.328281423 | 0.150596062 |
| ENSMUSG00000095217 | H2bc15 | 319187 | protein_coding | 1430.256471 | 0.150628105 | 0.14183297 | 1.062010509 | 0.288230915 | 0.53068843 | 0.150628105 |
| ENSMUSG00000028106 | Rprd2 | 75137 | protein_coding | 924.0184887 | -0.150637406 | 0.053867215 | -2.796458048 | 0.005166612 | 0.036592863 | 0.150637406 |
| ENSMUSG00000048100 | Taf13 | 99730 | protein_coding | 364.2167092 | 0.150672321 | 0.0910525 | 1.654785103 | 0.097968097 | 0.284518548 | 0.150672321 |
| ENSMUSG00000028683 | Eif2b3 | 108067 | protein_coding | 188.9907047 | -0.150690757 | 0.110096848 | -1.368710917 | 0.171089657 | 0.395692369 | 0.150690757 |
| ENSMUSG00000071369 | Map3k5 | 26408 | protein_coding | 1145.001222 | 0.150708179 | 0.052366987 | 2.877923462 | 0.004003022 | 0.030313495 | 0.150708179 |
| ENSMUSG00000028053 | Ash1l | 192195 | protein_coding | 2095.013473 | -0.150721296 | 0.047392362 | -3.180286642 | 0.001471294 | 0.013763089 | 0.150721296 |
| ENSMUSG00000024201 | Kdm4b | 193796 | protein_coding | 1047.753423 | 0.15073448 | 0.06604666 | 2.282242287 | 0.02247504 | 0.106731485 | 0.15073448 |

| ensembl_gene_id | external_gene_name | entrezgene_id | gene_biotype | baseMean | log2FoldChange | lfcSE | stat | pvalue | padj | absLog2FC |
|---|---|---|---|---|---|---|---|---|---|---|
| ENSMUSG00000041565 | L3mbtl4 | 320858 | protein_coding | 7.644783982 | -0.150740792 | 0.455753518 | -0.330750693 | 0.740832807 | 0.875937104 | 0.150740792 |
| ENSMUSG00000028261 | Ndufaf4 | 68493 | protein_coding | 587.7451093 | 0.150825315 | 0.078261554 | 1.927195494 | 0.053955271 | 0.194579678 | 0.150825315 |
| ENSMUSG00000062729 | Ppox | 19044 | protein_coding | 323.8126477 | 0.150828654 | 0.084408282 | 1.786894018 | 0.07395462 | 0.238294935 | 0.150828654 |
| ENSMUSG00000024170 | Telo2 | 71718 | protein_coding | 262.5563276 | -0.150836129 | 0.095044699 | -1.587002021 | 0.112512187 | 0.30863821 | 0.150836129 |
| ENSMUSG00000033813 | Tcea1 | 21399 | protein_coding | 1030.641307 | 0.150839211 | 0.065897554 | 2.288995609 | 0.022079608 | 0.105379648 | 0.150839211 |
| ENSMUSG00000025647 | Shisa5 | 66940 | protein_coding | 3411.320921 | -0.15084544 | 0.066769826 | -2.259185754 | 0.023871833 | 0.111137715 | 0.15084544 |
| ENSMUSG00000018411 | Mapt | 17762 | protein_coding | 112.7836444 | 0.150872115 | 0.146142511 | 1.032362959 | 0.301902118 | 0.545539031 | 0.150872115 |
| ENSMUSG00000037005 | Xpnpep2 | 170745 | protein_coding | 287.8395581 | -0.150879853 | 0.161311361 | -0.935333081 | 0.349616672 | 0.592006307 | 0.150879853 |
| ENSMUSG00000030002 | Dusp11 | 72102 | protein_coding | 2340.362572 | -0.150910896 | 0.040716379 | -3.706392862 | 0.000210232 | 0.002747673 | 0.150910896 |
| ENSMUSG00000021712 | Trim23 | 81003 | protein_coding | 259.7698114 | -0.150934981 | 0.105052192 | -1.436761842 | 0.150785676 | 0.366529497 | 0.150934981 |
| ENSMUSG00000015944 | Castor2 | 80909 | protein_coding | 138.9198859 | -0.15094345 | 0.1681685 | -0.897572676 | 0.369413411 | 0.611593275 | 0.15094345 |
| ENSMUSG00000013766 | Ly6g6e | 70274 | protein_coding | 190.2314492 | 0.150955713 | 0.146466124 | 1.030652748 | 0.30270369 | 0.546479617 | 0.150955713 |
| ENSMUSG00000039126 | Prune2 | 353211 | protein_coding | 21.73305689 | 0.150964956 | 0.269726224 | 0.559696991 | 0.575686135 | 0.76862707 | 0.150964956 |
| ENSMUSG00000041617 | Ccdc74a | 72315 | protein_coding | 18.16940681 | -0.150992702 | 0.293081324 | -0.51519046 | 0.606419935 | 0.789753 | 0.150992702 |
| ENSMUSG00000071076 | Jund | 16478 | protein_coding | 5503.124654 | 0.151066047 | 0.11135227 | 1.356649909 | 0.174892475 | 0.401207817 | 0.151066047 |
| ENSMUSG00000029020 | Mfn2 | 170731 | protein_coding | 2441.338271 | 0.151125257 | 0.060132386 | 2.513209072 | 0.011963841 | 0.067936924 | 0.151125257 |
| ENSMUSG00000026349 | Ccnt2 | 72949 | protein_coding | 1274.69483 | -0.151131462 | 0.047985669 | -3.149512451 | 0.001635431 | 0.01497706 | 0.151131462 |
| ENSMUSG00000068245 | Phf11d | 219132 | protein_coding | 104.8186231 | -0.151137568 | 0.173068227 | -0.873283161 | 0.382508757 | 0.623228806 | 0.151137568 |
| ENSMUSG00000042228 | Lyn | 17096 | protein_coding | 591.8317919 | 0.151141198 | 0.073018052 | 2.069915495 | 0.038460259 | 0.154385176 | 0.151141198 |
| ENSMUSG00000082361 | Btc | 12223 | protein_coding | 169.9628819 | 0.151142521 | 0.129823667 | 1.164213924 | 0.244337329 | 0.485573395 | 0.151142521 |
| ENSMUSG00000036672 | Cenpt | 320394 | protein_coding | 135.464013 | 0.151147543 | 0.129765138 | 1.164777733 | 0.244108974 | 0.485418672 | 0.151147543 |
| ENSMUSG00000095253 | Zfp799 | 240064 | protein_coding | 376.9700667 | -0.151159164 | 0.094210025 | -1.604491281 | 0.108605805 | 0.302001032 | 0.151159164 |
| ENSMUSG00000029640 | Usp12 | 22217 | protein_coding | 1744.424168 | -0.151179384 | 0.051709745 | -2.923614978 | 0.003459924 | 0.027024167 | 0.151179384 |
| ENSMUSG00000031174 | Rpgr | 19893 | protein_coding | 148.0749589 | -0.151180025 | 0.103272392 | -1.463895836 | 0.143222411 | 0.355127279 | 0.151180025 |
| ENSMUSG00000002948 | Map2k7 | 26400 | protein_coding | 435.6827551 | -0.151256066 | 0.081962204 | -1.845436793 | 0.064974025 | 0.219350242 | 0.151256066 |
| ENSMUSG00000026335 | Pam | 18484 | protein_coding | 6080.78666 | -0.151268806 | 0.054156592 | -2.793174378 | 0.005219355 | 0.036869266 | 0.151268806 |
| ENSMUSG00000044066 | Cep68 | 216543 | protein_coding | 532.1253352 | -0.151352502 | 0.070594833 | -2.143960068 | 0.032036087 | 0.136174477 | 0.151352502 |
| ENSMUSG00000030966 | Trim21 | 20821 | protein_coding | 299.2324462 | -0.151461429 | 0.104451703 | -1.450061837 | 0.147041276 | 0.360385206 | 0.151461429 |
| ENSMUSG00000086213 | A330040F15Rik | 74333 | lncRNA | 16.84003635 | 0.151473278 | 0.30683271 | 0.493667307 | 0.621541158 | 0.800250831 | 0.151473278 |
| ENSMUSG00000094338 | H2bc13 | 319185 | protein_coding | 598.6496911 | 0.151518504 | 0.130263496 | 1.163169337 | 0.244760805 | 0.486003975 | 0.151518504 |
| ENSMUSG00000030321 | Efcab12 | 212516 | protein_coding | 10.25033042 | 0.1515285 | 0.397487187 | 0.38121606 | 0.703042933 | 0.852351028 | 0.1515285 |
| ENSMUSG00000041396 | Mettl18 | 69962 | protein_coding | 81.95732947 | 0.151532392 | 0.145003452 | 1.045026101 | 0.296010906 | 0.538897588 | 0.151532392 |
| ENSMUSG00000071424 | Grid2 | 14804 | protein_coding | 56.11830631 | 0.151585276 | 0.167585215 | 0.904526547 | 0.365716264 | 0.60771831 | 0.151585276 |
| ENSMUSG00000039197 | Adk | 11534 | protein_coding | 3671.928562 | -0.151585876 | 0.049401985 | -3.068416692 | 0.002151963 | 0.01858849 | 0.151585876 |
| ENSMUSG00000032405 | Pias1 | 56469 | protein_coding | 554.7604383 | -0.1516466 | 0.059596634 | -2.544549772 | 0.010941872 | 0.063224495 | 0.1516466 |
| ENSMUSG00000030316 | Tamm41 | 68971 | protein_coding | 294.7683831 | -0.151655397 | 0.080260231 | -1.889545983 | 0.058818707 | 0.205867649 | 0.151655397 |
| ENSMUSG00000038807 | Rap1gap2 | 380711 | protein_coding | 3425.936 | -0.151686639 | 0.059884544 | -2.532984769 | 0.011309587 | 0.064907368 | 0.151686639 |
| ENSMUSG00000038368 | Focad | 230393 | protein_coding | 382.9973426 | 0.151705784 | 0.089455204 | 1.695885511 | 0.089907564 | 0.269655809 | 0.151705784 |
| ENSMUSG00000030103 | Bhlhe40 | 20893 | protein_coding | 3831.783123 | -0.15174677 | 0.072292807 | -2.09905765 | 0.035811819 | 0.146771529 | 0.15174677 |

| ensembl_gene_id | external_gene_name | entrezgene_id | gene_biotype | baseMean | log2FoldChange | lfcSE | stat | pvalue | padj | absLog2FC |
|---|---|---|---|---|---|---|---|---|---|---|
| ENSMUSG00000024580 | Grpel2 | 17714 | protein_coding | 1535.162614 | -0.15178912 | 0.070761098 | -2.145092784 | 0.031945429 | 0.135932437 | 0.15178912 |
| ENSMUSG00000044139 | Prss53 | 330657 | protein_coding | 8.274481969 | 0.151793082 | 0.415092038 | 0.365685363 | 0.71459987 | 0.859633975 | 0.151793082 |
| ENSMUSG00000098789 | Jmjd7 | 433466 | protein_coding | 28.39397962 | -0.151807767 | 0.243967206 | -0.622246611 | 0.533779721 | 0.741516545 | 0.151807767 |
| ENSMUSG00000036698 | Ago2 | 239528 | protein_coding | 1802.072388 | -0.151812466 | 0.085846779 | -1.768411903 | 0.076992068 | 0.24470853 | 0.151812466 |
| ENSMUSG00000035762 | Tmem161b | 72745 | protein_coding | 426.1859471 | 0.151858162 | 0.060853103 | 2.495487584 | 0.012578416 | 0.070557256 | 0.151858162 |
| ENSMUSG00000070704 | Ugt2b36 | 231396 | protein_coding | 112.2092778 | -0.151948439 | 0.125079866 | -1.214811334 | 0.224438057 | 0.462439318 | 0.151948439 |
| ENSMUSG00000091811 | Inafm1 | 66300 | protein_coding | 561.4923298 | -0.151965168 | 0.096943634 | -1.567562108 | 0.116983361 | 0.31618145 | 0.151965168 |
| ENSMUSG00000062794 | Zfp599 | 235048 | protein_coding | 124.4904094 | -0.151982873 | 0.111710581 | -1.360505618 | 0.173669976 | 0.399424529 | 0.151982873 |
| ENSMUSG00000003644 | Rps6ka1 | 20111 | protein_coding | 6098.125466 | 0.151986273 | 0.067120782 | 2.26436981 | 0.023551379 | 0.110048838 | 0.151986273 |
| ENSMUSG00000025132 | Arhgdia | 192662 | protein_coding | 5369.412411 | 0.15203892 | 0.059425536 | 2.558477867 | 0.010513151 | 0.061541049 | 0.15203892 |
| ENSMUSG00000002658 | Gtf2f1 | 98053 | protein_coding | 1102.748707 | 0.152060694 | 0.054760927 | 2.776810073 | 0.005489525 | 0.038300296 | 0.152060694 |
| ENSMUSG00000051335 | Gfod1 | 328232 | protein_coding | 183.2603816 | -0.152145047 | 0.113570989 | -1.339647111 | 0.1803601 | 0.407326553 | 0.152145047 |
| ENSMUSG00000040097 | Flywch1 | 224613 | protein_coding | 528.3594419 | 0.152150301 | 0.06951257 | 2.188817077 | 0.028610138 | 0.125858073 | 0.152150301 |
| ENSMUSG00000056445 | Hoxaas2 | 105242657 | lncRNA | 16.05051467 | -0.152175627 | 0.311781492 | -0.488084223 | 0.625490186 | 0.80236082 | 0.152175627 |
| ENSMUSG00000070291 | Ddx43 | 100048658 | protein_coding | 37.08666781 | -0.152194114 | 0.265138801 | -0.574016753 | 0.565956465 | 0.76212257 | 0.152194114 |
| ENSMUSG00000027775 | Mfsd1 | 66868 | protein_coding | 1500.448218 | -0.152204537 | 0.045502989 | -3.344934936 | 0.000823019 | 0.008563112 | 0.152204537 |
| ENSMUSG00000047246 | H2bc6 | 319179 | protein_coding | 1306.597228 | 0.152209397 | 0.126661746 | 1.201699818 | 0.229479852 | 0.468848355 | 0.152209397 |
| ENSMUSG00000056413 | Adap1 | 231821 | protein_coding | 1208.895971 | -0.152262729 | 0.07014901 | -2.170561338 | 0.029964347 | 0.130076724 | 0.152262729 |
| ENSMUSG00000001228 | Uhrf1 | 18140 | protein_coding | 709.8074092 | 0.152302193 | 0.170673519 | 0.892359833 | 0.372200093 | 0.61393518 | 0.152302193 |
| ENSMUSG00000027163 | Commd9 | 76501 | protein_coding | 521.4329305 | 0.152319021 | 0.076183817 | 1.999361901 | 0.045569211 | 0.173857267 | 0.152319021 |
| ENSMUSG00000030887 | Pdzd9 | 67983 | protein_coding | 8.410683808 | 0.152447275 | 0.423683523 | 0.359814028 | 0.718986212 | 0.862409121 | 0.152447275 |
| ENSMUSG00000090112 | Shprh | 268281 | protein_coding | 720.1465691 | -0.15245756 | 0.067035307 | -2.274287485 | 0.022948707 | 0.10799004 | 0.15245756 |
| ENSMUSG00000058979 | Hdhd5 | 214932 | protein_coding | 396.1723935 | 0.152530578 | 0.093127446 | 1.637869226 | 0.101448977 | 0.290288795 | 0.152530578 |
| ENSMUSG00000021749 | Fam3d | 18300 | protein_coding | 19095.88922 | 0.152598532 | 0.062770055 | 2.43107214 | 0.015054217 | 0.08032198 | 0.152598532 |
| ENSMUSG00000041308 | Sntb2 | 20650 | protein_coding | 354.2784856 | -0.152658543 | 0.071121892 | -2.146435348 | 0.031838262 | 0.135619558 | 0.152658543 |
| ENSMUSG00000007837 | Prrg2 | 65116 | protein_coding | 777.8287436 | 0.152739881 | 0.087587921 | 1.743846393 | 0.081185878 | 0.252562627 | 0.152739881 |
| ENSMUSG00000051499 | Zfp786 | 330301 | protein_coding | 38.41277466 | 0.152745816 | 0.211343047 | 0.722738781 | 0.469840385 | 0.694009261 | 0.152745816 |
| ENSMUSG00000000976 | Heatr6 | 217026 | protein_coding | 488.350577 | 0.152787161 | 0.069933557 | 2.184747445 | 0.028907369 | 0.126695316 | 0.152787161 |
| ENSMUSG00000110195 | Pde2a | 207728 | protein_coding | 32.36666922 | 0.152787653 | 0.245364806 | 0.622695875 | 0.533484394 | 0.741375537 | 0.152787653 |
| ENSMUSG00000117698 | BC037704 | 100502982 | lncRNA | 23.27659817 | 0.152824826 | 0.259560356 | 0.588783387 | 0.556006592 | 0.755288004 | 0.152824826 |
| ENSMUSG00000058392 | Rrp1b | 72462 | protein_coding | 196.2689048 | -0.152835919 | 0.103635006 | -1.474751872 | 0.140279265 | 0.351340845 | 0.152835919 |
| ENSMUSG00000022883 | Robo1 | 19876 | protein_coding | 27.88596176 | 0.152838072 | 0.231332789 | 0.660684865 | 0.508814432 | 0.724049267 | 0.152838072 |
| ENSMUSG00000051989 | Smim11 | 68936 | protein_coding | 329.7369183 | 0.152878015 | 0.068210846 | 2.241256693 | 0.025009452 | 0.114712014 | 0.152878015 |
| ENSMUSG00000030986 | Dhx32 | 101437 | protein_coding | 3432.097049 | 0.152882584 | 0.047940443 | 3.189010673 | 0.001427606 | 0.013434123 | 0.152882584 |
| ENSMUSG00000038637 | Lrrc56 | 70552 | protein_coding | 190.2562512 | 0.152917437 | 0.097184776 | 1.573471103 | 0.115609774 | 0.314143021 | 0.152917437 |
| ENSMUSG00000022754 | Tmem45a | 56277 | protein_coding | 7.482866174 | -0.152946938 | 0.569762074 | -0.268440011 | 0.788360645 | 0.90131094 | 0.152946938 |
| ENSMUSG00000021951 | Eef1akmt1 | 68043 | protein_coding | 287.1756997 | 0.15297159 | 0.079374047 | 1.927224272 | 0.053951686 | 0.194579678 | 0.15297159 |
| ENSMUSG00000031090 | Nadsyn1 | 78914 | protein_coding | 1280.109429 | -0.153035181 | 0.049754521 | -3.07580451 | 0.002099354 | 0.018231711 | 0.153035181 |
| ENSMUSG00000016933 | Plcg1 | 18803 | protein_coding | 541.3378657 | -0.153036226 | 0.059693965 | -2.56368001 | 0.010356898 | 0.060891613 | 0.153036226 |

| ensembl_gene_id | external_gene_name | entrezgene_id | gene_biotype | baseMean | log2FoldChange | lfcSE | stat | pvalue | padj | absLog2FC |
|---|---|---|---|---|---|---|---|---|---|---|
| ENSMUSG00000067006 | Serpinb5 | 20724 | protein_coding | 1899.621595 | -0.153044941 | 0.055492707 | -2.757928902 | 0.005816885 | 0.040151712 | 0.153044941 |
| ENSMUSG00000031684 | Slc10a7 | 76775 | protein_coding | 649.7313096 | 0.153050102 | 0.070249758 | 2.178656639 | 0.029357185 | 0.128165492 | 0.153050102 |
| ENSMUSG00000059991 | Nptx2 | 53324 | protein_coding | 11.42087184 | -0.15305841 | 0.368517795 | -0.415335194 | 0.67789654 | 0.836585484 | 0.15305841 |
| ENSMUSG00000044145 | 1810024B03Rik | 329509 | protein_coding | 25.13434283 | 0.153063456 | 0.246699079 | 0.620445998 | 0.534964197 | 0.741885541 | 0.153063456 |
| ENSMUSG00000050295 | Foxc1 | 17300 | protein_coding | 19.79350262 | -0.153108315 | 0.355997513 | -0.430082543 | 0.667135598 | 0.830161713 | 0.153108315 |
| ENSMUSG00000032068 | Plet1 | 76509 | protein_coding | 936.4121934 | -0.153121638 | 0.141184185 | -1.084552334 | 0.278119976 | 0.521009858 | 0.153121638 |
| ENSMUSG00000027665 | Pik3ca | 18706 | protein_coding | 1537.498709 | -0.153157117 | 0.069266076 | -2.211141808 | 0.027026019 | 0.120633435 | 0.153157117 |
| ENSMUSG00000030879 | Mrpl17 | 27397 | protein_coding | 848.4708342 | 0.153194217 | 0.055775632 | 2.74661555 | 0.006021367 | 0.041146859 | 0.153194217 |
| ENSMUSG00000043020 | Dnai3 | 242253 | protein_coding | 19.13548929 | 0.153260531 | 0.302939908 | 0.505910667 | 0.612919364 | 0.793748541 | 0.153260531 |
| ENSMUSG00000020142 | Slc1a4 | 55963 | protein_coding | 771.5973436 | -0.15326256 | 0.0839939 | -1.824686786 | 0.068048309 | 0.225933166 | 0.15326256 |
| ENSMUSG00000055692 | Tmem191 | 224019 | protein_coding | 65.67989636 | -0.153287511 | 0.191238363 | -0.801552096 | 0.422812098 | 0.658779673 | 0.153287511 |
| ENSMUSG00000043592 | Unc5cl | 76589 | protein_coding | 178.7237565 | 0.153359425 | 0.104290759 | 1.470498701 | 0.141426737 | 0.352354239 | 0.153359425 |
| ENSMUSG00000068323 | Slc4a5 | 232156 | protein_coding | 10.71054535 | -0.153373365 | 0.403631883 | -0.379983276 | 0.703957829 | 0.853011339 | 0.153373365 |
| ENSMUSG00000035722 | Abca7 | 27403 | protein_coding | 1764.922452 | 0.153414401 | 0.065521192 | 2.341447053 | 0.01920915 | 0.09546563 | 0.153414401 |
| ENSMUSG00000094091 | Ankrd40cl | 102639132 | protein_coding | 929.9471125 | 0.153422792 | 0.116431603 | 1.317707465 | 0.187601594 | 0.417346835 | 0.153422792 |
| ENSMUSG00000038375 | Trp53inp2 | 68728 | protein_coding | 2455.466194 | 0.153434009 | 0.067378664 | 2.277189833 | 0.022774892 | 0.107552177 | 0.153434009 |
| ENSMUSG00000032358 | Fam83b | 208994 | protein_coding | 1182.713842 | -0.153452718 | 0.048432356 | -3.168392613 | 0.001532844 | 0.014247936 | 0.153452718 |
| ENSMUSG00000033488 | Cryzl2 | 226527 | protein_coding | 252.1486871 | 0.153461124 | 0.110761063 | 1.38551509 | 0.165895022 | 0.389317377 | 0.153461124 |
| ENSMUSG00000030276 | Ttll3 | 101100 | protein_coding | 89.51720685 | 0.153485511 | 0.202482323 | 0.758019313 | 0.448439421 | 0.678988684 | 0.153485511 |
| ENSMUSG00000027692 | Tnik | 665113 | protein_coding | 822.6687934 | -0.153500273 | 0.062810882 | -2.443848394 | 0.014531532 | 0.078378495 | 0.153500273 |
| ENSMUSG00000045100 | Slc25a26 | 67582 | protein_coding | 170.2651539 | -0.153550562 | 0.100331134 | -1.530437824 | 0.125908393 | 0.329416718 | 0.153550562 |
| ENSMUSG00000050288 | Fzd2 | 57265 | protein_coding | 39.61368619 | 0.153576659 | 0.190558401 | 0.805929618 | 0.420283432 | 0.657347582 | 0.153576659 |
| ENSMUSG00000106847 | Peg13 | 353342 | lncRNA | 309.467718 | 0.153580102 | 0.113705201 | 1.350686691 | 0.176795817 | 0.403679194 | 0.153580102 |
| ENSMUSG00000001151 | Pcnt | 18541 | protein_coding | 1530.515878 | 0.153596367 | 0.059936623 | 2.562646319 | 0.010387781 | 0.061028685 | 0.153596367 |
| ENSMUSG00000038280 | Ostm1 | 14628 | protein_coding | 325.2018139 | -0.153603835 | 0.07866028 | -1.952749653 | 0.050849267 | 0.187225144 | 0.153603835 |
| ENSMUSG00000036968 | Cnpy4 | 66455 | protein_coding | 245.8250369 | 0.153635156 | 0.081248023 | 1.890940234 | 0.058632323 | 0.205334087 | 0.153635156 |
| ENSMUSG00000030987 | Stim1 | 20866 | protein_coding | 3460.704726 | -0.153643544 | 0.074708233 | -2.056581144 | 0.039726534 | 0.157761587 | 0.153643544 |
| ENSMUSG00000058056 | Palld | 72333 | protein_coding | 345.3278735 | -0.15367435 | 0.121280195 | -1.267101765 | 0.205118905 | 0.439595026 | 0.15367435 |
| ENSMUSG00000086247 | Gm15787 | 100504007 | lncRNA | 46.19571788 | -0.153676976 | 0.226090699 | -0.679713835 | 0.496685674 | 0.714991508 | 0.153676976 |
| ENSMUSG00000050846 | Zfp623 | 78834 | protein_coding | 407.287262 | -0.153700811 | 0.069172912 | -2.221979775 | 0.026284672 | 0.118778578 | 0.153700811 |
| ENSMUSG00000027002 | Nckap1 | 50884 | protein_coding | 5182.608314 | 0.153743959 | 0.044336661 | 3.467648609 | 0.000525033 | 0.005904611 | 0.153743959 |
| ENSMUSG00000032528 | Vipr1 | 22354 | protein_coding | 7014.362101 | -0.153765097 | 0.054632773 | -2.814521187 | 0.004884996 | 0.034966856 | 0.153765097 |
| ENSMUSG00000034575 | Tent4a | 210106 | protein_coding | 493.0452078 | -0.153779501 | 0.059683168 | -2.576597485 | 0.009977807 | 0.059145001 | 0.153779501 |
| ENSMUSG00000018554 | Ybx2 | 53422 | protein_coding | 169.2414737 | -0.153793231 | 0.104332438 | -1.474069182 | 0.140462965 | 0.351500309 | 0.153793231 |
| ENSMUSG00000062012 | Zfp13 | 22654 | protein_coding | 76.31216762 | -0.154000241 | 0.13846258 | -1.112215595 | 0.266045464 | 0.508053902 | 0.154000241 |
| ENSMUSG00000029484 | Anxa3 | 11745 | protein_coding | 2151.250281 | 0.154047192 | 0.104610278 | 1.472581814 | 0.140863831 | 0.35197344 | 0.154047192 |
| ENSMUSG00000006611 | Hfe | 15216 | protein_coding | 189.4924621 | -0.154085282 | 0.092834429 | -1.659785965 | 0.096957518 | 0.282652548 | 0.154085282 |
| ENSMUSG00000036333 | Kidins220 | 77480 | protein_coding | 3126.657114 | -0.154182626 | 0.042033752 | -3.66806721 | 0.000244391 | 0.003113373 | 0.154182626 |
| ENSMUSG00000000394 | Gcg | 14526 | protein_coding | 1244.777209 | -0.154213396 | 0.089091214 | -1.730960767 | 0.083458762 | 0.257239637 | 0.154213396 |

Supplementary Table 1

| ensembl_gene_id | external_gene_name | entrezgene_id | gene_biotype | baseMean | log2FoldChange | lfcSE | stat | pvalue | padj | absLog2FC |
|---|---|---|---|---|---|---|---|---|---|---|
| ENSMUSG00000024937 | Ehbp1l1 | 114601 | protein_coding | 1096.296729 | 0.154228284 | 0.096807235 | 1.593148318 | 0.111126922 | 0.306506563 | 0.154228284 |
| ENSMUSG00000044716 | Dok7 | 231134 | protein_coding | 132.6131067 | -0.15433492 | 0.139645625 | -1.105189793 | 0.269077359 | 0.511602283 | 0.15433492 |
| ENSMUSG00000053141 | Ptprt | 19281 | protein_coding | 54.53913452 | -0.154349165 | 0.201636404 | -0.765482629 | 0.443984206 | 0.674900387 | 0.154349165 |
| ENSMUSG00000031360 | Ctps2 | 55936 | protein_coding | 577.3558795 | -0.154355554 | 0.065058216 | -2.372575879 | 0.017664534 | 0.090113624 | 0.154355554 |
| ENSMUSG00000040952 | Rps19 | 20085 | protein_coding | 10118.04715 | 0.154359216 | 0.048013233 | 3.214930706 | 0.00130476 | 0.012548825 | 0.154359216 |
| ENSMUSG00000046082 | Tmem174 | 68344 | protein_coding | 28.39715562 | 0.154462286 | 0.320006237 | 0.482685235 | 0.629319249 | 0.804903761 | 0.154462286 |
| ENSMUSG00000027225 | Duoxa2 | 66811 | protein_coding | 60.79215394 | 0.154499633 | 0.17573557 | 0.879159708 | 0.379314688 | 0.620491412 | 0.154499633 |
| ENSMUSG00000070639 | Lrrc8b | 433926 | protein_coding | 653.6769062 | -0.154533482 | 0.083117897 | -1.859208284 | 0.062997621 | 0.21465511 | 0.154533482 |
| ENSMUSG00000050957 | Insl6 | 27356 | protein_coding | 91.14993614 | 0.154588976 | 0.135797718 | 1.138376832 | 0.254963164 | 0.496173639 | 0.154588976 |
| ENSMUSG00000019487 | Trip10 | 106628 | protein_coding | 1140.409603 | -0.154594699 | 0.064818341 | -2.38504558 | 0.017077005 | 0.088092517 | 0.154594699 |
| ENSMUSG00000013089 | Etv5 | 104156 | protein_coding | 516.5096674 | -0.154619404 | 0.121116344 | -1.276618816 | 0.201736856 | 0.435615504 | 0.154619404 |
| ENSMUSG00000031805 | Jak3 | 16453 | protein_coding | 192.6183276 | 0.1546395 | 0.121827589 | 1.269330704 | 0.20432314 | 0.438355631 | 0.1546395 |
| ENSMUSG00000029208 | Guf1 | 231279 | protein_coding | 624.0444026 | -0.154651316 | 0.081880635 | -1.88874105 | 0.058926535 | 0.205917277 | 0.154651316 |
| ENSMUSG00000039178 | Tbc1d19 | 67249 | protein_coding | 144.7703104 | 0.154668886 | 0.11966789 | 1.292484445 | 0.196189429 | 0.428764731 | 0.154668886 |
| ENSMUSG00000032306 | Mpi | 110119 | protein_coding | 1671.313905 | 0.154695932 | 0.053045971 | 2.916261678 | 0.003542532 | 0.027572597 | 0.154695932 |
| ENSMUSG00000028864 | Hgf | 15234 | protein_coding | 6.580847793 | 0.15470141 | 0.463832521 | 0.333528597 | 0.738735307 | 0.87479925 | 0.15470141 |
| ENSMUSG00000035266 | Helq | 191578 | protein_coding | 163.4380269 | 0.154721698 | 0.118784968 | 1.302536012 | 0.192733216 | 0.42444695 | 0.154721698 |
| ENSMUSG00000035851 | Ythdc1 | 231386 | protein_coding | 1067.974913 | -0.154778108 | 0.085452443 | -1.811277733 | 0.070097871 | 0.230050541 | 0.154778108 |
| ENSMUSG00000035208 | Slfn8 | 276950 | protein_coding | 7.62187677 | 0.154819194 | 0.435002165 | 0.355904422 | 0.721912144 | 0.864080715 | 0.154819194 |
| ENSMUSG00000028851 | Nudc | 18221 | protein_coding | 1047.305117 | 0.154852178 | 0.077043305 | 2.009936851 | 0.044437873 | 0.170874958 | 0.154852178 |
| ENSMUSG00000034813 | Grip1 | 74053 | protein_coding | 39.24437621 | -0.15486918 | 0.208702169 | -0.742058313 | 0.458052006 | 0.685606255 | 0.15486918 |
| ENSMUSG00000061306 | Slc38a10 | 72055 | protein_coding | 3016.333909 | 0.154935404 | 0.047933134 | 3.232323672 | 0.001227879 | 0.011943307 | 0.154935404 |
| ENSMUSG00000027806 | Tsc22d2 | 72033 | protein_coding | 742.8098874 | -0.154943899 | 0.098501572 | -1.573009407 | 0.11571664 | 0.314327481 | 0.154943899 |
| ENSMUSG00000021385 | Ippk | 75678 | protein_coding | 786.0564412 | 0.15497281 | 0.099080533 | 1.564109574 | 0.117791838 | 0.317250963 | 0.15497281 |
| ENSMUSG00000013973 | Dedd | 21945 | protein_coding | 170.1362583 | -0.154996432 | 0.115667884 | -1.340012688 | 0.18024122 | 0.407223247 | 0.154996432 |
| ENSMUSG00000022892 | App | 11820 | protein_coding | 8814.236578 | -0.155129112 | 0.036126094 | -4.294101416 | 1.75E-05 | 0.000320903 | 0.155129112 |
| ENSMUSG00000002835 | Chaf1a | 27221 | protein_coding | 240.4835874 | 0.155148699 | 0.157759404 | 0.983451347 | 0.32538535 | 0.569142032 | 0.155148699 |
| ENSMUSG00000035356 | Nfkbiz | 80859 | protein_coding | 1389.393079 | -0.155172198 | 0.106692526 | -1.45438677 | 0.145839106 | 0.358583208 | 0.155172198 |
| ENSMUSG00000026361 | Cdc73 | 214498 | protein_coding | 674.0278168 | 0.155180779 | 0.077074805 | 2.013378811 | 0.044074793 | 0.170111172 | 0.155180779 |
| ENSMUSG00000021379 | Id4 | 15904 | protein_coding | 85.82660022 | 0.155187282 | 0.136321312 | 1.138393399 | 0.254956249 | 0.496173639 | 0.155187282 |
| ENSMUSG00000029209 | Gnpda2 | 67980 | protein_coding | 137.0191513 | -0.155203949 | 0.135440397 | -1.145920659 | 0.251827982 | 0.492807896 | 0.155203949 |
| ENSMUSG00000034472 | Rasd2 | 75141 | protein_coding | 712.5170797 | -0.155233911 | 0.105313554 | -1.474016445 | 0.140477163 | 0.351500309 | 0.155233911 |
| ENSMUSG00000027765 | P2ry1 | 18441 | protein_coding | 679.6964146 | -0.155322266 | 0.077333092 | -2.008483848 | 0.044591901 | 0.171324583 | 0.155322266 |
| ENSMUSG00000013539 | Tango2 | 27883 | protein_coding | 378.7180717 | 0.155333486 | 0.07092666 | 2.190057831 | 0.028520043 | 0.125530222 | 0.155333486 |
| ENSMUSG00000021646 | Mccc2 | 78038 | protein_coding | 724.2412278 | 0.15533369 | 0.054037796 | 2.87453783 | 0.004046193 | 0.03050629 | 0.15533369 |
| ENSMUSG00000028719 | Cmpk1 | 66588 | protein_coding | 10029.43738 | 0.155359309 | 0.04109467 | 3.780522095 | 0.0001565 | 0.002142507 | 0.155359309 |
| ENSMUSG00000077714 | Snord17 | 100313519 | snoRNA | 1784.764122 | 0.15537936 | 0.248040062 | 0.626428482 | 0.531033933 | 0.73971984 | 0.15537936 |
| ENSMUSG00000084786 | Ubl5 | 66177 | protein_coding | 1955.365677 | 0.15541962 | 0.059971899 | 2.591540758 | 0.009554722 | 0.05751736 | 0.15541962 |
| ENSMUSG00000009894 | Snap47 | 67826 | protein_coding | 358.3664629 | 0.155455676 | 0.071905433 | 2.161946173 | 0.030622325 | 0.132046349 | 0.155455676 |

| ensembl_gene_id | external_gene_name | entrezgene_id | gene_biotype | baseMean | log2FoldChange | lfcSE | stat | pvalue | padj | absLog2FC |
|---|---|---|---|---|---|---|---|---|---|---|
| ENSMUSG00000005952 | Trpv1 | 193034 | protein_coding | 10.91501967 | -0.155536873 | 0.382400483 | -0.406738172 | 0.684200301 | 0.84094949 | 0.155536873 |
| ENSMUSG00000020115 | Tbk1 | 56480 | protein_coding | 949.8571228 | -0.155562727 | 0.064187122 | -2.423581595 | 0.0153683 | 0.081578678 | 0.155562727 |
| ENSMUSG00000028936 | Rpl22 | 19934 | protein_coding | 4654.072179 | -0.155568899 | 0.035794109 | -4.346215138 | 1.39E-05 | 0.000261864 | 0.155568899 |
| ENSMUSG00000055334 | Snupn | 66069 | protein_coding | 177.1073305 | 0.155577555 | 0.093349893 | 1.666606672 | 0.095592641 | 0.280216348 | 0.155577555 |
| ENSMUSG00000034311 | Kif4 | 16571 | protein_coding | 543.5073968 | 0.155581567 | 0.11917508 | 1.305487414 | 0.19172693 | 0.423163427 | 0.155581567 |
| ENSMUSG00000070473 | Cldn3 | 12739 | protein_coding | 15013.89998 | 0.155588037 | 0.075491872 | 2.060990573 | 0.039303939 | 0.156629283 | 0.155588037 |
| ENSMUSG00000038290 | Smg6 | 103677 | protein_coding | 976.6085545 | -0.155605946 | 0.052044877 | -2.989841751 | 0.00279122 | 0.022791902 | 0.155605946 |
| ENSMUSG00000040525 | Cblc | 80794 | protein_coding | 1585.139466 | 0.155615078 | 0.077501574 | 2.007895706 | 0.044654376 | 0.171421356 | 0.155615078 |
| ENSMUSG00000036501 | Fam13b | 225358 | protein_coding | 1199.139726 | -0.155616231 | 0.082302361 | -1.890786965 | 0.058652788 | 0.205361161 | 0.155616231 |
| ENSMUSG00000057531 | Dtnbp1 | 94245 | protein_coding | 336.7171547 | -0.155651034 | 0.068720944 | -2.264972297 | 0.02351438 | 0.109949666 | 0.155651034 |
| ENSMUSG00000027007 | Itprid2 | 70599 | protein_coding | 3114.731301 | -0.155685755 | 0.054541863 | -2.854426801 | 0.004311457 | 0.031970135 | 0.155685755 |
| ENSMUSG00000021048 | Mthfd1 | 108156 | protein_coding | 1976.284949 | -0.155700083 | 0.088855468 | -1.752284776 | 0.079724853 | 0.249745186 | 0.155700083 |
| ENSMUSG00000069520 | Tmem19 | 67226 | protein_coding | 976.6015659 | 0.155727468 | 0.055509009 | 2.805444932 | 0.005024717 | 0.035808045 | 0.155727468 |
| ENSMUSG00000022119 | Rbm26 | 74213 | protein_coding | 721.062899 | -0.155749595 | 0.082675769 | -1.883860259 | 0.05958388 | 0.207235951 | 0.155749595 |
| ENSMUSG00000032232 | Cgnl1 | 68178 | protein_coding | 1070.353967 | 0.155816668 | 0.047750551 | 3.26313864 | 0.001101856 | 0.010921716 | 0.155816668 |
| ENSMUSG00000085184 | 4933439K11Rik | 71319 | lncRNA | 54.61450134 | 0.155830076 | 0.198742169 | 0.784081592 | 0.432992236 | 0.667195279 | 0.155830076 |
| ENSMUSG00000017707 | Serinc3 | 26943 | protein_coding | 9001.232553 | -0.155947132 | 0.048759442 | -3.198296052 | 0.001382423 | 0.01306819 | 0.155947132 |
| ENSMUSG00000019891 | Dcbld1 | 66686 | protein_coding | 757.8626928 | 0.155972137 | 0.064358891 | 2.423474599 | 0.015372828 | 0.081578678 | 0.155972137 |
| ENSMUSG00000039463 | Slc9a8 | 77031 | protein_coding | 1089.442821 | -0.15600237 | 0.049126346 | -3.175533763 | 0.001495611 | 0.01394449 | 0.15600237 |
| ENSMUSG00000028423 | Nfx1 | 74164 | protein_coding | 1745.162513 | -0.156021336 | 0.077120016 | -2.023097808 | 0.043063065 | 0.167263497 | 0.156021336 |
| ENSMUSG00000034574 | Daam1 | 208846 | protein_coding | 1739.073348 | -0.156127547 | 0.048114703 | -3.24490304 | 0.001174906 | 0.011511368 | 0.156127547 |
| ENSMUSG00000039187 | Fanci | 208836 | protein_coding | 101.8018175 | 0.156160179 | 0.171684152 | 0.909578302 | 0.363044946 | 0.605769124 | 0.156160179 |
| ENSMUSG00000039826 | Trub2 | 227682 | protein_coding | 263.25372 | -0.156163949 | 0.088903413 | -1.756557408 | 0.078993278 | 0.248328382 | 0.156163949 |
| ENSMUSG00000054836 | Elp6 | 72341 | protein_coding | 125.4703928 | 0.156172653 | 0.116631188 | 1.33902994 | 0.180560926 | 0.407535035 | 0.156172653 |
| ENSMUSG00000027457 | Snph | 241727 | protein_coding | 121.1789399 | -0.156178049 | 0.150914659 | -1.034876593 | 0.300726547 | 0.544250592 | 0.156178049 |
| ENSMUSG00000032316 | Clk3 | 102414 | protein_coding | 1020.154795 | -0.156216257 | 0.06247388 | -2.500505109 | 0.012401634 | 0.069808432 | 0.156216257 |
| ENSMUSG00000002319 | Ipo4 | 75751 | protein_coding | 216.229356 | -0.156223129 | 0.13022773 | -1.199614925 | 0.230288927 | 0.469307431 | 0.156223129 |
| ENSMUSG00000021374 | Nup153 | 218210 | protein_coding | 1301.440116 | -0.156230923 | 0.0396189 | -3.943343314 | 8.04E-05 | 0.001224169 | 0.156230923 |
| ENSMUSG00000024870 | Rab1b | 76308 | protein_coding | 2862.095125 | 0.156286724 | 0.095382574 | 1.638524924 | 0.101312239 | 0.290200492 | 0.156286724 |
| ENSMUSG00000028410 | Dnaja1 | 15502 | protein_coding | 1489.387609 | -0.156301914 | 0.095615495 | -1.634692301 | 0.102113567 | 0.291609013 | 0.156301914 |
| ENSMUSG00000031803 | B3gnt3 | 72297 | protein_coding | 5355.543285 | 0.156316159 | 0.081162636 | 1.925962061 | 0.054109111 | 0.19482421 | 0.156316159 |
| ENSMUSG00000034427 | Myo15b | 217328 | protein_coding | 13030.61026 | -0.156323289 | 0.083539046 | -1.87126018 | 0.061309027 | 0.210995961 | 0.156323289 |
| ENSMUSG00000032673 | Prorsd1 | 67939 | protein_coding | 213.856177 | 0.156391438 | 0.095156318 | 1.643521315 | 0.100275119 | 0.288516832 | 0.156391438 |
| ENSMUSG00000040785 | Ttc3 | 22129 | protein_coding | 1939.081579 | -0.156419905 | 0.057951625 | -2.69914613 | 0.006951764 | 0.045592153 | 0.156419905 |
| ENSMUSG00000051396 | Gm45902 | 119065974 | protein_coding | 75.44508932 | -0.156461108 | 0.203242535 | -0.769824623 | 0.441403931 | 0.672888818 | 0.156461108 |
| ENSMUSG00000042063 | Zfp386 | 56220 | protein_coding | 628.6403063 | -0.156467498 | 0.104706461 | -1.494344258 | 0.135085663 | 0.343075037 | 0.156467498 |
| ENSMUSG00000037652 | Phc3 | 241915 | protein_coding | 765.3407456 | -0.156497392 | 0.067231832 | -2.327727607 | 0.019926572 | 0.097944477 | 0.156497392 |
| ENSMUSG00000029377 | Ereg | 13874 | protein_coding | 34.17401832 | 0.156508239 | 0.229108729 | 0.68311775 | 0.49453244 | 0.713491312 | 0.156508239 |
| ENSMUSG00000028664 | Ephb2 | 13844 | protein_coding | 2049.970253 | -0.156558759 | 0.055318896 | -2.830113598 | 0.004653148 | 0.033802395 | 0.156558759 |

| ensembl_gene_id | external_gene_name | entrezgene_id | gene_biotype | baseMean | log2FoldChange | lfcSE | stat | pvalue | padj | absLog2FC |
|---|---|---|---|---|---|---|---|---|---|---|
| ENSMUSG00000024026 | Glo1 | 109801 | protein_coding | 1708.881844 | 0.156608713 | 0.048448732 | 3.232462609 | 0.001227282 | 0.011943307 | 0.156608713 |
| ENSMUSG00000028447 | Dctn3 | 53598 | protein_coding | 1038.659396 | 0.15663653 | 0.064835565 | 2.415904451 | 0.01569618 | 0.082768143 | 0.15663653 |
| ENSMUSG00000019852 | Arfgef3 | 215821 | protein_coding | 1878.803377 | 0.156696689 | 0.066944661 | 2.340689866 | 0.019248149 | 0.095479458 | 0.156696689 |
| ENSMUSG00000021361 | Tmem14c | 66154 | protein_coding | 2116.768358 | 0.156749483 | 0.105470807 | 1.486188329 | 0.137229314 | 0.346747943 | 0.156749483 |
| ENSMUSG00000028842 | Ago3 | 214150 | protein_coding | 344.7860246 | -0.156763417 | 0.070405863 | -2.226567644 | 0.025976183 | 0.117839075 | 0.156763417 |
| ENSMUSG00000037991 | Rmi2 | 223970 | protein_coding | 60.09116965 | 0.156764416 | 0.235732951 | 0.665008503 | 0.506045055 | 0.721828109 | 0.156764416 |
| ENSMUSG00000032621 | Srek1 | 218543 | protein_coding | 738.2023925 | -0.156785451 | 0.114993452 | -1.363429383 | 0.172747226 | 0.397984931 | 0.156785451 |
| ENSMUSG00000050891 | Tatdn1 | 69694 | protein_coding | 169.6272145 | -0.156851581 | 0.110998706 | -1.413093777 | 0.157628153 | 0.376993804 | 0.156851581 |
| ENSMUSG00000022791 | Tnk2 | 51789 | protein_coding | 1072.546507 | 0.156999535 | 0.078543167 | 1.998894872 | 0.04561973 | 0.174003547 | 0.156999535 |
| ENSMUSG00000028392 | Bspry | 192120 | protein_coding | 4689.831983 | 0.157028987 | 0.065354042 | 2.402743311 | 0.016272605 | 0.085121085 | 0.157028987 |
| ENSMUSG00000031728 | Zfp821 | 75871 | protein_coding | 268.4348496 | -0.157033775 | 0.084127238 | -1.866622273 | 0.061954355 | 0.212446924 | 0.157033775 |
| ENSMUSG00000035811 | Ugt2b35 | 243085 | protein_coding | 975.0445474 | 0.15706034 | 0.089423126 | 1.756372737 | 0.079024785 | 0.248330613 | 0.15706034 |
| ENSMUSG00000116802 | Gm5165 | 102640673 | lncRNA | 156.4403345 | -0.157071922 | 0.101714659 | -1.544240758 | 0.122530018 | 0.32447317 | 0.157071922 |
| ENSMUSG00000000561 | Wdr77 | 70465 | protein_coding | 244.310787 | -0.157133303 | 0.099821448 | -1.574143693 | 0.115454233 | 0.313826128 | 0.157133303 |
| ENSMUSG00000026767 | Mindy3 | 66960 | protein_coding | 754.3942303 | 0.15713981 | 0.050604575 | 3.105249067 | 0.001901189 | 0.016874226 | 0.15713981 |
| ENSMUSG00000027201 | Myef2 | 17876 | protein_coding | 171.0537383 | 0.157154227 | 0.129715664 | 1.211528525 | 0.225692909 | 0.463780523 | 0.157154227 |
| ENSMUSG00000101972 | H3c11 | 319153 | protein_coding | 683.3011567 | 0.157155746 | 0.1720077 | 0.913655296 | 0.360898001 | 0.604050305 | 0.157155746 |
| ENSMUSG00000032058 | Ppp2r1b | 73699 | protein_coding | 666.0222774 | 0.157195558 | 0.074244475 | 2.117269442 | 0.034236987 | 0.142450951 | 0.157195558 |
| ENSMUSG00000069662 | Marcks | 17118 | protein_coding | 1470.660996 | 0.157205437 | 0.078136383 | 2.011936436 | 0.044226638 | 0.170480837 | 0.157205437 |
| ENSMUSG00000041057 | Wdr43 | 72515 | protein_coding | 980.6685205 | -0.157254595 | 0.067566548 | -2.327403123 | 0.01994382 | 0.097992668 | 0.157254595 |
| ENSMUSG00000038502 | Ptov1 | 84113 | protein_coding | 1139.93719 | 0.157266798 | 0.069342863 | 2.267959385 | 0.023331682 | 0.109264501 | 0.157266798 |
| ENSMUSG00000106864 | Gtf3c2 | 71752 | protein_coding | 397.668069 | -0.157300869 | 0.081074716 | -1.940196353 | 0.052355832 | 0.190915661 | 0.157300869 |
| ENSMUSG00000025169 | Ogfod3 | 66179 | protein_coding | 243.7723033 | -0.15730325 | 0.085232435 | -1.845579693 | 0.064953257 | 0.219350242 | 0.15730325 |
| ENSMUSG00000024292 | Cyp4f14 | 64385 | protein_coding | 5941.797691 | -0.157338514 | 0.075633739 | -2.080268889 | 0.037500877 | 0.15130764 | 0.157338514 |
| ENSMUSG00000018470 | Kcnab3 | 16499 | protein_coding | 8.046973257 | 0.157340694 | 0.461652182 | 0.340820861 | 0.733238445 | 0.871338619 | 0.157340694 |
| ENSMUSG00000005575 | Ube2m | 22192 | protein_coding | 1481.248251 | 0.157362489 | 0.100912059 | 1.559402231 | 0.118901209 | 0.319001795 | 0.157362489 |
| ENSMUSG00000056536 | Pign | 27392 | protein_coding | 1062.465503 | -0.157381287 | 0.060567638 | -2.59843859 | 0.009364879 | 0.056752643 | 0.157381287 |
| ENSMUSG00000025184 | R3hcc1l | 52013 | protein_coding | 510.7458345 | -0.157383873 | 0.060391534 | -2.606058558 | 0.00915908 | 0.055950184 | 0.157383873 |
| ENSMUSG00000091183 | Gm5141 | 380850 | protein_coding | 33.03288644 | -0.157395824 | 0.226063463 | -0.696246187 | 0.486274667 | 0.707337899 | 0.157395824 |
| ENSMUSG00000059586 | Nsmce2 | 68501 | protein_coding | 280.1656657 | -0.15741277 | 0.094435798 | -1.666876052 | 0.095539054 | 0.28018882 | 0.15741277 |
| ENSMUSG00000037318 | Traf3ip3 | 215243 | protein_coding | 12.68528693 | 0.157488108 | 0.447955143 | 0.351571158 | 0.725159899 | 0.865987203 | 0.157488108 |
| ENSMUSG00000040888 | Gfer | 11692 | protein_coding | 914.4168208 | 0.157513565 | 0.065497551 | 2.404877173 | 0.016177902 | 0.084790715 | 0.157513565 |
| ENSMUSG00000055188 | Rbm3os | 70226 | lncRNA | 25.12941024 | -0.157521381 | 0.248381695 | -0.634190781 | 0.525956324 | 0.736439527 | 0.157521381 |
| ENSMUSG00000001566 | Mnx1 | 15285 | protein_coding | 41.92372487 | -0.157592126 | 0.237160322 | -0.664496171 | 0.506372799 | 0.722065604 | 0.157592126 |
| ENSMUSG00000010095 | Slc3a2 | 17254 | protein_coding | 1025.034999 | -0.157658774 | 0.046737999 | -3.37324615 | 0.000742875 | 0.007881807 | 0.157658774 |
| ENSMUSG00000055866 | Per2 | 18627 | protein_coding | 450.2749817 | 0.157664167 | 0.128133536 | 1.230467619 | 0.218522046 | 0.454666616 | 0.157664167 |
| ENSMUSG00000050930 | Map10 | 74393 | protein_coding | 79.13556334 | -0.157736785 | 0.131569194 | -1.19888843 | 0.230571331 | 0.469603637 | 0.157736785 |
| ENSMUSG00000047205 | Dusp18 | 75219 | protein_coding | 357.012933 | 0.157772181 | 0.088886278 | 1.77498917 | 0.075899675 | 0.242575122 | 0.157772181 |
| ENSMUSG00000032500 | Dclk3 | 245038 | protein_coding | 119.9048225 | -0.157791909 | 0.171701797 | -0.918988101 | 0.358101796 | 0.601184005 | 0.157791909 |

| ensembl_gene_id | external_gene_name | entrezgene_id | gene_biotype | baseMean | log2FoldChange | lfcSE | stat | pvalue | padj | absLog2FC |
|---|---|---|---|---|---|---|---|---|---|---|
| ENSMUSG00000058706 | 0610030E20Rik | 68364 | protein_coding | 411.6306048 | -0.15781434 | 0.070738116 | -2.230966106 | 0.025683374 | 0.116930286 | 0.15781434 |
| ENSMUSG00000017493 | Igfbp4 | 16010 | protein_coding | 22.31199458 | 0.157873083 | 0.269433726 | 0.585944027 | 0.55791313 | 0.756598137 | 0.157873083 |
| ENSMUSG00000039701 | Usp53 | 99526 | protein_coding | 2888.101439 | -0.157875961 | 0.081595971 | -1.934849968 | 0.053008709 | 0.192495261 | 0.157875961 |
| ENSMUSG00000030042 | Pole4 | 66979 | protein_coding | 508.4817156 | 0.157910211 | 0.082822214 | 1.906616642 | 0.056570248 | 0.200957912 | 0.157910211 |
| ENSMUSG00000040296 | Rigi | 230073 | protein_coding | 456.0573685 | -0.157912105 | 0.097452673 | -1.620397888 | 0.105146833 | 0.297231022 | 0.157912105 |
| ENSMUSG00000015092 | Edf1 | 59022 | protein_coding | 4540.428327 | 0.157946581 | 0.061656917 | 2.561700936 | 0.010416097 | 0.061150489 | 0.157946581 |
| ENSMUSG00000021819 | Zswim8 | 268721 | protein_coding | 1026.490705 | -0.157983007 | 0.077187033 | -2.046755785 | 0.040682069 | 0.16048917 | 0.157983007 |
| ENSMUSG00000040520 | Manea | 242362 | protein_coding | 604.4322879 | 0.158034937 | 0.068553409 | 2.305281953 | 0.021150788 | 0.102131762 | 0.158034937 |
| ENSMUSG00000045273 | Cenph | 26886 | protein_coding | 58.68901351 | 0.1580372 | 0.167274375 | 0.944778301 | 0.344772072 | 0.58700794 | 0.1580372 |
| ENSMUSG00000080712 | H2bc26-ps | 382522 | protein_coding | 38.74876905 | 0.158069512 | 0.256961618 | 0.615148339 | 0.538456761 | 0.744805917 | 0.158069512 |
| ENSMUSG00000030386 | Zfp606 | 67370 | protein_coding | 287.0257172 | -0.158075083 | 0.08606695 | -1.836652541 | 0.066261206 | 0.222022535 | 0.158075083 |
| ENSMUSG00000038733 | Wdr26 | 226757 | protein_coding | 2557.370872 | -0.158101074 | 0.032493845 | -4.865569936 | 1.14E-06 | 2.76E-05 | 0.158101074 |
| ENSMUSG00000020988 | L2hgdh | 217666 | protein_coding | 590.2556627 | 0.158116344 | 0.063542601 | 2.488351784 | 0.012833672 | 0.071467067 | 0.158116344 |
| ENSMUSG00000038957 | Edc3 | 353190 | protein_coding | 389.2990229 | -0.158142474 | 0.066072943 | -2.393452837 | 0.016690627 | 0.086633325 | 0.158142474 |
| ENSMUSG00000038991 | Txndc5 | 105245 | protein_coding | 7038.780405 | 0.158143266 | 0.050995448 | 3.101125175 | 0.001927868 | 0.017026683 | 0.158143266 |
| ENSMUSG00000047342 | Zfp286 | 192651 | protein_coding | 12.98514269 | 0.158161603 | 0.35869869 | 0.440931644 | 0.659262484 | 0.825741594 | 0.158161603 |
| ENSMUSG00000090264 | Eif4ebp3 | 108112 | protein_coding | 596.1272452 | 0.158171359 | 0.128791638 | 1.228118239 | 0.219402585 | 0.455582357 | 0.158171359 |
| ENSMUSG00000025323 | Sp4 | 20688 | protein_coding | 249.4404109 | -0.158296535 | 0.102329316 | -1.546932399 | 0.121879548 | 0.323423905 | 0.158296535 |
| ENSMUSG00000042671 | Rgs8 | 67792 | protein_coding | 17.32728364 | 0.158297318 | 0.321886247 | 0.491780309 | 0.622874658 | 0.800888833 | 0.158297318 |
| ENSMUSG00000046794 | Ppp1r3b | 244416 | protein_coding | 981.4942047 | 0.158304104 | 0.070855676 | 2.234176757 | 0.025471445 | 0.116216716 | 0.158304104 |
| ENSMUSG00000026670 | Uap1 | 107652 | protein_coding | 5288.990968 | 0.158353033 | 0.042234324 | 3.749391934 | 0.000177264 | 0.002370426 | 0.158353033 |
| ENSMUSG00000029769 | Ccdc136 | 232664 | protein_coding | 7.47248003 | -0.15846539 | 0.449866217 | -0.352250033 | 0.724650758 | 0.865841503 | 0.15846539 |
| ENSMUSG00000029103 | Lrpap1 | 16976 | protein_coding | 1302.172315 | -0.158466545 | 0.052565276 | -3.014662104 | 0.002572657 | 0.021298377 | 0.158466545 |
| ENSMUSG00000051726 | Kcnf1 | 382571 | protein_coding | 531.045954 | -0.158476245 | 0.07738867 | -2.047796476 | 0.040579946 | 0.160203864 | 0.158476245 |
| ENSMUSG00000030105 | Arl8b | 67166 | protein_coding | 1766.450696 | 0.158491422 | 0.037498159 | 4.226645403 | 2.37E-05 | 0.000421293 | 0.158491422 |
| ENSMUSG00000042460 | C1galt1 | 94192 | protein_coding | 6010.740861 | 0.158512737 | 0.042743038 | 3.708504267 | 0.000208487 | 0.002731844 | 0.158512737 |
| ENSMUSG00000021357 | Exoc2 | 66482 | protein_coding | 727.8996869 | -0.158543466 | 0.052317749 | -3.030395386 | 0.002442338 | 0.020439845 | 0.158543466 |
| ENSMUSG00000027447 | Cst3 | 13010 | protein_coding | 1780.875389 | -0.158547147 | 0.063194227 | -2.508886584 | 0.012111236 | 0.068460536 | 0.158547147 |
| ENSMUSG00000045983 | Eif4g1 | 208643 | protein_coding | 10583.15378 | 0.158600525 | 0.061003934 | 2.599840939 | 0.009326698 | 0.056624534 | 0.158600525 |
| ENSMUSG00000040865 | Ino80d | 227195 | protein_coding | 1090.442054 | -0.15862338 | 0.062970816 | -2.518998316 | 0.011768922 | 0.067089927 | 0.15862338 |
| ENSMUSG00000055319 | Sec23ip | 207352 | protein_coding | 978.0336582 | 0.158630525 | 0.058433533 | 2.714717349 | 0.006633236 | 0.044045718 | 0.158630525 |
| ENSMUSG00000091119 | Ccdc152 | 100039139 | protein_coding | 6.28848199 | -0.158636229 | 0.546507106 | -0.290272949 | 0.771607432 | 0.892439856 | 0.158636229 |
| ENSMUSG00000032372 | Plscr2 | 18828 | protein_coding | 41.47202183 | -0.158671318 | 0.199275656 | -0.796240352 | 0.425892348 | 0.661460139 | 0.158671318 |
| ENSMUSG00000061356 | Nuggc | 100503545 | protein_coding | 8.081458184 | 0.15869908 | 0.497292369 | 0.319126313 | 0.749630731 | 0.880475742 | 0.15869908 |
| ENSMUSG00000029718 | Pcolce | 18542 | protein_coding | 58.41787136 | 0.15870318 | 0.163555218 | 0.970333944 | 0.331880063 | 0.575141012 | 0.15870318 |
| ENSMUSG00000044348 | Slc25a53 | 67062 | protein_coding | 52.14536605 | -0.158704762 | 0.193870244 | -0.818613313 | 0.413007069 | 0.650699981 | 0.158704762 |
| ENSMUSG00000038335 | Tsr1 | 104662 | protein_coding | 815.6827491 | -0.158718436 | 0.058569676 | -2.709908017 | 0.006730187 | 0.044482674 | 0.158718436 |
| ENSMUSG00000026415 | Fcamr | 64435 | protein_coding | 71.55495421 | 0.158745744 | 0.164233839 | 0.966583651 | 0.333752211 | 0.577079651 | 0.158745744 |
| ENSMUSG00000064036 | Mro | 71263 | protein_coding | 9.717417381 | 0.158806312 | 0.38272363 | 0.414937306 | 0.678187799 | 0.836812138 | 0.158806312 |

| ensembl_gene_id | external_gene_name | entrezgene_id | gene_biotype | baseMean | log2FoldChange | lfcSE | stat | pvalue | padj | absLog2FC |
|---|---|---|---|---|---|---|---|---|---|---|
| ENSMUSG00000022033 | Pbk | 52033 | protein_coding | 343.6359357 | 0.158826872 | 0.14421751 | 1.101300888 | 0.270765725 | 0.513373374 | 0.158826872 |
| ENSMUSG00000024276 | Zfp397 | 69256 | protein_coding | 714.7373892 | -0.158829004 | 0.081657922 | -1.945053219 | 0.051768573 | 0.189399225 | 0.158829004 |
| ENSMUSG00000064368 | mt-Nd6 | 17722 | protein_coding | 3615.85358 | 0.158880978 | 0.083483847 | 1.90313436 | 0.057023016 | 0.201934601 | 0.158880978 |
| ENSMUSG00000034187 | Nsf | 18195 | protein_coding | 1351.816414 | 0.158951554 | 0.055367141 | 2.87086439 | 0.004093511 | 0.030762372 | 0.158951554 |
| ENSMUSG00000017721 | Pigt | 78928 | protein_coding | 2179.438067 | 0.15895715 | 0.089920354 | 1.767754942 | 0.077101881 | 0.244881547 | 0.15895715 |
| ENSMUSG00000038393 | Txnip | 56338 | protein_coding | 20314.71186 | -0.158982697 | 0.120162961 | -1.323059077 | 0.185815732 | 0.415106014 | 0.158982697 |
| ENSMUSG00000070337 | Gpr179 | 217143 | protein_coding | 6.406967767 | 0.15900636 | 0.496210842 | 0.320441124 | 0.748633955 | 0.880475742 | 0.15900636 |
| ENSMUSG00000066735 | Vkorc1l1 | 69568 | protein_coding | 875.1425024 | 0.159020332 | 0.051643304 | 3.079205225 | 0.002075537 | 0.01807353 | 0.159020332 |
| ENSMUSG00000074166 | AW146154 | 101835 | protein_coding | 178.7028243 | -0.159080494 | 0.108706584 | -1.46339336 | 0.143359775 | 0.355303993 | 0.159080494 |
| ENSMUSG00000031447 | Lamp1 | 16783 | protein_coding | 9181.463669 | -0.159208764 | 0.033147177 | -4.803086723 | 1.56E-06 | 3.66E-05 | 0.159208764 |
| ENSMUSG00000026634 | Angel2 | 52477 | protein_coding | 1280.344835 | -0.159209621 | 0.052744612 | -3.018500177 | 0.002540293 | 0.021073713 | 0.159209621 |
| ENSMUSG00000031840 | Rab3a | 19339 | protein_coding | 180.3566993 | 0.159289202 | 0.096078507 | 1.657906704 | 0.097336299 | 0.283295883 | 0.159289202 |
| ENSMUSG00000026405 | C4bp | 12269 | protein_coding | 227.7152256 | 0.159294879 | 0.103551429 | 1.538316566 | 0.123971231 | 0.326892409 | 0.159294879 |
| ENSMUSG00000079469 | Pigb | 55981 | protein_coding | 188.6646405 | 0.159332553 | 0.094891828 | 1.679096666 | 0.093133206 | 0.275588846 | 0.159332553 |
| ENSMUSG00000001909 | Trmt1 | 212528 | protein_coding | 1058.57633 | -0.159345155 | 0.051279285 | -3.107398148 | 0.00188742 | 0.016779727 | 0.159345155 |
| ENSMUSG00000024943 | Smc5 | 226026 | protein_coding | 529.0161102 | -0.15936599 | 0.08312535 | -1.917176777 | 0.055215477 | 0.197660378 | 0.15936599 |
| ENSMUSG00000039103 | Nexn | 68810 | protein_coding | 24.69135198 | -0.159368281 | 0.284747502 | -0.559682805 | 0.575695813 | 0.76862707 | 0.159368281 |
| ENSMUSG00000071267 | Zfp942 | 73233 | protein_coding | 457.0097736 | -0.159376991 | 0.101428358 | -1.571325761 | 0.116107 | 0.314900319 | 0.159376991 |
| ENSMUSG00000005267 | Zfp287 | 170740 | protein_coding | 57.39376813 | 0.15941529 | 0.165945738 | 0.960647085 | 0.336729646 | 0.580296438 | 0.15941529 |
| ENSMUSG00000069727 | Zfp975 | 434179 | protein_coding | 130.5889737 | -0.15947069 | 0.172111912 | -0.926552315 | 0.354159023 | 0.596379092 | 0.15947069 |
| ENSMUSG00000002486 | Tchp | 77832 | protein_coding | 111.4012873 | -0.159480492 | 0.14467336 | -1.102348706 | 0.270310102 | 0.512797437 | 0.159480492 |
| ENSMUSG00000029762 | Akr1b8 | 14187 | protein_coding | 702.8094224 | -0.159491219 | 0.096575484 | -1.651466944 | 0.098643265 | 0.285758925 | 0.159491219 |
| ENSMUSG00000034259 | Exosc4 | 109075 | protein_coding | 347.3340068 | 0.15952505 | 0.087762414 | 1.817692133 | 0.069111201 | 0.228065548 | 0.15952505 |
| ENSMUSG00000046603 | Tcaim | 382117 | protein_coding | 207.4024111 | 0.159557579 | 0.092574149 | 1.723565169 | 0.084786371 | 0.259710803 | 0.159557579 |
| ENSMUSG00000001946 | Esam | 69524 | protein_coding | 21.2489967 | -0.159582783 | 0.26145694 | -0.610359713 | 0.541623549 | 0.74719957 | 0.159582783 |
| ENSMUSG00000024349 | Sting1 | 72512 | protein_coding | 111.7284201 | 0.159606424 | 0.120734267 | 1.32196458 | 0.186179946 | 0.415459248 | 0.159606424 |
| ENSMUSG00000069170 | Adgrv1 | 110789 | protein_coding | 19.64866914 | -0.159624583 | 0.289706723 | -0.550986809 | 0.581642717 | 0.773365661 | 0.159624583 |
| ENSMUSG00000047786 | Lix1 | 66643 | protein_coding | 15.06634351 | 0.159667886 | 0.398071681 | 0.401103354 | 0.688344032 | 0.843368224 | 0.159667886 |
| ENSMUSG00000029629 | Phf14 | 75725 | protein_coding | 738.0496707 | -0.159668441 | 0.06387785 | -2.499590087 | 0.012433708 | 0.069915764 | 0.159668441 |
| ENSMUSG00000028873 | Cdca8 | 52276 | protein_coding | 318.4662613 | 0.159675374 | 0.087534475 | 1.824142698 | 0.068130502 | 0.226076106 | 0.159675374 |
| ENSMUSG00000032086 | Bace1 | 23821 | protein_coding | 456.2016286 | -0.159682297 | 0.065738524 | -2.42905207 | 0.015138358 | 0.080626255 | 0.159682297 |
| ENSMUSG00000033713 | Foxn3 | 71375 | protein_coding | 1883.772691 | -0.159871605 | 0.070860186 | -2.25615561 | 0.024060887 | 0.111759773 | 0.159871605 |
| ENSMUSG00000023911 | Flywch2 | 76917 | protein_coding | 118.9231296 | 0.159918175 | 0.119365563 | 1.339734606 | 0.180331643 | 0.407326553 | 0.159918175 |
| ENSMUSG00000022550 | Adck5 | 268822 | protein_coding | 1823.56164 | -0.159926309 | 0.063759144 | -2.508288206 | 0.012131767 | 0.068515133 | 0.159926309 |
| ENSMUSG00000020420 | Zfp607a | 545938 | protein_coding | 139.3357295 | -0.159931673 | 0.134936297 | -1.185238341 | 0.235923224 | 0.476068282 | 0.159931673 |
| ENSMUSG00000031887 | Tradd | 71609 | protein_coding | 1642.962365 | 0.159943646 | 0.098836172 | 1.618270342 | 0.105604349 | 0.297761537 | 0.159943646 |
| ENSMUSG00000028173 | Wls | 68151 | protein_coding | 2951.108562 | -0.159959866 | 0.057629212 | -2.775673334 | 0.005508753 | 0.03837566 | 0.159959866 |
| ENSMUSG00000028933 | Xrcc2 | 57434 | protein_coding | 60.23951707 | 0.160012011 | 0.167361825 | 0.956084287 | 0.339029652 | 0.58240762 | 0.160012011 |
| ENSMUSG00000038774 | Ascc3 | 77987 | protein_coding | 1644.665538 | 0.160034781 | 0.069157099 | 2.314075953 | 0.020663556 | 0.100534642 | 0.160034781 |

| ensembl_gene_id | external_gene_name | entrezgene_id | gene_biotype | baseMean | log2FoldChange | lfcSE | stat | pvalue | padj | absLog2FC |
|---|---|---|---|---|---|---|---|---|---|---|
| ENSMUSG00000049488 | Tmem67 | 329795 | protein_coding | 101.9859328 | 0.16004539 | 0.136918751 | 1.168907754 | 0.242440799 | 0.483592179 | 0.16004539 |
| ENSMUSG00000007721 | Ccdc124 | 234388 | protein_coding | 762.811068 | 0.160066391 | 0.114160646 | 1.402115325 | 0.160880811 | 0.381379516 | 0.160066391 |
| ENSMUSG00000029378 | Areg | 11839 | protein_coding | 1520.637903 | 0.160133868 | 0.12343472 | 1.297316252 | 0.194522395 | 0.426521096 | 0.160133868 |
| ENSMUSG00000027506 | Tpd52 | 21985 | protein_coding | 8718.723047 | 0.160135286 | 0.032602224 | 4.911790227 | 9.02E-07 | 2.23E-05 | 0.160135286 |
| ENSMUSG00000054951 | 9130008F23Rik | 71583 | protein_coding | 341.8822386 | 0.16013774 | 0.112509861 | 1.423321824 | 0.154642883 | 0.372364958 | 0.16013774 |
| ENSMUSG00000058351 | Smim4 | 66487 | protein_coding | 171.2337082 | -0.160282671 | 0.114112834 | -1.404598102 | 0.160140821 | 0.380520263 | 0.160282671 |
| ENSMUSG00000037174 | Elf2 | 69257 | protein_coding | 1403.656006 | -0.160295776 | 0.052764746 | -3.037933221 | 0.002382067 | 0.020070847 | 0.160295776 |
| ENSMUSG00000020422 | Tns3 | 319939 | protein_coding | 3256.200114 | -0.160307274 | 0.038015031 | -4.216944433 | 2.48E-05 | 0.000437896 | 0.160307274 |
| ENSMUSG00000044367 | Slc16a13 | 69309 | protein_coding | 163.3469157 | 0.160318162 | 0.103955709 | 1.542177557 | 0.123030452 | 0.32531761 | 0.160318162 |
| ENSMUSG00000039910 | Cited2 | 17684 | protein_coding | 877.1526441 | -0.160351417 | 0.109933079 | -1.458627543 | 0.144667649 | 0.357173174 | 0.160351417 |
| ENSMUSG00000040502 | Marchf9 | 216438 | protein_coding | 88.87946504 | -0.160444002 | 0.149204377 | -1.075330395 | 0.282226833 | 0.525081686 | 0.160444002 |
| ENSMUSG00000073792 | Alg6 | 320438 | protein_coding | 376.7796277 | 0.160459808 | 0.076364686 | 2.101230505 | 0.035620737 | 0.14616681 | 0.160459808 |
| ENSMUSG00000019861 | Gopc | 94221 | protein_coding | 888.559379 | 0.160482615 | 0.053473093 | 3.001184451 | 0.002689316 | 0.022071552 | 0.160482615 |
| ENSMUSG00000033763 | Mtss2 | 244654 | protein_coding | 9.878968364 | -0.160523767 | 0.454812885 | -0.352944634 | 0.724129948 | 0.865580106 | 0.160523767 |
| ENSMUSG00000026385 | Dbi | 13167 | protein_coding | 4943.69091 | 0.160535521 | 0.04488884 | 3.576290271 | 0.000348505 | 0.004178688 | 0.160535521 |
| ENSMUSG00000036995 | Asap3 | 230837 | protein_coding | 34.68450008 | -0.160546275 | 0.225234509 | -0.712796081 | 0.475971951 | 0.699152897 | 0.160546275 |
| ENSMUSG00000033590 | Myo5c | 208943 | protein_coding | 4780.294417 | 0.160576339 | 0.05944107 | 2.701437573 | 0.006904044 | 0.045316043 | 0.160576339 |
| ENSMUSG00000021037 | Ahsa1 | 217737 | protein_coding | 1348.756142 | 0.160645172 | 0.094188098 | 1.705578261 | 0.08808663 | 0.266124593 | 0.160645172 |
| ENSMUSG00000072704 | Smim10l1 | 381820 | protein_coding | 2396.507145 | 0.160678355 | 0.033356343 | 4.817025452 | 1.46E-06 | 3.44E-05 | 0.160678355 |
| ENSMUSG00000051256 | Jagn1 | 67767 | protein_coding | 549.0292865 | 0.160697024 | 0.066073799 | 2.432083928 | 0.015012228 | 0.080166293 | 0.160697024 |
| ENSMUSG00000022765 | Snap29 | 67474 | protein_coding | 920.0761788 | 0.160705279 | 0.059043198 | 2.72182547 | 0.006492242 | 0.043336253 | 0.160705279 |
| ENSMUSG00000046991 | Wdr27 | 71682 | protein_coding | 24.13740926 | 0.160705809 | 0.252455898 | 0.636569832 | 0.524405081 | 0.735184911 | 0.160705809 |
| ENSMUSG00000025019 | Lcor | 212391 | protein_coding | 1747.032479 | -0.160725446 | 0.042711266 | -3.763069145 | 0.000167841 | 0.002268875 | 0.160725446 |
| ENSMUSG00000096215 | Smim22 | 432995 | protein_coding | 690.9125654 | -0.160776005 | 0.083842497 | -1.917595612 | 0.055162307 | 0.197645528 | 0.160776005 |
| ENSMUSG00000031196 | F8 | 14069 | protein_coding | 24.17229504 | 0.160792689 | 0.27386368 | 0.587126737 | 0.557118592 | 0.755953522 | 0.160792689 |
| ENSMUSG00000015133 | Lrrk1 | 233328 | protein_coding | 1084.794574 | 0.160886544 | 0.061264289 | 2.62610645 | 0.00863678 | 0.053509549 | 0.160886544 |
| ENSMUSG00000039195 | Bbln | 73737 | protein_coding | 266.8170374 | 0.160953645 | 0.122621428 | 1.312606181 | 0.189315686 | 0.419784692 | 0.160953645 |
| ENSMUSG00000020440 | Arf5 | 11844 | protein_coding | 2224.063459 | 0.160985424 | 0.076703866 | 2.098791522 | 0.035835283 | 0.146790856 | 0.160985424 |
| ENSMUSG00000048440 | Cyp4f16 | 70101 | protein_coding | 1128.618941 | -0.161022708 | 0.083512357 | -1.928130329 | 0.053838917 | 0.194372111 | 0.161022708 |
| ENSMUSG00000024007 | Ppil1 | 68816 | protein_coding | 630.6664772 | 0.161038923 | 0.074808491 | 2.152682421 | 0.031343646 | 0.134008213 | 0.161038923 |
| ENSMUSG00000020792 | Exoc7 | 53413 | protein_coding | 1045.820283 | 0.16105953 | 0.042176446 | 3.818707914 | 0.000134152 | 0.001896124 | 0.16105953 |
| ENSMUSG00000029580 | Actb | 11461 | protein_coding | 97992.59825 | 0.161059554 | 0.076053764 | 2.117706552 | 0.034199928 | 0.14237022 | 0.161059554 |
| ENSMUSG00000045409 | Trim39 | 79263 | protein_coding | 370.5140225 | -0.161066593 | 0.100214817 | -1.607213358 | 0.108007573 | 0.301043575 | 0.161066593 |
| ENSMUSG00000032913 | Lrig2 | 269473 | protein_coding | 660.0745482 | -0.161099361 | 0.068814647 | -2.341062076 | 0.01922897 | 0.09547586 | 0.161099361 |
| ENSMUSG00000051344 | Plekhm3 | 241075 | protein_coding | 84.37803109 | -0.161138221 | 0.139362829 | -1.156249642 | 0.24757906 | 0.488580666 | 0.161138221 |
| ENSMUSG00000002102 | Psmc3 | 19182 | protein_coding | 3095.136225 | 0.16116143 | 0.065878267 | 2.446351987 | 0.014431004 | 0.077980613 | 0.16116143 |
| ENSMUSG00000034055 | Phka1 | 18679 | protein_coding | 433.7068138 | 0.161162279 | 0.063869833 | 2.523292648 | 0.01162616 | 0.066299536 | 0.161162279 |
| ENSMUSG00000024789 | Jak2 | 16452 | protein_coding | 2078.80207 | -0.161236608 | 0.048906497 | -3.296834105 | 0.000977812 | 0.009861901 | 0.161236608 |
| ENSMUSG00000040463 | Mybbp1a | 18432 | protein_coding | 1330.386171 | -0.161252935 | 0.072173573 | -2.234237943 | 0.025467421 | 0.116216716 | 0.161252935 |

| ensembl_gene_id | external_gene_name | entrezgene_id | gene_biotype | baseMean | log2FoldChange | lfcSE | stat | pvalue | padj | absLog2FC |
|---|---|---|---|---|---|---|---|---|---|---|
| ENSMUSG00000031451 | Gas6 | 14456 | protein_coding | 10446.55071 | -0.161280213 | 0.067430086 | -2.39181383 | 0.016765344 | 0.086937204 | 0.161280213 |
| ENSMUSG00000019761 | Krt10 | 16661 | protein_coding | 41.23491149 | -0.161299059 | 0.222409373 | -0.725234987 | 0.468307882 | 0.693047027 | 0.161299059 |
| ENSMUSG00000032583 | Mon1a | 72825 | protein_coding | 878.2279089 | 0.161309185 | 0.073010747 | 2.2093896 | 0.027147554 | 0.121033168 | 0.161309185 |
| ENSMUSG00000028245 | Nsmaf | 18201 | protein_coding | 394.1198876 | -0.161329692 | 0.08346169 | -1.932978979 | 0.053238787 | 0.192853083 | 0.161329692 |
| ENSMUSG00000040048 | Ndufb10 | 68342 | protein_coding | 3126.65841 | 0.161349582 | 0.056230159 | 2.869449158 | 0.004111874 | 0.030837772 | 0.161349582 |
| ENSMUSG00000050605 | Zfp61 | 22719 | protein_coding | 105.1000861 | 0.161356277 | 0.134816436 | 1.196859088 | 0.231361483 | 0.47045349 | 0.161356277 |
| ENSMUSG00000034023 | Fancd2 | 211651 | protein_coding | 177.4990287 | 0.16137767 | 0.165091979 | 0.977501579 | 0.328320896 | 0.571937821 | 0.16137767 |
| ENSMUSG00000102543 | Pcdhgc5 | 93708 | protein_coding | 23.97316375 | 0.161399293 | 0.258871195 | 0.623473359 | 0.532973504 | 0.741098784 | 0.161399293 |
| ENSMUSG00000004207 | Psap | 19156 | protein_coding | 15451.67249 | -0.161425785 | 0.036021346 | -4.481392324 | 7.42E-06 | 0.000151003 | 0.161425785 |
| ENSMUSG00000043415 | Otud1 | 71198 | protein_coding | 86.18208738 | -0.161438816 | 0.1334645 | -1.209601175 | 0.226431967 | 0.464862544 | 0.161438816 |
| ENSMUSG00000007564 | Ppp2r1a | 51792 | protein_coding | 5377.818574 | 0.161475143 | 0.047666912 | 3.387572951 | 0.00070514 | 0.007586249 | 0.161475143 |
| ENSMUSG00000014444 | Piezo1 | 234839 | protein_coding | 297.7320021 | -0.161542683 | 0.104416287 | -1.547102359 | 0.121838565 | 0.32338554 | 0.161542683 |
| ENSMUSG00000015943 | Bola1 | 69168 | protein_coding | 311.2543752 | 0.161562469 | 0.094032371 | 1.718157983 | 0.085767803 | 0.261618566 | 0.161562469 |
| ENSMUSG00000078938 | Syce3 | 75459 | protein_coding | 16.66373726 | -0.161565999 | 0.296063125 | -0.545714699 | 0.585262067 | 0.775875162 | 0.161565999 |
| ENSMUSG00000021814 | Anxa7 | 11750 | protein_coding | 4356.693492 | -0.161568751 | 0.043049938 | -3.753054255 | 0.000174693 | 0.002345091 | 0.161568751 |
| ENSMUSG00000051339 | 2900026A02Rik | 243219 | protein_coding | 6312.433566 | -0.161611929 | 0.064916356 | -2.489541005 | 0.012790816 | 0.071277641 | 0.161611929 |
| ENSMUSG00000024369 | Nelfe | 27632 | protein_coding | 615.3778071 | 0.161757358 | 0.054839821 | 2.949633206 | 0.003181514 | 0.02531242 | 0.161757358 |
| ENSMUSG00000055044 | Pdlim1 | 54132 | protein_coding | 5391.539962 | 0.161775166 | 0.069778224 | 2.318419067 | 0.020426557 | 0.099774968 | 0.161775166 |
| ENSMUSG00000062157 | Ifnlr1 | 242700 | protein_coding | 340.5703523 | -0.161786407 | 0.095201963 | -1.69940201 | 0.089243464 | 0.268368093 | 0.161786407 |
| ENSMUSG00000022760 | Thap7 | 69009 | protein_coding | 334.9595311 | 0.161816888 | 0.0830033 | 1.94952354 | 0.051232934 | 0.188241916 | 0.161816888 |
| ENSMUSG00000036880 | Acaa2 | 52538 | protein_coding | 3485.17606 | -0.1618255 | 0.044305002 | -3.652533374 | 0.000259666 | 0.003276707 | 0.1618255 |
| ENSMUSG00000034765 | Dusp5 | 240672 | protein_coding | 465.7797136 | -0.161889101 | 0.170831154 | -0.947655612 | 0.343304805 | 0.585870539 | 0.161889101 |
| ENSMUSG00000003824 | Syce2 | 71846 | protein_coding | 73.23787986 | -0.161901015 | 0.189618823 | -0.85382354 | 0.393202769 | 0.632852401 | 0.161901015 |
| ENSMUSG00000060336 | Zfp937 | 245174 | protein_coding | 61.04968503 | -0.161959455 | 0.174244453 | -0.929495614 | 0.352632297 | 0.595229357 | 0.161959455 |
| ENSMUSG00000043219 | Hoxa6 | 15403 | protein_coding | 31.68228671 | -0.161963865 | 0.255540548 | -0.633808868 | 0.526205565 | 0.736578175 | 0.161963865 |
| ENSMUSG00000037257 | Aagab | 66939 | protein_coding | 1536.512549 | 0.162001209 | 0.053141998 | 3.048459159 | 0.002300182 | 0.019554576 | 0.162001209 |
| ENSMUSG00000034156 | Tspoap1 | 207777 | protein_coding | 7.330638994 | 0.16203176 | 0.547941094 | 0.295710181 | 0.767451421 | 0.890729692 | 0.16203176 |
| ENSMUSG00000018752 | Tnfsfm13 | 619441 | protein_coding | 10.55730809 | -0.162032718 | 0.365751285 | -0.443013393 | 0.657756048 | 0.824854726 | 0.162032718 |
| ENSMUSG00000020589 | Cyria | 76820 | protein_coding | 409.495975 | -0.162060904 | 0.141477752 | -1.145486846 | 0.252007541 | 0.493099444 | 0.162060904 |
| ENSMUSG00000024538 | Ppic | 19038 | protein_coding | 29.07318244 | -0.162113678 | 0.281954417 | -0.574964136 | 0.565315553 | 0.761818669 | 0.162113678 |
| ENSMUSG00000057103 | Nat8f1 | 66116 | protein_coding | 43.5301144 | -0.162282431 | 0.205418435 | -0.790009089 | 0.42952246 | 0.664591383 | 0.162282431 |
| ENSMUSG00000039270 | Megf9 | 230316 | protein_coding | 337.5979362 | 0.162298896 | 0.09907954 | 1.638066702 | 0.10140778 | 0.290238707 | 0.162298896 |
| ENSMUSG00000044393 | Dsg2 | 13511 | protein_coding | 6684.878334 | -0.16232286 | 0.045656906 | -3.555275072 | 0.000377584 | 0.004480717 | 0.16232286 |
| ENSMUSG00000074647 | Fam83c | 71405 | protein_coding | 217.7798381 | -0.162337305 | 0.116423859 | -1.394364573 | 0.163207541 | 0.38480234 | 0.162337305 |
| ENSMUSG00000034040 | Galnt17 | 212996 | protein_coding | 12.20862439 | 0.162358033 | 0.480543232 | 0.33786353 | 0.735466032 | 0.872889311 | 0.162358033 |
| ENSMUSG00000079553 | Kifc1 | 100502766 | protein_coding | 276.4618547 | 0.162405343 | 0.13932998 | 1.165616637 | 0.243769478 | 0.484922952 | 0.162405343 |
| ENSMUSG00000013698 | Pea15a | 18611 | protein_coding | 300.4728626 | 0.162513127 | 0.082720319 | 1.964609528 | 0.049459448 | 0.183913425 | 0.162513127 |
| ENSMUSG00000023009 | Nckap5l | 380969 | protein_coding | 6.560739152 | 0.162525992 | 0.476734561 | 0.340915062 | 0.733167526 | 0.871318547 | 0.162525992 |
| ENSMUSG00000010608 | Rbm25 | 67039 | protein_coding | 1621.702749 | -0.162559053 | 0.076919619 | -2.113362683 | 0.034569737 | 0.1434653 | 0.162559053 |

| ensembl_gene_id | external_gene_name | entrezgene_id | gene_biotype | baseMean | log2FoldChange | lfcSE | stat | pvalue | padj | absLog2FC |
|---|---|---|---|---|---|---|---|---|---|---|
| ENSMUSG00000006998 | Psmd2 | 21762 | protein_coding | 3826.463355 | 0.162561132 | 0.07037876 | 2.309803856 | 0.020899016 | 0.101348119 | 0.162561132 |
| ENSMUSG00000018537 | Pcgf2 | 22658 | protein_coding | 130.4783609 | -0.162593593 | 0.114103664 | -1.424963819 | 0.15416766 | 0.371583 | 0.162593593 |
| ENSMUSG00000078429 | Ctdsp2 | 52468 | protein_coding | 3602.642621 | -0.162596491 | 0.071947987 | -2.25991717 | 0.023826392 | 0.110958194 | 0.162596491 |
| ENSMUSG00000064901 | Snora21 | 100302498 | snoRNA | 43.65281802 | 0.162599902 | 0.220593676 | 0.737101376 | 0.461060703 | 0.687717915 | 0.162599902 |
| ENSMUSG00000020937 | Plcd3 | 72469 | protein_coding | 107.9494213 | 0.162601218 | 0.294877972 | 0.551418666 | 0.581346708 | 0.773163315 | 0.162601218 |
| ENSMUSG00000021666 | Gfm2 | 320806 | protein_coding | 1278.897399 | 0.162606104 | 0.053386299 | 3.045839624 | 0.002320316 | 0.019673885 | 0.162606104 |
| ENSMUSG00000030380 | Mzf1 | 109889 | protein_coding | 20.86654344 | 0.162616827 | 0.282072261 | 0.576507688 | 0.564272075 | 0.761252782 | 0.162616827 |
| ENSMUSG00000008036 | Ap2s1 | 232910 | protein_coding | 1957.952944 | 0.162620207 | 0.087950087 | 1.84900565 | 0.064456997 | 0.218015518 | 0.162620207 |
| ENSMUSG00000038473 | Nos1ap | 70729 | protein_coding | 106.3226691 | -0.162633743 | 0.123796371 | -1.313719797 | 0.188940516 | 0.419340469 | 0.162633743 |
| ENSMUSG00000005320 | Fgfr4 | 14186 | protein_coding | 431.6831693 | 0.162663628 | 0.062173027 | 2.61630543 | 0.008888701 | 0.054692132 | 0.162663628 |
| ENSMUSG00000070493 | Chchd2 | 14004 | protein_coding | 5630.227898 | 0.16268813 | 0.055391218 | 2.937074422 | 0.003313246 | 0.026166854 | 0.16268813 |
| ENSMUSG00000097726 | 9530036O11Rik | 654796 | lncRNA | 25.99101174 | -0.162690993 | 0.225504924 | -0.721452063 | 0.470631424 | 0.694916039 | 0.162690993 |
| ENSMUSG00000043154 | Ppp2r3a | 235542 | protein_coding | 536.2634992 | 0.1627355 | 0.05372909 | 3.028815497 | 0.002455146 | 0.020525731 | 0.1627355 |
| ENSMUSG00000037499 | Nenf | 66208 | protein_coding | 402.7286293 | -0.162837164 | 0.100505197 | -1.620186502 | 0.10519222 | 0.297255113 | 0.162837164 |
| ENSMUSG00000024486 | Hbegf | 15200 | protein_coding | 7666.99122 | -0.162838396 | 0.104698623 | -1.55530599 | 0.119873211 | 0.320277548 | 0.162838396 |
| ENSMUSG00000030796 | Tead2 | 21677 | protein_coding | 62.50106884 | -0.162839735 | 0.20042506 | -0.812471928 | 0.416520888 | 0.654445865 | 0.162839735 |
| ENSMUSG00000029919 | Hpgds | 54486 | protein_coding | 152.0902851 | -0.162844202 | 0.153953873 | -1.057746707 | 0.290170935 | 0.532649825 | 0.162844202 |
| ENSMUSG00000015759 | Cnih1 | 12793 | protein_coding | 984.7293332 | 0.16285009 | 0.054642552 | 2.980279704 | 0.002879853 | 0.023338386 | 0.16285009 |
| ENSMUSG00000109324 | Prmt1 | 15469 | protein_coding | 470.4528919 | 0.162897797 | 0.075090918 | 2.169340866 | 0.030056815 | 0.130372849 | 0.162897797 |
| ENSMUSG00000042423 | Fbrs | 14123 | protein_coding | 488.8013997 | -0.162905598 | 0.09563908 | -1.703337149 | 0.088504992 | 0.266988405 | 0.162905598 |
| ENSMUSG00000066042 | Med18 | 67219 | protein_coding | 155.7794357 | -0.162918187 | 0.096739901 | -1.684084708 | 0.092165299 | 0.273594537 | 0.162918187 |
| ENSMUSG00000031609 | Sap30 | 60406 | protein_coding | 211.7854896 | -0.162919976 | 0.111134257 | -1.4659744 | 0.142655261 | 0.354359599 | 0.162919976 |
| ENSMUSG00000044098 | Rsbn1 | 229675 | protein_coding | 509.6037827 | -0.162937317 | 0.057886856 | -2.814754982 | 0.004881444 | 0.034965035 | 0.162937317 |
| ENSMUSG00000028607 | Cpt2 | 12896 | protein_coding | 1933.134107 | 0.162958551 | 0.056019129 | 2.908980454 | 0.003626095 | 0.02807395 | 0.162958551 |
| ENSMUSG00000027634 | Ndrg3 | 29812 | protein_coding | 1497.118007 | -0.163061372 | 0.050159626 | -3.250849045 | 0.001150609 | 0.011328374 | 0.163061372 |
| ENSMUSG00000046573 | Lyrm4 | 380840 | protein_coding | 179.352824 | 0.163074673 | 0.106947331 | 1.524812926 | 0.127305765 | 0.33172727 | 0.163074673 |
| ENSMUSG00000036769 | Wdr44 | 72404 | protein_coding | 664.4901538 | -0.16308736 | 0.053858061 | -3.028095639 | 0.002461002 | 0.02056403 | 0.16308736 |
| ENSMUSG00000020788 | Atp2a3 | 53313 | protein_coding | 22215.39972 | 0.163146118 | 0.067332212 | 2.423002485 | 0.015392821 | 0.081631052 | 0.163146118 |
| ENSMUSG00000030007 | Cct7 | 12468 | protein_coding | 3359.978889 | 0.1631668 | 0.064436055 | 2.532228278 | 0.011334018 | 0.064978209 | 0.1631668 |
| ENSMUSG00000034121 | Mks1 | 380718 | protein_coding | 185.2046945 | -0.163231567 | 0.094806697 | -1.721730351 | 0.085118377 | 0.260228257 | 0.163231567 |
| ENSMUSG00000015247 | Nipsnap3b | 66536 | protein_coding | 1405.010756 | 0.163243834 | 0.048852884 | 3.34153937 | 0.000833152 | 0.008657373 | 0.163243834 |
| ENSMUSG00000037236 | Matr3 | 17184 | protein_coding | 1562.33645 | -0.163288202 | 0.065788691 | -2.482010198 | 0.013064354 | 0.072501318 | 0.163288202 |
| ENSMUSG00000026135 | Zfp142 | 77264 | protein_coding | 218.1235592 | 0.163294641 | 0.085943676 | 1.900019281 | 0.057430589 | 0.202887868 | 0.163294641 |
| ENSMUSG00000026705 | Klhl20 | 226541 | protein_coding | 242.923872 | -0.163372887 | 0.086723861 | -1.88382858 | 0.059588166 | 0.207235951 | 0.163372887 |
| ENSMUSG00000026358 | Rgs1 | 50778 | protein_coding | 136.9949989 | 0.163541634 | 0.116976496 | 1.3980726 | 0.162091267 | 0.382898178 | 0.163541634 |
| ENSMUSG00000053293 | Pom121 | 107939 | protein_coding | 279.1584783 | -0.163562279 | 0.116061938 | -1.409267167 | 0.158756183 | 0.378904983 | 0.163562279 |
| ENSMUSG00000025002 | Cyp2c55 | 72082 | protein_coding | 8129.768878 | -0.163618234 | 0.11763611 | -1.390884434 | 0.164260474 | 0.386381076 | 0.163618234 |
| ENSMUSG00000037315 | Jade3 | 382207 | protein_coding | 285.4554594 | -0.163719245 | 0.097102812 | -1.686040207 | 0.091788055 | 0.273111802 | 0.163719245 |
| ENSMUSG00000053094 | Tmem248 | 71667 | protein_coding | 1034.612082 | 0.163752013 | 0.051938857 | 3.152784323 | 0.001617212 | 0.014860846 | 0.163752013 |

| ensembl_gene_id | external_gene_name | entrezgene_id | gene_biotype | baseMean | log2FoldChange | lfcSE | stat | pvalue | padj | absLog2FC |
|---|---|---|---|---|---|---|---|---|---|---|
| ENSMUSG00000021144 | Mta1 | 116870 | protein_coding | 437.48639 | -0.163759826 | 0.081554114 | -2.007989748 | 0.044644381 | 0.171421356 | 0.163759826 |
| ENSMUSG00000002668 | Dennd1c | 70785 | protein_coding | 464.870019 | -0.163784267 | 0.071317043 | -2.296565606 | 0.021643563 | 0.104068496 | 0.163784267 |
| ENSMUSG00000005667 | Mthfd2 | 17768 | protein_coding | 130.5398795 | 0.163813952 | 0.171100532 | 0.957413456 | 0.338358609 | 0.582029301 | 0.163813952 |
| ENSMUSG00000038545 | Cul7 | 66515 | protein_coding | 385.5851183 | 0.163829174 | 0.101830612 | 1.608840124 | 0.107651305 | 0.300613293 | 0.163829174 |
| ENSMUSG00000023284 | Zfp605 | 675812 | protein_coding | 62.76689549 | -0.163829243 | 0.156726723 | -1.045317857 | 0.295876087 | 0.538897588 | 0.163829243 |
| ENSMUSG00000026987 | Baz2b | 407823 | protein_coding | 1364.77432 | -0.1638317 | 0.063730782 | -2.570683974 | 0.01014979 | 0.059914226 | 0.1638317 |
| ENSMUSG00000096914 | Galntl6 | 270049 | protein_coding | 146.6933452 | 0.163930913 | 0.107270933 | 1.528195085 | 0.126464109 | 0.330431117 | 0.163930913 |
| ENSMUSG00000037921 | Ddx60 | 234311 | protein_coding | 537.2657774 | -0.163932583 | 0.193705962 | -0.846296011 | 0.397387633 | 0.636981448 | 0.163932583 |
| ENSMUSG00000019256 | Ahr | 11622 | protein_coding | 444.6541898 | -0.163956929 | 0.070047837 | -2.340642302 | 0.019250601 | 0.095479458 | 0.163956929 |
| ENSMUSG00000005237 | Dnah2 | 327954 | protein_coding | 21.95273621 | 0.163968727 | 0.350539572 | 0.467760959 | 0.639955542 | 0.812603741 | 0.163968727 |
| ENSMUSG00000038203 | Hoxa13 | 15398 | protein_coding | 804.2789886 | -0.163981798 | 0.124350682 | -1.318704451 | 0.187267938 | 0.417073819 | 0.163981798 |
| ENSMUSG00000006586 | Runx1t1 | 12395 | protein_coding | 12.27718569 | -0.163994614 | 0.355816498 | -0.4608966 | 0.644872791 | 0.81580845 | 0.163994614 |
| ENSMUSG00000040651 | Tasor | 218850 | protein_coding | 1059.82843 | -0.164048553 | 0.067400476 | -2.433937612 | 0.014935569 | 0.079926167 | 0.164048553 |
| ENSMUSG00000045019 | Acer1 | 171168 | protein_coding | 1372.583148 | 0.16407254 | 0.063690643 | 2.57608547 | 0.009992594 | 0.059203 | 0.16407254 |
| ENSMUSG00000025465 | Echs1 | 93747 | protein_coding | 3668.911315 | -0.164106167 | 0.050248711 | -3.265878155 | 0.001091252 | 0.010823259 | 0.164106167 |
| ENSMUSG00000037295 | Ldlrap1 | 100017 | protein_coding | 441.4270337 | -0.164117841 | 0.077396058 | -2.120493551 | 0.033964445 | 0.141755849 | 0.164117841 |
| ENSMUSG00000030739 | Myh14 | 71960 | protein_coding | 19837.34827 | 0.164139224 | 0.091148693 | 1.800785269 | 0.071736732 | 0.23362243 | 0.164139224 |
| ENSMUSG00000035298 | Klhl35 | 72184 | protein_coding | 9.262481712 | -0.164183891 | 0.462929656 | -0.354662721 | 0.722842282 | 0.864654909 | 0.164183891 |
| ENSMUSG00000010651 | Acaa1b | 235674 | protein_coding | 185.7176604 | -0.1641975 | 0.171900682 | -0.955188187 | 0.339482539 | 0.582615186 | 0.1641975 |
| ENSMUSG00000020865 | Abcc3 | 76408 | protein_coding | 8226.122339 | -0.164247784 | 0.069998025 | -2.34646312 | 0.018952542 | 0.094722877 | 0.164247784 |
| ENSMUSG00000028035 | Dnajb4 | 67035 | protein_coding | 346.890219 | -0.164257452 | 0.086348967 | -1.902251486 | 0.057138285 | 0.202129099 | 0.164257452 |
| ENSMUSG00000058446 | Znrf2 | 387524 | protein_coding | 2381.226926 | -0.164307588 | 0.047451703 | -3.462627819 | 0.000534928 | 0.005996667 | 0.164307588 |
| ENSMUSG00000063108 | Zfp26 | 22688 | protein_coding | 688.6767465 | -0.16430891 | 0.065804167 | -2.496937764 | 0.012527094 | 0.070364658 | 0.16430891 |
| ENSMUSG00000027823 | Gmps | 229363 | protein_coding | 1220.355508 | 0.16432811 | 0.072400272 | 2.269716746 | 0.023224775 | 0.108932668 | 0.16432811 |
| ENSMUSG00000052253 | Zfp622 | 52521 | protein_coding | 1482.815768 | -0.164360016 | 0.046270125 | -3.552184402 | 0.000382047 | 0.004517062 | 0.164360016 |
| ENSMUSG00000015942 | Gtf2ird2 | 114674 | protein_coding | 177.5122245 | 0.164398015 | 0.097705866 | 1.68258081 | 0.092456269 | 0.274189454 | 0.164398015 |
| ENSMUSG00000096458 | Moap1 | 64113 | protein_coding | 7.425152565 | -0.164398821 | 0.442479663 | -0.371539835 | 0.710235492 | 0.856611363 | 0.164398821 |
| ENSMUSG00000000682 | Cd52 | 23833 | protein_coding | 41.46691647 | 0.164427122 | 0.190084075 | 0.865023136 | 0.38702609 | 0.627104366 | 0.164427122 |
| ENSMUSG00000031822 | Gse1 | 382034 | protein_coding | 779.5713699 | -0.164462831 | 0.087199078 | -1.886061585 | 0.059286656 | 0.206735531 | 0.164462831 |
| ENSMUSG00000042308 | Setd1a | 233904 | protein_coding | 165.1996478 | -0.16450002 | 0.10452364 | -1.573806844 | 0.115532111 | 0.313984894 | 0.16450002 |
| ENSMUSG00000040969 | Arhgef38 | 77669 | protein_coding | 1451.922572 | 0.164522465 | 0.077576754 | 2.120770169 | 0.033941148 | 0.141755849 | 0.164522465 |
| ENSMUSG00000055926 | Gm14137 | 623781 | protein_coding | 16.86648998 | 0.164535095 | 0.340045027 | 0.483862671 | 0.628483334 | 0.804287872 | 0.164535095 |
| ENSMUSG00000074749 | Kiz | 228730 | protein_coding | 502.1339003 | -0.164569212 | 0.081425551 | -2.021100382 | 0.043269375 | 0.167845772 | 0.164569212 |
| ENSMUSG00000021062 | Rab15 | 104886 | protein_coding | 2388.238288 | 0.164615381 | 0.060335165 | 2.728348912 | 0.006365224 | 0.042790188 | 0.164615381 |
| ENSMUSG00000026418 | Tnni1 | 21952 | protein_coding | 701.1527054 | -0.164636925 | 0.165204749 | -0.996562907 | 0.318976718 | 0.562649386 | 0.164636925 |
| ENSMUSG00000022831 | Hcls1 | 15163 | protein_coding | 36.87855408 | 0.164651397 | 0.218322884 | 0.754164628 | 0.450750374 | 0.679951459 | 0.164651397 |
| ENSMUSG00000029466 | Anapc7 | 56317 | protein_coding | 610.4100672 | -0.164656278 | 0.066601862 | -2.472247387 | 0.013426657 | 0.074131168 | 0.164656278 |
| ENSMUSG00000064351 | mt-Co1 | 17708 | protein_coding | 692323.5131 | 0.164689718 | 0.048544699 | 3.392537625 | 0.000692484 | 0.007465035 | 0.164689718 |
| ENSMUSG00000073147 | 5031425E22Rik | 269630 | lncRNA | 323.6690284 | -0.164701314 | 0.092898267 | -1.772921278 | 0.076241752 | 0.243234568 | 0.164701314 |

| ensembl_gene_id | external_gene_name | entrezgene_id | gene_biotype | baseMean | log2FoldChange | lfcSE | stat | pvalue | padj | absLog2FC |
|---|---|---|---|---|---|---|---|---|---|---|
| ENSMUSG00000015882 | Lcorl | 209707 | protein_coding | 1132.793536 | -0.16473459 | 0.075988009 | -2.16790245 | 0.030166109 | 0.130636103 | 0.16473459 |
| ENSMUSG00000052917 | Senp7 | 66315 | protein_coding | 348.6969839 | -0.164756467 | 0.07415 | -2.221934832 | 0.02628771 | 0.118778578 | 0.164756467 |
| ENSMUSG00000027281 | Slx4ip | 74243 | protein_coding | 280.1181563 | 0.164767134 | 0.095388038 | 1.727335388 | 0.084107447 | 0.258509777 | 0.164767134 |
| ENSMUSG00000073002 | Vamp5 | 53620 | protein_coding | 17.0957622 | 0.164779601 | 0.342507503 | 0.481097786 | 0.630447002 | 0.805900922 | 0.164779601 |
| ENSMUSG00000056498 | Tmem154 | 320782 | protein_coding | 18.30015518 | -0.164802483 | 0.291785037 | -0.564807864 | 0.572204457 | 0.767059504 | 0.164802483 |
| ENSMUSG00000004610 | Etfb | 110826 | protein_coding | 688.9533112 | 0.164806963 | 0.070306999 | 2.344104638 | 0.01907282 | 0.095081101 | 0.164806963 |
| ENSMUSG00000021993 | Mipep | 70478 | protein_coding | 562.1564205 | 0.164824883 | 0.074655252 | 2.207813646 | 0.027257266 | 0.121280976 | 0.164824883 |
| ENSMUSG00000038718 | Pbx3 | 18516 | protein_coding | 62.33905005 | 0.164894272 | 0.16186257 | 1.018730101 | 0.308331118 | 0.552028135 | 0.164894272 |
| ENSMUSG00000030865 | Chp2 | 70261 | protein_coding | 2604.495839 | -0.16490058 | 0.080111308 | -2.058393291 | 0.039552395 | 0.157263678 | 0.16490058 |
| ENSMUSG00000033970 | Rfc3 | 69263 | protein_coding | 308.7492401 | 0.164923088 | 0.120531608 | 1.368297416 | 0.171218999 | 0.395821216 | 0.164923088 |
| ENSMUSG00000037808 | Fam76b | 72826 | protein_coding | 544.2595051 | -0.165002282 | 0.061554788 | -2.680575897 | 0.00734956 | 0.0474675 | 0.165002282 |
| ENSMUSG00000038379 | Ttk | 22137 | protein_coding | 190.7853104 | 0.165012825 | 0.133222669 | 1.238624222 | 0.215484693 | 0.451138732 | 0.165012825 |
| ENSMUSG00000044676 | Zfp612 | 234725 | protein_coding | 231.2679551 | -0.165034822 | 0.11193386 | -1.474395886 | 0.140375032 | 0.351472935 | 0.165034822 |
| ENSMUSG00000004665 | Cnn2 | 12798 | protein_coding | 135.3295723 | -0.165048064 | 0.109300608 | -1.510037934 | 0.131033745 | 0.336976751 | 0.165048064 |
| ENSMUSG00000050777 | Tmem37 | 170706 | protein_coding | 171.4886258 | 0.165049643 | 0.170492253 | 0.96807708 | 0.333005876 | 0.576345327 | 0.165049643 |
| ENSMUSG00000027763 | Mbnl1 | 56758 | protein_coding | 2923.854039 | -0.165110605 | 0.048221262 | -3.424020842 | 0.000617019 | 0.006801547 | 0.165110605 |
| ENSMUSG00000044148 | 1810030O07Rik | 69155 | protein_coding | 692.1006838 | 0.165159008 | 0.064229625 | 2.571383651 | 0.010129304 | 0.059837104 | 0.165159008 |
| ENSMUSG00000068856 | Sf3b4 | 107701 | protein_coding | 257.4365883 | 0.165188895 | 0.094462145 | 1.748731144 | 0.080337504 | 0.250891349 | 0.165188895 |
| ENSMUSG00000047710 | Champ1 | 101994 | protein_coding | 482.67393 | -0.165235598 | 0.061294297 | -2.695774421 | 0.007022521 | 0.045906846 | 0.165235598 |
| ENSMUSG00000053801 | Grwd1 | 101612 | protein_coding | 299.1364149 | -0.165247682 | 0.083058253 | -1.989539597 | 0.046641676 | 0.176322153 | 0.165247682 |
| ENSMUSG00000041491 | Cep78 | 208518 | protein_coding | 163.39803 | 0.165248933 | 0.110151584 | 1.500195706 | 0.133563715 | 0.340981801 | 0.165248933 |
| ENSMUSG00000037824 | Tspan14 | 52588 | protein_coding | 973.7938288 | 0.165284986 | 0.059303585 | 2.787099389 | 0.005318217 | 0.037338649 | 0.165284986 |
| ENSMUSG00000061461 | Smim20 | 66278 | protein_coding | 848.7345519 | 0.165300057 | 0.051120606 | 3.233530863 | 0.001222701 | 0.011907309 | 0.165300057 |
| ENSMUSG00000028633 | Ctps | 51797 | protein_coding | 265.4951191 | -0.165314039 | 0.126849541 | -1.30322931 | 0.192496486 | 0.42427099 | 0.165314039 |
| ENSMUSG00000058756 | Thra | 21833 | protein_coding | 1237.285819 | -0.165356252 | 0.064270355 | -2.57282306 | 0.010087276 | 0.059625892 | 0.165356252 |
| ENSMUSG00000037344 | Slc12a9 | 83704 | protein_coding | 369.8928147 | -0.165356475 | 0.118716517 | -1.392868315 | 0.163659615 | 0.385451042 | 0.165356475 |
| ENSMUSG00000087260 | Lamtor5 | 68576 | protein_coding | 722.9742738 | 0.165374814 | 0.066911132 | 2.471559043 | 0.013452533 | 0.074195967 | 0.165374814 |
| ENSMUSG00000024052 | Lpin2 | 64898 | protein_coding | 1299.385512 | -0.165452948 | 0.078963461 | -2.095310241 | 0.036143423 | 0.147677978 | 0.165452948 |
| ENSMUSG00000052373 | Mpp3 | 13384 | protein_coding | 134.4332895 | -0.165457293 | 0.125326833 | -1.320206448 | 0.1867661 | 0.416363961 | 0.165457293 |
| ENSMUSG00000026201 | Stk16 | 20872 | protein_coding | 1230.825472 | 0.165576177 | 0.064730583 | 2.557928087 | 0.010529786 | 0.061552292 | 0.165576177 |
| ENSMUSG00000004996 | Mri1 | 67873 | protein_coding | 419.8678412 | -0.165629631 | 0.074618681 | -2.219680488 | 0.026440464 | 0.118917792 | 0.165629631 |
| ENSMUSG00000025224 | Gbf1 | 107338 | protein_coding | 1918.301194 | 0.165635603 | 0.057346902 | 2.888309499 | 0.003873186 | 0.0294918 | 0.165635603 |
| ENSMUSG00000038388 | Pals2 | 56524 | protein_coding | 1959.440456 | 0.165664456 | 0.105411225 | 1.571601653 | 0.116042963 | 0.314789716 | 0.165664456 |
| ENSMUSG00000039682 | Lap3 | 66988 | protein_coding | 1586.579786 | 0.165707711 | 0.109423252 | 1.514373846 | 0.129931049 | 0.335854788 | 0.165707711 |
| ENSMUSG00000025915 | Sgk3 | 170755 | protein_coding | 103.6728179 | 0.165730395 | 0.127638156 | 1.298439276 | 0.19413643 | 0.426123143 | 0.165730395 |
| ENSMUSG00000029600 | Rita1 | 100764 | protein_coding | 295.7262906 | 0.165762224 | 0.083905701 | 1.975577612 | 0.048202636 | 0.180362858 | 0.165762224 |
| ENSMUSG00000041135 | Ripk2 | 192656 | protein_coding | 230.2948017 | -0.165773317 | 0.102904486 | -1.610943546 | 0.107192028 | 0.29959893 | 0.165773317 |
| ENSMUSG00000031229 | Atrx | 22589 | protein_coding | 2568.525259 | 0.165789186 | 0.08319324 | 1.992820399 | 0.046281117 | 0.175601538 | 0.165789186 |
| ENSMUSG00000041777 | Cir1 | 66935 | protein_coding | 167.1595808 | -0.165819163 | 0.110578077 | -1.499566337 | 0.133726773 | 0.34107542 | 0.165819163 |

| ensembl_gene_id | external_gene_name | entrezgene_id | gene_biotype | baseMean | log2FoldChange | lfcSE | stat | pvalue | padj | absLog2FC |
|---|---|---|---|---|---|---|---|---|---|---|
| ENSMUSG00000030779 | Rbbp6 | 19647 | protein_coding | 1018.766526 | -0.165846694 | 0.066705955 | -2.486235227 | 0.01291026 | 0.071843948 | 0.165846694 |
| ENSMUSG00000036916 | Zfp280c | 208968 | protein_coding | 200.6167607 | -0.165897802 | 0.086920842 | -1.908607862 | 0.056312697 | 0.200512459 | 0.165897802 |
| ENSMUSG00000038170 | Pde4dip | 83679 | protein_coding | 3522.061209 | 0.16592176 | 0.035770937 | 4.638451587 | 3.51E-06 | 7.60E-05 | 0.16592176 |
| ENSMUSG00000017376 | Nlk | 18099 | protein_coding | 371.8235564 | 0.165934388 | 0.088628605 | 1.872244156 | 0.061172833 | 0.210759777 | 0.165934388 |
| ENSMUSG00000029920 | Smarcad1 | 13990 | protein_coding | 690.4515038 | -0.165967376 | 0.069715511 | -2.380637727 | 0.017282699 | 0.08873319 | 0.165967376 |
| ENSMUSG00000049960 | Mrps16 | 66242 | protein_coding | 783.423489 | 0.165972204 | 0.074052096 | 2.241289741 | 0.025007313 | 0.114712014 | 0.165972204 |
| ENSMUSG00000027107 | Chrna1 | 11435 | protein_coding | 31.64512673 | 0.166047942 | 0.274366694 | 0.605204441 | 0.545043165 | 0.749012844 | 0.166047942 |
| ENSMUSG00000038569 | Rad9b | 231724 | protein_coding | 55.52950857 | -0.166080229 | 0.232547355 | -0.714178102 | 0.475117056 | 0.69872449 | 0.166080229 |
| ENSMUSG00000022216 | Psme1 | 19186 | protein_coding | 2183.257622 | -0.166086155 | 0.095558987 | -1.738048507 | 0.082202267 | 0.254568403 | 0.166086155 |
| ENSMUSG00000030491 | Tdrd12 | 71981 | protein_coding | 19.95538948 | -0.166126069 | 0.284048367 | -0.584851344 | 0.558647677 | 0.757084966 | 0.166126069 |
| ENSMUSG00000063646 | Jakmip1 | 76071 | protein_coding | 196.5772735 | 0.166152522 | 0.090390583 | 1.838161861 | 0.066038558 | 0.221624129 | 0.166152522 |
| ENSMUSG00000001627 | Ifrd1 | 15982 | protein_coding | 1683.942455 | -0.166167441 | 0.111136748 | -1.495161991 | 0.134872171 | 0.342694737 | 0.166167441 |
| ENSMUSG00000040446 | Rprd1a | 225283 | protein_coding | 189.0230702 | -0.166173915 | 0.091364926 | -1.818793295 | 0.068942972 | 0.227836743 | 0.166173915 |
| ENSMUSG00000043671 | Dpy19l3 | 233115 | protein_coding | 146.5051929 | -0.166224263 | 0.102105303 | -1.627968955 | 0.103531469 | 0.294056357 | 0.166224263 |
| ENSMUSG00000021607 | Mrpl36 | 94066 | protein_coding | 801.4839288 | 0.166242654 | 0.080561076 | 2.063560496 | 0.039059406 | 0.155879992 | 0.166242654 |
| ENSMUSG00000048534 | Jaml | 270152 | protein_coding | 35.4952348 | -0.166292935 | 0.200099408 | -0.831051606 | 0.405944477 | 0.645118898 | 0.166292935 |
| ENSMUSG00000111731 | Gm48727 | 118567821 | lncRNA | 6.338460205 | 0.166309642 | 0.478228839 | 0.347761634 | 0.728019205 | 0.867832282 | 0.166309642 |
| ENSMUSG00000020514 | Mrpl22 | 216767 | protein_coding | 326.1198932 | 0.166317098 | 0.083554586 | 1.990520276 | 0.046533653 | 0.176202916 | 0.166317098 |
| ENSMUSG00000026739 | Bmi1 | 12151 | protein_coding | 424.9665461 | -0.166360433 | 0.070480164 | -2.360386591 | 0.018255899 | 0.092350341 | 0.166360433 |
| ENSMUSG00000029596 | Sdsl | 257635 | protein_coding | 399.689868 | 0.166385308 | 0.094714708 | 1.756699794 | 0.078968993 | 0.248300541 | 0.166385308 |
| ENSMUSG00000035171 | 1110059E24Rik | 66206 | protein_coding | 943.6105342 | 0.166502053 | 0.079711739 | 2.088802152 | 0.036725537 | 0.149187085 | 0.166502053 |
| ENSMUSG00000057766 | Ankrd29 | 225187 | protein_coding | 14.63007459 | -0.166570907 | 0.391357265 | -0.425623648 | 0.670382106 | 0.831826746 | 0.166570907 |
| ENSMUSG00000038085 | Cnbd2 | 70873 | protein_coding | 52.99810279 | -0.166576383 | 0.188035666 | -0.885876526 | 0.375684067 | 0.617285476 | 0.166576383 |
| ENSMUSG00000114456 | H2bc9 | 319182 | protein_coding | 835.1912401 | 0.166584625 | 0.135208646 | 1.232056008 | 0.217928163 | 0.454144426 | 0.166584625 |
| ENSMUSG00000000738 | Spg7 | 234847 | protein_coding | 2206.416392 | -0.166627963 | 0.039046541 | -4.267419297 | 1.98E-05 | 0.00035792 | 0.166627963 |
| ENSMUSG00000026303 | Mlph | 171531 | protein_coding | 1206.586114 | 0.166640702 | 0.069592696 | 2.394514256 | 0.016642396 | 0.086466472 | 0.166640702 |
| ENSMUSG00000020238 | Ncln | 103425 | protein_coding | 1993.12578 | 0.166695786 | 0.077882045 | 2.140362194 | 0.032325507 | 0.137151659 | 0.166695786 |
| ENSMUSG00000046311 | Zfp62 | 22720 | protein_coding | 711.6262586 | -0.166740566 | 0.072751751 | -2.29191138 | 0.021910757 | 0.104853052 | 0.166740566 |
| ENSMUSG00000041477 | Dcp1b | 319618 | protein_coding | 334.2478833 | -0.166761202 | 0.077455432 | -2.152995578 | 0.031319026 | 0.133956365 | 0.166761202 |
| ENSMUSG00000026565 | Pou2f1 | 18986 | protein_coding | 452.6948895 | -0.166776267 | 0.079125301 | -2.107748899 | 0.035052715 | 0.144724817 | 0.166776267 |
| ENSMUSG00000005481 | Ddx39a | 68278 | protein_coding | 1194.178152 | 0.166811092 | 0.092305954 | 1.807154189 | 0.070738241 | 0.231492618 | 0.166811092 |
| ENSMUSG00000070985 | Acnat1 | 230161 | protein_coding | 718.6401075 | 0.166812796 | 0.087170871 | 1.913630031 | 0.055667445 | 0.198761565 | 0.166812796 |
| ENSMUSG00000042699 | Dhx9 | 13211 | protein_coding | 1609.213242 | -0.166858881 | 0.083276465 | -2.003673916 | 0.045105002 | 0.17246929 | 0.166858881 |
| ENSMUSG00000022452 | Smdt1 | 69029 | protein_coding | 2466.450931 | 0.166871259 | 0.083756391 | 1.992340609 | 0.046333699 | 0.17573461 | 0.166871259 |
| ENSMUSG00000097471 | 5830432E09Rik | 67765 | lncRNA | 7.277056556 | 0.16692561 | 0.440434841 | 0.379001828 | 0.704686506 | 0.853157579 | 0.16692561 |
| ENSMUSG00000015950 | Ncf1 | 17969 | protein_coding | 18.74994807 | 0.167033511 | 0.264916596 | 0.630513577 | 0.528358623 | 0.737932232 | 0.167033511 |
| ENSMUSG00000090946 | Ccdc71l | 72123 | protein_coding | 1624.244976 | 0.16703593 | 0.087636104 | 1.906017292 | 0.056647962 | 0.201136435 | 0.16703593 |
| ENSMUSG00000025969 | Nrp2 | 18187 | protein_coding | 2390.245933 | -0.167056935 | 0.097765403 | -1.708753104 | 0.087496693 | 0.264813247 | 0.167056935 |
| ENSMUSG00000063652 | Slc22a21 | 56517 | protein_coding | 258.0932258 | -0.167083144 | 0.110476147 | -1.512391131 | 0.13043439 | 0.336485813 | 0.167083144 |

| ensembl_gene_id | external_gene_name | entrezgene_id | gene_biotype | baseMean | log2FoldChange | lfcSE | stat | pvalue | padj | absLog2FC |
|---|---|---|---|---|---|---|---|---|---|---|
| ENSMUSG00000026617 | Bpnt1 | 23827 | protein_coding | 3000.919362 | 0.16710602 | 0.076236998 | 2.19192812 | 0.028384697 | 0.125036879 | 0.16710602 |
| ENSMUSG00000078716 | Tmem8b | 242409 | protein_coding | 65.19023951 | -0.167135215 | 0.15560594 | -1.074092769 | 0.282781105 | 0.525455543 | 0.167135215 |
| ENSMUSG00000047528 | Flacc1 | 108812 | protein_coding | 54.16483846 | -0.167148139 | 0.214593535 | -0.77890575 | 0.436035236 | 0.669302737 | 0.167148139 |
| ENSMUSG00000096696 | Zfp960 | 449000 | protein_coding | 53.76358697 | -0.167173461 | 0.169382923 | -0.986955815 | 0.3236643 | 0.567732669 | 0.167173461 |
| ENSMUSG00000015290 | Ubl4a | 27643 | protein_coding | 951.3472204 | 0.167199459 | 0.049541235 | 3.374955372 | 0.000738277 | 0.007848508 | 0.167199459 |
| ENSMUSG00000041671 | Pyroxd1 | 232491 | protein_coding | 394.4099378 | -0.167318706 | 0.078500199 | -2.131443069 | 0.033052658 | 0.139211338 | 0.167318706 |
| ENSMUSG00000003623 | Crot | 74114 | protein_coding | 1560.176488 | 0.167409696 | 0.051157226 | 3.272454514 | 0.00106618 | 0.010613759 | 0.167409696 |
| ENSMUSG00000031398 | Plxna3 | 18846 | protein_coding | 113.6593321 | -0.167430965 | 0.133197492 | -1.257012891 | 0.208748968 | 0.444363068 | 0.167430965 |
| ENSMUSG00000024772 | Ehd1 | 13660 | protein_coding | 1454.090642 | 0.167450743 | 0.06551643 | 2.555858773 | 0.01059261 | 0.061835229 | 0.167450743 |
| ENSMUSG00000025137 | Pcyt2 | 68671 | protein_coding | 1762.534992 | -0.167472148 | 0.058789058 | -2.848695894 | 0.004389882 | 0.032385922 | 0.167472148 |
| ENSMUSG00000061458 | Nol10 | 217431 | protein_coding | 115.8312271 | 0.167515726 | 0.136822215 | 1.224331338 | 0.22082726 | 0.457336743 | 0.167515726 |
| ENSMUSG00000034664 | Itga2b | 16399 | protein_coding | 20.90622874 | 0.167522265 | 0.274022651 | 0.61134459 | 0.540971477 | 0.746550877 | 0.167522265 |
| ENSMUSG00000017210 | Med24 | 23989 | protein_coding | 823.1244437 | 0.167549979 | 0.081280776 | 2.061372777 | 0.03926749 | 0.156571643 | 0.167549979 |
| ENSMUSG00000028848 | Gpn2 | 100210 | protein_coding | 308.7463068 | 0.167551768 | 0.080403636 | 2.083882968 | 0.037170814 | 0.150503067 | 0.167551768 |
| ENSMUSG00000029439 | Sfswap | 231769 | protein_coding | 826.7182734 | -0.167568726 | 0.072755611 | -2.303172545 | 0.021269137 | 0.102482035 | 0.167568726 |
| ENSMUSG00000002489 | Tiam1 | 21844 | protein_coding | 324.34979 | -0.167578333 | 0.112702448 | -1.486909431 | 0.137038733 | 0.346479729 | 0.167578333 |
| ENSMUSG00000035953 | Pip4p1 | 219024 | protein_coding | 730.3205716 | -0.167583844 | 0.058336252 | -2.872722161 | 0.004069518 | 0.03065349 | 0.167583844 |
| ENSMUSG00000089804 | Gm16136 | 105245924 | lncRNA | 172.559078 | -0.167583885 | 0.099533487 | -1.683693493 | 0.092240919 | 0.273701805 | 0.167583885 |
| ENSMUSG00000042901 | Aida | 108909 | protein_coding | 386.3917542 | -0.167699976 | 0.079003753 | -2.122683655 | 0.03378037 | 0.141426796 | 0.167699976 |
| ENSMUSG00000061273 | Mmgt1 | 236792 | protein_coding | 865.5618068 | 0.167726968 | 0.060913012 | 2.753549066 | 0.005895293 | 0.040439555 | 0.167726968 |
| ENSMUSG00000030604 | Zfp626 | 71163 | protein_coding | 324.6830582 | -0.167745746 | 0.091875408 | -1.825795921 | 0.067881008 | 0.225575317 | 0.167745746 |
| ENSMUSG00000026600 | Soat1 | 20652 | protein_coding | 1533.848817 | -0.16775353 | 0.068713329 | -2.441353536 | 0.014632322 | 0.07881612 | 0.16775353 |
| ENSMUSG00000031683 | Lsm6 | 78651 | protein_coding | 485.0548023 | 0.167755651 | 0.087765662 | 1.911404154 | 0.055952663 | 0.199502233 | 0.167755651 |
| ENSMUSG00000001819 | Hoxd13 | 15433 | protein_coding | 306.893066 | 0.167799649 | 0.351034187 | 0.47801512 | 0.632639443 | 0.807820141 | 0.167799649 |
| ENSMUSG00000013483 | Card14 | 170720 | protein_coding | 448.9090063 | 0.167801942 | 0.085949376 | 1.952334611 | 0.050898491 | 0.187320849 | 0.167801942 |
| ENSMUSG00000006471 | Ndor1 | 78797 | protein_coding | 108.3978387 | -0.167806186 | 0.127228433 | -1.318936204 | 0.187190441 | 0.416996968 | 0.167806186 |
| ENSMUSG00000002944 | Cd36 | 12491 | protein_coding | 7.596273546 | 0.16786977 | 0.577687324 | 0.290589326 | 0.771365425 | 0.892332662 | 0.16786977 |
| ENSMUSG00000063439 | B9d2 | 232987 | protein_coding | 29.82322742 | 0.167895294 | 0.231029906 | 0.726725372 | 0.467394207 | 0.692926016 | 0.167895294 |
| ENSMUSG00000042389 | Tsen2 | 381802 | protein_coding | 97.60283456 | 0.167952452 | 0.135452888 | 1.239932602 | 0.215000324 | 0.45050036 | 0.167952452 |
| ENSMUSG00000037369 | Kdm6a | 22289 | protein_coding | 1728.277472 | 0.167995903 | 0.067837029 | 2.47646317 | 0.013269131 | 0.073410389 | 0.167995903 |
| ENSMUSG00000039367 | Sec24c | 218811 | protein_coding | 2895.017745 | 0.168009364 | 0.042409241 | 3.961621616 | 7.44E-05 | 0.001152143 | 0.168009364 |
| ENSMUSG00000063410 | Stk24 | 223255 | protein_coding | 1502.638185 | 0.168011675 | 0.047507529 | 3.536527334 | 0.000405425 | 0.004734454 | 0.168011675 |
| ENSMUSG00000021418 | Rpp40 | 208366 | protein_coding | 62.30998928 | -0.168013601 | 0.156387194 | -1.074343724 | 0.282668655 | 0.52536561 | 0.168013601 |
| ENSMUSG00000025429 | Pstpip2 | 19201 | protein_coding | 316.7461823 | -0.168030221 | 0.114417653 | -1.468569034 | 0.141949719 | 0.353057236 | 0.168030221 |
| ENSMUSG00000068959 | Zfp619 | 70227 | protein_coding | 73.46761734 | -0.168034082 | 0.164450499 | -1.021791251 | 0.306879711 | 0.550870027 | 0.168034082 |
| ENSMUSG00000039629 | Strip2 | 320609 | protein_coding | 234.8812871 | 0.16806405 | 0.116635155 | 1.440938196 | 0.149602144 | 0.36472166 | 0.16806405 |
| ENSMUSG00000032854 | Ugt8a | 22239 | protein_coding | 316.300783 | -0.168097537 | 0.143426428 | -1.172012294 | 0.24119212 | 0.481816589 | 0.168097537 |
| ENSMUSG00000006412 | Pfdn2 | 18637 | protein_coding | 193.876471 | 0.168114713 | 0.091389711 | 1.839536537 | 0.065836309 | 0.221242374 | 0.168114713 |
| ENSMUSG00000038236 | Hoxa7 | 15404 | protein_coding | 409.0334646 | 0.168161897 | 0.089870046 | 1.871167365 | 0.061321887 | 0.210995961 | 0.168161897 |

Supplementary Table 1

| ensembl_gene_id | external_gene_name | entrezgene_id | gene_biotype | baseMean | log2FoldChange | lfcSE | stat | pvalue | padj | absLog2FC |
|---|---|---|---|---|---|---|---|---|---|---|
| ENSMUSG00000075028 | Prdm11 | 100042784 | protein_coding | 42.50277722 | 0.168162066 | 0.197888311 | 0.849782714 | 0.395445902 | 0.63475227 | 0.168162066 |
| ENSMUSG00000025402 | Nab2 | 17937 | protein_coding | 14.49714278 | 0.168164448 | 0.318557057 | 0.527894279 | 0.597572716 | 0.78394657 | 0.168164448 |
| ENSMUSG00000020330 | Hmmr | 15366 | protein_coding | 538.357118 | 0.168182908 | 0.097355038 | 1.727521351 | 0.084074074 | 0.258505737 | 0.168182908 |
| ENSMUSG00000090115 | Usp49 | 224836 | protein_coding | 205.0748265 | -0.168202586 | 0.088689308 | -1.896537349 | 0.057889026 | 0.203825108 | 0.168202586 |
| ENSMUSG00000039789 | Zfp597 | 71063 | protein_coding | 255.0118693 | -0.168237031 | 0.105578656 | -1.593475782 | 0.111053497 | 0.30635644 | 0.168237031 |
| ENSMUSG00000004748 | Mtfp1 | 67900 | protein_coding | 292.2806522 | 0.168316368 | 0.094043521 | 1.789771006 | 0.073490732 | 0.237226789 | 0.168316368 |
| ENSMUSG00000020857 | Nme2 | 18103 | protein_coding | 205.3741299 | 0.168389687 | 0.152852337 | 1.101649408 | 0.27061412 | 0.513253429 | 0.168389687 |
| ENSMUSG00000026765 | Lypd6b | 71897 | protein_coding | 130.0169727 | 0.168425675 | 0.14230143 | 1.183583849 | 0.236577837 | 0.476792149 | 0.168425675 |
| ENSMUSG00000028277 | Ube2j1 | 56228 | protein_coding | 1375.595576 | 0.1684846 | 0.04389451 | 3.838398003 | 0.00012384 | 0.001773678 | 0.1684846 |
| ENSMUSG00000029165 | Agbl5 | 231093 | protein_coding | 211.4667963 | -0.1684963 | 0.112066518 | -1.503538278 | 0.132700294 | 0.339503983 | 0.1684963 |
| ENSMUSG00000016520 | Lnx2 | 140887 | protein_coding | 776.6953012 | -0.168508016 | 0.070355769 | -2.395084563 | 0.016616532 | 0.086415614 | 0.168508016 |
| ENSMUSG00000032386 | Trip4 | 56404 | protein_coding | 645.1485024 | 0.168513004 | 0.063074515 | 2.671649631 | 0.007547941 | 0.048515603 | 0.168513004 |
| ENSMUSG00000041417 | Pik3r1 | 18708 | protein_coding | 3776.430414 | -0.168529368 | 0.045446229 | -3.708324575 | 0.000208635 | 0.002731844 | 0.168529368 |
| ENSMUSG00000026383 | Epb41l5 | 226352 | protein_coding | 431.4973563 | -0.168621481 | 0.065271649 | -2.58338015 | 0.009783745 | 0.058437948 | 0.168621481 |
| ENSMUSG00000003119 | Cdk12 | 69131 | protein_coding | 1404.367722 | -0.168643649 | 0.040051745 | -4.210644238 | 2.55E-05 | 0.000448813 | 0.168643649 |
| ENSMUSG00000064061 | Dzip3 | 224170 | protein_coding | 141.6220733 | 0.168695222 | 0.110998879 | 1.519792124 | 0.128563229 | 0.333871046 | 0.168695222 |
| ENSMUSG00000088252 | Snord13 | 100217422 | snoRNA | 233.7677557 | -0.168740553 | 0.204069825 | -0.82687655 | 0.408307036 | 0.646860933 | 0.168740553 |
| ENSMUSG00000062007 | Hsh2d | 209488 | protein_coding | 49.27590091 | 0.168800417 | 0.174478718 | 0.967455625 | 0.333316314 | 0.576562634 | 0.168800417 |
| ENSMUSG00000055865 | Tafa3 | 329731 | protein_coding | 9.842479037 | -0.168803919 | 0.414036222 | -0.407703264 | 0.683491541 | 0.840526771 | 0.168803919 |
| ENSMUSG00000101174 | Hoxd4 | 15436 | protein_coding | 30.7808312 | -0.168817976 | 0.295150588 | -0.571972352 | 0.567340706 | 0.762827428 | 0.168817976 |
| ENSMUSG00000029504 | Ddx51 | 69663 | protein_coding | 240.723484 | -0.168817987 | 0.093300055 | -1.809409298 | 0.07038744 | 0.23074217 | 0.168817987 |
| ENSMUSG00000028636 | Ppcs | 106564 | protein_coding | 482.0493183 | 0.168841224 | 0.069396191 | 2.433004193 | 0.014974127 | 0.080047962 | 0.168841224 |
| ENSMUSG00000027165 | Iftap | 68170 | protein_coding | 111.8494116 | 0.168865936 | 0.134010393 | 1.26009582 | 0.207634798 | 0.442926771 | 0.168865936 |
| ENSMUSG00000025020 | Slit1 | 20562 | protein_coding | 5.934507614 | 0.168900567 | 0.526786961 | 0.320624047 | 0.748495312 | 0.880475742 | 0.168900567 |
| ENSMUSG00000036646 | Man1b1 | 227619 | protein_coding | 3137.041953 | 0.168940649 | 0.041953708 | 4.026834737 | 5.65E-05 | 0.000910793 | 0.168940649 |
| ENSMUSG00000097383 | 1500026H17Rik | 69002 | lncRNA | 27.61502588 | 0.168964531 | 0.242682871 | 0.696235915 | 0.486281099 | 0.707337899 | 0.168964531 |
| ENSMUSG00000071302 | 2610044O15Rik8 | 72139 | protein_coding | 190.5995167 | -0.168977825 | 0.126926751 | -1.331301903 | 0.183089692 | 0.411180542 | 0.168977825 |
| ENSMUSG00000052214 | Opa3 | 403187 | protein_coding | 1777.18698 | 0.168991741 | 0.058265268 | 2.900385522 | 0.003727039 | 0.028675443 | 0.168991741 |
| ENSMUSG00000023791 | Pigx | 72084 | protein_coding | 576.8002518 | -0.169031907 | 0.068726756 | -2.459477462 | 0.013913943 | 0.076012928 | 0.169031907 |
| ENSMUSG00000046269 | Usp27x | 54651 | protein_coding | 46.10082879 | 0.169040635 | 0.199617048 | 0.846824639 | 0.397092872 | 0.636592707 | 0.169040635 |
| ENSMUSG00000042363 | Lgalsl | 216551 | protein_coding | 929.1703532 | 0.169048482 | 0.063524059 | 2.661172572 | 0.007786904 | 0.049660074 | 0.169048482 |
| ENSMUSG00000019866 | Crybg1 | 11630 | protein_coding | 3020.940367 | 0.169087539 | 0.062094955 | 2.723047937 | 0.006468267 | 0.04322388 | 0.169087539 |
| ENSMUSG00000049580 | Tsku | 244152 | protein_coding | 707.8317033 | -0.16912768 | 0.136480593 | -1.23920681 | 0.215268919 | 0.450745503 | 0.16912768 |
| ENSMUSG00000034708 | Grn | 14824 | protein_coding | 1797.973969 | -0.169129898 | 0.066099893 | -2.558701537 | 0.01050639 | 0.061523803 | 0.169129898 |
| ENSMUSG00000031958 | Ldhd | 52815 | protein_coding | 183.97528 | 0.169215557 | 0.088089428 | 1.920951921 | 0.054737769 | 0.196492651 | 0.169215557 |
| ENSMUSG00000072494 | Ppp1r3e | 105651 | protein_coding | 88.71519231 | -0.169262803 | 0.152281764 | -1.111510655 | 0.266348606 | 0.508211544 | 0.169262803 |
| ENSMUSG00000035478 | Mbd3 | 17192 | protein_coding | 778.9645041 | 0.169286686 | 0.075245337 | 2.249796378 | 0.024461874 | 0.113101102 | 0.169286686 |
| ENSMUSG00000033065 | Pfkm | 18642 | protein_coding | 309.6788262 | -0.169343207 | 0.083769551 | -2.021536527 | 0.043224255 | 0.167727999 | 0.169343207 |
| ENSMUSG00000063142 | Kcnma1 | 16531 | protein_coding | 559.4272364 | 0.169343947 | 0.103912929 | 1.629671582 | 0.103170927 | 0.293497536 | 0.169343947 |

| ensembl_gene_id | external_gene_name | entrezgene_id | gene_biotype | baseMean | log2FoldChange | lfcSE | stat | pvalue | padj | absLog2FC |
|---|---|---|---|---|---|---|---|---|---|---|
| ENSMUSG00000026154 | Sdhaf4 | 68002 | protein_coding | 208.233737 | 0.169367559 | 0.084435853 | 2.005872535 | 0.044869849 | 0.171848571 | 0.169367559 |
| ENSMUSG00000021079 | Timm9 | 30056 | protein_coding | 233.3015378 | -0.169375071 | 0.091612779 | -1.848814901 | 0.064484545 | 0.21806296 | 0.169375071 |
| ENSMUSG00000068134 | Zfp120 | 104348 | protein_coding | 434.1675761 | -0.169427496 | 0.076303399 | -2.220444952 | 0.026388579 | 0.118807539 | 0.169427496 |
| ENSMUSG00000032363 | Adamts7 | 108153 | protein_coding | 9.248130029 | 0.169452592 | 0.384813895 | 0.440349463 | 0.659684021 | 0.82592378 | 0.169452592 |
| ENSMUSG00000029029 | Wrap73 | 59002 | protein_coding | 250.7188867 | 0.169460918 | 0.104671222 | 1.618982903 | 0.105450942 | 0.297761537 | 0.169460918 |
| ENSMUSG00000021811 | Dnajc9 | 108671 | protein_coding | 336.2272681 | 0.169470033 | 0.121328598 | 1.396785552 | 0.162478068 | 0.383474873 | 0.169470033 |
| ENSMUSG00000021069 | Pygl | 110095 | protein_coding | 188.7298569 | 0.169553697 | 0.105942467 | 1.600431835 | 0.109502818 | 0.303323933 | 0.169553697 |
| ENSMUSG00000005534 | Insr | 16337 | protein_coding | 3121.900189 | -0.169563954 | 0.05374608 | -3.15490832 | 0.001605485 | 0.014803696 | 0.169563954 |
| ENSMUSG00000099632 | 2900093K20Rik | 108168734 | lncRNA | 25.46448178 | -0.169610661 | 0.252156498 | -0.672640453 | 0.501176059 | 0.718355879 | 0.169610661 |
| ENSMUSG00000036377 | Cracd | 320827 | protein_coding | 1798.396276 | -0.169620676 | 0.072042365 | -2.35445736 | 0.018549772 | 0.093455848 | 0.169620676 |
| ENSMUSG00000007777 | 0610009B22Rik | 66050 | protein_coding | 308.1654574 | 0.169642304 | 0.074749629 | 2.269473505 | 0.023239547 | 0.108948886 | 0.169642304 |
| ENSMUSG00000049411 | Tmem241 | 338363 | protein_coding | 226.9689686 | 0.169653477 | 0.092345944 | 1.837151372 | 0.066187552 | 0.221867939 | 0.169653477 |
| ENSMUSG00000035521 | Gnptg | 214505 | protein_coding | 456.5315409 | -0.169674506 | 0.063232831 | -2.683329262 | 0.007289318 | 0.047172889 | 0.169674506 |
| ENSMUSG00000021606 | Ndufs6 | 407785 | protein_coding | 1923.785245 | 0.169675394 | 0.084090643 | 2.017767823 | 0.043615447 | 0.168799213 | 0.169675394 |
| ENSMUSG00000025374 | Nabp2 | 69917 | protein_coding | 542.3347767 | 0.169701617 | 0.067397156 | 2.517934385 | 0.011804531 | 0.067221635 | 0.169701617 |
| ENSMUSG00000017188 | Coa3 | 52469 | protein_coding | 2134.563101 | 0.169718306 | 0.076405748 | 2.221276678 | 0.026332227 | 0.118795321 | 0.169718306 |
| ENSMUSG00000044770 | Scml4 | 268297 | protein_coding | 1604.552656 | -0.169769493 | 0.057232927 | -2.966290585 | 0.003014156 | 0.024207812 | 0.169769493 |
| ENSMUSG00000073399 | Trim40 | 195359 | protein_coding | 187.1245877 | -0.169789862 | 0.142946399 | -1.187786909 | 0.234917373 | 0.474717773 | 0.169789862 |
| ENSMUSG00000035437 | Rabgap1 | 227800 | protein_coding | 1055.312991 | -0.16981538 | 0.055742318 | -3.04643555 | 0.002315721 | 0.019655598 | 0.16981538 |
| ENSMUSG00000041035 | Dpcd | 226162 | protein_coding | 346.7073715 | 0.169873009 | 0.088712861 | 1.914863383 | 0.055509929 | 0.198450148 | 0.169873009 |
| ENSMUSG00000076441 | Ass1 | 11898 | protein_coding | 391.1861304 | -0.169917129 | 0.135459033 | -1.254380199 | 0.209703846 | 0.44497561 | 0.169917129 |
| ENSMUSG00000021578 | Ccdc127 | 67433 | protein_coding | 605.9776856 | 0.169920625 | 0.058694003 | 2.895025327 | 0.003791279 | 0.029032271 | 0.169920625 |
| ENSMUSG00000005087 | Cd44 | 12505 | protein_coding | 856.1538583 | -0.169931071 | 0.075967024 | -2.236905735 | 0.025292502 | 0.115681275 | 0.169931071 |
| ENSMUSG00000026404 | Ddx59 | 67997 | protein_coding | 98.25981508 | 0.169935347 | 0.118786503 | 1.430594751 | 0.152546393 | 0.369310836 | 0.169935347 |
| ENSMUSG00000006720 | Zfp184 | 193452 | protein_coding | 19.55086365 | -0.169986338 | 0.270232007 | -0.629038505 | 0.529323848 | 0.73857118 | 0.169986338 |
| ENSMUSG00000046138 | 9930021J03Rik | 240613 | protein_coding | 1014.687488 | -0.17003397 | 0.056509536 | -3.008942922 | 0.002621584 | 0.021628492 | 0.17003397 |
| ENSMUSG00000031691 | Tnpo2 | 212999 | protein_coding | 1813.789766 | -0.170197388 | 0.047886142 | -3.55420965 | 0.000379117 | 0.004495601 | 0.170197388 |
| ENSMUSG00000063888 | Rpl7l1 | 66229 | protein_coding | 1331.965753 | 0.170354087 | 0.047453251 | 3.58993502 | 0.00033076 | 0.00399863 | 0.170354087 |
| ENSMUSG00000069743 | Zfp820 | 75424 | protein_coding | 147.8490885 | -0.17044788 | 0.16813003 | -1.013786053 | 0.310684852 | 0.554568301 | 0.17044788 |
| ENSMUSG00000024646 | Cyb5a | 109672 | protein_coding | 1948.099449 | 0.170483321 | 0.037532966 | 4.542228822 | 5.57E-06 | 0.000116732 | 0.170483321 |
| ENSMUSG00000031723 | Txnl4b | 234723 | protein_coding | 189.7130337 | 0.170492628 | 0.093049339 | 1.832281992 | 0.066909419 | 0.223405423 | 0.170492628 |
| ENSMUSG00000039219 | Arid4b | 94246 | protein_coding | 1161.059355 | -0.17052586 | 0.069095067 | -2.467988918 | 0.013587453 | 0.074684679 | 0.17052586 |
| ENSMUSG00000007880 | Arid1a | 93760 | protein_coding | 1098.487957 | -0.170566199 | 0.105824621 | -1.611781813 | 0.107009427 | 0.299347968 | 0.170566199 |
| ENSMUSG00000052298 | Cdc42se2 | 72729 | protein_coding | 1727.269167 | 0.170573623 | 0.059849331 | 2.850050604 | 0.004371227 | 0.032307417 | 0.170573623 |
| ENSMUSG00000027715 | Ccna2 | 12428 | protein_coding | 1273.856419 | 0.170640888 | 0.107018929 | 1.594492576 | 0.110825752 | 0.305832804 | 0.170640888 |
| ENSMUSG00000051586 | Mical3 | 194401 | protein_coding | 550.7757733 | 0.170645653 | 0.075283396 | 2.266710355 | 0.023407925 | 0.109547189 | 0.170645653 |
| ENSMUSG00000011114 | Tbrg1 | 21376 | protein_coding | 2809.239278 | 0.17065668 | 0.042854981 | 3.982190103 | 6.83E-05 | 0.001073278 | 0.17065668 |
| ENSMUSG00000042498 | Radx | 102871 | protein_coding | 125.2148498 | -0.170659998 | 0.14863237 | -1.148202088 | 0.250885148 | 0.491858105 | 0.170659998 |
| ENSMUSG00000003051 | Elf3 | 13710 | protein_coding | 10249.87384 | 0.170751903 | 0.056115276 | 3.042877386 | 0.002343278 | 0.019816488 | 0.170751903 |

| ensembl_gene_id | external_gene_name | entrezgene_id | gene_biotype | baseMean | log2FoldChange | lfcSE | stat | pvalue | padj | absLog2FC |
|---|---|---|---|---|---|---|---|---|---|---|
| ENSMUSG00000022965 | Ifngr2 | 15980 | protein_coding | 3494.36975 | -0.17079475 | 0.056563825 | -3.019504979 | 0.002531881 | 0.021014745 | 0.17079475 |
| ENSMUSG00000065738 | Gm24494 | 115489817 | snoRNA | 9.194689029 | -0.170824032 | 0.416570489 | -0.410072332 | 0.681752888 | 0.839028452 | 0.170824032 |
| ENSMUSG00000071291 | Zfp58 | 238693 | protein_coding | 69.55652271 | -0.170837881 | 0.149035063 | -1.14629321 | 0.251673852 | 0.492727559 | 0.170837881 |
| ENSMUSG00000030161 | Gabarapl1 | 57436 | protein_coding | 647.0589908 | -0.170876596 | 0.090032897 | -1.897935112 | 0.05770463 | 0.203588399 | 0.170876596 |
| ENSMUSG00000021704 | Mtx3 | 382793 | protein_coding | 444.7237154 | -0.170897911 | 0.08606141 | -1.985767032 | 0.047059198 | 0.177733885 | 0.170897911 |
| ENSMUSG00000003808 | Farsa | 66590 | protein_coding | 616.69836 | 0.170898972 | 0.064432208 | 2.652384229 | 0.007992553 | 0.050686553 | 0.170898972 |
| ENSMUSG00000022521 | Crebbp | 12914 | protein_coding | 1650.333437 | -0.170944729 | 0.099818408 | -1.712557167 | 0.086794039 | 0.263701482 | 0.170944729 |
| ENSMUSG00000030493 | Faap24 | 101831 | protein_coding | 124.9756384 | 0.170945404 | 0.119789654 | 1.427046474 | 0.153566499 | 0.370743553 | 0.170945404 |
| ENSMUSG00000060314 | Zfp941 | 407812 | protein_coding | 18.91044803 | -0.170976575 | 0.308986933 | -0.553345651 | 0.580026747 | 0.772081254 | 0.170976575 |
| ENSMUSG00000051169 | Rpusd3 | 101122 | protein_coding | 89.67969562 | 0.171021325 | 0.129056305 | 1.32516831 | 0.185115333 | 0.414061984 | 0.171021325 |
| ENSMUSG00000021028 | Mbip | 217588 | protein_coding | 190.0389336 | -0.171083616 | 0.11296853 | -1.514436061 | 0.129915279 | 0.335854788 | 0.171083616 |
| ENSMUSG00000028962 | Slc4a2 | 20535 | protein_coding | 5294.318308 | -0.171124513 | 0.074412207 | -2.299683337 | 0.021466167 | 0.103384952 | 0.171124513 |
| ENSMUSG00000030353 | Tead4 | 21679 | protein_coding | 74.19762794 | -0.171139646 | 0.147830814 | -1.157672351 | 0.246997771 | 0.487958829 | 0.171139646 |
| ENSMUSG00000020621 | Rdh14 | 105014 | protein_coding | 743.9977716 | 0.171164583 | 0.068780647 | 2.488557334 | 0.012826255 | 0.071450439 | 0.171164583 |
| ENSMUSG00000097842 | 9330104G04Rik | 320950 | lncRNA | 10.63078784 | 0.171180062 | 0.389183931 | 0.439843602 | 0.660050385 | 0.826041967 | 0.171180062 |
| ENSMUSG00000025545 | Clybl | 69634 | protein_coding | 791.9782033 | -0.171188712 | 0.069855742 | -2.450603288 | 0.014261704 | 0.07724278 | 0.171188712 |
| ENSMUSG00000028221 | Pip4p2 | 72519 | protein_coding | 481.6544594 | 0.171227782 | 0.074956365 | 2.284366139 | 0.022350022 | 0.106296214 | 0.171227782 |
| ENSMUSG00000059772 | Slx1b | 75764 | protein_coding | 346.446729 | -0.171279103 | 0.076878951 | -2.227906358 | 0.02588676 | 0.117532596 | 0.171279103 |
| ENSMUSG00000011752 | Pgam1 | 18648 | protein_coding | 998.8322076 | 0.171294486 | 0.086674652 | 1.976292742 | 0.048121631 | 0.180143349 | 0.171294486 |
| ENSMUSG00000026028 | Trak2 | 70827 | protein_coding | 1700.022942 | -0.171324383 | 0.054576077 | -3.139184663 | 0.001694186 | 0.015372629 | 0.171324383 |
| ENSMUSG00000014177 | Tvp23b | 67510 | protein_coding | 1224.234448 | 0.171335213 | 0.045135475 | 3.796021001 | 0.000147037 | 0.002037171 | 0.171335213 |
| ENSMUSG00000038721 | Hoxb7 | 15415 | protein_coding | 193.5672356 | -0.17135881 | 0.131203161 | -1.306057021 | 0.191533167 | 0.423020709 | 0.17135881 |
| ENSMUSG00000032867 | Fbxw8 | 231672 | protein_coding | 1244.247502 | 0.171381687 | 0.060681364 | 2.824288639 | 0.004738569 | 0.034222527 | 0.171381687 |
| ENSMUSG00000030335 | Mrpl51 | 66493 | protein_coding | 1608.902011 | 0.171408528 | 0.080968912 | 2.116967151 | 0.034262636 | 0.142520899 | 0.171408528 |
| ENSMUSG00000060373 | Hnrnpc | 15381 | protein_coding | 2877.653595 | 0.171418265 | 0.055248913 | 3.102654108 | 0.001917937 | 0.016985483 | 0.171418265 |
| ENSMUSG00000048497 | Mmgt2 | 216829 | protein_coding | 423.7056684 | 0.171428586 | 0.064858271 | 2.643126075 | 0.008214444 | 0.051626789 | 0.171428586 |
| ENSMUSG00000032340 | Neo1 | 18007 | protein_coding | 2624.064588 | -0.171440494 | 0.052039494 | -3.294430422 | 0.000986214 | 0.009934207 | 0.171440494 |
| ENSMUSG00000072620 | Slfn2 | 20556 | protein_coding | 75.8757043 | 0.171466665 | 0.254957248 | 0.672531047 | 0.501245682 | 0.718355879 | 0.171466665 |
| ENSMUSG00000027111 | Itga6 | 16403 | protein_coding | 6487.282355 | -0.171479006 | 0.047299981 | -3.625350417 | 0.00028857 | 0.003566105 | 0.171479006 |
| ENSMUSG00000060862 | Zbtb40 | 230848 | protein_coding | 392.7362982 | 0.171558563 | 0.080214773 | 2.138740228 | 0.032456713 | 0.137419114 | 0.171558563 |
| ENSMUSG00000029415 | Sdad1 | 231452 | protein_coding | 479.8783115 | -0.171563269 | 0.067260865 | -2.550714574 | 0.010750233 | 0.062318117 | 0.171563269 |
| ENSMUSG00000044167 | Foxo1 | 56458 | protein_coding | 344.8285559 | -0.171623128 | 0.070644643 | -2.429386293 | 0.015124408 | 0.08057857 | 0.171623128 |
| ENSMUSG00000060961 | Slc4a4 | 54403 | protein_coding | 3447.296138 | -0.171696782 | 0.075231798 | -2.282236869 | 0.022475359 | 0.106731485 | 0.171696782 |
| ENSMUSG00000026107 | Nabp1 | 109019 | protein_coding | 341.6925097 | 0.171699513 | 0.079584749 | 2.1574424 | 0.030971211 | 0.13301537 | 0.171699513 |
| ENSMUSG00000060538 | Tmem219 | 68742 | protein_coding | 401.8040797 | -0.171728297 | 0.071862236 | -2.38968764 | 0.016862708 | 0.087301729 | 0.171728297 |
| ENSMUSG00000036854 | Hspb6 | 243912 | protein_coding | 32.21159456 | 0.171940814 | 0.198832918 | 0.864750241 | 0.387175888 | 0.627221049 | 0.171940814 |
| ENSMUSG00000027375 | Mal | 17153 | protein_coding | 1455.360953 | -0.171984782 | 0.119456137 | -1.439731654 | 0.149943334 | 0.365277364 | 0.171984782 |
| ENSMUSG00000070806 | Zmynd12 | 332934 | protein_coding | 14.27203808 | 0.172012978 | 0.363699406 | 0.472953697 | 0.636246214 | 0.809845517 | 0.172012978 |
| ENSMUSG00000032827 | Ppp1r9a | 243725 | protein_coding | 1038.202445 | -0.172029403 | 0.070766603 | -2.430940508 | 0.015059687 | 0.08032198 | 0.172029403 |

Supplementary Table 1

| ensembl_gene_id | external_gene_name | entrezgene_id | gene_biotype | baseMean | log2FoldChange | lfcSE | stat | pvalue | padj | absLog2FC |
|---|---|---|---|---|---|---|---|---|---|---|
| ENSMUSG00000049717 | Lig4 | 319583 | protein_coding | 343.4969394 | 0.172031023 | 0.084201719 | 2.04308207 | 0.041044315 | 0.161483694 | 0.172031023 |
| ENSMUSG00000026393 | Nek7 | 59125 | protein_coding | 1167.947571 | 0.172046587 | 0.056977561 | 3.019549884 | 0.002531506 | 0.021014745 | 0.172046587 |
| ENSMUSG00000029621 | Arpc1a | 56443 | protein_coding | 1845.580225 | 0.172079057 | 0.044423388 | 3.873613991 | 0.000107233 | 0.001579682 | 0.172079057 |
| ENSMUSG00000038665 | Dgki | 320127 | protein_coding | 31.58716281 | 0.1721224 | 0.246478465 | 0.698326323 | 0.484973141 | 0.706263486 | 0.1721224 |
| ENSMUSG00000074247 | Dda1 | 66498 | protein_coding | 835.3354205 | 0.17214613 | 0.073601772 | 2.33888569 | 0.01934135 | 0.09573909 | 0.17214613 |
| ENSMUSG00000039747 | Orai2 | 269717 | protein_coding | 545.3541741 | 0.172161871 | 0.06670408 | 2.580979617 | 0.00985204 | 0.058661261 | 0.172161871 |
| ENSMUSG00000027379 | Bub1 | 12235 | protein_coding | 237.3234147 | 0.172162672 | 0.122330019 | 1.407362426 | 0.159319947 | 0.379408267 | 0.172162672 |
| ENSMUSG00000011832 | Evi5l | 213027 | protein_coding | 309.6324518 | 0.172167129 | 0.07337753 | 2.346319509 | 0.018959847 | 0.094722877 | 0.172167129 |
| ENSMUSG00000044231 | Nhlrc1 | 105193 | protein_coding | 80.37731627 | -0.172174278 | 0.153018822 | -1.125183654 | 0.260511218 | 0.502363316 | 0.172174278 |
| ENSMUSG00000028944 | Prkag2 | 108099 | protein_coding | 749.4226195 | -0.172186198 | 0.085196746 | -2.021041964 | 0.043275421 | 0.167845772 | 0.172186198 |
| ENSMUSG00000040322 | Slc25a24 | 229731 | protein_coding | 7995.846079 | 0.172246237 | 0.050140756 | 3.435254098 | 0.000591998 | 0.006575178 | 0.172246237 |
| ENSMUSG00000004040 | Stat3 | 20848 | protein_coding | 2537.089799 | 0.172285705 | 0.061209682 | 2.814680615 | 0.004882573 | 0.034965035 | 0.172285705 |
| ENSMUSG00000071516 | H2ac13 | 319191 | protein_coding | 249.853918 | 0.172291305 | 0.196941218 | 0.874836193 | 0.381663041 | 0.622454024 | 0.172291305 |
| ENSMUSG00000032440 | Tgfbr2 | 21813 | protein_coding | 1483.59197 | -0.17231239 | 0.045403234 | -3.795156774 | 0.00014755 | 0.002040458 | 0.17231239 |
| ENSMUSG00000044934 | Zfp367 | 238673 | protein_coding | 298.6600414 | 0.172321072 | 0.131337017 | 1.312052585 | 0.189502393 | 0.420140925 | 0.172321072 |
| ENSMUSG00000038150 | Ormdl3 | 66612 | protein_coding | 553.4288769 | 0.17232696 | 0.079145917 | 2.177332279 | 0.029455784 | 0.128491598 | 0.17232696 |
| ENSMUSG00000023886 | Smoc2 | 64074 | protein_coding | 710.3753738 | 0.172365515 | 0.101886709 | 1.691736997 | 0.090696131 | 0.271060828 | 0.172365515 |
| ENSMUSG00000028999 | Rint1 | 72772 | protein_coding | 577.8569106 | 0.172374221 | 0.056784464 | 3.03558772 | 0.002400674 | 0.020174917 | 0.172374221 |
| ENSMUSG00000040913 | Fbxw4 | 30838 | protein_coding | 176.0418095 | 0.172391227 | 0.092693795 | 1.859792522 | 0.062914886 | 0.214463826 | 0.172391227 |
| ENSMUSG00000024238 | Zeb1 | 21417 | protein_coding | 8.849971588 | 0.172392762 | 0.398118105 | 0.433019147 | 0.66500086 | 0.828976491 | 0.172392762 |
| ENSMUSG00000060147 | Serpinb6a | 20719 | protein_coding | 10328.49912 | -0.172397352 | 0.053652231 | -3.213237334 | 0.001312478 | 0.012599006 | 0.172397352 |
| ENSMUSG00000030894 | Tpp1 | 12751 | protein_coding | 672.5827857 | -0.172417351 | 0.052540854 | -3.281586403 | 0.001032249 | 0.010333382 | 0.172417351 |
| ENSMUSG00000027010 | Slc25a12 | 78830 | protein_coding | 678.3955318 | 0.172509378 | 0.125084818 | 1.379139221 | 0.167851833 | 0.391856762 | 0.172509378 |
| ENSMUSG00000016526 | Dyrk3 | 226419 | protein_coding | 19.58610777 | 0.172581985 | 0.280233752 | 0.615850102 | 0.537993456 | 0.744356595 | 0.172581985 |
| ENSMUSG00000042348 | Arl15 | 218639 | protein_coding | 189.81357 | 0.172605041 | 0.094887567 | 1.819048014 | 0.068904105 | 0.227801662 | 0.172605041 |
| ENSMUSG00000020211 | Sf3a2 | 20222 | protein_coding | 347.9768361 | 0.17263051 | 0.107720481 | 1.602578333 | 0.109027781 | 0.30241162 | 0.17263051 |
| ENSMUSG00000034714 | Ttyh2 | 117160 | protein_coding | 650.4945016 | -0.172641317 | 0.064671711 | -2.669502863 | 0.007596363 | 0.048768527 | 0.172641317 |
| ENSMUSG00000019699 | Akt3 | 23797 | protein_coding | 17.73856927 | -0.1726479 | 0.348096224 | -0.495977514 | 0.619910276 | 0.799032164 | 0.1726479 |
| ENSMUSG00000026283 | Ing5 | 66262 | protein_coding | 684.8483104 | -0.172650602 | 0.070417335 | -2.451819612 | 0.01421359 | 0.077108863 | 0.172650602 |
| ENSMUSG00000056515 | Rab31 | 106572 | protein_coding | 126.9916293 | 0.172727265 | 0.148074827 | 1.166486358 | 0.243417861 | 0.484522321 | 0.172727265 |
| ENSMUSG00000008193 | Spib | 272382 | protein_coding | 175.3469583 | 0.172798042 | 0.118561642 | 1.457453176 | 0.144991328 | 0.35758991 | 0.172798042 |
| ENSMUSG00000073787 | Gm10575 | 100126795 | lncRNA | 22.32591254 | -0.172850899 | 0.268482358 | -0.64380729 | 0.519700406 | 0.7315999 | 0.172850899 |
| ENSMUSG00000031788 | Kifc3 | 16582 | protein_coding | 577.0761897 | 0.172855222 | 0.077891101 | 2.219190885 | 0.026473741 | 0.118917792 | 0.172855222 |
| ENSMUSG00000035798 | Zdhhc17 | 320150 | protein_coding | 544.1686757 | -0.172858192 | 0.056958831 | -3.034791784 | 0.002407018 | 0.020200409 | 0.172858192 |
| ENSMUSG00000031934 | Panx1 | 55991 | protein_coding | 574.3606938 | -0.172873908 | 0.075354818 | -2.294132119 | 0.021782911 | 0.10455009 | 0.172873908 |
| ENSMUSG00000010505 | Myt1 | 17932 | protein_coding | 32.34930057 | -0.172930858 | 0.221095193 | -0.782155665 | 0.434123096 | 0.668066015 | 0.172930858 |
| ENSMUSG00000087424 | 5730405O15Rik | 70489 | lncRNA | 14.08818884 | -0.173045005 | 0.310974098 | -0.556461153 | 0.577895644 | 0.770352376 | 0.173045005 |
| ENSMUSG00000022555 | Dgat1 | 13350 | protein_coding | 994.9230191 | -0.173100608 | 0.062030082 | -2.7905913 | 0.005261186 | 0.037056142 | 0.173100608 |
| ENSMUSG00000022364 | Tbc1d31 | 210544 | protein_coding | 138.2320988 | 0.173114636 | 0.110822005 | 1.56209623 | 0.118265321 | 0.317992618 | 0.173114636 |

Supplementary Table 1

| ensembl_gene_id | external_gene_name | entrezgene_id | gene_biotype | baseMean | log2FoldChange | lfcSE | stat | pvalue | padj | absLog2FC |
|---|---|---|---|---|---|---|---|---|---|---|
| ENSMUSG00000059486 | Kbtbd2 | 210973 | protein_coding | 1022.216463 | -0.173198131 | 0.047799267 | -3.623447444 | 0.000290702 | 0.003581948 | 0.173198131 |
| ENSMUSG00000022010 | Tsc22d1 | 21807 | protein_coding | 9526.593728 | 0.173254877 | 0.044924035 | 3.856618755 | 0.000114966 | 0.001671239 | 0.173254877 |
| ENSMUSG00000062761 | Zfp512 | 269639 | protein_coding | 360.4390667 | -0.173264958 | 0.075805811 | -2.28564217 | 0.022275201 | 0.106124717 | 0.173264958 |
| ENSMUSG00000029847 | Slc23a4 | 243753 | protein_coding | 102.1233201 | -0.173289689 | 0.160428922 | -1.080164896 | 0.280068757 | 0.523118689 | 0.173289689 |
| ENSMUSG00000031730 | Dhodh | 56749 | protein_coding | 255.5108159 | 0.173320152 | 0.083256064 | 2.081772113 | 0.03736329 | 0.150979148 | 0.173320152 |
| ENSMUSG00000027324 | Rpusd2 | 271842 | protein_coding | 95.07098491 | -0.173356117 | 0.148463125 | -1.167671209 | 0.242939414 | 0.484047929 | 0.173356117 |
| ENSMUSG00000041341 | Atg2b | 76559 | protein_coding | 1089.77223 | -0.173397568 | 0.051745741 | -3.35095341 | 0.000805339 | 0.008411719 | 0.173397568 |
| ENSMUSG00000050854 | Tmem125 | 230678 | protein_coding | 568.2219718 | 0.173458679 | 0.068248639 | 2.541569788 | 0.011035591 | 0.063606498 | 0.173458679 |
| ENSMUSG00000030282 | Cmas | 12764 | protein_coding | 4455.780274 | 0.173481331 | 0.052870765 | 3.281233624 | 0.001033541 | 0.010339898 | 0.173481331 |
| ENSMUSG00000006289 | Osgep | 66246 | protein_coding | 626.6028713 | 0.173530169 | 0.067138534 | 2.584658305 | 0.009747555 | 0.058308165 | 0.173530169 |
| ENSMUSG00000024829 | Mrpl21 | 353242 | protein_coding | 807.9917669 | 0.173532976 | 0.096740685 | 1.793795195 | 0.072845864 | 0.236142883 | 0.173532976 |
| ENSMUSG00000041346 | Wrap53 | 216853 | protein_coding | 291.3109708 | 0.173538349 | 0.08450748 | 2.053526478 | 0.040021545 | 0.158581683 | 0.173538349 |
| ENSMUSG00000003309 | Ap1m2 | 11768 | protein_coding | 3575.307569 | 0.173588175 | 0.049449758 | 3.510394822 | 0.000447442 | 0.00515421 | 0.173588175 |
| ENSMUSG00000027667 | Zfp639 | 67778 | protein_coding | 334.2830667 | -0.173639443 | 0.080243946 | -2.163894622 | 0.030472438 | 0.131679798 | 0.173639443 |
| ENSMUSG00000023990 | Tfeb | 21425 | protein_coding | 534.3675714 | -0.173705366 | 0.111151107 | -1.562785748 | 0.118102997 | 0.31772177 | 0.173705366 |
| ENSMUSG00000038127 | Ccdc50 | 67501 | protein_coding | 1147.070227 | -0.173708458 | 0.062313357 | -2.7876601 | 0.005309021 | 0.037290326 | 0.173708458 |
| ENSMUSG00000025176 | Hoga1 | 67432 | protein_coding | 66.22254253 | 0.17371071 | 0.169744 | 1.023368776 | 0.306133517 | 0.550330534 | 0.17371071 |
| ENSMUSG00000022090 | Pdlim2 | 213019 | protein_coding | 515.8910339 | 0.17371458 | 0.104043835 | 1.669628767 | 0.094992835 | 0.279145308 | 0.17371458 |
| ENSMUSG00000036513 | Commd2 | 52245 | protein_coding | 313.2680604 | 0.173738186 | 0.067163132 | 2.586808876 | 0.009686931 | 0.058053189 | 0.173738186 |
| ENSMUSG00000002983 | Relb | 19698 | protein_coding | 1189.891327 | -0.173771675 | 0.060401952 | -2.87692149 | 0.004015755 | 0.03037621 | 0.173771675 |
| ENSMUSG00000050721 | Plekho2 | 102595 | protein_coding | 23.70380198 | 0.173911005 | 0.24461958 | 0.710944746 | 0.477118476 | 0.699880814 | 0.173911005 |
| ENSMUSG00000019850 | Tnfaip3 | 21929 | protein_coding | 856.4500968 | -0.173956428 | 0.07606102 | -2.28706409 | 0.022192083 | 0.105822506 | 0.173956428 |
| ENSMUSG00000085207 | Gm11767 | 102640451 | lncRNA | 10.14749553 | -0.174096247 | 0.365049605 | -0.476911205 | 0.633425352 | 0.808283189 | 0.174096247 |
| ENSMUSG00000006711 | D130043K22Rik | 210108 | protein_coding | 43.79324252 | -0.174139301 | 0.217180961 | -0.801816607 | 0.422659051 | 0.658779673 | 0.174139301 |
| ENSMUSG00000072763 | 5430403G16Rik | 77200 | protein_coding | 55.70958162 | -0.174253612 | 0.176293796 | -0.988427363 | 0.322943392 | 0.566904861 | 0.174253612 |
| ENSMUSG00000023266 | Frs3 | 107971 | protein_coding | 65.41623938 | -0.174334152 | 0.163962756 | -1.063254588 | 0.287666513 | 0.530090639 | 0.174334152 |
| ENSMUSG00000036523 | Greb1 | 268527 | protein_coding | 276.8591812 | -0.174344084 | 0.093065038 | -1.873357467 | 0.061019039 | 0.210492844 | 0.174344084 |
| ENSMUSG00000022673 | Mcm4 | 17217 | protein_coding | 581.3932138 | 0.17436138 | 0.124000148 | 1.406138487 | 0.159683006 | 0.379712155 | 0.17436138 |
| ENSMUSG00000050188 | Lsm10 | 116748 | protein_coding | 179.8587013 | 0.17436373 | 0.120137622 | 1.451366587 | 0.146677809 | 0.359905151 | 0.17436373 |
| ENSMUSG00000062638 | Btnl1 | 100038862 | protein_coding | 2352.743103 | -0.174467013 | 0.088213734 | -1.977776083 | 0.047953973 | 0.179737215 | 0.174467013 |
| ENSMUSG00000033845 | Mrpl15 | 27395 | protein_coding | 1027.024125 | 0.174505602 | 0.050085762 | 3.484135889 | 0.000493729 | 0.005591547 | 0.174505602 |
| ENSMUSG00000048022 | Tmem229a | 319832 | protein_coding | 15.69393033 | 0.174510217 | 0.287988338 | 0.60596279 | 0.544539462 | 0.749012844 | 0.174510217 |
| ENSMUSG00000094483 | Purb | 19291 | protein_coding | 4950.976847 | 0.174521968 | 0.052490287 | 3.324843066 | 0.000884683 | 0.009099034 | 0.174521968 |
| ENSMUSG00000031608 | Galnt7 | 108150 | protein_coding | 5814.471241 | 0.174549616 | 0.040152779 | 4.34713664 | 1.38E-05 | 0.000261072 | 0.174549616 |
| ENSMUSG00000006651 | Aplp1 | 11803 | protein_coding | 304.1067939 | 0.174624882 | 0.107602029 | 1.622877223 | 0.104615653 | 0.29617307 | 0.174624882 |
| ENSMUSG00000034263 | Ints14 | 69882 | protein_coding | 712.8842708 | 0.17462797 | 0.065771619 | 2.655065714 | 0.007929296 | 0.050384459 | 0.17462797 |
| ENSMUSG00000004936 | Map2k1 | 26395 | protein_coding | 5053.289585 | 0.174644372 | 0.062584262 | 2.790547739 | 0.005261894 | 0.037056142 | 0.174644372 |
| ENSMUSG00000022178 | Ajuba | 16475 | protein_coding | 79.00258585 | 0.174763717 | 0.147263902 | 1.186738333 | 0.23533085 | 0.475407818 | 0.174763717 |
| ENSMUSG00000025940 | Tmem70 | 70397 | protein_coding | 762.4322178 | 0.174826836 | 0.062259471 | 2.808036 | 0.004984465 | 0.035599856 | 0.174826836 |

| ensembl_gene_id | external_gene_name | entrezgene_id | gene_biotype | baseMean | log2FoldChange | lfcSE | stat | pvalue | padj | absLog2FC |
|---|---|---|---|---|---|---|---|---|---|---|
| ENSMUSG00000085551 | 1810063I02Rik | 69854 | lncRNA | 16.77235944 | -0.174827199 | 0.35306819 | -0.495165536 | 0.620483276 | 0.799499344 | 0.174827199 |
| ENSMUSG00000053769 | Lysmd1 | 217779 | protein_coding | 40.17902988 | -0.174855038 | 0.205517574 | -0.850803338 | 0.394878607 | 0.634346777 | 0.174855038 |
| ENSMUSG00000016252 | Atp5e | 67126 | protein_coding | 3733.343762 | 0.174879474 | 0.078908659 | 2.21622665 | 0.026675984 | 0.119434644 | 0.174879474 |
| ENSMUSG00000075543 | Urad | 231903 | protein_coding | 24.80537269 | 0.174898572 | 0.26467207 | 0.660812348 | 0.508732663 | 0.724049267 | 0.174898572 |
| ENSMUSG00000017713 | Tha1 | 71776 | protein_coding | 238.3827925 | -0.174910978 | 0.092451364 | -1.891924261 | 0.058501073 | 0.205052554 | 0.174910978 |
| ENSMUSG00000034345 | Gtf2h5 | 66467 | protein_coding | 529.3673301 | 0.174922931 | 0.076698874 | 2.280645361 | 0.022569441 | 0.106976995 | 0.174922931 |
| ENSMUSG00000005362 | Crbn | 58799 | protein_coding | 629.747013 | -0.174946283 | 0.066558211 | -2.628470338 | 0.008576984 | 0.053241345 | 0.174946283 |
| ENSMUSG00000002108 | Nr1h3 | 22259 | protein_coding | 606.8158116 | -0.174967911 | 0.074775363 | -2.339913893 | 0.019288186 | 0.095563927 | 0.174967911 |
| ENSMUSG00000080717 | B230307C23Rik | 245305 | protein_coding | 112.713895 | -0.174980492 | 0.142904437 | -1.22445808 | 0.220779471 | 0.457336743 | 0.174980492 |
| ENSMUSG00000026270 | Capn10 | 23830 | protein_coding | 571.1041789 | 0.175016598 | 0.070941058 | 2.467070592 | 0.01362235 | 0.074800013 | 0.175016598 |
| ENSMUSG00000032114 | Slc37a4 | 14385 | protein_coding | 1051.438902 | -0.175056461 | 0.0597636 | -2.929148534 | 0.00339892 | 0.026712659 | 0.175056461 |
| ENSMUSG00000045502 | Hcar2 | 80885 | protein_coding | 11.42466611 | -0.175064492 | 0.352166655 | -0.49710695 | 0.619113634 | 0.798277593 | 0.175064492 |
| ENSMUSG00000076435 | Acsf2 | 264895 | protein_coding | 2228.322543 | -0.175077148 | 0.070195633 | -2.494131626 | 0.012626571 | 0.070728973 | 0.175077148 |
| ENSMUSG00000103567 | Pcdhga5 | 93713 | protein_coding | 10.64607624 | 0.175085812 | 0.348913558 | 0.501802833 | 0.61580622 | 0.795599668 | 0.175085812 |
| ENSMUSG00000048728 | Zfp454 | 237758 | protein_coding | 11.1648259 | 0.175143802 | 0.452727838 | 0.386863337 | 0.698857385 | 0.849703939 | 0.175143802 |
| ENSMUSG00000002227 | Mov10 | 17454 | protein_coding | 827.1302334 | -0.175184017 | 0.12291915 | -1.425197103 | 0.154100234 | 0.371487597 | 0.175184017 |
| ENSMUSG00000004100 | Ppan | 235036 | protein_coding | 389.8590434 | -0.175222259 | 0.077087065 | -2.273043597 | 0.023023553 | 0.108250973 | 0.175222259 |
| ENSMUSG00000047242 | Taf9b | 407786 | protein_coding | 129.2829383 | 0.175280283 | 0.119059722 | 1.472204709 | 0.140965606 | 0.352075628 | 0.175280283 |
| ENSMUSG00000028750 | Pla2g2c | 18781 | protein_coding | 40.70667376 | 0.175339279 | 0.201915485 | 0.868379557 | 0.385186582 | 0.625632391 | 0.175339279 |
| ENSMUSG00000010110 | Stx5a | 56389 | protein_coding | 1601.617294 | 0.1754907 | 0.051305134 | 3.420529038 | 0.000624995 | 0.006875367 | 0.1754907 |
| ENSMUSG00000032067 | Pts | 19286 | protein_coding | 903.9372252 | 0.175498626 | 0.058452445 | 3.002417175 | 0.002678448 | 0.022015973 | 0.175498626 |
| ENSMUSG00000031391 | L1cam | 16728 | protein_coding | 2876.003581 | -0.175552161 | 0.045710186 | -3.840547924 | 0.00012276 | 0.001761238 | 0.175552161 |
| ENSMUSG00000009145 | Dqx1 | 93838 | protein_coding | 1132.890474 | -0.175552961 | 0.059706157 | -2.940282388 | 0.003279132 | 0.025948268 | 0.175552961 |
| ENSMUSG00000003929 | Zfp81 | 224694 | protein_coding | 238.3529422 | -0.175558589 | 0.094794841 | -1.851984637 | 0.064028029 | 0.217038598 | 0.175558589 |
| ENSMUSG00000003308 | Keap1 | 50868 | protein_coding | 1076.920013 | 0.175568951 | 0.096522486 | 1.818943515 | 0.068920048 | 0.227807669 | 0.175568951 |
| ENSMUSG00000038578 | Susd1 | 634731 | protein_coding | 25.25306601 | -0.175578781 | 0.253811525 | -0.691768354 | 0.489082813 | 0.709236362 | 0.175578781 |
| ENSMUSG00000029752 | Asns | 27053 | protein_coding | 621.599465 | 0.175615302 | 0.083873142 | 2.093820487 | 0.036275974 | 0.148069512 | 0.175615302 |
| ENSMUSG00000041577 | Prelp | 116847 | protein_coding | 117.3921538 | 0.175622839 | 0.171045065 | 1.026763555 | 0.304531814 | 0.548674402 | 0.175622839 |
| ENSMUSG00000072889 | Nfxl1 | 100978 | protein_coding | 339.7976362 | 0.175682876 | 0.074053087 | 2.372390973 | 0.017673378 | 0.090113624 | 0.175682876 |
| ENSMUSG00000051855 | Mest | 17294 | protein_coding | 14.72242811 | -0.175688658 | 0.312869247 | -0.561540197 | 0.574429336 | 0.768186995 | 0.175688658 |
| ENSMUSG00000039835 | Nhsl1 | 215819 | protein_coding | 1626.110241 | 0.175710923 | 0.057601299 | 3.050468042 | 0.00228485 | 0.019485867 | 0.175710923 |
| ENSMUSG00000043923 | Ccdc84 | 382073 | protein_coding | 183.9161854 | -0.175740061 | 0.159846441 | -1.099430551 | 0.271580311 | 0.51411852 | 0.175740061 |
| ENSMUSG00000034832 | Tet3 | 194388 | protein_coding | 2338.4114 | -0.175801011 | 0.071621701 | -2.454577446 | 0.014105028 | 0.076667268 | 0.175801011 |
| ENSMUSG00000025575 | Cant1 | 76025 | protein_coding | 6596.941533 | 0.175806304 | 0.0558052 | 3.150357041 | 0.00163071 | 0.014954671 | 0.175806304 |
| ENSMUSG00000002343 | Armc6 | 76813 | protein_coding | 178.5496737 | 0.175827139 | 0.113967456 | 1.542783754 | 0.122883253 | 0.325080339 | 0.175827139 |
| ENSMUSG00000005514 | Por | 18984 | protein_coding | 1855.899685 | -0.175844527 | 0.06736054 | -2.610497561 | 0.009041062 | 0.055418169 | 0.175844527 |
| ENSMUSG00000038775 | Vill | 22351 | protein_coding | 61.76530003 | -0.175874702 | 0.207190091 | -0.848856725 | 0.395961021 | 0.635262971 | 0.175874702 |
| ENSMUSG00000029787 | Avl9 | 78937 | protein_coding | 2943.413011 | -0.17596405 | 0.056420464 | -3.118798365 | 0.001815902 | 0.016260438 | 0.17596405 |
| ENSMUSG00000042745 | Id1 | 15901 | protein_coding | 6502.490541 | -0.176004903 | 0.097851407 | -1.798695686 | 0.072066832 | 0.234508029 | 0.176004903 |

Supplementary Table 1

| ensembl_gene_id | external_gene_name | entrezgene_id | gene_biotype | baseMean | log2FoldChange | lfcSE | stat | pvalue | padj | absLog2FC |
|---|---|---|---|---|---|---|---|---|---|---|
| ENSMUSG00000026675 | Hsd17b7 | 15490 | protein_coding | 738.4582085 | -0.176038137 | 0.103888503 | -1.694491046 | 0.090172012 | 0.270226918 | 0.176038137 |
| ENSMUSG00000071265 | 1700086L19Rik | 74284 | transcribed_unitary_pseudogene | 6.098942902 | 0.17606103 | 0.479217328 | 0.367392872 | 0.713325988 | 0.858813602 | 0.17606103 |
| ENSMUSG00000045314 | Sowahb | 78088 | protein_coding | 969.1758427 | -0.176099388 | 0.056047391 | -3.14197299 | 0.001678135 | 0.015264121 | 0.176099388 |
| ENSMUSG00000068732 | Tmem167b | 67495 | protein_coding | 1468.144702 | 0.176111687 | 0.056310296 | 3.127521965 | 0.001762866 | 0.01586756 | 0.176111687 |
| ENSMUSG00000027793 | Ccna1 | 12427 | protein_coding | 9.312629878 | -0.176140294 | 0.377678693 | -0.466376042 | 0.640946357 | 0.813324566 | 0.176140294 |
| ENSMUSG00000030471 | Zdhhc13 | 243983 | protein_coding | 2453.610333 | 0.17621839 | 0.040204508 | 4.383050543 | 1.17E-05 | 0.000225487 | 0.17621839 |
| ENSMUSG00000024028 | Tff2 | 21785 | protein_coding | 9.576691864 | 0.176271495 | 0.676302273 | 0.260640104 | 0.79437006 | 0.904717934 | 0.176271495 |
| ENSMUSG00000024472 | Dcp2 | 70640 | protein_coding | 1276.574576 | -0.176273734 | 0.056066675 | -3.144001945 | 0.001666543 | 0.015184375 | 0.176273734 |
| ENSMUSG00000054556 | Gm4876 | 232599 | lncRNA | 60.15391064 | -0.176303313 | 0.267840656 | -0.65823955 | 0.510384215 | 0.724852046 | 0.176303313 |
| ENSMUSG00000026775 | Yme1l1 | 27377 | protein_coding | 3705.097435 | 0.176590183 | 0.049794924 | 3.546349069 | 0.000390608 | 0.00459806 | 0.176590183 |
| ENSMUSG00000035226 | Rims4 | 241770 | protein_coding | 329.8366034 | -0.176590474 | 0.142431766 | -1.239825066 | 0.215040105 | 0.45050036 | 0.176590474 |
| ENSMUSG00000021594 | Srd5a1 | 78925 | protein_coding | 213.210783 | 0.176711215 | 0.090785972 | 1.946459463 | 0.051599573 | 0.189167154 | 0.176711215 |
| ENSMUSG00000038967 | Pdk2 | 18604 | protein_coding | 1103.790258 | 0.176768785 | 0.068751251 | 2.571135526 | 0.010136565 | 0.059858068 | 0.176768785 |
| ENSMUSG00000091994 | E130317F20Rik | 414101 | lncRNA | 48.43210207 | -0.176809292 | 0.220544547 | -0.801694236 | 0.422729851 | 0.658779673 | 0.176809292 |
| ENSMUSG00000053553 | 3110082I17Rik | 73212 | protein_coding | 87.52279956 | -0.176885925 | 0.159910899 | -1.106153025 | 0.268660291 | 0.511015761 | 0.176885925 |
| ENSMUSG00000024038 | Ndufv3 | 78330 | protein_coding | 3069.831806 | 0.176911703 | 0.053640549 | 3.298096438 | 0.000973427 | 0.009823815 | 0.176911703 |
| ENSMUSG00000022856 | Tmem41a | 66664 | protein_coding | 447.9742837 | -0.176999637 | 0.060622072 | -2.919722645 | 0.00350343 | 0.0272946 | 0.176999637 |
| ENSMUSG00000048550 | Thnsl1 | 208967 | protein_coding | 195.9386067 | 0.177003705 | 0.099430303 | 1.780178683 | 0.075046723 | 0.240742704 | 0.177003705 |
| ENSMUSG00000075700 | Selenot | 69227 | protein_coding | 2755.387697 | 0.177023133 | 0.041014417 | 4.316119644 | 1.59E-05 | 0.000296401 | 0.177023133 |
| ENSMUSG00000041658 | Rragb | 245670 | protein_coding | 18.41002989 | 0.177024762 | 0.322851987 | 0.548315542 | 0.583475257 | 0.774907805 | 0.177024762 |
| ENSMUSG00000015094 | Npdc1 | 18146 | protein_coding | 802.2264349 | -0.177119501 | 0.084001164 | -2.108536259 | 0.03498463 | 0.144554734 | 0.177119501 |
| ENSMUSG00000022241 | Tars | 110960 | protein_coding | 2155.159468 | 0.177140007 | 0.054906105 | 3.226235157 | 0.001254303 | 0.012163642 | 0.177140007 |
| ENSMUSG00000085287 | 4833418N02Rik | 74597 | lncRNA | 85.25816091 | 0.177169373 | 0.15512844 | 1.142081832 | 0.253420007 | 0.494722727 | 0.177169373 |
| ENSMUSG00000097061 | 9330151L19Rik | 414085 | lncRNA | 41.04907677 | -0.177179258 | 0.23852814 | -0.742802331 | 0.457601364 | 0.685405128 | 0.177179258 |
| ENSMUSG00000061731 | Ext1 | 14042 | protein_coding | 792.3535278 | 0.177229731 | 0.064837672 | 2.733437605 | 0.0062677 | 0.042345703 | 0.177229731 |
| ENSMUSG00000048911 | Rnf24 | 51902 | protein_coding | 60.10172457 | 0.177240204 | 0.149912244 | 1.18229305 | 0.237089442 | 0.477153008 | 0.177240204 |
| ENSMUSG00000045287 | Rtn4rl1 | 237847 | protein_coding | 137.9869477 | -0.177251225 | 0.117222071 | -1.512097711 | 0.130509007 | 0.336485813 | 0.177251225 |
| ENSMUSG00000029330 | Cds1 | 74596 | protein_coding | 4684.236568 | -0.177320048 | 0.062872229 | -2.820323878 | 0.00479752 | 0.034530138 | 0.177320048 |
| ENSMUSG00000028568 | Btf3l4 | 70533 | protein_coding | 336.1161528 | -0.177396731 | 0.069610393 | -2.54842308 | 0.010821115 | 0.062638951 | 0.177396731 |
| ENSMUSG00000030287 | Itpr2 | 16439 | protein_coding | 1647.803408 | -0.177421512 | 0.042748175 | -4.150387995 | 3.32E-05 | 0.000568836 | 0.177421512 |
| ENSMUSG00000027669 | Gnb4 | 14696 | protein_coding | 23.44825968 | -0.177456683 | 0.259611629 | -0.683546742 | 0.494261425 | 0.713491312 | 0.177456683 |
| ENSMUSG00000026203 | Dnajb2 | 56812 | protein_coding | 477.5849252 | 0.177528598 | 0.087443998 | 2.030197624 | 0.042336455 | 0.165140974 | 0.177528598 |
| ENSMUSG00000033629 | Hacd3 | 57874 | protein_coding | 1598.263141 | -0.17758342 | 0.053840643 | -3.298315368 | 0.000972668 | 0.009823815 | 0.17758342 |
| ENSMUSG00000046179 | E2f8 | 108961 | protein_coding | 244.9454057 | 0.177606965 | 0.176079614 | 1.008674207 | 0.313130905 | 0.557195422 | 0.177606965 |
| ENSMUSG00000046329 | Slc25a23 | 66972 | protein_coding | 173.6595128 | -0.177694201 | 0.09798117 | -1.813554591 | 0.069746327 | 0.229360057 | 0.177694201 |
| ENSMUSG00000041890 | Git2 | 26431 | protein_coding | 427.0166033 | -0.177694551 | 0.087243981 | -2.036754272 | 0.041674673 | 0.163167625 | 0.177694551 |
| ENSMUSG00000030337 | Vamp1 | 22317 | protein_coding | 185.2721368 | -0.177724429 | 0.098945687 | -1.796181666 | 0.07246563 | 0.235215876 | 0.177724429 |
| ENSMUSG00000012117 | Dhdds | 67422 | protein_coding | 573.3560991 | 0.177801521 | 0.056559835 | 3.143600402 | 0.001668832 | 0.015196638 | 0.177801521 |
| ENSMUSG00000025318 | Jph3 | 57340 | protein_coding | 6.025766583 | -0.177818592 | 0.571411145 | -0.311192026 | 0.755654644 | 0.884036449 | 0.177818592 |

| ensembl_gene_id | external_gene_name | entrezgene_id | gene_biotype | baseMean | log2FoldChange | lfcSE | stat | pvalue | padj | absLog2FC |
|---|---|---|---|---|---|---|---|---|---|---|
| ENSMUSG00000038895 | Zfp653 | 319601 | protein_coding | 106.8527621 | -0.177863235 | 0.137811166 | -1.29063007 | 0.196831984 | 0.429250765 | 0.177863235 |
| ENSMUSG00000022009 | Nufip1 | 27275 | protein_coding | 289.8847784 | -0.177873774 | 0.075137295 | -2.367316717 | 0.017917592 | 0.091124885 | 0.177873774 |
| ENSMUSG00000051124 | Gimap9 | 317758 | protein_coding | 109.6971287 | 0.177876296 | 0.130128903 | 1.366923811 | 0.171649188 | 0.396321966 | 0.177876296 |
| ENSMUSG00000030095 | Tmem43 | 74122 | protein_coding | 795.5376868 | 0.177934322 | 0.085517815 | 2.080669633 | 0.037464155 | 0.151214294 | 0.177934322 |
| ENSMUSG00000024194 | Cuta | 67675 | protein_coding | 831.3461818 | 0.17802425 | 0.055824965 | 3.188971966 | 0.001427797 | 0.013434123 | 0.17802425 |
| ENSMUSG00000028015 | Ctso | 229445 | protein_coding | 529.1551896 | -0.178100564 | 0.082453095 | -2.160022775 | 0.030770907 | 0.132451326 | 0.178100564 |
| ENSMUSG00000091844 | Gm8251 | 102636082 | protein_coding | 7.81679281 | 0.178139208 | 0.44644716 | 0.399015212 | 0.689881996 | 0.844374344 | 0.178139208 |
| ENSMUSG00000026856 | Dolpp1 | 57170 | protein_coding | 1261.227098 | 0.178175927 | 0.054021862 | 3.298218919 | 0.000973002 | 0.009823815 | 0.178175927 |
| ENSMUSG00000030714 | Sgf29 | 75565 | protein_coding | 608.8035261 | -0.17820632 | 0.067286439 | -2.648473045 | 0.00808563 | 0.051055971 | 0.17820632 |
| ENSMUSG00000024601 | Isoc1 | 66307 | protein_coding | 1216.529863 | -0.178235071 | 0.050126002 | -3.555740784 | 0.000376916 | 0.00447608 | 0.178235071 |
| ENSMUSG00000020528 | Prpsap2 | 212627 | protein_coding | 556.593544 | 0.17830119 | 0.064696766 | 2.755952145 | 0.005852156 | 0.040239329 | 0.17830119 |
| ENSMUSG00000052146 | Rps10 | 67097 | protein_coding | 164.8955294 | -0.178311553 | 0.112511232 | -1.584833351 | 0.113004203 | 0.309461501 | 0.178311553 |
| ENSMUSG00000000579 | Dynlt1c | 100040563 | protein_coding | 38.83184095 | -0.178318137 | 0.182269019 | -0.978323896 | 0.327914155 | 0.571645972 | 0.178318137 |
| ENSMUSG00000031429 | Psmd10 | 53380 | protein_coding | 465.4930296 | 0.178378233 | 0.059829858 | 2.981424953 | 0.002869104 | 0.023286378 | 0.178378233 |
| ENSMUSG00000043542 | Zc2hc1a | 67306 | protein_coding | 170.6853782 | 0.178394663 | 0.101984929 | 1.749225741 | 0.080252006 | 0.250770026 | 0.178394663 |
| ENSMUSG00000073131 | Vma21 | 67048 | protein_coding | 343.734539 | 0.178435411 | 0.076699548 | 2.326420619 | 0.019996123 | 0.09807709 | 0.178435411 |
| ENSMUSG00000015568 | Lpl | 16956 | protein_coding | 365.5062607 | -0.178466578 | 0.078576179 | -2.271255471 | 0.023131517 | 0.108600285 | 0.178466578 |
| ENSMUSG00000029093 | Sorcs2 | 81840 | protein_coding | 491.3563236 | -0.178487181 | 0.127153343 | -1.40371599 | 0.160403439 | 0.380799766 | 0.178487181 |
| ENSMUSG00000052137 | Rbm12b2 | 77604 | protein_coding | 182.957944 | -0.178530852 | 0.120969356 | -1.47583535 | 0.139988098 | 0.350881819 | 0.178530852 |
| ENSMUSG00000003549 | Ercc1 | 13870 | protein_coding | 188.9836046 | 0.178574748 | 0.094751766 | 1.884658786 | 0.05947592 | 0.207110275 | 0.178574748 |
| ENSMUSG00000031292 | Cdkl5 | 382253 | protein_coding | 103.3926733 | 0.178590337 | 0.119925378 | 1.489178856 | 0.136440277 | 0.345432975 | 0.178590337 |
| ENSMUSG00000069678 | Pcgf1 | 69837 | protein_coding | 131.6740113 | -0.178595638 | 0.105924742 | -1.686061581 | 0.091783939 | 0.273111802 | 0.178595638 |
| ENSMUSG00000020163 | Uqcr11 | 66594 | protein_coding | 5338.463357 | 0.178645542 | 0.081095884 | 2.202892826 | 0.027602303 | 0.122594973 | 0.178645542 |
| ENSMUSG00000037580 | Gch1 | 14528 | protein_coding | 297.8526644 | 0.178713889 | 0.109026134 | 1.63918395 | 0.101174955 | 0.290022548 | 0.178713889 |
| ENSMUSG00000034591 | Slc41a2 | 338365 | protein_coding | 711.7524403 | 0.178719261 | 0.0616551 | 2.89869389 | 0.003747205 | 0.028790463 | 0.178719261 |
| ENSMUSG00000020227 | Irak3 | 73914 | protein_coding | 103.5108905 | 0.178729715 | 0.142831675 | 1.251331091 | 0.210813707 | 0.445849236 | 0.178729715 |
| ENSMUSG00000024354 | Slc23a1 | 20522 | protein_coding | 14.51625215 | -0.178807953 | 0.335721355 | -0.53260822 | 0.594304811 | 0.781566802 | 0.178807953 |
| ENSMUSG00000001158 | Snrnp27 | 66618 | protein_coding | 433.1325119 | 0.178856008 | 0.070358998 | 2.542048828 | 0.011020478 | 0.063564822 | 0.178856008 |
| ENSMUSG00000026511 | Srp9 | 27058 | protein_coding | 1084.83479 | 0.178907164 | 0.05616444 | 3.185417015 | 0.001445456 | 0.013552826 | 0.178907164 |
| ENSMUSG00000021884 | Hacl1 | 56794 | protein_coding | 34.96696629 | -0.17892832 | 0.224603246 | -0.796641735 | 0.425659132 | 0.661307016 | 0.17892832 |
| ENSMUSG00000057137 | Tmem140 | 68487 | protein_coding | 66.98123753 | -0.178980469 | 0.165999848 | -1.078196583 | 0.280946037 | 0.523508209 | 0.178980469 |
| ENSMUSG00000028018 | Gstcd | 67553 | protein_coding | 180.3609716 | 0.179033223 | 0.104285512 | 1.716760261 | 0.086022985 | 0.262248142 | 0.179033223 |
| ENSMUSG00000027612 | Mmp24 | 17391 | protein_coding | 9.794013964 | -0.179045554 | 0.388403495 | -0.460978225 | 0.644814228 | 0.81580845 | 0.179045554 |
| ENSMUSG00000037681 | Esyt3 | 272636 | protein_coding | 11.03721934 | -0.179060418 | 0.346543428 | -0.516704123 | 0.605362718 | 0.789158883 | 0.179060418 |
| ENSMUSG00000035992 | Fnip1 | 216742 | protein_coding | 916.5633239 | -0.179083388 | 0.082428647 | -2.172586762 | 0.029811433 | 0.129657223 | 0.179083388 |
| ENSMUSG00000020650 | Bcap29 | 12033 | protein_coding | 141.7695973 | -0.179141058 | 0.105598156 | -1.696441165 | 0.089802363 | 0.269540799 | 0.179141058 |
| ENSMUSG00000040414 | Slc25a28 | 246696 | protein_coding | 421.4922718 | -0.179161091 | 0.070951421 | -2.525123364 | 0.011565768 | 0.066118803 | 0.179161091 |
| ENSMUSG00000013643 | Lypd8 | 70163 | protein_coding | 88211.82975 | -0.179178458 | 0.086644727 | -2.067967258 | 0.038643102 | 0.15463953 | 0.179178458 |
| ENSMUSG00000031262 | Cenpi | 102920 | protein_coding | 107.7300471 | 0.179207675 | 0.166352209 | 1.077278599 | 0.28135582 | 0.523831138 | 0.179207675 |

| ensembl_gene_id | external_gene_name | entrezgene_id | gene_biotype | baseMean | log2FoldChange | lfcSE | stat | pvalue | padj | absLog2FC |
|---|---|---|---|---|---|---|---|---|---|---|
| ENSMUSG00000073427 | Gm4924 | 237412 | protein_coding | 66.48978356 | 0.1792109 | 0.183534672 | 0.97644166 | 0.328845645 | 0.572498536 | 0.1792109 |
| ENSMUSG00000024487 | Yipf5 | 67180 | protein_coding | 758.9849492 | 0.179267496 | 0.052144588 | 3.437892621 | 0.00058626 | 0.006515932 | 0.179267496 |
| ENSMUSG00000021208 | Ifi27l2b | 217845 | protein_coding | 2144.117691 | -0.179363878 | 0.071446902 | -2.510450022 | 0.012057739 | 0.068277793 | 0.179363878 |
| ENSMUSG00000042116 | Vwa1 | 246228 | protein_coding | 23.47199207 | -0.179427624 | 0.324934392 | -0.552196471 | 0.580813752 | 0.772645663 | 0.179427624 |
| ENSMUSG00000029369 | Afm | 280662 | protein_coding | 72.88679348 | 0.179516891 | 0.155853463 | 1.151831262 | 0.249390419 | 0.490195121 | 0.179516891 |
| ENSMUSG00000042312 | S100a13 | 20196 | protein_coding | 648.2926591 | 0.179576883 | 0.076882784 | 2.335722953 | 0.019505685 | 0.096404593 | 0.179576883 |
| ENSMUSG00000024958 | Gpr137 | 107173 | protein_coding | 583.9030177 | 0.179592022 | 0.102377413 | 1.754215278 | 0.079393626 | 0.24905109 | 0.179592022 |
| ENSMUSG00000047180 | Neurl3 | 214854 | protein_coding | 3239.648444 | -0.179642278 | 0.061376302 | -2.926899652 | 0.003423593 | 0.026841171 | 0.179642278 |
| ENSMUSG00000094622 | Gm3055 | 100040944 | protein_coding | 9.371062849 | 0.179661908 | 0.426662379 | 0.421086829 | 0.673691679 | 0.834153989 | 0.179661908 |
| ENSMUSG00000015335 | Zdhhc12 | 66220 | protein_coding | 376.8682889 | 0.179815487 | 0.112114022 | 1.603862605 | 0.108744341 | 0.302001032 | 0.179815487 |
| ENSMUSG00000021696 | Elovl7 | 74559 | protein_coding | 2225.805738 | 0.17985223 | 0.069284056 | 2.59586751 | 0.009435244 | 0.057010933 | 0.17985223 |
| ENSMUSG00000025231 | Sufu | 24069 | protein_coding | 171.5495395 | -0.17986599 | 0.101429469 | -1.773310969 | 0.076177193 | 0.243124794 | 0.17986599 |
| ENSMUSG00000021725 | Parp8 | 52552 | protein_coding | 9.783762243 | 0.179892452 | 0.377448527 | 0.476601282 | 0.63364607 | 0.808283189 | 0.179892452 |
| ENSMUSG00000009030 | Pdcl | 67466 | protein_coding | 1150.491418 | 0.179924999 | 0.049645223 | 3.624215776 | 0.00028984 | 0.003576314 | 0.179924999 |
| ENSMUSG00000086549 | Gm13648 | 100415901 | lncRNA | 121.9686954 | -0.179986133 | 0.150179279 | -1.198475146 | 0.230732093 | 0.469647415 | 0.179986133 |
| ENSMUSG00000006678 | Pola1 | 18968 | protein_coding | 455.349012 | 0.179986421 | 0.165014605 | 1.090730246 | 0.275391597 | 0.518002859 | 0.179986421 |
| ENSMUSG00000029414 | Kntc1 | 208628 | protein_coding | 304.1155107 | 0.179994088 | 0.140804649 | 1.278324892 | 0.201134895 | 0.434870954 | 0.179994088 |
| ENSMUSG00000019659 | Ccdc12 | 72654 | protein_coding | 1287.757464 | 0.180040059 | 0.056190063 | 3.20412631 | 0.00135473 | 0.012866746 | 0.180040059 |
| ENSMUSG00000035948 | Acss3 | 380660 | protein_coding | 13.12591474 | 0.180128867 | 0.357262798 | 0.504191503 | 0.614126811 | 0.79448284 | 0.180128867 |
| ENSMUSG00000095432 | Zfp748 | 212276 | protein_coding | 376.6350517 | -0.18018012 | 0.072119236 | -2.498364244 | 0.012476792 | 0.070109138 | 0.18018012 |
| ENSMUSG00000021135 | Slc10a1 | 20493 | protein_coding | 10.3068473 | -0.180223746 | 0.392344659 | -0.459350578 | 0.645982433 | 0.816513895 | 0.180223746 |
| ENSMUSG00000030545 | Pex11a | 18631 | protein_coding | 371.7844574 | -0.180303728 | 0.084032116 | -2.145652598 | 0.031900706 | 0.135849665 | 0.180303728 |
| ENSMUSG00000016758 | Bik | 12124 | protein_coding | 69.01045617 | -0.180316175 | 0.145735564 | -1.237283269 | 0.215981937 | 0.451769221 | 0.180316175 |
| ENSMUSG00000022433 | Csnk1e | 27373 | protein_coding | 288.1159324 | -0.18036464 | 0.079252487 | -2.275823079 | 0.022856601 | 0.107689014 | 0.18036464 |
| ENSMUSG00000035757 | Selenoo | 223776 | protein_coding | 742.3748292 | -0.180437368 | 0.057214862 | -3.153680004 | 0.001612258 | 0.014823762 | 0.180437368 |
| ENSMUSG00000030602 | Pak4 | 70584 | protein_coding | 2400.791928 | 0.180461789 | 0.069138294 | 2.610156811 | 0.009050073 | 0.055431266 | 0.180461789 |
| ENSMUSG00000022280 | Rnf19a | 30945 | protein_coding | 897.1164973 | -0.180491999 | 0.053475816 | -3.37520794 | 0.000737599 | 0.007846481 | 0.180491999 |
| ENSMUSG00000018217 | Pmp22 | 18858 | protein_coding | 4230.417334 | 0.180506866 | 0.096182906 | 1.876704222 | 0.060558644 | 0.209217919 | 0.180506866 |
| ENSMUSG00000028649 | Macf1 | 11426 | protein_coding | 3714.262676 | 0.180521931 | 0.057836157 | 3.121264299 | 0.001800763 | 0.016160774 | 0.180521931 |
| ENSMUSG00000034303 | Ccdc15 | 245902 | protein_coding | 53.66258268 | -0.180524734 | 0.192506358 | -0.937759855 | 0.348367836 | 0.590760052 | 0.180524734 |
| ENSMUSG00000028977 | Casz1 | 69743 | protein_coding | 1810.233018 | -0.180601025 | 0.073451889 | -2.458766251 | 0.013941536 | 0.076112101 | 0.180601025 |
| ENSMUSG00000066637 | Ttc32 | 75516 | protein_coding | 177.2819026 | 0.18062936 | 0.106306494 | 1.69913758 | 0.089293264 | 0.268368093 | 0.18062936 |
| ENSMUSG00000032180 | Tmed1 | 17083 | protein_coding | 205.6559197 | -0.180635036 | 0.093182196 | -1.938514485 | 0.052560485 | 0.191471188 | 0.180635036 |
| ENSMUSG00000049013 | Prss48 | 368202 | protein_coding | 12.09622305 | -0.180667428 | 0.363756089 | -0.496671901 | 0.61942044 | 0.798512665 | 0.180667428 |
| ENSMUSG00000056671 | Prelid2 | 77619 | protein_coding | 704.415251 | -0.180704572 | 0.055863182 | -3.234770451 | 0.001217406 | 0.011862901 | 0.180704572 |
| ENSMUSG00000032264 | Zw10 | 26951 | protein_coding | 576.7191269 | 0.180726297 | 0.053863511 | 3.355263962 | 0.000792893 | 0.008303236 | 0.180726297 |
| ENSMUSG00000025161 | Slc16a3 | 80879 | protein_coding | 75.95094436 | 0.180786544 | 0.180614724 | 1.000951305 | 0.316850351 | 0.560658943 | 0.180786544 |
| ENSMUSG00000093904 | Tomm20 | 67952 | protein_coding | 1313.191512 | -0.180804285 | 0.05630052 | -3.211414141 | 0.001320834 | 0.012664147 | 0.180804285 |
| ENSMUSG00000092416 | Zfp141 | 434178 | protein_coding | 233.9392804 | -0.180828235 | 0.11835835 | -1.52780294 | 0.126561472 | 0.330535524 | 0.180828235 |

| ensembl_gene_id | external_gene_name | entrezgene_id | gene_biotype | baseMean | log2FoldChange | lfcSE | stat | pvalue | padj | absLog2FC |
|---|---|---|---|---|---|---|---|---|---|---|
| ENSMUSG00000036864 | Proser3 | 333193 | protein_coding | 55.13343448 | 0.180850358 | 0.157456283 | 1.148575051 | 0.25073125 | 0.491735724 | 0.180850358 |
| ENSMUSG00000068735 | Trp53i11 | 277414 | protein_coding | 2714.02458 | 0.180856585 | 0.086245643 | 2.09699386 | 0.035994119 | 0.147231287 | 0.180856585 |
| ENSMUSG00000041921 | Metap1d | 66559 | protein_coding | 266.7149838 | -0.180880057 | 0.101824951 | -1.776382464 | 0.075669899 | 0.24203262 | 0.180880057 |
| ENSMUSG00000074635 | 3110070M22Rik | 67304 | lncRNA | 7.63029112 | -0.180919722 | 0.528453669 | -0.342356829 | 0.732082373 | 0.870787945 | 0.180919722 |
| ENSMUSG00000029701 | Rbm28 | 68272 | protein_coding | 572.7848896 | 0.18092022 | 0.074030948 | 2.443845791 | 0.014531636 | 0.078378495 | 0.18092022 |
| ENSMUSG00000055799 | Tcf7l1 | 21415 | protein_coding | 170.7455906 | 0.180924821 | 0.160162597 | 1.129632161 | 0.258631254 | 0.500565425 | 0.180924821 |
| ENSMUSG00000029804 | Herc3 | 73998 | protein_coding | 580.6790792 | -0.180999036 | 0.078217741 | -2.314040715 | 0.020665489 | 0.100534642 | 0.180999036 |
| ENSMUSG00000018042 | Cyb5r3 | 109754 | protein_coding | 2105.395155 | 0.181024377 | 0.053761142 | 3.367197378 | 0.000759363 | 0.00801981 | 0.181024377 |
| ENSMUSG00000075704 | Txnrd2 | 26462 | protein_coding | 274.8032236 | -0.181054734 | 0.091772879 | -1.972856647 | 0.048511897 | 0.181275208 | 0.181054734 |
| ENSMUSG00000002871 | Tpra1 | 24100 | protein_coding | 686.6383531 | -0.181101086 | 0.07473181 | -2.423346716 | 0.015378242 | 0.08158056 | 0.181101086 |
| ENSMUSG00000032295 | Man2c1 | 73744 | protein_coding | 862.0764476 | -0.181217312 | 0.061151763 | -2.963402931 | 0.00304258 | 0.024375404 | 0.181217312 |
| ENSMUSG00000028980 | H6pd | 100198 | protein_coding | 644.8014965 | 0.181223968 | 0.069354407 | 2.613013008 | 0.008974789 | 0.055053791 | 0.181223968 |
| ENSMUSG00000045268 | Zfp691 | 195522 | protein_coding | 146.0657113 | 0.181378584 | 0.108263838 | 1.675338575 | 0.093867821 | 0.276898911 | 0.181378584 |
| ENSMUSG00000051149 | Adnp | 11538 | protein_coding | 38.6417 | -0.181448969 | 0.244812932 | -0.741173953 | 0.458587974 | 0.685788934 | 0.181448969 |
| ENSMUSG00000033209 | Ttc28 | 209683 | protein_coding | 7.812779968 | -0.181450331 | 0.456444016 | -0.397530311 | 0.69097644 | 0.844835257 | 0.181450331 |
| ENSMUSG00000100210 | H3c7 | 260423 | protein_coding | 2012.393955 | 0.181515235 | 0.123246664 | 1.472780105 | 0.140810339 | 0.351955161 | 0.181515235 |
| ENSMUSG00000013736 | Trnt1 | 70047 | protein_coding | 589.6560765 | -0.181523428 | 0.084823551 | -2.140012132 | 0.032353786 | 0.137199451 | 0.181523428 |
| ENSMUSG00000054312 | Mrps21 | 66292 | protein_coding | 922.923098 | 0.18153904 | 0.081618725 | 2.224232743 | 0.026132789 | 0.118377705 | 0.18153904 |
| ENSMUSG00000025377 | Tepsin | 78777 | protein_coding | 619.2413908 | 0.181557932 | 0.083400662 | 2.176936349 | 0.029485317 | 0.12855088 | 0.181557932 |
| ENSMUSG00000073775 | Kti12 | 100087 | protein_coding | 328.8998484 | 0.181624054 | 0.068809507 | 2.639519765 | 0.008302358 | 0.052037361 | 0.181624054 |
| ENSMUSG00000024587 | Nars | 70223 | protein_coding | 3675.674584 | 0.181688541 | 0.057931415 | 3.136269696 | 0.001711118 | 0.015476834 | 0.181688541 |
| ENSMUSG00000060224 | Pyroxd2 | 74580 | protein_coding | 176.5309106 | -0.181721298 | 0.128719942 | -1.411757149 | 0.15802148 | 0.377781372 | 0.181721298 |
| ENSMUSG00000029516 | Cit | 12704 | protein_coding | 299.7428289 | 0.181736139 | 0.090590716 | 2.006123226 | 0.044843102 | 0.171848571 | 0.181736139 |
| ENSMUSG00000039004 | Bmp6 | 12161 | protein_coding | 10.96651924 | -0.181769995 | 0.413413087 | -0.43968128 | 0.660167963 | 0.826041967 | 0.181769995 |
| ENSMUSG00000026728 | Vim | 22352 | protein_coding | 130.09174 | 0.181823227 | 0.112509615 | 1.616068337 | 0.106079539 | 0.298195002 | 0.181823227 |
| ENSMUSG00000039481 | Nrtn | 18188 | protein_coding | 28.38518444 | 0.181832823 | 0.25697629 | 0.707585992 | 0.479202395 | 0.701661251 | 0.181832823 |
| ENSMUSG00000079164 | Tlr5 | 53791 | protein_coding | 95.41314123 | -0.181933622 | 0.154264312 | -1.179363001 | 0.238253661 | 0.478614449 | 0.181933622 |
| ENSMUSG00000025962 | Fastkd2 | 75619 | protein_coding | 500.9273332 | 0.181991482 | 0.093586286 | 1.944638355 | 0.05181852 | 0.189495979 | 0.181991482 |
| ENSMUSG00000021109 | Hif1a | 15251 | protein_coding | 1917.119748 | -0.181999865 | 0.124512061 | -1.461704705 | 0.143822147 | 0.355902988 | 0.181999865 |
| ENSMUSG00000094870 | Zfp131 | 72465 | protein_coding | 535.2761777 | -0.182025591 | 0.080445221 | -2.262727223 | 0.02365251 | 0.110275809 | 0.182025591 |
| ENSMUSG00000056708 | Ier5 | 15939 | protein_coding | 1222.02253 | 0.182069805 | 0.089151052 | 2.042261994 | 0.04112555 | 0.161710035 | 0.182069805 |
| ENSMUSG00000001924 | Uba1 | 22201 | protein_coding | 7606.409172 | 0.182085708 | 0.063948902 | 2.847362525 | 0.004408313 | 0.032477324 | 0.182085708 |
| ENSMUSG00000000711 | Rab5b | 19344 | protein_coding | 1723.879481 | 0.182128865 | 0.050746837 | 3.588969827 | 0.000331987 | 0.004009493 | 0.182128865 |
| ENSMUSG00000053985 | Zfp14 | 243906 | protein_coding | 41.90839487 | -0.182152054 | 0.180157809 | -1.011069432 | 0.3119832 | 0.555723255 | 0.182152054 |
| ENSMUSG00000028698 | Pik3r3 | 18710 | protein_coding | 490.8849195 | 0.182158187 | 0.08588901 | 2.120855579 | 0.033933958 | 0.141755849 | 0.182158187 |
| ENSMUSG00000026166 | Ccl20 | 20297 | protein_coding | 18.72229477 | 0.182181117 | 0.281148908 | 0.647987993 | 0.516992716 | 0.729699066 | 0.182181117 |
| ENSMUSG00000033703 | Fcsk | 234730 | protein_coding | 1375.267998 | 0.182191598 | 0.082646958 | 2.204456183 | 0.027492278 | 0.122173593 | 0.182191598 |
| ENSMUSG00000048186 | Bend7 | 209645 | protein_coding | 226.0351319 | 0.182204843 | 0.091868091 | 1.983331116 | 0.047330456 | 0.178465808 | 0.182204843 |
| ENSMUSG00000034459 | Ifit1 | 15957 | protein_coding | 228.2282851 | -0.182289905 | 0.228455983 | -0.797921344 | 0.424916136 | 0.660875931 | 0.182289905 |

Supplementary Table 1

| ensembl_gene_id | external_gene_name | entrezgene_id | gene_biotype | baseMean | log2FoldChange | lfcSE | stat | pvalue | padj | absLog2FC |
|---|---|---|---|---|---|---|---|---|---|---|
| ENSMUSG00000022130 | Tgds | 76355 | protein_coding | 321.0712763 | 0.182291308 | 0.075271927 | 2.421770183 | 0.015445115 | 0.081881448 | 0.182291308 |
| ENSMUSG00000053675 | Tgm5 | 74176 | protein_coding | 18.50072089 | -0.182425461 | 0.271834784 | -0.67108947 | 0.502163535 | 0.718950524 | 0.182425461 |
| ENSMUSG00000001964 | Emd | 13726 | protein_coding | 471.586213 | 0.182436335 | 0.061873797 | 2.948523353 | 0.00319296 | 0.025378447 | 0.182436335 |
| ENSMUSG00000022184 | Fbxo4 | 106052 | protein_coding | 318.0350658 | 0.18244901 | 0.092050241 | 1.982059015 | 0.047472636 | 0.178786609 | 0.18244901 |
| ENSMUSG00000003299 | Mrpl4 | 66163 | protein_coding | 911.7980786 | 0.182459274 | 0.099146843 | 1.840293335 | 0.065725183 | 0.221007096 | 0.182459274 |
| ENSMUSG00000047654 | Tssk6 | 83984 | protein_coding | 28.28002757 | 0.182461362 | 0.219789075 | 0.830165748 | 0.406445078 | 0.645466424 | 0.182461362 |
| ENSMUSG00000078970 | Dnaaf10 | 103784 | protein_coding | 415.0197146 | 0.182481534 | 0.077231567 | 2.362784293 | 0.018138224 | 0.091928074 | 0.182481534 |
| ENSMUSG00000054619 | Mettl7a1 | 70152 | protein_coding | 4110.9115 | -0.182485404 | 0.079877452 | -2.284567157 | 0.022338221 | 0.106296214 | 0.182485404 |
| ENSMUSG00000032177 | Pde4a | 18577 | protein_coding | 40.9598898 | -0.182497283 | 0.204708578 | -0.891497976 | 0.372662077 | 0.61438574 | 0.182497283 |
| ENSMUSG00000074829 | 2010315B03Rik | 630836 | protein_coding | 790.1997898 | -0.182510488 | 0.076703293 | -2.379434859 | 0.017339207 | 0.088855862 | 0.182510488 |
| ENSMUSG00000054051 | Ercc6 | 319955 | protein_coding | 567.0513057 | -0.182518252 | 0.087477117 | -2.086468541 | 0.036936202 | 0.149703476 | 0.182518252 |
| ENSMUSG00000030588 | Yif1b | 77254 | protein_coding | 506.014456 | -0.182553109 | 0.117769771 | -1.550084613 | 0.121121209 | 0.322370555 | 0.182553109 |
| ENSMUSG00000003948 | Mmd | 67468 | protein_coding | 178.3292512 | -0.182598538 | 0.120127494 | -1.520039523 | 0.128501043 | 0.333871046 | 0.182598538 |
| ENSMUSG00000035566 | Pcdh17 | 219228 | protein_coding | 718.7259128 | 0.182615229 | 0.065866955 | 2.772486271 | 0.005562987 | 0.038720019 | 0.182615229 |
| ENSMUSG00000005949 | Ctns | 83429 | protein_coding | 449.1423645 | -0.182801724 | 0.08959232 | -2.04037271 | 0.041313218 | 0.162264555 | 0.182801724 |
| ENSMUSG00000024431 | Nr3c1 | 14815 | protein_coding | 3245.358808 | -0.182818673 | 0.045102838 | -4.053374022 | 5.05E-05 | 0.000827981 | 0.182818673 |
| ENSMUSG00000008999 | Bmp7 | 12162 | protein_coding | 68.89000007 | -0.182872901 | 0.200776207 | -0.910829544 | 0.362385193 | 0.605301503 | 0.182872901 |
| ENSMUSG00000004110 | Cacna1e | 12290 | protein_coding | 61.6556192 | -0.182917088 | 0.275498848 | -0.663948649 | 0.506723178 | 0.722412013 | 0.182917088 |
| ENSMUSG00000034353 | Ramp1 | 51801 | protein_coding | 1560.281219 | 0.183062075 | 0.086839519 | 2.108050302 | 0.035026639 | 0.144654187 | 0.183062075 |
| ENSMUSG00000062931 | Zfp938 | 237411 | protein_coding | 115.8652529 | -0.183192275 | 0.121164922 | -1.511924996 | 0.130552944 | 0.336509736 | 0.183192275 |
| ENSMUSG00000031095 | Cul4b | 72584 | protein_coding | 620.7168697 | -0.183210457 | 0.063070483 | -2.904852617 | 0.003674261 | 0.02835158 | 0.183210457 |
| ENSMUSG00000071659 | Hnrnpul2 | 68693 | protein_coding | 3252.151978 | 0.183226618 | 0.035959656 | 5.095338414 | 3.48E-07 | 9.36E-06 | 0.183226618 |
| ENSMUSG00000026686 | Lmx1a | 110648 | protein_coding | 37.54692348 | -0.183232932 | 0.215582914 | -0.849941808 | 0.39535744 | 0.634673446 | 0.183232932 |
| ENSMUSG00000060803 | Gstp1 | 14870 | protein_coding | 2375.242842 | -0.183245239 | 0.043077119 | -4.253887994 | 2.10E-05 | 0.000378562 | 0.183245239 |
| ENSMUSG00000038831 | Ralgps1 | 241308 | protein_coding | 283.7501608 | 0.183252338 | 0.08873453 | 2.065175045 | 0.038906441 | 0.155443076 | 0.183252338 |
| ENSMUSG00000021645 | Smn1 | 20595 | protein_coding | 125.7500106 | 0.183270592 | 0.119487223 | 1.533809119 | 0.125076619 | 0.328198932 | 0.183270592 |
| ENSMUSG00000056632 | Dsg3 | 13512 | protein_coding | 19.85877567 | 0.183415441 | 0.320272258 | 0.572686009 | 0.566857312 | 0.762762443 | 0.183415441 |
| ENSMUSG00000046523 | Kctd4 | 67516 | protein_coding | 10.84474875 | 0.183428289 | 0.360143806 | 0.509319571 | 0.610528242 | 0.792525305 | 0.183428289 |
| ENSMUSG00000015932 | Dstn | 56431 | protein_coding | 23862.98583 | 0.183430552 | 0.052377704 | 3.502073147 | 0.000461653 | 0.005291454 | 0.183430552 |
| ENSMUSG00000039886 | Tmem120a | 215210 | protein_coding | 716.3769128 | 0.183500539 | 0.082299708 | 2.229662098 | 0.025769884 | 0.11719992 | 0.183500539 |
| ENSMUSG00000041859 | Mcm3 | 17215 | protein_coding | 537.3314545 | 0.183535073 | 0.186452809 | 0.984351345 | 0.324942792 | 0.568614628 | 0.183535073 |
| ENSMUSG00000003848 | Nob1 | 67619 | protein_coding | 290.684153 | -0.183610057 | 0.084233316 | -2.179779524 | 0.029273808 | 0.127940029 | 0.183610057 |
| ENSMUSG00000025027 | Xpnpep1 | 170750 | protein_coding | 1674.296267 | 0.183710441 | 0.051651725 | 3.556714571 | 0.000375522 | 0.004462815 | 0.183710441 |
| ENSMUSG00000117232 | 1600002D24Rik | 69776 | lncRNA | 7.032963256 | -0.183713309 | 0.462919447 | -0.39685805 | 0.691472141 | 0.845249088 | 0.183713309 |
| ENSMUSG00000049538 | Adamts16 | 271127 | protein_coding | 24.38520429 | 0.183740583 | 0.269910308 | 0.680746818 | 0.496031706 | 0.714476238 | 0.183740583 |
| ENSMUSG00000067370 | B3galt4 | 54218 | protein_coding | 207.7010224 | 0.183787469 | 0.118424758 | 1.55193452 | 0.120677892 | 0.321468841 | 0.183787469 |
| ENSMUSG00000036136 | Fam110c | 104943 | protein_coding | 955.9810633 | -0.183926999 | 0.055738382 | -3.299826643 | 0.000967446 | 0.009794097 | 0.183926999 |
| ENSMUSG00000037979 | Ccdc92 | 215707 | protein_coding | 36.75162558 | 0.183974696 | 0.189831884 | 0.969145393 | 0.332472652 | 0.575843468 | 0.183974696 |
| ENSMUSG00000026158 | Ogfrl1 | 70155 | protein_coding | 95.86205039 | -0.183999459 | 0.129939124 | -1.416043556 | 0.156762752 | 0.375480655 | 0.183999459 |

| ensembl_gene_id | external_gene_name | entrezgene_id | gene_biotype | baseMean | log2FoldChange | lfcSE | stat | pvalue | padj | absLog2FC |
|---|---|---|---|---|---|---|---|---|---|---|
| ENSMUSG00000061536 | Sec22c | 215474 | protein_coding | 138.9407422 | -0.18400411 | 0.131906943 | -1.394953933 | 0.163029732 | 0.384495537 | 0.18400411 |
| ENSMUSG00000026696 | Vamp4 | 53330 | protein_coding | 464.4134182 | -0.184009844 | 0.083869464 | -2.194002847 | 0.028235205 | 0.124616625 | 0.184009844 |
| ENSMUSG00000020083 | Fam241b | 69894 | protein_coding | 417.1255198 | -0.184074071 | 0.073853778 | -2.492412386 | 0.012687863 | 0.070949086 | 0.184074071 |
| ENSMUSG00000003528 | Slc25a1 | 13358 | protein_coding | 2066.597525 | -0.184078702 | 0.061786568 | -2.979267306 | 0.002889386 | 0.023392132 | 0.184078702 |
| ENSMUSG00000061104 | Sap18b | 100041953 | protein_coding | 10.8360391 | -0.184176559 | 0.393066993 | -0.46856277 | 0.639382194 | 0.812170498 | 0.184176559 |
| ENSMUSG00000034557 | Zfyve9 | 230597 | protein_coding | 1034.698187 | -0.18422642 | 0.054024156 | -3.410074913 | 0.00064945 | 0.007100804 | 0.18422642 |
| ENSMUSG00000025494 | Sigirr | 24058 | protein_coding | 836.8901836 | -0.184244302 | 0.085045831 | -2.166411925 | 0.030279724 | 0.130987701 | 0.184244302 |
| ENSMUSG00000044468 | Tent5c | 74645 | protein_coding | 325.3274649 | 0.184310848 | 0.145948486 | 1.262848645 | 0.206643579 | 0.441566958 | 0.184310848 |
| ENSMUSG00000030451 | Herc2 | 15204 | protein_coding | 3876.101012 | 0.184328388 | 0.052803914 | 3.490809216 | 0.00048156 | 0.005472951 | 0.184328388 |
| ENSMUSG00000090236 | Car15 | 80733 | protein_coding | 11.41415805 | -0.184357437 | 0.340488076 | -0.541450493 | 0.588197112 | 0.778103095 | 0.184357437 |
| ENSMUSG00000050315 | Synpo2 | 118449 | protein_coding | 18.87752962 | -0.184362186 | 0.345585059 | -0.533478462 | 0.593702417 | 0.781156798 | 0.184362186 |
| ENSMUSG00000028189 | Ctbs | 74245 | protein_coding | 226.4865298 | -0.184427663 | 0.10541031 | -1.749616939 | 0.080184435 | 0.250607438 | 0.184427663 |
| ENSMUSG00000013523 | Bcas1 | 76960 | protein_coding | 2528.186884 | 0.184453903 | 0.06921063 | 2.665109426 | 0.007696328 | 0.04918511 | 0.184453903 |
| ENSMUSG00000044986 | Tst | 22117 | protein_coding | 3681.294314 | 0.184504547 | 0.072243458 | 2.553927397 | 0.010651546 | 0.061991154 | 0.184504547 |
| ENSMUSG00000077391 | Gm24336 | 115489055 | snoRNA | 719.5737901 | -0.184554048 | 0.069001982 | -2.674619534 | 0.00748141 | 0.048137725 | 0.184554048 |
| ENSMUSG00000037965 | Zc3h7a | 106205 | protein_coding | 1502.215078 | 0.184589653 | 0.080302307 | 2.298684322 | 0.021522871 | 0.103580855 | 0.184589653 |
| ENSMUSG00000052955 | Cpvl | 71287 | protein_coding | 9.038814395 | 0.184642645 | 0.406730498 | 0.453968036 | 0.649851833 | 0.819466769 | 0.184642645 |
| ENSMUSG00000059555 | Tor4a | 227612 | protein_coding | 1302.047533 | 0.184670029 | 0.071089125 | 2.597725455 | 0.009384349 | 0.056796115 | 0.184670029 |
| ENSMUSG00000042616 | Oscp1 | 230751 | protein_coding | 27.09053125 | 0.184688998 | 0.29722464 | 0.62137849 | 0.534350621 | 0.741606925 | 0.184688998 |
| ENSMUSG00000014837 | 4931428F04Rik | 74356 | protein_coding | 41.86826669 | 0.184718838 | 0.197625149 | 0.934692974 | 0.349946549 | 0.592368513 | 0.184718838 |
| ENSMUSG00000003518 | Dusp3 | 72349 | protein_coding | 663.5371315 | -0.184731832 | 0.054680522 | -3.378384569 | 0.00072913 | 0.0077847 | 0.184731832 |
| ENSMUSG00000063576 | Klhdc3 | 71765 | protein_coding | 501.5449157 | -0.184756288 | 0.078116142 | -2.365148648 | 0.018022835 | 0.091458234 | 0.184756288 |
| ENSMUSG00000038702 | Dsel | 319901 | protein_coding | 187.0371175 | 0.184759916 | 0.121464797 | 1.521098461 | 0.128235131 | 0.333379077 | 0.184759916 |
| ENSMUSG00000030890 | Ilk | 16202 | protein_coding | 17.19120837 | -0.184820323 | 0.294969365 | -0.626574638 | 0.530938098 | 0.73971984 | 0.184820323 |
| ENSMUSG00000034610 | Tut4 | 230594 | protein_coding | 403.9054661 | -0.184891666 | 0.118776987 | -1.556628685 | 0.119558669 | 0.319914163 | 0.184891666 |
| ENSMUSG00000067199 | Frat1 | 14296 | protein_coding | 597.729979 | -0.184915555 | 0.10017171 | -1.845985811 | 0.064894266 | 0.219231568 | 0.184915555 |
| ENSMUSG00000026816 | Gtf3c5 | 70239 | protein_coding | 437.1006629 | 0.184957888 | 0.062796609 | 2.945348339 | 0.003225913 | 0.025589912 | 0.184957888 |
| ENSMUSG00000055639 | Dach1 | 13134 | protein_coding | 87.75473992 | -0.184971009 | 0.145311656 | -1.272926171 | 0.203044241 | 0.437065467 | 0.184971009 |
| ENSMUSG00000022702 | Hira | 15260 | protein_coding | 189.0996436 | -0.184980922 | 0.096975997 | -1.907491834 | 0.056456927 | 0.200767557 | 0.184980922 |
| ENSMUSG00000018740 | Slc25a35 | 71998 | protein_coding | 425.9728647 | 0.18501278 | 0.074295744 | 2.490220433 | 0.012766389 | 0.071215342 | 0.18501278 |
| ENSMUSG00000020577 | Tspan13 | 66109 | protein_coding | 2362.93591 | 0.185033324 | 0.082618583 | 2.239609012 | 0.025116317 | 0.115038582 | 0.185033324 |
| ENSMUSG00000034918 | Cdhr2 | 268663 | protein_coding | 15544.11759 | -0.18504019 | 0.060155039 | -3.076054715 | 0.002097593 | 0.018226234 | 0.18504019 |
| ENSMUSG00000032009 | Sesn3 | 75747 | protein_coding | 796.5386141 | 0.185043375 | 0.056103782 | 3.298233499 | 0.000972952 | 0.009823815 | 0.185043375 |
| ENSMUSG00000051098 | Mblac2 | 72852 | protein_coding | 119.8099628 | -0.185079613 | 0.134681514 | -1.374202046 | 0.169378966 | 0.393655366 | 0.185079613 |
| ENSMUSG00000033781 | Asb13 | 142688 | protein_coding | 350.6534557 | -0.185191528 | 0.08393092 | -2.206475609 | 0.027350716 | 0.121645063 | 0.185191528 |
| ENSMUSG00000025351 | Cd63 | 12512 | protein_coding | 4728.222563 | -0.185194583 | 0.043562215 | -4.251266423 | 2.13E-05 | 0.000381741 | 0.185194583 |
| ENSMUSG00000032580 | Rbm5 | 83486 | protein_coding | 1977.939381 | -0.185338596 | 0.077513636 | -2.391045049 | 0.016800491 | 0.087063472 | 0.185338596 |
| ENSMUSG00000028347 | Tmeff1 | 230157 | protein_coding | 244.6005541 | -0.185369063 | 0.121969093 | -1.519803561 | 0.128560354 | 0.333871046 | 0.185369063 |
| ENSMUSG00000022604 | Cep97 | 74201 | protein_coding | 108.3185144 | -0.185407786 | 0.11475177 | -1.615729213 | 0.106152872 | 0.298245186 | 0.185407786 |

| ensembl_gene_id | external_gene_name | entrezgene_id | gene_biotype | baseMean | log2FoldChange | lfcSE | stat | pvalue | padj | absLog2FC |
|---|---|---|---|---|---|---|---|---|---|---|
| ENSMUSG00000035399 | Oser1 | 66680 | protein_coding | 281.1136565 | -0.185408056 | 0.082527779 | -2.246613916 | 0.024664712 | 0.113777978 | 0.185408056 |
| ENSMUSG00000059713 | Rcan3 | 53902 | protein_coding | 872.2866519 | 0.185410831 | 0.051423721 | 3.605550654 | 0.000311492 | 0.003796997 | 0.185410831 |
| ENSMUSG00000027599 | Armc1 | 74252 | protein_coding | 684.9996689 | 0.185564918 | 0.058887708 | 3.151165594 | 0.001626203 | 0.014934949 | 0.185564918 |
| ENSMUSG00000029570 | Lfng | 16848 | protein_coding | 403.222071 | -0.185657109 | 0.113525395 | -1.63537955 | 0.101969507 | 0.291364677 | 0.185657109 |
| ENSMUSG00000025241 | Fyco1 | 17281 | protein_coding | 974.637752 | -0.185703656 | 0.053249476 | -3.487426883 | 0.000487692 | 0.005527065 | 0.185703656 |
| ENSMUSG00000048478 | Spata33 | 320869 | protein_coding | 10.85629881 | 0.18571928 | 0.386338537 | 0.480716425 | 0.630718056 | 0.805989706 | 0.18571928 |
| ENSMUSG00000046808 | Atp10d | 231287 | protein_coding | 16.43882788 | 0.185736985 | 0.346295233 | 0.536354436 | 0.591713616 | 0.780262005 | 0.185736985 |
| ENSMUSG00000032840 | Spring1 | 76792 | protein_coding | 302.743985 | 0.18574915 | 0.075798792 | 2.450555553 | 0.014263595 | 0.07724278 | 0.18574915 |
| ENSMUSG00000024742 | Fen1 | 14156 | protein_coding | 40.32429103 | 0.185801247 | 0.228558122 | 0.812927781 | 0.416259465 | 0.654207378 | 0.185801247 |
| ENSMUSG00000069267 | H3c2 | 319150 | protein_coding | 952.3339177 | 0.185831023 | 0.131070442 | 1.417795041 | 0.156250612 | 0.374715872 | 0.185831023 |
| ENSMUSG00000079173 | Zan | 22635 | protein_coding | 1568.735473 | -0.185836575 | 0.086102111 | -2.158327743 | 0.030902359 | 0.132825785 | 0.185836575 |
| ENSMUSG00000114613 | Gm10732 | 100038690 | lncRNA | 31.19865845 | 0.185856015 | 0.234459159 | 0.792701019 | 0.427952036 | 0.663358563 | 0.185856015 |
| ENSMUSG00000022472 | Desi1 | 28075 | protein_coding | 387.953533 | -0.185878643 | 0.069773746 | -2.664019821 | 0.007721301 | 0.049315416 | 0.185878643 |
| ENSMUSG00000038349 | Plcl1 | 227120 | protein_coding | 26.65116555 | -0.185912168 | 0.235721737 | -0.788693355 | 0.430291252 | 0.665280626 | 0.185912168 |
| ENSMUSG00000061371 | Zfp873 | 408062 | protein_coding | 117.1993004 | -0.185926033 | 0.130160381 | -1.428437992 | 0.153165831 | 0.370189171 | 0.185926033 |
| ENSMUSG00000028127 | Abcd3 | 19299 | protein_coding | 5131.668286 | 0.186014673 | 0.050433421 | 3.688321545 | 0.000225738 | 0.002907732 | 0.186014673 |
| ENSMUSG00000020471 | Pold2 | 18972 | protein_coding | 720.9834012 | 0.186108987 | 0.104259004 | 1.785063934 | 0.074250947 | 0.239010983 | 0.186108987 |
| ENSMUSG00000058486 | Wdr91 | 101240 | protein_coding | 98.52322447 | -0.186282814 | 0.148029153 | -1.258419776 | 0.208239984 | 0.443565039 | 0.186282814 |
| ENSMUSG00000086158 | Pierce2 | 546143 | protein_coding | 118.4524814 | 0.18640601 | 0.10894481 | 1.711013208 | 0.087078675 | 0.264317294 | 0.18640601 |
| ENSMUSG00000050270 | Tmem220 | 338369 | protein_coding | 108.2063259 | 0.186409597 | 0.116799672 | 1.595977061 | 0.110493915 | 0.305178175 | 0.186409597 |
| ENSMUSG00000042831 | Alkbh6 | 233065 | protein_coding | 402.1267518 | 0.186498842 | 0.083776284 | 2.226153178 | 0.026003923 | 0.117931739 | 0.186498842 |
| ENSMUSG00000021590 | Spata9 | 75571 | protein_coding | 7.674439807 | 0.186567614 | 0.475009367 | 0.392766179 | 0.694492187 | 0.847142841 | 0.186567614 |
| ENSMUSG00000036249 | Rbm43 | 71684 | protein_coding | 162.0286396 | 0.186635103 | 0.094337919 | 1.978367824 | 0.047887227 | 0.179682947 | 0.186635103 |
| ENSMUSG00000000957 | Mmp14 | 17387 | protein_coding | 878.3679135 | 0.186639513 | 0.07730561 | 2.414307479 | 0.015765152 | 0.082978005 | 0.186639513 |
| ENSMUSG00000018286 | Psmb6 | 19175 | protein_coding | 2406.123309 | 0.186814268 | 0.067463053 | 2.769134507 | 0.005620543 | 0.039036387 | 0.186814268 |
| ENSMUSG00000033906 | Zdhhc15 | 108672 | protein_coding | 130.6294139 | -0.18692468 | 0.134460987 | -1.390177802 | 0.164474892 | 0.386772614 | 0.18692468 |
| ENSMUSG00000027274 | Mkks | 59030 | protein_coding | 461.3533218 | 0.186945808 | 0.080063451 | 2.334970632 | 0.019544955 | 0.096539507 | 0.186945808 |
| ENSMUSG00000031214 | Ophn1 | 94190 | protein_coding | 551.2228004 | -0.186981469 | 0.064732361 | -2.888531572 | 0.003870452 | 0.0294918 | 0.186981469 |
| ENSMUSG00000030330 | Ing4 | 28019 | protein_coding | 25.55250493 | -0.186998974 | 0.289406413 | -0.646146615 | 0.518184409 | 0.730504162 | 0.186998974 |
| ENSMUSG00000034801 | Sos2 | 20663 | protein_coding | 1036.693534 | -0.187014611 | 0.048297042 | -3.872175208 | 0.000107868 | 0.00158722 | 0.187014611 |
| ENSMUSG00000010025 | Aldh3a2 | 11671 | protein_coding | 4009.610709 | 0.187020738 | 0.042254602 | 4.426044227 | 9.60E-06 | 0.000189451 | 0.187020738 |
| ENSMUSG00000020946 | Gosr2 | 56494 | protein_coding | 1358.0028 | 0.187083678 | 0.065224856 | 2.868288092 | 0.004126996 | 0.030904873 | 0.187083678 |
| ENSMUSG00000055723 | Rras2 | 66922 | protein_coding | 1250.390944 | -0.187086649 | 0.045540222 | -4.108162872 | 3.99E-05 | 0.000666808 | 0.187086649 |
| ENSMUSG00000055137 | Sugct | 192136 | protein_coding | 59.05086944 | -0.187295365 | 0.168522341 | -1.111397841 | 0.266397141 | 0.50824402 | 0.187295365 |
| ENSMUSG00000024124 | Prss30 | 30943 | protein_coding | 2991.788975 | -0.187306898 | 0.123743056 | -1.51367603 | 0.130108027 | 0.336096773 | 0.187306898 |
| ENSMUSG00000020657 | Dnajc27 | 217378 | protein_coding | 37.27999007 | -0.187327392 | 0.206832949 | -0.905694151 | 0.365097759 | 0.607253668 | 0.187327392 |
| ENSMUSG00000030498 | Gas2 | 14453 | protein_coding | 66.98513949 | -0.187342356 | 0.171204629 | -1.094259878 | 0.273841017 | 0.516458201 | 0.187342356 |
| ENSMUSG00000019804 | Snx3 | 54198 | protein_coding | 1687.646979 | 0.187355755 | 0.082214508 | 2.278864874 | 0.022675099 | 0.107237924 | 0.187355755 |
| ENSMUSG00000041220 | Elovl6 | 170439 | protein_coding | 1666.357618 | 0.187374338 | 0.058575458 | 3.198854004 | 0.00137975 | 0.013050576 | 0.187374338 |

| ensembl_gene_id | external_gene_name | entrezgene_id | gene_biotype | baseMean | log2FoldChange | lfcSE | stat | pvalue | padj | absLog2FC |
|---|---|---|---|---|---|---|---|---|---|---|
| ENSMUSG00000052131 | Akr1b7 | 11997 | protein_coding | 51.33730697 | -0.187433613 | 0.231857777 | -0.808399079 | 0.418860882 | 0.656326613 | 0.187433613 |
| ENSMUSG00000042489 | Clspn | 269582 | protein_coding | 227.0500725 | 0.187493967 | 0.207915709 | 0.901778743 | 0.367174414 | 0.609576053 | 0.187493967 |
| ENSMUSG00000052516 | Robo2 | 268902 | protein_coding | 645.6881337 | 0.187515051 | 0.117467151 | 1.596319051 | 0.110417579 | 0.305071833 | 0.187515051 |
| ENSMUSG00000022322 | Shcbp1 | 20419 | protein_coding | 210.4038862 | 0.187541191 | 0.136817 | 1.370744801 | 0.170454523 | 0.394876088 | 0.187541191 |
| ENSMUSG00000019689 | Fmc1 | 66117 | protein_coding | 770.8254789 | 0.187576995 | 0.100325287 | 1.869688096 | 0.061527144 | 0.211521689 | 0.187576995 |
| ENSMUSG00000005198 | Polr2a | 20020 | protein_coding | 842.7245449 | -0.187687512 | 0.167915657 | -1.117748725 | 0.263674326 | 0.504916665 | 0.187687512 |
| ENSMUSG00000027130 | Slc12a6 | 107723 | protein_coding | 1015.228986 | -0.187687746 | 0.053084522 | -3.535639753 | 0.000406789 | 0.004743519 | 0.187687746 |
| ENSMUSG00000061544 | Zfp229 | 381067 | protein_coding | 134.5232987 | 0.187693968 | 0.113688855 | 1.650944309 | 0.098749947 | 0.285896663 | 0.187693968 |
| ENSMUSG00000036622 | Atp13a2 | 74772 | protein_coding | 893.5130354 | 0.187699967 | 0.095538399 | 1.964654739 | 0.049454212 | 0.183913425 | 0.187699967 |
| ENSMUSG00000054662 | Ano9 | 71345 | protein_coding | 1318.948329 | -0.187727393 | 0.103159654 | -1.819775333 | 0.068793226 | 0.227621738 | 0.187727393 |
| ENSMUSG00000046096 | Mosmo | 233812 | protein_coding | 135.5819882 | -0.187816589 | 0.101893687 | -1.843260304 | 0.065291014 | 0.220005888 | 0.187816589 |
| ENSMUSG00000046572 | Zfp518b | 100515 | protein_coding | 99.9922653 | 0.187828107 | 0.14566442 | 1.289457693 | 0.197239015 | 0.42973164 | 0.187828107 |
| ENSMUSG00000010054 | Tusc2 | 80385 | protein_coding | 716.0361994 | 0.187866736 | 0.052910861 | 3.550627103 | 0.000384315 | 0.004537457 | 0.187866736 |
| ENSMUSG00000025026 | Add3 | 27360 | protein_coding | 3764.218286 | -0.187868404 | 0.050849494 | -3.694597298 | 0.000220236 | 0.002859216 | 0.187868404 |
| ENSMUSG00000038860 | Garnl3 | 99326 | protein_coding | 15.49175926 | -0.187963974 | 0.315728271 | -0.595334631 | 0.551619811 | 0.752493038 | 0.187963974 |
| ENSMUSG00000036208 | Nepro | 212547 | protein_coding | 330.7696621 | -0.187981604 | 0.088906839 | -2.114366074 | 0.034484013 | 0.143220107 | 0.187981604 |
| ENSMUSG00000042677 | Zc3h12a | 230738 | protein_coding | 408.9217457 | -0.187999801 | 0.123051723 | -1.5278112 | 0.126559421 | 0.330535524 | 0.187999801 |
| ENSMUSG00000051184 | Zfp524 | 66056 | protein_coding | 130.1778436 | -0.188036279 | 0.129345533 | -1.453751622 | 0.14601518 | 0.358961405 | 0.188036279 |
| ENSMUSG00000036585 | Fgf1 | 14164 | protein_coding | 868.9949234 | 0.188039512 | 0.049396111 | 3.806767499 | 0.000140795 | 0.00197128 | 0.188039512 |
| ENSMUSG00000015702 | Anxa9 | 71790 | protein_coding | 14.67386897 | 0.1880567 | 0.349369931 | 0.538273857 | 0.590387997 | 0.779594467 | 0.1880567 |
| ENSMUSG00000087881 | Gm22442 | 115487906 | scaRNA | 1232.569508 | -0.188097249 | 0.059284045 | -3.17281404 | 0.001509692 | 0.014057044 | 0.188097249 |
| ENSMUSG00000038917 | 3930402G23Rik | 665306 | lncRNA | 13.255088 | 0.188105998 | 0.323746628 | 0.58102844 | 0.561221286 | 0.759206347 | 0.188105998 |
| ENSMUSG00000005681 | Apoa2 | 11807 | protein_coding | 24.57446243 | -0.188114382 | 0.249113536 | -0.755135129 | 0.450167907 | 0.679535938 | 0.188114382 |
| ENSMUSG00000032698 | Lmo2 | 16909 | protein_coding | 24.11884509 | -0.188161779 | 0.250199008 | -0.752048461 | 0.452021916 | 0.680967534 | 0.188161779 |
| ENSMUSG00000095930 | Nim1k | 245269 | protein_coding | 36.17687896 | 0.188341532 | 0.281667253 | 0.66866677 | 0.50370807 | 0.720023049 | 0.188341532 |
| ENSMUSG00000030839 | Sergef | 27414 | protein_coding | 123.9745215 | -0.188370238 | 0.14792186 | -1.273444224 | 0.202860453 | 0.436976894 | 0.188370238 |
| ENSMUSG00000075225 | Ccdc162 | 75973 | protein_coding | 19.81773235 | -0.188407091 | 0.432288063 | -0.435836905 | 0.662955066 | 0.827707324 | 0.188407091 |
| ENSMUSG00000039128 | Cdc123 | 98828 | protein_coding | 1090.478893 | 0.188429921 | 0.059004888 | 3.193462883 | 0.001405774 | 0.013257847 | 0.188429921 |
| ENSMUSG00000029068 | Ccnl2 | 56036 | protein_coding | 1873.399146 | -0.188440998 | 0.095861548 | -1.965762102 | 0.049326097 | 0.183629264 | 0.188440998 |
| ENSMUSG00000031093 | Dock11 | 75974 | protein_coding | 299.5181938 | -0.188587615 | 0.098552783 | -1.913569662 | 0.055675165 | 0.198761565 | 0.188587615 |
| ENSMUSG00000020309 | Chac2 | 68044 | protein_coding | 193.3885181 | 0.188609476 | 0.096656445 | 1.951338846 | 0.051016751 | 0.187670425 | 0.188609476 |
| ENSMUSG00000090698 | Apold1 | 381823 | protein_coding | 8.720874526 | -0.188625806 | 0.403818099 | -0.467105875 | 0.640424131 | 0.812950957 | 0.188625806 |
| ENSMUSG00000031666 | Rbl2 | 19651 | protein_coding | 1055.195158 | -0.188640905 | 0.056607131 | -3.332458314 | 0.000860824 | 0.00888196 | 0.188640905 |
| ENSMUSG00000025040 | Fundc1 | 72018 | protein_coding | 423.0423673 | 0.188651437 | 0.064762937 | 2.91295372 | 0.003580277 | 0.027799288 | 0.188651437 |
| ENSMUSG00000046417 | Lrrc75a | 192976 | protein_coding | 1060.835982 | 0.188746684 | 0.057528834 | 3.280905786 | 0.001034743 | 0.010345506 | 0.188746684 |
| ENSMUSG00000029864 | Gstk1 | 76263 | protein_coding | 963.897113 | 0.188837122 | 0.054035554 | 3.49468282 | 0.000474625 | 0.005414954 | 0.188837122 |
| ENSMUSG00000069094 | Pde7a | 18583 | protein_coding | 1873.018451 | -0.188861657 | 0.090651378 | -2.08338429 | 0.037216209 | 0.150573458 | 0.188861657 |
| ENSMUSG00000074733 | Zfp950 | 414758 | protein_coding | 317.061344 | -0.188896575 | 0.122031402 | -1.547934158 | 0.12163815 | 0.323066278 | 0.188896575 |
| ENSMUSG00000033857 | Engase | 217364 | protein_coding | 978.8434202 | 0.189025298 | 0.053588067 | 3.527376711 | 0.000419699 | 0.004869416 | 0.189025298 |

| ensembl_gene_id | external_gene_name | entrezgene_id | gene_biotype | baseMean | log2FoldChange | lfcSE | stat | pvalue | padj | absLog2FC |
|---|---|---|---|---|---|---|---|---|---|---|
| ENSMUSG00000022705 | Drd3 | 13490 | protein_coding | 8.577133292 | -0.189110961 | 0.437453088 | -0.432299979 | 0.665523402 | 0.82924329 | 0.189110961 |
| ENSMUSG00000031558 | Slit2 | 20563 | protein_coding | 114.6128618 | -0.189139956 | 0.130827177 | -1.445723738 | 0.148254703 | 0.362422859 | 0.189139956 |
| ENSMUSG00000018379 | Srsf1 | 110809 | protein_coding | 3785.45653 | 0.189148231 | 0.049839928 | 3.795114474 | 0.000147575 | 0.002040458 | 0.189148231 |
| ENSMUSG00000034586 | Hid1 | 217310 | protein_coding | 1883.177289 | 0.189216768 | 0.089099642 | 2.123653513 | 0.033699127 | 0.141233323 | 0.189216768 |
| ENSMUSG00000032688 | Malt1 | 240354 | protein_coding | 744.0464504 | -0.189230852 | 0.073811734 | -2.563696062 | 0.010356419 | 0.060891613 | 0.189230852 |
| ENSMUSG00000021108 | Prkch | 18755 | protein_coding | 198.3445271 | 0.189257224 | 0.118598369 | 1.595782685 | 0.11053732 | 0.305193523 | 0.189257224 |
| ENSMUSG00000049115 | Agtr1a | 11607 | protein_coding | 10.89936939 | 0.189271411 | 0.345397631 | 0.547981208 | 0.583704806 | 0.775075241 | 0.189271411 |
| ENSMUSG00000019066 | Rab3d | 19340 | protein_coding | 3648.396767 | 0.189331038 | 0.055371275 | 3.419300656 | 0.000627823 | 0.006901775 | 0.189331038 |
| ENSMUSG00000020230 | Prmt2 | 15468 | protein_coding | 348.2526281 | 0.189342821 | 0.116352026 | 1.627327239 | 0.103667616 | 0.294235769 | 0.189342821 |
| ENSMUSG00000015451 | C4a | 625018 | lncRNA | 6.976887367 | -0.189382151 | 0.532865719 | -0.355403142 | 0.722287595 | 0.864247815 | 0.189382151 |
| ENSMUSG00000032591 | Mst1 | 15235 | protein_coding | 15.36663562 | -0.189535855 | 0.440667577 | -0.430110734 | 0.667115092 | 0.830161713 | 0.189535855 |
| ENSMUSG00000050677 | Ccdc96 | 66717 | protein_coding | 5.919559832 | -0.189601419 | 0.51284904 | -0.3697022 | 0.711604392 | 0.857573161 | 0.189601419 |
| ENSMUSG00000021608 | Lpcat1 | 210992 | protein_coding | 279.5177712 | -0.189661727 | 0.089750639 | -2.113207533 | 0.034583009 | 0.143483454 | 0.189661727 |
| ENSMUSG00000070509 | Rgma | 244058 | protein_coding | 30.07694116 | 0.189675872 | 0.246624314 | 0.769088292 | 0.441840898 | 0.673006845 | 0.189675872 |
| ENSMUSG00000026437 | Cdk18 | 18557 | protein_coding | 455.2022805 | 0.189697867 | 0.09224513 | 2.056453997 | 0.039738776 | 0.157771355 | 0.189697867 |
| ENSMUSG00000018909 | Arrb1 | 109689 | protein_coding | 2616.600403 | 0.189698469 | 0.044376389 | 4.274761227 | 1.91E-05 | 0.000347889 | 0.189698469 |
| ENSMUSG00000114442 | F630042J09Rik | 100038627 | lncRNA | 17.19203657 | -0.189714682 | 0.340006413 | -0.557973835 | 0.576862251 | 0.769561988 | 0.189714682 |
| ENSMUSG00000021203 | Otub2 | 68149 | protein_coding | 38.67666361 | 0.189744729 | 0.200283236 | 0.94738198 | 0.34344417 | 0.586046358 | 0.189744729 |
| ENSMUSG00000011158 | Brf1 | 72308 | protein_coding | 604.6348567 | -0.189750292 | 0.081324566 | -2.3332469 | 0.019635191 | 0.096777726 | 0.189750292 |
| ENSMUSG00000030075 | Cntn3 | 18488 | protein_coding | 50.54339183 | -0.189793594 | 0.201601249 | -0.941430647 | 0.346484214 | 0.588680038 | 0.189793594 |
| ENSMUSG00000020198 | Ap3d1 | 11776 | protein_coding | 3531.288262 | 0.189879945 | 0.03508993 | 5.411237485 | 6.26E-08 | 1.98E-06 | 0.189879945 |
| ENSMUSG00000033149 | Phldb2 | 208177 | protein_coding | 207.1172866 | -0.189899734 | 0.10843913 | -1.751210423 | 0.079909671 | 0.249990933 | 0.189899734 |
| ENSMUSG00000006782 | Cnp | 12799 | protein_coding | 791.9807027 | 0.189907 | 0.091668073 | 2.071680952 | 0.038295205 | 0.153937911 | 0.189907 |
| ENSMUSG00000029162 | Khk | 16548 | protein_coding | 387.0017902 | -0.190023593 | 0.06733831 | -2.821923997 | 0.004773649 | 0.034445027 | 0.190023593 |
| ENSMUSG00000059326 | Csf2ra | 12982 | protein_coding | 234.1650596 | 0.190057315 | 0.130403826 | 1.457451989 | 0.144991655 | 0.35758991 | 0.190057315 |
| ENSMUSG00000026411 | Tmem9 | 66241 | protein_coding | 234.1472124 | -0.190058346 | 0.10883527 | -1.746293687 | 0.080759932 | 0.251771782 | 0.190058346 |
| ENSMUSG00000025170 | Rab40b | 217371 | protein_coding | 21.10773383 | -0.190184044 | 0.265857734 | -0.715360209 | 0.474386493 | 0.698223143 | 0.190184044 |
| ENSMUSG00000066621 | Tecpr1 | 70381 | protein_coding | 1604.336619 | 0.190273611 | 0.05927564 | 3.209979881 | 0.001327443 | 0.012691889 | 0.190273611 |
| ENSMUSG00000036216 | Leap2 | 259301 | protein_coding | 6.320753612 | -0.190314453 | 0.62144173 | -0.306246658 | 0.75941685 | 0.886076667 | 0.190314453 |
| ENSMUSG00000032415 | Ube2cbp | 70348 | protein_coding | 71.36746819 | -0.190317611 | 0.144423775 | -1.317772029 | 0.187579974 | 0.417346835 | 0.190317611 |
| ENSMUSG00000024019 | Cmtr1 | 74157 | protein_coding | 104.9272529 | -0.190360717 | 0.136485394 | -1.394733244 | 0.163096296 | 0.384596281 | 0.190360717 |
| ENSMUSG00000032593 | Amigo3 | 320844 | protein_coding | 202.3055102 | 0.190361236 | 0.134605317 | 1.414217807 | 0.157297961 | 0.376539145 | 0.190361236 |
| ENSMUSG00000090019 | Gimap1 | 16205 | protein_coding | 56.66881594 | 0.190364252 | 0.156555556 | 1.215953342 | 0.224002696 | 0.461733499 | 0.190364252 |
| ENSMUSG00000033184 | Tmed7 | 66676 | protein_coding | 4032.634277 | 0.190378353 | 0.03836179 | 4.962707714 | 6.95E-07 | 1.77E-05 | 0.190378353 |
| ENSMUSG00000030603 | Psmc4 | 23996 | protein_coding | 1623.554686 | 0.190407983 | 0.069367929 | 2.744899338 | 0.006052946 | 0.041292666 | 0.190407983 |
| ENSMUSG00000033792 | Atp7a | 11977 | protein_coding | 786.1977868 | -0.19050609 | 0.061429177 | -3.101231348 | 0.001927177 | 0.017026683 | 0.19050609 |
| ENSMUSG00000040028 | Elavl1 | 15568 | protein_coding | 1362.775434 | 0.190517682 | 0.048339958 | 3.941204935 | 8.11E-05 | 0.001232299 | 0.190517682 |
| ENSMUSG00000021708 | Rasgrf2 | 19418 | protein_coding | 271.4989895 | 0.190541887 | 0.123093025 | 1.54795032 | 0.121634258 | 0.323066278 | 0.190541887 |
| ENSMUSG00000024331 | Dsc2 | 13506 | protein_coding | 5996.959536 | 0.190543935 | 0.050961969 | 3.738943702 | 0.000184795 | 0.002454852 | 0.190543935 |

| ensembl_gene_id | external_gene_name | entrezgene_id | gene_biotype | baseMean | log2FoldChange | lfcSE | stat | pvalue | padj | absLog2FC |
|---|---|---|---|---|---|---|---|---|---|---|
| ENSMUSG00000071181 | 3830408C21Rik | 100040322 | lncRNA | 7.1201104 | 0.190549553 | 0.499497567 | 0.381482445 | 0.702845295 | 0.852303638 | 0.190549553 |
| ENSMUSG00000019987 | Arg1 | 11846 | protein_coding | 17.72586422 | -0.190706811 | 0.313456081 | -0.60840042 | 0.542921933 | 0.747967026 | 0.190706811 |
| ENSMUSG00000074088 | Snrnp40 | 66585 | protein_coding | 707.3591039 | 0.190721 | 0.067448451 | 2.827655751 | 0.00468902 | 0.033986438 | 0.190721 |
| ENSMUSG00000019179 | Mdh2 | 17448 | protein_coding | 5480.175635 | 0.190827875 | 0.071736624 | 2.660117882 | 0.007811331 | 0.049772157 | 0.190827875 |
| ENSMUSG00000030538 | Cib1 | 23991 | protein_coding | 3288.193033 | 0.190865091 | 0.066161469 | 2.884837558 | 0.003916157 | 0.029748403 | 0.190865091 |
| ENSMUSG00000030629 | Zfand6 | 65098 | protein_coding | 1674.88577 | 0.190885383 | 0.047887858 | 3.986091446 | 6.72E-05 | 0.001059942 | 0.190885383 |
| ENSMUSG00000021843 | Ktn1 | 16709 | protein_coding | 1635.586768 | -0.190900777 | 0.080100759 | -2.383258014 | 0.017160162 | 0.088359495 | 0.190900777 |
| ENSMUSG00000019188 | H13 | 14950 | protein_coding | 2110.664026 | 0.190931365 | 0.060994289 | 3.130315451 | 0.001746187 | 0.015741059 | 0.190931365 |
| ENSMUSG00000025081 | Tdrd1 | 83561 | protein_coding | 66.38135773 | -0.190948743 | 0.279496852 | -0.683187455 | 0.494488399 | 0.713491312 | 0.190948743 |
| ENSMUSG00000020423 | Btg2 | 12227 | protein_coding | 15959.37431 | 0.191052475 | 0.078470214 | 2.434713329 | 0.014903592 | 0.079850573 | 0.191052475 |
| ENSMUSG00000009418 | Nav1 | 215690 | protein_coding | 129.739817 | 0.191079603 | 0.152946036 | 1.249326938 | 0.211545519 | 0.446836226 | 0.191079603 |
| ENSMUSG00000024462 | Gabbr1 | 54393 | protein_coding | 87.61455424 | -0.191116722 | 0.151622906 | -1.260473941 | 0.207498443 | 0.442811616 | 0.191116722 |
| ENSMUSG00000069793 | Slfn9 | 237886 | protein_coding | 606.3885761 | 0.191129184 | 0.153159038 | 1.24791319 | 0.21206285 | 0.447440166 | 0.191129184 |
| ENSMUSG00000100738 | 2010106C02Rik | 72104 | lncRNA | 11.74786385 | -0.191223212 | 0.402339886 | -0.475277791 | 0.634588987 | 0.80894922 | 0.191223212 |
| ENSMUSG00000056492 | Adgrf5 | 224792 | protein_coding | 29.06576709 | -0.191304848 | 0.246744991 | -0.775314007 | 0.438154134 | 0.671081619 | 0.191304848 |
| ENSMUSG00000027323 | Rad51 | 19361 | protein_coding | 80.14563458 | 0.191338929 | 0.184832366 | 1.035202509 | 0.300574347 | 0.544111073 | 0.191338929 |
| ENSMUSG00000026914 | Psmd14 | 59029 | protein_coding | 1227.226747 | 0.191408897 | 0.090624461 | 2.112110749 | 0.034676951 | 0.143725313 | 0.191408897 |
| ENSMUSG00000034175 | Rhbdd3 | 279766 | protein_coding | 232.8850981 | -0.191429088 | 0.09136578 | -2.095194596 | 0.036153697 | 0.147682543 | 0.191429088 |
| ENSMUSG00000032265 | Tent5a | 212943 | protein_coding | 2490.852463 | -0.191544651 | 0.053789451 | -3.56100772 | 0.000369434 | 0.004400196 | 0.191544651 |
| ENSMUSG00000062753 | Smim29 | 106672 | protein_coding | 166.5969953 | -0.191558813 | 0.114433794 | -1.673970648 | 0.094136368 | 0.277386663 | 0.191558813 |
| ENSMUSG00000044024 | Rell2 | 225392 | protein_coding | 8.814409859 | 0.191573142 | 0.389278534 | 0.492123569 | 0.622631991 | 0.800860275 | 0.191573142 |
| ENSMUSG00000031628 | Casp3 | 12367 | protein_coding | 1579.404128 | -0.191637367 | 0.051202552 | -3.742730772 | 0.000182031 | 0.002422127 | 0.191637367 |
| ENSMUSG00000018548 | Trim37 | 68729 | protein_coding | 653.5875861 | 0.191703496 | 0.095323835 | 2.0110762 | 0.044317409 | 0.170737422 | 0.191703496 |
| ENSMUSG00000031838 | Ifi30 | 65972 | protein_coding | 1958.238127 | -0.191750218 | 0.062795085 | -3.053586405 | 0.002261236 | 0.019356132 | 0.191750218 |
| ENSMUSG00000033282 | Rpgrip1l | 244585 | protein_coding | 192.7849554 | 0.191760379 | 0.094471382 | 2.029825053 | 0.042374325 | 0.165145178 | 0.191760379 |
| ENSMUSG00000027804 | Ppid | 67738 | protein_coding | 542.1389578 | -0.19179069 | 0.084097443 | -2.280576952 | 0.022573492 | 0.106976995 | 0.19179069 |
| ENSMUSG00000042367 | Gjb3 | 14620 | protein_coding | 1205.296763 | -0.191830773 | 0.08192582 | -2.34151788 | 0.019205506 | 0.09546563 | 0.191830773 |
| ENSMUSG00000054676 | 1600014C10Rik | 72244 | protein_coding | 274.7525931 | 0.191898709 | 0.075701928 | 2.534924974 | 0.011247141 | 0.064641 | 0.191898709 |
| ENSMUSG00000032041 | Tirap | 117149 | protein_coding | 371.0904439 | 0.19196223 | 0.065594366 | 2.926504826 | 0.003427942 | 0.026849159 | 0.19196223 |
| ENSMUSG00000025278 | Flnb | 286940 | protein_coding | 15382.7946 | 0.191996195 | 0.084340885 | 2.276430876 | 0.022820233 | 0.10767171 | 0.191996195 |
| ENSMUSG00000032815 | Fanca | 14087 | protein_coding | 176.5675793 | 0.192050574 | 0.150691313 | 1.274463473 | 0.20249921 | 0.436583513 | 0.192050574 |
| ENSMUSG00000031242 | Chmp1b2 | 67028 | protein_coding | 755.5758751 | 0.192067754 | 0.055959966 | 3.432234969 | 0.000598629 | 0.006630553 | 0.192067754 |
| ENSMUSG00000053841 | Txlna | 109658 | protein_coding | 1629.899788 | 0.192121184 | 0.047883285 | 4.012280778 | 6.01E-05 | 0.000962527 | 0.192121184 |
| ENSMUSG00000039217 | Il18 | 16173 | protein_coding | 1143.621405 | -0.19215797 | 0.137218319 | -1.400381312 | 0.161399163 | 0.381977497 | 0.19215797 |
| ENSMUSG00000016624 | Phf21b | 271305 | protein_coding | 137.0177766 | -0.192167563 | 0.131690507 | -1.459236253 | 0.144500094 | 0.357032789 | 0.192167563 |
| ENSMUSG00000028911 | Srsf4 | 57317 | protein_coding | 219.5776417 | -0.192199528 | 0.087327263 | -2.200910939 | 0.027742329 | 0.122979809 | 0.192199528 |
| ENSMUSG00000046404 | Yod1 | 226418 | protein_coding | 305.8068775 | -0.192294898 | 0.093121555 | -2.064988062 | 0.03892413 | 0.155455533 | 0.192294898 |
| ENSMUSG00000025498 | Irf7 | 54123 | protein_coding | 1246.92812 | -0.192313131 | 0.123342213 | -1.5591834 | 0.118952979 | 0.31903454 | 0.192313131 |
| ENSMUSG00000031010 | Usp9x | 22284 | protein_coding | 5739.342021 | 0.192340777 | 0.039721665 | 4.842213313 | 1.28E-06 | 3.07E-05 | 0.192340777 |

| ensembl_gene_id | external_gene_name | entrezgene_id | gene_biotype | baseMean | log2FoldChange | lfcSE | stat | pvalue | padj | absLog2FC |
|---|---|---|---|---|---|---|---|---|---|---|
| ENSMUSG00000047193 | Dync2h1 | 110350 | protein_coding | 206.1786322 | -0.192381468 | 0.10732353 | -1.79253765 | 0.073046884 | 0.236530829 | 0.192381468 |
| ENSMUSG00000044252 | Osbpl1a | 64291 | protein_coding | 388.7072217 | -0.192462949 | 0.08207866 | -2.344859793 | 0.019034236 | 0.094932209 | 0.192462949 |
| ENSMUSG00000059436 | Max | 17187 | protein_coding | 2160.553781 | -0.192571858 | 0.062174996 | -3.097255657 | 0.001953213 | 0.017222238 | 0.192571858 |
| ENSMUSG00000027259 | Adal | 75894 | protein_coding | 159.0301054 | -0.19258912 | 0.116743058 | -1.649683702 | 0.099007645 | 0.28619758 | 0.19258912 |
| ENSMUSG00000047040 | Prr15l | 217138 | protein_coding | 6112.122912 | 0.192608189 | 0.049873056 | 3.861968832 | 0.000112477 | 0.001644529 | 0.192608189 |
| ENSMUSG00000072762 | 4930522L14Rik | 100041734 | protein_coding | 85.62875708 | -0.192713392 | 0.157377792 | -1.224527236 | 0.220753398 | 0.457336743 | 0.192713392 |
| ENSMUSG00000063535 | Zfp773 | 76373 | protein_coding | 28.91971708 | 0.192788941 | 0.24548281 | 0.785345991 | 0.432250738 | 0.666753481 | 0.192788941 |
| ENSMUSG00000046753 | Ccdc66 | 320234 | protein_coding | 209.7292485 | 0.19286477 | 0.109259102 | 1.765205517 | 0.077529232 | 0.245659762 | 0.19286477 |
| ENSMUSG00000024799 | Tm7sf2 | 73166 | protein_coding | 815.7875525 | 0.192866892 | 0.082239766 | 2.345178032 | 0.019017996 | 0.094931644 | 0.192866892 |
| ENSMUSG00000029320 | 1700016H13Rik | 74218 | protein_coding | 16.56732596 | -0.193055005 | 0.326614825 | -0.591078513 | 0.554467816 | 0.75465461 | 0.193055005 |
| ENSMUSG00000020308 | Tpgs1 | 110012 | protein_coding | 376.721151 | 0.193083774 | 0.122367324 | 1.577903051 | 0.114587884 | 0.312196844 | 0.193083774 |
| ENSMUSG00000006464 | Bbs1 | 52028 | protein_coding | 56.11787613 | 0.193115547 | 0.170786816 | 1.130740368 | 0.258164385 | 0.500132153 | 0.193115547 |
| ENSMUSG00000029234 | Tmem165 | 21982 | protein_coding | 956.5179098 | 0.193131788 | 0.045171109 | 4.275560015 | 1.91E-05 | 0.000347035 | 0.193131788 |
| ENSMUSG00000022105 | Rb1 | 19645 | protein_coding | 1257.807388 | -0.193139795 | 0.065044165 | -2.969363882 | 0.00298417 | 0.024014824 | 0.193139795 |
| ENSMUSG00000059588 | Calcrl | 54598 | protein_coding | 89.57972865 | -0.193236217 | 0.141666723 | -1.364019812 | 0.17256133 | 0.397669477 | 0.193236217 |
| ENSMUSG00000020538 | Srebf1 | 20787 | protein_coding | 3201.565353 | 0.193254369 | 0.088373969 | 2.186779325 | 0.028758637 | 0.126236996 | 0.193254369 |
| ENSMUSG00000017221 | Psmd3 | 22123 | protein_coding | 2024.06797 | 0.193355528 | 0.078371317 | 2.467172098 | 0.013618489 | 0.074800013 | 0.193355528 |
| ENSMUSG00000006411 | Nectin4 | 71740 | protein_coding | 103.9172893 | -0.193369084 | 0.13062364 | -1.4803529 | 0.138779093 | 0.348921317 | 0.193369084 |
| ENSMUSG00000025544 | Tm9sf2 | 68059 | protein_coding | 13366.86893 | 0.193383147 | 0.03321033 | 5.822981799 | 5.78E-09 | 2.12E-07 | 0.193383147 |
| ENSMUSG00000030611 | Mrps11 | 67994 | protein_coding | 325.8728421 | 0.193390686 | 0.080404991 | 2.405207479 | 0.016163286 | 0.084790715 | 0.193390686 |
| ENSMUSG00000057706 | Mex3b | 108797 | protein_coding | 35.99624303 | 0.19346841 | 0.220204137 | 0.878586627 | 0.379625449 | 0.62053716 | 0.19346841 |
| ENSMUSG00000067206 | Lrrc66 | 231296 | protein_coding | 1349.771319 | 0.193585889 | 0.067615332 | 2.863047233 | 0.00419588 | 0.031312799 | 0.193585889 |
| ENSMUSG00000029699 | Ssc4d | 109267 | protein_coding | 96.31470778 | -0.193587159 | 0.135810388 | -1.425422323 | 0.154035159 | 0.371430175 | 0.193587159 |
| ENSMUSG00000027459 | Fam110a | 73847 | protein_coding | 214.9636119 | 0.193590416 | 0.106264752 | 1.821774502 | 0.068489208 | 0.226895123 | 0.193590416 |
| ENSMUSG00000074922 | Fam122a | 68034 | protein_coding | 134.8505651 | 0.193597694 | 0.107972883 | 1.793021435 | 0.072969497 | 0.236348478 | 0.193597694 |
| ENSMUSG00000031227 | Magee1 | 107528 | protein_coding | 22.044496 | -0.193722729 | 0.324190392 | -0.597558514 | 0.550134552 | 0.751671802 | 0.193722729 |
| ENSMUSG00000026799 | Med27 | 68975 | protein_coding | 190.0099202 | 0.193735676 | 0.097462298 | 1.987801236 | 0.046833677 | 0.177006494 | 0.193735676 |
| ENSMUSG00000022677 | Cep20 | 66086 | protein_coding | 805.5601483 | 0.19378353 | 0.055933323 | 3.464545298 | 0.000531129 | 0.00596069 | 0.19378353 |
| ENSMUSG00000075227 | Znhit2 | 29805 | protein_coding | 359.8308531 | 0.193800488 | 0.103724913 | 1.868408298 | 0.061705181 | 0.211953026 | 0.193800488 |
| ENSMUSG00000072774 | Zfp951 | 626391 | protein_coding | 44.55057362 | -0.193813169 | 0.207113106 | -0.935784183 | 0.349384317 | 0.591799273 | 0.193813169 |
| ENSMUSG00000116138 | C030006K11Rik | 223665 | protein_coding | 176.775916 | -0.193883421 | 0.118169863 | -1.640717991 | 0.100855969 | 0.289568133 | 0.193883421 |
| ENSMUSG00000030737 | Slco2b1 | 101488 | protein_coding | 391.6827667 | -0.194026302 | 0.107224902 | -1.809526501 | 0.070369247 | 0.23074217 | 0.194026302 |
| ENSMUSG00000059149 | Mfsd4a | 213006 | protein_coding | 3376.987238 | 0.194027784 | 0.094438761 | 2.05453547 | 0.039923894 | 0.15823363 | 0.194027784 |
| ENSMUSG00000025579 | Gaa | 14387 | protein_coding | 1089.748116 | -0.194097607 | 0.071493898 | -2.714883519 | 0.006629908 | 0.044045718 | 0.194097607 |
| ENSMUSG00000026836 | Acvr1 | 11477 | protein_coding | 213.6490524 | 0.194222804 | 0.095083697 | 2.042650949 | 0.041087004 | 0.161612222 | 0.194222804 |
| ENSMUSG00000024164 | C3 | 12266 | protein_coding | 53.90496203 | 0.194286938 | 0.377415474 | 0.51478265 | 0.60670491 | 0.789765867 | 0.194286938 |
| ENSMUSG00000028692 | Akr1a1 | 58810 | protein_coding | 10222.85285 | 0.194342958 | 0.052241158 | 3.720112009 | 0.000199134 | 0.002630173 | 0.194342958 |
| ENSMUSG00000024664 | Fads3 | 60527 | protein_coding | 288.5340158 | -0.194375834 | 0.100982808 | -1.924840862 | 0.05424927 | 0.195285263 | 0.194375834 |
| ENSMUSG00000053783 | 1700016K19Rik | 74230 | protein_coding | 23.48247707 | 0.19438391 | 0.24605696 | 0.789995576 | 0.429530352 | 0.664591383 | 0.19438391 |

| ensembl_gene_id | external_gene_name | entrezgene_id | gene_biotype | baseMean | log2FoldChange | lfcSE | stat | pvalue | padj | absLog2FC |
|---|---|---|---|---|---|---|---|---|---|---|
| ENSMUSG00000000782 | Tcf7 | 21414 | protein_coding | 67.44429699 | -0.194403195 | 0.160194583 | -1.21354413 | 0.224921853 | 0.463022048 | 0.194403195 |
| ENSMUSG00000027456 | Sdcbp2 | 228765 | protein_coding | 3832.217489 | 0.194416738 | 0.056325491 | 3.451665187 | 0.000557139 | 0.006217975 | 0.194416738 |
| ENSMUSG00000039233 | Tbce | 70430 | protein_coding | 457.9697796 | -0.194440684 | 0.083291764 | -2.334452717 | 0.019572029 | 0.09661718 | 0.194440684 |
| ENSMUSG00000038518 | Jarid2 | 16468 | protein_coding | 1335.219977 | -0.194443587 | 0.03795919 | -5.122437716 | 3.02E-07 | 8.22E-06 | 0.194443587 |
| ENSMUSG00000031864 | Ints10 | 70885 | protein_coding | 861.7221689 | 0.194470829 | 0.095647067 | 2.033212686 | 0.04203104 | 0.164204113 | 0.194470829 |
| ENSMUSG00000066571 | Garre1 | 233103 | protein_coding | 455.9732183 | -0.194501676 | 0.060719876 | -3.203262096 | 0.001358803 | 0.012897827 | 0.194501676 |
| ENSMUSG00000028690 | Mmachc | 67096 | protein_coding | 319.1836462 | 0.194502868 | 0.069270985 | 2.807854794 | 0.004987271 | 0.035604125 | 0.194502868 |
| ENSMUSG00000031371 | Haus7 | 73738 | protein_coding | 171.3843826 | 0.194514641 | 0.106417621 | 1.827842413 | 0.067573206 | 0.224876824 | 0.194514641 |
| ENSMUSG00000024660 | Incenp | 16319 | protein_coding | 540.6894517 | 0.194632352 | 0.104034614 | 1.870842258 | 0.061366948 | 0.211105968 | 0.194632352 |
| ENSMUSG00000057132 | Rpgrip1 | 77945 | protein_coding | 15.41667912 | 0.194661617 | 0.342684657 | 0.568048826 | 0.570001814 | 0.765233601 | 0.194661617 |
| ENSMUSG00000070661 | Rnf186 | 66825 | protein_coding | 3608.166102 | -0.194866314 | 0.090269157 | -2.158725302 | 0.030871485 | 0.132728454 | 0.194866314 |
| ENSMUSG00000039713 | Plekhg5 | 269608 | protein_coding | 256.7548244 | -0.194928563 | 0.085815988 | -2.271471414 | 0.023118456 | 0.108600285 | 0.194928563 |
| ENSMUSG00000004043 | Stat5a | 20850 | protein_coding | 281.0010023 | -0.194935588 | 0.077693804 | -2.509023618 | 0.012106539 | 0.068457978 | 0.194935588 |
| ENSMUSG00000042870 | Tom1 | 21968 | protein_coding | 333.6789881 | -0.194953195 | 0.074883921 | -2.603405274 | 0.009230277 | 0.056214758 | 0.194953195 |
| ENSMUSG00000031523 | Dlc1 | 50768 | protein_coding | 12.11426377 | -0.195186166 | 0.393552104 | -0.495960164 | 0.619922517 | 0.799032164 | 0.195186166 |
| ENSMUSG00000047003 | Zfp41 | 22701 | protein_coding | 151.6962395 | 0.19519722 | 0.107881672 | 1.809364071 | 0.070394461 | 0.23074217 | 0.19519722 |
| ENSMUSG00000032458 | Copb2 | 50797 | protein_coding | 5970.238703 | 0.195235698 | 0.04691747 | 4.161258055 | 3.16E-05 | 0.000544156 | 0.195235698 |
| ENSMUSG00000030347 | D6Wsu163e | 28040 | protein_coding | 310.1064272 | 0.195406553 | 0.081568158 | 2.395622963 | 0.016592148 | 0.086330552 | 0.195406553 |
| ENSMUSG00000040249 | Lrp1 | 16971 | protein_coding | 5654.284489 | -0.195434491 | 0.084212074 | -2.320741917 | 0.020300777 | 0.099299554 | 0.195434491 |
| ENSMUSG00000039914 | Coq10a | 210582 | protein_coding | 267.2365071 | -0.195498725 | 0.081025149 | -2.412815361 | 0.015829836 | 0.083182722 | 0.195498725 |
| ENSMUSG00000021662 | Arhgef28 | 110596 | protein_coding | 2212.600693 | -0.195527264 | 0.04883043 | -4.004209319 | 6.22E-05 | 0.000988977 | 0.195527264 |
| ENSMUSG00000039929 | Urb1 | 207932 | protein_coding | 301.1971675 | -0.195556933 | 0.07686657 | -2.544109008 | 0.010955689 | 0.063259266 | 0.195556933 |
| ENSMUSG00000056228 | Cars2 | 71941 | protein_coding | 480.1396471 | 0.195618799 | 0.071725055 | 2.727342616 | 0.006384671 | 0.042881783 | 0.195618799 |
| ENSMUSG00000055408 | Hottip | 791364 | lncRNA | 300.1882144 | 0.195621854 | 0.122139988 | 1.601620049 | 0.109239655 | 0.302750973 | 0.195621854 |
| ENSMUSG00000002897 | Il17ra | 16172 | protein_coding | 490.8767527 | -0.195659896 | 0.067126417 | -2.914797241 | 0.003559197 | 0.027655107 | 0.195659896 |
| ENSMUSG00000061589 | Dot1l | 208266 | protein_coding | 623.2605776 | -0.195751908 | 0.091093995 | -2.148900237 | 0.031642309 | 0.134998814 | 0.195751908 |
| ENSMUSG00000037762 | Slc16a9 | 66859 | protein_coding | 447.6194358 | -0.195755898 | 0.082220174 | -2.380874278 | 0.017271605 | 0.08873319 | 0.195755898 |
| ENSMUSG00000038612 | Mcl1 | 17210 | protein_coding | 8506.792581 | -0.195857306 | 0.068975906 | -2.839503198 | 0.004518384 | 0.033016757 | 0.195857306 |
| ENSMUSG00000022108 | Itm2b | 16432 | protein_coding | 15005.8119 | -0.195923719 | 0.056307082 | -3.479557292 | 0.000502243 | 0.005665866 | 0.195923719 |
| ENSMUSG00000037868 | Egr2 | 13654 | protein_coding | 108.0058775 | -0.195934411 | 0.206629427 | -0.948240594 | 0.343006986 | 0.585633643 | 0.195934411 |
| ENSMUSG00000035441 | Myo1d | 338367 | protein_coding | 7652.492002 | 0.196032922 | 0.060823642 | 3.222972457 | 0.001268678 | 0.012273523 | 0.196032922 |
| ENSMUSG00000063406 | Tmed5 | 73130 | protein_coding | 775.68303 | 0.19605165 | 0.067058816 | 2.923577553 | 0.00346034 | 0.027024167 | 0.19605165 |
| ENSMUSG00000043831 | Lysmd4 | 75099 | protein_coding | 289.7174039 | 0.196074205 | 0.085623708 | 2.289952283 | 0.022024083 | 0.105208053 | 0.196074205 |
| ENSMUSG00000005947 | Itgae | 16407 | protein_coding | 77.19850913 | 0.196082021 | 0.175395807 | 1.11794019 | 0.263592539 | 0.504916665 | 0.196082021 |
| ENSMUSG00000021493 | Pdlim7 | 67399 | protein_coding | 85.36013279 | 0.196108739 | 0.164089403 | 1.195133479 | 0.232034883 | 0.471169298 | 0.196108739 |
| ENSMUSG00000031167 | Rbm3 | 19652 | protein_coding | 2156.777059 | -0.19624122 | 0.103878997 | -1.889132795 | 0.058874037 | 0.205867649 | 0.19624122 |
| ENSMUSG00000047638 | Nr1h4 | 20186 | protein_coding | 2113.959776 | -0.196276499 | 0.05757903 | -3.408819117 | 0.000652447 | 0.007123911 | 0.196276499 |
| ENSMUSG00000018927 | Ccl6 | 20305 | protein_coding | 6051.467573 | 0.196347158 | 0.051113116 | 3.841424141 | 0.000122323 | 0.00175663 | 0.196347158 |
| ENSMUSG00000053846 | Lipg | 16891 | protein_coding | 1469.434677 | -0.196364741 | 0.12128084 | -1.619091203 | 0.105427641 | 0.297761537 | 0.196364741 |

| ensembl_gene_id | external_gene_name | entrezgene_id | gene_biotype | baseMean | log2FoldChange | lfcSE | stat | pvalue | padj | absLog2FC |
|---|---|---|---|---|---|---|---|---|---|---|
| ENSMUSG00000008129 | Brme1 | 74666 | protein_coding | 17.92237859 | -0.196365661 | 0.332739463 | -0.590148397 | 0.555091165 | 0.755056951 | 0.196365661 |
| ENSMUSG00000000934 | Top1mt | 72960 | protein_coding | 143.3786985 | -0.196410662 | 0.124448071 | -1.578253963 | 0.114507278 | 0.31208616 | 0.196410662 |
| ENSMUSG00000034686 | Prr7 | 432763 | protein_coding | 9.60610759 | -0.196440067 | 0.387698886 | -0.506682052 | 0.612377926 | 0.793255831 | 0.196440067 |
| ENSMUSG00000025880 | Smad7 | 17131 | protein_coding | 1369.669616 | -0.196441736 | 0.085458339 | -2.298684214 | 0.021522878 | 0.103580855 | 0.196441736 |
| ENSMUSG00000038069 | Cdkn2aip | 70925 | protein_coding | 457.1579572 | -0.19649353 | 0.071454094 | -2.749926831 | 0.005960857 | 0.04081985 | 0.19649353 |
| ENSMUSG00000030361 | Klrb1a | 17057 | protein_coding | 8.419114822 | 0.196495386 | 0.608212882 | 0.323070082 | 0.746642167 | 0.879811357 | 0.196495386 |
| ENSMUSG00000033068 | Entpd6 | 12497 | protein_coding | 1576.246018 | 0.196550825 | 0.055601966 | 3.534961789 | 0.000407834 | 0.00475227 | 0.196550825 |
| ENSMUSG00000021906 | Oxnad1 | 218885 | protein_coding | 639.4874438 | 0.196586891 | 0.054017579 | 3.639313238 | 0.000273366 | 0.003425466 | 0.196586891 |
| ENSMUSG00000044894 | Uqcrq | 22272 | protein_coding | 3603.229232 | 0.19661408 | 0.100102504 | 1.964127487 | 0.049515309 | 0.184036274 | 0.19661408 |
| ENSMUSG00000032596 | Uba7 | 74153 | protein_coding | 1008.682098 | -0.196666552 | 0.092835994 | -2.118429966 | 0.03413867 | 0.142262102 | 0.196666552 |
| ENSMUSG00000032066 | Bco2 | 170752 | protein_coding | 346.8163653 | -0.196669329 | 0.114219862 | -1.721848777 | 0.085096916 | 0.260220234 | 0.196669329 |
| ENSMUSG00000015627 | Gata5 | 14464 | protein_coding | 34.98081214 | 0.196685923 | 0.320976876 | 0.612772875 | 0.540026529 | 0.745761931 | 0.196685923 |
| ENSMUSG00000042473 | Tbc1d8b | 245638 | protein_coding | 1022.734235 | 0.196900145 | 0.050732602 | 3.881136343 | 0.00010397 | 0.001539685 | 0.196900145 |
| ENSMUSG00000031904 | Slc7a6 | 330836 | protein_coding | 198.0690694 | -0.196902481 | 0.114425616 | -1.720790228 | 0.085288896 | 0.260650754 | 0.196902481 |
| ENSMUSG00000034951 | Cog7 | 233824 | protein_coding | 928.8981729 | 0.197076916 | 0.051162913 | 3.851948729 | 0.000117182 | 0.001697944 | 0.197076916 |
| ENSMUSG00000097415 | AU020206 | 108167440 | lncRNA | 133.7859849 | 0.197111583 | 0.139858743 | 1.409361893 | 0.158728186 | 0.378894236 | 0.197111583 |
| ENSMUSG00000022131 | Gpr180 | 58245 | protein_coding | 624.6697967 | 0.197142211 | 0.063970854 | 3.081750483 | 0.002057873 | 0.017948789 | 0.197142211 |
| ENSMUSG00000030924 | Rexo5 | 434234 | protein_coding | 121.0096901 | -0.197236774 | 0.117117863 | -1.68408788 | 0.092164686 | 0.273594537 | 0.197236774 |
| ENSMUSG00000075486 | Commd6 | 66200 | protein_coding | 642.4124102 | 0.197248794 | 0.056908128 | 3.466091784 | 0.000528083 | 0.005930637 | 0.197248794 |
| ENSMUSG00000061232 | H2-K1 | 14972 | protein_coding | 11666.18048 | -0.197301954 | 0.105866555 | -1.863685416 | 0.062365895 | 0.213194897 | 0.197301954 |
| ENSMUSG00000063952 | Brpf3 | 268936 | protein_coding | 788.6381372 | -0.197394306 | 0.086608474 | -2.279156956 | 0.022657737 | 0.107187246 | 0.197394306 |
| ENSMUSG00000011884 | Gltp | 56356 | protein_coding | 1134.079117 | 0.197469536 | 0.060018772 | 3.290129529 | 0.001001413 | 0.010043397 | 0.197469536 |
| ENSMUSG00000022523 | Fgf12 | 14167 | protein_coding | 40.54118343 | -0.197610777 | 0.210913878 | -0.936926386 | 0.348796425 | 0.59123817 | 0.197610777 |
| ENSMUSG00000030685 | Kctd13 | 233877 | protein_coding | 100.5420376 | -0.197635321 | 0.163607056 | -1.20798776 | 0.22705197 | 0.465615713 | 0.197635321 |
| ENSMUSG00000049799 | Lrrc19 | 100061 | protein_coding | 2095.411026 | -0.197638145 | 0.082727838 | -2.389016189 | 0.016893558 | 0.087430799 | 0.197638145 |
| ENSMUSG00000044876 | Zfp444 | 72667 | protein_coding | 393.2260583 | 0.19767565 | 0.100767012 | 1.961709957 | 0.049796262 | 0.184654936 | 0.19767565 |
| ENSMUSG00000053581 | Zfand2a | 100494 | protein_coding | 524.8912095 | -0.197678068 | 0.10723727 | -1.84337094 | 0.06527487 | 0.219997456 | 0.197678068 |
| ENSMUSG00000026788 | Zbtb43 | 71834 | protein_coding | 969.3180973 | -0.197705866 | 0.052869364 | -3.73951663 | 0.000184374 | 0.002451283 | 0.197705866 |
| ENSMUSG00000070709 | Zfp974 | 73430 | protein_coding | 172.449678 | -0.197903229 | 0.117581931 | -1.683109189 | 0.092353956 | 0.2739868 | 0.197903229 |
| ENSMUSG00000022940 | Pigp | 56176 | protein_coding | 453.3174609 | -0.197947801 | 0.089195075 | -2.219268278 | 0.026468478 | 0.118917792 | 0.197947801 |
| ENSMUSG00000024846 | Cst6 | 73720 | protein_coding | 67.41663713 | -0.197975453 | 0.187126029 | -1.057979236 | 0.290064909 | 0.532545171 | 0.197975453 |
| ENSMUSG00000062382 | Ftl1-ps1 | 434624 | protein_coding | 7.069868732 | 0.19800089 | 0.452414305 | 0.437653911 | 0.661637184 | 0.827034368 | 0.19800089 |
| ENSMUSG00000031907 | Zfp90 | 22751 | protein_coding | 113.3978062 | -0.19804472 | 0.120755948 | -1.640041115 | 0.100996619 | 0.289868743 | 0.19804472 |
| ENSMUSG00000051735 | Rinl | 320435 | protein_coding | 32.56816478 | 0.198086184 | 0.249710988 | 0.793261785 | 0.427625317 | 0.663107066 | 0.198086184 |
| ENSMUSG00000016319 | Slc25a5 | 11740 | protein_coding | 27822.50834 | 0.198096069 | 0.058410101 | 3.39146937 | 0.000695189 | 0.00748652 | 0.198096069 |
| ENSMUSG00000070544 | Top1 | 21969 | protein_coding | 2116.486692 | 0.198173292 | 0.053217815 | 3.723814896 | 0.000196235 | 0.002593999 | 0.198173292 |
| ENSMUSG00000040447 | Spns2 | 216892 | protein_coding | 41.32380578 | -0.198186801 | 0.19933778 | -0.99422599 | 0.320112858 | 0.563525822 | 0.198186801 |
| ENSMUSG00000004268 | Emg1 | 14791 | protein_coding | 1120.409489 | 0.198216986 | 0.05520496 | 3.590564771 | 0.000329962 | 0.003991974 | 0.198216986 |
| ENSMUSG00000025867 | Cplx2 | 12890 | protein_coding | 134.3871842 | -0.198225441 | 0.104631502 | -1.894510139 | 0.05815733 | 0.204158306 | 0.198225441 |

| ensembl_gene_id | external_gene_name | entrezgene_id | gene_biotype | baseMean | log2FoldChange | lfcSE | stat | pvalue | padj | absLog2FC |
|---|---|---|---|---|---|---|---|---|---|---|
| ENSMUSG00000051079 | Rgs13 | 246709 | protein_coding | 195.9104865 | 0.198419163 | 0.118145624 | 1.679445731 | 0.093065207 | 0.275438171 | 0.198419163 |
| ENSMUSG00000021244 | Ylpm1 | 56531 | protein_coding | 489.498871 | -0.19844353 | 0.089114915 | -2.226827346 | 0.025958815 | 0.117793419 | 0.19844353 |
| ENSMUSG00000029004 | Kmt2e | 69188 | protein_coding | 773.3384965 | -0.198490714 | 0.086546562 | -2.293455794 | 0.021821778 | 0.104644605 | 0.198490714 |
| ENSMUSG00000031711 | Zfp330 | 30932 | protein_coding | 688.30218 | 0.198517516 | 0.062342773 | 3.184290775 | 0.001451092 | 0.013597766 | 0.198517516 |
| ENSMUSG00000061436 | Hipk2 | 15258 | protein_coding | 781.866499 | -0.198532806 | 0.09596725 | -2.068755802 | 0.038569008 | 0.154496371 | 0.198532806 |
| ENSMUSG00000068452 | Duox2 | 214593 | protein_coding | 1266.481949 | 0.198535019 | 0.097056211 | 2.045567373 | 0.040798954 | 0.160753661 | 0.198535019 |
| ENSMUSG00000044730 | 9930104L06Rik | 194268 | protein_coding | 154.8649861 | -0.198553011 | 0.114123005 | -1.739815836 | 0.081891361 | 0.254171925 | 0.198553011 |
| ENSMUSG00000041235 | Chd7 | 320790 | protein_coding | 2077.267601 | -0.198573248 | 0.038508083 | -5.156664095 | 2.51E-07 | 7.01E-06 | 0.198573248 |
| ENSMUSG00000017204 | Gsdma | 57911 | protein_coding | 12.94284033 | 0.198580949 | 0.344215893 | 0.576908135 | 0.564001515 | 0.761142463 | 0.198580949 |
| ENSMUSG00000048878 | Hexim1 | 192231 | protein_coding | 1150.037787 | 0.198639847 | 0.083474177 | 2.379656245 | 0.017328795 | 0.088855862 | 0.198639847 |
| ENSMUSG00000036372 | Tmem258 | 69038 | protein_coding | 1107.715003 | 0.198647542 | 0.070655211 | 2.811505901 | 0.004931019 | 0.035280629 | 0.198647542 |
| ENSMUSG00000054499 | Dedd2 | 67379 | protein_coding | 1030.463668 | -0.198698901 | 0.066843814 | -2.972584754 | 0.002953037 | 0.023776147 | 0.198698901 |
| ENSMUSG00000023961 | Enpp4 | 224794 | protein_coding | 323.7173631 | -0.198771838 | 0.07792211 | -2.550904205 | 0.010744385 | 0.062318117 | 0.198771838 |
| ENSMUSG00000009555 | Cdk9 | 107951 | protein_coding | 871.1584567 | -0.198853104 | 0.051659497 | -3.84930391 | 0.000118454 | 0.001710212 | 0.198853104 |
| ENSMUSG00000073758 | Sh3d21 | 66938 | protein_coding | 967.5019705 | 0.198908508 | 0.102295604 | 1.944448258 | 0.051841419 | 0.189536742 | 0.198908508 |
| ENSMUSG00000052557 | Gan | 209239 | protein_coding | 891.412756 | -0.198958008 | 0.055781578 | -3.566733247 | 0.000361459 | 0.00432437 | 0.198958008 |
| ENSMUSG00000032235 | Ice2 | 93697 | protein_coding | 376.1895483 | -0.198965839 | 0.088792231 | -2.240802346 | 0.025038881 | 0.11481434 | 0.198965839 |
| ENSMUSG00000025401 | Myo1a | 432516 | protein_coding | 16852.3418 | -0.199005596 | 0.056694432 | -3.510143561 | 0.000447865 | 0.0051554 | 0.199005596 |
| ENSMUSG00000097251 | 5033417F24Rik | 75995 | lncRNA | 12.66565164 | 0.199085306 | 0.35391578 | 0.562521702 | 0.573760622 | 0.76815045 | 0.199085306 |
| ENSMUSG00000006476 | Nsmf | 56876 | protein_coding | 1194.421548 | -0.199117398 | 0.045117282 | -4.41332879 | 1.02E-05 | 0.00019971 | 0.199117398 |
| ENSMUSG00000055980 | Irs1 | 16367 | protein_coding | 173.4785639 | 0.199129172 | 0.167441289 | 1.189247724 | 0.2343422 | 0.473944903 | 0.199129172 |
| ENSMUSG00000008206 | Cers4 | 67260 | protein_coding | 1061.6853 | -0.1991909 | 0.060254451 | -3.305828818 | 0.000946959 | 0.009610832 | 0.1991909 |
| ENSMUSG00000026339 | Ccdc93 | 70829 | protein_coding | 1866.766957 | -0.19936254 | 0.045225986 | -4.408141365 | 1.04E-05 | 0.000203809 | 0.19936254 |
| ENSMUSG00000028438 | Kif24 | 109242 | protein_coding | 125.6944362 | 0.199394599 | 0.128001357 | 1.557753793 | 0.119291623 | 0.31943494 | 0.199394599 |
| ENSMUSG00000030317 | Timp4 | 110595 | protein_coding | 95.25565265 | 0.199424411 | 0.191621371 | 1.040721141 | 0.298004989 | 0.541512839 | 0.199424411 |
| ENSMUSG00000033972 | Zfp944 | 319615 | protein_coding | 285.2643033 | -0.199451057 | 0.087941752 | -2.267990497 | 0.023329786 | 0.109264501 | 0.199451057 |
| ENSMUSG00000037211 | Spry1 | 24063 | protein_coding | 378.6217262 | -0.199648229 | 0.104701007 | -1.906841541 | 0.05654111 | 0.200957912 | 0.199648229 |
| ENSMUSG00000074653 | Lrrc31 | 320352 | protein_coding | 572.7348297 | -0.199978902 | 0.092121057 | -2.170827278 | 0.029944231 | 0.130024398 | 0.199978902 |
| ENSMUSG00000080316 | Spaca6 | 75202 | protein_coding | 11.72742305 | 0.199981833 | 0.460399517 | 0.434365862 | 0.664022784 | 0.828333496 | 0.199981833 |
| ENSMUSG00000042155 | Klhl23 | 277396 | protein_coding | 90.70777324 | 0.199990567 | 0.149240023 | 1.340059877 | 0.180225879 | 0.407223247 | 0.199990567 |
| ENSMUSG00000035638 | Muc20 | 224116 | protein_coding | 9.08715141 | -0.200009078 | 0.381201358 | -0.524680917 | 0.599805025 | 0.785533185 | 0.200009078 |
| ENSMUSG00000029422 | Rsrc2 | 208606 | protein_coding | 1607.729876 | -0.20001551 | 0.081278532 | -2.460865179 | 0.013860244 | 0.075770899 | 0.20001551 |
| ENSMUSG00000049577 | Zfpm1 | 22761 | protein_coding | 88.59081023 | 0.200119404 | 0.142093783 | 1.408361436 | 0.159024072 | 0.37913729 | 0.200119404 |
| ENSMUSG00000061898 | Rbak | 57782 | protein_coding | 214.9337191 | -0.200187939 | 0.116204641 | -1.72271896 | 0.08493936 | 0.260026017 | 0.200187939 |
| ENSMUSG00000050947 | Amigo1 | 229715 | protein_coding | 305.6467758 | -0.200193925 | 0.090578936 | -2.210159829 | 0.027094072 | 0.120870289 | 0.200193925 |
| ENSMUSG00000042515 | Pwwp3b | 245631 | protein_coding | 8.087770568 | 0.200231175 | 0.409656228 | 0.488778545 | 0.624998489 | 0.802084296 | 0.200231175 |
| ENSMUSG00000047632 | Fgfbp3 | 72514 | protein_coding | 29.46436426 | 0.200261851 | 0.224036412 | 0.893880815 | 0.371385663 | 0.613347321 | 0.200261851 |
| ENSMUSG00000031937 | Vstm5 | 69137 | protein_coding | 198.2843224 | -0.200267947 | 0.11706816 | -1.710695264 | 0.087137382 | 0.264392609 | 0.200267947 |
| ENSMUSG00000058093 | Zfp729b | 100416706 | protein_coding | 247.1786365 | -0.200289182 | 0.086961763 | -2.303186785 | 0.021268336 | 0.102482035 | 0.200289182 |

| ensembl_gene_id | external_gene_name | entrezgene_id | gene_biotype | baseMean | log2FoldChange | lfcSE | stat | pvalue | padj | absLog2FC |
|---|---|---|---|---|---|---|---|---|---|---|
| ENSMUSG00000015478 | Rnf5 | 54197 | protein_coding | 350.0659357 | 0.200382421 | 0.093342035 | 2.146754362 | 0.031812842 | 0.135547083 | 0.200382421 |
| ENSMUSG00000037001 | Zfp39 | 22698 | protein_coding | 95.36593575 | -0.200595919 | 0.151253072 | -1.326227069 | 0.184764496 | 0.413617021 | 0.200595919 |
| ENSMUSG00000071662 | Polr2g | 67710 | protein_coding | 450.7745105 | 0.200659434 | 0.064480166 | 3.111955914 | 0.001858523 | 0.016577651 | 0.200659434 |
| ENSMUSG00000070803 | Cited4 | 56222 | protein_coding | 174.7505589 | 0.200684671 | 0.123478321 | 1.625262388 | 0.104106662 | 0.295082324 | 0.200684671 |
| ENSMUSG00000036792 | Mbd5 | 109241 | protein_coding | 272.0802313 | -0.200705374 | 0.079514029 | -2.52415048 | 0.011597827 | 0.066254458 | 0.200705374 |
| ENSMUSG00000024269 | Tpgs2 | 66648 | protein_coding | 130.9067842 | 0.200774864 | 0.145085005 | 1.383842972 | 0.166406542 | 0.389893694 | 0.200774864 |
| ENSMUSG00000024055 | Cyp4f13 | 170716 | protein_coding | 1027.429843 | -0.200860886 | 0.071740775 | -2.799814812 | 0.005113193 | 0.036262298 | 0.200860886 |
| ENSMUSG00000031757 | Mt4 | 17752 | protein_coding | 6.61491523 | -0.200895723 | 0.48016414 | -0.418389685 | 0.675662227 | 0.835024575 | 0.200895723 |
| ENSMUSG00000005374 | Tbl2 | 27368 | protein_coding | 568.8970617 | 0.200913713 | 0.071918741 | 2.793621122 | 0.005212151 | 0.03683451 | 0.200913713 |
| ENSMUSG00000022228 | Zscan26 | 432731 | protein_coding | 815.9763847 | -0.201011959 | 0.093396151 | -2.152250992 | 0.031377591 | 0.134058533 | 0.201011959 |
| ENSMUSG00000097170 | Gm16982 | 100036523 | lncRNA | 12.48856277 | -0.20101606 | 0.39465856 | -0.509341695 | 0.610512737 | 0.792525305 | 0.20101606 |
| ENSMUSG00000028843 | Sh3bgrl3 | 73723 | protein_coding | 1333.976899 | 0.201043982 | 0.075547363 | 2.661164802 | 0.007787083 | 0.049660074 | 0.201043982 |
| ENSMUSG00000027286 | Lrrc57 | 66606 | protein_coding | 193.0802745 | -0.201069433 | 0.099375832 | -2.02332327 | 0.043039829 | 0.167213521 | 0.201069433 |
| ENSMUSG00000065778 | Gm22154 | 115490050 | snoRNA | 551.9628916 | 0.201113403 | 0.382453207 | 0.525851005 | 0.598991733 | 0.784827956 | 0.201113403 |
| ENSMUSG00000017802 | Retreg3 | 67998 | protein_coding | 2120.006603 | -0.201147751 | 0.044725982 | -4.497335595 | 6.88E-06 | 0.000141004 | 0.201147751 |
| ENSMUSG00000048696 | Mex3d | 237400 | protein_coding | 209.3130462 | -0.201150178 | 0.109639509 | -1.834650481 | 0.066557494 | 0.222414567 | 0.201150178 |
| ENSMUSG00000040183 | Ankrd6 | 140577 | protein_coding | 7.937313501 | -0.201159566 | 0.448234513 | -0.448781966 | 0.65358895 | 0.821663954 | 0.201159566 |
| ENSMUSG00000026766 | Mmadhc | 109129 | protein_coding | 1105.284848 | 0.201227773 | 0.052628187 | 3.823574073 | 0.000131531 | 0.001863973 | 0.201227773 |
| ENSMUSG00000040016 | Ptger3 | 19218 | protein_coding | 202.5706316 | -0.201385059 | 0.120677283 | -1.668790126 | 0.09515898 | 0.279429875 | 0.201385059 |
| ENSMUSG00000073471 | Rsph3a | 66832 | protein_coding | 284.4848849 | 0.201389216 | 0.076650499 | 2.627369913 | 0.008604774 | 0.053393299 | 0.201389216 |
| ENSMUSG00000034667 | Xpot | 73192 | protein_coding | 1021.306106 | 0.201405013 | 0.07223002 | 2.788383729 | 0.005297176 | 0.037252106 | 0.201405013 |
| ENSMUSG00000037364 | Srrt | 83701 | protein_coding | 1771.502128 | 0.201470086 | 0.056979199 | 3.535853226 | 0.00040646 | 0.004743117 | 0.201470086 |
| ENSMUSG00000025223 | Ldb1 | 16825 | protein_coding | 849.9844759 | -0.201472792 | 0.080589599 | -2.499985039 | 0.012419855 | 0.069862227 | 0.201472792 |
| ENSMUSG00000028609 | Magoh | 17149 | protein_coding | 581.3566955 | 0.201526085 | 0.072742748 | 2.770394174 | 0.005598849 | 0.038902473 | 0.201526085 |
| ENSMUSG00000041528 | Rnf123 | 84585 | protein_coding | 643.248255 | -0.20154997 | 0.073733994 | -2.733474187 | 0.006267004 | 0.042345703 | 0.20154997 |
| ENSMUSG00000053580 | Tanc2 | 77097 | protein_coding | 205.7974726 | -0.201555676 | 0.110468268 | -1.82455721 | 0.068067876 | 0.225933166 | 0.201555676 |
| ENSMUSG00000027219 | Slc28a2 | 269346 | protein_coding | 152.4859396 | -0.201619326 | 0.154371798 | -1.30606321 | 0.191531063 | 0.423020709 | 0.201619326 |
| ENSMUSG00000038963 | Slco4a1 | 108115 | protein_coding | 131.762701 | -0.201741769 | 0.108336689 | -1.862174031 | 0.062578564 | 0.213633466 | 0.201741769 |
| ENSMUSG00000055436 | Srsf11 | 69207 | protein_coding | 1389.512738 | -0.201821772 | 0.077457857 | -2.605568759 | 0.009172186 | 0.05598783 | 0.201821772 |
| ENSMUSG00000023010 | Tmbim6 | 110213 | protein_coding | 27041.36146 | 0.201915272 | 0.059096836 | 3.416684979 | 0.000633886 | 0.006954202 | 0.201915272 |
| ENSMUSG00000090191 | 9230105E05Rik | 320626 | lncRNA | 144.4330089 | 0.201960544 | 0.117164185 | 1.723739584 | 0.084754866 | 0.259707718 | 0.201960544 |
| ENSMUSG00000063021 | H2ac15 | 319169 | protein_coding | 589.6535452 | 0.201983255 | 0.142034715 | 1.422069637 | 0.155006036 | 0.372948208 | 0.201983255 |
| ENSMUSG00000058886 | Deaf1 | 54006 | protein_coding | 529.7693099 | -0.201985496 | 0.057791613 | -3.495065885 | 0.000473945 | 0.005413107 | 0.201985496 |
| ENSMUSG00000034206 | Polq | 77782 | protein_coding | 120.0579704 | 0.201994917 | 0.128701882 | 1.569479121 | 0.116536341 | 0.31570327 | 0.201994917 |
| ENSMUSG00000047766 | Lrrc49 | 102747 | protein_coding | 222.5348952 | 0.201999765 | 0.107499247 | 1.879080738 | 0.06023347 | 0.208675653 | 0.201999765 |
| ENSMUSG00000022994 | Adcy6 | 11512 | protein_coding | 6656.996144 | -0.202063122 | 0.050300879 | -4.017089296 | 5.89E-05 | 0.000944558 | 0.202063122 |
| ENSMUSG00000090038 | Gm16573 | 102635191 | lncRNA | 7.296874197 | -0.202065854 | 0.502892179 | -0.401807509 | 0.687825696 | 0.843220448 | 0.202065854 |
| ENSMUSG00000068917 | Clk2 | 12748 | protein_coding | 435.5697388 | -0.202069253 | 0.083645884 | -2.415770431 | 0.015701958 | 0.082768143 | 0.202069253 |
| ENSMUSG00000021994 | Wnt5a | 22418 | protein_coding | 48.78080285 | -0.202132142 | 0.212981831 | -0.949058147 | 0.342591039 | 0.585270749 | 0.202132142 |

| ensembl_gene_id | external_gene_name | entrezgene_id | gene_biotype | baseMean | log2FoldChange | lfcSE | stat | pvalue | padj | absLog2FC |
|---|---|---|---|---|---|---|---|---|---|---|
| ENSMUSG00000037172 | Dennd11 | 243780 | protein_coding | 1391.065256 | 0.202134546 | 0.04991995 | 4.049173673 | 5.14E-05 | 0.000838976 | 0.202134546 |
| ENSMUSG00000028076 | Cd1d1 | 12479 | protein_coding | 132.5896542 | 0.202136495 | 0.113142875 | 1.786559652 | 0.074008688 | 0.238421516 | 0.202136495 |
| ENSMUSG00000068040 | Tm9sf4 | 99237 | protein_coding | 3958.04887 | 0.202225899 | 0.068886038 | 2.935658722 | 0.003328403 | 0.026257023 | 0.202225899 |
| ENSMUSG00000006576 | Slc4a3 | 20536 | protein_coding | 12.57670082 | 0.202312739 | 0.408839751 | 0.494846058 | 0.62070879 | 0.799662139 | 0.202312739 |
| ENSMUSG00000021281 | Tnfaip2 | 21928 | protein_coding | 10.24741642 | 0.202333625 | 0.370460712 | 0.546167564 | 0.58495076 | 0.775781324 | 0.202333625 |
| ENSMUSG00000041681 | Iapp | 15874 | protein_coding | 12.45390618 | 0.20243775 | 0.3191174 | 0.634367636 | 0.525840926 | 0.73638734 | 0.20243775 |
| ENSMUSG00000031197 | Vbp1 | 22327 | protein_coding | 706.3154999 | 0.202465792 | 0.052618914 | 3.847775972 | 0.000119195 | 0.001717835 | 0.202465792 |
| ENSMUSG00000031546 | Gins4 | 109145 | protein_coding | 390.4371212 | 0.202529641 | 0.069928671 | 2.896231784 | 0.003776733 | 0.028948374 | 0.202529641 |
| ENSMUSG00000052974 | Cyp2f2 | 13107 | protein_coding | 238.0509619 | -0.202535538 | 0.125475649 | -1.614142192 | 0.106496588 | 0.298794445 | 0.202535538 |
| ENSMUSG00000020834 | Dhrs13 | 70451 | protein_coding | 151.2544711 | -0.202582787 | 0.114868567 | -1.763605075 | 0.077798493 | 0.246155835 | 0.202582787 |
| ENSMUSG00000029534 | St7 | 64213 | protein_coding | 224.3968476 | -0.202596461 | 0.117844242 | -1.719188464 | 0.085580061 | 0.261293002 | 0.202596461 |
| ENSMUSG00000055675 | Kbtbd11 | 74901 | protein_coding | 2111.563205 | 0.202599089 | 0.118500603 | 1.709688254 | 0.087323536 | 0.26456259 | 0.202599089 |
| ENSMUSG00000039653 | Baat | 12012 | protein_coding | 27.44632585 | -0.202823455 | 0.239049905 | -0.84845654 | 0.396183766 | 0.635493892 | 0.202823455 |
| ENSMUSG00000036944 | Tmem71 | 213068 | protein_coding | 14.06073707 | -0.202833548 | 0.310162243 | -0.65395951 | 0.513137892 | 0.726547391 | 0.202833548 |
| ENSMUSG00000094942 | Gm3604 | 100041979 | protein_coding | 46.85667805 | -0.202860937 | 0.193022419 | -1.050970853 | 0.293271977 | 0.536272425 | 0.202860937 |
| ENSMUSG00000029195 | Klb | 83379 | protein_coding | 324.7379809 | -0.202890798 | 0.154774684 | -1.310878456 | 0.189898831 | 0.420477604 | 0.202890798 |
| ENSMUSG00000020593 | Lpin1 | 14245 | protein_coding | 367.4266146 | 0.20290919 | 0.134973493 | 1.503326217 | 0.132754943 | 0.339551732 | 0.20290919 |
| ENSMUSG00000099474 | 1700097N02Rik | 67522 | lncRNA | 21.55598049 | -0.202991583 | 0.258171638 | -0.786266007 | 0.431711663 | 0.666254386 | 0.202991583 |
| ENSMUSG00000022543 | Dnaaf8 | 74684 | transcribed_unitary_pseudogene | 18.88041385 | 0.203023487 | 0.273687891 | 0.74180661 | 0.458204515 | 0.685606255 | 0.203023487 |
| ENSMUSG00000032606 | Nicn1 | 66257 | protein_coding | 151.0610455 | 0.203061421 | 0.128057012 | 1.585711064 | 0.112804868 | 0.309125592 | 0.203061421 |
| ENSMUSG00000064370 | mt-Cytb | 17711 | protein_coding | 99793.94118 | 0.203064213 | 0.051496733 | 3.943244557 | 8.04E-05 | 0.001224169 | 0.203064213 |
| ENSMUSG00000034245 | Hdac11 | 232232 | protein_coding | 914.8885577 | -0.203163184 | 0.076717975 | -2.648182299 | 0.008092587 | 0.051079904 | 0.203163184 |
| ENSMUSG00000055150 | Zfp78 | 330463 | protein_coding | 17.0054031 | -0.203345743 | 0.292563568 | -0.69504807 | 0.487025177 | 0.708037053 | 0.203345743 |
| ENSMUSG00000059409 | Ppp2r5d | 21770 | protein_coding | 1386.370997 | 0.203358022 | 0.050211984 | 4.049989747 | 5.12E-05 | 0.000838602 | 0.203358022 |
| ENSMUSG00000044337 | Ackr3 | 12778 | protein_coding | 18.7172983 | 0.203431248 | 0.2768814 | 0.734723417 | 0.462507956 | 0.688854258 | 0.203431248 |
| ENSMUSG00000039943 | Plcb4 | 18798 | protein_coding | 528.5301836 | 0.203486289 | 0.098332763 | 2.069364085 | 0.038511934 | 0.154396187 | 0.203486289 |
| ENSMUSG00000051557 | Pusl1 | 433813 | protein_coding | 126.0183461 | -0.20350221 | 0.121403864 | -1.676241625 | 0.093690874 | 0.276680594 | 0.20350221 |
| ENSMUSG00000002844 | Adprh | 11544 | protein_coding | 633.7123552 | 0.203508278 | 0.079378674 | 2.563765156 | 0.010354358 | 0.060891613 | 0.203508278 |
| ENSMUSG00000032409 | Atr | 245000 | protein_coding | 562.0997081 | -0.203550813 | 0.091370479 | -2.227752492 | 0.025897024 | 0.117546106 | 0.203550813 |
| ENSMUSG00000054405 | Dnajc8 | 68598 | protein_coding | 947.6961618 | -0.203653752 | 0.04632538 | -4.396159359 | 1.10E-05 | 0.000213747 | 0.203653752 |
| ENSMUSG00000026727 | Rsu1 | 20163 | protein_coding | 330.5038946 | 0.203672804 | 0.068417046 | 2.976930714 | 0.002911498 | 0.023512132 | 0.203672804 |
| ENSMUSG00000035165 | Kcne3 | 57442 | protein_coding | 3307.473787 | -0.203696027 | 0.077916552 | -2.614284403 | 0.008941457 | 0.054911988 | 0.203696027 |
| ENSMUSG00000029217 | Tec | 21682 | protein_coding | 340.9098974 | -0.203752612 | 0.077406587 | -2.632238654 | 0.008482426 | 0.052776264 | 0.203752612 |
| ENSMUSG00000025159 | Mms19 | 72199 | protein_coding | 658.1177057 | -0.203772491 | 0.051319291 | -3.970680194 | 7.17E-05 | 0.001113473 | 0.203772491 |
| ENSMUSG00000007812 | Zfp655 | 72611 | protein_coding | 920.3437152 | -0.203810422 | 0.088687752 | -2.298067297 | 0.021557959 | 0.103718737 | 0.203810422 |
| ENSMUSG00000032194 | Kank2 | 235041 | protein_coding | 636.0407081 | -0.203833612 | 0.105408534 | -1.93374866 | 0.053144038 | 0.192705082 | 0.203833612 |
| ENSMUSG00000041459 | Tardbp | 230908 | protein_coding | 1838.471724 | 0.203840254 | 0.067984309 | 2.998342674 | 0.002714523 | 0.022244465 | 0.203840254 |
| ENSMUSG00000116564 | Riok2 | 67045 | protein_coding | 443.367969 | -0.204134906 | 0.085385094 | -2.390755779 | 0.016813733 | 0.087104102 | 0.204134906 |
| ENSMUSG00000021569 | Trip13 | 69716 | protein_coding | 137.2991302 | 0.204165841 | 0.14408166 | 1.417014771 | 0.156478609 | 0.375078854 | 0.204165841 |

| ensembl_gene_id | external_gene_name | entrezgene_id | gene_biotype | baseMean | log2FoldChange | lfcSE | stat | pvalue | padj | absLog2FC |
|---|---|---|---|---|---|---|---|---|---|---|
| ENSMUSG00000029810 | Tmem176b | 65963 | protein_coding | 1030.30804 | 0.204180878 | 0.053040585 | 3.84952161 | 0.000118349 | 0.001710212 | 0.204180878 |
| ENSMUSG00000041773 | Enc1 | 13803 | protein_coding | 317.0451222 | -0.204208597 | 0.096201654 | -2.122713991 | 0.033777826 | 0.141426796 | 0.204208597 |
| ENSMUSG00000027332 | Ivd | 56357 | protein_coding | 1871.654289 | 0.204209551 | 0.050435578 | 4.048918611 | 5.15E-05 | 0.000839042 | 0.204209551 |
| ENSMUSG00000028536 | 2610528J11Rik | 66451 | protein_coding | 1027.022398 | 0.20425497 | 0.094041667 | 2.17196246 | 0.029858495 | 0.129791898 | 0.20425497 |
| ENSMUSG00000078877 | Gm14295 | 628147 | protein_coding | 63.64960738 | -0.204301852 | 0.151988907 | -1.344189233 | 0.178887202 | 0.405982819 | 0.204301852 |
| ENSMUSG00000042688 | Mapk6 | 50772 | protein_coding | 2480.446584 | -0.204325143 | 0.042416322 | -4.817134903 | 1.46E-06 | 3.44E-05 | 0.204325143 |
| ENSMUSG00000030177 | Ccdc77 | 67200 | protein_coding | 200.0432731 | -0.20434264 | 0.116447802 | -1.754800321 | 0.079293469 | 0.248883957 | 0.20434264 |
| ENSMUSG00000027371 | Fahd2a | 68126 | protein_coding | 264.0439786 | -0.204420881 | 0.077101219 | -2.651331366 | 0.008017514 | 0.050786438 | 0.204420881 |
| ENSMUSG00000021149 | Gtpbp4 | 69237 | protein_coding | 709.0670778 | -0.204480133 | 0.060901903 | -3.357532741 | 0.000786414 | 0.008251466 | 0.204480133 |
| ENSMUSG00000018427 | Ypel2 | 77864 | protein_coding | 649.0281615 | 0.204590934 | 0.090802215 | 2.253149158 | 0.024249746 | 0.112442683 | 0.204590934 |
| ENSMUSG00000033948 | Zswim5 | 74464 | protein_coding | 388.4984927 | -0.204648335 | 0.061095834 | -3.34962832 | 0.000809201 | 0.008441125 | 0.204648335 |
| ENSMUSG00000022475 | Hdac7 | 56233 | protein_coding | 113.1594503 | -0.204676748 | 0.140276375 | -1.459096357 | 0.144538589 | 0.357073195 | 0.204676748 |
| ENSMUSG00000040528 | Milr1 | 380732 | protein_coding | 7.170477055 | 0.204810084 | 0.505757456 | 0.404957122 | 0.685509029 | 0.841671981 | 0.204810084 |
| ENSMUSG00000071722 | Spin4 | 270624 | protein_coding | 21.06939226 | 0.20486304 | 0.252339795 | 0.81185387 | 0.416875487 | 0.654811627 | 0.20486304 |
| ENSMUSG00000035930 | Chst4 | 26887 | protein_coding | 1425.424112 | 0.204945562 | 0.069215893 | 2.960961033 | 0.003066807 | 0.024532938 | 0.204945562 |
| ENSMUSG00000027166 | Dnajc24 | 99349 | protein_coding | 154.9430234 | 0.204996488 | 0.117017441 | 1.751845587 | 0.079800363 | 0.249794343 | 0.204996488 |
| ENSMUSG00000019796 | Lrp11 | 237253 | protein_coding | 44.02421956 | 0.205013927 | 0.187734752 | 1.092040362 | 0.274815361 | 0.517267428 | 0.205013927 |
| ENSMUSG00000006360 | Crip1 | 12925 | protein_coding | 10668.04118 | -0.205029461 | 0.089753811 | -2.284353828 | 0.022350745 | 0.106296214 | 0.205029461 |
| ENSMUSG00000024313 | Mep1b | 17288 | protein_coding | 2825.690544 | -0.205099463 | 0.091866419 | -2.232583635 | 0.025576414 | 0.116583051 | 0.205099463 |
| ENSMUSG00000036459 | Wtip | 101543 | protein_coding | 188.741107 | -0.20510697 | 0.101861903 | -2.013578818 | 0.044053772 | 0.170087427 | 0.20510697 |
| ENSMUSG00000019505 | Ubb | 22187 | protein_coding | 7301.347916 | 0.205139998 | 0.093135465 | 2.2025981 | 0.027623088 | 0.122653508 | 0.205139998 |
| ENSMUSG00000041471 | Shld2 | 75698 | protein_coding | 151.4538964 | -0.205155742 | 0.09803632 | -2.092650384 | 0.036380375 | 0.148382983 | 0.205155742 |
| ENSMUSG00000071454 | Dtnb | 13528 | protein_coding | 134.6910229 | 0.205179958 | 0.108356178 | 1.893569536 | 0.05828217 | 0.204552026 | 0.205179958 |
| ENSMUSG00000116056 | Gm4544 | 100043600 | lncRNA | 17.38838261 | -0.20525956 | 0.307105042 | -0.668369227 | 0.503897934 | 0.720166783 | 0.20525956 |
| ENSMUSG00000044734 | Serpinb1a | 66222 | protein_coding | 9056.842738 | 0.205263357 | 0.061438439 | 3.340959816 | 0.000834893 | 0.008669878 | 0.205263357 |
| ENSMUSG00000001576 | Ergic1 | 67458 | protein_coding | 2429.65581 | 0.205277027 | 0.056800142 | 3.614023112 | 0.000301482 | 0.003688925 | 0.205277027 |
| ENSMUSG00000046387 | Pcdhb17 | 93888 | protein_coding | 8.989294018 | 0.205300951 | 0.430948932 | 0.47639276 | 0.633794592 | 0.8083434 | 0.205300951 |
| ENSMUSG00000062866 | Phactr2 | 215789 | protein_coding | 541.0278331 | -0.205304858 | 0.079382329 | -2.586279088 | 0.009701834 | 0.058099322 | 0.205304858 |
| ENSMUSG00000015652 | Steap1 | 70358 | protein_coding | 87.86215957 | 0.20542606 | 0.168554983 | 1.21874807 | 0.222939827 | 0.460352188 | 0.20542606 |
| ENSMUSG00000041261 | Car8 | 12319 | protein_coding | 1857.708857 | -0.205451197 | 0.106874646 | -1.92235676 | 0.054560883 | 0.196218366 | 0.205451197 |
| ENSMUSG00000087128 | Gm12655 | 115489886 | lncRNA | 7.458193561 | -0.205475765 | 0.470244888 | -0.436954808 | 0.662144123 | 0.827333871 | 0.205475765 |
| ENSMUSG00000041762 | Gpr155 | 68526 | protein_coding | 19.26988772 | -0.205482257 | 0.307068729 | -0.669173502 | 0.503384807 | 0.719943844 | 0.205482257 |
| ENSMUSG00000055978 | Fut2 | 14344 | protein_coding | 3123.458693 | 0.205499799 | 0.055374618 | 3.711082905 | 0.000206374 | 0.002710262 | 0.205499799 |
| ENSMUSG00000054519 | Zfp867 | 237775 | protein_coding | 112.8641393 | -0.205611927 | 0.137099684 | -1.499725754 | 0.133685457 | 0.34107542 | 0.205611927 |
| ENSMUSG00000047473 | Zfp30 | 22693 | protein_coding | 26.71288677 | 0.205695069 | 0.258003079 | 0.797258195 | 0.425301094 | 0.661079917 | 0.205695069 |
| ENSMUSG00000022052 | Ppp2r2a | 71978 | protein_coding | 341.9928036 | -0.205742929 | 0.11545012 | -1.782093681 | 0.074733956 | 0.240086556 | 0.205742929 |
| ENSMUSG00000037395 | Rcor3 | 214742 | protein_coding | 448.9413669 | -0.205798508 | 0.074964209 | -2.745290204 | 0.006045741 | 0.041278435 | 0.205798508 |
| ENSMUSG00000089542 | Gm25835 | 115487914 | snoRNA | 345.0509581 | 0.205815681 | 0.32373479 | 0.635753978 | 0.524936788 | 0.735696149 | 0.205815681 |
| ENSMUSG00000091345 | Col6a5 | 665033 | protein_coding | 5.932674447 | 0.205832848 | 0.629115435 | 0.327178188 | 0.743533118 | 0.877686912 | 0.205832848 |

| ensembl_gene_id | external_gene_name | entrezgene_id | gene_biotype | baseMean | log2FoldChange | lfcSE | stat | pvalue | padj | absLog2FC |
|---|---|---|---|---|---|---|---|---|---|---|
| ENSMUSG00000072082 | Ccnf | 12449 | protein_coding | 294.2143064 | 0.20589312 | 0.119122565 | 1.728414096 | 0.08391401 | 0.258112004 | 0.20589312 |
| ENSMUSG00000027978 | Prss12 | 19142 | protein_coding | 13.9477666 | -0.206008889 | 0.398284165 | -0.517240973 | 0.604987954 | 0.789053031 | 0.206008889 |
| ENSMUSG00000025497 | Cdhr5 | 72040 | protein_coding | 14480.80096 | -0.206100007 | 0.07036596 | -2.928973126 | 0.003400838 | 0.026714719 | 0.206100007 |
| ENSMUSG00000006386 | Tek | 21687 | protein_coding | 15.39271779 | 0.206113551 | 0.317361525 | 0.649459794 | 0.516041226 | 0.728738778 | 0.206113551 |
| ENSMUSG00000032598 | Nckipsd | 80987 | protein_coding | 148.2955167 | 0.206229641 | 0.113146319 | 1.822680948 | 0.068351727 | 0.226532739 | 0.206229641 |
| ENSMUSG00000024986 | Hhex | 15242 | protein_coding | 21.19337503 | -0.206275419 | 0.297446178 | -0.693488214 | 0.48800322 | 0.708500894 | 0.206275419 |
| ENSMUSG00000092083 | Kcnb2 | 98741 | protein_coding | 17.80189683 | -0.206378319 | 0.360162502 | -0.573014454 | 0.566634908 | 0.762590433 | 0.206378319 |
| ENSMUSG00000026189 | Pecr | 111175 | protein_coding | 142.6158123 | 0.206474196 | 0.111698367 | 1.848497888 | 0.06453035 | 0.218172107 | 0.206474196 |
| ENSMUSG00000038729 | Pakap | 677884 | protein_coding | 49.85550688 | -0.206496102 | 0.178767869 | -1.155107476 | 0.248046417 | 0.489027453 | 0.206496102 |
| ENSMUSG00000013701 | Timm23 | 53600 | protein_coding | 813.2550947 | 0.206660585 | 0.08555212 | 2.415610323 | 0.015708863 | 0.082768143 | 0.206660585 |
| ENSMUSG00000034818 | Celf5 | 319586 | protein_coding | 58.29840993 | -0.206681881 | 0.213281691 | -0.9690559 | 0.332517299 | 0.575843468 | 0.206681881 |
| ENSMUSG00000097277 | 2900076A07Rik | 100504421 | lncRNA | 46.51572887 | -0.206711804 | 0.216979863 | -0.952677367 | 0.340753564 | 0.583616475 | 0.206711804 |
| ENSMUSG00000021917 | Spcs1 | 69019 | protein_coding | 1466.440197 | 0.206725292 | 0.060503194 | 3.416766579 | 0.000633696 | 0.006954202 | 0.206725292 |
| ENSMUSG00000101031 | Ms4a12 | 381213 | protein_coding | 417.7071965 | -0.206741132 | 0.094984699 | -2.176573013 | 0.029512441 | 0.128599602 | 0.206741132 |
| ENSMUSG00000037593 | Rskr | 216976 | protein_coding | 6.694005554 | -0.206855666 | 0.521897132 | -0.39635333 | 0.69184439 | 0.845511858 | 0.206855666 |
| ENSMUSG00000022124 | Fbxl3 | 50789 | protein_coding | 3035.981018 | -0.206913474 | 0.066656489 | -3.104176008 | 0.001908098 | 0.016907634 | 0.206913474 |
| ENSMUSG00000038065 | Mturn | 68235 | protein_coding | 40.22381636 | 0.206949287 | 0.195894282 | 1.056433525 | 0.290770196 | 0.5332246 | 0.206949287 |
| ENSMUSG00000046574 | Prr12 | 233210 | protein_coding | 145.5024486 | -0.20695107 | 0.154368328 | -1.340631671 | 0.180040069 | 0.4070526 | 0.20695107 |
| ENSMUSG00000020847 | Rph3al | 380714 | protein_coding | 546.0920982 | 0.207013725 | 0.062088843 | 3.334153364 | 0.000855595 | 0.008850658 | 0.207013725 |
| ENSMUSG00000065523 | Mir192 | 387187 | miRNA | 20.42941418 | 0.207055634 | 0.293536835 | 0.705382115 | 0.480572471 | 0.702901527 | 0.207055634 |
| ENSMUSG00000033031 | Cip2a | 224171 | protein_coding | 475.9954453 | 0.20706694 | 0.092020025 | 2.250237813 | 0.024433853 | 0.11303636 | 0.20706694 |
| ENSMUSG00000039697 | Ncoa7 | 211329 | protein_coding | 745.282381 | -0.20709323 | 0.156748934 | -1.321177918 | 0.186442047 | 0.415699004 | 0.20709323 |
| ENSMUSG00000055407 | Map6 | 17760 | protein_coding | 13.38769802 | 0.207136788 | 0.341903802 | 0.605833533 | 0.5446253 | 0.749012844 | 0.207136788 |
| ENSMUSG00000004933 | Matk | 17179 | protein_coding | 123.9329465 | -0.207174519 | 0.117698981 | -1.760206571 | 0.078372788 | 0.247293672 | 0.207174519 |
| ENSMUSG00000063179 | Pstk | 214580 | protein_coding | 356.7999136 | 0.20721354 | 0.076320436 | 2.715046619 | 0.006626644 | 0.044045718 | 0.20721354 |
| ENSMUSG00000068335 | Dok1 | 13448 | protein_coding | 306.1698907 | -0.20722985 | 0.076504926 | -2.708712489 | 0.006754484 | 0.044588445 | 0.20722985 |
| ENSMUSG00000015377 | Dennd6b | 69440 | protein_coding | 156.7560423 | -0.207322788 | 0.102513526 | -2.022394475 | 0.043135615 | 0.167504953 | 0.207322788 |
| ENSMUSG00000024271 | Elp2 | 58523 | protein_coding | 537.9760858 | 0.207378693 | 0.066678218 | 3.110141501 | 0.001869977 | 0.016633825 | 0.207378693 |
| ENSMUSG00000050323 | Ndufaf6 | 76947 | protein_coding | 129.0870878 | -0.207427722 | 0.106582047 | -1.946178807 | 0.051633265 | 0.189222442 | 0.207427722 |
| ENSMUSG00000030849 | Fgfr2 | 14183 | protein_coding | 2324.874016 | 0.207434452 | 0.106019038 | 1.956577389 | 0.050397176 | 0.186070341 | 0.207434452 |
| ENSMUSG00000028756 | Pink1 | 68943 | protein_coding | 3530.119871 | 0.207441369 | 0.064382669 | 3.222006377 | 0.001272963 | 0.012300224 | 0.207441369 |
| ENSMUSG00000024299 | Adamts10 | 224697 | protein_coding | 56.44191017 | 0.207671378 | 0.213885064 | 0.970948483 | 0.331573933 | 0.574895496 | 0.207671378 |
| ENSMUSG00000025158 | Rfng | 19719 | protein_coding | 291.6091829 | 0.207675788 | 0.084190046 | 2.46674989 | 0.013634556 | 0.074841551 | 0.207675788 |
| ENSMUSG00000032849 | Abcc4 | 239273 | protein_coding | 248.7014657 | 0.207692795 | 0.076221148 | 2.724871 | 0.006432661 | 0.043063317 | 0.207692795 |
| ENSMUSG00000028744 | Slc66a1 | 212555 | protein_coding | 158.1793354 | -0.207718625 | 0.112947011 | -1.839080322 | 0.065903373 | 0.221421603 | 0.207718625 |
| ENSMUSG00000028188 | Spata1 | 70951 | protein_coding | 31.69265144 | -0.207725307 | 0.218284278 | -0.951627434 | 0.341285963 | 0.584131301 | 0.207725307 |
| ENSMUSG00000049751 | Rpl36al | 66483 | protein_coding | 305.1253836 | 0.207753029 | 0.118916201 | 1.747054036 | 0.080627965 | 0.251408971 | 0.207753029 |
| ENSMUSG00000020649 | Rrm2 | 20135 | protein_coding | 323.2742549 | 0.20779188 | 0.153232055 | 1.356060127 | 0.175080038 | 0.401466767 | 0.20779188 |
| ENSMUSG00000078435 | AU041133 | 216177 | protein_coding | 100.6003561 | -0.207872041 | 0.144968544 | -1.433911352 | 0.151597562 | 0.367724228 | 0.207872041 |

Supplementary Table 1

| ensembl_gene_id | external_gene_name | entrezgene_id | gene_biotype | baseMean | log2FoldChange | lfcSE | stat | pvalue | padj | absLog2FC |
|---|---|---|---|---|---|---|---|---|---|---|
| ENSMUSG00000024411 | Aqp4 | 11829 | protein_coding | 10128.72625 | 0.207936069 | 0.112226711 | 1.852821548 | 0.063907941 | 0.216756548 | 0.207936069 |
| ENSMUSG00000018648 | Dusp14 | 56405 | protein_coding | 36.59386178 | 0.207941687 | 0.237964395 | 0.873835293 | 0.382207957 | 0.622927511 | 0.207941687 |
| ENSMUSG00000022747 | St3gal6 | 54613 | protein_coding | 2319.686506 | -0.207960409 | 0.091288644 | -2.278053427 | 0.022723395 | 0.107434824 | 0.207960409 |
| ENSMUSG00000022687 | Boc | 117606 | protein_coding | 1428.653301 | -0.207991322 | 0.049670931 | -4.187385245 | 2.82E-05 | 0.000490919 | 0.207991322 |
| ENSMUSG00000038980 | Rbbp8nl | 271887 | protein_coding | 69.80433413 | 0.208004209 | 0.146312483 | 1.421643628 | 0.155129733 | 0.373121133 | 0.208004209 |
| ENSMUSG00000066258 | Trim12a | 76681 | protein_coding | 334.5824772 | -0.208127334 | 0.112078316 | -1.856981279 | 0.063313818 | 0.215277659 | 0.208127334 |
| ENSMUSG00000017144 | Rnd3 | 74194 | protein_coding | 1626.974468 | -0.208216732 | 0.075379443 | -2.762248225 | 0.005740482 | 0.039783734 | 0.208216732 |
| ENSMUSG00000030246 | Ldhb | 16832 | protein_coding | 178.3257962 | 0.208277117 | 0.112158394 | 1.856990898 | 0.063312449 | 0.215277659 | 0.208277117 |
| ENSMUSG00000027433 | Xrn2 | 24128 | protein_coding | 1823.156281 | -0.208285948 | 0.065746985 | -3.167992387 | 0.001534955 | 0.014251137 | 0.208285948 |
| ENSMUSG00000022285 | Ywhaz | 22631 | protein_coding | 5697.615901 | 0.208304842 | 0.041303413 | 5.043283946 | 4.58E-07 | 1.20E-05 | 0.208304842 |
| ENSMUSG00000001440 | Kpnb1 | 16211 | protein_coding | 5168.699965 | 0.208335077 | 0.057814708 | 3.603496162 | 0.000313965 | 0.00382426 | 0.208335077 |
| ENSMUSG00000030666 | Calcb | 116903 | protein_coding | 6.62910484 | 0.208360119 | 0.511861305 | 0.407063625 | 0.683961258 | 0.840783841 | 0.208360119 |
| ENSMUSG00000006362 | Cbfa2t3 | 12398 | protein_coding | 315.8948001 | -0.208432884 | 0.096780605 | -2.153663785 | 0.031266548 | 0.133784989 | 0.208432884 |
| ENSMUSG00000028184 | Adgrl2 | 99633 | protein_coding | 600.2264771 | -0.208548692 | 0.090972936 | -2.292425663 | 0.021881093 | 0.104773272 | 0.208548692 |
| ENSMUSG00000002379 | Ndufa11 | 69875 | protein_coding | 1654.702215 | 0.208605577 | 0.081362867 | 2.563891683 | 0.010350584 | 0.060891613 | 0.208605577 |
| ENSMUSG00000020015 | Cdk17 | 237459 | protein_coding | 268.1111082 | -0.20865078 | 0.073760138 | -2.828774277 | 0.004672664 | 0.033928886 | 0.20865078 |
| ENSMUSG00000046997 | Spsb4 | 211949 | protein_coding | 57.98786847 | -0.208684416 | 0.179016554 | -1.165726923 | 0.243724871 | 0.484894029 | 0.208684416 |
| ENSMUSG00000037601 | Nme1 | 18102 | protein_coding | 451.866652 | 0.208740412 | 0.132302398 | 1.57775229 | 0.114622528 | 0.312196844 | 0.208740412 |
| ENSMUSG00000024376 | Epb41l4a | 13824 | protein_coding | 247.1561664 | -0.208781258 | 0.082933233 | -2.517461939 | 0.011820374 | 0.067240721 | 0.208781258 |
| ENSMUSG00000026683 | Nuf2 | 66977 | protein_coding | 193.4462759 | 0.208862889 | 0.150850402 | 1.384569656 | 0.166184095 | 0.389715871 | 0.208862889 |
| ENSMUSG00000038000 | Acd | 497652 | protein_coding | 265.047387 | -0.208889281 | 0.087974748 | -2.374423174 | 0.017576393 | 0.089842947 | 0.208889281 |
| ENSMUSG00000044005 | Gls2 | 216456 | protein_coding | 11.39983815 | 0.209048551 | 0.372430087 | 0.561309514 | 0.574586557 | 0.768227276 | 0.209048551 |
| ENSMUSG00000074738 | Fndc10 | 230991 | protein_coding | 45.08267141 | -0.20907875 | 0.215341637 | -0.970916505 | 0.331589858 | 0.574895496 | 0.20907875 |
| ENSMUSG00000017734 | Dbndd2 | 52840 | protein_coding | 62.43195459 | -0.209377925 | 0.197546952 | -1.059889422 | 0.289194908 | 0.531418201 | 0.209377925 |
| ENSMUSG00000060487 | Samd5 | 320825 | protein_coding | 61.33619983 | -0.209378955 | 0.244310188 | -0.857020973 | 0.391433294 | 0.631323671 | 0.209378955 |
| ENSMUSG00000039903 | Eva1c | 70967 | protein_coding | 10.46283155 | -0.209389981 | 0.406471868 | -0.515140155 | 0.606455085 | 0.789753 | 0.209389981 |
| ENSMUSG00000011958 | Bnip2 | 12175 | protein_coding | 255.4759706 | -0.209418054 | 0.115012572 | -1.820827495 | 0.068633083 | 0.227231726 | 0.209418054 |
| ENSMUSG00000028772 | Zcchc17 | 619605 | protein_coding | 593.7561218 | 0.209469731 | 0.056914298 | 3.680441266 | 0.000232831 | 0.002991921 | 0.209469731 |
| ENSMUSG00000029283 | Cdc7 | 12545 | protein_coding | 72.28378699 | -0.209481075 | 0.195094938 | -1.07373916 | 0.282939605 | 0.525566345 | 0.209481075 |
| ENSMUSG00000055612 | Cdca7 | 66953 | protein_coding | 895.4869866 | -0.209535131 | 0.07830121 | -2.676013958 | 0.007450354 | 0.048003141 | 0.209535131 |
| ENSMUSG00000064264 | Zfp428 | 232969 | protein_coding | 34.33574774 | 0.209645714 | 0.255483574 | 0.820583924 | 0.411883308 | 0.649945412 | 0.209645714 |
| ENSMUSG00000042594 | Sh2b3 | 16923 | protein_coding | 346.8030924 | -0.209702308 | 0.074858736 | -2.801307094 | 0.005089606 | 0.036126794 | 0.209702308 |
| ENSMUSG00000027834 | Serpini1 | 20713 | protein_coding | 10.96869733 | 0.209771458 | 0.422074814 | 0.497000652 | 0.619188592 | 0.798277593 | 0.209771458 |
| ENSMUSG00000025196 | Cpn1 | 93721 | protein_coding | 183.2099123 | 0.209780752 | 0.174774689 | 1.200292519 | 0.230025754 | 0.46920123 | 0.209780752 |
| ENSMUSG00000039199 | Zdhhc1 | 70796 | protein_coding | 260.9164016 | -0.209899903 | 0.114094641 | -1.839699936 | 0.065812303 | 0.221242374 | 0.209899903 |
| ENSMUSG00000034826 | Nup54 | 269113 | protein_coding | 378.8264902 | -0.209904987 | 0.094749684 | -2.215363448 | 0.026735128 | 0.119666226 | 0.209904987 |
| ENSMUSG00000031264 | Btk | 12229 | protein_coding | 16.87981954 | 0.209956336 | 0.354134505 | 0.592871729 | 0.553266999 | 0.753561 | 0.209956336 |
| ENSMUSG00000028048 | Gba | 14466 | protein_coding | 1141.123252 | 0.209983026 | 0.06808966 | 3.083919465 | 0.002042929 | 0.017857081 | 0.209983026 |
| ENSMUSG00000030590 | Fam98c | 73833 | protein_coding | 811.0082956 | 0.210074056 | 0.092257879 | 2.27703106 | 0.022784371 | 0.107565442 | 0.210074056 |

Supplementary Table 1

| ensembl_gene_id | external_gene_name | entrezgene_id | gene_biotype | baseMean | log2FoldChange | lfcSE | stat | pvalue | padj | absLog2FC |
|---|---|---|---|---|---|---|---|---|---|---|
| ENSMUSG00000024959 | Bad | 12015 | protein_coding | 826.1382853 | 0.210078583 | 0.09223816 | 2.27756692 | 0.022752393 | 0.107508891 | 0.210078583 |
| ENSMUSG00000027296 | Itpka | 228550 | protein_coding | 293.7670066 | -0.210117777 | 0.102820942 | -2.043530949 | 0.040999907 | 0.16134834 | 0.210117777 |
| ENSMUSG00000096210 | H1f0 | 14958 | protein_coding | 6168.097885 | 0.210188896 | 0.071103752 | 2.956087276 | 0.003115689 | 0.024878349 | 0.210188896 |
| ENSMUSG00000019817 | Plagl1 | 22634 | protein_coding | 23.77891485 | 0.210206347 | 0.260933328 | 0.80559409 | 0.420476935 | 0.657586455 | 0.210206347 |
| ENSMUSG00000085958 | A930019D19Rik | 77806 | lncRNA | 7.593256584 | 0.210278603 | 0.538623691 | 0.39039984 | 0.696240906 | 0.84812124 | 0.210278603 |
| ENSMUSG00000030871 | Ears2 | 67417 | protein_coding | 405.332653 | 0.210451061 | 0.073986834 | 2.844439335 | 0.004448966 | 0.032627772 | 0.210451061 |
| ENSMUSG00000037260 | Hgsnat | 52120 | protein_coding | 794.2990276 | -0.210523574 | 0.056888623 | -3.700626992 | 0.000215067 | 0.002799348 | 0.210523574 |
| ENSMUSG00000101355 | H3c10 | 319152 | protein_coding | 394.911223 | 0.210546284 | 0.191808488 | 1.097690127 | 0.272339823 | 0.514779021 | 0.210546284 |
| ENSMUSG00000028601 | Echdc2 | 52430 | protein_coding | 240.1931339 | -0.210592201 | 0.101630102 | -2.07214395 | 0.038252019 | 0.153837982 | 0.210592201 |
| ENSMUSG00000062585 | Cnr2 | 12802 | protein_coding | 10.14749074 | 0.21059223 | 0.428442082 | 0.491530218 | 0.623051485 | 0.800888833 | 0.21059223 |
| ENSMUSG00000029478 | Ncor2 | 20602 | protein_coding | 411.443117 | -0.210608161 | 0.131344549 | -1.603478503 | 0.108829051 | 0.302010618 | 0.210608161 |
| ENSMUSG00000079442 | St6galnac4 | 20448 | protein_coding | 636.9306269 | 0.210705599 | 0.088764376 | 2.373763102 | 0.017607843 | 0.089975188 | 0.210705599 |
| ENSMUSG00000042747 | Krtcap2 | 66059 | protein_coding | 916.7625513 | 0.210720203 | 0.078742375 | 2.676071225 | 0.007449081 | 0.048003141 | 0.210720203 |
| ENSMUSG00000012422 | Tmem167 | 66074 | protein_coding | 1122.942224 | 0.210812567 | 0.058085064 | 3.629376538 | 0.000284107 | 0.003521742 | 0.210812567 |
| ENSMUSG00000020034 | Tcp11l2 | 216198 | protein_coding | 210.8394079 | -0.210837068 | 0.094445568 | -2.232365933 | 0.025590787 | 0.116615606 | 0.210837068 |
| ENSMUSG00000020656 | Grhl1 | 195733 | protein_coding | 70.5638095 | 0.210951174 | 0.146970583 | 1.435329233 | 0.151193301 | 0.367102434 | 0.210951174 |
| ENSMUSG00000027230 | Creb3l1 | 26427 | protein_coding | 1961.028287 | 0.210980245 | 0.112697392 | 1.87209518 | 0.061193437 | 0.210759777 | 0.210980245 |
| ENSMUSG00000052609 | Plekhg3 | 263406 | protein_coding | 1322.651682 | -0.21101824 | 0.051727999 | -4.079381482 | 4.52E-05 | 0.000746133 | 0.21101824 |
| ENSMUSG00000051341 | Zfp52 | 22710 | protein_coding | 93.22724064 | 0.211037374 | 0.147515471 | 1.430611808 | 0.152541501 | 0.369310836 | 0.211037374 |
| ENSMUSG00000013091 | Tmem190 | 78052 | protein_coding | 23.4029225 | 0.211140691 | 0.267243909 | 0.790067366 | 0.429488427 | 0.664591383 | 0.211140691 |
| ENSMUSG00000063802 | Hspbp1 | 66245 | protein_coding | 441.144406 | 0.211149781 | 0.122489833 | 1.723814752 | 0.084741291 | 0.259707718 | 0.211149781 |
| ENSMUSG00000002825 | Qtrt1 | 60507 | protein_coding | 265.5181126 | -0.211259509 | 0.084686924 | -2.4945942 | 0.012610125 | 0.070685953 | 0.211259509 |
| ENSMUSG00000021824 | Ap3m1 | 55946 | protein_coding | 1341.929307 | 0.211307744 | 0.051523297 | 4.101207738 | 4.11E-05 | 0.00068473 | 0.211307744 |
| ENSMUSG00000021916 | Glt8d1 | 76485 | protein_coding | 154.4681601 | 0.211388506 | 0.11291121 | 1.872165808 | 0.061183668 | 0.210759777 | 0.211388506 |
| ENSMUSG00000039936 | Pik3cd | 18707 | protein_coding | 72.02485666 | 0.21139323 | 0.17675212 | 1.195986954 | 0.23170165 | 0.470848351 | 0.21139323 |
| ENSMUSG00000090093 | Gm14399 | 100043761 | protein_coding | 50.93560881 | -0.211401329 | 0.188841253 | -1.119465826 | 0.262941462 | 0.504855347 | 0.211401329 |
| ENSMUSG00000038369 | Ncoa6 | 56406 | protein_coding | 584.3735933 | -0.211456977 | 0.111685868 | -1.893319018 | 0.058315458 | 0.204579809 | 0.211456977 |
| ENSMUSG00000021303 | Gng4 | 14706 | protein_coding | 23.13363914 | 0.211484262 | 0.276738307 | 0.764203063 | 0.444746242 | 0.675687484 | 0.211484262 |
| ENSMUSG00000068015 | Lrch1 | 380916 | protein_coding | 681.1167924 | -0.211490907 | 0.063935568 | -3.307875611 | 0.000940066 | 0.00955289 | 0.211490907 |
| ENSMUSG00000025078 | Nhlrc2 | 66866 | protein_coding | 1950.645243 | -0.211574517 | 0.046427958 | -4.557049768 | 5.19E-06 | 0.000109362 | 0.211574517 |
| ENSMUSG00000061451 | Tmem151a | 381199 | protein_coding | 30.31399317 | 0.211695972 | 0.222470015 | 0.951570807 | 0.341314693 | 0.584131301 | 0.211695972 |
| ENSMUSG00000031399 | Fam3a | 66294 | protein_coding | 800.228064 | 0.211839969 | 0.069856757 | 3.032490754 | 0.002425445 | 0.020319562 | 0.211839969 |
| ENSMUSG00000021902 | Phf7 | 71838 | protein_coding | 163.6902423 | -0.211874019 | 0.09555052 | -2.217403103 | 0.026595558 | 0.119191767 | 0.211874019 |
| ENSMUSG00000035357 | Pdzrn3 | 55983 | protein_coding | 143.5082923 | -0.211889471 | 0.129862009 | -1.631650962 | 0.102753036 | 0.29253061 | 0.211889471 |
| ENSMUSG00000032714 | Syde1 | 71709 | protein_coding | 20.21333871 | -0.211923926 | 0.328015594 | -0.646078814 | 0.518228315 | 0.730504162 | 0.211923926 |
| ENSMUSG00000022091 | Sorbs3 | 20410 | protein_coding | 41.18868012 | 0.211926722 | 0.198915881 | 1.065408756 | 0.286690996 | 0.529168268 | 0.211926722 |
| ENSMUSG00000024875 | Yif1a | 68090 | protein_coding | 1434.357034 | 0.211989483 | 0.071178948 | 2.978260981 | 0.00289889 | 0.023422046 | 0.211989483 |
| ENSMUSG00000065725 | Gm26165 | 115486130 | snoRNA | 59.93308147 | 0.212121934 | 0.239377419 | 0.886140113 | 0.37554203 | 0.617215876 | 0.212121934 |
| ENSMUSG00000021684 | Pde8b | 218461 | protein_coding | 824.7208947 | -0.212212417 | 0.098045613 | -2.164425415 | 0.030431715 | 0.13153907 | 0.212212417 |

| ensembl_gene_id | external_gene_name | entrezgene_id | gene_biotype | baseMean | log2FoldChange | lfcSE | stat | pvalue | padj | absLog2FC |
|---|---|---|---|---|---|---|---|---|---|---|
| ENSMUSG00000097321 | 1700028E10Rik | 625175 | lncRNA | 8.654159609 | -0.212320735 | 0.423127188 | -0.501789394 | 0.615815674 | 0.795599668 | 0.212320735 |
| ENSMUSG00000061981 | Flot2 | 14252 | protein_coding | 304.3609773 | -0.212334946 | 0.101613306 | -2.089637219 | 0.036650401 | 0.148994457 | 0.212334946 |
| ENSMUSG00000105402 | Gm3716 | 100042198 | lncRNA | 12.62739653 | 0.212344524 | 0.38282355 | 0.55467989 | 0.579113634 | 0.771529581 | 0.212344524 |
| ENSMUSG00000029910 | Mad2l1 | 56150 | protein_coding | 441.1407506 | 0.212367046 | 0.107169932 | 1.981591679 | 0.047524959 | 0.178864648 | 0.212367046 |
| ENSMUSG00000007379 | Dennd2c | 329727 | protein_coding | 59.22465371 | -0.212414578 | 0.163864195 | -1.296284268 | 0.194877566 | 0.426951571 | 0.212414578 |
| ENSMUSG00000026024 | Als2 | 74018 | protein_coding | 380.7678311 | -0.212472427 | 0.078358055 | -2.711558199 | 0.006696778 | 0.04431635 | 0.212472427 |
| ENSMUSG00000042388 | Dlgap3 | 242667 | protein_coding | 6.972355843 | -0.212484371 | 0.448096742 | -0.474193073 | 0.635362233 | 0.809423077 | 0.212484371 |
| ENSMUSG00000029469 | Ift81 | 12589 | protein_coding | 87.58373898 | -0.212508365 | 0.18594749 | -1.142840731 | 0.253104724 | 0.494286739 | 0.212508365 |
| ENSMUSG00000041323 | Ak7 | 78801 | protein_coding | 11.12522165 | -0.212623227 | 0.410270666 | -0.518251107 | 0.604283081 | 0.788707774 | 0.212623227 |
| ENSMUSG00000020661 | Dnmt3a | 13435 | protein_coding | 1151.092847 | -0.21265956 | 0.042036239 | -5.058957811 | 4.22E-07 | 1.11E-05 | 0.21265956 |
| ENSMUSG00000033411 | Ctdspl2 | 329506 | protein_coding | 696.9885097 | -0.212663667 | 0.077373686 | -2.74852704 | 0.00598637 | 0.040959773 | 0.212663667 |
| ENSMUSG00000037995 | Igsf9 | 93842 | protein_coding | 2890.771901 | 0.212762589 | 0.095168793 | 2.235633994 | 0.025375756 | 0.115930543 | 0.212762589 |
| ENSMUSG00000027074 | Slc43a3 | 58207 | protein_coding | 19.8183991 | 0.21277957 | 0.406533209 | 0.523400216 | 0.600695774 | 0.786125192 | 0.21277957 |
| ENSMUSG00000040429 | Mterf1a | 545725 | protein_coding | 39.7079257 | -0.212786445 | 0.182165297 | -1.168095399 | 0.242768286 | 0.483886312 | 0.212786445 |
| ENSMUSG00000000378 | Ccm2 | 216527 | protein_coding | 399.0626138 | 0.212810957 | 0.071810906 | 2.963490762 | 0.003041712 | 0.024375404 | 0.212810957 |
| ENSMUSG00000064380 | Gm26448 | 115487256 | snoRNA | 985.740141 | 0.212864968 | 0.260099909 | 0.818396934 | 0.413130572 | 0.650767409 | 0.212864968 |
| ENSMUSG00000100594 | 2810414N06Rik | 102640772 | lncRNA | 26.40722781 | 0.212884041 | 0.249836201 | 0.852094452 | 0.394161671 | 0.633763237 | 0.212884041 |
| ENSMUSG00000064341 | mt-Nd1 | 17716 | protein_coding | 62322.7501 | 0.212891232 | 0.065010981 | 3.274696492 | 0.001057755 | 0.010542907 | 0.212891232 |
| ENSMUSG00000043019 | Edem3 | 66967 | protein_coding | 3608.426564 | 0.212919974 | 0.043291929 | 4.918237154 | 8.73E-07 | 2.17E-05 | 0.212919974 |
| ENSMUSG00000064141 | Zfp69 | 381549 | protein_coding | 40.09279237 | 0.212967177 | 0.221923164 | 0.959643749 | 0.337234543 | 0.580796825 | 0.212967177 |
| ENSMUSG00000002617 | Zfp40 | 22700 | protein_coding | 100.7211762 | -0.213114278 | 0.171617354 | -1.241799112 | 0.21431069 | 0.450024545 | 0.213114278 |
| ENSMUSG00000020434 | 4921536K21Rik | 67430 | protein_coding | 46.73852391 | 0.21313952 | 0.189027216 | 1.127559962 | 0.259505804 | 0.501083584 | 0.21313952 |
| ENSMUSG00000037788 | Vopp1 | 232023 | protein_coding | 440.538721 | 0.213208801 | 0.084048041 | 2.536749206 | 0.011188708 | 0.064373988 | 0.213208801 |
| ENSMUSG00000029446 | Psph | 100678 | protein_coding | 128.0926854 | 0.213214799 | 0.1201046 | 1.775242569 | 0.075857844 | 0.242489484 | 0.213214799 |
| ENSMUSG00000026785 | Pkn3 | 263803 | protein_coding | 153.5568743 | -0.21330239 | 0.115434538 | -1.847821223 | 0.06462821 | 0.218457165 | 0.21330239 |
| ENSMUSG00000046062 | Ppp1r15b | 108954 | protein_coding | 4092.332482 | -0.213309372 | 0.05411822 | -3.941544475 | 8.10E-05 | 0.001231717 | 0.213309372 |
| ENSMUSG00000049321 | Zfp2 | 22678 | protein_coding | 91.70237388 | 0.213328189 | 0.130525855 | 1.634374959 | 0.102180143 | 0.291648396 | 0.213328189 |
| ENSMUSG00000027330 | Cdc25b | 12531 | protein_coding | 416.1925528 | -0.213353989 | 0.077727141 | -2.744909786 | 0.006052753 | 0.041292666 | 0.213353989 |
| ENSMUSG00000020806 | Rhbdf2 | 217344 | protein_coding | 402.176155 | -0.213363004 | 0.076001232 | -2.807362423 | 0.004994901 | 0.035627054 | 0.213363004 |
| ENSMUSG00000091264 | Smim13 | 108934 | protein_coding | 296.5494979 | -0.213364306 | 0.093195672 | -2.289422895 | 0.022054793 | 0.105292377 | 0.213364306 |
| ENSMUSG00000030760 | Acer3 | 66190 | protein_coding | 556.2515636 | -0.213366786 | 0.066213019 | -3.222429512 | 0.001271084 | 0.012289435 | 0.213366786 |
| ENSMUSG00000067212 | H2-T23 | 15040 | protein_coding | 4098.992639 | -0.213415969 | 0.086817911 | -2.458202064 | 0.013963458 | 0.076205988 | 0.213415969 |
| ENSMUSG00000039307 | Hexdc | 238023 | protein_coding | 375.0556081 | -0.213609836 | 0.081258175 | -2.628779635 | 0.008569187 | 0.05321343 | 0.213609836 |
| ENSMUSG00000021693 | Kif2a | 16563 | protein_coding | 557.432068 | -0.213652714 | 0.099952739 | -2.137537351 | 0.032554311 | 0.137687746 | 0.213652714 |
| ENSMUSG00000072980 | Oip5 | 70645 | protein_coding | 74.70479829 | 0.213655726 | 0.164579344 | 1.298192841 | 0.194221077 | 0.426123143 | 0.213655726 |
| ENSMUSG00000000340 | Dbt | 13171 | protein_coding | 967.0915348 | -0.213767226 | 0.055293539 | -3.866043451 | 0.000110615 | 0.001620247 | 0.213767226 |
| ENSMUSG00000028398 | Dmac1 | 66928 | protein_coding | 412.4998015 | 0.213802019 | 0.104382051 | 2.048264208 | 0.040534118 | 0.160101328 | 0.213802019 |
| ENSMUSG00000052534 | Pbx1 | 18514 | protein_coding | 1065.66097 | -0.213806894 | 0.075038858 | -2.849282358 | 0.004381797 | 0.032358233 | 0.213806894 |
| ENSMUSG00000024743 | Syt7 | 54525 | protein_coding | 1298.290616 | -0.213814964 | 0.0533753 | -4.005878436 | 6.18E-05 | 0.000985604 | 0.213814964 |

Supplementary Table 1

| ensembl_gene_id | external_gene_name | entrezgene_id | gene_biotype | baseMean | log2FoldChange | lfcSE | stat | pvalue | padj | absLog2FC |
|---|---|---|---|---|---|---|---|---|---|---|
| ENSMUSG00000039683 | Sdk1 | 330222 | protein_coding | 11.83897618 | 0.213919281 | 0.363662723 | 0.588235383 | 0.556374311 | 0.755428309 | 0.213919281 |
| ENSMUSG00000038855 | Itpkb | 320404 | protein_coding | 312.7719497 | 0.213997766 | 0.119328058 | 1.793356644 | 0.072915915 | 0.236269835 | 0.213997766 |
| ENSMUSG00000033965 | Slc16a2 | 20502 | protein_coding | 24.57192061 | -0.214187025 | 0.246702707 | -0.868198925 | 0.385285443 | 0.625729943 | 0.214187025 |
| ENSMUSG00000073599 | Ecscr | 68545 | protein_coding | 35.46634274 | -0.214216384 | 0.23807403 | -0.899788963 | 0.368232569 | 0.610334928 | 0.214216384 |
| ENSMUSG00000030539 | Sema4b | 20352 | protein_coding | 2372.995925 | 0.214224152 | 0.066734255 | 3.210107781 | 0.001326852 | 0.012691889 | 0.214224152 |
| ENSMUSG00000021779 | Thrb | 21834 | protein_coding | 884.0975818 | -0.214225834 | 0.063489273 | -3.374205167 | 0.000740292 | 0.007859568 | 0.214225834 |
| ENSMUSG00000055707 | Klhl26 | 234378 | protein_coding | 341.9002365 | -0.214311169 | 0.078594764 | -2.72678686 | 0.006395434 | 0.042920997 | 0.214311169 |
| ENSMUSG00000100017 | 2410022M11Rik | 100503842 | lncRNA | 88.45452938 | -0.214399961 | 0.139465402 | -1.537298552 | 0.124220216 | 0.327122701 | 0.214399961 |
| ENSMUSG00000037185 | Krt80 | 74127 | protein_coding | 6.571972563 | -0.214421899 | 0.539041029 | -0.397784004 | 0.690789409 | 0.844734668 | 0.214421899 |
| ENSMUSG00000025921 | Rdh10 | 98711 | protein_coding | 529.9662306 | -0.214437843 | 0.059751344 | -3.588837175 | 0.000332156 | 0.004009493 | 0.214437843 |
| ENSMUSG00000031520 | Vegfc | 22341 | protein_coding | 79.83447115 | 0.214493959 | 0.144638763 | 1.482963176 | 0.138084195 | 0.348113774 | 0.214493959 |
| ENSMUSG00000035504 | Reep6 | 70335 | protein_coding | 333.232328 | 0.214542502 | 0.108023496 | 1.986072567 | 0.047025267 | 0.177668978 | 0.214542502 |
| ENSMUSG00000024308 | Tapbp | 21356 | protein_coding | 3895.466934 | -0.214550882 | 0.120371079 | -1.782412216 | 0.074682035 | 0.239967558 | 0.214550882 |
| ENSMUSG00000108621 | Gm33989 | 102637087 | lncRNA | 502.830828 | -0.214577168 | 0.072379336 | -2.96461918 | 0.003030578 | 0.024315491 | 0.214577168 |
| ENSMUSG00000111709 | Gm3776 | 100042295 | protein_coding | 14.7835796 | 0.214704672 | 0.378476466 | 0.567286718 | 0.5705194 | 0.765709584 | 0.214704672 |
| ENSMUSG00000021699 | Pde4d | 238871 | protein_coding | 605.7978946 | -0.214722723 | 0.062261409 | -3.448728929 | 0.000563232 | 0.006281632 | 0.214722723 |
| ENSMUSG00000068270 | Shroom4 | 208431 | protein_coding | 42.1414037 | -0.214785957 | 0.249690295 | -0.860209473 | 0.389673584 | 0.629496733 | 0.214785957 |
| ENSMUSG00000033318 | Gstt2 | 14872 | protein_coding | 698.8000846 | 0.214811545 | 0.0620385 | 3.462552207 | 0.000535078 | 0.005996667 | 0.214811545 |
| ENSMUSG00000032553 | Srprb | 20818 | protein_coding | 873.2846876 | 0.214819758 | 0.065607094 | 3.274337331 | 0.001059101 | 0.010549797 | 0.214819758 |
| ENSMUSG00000027854 | Sike1 | 66641 | protein_coding | 1521.725217 | -0.21487386 | 0.063499349 | -3.38387499 | 0.000714705 | 0.007673804 | 0.21487386 |
| ENSMUSG00000051695 | Pcbp1 | 23983 | protein_coding | 2014.112261 | 0.214888111 | 0.109745146 | 1.958064838 | 0.050222405 | 0.185765304 | 0.214888111 |
| ENSMUSG00000030657 | Xylt1 | 233781 | protein_coding | 232.7638287 | 0.214963806 | 0.105453036 | 2.038479066 | 0.041502046 | 0.162728786 | 0.214963806 |
| ENSMUSG00000070462 | Tlnrd1 | 80889 | protein_coding | 2009.850077 | 0.214988248 | 0.095218855 | 2.257832729 | 0.02395609 | 0.111401344 | 0.214988248 |
| ENSMUSG00000025384 | Faap100 | 71885 | protein_coding | 627.6808677 | -0.215023691 | 0.071626432 | -3.002015948 | 0.002681981 | 0.022022562 | 0.215023691 |
| ENSMUSG00000026687 | Aldh9a1 | 56752 | protein_coding | 1516.334289 | 0.215024031 | 0.048849871 | 4.401731817 | 1.07E-05 | 0.000209417 | 0.215024031 |
| ENSMUSG00000045216 | Hs6st1 | 50785 | protein_coding | 1797.479985 | 0.215025241 | 0.076696604 | 2.803582307 | 0.005053833 | 0.035951992 | 0.215025241 |
| ENSMUSG00000008682 | Rpl10 | 110954 | protein_coding | 12619.15988 | 0.215044065 | 0.045127764 | 4.765227523 | 1.89E-06 | 4.35E-05 | 0.215044065 |
| ENSMUSG00000027641 | Rbl1 | 19650 | protein_coding | 858.5135838 | -0.215076582 | 0.06178444 | -3.481080031 | 0.000499396 | 0.005643843 | 0.215076582 |
| ENSMUSG00000098923 | Tmem185b | 226351 | protein_coding | 322.6234964 | -0.215091982 | 0.086071738 | -2.498984997 | 0.012454958 | 0.070010845 | 0.215091982 |
| ENSMUSG00000028017 | Egf | 13645 | protein_coding | 212.0941003 | 0.215103244 | 0.102072727 | 2.10735278 | 0.035087011 | 0.144829341 | 0.215103244 |
| ENSMUSG00000028654 | Mycl | 16918 | protein_coding | 417.065802 | -0.215109224 | 0.084282135 | -2.552251707 | 0.010702917 | 0.062200337 | 0.215109224 |
| ENSMUSG00000021752 | Kctd6 | 71393 | protein_coding | 188.6995944 | -0.215165785 | 0.095840702 | -2.245035575 | 0.024765849 | 0.114048785 | 0.215165785 |
| ENSMUSG00000043004 | Gng2 | 14702 | protein_coding | 16.77461214 | 0.215185162 | 0.291965298 | 0.737023076 | 0.461108316 | 0.687717915 | 0.215185162 |
| ENSMUSG00000027840 | Wnt2b | 22414 | protein_coding | 10.71885897 | -0.21528849 | 0.40097169 | -0.536916933 | 0.591324994 | 0.780129105 | 0.21528849 |
| ENSMUSG00000059602 | Syn3 | 27204 | protein_coding | 43.71839318 | -0.21533081 | 0.192697557 | -1.117454802 | 0.263799915 | 0.50508147 | 0.21533081 |
| ENSMUSG00000109179 | Gm35339 | 102638882 | protein_coding | 32.24297161 | 0.215355216 | 0.24668636 | 0.872991991 | 0.382667443 | 0.62329844 | 0.215355216 |
| ENSMUSG00000015027 | Galns | 50917 | protein_coding | 601.0049067 | -0.215447791 | 0.067667064 | -3.1839388 | 0.001452857 | 0.013606405 | 0.215447791 |
| ENSMUSG00000062526 | Mppe1 | 225651 | protein_coding | 388.326428 | 0.215502334 | 0.067178558 | 3.207903537 | 0.001337063 | 0.012743984 | 0.215502334 |
| ENSMUSG00000026162 | Nhej1 | 75570 | protein_coding | 97.72940609 | 0.215616417 | 0.140269347 | 1.537159908 | 0.124254156 | 0.327154117 | 0.215616417 |

| ensembl_gene_id | external_gene_name | entrezgene_id | gene_biotype | baseMean | log2FoldChange | lfcSE | stat | pvalue | padj | absLog2FC |
|---|---|---|---|---|---|---|---|---|---|---|
| ENSMUSG00000036225 | Kctd1 | 106931 | protein_coding | 116.0575174 | 0.215621352 | 0.122922209 | 1.754128512 | 0.079408489 | 0.24905109 | 0.215621352 |
| ENSMUSG00000057093 | Zfp607b | 112415 | protein_coding | 17.22022305 | -0.215854443 | 0.282006253 | -0.765424314 | 0.444018919 | 0.674900387 | 0.215854443 |
| ENSMUSG00000032083 | Apoa1 | 11806 | protein_coding | 376.3191895 | 0.21588956 | 0.764469436 | 0.282404436 | 0.777633414 | 0.89645106 | 0.21588956 |
| ENSMUSG00000089687 | Rab42 | 242681 | protein_coding | 9.550475212 | 0.21593002 | 0.394961241 | 0.546711925 | 0.58457666 | 0.775604097 | 0.21593002 |
| ENSMUSG00000020644 | Id2 | 15902 | protein_coding | 425.2446576 | -0.215931726 | 0.103557896 | -2.08513049 | 0.037057457 | 0.150119471 | 0.215931726 |
| ENSMUSG00000052485 | Tmem171 | 380863 | protein_coding | 2714.295303 | -0.21595905 | 0.058313755 | -3.703398106 | 0.000212731 | 0.002772663 | 0.21595905 |
| ENSMUSG00000053835 | H2-T24 | 15042 | protein_coding | 27.77112915 | -0.216108464 | 0.279484403 | -0.773239801 | 0.439380475 | 0.671700891 | 0.216108464 |
| ENSMUSG00000026123 | Plekhb2 | 226971 | protein_coding | 8413.7935 | 0.21616544 | 0.050153734 | 4.310056725 | 1.63E-05 | 0.000301734 | 0.21616544 |
| ENSMUSG00000024998 | Plce1 | 74055 | protein_coding | 2936.770106 | -0.216174491 | 0.049308025 | -4.384164453 | 1.16E-05 | 0.000224605 | 0.216174491 |
| ENSMUSG00000001741 | Il16 | 16170 | protein_coding | 21.84129206 | 0.216244918 | 0.288776727 | 0.748830837 | 0.453959171 | 0.682675952 | 0.216244918 |
| ENSMUSG00000055320 | Tead1 | 21676 | protein_coding | 318.373474 | -0.216252628 | 0.124004253 | -1.743912984 | 0.081174264 | 0.252562627 | 0.216252628 |
| ENSMUSG00000045503 | Sys1 | 66460 | protein_coding | 496.8027018 | 0.216326224 | 0.069938587 | 3.093088276 | 0.001980852 | 0.01741389 | 0.216326224 |
| ENSMUSG00000071637 | Cebpd | 12609 | protein_coding | 531.4327816 | -0.216370954 | 0.150305799 | -1.439538299 | 0.149998067 | 0.365355509 | 0.216370954 |
| ENSMUSG00000042613 | Pbxip1 | 229534 | protein_coding | 2335.566777 | -0.216403435 | 0.046133427 | -4.69081633 | 2.72E-06 | 6.05E-05 | 0.216403435 |
| ENSMUSG00000024774 | Ankrd22 | 52024 | protein_coding | 159.3349254 | 0.216418224 | 0.143716376 | 1.505870306 | 0.132100471 | 0.338693846 | 0.216418224 |
| ENSMUSG00000041124 | Msantd4 | 78100 | protein_coding | 842.1068979 | 0.216561317 | 0.059618321 | 3.632462539 | 0.000280729 | 0.003493304 | 0.216561317 |
| ENSMUSG00000037902 | Sirpa | 19261 | protein_coding | 26.05745473 | 0.21671288 | 0.290755986 | 0.745342799 | 0.456064514 | 0.684117981 | 0.21671288 |
| ENSMUSG00000018199 | Ro60 | 20822 | protein_coding | 598.1796343 | 0.216728186 | 0.063407842 | 3.418002882 | 0.000630824 | 0.006930044 | 0.216728186 |
| ENSMUSG00000081683 | Fzd10 | 93897 | protein_coding | 21.62459419 | -0.216729968 | 0.295544239 | -0.733324962 | 0.463360254 | 0.68941884 | 0.216729968 |
| ENSMUSG00000070605 | Zfp992 | 433791 | protein_coding | 52.56088907 | -0.216817658 | 0.316213485 | -0.685668602 | 0.492922111 | 0.712333599 | 0.216817658 |
| ENSMUSG00000038607 | Gng10 | 14700 | protein_coding | 15.31246097 | -0.216822572 | 0.32173716 | -0.673912121 | 0.500367185 | 0.717537044 | 0.216822572 |
| ENSMUSG00000003037 | Rab8a | 17274 | protein_coding | 2513.940225 | 0.216865006 | 0.061795878 | 3.50937656 | 0.000449158 | 0.005166604 | 0.216865006 |
| ENSMUSG00000009293 | Ube2g2 | 22213 | protein_coding | 611.5595919 | 0.216915806 | 0.06680491 | 3.247003935 | 0.001166268 | 0.011447596 | 0.216915806 |
| ENSMUSG00000028293 | Slc35a1 | 24060 | protein_coding | 3845.998235 | 0.216981775 | 0.044226981 | 4.906095128 | 9.29E-07 | 2.29E-05 | 0.216981775 |
| ENSMUSG00000047822 | Angptl8 | 624219 | protein_coding | 33.23496169 | -0.217027941 | 0.216100607 | -1.004291216 | 0.315238266 | 0.559439557 | 0.217027941 |
| ENSMUSG00000015790 | Surf1 | 20930 | protein_coding | 1432.780871 | -0.217028322 | 0.038271819 | -5.670708252 | 1.42E-08 | 5.00E-07 | 0.217028322 |
| ENSMUSG00000019732 | Calr3 | 73316 | protein_coding | 9.436248982 | -0.217071208 | 0.394098182 | -0.550804897 | 0.581767427 | 0.773403948 | 0.217071208 |
| ENSMUSG00000002741 | Ykt6 | 56418 | protein_coding | 2200.662889 | 0.217151363 | 0.049310549 | 4.403750696 | 1.06E-05 | 0.000207728 | 0.217151363 |
| ENSMUSG00000021510 | Zfp729a | 212281 | protein_coding | 359.0031956 | -0.217168375 | 0.09646913 | -2.25116961 | 0.024374797 | 0.112857553 | 0.217168375 |
| ENSMUSG00000021514 | Zfp369 | 170936 | protein_coding | 382.127657 | -0.217341414 | 0.08797974 | -2.470357537 | 0.013497807 | 0.074394782 | 0.217341414 |
| ENSMUSG00000113836 | Gm3325 | 100041420 | transcribed_processed_pseudogene | 68.29535827 | -0.217362098 | 0.177680637 | -1.223330249 | 0.221204987 | 0.457725814 | 0.217362098 |
| ENSMUSG00000001663 | Gstt1 | 14871 | protein_coding | 234.7328201 | -0.217486171 | 0.08150111 | -2.668505643 | 0.00761895 | 0.048855191 | 0.217486171 |
| ENSMUSG00000025049 | Taf5 | 226182 | protein_coding | 199.0781434 | -0.21757983 | 0.090825463 | -2.395581846 | 0.016594009 | 0.086330552 | 0.21757983 |
| ENSMUSG00000021287 | Xrcc3 | 74335 | protein_coding | 149.8670294 | 0.217759843 | 0.141464895 | 1.539320718 | 0.123726019 | 0.326407574 | 0.217759843 |
| ENSMUSG00000078923 | Ube2v1 | 66589 | protein_coding | 19.91350143 | -0.217780115 | 0.304235182 | -0.715828176 | 0.474097452 | 0.697951154 | 0.217780115 |
| ENSMUSG00000036473 | Tbc1d24 | 224617 | protein_coding | 619.9358281 | -0.217802478 | 0.082651714 | -2.635184046 | 0.008409167 | 0.052462144 | 0.217802478 |
| ENSMUSG00000020182 | Ddc | 13195 | protein_coding | 1745.531353 | 0.217851216 | 0.080771677 | 2.697123832 | 0.006994126 | 0.045795479 | 0.217851216 |
| ENSMUSG00000020023 | Tmcc3 | 319880 | protein_coding | 2144.972945 | 0.217891581 | 0.048879831 | 4.457699145 | 8.28E-06 | 0.000166379 | 0.217891581 |
| ENSMUSG00000005958 | Ephb3 | 13845 | protein_coding | 744.3948977 | -0.217893861 | 0.091519804 | -2.380838363 | 0.017273289 | 0.08873319 | 0.217893861 |

| ensembl_gene_id | external_gene_name | entrezgene_id | gene_biotype | baseMean | log2FoldChange | lfcSE | stat | pvalue | padj | absLog2FC |
|---|---|---|---|---|---|---|---|---|---|---|
| ENSMUSG00000038175 | Mylip | 218203 | protein_coding | 593.5172268 | -0.217933694 | 0.085308549 | -2.554652456 | 0.010629387 | 0.061938406 | 0.217933694 |
| ENSMUSG00000022574 | Naprt | 223646 | protein_coding | 1713.896024 | -0.218039163 | 0.072565006 | -3.00474258 | 0.002658057 | 0.021870656 | 0.218039163 |
| ENSMUSG00000041476 | Smpx | 66106 | protein_coding | 127.1606689 | 0.218170713 | 0.151434242 | 1.440696038 | 0.149670575 | 0.364802272 | 0.218170713 |
| ENSMUSG00000096918 | Gm16863 | 328646 | lncRNA | 16.31939865 | -0.218249295 | 0.29356142 | -0.743453602 | 0.457207103 | 0.685194586 | 0.218249295 |
| ENSMUSG00000059208 | Hnrnpm | 76936 | protein_coding | 3268.426592 | 0.218256544 | 0.07098142 | 3.074840478 | 0.002106152 | 0.018271064 | 0.218256544 |
| ENSMUSG00000022797 | Tfrc | 22042 | protein_coding | 11646.24633 | 0.218333524 | 0.178613866 | 1.222377235 | 0.221565004 | 0.458276109 | 0.218333524 |
| ENSMUSG00000022564 | Grina | 66168 | protein_coding | 704.1848657 | -0.218487483 | 0.084042067 | -2.599739519 | 0.009329455 | 0.056624534 | 0.218487483 |
| ENSMUSG00000031906 | Smpd3 | 58994 | protein_coding | 4036.899353 | 0.218508318 | 0.068125929 | 3.207417797 | 0.001339323 | 0.012757984 | 0.218508318 |
| ENSMUSG00000052565 | H1f3 | 14957 | protein_coding | 4933.713024 | 0.218512568 | 0.109165254 | 2.001667738 | 0.045320478 | 0.173037093 | 0.218512568 |
| ENSMUSG00000086427 | Hoxa11os | 15397 | lncRNA | 428.5087207 | -0.218552325 | 0.067679358 | -3.22923165 | 0.001241233 | 0.012065923 | 0.218552325 |
| ENSMUSG00000026553 | Copa | 12847 | protein_coding | 8313.321906 | 0.218556847 | 0.041793946 | 5.229390069 | 1.70E-07 | 4.92E-06 | 0.218556847 |
| ENSMUSG00000040270 | Bach2 | 12014 | protein_coding | 66.32781645 | -0.218573908 | 0.1720959 | -1.270070399 | 0.204059554 | 0.438306743 | 0.218573908 |
| ENSMUSG00000085457 | 1110046J04Rik | 68808 | lncRNA | 9.948079928 | -0.218595687 | 0.397722832 | -0.54961815 | 0.582581308 | 0.774230634 | 0.218595687 |
| ENSMUSG00000029072 | Tas1r3 | 83771 | protein_coding | 65.79281415 | -0.218601378 | 0.214592237 | -1.018682603 | 0.308353674 | 0.552028135 | 0.218601378 |
| ENSMUSG00000001632 | Brpf1 | 78783 | protein_coding | 796.7886228 | -0.218621976 | 0.056674872 | -3.857476293 | 0.000114564 | 0.001668988 | 0.218621976 |
| ENSMUSG00000066149 | Cdc26 | 66440 | protein_coding | 385.6388887 | 0.218651228 | 0.076853123 | 2.845053267 | 0.0044404 | 0.032609438 | 0.218651228 |
| ENSMUSG00000037353 | Letmd1 | 68614 | protein_coding | 535.4510063 | -0.218686228 | 0.06795958 | -3.217886678 | 0.001291388 | 0.012463327 | 0.218686228 |
| ENSMUSG00000040713 | Creg1 | 433375 | protein_coding | 989.9441637 | 0.218787522 | 0.059384614 | 3.684245926 | 0.000229381 | 0.002949938 | 0.218787522 |
| ENSMUSG00000034165 | Ccnd3 | 12445 | protein_coding | 895.8787887 | 0.218805512 | 0.07748879 | 2.823705359 | 0.0047472 | 0.034269516 | 0.218805512 |
| ENSMUSG00000030096 | Slc6a6 | 21366 | protein_coding | 3827.900281 | -0.218837508 | 0.053270319 | -4.108056996 | 3.99E-05 | 0.000666808 | 0.218837508 |
| ENSMUSG00000042766 | Trim46 | 360213 | protein_coding | 254.6940291 | 0.218862487 | 0.124424156 | 1.759003189 | 0.078576967 | 0.247744036 | 0.218862487 |
| ENSMUSG00000023048 | Prr13 | 66151 | protein_coding | 2617.079928 | 0.218871202 | 0.084302083 | 2.596272756 | 0.009424122 | 0.056965074 | 0.218871202 |
| ENSMUSG00000026222 | Sp100 | 20684 | protein_coding | 82.63539438 | -0.218908231 | 0.248573064 | -0.880659502 | 0.378502143 | 0.619709567 | 0.218908231 |
| ENSMUSG00000026005 | Rpe | 66646 | protein_coding | 979.4987139 | 0.218932409 | 0.060202584 | 3.63659491 | 0.000276266 | 0.003451077 | 0.218932409 |
| ENSMUSG00000037973 | Itprid1 | 232016 | protein_coding | 20.29383943 | -0.218982444 | 0.283835825 | -0.771510939 | 0.440404141 | 0.672114736 | 0.218982444 |
| ENSMUSG00000047250 | Ptgs1 | 19224 | protein_coding | 83.56597791 | 0.219005218 | 0.133673252 | 1.638362303 | 0.101346138 | 0.290200492 | 0.219005218 |
| ENSMUSG00000030854 | Ptpn5 | 19259 | protein_coding | 17.55911962 | -0.219016785 | 0.535137753 | -0.409271787 | 0.682340217 | 0.839430978 | 0.219016785 |
| ENSMUSG00000018770 | Atp5g3 | 228033 | protein_coding | 9525.63072 | 0.219062939 | 0.07512839 | 2.915847639 | 0.003547237 | 0.027595894 | 0.219062939 |
| ENSMUSG00000037474 | Dtl | 76843 | protein_coding | 228.5334526 | 0.219067523 | 0.169457071 | 1.29276118 | 0.196093671 | 0.428683669 | 0.219067523 |
| ENSMUSG00000025701 | Alox5 | 11689 | protein_coding | 130.0464214 | 0.219091489 | 0.144002842 | 1.521438653 | 0.128149795 | 0.333333335 | 0.219091489 |
| ENSMUSG00000019832 | Rab32 | 67844 | protein_coding | 365.7425565 | 0.219097708 | 0.10966066 | 1.997960882 | 0.045720901 | 0.174229909 | 0.219097708 |
| ENSMUSG00000035247 | Hectd1 | 207304 | protein_coding | 6224.668214 | 0.21911421 | 0.041121243 | 5.328491853 | 9.90E-08 | 3.03E-06 | 0.21911421 |
| ENSMUSG00000025500 | Lmntd2 | 72000 | protein_coding | 103.0399491 | -0.219149441 | 0.143213171 | -1.530232442 | 0.125959204 | 0.329442764 | 0.219149441 |
| ENSMUSG00000019478 | Rab4a | 19341 | protein_coding | 767.224118 | 0.219259406 | 0.071229722 | 3.078201052 | 0.002082544 | 0.018114984 | 0.219259406 |
| ENSMUSG00000055629 | B4galnt4 | 330671 | protein_coding | 87.30692214 | -0.219306843 | 0.156444234 | -1.401821199 | 0.160968646 | 0.381467566 | 0.219306843 |
| ENSMUSG00000039218 | Srrm2 | 75956 | protein_coding | 4251.471674 | -0.219316137 | 0.083367935 | -2.630701317 | 0.008520889 | 0.0529747 | 0.219316137 |
| ENSMUSG00000037685 | Atp8a1 | 11980 | protein_coding | 2841.554712 | 0.219364415 | 0.057824402 | 3.793630488 | 0.000148461 | 0.002048095 | 0.219364415 |
| ENSMUSG00000021720 | Rnf180 | 71816 | protein_coding | 166.2970032 | -0.219398938 | 0.115696057 | -1.896338939 | 0.05791524 | 0.203825108 | 0.219398938 |
| ENSMUSG00000056763 | Cspp1 | 211660 | protein_coding | 520.2340314 | -0.2194189 | 0.107744865 | -2.036467347 | 0.041703449 | 0.16319418 | 0.2194189 |

| ensembl_gene_id | external_gene_name | entrezgene_id | gene_biotype | baseMean | log2FoldChange | lfcSE | stat | pvalue | padj | absLog2FC |
|---|---|---|---|---|---|---|---|---|---|---|
| ENSMUSG00000042429 | Adora1 | 11539 | protein_coding | 22.07195254 | -0.219470161 | 0.261936684 | -0.837874853 | 0.402100993 | 0.641349295 | 0.219470161 |
| ENSMUSG00000028060 | Khdc4 | 74200 | protein_coding | 1002.123438 | -0.219475561 | 0.100873349 | -2.175753687 | 0.029573683 | 0.128824466 | 0.219475561 |
| ENSMUSG00000090015 | Gm15446 | 100642166 | protein_coding | 133.415549 | -0.219490683 | 0.144636375 | -1.51753446 | 0.1291318 | 0.334754628 | 0.219490683 |
| ENSMUSG00000045107 | Saysd1 | 67509 | protein_coding | 265.7107556 | -0.219565218 | 0.093109263 | -2.358145803 | 0.018366478 | 0.092761185 | 0.219565218 |
| ENSMUSG00000026389 | Steap3 | 68428 | protein_coding | 712.3819375 | 0.219606806 | 0.082399337 | 2.665152583 | 0.00769534 | 0.04918511 | 0.219606806 |
| ENSMUSG00000019883 | Echdc1 | 52665 | protein_coding | 317.0371945 | 0.21966896 | 0.113402062 | 1.937080829 | 0.052735464 | 0.191891882 | 0.21966896 |
| ENSMUSG00000032300 | 1700017B05Rik | 74211 | protein_coding | 205.6164005 | -0.219719441 | 0.0988923 | -2.221805361 | 0.026296462 | 0.118778578 | 0.219719441 |
| ENSMUSG00000034422 | Parp14 | 547253 | protein_coding | 1220.462957 | -0.219731337 | 0.123513086 | -1.779012611 | 0.075237695 | 0.241032638 | 0.219731337 |
| ENSMUSG00000040680 | Kremen2 | 73016 | protein_coding | 142.1043697 | -0.219916689 | 0.121046128 | -1.816800687 | 0.069247638 | 0.228375593 | 0.219916689 |
| ENSMUSG00000001604 | Tcea3 | 21401 | protein_coding | 969.2158949 | -0.219936089 | 0.082119006 | -2.678260469 | 0.007400564 | 0.047758664 | 0.219936089 |
| ENSMUSG00000051246 | Msantd1 | 403174 | protein_coding | 38.90834654 | -0.219976082 | 0.194360538 | -1.131793954 | 0.25772107 | 0.499713206 | 0.219976082 |
| ENSMUSG00000066894 | Vsig10 | 231668 | protein_coding | 4097.140051 | -0.220116992 | 0.044573633 | -4.938278045 | 7.88E-07 | 1.98E-05 | 0.220116992 |
| ENSMUSG00000026965 | Anapc2 | 99152 | protein_coding | 1106.404329 | -0.220168878 | 0.064896802 | -3.392599816 | 0.000692327 | 0.007465035 | 0.220168878 |
| ENSMUSG00000073406 | H2-Bl | 14963 | protein_coding | 32.55375565 | -0.22017982 | 0.244036121 | -0.902242746 | 0.366927932 | 0.609292294 | 0.22017982 |
| ENSMUSG00000042549 | Map2k3os | 24082 | lncRNA | 42.17051673 | -0.22020109 | 0.181855606 | -1.21085676 | 0.225950307 | 0.464143013 | 0.22020109 |
| ENSMUSG00000073700 | Klhl21 | 242785 | protein_coding | 587.4645993 | 0.22035765 | 0.078603863 | 2.803394663 | 0.005056775 | 0.035957058 | 0.22035765 |
| ENSMUSG00000048827 | Pkd1l3 | 244646 | protein_coding | 21.54043844 | -0.220365401 | 0.287190817 | -0.767313536 | 0.442895122 | 0.673954486 | 0.220365401 |
| ENSMUSG00000015966 | Il17rb | 50905 | protein_coding | 44.89615946 | -0.22036706 | 0.182446429 | -1.207845287 | 0.227106778 | 0.465617977 | 0.22036706 |
| ENSMUSG00000043557 | Mdga1 | 74762 | protein_coding | 12.46602155 | 0.220400118 | 0.363253521 | 0.60673911 | 0.544024063 | 0.748781775 | 0.220400118 |
| ENSMUSG00000034171 | Faah | 14073 | protein_coding | 746.0372488 | -0.220616972 | 0.053187021 | -4.147947549 | 3.35E-05 | 0.000573713 | 0.220616972 |
| ENSMUSG00000018395 | Kif3a | 16568 | protein_coding | 142.3775534 | 0.220627219 | 0.101490119 | 2.173878804 | 0.029714238 | 0.129304242 | 0.220627219 |
| ENSMUSG00000027104 | Atf2 | 11909 | protein_coding | 1082.522495 | 0.220642069 | 0.055996702 | 3.94026899 | 8.14E-05 | 0.001235952 | 0.220642069 |
| ENSMUSG00000038793 | Lefty1 | 13590 | protein_coding | 113.7516251 | -0.220650112 | 0.148995051 | -1.480922421 | 0.138627248 | 0.348719642 | 0.220650112 |
| ENSMUSG00000073888 | Ccl27a | 20301 | protein_coding | 37.14740299 | -0.220663833 | 0.221477492 | -0.996326226 | 0.319091665 | 0.562649386 | 0.220663833 |
| ENSMUSG00000086316 | Nbdy | 70123 | protein_coding | 124.7858668 | 0.220686046 | 0.103395179 | 2.134393966 | 0.032810545 | 0.138444705 | 0.220686046 |
| ENSMUSG00000026103 | Gls | 14660 | protein_coding | 2470.236868 | -0.220792358 | 0.066581743 | -3.316109626 | 0.0009128 | 0.009340563 | 0.220792358 |
| ENSMUSG00000040213 | Kyat3 | 229905 | protein_coding | 218.7733844 | 0.220869192 | 0.085346634 | 2.587907477 | 0.009656091 | 0.057911411 | 0.220869192 |
| ENSMUSG00000042632 | Pla2g6 | 53357 | protein_coding | 230.906847 | 0.220879358 | 0.109116301 | 2.024256278 | 0.04294379 | 0.166920824 | 0.220879358 |
| ENSMUSG00000071650 | Ganab | 14376 | protein_coding | 5047.894896 | 0.22094331 | 0.046569808 | 4.744346628 | 2.09E-06 | 4.79E-05 | 0.22094331 |
| ENSMUSG00000061947 | Serpina10 | 217847 | protein_coding | 15.61899764 | -0.220966348 | 0.436367324 | -0.506376935 | 0.612592063 | 0.793452108 | 0.220966348 |
| ENSMUSG00000089235 | Gm23119 | 115486069 | scaRNA | 120.6167252 | -0.220982632 | 0.194806506 | -1.134369876 | 0.256639429 | 0.498533375 | 0.220982632 |
| ENSMUSG00000034135 | Sik3 | 70661 | protein_coding | 618.2618945 | -0.220986455 | 0.098572114 | -2.241875988 | 0.024969388 | 0.114626053 | 0.220986455 |
| ENSMUSG00000027533 | Fabp5 | 16592 | protein_coding | 90.6500411 | -0.221070496 | 0.144277351 | -1.532260567 | 0.125458148 | 0.328615354 | 0.221070496 |
| ENSMUSG00000070390 | Nlrp1b | 637515 | protein_coding | 572.8103606 | -0.221117343 | 0.095194416 | -2.322797411 | 0.020190038 | 0.098877846 | 0.221117343 |
| ENSMUSG00000052456 | Get3 | 56495 | protein_coding | 1153.341302 | 0.221186233 | 0.058347311 | 3.790855646 | 0.000150129 | 0.002065814 | 0.221186233 |
| ENSMUSG00000025747 | Tyms | 22171 | protein_coding | 663.763277 | 0.221206095 | 0.10339387 | 2.139450767 | 0.032399179 | 0.137270396 | 0.221206095 |
| ENSMUSG00000002028 | Kmt2a | 214162 | protein_coding | 772.9287541 | -0.221261451 | 0.091929841 | -2.406851226 | 0.016090723 | 0.084471059 | 0.221261451 |
| ENSMUSG00000027944 | Hax1 | 23897 | protein_coding | 433.5835044 | 0.221262109 | 0.072337694 | 3.058738774 | 0.002222709 | 0.019066821 | 0.221262109 |
| ENSMUSG00000041794 | Myrip | 245049 | protein_coding | 143.4760875 | -0.221272862 | 0.158110894 | -1.399478911 | 0.161669418 | 0.382237605 | 0.221272862 |

| ensembl_gene_id | external_gene_name | entrezgene_id | gene_biotype | baseMean | log2FoldChange | lfcSE | stat | pvalue | padj | absLog2FC |
|---|---|---|---|---|---|---|---|---|---|---|
| ENSMUSG00000023057 | Fabp2 | 14079 | protein_coding | 7485.694777 | 0.221294012 | 0.077735009 | 2.846774106 | 0.004416469 | 0.032496922 | 0.221294012 |
| ENSMUSG00000023021 | Cers5 | 71949 | protein_coding | 735.7166809 | -0.22138991 | 0.080978319 | -2.733940539 | 0.006258134 | 0.042325937 | 0.22138991 |
| ENSMUSG00000036611 | Eepd1 | 67484 | protein_coding | 197.4988584 | -0.22142427 | 0.107562589 | -2.058562113 | 0.039536205 | 0.157238072 | 0.22142427 |
| ENSMUSG00000033439 | Trmt13 | 229780 | protein_coding | 79.99637929 | -0.221479859 | 0.168843163 | -1.311749057 | 0.189604819 | 0.420211186 | 0.221479859 |
| ENSMUSG00000004937 | Sgta | 52551 | protein_coding | 1656.600037 | 0.221504459 | 0.066509468 | 3.330419964 | 0.000867151 | 0.00893581 | 0.221504459 |
| ENSMUSG00000031570 | Plpp5 | 71910 | protein_coding | 1152.545022 | 0.221510715 | 0.063387541 | 3.494546604 | 0.000474868 | 0.005414954 | 0.221510715 |
| ENSMUSG00000019762 | Iyd | 70337 | protein_coding | 229.5610574 | -0.22155155 | 0.114183434 | -1.940312551 | 0.052341717 | 0.190915661 | 0.22155155 |
| ENSMUSG00000110206 | Flt3l | 14256 | protein_coding | 22.92476283 | -0.221623654 | 0.2610435 | -0.848991275 | 0.395886147 | 0.635262971 | 0.221623654 |
| ENSMUSG00000029136 | Rbks | 71336 | protein_coding | 187.9294933 | 0.221633914 | 0.092813602 | 2.387946475 | 0.016942809 | 0.087561384 | 0.221633914 |
| ENSMUSG00000004319 | Clcn3 | 12725 | protein_coding | 3129.805918 | 0.22168408 | 0.040358597 | 5.492858885 | 3.95E-08 | 1.30E-06 | 0.22168408 |
| ENSMUSG00000036620 | Mgat4b | 103534 | protein_coding | 5466.527747 | 0.221693983 | 0.062438223 | 3.550613275 | 0.000384335 | 0.004537457 | 0.221693983 |
| ENSMUSG00000032042 | Srpr | 67398 | protein_coding | 2550.999011 | 0.221713127 | 0.038035278 | 5.82914437 | 5.57E-09 | 2.06E-07 | 0.221713127 |
| ENSMUSG00000035397 | Klf16 | 118445 | protein_coding | 485.353279 | -0.221726392 | 0.113950316 | -1.945816388 | 0.051676799 | 0.189222442 | 0.221726392 |
| ENSMUSG00000054074 | Skida1 | 72668 | protein_coding | 43.92321751 | -0.221829109 | 0.219266773 | -1.011685926 | 0.311688247 | 0.555505998 | 0.221829109 |
| ENSMUSG00000059182 | Skap2 | 54353 | protein_coding | 1140.971711 | 0.221930621 | 0.085631727 | 2.591686853 | 0.009550666 | 0.057517249 | 0.221930621 |
| ENSMUSG00000042675 | Ypel3 | 66090 | protein_coding | 639.9209369 | -0.221964276 | 0.128057288 | -1.733320129 | 0.083038785 | 0.256348065 | 0.221964276 |
| ENSMUSG00000044442 | N6amt1 | 67768 | protein_coding | 227.3672764 | -0.222008507 | 0.106371507 | -2.087105021 | 0.036878642 | 0.149582964 | 0.222008507 |
| ENSMUSG00000091243 | Vgll3 | 73569 | protein_coding | 9.396133397 | 0.222021448 | 0.459088244 | 0.483613883 | 0.62865992 | 0.804315631 | 0.222021448 |
| ENSMUSG00000096146 | Kcnj11 | 16514 | protein_coding | 13.0445749 | -0.222113906 | 0.318535947 | -0.697296204 | 0.485617443 | 0.706867216 | 0.222113906 |
| ENSMUSG00000026281 | Dtymk | 21915 | protein_coding | 348.1630317 | 0.222127868 | 0.104066772 | 2.13447447 | 0.032803961 | 0.138444705 | 0.222127868 |
| ENSMUSG00000107456 | Gm10400 | 100093700 | TEC | 15.20489115 | -0.222146779 | 0.353122694 | -0.629092331 | 0.529288611 | 0.73857118 | 0.222146779 |
| ENSMUSG00000061482 | H4c4 | 319156 | protein_coding | 5531.109375 | 0.222165539 | 0.102704286 | 2.163157423 | 0.030529074 | 0.131853876 | 0.222165539 |
| ENSMUSG00000026955 | Sapcd2 | 72080 | protein_coding | 153.8877631 | 0.222177187 | 0.112268749 | 1.97897624 | 0.047818681 | 0.179562869 | 0.222177187 |
| ENSMUSG00000022463 | Srebf2 | 20788 | protein_coding | 3883.158333 | -0.222191538 | 0.07157122 | -3.104481636 | 0.001906128 | 0.016899461 | 0.222191538 |
| ENSMUSG00000024503 | Spink1 | 20730 | protein_coding | 5629.939879 | 0.22221153 | 0.113522838 | 1.957416978 | 0.050298464 | 0.185896496 | 0.22221153 |
| ENSMUSG00000024188 | Luc7l | 66978 | protein_coding | 760.5335234 | -0.222257825 | 0.105096276 | -2.114802097 | 0.034446818 | 0.143118696 | 0.222257825 |
| ENSMUSG00000012519 | Mlkl | 74568 | protein_coding | 380.7108342 | -0.222292389 | 0.096203525 | -2.310647017 | 0.02085236 | 0.101260467 | 0.222292389 |
| ENSMUSG00000056687 | Gm11696 | 100036768 | lncRNA | 53.21110188 | 0.222341246 | 0.188876073 | 1.177180584 | 0.239123437 | 0.479465829 | 0.222341246 |
| ENSMUSG00000085315 | A430018G15Rik | 115489452 | lncRNA | 21.30708031 | -0.222383728 | 0.268496468 | -0.82825569 | 0.407525711 | 0.646183716 | 0.222383728 |
| ENSMUSG00000023495 | Pcbp4 | 59092 | protein_coding | 71.36056112 | -0.222418462 | 0.167312377 | -1.329360477 | 0.183729071 | 0.412214625 | 0.222418462 |
| ENSMUSG00000022829 | Stxbp5l | 207227 | protein_coding | 6.114792493 | 0.222423469 | 0.474076055 | 0.469172545 | 0.638946308 | 0.81200056 | 0.222423469 |
| ENSMUSG00000027936 | Crtc2 | 74343 | protein_coding | 234.5567692 | -0.222427181 | 0.119314928 | -1.864202451 | 0.062293279 | 0.213110674 | 0.222427181 |
| ENSMUSG00000042414 | Prdm14 | 383491 | protein_coding | 38.39841211 | 0.222454905 | 0.222015975 | 1.00197702 | 0.316354692 | 0.560212157 | 0.222454905 |
| ENSMUSG00000021509 | Slc25a48 | 328258 | protein_coding | 13.89702622 | 0.222504868 | 0.321943276 | 0.691130655 | 0.489483437 | 0.70956399 | 0.222504868 |
| ENSMUSG00000028378 | Ptgr1 | 67103 | protein_coding | 438.6391908 | 0.222551426 | 0.084094079 | 2.646457743 | 0.008133967 | 0.051260837 | 0.222551426 |
| ENSMUSG00000071451 | Psmg4 | 69666 | protein_coding | 127.1735824 | -0.22256735 | 0.113531225 | -1.9604065 | 0.049948297 | 0.185048514 | 0.22256735 |
| ENSMUSG00000108171 | Gm43915 | 215458 | lncRNA | 6.04449081 | -0.222581945 | 0.477548341 | -0.466093012 | 0.641148925 | 0.813517602 | 0.222581945 |
| ENSMUSG00000045934 | Mtmr11 | 194126 | protein_coding | 932.6962301 | 0.222614235 | 0.077622907 | 2.867893579 | 0.004132145 | 0.030908674 | 0.222614235 |
| ENSMUSG00000062980 | Cped1 | 214642 | protein_coding | 19.5923927 | -0.2226663 | 0.311629067 | -0.714523527 | 0.474903514 | 0.698573273 | 0.2226663 |

| ensembl_gene_id | external_gene_name | entrezgene_id | gene_biotype | baseMean | log2FoldChange | lfcSE | stat | pvalue | padj | absLog2FC |
|---|---|---|---|---|---|---|---|---|---|---|
| ENSMUSG00000037492 | Zmat4 | 320158 | protein_coding | 49.88283932 | -0.222724213 | 0.278624392 | -0.799370836 | 0.424075416 | 0.659950229 | 0.222724213 |
| ENSMUSG00000060904 | Arl1 | 104303 | protein_coding | 3916.825685 | 0.222792705 | 0.060883471 | 3.659329868 | 0.000252876 | 0.003211123 | 0.222792705 |
| ENSMUSG00000030336 | Cd27 | 21940 | protein_coding | 9.323585755 | 0.222824935 | 0.404313367 | 0.551119387 | 0.581551836 | 0.773321692 | 0.222824935 |
| ENSMUSG00000037390 | Muc3 | 666339 | protein_coding | 12096.5337 | -0.222868563 | 0.08711809 | -2.558235186 | 0.010520491 | 0.061552292 | 0.222868563 |
| ENSMUSG00000084803 | 5830444B04Rik | 641454 | lncRNA | 90.62900427 | -0.22293807 | 0.329584168 | -0.676422268 | 0.498772583 | 0.716014372 | 0.22293807 |
| ENSMUSG00000022174 | Dad1 | 13135 | protein_coding | 1467.1035 | 0.222948513 | 0.073091644 | 3.050259949 | 0.002286434 | 0.019489068 | 0.222948513 |
| ENSMUSG00000028780 | Sema3c | 20348 | protein_coding | 3561.617573 | -0.222957482 | 0.048903095 | -4.559169161 | 5.14E-06 | 0.000108406 | 0.222957482 |
| ENSMUSG00000060981 | H4c8 | 69386 | protein_coding | 5146.361897 | 0.223017717 | 0.09069027 | 2.45911406 | 0.013928036 | 0.07606415 | 0.223017717 |
| ENSMUSG00000027384 | Ndufaf5 | 69487 | protein_coding | 294.2042734 | 0.22313302 | 0.081141363 | 2.749929406 | 0.005960811 | 0.04081985 | 0.22313302 |
| ENSMUSG00000002797 | Ggct | 110175 | protein_coding | 74.10797427 | 0.223180162 | 0.148833952 | 1.499524529 | 0.13373761 | 0.34107542 | 0.223180162 |
| ENSMUSG00000029084 | Cd38 | 12494 | protein_coding | 3555.385055 | -0.223183571 | 0.057748795 | -3.864731195 | 0.000111212 | 0.001627504 | 0.223183571 |
| ENSMUSG00000113949 | Scamp4 | 56214 | protein_coding | 9.291493479 | -0.223185651 | 0.394301769 | -0.566027517 | 0.571375075 | 0.766284061 | 0.223185651 |
| ENSMUSG00000025198 | Erlin1 | 226144 | protein_coding | 1256.040488 | -0.223218497 | 0.057920167 | -3.853899379 | 0.000116251 | 0.001687476 | 0.223218497 |
| ENSMUSG00000032113 | Chek1 | 12649 | protein_coding | 98.30533534 | 0.223297391 | 0.182202436 | 1.225545584 | 0.220369726 | 0.456836916 | 0.223297391 |
| ENSMUSG00000069274 | H4c6 | 319157 | protein_coding | 2299.613099 | 0.223372487 | 0.149297571 | 1.4961562 | 0.134612956 | 0.342321275 | 0.223372487 |
| ENSMUSG00000068417 | Pnp2 | 667034 | protein_coding | 422.1553902 | 0.223439768 | 0.087418473 | 2.555978854 | 0.010588955 | 0.061835229 | 0.223439768 |
| ENSMUSG00000020952 | Scfd1 | 76983 | protein_coding | 1261.171897 | 0.223525067 | 0.05186012 | 4.31015329 | 1.63E-05 | 0.000301734 | 0.223525067 |
| ENSMUSG00000025981 | Coq10b | 67876 | protein_coding | 660.397284 | -0.223629403 | 0.09792206 | -2.283748959 | 0.022386289 | 0.1064025 | 0.223629403 |
| ENSMUSG00000036943 | Rab8b | 235442 | protein_coding | 266.8766482 | 0.223709821 | 0.092035478 | 2.430691149 | 0.015070055 | 0.080342073 | 0.223709821 |
| ENSMUSG00000031732 | Phlpp2 | 244650 | protein_coding | 1108.864724 | -0.223790433 | 0.109508967 | -2.043580903 | 0.040994967 | 0.16134834 | 0.223790433 |
| ENSMUSG00000029433 | Diablo | 66593 | protein_coding | 113.1994178 | -0.2238795 | 0.129738386 | -1.725622664 | 0.084415319 | 0.258962499 | 0.2238795 |
| ENSMUSG00000022186 | Oxct1 | 67041 | protein_coding | 5468.845936 | -0.2239008 | 0.073710483 | -3.037570657 | 0.002384935 | 0.020074032 | 0.2239008 |
| ENSMUSG00000095990 | Zfp97 | 22759 | protein_coding | 107.5431992 | -0.224044943 | 0.132363819 | -1.692644907 | 0.090523078 | 0.270847064 | 0.224044943 |
| ENSMUSG00000074884 | Serf2 | 378702 | protein_coding | 2207.051605 | 0.224139471 | 0.057136335 | 3.922888502 | 8.75E-05 | 0.001316241 | 0.224139471 |
| ENSMUSG00000034998 | Foxn2 | 14236 | protein_coding | 591.3261944 | -0.224245952 | 0.07527728 | -2.978932718 | 0.002892543 | 0.023394202 | 0.224245952 |
| ENSMUSG00000051457 | Spn | 20737 | protein_coding | 28.81665821 | 0.224365608 | 0.297098965 | 0.755188118 | 0.450136117 | 0.679535938 | 0.224365608 |
| ENSMUSG00000049285 | Mblac1 | 330216 | protein_coding | 67.99468409 | 0.224367585 | 0.163999244 | 1.368101342 | 0.171280357 | 0.395906308 | 0.224367585 |
| ENSMUSG00000046805 | Mpeg1 | 17476 | protein_coding | 12.48620921 | -0.224418905 | 0.354660398 | -0.632771254 | 0.526883031 | 0.737165628 | 0.224418905 |
| ENSMUSG00000029402 | Snrnp35 | 76167 | protein_coding | 149.8439351 | 0.224447128 | 0.104027707 | 2.157570651 | 0.030961229 | 0.13300792 | 0.224447128 |
| ENSMUSG00000001089 | Luzp1 | 269593 | protein_coding | 4008.17933 | -0.224517545 | 0.03873613 | -5.796075731 | 6.79E-09 | 2.45E-07 | 0.224517545 |
| ENSMUSG00000064427 | Gm22748 | 115488064 | snoRNA | 142.949552 | -0.224532449 | 0.194576984 | -1.153951738 | 0.248519956 | 0.4893628 | 0.224532449 |
| ENSMUSG00000019461 | Plscr3 | 70310 | protein_coding | 211.5264814 | -0.224620075 | 0.112694245 | -1.993181412 | 0.046241586 | 0.175509073 | 0.224620075 |
| ENSMUSG00000055917 | Zfp277 | 246196 | protein_coding | 551.4268667 | -0.224622333 | 0.078697176 | -2.854261676 | 0.004313698 | 0.031970135 | 0.224622333 |
| ENSMUSG00000086013 | Gm15706 | 100039495 | lncRNA | 73.49924782 | -0.224637099 | 0.172066275 | -1.305526596 | 0.191713597 | 0.423163427 | 0.224637099 |
| ENSMUSG00000027865 | Gdap2 | 14547 | protein_coding | 440.3510967 | 0.224643552 | 0.070462294 | 3.188138493 | 0.001431919 | 0.013462227 | 0.224643552 |
| ENSMUSG00000026082 | Rev1 | 56210 | protein_coding | 218.5421714 | -0.224690559 | 0.086375601 | -2.601319764 | 0.009286585 | 0.056432441 | 0.224690559 |
| ENSMUSG00000048895 | Cdk5r1 | 12569 | protein_coding | 17.1346574 | -0.224810546 | 0.29099837 | -0.772549159 | 0.439789243 | 0.671740242 | 0.224810546 |
| ENSMUSG00000021071 | Trim9 | 94090 | protein_coding | 14.85773072 | -0.224953916 | 0.40166708 | -0.560050668 | 0.575444878 | 0.768546546 | 0.224953916 |
| ENSMUSG00000043091 | Tuba1c | 22146 | protein_coding | 2626.906851 | 0.224979755 | 0.082447889 | 2.728750934 | 0.00635747 | 0.042790188 | 0.224979755 |

Supplementary Table 1

| ensembl_gene_id | external_gene_name | entrezgene_id | gene_biotype | baseMean | log2FoldChange | lfcSE | stat | pvalue | padj | absLog2FC |
|---|---|---|---|---|---|---|---|---|---|---|
| ENSMUSG00000037129 | Tmprss13 | 214531 | protein_coding | 153.9203908 | -0.225193824 | 0.199812922 | -1.127023328 | 0.259732618 | 0.50123327 | 0.225193824 |
| ENSMUSG00000074305 | Peak1 | 244895 | protein_coding | 545.7457984 | -0.225255214 | 0.070471341 | -3.196408786 | 0.001391498 | 0.013138578 | 0.225255214 |
| ENSMUSG00000036192 | Rorb | 225998 | protein_coding | 14.18250552 | 0.225407099 | 0.369297083 | 0.610367939 | 0.541618101 | 0.74719957 | 0.225407099 |
| ENSMUSG00000024327 | Slc39a7 | 14977 | protein_coding | 352.4155045 | 0.225423134 | 0.112132635 | 2.010325845 | 0.044396713 | 0.170791501 | 0.225423134 |
| ENSMUSG00000056214 | Pard6g | 93737 | protein_coding | 21.60347046 | 0.225484376 | 0.333257301 | 0.67660746 | 0.498655044 | 0.716014372 | 0.225484376 |
| ENSMUSG00000040044 | Orc3 | 50793 | protein_coding | 609.108708 | -0.225508915 | 0.07129371 | -3.163096912 | 0.001561004 | 0.014456304 | 0.225508915 |
| ENSMUSG00000113722 | Snhg10 | 69434 | lncRNA | 10.99057031 | -0.225536466 | 0.356532484 | -0.632583219 | 0.527005848 | 0.737165628 | 0.225536466 |
| ENSMUSG00000045136 | Tubb2b | 73710 | protein_coding | 20.75921078 | -0.22555687 | 0.28832696 | -0.782295455 | 0.434040957 | 0.668045287 | 0.22555687 |
| ENSMUSG00000073096 | Lrrc61 | 243371 | protein_coding | 134.2280802 | -0.225568743 | 0.120521283 | -1.871609207 | 0.061260689 | 0.210875374 | 0.225568743 |
| ENSMUSG00000077192 | Snora17 | 100306954 | snoRNA | 48.07276185 | 0.225578272 | 0.26011234 | 0.867234027 | 0.385813796 | 0.625831733 | 0.225578272 |
| ENSMUSG00000025193 | Cutc | 66388 | protein_coding | 287.3618884 | -0.225596951 | 0.092143703 | -2.448316522 | 0.014352551 | 0.077646289 | 0.225596951 |
| ENSMUSG00000021738 | Atxn7 | 246103 | protein_coding | 902.5139677 | -0.22560864 | 0.057688852 | -3.910783973 | 9.20E-05 | 0.001381412 | 0.22560864 |
| ENSMUSG00000025511 | Tspan4 | 64540 | protein_coding | 20.89756951 | 0.225681583 | 0.272507653 | 0.828166038 | 0.407576475 | 0.646184212 | 0.225681583 |
| ENSMUSG00000029247 | Paics | 67054 | protein_coding | 1899.746535 | -0.225683598 | 0.074031212 | -3.048492556 | 0.002299926 | 0.019554576 | 0.225683598 |
| ENSMUSG00000036006 | Ripor2 | 193385 | protein_coding | 57.2107036 | 0.225726923 | 0.172311288 | 1.309994988 | 0.190197531 | 0.420872061 | 0.225726923 |
| ENSMUSG00000024298 | Zfp871 | 208292 | protein_coding | 1409.754772 | -0.225798239 | 0.06193674 | -3.645626777 | 0.000266741 | 0.003350257 | 0.225798239 |
| ENSMUSG00000027173 | Depdc7 | 211896 | protein_coding | 29.30145361 | -0.225804932 | 0.28085568 | -0.803989196 | 0.421403215 | 0.65807782 | 0.225804932 |
| ENSMUSG00000001768 | Rin2 | 74030 | protein_coding | 2151.999164 | -0.225855441 | 0.068213658 | -3.311000268 | 0.000929631 | 0.009476714 | 0.225855441 |
| ENSMUSG00000118087 | 4833438C02Rik | 100503392 | lncRNA | 176.2465959 | -0.225911537 | 0.093989052 | -2.403594152 | 0.016234786 | 0.084969221 | 0.225911537 |
| ENSMUSG00000004071 | Cdip1 | 66626 | protein_coding | 722.6371215 | -0.225947667 | 0.066896483 | -3.377571681 | 0.000731289 | 0.007789628 | 0.225947667 |
| ENSMUSG00000026768 | Itga8 | 241226 | protein_coding | 8.866711201 | -0.225964506 | 0.457366326 | -0.494055845 | 0.621266741 | 0.80018918 | 0.225964506 |
| ENSMUSG00000032530 | Lyzl4 | 69032 | protein_coding | 28.27343119 | -0.226004501 | 0.248906228 | -0.907990541 | 0.363883219 | 0.60618049 | 0.226004501 |
| ENSMUSG00000071203 | Naip5 | 17951 | protein_coding | 1197.380684 | -0.226068976 | 0.088302144 | -2.560175402 | 0.010461935 | 0.061397241 | 0.226068976 |
| ENSMUSG00000027999 | Pla2g12a | 66350 | protein_coding | 166.0409995 | -0.226243939 | 0.103662333 | -2.182508662 | 0.029072013 | 0.127299579 | 0.226243939 |
| ENSMUSG00000065400 | Mir200a | 387242 | miRNA | 14.66879009 | -0.226290599 | 0.383780714 | -0.589635151 | 0.55543528 | 0.75524772 | 0.226290599 |
| ENSMUSG00000034342 | Cbl | 12402 | protein_coding | 1458.528576 | -0.226293082 | 0.078932462 | -2.866920373 | 0.004144873 | 0.030989506 | 0.226293082 |
| ENSMUSG00000044505 | Lingo4 | 320747 | protein_coding | 9.903577774 | 0.226308464 | 0.366904881 | 0.616804178 | 0.537363896 | 0.743740778 | 0.226308464 |
| ENSMUSG00000005674 | Tomm40l | 641376 | protein_coding | 254.8710811 | 0.226313843 | 0.133180733 | 1.699298678 | 0.089262922 | 0.268368093 | 0.226313843 |
| ENSMUSG00000038530 | Rgs4 | 19736 | protein_coding | 6.793858366 | -0.226313913 | 0.517124666 | -0.437638983 | 0.661648008 | 0.827034368 | 0.226313913 |
| ENSMUSG00000020718 | Polg2 | 50776 | protein_coding | 281.1102334 | -0.226319055 | 0.079888524 | -2.832935754 | 0.004612265 | 0.033565887 | 0.226319055 |
| ENSMUSG00000040441 | Slc26a10 | 216441 | protein_coding | 31.49670801 | 0.226334457 | 0.245176507 | 0.923149039 | 0.355929542 | 0.598619525 | 0.226334457 |
| ENSMUSG00000020601 | Trib2 | 217410 | protein_coding | 74.24537714 | 0.226348963 | 0.161231663 | 1.403874145 | 0.16035633 | 0.380751845 | 0.226348963 |
| ENSMUSG00000020263 | Appl2 | 216190 | protein_coding | 2977.479762 | -0.226362148 | 0.081489824 | -2.777796495 | 0.005472888 | 0.038241451 | 0.226362148 |
| ENSMUSG00000003348 | Mob3a | 208228 | protein_coding | 692.8384182 | 0.226370724 | 0.072571443 | 3.119280992 | 0.00181293 | 0.016251873 | 0.226370724 |
| ENSMUSG00000037683 | Armc3 | 70882 | protein_coding | 11.7366086 | -0.226435945 | 0.414816337 | -0.545870365 | 0.58515505 | 0.775860848 | 0.226435945 |
| ENSMUSG00000085929 | Gm13421 | 105244102 | lncRNA | 26.21407075 | 0.226498146 | 0.242413056 | 0.934347967 | 0.350124429 | 0.592368513 | 0.226498146 |
| ENSMUSG00000091764 | Zfp964 | 636741 | protein_coding | 63.19555006 | -0.226502236 | 0.168326106 | -1.345615613 | 0.178426514 | 0.405619009 | 0.226502236 |
| ENSMUSG00000085396 | Firre | 103012 | lncRNA | 995.4316357 | -0.226541615 | 0.147384008 | -1.537084092 | 0.124272718 | 0.327154117 | 0.226541615 |
| ENSMUSG00000078612 | Fyb2 | 242594 | protein_coding | 77.55378189 | -0.226570098 | 0.155780323 | -1.454420519 | 0.145829754 | 0.358583208 | 0.226570098 |

| ensembl_gene_id | external_gene_name | entrezgene_id | gene_biotype | baseMean | log2FoldChange | lfcSE | stat | pvalue | padj | absLog2FC |
|---|---|---|---|---|---|---|---|---|---|---|
| ENSMUSG00000002233 | Rhoc | 11853 | protein_coding | 1813.999626 | -0.226667589 | 0.115427216 | -1.963727411 | 0.049561712 | 0.184123872 | 0.226667589 |
| ENSMUSG00000047238 | Mageh1 | 75625 | protein_coding | 110.321803 | 0.226788289 | 0.125159232 | 1.81199809 | 0.069986492 | 0.229825321 | 0.226788289 |
| ENSMUSG00000053198 | Prx | 19153 | protein_coding | 78.14149764 | 0.226808572 | 0.13263524 | 1.710017422 | 0.087262651 | 0.264527214 | 0.226808572 |
| ENSMUSG00000040283 | Btnl9 | 237754 | protein_coding | 6.618457714 | -0.2268161 | 0.460639374 | -0.492394078 | 0.622440785 | 0.800805946 | 0.2268161 |
| ENSMUSG00000036504 | Phpt1 | 75454 | protein_coding | 33.8778889 | 0.226817993 | 0.2126417 | 1.066667513 | 0.286122002 | 0.528797792 | 0.226817993 |
| ENSMUSG00000022941 | Ripply3 | 170765 | protein_coding | 6.759668435 | 0.226941819 | 0.446068982 | 0.50875947 | 0.610920832 | 0.792706869 | 0.226941819 |
| ENSMUSG00000017418 | Arl5b | 75869 | protein_coding | 846.6000463 | -0.227095492 | 0.066539604 | -3.41293725 | 0.000642667 | 0.007036183 | 0.227095492 |
| ENSMUSG00000030180 | Kdm5a | 214899 | protein_coding | 1636.311277 | -0.227111174 | 0.070937489 | -3.201567696 | 0.00136682 | 0.012951057 | 0.227111174 |
| ENSMUSG00000058748 | Zfp958 | 233987 | protein_coding | 238.7225731 | -0.227112089 | 0.087689171 | -2.589967347 | 0.009598504 | 0.05763033 | 0.227112089 |
| ENSMUSG00000048677 | Tpcn2 | 233979 | protein_coding | 257.5214011 | -0.227113096 | 0.088039212 | -2.579681156 | 0.009889157 | 0.058784535 | 0.227113096 |
| ENSMUSG00000000440 | Pparg | 19016 | protein_coding | 1718.325297 | -0.227188786 | 0.061787645 | -3.67692903 | 0.000236059 | 0.003024955 | 0.227188786 |
| ENSMUSG00000026637 | Traf5 | 22033 | protein_coding | 48.51232824 | -0.227193608 | 0.226706186 | -1.002150017 | 0.316271144 | 0.560212157 | 0.227193608 |
| ENSMUSG00000039994 | Timeless | 21853 | protein_coding | 165.1708452 | 0.227210272 | 0.192170091 | 1.182339407 | 0.237071055 | 0.477153008 | 0.227210272 |
| ENSMUSG00000069310 | H3c3 | 319148 | protein_coding | 1243.745756 | 0.227234845 | 0.136654806 | 1.662838297 | 0.096344808 | 0.28168106 | 0.227234845 |
| ENSMUSG00000024480 | Ap3s1 | 11777 | protein_coding | 682.3976819 | 0.22727805 | 0.067437609 | 3.370197306 | 0.000751144 | 0.007948619 | 0.22727805 |
| ENSMUSG00000037243 | Zfp692 | 103836 | protein_coding | 300.8594753 | -0.227338856 | 0.097433951 | -2.333261194 | 0.019634441 | 0.096777726 | 0.227338856 |
| ENSMUSG00000050310 | Rictor | 78757 | protein_coding | 1357.147504 | -0.227379667 | 0.060371537 | -3.766338878 | 0.000165659 | 0.002243141 | 0.227379667 |
| ENSMUSG00000022221 | Ripk3 | 56532 | protein_coding | 302.6684423 | -0.227399809 | 0.115636571 | -1.966504253 | 0.049240392 | 0.183499374 | 0.227399809 |
| ENSMUSG00000040570 | Rundc3b | 242819 | protein_coding | 592.6194396 | -0.227452939 | 0.056692358 | -4.012056402 | 6.02E-05 | 0.000962527 | 0.227452939 |
| ENSMUSG00000014704 | Hoxa2 | 15399 | protein_coding | 32.23096802 | -0.227565028 | 0.205290951 | -1.108500043 | 0.26764592 | 0.51014369 | 0.227565028 |
| ENSMUSG00000030245 | Golt1b | 66964 | protein_coding | 608.0153795 | 0.227590956 | 0.094787637 | 2.401061618 | 0.016347584 | 0.085418361 | 0.227590956 |
| ENSMUSG00000108714 | Gm21057 | 100859962 | lncRNA | 7.847289258 | -0.227592434 | 0.468493181 | -0.485796686 | 0.627111324 | 0.803226458 | 0.227592434 |
| ENSMUSG00000005410 | Mcm5 | 17218 | protein_coding | 880.0068629 | 0.227633673 | 0.16922746 | 1.345134375 | 0.178581844 | 0.405715197 | 0.227633673 |
| ENSMUSG00000020140 | Lgr5 | 14160 | protein_coding | 308.355566 | 0.2277701 | 0.099289796 | 2.293993022 | 0.0217909 | 0.10455009 | 0.2277701 |
| ENSMUSG00000032501 | Trib1 | 211770 | protein_coding | 2973.44657 | 0.227773127 | 0.051771064 | 4.399622288 | 1.08E-05 | 0.000210953 | 0.227773127 |
| ENSMUSG00000002057 | Foxn1 | 15218 | protein_coding | 20.9538999 | 0.227883614 | 0.262666074 | 0.86757917 | 0.385624754 | 0.625831733 | 0.227883614 |
| ENSMUSG00000028182 | Lrriq3 | 74435 | protein_coding | 30.9341856 | -0.227903643 | 0.219674942 | -1.037458535 | 0.299522211 | 0.543107116 | 0.227903643 |
| ENSMUSG00000025257 | Ribc1 | 66611 | protein_coding | 57.33340629 | -0.227926375 | 0.173819395 | -1.311282757 | 0.189762252 | 0.42043505 | 0.227926375 |
| ENSMUSG00000027016 | Zfp385b | 241494 | protein_coding | 57.52992262 | -0.227954447 | 0.251195729 | -0.907477399 | 0.364154396 | 0.606340896 | 0.227954447 |
| ENSMUSG00000029073 | Cptp | 79554 | protein_coding | 672.0049425 | 0.228018254 | 0.059259111 | 3.847817639 | 0.000119175 | 0.001717835 | 0.228018254 |
| ENSMUSG00000021676 | Iqgap2 | 544963 | protein_coding | 10553.78924 | 0.22815632 | 0.081394881 | 2.803079481 | 0.005061719 | 0.035960506 | 0.22815632 |
| ENSMUSG00000022793 | B4galt4 | 56375 | protein_coding | 1101.295913 | 0.228167922 | 0.052525825 | 4.343918819 | 1.40E-05 | 0.000264308 | 0.228167922 |
| ENSMUSG00000032565 | Nudt16 | 75686 | protein_coding | 518.7531842 | 0.228214767 | 0.067478363 | 3.382043641 | 0.000719487 | 0.007714771 | 0.228214767 |
| ENSMUSG00000049649 | Gpr3 | 14748 | protein_coding | 6.596493654 | -0.228219312 | 0.446143333 | -0.511538098 | 0.608974318 | 0.791308423 | 0.228219312 |
| ENSMUSG00000031226 | Pbdc1 | 67683 | protein_coding | 396.1224396 | 0.228269608 | 0.074037384 | 3.08316685 | 0.002048103 | 0.017882921 | 0.228269608 |
| ENSMUSG00000040586 | Ofd1 | 237222 | protein_coding | 264.4725001 | -0.228293745 | 0.116523948 | -1.959200232 | 0.050089342 | 0.185400693 | 0.228293745 |
| ENSMUSG00000026971 | Itgb6 | 16420 | protein_coding | 860.9548642 | 0.228321909 | 0.058395279 | 3.909937822 | 9.23E-05 | 0.00138497 | 0.228321909 |
| ENSMUSG00000039238 | Zfp750 | 319530 | protein_coding | 146.114528 | 0.228345639 | 0.158731027 | 1.438569656 | 0.150272489 | 0.365794955 | 0.228345639 |
| ENSMUSG00000026622 | Nek2 | 18005 | protein_coding | 661.0566589 | 0.228350998 | 0.117835639 | 1.9378772 | 0.052638206 | 0.191597869 | 0.228350998 |

Supplementary Table 1

| ensembl_gene_id | external_gene_name | entrezgene_id | gene_biotype | baseMean | log2FoldChange | lfcSE | stat | pvalue | padj | absLog2FC |
|---|---|---|---|---|---|---|---|---|---|---|
| ENSMUSG00000056131 | Pgm3 | 109785 | protein_coding | 1037.266684 | 0.228396137 | 0.07351156 | 3.106941767 | 0.001890336 | 0.016796393 | 0.228396137 |
| ENSMUSG00000054385 | Ceacam2 | 26367 | protein_coding | 126.367905 | -0.228435017 | 0.12536518 | -1.822156816 | 0.068431195 | 0.226749513 | 0.228435017 |
| ENSMUSG00000014294 | Ndufa2 | 17991 | protein_coding | 2078.98973 | 0.228517155 | 0.107673739 | 2.12231095 | 0.033811635 | 0.141484232 | 0.228517155 |
| ENSMUSG00000054115 | Skp2 | 27401 | protein_coding | 91.48682469 | 0.228624329 | 0.158430606 | 1.44305658 | 0.149004533 | 0.363539502 | 0.228624329 |
| ENSMUSG00000025871 | 4833439L19Rik | 97820 | protein_coding | 4050.575358 | 0.2286912 | 0.038995989 | 5.864480038 | 4.51E-09 | 1.68E-07 | 0.2286912 |
| ENSMUSG00000015747 | Vps45 | 22365 | protein_coding | 392.8594159 | 0.228691262 | 0.080350149 | 2.846183422 | 0.00442467 | 0.032538374 | 0.228691262 |
| ENSMUSG00000033061 | Resp18 | 19711 | protein_coding | 65.36683418 | -0.2287646 | 0.166935333 | -1.370378558 | 0.170568762 | 0.394997476 | 0.2287646 |
| ENSMUSG00000027761 | Aadac | 67758 | protein_coding | 1455.55565 | 0.2288404 | 0.080677238 | 2.836492744 | 0.004561201 | 0.033299455 | 0.2288404 |
| ENSMUSG00000032834 | Pwp2 | 110816 | protein_coding | 356.2902196 | -0.228933365 | 0.074135402 | -3.088043757 | 0.002014788 | 0.017639786 | 0.228933365 |
| ENSMUSG00000057113 | Npm1 | 18148 | protein_coding | 5543.057083 | -0.228946325 | 0.079026914 | -2.897067765 | 0.003766684 | 0.02891257 | 0.228946325 |
| ENSMUSG00000085427 | 6430710C18Rik | 76218 | lncRNA | 9.021038209 | -0.228984369 | 0.420674253 | -0.544327035 | 0.586216446 | 0.776502064 | 0.228984369 |
| ENSMUSG00000029049 | Morn1 | 76866 | protein_coding | 67.82770419 | -0.229059865 | 0.140611565 | -1.629025779 | 0.103307562 | 0.293679015 | 0.229059865 |
| ENSMUSG00000063568 | Jazf1 | 231986 | protein_coding | 11.60512732 | -0.229386752 | 0.411223892 | -0.557814748 | 0.576970891 | 0.769628582 | 0.229386752 |
| ENSMUSG00000043015 | Nemp2 | 227094 | protein_coding | 81.26579097 | -0.229397766 | 0.182631327 | -1.256070194 | 0.209090522 | 0.444592741 | 0.229397766 |
| ENSMUSG00000041132 | N4bp2l1 | 100637 | protein_coding | 38.23740509 | -0.22942416 | 0.215410967 | -1.065053294 | 0.286851814 | 0.529236838 | 0.22942416 |
| ENSMUSG00000024583 | Txnl1 | 53382 | protein_coding | 1498.486749 | 0.229451105 | 0.063401646 | 3.619008648 | 0.000295734 | 0.003635136 | 0.229451105 |
| ENSMUSG00000004815 | Dgkq | 110524 | protein_coding | 892.7160764 | -0.229466278 | 0.091268815 | -2.51418052 | 0.011930935 | 0.067773928 | 0.229466278 |
| ENSMUSG00000029174 | Tbc1d1 | 57915 | protein_coding | 899.959542 | 0.229516999 | 0.072980332 | 3.144915812 | 0.001661346 | 0.015154147 | 0.229516999 |
| ENSMUSG00000002845 | Tmem39a | 67846 | protein_coding | 508.1993679 | 0.229540044 | 0.071140544 | 3.226571381 | 0.00125283 | 0.01215667 | 0.229540044 |
| ENSMUSG00000038615 | Nfe2l1 | 18023 | protein_coding | 4753.103122 | 0.229591655 | 0.04396556 | 5.222079637 | 1.77E-07 | 5.10E-06 | 0.229591655 |
| ENSMUSG00000032883 | Acsl3 | 74205 | protein_coding | 3906.460145 | -0.229603927 | 0.054248495 | -4.232447839 | 2.31E-05 | 0.00041284 | 0.229603927 |
| ENSMUSG00000040537 | Adam22 | 11496 | protein_coding | 57.63402966 | -0.229604285 | 0.183702818 | -1.249868059 | 0.211347749 | 0.44665249 | 0.229604285 |
| ENSMUSG00000037363 | Letm2 | 270035 | protein_coding | 36.84056365 | -0.229611214 | 0.20613426 | -1.113891565 | 0.265325709 | 0.507219974 | 0.229611214 |
| ENSMUSG00000060149 | BC002059 | 213811 | lncRNA | 138.6845998 | -0.229618585 | 0.1017617 | -2.256434239 | 0.024043449 | 0.11171095 | 0.229618585 |
| ENSMUSG00000036002 | Atosb | 230088 | protein_coding | 1026.895949 | 0.229618671 | 0.097726126 | 2.349613972 | 0.01879289 | 0.094291987 | 0.229618671 |
| ENSMUSG00000037326 | Capn15 | 50817 | protein_coding | 484.9911285 | 0.229622262 | 0.079111834 | 2.902502066 | 0.003701947 | 0.028497042 | 0.229622262 |
| ENSMUSG00000069309 | H2ac22 | 319170 | protein_coding | 51.07772409 | 0.229629676 | 0.187014124 | 1.227873443 | 0.21949448 | 0.455688399 | 0.229629676 |
| ENSMUSG00000023206 | Il15ra | 16169 | protein_coding | 177.4244477 | -0.229736413 | 0.136352707 | -1.684868735 | 0.0920139 | 0.273475636 | 0.229736413 |
| ENSMUSG00000070287 | Slc35g2 | 245020 | protein_coding | 160.5064601 | 0.229758465 | 0.096336421 | 2.384959519 | 0.017081001 | 0.088092517 | 0.229758465 |
| ENSMUSG00000068854 | H2bc21 | 319190 | protein_coding | 262.2025612 | 0.229776082 | 0.092163188 | 2.493143804 | 0.012661755 | 0.070827655 | 0.229776082 |
| ENSMUSG00000086494 | 2210417A02Rik | 70138 | lncRNA | 11.60392021 | 0.22988537 | 0.425249249 | 0.540589714 | 0.588790409 | 0.778426904 | 0.22988537 |
| ENSMUSG00000024340 | Btnl2 | 547431 | protein_coding | 94.00273659 | -0.229916187 | 0.205653623 | -1.117977812 | 0.26357647 | 0.504916665 | 0.229916187 |
| ENSMUSG00000065653 | Gm23105 | 115487521 | snoRNA | 115.3689027 | 0.229949988 | 0.233997641 | 0.982702165 | 0.325754045 | 0.56923128 | 0.229949988 |
| ENSMUSG00000024905 | Tesmin | 17771 | protein_coding | 8.961468452 | -0.229978534 | 0.381897127 | -0.602200221 | 0.547040869 | 0.75018096 | 0.229978534 |
| ENSMUSG00000032231 | Anxa2 | 12306 | protein_coding | 12728.47736 | 0.230025354 | 0.077180811 | 2.980343828 | 0.00287925 | 0.023338386 | 0.230025354 |
| ENSMUSG00000072572 | Slc39a2 | 214922 | protein_coding | 8.274500684 | 0.230026802 | 0.485737943 | 0.473561527 | 0.635812616 | 0.80962603 | 0.230026802 |
| ENSMUSG00000043079 | Synpo | 104027 | protein_coding | 882.2224925 | -0.230086559 | 0.069181051 | -3.325860998 | 0.000881459 | 0.009077449 | 0.230086559 |
| ENSMUSG00000023972 | Ptk7 | 71461 | protein_coding | 466.4262126 | -0.230112667 | 0.082847238 | -2.777553862 | 0.005476976 | 0.038253441 | 0.230112667 |
| ENSMUSG00000022999 | Lmbr1l | 74775 | protein_coding | 438.2229743 | -0.230161731 | 0.097811317 | -2.353119627 | 0.018616644 | 0.093675699 | 0.230161731 |

| ensembl_gene_id | external_gene_name | entrezgene_id | gene_biotype | baseMean | log2FoldChange | lfcSE | stat | pvalue | padj | absLog2FC |
|---|---|---|---|---|---|---|---|---|---|---|
| ENSMUSG00000024236 | Svil | 225115 | protein_coding | 198.2942886 | 0.230257109 | 0.115338277 | 1.996363361 | 0.045894386 | 0.174478727 | 0.230257109 |
| ENSMUSG00000033576 | Apol6 | 71939 | protein_coding | 286.2554801 | 0.230582355 | 0.156608123 | 1.472352457 | 0.140925725 | 0.352030539 | 0.230582355 |
| ENSMUSG00000004473 | Clec11a | 20256 | protein_coding | 40.46294637 | 0.230628255 | 0.185360706 | 1.244213292 | 0.213421073 | 0.449133255 | 0.230628255 |
| ENSMUSG00000059883 | Irak4 | 266632 | protein_coding | 821.8593261 | 0.23063086 | 0.05483032 | 4.20626508 | 2.60E-05 | 0.000455107 | 0.23063086 |
| ENSMUSG00000044469 | Tnfaip8l1 | 66443 | protein_coding | 402.1998331 | -0.230637921 | 0.085050104 | -2.711788824 | 0.006692121 | 0.04430371 | 0.230637921 |
| ENSMUSG00000038965 | Ube2l3 | 22195 | protein_coding | 1701.597772 | 0.230710508 | 0.05921189 | 3.896354417 | 9.77E-05 | 0.001454132 | 0.230710508 |
| ENSMUSG00000030031 | Kbtbd8 | 243574 | protein_coding | 231.8663811 | 0.230791347 | 0.099633094 | 2.316412524 | 0.020535756 | 0.10005442 | 0.230791347 |
| ENSMUSG00000030228 | Pik3c2g | 18705 | protein_coding | 28.2363829 | -0.230834815 | 0.293014367 | -0.787793503 | 0.430817502 | 0.665561246 | 0.230834815 |
| ENSMUSG00000038208 | Pgap3 | 320655 | protein_coding | 709.3390718 | 0.230856425 | 0.06757848 | 3.416123368 | 0.000635195 | 0.006963823 | 0.230856425 |
| ENSMUSG00000028345 | Tex10 | 269536 | protein_coding | 381.6416831 | -0.230860626 | 0.090098485 | -2.562314184 | 0.010397721 | 0.061064839 | 0.230860626 |
| ENSMUSG00000034429 | Zfp707 | 69020 | protein_coding | 287.7357651 | -0.23097821 | 0.085013025 | -2.716974371 | 0.006588171 | 0.043849539 | 0.23097821 |
| ENSMUSG00000042010 | Acacb | 100705 | protein_coding | 361.8515334 | 0.231119732 | 0.096408963 | 2.397284705 | 0.016517084 | 0.086176324 | 0.231119732 |
| ENSMUSG00000026399 | Cd55 | 13136 | protein_coding | 387.96165 | -0.231150538 | 0.132165212 | -1.748951451 | 0.080299412 | 0.25082096 | 0.231150538 |
| ENSMUSG00000025731 | Mettl26 | 68347 | protein_coding | 1290.18419 | -0.2311553 | 0.052918049 | -4.368175019 | 1.25E-05 | 0.000239969 | 0.2311553 |
| ENSMUSG00000037110 | Ralgapa2 | 241694 | protein_coding | 3382.482291 | -0.231219196 | 0.045834791 | -5.044621986 | 4.54E-07 | 1.20E-05 | 0.231219196 |
| ENSMUSG00000032178 | Ilf3 | 16201 | protein_coding | 817.1364225 | -0.231291771 | 0.063395244 | -3.648408891 | 0.000263869 | 0.003319362 | 0.231291771 |
| ENSMUSG00000067377 | Tspan6 | 56496 | protein_coding | 190.5921615 | 0.231308018 | 0.100975123 | 2.290742614 | 0.021978304 | 0.105020479 | 0.231308018 |
| ENSMUSG00000041959 | S100a10 | 20194 | protein_coding | 6720.541375 | 0.231351427 | 0.073142052 | 3.163042618 | 0.001561295 | 0.014456304 | 0.231351427 |
| ENSMUSG00000010721 | Lmbr1 | 56873 | protein_coding | 459.9550342 | -0.231489659 | 0.075198953 | -3.078362799 | 0.002081413 | 0.018114925 | 0.231489659 |
| ENSMUSG00000007570 | Fance | 72775 | protein_coding | 309.9666665 | -0.231551608 | 0.075318596 | -3.074295333 | 0.002110004 | 0.018291696 | 0.231551608 |
| ENSMUSG00000026977 | Marchf7 | 57438 | protein_coding | 2296.067228 | -0.231590219 | 0.081217016 | -2.851498748 | 0.004351366 | 0.032175367 | 0.231590219 |
| ENSMUSG00000035595 | Fam174c | 69770 | protein_coding | 177.4086138 | 0.231619888 | 0.110382551 | 2.098337887 | 0.035875308 | 0.146880196 | 0.231619888 |
| ENSMUSG00000018239 | Zcchc10 | 67966 | protein_coding | 35.56350865 | -0.231651046 | 0.218176179 | -1.061761401 | 0.288344017 | 0.530775478 | 0.231651046 |
| ENSMUSG00000051316 | Taf7 | 24074 | protein_coding | 364.7586143 | -0.23175173 | 0.110932737 | -2.089119367 | 0.036696979 | 0.149108639 | 0.23175173 |
| ENSMUSG00000028459 | Cd72 | 12517 | protein_coding | 8.983342207 | -0.23178639 | 0.388955724 | -0.595919729 | 0.551228853 | 0.752144991 | 0.23178639 |
| ENSMUSG00000052155 | Acvr2a | 11480 | protein_coding | 1236.342688 | 0.231823448 | 0.044566718 | 5.201716815 | 1.97E-07 | 5.65E-06 | 0.231823448 |
| ENSMUSG00000030406 | Gipr | 381853 | protein_coding | 14.26259547 | -0.231882828 | 0.388956392 | -0.596166647 | 0.551063905 | 0.75210185 | 0.231882828 |
| ENSMUSG00000055945 | Prr18 | 320111 | protein_coding | 5.9416236 | 0.231890601 | 0.498579973 | 0.465102116 | 0.641858331 | 0.813747171 | 0.231890601 |
| ENSMUSG00000031286 | Glt28d2 | 320302 | protein_coding | 206.6229255 | 0.231894597 | 0.10860788 | 2.135154437 | 0.032748397 | 0.138291019 | 0.231894597 |
| ENSMUSG00000112188 | Gm47708 | 102641383 | lncRNA | 25.45389229 | -0.231946302 | 0.276350387 | -0.839319622 | 0.401289975 | 0.640499152 | 0.231946302 |
| ENSMUSG00000053205 | Styx | 56291 | protein_coding | 386.4027187 | -0.231980662 | 0.125597463 | -1.847017103 | 0.064744662 | 0.218804936 | 0.231980662 |
| ENSMUSG00000087593 | Gm16174 | 102638408 | lncRNA | 16.64798724 | 0.231981282 | 0.286033928 | 0.811027151 | 0.417350079 | 0.655174216 | 0.231981282 |
| ENSMUSG00000021950 | Anxa8 | 11752 | protein_coding | 430.6306266 | 0.232047318 | 0.159015891 | 1.459271249 | 0.144490465 | 0.357032789 | 0.232047318 |
| ENSMUSG00000025532 | Crcp | 12909 | protein_coding | 341.975941 | 0.232062431 | 0.067632885 | 3.431207026 | 0.000600902 | 0.006643885 | 0.232062431 |
| ENSMUSG00000046727 | Cystm1 | 66060 | protein_coding | 4614.350431 | 0.232109978 | 0.083085087 | 2.79364187 | 0.005211816 | 0.03683451 | 0.232109978 |
| ENSMUSG00000074165 | Zfp788 | 67607 | protein_coding | 185.4422434 | -0.232119906 | 0.110187161 | -2.106596661 | 0.035152556 | 0.145025653 | 0.232119906 |
| ENSMUSG00000033276 | Stk36 | 269209 | protein_coding | 10.19914317 | 0.232138273 | 0.427181641 | 0.543418188 | 0.586841906 | 0.777011446 | 0.232138273 |
| ENSMUSG00000035944 | Ttc38 | 239570 | protein_coding | 2271.246556 | 0.232331054 | 0.043243448 | 5.372630244 | 7.76E-08 | 2.42E-06 | 0.232331054 |
| ENSMUSG00000028307 | Aldob | 230163 | protein_coding | 4812.965813 | 0.23233454 | 0.063258382 | 3.672786654 | 0.00023992 | 0.003068348 | 0.23233454 |

Supplementary Table 1

| ensembl_gene_id | external_gene_name | entrezgene_id | gene_biotype | baseMean | log2FoldChange | lfcSE | stat | pvalue | padj | absLog2FC |
|---|---|---|---|---|---|---|---|---|---|---|
| ENSMUSG00000098188 | Sowahc | 268301 | protein_coding | 2096.569418 | -0.232360506 | 0.070439932 | -3.298704286 | 0.000971322 | 0.009823815 | 0.232360506 |
| ENSMUSG00000022807 | Osbpl11 | 106326 | protein_coding | 626.7999084 | -0.232365939 | 0.06429192 | -3.614232362 | 0.000301239 | 0.003688746 | 0.232365939 |
| ENSMUSG00000029027 | Dffb | 13368 | protein_coding | 221.6274562 | -0.232512535 | 0.096972528 | -2.397715512 | 0.016497673 | 0.086102903 | 0.232512535 |
| ENSMUSG00000056411 | Gm12500 | 791415 | lncRNA | 12.10643996 | -0.232556295 | 0.390214662 | -0.595970162 | 0.55119516 | 0.752144991 | 0.232556295 |
| ENSMUSG00000034731 | Dgkh | 380921 | protein_coding | 1757.238366 | -0.232566876 | 0.076182606 | -3.052755579 | 0.002267505 | 0.019391822 | 0.232566876 |
| ENSMUSG00000024131 | Slc3a1 | 20532 | protein_coding | 780.319486 | -0.232645901 | 0.152418259 | -1.526365029 | 0.126918982 | 0.331093889 | 0.232645901 |
| ENSMUSG00000047613 | A430005L14Rik | 97159 | protein_coding | 405.5182174 | 0.232654656 | 0.081302372 | 2.86159741 | 0.00421512 | 0.031427292 | 0.232654656 |
| ENSMUSG00000028483 | Snapc3 | 77634 | protein_coding | 229.4016769 | -0.232682629 | 0.108001216 | -2.154444529 | 0.031205328 | 0.133664893 | 0.232682629 |
| ENSMUSG00000040703 | Cyp2s1 | 74134 | protein_coding | 2283.208191 | -0.232685544 | 0.068365493 | -3.403552502 | 0.000665156 | 0.007233294 | 0.232685544 |
| ENSMUSG00000106583 | 4930447N08Rik | 71647 | lncRNA | 16.09278706 | 0.232742876 | 0.324732903 | 0.716720952 | 0.473546295 | 0.697472342 | 0.232742876 |
| ENSMUSG00000047749 | Zc3hav1l | 209032 | protein_coding | 31.82825069 | -0.232847059 | 0.211474395 | -1.10106502 | 0.27086836 | 0.513373374 | 0.232847059 |
| ENSMUSG00000079559 | Pou2af3 | 100502940 | protein_coding | 22.30528784 | 0.232938387 | 0.346505208 | 0.672250754 | 0.501424075 | 0.718418124 | 0.232938387 |
| ENSMUSG00000038500 | Prr3 | 75210 | protein_coding | 70.96188738 | -0.232953246 | 0.138658519 | -1.680050007 | 0.092947587 | 0.275342713 | 0.232953246 |
| ENSMUSG00000001829 | Clpb | 20480 | protein_coding | 757.5808989 | 0.232962204 | 0.098172705 | 2.372983443 | 0.017645055 | 0.090105959 | 0.232962204 |
| ENSMUSG00000036186 | Dipk1b | 56279 | protein_coding | 75.98926541 | -0.232968664 | 0.142164464 | -1.638726432 | 0.101270247 | 0.29018928 | 0.232968664 |
| ENSMUSG00000067656 | Slc22a27 | 171405 | protein_coding | 9.377895755 | -0.232973193 | 0.398220033 | -0.585036345 | 0.558523279 | 0.756983295 | 0.232973193 |
| ENSMUSG00000079184 | Mphosph8 | 75339 | protein_coding | 254.9842578 | 0.233004722 | 0.097202671 | 2.397102057 | 0.01652532 | 0.086179968 | 0.233004722 |
| ENSMUSG00000047090 | Tmem198b | 73827 | protein_coding | 44.47845009 | -0.233069245 | 0.179081899 | -1.301467355 | 0.193098532 | 0.424923197 | 0.233069245 |
| ENSMUSG00000072244 | Trim6 | 94088 | protein_coding | 96.12981962 | 0.233125862 | 0.176803787 | 1.318556951 | 0.187317274 | 0.417073819 | 0.233125862 |
| ENSMUSG00000002458 | Rgs19 | 56470 | protein_coding | 53.33722924 | 0.233144603 | 0.189757093 | 1.228647635 | 0.219203947 | 0.455544189 | 0.233144603 |
| ENSMUSG00000052504 | Epha3 | 13837 | protein_coding | 8.403969055 | -0.233183976 | 0.507122365 | -0.459817969 | 0.645646884 | 0.816434847 | 0.233183976 |
| ENSMUSG00000048330 | Ric3 | 320360 | protein_coding | 37.53470488 | 0.233207405 | 0.252480637 | 0.923664516 | 0.355661012 | 0.598408466 | 0.233207405 |
| ENSMUSG00000025790 | Slco3a1 | 108116 | protein_coding | 22.58138576 | 0.233251194 | 0.236462269 | 0.986420353 | 0.323926882 | 0.567946164 | 0.233251194 |
| ENSMUSG00000087026 | A230103J11Rik | 320466 | lncRNA | 14.69058327 | -0.233323603 | 0.359225009 | -0.649519373 | 0.516002729 | 0.728738778 | 0.233323603 |
| ENSMUSG00000041319 | Thoc6 | 386612 | protein_coding | 210.2508024 | 0.233344291 | 0.121475917 | 1.920909885 | 0.054743069 | 0.196492651 | 0.233344291 |
| ENSMUSG00000025876 | Unc5a | 107448 | protein_coding | 121.9157009 | 0.233353618 | 0.180099203 | 1.295694891 | 0.195080622 | 0.427338385 | 0.233353618 |
| ENSMUSG00000032640 | Chsy1 | 269941 | protein_coding | 130.0697496 | 0.233382386 | 0.127993469 | 1.823392931 | 0.0682439 | 0.226268376 | 0.233382386 |
| ENSMUSG00000020088 | Sar1a | 20224 | protein_coding | 2199.603288 | 0.23338365 | 0.045523047 | 5.126714114 | 2.95E-07 | 8.08E-06 | 0.23338365 |
| ENSMUSG00000048832 | Vps37c | 107305 | protein_coding | 986.5254917 | 0.233404743 | 0.071774138 | 3.251933776 | 0.001146227 | 0.011299026 | 0.233404743 |
| ENSMUSG00000015202 | Cnksr3 | 215748 | protein_coding | 1062.755188 | -0.233518636 | 0.088534916 | -2.637588034 | 0.008349795 | 0.052220563 | 0.233518636 |
| ENSMUSG00000023122 | Sult1c2 | 69083 | protein_coding | 1365.203819 | -0.233583176 | 0.09830424 | -2.376125141 | 0.017495528 | 0.089457952 | 0.233583176 |
| ENSMUSG00000040891 | Foxa3 | 15377 | protein_coding | 1362.797528 | 0.233648484 | 0.091839075 | 2.544107555 | 0.010955734 | 0.063259266 | 0.233648484 |
| ENSMUSG00000095315 | Gm10130 | 102633750 | protein_coding | 65.23354615 | -0.2336706 | 0.149768141 | -1.56021567 | 0.118708924 | 0.318807809 | 0.2336706 |
| ENSMUSG00000040841 | Six5 | 20475 | protein_coding | 92.38733563 | 0.23370599 | 0.141353774 | 1.653340997 | 0.098261484 | 0.28511388 | 0.23370599 |
| ENSMUSG00000064345 | mt-Nd2 | 17717 | protein_coding | 13711.56301 | 0.233719439 | 0.073981281 | 3.159169943 | 0.001582192 | 0.014622356 | 0.233719439 |
| ENSMUSG00000023262 | Acy1 | 109652 | protein_coding | 965.4495195 | 0.233899717 | 0.095788674 | 2.44183061 | 0.014613001 | 0.078738347 | 0.233899717 |
| ENSMUSG00000040035 | Disp2 | 214240 | protein_coding | 76.30682794 | 0.233962928 | 0.14388406 | 1.626051752 | 0.103938647 | 0.29471814 | 0.233962928 |
| ENSMUSG00000006127 | Inpp5k | 19062 | protein_coding | 640.1138441 | -0.234069552 | 0.072961115 | -3.208141119 | 0.001335959 | 0.012740991 | 0.234069552 |
| ENSMUSG00000026649 | Cfap126 | 75472 | protein_coding | 17.25128323 | 0.234091961 | 0.404200365 | 0.579148317 | 0.5624891 | 0.76019675 | 0.234091961 |

| ensembl_gene_id | external_gene_name | entrezgene_id | gene_biotype | baseMean | log2FoldChange | lfcSE | stat | pvalue | padj | absLog2FC |
|---|---|---|---|---|---|---|---|---|---|---|
| ENSMUSG00000039967 | Zfp292 | 30046 | protein_coding | 1785.211007 | -0.234139916 | 0.064549821 | -3.627274425 | 0.000286429 | 0.00354236 | 0.234139916 |
| ENSMUSG00000020453 | Patz1 | 56218 | protein_coding | 208.3885928 | -0.234182722 | 0.104677554 | -2.237181835 | 0.025274458 | 0.115631542 | 0.234182722 |
| ENSMUSG00000038803 | Ost4 | 67695 | protein_coding | 713.4401036 | 0.234231282 | 0.096351461 | 2.431009138 | 0.015056835 | 0.08032198 | 0.234231282 |
| ENSMUSG00000039033 | Tasp1 | 75812 | protein_coding | 260.556106 | 0.234350613 | 0.107041472 | 2.189344073 | 0.028571841 | 0.125723897 | 0.234350613 |
| ENSMUSG00000026425 | Srgap2 | 14270 | protein_coding | 204.6811166 | 0.234415909 | 0.101223422 | 2.315826766 | 0.020567729 | 0.100149689 | 0.234415909 |
| ENSMUSG00000041737 | Tmem45b | 235135 | protein_coding | 9388.272977 | -0.234505059 | 0.063931774 | -3.668051827 | 0.000244406 | 0.003113373 | 0.234505059 |
| ENSMUSG00000033405 | Nudt15 | 214254 | protein_coding | 40.89387092 | 0.234556391 | 0.193492922 | 1.212222075 | 0.225427383 | 0.463412035 | 0.234556391 |
| ENSMUSG00000059654 | Reg1 | 19692 | protein_coding | 52.51308022 | -0.234574716 | 0.804189144 | -0.291690976 | 0.770522909 | 0.892047592 | 0.234574716 |
| ENSMUSG00000022360 | Atad2 | 70472 | protein_coding | 617.443705 | 0.234625775 | 0.145968079 | 1.607377285 | 0.10797163 | 0.300995415 | 0.234625775 |
| ENSMUSG00000051209 | Gpr119 | 236781 | protein_coding | 11.06171057 | -0.234689838 | 0.346644972 | -0.677032287 | 0.498385469 | 0.715904735 | 0.234689838 |
| ENSMUSG00000037151 | Lrrc20 | 216011 | protein_coding | 197.0131085 | 0.234783643 | 0.12918412 | 1.817434239 | 0.069150649 | 0.228149042 | 0.234783643 |
| ENSMUSG00000023909 | Paqr4 | 76498 | protein_coding | 3420.002733 | 0.234921657 | 0.065312119 | 3.596907582 | 0.000322023 | 0.003904708 | 0.234921657 |
| ENSMUSG00000044562 | Rasip1 | 69903 | protein_coding | 24.28297271 | 0.234978309 | 0.272744873 | 0.861531533 | 0.388945362 | 0.628768653 | 0.234978309 |
| ENSMUSG00000021427 | Ssr1 | 107513 | protein_coding | 3536.407777 | 0.235000573 | 0.054420777 | 4.318214234 | 1.57E-05 | 0.000293943 | 0.235000573 |
| ENSMUSG00000040263 | Klhdc4 | 234825 | protein_coding | 499.9678194 | 0.235082606 | 0.103067976 | 2.280850123 | 0.022557317 | 0.106976995 | 0.235082606 |
| ENSMUSG00000028789 | Azin2 | 242669 | protein_coding | 6.288340338 | -0.235199806 | 0.448560125 | -0.52434399 | 0.600039305 | 0.785648606 | 0.235199806 |
| ENSMUSG00000032221 | Mns1 | 17427 | protein_coding | 36.19531646 | 0.235236428 | 0.24716411 | 0.951741853 | 0.341227918 | 0.584118738 | 0.235236428 |
| ENSMUSG00000068205 | Macrod2 | 72899 | protein_coding | 176.1475149 | 0.235276555 | 0.120221421 | 1.957026899 | 0.050344306 | 0.186002892 | 0.235276555 |
| ENSMUSG00000031700 | Gpt2 | 108682 | protein_coding | 1086.285608 | -0.23528127 | 0.072673696 | -3.237502483 | 0.001205809 | 0.011764114 | 0.23528127 |
| ENSMUSG00000024197 | Plin3 | 66905 | protein_coding | 2411.651957 | 0.235296438 | 0.052956221 | 4.443225596 | 8.86E-06 | 0.000176007 | 0.235296438 |
| ENSMUSG00000059401 | Mamld1 | 333639 | protein_coding | 15.43779359 | 0.235360104 | 0.329446958 | 0.714409707 | 0.474973871 | 0.698577622 | 0.235360104 |
| ENSMUSG00000025804 | Ccr1 | 12768 | protein_coding | 20.39223153 | 0.235422377 | 0.27355093 | 0.860616255 | 0.38944943 | 0.629260691 | 0.235422377 |
| ENSMUSG00000041328 | Pcf11 | 74737 | protein_coding | 1916.958039 | -0.235426005 | 0.07153336 | -3.291135847 | 0.000997837 | 0.010029817 | 0.235426005 |
| ENSMUSG00000086712 | Mexis | 381524 | lncRNA | 36.21987874 | 0.235460016 | 0.224738594 | 1.047706191 | 0.294773994 | 0.537823306 | 0.235460016 |
| ENSMUSG00000053862 | Slc51b | 330962 | protein_coding | 886.6737284 | -0.235544001 | 0.0811221 | -2.903573778 | 0.0036893 | 0.028426826 | 0.235544001 |
| ENSMUSG00000095362 | Gm14325 | 329575 | protein_coding | 109.8674034 | -0.235625403 | 0.137943046 | -1.708135426 | 0.087611217 | 0.264986139 | 0.235625403 |
| ENSMUSG00000017307 | Acot8 | 170789 | protein_coding | 663.3277748 | 0.23563455 | 0.085855921 | 2.744534654 | 0.006059676 | 0.041321095 | 0.23563455 |
| ENSMUSG00000021025 | Nfkbia | 18035 | protein_coding | 1880.247172 | -0.235641136 | 0.066618627 | -3.537165905 | 0.000404446 | 0.004729872 | 0.235641136 |
| ENSMUSG00000048264 | Dip2c | 208440 | protein_coding | 508.1860109 | -0.235666546 | 0.058929539 | -3.999124192 | 6.36E-05 | 0.001008172 | 0.235666546 |
| ENSMUSG00000027249 | F2 | 14061 | protein_coding | 10.92916038 | -0.23567694 | 0.393152612 | -0.599454087 | 0.548870118 | 0.751266959 | 0.23567694 |
| ENSMUSG00000020886 | Dlg4 | 13385 | protein_coding | 13.16234725 | 0.235877681 | 0.373816734 | 0.630998186 | 0.528041711 | 0.737831675 | 0.235877681 |
| ENSMUSG00000058435 | Btnl4 | 632126 | protein_coding | 1780.726135 | -0.235942505 | 0.08160875 | -2.891142248 | 0.003838443 | 0.029309932 | 0.235942505 |
| ENSMUSG00000090394 | 4930523C07Rik | 67647 | protein_coding | 178.0684326 | -0.236062921 | 0.128939951 | -1.830797349 | 0.067130796 | 0.223773946 | 0.236062921 |
| ENSMUSG00000025911 | Adhfe1 | 76187 | protein_coding | 51.76218614 | -0.236114402 | 0.17793155 | -1.326995703 | 0.184510104 | 0.413206122 | 0.236114402 |
| ENSMUSG00000036834 | Plch1 | 269437 | protein_coding | 1052.26826 | 0.236129393 | 0.081012512 | 2.914727466 | 0.003559993 | 0.027655107 | 0.236129393 |
| ENSMUSG00000063694 | Cycs | 13063 | protein_coding | 3336.463812 | 0.236135583 | 0.093033277 | 2.538184086 | 0.011142935 | 0.064133519 | 0.236135583 |
| ENSMUSG00000025735 | Rhbdl1 | 214951 | protein_coding | 9.943129207 | -0.236139704 | 0.455020557 | -0.518964912 | 0.60378521 | 0.78841372 | 0.236139704 |
| ENSMUSG00000022575 | Gsdmd | 69146 | protein_coding | 1732.12139 | -0.236172091 | 0.095632267 | -2.469585832 | 0.013526956 | 0.07445366 | 0.236172091 |
| ENSMUSG00000030641 | Ddias | 74041 | protein_coding | 73.84274644 | 0.236282211 | 0.153781909 | 1.536475986 | 0.124421683 | 0.327353023 | 0.236282211 |

| ensembl_gene_id | external_gene_name | entrezgene_id | gene_biotype | baseMean | log2FoldChange | lfcSE | stat | pvalue | padj | absLog2FC |
|---|---|---|---|---|---|---|---|---|---|---|
| ENSMUSG00000005469 | Prkaca | 18747 | protein_coding | 1370.382933 | 0.236388499 | 0.057952312 | 4.079017598 | 4.52E-05 | 0.000746537 | 0.236388499 |
| ENSMUSG00000030401 | Rtn2 | 20167 | protein_coding | 44.66867286 | -0.23639784 | 0.309570575 | -0.763631491 | 0.445086877 | 0.676141303 | 0.23639784 |
| ENSMUSG00000074825 | Itpripl1 | 73338 | protein_coding | 79.72308659 | -0.236467503 | 0.160476272 | -1.473535621 | 0.140606666 | 0.351653742 | 0.236467503 |
| ENSMUSG00000054723 | Vmac | 106639 | protein_coding | 551.7505138 | 0.236471074 | 0.077534136 | 3.049896306 | 0.002289204 | 0.019496529 | 0.236471074 |
| ENSMUSG00000022265 | Ank | 11732 | protein_coding | 2337.708076 | -0.236509328 | 0.059750339 | -3.95829265 | 7.55E-05 | 0.001164963 | 0.236509328 |
| ENSMUSG00000069744 | Psmb3 | 26446 | protein_coding | 1618.178977 | 0.236530472 | 0.062091374 | 3.809393433 | 0.000139308 | 0.001955285 | 0.236530472 |
| ENSMUSG00000031930 | Wwp2 | 66894 | protein_coding | 3261.862615 | -0.236531974 | 0.054649671 | -4.328150048 | 1.50E-05 | 0.000281646 | 0.236531974 |
| ENSMUSG00000017561 | Crlf3 | 54394 | protein_coding | 278.3214032 | -0.236552811 | 0.089024953 | -2.657151774 | 0.007880396 | 0.050113226 | 0.236552811 |
| ENSMUSG00000020838 | Slc6a4 | 15567 | protein_coding | 453.6741054 | -0.23662805 | 0.099442166 | -2.379554457 | 0.017333582 | 0.088855862 | 0.23662805 |
| ENSMUSG00000064945 | Rny3 | 19874 | misc_RNA | 441.2149722 | 0.236680723 | 0.344714416 | 0.686599436 | 0.492335184 | 0.711842345 | 0.236680723 |
| ENSMUSG00000025529 | Zfp711 | 245595 | protein_coding | 41.14729083 | -0.236695419 | 0.202699232 | -1.167717393 | 0.242920778 | 0.484047929 | 0.236695419 |
| ENSMUSG00000029385 | Ccng2 | 12452 | protein_coding | 1783.789766 | -0.23679234 | 0.088119414 | -2.687175605 | 0.007205906 | 0.046820968 | 0.23679234 |
| ENSMUSG00000037622 | Wdtc1 | 230796 | protein_coding | 1148.075679 | -0.236821789 | 0.090406226 | -2.619529645 | 0.008805112 | 0.054260619 | 0.236821789 |
| ENSMUSG00000015653 | Steap2 | 74051 | protein_coding | 387.6836978 | 0.23693482 | 0.072234105 | 3.280096299 | 0.001037717 | 0.010368808 | 0.23693482 |
| ENSMUSG00000022048 | Dpysl2 | 12934 | protein_coding | 1819.003989 | 0.236970805 | 0.086740397 | 2.73195434 | 0.006295987 | 0.042447899 | 0.236970805 |
| ENSMUSG00000065118 | Gm23297 | 115486827 | snoRNA | 12.76449124 | 0.237003519 | 0.394767656 | 0.600362048 | 0.548264975 | 0.750864769 | 0.237003519 |
| ENSMUSG00000015488 | Cacfd1 | 381356 | protein_coding | 688.675277 | 0.237040638 | 0.078200724 | 3.031182143 | 0.002435983 | 0.020397243 | 0.237040638 |
| ENSMUSG00000038954 | Supt3 | 109115 | protein_coding | 59.03750656 | 0.237081623 | 0.159937961 | 1.482334913 | 0.138251203 | 0.348262598 | 0.237081623 |
| ENSMUSG00000020377 | Ltc4s | 17001 | protein_coding | 34.43884631 | 0.237165533 | 0.227438737 | 1.042766664 | 0.297056374 | 0.540093365 | 0.237165533 |
| ENSMUSG00000026170 | Cyp27a1 | 104086 | protein_coding | 424.5821831 | -0.237258826 | 0.16250844 | -1.45997848 | 0.144295988 | 0.356748847 | 0.237258826 |
| ENSMUSG00000002475 | Abhd3 | 106861 | protein_coding | 147.1412199 | 0.23735145 | 0.108518872 | 2.187190536 | 0.028728617 | 0.126172771 | 0.23735145 |
| ENSMUSG00000027257 | Pacsin3 | 80708 | protein_coding | 123.6205493 | 0.237396347 | 0.129868137 | 1.827979923 | 0.067552565 | 0.224854533 | 0.237396347 |
| ENSMUSG00000021147 | Wdr37 | 207615 | protein_coding | 482.5154662 | -0.237473545 | 0.077322034 | -3.071227364 | 0.002131808 | 0.018434134 | 0.237473545 |
| ENSMUSG00000007827 | Ankrd26 | 232339 | protein_coding | 156.7459923 | 0.237483361 | 0.113590856 | 2.090690826 | 0.036555788 | 0.148759075 | 0.237483361 |
| ENSMUSG00000090173 | Fbxw10 | 213980 | protein_coding | 7.062869685 | 0.237521366 | 0.490080768 | 0.484657594 | 0.627919252 | 0.803865895 | 0.237521366 |
| ENSMUSG00000021223 | Papln | 170721 | protein_coding | 33.3227711 | -0.237587778 | 0.215983384 | -1.100028037 | 0.271319906 | 0.513806497 | 0.237587778 |
| ENSMUSG00000022528 | Hes1 | 15205 | protein_coding | 2205.135555 | -0.237689577 | 0.083516715 | -2.846012058 | 0.004427052 | 0.032541051 | 0.237689577 |
| ENSMUSG00000026202 | Tuba4a | 22145 | protein_coding | 2448.085977 | 0.237827945 | 0.084842382 | 2.803173844 | 0.005060239 | 0.035960506 | 0.237827945 |
| ENSMUSG00000117431 | 8430430B14Rik | 78148 | lncRNA | 20.31360063 | 0.237861328 | 0.254660011 | 0.934034864 | 0.350285909 | 0.592517397 | 0.237861328 |
| ENSMUSG00000017756 | Slc12a7 | 20499 | protein_coding | 2286.223741 | -0.23794467 | 0.050813458 | -4.682709685 | 2.83E-06 | 6.28E-05 | 0.23794467 |
| ENSMUSG00000040033 | Stat2 | 20847 | protein_coding | 451.8445334 | 0.237975539 | 0.124674509 | 1.908774619 | 0.056291172 | 0.200487575 | 0.237975539 |
| ENSMUSG00000032093 | Cd3e | 12501 | protein_coding | 17.18809075 | 0.237996767 | 0.321391501 | 0.74051979 | 0.458984658 | 0.685946212 | 0.237996767 |
| ENSMUSG00000034936 | Arl4d | 80981 | protein_coding | 101.1426738 | 0.238021795 | 0.158485444 | 1.501852718 | 0.13313515 | 0.340264748 | 0.238021795 |
| ENSMUSG00000022763 | Aifm3 | 72168 | protein_coding | 42.58418759 | -0.238058158 | 0.185296885 | -1.284739124 | 0.198883462 | 0.432263286 | 0.238058158 |
| ENSMUSG00000061615 | H2ac4 | 319172 | protein_coding | 527.2129988 | 0.238067107 | 0.132701875 | 1.793999573 | 0.072813237 | 0.2361269 | 0.238067107 |
| ENSMUSG00000023473 | Celsr3 | 107934 | protein_coding | 19.22960284 | 0.238126787 | 0.279512279 | 0.851936766 | 0.394249189 | 0.633814677 | 0.238126787 |
| ENSMUSG00000033705 | Stard9 | 668880 | protein_coding | 9.505116187 | 0.238188987 | 0.419275247 | 0.568096945 | 0.569969142 | 0.765233601 | 0.238188987 |
| ENSMUSG00000059013 | Sh2d3c | 27387 | protein_coding | 90.27985805 | 0.238222233 | 0.14852436 | 1.603927009 | 0.108730143 | 0.302001032 | 0.238222233 |
| ENSMUSG00000017311 | Pyy | 217212 | protein_coding | 894.7255698 | -0.238328526 | 0.075514884 | -3.156047031 | 0.001599231 | 0.014762904 | 0.238328526 |

| ensembl_gene_id | external_gene_name | entrezgene_id | gene_biotype | baseMean | log2FoldChange | lfcSE | stat | pvalue | padj | absLog2FC |
|---|---|---|---|---|---|---|---|---|---|---|
| ENSMUSG00000038535 | Zfp280d | 235469 | protein_coding | 459.8667346 | -0.238383455 | 0.104563529 | -2.279795423 | 0.022619825 | 0.107070711 | 0.238383455 |
| ENSMUSG00000041301 | Cftr | 12638 | protein_coding | 5328.930403 | -0.238440338 | 0.086385877 | -2.760177325 | 0.005777 | 0.03993385 | 0.238440338 |
| ENSMUSG00000063931 | Pepd | 18624 | protein_coding | 1090.742601 | 0.23854853 | 0.060676256 | 3.931497195 | 8.44E-05 | 0.001274735 | 0.23854853 |
| ENSMUSG00000024084 | Qpct | 70536 | protein_coding | 12.77994808 | 0.238549696 | 0.321791691 | 0.74131714 | 0.458501172 | 0.685788934 | 0.238549696 |
| ENSMUSG00000021258 | Ccnk | 12454 | protein_coding | 188.0662662 | -0.238587753 | 0.101296783 | -2.355333972 | 0.018506065 | 0.093297675 | 0.238587753 |
| ENSMUSG00000022667 | Cd200r1 | 57781 | protein_coding | 6.861342736 | 0.238594759 | 0.452040715 | 0.527816966 | 0.59762638 | 0.783947174 | 0.238594759 |
| ENSMUSG00000031562 | Dctd | 320685 | protein_coding | 90.51942851 | 0.238595487 | 0.16286071 | 1.465027921 | 0.1429133 | 0.35463345 | 0.238595487 |
| ENSMUSG00000027555 | Car13 | 71934 | protein_coding | 811.1046703 | 0.238600537 | 0.059948089 | 3.98011918 | 6.89E-05 | 0.00107936 | 0.238600537 |
| ENSMUSG00000049971 | Glt1d1 | 319804 | protein_coding | 455.7019001 | -0.238647765 | 0.095786566 | -2.491453394 | 0.012722166 | 0.071066978 | 0.238647765 |
| ENSMUSG00000065089 | Gm26493 | 115487044 | snoRNA | 10.42994391 | -0.238706095 | 0.443893547 | -0.537755271 | 0.590746016 | 0.779660849 | 0.238706095 |
| ENSMUSG00000034880 | Mrpl34 | 94065 | protein_coding | 850.7973303 | 0.238814533 | 0.107521234 | 2.221091826 | 0.026344743 | 0.118795321 | 0.238814533 |
| ENSMUSG00000062124 | Defb45 | 433490 | protein_coding | 16.97051176 | -0.238867838 | 0.396330845 | -0.602698076 | 0.546709562 | 0.749854321 | 0.238867838 |
| ENSMUSG00000038244 | Mical2 | 320878 | protein_coding | 2607.324473 | -0.238965299 | 0.05404658 | -4.421469408 | 9.80E-06 | 0.0001928 | 0.238965299 |
| ENSMUSG00000030978 | Rrm1 | 20133 | protein_coding | 1003.134381 | 0.238988057 | 0.125235427 | 1.908310314 | 0.056351121 | 0.200523946 | 0.238988057 |
| ENSMUSG00000020659 | Cbll1 | 104836 | protein_coding | 146.9743811 | -0.238999596 | 0.129129573 | -1.850850973 | 0.064190996 | 0.21752641 | 0.238999596 |
| ENSMUSG00000069835 | Sat2 | 69215 | protein_coding | 21.474757 | 0.239003839 | 0.338991486 | 0.70504378 | 0.480782991 | 0.70308191 | 0.239003839 |
| ENSMUSG00000050379 | Septin6 | 56526 | protein_coding | 137.820841 | -0.239089565 | 0.120137019 | -1.990140651 | 0.046575444 | 0.176237021 | 0.239089565 |
| ENSMUSG00000070576 | Mn1 | 433938 | protein_coding | 13.64287064 | 0.239155636 | 0.31364274 | 0.762509713 | 0.445755848 | 0.67671134 | 0.239155636 |
| ENSMUSG00000053907 | Mat2a | 232087 | protein_coding | 4448.602792 | -0.239223078 | 0.104743754 | -2.283888728 | 0.022378071 | 0.106394798 | 0.239223078 |
| ENSMUSG00000000942 | Hoxa4 | 15401 | protein_coding | 12.44028721 | -0.239223571 | 0.337347652 | -0.709130684 | 0.478243383 | 0.700798986 | 0.239223571 |
| ENSMUSG00000042333 | Tnfrsf14 | 230979 | protein_coding | 302.7240986 | -0.239283461 | 0.089146055 | -2.684173314 | 0.00727094 | 0.047118814 | 0.239283461 |
| ENSMUSG00000054453 | Sytl5 | 236643 | protein_coding | 39.91773446 | 0.239300229 | 0.207124387 | 1.155345499 | 0.247948971 | 0.488954885 | 0.239300229 |
| ENSMUSG00000094651 | Gal3st2 | 381334 | protein_coding | 469.7639348 | 0.239337476 | 0.094764888 | 2.525592341 | 0.011550342 | 0.066100912 | 0.239337476 |
| ENSMUSG00000041355 | Ssr2 | 66256 | protein_coding | 2720.029549 | 0.239339718 | 0.073696012 | 3.247661738 | 0.001163575 | 0.011435086 | 0.239339718 |
| ENSMUSG00000033099 | Nol12 | 97961 | protein_coding | 261.4437718 | 0.239419818 | 0.115684586 | 2.06959135 | 0.038490629 | 0.154396187 | 0.239419818 |
| ENSMUSG00000003573 | Homer3 | 26558 | protein_coding | 21.66451118 | -0.239436887 | 0.243114127 | -0.98487443 | 0.324685755 | 0.568485113 | 0.239436887 |
| ENSMUSG00000087412 | Gm15501 | 100040298 | transcribed_processed_pseudogene | 7.980572199 | 0.239438063 | 0.488603708 | 0.490045529 | 0.624101682 | 0.801591615 | 0.239438063 |
| ENSMUSG00000103749 | Pcdhgb5 | 93702 | protein_coding | 13.945775 | 0.239627172 | 0.362939422 | 0.660240133 | 0.509099742 | 0.724201408 | 0.239627172 |
| ENSMUSG00000091956 | C2cd4b | 75697 | protein_coding | 107.4453088 | 0.239672365 | 0.169035558 | 1.417881352 | 0.156225408 | 0.374715872 | 0.239672365 |
| ENSMUSG00000053877 | Srcap | 100043597 | protein_coding | 160.1151351 | -0.239703363 | 0.115826968 | -2.069495284 | 0.038499633 | 0.154396187 | 0.239703363 |
| ENSMUSG00000063760 | Rnf217 | 268291 | protein_coding | 236.1169145 | 0.239750282 | 0.081801953 | 2.930862571 | 0.003380223 | 0.026604614 | 0.239750282 |
| ENSMUSG00000037820 | Tgm2 | 21817 | protein_coding | 610.9717142 | 0.23984519 | 0.084448778 | 2.840126249 | 0.004509568 | 0.03299719 | 0.23984519 |
| ENSMUSG00000005125 | Ndrg1 | 17988 | protein_coding | 3691.868634 | -0.239872992 | 0.059234035 | -4.04958047 | 5.13E-05 | 0.000838657 | 0.239872992 |
| ENSMUSG00000032123 | Dpagt1 | 13478 | protein_coding | 692.6583658 | 0.239961086 | 0.050404408 | 4.760716249 | 1.93E-06 | 4.44E-05 | 0.239961086 |
| ENSMUSG00000026701 | Prdx6 | 11758 | protein_coding | 19900.15742 | -0.239979463 | 0.182647018 | -1.313897516 | 0.188880695 | 0.419339065 | 0.239979463 |
| ENSMUSG00000028152 | Tspan5 | 56224 | protein_coding | 264.7268157 | 0.24001962 | 0.079774519 | 3.008725376 | 0.002623461 | 0.021630144 | 0.24001962 |
| ENSMUSG00000031503 | Col4a2 | 12827 | protein_coding | 24.20118224 | 0.240037829 | 0.320479912 | 0.748994929 | 0.453860262 | 0.682652904 | 0.240037829 |
| ENSMUSG00000022335 | Zfat | 380993 | protein_coding | 119.0695025 | 0.240106139 | 0.113856069 | 2.108856756 | 0.034956948 | 0.144514406 | 0.240106139 |
| ENSMUSG00000038762 | Abcf1 | 224742 | protein_coding | 2366.736884 | 0.240117473 | 0.040554803 | 5.920814717 | 3.20E-09 | 1.21E-07 | 0.240117473 |

| ensembl_gene_id | external_gene_name | entrezgene_id | gene_biotype | baseMean | log2FoldChange | lfcSE | stat | pvalue | padj | absLog2FC |
|---|---|---|---|---|---|---|---|---|---|---|
| ENSMUSG00000027035 | Cers6 | 241447 | protein_coding | 5197.434227 | -0.240125406 | 0.031041438 | -7.735640606 | 1.03E-14 | 6.33E-13 | 0.240125406 |
| ENSMUSG00000019843 | Fyn | 14360 | protein_coding | 56.1455451 | 0.240178461 | 0.176489017 | 1.360869163 | 0.17355504 | 0.399332591 | 0.240178461 |
| ENSMUSG00000042258 | Isl1 | 16392 | protein_coding | 20.87134016 | 0.240237013 | 0.269616234 | 0.891033189 | 0.372911366 | 0.614664445 | 0.240237013 |
| ENSMUSG00000036437 | Npy1r | 18166 | protein_coding | 103.3942518 | -0.240249693 | 0.18395146 | -1.306049393 | 0.191535761 | 0.423020709 | 0.240249693 |
| ENSMUSG00000028550 | Atg4c | 242557 | protein_coding | 256.7374514 | 0.240268149 | 0.083130907 | 2.890238496 | 0.003849497 | 0.029380422 | 0.240268149 |
| ENSMUSG00000054893 | Zfp667 | 384763 | protein_coding | 83.0157843 | 0.240307826 | 0.14323364 | 1.677733146 | 0.093399206 | 0.276072029 | 0.240307826 |
| ENSMUSG00000075502 | Kbtbd6 | 432879 | protein_coding | 9.274302397 | -0.240437886 | 0.46707318 | -0.51477562 | 0.606709824 | 0.789765867 | 0.240437886 |
| ENSMUSG00000021629 | Slc30a5 | 69048 | protein_coding | 1545.597653 | 0.240440873 | 0.043606694 | 5.513852416 | 3.51E-08 | 1.17E-06 | 0.240440873 |
| ENSMUSG00000028522 | Mier1 | 71148 | protein_coding | 3150.454412 | -0.240472149 | 0.104097688 | -2.310062339 | 0.020884703 | 0.101348119 | 0.240472149 |
| ENSMUSG00000024388 | Myo7b | 17922 | protein_coding | 6525.091763 | -0.240563722 | 0.065266707 | -3.685856604 | 0.000227935 | 0.002933681 | 0.240563722 |
| ENSMUSG00000029228 | Lnx1 | 16924 | protein_coding | 628.7097225 | -0.240584604 | 0.068396743 | -3.517486251 | 0.000435655 | 0.005032813 | 0.240584604 |
| ENSMUSG00000026204 | Ptprn | 19275 | protein_coding | 31.85822553 | -0.240650552 | 0.213422714 | -1.127577041 | 0.259498587 | 0.501083584 | 0.240650552 |
| ENSMUSG00000017764 | Zswim1 | 71971 | protein_coding | 193.4772098 | 0.240695172 | 0.111287667 | 2.162819816 | 0.030555041 | 0.131915576 | 0.240695172 |
| ENSMUSG00000036964 | Trim17 | 56631 | protein_coding | 6.339993166 | -0.240789649 | 0.469469277 | -0.512897565 | 0.608022977 | 0.790773109 | 0.240789649 |
| ENSMUSG00000087385 | Frg2f1 | 433752 | protein_coding | 89.36432758 | -0.24094383 | 0.135116264 | -1.783233361 | 0.074548324 | 0.239585656 | 0.24094383 |
| ENSMUSG00000016918 | Sulf1 | 240725 | protein_coding | 11.856648 | 0.240970905 | 0.398604235 | 0.604536741 | 0.54548685 | 0.749069007 | 0.240970905 |
| ENSMUSG00000038055 | Dexi | 58239 | protein_coding | 253.887451 | 0.241010508 | 0.114324579 | 2.108125037 | 0.035020175 | 0.144654187 | 0.241010508 |
| ENSMUSG00000041189 | Chrnb1 | 11443 | protein_coding | 43.24293147 | -0.241012028 | 0.186226844 | -1.294185216 | 0.195601452 | 0.428130377 | 0.241012028 |
| ENSMUSG00000022016 | Akap11 | 219181 | protein_coding | 1691.621996 | -0.241080434 | 0.060376207 | -3.992970819 | 6.53E-05 | 0.001032676 | 0.241080434 |
| ENSMUSG00000024158 | Hagh | 14651 | protein_coding | 1645.992412 | 0.241102829 | 0.068105285 | 3.540148611 | 0.000399902 | 0.004683526 | 0.241102829 |
| ENSMUSG00000022391 | Rangap1 | 19387 | protein_coding | 2342.391217 | 0.241158377 | 0.090313324 | 2.670241409 | 0.007579673 | 0.048680759 | 0.241158377 |
| ENSMUSG00000033220 | Rac2 | 19354 | protein_coding | 98.4579742 | 0.241358628 | 0.151293943 | 1.595296038 | 0.110646051 | 0.305389161 | 0.241358628 |
| ENSMUSG00000040562 | Gstm2 | 14863 | protein_coding | 1699.672485 | 0.241399477 | 0.08113828 | 2.975161392 | 0.002928344 | 0.023612705 | 0.241399477 |
| ENSMUSG00000001911 | Nfix | 18032 | protein_coding | 554.2778584 | -0.241497313 | 0.114487358 | -2.109379739 | 0.034911817 | 0.144401865 | 0.241497313 |
| ENSMUSG00000074867 | Zfp808 | 630579 | protein_coding | 36.3759229 | -0.241578202 | 0.21838437 | -1.106206466 | 0.268637165 | 0.511015761 | 0.241578202 |
| ENSMUSG00000029781 | Fkbp9 | 27055 | protein_coding | 1521.493347 | 0.241591792 | 0.072031382 | 3.353979665 | 0.000796582 | 0.008336457 | 0.241591792 |
| ENSMUSG00000031176 | Dynlt3 | 67117 | protein_coding | 1419.519766 | 0.241668199 | 0.050387383 | 4.79620465 | 1.62E-06 | 3.78E-05 | 0.241668199 |
| ENSMUSG00000051747 | Ttn | 22138 | protein_coding | 24.10487544 | 0.241690632 | 0.429717399 | 0.56244088 | 0.573815673 | 0.76815045 | 0.241690632 |
| ENSMUSG00000100199 | Gm20324 | 100504637 | lncRNA | 20.29106235 | -0.241767302 | 0.333366816 | -0.725229058 | 0.468311519 | 0.693047027 | 0.241767302 |
| ENSMUSG00000030088 | Aldh1l1 | 107747 | protein_coding | 3823.536821 | -0.24188554 | 0.099653909 | -2.42725592 | 0.01521352 | 0.080952136 | 0.24188554 |
| ENSMUSG00000001794 | Capns1 | 12336 | protein_coding | 4447.068593 | 0.241912 | 0.062972704 | 3.84153743 | 0.000122266 | 0.00175663 | 0.241912 |
| ENSMUSG00000025265 | Fgd1 | 14163 | protein_coding | 65.59569588 | 0.241950734 | 0.175016136 | 1.382448154 | 0.166834139 | 0.390611812 | 0.241950734 |
| ENSMUSG00000049396 | Gemin4 | 276919 | protein_coding | 10.83952932 | -0.241970744 | 0.371321568 | -0.651647427 | 0.514628645 | 0.727380907 | 0.241970744 |
| ENSMUSG00000027940 | Tpm3 | 59069 | protein_coding | 5937.53716 | 0.241996681 | 0.0527654 | 4.586275911 | 4.51E-06 | 9.65E-05 | 0.241996681 |
| ENSMUSG00000001249 | Hpn | 15451 | protein_coding | 144.0024698 | -0.242104026 | 0.122763426 | -1.97211853 | 0.048596077 | 0.181540175 | 0.242104026 |
| ENSMUSG00000046782 | Ttc6 | 70846 | protein_coding | 110.8244181 | -0.242156902 | 0.131934496 | -1.83543281 | 0.066441586 | 0.222380129 | 0.242156902 |
| ENSMUSG00000024002 | Brd4 | 57261 | protein_coding | 577.0463433 | -0.242319507 | 0.093270322 | -2.59803441 | 0.00937591 | 0.056780059 | 0.242319507 |
| ENSMUSG00000023043 | Krt18 | 16668 | protein_coding | 10734.52937 | 0.242444932 | 0.041066837 | 5.903667093 | 3.56E-09 | 1.33E-07 | 0.242444932 |
| ENSMUSG00000033581 | Igf2bp2 | 319765 | protein_coding | 938.104434 | -0.242542979 | 0.061862515 | -3.920677629 | 8.83E-05 | 0.001327138 | 0.242542979 |

| ensembl_gene_id | external_gene_name | entrezgene_id | gene_biotype | baseMean | log2FoldChange | lfcSE | stat | pvalue | padj | absLog2FC |
|---|---|---|---|---|---|---|---|---|---|---|
| ENSMUSG00000028801 | Stpg1 | 78806 | protein_coding | 16.46261041 | 0.242550165 | 0.333939463 | 0.726329744 | 0.467636649 | 0.692926016 | 0.242550165 |
| ENSMUSG00000025997 | Ikzf2 | 22779 | protein_coding | 560.1204481 | 0.242589017 | 0.083290232 | 2.912574614 | 0.003584626 | 0.027819661 | 0.242589017 |
| ENSMUSG00000048856 | Slc25a47 | 104910 | protein_coding | 134.3126469 | 0.242690801 | 0.136522639 | 1.777659756 | 0.075459753 | 0.241648022 | 0.242690801 |
| ENSMUSG00000034908 | Sidt2 | 214597 | protein_coding | 1568.191712 | -0.242789831 | 0.043112797 | -5.631502615 | 1.79E-08 | 6.20E-07 | 0.242789831 |
| ENSMUSG00000035325 | Sec31a | 69162 | protein_coding | 4030.896605 | 0.24279914 | 0.045288099 | 5.361212937 | 8.27E-08 | 2.56E-06 | 0.24279914 |
| ENSMUSG00000027843 | Ptpn22 | 19260 | protein_coding | 29.16266185 | 0.242886868 | 0.252767683 | 0.960909502 | 0.336597673 | 0.580252873 | 0.242886868 |
| ENSMUSG00000063382 | Bcl9l | 80288 | protein_coding | 259.0870186 | -0.243059209 | 0.12902977 | -1.88374519 | 0.059599451 | 0.207235951 | 0.243059209 |
| ENSMUSG00000063511 | Snrnp70 | 20637 | protein_coding | 2352.803521 | 0.243080115 | 0.090566995 | 2.683981226 | 0.007275119 | 0.047118814 | 0.243080115 |
| ENSMUSG00000033644 | Piwil2 | 57746 | protein_coding | 9.521511967 | -0.243102743 | 0.414628274 | -0.586314919 | 0.557663907 | 0.75632376 | 0.243102743 |
| ENSMUSG00000030058 | Copg1 | 54161 | protein_coding | 3392.89603 | 0.243167708 | 0.055753515 | 4.361477622 | 1.29E-05 | 0.000247141 | 0.243167708 |
| ENSMUSG00000087331 | 1810021B22Rik | 69120 | lncRNA | 43.9777751 | 0.243274772 | 0.173105039 | 1.405359271 | 0.159914471 | 0.380150454 | 0.243274772 |
| ENSMUSG00000028899 | Taf12 | 66464 | protein_coding | 488.8768555 | 0.243284231 | 0.065960068 | 3.688356299 | 0.000225707 | 0.002907732 | 0.243284231 |
| ENSMUSG00000026239 | Pde6d | 18582 | protein_coding | 184.7857763 | 0.243336627 | 0.108429905 | 2.244183727 | 0.024820583 | 0.114189231 | 0.243336627 |
| ENSMUSG00000020948 | Klhl28 | 66689 | protein_coding | 304.7552534 | -0.243486615 | 0.087688131 | -2.776733998 | 0.00549081 | 0.038300296 | 0.243486615 |
| ENSMUSG00000033308 | Dpyd | 99586 | protein_coding | 1141.999503 | -0.243581055 | 0.075972874 | -3.206158239 | 0.0013452 | 0.012791297 | 0.243581055 |
| ENSMUSG00000025060 | Slk | 20874 | protein_coding | 3864.854001 | -0.243728833 | 0.051038615 | -4.775381039 | 1.79E-06 | 4.16E-05 | 0.243728833 |
| ENSMUSG00000049904 | Tmem17 | 103765 | protein_coding | 25.0298571 | 0.243738292 | 0.226203584 | 1.077517373 | 0.281249194 | 0.523693078 | 0.243738292 |
| ENSMUSG00000026578 | Ccdc181 | 74895 | protein_coding | 208.8172745 | 0.243853982 | 0.092638243 | 2.632325213 | 0.008480265 | 0.052776264 | 0.243853982 |
| ENSMUSG00000015766 | Eps8 | 13860 | protein_coding | 3651.879501 | -0.243932014 | 0.077161111 | -3.16133361 | 0.001570485 | 0.014522485 | 0.243932014 |
| ENSMUSG00000042351 | Grap2 | 17444 | protein_coding | 45.26010877 | 0.243952778 | 0.195851085 | 1.245603404 | 0.212910032 | 0.448543447 | 0.243952778 |
| ENSMUSG00000030979 | Uros | 22276 | protein_coding | 351.4452924 | 0.243989526 | 0.072941803 | 3.344988952 | 0.000822858 | 0.008563112 | 0.243989526 |
| ENSMUSG00000029154 | Cwh43 | 231293 | protein_coding | 729.5473594 | -0.244089404 | 0.089187004 | -2.73682704 | 0.00620349 | 0.042017506 | 0.244089404 |
| ENSMUSG00000031903 | Pla2g15 | 192654 | protein_coding | 119.2813825 | 0.244104554 | 0.112676932 | 2.166411072 | 0.030279789 | 0.130987701 | 0.244104554 |
| ENSMUSG00000027200 | Sema6d | 214968 | protein_coding | 865.3359108 | 0.244105881 | 0.054774952 | 4.456523906 | 8.33E-06 | 0.000166878 | 0.244105881 |
| ENSMUSG00000001763 | Tspan33 | 232670 | protein_coding | 473.1549425 | 0.244188798 | 0.082808677 | 2.948831071 | 0.003189783 | 0.025365694 | 0.244188798 |
| ENSMUSG00000031748 | Gnao1 | 14681 | protein_coding | 40.99156136 | -0.244211293 | 0.19569487 | -1.247918727 | 0.212060822 | 0.447440166 | 0.244211293 |
| ENSMUSG00000041229 | Phf8 | 320595 | protein_coding | 736.7606634 | 0.244243746 | 0.056375074 | 4.332477581 | 1.47E-05 | 0.000276807 | 0.244243746 |
| ENSMUSG00000047155 | Cyp4x1 | 81906 | protein_coding | 9.424144656 | -0.244283631 | 0.386469008 | -0.632091129 | 0.527327332 | 0.737316445 | 0.244283631 |
| ENSMUSG00000029094 | Afap1 | 70292 | protein_coding | 286.3148484 | 0.24434382 | 0.094922254 | 2.574146838 | 0.010048762 | 0.059470231 | 0.24434382 |
| ENSMUSG00000063145 | Bbs5 | 72569 | protein_coding | 70.12088198 | 0.244380403 | 0.145469013 | 1.679948172 | 0.0929674 | 0.275350828 | 0.244380403 |
| ENSMUSG00000032589 | Bsn | 12217 | protein_coding | 292.9248841 | -0.244548315 | 0.076032919 | -3.216347831 | 0.001298334 | 0.012507901 | 0.244548315 |
| ENSMUSG00000032080 | Apoa4 | 11808 | protein_coding | 99.71291572 | 0.244559666 | 1.022469037 | 0.239185401 | 0.810961823 | 0.914838553 | 0.244559666 |
| ENSMUSG00000021270 | Hsp90aa1 | 15519 | protein_coding | 4695.870467 | 0.244574887 | 0.116199803 | 2.104778849 | 0.035310563 | 0.145383562 | 0.244574887 |
| ENSMUSG00000030157 | Clec2d | 93694 | protein_coding | 1574.113267 | -0.244701093 | 0.076781036 | -3.186999095 | 0.001437572 | 0.013494603 | 0.244701093 |
| ENSMUSG00000020087 | Tysnd1 | 71767 | protein_coding | 717.655049 | 0.244709112 | 0.088618249 | 2.761385091 | 0.005755677 | 0.039815194 | 0.244709112 |
| ENSMUSG00000076432 | Ywhaq | 22630 | protein_coding | 2129.712774 | 0.244709801 | 0.061980177 | 3.948194645 | 7.87E-05 | 0.001205968 | 0.244709801 |
| ENSMUSG00000031493 | Ggn | 243897 | protein_coding | 7.878605215 | -0.24477497 | 0.459756492 | -0.532401334 | 0.594448061 | 0.781691445 | 0.24477497 |
| ENSMUSG00000020910 | Adprm | 66358 | protein_coding | 308.3873683 | -0.244806549 | 0.101207414 | -2.418859838 | 0.015569238 | 0.082404037 | 0.244806549 |
| ENSMUSG00000073197 | 5730507C01Rik | 236366 | protein_coding | 28.70355064 | -0.244832084 | 0.227656799 | -1.075443757 | 0.2821761 | 0.525055263 | 0.244832084 |

| ensembl_gene_id | external_gene_name | entrezgene_id | gene_biotype | baseMean | log2FoldChange | lfcSE | stat | pvalue | padj | absLog2FC |
|---|---|---|---|---|---|---|---|---|---|---|
| ENSMUSG00000073600 | Prob1 | 381148 | protein_coding | 16.63985226 | 0.244896132 | 0.311140714 | 0.787091246 | 0.431228454 | 0.666004719 | 0.244896132 |
| ENSMUSG00000022123 | Scel | 64929 | protein_coding | 109.5421326 | -0.244900598 | 0.14883489 | -1.645451532 | 0.09987673 | 0.287678519 | 0.244900598 |
| ENSMUSG00000031613 | Hpgd | 15446 | protein_coding | 5232.178242 | -0.244907866 | 0.08038255 | -3.046779005 | 0.002313077 | 0.019643493 | 0.244907866 |
| ENSMUSG00000019888 | Mgat4c | 67569 | protein_coding | 10110.41645 | 0.244921744 | 0.078558525 | 3.117697846 | 0.001822696 | 0.016312216 | 0.244921744 |
| ENSMUSG00000037541 | Shank2 | 210274 | protein_coding | 609.0863494 | -0.244927917 | 0.057706095 | -4.244402865 | 2.19E-05 | 0.00039274 | 0.244927917 |
| ENSMUSG00000079036 | Alkbh1 | 211064 | protein_coding | 315.4465001 | -0.244991123 | 0.120686984 | -2.029971382 | 0.042359448 | 0.165140974 | 0.244991123 |
| ENSMUSG00000094936 | Rbm4 | 19653 | protein_coding | 81.37938026 | -0.245054291 | 0.166381255 | -1.47284796 | 0.140792037 | 0.351955161 | 0.245054291 |
| ENSMUSG00000036106 | Prr5 | 109270 | protein_coding | 340.1819133 | 0.245113868 | 0.083060885 | 2.9510144 | 0.003167322 | 0.02524933 | 0.245113868 |
| ENSMUSG00000058063 | Trim31 | 224762 | protein_coding | 1548.936071 | 0.245201174 | 0.076466667 | 3.206641313 | 0.001342943 | 0.012784915 | 0.245201174 |
| ENSMUSG00000058267 | Mrps14 | 64659 | protein_coding | 699.3885651 | 0.245211013 | 0.064708362 | 3.789479521 | 0.000150963 | 0.00207552 | 0.245211013 |
| ENSMUSG00000080797 | Gm15760 | 100042948 | transcribed_processed_pseudogene | 25.29439057 | -0.245275596 | 0.248718814 | -0.986156182 | 0.324056478 | 0.567992068 | 0.245275596 |
| ENSMUSG00000031015 | Swap70 | 20947 | protein_coding | 945.1135919 | -0.245324649 | 0.054896534 | -4.468854993 | 7.86E-06 | 0.000158925 | 0.245324649 |
| ENSMUSG00000030207 | Fam234b | 74525 | protein_coding | 16.45687969 | 0.245337745 | 0.40501639 | 0.605747696 | 0.544682307 | 0.749012844 | 0.245337745 |
| ENSMUSG00000041571 | Selenow | 20364 | protein_coding | 1039.943078 | -0.245341169 | 0.04185445 | -5.861770296 | 4.58E-09 | 1.71E-07 | 0.245341169 |
| ENSMUSG00000022559 | Fbxl6 | 30840 | protein_coding | 367.6530648 | -0.245426727 | 0.090433194 | -2.713900911 | 0.006649605 | 0.04413094 | 0.245426727 |
| ENSMUSG00000086040 | Wipf3 | 330319 | protein_coding | 26.01780645 | -0.245442762 | 0.251967509 | -0.974104807 | 0.330004503 | 0.573705524 | 0.245442762 |
| ENSMUSG00000039748 | Exo1 | 26909 | protein_coding | 128.7167102 | 0.245473857 | 0.19666801 | 1.248163627 | 0.211971141 | 0.447440166 | 0.245473857 |
| ENSMUSG00000022562 | Oplah | 75475 | protein_coding | 1094.042496 | -0.245548051 | 0.076734322 | -3.199976791 | 0.001374387 | 0.013007472 | 0.245548051 |
| ENSMUSG00000049092 | Gpr137c | 70713 | protein_coding | 16.1315098 | -0.245613855 | 0.317970089 | -0.77244327 | 0.439851934 | 0.671740242 | 0.245613855 |
| ENSMUSG00000028381 | Ugcg | 22234 | protein_coding | 1822.553471 | -0.245772232 | 0.051658189 | -4.757662561 | 1.96E-06 | 4.50E-05 | 0.245772232 |
| ENSMUSG00000033863 | Klf9 | 16601 | protein_coding | 2545.688218 | -0.245819607 | 0.073448862 | -3.346812992 | 0.000817463 | 0.0085163 | 0.245819607 |
| ENSMUSG00000037669 | Ldah | 68832 | protein_coding | 1107.758288 | 0.245893276 | 0.052818101 | 4.655473654 | 3.23E-06 | 7.06E-05 | 0.245893276 |
| ENSMUSG00000030357 | Fkbp4 | 14228 | protein_coding | 6386.910936 | 0.245921555 | 0.101327139 | 2.427005798 | 0.015224013 | 0.080975478 | 0.245921555 |
| ENSMUSG00000036885 | Arhgef26 | 622434 | protein_coding | 198.6713053 | -0.245946451 | 0.103399632 | -2.378600832 | 0.017378483 | 0.088972317 | 0.245946451 |
| ENSMUSG00000021671 | Poc5 | 67463 | protein_coding | 317.1190114 | 0.245981597 | 0.086200911 | 2.853584648 | 0.004322901 | 0.032023621 | 0.245981597 |
| ENSMUSG00000085037 | 4933421O10Rik | 100504166 | lncRNA | 125.2923789 | 0.245998575 | 0.109943039 | 2.237509315 | 0.025253071 | 0.115566481 | 0.245998575 |
| ENSMUSG00000021589 | Rhobtb3 | 73296 | protein_coding | 220.1544184 | 0.246083513 | 0.101768521 | 2.418071038 | 0.01560303 | 0.082474622 | 0.246083513 |
| ENSMUSG00000020173 | Cobl | 12808 | protein_coding | 1818.055049 | -0.246233567 | 0.051894735 | -4.744866036 | 2.09E-06 | 4.78E-05 | 0.246233567 |
| ENSMUSG00000031173 | Otc | 18416 | protein_coding | 683.0521951 | -0.24623553 | 0.097762481 | -2.518711952 | 0.011778497 | 0.067120535 | 0.24623553 |
| ENSMUSG00000074219 | Gm10644 | 100126034 | lncRNA | 7.574773065 | -0.246332131 | 0.485275753 | -0.507612692 | 0.611724988 | 0.79296591 | 0.246332131 |
| ENSMUSG00000032059 | Alg9 | 102580 | protein_coding | 984.1670426 | 0.246482873 | 0.04986625 | 4.942879711 | 7.70E-07 | 1.94E-05 | 0.246482873 |
| ENSMUSG00000038650 | Rnh1 | 107702 | protein_coding | 2805.794838 | 0.24653169 | 0.049859543 | 4.944523656 | 7.63E-07 | 1.93E-05 | 0.24653169 |
| ENSMUSG00000052105 | Mtcl1 | 68617 | protein_coding | 127.7700709 | -0.246777595 | 0.123837083 | -1.992760081 | 0.046287725 | 0.175601538 | 0.246777595 |
| ENSMUSG00000021236 | Entpd5 | 12499 | protein_coding | 7132.236481 | -0.247082221 | 0.063457171 | -3.893684787 | 9.87E-05 | 0.001467526 | 0.247082221 |
| ENSMUSG00000027078 | Ube2l6 | 56791 | protein_coding | 156.7619215 | -0.247235313 | 0.154699486 | -1.598165059 | 0.110006248 | 0.304322033 | 0.247235313 |
| ENSMUSG00000021194 | Chga | 12652 | protein_coding | 1121.316274 | -0.247275869 | 0.095128023 | -2.599400894 | 0.009338664 | 0.056639577 | 0.247275869 |
| ENSMUSG00000028525 | Pde4b | 18578 | protein_coding | 37.93528032 | 0.247300422 | 0.188239745 | 1.313752428 | 0.188929531 | 0.419340469 | 0.247300422 |
| ENSMUSG00000054752 | Fsd1l | 319636 | protein_coding | 285.8796382 | -0.247307849 | 0.076199119 | -3.245547346 | 0.001172251 | 0.011492333 | 0.247307849 |
| ENSMUSG00000095620 | Csta2 | 76770 | protein_coding | 10.99784891 | 0.247310836 | 0.463711168 | 0.53332948 | 0.593805524 | 0.781164996 | 0.247310836 |

| ensembl_gene_id | external_gene_name | entrezgene_id | gene_biotype | baseMean | log2FoldChange | lfcSE | stat | pvalue | padj | absLog2FC |
|---|---|---|---|---|---|---|---|---|---|---|
| ENSMUSG00000075271 | Ttc30a1 | 78802 | protein_coding | 87.48409217 | 0.247365409 | 0.141579104 | 1.747188692 | 0.080604613 | 0.251384759 | 0.247365409 |
| ENSMUSG00000079017 | Ifi27l2a | 76933 | protein_coding | 8.142432645 | -0.247402121 | 0.473562026 | -0.522428125 | 0.601372278 | 0.786583124 | 0.247402121 |
| ENSMUSG00000015749 | Anp32e | 66471 | protein_coding | 898.0058197 | 0.247412837 | 0.092746348 | 2.667628897 | 0.007638858 | 0.04892449 | 0.247412837 |
| ENSMUSG00000039887 | Alg14 | 66789 | protein_coding | 316.5479129 | 0.24742876 | 0.083578243 | 2.960444624 | 0.003071953 | 0.02456192 | 0.24742876 |
| ENSMUSG00000035637 | Grhpr | 76238 | protein_coding | 501.0684589 | 0.247446042 | 0.056992297 | 4.341745357 | 1.41E-05 | 0.000266624 | 0.247446042 |
| ENSMUSG00000049241 | Hcar1 | 243270 | protein_coding | 8.456643657 | -0.24744888 | 0.440092122 | -0.562266098 | 0.573934733 | 0.76815045 | 0.24744888 |
| ENSMUSG00000029512 | Ulk1 | 22241 | protein_coding | 466.0410875 | -0.247478014 | 0.079947286 | -3.095514889 | 0.001964715 | 0.017314186 | 0.247478014 |
| ENSMUSG00000054414 | Slc30a7 | 66500 | protein_coding | 1493.30176 | 0.247581481 | 0.045547047 | 5.43573063 | 5.46E-08 | 1.75E-06 | 0.247581481 |
| ENSMUSG00000032109 | Nlrx1 | 270151 | protein_coding | 707.9824572 | 0.247634825 | 0.108601677 | 2.280211792 | 0.02259513 | 0.107008585 | 0.247634825 |
| ENSMUSG00000032786 | Alas1 | 11655 | protein_coding | 3232.677512 | 0.24781948 | 0.076034032 | 3.259323133 | 0.001116784 | 0.011052553 | 0.24781948 |
| ENSMUSG00000022324 | Matn2 | 17181 | protein_coding | 179.5843901 | 0.247897565 | 0.148063775 | 1.674262085 | 0.094079102 | 0.277319263 | 0.247897565 |
| ENSMUSG00000002833 | Hdgfl2 | 15193 | protein_coding | 627.1074637 | 0.24792376 | 0.062656903 | 3.956846689 | 7.59E-05 | 0.001169795 | 0.24792376 |
| ENSMUSG00000049588 | Ccdc69 | 52570 | protein_coding | 10.05499794 | 0.247977493 | 0.397383666 | 0.624025379 | 0.532610918 | 0.740799392 | 0.247977493 |
| ENSMUSG00000052584 | Serp2 | 72661 | protein_coding | 8.23221529 | 0.247979812 | 0.406583301 | 0.609911453 | 0.541920465 | 0.747417373 | 0.247979812 |
| ENSMUSG00000035852 | Misp | 78906 | protein_coding | 12329.04393 | 0.248005693 | 0.07648509 | 3.242536464 | 0.001184708 | 0.011600354 | 0.248005693 |
| ENSMUSG00000027253 | Lrp4 | 228357 | protein_coding | 1549.146829 | -0.248061811 | 0.057548097 | -4.310512877 | 1.63E-05 | 0.000301734 | 0.248061811 |
| ENSMUSG00000024897 | Apba1 | 319924 | protein_coding | 11.69881634 | -0.248270148 | 0.404653282 | -0.613537958 | 0.539520693 | 0.745510257 | 0.248270148 |
| ENSMUSG00000055024 | Ep300 | 328572 | protein_coding | 1381.895748 | -0.248292531 | 0.084701796 | -2.93137268 | 0.003374677 | 0.026573931 | 0.248292531 |
| ENSMUSG00000053950 | Adnp2 | 240442 | protein_coding | 738.1726919 | 0.24846646 | 0.068720528 | 3.615607552 | 0.000299644 | 0.003677595 | 0.24846646 |
| ENSMUSG00000010911 | Apip | 56369 | protein_coding | 131.0491503 | 0.24846911 | 0.110431473 | 2.249984576 | 0.024449924 | 0.113078271 | 0.24846911 |
| ENSMUSG00000041642 | Kif21b | 16565 | protein_coding | 1217.578729 | -0.248589799 | 0.090228445 | -2.75511564 | 0.005867139 | 0.040297853 | 0.248589799 |
| ENSMUSG00000027297 | Ltk | 17005 | protein_coding | 22.27784336 | 0.248688509 | 0.323584261 | 0.768543278 | 0.442164489 | 0.673287388 | 0.248688509 |
| ENSMUSG00000025665 | Rps6ka6 | 67071 | protein_coding | 70.18133962 | 0.248698347 | 0.180943992 | 1.37444932 | 0.169302234 | 0.393611906 | 0.248698347 |
| ENSMUSG00000031672 | Got2 | 14719 | protein_coding | 1436.032235 | 0.248877925 | 0.062522975 | 3.980583555 | 6.87E-05 | 0.001078474 | 0.248877925 |
| ENSMUSG00000037664 | Cdkn1c | 12577 | protein_coding | 194.7759665 | 0.24894274 | 0.119544157 | 2.082433362 | 0.037302904 | 0.150848528 | 0.24894274 |
| ENSMUSG00000022974 | Paxbp1 | 67367 | protein_coding | 613.7062971 | -0.248992145 | 0.102872397 | -2.420398006 | 0.015503528 | 0.08213712 | 0.248992145 |
| ENSMUSG00000029188 | Slc34a2 | 20531 | protein_coding | 880.2188133 | 0.249147356 | 0.222780214 | 1.118354953 | 0.263415425 | 0.504916665 | 0.249147356 |
| ENSMUSG00000025102 | 3110040N11Rik | 67290 | protein_coding | 272.8729604 | 0.249152522 | 0.091128632 | 2.734075085 | 0.006255578 | 0.042325937 | 0.249152522 |
| ENSMUSG00000023988 | Bysl | 53414 | protein_coding | 342.0960775 | 0.249155809 | 0.085098588 | 2.927848899 | 0.003413159 | 0.026787965 | 0.249155809 |
| ENSMUSG00000033569 | Adgrb3 | 210933 | protein_coding | 20.45307023 | -0.24918385 | 0.318814086 | -0.781596112 | 0.434451973 | 0.668221346 | 0.24918385 |
| ENSMUSG00000024246 | Thumpd2 | 72167 | protein_coding | 66.30931518 | -0.249189769 | 0.155828444 | -1.59912891 | 0.109791962 | 0.303863905 | 0.249189769 |
| ENSMUSG00000045948 | Mrps12 | 24030 | protein_coding | 703.4433994 | 0.249200651 | 0.091342314 | 2.728206017 | 0.006367982 | 0.042790188 | 0.249200651 |
| ENSMUSG00000092680 | Snord55 | 100216533 | snoRNA | 16.53321407 | -0.249243152 | 0.395275923 | -0.630554855 | 0.528331625 | 0.737932232 | 0.249243152 |
| ENSMUSG00000031662 | Snx20 | 71607 | protein_coding | 334.6688024 | 0.249482589 | 0.094771657 | 2.632459913 | 0.008476903 | 0.052776264 | 0.249482589 |
| ENSMUSG00000085057 | Gm13415 | 102638285 | lncRNA | 13.92043464 | -0.249483963 | 0.320361476 | -0.778757691 | 0.436122465 | 0.669302737 | 0.249483963 |
| ENSMUSG00000031209 | Heph | 15203 | protein_coding | 5199.192145 | 0.249505241 | 0.056401468 | 4.423736646 | 9.70E-06 | 0.000191253 | 0.249505241 |
| ENSMUSG00000031379 | Pir | 69656 | protein_coding | 91.62230999 | -0.249569715 | 0.133719686 | -1.866364804 | 0.061990344 | 0.212479973 | 0.249569715 |
| ENSMUSG00000030465 | Psd3 | 234353 | protein_coding | 1570.58167 | -0.249598355 | 0.052676664 | -4.738309836 | 2.16E-06 | 4.90E-05 | 0.249598355 |
| ENSMUSG00000021506 | Pitx1 | 18740 | protein_coding | 20.14786602 | 0.249620452 | 0.301715288 | 0.827337762 | 0.408045646 | 0.646554316 | 0.249620452 |

| ensembl_gene_id | external_gene_name | entrezgene_id | gene_biotype | baseMean | log2FoldChange | lfcSE | stat | pvalue | padj | absLog2FC |
|---|---|---|---|---|---|---|---|---|---|---|
| ENSMUSG00000018169 | Mfng | 17305 | protein_coding | 7.413854102 | 0.249671777 | 0.472411927 | 0.52850439 | 0.597149301 | 0.783469018 | 0.249671777 |
| ENSMUSG00000036752 | Tubb4b | 227613 | protein_coding | 11460.27478 | 0.249739005 | 0.085884896 | 2.907833811 | 0.003639417 | 0.028136566 | 0.249739005 |
| ENSMUSG00000054598 | 9130230L23Rik | 231253 | lncRNA | 71.7081903 | -0.249848851 | 0.170049899 | -1.469267858 | 0.141760151 | 0.352748952 | 0.249848851 |
| ENSMUSG00000034361 | Cpne2 | 234577 | protein_coding | 319.1488224 | -0.249876249 | 0.094884134 | -2.633488214 | 0.008451277 | 0.052651813 | 0.249876249 |
| ENSMUSG00000074513 | Arfip1 | 99889 | protein_coding | 435.5064479 | 0.249918531 | 0.069180342 | 3.612565701 | 0.000303182 | 0.003706914 | 0.249918531 |
| ENSMUSG00000034066 | Farp2 | 227377 | protein_coding | 931.0763199 | -0.249997252 | 0.063222483 | -3.954246024 | 7.68E-05 | 0.001181462 | 0.249997252 |
| ENSMUSG00000015342 | Xk | 22439 | protein_coding | 465.2676178 | 0.250231903 | 0.074961576 | 3.338135577 | 0.000843426 | 0.008741599 | 0.250231903 |
| ENSMUSG00000017146 | Brca1 | 12189 | protein_coding | 261.6272723 | 0.25025968 | 0.17633266 | 1.419247458 | 0.155826884 | 0.374239786 | 0.25025968 |
| ENSMUSG00000043088 | Il17re | 57890 | protein_coding | 676.5857234 | 0.250312853 | 0.076893515 | 3.255318116 | 0.001132654 | 0.01118574 | 0.250312853 |
| ENSMUSG00000039997 | Ifi203 | 15950 | protein_coding | 9.565228428 | 0.250364423 | 0.43054407 | 0.581507074 | 0.560898752 | 0.759175676 | 0.250364423 |
| ENSMUSG00000032410 | Xrn1 | 24127 | protein_coding | 1334.432863 | -0.250580164 | 0.068609839 | -3.652248231 | 0.000259954 | 0.003277783 | 0.250580164 |
| ENSMUSG00000035632 | Cnot3 | 232791 | protein_coding | 457.0409065 | 0.250653035 | 0.070245604 | 3.568238041 | 0.00035939 | 0.004302807 | 0.250653035 |
| ENSMUSG00000046826 | Fam187b | 76415 | protein_coding | 71.64164487 | 0.25071099 | 0.177873229 | 1.409492545 | 0.158689576 | 0.378894236 | 0.25071099 |
| ENSMUSG00000030410 | Dmwd | 13401 | protein_coding | 26.56257986 | 0.25081479 | 0.251127382 | 0.998755246 | 0.317913271 | 0.561627805 | 0.25081479 |
| ENSMUSG00000033684 | Qsox1 | 104009 | protein_coding | 3400.338627 | 0.251104397 | 0.063052991 | 3.982434362 | 6.82E-05 | 0.00107324 | 0.251104397 |
| ENSMUSG00000022383 | Ppara | 19013 | protein_coding | 836.4169498 | -0.251203891 | 0.115239425 | -2.17984332 | 0.029269078 | 0.127940029 | 0.251203891 |
| ENSMUSG00000008035 | Mid1ip1 | 68041 | protein_coding | 1649.039709 | 0.251240301 | 0.113681966 | 2.210027753 | 0.027103236 | 0.120877735 | 0.251240301 |
| ENSMUSG00000025271 | Pfkfb1 | 18639 | protein_coding | 30.8960292 | -0.251345611 | 0.233903112 | -1.074571472 | 0.28256663 | 0.525297065 | 0.251345611 |
| ENSMUSG00000026771 | Spopl | 76857 | protein_coding | 482.8765014 | -0.251568738 | 0.06693943 | -3.75815476 | 0.000171171 | 0.002308089 | 0.251568738 |
| ENSMUSG00000020527 | Myo19 | 66196 | protein_coding | 1237.93802 | -0.251638534 | 0.081782157 | -3.076936864 | 0.002091396 | 0.018182179 | 0.251638534 |
| ENSMUSG00000078157 | Fem1al | 216622 | protein_coding | 72.08121093 | -0.251810613 | 0.151863206 | -1.658141034 | 0.097289003 | 0.283264621 | 0.251810613 |
| ENSMUSG00000063524 | Eno1 | 13806 | protein_coding | 5458.90183 | 0.251876274 | 0.076538122 | 3.290860373 | 0.000998815 | 0.010029817 | 0.251876274 |
| ENSMUSG00000018822 | Sfrp5 | 54612 | protein_coding | 8.101030205 | 0.251928519 | 0.427483071 | 0.589329815 | 0.555640048 | 0.75524772 | 0.251928519 |
| ENSMUSG00000040699 | Limd2 | 67803 | protein_coding | 136.8330688 | 0.251945719 | 0.121491813 | 2.073767055 | 0.03810095 | 0.153453234 | 0.251945719 |
| ENSMUSG00000025855 | Prkar1b | 19085 | protein_coding | 14.27300954 | 0.252021921 | 0.335278983 | 0.751678254 | 0.452244571 | 0.68117317 | 0.252021921 |
| ENSMUSG00000021495 | Fam193b | 212483 | protein_coding | 567.6061333 | -0.25206408 | 0.119821288 | -2.103666931 | 0.035407511 | 0.145593302 | 0.25206408 |
| ENSMUSG00000022244 | Amacr | 17117 | protein_coding | 322.943919 | 0.25214175 | 0.079061637 | 3.189179482 | 0.001426773 | 0.013434123 | 0.25214175 |
| ENSMUSG00000021061 | Sptb | 20741 | protein_coding | 79.12966793 | -0.252163949 | 0.145123762 | -1.737578643 | 0.082285086 | 0.254690744 | 0.252163949 |
| ENSMUSG00000032599 | Ip6k2 | 76500 | protein_coding | 418.7042094 | -0.252361722 | 0.092918356 | -2.715951211 | 0.006608566 | 0.043967136 | 0.252361722 |
| ENSMUSG00000046312 | Myorg | 329828 | protein_coding | 606.9202706 | 0.252372069 | 0.060936532 | 4.141556148 | 3.45E-05 | 0.000588069 | 0.252372069 |
| ENSMUSG00000065444 | Mir27a | 387220 | miRNA | 31.46566285 | -0.252483488 | 0.329631945 | -0.765955763 | 0.443702624 | 0.674737597 | 0.252483488 |
| ENSMUSG00000024594 | Prrc1 | 73137 | protein_coding | 1083.608065 | 0.252500251 | 0.075219509 | 3.356845245 | 0.000788372 | 0.008266632 | 0.252500251 |
| ENSMUSG00000036931 | Nfkbid | 243910 | protein_coding | 81.19443587 | -0.252609834 | 0.179533224 | -1.407036696 | 0.159416508 | 0.379526132 | 0.252609834 |
| ENSMUSG00000048076 | Arf1 | 11840 | protein_coding | 8759.725763 | 0.252697096 | 0.061187536 | 4.129878632 | 3.63E-05 | 0.000615497 | 0.252697096 |
| ENSMUSG00000021556 | Golm1 | 105348 | protein_coding | 13970.74043 | 0.252714814 | 0.05770887 | 4.379132935 | 1.19E-05 | 0.000228758 | 0.252714814 |
| ENSMUSG00000069268 | H2bc7 | 319180 | protein_coding | 320.2790324 | 0.252748983 | 0.142824652 | 1.769645357 | 0.076786238 | 0.24422641 | 0.252748983 |
| ENSMUSG00000028600 | Podn | 242608 | protein_coding | 8.288210731 | -0.252884483 | 0.415172806 | -0.609106568 | 0.542453804 | 0.747864433 | 0.252884483 |
| ENSMUSG00000072115 | Ang | 11727 | protein_coding | 513.2323125 | -0.25292446 | 0.069155122 | -3.657349602 | 0.000254837 | 0.003228399 | 0.25292446 |
| ENSMUSG00000014504 | Srp19 | 66384 | protein_coding | 478.3244323 | 0.252930226 | 0.067027738 | 3.773515742 | 0.000160963 | 0.002188745 | 0.252930226 |

| ensembl_gene_id | external_gene_name | entrezgene_id | gene_biotype | baseMean | log2FoldChange | lfcSE | stat | pvalue | padj | absLog2FC |
|---|---|---|---|---|---|---|---|---|---|---|
| ENSMUSG00000007817 | Zmiz1 | 328365 | protein_coding | 1530.742433 | -0.252984105 | 0.075092586 | -3.368962466 | 0.000754517 | 0.007978132 | 0.252984105 |
| ENSMUSG00000020826 | Nos2 | 18126 | protein_coding | 11.71446699 | -0.252988098 | 0.487850877 | -0.5185767 | 0.604055961 | 0.788602807 | 0.252988098 |
| ENSMUSG00000035790 | Cep19 | 66994 | protein_coding | 506.9136902 | 0.253274941 | 0.060880221 | 4.160217171 | 3.18E-05 | 0.00054606 | 0.253274941 |
| ENSMUSG00000087968 | Gm25395 | 115485773 | scaRNA | 795.8391185 | 0.253353078 | 0.233451285 | 1.085250303 | 0.27781081 | 0.520660159 | 0.253353078 |
| ENSMUSG00000033767 | Tmem131l | 229473 | protein_coding | 432.4790687 | -0.253378783 | 0.083987539 | -3.016861611 | 0.002554064 | 0.021166181 | 0.253378783 |
| ENSMUSG00000031434 | Morc4 | 75746 | protein_coding | 160.2185227 | 0.253482926 | 0.122541783 | 2.068542832 | 0.038589007 | 0.154536441 | 0.253482926 |
| ENSMUSG00000042759 | Apobr | 171504 | protein_coding | 15.27318688 | 0.253542196 | 0.29646764 | 0.855210353 | 0.392434706 | 0.632120905 | 0.253542196 |
| ENSMUSG00000097073 | 9430037G07Rik | 320692 | lncRNA | 17.25485863 | 0.253592507 | 0.341432914 | 0.742730115 | 0.457645093 | 0.685405128 | 0.253592507 |
| ENSMUSG00000025736 | Jmjd8 | 72106 | protein_coding | 408.3396834 | 0.253603152 | 0.08903418 | 2.848379696 | 0.004394246 | 0.032388488 | 0.253603152 |
| ENSMUSG00000027423 | Snx5 | 69178 | protein_coding | 2795.931502 | -0.25365085 | 0.070432034 | -3.60135634 | 0.000316561 | 0.003847162 | 0.25365085 |
| ENSMUSG00000050762 | Prss27 | 213171 | protein_coding | 9.136570992 | -0.253695737 | 0.440494146 | -0.575934412 | 0.564659515 | 0.761458074 | 0.253695737 |
| ENSMUSG00000104960 | Snhg8 | 69895 | lncRNA | 264.8608635 | -0.253718494 | 0.099391621 | -2.552715124 | 0.010688688 | 0.06217684 | 0.253718494 |
| ENSMUSG00000056870 | Gulp1 | 70676 | protein_coding | 119.6988145 | 0.254095952 | 0.15607375 | 1.628050533 | 0.103514171 | 0.294056357 | 0.254095952 |
| ENSMUSG00000045009 | Prrt3 | 210673 | protein_coding | 12.98442663 | 0.254209096 | 0.356130363 | 0.713809105 | 0.475345229 | 0.69880515 | 0.254209096 |
| ENSMUSG00000062861 | Zfp28 | 22690 | protein_coding | 55.09358839 | -0.254211312 | 0.165638099 | -1.534739372 | 0.12484786 | 0.32795219 | 0.254211312 |
| ENSMUSG00000052605 | Isoc2b | 67441 | protein_coding | 339.6720638 | 0.254316923 | 0.075239699 | 3.380089598 | 0.000724622 | 0.007744188 | 0.254316923 |
| ENSMUSG00000044037 | Als2cl | 235633 | protein_coding | 664.8330498 | -0.254401658 | 0.098423852 | -2.584756165 | 0.009744789 | 0.058308165 | 0.254401658 |
| ENSMUSG00000048965 | Mrgpre | 244238 | protein_coding | 14.91404442 | -0.25443315 | 0.316927648 | -0.802811467 | 0.422083713 | 0.658502858 | 0.25443315 |
| ENSMUSG00000026500 | Cox20 | 66359 | protein_coding | 16.86386254 | 0.254459765 | 0.324377 | 0.784456867 | 0.432772082 | 0.666983502 | 0.254459765 |
| ENSMUSG00000044626 | Liph | 239759 | protein_coding | 1360.650337 | 0.254537922 | 0.043722645 | 5.82164966 | 5.83E-09 | 2.13E-07 | 0.254537922 |
| ENSMUSG00000028687 | Mutyh | 70603 | protein_coding | 29.99068811 | 0.254638361 | 0.216763389 | 1.174729562 | 0.240102928 | 0.480533622 | 0.254638361 |
| ENSMUSG00000020151 | Ptprr | 19279 | protein_coding | 660.3514533 | -0.25474753 | 0.132636791 | -1.920640025 | 0.054777106 | 0.196571068 | 0.25474753 |
| ENSMUSG00000024691 | Fam111a | 107373 | protein_coding | 230.9156002 | 0.254884888 | 0.164356716 | 1.550802999 | 0.120948902 | 0.322031193 | 0.254884888 |
| ENSMUSG00000027224 | Duoxa1 | 213696 | protein_coding | 7.385770634 | 0.255129161 | 0.433741917 | 0.588204991 | 0.556394708 | 0.755428309 | 0.255129161 |
| ENSMUSG00000061808 | Ttr | 22139 | protein_coding | 468.7957732 | 0.255199477 | 0.129802593 | 1.966058388 | 0.049291866 | 0.1835442 | 0.255199477 |
| ENSMUSG00000021337 | Scgn | 214189 | protein_coding | 35.94253577 | -0.255322102 | 0.236709096 | -1.078632411 | 0.280751627 | 0.523508209 | 0.255322102 |
| ENSMUSG00000040247 | Tbc1d10c | 108995 | protein_coding | 11.00639094 | -0.255331676 | 0.368739918 | -0.692443815 | 0.488658658 | 0.708970987 | 0.255331676 |
| ENSMUSG00000025178 | Pi4k2a | 84095 | protein_coding | 620.6202475 | 0.255383592 | 0.070802671 | 3.606976792 | 0.000309785 | 0.003779053 | 0.255383592 |
| ENSMUSG00000045751 | Mms22l | 212377 | protein_coding | 208.9346728 | 0.255423707 | 0.127904646 | 1.996985364 | 0.045826773 | 0.174452968 | 0.255423707 |
| ENSMUSG00000075324 | Fign | 60344 | protein_coding | 8.197467234 | 0.255483094 | 0.62756544 | 0.407101918 | 0.683933135 | 0.840783841 | 0.255483094 |
| ENSMUSG00000048058 | Ldlrad3 | 241576 | protein_coding | 612.3457213 | -0.255504413 | 0.093833435 | -2.72295705 | 0.006470047 | 0.04322388 | 0.255504413 |
| ENSMUSG00000117679 | Apbb3 | 225372 | protein_coding | 223.3095036 | -0.255505646 | 0.107677631 | -2.372875821 | 0.017650197 | 0.090105959 | 0.255505646 |
| ENSMUSG00000037235 | Mxd4 | 17122 | protein_coding | 3199.33637 | 0.255536784 | 0.056355156 | 4.534399414 | 5.78E-06 | 0.000120676 | 0.255536784 |
| ENSMUSG00000114828 | AI463229 | 100042802 | lncRNA | 44.78895352 | 0.255576308 | 0.185484471 | 1.377885207 | 0.168238733 | 0.392248958 | 0.255576308 |
| ENSMUSG00000020660 | Pomc | 18976 | protein_coding | 6.061238168 | -0.255688924 | 0.518438512 | -0.493190452 | 0.621878024 | 0.800401189 | 0.255688924 |
| ENSMUSG00000016487 | Ppfibp1 | 67533 | protein_coding | 551.2286249 | 0.255693944 | 0.05799134 | 4.409174618 | 1.04E-05 | 0.000203085 | 0.255693944 |
| ENSMUSG00000025499 | Hras | 15461 | protein_coding | 1386.119992 | 0.25579034 | 0.062489459 | 4.093335837 | 4.25E-05 | 0.000706218 | 0.25579034 |
| ENSMUSG00000039765 | Cc2d2a | 231214 | protein_coding | 566.8791521 | -0.255922776 | 0.08180022 | -3.128631863 | 0.001756222 | 0.015822676 | 0.255922776 |
| ENSMUSG00000000149 | Gna12 | 14673 | protein_coding | 639.1828067 | 0.25595078 | 0.056353839 | 4.541851705 | 5.58E-06 | 0.000116789 | 0.25595078 |

| ensembl_gene_id | external_gene_name | entrezgene_id | gene_biotype | baseMean | log2FoldChange | lfcSE | stat | pvalue | padj | absLog2FC |
|---|---|---|---|---|---|---|---|---|---|---|
| ENSMUSG00000028339 | Col15a1 | 12819 | protein_coding | 39.843964 | 0.25606664 | 0.199562482 | 1.283140187 | 0.199442967 | 0.432954196 | 0.25606664 |
| ENSMUSG00000080727 | C920021L13Rik | 100042889 | lncRNA | 18.37218659 | -0.256138951 | 0.288310668 | -0.888413021 | 0.374318624 | 0.616171935 | 0.256138951 |
| ENSMUSG00000027496 | Aurka | 20878 | protein_coding | 354.9073019 | 0.256207321 | 0.098555637 | 2.599621164 | 0.009332673 | 0.056624534 | 0.256207321 |
| ENSMUSG00000038695 | Josd2 | 66124 | protein_coding | 345.0240791 | 0.256214671 | 0.098451056 | 2.602457305 | 0.009255834 | 0.056349126 | 0.256214671 |
| ENSMUSG00000034761 | Map4k5 | 399510 | protein_coding | 703.2872662 | -0.256286222 | 0.05751391 | -4.456073705 | 8.35E-06 | 0.000167021 | 0.256286222 |
| ENSMUSG00000020744 | Slc25a19 | 67283 | protein_coding | 381.5531153 | 0.256287661 | 0.064970844 | 3.944656455 | 7.99E-05 | 0.001219282 | 0.256287661 |
| ENSMUSG00000055681 | Cope | 59042 | protein_coding | 2727.941407 | 0.256288462 | 0.067249874 | 3.81098801 | 0.000138412 | 0.001947167 | 0.256288462 |
| ENSMUSG00000074656 | Eif2s2 | 67204 | protein_coding | 5461.08608 | -0.256341013 | 0.069218921 | -3.703337287 | 0.000212782 | 0.002772663 | 0.256341013 |
| ENSMUSG00000028145 | Them4 | 75778 | protein_coding | 223.6735548 | -0.256487933 | 0.087049007 | -2.946477416 | 0.003214159 | 0.025521783 | 0.256487933 |
| ENSMUSG00000050100 | Hmx2 | 15372 | protein_coding | 22.48828755 | 0.25657049 | 0.308050512 | 0.832884479 | 0.404909884 | 0.644152928 | 0.25657049 |
| ENSMUSG00000036390 | Gadd45a | 13197 | protein_coding | 476.7196938 | -0.256806761 | 0.085011492 | -3.020847575 | 0.002520682 | 0.020943349 | 0.256806761 |
| ENSMUSG00000065176 | Rnu12 | 104307 | snRNA | 41.21851016 | 0.257008848 | 0.233974626 | 1.098447521 | 0.272009121 | 0.514567871 | 0.257008848 |
| ENSMUSG00000035704 | Alg8 | 381903 | protein_coding | 205.9682761 | 0.257053093 | 0.136363579 | 1.885056812 | 0.059422168 | 0.206976523 | 0.257053093 |
| ENSMUSG00000018387 | Shroom1 | 71774 | protein_coding | 18.34503065 | 0.257152528 | 0.269416163 | 0.9544807 | 0.339840374 | 0.583105193 | 0.257152528 |
| ENSMUSG00000042644 | Itpr3 | 16440 | protein_coding | 5224.19074 | -0.257319238 | 0.061561298 | -4.179886489 | 2.92E-05 | 0.000505211 | 0.257319238 |
| ENSMUSG00000035126 | Dnai4 | 242584 | protein_coding | 45.05232355 | 0.257372749 | 0.181543487 | 1.417692001 | 0.156280706 | 0.374715872 | 0.257372749 |
| ENSMUSG00000040557 | Mettl27 | 79565 | protein_coding | 343.1892869 | -0.257408363 | 0.102009479 | -2.523376913 | 0.011623374 | 0.066299536 | 0.257408363 |
| ENSMUSG00000032449 | Slc25a36 | 192287 | protein_coding | 1357.567228 | -0.257469359 | 0.081266211 | -3.168221537 | 0.001533746 | 0.014248111 | 0.257469359 |
| ENSMUSG00000022070 | Bora | 77744 | protein_coding | 291.4551109 | -0.257568797 | 0.110339578 | -2.334328277 | 0.019578539 | 0.09661718 | 0.257568797 |
| ENSMUSG00000038871 | Bpgm | 12183 | protein_coding | 835.0226583 | 0.257613529 | 0.047043571 | 5.476062335 | 4.35E-08 | 1.41E-06 | 0.257613529 |
| ENSMUSG00000001211 | Agpat3 | 28169 | protein_coding | 1245.56504 | 0.257774308 | 0.066407904 | 3.881681119 | 0.000103737 | 0.001537651 | 0.257774308 |
| ENSMUSG00000057666 | Gapdh | 14433 | protein_coding | 12187.1148 | 0.25777664 | 0.052347843 | 4.924303011 | 8.47E-07 | 2.11E-05 | 0.25777664 |
| ENSMUSG00000064210 | Ano6 | 105722 | protein_coding | 632.0337939 | 0.257796238 | 0.072704696 | 3.545799 | 0.000391425 | 0.004604307 | 0.257796238 |
| ENSMUSG00000010362 | Rdm1 | 66599 | protein_coding | 298.3039627 | -0.257898094 | 0.079924712 | -3.22676287 | 0.001251992 | 0.012155852 | 0.257898094 |
| ENSMUSG00000046697 | Enpp7 | 238011 | protein_coding | 38.34790294 | 0.257948812 | 0.942073889 | 0.273809534 | 0.784230997 | 0.899522993 | 0.257948812 |
| ENSMUSG00000038160 | Atg5 | 11793 | protein_coding | 977.0147246 | 0.257951024 | 0.055881426 | 4.616042238 | 3.91E-06 | 8.39E-05 | 0.257951024 |
| ENSMUSG00000067229 | Cyp2c66 | 69888 | protein_coding | 92.16940004 | -0.257990837 | 0.140143495 | -1.840904837 | 0.065635505 | 0.220797621 | 0.257990837 |
| ENSMUSG00000072770 | Acrbp | 54137 | protein_coding | 37.87356147 | -0.258009068 | 0.213134083 | -1.210548142 | 0.226068629 | 0.464315218 | 0.258009068 |
| ENSMUSG00000027071 | P2rx3 | 228139 | protein_coding | 11.38000581 | -0.258217284 | 0.403721748 | -0.639592207 | 0.52243775 | 0.733404735 | 0.258217284 |
| ENSMUSG00000062822 | 4833420G17Rik | 67392 | protein_coding | 596.5771618 | -0.258236855 | 0.132461418 | -1.949525078 | 0.051232751 | 0.188241916 | 0.258236855 |
| ENSMUSG00000006445 | Epha2 | 13836 | protein_coding | 1013.400987 | -0.258237184 | 0.121396222 | -2.127225865 | 0.033401321 | 0.140386527 | 0.258237184 |
| ENSMUSG00000024474 | Ik | 24010 | protein_coding | 1778.60928 | 0.258365411 | 0.051556697 | 5.011287135 | 5.41E-07 | 1.40E-05 | 0.258365411 |
| ENSMUSG00000026715 | Serpinc1 | 11905 | protein_coding | 8.446846447 | -0.258369892 | 0.468233333 | -0.551797306 | 0.581087233 | 0.772887622 | 0.258369892 |
| ENSMUSG00000069272 | H2ac8 | 319166 | protein_coding | 314.2581076 | 0.258458906 | 0.171113463 | 1.510453363 | 0.130927781 | 0.336836503 | 0.258458906 |
| ENSMUSG00000044966 | Fbxo48 | 319701 | protein_coding | 11.98273956 | 0.258504709 | 0.39415259 | 0.655849322 | 0.511921078 | 0.725588963 | 0.258504709 |
| ENSMUSG00000024696 | Lpxn | 107321 | protein_coding | 29.64156722 | 0.258552144 | 0.219271279 | 1.179142771 | 0.23834133 | 0.478679126 | 0.258552144 |
| ENSMUSG00000004113 | Cacna1b | 12287 | protein_coding | 164.1847666 | 0.258555083 | 0.098081004 | 2.636138206 | 0.008385557 | 0.052335092 | 0.258555083 |
| ENSMUSG00000020387 | Jade2 | 76901 | protein_coding | 478.7271808 | -0.258566132 | 0.081012083 | -3.191698335 | 0.00141439 | 0.013331305 | 0.258566132 |
| ENSMUSG00000043998 | Mgat2 | 217664 | protein_coding | 1125.472453 | 0.258573343 | 0.055454603 | 4.662793149 | 3.12E-06 | 6.87E-05 | 0.258573343 |

Supplementary Table 1

| ensembl_gene_id | external_gene_name | entrezgene_id | gene_biotype | baseMean | log2FoldChange | lfcSE | stat | pvalue | padj | absLog2FC |
|---|---|---|---|---|---|---|---|---|---|---|
| ENSMUSG00000045176 | Borcs6 | 71923 | protein_coding | 342.0148216 | 0.258670202 | 0.080676963 | 3.206246137 | 0.001344789 | 0.012791297 | 0.258670202 |
| ENSMUSG00000099517 | H3c8 | 97908 | protein_coding | 313.0725347 | 0.258797557 | 0.172661229 | 1.498874748 | 0.133906129 | 0.341261715 | 0.258797557 |
| ENSMUSG00000109564 | Muc16 | 73732 | protein_coding | 26.52545791 | -0.258817247 | 0.49248571 | -0.525532501 | 0.599213066 | 0.78494916 | 0.258817247 |
| ENSMUSG00000033216 | Eefsec | 65967 | protein_coding | 329.6660686 | 0.25883264 | 0.086156673 | 3.004208851 | 0.002662725 | 0.02189789 | 0.25883264 |
| ENSMUSG00000028278 | Rragd | 52187 | protein_coding | 18.75752976 | -0.258873006 | 0.359588129 | -0.719915329 | 0.471577129 | 0.696004848 | 0.258873006 |
| ENSMUSG00000016024 | Lbp | 16803 | protein_coding | 15.69162451 | 0.258926062 | 0.346513901 | 0.747231385 | 0.454923903 | 0.682988492 | 0.258926062 |
| ENSMUSG00000035198 | Tubg1 | 103733 | protein_coding | 288.9166886 | 0.258939078 | 0.103503674 | 2.501738023 | 0.012358534 | 0.069590109 | 0.258939078 |
| ENSMUSG00000030551 | Nr2f2 | 11819 | protein_coding | 248.8948825 | -0.258939733 | 0.090682816 | -2.855444324 | 0.004297666 | 0.031924668 | 0.258939733 |
| ENSMUSG00000071537 | Klrg2 | 74253 | protein_coding | 7.103536201 | 0.258956817 | 0.463984563 | 0.558115156 | 0.576765752 | 0.769561988 | 0.258956817 |
| ENSMUSG00000020534 | Shmt1 | 20425 | protein_coding | 1907.860846 | 0.258967048 | 0.06891765 | 3.757630301 | 0.00017153 | 0.00230963 | 0.258967048 |
| ENSMUSG00000040111 | Gramd1b | 235283 | protein_coding | 350.4120495 | -0.259022098 | 0.096266593 | -2.690674826 | 0.007130766 | 0.046463784 | 0.259022098 |
| ENSMUSG00000024597 | Slc12a2 | 20496 | protein_coding | 12316.71415 | -0.259061626 | 0.05476636 | -4.730305759 | 2.24E-06 | 5.08E-05 | 0.259061626 |
| ENSMUSG00000032514 | Ttc21a | 74052 | protein_coding | 6.388484046 | 0.259094597 | 0.506417749 | 0.511622267 | 0.608915398 | 0.791308423 | 0.259094597 |
| ENSMUSG00000095098 | Ccdc85b | 240514 | protein_coding | 321.2252459 | -0.259133636 | 0.098923188 | -2.619543915 | 0.008804744 | 0.054260619 | 0.259133636 |
| ENSMUSG00000015659 | Serac1 | 321007 | protein_coding | 139.3895151 | -0.259199744 | 0.133993516 | -1.934420047 | 0.053061504 | 0.192513581 | 0.259199744 |
| ENSMUSG00000028803 | Nipal3 | 74552 | protein_coding | 2424.391816 | 0.259213935 | 0.066429339 | 3.902100171 | 9.54E-05 | 0.001424611 | 0.259213935 |
| ENSMUSG00000039531 | Zup1 | 72580 | protein_coding | 340.875373 | -0.259386991 | 0.094353412 | -2.749100261 | 0.00597591 | 0.040905563 | 0.259386991 |
| ENSMUSG00000031373 | Car5b | 56078 | protein_coding | 15.18134439 | 0.259440703 | 0.314066254 | 0.826069975 | 0.408764399 | 0.647240398 | 0.259440703 |
| ENSMUSG00000035597 | Prpf39 | 328110 | protein_coding | 509.1844562 | -0.259470649 | 0.142956809 | -1.81502826 | 0.069519567 | 0.228944671 | 0.259470649 |
| ENSMUSG00000026062 | Slc9a2 | 226999 | protein_coding | 8759.148859 | -0.259483954 | 0.070268297 | -3.692759935 | 0.000221833 | 0.002870449 | 0.259483954 |
| ENSMUSG00000052337 | Immt | 76614 | protein_coding | 6372.113491 | 0.259625923 | 0.048912684 | 5.307946787 | 1.11E-07 | 3.35E-06 | 0.259625923 |
| ENSMUSG00000026004 | Kansl1l | 68691 | protein_coding | 280.7621809 | -0.259778473 | 0.0921041 | -2.820487621 | 0.004795072 | 0.034530138 | 0.259778473 |
| ENSMUSG00000030541 | Idh2 | 269951 | protein_coding | 2761.346375 | -0.25983802 | 0.048090964 | -5.403052888 | 6.55E-08 | 2.06E-06 | 0.25983802 |
| ENSMUSG00000035818 | Plekhs1 | 226245 | protein_coding | 21.18894595 | -0.259892896 | 0.330787026 | -0.785680438 | 0.432054726 | 0.666514881 | 0.259892896 |
| ENSMUSG00000118376 | Gm9926 | 100503565 | lncRNA | 278.7354062 | -0.260010812 | 0.113435766 | -2.292141367 | 0.021897487 | 0.104820651 | 0.260010812 |
| ENSMUSG00000044122 | Proca1 | 216974 | protein_coding | 6.414647188 | -0.260101764 | 0.559144191 | -0.465178336 | 0.641803751 | 0.813747171 | 0.260101764 |
| ENSMUSG00000029822 | Osbpl3 | 71720 | protein_coding | 486.1366593 | -0.260499893 | 0.071176704 | -3.659903831 | 0.00025231 | 0.003206463 | 0.260499893 |
| ENSMUSG00000064288 | H4c12 | 319160 | protein_coding | 1483.40441 | 0.260621816 | 0.136276499 | 1.912448726 | 0.055818663 | 0.199156545 | 0.260621816 |
| ENSMUSG00000025578 | Cbx8 | 30951 | protein_coding | 206.4848794 | 0.26062552 | 0.092518318 | 2.817015317 | 0.004847221 | 0.034804603 | 0.26062552 |
| ENSMUSG00000098557 | Kctd12 | 239217 | protein_coding | 1060.729098 | 0.260736375 | 0.100810873 | 2.586391399 | 0.009698673 | 0.058099322 | 0.260736375 |
| ENSMUSG00000041842 | Fhdc1 | 229474 | protein_coding | 307.6284085 | -0.260754741 | 0.085650775 | -3.044394415 | 0.002331493 | 0.019737522 | 0.260754741 |
| ENSMUSG00000061046 | Haghl | 68977 | protein_coding | 165.5001389 | -0.26076974 | 0.116585055 | -2.236733867 | 0.025303739 | 0.115699858 | 0.26076974 |
| ENSMUSG00000026956 | Uap1l1 | 227620 | protein_coding | 501.0061129 | -0.260858134 | 0.075402819 | -3.459527617 | 0.000541124 | 0.006056005 | 0.260858134 |
| ENSMUSG00000047649 | Cd3eap | 70333 | protein_coding | 157.5613882 | -0.260956676 | 0.095924256 | -2.720445154 | 0.006519409 | 0.043445662 | 0.260956676 |
| ENSMUSG00000057110 | Cntrl | 26920 | protein_coding | 1473.717 | -0.260974687 | 0.073570139 | -3.54729093 | 0.000389215 | 0.004587415 | 0.260974687 |
| ENSMUSG00000032116 | Stt3a | 16430 | protein_coding | 4043.601333 | 0.260986371 | 0.050095004 | 5.209828328 | 1.89E-07 | 5.44E-06 | 0.260986371 |
| ENSMUSG00000023070 | Rgn | 19733 | protein_coding | 35.88727643 | 0.261013955 | 0.19383793 | 1.346557684 | 0.178122731 | 0.405350432 | 0.261013955 |
| ENSMUSG00000031618 | Nr3c2 | 110784 | protein_coding | 5507.689413 | -0.261026102 | 0.054041851 | -4.830073328 | 1.36E-06 | 3.25E-05 | 0.261026102 |
| ENSMUSG00000059659 | Gm10069 | 791299 | lncRNA | 8.414234889 | 0.26107161 | 0.397806873 | 0.656277274 | 0.511645737 | 0.725473655 | 0.26107161 |

Supplementary Table 1

| ensembl_gene_id | external_gene_name | entrezgene_id | gene_biotype | baseMean | log2FoldChange | lfcSE | stat | pvalue | padj | absLog2FC |
|---|---|---|---|---|---|---|---|---|---|---|
| ENSMUSG00000039852 | Rere | 68703 | protein_coding | 1251.013893 | -0.261078387 | 0.088681508 | -2.944000302 | 0.003239997 | 0.025676378 | 0.261078387 |
| ENSMUSG00000008090 | Fgfrl1 | 116701 | protein_coding | 456.3947813 | -0.261108758 | 0.080254013 | -3.253528999 | 0.001139811 | 0.011242649 | 0.261108758 |
| ENSMUSG00000032661 | Oas3 | 246727 | protein_coding | 101.2072404 | -0.261428735 | 0.310307964 | -0.842481553 | 0.399518461 | 0.638873001 | 0.261428735 |
| ENSMUSG00000026110 | Mgat4a | 269181 | protein_coding | 10198.98536 | -0.261438207 | 0.084302422 | -3.101194481 | 0.001927417 | 0.017026683 | 0.261438207 |
| ENSMUSG00000055850 | Rnf181 | 66510 | protein_coding | 1265.244846 | 0.261472126 | 0.066276966 | 3.945143246 | 7.98E-05 | 0.001219114 | 0.261472126 |
| ENSMUSG00000029828 | 4921507P07Rik | 70821 | protein_coding | 22.51806569 | -0.261536538 | 0.267380425 | -0.978143925 | 0.328003146 | 0.571645972 | 0.261536538 |
| ENSMUSG00000046876 | Atxn1 | 20238 | protein_coding | 1404.179528 | -0.26163222 | 0.067285129 | -3.888410771 | 0.000100903 | 0.001498397 | 0.26163222 |
| ENSMUSG00000117602 | 9130209A04Rik | 77668 | lncRNA | 27.99301682 | -0.261672266 | 0.278672981 | -0.938994031 | 0.347733809 | 0.589933004 | 0.261672266 |
| ENSMUSG00000073437 | D330041H03Rik | 654822 | lncRNA | 32.65659286 | -0.261700015 | 0.223916843 | -1.168737513 | 0.242509403 | 0.483669199 | 0.261700015 |
| ENSMUSG00000033594 | Spata2l | 78779 | protein_coding | 215.7982929 | -0.261711355 | 0.090924994 | -2.878321384 | 0.003997976 | 0.030298572 | 0.261711355 |
| ENSMUSG00000003545 | Fosb | 14282 | protein_coding | 8511.339528 | -0.261761307 | 0.191915198 | -1.363942563 | 0.172585644 | 0.397669477 | 0.261761307 |
| ENSMUSG00000079110 | Capn3 | 12335 | protein_coding | 18.25170125 | -0.262020879 | 0.335900668 | -0.780054652 | 0.435358707 | 0.668860614 | 0.262020879 |
| ENSMUSG00000052512 | Nav2 | 78286 | protein_coding | 309.6340617 | -0.262033027 | 0.105713674 | -2.478705137 | 0.013186026 | 0.073050857 | 0.262033027 |
| ENSMUSG00000057156 | Homez | 239099 | protein_coding | 205.6246221 | 0.262067155 | 0.091369071 | 2.868226116 | 0.004127804 | 0.030904873 | 0.262067155 |
| ENSMUSG00000038580 | Sct | 20287 | protein_coding | 383.2444093 | -0.262122799 | 0.099100609 | -2.645017035 | 0.00816868 | 0.051459496 | 0.262122799 |
| ENSMUSG00000055730 | Ces2a | 102022 | protein_coding | 1404.587618 | 0.262268985 | 0.098037504 | 2.67519035 | 0.007468683 | 0.048102018 | 0.262268985 |
| ENSMUSG00000034612 | Chst11 | 58250 | protein_coding | 191.9310881 | 0.262320697 | 0.105459817 | 2.487399508 | 0.01286808 | 0.071633942 | 0.262320697 |
| ENSMUSG00000078851 | H2ac25 | 319162 | protein_coding | 83.40132631 | 0.262504744 | 0.134924251 | 1.945571253 | 0.051706263 | 0.189254514 | 0.262504744 |
| ENSMUSG00000039461 | Tcta | 102791 | protein_coding | 969.4419468 | 0.262506672 | 0.06611522 | 3.970442396 | 7.17E-05 | 0.001113513 | 0.262506672 |
| ENSMUSG00000060935 | Tmem263 | 103266 | protein_coding | 546.9590881 | 0.262513183 | 0.067996313 | 3.860697321 | 0.000113064 | 0.001648627 | 0.262513183 |
| ENSMUSG00000024339 | Tap2 | 21355 | protein_coding | 409.1717589 | -0.262556243 | 0.139143063 | -1.886951725 | 0.059166819 | 0.206435622 | 0.262556243 |
| ENSMUSG00000063253 | Scoc | 56367 | protein_coding | 107.9601754 | 0.262593315 | 0.116652749 | 2.251068386 | 0.024381206 | 0.112857553 | 0.262593315 |
| ENSMUSG00000013822 | Elof1 | 66126 | protein_coding | 1226.21924 | 0.262624177 | 0.074246662 | 3.537184972 | 0.000404416 | 0.004729872 | 0.262624177 |
| ENSMUSG00000036882 | Arhgap33 | 233071 | protein_coding | 39.80956416 | -0.262761998 | 0.181977154 | -1.443928496 | 0.14875909 | 0.36310547 | 0.262761998 |
| ENSMUSG00000029859 | Epha1 | 13835 | protein_coding | 729.6005954 | -0.262827171 | 0.07726186 | -3.40177122 | 0.000669507 | 0.007270796 | 0.262827171 |
| ENSMUSG00000021877 | Arf4 | 11843 | protein_coding | 4268.577787 | 0.262869049 | 0.039421544 | 6.668157187 | 2.59E-11 | 1.20E-09 | 0.262869049 |
| ENSMUSG00000031432 | Prps1 | 19139 | protein_coding | 301.8008211 | 0.262873479 | 0.072334656 | 3.634129135 | 0.000278921 | 0.003473484 | 0.262873479 |
| ENSMUSG00000061393 | Acvr2b | 11481 | protein_coding | 112.9936022 | -0.262881069 | 0.151389006 | -1.73646076 | 0.082482397 | 0.254924035 | 0.262881069 |
| ENSMUSG00000035697 | Arhgap45 | 70719 | protein_coding | 85.15094646 | 0.262909538 | 0.169571285 | 1.55043667 | 0.121036743 | 0.322211877 | 0.262909538 |
| ENSMUSG00000046711 | Hmga1 | 15361 | protein_coding | 560.4661845 | -0.262927128 | 0.068996983 | -3.810704712 | 0.000138571 | 0.001947167 | 0.262927128 |
| ENSMUSG00000086968 | 4933431E20Rik | 329735 | lncRNA | 505.2773484 | -0.262929778 | 0.156541973 | -1.679612008 | 0.09303283 | 0.275392886 | 0.262929778 |
| ENSMUSG00000031016 | Wee1 | 22390 | protein_coding | 347.988224 | -0.262993296 | 0.089128719 | -2.95071328 | 0.003170411 | 0.025261469 | 0.262993296 |
| ENSMUSG00000021553 | Slc28a3 | 114304 | protein_coding | 111.9649701 | 0.263001672 | 0.196221175 | 1.340332775 | 0.180137181 | 0.407157997 | 0.263001672 |
| ENSMUSG00000109408 | A930037H05Rik | 78439 | lncRNA | 42.59419398 | -0.263093054 | 0.246970689 | -1.065280481 | 0.286749023 | 0.529168268 | 0.263093054 |
| ENSMUSG00000031158 | Timm17b | 21855 | protein_coding | 601.4996207 | 0.263196118 | 0.095940472 | 2.743327313 | 0.006082003 | 0.04141158 | 0.263196118 |
| ENSMUSG00000040600 | Eps8l3 | 99662 | protein_coding | 3568.998381 | -0.263217117 | 0.056303309 | -4.674984822 | 2.94E-06 | 6.49E-05 | 0.263217117 |
| ENSMUSG00000041609 | Bicdl1 | 75665 | protein_coding | 450.8689881 | 0.263326572 | 0.060398211 | 4.359840584 | 1.30E-05 | 0.000248702 | 0.263326572 |
| ENSMUSG00000102224 | 4930447F24Rik | 76873 | lncRNA | 15.44600115 | -0.26342469 | 0.355654751 | -0.740675303 | 0.458890338 | 0.685946212 | 0.26342469 |
| ENSMUSG00000006014 | Prg4 | 96875 | protein_coding | 6.231169352 | -0.263571046 | 0.520209836 | -0.506662941 | 0.612391337 | 0.793255831 | 0.263571046 |

Supplementary Table 1

| ensembl_gene_id | external_gene_name | entrezgene_id | gene_biotype | baseMean | log2FoldChange | lfcSE | stat | pvalue | padj | absLog2FC |
|---|---|---|---|---|---|---|---|---|---|---|
| ENSMUSG00000078870 | Gm14410 | 100043403 | protein_coding | 54.21537869 | -0.263639205 | 0.234669487 | -1.123449019 | 0.261246841 | 0.503084198 | 0.263639205 |
| ENSMUSG00000050627 | Gpd1l | 333433 | protein_coding | 3657.633632 | 0.263642275 | 0.032036654 | 8.229394772 | 1.88E-16 | 1.29E-14 | 0.263642275 |
| ENSMUSG00000090210 | Itga10 | 213119 | protein_coding | 13.64797306 | 0.263689057 | 0.372713914 | 0.707483801 | 0.479265876 | 0.701690493 | 0.263689057 |
| ENSMUSG00000023800 | Tiam2 | 24001 | protein_coding | 648.8248771 | 0.263690197 | 0.084595012 | 3.117089202 | 0.001826463 | 0.016336866 | 0.263690197 |
| ENSMUSG00000022900 | Ildr1 | 106347 | protein_coding | 5863.834629 | 0.263759406 | 0.058830726 | 4.483361431 | 7.35E-06 | 0.000149804 | 0.263759406 |
| ENSMUSG00000044067 | Gpr22 | 73010 | protein_coding | 14.06755235 | -0.26389456 | 0.320135604 | -0.824321181 | 0.409757086 | 0.648303005 | 0.26389456 |
| ENSMUSG00000026628 | Atf3 | 11910 | protein_coding | 15436.93399 | 0.263933213 | 0.065907251 | 4.004615736 | 6.21E-05 | 0.000988927 | 0.263933213 |
| ENSMUSG00000049922 | Slc35c1 | 228368 | protein_coding | 4068.588708 | 0.26393922 | 0.086495098 | 3.051493404 | 0.002277061 | 0.019450294 | 0.26393922 |
| ENSMUSG00000087963 | Gm25394 | 115488237 | snoRNA | 59.16753106 | 0.264054236 | 0.309086394 | 0.854305595 | 0.392935688 | 0.632548796 | 0.264054236 |
| ENSMUSG00000028419 | Chmp5 | 76959 | protein_coding | 2562.412964 | 0.264185626 | 0.04180379 | 6.319657352 | 2.62E-10 | 1.11E-08 | 0.264185626 |
| ENSMUSG00000031059 | Ndufb11 | 104130 | protein_coding | 2717.753553 | 0.264228658 | 0.05025605 | 5.257648682 | 1.46E-07 | 4.32E-06 | 0.264228658 |
| ENSMUSG00000099583 | H3c4 | 319149 | protein_coding | 507.8071979 | 0.264286206 | 0.115255054 | 2.293055245 | 0.021844825 | 0.104692864 | 0.264286206 |
| ENSMUSG00000045639 | Zfp629 | 320683 | protein_coding | 639.434642 | -0.264312659 | 0.070192906 | -3.765518081 | 0.000166204 | 0.002248635 | 0.264312659 |
| ENSMUSG00000002017 | Fam98a | 72722 | protein_coding | 558.4690025 | 0.264469108 | 0.077864711 | 3.396520773 | 0.000682484 | 0.007381902 | 0.264469108 |
| ENSMUSG00000050222 | Il17d | 239114 | protein_coding | 69.50276842 | -0.264732356 | 0.156094144 | -1.695978779 | 0.089889899 | 0.269652976 | 0.264732356 |
| ENSMUSG00000048445 | Ccdc57 | 71276 | protein_coding | 87.20956234 | -0.264819374 | 0.149512883 | -1.771214414 | 0.076525053 | 0.243800777 | 0.264819374 |
| ENSMUSG00000043461 | Sptssb | 66183 | protein_coding | 710.9696801 | 0.264921463 | 0.110088551 | 2.406439729 | 0.016108862 | 0.084538762 | 0.264921463 |
| ENSMUSG00000025473 | Adam8 | 11501 | protein_coding | 12.17282046 | 0.265093588 | 0.369122965 | 0.718171484 | 0.472651558 | 0.696814304 | 0.265093588 |
| ENSMUSG00000033917 | Gde1 | 56209 | protein_coding | 1214.547427 | 0.265115799 | 0.094519466 | 2.804880409 | 0.005033525 | 0.035839144 | 0.265115799 |
| ENSMUSG00000021185 | Dglucy | 217830 | protein_coding | 720.0876543 | 0.265151638 | 0.05579667 | 4.752105064 | 2.01E-06 | 4.62E-05 | 0.265151638 |
| ENSMUSG00000005268 | Prlr | 19116 | protein_coding | 1006.377044 | -0.265240753 | 0.073032353 | -3.63182536 | 0.000281424 | 0.003496865 | 0.265240753 |
| ENSMUSG00000024622 | Hmgxb3 | 106894 | protein_coding | 1027.326634 | -0.265249549 | 0.056506874 | -4.694111149 | 2.68E-06 | 5.97E-05 | 0.265249549 |
| ENSMUSG00000001525 | Tubb5 | 22154 | protein_coding | 4776.240854 | 0.26533969 | 0.104102746 | 2.548825071 | 0.01080865 | 0.062589264 | 0.26533969 |
| ENSMUSG00000002083 | Bbc3 | 170770 | protein_coding | 241.6988746 | -0.265449444 | 0.144126171 | -1.841785164 | 0.065506581 | 0.220501906 | 0.265449444 |
| ENSMUSG00000001552 | Jup | 16480 | protein_coding | 6725.745475 | 0.265620437 | 0.089265885 | 2.975609758 | 0.002924067 | 0.023596044 | 0.265620437 |
| ENSMUSG00000007035 | Msh5 | 17687 | protein_coding | 24.80433912 | -0.265775989 | 0.278152653 | -0.955504059 | 0.339322854 | 0.582473517 | 0.265775989 |
| ENSMUSG00000023959 | Clic5 | 224796 | protein_coding | 4991.224722 | -0.265855113 | 0.116906589 | -2.274081512 | 0.022961086 | 0.10799004 | 0.265855113 |
| ENSMUSG00000032482 | Cspg5 | 29873 | protein_coding | 23.76497456 | -0.265897018 | 0.262864238 | -1.01153744 | 0.311759271 | 0.555505998 | 0.265897018 |
| ENSMUSG00000029334 | Prkg2 | 19092 | protein_coding | 1652.301873 | -0.265961639 | 0.142511999 | -1.866240324 | 0.06200775 | 0.212494471 | 0.265961639 |
| ENSMUSG00000008855 | Hdac5 | 15184 | protein_coding | 1326.170925 | -0.266227855 | 0.097355091 | -2.734606417 | 0.00624549 | 0.042284225 | 0.266227855 |
| ENSMUSG00000039345 | Mettl22 | 239706 | protein_coding | 100.2476366 | -0.266239262 | 0.124817626 | -2.133026175 | 0.032922579 | 0.138772199 | 0.266239262 |
| ENSMUSG00000002625 | Akap8l | 54194 | protein_coding | 633.0923843 | -0.266243167 | 0.112458464 | -2.367480015 | 0.017909687 | 0.091124885 | 0.266243167 |
| ENSMUSG00000064202 | Spata6l | 381218 | protein_coding | 31.8026094 | 0.266317468 | 0.201538581 | 1.321421771 | 0.186360771 | 0.415618234 | 0.266317468 |
| ENSMUSG00000029130 | Rnf32 | 56874 | protein_coding | 278.4500001 | -0.266320861 | 0.132966199 | -2.002921522 | 0.045185713 | 0.172680092 | 0.266320861 |
| ENSMUSG00000036745 | Ttll7 | 70892 | protein_coding | 213.6292283 | 0.266323019 | 0.125047719 | 2.129771117 | 0.033190514 | 0.13964608 | 0.266323019 |
| ENSMUSG00000042446 | Zmym4 | 67785 | protein_coding | 495.766588 | -0.266356297 | 0.093224328 | -2.857154385 | 0.004274579 | 0.031797108 | 0.266356297 |
| ENSMUSG00000029811 | Aoc1 | 76507 | protein_coding | 9944.311559 | -0.266367581 | 0.069070043 | -3.856484941 | 0.000115029 | 0.001671239 | 0.266367581 |
| ENSMUSG00000071856 | Mcc | 328949 | protein_coding | 583.9144374 | -0.266463186 | 0.076260016 | -3.494140185 | 0.000475591 | 0.005416567 | 0.266463186 |
| ENSMUSG00000031925 | Maml2 | 270118 | protein_coding | 8.385737191 | 0.266568354 | 0.427996432 | 0.62282845 | 0.533397261 | 0.741375537 | 0.266568354 |

| ensembl_gene_id | external_gene_name | entrezgene_id | gene_biotype | baseMean | log2FoldChange | lfcSE | stat | pvalue | padj | absLog2FC |
|---|---|---|---|---|---|---|---|---|---|---|
| ENSMUSG00000031725 | Ces1f | 234564 | protein_coding | 2096.41761 | -0.266688144 | 0.074851123 | -3.56291441 | 0.00036676 | 0.004378048 | 0.266688144 |
| ENSMUSG00000002984 | Tomm40 | 53333 | protein_coding | 1267.551104 | 0.266832327 | 0.084869915 | 3.144015502 | 0.001666466 | 0.015184375 | 0.266832327 |
| ENSMUSG00000044456 | Rin3 | 217835 | protein_coding | 74.86061627 | 0.266871972 | 0.149134633 | 1.789470133 | 0.073539133 | 0.23731003 | 0.266871972 |
| ENSMUSG00000039089 | L3mbtl3 | 237339 | protein_coding | 423.4660565 | 0.266895248 | 0.064816522 | 4.117703937 | 3.83E-05 | 0.000644181 | 0.266895248 |
| ENSMUSG00000028749 | Pla2g2f | 26971 | protein_coding | 1081.114802 | 0.266904915 | 0.107052539 | 2.493214247 | 0.012659244 | 0.070827655 | 0.266904915 |
| ENSMUSG00000047022 | Mipol1 | 73490 | protein_coding | 85.19920268 | -0.267065367 | 0.147119023 | -1.815301393 | 0.069477605 | 0.228899966 | 0.267065367 |
| ENSMUSG00000040591 | Cstpp1 | 228356 | protein_coding | 75.66130896 | -0.267158575 | 0.152861903 | -1.747711954 | 0.080513918 | 0.251296295 | 0.267158575 |
| ENSMUSG00000002043 | Trappc6a | 67091 | protein_coding | 1562.834852 | 0.267211774 | 0.088714568 | 3.012039411 | 0.002594989 | 0.021439236 | 0.267211774 |
| ENSMUSG00000021701 | Plk2 | 20620 | protein_coding | 861.3554901 | -0.267288193 | 0.073268413 | -3.648068571 | 0.000264219 | 0.003321171 | 0.267288193 |
| ENSMUSG00000096145 | Vkorc1 | 27973 | protein_coding | 166.54682 | 0.267328434 | 0.093500646 | 2.859107891 | 0.004248343 | 0.031631128 | 0.267328434 |
| ENSMUSG00000030047 | Arhgap25 | 232201 | protein_coding | 10.36012422 | 0.267328438 | 0.378115676 | 0.707001733 | 0.479565401 | 0.702001563 | 0.267328438 |
| ENSMUSG00000025420 | Katnal2 | 71206 | protein_coding | 5.994741602 | -0.267605219 | 0.533021992 | -0.502052866 | 0.615630334 | 0.795599668 | 0.267605219 |
| ENSMUSG00000031216 | Stard8 | 236920 | protein_coding | 94.56096808 | -0.267608265 | 0.141640004 | -1.889355116 | 0.058844261 | 0.205867649 | 0.267608265 |
| ENSMUSG00000026509 | Capn2 | 12334 | protein_coding | 1591.176338 | 0.267679603 | 0.087685743 | 3.052715243 | 0.00226781 | 0.019391822 | 0.267679603 |
| ENSMUSG00000071478 | H2ac7 | 319165 | protein_coding | 100.4351411 | 0.267688036 | 0.213575973 | 1.253362129 | 0.210073947 | 0.445219276 | 0.267688036 |
| ENSMUSG00000039976 | Tbc1d16 | 207592 | protein_coding | 411.0912993 | -0.267894169 | 0.095502644 | -2.805096896 | 0.005030146 | 0.0358309 | 0.267894169 |
| ENSMUSG00000028795 | Ccdc28b | 66264 | protein_coding | 295.7789123 | -0.267894979 | 0.076933053 | -3.48218311 | 0.000497343 | 0.005628532 | 0.267894979 |
| ENSMUSG00000037568 | Vash2 | 226841 | protein_coding | 6.056098924 | -0.267900059 | 0.50050891 | -0.535255325 | 0.592473314 | 0.780262005 | 0.267900059 |
| ENSMUSG00000071636 | Rimbp3 | 239731 | protein_coding | 24.57786508 | 0.267965942 | 0.250015895 | 1.071795623 | 0.28381184 | 0.526458884 | 0.267965942 |
| ENSMUSG00000065637 | Gm26397 | 115487043 | snoRNA | 411.3201982 | -0.268157959 | 0.261907751 | -1.023864157 | 0.305899442 | 0.55012849 | 0.268157959 |
| ENSMUSG00000031844 | Hsd17b2 | 15486 | protein_coding | 3165.714514 | -0.268190349 | 0.059710943 | -4.491477365 | 7.07E-06 | 0.000144572 | 0.268190349 |
| ENSMUSG00000000295 | Hddc2 | 69692 | protein_coding | 265.9236185 | 0.268359806 | 0.087776688 | 3.057301565 | 0.002233395 | 0.019138127 | 0.268359806 |
| ENSMUSG00000043931 | Gimap7 | 231932 | protein_coding | 13.43622117 | 0.268415645 | 0.340838121 | 0.787516505 | 0.430979571 | 0.665684085 | 0.268415645 |
| ENSMUSG00000113035 | 5830428M24Rik | 76062 | lncRNA | 34.84672241 | -0.26848642 | 0.227075577 | -1.182365904 | 0.237060546 | 0.477153008 | 0.26848642 |
| ENSMUSG00000048249 | Crebrf | 77128 | protein_coding | 1575.089894 | -0.268506476 | 0.114643043 | -2.342108773 | 0.019175126 | 0.095444865 | 0.268506476 |
| ENSMUSG00000027171 | Prrg4 | 228413 | protein_coding | 21.01057904 | 0.268514355 | 0.284067873 | 0.945247174 | 0.344532701 | 0.58688999 | 0.268514355 |
| ENSMUSG00000034768 | Asb16 | 217217 | protein_coding | 11.82840425 | 0.268561879 | 0.356644927 | 0.753023129 | 0.451436015 | 0.680529877 | 0.268561879 |
| ENSMUSG00000083282 | Ctsf | 56464 | protein_coding | 285.5960932 | -0.268607349 | 0.105006073 | -2.558017291 | 0.010527086 | 0.061552292 | 0.268607349 |
| ENSMUSG00000059791 | Nrm | 106582 | protein_coding | 162.2180881 | -0.268852054 | 0.127925526 | -2.101629466 | 0.035585747 | 0.146102682 | 0.268852054 |
| ENSMUSG00000028909 | Ptpru | 19273 | protein_coding | 144.5435831 | -0.268864462 | 0.168446479 | -1.596141775 | 0.110457144 | 0.305128872 | 0.268864462 |
| ENSMUSG00000009378 | Slc16a12 | 240638 | protein_coding | 151.6762509 | 0.268880912 | 0.162891026 | 1.650679709 | 0.098803993 | 0.285916382 | 0.268880912 |
| ENSMUSG00000047115 | Fam221a | 231946 | protein_coding | 115.8896384 | 0.26888866 | 0.157407244 | 1.70823561 | 0.087592633 | 0.264979628 | 0.26888866 |
| ENSMUSG00000021306 | Gpr137b | 83924 | protein_coding | 196.0346399 | -0.269043636 | 0.146917573 | -1.831255655 | 0.067062393 | 0.223684636 | 0.269043636 |
| ENSMUSG00000000531 | Tamalin | 56149 | protein_coding | 118.7925532 | -0.269087503 | 0.153040358 | -1.758278056 | 0.07870021 | 0.247934382 | 0.269087503 |
| ENSMUSG00000040722 | Scamp5 | 56807 | protein_coding | 1180.421653 | -0.269191604 | 0.04198438 | -6.411708488 | 1.44E-10 | 6.22E-09 | 0.269191604 |
| ENSMUSG00000026873 | Phf19 | 74016 | protein_coding | 125.0683493 | -0.269218961 | 0.136190779 | -1.976778186 | 0.048066708 | 0.179979523 | 0.269218961 |
| ENSMUSG00000052040 | Klf13 | 50794 | protein_coding | 2068.709822 | -0.269254844 | 0.051396069 | -5.238821709 | 1.62E-07 | 4.70E-06 | 0.269254844 |
| ENSMUSG00000027956 | Tmem144 | 70652 | protein_coding | 247.6057598 | -0.269354211 | 0.083812469 | -3.213772524 | 0.001310034 | 0.012583038 | 0.269354211 |
| ENSMUSG00000065258 | Gm23969 | 115489598 | snoRNA | 12.00176987 | -0.269398524 | 0.369162434 | -0.729756062 | 0.465539306 | 0.691532821 | 0.269398524 |

Supplementary Table 1

| ensembl_gene_id | external_gene_name | entrezgene_id | gene_biotype | baseMean | log2FoldChange | lfcSE | stat | pvalue | padj | absLog2FC |
|---|---|---|---|---|---|---|---|---|---|---|
| ENSMUSG00000034154 | Ino80 | 68142 | protein_coding | 923.091203 | -0.269483385 | 0.060160608 | -4.479399335 | 7.49E-06 | 0.000152227 | 0.269483385 |
| ENSMUSG00000031960 | Aars | 234734 | protein_coding | 2057.445819 | 0.269698318 | 0.077058888 | 3.499898907 | 0.000465435 | 0.005319677 | 0.269698318 |
| ENSMUSG00000022947 | Cbr3 | 109857 | protein_coding | 351.5214268 | 0.269698538 | 0.163024099 | 1.654347663 | 0.098056894 | 0.284673902 | 0.269698538 |
| ENSMUSG00000026974 | Zmynd19 | 67187 | protein_coding | 340.397109 | 0.269807448 | 0.078593099 | 3.432966131 | 0.000597017 | 0.006617243 | 0.269807448 |
| ENSMUSG00000097327 | E030030I06Rik | 319887 | transcribed_unprocessed_pseudogene | 113.5174332 | 0.269888517 | 0.162361958 | 1.662264491 | 0.096459754 | 0.281863826 | 0.269888517 |
| ENSMUSG00000051037 | Zfp455 | 218311 | protein_coding | 34.8611751 | -0.269968882 | 0.221071406 | -1.221184079 | 0.222016331 | 0.459085065 | 0.269968882 |
| ENSMUSG00000030134 | Rasgef1a | 70727 | protein_coding | 10.3033821 | -0.26998779 | 0.431011958 | -0.626404407 | 0.53104972 | 0.73971984 | 0.26998779 |
| ENSMUSG00000028937 | Acot7 | 70025 | protein_coding | 1595.387477 | 0.269994083 | 0.114656526 | 2.354807821 | 0.018532287 | 0.093396937 | 0.269994083 |
| ENSMUSG00000067889 | Sptbn2 | 20743 | protein_coding | 510.7586274 | -0.270204595 | 0.084422182 | -3.200635052 | 0.001371251 | 0.012985416 | 0.270204595 |
| ENSMUSG00000059742 | Kcnh7 | 170738 | protein_coding | 32.88799784 | -0.270247284 | 0.248618981 | -1.086993771 | 0.277039562 | 0.519815826 | 0.270247284 |
| ENSMUSG00000037936 | Scarb1 | 20778 | protein_coding | 318.7954669 | -0.270302944 | 0.093199951 | -2.900247697 | 0.003728679 | 0.028675443 | 0.270302944 |
| ENSMUSG00000030946 | Lhpp | 76429 | protein_coding | 411.0052521 | -0.270410559 | 0.078827586 | -3.430405181 | 0.000602681 | 0.006657143 | 0.270410559 |
| ENSMUSG00000021754 | Map3k1 | 26401 | protein_coding | 1245.091826 | -0.270413796 | 0.056141666 | -4.816632919 | 1.46E-06 | 3.44E-05 | 0.270413796 |
| ENSMUSG00000093989 | Rnasek | 52898 | protein_coding | 7.901010058 | -0.270609522 | 0.427364419 | -0.633205548 | 0.526599423 | 0.737001553 | 0.270609522 |
| ENSMUSG00000024018 | Ccdc167 | 68597 | protein_coding | 77.8730379 | 0.27071475 | 0.169971484 | 1.592706867 | 0.111225966 | 0.306582205 | 0.27071475 |
| ENSMUSG00000016496 | Cd274 | 60533 | protein_coding | 40.68940791 | -0.270761083 | 0.327114319 | -0.827726172 | 0.407825594 | 0.646388536 | 0.270761083 |
| ENSMUSG00000026034 | Clk1 | 12747 | protein_coding | 2340.681166 | -0.270784691 | 0.115788405 | -2.338616652 | 0.019355282 | 0.095778655 | 0.270784691 |
| ENSMUSG00000060508 | Nlrp9b | 243874 | protein_coding | 184.5566861 | -0.270937818 | 0.190564981 | -1.421760793 | 0.155095705 | 0.373094934 | 0.270937818 |
| ENSMUSG00000029201 | Ugdh | 22235 | protein_coding | 12958.28879 | -0.271067316 | 0.086294129 | -3.141202302 | 0.001682558 | 0.015287103 | 0.271067316 |
| ENSMUSG00000021714 | Cenpk | 60411 | protein_coding | 79.43961227 | 0.271071643 | 0.144948057 | 1.870129543 | 0.061465831 | 0.21135596 | 0.271071643 |
| ENSMUSG00000020474 | Polm | 54125 | protein_coding | 221.7587993 | -0.271237207 | 0.093277066 | -2.907865996 | 0.003639042 | 0.028136566 | 0.271237207 |
| ENSMUSG00000070867 | Trabd2b | 666048 | protein_coding | 22.9218051 | -0.271439967 | 0.283998865 | -0.955778349 | 0.339184229 | 0.582413124 | 0.271439967 |
| ENSMUSG00000052713 | Zfp608 | 269023 | protein_coding | 429.8009217 | -0.271476949 | 0.134706119 | -2.015327519 | 0.043870343 | 0.169495017 | 0.271476949 |
| ENSMUSG00000053510 | Nrd1 | 230598 | protein_coding | 2040.651212 | 0.271555184 | 0.053349552 | 5.090111774 | 3.58E-07 | 9.60E-06 | 0.271555184 |
| ENSMUSG00000032081 | Apoc3 | 11814 | protein_coding | 25.03195104 | 0.271760019 | 0.558710212 | 0.486406035 | 0.626679313 | 0.803180854 | 0.271760019 |
| ENSMUSG00000025574 | Tk1 | 21877 | protein_coding | 561.9888184 | 0.271810663 | 0.115506407 | 2.353208536 | 0.018612193 | 0.093675699 | 0.271810663 |
| ENSMUSG00000025738 | Fbxl16 | 214931 | protein_coding | 14.19262846 | 0.271895242 | 0.309774922 | 0.877718702 | 0.380096392 | 0.621026335 | 0.271895242 |
| ENSMUSG00000086241 | 4930483K19Rik | 74953 | lncRNA | 6.17003764 | -0.272225019 | 0.489920363 | -0.55565157 | 0.57844907 | 0.770962657 | 0.272225019 |
| ENSMUSG00000097691 | 9030616G12Rik | 77699 | lncRNA | 31.42745234 | -0.272280437 | 0.265361768 | -1.026072591 | 0.304857369 | 0.549138254 | 0.272280437 |
| ENSMUSG00000086550 | Pla2g10os | 76684 | lncRNA | 166.8229527 | -0.272303895 | 0.095725764 | -2.844624944 | 0.004446374 | 0.032623603 | 0.272303895 |
| ENSMUSG00000034773 | Hrob | 217216 | protein_coding | 43.09580848 | 0.272341211 | 0.202962018 | 1.341833381 | 0.179650026 | 0.406744622 | 0.272341211 |
| ENSMUSG00000052707 | Tnrc6a | 233833 | protein_coding | 1153.513991 | -0.27288113 | 0.071805249 | -3.800295007 | 0.000144524 | 0.0020054 | 0.27288113 |
| ENSMUSG00000021665 | Hexb | 15212 | protein_coding | 2757.507238 | -0.272951896 | 0.102101868 | -2.673329108 | 0.007510253 | 0.048292604 | 0.272951896 |
| ENSMUSG00000040269 | Mrps28 | 66230 | protein_coding | 493.6229012 | 0.272953209 | 0.100981363 | 2.703005793 | 0.006871555 | 0.045157932 | 0.272953209 |
| ENSMUSG00000054434 | Tmem120b | 330189 | protein_coding | 56.92854395 | -0.273067945 | 0.171522583 | -1.592023277 | 0.111379474 | 0.306783396 | 0.273067945 |
| ENSMUSG00000021477 | Ctsl | 13039 | protein_coding | 1246.329229 | -0.273084124 | 0.061066566 | -4.471908929 | 7.75E-06 | 0.000156868 | 0.273084124 |
| ENSMUSG00000032489 | Kif9 | 16578 | protein_coding | 41.69382378 | 0.273120825 | 0.187411393 | 1.457333094 | 0.145024456 | 0.357616116 | 0.273120825 |
| ENSMUSG00000035314 | Gdpd5 | 233552 | protein_coding | 168.4557057 | -0.273315304 | 0.114882827 | -2.379078849 | 0.017355963 | 0.088913474 | 0.273315304 |
| ENSMUSG00000035954 | Dock4 | 238130 | protein_coding | 29.69917197 | -0.273325057 | 0.294672813 | -0.927554378 | 0.353638772 | 0.595956629 | 0.273325057 |

| ensembl_gene_id | external_gene_name | entrezgene_id | gene_biotype | baseMean | log2FoldChange | lfcSE | stat | pvalue | padj | absLog2FC |
|---|---|---|---|---|---|---|---|---|---|---|
| ENSMUSG00000006587 | Snai3 | 30927 | protein_coding | 13.83697492 | 0.27341132 | 0.326953658 | 0.836238755 | 0.4030206 | 0.642359644 | 0.27341132 |
| ENSMUSG00000033268 | Duox1 | 99439 | protein_coding | 7.962636428 | 0.27350477 | 0.438791092 | 0.623314318 | 0.533077991 | 0.741142191 | 0.27350477 |
| ENSMUSG00000036992 | Nxt1 | 56488 | protein_coding | 200.7237269 | 0.273579582 | 0.114407303 | 2.391277258 | 0.016789868 | 0.087036389 | 0.273579582 |
| ENSMUSG00000038412 | Higd1a | 56295 | protein_coding | 3540.484508 | 0.273672977 | 0.082597754 | 3.313322268 | 0.000921947 | 0.009422203 | 0.273672977 |
| ENSMUSG00000040329 | Il7 | 16196 | protein_coding | 187.9911554 | 0.273755151 | 0.097086643 | 2.819699431 | 0.004806865 | 0.034576411 | 0.273755151 |
| ENSMUSG00000074398 | Gm15441 | 100038464 | lncRNA | 11.02924531 | -0.273758919 | 0.36646872 | -0.747018514 | 0.455052386 | 0.682988492 | 0.273758919 |
| ENSMUSG00000022479 | Vdr | 22337 | protein_coding | 7580.354679 | -0.273789596 | 0.041805323 | -6.549156345 | 5.79E-11 | 2.60E-09 | 0.273789596 |
| ENSMUSG00000049422 | Chchd10 | 103172 | protein_coding | 5225.818978 | 0.273812405 | 0.088553807 | 3.092045572 | 0.001987823 | 0.017432097 | 0.273812405 |
| ENSMUSG00000085227 | 6330418K02Rik | 69004 | lncRNA | 17.83920821 | 0.273822562 | 0.293114597 | 0.934182618 | 0.3502097 | 0.592450628 | 0.273822562 |
| ENSMUSG00000020882 | Cacnb1 | 12295 | protein_coding | 55.92225272 | 0.273843333 | 0.155612205 | 1.759780562 | 0.07844502 | 0.247473168 | 0.273843333 |
| ENSMUSG00000070802 | Pnma8b | 434128 | protein_coding | 56.65469494 | -0.273930299 | 0.16926776 | -1.618325298 | 0.105592511 | 0.297761537 | 0.273930299 |
| ENSMUSG00000024424 | Ttc39c | 72747 | protein_coding | 36.05072936 | 0.273951063 | 0.282352039 | 0.970246447 | 0.331923664 | 0.575141012 | 0.273951063 |
| ENSMUSG00000020651 | Slc26a4 | 23985 | protein_coding | 7.391931585 | -0.274107551 | 0.425232012 | -0.644607046 | 0.519181868 | 0.730997554 | 0.274107551 |
| ENSMUSG00000075520 | Malrd1 | 102635496 | protein_coding | 1615.936714 | -0.274198786 | 0.136577784 | -2.007638275 | 0.044681744 | 0.171444799 | 0.274198786 |
| ENSMUSG00000035929 | H2-Q4 | 15015 | protein_coding | 393.0008591 | -0.274246139 | 0.129647897 | -2.115314977 | 0.034403111 | 0.14299458 | 0.274246139 |
| ENSMUSG00000025492 | Ifitm3 | 66141 | protein_coding | 132.6691681 | 0.274388637 | 0.16254598 | 1.688067814 | 0.091398212 | 0.272408741 | 0.274388637 |
| ENSMUSG00000001175 | Calm1 | 12313 | protein_coding | 28252.28147 | 0.27451526 | 0.043456881 | 6.316957311 | 2.67E-10 | 1.12E-08 | 0.27451526 |
| ENSMUSG00000034159 | Mab21l4 | 71874 | protein_coding | 386.2714744 | 0.274684291 | 0.087419444 | 3.14214182 | 0.001677168 | 0.01526393 | 0.274684291 |
| ENSMUSG00000028218 | Cibar1 | 68099 | protein_coding | 85.85389543 | 0.274871277 | 0.141004703 | 1.949376658 | 0.05125046 | 0.188241916 | 0.274871277 |
| ENSMUSG00000034110 | Kctd7 | 212919 | protein_coding | 144.6339475 | 0.274885524 | 0.150431124 | 1.827318154 | 0.067651947 | 0.225092418 | 0.274885524 |
| ENSMUSG00000001751 | Naglu | 27419 | protein_coding | 699.7477329 | 0.275247677 | 0.09570022 | 2.87614465 | 0.004025652 | 0.030422534 | 0.275247677 |
| ENSMUSG00000032656 | Marchf3 | 320253 | protein_coding | 115.4305845 | 0.275291444 | 0.142257939 | 1.935156982 | 0.052971035 | 0.192401774 | 0.275291444 |
| ENSMUSG00000072235 | Tuba1a | 22142 | protein_coding | 165.6168077 | 0.275353605 | 0.109966136 | 2.503985458 | 0.012280308 | 0.069246339 | 0.275353605 |
| ENSMUSG00000069805 | Fbp1 | 14121 | protein_coding | 47.7500143 | -0.275374544 | 0.211592523 | -1.301437974 | 0.193108583 | 0.424923197 | 0.275374544 |
| ENSMUSG00000064215 | Ifi27 | 52668 | protein_coding | 682.115521 | -0.275421378 | 0.064740842 | -4.254213747 | 2.10E-05 | 0.000378434 | 0.275421378 |
| ENSMUSG00000022799 | Arhgap31 | 12549 | protein_coding | 251.5519499 | 0.275444389 | 0.099752094 | 2.761289288 | 0.005757365 | 0.039815194 | 0.275444389 |
| ENSMUSG00000055760 | Gemin6 | 67242 | protein_coding | 84.67158435 | 0.275584275 | 0.160558563 | 1.716409699 | 0.086087083 | 0.262393948 | 0.275584275 |
| ENSMUSG00000040747 | Cd53 | 12508 | protein_coding | 24.84067234 | 0.275734472 | 0.252303952 | 1.092866243 | 0.274452532 | 0.5169296 | 0.275734472 |
| ENSMUSG00000005360 | Slc1a3 | 20512 | protein_coding | 28.30894583 | -0.276160557 | 0.24500006 | -1.127185673 | 0.259663987 | 0.50123327 | 0.276160557 |
| ENSMUSG00000021252 | Erg28 | 58520 | protein_coding | 898.7074825 | 0.276205322 | 0.067084117 | 4.117298283 | 3.83E-05 | 0.000644642 | 0.276205322 |
| ENSMUSG00000027784 | Ppm1l | 242083 | protein_coding | 351.6093716 | 0.276206204 | 0.071184286 | 3.880156975 | 0.000104389 | 0.001544479 | 0.276206204 |
| ENSMUSG00000021876 | Rnase4 | 58809 | protein_coding | 3352.864099 | -0.276215449 | 0.067056705 | -4.119132473 | 3.80E-05 | 0.00064154 | 0.276215449 |
| ENSMUSG00000097583 | 6430590A07Rik | 619719 | lncRNA | 18.49361825 | -0.276217074 | 0.328837575 | -0.839980268 | 0.40091945 | 0.640224574 | 0.276217074 |
| ENSMUSG00000035200 | Chrnb4 | 108015 | protein_coding | 84.5009746 | -0.27628645 | 0.159068579 | -1.736901477 | 0.082404563 | 0.254878861 | 0.27628645 |
| ENSMUSG00000029464 | Gpn3 | 68080 | protein_coding | 336.0191504 | 0.276303879 | 0.083259332 | 3.318593513 | 0.00090472 | 0.009275539 | 0.276303879 |
| ENSMUSG00000023367 | Tmem176a | 66058 | protein_coding | 687.2748674 | 0.276316415 | 0.065829718 | 4.197441879 | 2.70E-05 | 0.000471151 | 0.276316415 |
| ENSMUSG00000053054 | Adh6a | 69117 | protein_coding | 34.1601917 | -0.276368517 | 0.506367756 | -0.545786168 | 0.585212932 | 0.775873805 | 0.276368517 |
| ENSMUSG00000028369 | Svep1 | 64817 | protein_coding | 13.37940321 | -0.276471914 | 0.393759704 | -0.70213359 | 0.482595857 | 0.70472411 | 0.276471914 |
| ENSMUSG00000033998 | Kcnk1 | 16525 | protein_coding | 1625.375897 | 0.276480184 | 0.050950535 | 5.426443196 | 5.75E-08 | 1.83E-06 | 0.276480184 |

| ensembl_gene_id | external_gene_name | entrezgene_id | gene_biotype | baseMean | log2FoldChange | lfcSE | stat | pvalue | padj | absLog2FC |
|---|---|---|---|---|---|---|---|---|---|---|
| ENSMUSG00000030203 | Dusp16 | 70686 | protein_coding | 743.3188299 | -0.276534286 | 0.096189886 | -2.874879017 | 0.004041823 | 0.030487599 | 0.276534286 |
| ENSMUSG00000079339 | Ifit1bl1 | 667373 | protein_coding | 1367.883812 | -0.276589809 | 0.318089216 | -0.869535323 | 0.384554396 | 0.624972092 | 0.276589809 |
| ENSMUSG00000022967 | Ifnar1 | 15975 | protein_coding | 1074.880197 | 0.276924055 | 0.04553046 | 6.082171232 | 1.19E-09 | 4.68E-08 | 0.276924055 |
| ENSMUSG00000078664 | Sprr2a1 | 20755 | protein_coding | 383.9843404 | -0.277004941 | 0.726125407 | -0.381483609 | 0.702844431 | 0.852303638 | 0.277004941 |
| ENSMUSG00000030149 | Klrk1 | 27007 | protein_coding | 18.86366222 | 0.277209046 | 0.31548882 | 0.878665197 | 0.379582834 | 0.62053716 | 0.277209046 |
| ENSMUSG00000042195 | Slc35f2 | 72022 | protein_coding | 348.3094977 | 0.277312105 | 0.081638562 | 3.396827406 | 0.00068172 | 0.007378584 | 0.277312105 |
| ENSMUSG00000015647 | Lama5 | 16776 | protein_coding | 90.10026761 | 0.277497696 | 0.172499807 | 1.608684101 | 0.107685434 | 0.300613293 | 0.277497696 |
| ENSMUSG00000020359 | Phykpl | 72947 | protein_coding | 474.3081394 | -0.277502881 | 0.122603405 | -2.263419047 | 0.02360987 | 0.110140691 | 0.277502881 |
| ENSMUSG00000098332 | Pigbos1 | 66364 | protein_coding | 165.3812391 | 0.277572218 | 0.098984606 | 2.804195812 | 0.005044226 | 0.035899485 | 0.277572218 |
| ENSMUSG00000049037 | Clec4a1 | 269799 | protein_coding | 9.218604975 | 0.277607633 | 0.430267594 | 0.645197631 | 0.518799121 | 0.730713836 | 0.277607633 |
| ENSMUSG00000034160 | Ogt | 108155 | protein_coding | 1890.55841 | -0.277667972 | 0.09977183 | -2.783029776 | 0.005385386 | 0.037711735 | 0.277667972 |
| ENSMUSG00000005233 | Spc25 | 66442 | protein_coding | 249.1166144 | 0.277769858 | 0.113940173 | 2.437857097 | 0.014774613 | 0.079396931 | 0.277769858 |
| ENSMUSG00000006567 | Atp7b | 11979 | protein_coding | 54.96746471 | 0.277780775 | 0.223898173 | 1.240656732 | 0.214732585 | 0.450382676 | 0.277780775 |
| ENSMUSG00000035901 | Dennd5a | 19347 | protein_coding | 50.06576895 | 0.277933459 | 0.174803044 | 1.589980659 | 0.111839165 | 0.3074195 | 0.277933459 |
| ENSMUSG00000035671 | Zswim4 | 212168 | protein_coding | 417.9395745 | -0.277972733 | 0.071882193 | -3.867059712 | 0.000110155 | 0.001614979 | 0.277972733 |
| ENSMUSG00000063894 | Zkscan8 | 93681 | protein_coding | 395.431613 | -0.278100488 | 0.08533095 | -3.259081132 | 0.001117737 | 0.011052553 | 0.278100488 |
| ENSMUSG00000027746 | Ufm1 | 67890 | protein_coding | 1246.561719 | 0.278137337 | 0.05986723 | 4.645902855 | 3.39E-06 | 7.37E-05 | 0.278137337 |
| ENSMUSG00000037053 | Azgp1 | 12007 | protein_coding | 48.41971616 | -0.278199396 | 0.22420204 | -1.240842396 | 0.214663976 | 0.450343118 | 0.278199396 |
| ENSMUSG00000003282 | Plag1 | 56711 | protein_coding | 85.39375357 | -0.278202361 | 0.16732874 | -1.662609548 | 0.096390619 | 0.281763913 | 0.278202361 |
| ENSMUSG00000021965 | Ska3 | 219114 | protein_coding | 103.7536898 | 0.278334411 | 0.135828137 | 2.049166078 | 0.040445878 | 0.159861993 | 0.278334411 |
| ENSMUSG00000079659 | Tmem243 | 652925 | protein_coding | 483.0528659 | -0.278388089 | 0.082636811 | -3.368814514 | 0.000754922 | 0.007978132 | 0.278388089 |
| ENSMUSG00000057265 | Bbof1 | 72873 | protein_coding | 28.77162518 | -0.278419739 | 0.273137316 | -1.019339805 | 0.308041673 | 0.551915127 | 0.278419739 |
| ENSMUSG00000026547 | Tagln2 | 21346 | protein_coding | 3626.10298 | 0.278547302 | 0.096845909 | 2.876190687 | 0.004025065 | 0.030422534 | 0.278547302 |
| ENSMUSG00000015668 | Pdzd11 | 72621 | protein_coding | 654.6808617 | 0.278765619 | 0.082031141 | 3.398290177 | 0.000678085 | 0.007346903 | 0.278765619 |
| ENSMUSG00000041936 | Agrn | 11603 | protein_coding | 1516.566715 | -0.278807134 | 0.072022783 | -3.871096401 | 0.000108347 | 0.001591358 | 0.278807134 |
| ENSMUSG00000116895 | Gm3435 | 100041621 | protein_coding | 12.26337259 | 0.278847771 | 0.336262821 | 0.829255434 | 0.406959883 | 0.645650963 | 0.278847771 |
| ENSMUSG00000040146 | Rgl3 | 71746 | protein_coding | 68.16420726 | 0.278945507 | 0.163174726 | 1.709489662 | 0.087360285 | 0.26457452 | 0.278945507 |
| ENSMUSG00000035547 | Capn5 | 12337 | protein_coding | 2382.491546 | 0.27898976 | 0.070390966 | 3.963431324 | 7.39E-05 | 0.001144539 | 0.27898976 |
| ENSMUSG00000064326 | Siva1 | 30954 | protein_coding | 225.6201251 | -0.278995171 | 0.095043639 | -2.935442862 | 0.003330719 | 0.026257023 | 0.278995171 |
| ENSMUSG00000013076 | Amotl1 | 75723 | protein_coding | 196.7064093 | 0.279106755 | 0.121672855 | 2.293911456 | 0.021795585 | 0.10455009 | 0.279106755 |
| ENSMUSG00000009621 | Vav2 | 22325 | protein_coding | 605.8041291 | -0.279110849 | 0.097237197 | -2.870412344 | 0.004099368 | 0.030792042 | 0.279110849 |
| ENSMUSG00000032586 | Traip | 22036 | protein_coding | 80.22376596 | 0.279218923 | 0.174072274 | 1.604040185 | 0.108705195 | 0.302001032 | 0.279218923 |
| ENSMUSG00000040729 | Cep126 | 234915 | protein_coding | 77.85269762 | 0.279273182 | 0.158128904 | 1.766110904 | 0.077377244 | 0.245400751 | 0.279273182 |
| ENSMUSG00000006456 | Rbm14 | 56275 | protein_coding | 678.4442636 | 0.279314753 | 0.074799253 | 3.734191766 | 0.000188319 | 0.002499605 | 0.279314753 |
| ENSMUSG00000032050 | Rdx | 19684 | protein_coding | 348.141283 | 0.279353041 | 0.170388761 | 1.639503912 | 0.101108357 | 0.290022548 | 0.279353041 |
| ENSMUSG00000036026 | Tmem63b | 224807 | protein_coding | 1660.233386 | -0.279389286 | 0.057924003 | -4.823376677 | 1.41E-06 | 3.34E-05 | 0.279389286 |
| ENSMUSG00000022704 | Qtrt2 | 106248 | protein_coding | 294.8904252 | -0.279514439 | 0.110729696 | -2.524295181 | 0.011593054 | 0.066251304 | 0.279514439 |
| ENSMUSG00000022339 | Ebag9 | 55960 | protein_coding | 524.2716529 | 0.279605805 | 0.062699839 | 4.459434191 | 8.22E-06 | 0.000165657 | 0.279605805 |
| ENSMUSG00000070305 | Mpzl3 | 319742 | protein_coding | 685.2378236 | -0.279811004 | 0.059004856 | -4.74216908 | 2.11E-06 | 4.83E-05 | 0.279811004 |

| ensembl_gene_id | external_gene_name | entrezgene_id | gene_biotype | baseMean | log2FoldChange | lfcSE | stat | pvalue | padj | absLog2FC |
|---|---|---|---|---|---|---|---|---|---|---|
| ENSMUSG00000038936 | Sccpdh | 109232 | protein_coding | 387.341777 | 0.279815423 | 0.077480788 | 3.611416842 | 0.000304529 | 0.003720557 | 0.279815423 |
| ENSMUSG00000026811 | St6galnac6 | 50935 | protein_coding | 10962.58784 | 0.279841914 | 0.084465748 | 3.313081585 | 0.000922741 | 0.009424343 | 0.279841914 |
| ENSMUSG00000028702 | Rad54l | 19366 | protein_coding | 130.7314697 | 0.279906907 | 0.162119352 | 1.726548399 | 0.0842488 | 0.25884557 | 0.279906907 |
| ENSMUSG00000008461 | Fut1 | 14343 | protein_coding | 32.11457325 | 0.280113247 | 0.21258557 | 1.317649394 | 0.187621042 | 0.417346835 | 0.280113247 |
| ENSMUSG00000032988 | Slc16a8 | 57274 | protein_coding | 17.98756011 | -0.28046968 | 0.393541288 | -0.712681717 | 0.476042733 | 0.699193183 | 0.28046968 |
| ENSMUSG00000039646 | Vasn | 246154 | protein_coding | 156.6888059 | -0.280519122 | 0.116081214 | -2.416576389 | 0.015667238 | 0.082759762 | 0.280519122 |
| ENSMUSG00000027961 | Lrrc39 | 109245 | protein_coding | 6.568012145 | -0.280521935 | 0.464445803 | -0.603992831 | 0.545848409 | 0.749242602 | 0.280521935 |
| ENSMUSG00000034007 | Scaper | 244891 | protein_coding | 583.6249041 | -0.28059096 | 0.081766877 | -3.431596883 | 0.000600039 | 0.00664161 | 0.28059096 |
| ENSMUSG00000046841 | Ckap4 | 216197 | protein_coding | 656.6215964 | 0.280664692 | 0.107357404 | 2.614302148 | 0.008940993 | 0.054911988 | 0.280664692 |
| ENSMUSG00000085719 | 4933424L21Rik | 78762 | lncRNA | 13.99613634 | -0.280673954 | 0.36116404 | -0.777137044 | 0.43707792 | 0.670224933 | 0.280673954 |
| ENSMUSG00000070324 | Fbxw22 | 382156 | protein_coding | 16.69041053 | -0.280714979 | 0.357576853 | -0.78504796 | 0.43242545 | 0.666779423 | 0.280714979 |
| ENSMUSG00000073987 | Ggh | 14590 | protein_coding | 4266.523131 | -0.280736346 | 0.10606205 | -2.646906659 | 0.008123177 | 0.051214509 | 0.280736346 |
| ENSMUSG00000084989 | Crocc2 | 381284 | protein_coding | 7.566877169 | 0.280805616 | 0.432172322 | 0.649753816 | 0.515851256 | 0.728649792 | 0.280805616 |
| ENSMUSG00000021391 | Cenpp | 66336 | protein_coding | 64.95414514 | -0.280964033 | 0.165167123 | -1.701089347 | 0.088926211 | 0.267808218 | 0.280964033 |
| ENSMUSG00000079298 | Klrb1b | 80782 | protein_coding | 6.247064425 | 0.280987618 | 0.524052803 | 0.53618188 | 0.591832855 | 0.780262005 | 0.280987618 |
| ENSMUSG00000062901 | Klhl24 | 75785 | protein_coding | 2330.512563 | -0.28106681 | 0.096119192 | -2.924148691 | 0.003453997 | 0.027013876 | 0.28106681 |
| ENSMUSG00000042104 | Uggt2 | 66435 | protein_coding | 26.21948441 | 0.281148128 | 0.247059339 | 1.137978142 | 0.255129609 | 0.496437646 | 0.281148128 |
| ENSMUSG00000056004 | Elapor2 | 231014 | protein_coding | 26.16721794 | -0.28115938 | 0.233774567 | -1.202694473 | 0.229094573 | 0.468321044 | 0.28115938 |
| ENSMUSG00000020027 | Socs2 | 216233 | protein_coding | 960.5698857 | -0.281174766 | 0.063184018 | -4.450093136 | 8.58E-06 | 0.000170893 | 0.281174766 |
| ENSMUSG00000018449 | Rpain | 69723 | protein_coding | 175.0738753 | 0.281199311 | 0.092969869 | 3.024628456 | 0.002489387 | 0.020731319 | 0.281199311 |
| ENSMUSG00000036466 | Megf11 | 214058 | protein_coding | 46.22061425 | -0.28130463 | 0.21953629 | -1.281358218 | 0.200067873 | 0.433483049 | 0.28130463 |
| ENSMUSG00000074207 | Adh1 | 11522 | protein_coding | 15541.32168 | -0.281366532 | 0.109934443 | -2.559402889 | 0.010485215 | 0.061486489 | 0.281366532 |
| ENSMUSG00000007591 | Tssk4 | 71099 | protein_coding | 31.55269789 | 0.281517626 | 0.211380451 | 1.331805398 | 0.182924144 | 0.410980449 | 0.281517626 |
| ENSMUSG00000050335 | Lgals3 | 16854 | protein_coding | 29580.27746 | 0.281611276 | 0.095266364 | 2.956040963 | 0.003116157 | 0.024878349 | 0.281611276 |
| ENSMUSG00000001095 | Slc13a2 | 20500 | protein_coding | 1448.681313 | -0.281683614 | 0.111946662 | -2.51623058 | 0.011861755 | 0.06745223 | 0.281683614 |
| ENSMUSG00000037432 | Fer1l5 | 100534273 | protein_coding | 50.40802284 | 0.28175621 | 0.180058184 | 1.564806466 | 0.117628295 | 0.317145235 | 0.28175621 |
| ENSMUSG00000021187 | Tc2n | 74413 | protein_coding | 917.7032404 | 0.281800934 | 0.05563823 | 5.064879585 | 4.09E-07 | 1.08E-05 | 0.281800934 |
| ENSMUSG00000053702 | Nebl | 74103 | protein_coding | 139.28645 | 0.28180367 | 0.121649885 | 2.316514065 | 0.020530217 | 0.10005442 | 0.28180367 |
| ENSMUSG00000046093 | Hpcal4 | 170638 | protein_coding | 9.81447985 | 0.281921799 | 0.442335645 | 0.637348137 | 0.523898102 | 0.734827388 | 0.281921799 |
| ENSMUSG00000107768 | Gm36582 | 102640545 | lncRNA | 7.301727363 | 0.281969448 | 0.429587925 | 0.656371914 | 0.511584857 | 0.725473655 | 0.281969448 |
| ENSMUSG00000021990 | Spata13 | 219140 | protein_coding | 1480.511721 | -0.281981482 | 0.047449731 | -5.942741381 | 2.80E-09 | 1.06E-07 | 0.281981482 |
| ENSMUSG00000042426 | Dhx29 | 218629 | protein_coding | 363.8896649 | 0.282056096 | 0.089642415 | 3.146458016 | 0.00165261 | 0.015091528 | 0.282056096 |
| ENSMUSG00000032724 | Abtb2 | 99382 | protein_coding | 225.5164215 | 0.282141586 | 0.136291863 | 2.070127885 | 0.03844037 | 0.154385176 | 0.282141586 |
| ENSMUSG00000084798 | 4930533B01Rik | 75166 | lncRNA | 6.228381105 | 0.282266321 | 0.509822228 | 0.553656364 | 0.579814044 | 0.771934807 | 0.282266321 |
| ENSMUSG00000037905 | Bri3bp | 76809 | protein_coding | 544.19077 | -0.282395282 | 0.068753347 | -4.10736776 | 4.00E-05 | 0.000668108 | 0.282395282 |
| ENSMUSG00000032513 | Gorasp1 | 74498 | protein_coding | 842.9976535 | 0.282405009 | 0.077695779 | 3.634753575 | 0.000278246 | 0.003467759 | 0.282405009 |
| ENSMUSG00000024511 | Rab27b | 80718 | protein_coding | 2209.235976 | 0.282414223 | 0.043523109 | 6.488833863 | 8.65E-11 | 3.83E-09 | 0.282414223 |
| ENSMUSG00000030222 | Rerg | 232441 | protein_coding | 5.916751181 | -0.282523625 | 0.564891978 | -0.500137435 | 0.616978309 | 0.796574268 | 0.282523625 |
| ENSMUSG00000086682 | Gm16023 | 100502734 | lncRNA | 30.78580146 | -0.28255469 | 0.245313331 | -1.151811395 | 0.249398584 | 0.490195121 | 0.28255469 |

| ensembl_gene_id | external_gene_name | entrezgene_id | gene_biotype | baseMean | log2FoldChange | lfcSE | stat | pvalue | padj | absLog2FC |
|---|---|---|---|---|---|---|---|---|---|---|
| ENSMUSG00000078247 | Airn | 104103 | lncRNA | 74.85110525 | -0.28258324 | 0.156384011 | -1.806982943 | 0.070764939 | 0.231533002 | 0.28258324 |
| ENSMUSG00000085502 | Gm12320 | 102635935 | lncRNA | 82.80046224 | 0.282685875 | 0.157924852 | 1.790002466 | 0.073453515 | 0.237154121 | 0.282685875 |
| ENSMUSG00000039114 | Nrn1 | 68404 | protein_coding | 117.7632417 | 0.282776763 | 0.117046018 | 2.41594517 | 0.015694424 | 0.082768143 | 0.282776763 |
| ENSMUSG00000041141 | Pnma8a | 71691 | protein_coding | 18.7142517 | -0.282959632 | 0.272882386 | -1.036928899 | 0.299768994 | 0.543493488 | 0.282959632 |
| ENSMUSG00000028327 | Stra6l | 74152 | protein_coding | 604.6543405 | -0.283040759 | 0.134693888 | -2.101363051 | 0.035609109 | 0.146161389 | 0.283040759 |
| ENSMUSG00000040987 | Mill2 | 243864 | protein_coding | 134.6053416 | -0.283243134 | 0.133692816 | -2.11861148 | 0.034123315 | 0.142262102 | 0.283243134 |
| ENSMUSG00000053460 | Ggcx | 56316 | protein_coding | 540.9267252 | 0.28333947 | 0.083495915 | 3.393453082 | 0.000690174 | 0.007453027 | 0.28333947 |
| ENSMUSG00000024427 | Spry4 | 24066 | protein_coding | 58.81837033 | -0.283714753 | 0.217344805 | -1.305367079 | 0.191767883 | 0.423163427 | 0.283714753 |
| ENSMUSG00000067430 | Zfp763 | 73451 | protein_coding | 179.7444216 | -0.284221286 | 0.109585127 | -2.593611881 | 0.009497363 | 0.057278974 | 0.284221286 |
| ENSMUSG00000038045 | Sult6b1 | 73671 | protein_coding | 11.99334409 | -0.284274943 | 0.328143621 | -0.866312569 | 0.386318775 | 0.626292308 | 0.284274943 |
| ENSMUSG00000029016 | Clcn6 | 26372 | protein_coding | 69.5382682 | -0.284382613 | 0.142850069 | -1.990776871 | 0.046505423 | 0.176137379 | 0.284382613 |
| ENSMUSG00000026994 | Galnt3 | 14425 | protein_coding | 1301.680495 | 0.284548948 | 0.058302632 | 4.880550608 | 1.06E-06 | 2.58E-05 | 0.284548948 |
| ENSMUSG00000049897 | Stkld1 | 279029 | protein_coding | 7.360199135 | -0.284689182 | 0.523869681 | -0.543435118 | 0.586830252 | 0.777011446 | 0.284689182 |
| ENSMUSG00000032181 | Scg3 | 20255 | protein_coding | 29.08275019 | -0.284704457 | 0.266345157 | -1.068930482 | 0.285100995 | 0.527999972 | 0.284704457 |
| ENSMUSG00000056305 | Usp39 | 28035 | protein_coding | 462.4642521 | 0.284737537 | 0.072876728 | 3.90711199 | 9.34E-05 | 0.00139866 | 0.284737537 |
| ENSMUSG00000020812 | Snhg16 | 66293 | lncRNA | 114.882207 | -0.284765584 | 0.120660071 | -2.360064793 | 0.018271743 | 0.092401507 | 0.284765584 |
| ENSMUSG00000097392 | Thoc2l | 100042165 | protein_coding | 140.9442773 | -0.284808864 | 0.137597567 | -2.069868463 | 0.038464664 | 0.154385176 | 0.284808864 |
| ENSMUSG00000037455 | Slc18b1 | 76306 | protein_coding | 538.5881787 | 0.284814124 | 0.075398382 | 3.777456712 | 0.000158438 | 0.002161701 | 0.284814124 |
| ENSMUSG00000110151 | Gm38416 | 330850 | lncRNA | 34.31093559 | 0.28481671 | 0.213847267 | 1.331869765 | 0.182902988 | 0.410980449 | 0.28481671 |
| ENSMUSG00000057130 | Txnl4a | 27366 | protein_coding | 422.1628887 | 0.284867205 | 0.092668006 | 3.07406211 | 0.002111655 | 0.018291696 | 0.284867205 |
| ENSMUSG00000058586 | Serhl | 68607 | protein_coding | 958.1572758 | -0.284901115 | 0.062424683 | -4.56391772 | 5.02E-06 | 0.00010626 | 0.284901115 |
| ENSMUSG00000031104 | Rab33a | 19337 | protein_coding | 12.09110864 | -0.285141778 | 0.376341359 | -0.757667929 | 0.448649803 | 0.678988684 | 0.285141778 |
| ENSMUSG00000075415 | Fnbp1 | 14269 | protein_coding | 354.3858442 | -0.28518105 | 0.082541745 | -3.454991793 | 0.00055031 | 0.006150272 | 0.28518105 |
| ENSMUSG00000024485 | Slc4a9 | 240215 | protein_coding | 10.29996063 | -0.285211554 | 0.521911181 | -0.546475271 | 0.584739282 | 0.775689464 | 0.285211554 |
| ENSMUSG00000051510 | Mafg | 17134 | protein_coding | 617.3300975 | -0.285451176 | 0.08915347 | -3.201795468 | 0.001365739 | 0.012948429 | 0.285451176 |
| ENSMUSG00000038608 | Dock10 | 210293 | protein_coding | 54.91146273 | 0.285459572 | 0.17708545 | 1.611987726 | 0.106964611 | 0.299347968 | 0.285459572 |
| ENSMUSG00000094840 | Muc3a | 619309 | protein_coding | 306.3705729 | -0.285557247 | 0.137951569 | -2.069981865 | 0.038454043 | 0.154385176 | 0.285557247 |
| ENSMUSG00000003123 | Lipe | 16890 | protein_coding | 235.9676065 | -0.285590838 | 0.100599377 | -2.838892711 | 0.004527038 | 0.033065007 | 0.285590838 |
| ENSMUSG00000026490 | Cdc42bpa | 226751 | protein_coding | 1177.761326 | 0.285777849 | 0.065659241 | 4.35243913 | 1.35E-05 | 0.0002563 | 0.285777849 |
| ENSMUSG00000032776 | Mctp2 | 244049 | protein_coding | 1304.187582 | 0.2857995 | 0.057116037 | 5.00383979 | 5.62E-07 | 1.45E-05 | 0.2857995 |
| ENSMUSG00000022948 | Setd4 | 224440 | protein_coding | 158.6720667 | -0.285833765 | 0.120667775 | -2.36876635 | 0.017847524 | 0.090854491 | 0.285833765 |
| ENSMUSG00000022092 | Ppp3cc | 19057 | protein_coding | 154.9324524 | -0.285853668 | 0.152288128 | -1.877058117 | 0.060510129 | 0.209095104 | 0.285853668 |
| ENSMUSG00000028518 | Prkaa2 | 108079 | protein_coding | 3570.930234 | -0.285860977 | 0.037767347 | -7.568997985 | 3.76E-14 | 2.20E-12 | 0.285860977 |
| ENSMUSG00000068794 | Col28a1 | 213945 | protein_coding | 25.04104816 | 0.285947481 | 0.32530509 | 0.879013241 | 0.379394096 | 0.620491412 | 0.285947481 |
| ENSMUSG00000036820 | Amdhd2 | 245847 | protein_coding | 949.5314086 | -0.28601875 | 0.069584088 | -4.110404504 | 3.95E-05 | 0.000661436 | 0.28601875 |
| ENSMUSG00000021179 | Nrde2 | 217827 | protein_coding | 498.435919 | 0.286043561 | 0.066005738 | 4.333616567 | 1.47E-05 | 0.000275699 | 0.286043561 |
| ENSMUSG00000041406 | BC055324 | 381306 | protein_coding | 83.2975748 | 0.286064646 | 0.184791157 | 1.54804294 | 0.121611958 | 0.323066278 | 0.286064646 |
| ENSMUSG00000027188 | Pamr1 | 210622 | protein_coding | 84.65332899 | 0.286136735 | 0.24534551 | 1.166260326 | 0.243509209 | 0.484584527 | 0.286136735 |
| ENSMUSG00000042118 | Bhmt2 | 64918 | protein_coding | 15.51322155 | -0.286266689 | 0.295579566 | -0.968492823 | 0.332798302 | 0.576171574 | 0.286266689 |

| ensembl_gene_id | external_gene_name | entrezgene_id | gene_biotype | baseMean | log2FoldChange | lfcSE | stat | pvalue | padj | absLog2FC |
|---|---|---|---|---|---|---|---|---|---|---|
| ENSMUSG00000112640 | Gm32687 | 102635315 | protein_coding | 23.12807057 | -0.286270223 | 0.291604698 | -0.981706485 | 0.32624447 | 0.569717873 | 0.286270223 |
| ENSMUSG00000042035 | Igsf3 | 78908 | protein_coding | 1024.160051 | -0.286350178 | 0.059521487 | -4.810870724 | 1.50E-06 | 3.53E-05 | 0.286350178 |
| ENSMUSG00000087141 | Plcxd2 | 433022 | protein_coding | 361.3179972 | 0.286444462 | 0.066413358 | 4.313054977 | 1.61E-05 | 0.000298812 | 0.286444462 |
| ENSMUSG00000043621 | Ubxn10 | 212190 | protein_coding | 170.2949964 | 0.286619681 | 0.097336344 | 2.944631671 | 0.003233393 | 0.025636644 | 0.286619681 |
| ENSMUSG00000032718 | Mansc1 | 67729 | protein_coding | 1355.022455 | 0.286708669 | 0.061048699 | 4.696392753 | 2.65E-06 | 5.91E-05 | 0.286708669 |
| ENSMUSG00000051844 | B230319C09Rik | 320775 | lncRNA | 28.38542722 | -0.28671703 | 0.292710802 | -0.979523226 | 0.327321518 | 0.571042203 | 0.28671703 |
| ENSMUSG00000028093 | Acp6 | 66659 | protein_coding | 837.7697885 | -0.2868626 | 0.06154036 | -4.661373408 | 3.14E-06 | 6.90E-05 | 0.2868626 |
| ENSMUSG00000029190 | D5Ertd579e | 320661 | protein_coding | 869.874272 | -0.286889589 | 0.070886209 | -4.047184804 | 5.18E-05 | 0.000844425 | 0.286889589 |
| ENSMUSG00000028571 | Cyp2j13 | 230459 | protein_coding | 10.04193711 | 0.287048199 | 0.418657141 | 0.68564028 | 0.492939974 | 0.712333599 | 0.287048199 |
| ENSMUSG00000093805 | Gal3st2b | 100041596 | protein_coding | 318.3627042 | 0.287053018 | 0.143192482 | 2.004665417 | 0.044998828 | 0.17221075 | 0.287053018 |
| ENSMUSG00000019970 | Sgk1 | 20393 | protein_coding | 3770.053672 | 0.287061353 | 0.259134604 | 1.107769278 | 0.267961471 | 0.510503797 | 0.287061353 |
| ENSMUSG00000053774 | Ubxn7 | 224111 | protein_coding | 840.8930134 | -0.287132265 | 0.063275357 | -4.537821341 | 5.68E-06 | 0.000118889 | 0.287132265 |
| ENSMUSG00000038225 | Primpol | 408022 | protein_coding | 213.7069297 | -0.287133001 | 0.117972865 | -2.433890212 | 0.014937525 | 0.079926167 | 0.287133001 |
| ENSMUSG00000038252 | Ncapd2 | 68298 | protein_coding | 1164.812809 | 0.287145508 | 0.094031263 | 3.053723831 | 0.0022602 | 0.019356132 | 0.287145508 |
| ENSMUSG00000040105 | Plpp6 | 74411 | protein_coding | 1135.146212 | 0.287145917 | 0.045592538 | 6.298090241 | 3.01E-10 | 1.26E-08 | 0.287145917 |
| ENSMUSG00000046085 | 4931422A03Rik | 70986 | lncRNA | 6.972980104 | -0.287238007 | 0.532233479 | -0.539684214 | 0.589414828 | 0.77875577 | 0.287238007 |
| ENSMUSG00000024922 | Ovol1 | 18426 | protein_coding | 791.2467054 | -0.287259866 | 0.069269199 | -4.147007201 | 3.37E-05 | 0.000575463 | 0.287259866 |
| ENSMUSG00000025993 | Slc40a1 | 53945 | protein_coding | 5928.895197 | -0.287275047 | 0.123202632 | -2.331728156 | 0.019714997 | 0.097082061 | 0.287275047 |
| ENSMUSG00000069910 | Spdl1 | 70385 | protein_coding | 143.5695079 | 0.287514313 | 0.158476119 | 1.814243773 | 0.069640204 | 0.229201544 | 0.287514313 |
| ENSMUSG00000032425 | Zfp949 | 71640 | protein_coding | 231.9751667 | -0.287623896 | 0.129642059 | -2.218600179 | 0.026513937 | 0.119015143 | 0.287623896 |
| ENSMUSG00000041961 | Znrf3 | 407821 | protein_coding | 386.2756673 | -0.287815276 | 0.100315725 | -2.869094318 | 0.00411649 | 0.03084881 | 0.287815276 |
| ENSMUSG00000062727 | H2bc12 | 319184 | protein_coding | 648.0888046 | 0.28785902 | 0.163178111 | 1.764078637 | 0.077718741 | 0.246000026 | 0.28785902 |
| ENSMUSG00000097571 | Jpx | 70252 | lncRNA | 44.50719104 | -0.287999148 | 0.196446663 | -1.466042455 | 0.142636721 | 0.354359599 | 0.287999148 |
| ENSMUSG00000050368 | Hoxd10 | 15430 | protein_coding | 15.24767519 | 0.288039538 | 0.422207939 | 0.682221985 | 0.495098595 | 0.713812781 | 0.288039538 |
| ENSMUSG00000028251 | Tstd3 | 77032 | protein_coding | 117.9508251 | 0.28806891 | 0.122660635 | 2.348503337 | 0.01884903 | 0.094432529 | 0.28806891 |
| ENSMUSG00000097281 | Gm26685 | 102635083 | lncRNA | 25.72017111 | 0.28809143 | 0.249648066 | 1.153990236 | 0.248504172 | 0.4893628 | 0.28809143 |
| ENSMUSG00000085696 | Hoxaas3 | 72628 | lncRNA | 224.3176506 | 0.2881074 | 0.097645964 | 2.950530545 | 0.003172287 | 0.025263935 | 0.2881074 |
| ENSMUSG00000031766 | Slc12a3 | 20497 | protein_coding | 7.442413411 | 0.288217248 | 0.452845312 | 0.636458499 | 0.524477622 | 0.735184911 | 0.288217248 |
| ENSMUSG00000028763 | Hspg2 | 15530 | protein_coding | 450.3976969 | -0.288510036 | 0.12650276 | -2.280661982 | 0.022568456 | 0.106976995 | 0.288510036 |
| ENSMUSG00000029328 | Hnrnpdl | 50926 | protein_coding | 2297.042611 | -0.28853576 | 0.077393195 | -3.728179969 | 0.000192868 | 0.002555773 | 0.28853576 |
| ENSMUSG00000037071 | Scd1 | 20249 | protein_coding | 1196.651096 | 0.288635492 | 0.123490348 | 2.337312156 | 0.019422959 | 0.096084067 | 0.288635492 |
| ENSMUSG00000028034 | Fubp1 | 51886 | protein_coding | 1058.192163 | -0.288740066 | 0.078925743 | -3.658376271 | 0.000253818 | 0.003218024 | 0.288740066 |
| ENSMUSG00000054435 | Gimap4 | 107526 | protein_coding | 78.78390967 | 0.28880604 | 0.180013683 | 1.604356046 | 0.108635594 | 0.302001032 | 0.28880604 |
| ENSMUSG00000050410 | Tcf19 | 106795 | protein_coding | 255.5986017 | 0.288814124 | 0.166746901 | 1.732050926 | 0.083264496 | 0.256750769 | 0.288814124 |
| ENSMUSG00000087230 | Mroh3 | 76422 | protein_coding | 90.50975314 | -0.288926687 | 0.209294851 | -1.380476805 | 0.167439886 | 0.391461735 | 0.288926687 |
| ENSMUSG00000043614 | Vps37d | 194309 | protein_coding | 13.27022387 | 0.288973549 | 0.362035737 | 0.798190675 | 0.424759848 | 0.660696572 | 0.288973549 |
| ENSMUSG00000037847 | Nmrk1 | 225994 | protein_coding | 70.38062103 | 0.289057921 | 0.15310001 | 1.888033331 | 0.059021476 | 0.206112189 | 0.289057921 |
| ENSMUSG00000046733 | Gprc5a | 232431 | protein_coding | 1388.203942 | 0.289150064 | 0.124281591 | 2.326571963 | 0.019988058 | 0.098067361 | 0.289150064 |
| ENSMUSG00000034412 | Tbc1d10a | 103724 | protein_coding | 363.7970618 | 0.289242644 | 0.072742149 | 3.97627301 | 7.00E-05 | 0.001092885 | 0.289242644 |

| ensembl_gene_id | external_gene_name | entrezgene_id | gene_biotype | baseMean | log2FoldChange | lfcSE | stat | pvalue | padj | absLog2FC |
|---|---|---|---|---|---|---|---|---|---|---|
| ENSMUSG00000027742 | Cog6 | 67542 | protein_coding | 1385.253566 | 0.289313079 | 0.052239999 | 5.538152483 | 3.06E-08 | 1.03E-06 | 0.289313079 |
| ENSMUSG00000089762 | Ier5l | 72500 | protein_coding | 25.01806851 | -0.28935943 | 0.234636686 | -1.23322331 | 0.217492461 | 0.453693043 | 0.28935943 |
| ENSMUSG00000040415 | Dtx3 | 80904 | protein_coding | 41.00644561 | -0.289364292 | 0.216249675 | -1.338102782 | 0.180862934 | 0.407746288 | 0.289364292 |
| ENSMUSG00000032726 | Bmp8a | 12163 | protein_coding | 152.9944043 | 0.289379805 | 0.127113758 | 2.276541973 | 0.022813591 | 0.10767171 | 0.289379805 |
| ENSMUSG00000087088 | Gm16638 | 102631889 | lncRNA | 16.79942307 | -0.289380151 | 0.311574329 | -0.92876763 | 0.353009523 | 0.59550048 | 0.289380151 |
| ENSMUSG00000029059 | Prxl2b | 66469 | protein_coding | 254.6312907 | -0.289410094 | 0.111700133 | -2.590955675 | 0.009570982 | 0.057550792 | 0.289410094 |
| ENSMUSG00000032350 | Gclc | 14629 | protein_coding | 2029.498868 | -0.289491527 | 0.062350902 | -4.642940504 | 3.43E-06 | 7.46E-05 | 0.289491527 |
| ENSMUSG00000017639 | Rab11fip4 | 268451 | protein_coding | 1174.239892 | -0.289554551 | 0.05256812 | -5.508177826 | 3.63E-08 | 1.20E-06 | 0.289554551 |
| ENSMUSG00000029632 | Ndufa4 | 17992 | protein_coding | 3178.140305 | 0.289563791 | 0.05322291 | 5.440585502 | 5.31E-08 | 1.71E-06 | 0.289563791 |
| ENSMUSG00000033458 | Fan1 | 330554 | protein_coding | 198.5528363 | -0.289812791 | 0.100906621 | -2.872088942 | 0.004077682 | 0.030686316 | 0.289812791 |
| ENSMUSG00000033721 | Vav3 | 57257 | protein_coding | 546.0903384 | -0.290234537 | 0.099826679 | -2.907384474 | 0.003644649 | 0.028150029 | 0.290234537 |
| ENSMUSG00000079445 | B3gnt7 | 227327 | protein_coding | 8510.360107 | -0.290293346 | 0.179750507 | -1.614979289 | 0.10631518 | 0.298545169 | 0.290293346 |
| ENSMUSG00000058571 | Gpc6 | 23888 | protein_coding | 17.23317338 | 0.290318421 | 0.354702513 | 0.818484251 | 0.413080731 | 0.650752462 | 0.290318421 |
| ENSMUSG00000067613 | Krt83 | 100126226 | protein_coding | 11.28884768 | -0.290350788 | 0.394138662 | -0.736671673 | 0.461322037 | 0.687894338 | 0.290350788 |
| ENSMUSG00000026447 | Pik3c2b | 240752 | protein_coding | 2224.744139 | -0.290483197 | 0.060635338 | -4.790658469 | 1.66E-06 | 3.88E-05 | 0.290483197 |
| ENSMUSG00000033849 | B3galt2 | 26878 | protein_coding | 194.2524692 | -0.29049286 | 0.121314843 | -2.394536839 | 0.016641372 | 0.086466472 | 0.29049286 |
| ENSMUSG00000026065 | Slc9a4 | 110895 | protein_coding | 14.0464521 | -0.290560223 | 0.306805773 | -0.947049402 | 0.343613606 | 0.586128211 | 0.290560223 |
| ENSMUSG00000027357 | Crls1 | 66586 | protein_coding | 853.3839213 | 0.290666255 | 0.061055137 | 4.760717466 | 1.93E-06 | 4.44E-05 | 0.290666255 |
| ENSMUSG00000002228 | Ppm1j | 71887 | protein_coding | 137.6790659 | -0.290668606 | 0.129418847 | -2.245952684 | 0.024707039 | 0.113875511 | 0.290668606 |
| ENSMUSG00000028128 | F3 | 14066 | protein_coding | 566.1729212 | -0.29076401 | 0.092180775 | -3.154280393 | 0.001608944 | 0.014818642 | 0.29076401 |
| ENSMUSG00000026200 | Glb1l | 74577 | protein_coding | 47.69308761 | -0.290830063 | 0.174077186 | -1.670696026 | 0.094781734 | 0.278626509 | 0.290830063 |
| ENSMUSG00000059474 | Mbtd1 | 103537 | protein_coding | 595.2933044 | -0.290923263 | 0.095061755 | -3.060360733 | 0.002210705 | 0.018994165 | 0.290923263 |
| ENSMUSG00000024378 | Stard4 | 170459 | protein_coding | 5284.89687 | 0.290972459 | 0.048899426 | 5.950426931 | 2.67E-09 | 1.01E-07 | 0.290972459 |
| ENSMUSG00000027879 | Sec22b | 20333 | protein_coding | 1081.462833 | 0.291097584 | 0.069809121 | 4.169907613 | 3.05E-05 | 0.000526153 | 0.291097584 |
| ENSMUSG00000026388 | 3110009E18Rik | 73103 | protein_coding | 23.39099103 | -0.291102658 | 0.241570061 | -1.205044437 | 0.228186148 | 0.46714214 | 0.291102658 |
| ENSMUSG00000028248 | Pnisr | 66625 | protein_coding | 1075.91873 | -0.291114345 | 0.114465168 | -2.543257035 | 0.010982441 | 0.063390773 | 0.291114345 |
| ENSMUSG00000028138 | Adh5 | 11532 | protein_coding | 2944.080586 | -0.291274217 | 0.055515 | -5.246766021 | 1.55E-07 | 4.56E-06 | 0.291274217 |
| ENSMUSG00000041025 | Iffo2 | 212632 | protein_coding | 428.0625582 | -0.291338437 | 0.082630266 | -3.525807795 | 0.000422193 | 0.004894832 | 0.291338437 |
| ENSMUSG00000027082 | Tfpi | 21788 | protein_coding | 83.52022135 | -0.29160623 | 0.204392047 | -1.426700475 | 0.153666249 | 0.37087333 | 0.29160623 |
| ENSMUSG00000020282 | Rhbdf1 | 13650 | protein_coding | 541.0480105 | -0.291641066 | 0.114139357 | -2.555131506 | 0.010614769 | 0.061911166 | 0.291641066 |
| ENSMUSG00000033170 | Card10 | 105844 | protein_coding | 691.7201176 | -0.291737539 | 0.077271367 | -3.775493433 | 0.000159691 | 0.002176957 | 0.291737539 |
| ENSMUSG00000058952 | Cfi | 12630 | protein_coding | 7.794020238 | 0.292116689 | 0.418267536 | 0.698396753 | 0.484929107 | 0.706263486 | 0.292116689 |
| ENSMUSG00000022773 | Ypel1 | 106369 | protein_coding | 18.36197046 | -0.292149784 | 0.279537474 | -1.045118492 | 0.295968208 | 0.538897588 | 0.292149784 |
| ENSMUSG00000023277 | Twf2 | 23999 | protein_coding | 351.6312461 | 0.292246894 | 0.081459537 | 3.587632645 | 0.000333694 | 0.004025043 | 0.292246894 |
| ENSMUSG00000052684 | Jun | 16476 | protein_coding | 21704.71589 | 0.292267112 | 0.087259704 | 3.349393803 | 0.000809886 | 0.008442812 | 0.292267112 |
| ENSMUSG00000053080 | Zfta | 108899 | protein_coding | 113.4612206 | -0.292428518 | 0.13315838 | -2.196095495 | 0.028085109 | 0.12424678 | 0.292428518 |
| ENSMUSG00000074797 | Itpa | 16434 | protein_coding | 678.4609679 | 0.292433437 | 0.079488038 | 3.678961554 | 0.000234186 | 0.003006935 | 0.292433437 |
| ENSMUSG00000027358 | Bmp2 | 12156 | protein_coding | 978.297198 | -0.292454913 | 0.141434101 | -2.067782183 | 0.03866051 | 0.154660479 | 0.292454913 |
| ENSMUSG00000099083 | Atf7 | 223922 | protein_coding | 271.814223 | -0.292599528 | 0.136452529 | -2.144332024 | 0.032006293 | 0.136119589 | 0.292599528 |

| ensembl_gene_id | external_gene_name | entrezgene_id | gene_biotype | baseMean | log2FoldChange | lfcSE | stat | pvalue | padj | absLog2FC |
|---|---|---|---|---|---|---|---|---|---|---|
| ENSMUSG00000026450 | Chit1 | 71884 | protein_coding | 11.59105782 | -0.292743214 | 0.453689963 | -0.645249484 | 0.518765523 | 0.730713836 | 0.292743214 |
| ENSMUSG00000050824 | Sstr5 | 20609 | protein_coding | 13.47399518 | -0.292809021 | 0.410063775 | -0.714057273 | 0.475191765 | 0.698770639 | 0.292809021 |
| ENSMUSG00000000811 | Txnrd3 | 232223 | protein_coding | 331.6272036 | -0.292863668 | 0.088285174 | -3.317246304 | 0.000909094 | 0.009314463 | 0.292863668 |
| ENSMUSG00000041954 | Tnfrsf18 | 21936 | protein_coding | 27.78324328 | 0.292908277 | 0.224378892 | 1.305418143 | 0.191750504 | 0.423163427 | 0.292908277 |
| ENSMUSG00000042523 | Dnal1 | 105000 | protein_coding | 28.07264661 | 0.293044456 | 0.242788931 | 1.206992653 | 0.227434973 | 0.466053852 | 0.293044456 |
| ENSMUSG00000001229 | Dpp9 | 224897 | protein_coding | 1016.052587 | 0.29310449 | 0.100422927 | 2.918700927 | 0.003514933 | 0.02737099 | 0.29310449 |
| ENSMUSG00000037887 | Dusp8 | 18218 | protein_coding | 427.8834243 | 0.293360132 | 0.114833696 | 2.554651999 | 0.010629401 | 0.061938406 | 0.293360132 |
| ENSMUSG00000027422 | Rrbp1 | 81910 | protein_coding | 10453.23199 | 0.293780435 | 0.043143213 | 6.809424071 | 9.80E-12 | 4.70E-10 | 0.293780435 |
| ENSMUSG00000022330 | Osr2 | 107587 | protein_coding | 799.5700867 | -0.293982297 | 0.141676021 | -2.075032142 | 0.037983556 | 0.153140202 | 0.293982297 |
| ENSMUSG00000022428 | Cby1 | 73739 | protein_coding | 255.7233469 | 0.294023559 | 0.081505882 | 3.607390698 | 0.000309292 | 0.003775888 | 0.294023559 |
| ENSMUSG00000027429 | Sec23b | 27054 | protein_coding | 1671.08038 | 0.294032637 | 0.068876392 | 4.268990101 | 1.96E-05 | 0.000355809 | 0.294032637 |
| ENSMUSG00000020255 | Nopchap1 | 28109 | protein_coding | 919.4962444 | 0.294067476 | 0.05394517 | 5.451229045 | 5.00E-08 | 1.62E-06 | 0.294067476 |
| ENSMUSG00000115149 | 9330188P03Rik | 380930 | lncRNA | 15.98622933 | -0.294067669 | 0.292394662 | -1.005721742 | 0.314549442 | 0.558770915 | 0.294067669 |
| ENSMUSG00000026179 | Pnkd | 56695 | protein_coding | 413.3837975 | 0.294151321 | 0.071759653 | 4.099118474 | 4.15E-05 | 0.00068978 | 0.294151321 |
| ENSMUSG00000028150 | Rorc | 19885 | protein_coding | 1191.027163 | 0.294330217 | 0.100799808 | 2.9199482 | 0.003500896 | 0.027288035 | 0.294330217 |
| ENSMUSG00000068329 | Htra2 | 64704 | protein_coding | 297.8921275 | -0.294411717 | 0.072219973 | -4.076596872 | 4.57E-05 | 0.000753576 | 0.294411717 |
| ENSMUSG00000004383 | Large1 | 16795 | protein_coding | 394.5604355 | -0.294602514 | 0.095197362 | -3.094649969 | 0.001970453 | 0.017345794 | 0.294602514 |
| ENSMUSG00000055184 | Fam72a | 108900 | protein_coding | 19.73659001 | 0.294606315 | 0.265125996 | 1.111193621 | 0.266485016 | 0.508351533 | 0.294606315 |
| ENSMUSG00000072972 | Adam4 | 11498 | protein_coding | 16.75568153 | -0.295053707 | 0.35070942 | -0.841305338 | 0.400176901 | 0.639482053 | 0.295053707 |
| ENSMUSG00000025324 | Atp10a | 11982 | protein_coding | 49.93870444 | -0.295055709 | 0.218626031 | -1.349590931 | 0.177147235 | 0.404081111 | 0.295055709 |
| ENSMUSG00000040877 | Wdr25 | 212198 | protein_coding | 121.7517777 | -0.295117206 | 0.109951876 | -2.68405795 | 0.00727345 | 0.047118814 | 0.295117206 |
| ENSMUSG00000079507 | H2-Q1 | 15006 | protein_coding | 258.7255595 | -0.295175506 | 0.180893786 | -1.631761449 | 0.10272975 | 0.29253061 | 0.295175506 |
| ENSMUSG00000063146 | Clip2 | 269713 | protein_coding | 2144.983834 | 0.295319489 | 0.040975922 | 7.207146834 | 5.71E-13 | 3.01E-11 | 0.295319489 |
| ENSMUSG00000058794 | Nfe2 | 18022 | protein_coding | 9.822118229 | 0.295382234 | 0.404656026 | 0.729958817 | 0.465415358 | 0.691455871 | 0.295382234 |
| ENSMUSG00000028949 | Smarcd3 | 66993 | protein_coding | 15.10276211 | -0.29544951 | 0.291759844 | -1.012646241 | 0.311229164 | 0.555097626 | 0.29544951 |
| ENSMUSG00000061825 | Ces2c | 234671 | protein_coding | 2577.970355 | -0.295477726 | 0.108112238 | -2.733064548 | 0.006274803 | 0.042375944 | 0.295477726 |
| ENSMUSG00000020897 | Aurkb | 20877 | protein_coding | 420.3874355 | 0.295570243 | 0.130869897 | 2.258504434 | 0.023914228 | 0.111302962 | 0.295570243 |
| ENSMUSG00000060548 | Tnfrsf19 | 29820 | protein_coding | 41.37821577 | 0.295635793 | 0.262390861 | 1.126700037 | 0.259869327 | 0.501305339 | 0.295635793 |
| ENSMUSG00000039021 | Ttc16 | 338348 | protein_coding | 9.846212184 | -0.29569947 | 0.405261068 | -0.729651806 | 0.465603047 | 0.691532821 | 0.29569947 |
| ENSMUSG00000072618 | Gm10384 | 100038735 | lncRNA | 67.47293317 | -0.296016753 | 0.204854166 | -1.445012119 | 0.148454482 | 0.362651904 | 0.296016753 |
| ENSMUSG00000038828 | Tmem214 | 68796 | protein_coding | 1346.351002 | 0.296138389 | 0.06127844 | 4.832668555 | 1.35E-06 | 3.21E-05 | 0.296138389 |
| ENSMUSG00000033855 | Ston1 | 77057 | protein_coding | 110.4921082 | -0.296254681 | 0.157300462 | -1.883368156 | 0.059650493 | 0.207368721 | 0.296254681 |
| ENSMUSG00000005225 | Plekha8 | 231999 | protein_coding | 2318.994388 | -0.296283703 | 0.053753853 | -5.511859815 | 3.55E-08 | 1.18E-06 | 0.296283703 |
| ENSMUSG00000072915 | Gm12258 | 108167848 | protein_coding | 31.83298382 | 0.29641849 | 0.215051662 | 1.378359448 | 0.168092337 | 0.391964304 | 0.29641849 |
| ENSMUSG00000022983 | Scaf4 | 224432 | protein_coding | 395.4641372 | -0.296630219 | 0.090814223 | -3.266340997 | 0.00108947 | 0.010812233 | 0.296630219 |
| ENSMUSG00000057596 | Trim30d | 209387 | protein_coding | 376.8144409 | -0.296697509 | 0.134148189 | -2.211714612 | 0.02698639 | 0.120556652 | 0.296697509 |
| ENSMUSG00000021728 | Emb | 13723 | protein_coding | 29.02615246 | 0.297057797 | 0.276954236 | 1.072588022 | 0.283456001 | 0.526101385 | 0.297057797 |
| ENSMUSG00000005983 | 1700037C18Rik | 73261 | protein_coding | 101.7691568 | 0.297094885 | 0.118361534 | 2.510062819 | 0.012070969 | 0.068328716 | 0.297094885 |
| ENSMUSG00000020936 | Nmt1 | 18107 | protein_coding | 2248.121675 | 0.297106518 | 0.054120527 | 5.489719609 | 4.03E-08 | 1.32E-06 | 0.297106518 |

| ensembl_gene_id | external_gene_name | entrezgene_id | gene_biotype | baseMean | log2FoldChange | lfcSE | stat | pvalue | padj | absLog2FC |
|---|---|---|---|---|---|---|---|---|---|---|
| ENSMUSG00000097397 | Gm16861 | 100503704 | lncRNA | 11.71657689 | -0.297134582 | 0.407048611 | -0.729973213 | 0.465406559 | 0.691455871 | 0.297134582 |
| ENSMUSG00000037072 | Selenof | 93684 | protein_coding | 3354.515065 | 0.297221577 | 0.047548 | 6.25097963 | 4.08E-10 | 1.68E-08 | 0.297221577 |
| ENSMUSG00000055491 | Pprc1 | 226169 | protein_coding | 283.9706804 | -0.297227705 | 0.123611888 | -2.404523633 | 0.016193559 | 0.084845201 | 0.297227705 |
| ENSMUSG00000056895 | H2bc27 | 78303 | protein_coding | 27.7939217 | 0.297366078 | 0.291045505 | 1.021716785 | 0.306914965 | 0.550870027 | 0.297366078 |
| ENSMUSG00000033487 | Fndc3a | 319448 | protein_coding | 3579.339291 | 0.297371061 | 0.039863487 | 7.459735378 | 8.67E-14 | 4.87E-12 | 0.297371061 |
| ENSMUSG00000069300 | H2bc11 | 319183 | protein_coding | 492.8998322 | 0.297442829 | 0.150288452 | 1.979146261 | 0.047799541 | 0.179562869 | 0.297442829 |
| ENSMUSG00000046295 | Ankle1 | 234396 | protein_coding | 80.49944985 | -0.297558388 | 0.153692793 | -1.936059466 | 0.052860418 | 0.192216452 | 0.297558388 |
| ENSMUSG00000092618 | Btnl6 | 624681 | protein_coding | 2308.004525 | -0.297800789 | 0.121234704 | -2.456398858 | 0.01403373 | 0.076460123 | 0.297800789 |
| ENSMUSG00000037419 | Endod1 | 71946 | protein_coding | 6376.960379 | 0.297831253 | 0.089575072 | 3.324934548 | 0.000884393 | 0.009099034 | 0.297831253 |
| ENSMUSG00000027569 | Mrgbp | 73247 | protein_coding | 395.0091337 | -0.29799767 | 0.079088168 | -3.767917196 | 0.000164615 | 0.002230883 | 0.29799767 |
| ENSMUSG00000026149 | Tm4sf20 | 66261 | protein_coding | 3297.058605 | -0.298074757 | 0.055854722 | -5.336608048 | 9.47E-08 | 2.91E-06 | 0.298074757 |
| ENSMUSG00000021368 | Tbc1d7 | 67046 | protein_coding | 118.8309274 | 0.298168469 | 0.13254077 | 2.249635853 | 0.024472071 | 0.113115816 | 0.298168469 |
| ENSMUSG00000031465 | Angpt2 | 11601 | protein_coding | 77.13043146 | -0.298176928 | 0.144833626 | -2.05875484 | 0.03951773 | 0.157203362 | 0.298176928 |
| ENSMUSG00000078317 | F8a | 14070 | protein_coding | 443.4936534 | 0.298233839 | 0.084700703 | 3.521031464 | 0.000429872 | 0.004976697 | 0.298233839 |
| ENSMUSG00000073739 | Gm16287 | 100038595 | lncRNA | 18.00261538 | -0.298333179 | 0.327027462 | -0.912257268 | 0.361633304 | 0.604649053 | 0.298333179 |
| ENSMUSG00000100632 | 9430060I03Rik | 100037260 | lncRNA | 18.69140187 | 0.298411564 | 0.268272846 | 1.112343526 | 0.265990476 | 0.508009049 | 0.298411564 |
| ENSMUSG00000030352 | Tspan9 | 109246 | protein_coding | 184.4185423 | 0.298467772 | 0.130094173 | 2.294243978 | 0.021776488 | 0.10455009 | 0.298467772 |
| ENSMUSG00000023460 | Rab12 | 19328 | protein_coding | 137.5980842 | -0.298496611 | 0.108241159 | -2.75769968 | 0.005820965 | 0.040151712 | 0.298496611 |
| ENSMUSG00000031970 | Dbndd1 | 72185 | protein_coding | 23.74731444 | -0.298683897 | 0.257413904 | -1.160325422 | 0.245916338 | 0.487025714 | 0.298683897 |
| ENSMUSG00000020328 | Nudcd2 | 52653 | protein_coding | 486.3387816 | 0.298731766 | 0.069313108 | 4.309888498 | 1.63E-05 | 0.000301734 | 0.298731766 |
| ENSMUSG00000047875 | Gpr157 | 269604 | protein_coding | 47.36693259 | -0.29874254 | 0.205622361 | -1.452869907 | 0.146259878 | 0.359247178 | 0.29874254 |
| ENSMUSG00000040459 | Arglu1 | 234023 | protein_coding | 1020.79573 | -0.298774301 | 0.111535937 | -2.678726767 | 0.007390267 | 0.047711305 | 0.298774301 |
| ENSMUSG00000000078 | Klf6 | 23849 | protein_coding | 14072.43117 | 0.298857665 | 0.067386312 | 4.434990644 | 9.21E-06 | 0.000182422 | 0.298857665 |
| ENSMUSG00000024871 | Doc2g | 60425 | protein_coding | 14.86350164 | -0.298878665 | 0.309476297 | -0.965756242 | 0.334166169 | 0.577547724 | 0.298878665 |
| ENSMUSG00000022193 | Psmb5 | 19173 | protein_coding | 1119.080351 | 0.298932273 | 0.078663674 | 3.800131087 | 0.00014462 | 0.0020054 | 0.298932273 |
| ENSMUSG00000075590 | Nrbp2 | 223649 | protein_coding | 497.6501723 | -0.299005423 | 0.121669059 | -2.457530495 | 0.013989593 | 0.076271187 | 0.299005423 |
| ENSMUSG00000070283 | Ndufaf3 | 66706 | protein_coding | 485.7871827 | 0.299050764 | 0.09611438 | 3.111405011 | 0.001861994 | 0.016590261 | 0.299050764 |
| ENSMUSG00000038587 | Akap12 | 83397 | protein_coding | 7.78769633 | 0.299175212 | 0.465821107 | 0.642253447 | 0.520708635 | 0.732316051 | 0.299175212 |
| ENSMUSG00000097080 | 1700086O06Rik | 73516 | lncRNA | 38.73086542 | -0.299193497 | 0.19690574 | -1.519475749 | 0.128642788 | 0.334023867 | 0.299193497 |
| ENSMUSG00000070304 | Scn2b | 72821 | protein_coding | 126.0827263 | -0.299344606 | 0.171063238 | -1.749906114 | 0.080134516 | 0.250536319 | 0.299344606 |
| ENSMUSG00000025050 | Pcgf6 | 71041 | protein_coding | 201.5634653 | -0.299376187 | 0.085812462 | -3.488726212 | 0.000485328 | 0.005508011 | 0.299376187 |
| ENSMUSG00000004263 | Atn1 | 13498 | protein_coding | 25.50390206 | -0.299566889 | 0.293905459 | -1.019262759 | 0.308078239 | 0.551919325 | 0.299566889 |
| ENSMUSG00000075463 | 4930594M22Rik | 105245866 | lncRNA | 14.70297876 | -0.299678575 | 0.357792808 | -0.837575737 | 0.402269024 | 0.641553852 | 0.299678575 |
| ENSMUSG00000031556 | Tm2d2 | 69742 | protein_coding | 1888.047715 | 0.299823497 | 0.056824566 | 5.276300666 | 1.32E-07 | 3.94E-06 | 0.299823497 |
| ENSMUSG00000073144 | 4930599N23Rik | 75379 | lncRNA | 6.886286396 | 0.299923751 | 0.520761343 | 0.575933209 | 0.564660328 | 0.761458074 | 0.299923751 |
| ENSMUSG00000037111 | Setd7 | 73251 | protein_coding | 955.3891201 | -0.30004 | 0.10506618 | -2.855723892 | 0.004293884 | 0.031911273 | 0.30004 |
| ENSMUSG00000055296 | Tmem245 | 242474 | protein_coding | 927.4222546 | -0.300169738 | 0.076421846 | -3.927800151 | 8.57E-05 | 0.001292065 | 0.300169738 |
| ENSMUSG00000030720 | Cln3 | 12752 | protein_coding | 560.0183541 | -0.300491529 | 0.074899274 | -4.011941788 | 6.02E-05 | 0.000962527 | 0.300491529 |
| ENSMUSG00000008200 | Fnbp4 | 55935 | protein_coding | 773.3285931 | -0.300617699 | 0.09779293 | -3.074022837 | 0.002111933 | 0.018291696 | 0.300617699 |

| ensembl_gene_id | external_gene_name | entrezgene_id | gene_biotype | baseMean | log2FoldChange | lfcSE | stat | pvalue | padj | absLog2FC |
|---|---|---|---|---|---|---|---|---|---|---|
| ENSMUSG00000062078 | Qki | 19317 | protein_coding | 14.27623229 | 0.300886972 | 0.339579383 | 0.886057833 | 0.375586364 | 0.617215876 | 0.300886972 |
| ENSMUSG00000024785 | Rcl1 | 59028 | protein_coding | 289.4258902 | 0.300964723 | 0.08399555 | 3.583103173 | 0.000339536 | 0.004083299 | 0.300964723 |
| ENSMUSG00000026248 | Mrpl44 | 69163 | protein_coding | 645.3417283 | 0.300979512 | 0.067457778 | 4.461746604 | 8.13E-06 | 0.000164084 | 0.300979512 |
| ENSMUSG00000019916 | P4ha1 | 18451 | protein_coding | 349.0304567 | 0.301108985 | 0.122325499 | 2.461538993 | 0.013834236 | 0.075680028 | 0.301108985 |
| ENSMUSG00000021466 | Ptch1 | 19206 | protein_coding | 585.0296097 | -0.301294805 | 0.073503934 | -4.09902963 | 4.15E-05 | 0.00068978 | 0.301294805 |
| ENSMUSG00000054708 | Ankrd24 | 70615 | protein_coding | 169.2094211 | -0.301524159 | 0.140376982 | -2.147960114 | 0.031716924 | 0.135245594 | 0.301524159 |
| ENSMUSG00000074500 | Zfp558 | 72230 | protein_coding | 39.16377808 | -0.302018162 | 0.204614169 | -1.476037377 | 0.139933858 | 0.350881819 | 0.302018162 |
| ENSMUSG00000033669 | Zfp7 | 223669 | protein_coding | 577.3073274 | -0.3020449 | 0.068767676 | -4.392251071 | 1.12E-05 | 0.000216928 | 0.3020449 |
| ENSMUSG00000024644 | Cndp2 | 66054 | protein_coding | 2326.104584 | -0.302091774 | 0.080044253 | -3.774059517 | 0.000160612 | 0.002187666 | 0.302091774 |
| ENSMUSG00000029860 | Zyx | 22793 | protein_coding | 802.6401604 | -0.302439055 | 0.107237536 | -2.820272323 | 0.004798291 | 0.034530138 | 0.302439055 |
| ENSMUSG00000064647 | Gm23301 | 115488997 | snoRNA | 10.2734888 | -0.302537776 | 0.421964744 | -0.71697406 | 0.473390103 | 0.697472342 | 0.302537776 |
| ENSMUSG00000091405 | H4c14 | 97122 | protein_coding | 1071.334327 | 0.302609975 | 0.14238552 | 2.125286158 | 0.033562743 | 0.140808103 | 0.302609975 |
| ENSMUSG00000040350 | Trim7 | 94089 | protein_coding | 74.05738894 | -0.302711129 | 0.173494233 | -1.744790727 | 0.081021304 | 0.252369884 | 0.302711129 |
| ENSMUSG00000053332 | Gas5 | 14455 | lncRNA | 1690.64575 | -0.302755677 | 0.071541039 | -4.231916141 | 2.32E-05 | 0.000413359 | 0.302755677 |
| ENSMUSG00000011256 | Adam19 | 11492 | protein_coding | 13.07065118 | 0.302852421 | 0.3905781 | 0.775395294 | 0.438106115 | 0.671081619 | 0.302852421 |
| ENSMUSG00000053647 | Gper1 | 76854 | protein_coding | 18.13356378 | -0.302862646 | 0.290007092 | -1.044328414 | 0.296333471 | 0.539082898 | 0.302862646 |
| ENSMUSG00000110062 | B430319F04Rik | 619306 | TEC | 31.7477883 | 0.303063087 | 0.233133613 | 1.299954493 | 0.193616567 | 0.425472739 | 0.303063087 |
| ENSMUSG00000058248 | Kcnh1 | 16510 | protein_coding | 50.54431114 | -0.303119788 | 0.225690718 | -1.343076005 | 0.179247364 | 0.406457009 | 0.303119788 |
| ENSMUSG00000054252 | Fgfr3 | 14184 | protein_coding | 1259.748485 | -0.303176762 | 0.071958489 | -4.213217468 | 2.52E-05 | 0.000444213 | 0.303176762 |
| ENSMUSG00000047798 | Cd300lf | 246746 | protein_coding | 33.20588348 | -0.303249082 | 0.230795466 | -1.313929983 | 0.188869768 | 0.419339065 | 0.303249082 |
| ENSMUSG00000030407 | Qpctl | 67369 | protein_coding | 300.0785899 | 0.303508625 | 0.082800865 | 3.665524806 | 0.000246832 | 0.003141798 | 0.303508625 |
| ENSMUSG00000020396 | Nefh | 380684 | protein_coding | 16.3425373 | 0.303509554 | 0.304266155 | 0.997513358 | 0.318515393 | 0.562309551 | 0.303509554 |
| ENSMUSG00000089704 | Galnt2 | 108148 | protein_coding | 20.5743112 | -0.303594698 | 0.304403281 | -0.997343712 | 0.318597703 | 0.562331745 | 0.303594698 |
| ENSMUSG00000086914 | Gm16124 | 102636154 | lncRNA | 22.69292447 | -0.303618582 | 0.290041605 | -1.046810447 | 0.295187011 | 0.538150682 | 0.303618582 |
| ENSMUSG00000026924 | Sec16a | 227648 | protein_coding | 3690.960209 | 0.30369936 | 0.054242935 | 5.598873993 | 2.16E-08 | 7.42E-07 | 0.30369936 |
| ENSMUSG00000001281 | Itgb7 | 16421 | protein_coding | 46.89322576 | 0.303730593 | 0.216927728 | 1.400146474 | 0.161469461 | 0.382044893 | 0.303730593 |
| ENSMUSG00000051498 | Tlr6 | 21899 | protein_coding | 20.01685487 | 0.303858482 | 0.276203171 | 1.100126698 | 0.271276923 | 0.513806497 | 0.303858482 |
| ENSMUSG00000118384 | 9030625G05Rik | 71545 | lncRNA | 10.47815841 | 0.30399798 | 0.377858404 | 0.804528831 | 0.421091626 | 0.657973515 | 0.30399798 |
| ENSMUSG00000090534 | Gm4675 | 105245869 | lncRNA | 19.61201397 | -0.304114781 | 0.30456395 | -0.998525206 | 0.318024748 | 0.561627805 | 0.304114781 |
| ENSMUSG00000013160 | Atp6v0d1 | 11972 | protein_coding | 2042.224111 | 0.304316611 | 0.060847169 | 5.001327346 | 5.69E-07 | 1.47E-05 | 0.304316611 |
| ENSMUSG00000071226 | Cecr2 | 330409 | protein_coding | 20.74805956 | -0.304349432 | 0.401689565 | -0.757673235 | 0.448646626 | 0.678988684 | 0.304349432 |
| ENSMUSG00000027244 | Atg13 | 51897 | protein_coding | 656.7659122 | 0.304371483 | 0.073561487 | 4.137647227 | 3.51E-05 | 0.000596283 | 0.304371483 |
| ENSMUSG00000068874 | Selenbp1 | 20341 | protein_coding | 43132.17479 | -0.304472918 | 0.074528048 | -4.085346724 | 4.40E-05 | 0.000727965 | 0.304472918 |
| ENSMUSG00000027239 | Mdk | 17242 | protein_coding | 7.61535576 | -0.304542204 | 0.427111572 | -0.713027282 | 0.475828875 | 0.699133771 | 0.304542204 |
| ENSMUSG00000031157 | Pqbp1 | 54633 | protein_coding | 551.5198522 | 0.30461131 | 0.062440828 | 4.878399582 | 1.07E-06 | 2.60E-05 | 0.30461131 |
| ENSMUSG00000040297 | Suco | 226551 | protein_coding | 1354.58485 | 0.304623527 | 0.047597417 | 6.400001209 | 1.55E-10 | 6.66E-09 | 0.304623527 |
| ENSMUSG00000034584 | Exph5 | 320051 | protein_coding | 2061.75537 | -0.304886059 | 0.087068449 | -3.501682462 | 0.00046233 | 0.005293007 | 0.304886059 |
| ENSMUSG00000012123 | Crybg2 | 230806 | protein_coding | 1222.957291 | -0.304899467 | 0.111341645 | -2.738413517 | 0.00617364 | 0.041868076 | 0.304899467 |
| ENSMUSG00000054942 | Miga1 | 215708 | protein_coding | 273.7587421 | -0.304940999 | 0.116611574 | -2.615014856 | 0.008922357 | 0.054836455 | 0.304940999 |

| ensembl_gene_id | external_gene_name | entrezgene_id | gene_biotype | baseMean | log2FoldChange | lfcSE | stat | pvalue | padj | absLog2FC |
|---|---|---|---|---|---|---|---|---|---|---|
| ENSMUSG00000029727 | Cyp3a13 | 13113 | protein_coding | 3828.472836 | -0.305200553 | 0.089915227 | -3.394314436 | 0.000688006 | 0.007436649 | 0.305200553 |
| ENSMUSG00000022679 | Mpv17l | 93734 | protein_coding | 204.2236508 | 0.305270308 | 0.093668039 | 3.259065862 | 0.001117797 | 0.011052553 | 0.305270308 |
| ENSMUSG00000036553 | Sh3tc1 | 231147 | protein_coding | 857.295784 | 0.305276227 | 0.091413273 | 3.33951753 | 0.000839241 | 0.008703815 | 0.305276227 |
| ENSMUSG00000042784 | Muc1 | 17829 | protein_coding | 114.8392016 | -0.305299102 | 0.137009337 | -2.22830874 | 0.025859934 | 0.117491746 | 0.305299102 |
| ENSMUSG00000078862 | Gm14326 | 628147 | protein_coding | 94.54285263 | -0.305336611 | 0.170633661 | -1.789427774 | 0.07354595 | 0.23731003 | 0.305336611 |
| ENSMUSG00000041831 | Sytl3 | 83672 | protein_coding | 68.11006362 | -0.305433823 | 0.200081757 | -1.526545082 | 0.126874172 | 0.331056319 | 0.305433823 |
| ENSMUSG00000028399 | Ptprd | 19266 | protein_coding | 2767.276395 | -0.305530197 | 0.057308014 | -5.331369478 | 9.75E-08 | 2.99E-06 | 0.305530197 |
| ENSMUSG00000022176 | Rem2 | 140743 | protein_coding | 9.149382111 | -0.305560701 | 0.409475755 | -0.746224159 | 0.455532017 | 0.683446351 | 0.305560701 |
| ENSMUSG00000024049 | Myom1 | 17929 | protein_coding | 30.40478542 | -0.305647184 | 0.267093987 | -1.144343186 | 0.25248134 | 0.493560011 | 0.305647184 |
| ENSMUSG00000042726 | Trafd1 | 231712 | protein_coding | 767.3033779 | -0.305870102 | 0.097535697 | -3.135981102 | 0.001712802 | 0.015483388 | 0.305870102 |
| ENSMUSG00000001943 | Vsig2 | 57276 | protein_coding | 211.0460148 | 0.305913187 | 0.14489477 | 2.111278331 | 0.034748395 | 0.143910471 | 0.305913187 |
| ENSMUSG00000048218 | Amigo2 | 105827 | protein_coding | 41.34473318 | -0.306131164 | 0.263707635 | -1.160873344 | 0.24569341 | 0.486948216 | 0.306131164 |
| ENSMUSG00000028849 | Map7d1 | 245877 | protein_coding | 159.1299808 | 0.306177999 | 0.140146119 | 2.18470551 | 0.028910446 | 0.126695316 | 0.306177999 |
| ENSMUSG00000097768 | 2310043M15Rik | 102633000 | lncRNA | 9.660487499 | -0.306339438 | 0.397562261 | -0.770544562 | 0.440976932 | 0.672622244 | 0.306339438 |
| ENSMUSG00000050103 | Agmo | 319660 | protein_coding | 225.7972383 | -0.306384955 | 0.1167774 | -2.623666537 | 0.008698891 | 0.053791031 | 0.306384955 |
| ENSMUSG00000031220 | Awat2 | 245532 | protein_coding | 15.47000533 | 0.306456276 | 0.348109186 | 0.880345271 | 0.378672295 | 0.619821175 | 0.306456276 |
| ENSMUSG00000022040 | Ephx2 | 13850 | protein_coding | 860.1869099 | -0.306505019 | 0.089034538 | -3.442540664 | 0.000576277 | 0.006413818 | 0.306505019 |
| ENSMUSG00000047415 | Gpr68 | 238377 | protein_coding | 24.50564499 | -0.306655537 | 0.279846838 | -1.095797756 | 0.273167291 | 0.515530122 | 0.306655537 |
| ENSMUSG00000026694 | Eef1aknmt | 71449 | protein_coding | 249.6632142 | -0.306719689 | 0.087499819 | -3.505375143 | 0.000455964 | 0.005241152 | 0.306719689 |
| ENSMUSG00000074218 | Cox7a1 | 12865 | protein_coding | 1114.681187 | 0.306857507 | 0.091330781 | 3.359847619 | 0.000779855 | 0.008193301 | 0.306857507 |
| ENSMUSG00000079243 | Xirp1 | 22437 | protein_coding | 9.133206896 | -0.306972553 | 0.447784861 | -0.685535799 | 0.493005878 | 0.712365003 | 0.306972553 |
| ENSMUSG00000026921 | Egfl7 | 353156 | protein_coding | 20.65131176 | -0.307023252 | 0.287319793 | -1.068576756 | 0.285260427 | 0.528173927 | 0.307023252 |
| ENSMUSG00000020178 | Adora2a | 11540 | protein_coding | 10.79931143 | 0.307028917 | 0.398367089 | 0.77071858 | 0.440873756 | 0.672592098 | 0.307028917 |
| ENSMUSG00000017692 | Rhbdl3 | 246104 | protein_coding | 106.3847869 | -0.307069664 | 0.160338792 | -1.915130212 | 0.0554759 | 0.198372469 | 0.307069664 |
| ENSMUSG00000002908 | Kcnn1 | 84036 | protein_coding | 10.17591742 | -0.3070875 | 0.403508469 | -0.761043508 | 0.446631081 | 0.677580465 | 0.3070875 |
| ENSMUSG00000099032 | Tcf24 | 100039596 | protein_coding | 30.66877839 | 0.307117744 | 0.246257001 | 1.247143199 | 0.212344996 | 0.447821073 | 0.307117744 |
| ENSMUSG00000019124 | Scrn1 | 69938 | protein_coding | 8.031908052 | -0.307162199 | 0.416915658 | -0.736749011 | 0.461274996 | 0.687894338 | 0.307162199 |
| ENSMUSG00000020869 | Lrrc59 | 98238 | protein_coding | 3280.250029 | 0.307521002 | 0.081039771 | 3.7946924 | 0.000147827 | 0.002041096 | 0.307521002 |
| ENSMUSG00000020807 | 4933427D14Rik | 74477 | protein_coding | 10.65563183 | 0.307862281 | 0.37211579 | 0.82732926 | 0.408050463 | 0.646554316 | 0.307862281 |
| ENSMUSG00000033278 | Ptprm | 19274 | protein_coding | 118.7586694 | 0.307916861 | 0.140368929 | 2.193625497 | 0.028262344 | 0.124668167 | 0.307916861 |
| ENSMUSG00000071551 | Akr1c19 | 432720 | protein_coding | 8445.161321 | -0.307933522 | 0.087230229 | -3.530123956 | 0.000415365 | 0.0048226 | 0.307933522 |
| ENSMUSG00000020063 | Sirt1 | 93759 | protein_coding | 573.2103267 | -0.308096749 | 0.067215739 | -4.583699503 | 4.57E-06 | 9.76E-05 | 0.308096749 |
| ENSMUSG00000036492 | Rnf39 | 386454 | protein_coding | 231.3998992 | -0.308104615 | 0.1034421 | -2.97852244 | 0.002896418 | 0.023413803 | 0.308104615 |
| ENSMUSG00000101970 | Chaserr | 69170 | lncRNA | 132.6726259 | -0.308169917 | 0.148535148 | -2.074727233 | 0.038011822 | 0.153215859 | 0.308169917 |
| ENSMUSG00000021176 | Efcab11 | 78767 | protein_coding | 44.18833356 | 0.308289852 | 0.180194289 | 1.710874712 | 0.087104244 | 0.264345152 | 0.308289852 |
| ENSMUSG00000031749 | St3gal2 | 20444 | protein_coding | 21.49419145 | 0.308364214 | 0.286343884 | 1.076901697 | 0.281524185 | 0.524084097 | 0.308364214 |
| ENSMUSG00000030703 | Gdpd3 | 68616 | protein_coding | 30.66785291 | -0.308370694 | 0.223375153 | -1.380505802 | 0.167430965 | 0.391461735 | 0.308370694 |
| ENSMUSG00000017697 | Ada | 11486 | protein_coding | 163.666552 | 0.308451967 | 0.118711658 | 2.598329201 | 0.009367864 | 0.056752643 | 0.308451967 |
| ENSMUSG00000070808 | Bicra | 243842 | protein_coding | 148.6153834 | -0.308502135 | 0.140686359 | -2.192836158 | 0.028319186 | 0.124816482 | 0.308502135 |

| ensembl_gene_id | external_gene_name | entrezgene_id | gene_biotype | baseMean | log2FoldChange | lfcSE | stat | pvalue | padj | absLog2FC |
|---|---|---|---|---|---|---|---|---|---|---|
| ENSMUSG00000053164 | Gpr21 | 338346 | protein_coding | 13.02501714 | -0.308553275 | 0.394944371 | -0.781257558 | 0.434651027 | 0.668304413 | 0.308553275 |
| ENSMUSG00000079588 | Tmem182 | 381339 | protein_coding | 42.11133296 | 0.308560799 | 0.248816506 | 1.240113865 | 0.214933281 | 0.45050036 | 0.308560799 |
| ENSMUSG00000026435 | Slc45a3 | 212980 | protein_coding | 730.0227442 | 0.308561135 | 0.111988834 | 2.755284826 | 0.005864106 | 0.04029418 | 0.308561135 |
| ENSMUSG00000025817 | Nudt5 | 53893 | protein_coding | 510.1565057 | 0.308673983 | 0.086158417 | 3.58263295 | 0.000340148 | 0.00408761 | 0.308673983 |
| ENSMUSG00000024730 | Ms4a8a | 64381 | protein_coding | 7246.054361 | -0.308743399 | 0.05748309 | -5.371030011 | 7.83E-08 | 2.43E-06 | 0.308743399 |
| ENSMUSG00000039304 | Tnfsf10 | 22035 | protein_coding | 373.2286506 | -0.308781437 | 0.173489608 | -1.779826707 | 0.075104325 | 0.240748848 | 0.308781437 |
| ENSMUSG00000017390 | Aldoc | 11676 | protein_coding | 461.3806199 | -0.308800215 | 0.082881477 | -3.725804946 | 0.000194693 | 0.002575728 | 0.308800215 |
| ENSMUSG00000022837 | Iqcb1 | 320299 | protein_coding | 51.18323584 | 0.308831902 | 0.250398998 | 1.233359178 | 0.217441788 | 0.453693043 | 0.308831902 |
| ENSMUSG00000032311 | Nrg4 | 83961 | protein_coding | 23.34362444 | -0.308960348 | 0.24461634 | -1.263040518 | 0.206574619 | 0.441502442 | 0.308960348 |
| ENSMUSG00000036983 | Tfb1m | 224481 | protein_coding | 183.7852047 | 0.308994916 | 0.096252111 | 3.210266389 | 0.00132612 | 0.012691889 | 0.308994916 |
| ENSMUSG00000061758 | Akr1b10 | 67861 | protein_coding | 105.9473693 | 0.309071572 | 0.130622279 | 2.366147445 | 0.017974284 | 0.091350728 | 0.309071572 |
| ENSMUSG00000021614 | Vcan | 13003 | protein_coding | 142.0514269 | 0.309118423 | 0.15690959 | 1.970041613 | 0.048833602 | 0.182174298 | 0.309118423 |
| ENSMUSG00000024109 | Nrxn1 | 18189 | protein_coding | 15.64651601 | 0.309124555 | 0.337440576 | 0.916085904 | 0.35962184 | 0.603010752 | 0.309124555 |
| ENSMUSG00000020032 | Nuak1 | 77976 | protein_coding | 26.58795095 | 0.309278033 | 0.280320632 | 1.103300997 | 0.269896474 | 0.512314081 | 0.309278033 |
| ENSMUSG00000091575 | 2010016I18Rik | 69206 | lncRNA | 123.3709447 | -0.309299955 | 0.181875466 | -1.700613949 | 0.089015503 | 0.267960217 | 0.309299955 |
| ENSMUSG00000041674 | BC006965 | 217294 | lncRNA | 6.945609638 | 0.309354285 | 0.450628332 | 0.686495418 | 0.492400753 | 0.711873325 | 0.309354285 |
| ENSMUSG00000038884 | Shfl | 319278 | protein_coding | 26.2485292 | -0.309451364 | 0.228705285 | -1.353057339 | 0.176037313 | 0.402521936 | 0.309451364 |
| ENSMUSG00000042851 | Zc3h6 | 78751 | protein_coding | 157.5082477 | -0.309480935 | 0.141003295 | -2.194848967 | 0.028174434 | 0.124416509 | 0.309480935 |
| ENSMUSG00000094441 | Zfp955a | 77652 | protein_coding | 273.7465687 | -0.30948654 | 0.098321026 | -3.147714713 | 0.001645522 | 0.015035317 | 0.30948654 |
| ENSMUSG00000054083 | Capn12 | 60594 | protein_coding | 7.892609938 | -0.309617061 | 0.432789629 | -0.715398521 | 0.474362826 | 0.698223143 | 0.309617061 |
| ENSMUSG00000020149 | Rab1a | 19324 | protein_coding | 5229.360865 | 0.309672515 | 0.045260548 | 6.841996555 | 7.81E-12 | 3.79E-10 | 0.309672515 |
| ENSMUSG00000043953 | Ccrl2 | 54199 | protein_coding | 24.53954202 | -0.309790897 | 0.28228709 | -1.09743204 | 0.272452574 | 0.514863213 | 0.309790897 |
| ENSMUSG00000029438 | Bcl7a | 77045 | protein_coding | 196.9072058 | -0.309829406 | 0.100096063 | -3.095320602 | 0.001966003 | 0.01731607 | 0.309829406 |
| ENSMUSG00000040380 | Cbln3 | 56410 | protein_coding | 90.82937205 | 0.309902693 | 0.153416643 | 2.02000701 | 0.04338266 | 0.168180809 | 0.309902693 |
| ENSMUSG00000090489 | Gm17415 | 108167670 | processed_pseudogene | 23.00900224 | 0.310008001 | 0.240284445 | 1.290170912 | 0.196991323 | 0.429482097 | 0.310008001 |
| ENSMUSG00000040740 | Slc25a34 | 384071 | protein_coding | 897.5287464 | -0.310323181 | 0.094311002 | -3.290423954 | 0.001000365 | 0.010039137 | 0.310323181 |
| ENSMUSG00000033715 | Akr1c14 | 105387 | protein_coding | 6517.228684 | -0.310403603 | 0.069680318 | -4.45468121 | 8.40E-06 | 0.000167693 | 0.310403603 |
| ENSMUSG00000027534 | Snx16 | 74718 | protein_coding | 173.9784952 | -0.310458599 | 0.118400246 | -2.622111089 | 0.008738694 | 0.053942191 | 0.310458599 |
| ENSMUSG00000117975 | Itprip | 414801 | protein_coding | 59.12571187 | 0.310584219 | 0.151099749 | 2.055491295 | 0.039831576 | 0.158042883 | 0.310584219 |
| ENSMUSG00000024924 | Vldlr | 22359 | protein_coding | 460.4833371 | -0.31066582 | 0.079617218 | -3.901992896 | 9.54E-05 | 0.001424611 | 0.31066582 |
| ENSMUSG00000048371 | Pdp2 | 382051 | protein_coding | 1055.498335 | -0.310694658 | 0.107267801 | -2.896439152 | 0.003774238 | 0.028948374 | 0.310694658 |
| ENSMUSG00000079260 | Tmppe | 100504715 | protein_coding | 273.0570549 | -0.310755137 | 0.103567377 | -3.000511817 | 0.002695263 | 0.022109108 | 0.310755137 |
| ENSMUSG00000044072 | Eml6 | 237711 | protein_coding | 120.8721659 | 0.31112326 | 0.114354726 | 2.720685634 | 0.006514668 | 0.043445662 | 0.31112326 |
| ENSMUSG00000019900 | Rfx6 | 320995 | protein_coding | 18.21995772 | -0.311444015 | 0.279467999 | -1.114417449 | 0.265100141 | 0.506909435 | 0.311444015 |
| ENSMUSG00000034647 | Ankrd12 | 106585 | protein_coding | 597.8641854 | -0.311849367 | 0.109531391 | -2.847123235 | 0.004411628 | 0.032486907 | 0.311849367 |
| ENSMUSG00000039264 | Gimap3 | 83408 | protein_coding | 69.72307064 | 0.31194294 | 0.185997267 | 1.677137223 | 0.093515651 | 0.276314934 | 0.31194294 |
| ENSMUSG00000030325 | Klrb1c | 17059 | protein_coding | 7.583230071 | 0.311999204 | 0.418360127 | 0.745767065 | 0.455808138 | 0.683797009 | 0.311999204 |
| ENSMUSG00000036613 | Eipr1 | 380752 | protein_coding | 500.203849 | 0.312094783 | 0.070575468 | 4.422142567 | 9.77E-06 | 0.000192435 | 0.312094783 |
| ENSMUSG00000051331 | Cacna1c | 12288 | protein_coding | 60.47797931 | 0.312338795 | 0.155901257 | 2.003439872 | 0.045130095 | 0.172508426 | 0.312338795 |

Supplementary Table 1

| ensembl_gene_id | external_gene_name | entrezgene_id | gene_biotype | baseMean | log2FoldChange | lfcSE | stat | pvalue | padj | absLog2FC |
|---|---|---|---|---|---|---|---|---|---|---|
| ENSMUSG00000015452 | Ager | 11596 | protein_coding | 11.21811919 | -0.312347935 | 0.355508693 | -0.878594366 | 0.379621251 | 0.62053716 | 0.312347935 |
| ENSMUSG00000097789 | Ctxnd1 | 100039239 | protein_coding | 294.2601767 | -0.31253385 | 0.129225014 | -2.4185244 | 0.0155836 | 0.082452993 | 0.31253385 |
| ENSMUSG00000004446 | Bid | 12122 | protein_coding | 290.3511399 | -0.312546481 | 0.108925712 | -2.869354501 | 0.004113105 | 0.030837772 | 0.312546481 |
| ENSMUSG00000070942 | Il1rl2 | 107527 | protein_coding | 31.77328673 | 0.31283987 | 0.274262082 | 1.140660304 | 0.254011312 | 0.495031103 | 0.31283987 |
| ENSMUSG00000036398 | Ppp1r11 | 76497 | protein_coding | 846.6451569 | 0.313066212 | 0.067240713 | 4.655902609 | 3.23E-06 | 7.06E-05 | 0.313066212 |
| ENSMUSG00000073733 | Cplane2 | 76166 | protein_coding | 9.506343053 | -0.31318212 | 0.415737757 | -0.753316519 | 0.451259734 | 0.680391336 | 0.31318212 |
| ENSMUSG00000031347 | Cetn2 | 26370 | protein_coding | 330.3128681 | 0.313349884 | 0.080263472 | 3.904016091 | 9.46E-05 | 0.001415368 | 0.313349884 |
| ENSMUSG00000114279 | H2bc14 | 319186 | protein_coding | 772.648363 | 0.313524612 | 0.157402961 | 1.991859675 | 0.046386456 | 0.175769357 | 0.313524612 |
| ENSMUSG00000007039 | Ddah2 | 51793 | protein_coding | 23.40334792 | -0.313584245 | 0.241884756 | -1.296420037 | 0.194830813 | 0.426907135 | 0.313584245 |
| ENSMUSG00000020573 | Pik3cg | 30955 | protein_coding | 249.8972276 | 0.313705962 | 0.082545376 | 3.800406217 | 0.000144459 | 0.0020054 | 0.313705962 |
| ENSMUSG00000025612 | Bach1 | 12013 | protein_coding | 1657.645707 | -0.313732244 | 0.061047532 | -5.139147019 | 2.76E-07 | 7.63E-06 | 0.313732244 |
| ENSMUSG00000021286 | Zfyve21 | 68520 | protein_coding | 755.8373394 | 0.313741928 | 0.074859055 | 4.191101875 | 2.78E-05 | 0.000483467 | 0.313741928 |
| ENSMUSG00000048240 | Gng7 | 14708 | protein_coding | 10.53030763 | -0.31404914 | 0.503529331 | -0.623695823 | 0.532827367 | 0.741023365 | 0.31404914 |
| ENSMUSG00000026712 | Mrc1 | 17533 | protein_coding | 11.82402523 | 0.314093418 | 0.389202935 | 0.807017084 | 0.419656641 | 0.657004431 | 0.314093418 |
| ENSMUSG00000014633 | Cmc2 | 66531 | protein_coding | 280.4991765 | 0.31411997 | 0.088730005 | 3.540177516 | 0.000399858 | 0.004683526 | 0.31411997 |
| ENSMUSG00000071350 | Setdb2 | 239122 | protein_coding | 202.0031557 | 0.314200237 | 0.110305292 | 2.848460223 | 0.004393134 | 0.032388488 | 0.314200237 |
| ENSMUSG00000036256 | Igfbp7 | 29817 | protein_coding | 6.559839965 | 0.314401085 | 0.488926728 | 0.643043358 | 0.520195966 | 0.731850331 | 0.314401085 |
| ENSMUSG00000031853 | Map3k21 | 234878 | protein_coding | 2307.050166 | -0.314412739 | 0.059433926 | -5.290122351 | 1.22E-07 | 3.67E-06 | 0.314412739 |
| ENSMUSG00000020447 | Npc1l1 | 237636 | protein_coding | 78.67691702 | -0.314461957 | 0.503768865 | -0.624218722 | 0.532483953 | 0.740799392 | 0.314461957 |
| ENSMUSG00000020904 | Cfap52 | 71860 | protein_coding | 27.47340253 | -0.314466878 | 0.255902261 | -1.228855409 | 0.219126023 | 0.455509844 | 0.314466878 |
| ENSMUSG00000062929 | Cfl2 | 12632 | protein_coding | 751.2824121 | -0.314641533 | 0.123554546 | -2.546579983 | 0.010878428 | 0.062902978 | 0.314641533 |
| ENSMUSG00000001143 | Lman2l | 214895 | protein_coding | 264.2765487 | 0.314701819 | 0.091646916 | 3.433850611 | 0.000595072 | 0.006603514 | 0.314701819 |
| ENSMUSG00000027420 | Bfsp1 | 12075 | protein_coding | 165.6299775 | -0.314730045 | 0.105818698 | -2.9742385 | 0.002937167 | 0.023672009 | 0.314730045 |
| ENSMUSG00000080365 | Gm25776 | 115488078 | snoRNA | 170.1568196 | 0.314736145 | 0.462214355 | 0.680931135 | 0.495915066 | 0.714456162 | 0.314736145 |
| ENSMUSG00000025082 | Vwa2 | 240675 | protein_coding | 23.19081922 | 0.314878681 | 0.27063558 | 1.163478509 | 0.244635413 | 0.485926259 | 0.314878681 |
| ENSMUSG00000034579 | Pla2g3 | 237625 | protein_coding | 264.2597436 | -0.314939463 | 0.127916472 | -2.462071204 | 0.013813724 | 0.07559346 | 0.314939463 |
| ENSMUSG00000027317 | Ppp1r14d | 72112 | protein_coding | 2235.863913 | 0.315083069 | 0.082879942 | 3.801680611 | 0.000143718 | 0.001999775 | 0.315083069 |
| ENSMUSG00000018476 | Kdm6b | 216850 | protein_coding | 513.7406495 | -0.315152489 | 0.129220578 | -2.438872308 | 0.014733173 | 0.079253464 | 0.315152489 |
| ENSMUSG00000093445 | Lrch4 | 231798 | protein_coding | 6.355231796 | -0.315155862 | 0.475426794 | -0.66289041 | 0.507400744 | 0.722866767 | 0.315155862 |
| ENSMUSG00000028133 | Rwdd3 | 66568 | protein_coding | 52.91859576 | -0.315497216 | 0.219804193 | -1.435355772 | 0.151185742 | 0.367102434 | 0.315497216 |
| ENSMUSG00000020385 | Clk4 | 12750 | protein_coding | 678.0962722 | -0.315720096 | 0.102666381 | -3.075204301 | 0.002103584 | 0.018258611 | 0.315720096 |
| ENSMUSG00000064247 | Plcxd1 | 403178 | protein_coding | 18.72141968 | -0.316003514 | 0.307550974 | -1.027483376 | 0.304192909 | 0.548247546 | 0.316003514 |
| ENSMUSG00000025314 | Ptprj | 19271 | protein_coding | 3154.646151 | -0.316007137 | 0.069211161 | -4.565840748 | 4.97E-06 | 0.000105429 | 0.316007137 |
| ENSMUSG00000092074 | Dynlt1a | 100310872 | protein_coding | 83.88195788 | -0.316042126 | 0.161860591 | -1.95255759 | 0.050872041 | 0.18726624 | 0.316042126 |
| ENSMUSG00000009585 | Apobec3 | 80287 | protein_coding | 283.1033258 | -0.316272971 | 0.085611402 | -3.694285626 | 0.000220506 | 0.002859216 | 0.316272971 |
| ENSMUSG00000004317 | Clcn5 | 12728 | protein_coding | 1190.175068 | -0.316297645 | 0.067205182 | -4.706447273 | 2.52E-06 | 5.65E-05 | 0.316297645 |
| ENSMUSG00000078866 | Zfp970 | 628308 | protein_coding | 70.87373409 | -0.316358249 | 0.185417264 | -1.70619629 | 0.087971539 | 0.265849589 | 0.316358249 |
| ENSMUSG00000072612 | Gm10382 | 639281 | lncRNA | 329.5024325 | -0.316459754 | 0.204183511 | -1.549879088 | 0.12117054 | 0.322370555 | 0.316459754 |
| ENSMUSG00000029554 | Mad1l1 | 17120 | protein_coding | 340.9275502 | 0.316489999 | 0.108248064 | 2.923747444 | 0.003458452 | 0.027024167 | 0.316489999 |

| ensembl_gene_id | external_gene_name | entrezgene_id | gene_biotype | baseMean | log2FoldChange | lfcSE | stat | pvalue | padj | absLog2FC |
|---|---|---|---|---|---|---|---|---|---|---|
| ENSMUSG00000027950 | Chrnb2 | 11444 | protein_coding | 78.95273832 | -0.316655214 | 0.14320363 | -2.211223372 | 0.027020373 | 0.120633435 | 0.316655214 |
| ENSMUSG00000049119 | Fam110b | 242297 | protein_coding | 36.99361755 | -0.317108628 | 0.194284863 | -1.63218391 | 0.102640749 | 0.292453598 | 0.317108628 |
| ENSMUSG00000024579 | Pcyox1l | 240334 | protein_coding | 158.7414088 | 0.317149003 | 0.111189848 | 2.852319793 | 0.004340141 | 0.032121826 | 0.317149003 |
| ENSMUSG00000077394 | Gm24339 | 115487283 | snoRNA | 273.4590469 | -0.31721892 | 0.163914271 | -1.935273353 | 0.052956761 | 0.192401774 | 0.31721892 |
| ENSMUSG00000020472 | Zkscan17 | 268417 | protein_coding | 362.7926074 | -0.317283429 | 0.084543114 | -3.752918667 | 0.000174788 | 0.002345091 | 0.317283429 |
| ENSMUSG00000034771 | Tle2 | 21886 | protein_coding | 45.01589121 | -0.317328055 | 0.21349916 | -1.486319923 | 0.137194519 | 0.346738131 | 0.317328055 |
| ENSMUSG00000033386 | Frrs1 | 20321 | protein_coding | 921.7702149 | -0.317507435 | 0.084499372 | -3.757512345 | 0.000171611 | 0.00230963 | 0.317507435 |
| ENSMUSG00000039760 | Il22ra2 | 237310 | protein_coding | 5.875146695 | 0.317658644 | 0.485024924 | 0.654932621 | 0.512511136 | 0.726170014 | 0.317658644 |
| ENSMUSG00000032436 | Cmtm7 | 102545 | protein_coding | 144.1035992 | 0.31781268 | 0.125942217 | 2.523480117 | 0.011619963 | 0.066299536 | 0.31781268 |
| ENSMUSG00000028528 | Dnajc6 | 72685 | protein_coding | 7.95677447 | 0.318246664 | 0.501109168 | 0.635084497 | 0.525373307 | 0.735943093 | 0.318246664 |
| ENSMUSG00000040287 | Stac3 | 237611 | protein_coding | 6.579495702 | -0.318282355 | 0.53249221 | -0.597722086 | 0.550025386 | 0.751653199 | 0.318282355 |
| ENSMUSG00000057378 | Ryr3 | 20192 | protein_coding | 20.37958447 | 0.318320076 | 0.323610518 | 0.983651822 | 0.325286736 | 0.56903126 | 0.318320076 |
| ENSMUSG00000038437 | Mllt6 | 246198 | protein_coding | 1685.195771 | -0.318533172 | 0.056579229 | -5.629860551 | 1.80E-08 | 6.24E-07 | 0.318533172 |
| ENSMUSG00000040511 | Pvr | 52118 | protein_coding | 314.4967785 | -0.318942259 | 0.116324483 | -2.741832584 | 0.006109747 | 0.041539585 | 0.318942259 |
| ENSMUSG00000021690 | Jmy | 57748 | protein_coding | 745.651204 | -0.319073435 | 0.085058147 | -3.751238967 | 0.000175963 | 0.002356938 | 0.319073435 |
| ENSMUSG00000057967 | Fgf18 | 14172 | protein_coding | 8.296770133 | -0.319137828 | 0.536678951 | -0.594653149 | 0.552075345 | 0.752630378 | 0.319137828 |
| ENSMUSG00000042487 | Leo1 | 235497 | protein_coding | 574.5810189 | 0.319194655 | 0.074898144 | 4.261716497 | 2.03E-05 | 0.000366768 | 0.319194655 |
| ENSMUSG00000055198 | Gm12830 | 433746 | lncRNA | 12.99766815 | 0.319227753 | 0.34608187 | 0.922405306 | 0.356317203 | 0.599075014 | 0.319227753 |
| ENSMUSG00000029074 | Ttll10 | 330010 | protein_coding | 213.3763355 | -0.319240936 | 0.14289264 | -2.234131416 | 0.025474427 | 0.116216716 | 0.319240936 |
| ENSMUSG00000006356 | Crip2 | 68337 | protein_coding | 18.48909818 | -0.319392156 | 0.293204693 | -1.08931461 | 0.276015172 | 0.518738688 | 0.319392156 |
| ENSMUSG00000041779 | Tram2 | 170829 | protein_coding | 45.07598629 | -0.319395099 | 0.186460205 | -1.712939758 | 0.086723623 | 0.263676126 | 0.319395099 |
| ENSMUSG00000048970 | C1galt1c1 | 59048 | protein_coding | 1532.25911 | 0.319555097 | 0.055654405 | 5.741775405 | 9.37E-09 | 3.35E-07 | 0.319555097 |
| ENSMUSG00000049044 | Rapgef4 | 56508 | protein_coding | 131.4006923 | -0.319615783 | 0.150654206 | -2.121519149 | 0.033878138 | 0.141652252 | 0.319615783 |
| ENSMUSG00000019368 | Sec14l4 | 103655 | protein_coding | 14.41478061 | -0.319706862 | 0.40834402 | -0.78293509 | 0.43366523 | 0.667785655 | 0.319706862 |
| ENSMUSG00000020883 | Fbxl20 | 72194 | protein_coding | 960.1100334 | -0.319803005 | 0.0860124 | -3.718103502 | 0.000200724 | 0.002648999 | 0.319803005 |
| ENSMUSG00000028785 | Hpca | 15444 | protein_coding | 24.04233098 | 0.32012814 | 0.277770427 | 1.152491802 | 0.249119036 | 0.489920593 | 0.32012814 |
| ENSMUSG00000109771 | Gm35315 | 102638847 | protein_coding | 42.27051872 | 0.320158257 | 0.198522347 | 1.61270639 | 0.106808312 | 0.299347968 | 0.320158257 |
| ENSMUSG00000020232 | Hmg20b | 15353 | protein_coding | 1002.923266 | 0.320218845 | 0.074698196 | 4.286835067 | 1.81E-05 | 0.000330634 | 0.320218845 |
| ENSMUSG00000065611 | Mir23a | 387216 | miRNA | 29.26461905 | -0.32025069 | 0.368187586 | -0.869803063 | 0.384408037 | 0.624972092 | 0.32025069 |
| ENSMUSG00000046667 | Rbm12b1 | 72397 | protein_coding | 26.24855982 | 0.320423117 | 0.283679661 | 1.129524464 | 0.258676655 | 0.500565425 | 0.320423117 |
| ENSMUSG00000024731 | Ms4a10 | 69826 | protein_coding | 40.98088154 | 0.320437473 | 0.258783642 | 1.238244698 | 0.215625342 | 0.451374597 | 0.320437473 |
| ENSMUSG00000009406 | Elk1 | 13712 | protein_coding | 118.7731108 | 0.320439392 | 0.1346081 | 2.38053573 | 0.017287484 | 0.08873319 | 0.320439392 |
| ENSMUSG00000078771 | Evi2a | 14017 | protein_coding | 6.559901581 | -0.320573959 | 0.503046721 | -0.637264783 | 0.523952386 | 0.734827388 | 0.320573959 |
| ENSMUSG00000029714 | Gigyf1 | 57330 | protein_coding | 390.7121173 | -0.320688408 | 0.095457321 | -3.35949515 | 0.00078085 | 0.008198419 | 0.320688408 |
| ENSMUSG00000038155 | Gstp2 | 14869 | protein_coding | 572.7572303 | -0.320690989 | 0.110005306 | -2.915232017 | 0.003554242 | 0.027637059 | 0.320690989 |
| ENSMUSG00000022292 | Rrm2b | 382985 | protein_coding | 161.3146883 | -0.320696255 | 0.117511658 | -2.729059059 | 0.006351533 | 0.04278662 | 0.320696255 |
| ENSMUSG00000052727 | Map1b | 17755 | protein_coding | 91.3427721 | 0.320708354 | 0.146669004 | 2.186613017 | 0.028770786 | 0.126254822 | 0.320708354 |
| ENSMUSG00000027359 | Slc27a2 | 26458 | protein_coding | 196.5684205 | 0.320713149 | 0.113911494 | 2.815459066 | 0.00487076 | 0.034926967 | 0.320713149 |
| ENSMUSG00000027639 | Samhd1 | 56045 | protein_coding | 899.7198516 | -0.320881394 | 0.124311922 | -2.581260026 | 0.00984404 | 0.058646044 | 0.320881394 |

| ensembl_gene_id | external_gene_name | entrezgene_id | gene_biotype | baseMean | log2FoldChange | lfcSE | stat | pvalue | padj | absLog2FC |
|---|---|---|---|---|---|---|---|---|---|---|
| ENSMUSG00000042216 | Sgsm1 | 52850 | protein_coding | 565.4603351 | 0.320948048 | 0.084835778 | 3.783168557 | 0.000154844 | 0.002123669 | 0.320948048 |
| ENSMUSG00000052135 | Foxo6 | 329934 | protein_coding | 41.30441707 | -0.321000156 | 0.199018509 | -1.612916097 | 0.106762737 | 0.299280839 | 0.321000156 |
| ENSMUSG00000025791 | Pgm1 | 72157 | protein_coding | 2171.526449 | 0.321067501 | 0.04004907 | 8.016852941 | 1.08E-15 | 7.14E-14 | 0.321067501 |
| ENSMUSG00000031099 | Smarca1 | 93761 | protein_coding | 9.626838877 | -0.321083874 | 0.375011043 | -0.85619845 | 0.39188802 | 0.631745112 | 0.321083874 |
| ENSMUSG00000002633 | Shh | 20423 | protein_coding | 23.59411211 | -0.321094522 | 0.288552112 | -1.112778275 | 0.265803668 | 0.507758471 | 0.321094522 |
| ENSMUSG00000037904 | Ankrd9 | 74251 | protein_coding | 819.9091801 | -0.321123587 | 0.09429425 | -3.405547915 | 0.000660314 | 0.007204931 | 0.321123587 |
| ENSMUSG00000032965 | Ift57 | 73916 | protein_coding | 7.852048704 | -0.321405017 | 0.437967333 | -0.733856142 | 0.46303642 | 0.689357631 | 0.321405017 |
| ENSMUSG00000029798 | Herc6 | 67138 | protein_coding | 84.28221421 | -0.321697332 | 0.281403256 | -1.143189801 | 0.252959796 | 0.494063538 | 0.321697332 |
| ENSMUSG00000031901 | Dus2 | 66369 | protein_coding | 148.5351143 | 0.321806389 | 0.106402887 | 3.024414063 | 0.002491152 | 0.020731319 | 0.321806389 |
| ENSMUSG00000049791 | Fzd4 | 14366 | protein_coding | 192.3275988 | 0.321847117 | 0.097689141 | 3.294604835 | 0.000985602 | 0.009934207 | 0.321847117 |
| ENSMUSG00000054640 | Slc8a1 | 20541 | protein_coding | 2571.141234 | -0.321860981 | 0.112223389 | -2.868038319 | 0.004130255 | 0.030908674 | 0.321860981 |
| ENSMUSG00000020444 | Guk1 | 14923 | protein_coding | 1356.142085 | 0.321897956 | 0.059878895 | 5.37581659 | 7.62E-08 | 2.38E-06 | 0.321897956 |
| ENSMUSG00000026430 | Rassf5 | 54354 | protein_coding | 26.42008535 | 0.321936771 | 0.277831043 | 1.158750181 | 0.24655803 | 0.48740394 | 0.321936771 |
| ENSMUSG00000041343 | Ankrd42 | 73845 | protein_coding | 52.99196929 | 0.321973032 | 0.199800555 | 1.611472161 | 0.10707685 | 0.299484627 | 0.321973032 |
| ENSMUSG00000050050 | Ccdc158 | 320696 | protein_coding | 6.248390538 | -0.321981736 | 0.504885941 | -0.637731634 | 0.523648389 | 0.734592691 | 0.321981736 |
| ENSMUSG00000026930 | Gpsm1 | 67839 | protein_coding | 149.1788397 | -0.322025432 | 0.128794881 | -2.500296828 | 0.012408929 | 0.06982512 | 0.322025432 |
| ENSMUSG00000067455 | H4c11 | 319159 | protein_coding | 1123.142838 | 0.322122178 | 0.153969894 | 2.092111445 | 0.036428548 | 0.148466815 | 0.322122178 |
| ENSMUSG00000078974 | Sec61g | 20335 | protein_coding | 579.9602792 | 0.322375474 | 0.095508155 | 3.375371182 | 0.000737162 | 0.007846481 | 0.322375474 |
| ENSMUSG00000021846 | Peli2 | 93834 | protein_coding | 407.3654632 | -0.322440174 | 0.071601135 | -4.503282998 | 6.69E-06 | 0.000137814 | 0.322440174 |
| ENSMUSG00000044408 | Sptssa | 104725 | protein_coding | 1202.95962 | 0.322451171 | 0.063766315 | 5.056763467 | 4.26E-07 | 1.13E-05 | 0.322451171 |
| ENSMUSG00000051008 | 4930412M03Rik | 100504140 | lncRNA | 17.82984236 | -0.322484319 | 0.359658254 | -0.896640951 | 0.369910539 | 0.612036907 | 0.322484319 |
| ENSMUSG00000086629 | 2810403D21Rik | 69964 | lncRNA | 56.70280411 | -0.322507359 | 0.200875111 | -1.605511788 | 0.108381221 | 0.301615868 | 0.322507359 |
| ENSMUSG00000046108 | Il17c | 234836 | protein_coding | 6.526514409 | -0.322567877 | 0.502794555 | -0.641550061 | 0.521165367 | 0.732548832 | 0.322567877 |
| ENSMUSG00000005973 | Rcn1 | 19672 | protein_coding | 126.3651387 | -0.322573791 | 0.168655459 | -1.912619924 | 0.055796727 | 0.199122332 | 0.322573791 |
| ENSMUSG00000035181 | Heatr5a | 320487 | protein_coding | 1929.870384 | 0.322671309 | 0.080250916 | 4.020780384 | 5.80E-05 | 0.00093173 | 0.322671309 |
| ENSMUSG00000045838 | Ccdc9b | 214239 | protein_coding | 388.6948647 | -0.322676987 | 0.185762921 | -1.737036564 | 0.082380717 | 0.254853985 | 0.322676987 |
| ENSMUSG00000070336 | Fbxo47 | 72973 | protein_coding | 17.16077683 | -0.322720912 | 0.307979153 | -1.047866092 | 0.294700306 | 0.537810551 | 0.322720912 |
| ENSMUSG00000042249 | Grk3 | 320129 | protein_coding | 99.54762502 | -0.322880946 | 0.170912035 | -1.889164478 | 0.058869793 | 0.205867649 | 0.322880946 |
| ENSMUSG00000050022 | Amz1 | 231842 | protein_coding | 6.095115544 | 0.322952832 | 0.486802864 | 0.663416048 | 0.50706413 | 0.722697656 | 0.322952832 |
| ENSMUSG00000032532 | Cck | 12424 | protein_coding | 37.81951111 | -0.323257287 | 0.259314366 | -1.246584571 | 0.212549862 | 0.448053195 | 0.323257287 |
| ENSMUSG00000027217 | Tspan18 | 241556 | protein_coding | 6.766073375 | -0.3234257 | 0.545223333 | -0.593198567 | 0.553048272 | 0.753421987 | 0.3234257 |
| ENSMUSG00000039062 | Anpep | 16790 | protein_coding | 971.7628183 | -0.323698877 | 0.377473498 | -0.857540674 | 0.391146146 | 0.631053811 | 0.323698877 |
| ENSMUSG00000070822 | Zscan18 | 232875 | protein_coding | 7.304793562 | 0.32371056 | 0.463297836 | 0.698709416 | 0.484733648 | 0.706263486 | 0.32371056 |
| ENSMUSG00000049086 | Bmyc | 107771 | protein_coding | 31.96583031 | -0.323767048 | 0.283510626 | -1.141992636 | 0.253457081 | 0.494735214 | 0.323767048 |
| ENSMUSG00000026360 | Rgs2 | 19735 | protein_coding | 568.4776892 | 0.32388856 | 0.120486025 | 2.688183638 | 0.007184188 | 0.046747707 | 0.32388856 |
| ENSMUSG00000046743 | Fat4 | 329628 | protein_coding | 20.92825347 | -0.323958972 | 0.303355143 | -1.067919828 | 0.285556678 | 0.528457779 | 0.323958972 |
| ENSMUSG00000020895 | Tmem107 | 66910 | protein_coding | 133.4080322 | 0.324090046 | 0.144417419 | 2.244120188 | 0.024824669 | 0.114189231 | 0.324090046 |
| ENSMUSG00000049436 | Upk1b | 22268 | protein_coding | 37.00269725 | 0.324124861 | 0.240587398 | 1.347222935 | 0.177908444 | 0.405210667 | 0.324124861 |
| ENSMUSG00000092570 | Plut | 70147 | lncRNA | 7.340534595 | 0.324155868 | 0.447030619 | 0.725131242 | 0.46837152 | 0.693047027 | 0.324155868 |

| ensembl_gene_id | external_gene_name | entrezgene_id | gene_biotype | baseMean | log2FoldChange | lfcSE | stat | pvalue | padj | absLog2FC |
|---|---|---|---|---|---|---|---|---|---|---|
| ENSMUSG00000001891 | Ugp2 | 216558 | protein_coding | 4600.478639 | 0.324185319 | 0.039087176 | 8.29390491 | 1.10E-16 | 7.62E-15 | 0.324185319 |
| ENSMUSG00000031982 | Arv1 | 68865 | protein_coding | 48.93597371 | -0.324274374 | 0.176433278 | -1.837943373 | 0.06607075 | 0.2216607 | 0.324274374 |
| ENSMUSG00000041774 | Ydjc | 69101 | protein_coding | 575.0678166 | 0.32428889 | 0.062990618 | 5.148209342 | 2.63E-07 | 7.30E-06 | 0.32428889 |
| ENSMUSG00000090231 | Cfb | 14962 | protein_coding | 278.0466779 | -0.324432948 | 0.08943076 | -3.627755681 | 0.000285896 | 0.003538479 | 0.324432948 |
| ENSMUSG00000005131 | 4930550C14Rik | 75311 | protein_coding | 5.952442829 | -0.324452386 | 0.502465729 | -0.645720428 | 0.518460429 | 0.730599032 | 0.324452386 |
| ENSMUSG00000021280 | Exoc3l4 | 74190 | protein_coding | 38.3685534 | 0.324714472 | 0.27975908 | 1.160693235 | 0.245766674 | 0.486955333 | 0.324714472 |
| ENSMUSG00000031024 | Denn2b | 76954 | protein_coding | 1004.420887 | 0.324738826 | 0.075626681 | 4.293971678 | 1.76E-05 | 0.000320903 | 0.324738826 |
| ENSMUSG00000052676 | Zmat1 | 215693 | protein_coding | 21.19169905 | 0.324822099 | 0.277577661 | 1.170202594 | 0.241919449 | 0.482850861 | 0.324822099 |
| ENSMUSG00000111761 | 9230112J17Rik | 100504561 | lncRNA | 17.24706445 | -0.324861402 | 0.313032196 | -1.037789103 | 0.299368251 | 0.543015856 | 0.324861402 |
| ENSMUSG00000069917 | Hba-a2 | 110257 | protein_coding | 13.82535993 | 0.324987898 | 0.572321021 | 0.567841974 | 0.570142276 | 0.765267123 | 0.324987898 |
| ENSMUSG00000035109 | Shc4 | 271849 | protein_coding | 6.624459269 | 0.325249895 | 0.515287183 | 0.631201213 | 0.527908969 | 0.737831675 | 0.325249895 |
| ENSMUSG00000047878 | A4galt | 239559 | protein_coding | 268.0725451 | 0.325336051 | 0.135889575 | 2.394120748 | 0.016660263 | 0.086531421 | 0.325336051 |
| ENSMUSG00000029878 | Dbpht2 | 386753 | lncRNA | 10.25523023 | 0.325566824 | 0.361350016 | 0.900973598 | 0.367602358 | 0.609972551 | 0.325566824 |
| ENSMUSG00000068220 | Lgals1 | 16852 | protein_coding | 10.9432402 | 0.325613498 | 0.36545971 | 0.890969618 | 0.37294547 | 0.614664445 | 0.325613498 |
| ENSMUSG00000024225 | Clps | 109791 | protein_coding | 192.5626685 | 0.325676095 | 0.123376428 | 2.639694631 | 0.008298076 | 0.052030743 | 0.325676095 |
| ENSMUSG00000027636 | Sla2 | 77799 | protein_coding | 15.96331609 | 0.325680156 | 0.332932909 | 0.978215572 | 0.327967716 | 0.571645972 | 0.325680156 |
| ENSMUSG00000024713 | Pcsk5 | 18552 | protein_coding | 1193.255735 | -0.325694497 | 0.080331764 | -4.054367547 | 5.03E-05 | 0.000825566 | 0.325694497 |
| ENSMUSG00000039672 | Kcne2 | 246133 | protein_coding | 7.143314743 | 0.325804032 | 0.532092281 | 0.612307382 | 0.540334406 | 0.74605933 | 0.325804032 |
| ENSMUSG00000005107 | Slc2a9 | 117591 | protein_coding | 233.9900697 | -0.326004528 | 0.094942205 | -3.433715575 | 0.000595369 | 0.006603514 | 0.326004528 |
| ENSMUSG00000078942 | Naip6 | 17952 | protein_coding | 2520.886116 | -0.326090252 | 0.084554921 | -3.856549646 | 0.000114999 | 0.001671239 | 0.326090252 |
| ENSMUSG00000024371 | C2 | 12263 | protein_coding | 22.20094545 | -0.326115541 | 0.285815419 | -1.141000516 | 0.253869708 | 0.495031103 | 0.326115541 |
| ENSMUSG00000063286 | Gvin-ps7 | 668139 | transcribed_unprocessed_pseudogene | 38.12880283 | 0.326159878 | 0.201704025 | 1.617022159 | 0.105873498 | 0.297926072 | 0.326159878 |
| ENSMUSG00000042808 | Gpx2 | 14776 | protein_coding | 5634.230272 | 0.326214483 | 0.067479063 | 4.834306682 | 1.34E-06 | 3.19E-05 | 0.326214483 |
| ENSMUSG00000074766 | Ism1 | 319909 | protein_coding | 9.976147822 | 0.326320601 | 0.622321443 | 0.524360207 | 0.600028028 | 0.785648606 | 0.326320601 |
| ENSMUSG00000029757 | Dync1i1 | 13426 | protein_coding | 63.58726822 | 0.326335782 | 0.148587639 | 2.196251212 | 0.028073968 | 0.12424678 | 0.326335782 |
| ENSMUSG00000035429 | Ptprh | 545902 | protein_coding | 1332.421388 | -0.326442627 | 0.088075942 | -3.706376789 | 0.000210245 | 0.002747673 | 0.326442627 |
| ENSMUSG00000032051 | Fdx1 | 14148 | protein_coding | 780.857224 | 0.326466795 | 0.065528753 | 4.982038873 | 6.29E-07 | 1.61E-05 | 0.326466795 |
| ENSMUSG00000002580 | Mien1 | 103742 | protein_coding | 858.6567546 | 0.326475074 | 0.07843528 | 4.162349804 | 3.15E-05 | 0.000543216 | 0.326475074 |
| ENSMUSG00000015085 | Entpd2 | 12496 | protein_coding | 229.8040201 | -0.326678131 | 0.088320671 | -3.698773195 | 0.000216644 | 0.002817596 | 0.326678131 |
| ENSMUSG00000021565 | Slc6a19 | 74338 | protein_coding | 137.5824146 | -0.326704031 | 0.287228784 | -1.13743486 | 0.255356541 | 0.496639525 | 0.326704031 |
| ENSMUSG00000056293 | Gsdmc2 | 331063 | protein_coding | 5925.286041 | 0.326950824 | 0.124563307 | 2.624776361 | 0.00867059 | 0.053657175 | 0.326950824 |
| ENSMUSG00000055912 | Tmem150a | 232086 | protein_coding | 14.4983129 | -0.326953854 | 0.341706913 | -0.956825402 | 0.338655389 | 0.582186916 | 0.326953854 |
| ENSMUSG00000034480 | Diaph2 | 54004 | protein_coding | 375.2036384 | -0.327074852 | 0.081072312 | -4.0343595 | 5.48E-05 | 0.000887414 | 0.327074852 |
| ENSMUSG00000042622 | Maff | 17133 | protein_coding | 870.5831546 | -0.327120824 | 0.13068325 | -2.503158011 | 0.012309058 | 0.069359948 | 0.327120824 |
| ENSMUSG00000026278 | Bok | 51800 | protein_coding | 288.5172156 | 0.327132827 | 0.091889509 | 3.560067214 | 0.00037076 | 0.004412727 | 0.327132827 |
| ENSMUSG00000061577 | Adgrg5 | 382045 | protein_coding | 9.154953878 | -0.327547524 | 0.430711995 | -0.760479223 | 0.446968184 | 0.677849907 | 0.327547524 |
| ENSMUSG00000112160 | BC024063 | 666584 | protein_coding | 36.52425252 | -0.327606186 | 0.217139023 | -1.508739334 | 0.13136541 | 0.337453004 | 0.327606186 |
| ENSMUSG00000059336 | Slc14a1 | 108052 | protein_coding | 5.928153436 | -0.327621267 | 0.551694678 | -0.593845256 | 0.552615617 | 0.753150685 | 0.327621267 |
| ENSMUSG00000031207 | Msn | 17698 | protein_coding | 51.95460154 | 0.327713048 | 0.179538372 | 1.825309236 | 0.067954377 | 0.225772608 | 0.327713048 |

| ensembl_gene_id | external_gene_name | entrezgene_id | gene_biotype | baseMean | log2FoldChange | lfcSE | stat | pvalue | padj | absLog2FC |
|---|---|---|---|---|---|---|---|---|---|---|
| ENSMUSG00000026395 | Ptprc | 19264 | protein_coding | 298.0909179 | 0.32779864 | 0.128702957 | 2.546939459 | 0.010867228 | 0.062860758 | 0.32779864 |
| ENSMUSG00000026679 | Enkur | 71233 | protein_coding | 14.71258156 | 0.327830205 | 0.326714146 | 1.003416012 | 0.315660182 | 0.559864561 | 0.327830205 |
| ENSMUSG00000029449 | Rhof | 23912 | protein_coding | 384.3527378 | 0.327851327 | 0.091470795 | 3.584218631 | 0.000338089 | 0.004068924 | 0.327851327 |
| ENSMUSG00000043029 | Trpv3 | 246788 | protein_coding | 374.2387908 | -0.328359139 | 0.139925543 | -2.346670471 | 0.018941999 | 0.094722877 | 0.328359139 |
| ENSMUSG00000036281 | Snapc4 | 227644 | protein_coding | 240.5331566 | -0.328506907 | 0.101764077 | -3.228122511 | 0.001246056 | 0.012105509 | 0.328506907 |
| ENSMUSG00000017057 | Il13ra1 | 16164 | protein_coding | 3114.066827 | 0.328585161 | 0.054452599 | 6.034333799 | 1.60E-09 | 6.22E-08 | 0.328585161 |
| ENSMUSG00000053192 | Mllt11 | 56772 | protein_coding | 126.3063627 | -0.328821661 | 0.166670701 | -1.972882217 | 0.048508983 | 0.181275208 | 0.328821661 |
| ENSMUSG00000069265 | H3c1 | 360198 | protein_coding | 886.5474536 | 0.329238618 | 0.116455903 | 2.827152678 | 0.004696393 | 0.034024587 | 0.329238618 |
| ENSMUSG00000055480 | Zfp458 | 238690 | protein_coding | 71.35970101 | -0.329568049 | 0.177662057 | -1.855027765 | 0.063592262 | 0.215996716 | 0.329568049 |
| ENSMUSG00000025464 | Paox | 212503 | protein_coding | 202.1674158 | -0.329609877 | 0.083467337 | -3.948968417 | 7.85E-05 | 0.00120322 | 0.329609877 |
| ENSMUSG00000037300 | Ttc13 | 234875 | protein_coding | 980.6227766 | 0.329627763 | 0.054823083 | 6.012572489 | 1.83E-09 | 7.06E-08 | 0.329627763 |
| ENSMUSG00000028701 | Lurap1 | 68075 | protein_coding | 42.95739468 | -0.329766611 | 0.195438432 | -1.687317111 | 0.091542392 | 0.27253169 | 0.329766611 |
| ENSMUSG00000044566 | Cage1 | 71213 | protein_coding | 73.54760706 | -0.329877285 | 0.152246849 | -2.166726527 | 0.030255713 | 0.130971062 | 0.329877285 |
| ENSMUSG00000078779 | Zfp59 | 22717 | protein_coding | 165.1234202 | -0.330053377 | 0.128146355 | -2.575597073 | 0.010006718 | 0.059264907 | 0.330053377 |
| ENSMUSG00000038598 | Shoc1 | 100155 | protein_coding | 19.80322275 | -0.330173258 | 0.319443405 | -1.033589214 | 0.301328244 | 0.544911481 | 0.330173258 |
| ENSMUSG00000055067 | Smyd3 | 69726 | protein_coding | 212.8718873 | -0.330189282 | 0.087082651 | -3.791676956 | 0.000149633 | 0.002060751 | 0.330189282 |
| ENSMUSG00000018899 | Irf1 | 16362 | protein_coding | 2170.968282 | -0.330205076 | 0.139733736 | -2.363102046 | 0.018122679 | 0.091896763 | 0.330205076 |
| ENSMUSG00000029605 | Oas1b | 23961 | protein_coding | 22.80110348 | -0.330344005 | 0.258444665 | -1.278200132 | 0.20117887 | 0.434907725 | 0.330344005 |
| ENSMUSG00000078630 | Tomt | 791260 | protein_coding | 7.022205853 | -0.330437018 | 0.447684642 | -0.738102197 | 0.460452349 | 0.687265524 | 0.330437018 |
| ENSMUSG00000002699 | Lcp2 | 16822 | protein_coding | 26.74892164 | 0.330469363 | 0.240758555 | 1.372617325 | 0.169871343 | 0.394231565 | 0.330469363 |
| ENSMUSG00000019971 | Cep290 | 216274 | protein_coding | 358.1025025 | 0.330530885 | 0.106590404 | 3.100944101 | 0.001929047 | 0.017027773 | 0.330530885 |
| ENSMUSG00000029022 | Miip | 28010 | protein_coding | 339.9875777 | -0.330601964 | 0.080277279 | -4.118250734 | 3.82E-05 | 0.000643326 | 0.330601964 |
| ENSMUSG00000020072 | Pbld2 | 67307 | protein_coding | 618.5798967 | -0.330881836 | 0.119011714 | -2.780245951 | 0.005431774 | 0.037987087 | 0.330881836 |
| ENSMUSG00000016206 | H2-M3 | 14991 | protein_coding | 103.4381262 | 0.330892515 | 0.132837148 | 2.490963702 | 0.012739713 | 0.071115735 | 0.330892515 |
| ENSMUSG00000089960 | Ugt1a1 | 394436 | protein_coding | 832.1238187 | -0.330912154 | 0.115006994 | -2.87732199 | 0.004010661 | 0.030351916 | 0.330912154 |
| ENSMUSG00000097577 | 6230400D17Rik | 76133 | lncRNA | 26.0279133 | -0.330929193 | 0.2841 | -1.164833488 | 0.244086401 | 0.485418672 | 0.330929193 |
| ENSMUSG00000024558 | Mapk4 | 225724 | protein_coding | 5.930053562 | 0.330951836 | 0.506897772 | 0.652896609 | 0.513822935 | 0.727070499 | 0.330951836 |
| ENSMUSG00000002033 | Cd3g | 12502 | protein_coding | 49.22818354 | 0.330958321 | 0.18778906 | 1.762394041 | 0.078002745 | 0.246560224 | 0.330958321 |
| ENSMUSG00000054169 | Ceacam10 | 26366 | protein_coding | 344.224701 | 0.331060356 | 0.14141301 | 2.341088399 | 0.019227614 | 0.09547586 | 0.331060356 |
| ENSMUSG00000018844 | Fndc8 | 78919 | protein_coding | 6.968932409 | -0.331135116 | 0.525188217 | -0.630507512 | 0.52836259 | 0.737932232 | 0.331135116 |
| ENSMUSG00000018405 | Mrm1 | 217038 | protein_coding | 252.6974776 | -0.331147625 | 0.111540552 | -2.968854108 | 0.002989125 | 0.024042701 | 0.331147625 |
| ENSMUSG00000027778 | Ift80 | 68259 | protein_coding | 128.592905 | 0.331157886 | 0.108982736 | 3.038627025 | 0.002376589 | 0.020035154 | 0.331157886 |
| ENSMUSG00000075389 | Reno1 | 100042332 | lncRNA | 57.39347428 | -0.331214404 | 0.167859525 | -1.973164184 | 0.04847686 | 0.181262767 | 0.331214404 |
| ENSMUSG00000057777 | Mab21l2 | 23937 | protein_coding | 6.572980835 | 0.331461521 | 0.513108784 | 0.645986839 | 0.518287879 | 0.730504162 | 0.331461521 |
| ENSMUSG00000078776 | 9530053A07Rik | 319482 | protein_coding | 16.70446084 | 0.33163034 | 0.313109928 | 1.059149868 | 0.289531532 | 0.531747525 | 0.33163034 |
| ENSMUSG00000027187 | Cat | 12359 | protein_coding | 3437.198298 | 0.331696872 | 0.046715733 | 7.100324713 | 1.24E-12 | 6.43E-11 | 0.331696872 |
| ENSMUSG00000025656 | Arhgef9 | 236915 | protein_coding | 24.66647299 | -0.331750764 | 0.242510138 | -1.367987193 | 0.171316085 | 0.395932144 | 0.331750764 |
| ENSMUSG00000041763 | Tpp2 | 22019 | protein_coding | 1960.708958 | 0.331813599 | 0.047332369 | 7.010289345 | 2.38E-12 | 1.21E-10 | 0.331813599 |
| ENSMUSG00000048174 | Tmem81 | 74626 | protein_coding | 46.26775161 | -0.331920218 | 0.193933435 | -1.711516209 | 0.086985862 | 0.264135001 | 0.331920218 |

| ensembl_gene_id | external_gene_name | entrezgene_id | gene_biotype | baseMean | log2FoldChange | lfcSE | stat | pvalue | padj | absLog2FC |
|---|---|---|---|---|---|---|---|---|---|---|
| ENSMUSG00000117322 | 6330415G19Rik | 320020 | lncRNA | 15.01248387 | 0.331941641 | 0.330033888 | 1.005780476 | 0.314521181 | 0.558770915 | 0.331941641 |
| ENSMUSG00000021457 | Syk | 20963 | protein_coding | 1580.865673 | 0.332049256 | 0.125258396 | 2.650914162 | 0.008027424 | 0.050786438 | 0.332049256 |
| ENSMUSG00000007670 | Khsrp | 16549 | protein_coding | 2007.754272 | 0.332061094 | 0.079791328 | 4.161618844 | 3.16E-05 | 0.000543877 | 0.332061094 |
| ENSMUSG00000021751 | Acox2 | 93732 | protein_coding | 54.92168139 | 0.33206867 | 0.192235908 | 1.727401886 | 0.084095512 | 0.258509777 | 0.33206867 |
| ENSMUSG00000044461 | Shisa2 | 219134 | protein_coding | 1649.418559 | -0.332109187 | 0.078768992 | -4.216242699 | 2.48E-05 | 0.000438779 | 0.332109187 |
| ENSMUSG00000002997 | Prkar2b | 19088 | protein_coding | 121.0526969 | 0.332258979 | 0.139290107 | 2.385373847 | 0.017061773 | 0.088049668 | 0.332258979 |
| ENSMUSG00000015312 | Gadd45b | 17873 | protein_coding | 231.2852098 | 0.33241735 | 0.104707669 | 3.174718265 | 0.00149982 | 0.013973197 | 0.33241735 |
| ENSMUSG00000104063 | Pcdhgb7 | 93704 | protein_coding | 10.95882673 | 0.332479747 | 0.366175132 | 0.907980137 | 0.363888716 | 0.60618049 | 0.332479747 |
| ENSMUSG00000021591 | Glrx | 93692 | protein_coding | 1860.230935 | -0.332547574 | 0.091740331 | -3.624878725 | 0.000289097 | 0.003569884 | 0.332547574 |
| ENSMUSG00000009646 | Pla2g12b | 69836 | protein_coding | 70.3875297 | 0.332560741 | 0.247747418 | 1.342337868 | 0.17948647 | 0.406691669 | 0.332560741 |
| ENSMUSG00000090083 | Rnf8 | 58230 | protein_coding | 166.2377397 | -0.332584701 | 0.108985459 | -3.051642902 | 0.002275927 | 0.019450294 | 0.332584701 |
| ENSMUSG00000014959 | Gorasp2 | 70231 | protein_coding | 2216.030891 | 0.332630845 | 0.063454128 | 5.242067869 | 1.59E-07 | 4.66E-06 | 0.332630845 |
| ENSMUSG00000054720 | Lrrc8c | 100604 | protein_coding | 13.8894503 | 0.332708782 | 0.311029889 | 1.069700352 | 0.284754208 | 0.527617708 | 0.332708782 |
| ENSMUSG00000004668 | Abca13 | 268379 | protein_coding | 23.10176547 | -0.332850218 | 0.413623393 | -0.804718069 | 0.42098239 | 0.657926757 | 0.332850218 |
| ENSMUSG00000030269 | Mtmr14 | 97287 | protein_coding | 950.6167673 | 0.332925632 | 0.051205806 | 6.501716418 | 7.94E-11 | 3.55E-09 | 0.332925632 |
| ENSMUSG00000031377 | Bmx | 12169 | protein_coding | 69.50561432 | 0.332945595 | 0.172608555 | 1.928905518 | 0.053742593 | 0.194173579 | 0.332945595 |
| ENSMUSG00000044052 | Ccr10 | 12777 | protein_coding | 13.14041363 | -0.332950509 | 0.325053998 | -1.024292918 | 0.305696942 | 0.54999264 | 0.332950509 |
| ENSMUSG00000031506 | Ptpn7 | 320139 | protein_coding | 9.571954949 | 0.33304028 | 0.541697382 | 0.614808731 | 0.538681042 | 0.74493282 | 0.33304028 |
| ENSMUSG00000036890 | Gtdc1 | 227835 | protein_coding | 121.3662903 | -0.333136143 | 0.119319358 | -2.791970632 | 0.005238811 | 0.036958139 | 0.333136143 |
| ENSMUSG00000075420 | Smim6 | 68528 | protein_coding | 2036.109196 | 0.333165862 | 0.055508761 | 6.002041062 | 1.95E-09 | 7.50E-08 | 0.333165862 |
| ENSMUSG00000025422 | Agap2 | 216439 | protein_coding | 22.16942488 | 0.333171053 | 0.326240485 | 1.021243741 | 0.307138974 | 0.551093706 | 0.333171053 |
| ENSMUSG00000030257 | Srgap3 | 259302 | protein_coding | 266.2623751 | -0.333262663 | 0.09554273 | -3.488100706 | 0.000486465 | 0.005517031 | 0.333262663 |
| ENSMUSG00000006958 | Chrd | 12667 | protein_coding | 10.18313631 | 0.333280517 | 0.386378793 | 0.862574558 | 0.388371424 | 0.628274245 | 0.333280517 |
| ENSMUSG00000061578 | Ksr2 | 333050 | protein_coding | 10.57975283 | 0.333377491 | 0.433921686 | 0.768289538 | 0.442315189 | 0.673453271 | 0.333377491 |
| ENSMUSG00000047369 | Dnah14 | 240960 | protein_coding | 9.31201402 | -0.333484987 | 0.396656887 | -0.840739183 | 0.400494064 | 0.639735319 | 0.333484987 |
| ENSMUSG00000075296 | Aldh3b2 | 621603 | protein_coding | 190.4115176 | 0.333519312 | 0.115978581 | 2.875697466 | 0.004031359 | 0.030437138 | 0.333519312 |
| ENSMUSG00000015468 | Notch4 | 18132 | protein_coding | 149.1297934 | 0.333651581 | 0.117463967 | 2.840458997 | 0.004504866 | 0.032977749 | 0.333651581 |
| ENSMUSG00000057561 | Eif1a | 13664 | protein_coding | 1764.426692 | 0.333819344 | 0.057548012 | 5.800710269 | 6.60E-09 | 2.40E-07 | 0.333819344 |
| ENSMUSG00000037336 | Mfsd2b | 432628 | protein_coding | 15.59080255 | -0.334104433 | 0.32520078 | -1.027378942 | 0.304242063 | 0.548274864 | 0.334104433 |
| ENSMUSG00000072761 | Gm6712 | 100862203 | lncRNA | 66.38998814 | -0.334397706 | 0.159629144 | -2.09484119 | 0.036185112 | 0.147736574 | 0.334397706 |
| ENSMUSG00000001053 | N4bp3 | 212706 | protein_coding | 169.7210457 | -0.334446642 | 0.107083588 | -3.1232297 | 0.001788781 | 0.016080083 | 0.334446642 |
| ENSMUSG00000039568 | Ubald1 | 207740 | protein_coding | 596.1909327 | -0.334550106 | 0.122713343 | -2.726273251 | 0.006405395 | 0.042967024 | 0.334550106 |
| ENSMUSG00000032679 | Cd59a | 12509 | protein_coding | 59.70944496 | 0.334596714 | 0.257812693 | 1.297828707 | 0.194346203 | 0.426250675 | 0.334596714 |
| ENSMUSG00000032293 | Ireb2 | 64602 | protein_coding | 1256.811457 | -0.334658753 | 0.124701787 | -2.68367248 | 0.00728184 | 0.047143412 | 0.334658753 |
| ENSMUSG00000064358 | mt-Co3 | 17710 | protein_coding | 6.655980121 | 0.334725204 | 0.583831047 | 0.573325461 | 0.566424349 | 0.762497953 | 0.334725204 |
| ENSMUSG00000001918 | Slc1a5 | 20514 | protein_coding | 3309.369402 | -0.334897828 | 0.054613303 | -6.13216574 | 8.67E-10 | 3.46E-08 | 0.334897828 |
| ENSMUSG00000093803 | Ppp2r3d | 19054 | protein_coding | 576.4983556 | -0.33491635 | 0.129678075 | -2.582675224 | 0.009803756 | 0.058492482 | 0.33491635 |
| ENSMUSG00000026097 | Ormdl1 | 227102 | protein_coding | 351.3680746 | 0.334921764 | 0.068834225 | 4.865628466 | 1.14E-06 | 2.76E-05 | 0.334921764 |
| ENSMUSG00000039168 | Dap | 223453 | protein_coding | 754.2439218 | 0.334994437 | 0.07122873 | 4.703080315 | 2.56E-06 | 5.74E-05 | 0.334994437 |

Supplementary Table 1

| ensembl_gene_id | external_gene_name | entrezgene_id | gene_biotype | baseMean | log2FoldChange | lfcSE | stat | pvalue | padj | absLog2FC |
|---|---|---|---|---|---|---|---|---|---|---|
| ENSMUSG00000031883 | Car7 | 12354 | protein_coding | 9.46319893 | -0.335097713 | 0.399815978 | -0.838129869 | 0.401957769 | 0.641247695 | 0.335097713 |
| ENSMUSG00000087444 | Gm5475 | 432982 | lncRNA | 6.836351941 | -0.33514257 | 0.48601343 | -0.68957471 | 0.490461676 | 0.710394783 | 0.33514257 |
| ENSMUSG00000025854 | Fam20c | 80752 | protein_coding | 8.411828136 | 0.335192488 | 0.402733388 | 0.832293766 | 0.405243148 | 0.644334723 | 0.335192488 |
| ENSMUSG00000102752 | Gm7694 | 665574 | protein_coding | 145.1730601 | -0.335203749 | 0.143301824 | -2.339145034 | 0.019327929 | 0.095731421 | 0.335203749 |
| ENSMUSG00000027199 | Gatm | 67092 | protein_coding | 321.6016539 | 0.335221815 | 0.141923254 | 2.361993574 | 0.018176957 | 0.092025153 | 0.335221815 |
| ENSMUSG00000049932 | H2ax | 15270 | protein_coding | 988.879126 | 0.335241227 | 0.106871613 | 3.136859429 | 0.00170768 | 0.015463084 | 0.335241227 |
| ENSMUSG00000028453 | Fancg | 60534 | protein_coding | 152.7159132 | 0.335243714 | 0.100640673 | 3.331095704 | 0.000865049 | 0.008919845 | 0.335243714 |
| ENSMUSG00000015709 | Arnt2 | 11864 | protein_coding | 58.83439811 | -0.335261299 | 0.189386725 | -1.770247087 | 0.076685988 | 0.244031351 | 0.335261299 |
| ENSMUSG00000020839 | Tmigd1 | 66601 | protein_coding | 2371.999404 | -0.335400296 | 0.145319819 | -2.308014819 | 0.020998312 | 0.101663099 | 0.335400296 |
| ENSMUSG00000026494 | Kif26b | 269152 | protein_coding | 28.12143221 | -0.335577312 | 0.273193716 | -1.228349308 | 0.219315869 | 0.455552165 | 0.335577312 |
| ENSMUSG00000047642 | D930020B18Rik | 216393 | protein_coding | 27.54714563 | 0.335619008 | 0.258753561 | 1.297060441 | 0.194610391 | 0.426540063 | 0.335619008 |
| ENSMUSG00000091649 | Phf11b | 236451 | protein_coding | 46.4660853 | -0.335638839 | 0.240171774 | -1.39749494 | 0.162264787 | 0.383043628 | 0.335638839 |
| ENSMUSG00000036764 | Dnajc12 | 30045 | protein_coding | 110.5943538 | 0.335708919 | 0.120080127 | 2.79570756 | 0.005178624 | 0.036661836 | 0.335708919 |
| ENSMUSG00000050711 | Scg2 | 20254 | protein_coding | 36.00125556 | 0.335816281 | 0.264468932 | 1.26977592 | 0.20416446 | 0.438306743 | 0.335816281 |
| ENSMUSG00000078700 | D030028A08Rik | 319371 | lncRNA | 149.1901934 | 0.335883694 | 0.105731009 | 3.176775643 | 0.001489222 | 0.013906591 | 0.335883694 |
| ENSMUSG00000029993 | Nfu1 | 56748 | protein_coding | 526.9635824 | -0.336114265 | 0.083443092 | -4.028065814 | 5.62E-05 | 0.000907852 | 0.336114265 |
| ENSMUSG00000028032 | Papss1 | 23971 | protein_coding | 4994.541468 | 0.336159121 | 0.044452786 | 7.562160938 | 3.96E-14 | 2.31E-12 | 0.336159121 |
| ENSMUSG00000035104 | Eva1a | 232146 | protein_coding | 32.32932801 | -0.336423033 | 0.209361855 | -1.606897461 | 0.108076863 | 0.301132615 | 0.336423033 |
| ENSMUSG00000034587 | 8430429K09Rik | 71523 | lncRNA | 78.3006164 | -0.336488672 | 0.148308502 | -2.26884277 | 0.023277889 | 0.109096924 | 0.336488672 |
| ENSMUSG00000051703 | Tmem198 | 319998 | protein_coding | 23.22395383 | 0.336493923 | 0.247557918 | 1.359253321 | 0.17406633 | 0.399938411 | 0.336493923 |
| ENSMUSG00000109186 | Gm34821 | 102638204 | lncRNA | 17.85652738 | -0.336569712 | 0.297760254 | -1.130337941 | 0.258333853 | 0.50034619 | 0.336569712 |
| ENSMUSG00000037709 | Fam13a | 58909 | protein_coding | 944.3355427 | 0.336572116 | 0.083567027 | 4.027570768 | 5.64E-05 | 0.000908855 | 0.336572116 |
| ENSMUSG00000049526 | Tmem202 | 73893 | protein_coding | 13.70033116 | -0.336661976 | 0.355817941 | -0.946163579 | 0.34406516 | 0.586511041 | 0.336661976 |
| ENSMUSG00000034413 | Neurl1b | 240055 | protein_coding | 2452.813973 | -0.336667691 | 0.082326073 | -4.089441882 | 4.32E-05 | 0.000716702 | 0.336667691 |
| ENSMUSG00000020331 | Hcn2 | 15166 | protein_coding | 199.1346105 | -0.337243742 | 0.122580597 | -2.751200037 | 0.005937737 | 0.040696085 | 0.337243742 |
| ENSMUSG00000068299 | Nat8f4 | 75541 | protein_coding | 446.0712515 | -0.33750161 | 0.108331961 | -3.115438953 | 0.001836714 | 0.016419446 | 0.33750161 |
| ENSMUSG00000045045 | Lrfn4 | 225875 | protein_coding | 585.6264301 | -0.337691905 | 0.079852296 | -4.22895674 | 2.35E-05 | 0.000417448 | 0.337691905 |
| ENSMUSG00000029603 | Dtx1 | 14357 | protein_coding | 28.80414666 | 0.337911496 | 0.299117536 | 1.129694703 | 0.25860489 | 0.500565425 | 0.337911496 |
| ENSMUSG00000041679 | Fbxl9 | 234684 | protein_coding | 98.70895537 | -0.337966243 | 0.130469764 | -2.590379809 | 0.009587009 | 0.05760421 | 0.337966243 |
| ENSMUSG00000020154 | Ptprb | 19263 | protein_coding | 12.9166064 | -0.338063324 | 0.412340197 | -0.819865068 | 0.412293033 | 0.649970343 | 0.338063324 |
| ENSMUSG00000027347 | Rasgrp1 | 19419 | protein_coding | 24.69101082 | 0.338146139 | 0.234012868 | 1.444989504 | 0.148460834 | 0.362651904 | 0.338146139 |
| ENSMUSG00000028555 | Ttc39a | 230603 | protein_coding | 2438.741832 | 0.338353102 | 0.067241283 | 5.031925147 | 4.86E-07 | 1.27E-05 | 0.338353102 |
| ENSMUSG00000036634 | Mag | 17136 | protein_coding | 6.86151516 | -0.338412714 | 0.513133776 | -0.659501926 | 0.50957351 | 0.72436245 | 0.338412714 |
| ENSMUSG00000078502 | Zfp268 | 433801 | protein_coding | 51.50549486 | -0.338429859 | 0.220975341 | -1.531527713 | 0.125639023 | 0.328925409 | 0.338429859 |
| ENSMUSG00000022445 | Cyp2d26 | 76279 | protein_coding | 1650.054754 | -0.338454727 | 0.141205591 | -2.39689324 | 0.016534741 | 0.086179968 | 0.338454727 |
| ENSMUSG00000085795 | Zfp703 | 353310 | protein_coding | 276.2182943 | -0.338564115 | 0.128631079 | -2.632055318 | 0.008487004 | 0.052784388 | 0.338564115 |
| ENSMUSG00000031627 | Irf2 | 16363 | protein_coding | 568.0328266 | -0.338613189 | 0.102356554 | -3.308173021 | 0.000939068 | 0.009548768 | 0.338613189 |
| ENSMUSG00000029865 | Sval1 | 71578 | protein_coding | 4053.03161 | 0.338671359 | 0.121354523 | 2.790760096 | 0.005258443 | 0.037056142 | 0.338671359 |
| ENSMUSG00000033021 | Gmppa | 69080 | protein_coding | 1333.963263 | 0.338707029 | 0.070873498 | 4.779036451 | 1.76E-06 | 4.09E-05 | 0.338707029 |

| ensembl_gene_id | external_gene_name | entrezgene_id | gene_biotype | baseMean | log2FoldChange | lfcSE | stat | pvalue | padj | absLog2FC |
|---|---|---|---|---|---|---|---|---|---|---|
| ENSMUSG00000053695 | Defb37 | 353320 | protein_coding | 39.80938159 | -0.33871156 | 0.206684118 | -1.638788523 | 0.10125731 | 0.29018928 | 0.33871156 |
| ENSMUSG00000035835 | Plppr3 | 216152 | protein_coding | 79.28061407 | -0.338738676 | 0.209538915 | -1.616590775 | 0.105966645 | 0.298033498 | 0.338738676 |
| ENSMUSG00000074578 | Zfas1 | 68949 | lncRNA | 641.7351323 | -0.338868013 | 0.08562746 | -3.957468948 | 7.57E-05 | 0.001167868 | 0.338868013 |
| ENSMUSG00000021416 | Eci3 | 69123 | protein_coding | 367.8294578 | 0.33898992 | 0.0657939 | 5.152300153 | 2.57E-07 | 7.17E-06 | 0.33898992 |
| ENSMUSG00000029771 | Irf5 | 27056 | protein_coding | 14.48035749 | 0.339061587 | 0.360348237 | 0.940927557 | 0.346741985 | 0.588931858 | 0.339061587 |
| ENSMUSG00000036223 | Ska1 | 66468 | protein_coding | 55.91260659 | 0.339138371 | 0.243490253 | 1.392821138 | 0.163673885 | 0.385451042 | 0.339138371 |
| ENSMUSG00000037962 | Rflna | 73121 | protein_coding | 13.90794505 | -0.339174463 | 0.408595734 | -0.830097903 | 0.406483432 | 0.645466424 | 0.339174463 |
| ENSMUSG00000029034 | Ints11 | 71957 | protein_coding | 571.0917032 | 0.339385247 | 0.069520903 | 4.881772727 | 1.05E-06 | 2.57E-05 | 0.339385247 |
| ENSMUSG00000118401 | Gpr52 | 620246 | protein_coding | 8.583281261 | -0.339541375 | 0.413083363 | -0.821968168 | 0.411095014 | 0.649146117 | 0.339541375 |
| ENSMUSG00000102018 | Iqank1 | 432964 | protein_coding | 62.72301061 | -0.339742479 | 0.152896942 | -2.222035811 | 0.026280886 | 0.118778578 | 0.339742479 |
| ENSMUSG00000057246 | BC051142 | 407788 | protein_coding | 18.52834441 | -0.339763076 | 0.299742771 | -1.133515497 | 0.256997837 | 0.498869056 | 0.339763076 |
| ENSMUSG00000042734 | Ttc9 | 69480 | protein_coding | 6.183449763 | 0.339978472 | 0.527099695 | 0.644998423 | 0.518928208 | 0.730831823 | 0.339978472 |
| ENSMUSG00000021919 | Chat | 12647 | protein_coding | 25.65198534 | -0.340133508 | 0.275128151 | -1.236273012 | 0.216357099 | 0.452297019 | 0.340133508 |
| ENSMUSG00000003762 | Coq8b | 76889 | protein_coding | 494.019095 | 0.340174921 | 0.077426916 | 4.393496955 | 1.12E-05 | 0.000215946 | 0.340174921 |
| ENSMUSG00000029560 | Snx8 | 231834 | protein_coding | 123.4847901 | 0.340585107 | 0.108674592 | 3.133990216 | 0.001724466 | 0.015571368 | 0.340585107 |
| ENSMUSG00000005089 | Slc1a2 | 20511 | protein_coding | 17.17061031 | -0.340766334 | 0.302379606 | -1.126948801 | 0.259764129 | 0.50123327 | 0.340766334 |
| ENSMUSG00000028100 | Nudt17 | 78373 | protein_coding | 57.7912225 | -0.340771149 | 0.160224433 | -2.126836355 | 0.033433683 | 0.140437951 | 0.340771149 |
| ENSMUSG00000004661 | Arid3b | 56380 | protein_coding | 122.6325288 | -0.340962158 | 0.110966259 | -3.072665174 | 0.002121564 | 0.018357166 | 0.340962158 |
| ENSMUSG00000032094 | Cd3d | 12500 | protein_coding | 23.06691659 | 0.341067131 | 0.384324645 | 0.887445382 | 0.374839159 | 0.616422248 | 0.341067131 |
| ENSMUSG00000001424 | Snd1 | 56463 | protein_coding | 3304.807771 | 0.341134327 | 0.070260579 | 4.855273498 | 1.20E-06 | 2.89E-05 | 0.341134327 |
| ENSMUSG00000021278 | Amn | 93835 | protein_coding | 220.9248607 | -0.341157895 | 0.128732191 | -2.650136626 | 0.008045923 | 0.050864992 | 0.341157895 |
| ENSMUSG00000049154 | Fam183b | 75429 | protein_coding | 14.10636195 | 0.34116675 | 0.339240666 | 1.005677632 | 0.314570666 | 0.558770915 | 0.34116675 |
| ENSMUSG00000042179 | Pnliprp1 | 18946 | protein_coding | 8.34208 | -0.341173101 | 0.425011698 | -0.802738141 | 0.422126103 | 0.658505287 | 0.341173101 |
| ENSMUSG00000090700 | Cyp4f40 | 631304 | protein_coding | 2184.655544 | 0.341362285 | 0.067950536 | 5.023687925 | 5.07E-07 | 1.32E-05 | 0.341362285 |
| ENSMUSG00000030298 | Sec13 | 110379 | protein_coding | 2544.205311 | 0.341789559 | 0.073019872 | 4.680774541 | 2.86E-06 | 6.33E-05 | 0.341789559 |
| ENSMUSG00000041654 | Slc39a11 | 69806 | protein_coding | 2545.090953 | 0.341801313 | 0.084072882 | 4.065535816 | 4.79E-05 | 0.00078781 | 0.341801313 |
| ENSMUSG00000042500 | Ago4 | 76850 | protein_coding | 213.3149604 | -0.341974304 | 0.095042191 | -3.598131523 | 0.000320512 | 0.003889307 | 0.341974304 |
| ENSMUSG00000097075 | Cdiptos | 381922 | protein_coding | 8.875143761 | -0.342104633 | 0.409223727 | -0.835984353 | 0.403163705 | 0.642408959 | 0.342104633 |
| ENSMUSG00000056185 | Snx32 | 225861 | protein_coding | 59.02764041 | -0.342182873 | 0.273481094 | -1.25121217 | 0.210857079 | 0.445849236 | 0.342182873 |
| ENSMUSG00000039693 | Msantd3 | 66665 | protein_coding | 49.14978253 | 0.342293447 | 0.19229744 | 1.780020825 | 0.075072553 | 0.240742704 | 0.342293447 |
| ENSMUSG00000079084 | Ccdc82 | 66396 | protein_coding | 365.390679 | -0.34235186 | 0.125658359 | -2.724465471 | 0.006440566 | 0.043098347 | 0.34235186 |
| ENSMUSG00000024677 | Ms4a6b | 69774 | protein_coding | 10.35290088 | 0.342383065 | 0.37907311 | 0.903211166 | 0.366413833 | 0.608689316 | 0.342383065 |
| ENSMUSG00000015806 | Qdpr | 110391 | protein_coding | 2033.741555 | 0.342551808 | 0.037840527 | 9.052511542 | 1.40E-19 | 1.16E-17 | 0.342551808 |
| ENSMUSG00000028039 | Efna3 | 13638 | protein_coding | 59.76734964 | -0.342697274 | 0.178284534 | -1.922192948 | 0.054581484 | 0.196218366 | 0.342697274 |
| ENSMUSG00000025268 | Maged2 | 80884 | protein_coding | 377.0292348 | 0.342845198 | 0.070137086 | 4.888215635 | 1.02E-06 | 2.49E-05 | 0.342845198 |
| ENSMUSG00000055561 | Spink5 | 72432 | protein_coding | 7.210882556 | 0.342869836 | 0.566838975 | 0.604880489 | 0.545258408 | 0.749029274 | 0.342869836 |
| ENSMUSG00000020654 | Adcy3 | 104111 | protein_coding | 17.72652607 | -0.343047709 | 0.344130892 | -0.996852411 | 0.318836154 | 0.562506362 | 0.343047709 |
| ENSMUSG00000097162 | 2310010J17Rik | 78329 | lncRNA | 54.94150395 | -0.343228332 | 0.186724658 | -1.83815215 | 0.066039989 | 0.221624129 | 0.343228332 |
| ENSMUSG00000020366 | Mapk9 | 26420 | protein_coding | 1165.582862 | 0.343349678 | 0.046896246 | 7.321474748 | 2.45E-13 | 1.34E-11 | 0.343349678 |

| ensembl_gene_id | external_gene_name | entrezgene_id | gene_biotype | baseMean | log2FoldChange | lfcSE | stat | pvalue | padj | absLog2FC |
|---|---|---|---|---|---|---|---|---|---|---|
| ENSMUSG00000025512 | Chid1 | 68038 | protein_coding | 1156.347586 | 0.343381295 | 0.068604614 | 5.00522162 | 5.58E-07 | 1.44E-05 | 0.343381295 |
| ENSMUSG00000019872 | Smpdl3a | 57319 | protein_coding | 4088.182368 | -0.343628789 | 0.102233458 | -3.361216545 | 0.000776 | 0.008163435 | 0.343628789 |
| ENSMUSG00000105975 | Gm9831 | 101056032 | lncRNA | 5.921340332 | -0.343777153 | 0.565124129 | -0.608321492 | 0.54297427 | 0.747975232 | 0.343777153 |
| ENSMUSG00000085995 | Gm2788 | 108167326 | lncRNA | 13.32053108 | 0.344004595 | 0.337139378 | 1.020363143 | 0.307556267 | 0.551334926 | 0.344004595 |
| ENSMUSG00000033697 | Arhgap39 | 223666 | protein_coding | 857.759862 | 0.344086782 | 0.064118909 | 5.366385458 | 8.03E-08 | 2.49E-06 | 0.344086782 |
| ENSMUSG00000084834 | 4930565N06Rik | 68306 | lncRNA | 39.8111646 | -0.344134857 | 0.274702544 | -1.252754533 | 0.210295053 | 0.445347562 | 0.344134857 |
| ENSMUSG00000030428 | Ttyh1 | 57776 | protein_coding | 36.42176053 | 0.344142588 | 0.227659999 | 1.511651543 | 0.130622532 | 0.336509736 | 0.344142588 |
| ENSMUSG00000000266 | Mid2 | 23947 | protein_coding | 1501.457048 | 0.344249561 | 0.046581783 | 7.39021871 | 1.47E-13 | 8.18E-12 | 0.344249561 |
| ENSMUSG00000025608 | Podxl | 27205 | protein_coding | 6.850971677 | 0.34425671 | 0.4924848 | 0.699019969 | 0.484539551 | 0.706263486 | 0.34425671 |
| ENSMUSG00000020431 | Adcy1 | 432530 | protein_coding | 13.95013932 | -0.344416219 | 0.33251424 | -1.035793894 | 0.300298307 | 0.543841751 | 0.344416219 |
| ENSMUSG00000030214 | Plbd1 | 66857 | protein_coding | 1623.522275 | -0.344458354 | 0.076501255 | -4.502649733 | 6.71E-06 | 0.000138049 | 0.344458354 |
| ENSMUSG00000033177 | Tmprss7 | 208171 | protein_coding | 27.80411216 | 0.34461168 | 0.232340723 | 1.48321687 | 0.1380168 | 0.347998271 | 0.34461168 |
| ENSMUSG00000003402 | Prkcsh | 19089 | protein_coding | 3027.479662 | 0.344746975 | 0.06977831 | 4.940603655 | 7.79E-07 | 1.96E-05 | 0.344746975 |
| ENSMUSG00000031831 | Dnaaf1 | 68270 | protein_coding | 35.49954807 | 0.344891489 | 0.220613911 | 1.56332612 | 0.117975908 | 0.317522941 | 0.344891489 |
| ENSMUSG00000079057 | Cyp4v3 | 102294 | protein_coding | 393.2665959 | -0.344908533 | 0.141787882 | -2.432567073 | 0.014992214 | 0.080085937 | 0.344908533 |
| ENSMUSG00000031671 | Setd6 | 66083 | protein_coding | 163.9282863 | -0.344933423 | 0.135051738 | -2.554083561 | 0.01064677 | 0.061985727 | 0.344933423 |
| ENSMUSG00000029335 | Bmp3 | 110075 | protein_coding | 2858.484348 | -0.344997762 | 0.144152301 | -2.393286536 | 0.016698195 | 0.086644719 | 0.344997762 |
| ENSMUSG00000028777 | Gnat3 | 242851 | protein_coding | 71.51650328 | 0.345108804 | 0.145802568 | 2.366959716 | 0.017934885 | 0.091184076 | 0.345108804 |
| ENSMUSG00000028023 | Pitx2 | 18741 | protein_coding | 471.9560051 | 0.345118499 | 0.191200414 | 1.805009166 | 0.071073248 | 0.232315871 | 0.345118499 |
| ENSMUSG00000057406 | Nsd2 | 107823 | protein_coding | 1406.655641 | 0.345288995 | 0.069295902 | 4.982819843 | 6.27E-07 | 1.61E-05 | 0.345288995 |
| ENSMUSG00000024553 | Galr1 | 14427 | protein_coding | 8.722438168 | 0.345347576 | 0.460858166 | 0.749357615 | 0.45364169 | 0.682451449 | 0.345347576 |
| ENSMUSG00000110754 | 4930405J17Rik | 73811 | lncRNA | 15.67026319 | 0.345496093 | 0.300632845 | 1.149229364 | 0.250461416 | 0.491535798 | 0.345496093 |
| ENSMUSG00000005682 | Pan2 | 103135 | protein_coding | 730.6645287 | 0.345624222 | 0.069999767 | 4.937505318 | 7.91E-07 | 1.98E-05 | 0.345624222 |
| ENSMUSG00000004187 | Kifc2 | 16581 | protein_coding | 194.6001297 | -0.345711216 | 0.155098028 | -2.228985237 | 0.025814887 | 0.117338412 | 0.345711216 |
| ENSMUSG00000049858 | Suox | 211389 | protein_coding | 835.2030891 | 0.345725888 | 0.115735351 | 2.98721077 | 0.002815356 | 0.022942515 | 0.345725888 |
| ENSMUSG00000031919 | Tmed6 | 66269 | protein_coding | 48.14205404 | -0.345957973 | 0.207190584 | -1.669757219 | 0.094967408 | 0.279121448 | 0.345957973 |
| ENSMUSG00000037355 | Uvssa | 71101 | protein_coding | 369.315784 | -0.346087316 | 0.096806812 | -3.575030609 | 0.000350187 | 0.004195739 | 0.346087316 |
| ENSMUSG00000042371 | Slc5a10 | 109342 | protein_coding | 5.996439452 | -0.346132362 | 0.47187835 | -0.733520328 | 0.463241134 | 0.689357631 | 0.346132362 |
| ENSMUSG00000042354 | Gnl3 | 30877 | protein_coding | 470.7429566 | -0.346258398 | 0.132980834 | -2.603821831 | 0.009219067 | 0.056167696 | 0.346258398 |
| ENSMUSG00000022451 | Twf1 | 19230 | protein_coding | 2783.943223 | 0.346397262 | 0.046135778 | 7.508213237 | 5.99E-14 | 3.43E-12 | 0.346397262 |
| ENSMUSG00000026288 | Inpp5d | 16331 | protein_coding | 121.3098013 | 0.346442988 | 0.116089915 | 2.984264296 | 0.002842611 | 0.023082976 | 0.346442988 |
| ENSMUSG00000055053 | Nfic | 18029 | protein_coding | 256.5038037 | -0.346610455 | 0.089778863 | -3.860713368 | 0.000113056 | 0.001648627 | 0.346610455 |
| ENSMUSG00000019944 | Rhobtb1 | 69288 | protein_coding | 72.21570887 | -0.346630604 | 0.19411362 | -1.785709849 | 0.07414625 | 0.238816971 | 0.346630604 |
| ENSMUSG00000021209 | Ppp4r4 | 74521 | protein_coding | 11.5461565 | 0.346637344 | 0.357285047 | 0.970198297 | 0.331947659 | 0.575141012 | 0.346637344 |
| ENSMUSG00000086290 | Snhg12 | 100039864 | lncRNA | 230.6120973 | -0.346673168 | 0.092481931 | -3.74855026 | 0.00017786 | 0.002374456 | 0.346673168 |
| ENSMUSG00000021756 | Il6st | 16195 | protein_coding | 1079.068985 | 0.346676251 | 0.052862926 | 6.558022348 | 5.45E-11 | 2.46E-09 | 0.346676251 |
| ENSMUSG00000002808 | Epdr1 | 105298 | protein_coding | 20.51944081 | -0.346714979 | 0.276387629 | -1.254451875 | 0.209677807 | 0.44497561 | 0.346714979 |
| ENSMUSG00000041935 | Rimoc1 | 106064 | protein_coding | 696.9659693 | -0.346735137 | 0.090340907 | -3.83807456 | 0.000124003 | 0.00177444 | 0.346735137 |
| ENSMUSG00000043445 | Pgp | 67078 | protein_coding | 2114.163409 | 0.346778099 | 0.08453297 | 4.102282214 | 4.09E-05 | 0.000682262 | 0.346778099 |

| ensembl_gene_id | external_gene_name | entrezgene_id | gene_biotype | baseMean | log2FoldChange | lfcSE | stat | pvalue | padj | absLog2FC |
|---|---|---|---|---|---|---|---|---|---|---|
| ENSMUSG00000102918 | Pcdhgc3 | 93706 | protein_coding | 98.3281417 | 0.346820662 | 0.142887428 | 2.427230061 | 0.015214604 | 0.080952136 | 0.346820662 |
| ENSMUSG00000035277 | Arx | 11878 | protein_coding | 7.578787392 | 0.346866413 | 0.476611552 | 0.727775925 | 0.466750764 | 0.692404394 | 0.346866413 |
| ENSMUSG00000006057 | Atp5g1 | 11951 | protein_coding | 5819.776798 | 0.346988617 | 0.084884398 | 4.087778482 | 4.36E-05 | 0.000721118 | 0.346988617 |
| ENSMUSG00000024177 | Nme4 | 56520 | protein_coding | 83.55941278 | -0.347056267 | 0.127970122 | -2.71201013 | 0.006687655 | 0.044292325 | 0.347056267 |
| ENSMUSG00000057667 | Bloc1s3 | 232946 | protein_coding | 459.6479487 | 0.34720506 | 0.105927943 | 3.277747572 | 0.001046389 | 0.010442524 | 0.34720506 |
| ENSMUSG00000014776 | Nol3 | 78688 | protein_coding | 48.97064317 | 0.347292241 | 0.194216859 | 1.788167325 | 0.073749016 | 0.237870077 | 0.347292241 |
| ENSMUSG00000048603 | Gm9828 | 105246056 | lncRNA | 18.7166644 | -0.347509437 | 0.278702307 | -1.24688396 | 0.212440049 | 0.447904389 | 0.347509437 |
| ENSMUSG00000085990 | Gm16731 | 105247126 | lncRNA | 17.42643748 | 0.34769248 | 0.280410485 | 1.239941081 | 0.214997188 | 0.45050036 | 0.34769248 |
| ENSMUSG00000023966 | Rsph9 | 75564 | protein_coding | 18.56126058 | -0.348063787 | 0.291348567 | -1.19466449 | 0.232218142 | 0.471304049 | 0.348063787 |
| ENSMUSG00000106464 | C130083M11Rik | 414111 | lncRNA | 22.07637728 | -0.348076842 | 0.259131003 | -1.343246614 | 0.179192132 | 0.406457009 | 0.348076842 |
| ENSMUSG00000061531 | Tmem236 | 625286 | protein_coding | 2925.040264 | -0.348092795 | 0.077390039 | -4.497901794 | 6.86E-06 | 0.000140808 | 0.348092795 |
| ENSMUSG00000025069 | Gsto2 | 68214 | protein_coding | 18.03195634 | 0.348342089 | 0.333503947 | 1.044491655 | 0.296257978 | 0.53906718 | 0.348342089 |
| ENSMUSG00000013846 | St3gal1 | 20442 | protein_coding | 242.9165612 | 0.348391383 | 0.095311067 | 3.655308822 | 0.000256873 | 0.003249085 | 0.348391383 |
| ENSMUSG00000022885 | St6gal1 | 20440 | protein_coding | 2555.486142 | 0.348408419 | 0.067059995 | 5.195473378 | 2.04E-07 | 5.82E-06 | 0.348408419 |
| ENSMUSG00000056822 | Or2l13 | 259071 | protein_coding | 5.874628185 | 0.348497662 | 0.503827466 | 0.691700405 | 0.489125493 | 0.709236362 | 0.348497662 |
| ENSMUSG00000051652 | Lrrc3 | 237387 | protein_coding | 12.49756211 | -0.348779408 | 0.323533841 | -1.078030683 | 0.281020064 | 0.523508209 | 0.348779408 |
| ENSMUSG00000074444 | Defa30 | 100038927 | protein_coding | 76.86447198 | 0.348786413 | 1.157969505 | 0.30120518 | 0.76325804 | 0.888612649 | 0.348786413 |
| ENSMUSG00000021273 | Fdft1 | 14137 | protein_coding | 3186.725253 | 0.348945939 | 0.115639123 | 3.017542255 | 0.002548335 | 0.021129563 | 0.348945939 |
| ENSMUSG00000079111 | Kdelr2 | 66913 | protein_coding | 4214.44071 | 0.349224024 | 0.058202939 | 6.000109774 | 1.97E-09 | 7.55E-08 | 0.349224024 |
| ENSMUSG00000036533 | Cdc42ep3 | 260409 | protein_coding | 389.7226931 | 0.34937615 | 0.098494831 | 3.547152116 | 0.00038942 | 0.004587415 | 0.34937615 |
| ENSMUSG00000053182 | Cd200l1 | 208166 | protein_coding | 326.6340268 | -0.349646307 | 0.125961376 | -2.775821587 | 0.005506241 | 0.03837566 | 0.349646307 |
| ENSMUSG00000022043 | Trim35 | 66854 | protein_coding | 345.0036338 | 0.349703978 | 0.095843203 | 3.648709215 | 0.000263561 | 0.003318073 | 0.349703978 |
| ENSMUSG00000028334 | Nans | 94181 | protein_coding | 9822.111664 | 0.349867164 | 0.089705672 | 3.900167694 | 9.61E-05 | 0.001434066 | 0.349867164 |
| ENSMUSG00000028088 | Fmo5 | 14263 | protein_coding | 632.7452018 | -0.350152374 | 0.16002615 | -2.188094725 | 0.028662703 | 0.125951884 | 0.350152374 |
| ENSMUSG00000036814 | Slc6a20a | 102680 | protein_coding | 195.310011 | -0.350266522 | 0.173689804 | -2.016621087 | 0.043735071 | 0.169099853 | 0.350266522 |
| ENSMUSG00000043535 | Setx | 269254 | protein_coding | 2179.642697 | 0.350373884 | 0.049131015 | 7.131419673 | 9.93E-13 | 5.17E-11 | 0.350373884 |
| ENSMUSG00000049382 | Krt8 | 16691 | protein_coding | 113769.281 | 0.350434263 | 0.063282215 | 5.537642162 | 3.07E-08 | 1.03E-06 | 0.350434263 |
| ENSMUSG00000074457 | S100a16 | 67860 | protein_coding | 2497.100208 | 0.350482393 | 0.092096332 | 3.805606418 | 0.000141457 | 0.001976135 | 0.350482393 |
| ENSMUSG00000040794 | C1qtnf4 | 67445 | protein_coding | 48.55232761 | -0.350540421 | 0.175125457 | -2.001653147 | 0.045322049 | 0.173037093 | 0.350540421 |
| ENSMUSG00000028134 | Ptbp2 | 56195 | protein_coding | 177.4339336 | -0.350664211 | 0.149527246 | -2.345152604 | 0.019019293 | 0.094931644 | 0.350664211 |
| ENSMUSG00000078521 | Aunip | 69885 | protein_coding | 39.81562892 | 0.350745935 | 0.228771635 | 1.533170557 | 0.125233837 | 0.328321636 | 0.350745935 |
| ENSMUSG00000022253 | Nadk2 | 68646 | protein_coding | 814.2278451 | 0.350774362 | 0.061928763 | 5.66415904 | 1.48E-08 | 5.17E-07 | 0.350774362 |
| ENSMUSG00000041920 | Slc16a6 | 104681 | protein_coding | 143.2491733 | 0.350904967 | 0.123910044 | 2.831933183 | 0.004626752 | 0.033640948 | 0.350904967 |
| ENSMUSG00000078864 | Gm14322 | 626802 | protein_coding | 9.204032964 | -0.351222584 | 0.443625788 | -0.791709124 | 0.428530301 | 0.663553352 | 0.351222584 |
| ENSMUSG00000039116 | Adgrg6 | 215798 | protein_coding | 38.26550759 | 0.351236047 | 0.189852157 | 1.850050335 | 0.064306295 | 0.217779845 | 0.351236047 |
| ENSMUSG00000047344 | Lancl3 | 236285 | protein_coding | 95.73485695 | 0.351329784 | 0.137634191 | 2.55263449 | 0.010691163 | 0.06217684 | 0.351329784 |
| ENSMUSG00000032192 | Gnb5 | 14697 | protein_coding | 144.1863848 | -0.351371098 | 0.097074635 | -3.619597403 | 0.000295062 | 0.003632413 | 0.351371098 |
| ENSMUSG00000078570 | 1110065P20Rik | 68920 | protein_coding | 140.9752769 | 0.351373149 | 0.128037472 | 2.744299329 | 0.006064022 | 0.04133159 | 0.351373149 |
| ENSMUSG00000015405 | Ace2 | 70008 | protein_coding | 582.7888472 | -0.35154217 | 0.10675461 | -3.292992884 | 0.00099127 | 0.009978907 | 0.35154217 |

| ensembl_gene_id | external_gene_name | entrezgene_id | gene_biotype | baseMean | log2FoldChange | lfcSE | stat | pvalue | padj | absLog2FC |
|---|---|---|---|---|---|---|---|---|---|---|
| ENSMUSG00000037266 | Rsrp1 | 27981 | protein_coding | 3225.47454 | -0.35159078 | 0.128347763 | -2.739360403 | 0.006155885 | 0.041765233 | 0.35159078 |
| ENSMUSG00000026080 | Chst10 | 98388 | protein_coding | 17.73524617 | 0.351621834 | 0.414903279 | 0.847479043 | 0.396728163 | 0.636177298 | 0.351621834 |
| ENSMUSG00000029368 | Alb | 11657 | protein_coding | 26.26826958 | 0.351651789 | 0.282924955 | 1.242915416 | 0.213899004 | 0.449687034 | 0.351651789 |
| ENSMUSG00000022621 | Rabl2 | 68708 | protein_coding | 78.05720285 | 0.351715329 | 0.154662896 | 2.274076962 | 0.02296136 | 0.10799004 | 0.351715329 |
| ENSMUSG00000022993 | Tex49 | 73863 | protein_coding | 31.61800469 | -0.351813795 | 0.220880074 | -1.592781952 | 0.111209115 | 0.306582205 | 0.351813795 |
| ENSMUSG00000038205 | Prkab2 | 108097 | protein_coding | 81.89527889 | -0.351875639 | 0.168936563 | -2.082886215 | 0.037261596 | 0.150719279 | 0.351875639 |
| ENSMUSG00000068617 | Efcab1 | 66793 | protein_coding | 10.76870547 | 0.351930032 | 0.41945191 | 0.839023552 | 0.401456093 | 0.640672993 | 0.351930032 |
| ENSMUSG00000100826 | Snhg14 | 16353 | lncRNA | 51.33312537 | -0.352233119 | 0.200452841 | -1.757186965 | 0.078885946 | 0.248300541 | 0.352233119 |
| ENSMUSG00000025213 | Kazald1 | 107250 | protein_coding | 49.0865721 | -0.352325998 | 0.194173511 | -1.814490538 | 0.069602238 | 0.229123351 | 0.352325998 |
| ENSMUSG00000022385 | Gtse1 | 29870 | protein_coding | 185.429346 | 0.35244567 | 0.136712505 | 2.578006087 | 0.009937225 | 0.059005016 | 0.35244567 |
| ENSMUSG00000049047 | Armcx3 | 71703 | protein_coding | 239.7910915 | 0.352585051 | 0.104726041 | 3.366737126 | 0.000760632 | 0.008027948 | 0.352585051 |
| ENSMUSG00000054850 | Smim10l2a | 320237 | protein_coding | 58.20625114 | 0.352693137 | 0.187816375 | 1.877861485 | 0.060400116 | 0.20889399 | 0.352693137 |
| ENSMUSG00000030492 | Slc7a9 | 30962 | protein_coding | 375.3531044 | -0.352725335 | 0.150611238 | -2.341958938 | 0.019182825 | 0.095444865 | 0.352725335 |
| ENSMUSG00000041891 | Lman1 | 70361 | protein_coding | 1866.009937 | 0.353240752 | 0.074590853 | 4.735711408 | 2.18E-06 | 4.96E-05 | 0.353240752 |
| ENSMUSG00000024734 | Zp1 | 22786 | protein_coding | 10.06743462 | 0.353287648 | 0.4132162 | 0.854970467 | 0.392567498 | 0.632267311 | 0.353287648 |
| ENSMUSG00000029428 | Stx2 | 13852 | protein_coding | 12.41314399 | 0.353322508 | 0.363229339 | 0.972725685 | 0.330689656 | 0.574280957 | 0.353322508 |
| ENSMUSG00000037907 | Ankrd13b | 268445 | protein_coding | 134.6052915 | -0.353539964 | 0.122021658 | -2.897354196 | 0.003763246 | 0.028899939 | 0.353539964 |
| ENSMUSG00000074896 | Ifit3 | 15959 | protein_coding | 65.29208858 | -0.353660481 | 0.26256117 | -1.346964144 | 0.177991781 | 0.405224566 | 0.353660481 |
| ENSMUSG00000070644 | Etnk2 | 214253 | protein_coding | 17.41604679 | 0.354054461 | 0.276903377 | 1.278620956 | 0.201030567 | 0.434761962 | 0.354054461 |
| ENSMUSG00000018654 | Ikzf1 | 22778 | protein_coding | 31.10262001 | 0.354084275 | 0.267191292 | 1.325208889 | 0.185101878 | 0.414061984 | 0.354084275 |
| ENSMUSG00000069266 | H4c2 | 326620 | protein_coding | 511.7105674 | 0.354235311 | 0.145721783 | 2.43090158 | 0.015061305 | 0.08032198 | 0.354235311 |
| ENSMUSG00000036299 | BC031181 | 407819 | protein_coding | 1844.457173 | 0.354418495 | 0.066494549 | 5.330038333 | 9.82E-08 | 3.01E-06 | 0.354418495 |
| ENSMUSG00000075270 | Pde11a | 241489 | protein_coding | 53.65401427 | 0.354479023 | 0.168842745 | 2.099462566 | 0.035776144 | 0.146697656 | 0.354479023 |
| ENSMUSG00000038037 | Socs1 | 12703 | protein_coding | 50.22890632 | -0.354563071 | 0.253009551 | -1.401382157 | 0.161099825 | 0.381618226 | 0.354563071 |
| ENSMUSG00000055333 | Fat2 | 245827 | protein_coding | 62.04199448 | -0.354680722 | 0.164430286 | -2.157027944 | 0.031003487 | 0.133047695 | 0.354680722 |
| ENSMUSG00000038028 | Tigar | 319801 | protein_coding | 428.3423969 | 0.354714295 | 0.070601211 | 5.024195641 | 5.06E-07 | 1.32E-05 | 0.354714295 |
| ENSMUSG00000097368 | Gm10390 | 100503427 | lncRNA | 13.1191688 | -0.354788412 | 0.360974591 | -0.982862563 | 0.325675085 | 0.56923128 | 0.354788412 |
| ENSMUSG00000029630 | Cyp3a25 | 56388 | protein_coding | 11.73358411 | 0.354939152 | 0.63606619 | 0.558022352 | 0.576829121 | 0.769561988 | 0.354939152 |
| ENSMUSG00000045281 | Gpr20 | 239530 | protein_coding | 373.8518534 | 0.35494044 | 0.132162901 | 2.685628397 | 0.007239355 | 0.046962624 | 0.35494044 |
| ENSMUSG00000029053 | Prkcz | 18762 | protein_coding | 2739.949384 | 0.354951898 | 0.040265135 | 8.815365925 | 1.19E-18 | 9.43E-17 | 0.354951898 |
| ENSMUSG00000034974 | Dapk3 | 13144 | protein_coding | 753.0507197 | 0.354968594 | 0.054887196 | 6.467238614 | 9.98E-11 | 4.41E-09 | 0.354968594 |
| ENSMUSG00000072294 | Klf12 | 16597 | protein_coding | 263.8187017 | 0.355033085 | 0.077753522 | 4.566135076 | 4.97E-06 | 0.000105419 | 0.355033085 |
| ENSMUSG00000050592 | Fam78a | 241303 | protein_coding | 99.07475518 | -0.355113493 | 0.168718519 | -2.104768908 | 0.035311429 | 0.145383562 | 0.355113493 |
| ENSMUSG00000084925 | 1810062O18Rik | 75602 | lncRNA | 19.31310927 | -0.355129363 | 0.289519504 | -1.22661637 | 0.219966814 | 0.456081872 | 0.355129363 |
| ENSMUSG00000017314 | Mpp2 | 50997 | protein_coding | 22.89736098 | 0.355138446 | 0.267557489 | 1.3273351 | 0.184397858 | 0.413082963 | 0.355138446 |
| ENSMUSG00000062232 | Rapgef2 | 76089 | protein_coding | 820.9403025 | 0.355142636 | 0.051006679 | 6.962669233 | 3.34E-12 | 1.66E-10 | 0.355142636 |
| ENSMUSG00000027514 | Zbp1 | 58203 | protein_coding | 176.1712441 | -0.355186354 | 0.286978987 | -1.237673735 | 0.215837062 | 0.451641923 | 0.355186354 |
| ENSMUSG00000020911 | Krt19 | 16669 | protein_coding | 68956.10272 | 0.355428024 | 0.096211645 | 3.694230812 | 0.000220553 | 0.002859216 | 0.355428024 |
| ENSMUSG00000018924 | Alox15 | 11687 | protein_coding | 8.347421045 | 0.35547943 | 0.58868112 | 0.603857365 | 0.545938477 | 0.749242602 | 0.35547943 |

| ensembl_gene_id | external_gene_name | entrezgene_id | gene_biotype | baseMean | log2FoldChange | lfcSE | stat | pvalue | padj | absLog2FC |
|---|---|---|---|---|---|---|---|---|---|---|
| ENSMUSG00000069301 | H2ac11 | 319167 | protein_coding | 74.97887929 | 0.355615529 | 0.180354933 | 1.971753821 | 0.048637716 | 0.181569549 | 0.355615529 |
| ENSMUSG00000004896 | Rrnad1 | 229503 | protein_coding | 186.1959905 | -0.355697398 | 0.126635284 | -2.808833268 | 0.004972139 | 0.035527552 | 0.355697398 |
| ENSMUSG00000000058 | Cav2 | 12390 | protein_coding | 135.9640279 | 0.355803709 | 0.130514678 | 2.726158588 | 0.00640762 | 0.042967024 | 0.355803709 |
| ENSMUSG00000085589 | A430078I02Rik | 320486 | lncRNA | 40.61065755 | 0.355961441 | 0.184359219 | 1.930803586 | 0.053507347 | 0.193651926 | 0.355961441 |
| ENSMUSG00000028919 | Arhgef19 | 213649 | protein_coding | 652.9912322 | -0.356175957 | 0.08279263 | -4.302024885 | 1.69E-05 | 0.000311576 | 0.356175957 |
| ENSMUSG00000035275 | Raver2 | 242570 | protein_coding | 9.435094168 | 0.356412037 | 0.41082951 | 0.867542445 | 0.385644866 | 0.625831733 | 0.356412037 |
| ENSMUSG00000017929 | B4galt5 | 56336 | protein_coding | 1267.671593 | -0.356416847 | 0.074763615 | -4.767250019 | 1.87E-06 | 4.32E-05 | 0.356416847 |
| ENSMUSG00000024423 | Impact | 16210 | protein_coding | 159.0970626 | -0.356860492 | 0.136237893 | -2.619392331 | 0.008808657 | 0.054261734 | 0.356860492 |
| ENSMUSG00000024270 | Slc39a6 | 106957 | protein_coding | 61.35437263 | 0.357098455 | 0.150844341 | 2.367330803 | 0.01791691 | 0.091124885 | 0.357098455 |
| ENSMUSG00000042138 | Msantd2 | 235184 | protein_coding | 132.05148 | -0.35712618 | 0.13754747 | -2.596384945 | 0.009421045 | 0.056965074 | 0.35712618 |
| ENSMUSG00000022723 | Crybg3 | 224273 | protein_coding | 578.0587753 | 0.357249953 | 0.150587348 | 2.372376947 | 0.017674049 | 0.090113624 | 0.357249953 |
| ENSMUSG00000024268 | Celf4 | 108013 | protein_coding | 38.80936666 | -0.357442107 | 0.22378644 | -1.597246494 | 0.110210773 | 0.304713258 | 0.357442107 |
| ENSMUSG00000005470 | Asf1b | 66929 | protein_coding | 182.5176608 | 0.357510143 | 0.149718046 | 2.387889451 | 0.016945438 | 0.087561384 | 0.357510143 |
| ENSMUSG00000104900 | 4930596I21Rik | 78229 | lncRNA | 8.541209697 | -0.357533925 | 0.50249655 | -0.711515183 | 0.476765045 | 0.699625498 | 0.357533925 |
| ENSMUSG00000051615 | Rap2a | 76108 | protein_coding | 1650.672437 | 0.357570236 | 0.063101063 | 5.666627788 | 1.46E-08 | 5.11E-07 | 0.357570236 |
| ENSMUSG00000086358 | Gm13270 | 105244084 | lncRNA | 7.647720315 | -0.357651169 | 0.439687475 | -0.813421324 | 0.415976537 | 0.653976386 | 0.357651169 |
| ENSMUSG00000093930 | Hmgcs1 | 208715 | protein_coding | 10373.34746 | 0.357771819 | 0.179183791 | 1.996675127 | 0.045860486 | 0.174452968 | 0.357771819 |
| ENSMUSG00000031821 | Gins2 | 272551 | protein_coding | 149.0758753 | 0.357828008 | 0.163109708 | 2.193787313 | 0.028250703 | 0.124650914 | 0.357828008 |
| ENSMUSG00000017677 | Wsb1 | 78889 | protein_coding | 2882.719549 | -0.357960413 | 0.104522536 | -3.42471993 | 0.000615433 | 0.006788711 | 0.357960413 |
| ENSMUSG00000067158 | Col4a4 | 12829 | protein_coding | 6.107021606 | -0.357962581 | 0.552566806 | -0.647817743 | 0.517102838 | 0.729722934 | 0.357962581 |
| ENSMUSG00000037379 | Spon2 | 100689 | protein_coding | 49.81548653 | 0.357983897 | 0.180619795 | 1.981974886 | 0.047482051 | 0.178786609 | 0.357983897 |
| ENSMUSG00000108356 | 9130221F21Rik | 330350 | lncRNA | 54.73828198 | 0.358024608 | 0.248787718 | 1.4390767 | 0.150128793 | 0.365508309 | 0.358024608 |
| ENSMUSG00000027168 | Pax6 | 18508 | protein_coding | 19.09835283 | 0.358389476 | 0.29853446 | 1.20049617 | 0.229946699 | 0.469173889 | 0.358389476 |
| ENSMUSG00000110803 | Gm20275 | 100504539 | lncRNA | 71.35391472 | -0.358433493 | 0.153054165 | -2.341873495 | 0.019187217 | 0.095444865 | 0.358433493 |
| ENSMUSG00000049709 | Nlrp10 | 244202 | protein_coding | 106.2493006 | -0.358485177 | 0.166730631 | -2.150085888 | 0.031548422 | 0.134705162 | 0.358485177 |
| ENSMUSG00000036500 | Akp3 | 11648 | protein_coding | 80.02031793 | -0.358508715 | 1.029724531 | -0.348159828 | 0.727720155 | 0.867614603 | 0.358508715 |
| ENSMUSG00000038811 | Gngt2 | 14710 | protein_coding | 68.47383854 | -0.358511401 | 0.182390753 | -1.965622685 | 0.049342212 | 0.183646861 | 0.358511401 |
| ENSMUSG00000050786 | Ccdc126 | 57895 | protein_coding | 62.15283452 | 0.358822081 | 0.149200815 | 2.404960595 | 0.01617421 | 0.084790715 | 0.358822081 |
| ENSMUSG00000026959 | Grin1 | 14810 | protein_coding | 46.48591652 | 0.35911159 | 0.188161566 | 1.908527859 | 0.056323026 | 0.200512459 | 0.35911159 |
| ENSMUSG00000026571 | Dcaf6 | 74106 | protein_coding | 652.1917856 | -0.359135424 | 0.068947523 | -5.208822698 | 1.90E-07 | 5.45E-06 | 0.359135424 |
| ENSMUSG00000035699 | Slc51a | 106407 | protein_coding | 2504.374873 | -0.359369355 | 0.12989446 | -2.766625731 | 0.005663974 | 0.039304177 | 0.359369355 |
| ENSMUSG00000025931 | Paqr8 | 74229 | protein_coding | 381.0266697 | 0.359692174 | 0.093738234 | 3.837198089 | 0.000124446 | 0.001776055 | 0.359692174 |
| ENSMUSG00000064065 | Ipcef1 | 320495 | protein_coding | 36.16323515 | 0.359702007 | 0.233203222 | 1.542440126 | 0.122966677 | 0.325202296 | 0.359702007 |
| ENSMUSG00000071855 | Ccdc112 | 240261 | protein_coding | 58.34979381 | 0.359751031 | 0.152995863 | 2.351377512 | 0.018704047 | 0.093968897 | 0.359751031 |
| ENSMUSG00000086541 | Has2os | 594843 | lncRNA | 24.30130743 | -0.35983492 | 0.26176455 | -1.374651077 | 0.169239645 | 0.393611906 | 0.35983492 |
| ENSMUSG00000035983 | Gm7008 | 629820 | lncRNA | 26.59341161 | -0.359856937 | 0.25993587 | -1.384406613 | 0.166233985 | 0.389715871 | 0.359856937 |
| ENSMUSG00000024966 | Stip1 | 20867 | protein_coding | 1599.036434 | 0.359864975 | 0.101045725 | 3.561407228 | 0.000368872 | 0.004396752 | 0.359864975 |
| ENSMUSG00000029088 | Kcnip4 | 80334 | protein_coding | 7.316717543 | -0.359922276 | 0.453138273 | -0.794287964 | 0.427027809 | 0.662499036 | 0.359922276 |
| ENSMUSG00000032527 | Pccb | 66904 | protein_coding | 4900.770354 | 0.359922441 | 0.049730642 | 7.237438013 | 4.57E-13 | 2.44E-11 | 0.359922441 |

| ensembl_gene_id | external_gene_name | entrezgene_id | gene_biotype | baseMean | log2FoldChange | lfcSE | stat | pvalue | padj | absLog2FC |
|---|---|---|---|---|---|---|---|---|---|---|
| ENSMUSG00000055818 | A230083G16Rik | 442825 | lncRNA | 24.15853996 | -0.36000669 | 0.331734913 | -1.085224004 | 0.277822455 | 0.520660159 | 0.36000669 |
| ENSMUSG00000039716 | Dock3 | 208869 | protein_coding | 20.97238418 | 0.360013803 | 0.278834738 | 1.291136841 | 0.196656231 | 0.429095149 | 0.360013803 |
| ENSMUSG00000051439 | Cd14 | 12475 | protein_coding | 608.093382 | 0.360270763 | 0.093007725 | 3.873557424 | 0.000107258 | 0.001579682 | 0.360270763 |
| ENSMUSG00000022790 | Igsf11 | 207683 | protein_coding | 24.55062604 | 0.360276914 | 0.256812442 | 1.402879516 | 0.16065277 | 0.381119039 | 0.360276914 |
| ENSMUSG00000028461 | Ccdc107 | 622404 | protein_coding | 715.6283355 | -0.360365713 | 0.068762956 | -5.240695494 | 1.60E-07 | 4.67E-06 | 0.360365713 |
| ENSMUSG00000026311 | Asb1 | 65247 | protein_coding | 128.9598435 | -0.36039644 | 0.109167579 | -3.301313842 | 0.000962332 | 0.009748444 | 0.36039644 |
| ENSMUSG00000038301 | Snx10 | 71982 | protein_coding | 248.0681507 | -0.360447126 | 0.089988205 | -4.005493006 | 6.19E-05 | 0.000986237 | 0.360447126 |
| ENSMUSG00000032065 | Tex12 | 66654 | protein_coding | 18.44635026 | -0.360603366 | 0.282401854 | -1.27691572 | 0.201632004 | 0.435478682 | 0.360603366 |
| ENSMUSG00000050931 | Sgms2 | 74442 | protein_coding | 1952.234658 | -0.360993962 | 0.047544208 | -7.592806298 | 3.13E-14 | 1.84E-12 | 0.360993962 |
| ENSMUSG00000024426 | Atat1 | 73242 | protein_coding | 80.48082188 | -0.361020575 | 0.164424729 | -2.19565863 | 0.028116386 | 0.12424678 | 0.361020575 |
| ENSMUSG00000000204 | Slfn4 | 20558 | protein_coding | 488.2932524 | -0.361232236 | 0.15296743 | -2.36149771 | 0.018201284 | 0.092102953 | 0.361232236 |
| ENSMUSG00000076434 | Wfdc3 | 71856 | protein_coding | 7.295006212 | -0.361291333 | 0.451219434 | -0.800699851 | 0.423305429 | 0.659294819 | 0.361291333 |
| ENSMUSG00000100182 | 1810006J02Rik | 69030 | lncRNA | 9.943734902 | -0.361333476 | 0.418059111 | -0.864311928 | 0.38741656 | 0.627358857 | 0.361333476 |
| ENSMUSG00000032959 | Pebp1 | 23980 | protein_coding | 1904.731283 | 0.361523514 | 0.068816949 | 5.25340803 | 1.49E-07 | 4.41E-06 | 0.361523514 |
| ENSMUSG00000023094 | Msrb2 | 76467 | protein_coding | 148.3444027 | 0.361796691 | 0.103921847 | 3.481430517 | 0.000498743 | 0.005640414 | 0.361796691 |
| ENSMUSG00000071547 | Nt5dc2 | 70021 | protein_coding | 41.047247 | 0.361801791 | 0.18557227 | 1.94965439 | 0.051217326 | 0.188241916 | 0.361801791 |
| ENSMUSG00000064373 | Selenop | 20363 | protein_coding | 10534.26637 | -0.362122583 | 0.134284868 | -2.696674517 | 0.007003569 | 0.045838699 | 0.362122583 |
| ENSMUSG00000021768 | Dusp13 | 27389 | protein_coding | 7.969136445 | -0.362321573 | 0.427731333 | -0.847077464 | 0.396951946 | 0.63647286 | 0.362321573 |
| ENSMUSG00000049103 | Ccr2 | 12772 | protein_coding | 6.032357498 | 0.362472963 | 0.601227295 | 0.602888401 | 0.546582933 | 0.749829352 | 0.362472963 |
| ENSMUSG00000053714 | 4732471J01Rik | 654804 | lncRNA | 30.16693587 | -0.362552457 | 0.232639877 | -1.558427822 | 0.119131866 | 0.319248853 | 0.362552457 |
| ENSMUSG00000006378 | Gcat | 26912 | protein_coding | 118.8498789 | 0.36298802 | 0.135921411 | 2.67057277 | 0.007572196 | 0.048652111 | 0.36298802 |
| ENSMUSG00000096910 | Zfp955b | 100043468 | protein_coding | 328.0083626 | -0.363154958 | 0.086081713 | -4.218723664 | 2.46E-05 | 0.000435306 | 0.363154958 |
| ENSMUSG00000022617 | Chkb | 12651 | protein_coding | 178.5908653 | -0.363160594 | 0.169553926 | -2.141858952 | 0.032204834 | 0.136747592 | 0.363160594 |
| ENSMUSG00000030082 | Sec61a1 | 53421 | protein_coding | 4729.896433 | 0.363277986 | 0.059249032 | 6.131374155 | 8.71E-10 | 3.47E-08 | 0.363277986 |
| ENSMUSG00000087177 | E130307A14Rik | 327744 | lncRNA | 57.19987165 | -0.363287732 | 0.203220895 | -1.787649503 | 0.073832573 | 0.238091964 | 0.363287732 |
| ENSMUSG00000041649 | Klf8 | 245671 | protein_coding | 31.88827074 | -0.363291613 | 0.231554312 | -1.568926143 | 0.116665149 | 0.315962272 | 0.363291613 |
| ENSMUSG00000021248 | Tmed10 | 68581 | protein_coding | 2939.814961 | 0.363647629 | 0.041800767 | 8.699544376 | 3.33E-18 | 2.57E-16 | 0.363647629 |
| ENSMUSG00000034173 | Chchd2l | 71970 | protein_coding | 11.96121667 | -0.363747758 | 0.368739887 | -0.986461652 | 0.323906624 | 0.567946164 | 0.363747758 |
| ENSMUSG00000108034 | 4930557K07Rik | 78202 | lncRNA | 5.921140879 | -0.363754751 | 0.532617832 | -0.682956387 | 0.494634401 | 0.713491312 | 0.363754751 |
| ENSMUSG00000053693 | Mast1 | 56527 | protein_coding | 14.91911065 | -0.363849567 | 0.335425276 | -1.084741053 | 0.27803636 | 0.520958798 | 0.363849567 |
| ENSMUSG00000064043 | Trerf1 | 224829 | protein_coding | 6.471576944 | 0.36421211 | 0.491633558 | 0.740820279 | 0.458802418 | 0.685928116 | 0.36421211 |
| ENSMUSG00000032135 | Mcam | 84004 | protein_coding | 6.579230258 | -0.364245117 | 0.567691569 | -0.641625025 | 0.52111668 | 0.732548832 | 0.364245117 |
| ENSMUSG00000022150 | Dab2 | 13132 | protein_coding | 11.18758467 | -0.364327722 | 0.368940659 | -0.987496806 | 0.323399148 | 0.567390999 | 0.364327722 |
| ENSMUSG00000019960 | Dusp6 | 67603 | protein_coding | 1322.338806 | 0.364372027 | 0.07704834 | 4.729135339 | 2.25E-06 | 5.11E-05 | 0.364372027 |
| ENSMUSG00000021604 | Irx4 | 50916 | protein_coding | 27.52157369 | 0.364558985 | 0.24809585 | 1.469427983 | 0.141716742 | 0.352697729 | 0.364558985 |
| ENSMUSG00000026786 | Apbb1ip | 54519 | protein_coding | 17.7992217 | 0.364566547 | 0.300174545 | 1.214515197 | 0.22455105 | 0.462553939 | 0.364566547 |
| ENSMUSG00000069515 | Lyz1 | 17110 | protein_coding | 182.455633 | 0.36475145 | 1.038006425 | 0.351396139 | 0.725291179 | 0.865987203 | 0.36475145 |
| ENSMUSG00000004462 | Tbccd1 | 70573 | protein_coding | 878.9789314 | 0.364936097 | 0.070973845 | 5.141839142 | 2.72E-07 | 7.54E-06 | 0.364936097 |
| ENSMUSG00000068966 | Zbtb34 | 241311 | protein_coding | 122.3439468 | -0.364957969 | 0.135781351 | -2.687835741 | 0.007191676 | 0.046766196 | 0.364957969 |

Supplementary Table 1

| ensembl_gene_id | external_gene_name | entrezgene_id | gene_biotype | baseMean | log2FoldChange | lfcSE | stat | pvalue | padj | absLog2FC |
|---|---|---|---|---|---|---|---|---|---|---|
| ENSMUSG00000096981 | Gm16845 | 100503652 | lncRNA | 16.99195191 | -0.365143005 | 0.347621817 | -1.050403015 | 0.29353286 | 0.536379174 | 0.365143005 |
| ENSMUSG00000044062 | Plekhd1os | 68354 | lncRNA | 32.98969737 | 0.365511319 | 0.251458953 | 1.453562557 | 0.146067624 | 0.3590356 | 0.365511319 |
| ENSMUSG00000066595 | Flvcr1 | 226844 | protein_coding | 315.7753215 | 0.365607632 | 0.096643304 | 3.783062227 | 0.000154911 | 0.002123669 | 0.365607632 |
| ENSMUSG00000009687 | Fxyd5 | 18301 | protein_coding | 20.46730181 | 0.365821826 | 0.297876005 | 1.228101023 | 0.219409047 | 0.455582357 | 0.365821826 |
| ENSMUSG00000059142 | Zfp945 | 240041 | protein_coding | 243.6750684 | -0.365861819 | 0.122498981 | -2.986651949 | 0.002820506 | 0.022970241 | 0.365861819 |
| ENSMUSG00000029672 | Fam3c | 27999 | protein_coding | 340.1224063 | 0.36587672 | 0.096152102 | 3.805186904 | 0.000141697 | 0.001976773 | 0.36587672 |
| ENSMUSG00000049001 | Ndnf | 68169 | protein_coding | 98.53553308 | 0.366018855 | 0.156284289 | 2.342006729 | 0.019180369 | 0.095444865 | 0.366018855 |
| ENSMUSG00000032377 | Plscr4 | 235527 | protein_coding | 68.00799292 | -0.366143951 | 0.235874132 | -1.552285311 | 0.120593971 | 0.321404557 | 0.366143951 |
| ENSMUSG00000064823 | Snord82 | 80828 | snoRNA | 13.11190405 | -0.366159149 | 0.449603683 | -0.814404246 | 0.415413407 | 0.653406029 | 0.366159149 |
| ENSMUSG00000044365 | Cxxc4 | 319478 | protein_coding | 31.38989659 | 0.366198704 | 0.207195853 | 1.767403636 | 0.077160654 | 0.24490649 | 0.366198704 |
| ENSMUSG00000083674 | Zfp133-ps | 668917 | transcribed_unitary_pseudogene | 11.6463539 | 0.366340986 | 0.393239681 | 0.931597202 | 0.35154473 | 0.593903542 | 0.366340986 |
| ENSMUSG00000045331 | 2310079G19Rik | 69699 | protein_coding | 32.57767758 | -0.36634963 | 0.344548562 | -1.063274297 | 0.287657578 | 0.530090639 | 0.36634963 |
| ENSMUSG00000042787 | Exog | 208194 | protein_coding | 149.4897555 | 0.366374103 | 0.131064306 | 2.795376674 | 0.005183928 | 0.036667193 | 0.366374103 |
| ENSMUSG00000011171 | Vipr2 | 22355 | protein_coding | 64.84665905 | -0.366420192 | 0.215359494 | -1.701435058 | 0.088861323 | 0.267762809 | 0.366420192 |
| ENSMUSG00000005373 | Mlxipl | 58805 | protein_coding | 265.9526632 | -0.366706319 | 0.115942444 | -3.162830664 | 0.001562432 | 0.014456304 | 0.366706319 |
| ENSMUSG00000073771 | Btbd19 | 78611 | protein_coding | 73.24684917 | -0.366848866 | 0.244768592 | -1.498757918 | 0.133936445 | 0.341285046 | 0.366848866 |
| ENSMUSG00000102101 | Zbtb11os1 | 66391 | lncRNA | 23.95288125 | -0.366924721 | 0.276279478 | -1.328092566 | 0.184147529 | 0.412866286 | 0.366924721 |
| ENSMUSG00000051984 | Sec31b | 240667 | protein_coding | 23.74100095 | -0.366995888 | 0.278423079 | -1.318123086 | 0.187462448 | 0.417281835 | 0.366995888 |
| ENSMUSG00000000392 | Fap | 14089 | protein_coding | 77.92143797 | 0.367073314 | 0.153657294 | 2.388909143 | 0.016898481 | 0.087430799 | 0.367073314 |
| ENSMUSG00000038894 | Irs2 | 384783 | protein_coding | 1475.860775 | -0.367248305 | 0.104019461 | -3.530573053 | 0.00041466 | 0.004817897 | 0.367248305 |
| ENSMUSG00000051650 | B3gnt2 | 53625 | protein_coding | 139.3286431 | -0.36733913 | 0.139259777 | -2.637797768 | 0.008344633 | 0.052220563 | 0.36733913 |
| ENSMUSG00000060771 | Tsga10 | 211484 | protein_coding | 119.4525149 | 0.367428706 | 0.107266925 | 3.425368121 | 0.000613967 | 0.006777169 | 0.367428706 |
| ENSMUSG00000001802 | Lrp3 | 435965 | protein_coding | 15.38948659 | -0.367473912 | 0.330548273 | -1.111710277 | 0.26626274 | 0.508211544 | 0.367473912 |
| ENSMUSG00000028680 | Plk3 | 12795 | protein_coding | 1814.316293 | -0.367480182 | 0.150799399 | -2.436880953 | 0.014814556 | 0.07953207 | 0.367480182 |
| ENSMUSG00000007872 | Id3 | 15903 | protein_coding | 3710.706004 | 0.367568203 | 0.087131831 | 4.218529529 | 2.46E-05 | 0.000435306 | 0.367568203 |
| ENSMUSG00000040594 | Ranbp17 | 66011 | protein_coding | 25.32766529 | -0.367821819 | 0.236894489 | -1.552682044 | 0.120499114 | 0.321263055 | 0.367821819 |
| ENSMUSG00000043629 | 1700019D03Rik | 67080 | protein_coding | 446.6599987 | 0.36791998 | 0.069110717 | 5.323631373 | 1.02E-07 | 3.10E-06 | 0.36791998 |
| ENSMUSG00000001666 | Ddt | 13202 | protein_coding | 425.8963745 | 0.367984182 | 0.089067948 | 4.131499506 | 3.60E-05 | 0.000611815 | 0.367984182 |
| ENSMUSG00000050463 | Krt78 | 332131 | protein_coding | 7.68330065 | 0.368173584 | 0.42671863 | 0.862801757 | 0.388246473 | 0.628261175 | 0.368173584 |
| ENSMUSG00000032375 | Aph1b | 208117 | protein_coding | 365.5900676 | 0.368198023 | 0.070216185 | 5.243777121 | 1.57E-07 | 4.62E-06 | 0.368198023 |
| ENSMUSG00000049690 | Nckap5 | 210356 | protein_coding | 107.3082316 | -0.368226745 | 0.179425223 | -2.052257412 | 0.040144653 | 0.158991358 | 0.368226745 |
| ENSMUSG00000045007 | Tubg2 | 103768 | protein_coding | 12.19249386 | -0.368406363 | 0.423713618 | -0.869470197 | 0.384590002 | 0.624972092 | 0.368406363 |
| ENSMUSG00000026923 | Notch1 | 18128 | protein_coding | 1047.864234 | -0.368466769 | 0.081401364 | -4.526542923 | 6.00E-06 | 0.000124764 | 0.368466769 |
| ENSMUSG00000038541 | Srd5a2 | 94224 | protein_coding | 81.00832888 | 0.368482327 | 0.179939886 | 2.047807943 | 0.040578822 | 0.160203864 | 0.368482327 |
| ENSMUSG00000062488 | Ifit3b | 667370 | protein_coding | 22.83984616 | -0.368505109 | 0.269591939 | -1.366899583 | 0.171656783 | 0.396321966 | 0.368505109 |
| ENSMUSG00000033949 | Trim36 | 28105 | protein_coding | 549.8516245 | -0.368700507 | 0.062762265 | -5.874557069 | 4.24E-09 | 1.59E-07 | 0.368700507 |
| ENSMUSG00000078486 | Perm1 | 74183 | protein_coding | 14.97467333 | -0.368799355 | 0.317207882 | -1.162642468 | 0.244974593 | 0.486120986 | 0.368799355 |
| ENSMUSG00000074158 | Zfp976 | 208111 | protein_coding | 116.758629 | -0.368935827 | 0.14977263 | -2.463306052 | 0.013766235 | 0.075435975 | 0.368935827 |
| ENSMUSG00000022436 | Sh3bp1 | 20401 | protein_coding | 8.725776211 | -0.369001569 | 0.404512965 | -0.912211969 | 0.361657144 | 0.604649053 | 0.369001569 |

| ensembl_gene_id | external_gene_name | entrezgene_id | gene_biotype | baseMean | log2FoldChange | lfcSE | stat | pvalue | padj | absLog2FC |
|---|---|---|---|---|---|---|---|---|---|---|
| ENSMUSG00000019437 | Tlcd1 | 68385 | protein_coding | 282.6127384 | -0.370143031 | 0.096534434 | -3.834310881 | 0.000125917 | 0.00179387 | 0.370143031 |
| ENSMUSG00000031972 | Acta1 | 11459 | protein_coding | 375.4384121 | 0.370181756 | 0.123057687 | 3.008196933 | 0.002628028 | 0.021656722 | 0.370181756 |
| ENSMUSG00000022094 | Slc39a14 | 213053 | protein_coding | 1259.967308 | 0.370594038 | 0.048251296 | 7.680499224 | 1.58E-14 | 9.57E-13 | 0.370594038 |
| ENSMUSG00000026815 | Gfi1b | 14582 | protein_coding | 14.03992549 | 0.370640127 | 0.315865525 | 1.173411143 | 0.240630971 | 0.481172433 | 0.370640127 |
| ENSMUSG00000055193 | Klk15 | 317652 | protein_coding | 17.41586538 | 0.370742148 | 0.478740043 | 0.774412238 | 0.438687049 | 0.671427662 | 0.370742148 |
| ENSMUSG00000021136 | Smoc1 | 64075 | protein_coding | 9.966297984 | -0.370952089 | 0.605508946 | -0.612628585 | 0.540121953 | 0.745829843 | 0.370952089 |
| ENSMUSG00000028443 | Nudt2 | 66401 | protein_coding | 367.2731908 | 0.371026662 | 0.086225915 | 4.302959986 | 1.69E-05 | 0.000310618 | 0.371026662 |
| ENSMUSG00000074802 | Gas2l3 | 237436 | protein_coding | 236.1009731 | 0.371112485 | 0.096635209 | 3.840344398 | 0.000122862 | 0.001761238 | 0.371112485 |
| ENSMUSG00000031383 | Dusp9 | 75590 | protein_coding | 19.45829713 | -0.371731012 | 0.279334244 | -1.330774941 | 0.183263076 | 0.411471688 | 0.371731012 |
| ENSMUSG00000041930 | Fam222a | 433940 | protein_coding | 102.0654778 | 0.3721769 | 0.138453391 | 2.688102455 | 0.007185934 | 0.046747707 | 0.3721769 |
| ENSMUSG00000078490 | Cfap74 | 544678 | protein_coding | 33.62522951 | -0.372219391 | 0.358685872 | -1.03773084 | 0.299395383 | 0.543015856 | 0.372219391 |
| ENSMUSG00000031485 | Plpbp | 114863 | protein_coding | 1099.269054 | 0.37234129 | 0.048954432 | 7.605875061 | 2.83E-14 | 1.67E-12 | 0.37234129 |
| ENSMUSG00000073705 | Cenps | 69928 | protein_coding | 41.26298208 | 0.372522398 | 0.217571876 | 1.712180846 | 0.086863346 | 0.263862344 | 0.372522398 |
| ENSMUSG00000054509 | Parp4 | 328417 | protein_coding | 543.1646331 | -0.372550354 | 0.128783746 | -2.892836743 | 0.003817797 | 0.02917991 | 0.372550354 |
| ENSMUSG00000020681 | Ace | 11421 | protein_coding | 783.0301926 | -0.372785252 | 0.169785805 | -2.195620849 | 0.028119093 | 0.12424678 | 0.372785252 |
| ENSMUSG00000078451 | Ppil6 | 73075 | protein_coding | 16.46662578 | -0.372817534 | 0.335001972 | -1.112881609 | 0.26575928 | 0.507758471 | 0.372817534 |
| ENSMUSG00000051379 | Flrt3 | 71436 | protein_coding | 1115.217077 | 0.372955928 | 0.0847998 | 4.398075589 | 1.09E-05 | 0.000212206 | 0.372955928 |
| ENSMUSG00000067642 | Adgrf3 | 381628 | protein_coding | 6.370652217 | -0.373003298 | 0.611249057 | -0.610231286 | 0.541708608 | 0.747252991 | 0.373003298 |
| ENSMUSG00000018008 | Cyth4 | 72318 | protein_coding | 24.916743 | 0.373222283 | 0.275348071 | 1.35545632 | 0.175272216 | 0.401621914 | 0.373222283 |
| ENSMUSG00000056501 | Cebpb | 12608 | protein_coding | 257.3507649 | -0.373238417 | 0.188695583 | -1.977992334 | 0.047929572 | 0.179737215 | 0.373238417 |
| ENSMUSG00000045004 | Spata21 | 329972 | protein_coding | 6.105393263 | -0.373321596 | 0.514839748 | -0.725121937 | 0.468377227 | 0.693047027 | 0.373321596 |
| ENSMUSG00000026223 | Itm2c | 64294 | protein_coding | 1199.374159 | 0.373336532 | 0.051184402 | 7.293951253 | 3.01E-13 | 1.63E-11 | 0.373336532 |
| ENSMUSG00000098112 | Bin2 | 668218 | protein_coding | 29.89253616 | 0.373385278 | 0.256975215 | 1.453001129 | 0.14622344 | 0.359247178 | 0.373385278 |
| ENSMUSG00000040928 | S100pbp | 74648 | protein_coding | 276.1541895 | -0.373638158 | 0.096507928 | -3.871579931 | 0.000108132 | 0.001589652 | 0.373638158 |
| ENSMUSG00000040231 | Syngr4 | 58867 | protein_coding | 5.991255993 | 0.373700818 | 0.508023069 | 0.735598126 | 0.461975303 | 0.688426884 | 0.373700818 |
| ENSMUSG00000054387 | Mdm4 | 17248 | protein_coding | 1256.58991 | -0.373747761 | 0.106306408 | -3.515759467 | 0.000438498 | 0.005058411 | 0.373747761 |
| ENSMUSG00000028581 | Laptm5 | 16792 | protein_coding | 52.02109515 | 0.373756919 | 0.210311685 | 1.777157174 | 0.075542383 | 0.241780511 | 0.373756919 |
| ENSMUSG00000048285 | Frmd6 | 319710 | protein_coding | 73.83153355 | 0.373794563 | 0.150356176 | 2.48606059 | 0.012916597 | 0.071852934 | 0.373794563 |
| ENSMUSG00000069874 | Irgm2 | 54396 | protein_coding | 295.1352117 | -0.37381946 | 0.383584864 | -0.974541738 | 0.329787626 | 0.573541103 | 0.37381946 |
| ENSMUSG00000045193 | Cirbp | 12696 | protein_coding | 600.4978059 | -0.37406941 | 0.110656415 | -3.380458431 | 0.00072365 | 0.00773893 | 0.37406941 |
| ENSMUSG00000025240 | Sacm1l | 83493 | protein_coding | 1381.422182 | 0.374248835 | 0.050699008 | 7.381778236 | 1.56E-13 | 8.69E-12 | 0.374248835 |
| ENSMUSG00000015879 | Fam184b | 58227 | protein_coding | 31.3551322 | -0.374280042 | 0.275141488 | -1.36031845 | 0.173729172 | 0.399424529 | 0.374280042 |
| ENSMUSG00000118117 | Gm30961 | 102632967 | lncRNA | 7.43752108 | -0.37452103 | 0.483628056 | -0.774398892 | 0.438694939 | 0.671427662 | 0.37452103 |
| ENSMUSG00000026064 | Ptp4a1 | 19243 | protein_coding | 334.373886 | -0.374603041 | 0.128929905 | -2.905478301 | 0.003666923 | 0.028308503 | 0.374603041 |
| ENSMUSG00000029644 | Pdx1 | 18609 | protein_coding | 8.494067138 | 0.374835945 | 0.39232074 | 0.955432395 | 0.339359078 | 0.582473517 | 0.374835945 |
| ENSMUSG00000097743 | Gm16973 | 100503823 | lncRNA | 38.74872973 | -0.375018905 | 0.214141422 | -1.751267464 | 0.079899849 | 0.249990933 | 0.375018905 |
| ENSMUSG00000036091 | Hyal3 | 109685 | protein_coding | 9.561662423 | -0.375310898 | 0.404625424 | -0.927551447 | 0.353640293 | 0.595956629 | 0.375310898 |
| ENSMUSG00000028051 | Hcn3 | 15168 | protein_coding | 14.25739367 | 0.375397997 | 0.331068825 | 1.13389715 | 0.256837693 | 0.498650401 | 0.375397997 |
| ENSMUSG00000058624 | Gda | 14544 | protein_coding | 1354.416089 | -0.375660102 | 0.094691102 | -3.967216504 | 7.27E-05 | 0.001127602 | 0.375660102 |

| ensembl_gene_id | external_gene_name | entrezgene_id | gene_biotype | baseMean | log2FoldChange | lfcSE | stat | pvalue | padj | absLog2FC |
|---|---|---|---|---|---|---|---|---|---|---|
| ENSMUSG00000010592 | Dazl | 13164 | protein_coding | 7.089282718 | -0.375814693 | 0.464104622 | -0.809762875 | 0.418076474 | 0.655860109 | 0.375814693 |
| ENSMUSG00000023031 | Cela1 | 109901 | protein_coding | 3342.006208 | -0.376052101 | 0.120721641 | -3.115034705 | 0.001839233 | 0.016432856 | 0.376052101 |
| ENSMUSG00000040274 | Cdk6 | 12571 | protein_coding | 1407.488443 | -0.376156426 | 0.083240888 | -4.518890137 | 6.22E-06 | 0.00012886 | 0.376156426 |
| ENSMUSG00000078495 | Zfp984 | 100041677 | protein_coding | 228.0986586 | -0.376487579 | 0.107518222 | -3.501616482 | 0.000462445 | 0.005293007 | 0.376487579 |
| ENSMUSG00000066705 | Fxyd6 | 59095 | protein_coding | 17.89226508 | 0.376591798 | 0.324142103 | 1.161810805 | 0.245312324 | 0.486611544 | 0.376591798 |
| ENSMUSG00000024253 | Dync2li1 | 213575 | protein_coding | 31.73739465 | 0.376818266 | 0.223056223 | 1.689342089 | 0.091153892 | 0.272039484 | 0.376818266 |
| ENSMUSG00000085424 | Fhad1os1 | 545691 | lncRNA | 6.670935907 | 0.376969232 | 0.45813449 | 0.822835303 | 0.410601658 | 0.648748206 | 0.376969232 |
| ENSMUSG00000032420 | Nt5e | 23959 | protein_coding | 723.93345 | 0.377398321 | 0.270935185 | 1.392946882 | 0.163635854 | 0.385451042 | 0.377398321 |
| ENSMUSG00000028884 | Rpa2 | 19891 | protein_coding | 275.3817875 | 0.377875456 | 0.141652032 | 2.66763174 | 0.007638793 | 0.04892449 | 0.377875456 |
| ENSMUSG00000025153 | Fasn | 14104 | protein_coding | 9561.080339 | 0.377901199 | 0.120128906 | 3.145797397 | 0.001656347 | 0.015117096 | 0.377901199 |
| ENSMUSG00000022519 | Srl | 106393 | protein_coding | 184.3203605 | 0.378177077 | 0.103824974 | 3.642448064 | 0.000270057 | 0.003386638 | 0.378177077 |
| ENSMUSG00000058145 | Adamts17 | 233332 | protein_coding | 85.11590801 | 0.378186925 | 0.18362703 | 2.059538426 | 0.039442688 | 0.156982287 | 0.378186925 |
| ENSMUSG00000024666 | Tmem138 | 72982 | protein_coding | 46.26286948 | 0.378363293 | 0.180992872 | 2.090487259 | 0.036574052 | 0.148759075 | 0.378363293 |
| ENSMUSG00000050332 | Amer1 | 72345 | protein_coding | 304.0111303 | -0.378436425 | 0.093840079 | -4.032780345 | 5.51E-05 | 0.000892503 | 0.378436425 |
| ENSMUSG00000034324 | Tmem132c | 208213 | protein_coding | 91.26620147 | -0.378481843 | 0.201716125 | -1.876309309 | 0.06061282 | 0.209360236 | 0.378481843 |
| ENSMUSG00000087543 | Gm16576 | 100504191 | lncRNA | 51.28776857 | -0.378693157 | 0.181655446 | -2.084678246 | 0.037098517 | 0.15024806 | 0.378693157 |
| ENSMUSG00000018068 | Ints2 | 70422 | protein_coding | 493.7794538 | 0.378895052 | 0.109095186 | 3.473068474 | 0.000514544 | 0.005794729 | 0.378895052 |
| ENSMUSG00000072066 | 6720489N17Rik | 211378 | protein_coding | 29.70853149 | 0.378964954 | 0.240475047 | 1.57590136 | 0.115048538 | 0.313039947 | 0.378964954 |
| ENSMUSG00000040606 | Kazn | 71529 | protein_coding | 190.1914987 | -0.379156089 | 0.131549832 | -2.882224048 | 0.003948788 | 0.029968052 | 0.379156089 |
| ENSMUSG00000113585 | Gm19605 | 102638940 | lncRNA | 83.4180202 | 0.379184588 | 0.151934543 | 2.495710191 | 0.012570526 | 0.070537532 | 0.379184588 |
| ENSMUSG00000054958 | Nt5c1a | 230718 | protein_coding | 47.05938952 | 0.379226497 | 0.293102349 | 1.29383643 | 0.195721926 | 0.428222446 | 0.379226497 |
| ENSMUSG00000018570 | 2810408A11Rik | 70419 | protein_coding | 117.8096915 | -0.379226659 | 0.143056636 | -2.650884785 | 0.008028122 | 0.050786438 | 0.379226659 |
| ENSMUSG00000023236 | Scg5 | 20394 | protein_coding | 31.84216704 | -0.379603787 | 0.215858496 | -1.758577003 | 0.078649382 | 0.247826999 | 0.379603787 |
| ENSMUSG00000062743 | Zfp677 | 210503 | protein_coding | 87.87353182 | 0.379972769 | 0.136709835 | 2.779410626 | 0.005445764 | 0.038068414 | 0.379972769 |
| ENSMUSG00000032013 | Trim29 | 72169 | protein_coding | 8.919873035 | -0.380132726 | 0.389860994 | -0.97504683 | 0.329537031 | 0.573356749 | 0.380132726 |
| ENSMUSG00000070604 | Vsig10l | 75690 | protein_coding | 31.96173448 | -0.38022796 | 0.251281822 | -1.51315347 | 0.13024068 | 0.336231563 | 0.38022796 |
| ENSMUSG00000091476 | Catspere2 | 545391 | protein_coding | 30.66440035 | 0.380394884 | 0.212634141 | 1.788964283 | 0.073620568 | 0.23750328 | 0.380394884 |
| ENSMUSG00000105504 | Gbp5 | 229898 | protein_coding | 15.28827467 | -0.380492553 | 0.531421104 | -0.71599067 | 0.47399711 | 0.69790481 | 0.380492553 |
| ENSMUSG00000068740 | Celsr2 | 53883 | protein_coding | 91.17507379 | -0.380660539 | 0.166351566 | -2.288289486 | 0.022120668 | 0.105513168 | 0.380660539 |
| ENSMUSG00000022455 | Wbp2nl | 74716 | protein_coding | 10.38868231 | 0.380717247 | 0.373916261 | 1.018188528 | 0.30858837 | 0.552278995 | 0.380717247 |
| ENSMUSG00000024793 | Tnfrsf25 | 85030 | protein_coding | 12.43453788 | -0.380870999 | 0.349385048 | -1.090118198 | 0.275661081 | 0.518314825 | 0.380870999 |
| ENSMUSG00000027902 | Chil6 | 229688 | protein_coding | 17.47056633 | -0.381012968 | 0.307842619 | -1.237687523 | 0.215831947 | 0.451641923 | 0.381012968 |
| ENSMUSG00000051910 | Sox6 | 20679 | protein_coding | 465.9352375 | -0.381022342 | 0.156308056 | -2.437637265 | 0.0147836 | 0.079418761 | 0.381022342 |
| ENSMUSG00000039137 | Whrn | 73750 | protein_coding | 200.9166711 | -0.381072972 | 0.127681076 | -2.984568921 | 0.002839782 | 0.023082976 | 0.381072972 |
| ENSMUSG00000034157 | Cipc | 217732 | protein_coding | 3448.16872 | -0.381074922 | 0.065374151 | -5.829137607 | 5.57E-09 | 2.06E-07 | 0.381074922 |
| ENSMUSG00000061048 | Cdh3 | 12560 | protein_coding | 7.264271644 | 0.381092042 | 0.539304903 | 0.706635596 | 0.479792962 | 0.702150736 | 0.381092042 |
| ENSMUSG00000029392 | Rilpl1 | 75695 | protein_coding | 14.89931825 | -0.381178835 | 0.311543335 | -1.223517863 | 0.221134161 | 0.457721775 | 0.381178835 |
| ENSMUSG00000044229 | Nxpe4 | 244853 | protein_coding | 1124.903932 | -0.381474144 | 0.151168847 | -2.523497077 | 0.011619403 | 0.066299536 | 0.381474144 |
| ENSMUSG00000030110 | Ret | 19713 | protein_coding | 65.8468312 | 0.382141193 | 0.167800751 | 2.277350906 | 0.022765279 | 0.107538272 | 0.382141193 |

| ensembl_gene_id | external_gene_name | entrezgene_id | gene_biotype | baseMean | log2FoldChange | lfcSE | stat | pvalue | padj | absLog2FC |
|---|---|---|---|---|---|---|---|---|---|---|
| ENSMUSG00000075334 | Rprm | 67874 | protein_coding | 56.96603721 | 0.382290023 | 0.212081962 | 1.802557938 | 0.071457668 | 0.233184502 | 0.382290023 |
| ENSMUSG00000021959 | Lats2 | 50523 | protein_coding | 123.5747244 | -0.382315176 | 0.144015854 | -2.654674228 | 0.007938503 | 0.050423097 | 0.382315176 |
| ENSMUSG00000033967 | Rnf225 | 381845 | protein_coding | 9.236697866 | -0.382324761 | 0.426376487 | -0.896683503 | 0.369887826 | 0.612036907 | 0.382324761 |
| ENSMUSG00000035184 | Fam124a | 629059 | protein_coding | 159.2222042 | 0.382408075 | 0.100766758 | 3.794982436 | 0.000147654 | 0.002040458 | 0.382408075 |
| ENSMUSG00000033059 | Pygb | 110078 | protein_coding | 4738.343955 | 0.382500327 | 0.065366616 | 5.851615848 | 4.87E-09 | 1.81E-07 | 0.382500327 |
| ENSMUSG00000070348 | Ccnd1 | 12443 | protein_coding | 1339.331801 | 0.382577032 | 0.059207302 | 6.461652904 | 1.04E-10 | 4.55E-09 | 0.382577032 |
| ENSMUSG00000020469 | Myl7 | 17898 | protein_coding | 37.84057573 | -0.382616376 | 0.281188223 | -1.360712662 | 0.173604512 | 0.399389438 | 0.382616376 |
| ENSMUSG00000027712 | Anxa5 | 11747 | protein_coding | 914.3019457 | 0.382674468 | 0.068785608 | 5.563292662 | 2.65E-08 | 9.01E-07 | 0.382674468 |
| ENSMUSG00000027381 | Bcl2l11 | 12125 | protein_coding | 282.1074286 | -0.382742555 | 0.104453582 | -3.664235788 | 0.000248078 | 0.003155171 | 0.382742555 |
| ENSMUSG00000016552 | Foxred2 | 239554 | protein_coding | 87.98065629 | -0.382928971 | 0.153008929 | -2.502657672 | 0.012326471 | 0.069433808 | 0.382928971 |
| ENSMUSG00000027356 | Fermt1 | 241639 | protein_coding | 3742.929232 | 0.382954006 | 0.035710879 | 10.72373493 | 7.88E-27 | 9.27E-25 | 0.382954006 |
| ENSMUSG00000040339 | Fam102b | 329739 | protein_coding | 1027.527756 | 0.383146537 | 0.127625043 | 3.002126601 | 0.002681007 | 0.022022562 | 0.383146537 |
| ENSMUSG00000023927 | Satb1 | 20230 | protein_coding | 12.50640575 | 0.383299607 | 0.319528626 | 1.199578304 | 0.230303157 | 0.469307431 | 0.383299607 |
| ENSMUSG00000064994 | Gm22422 | 115486004 | snoRNA | 157.19577 | 0.383358129 | 0.221179631 | 1.733243366 | 0.083052422 | 0.256348065 | 0.383358129 |
| ENSMUSG00000033467 | Crlf2 | 57914 | protein_coding | 69.70489743 | -0.383676418 | 0.165929255 | -2.312289164 | 0.020761753 | 0.100942055 | 0.383676418 |
| ENSMUSG00000056043 | Rgs9bp | 243923 | protein_coding | 10.89129654 | 0.38387656 | 0.463874552 | 0.827543908 | 0.407928846 | 0.646488649 | 0.38387656 |
| ENSMUSG00000044471 | Lncpint | 232685 | lncRNA | 137.419221 | -0.383979277 | 0.195652194 | -1.962560554 | 0.049697258 | 0.184385584 | 0.383979277 |
| ENSMUSG00000039110 | Mycbpap | 104601 | protein_coding | 103.5213124 | -0.384075301 | 0.15115265 | -2.540976297 | 0.011054341 | 0.063669058 | 0.384075301 |
| ENSMUSG00000036959 | Bcorl1 | 320376 | protein_coding | 223.7976197 | -0.384163717 | 0.12857125 | -2.98794418 | 0.002808608 | 0.022899104 | 0.384163717 |
| ENSMUSG00000044948 | Cfap43 | 100048534 | protein_coding | 7.803046652 | -0.384355569 | 0.476424464 | -0.80675028 | 0.41981037 | 0.657014281 | 0.384355569 |
| ENSMUSG00000034919 | Ttc22 | 230576 | protein_coding | 1443.727497 | 0.384381353 | 0.078789575 | 4.878581384 | 1.07E-06 | 2.60E-05 | 0.384381353 |
| ENSMUSG00000022951 | Rcan1 | 54720 | protein_coding | 718.2508259 | 0.384390035 | 0.079028481 | 4.863943119 | 1.15E-06 | 2.77E-05 | 0.384390035 |
| ENSMUSG00000069273 | H3c6 | 319151 | protein_coding | 2369.466531 | 0.384405332 | 0.130855017 | 2.937643064 | 0.003307175 | 0.026131709 | 0.384405332 |
| ENSMUSG00000053279 | Aldh1a1 | 11668 | protein_coding | 4938.944422 | -0.384515395 | 0.137885839 | -2.788650357 | 0.005292817 | 0.037252106 | 0.384515395 |
| ENSMUSG00000022180 | Slc7a8 | 50934 | protein_coding | 184.4406042 | -0.384570778 | 0.161749016 | -2.37757723 | 0.017426793 | 0.089163038 | 0.384570778 |
| ENSMUSG00000086844 | B230206H07Rik | 320871 | lncRNA | 25.94483313 | -0.384683249 | 0.305184299 | -1.260494887 | 0.207490891 | 0.442811616 | 0.384683249 |
| ENSMUSG00000072941 | Sod3 | 20657 | protein_coding | 8.596737857 | 0.38485523 | 0.421057802 | 0.914019947 | 0.360706365 | 0.603998706 | 0.38485523 |
| ENSMUSG00000049866 | Arl4c | 320982 | protein_coding | 25.17944128 | 0.385000887 | 0.23372942 | 1.647207641 | 0.099515374 | 0.286945187 | 0.385000887 |
| ENSMUSG00000073079 | Srp54a | 24067 | protein_coding | 119.2184758 | 0.385028309 | 0.118623364 | 3.245805014 | 0.00117119 | 0.011488921 | 0.385028309 |
| ENSMUSG00000097727 | F630040K05Rik | 433111 | lncRNA | 6.339017103 | 0.385102792 | 0.574636467 | 0.670167688 | 0.5027509 | 0.719356203 | 0.385102792 |
| ENSMUSG00000002032 | Tmem25 | 71687 | protein_coding | 172.0035627 | -0.385310544 | 0.119817674 | -3.215807252 | 0.001300782 | 0.012524004 | 0.385310544 |
| ENSMUSG00000042042 | Csgalnact2 | 78752 | protein_coding | 272.8393183 | 0.385415359 | 0.074011218 | 5.207526247 | 1.91E-07 | 5.48E-06 | 0.385415359 |
| ENSMUSG00000038146 | Notch3 | 18131 | protein_coding | 8.456810907 | -0.385493154 | 0.512466126 | -0.752231484 | 0.451911863 | 0.680967534 | 0.385493154 |
| ENSMUSG00000022488 | Nckap1l | 105855 | protein_coding | 25.45123939 | 0.385612622 | 0.236088918 | 1.633336391 | 0.102398267 | 0.292072312 | 0.385612622 |
| ENSMUSG00000097615 | Gm2061 | 100039138 | lncRNA | 88.76367567 | 0.385641928 | 0.147172711 | 2.620335821 | 0.008784322 | 0.054173902 | 0.385641928 |
| ENSMUSG00000035413 | Tmem98 | 103743 | protein_coding | 3303.249761 | 0.385929536 | 0.041598071 | 9.277582481 | 1.73E-20 | 1.47E-18 | 0.385929536 |
| ENSMUSG00000073678 | Pgap1 | 241062 | protein_coding | 1024.954477 | 0.386157013 | 0.057594613 | 6.704741837 | 2.02E-11 | 9.40E-10 | 0.386157013 |
| ENSMUSG00000045441 | Gprin3 | 243385 | protein_coding | 1008.184803 | -0.386241533 | 0.067033831 | -5.761889584 | 8.32E-09 | 2.98E-07 | 0.386241533 |
| ENSMUSG00000058070 | Eml1 | 68519 | protein_coding | 71.02880332 | 0.386297897 | 0.150043072 | 2.574580029 | 0.010036187 | 0.059417614 | 0.386297897 |

| ensembl_gene_id | external_gene_name | entrezgene_id | gene_biotype | baseMean | log2FoldChange | lfcSE | stat | pvalue | padj | absLog2FC |
|---|---|---|---|---|---|---|---|---|---|---|
| ENSMUSG00000012777 | Acp4 | 100503991 | protein_coding | 6.061756344 | -0.386372913 | 0.557921942 | -0.692521451 | 0.48860992 | 0.708970987 | 0.386372913 |
| ENSMUSG00000037362 | Ccn3 | 18133 | protein_coding | 3235.675009 | 0.386570974 | 0.257023606 | 1.504029065 | 0.132573883 | 0.339503983 | 0.386570974 |
| ENSMUSG00000116953 | 9030025P20Rik | 100041574 | protein_coding | 17.91035679 | -0.386619068 | 0.276213885 | -1.399709024 | 0.16160047 | 0.382186653 | 0.386619068 |
| ENSMUSG00000043099 | Hic1 | 15248 | protein_coding | 20.01509609 | 0.386975619 | 0.334990142 | 1.155185095 | 0.248014637 | 0.489024582 | 0.386975619 |
| ENSMUSG00000036934 | 4921524J17Rik | 66714 | protein_coding | 153.3504242 | -0.387124434 | 0.120863486 | -3.202989156 | 0.001360091 | 0.012902464 | 0.387124434 |
| ENSMUSG00000024970 | Spindoc | 68229 | protein_coding | 322.696375 | 0.387356958 | 0.072308954 | 5.356970827 | 8.46E-08 | 2.61E-06 | 0.387356958 |
| ENSMUSG00000066057 | Gm1976 | 100038746 | lncRNA | 45.52402006 | -0.387401816 | 0.239165596 | -1.619805786 | 0.105274003 | 0.2974341 | 0.387401816 |
| ENSMUSG00000028207 | Asph | 65973 | protein_coding | 2699.628189 | 0.387575675 | 0.040636395 | 9.537649094 | 1.46E-21 | 1.32E-19 | 0.387575675 |
| ENSMUSG00000098488 | Pla2g4b | 211429 | protein_coding | 7.890444344 | -0.388270285 | 0.412208748 | -0.941926359 | 0.346230344 | 0.588620784 | 0.388270285 |
| ENSMUSG00000035713 | Usp35 | 244144 | protein_coding | 33.34037205 | -0.3883161 | 0.224973438 | -1.726053101 | 0.08433786 | 0.258921887 | 0.3883161 |
| ENSMUSG00000020828 | Pld2 | 18806 | protein_coding | 73.33468229 | 0.388489146 | 0.191418771 | 2.029524818 | 0.042404864 | 0.165184357 | 0.388489146 |
| ENSMUSG00000036053 | Fmnl2 | 71409 | protein_coding | 57.05227655 | 0.388616226 | 0.16116613 | 2.411277272 | 0.015896758 | 0.083507172 | 0.388616226 |
| ENSMUSG00000000594 | Gm2a | 14667 | protein_coding | 340.293181 | -0.389088993 | 0.108303975 | -3.592564306 | 0.00032744 | 0.003964431 | 0.389088993 |
| ENSMUSG00000047298 | Kcnv2 | 240595 | protein_coding | 11.79041502 | 0.389193761 | 0.401054768 | 0.970425468 | 0.331834459 | 0.575141012 | 0.389193761 |
| ENSMUSG00000038843 | Gcnt1 | 14537 | protein_coding | 971.2061153 | -0.389207433 | 0.086605673 | -4.494017745 | 6.99E-06 | 0.000143039 | 0.389207433 |
| ENSMUSG00000111356 | Rpl32l | 621697 | processed_pseudogene | 12.39963634 | -0.389244096 | 0.337626922 | -1.152882282 | 0.248958705 | 0.489898763 | 0.389244096 |
| ENSMUSG00000027796 | Smad9 | 55994 | protein_coding | 17.46673503 | -0.389284154 | 0.310282693 | -1.25461124 | 0.209619921 | 0.444925042 | 0.389284154 |
| ENSMUSG00000004032 | Gstm5 | 14866 | protein_coding | 507.5165709 | 0.389420549 | 0.072363335 | 5.381462152 | 7.39E-08 | 2.31E-06 | 0.389420549 |
| ENSMUSG00000020131 | Pcsk4 | 18551 | protein_coding | 48.35422844 | -0.38960166 | 0.166452439 | -2.340618509 | 0.019251827 | 0.095479458 | 0.38960166 |
| ENSMUSG00000097534 | Gm16675 | 100503498 | lncRNA | 40.71582035 | -0.389941524 | 0.1945003 | -2.004837652 | 0.044980406 | 0.172181107 | 0.389941524 |
| ENSMUSG00000036098 | Myrf | 225908 | protein_coding | 6.01186354 | 0.390204096 | 0.529723677 | 0.736618189 | 0.461354571 | 0.687894338 | 0.390204096 |
| ENSMUSG00000031981 | Capn9 | 73647 | protein_coding | 1175.836338 | 0.39027658 | 0.077384819 | 5.04332225 | 4.58E-07 | 1.20E-05 | 0.39027658 |
| ENSMUSG00000048865 | Arhgap30 | 226652 | protein_coding | 38.03238191 | 0.390408186 | 0.260856402 | 1.496640233 | 0.134486895 | 0.342309054 | 0.390408186 |
| ENSMUSG00000020042 | Btbd11 | 74007 | protein_coding | 25.31210643 | 0.390588547 | 0.271827446 | 1.43689886 | 0.150746734 | 0.366514786 | 0.390588547 |
| ENSMUSG00000032757 | Bet1 | 12068 | protein_coding | 656.7432392 | 0.390983686 | 0.074048649 | 5.280092118 | 1.29E-07 | 3.86E-06 | 0.390983686 |
| ENSMUSG00000066026 | Dhrs3 | 20148 | protein_coding | 830.1109983 | -0.391287625 | 0.076532957 | -5.112668329 | 3.18E-07 | 8.62E-06 | 0.391287625 |
| ENSMUSG00000001403 | Ube2c | 68612 | protein_coding | 632.4554257 | 0.391673944 | 0.090354924 | 4.334837829 | 1.46E-05 | 0.000274493 | 0.391673944 |
| ENSMUSG00000016356 | Col20a1 | 73368 | protein_coding | 59.9101278 | 0.391697859 | 0.191157347 | 2.049086081 | 0.040453698 | 0.159861993 | 0.391697859 |
| ENSMUSG00000021613 | Hapln1 | 12950 | protein_coding | 11.59794032 | 0.39177845 | 0.364070944 | 1.076104688 | 0.281880439 | 0.524623111 | 0.39177845 |
| ENSMUSG00000020752 | Recql5 | 170472 | protein_coding | 985.6976913 | -0.39187455 | 0.125564202 | -3.120909801 | 0.001802932 | 0.016171241 | 0.39187455 |
| ENSMUSG00000000216 | Scnn1g | 20278 | protein_coding | 169.0305024 | -0.391881837 | 0.284756255 | -1.376200979 | 0.168759419 | 0.392724841 | 0.391881837 |
| ENSMUSG00000021537 | Cetn3 | 12626 | protein_coding | 715.6470739 | 0.391972491 | 0.08907759 | 4.400349096 | 1.08E-05 | 0.000210502 | 0.391972491 |
| ENSMUSG00000064600 | Gm25636 | 115486823 | snoRNA | 17.00703737 | -0.392072967 | 0.382319383 | -1.025511615 | 0.305121848 | 0.549532801 | 0.392072967 |
| ENSMUSG00000097011 | Gm4651 | 100043793 | lncRNA | 22.06504936 | -0.392164612 | 0.272143389 | -1.441022006 | 0.149578466 | 0.364719069 | 0.392164612 |
| ENSMUSG00000054582 | Pabpc1l | 381404 | protein_coding | 6.879115571 | 0.392206334 | 0.467087854 | 0.839684293 | 0.401085423 | 0.640407651 | 0.392206334 |
| ENSMUSG00000085408 | C530005A16Rik | 654318 | lncRNA | 12.13710519 | -0.392244136 | 0.44561771 | -0.880225645 | 0.378737083 | 0.619821175 | 0.392244136 |
| ENSMUSG00000031709 | Tbc1d9 | 71310 | protein_coding | 190.1779401 | 0.392349095 | 0.125452508 | 3.127471107 | 0.001763171 | 0.01586756 | 0.392349095 |
| ENSMUSG00000034684 | Sema3f | 20350 | protein_coding | 32.48030741 | -0.392417221 | 0.22679033 | -1.730308435 | 0.083575183 | 0.25741348 | 0.392417221 |
| ENSMUSG00000029657 | Hsph1 | 15505 | protein_coding | 684.2874938 | 0.392832481 | 0.131142757 | 2.995456936 | 0.00274034 | 0.022421848 | 0.392832481 |

| ensembl_gene_id | external_gene_name | entrezgene_id | gene_biotype | baseMean | log2FoldChange | lfcSE | stat | pvalue | padj | absLog2FC |
|---|---|---|---|---|---|---|---|---|---|---|
| ENSMUSG00000031712 | Il15 | 16168 | protein_coding | 322.0852762 | -0.393034275 | 0.094075167 | -4.177874847 | 2.94E-05 | 0.000509152 | 0.393034275 |
| ENSMUSG00000038210 | Hoxa11 | 15396 | protein_coding | 34.63556246 | 0.393500647 | 0.253025118 | 1.555184124 | 0.119902224 | 0.320286898 | 0.393500647 |
| ENSMUSG00000115526 | 4930413F20Rik | 71637 | lncRNA | 6.598469422 | -0.393750689 | 0.547518709 | -0.719154767 | 0.472045566 | 0.696557676 | 0.393750689 |
| ENSMUSG00000034028 | Cd226 | 225825 | protein_coding | 18.7080388 | 0.393786295 | 0.281741859 | 1.397684733 | 0.162207761 | 0.38300506 | 0.393786295 |
| ENSMUSG00000003934 | Efnb3 | 13643 | protein_coding | 7.34021404 | 0.393819308 | 0.493890678 | 0.797381537 | 0.425229479 | 0.661044515 | 0.393819308 |
| ENSMUSG00000037813 | D630003M21Rik | 228846 | protein_coding | 9.20751022 | 0.39394917 | 0.405517363 | 0.971473003 | 0.33131279 | 0.574833929 | 0.39394917 |
| ENSMUSG00000042408 | Zmym6 | 100177 | protein_coding | 297.4706813 | -0.394143501 | 0.111544785 | -3.533500045 | 0.000410096 | 0.004775176 | 0.394143501 |
| ENSMUSG00000040943 | Tet2 | 214133 | protein_coding | 771.8347121 | -0.39422019 | 0.085209939 | -4.626457855 | 3.72E-06 | 8.01E-05 | 0.39422019 |
| ENSMUSG00000026737 | Pip4k2a | 18718 | protein_coding | 34.00273742 | 0.394467015 | 0.221399852 | 1.781695027 | 0.074798978 | 0.240247585 | 0.394467015 |
| ENSMUSG00000050908 | Tvp23a | 383103 | protein_coding | 8.843478889 | -0.394501096 | 0.533399636 | -0.739597611 | 0.459544192 | 0.68653709 | 0.394501096 |
| ENSMUSG00000033307 | Mif | 17319 | protein_coding | 1143.81302 | 0.394671369 | 0.111408932 | 3.542546925 | 0.000396283 | 0.004647894 | 0.394671369 |
| ENSMUSG00000078485 | Plekhn1 | 231002 | protein_coding | 341.7952761 | -0.395017819 | 0.075268329 | -5.248127908 | 1.54E-07 | 4.53E-06 | 0.395017819 |
| ENSMUSG00000074261 | Erich4 | 632778 | protein_coding | 59.13533022 | -0.395204498 | 0.16875512 | -2.34188153 | 0.019186804 | 0.095444865 | 0.395204498 |
| ENSMUSG00000040667 | Nup88 | 19069 | protein_coding | 1427.370882 | 0.395237706 | 0.051251501 | 7.711729357 | 1.24E-14 | 7.58E-13 | 0.395237706 |
| ENSMUSG00000022629 | Kif21a | 16564 | protein_coding | 2010.92102 | 0.395257421 | 0.082485685 | 4.791830472 | 1.65E-06 | 3.86E-05 | 0.395257421 |
| ENSMUSG00000032327 | Stra6 | 20897 | protein_coding | 6.007607278 | 0.395358366 | 0.539188664 | 0.733246807 | 0.463407912 | 0.689426144 | 0.395358366 |
| ENSMUSG00000050860 | Phospho1 | 237928 | protein_coding | 26.02355708 | -0.395455969 | 0.265447283 | -1.489772151 | 0.136284156 | 0.345248914 | 0.395455969 |
| ENSMUSG00000030793 | Pycard | 66824 | protein_coding | 3315.217419 | 0.395571152 | 0.066282756 | 5.967934604 | 2.40E-09 | 9.16E-08 | 0.395571152 |
| ENSMUSG00000085175 | Gm11423 | 100503272 | lncRNA | 44.76106406 | -0.395705065 | 0.212972503 | -1.85801012 | 0.063167577 | 0.214961704 | 0.395705065 |
| ENSMUSG00000008540 | Mgst1 | 56615 | protein_coding | 2194.529722 | -0.395795739 | 0.080072573 | -4.942962646 | 7.69E-07 | 1.94E-05 | 0.395795739 |
| ENSMUSG00000024749 | Tmc1 | 13409 | protein_coding | 33.06578892 | -0.395905707 | 0.240316443 | -1.647434949 | 0.099468677 | 0.286913137 | 0.395905707 |
| ENSMUSG00000010342 | Tex14 | 83560 | protein_coding | 7.661970567 | -0.3961195 | 0.44341548 | -0.893337102 | 0.371676675 | 0.613533062 | 0.3961195 |
| ENSMUSG00000040432 | Ltb4r2 | 57260 | protein_coding | 24.57307645 | 0.396396606 | 0.287859209 | 1.377050287 | 0.168496701 | 0.392565467 | 0.396396606 |
| ENSMUSG00000041695 | Kcnj2 | 16518 | protein_coding | 70.1098707 | -0.39671481 | 0.216234491 | -1.834650935 | 0.066557426 | 0.222414567 | 0.39671481 |
| ENSMUSG00000023826 | Prkn | 50873 | protein_coding | 30.54496932 | -0.39675334 | 0.227344876 | -1.745160689 | 0.080956902 | 0.252239559 | 0.39675334 |
| ENSMUSG00000014329 | Bicc1 | 83675 | protein_coding | 14.94637856 | -0.396780853 | 0.354340428 | -1.119773025 | 0.262810497 | 0.504745134 | 0.396780853 |
| ENSMUSG00000070574 | 2310016G11Rik | 69578 | lncRNA | 115.8939436 | -0.396803309 | 0.185581073 | -2.138166912 | 0.032503199 | 0.137579813 | 0.396803309 |
| ENSMUSG00000025350 | Rdh5 | 19682 | protein_coding | 62.84688037 | -0.396891872 | 0.191593021 | -2.071536167 | 0.038308718 | 0.153950835 | 0.396891872 |
| ENSMUSG00000034320 | Slc26a2 | 13521 | protein_coding | 22064.16424 | -0.396973754 | 0.078258602 | -5.072589363 | 3.92E-07 | 1.04E-05 | 0.396973754 |
| ENSMUSG00000080268 | Brms1 | 107392 | protein_coding | 8.035694996 | -0.397076148 | 0.42659917 | -0.93079447 | 0.351959891 | 0.594351535 | 0.397076148 |
| ENSMUSG00000035062 | Zc4h2 | 245522 | protein_coding | 15.31116924 | 0.397283257 | 0.321042018 | 1.237480561 | 0.215908727 | 0.451733269 | 0.397283257 |
| ENSMUSG00000044726 | Erich5 | 239368 | protein_coding | 13.48565565 | -0.397472291 | 0.327760157 | -1.212692521 | 0.2252474 | 0.463289311 | 0.397472291 |
| ENSMUSG00000033227 | Wnt6 | 22420 | protein_coding | 12.4244305 | -0.397488799 | 0.354200594 | -1.122213811 | 0.261771542 | 0.50370954 | 0.397488799 |
| ENSMUSG00000031508 | Ankrd10 | 102334 | protein_coding | 1221.694514 | -0.397577629 | 0.077557113 | -5.126256186 | 2.96E-07 | 8.08E-06 | 0.397577629 |
| ENSMUSG00000055809 | Dnaaf3 | 436022 | protein_coding | 8.007643118 | 0.397601792 | 0.420781621 | 0.944912448 | 0.344703576 | 0.586953285 | 0.397601792 |
| ENSMUSG00000020728 | Cep112 | 76380 | protein_coding | 21.34360428 | 0.39776036 | 0.256897613 | 1.548322524 | 0.121544664 | 0.323030784 | 0.39776036 |
| ENSMUSG00000029403 | Cdkl2 | 53886 | protein_coding | 300.8122448 | 0.398121696 | 0.076929918 | 5.175121824 | 2.28E-07 | 6.43E-06 | 0.398121696 |
| ENSMUSG00000098912 | 1500004A13Rik | 319830 | lncRNA | 32.56900605 | -0.398216573 | 0.214908699 | -1.852956979 | 0.063888525 | 0.216756548 | 0.398216573 |
| ENSMUSG00000024232 | Bambi | 68010 | protein_coding | 155.2419305 | -0.398275484 | 0.094850542 | -4.198979502 | 2.68E-05 | 0.00046847 | 0.398275484 |

| ensembl_gene_id | external_gene_name | entrezgene_id | gene_biotype | baseMean | log2FoldChange | lfcSE | stat | pvalue | padj | absLog2FC |
|---|---|---|---|---|---|---|---|---|---|---|
| ENSMUSG00000041633 | Kctd12b | 207474 | protein_coding | 5.892908326 | 0.398472642 | 0.563701684 | 0.706885669 | 0.47963753 | 0.702043425 | 0.398472642 |
| ENSMUSG00000032338 | Hcn4 | 330953 | protein_coding | 9.081229481 | -0.398514656 | 0.41025345 | -0.971386483 | 0.331355857 | 0.574846806 | 0.398514656 |
| ENSMUSG00000027983 | Cyp2u1 | 71519 | protein_coding | 65.85290333 | -0.398865471 | 0.161052286 | -2.476620978 | 0.013263267 | 0.073403124 | 0.398865471 |
| ENSMUSG00000041046 | Ramp3 | 56089 | protein_coding | 8.412962501 | 0.398921914 | 0.46234491 | 0.862823198 | 0.388234682 | 0.628261175 | 0.398921914 |
| ENSMUSG00000003585 | Sec14l2 | 67815 | protein_coding | 77.76285255 | 0.399171746 | 0.168892674 | 2.363463949 | 0.018104988 | 0.091846223 | 0.399171746 |
| ENSMUSG00000028037 | Ifi44 | 99899 | protein_coding | 36.31167476 | 0.399188966 | 0.330348087 | 1.208388914 | 0.226897702 | 0.465544432 | 0.399188966 |
| ENSMUSG00000063889 | Crem | 12916 | protein_coding | 94.3257952 | 0.399257658 | 0.121740106 | 3.279590201 | 0.00103958 | 0.010380992 | 0.399257658 |
| ENSMUSG00000040187 | Arntl2 | 272322 | protein_coding | 16.71660754 | -0.399403686 | 0.301956149 | -1.322720822 | 0.185928236 | 0.415160239 | 0.399403686 |
| ENSMUSG00000042810 | Krba1 | 77827 | protein_coding | 548.8681159 | 0.399456863 | 0.072773182 | 5.489066882 | 4.04E-08 | 1.32E-06 | 0.399456863 |
| ENSMUSG00000092517 | Art2a | 11871 | protein_coding | 62.26123415 | -0.399776252 | 0.220371672 | -1.814100013 | 0.06966233 | 0.229227584 | 0.399776252 |
| ENSMUSG00000079470 | Utp14b | 195434 | protein_coding | 1990.931098 | -0.399937794 | 0.119590493 | -3.344227322 | 0.000825121 | 0.008579449 | 0.399937794 |
| ENSMUSG00000016664 | Pacsin2 | 23970 | protein_coding | 4122.404553 | 0.400064428 | 0.031797951 | 12.58145287 | 2.67E-36 | 4.35E-34 | 0.400064428 |
| ENSMUSG00000038422 | Hdhd3 | 72748 | protein_coding | 374.6561109 | 0.40020501 | 0.078395018 | 5.104980152 | 3.31E-07 | 8.95E-06 | 0.40020501 |
| ENSMUSG00000080440 | Gm25848 | 115488713 | snoRNA | 33.34961042 | 0.400207797 | 0.44851669 | 0.892291872 | 0.372236509 | 0.61393518 | 0.400207797 |
| ENSMUSG00000046618 | Olfml2a | 241327 | protein_coding | 10.52299385 | -0.400255972 | 0.402966089 | -0.993274578 | 0.320576163 | 0.56409512 | 0.400255972 |
| ENSMUSG00000028836 | Slc30a2 | 230810 | protein_coding | 16.11378016 | -0.400462976 | 0.460323217 | -0.869960413 | 0.384322039 | 0.624972092 | 0.400462976 |
| ENSMUSG00000065434 | Mir7-1 | 723902 | miRNA | 12.9069616 | -0.400689792 | 0.358446019 | -1.117852537 | 0.263629979 | 0.504916665 | 0.400689792 |
| ENSMUSG00000026564 | Dusp27 | 240892 | protein_coding | 6.561083811 | 0.400903483 | 0.582398001 | 0.688366859 | 0.491221788 | 0.710862177 | 0.400903483 |
| ENSMUSG00000040990 | Sh3kbp1 | 58194 | protein_coding | 299.6486917 | 0.400940096 | 0.100689065 | 3.981962647 | 6.83E-05 | 0.001073278 | 0.400940096 |
| ENSMUSG00000027513 | Pck1 | 18534 | protein_coding | 7104.358333 | -0.401009787 | 0.080965425 | -4.952852246 | 7.31E-07 | 1.85E-05 | 0.401009787 |
| ENSMUSG00000042942 | Greb1l | 381157 | protein_coding | 175.4625523 | -0.401535767 | 0.124898508 | -3.21489642 | 0.001304916 | 0.012548825 | 0.401535767 |
| ENSMUSG00000039585 | Myo9a | 270163 | protein_coding | 404.9004815 | -0.401627119 | 0.114998042 | -3.492469203 | 0.000478577 | 0.005446724 | 0.401627119 |
| ENSMUSG00000069227 | Gprin1 | 26913 | protein_coding | 22.08143026 | -0.401844146 | 0.262964808 | -1.528128986 | 0.126480516 | 0.330431117 | 0.401844146 |
| ENSMUSG00000044854 | 1700056E22Rik | 73363 | protein_coding | 19.60386806 | 0.401978241 | 0.306082318 | 1.313301086 | 0.189081513 | 0.419461348 | 0.401978241 |
| ENSMUSG00000042770 | Hebp1 | 15199 | protein_coding | 8.911452333 | -0.401987733 | 0.418484988 | -0.960578621 | 0.336764083 | 0.580296438 | 0.401987733 |
| ENSMUSG00000022270 | Retreg1 | 66270 | protein_coding | 3710.832228 | -0.402095492 | 0.06281755 | -6.401005599 | 1.54E-10 | 6.64E-09 | 0.402095492 |
| ENSMUSG00000035775 | Krt20 | 66809 | protein_coding | 11721.72146 | 0.402240332 | 0.105557233 | 3.810637314 | 0.000138609 | 0.001947167 | 0.402240332 |
| ENSMUSG00000038773 | Kdm3b | 277250 | protein_coding | 2278.516338 | 0.402366757 | 0.038935816 | 10.33410356 | 4.94E-25 | 5.38E-23 | 0.402366757 |
| ENSMUSG00000064037 | Gpn1 | 74254 | protein_coding | 338.2693494 | 0.402539865 | 0.082240731 | 4.894653313 | 9.85E-07 | 2.42E-05 | 0.402539865 |
| ENSMUSG00000045349 | Sh2d5 | 230863 | protein_coding | 6.007546803 | 0.402607351 | 0.483070711 | 0.833433578 | 0.404600244 | 0.644017631 | 0.402607351 |
| ENSMUSG00000027245 | Hypk | 67693 | protein_coding | 28.94211331 | -0.402625165 | 0.329504463 | -1.221911114 | 0.221741242 | 0.458581818 | 0.402625165 |
| ENSMUSG00000028597 | Gpx7 | 67305 | protein_coding | 26.40758836 | 0.402681177 | 0.244752128 | 1.645261188 | 0.09991596 | 0.28770764 | 0.402681177 |
| ENSMUSG00000038740 | Mvb12b | 72543 | protein_coding | 120.6055025 | -0.402760868 | 0.179484652 | -2.243985001 | 0.024833366 | 0.114196664 | 0.402760868 |
| ENSMUSG00000031530 | Dusp4 | 319520 | protein_coding | 129.8242308 | -0.402969558 | 0.134845345 | -2.98838315 | 0.002804577 | 0.022889381 | 0.402969558 |
| ENSMUSG00000006736 | Tspan31 | 67125 | protein_coding | 796.680028 | 0.403029139 | 0.051409814 | 7.839537071 | 4.52E-15 | 2.89E-13 | 0.403029139 |
| ENSMUSG00000003378 | Grik5 | 14809 | protein_coding | 42.94712152 | -0.403242526 | 0.255502214 | -1.578234961 | 0.114511641 | 0.31208616 | 0.403242526 |
| ENSMUSG00000048707 | Tprn | 97031 | protein_coding | 2576.344621 | 0.40335853 | 0.081433961 | 4.953197994 | 7.30E-07 | 1.85E-05 | 0.40335853 |
| ENSMUSG00000067813 | Xkr9 | 381246 | protein_coding | 83.80301401 | -0.403366949 | 0.188540443 | -2.139418695 | 0.032401774 | 0.137270396 | 0.403366949 |
| ENSMUSG00000026614 | Slc30a10 | 226781 | protein_coding | 881.635459 | 0.40352603 | 0.143544671 | 2.811152986 | 0.004936431 | 0.035303689 | 0.40352603 |

| ensembl_gene_id | external_gene_name | entrezgene_id | gene_biotype | baseMean | log2FoldChange | lfcSE | stat | pvalue | padj | absLog2FC |
|---|---|---|---|---|---|---|---|---|---|---|
| ENSMUSG00000086287 | Gm15972 | 100502834 | lncRNA | 33.47094995 | -0.403560413 | 0.211888688 | -1.90458687 | 0.056833795 | 0.201530037 | 0.403560413 |
| ENSMUSG00000029204 | Rhoh | 74734 | protein_coding | 16.27072534 | 0.403591372 | 0.309725371 | 1.303062033 | 0.192553584 | 0.424308967 | 0.403591372 |
| ENSMUSG00000033987 | Dnah17 | 69926 | protein_coding | 19.60385552 | -0.403840489 | 0.317163156 | -1.273289414 | 0.202915362 | 0.437021373 | 0.403840489 |
| ENSMUSG00000024873 | Cnih2 | 12794 | protein_coding | 12.11468908 | -0.403983605 | 0.396971611 | -1.017663717 | 0.308837796 | 0.552491851 | 0.403983605 |
| ENSMUSG00000030707 | Coro1a | 12721 | protein_coding | 108.402092 | 0.404201696 | 0.171812903 | 2.35256892 | 0.018644234 | 0.093749523 | 0.404201696 |
| ENSMUSG00000028024 | Enpep | 13809 | protein_coding | 1223.996532 | -0.404433172 | 0.248652188 | -1.62650156 | 0.103843002 | 0.294578028 | 0.404433172 |
| ENSMUSG00000036966 | Spryd3 | 223918 | protein_coding | 929.8311388 | 0.404991128 | 0.06464265 | 6.265076206 | 3.73E-10 | 1.54E-08 | 0.404991128 |
| ENSMUSG00000050503 | Fbxl22 | 74165 | protein_coding | 32.60404714 | -0.405005722 | 0.215667511 | -1.877917168 | 0.060392497 | 0.20889399 | 0.405005722 |
| ENSMUSG00000040189 | Odad1 | 211535 | protein_coding | 86.69263309 | 0.40523217 | 0.128989652 | 3.141586672 | 0.001680351 | 0.015275657 | 0.40523217 |
| ENSMUSG00000030472 | Ceacam18 | 72431 | protein_coding | 218.7142334 | -0.405270029 | 0.090968233 | -4.455072002 | 8.39E-06 | 0.000167595 | 0.405270029 |
| ENSMUSG00000022973 | Synj1 | 104015 | protein_coding | 618.8371136 | -0.405274515 | 0.059950919 | -6.760105092 | 1.38E-11 | 6.50E-10 | 0.405274515 |
| ENSMUSG00000049295 | Zfp219 | 69890 | protein_coding | 1139.762427 | 0.405333063 | 0.077333813 | 5.241343275 | 1.59E-07 | 4.67E-06 | 0.405333063 |
| ENSMUSG00000029054 | Gabrd | 14403 | protein_coding | 20.56040753 | -0.405539023 | 0.417598773 | -0.971121205 | 0.331487925 | 0.574895496 | 0.405539023 |
| ENSMUSG00000072571 | Tmem253 | 619301 | protein_coding | 155.5981619 | -0.405551339 | 0.102962992 | -3.938806852 | 8.19E-05 | 0.00124117 | 0.405551339 |
| ENSMUSG00000100680 | 1810044D09Rik | 69798 | lncRNA | 36.19219038 | -0.405841422 | 0.227501495 | -1.783906614 | 0.074438841 | 0.2394347 | 0.405841422 |
| ENSMUSG00000028931 | Kcnab2 | 16498 | protein_coding | 17.48888093 | 0.40604036 | 0.307109043 | 1.322137425 | 0.186122394 | 0.415388298 | 0.40604036 |
| ENSMUSG00000039246 | Lyplal1 | 226791 | protein_coding | 171.2529829 | 0.406219772 | 0.109359774 | 3.714526429 | 0.000203585 | 0.00268017 | 0.406219772 |
| ENSMUSG00000030823 | 9130019O22Rik | 78921 | protein_coding | 143.1634947 | -0.406351881 | 0.114894883 | -3.536727389 | 0.000405118 | 0.004734298 | 0.406351881 |
| ENSMUSG00000051043 | Gprc5c | 70355 | protein_coding | 24.01030221 | -0.40648321 | 0.277061467 | -1.467122855 | 0.142342638 | 0.353816233 | 0.40648321 |
| ENSMUSG00000028357 | Kif12 | 16552 | protein_coding | 435.5807576 | -0.406568742 | 0.106274963 | -3.825630535 | 0.000130438 | 0.001851734 | 0.406568742 |
| ENSMUSG00000006777 | Krt23 | 94179 | protein_coding | 195.5795993 | 0.406745999 | 0.125488845 | 3.241292078 | 0.001189892 | 0.011636985 | 0.406745999 |
| ENSMUSG00000032727 | Mier3 | 218613 | protein_coding | 2412.819582 | -0.406801966 | 0.092119643 | -4.416017627 | 1.01E-05 | 0.000197484 | 0.406801966 |
| ENSMUSG00000086370 | Ftx | 78878 | lncRNA | 98.78418288 | -0.406894424 | 0.194017714 | -2.097202442 | 0.035975658 | 0.147216301 | 0.406894424 |
| ENSMUSG00000006442 | Srm | 20810 | protein_coding | 307.8415057 | 0.406936711 | 0.092570152 | 4.395981881 | 1.10E-05 | 0.000213747 | 0.406936711 |
| ENSMUSG00000021610 | Clptm1l | 218335 | protein_coding | 3309.156269 | 0.407258442 | 0.056281335 | 7.236119095 | 4.62E-13 | 2.46E-11 | 0.407258442 |
| ENSMUSG00000030036 | Mogs | 57377 | protein_coding | 1933.427647 | 0.407259571 | 0.079283491 | 5.136751252 | 2.80E-07 | 7.72E-06 | 0.407259571 |
| ENSMUSG00000052396 | Mogat2 | 233549 | protein_coding | 124.2810644 | 0.407387758 | 0.165336987 | 2.463984411 | 0.013740209 | 0.075318948 | 0.407387758 |
| ENSMUSG00000024810 | Il33 | 77125 | protein_coding | 15.29815262 | -0.40748639 | 0.368858158 | -1.10472381 | 0.269279283 | 0.511744874 | 0.40748639 |
| ENSMUSG00000117655 | 6720468P15Rik | 78073 | lncRNA | 21.50336632 | 0.407508941 | 0.270988061 | 1.503789279 | 0.132635633 | 0.339503983 | 0.407508941 |
| ENSMUSG00000039396 | Neil3 | 234258 | protein_coding | 91.42451458 | -0.407566315 | 0.185109816 | -2.201754209 | 0.027682675 | 0.122794781 | 0.407566315 |
| ENSMUSG00000115518 | Gm10791 | 100038614 | lncRNA | 14.98428494 | -0.40757302 | 0.309309574 | -1.317686405 | 0.187608647 | 0.417346835 | 0.40757302 |
| ENSMUSG00000041670 | Rims1 | 116837 | protein_coding | 7.925321025 | 0.407772191 | 0.457734441 | 0.890848829 | 0.373010275 | 0.614708431 | 0.407772191 |
| ENSMUSG00000078496 | Zfp982 | 195531 | protein_coding | 8.550701913 | -0.408383774 | 0.417881671 | -0.977271323 | 0.328434846 | 0.572028827 | 0.408383774 |
| ENSMUSG00000039145 | Camk1d | 227541 | protein_coding | 2628.196672 | -0.408659763 | 0.078218563 | -5.224588002 | 1.75E-07 | 5.04E-06 | 0.408659763 |
| ENSMUSG00000039364 | Sectm1b | 58210 | protein_coding | 2290.521889 | -0.408726306 | 0.134629581 | -3.035932395 | 0.002397931 | 0.020162376 | 0.408726306 |
| ENSMUSG00000028635 | Edn2 | 13615 | protein_coding | 513.1446583 | -0.408891674 | 0.161349675 | -2.534195828 | 0.011270573 | 0.064744392 | 0.408891674 |
| ENSMUSG00000045969 | Ing1 | 26356 | protein_coding | 705.0958776 | 0.408988183 | 0.049154384 | 8.320482353 | 8.76E-17 | 6.17E-15 | 0.408988183 |
| ENSMUSG00000108030 | 9530062K07Rik | 78739 | lncRNA | 8.655056622 | 0.409058265 | 0.449643545 | 0.90973899 | 0.362960176 | 0.605769124 | 0.409058265 |
| ENSMUSG00000030579 | Tyrobp | 22177 | protein_coding | 16.37778486 | 0.409113816 | 0.362384364 | 1.128949967 | 0.258918941 | 0.500729794 | 0.409113816 |

| ensembl_gene_id | external_gene_name | entrezgene_id | gene_biotype | baseMean | log2FoldChange | lfcSE | stat | pvalue | padj | absLog2FC |
|---|---|---|---|---|---|---|---|---|---|---|
| ENSMUSG00000031668 | Eif2ak3 | 13666 | protein_coding | 1154.062063 | 0.409258965 | 0.046619816 | 8.778648149 | 1.65E-18 | 1.30E-16 | 0.409258965 |
| ENSMUSG00000097855 | A930007I19Rik | 77779 | lncRNA | 12.66546799 | 0.409273327 | 0.383927008 | 1.066018588 | 0.286415239 | 0.529013894 | 0.409273327 |
| ENSMUSG00000025190 | Got1 | 14718 | protein_coding | 1412.342167 | 0.409313754 | 0.075050883 | 5.453816663 | 4.93E-08 | 1.60E-06 | 0.409313754 |
| ENSMUSG00000030589 | Rasgrp4 | 233046 | protein_coding | 13.51946525 | -0.409567598 | 0.322303269 | -1.270752233 | 0.203816807 | 0.437979805 | 0.409567598 |
| ENSMUSG00000026601 | Axdnd1 | 77352 | protein_coding | 5.895027325 | 0.409571576 | 0.527676758 | 0.776178919 | 0.437643351 | 0.670771892 | 0.409571576 |
| ENSMUSG00000006546 | Cryba2 | 12958 | protein_coding | 6.880963049 | -0.409728407 | 0.473512361 | -0.865296117 | 0.386876281 | 0.626924617 | 0.409728407 |
| ENSMUSG00000028671 | Gale | 74246 | protein_coding | 5946.302226 | 0.409747869 | 0.097370734 | 4.208121391 | 2.58E-05 | 0.000453356 | 0.409747869 |
| ENSMUSG00000037594 | Clba1 | 217887 | protein_coding | 46.6685703 | 0.409755911 | 0.178675612 | 2.293295127 | 0.02183102 | 0.104657804 | 0.409755911 |
| ENSMUSG00000038642 | Ctss | 13040 | protein_coding | 4501.965122 | -0.409927723 | 0.07207982 | -5.6871358 | 1.29E-08 | 4.57E-07 | 0.409927723 |
| ENSMUSG00000035545 | Leng8 | 232798 | protein_coding | 1149.456316 | -0.410056463 | 0.113027138 | -3.627946966 | 0.000285684 | 0.003538479 | 0.410056463 |
| ENSMUSG00000020435 | Osbp2 | 74309 | protein_coding | 7.665496238 | -0.410236303 | 0.427141837 | -0.960421731 | 0.336843006 | 0.580370422 | 0.410236303 |
| ENSMUSG00000074183 | Gsta1 | 14857 | protein_coding | 129.5659582 | 0.410246301 | 0.127198949 | 3.225233418 | 0.0012587 | 0.012198951 | 0.410246301 |
| ENSMUSG00000090290 | Tarbp1 | 212728 | protein_coding | 400.2758773 | -0.410639869 | 0.109443954 | -3.752056234 | 0.00017539 | 0.002351219 | 0.410639869 |
| ENSMUSG00000032690 | Oas2 | 246728 | protein_coding | 35.02865165 | -0.41065108 | 0.25433308 | -1.614619221 | 0.10639318 | 0.298608218 | 0.41065108 |
| ENSMUSG00000031502 | Col4a1 | 12826 | protein_coding | 33.7670135 | 0.410756956 | 0.320471681 | 1.281726218 | 0.199938704 | 0.433436829 | 0.410756956 |
| ENSMUSG00000039084 | Chad | 12643 | protein_coding | 32.34791518 | -0.411074602 | 0.242098349 | -1.697965321 | 0.089514307 | 0.268926459 | 0.411074602 |
| ENSMUSG00000024687 | Osbp | 76303 | protein_coding | 2794.264171 | 0.41124626 | 0.038608656 | 10.65165962 | 1.71E-26 | 1.95E-24 | 0.41124626 |
| ENSMUSG00000028976 | Slc2a5 | 56485 | protein_coding | 112.350709 | -0.411380131 | 0.185488628 | -2.217818614 | 0.026567202 | 0.119146072 | 0.411380131 |
| ENSMUSG00000074818 | Pdzd7 | 100503041 | protein_coding | 167.3290092 | -0.411444046 | 0.119199449 | -3.451727741 | 0.000557009 | 0.006217975 | 0.411444046 |
| ENSMUSG00000002980 | Bcam | 57278 | protein_coding | 97.62422621 | -0.411510067 | 0.122785912 | -3.35144366 | 0.000803914 | 0.008402281 | 0.411510067 |
| ENSMUSG00000087590 | Epb41l4aos | 69749 | lncRNA | 249.9570465 | -0.411568823 | 0.086845953 | -4.739067388 | 2.15E-06 | 4.89E-05 | 0.411568823 |
| ENSMUSG00000039055 | Eme1 | 268465 | protein_coding | 70.22573437 | 0.411582593 | 0.171130933 | 2.405074204 | 0.016169182 | 0.084790715 | 0.411582593 |
| ENSMUSG00000054146 | Krt15 | 16665 | protein_coding | 12.86842565 | -0.41171488 | 0.496378274 | -0.829437752 | 0.406856747 | 0.645614361 | 0.41171488 |
| ENSMUSG00000074024 | 4632427E13Rik | 666737 | lncRNA | 169.4576785 | -0.411916779 | 0.207408072 | -1.986020968 | 0.047030996 | 0.177668978 | 0.411916779 |
| ENSMUSG00000102428 | Pcdhga12 | 93724 | protein_coding | 6.741804936 | -0.412433899 | 0.482046675 | -0.855589136 | 0.392225081 | 0.63203576 | 0.412433899 |
| ENSMUSG00000027660 | Skil | 20482 | protein_coding | 1752.113781 | 0.412718107 | 0.074966054 | 5.505399923 | 3.68E-08 | 1.22E-06 | 0.412718107 |
| ENSMUSG00000046447 | Camk2n1 | 66259 | protein_coding | 1091.634256 | 0.412973528 | 0.094283758 | 4.380113156 | 1.19E-05 | 0.000228003 | 0.412973528 |
| ENSMUSG00000020102 | Slc16a7 | 20503 | protein_coding | 111.6062445 | -0.413118649 | 0.149839424 | -2.757075805 | 0.005832083 | 0.040194018 | 0.413118649 |
| ENSMUSG00000062110 | Scfd2 | 212986 | protein_coding | 355.1077421 | 0.413297125 | 0.082258254 | 5.024384851 | 5.05E-07 | 1.32E-05 | 0.413297125 |
| ENSMUSG00000062093 | Gm10110 | 328451 | transcribed_processed_pseudogene | 10.06045575 | -0.413436894 | 0.380823879 | -1.085638049 | 0.277639159 | 0.52063799 | 0.413436894 |
| ENSMUSG00000048489 | Depp1 | 213393 | protein_coding | 134.4406756 | -0.41372766 | 0.212238934 | -1.949348555 | 0.051253814 | 0.188241916 | 0.41372766 |
| ENSMUSG00000046589 | Lrrc8e | 72267 | protein_coding | 435.1480738 | -0.413868643 | 0.078711425 | -5.258050428 | 1.46E-07 | 4.32E-06 | 0.413868643 |
| ENSMUSG00000086503 | Xist | 213742 | lncRNA | 5865.464348 | -0.414075775 | 0.117571711 | -3.521899718 | 0.000428466 | 0.00496399 | 0.414075775 |
| ENSMUSG00000012889 | Podnl1 | 244550 | protein_coding | 6.193452899 | -0.414137239 | 0.51018251 | -0.811743308 | 0.416938938 | 0.654847512 | 0.414137239 |
| ENSMUSG00000019055 | Plod1 | 18822 | protein_coding | 234.2804898 | 0.414196689 | 0.130197916 | 3.181285085 | 0.001466233 | 0.013723699 | 0.414196689 |
| ENSMUSG00000079554 | Aox2 | 213043 | protein_coding | 6.604149197 | -0.41444636 | 0.46334062 | -0.89447448 | 0.371068078 | 0.613223192 | 0.41444636 |
| ENSMUSG00000026692 | Fmo4 | 226564 | protein_coding | 97.19138788 | -0.414465149 | 0.145001007 | -2.858360489 | 0.004258363 | 0.031691104 | 0.414465149 |
| ENSMUSG00000005677 | Nr1i3 | 12355 | protein_coding | 25.11118837 | -0.414485348 | 0.273329884 | -1.516428949 | 0.129410924 | 0.335208796 | 0.414485348 |
| ENSMUSG00000051675 | Trim32 | 69807 | protein_coding | 414.5266016 | 0.414787574 | 0.072661045 | 5.708527482 | 1.14E-08 | 4.05E-07 | 0.414787574 |

| ensembl_gene_id | external_gene_name | entrezgene_id | gene_biotype | baseMean | log2FoldChange | lfcSE | stat | pvalue | padj | absLog2FC |
|---|---|---|---|---|---|---|---|---|---|---|
| ENSMUSG00000029167 | Ppargc1a | 19017 | protein_coding | 1041.093351 | -0.414887577 | 0.07813958 | -5.309570066 | 1.10E-07 | 3.34E-06 | 0.414887577 |
| ENSMUSG00000090891 | D6Ertd527e | 52372 | lncRNA | 7.931634829 | -0.414904948 | 0.477079161 | -0.869677365 | 0.384476746 | 0.624972092 | 0.414904948 |
| ENSMUSG00000080059 | Rps19-ps3 | 277692 | processed_pseudogene | 16.06364565 | -0.415112007 | 0.3261878 | -1.272616594 | 0.203154127 | 0.437185296 | 0.415112007 |
| ENSMUSG00000039985 | Sinhcaf | 56306 | protein_coding | 290.8819012 | -0.415136699 | 0.114857387 | -3.614366551 | 0.000301083 | 0.003688746 | 0.415136699 |
| ENSMUSG00000047793 | Sned1 | 208777 | protein_coding | 432.6462232 | -0.4152284 | 0.09869623 | -4.207135375 | 2.59E-05 | 0.000454338 | 0.4152284 |
| ENSMUSG00000028344 | Invs | 16348 | protein_coding | 102.1653068 | 0.415247656 | 0.133135717 | 3.118980124 | 0.001814782 | 0.016259439 | 0.415247656 |
| ENSMUSG00000032280 | Tle3 | 21887 | protein_coding | 1182.561777 | -0.415253321 | 0.124298842 | -3.340765808 | 0.000835477 | 0.008670355 | 0.415253321 |
| ENSMUSG00000028044 | Cks1b | 54124 | protein_coding | 155.9268964 | 0.415254129 | 0.151693653 | 2.737452227 | 0.006191711 | 0.041961584 | 0.415254129 |
| ENSMUSG00000039620 | Trmt9b | 319582 | protein_coding | 39.93988408 | -0.415300402 | 0.218779712 | -1.898258286 | 0.057662066 | 0.20352728 | 0.415300402 |
| ENSMUSG00000028631 | Kcnq4 | 60613 | protein_coding | 10.60079713 | -0.415418645 | 0.376449452 | -1.10351773 | 0.269802396 | 0.512256092 | 0.415418645 |
| ENSMUSG00000087478 | 4930506C21Rik | 75060 | lncRNA | 9.056892336 | -0.415492728 | 0.471776585 | -0.88069807 | 0.378481261 | 0.619709567 | 0.415492728 |
| ENSMUSG00000033538 | Casp4 | 12363 | protein_coding | 410.3395417 | -0.41567636 | 0.09114551 | -4.560579685 | 5.10E-06 | 0.000107822 | 0.41567636 |
| ENSMUSG00000051166 | Eml5 | 319670 | protein_coding | 35.64854193 | -0.415711653 | 0.222863939 | -1.865315915 | 0.062137136 | 0.212757025 | 0.415711653 |
| ENSMUSG00000038930 | Rccd1 | 269955 | protein_coding | 131.3727395 | -0.415957965 | 0.137573572 | -3.023531039 | 0.002498434 | 0.02077991 | 0.415957965 |
| ENSMUSG00000056832 | Ttc26 | 264134 | protein_coding | 35.46160532 | 0.416127815 | 0.213680567 | 1.947429381 | 0.051483278 | 0.188869616 | 0.416127815 |
| ENSMUSG00000021733 | Slc4a7 | 218756 | protein_coding | 5183.967678 | 0.416207243 | 0.055136336 | 7.548692502 | 4.40E-14 | 2.55E-12 | 0.416207243 |
| ENSMUSG00000066800 | Rnasel | 24014 | protein_coding | 9653.834141 | -0.416233427 | 0.078402832 | -5.308908064 | 1.10E-07 | 3.34E-06 | 0.416233427 |
| ENSMUSG00000026482 | Rgl1 | 19731 | protein_coding | 13.69653023 | 0.416440068 | 0.349151714 | 1.19271953 | 0.232979234 | 0.472500517 | 0.416440068 |
| ENSMUSG00000026839 | Upp2 | 76654 | protein_coding | 78.40533309 | 0.416566907 | 0.173440231 | 2.401789389 | 0.016315099 | 0.085315693 | 0.416566907 |
| ENSMUSG00000030795 | Fus | 233908 | protein_coding | 636.7526715 | -0.416772926 | 0.088367171 | -4.716377379 | 2.40E-06 | 5.40E-05 | 0.416772926 |
| ENSMUSG00000043505 | Gimap5 | 317757 | protein_coding | 20.35031555 | 0.416869881 | 0.303793917 | 1.372212732 | 0.169997223 | 0.394410188 | 0.416869881 |
| ENSMUSG00000017316 | Ppy | 19064 | protein_coding | 21.63100479 | -0.416982961 | 0.343571601 | -1.213671211 | 0.224873302 | 0.4629812 | 0.416982961 |
| ENSMUSG00000027792 | Bche | 12038 | protein_coding | 1979.478184 | -0.416986176 | 0.122471726 | -3.404754621 | 0.000662235 | 0.007209695 | 0.416986176 |
| ENSMUSG00000027820 | Mme | 17380 | protein_coding | 113.6039377 | -0.417790627 | 0.284188379 | -1.470118619 | 0.141529631 | 0.352392829 | 0.417790627 |
| ENSMUSG00000026573 | Xcl1 | 16963 | protein_coding | 26.63942219 | 0.418183448 | 0.309663714 | 1.350443818 | 0.176873664 | 0.40378736 | 0.418183448 |
| ENSMUSG00000021747 | Cfap20dc | 74430 | protein_coding | 31.63901274 | 0.418540957 | 0.222837038 | 1.878237839 | 0.060348636 | 0.208850312 | 0.418540957 |
| ENSMUSG00000027490 | E2f1 | 13555 | protein_coding | 105.0118874 | 0.419305369 | 0.198455096 | 2.112847577 | 0.034613816 | 0.143574334 | 0.419305369 |
| ENSMUSG00000055862 | Izumo4 | 71564 | protein_coding | 189.1476547 | -0.419319492 | 0.113229783 | -3.703261477 | 0.000212845 | 0.002772663 | 0.419319492 |
| ENSMUSG00000002885 | Adgre5 | 26364 | protein_coding | 630.6773611 | 0.419329585 | 0.06824358 | 6.144601198 | 8.02E-10 | 3.22E-08 | 0.419329585 |
| ENSMUSG00000032175 | Tyk2 | 54721 | protein_coding | 1576.615202 | 0.419339504 | 0.048462138 | 8.652930396 | 5.02E-18 | 3.75E-16 | 0.419339504 |
| ENSMUSG00000027962 | Vcam1 | 22329 | protein_coding | 7.443589973 | -0.419366293 | 0.477137091 | -0.878922015 | 0.379443561 | 0.620491412 | 0.419366293 |
| ENSMUSG00000034593 | Myo5a | 17918 | protein_coding | 36.77181585 | 0.419440653 | 0.264005948 | 1.588754561 | 0.112115816 | 0.308074929 | 0.419440653 |
| ENSMUSG00000106951 | 5930430L01Rik | 319982 | lncRNA | 28.25777247 | -0.419464477 | 0.255281093 | -1.643147451 | 0.100352429 | 0.288636281 | 0.419464477 |
| ENSMUSG00000056476 | Med12l | 329650 | protein_coding | 51.9695598 | 0.419481986 | 0.219811167 | 1.908374322 | 0.056342853 | 0.200523946 | 0.419481986 |
| ENSMUSG00000045382 | Cxcr4 | 12767 | protein_coding | 29.9786344 | 0.419532491 | 0.291696108 | 1.438251932 | 0.150362585 | 0.365794955 | 0.419532491 |
| ENSMUSG00000097767 | Miat | 330166 | lncRNA | 6.816124847 | -0.419836108 | 0.520703574 | -0.806286204 | 0.420077843 | 0.657152071 | 0.419836108 |
| ENSMUSG00000029706 | Pax4 | 18506 | protein_coding | 14.98748466 | -0.420081828 | 0.317399738 | -1.323510318 | 0.185665727 | 0.414943346 | 0.420081828 |
| ENSMUSG00000034275 | Igsf9b | 235086 | protein_coding | 312.5901149 | 0.42021769 | 0.136283666 | 3.083404645 | 0.002046467 | 0.017878316 | 0.42021769 |
| ENSMUSG00000017724 | Etv4 | 18612 | protein_coding | 311.8856038 | 0.420257887 | 0.161996467 | 2.594241073 | 0.009479999 | 0.057217045 | 0.420257887 |

| ensembl_gene_id | external_gene_name | entrezgene_id | gene_biotype | baseMean | log2FoldChange | lfcSE | stat | pvalue | padj | absLog2FC |
|---|---|---|---|---|---|---|---|---|---|---|
| ENSMUSG00000112778 | Gm33677 | 102636674 | lncRNA | 12.30567322 | -0.420258735 | 0.371374169 | -1.131631572 | 0.257789361 | 0.499743842 | 0.420258735 |
| ENSMUSG00000026228 | Htr2b | 15559 | protein_coding | 6.473560711 | -0.420329989 | 0.741895106 | -0.566562558 | 0.571011419 | 0.765987453 | 0.420329989 |
| ENSMUSG00000069919 | Hba-a1 | 15122 | protein_coding | 19.6697463 | 0.420518492 | 0.441462644 | 0.952557362 | 0.340814389 | 0.583658666 | 0.420518492 |
| ENSMUSG00000038387 | Rras | 20130 | protein_coding | 82.46404699 | -0.42061959 | 0.148842651 | -2.825934547 | 0.00471429 | 0.034123585 | 0.42061959 |
| ENSMUSG00000069581 | Tspear | 252974 | protein_coding | 10.57537334 | -0.420657567 | 0.421584896 | -0.997800374 | 0.318376169 | 0.562186847 | 0.420657567 |
| ENSMUSG00000078919 | Dpm1 | 13480 | protein_coding | 72.92705896 | -0.420740308 | 0.211220791 | -1.991945519 | 0.046377036 | 0.175769357 | 0.420740308 |
| ENSMUSG00000040282 | Cdin1 | 399568 | protein_coding | 86.69587685 | 0.421120581 | 0.152622705 | 2.759226291 | 0.00579384 | 0.040015956 | 0.421120581 |
| ENSMUSG00000110080 | Gm6145 | 620376 | lncRNA | 7.901130086 | 0.421163896 | 0.477122712 | 0.882716093 | 0.377389685 | 0.61867166 | 0.421163896 |
| ENSMUSG00000031939 | Taf1d | 75316 | protein_coding | 1927.258032 | -0.421388032 | 0.114820903 | -3.669959214 | 0.000242589 | 0.003096681 | 0.421388032 |
| ENSMUSG00000053317 | Sec61b | 66212 | protein_coding | 1802.132085 | 0.421482422 | 0.107175386 | 3.932641996 | 8.40E-05 | 0.001271058 | 0.421482422 |
| ENSMUSG00000003813 | Rad23a | 19358 | protein_coding | 734.1607574 | -0.421720212 | 0.096125877 | -4.387166326 | 1.15E-05 | 0.000221794 | 0.421720212 |
| ENSMUSG00000063765 | Chadl | 214685 | protein_coding | 7.247288291 | -0.421819596 | 0.481187995 | -0.876621195 | 0.380692423 | 0.621615656 | 0.421819596 |
| ENSMUSG00000032308 | Ulk3 | 71742 | protein_coding | 152.4082236 | -0.422108981 | 0.118469647 | -3.563013758 | 0.000366621 | 0.004378048 | 0.422108981 |
| ENSMUSG00000019122 | Ccl9 | 20308 | protein_coding | 421.0379334 | 0.422127024 | 0.106279498 | 3.971857516 | 7.13E-05 | 0.001110122 | 0.422127024 |
| ENSMUSG00000024659 | Anxa1 | 16952 | protein_coding | 2876.779371 | 0.422185488 | 0.107454327 | 3.928976129 | 8.53E-05 | 0.001286966 | 0.422185488 |
| ENSMUSG00000099689 | Zfp383 | 73729 | protein_coding | 39.11406322 | 0.422354557 | 0.21017496 | 2.009537941 | 0.044480115 | 0.170996619 | 0.422354557 |
| ENSMUSG00000040374 | Pex2 | 19302 | protein_coding | 1209.620508 | 0.422519221 | 0.048590728 | 8.695470099 | 3.45E-18 | 2.64E-16 | 0.422519221 |
| ENSMUSG00000001039 | B9d1 | 27078 | protein_coding | 63.22246345 | 0.422604942 | 0.170230883 | 2.482539793 | 0.01304495 | 0.072418556 | 0.422604942 |
| ENSMUSG00000022204 | Ngdn | 68966 | protein_coding | 788.6510091 | 0.423124825 | 0.053008067 | 7.982272228 | 1.44E-15 | 9.38E-14 | 0.423124825 |
| ENSMUSG00000030346 | Rad51ap1 | 19362 | protein_coding | 93.57528293 | 0.423153885 | 0.186896777 | 2.264104771 | 0.023567672 | 0.110048838 | 0.423153885 |
| ENSMUSG00000046806 | Cyren | 78412 | protein_coding | 37.69287353 | 0.423311773 | 0.232879448 | 1.817729203 | 0.069105532 | 0.228065548 | 0.423311773 |
| ENSMUSG00000003847 | Nfat5 | 54446 | protein_coding | 610.7404064 | -0.423322846 | 0.106929827 | -3.958884593 | 7.53E-05 | 0.001163192 | 0.423322846 |
| ENSMUSG00000110344 | Smim36 | 121811722 | protein_coding | 31.55298442 | -0.423416449 | 0.210411231 | -2.012328175 | 0.044185354 | 0.170432243 | 0.423416449 |
| ENSMUSG00000096992 | Gm26788 | 108167617 | lncRNA | 21.7786853 | -0.423615033 | 0.323840144 | -1.308099201 | 0.190839664 | 0.421829944 | 0.423615033 |
| ENSMUSG00000069540 | Gm4925 | 237433 | transcribed_processed_pseudogene | 9.779571398 | 0.424099065 | 0.36560791 | 1.159983287 | 0.246055611 | 0.487067489 | 0.424099065 |
| ENSMUSG00000068855 | H2ac20 | 319176 | protein_coding | 352.9620572 | 0.424169567 | 0.107792885 | 3.935042361 | 8.32E-05 | 0.001259602 | 0.424169567 |
| ENSMUSG00000035164 | Zc3h12c | 244871 | protein_coding | 41.4282265 | 0.424207144 | 0.197340186 | 2.14962372 | 0.031584991 | 0.134825607 | 0.424207144 |
| ENSMUSG00000087231 | E230016M11Rik | 320172 | lncRNA | 21.85987683 | -0.424285283 | 0.343210073 | -1.236226196 | 0.216374496 | 0.452297019 | 0.424285283 |
| ENSMUSG00000030699 | Tbx6 | 21389 | protein_coding | 12.37657267 | -0.424292766 | 0.392079161 | -1.082160971 | 0.279181007 | 0.522078519 | 0.424292766 |
| ENSMUSG00000049608 | Gpr55 | 227326 | protein_coding | 27.87965284 | 0.424418056 | 0.283012977 | 1.49964168 | 0.133707245 | 0.34107542 | 0.424418056 |
| ENSMUSG00000003526 | Prodh | 19125 | protein_coding | 506.6296521 | -0.424550477 | 0.098385047 | -4.31519313 | 1.59E-05 | 0.000297303 | 0.424550477 |
| ENSMUSG00000025986 | Slc39a10 | 227059 | protein_coding | 201.8576314 | 0.42461798 | 0.124482458 | 3.411066806 | 0.000647092 | 0.007079822 | 0.42461798 |
| ENSMUSG00000026782 | Abi2 | 329165 | protein_coding | 114.0488304 | 0.424737988 | 0.121262102 | 3.50264411 | 0.000460665 | 0.005283881 | 0.424737988 |
| ENSMUSG00000052560 | Cpne8 | 66871 | protein_coding | 416.7648414 | -0.424938705 | 0.087188265 | -4.873806153 | 1.09E-06 | 2.66E-05 | 0.424938705 |
| ENSMUSG00000112925 | Gm31763 | 102634101 | lncRNA | 7.084057169 | -0.425099179 | 0.568960172 | -0.747151031 | 0.4549724 | 0.682988492 | 0.425099179 |
| ENSMUSG00000020261 | Slc36a1 | 215335 | protein_coding | 1675.947451 | -0.425149152 | 0.150337351 | -2.827967567 | 0.004684455 | 0.033968619 | 0.425149152 |
| ENSMUSG00000068227 | Il2rb | 16185 | protein_coding | 88.54210165 | 0.425311402 | 0.137870652 | 3.084858118 | 0.002036493 | 0.017810477 | 0.425311402 |
| ENSMUSG00000023951 | Vegfa | 22339 | protein_coding | 953.1432815 | -0.425468302 | 0.091248561 | -4.662739858 | 3.12E-06 | 6.87E-05 | 0.425468302 |
| ENSMUSG00000038224 | Serpinf2 | 18816 | protein_coding | 22.58042065 | 0.425517796 | 0.284697856 | 1.494629436 | 0.135011179 | 0.342958039 | 0.425517796 |

| ensembl_gene_id | external_gene_name | entrezgene_id | gene_biotype | baseMean | log2FoldChange | lfcSE | stat | pvalue | padj | absLog2FC |
|---|---|---|---|---|---|---|---|---|---|---|
| ENSMUSG00000095789 | Nupr1l | 69034 | protein_coding | 13.32070009 | -0.425571286 | 0.346433752 | -1.228434826 | 0.219283781 | 0.455544189 | 0.425571286 |
| ENSMUSG00000021190 | Lgmn | 19141 | protein_coding | 618.3714793 | -0.42560005 | 0.087922282 | -4.840639231 | 1.29E-06 | 3.09E-05 | 0.42560005 |
| ENSMUSG00000027006 | Dnajc10 | 66861 | protein_coding | 3055.39065 | 0.425769789 | 0.067827714 | 6.277224518 | 3.45E-10 | 1.43E-08 | 0.425769789 |
| ENSMUSG00000036833 | Pnpla7 | 241274 | protein_coding | 302.4830105 | -0.425921878 | 0.085967331 | -4.954462047 | 7.25E-07 | 1.84E-05 | 0.425921878 |
| ENSMUSG00000024978 | Gpam | 14732 | protein_coding | 1418.209002 | -0.425954956 | 0.090126863 | -4.726170883 | 2.29E-06 | 5.17E-05 | 0.425954956 |
| ENSMUSG00000054523 | Ms4a5 | 269063 | protein_coding | 10.39754564 | -0.426021976 | 0.403756088 | -1.055146879 | 0.291358155 | 0.533764962 | 0.426021976 |
| ENSMUSG00000077274 | Gm22786 | 115485822 | snoRNA | 11.34636466 | 0.426403866 | 0.396632435 | 1.075060504 | 0.282347641 | 0.525132096 | 0.426403866 |
| ENSMUSG00000030543 | Mesp2 | 17293 | protein_coding | 8.146498411 | 0.4265031 | 0.451844077 | 0.943916546 | 0.345212296 | 0.587571366 | 0.4265031 |
| ENSMUSG00000022332 | Khdrbs3 | 13992 | protein_coding | 633.7532019 | 0.42658384 | 0.091528299 | 4.660677018 | 3.15E-06 | 6.92E-05 | 0.42658384 |
| ENSMUSG00000117042 | 2700054A10Rik | 72578 | lncRNA | 13.78316662 | -0.426605973 | 0.314918972 | -1.354653134 | 0.175528096 | 0.401865646 | 0.426605973 |
| ENSMUSG00000017167 | Cntnap1 | 53321 | protein_coding | 149.7718855 | -0.426685216 | 0.114779254 | -3.717441964 | 0.00020125 | 0.002651838 | 0.426685216 |
| ENSMUSG00000003809 | Gcdh | 270076 | protein_coding | 1070.717941 | 0.427010277 | 0.061158921 | 6.981978651 | 2.91E-12 | 1.47E-10 | 0.427010277 |
| ENSMUSG00000031504 | Rab20 | 19332 | protein_coding | 503.9149484 | 0.42702585 | 0.075880874 | 5.627582154 | 1.83E-08 | 6.31E-07 | 0.42702585 |
| ENSMUSG00000071723 | Gspt2 | 14853 | protein_coding | 57.05912523 | 0.427143679 | 0.189423335 | 2.254968637 | 0.024135297 | 0.112008616 | 0.427143679 |
| ENSMUSG00000067231 | Cyp2c65 | 72303 | protein_coding | 4597.316468 | -0.427194064 | 0.084090354 | -5.080179182 | 3.77E-07 | 1.01E-05 | 0.427194064 |
| ENSMUSG00000044177 | Wfikkn2 | 278507 | protein_coding | 43.24733459 | 0.427553822 | 0.183285374 | 2.332721988 | 0.019662741 | 0.096883908 | 0.427553822 |
| ENSMUSG00000028553 | Angptl3 | 30924 | protein_coding | 7.801104058 | 0.42763009 | 0.551919973 | 0.77480452 | 0.438455178 | 0.671251818 | 0.42763009 |
| ENSMUSG00000020326 | Ccng1 | 12450 | protein_coding | 3387.846373 | 0.427645846 | 0.046584329 | 9.180036557 | 4.31E-20 | 3.58E-18 | 0.427645846 |
| ENSMUSG00000028159 | Dapp1 | 26377 | protein_coding | 291.7438811 | -0.427764667 | 0.15044603 | -2.843309775 | 0.004464765 | 0.032713889 | 0.427764667 |
| ENSMUSG00000019802 | Sec63 | 140740 | protein_coding | 1720.305985 | 0.428012865 | 0.047392475 | 9.031241139 | 1.70E-19 | 1.40E-17 | 0.428012865 |
| ENSMUSG00000058046 | 4933430I17Rik | 214106 | protein_coding | 6.523081572 | -0.428141311 | 0.47060824 | -0.909761613 | 0.362948243 | 0.605769124 | 0.428141311 |
| ENSMUSG00000001999 | Blvra | 109778 | protein_coding | 377.8860517 | 0.428413822 | 0.077068927 | 5.558839854 | 2.72E-08 | 9.22E-07 | 0.428413822 |
| ENSMUSG00000038271 | Iffo1 | 320678 | protein_coding | 18.54436329 | -0.4285303 | 0.316325151 | -1.354714596 | 0.175508506 | 0.401865646 | 0.4285303 |
| ENSMUSG00000022657 | Cd96 | 84544 | protein_coding | 29.43426614 | 0.428713155 | 0.268938997 | 1.5940907 | 0.110915722 | 0.306028716 | 0.428713155 |
| ENSMUSG00000025144 | Cenpx | 20892 | protein_coding | 470.7358393 | 0.428780353 | 0.073647409 | 5.822069744 | 5.81E-09 | 2.13E-07 | 0.428780353 |
| ENSMUSG00000006800 | Sulf2 | 72043 | protein_coding | 241.0951353 | -0.42886031 | 0.159357581 | -2.691182345 | 0.007119927 | 0.046411907 | 0.42886031 |
| ENSMUSG00000043424 | Eif3j2 | 100042807 | protein_coding | 6.389834951 | -0.428866145 | 0.465484567 | -0.921332683 | 0.356876762 | 0.599765689 | 0.428866145 |
| ENSMUSG00000002289 | Angptl4 | 57875 | protein_coding | 39.88751085 | -0.428910897 | 0.227084238 | -1.888774412 | 0.058922063 | 0.205917277 | 0.428910897 |
| ENSMUSG00000039607 | Rbms3 | 207181 | protein_coding | 79.72851227 | -0.428982719 | 0.17804893 | -2.409352969 | 0.015980834 | 0.083921494 | 0.428982719 |
| ENSMUSG00000020914 | Top2a | 21973 | protein_coding | 3741.77367 | 0.428992527 | 0.125029449 | 3.43113188 | 0.000601068 | 0.006643885 | 0.428992527 |
| ENSMUSG00000032014 | Oaf | 102644 | protein_coding | 355.1342276 | -0.429254664 | 0.118748378 | -3.614825488 | 0.00030055 | 0.003685909 | 0.429254664 |
| ENSMUSG00000026941 | Mamdc4 | 381352 | protein_coding | 84.38987332 | -0.4294887 | 0.187163003 | -2.294730762 | 0.021748559 | 0.104486659 | 0.4294887 |
| ENSMUSG00000032946 | Rasgrp2 | 19395 | protein_coding | 9.604431977 | 0.429532194 | 0.424162195 | 1.01266025 | 0.311222471 | 0.555097626 | 0.429532194 |
| ENSMUSG00000057400 | Ces1c | 13884 | protein_coding | 8.027832709 | 0.429647106 | 0.560275296 | 0.766849991 | 0.44317071 | 0.674226502 | 0.429647106 |
| ENSMUSG00000039234 | Sec24d | 69608 | protein_coding | 1438.999108 | 0.42985717 | 0.087616474 | 4.906122652 | 9.29E-07 | 2.29E-05 | 0.42985717 |
| ENSMUSG00000000690 | Hoxb6 | 15414 | protein_coding | 459.9435643 | -0.429935483 | 0.133332602 | -3.224533804 | 0.00126178 | 0.012221453 | 0.429935483 |
| ENSMUSG00000025212 | Sfxn3 | 94280 | protein_coding | 351.6666319 | -0.430283315 | 0.115935559 | -3.711400687 | 0.000206116 | 0.002709067 | 0.430283315 |
| ENSMUSG00000003992 | Ssbp2 | 66970 | protein_coding | 80.04368149 | -0.430686817 | 0.15036862 | -2.86420675 | 0.00418055 | 0.031212841 | 0.430686817 |
| ENSMUSG00000023328 | Ache | 11423 | protein_coding | 46.59156496 | -0.430782192 | 0.20590151 | -2.092175972 | 0.036422777 | 0.148466815 | 0.430782192 |

| ensembl_gene_id | external_gene_name | entrezgene_id | gene_biotype | baseMean | log2FoldChange | lfcSE | stat | pvalue | padj | absLog2FC |
|---|---|---|---|---|---|---|---|---|---|---|
| ENSMUSG00000029471 | Camkk2 | 207565 | protein_coding | 76.336045 | -0.430822411 | 0.162104272 | -2.657686961 | 0.007867894 | 0.0501028 | 0.430822411 |
| ENSMUSG00000003865 | Gys1 | 14936 | protein_coding | 693.0646259 | 0.431024605 | 0.066753677 | 6.456941718 | 1.07E-10 | 4.68E-09 | 0.431024605 |
| ENSMUSG00000014873 | Surf2 | 20931 | protein_coding | 269.3797548 | -0.43109601 | 0.095113506 | -4.53243737 | 5.83E-06 | 0.000121645 | 0.43109601 |
| ENSMUSG00000025912 | Mybl1 | 17864 | protein_coding | 39.7079783 | 0.431178027 | 0.219560818 | 1.963820465 | 0.049550916 | 0.184123872 | 0.431178027 |
| ENSMUSG00000075702 | Selenom | 114679 | protein_coding | 1389.751007 | 0.431283279 | 0.081962673 | 5.261947438 | 1.43E-07 | 4.23E-06 | 0.431283279 |
| ENSMUSG00000050921 | P2ry10 | 78826 | protein_coding | 38.33096607 | 0.431327561 | 0.197960175 | 2.178860278 | 0.029342049 | 0.128165492 | 0.431327561 |
| ENSMUSG00000026885 | Ttll11 | 74410 | protein_coding | 25.98612036 | -0.431355667 | 0.250432043 | -1.722445983 | 0.08498876 | 0.260078507 | 0.431355667 |
| ENSMUSG00000098973 | Mir6236 | 102465959 | miRNA | 14567.10105 | 0.431491452 | 0.283853387 | 1.52012085 | 0.128480605 | 0.333871046 | 0.431491452 |
| ENSMUSG00000074170 | Plekhf1 | 72287 | protein_coding | 42.43646018 | 0.431505303 | 0.20222288 | 2.133810488 | 0.032858297 | 0.138573681 | 0.431505303 |
| ENSMUSG00000030934 | Oat | 18242 | protein_coding | 7114.218936 | 0.431619577 | 0.056549155 | 7.632644096 | 2.30E-14 | 1.37E-12 | 0.431619577 |
| ENSMUSG00000038628 | Polr3k | 67005 | protein_coding | 819.7331098 | 0.431700789 | 0.059882975 | 7.209073807 | 5.63E-13 | 2.98E-11 | 0.431700789 |
| ENSMUSG00000023931 | Efhb | 211482 | protein_coding | 21.81166348 | 0.431860398 | 0.267285082 | 1.61572952 | 0.106152806 | 0.298245186 | 0.431860398 |
| ENSMUSG00000056648 | Hoxb8 | 15416 | protein_coding | 95.92933257 | -0.432098672 | 0.131299394 | -3.290941862 | 0.000998525 | 0.010029817 | 0.432098672 |
| ENSMUSG00000040253 | Gbp7 | 229900 | protein_coding | 173.7306102 | -0.432393739 | 0.367592634 | -1.176285103 | 0.239480967 | 0.480032716 | 0.432393739 |
| ENSMUSG00000037722 | Gnpnat1 | 54342 | protein_coding | 129.2239823 | -0.432499618 | 0.156196198 | -2.768950989 | 0.005623709 | 0.039041568 | 0.432499618 |
| ENSMUSG00000087107 | AI662270 | 100043636 | lncRNA | 7.437766974 | 0.432572461 | 0.491878935 | 0.879428718 | 0.379168867 | 0.620356733 | 0.432572461 |
| ENSMUSG00000021087 | Rtn1 | 104001 | protein_coding | 31.45178792 | 0.432698197 | 0.379035892 | 1.141575787 | 0.253630393 | 0.494893811 | 0.432698197 |
| ENSMUSG00000059395 | Nkapl | 66707 | protein_coding | 8.99667188 | -0.432793103 | 0.426081692 | -1.015751467 | 0.309747753 | 0.553189591 | 0.432793103 |
| ENSMUSG00000029134 | Plb1 | 665270 | protein_coding | 18.88693474 | 0.432828156 | 0.6142947 | 0.704593668 | 0.48106314 | 0.703236518 | 0.432828156 |
| ENSMUSG00000045411 | 2410002F23Rik | 668661 | protein_coding | 616.8131567 | -0.43293499 | 0.111186405 | -3.893776298 | 9.87E-05 | 0.001467526 | 0.43293499 |
| ENSMUSG00000009013 | Dynll1 | 56455 | protein_coding | 1732.814981 | 0.432961762 | 0.084086482 | 5.149005531 | 2.62E-07 | 7.28E-06 | 0.432961762 |
| ENSMUSG00000069922 | Ces3a | 382053 | protein_coding | 9.495757498 | 0.433563779 | 0.484211472 | 0.895401707 | 0.370572388 | 0.612568768 | 0.433563779 |
| ENSMUSG00000058589 | Anks1b | 77531 | protein_coding | 7.179910108 | 0.43370069 | 0.489832456 | 0.885406194 | 0.375937593 | 0.617324668 | 0.43370069 |
| ENSMUSG00000038352 | Arl5c | 217151 | protein_coding | 23.61895584 | 0.434526859 | 0.25741307 | 1.688052823 | 0.091401089 | 0.272408741 | 0.434526859 |
| ENSMUSG00000039908 | Slc26a11 | 268512 | protein_coding | 136.1965343 | -0.434843814 | 0.127896416 | -3.39996871 | 0.000673936 | 0.007313971 | 0.434843814 |
| ENSMUSG00000030598 | Fbxo17 | 50760 | protein_coding | 24.65646983 | -0.435065165 | 0.238187978 | -1.826562228 | 0.067765617 | 0.225377625 | 0.435065165 |
| ENSMUSG00000066512 | Klk1b5 | 16622 | protein_coding | 23.62092172 | 0.43508624 | 0.28695169 | 1.516235157 | 0.129459901 | 0.335281809 | 0.43508624 |
| ENSMUSG00000001334 | Fndc5 | 384061 | protein_coding | 279.4183687 | 0.435091605 | 0.107555434 | 4.04527776 | 5.23E-05 | 0.000849612 | 0.435091605 |
| ENSMUSG00000058569 | Tmed9 | 67511 | protein_coding | 2014.105284 | 0.435808745 | 0.061794317 | 7.052569948 | 1.76E-12 | 9.05E-11 | 0.435808745 |
| ENSMUSG00000040550 | Otud6b | 72201 | protein_coding | 805.9571125 | 0.435871534 | 0.057001026 | 7.646731404 | 2.06E-14 | 1.24E-12 | 0.435871534 |
| ENSMUSG00000023259 | Slc26a6 | 171429 | protein_coding | 108.4547703 | 0.435949929 | 0.229997418 | 1.895455756 | 0.058032048 | 0.203984926 | 0.435949929 |
| ENSMUSG00000028464 | Tpm2 | 22004 | protein_coding | 13.89738619 | -0.436056586 | 0.341074292 | -1.278479781 | 0.201080309 | 0.43481123 | 0.436056586 |
| ENSMUSG00000094410 | Zbed6 | 667118 | protein_coding | 101.1149575 | -0.436103317 | 0.145765815 | -2.991807904 | 0.002773307 | 0.022657104 | 0.436103317 |
| ENSMUSG00000042328 | Hps4 | 192232 | protein_coding | 682.4496181 | 0.436168041 | 0.068868819 | 6.333316727 | 2.40E-10 | 1.02E-08 | 0.436168041 |
| ENSMUSG00000022861 | Dgkg | 110197 | protein_coding | 186.3615815 | -0.43659293 | 0.131391133 | -3.322849256 | 0.000891031 | 0.009146817 | 0.43659293 |
| ENSMUSG00000065232 | Gm22973 | 115488085 | snRNA | 103.3195266 | 0.436712166 | 0.32464712 | 1.345190328 | 0.178563779 | 0.405715197 | 0.436712166 |
| ENSMUSG00000023039 | Krt7 | 110310 | protein_coding | 3266.296674 | 0.436762333 | 0.050471535 | 8.653636872 | 4.99E-18 | 3.75E-16 | 0.436762333 |
| ENSMUSG00000017723 | Wfdc2 | 67701 | protein_coding | 3388.076818 | -0.436870988 | 0.110747756 | -3.944738977 | 7.99E-05 | 0.001219282 | 0.436870988 |
| ENSMUSG00000009394 | Syn2 | 20965 | protein_coding | 124.7169956 | -0.437051926 | 0.162038082 | -2.697217346 | 0.006992162 | 0.045795479 | 0.437051926 |

| ensembl_gene_id | external_gene_name | entrezgene_id | gene_biotype | baseMean | log2FoldChange | lfcSE | stat | pvalue | padj | absLog2FC |
|---|---|---|---|---|---|---|---|---|---|---|
| ENSMUSG00000028456 | Unc13b | 22249 | protein_coding | 1644.659814 | 0.437135504 | 0.060555757 | 7.218727374 | 5.25E-13 | 2.78E-11 | 0.437135504 |
| ENSMUSG00000021066 | Atl1 | 73991 | protein_coding | 41.5082832 | -0.437374125 | 0.203874075 | -2.145315075 | 0.031927664 | 0.135892699 | 0.437374125 |
| ENSMUSG00000034220 | Gpc1 | 14733 | protein_coding | 79.57330973 | -0.437428256 | 0.169292192 | -2.583865505 | 0.009769988 | 0.058377401 | 0.437428256 |
| ENSMUSG00000031342 | Gpm6b | 14758 | protein_coding | 14.43566953 | -0.437531855 | 0.399505555 | -1.095183407 | 0.273436294 | 0.515936247 | 0.437531855 |
| ENSMUSG00000001225 | Slc26a3 | 13487 | protein_coding | 12488.32639 | -0.437831788 | 0.079952176 | -5.476170999 | 4.35E-08 | 1.41E-06 | 0.437831788 |
| ENSMUSG00000030309 | Caprin2 | 232560 | protein_coding | 125.4001275 | -0.437877641 | 0.141850748 | -3.086889887 | 0.002022625 | 0.017698792 | 0.437877641 |
| ENSMUSG00000025352 | Gdf11 | 14561 | protein_coding | 15.81021376 | -0.438129054 | 0.299641807 | -1.462175982 | 0.143692992 | 0.355746355 | 0.438129054 |
| ENSMUSG00000031355 | Arhgap6 | 11856 | protein_coding | 52.49466659 | 0.438274963 | 0.176334456 | 2.485475461 | 0.012937851 | 0.071923033 | 0.438274963 |
| ENSMUSG00000087368 | BC065397 | 436230 | lncRNA | 19.40987451 | -0.438520896 | 0.298530424 | -1.468932012 | 0.141851231 | 0.352866698 | 0.438520896 |
| ENSMUSG00000050982 | Apol10a | 245282 | protein_coding | 4036.018303 | -0.438780743 | 0.208674818 | -2.102700972 | 0.035491917 | 0.145828827 | 0.438780743 |
| ENSMUSG00000071192 | Wfikkn1 | 215001 | protein_coding | 24.84346805 | -0.43913127 | 0.319729378 | -1.373446733 | 0.16961351 | 0.39391111 | 0.43913127 |
| ENSMUSG00000037032 | Apbb1 | 11785 | protein_coding | 21.69765108 | 0.439192895 | 0.271483876 | 1.617749465 | 0.105716601 | 0.29784969 | 0.439192895 |
| ENSMUSG00000004562 | Arhgef40 | 268739 | protein_coding | 419.267208 | 0.439259351 | 0.080619959 | 5.448518674 | 5.08E-08 | 1.64E-06 | 0.439259351 |
| ENSMUSG00000029161 | Cgref1 | 68567 | protein_coding | 1728.656296 | 0.439269439 | 0.059554581 | 7.375913501 | 1.63E-13 | 9.05E-12 | 0.439269439 |
| ENSMUSG00000097276 | 4930525G20Rik | 320163 | lncRNA | 11.85818644 | 0.439439747 | 0.373742605 | 1.175781784 | 0.239682087 | 0.480227733 | 0.439439747 |
| ENSMUSG00000031669 | Gins3 | 78833 | protein_coding | 60.25590342 | 0.439520438 | 0.197017594 | 2.230868972 | 0.02568981 | 0.116930286 | 0.439520438 |
| ENSMUSG00000051452 | Gm11437 | 628813 | protein_coding | 179.0447221 | -0.439546812 | 0.142117525 | -3.092840327 | 0.001982508 | 0.01741389 | 0.439546812 |
| ENSMUSG00000053178 | Mterf1b | 208595 | protein_coding | 17.4232804 | 0.439578546 | 0.282261903 | 1.557342811 | 0.119389116 | 0.319566519 | 0.439578546 |
| ENSMUSG00000053746 | Ptrh1 | 329384 | protein_coding | 256.8413082 | 0.439816577 | 0.119618465 | 3.676828469 | 0.000236152 | 0.003024955 | 0.439816577 |
| ENSMUSG00000040563 | Plppr2 | 235044 | protein_coding | 15.72388743 | -0.439989199 | 0.326658909 | -1.346937697 | 0.178000299 | 0.405224566 | 0.439989199 |
| ENSMUSG00000038304 | Cd160 | 54215 | protein_coding | 52.62118863 | 0.440045849 | 0.207817712 | 2.117460748 | 0.034220763 | 0.142420193 | 0.440045849 |
| ENSMUSG00000021411 | Pxdc1 | 66895 | protein_coding | 49.39485646 | 0.440124438 | 0.188599864 | 2.333641337 | 0.01961451 | 0.096764517 | 0.440124438 |
| ENSMUSG00000053199 | Arhgap20 | 244867 | protein_coding | 889.7501568 | -0.440165725 | 0.103005614 | -4.273220745 | 1.93E-05 | 0.000349806 | 0.440165725 |
| ENSMUSG00000022211 | Carmil3 | 268747 | protein_coding | 9.33591298 | -0.440317272 | 0.387982043 | -1.134890854 | 0.256421053 | 0.498229195 | 0.440317272 |
| ENSMUSG00000030872 | Gga2 | 74105 | protein_coding | 207.6443146 | 0.440720234 | 0.101752167 | 4.331310557 | 1.48E-05 | 0.000277955 | 0.440720234 |
| ENSMUSG00000072974 | Gm4787 | 214321 | protein_coding | 12.90439475 | -0.440805045 | 0.342442597 | -1.287237769 | 0.198011426 | 0.430948755 | 0.440805045 |
| ENSMUSG00000023050 | Map3k12 | 26404 | protein_coding | 45.4471612 | -0.440848666 | 0.240052958 | -1.836464208 | 0.066289031 | 0.22202351 | 0.440848666 |
| ENSMUSG00000020226 | Slc5a4b | 64454 | protein_coding | 16.44332933 | 0.440976494 | 0.497230506 | 0.886865324 | 0.375151413 | 0.616661588 | 0.440976494 |
| ENSMUSG00000021298 | Gpr132 | 56696 | protein_coding | 9.289239934 | 0.441075613 | 0.449572422 | 0.981100245 | 0.32654331 | 0.57003615 | 0.441075613 |
| ENSMUSG00000069302 | H2ac12 | 319168 | protein_coding | 75.24618833 | 0.44172674 | 0.213463813 | 2.069328444 | 0.038515276 | 0.154396187 | 0.44172674 |
| ENSMUSG00000000028 | Cdc45 | 12544 | protein_coding | 170.4441486 | 0.442011734 | 0.191624159 | 2.30665975 | 0.021073796 | 0.101895312 | 0.442011734 |
| ENSMUSG00000042607 | Asb4 | 65255 | protein_coding | 43.95349625 | -0.442014921 | 0.226588005 | -1.950742812 | 0.051087648 | 0.187888369 | 0.442014921 |
| ENSMUSG00000047117 | Ankdd1b | 271144 | protein_coding | 57.44919088 | 0.442118214 | 0.189987903 | 2.327086128 | 0.019960682 | 0.097992668 | 0.442118214 |
| ENSMUSG00000063434 | Sorcs3 | 66673 | protein_coding | 19.83006522 | -0.442148657 | 0.255694074 | -1.729209638 | 0.083771582 | 0.257723064 | 0.442148657 |
| ENSMUSG00000022822 | Abcc5 | 27416 | protein_coding | 202.1072319 | -0.442157325 | 0.167012641 | -2.647448261 | 0.008110177 | 0.051170901 | 0.442157325 |
| ENSMUSG00000029814 | Igf2bp3 | 140488 | protein_coding | 14.3995252 | 0.44251195 | 0.32599222 | 1.357431014 | 0.174644299 | 0.40092091 | 0.44251195 |
| ENSMUSG00000061479 | Snrpa | 53607 | protein_coding | 709.4273085 | 0.442550476 | 0.0892292 | 4.959704604 | 7.06E-07 | 1.80E-05 | 0.442550476 |
| ENSMUSG00000038312 | Edem2 | 108687 | protein_coding | 1288.639447 | 0.442778296 | 0.060808908 | 7.281470963 | 3.30E-13 | 1.78E-11 | 0.442778296 |
| ENSMUSG00000004347 | Pde1c | 18575 | protein_coding | 24.12957848 | 0.442831424 | 0.271518459 | 1.63094408 | 0.102902119 | 0.292836153 | 0.442831424 |

| ensembl_gene_id | external_gene_name | entrezgene_id | gene_biotype | baseMean | log2FoldChange | lfcSE | stat | pvalue | padj | absLog2FC |
|---|---|---|---|---|---|---|---|---|---|---|
| ENSMUSG00000048521 | Cxcr6 | 80901 | protein_coding | 22.58700471 | 0.443081161 | 0.27642818 | 1.602879858 | 0.108961181 | 0.302290895 | 0.443081161 |
| ENSMUSG00000055200 | Sertad3 | 170742 | protein_coding | 407.7976443 | 0.443123576 | 0.083692581 | 5.294657779 | 1.19E-07 | 3.59E-06 | 0.443123576 |
| ENSMUSG00000028078 | Dclk2 | 70762 | protein_coding | 54.45774579 | 0.443231287 | 0.171661529 | 2.582007094 | 0.009822757 | 0.05858417 | 0.443231287 |
| ENSMUSG00000050567 | Maml1 | 103806 | protein_coding | 416.7854673 | -0.443430672 | 0.08908412 | -4.977662377 | 6.44E-07 | 1.65E-05 | 0.443430672 |
| ENSMUSG00000067925 | Rtl8a | 66158 | protein_coding | 22.18539387 | 0.443554195 | 0.244303176 | 1.815589144 | 0.06943342 | 0.228801137 | 0.443554195 |
| ENSMUSG00000047394 | Odf3b | 70113 | protein_coding | 122.9927361 | -0.443642078 | 0.171044272 | -2.593726598 | 0.009494195 | 0.057278974 | 0.443642078 |
| ENSMUSG00000052305 | Hbb-bs | 15129 | protein_coding | 97.78522145 | 0.443885017 | 0.497812459 | 0.891671168 | 0.372569212 | 0.61438574 | 0.443885017 |
| ENSMUSG00000036908 | Unc93b1 | 54445 | protein_coding | 662.9699632 | -0.444041235 | 0.092325587 | -4.809514319 | 1.51E-06 | 3.55E-05 | 0.444041235 |
| ENSMUSG00000090564 | A430057M04Rik | 319486 | lncRNA | 11.95691691 | -0.444097271 | 0.336843008 | -1.318410241 | 0.187366354 | 0.417125511 | 0.444097271 |
| ENSMUSG00000060090 | Rp2 | 19889 | protein_coding | 1528.72384 | 0.444286358 | 0.072214553 | 6.152310613 | 7.64E-10 | 3.08E-08 | 0.444286358 |
| ENSMUSG00000087440 | 4930577N17Rik | 67746 | TEC | 23.12334211 | -0.444515483 | 0.248000919 | -1.792394499 | 0.073069795 | 0.236530829 | 0.444515483 |
| ENSMUSG00000025185 | Loxl4 | 67573 | protein_coding | 6.15023009 | 0.444661581 | 0.539751444 | 0.823826572 | 0.410038107 | 0.64854129 | 0.444661581 |
| ENSMUSG00000052572 | Dlg2 | 23859 | protein_coding | 13.070464 | 0.444707649 | 0.324961917 | 1.368491587 | 0.171158254 | 0.395794258 | 0.444707649 |
| ENSMUSG00000097325 | Gm16897 | 320400 | lncRNA | 7.441152453 | 0.44477366 | 0.423822593 | 1.049433578 | 0.293978611 | 0.536918806 | 0.44477366 |
| ENSMUSG00000085355 | 3010003L21Rik | 109163 | lncRNA | 94.13711115 | -0.444959042 | 0.171024054 | -2.601733685 | 0.009275385 | 0.056413087 | 0.444959042 |
| ENSMUSG00000032020 | Ubash3b | 72828 | protein_coding | 52.11592741 | 0.445070692 | 0.185117695 | 2.404257963 | 0.016205333 | 0.084879315 | 0.445070692 |
| ENSMUSG00000030556 | Lrrc28 | 67867 | protein_coding | 163.9262523 | 0.445471694 | 0.118923097 | 3.745880351 | 0.000179762 | 0.002397871 | 0.445471694 |
| ENSMUSG00000009545 | Kcnq1 | 16535 | protein_coding | 1953.776534 | -0.445671048 | 0.065747629 | -6.778511281 | 1.21E-11 | 5.81E-10 | 0.445671048 |
| ENSMUSG00000063972 | Nr6a1 | 14536 | protein_coding | 223.9051819 | -0.445760338 | 0.11615674 | -3.837576166 | 0.000124255 | 0.001774894 | 0.445760338 |
| ENSMUSG00000041632 | Mrps27 | 218506 | protein_coding | 512.5367646 | 0.445863333 | 0.071826825 | 6.207476563 | 5.38E-10 | 2.19E-08 | 0.445863333 |
| ENSMUSG00000031364 | Grpr | 14829 | protein_coding | 65.22657467 | 0.445875771 | 0.171352205 | 2.602101164 | 0.009265452 | 0.056386392 | 0.445875771 |
| ENSMUSG00000026664 | Phyh | 16922 | protein_coding | 3802.669065 | 0.445988645 | 0.049766476 | 8.961627924 | 3.20E-19 | 2.58E-17 | 0.445988645 |
| ENSMUSG00000053111 | Fank1 | 66930 | protein_coding | 12.32617132 | -0.446301913 | 0.331664921 | -1.345640991 | 0.178418325 | 0.405619009 | 0.446301913 |
| ENSMUSG00000007038 | Neu1 | 18010 | protein_coding | 4661.185824 | -0.446332812 | 0.108421784 | -4.1166341 | 3.84E-05 | 0.000645829 | 0.446332812 |
| ENSMUSG00000044788 | Fads6 | 328035 | protein_coding | 16.97903208 | -0.446357627 | 0.331200911 | -1.347694443 | 0.17775668 | 0.404955782 | 0.446357627 |
| ENSMUSG00000026723 | Trdmt1 | 13434 | protein_coding | 115.6292649 | -0.446992154 | 0.13393697 | -3.337332137 | 0.000845868 | 0.00876128 | 0.446992154 |
| ENSMUSG00000028608 | Czib | 74098 | protein_coding | 500.5880372 | 0.447534571 | 0.064435355 | 6.945481537 | 3.77E-12 | 1.86E-10 | 0.447534571 |
| ENSMUSG00000051238 | Swsap1 | 66962 | protein_coding | 247.3596992 | 0.447591253 | 0.081674545 | 5.48018054 | 4.25E-08 | 1.38E-06 | 0.447591253 |
| ENSMUSG00000077565 | Gm23201 | 115486177 | snoRNA | 8.35858609 | -0.447798159 | 0.449141648 | -0.997008763 | 0.318760256 | 0.562495531 | 0.447798159 |
| ENSMUSG00000062995 | Ica1 | 15893 | protein_coding | 850.5296503 | 0.44784988 | 0.072640283 | 6.165310222 | 7.03E-10 | 2.84E-08 | 0.44784988 |
| ENSMUSG00000020576 | Nbas | 71169 | protein_coding | 1496.99354 | 0.447910195 | 0.071837894 | 6.235012869 | 4.52E-10 | 1.85E-08 | 0.447910195 |
| ENSMUSG00000002732 | Fkbp7 | 14231 | protein_coding | 33.06604734 | 0.44799406 | 0.209114816 | 2.142335338 | 0.032166507 | 0.136656813 | 0.44799406 |
| ENSMUSG00000032006 | Pdgfd | 71785 | protein_coding | 71.6693471 | -0.448141554 | 0.18076289 | -2.479167901 | 0.01316893 | 0.072981214 | 0.448141554 |
| ENSMUSG00000066513 | Klk1b4 | 18048 | protein_coding | 16.55111384 | -0.449350645 | 0.294466549 | -1.525981973 | 0.127014354 | 0.33118197 | 0.449350645 |
| ENSMUSG00000118339 | 4930592I03Rik | 75853 | lncRNA | 6.294702708 | 0.449712076 | 0.495556693 | 0.907488652 | 0.364148448 | 0.606340896 | 0.449712076 |
| ENSMUSG00000037016 | Frem2 | 242022 | protein_coding | 46.78397065 | 0.449917421 | 0.177661822 | 2.532437284 | 0.011327263 | 0.064962579 | 0.449917421 |
| ENSMUSG00000034675 | Dbn1 | 56320 | protein_coding | 32.9440353 | 0.450270378 | 0.233739795 | 1.926374489 | 0.05405763 | 0.1947258 | 0.450270378 |
| ENSMUSG00000048763 | Hoxb3 | 15410 | protein_coding | 100.9052814 | -0.45031309 | 0.142652642 | -3.156710487 | 0.001595597 | 0.014737795 | 0.45031309 |
| ENSMUSG00000071506 | Tmem139 | 109218 | protein_coding | 651.4054637 | 0.450880647 | 0.069839475 | 6.455956961 | 1.08E-10 | 4.70E-09 | 0.450880647 |

Supplementary Table 1

| ensembl_gene_id | external_gene_name | entrezgene_id | gene_biotype | baseMean | log2FoldChange | lfcSE | stat | pvalue | padj | absLog2FC |
|---|---|---|---|---|---|---|---|---|---|---|
| ENSMUSG00000070953 | Rabepk | 227746 | protein_coding | 525.7999654 | 0.451518594 | 0.088548341 | 5.099119751 | 3.41E-07 | 9.20E-06 | 0.451518594 |
| ENSMUSG00000063445 | Nmral1 | 67824 | protein_coding | 99.11451741 | 0.451976854 | 0.189270776 | 2.38799071 | 0.01694077 | 0.087561384 | 0.451976854 |
| ENSMUSG00000030223 | Ptpro | 19277 | protein_coding | 67.04959841 | 0.452051171 | 0.15992228 | 2.826692877 | 0.004703141 | 0.034058175 | 0.452051171 |
| ENSMUSG00000032062 | 2310030G06Rik | 66952 | protein_coding | 381.4797102 | 0.4520681 | 0.064385554 | 7.021266 | 2.20E-12 | 1.13E-10 | 0.4520681 |
| ENSMUSG00000054892 | Txk | 22165 | protein_coding | 16.80266262 | 0.452147986 | 0.307411019 | 1.470825565 | 0.141338297 | 0.352354239 | 0.452147986 |
| ENSMUSG00000029009 | Mthfr | 17769 | protein_coding | 527.9476683 | -0.452305874 | 0.088476717 | -5.112145792 | 3.19E-07 | 8.63E-06 | 0.452305874 |
| ENSMUSG00000029406 | Pitpnm2 | 19679 | protein_coding | 78.95021553 | -0.452530303 | 0.148751256 | -3.042194835 | 0.002348598 | 0.019840672 | 0.452530303 |
| ENSMUSG00000030220 | Arhgdib | 11857 | protein_coding | 36.29321563 | 0.452652285 | 0.286895138 | 1.577762134 | 0.114620266 | 0.312196844 | 0.452652285 |
| ENSMUSG00000025787 | Tgm4 | 331046 | protein_coding | 11.39364656 | -0.453051697 | 0.364641314 | -1.242458494 | 0.214067445 | 0.449773548 | 0.453051697 |
| ENSMUSG00000026238 | Ptma | 19231 | protein_coding | 9417.614364 | 0.45311205 | 0.067614933 | 6.70136058 | 2.06E-11 | 9.60E-10 | 0.45311205 |
| ENSMUSG00000039704 | Lmbrd2 | 320506 | protein_coding | 1567.349639 | 0.453162185 | 0.064712999 | 7.002645389 | 2.51E-12 | 1.28E-10 | 0.453162185 |
| ENSMUSG00000027499 | Pkia | 18767 | protein_coding | 165.3114616 | 0.45324354 | 0.1041947 | 4.349967331 | 1.36E-05 | 0.000258331 | 0.45324354 |
| ENSMUSG00000002109 | Ddb2 | 107986 | protein_coding | 156.1364392 | -0.453306693 | 0.138757923 | -3.266888713 | 0.001087364 | 0.010812233 | 0.453306693 |
| ENSMUSG00000028982 | Slc25a33 | 70556 | protein_coding | 128.0666263 | -0.453905387 | 0.133763403 | -3.393345093 | 0.000690446 | 0.007453027 | 0.453905387 |
| ENSMUSG00000053399 | Adamts18 | 208936 | protein_coding | 25.19154282 | 0.454089944 | 0.4123758 | 1.101155655 | 0.270828918 | 0.513373374 | 0.454089944 |
| ENSMUSG00000031552 | Adam18 | 13524 | protein_coding | 9.290661961 | -0.454937693 | 0.429030586 | -1.060385221 | 0.288969383 | 0.531357026 | 0.454937693 |
| ENSMUSG00000020335 | Zfp354b | 27274 | protein_coding | 6.249112711 | -0.454977432 | 0.476872902 | -0.954085313 | 0.34004046 | 0.583162735 | 0.454977432 |
| ENSMUSG00000035505 | Cox18 | 231430 | protein_coding | 149.6219188 | 0.455110801 | 0.120478334 | 3.777532335 | 0.00015839 | 0.002161701 | 0.455110801 |
| ENSMUSG00000025650 | Col7a1 | 12836 | protein_coding | 31.60431162 | 0.455306717 | 0.276322992 | 1.647733736 | 0.099407322 | 0.286840423 | 0.455306717 |
| ENSMUSG00000026773 | Pfkfb3 | 170768 | protein_coding | 48.01745898 | 0.455374646 | 0.200881629 | 2.266880497 | 0.023397527 | 0.10953031 | 0.455374646 |
| ENSMUSG00000035692 | Isg15 | 100038882 | protein_coding | 77.16976561 | 0.455391428 | 0.272364102 | 1.67199504 | 0.0945253 | 0.27822769 | 0.455391428 |
| ENSMUSG00000028071 | Sh2d2a | 27371 | protein_coding | 20.69461096 | 0.4554032 | 0.288075315 | 1.580847704 | 0.113912864 | 0.311104616 | 0.4554032 |
| ENSMUSG00000075602 | Ly6a | 110454 | protein_coding | 905.1699943 | -0.455431398 | 0.380826851 | -1.195901487 | 0.231735005 | 0.4708568 | 0.455431398 |
| ENSMUSG00000032557 | Uba5 | 66663 | protein_coding | 771.5384361 | 0.455742859 | 0.088902443 | 5.126325464 | 2.95E-07 | 8.08E-06 | 0.455742859 |
| ENSMUSG00000042129 | Rassf4 | 213391 | protein_coding | 262.3061417 | -0.455870908 | 0.115321062 | -3.953058517 | 7.72E-05 | 0.001186212 | 0.455870908 |
| ENSMUSG00000032011 | Thy1 | 21838 | protein_coding | 15.05994854 | 0.456083798 | 0.382244403 | 1.193173255 | 0.232801527 | 0.472309752 | 0.456083798 |
| ENSMUSG00000030263 | Irag2 | 16970 | protein_coding | 618.7687993 | 0.456177388 | 0.12094718 | 3.771707533 | 0.000162134 | 0.002199107 | 0.456177388 |
| ENSMUSG00000039286 | Fndc3b | 72007 | protein_coding | 950.4486737 | 0.456494633 | 0.062194791 | 7.33975669 | 2.14E-13 | 1.17E-11 | 0.456494633 |
| ENSMUSG00000058715 | Fcer1g | 14127 | protein_coding | 15.51595017 | 0.456705905 | 0.369103656 | 1.237337799 | 0.215961701 | 0.451769221 | 0.456705905 |
| ENSMUSG00000091086 | Rpl6l | 432502 | processed_pseudogene | 6.95750593 | -0.457151567 | 0.462452529 | -0.988537282 | 0.322889585 | 0.566904861 | 0.457151567 |
| ENSMUSG00000030322 | Mbd4 | 17193 | protein_coding | 55.06094131 | 0.457267868 | 0.184907308 | 2.472957254 | 0.013400017 | 0.074032913 | 0.457267868 |
| ENSMUSG00000029463 | Fam216a | 68948 | protein_coding | 20.8085222 | 0.457470115 | 0.31427199 | 1.455650295 | 0.145489317 | 0.358198217 | 0.457470115 |
| ENSMUSG00000033029 | 1700088E04Rik | 27660 | protein_coding | 17.5260735 | 0.45762153 | 0.296220791 | 1.544866342 | 0.122378596 | 0.324285182 | 0.45762153 |
| ENSMUSG00000045752 | Tssc4 | 56844 | protein_coding | 388.9334613 | 0.457655797 | 0.083018676 | 5.51268484 | 3.53E-08 | 1.18E-06 | 0.457655797 |
| ENSMUSG00000035486 | Plk5 | 216166 | protein_coding | 5.886893579 | -0.457703732 | 0.541195963 | -0.845726433 | 0.397705374 | 0.637110813 | 0.457703732 |
| ENSMUSG00000056290 | Ms4a4b | 60361 | protein_coding | 15.53503893 | 0.457967524 | 0.301724063 | 1.517835597 | 0.129055849 | 0.334611523 | 0.457967524 |
| ENSMUSG00000025321 | Itgb8 | 320910 | protein_coding | 36.40832208 | 0.458084298 | 0.318569016 | 1.437943663 | 0.15045004 | 0.365903753 | 0.458084298 |
| ENSMUSG00000085385 | Snhg17 | 68108 | lncRNA | 209.5484856 | -0.458092313 | 0.162196139 | -2.824310835 | 0.004738241 | 0.034222527 | 0.458092313 |
| ENSMUSG00000030718 | Ppme1 | 72590 | protein_coding | 943.4645121 | 0.45836928 | 0.065572226 | 6.990296135 | 2.74E-12 | 1.39E-10 | 0.45836928 |

| ensembl_gene_id | external_gene_name | entrezgene_id | gene_biotype | baseMean | log2FoldChange | lfcSE | stat | pvalue | padj | absLog2FC |
|---|---|---|---|---|---|---|---|---|---|---|
| ENSMUSG00000027863 | Cd2 | 12481 | protein_coding | 8.934589135 | 0.458423949 | 0.384781444 | 1.19138788 | 0.233501349 | 0.472837299 | 0.458423949 |
| ENSMUSG00000093865 | Lrit3 | 242235 | protein_coding | 10.01263751 | 0.458605489 | 0.419207366 | 1.093982421 | 0.273962688 | 0.516506461 | 0.458605489 |
| ENSMUSG00000025810 | Nrp1 | 18186 | protein_coding | 113.2687768 | 0.459916643 | 0.142045485 | 3.237812466 | 0.0012045 | 0.011764114 | 0.459916643 |
| ENSMUSG00000073158 | 9030624G23Rik | 66808 | protein_coding | 7.350398062 | -0.460138348 | 0.448819671 | -1.025218763 | 0.305259977 | 0.549639002 | 0.460138348 |
| ENSMUSG00000027889 | Ampd2 | 109674 | protein_coding | 709.6255549 | 0.460211159 | 0.066683068 | 6.901469476 | 5.15E-12 | 2.52E-10 | 0.460211159 |
| ENSMUSG00000022844 | Pdia5 | 72599 | protein_coding | 1451.161031 | 0.460504098 | 0.069754172 | 6.601814407 | 4.06E-11 | 1.84E-09 | 0.460504098 |
| ENSMUSG00000074340 | Ovgp1 | 12659 | protein_coding | 6.843896481 | -0.460661874 | 0.584153095 | -0.788597849 | 0.430347088 | 0.665280626 | 0.460661874 |
| ENSMUSG00000065273 | Gm25128 | 115488831 | snoRNA | 7.125072057 | -0.460984581 | 0.478807659 | -0.96277612 | 0.335659887 | 0.579221524 | 0.460984581 |
| ENSMUSG00000046718 | Bst2 | 69550 | protein_coding | 155.8373124 | 0.461238858 | 0.198553341 | 2.322997212 | 0.020179302 | 0.098855288 | 0.461238858 |
| ENSMUSG00000022912 | Pros1 | 19128 | protein_coding | 147.632499 | 0.461251586 | 0.12084228 | 3.816971893 | 0.0001351 | 0.001906168 | 0.461251586 |
| ENSMUSG00000032418 | Me1 | 17436 | protein_coding | 107.5364948 | 0.461359034 | 0.158002129 | 2.919954545 | 0.003500824 | 0.027288035 | 0.461359034 |
| ENSMUSG00000065979 | Cpped1 | 223978 | protein_coding | 1880.321352 | 0.461396597 | 0.040882119 | 11.28602457 | 1.54E-29 | 2.00E-27 | 0.461396597 |
| ENSMUSG00000054589 | Gm9949 | 225609 | lncRNA | 9.405571187 | 0.461620457 | 0.4224146 | 1.092813689 | 0.27447561 | 0.5169296 | 0.461620457 |
| ENSMUSG00000029344 | Tpst2 | 22022 | protein_coding | 350.3129013 | 0.461654956 | 0.090035574 | 5.127472784 | 2.94E-07 | 8.07E-06 | 0.461654956 |
| ENSMUSG00000079440 | Alpi | 76768 | protein_coding | 506.6698879 | -0.461815176 | 0.135900403 | -3.398188425 | 0.000678337 | 0.007346903 | 0.461815176 |
| ENSMUSG00000039824 | Myl6b | 216459 | protein_coding | 24.02576169 | -0.462297917 | 0.27669966 | -1.670757082 | 0.094769669 | 0.278626509 | 0.462297917 |
| ENSMUSG00000052889 | Prkcb | 18751 | protein_coding | 12.75076436 | 0.462510429 | 0.357703487 | 1.292999498 | 0.196011233 | 0.428620088 | 0.462510429 |
| ENSMUSG00000052496 | Pkdrej | 18766 | protein_coding | 58.41353978 | 0.462692711 | 0.156846672 | 2.949968293 | 0.003178065 | 0.025297464 | 0.462692711 |
| ENSMUSG00000117869 | Snhg4 | 100503380 | lncRNA | 162.5652241 | -0.462754727 | 0.163316476 | -2.833484647 | 0.004604352 | 0.033538566 | 0.462754727 |
| ENSMUSG00000031410 | Nxf7 | 170722 | protein_coding | 14.53383951 | -0.462992367 | 0.399128371 | -1.160008661 | 0.24604528 | 0.487067489 | 0.462992367 |
| ENSMUSG00000097537 | 2610020C07Rik | 69918 | lncRNA | 15.40660406 | -0.463441729 | 0.317316993 | -1.460500819 | 0.144152483 | 0.356610996 | 0.463441729 |
| ENSMUSG00000029601 | Iqcd | 75732 | protein_coding | 24.83072374 | 0.463611691 | 0.227134024 | 2.041137134 | 0.041237198 | 0.162044907 | 0.463611691 |
| ENSMUSG00000071711 | Mpst | 246221 | protein_coding | 5400.999638 | 0.463899131 | 0.107517009 | 4.314658061 | 1.60E-05 | 0.000297679 | 0.463899131 |
| ENSMUSG00000027293 | Ehd4 | 98878 | protein_coding | 1180.387827 | 0.463958075 | 0.083580559 | 5.551028617 | 2.84E-08 | 9.62E-07 | 0.463958075 |
| ENSMUSG00000041193 | Pla2g5 | 18784 | protein_coding | 135.3166028 | 0.464067845 | 0.201065568 | 2.308042344 | 0.020996781 | 0.101663099 | 0.464067845 |
| ENSMUSG00000001128 | Cfp | 18636 | protein_coding | 24.49439423 | 0.464096846 | 0.266269933 | 1.742956255 | 0.081341256 | 0.252899642 | 0.464096846 |
| ENSMUSG00000019890 | Nts | 67405 | protein_coding | 114.8050747 | -0.464102121 | 0.201376288 | -2.304651288 | 0.021186111 | 0.102159742 | 0.464102121 |
| ENSMUSG00000027829 | Ccnl1 | 56706 | protein_coding | 2213.156519 | -0.464197989 | 0.150099154 | -3.092608968 | 0.001984054 | 0.017417983 | 0.464197989 |
| ENSMUSG00000079436 | Kcnj13 | 100040591 | protein_coding | 9.712393818 | -0.46486282 | 0.42378693 | -1.096925806 | 0.272673827 | 0.515160591 | 0.46486282 |
| ENSMUSG00000021238 | Aldh6a1 | 104776 | protein_coding | 1488.407235 | 0.465165105 | 0.06097876 | 7.628313656 | 2.38E-14 | 1.41E-12 | 0.465165105 |
| ENSMUSG00000062937 | Mtap | 66902 | protein_coding | 80.38515349 | 0.465178172 | 0.182094264 | 2.554600912 | 0.010630961 | 0.061938406 | 0.465178172 |
| ENSMUSG00000073293 | Nudt10 | 102954 | protein_coding | 29.20681956 | 0.4652908 | 0.240673282 | 1.933288135 | 0.053200713 | 0.192801774 | 0.4652908 |
| ENSMUSG00000045008 | 9030612E09Rik | 74530 | lncRNA | 8.048734942 | -0.465336335 | 0.402100649 | -1.157263325 | 0.247164792 | 0.488015969 | 0.465336335 |
| ENSMUSG00000103613 | Gm34882 | 102638281 | lncRNA | 6.365415832 | -0.465337324 | 0.477684404 | -0.974152223 | 0.329980963 | 0.573705524 | 0.465337324 |
| ENSMUSG00000097814 | Panct2 | 329217 | lncRNA | 59.52608799 | 0.465385687 | 0.151714893 | 3.067501667 | 0.002158563 | 0.018625542 | 0.465385687 |
| ENSMUSG00000013236 | Ptprs | 19280 | protein_coding | 770.1821621 | -0.465397552 | 0.061935383 | -7.514243521 | 5.72E-14 | 3.29E-12 | 0.465397552 |
| ENSMUSG00000050097 | Ces2b | 234669 | protein_coding | 328.2826535 | -0.46554205 | 0.124339296 | -3.744126485 | 0.000181022 | 0.002410693 | 0.46554205 |
| ENSMUSG00000044244 | Il20rb | 213208 | protein_coding | 24.91003176 | -0.465869815 | 0.245161769 | -1.900254743 | 0.057399697 | 0.202823164 | 0.465869815 |
| ENSMUSG00000055745 | Rtl6 | 223732 | protein_coding | 14.41806196 | 0.466337762 | 0.350504794 | 1.330474704 | 0.183361915 | 0.411619934 | 0.466337762 |

| ensembl_gene_id | external_gene_name | entrezgene_id | gene_biotype | baseMean | log2FoldChange | lfcSE | stat | pvalue | padj | absLog2FC |
|---|---|---|---|---|---|---|---|---|---|---|
| ENSMUSG00000006369 | Fbln1 | 14114 | protein_coding | 50.51251641 | 0.466431455 | 0.247306635 | 1.886045053 | 0.059288884 | 0.206735531 | 0.466431455 |
| ENSMUSG00000031089 | Slc6a14 | 56774 | protein_coding | 19852.56745 | -0.466789681 | 0.068933957 | -6.771549197 | 1.27E-11 | 6.04E-10 | 0.466789681 |
| ENSMUSG00000034177 | Rnf43 | 207742 | protein_coding | 986.3765009 | -0.466906889 | 0.066444218 | -7.027050689 | 2.11E-12 | 1.08E-10 | 0.466906889 |
| ENSMUSG00000060802 | B2m | 12010 | protein_coding | 5975.632171 | -0.466943719 | 0.210609471 | -2.217106938 | 0.026615785 | 0.119231323 | 0.466943719 |
| ENSMUSG00000002020 | Ltbp2 | 16997 | protein_coding | 6.294316351 | -0.466959229 | 0.529140709 | -0.882485926 | 0.377514087 | 0.61867166 | 0.466959229 |
| ENSMUSG00000086189 | Gm15462 | 108168861 | lncRNA | 39.07899864 | -0.467002148 | 0.211126545 | -2.211953722 | 0.026969863 | 0.120516203 | 0.467002148 |
| ENSMUSG00000040327 | Cul9 | 78309 | protein_coding | 136.7235017 | -0.46706758 | 0.193141922 | -2.41826101 | 0.015594886 | 0.082474622 | 0.46706758 |
| ENSMUSG00000040121 | Rep15 | 66532 | protein_coding | 2419.220082 | 0.467075603 | 0.083040095 | 5.624699758 | 1.86E-08 | 6.40E-07 | 0.467075603 |
| ENSMUSG00000059493 | Nhs | 195727 | protein_coding | 38.67705734 | -0.46719547 | 0.199526503 | -2.341520866 | 0.019205352 | 0.09546563 | 0.46719547 |
| ENSMUSG00000020646 | Mboat2 | 67216 | protein_coding | 236.4478508 | 0.467254596 | 0.085122365 | 5.489210668 | 4.04E-08 | 1.32E-06 | 0.467254596 |
| ENSMUSG00000000817 | Fasl | 14103 | protein_coding | 10.35128096 | 0.467358777 | 0.455937784 | 1.025049454 | 0.305339854 | 0.549639002 | 0.467358777 |
| ENSMUSG00000064672 | Gm22806 | 115488214 | snoRNA | 6.145601504 | -0.467387701 | 0.606835929 | -0.770204398 | 0.441178656 | 0.672739049 | 0.467387701 |
| ENSMUSG00000006638 | Abhd1 | 57742 | protein_coding | 7.752750861 | -0.467443365 | 0.427876077 | -1.092473708 | 0.274624941 | 0.517029701 | 0.467443365 |
| ENSMUSG00000046434 | Hnrnpa1 | 15382 | protein_coding | 3385.807183 | -0.467445405 | 0.087971448 | -5.313603629 | 1.07E-07 | 3.27E-06 | 0.467445405 |
| ENSMUSG00000047996 | Prrg1 | 546336 | protein_coding | 65.41778973 | -0.467568427 | 0.172718207 | -2.707117183 | 0.00678703 | 0.044730048 | 0.467568427 |
| ENSMUSG00000019738 | Polr2i | 69920 | protein_coding | 457.2620478 | 0.467706388 | 0.085144125 | 5.493114061 | 3.95E-08 | 1.30E-06 | 0.467706388 |
| ENSMUSG00000067924 | Rtl8b | 553127 | protein_coding | 13.18080124 | 0.467842174 | 0.328428242 | 1.424488258 | 0.154305182 | 0.371849022 | 0.467842174 |
| ENSMUSG00000030323 | Ift122 | 81896 | protein_coding | 164.6513602 | 0.467855963 | 0.102231347 | 4.576443291 | 4.73E-06 | 0.000100623 | 0.467855963 |
| ENSMUSG00000019947 | Arid5b | 71371 | protein_coding | 770.5190033 | -0.467984072 | 0.096405939 | -4.8543075 | 1.21E-06 | 2.90E-05 | 0.467984072 |
| ENSMUSG00000063522 | Ly6m | 67038 | protein_coding | 408.3751647 | 0.468185365 | 0.193152011 | 2.423921777 | 0.015353912 | 0.081558809 | 0.468185365 |
| ENSMUSG00000054364 | Rhob | 11852 | protein_coding | 10163.66925 | 0.468296353 | 0.116828827 | 4.008397295 | 6.11E-05 | 0.000976118 | 0.468296353 |
| ENSMUSG00000054793 | Cadm4 | 260299 | protein_coding | 31.79037099 | 0.468545727 | 0.22221254 | 2.108547642 | 0.034983646 | 0.144554734 | 0.468545727 |
| ENSMUSG00000022686 | B3gnt5 | 108105 | protein_coding | 1823.288064 | 0.468830074 | 0.092403525 | 5.073725 | 3.90E-07 | 1.04E-05 | 0.468830074 |
| ENSMUSG00000029380 | Cxcl1 | 14825 | protein_coding | 25.52633001 | -0.468917873 | 0.357725742 | -1.310830667 | 0.189914979 | 0.420477604 | 0.468917873 |
| ENSMUSG00000028186 | Uox | 22262 | protein_coding | 34.51197133 | 0.46899627 | 0.213959814 | 2.191982978 | 0.028380736 | 0.125036879 | 0.46899627 |
| ENSMUSG00000097350 | 4732491K20Rik | 224523 | lncRNA | 33.80675533 | 0.469137817 | 0.203541816 | 2.304871922 | 0.021173748 | 0.102159742 | 0.469137817 |
| ENSMUSG00000018599 | Mief2 | 237781 | protein_coding | 305.4821367 | 0.46917373 | 0.105263803 | 4.457123138 | 8.31E-06 | 0.000166619 | 0.46917373 |
| ENSMUSG00000030830 | Itgal | 16408 | protein_coding | 17.52737774 | 0.469232044 | 0.358365086 | 1.309368747 | 0.190409473 | 0.421109924 | 0.469232044 |
| ENSMUSG00000019768 | Esr1 | 13982 | protein_coding | 52.23333446 | 0.469794006 | 0.253596253 | 1.852527396 | 0.063950127 | 0.216846868 | 0.469794006 |
| ENSMUSG00000028542 | Slc6a9 | 14664 | protein_coding | 199.1494833 | 0.470209218 | 0.089886946 | 5.231117985 | 1.68E-07 | 4.89E-06 | 0.470209218 |
| ENSMUSG00000020108 | Ddit4 | 74747 | protein_coding | 1371.061673 | 0.470247076 | 0.118164119 | 3.979609725 | 6.90E-05 | 0.00107975 | 0.470247076 |
| ENSMUSG00000022365 | Derl1 | 67819 | protein_coding | 2277.945277 | 0.470562721 | 0.058614439 | 8.028102386 | 9.90E-16 | 6.54E-14 | 0.470562721 |
| ENSMUSG00000086938 | 4930481A15Rik | 74931 | lncRNA | 165.2941002 | 0.470634886 | 0.098587958 | 4.773756316 | 1.81E-06 | 4.19E-05 | 0.470634886 |
| ENSMUSG00000050511 | Oprd1 | 18386 | protein_coding | 23.70950462 | 0.470665819 | 0.261162022 | 1.80219856 | 0.071514171 | 0.233227308 | 0.470665819 |
| ENSMUSG00000042097 | Zfp239 | 22685 | protein_coding | 200.3683564 | 0.470780609 | 0.08732431 | 5.391174697 | 7.00E-08 | 2.19E-06 | 0.470780609 |
| ENSMUSG00000038354 | Ankrd35 | 213121 | protein_coding | 56.70536788 | 0.471121565 | 0.169731077 | 2.775694193 | 0.005508399 | 0.03837566 | 0.471121565 |
| ENSMUSG00000048612 | Myof | 226101 | protein_coding | 1027.075255 | 0.471138945 | 0.073178327 | 6.43823065 | 1.21E-10 | 5.24E-09 | 0.471138945 |
| ENSMUSG00000034161 | Scx | 20289 | protein_coding | 7.770781074 | -0.471184975 | 0.520312518 | -0.905580701 | 0.365157827 | 0.607290944 | 0.471184975 |
| ENSMUSG00000019232 | Etnppl | 71760 | protein_coding | 117.6670157 | 0.471980094 | 0.122988799 | 3.837586003 | 0.00012425 | 0.001774894 | 0.471980094 |

| ensembl_gene_id | external_gene_name | entrezgene_id | gene_biotype | baseMean | log2FoldChange | lfcSE | stat | pvalue | padj | absLog2FC |
|---|---|---|---|---|---|---|---|---|---|---|
| ENSMUSG00000040978 | Gm11992 | 626870 | protein_coding | 53.31919635 | 0.472203305 | 0.172211153 | 2.742001874 | 0.006106599 | 0.041535693 | 0.472203305 |
| ENSMUSG00000109644 | 0610005C13Rik | 71661 | lncRNA | 565.9222448 | 0.472272074 | 0.090667207 | 5.20885214 | 1.90E-07 | 5.45E-06 | 0.472272074 |
| ENSMUSG00000074354 | Arhgap20os | 408028 | lncRNA | 11.72081704 | -0.472831621 | 0.384265197 | -1.230482556 | 0.218516455 | 0.454666616 | 0.472831621 |
| ENSMUSG00000067297 | Ifit1bl2 | 112419 | protein_coding | 192.4362515 | -0.473213503 | 0.177778147 | -2.661820428 | 0.007771933 | 0.049619145 | 0.473213503 |
| ENSMUSG00000032633 | Flcn | 216805 | protein_coding | 770.7743427 | -0.473352507 | 0.083925433 | -5.640155694 | 1.70E-08 | 5.91E-07 | 0.473352507 |
| ENSMUSG00000044339 | Alkbh2 | 231642 | protein_coding | 62.90399538 | 0.473821847 | 0.196302345 | 2.413735027 | 0.015789941 | 0.083081362 | 0.473821847 |
| ENSMUSG00000096917 | 2500002B13Rik | 78353 | lncRNA | 32.86839696 | -0.473833416 | 0.292823924 | -1.618151306 | 0.105629994 | 0.297761739 | 0.473833416 |
| ENSMUSG00000028457 | Atp8b5 | 320571 | protein_coding | 54.82282054 | -0.473847117 | 0.172213098 | -2.75151613 | 0.00593201 | 0.040674117 | 0.473847117 |
| ENSMUSG00000032289 | Thsd4 | 207596 | protein_coding | 614.1251644 | -0.473974632 | 0.08174575 | -5.798156225 | 6.70E-09 | 2.43E-07 | 0.473974632 |
| ENSMUSG00000028519 | Dab1 | 13131 | protein_coding | 44.97306226 | 0.474183666 | 0.301664946 | 1.571888522 | 0.115976407 | 0.314662099 | 0.474183666 |
| ENSMUSG00000097167 | Gm16740 | 100504029 | lncRNA | 7.220400465 | -0.474226202 | 0.457024319 | -1.03763888 | 0.29943821 | 0.543015856 | 0.474226202 |
| ENSMUSG00000052783 | Grk4 | 14772 | protein_coding | 11.97658318 | 0.474271609 | 0.371133064 | 1.277901794 | 0.201284055 | 0.435076794 | 0.474271609 |
| ENSMUSG00000032121 | Tmem218 | 66279 | protein_coding | 182.5846863 | 0.474348427 | 0.112755037 | 4.206893453 | 2.59E-05 | 0.000454338 | 0.474348427 |
| ENSMUSG00000001823 | Hoxd12 | 15432 | protein_coding | 20.26392839 | 0.474405987 | 0.467018963 | 1.015817395 | 0.309716351 | 0.553189591 | 0.474405987 |
| ENSMUSG00000041012 | Cmtm8 | 70031 | protein_coding | 113.5416081 | 0.474548258 | 0.145790381 | 3.255003903 | 0.001133908 | 0.01119127 | 0.474548258 |
| ENSMUSG00000004105 | Angptl2 | 26360 | protein_coding | 13.5343013 | -0.474842556 | 0.396421968 | -1.197820993 | 0.230986713 | 0.469953022 | 0.474842556 |
| ENSMUSG00000027907 | S100a11 | 20195 | protein_coding | 394.4834312 | 0.475881515 | 0.076848199 | 6.192487548 | 5.92E-10 | 2.40E-08 | 0.475881515 |
| ENSMUSG00000019853 | Hebp2 | 56016 | protein_coding | 124.5236555 | 0.475961772 | 0.124413202 | 3.825653254 | 0.000130426 | 0.001851734 | 0.475961772 |
| ENSMUSG00000021200 | Asb2 | 65256 | protein_coding | 16.88539768 | 0.476002446 | 0.332572526 | 1.431274108 | 0.152351671 | 0.36913693 | 0.476002446 |
| ENSMUSG00000001313 | Rnd2 | 11858 | protein_coding | 24.42999262 | -0.476229122 | 0.255505931 | -1.863867188 | 0.062340358 | 0.213181286 | 0.476229122 |
| ENSMUSG00000029607 | Ankrd61 | 66729 | protein_coding | 12.57502154 | -0.476575789 | 0.480105317 | -0.992648429 | 0.320881315 | 0.564467652 | 0.476575789 |
| ENSMUSG00000068762 | Gstm6 | 14867 | protein_coding | 214.1882747 | 0.476601727 | 0.096173748 | 4.955632242 | 7.21E-07 | 1.83E-05 | 0.476601727 |
| ENSMUSG00000011751 | Sptbn4 | 80297 | protein_coding | 30.09540084 | 0.476632381 | 0.214853728 | 2.218404055 | 0.026527295 | 0.119015238 | 0.476632381 |
| ENSMUSG00000000142 | Axin2 | 12006 | protein_coding | 850.8222611 | -0.476782359 | 0.085433063 | -5.580770981 | 2.39E-08 | 8.16E-07 | 0.476782359 |
| ENSMUSG00000085793 | Lin52 | 217708 | protein_coding | 191.4599693 | 0.476922489 | 0.092047785 | 5.181248935 | 2.20E-07 | 6.25E-06 | 0.476922489 |
| ENSMUSG00000033436 | Armcx2 | 67416 | protein_coding | 10.02264972 | 0.476969214 | 0.406409863 | 1.173616236 | 0.240548775 | 0.481155234 | 0.476969214 |
| ENSMUSG00000020653 | Klf11 | 194655 | protein_coding | 581.0415897 | -0.47718221 | 0.18750077 | -2.544961331 | 0.010928984 | 0.063172661 | 0.47718221 |
| ENSMUSG00000074796 | Slc4a11 | 269356 | protein_coding | 7.751435618 | 0.477329753 | 0.409925492 | 1.164430517 | 0.244249587 | 0.485573395 | 0.477329753 |
| ENSMUSG00000032997 | Chpf | 74241 | protein_coding | 355.1754268 | 0.477390776 | 0.128653315 | 3.710676075 | 0.000206706 | 0.002712413 | 0.477390776 |
| ENSMUSG00000029223 | Uchl1 | 22223 | protein_coding | 13.1737546 | -0.477512372 | 0.397482398 | -1.201342182 | 0.229618494 | 0.46891952 | 0.477512372 |
| ENSMUSG00000084880 | Tomm6os | 100415787 | lncRNA | 9.70772458 | -0.477693416 | 0.436951688 | -1.093240808 | 0.274288084 | 0.516878235 | 0.477693416 |
| ENSMUSG00000014592 | Camta1 | 100072 | protein_coding | 129.7176804 | 0.477896998 | 0.118772593 | 4.023630237 | 5.73E-05 | 0.00092144 | 0.477896998 |
| ENSMUSG00000021294 | Kif26a | 668303 | protein_coding | 8.90449128 | -0.478300754 | 0.500206717 | -0.956206179 | 0.338968079 | 0.58240762 | 0.478300754 |
| ENSMUSG00000000037 | Scml2 | 107815 | protein_coding | 39.31990089 | 0.478527803 | 0.185354044 | 2.581696047 | 0.009831613 | 0.058615315 | 0.478527803 |
| ENSMUSG00000054545 | Ugt1a6a | 94284 | protein_coding | 104.3263731 | -0.478706397 | 0.20109237 | -2.380529887 | 0.017287759 | 0.08873319 | 0.478706397 |
| ENSMUSG00000117286 | Gm1043 | 381634 | protein_coding | 11.82004695 | 0.479955469 | 0.362186167 | 1.325162341 | 0.185117313 | 0.414061984 | 0.479955469 |
| ENSMUSG00000025935 | Tram1 | 72265 | protein_coding | 3926.813425 | 0.480227942 | 0.035127751 | 13.6708992 | 1.52E-42 | 2.81E-40 | 0.480227942 |
| ENSMUSG00000043286 | Pnpla1 | 433091 | protein_coding | 43.1888348 | 0.480747839 | 0.215007495 | 2.235958512 | 0.025354489 | 0.115866207 | 0.480747839 |
| ENSMUSG00000047123 | Ticam1 | 106759 | protein_coding | 711.3646458 | 0.480915128 | 0.075433785 | 6.37532807 | 1.83E-10 | 7.81E-09 | 0.480915128 |

| ensembl_gene_id | external_gene_name | entrezgene_id | gene_biotype | baseMean | log2FoldChange | lfcSE | stat | pvalue | padj | absLog2FC |
|---|---|---|---|---|---|---|---|---|---|---|
| ENSMUSG00000063838 | Cdc42ep5 | 58804 | protein_coding | 2781.553248 | 0.481165856 | 0.080797597 | 5.955200107 | 2.60E-09 | 9.88E-08 | 0.481165856 |
| ENSMUSG00000065952 | Rps23rg1 | 546049 | protein_coding | 25.69932574 | 0.481252648 | 0.244859869 | 1.965420669 | 0.049365569 | 0.183691401 | 0.481252648 |
| ENSMUSG00000104945 | Gm35715 | 102639389 | lncRNA | 7.201379677 | -0.481374301 | 0.506090828 | -0.951161875 | 0.341522209 | 0.584243096 | 0.481374301 |
| ENSMUSG00000028341 | Nr4a3 | 18124 | protein_coding | 16.35163531 | -0.481392468 | 0.350220721 | -1.374540223 | 0.169274032 | 0.393611906 | 0.481392468 |
| ENSMUSG00000025037 | Maoa | 17161 | protein_coding | 24141.66084 | -0.482004682 | 0.067077564 | -7.185780966 | 6.68E-13 | 3.50E-11 | 0.482004682 |
| ENSMUSG00000087138 | Gm15545 | 100502630 | lncRNA | 39.36106275 | -0.482012995 | 0.190709742 | -2.527469184 | 0.011488791 | 0.065795357 | 0.482012995 |
| ENSMUSG00000029189 | Sel1l3 | 231238 | protein_coding | 4765.689774 | 0.482067322 | 0.045094981 | 10.69004385 | 1.13E-26 | 1.31E-24 | 0.482067322 |
| ENSMUSG00000011305 | Plin5 | 66968 | protein_coding | 6.378207671 | -0.482107873 | 0.463774145 | -1.039531587 | 0.298557576 | 0.542138138 | 0.482107873 |
| ENSMUSG00000087020 | Gm16283 | 102635853 | lncRNA | 6.121158326 | -0.482824086 | 0.520995215 | -0.926734204 | 0.354064554 | 0.596344549 | 0.482824086 |
| ENSMUSG00000038742 | Angptl6 | 70726 | protein_coding | 25.77359742 | 0.482929464 | 0.262958568 | 1.836523026 | 0.06628034 | 0.22202351 | 0.482929464 |
| ENSMUSG00000020334 | Slc22a4 | 30805 | protein_coding | 80.16466169 | -0.483097914 | 0.204966244 | -2.356963303 | 0.018425068 | 0.09298871 | 0.483097914 |
| ENSMUSG00000028943 | Espn | 56226 | protein_coding | 674.0540588 | -0.483102022 | 0.10137637 | -4.765430254 | 1.88E-06 | 4.35E-05 | 0.483102022 |
| ENSMUSG00000022623 | Shank3 | 58234 | protein_coding | 8.088848051 | 0.483118098 | 0.39826631 | 1.21305289 | 0.2251096 | 0.463123925 | 0.483118098 |
| ENSMUSG00000041644 | Slc5a12 | 241612 | protein_coding | 13.57715696 | 0.483144436 | 0.819175394 | 0.589793639 | 0.555329008 | 0.755247648 | 0.483144436 |
| ENSMUSG00000024399 | Ltb | 16994 | protein_coding | 15.88472362 | 0.483340314 | 0.354942841 | 1.36174127 | 0.173279552 | 0.398908703 | 0.483340314 |
| ENSMUSG00000020674 | Pxdn | 69675 | protein_coding | 6.545058458 | 0.483484837 | 0.4751845 | 1.017467608 | 0.308931034 | 0.552526426 | 0.483484837 |
| ENSMUSG00000038070 | Cntln | 338349 | protein_coding | 56.6197916 | 0.483655325 | 0.212483163 | 2.276205413 | 0.022833718 | 0.107689014 | 0.483655325 |
| ENSMUSG00000089371 | Mir1938 | 100316693 | miRNA | 90.82521865 | -0.483688288 | 0.161556875 | -2.993919555 | 0.002754186 | 0.022512291 | 0.483688288 |
| ENSMUSG00000027194 | Ttc17 | 74569 | protein_coding | 1740.908158 | 0.483696787 | 0.037260175 | 12.98160269 | 1.56E-38 | 2.64E-36 | 0.483696787 |
| ENSMUSG00000047909 | Ankrd16 | 320816 | protein_coding | 319.9291227 | -0.483704444 | 0.158861494 | -3.044818676 | 0.002328206 | 0.019720054 | 0.483704444 |
| ENSMUSG00000074824 | Rslcan18 | 432770 | protein_coding | 10.59909732 | 0.48410897 | 0.375077418 | 1.290690793 | 0.196810919 | 0.429250765 | 0.48410897 |
| ENSMUSG00000041298 | Katnal1 | 231912 | protein_coding | 72.84233681 | 0.4844129 | 0.173206141 | 2.796742059 | 0.005162073 | 0.036576778 | 0.4844129 |
| ENSMUSG00000071103 | 1700029J07Rik | 69479 | protein_coding | 15.79689904 | -0.484902443 | 0.332130437 | -1.459975926 | 0.14429669 | 0.356748847 | 0.484902443 |
| ENSMUSG00000055301 | Adh7 | 11529 | protein_coding | 6.027619153 | -0.485158029 | 0.586121398 | -0.827743246 | 0.407815922 | 0.646388536 | 0.485158029 |
| ENSMUSG00000028560 | Usp1 | 230484 | protein_coding | 1072.568819 | 0.485163864 | 0.080406846 | 6.033862629 | 1.60E-09 | 6.22E-08 | 0.485163864 |
| ENSMUSG00000025163 | Cd7 | 12516 | protein_coding | 91.72322127 | 0.485530777 | 0.171909668 | 2.824336657 | 0.004737859 | 0.034222527 | 0.485530777 |
| ENSMUSG00000112129 | Pbld1 | 68371 | protein_coding | 301.5719316 | -0.486234778 | 0.114473346 | -4.247580713 | 2.16E-05 | 0.000387642 | 0.486234778 |
| ENSMUSG00000097177 | 9330159M07Rik | 319673 | lncRNA | 31.91129851 | 0.486283565 | 0.209603812 | 2.320012979 | 0.020340175 | 0.099431951 | 0.486283565 |
| ENSMUSG00000022159 | Rab2b | 76338 | protein_coding | 14.31962748 | -0.486467585 | 0.355086277 | -1.369998269 | 0.170687442 | 0.395158825 | 0.486467585 |
| ENSMUSG00000057335 | Cep170 | 545389 | protein_coding | 23.73365378 | 0.486490662 | 0.243672508 | 1.996493843 | 0.045880195 | 0.174465906 | 0.486490662 |
| ENSMUSG00000085007 | Smim43 | 100503068 | protein_coding | 8.109820963 | 0.486774594 | 0.437326015 | 1.113070287 | 0.265678243 | 0.507713359 | 0.486774594 |
| ENSMUSG00000068758 | Il3ra | 16188 | protein_coding | 16.93249528 | 0.486807694 | 0.307245905 | 1.584423702 | 0.113097332 | 0.309576536 | 0.486807694 |
| ENSMUSG00000115548 | 5430440P10Rik | 71362 | lncRNA | 7.067450052 | -0.486842057 | 0.465166427 | -1.04659758 | 0.295285218 | 0.538208038 | 0.486842057 |
| ENSMUSG00000027867 | Spag17 | 74362 | protein_coding | 14.19074575 | 0.486842434 | 0.320464356 | 1.519178108 | 0.12871767 | 0.334110714 | 0.486842434 |
| ENSMUSG00000038188 | Scarf1 | 380713 | protein_coding | 83.94458144 | -0.487068924 | 0.165929298 | -2.935400375 | 0.003331176 | 0.026257023 | 0.487068924 |
| ENSMUSG00000020395 | Itk | 16428 | protein_coding | 23.35083391 | 0.487944125 | 0.322887726 | 1.511188211 | 0.130740505 | 0.336624075 | 0.487944125 |
| ENSMUSG00000019429 | Ffar3 | 233080 | protein_coding | 28.42005061 | 0.488183187 | 0.2430667 | 2.008433026 | 0.044597296 | 0.171324583 | 0.488183187 |
| ENSMUSG00000073565 | Prr16 | 71373 | protein_coding | 10.33018745 | 0.48831132 | 0.415850147 | 1.174248282 | 0.240295592 | 0.480740232 | 0.48831132 |
| ENSMUSG00000040276 | Pacsin1 | 23969 | protein_coding | 58.14818631 | -0.488535194 | 0.282985476 | -1.726361371 | 0.084282421 | 0.258850238 | 0.488535194 |

| ensembl_gene_id | external_gene_name | entrezgene_id | gene_biotype | baseMean | log2FoldChange | lfcSE | stat | pvalue | padj | absLog2FC |
|---|---|---|---|---|---|---|---|---|---|---|
| ENSMUSG00000031112 | Stk26 | 70415 | protein_coding | 231.8877961 | 0.488573201 | 0.109217416 | 4.473400103 | 7.70E-06 | 0.000155973 | 0.488573201 |
| ENSMUSG00000021003 | Galc | 14420 | protein_coding | 125.8910319 | 0.488761087 | 0.110276536 | 4.432140362 | 9.33E-06 | 0.000184398 | 0.488761087 |
| ENSMUSG00000030747 | Dgat2 | 67800 | protein_coding | 793.2360894 | 0.488778813 | 0.085501874 | 5.716585996 | 1.09E-08 | 3.87E-07 | 0.488778813 |
| ENSMUSG00000022372 | Sla | 20491 | protein_coding | 21.22984602 | 0.488896246 | 0.271903431 | 1.79805104 | 0.07216892 | 0.234551147 | 0.488896246 |
| ENSMUSG00000114469 | C730002L08Rik | 320155 | lncRNA | 11.89754733 | -0.489292671 | 0.387772113 | -1.26180469 | 0.207019074 | 0.442081119 | 0.489292671 |
| ENSMUSG00000015812 | Gnrh1 | 14714 | protein_coding | 6.812596161 | -0.48934917 | 0.481161066 | -1.017017387 | 0.309145158 | 0.552606516 | 0.48934917 |
| ENSMUSG00000043432 | Leng9 | 243813 | protein_coding | 1083.789773 | 0.48935164 | 0.071751906 | 6.820050709 | 9.10E-12 | 4.41E-10 | 0.48935164 |
| ENSMUSG00000101581 | C430002N11Rik | 319707 | lncRNA | 8.300107204 | -0.490290437 | 0.438068544 | -1.119209413 | 0.263050811 | 0.504886203 | 0.490290437 |
| ENSMUSG00000024029 | Tff3 | 21786 | protein_coding | 27281.73506 | 0.49045038 | 0.073613409 | 6.662514127 | 2.69E-11 | 1.24E-09 | 0.49045038 |
| ENSMUSG00000030790 | Adm | 11535 | protein_coding | 64.511179 | 0.490597736 | 0.197059932 | 2.489586453 | 0.012789181 | 0.071277641 | 0.490597736 |
| ENSMUSG00000108622 | Gm36864 | 102640913 | protein_coding | 9.161964426 | 0.490708787 | 0.469020497 | 1.046241667 | 0.29544947 | 0.538446559 | 0.490708787 |
| ENSMUSG00000002014 | Ssr4 | 20832 | protein_coding | 2999.63687 | 0.491512469 | 0.051885712 | 9.472983063 | 2.72E-21 | 2.38E-19 | 0.491512469 |
| ENSMUSG00000031486 | Adgra2 | 78560 | protein_coding | 33.26078229 | -0.491594762 | 0.277453348 | -1.771810523 | 0.076426015 | 0.243533362 | 0.491594762 |
| ENSMUSG00000098184 | Gm27011 | 115486886 | lncRNA | 7.807515728 | 0.491773857 | 0.409476968 | 1.200980508 | 0.229758762 | 0.469064473 | 0.491773857 |
| ENSMUSG00000078878 | Gm14305 | 100043387 | protein_coding | 64.50919216 | -0.492132825 | 0.194235701 | -2.533688833 | 0.011286891 | 0.064800174 | 0.492132825 |
| ENSMUSG00000032978 | Guca2b | 14916 | protein_coding | 350.6798246 | -0.492361852 | 0.137472938 | -3.581518372 | 0.000341603 | 0.004102035 | 0.492361852 |
| ENSMUSG00000077222 | Gm22270 | 115490266 | snoRNA | 18.33183252 | 0.492975744 | 0.284100405 | 1.735216615 | 0.082702445 | 0.255506193 | 0.492975744 |
| ENSMUSG00000021453 | Gadd45g | 23882 | protein_coding | 209.6726183 | 0.493233336 | 0.155923075 | 3.163312012 | 0.001559851 | 0.014456304 | 0.493233336 |
| ENSMUSG00000044033 | Ccdc141 | 545428 | protein_coding | 39.73877602 | -0.493806763 | 0.236554543 | -2.087496431 | 0.036843283 | 0.149514754 | 0.493806763 |
| ENSMUSG00000054404 | Slfn5 | 327978 | protein_coding | 10.13960893 | -0.493903919 | 0.647622697 | -0.762641459 | 0.445677252 | 0.67671134 | 0.493903919 |
| ENSMUSG00000036712 | Cyld | 74256 | protein_coding | 1643.646141 | 0.494145753 | 0.051487851 | 9.597327132 | 8.20E-22 | 7.52E-20 | 0.494145753 |
| ENSMUSG00000047904 | Sstr2 | 20606 | protein_coding | 75.09287898 | -0.494774799 | 0.162332591 | -3.047907967 | 0.002304405 | 0.019580157 | 0.494774799 |
| ENSMUSG00000060600 | Eno3 | 13808 | protein_coding | 1691.38111 | -0.494834384 | 0.20283163 | -2.439631258 | 0.01470226 | 0.079113565 | 0.494834384 |
| ENSMUSG00000060063 | Alox5ap | 11690 | protein_coding | 45.22077955 | 0.495028226 | 0.20105193 | 2.462190868 | 0.013809116 | 0.07559346 | 0.495028226 |
| ENSMUSG00000060791 | Gmfg | 63986 | protein_coding | 285.1167166 | 0.495093842 | 0.105672505 | 4.685171808 | 2.80E-06 | 6.21E-05 | 0.495093842 |
| ENSMUSG00000044707 | Ccnjl | 380694 | protein_coding | 94.30517978 | 0.49514302 | 0.142529309 | 3.473973332 | 0.000512812 | 0.005779259 | 0.49514302 |
| ENSMUSG00000069516 | Lyz2 | 17105 | protein_coding | 27.77022872 | 0.495205854 | 0.262679858 | 1.885206795 | 0.059401924 | 0.206976523 | 0.495205854 |
| ENSMUSG00000001642 | Akr1b3 | 11677 | protein_coding | 1056.490783 | 0.495726516 | 0.080882025 | 6.129007222 | 8.84E-10 | 3.50E-08 | 0.495726516 |
| ENSMUSG00000022659 | Gcsam | 14525 | protein_coding | 8.623472745 | 0.496037549 | 0.4054375 | 1.22346243 | 0.221155086 | 0.457721775 | 0.496037549 |
| ENSMUSG00000025875 | Tspan17 | 74257 | protein_coding | 231.6530775 | -0.496043884 | 0.088855375 | -5.5825985 | 2.37E-08 | 8.10E-07 | 0.496043884 |
| ENSMUSG00000039096 | Rsad1 | 237926 | protein_coding | 87.19259873 | -0.496467052 | 0.161678705 | -3.07070157 | 0.002135565 | 0.018456728 | 0.496467052 |
| ENSMUSG00000079497 | Gm13420 | 433415 | protein_coding | 6.69035571 | 0.496519671 | 0.471432316 | 1.053215179 | 0.292242383 | 0.534838052 | 0.496519671 |
| ENSMUSG00000047669 | Msl3l2 | 73390 | protein_coding | 19.34347713 | 0.496908829 | 0.281018812 | 1.76824044 | 0.077020716 | 0.244751348 | 0.496908829 |
| ENSMUSG00000033530 | Ttc7b | 104718 | protein_coding | 91.80071821 | 0.49713459 | 0.145255212 | 3.422490543 | 0.000620503 | 0.006830611 | 0.49713459 |
| ENSMUSG00000047777 | Phf13 | 230936 | protein_coding | 343.4625443 | 0.497837011 | 0.09647571 | 5.160231666 | 2.47E-07 | 6.91E-06 | 0.497837011 |
| ENSMUSG00000027287 | Snap23 | 20619 | protein_coding | 1161.883477 | 0.498254997 | 0.055574103 | 8.965596736 | 3.09E-19 | 2.50E-17 | 0.498254997 |
| ENSMUSG00000026610 | Esrrg | 26381 | protein_coding | 160.9376498 | -0.498332744 | 0.178949499 | -2.784767473 | 0.005356612 | 0.037542844 | 0.498332744 |
| ENSMUSG00000028940 | Hes2 | 15206 | protein_coding | 102.0219613 | 0.498337601 | 0.147687324 | 3.374274697 | 0.000740105 | 0.007859568 | 0.498337601 |
| ENSMUSG00000022026 | Olfm4 | 380924 | protein_coding | 93.03544583 | 0.498372683 | 0.966280257 | 0.515764117 | 0.606019167 | 0.789753 | 0.498372683 |

| ensembl_gene_id | external_gene_name | entrezgene_id | gene_biotype | baseMean | log2FoldChange | lfcSE | stat | pvalue | padj | absLog2FC |
|---|---|---|---|---|---|---|---|---|---|---|
| ENSMUSG00000045827 | Serpinb9 | 20723 | protein_coding | 74.88634499 | -0.499534036 | 0.165818462 | -3.012535694 | 0.00259075 | 0.021415182 | 0.499534036 |
| ENSMUSG00000055994 | Nod2 | 257632 | protein_coding | 73.09977776 | -0.499725062 | 0.161716234 | -3.090135423 | 0.002000653 | 0.017525542 | 0.499725062 |
| ENSMUSG00000008307 | 1700109H08Rik | 77036 | protein_coding | 20.79051373 | -0.499726805 | 0.418081154 | -1.19528661 | 0.23197507 | 0.471166492 | 0.499726805 |
| ENSMUSG00000017868 | Sgk2 | 27219 | protein_coding | 668.0374811 | -0.500171961 | 0.130756348 | -3.825221262 | 0.000130655 | 0.001853182 | 0.500171961 |
| ENSMUSG00000092335 | Zfp977 | 637776 | protein_coding | 8.120272779 | -0.500263844 | 0.463046408 | -1.080375176 | 0.279975146 | 0.523118689 | 0.500263844 |
| ENSMUSG00000002384 | Bmp8b | 12164 | protein_coding | 14.47763288 | 0.500641778 | 0.35751936 | 1.400320748 | 0.16141729 | 0.381977497 | 0.500641778 |
| ENSMUSG00000096727 | Psmb9 | 16912 | protein_coding | 305.5364194 | -0.500652532 | 0.321624086 | -1.556638803 | 0.119556266 | 0.319914163 | 0.500652532 |
| ENSMUSG00000028906 | Epb41 | 269587 | protein_coding | 2406.051645 | 0.500736116 | 0.047437008 | 10.55581156 | 4.77E-26 | 5.31E-24 | 0.500736116 |
| ENSMUSG00000056888 | Glipr1 | 73690 | protein_coding | 10.48388527 | 0.500773659 | 0.378978457 | 1.321377639 | 0.186375478 | 0.415618234 | 0.500773659 |
| ENSMUSG00000006463 | Zdhhc24 | 70605 | protein_coding | 308.7055254 | 0.500842846 | 0.096586442 | 5.185436356 | 2.16E-07 | 6.12E-06 | 0.500842846 |
| ENSMUSG00000021013 | Ttc8 | 76260 | protein_coding | 26.41140835 | 0.500865751 | 0.250809677 | 1.996995321 | 0.045825691 | 0.174452968 | 0.500865751 |
| ENSMUSG00000074899 | Sptbn5 | 640524 | protein_coding | 26.53575933 | 0.501046686 | 0.243534914 | 2.057391602 | 0.039648572 | 0.15749077 | 0.501046686 |
| ENSMUSG00000035202 | Lars2 | 102436 | protein_coding | 14775.47071 | 0.501152143 | 0.242299268 | 2.06831885 | 0.03861005 | 0.154545615 | 0.501152143 |
| ENSMUSG00000023084 | Lrrc71 | 74485 | protein_coding | 10.40151646 | -0.501207438 | 0.394829093 | -1.269428843 | 0.204288155 | 0.438338886 | 0.501207438 |
| ENSMUSG00000078552 | Dcdc2b | 100504491 | protein_coding | 18.25956151 | -0.501808103 | 0.311257039 | -1.612198408 | 0.106918772 | 0.299347968 | 0.501808103 |
| ENSMUSG00000019464 | Ptger1 | 19216 | protein_coding | 19.84316689 | -0.502386054 | 0.311692103 | -1.611802319 | 0.107004963 | 0.299347968 | 0.502386054 |
| ENSMUSG00000027896 | Slc16a4 | 229699 | protein_coding | 9.927832257 | -0.502402792 | 0.39449985 | -1.273518333 | 0.202834171 | 0.436976894 | 0.502402792 |
| ENSMUSG00000037621 | Atoh8 | 71093 | protein_coding | 68.64070708 | -0.5027181 | 0.177760324 | -2.828066974 | 0.004683001 | 0.033968619 | 0.5027181 |
| ENSMUSG00000050520 | Cldn8 | 54420 | protein_coding | 2286.714093 | 0.503611324 | 0.103978993 | 4.843394899 | 1.28E-06 | 3.06E-05 | 0.503611324 |
| ENSMUSG00000038539 | Atf5 | 107503 | protein_coding | 169.1765023 | 0.503954929 | 0.130006259 | 3.876389757 | 0.000106018 | 0.001565704 | 0.503954929 |
| ENSMUSG00000039478 | Micu3 | 78506 | protein_coding | 37.28235005 | -0.504220009 | 0.224296824 | -2.248003335 | 0.024575977 | 0.113433539 | 0.504220009 |
| ENSMUSG00000034730 | Adgrb1 | 107831 | protein_coding | 11.57892141 | -0.504569868 | 0.403262617 | -1.251219049 | 0.21085457 | 0.445849236 | 0.504569868 |
| ENSMUSG00000026173 | Plcd4 | 18802 | protein_coding | 21.82181465 | -0.504584649 | 0.257850805 | -1.956886072 | 0.050360864 | 0.186021452 | 0.504584649 |
| ENSMUSG00000051811 | Cox6b2 | 333182 | protein_coding | 17.73394475 | -0.504669673 | 0.336880075 | -1.498069223 | 0.134115266 | 0.34157879 | 0.504669673 |
| ENSMUSG00000054206 | Gzmm | 16904 | protein_coding | 7.921429789 | -0.505015736 | 0.416289125 | -1.21313699 | 0.22507745 | 0.463123925 | 0.505015736 |
| ENSMUSG00000028011 | Tdo2 | 56720 | protein_coding | 8.51969139 | -0.505032655 | 0.456134501 | -1.107201174 | 0.26820696 | 0.510669852 | 0.505032655 |
| ENSMUSG00000055805 | Fmnl1 | 57778 | protein_coding | 18.28424377 | 0.505566447 | 0.307223259 | 1.645599517 | 0.099846239 | 0.287678519 | 0.505566447 |
| ENSMUSG00000074004 | B3gnt6 | 272411 | protein_coding | 383.2933075 | -0.505850345 | 0.123039876 | -4.111271575 | 3.93E-05 | 0.000659641 | 0.505850345 |
| ENSMUSG00000038094 | Atp13a4 | 224079 | protein_coding | 10.94054017 | 0.506108196 | 0.417558156 | 1.212066363 | 0.225486979 | 0.463452522 | 0.506108196 |
| ENSMUSG00000023341 | Mx2 | 17858 | protein_coding | 12.88299047 | -0.506531147 | 0.488870287 | -1.036125861 | 0.300143429 | 0.543683373 | 0.506531147 |
| ENSMUSG00000022537 | Tmem44 | 224090 | protein_coding | 177.8343708 | -0.506645017 | 0.14155642 | -3.579103056 | 0.000344775 | 0.004137049 | 0.506645017 |
| ENSMUSG00000089900 | Tbc1d22bos | 74092 | lncRNA | 6.353277358 | -0.507115413 | 0.500239503 | -1.013745237 | 0.310704333 | 0.554568301 | 0.507115413 |
| ENSMUSG00000032519 | Slc25a38 | 208638 | protein_coding | 569.6126423 | 0.50758792 | 0.083693424 | 6.064848273 | 1.32E-09 | 5.16E-08 | 0.50758792 |
| ENSMUSG00000023008 | Fmnl3 | 22379 | protein_coding | 35.97897285 | 0.507664008 | 0.221442664 | 2.292530259 | 0.021875064 | 0.104773272 | 0.507664008 |
| ENSMUSG00000056758 | Hmga2 | 15364 | protein_coding | 272.4172016 | -0.50869192 | 0.123183435 | -4.129548093 | 3.63E-05 | 0.000615735 | 0.50869192 |
| ENSMUSG00000086728 | Man2c1os | 76794 | lncRNA | 20.51001535 | -0.50876732 | 0.29405777 | -1.730161117 | 0.083601493 | 0.257445347 | 0.50876732 |
| ENSMUSG00000034248 | Slc25a37 | 67712 | protein_coding | 581.4784137 | -0.508879352 | 0.116109773 | -4.382743482 | 1.17E-05 | 0.000225536 | 0.508879352 |
| ENSMUSG00000032932 | Hspa13 | 110920 | protein_coding | 767.5647466 | 0.508964014 | 0.078856038 | 6.454344271 | 1.09E-10 | 4.74E-09 | 0.508964014 |
| ENSMUSG00000053310 | Nrgn | 64011 | protein_coding | 14.57123666 | 0.508989935 | 0.354446995 | 1.436011428 | 0.15099909 | 0.36685181 | 0.508989935 |

| ensembl_gene_id | external_gene_name | entrezgene_id | gene_biotype | baseMean | log2FoldChange | lfcSE | stat | pvalue | padj | absLog2FC |
|---|---|---|---|---|---|---|---|---|---|---|
| ENSMUSG00000037287 | Tbcel | 272589 | protein_coding | 498.6792492 | -0.509151917 | 0.100616251 | -5.06033481 | 4.19E-07 | 1.11E-05 | 0.509151917 |
| ENSMUSG00000022838 | Eaf2 | 106389 | protein_coding | 9.536605378 | 0.509195179 | 0.375633241 | 1.355564746 | 0.175237695 | 0.401599873 | 0.509195179 |
| ENSMUSG00000037579 | Kcnh3 | 16512 | protein_coding | 539.6592444 | -0.509217299 | 0.107857466 | -4.721205842 | 2.34E-06 | 5.30E-05 | 0.509217299 |
| ENSMUSG00000047281 | Sfn | 55948 | protein_coding | 5225.19621 | 0.509294437 | 0.079106269 | 6.438104602 | 1.21E-10 | 5.24E-09 | 0.509294437 |
| ENSMUSG00000030662 | Ipo5 | 70572 | protein_coding | 3285.184708 | 0.509698445 | 0.071253208 | 7.153340297 | 8.47E-13 | 4.42E-11 | 0.509698445 |
| ENSMUSG00000049807 | Arhgap23 | 58996 | protein_coding | 12.80802273 | -0.509799833 | 0.39265551 | -1.29833867 | 0.194170984 | 0.426123143 | 0.509799833 |
| ENSMUSG00000048217 | Nags | 217214 | protein_coding | 233.7784973 | -0.509896245 | 0.131246714 | -3.885021036 | 0.000102321 | 0.00151806 | 0.509896245 |
| ENSMUSG00000039485 | Tspyl4 | 72480 | protein_coding | 19.09700578 | 0.510824345 | 0.300969019 | 1.697265545 | 0.089646468 | 0.269173076 | 0.510824345 |
| ENSMUSG00000053128 | Rnf26 | 213211 | protein_coding | 59.58137798 | 0.511041837 | 0.166320309 | 3.072636414 | 0.002121768 | 0.018357166 | 0.511041837 |
| ENSMUSG00000046814 | Gchfr | 320415 | protein_coding | 45.15113988 | 0.511049819 | 0.201269336 | 2.539134025 | 0.011112724 | 0.063982472 | 0.511049819 |
| ENSMUSG00000101605 | Ace3 | 217246 | protein_coding | 7.997215279 | 0.511177679 | 0.451756249 | 1.131534273 | 0.257830287 | 0.499763105 | 0.511177679 |
| ENSMUSG00000026791 | Slc2a8 | 56017 | protein_coding | 279.7497596 | 0.51119962 | 0.083136319 | 6.148932604 | 7.80E-10 | 3.14E-08 | 0.51119962 |
| ENSMUSG00000026077 | Npas2 | 18143 | protein_coding | 360.064052 | 0.511410826 | 0.101228861 | 5.052025857 | 4.37E-07 | 1.15E-05 | 0.511410826 |
| ENSMUSG00000040428 | Plekha4 | 69217 | protein_coding | 16.08770035 | -0.511523407 | 0.403072957 | -1.269059108 | 0.204419983 | 0.438439283 | 0.511523407 |
| ENSMUSG00000030824 | Nucb1 | 18220 | protein_coding | 9275.35849 | 0.511672766 | 0.066308158 | 7.716588386 | 1.19E-14 | 7.33E-13 | 0.511672766 |
| ENSMUSG00000024352 | Spata24 | 71242 | protein_coding | 15.5444343 | 0.51186825 | 0.291221892 | 1.757657181 | 0.078805858 | 0.248174588 | 0.51186825 |
| ENSMUSG00000004655 | Aqp1 | 11826 | protein_coding | 2270.625567 | -0.511886096 | 0.057333721 | -8.928185441 | 4.33E-19 | 3.47E-17 | 0.511886096 |
| ENSMUSG00000032258 | Lca5 | 75782 | protein_coding | 36.83046733 | 0.511985096 | 0.214923656 | 2.382171913 | 0.017210861 | 0.088535742 | 0.511985096 |
| ENSMUSG00000065818 | Snord35a | 27211 | snoRNA | 6.62019537 | -0.512390994 | 0.535994456 | -0.955963235 | 0.339090809 | 0.58240762 | 0.512390994 |
| ENSMUSG00000028541 | B4galt2 | 53418 | protein_coding | 5.868604585 | 0.512596281 | 0.497640008 | 1.030054402 | 0.302984468 | 0.546864076 | 0.512596281 |
| ENSMUSG00000071716 | Apol7e | 666348 | protein_coding | 10.14754765 | 0.512942284 | 0.427168972 | 1.200794809 | 0.229830806 | 0.469064473 | 0.512942284 |
| ENSMUSG00000040599 | Mis12 | 67139 | protein_coding | 456.7007254 | 0.51301626 | 0.085461724 | 6.002877525 | 1.94E-09 | 7.48E-08 | 0.51301626 |
| ENSMUSG00000087476 | Rap1gapos | 69808 | lncRNA | 49.78341384 | 0.513334607 | 0.179448534 | 2.860623021 | 0.004228095 | 0.031494912 | 0.513334607 |
| ENSMUSG00000064262 | Gimap8 | 243374 | protein_coding | 57.76647306 | 0.513802724 | 0.161586734 | 3.179733343 | 0.001474106 | 0.013781399 | 0.513802724 |
| ENSMUSG00000015980 | Lrrc27 | 76612 | protein_coding | 7.015644268 | -0.513870061 | 0.477900175 | -1.075266526 | 0.282255418 | 0.525081686 | 0.513870061 |
| ENSMUSG00000074627 | Mroh8 | 629499 | protein_coding | 16.19086515 | -0.514041632 | 0.291260229 | -1.764887827 | 0.07758262 | 0.245713848 | 0.514041632 |
| ENSMUSG00000024697 | Gna14 | 14675 | protein_coding | 1267.658518 | 0.514287889 | 0.077666849 | 6.621716924 | 3.55E-11 | 1.62E-09 | 0.514287889 |
| ENSMUSG00000000290 | Itgb2 | 16414 | protein_coding | 32.53335747 | 0.515137484 | 0.226337835 | 2.275967186 | 0.022847974 | 0.107689014 | 0.515137484 |
| ENSMUSG00000074671 | Tspyl3 | 241732 | protein_coding | 8.880793338 | 0.51528075 | 0.405062778 | 1.272100963 | 0.203337249 | 0.437462622 | 0.51528075 |
| ENSMUSG00000038599 | Capn8 | 170725 | protein_coding | 487.826381 | 0.516601036 | 0.064553322 | 8.00270251 | 1.22E-15 | 7.98E-14 | 0.516601036 |
| ENSMUSG00000045064 | Zc2hc1c | 72350 | protein_coding | 27.0518807 | -0.517173948 | 0.23002508 | -2.248337218 | 0.024554695 | 0.113432704 | 0.517173948 |
| ENSMUSG00000098449 | Gm7467 | 665055 | lncRNA | 34.18643458 | -0.517196738 | 0.281344723 | -1.83830261 | 0.066017827 | 0.221624129 | 0.517196738 |
| ENSMUSG00000102748 | Pcdhgb2 | 93700 | protein_coding | 8.586033757 | 0.51744635 | 0.462707784 | 1.118300507 | 0.26343867 | 0.504916665 | 0.51744635 |
| ENSMUSG00000020143 | Dock2 | 94176 | protein_coding | 43.83990236 | 0.517745751 | 0.191264453 | 2.706962756 | 0.006790187 | 0.044732579 | 0.517745751 |
| ENSMUSG00000036782 | Klhl13 | 67455 | protein_coding | 73.64064608 | 0.517811214 | 0.187200202 | 2.766082552 | 0.005673417 | 0.039352773 | 0.517811214 |
| ENSMUSG00000031932 | Gpr83 | 14608 | protein_coding | 25.07151182 | 0.517948678 | 0.307130194 | 1.68641406 | 0.091716075 | 0.272998366 | 0.517948678 |
| ENSMUSG00000098702 | 1500015A07Rik | 68982 | lncRNA | 49.11210742 | -0.518310765 | 0.222474811 | -2.329750334 | 0.019819351 | 0.097536364 | 0.518310765 |
| ENSMUSG00000037795 | N4bp2 | 333789 | protein_coding | 994.1757232 | 0.51860713 | 0.066473698 | 7.801689225 | 6.11E-15 | 3.83E-13 | 0.51860713 |
| ENSMUSG00000020099 | Unc5b | 107449 | protein_coding | 2645.278966 | 0.518617937 | 0.063661831 | 8.146450221 | 3.75E-16 | 2.52E-14 | 0.518617937 |

| ensembl_gene_id | external_gene_name | entrezgene_id | gene_biotype | baseMean | log2FoldChange | lfcSE | stat | pvalue | padj | absLog2FC |
|---|---|---|---|---|---|---|---|---|---|---|
| ENSMUSG00000031842 | Pde4c | 110385 | protein_coding | 6.156014855 | 0.519092192 | 0.492402046 | 1.054203971 | 0.291789543 | 0.534312473 | 0.519092192 |
| ENSMUSG00000041096 | Tspyl2 | 52808 | protein_coding | 147.6464927 | 0.519347296 | 0.119144005 | 4.358988064 | 1.31E-05 | 0.000249377 | 0.519347296 |
| ENSMUSG00000043243 | Niban3 | 100037278 | protein_coding | 11.49136007 | 0.519378942 | 0.38150371 | 1.361399453 | 0.173387489 | 0.399060944 | 0.519378942 |
| ENSMUSG00000025716 | Myo3a | 667663 | protein_coding | 154.1529688 | -0.519398003 | 0.162472494 | -3.196836532 | 0.001389436 | 0.013126796 | 0.519398003 |
| ENSMUSG00000028019 | Pdgfc | 54635 | protein_coding | 53.27566546 | 0.519517274 | 0.219277456 | 2.369223374 | 0.017825484 | 0.090770944 | 0.519517274 |
| ENSMUSG00000018604 | Tbx3 | 21386 | protein_coding | 353.0950349 | -0.519617742 | 0.120330376 | -4.318259105 | 1.57E-05 | 0.000293943 | 0.519617742 |
| ENSMUSG00000031101 | Sash3 | 74131 | protein_coding | 30.17271851 | 0.51981141 | 0.23400005 | 2.221415813 | 0.026322811 | 0.118795321 | 0.51981141 |
| ENSMUSG00000040836 | Gpr161 | 240888 | protein_coding | 7.626514671 | 0.51984161 | 0.691140941 | 0.752149929 | 0.451960901 | 0.680967534 | 0.51984161 |
| ENSMUSG00000031896 | Ctrl | 109660 | protein_coding | 9.795641198 | -0.51989747 | 0.432498592 | -1.20207899 | 0.229332926 | 0.468692446 | 0.51989747 |
| ENSMUSG00000085852 | Gm13807 | 639658 | lncRNA | 24.39482738 | -0.520471757 | 0.28104047 | -1.851945938 | 0.064033586 | 0.217038598 | 0.520471757 |
| ENSMUSG00000086894 | Gm15708 | 100504231 | lncRNA | 31.0762416 | -0.520625297 | 0.278757319 | -1.867665032 | 0.061808776 | 0.212128139 | 0.520625297 |
| ENSMUSG00000020268 | Lyrm7 | 75530 | protein_coding | 93.33074679 | 0.520759654 | 0.130234223 | 3.998639065 | 6.37E-05 | 0.001009248 | 0.520759654 |
| ENSMUSG00000023132 | Gzma | 14938 | protein_coding | 30.00841206 | -0.520884242 | 0.491038 | -1.060781939 | 0.288789012 | 0.531170873 | 0.520884242 |
| ENSMUSG00000064284 | Cdpf1 | 72355 | protein_coding | 560.9924941 | 0.520923927 | 0.08997828 | 5.789440792 | 7.06E-09 | 2.54E-07 | 0.520923927 |
| ENSMUSG00000026980 | Ly75 | 17076 | protein_coding | 966.7287449 | -0.521063306 | 0.115838295 | -4.49819558 | 6.85E-06 | 0.000140793 | 0.521063306 |
| ENSMUSG00000029390 | Tmed2 | 56334 | protein_coding | 2130.162874 | 0.521214571 | 0.048602604 | 10.72400514 | 7.85E-27 | 9.27E-25 | 0.521214571 |
| ENSMUSG00000058975 | Kcnc1 | 16502 | protein_coding | 7.023825494 | 0.521330091 | 0.518118393 | 1.006198773 | 0.314319962 | 0.558571684 | 0.521330091 |
| ENSMUSG00000116660 | 9030404E10Rik | 71528 | lncRNA | 23.64743739 | -0.52157447 | 0.265404674 | -1.965204536 | 0.049390569 | 0.183742032 | 0.52157447 |
| ENSMUSG00000035828 | Pim3 | 223775 | protein_coding | 3022.991785 | -0.521636408 | 0.141264504 | -3.692621954 | 0.000221954 | 0.002870449 | 0.521636408 |
| ENSMUSG00000061474 | Mrps36 | 66128 | protein_coding | 431.6752496 | 0.521658913 | 0.119860569 | 4.352214549 | 1.35E-05 | 0.0002563 | 0.521658913 |
| ENSMUSG00000038179 | Slamf7 | 75345 | protein_coding | 8.847302038 | 0.521979561 | 0.461857224 | 1.130175158 | 0.258402425 | 0.500383507 | 0.521979561 |
| ENSMUSG00000070331 | Qrich2 | 217341 | protein_coding | 10.92804309 | 0.521991759 | 0.419546416 | 1.244181188 | 0.213432886 | 0.449133255 | 0.521991759 |
| ENSMUSG00000097637 | 4933417D19Rik | 71186 | lncRNA | 9.615083262 | -0.522200249 | 0.390293714 | -1.337967358 | 0.180907078 | 0.407747138 | 0.522200249 |
| ENSMUSG00000010051 | Hyal1 | 15586 | protein_coding | 29.9766944 | -0.522214756 | 0.238843982 | -2.186426268 | 0.028784434 | 0.126280348 | 0.522214756 |
| ENSMUSG00000027227 | Sord | 20322 | protein_coding | 3851.297335 | -0.522307696 | 0.050992855 | -10.24276239 | 1.28E-24 | 1.34E-22 | 0.522307696 |
| ENSMUSG00000029674 | Limk1 | 16885 | protein_coding | 68.34436878 | 0.52231796 | 0.180973358 | 2.886159405 | 0.003899746 | 0.029651674 | 0.52231796 |
| ENSMUSG00000032355 | Mlip | 69642 | protein_coding | 8.996461554 | 0.52243434 | 0.476366941 | 1.0967057 | 0.272770064 | 0.515282046 | 0.52243434 |
| ENSMUSG00000025007 | Aldh18a1 | 56454 | protein_coding | 1175.676182 | 0.523005213 | 0.077222744 | 6.772683655 | 1.26E-11 | 6.01E-10 | 0.523005213 |
| ENSMUSG00000044197 | Gpr146 | 80290 | protein_coding | 15.81100754 | -0.523191442 | 0.331634961 | -1.577612447 | 0.114654671 | 0.312220666 | 0.523191442 |
| ENSMUSG00000025175 | Fn3k | 63828 | protein_coding | 27.54479298 | -0.523240088 | 0.282761449 | -1.850464731 | 0.064246597 | 0.21766909 | 0.523240088 |
| ENSMUSG00000026817 | Ak1 | 11636 | protein_coding | 18.48529971 | -0.523248844 | 0.334538589 | -1.564091142 | 0.117796166 | 0.317250963 | 0.523248844 |
| ENSMUSG00000097006 | 9530082P21Rik | 638247 | lncRNA | 128.9320559 | -0.523286664 | 0.176497068 | -2.964846218 | 0.003028343 | 0.024309648 | 0.523286664 |
| ENSMUSG00000029408 | Abcb9 | 56325 | protein_coding | 264.1422187 | 0.523330475 | 0.086094923 | 6.078528888 | 1.21E-09 | 4.76E-08 | 0.523330475 |
| ENSMUSG00000073940 | Hbb-bt | 101488143 | protein_coding | 10.32729345 | 0.523702693 | 0.491459186 | 1.0656077 | 0.286601017 | 0.529076763 | 0.523702693 |
| ENSMUSG00000032292 | Nr2e3 | 23958 | protein_coding | 81.51390592 | 0.523842153 | 0.166877964 | 3.13907326 | 0.001694831 | 0.015372629 | 0.523842153 |
| ENSMUSG00000001270 | Ckb | 12709 | protein_coding | 147.5028978 | 0.524167194 | 0.157136257 | 3.335749532 | 0.000850698 | 0.008805649 | 0.524167194 |
| ENSMUSG00000073598 | Smim33 | 73449 | protein_coding | 358.0063726 | 0.524184355 | 0.11602884 | 4.517707437 | 6.25E-06 | 0.000129415 | 0.524184355 |
| ENSMUSG00000056458 | Mok | 26448 | protein_coding | 10.6179542 | -0.524877732 | 0.430786485 | -1.21841736 | 0.223065412 | 0.460551754 | 0.524877732 |
| ENSMUSG00000074364 | Ehd2 | 259300 | protein_coding | 6.866746323 | 0.525625139 | 0.464658423 | 1.1312076 | 0.257967724 | 0.499952265 | 0.525625139 |

Supplementary Table 1

| ensembl_gene_id | external_gene_name | entrezgene_id | gene_biotype | baseMean | log2FoldChange | lfcSE | stat | pvalue | padj | absLog2FC |
|---|---|---|---|---|---|---|---|---|---|---|
| ENSMUSG00000001750 | Tcirg1 | 27060 | protein_coding | 509.8524762 | -0.525663581 | 0.068199138 | -7.707774559 | 1.28E-14 | 7.79E-13 | 0.525663581 |
| ENSMUSG00000040855 | Reps2 | 194590 | protein_coding | 1154.010203 | 0.526001824 | 0.055322434 | 9.50792985 | 1.94E-21 | 1.74E-19 | 0.526001824 |
| ENSMUSG00000031951 | Tmem231 | 234740 | protein_coding | 62.11948484 | 0.526043422 | 0.150122409 | 3.504096599 | 0.000458159 | 0.00526263 | 0.526043422 |
| ENSMUSG00000005413 | Hmox1 | 15368 | protein_coding | 49.25173591 | 0.526124396 | 0.209420053 | 2.512292348 | 0.011994967 | 0.068017876 | 0.526124396 |
| ENSMUSG00000041293 | Adgrf1 | 77596 | protein_coding | 100.8989715 | 0.52731801 | 0.185612226 | 2.840965922 | 0.004497712 | 0.032940326 | 0.52731801 |
| ENSMUSG00000073607 | Gm10548 | 633395 | transcribed_processed_pseudogene | 16.37544458 | -0.527625084 | 0.305053045 | -1.729617496 | 0.083698638 | 0.257588275 | 0.527625084 |
| ENSMUSG00000035875 | AI182371 | 98870 | protein_coding | 15.28826307 | -0.527654012 | 0.319104214 | -1.653547617 | 0.098219463 | 0.285043241 | 0.527654012 |
| ENSMUSG00000043102 | Qrfp | 227717 | protein_coding | 6.763093115 | -0.527801606 | 0.507483245 | -1.0400375 | 0.298322479 | 0.541946771 | 0.527801606 |
| ENSMUSG00000025433 | Crisp3 | 11572 | protein_coding | 38.55651378 | 0.528239954 | 0.200202815 | 2.638524108 | 0.008326778 | 0.052170143 | 0.528239954 |
| ENSMUSG00000092981 | Mir5125 | 100628593 | miRNA | 13.34009568 | -0.529100993 | 0.357773005 | -1.478873433 | 0.139174145 | 0.349495708 | 0.529100993 |
| ENSMUSG00000066687 | Zbtb16 | 235320 | protein_coding | 58.76218326 | -0.529149831 | 0.210848258 | -2.509623917 | 0.01208598 | 0.068389686 | 0.529149831 |
| ENSMUSG00000029638 | Glcci1 | 170772 | protein_coding | 48.5119125 | -0.529857887 | 0.250925486 | -2.111614469 | 0.03471953 | 0.143840743 | 0.529857887 |
| ENSMUSG00000031133 | Arhgef6 | 73341 | protein_coding | 15.18287624 | 0.530010019 | 0.321992204 | 1.646033702 | 0.099756821 | 0.287589962 | 0.530010019 |
| ENSMUSG00000039275 | Foxk2 | 68837 | protein_coding | 1038.350379 | 0.530055888 | 0.072738656 | 7.287127911 | 3.17E-13 | 1.71E-11 | 0.530055888 |
| ENSMUSG00000084771 | A230072E10Rik | 331547 | lncRNA | 10.02661123 | 0.530273181 | 0.389306912 | 1.362095472 | 0.173167757 | 0.398725931 | 0.530273181 |
| ENSMUSG00000058420 | Syt17 | 110058 | protein_coding | 13.2360767 | 0.530452547 | 0.327917737 | 1.617639082 | 0.105740401 | 0.297864708 | 0.530452547 |
| ENSMUSG00000024642 | Tle4 | 21888 | protein_coding | 1387.748456 | -0.53157348 | 0.104701467 | -5.077039448 | 3.83E-07 | 1.02E-05 | 0.53157348 |
| ENSMUSG00000026209 | Dnpep | 13437 | protein_coding | 2532.128334 | 0.531655408 | 0.061512994 | 8.642977188 | 5.48E-18 | 4.05E-16 | 0.531655408 |
| ENSMUSG00000089661 | Mia | 12587 | protein_coding | 8.207956096 | 0.531707659 | 0.460483449 | 1.154672683 | 0.24822449 | 0.489252203 | 0.531707659 |
| ENSMUSG00000070031 | Sp140 | 434484 | protein_coding | 37.57464409 | -0.532140381 | 0.334384676 | -1.591401819 | 0.111519175 | 0.306958481 | 0.532140381 |
| ENSMUSG00000047757 | Fancb | 237211 | protein_coding | 46.23644411 | 0.532419261 | 0.187496772 | 2.839618276 | 0.004516755 | 0.033016757 | 0.532419261 |
| ENSMUSG00000085169 | Gm10785 | 100038392 | lncRNA | 21.74688426 | -0.533594655 | 0.330510533 | -1.614455826 | 0.106428591 | 0.298655627 | 0.533594655 |
| ENSMUSG00000035849 | Krt222 | 268481 | protein_coding | 39.58844253 | -0.533638784 | 0.234630104 | -2.274383274 | 0.022942952 | 0.10799004 | 0.533638784 |
| ENSMUSG00000032202 | Rab27a | 11891 | protein_coding | 1086.52426 | 0.533813468 | 0.07645411 | 6.982142204 | 2.91E-12 | 1.47E-10 | 0.533813468 |
| ENSMUSG00000050822 | Slc29a4 | 243328 | protein_coding | 23.93447597 | -0.533837586 | 0.254579589 | -2.09693789 | 0.035999074 | 0.147231287 | 0.533837586 |
| ENSMUSG00000031422 | Morf4l2 | 56397 | protein_coding | 2793.671698 | 0.533842318 | 0.068665628 | 7.774520285 | 7.57E-15 | 4.70E-13 | 0.533842318 |
| ENSMUSG00000037145 | Lypd8l | 78354 | protein_coding | 367.3575229 | -0.534023791 | 0.194667454 | -2.7432618 | 0.006083216 | 0.04141158 | 0.534023791 |
| ENSMUSG00000039781 | Cep131 | 12009 | protein_coding | 281.6624398 | 0.534941229 | 0.079841956 | 6.700001541 | 2.08E-11 | 9.66E-10 | 0.534941229 |
| ENSMUSG00000037341 | Slc9a7 | 236727 | protein_coding | 18.40866983 | 0.535035871 | 0.305356196 | 1.752169691 | 0.079744634 | 0.249745186 | 0.535035871 |
| ENSMUSG00000020439 | Smtn | 29856 | protein_coding | 206.5051634 | 0.535091391 | 0.116558927 | 4.590737092 | 4.42E-06 | 9.46E-05 | 0.535091391 |
| ENSMUSG00000087396 | 4933407K13Rik | 74396 | lncRNA | 56.29722573 | -0.535131436 | 0.20402053 | -2.62292935 | 0.008717735 | 0.053867579 | 0.535131436 |
| ENSMUSG00000024300 | Myo1f | 17916 | protein_coding | 32.44049592 | 0.535258354 | 0.248492537 | 2.154021846 | 0.031238459 | 0.133735764 | 0.535258354 |
| ENSMUSG00000074006 | Omp | 18378 | protein_coding | 118.2550861 | 0.535341705 | 0.19171149 | 2.792434121 | 0.005231312 | 0.036921386 | 0.535341705 |
| ENSMUSG00000035042 | Ccl5 | 20304 | protein_coding | 47.31259015 | 0.536224757 | 0.218719046 | 2.451660101 | 0.014219891 | 0.077109679 | 0.536224757 |
| ENSMUSG00000037405 | Icam1 | 15894 | protein_coding | 23.17169148 | -0.536352917 | 0.397542152 | -1.349172442 | 0.177281584 | 0.404273205 | 0.536352917 |
| ENSMUSG00000024451 | Arap3 | 106952 | protein_coding | 19.15261565 | 0.536453673 | 0.356726297 | 1.5038243 | 0.132626613 | 0.339503983 | 0.536453673 |
| ENSMUSG00000019235 | Rps6kl1 | 238323 | protein_coding | 36.51117833 | -0.536655239 | 0.229749322 | -2.335829475 | 0.019500131 | 0.096404593 | 0.536655239 |
| ENSMUSG00000052852 | Reep1 | 52250 | protein_coding | 11.4964372 | 0.536935547 | 0.338199449 | 1.587629868 | 0.11237006 | 0.308615797 | 0.536935547 |
| ENSMUSG00000059005 | Hnrnpa3 | 229279 | protein_coding | 2441.928408 | 0.537018348 | 0.085086789 | 6.31141867 | 2.76E-10 | 1.16E-08 | 0.537018348 |

| ensembl_gene_id | external_gene_name | entrezgene_id | gene_biotype | baseMean | log2FoldChange | lfcSE | stat | pvalue | padj | absLog2FC |
|---|---|---|---|---|---|---|---|---|---|---|
| ENSMUSG00000037254 | Itih2 | 16425 | protein_coding | 311.9754847 | 0.537023066 | 0.115649169 | 4.643553176 | 3.42E-06 | 7.44E-05 | 0.537023066 |
| ENSMUSG00000020900 | Myh10 | 77579 | protein_coding | 38.30621883 | -0.537354059 | 0.298302213 | -1.801374698 | 0.071643841 | 0.233461352 | 0.537354059 |
| ENSMUSG00000042828 | Trim72 | 434246 | protein_coding | 101.0786727 | 0.537502518 | 0.133324051 | 4.031549562 | 5.54E-05 | 0.000896291 | 0.537502518 |
| ENSMUSG00000026496 | Parp1 | 11545 | protein_coding | 1513.802657 | 0.537639882 | 0.081411981 | 6.603940518 | 4.00E-11 | 1.82E-09 | 0.537639882 |
| ENSMUSG00000055022 | Cntn1 | 12805 | protein_coding | 6.798311515 | 0.53800632 | 0.498264489 | 1.079760512 | 0.28024884 | 0.523280426 | 0.53800632 |
| ENSMUSG00000027524 | Edn3 | 13616 | protein_coding | 145.4839994 | 0.53878033 | 0.133829125 | 4.025882496 | 5.68E-05 | 0.000913574 | 0.53878033 |
| ENSMUSG00000031594 | Fgl1 | 234199 | protein_coding | 25.14586409 | -0.538850831 | 0.240206304 | -2.24328347 | 0.024878541 | 0.114352201 | 0.538850831 |
| ENSMUSG00000023393 | Slc17a9 | 228993 | protein_coding | 1068.12706 | 0.539141565 | 0.072204601 | 7.466858846 | 8.21E-14 | 4.63E-12 | 0.539141565 |
| ENSMUSG00000031891 | Hsd11b2 | 15484 | protein_coding | 6774.782371 | 0.539172888 | 0.116588767 | 4.624569758 | 3.75E-06 | 8.07E-05 | 0.539172888 |
| ENSMUSG00000106943 | Dancr | 70036 | lncRNA | 49.36810772 | -0.539218631 | 0.189887358 | -2.839676306 | 0.004515933 | 0.033016757 | 0.539218631 |
| ENSMUSG00000055214 | Pld5 | 319455 | protein_coding | 11.42248062 | 0.539551817 | 0.351322937 | 1.53577168 | 0.124594388 | 0.327433832 | 0.539551817 |
| ENSMUSG00000078920 | Ifi47 | 15953 | protein_coding | 101.1678668 | -0.539605598 | 0.439227351 | -1.228533691 | 0.21924669 | 0.455544189 | 0.539605598 |
| ENSMUSG00000029359 | Tesc | 57816 | protein_coding | 80.57826733 | 0.539769122 | 0.1652492 | 3.266394764 | 0.001089263 | 0.010812233 | 0.539769122 |
| ENSMUSG00000019989 | Enpp3 | 209558 | protein_coding | 58.53568385 | 0.539966916 | 0.19870326 | 2.717453729 | 0.006578635 | 0.04380415 | 0.539966916 |
| ENSMUSG00000026117 | Zap70 | 22637 | protein_coding | 24.71246862 | 0.540395644 | 0.27292232 | 1.98003463 | 0.047699638 | 0.179396468 | 0.540395644 |
| ENSMUSG00000022371 | Col14a1 | 12818 | protein_coding | 7.01727359 | 0.540639459 | 0.566969665 | 0.953559763 | 0.340306532 | 0.583213766 | 0.540639459 |
| ENSMUSG00000020086 | Macroh2a2 | 404634 | protein_coding | 11.34089251 | 0.540752789 | 0.395031992 | 1.368883535 | 0.171035683 | 0.395624278 | 0.540752789 |
| ENSMUSG00000059900 | Tmem40 | 94346 | protein_coding | 21.16874985 | 0.540865974 | 0.250093042 | 2.162659025 | 0.030567415 | 0.131915576 | 0.540865974 |
| ENSMUSG00000020717 | Pecam1 | 18613 | protein_coding | 27.77668567 | -0.541286141 | 0.26466919 | -2.045142241 | 0.040840837 | 0.160879379 | 0.541286141 |
| ENSMUSG00000027246 | Ell3 | 269344 | protein_coding | 12.16744363 | -0.541306413 | 0.383587796 | -1.411166933 | 0.158195398 | 0.378014103 | 0.541306413 |
| ENSMUSG00000035836 | Ugt2b1 | 71773 | protein_coding | 6.74381785 | -0.541370889 | 0.500385765 | -1.081907054 | 0.27929383 | 0.522167763 | 0.541370889 |
| ENSMUSG00000032040 | Dcps | 69305 | protein_coding | 430.5025196 | 0.541823726 | 0.077890496 | 6.956223813 | 3.50E-12 | 1.73E-10 | 0.541823726 |
| ENSMUSG00000029452 | Tmem116 | 77462 | protein_coding | 39.02749595 | 0.542114406 | 0.181653135 | 2.984338294 | 0.002841924 | 0.023082976 | 0.542114406 |
| ENSMUSG00000056025 | Clca3a1 | 12722 | protein_coding | 40.66667819 | -0.542778239 | 0.249252488 | -2.177624153 | 0.02943403 | 0.128431439 | 0.542778239 |
| ENSMUSG00000030711 | Sult1a1 | 20887 | protein_coding | 9057.2022 | -0.543122377 | 0.041242881 | -13.16887565 | 1.33E-39 | 2.32E-37 | 0.543122377 |
| ENSMUSG00000089789 | Rdh1 | 107605 | protein_coding | 80.91611035 | -0.543335972 | 0.188103096 | -2.888500955 | 0.003870829 | 0.0294918 | 0.543335972 |
| ENSMUSG00000040003 | Magi2 | 50791 | protein_coding | 9.358281218 | 0.543943535 | 0.405028524 | 1.34297587 | 0.179279788 | 0.406473377 | 0.543943535 |
| ENSMUSG00000068196 | Col8a1 | 12837 | protein_coding | 538.804752 | -0.54394367 | 0.185219017 | -2.936759298 | 0.003316614 | 0.026180635 | 0.54394367 |
| ENSMUSG00000039384 | Dusp10 | 63953 | protein_coding | 96.0303916 | 0.544027258 | 0.129961587 | 4.18606199 | 2.84E-05 | 0.000492726 | 0.544027258 |
| ENSMUSG00000026986 | Hnmt | 140483 | protein_coding | 23.41321621 | 0.544107663 | 0.261161329 | 2.083415889 | 0.037213331 | 0.150573458 | 0.544107663 |
| ENSMUSG00000030671 | Pde3b | 18576 | protein_coding | 332.3080592 | 0.544387232 | 0.100437934 | 5.420135726 | 5.96E-08 | 1.89E-06 | 0.544387232 |
| ENSMUSG00000063239 | Grm4 | 268934 | protein_coding | 9.164365399 | -0.5445822 | 0.421539331 | -1.291889417 | 0.196395443 | 0.428914823 | 0.5445822 |
| ENSMUSG00000023912 | Slc25a27 | 74011 | protein_coding | 22.72533806 | -0.545081339 | 0.288716861 | -1.887944253 | 0.059033435 | 0.206112189 | 0.545081339 |
| ENSMUSG00000020076 | Ddx50 | 94213 | protein_coding | 1508.649185 | 0.545108748 | 0.049256111 | 11.06682474 | 1.82E-28 | 2.22E-26 | 0.545108748 |
| ENSMUSG00000072214 | Septin5 | 18951 | protein_coding | 683.1555513 | 0.545618683 | 0.078722483 | 6.930913037 | 4.18E-12 | 2.06E-10 | 0.545618683 |
| ENSMUSG00000025739 | Gng13 | 64337 | protein_coding | 6.023997145 | 0.545868018 | 0.520673327 | 1.048388673 | 0.29445957 | 0.537492868 | 0.545868018 |
| ENSMUSG00000050675 | Gp1ba | 14723 | protein_coding | 9.607866127 | -0.545943482 | 0.375784498 | -1.452810016 | 0.14627651 | 0.359247178 | 0.545943482 |
| ENSMUSG00000040345 | Arhgap9 | 216445 | protein_coding | 73.25890301 | -0.546210319 | 0.14316398 | -3.815277547 | 0.00013603 | 0.001917618 | 0.546210319 |
| ENSMUSG00000046351 | Zfp322a | 218100 | protein_coding | 437.9769331 | 0.546351019 | 0.084515717 | 6.464490137 | 1.02E-10 | 4.48E-09 | 0.546351019 |

| ensembl_gene_id | external_gene_name | entrezgene_id | gene_biotype | baseMean | log2FoldChange | lfcSE | stat | pvalue | padj | absLog2FC |
|---|---|---|---|---|---|---|---|---|---|---|
| ENSMUSG00000000983 | Wfdc18 | 14038 | protein_coding | 51.45178247 | 0.54693011 | 0.183034707 | 2.988122404 | 0.002806971 | 0.022897331 | 0.54693011 |
| ENSMUSG00000040621 | Gemin8 | 237221 | protein_coding | 38.63276458 | 0.547033321 | 0.188393773 | 2.903669867 | 0.003688168 | 0.028426826 | 0.547033321 |
| ENSMUSG00000025592 | Dach2 | 93837 | protein_coding | 16.53143273 | -0.547425592 | 0.290502778 | -1.884407426 | 0.059509886 | 0.207110275 | 0.547425592 |
| ENSMUSG00000040985 | Sun3 | 194974 | protein_coding | 20.15125396 | 0.547650903 | 0.341344635 | 1.604392881 | 0.108627479 | 0.302001032 | 0.547650903 |
| ENSMUSG00000041481 | Serpina3g | 20715 | protein_coding | 6.49253551 | 0.547683207 | 0.490461745 | 1.116668552 | 0.264136069 | 0.505545025 | 0.547683207 |
| ENSMUSG00000075701 | Selenos | 109815 | protein_coding | 1576.118824 | 0.547759534 | 0.039634467 | 13.82028245 | 1.92E-43 | 3.78E-41 | 0.547759534 |
| ENSMUSG00000054967 | Zfp647 | 239546 | protein_coding | 16.54719629 | -0.548099443 | 0.303096609 | -1.808332479 | 0.07055477 | 0.23105592 | 0.548099443 |
| ENSMUSG00000073067 | 9130019P16Rik | 100042056 | lncRNA | 9.184097178 | -0.548187737 | 0.428051992 | -1.280656901 | 0.200314204 | 0.433794437 | 0.548187737 |
| ENSMUSG00000043251 | Exoc3l | 277978 | protein_coding | 9.268797918 | 0.548250758 | 0.367778825 | 1.490707787 | 0.136038231 | 0.344951029 | 0.548250758 |
| ENSMUSG00000024885 | Aldh3b1 | 67689 | protein_coding | 136.0492168 | -0.54932865 | 0.132407974 | -4.148758078 | 3.34E-05 | 0.000572293 | 0.54932865 |
| ENSMUSG00000061126 | Cyp4f39 | 320997 | protein_coding | 8.236371109 | -0.550037895 | 0.490465045 | -1.121461967 | 0.262091272 | 0.503963985 | 0.550037895 |
| ENSMUSG00000044254 | Pcsk9 | 100102 | protein_coding | 1080.946606 | -0.55008094 | 0.249057919 | -2.208646659 | 0.027199227 | 0.121104897 | 0.55008094 |
| ENSMUSG00000092341 | Malat1 | 72289 | lncRNA | 52368.84961 | -0.550417878 | 0.283777537 | -1.939610454 | 0.052427049 | 0.191071418 | 0.550417878 |
| ENSMUSG00000032353 | Tmed3 | 66111 | protein_coding | 2095.516928 | 0.550822535 | 0.065963365 | 8.350431124 | 6.80E-17 | 4.81E-15 | 0.550822535 |
| ENSMUSG00000031340 | Gabre | 14404 | protein_coding | 24.1451571 | -0.551122952 | 0.350579009 | -1.572036368 | 0.115942117 | 0.314662099 | 0.551122952 |
| ENSMUSG00000029082 | Bst1 | 12182 | protein_coding | 9.804003297 | 0.551295621 | 0.58838374 | 0.936966105 | 0.348775993 | 0.59123817 | 0.551295621 |
| ENSMUSG00000025203 | Scd2 | 20250 | protein_coding | 24146.94792 | 0.551370953 | 0.17991306 | 3.064652198 | 0.002179233 | 0.018753727 | 0.551370953 |
| ENSMUSG00000043833 | 2900005J15Rik | 67261 | lncRNA | 14.81441342 | -0.551425723 | 0.340325752 | -1.620287976 | 0.105170431 | 0.297245625 | 0.551425723 |
| ENSMUSG00000001948 | Spa17 | 20686 | protein_coding | 17.37024902 | 0.551559462 | 0.282417733 | 1.952991604 | 0.05082059 | 0.187205038 | 0.551559462 |
| ENSMUSG00000020641 | Rsad2 | 58185 | protein_coding | 87.1793321 | -0.551583529 | 0.221964218 | -2.48501103 | 0.012954742 | 0.071992118 | 0.551583529 |
| ENSMUSG00000043753 | Dmrta1 | 242523 | protein_coding | 9.185220357 | 0.552407391 | 0.427869223 | 1.291065964 | 0.196680805 | 0.429095149 | 0.552407391 |
| ENSMUSG00000064899 | Snord118 | 100216530 | snoRNA | 77.06040607 | 0.552489751 | 0.180249082 | 3.065145989 | 0.002175638 | 0.018732787 | 0.552489751 |
| ENSMUSG00000028521 | Slc35d1 | 242585 | protein_coding | 667.1108809 | -0.552799795 | 0.117616717 | -4.700010398 | 2.60E-06 | 5.82E-05 | 0.552799795 |
| ENSMUSG00000036676 | Tmtc3 | 237500 | protein_coding | 559.752325 | 0.553136451 | 0.067702311 | 8.170126637 | 3.08E-16 | 2.09E-14 | 0.553136451 |
| ENSMUSG00000004988 | Fxyd4 | 108017 | protein_coding | 668.9197949 | -0.553210221 | 0.307400177 | -1.799641846 | 0.071917209 | 0.234068382 | 0.553210221 |
| ENSMUSG00000109372 | Gm19410 | 100502846 | protein_coding | 7.919925828 | 0.553446267 | 0.437713011 | 1.264404421 | 0.206084907 | 0.440819483 | 0.553446267 |
| ENSMUSG00000075705 | Msrb1 | 27361 | protein_coding | 1198.633109 | 0.553546126 | 0.054387136 | 10.1778871 | 2.49E-24 | 2.56E-22 | 0.553546126 |
| ENSMUSG00000021675 | F2rl2 | 14064 | protein_coding | 13.23908501 | -0.554105318 | 0.370594092 | -1.495181196 | 0.13486716 | 0.342694737 | 0.554105318 |
| ENSMUSG00000039474 | Wfs1 | 22393 | protein_coding | 292.3829718 | 0.554278432 | 0.083472879 | 6.640221848 | 3.13E-11 | 1.43E-09 | 0.554278432 |
| ENSMUSG00000055172 | C1ra | 50909 | protein_coding | 26.46856775 | -0.554457376 | 0.280243815 | -1.978482116 | 0.047874344 | 0.179676413 | 0.554457376 |
| ENSMUSG00000092274 | Neat1 | 66961 | lncRNA | 11334.5788 | -0.555132789 | 0.312019055 | -1.779163096 | 0.075213027 | 0.241001487 | 0.555132789 |
| ENSMUSG00000037801 | Iqch | 78250 | protein_coding | 29.70139016 | -0.555841215 | 0.283434481 | -1.96109243 | 0.049868242 | 0.18483685 | 0.555841215 |
| ENSMUSG00000060992 | Copz1 | 56447 | protein_coding | 2551.354768 | 0.555900939 | 0.061717735 | 9.007150627 | 2.11E-19 | 1.73E-17 | 0.555900939 |
| ENSMUSG00000020080 | Hkdc1 | 216019 | protein_coding | 1216.424505 | -0.555932287 | 0.142250344 | -3.90812614 | 9.30E-05 | 0.001394097 | 0.555932287 |
| ENSMUSG00000037686 | Aspg | 104816 | protein_coding | 1347.926434 | -0.556030074 | 0.129735765 | -4.285865786 | 1.82E-05 | 0.000331704 | 0.556030074 |
| ENSMUSG00000033453 | Adamts15 | 235130 | protein_coding | 573.0760636 | 0.556150777 | 0.158014989 | 3.519607739 | 0.000432185 | 0.004999896 | 0.556150777 |
| ENSMUSG00000060176 | Kif27 | 75050 | protein_coding | 52.8544311 | -0.556596446 | 0.240906795 | -2.310422361 | 0.020864782 | 0.10129029 | 0.556596446 |
| ENSMUSG00000049350 | Zg16 | 69036 | protein_coding | 90928.39472 | 0.556720745 | 0.083796487 | 6.643724167 | 3.06E-11 | 1.40E-09 | 0.556720745 |
| ENSMUSG00000037447 | Arid5a | 214855 | protein_coding | 154.1950011 | -0.556849701 | 0.140150585 | -3.97322424 | 7.09E-05 | 0.001105904 | 0.556849701 |

| ensembl_gene_id | external_gene_name | entrezgene_id | gene_biotype | baseMean | log2FoldChange | lfcSE | stat | pvalue | padj | absLog2FC |
|---|---|---|---|---|---|---|---|---|---|---|
| ENSMUSG00000096856 | Gm10778 | 100233208 | protein_coding | 7.456877323 | -0.556949351 | 0.428182127 | -1.300730029 | 0.19335088 | 0.425279965 | 0.556949351 |
| ENSMUSG00000019935 | Slc17a8 | 216227 | protein_coding | 10.97212936 | 0.557687196 | 0.345366677 | 1.614768401 | 0.106360859 | 0.298608218 | 0.557687196 |
| ENSMUSG00000073102 | Drc1 | 381738 | protein_coding | 21.87201045 | 0.558045268 | 0.280264823 | 1.991135607 | 0.046465981 | 0.176029333 | 0.558045268 |
| ENSMUSG00000041809 | Efhc1 | 71877 | protein_coding | 8.442235562 | 0.558707607 | 0.468617937 | 1.192245459 | 0.233165013 | 0.472624422 | 0.558707607 |
| ENSMUSG00000032035 | Ets1 | 23871 | protein_coding | 118.5696409 | 0.559511655 | 0.171806826 | 3.256632275 | 0.001127424 | 0.01114091 | 0.559511655 |
| ENSMUSG00000069633 | Pex11g | 69129 | protein_coding | 207.946841 | 0.559910793 | 0.089761436 | 6.237765554 | 4.44E-10 | 1.83E-08 | 0.559910793 |
| ENSMUSG00000047819 | Tigd4 | 403175 | protein_coding | 7.317776514 | 0.560728547 | 0.488598778 | 1.147625767 | 0.251123088 | 0.492145102 | 0.560728547 |
| ENSMUSG00000070284 | Gmppb | 331026 | protein_coding | 1219.750945 | 0.560767331 | 0.064910127 | 8.639134763 | 5.66E-18 | 4.17E-16 | 0.560767331 |
| ENSMUSG00000009739 | Pou6f1 | 19009 | protein_coding | 56.27396319 | -0.56145203 | 0.181013578 | -3.101712241 | 0.001924049 | 0.01702092 | 0.56145203 |
| ENSMUSG00000029651 | Mtus2 | 77521 | protein_coding | 89.12185876 | -0.561517954 | 0.136030402 | -4.127885713 | 3.66E-05 | 0.000619329 | 0.561517954 |
| ENSMUSG00000000869 | Il4 | 16189 | protein_coding | 5.948993319 | 0.561528911 | 0.611066394 | 0.918932732 | 0.358130758 | 0.601184005 | 0.561528911 |
| ENSMUSG00000036853 | Mcoln3 | 171166 | protein_coding | 7.722377487 | 0.561561504 | 0.420557966 | 1.335277295 | 0.181785607 | 0.409163501 | 0.561561504 |
| ENSMUSG00000020135 | Apc2 | 23805 | protein_coding | 14.30982543 | -0.561563127 | 0.407002031 | -1.379755099 | 0.167662062 | 0.391604912 | 0.561563127 |
| ENSMUSG00000048572 | Tmem252 | 226040 | protein_coding | 441.6479423 | -0.562565506 | 0.121342212 | -4.636189649 | 3.55E-06 | 7.66E-05 | 0.562565506 |
| ENSMUSG00000038372 | Gmds | 218138 | protein_coding | 5566.419001 | 0.562719738 | 0.055202161 | 10.19379911 | 2.11E-24 | 2.18E-22 | 0.562719738 |
| ENSMUSG00000070563 | Spaca4 | 69363 | protein_coding | 193.6138846 | -0.563976344 | 0.147891376 | -3.813449825 | 0.00013704 | 0.001930175 | 0.563976344 |
| ENSMUSG00000035640 | Cbarp | 100503659 | protein_coding | 43.59266212 | -0.564032306 | 0.2085893 | -2.704032777 | 0.006850353 | 0.04505532 | 0.564032306 |
| ENSMUSG00000031367 | Ap1s2 | 108012 | protein_coding | 39.20066845 | 0.5642638 | 0.230137282 | 2.45185741 | 0.014212097 | 0.077108863 | 0.5642638 |
| ENSMUSG00000028996 | Rbp7 | 63954 | protein_coding | 13.11148393 | -0.565092302 | 0.383997872 | -1.471602692 | 0.141128198 | 0.352173719 | 0.565092302 |
| ENSMUSG00000056148 | Rdh9 | 103142 | protein_coding | 31.17385618 | -0.565098116 | 0.248701412 | -2.272195043 | 0.023074733 | 0.108428383 | 0.565098116 |
| ENSMUSG00000072875 | Gpr27 | 14761 | protein_coding | 15.81357555 | -0.565171692 | 0.309872424 | -1.823885083 | 0.068169447 | 0.226114495 | 0.565171692 |
| ENSMUSG00000025528 | 2010106E10Rik | 67715 | protein_coding | 119.1354735 | -0.565283568 | 0.22535359 | -2.508429385 | 0.01212692 | 0.068515133 | 0.565283568 |
| ENSMUSG00000021838 | Samd4 | 74480 | protein_coding | 6.564366976 | 0.565311581 | 0.562687804 | 1.004662936 | 0.315059181 | 0.559370353 | 0.565311581 |
| ENSMUSG00000026874 | Hc | 15139 | protein_coding | 21.17623912 | -0.565344067 | 0.29984985 | -1.885423878 | 0.059372633 | 0.20693807 | 0.565344067 |
| ENSMUSG00000047228 | A2ml1 | 232400 | protein_coding | 14.23157706 | -0.565364531 | 0.320930723 | -1.76164041 | 0.078130072 | 0.246748312 | 0.565364531 |
| ENSMUSG00000040152 | Thbs1 | 21825 | protein_coding | 4869.781055 | -0.565387198 | 0.121906867 | -4.637861784 | 3.52E-06 | 7.61E-05 | 0.565387198 |
| ENSMUSG00000020089 | Ppa1 | 67895 | protein_coding | 5697.784315 | 0.565446373 | 0.110366506 | 5.123351241 | 3.00E-07 | 8.19E-06 | 0.565446373 |
| ENSMUSG00000100291 | 2310069B03Rik | 69652 | lncRNA | 9.051525223 | -0.566217012 | 0.506577843 | -1.117729524 | 0.263682529 | 0.504916665 | 0.566217012 |
| ENSMUSG00000037784 | Dzip1l | 72507 | protein_coding | 31.15329044 | -0.566280799 | 0.229789506 | -2.464345785 | 0.013726362 | 0.07529423 | 0.566280799 |
| ENSMUSG00000064267 | Hvcn1 | 74096 | protein_coding | 8.8645545 | 0.566492881 | 0.498006925 | 1.137520088 | 0.255320932 | 0.496630161 | 0.566492881 |
| ENSMUSG00000032548 | Slco2a1 | 24059 | protein_coding | 784.9416083 | -0.566565268 | 0.116799774 | -4.850739429 | 1.23E-06 | 2.95E-05 | 0.566565268 |
| ENSMUSG00000042306 | S100a14 | 66166 | protein_coding | 1414.419977 | 0.567451316 | 0.14061284 | 4.035558301 | 5.45E-05 | 0.000883782 | 0.567451316 |
| ENSMUSG00000110622 | Iqcn | 637079 | protein_coding | 16.99224454 | -0.567744225 | 0.415284096 | -1.367122487 | 0.171586916 | 0.396321966 | 0.567744225 |
| ENSMUSG00000103585 | Pcdhgb4 | 93701 | protein_coding | 10.65943197 | -0.568589803 | 0.367787359 | -1.545974297 | 0.122110775 | 0.323870851 | 0.568589803 |
| ENSMUSG00000041696 | Rasl12 | 70784 | protein_coding | 22.72111231 | -0.569650518 | 0.319759959 | -1.781494219 | 0.074831748 | 0.240298679 | 0.569650518 |
| ENSMUSG00000085148 | Mir22hg | 100042498 | lncRNA | 362.8808902 | -0.569913174 | 0.137722156 | -4.138137185 | 3.50E-05 | 0.000596268 | 0.569913174 |
| ENSMUSG00000034786 | Gpsm3 | 106512 | protein_coding | 10.1776343 | 0.569921892 | 0.380711598 | 1.496991148 | 0.134395561 | 0.342130579 | 0.569921892 |
| ENSMUSG00000039316 | Rftn1 | 76438 | protein_coding | 10.18861137 | 0.570109435 | 0.393562545 | 1.448586615 | 0.147453062 | 0.361065218 | 0.570109435 |
| ENSMUSG00000045629 | Sh3tc2 | 225608 | protein_coding | 42.19639553 | 0.570145496 | 0.203565676 | 2.800793859 | 0.005097707 | 0.036168378 | 0.570145496 |

Supplementary Table 1

| ensembl_gene_id | external_gene_name | entrezgene_id | gene_biotype | baseMean | log2FoldChange | lfcSE | stat | pvalue | padj | absLog2FC |
|---|---|---|---|---|---|---|---|---|---|---|
| ENSMUSG00000041845 | Rhod | 11854 | protein_coding | 129.8533372 | 0.570147481 | 0.163831724 | 3.480079851 | 0.000501264 | 0.005660988 | 0.570147481 |
| ENSMUSG00000038276 | Asic3 | 171209 | protein_coding | 9.900631063 | -0.57015495 | 0.430053076 | -1.325778103 | 0.184913208 | 0.413835041 | 0.57015495 |
| ENSMUSG00000078234 | Klhdc7a | 242721 | protein_coding | 489.2341991 | 0.570171318 | 0.104903946 | 5.435175149 | 5.47E-08 | 1.75E-06 | 0.570171318 |
| ENSMUSG00000016493 | Cd46 | 17221 | protein_coding | 39.11822437 | -0.570241943 | 0.238295666 | -2.393001737 | 0.016711162 | 0.086684114 | 0.570241943 |
| ENSMUSG00000078492 | 1700045H11Rik | 73348 | lncRNA | 14.42859926 | -0.570350529 | 0.400728423 | -1.423284437 | 0.154653717 | 0.372364958 | 0.570350529 |
| ENSMUSG00000091387 | Gcnt4 | 218476 | protein_coding | 295.3911517 | -0.570893469 | 0.150912986 | -3.78293138 | 0.000154992 | 0.002123669 | 0.570893469 |
| ENSMUSG00000031431 | Tsc22d3 | 14605 | protein_coding | 1504.364662 | 0.571145826 | 0.085751018 | 6.66051364 | 2.73E-11 | 1.25E-09 | 0.571145826 |
| ENSMUSG00000036923 | Stox1 | 216021 | protein_coding | 5.977161413 | 0.571181662 | 0.497186002 | 1.148828928 | 0.250626528 | 0.491645551 | 0.571181662 |
| ENSMUSG00000026070 | Il18r1 | 16182 | protein_coding | 17.11762603 | 0.571306942 | 0.329781031 | 1.732382665 | 0.083205452 | 0.256715961 | 0.571306942 |
| ENSMUSG00000094595 | Fsbp | 100503583 | protein_coding | 5.868379642 | 0.571315975 | 0.53483365 | 1.068212471 | 0.285424681 | 0.528356729 | 0.571315975 |
| ENSMUSG00000062296 | Trank1 | 320429 | protein_coding | 7.942663806 | 0.571317181 | 0.412417018 | 1.385290022 | 0.165963803 | 0.389376941 | 0.571317181 |
| ENSMUSG00000025467 | Prap1 | 22264 | protein_coding | 217.6073185 | -0.571335532 | 0.189881612 | -3.008903949 | 0.00262192 | 0.021628492 | 0.571335532 |
| ENSMUSG00000029156 | Sgcb | 24051 | protein_coding | 58.17256731 | -0.57153775 | 0.17632037 | -3.241473168 | 0.001189136 | 0.011636651 | 0.57153775 |
| ENSMUSG00000058396 | Gpr182 | 11536 | protein_coding | 7.961983773 | 0.571710181 | 0.472158874 | 1.210842818 | 0.225955652 | 0.464143013 | 0.571710181 |
| ENSMUSG00000072844 | G530011O06Rik | 654820 | lncRNA | 6.495490822 | -0.572308361 | 0.519390746 | -1.101884016 | 0.270512098 | 0.513120279 | 0.572308361 |
| ENSMUSG00000052415 | Tchh | 99681 | protein_coding | 35.53604869 | 0.572346629 | 0.223661445 | 2.558986543 | 0.010497781 | 0.061495716 | 0.572346629 |
| ENSMUSG00000017607 | Tns4 | 217169 | protein_coding | 2758.041715 | -0.572379334 | 0.055527133 | -10.30810178 | 6.48E-25 | 6.96E-23 | 0.572379334 |
| ENSMUSG00000033634 | Nat8f2 | 93673 | protein_coding | 8.369090638 | -0.572891783 | 0.45313617 | -1.264281734 | 0.206128923 | 0.440822324 | 0.572891783 |
| ENSMUSG00000002068 | Ccne1 | 12447 | protein_coding | 94.57936824 | 0.573412677 | 0.195934255 | 2.926556543 | 0.003427372 | 0.026849159 | 0.573412677 |
| ENSMUSG00000037736 | Limch1 | 77569 | protein_coding | 76.87284281 | 0.574126442 | 0.21400305 | 2.682795604 | 0.00730096 | 0.047191414 | 0.574126442 |
| ENSMUSG00000078566 | Bnip3 | 12176 | protein_coding | 147.216942 | -0.574636342 | 0.125556755 | -4.576705902 | 4.72E-06 | 0.000100623 | 0.574636342 |
| ENSMUSG00000020101 | Vsir | 74048 | protein_coding | 35.18858951 | 0.575453467 | 0.223988093 | 2.569125259 | 0.01019556 | 0.060118388 | 0.575453467 |
| ENSMUSG00000085196 | Gm14963 | 105246847 | lncRNA | 13.21685753 | -0.575599875 | 0.417446323 | -1.378859612 | 0.167938043 | 0.391953183 | 0.575599875 |
| ENSMUSG00000069920 | B3gnt9 | 97440 | protein_coding | 77.69292257 | -0.575602285 | 0.180551054 | -3.188030602 | 0.001432454 | 0.013462227 | 0.575602285 |
| ENSMUSG00000040658 | Dnph1 | 381101 | protein_coding | 67.98665954 | -0.57570903 | 0.151264907 | -3.805965591 | 0.000141252 | 0.001975691 | 0.57570903 |
| ENSMUSG00000052142 | Rasal3 | 320484 | protein_coding | 23.9489261 | 0.576762843 | 0.302548172 | 1.906350449 | 0.056604753 | 0.201027274 | 0.576762843 |
| ENSMUSG00000042589 | Cux2 | 13048 | protein_coding | 34.46997342 | -0.577020041 | 0.267577175 | -2.156462118 | 0.031047599 | 0.133152759 | 0.577020041 |
| ENSMUSG00000037649 | H2-DMa | 14998 | protein_coding | 61.16767188 | -0.577096013 | 0.378901834 | -1.523075271 | 0.127739875 | 0.332543725 | 0.577096013 |
| ENSMUSG00000056724 | Nbeal2 | 235627 | protein_coding | 1800.527593 | 0.577245912 | 0.061574759 | 9.374716555 | 6.94E-21 | 5.98E-19 | 0.577245912 |
| ENSMUSG00000078768 | Zfp566 | 72556 | protein_coding | 21.18972739 | -0.577560029 | 0.287209563 | -2.010935926 | 0.044332225 | 0.170740692 | 0.577560029 |
| ENSMUSG00000029346 | Srrd | 70118 | protein_coding | 136.8290754 | 0.577717239 | 0.109509446 | 5.275501422 | 1.32E-07 | 3.95E-06 | 0.577717239 |
| ENSMUSG00000025557 | Slc15a1 | 56643 | protein_coding | 1831.653878 | -0.578689905 | 0.23649103 | -2.44698459 | 0.0144057 | 0.077881569 | 0.578689905 |
| ENSMUSG00000034614 | Pik3ip1 | 216505 | protein_coding | 83.90717719 | -0.579072268 | 0.157764778 | -3.670478758 | 0.000242097 | 0.003093734 | 0.579072268 |
| ENSMUSG00000085154 | C130046K22Rik | 399609 | lncRNA | 12.88189003 | 0.57912283 | 0.337461531 | 1.716115102 | 0.086140978 | 0.262471712 | 0.57912283 |
| ENSMUSG00000097908 | 4933404O12Rik | 66752 | lncRNA | 48.9191053 | -0.579511086 | 0.263890954 | -2.19602482 | 0.028090167 | 0.12424678 | 0.579511086 |
| ENSMUSG00000071540 | 3425401B19Rik | 100504518 | protein_coding | 16.04526376 | 0.579520641 | 0.368469745 | 1.572776732 | 0.115770525 | 0.314420893 | 0.579520641 |
| ENSMUSG00000026830 | Ermn | 77767 | protein_coding | 12.66788013 | 0.580668081 | 0.353838415 | 1.641054383 | 0.100786127 | 0.289522242 | 0.580668081 |
| ENSMUSG00000029767 | Calu | 12321 | protein_coding | 3586.19217 | 0.580918298 | 0.051213793 | 11.34300481 | 8.03E-30 | 1.06E-27 | 0.580918298 |
| ENSMUSG00000022678 | Nde1 | 67203 | protein_coding | 542.2161311 | 0.581377422 | 0.084351033 | 6.892356878 | 5.49E-12 | 2.67E-10 | 0.581377422 |

| ensembl_gene_id | external_gene_name | entrezgene_id | gene_biotype | baseMean | log2FoldChange | lfcSE | stat | pvalue | padj | absLog2FC |
|---|---|---|---|---|---|---|---|---|---|---|
| ENSMUSG00000003411 | Rab3b | 69908 | protein_coding | 28.89699264 | 0.581462521 | 0.258530656 | 2.249104732 | 0.024505834 | 0.113239421 | 0.581462521 |
| ENSMUSG00000028758 | Kif17 | 16559 | protein_coding | 8.815786529 | 0.581979308 | 0.520498333 | 1.118119446 | 0.263515982 | 0.504916665 | 0.581979308 |
| ENSMUSG00000045555 | Mettl24 | 327747 | protein_coding | 37.39309695 | 0.58209563 | 0.206786094 | 2.814965062 | 0.004878254 | 0.034965035 | 0.58209563 |
| ENSMUSG00000055748 | Gsdmc4 | 74548 | protein_coding | 16683.12875 | 0.582214936 | 0.129047377 | 4.511637107 | 6.43E-06 | 0.000132834 | 0.582214936 |
| ENSMUSG00000030670 | Cyp2r1 | 244209 | protein_coding | 68.48741378 | 0.583325194 | 0.146235061 | 3.988955791 | 6.64E-05 | 0.001048252 | 0.583325194 |
| ENSMUSG00000034842 | Art3 | 109979 | protein_coding | 6.675408913 | 0.583466734 | 0.460418693 | 1.267252489 | 0.205065023 | 0.439537962 | 0.583466734 |
| ENSMUSG00000084799 | Ino80dos | 100380944 | lncRNA | 26.47262788 | -0.583690836 | 0.252962005 | -2.307424928 | 0.021031143 | 0.101791489 | 0.583690836 |
| ENSMUSG00000049999 | Ppp1r3d | 228966 | protein_coding | 98.13721906 | 0.584268834 | 0.129261874 | 4.520039929 | 6.18E-06 | 0.000128327 | 0.584268834 |
| ENSMUSG00000003352 | Cacnb3 | 12297 | protein_coding | 965.23093 | -0.584483487 | 0.080100363 | -7.296889381 | 2.94E-13 | 1.60E-11 | 0.584483487 |
| ENSMUSG00000024771 | Lipk | 240633 | protein_coding | 22.68982742 | 0.584877958 | 0.300587499 | 1.945782708 | 0.051680847 | 0.189222442 | 0.584877958 |
| ENSMUSG00000026969 | Fam166a | 68222 | protein_coding | 19.91078943 | -0.584941921 | 0.408926599 | -1.430432557 | 0.15259291 | 0.369310836 | 0.584941921 |
| ENSMUSG00000047867 | Gimap6 | 231931 | protein_coding | 65.65045635 | 0.585756026 | 0.199246692 | 2.939853202 | 0.003283678 | 0.025971492 | 0.585756026 |
| ENSMUSG00000031060 | Rbm10 | 236732 | protein_coding | 1653.196691 | 0.585773109 | 0.060309574 | 9.712771421 | 2.66E-22 | 2.55E-20 | 0.585773109 |
| ENSMUSG00000027827 | Kcnab1 | 16497 | protein_coding | 8.409876036 | 0.586018203 | 0.435493966 | 1.345640236 | 0.178418569 | 0.405619009 | 0.586018203 |
| ENSMUSG00000025515 | Muc2 | 17831 | protein_coding | 56327.11605 | -0.586626949 | 0.1200515 | -4.88646079 | 1.03E-06 | 2.51E-05 | 0.586626949 |
| ENSMUSG00000037752 | Xkr8 | 381560 | protein_coding | 34.32178592 | -0.586914054 | 0.26737929 | -2.195061761 | 0.028159168 | 0.124383155 | 0.586914054 |
| ENSMUSG00000042213 | Zfand4 | 67492 | protein_coding | 10.64325312 | 0.586954119 | 0.379622372 | 1.546152604 | 0.122067717 | 0.323833866 | 0.586954119 |
| ENSMUSG00000040584 | Abcb1a | 18671 | protein_coding | 9831.363996 | -0.587335577 | 0.072941513 | -8.052144169 | 8.14E-16 | 5.40E-14 | 0.587335577 |
| ENSMUSG00000079299 | Klrb1 | 100043861 | protein_coding | 7.609031059 | 0.587437666 | 0.43140425 | 1.36168725 | 0.173296607 | 0.398908703 | 0.587437666 |
| ENSMUSG00000004451 | Ralb | 64143 | protein_coding | 2359.206752 | 0.587530481 | 0.061820856 | 9.503758349 | 2.02E-21 | 1.80E-19 | 0.587530481 |
| ENSMUSG00000096957 | E230013L22Rik | 102132 | lncRNA | 60.25798143 | 0.587546742 | 0.174615818 | 3.364796778 | 0.000766001 | 0.008068773 | 0.587546742 |
| ENSMUSG00000032033 | Barx2 | 12023 | protein_coding | 42.6628051 | 0.587731332 | 0.239376568 | 2.455258416 | 0.014078334 | 0.076614782 | 0.587731332 |
| ENSMUSG00000048216 | Gpr85 | 64450 | protein_coding | 8.854055129 | 0.588024423 | 0.434186685 | 1.354312426 | 0.175636723 | 0.402057266 | 0.588024423 |
| ENSMUSG00000040852 | Plekhh2 | 213556 | protein_coding | 30.61770584 | 0.588097022 | 0.222183676 | 2.646895724 | 0.00812344 | 0.051214509 | 0.588097022 |
| ENSMUSG00000088990 | Gm22767 | 115490051 | scaRNA | 13.62318294 | -0.588224847 | 0.416739437 | -1.411493117 | 0.158099264 | 0.377840372 | 0.588224847 |
| ENSMUSG00000029591 | Ung | 22256 | protein_coding | 337.1188214 | 0.588584386 | 0.111220504 | 5.292049279 | 1.21E-07 | 3.64E-06 | 0.588584386 |
| ENSMUSG00000033389 | Arhgap44 | 216831 | protein_coding | 11.71951504 | -0.588592827 | 0.403104348 | -1.460150031 | 0.144248845 | 0.356739936 | 0.588592827 |
| ENSMUSG00000110086 | Smim32 | 115488204 | protein_coding | 13.13218533 | 0.589012338 | 0.332691755 | 1.770444653 | 0.076653097 | 0.244015888 | 0.589012338 |
| ENSMUSG00000026880 | Stom | 13830 | protein_coding | 635.0133498 | -0.589124985 | 0.124900731 | -4.716745692 | 2.40E-06 | 5.40E-05 | 0.589124985 |
| ENSMUSG00000033900 | Map9 | 213582 | protein_coding | 36.24886026 | 0.589229966 | 0.215673323 | 2.732048439 | 0.006294189 | 0.042447899 | 0.589229966 |
| ENSMUSG00000043687 | 1190005I06Rik | 68918 | protein_coding | 7.795580766 | -0.589479047 | 0.481463857 | -1.224347452 | 0.220821183 | 0.457336743 | 0.589479047 |
| ENSMUSG00000004698 | Hdac9 | 79221 | protein_coding | 62.63947519 | -0.589934282 | 0.178523568 | -3.304517649 | 0.0009514 | 0.009649827 | 0.589934282 |
| ENSMUSG00000049744 | Arhgap15 | 76117 | protein_coding | 8.047292626 | 0.59035978 | 0.522551303 | 1.129764247 | 0.258575577 | 0.500565425 | 0.59035978 |
| ENSMUSG00000073856 | Iqck | 434232 | protein_coding | 7.862416504 | 0.590406304 | 0.421108734 | 1.40202816 | 0.160906838 | 0.381385152 | 0.590406304 |
| ENSMUSG00000032207 | Lipc | 15450 | protein_coding | 7.972909174 | 0.590657742 | 0.417097508 | 1.416114292 | 0.156742044 | 0.375480655 | 0.590657742 |
| ENSMUSG00000022805 | Cfap91 | 320214 | protein_coding | 17.35441184 | 0.590756363 | 0.32139326 | 1.838110618 | 0.066046107 | 0.221624129 | 0.590756363 |
| ENSMUSG00000006216 | Clcnkb | 56365 | protein_coding | 15.92772636 | 0.59083945 | 0.322267043 | 1.833384652 | 0.066745388 | 0.222950058 | 0.59083945 |
| ENSMUSG00000054909 | Wbscr25 | 71304 | lncRNA | 21.03252694 | -0.591316795 | 0.282726457 | -2.091480229 | 0.036485037 | 0.148621922 | 0.591316795 |
| ENSMUSG00000044906 | 4930503L19Rik | 269033 | protein_coding | 6.427208229 | 0.592109775 | 0.496290763 | 1.193070312 | 0.232841837 | 0.472332114 | 0.592109775 |

Supplementary Table 1

| ensembl_gene_id | external_gene_name | entrezgene_id | gene_biotype | baseMean | log2FoldChange | lfcSE | stat | pvalue | padj | absLog2FC |
|---|---|---|---|---|---|---|---|---|---|---|
| ENSMUSG00000004707 | Ly9 | 17085 | protein_coding | 11.5584429 | 0.592158035 | 0.35359383 | 1.674684299 | 0.09399619 | 0.277226874 | 0.592158035 |
| ENSMUSG00000041377 | Ninj2 | 29862 | protein_coding | 15.84500957 | 0.592220784 | 0.378385711 | 1.565124601 | 0.117553696 | 0.317074504 | 0.592220784 |
| ENSMUSG00000030621 | Me3 | 109264 | protein_coding | 98.13839173 | -0.592502191 | 0.182349213 | -3.249271988 | 0.001157008 | 0.01137748 | 0.592502191 |
| ENSMUSG00000054901 | Arhgef33 | 381112 | protein_coding | 7.884510541 | 0.593297457 | 0.405785395 | 1.462096627 | 0.143714733 | 0.355746355 | 0.593297457 |
| ENSMUSG00000066952 | Myo1h | 231646 | protein_coding | 16.1856479 | -0.5939043 | 0.300327242 | -1.977523905 | 0.047982441 | 0.179789226 | 0.5939043 |
| ENSMUSG00000117226 | Gm31235 | 102633401 | lncRNA | 63.71708448 | 0.594071872 | 0.168157422 | 3.532831695 | 0.000411134 | 0.004783809 | 0.594071872 |
| ENSMUSG00000052563 | D930048N14Rik | 97775 | lncRNA | 34.76355828 | -0.594880266 | 0.206708547 | -2.877869712 | 0.004003705 | 0.030313495 | 0.594880266 |
| ENSMUSG00000047787 | Flrt1 | 396184 | protein_coding | 28.90817871 | -0.595353431 | 0.320286688 | -1.858814159 | 0.063053485 | 0.21475471 | 0.595353431 |
| ENSMUSG00000070044 | Fam149a | 212326 | protein_coding | 119.192882 | 0.595355991 | 0.148685204 | 4.004137431 | 6.22E-05 | 0.000988977 | 0.595355991 |
| ENSMUSG00000024421 | Lama3 | 16774 | protein_coding | 1223.652663 | 0.595356347 | 0.140021844 | 4.251881909 | 2.12E-05 | 0.000381535 | 0.595356347 |
| ENSMUSG00000086468 | Etaa1os | 102633673 | lncRNA | 16.79942031 | 0.595397516 | 0.315155485 | 1.889218319 | 0.058862581 | 0.205867649 | 0.595397516 |
| ENSMUSG00000016239 | Lonrf3 | 74365 | protein_coding | 161.1900843 | 0.595731144 | 0.107870116 | 5.522670822 | 3.34E-08 | 1.12E-06 | 0.595731144 |
| ENSMUSG00000048731 | Ggnbp1 | 70772 | protein_coding | 26.10628456 | -0.59577239 | 0.26644189 | -2.236031243 | 0.025349725 | 0.115866207 | 0.59577239 |
| ENSMUSG00000060208 | Defa17 | 23855 | protein_coding | 31.73352136 | 0.595857896 | 0.998273023 | 0.596888709 | 0.550581686 | 0.752000715 | 0.595857896 |
| ENSMUSG00000045259 | Klhdc9 | 68874 | protein_coding | 21.55040051 | -0.596048778 | 0.297474167 | -2.00369929 | 0.045102282 | 0.17246929 | 0.596048778 |
| ENSMUSG00000029664 | Tfpi2 | 21789 | protein_coding | 271.6476291 | 0.596169966 | 0.075730443 | 7.872263007 | 3.48E-15 | 2.24E-13 | 0.596169966 |
| ENSMUSG00000013083 | 2200002J24Rik | 69147 | protein_coding | 10.64250648 | -0.597302851 | 0.361553003 | -1.652047822 | 0.098524802 | 0.285589823 | 0.597302851 |
| ENSMUSG00000049907 | Rasl11b | 68939 | protein_coding | 28.52921826 | 0.597790342 | 0.227964335 | 2.622297659 | 0.008733911 | 0.053942191 | 0.597790342 |
| ENSMUSG00000006675 | P4htm | 74443 | protein_coding | 6.450158419 | 0.598043692 | 0.48241584 | 1.239685024 | 0.215091919 | 0.45055038 | 0.598043692 |
| ENSMUSG00000024083 | Pja2 | 224938 | protein_coding | 1548.609018 | 0.598622151 | 0.055997688 | 10.69012259 | 1.13E-26 | 1.31E-24 | 0.598622151 |
| ENSMUSG00000039452 | Snx22 | 382083 | protein_coding | 6.864694036 | 0.598902756 | 0.482043184 | 1.242425525 | 0.214079603 | 0.449773548 | 0.598902756 |
| ENSMUSG00000023979 | Guca1b | 107477 | protein_coding | 5.928084845 | -0.598936022 | 0.538618927 | -1.11198473 | 0.266144716 | 0.508183262 | 0.598936022 |
| ENSMUSG00000017740 | Slc12a5 | 57138 | protein_coding | 6.19843528 | 0.599091201 | 0.628562409 | 0.953113315 | 0.340532661 | 0.583344264 | 0.599091201 |
| ENSMUSG00000018774 | Cd68 | 12514 | protein_coding | 13.71489462 | -0.599152433 | 0.533197913 | -1.123696133 | 0.261141958 | 0.503084198 | 0.599152433 |
| ENSMUSG00000113634 | Gm36756 | 102640764 | lncRNA | 6.825242895 | -0.599351234 | 0.617199622 | -0.97108166 | 0.331507616 | 0.574895496 | 0.599351234 |
| ENSMUSG00000021534 | 1700001L19Rik | 69315 | protein_coding | 6.676136705 | -0.599474408 | 0.454194946 | -1.319861468 | 0.186881274 | 0.416448019 | 0.599474408 |
| ENSMUSG00000069804 | Gm10277 | 791303 | protein_coding | 38.48326508 | -0.59966944 | 0.311072495 | -1.927748194 | 0.053886454 | 0.194492779 | 0.59966944 |
| ENSMUSG00000021262 | Evl | 14026 | protein_coding | 131.4025626 | -0.599700042 | 0.177856751 | -3.37181489 | 0.000746746 | 0.00791767 | 0.599700042 |
| ENSMUSG00000014867 | Surf4 | 20932 | protein_coding | 6314.251468 | 0.599996744 | 0.056752036 | 10.57225061 | 4.01E-26 | 4.49E-24 | 0.599996744 |
| ENSMUSG00000022415 | Syngr1 | 20972 | protein_coding | 37.57766864 | -0.600366923 | 0.191382009 | -3.137008162 | 0.001706814 | 0.015463084 | 0.600366923 |
| ENSMUSG00000055435 | Maf | 17132 | protein_coding | 236.6688781 | -0.60070592 | 0.14941006 | -4.020518569 | 5.81E-05 | 0.000931838 | 0.60070592 |
| ENSMUSG00000038526 | Car14 | 23831 | protein_coding | 7.746109422 | -0.601863999 | 0.522770113 | -1.151297642 | 0.249609807 | 0.490462051 | 0.601863999 |
| ENSMUSG00000004612 | Nkg7 | 72310 | protein_coding | 23.31762729 | 0.602483136 | 0.284894826 | 2.114756326 | 0.034450721 | 0.143118696 | 0.602483136 |
| ENSMUSG00000026875 | Traf1 | 22029 | protein_coding | 268.628036 | -0.602929656 | 0.120881111 | -4.987790507 | 6.11E-07 | 1.57E-05 | 0.602929656 |
| ENSMUSG00000036278 | Macrod1 | 107227 | protein_coding | 239.7036549 | -0.602961 | 0.098516537 | -6.120403915 | 9.33E-10 | 3.69E-08 | 0.602961 |
| ENSMUSG00000039081 | Zfp503 | 218820 | protein_coding | 56.10621545 | -0.603188456 | 0.208376173 | -2.894709345 | 0.003795098 | 0.029047716 | 0.603188456 |
| ENSMUSG00000030513 | Pcsk6 | 18553 | protein_coding | 10.0957653 | 0.603225609 | 0.409940391 | 1.47149591 | 0.141157053 | 0.352173719 | 0.603225609 |
| ENSMUSG00000096768 | Gm47283 | 170942 | lncRNA | 17.26111088 | -0.603944789 | 0.647092149 | -0.933321151 | 0.350654178 | 0.592953751 | 0.603944789 |
| ENSMUSG00000021700 | Rab3c | 67295 | protein_coding | 35.29942023 | 0.604308231 | 0.234144911 | 2.580915501 | 0.00985387 | 0.058661261 | 0.604308231 |

| ensembl_gene_id | external_gene_name | entrezgene_id | gene_biotype | baseMean | log2FoldChange | lfcSE | stat | pvalue | padj | absLog2FC |
|---|---|---|---|---|---|---|---|---|---|---|
| ENSMUSG00000038060 | Dlec1 | 320256 | protein_coding | 11.24836381 | 0.604333272 | 0.411272092 | 1.469424463 | 0.141717696 | 0.352697729 | 0.604333272 |
| ENSMUSG00000044162 | Tnip3 | 414084 | protein_coding | 393.4358115 | 0.604750386 | 0.16627204 | 3.637114117 | 0.00027571 | 0.0034468 | 0.604750386 |
| ENSMUSG00000056735 | A930024E05Rik | 109202 | lncRNA | 10.54167373 | -0.604778485 | 0.373704559 | -1.618333173 | 0.105590815 | 0.297761537 | 0.604778485 |
| ENSMUSG00000031740 | Mmp2 | 17390 | protein_coding | 9.640770842 | 0.605471647 | 0.403119226 | 1.501966685 | 0.133105713 | 0.340243435 | 0.605471647 |
| ENSMUSG00000064390 | Snord73b | 19871 | snoRNA | 8.956266618 | -0.606243572 | 0.456335069 | -1.328505333 | 0.184011223 | 0.412618047 | 0.606243572 |
| ENSMUSG00000074580 | 4931440P22Rik | 71004 | transcribed_processed_pseudogene | 43.31336736 | -0.606874397 | 0.237066723 | -2.559930767 | 0.010469302 | 0.061418126 | 0.606874397 |
| ENSMUSG00000028641 | P3h1 | 56401 | protein_coding | 27.49181492 | 0.607109717 | 0.231539408 | 2.622057831 | 0.00874006 | 0.053942191 | 0.607109717 |
| ENSMUSG00000059060 | Rad51b | 19363 | protein_coding | 38.31430659 | -0.607138263 | 0.232878054 | -2.607108108 | 0.009131053 | 0.055842429 | 0.607138263 |
| ENSMUSG00000097527 | 1700112J16Rik | 76628 | lncRNA | 11.81187849 | 0.607269287 | 0.356292998 | 1.704409829 | 0.088304549 | 0.266583201 | 0.607269287 |
| ENSMUSG00000109305 | Smim38 | 100040049 | protein_coding | 109.0696302 | -0.607774177 | 0.209273411 | -2.904211164 | 0.003681797 | 0.028396149 | 0.607774177 |
| ENSMUSG00000118506 | Cfap141 | 73545 | protein_coding | 6.166810286 | -0.608223301 | 0.535714004 | -1.135350759 | 0.256228383 | 0.497914824 | 0.608223301 |
| ENSMUSG00000098286 | Mir8091 | 102465884 | miRNA | 9.675028545 | -0.608395187 | 0.450692054 | -1.349913275 | 0.177043803 | 0.403959453 | 0.608395187 |
| ENSMUSG00000081058 | H3c15 | 97114 | protein_coding | 14.16768985 | 0.608771304 | 0.378693688 | 1.607555981 | 0.10793246 | 0.300944072 | 0.608771304 |
| ENSMUSG00000023348 | Trip6 | 22051 | protein_coding | 156.2798295 | -0.609077054 | 0.115486748 | -5.273999543 | 1.33E-07 | 3.97E-06 | 0.609077054 |
| ENSMUSG00000022498 | Txndc11 | 106200 | protein_coding | 1110.102335 | 0.60966812 | 0.053124602 | 11.47619186 | 1.74E-30 | 2.34E-28 | 0.60966812 |
| ENSMUSG00000022376 | Adcy8 | 11514 | protein_coding | 26.39916122 | -0.609915389 | 0.27491352 | -2.21857182 | 0.026515868 | 0.119015143 | 0.609915389 |
| ENSMUSG00000048578 | Mlec | 109154 | protein_coding | 12019.00405 | 0.609983246 | 0.069313141 | 8.80039829 | 1.36E-18 | 1.07E-16 | 0.609983246 |
| ENSMUSG00000086784 | Isoc2a | 664994 | protein_coding | 624.7028379 | 0.610026203 | 0.087521852 | 6.969987385 | 3.17E-12 | 1.58E-10 | 0.610026203 |
| ENSMUSG00000031387 | Renbp | 19703 | protein_coding | 20.8678027 | -0.611742459 | 0.293893439 | -2.081511111 | 0.037387148 | 0.15103771 | 0.611742459 |
| ENSMUSG00000042524 | Sun2 | 223697 | protein_coding | 1996.315626 | -0.612262864 | 0.126846299 | -4.826809041 | 1.39E-06 | 3.30E-05 | 0.612262864 |
| ENSMUSG00000026475 | Rgs16 | 19734 | protein_coding | 98.96701551 | 0.612641484 | 0.217118006 | 2.821698191 | 0.004777011 | 0.034453872 | 0.612641484 |
| ENSMUSG00000072623 | Zfp9 | 22750 | protein_coding | 116.0602564 | 0.613730402 | 0.136692695 | 4.489855162 | 7.13E-06 | 0.000145493 | 0.613730402 |
| ENSMUSG00000046442 | Ppm1e | 320472 | protein_coding | 15.48299227 | 0.614166735 | 0.295405968 | 2.079060009 | 0.037611835 | 0.151717375 | 0.614166735 |
| ENSMUSG00000035493 | Tgfbi | 21810 | protein_coding | 639.1914594 | -0.614798755 | 0.253442796 | -2.425789037 | 0.015275147 | 0.081167147 | 0.614798755 |
| ENSMUSG00000025189 | Cnnm1 | 83674 | protein_coding | 30.64371978 | -0.61485328 | 0.217209936 | -2.830686714 | 0.004644819 | 0.033757097 | 0.61485328 |
| ENSMUSG00000062070 | Pgk1 | 18655 | protein_coding | 4094.789843 | 0.61514463 | 0.075410121 | 8.157321897 | 3.43E-16 | 2.31E-14 | 0.61514463 |
| ENSMUSG00000028343 | Erp44 | 76299 | protein_coding | 1931.029703 | 0.615158137 | 0.055305966 | 11.12281704 | 9.72E-29 | 1.21E-26 | 0.615158137 |
| ENSMUSG00000010841 | Cfap97d1 | 75437 | protein_coding | 9.208048445 | -0.615364528 | 0.444818967 | -1.383404428 | 0.166540894 | 0.3901518 | 0.615364528 |
| ENSMUSG00000000167 | Pih1d2 | 72614 | protein_coding | 23.61628964 | 0.615842221 | 0.237545037 | 2.592528253 | 0.009527336 | 0.057416794 | 0.615842221 |
| ENSMUSG00000074102 | Rbm15b | 109095 | protein_coding | 2218.804615 | 0.616768306 | 0.048562867 | 12.70040973 | 5.88E-37 | 9.68E-35 | 0.616768306 |
| ENSMUSG00000031430 | Vsig1 | 78789 | protein_coding | 155.7216451 | 0.616864311 | 0.142102049 | 4.340995185 | 1.42E-05 | 0.000267224 | 0.616864311 |
| ENSMUSG00000036306 | Lzts1 | 211134 | protein_coding | 7.969253383 | 0.617019133 | 0.449257639 | 1.373419345 | 0.169622019 | 0.39391111 | 0.617019133 |
| ENSMUSG00000097032 | 4930539J05Rik | 319587 | lncRNA | 11.16048762 | -0.617590067 | 0.429353389 | -1.438418986 | 0.150315209 | 0.365794955 | 0.617590067 |
| ENSMUSG00000038209 | Itln1 | 16429 | protein_coding | 143.1539032 | 0.617762014 | 0.469379649 | 1.316124411 | 0.188132289 | 0.418302123 | 0.617762014 |
| ENSMUSG00000027849 | Syt6 | 54524 | protein_coding | 19.86356762 | -0.618250729 | 0.400246791 | -1.544673791 | 0.122425187 | 0.324355347 | 0.618250729 |
| ENSMUSG00000089706 | B230216N24Rik | 78603 | lncRNA | 14.29862719 | -0.618832095 | 0.416071434 | -1.487321753 | 0.136929852 | 0.346340609 | 0.618832095 |
| ENSMUSG00000038700 | Hoxb5 | 15413 | protein_coding | 69.99154333 | -0.619377037 | 0.229499615 | -2.698815148 | 0.006958682 | 0.04560043 | 0.619377037 |
| ENSMUSG00000074195 | Clca4b | 99709 | protein_coding | 218.1329515 | 0.619967466 | 0.262098608 | 2.365397785 | 0.018010714 | 0.091425492 | 0.619967466 |
| ENSMUSG00000051998 | Lax1 | 240754 | protein_coding | 10.57081987 | 0.620282279 | 0.398128499 | 1.557995171 | 0.119234393 | 0.319397925 | 0.620282279 |

| ensembl_gene_id | external_gene_name | entrezgene_id | gene_biotype | baseMean | log2FoldChange | lfcSE | stat | pvalue | padj | absLog2FC |
|---|---|---|---|---|---|---|---|---|---|---|
| ENSMUSG00000018669 | Cdk5rap3 | 80280 | protein_coding | 875.3642751 | 0.620307586 | 0.064547829 | 9.610045726 | 7.25E-22 | 6.68E-20 | 0.620307586 |
| ENSMUSG00000000296 | Tpd52l1 | 21987 | protein_coding | 129.1413028 | 0.621435125 | 0.109477609 | 5.676367355 | 1.38E-08 | 4.84E-07 | 0.621435125 |
| ENSMUSG00000039236 | Isg20 | 57444 | protein_coding | 365.7158759 | 0.622800559 | 0.095970997 | 6.489466406 | 8.61E-11 | 3.83E-09 | 0.622800559 |
| ENSMUSG00000035711 | Dok3 | 27261 | protein_coding | 8.304162976 | 0.623471014 | 0.506230753 | 1.231594506 | 0.218100594 | 0.45437389 | 0.623471014 |
| ENSMUSG00000060441 | Trim5 | 667823 | protein_coding | 38.21351748 | -0.624249455 | 0.261178345 | -2.390127157 | 0.01684254 | 0.087225321 | 0.624249455 |
| ENSMUSG00000008398 | Elk3 | 13713 | protein_coding | 45.31637711 | 0.62518771 | 0.179036678 | 3.491953251 | 0.000479502 | 0.005453407 | 0.62518771 |
| ENSMUSG00000097039 | Pvt1 | 19296 | lncRNA | 18.96183379 | 0.625501011 | 0.27647544 | 2.262410764 | 0.023672036 | 0.110334952 | 0.625501011 |
| ENSMUSG00000026354 | Lct | 226413 | protein_coding | 209.5377791 | 0.625508877 | 0.890742662 | 0.702232983 | 0.482533879 | 0.70472411 | 0.625508877 |
| ENSMUSG00000030247 | Kcnj8 | 16523 | protein_coding | 23.73100667 | 0.625631054 | 0.304987868 | 2.051330952 | 0.040234728 | 0.159269873 | 0.625631054 |
| ENSMUSG00000029345 | Tfip11 | 54723 | protein_coding | 927.9206191 | 0.62566661 | 0.05812414 | 10.76431602 | 5.07E-27 | 6.11E-25 | 0.62566661 |
| ENSMUSG00000056737 | Capg | 12332 | protein_coding | 272.910749 | 0.62600049 | 0.150403514 | 4.162140064 | 3.15E-05 | 0.000543216 | 0.62600049 |
| ENSMUSG00000040488 | Ltbp4 | 108075 | protein_coding | 14.13410428 | 0.626023395 | 0.3111591 | 2.011907717 | 0.044229666 | 0.170480837 | 0.626023395 |
| ENSMUSG00000036052 | Dnajb5 | 56323 | protein_coding | 104.6135577 | 0.626069493 | 0.121328918 | 5.160101166 | 2.47E-07 | 6.91E-06 | 0.626069493 |
| ENSMUSG00000031847 | 1700030J22Rik | 69528 | protein_coding | 16.92270239 | 0.626233202 | 0.288841939 | 2.168082667 | 0.030152397 | 0.130611795 | 0.626233202 |
| ENSMUSG00000045896 | Paip2b | 232164 | protein_coding | 929.0802674 | 0.626717533 | 0.069724481 | 8.988486119 | 2.51E-19 | 2.04E-17 | 0.626717533 |
| ENSMUSG00000021367 | Edn1 | 13614 | protein_coding | 467.9894655 | -0.626992195 | 0.1868019 | -3.35645513 | 0.000789485 | 0.008272924 | 0.626992195 |
| ENSMUSG00000045312 | Lhfpl2 | 218454 | protein_coding | 1393.343277 | -0.627011878 | 0.10774898 | -5.819190834 | 5.91E-09 | 2.16E-07 | 0.627011878 |
| ENSMUSG00000051236 | Msrb3 | 320183 | protein_coding | 8.491724637 | -0.627576132 | 0.502834104 | -1.248077899 | 0.212002531 | 0.447440166 | 0.627576132 |
| ENSMUSG00000020604 | Arsg | 74008 | protein_coding | 233.7983087 | -0.627980736 | 0.105960555 | -5.926551941 | 3.09E-09 | 1.17E-07 | 0.627980736 |
| ENSMUSG00000038692 | Hoxb4 | 15412 | protein_coding | 97.22350242 | -0.62805188 | 0.269877036 | -2.327177919 | 0.019955798 | 0.097992668 | 0.62805188 |
| ENSMUSG00000010492 | Uckl1os | 100271841 | lncRNA | 35.33469764 | -0.628311552 | 0.288911456 | -2.17475472 | 0.029648502 | 0.129053007 | 0.628311552 |
| ENSMUSG00000031125 | 3830403N18Rik | 70691 | protein_coding | 9.659395668 | 0.6288718 | 1.319456597 | 0.476614238 | 0.633636843 | 0.808283189 | 0.6288718 |
| ENSMUSG00000022987 | Zfp641 | 239652 | protein_coding | 11.55570537 | 0.629388782 | 0.45396135 | 1.386436932 | 0.165613526 | 0.388863072 | 0.629388782 |
| ENSMUSG00000034457 | Eda2r | 245527 | protein_coding | 33.71264633 | 0.629493716 | 0.283334121 | 2.221736349 | 0.026301129 | 0.118778578 | 0.629493716 |
| ENSMUSG00000035239 | Neu3 | 50877 | protein_coding | 173.089881 | -0.629534203 | 0.206732239 | -3.045167052 | 0.002325511 | 0.019707576 | 0.629534203 |
| ENSMUSG00000050075 | Gpr171 | 229323 | protein_coding | 11.45822363 | 0.630202883 | 0.398347883 | 1.582041503 | 0.113640096 | 0.31067534 | 0.630202883 |
| ENSMUSG00000047854 | Stx19 | 68159 | protein_coding | 38.41686534 | 0.630357897 | 0.199879713 | 3.153686221 | 0.001612223 | 0.014823762 | 0.630357897 |
| ENSMUSG00000019194 | Scn1b | 20266 | protein_coding | 12.44013917 | 0.631790732 | 0.409110169 | 1.544304639 | 0.122514549 | 0.32447317 | 0.631790732 |
| ENSMUSG00000087341 | 0610040F04Rik | 75394 | lncRNA | 10.35551199 | 0.632075974 | 0.428721956 | 1.47432611 | 0.140393808 | 0.351472935 | 0.632075974 |
| ENSMUSG00000026958 | Dpp7 | 83768 | protein_coding | 282.2346814 | -0.63213071 | 0.099216902 | -6.371199823 | 1.88E-10 | 8.00E-09 | 0.63213071 |
| ENSMUSG00000030319 | Cand2 | 67088 | protein_coding | 73.1030086 | 0.632903258 | 0.172979555 | 3.658832729 | 0.000253367 | 0.003214826 | 0.632903258 |
| ENSMUSG00000027876 | Reg4 | 67709 | protein_coding | 9460.489261 | 0.633419424 | 0.13585902 | 4.662328834 | 3.13E-06 | 6.88E-05 | 0.633419424 |
| ENSMUSG00000058163 | Gm5431 | 432555 | protein_coding | 117.045603 | -0.633649655 | 0.250047672 | -2.534115395 | 0.01127316 | 0.064744392 | 0.633649655 |
| ENSMUSG00000036960 | Clca2 | 229933 | protein_coding | 10.67553715 | 0.633976099 | 0.368242895 | 1.721624796 | 0.085137508 | 0.260237414 | 0.633976099 |
| ENSMUSG00000040828 | Catsperd | 106757 | protein_coding | 10.91631785 | -0.634168424 | 0.382015765 | -1.660058255 | 0.096902734 | 0.282646118 | 0.634168424 |
| ENSMUSG00000040136 | Abcc8 | 20927 | protein_coding | 63.75434408 | -0.634173859 | 0.197153056 | -3.216657507 | 0.001296933 | 0.012507901 | 0.634173859 |
| ENSMUSG00000097482 | Gm17634 | 100504568 | lncRNA | 12.44280288 | -0.634435496 | 0.341640434 | -1.857026957 | 0.063307319 | 0.215277659 | 0.634435496 |
| ENSMUSG00000041798 | Gck | 103988 | protein_coding | 10.74939565 | -0.634764438 | 0.412265009 | -1.539700009 | 0.123633495 | 0.326323628 | 0.634764438 |
| ENSMUSG00000030870 | Ubfd1 | 28018 | protein_coding | 1379.747058 | 0.635813001 | 0.055601083 | 11.43526282 | 2.79E-30 | 3.71E-28 | 0.635813001 |

| ensembl_gene_id | external_gene_name | entrezgene_id | gene_biotype | baseMean | log2FoldChange | lfcSE | stat | pvalue | padj | absLog2FC |
|---|---|---|---|---|---|---|---|---|---|---|
| ENSMUSG00000118522 | Gm31160 | 102633301 | protein_coding | 27.39979201 | -0.635929763 | 0.248829522 | -2.555684542 | 0.010597915 | 0.061835229 | 0.635929763 |
| ENSMUSG00000071633 | Gm4952 | 240549 | protein_coding | 8.710017777 | 0.636306458 | 0.413013709 | 1.540642464 | 0.123403827 | 0.326064971 | 0.636306458 |
| ENSMUSG00000053931 | Cnn3 | 71994 | protein_coding | 480.936586 | -0.636322949 | 0.099329917 | -6.406156038 | 1.49E-10 | 6.44E-09 | 0.636322949 |
| ENSMUSG00000025726 | Slc28a1 | 434203 | protein_coding | 7.748184702 | -0.637911001 | 0.852863474 | -0.747963795 | 0.454481996 | 0.682696642 | 0.637911001 |
| ENSMUSG00000043644 | 0610009L18Rik | 66838 | lncRNA | 80.44210233 | -0.638136414 | 0.152245438 | -4.191497761 | 2.77E-05 | 0.000483146 | 0.638136414 |
| ENSMUSG00000022505 | Emp2 | 13731 | protein_coding | 709.7920153 | -0.638451778 | 0.077857993 | -8.200208551 | 2.40E-16 | 1.63E-14 | 0.638451778 |
| ENSMUSG00000051674 | Dcun1d4 | 100737 | protein_coding | 272.4335686 | 0.638732258 | 0.077846459 | 8.205026498 | 2.31E-16 | 1.58E-14 | 0.638732258 |
| ENSMUSG00000053040 | Aph1c | 68318 | protein_coding | 11.56822678 | 0.638815483 | 0.35452932 | 1.801869259 | 0.071565978 | 0.233254754 | 0.638815483 |
| ENSMUSG00000062939 | Stat4 | 20849 | protein_coding | 8.997296589 | 0.638924376 | 0.487069093 | 1.311773597 | 0.189596536 | 0.420211186 | 0.638924376 |
| ENSMUSG00000046908 | Ltb4r1 | 16995 | protein_coding | 6.264971947 | 0.639097404 | 0.497362181 | 1.284973865 | 0.198801417 | 0.432143207 | 0.639097404 |
| ENSMUSG00000015222 | Map2 | 17756 | protein_coding | 160.9992578 | 0.639423897 | 0.122208578 | 5.232234162 | 1.67E-07 | 4.87E-06 | 0.639423897 |
| ENSMUSG00000047935 | Sox1ot | 434280 | lncRNA | 11.03400483 | 0.639833772 | 0.375938712 | 1.701962985 | 0.088762307 | 0.267564435 | 0.639833772 |
| ENSMUSG00000023068 | Nus1 | 52014 | protein_coding | 2496.482754 | 0.639849044 | 0.036237864 | 17.65691951 | 9.00E-70 | 2.59E-67 | 0.639849044 |
| ENSMUSG00000027808 | Serp1 | 28146 | protein_coding | 5680.168013 | 0.640547628 | 0.066280642 | 9.664173503 | 4.28E-22 | 4.06E-20 | 0.640547628 |
| ENSMUSG00000073198 | Bnip3l-ps | 100043324 | transcribed_processed_pseudogene | 6.650212967 | 0.640770694 | 0.6111261 | 1.048508146 | 0.294404551 | 0.537453271 | 0.640770694 |
| ENSMUSG00000016087 | Fli1 | 14247 | protein_coding | 15.47770932 | 0.64158 | 0.363189397 | 1.766516328 | 0.077309265 | 0.245233381 | 0.64158 |
| ENSMUSG00000061175 | Fnip2 | 329679 | protein_coding | 903.2630556 | -0.642045314 | 0.085437895 | -7.514760473 | 5.70E-14 | 3.28E-12 | 0.642045314 |
| ENSMUSG00000071356 | Reg3b | 18489 | protein_coding | 1010.275826 | -0.642085532 | 0.3392874 | -1.892453218 | 0.058430621 | 0.204850136 | 0.642085532 |
| ENSMUSG00000034833 | Tespa1 | 67596 | protein_coding | 6.227270291 | 0.642397532 | 0.487193228 | 1.31856827 | 0.187313487 | 0.417073819 | 0.642397532 |
| ENSMUSG00000027822 | Slc33a1 | 11416 | protein_coding | 1002.801027 | 0.643080849 | 0.064022817 | 10.04455727 | 9.71E-24 | 9.84E-22 | 0.643080849 |
| ENSMUSG00000048163 | Selplg | 20345 | protein_coding | 50.80559905 | 0.643088574 | 0.223648222 | 2.875446847 | 0.004034561 | 0.030447057 | 0.643088574 |
| ENSMUSG00000091971 | Hspa1a | 193740 | protein_coding | 393.455508 | 0.643136095 | 0.392611257 | 1.638098967 | 0.10140105 | 0.290238707 | 0.643136095 |
| ENSMUSG00000087281 | Gm16015 | 115490183 | lncRNA | 11.45930772 | -0.644011867 | 0.426387564 | -1.510390829 | 0.130943727 | 0.336836503 | 0.644011867 |
| ENSMUSG00000035969 | Rusc2 | 100213 | protein_coding | 202.2411817 | -0.644117408 | 0.115087871 | -5.596744502 | 2.18E-08 | 7.49E-07 | 0.644117408 |
| ENSMUSG00000009614 | Sardh | 192166 | protein_coding | 150.0573569 | 0.644220929 | 0.112378673 | 5.732590651 | 9.89E-09 | 3.53E-07 | 0.644220929 |
| ENSMUSG00000045991 | Onecut2 | 225631 | protein_coding | 6.280860592 | 0.644563334 | 0.776934106 | 0.829624197 | 0.406751291 | 0.645607651 | 0.644563334 |
| ENSMUSG00000086859 | Snhg20 | 76972 | lncRNA | 58.69714184 | -0.644961563 | 0.205620363 | -3.136661913 | 0.001708831 | 0.01546482 | 0.644961563 |
| ENSMUSG00000093042 | Mir3109 | 100526560 | miRNA | 29.9807374 | -0.645415552 | 0.325535997 | -1.982624221 | 0.04740942 | 0.178679998 | 0.645415552 |
| ENSMUSG00000039563 | 2210406O10Rik | 76710 | lncRNA | 8.794263709 | 0.645602616 | 0.476968355 | 1.353554402 | 0.175878583 | 0.402268317 | 0.645602616 |
| ENSMUSG00000044933 | Sstr3 | 20607 | protein_coding | 22.08416523 | 0.645661326 | 0.264649579 | 2.439683932 | 0.014700117 | 0.079113565 | 0.645661326 |
| ENSMUSG00000051790 | Nlgn2 | 216856 | protein_coding | 21.56829708 | -0.64573411 | 0.287243582 | -2.248036689 | 0.024573851 | 0.113433539 | 0.64573411 |
| ENSMUSG00000058216 | Gstp3 | 225884 | protein_coding | 52.7101853 | -0.645890258 | 0.228029777 | -2.832482086 | 0.004618815 | 0.033598393 | 0.645890258 |
| ENSMUSG00000029384 | 2010109A12Rik | 75610 | protein_coding | 9.38029048 | -0.646435323 | 0.441792349 | -1.463210769 | 0.143409715 | 0.355373152 | 0.646435323 |
| ENSMUSG00000025150 | Cbr2 | 12409 | protein_coding | 373.9716049 | 0.646466725 | 0.145253046 | 4.450624228 | 8.56E-06 | 0.000170681 | 0.646466725 |
| ENSMUSG00000054999 | Naaladl1 | 381204 | protein_coding | 3459.824706 | 0.647372364 | 0.1413541 | 4.579791923 | 4.65E-06 | 9.93E-05 | 0.647372364 |
| ENSMUSG00000085421 | 4732490B19Rik | 319871 | lncRNA | 30.82370355 | -0.647504279 | 0.261419103 | -2.476882032 | 0.01325357 | 0.073374639 | 0.647504279 |
| ENSMUSG00000038508 | Gdf15 | 23886 | protein_coding | 316.7849623 | 0.6475624 | 0.150638822 | 4.298774998 | 1.72E-05 | 0.0003151 | 0.6475624 |
| ENSMUSG00000086254 | 3100003L05Rik | 73076 | lncRNA | 10.97896257 | -0.647774232 | 0.431760094 | -1.500310568 | 0.133533973 | 0.340959846 | 0.647774232 |
| ENSMUSG00000021484 | Lman2 | 66890 | protein_coding | 6538.788451 | 0.648517745 | 0.063577068 | 10.20049781 | 1.97E-24 | 2.05E-22 | 0.648517745 |

| ensembl_gene_id | external_gene_name | entrezgene_id | gene_biotype | baseMean | log2FoldChange | lfcSE | stat | pvalue | padj | absLog2FC |
|---|---|---|---|---|---|---|---|---|---|---|
| ENSMUSG00000044646 | Zbtb7c | 207259 | protein_coding | 410.2963474 | 0.64884705 | 0.135499686 | 4.788550214 | 1.68E-06 | 3.92E-05 | 0.64884705 |
| ENSMUSG00000026639 | Lamb3 | 16780 | protein_coding | 4689.224431 | 0.649381347 | 0.07692574 | 8.441665221 | 3.13E-17 | 2.24E-15 | 0.649381347 |
| ENSMUSG00000034088 | Hdlbp | 110611 | protein_coding | 12740.26249 | 0.649408847 | 0.047299979 | 13.72958015 | 6.75E-43 | 1.30E-40 | 0.649408847 |
| ENSMUSG00000031543 | Ank1 | 11733 | protein_coding | 7.417218921 | 0.650672966 | 0.555884108 | 1.170519101 | 0.241792132 | 0.482716231 | 0.650672966 |
| ENSMUSG00000090877 | Hspa1b | 15511 | protein_coding | 1155.854783 | 0.651086924 | 0.309283226 | 2.105147869 | 0.035278439 | 0.145359064 | 0.651086924 |
| ENSMUSG00000031149 | Praf2 | 54637 | protein_coding | 148.9989763 | 0.65115369 | 0.137400827 | 4.739081315 | 2.15E-06 | 4.89E-05 | 0.65115369 |
| ENSMUSG00000022722 | Arl6 | 56297 | protein_coding | 32.59409045 | 0.651644941 | 0.22824174 | 2.855064722 | 0.004302806 | 0.031948136 | 0.651644941 |
| ENSMUSG00000020234 | Tektip1 | 432479 | protein_coding | 13.36576729 | 0.651863583 | 0.363030671 | 1.795615729 | 0.072555653 | 0.235386242 | 0.651863583 |
| ENSMUSG00000044390 | Tigd3 | 332359 | protein_coding | 6.421181724 | 0.651891234 | 0.495743754 | 1.314976191 | 0.188517904 | 0.418937335 | 0.651891234 |
| ENSMUSG00000079103 | Tgm7 | 640543 | protein_coding | 13.59891634 | -0.652128534 | 0.408429355 | -1.596674005 | 0.110338393 | 0.304905289 | 0.652128534 |
| ENSMUSG00000024042 | Sik1 | 17691 | protein_coding | 747.8727937 | -0.652304682 | 0.118344087 | -5.511933023 | 3.55E-08 | 1.18E-06 | 0.652304682 |
| ENSMUSG00000030259 | Rassf8 | 71323 | protein_coding | 44.35356403 | 0.652639522 | 0.196491893 | 3.321457749 | 0.000895485 | 0.0091867 | 0.652639522 |
| ENSMUSG00000081769 | Gm12216 | 622459 | lncRNA | 32.86633396 | -0.6532779 | 0.234669292 | -2.783823539 | 0.005372225 | 0.037635915 | 0.6532779 |
| ENSMUSG00000047730 | Fcgbp | 215384 | protein_coding | 243271.9589 | 0.653393316 | 0.112374797 | 5.814411529 | 6.08E-09 | 2.22E-07 | 0.653393316 |
| ENSMUSG00000052819 | Best2 | 212989 | protein_coding | 1407.183679 | 0.653472359 | 0.138697261 | 4.711501537 | 2.46E-06 | 5.52E-05 | 0.653472359 |
| ENSMUSG00000018819 | Lsp1 | 16985 | protein_coding | 36.59948129 | 0.654086867 | 0.238944279 | 2.73740334 | 0.006192632 | 0.041961584 | 0.654086867 |
| ENSMUSG00000020424 | Castor1 | 71962 | protein_coding | 25.79433395 | 0.654102749 | 0.243371934 | 2.687667136 | 0.007195308 | 0.046770954 | 0.654102749 |
| ENSMUSG00000001435 | Col18a1 | 12822 | protein_coding | 13.71422329 | 0.654772388 | 0.343699368 | 1.905073006 | 0.056770582 | 0.201440865 | 0.654772388 |
| ENSMUSG00000085957 | Syna | 214292 | protein_coding | 12.27111648 | 0.654873404 | 0.326294713 | 2.006999743 | 0.04474969 | 0.171664666 | 0.654873404 |
| ENSMUSG00000019992 | Mtfr2 | 71804 | protein_coding | 26.72951573 | 0.656092121 | 0.26157536 | 2.508233653 | 0.01213364 | 0.068515133 | 0.656092121 |
| ENSMUSG00000043635 | Adamts3 | 330119 | protein_coding | 14.11988471 | 0.656484665 | 0.323401596 | 2.029936379 | 0.042363007 | 0.165140974 | 0.656484665 |
| ENSMUSG00000022518 | 4930562C15Rik | 78809 | protein_coding | 9.592285118 | -0.657003284 | 0.458227948 | -1.433791383 | 0.151631805 | 0.367724228 | 0.657003284 |
| ENSMUSG00000078810 | Gp6 | 243816 | protein_coding | 18.73358433 | 0.657376957 | 0.34930564 | 1.881953458 | 0.059842335 | 0.2077222 | 0.657376957 |
| ENSMUSG00000038503 | Mesd | 67943 | protein_coding | 678.7265827 | 0.657399204 | 0.071097167 | 9.246489415 | 2.32E-20 | 1.96E-18 | 0.657399204 |
| ENSMUSG00000024112 | Cacna1h | 58226 | protein_coding | 306.8363891 | -0.657572563 | 0.177491948 | -3.704802218 | 0.000211556 | 0.002762558 | 0.657572563 |
| ENSMUSG00000024330 | Col11a2 | 12815 | protein_coding | 28.48693649 | 0.658111162 | 0.244688178 | 2.689591171 | 0.00715396 | 0.046596089 | 0.658111162 |
| ENSMUSG00000025724 | Sec11a | 56529 | protein_coding | 1425.535289 | 0.65843233 | 0.046427567 | 14.18192633 | 1.19E-45 | 2.52E-43 | 0.65843233 |
| ENSMUSG00000057176 | Ccdc189 | 233899 | protein_coding | 36.09678705 | -0.658535521 | 0.249319383 | -2.641333025 | 0.00825805 | 0.05184024 | 0.658535521 |
| ENSMUSG00000097990 | Gm19557 | 100503104 | lncRNA | 11.12550774 | -0.658639966 | 0.385446325 | -1.708772201 | 0.087493154 | 0.264813247 | 0.658639966 |
| ENSMUSG00000048647 | Exd1 | 241624 | protein_coding | 33.1870616 | 0.658807984 | 0.224865249 | 2.92979012 | 0.00339191 | 0.026670568 | 0.658807984 |
| ENSMUSG00000031861 | Lpar2 | 53978 | protein_coding | 34.596924 | -0.660146379 | 0.230628329 | -2.862382011 | 0.004204698 | 0.03136409 | 0.660146379 |
| ENSMUSG00000085028 | Slc2a4rg-ps | 329584 | transcribed_unitary_pseudogene | 71.80957008 | -0.660434057 | 0.244188952 | -2.704602535 | 0.006838616 | 0.044996476 | 0.660434057 |
| ENSMUSG00000029407 | Uso1 | 56041 | protein_coding | 1426.297351 | 0.660468348 | 0.08591163 | 7.687764151 | 1.50E-14 | 9.08E-13 | 0.660468348 |
| ENSMUSG00000070883 | Ccdc173 | 75051 | protein_coding | 9.207664306 | -0.660623368 | 0.40092915 | -1.647730945 | 0.099407895 | 0.286840423 | 0.660623368 |
| ENSMUSG00000032525 | Nktr | 18087 | protein_coding | 2128.338399 | 0.661556486 | 0.086457515 | 7.651810064 | 1.98E-14 | 1.19E-12 | 0.661556486 |
| ENSMUSG00000050394 | Armcx6 | 278097 | protein_coding | 28.69986217 | 0.662102671 | 0.224361901 | 2.951047698 | 0.00316698 | 0.02524933 | 0.662102671 |
| ENSMUSG00000004035 | Gstm7 | 68312 | protein_coding | 297.1625149 | 0.662362005 | 0.098410819 | 6.73058118 | 1.69E-11 | 7.90E-10 | 0.662362005 |
| ENSMUSG00000028255 | Clca1 | 23844 | protein_coding | 101155.2319 | 0.662444784 | 0.189071253 | 3.503677976 | 0.00045888 | 0.005267157 | 0.662444784 |
| ENSMUSG00000040812 | Agbl2 | 271813 | protein_coding | 18.98226689 | 0.662669969 | 0.286275851 | 2.314795209 | 0.020624142 | 0.100394066 | 0.662669969 |

Supplementary Table 1

| ensembl_gene_id | external_gene_name | entrezgene_id | gene_biotype | baseMean | log2FoldChange | lfcSE | stat | pvalue | padj | absLog2FC |
|---|---|---|---|---|---|---|---|---|---|---|
| ENSMUSG00000011267 | Zfp296 | 63872 | protein_coding | 11.47199906 | -0.662765166 | 0.36278653 | -1.826873687 | 0.067718763 | 0.225268254 | 0.662765166 |
| ENSMUSG00000002664 | Pspn | 19197 | protein_coding | 8.996628125 | 0.663215974 | 0.377776154 | 1.755579239 | 0.07916028 | 0.248514275 | 0.663215974 |
| ENSMUSG00000026579 | F5 | 14067 | protein_coding | 11.51762146 | 0.663662259 | 0.359660243 | 1.845247763 | 0.065001505 | 0.219397086 | 0.663662259 |
| ENSMUSG00000051373 | Plpp7 | 227721 | protein_coding | 6.548158394 | -0.663815685 | 0.458642977 | -1.447347321 | 0.147799672 | 0.361643489 | 0.663815685 |
| ENSMUSG00000024165 | Jpt2 | 52009 | protein_coding | 2077.05867 | 0.663832109 | 0.065967707 | 10.06298592 | 8.05E-24 | 8.22E-22 | 0.663832109 |
| ENSMUSG00000030155 | Clec2e | 232409 | protein_coding | 548.5378337 | -0.663917249 | 0.166357551 | -3.990905399 | 6.58E-05 | 0.00104069 | 0.663917249 |
| ENSMUSG00000079396 | Gm3411 | 100861615 | protein_coding | 6.077734908 | 0.66392614 | 0.484710243 | 1.36973821 | 0.170768637 | 0.395233324 | 0.66392614 |
| ENSMUSG00000099825 | 1810012K16Rik | 69108 | lncRNA | 6.767314347 | -0.664097145 | 0.517114065 | -1.28423725 | 0.199058955 | 0.432411607 | 0.664097145 |
| ENSMUSG00000032940 | Rbm11 | 224344 | protein_coding | 10.37305279 | -0.665457917 | 0.448791866 | -1.482776243 | 0.13813387 | 0.348161805 | 0.665457917 |
| ENSMUSG00000075025 | Gm10804 | 100038525 | lncRNA | 24.91616469 | -0.665494647 | 0.30649302 | -2.171320726 | 0.029906937 | 0.129900331 | 0.665494647 |
| ENSMUSG00000086782 | E130102H24Rik | 77866 | lncRNA | 13.99433833 | -0.665574437 | 0.397569391 | -1.674108854 | 0.094109208 | 0.27735731 | 0.665574437 |
| ENSMUSG00000053411 | Cbx7 | 52609 | protein_coding | 480.4027508 | -0.666461873 | 0.104970626 | -6.349032066 | 2.17E-10 | 9.22E-09 | 0.666461873 |
| ENSMUSG00000038022 | Mindy4 | 330323 | protein_coding | 13.67025861 | 0.667740053 | 0.33838586 | 1.973309559 | 0.048460305 | 0.181262767 | 0.667740053 |
| ENSMUSG00000029163 | Emilin1 | 100952 | protein_coding | 38.52697203 | 0.668032696 | 0.193056592 | 3.460294661 | 0.000539585 | 0.006042973 | 0.668032696 |
| ENSMUSG00000069255 | Dusp22 | 105352 | protein_coding | 67.45098843 | 0.668174132 | 0.148066993 | 4.512647419 | 6.40E-06 | 0.000132372 | 0.668174132 |
| ENSMUSG00000033526 | Ppip5k1 | 327655 | protein_coding | 2751.775579 | 0.668174954 | 0.038011795 | 17.57809529 | 3.63E-69 | 1.01E-66 | 0.668174954 |
| ENSMUSG00000020546 | Stxbp4 | 20913 | protein_coding | 45.12384292 | 0.668895997 | 0.177049446 | 3.778018017 | 0.000158081 | 0.002160491 | 0.668895997 |
| ENSMUSG00000023828 | Slc22a3 | 20519 | protein_coding | 33.28697322 | -0.669105194 | 0.230797681 | -2.89909843 | 0.003742374 | 0.028767047 | 0.669105194 |
| ENSMUSG00000085004 | 5430427M07Rik | 71349 | lncRNA | 38.8752921 | -0.669459836 | 0.230223162 | -2.907873516 | 0.003638955 | 0.028136566 | 0.669459836 |
| ENSMUSG00000026315 | Serpinb8 | 20725 | protein_coding | 9.515804212 | -0.669706126 | 0.422065173 | -1.586736287 | 0.112572384 | 0.308750823 | 0.669706126 |
| ENSMUSG00000032549 | Rab6b | 270192 | protein_coding | 25.50806167 | 0.671165661 | 0.245986612 | 2.728464188 | 0.006363 | 0.042790188 | 0.671165661 |
| ENSMUSG00000041084 | Ostc | 66357 | protein_coding | 1716.92428 | 0.671386763 | 0.070938167 | 9.464393961 | 2.95E-21 | 2.57E-19 | 0.671386763 |
| ENSMUSG00000074882 | Cyp2c68 | 433247 | protein_coding | 1373.090441 | -0.671574065 | 0.123398989 | -5.442297943 | 5.26E-08 | 1.69E-06 | 0.671574065 |
| ENSMUSG00000032437 | Stt3b | 68292 | protein_coding | 7810.359221 | 0.671671372 | 0.048191011 | 13.93769 | 3.74E-44 | 7.63E-42 | 0.671671372 |
| ENSMUSG00000061740 | Cyp2d22 | 56448 | protein_coding | 178.0546249 | -0.67173772 | 0.173313328 | -3.875857258 | 0.00010625 | 0.001567697 | 0.67173772 |
| ENSMUSG00000021360 | Gcnt2 | 14538 | protein_coding | 260.3083167 | -0.6727486 | 0.110739879 | -6.075034643 | 1.24E-09 | 4.85E-08 | 0.6727486 |
| ENSMUSG00000039676 | Capsl | 75568 | protein_coding | 104.3781517 | 0.672795838 | 0.154459717 | 4.355801325 | 1.33E-05 | 0.000252734 | 0.672795838 |
| ENSMUSG00000071661 | Zbtb3 | 75291 | protein_coding | 53.50189432 | 0.672887021 | 0.166013398 | 4.05320913 | 5.05E-05 | 0.000827981 | 0.672887021 |
| ENSMUSG00000085333 | 1700030A11Rik | 74179 | lncRNA | 23.3424713 | -0.67358654 | 0.353023699 | -1.90804907 | 0.056384874 | 0.200599795 | 0.67358654 |
| ENSMUSG00000100372 | 1700003I22Rik | 69356 | lncRNA | 9.453306711 | 0.674330366 | 0.417450554 | 1.615353864 | 0.106234085 | 0.298369399 | 0.674330366 |
| ENSMUSG00000036097 | Slf2 | 226151 | protein_coding | 1060.974981 | 0.674371888 | 0.068614385 | 9.82843309 | 8.49E-23 | 8.40E-21 | 0.674371888 |
| ENSMUSG00000022225 | Cma1 | 17228 | protein_coding | 19.21020944 | 0.674433135 | 0.355834958 | 1.89535379 | 0.058045546 | 0.203987912 | 0.674433135 |
| ENSMUSG00000055653 | Gpc3 | 14734 | protein_coding | 17.2958385 | 0.674968053 | 0.300551726 | 2.245763359 | 0.024719169 | 0.11389887 | 0.674968053 |
| ENSMUSG00000024672 | Ms4a7 | 109225 | protein_coding | 13.96630844 | -0.674986161 | 0.349172951 | -1.933099797 | 0.053223905 | 0.192842489 | 0.674986161 |
| ENSMUSG00000019579 | Mydgf | 28106 | protein_coding | 1840.803843 | 0.675051347 | 0.088319196 | 7.643314015 | 2.12E-14 | 1.26E-12 | 0.675051347 |
| ENSMUSG00000052949 | Rnf157 | 217340 | protein_coding | 31.01274173 | 0.675138398 | 0.212493656 | 3.177216732 | 0.001486959 | 0.013893501 | 0.675138398 |
| ENSMUSG00000049804 | Armcx4 | 100503043 | protein_coding | 10.89465535 | 0.675667675 | 0.431557035 | 1.565650935 | 0.117430359 | 0.316953874 | 0.675667675 |
| ENSMUSG00000032417 | Rwdd2a | 69519 | protein_coding | 11.53388134 | -0.675751719 | 0.378641164 | -1.784675789 | 0.07431392 | 0.239165953 | 0.675751719 |
| ENSMUSG00000030745 | Il21r | 60504 | protein_coding | 17.47418068 | 0.676368237 | 0.359972307 | 1.878945197 | 0.060251977 | 0.208682378 | 0.676368237 |

| ensembl_gene_id | external_gene_name | entrezgene_id | gene_biotype | baseMean | log2FoldChange | lfcSE | stat | pvalue | padj | absLog2FC |
|---|---|---|---|---|---|---|---|---|---|---|
| ENSMUSG00000035158 | Mitf | 17342 | protein_coding | 43.66264487 | 0.677743371 | 0.182766298 | 3.708251342 | 0.000208695 | 0.002731844 | 0.677743371 |
| ENSMUSG00000073295 | Nudt11 | 58242 | protein_coding | 27.36606285 | 0.677758339 | 0.242839243 | 2.79097534 | 0.005254948 | 0.037055767 | 0.677758339 |
| ENSMUSG00000035227 | Spcs2 | 66624 | protein_coding | 1960.418095 | 0.678327726 | 0.054916805 | 12.35191538 | 4.76E-35 | 7.04E-33 | 0.678327726 |
| ENSMUSG00000027777 | Schip1 | 30953 | protein_coding | 7.982697829 | -0.678819457 | 0.44481628 | -1.526067026 | 0.126993173 | 0.331180288 | 0.678819457 |
| ENSMUSG00000014846 | Tppp3 | 67971 | protein_coding | 886.2264746 | 0.678828501 | 0.153140597 | 4.432714208 | 9.31E-06 | 0.000184133 | 0.678828501 |
| ENSMUSG00000094806 | Cyp2d10 | 13101 | protein_coding | 469.9470796 | -0.678832482 | 0.125569005 | -5.406051307 | 6.44E-08 | 2.03E-06 | 0.678832482 |
| ENSMUSG00000041180 | Hectd2 | 226098 | protein_coding | 14.29735349 | 0.678959103 | 0.373506936 | 1.817795169 | 0.069095445 | 0.228065548 | 0.678959103 |
| ENSMUSG00000074445 | Sprr2a3 | 100042514 | protein_coding | 2373.266675 | 0.679163961 | 0.207907927 | 3.266657362 | 0.001088253 | 0.010812233 | 0.679163961 |
| ENSMUSG00000051506 | Wdfy4 | 545030 | protein_coding | 11.13253388 | 0.679382372 | 0.399298378 | 1.701440349 | 0.08886033 | 0.267762809 | 0.679382372 |
| ENSMUSG00000030035 | Wbp1 | 22377 | protein_coding | 1893.766168 | 0.679571199 | 0.061217614 | 11.10090962 | 1.24E-28 | 1.54E-26 | 0.679571199 |
| ENSMUSG00000095562 | Gm55594 | 170942 | lncRNA | 46.792848 | -0.679619435 | 0.780545585 | -0.870697942 | 0.383919102 | 0.62483231 | 0.679619435 |
| ENSMUSG00000085980 | Gm12408 | 100040557 | lncRNA | 33.12013733 | 0.680069183 | 0.252237592 | 2.696145243 | 0.007014707 | 0.045874365 | 0.680069183 |
| ENSMUSG00000086915 | Gm16364 | 100504626 | lncRNA | 10.69383481 | -0.680468407 | 0.431623797 | -1.576531255 | 0.114903422 | 0.312750629 | 0.680468407 |
| ENSMUSG00000002688 | Prkd1 | 18760 | protein_coding | 19.43008275 | 0.680513429 | 0.299672872 | 2.270854294 | 0.0231558 | 0.10868265 | 0.680513429 |
| ENSMUSG00000031574 | Star | 20845 | protein_coding | 46.20528898 | -0.680640748 | 0.189889742 | -3.584399767 | 0.000337854 | 0.004068924 | 0.680640748 |
| ENSMUSG00000050357 | Carmil2 | 234695 | protein_coding | 10.51689911 | 0.680876308 | 0.415365875 | 1.639220621 | 0.101167321 | 0.290022548 | 0.680876308 |
| ENSMUSG00000015467 | Egfl8 | 81701 | protein_coding | 14.33768259 | -0.681698652 | 0.336031719 | -2.028673526 | 0.042491554 | 0.165460865 | 0.681698652 |
| ENSMUSG00000073414 | Mpig6b | 106722 | protein_coding | 11.01578357 | -0.683222464 | 0.393055592 | -1.73823367 | 0.082169649 | 0.25454282 | 0.683222464 |
| ENSMUSG00000031673 | Cdh11 | 12552 | protein_coding | 7.810420068 | 0.684380385 | 0.42477966 | 1.61114208 | 0.107148758 | 0.299598598 | 0.684380385 |
| ENSMUSG00000064354 | mt-Co2 | 17709 | protein_coding | 7.146548639 | 0.684561557 | 0.514920662 | 1.329450548 | 0.18369937 | 0.412210695 | 0.684561557 |
| ENSMUSG00000037321 | Tap1 | 21354 | protein_coding | 505.2984451 | -0.685103245 | 0.333399562 | -2.054901455 | 0.039888524 | 0.158163814 | 0.685103245 |
| ENSMUSG00000019039 | Dalrd3 | 67789 | protein_coding | 867.9113576 | 0.685634837 | 0.052682305 | 13.01451866 | 1.01E-38 | 1.74E-36 | 0.685634837 |
| ENSMUSG00000035274 | Tpbg | 21983 | protein_coding | 143.4002641 | -0.685641205 | 0.13711323 | -5.000547403 | 5.72E-07 | 1.47E-05 | 0.685641205 |
| ENSMUSG00000036123 | Slc9a3 | 105243 | protein_coding | 3484.50909 | -0.685985947 | 0.129828677 | -5.283778325 | 1.27E-07 | 3.79E-06 | 0.685985947 |
| ENSMUSG00000032740 | Ccdc88a | 108686 | protein_coding | 21.29424108 | 0.686158453 | 0.252424961 | 2.718267046 | 0.006562485 | 0.043714661 | 0.686158453 |
| ENSMUSG00000087626 | Gm15050 | 100504154 | lncRNA | 6.114187657 | -0.686557871 | 0.50039108 | -1.372042585 | 0.170050181 | 0.394462093 | 0.686557871 |
| ENSMUSG00000031576 | Kcnu1 | 16532 | protein_coding | 12.63658056 | -0.687142161 | 0.411810773 | -1.668587142 | 0.095199229 | 0.279497171 | 0.687142161 |
| ENSMUSG00000059540 | Tcea2 | 21400 | protein_coding | 27.44372271 | -0.688055879 | 0.234693587 | -2.931719976 | 0.003370905 | 0.026557202 | 0.688055879 |
| ENSMUSG00000044595 | Dnd1 | 213236 | protein_coding | 7.322683822 | 0.688225986 | 0.445446156 | 1.545026212 | 0.122339923 | 0.324281511 | 0.688225986 |
| ENSMUSG00000097487 | Ptges3l | 73635 | protein_coding | 31.5950611 | 0.690840626 | 0.211489184 | 3.266552994 | 0.001088654 | 0.010812233 | 0.690840626 |
| ENSMUSG00000042834 | Nrep | 27528 | protein_coding | 483.433496 | -0.690906986 | 0.134546058 | -5.135096441 | 2.82E-07 | 7.77E-06 | 0.690906986 |
| ENSMUSG00000001827 | Folr1 | 14275 | protein_coding | 37.13204534 | -0.690941973 | 0.339809564 | -2.033321149 | 0.042020088 | 0.164201104 | 0.690941973 |
| ENSMUSG00000003273 | Car11 | 12348 | protein_coding | 11.68426037 | -0.691192674 | 0.398044084 | -1.73647267 | 0.082480293 | 0.254924035 | 0.691192674 |
| ENSMUSG00000050304 | Slc25a2 | 83885 | protein_coding | 6.486412209 | -0.691442424 | 0.471449726 | -1.466630239 | 0.14247667 | 0.354094814 | 0.691442424 |
| ENSMUSG00000024516 | Sec11c | 66286 | protein_coding | 2202.555909 | 0.691903251 | 0.037307985 | 18.54571503 | 8.83E-77 | 3.10E-74 | 0.691903251 |
| ENSMUSG00000026676 | Ccdc3 | 74186 | protein_coding | 30.56310327 | 0.692887692 | 0.232595501 | 2.978938503 | 0.002892488 | 0.023394202 | 0.692887692 |
| ENSMUSG00000100281 | 4930422M22Rik | 73774 | lncRNA | 6.554182988 | -0.692996269 | 0.512518716 | -1.352138463 | 0.176331025 | 0.402927855 | 0.692996269 |
| ENSMUSG00000040723 | Rcsd1 | 226594 | protein_coding | 20.65140298 | 0.69325309 | 0.295258982 | 2.347949196 | 0.018877096 | 0.094455763 | 0.69325309 |
| ENSMUSG00000053137 | Mapk11 | 19094 | protein_coding | 15.47568005 | -0.695746105 | 0.312866797 | -2.223777377 | 0.026163426 | 0.118455243 | 0.695746105 |

| ensembl_gene_id | external_gene_name | entrezgene_id | gene_biotype | baseMean | log2FoldChange | lfcSE | stat | pvalue | padj | absLog2FC |
|---|---|---|---|---|---|---|---|---|---|---|
| ENSMUSG00000032554 | Trf | 22041 | protein_coding | 350.4934878 | 0.696389469 | 0.209962191 | 3.316737479 | 0.000910751 | 0.009325517 | 0.696389469 |
| ENSMUSG00000087366 | Junos | 230451 | lncRNA | 47.77863385 | -0.698578487 | 0.256149852 | -2.72722581 | 0.006386931 | 0.042881783 | 0.698578487 |
| ENSMUSG00000045790 | Ccdc149 | 100503884 | protein_coding | 307.889611 | 0.69947706 | 0.084293238 | 8.298139687 | 1.06E-16 | 7.38E-15 | 0.69947706 |
| ENSMUSG00000036411 | 9530077C05Rik | 68283 | protein_coding | 7.941114541 | -0.701574014 | 0.549640273 | -1.276423961 | 0.201805691 | 0.435678765 | 0.701574014 |
| ENSMUSG00000017002 | Slpi | 20568 | protein_coding | 27.92587737 | 0.702855516 | 0.291108153 | 2.414413713 | 0.015760556 | 0.082978005 | 0.702855516 |
| ENSMUSG00000064451 | Snora23 | 100379145 | snoRNA | 17.5065884 | 0.703590679 | 0.382119171 | 1.841285998 | 0.065579659 | 0.220701827 | 0.703590679 |
| ENSMUSG00000060550 | H2-Q7 | 15018 | protein_coding | 58.39057626 | -0.703791558 | 0.421586712 | -1.66938743 | 0.095040623 | 0.27921863 | 0.703791558 |
| ENSMUSG00000023073 | Slc10a2 | 20494 | protein_coding | 3130.108561 | -0.704312901 | 0.250872105 | -2.807458009 | 0.004993419 | 0.035627054 | 0.704312901 |
| ENSMUSG00000056468 | 5730596B20Rik | 77580 | TEC | 17.06416118 | -0.704346182 | 0.390541682 | -1.803510904 | 0.071308015 | 0.23288464 | 0.704346182 |
| ENSMUSG00000049928 | Glp2r | 93896 | protein_coding | 7.821747299 | -0.704458981 | 0.496323583 | -1.419354237 | 0.155795767 | 0.374220783 | 0.704458981 |
| ENSMUSG00000060780 | Lrrtm1 | 74342 | protein_coding | 49.58281294 | 0.706331021 | 0.300444011 | 2.35095723 | 0.018725186 | 0.094045804 | 0.706331021 |
| ENSMUSG00000026532 | Spta1 | 20739 | protein_coding | 30.51492245 | 0.707733167 | 0.387529983 | 1.826266863 | 0.067810074 | 0.225459333 | 0.707733167 |
| ENSMUSG00000043811 | Rtn4r | 65079 | protein_coding | 132.7786149 | -0.708202461 | 0.127710257 | -5.545384341 | 2.93E-08 | 9.90E-07 | 0.708202461 |
| ENSMUSG00000020019 | Ntn4 | 57764 | protein_coding | 519.1718691 | -0.708393565 | 0.121311692 | -5.839450053 | 5.24E-09 | 1.94E-07 | 0.708393565 |
| ENSMUSG00000077611 | Gm23946 | 115488402 | snoRNA | 11.99335276 | 0.708401298 | 0.37914219 | 1.86843173 | 0.061701918 | 0.211953026 | 0.708401298 |
| ENSMUSG00000029552 | Tes | 21753 | protein_coding | 4469.918977 | 0.708900426 | 0.038988626 | 18.18223686 | 7.14E-74 | 2.35E-71 | 0.708900426 |
| ENSMUSG00000070369 | Itgad | 381924 | protein_coding | 6.24251845 | 0.709592894 | 0.544123119 | 1.304103555 | 0.192198277 | 0.423671625 | 0.709592894 |
| ENSMUSG00000039405 | Prss23 | 76453 | protein_coding | 679.6704244 | -0.710520068 | 0.161022346 | -4.412555688 | 1.02E-05 | 0.000200182 | 0.710520068 |
| ENSMUSG00000048652 | Samd13 | 75015 | protein_coding | 17.45673586 | -0.71164073 | 0.381683721 | -1.864477552 | 0.062254671 | 0.21306899 | 0.71164073 |
| ENSMUSG00000035778 | Ggta1 | 14594 | protein_coding | 9.720958757 | 0.712176387 | 0.394791167 | 1.803931917 | 0.071241981 | 0.232716108 | 0.712176387 |
| ENSMUSG00000037370 | Enpp1 | 18605 | protein_coding | 460.4713698 | 0.712618678 | 0.07401536 | 9.627983661 | 6.09E-22 | 5.68E-20 | 0.712618678 |
| ENSMUSG00000026778 | Prkcq | 18761 | protein_coding | 13.35384523 | 0.712650934 | 0.360406797 | 1.977351536 | 0.048001908 | 0.179820388 | 0.712650934 |
| ENSMUSG00000059854 | Hydin | 244653 | protein_coding | 7.115653208 | 0.714211571 | 0.632441213 | 1.129293214 | 0.258774162 | 0.500569859 | 0.714211571 |
| ENSMUSG00000032826 | Ank2 | 109676 | protein_coding | 12.50622779 | 0.714870217 | 0.414764788 | 1.723555705 | 0.084788081 | 0.259710803 | 0.714870217 |
| ENSMUSG00000020262 | Adarb1 | 110532 | protein_coding | 8.535405745 | 0.715090764 | 0.550388983 | 1.299246145 | 0.19385947 | 0.425646245 | 0.715090764 |
| ENSMUSG00000051111 | Sv2c | 75209 | protein_coding | 151.7278871 | -0.715094364 | 0.179735386 | -3.978595306 | 6.93E-05 | 0.001083316 | 0.715094364 |
| ENSMUSG00000032122 | Slc37a2 | 56857 | protein_coding | 6117.109007 | -0.715651695 | 0.207806615 | -3.443834998 | 0.000573526 | 0.006387602 | 0.715651695 |
| ENSMUSG00000026156 | B3gat2 | 280645 | protein_coding | 23.17867875 | -0.715664919 | 0.277744099 | -2.576706117 | 0.009974672 | 0.059145001 | 0.715664919 |
| ENSMUSG00000036432 | Siah2 | 20439 | protein_coding | 507.4311232 | 0.716758722 | 0.072203209 | 9.926964915 | 3.18E-23 | 3.18E-21 | 0.716758722 |
| ENSMUSG00000061958 | Defa38 | 634825 | protein_coding | 52.8781969 | 0.716855891 | 1.057953279 | 0.677587475 | 0.49803329 | 0.715717596 | 0.716855891 |
| ENSMUSG00000031647 | Mfap3l | 71306 | protein_coding | 1560.874359 | -0.717313548 | 0.116996162 | -6.131086137 | 8.73E-10 | 3.47E-08 | 0.717313548 |
| ENSMUSG00000034595 | Ppp1r18 | 76448 | protein_coding | 15.25919542 | 0.717755775 | 0.335590793 | 2.138782677 | 0.032453273 | 0.137419114 | 0.717755775 |
| ENSMUSG00000027985 | Lef1 | 16842 | protein_coding | 6.275608145 | 0.718865306 | 0.546929312 | 1.314366026 | 0.188723058 | 0.419197611 | 0.718865306 |
| ENSMUSG00000092008 | Cyp2c69 | 100043108 | protein_coding | 88.28475773 | -0.721027539 | 0.146557799 | -4.919748691 | 8.67E-07 | 2.16E-05 | 0.721027539 |
| ENSMUSG00000090272 | Mndal | 100040462 | protein_coding | 10.57811664 | 0.721262831 | 0.39931742 | 1.806239336 | 0.070880963 | 0.231865576 | 0.721262831 |
| ENSMUSG00000040706 | Agmat | 75986 | protein_coding | 7.322039257 | 0.721266071 | 0.415222648 | 1.737058597 | 0.082376828 | 0.254853985 | 0.721266071 |
| ENSMUSG00000085498 | Gm14023 | 100503468 | lncRNA | 6.471998541 | -0.722103045 | 0.48753813 | -1.481121167 | 0.138574288 | 0.34868951 | 0.722103045 |
| ENSMUSG00000035032 | Nek11 | 208583 | protein_coding | 7.329226261 | 0.722524784 | 0.475573611 | 1.519270136 | 0.128694514 | 0.334104382 | 0.722524784 |
| ENSMUSG00000032788 | Pdxk | 216134 | protein_coding | 840.7042827 | 0.722817315 | 0.093067274 | 7.766611018 | 8.06E-15 | 4.98E-13 | 0.722817315 |

| ensembl_gene_id | external_gene_name | entrezgene_id | gene_biotype | baseMean | log2FoldChange | lfcSE | stat | pvalue | padj | absLog2FC |
|---|---|---|---|---|---|---|---|---|---|---|
| ENSMUSG00000025151 | Maged1 | 94275 | protein_coding | 1759.506047 | 0.723633339 | 0.1036483 | 6.981622824 | 2.92E-12 | 1.47E-10 | 0.723633339 |
| ENSMUSG00000026018 | Ica1l | 70375 | protein_coding | 13.17933356 | -0.72431428 | 0.428169566 | -1.691652879 | 0.090712178 | 0.271060828 | 0.72431428 |
| ENSMUSG00000090622 | A930033H14Rik | 320700 | lncRNA | 6.895360218 | -0.724384325 | 0.5398487 | -1.341828415 | 0.179651637 | 0.406744622 | 0.724384325 |
| ENSMUSG00000104368 | 2010103J01Rik | 69889 | lncRNA | 8.312584441 | -0.725061904 | 0.426724049 | -1.699135323 | 0.089293689 | 0.268368093 | 0.725061904 |
| ENSMUSG00000107455 | Gm43868 | 115490363 | TEC | 18.14209261 | -0.725084435 | 0.354599842 | -2.044796273 | 0.040874947 | 0.160974426 | 0.725084435 |
| ENSMUSG00000050628 | Ubald2 | 319370 | protein_coding | 1305.506137 | 0.725125543 | 0.104344899 | 6.949314748 | 3.67E-12 | 1.82E-10 | 0.725125543 |
| ENSMUSG00000022102 | Dok2 | 13449 | protein_coding | 14.62059634 | 0.725139862 | 0.395145322 | 1.83512197 | 0.066487619 | 0.222411499 | 0.725139862 |
| ENSMUSG00000047150 | 1700001C19Rik | 75462 | protein_coding | 12.91047783 | -0.725243319 | 0.361021686 | -2.008863586 | 0.044551603 | 0.171230623 | 0.725243319 |
| ENSMUSG00000029007 | Agtrap | 11610 | protein_coding | 918.6817753 | 0.725942266 | 0.064686555 | 11.22245988 | 3.16E-29 | 4.05E-27 | 0.725942266 |
| ENSMUSG00000074682 | Zcchc3 | 67917 | protein_coding | 39.32449792 | 0.726252446 | 0.240115119 | 3.024601066 | 0.002489613 | 0.020731319 | 0.726252446 |
| ENSMUSG00000018401 | Mtmr4 | 170749 | protein_coding | 2537.140618 | 0.726255528 | 0.0552597 | 13.14258897 | 1.88E-39 | 3.25E-37 | 0.726255528 |
| ENSMUSG00000073409 | H2-Q6 | 110557 | protein_coding | 50.26845193 | -0.726501754 | 0.357725442 | -2.030892046 | 0.042265947 | 0.164961985 | 0.726501754 |
| ENSMUSG00000027452 | Acss1 | 68738 | protein_coding | 3608.52268 | 0.727079094 | 0.039962529 | 18.19402109 | 5.76E-74 | 1.93E-71 | 0.727079094 |
| ENSMUSG00000029108 | Pcdh7 | 54216 | protein_coding | 8.080161691 | 0.727992659 | 0.460373202 | 1.581309806 | 0.113807219 | 0.310868736 | 0.727992659 |
| ENSMUSG00000054871 | Tmem158 | 72309 | protein_coding | 15.77184832 | 0.728002725 | 0.342869691 | 2.123263572 | 0.033731771 | 0.141296696 | 0.728002725 |
| ENSMUSG00000048279 | Sacs | 50720 | protein_coding | 24.22565302 | 0.728160003 | 0.279353092 | 2.606593674 | 0.009144781 | 0.055884001 | 0.728160003 |
| ENSMUSG00000087330 | Bloodlinc | 105463053 | lncRNA | 10.32998733 | 0.728405363 | 0.424609962 | 1.71546932 | 0.086259216 | 0.262627924 | 0.728405363 |
| ENSMUSG00000045932 | Ifit2 | 15958 | protein_coding | 66.79827451 | -0.729955233 | 0.435593984 | -1.675769777 | 0.093783296 | 0.276801468 | 0.729955233 |
| ENSMUSG00000037892 | Pcdh18 | 73173 | protein_coding | 50.20433935 | 0.730499593 | 0.197962626 | 3.690088421 | 0.000224176 | 0.002892231 | 0.730499593 |
| ENSMUSG00000020185 | E2f7 | 52679 | protein_coding | 57.40557553 | 0.730982822 | 0.240776793 | 3.035935531 | 0.002397906 | 0.020162376 | 0.730982822 |
| ENSMUSG00000037139 | Myom3 | 242702 | protein_coding | 49.77479844 | -0.732527013 | 0.21863655 | -3.350432552 | 0.000806855 | 0.008422099 | 0.732527013 |
| ENSMUSG00000045091 | Aqp12 | 208760 | protein_coding | 9.574055887 | 0.733634209 | 0.374598408 | 1.958455224 | 0.05017662 | 0.185651138 | 0.733634209 |
| ENSMUSG00000010476 | Ebf3 | 13593 | protein_coding | 6.730749905 | 0.734437809 | 0.52834279 | 1.390078227 | 0.164505124 | 0.386787306 | 0.734437809 |
| ENSMUSG00000047759 | Hs3st3a1 | 15478 | protein_coding | 18.74267475 | 0.734455615 | 0.299727245 | 2.450413256 | 0.014269234 | 0.077247377 | 0.734455615 |
| ENSMUSG00000095385 | D630033O11Rik | 235302 | lncRNA | 59.82155426 | -0.735668674 | 0.160830712 | -4.574180295 | 4.78E-06 | 0.000101583 | 0.735668674 |
| ENSMUSG00000015829 | Tnr | 21960 | protein_coding | 8.126103933 | 0.736854481 | 0.524358111 | 1.405250467 | 0.159946812 | 0.380171295 | 0.736854481 |
| ENSMUSG00000074115 | Saa1 | 20208 | protein_coding | 7548.107443 | -0.736891251 | 0.193002813 | -3.818033727 | 0.00013452 | 0.001899647 | 0.736891251 |
| ENSMUSG00000019433 | Gipc1 | 67903 | protein_coding | 1806.9384 | 0.737064706 | 0.079277622 | 9.2972605 | 1.44E-20 | 1.23E-18 | 0.737064706 |
| ENSMUSG00000085105 | Gm12758 | 100126229 | lncRNA | 8.358426909 | -0.737440654 | 0.474019525 | -1.555717888 | 0.119775191 | 0.320093959 | 0.737440654 |
| ENSMUSG00000025197 | Cyp2c23 | 226143 | protein_coding | 7.777762166 | -0.737552944 | 0.51984872 | -1.418783803 | 0.155962056 | 0.374452893 | 0.737552944 |
| ENSMUSG00000027339 | Rassf2 | 215653 | protein_coding | 21.04204853 | 0.737812989 | 0.292166646 | 2.525315604 | 0.011559443 | 0.066117863 | 0.737812989 |
| ENSMUSG00000019564 | Arid3a | 13496 | protein_coding | 119.1774444 | 0.737907611 | 0.118508026 | 6.226646714 | 4.77E-10 | 1.95E-08 | 0.737907611 |
| ENSMUSG00000094559 | Cyp2d34 | 223706 | protein_coding | 2418.064165 | -0.738374145 | 0.09787601 | -7.543974736 | 4.56E-14 | 2.63E-12 | 0.738374145 |
| ENSMUSG00000033854 | Kcnk10 | 72258 | protein_coding | 9.407787357 | 0.739917 | 0.505214797 | 1.46455924 | 0.143041209 | 0.354896244 | 0.739917 |
| ENSMUSG00000026825 | Dnm1 | 13429 | protein_coding | 26.80706569 | -0.742164037 | 0.314889946 | -2.356899756 | 0.018428221 | 0.09298871 | 0.742164037 |
| ENSMUSG00000030825 | Hsd17b14 | 66065 | protein_coding | 23.47321121 | -0.742503335 | 0.250555557 | -2.963427932 | 0.003042333 | 0.024375404 | 0.742503335 |
| ENSMUSG00000022582 | Ly6g | 546644 | protein_coding | 316.9547525 | -0.742693891 | 0.230770131 | -3.21832764 | 0.001289404 | 0.012451632 | 0.742693891 |
| ENSMUSG00000026581 | Sell | 20343 | protein_coding | 6.011233689 | 0.744055847 | 0.625435177 | 1.189661014 | 0.234179654 | 0.473794416 | 0.744055847 |
| ENSMUSG00000026950 | Neb | 17996 | protein_coding | 9.65865651 | 0.744115051 | 0.647523277 | 1.149171122 | 0.250485426 | 0.491535798 | 0.744115051 |

| ensembl_gene_id | external_gene_name | entrezgene_id | gene_biotype | baseMean | log2FoldChange | lfcSE | stat | pvalue | padj | absLog2FC |
|---|---|---|---|---|---|---|---|---|---|---|
| ENSMUSG00000001588 | Acap1 | 216859 | protein_coding | 40.37016978 | 0.746416294 | 0.225590118 | 3.308727796 | 0.000937209 | 0.009541901 | 0.746416294 |
| ENSMUSG00000070476 | Fam217b | 71532 | protein_coding | 58.06183065 | 0.746463232 | 0.173968002 | 4.290807647 | 1.78E-05 | 0.000325142 | 0.746463232 |
| ENSMUSG00000041073 | Nacad | 192950 | protein_coding | 63.18669415 | 0.746707406 | 0.187968022 | 3.972523612 | 7.11E-05 | 0.001108091 | 0.746707406 |
| ENSMUSG00000026520 | Pycr2 | 69051 | protein_coding | 630.5788895 | 0.747454586 | 0.084751719 | 8.819344201 | 1.15E-18 | 9.15E-17 | 0.747454586 |
| ENSMUSG00000020774 | Aspa | 11484 | protein_coding | 143.8563514 | -0.74765701 | 0.145682504 | -5.132098834 | 2.87E-07 | 7.89E-06 | 0.74765701 |
| ENSMUSG00000020212 | Mdm1 | 17245 | protein_coding | 113.9397332 | 0.747956935 | 0.146280834 | 5.113157426 | 3.17E-07 | 8.62E-06 | 0.747956935 |
| ENSMUSG00000017499 | Cdc6 | 23834 | protein_coding | 151.6885242 | 0.748214854 | 0.185371929 | 4.036289947 | 5.43E-05 | 0.00088192 | 0.748214854 |
| ENSMUSG00000020319 | Wdpcp | 216560 | protein_coding | 65.54154728 | 0.748771803 | 0.198129155 | 3.779210595 | 0.000157326 | 0.002151995 | 0.748771803 |
| ENSMUSG00000103409 | Lsmem2 | 434436 | protein_coding | 17.3351815 | -0.749341484 | 0.385053513 | -1.946071022 | 0.051646209 | 0.189222442 | 0.749341484 |
| ENSMUSG00000111329 | A830035O19Rik | 100101457 | lncRNA | 19.07777017 | -0.749907575 | 0.333132142 | -2.251081421 | 0.024380381 | 0.112857553 | 0.749907575 |
| ENSMUSG00000035232 | Pdk3 | 236900 | protein_coding | 673.011334 | 0.750425244 | 0.085506234 | 8.7762635 | 1.69E-18 | 1.32E-16 | 0.750425244 |
| ENSMUSG00000031532 | Saraf | 67887 | protein_coding | 3321.996148 | 0.750635435 | 0.056362547 | 13.31798295 | 1.82E-40 | 3.26E-38 | 0.750635435 |
| ENSMUSG00000040653 | Ppp1r14c | 76142 | protein_coding | 31.83219194 | 0.750650559 | 0.203117175 | 3.695652815 | 0.000219323 | 0.002850132 | 0.750650559 |
| ENSMUSG00000053219 | Raet1e | 379043 | protein_coding | 143.0834986 | -0.750743538 | 0.153240861 | -4.89910806 | 9.63E-07 | 2.37E-05 | 0.750743538 |
| ENSMUSG00000045802 | Hsf3 | 245525 | protein_coding | 19.80042128 | -0.75175231 | 0.283401136 | -2.652608669 | 0.007987241 | 0.050672793 | 0.75175231 |
| ENSMUSG00000038295 | Atg9b | 213948 | protein_coding | 6.665180992 | 0.751813956 | 0.561301995 | 1.339410803 | 0.180436974 | 0.40741624 | 0.751813956 |
| ENSMUSG00000089768 | Tmsb15b1 | 666244 | protein_coding | 6.193148771 | 0.752500296 | 0.504865517 | 1.490496518 | 0.136093731 | 0.345013444 | 0.752500296 |
| ENSMUSG00000031165 | Was | 22376 | protein_coding | 14.10453044 | 0.752740417 | 0.376338369 | 2.000169209 | 0.045481995 | 0.17356558 | 0.752740417 |
| ENSMUSG00000038147 | Cd84 | 12523 | protein_coding | 7.980121453 | 0.754081669 | 0.421960852 | 1.787089171 | 0.073923077 | 0.238288521 | 0.754081669 |
| ENSMUSG00000020773 | Trim47 | 217333 | protein_coding | 41.06685503 | 0.757602756 | 0.230694654 | 3.28400656 | 0.001023425 | 0.010257788 | 0.757602756 |
| ENSMUSG00000029095 | Ablim2 | 231148 | protein_coding | 180.8161316 | -0.759215299 | 0.140547944 | -5.401824286 | 6.60E-08 | 2.07E-06 | 0.759215299 |
| ENSMUSG00000020364 | Zfp354a | 21408 | protein_coding | 26.08920413 | -0.760029909 | 0.282329673 | -2.691994435 | 0.007102613 | 0.046336505 | 0.760029909 |
| ENSMUSG00000025017 | Pik3ap1 | 83490 | protein_coding | 38.38041171 | 0.760186359 | 0.220023972 | 3.455016074 | 0.00055026 | 0.006150272 | 0.760186359 |
| ENSMUSG00000073968 | Trim68 | 101700 | protein_coding | 280.3740266 | 0.761147438 | 0.0875813 | 8.690752944 | 3.60E-18 | 2.74E-16 | 0.761147438 |
| ENSMUSG00000040026 | Saa3 | 20210 | protein_coding | 235.4919048 | 0.761698632 | 0.191158336 | 3.984647749 | 6.76E-05 | 0.001065364 | 0.761698632 |
| ENSMUSG00000056130 | Ticam2 | 225471 | protein_coding | 7.212148749 | -0.762125339 | 0.514495264 | -1.481306812 | 0.138524834 | 0.348679833 | 0.762125339 |
| ENSMUSG00000000902 | Smarcb1 | 20587 | protein_coding | 1249.492001 | 0.762645145 | 0.088170014 | 8.649711078 | 5.16E-18 | 3.84E-16 | 0.762645145 |
| ENSMUSG00000038068 | Rnf144b | 218215 | protein_coding | 7.571924309 | 0.762836462 | 0.469693259 | 1.624116266 | 0.104350997 | 0.295551199 | 0.762836462 |
| ENSMUSG00000027800 | Tm4sf1 | 17112 | protein_coding | 44.93945995 | 0.762854454 | 0.210070865 | 3.631414832 | 0.000281872 | 0.003496865 | 0.762854454 |
| ENSMUSG00000034449 | Dhrs11 | 192970 | protein_coding | 3095.427884 | 0.763646259 | 0.078567334 | 9.71964069 | 2.49E-22 | 2.42E-20 | 0.763646259 |
| ENSMUSG00000042156 | Dzip1 | 66573 | protein_coding | 7.254055672 | 0.764483934 | 0.490011391 | 1.560135025 | 0.118727976 | 0.318807809 | 0.764483934 |
| ENSMUSG00000023247 | Guca2a | 14915 | protein_coding | 36695.03862 | 0.764510424 | 0.094210506 | 8.114916891 | 4.86E-16 | 3.25E-14 | 0.764510424 |
| ENSMUSG00000085720 | Gm7854 | 665934 | lncRNA | 17.92922833 | -0.764567639 | 0.288385133 | -2.651203378 | 0.008020553 | 0.050786438 | 0.764567639 |
| ENSMUSG00000113918 | Gm6566 | 625237 | lncRNA | 9.358772073 | -0.764677208 | 0.403368046 | -1.895730748 | 0.057995657 | 0.203901453 | 0.764677208 |
| ENSMUSG00000034057 | Myrfl | 237558 | protein_coding | 30.5922117 | -0.764805379 | 0.311485876 | -2.455345292 | 0.014074932 | 0.076614782 | 0.764805379 |
| ENSMUSG00000025091 | Pnliprp2 | 18947 | protein_coding | 29.59958956 | 0.765328728 | 0.277134963 | 2.761574072 | 0.005752347 | 0.039815194 | 0.765328728 |
| ENSMUSG00000031351 | Zfp185 | 22673 | protein_coding | 14.75202657 | 0.766419507 | 0.344445647 | 2.225081126 | 0.026075793 | 0.118191205 | 0.766419507 |
| ENSMUSG00000028356 | Ambp | 11699 | protein_coding | 39.53166011 | -0.767046189 | 0.222948176 | -3.440468551 | 0.000580708 | 0.006458674 | 0.767046189 |
| ENSMUSG00000028527 | Ak4 | 11639 | protein_coding | 150.4891388 | -0.768163938 | 0.163650013 | -4.693943648 | 2.68E-06 | 5.97E-05 | 0.768163938 |

Supplementary Table 1

| ensembl_gene_id | external_gene_name | entrezgene_id | gene_biotype | baseMean | log2FoldChange | lfcSE | stat | pvalue | padj | absLog2FC |
|---|---|---|---|---|---|---|---|---|---|---|
| ENSMUSG00000038151 | Prdm1 | 12142 | protein_coding | 39.13258228 | 0.768749162 | 0.216596152 | 3.549228162 | 0.000386362 | 0.004558056 | 0.768749162 |
| ENSMUSG00000022661 | Cd200 | 17470 | protein_coding | 62.80498986 | -0.769130829 | 0.169156265 | -4.546865745 | 5.45E-06 | 0.000114488 | 0.769130829 |
| ENSMUSG00000055116 | Arntl | 11865 | protein_coding | 1034.375254 | 0.769862824 | 0.088874586 | 8.662350559 | 4.62E-18 | 3.50E-16 | 0.769862824 |
| ENSMUSG00000052212 | Cd177 | 68891 | protein_coding | 2831.856225 | 0.769866233 | 0.103068465 | 7.46946444 | 8.05E-14 | 4.56E-12 | 0.769866233 |
| ENSMUSG00000004709 | Cd244a | 18106 | protein_coding | 29.78202741 | 0.770412165 | 0.226280395 | 3.40467925 | 0.000662418 | 0.007209695 | 0.770412165 |
| ENSMUSG00000034854 | Mfsd12 | 73822 | protein_coding | 445.7107838 | 0.770761199 | 0.079831765 | 9.654818491 | 4.69E-22 | 4.40E-20 | 0.770761199 |
| ENSMUSG00000040134 | Rdh7 | 54150 | protein_coding | 16.53551745 | 0.771096716 | 0.487554539 | 1.581559918 | 0.11375007 | 0.310817924 | 0.771096716 |
| ENSMUSG00000073805 | Insyn2a | 627214 | protein_coding | 32.28366353 | -0.771289323 | 0.284270762 | -2.713220723 | 0.006663271 | 0.044203442 | 0.771289323 |
| ENSMUSG00000046240 | Hepacam | 72927 | protein_coding | 11.59848885 | -0.774775033 | 0.466958787 | -1.659193603 | 0.097076786 | 0.282795761 | 0.774775033 |
| ENSMUSG00000020312 | Shc2 | 216148 | protein_coding | 14.06617392 | 0.774918985 | 0.319440015 | 2.425866983 | 0.015271867 | 0.081167147 | 0.774918985 |
| ENSMUSG00000000157 | Itgb2l | 16415 | protein_coding | 152.8781349 | 0.775565162 | 0.180392233 | 4.299326796 | 1.71E-05 | 0.000314674 | 0.775565162 |
| ENSMUSG00000032478 | Nme6 | 54369 | protein_coding | 309.6219886 | 0.776224635 | 0.104359917 | 7.437957558 | 1.02E-13 | 5.73E-12 | 0.776224635 |
| ENSMUSG00000029075 | Tnfrsf4 | 22163 | protein_coding | 5.915205682 | -0.77645237 | 0.553983954 | -1.401579169 | 0.161040951 | 0.381534805 | 0.77645237 |
| ENSMUSG00000035337 | Uchl4 | 93841 | protein_coding | 9.567784249 | -0.776668644 | 0.483134407 | -1.607562271 | 0.107931081 | 0.300944072 | 0.776668644 |
| ENSMUSG00000007207 | Stx1a | 20907 | protein_coding | 18.75867936 | -0.778944061 | 0.314100056 | -2.479923341 | 0.013141063 | 0.072876866 | 0.778944061 |
| ENSMUSG00000033174 | Mgll | 23945 | protein_coding | 694.7559117 | 0.77900192 | 0.114968939 | 6.775759842 | 1.24E-11 | 5.90E-10 | 0.77900192 |
| ENSMUSG00000086645 | Gm15743 | 100047123 | lncRNA | 23.64690965 | -0.779281656 | 0.298134416 | -2.613860104 | 0.008952569 | 0.054959297 | 0.779281656 |
| ENSMUSG00000087213 | 2810408I11Rik | 69941 | lncRNA | 17.58992689 | 0.780606482 | 0.326130867 | 2.393537569 | 0.016686772 | 0.086633325 | 0.780606482 |
| ENSMUSG00000078439 | Smim24 | 72273 | protein_coding | 4170.680668 | -0.780749891 | 0.093160434 | -8.380702616 | 5.26E-17 | 3.74E-15 | 0.780749891 |
| ENSMUSG00000034687 | Fras1 | 231470 | protein_coding | 15.352648 | 0.781190209 | 0.426995578 | 1.829504214 | 0.067324109 | 0.224318083 | 0.781190209 |
| ENSMUSG00000020077 | Srgn | 19073 | protein_coding | 35.61796728 | 0.781945801 | 0.246764145 | 3.168798287 | 0.001530706 | 0.014236271 | 0.781945801 |
| ENSMUSG00000085440 | Sorbs2os | 319940 | lncRNA | 8.453624531 | -0.782074639 | 0.451522717 | -1.732082593 | 0.083258858 | 0.256750769 | 0.782074639 |
| ENSMUSG00000044453 | Ffar1 | 233081 | protein_coding | 8.478390842 | -0.783017499 | 0.445925687 | -1.755937192 | 0.079099133 | 0.248419031 | 0.783017499 |
| ENSMUSG00000041801 | Phlda3 | 27280 | protein_coding | 22.31495082 | 0.783098861 | 0.275270538 | 2.844833547 | 0.004443464 | 0.032617086 | 0.783098861 |
| ENSMUSG00000050148 | Ubqln2 | 54609 | protein_coding | 445.041983 | 0.783170458 | 0.114905012 | 6.815807646 | 9.37E-12 | 4.53E-10 | 0.783170458 |
| ENSMUSG00000031232 | Magt1 | 67075 | protein_coding | 4282.491832 | 0.783885488 | 0.039172234 | 20.01125313 | 4.39E-89 | 1.73E-86 | 0.783885488 |
| ENSMUSG00000018500 | Adora2b | 11541 | protein_coding | 18.21352813 | -0.784118486 | 0.31986475 | -2.451406375 | 0.01422992 | 0.077138124 | 0.784118486 |
| ENSMUSG00000109864 | Eid3 | 66341 | protein_coding | 32.80980434 | -0.784459749 | 0.275334668 | -2.849113609 | 0.004384122 | 0.032358233 | 0.784459749 |
| ENSMUSG00000028957 | Per3 | 18628 | protein_coding | 459.0219079 | -0.785342591 | 0.125957655 | -6.234973114 | 4.52E-10 | 1.85E-08 | 0.785342591 |
| ENSMUSG00000064844 | Gm26202 | 115488998 | snoRNA | 14.09034471 | -0.787299501 | 0.343546293 | -2.29168388 | 0.021923891 | 0.104860209 | 0.787299501 |
| ENSMUSG00000058626 | Capn11 | 268958 | protein_coding | 16.20050113 | -0.787642504 | 1.652461199 | -0.476648108 | 0.63361272 | 0.808283189 | 0.787642504 |
| ENSMUSG00000115667 | Gm10389 | 100038608 | lncRNA | 11.29139337 | -0.787820748 | 0.456513465 | -1.72573387 | 0.084395302 | 0.258950348 | 0.787820748 |
| ENSMUSG00000027674 | Pex5l | 58869 | protein_coding | 60.97098731 | -0.788697225 | 0.169532219 | -4.652196666 | 3.28E-06 | 7.17E-05 | 0.788697225 |
| ENSMUSG00000040875 | Osbpl10 | 74486 | protein_coding | 531.1269012 | 0.790673539 | 0.07651727 | 10.333269 | 4.98E-25 | 5.39E-23 | 0.790673539 |
| ENSMUSG00000028926 | Cdk14 | 18647 | protein_coding | 89.47042034 | 0.790711244 | 0.141587495 | 5.584612157 | 2.34E-08 | 8.02E-07 | 0.790711244 |
| ENSMUSG00000022650 | Retnlb | 57263 | protein_coding | 1565.309701 | 0.791126518 | 0.291648087 | 2.712606571 | 0.006675631 | 0.044230856 | 0.791126518 |
| ENSMUSG00000030873 | Scnn1b | 20277 | protein_coding | 359.9455415 | 0.791817034 | 0.192569471 | 4.11185131 | 3.92E-05 | 0.000658671 | 0.791817034 |
| ENSMUSG00000031374 | Zfp92 | 22754 | protein_coding | 7.883092208 | 0.792897813 | 0.519420255 | 1.526505378 | 0.126884052 | 0.331056319 | 0.792897813 |
| ENSMUSG00000096887 | Gm20594 | 100463512 | protein_coding | 6.759215339 | -0.795854832 | 0.576556551 | -1.380358667 | 0.16747624 | 0.391489973 | 0.795854832 |

| ensembl_gene_id | external_gene_name | entrezgene_id | gene_biotype | baseMean | log2FoldChange | lfcSE | stat | pvalue | padj | absLog2FC |
|---|---|---|---|---|---|---|---|---|---|---|
| ENSMUSG00000023973 | Cnpy3 | 72029 | protein_coding | 845.7191922 | 0.795960111 | 0.058175041 | 13.68215815 | 1.30E-42 | 2.46E-40 | 0.795960111 |
| ENSMUSG00000040118 | Cacna2d1 | 12293 | protein_coding | 89.54871502 | 0.797414838 | 0.137616043 | 5.794490402 | 6.85E-09 | 2.47E-07 | 0.797414838 |
| ENSMUSG00000014905 | Dnajb9 | 27362 | protein_coding | 1216.428398 | 0.798429408 | 0.076494699 | 10.43770903 | 1.67E-25 | 1.83E-23 | 0.798429408 |
| ENSMUSG00000106445 | Gm21190 | 100861755 | protein_coding | 8.544459219 | -0.798541202 | 0.408188923 | -1.956302968 | 0.050429475 | 0.18610438 | 0.798541202 |
| ENSMUSG00000026246 | Alppl2 | 11650 | protein_coding | 6.237262583 | -0.799021556 | 0.497228423 | -1.606950689 | 0.108065186 | 0.301132615 | 0.799021556 |
| ENSMUSG00000023224 | Serping1 | 12258 | protein_coding | 6.530581715 | 0.799372613 | 0.616223049 | 1.297213102 | 0.194557874 | 0.426539942 | 0.799372613 |
| ENSMUSG00000050192 | Eif5a2 | 208691 | protein_coding | 15.33434239 | 0.799457449 | 0.30310741 | 2.637538447 | 0.008351016 | 0.052220563 | 0.799457449 |
| ENSMUSG00000047992 | Dipk1c | 240479 | protein_coding | 13.68910629 | 0.800392117 | 0.343158758 | 2.332425151 | 0.019678336 | 0.096931133 | 0.800392117 |
| ENSMUSG00000028838 | Extl1 | 56219 | protein_coding | 24.41709397 | -0.800518894 | 0.309742312 | -2.584467352 | 0.009752954 | 0.05831883 | 0.800518894 |
| ENSMUSG00000030731 | Syt3 | 20981 | protein_coding | 12.45692785 | -0.800541629 | 0.358565686 | -2.23262197 | 0.025573884 | 0.116583051 | 0.800541629 |
| ENSMUSG00000027559 | Car3 | 12350 | protein_coding | 9.081862941 | 0.801381086 | 0.38805491 | 2.065122913 | 0.038911372 | 0.155443076 | 0.801381086 |
| ENSMUSG00000083012 | Fam220a | 67238 | protein_coding | 6.596985991 | -0.801509833 | 0.568682 | -1.40941657 | 0.158712027 | 0.378894236 | 0.801509833 |
| ENSMUSG00000024030 | Abcg1 | 11307 | protein_coding | 94.81824746 | 0.801900105 | 0.184296582 | 4.35113932 | 1.35E-05 | 0.000257257 | 0.801900105 |
| ENSMUSG00000018581 | Dnah11 | 13411 | protein_coding | 8.059079405 | 0.802300269 | 0.522508883 | 1.535476801 | 0.124666751 | 0.327549804 | 0.802300269 |
| ENSMUSG00000022877 | Hrg | 94175 | protein_coding | 7.386637496 | -0.802658561 | 0.486224034 | -1.650799849 | 0.098779451 | 0.285896663 | 0.802658561 |
| ENSMUSG00000064105 | Cnnm2 | 94219 | protein_coding | 206.5418226 | 0.80287406 | 0.092064501 | 8.72077788 | 2.76E-18 | 2.14E-16 | 0.80287406 |
| ENSMUSG00000015363 | Trabd | 67976 | protein_coding | 2277.930698 | 0.803371134 | 0.053778924 | 14.93840107 | 1.85E-50 | 4.27E-48 | 0.803371134 |
| ENSMUSG00000033486 | Catsper2 | 212670 | protein_coding | 37.28717461 | 0.805015419 | 0.196807859 | 4.090362165 | 4.31E-05 | 0.000714599 | 0.805015419 |
| ENSMUSG00000084984 | Far1os | 69295 | lncRNA | 36.81663461 | 0.805455036 | 0.243452149 | 3.308473712 | 0.00093806 | 0.009544538 | 0.805455036 |
| ENSMUSG00000020407 | Upp1 | 22271 | protein_coding | 875.718231 | -0.805738977 | 0.401559613 | -2.006523939 | 0.044800377 | 0.171777386 | 0.805738977 |
| ENSMUSG00000024640 | Psat1 | 107272 | protein_coding | 168.111349 | 0.806477151 | 0.181484762 | 4.443773355 | 8.84E-06 | 0.000175776 | 0.806477151 |
| ENSMUSG00000040710 | St8sia4 | 20452 | protein_coding | 29.73297526 | 0.808018699 | 0.251816569 | 3.208759066 | 0.001333092 | 0.012728696 | 0.808018699 |
| ENSMUSG00000048794 | Cfap100 | 243538 | protein_coding | 6.130290401 | -0.808311994 | 0.54772338 | -1.475766826 | 0.140006499 | 0.350881819 | 0.808311994 |
| ENSMUSG00000067599 | Klra7 | 16638 | protein_coding | 15.38868585 | 0.808529445 | 0.298200674 | 2.711360221 | 0.006700779 | 0.044324634 | 0.808529445 |
| ENSMUSG00000018920 | Cxcl16 | 66102 | protein_coding | 61.73485941 | -0.808683356 | 0.22239622 | -3.63622797 | 0.00027666 | 0.003453319 | 0.808683356 |
| ENSMUSG00000046959 | Slc26a1 | 231583 | protein_coding | 100.974938 | 0.808909603 | 0.218981549 | 3.693962368 | 0.000220786 | 0.002859939 | 0.808909603 |
| ENSMUSG00000032171 | Pin1 | 23988 | protein_coding | 642.5425943 | 0.810346773 | 0.075626462 | 10.71512211 | 8.64E-27 | 1.01E-24 | 0.810346773 |
| ENSMUSG00000020176 | Grb10 | 14783 | protein_coding | 75.62240451 | 0.811776491 | 0.14342439 | 5.659961263 | 1.51E-08 | 5.29E-07 | 0.811776491 |
| ENSMUSG00000013415 | Igf2bp1 | 140486 | protein_coding | 21.32979272 | -0.812136165 | 0.284687451 | -2.85272906 | 0.004334556 | 0.032095218 | 0.812136165 |
| ENSMUSG00000020484 | Xbp1 | 22433 | protein_coding | 4903.650164 | 0.81253845 | 0.027623565 | 29.4146845 | 3.56E-190 | 4.42E-187 | 0.81253845 |
| ENSMUSG00000017453 | Pipox | 19193 | protein_coding | 87.01219963 | -0.814172605 | 0.165741896 | -4.9122921 | 9.00E-07 | 2.23E-05 | 0.814172605 |
| ENSMUSG00000118516 | Cyp2d13 | 68444 | protein_coding | 248.5210638 | -0.814412191 | 0.160144845 | -5.085472409 | 3.67E-07 | 9.82E-06 | 0.814412191 |
| ENSMUSG00000064213 | Defa24 | 503491 | protein_coding | 210.5465189 | 0.815207808 | 1.117151269 | 0.729720165 | 0.465561253 | 0.691532821 | 0.815207808 |
| ENSMUSG00000090171 | Ugt1a2 | 22236 | protein_coding | 8.105851444 | -0.816208766 | 0.489754594 | -1.66656684 | 0.095600567 | 0.280216348 | 0.816208766 |
| ENSMUSG00000026764 | Kif5c | 16574 | protein_coding | 54.57561426 | 0.816721499 | 0.222125214 | 3.676851833 | 0.00023613 | 0.003024955 | 0.816721499 |
| ENSMUSG00000030605 | Mfge8 | 17304 | protein_coding | 327.6640883 | 0.816797808 | 0.096355678 | 8.476903734 | 2.31E-17 | 1.67E-15 | 0.816797808 |
| ENSMUSG00000093979 | Gm2237 | 100039441 | protein_coding | 8.112688158 | -0.817187112 | 0.410730679 | -1.989593554 | 0.046635727 | 0.176322153 | 0.817187112 |
| ENSMUSG00000045466 | Zfp956 | 101197 | protein_coding | 214.0300415 | 0.817905205 | 0.094522528 | 8.653018763 | 5.02E-18 | 3.75E-16 | 0.817905205 |
| ENSMUSG00000030789 | Itgax | 16411 | protein_coding | 21.25979419 | 0.818242095 | 0.297735284 | 2.748220106 | 0.005991977 | 0.040980752 | 0.818242095 |

| ensembl_gene_id | external_gene_name | entrezgene_id | gene_biotype | baseMean | log2FoldChange | lfcSE | stat | pvalue | padj | absLog2FC |
|---|---|---|---|---|---|---|---|---|---|---|
| ENSMUSG00000025044 | Msr1 | 20288 | protein_coding | 6.577992759 | -0.818435898 | 0.47079741 | -1.738403568 | 0.082139728 | 0.254517969 | 0.818435898 |
| ENSMUSG00000050395 | Tnfsf15 | 326623 | protein_coding | 42.6526174 | -0.819707508 | 0.190563947 | -4.301482632 | 1.70E-05 | 0.000311983 | 0.819707508 |
| ENSMUSG00000043313 | Pcdhb19 | 93890 | protein_coding | 7.780591124 | 0.819884562 | 0.431312752 | 1.90090499 | 0.057314458 | 0.202566353 | 0.819884562 |
| ENSMUSG00000101360 | 4933417O13Rik | 71153 | lncRNA | 15.50115719 | -0.825197834 | 0.419657614 | -1.966359734 | 0.049257071 | 0.183499374 | 0.825197834 |
| ENSMUSG00000024912 | Fosl1 | 14283 | protein_coding | 57.84393753 | -0.825342224 | 0.310742918 | -2.65602907 | 0.00790668 | 0.050260554 | 0.825342224 |
| ENSMUSG00000008318 | Relt | 320100 | protein_coding | 7.906914668 | 0.826060742 | 0.435724844 | 1.895831174 | 0.057982372 | 0.203899197 | 0.826060742 |
| ENSMUSG00000004837 | Grap | 71520 | protein_coding | 7.875505925 | 0.826389627 | 0.500706979 | 1.650445594 | 0.098851832 | 0.286003499 | 0.826389627 |
| ENSMUSG00000024575 | Pde6a | 225600 | protein_coding | 15.9553441 | -0.826985515 | 0.306718541 | -2.696235816 | 0.0070128 | 0.045874365 | 0.826985515 |
| ENSMUSG00000094584 | Ms4a18 | 76002 | protein_coding | 21.26576785 | -0.827101397 | 0.422005613 | -1.959929849 | 0.05000399 | 0.18516977 | 0.827101397 |
| ENSMUSG00000079363 | Gbp4 | 17472 | protein_coding | 17.40950593 | -0.827235184 | 0.54594083 | -1.515246963 | 0.129709874 | 0.335781899 | 0.827235184 |
| ENSMUSG00000086446 | Prkag2os1 | 71161 | lncRNA | 6.371227563 | -0.827819279 | 0.506984683 | -1.632828973 | 0.102504972 | 0.292221615 | 0.827819279 |
| ENSMUSG00000090145 | Ugt1a6b | 394435 | protein_coding | 19.11947176 | -0.829599323 | 0.330599284 | -2.509380276 | 0.012094321 | 0.068412876 | 0.829599323 |
| ENSMUSG00000037849 | Ifi206 | 102639543 | protein_coding | 5.897347197 | -0.830178619 | 0.559297607 | -1.484323568 | 0.137723101 | 0.347583794 | 0.830178619 |
| ENSMUSG00000024901 | Peli3 | 240518 | protein_coding | 48.58830564 | -0.830497227 | 0.179432998 | -4.628453153 | 3.68E-06 | 7.94E-05 | 0.830497227 |
| ENSMUSG00000051985 | Igfn1 | 226438 | protein_coding | 6.002490663 | 0.831465509 | 0.530281029 | 1.567971442 | 0.116887797 | 0.316071346 | 0.831465509 |
| ENSMUSG00000052229 | Gpr17 | 574402 | protein_coding | 15.4665632 | -0.832012128 | 0.361803047 | -2.2996272 | 0.02146935 | 0.103384952 | 0.832012128 |
| ENSMUSG00000021123 | Rdh12 | 77974 | protein_coding | 8.728293464 | -0.832673498 | 0.438470478 | -1.89904119 | 0.05755906 | 0.20320818 | 0.832673498 |
| ENSMUSG00000006179 | Prss16 | 54373 | protein_coding | 25.1430292 | 0.832948166 | 0.300123312 | 2.77535311 | 0.00551418 | 0.038396884 | 0.832948166 |
| ENSMUSG00000097587 | 4930578M01Rik | 75951 | lncRNA | 10.95483314 | -0.833984848 | 0.399443631 | -2.08786618 | 0.036809908 | 0.149416909 | 0.833984848 |
| ENSMUSG00000107722 | 2900060B14Rik | 68204 | lncRNA | 11.64480517 | -0.83482119 | 0.432404915 | -1.930646858 | 0.053526739 | 0.193678645 | 0.83482119 |
| ENSMUSG00000002799 | Jag2 | 16450 | protein_coding | 102.087395 | -0.836181287 | 0.150662954 | -5.550012567 | 2.86E-08 | 9.66E-07 | 0.836181287 |
| ENSMUSG00000032038 | St3gal4 | 20443 | protein_coding | 6452.442716 | -0.83850901 | 0.115301553 | -7.272313234 | 3.53E-13 | 1.90E-11 | 0.83850901 |
| ENSMUSG00000027690 | Slc2a2 | 20526 | protein_coding | 24.35822078 | 0.839599584 | 0.670697176 | 1.251831101 | 0.210631415 | 0.445605776 | 0.839599584 |
| ENSMUSG00000031910 | Has3 | 15118 | protein_coding | 15.3400781 | -0.840144806 | 0.417571089 | -2.011980305 | 0.044222013 | 0.170480837 | 0.840144806 |
| ENSMUSG00000027828 | Ssr3 | 67437 | protein_coding | 5797.639179 | 0.841151528 | 0.053636246 | 15.6825204 | 1.99E-55 | 5.02E-53 | 0.841151528 |
| ENSMUSG00000053334 | Ficd | 231630 | protein_coding | 248.0613024 | 0.841584587 | 0.082324914 | 10.22272057 | 1.57E-24 | 1.64E-22 | 0.841584587 |
| ENSMUSG00000078427 | Sarnp | 66118 | protein_coding | 610.0104405 | 0.841990225 | 0.086663464 | 9.715630835 | 2.59E-22 | 2.50E-20 | 0.841990225 |
| ENSMUSG00000021266 | Wars1 | 22375 | protein_coding | 3679.648641 | 0.842601508 | 0.101462846 | 8.304532516 | 1.00E-16 | 7.03E-15 | 0.842601508 |
| ENSMUSG00000027642 | Rpn2 | 20014 | protein_coding | 9808.460069 | 0.843455592 | 0.067094886 | 12.57108622 | 3.05E-36 | 4.91E-34 | 0.843455592 |
| ENSMUSG00000070423 | Or51e1 | 259097 | protein_coding | 10.39955641 | -0.847797546 | 0.407421808 | -2.080884062 | 0.037444519 | 0.151214294 | 0.847797546 |
| ENSMUSG00000024357 | Sil1 | 81500 | protein_coding | 1184.761224 | 0.848381875 | 0.057001003 | 14.88363063 | 4.21E-50 | 9.43E-48 | 0.848381875 |
| ENSMUSG00000073125 | Xlr3b | 574437 | protein_coding | 27.88550773 | 0.849400941 | 0.839597337 | 1.011676555 | 0.311692729 | 0.555505998 | 0.849400941 |
| ENSMUSG00000026193 | Fn1 | 14268 | protein_coding | 17.16382724 | 0.85008131 | 0.294318337 | 2.888305636 | 0.003873233 | 0.0294918 | 0.85008131 |
| ENSMUSG00000047910 | Pcdhb16 | 93887 | protein_coding | 6.224075045 | 0.852220203 | 0.477327745 | 1.785398423 | 0.074196714 | 0.238931791 | 0.852220203 |
| ENSMUSG00000089736 | Tgfbr3l | 100044509 | protein_coding | 14.11431557 | -0.852402785 | 0.383794801 | -2.220985752 | 0.026351926 | 0.118795321 | 0.852402785 |
| ENSMUSG00000047990 | C2cd4a | 244911 | protein_coding | 133.1477844 | 0.853823656 | 0.253338817 | 3.370283579 | 0.000750909 | 0.007948619 | 0.853823656 |
| ENSMUSG00000043648 | Pld6 | 194908 | protein_coding | 6.100429068 | -0.855897979 | 0.497031487 | -1.722019634 | 0.085065962 | 0.260216003 | 0.855897979 |
| ENSMUSG00000033676 | Gabrb3 | 14402 | protein_coding | 6.0166464 | -0.856016339 | 0.53249867 | -1.607546437 | 0.107934551 | 0.300944072 | 0.856016339 |
| ENSMUSG00000051727 | Kctd14 | 233529 | protein_coding | 220.0874039 | 0.85650526 | 0.103479968 | 8.2770151 | 1.26E-16 | 8.74E-15 | 0.85650526 |

Supplementary Table 1

| ensembl_gene_id | external_gene_name | entrezgene_id | gene_biotype | baseMean | log2FoldChange | lfcSE | stat | pvalue | padj | absLog2FC |
|---|---|---|---|---|---|---|---|---|---|---|
| ENSMUSG00000028452 | Vcp | 269523 | protein_coding | 5896.476063 | 0.857255859 | 0.068964523 | 12.43038913 | 1.79E-35 | 2.72E-33 | 0.857255859 |
| ENSMUSG00000080115 | Eef1akmt3 | 100504608 | protein_coding | 21.30782961 | 0.858207639 | 0.284574952 | 3.015752561 | 0.002563424 | 0.021232836 | 0.858207639 |
| ENSMUSG00000020044 | Timp3 | 21859 | protein_coding | 123.4119803 | -0.859578261 | 0.188891496 | -4.550645639 | 5.35E-06 | 0.000112598 | 0.859578261 |
| ENSMUSG00000030041 | M1ap | 110958 | protein_coding | 10.9869072 | 0.860238675 | 0.457918575 | 1.878584365 | 0.060301268 | 0.208758782 | 0.860238675 |
| ENSMUSG00000021490 | Slc34a1 | 20505 | protein_coding | 19.32125064 | 0.860707767 | 0.295766855 | 2.910088644 | 0.003613263 | 0.028014945 | 0.860707767 |
| ENSMUSG00000040466 | Blvrb | 233016 | protein_coding | 724.8050051 | 0.860889144 | 0.068845519 | 12.50465033 | 7.04E-36 | 1.08E-33 | 0.860889144 |
| ENSMUSG00000020321 | Mdh1 | 17449 | protein_coding | 11488.56524 | 0.862055853 | 0.058852767 | 14.6476691 | 1.39E-48 | 3.00E-46 | 0.862055853 |
| ENSMUSG00000042682 | Selenok | 80795 | protein_coding | 1305.27816 | 0.86434525 | 0.058265131 | 14.83469167 | 8.74E-50 | 1.90E-47 | 0.86434525 |
| ENSMUSG00000114362 | 4933433G19Rik | 71226 | lncRNA | 16.48837486 | -0.86439098 | 0.367438378 | -2.352478759 | 0.018648755 | 0.093749523 | 0.86439098 |
| ENSMUSG00000001444 | Tbx21 | 57765 | protein_coding | 10.56033542 | 0.866597955 | 0.41021536 | 2.112543898 | 0.034639825 | 0.143636725 | 0.866597955 |
| ENSMUSG00000046546 | Fam43a | 224093 | protein_coding | 115.2211465 | -0.866790727 | 0.255586126 | -3.391384109 | 0.000695406 | 0.00748652 | 0.866790727 |
| ENSMUSG00000031539 | Ap3m2 | 64933 | protein_coding | 23.87619682 | 0.868574445 | 0.257132361 | 3.377927392 | 0.000730344 | 0.0077847 | 0.868574445 |
| ENSMUSG00000027204 | Fbn1 | 14118 | protein_coding | 58.6417167 | -0.870965938 | 0.219314278 | -3.971314352 | 7.15E-05 | 0.001111584 | 0.870965938 |
| ENSMUSG00000037386 | Rims2 | 116838 | protein_coding | 99.2406864 | -0.872494319 | 0.589645465 | -1.479693088 | 0.138955171 | 0.349163297 | 0.872494319 |
| ENSMUSG00000023571 | C1qtnf12 | 67389 | protein_coding | 208.2034995 | -0.872962618 | 0.094786678 | -9.209760625 | 3.27E-20 | 2.75E-18 | 0.872962618 |
| ENSMUSG00000037003 | Tns2 | 209039 | protein_coding | 391.7743418 | 0.877372109 | 0.063348939 | 13.84983126 | 1.28E-43 | 2.54E-41 | 0.877372109 |
| ENSMUSG00000003153 | Slc2a3 | 20527 | protein_coding | 6.54829665 | 0.878368555 | 0.479719803 | 1.831003325 | 0.067100047 | 0.223743052 | 0.878368555 |
| ENSMUSG00000020175 | Rab36 | 76877 | protein_coding | 9.046101423 | 0.881029546 | 0.449465329 | 1.960172431 | 0.04997564 | 0.18510729 | 0.881029546 |
| ENSMUSG00000024114 | Prss41 | 71003 | protein_coding | 15.59815091 | -0.881032962 | 0.386810204 | -2.277688007 | 0.022745173 | 0.107506272 | 0.881032962 |
| ENSMUSG00000045092 | S1pr1 | 13609 | protein_coding | 10.00707741 | 0.882561943 | 0.431733893 | 2.044226678 | 0.040931159 | 0.161156445 | 0.882561943 |
| ENSMUSG00000032625 | Thsd7a | 330267 | protein_coding | 10.86352259 | -0.883563819 | 0.456936989 | -1.933666653 | 0.053154126 | 0.192705082 | 0.883563819 |
| ENSMUSG00000037060 | Cavin3 | 109042 | protein_coding | 19.44208065 | 0.884107082 | 0.273215213 | 3.235936503 | 0.001212444 | 0.011821691 | 0.884107082 |
| ENSMUSG00000029373 | Pf4 | 56744 | protein_coding | 7.506333973 | 0.884968097 | 0.425721208 | 2.07875032 | 0.037640305 | 0.151794249 | 0.884968097 |
| ENSMUSG00000029503 | P2rx2 | 231602 | protein_coding | 9.555640077 | -0.885496074 | 0.412309117 | -2.147650966 | 0.031741493 | 0.135278822 | 0.885496074 |
| ENSMUSG00000038456 | Dennd2a | 209773 | protein_coding | 6.048484391 | 0.885838821 | 0.471472493 | 1.878876996 | 0.060261291 | 0.208682378 | 0.885838821 |
| ENSMUSG00000028415 | Spink4 | 20731 | protein_coding | 11044.3378 | 0.885962386 | 0.084722586 | 10.45721607 | 1.36E-25 | 1.50E-23 | 0.885962386 |
| ENSMUSG00000042763 | Maneal | 215090 | protein_coding | 7.057343097 | -0.88732883 | 0.518886529 | -1.710063336 | 0.087254162 | 0.264527214 | 0.88732883 |
| ENSMUSG00000056749 | Nfil3 | 18030 | protein_coding | 471.0996866 | 0.888827191 | 0.163486393 | 5.436704388 | 5.43E-08 | 1.74E-06 | 0.888827191 |
| ENSMUSG00000020340 | Cyfip2 | 76884 | protein_coding | 30.52996872 | 0.889600972 | 0.234091662 | 3.800224937 | 0.000144565 | 0.0020054 | 0.889600972 |
| ENSMUSG00000022064 | Pibf1 | 52023 | protein_coding | 463.6928835 | 0.889712186 | 0.092972633 | 9.569613781 | 1.07E-21 | 9.72E-20 | 0.889712186 |
| ENSMUSG00000026365 | Cfh | 12628 | protein_coding | 9.591176659 | -0.890186212 | 0.52614302 | -1.691909191 | 0.090663289 | 0.271060828 | 0.890186212 |
| ENSMUSG00000028757 | Ddost | 13200 | protein_coding | 7175.439687 | 0.89030421 | 0.075567678 | 11.78154776 | 4.86E-32 | 6.64E-30 | 0.89030421 |
| ENSMUSG00000009628 | Tex15 | 104271 | protein_coding | 6.906641953 | 0.890327131 | 0.541156845 | 1.645229361 | 0.099922521 | 0.28770764 | 0.890327131 |
| ENSMUSG00000024186 | Rgs11 | 50782 | protein_coding | 48.07230093 | -0.890913257 | 0.253398599 | -3.515857071 | 0.000438337 | 0.005058411 | 0.890913257 |
| ENSMUSG00000050195 | Scd4 | 329065 | protein_coding | 7.629780873 | -0.891476378 | 0.492568645 | -1.809852063 | 0.070318732 | 0.230634571 | 0.891476378 |
| ENSMUSG00000022032 | Scara5 | 71145 | protein_coding | 35.39930472 | 0.892347672 | 0.250494907 | 3.562338577 | 0.000367566 | 0.00438442 | 0.892347672 |
| ENSMUSG00000027932 | Slc27a3 | 26568 | protein_coding | 38.74383944 | 0.893018937 | 0.2053094 | 4.34962519 | 1.36E-05 | 0.000258431 | 0.893018937 |
| ENSMUSG00000002900 | Lamb1 | 16777 | protein_coding | 57.06409792 | -0.893552741 | 0.185229937 | -4.824019018 | 1.41E-06 | 3.33E-05 | 0.893552741 |
| ENSMUSG00000035780 | Ugt2a3 | 72094 | protein_coding | 12.40509513 | 0.89412645 | 0.805644115 | 1.109828066 | 0.267073123 | 0.509292687 | 0.89412645 |

| ensembl_gene_id | external_gene_name | entrezgene_id | gene_biotype | baseMean | log2FoldChange | lfcSE | stat | pvalue | padj | absLog2FC |
|---|---|---|---|---|---|---|---|---|---|---|
| ENSMUSG00000071847 | Apcdd1 | 494504 | protein_coding | 6.914005875 | 0.894810273 | 0.502586732 | 1.780409663 | 0.075008942 | 0.240742704 | 0.894810273 |
| ENSMUSG00000042357 | Gjb5 | 14622 | protein_coding | 32.00405277 | -0.896013863 | 0.284460617 | -3.14986965 | 0.001633433 | 0.01496726 | 0.896013863 |
| ENSMUSG00000086992 | Gm15941 | 100503538 | lncRNA | 9.745737934 | -0.897449526 | 0.458497637 | -1.957370012 | 0.050303982 | 0.185896496 | 0.897449526 |
| ENSMUSG00000026421 | Csrp1 | 13007 | protein_coding | 910.8110901 | 0.898286595 | 0.06769653 | 13.26931524 | 3.49E-40 | 6.18E-38 | 0.898286595 |
| ENSMUSG00000097403 | 9230116N13Rik | 320054 | lncRNA | 6.65370747 | -0.898376659 | 0.480855741 | -1.86828727 | 0.06172204 | 0.211965788 | 0.898376659 |
| ENSMUSG00000063751 | Platr8 | 100038454 | lncRNA | 7.032036318 | -0.900070647 | 0.523609157 | -1.718974229 | 0.085619065 | 0.261362607 | 0.900070647 |
| ENSMUSG00000040462 | Os9 | 216440 | protein_coding | 5323.927385 | 0.901747805 | 0.049670859 | 18.15446373 | 1.18E-73 | 3.82E-71 | 0.901747805 |
| ENSMUSG00000085944 | 1700003D09Rik | 69338 | lncRNA | 41.12524174 | -0.903424651 | 0.324862452 | -2.780945125 | 0.00542009 | 0.037921817 | 0.903424651 |
| ENSMUSG00000035941 | Ibtk | 108837 | protein_coding | 2890.261098 | 0.903541811 | 0.060785615 | 14.86440179 | 5.61E-50 | 1.24E-47 | 0.903541811 |
| ENSMUSG00000062421 | Arf2 | 11841 | protein_coding | 376.4324204 | 0.905808699 | 0.079943019 | 11.33067909 | 9.25E-30 | 1.21E-27 | 0.905808699 |
| ENSMUSG00000026258 | Snorc | 73234 | protein_coding | 6.411684949 | -0.906456742 | 0.611075098 | -1.483380267 | 0.137973407 | 0.347997676 | 0.906456742 |
| ENSMUSG00000058618 | Defa39 | 382000 | protein_coding | 34.3954969 | 0.906848151 | 1.101453448 | 0.823319544 | 0.410326302 | 0.648598559 | 0.906848151 |
| ENSMUSG00000085887 | Arhgap27os3 | 653030 | lncRNA | 35.19984849 | -0.907053794 | 0.381199995 | -2.379469588 | 0.017337574 | 0.088855862 | 0.907053794 |
| ENSMUSG00000012017 | Scarf2 | 224024 | protein_coding | 50.82077469 | 0.907669209 | 0.195336952 | 4.646684615 | 3.37E-06 | 7.35E-05 | 0.907669209 |
| ENSMUSG00000030062 | Rpn1 | 103963 | protein_coding | 8445.262172 | 0.909090167 | 0.071506664 | 12.71336279 | 4.98E-37 | 8.29E-35 | 0.909090167 |
| ENSMUSG00000031661 | Nkd1 | 93960 | protein_coding | 20.3199828 | -0.909664294 | 0.273686451 | -3.323746176 | 0.00088817 | 0.009129074 | 0.909664294 |
| ENSMUSG00000061186 | Sfmbt2 | 353282 | protein_coding | 9.253421656 | 0.910053964 | 0.421288585 | 2.16016763 | 0.030759695 | 0.132451326 | 0.910053964 |
| ENSMUSG00000095061 | E030018B13Rik | 381994 | protein_coding | 23.78892734 | -0.910742583 | 0.422714871 | -2.154508028 | 0.031200353 | 0.133664893 | 0.910742583 |
| ENSMUSG00000027238 | Frmd5 | 228564 | protein_coding | 12.68018002 | 0.912087597 | 0.349536167 | 2.609422665 | 0.009069514 | 0.055508181 | 0.912087597 |
| ENSMUSG00000013629 | Cad | 69719 | protein_coding | 1407.812079 | 0.91278654 | 0.115584352 | 7.897146337 | 2.85E-15 | 1.85E-13 | 0.91278654 |
| ENSMUSG00000043165 | Lor | 16939 | protein_coding | 9.178272356 | 0.913073698 | 0.515792744 | 1.770233699 | 0.076688218 | 0.244031351 | 0.913073698 |
| ENSMUSG00000066456 | Hmgn3 | 94353 | protein_coding | 12.29517705 | -0.914715247 | 0.346719696 | -2.638198108 | 0.008334787 | 0.052179782 | 0.914715247 |
| ENSMUSG00000073422 | H2-Ke6 | 14979 | protein_coding | 9.923200712 | -0.915739293 | 0.428377338 | -2.137693128 | 0.032541658 | 0.137670334 | 0.915739293 |
| ENSMUSG00000055385 | Rnf212 | 671564 | protein_coding | 54.26161913 | -0.918880948 | 0.259188553 | -3.545221958 | 0.000392283 | 0.004607678 | 0.918880948 |
| ENSMUSG00000026544 | Dusp23 | 68440 | protein_coding | 266.7917002 | 0.921834962 | 0.09705102 | 9.498457239 | 2.13E-21 | 1.89E-19 | 0.921834962 |
| ENSMUSG00000022240 | Ctnnd2 | 18163 | protein_coding | 9.64318 | 0.921894989 | 0.52838717 | 1.744733863 | 0.081031206 | 0.252369884 | 0.921894989 |
| ENSMUSG00000070427 | Il18bp | 16068 | protein_coding | 105.4755935 | -0.923177632 | 0.311770682 | -2.961079042 | 0.003065633 | 0.024532938 | 0.923177632 |
| ENSMUSG00000023176 | Cpn2 | 71756 | protein_coding | 27.34214218 | 0.924115867 | 0.350468951 | 2.636798111 | 0.008369263 | 0.052253619 | 0.924115867 |
| ENSMUSG00000052026 | Slc6a7 | 240332 | protein_coding | 70.85949493 | -0.924632033 | 0.222532449 | -4.155043622 | 3.25E-05 | 0.000557969 | 0.924632033 |
| ENSMUSG00000086315 | Arl4aos | 100043606 | lncRNA | 8.459818596 | -0.925159825 | 0.410820997 | -2.251977946 | 0.024323666 | 0.11275302 | 0.925159825 |
| ENSMUSG00000090799 | Klhl33 | 546611 | protein_coding | 6.152998339 | 0.92528482 | 0.529390301 | 1.747831077 | 0.080493282 | 0.251280519 | 0.92528482 |
| ENSMUSG00000035041 | Creb3l3 | 208677 | protein_coding | 62.45951975 | 0.927367398 | 0.306352282 | 3.027127437 | 0.002468898 | 0.02061933 | 0.927367398 |
| ENSMUSG00000028613 | Lrp8 | 16975 | protein_coding | 84.06786162 | 0.927915795 | 0.204912782 | 4.528345123 | 5.94E-06 | 0.000123864 | 0.927915795 |
| ENSMUSG00000050600 | Zfp831 | 100043757 | protein_coding | 11.52247278 | 0.928314332 | 0.367551963 | 2.525668274 | 0.011547846 | 0.066100912 | 0.928314332 |
| ENSMUSG00000026934 | Lhx3 | 16871 | protein_coding | 29.06425425 | 0.92889168 | 0.336884222 | 2.757302414 | 0.005828042 | 0.040183343 | 0.92889168 |
| ENSMUSG00000000305 | Cdh4 | 12561 | protein_coding | 6.569219516 | 0.929641998 | 0.562852345 | 1.651662298 | 0.098603413 | 0.285746134 | 0.929641998 |
| ENSMUSG00000031924 | Cyb5b | 66427 | protein_coding | 11942.05271 | 0.929782532 | 0.065898958 | 14.10921445 | 3.33E-45 | 6.98E-43 | 0.929782532 |
| ENSMUSG00000063903 | Klk1 | 16612 | protein_coding | 33014.69042 | 0.932773909 | 0.086673768 | 10.76189404 | 5.21E-27 | 6.22E-25 | 0.932773909 |
| ENSMUSG00000029913 | Prdm5 | 70779 | protein_coding | 28.72859366 | 0.935622578 | 0.243510256 | 3.842230689 | 0.000121921 | 0.00175399 | 0.935622578 |

| ensembl_gene_id | external_gene_name | entrezgene_id | gene_biotype | baseMean | log2FoldChange | lfcSE | stat | pvalue | padj | absLog2FC |
|---|---|---|---|---|---|---|---|---|---|---|
| ENSMUSG00000118314 | 1700001K23Rik | 69319 | lncRNA | 7.651783068 | -0.937638813 | 0.44461196 | -2.108892464 | 0.034953865 | 0.144514406 | 0.937638813 |
| ENSMUSG00000085645 | Hoxb5os | 75395 | lncRNA | 40.9378206 | -0.941485397 | 0.24335406 | -3.868788538 | 0.000109377 | 0.001605032 | 0.941485397 |
| ENSMUSG00000097328 | Tnfsf12 | 21944 | protein_coding | 6.951526677 | -0.946326827 | 0.475500389 | -1.990170458 | 0.046572161 | 0.176237021 | 0.946326827 |
| ENSMUSG00000079845 | Xlr4a | 434794 | protein_coding | 11.57146914 | 0.947245337 | 0.974167574 | 0.972363854 | 0.330869567 | 0.574373898 | 0.947245337 |
| ENSMUSG00000031381 | Piga | 18700 | protein_coding | 678.0769136 | 0.947589259 | 0.07705653 | 12.29732592 | 9.36E-35 | 1.36E-32 | 0.947589259 |
| ENSMUSG00000004902 | Slc25a18 | 71803 | protein_coding | 8.510169253 | -0.948752326 | 0.514793582 | -1.842976212 | 0.065332484 | 0.220053669 | 0.948752326 |
| ENSMUSG00000006313 | Upk1a | 109637 | protein_coding | 288.2802971 | 0.949739724 | 0.098795849 | 9.613154127 | 7.04E-22 | 6.52E-20 | 0.949739724 |
| ENSMUSG00000041895 | Wipi1 | 52639 | protein_coding | 583.1456136 | 0.951220255 | 0.07803544 | 12.18959305 | 3.53E-34 | 5.00E-32 | 0.951220255 |
| ENSMUSG00000010307 | Tmem86a | 67893 | protein_coding | 114.1038979 | -0.952091477 | 0.239223384 | -3.979926458 | 6.89E-05 | 0.00107936 | 0.952091477 |
| ENSMUSG00000018442 | Derl2 | 116891 | protein_coding | 1220.01849 | 0.952409784 | 0.063967145 | 14.88904633 | 3.88E-50 | 8.82E-48 | 0.952409784 |
| ENSMUSG00000106514 | Mir6976 | 102466778 | miRNA | 6.197990718 | -0.952465538 | 0.732031761 | -1.301125975 | 0.193215338 | 0.425042116 | 0.952465538 |
| ENSMUSG00000043008 | Klhl6 | 239743 | protein_coding | 8.507261295 | 0.952651089 | 0.427531416 | 2.228259849 | 0.025863192 | 0.117491746 | 0.952651089 |
| ENSMUSG00000023467 | Tulp2 | 56734 | protein_coding | 27.41578282 | -0.9526704 | 0.285748694 | -3.333944898 | 0.000856236 | 0.008851616 | 0.9526704 |
| ENSMUSG00000023235 | Ccl25 | 20300 | protein_coding | 237.506863 | -0.954425004 | 0.211925153 | -4.503594733 | 6.68E-06 | 0.000137788 | 0.954425004 |
| ENSMUSG00000068086 | Cyp2d9 | 13105 | protein_coding | 203.8927247 | -0.956213282 | 0.185343883 | -5.159130509 | 2.48E-07 | 6.93E-06 | 0.956213282 |
| ENSMUSG00000022758 | P2rx6 | 18440 | protein_coding | 11.51651342 | -0.957311726 | 0.354241446 | -2.702427217 | 0.006883525 | 0.045218172 | 0.957311726 |
| ENSMUSG00000085024 | C230035I16Rik | 320842 | lncRNA | 9.528403544 | -0.958421491 | 0.389393206 | -2.461320524 | 0.013842664 | 0.075700453 | 0.958421491 |
| ENSMUSG00000003279 | Dlgap1 | 224997 | protein_coding | 11.88713117 | 0.958835937 | 0.391343185 | 2.450115332 | 0.014281046 | 0.077285381 | 0.958835937 |
| ENSMUSG00000036599 | Chst12 | 59031 | protein_coding | 13.67641645 | 0.962582332 | 0.34921666 | 2.756404383 | 0.00584407 | 0.040237686 | 0.962582332 |
| ENSMUSG00000036832 | Lpar3 | 65086 | protein_coding | 8.484969549 | 0.968080792 | 0.448297749 | 2.159459407 | 0.030814543 | 0.132518971 | 0.968080792 |
| ENSMUSG00000021679 | S100z | 268686 | protein_coding | 6.476975076 | 0.968767922 | 0.508522377 | 1.905064489 | 0.056771689 | 0.201440865 | 0.968767922 |
| ENSMUSG00000047108 | Dnajb7 | 57755 | protein_coding | 5.884289122 | -0.969715892 | 0.515213637 | -1.882162705 | 0.059813928 | 0.207699031 | 0.969715892 |
| ENSMUSG00000030087 | Klf15 | 66277 | protein_coding | 23.10963782 | -0.969765632 | 0.251768399 | -3.85181634 | 0.000117245 | 0.001697944 | 0.969765632 |
| ENSMUSG00000091721 | Gimd1 | 433653 | protein_coding | 7.159651137 | 0.969924571 | 0.743568281 | 1.30441897 | 0.192090771 | 0.423550432 | 0.969924571 |
| ENSMUSG00000085747 | Slc13a2os | 790911 | lncRNA | 84.92055983 | -0.971529283 | 0.284657888 | -3.41297159 | 0.000642586 | 0.007036183 | 0.971529283 |
| ENSMUSG00000047671 | Dynlt4 | 242646 | protein_coding | 45.85313446 | -0.972448041 | 0.350988075 | -2.77060137 | 0.005595288 | 0.038894487 | 0.972448041 |
| ENSMUSG00000047619 | Ddi1 | 71829 | protein_coding | 6.292478614 | -0.973217822 | 0.531979775 | -1.829426356 | 0.067335762 | 0.224318083 | 0.973217822 |
| ENSMUSG00000040204 | Pclaf | 68026 | protein_coding | 789.8774603 | 0.976259335 | 0.155332773 | 6.284954031 | 3.28E-10 | 1.37E-08 | 0.976259335 |
| ENSMUSG00000057074 | Ces1g | 12623 | protein_coding | 1327.935083 | -0.977259409 | 0.155623473 | -6.279640161 | 3.39E-10 | 1.41E-08 | 0.977259409 |
| ENSMUSG00000026307 | Scly | 50880 | protein_coding | 390.2309345 | 0.977340679 | 0.084203285 | 11.60691863 | 3.80E-31 | 5.15E-29 | 0.977340679 |
| ENSMUSG00000030483 | Cyp2b10 | 13088 | protein_coding | 9.554681907 | 0.977477124 | 0.738723802 | 1.323197008 | 0.185769871 | 0.415061056 | 0.977477124 |
| ENSMUSG00000062017 | Abca14 | 67928 | protein_coding | 6.759075589 | -0.978544253 | 0.543292884 | -1.801135781 | 0.071681481 | 0.233536818 | 0.978544253 |
| ENSMUSG00000087203 | Gm13986 | 329496 | lncRNA | 20.75130914 | -0.978606377 | 0.298629819 | -3.276988146 | 0.001049208 | 0.010464176 | 0.978606377 |
| ENSMUSG00000078670 | Fam174b | 100038347 | protein_coding | 2033.161744 | 0.979147659 | 0.048831494 | 20.05156067 | 1.96E-89 | 7.89E-87 | 0.979147659 |
| ENSMUSG00000023153 | Tmem52 | 69671 | protein_coding | 7.514572223 | -0.980752254 | 0.485603271 | -2.019657428 | 0.043418934 | 0.168280977 | 0.980752254 |
| ENSMUSG00000048349 | Pou4f1 | 18996 | protein_coding | 9.292924992 | -0.9823877 | 0.460542674 | -2.13310895 | 0.03291579 | 0.138772199 | 0.9823877 |
| ENSMUSG00000022757 | Tfg | 21787 | protein_coding | 1706.372242 | 0.982491207 | 0.043641794 | 22.51262202 | 3.12E-112 | 1.53E-109 | 0.982491207 |
| ENSMUSG00000013338 | Fer1l4 | 74562 | protein_coding | 62.5423182 | 0.983240243 | 0.199953606 | 4.917341886 | 8.77E-07 | 2.18E-05 | 0.983240243 |
| ENSMUSG00000097331 | F420014N23Rik | 100038591 | lncRNA | 11.8828019 | -0.984159465 | 0.450559273 | -2.184306312 | 0.028939747 | 0.126769891 | 0.984159465 |

Supplementary Table 1

| ensembl_gene_id | external_gene_name | entrezgene_id | gene_biotype | baseMean | log2FoldChange | lfcSE | stat | pvalue | padj | absLog2FC |
|---|---|---|---|---|---|---|---|---|---|---|
| ENSMUSG00000030017 | Reg3g | 19695 | protein_coding | 95.62246605 | -0.984769695 | 0.258691541 | -3.806733271 | 0.000140814 | 0.00197128 | 0.984769695 |
| ENSMUSG00000097267 | 1700109K24Rik | 74303 | lncRNA | 11.45425794 | -0.984987948 | 0.41319947 | -2.383807388 | 0.017134568 | 0.088255885 | 0.984987948 |
| ENSMUSG00000090386 | Mir99ahg | 77994 | lncRNA | 9.667933463 | 0.987916201 | 0.391404395 | 2.524029405 | 0.011601822 | 0.066254458 | 0.987916201 |
| ENSMUSG00000050821 | Fam131a | 78408 | protein_coding | 12.61305033 | -0.988018934 | 0.359605587 | -2.747507181 | 0.006005019 | 0.041052542 | 0.988018934 |
| ENSMUSG00000057751 | Megf6 | 230971 | protein_coding | 17.52710168 | -0.9913776 | 0.325921282 | -3.041770066 | 0.002351915 | 0.019858288 | 0.9913776 |
| ENSMUSG00000035407 | Kank4 | 242553 | protein_coding | 6.091286687 | -0.991689548 | 0.710509747 | -1.395743763 | 0.162791671 | 0.384046412 | 0.991689548 |
| ENSMUSG00000097021 | 4933433G15Rik | 71274 | lncRNA | 12.96594858 | -0.992287682 | 0.403715832 | -2.457886471 | 0.013975735 | 0.076221398 | 0.992287682 |
| ENSMUSG00000078931 | Pdf | 68023 | protein_coding | 8.235715573 | -0.994822713 | 0.472366009 | -2.106042125 | 0.035200693 | 0.145122272 | 0.994822713 |
| ENSMUSG00000030098 | Grip2 | 243547 | protein_coding | 6.433188531 | -0.997364513 | 0.640180619 | -1.557942372 | 0.119246909 | 0.319397925 | 0.997364513 |
| ENSMUSG00000074139 | 1700057G04Rik | 78459 | protein_coding | 11.32099805 | -0.998768008 | 0.403446041 | -2.475592535 | 0.013301529 | 0.073539168 | 0.998768008 |
| ENSMUSG00000052928 | Ctif | 269037 | protein_coding | 212.3638602 | 1.000065463 | 0.094083443 | 10.62955854 | 2.17E-26 | 2.45E-24 | 1.000065463 |
| ENSMUSG00000032383 | Ppib | 19035 | protein_coding | 2921.754942 | 1.002195685 | 0.072274093 | 13.86659655 | 1.01E-43 | 2.04E-41 | 1.002195685 |
| ENSMUSG00000030815 | Phkg2 | 68961 | protein_coding | 316.8884868 | 1.002930932 | 0.08083033 | 12.40785392 | 2.37E-35 | 3.57E-33 | 1.002930932 |
| ENSMUSG00000044294 | Krt84 | 16680 | protein_coding | 93.60988729 | 1.004058676 | 0.237710994 | 4.223863016 | 2.40E-05 | 0.000426062 | 1.004058676 |
| ENSMUSG00000104667 | Gm4961 | 242914 | lncRNA | 11.99697106 | 1.005344706 | 0.366836395 | 2.740580594 | 0.006133074 | 0.041645508 | 1.005344706 |
| ENSMUSG00000058057 | Mettl7a3 | 668178 | protein_coding | 19.83855953 | -1.006616906 | 0.424676161 | -2.370316488 | 0.017772864 | 0.090551523 | 1.006616906 |
| ENSMUSG00000073405 | H2-T-ps | 667803 | unprocessed_pseudogene | 25.26658695 | 1.006782746 | 0.297748337 | 3.381321139 | 0.000721382 | 0.007724916 | 1.006782746 |
| ENSMUSG00000035845 | Alg12 | 223774 | protein_coding | 445.0950267 | 1.009187288 | 0.080521667 | 12.53311469 | 4.92E-36 | 7.63E-34 | 1.009187288 |
| ENSMUSG00000102737 | 1700016A09Rik | 74227 | TEC | 17.16456971 | -1.009260673 | 0.343138891 | -2.941259935 | 0.003268801 | 0.025879212 | 1.009260673 |
| ENSMUSG00000029616 | Erp29 | 67397 | protein_coding | 2279.489281 | 1.010528251 | 0.051926578 | 19.46071327 | 2.36E-84 | 8.87E-82 | 1.010528251 |
| ENSMUSG00000021268 | Meg3 | 17263 | lncRNA | 27.45686548 | -1.013107181 | 0.412584347 | -2.455515309 | 0.014068276 | 0.076614782 | 1.013107181 |
| ENSMUSG00000075304 | Sp5 | 64406 | protein_coding | 13.29547801 | 1.013249493 | 0.361995181 | 2.799069009 | 0.005125018 | 0.036330185 | 1.013249493 |
| ENSMUSG00000019906 | Lin7a | 108030 | protein_coding | 9.980774366 | -1.015071145 | 0.416989804 | -2.434282892 | 0.014921328 | 0.07991905 | 1.015071145 |
| ENSMUSG00000056629 | Fkbp2 | 14227 | protein_coding | 984.1928321 | 1.015206575 | 0.067102426 | 15.1292082 | 1.04E-51 | 2.47E-49 | 1.015206575 |
| ENSMUSG00000026442 | Nfasc | 269116 | protein_coding | 25.06142866 | 1.015261685 | 0.257077791 | 3.949239176 | 7.84E-05 | 0.001203003 | 1.015261685 |
| ENSMUSG00000022610 | Mapk12 | 29857 | protein_coding | 9.092885427 | -1.016198849 | 0.456784097 | -2.224680884 | 0.026102668 | 0.118279779 | 1.016198849 |
| ENSMUSG00000031824 | 6430548M08Rik | 234797 | protein_coding | 1759.993104 | 1.017815069 | 0.056439467 | 18.03374712 | 1.06E-72 | 3.28E-70 | 1.017815069 |
| ENSMUSG00000032558 | Nphp3 | 74025 | protein_coding | 224.6790501 | 1.018230837 | 0.084094336 | 12.10819754 | 9.56E-34 | 1.33E-31 | 1.018230837 |
| ENSMUSG00000096878 | Gm21083 | 105242399 | protein_coding | 9.470483392 | -1.021245202 | 0.451778373 | -2.26050042 | 0.02379021 | 0.110821699 | 1.021245202 |
| ENSMUSG00000045942 | BC049762 | 193286 | protein_coding | 7.049668009 | -1.021543748 | 0.472983507 | -2.159787251 | 0.030789143 | 0.132480392 | 1.021543748 |
| ENSMUSG00000073529 | F830208F22Rik | 619308 | lncRNA | 8.898781811 | -1.021915466 | 0.530407664 | -1.926660447 | 0.05402196 | 0.194647414 | 1.021915466 |
| ENSMUSG00000027995 | Tlr2 | 24088 | protein_coding | 449.90268 | -1.022088918 | 0.130847641 | -7.811290371 | 5.66E-15 | 3.58E-13 | 1.022088918 |
| ENSMUSG00000043067 | Dpy19l1 | 244745 | protein_coding | 170.4959127 | 1.022635063 | 0.099748323 | 10.25215294 | 1.16E-24 | 1.23E-22 | 1.022635063 |
| ENSMUSG00000026109 | Tmeff2 | 56363 | protein_coding | 7.638161721 | -1.024086773 | 0.448111633 | -2.285338512 | 0.022292986 | 0.106146735 | 1.024086773 |
| ENSMUSG00000084781 | D930015M05Rik | 320507 | lncRNA | 16.79419756 | -1.025889229 | 0.387216948 | -2.64939134 | 0.00806369 | 0.050937378 | 1.025889229 |
| ENSMUSG00000045672 | Col27a1 | 373864 | protein_coding | 7.339960082 | 1.026415379 | 0.608040313 | 1.688071263 | 0.09139755 | 0.272408741 | 1.026415379 |
| ENSMUSG00000028360 | Slc44a5 | 242259 | protein_coding | 7.43922578 | -1.027579096 | 0.498407416 | -2.061725131 | 0.039233913 | 0.156498963 | 1.027579096 |
| ENSMUSG00000048458 | Inka2 | 109050 | protein_coding | 6.719940498 | 1.028071465 | 0.461226562 | 2.228994489 | 0.025814271 | 0.117338412 | 1.028071465 |
| ENSMUSG00000005503 | Evx1 | 14028 | protein_coding | 15.87956666 | 1.028668911 | 0.327616626 | 3.139855646 | 0.001690311 | 0.0153489 | 1.028668911 |

| ensembl_gene_id | external_gene_name | entrezgene_id | gene_biotype | baseMean | log2FoldChange | lfcSE | stat | pvalue | padj | absLog2FC |
|---|---|---|---|---|---|---|---|---|---|---|
| ENSMUSG00000022864 | D16Ertd472e | 67102 | protein_coding | 29.44854661 | 1.029160445 | 0.259842273 | 3.960712133 | 7.47E-05 | 0.001155432 | 1.029160445 |
| ENSMUSG00000026452 | Syt2 | 20980 | protein_coding | 169.2129879 | -1.030069771 | 0.102555659 | -10.04400709 | 9.76E-24 | 9.84E-22 | 1.030069771 |
| ENSMUSG00000050232 | Cxcr3 | 12766 | protein_coding | 11.05882788 | 1.030513489 | 0.46409462 | 2.220481436 | 0.026386104 | 0.118807539 | 1.030513489 |
| ENSMUSG00000044285 | Ubb-ps | 218963 | transcribed_processed_pseudogene | 24.09039178 | 1.031599414 | 0.677676676 | 1.522258993 | 0.127944199 | 0.33296048 | 1.031599414 |
| ENSMUSG00000104713 | Gbp6 | 100702 | protein_coding | 16.42883908 | -1.033719306 | 0.48182595 | -2.145420576 | 0.031919235 | 0.13589269 | 1.033719306 |
| ENSMUSG00000052353 | Cemip | 80982 | protein_coding | 8.276914984 | 1.0348062 | 0.529379524 | 1.954752978 | 0.050612234 | 0.186591593 | 1.0348062 |
| ENSMUSG00000059708 | Akap17b | 338351 | protein_coding | 38.57215359 | 1.035584286 | 0.200136278 | 5.174395649 | 2.29E-07 | 6.44E-06 | 1.035584286 |
| ENSMUSG00000022439 | Parvg | 64099 | protein_coding | 10.54975487 | 1.035693967 | 0.387240298 | 2.674551104 | 0.007482937 | 0.048137725 | 1.035693967 |
| ENSMUSG00000078606 | Gvin2 | 100042856 | protein_coding | 6.37663073 | -1.035984357 | 0.612407429 | -1.691658703 | 0.090711067 | 0.271060828 | 1.035984357 |
| ENSMUSG00000031765 | Mt1 | 17748 | protein_coding | 8976.327096 | -1.036381574 | 0.093443649 | -11.09097928 | 1.39E-28 | 1.71E-26 | 1.036381574 |
| ENSMUSG00000054408 | Spcs3 | 76687 | protein_coding | 3000.570582 | 1.03870666 | 0.053848284 | 19.28950341 | 6.58E-83 | 2.41E-80 | 1.03870666 |
| ENSMUSG00000000489 | Pdgfb | 18591 | protein_coding | 21.33851446 | -1.039743405 | 0.309377398 | -3.360760714 | 0.000777281 | 0.008171588 | 1.039743405 |
| ENSMUSG00000031168 | Ebp | 13595 | protein_coding | 2942.204565 | 1.040406779 | 0.085695931 | 12.14067889 | 6.43E-34 | 9.02E-32 | 1.040406779 |
| ENSMUSG00000018906 | P4ha2 | 18452 | protein_coding | 114.8896998 | 1.040408424 | 0.178605595 | 5.825172643 | 5.71E-09 | 2.10E-07 | 1.040408424 |
| ENSMUSG00000077505 | Gm24233 | 115485895 | snoRNA | 10.91699981 | 1.041735413 | 0.444687004 | 2.342626168 | 0.019148559 | 0.095399692 | 1.041735413 |
| ENSMUSG00000028655 | Mfsd2a | 76574 | protein_coding | 52.6351428 | 1.041736767 | 0.27463467 | 3.793172827 | 0.000148735 | 0.002050122 | 1.041736767 |
| ENSMUSG00000019893 | Ros1 | 19886 | protein_coding | 62.70105093 | 1.044843174 | 0.171876454 | 6.079036141 | 1.21E-09 | 4.76E-08 | 1.044843174 |
| ENSMUSG00000034118 | Tpst1 | 22021 | protein_coding | 53.91903072 | 1.044964886 | 0.212482022 | 4.917897891 | 8.75E-07 | 2.17E-05 | 1.044964886 |
| ENSMUSG00000006310 | Zbtb32 | 58206 | protein_coding | 143.4574765 | 1.047293763 | 0.101680229 | 10.29987609 | 7.06E-25 | 7.54E-23 | 1.047293763 |
| ENSMUSG00000063689 | H2ac21 | 621893 | protein_coding | 41.01475204 | 1.052449372 | 0.194617858 | 5.407773889 | 6.38E-08 | 2.02E-06 | 1.052449372 |
| ENSMUSG00000033508 | Asprv1 | 67855 | protein_coding | 7.741230001 | -1.053346578 | 0.542068682 | -1.943197631 | 0.051992284 | 0.189926522 | 1.053346578 |
| ENSMUSG00000050953 | Gja1 | 14609 | protein_coding | 188.7529233 | -1.054809867 | 0.195224764 | -5.403053619 | 6.55E-08 | 2.06E-06 | 1.054809867 |
| ENSMUSG00000048040 | Arxes2 | 76976 | protein_coding | 6.188295837 | 1.060838579 | 0.479732051 | 2.211314788 | 0.027014046 | 0.120633435 | 1.060838579 |
| ENSMUSG00000103948 | 4930594C11Rik | 77633 | TEC | 16.875875 | -1.062040987 | 0.367275409 | -2.891674644 | 0.003831946 | 0.029274176 | 1.062040987 |
| ENSMUSG00000039873 | Neurl2 | 415115 | protein_coding | 5.8646324 | -1.063514728 | 0.552196328 | -1.925972113 | 0.054107856 | 0.19482421 | 1.063514728 |
| ENSMUSG00000014351 | Gip | 14607 | protein_coding | 5.895697169 | 1.063738252 | 0.755502559 | 1.407987623 | 0.159134735 | 0.37924721 | 1.063738252 |
| ENSMUSG00000067653 | Ankrd23 | 78321 | protein_coding | 5.911708096 | -1.065889749 | 0.561763698 | -1.89739877 | 0.057775328 | 0.20374868 | 1.065889749 |
| ENSMUSG00000004500 | Zfp324 | 243834 | protein_coding | 380.0717676 | 1.069683061 | 0.087175588 | 12.2704428 | 1.31E-34 | 1.86E-32 | 1.069683061 |
| ENSMUSG00000027068 | Dhrs9 | 241452 | protein_coding | 257.4197806 | 1.076826809 | 0.15138146 | 7.113333506 | 1.13E-12 | 5.87E-11 | 1.076826809 |
| ENSMUSG00000085975 | Gm13572 | 102636022 | lncRNA | 8.894443441 | -1.077387311 | 0.535931814 | -2.010306691 | 0.044398739 | 0.170791501 | 1.077387311 |
| ENSMUSG00000038648 | Creb3l2 | 208647 | protein_coding | 245.0924924 | 1.078378481 | 0.096629981 | 11.1598747 | 6.41E-29 | 8.07E-27 | 1.078378481 |
| ENSMUSG00000062785 | Kcnc3 | 16504 | protein_coding | 20.0471537 | 1.078932153 | 0.342020252 | 3.154585574 | 0.001607262 | 0.014811611 | 1.078932153 |
| ENSMUSG00000018983 | E2f2 | 242705 | protein_coding | 699.7057039 | 1.079591077 | 0.135791346 | 7.950367286 | 1.86E-15 | 1.21E-13 | 1.079591077 |
| ENSMUSG00000041120 | Nbl1 | 17965 | protein_coding | 246.329869 | -1.081393699 | 0.203975124 | -5.301595992 | 1.15E-07 | 3.47E-06 | 1.081393699 |
| ENSMUSG00000075588 | Hoxb2 | 103889 | protein_coding | 5.957570861 | -1.081870463 | 0.627419286 | -1.724318152 | 0.084650424 | 0.259486294 | 1.081870463 |
| ENSMUSG00000035107 | Dcbld2 | 73379 | protein_coding | 238.6162385 | 1.084738809 | 0.144919003 | 7.485138504 | 7.15E-14 | 4.07E-12 | 1.084738809 |
| ENSMUSG00000042641 | Rgsl1 | 240816 | protein_coding | 8.274877415 | -1.085257437 | 0.540478408 | -2.007957062 | 0.044647855 | 0.171421356 | 1.085257437 |
| ENSMUSG00000030847 | Bag3 | 29810 | protein_coding | 825.8150301 | 1.085765681 | 0.11607064 | 9.354352497 | 8.41E-21 | 7.22E-19 | 1.085765681 |
| ENSMUSG00000022389 | Tef | 21685 | protein_coding | 1085.97329 | -1.086142862 | 0.124866184 | -8.698454811 | 3.36E-18 | 2.58E-16 | 1.086142862 |

| ensembl_gene_id | external_gene_name | entrezgene_id | gene_biotype | baseMean | log2FoldChange | lfcSE | stat | pvalue | padj | absLog2FC |
|---|---|---|---|---|---|---|---|---|---|---|
| ENSMUSG00000009108 | Gnat2 | 14686 | protein_coding | 29.27325303 | 1.087263541 | 0.255234601 | 4.259859499 | 2.05E-05 | 0.000369414 | 1.087263541 |
| ENSMUSG00000020607 | Lratd1 | 105005 | protein_coding | 207.3044666 | 1.091298631 | 0.111355727 | 9.800112285 | 1.12E-22 | 1.11E-20 | 1.091298631 |
| ENSMUSG00000020427 | Igfbp3 | 16009 | protein_coding | 14.58833562 | 1.091532866 | 0.383864303 | 2.843538351 | 0.004461564 | 0.032705292 | 1.091532866 |
| ENSMUSG00000000159 | Igsf5 | 72058 | protein_coding | 606.6430453 | 1.091935232 | 0.087110813 | 12.53501373 | 4.80E-36 | 7.52E-34 | 1.091935232 |
| ENSMUSG00000034645 | Zyg11a | 230590 | protein_coding | 12.02743039 | -1.09199946 | 0.414085606 | -2.637134553 | 0.008360966 | 0.052253619 | 1.09199946 |
| ENSMUSG00000007594 | Hapln4 | 330790 | protein_coding | 12.77578669 | 1.092222657 | 0.410433618 | 2.66114326 | 0.007787582 | 0.049660074 | 1.092222657 |
| ENSMUSG00000040564 | Apoc1 | 11812 | protein_coding | 14.35013893 | -1.095443929 | 0.380480537 | -2.879106342 | 0.003988038 | 0.030237468 | 1.095443929 |
| ENSMUSG00000074151 | Nlrc5 | 434341 | protein_coding | 51.14862369 | -1.097131997 | 0.603055068 | -1.819289905 | 0.068867213 | 0.22777308 | 1.097131997 |
| ENSMUSG00000044937 | Ttc41 | 103220 | protein_coding | 47.11211027 | 1.097583139 | 0.18546741 | 5.91792995 | 3.26E-09 | 1.23E-07 | 1.097583139 |
| ENSMUSG00000075266 | Cenpw | 66311 | protein_coding | 131.6441258 | 1.097951315 | 0.205276781 | 5.348638591 | 8.86E-08 | 2.73E-06 | 1.097951315 |
| ENSMUSG00000050359 | Sprr1a | 20753 | protein_coding | 43.75311573 | 1.101001936 | 0.235398616 | 4.677181013 | 2.91E-06 | 6.43E-05 | 1.101001936 |
| ENSMUSG00000091017 | Garin4 | 619288 | protein_coding | 44.12851653 | -1.105314947 | 0.653799159 | -1.690603195 | 0.090912614 | 0.271508838 | 1.105314947 |
| ENSMUSG00000091996 | Smim35 | 408059 | protein_coding | 8.029304375 | -1.106937085 | 0.464662967 | -2.382236509 | 0.017207842 | 0.088535742 | 1.106937085 |
| ENSMUSG00000067235 | H2-Q10 | 15007 | protein_coding | 80.6028603 | -1.107869849 | 0.180548126 | -6.136147037 | 8.45E-10 | 3.38E-08 | 1.107869849 |
| ENSMUSG00000017754 | Pltp | 18830 | protein_coding | 11.059349 | 1.110110441 | 0.435156684 | 2.551059152 | 0.01073961 | 0.062318117 | 1.110110441 |
| ENSMUSG00000036110 | Slc17a2 | 218103 | protein_coding | 56.54105612 | -1.11111884 | 0.295462789 | -3.760604994 | 0.000169503 | 0.002287509 | 1.11111884 |
| ENSMUSG00000071178 | Serpina1b | 20701 | protein_coding | 10.25475651 | 1.111205396 | 0.431853017 | 2.573110184 | 0.010078911 | 0.059604916 | 1.111205396 |
| ENSMUSG00000025194 | Abcc2 | 12780 | protein_coding | 70.33322215 | -1.112744072 | 0.357469213 | -3.112838898 | 0.001852971 | 0.016537282 | 1.112744072 |
| ENSMUSG00000039579 | Grin3a | 242443 | protein_coding | 383.4681901 | -1.113880863 | 0.299638192 | -3.717419514 | 0.000201268 | 0.002651838 | 1.113880863 |
| ENSMUSG00000091618 | H60c | 670558 | protein_coding | 40.64124053 | 1.114013727 | 0.246235595 | 4.52417826 | 6.06E-06 | 0.000126004 | 1.114013727 |
| ENSMUSG00000020889 | Nr1d1 | 217166 | protein_coding | 1322.483995 | -1.114923967 | 0.130995342 | -8.51117261 | 1.72E-17 | 1.25E-15 | 1.114923967 |
| ENSMUSG00000021775 | Nr1d2 | 353187 | protein_coding | 1239.494348 | -1.115898637 | 0.125521512 | -8.890098767 | 6.11E-19 | 4.87E-17 | 1.115898637 |
| ENSMUSG00000111296 | 2010001M07Rik | 115487174 | lncRNA | 21.56338625 | -1.116301443 | 0.302370851 | -3.691828887 | 0.000222647 | 0.002877109 | 1.116301443 |
| ENSMUSG00000030109 | Slc6a12 | 14411 | protein_coding | 17.21173055 | -1.116643893 | 0.300690525 | -3.71359854 | 0.000204333 | 0.002687827 | 1.116643893 |
| ENSMUSG00000035186 | Ubd | 24108 | protein_coding | 8.686482408 | -1.117286552 | 0.699580558 | -1.597080621 | 0.110247737 | 0.304759215 | 1.117286552 |
| ENSMUSG00000053398 | Phgdh | 236539 | protein_coding | 281.6185455 | 1.119851407 | 0.143864471 | 7.784072053 | 7.02E-15 | 4.39E-13 | 1.119851407 |
| ENSMUSG00000037418 | Best1 | 24115 | protein_coding | 12.36296176 | -1.123919031 | 0.433478714 | -2.592789437 | 0.009520104 | 0.057394659 | 1.123919031 |
| ENSMUSG00000016262 | Sertad4 | 214791 | protein_coding | 7.60790174 | -1.126904044 | 0.452177697 | -2.492170783 | 0.012696497 | 0.070972759 | 1.126904044 |
| ENSMUSG00000039759 | Thap3 | 69876 | protein_coding | 420.1459099 | 1.127042101 | 0.087737289 | 12.84564534 | 9.10E-38 | 1.53E-35 | 1.127042101 |
| ENSMUSG00000015134 | Aldh1a3 | 56847 | protein_coding | 31.20916403 | 1.127798572 | 0.296475527 | 3.804019109 | 0.000142367 | 0.001982691 | 1.127798572 |
| ENSMUSG00000001493 | Meox1 | 17285 | protein_coding | 13.78804252 | 1.128143322 | 0.331355692 | 3.40462937 | 0.000662539 | 0.007209695 | 1.128143322 |
| ENSMUSG00000056035 | Cyp3a11 | 13112 | protein_coding | 9.880893551 | 1.128729172 | 0.671187407 | 1.681690032 | 0.092628962 | 0.274499682 | 1.128729172 |
| ENSMUSG00000026414 | Tnnt2 | 21956 | protein_coding | 46.09089524 | -1.129306584 | 0.334966445 | -3.371402123 | 0.000747866 | 0.007924334 | 1.129306584 |
| ENSMUSG00000024150 | Mcfd2 | 193813 | protein_coding | 2065.727851 | 1.132428002 | 0.063988997 | 17.69723006 | 4.40E-70 | 1.29E-67 | 1.132428002 |
| ENSMUSG00000019846 | Lama4 | 16775 | protein_coding | 12.9475513 | 1.133267934 | 0.360017622 | 3.147812398 | 0.001644972 | 0.015035317 | 1.133267934 |
| ENSMUSG00000078657 | Crnn | 381457 | protein_coding | 7.234893983 | 1.135019821 | 0.473557251 | 2.396795356 | 0.016539158 | 0.086179968 | 1.135019821 |
| ENSMUSG00000020581 | Agr2 | 23795 | protein_coding | 31377.13531 | 1.137422568 | 0.083194242 | 13.67189051 | 1.49E-42 | 2.80E-40 | 1.137422568 |
| ENSMUSG00000016494 | Cd34 | 12490 | protein_coding | 8.95241916 | 1.139000374 | 0.442063908 | 2.576551383 | 0.009979137 | 0.059145001 | 1.139000374 |
| ENSMUSG00000030775 | Trat1 | 77647 | protein_coding | 7.181267963 | 1.139422986 | 0.508192986 | 2.242106871 | 0.024954466 | 0.114590169 | 1.139422986 |

| ensembl_gene_id | external_gene_name | entrezgene_id | gene_biotype | baseMean | log2FoldChange | lfcSE | stat | pvalue | padj | absLog2FC |
|---|---|---|---|---|---|---|---|---|---|---|
| ENSMUSG00000079516 | Reg3a | 19694 | protein_coding | 24.22572268 | -1.14168893 | 0.462726869 | -2.467306323 | 0.013613384 | 0.074800013 | 1.14168893 |
| ENSMUSG00000024338 | Psmb8 | 16913 | protein_coding | 306.5507725 | -1.143660494 | 0.454785275 | -2.514726307 | 0.011912482 | 0.067692953 | 1.143660494 |
| ENSMUSG00000002007 | Srpk3 | 56504 | protein_coding | 10.26390249 | -1.143766368 | 0.441848261 | -2.588595383 | 0.009636825 | 0.057838885 | 1.143766368 |
| ENSMUSG00000052736 | Klrc2 | 16642 | protein_coding | 7.010612226 | 1.145870065 | 0.469992242 | 2.438061661 | 0.014766255 | 0.079378465 | 1.145870065 |
| ENSMUSG00000031617 | Tmem184c | 234463 | protein_coding | 227.0897105 | 1.146070644 | 0.139161431 | 8.235548005 | 1.79E-16 | 1.23E-14 | 1.146070644 |
| ENSMUSG00000020368 | Canx | 12330 | protein_coding | 20207.86079 | 1.146977091 | 0.045163805 | 25.39593549 | 2.80E-142 | 1.73E-139 | 1.146977091 |
| ENSMUSG00000021760 | Gpx8 | 69590 | protein_coding | 5.906001983 | 1.152213744 | 0.511675355 | 2.2518453 | 0.02433205 | 0.112759473 | 1.152213744 |
| ENSMUSG00000059743 | Fdps | 110196 | protein_coding | 804.0400161 | 1.152371936 | 0.134067284 | 8.595474636 | 8.29E-18 | 6.08E-16 | 1.152371936 |
| ENSMUSG00000027368 | Dusp2 | 13537 | protein_coding | 15.08419995 | 1.153137274 | 0.411617061 | 2.801480754 | 0.005086868 | 0.036123255 | 1.153137274 |
| ENSMUSG00000032491 | Nradd | 67169 | protein_coding | 35.29816635 | -1.157211276 | 0.241880653 | -4.78422422 | 1.72E-06 | 3.99E-05 | 1.157211276 |
| ENSMUSG00000004929 | Thop1 | 50492 | protein_coding | 1137.589045 | 1.158085548 | 0.104820144 | 11.04831097 | 2.23E-28 | 2.71E-26 | 1.158085548 |
| ENSMUSG00000085034 | Gm11240 | 667768 | lncRNA | 10.60386966 | 1.160975505 | 0.381430488 | 3.043740713 | 0.002336564 | 0.01977008 | 1.160975505 |
| ENSMUSG00000044628 | Rnf208 | 68846 | protein_coding | 11.30639054 | -1.162431206 | 0.412725517 | -2.816475257 | 0.004855378 | 0.034832155 | 1.162431206 |
| ENSMUSG00000026726 | Cubn | 65969 | protein_coding | 662.1689195 | -1.164119716 | 0.235167555 | -4.950171455 | 7.41E-07 | 1.87E-05 | 1.164119716 |
| ENSMUSG00000020038 | Cry1 | 12952 | protein_coding | 357.8188181 | 1.164168101 | 0.160214537 | 7.266307576 | 3.69E-13 | 1.98E-11 | 1.164168101 |
| ENSMUSG00000101009 | 1700108F19Rik | 73272 | lncRNA | 9.963073706 | 1.164582827 | 0.388617101 | 2.996735925 | 0.00272887 | 0.022339332 | 1.164582827 |
| ENSMUSG00000016179 | Camk1g | 215303 | protein_coding | 6.287575332 | -1.165674828 | 0.547055791 | -2.130815261 | 0.033104365 | 0.139356327 | 1.165674828 |
| ENSMUSG00000024304 | Cdh2 | 12558 | protein_coding | 11.89548649 | 1.167361649 | 0.418014784 | 2.792632444 | 0.005228106 | 0.036914915 | 1.167361649 |
| ENSMUSG00000026463 | Atp2b4 | 381290 | protein_coding | 15.18917821 | 1.1673807 | 0.345188472 | 3.381864674 | 0.000719956 | 0.007714771 | 1.1673807 |
| ENSMUSG00000002279 | Lmf1 | 76483 | protein_coding | 1058.069924 | 1.17083814 | 0.071129695 | 16.46060953 | 7.04E-61 | 1.86E-58 | 1.17083814 |
| ENSMUSG00000000805 | Car4 | 12351 | protein_coding | 3648.010082 | -1.172661042 | 0.177727451 | -6.598086211 | 4.16E-11 | 1.88E-09 | 1.172661042 |
| ENSMUSG00000040694 | Apobec2 | 11811 | protein_coding | 47.03194472 | 1.175551295 | 0.243605004 | 4.825645113 | 1.40E-06 | 3.31E-05 | 1.175551295 |
| ENSMUSG00000040612 | Ildr2 | 100039795 | protein_coding | 6.323397162 | 1.180814152 | 0.53172571 | 2.220720439 | 0.026369902 | 0.118795321 | 1.180814152 |
| ENSMUSG00000070424 | Art5 | 11875 | protein_coding | 11.18227271 | 1.182763126 | 0.484642895 | 2.440483784 | 0.014667604 | 0.078979785 | 1.182763126 |
| ENSMUSG00000032126 | Hmbs | 15288 | protein_coding | 1046.719502 | 1.184186294 | 0.067170896 | 17.629455 | 1.46E-69 | 4.14E-67 | 1.184186294 |
| ENSMUSG00000050241 | Klre1 | 243655 | protein_coding | 6.256056336 | 1.189282631 | 0.555813536 | 2.139715126 | 0.032377796 | 0.137265172 | 1.189282631 |
| ENSMUSG00000026211 | Obsl1 | 98733 | protein_coding | 10.80711052 | 1.1976188 | 0.432011025 | 2.772194994 | 0.005567967 | 0.038737967 | 1.1976188 |
| ENSMUSG00000029700 | Slc13a1 | 55961 | protein_coding | 738.9502054 | -1.19833102 | 0.290312562 | -4.127727068 | 3.66E-05 | 0.000619329 | 1.19833102 |
| ENSMUSG00000045053 | Kcng3 | 225030 | protein_coding | 6.015120471 | -1.198759677 | 0.534144383 | -2.244261505 | 0.024815581 | 0.114189231 | 1.198759677 |
| ENSMUSG00000042349 | Ikbke | 56489 | protein_coding | 1421.272756 | 1.198889768 | 0.066593808 | 18.00302168 | 1.84E-72 | 5.61E-70 | 1.198889768 |
| ENSMUSG00000030844 | Rgs10 | 67865 | protein_coding | 18.27930203 | 1.200601963 | 0.286753042 | 4.186884833 | 2.83E-05 | 0.000491472 | 1.200601963 |
| ENSMUSG00000040808 | S100g | 12309 | protein_coding | 906.8947836 | -1.201748332 | 0.283401448 | -4.24044528 | 2.23E-05 | 0.000399285 | 1.201748332 |
| ENSMUSG00000086564 | Cd101 | 630146 | protein_coding | 12.92226477 | 1.20342044 | 0.374544034 | 3.213027922 | 0.001313435 | 0.012600696 | 1.20342044 |
| ENSMUSG00000024781 | Lipa | 16889 | protein_coding | 1005.398315 | 1.204335576 | 0.059302589 | 20.30831363 | 1.09E-91 | 4.49E-89 | 1.204335576 |
| ENSMUSG00000025130 | P4hb | 18453 | protein_coding | 26358.21941 | 1.210306569 | 0.048352114 | 25.03109945 | 2.80E-138 | 1.68E-135 | 1.210306569 |
| ENSMUSG00000087658 | Hotairm1 | 105242658 | lncRNA | 21.81886597 | 1.217770458 | 0.267986591 | 4.544146973 | 5.52E-06 | 0.000115825 | 1.217770458 |
| ENSMUSG00000078650 | G6pc | 14377 | protein_coding | 10.05770029 | 1.218339861 | 0.463741042 | 2.627198697 | 0.008609105 | 0.053399627 | 1.218339861 |
| ENSMUSG00000046550 | Spin2c | 278240 | protein_coding | 19.72623621 | 1.220965099 | 0.324600428 | 3.761440202 | 0.000168938 | 0.002281792 | 1.220965099 |
| ENSMUSG00000062833 | Sval3 | 387564 | protein_coding | 9.442836979 | 1.221362017 | 0.459136113 | 2.660130583 | 0.007811036 | 0.049772157 | 1.221362017 |

| ensembl_gene_id | external_gene_name | entrezgene_id | gene_biotype | baseMean | log2FoldChange | lfcSE | stat | pvalue | padj | absLog2FC |
|---|---|---|---|---|---|---|---|---|---|---|
| ENSMUSG00000027313 | Chac1 | 69065 | protein_coding | 9.735083749 | 1.227354272 | 0.391043056 | 3.138667864 | 0.001697177 | 0.015385257 | 1.227354272 |
| ENSMUSG00000022558 | Mroh1 | 223658 | protein_coding | 2393.764368 | 1.229443226 | 0.047948251 | 25.64104446 | 5.32E-145 | 3.58E-142 | 1.229443226 |
| ENSMUSG00000058153 | Sez6l | 56747 | protein_coding | 6.509133361 | 1.229570313 | 0.500834717 | 2.455042095 | 0.014086809 | 0.076614782 | 1.229570313 |
| ENSMUSG00000046768 | Rhoj | 80837 | protein_coding | 6.597929113 | 1.229964717 | 0.541261375 | 2.272404378 | 0.023062098 | 0.108400597 | 1.229964717 |
| ENSMUSG00000079227 | Ccr5 | 12774 | protein_coding | 6.825095614 | 1.230396716 | 0.560981343 | 2.19329347 | 0.028286242 | 0.124739465 | 1.230396716 |
| ENSMUSG00000063851 | Rnf183 | 76072 | protein_coding | 8.805556219 | 1.230470807 | 0.443104268 | 2.776932871 | 0.005487451 | 0.038300296 | 1.230470807 |
| ENSMUSG00000074569 | Gcnt7 | 654821 | protein_coding | 19.99327284 | -1.231655255 | 0.371840389 | -3.312322414 | 0.000925248 | 0.009438003 | 1.231655255 |
| ENSMUSG00000006342 | Susd2 | 71733 | protein_coding | 9.134557854 | 1.232224883 | 0.470864333 | 2.616942494 | 0.008872129 | 0.054611 | 1.232224883 |
| ENSMUSG00000080626 | Mir1192 | 100316672 | miRNA | 10.18968311 | -1.233230265 | 0.506296858 | -2.435784947 | 0.014859516 | 0.079719751 | 1.233230265 |
| ENSMUSG00000067276 | Capn6 | 12338 | protein_coding | 16.06682493 | 1.235177901 | 0.305019868 | 4.049499821 | 5.13E-05 | 0.000838657 | 1.235177901 |
| ENSMUSG00000030708 | Dnajb13 | 69387 | protein_coding | 122.1963777 | -1.236355788 | 0.2995698 | -4.127104227 | 3.67E-05 | 0.000620357 | 1.236355788 |
| ENSMUSG00000006517 | Mvd | 192156 | protein_coding | 1998.245621 | 1.237639918 | 0.128044045 | 9.665735901 | 4.22E-22 | 4.02E-20 | 1.237639918 |
| ENSMUSG00000030022 | Adamts9 | 101401 | protein_coding | 28.54446602 | -1.245552646 | 0.342599333 | -3.635595649 | 0.000277339 | 0.003459124 | 1.245552646 |
| ENSMUSG00000029467 | Atp2a2 | 11938 | protein_coding | 10013.64543 | 1.249491485 | 0.040079821 | 31.1750768 | 2.32E-213 | 3.40E-210 | 1.249491485 |
| ENSMUSG00000033730 | Egr3 | 13655 | protein_coding | 17.33207683 | -1.251913483 | 0.404770738 | -3.092895227 | 0.001982141 | 0.01741389 | 1.251913483 |
| ENSMUSG00000030762 | Aqp8 | 11833 | protein_coding | 1812.669483 | -1.252509679 | 0.299553949 | -4.181249103 | 2.90E-05 | 0.000502732 | 1.252509679 |
| ENSMUSG00000044034 | Npb | 208990 | protein_coding | 7.815770308 | -1.257527013 | 0.480356357 | -2.617904385 | 0.008847159 | 0.054478097 | 1.257527013 |
| ENSMUSG00000047554 | Tmem41b | 233724 | protein_coding | 1369.683422 | 1.258997098 | 0.055720335 | 22.5949305 | 4.86E-113 | 2.45E-110 | 1.258997098 |
| ENSMUSG00000042567 | Nek10 | 674895 | protein_coding | 7.447567853 | 1.259990946 | 0.502429914 | 2.50779444 | 0.012148732 | 0.068576338 | 1.259990946 |
| ENSMUSG00000096852 | Cyp2d12 | 380997 | protein_coding | 181.5442138 | -1.26910643 | 0.224422034 | -5.654999223 | 1.56E-08 | 5.43E-07 | 1.26910643 |
| ENSMUSG00000055413 | H2-Q5 | 15016 | protein_coding | 8.894733101 | -1.272689196 | 0.56811973 | -2.24017778 | 0.025079384 | 0.114934703 | 1.272689196 |
| ENSMUSG00000013419 | Zfp651 | 270210 | protein_coding | 6.710064225 | 1.278335492 | 0.452471173 | 2.825230794 | 0.004724657 | 0.034183286 | 1.278335492 |
| ENSMUSG00000061039 | Or8b53 | 258783 | protein_coding | 6.555005391 | -1.282319756 | 0.550623505 | -2.328850374 | 0.019866994 | 0.097711195 | 1.282319756 |
| ENSMUSG00000033847 | Pla2g4c | 232889 | protein_coding | 64.83051064 | 1.282856295 | 0.545859579 | 2.35015807 | 0.01876544 | 0.09418931 | 1.282856295 |
| ENSMUSG00000086962 | Gm12248 | 676527 | protein_coding | 7.970800464 | -1.285363896 | 0.51052703 | -2.517719574 | 0.011811732 | 0.0672389 | 1.285363896 |
| ENSMUSG00000027933 | Ints3 | 229543 | protein_coding | 2455.536259 | 1.293673578 | 0.081555701 | 15.86245423 | 1.15E-56 | 2.95E-54 | 1.293673578 |
| ENSMUSG00000044362 | Ccdc89 | 70054 | protein_coding | 21.26597175 | 1.296561065 | 0.274822468 | 4.717813185 | 2.38E-06 | 5.38E-05 | 1.296561065 |
| ENSMUSG00000090254 | Gm1965 | 434065 | lncRNA | 40.04749715 | -1.296888057 | 0.46964797 | -2.761404583 | 0.005755333 | 0.039815194 | 1.296888057 |
| ENSMUSG00000091648 | C2cd4d | 271944 | protein_coding | 6.638738868 | 1.302580274 | 0.49801276 | 2.615556021 | 0.008908231 | 0.054791394 | 1.302580274 |
| ENSMUSG00000022816 | Fstl1 | 14314 | protein_coding | 6.323182133 | 1.305407722 | 0.513707471 | 2.541149964 | 0.011048852 | 0.063660176 | 1.305407722 |
| ENSMUSG00000074743 | Thbd | 21824 | protein_coding | 13.19956907 | -1.307482653 | 0.459550089 | -2.845136329 | 0.004439242 | 0.032609438 | 1.307482653 |
| ENSMUSG00000030104 | Edem1 | 192193 | protein_coding | 5531.155739 | 1.307488601 | 0.050674037 | 25.80194288 | 8.43E-147 | 5.91E-144 | 1.307488601 |
| ENSMUSG00000051817 | Sox12 | 20667 | protein_coding | 28.11703572 | 1.309800718 | 0.259974112 | 5.038196716 | 4.70E-07 | 1.23E-05 | 1.309800718 |
| ENSMUSG00000020363 | Gfpt2 | 14584 | protein_coding | 6.052668234 | 1.320492587 | 0.670708639 | 1.968802113 | 0.04897582 | 0.182662591 | 1.320492587 |
| ENSMUSG00000022229 | Atp12a | 192113 | protein_coding | 5150.40604 | -1.32713853 | 0.307680196 | -4.313370013 | 1.61E-05 | 0.00029873 | 1.32713853 |
| ENSMUSG00000024856 | Cdk2ap2 | 52004 | protein_coding | 2365.742441 | 1.329351059 | 0.087068549 | 15.26786744 | 1.25E-52 | 3.01E-50 | 1.329351059 |
| ENSMUSG00000031098 | Syt8 | 55925 | protein_coding | 22.34152606 | 1.331421501 | 0.321449998 | 4.141924129 | 3.44E-05 | 0.000587747 | 1.331421501 |
| ENSMUSG00000042842 | Serpinb6b | 20708 | protein_coding | 8.37732313 | 1.331733503 | 0.432289387 | 3.080652781 | 0.002065474 | 0.018005347 | 1.331733503 |
| ENSMUSG00000035431 | Sstr1 | 20605 | protein_coding | 498.1161821 | -1.332002467 | 0.106176104 | -12.54521889 | 4.22E-36 | 6.68E-34 | 1.332002467 |

| ensembl_gene_id | external_gene_name | entrezgene_id | gene_biotype | baseMean | log2FoldChange | lfcSE | stat | pvalue | padj | absLog2FC |
|---|---|---|---|---|---|---|---|---|---|---|
| ENSMUSG00000030759 | Far1 | 67420 | protein_coding | 7814.276199 | 1.334709204 | 0.064369941 | 20.73497621 | 1.68E-95 | 7.11E-93 | 1.334709204 |
| ENSMUSG00000068114 | Ccdc134 | 76457 | protein_coding | 348.9417474 | 1.343611603 | 0.107098042 | 12.54562246 | 4.20E-36 | 6.68E-34 | 1.343611603 |
| ENSMUSG00000030826 | Bcat2 | 12036 | protein_coding | 1052.322799 | 1.344101164 | 0.059181225 | 22.7116145 | 3.44E-114 | 1.79E-111 | 1.344101164 |
| ENSMUSG00000049109 | Themis | 210757 | protein_coding | 6.139543748 | 1.356400458 | 0.583398976 | 2.324996294 | 0.020072158 | 0.098390182 | 1.356400458 |
| ENSMUSG00000031762 | Mt2 | 17750 | protein_coding | 1435.791617 | -1.366416311 | 0.11401196 | -11.98485061 | 4.27E-33 | 5.88E-31 | 1.366416311 |
| ENSMUSG00000005397 | Nid1 | 18073 | protein_coding | 8.561981677 | 1.369228283 | 0.447483956 | 3.059837712 | 0.00221457 | 0.019007112 | 1.369228283 |
| ENSMUSG00000003355 | Fkbp11 | 66120 | protein_coding | 177.2937825 | 1.386921573 | 0.144604142 | 9.591160762 | 8.71E-22 | 7.94E-20 | 1.386921573 |
| ENSMUSG00000032323 | Cyp11a1 | 13070 | protein_coding | 12.89732047 | -1.39301254 | 0.528818276 | -2.634198937 | 0.008433606 | 0.05258644 | 1.39301254 |
| ENSMUSG00000038264 | Sema7a | 20361 | protein_coding | 56.582478 | -1.395347074 | 0.189346198 | -7.369290145 | 1.72E-13 | 9.47E-12 | 1.395347074 |
| ENSMUSG00000073988 | Ttpa | 50500 | protein_coding | 41.85159069 | 1.39550315 | 0.262074447 | 5.324834851 | 1.01E-07 | 3.09E-06 | 1.39550315 |
| ENSMUSG00000030366 | Ceacam12 | 67315 | protein_coding | 25.59760887 | -1.40179365 | 0.351847075 | -3.9840992 | 6.77E-05 | 0.001066785 | 1.40179365 |
| ENSMUSG00000078921 | Tgtp2 | 100039796 | protein_coding | 13.41895768 | -1.402059311 | 0.814282569 | -1.721833874 | 0.085099616 | 0.260220234 | 1.402059311 |
| ENSMUSG00000019929 | Dcn | 13179 | protein_coding | 14.29610265 | 1.405947163 | 0.539939672 | 2.603896759 | 0.009217052 | 0.056167696 | 1.405947163 |
| ENSMUSG00000089943 | Ugt1a5 | 394433 | protein_coding | 8.559026373 | -1.407592165 | 0.454426628 | -3.09751251 | 0.001951522 | 0.017216733 | 1.407592165 |
| ENSMUSG00000074489 | Bglap3 | 12095 | protein_coding | 34.43191774 | -1.416174944 | 0.257363127 | -5.502633419 | 3.74E-08 | 1.24E-06 | 1.416174944 |
| ENSMUSG00000025425 | St8sia5 | 225742 | protein_coding | 47.4740242 | -1.416627045 | 0.372259652 | -3.805481031 | 0.000141529 | 0.001976135 | 1.416627045 |
| ENSMUSG00000017897 | Eya2 | 14049 | protein_coding | 14.68135879 | 1.417391061 | 0.333375373 | 4.251636975 | 2.12E-05 | 0.000381535 | 1.417391061 |
| ENSMUSG00000038143 | Stox2 | 71069 | protein_coding | 305.0061851 | 1.425593182 | 0.102180768 | 13.95167804 | 3.07E-44 | 6.35E-42 | 1.425593182 |
| ENSMUSG00000030373 | Psg28 | 114871 | protein_coding | 6.068686262 | -1.426647173 | 0.80861049 | -1.764319398 | 0.07767822 | 0.245920034 | 1.426647173 |
| ENSMUSG00000034855 | Cxcl10 | 15945 | protein_coding | 17.97567434 | -1.43786284 | 0.447941807 | -3.20993222 | 0.001327663 | 0.012691889 | 1.43786284 |
| ENSMUSG00000038738 | Shank1 | 243961 | protein_coding | 15.41991031 | 1.448236853 | 0.417723865 | 3.466971787 | 0.000526357 | 0.005915374 | 1.448236853 |
| ENSMUSG00000038011 | Dnah10 | 56087 | protein_coding | 8.718076936 | 1.450480348 | 0.511786254 | 2.834152609 | 0.004594738 | 0.0334988 | 1.450480348 |
| ENSMUSG00000020873 | Slc35b1 | 110172 | protein_coding | 2040.641185 | 1.453083513 | 0.081615887 | 17.80392977 | 6.59E-71 | 1.97E-68 | 1.453083513 |
| ENSMUSG00000071176 | Arhgef10 | 234094 | protein_coding | 100.302571 | 1.462798212 | 0.181463735 | 8.061104943 | 7.56E-16 | 5.04E-14 | 1.462798212 |
| ENSMUSG00000034532 | Fbxo16 | 50759 | protein_coding | 6.422994924 | 1.465452376 | 0.486185179 | 3.014185624 | 0.002576701 | 0.021320912 | 1.465452376 |
| ENSMUSG00000024807 | Syvn1 | 74126 | protein_coding | 1002.133356 | 1.480926223 | 0.068709363 | 21.55348506 | 4.91E-103 | 2.26E-100 | 1.480926223 |
| ENSMUSG00000097462 | 9530026P05Rik | 330385 | lncRNA | 10.20223611 | -1.485320763 | 0.484123931 | -3.068058956 | 0.002154541 | 0.018600794 | 1.485320763 |
| ENSMUSG00000052271 | Bhlha15 | 17341 | protein_coding | 59.5811162 | 1.485474475 | 0.22054759 | 6.735392003 | 1.63E-11 | 7.66E-10 | 1.485474475 |
| ENSMUSG00000035896 | Rnase1 | 19752 | protein_coding | 7.365279801 | -1.487136629 | 0.507964026 | -2.927641632 | 0.003415435 | 0.026790231 | 1.487136629 |
| ENSMUSG00000020241 | Col6a2 | 12834 | protein_coding | 7.303505517 | 1.491154053 | 0.51537101 | 2.893360365 | 0.003811437 | 0.029145115 | 1.491154053 |
| ENSMUSG00000043439 | Epop | 103551 | protein_coding | 30.3693649 | -1.494104251 | 0.248977319 | -6.000965286 | 1.96E-09 | 7.53E-08 | 1.494104251 |
| ENSMUSG00000023809 | Rps6ka2 | 20112 | protein_coding | 51.48567857 | 1.494799459 | 0.207771721 | 7.194431722 | 6.27E-13 | 3.29E-11 | 1.494799459 |
| ENSMUSG00000006095 | Tbcb | 66411 | protein_coding | 1718.913457 | 1.498413291 | 0.075465124 | 19.85570578 | 9.84E-88 | 3.78E-85 | 1.498413291 |
| ENSMUSG00000038576 | Susd4 | 96935 | protein_coding | 38.42318851 | 1.501180432 | 0.242121641 | 6.200108439 | 5.64E-10 | 2.29E-08 | 1.501180432 |
| ENSMUSG00000074447 | Defa21 | 66298 | protein_coding | 13.15579058 | 1.505312643 | 1.179173003 | 1.276583368 | 0.201749377 | 0.435615504 | 1.505312643 |
| ENSMUSG00000031382 | Asb11 | 68854 | protein_coding | 19.86883749 | -1.506158091 | 0.305619518 | -4.928213029 | 8.30E-07 | 2.07E-05 | 1.506158091 |
| ENSMUSG00000006519 | Cyba | 13057 | protein_coding | 693.1134664 | -1.506440932 | 0.069321948 | -21.73108195 | 1.04E-104 | 4.95E-102 | 1.506440932 |
| ENSMUSG00000001119 | Col6a1 | 12833 | protein_coding | 6.53111587 | 1.508388513 | 0.52940246 | 2.849228378 | 0.004382541 | 0.032358233 | 1.508388513 |
| ENSMUSG00000110404 | Rnf223 | 100045778 | protein_coding | 7.344415219 | 1.516161285 | 0.508776906 | 2.980012001 | 0.002882371 | 0.02334706 | 1.516161285 |

| ensembl_gene_id | external_gene_name | entrezgene_id | gene_biotype | baseMean | log2FoldChange | lfcSE | stat | pvalue | padj | absLog2FC |
|---|---|---|---|---|---|---|---|---|---|---|
| ENSMUSG00000021719 | Rgs7bp | 52882 | protein_coding | 9.771173434 | 1.519193191 | 0.538854317 | 2.819302255 | 0.004812818 | 0.034603793 | 1.519193191 |
| ENSMUSG00000027931 | Npr1 | 18160 | protein_coding | 8.536847011 | 1.520057695 | 0.434997162 | 3.494408305 | 0.000475114 | 0.005414954 | 1.520057695 |
| ENSMUSG00000020990 | Cdkl1 | 71091 | protein_coding | 116.1885288 | -1.524893095 | 0.234970731 | -6.489715062 | 8.60E-11 | 3.83E-09 | 1.524893095 |
| ENSMUSG00000084174 | Sycn | 68416 | protein_coding | 3923.238788 | -1.537342679 | 0.194726863 | -7.894866962 | 2.91E-15 | 1.87E-13 | 1.537342679 |
| ENSMUSG00000032374 | Plod2 | 26432 | protein_coding | 56.18802782 | 1.538320601 | 0.22583067 | 6.811832073 | 9.64E-12 | 4.64E-10 | 1.538320601 |
| ENSMUSG00000026043 | Col3a1 | 12825 | protein_coding | 54.53488507 | 1.544740608 | 0.40771431 | 3.788781922 | 0.000151388 | 0.002079583 | 1.544740608 |
| ENSMUSG00000055322 | Tns1 | 21961 | protein_coding | 7.623433378 | 1.549104524 | 0.601912671 | 2.573636672 | 0.010063589 | 0.059536134 | 1.549104524 |
| ENSMUSG00000066279 | Chrna10 | 504186 | protein_coding | 9.215911978 | 1.549624019 | 0.525977469 | 2.946179466 | 0.003217257 | 0.025533809 | 1.549624019 |
| ENSMUSG00000000791 | Il12rb1 | 16161 | protein_coding | 8.468789045 | 1.555579487 | 0.43783497 | 3.552890002 | 0.000381024 | 0.004508269 | 1.555579487 |
| ENSMUSG00000037470 | Uggt1 | 320011 | protein_coding | 5447.286712 | 1.558238267 | 0.074687073 | 20.86356052 | 1.15E-96 | 5.00E-94 | 1.558238267 |
| ENSMUSG00000031551 | Ido1 | 15930 | protein_coding | 24.27929373 | -1.575695474 | 0.607984639 | -2.591669877 | 0.009551137 | 0.057517249 | 1.575695474 |
| ENSMUSG00000042686 | Jph1 | 57339 | protein_coding | 15.36826672 | 1.579852265 | 0.375454635 | 4.207837955 | 2.58E-05 | 0.00045343 | 1.579852265 |
| ENSMUSG00000028354 | Fmn2 | 54418 | protein_coding | 14.89426149 | 1.593788658 | 0.343535569 | 4.639370132 | 3.49E-06 | 7.58E-05 | 1.593788658 |
| ENSMUSG00000039438 | Ttc36 | 192653 | protein_coding | 28.74017691 | -1.595910856 | 0.290677637 | -5.490311783 | 4.01E-08 | 1.32E-06 | 1.595910856 |
| ENSMUSG00000028402 | Mpdz | 17475 | protein_coding | 30.20815246 | 1.609526909 | 0.316677568 | 5.082541591 | 3.72E-07 | 9.96E-06 | 1.609526909 |
| ENSMUSG00000079018 | Ly6c1 | 17067 | protein_coding | 22.87619181 | -1.613062711 | 0.698364474 | -2.309772005 | 0.02090078 | 0.101348119 | 1.613062711 |
| ENSMUSG00000050737 | Ptges | 64292 | protein_coding | 29.81559657 | 1.634781032 | 0.301296342 | 5.42582437 | 5.77E-08 | 1.83E-06 | 1.634781032 |
| ENSMUSG00000022964 | Tmem50b | 77975 | protein_coding | 1867.772265 | 1.635173594 | 0.095756135 | 17.07643685 | 2.22E-65 | 6.08E-63 | 1.635173594 |
| ENSMUSG00000062181 | Ces3b | 13909 | protein_coding | 6.261793052 | 1.643187252 | 0.57985673 | 2.833781462 | 0.004600078 | 0.033522574 | 1.643187252 |
| ENSMUSG00000028713 | Cyp4b1 | 13120 | protein_coding | 1009.307873 | -1.654325263 | 0.291189939 | -5.681258323 | 1.34E-08 | 4.72E-07 | 1.654325263 |
| ENSMUSG00000019139 | Isyna1 | 71780 | protein_coding | 2277.271324 | 1.683320107 | 0.10845641 | 15.52070648 | 2.51E-54 | 6.23E-52 | 1.683320107 |
| ENSMUSG00000027790 | Sis | 69983 | protein_coding | 1014.198188 | -1.702821781 | 0.333941889 | -5.099155982 | 3.41E-07 | 9.20E-06 | 1.702821781 |
| ENSMUSG00000022226 | Mcpt2 | 17225 | protein_coding | 25.54690871 | 1.741569811 | 0.522493071 | 3.333192168 | 0.000858556 | 0.008864236 | 1.741569811 |
| ENSMUSG00000019876 | Pkib | 18768 | protein_coding | 38.49085669 | 1.764184124 | 0.231369013 | 7.624980106 | 2.44E-14 | 1.44E-12 | 1.764184124 |
| ENSMUSG00000022227 | Mcpt1 | 17224 | protein_coding | 62.8478353 | 1.777408925 | 0.652172854 | 2.725364779 | 0.006423048 | 0.043034689 | 1.777408925 |
| ENSMUSG00000079547 | H2-DMb1 | 14999 | protein_coding | 34.64760903 | -1.78284053 | 0.718623595 | -2.480910091 | 0.013104742 | 0.07270044 | 1.78284053 |
| ENSMUSG00000030284 | Creld1 | 171508 | protein_coding | 157.3406272 | 1.79899301 | 0.117121215 | 15.36009517 | 3.03E-53 | 7.41E-51 | 1.79899301 |
| ENSMUSG00000033327 | Tnxb | 81877 | protein_coding | 7.019768249 | 1.853324924 | 0.548635636 | 3.378061506 | 0.000729988 | 0.0077847 | 1.853324924 |
| ENSMUSG00000026249 | Serpine2 | 20720 | protein_coding | 39.47523742 | 1.861707494 | 0.238187653 | 7.816137705 | 5.45E-15 | 3.46E-13 | 1.861707494 |
| ENSMUSG00000015243 | Abca1 | 11303 | protein_coding | 541.3169369 | 1.900838571 | 0.104405415 | 18.20632169 | 4.60E-74 | 1.58E-71 | 1.900838571 |
| ENSMUSG00000001506 | Col1a1 | 12842 | protein_coding | 19.88482434 | 1.904023601 | 0.366157389 | 5.200014142 | 1.99E-07 | 5.69E-06 | 1.904023601 |
| ENSMUSG00000031155 | Pim2 | 18715 | protein_coding | 120.7413441 | 1.911631424 | 0.118095821 | 16.18712161 | 6.22E-59 | 1.62E-56 | 1.911631424 |
| ENSMUSG00000061353 | Cxcl12 | 20315 | protein_coding | 30.16424976 | -1.929794299 | 0.371686452 | -5.191995264 | 2.08E-07 | 5.92E-06 | 1.929794299 |
| ENSMUSG00000020405 | Fabp6 | 16204 | protein_coding | 597.2061998 | -1.93679198 | 0.389781808 | -4.968913219 | 6.73E-07 | 1.72E-05 | 1.93679198 |
| ENSMUSG00000032098 | Treh | 58866 | protein_coding | 111.058062 | 1.943586016 | 0.248668668 | 7.815966637 | 5.45E-15 | 3.46E-13 | 1.943586016 |
| ENSMUSG00000070436 | Serpinh1 | 12406 | protein_coding | 10.46380408 | 1.950538558 | 0.519124655 | 3.757360661 | 0.000171715 | 0.00230963 | 1.950538558 |
| ENSMUSG00000025353 | Ormdl2 | 66844 | protein_coding | 1908.059892 | 1.953378669 | 0.071010146 | 27.50844454 | 1.39E-166 | 1.12E-163 | 1.953378669 |
| ENSMUSG00000059824 | Dbp | 13170 | protein_coding | 354.5395666 | -1.961187426 | 0.201464477 | -9.734656209 | 2.15E-22 | 2.10E-20 | 1.961187426 |
| ENSMUSG00000026837 | Col5a1 | 12831 | protein_coding | 6.857000693 | 1.973670008 | 0.634532059 | 3.110433868 | 0.001868127 | 0.016626538 | 1.973670008 |

Supplementary Table 1

| ensembl_gene_id | external_gene_name | entrezgene_id | gene_biotype | baseMean | log2FoldChange | lfcSE | stat | pvalue | padj | absLog2FC |
|---|---|---|---|---|---|---|---|---|---|---|
| ENSMUSG00000020388 | Pdlim4 | 30794 | protein_coding | 21.396834 | -1.976495617 | 0.328182096 | -6.022557729 | 1.72E-09 | 6.66E-08 | 1.976495617 |
| ENSMUSG00000022136 | Dnajc3 | 100037258 | protein_coding | 7301.315443 | 1.981806734 | 0.054477638 | 36.37835279 | 9.36E-290 | 2.16E-286 | 1.981806734 |
| ENSMUSG00000025408 | Ddit3 | 13198 | protein_coding | 832.3552332 | 1.985240671 | 0.087136819 | 22.78302896 | 6.76E-115 | 3.63E-112 | 1.985240671 |
| ENSMUSG00000078349 | AW011738 | 100382 | lncRNA | 52.64982354 | 1.997813188 | 0.206797074 | 9.660742029 | 4.43E-22 | 4.17E-20 | 1.997813188 |
| ENSMUSG00000026833 | Olfm1 | 56177 | protein_coding | 25.68995464 | 1.999390511 | 0.271829931 | 7.355299338 | 1.90E-13 | 1.05E-11 | 1.999390511 |
| ENSMUSG00000056487 | Mettl7a2 | 393082 | protein_coding | 37.88212801 | -2.009542656 | 0.450761226 | -4.458108949 | 8.27E-06 | 0.000166269 | 2.009542656 |
| ENSMUSG00000031770 | Herpud1 | 64209 | protein_coding | 9038.054133 | 2.032888636 | 0.051594418 | 39.40132873 | 0 | 0 | 2.032888636 |
| ENSMUSG00000026663 | Atf6 | 226641 | protein_coding | 4834.247381 | 2.043895542 | 0.069996528 | 29.19995615 | 1.94E-187 | 2.09E-184 | 2.043895542 |
| ENSMUSG00000021214 | Akr1c18 | 105349 | protein_coding | 48.19981741 | -2.068027368 | 0.265961397 | -7.775667399 | 7.51E-15 | 4.67E-13 | 2.068027368 |
| ENSMUSG00000094430 | Smbd1 | 381043 | protein_coding | 37.40545007 | 2.069597345 | 0.242113391 | 8.548049892 | 1.25E-17 | 9.09E-16 | 2.069597345 |
| ENSMUSG00000007655 | Cav1 | 12389 | protein_coding | 6.319000565 | 2.079235365 | 0.609801228 | 3.409693637 | 0.000650359 | 0.00710592 | 2.079235365 |
| ENSMUSG00000027737 | Slc7a11 | 26570 | protein_coding | 100.5109837 | 2.101296989 | 0.17093816 | 12.29273199 | 9.91E-35 | 1.43E-32 | 2.101296989 |
| ENSMUSG00000004460 | Dnajb11 | 67838 | protein_coding | 7049.594719 | 2.111108898 | 0.082415049 | 25.61557524 | 1.02E-144 | 6.60E-142 | 2.111108898 |
| ENSMUSG00000029661 | Col1a2 | 12843 | protein_coding | 19.89290386 | 2.124014945 | 0.433815757 | 4.896122164 | 9.77E-07 | 2.40E-05 | 2.124014945 |
| ENSMUSG00000030559 | Rab38 | 72433 | protein_coding | 123.3437697 | 2.14558157 | 0.143099486 | 14.99363578 | 8.08E-51 | 1.89E-48 | 2.14558157 |
| ENSMUSG00000018593 | Sparc | 20692 | protein_coding | 12.65456989 | 2.149235893 | 0.465060237 | 4.62141401 | 3.81E-06 | 8.18E-05 | 2.149235893 |
| ENSMUSG00000060615 | Ang4 | 219033 | protein_coding | 1511.062456 | 2.211595027 | 0.522763193 | 4.23058673 | 2.33E-05 | 0.000414892 | 2.211595027 |
| ENSMUSG00000038550 | Ciart | 229599 | protein_coding | 75.37722362 | -2.250721792 | 0.345068298 | -6.522540039 | 6.91E-11 | 3.10E-09 | 2.250721792 |
| ENSMUSG00000007783 | Cpt1c | 78070 | protein_coding | 11.99372343 | 2.256879984 | 0.378021067 | 5.970249228 | 2.37E-09 | 9.05E-08 | 2.256879984 |
| ENSMUSG00000027248 | Pdia3 | 14827 | protein_coding | 22916.4404 | 2.273402674 | 0.066279342 | 34.30032035 | 7.76E-258 | 1.56E-254 | 2.273402674 |
| ENSMUSG00000030145 | Zfp248 | 72720 | protein_coding | 6.67475299 | 2.338268948 | 0.544029373 | 4.298056433 | 1.72E-05 | 0.000315764 | 2.338268948 |
| ENSMUSG00000057069 | Ero1b | 67475 | protein_coding | 445.4876904 | 2.354847291 | 0.086267998 | 27.29688126 | 4.62E-164 | 3.55E-161 | 2.354847291 |
| ENSMUSG00000020964 | Sel1l | 20338 | protein_coding | 6878.299058 | 2.373731915 | 0.058020648 | 40.91184765 | 0 | 0 | 2.373731915 |
| ENSMUSG00000074443 | Defa22 | 382059 | protein_coding | 14.10170136 | 2.442262227 | 0.769099525 | 3.175482685 | 0.001495874 | 0.01394449 | 2.442262227 |
| ENSMUSG00000099413 | Gm17767 | 100045066 | lncRNA | 17.6056137 | 2.450121289 | 0.351039658 | 6.979613934 | 2.96E-12 | 1.48E-10 | 2.450121289 |
| ENSMUSG00000032875 | Arhgef17 | 207212 | protein_coding | 111.6770254 | 2.461132345 | 0.145625453 | 16.90042709 | 4.47E-64 | 1.20E-61 | 2.461132345 |
| ENSMUSG00000053550 | Shisa7 | 232813 | protein_coding | 74.7391542 | 2.469090316 | 0.182394594 | 13.53708057 | 9.45E-42 | 1.73E-39 | 2.469090316 |
| ENSMUSG00000095105 | Edaradd | 171211 | protein_coding | 31.15723183 | 2.469645319 | 0.260247328 | 9.489608751 | 2.32E-21 | 2.04E-19 | 2.469645319 |
| ENSMUSG00000024184 | Pdia2 | 69191 | protein_coding | 8.320568763 | 2.497550031 | 0.482616832 | 5.175016426 | 2.28E-07 | 6.43E-06 | 2.497550031 |
| ENSMUSG00000026185 | Igfbp5 | 16011 | protein_coding | 6.462237391 | 2.534137136 | 0.653660562 | 3.876839575 | 0.000105822 | 0.001564246 | 2.534137136 |
| ENSMUSG00000003814 | Calr | 12317 | protein_coding | 38500.50291 | 2.59802443 | 0.107449798 | 24.17896049 | 3.70E-129 | 2.06E-126 | 2.59802443 |
| ENSMUSG00000057836 | Xlr3a | 22445 | protein_coding | 158.0192915 | 2.625442999 | 0.277595935 | 9.457786178 | 3.15E-21 | 2.73E-19 | 2.625442999 |
| ENSMUSG00000020571 | Pdia6 | 71853 | protein_coding | 12343.51873 | 2.649079889 | 0.090408432 | 29.30124801 | 1.00E-188 | 1.15E-185 | 2.649079889 |
| ENSMUSG00000018623 | Mmp7 | 17393 | protein_coding | 153.726548 | 2.661532419 | 0.198868105 | 13.38340512 | 7.56E-41 | 1.37E-38 | 2.661532419 |
| ENSMUSG00000018340 | Anxa6 | 11749 | protein_coding | 459.8262722 | 2.679531473 | 0.125981867 | 21.26918367 | 2.19E-100 | 9.81E-98 | 2.679531473 |
| ENSMUSG00000058013 | Septin11 | 52398 | protein_coding | 1937.360891 | 2.730335346 | 0.080761986 | 33.80718418 | 1.55E-250 | 2.49E-247 | 2.730335346 |
| ENSMUSG00000029844 | Hoxa1 | 15394 | protein_coding | 18.58195171 | 2.783520407 | 0.356507724 | 7.807742216 | 5.82E-15 | 3.67E-13 | 2.783520407 |
| ENSMUSG00000070407 | Hs3st3b1 | 54710 | protein_coding | 799.3707605 | 2.835787146 | 0.093271673 | 30.40351982 | 4.93E-203 | 6.63E-200 | 2.835787146 |
| ENSMUSG00000030659 | Nucb2 | 53322 | protein_coding | 456.5347818 | 3.187658154 | 0.115041286 | 27.70881896 | 5.47E-169 | 4.64E-166 | 3.187658154 |

Supplementary Table 1

| ensembl_gene_id | external_gene_name | entrezgene_id | gene_biotype | baseMean | log2FoldChange | lfcSE | stat | pvalue | padj | absLog2FC |
|---|---|---|---|---|---|---|---|---|---|---|
| ENSMUSG00000023272 | Creld2 | 76737 | protein_coding | 2656.538137 | 3.200577546 | 0.121004734 | 26.45001919 | 3.65E-154 | 2.67E-151 | 3.200577546 |
| ENSMUSG00000032115 | Hyou1 | 12282 | protein_coding | 13345.86755 | 3.211051015 | 0.114760896 | 27.98035841 | 2.82E-172 | 2.67E-169 | 3.211051015 |
| ENSMUSG00000060671 | Atp8b2 | 54667 | protein_coding | 338.1305663 | 3.307401039 | 0.09691073 | 34.12832639 | 2.80E-255 | 5.02E-252 | 3.307401039 |
| ENSMUSG00000032715 | Trib3 | 228775 | protein_coding | 30.11750252 | 3.339878378 | 0.29926184 | 11.16038845 | 6.37E-29 | 8.07E-27 | 3.339878378 |
| ENSMUSG00000020048 | Hsp90b1 | 22027 | protein_coding | 45079.81948 | 3.578200638 | 0.089927167 | 39.78998522 | 0 | 0 | 3.578200638 |
| ENSMUSG00000032575 | Manf | 74840 | protein_coding | 7961.356644 | 3.585189138 | 0.085650945 | 41.85813881 | 0 | 0 | 3.585189138 |
| ENSMUSG00000026864 | Hspa5 | 14828 | protein_coding | 71086.24427 | 3.735873883 | 0.082223324 | 45.43569493 | 0 | 0 | 3.735873883 |
| ENSMUSG00000025823 | Pdia4 | 12304 | protein_coding | 24458.72756 | 3.74956984 | 0.09553845 | 39.24670995 | 0 | 0 | 3.74956984 |
| ENSMUSG00000036067 | Slc2a6 | 227659 | protein_coding | 408.8146186 | 3.751742178 | 0.133803476 | 28.03919821 | 5.41E-173 | 5.45E-170 | 3.751742178 |
| ENSMUSG00000022769 | Sdf2l1 | 64136 | protein_coding | 4335.772513 | 3.820220329 | 0.200523814 | 19.05120516 | 6.42E-81 | 2.30E-78 | 3.820220329 |
| ENSMUSG00000021622 | Ckmt2 | 76722 | protein_coding | 808.0339639 | 3.853417169 | 0.155445601 | 24.78949002 | 1.16E-135 | 6.70E-133 | 3.853417169 |
| ENSMUSG00000033768 | Nrxn2 | 18190 | protein_coding | 31.35696658 | 4.087217857 | 0.362545755 | 11.27366078 | 1.77E-29 | 2.28E-27 | 4.087217857 |
| ENSMUSG00000001120 | Pcbp3 | 59093 | protein_coding | 12.84309949 | 4.238911134 | 0.566561825 | 7.481815661 | 7.33E-14 | 4.16E-12 | 4.238911134 |
| ENSMUSG00000028558 | Calr4 | 108802 | protein_coding | 8.388791774 | 4.537471887 | 0.865778601 | 5.240914804 | 1.60E-07 | 4.67E-06 | 4.537471887 |
| ENSMUSG00000029151 | Slc30a3 | 22784 | protein_coding | 45.49460144 | 5.565410262 | 0.450628341 | 12.35033342 | 4.85E-35 | 7.11E-33 | 5.565410262 |
| ENSMUSG00000072720 | Myo18b | 74376 | protein_coding | 49.09215487 | 6.239168327 | 0.504739832 | 12.36115703 | 4.24E-35 | 6.33E-33 | 6.239168327 |
| ENSMUSG00000019789 | Hey2 | 15214 | protein_coding | 26.02856265 | 6.485109446 | 0.755188249 | 8.587407782 | 8.90E-18 | 6.49E-16 | 6.485109446 |
| ENSMUSG00000017446 | C1qtnf1 | 56745 | protein_coding | 936.9024332 | 6.511563321 | 0.359987095 | 18.0883243 | 3.94E-73 | 1.25E-70 | 6.511563321 |
| ENSMUSG00000036036 | Zfp57 | 22715 | protein_coding | 13.78597939 | 7.440368588 | 0.887478446 | 8.383717508 | 5.13E-17 | 3.66E-15 | 7.440368588 |
| ENSMUSG00000009092 | Derl3 | 70377 | protein_coding | 4133.194737 | 7.442074117 | 0.697866778 | 10.66403267 | 1.50E-26 | 1.72E-24 | 7.442074117 |
| ENSMUSG00000098682 | Otx2os1 | 606497 | lncRNA | 28.10156827 | 8.460384981 | 0.857642454 | 9.864699379 | 5.92E-23 | 5.89E-21 | 8.460384981 |

| ensembl_gene_id | external_gene_name | entrezgene_id | gene_biotype | baseMean | log2FoldChange | lfcSE | stat | pvalue | padj | absLog2FC |
|---|---|---|---|---|---|---|---|---|---|---|
| ENSMUSG00000040151 | Hs2st1 | 23908 | protein_coding | 346.3454006 | -2.96E-05 | 0.0935685 | -0.000316761 | 0.999747262 | 0.999820998 | 2.96E-05 |
| ENSMUSG00000020516 | Rps6kb1 | 72508 | protein_coding | 692.2160693 | 4.51E-05 | 0.073819131 | 0.000610581 | 0.999512827 | 0.999815293 | 4.51E-05 |
| ENSMUSG00000034071 | Zfp551 | 619331 | protein_coding | 40.1961098 | 6.14E-05 | 0.229339111 | 0.000267639 | 0.999786455 | 0.999820998 | 6.14E-05 |
| ENSMUSG00000072761 | Gm6712 | 100862203 | lncRNA | 62.9272091 | 7.73E-05 | 0.165438938 | 0.000467512 | 0.99962698 | 0.999815293 | 7.73E-05 |
| ENSMUSG00000041840 | Haus1 | 225745 | protein_coding | 116.4161239 | 7.97E-05 | 0.140581215 | 0.000566834 | 0.999547732 | 0.999815293 | 7.97E-05 |
| ENSMUSG00000029802 | Abcg2 | 26357 | protein_coding | 1493.311754 | -8.22E-05 | 0.163427988 | -0.000503166 | 0.999598531 | 0.999815293 | 8.22E-05 |
| ENSMUSG00000055720 | Ubl7 | 69459 | protein_coding | 249.9219297 | 0.000100499 | 0.091183446 | 0.001102164 | 0.999120601 | 0.999559885 | 0.000100499 |
| ENSMUSG00000033111 | 3830406C13Rik | 218734 | protein_coding | 253.7719071 | 0.00011661 | 0.084473369 | 0.001380437 | 0.998898571 | 0.999559885 | 0.00011661 |
| ENSMUSG00000033253 | Szt2 | 230676 | protein_coding | 512.6756741 | 0.000166325 | 0.148844108 | 0.001117447 | 0.999108407 | 0.999559885 | 0.000166325 |
| ENSMUSG00000006392 | Med8 | 80509 | protein_coding | 488.0024346 | -0.000168189 | 0.093160033 | -0.001805381 | 0.998559515 | 0.999437977 | 0.000168189 |
| ENSMUSG00000058093 | Zfp729b | 100416706 | protein_coding | 260.1048869 | -0.000218276 | 0.106135571 | -0.00205658 | 0.998359088 | 0.999365516 | 0.000218276 |
| ENSMUSG00000068823 | Csde1 | 229663 | protein_coding | 6592.485948 | 0.000263986 | 0.045100398 | 0.005853303 | 0.995329766 | 0.998240742 | 0.000263986 |
| ENSMUSG00000087174 | 5530601H04Rik | 71445 | lncRNA | 101.5974485 | -0.000267528 | 0.135474376 | -0.001974753 | 0.998424376 | 0.999365516 | 0.000267528 |
| ENSMUSG00000046229 | Scand1 | 19018 | protein_coding | 157.8620293 | 0.000268285 | 0.225262356 | 0.001190989 | 0.999049729 | 0.999559885 | 0.000268285 |
| ENSMUSG00000034926 | Dhcr24 | 74754 | protein_coding | 1861.566801 | 0.000271554 | 0.117554063 | 0.002310035 | 0.99815686 | 0.999365516 | 0.000271554 |
| ENSMUSG00000040548 | Tex2 | 21763 | protein_coding | 764.2503306 | -0.000283295 | 0.054786083 | -0.005170935 | 0.995874209 | 0.998558825 | 0.000283295 |
| ENSMUSG00000029148 | Nrbp1 | 192292 | protein_coding | 1073.738314 | -0.000286947 | 0.077878581 | -0.003684549 | 0.997060162 | 0.9987142 | 0.000286947 |
| ENSMUSG00000046456 | Tmem150b | 330460 | protein_coding | 224.6678009 | -0.000293502 | 0.170212811 | -0.001724326 | 0.998624188 | 0.999439903 | 0.000293502 |
| ENSMUSG00000024180 | Pgap6 | 60455 | protein_coding | 1222.207399 | 0.00037415 | 0.079080561 | 0.004731246 | 0.996225026 | 0.998642884 | 0.00037415 |
| ENSMUSG00000034501 | Pcnx4 | 67708 | protein_coding | 86.62653796 | 0.00037887 | 0.171761299 | 0.002205796 | 0.998240031 | 0.999365516 | 0.00037887 |
| ENSMUSG00000055652 | Klhl25 | 207952 | protein_coding | 200.2628894 | -0.000392347 | 0.092929288 | -0.004221999 | 0.996631342 | 0.9987142 | 0.000392347 |
| ENSMUSG00000047534 | Mis18bp1 | 217653 | protein_coding | 262.4875437 | -0.000407235 | 0.118833223 | -0.003426942 | 0.997265702 | 0.998781261 | 0.000407235 |
| ENSMUSG00000033964 | Zbtb41 | 226470 | protein_coding | 705.0080311 | 0.000415242 | 0.101602816 | 0.004086918 | 0.99673912 | 0.9987142 | 0.000415242 |
| ENSMUSG00000030983 | Bccip | 66165 | protein_coding | 600.4896404 | 0.000522586 | 0.102340915 | 0.005106325 | 0.99592576 | 0.998558825 | 0.000522586 |
| ENSMUSG00000030045 | Mrpl19 | 56284 | protein_coding | 339.0595156 | -0.000524128 | 0.117750177 | -0.004451187 | 0.996448478 | 0.998642884 | 0.000524128 |
| ENSMUSG00000062762 | Ei24 | 13663 | protein_coding | 1383.473333 | -0.000536334 | 0.056661339 | -0.009465613 | 0.992447646 | 0.996654803 | 0.000536334 |
| ENSMUSG00000021747 | Cfap20dc | 74430 | protein_coding | 21.93128798 | 0.000604652 | 0.298536687 | 0.002025385 | 0.998383978 | 0.999365516 | 0.000604652 |
| ENSMUSG00000001707 | Eef1e1 | 66143 | protein_coding | 229.1364033 | 0.000623737 | 0.153016017 | 0.004076286 | 0.996747603 | 0.9987142 | 0.000623737 |
| ENSMUSG00000079557 | Marchf2 | 224703 | protein_coding | 566.6524741 | -0.000651476 | 0.083506529 | -0.007801502 | 0.993775365 | 0.997470163 | 0.000651476 |
| ENSMUSG00000017299 | Dnttip1 | 76233 | protein_coding | 220.2280229 | -0.000658206 | 0.109498806 | -0.006011079 | 0.995203881 | 0.998240742 | 0.000658206 |
| ENSMUSG00000035206 | Sppl2b | 73218 | protein_coding | 409.9265934 | -0.000666861 | 0.084110482 | -0.007928389 | 0.993674127 | 0.997431403 | 0.000666861 |
| ENSMUSG00000037007 | Zfp113 | 56314 | protein_coding | 228.8469577 | -0.000683426 | 0.168883454 | -0.004046731 | 0.996771185 | 0.9987142 | 0.000683426 |
| ENSMUSG00000020491 | 2810021J22Rik | 69944 | protein_coding | 115.7935246 | -0.000697942 | 0.153254676 | -0.004554134 | 0.996366339 | 0.998642884 | 0.000697942 |
| ENSMUSG00000033565 | Rbfox2 | 93686 | protein_coding | 387.1339019 | -0.000720753 | 0.097615084 | -0.007383626 | 0.994108773 | 0.997679071 | 0.000720753 |
| ENSMUSG00000046034 | Otulin | 432940 | protein_coding | 248.6109641 | 0.000857273 | 0.091550546 | 0.009363935 | 0.99252877 | 0.996654803 | 0.000857273 |
| ENSMUSG00000044477 | Zfand3 | 21769 | protein_coding | 722.5690517 | 0.00086867 | 0.096061059 | 0.009042891 | 0.992784915 | 0.996790099 | 0.00086867 |
| ENSMUSG00000046985 | Tapt1 | 231225 | protein_coding | 790.1481193 | -0.000883105 | 0.078832534 | -0.011202286 | 0.991062056 | 0.995813552 | 0.000883105 |
| ENSMUSG00000028797 | Tmem234 | 76799 | protein_coding | 1114.660247 | -0.000907351 | 0.089691172 | -0.01011639 | 0.991928426 | 0.996558343 | 0.000907351 |

| ensembl_gene_id | external_gene_name | entrezgene_id | gene_biotype | baseMean | log2FoldChange | lfcSE | stat | pvalue | padj | absLog2FC |
|---|---|---|---|---|---|---|---|---|---|---|
| ENSMUSG00000053886 | Sh2d4a | 72281 | protein_coding | 1692.232429 | -0.000909606 | 0.096960282 | -0.009381224 | 0.992514976 | 0.996654803 | 0.000909606 |
| ENSMUSG00000004552 | Ctse | 13034 | protein_coding | 707.7326894 | -0.001005598 | 0.272661517 | -0.003688084 | 0.997057342 | 0.9987142 | 0.001005598 |
| ENSMUSG00000041133 | Smc1a | 24061 | protein_coding | 1431.077878 | -0.001016247 | 0.070520972 | -0.014410569 | 0.988502427 | 0.994441098 | 0.001016247 |
| ENSMUSG00000020612 | Prkar1a | 19084 | protein_coding | 1173.340095 | 0.001062942 | 0.110867685 | 0.009587487 | 0.99235041 | 0.996654803 | 0.001062942 |
| ENSMUSG00000067297 | Ifit1bl2 | 112419 | protein_coding | 152.9144972 | 0.001065561 | 0.319504033 | 0.003335047 | 0.997339023 | 0.998781261 | 0.001065561 |
| ENSMUSG00000063522 | Ly6m | 67038 | protein_coding | 592.9325374 | -0.00113344 | 0.508360709 | -0.002229598 | 0.99822104 | 0.999365516 | 0.00113344 |
| ENSMUSG00000030374 | Strn4 | 97387 | protein_coding | 1049.555625 | 0.001133715 | 0.08607355 | 0.013171463 | 0.989490997 | 0.99479976 | 0.001133715 |
| ENSMUSG00000039478 | Micu3 | 78506 | protein_coding | 34.97827321 | -0.001138727 | 0.246869751 | -0.004612663 | 0.99631964 | 0.998642884 | 0.001138727 |
| ENSMUSG00000051390 | Zbtb22 | 81630 | protein_coding | 440.4768191 | 0.001185246 | 0.078767664 | 0.015047361 | 0.987994396 | 0.994255508 | 0.001185246 |
| ENSMUSG00000031534 | Smim19 | 102032 | protein_coding | 139.8030316 | 0.001195958 | 0.160639019 | 0.007445005 | 0.994059801 | 0.997679071 | 0.001195958 |
| ENSMUSG00000013662 | Atad1 | 67979 | protein_coding | 1861.573756 | 0.001227262 | 0.064744523 | 0.018955452 | 0.984876644 | 0.993212987 | 0.001227262 |
| ENSMUSG00000110424 | Zbed5-ps | 75479 | lncRNA | 13.96972332 | 0.001240776 | 0.33949864 | 0.003654731 | 0.997083953 | 0.9987142 | 0.001240776 |
| ENSMUSG00000096001 | 2610528A11Rik | 70045 | protein_coding | 4304.803913 | 0.001271204 | 0.092973886 | 0.013672704 | 0.989091101 | 0.994743066 | 0.001271204 |
| ENSMUSG00000032369 | Plscr1 | 22038 | protein_coding | 2127.805055 | -0.001273466 | 0.1907294 | -0.006676821 | 0.994672707 | 0.997993504 | 0.001273466 |
| ENSMUSG00000032399 | Rpl4 | 67891 | protein_coding | 15903.39693 | -0.001273844 | 0.077522015 | -0.016432024 | 0.986889732 | 0.993752664 | 0.001273844 |
| ENSMUSG00000118117 | Gm30961 | 102632967 | lncRNA | 8.497096786 | 0.001312496 | 0.451618788 | 0.002906204 | 0.997681188 | 0.998998301 | 0.001312496 |
| ENSMUSG00000074882 | Cyp2c68 | 433247 | protein_coding | 1026.808834 | -0.00131417 | 0.343435713 | -0.003826539 | 0.996946871 | 0.9987142 | 0.00131417 |
| ENSMUSG00000095362 | Gm14325 | 329575 | protein_coding | 136.4202965 | -0.001326365 | 0.188563057 | -0.007034068 | 0.994387672 | 0.997771772 | 0.001326365 |
| ENSMUSG00000052738 | Suclg1 | 56451 | protein_coding | 3708.650705 | -0.001352936 | 0.068867583 | -0.019645466 | 0.984326194 | 0.99313545 | 0.001352936 |
| ENSMUSG00000026062 | Slc9a2 | 226999 | protein_coding | 3895.354087 | 0.00139175 | 0.14599834 | 0.009532644 | 0.992394166 | 0.996654803 | 0.00139175 |
| ENSMUSG00000032667 | Pon2 | 330260 | protein_coding | 2985.36518 | -0.001408709 | 0.088201244 | -0.015971535 | 0.9872571 | 0.993846083 | 0.001408709 |
| ENSMUSG00000022142 | Nup155 | 170762 | protein_coding | 411.1034915 | 0.001424147 | 0.081664352 | 0.017439033 | 0.98608637 | 0.993716285 | 0.001424147 |
| ENSMUSG00000068794 | Col28a1 | 213945 | protein_coding | 37.6083111 | 0.00144208 | 0.22875663 | 0.006303992 | 0.994970175 | 0.998166212 | 0.00144208 |
| ENSMUSG00000053714 | 4732471J01Rik | 654804 | lncRNA | 34.50674261 | 0.001509654 | 0.270271713 | 0.005585689 | 0.995543288 | 0.998301026 | 0.001509654 |
| ENSMUSG00000042642 | Flad1 | 319945 | protein_coding | 396.0769754 | 0.001537522 | 0.09080229 | 0.016932635 | 0.986490357 | 0.993752664 | 0.001537522 |
| ENSMUSG00000079144 | A130010J15Rik | 319266 | protein_coding | 216.7701631 | 0.001568326 | 0.091844023 | 0.017075972 | 0.986376008 | 0.993752664 | 0.001568326 |
| ENSMUSG00000025266 | Gnl3l | 237107 | protein_coding | 939.0645156 | -0.001583817 | 0.071382407 | -0.022187782 | 0.982298164 | 0.992454497 | 0.001583817 |
| ENSMUSG00000023104 | Rfc2 | 19718 | protein_coding | 568.1489299 | 0.001589576 | 0.139762186 | 0.011373435 | 0.990925507 | 0.995739164 | 0.001589576 |
| ENSMUSG00000029169 | Dhx15 | 13204 | protein_coding | 2413.825731 | 0.001615859 | 0.10456699 | 0.015452857 | 0.987670895 | 0.994036523 | 0.001615859 |
| ENSMUSG00000063646 | Jakmip1 | 76071 | protein_coding | 139.1595852 | 0.001616933 | 0.130055726 | 0.012432618 | 0.990080462 | 0.995203938 | 0.001616933 |
| ENSMUSG00000029086 | Prom1 | 19126 | protein_coding | 7117.472476 | 0.00164941 | 0.192902165 | 0.008550499 | 0.993177772 | 0.996996063 | 0.00164941 |
| ENSMUSG00000085772 | D630024D03Rik | 414116 | lncRNA | 11.05727046 | -0.001649441 | 0.35697586 | -0.004620596 | 0.996313311 | 0.998642884 | 0.001649441 |
| ENSMUSG00000017707 | Serinc3 | 26943 | protein_coding | 7828.763151 | 0.001655078 | 0.090407149 | 0.018306939 | 0.985393992 | 0.993376931 | 0.001655078 |
| ENSMUSG00000024791 | Cdca5 | 67849 | protein_coding | 138.566731 | -0.001663581 | 0.144695726 | -0.011497096 | 0.990826847 | 0.99570284 | 0.001663581 |
| ENSMUSG00000027678 | Ncoa3 | 17979 | protein_coding | 843.0369836 | -0.001695062 | 0.094969032 | -0.017848574 | 0.985759654 | 0.99361978 | 0.001695062 |
| ENSMUSG00000024308 | Tapbp | 21356 | protein_coding | 1896.039448 | -0.001705677 | 0.096516416 | -0.017672404 | 0.985900196 | 0.993698558 | 0.001705677 |
| ENSMUSG00000039640 | Mrpl12 | 56282 | protein_coding | 1532.913065 | -0.001762239 | 0.095956522 | -0.018364972 | 0.985347696 | 0.993376931 | 0.001762239 |
| ENSMUSG00000021215 | Net1 | 56349 | protein_coding | 3208.00656 | 0.001768166 | 0.108525085 | 0.01629269 | 0.987000889 | 0.993799642 | 0.001768166 |

| ensembl_gene_id | external_gene_name | entrezgene_id | gene_biotype | baseMean | log2FoldChange | lfcSE | stat | pvalue | padj | absLog2FC |
|---|---|---|---|---|---|---|---|---|---|---|
| ENSMUSG00000028149 | Rap1gds1 | 229877 | protein_coding | 706.3028152 | 0.001777364 | 0.070146514 | 0.025337881 | 0.979785459 | 0.990929528 | 0.001777364 |
| ENSMUSG00000000078 | Klf6 | 23849 | protein_coding | 7991.394789 | -0.001794515 | 0.209692545 | -0.008557841 | 0.993171914 | 0.996996063 | 0.001794515 |
| ENSMUSG00000052911 | Lamb2 | 16779 | protein_coding | 346.4565582 | -0.001834208 | 0.098947488 | -0.018537187 | 0.985210312 | 0.993376931 | 0.001834208 |
| ENSMUSG00000045095 | Magi1 | 14924 | protein_coding | 904.3438191 | -0.001838739 | 0.088239347 | -0.020838087 | 0.983374815 | 0.992816672 | 0.001838739 |
| ENSMUSG00000057375 | Yipf1 | 230584 | protein_coding | 1744.653046 | -0.001865392 | 0.088635093 | -0.021045753 | 0.983209158 | 0.992816672 | 0.001865392 |
| ENSMUSG00000034121 | Mks1 | 380718 | protein_coding | 102.1250836 | -0.001886526 | 0.140799121 | -0.013398704 | 0.989309701 | 0.994743066 | 0.001886526 |
| ENSMUSG00000034377 | Tulp4 | 68842 | protein_coding | 1140.071309 | -0.001977109 | 0.065354717 | -0.03025197 | 0.975866101 | 0.989281776 | 0.001977109 |
| ENSMUSG00000075467 | Dnlz | 52838 | protein_coding | 274.6645698 | 0.002007583 | 0.119705816 | 0.016770973 | 0.986619327 | 0.993752664 | 0.002007583 |
| ENSMUSG00000044393 | Dsg2 | 13511 | protein_coding | 4299.988899 | -0.002047781 | 0.124556118 | -0.016440631 | 0.986882865 | 0.993752664 | 0.002047781 |
| ENSMUSG00000025486 | Sirt3 | 64384 | protein_coding | 176.7906596 | 0.00205883 | 0.128689659 | 0.015998413 | 0.987235658 | 0.993846083 | 0.00205883 |
| ENSMUSG00000019851 | Perp | 64058 | protein_coding | 3987.681356 | -0.00207449 | 0.120860654 | -0.017164316 | 0.98630553 | 0.993752664 | 0.00207449 |
| ENSMUSG00000033335 | Dnm2 | 13430 | protein_coding | 2837.852851 | 0.002086971 | 0.088113003 | 0.023685162 | 0.981103742 | 0.991688057 | 0.002086971 |
| ENSMUSG00000017314 | Mpp2 | 50997 | protein_coding | 17.26439943 | -0.002089387 | 0.359116315 | -0.005818134 | 0.995357827 | 0.998240742 | 0.002089387 |
| ENSMUSG00000002733 | Plekha3 | 83435 | protein_coding | 625.0150956 | -0.002095279 | 0.082911203 | -0.025271358 | 0.97983852 | 0.990929528 | 0.002095279 |
| ENSMUSG00000032396 | Dis3l | 213550 | protein_coding | 363.7035602 | 0.002105812 | 0.083609878 | 0.025186168 | 0.97990647 | 0.990929528 | 0.002105812 |
| ENSMUSG00000024084 | Qpct | 70536 | protein_coding | 12.62089845 | -0.002138008 | 0.324241179 | -0.006593881 | 0.994738882 | 0.997997034 | 0.002138008 |
| ENSMUSG00000026578 | Ccdc181 | 74895 | protein_coding | 155.2500995 | -0.002148773 | 0.179718165 | -0.011956346 | 0.990460443 | 0.995397435 | 0.002148773 |
| ENSMUSG00000029401 | Rilpl2 | 80291 | protein_coding | 124.2338528 | 0.002154033 | 0.148069823 | 0.014547415 | 0.988393252 | 0.994441098 | 0.002154033 |
| ENSMUSG00000019791 | Hint3 | 66847 | protein_coding | 447.0771694 | 0.00215662 | 0.122819457 | 0.017559271 | 0.985990449 | 0.993716285 | 0.00215662 |
| ENSMUSG00000036104 | Rab3gap1 | 226407 | protein_coding | 1206.729099 | -0.002171527 | 0.085099452 | -0.025517516 | 0.979642177 | 0.990929528 | 0.002171527 |
| ENSMUSG00000026499 | Acbd3 | 170760 | protein_coding | 1453.218246 | 0.002196291 | 0.057338093 | 0.038304223 | 0.969445124 | 0.986024667 | 0.002196291 |
| ENSMUSG00000018921 | Pelp1 | 75273 | protein_coding | 227.7687407 | 0.002211663 | 0.113994718 | 0.019401449 | 0.984520854 | 0.993188179 | 0.002211663 |
| ENSMUSG00000024122 | Pdpk1 | 18607 | protein_coding | 1594.998034 | -0.002255061 | 0.063255394 | -0.035650097 | 0.971561362 | 0.987242703 | 0.002255061 |
| ENSMUSG00000006335 | Tfpt | 69714 | protein_coding | 121.6145634 | 0.002257198 | 0.118223368 | 0.019092651 | 0.984767194 | 0.993212987 | 0.002257198 |
| ENSMUSG00000030231 | Plekha5 | 109135 | protein_coding | 1242.256852 | 0.002325445 | 0.10104936 | 0.023012961 | 0.981639935 | 0.991982127 | 0.002325445 |
| ENSMUSG00000031479 | Vps36 | 70160 | protein_coding | 656.1378783 | 0.00236426 | 0.064702528 | 0.03654046 | 0.970851418 | 0.986753935 | 0.00236426 |
| ENSMUSG00000002365 | Snx9 | 66616 | protein_coding | 937.8410934 | -0.002390264 | 0.086820514 | -0.027531093 | 0.978036141 | 0.99028413 | 0.002390264 |
| ENSMUSG00000042492 | Tbc1d10b | 68449 | protein_coding | 605.4767302 | -0.002406082 | 0.089344481 | -0.026930393 | 0.978515252 | 0.990328564 | 0.002406082 |
| ENSMUSG00000068523 | Gng5 | 14707 | protein_coding | 1013.994569 | 0.002421452 | 0.093285117 | 0.02595754 | 0.979291205 | 0.990902766 | 0.002421452 |
| ENSMUSG00000027580 | Helz2 | 229003 | protein_coding | 1207.557276 | -0.002433264 | 0.123066533 | -0.01977194 | 0.984225302 | 0.99313545 | 0.002433264 |
| ENSMUSG00000039740 | Alg2 | 56737 | protein_coding | 266.5200458 | 0.002446603 | 0.1416814 | 0.017268339 | 0.986222543 | 0.993752664 | 0.002446603 |
| ENSMUSG00000042496 | Prdm10 | 382066 | protein_coding | 259.255971 | -0.002497731 | 0.126215708 | -0.019789381 | 0.984211389 | 0.99313545 | 0.002497731 |
| ENSMUSG00000020375 | Rufy1 | 216724 | protein_coding | 381.1987278 | 0.002500318 | 0.083882146 | 0.029807508 | 0.976220571 | 0.989389237 | 0.002500318 |
| ENSMUSG00000033454 | Zbtb1 | 268564 | protein_coding | 486.340194 | 0.002509814 | 0.077214183 | 0.032504568 | 0.974069673 | 0.9886562 | 0.002509814 |
| ENSMUSG00000027952 | Pmvk | 68603 | protein_coding | 355.975483 | -0.002519558 | 0.132692092 | -0.018988005 | 0.984850674 | 0.993212987 | 0.002519558 |
| ENSMUSG00000033809 | Alg3 | 208624 | protein_coding | 81.07599015 | -0.002550768 | 0.202025587 | -0.012625966 | 0.989926204 | 0.99511168 | 0.002550768 |
| ENSMUSG00000019809 | Pex3 | 56535 | protein_coding | 292.7701744 | 0.002563344 | 0.08367229 | 0.030635515 | 0.975560218 | 0.989201611 | 0.002563344 |
| ENSMUSG00000022110 | Sucla2 | 20916 | protein_coding | 1686.085172 | 0.002578916 | 0.072915845 | 0.035368384 | 0.971785995 | 0.987344985 | 0.002578916 |

| ensembl_gene_id | external_gene_name | entrezgene_id | gene_biotype | baseMean | log2FoldChange | lfcSE | stat | pvalue | padj | absLog2FC |
|---|---|---|---|---|---|---|---|---|---|---|
| ENSMUSG00000042790 | Rnf214 | 235315 | protein_coding | 318.41418 | -0.002596001 | 0.103130751 | -0.025171941 | 0.979917817 | 0.990929528 | 0.002596001 |
| ENSMUSG00000071604 | Entrep1 | 381217 | protein_coding | 589.6292508 | -0.00260265 | 0.188299949 | -0.013821831 | 0.988972126 | 0.994743066 | 0.00260265 |
| ENSMUSG00000019710 | Mrpl24 | 67707 | protein_coding | 648.3454797 | 0.002684707 | 0.07199826 | 0.037288493 | 0.970254981 | 0.986376176 | 0.002684707 |
| ENSMUSG00000018677 | Slc25a39 | 68066 | protein_coding | 1847.927241 | -0.00270198 | 0.086238192 | -0.031331592 | 0.975005096 | 0.989003404 | 0.00270198 |
| ENSMUSG00000028080 | Lrba | 80877 | protein_coding | 3245.765188 | 0.002731762 | 0.091386404 | 0.029892431 | 0.976152842 | 0.989389237 | 0.002731762 |
| ENSMUSG00000025792 | Slc25a10 | 27376 | protein_coding | 1801.752052 | -0.002734089 | 0.133675783 | -0.020453138 | 0.983681895 | 0.992816672 | 0.002734089 |
| ENSMUSG00000070056 | Mfhas1 | 52065 | protein_coding | 526.3321002 | 0.00273526 | 0.088097722 | 0.031048018 | 0.975231245 | 0.989135397 | 0.00273526 |
| ENSMUSG00000068290 | Ddrgk1 | 77006 | protein_coding | 942.8414693 | 0.002764313 | 0.095517947 | 0.028940243 | 0.97691225 | 0.989550194 | 0.002764313 |
| ENSMUSG00000024535 | Snx24 | 69226 | protein_coding | 312.3833831 | 0.002793196 | 0.110320542 | 0.025318914 | 0.979800588 | 0.990929528 | 0.002793196 |
| ENSMUSG00000025332 | Kdm5c | 20591 | protein_coding | 1798.777611 | -0.002835776 | 0.108325252 | -0.026178351 | 0.979115083 | 0.990809723 | 0.002835776 |
| ENSMUSG00000034932 | Mrpl54 | 66047 | protein_coding | 533.7881204 | -0.002851338 | 0.107419691 | -0.026543903 | 0.978823516 | 0.990577608 | 0.002851338 |
| ENSMUSG00000038708 | Golga4 | 54214 | protein_coding | 3943.283775 | -0.002858395 | 0.076660599 | -0.03728637 | 0.970256673 | 0.986376176 | 0.002858395 |
| ENSMUSG00000048109 | Rbm15 | 229700 | protein_coding | 790.8119693 | -0.002865044 | 0.08579182 | -0.033395304 | 0.973359355 | 0.988124143 | 0.002865044 |
| ENSMUSG00000028910 | Mecr | 26922 | protein_coding | 806.5508858 | 0.002879673 | 0.137871797 | 0.020886603 | 0.983336113 | 0.992816672 | 0.002879673 |
| ENSMUSG00000052962 | Mrpl35 | 66223 | protein_coding | 395.1805988 | 0.00295671 | 0.114799435 | 0.025755438 | 0.979452405 | 0.990922997 | 0.00295671 |
| ENSMUSG00000027215 | Cd82 | 12521 | protein_coding | 1190.120493 | -0.003002511 | 0.057427727 | -0.052283294 | 0.958302964 | 0.979330785 | 0.003002511 |
| ENSMUSG00000074867 | Zfp808 | 630579 | protein_coding | 63.16371034 | -0.003013237 | 0.178302678 | -0.016899559 | 0.986516745 | 0.993752664 | 0.003013237 |
| ENSMUSG00000022100 | Xpo7 | 65246 | protein_coding | 867.3968538 | -0.003014906 | 0.073979927 | -0.04075303 | 0.967492785 | 0.9851917 | 0.003014906 |
| ENSMUSG00000038005 | Hpf1 | 72612 | protein_coding | 225.9725826 | 0.00303196 | 0.117032085 | 0.025907083 | 0.979331451 | 0.990902766 | 0.00303196 |
| ENSMUSG00000026893 | Gca | 227960 | protein_coding | 219.7863465 | 0.003111851 | 0.106848308 | 0.029124009 | 0.976765687 | 0.989500977 | 0.003111851 |
| ENSMUSG00000032393 | Dpp8 | 74388 | protein_coding | 2313.541425 | 0.003133887 | 0.043119084 | 0.072679822 | 0.942060906 | 0.971992316 | 0.003133887 |
| ENSMUSG00000004056 | Akt2 | 11652 | protein_coding | 484.9184081 | -0.003228479 | 0.081713216 | -0.039509873 | 0.968483882 | 0.985435753 | 0.003228479 |
| ENSMUSG00000039016 | Timm8b | 30057 | protein_coding | 501.528027 | -0.003236949 | 0.118551106 | -0.027304251 | 0.978217066 | 0.990328564 | 0.003236949 |
| ENSMUSG00000035439 | Haus8 | 76478 | protein_coding | 178.2725783 | 0.003259343 | 0.112751553 | 0.0289073 | 0.976938523 | 0.989550194 | 0.003259343 |
| ENSMUSG00000023143 | Nagpa | 27426 | protein_coding | 189.7916433 | 0.003360736 | 0.103914181 | 0.032341458 | 0.974199747 | 0.988725218 | 0.003360736 |
| ENSMUSG00000037031 | Tspan15 | 70423 | protein_coding | 1576.967247 | 0.00337128 | 0.102054048 | 0.033034258 | 0.973647269 | 0.988353432 | 0.00337128 |
| ENSMUSG00000022672 | Prkdc | 19090 | protein_coding | 496.2804631 | 0.00337628 | 0.132258108 | 0.025527964 | 0.979633844 | 0.990929528 | 0.00337628 |
| ENSMUSG00000021096 | Ppm1a | 19042 | protein_coding | 1758.959589 | 0.003379085 | 0.07018649 | 0.048144378 | 0.961601179 | 0.981316221 | 0.003379085 |
| ENSMUSG00000022336 | Eif3e | 16341 | protein_coding | 2711.898246 | 0.003420283 | 0.089548501 | 0.038194756 | 0.969532402 | 0.986024667 | 0.003420283 |
| ENSMUSG00000020402 | Vdac1 | 22333 | protein_coding | 4408.729771 | 0.003493638 | 0.060089846 | 0.058140234 | 0.953636927 | 0.976611016 | 0.003493638 |
| ENSMUSG00000038893 | Fam117a | 215512 | protein_coding | 111.0903879 | 0.003502079 | 0.206172737 | 0.016986142 | 0.986447671 | 0.993752664 | 0.003502079 |
| ENSMUSG00000039623 | Ap5z1 | 231855 | protein_coding | 195.9606433 | -0.003535091 | 0.160090111 | -0.022081885 | 0.982382637 | 0.992476889 | 0.003535091 |
| ENSMUSG00000092417 | Gpank1 | 81845 | protein_coding | 154.1167458 | -0.003549674 | 0.112466764 | -0.031561982 | 0.974821362 | 0.988988898 | 0.003549674 |
| ENSMUSG00000024896 | Minpp1 | 17330 | protein_coding | 526.3440936 | -0.003549915 | 0.07968288 | -0.044550534 | 0.964465572 | 0.983357293 | 0.003549915 |
| ENSMUSG00000070031 | Sp140 | 434484 | protein_coding | 41.6608607 | 0.003588105 | 0.358655559 | 0.010004319 | 0.992017841 | 0.996583807 | 0.003588105 |
| ENSMUSG00000039450 | Dcxr | 67880 | protein_coding | 595.148061 | -0.003634993 | 0.114203276 | -0.031829152 | 0.974608298 | 0.988950817 | 0.003634993 |
| ENSMUSG00000067430 | Zfp763 | 73451 | protein_coding | 124.1855358 | -0.003636182 | 0.229033098 | -0.01587623 | 0.987333133 | 0.993846083 | 0.003636182 |
| ENSMUSG00000090553 | Snrpe | 20643 | protein_coding | 724.3038457 | 0.003675328 | 0.120987611 | 0.03037772 | 0.975765814 | 0.989243071 | 0.003675328 |

Supplementary Table 1

| ensembl_gene_id | external_gene_name | entrezgene_id | gene_biotype | baseMean | log2FoldChange | lfcSE | stat | pvalue | padj | absLog2FC |
|---|---|---|---|---|---|---|---|---|---|---|
| ENSMUSG00000021222 | Dcaf4 | 73828 | protein_coding | 162.9277229 | 0.003681296 | 0.176496531 | 0.020857612 | 0.98335924 | 0.992816672 | 0.003681296 |
| ENSMUSG00000021023 | Prorp | 66132 | protein_coding | 283.2962219 | -0.003695457 | 0.092628816 | -0.039895325 | 0.968176578 | 0.985364183 | 0.003695457 |
| ENSMUSG00000048706 | Lurap1l | 52829 | protein_coding | 1871.341563 | -0.003738816 | 0.100178563 | -0.037321515 | 0.970228651 | 0.986376176 | 0.003738816 |
| ENSMUSG00000044933 | Sstr3 | 20607 | protein_coding | 12.71699373 | 0.003815005 | 0.410647847 | 0.009290211 | 0.992587591 | 0.996654803 | 0.003815005 |
| ENSMUSG00000026816 | Gtf3c5 | 70239 | protein_coding | 259.8950809 | -0.003948366 | 0.094923607 | -0.041595193 | 0.966821405 | 0.984812862 | 0.003948366 |
| ENSMUSG00000022772 | Senp5 | 320213 | protein_coding | 579.4513163 | -0.003992003 | 0.073838741 | -0.054063796 | 0.956884336 | 0.978257619 | 0.003992003 |
| ENSMUSG00000036500 | Akp3 | 11648 | protein_coding | 13.49511555 | 0.004077877 | 1.051711383 | 0.003877373 | 0.996906312 | 0.9987142 | 0.004077877 |
| ENSMUSG00000028494 | Plin2 | 11520 | protein_coding | 1280.797349 | -0.004108691 | 0.130439162 | -0.031498909 | 0.974871662 | 0.988988898 | 0.004108691 |
| ENSMUSG00000039982 | Dtx4 | 207521 | protein_coding | 313.0200507 | -0.00415927 | 0.084685821 | -0.049114123 | 0.960828349 | 0.980957331 | 0.00415927 |
| ENSMUSG00000052906 | Ubxn8 | 108159 | protein_coding | 427.2784572 | -0.004173936 | 0.122085536 | -0.034188616 | 0.972726744 | 0.987733749 | 0.004173936 |
| ENSMUSG00000089872 | Rps6kc1 | 320119 | protein_coding | 360.6303025 | 0.004235935 | 0.077157942 | 0.05489954 | 0.956218499 | 0.977890738 | 0.004235935 |
| ENSMUSG00000079559 | Pou2af3 | 100502940 | protein_coding | 19.34740833 | -0.004249432 | 0.2681628 | -0.015846462 | 0.987356882 | 0.993846083 | 0.004249432 |
| ENSMUSG00000095930 | Nim1k | 245269 | protein_coding | 26.11274528 | -0.004252015 | 0.355318283 | -0.011966778 | 0.990452121 | 0.995397435 | 0.004252015 |
| ENSMUSG00000028140 | Mrpl9 | 78523 | protein_coding | 926.6177364 | -0.004260865 | 0.087612244 | -0.048633213 | 0.961211601 | 0.981044366 | 0.004260865 |
| ENSMUSG00000029145 | Eif2b4 | 13667 | protein_coding | 394.3277578 | 0.004285771 | 0.113112805 | 0.037889352 | 0.969775903 | 0.98612044 | 0.004285771 |
| ENSMUSG00000053333 | Dis3l2 | 208718 | protein_coding | 274.6731474 | -0.004295744 | 0.078600569 | -0.054652832 | 0.956415048 | 0.977981234 | 0.004295744 |
| ENSMUSG00000030738 | Eif3c | 56347 | protein_coding | 3877.607792 | 0.004309888 | 0.066114327 | 0.065188412 | 0.948023988 | 0.974453154 | 0.004309888 |
| ENSMUSG00000001741 | Il16 | 16170 | protein_coding | 22.29599411 | 0.004320423 | 0.317978508 | 0.013587155 | 0.989159352 | 0.994743066 | 0.004320423 |
| ENSMUSG00000035506 | Slc12a8 | 171286 | protein_coding | 1695.596735 | -0.004371757 | 0.127800087 | -0.034207777 | 0.972711465 | 0.987733749 | 0.004371757 |
| ENSMUSG00000074890 | Lcmt2 | 329504 | protein_coding | 152.4877549 | -0.004381085 | 0.181547431 | -0.024131906 | 0.980747394 | 0.991453699 | 0.004381085 |
| ENSMUSG00000026240 | Cops7b | 26895 | protein_coding | 145.4449722 | 0.004387209 | 0.096280511 | 0.045566951 | 0.963655411 | 0.982594161 | 0.004387209 |
| ENSMUSG00000021731 | Mrps30 | 59054 | protein_coding | 448.3960296 | -0.004396844 | 0.088545698 | -0.04965621 | 0.960396353 | 0.980840799 | 0.004396844 |
| ENSMUSG00000068566 | Myadm | 50918 | protein_coding | 796.1272693 | -0.004399601 | 0.109524138 | -0.040170152 | 0.967957473 | 0.985364183 | 0.004399601 |
| ENSMUSG00000031221 | Igbp1 | 18518 | protein_coding | 1190.261998 | 0.004408621 | 0.096416358 | 0.04572482 | 0.963529581 | 0.982591662 | 0.004408621 |
| ENSMUSG00000026718 | Stam | 20844 | protein_coding | 834.6874991 | -0.004418466 | 0.073651282 | -0.059991705 | 0.952162241 | 0.975872864 | 0.004418466 |
| ENSMUSG00000032480 | Dhx30 | 72831 | protein_coding | 579.9507474 | -0.00441893 | 0.126867122 | -0.034831165 | 0.97221437 | 0.987518805 | 0.00441893 |
| ENSMUSG00000027171 | Prrg4 | 228413 | protein_coding | 34.36355167 | 0.004438815 | 0.284113472 | 0.015623387 | 0.987534848 | 0.993962405 | 0.004438815 |
| ENSMUSG00000026176 | Ctdsp1 | 227292 | protein_coding | 689.2648543 | -0.004454652 | 0.066827057 | -0.066659407 | 0.946852851 | 0.973877839 | 0.004454652 |
| ENSMUSG00000027782 | Kpna4 | 16649 | protein_coding | 1319.70287 | 0.004465169 | 0.108427412 | 0.041181186 | 0.967151452 | 0.985023872 | 0.004465169 |
| ENSMUSG00000059119 | Nap1l4 | 17955 | protein_coding | 990.2877053 | -0.004498025 | 0.072931627 | -0.061674548 | 0.950822009 | 0.975614007 | 0.004498025 |
| ENSMUSG00000097487 | Ptges3l | 73635 | protein_coding | 23.0011368 | -0.004522726 | 0.231024324 | -0.019576839 | 0.98438094 | 0.99313545 | 0.004522726 |
| ENSMUSG00000026779 | Mastl | 67121 | protein_coding | 304.0020136 | 0.004532914 | 0.167434583 | 0.027072747 | 0.978401712 | 0.990328564 | 0.004532914 |
| ENSMUSG00000021123 | Rdh12 | 77974 | protein_coding | 7.880645863 | 0.004571864 | 0.449098707 | 0.010180088 | 0.991877605 | 0.996558343 | 0.004571864 |
| ENSMUSG00000055407 | Map6 | 17760 | protein_coding | 11.82827812 | 0.004624969 | 0.381943501 | 0.012109041 | 0.990338619 | 0.995397435 | 0.004624969 |
| ENSMUSG00000003660 | Snrnp200 | 320632 | protein_coding | 1778.65222 | 0.004634297 | 0.068870378 | 0.067290136 | 0.946350729 | 0.973812302 | 0.004634297 |
| ENSMUSG00000012429 | Mplkip | 66308 | protein_coding | 195.1861159 | -0.004659413 | 0.116648718 | -0.039943968 | 0.968137797 | 0.985364183 | 0.004659413 |
| ENSMUSG00000020460 | Rps27a | 78294 | protein_coding | 5558.315257 | 0.004661824 | 0.098899583 | 0.047136947 | 0.96240408 | 0.98187404 | 0.004661824 |
| ENSMUSG00000028392 | Bspry | 192120 | protein_coding | 3000.30785 | -0.004749281 | 0.132190209 | -0.03592763 | 0.971340065 | 0.987080806 | 0.004749281 |

Supplementary Table 1

| ensembl_gene_id | external_gene_name | entrezgene_id | gene_biotype | baseMean | log2FoldChange | lfcSE | stat | pvalue | padj | absLog2FC |
|---|---|---|---|---|---|---|---|---|---|---|
| ENSMUSG00000071415 | Rpl23 | 65019 | protein_coding | 6810.724613 | -0.004801106 | 0.083233088 | -0.057682657 | 0.954001408 | 0.976611016 | 0.004801106 |
| ENSMUSG00000042694 | Stn1 | 108689 | protein_coding | 146.6957912 | 0.004816065 | 0.20230425 | 0.023806049 | 0.981007315 | 0.99165352 | 0.004816065 |
| ENSMUSG00000025997 | Ikzf2 | 22779 | protein_coding | 321.6501757 | 0.004836859 | 0.163888406 | 0.029513126 | 0.976455351 | 0.989434359 | 0.004836859 |
| ENSMUSG00000036427 | Gpi1 | 14751 | protein_coding | 6303.507519 | 0.004878063 | 0.071901796 | 0.067843405 | 0.945910291 | 0.973769693 | 0.004878063 |
| ENSMUSG00000087060 | Eldr | 72735 | lncRNA | 70.82330597 | -0.004879752 | 0.198351364 | -0.024601554 | 0.98037278 | 0.991275304 | 0.004879752 |
| ENSMUSG00000032705 | Exd2 | 97827 | protein_coding | 340.5072826 | -0.00490084 | 0.079040791 | -0.062003938 | 0.950559696 | 0.975605911 | 0.00490084 |
| ENSMUSG00000005774 | Rfx5 | 53970 | protein_coding | 126.0791903 | 0.004929583 | 0.155436408 | 0.031714465 | 0.974699759 | 0.98898062 | 0.004929583 |
| ENSMUSG00000024660 | Incenp | 16319 | protein_coding | 428.9093342 | 0.004957046 | 0.09305766 | 0.053268545 | 0.957517942 | 0.978639692 | 0.004957046 |
| ENSMUSG00000028071 | Sh2d2a | 27371 | protein_coding | 13.630432 | -0.004960369 | 0.504155897 | -0.009838959 | 0.992149773 | 0.996592141 | 0.004960369 |
| ENSMUSG00000031641 | Cbr4 | 234309 | protein_coding | 325.5731116 | -0.004988465 | 0.12370148 | -0.040326639 | 0.967832717 | 0.985339694 | 0.004988465 |
| ENSMUSG00000026976 | Pax8 | 18510 | protein_coding | 45.61756908 | 0.005020031 | 0.170719089 | 0.029405215 | 0.976541414 | 0.989434359 | 0.005020031 |
| ENSMUSG00000058435 | Btnl4 | 632126 | protein_coding | 902.0001546 | -0.005030369 | 0.143497505 | -0.035055446 | 0.972035529 | 0.987488745 | 0.005030369 |
| ENSMUSG00000042502 | Cd2bp2 | 70233 | protein_coding | 619.842778 | -0.005065327 | 0.078208992 | -0.064766561 | 0.948359866 | 0.974735493 | 0.005065327 |
| ENSMUSG00000046808 | Atp10d | 231287 | protein_coding | 12.38367004 | -0.005078318 | 0.511525997 | -0.00992778 | 0.992078908 | 0.996583807 | 0.005078318 |
| ENSMUSG00000054309 | Cpsf3 | 54451 | protein_coding | 678.3793936 | -0.005084609 | 0.065220915 | -0.077959795 | 0.937860034 | 0.969385764 | 0.005084609 |
| ENSMUSG00000042472 | Zfp410 | 52708 | protein_coding | 461.5524402 | 0.005098097 | 0.068248993 | 0.074698495 | 0.940454605 | 0.971059312 | 0.005098097 |
| ENSMUSG00000028134 | Ptbp2 | 56195 | protein_coding | 179.1885438 | 0.005114437 | 0.174364676 | 0.029331839 | 0.976599934 | 0.989434359 | 0.005114437 |
| ENSMUSG00000032571 | Pik3r4 | 75669 | protein_coding | 486.0831786 | 0.005124036 | 0.087005593 | 0.05889318 | 0.95303719 | 0.976295359 | 0.005124036 |
| ENSMUSG00000023861 | Mpc1 | 55951 | protein_coding | 601.4363274 | -0.005154936 | 0.104911944 | -0.049135837 | 0.960811044 | 0.980957331 | 0.005154936 |
| ENSMUSG00000022231 | Sema5a | 20356 | protein_coding | 1359.083892 | 0.005155887 | 0.18226159 | 0.028288389 | 0.977432141 | 0.98986134 | 0.005155887 |
| ENSMUSG00000034442 | Trmt5 | 76357 | protein_coding | 141.9977842 | -0.005192518 | 0.13041134 | -0.039816458 | 0.968239455 | 0.985364183 | 0.005192518 |
| ENSMUSG00000037033 | Clca3b | 229927 | protein_coding | 4134.769511 | -0.005193638 | 0.122280789 | -0.042473052 | 0.966121593 | 0.984289673 | 0.005193638 |
| ENSMUSG00000024858 | Grk2 | 110355 | protein_coding | 1139.254577 | 0.005217825 | 0.094753604 | 0.055067297 | 0.95608485 | 0.977816842 | 0.005217825 |
| ENSMUSG00000044224 | Dnajc21 | 78244 | protein_coding | 420.3866021 | 0.005221159 | 0.090689239 | 0.057571981 | 0.954089569 | 0.976611016 | 0.005221159 |
| ENSMUSG00000060512 | 0610040J01Rik | 76261 | protein_coding | 1000.353934 | -0.005239854 | 0.09595862 | -0.054605353 | 0.956452874 | 0.977981234 | 0.005239854 |
| ENSMUSG00000109179 | Gm35339 | 102638882 | protein_coding | 33.64367648 | 0.005275897 | 0.276253725 | 0.019098011 | 0.984762919 | 0.993212987 | 0.005275897 |
| ENSMUSG00000061544 | Zfp229 | 381067 | protein_coding | 122.0409404 | -0.005421866 | 0.121992716 | -0.044444181 | 0.964550345 | 0.983380778 | 0.005421866 |
| ENSMUSG00000078185 | Chml | 12663 | protein_coding | 471.0262461 | 0.005441057 | 0.099763815 | 0.054539384 | 0.956505431 | 0.977981234 | 0.005441057 |
| ENSMUSG00000026509 | Capn2 | 12334 | protein_coding | 983.0917394 | -0.005463532 | 0.176367554 | -0.030978101 | 0.975287004 | 0.989135397 | 0.005463532 |
| ENSMUSG00000030002 | Dusp11 | 72102 | protein_coding | 1928.896378 | -0.005547973 | 0.052901847 | -0.104872962 | 0.916476614 | 0.958597942 | 0.005547973 |
| ENSMUSG00000022724 | Riox2 | 67014 | protein_coding | 305.3840222 | 0.005559747 | 0.096419193 | 0.057662246 | 0.954017667 | 0.976611016 | 0.005559747 |
| ENSMUSG00000024985 | Tcf7l2 | 21416 | protein_coding | 464.2611038 | -0.005571571 | 0.189314658 | -0.029430213 | 0.976521477 | 0.989434359 | 0.005571571 |
| ENSMUSG00000018474 | Chd3 | 216848 | protein_coding | 1065.332576 | 0.00558069 | 0.099223414 | 0.056243678 | 0.955147686 | 0.977172095 | 0.00558069 |
| ENSMUSG00000078201 | Tmem203 | 227615 | protein_coding | 39.00041605 | 0.005592932 | 0.186162242 | 0.03004332 | 0.976032504 | 0.989387496 | 0.005592932 |
| ENSMUSG00000022035 | Ccdc25 | 67179 | protein_coding | 800.2495704 | 0.00560003 | 0.091818506 | 0.060990209 | 0.951367007 | 0.975614007 | 0.00560003 |
| ENSMUSG00000043411 | Usp48 | 170707 | protein_coding | 1030.523821 | 0.005690218 | 0.05062758 | 0.112393633 | 0.910511304 | 0.955759393 | 0.005690218 |
| ENSMUSG00000039754 | Alkbh4 | 72041 | protein_coding | 107.9202457 | 0.005697652 | 0.135554631 | 0.042032144 | 0.966473074 | 0.984521816 | 0.005697652 |
| ENSMUSG00000030753 | Thap12 | 72981 | protein_coding | 918.0507968 | 0.005699939 | 0.106138479 | 0.053702854 | 0.957171909 | 0.978488811 | 0.005699939 |

| ensembl_gene_id | external_gene_name | entrezgene_id | gene_biotype | baseMean | log2FoldChange | lfcSE | stat | pvalue | padj | absLog2FC |
|---|---|---|---|---|---|---|---|---|---|---|
| ENSMUSG00000053799 | Exoc6 | 107371 | protein_coding | 570.6252872 | 0.005729023 | 0.099685561 | 0.057470939 | 0.954170055 | 0.976611016 | 0.005729023 |
| ENSMUSG00000020829 | Slc46a1 | 52466 | protein_coding | 109.1762701 | 0.005791601 | 0.203702709 | 0.028431633 | 0.977317895 | 0.989853998 | 0.005791601 |
| ENSMUSG00000089876 | Tmem102 | 380705 | protein_coding | 323.4433929 | -0.005808521 | 0.119377834 | -0.048656609 | 0.961192956 | 0.981044366 | 0.005808521 |
| ENSMUSG00000079477 | Rab7 | 19349 | protein_coding | 1440.235596 | -0.00585594 | 0.085506829 | -0.068485055 | 0.945399516 | 0.97338872 | 0.00585594 |
| ENSMUSG00000029416 | Slc15a4 | 100561 | protein_coding | 331.3578625 | -0.005859142 | 0.095706506 | -0.061219894 | 0.951184086 | 0.975614007 | 0.005859142 |
| ENSMUSG00000020358 | Hnrnpab | 15384 | protein_coding | 2912.159942 | 0.005859778 | 0.148067536 | 0.039575039 | 0.968431928 | 0.985435753 | 0.005859778 |
| ENSMUSG00000015126 | Tsr3 | 68327 | protein_coding | 116.5434555 | 0.005909346 | 0.115490477 | 0.05116739 | 0.959192137 | 0.980006131 | 0.005909346 |
| ENSMUSG00000026349 | Ccnt2 | 72949 | protein_coding | 951.8224621 | -0.00594626 | 0.096920966 | -0.061351635 | 0.951079169 | 0.975614007 | 0.00594626 |
| ENSMUSG00000021686 | Ap3b1 | 11774 | protein_coding | 1964.713368 | -0.005998521 | 0.065367223 | -0.091766495 | 0.926883565 | 0.963794322 | 0.005998521 |
| ENSMUSG00000020305 | Asb3 | 65257 | protein_coding | 177.1992043 | 0.005999708 | 0.122825937 | 0.048847241 | 0.961041034 | 0.981044366 | 0.005999708 |
| ENSMUSG00000037523 | Mavs | 228607 | protein_coding | 632.0823104 | -0.006019941 | 0.091140609 | -0.066051141 | 0.94733711 | 0.974114993 | 0.006019941 |
| ENSMUSG00000041879 | Ipo9 | 226432 | protein_coding | 478.2832179 | 0.006022648 | 0.09106266 | 0.06613741 | 0.947268427 | 0.974114993 | 0.006022648 |
| ENSMUSG00000015013 | Trappc2l | 59005 | protein_coding | 246.8029334 | -0.006024423 | 0.122823736 | -0.049049338 | 0.960879977 | 0.980957331 | 0.006024423 |
| ENSMUSG00000025917 | Cops5 | 26754 | protein_coding | 769.5960077 | 0.006073171 | 0.082613846 | 0.073512745 | 0.941398102 | 0.97146673 | 0.006073171 |
| ENSMUSG00000024258 | Polr2d | 69241 | protein_coding | 201.4268924 | -0.006088262 | 0.137462368 | -0.044290389 | 0.964672933 | 0.983402378 | 0.006088262 |
| ENSMUSG00000022359 | Wdyhv1 | 76773 | protein_coding | 168.5887001 | 0.006092327 | 0.1379896 | 0.044150624 | 0.96478434 | 0.983422954 | 0.006092327 |
| ENSMUSG00000033752 | Mnd1 | 76915 | protein_coding | 28.03870003 | -0.006110661 | 0.226596792 | -0.026967111 | 0.978485966 | 0.990328564 | 0.006110661 |
| ENSMUSG00000046671 | Mtfr1l | 76824 | protein_coding | 472.8702515 | -0.006173436 | 0.070580109 | -0.087467077 | 0.930300254 | 0.965393344 | 0.006173436 |
| ENSMUSG00000056498 | Tmem154 | 320782 | protein_coding | 17.94378523 | 0.006188235 | 0.301719917 | 0.020509866 | 0.983636641 | 0.992816672 | 0.006188235 |
| ENSMUSG00000033793 | Atp6v1h | 108664 | protein_coding | 925.878293 | -0.006193823 | 0.07743645 | -0.079985889 | 0.936248479 | 0.968704702 | 0.006193823 |
| ENSMUSG00000035983 | Gm7008 | 629820 | lncRNA | 8.434903296 | -0.006211843 | 0.460175861 | -0.013498846 | 0.989229806 | 0.994743066 | 0.006211843 |
| ENSMUSG00000015314 | Slamf6 | 30925 | protein_coding | 10.50522593 | 0.006220809 | 1.000373031 | 0.006218489 | 0.995038396 | 0.998171783 | 0.006220809 |
| ENSMUSG00000015214 | Mtmr1 | 53332 | protein_coding | 541.0383345 | -0.00624759 | 0.087518641 | -0.071385818 | 0.943090697 | 0.972533548 | 0.00624759 |
| ENSMUSG00000008384 | Sertad1 | 55942 | protein_coding | 724.6328973 | -0.006248619 | 0.242891002 | -0.025726019 | 0.97947587 | 0.990922997 | 0.006248619 |
| ENSMUSG00000077192 | Snora17 | 100306954 | snoRNA | 15.67490729 | 0.006281761 | 0.303297562 | 0.020711546 | 0.983475759 | 0.992816672 | 0.006281761 |
| ENSMUSG00000020955 | Ap4s1 | 11782 | protein_coding | 152.0470839 | -0.00631868 | 0.106515783 | -0.059321538 | 0.952696007 | 0.976168671 | 0.00631868 |
| ENSMUSG00000027668 | Mfn1 | 67414 | protein_coding | 1216.858308 | -0.00636407 | 0.071629041 | -0.088847619 | 0.929203012 | 0.965135023 | 0.00636407 |
| ENSMUSG00000009630 | Ppp2cb | 19053 | protein_coding | 1097.258281 | -0.006390402 | 0.10038111 | -0.063661398 | 0.949239842 | 0.975117538 | 0.006390402 |
| ENSMUSG00000034903 | Cobll1 | 319876 | protein_coding | 319.4661574 | -0.006393561 | 0.110834422 | -0.057685695 | 0.953998989 | 0.976611016 | 0.006393561 |
| ENSMUSG00000062169 | Cnih4 | 98417 | protein_coding | 638.3401015 | -0.006405893 | 0.075312505 | -0.085057493 | 0.932215683 | 0.96605218 | 0.006405893 |
| ENSMUSG00000025533 | Asl | 109900 | protein_coding | 396.6130169 | -0.006416154 | 0.105688026 | -0.060708428 | 0.951591419 | 0.975632393 | 0.006416154 |
| ENSMUSG00000019849 | Prep | 19072 | protein_coding | 606.6350319 | -0.006439778 | 0.09697051 | -0.066409659 | 0.94705168 | 0.974019446 | 0.006439778 |
| ENSMUSG00000038195 | Rilp | 280408 | protein_coding | 137.1636573 | -0.006444071 | 0.141199693 | -0.045637996 | 0.963598785 | 0.982594161 | 0.006444071 |
| ENSMUSG00000052974 | Cyp2f2 | 13107 | protein_coding | 132.241536 | -0.00650963 | 0.300905469 | -0.021633472 | 0.982740333 | 0.992775292 | 0.00650963 |
| ENSMUSG00000024797 | Vps51 | 68505 | protein_coding | 426.350569 | -0.006529045 | 0.112933538 | -0.05781316 | 0.953897455 | 0.976611016 | 0.006529045 |
| ENSMUSG00000060657 | Marf1 | 223989 | protein_coding | 1250.305124 | -0.006565765 | 0.092603298 | -0.070902063 | 0.943475701 | 0.97253954 | 0.006565765 |
| ENSMUSG00000069516 | Lyz2 | 17105 | protein_coding | 30.39176052 | -0.00656915 | 0.456153395 | -0.014401186 | 0.988509913 | 0.994441098 | 0.00656915 |
| ENSMUSG00000030432 | Rpl28 | 19943 | protein_coding | 4039.223686 | 0.006570219 | 0.089172659 | 0.073679752 | 0.941265211 | 0.971456555 | 0.006570219 |

Supplementary Table 1

| ensembl_gene_id | external_gene_name | entrezgene_id | gene_biotype | baseMean | log2FoldChange | lfcSE | stat | pvalue | padj | absLog2FC |
|---|---|---|---|---|---|---|---|---|---|---|
| ENSMUSG00000055917 | Zfp277 | 246196 | protein_coding | 422.9549635 | 0.006584921 | 0.093779483 | 0.070217076 | 0.944020883 | 0.972723808 | 0.006584921 |
| ENSMUSG00000046722 | Cdc42se1 | 57912 | protein_coding | 1123.350787 | 0.006585045 | 0.105345173 | 0.062509221 | 0.950157319 | 0.975404564 | 0.006585045 |
| ENSMUSG00000036435 | Exoc1 | 69940 | protein_coding | 565.7417904 | 0.006607611 | 0.066386023 | 0.099533165 | 0.920714957 | 0.96082997 | 0.006607611 |
| ENSMUSG00000035878 | Hykk | 235386 | protein_coding | 175.7462239 | 0.006644597 | 0.284017662 | 0.023395015 | 0.981335182 | 0.99185905 | 0.006644597 |
| ENSMUSG00000056608 | Chd9 | 109151 | protein_coding | 1079.204277 | -0.006655583 | 0.124254223 | -0.053564238 | 0.95728235 | 0.978538908 | 0.006655583 |
| ENSMUSG00000045273 | Cenph | 26886 | protein_coding | 59.43974972 | -0.00666581 | 0.191878299 | -0.034739781 | 0.972287239 | 0.987518805 | 0.00666581 |
| ENSMUSG00000026227 | 2810459M11Rik | 72792 | protein_coding | 490.246188 | 0.006793197 | 0.176505756 | 0.038487112 | 0.969299307 | 0.985985438 | 0.006793197 |
| ENSMUSG00000026466 | Tor1aip1 | 208263 | protein_coding | 363.7694866 | 0.006820751 | 0.076112881 | 0.089613628 | 0.928594254 | 0.964808485 | 0.006820751 |
| ENSMUSG00000021000 | Mia2 | 338320 | protein_coding | 1927.814073 | 0.006823711 | 0.052611708 | 0.129699484 | 0.896804191 | 0.948387789 | 0.006823711 |
| ENSMUSG00000090733 | Rps27 | 57294 | protein_coding | 8024.65671 | -0.006845263 | 0.09967576 | -0.068675301 | 0.945248078 | 0.973358637 | 0.006845263 |
| ENSMUSG00000031154 | Otud5 | 54644 | protein_coding | 836.6113277 | -0.006875232 | 0.087321557 | -0.078734651 | 0.937243683 | 0.969063122 | 0.006875232 |
| ENSMUSG00000024491 | Rbm27 | 225432 | protein_coding | 829.2000561 | 0.006909991 | 0.091839248 | 0.075240069 | 0.940023704 | 0.970803187 | 0.006909991 |
| ENSMUSG00000047264 | Zfp358 | 140482 | protein_coding | 161.0107197 | -0.007010059 | 0.148706222 | -0.047140318 | 0.962401394 | 0.98187404 | 0.007010059 |
| ENSMUSG00000037772 | Mrpl23 | 19935 | protein_coding | 123.3102192 | -0.00701335 | 0.125267312 | -0.055987073 | 0.955352105 | 0.977318454 | 0.00701335 |
| ENSMUSG00000034165 | Ccnd3 | 12445 | protein_coding | 380.3151033 | 0.007021439 | 0.088732044 | 0.07913082 | 0.936928569 | 0.968800198 | 0.007021439 |
| ENSMUSG00000028868 | Wasf2 | 242687 | protein_coding | 1074.241803 | -0.007074881 | 0.10532012 | -0.067175022 | 0.94644237 | 0.973812302 | 0.007074881 |
| ENSMUSG00000060427 | Zfp868 | 234362 | protein_coding | 255.5342714 | 0.007082312 | 0.104634169 | 0.067686418 | 0.946035262 | 0.973769693 | 0.007082312 |
| ENSMUSG00000071657 | Bscl2 | 14705 | protein_coding | 450.6688598 | -0.0071275 | 0.100195744 | -0.071135754 | 0.943289713 | 0.972533548 | 0.0071275 |
| ENSMUSG00000041645 | Ddx24 | 27225 | protein_coding | 945.942092 | -0.00713007 | 0.113227985 | -0.062970914 | 0.949789666 | 0.975281565 | 0.00713007 |
| ENSMUSG00000038274 | Fau | 14109 | protein_coding | 4558.420055 | 0.007132082 | 0.100249261 | 0.071143484 | 0.94328356 | 0.972533548 | 0.007132082 |
| ENSMUSG00000035649 | Zcchc7 | 319885 | protein_coding | 597.453271 | -0.007137782 | 0.098626909 | -0.072371547 | 0.942306228 | 0.972026009 | 0.007137782 |
| ENSMUSG00000021264 | Yy1 | 22632 | protein_coding | 1735.932343 | -0.007207324 | 0.041265362 | -0.174657963 | 0.861348401 | 0.927926916 | 0.007207324 |
| ENSMUSG00000028796 | Phc2 | 54383 | protein_coding | 528.9178485 | 0.007212436 | 0.091795682 | 0.078570541 | 0.93737422 | 0.96913518 | 0.007212436 |
| ENSMUSG00000031487 | Brf2 | 66653 | protein_coding | 87.96497134 | -0.007251089 | 0.141984365 | -0.05106963 | 0.959270036 | 0.980006131 | 0.007251089 |
| ENSMUSG00000117602 | 9130209A04Rik | 77668 | lncRNA | 42.18225475 | -0.007258524 | 0.227651255 | -0.031884399 | 0.97456424 | 0.988950817 | 0.007258524 |
| ENSMUSG00000053134 | Supt7l | 72195 | protein_coding | 203.6671372 | 0.007276687 | 0.100539899 | 0.072376109 | 0.942302597 | 0.972026009 | 0.007276687 |
| ENSMUSG00000030126 | Tmcc1 | 330401 | protein_coding | 511.9658661 | 0.007326526 | 0.085229432 | 0.085962393 | 0.931496312 | 0.965879387 | 0.007326526 |
| ENSMUSG00000047824 | Pygo2 | 68911 | protein_coding | 533.4044312 | -0.007345731 | 0.097595896 | -0.075266802 | 0.940002434 | 0.970803187 | 0.007345731 |
| ENSMUSG00000038732 | Mboat1 | 218121 | protein_coding | 4243.335638 | -0.007357469 | 0.192791811 | -0.038162768 | 0.969557906 | 0.986024667 | 0.007357469 |
| ENSMUSG00000024960 | Plcb3 | 18797 | protein_coding | 4733.938865 | -0.007376389 | 0.051574864 | -0.143022955 | 0.886272052 | 0.94220607 | 0.007376389 |
| ENSMUSG00000027752 | Exosc8 | 69639 | protein_coding | 195.4403345 | -0.007376885 | 0.124088583 | -0.059448543 | 0.95259485 | 0.976127816 | 0.007376885 |
| ENSMUSG00000040591 | Cstpp1 | 228356 | protein_coding | 61.73951589 | -0.007441662 | 0.161378016 | -0.046113234 | 0.963219998 | 0.98246466 | 0.007441662 |
| ENSMUSG00000034807 | Colgalt1 | 234407 | protein_coding | 655.6755448 | -0.007482187 | 0.072427057 | -0.103306519 | 0.917719702 | 0.959208789 | 0.007482187 |
| ENSMUSG00000054523 | Ms4a5 | 269063 | protein_coding | 15.80136107 | -0.007515192 | 0.353413433 | -0.02126459 | 0.98303459 | 0.992816672 | 0.007515192 |
| ENSMUSG00000022762 | Ncam2 | 17968 | protein_coding | 7.377938283 | 0.007522973 | 1.069771751 | 0.007032316 | 0.99438907 | 0.997771772 | 0.007522973 |
| ENSMUSG00000078652 | Psme3 | 19192 | protein_coding | 851.5991591 | -0.007534421 | 0.127273852 | -0.059198498 | 0.952794006 | 0.976206287 | 0.007534421 |
| ENSMUSG00000030689 | Ino80e | 233875 | protein_coding | 179.9667271 | -0.007563038 | 0.11846469 | -0.063842127 | 0.949095935 | 0.975117538 | 0.007563038 |
| ENSMUSG00000031988 | Vps26b | 69091 | protein_coding | 725.3136896 | -0.007573072 | 0.09490271 | -0.079798267 | 0.936397703 | 0.968704702 | 0.007573072 |

| ensembl_gene_id | external_gene_name | entrezgene_id | gene_biotype | baseMean | log2FoldChange | lfcSE | stat | pvalue | padj | absLog2FC |
|---|---|---|---|---|---|---|---|---|---|---|
| ENSMUSG00000003534 | Ddr1 | 12305 | protein_coding | 1258.830434 | 0.007573152 | 0.082796558 | 0.091466992 | 0.927121533 | 0.963947469 | 0.007573152 |
| ENSMUSG00000022507 | 1810013L24Rik | 69053 | protein_coding | 1097.289099 | -0.007630407 | 0.093985805 | -0.081186804 | 0.935293393 | 0.967926267 | 0.007630407 |
| ENSMUSG00000037572 | Wdhd1 | 218973 | protein_coding | 219.6408335 | -0.007636545 | 0.109060418 | -0.070021232 | 0.94417676 | 0.972731213 | 0.007636545 |
| ENSMUSG00000026889 | Rbm18 | 67889 | protein_coding | 420.5185068 | -0.007683079 | 0.089284778 | -0.086051389 | 0.931425566 | 0.965868908 | 0.007683079 |
| ENSMUSG00000060149 | BC002059 | 213811 | lncRNA | 171.3352672 | -0.007716303 | 0.11549589 | -0.066810192 | 0.94673281 | 0.973817569 | 0.007716303 |
| ENSMUSG00000028843 | Sh3bgrl3 | 73723 | protein_coding | 546.1605232 | -0.007744006 | 0.128018158 | -0.060491464 | 0.951764214 | 0.975678597 | 0.007744006 |
| ENSMUSG00000040594 | Ranbp17 | 66011 | protein_coding | 29.81891227 | 0.007746915 | 0.278948391 | 0.027771856 | 0.977844113 | 0.99015264 | 0.007746915 |
| ENSMUSG00000077565 | Gm23201 | 115486177 | snoRNA | 7.592092813 | 0.007762039 | 0.470194844 | 0.016508133 | 0.986829014 | 0.993752664 | 0.007762039 |
| ENSMUSG00000035173 | Ccdc186 | 213993 | protein_coding | 1218.373854 | 0.007768552 | 0.11773085 | 0.065985699 | 0.947389211 | 0.974114993 | 0.007768552 |
| ENSMUSG00000024260 | Sap130 | 269003 | protein_coding | 390.0130473 | 0.007792683 | 0.119005406 | 0.065481756 | 0.947790432 | 0.974338841 | 0.007792683 |
| ENSMUSG00000050549 | Fam241a | 70617 | protein_coding | 204.8849988 | -0.007792873 | 0.127622543 | -0.061061883 | 0.951309925 | 0.975614007 | 0.007792873 |
| ENSMUSG00000002342 | Tmem161a | 234371 | protein_coding | 356.9281485 | 0.007796288 | 0.079240306 | 0.098387912 | 0.921624273 | 0.961086252 | 0.007796288 |
| ENSMUSG00000039768 | Dnajc11 | 230935 | protein_coding | 1044.283442 | -0.007796464 | 0.084731598 | -0.092013655 | 0.92668719 | 0.963740176 | 0.007796464 |
| ENSMUSG00000027180 | Fbxo3 | 57443 | protein_coding | 965.5014414 | -0.007800553 | 0.047858003 | -0.162993702 | 0.870523393 | 0.933277015 | 0.007800553 |
| ENSMUSG00000031552 | Adam18 | 13524 | protein_coding | 14.7072973 | -0.007816464 | 0.380795421 | -0.020526675 | 0.983623233 | 0.992816672 | 0.007816464 |
| ENSMUSG00000032763 | Ilvbl | 216136 | protein_coding | 733.2439665 | 0.007858511 | 0.10363639 | 0.075827716 | 0.939556166 | 0.970446184 | 0.007858511 |
| ENSMUSG00000040018 | Cox15 | 226139 | protein_coding | 792.9742744 | -0.007878518 | 0.098798393 | -0.079743381 | 0.936441356 | 0.968704702 | 0.007878518 |
| ENSMUSG00000031314 | Taf1 | 270627 | protein_coding | 1223.310565 | -0.00789558 | 0.074524116 | -0.10594664 | 0.915624689 | 0.95821748 | 0.00789558 |
| ENSMUSG00000025316 | Banp | 53325 | protein_coding | 227.0691426 | -0.007924558 | 0.181072319 | -0.043764603 | 0.965092043 | 0.983535029 | 0.007924558 |
| ENSMUSG00000022969 | Il10rb | 16155 | protein_coding | 585.9259831 | 0.007982453 | 0.066064609 | 0.120827975 | 0.903827291 | 0.952525195 | 0.007982453 |
| ENSMUSG00000024953 | Prdx5 | 54683 | protein_coding | 1732.724986 | -0.007988947 | 0.112216666 | -0.071192164 | 0.943244818 | 0.972533548 | 0.007988947 |
| ENSMUSG00000020495 | Smg8 | 74133 | protein_coding | 234.7348452 | -0.008009462 | 0.114301319 | -0.070073224 | 0.944135379 | 0.972731213 | 0.008009462 |
| ENSMUSG00000002129 | Sf3a1 | 67465 | protein_coding | 884.16967 | 0.008012279 | 0.085803833 | 0.093379035 | 0.925602446 | 0.963531518 | 0.008012279 |
| ENSMUSG00000063281 | Zfp35 | 22694 | protein_coding | 292.7666051 | 0.008019846 | 0.111548689 | 0.071895473 | 0.942685093 | 0.972290887 | 0.008019846 |
| ENSMUSG00000015217 | Hmgb3 | 15354 | protein_coding | 348.1387478 | 0.008049357 | 0.092187812 | 0.087314763 | 0.93042132 | 0.965456077 | 0.008049357 |
| ENSMUSG00000030770 | Parva | 57342 | protein_coding | 1512.602349 | -0.008051872 | 0.083094962 | -0.096899635 | 0.922806098 | 0.961877857 | 0.008051872 |
| ENSMUSG00000006589 | Aprt | 11821 | protein_coding | 801.2879941 | 0.008061568 | 0.125687147 | 0.064139959 | 0.948858785 | 0.975059531 | 0.008061568 |
| ENSMUSG00000074582 | Arfgef2 | 99371 | protein_coding | 1280.622206 | 0.00811266 | 0.086316788 | 0.09398705 | 0.925119444 | 0.963197355 | 0.00811266 |
| ENSMUSG00000042462 | Dctpp1 | 66422 | protein_coding | 239.9019761 | -0.008124083 | 0.097468913 | -0.083350502 | 0.933572845 | 0.966899556 | 0.008124083 |
| ENSMUSG00000024576 | Csnk1a1 | 93687 | protein_coding | 7024.934601 | 0.008139359 | 0.085588256 | 0.095099015 | 0.92423618 | 0.962738466 | 0.008139359 |
| ENSMUSG00000023947 | Nfkbie | 18037 | protein_coding | 32.35152319 | 0.008152534 | 0.212062192 | 0.038444069 | 0.969333625 | 0.985985438 | 0.008152534 |
| ENSMUSG00000029032 | Arhgef16 | 230972 | protein_coding | 1453.666508 | 0.008204395 | 0.085247958 | 0.09624154 | 0.92332874 | 0.962296681 | 0.008204395 |
| ENSMUSG00000032342 | Mto1 | 68291 | protein_coding | 268.1166445 | 0.008205467 | 0.093651166 | 0.08761735 | 0.930180812 | 0.965369999 | 0.008205467 |
| ENSMUSG00000097333 | Zfp87 | 170763 | protein_coding | 198.1957183 | 0.008208547 | 0.101010173 | 0.081264553 | 0.935231563 | 0.967926267 | 0.008208547 |
| ENSMUSG00000048911 | Rnf24 | 51902 | protein_coding | 54.84052611 | 0.008264772 | 0.269029832 | 0.030720652 | 0.975492321 | 0.989201611 | 0.008264772 |
| ENSMUSG00000034729 | Mrps10 | 64657 | protein_coding | 295.5222646 | 0.008289903 | 0.09261532 | 0.089508986 | 0.928677413 | 0.964808485 | 0.008289903 |
| ENSMUSG00000020115 | Tbk1 | 56480 | protein_coding | 652.2168984 | 0.008330952 | 0.07034959 | 0.118422188 | 0.905733146 | 0.953387557 | 0.008330952 |
| ENSMUSG00000068747 | Sort1 | 20661 | protein_coding | 6434.179703 | 0.008339517 | 0.076238688 | 0.109386943 | 0.91289559 | 0.957122232 | 0.008339517 |

| ensembl_gene_id | external_gene_name | entrezgene_id | gene_biotype | baseMean | log2FoldChange | lfcSE | stat | pvalue | padj | absLog2FC |
|---|---|---|---|---|---|---|---|---|---|---|
| ENSMUSG00000068267 | Cenpb | 12616 | protein_coding | 2370.023127 | 0.008343573 | 0.084706758 | 0.098499492 | 0.921535676 | 0.961086252 | 0.008343573 |
| ENSMUSG00000062203 | Gspt1 | 14852 | protein_coding | 1338.248887 | 0.008397826 | 0.125397136 | 0.066969835 | 0.946605718 | 0.973812302 | 0.008397826 |
| ENSMUSG00000036960 | Clca2 | 229933 | protein_coding | 9.214924703 | -0.008400482 | 0.469418417 | -0.017895509 | 0.985722211 | 0.99361978 | 0.008400482 |
| ENSMUSG00000053716 | Dusp7 | 235584 | protein_coding | 103.4045569 | 0.00844403 | 0.283133062 | 0.029823538 | 0.976207787 | 0.989389237 | 0.00844403 |
| ENSMUSG00000039456 | Morc3 | 338467 | protein_coding | 817.9782688 | -0.008447835 | 0.051757377 | -0.163219927 | 0.870345277 | 0.933211685 | 0.008447835 |
| ENSMUSG00000022248 | Rad1 | 19355 | protein_coding | 124.9627563 | 0.00850971 | 0.139315684 | 0.061082208 | 0.951293739 | 0.975614007 | 0.00850971 |
| ENSMUSG00000071103 | 1700029J07Rik | 69479 | protein_coding | 12.93417284 | -0.008516081 | 0.39780119 | -0.021407882 | 0.982920286 | 0.992816672 | 0.008516081 |
| ENSMUSG00000025920 | Stau2 | 29819 | protein_coding | 265.9205265 | -0.008527305 | 0.081095796 | -0.105151011 | 0.916255983 | 0.958563238 | 0.008527305 |
| ENSMUSG00000027206 | Cops2 | 12848 | protein_coding | 1375.409243 | 0.008547197 | 0.059725291 | 0.143108499 | 0.886204493 | 0.94220607 | 0.008547197 |
| ENSMUSG00000038489 | Polr2l | 66491 | protein_coding | 93.85653703 | 0.008555795 | 0.151777809 | 0.056370526 | 0.955046636 | 0.977131476 | 0.008555795 |
| ENSMUSG00000020448 | Rnf185 | 193670 | protein_coding | 398.1135636 | 0.008570951 | 0.066134462 | 0.129598858 | 0.896883807 | 0.948387789 | 0.008570951 |
| ENSMUSG00000024045 | Akap8 | 56399 | protein_coding | 1156.022421 | -0.008593881 | 0.062188214 | -0.138191467 | 0.890089099 | 0.94465213 | 0.008593881 |
| ENSMUSG00000029414 | Kntc1 | 208628 | protein_coding | 284.7478247 | 0.008667176 | 0.139657967 | 0.062060016 | 0.950515038 | 0.975605911 | 0.008667176 |
| ENSMUSG00000068206 | Pick1 | 18693 | protein_coding | 209.5251989 | -0.008677411 | 0.112274318 | -0.077287584 | 0.938394768 | 0.969812604 | 0.008677411 |
| ENSMUSG00000040035 | Disp2 | 214240 | protein_coding | 63.38923967 | -0.008677709 | 0.247683766 | -0.035035437 | 0.972051483 | 0.987488745 | 0.008677709 |
| ENSMUSG00000027412 | Lpin3 | 64899 | protein_coding | 503.3169611 | -0.008708749 | 0.087599285 | -0.099415754 | 0.920808175 | 0.960864296 | 0.008708749 |
| ENSMUSG00000037279 | Ovol2 | 107586 | protein_coding | 252.7641098 | 0.008710159 | 0.149929317 | 0.058095102 | 0.953672876 | 0.976611016 | 0.008710159 |
| ENSMUSG00000026867 | Gapvd1 | 66691 | protein_coding | 1255.936043 | 0.008722015 | 0.067134107 | 0.129919277 | 0.896630293 | 0.948360442 | 0.008722015 |
| ENSMUSG00000027312 | Atrn | 11990 | protein_coding | 838.1047198 | 0.008738716 | 0.078973663 | 0.110653545 | 0.911891085 | 0.956746934 | 0.008738716 |
| ENSMUSG00000031410 | Nxf7 | 170722 | protein_coding | 8.015758876 | -0.008749262 | 0.525529458 | -0.016648471 | 0.986717056 | 0.993752664 | 0.008749262 |
| ENSMUSG00000021180 | Rps6ka5 | 73086 | protein_coding | 58.64367184 | -0.00876214 | 0.173198852 | -0.050590059 | 0.959652184 | 0.980212886 | 0.00876214 |
| ENSMUSG00000024949 | Sf1 | 22668 | protein_coding | 1221.240189 | -0.008888492 | 0.100180706 | -0.08872459 | 0.929300789 | 0.965173642 | 0.008888492 |
| ENSMUSG00000001844 | Zdhhc4 | 72881 | protein_coding | 218.8471948 | -0.008921796 | 0.083662056 | -0.106640885 | 0.915073882 | 0.958081692 | 0.008921796 |
| ENSMUSG00000021311 | Mtr | 238505 | protein_coding | 160.771654 | -0.008927003 | 0.099901074 | -0.089358431 | 0.928797059 | 0.964808485 | 0.008927003 |
| ENSMUSG00000075327 | Zbtb2 | 381990 | protein_coding | 215.7725084 | 0.008957854 | 0.090901367 | 0.098544764 | 0.921499729 | 0.961086252 | 0.008957854 |
| ENSMUSG00000092103 | Gm17039 | 102636903 | lncRNA | 8.41808897 | -0.00895873 | 0.458361209 | -0.019545132 | 0.984406234 | 0.99313545 | 0.00895873 |
| ENSMUSG00000027536 | Chmp4c | 66371 | protein_coding | 2358.777532 | 0.008966043 | 0.078267586 | 0.114556271 | 0.908796842 | 0.954836155 | 0.008966043 |
| ENSMUSG00000014243 | Zswim7 | 69747 | protein_coding | 71.33144831 | -0.008984137 | 0.142728017 | -0.062945854 | 0.949809621 | 0.975281565 | 0.008984137 |
| ENSMUSG00000050600 | Zfp831 | 100043757 | protein_coding | 7.454545103 | -0.0089932 | 0.668960046 | -0.013443554 | 0.989273919 | 0.994743066 | 0.0089932 |
| ENSMUSG00000024776 | Stambpl1 | 76630 | protein_coding | 231.0937521 | -0.008999879 | 0.128086503 | -0.070264072 | 0.943983478 | 0.972723808 | 0.008999879 |
| ENSMUSG00000005951 | Shpk | 74637 | protein_coding | 238.6431852 | 0.009031355 | 0.149449509 | 0.060430811 | 0.95181252 | 0.975678597 | 0.009031355 |
| ENSMUSG00000026470 | Stx6 | 58244 | protein_coding | 375.9542996 | 0.009038373 | 0.08768023 | 0.103083365 | 0.917896808 | 0.959208789 | 0.009038373 |
| ENSMUSG00000058897 | Col25a1 | 77018 | protein_coding | 17.80253815 | 0.00910302 | 0.481599887 | 0.018901625 | 0.984919584 | 0.993212987 | 0.00910302 |
| ENSMUSG00000024515 | Smad4 | 17128 | protein_coding | 1641.295049 | -0.009115404 | 0.107075761 | -0.085130418 | 0.932157708 | 0.96605218 | 0.009115404 |
| ENSMUSG00000009647 | Mcu | 215999 | protein_coding | 1766.200397 | -0.009133837 | 0.105399647 | -0.086659086 | 0.930942499 | 0.96561944 | 0.009133837 |
| ENSMUSG00000029592 | Usp30 | 100756 | protein_coding | 258.4404299 | -0.009173659 | 0.085292605 | -0.107555151 | 0.914348574 | 0.957700013 | 0.009173659 |
| ENSMUSG00000034931 | Dhx8 | 217207 | protein_coding | 639.5029998 | -0.009197991 | 0.086929597 | -0.105809655 | 0.915733376 | 0.958268262 | 0.009197991 |
| ENSMUSG00000018774 | Cd68 | 12514 | protein_coding | 21.07018009 | 0.009204673 | 0.324389235 | 0.028375397 | 0.977362746 | 0.989853998 | 0.009204673 |

| ensembl_gene_id | external_gene_name | entrezgene_id | gene_biotype | baseMean | log2FoldChange | lfcSE | stat | pvalue | padj | absLog2FC |
|---|---|---|---|---|---|---|---|---|---|---|
| ENSMUSG00000027079 | Clp1 | 98985 | protein_coding | 250.1345086 | -0.009226568 | 0.096633622 | -0.095479899 | 0.923933655 | 0.962592167 | 0.009226568 |
| ENSMUSG00000022443 | Myh9 | 17886 | protein_coding | 10386.98876 | 0.009251114 | 0.112578487 | 0.082174795 | 0.934507716 | 0.967553497 | 0.009251114 |
| ENSMUSG00000025860 | Xiap | 11798 | protein_coding | 1968.27254 | -0.009260442 | 0.137765176 | -0.067219036 | 0.946407331 | 0.973812302 | 0.009260442 |
| ENSMUSG00000028514 | Usp24 | 329908 | protein_coding | 1426.101375 | -0.009281046 | 0.089688518 | -0.103480872 | 0.91758133 | 0.95913082 | 0.009281046 |
| ENSMUSG00000031134 | Rbmx | 19655 | protein_coding | 463.9023717 | 0.009288941 | 0.067611453 | 0.137387092 | 0.890724834 | 0.944949058 | 0.009288941 |
| ENSMUSG00000026927 | Entr1 | 68112 | protein_coding | 447.2903351 | 0.009295584 | 0.144876912 | 0.064161942 | 0.948841281 | 0.975059531 | 0.009295584 |
| ENSMUSG00000002963 | Pnkp | 59047 | protein_coding | 341.5756971 | -0.009296105 | 0.100061149 | -0.092904242 | 0.925979636 | 0.963686666 | 0.009296105 |
| ENSMUSG00000015882 | Lcorl | 209707 | protein_coding | 939.8663254 | -0.00930474 | 0.092124828 | -0.101001433 | 0.919549323 | 0.960107027 | 0.00930474 |
| ENSMUSG00000020009 | Ifngr1 | 15979 | protein_coding | 2805.12609 | 0.009316276 | 0.109228803 | 0.085291391 | 0.932029736 | 0.96605218 | 0.009316276 |
| ENSMUSG00000035495 | Tstd2 | 272027 | protein_coding | 454.7436114 | 0.009337148 | 0.065696341 | 0.142125842 | 0.886980606 | 0.942483462 | 0.009337148 |
| ENSMUSG00000058729 | Lin9 | 72568 | protein_coding | 174.6157361 | 0.009371334 | 0.117852147 | 0.07951772 | 0.936620837 | 0.968704702 | 0.009371334 |
| ENSMUSG00000028099 | Polr3c | 74414 | protein_coding | 582.6531347 | 0.009485406 | 0.0559362 | 0.169575446 | 0.865344034 | 0.930165999 | 0.009485406 |
| ENSMUSG00000045410 | Akr1e1 | 56043 | protein_coding | 665.2928861 | 0.009491143 | 0.130413833 | 0.072777117 | 0.941983481 | 0.971992316 | 0.009491143 |
| ENSMUSG00000102918 | Pcdhgc3 | 93706 | protein_coding | 77.42745494 | 0.009528791 | 0.234710877 | 0.040597997 | 0.967616381 | 0.985245395 | 0.009528791 |
| ENSMUSG00000028617 | Lrrc42 | 77809 | protein_coding | 787.5925163 | -0.009551384 | 0.135357809 | -0.070563967 | 0.943744789 | 0.972663741 | 0.009551384 |
| ENSMUSG00000032293 | Ireb2 | 64602 | protein_coding | 980.1339635 | 0.009554013 | 0.208751174 | 0.045767471 | 0.963495586 | 0.982591662 | 0.009554013 |
| ENSMUSG00000016940 | Kctd2 | 70382 | protein_coding | 258.9347104 | 0.009581979 | 0.138141574 | 0.069363471 | 0.944700305 | 0.973109129 | 0.009581979 |
| ENSMUSG00000017670 | Elmo2 | 140579 | protein_coding | 200.6272239 | 0.009597342 | 0.096766734 | 0.099180181 | 0.92099521 | 0.960868211 | 0.009597342 |
| ENSMUSG00000022779 | Top3b | 21976 | protein_coding | 529.2866103 | 0.009611036 | 0.07826433 | 0.122802254 | 0.90226369 | 0.951737121 | 0.009611036 |
| ENSMUSG00000041343 | Ankrd42 | 73845 | protein_coding | 29.8832234 | -0.00964597 | 0.242645834 | -0.039753289 | 0.968289817 | 0.985364183 | 0.00964597 |
| ENSMUSG00000020311 | Erlec1 | 66753 | protein_coding | 433.6824476 | 0.009647782 | 0.128740845 | 0.074939558 | 0.940262802 | 0.970975969 | 0.009647782 |
| ENSMUSG00000007589 | Tinf2 | 28113 | protein_coding | 297.5245047 | -0.009758584 | 0.089182103 | -0.109423117 | 0.912866899 | 0.957122232 | 0.009758584 |
| ENSMUSG00000042810 | Krba1 | 77827 | protein_coding | 280.8146819 | 0.009787351 | 0.176754848 | 0.055372459 | 0.955841737 | 0.977693757 | 0.009787351 |
| ENSMUSG00000031072 | LTO1 | 72284 | protein_coding | 223.7672798 | 0.00978803 | 0.083659673 | 0.116998185 | 0.906861492 | 0.954080995 | 0.00978803 |
| ENSMUSG00000053080 | Zfta | 108899 | protein_coding | 57.40803787 | 0.009803144 | 0.24646472 | 0.039775042 | 0.968272474 | 0.985364183 | 0.009803144 |
| ENSMUSG00000029613 | Eif2ak1 | 15467 | protein_coding | 668.8735774 | -0.009823757 | 0.06814805 | -0.144153165 | 0.885379523 | 0.94220607 | 0.009823757 |
| ENSMUSG00000025870 | Arl10 | 56795 | protein_coding | 123.6603333 | 0.009843458 | 0.133454498 | 0.073758905 | 0.941202227 | 0.971456555 | 0.009843458 |
| ENSMUSG00000022680 | Pdxdc1 | 94184 | protein_coding | 8878.941852 | -0.009884886 | 0.097789676 | -0.101083122 | 0.919484476 | 0.960107027 | 0.009884886 |
| ENSMUSG00000034538 | Zfp418 | 232854 | protein_coding | 62.60643474 | 0.009903747 | 0.167661044 | 0.059070052 | 0.952896312 | 0.976248309 | 0.009903747 |
| ENSMUSG00000036438 | Calm2 | 12314 | protein_coding | 5842.635511 | -0.009906435 | 0.076764068 | -0.129050422 | 0.897317751 | 0.948703875 | 0.009906435 |
| ENSMUSG00000046364 | Rpl27a | 26451 | protein_coding | 3207.664525 | 0.00991788 | 0.087981829 | 0.112726463 | 0.910247421 | 0.955759393 | 0.00991788 |
| ENSMUSG00000024841 | Eif1ad | 69860 | protein_coding | 422.318851 | 0.009982861 | 0.105555594 | 0.094574436 | 0.924652856 | 0.963036092 | 0.009982861 |
| ENSMUSG00000038685 | Rtel1 | 269400 | protein_coding | 316.6650726 | -0.010010424 | 0.093069463 | -0.107558633 | 0.914345813 | 0.957700013 | 0.010010424 |
| ENSMUSG00000024758 | Rtn3 | 20168 | protein_coding | 1989.159059 | -0.010028628 | 0.069648421 | -0.143989301 | 0.885508918 | 0.94220607 | 0.010028628 |
| ENSMUSG00000039518 | Cdsn | 386463 | protein_coding | 13.24624882 | -0.010046922 | 0.413969195 | -0.024269734 | 0.980637455 | 0.991405483 | 0.010046922 |
| ENSMUSG00000019718 | L3hypdh | 67217 | protein_coding | 11.70508733 | 0.010086233 | 0.481538951 | 0.02094583 | 0.983288868 | 0.992816672 | 0.010086233 |
| ENSMUSG00000026688 | Mgst3 | 66447 | protein_coding | 5235.40463 | -0.010113836 | 0.140825182 | -0.071818379 | 0.942746446 | 0.972291206 | 0.010113836 |
| ENSMUSG00000010047 | Hyal2 | 15587 | protein_coding | 526.1080675 | 0.01012809 | 0.111104435 | 0.091158283 | 0.927366822 | 0.964045082 | 0.01012809 |

| ensembl_gene_id | external_gene_name | entrezgene_id | gene_biotype | baseMean | log2FoldChange | lfcSE | stat | pvalue | padj | absLog2FC |
|---|---|---|---|---|---|---|---|---|---|---|
| ENSMUSG00000039354 | Smarcal1 | 54380 | protein_coding | 278.5141563 | -0.010134729 | 0.107268376 | -0.094480122 | 0.924727773 | 0.963036092 | 0.010134729 |
| ENSMUSG00000030589 | Rasgrp4 | 233046 | protein_coding | 12.00033513 | -0.010144937 | 0.43744625 | -0.023191276 | 0.981497697 | 0.991960363 | 0.010144937 |
| ENSMUSG00000001786 | Fbxo7 | 69754 | protein_coding | 226.6528563 | -0.010177697 | 0.089619017 | -0.113566263 | 0.909581633 | 0.955345615 | 0.010177697 |
| ENSMUSG00000031767 | Nudt7 | 67528 | protein_coding | 294.9342533 | -0.010202964 | 0.161711466 | -0.063093635 | 0.949691942 | 0.975281565 | 0.010202964 |
| ENSMUSG00000094942 | Gm3604 | 100041979 | protein_coding | 70.47314907 | -0.010206115 | 0.211189107 | -0.048326902 | 0.961455715 | 0.981230642 | 0.010206115 |
| ENSMUSG00000057691 | Zfp746 | 69228 | protein_coding | 431.4406496 | -0.010283571 | 0.112528369 | -0.091386479 | 0.927185505 | 0.963947469 | 0.010283571 |
| ENSMUSG00000011831 | Evi5 | 14020 | protein_coding | 629.8631958 | -0.010287857 | 0.059798667 | -0.172041576 | 0.863404843 | 0.929084741 | 0.010287857 |
| ENSMUSG00000015189 | Casd1 | 213819 | protein_coding | 1660.91855 | 0.010296303 | 0.141780168 | 0.0726216 | 0.942107238 | 0.971992316 | 0.010296303 |
| ENSMUSG00000027365 | Trpm7 | 58800 | protein_coding | 1663.432584 | 0.010302588 | 0.088132304 | 0.116899112 | 0.906940002 | 0.954080995 | 0.010302588 |
| ENSMUSG00000024048 | Myl12a | 67268 | protein_coding | 705.8686688 | -0.010324911 | 0.150660635 | -0.068530914 | 0.945363012 | 0.97338872 | 0.010324911 |
| ENSMUSG00000062638 | Btnl1 | 100038862 | protein_coding | 1298.000123 | -0.01033164 | 0.146402155 | -0.070570275 | 0.943739769 | 0.972663741 | 0.01033164 |
| ENSMUSG00000060126 | Tpt1 | 22070 | protein_coding | 11213.43522 | -0.010342112 | 0.066104455 | -0.156451066 | 0.875677488 | 0.936748192 | 0.010342112 |
| ENSMUSG00000022561 | Gpaa1 | 14731 | protein_coding | 515.1764687 | -0.010359489 | 0.102361731 | -0.10120471 | 0.919387958 | 0.960107027 | 0.010359489 |
| ENSMUSG00000004233 | Wars2 | 70560 | protein_coding | 269.9317369 | 0.010381731 | 0.08859591 | 0.117180701 | 0.90671686 | 0.954080995 | 0.010381731 |
| ENSMUSG00000029624 | Ptcd1 | 71799 | protein_coding | 293.6311616 | -0.010469889 | 0.109100276 | -0.095965744 | 0.923547779 | 0.96235224 | 0.010469889 |
| ENSMUSG00000021905 | Dph3 | 105638 | protein_coding | 439.2217711 | 0.010502352 | 0.1264317 | 0.083067396 | 0.933797951 | 0.967069815 | 0.010502352 |
| ENSMUSG00000036114 | Rpp25l | 69961 | protein_coding | 208.3418027 | -0.010516519 | 0.123559947 | -0.08511269 | 0.932171802 | 0.96605218 | 0.010516519 |
| ENSMUSG00000028413 | B4galt1 | 14595 | protein_coding | 1227.396624 | -0.010522452 | 0.141175097 | -0.074534762 | 0.940584882 | 0.971067928 | 0.010522452 |
| ENSMUSG00000062997 | Rpl35 | 66489 | protein_coding | 1953.050586 | 0.010536195 | 0.096612611 | 0.109056107 | 0.913157989 | 0.95727132 | 0.010536195 |
| ENSMUSG00000027751 | Supt20 | 56790 | protein_coding | 653.7925283 | 0.010562599 | 0.066160181 | 0.159651902 | 0.873155291 | 0.93490303 | 0.010562599 |
| ENSMUSG00000048661 | Lemd3 | 380664 | protein_coding | 384.1721527 | 0.010590917 | 0.120451052 | 0.087927144 | 0.929934582 | 0.965328339 | 0.010590917 |
| ENSMUSG00000022799 | Arhgap31 | 12549 | protein_coding | 143.7329704 | -0.010599539 | 0.166653524 | -0.063602252 | 0.949286939 | 0.975117538 | 0.010599539 |
| ENSMUSG00000030168 | Adipor2 | 68465 | protein_coding | 3188.063002 | 0.0106385 | 0.097920958 | 0.108643751 | 0.913485058 | 0.957282345 | 0.0106385 |
| ENSMUSG00000037805 | Rpl10a | 19896 | protein_coding | 3520.713659 | 0.010674035 | 0.097754855 | 0.109191865 | 0.913050312 | 0.957221442 | 0.010674035 |
| ENSMUSG00000003438 | Timm50 | 66525 | protein_coding | 430.7984282 | -0.010767981 | 0.14838888 | -0.072565957 | 0.942151517 | 0.971992316 | 0.010767981 |
| ENSMUSG00000027598 | Itch | 16396 | protein_coding | 1887.270864 | 0.010791091 | 0.074588735 | 0.144674547 | 0.884967837 | 0.942104511 | 0.010791091 |
| ENSMUSG00000099826 | Scgb2b10 | 624584 | unprocessed_pseudogene | 7.435294657 | 0.010792367 | 0.469067623 | 0.023008127 | 0.98164379 | 0.991982127 | 0.010792367 |
| ENSMUSG00000047414 | Flrt2 | 399558 | protein_coding | 32.60327934 | -0.010814155 | 0.296388483 | -0.036486422 | 0.970894505 | 0.986753935 | 0.010814155 |
| ENSMUSG00000025354 | Dnajc14 | 74330 | protein_coding | 1200.807277 | 0.010892951 | 0.068188217 | 0.159748289 | 0.87307936 | 0.934884575 | 0.010892951 |
| ENSMUSG00000036225 | Kctd1 | 106931 | protein_coding | 58.61851664 | 0.010894093 | 0.282108094 | 0.038616732 | 0.969195962 | 0.985971341 | 0.010894093 |
| ENSMUSG00000020580 | Rock2 | 19878 | protein_coding | 2687.648509 | -0.010942807 | 0.178058359 | -0.061456292 | 0.950995822 | 0.975614007 | 0.010942807 |
| ENSMUSG00000043510 | Hscb | 100900 | protein_coding | 158.9024353 | 0.010944963 | 0.138061736 | 0.07927586 | 0.936813207 | 0.968743801 | 0.010944963 |
| ENSMUSG00000069793 | Slfn9 | 237886 | protein_coding | 641.7124054 | 0.010951923 | 0.126782234 | 0.086383736 | 0.931161376 | 0.96574404 | 0.010951923 |
| ENSMUSG00000047248 | C2cd3 | 277939 | protein_coding | 428.3843874 | 0.010983417 | 0.129999838 | 0.084487929 | 0.9326685 | 0.966214235 | 0.010983417 |
| ENSMUSG00000001138 | Cnnm3 | 94218 | protein_coding | 279.7931505 | 0.01099367 | 0.093882597 | 0.117100196 | 0.906780654 | 0.954080995 | 0.01099367 |
| ENSMUSG00000016481 | Cr1l | 12946 | protein_coding | 1139.138178 | -0.011008743 | 0.087308275 | -0.12609049 | 0.899660295 | 0.950188262 | 0.011008743 |
| ENSMUSG00000032077 | Bud13 | 215051 | protein_coding | 145.9452256 | -0.011028856 | 0.097794369 | -0.112775981 | 0.910208162 | 0.955759393 | 0.011028856 |
| ENSMUSG00000021051 | Ppp2r5e | 26932 | protein_coding | 609.14956 | -0.011035069 | 0.091039805 | -0.121211474 | 0.903523536 | 0.952386474 | 0.011035069 |

| ensembl_gene_id | external_gene_name | entrezgene_id | gene_biotype | baseMean | log2FoldChange | lfcSE | stat | pvalue | padj | absLog2FC |
|---|---|---|---|---|---|---|---|---|---|---|
| ENSMUSG00000027509 | Rae1 | 66679 | protein_coding | 455.5029221 | -0.011081563 | 0.092763467 | -0.119460427 | 0.90491059 | 0.953079143 | 0.011081563 |
| ENSMUSG00000029599 | Ddx54 | 71990 | protein_coding | 894.0613695 | -0.011090017 | 0.095914071 | -0.115624509 | 0.907950138 | 0.954450224 | 0.011090017 |
| ENSMUSG00000003208 | Yju2 | 72886 | protein_coding | 114.4535951 | 0.011105192 | 0.131597136 | 0.084387797 | 0.932748109 | 0.966233864 | 0.011105192 |
| ENSMUSG00000066442 | Mthfs | 107885 | protein_coding | 41.00463776 | -0.011137905 | 0.204439535 | -0.054480192 | 0.95655259 | 0.977981234 | 0.011137905 |
| ENSMUSG00000028702 | Rad54l | 19366 | protein_coding | 122.4685013 | -0.011243926 | 0.191630681 | -0.058674978 | 0.95321099 | 0.976319506 | 0.011243926 |
| ENSMUSG00000061104 | Sap18b | 100041953 | protein_coding | 9.148019423 | 0.011258414 | 0.365171738 | 0.030830464 | 0.975404745 | 0.989191824 | 0.011258414 |
| ENSMUSG00000021669 | Cert1 | 68018 | protein_coding | 1199.955518 | 0.011270515 | 0.063647181 | 0.177077997 | 0.859447125 | 0.926766895 | 0.011270515 |
| ENSMUSG00000019863 | Qrsl1 | 76563 | protein_coding | 237.1576517 | 0.011276029 | 0.126029296 | 0.089471492 | 0.928707209 | 0.964808485 | 0.011276029 |
| ENSMUSG00000031901 | Dus2 | 66369 | protein_coding | 92.4126519 | 0.011415482 | 0.131445739 | 0.086845583 | 0.930794254 | 0.96552855 | 0.011415482 |
| ENSMUSG00000030768 | Disp1 | 68897 | protein_coding | 108.3316975 | 0.011442043 | 0.134969899 | 0.08477478 | 0.932440444 | 0.966103649 | 0.011442043 |
| ENSMUSG00000028479 | Gne | 50798 | protein_coding | 7830.557424 | -0.011460787 | 0.146294518 | -0.07834051 | 0.937557194 | 0.969210087 | 0.011460787 |
| ENSMUSG00000101892 | 9130401M01Rik | 75758 | protein_coding | 78.77395413 | 0.011490107 | 0.143379001 | 0.080138002 | 0.936127499 | 0.968663601 | 0.011490107 |
| ENSMUSG00000032459 | Mrps22 | 64655 | protein_coding | 355.5227563 | 0.011491819 | 0.109955645 | 0.104513222 | 0.916762076 | 0.95865194 | 0.011491819 |
| ENSMUSG00000022897 | Dyrk1a | 13548 | protein_coding | 1063.93648 | 0.011510857 | 0.066407977 | 0.173335454 | 0.862387757 | 0.928297339 | 0.011510857 |
| ENSMUSG00000036977 | Anapc10 | 68999 | protein_coding | 156.8737979 | 0.011521137 | 0.104212869 | 0.110553878 | 0.911970123 | 0.956746934 | 0.011521137 |
| ENSMUSG00000037788 | Vopp1 | 232023 | protein_coding | 331.2511072 | 0.011540653 | 0.125463236 | 0.091984342 | 0.92671048 | 0.963740176 | 0.011540653 |
| ENSMUSG00000029199 | Lias | 79464 | protein_coding | 410.1819305 | 0.011571688 | 0.096828579 | 0.119506946 | 0.904873737 | 0.953079143 | 0.011571688 |
| ENSMUSG00000028798 | Eif3i | 54709 | protein_coding | 2099.163801 | 0.011579412 | 0.093754003 | 0.123508458 | 0.901704477 | 0.951274931 | 0.011579412 |
| ENSMUSG00000003131 | Pafah1b2 | 18475 | protein_coding | 1113.31015 | 0.011599891 | 0.112977824 | 0.102674053 | 0.918221668 | 0.959359444 | 0.011599891 |
| ENSMUSG00000019777 | Hdac2 | 15182 | protein_coding | 839.3295899 | 0.011628207 | 0.101296737 | 0.114793499 | 0.908608802 | 0.954764547 | 0.011628207 |
| ENSMUSG00000038822 | Hace1 | 209462 | protein_coding | 334.7835132 | 0.011673712 | 0.094152235 | 0.123987623 | 0.901325075 | 0.951000649 | 0.011673712 |
| ENSMUSG00000031791 | Tmem38a | 74166 | protein_coding | 50.37054129 | -0.011680902 | 0.171643388 | -0.06805332 | 0.94574319 | 0.973679628 | 0.011680902 |
| ENSMUSG00000020608 | Smc6 | 67241 | protein_coding | 1113.108453 | -0.011695219 | 0.068056887 | -0.171844759 | 0.863559577 | 0.929125705 | 0.011695219 |
| ENSMUSG00000009291 | Pttg1ip | 108705 | protein_coding | 2635.439607 | 0.011708439 | 0.09706983 | 0.120618727 | 0.903993035 | 0.952525195 | 0.011708439 |
| ENSMUSG00000029787 | Avl9 | 78937 | protein_coding | 1950.442836 | 0.011760369 | 0.055828825 | 0.210650489 | 0.833160012 | 0.911190103 | 0.011760369 |
| ENSMUSG00000061032 | Rrp1 | 18114 | protein_coding | 1253.093051 | -0.01177024 | 0.099401697 | -0.11841086 | 0.905742121 | 0.953387557 | 0.01177024 |
| ENSMUSG00000021870 | Slmap | 83997 | protein_coding | 1469.077537 | -0.011796501 | 0.06737961 | -0.175075235 | 0.861020518 | 0.927835384 | 0.011796501 |
| ENSMUSG00000024286 | Ccny | 67974 | protein_coding | 1617.934879 | -0.011825886 | 0.073990671 | -0.159829425 | 0.873015445 | 0.934878984 | 0.011825886 |
| ENSMUSG00000022843 | Clcn2 | 12724 | protein_coding | 2501.736118 | 0.011849699 | 0.132542952 | 0.089402706 | 0.928761873 | 0.964808485 | 0.011849699 |
| ENSMUSG00000095325 | Zfp870 | 240066 | protein_coding | 127.4220553 | -0.011868944 | 0.171770002 | -0.069097887 | 0.944911703 | 0.973212655 | 0.011868944 |
| ENSMUSG00000024283 | Wac | 225131 | protein_coding | 1339.671504 | -0.011894224 | 0.063059493 | -0.188619084 | 0.850391375 | 0.920992304 | 0.011894224 |
| ENSMUSG00000022022 | Mtrf1 | 211253 | protein_coding | 100.7480032 | 0.011903689 | 0.13291712 | 0.089557229 | 0.928639074 | 0.964808485 | 0.011903689 |
| ENSMUSG00000050052 | Tdrp | 72148 | protein_coding | 7.202157158 | -0.011944506 | 0.485801566 | -0.024587211 | 0.98038422 | 0.991275304 | 0.011944506 |
| ENSMUSG00000041313 | Slc7a1 | 11987 | protein_coding | 661.2710651 | 0.011947571 | 0.245330503 | 0.048699899 | 0.961158456 | 0.981044366 | 0.011947571 |
| ENSMUSG00000029028 | Lrrc47 | 72946 | protein_coding | 346.4130008 | 0.011953254 | 0.120735803 | 0.099003392 | 0.921135577 | 0.960891183 | 0.011953254 |
| ENSMUSG00000110185 | Igip | 109169 | protein_coding | 177.574654 | 0.011977615 | 0.215067226 | 0.055692422 | 0.955586837 | 0.977495802 | 0.011977615 |
| ENSMUSG00000028412 | Slc44a1 | 100434 | protein_coding | 6723.521446 | -0.012054093 | 0.070535778 | -0.170893311 | 0.864307657 | 0.929553839 | 0.012054093 |
| ENSMUSG00000034917 | Tjp3 | 27375 | protein_coding | 3468.735765 | 0.012059028 | 0.088837248 | 0.135742927 | 0.892024511 | 0.945409888 | 0.012059028 |

Supplementary Table 1

| ensembl_gene_id | external_gene_name | entrezgene_id | gene_biotype | baseMean | log2FoldChange | lfcSE | stat | pvalue | padj | absLog2FC |
|---|---|---|---|---|---|---|---|---|---|---|
| ENSMUSG00000022808 | Snx4 | 69150 | protein_coding | 969.7104987 | -0.012110652 | 0.060079749 | -0.201576266 | 0.840248001 | 0.915836464 | 0.012110652 |
| ENSMUSG00000033128 | Gga1 | 106039 | protein_coding | 590.4566348 | -0.012111926 | 0.09061168 | -0.133668482 | 0.893664729 | 0.946554848 | 0.012111926 |
| ENSMUSG00000031590 | Frg1 | 14300 | protein_coding | 750.2303218 | 0.012122265 | 0.101686324 | 0.119212343 | 0.905107128 | 0.953132467 | 0.012122265 |
| ENSMUSG00000043085 | Tmem82 | 213989 | protein_coding | 156.8596502 | -0.012132573 | 0.139156992 | -0.087186227 | 0.930523487 | 0.965468682 | 0.012132573 |
| ENSMUSG00000010277 | Bltp2 | 72503 | protein_coding | 5087.430184 | -0.012151653 | 0.061017116 | -0.199151547 | 0.8421442 | 0.916587498 | 0.012151653 |
| ENSMUSG00000060036 | Rpl3 | 27367 | protein_coding | 9414.559125 | 0.012157461 | 0.092693019 | 0.131158328 | 0.895650061 | 0.947964261 | 0.012157461 |
| ENSMUSG00000021661 | Ankra2 | 68558 | protein_coding | 241.8551557 | -0.012163225 | 0.112055642 | -0.108546294 | 0.91356236 | 0.957282345 | 0.012163225 |
| ENSMUSG00000055560 | Zfp459 | 328274 | protein_coding | 57.59771762 | 0.012177016 | 0.159789607 | 0.076206559 | 0.939254764 | 0.970326602 | 0.012177016 |
| ENSMUSG00000034674 | Tdg | 21665 | protein_coding | 326.2730131 | 0.012212961 | 0.104573335 | 0.116788485 | 0.907027669 | 0.954095762 | 0.012212961 |
| ENSMUSG00000066232 | Ipo7 | 233726 | protein_coding | 3295.621222 | -0.012232892 | 0.065708224 | -0.186169881 | 0.852311543 | 0.921945066 | 0.012232892 |
| ENSMUSG00000036061 | Smug1 | 71726 | protein_coding | 295.07091 | 0.012245397 | 0.091509506 | 0.133815571 | 0.893548414 | 0.94649459 | 0.012245397 |
| ENSMUSG00000062054 | Iah1 | 67732 | protein_coding | 411.6388018 | 0.012264822 | 0.113026874 | 0.108512444 | 0.91358921 | 0.957282345 | 0.012264822 |
| ENSMUSG00000044707 | Ccnjl | 380694 | protein_coding | 57.20587255 | 0.012267351 | 0.237323616 | 0.051690394 | 0.958775391 | 0.979687864 | 0.012267351 |
| ENSMUSG00000032288 | Imp3 | 102462 | protein_coding | 516.0183045 | -0.012333713 | 0.106003175 | -0.116352294 | 0.907373342 | 0.954095762 | 0.012333713 |
| ENSMUSG00000025898 | Cwf19l2 | 244672 | protein_coding | 401.0069136 | -0.012341338 | 0.088041078 | -0.140177048 | 0.888520106 | 0.943426988 | 0.012341338 |
| ENSMUSG00000030177 | Ccdc77 | 67200 | protein_coding | 183.716387 | 0.012343896 | 0.107204027 | 0.115143958 | 0.908331018 | 0.954575761 | 0.012343896 |
| ENSMUSG00000031442 | Mcf2l | 17207 | protein_coding | 1364.581019 | 0.012415587 | 0.112866345 | 0.110002565 | 0.91240734 | 0.956925273 | 0.012415587 |
| ENSMUSG00000042102 | Dmgdh | 74129 | protein_coding | 12.46247902 | -0.012498347 | 0.379295226 | -0.0329515 | 0.973713264 | 0.988357435 | 0.012498347 |
| ENSMUSG00000031584 | Gsr | 14782 | protein_coding | 2031.796444 | -0.012524065 | 0.064301469 | -0.194771057 | 0.845572182 | 0.918484679 | 0.012524065 |
| ENSMUSG00000059923 | Grb2 | 14784 | protein_coding | 575.9256271 | 0.012525175 | 0.074600965 | 0.16789562 | 0.866665396 | 0.930916197 | 0.012525175 |
| ENSMUSG00000031751 | Amfr | 23802 | protein_coding | 2456.625778 | 0.01257911 | 0.096304954 | 0.130617479 | 0.896077915 | 0.948165086 | 0.01257911 |
| ENSMUSG00000038366 | Lasp1 | 16796 | protein_coding | 3590.311123 | 0.01259559 | 0.110155052 | 0.114344186 | 0.908964957 | 0.954949794 | 0.01259559 |
| ENSMUSG00000073139 | Tmem185a | 236848 | protein_coding | 185.417485 | -0.012629373 | 0.113148845 | -0.111617339 | 0.911126823 | 0.95627491 | 0.012629373 |
| ENSMUSG00000021611 | Tert | 21752 | protein_coding | 43.51752455 | 0.012633903 | 0.252533299 | 0.050028661 | 0.960099549 | 0.980600552 | 0.012633903 |
| ENSMUSG00000022340 | Sybu | 319613 | protein_coding | 160.3323522 | -0.012669637 | 0.296990136 | -0.042660127 | 0.965972464 | 0.984263652 | 0.012669637 |
| ENSMUSG00000026356 | Dars | 226414 | protein_coding | 1113.603052 | 0.01267481 | 0.087359103 | 0.145088596 | 0.884640922 | 0.941945358 | 0.01267481 |
| ENSMUSG00000008429 | Herpud2 | 80517 | protein_coding | 386.5346984 | -0.012708267 | 0.080132251 | -0.158591167 | 0.873990988 | 0.935546268 | 0.012708267 |
| ENSMUSG00000054302 | Eapp | 66266 | protein_coding | 494.6380817 | 0.012751212 | 0.099387224 | 0.128298302 | 0.897912909 | 0.949034957 | 0.012751212 |
| ENSMUSG00000058174 | Gm5148 | 381438 | protein_coding | 20.83051931 | 0.01276045 | 0.342440631 | 0.037263249 | 0.970275109 | 0.986376176 | 0.01276045 |
| ENSMUSG00000002031 | Ift46 | 76568 | protein_coding | 358.3074707 | 0.01276065 | 0.109326373 | 0.116720696 | 0.907081389 | 0.954095762 | 0.01276065 |
| ENSMUSG00000043733 | Ptpn11 | 19247 | protein_coding | 1188.820602 | -0.012795508 | 0.101077197 | -0.126591443 | 0.89926377 | 0.950084461 | 0.012795508 |
| ENSMUSG00000028124 | Gclm | 14630 | protein_coding | 1303.506152 | -0.012853775 | 0.107077887 | -0.120041362 | 0.904450383 | 0.952783446 | 0.012853775 |
| ENSMUSG00000028436 | Dcaf12 | 68970 | protein_coding | 767.6118612 | -0.012860086 | 0.077616647 | -0.165687212 | 0.86840311 | 0.932078223 | 0.012860086 |
| ENSMUSG00000015488 | Cacfd1 | 381356 | protein_coding | 418.3659582 | -0.012887117 | 0.083424806 | -0.154475836 | 0.877234562 | 0.937796695 | 0.012887117 |
| ENSMUSG00000027710 | Acad9 | 229211 | protein_coding | 662.4988065 | -0.012951549 | 0.081970887 | -0.158001811 | 0.874455372 | 0.935867429 | 0.012951549 |
| ENSMUSG00000026915 | Strbp | 20744 | protein_coding | 2614.660369 | 0.012986837 | 0.06218884 | 0.208829065 | 0.834581681 | 0.911868364 | 0.012986837 |
| ENSMUSG00000030264 | Thumpd3 | 14911 | protein_coding | 1100.342422 | 0.012999916 | 0.1021325 | 0.127284815 | 0.898714978 | 0.949630634 | 0.012999916 |
| ENSMUSG00000003363 | Pld3 | 18807 | protein_coding | 349.1104916 | 0.013005848 | 0.095237839 | 0.136561773 | 0.891377194 | 0.945137528 | 0.013005848 |

| ensembl_gene_id | external_gene_name | entrezgene_id | gene_biotype | baseMean | log2FoldChange | lfcSE | stat | pvalue | padj | absLog2FC |
|---|---|---|---|---|---|---|---|---|---|---|
| ENSMUSG00000114797 | Gm49336 | 71844 | protein_coding | 1149.767845 | 0.013008721 | 0.08181665 | 0.158998461 | 0.873670087 | 0.935328481 | 0.013008721 |
| ENSMUSG00000023953 | Polh | 80905 | protein_coding | 224.9955996 | -0.013038633 | 0.091960382 | -0.141785332 | 0.887249571 | 0.942706368 | 0.013038633 |
| ENSMUSG00000026988 | Wdsub1 | 72137 | protein_coding | 240.9608279 | 0.013053114 | 0.094772848 | 0.137730528 | 0.890453392 | 0.944864393 | 0.013053114 |
| ENSMUSG00000063320 | 1190007I07Rik | 544717 | protein_coding | 86.01998856 | 0.013059051 | 0.141526998 | 0.092272505 | 0.926481533 | 0.963740176 | 0.013059051 |
| ENSMUSG00000032601 | Prkar2a | 19087 | protein_coding | 2155.632686 | -0.013119993 | 0.069032885 | -0.190054242 | 0.849266626 | 0.92046263 | 0.013119993 |
| ENSMUSG00000026319 | Relch | 227446 | protein_coding | 1655.710836 | -0.013125101 | 0.060646934 | -0.216418209 | 0.828661774 | 0.909295436 | 0.013125101 |
| ENSMUSG00000034445 | Cyb561a3 | 225912 | protein_coding | 249.582866 | 0.013127499 | 0.104082579 | 0.126125804 | 0.899632342 | 0.950188262 | 0.013127499 |
| ENSMUSG00000051285 | Pcmtd1 | 319263 | protein_coding | 873.346132 | -0.01313416 | 0.100812061 | -0.130283614 | 0.896342043 | 0.948192606 | 0.01313416 |
| ENSMUSG00000066892 | Fbxl12 | 30843 | protein_coding | 91.72955661 | -0.013144657 | 0.175551029 | -0.074876558 | 0.940312928 | 0.970975969 | 0.013144657 |
| ENSMUSG00000001672 | Marveld3 | 73608 | protein_coding | 787.8629712 | -0.013151935 | 0.095009344 | -0.138427803 | 0.889902325 | 0.944516841 | 0.013151935 |
| ENSMUSG00000003421 | Nosip | 66394 | protein_coding | 378.0093101 | 0.013164122 | 0.096487403 | 0.136433587 | 0.891478523 | 0.945182049 | 0.013164122 |
| ENSMUSG00000025816 | Sec61a2 | 57743 | protein_coding | 132.1432606 | 0.013177484 | 0.126280931 | 0.104350548 | 0.916891166 | 0.958723986 | 0.013177484 |
| ENSMUSG00000047045 | Tmem164 | 209497 | protein_coding | 2017.108087 | -0.013195262 | 0.059281532 | -0.22258638 | 0.823857438 | 0.906994834 | 0.013195262 |
| ENSMUSG00000017861 | Mybl2 | 17865 | protein_coding | 74.57335199 | 0.013195451 | 0.17058114 | 0.077355863 | 0.938340452 | 0.969812604 | 0.013195451 |
| ENSMUSG00000070733 | Fryl | 72313 | protein_coding | 3235.546123 | 0.013196584 | 0.102768778 | 0.128410442 | 0.897824168 | 0.949004137 | 0.013196584 |
| ENSMUSG00000022756 | Slc7a4 | 224022 | protein_coding | 270.6617244 | -0.013204345 | 0.125138602 | -0.10551776 | 0.915964979 | 0.958406652 | 0.013204345 |
| ENSMUSG00000040158 | Tax1bp3 | 76281 | protein_coding | 641.7096057 | -0.013310311 | 0.107250728 | -0.124104626 | 0.901232436 | 0.950997373 | 0.013310311 |
| ENSMUSG00000000168 | Dlat | 235339 | protein_coding | 1324.470998 | 0.013328263 | 0.091322868 | 0.145946612 | 0.883963535 | 0.941599221 | 0.013328263 |
| ENSMUSG00000028647 | Mycbp | 56309 | protein_coding | 273.4130789 | 0.013331307 | 0.104012583 | 0.128170141 | 0.89801433 | 0.949079175 | 0.013331307 |
| ENSMUSG00000009687 | Fxyd5 | 18301 | protein_coding | 15.10916439 | -0.013360647 | 0.495958097 | -0.026939065 | 0.978508336 | 0.990328564 | 0.013360647 |
| ENSMUSG00000023873 | 1700010I14Rik | 66931 | protein_coding | 55.84713371 | 0.013376249 | 0.214983039 | 0.062220018 | 0.950387621 | 0.975560613 | 0.013376249 |
| ENSMUSG00000025728 | Pigq | 14755 | protein_coding | 741.4849814 | -0.013424659 | 0.064559297 | -0.207943084 | 0.835273408 | 0.912561551 | 0.013424659 |
| ENSMUSG00000032013 | Trim29 | 72169 | protein_coding | 29.82997719 | -0.013449051 | 0.595283373 | -0.022592688 | 0.981975177 | 0.992254051 | 0.013449051 |
| ENSMUSG00000038644 | Pold1 | 18971 | protein_coding | 278.3043331 | -0.013466731 | 0.152557439 | -0.088273188 | 0.929659549 | 0.965328339 | 0.013466731 |
| ENSMUSG00000040325 | Dcaf1 | 321006 | protein_coding | 946.8997568 | -0.013472633 | 0.07228443 | -0.186383607 | 0.852143947 | 0.921945066 | 0.013472633 |
| ENSMUSG00000029701 | Rbm28 | 68272 | protein_coding | 423.5576987 | 0.013473741 | 0.090383523 | 0.149072976 | 0.88149605 | 0.939979185 | 0.013473741 |
| ENSMUSG00000025362 | Rps26 | 27370 | protein_coding | 4781.379169 | -0.013496678 | 0.080727436 | -0.167188236 | 0.86722194 | 0.931241897 | 0.013496678 |
| ENSMUSG00000056888 | Glipr1 | 73690 | protein_coding | 12.32041624 | 0.013519629 | 0.461667359 | 0.029284351 | 0.976637807 | 0.989434359 | 0.013519629 |
| ENSMUSG00000053897 | Slc39a8 | 67547 | protein_coding | 936.3389036 | -0.013579446 | 0.111135799 | -0.122187866 | 0.902750236 | 0.951936826 | 0.013579446 |
| ENSMUSG00000049672 | Zbtb14 | 22666 | protein_coding | 219.8795193 | -0.01358063 | 0.103715764 | -0.130940849 | 0.8958221 | 0.948071543 | 0.01358063 |
| ENSMUSG00000044502 | Bod1 | 69556 | protein_coding | 478.5702896 | -0.013616754 | 0.094638021 | -0.143882484 | 0.885593268 | 0.94220607 | 0.013616754 |
| ENSMUSG00000039298 | Cdk5rap2 | 214444 | protein_coding | 380.84178 | 0.013624124 | 0.081869127 | 0.166413446 | 0.867831593 | 0.931770921 | 0.013624124 |
| ENSMUSG00000003527 | Ess2 | 27886 | protein_coding | 200.6133506 | 0.013658944 | 0.089040066 | 0.153402219 | 0.878081095 | 0.938098339 | 0.013658944 |
| ENSMUSG00000025939 | Ube2w | 66799 | protein_coding | 431.6634314 | 0.013661145 | 0.087000774 | 0.157023259 | 0.875226517 | 0.936491198 | 0.013661145 |
| ENSMUSG00000063236 | 1110038F14Rik | 117171 | protein_coding | 163.4532503 | 0.013684975 | 0.104114589 | 0.131441475 | 0.895426081 | 0.94788821 | 0.013684975 |
| ENSMUSG00000018417 | Myo1b | 17912 | protein_coding | 1266.219472 | -0.01369271 | 0.082484374 | -0.166003679 | 0.868154055 | 0.932000849 | 0.01369271 |
| ENSMUSG00000021188 | Trip11 | 109181 | protein_coding | 1606.683313 | -0.013706804 | 0.083772271 | -0.163619821 | 0.87003044 | 0.933188205 | 0.013706804 |
| ENSMUSG00000020716 | Nf1 | 18015 | protein_coding | 879.2852496 | -0.013756107 | 0.149227328 | -0.092182223 | 0.926553262 | 0.963740176 | 0.013756107 |

| ensembl_gene_id | external_gene_name | entrezgene_id | gene_biotype | baseMean | log2FoldChange | lfcSE | stat | pvalue | padj | absLog2FC |
|---|---|---|---|---|---|---|---|---|---|---|
| ENSMUSG00000039795 | Zfand1 | 66361 | protein_coding | 296.5884297 | -0.013802994 | 0.1278439 | -0.107967557 | 0.914021428 | 0.957546258 | 0.013802994 |
| ENSMUSG00000068580 | Zfyve19 | 72008 | protein_coding | 321.8304669 | -0.01380895 | 0.094897293 | -0.145514693 | 0.884304516 | 0.94174395 | 0.01380895 |
| ENSMUSG00000043131 | Mob1a | 232157 | protein_coding | 2439.234133 | -0.013809122 | 0.079270462 | -0.174202622 | 0.861706225 | 0.928009246 | 0.013809122 |
| ENSMUSG00000042178 | Armc5 | 233912 | protein_coding | 269.1669137 | -0.013809535 | 0.135447039 | -0.101955239 | 0.918792204 | 0.959766673 | 0.013809535 |
| ENSMUSG00000075254 | Heg1 | 77446 | protein_coding | 136.0953145 | 0.013848307 | 0.14703166 | 0.094185888 | 0.924961495 | 0.963179046 | 0.013848307 |
| ENSMUSG00000014932 | Yes1 | 22612 | protein_coding | 1392.313685 | -0.013871808 | 0.085461962 | -0.162315585 | 0.871057343 | 0.933600906 | 0.013871808 |
| ENSMUSG00000041040 | Fam117b | 72750 | protein_coding | 911.4378399 | -0.013920947 | 0.062964139 | -0.221093266 | 0.825019813 | 0.907748917 | 0.013920947 |
| ENSMUSG00000031434 | Morc4 | 75746 | protein_coding | 148.3718825 | 0.013955771 | 0.129862673 | 0.107465607 | 0.914419609 | 0.957711437 | 0.013955771 |
| ENSMUSG00000031066 | Usp11 | 236733 | protein_coding | 43.05394235 | -0.013961909 | 0.241102144 | -0.057908689 | 0.953821362 | 0.976611016 | 0.013961909 |
| ENSMUSG00000004895 | Prcc | 94315 | protein_coding | 572.1644986 | 0.013991508 | 0.07474869 | 0.187180643 | 0.851519001 | 0.921798204 | 0.013991508 |
| ENSMUSG00000079555 | Haus3 | 231123 | protein_coding | 179.9322496 | -0.014016721 | 0.091373885 | -0.153399635 | 0.878083132 | 0.938098339 | 0.014016721 |
| ENSMUSG00000045140 | Pigw | 70325 | protein_coding | 108.9197455 | 0.014117206 | 0.168087243 | 0.083987376 | 0.933066469 | 0.966437945 | 0.014117206 |
| ENSMUSG00000034484 | Snx2 | 67804 | protein_coding | 1251.392314 | -0.014196093 | 0.079436668 | -0.17870957 | 0.858165755 | 0.926176542 | 0.014196093 |
| ENSMUSG00000058600 | Rpl30 | 19946 | protein_coding | 3006.536251 | -0.014215002 | 0.090968704 | -0.156262554 | 0.875826071 | 0.936755148 | 0.014215002 |
| ENSMUSG00000033540 | Idua | 15932 | protein_coding | 121.3002733 | -0.014269048 | 0.132293903 | -0.107858697 | 0.914107781 | 0.957573741 | 0.014269048 |
| ENSMUSG00000044626 | Liph | 239759 | protein_coding | 919.1211544 | -0.014293061 | 0.133526034 | -0.107043255 | 0.914754664 | 0.957833724 | 0.014293061 |
| ENSMUSG00000058388 | Phtf1 | 18685 | protein_coding | 211.2880267 | -0.014334542 | 0.087809019 | -0.163246803 | 0.870324117 | 0.933211685 | 0.014334542 |
| ENSMUSG00000024887 | Asah2 | 54447 | protein_coding | 464.3839681 | -0.014384505 | 0.086657505 | -0.165992605 | 0.86816277 | 0.932000849 | 0.014384505 |
| ENSMUSG00000071281 | Zfp65 | 235907 | protein_coding | 325.4772216 | 0.014392321 | 0.117516256 | 0.122470892 | 0.902526097 | 0.951793254 | 0.014392321 |
| ENSMUSG00000031778 | Cx3cl1 | 20312 | protein_coding | 826.1459652 | -0.014408511 | 0.10522816 | -0.13692638 | 0.891088987 | 0.945137528 | 0.014408511 |
| ENSMUSG00000015468 | Notch4 | 18132 | protein_coding | 88.5005302 | -0.014409546 | 0.222869886 | -0.064654521 | 0.948449074 | 0.974764282 | 0.014409546 |
| ENSMUSG00000043241 | Upf2 | 326622 | protein_coding | 553.1192669 | 0.01444821 | 0.100382645 | 0.143931356 | 0.885554675 | 0.94220607 | 0.01444821 |
| ENSMUSG00000049950 | Rpp38 | 227522 | protein_coding | 72.76452397 | 0.014457685 | 0.168388236 | 0.085859234 | 0.931578318 | 0.965901539 | 0.014457685 |
| ENSMUSG00000021003 | Galc | 14420 | protein_coding | 111.9436823 | -0.014461642 | 0.155846975 | -0.092793856 | 0.926067332 | 0.963686666 | 0.014461642 |
| ENSMUSG00000039968 | Rsbn1l | 242860 | protein_coding | 347.6979224 | -0.014485516 | 0.150780599 | -0.096070158 | 0.923464852 | 0.96235224 | 0.014485516 |
| ENSMUSG00000025899 | Alkbh8 | 67667 | protein_coding | 318.3099454 | -0.014507303 | 0.106470177 | -0.136256961 | 0.891618147 | 0.945204249 | 0.014507303 |
| ENSMUSG00000032373 | Car12 | 76459 | protein_coding | 1334.912815 | 0.014511444 | 0.246557166 | 0.058856305 | 0.953066562 | 0.976295359 | 0.014511444 |
| ENSMUSG00000027519 | Rab22a | 19334 | protein_coding | 856.3726526 | -0.01451187 | 0.065953202 | -0.22003283 | 0.825845587 | 0.908116569 | 0.01451187 |
| ENSMUSG00000002210 | Smg9 | 71997 | protein_coding | 298.2353674 | -0.014543206 | 0.100295249 | -0.145003939 | 0.884707762 | 0.941953558 | 0.014543206 |
| ENSMUSG00000063849 | Ppcdc | 66812 | protein_coding | 220.3402811 | -0.014612571 | 0.097207066 | -0.150324169 | 0.880508866 | 0.939399502 | 0.014612571 |
| ENSMUSG00000024014 | Pim1 | 18712 | protein_coding | 190.5780458 | -0.014637227 | 0.117121611 | -0.124974605 | 0.900543655 | 0.95050289 | 0.014637227 |
| ENSMUSG00000035027 | Map2k2 | 26396 | protein_coding | 1147.213344 | 0.014653266 | 0.121535981 | 0.120567305 | 0.904033767 | 0.952525195 | 0.014653266 |
| ENSMUSG00000049354 | Dcaf7 | 71833 | protein_coding | 1363.079161 | 0.014682809 | 0.07002738 | 0.209672398 | 0.83392337 | 0.911286376 | 0.014682809 |
| ENSMUSG00000062647 | Rpl7a | 27176 | protein_coding | 10677.23386 | 0.014690329 | 0.086506571 | 0.169817492 | 0.86515367 | 0.930024138 | 0.014690329 |
| ENSMUSG00000062646 | Ganc | 76051 | protein_coding | 249.2875791 | -0.014699686 | 0.118161974 | -0.12440285 | 0.900996317 | 0.950842731 | 0.014699686 |
| ENSMUSG00000030835 | Nomo1 | 211548 | protein_coding | 1470.724698 | 0.014705922 | 0.117588066 | 0.125063052 | 0.900473634 | 0.95050289 | 0.014705922 |
| ENSMUSG00000015869 | Prpsap1 | 67763 | protein_coding | 1347.435164 | -0.014706538 | 0.156551283 | -0.093940708 | 0.925156256 | 0.963197355 | 0.014706538 |
| ENSMUSG00000026014 | Raph1 | 77300 | protein_coding | 717.9204875 | -0.014785101 | 0.160700859 | -0.092003869 | 0.926694966 | 0.963740176 | 0.014785101 |

| ensembl_gene_id | external_gene_name | entrezgene_id | gene_biotype | baseMean | log2FoldChange | lfcSE | stat | pvalue | padj | absLog2FC |
|---|---|---|---|---|---|---|---|---|---|---|
| ENSMUSG00000025795 | Rassf3 | 192678 | protein_coding | 810.6350122 | -0.014846496 | 0.074370043 | -0.199630063 | 0.841769914 | 0.916390622 | 0.014846496 |
| ENSMUSG00000019528 | Gyg | 27357 | protein_coding | 338.8052406 | -0.014860601 | 0.121100238 | -0.122713224 | 0.902334192 | 0.951737121 | 0.014860601 |
| ENSMUSG00000059939 | 9430015G10Rik | 230996 | protein_coding | 126.5213939 | 0.014902052 | 0.137291133 | 0.108543443 | 0.913564622 | 0.957282345 | 0.014902052 |
| ENSMUSG00000039244 | Ints15 | 231868 | protein_coding | 229.849179 | 0.014904179 | 0.140550171 | 0.106041699 | 0.915549268 | 0.95821748 | 0.014904179 |
| ENSMUSG00000029311 | Hsd17b11 | 114664 | protein_coding | 1107.415955 | 0.014954569 | 0.100853556 | 0.148280033 | 0.882121772 | 0.940394719 | 0.014954569 |
| ENSMUSG00000032193 | Ldlr | 16835 | protein_coding | 3146.501418 | -0.014969969 | 0.182472834 | -0.082039442 | 0.934615349 | 0.96756277 | 0.014969969 |
| ENSMUSG00000053286 | Trmt1l | 98685 | protein_coding | 461.0191739 | 0.014991152 | 0.089301677 | 0.167870892 | 0.86668485 | 0.930916197 | 0.014991152 |
| ENSMUSG00000050199 | Lgr4 | 107515 | protein_coding | 4096.321246 | -0.015047933 | 0.120942074 | -0.124422644 | 0.900980646 | 0.950842731 | 0.015047933 |
| ENSMUSG00000021532 | Fastkd3 | 69577 | protein_coding | 298.0492006 | -0.015051522 | 0.146006787 | -0.103087825 | 0.917893268 | 0.959208789 | 0.015051522 |
| ENSMUSG00000050708 | Ftl1 | 14325 | protein_coding | 6756.625343 | 0.015066427 | 0.136584177 | 0.110308732 | 0.912164533 | 0.956746934 | 0.015066427 |
| ENSMUSG00000038174 | Hycc2 | 213056 | protein_coding | 652.5391585 | -0.015087155 | 0.107166303 | -0.140782641 | 0.888041657 | 0.943170468 | 0.015087155 |
| ENSMUSG00000032841 | Prr5l | 72446 | protein_coding | 41.07083904 | -0.015116618 | 0.251616469 | -0.060078014 | 0.9520935 | 0.975865204 | 0.015116618 |
| ENSMUSG00000032297 | Celf6 | 76183 | protein_coding | 21.90268423 | 0.015162182 | 0.283600568 | 0.053463159 | 0.957362883 | 0.978558429 | 0.015162182 |
| ENSMUSG00000041658 | Rragb | 245670 | protein_coding | 9.076926523 | -0.01518225 | 0.497666596 | -0.03050687 | 0.975662814 | 0.989201611 | 0.01518225 |
| ENSMUSG00000028180 | Zranb2 | 53861 | protein_coding | 1015.464103 | 0.015196616 | 0.089091638 | 0.170572871 | 0.864559631 | 0.929652738 | 0.015196616 |
| ENSMUSG00000043059 | Zfp513 | 101023 | protein_coding | 179.874329 | -0.015202083 | 0.103993801 | -0.146182589 | 0.88377725 | 0.941592243 | 0.015202083 |
| ENSMUSG00000004846 | Plod3 | 26433 | protein_coding | 312.7085153 | -0.015244228 | 0.090604543 | -0.168250146 | 0.866386492 | 0.930721255 | 0.015244228 |
| ENSMUSG00000021694 | Ercc8 | 71991 | protein_coding | 122.6710212 | 0.015248793 | 0.143358989 | 0.106367888 | 0.91529047 | 0.95821748 | 0.015248793 |
| ENSMUSG00000024068 | Spast | 50850 | protein_coding | 686.4419918 | -0.015340612 | 0.083817159 | -0.183024722 | 0.854778621 | 0.923733895 | 0.015340612 |
| ENSMUSG00000003435 | Supt5 | 20924 | protein_coding | 1076.553157 | -0.015355133 | 0.069695728 | -0.220316702 | 0.825624514 | 0.907998844 | 0.015355133 |
| ENSMUSG00000063268 | Parp10 | 671535 | protein_coding | 404.8802391 | 0.015357255 | 0.093764646 | 0.163785131 | 0.869900297 | 0.933174295 | 0.015357255 |
| ENSMUSG00000023911 | Flywch2 | 76917 | protein_coding | 98.25772394 | -0.01535829 | 0.170242719 | -0.090214078 | 0.928117097 | 0.964521871 | 0.01535829 |
| ENSMUSG00000039217 | Il18 | 16173 | protein_coding | 1064.784217 | 0.015414312 | 0.223622177 | 0.06893016 | 0.945045212 | 0.973212655 | 0.015414312 |
| ENSMUSG00000028552 | Eps15 | 13858 | protein_coding | 763.6772973 | -0.015457084 | 0.055457022 | -0.27872185 | 0.780458293 | 0.880641803 | 0.015457084 |
| ENSMUSG00000113918 | Gm6566 | 625237 | lncRNA | 8.839582564 | 0.015475578 | 0.443287659 | 0.034910915 | 0.972150777 | 0.987518805 | 0.015475578 |
| ENSMUSG00000025066 | Sfr1 | 67788 | protein_coding | 711.4978091 | 0.015487725 | 0.101897439 | 0.151993269 | 0.879192245 | 0.938779518 | 0.015487725 |
| ENSMUSG00000021552 | Gkap1 | 56278 | protein_coding | 52.83731658 | -0.015499012 | 0.200805525 | -0.077184193 | 0.938477017 | 0.96983468 | 0.015499012 |
| ENSMUSG00000060477 | Irak2 | 108960 | protein_coding | 387.4356227 | -0.01552467 | 0.095952816 | -0.16179483 | 0.871467424 | 0.933849107 | 0.01552467 |
| ENSMUSG00000022111 | Uchl3 | 50933 | protein_coding | 322.7007318 | 0.015530441 | 0.121390559 | 0.127937796 | 0.8981982 | 0.949210518 | 0.015530441 |
| ENSMUSG00000077450 | Rab11b | 19326 | protein_coding | 655.2540601 | 0.015553404 | 0.072547356 | 0.214389667 | 0.830243199 | 0.9095933 | 0.015553404 |
| ENSMUSG00000035085 | 1700020L24Rik | 66330 | protein_coding | 24.32286463 | -0.015553686 | 0.355820977 | -0.043712112 | 0.965133884 | 0.983535029 | 0.015553686 |
| ENSMUSG00000038970 | Lmtk2 | 231876 | protein_coding | 1429.501548 | 0.015574599 | 0.11476971 | 0.135703044 | 0.892056041 | 0.945409888 | 0.015574599 |
| ENSMUSG00000032698 | Lmo2 | 16909 | protein_coding | 8.221749052 | -0.015588737 | 0.423580382 | -0.03680231 | 0.970642632 | 0.98667093 | 0.015588737 |
| ENSMUSG00000041859 | Mcm3 | 17215 | protein_coding | 356.74188 | 0.015617081 | 0.177607593 | 0.087930256 | 0.929932109 | 0.965328339 | 0.015617081 |
| ENSMUSG00000031819 | Emc8 | 18117 | protein_coding | 305.2024096 | -0.015653476 | 0.093002636 | -0.16831217 | 0.8663377 | 0.930721255 | 0.015653476 |
| ENSMUSG00000051232 | Tmem199 | 195040 | protein_coding | 159.9956035 | 0.015728124 | 0.101846455 | 0.154429759 | 0.877270891 | 0.937796695 | 0.015728124 |
| ENSMUSG00000030672 | Mylpf | 17907 | protein_coding | 17.87407845 | 0.015748192 | 0.27795195 | 0.056657964 | 0.95481766 | 0.977022722 | 0.015748192 |
| ENSMUSG00000025261 | Huwe1 | 59026 | protein_coding | 3522.211056 | 0.015796634 | 0.157533883 | 0.100274518 | 0.920126388 | 0.96046285 | 0.015796634 |

| ensembl_gene_id | external_gene_name | entrezgene_id | gene_biotype | baseMean | log2FoldChange | lfcSE | stat | pvalue | padj | absLog2FC |
|---|---|---|---|---|---|---|---|---|---|---|
| ENSMUSG00000027669 | Gnb4 | 14696 | protein_coding | 24.1953286 | -0.015806725 | 0.330669657 | -0.047802164 | 0.961873912 | 0.981468781 | 0.015806725 |
| ENSMUSG00000036879 | Phkb | 102093 | protein_coding | 949.8247982 | 0.015815953 | 0.079464766 | 0.199031011 | 0.842238487 | 0.916587498 | 0.015815953 |
| ENSMUSG00000024639 | Gnaq | 14682 | protein_coding | 600.5742232 | -0.015830135 | 0.083848219 | -0.188795121 | 0.850253396 | 0.920905487 | 0.015830135 |
| ENSMUSG00000015357 | Clpx | 270166 | protein_coding | 789.152537 | -0.015842892 | 0.083544128 | -0.18963501 | 0.849595151 | 0.920572697 | 0.015842892 |
| ENSMUSG00000025745 | Hadha | 97212 | protein_coding | 4116.213822 | -0.015865579 | 0.080429769 | -0.197260031 | 0.843624056 | 0.917176415 | 0.015865579 |
| ENSMUSG00000035620 | Ric8b | 237422 | protein_coding | 376.6913596 | 0.015873388 | 0.083729821 | 0.189578666 | 0.849639306 | 0.920572697 | 0.015873388 |
| ENSMUSG00000032097 | Ddx6 | 13209 | protein_coding | 2253.231152 | -0.015901509 | 0.13246497 | -0.120043126 | 0.904448985 | 0.952783446 | 0.015901509 |
| ENSMUSG00000031255 | Sytl4 | 27359 | protein_coding | 457.1567788 | 0.015943945 | 0.11160548 | 0.142859879 | 0.886400845 | 0.94220607 | 0.015943945 |
| ENSMUSG00000026456 | Cyb5r1 | 72017 | protein_coding | 162.8239175 | -0.015946619 | 0.1159222 | -0.137563117 | 0.890585706 | 0.944864393 | 0.015946619 |
| ENSMUSG00000030560 | Ctsc | 13032 | protein_coding | 1258.434005 | 0.015964098 | 0.134074067 | 0.119069245 | 0.905220497 | 0.953132467 | 0.015964098 |
| ENSMUSG00000023004 | Tuba1b | 22143 | protein_coding | 545.5799429 | 0.016040115 | 0.13242227 | 0.121128529 | 0.903589233 | 0.952386474 | 0.016040115 |
| ENSMUSG00000063488 | Zkscan7 | 382118 | protein_coding | 235.2685935 | 0.016165828 | 0.145551116 | 0.111066327 | 0.911563751 | 0.956544398 | 0.016165828 |
| ENSMUSG00000051768 | Xrcc1 | 22594 | protein_coding | 296.7952543 | 0.01617012 | 0.094834108 | 0.170509541 | 0.864609431 | 0.929652738 | 0.01617012 |
| ENSMUSG00000020349 | Ppp2ca | 19052 | protein_coding | 4284.438111 | 0.016183643 | 0.092770494 | 0.174448169 | 0.861513262 | 0.927926916 | 0.016183643 |
| ENSMUSG00000038227 | Hoxa9 | 15405 | protein_coding | 378.9723395 | 0.016195725 | 0.232798817 | 0.069569619 | 0.944536219 | 0.973003033 | 0.016195725 |
| ENSMUSG00000039774 | Galnt12 | 230145 | protein_coding | 1912.87031 | 0.016276678 | 0.102327236 | 0.159064959 | 0.873617695 | 0.935328481 | 0.016276678 |
| ENSMUSG00000020648 | Dus4l | 71916 | protein_coding | 56.39256544 | -0.016323669 | 0.171925015 | -0.094946447 | 0.924357364 | 0.962801732 | 0.016323669 |
| ENSMUSG00000065089 | Gm26493 | 115487044 | snoRNA | 10.15823673 | 0.016356276 | 0.38495091 | 0.042489251 | 0.96610868 | 0.984289673 | 0.016356276 |
| ENSMUSG00000070730 | Rmdn3 | 67809 | protein_coding | 767.4196552 | 0.016359136 | 0.087569789 | 0.186812561 | 0.851807598 | 0.921803579 | 0.016359136 |
| ENSMUSG00000074238 | Ap1ar | 211556 | protein_coding | 1306.869612 | -0.016382839 | 0.071415283 | -0.229402423 | 0.818556153 | 0.903594317 | 0.016382839 |
| ENSMUSG00000019818 | Cd164 | 53599 | protein_coding | 4276.422585 | 0.016408948 | 0.056617593 | 0.289820655 | 0.771953445 | 0.876072282 | 0.016408948 |
| ENSMUSG00000036721 | Zscan12 | 22758 | protein_coding | 116.0394732 | -0.016415326 | 0.138218842 | -0.1187633 | 0.905462885 | 0.953282625 | 0.016415326 |
| ENSMUSG00000008398 | Elk3 | 13713 | protein_coding | 27.93787771 | 0.016512727 | 0.273464139 | 0.060383521 | 0.951850183 | 0.975678597 | 0.016512727 |
| ENSMUSG00000027498 | Cstf1 | 67337 | protein_coding | 328.4109081 | -0.016521336 | 0.088806755 | -0.186036929 | 0.852415801 | 0.921945066 | 0.016521336 |
| ENSMUSG00000042043 | Tbca | 21371 | protein_coding | 798.7569272 | 0.01652403 | 0.103996706 | 0.158889933 | 0.873755592 | 0.935357152 | 0.01652403 |
| ENSMUSG00000013766 | Ly6g6e | 70274 | protein_coding | 112.8916289 | 0.016527968 | 0.262094422 | 0.063061121 | 0.949717833 | 0.975281565 | 0.016527968 |
| ENSMUSG00000027770 | Dhx36 | 72162 | protein_coding | 914.7914697 | -0.016530208 | 0.058949618 | -0.280412466 | 0.779161074 | 0.879926091 | 0.016530208 |
| ENSMUSG00000022350 | Washc5 | 223593 | protein_coding | 763.4977663 | 0.016566196 | 0.068701811 | 0.241131862 | 0.809452919 | 0.898527152 | 0.016566196 |
| ENSMUSG00000038578 | Susd1 | 634731 | protein_coding | 23.80125018 | 0.016635488 | 0.253879751 | 0.065525068 | 0.947755948 | 0.974338841 | 0.016635488 |
| ENSMUSG00000037262 | Kin | 16588 | protein_coding | 193.5506616 | -0.016637447 | 0.110886119 | -0.150040842 | 0.880732393 | 0.939542265 | 0.016637447 |
| ENSMUSG00000035069 | Oma1 | 67013 | protein_coding | 666.7901107 | 0.016640948 | 0.13591632 | 0.122435246 | 0.902554327 | 0.951793254 | 0.016640948 |
| ENSMUSG00000025907 | Rb1cc1 | 12421 | protein_coding | 1681.777657 | 0.016645122 | 0.086615377 | 0.192172828 | 0.847606829 | 0.919539679 | 0.016645122 |
| ENSMUSG00000028488 | Sh3gl2 | 20404 | protein_coding | 168.5962526 | -0.016645182 | 0.171759487 | -0.096909828 | 0.922798004 | 0.961877857 | 0.016645182 |
| ENSMUSG00000031065 | Cdk16 | 18555 | protein_coding | 1301.467647 | 0.016645971 | 0.069868107 | 0.238248495 | 0.811688366 | 0.899754492 | 0.016645971 |
| ENSMUSG00000005362 | Crbn | 58799 | protein_coding | 454.8217538 | 0.016671393 | 0.094949627 | 0.175581442 | 0.860622783 | 0.927608253 | 0.016671393 |
| ENSMUSG00000055835 | Zfp1 | 22640 | protein_coding | 94.83711842 | 0.016762783 | 0.14889091 | 0.112584325 | 0.910360113 | 0.955759393 | 0.016762783 |
| ENSMUSG00000021156 | Zmynd11 | 66505 | protein_coding | 802.2056271 | 0.016792199 | 0.057977558 | 0.289632736 | 0.77209722 | 0.876172816 | 0.016792199 |
| ENSMUSG00000020737 | Jpt1 | 15374 | protein_coding | 2112.761388 | -0.016813919 | 0.094885459 | -0.177202269 | 0.859349514 | 0.926766895 | 0.016813919 |

Supplementary Table 1

| ensembl_gene_id | external_gene_name | entrezgene_id | gene_biotype | baseMean | log2FoldChange | lfcSE | stat | pvalue | padj | absLog2FC |
|---|---|---|---|---|---|---|---|---|---|---|
| ENSMUSG00000000732 | Icosl | 50723 | protein_coding | 81.39390056 | -0.016845175 | 0.277648587 | -0.060670847 | 0.95162135 | 0.975632393 | 0.016845175 |
| ENSMUSG00000033545 | Znrf1 | 170737 | protein_coding | 182.7642726 | -0.016856552 | 0.127503765 | -0.13220435 | 0.894822661 | 0.947466286 | 0.016856552 |
| ENSMUSG00000003033 | Ap1m1 | 11767 | protein_coding | 164.0537852 | -0.016884414 | 0.140785181 | -0.119930336 | 0.904538333 | 0.952813079 | 0.016884414 |
| ENSMUSG00000028703 | Lrrc41 | 230654 | protein_coding | 391.0434047 | 0.016895642 | 0.085659939 | 0.19724088 | 0.843639042 | 0.917176415 | 0.016895642 |
| ENSMUSG00000053477 | Tcf4 | 21413 | protein_coding | 440.3609216 | 0.0169014 | 0.16744458 | 0.100937277 | 0.919600252 | 0.960107027 | 0.0169014 |
| ENSMUSG00000025395 | Prim1 | 19075 | protein_coding | 224.5508366 | 0.016924038 | 0.132570737 | 0.127660434 | 0.898417703 | 0.949379498 | 0.016924038 |
| ENSMUSG00000074899 | Sptbn5 | 640524 | protein_coding | 34.60485605 | -0.016958938 | 0.414043518 | -0.040959314 | 0.967328331 | 0.985141027 | 0.016958938 |
| ENSMUSG00000027430 | Dtd1 | 66044 | protein_coding | 120.7521396 | 0.017023532 | 0.114822362 | 0.148259729 | 0.882137796 | 0.940394719 | 0.017023532 |
| ENSMUSG00000056014 | A430033K04Rik | 243308 | protein_coding | 172.9701165 | -0.017076553 | 0.164679462 | -0.103695704 | 0.917410836 | 0.959078485 | 0.017076553 |
| ENSMUSG00000117431 | 8430430B14Rik | 78148 | lncRNA | 15.63677579 | -0.017085443 | 0.276208965 | -0.061856946 | 0.950676754 | 0.975605911 | 0.017085443 |
| ENSMUSG00000036197 | Gxylt1 | 223827 | protein_coding | 640.9157997 | -0.017104497 | 0.136045624 | -0.125726185 | 0.899948674 | 0.950214744 | 0.017104497 |
| ENSMUSG00000020182 | Ddc | 13195 | protein_coding | 973.9420003 | 0.017135502 | 0.159566591 | 0.107387779 | 0.914481349 | 0.957713126 | 0.017135502 |
| ENSMUSG00000031505 | Naxd | 69225 | protein_coding | 709.7262284 | -0.017156563 | 0.076684747 | -0.223728497 | 0.822968571 | 0.906288656 | 0.017156563 |
| ENSMUSG00000067288 | Rps28 | 54127 | protein_coding | 4201.633182 | 0.017160505 | 0.110204383 | 0.15571527 | 0.87625746 | 0.937090696 | 0.017160505 |
| ENSMUSG00000025381 | Cnpy2 | 56530 | protein_coding | 932.018517 | 0.017169403 | 0.095738741 | 0.179335997 | 0.857673884 | 0.925781659 | 0.017169403 |
| ENSMUSG00000022770 | Dlg1 | 13383 | protein_coding | 1323.627703 | 0.017173812 | 0.057691016 | 0.297686071 | 0.765942775 | 0.872166845 | 0.017173812 |
| ENSMUSG00000006169 | Clint1 | 216705 | protein_coding | 3362.058645 | 0.017201455 | 0.060795293 | 0.282940576 | 0.77722239 | 0.878983118 | 0.017201455 |
| ENSMUSG00000027799 | Nbea | 26422 | protein_coding | 356.3109504 | -0.017226478 | 0.158138776 | -0.108932666 | 0.913255897 | 0.957282345 | 0.017226478 |
| ENSMUSG00000041236 | Vps41 | 218035 | protein_coding | 1104.17431 | -0.017234041 | 0.068352229 | -0.252135758 | 0.800936132 | 0.893243993 | 0.017234041 |
| ENSMUSG00000006941 | Eif1b | 68969 | protein_coding | 197.9397001 | 0.017238279 | 0.154150063 | 0.111827909 | 0.910959858 | 0.956162677 | 0.017238279 |
| ENSMUSG00000025085 | Ablim1 | 226251 | protein_coding | 1692.564772 | -0.017243945 | 0.091079886 | -0.189327691 | 0.849835993 | 0.920641209 | 0.017243945 |
| ENSMUSG00000055435 | Maf | 17132 | protein_coding | 88.90706767 | -0.017246609 | 0.312439071 | -0.05519991 | 0.9559792 | 0.977771573 | 0.017246609 |
| ENSMUSG00000023286 | Ube2j2 | 140499 | protein_coding | 671.3316731 | 0.017246715 | 0.107634507 | 0.16023407 | 0.872696692 | 0.934663321 | 0.017246715 |
| ENSMUSG00000040738 | Ints8 | 72656 | protein_coding | 690.5988168 | 0.017376084 | 0.088750612 | 0.195785515 | 0.844778048 | 0.918096667 | 0.017376084 |
| ENSMUSG00000074715 | Ccl28 | 56838 | protein_coding | 2052.031714 | -0.017386393 | 0.181878669 | -0.095593359 | 0.92384354 | 0.962581272 | 0.017386393 |
| ENSMUSG00000031578 | Mak16 | 67920 | protein_coding | 261.9541262 | 0.017386424 | 0.147863034 | 0.117584659 | 0.906396761 | 0.953887519 | 0.017386424 |
| ENSMUSG00000030337 | Vamp1 | 22317 | protein_coding | 122.9043909 | -0.017387076 | 0.175072491 | -0.099313583 | 0.920889294 | 0.960868211 | 0.017387076 |
| ENSMUSG00000024234 | Mtpap | 67440 | protein_coding | 410.2624232 | -0.017421585 | 0.07954843 | -0.219006026 | 0.826645354 | 0.908412958 | 0.017421585 |
| ENSMUSG00000038900 | Rpl12 | 269261 | protein_coding | 5882.305675 | -0.01745806 | 0.105697927 | -0.165169372 | 0.868810672 | 0.93225656 | 0.01745806 |
| ENSMUSG00000071649 | B3gat3 | 72727 | protein_coding | 346.6180567 | -0.01746425 | 0.105410829 | -0.165677951 | 0.868410398 | 0.932078223 | 0.01746425 |
| ENSMUSG00000022667 | Cd200r1 | 57781 | protein_coding | 9.746958877 | -0.017479898 | 0.451313181 | -0.038731193 | 0.969104703 | 0.985941474 | 0.017479898 |
| ENSMUSG00000034930 | Rtkn | 20166 | protein_coding | 192.319843 | 0.017495312 | 0.113410046 | 0.154265981 | 0.877400019 | 0.937871788 | 0.017495312 |
| ENSMUSG00000002661 | Alkbh7 | 66400 | protein_coding | 180.5325328 | 0.017500662 | 0.154044553 | 0.113607798 | 0.909548706 | 0.955345615 | 0.017500662 |
| ENSMUSG00000059851 | Kmt5c | 232811 | protein_coding | 430.3036908 | -0.017519547 | 0.102766315 | -0.170479472 | 0.864633077 | 0.929652738 | 0.017519547 |
| ENSMUSG00000039782 | Cpeb2 | 231207 | protein_coding | 304.9483525 | -0.017542618 | 0.151984526 | -0.11542371 | 0.908109286 | 0.954554525 | 0.017542618 |
| ENSMUSG00000038525 | Armc10 | 67211 | protein_coding | 599.4142004 | 0.017589099 | 0.07150487 | 0.245984627 | 0.805694133 | 0.895535555 | 0.017589099 |
| ENSMUSG00000044768 | Macir | 52392 | protein_coding | 930.1805509 | 0.017614684 | 0.084826297 | 0.20765593 | 0.83549763 | 0.912743913 | 0.017614684 |
| ENSMUSG00000041263 | Rusc1 | 72296 | protein_coding | 99.71714657 | -0.017629389 | 0.119478898 | -0.14755232 | 0.882696086 | 0.940754985 | 0.017629389 |

| ensembl_gene_id | external_gene_name | entrezgene_id | gene_biotype | baseMean | log2FoldChange | lfcSE | stat | pvalue | padj | absLog2FC |
|---|---|---|---|---|---|---|---|---|---|---|
| ENSMUSG00000026159 | Agfg1 | 15463 | protein_coding | 776.8752106 | -0.017635673 | 0.09230292 | -0.191062998 | 0.848476237 | 0.919988003 | 0.017635673 |
| ENSMUSG00000025656 | Arhgef9 | 236915 | protein_coding | 18.27352239 | -0.017648315 | 0.28214155 | -0.062551279 | 0.950123827 | 0.975404564 | 0.017648315 |
| ENSMUSG00000038845 | Phb | 18673 | protein_coding | 764.8472253 | -0.017674288 | 0.091822159 | -0.192483913 | 0.847363168 | 0.919400582 | 0.017674288 |
| ENSMUSG00000041688 | Amot | 27494 | protein_coding | 631.1131704 | 0.017729099 | 0.102760744 | 0.172527932 | 0.863022504 | 0.928838863 | 0.017729099 |
| ENSMUSG00000069184 | Zfp72 | 238722 | protein_coding | 85.64594912 | -0.017733955 | 0.214642074 | -0.082621059 | 0.934152856 | 0.967374468 | 0.017733955 |
| ENSMUSG00000023075 | Akirin1 | 68050 | protein_coding | 966.1682761 | -0.017747839 | 0.160543384 | -0.110548554 | 0.911974345 | 0.956746934 | 0.017747839 |
| ENSMUSG00000061950 | Ppp4r1 | 70351 | protein_coding | 1029.543228 | -0.017797041 | 0.14662576 | -0.121377318 | 0.903392182 | 0.952386474 | 0.017797041 |
| ENSMUSG00000001420 | Tmem79 | 71913 | protein_coding | 237.2406257 | -0.017858569 | 0.116812597 | -0.152882219 | 0.878491158 | 0.93834699 | 0.017858569 |
| ENSMUSG00000011254 | Thg1l | 66628 | protein_coding | 132.8073206 | 0.017880917 | 0.128038846 | 0.139652281 | 0.888934731 | 0.943741412 | 0.017880917 |
| ENSMUSG00000016757 | Ttll12 | 223723 | protein_coding | 294.9745514 | 0.017947632 | 0.101470712 | 0.176875003 | 0.859606574 | 0.926876084 | 0.017947632 |
| ENSMUSG00000020794 | Ube2g1 | 67128 | protein_coding | 811.8683051 | -0.017963337 | 0.060179388 | -0.298496506 | 0.765324242 | 0.871716571 | 0.017963337 |
| ENSMUSG00000062014 | Gmfb | 63985 | protein_coding | 1584.223801 | 0.017983955 | 0.089482032 | 0.200978393 | 0.840715469 | 0.915931326 | 0.017983955 |
| ENSMUSG00000036992 | Nxt1 | 56488 | protein_coding | 146.8922367 | -0.018008011 | 0.132597174 | -0.135809916 | 0.891971552 | 0.945409888 | 0.018008011 |
| ENSMUSG00000073725 | Lmbrd1 | 68421 | protein_coding | 901.1614583 | -0.018075185 | 0.061416756 | -0.294303811 | 0.768525761 | 0.873738638 | 0.018075185 |
| ENSMUSG00000001323 | Srr | 27364 | protein_coding | 242.5145691 | 0.01807918 | 0.101232832 | 0.178590081 | 0.858259584 | 0.926176542 | 0.01807918 |
| ENSMUSG00000061689 | Dlgap4 | 228836 | protein_coding | 297.3614735 | -0.018138835 | 0.100363251 | -0.180731839 | 0.856578066 | 0.92486276 | 0.018138835 |
| ENSMUSG00000041695 | Kcnj2 | 16518 | protein_coding | 58.1182018 | 0.01814532 | 0.296732086 | 0.061150516 | 0.951239338 | 0.975614007 | 0.01814532 |
| ENSMUSG00000030534 | Vps33b | 233405 | protein_coding | 266.3560688 | -0.018157183 | 0.084271814 | -0.215459743 | 0.829408895 | 0.909405742 | 0.018157183 |
| ENSMUSG00000020705 | Ddx42 | 72047 | protein_coding | 1097.143717 | -0.018163178 | 0.059338252 | -0.306095604 | 0.759531855 | 0.868410263 | 0.018163178 |
| ENSMUSG00000017288 | Vps53 | 68299 | protein_coding | 684.1698407 | -0.018209751 | 0.073128813 | -0.249009251 | 0.803353626 | 0.894249532 | 0.018209751 |
| ENSMUSG00000021270 | Hsp90aa1 | 15519 | protein_coding | 6026.322644 | 0.018209818 | 0.18979198 | 0.095946194 | 0.923563307 | 0.96235224 | 0.018209818 |
| ENSMUSG00000057863 | Rpl36 | 54217 | protein_coding | 2860.445016 | -0.018274758 | 0.102914267 | -0.177572638 | 0.859058615 | 0.92672439 | 0.018274758 |
| ENSMUSG00000019761 | Krt10 | 16661 | protein_coding | 27.54049927 | 0.018298037 | 0.248608611 | 0.073601783 | 0.941327252 | 0.971456555 | 0.018298037 |
| ENSMUSG00000026500 | Cox20 | 66359 | protein_coding | 9.066931032 | 0.018306503 | 0.48093851 | 0.038064123 | 0.969636556 | 0.986041695 | 0.018306503 |
| ENSMUSG00000056708 | Ier5 | 15939 | protein_coding | 654.6729378 | 0.018324177 | 0.094095535 | 0.194740133 | 0.845596392 | 0.918484679 | 0.018324177 |
| ENSMUSG00000079083 | Jrkl | 77532 | protein_coding | 185.4678435 | -0.018369103 | 0.123392406 | -0.148867368 | 0.881658291 | 0.940089253 | 0.018369103 |
| ENSMUSG00000042901 | Aida | 108909 | protein_coding | 321.6361156 | 0.018382436 | 0.073151561 | 0.251292465 | 0.801588001 | 0.893676478 | 0.018382436 |
| ENSMUSG00000022376 | Adcy8 | 11514 | protein_coding | 17.36482127 | 0.018388497 | 0.45557033 | 0.040363685 | 0.967803182 | 0.985339694 | 0.018388497 |
| ENSMUSG00000075054 | Yae1d1 | 67008 | protein_coding | 229.4870576 | -0.018460784 | 0.095706096 | -0.192890366 | 0.84704483 | 0.919400582 | 0.018460784 |
| ENSMUSG00000019795 | Pcmt1 | 18537 | protein_coding | 591.0692658 | 0.018491754 | 0.09579622 | 0.193032184 | 0.846933763 | 0.919400582 | 0.018491754 |
| ENSMUSG00000018481 | Appbp2 | 66884 | protein_coding | 1262.447411 | 0.018513567 | 0.068426371 | 0.270561875 | 0.786728022 | 0.884458731 | 0.018513567 |
| ENSMUSG00000055114 | Anxa13 | 69787 | protein_coding | 223.4047249 | -0.01852558 | 0.199951564 | -0.092650337 | 0.926181352 | 0.96369327 | 0.01852558 |
| ENSMUSG00000031592 | Pcm1 | 18536 | protein_coding | 1307.101307 | -0.018557593 | 0.069509911 | -0.266977652 | 0.789486368 | 0.885815145 | 0.018557593 |
| ENSMUSG00000009907 | Vps4b | 20479 | protein_coding | 2320.383015 | 0.018562757 | 0.062386973 | 0.297542199 | 0.766052596 | 0.872171962 | 0.018562757 |
| ENSMUSG00000001942 | Siae | 22619 | protein_coding | 430.8785504 | -0.018562876 | 0.155794919 | -0.119149434 | 0.905156967 | 0.953132467 | 0.018562876 |
| ENSMUSG00000029518 | Rab35 | 77407 | protein_coding | 808.2422318 | 0.018615539 | 0.110115155 | 0.169055197 | 0.865753225 | 0.930354701 | 0.018615539 |
| ENSMUSG00000024136 | Dnase1l2 | 66705 | protein_coding | 54.22496037 | 0.018686722 | 0.218042398 | 0.08570224 | 0.931703121 | 0.96596806 | 0.018686722 |
| ENSMUSG00000021930 | Sprydd7 | 66674 | protein_coding | 457.5691211 | 0.0186877 | 0.131353484 | 0.142270302 | 0.886866504 | 0.942483462 | 0.0186877 |

| ensembl_gene_id | external_gene_name | entrezgene_id | gene_biotype | baseMean | log2FoldChange | lfcSE | stat | pvalue | padj | absLog2FC |
|---|---|---|---|---|---|---|---|---|---|---|
| ENSMUSG00000025218 | Poll | 56626 | protein_coding | 122.0142476 | 0.018732001 | 0.126392445 | 0.148205068 | 0.882180933 | 0.940394719 | 0.018732001 |
| ENSMUSG00000039001 | Rps21 | 66481 | protein_coding | 4026.18803 | 0.018738061 | 0.1198504 | 0.15634542 | 0.875760757 | 0.936748192 | 0.018738061 |
| ENSMUSG00000004266 | Ptpn6 | 15170 | protein_coding | 352.6788672 | -0.01874438 | 0.082413072 | -0.227444262 | 0.820078306 | 0.904579103 | 0.01874438 |
| ENSMUSG00000028821 | Syf2 | 68592 | protein_coding | 619.7192718 | 0.01882162 | 0.087781793 | 0.214413708 | 0.830224453 | 0.9095933 | 0.01882162 |
| ENSMUSG00000051864 | Tbc1d22a | 223754 | protein_coding | 725.2085007 | -0.018824247 | 0.124992007 | -0.150603605 | 0.880288418 | 0.939320286 | 0.018824247 |
| ENSMUSG00000028653 | Trit1 | 66966 | protein_coding | 183.1929988 | 0.018914 | 0.108397039 | 0.174488158 | 0.861481837 | 0.927926916 | 0.018914 |
| ENSMUSG00000022323 | Rida | 15473 | protein_coding | 312.0216172 | 0.018938247 | 0.12109036 | 0.156397641 | 0.875719596 | 0.936748192 | 0.018938247 |
| ENSMUSG00000031146 | Plp2 | 18824 | protein_coding | 994.2568833 | 0.018943162 | 0.125147742 | 0.151366391 | 0.8796867 | 0.939055677 | 0.018943162 |
| ENSMUSG00000022307 | Oxr1 | 170719 | protein_coding | 668.5561279 | -0.018960429 | 0.076215966 | -0.248772407 | 0.803536836 | 0.89432847 | 0.018960429 |
| ENSMUSG00000001280 | Sp1 | 20683 | protein_coding | 1769.432828 | -0.018974869 | 0.066898756 | -0.2836356 | 0.776689653 | 0.878567771 | 0.018974869 |
| ENSMUSG00000060679 | Mrps9 | 69527 | protein_coding | 602.1595108 | 0.019062805 | 0.077178705 | 0.246995661 | 0.804911582 | 0.894920542 | 0.019062805 |
| ENSMUSG00000046756 | Mrps7 | 50529 | protein_coding | 545.6305658 | 0.019097164 | 0.116981156 | 0.163249919 | 0.870321664 | 0.933211685 | 0.019097164 |
| ENSMUSG00000025511 | Tspan4 | 64540 | protein_coding | 17.6852223 | 0.019125293 | 0.300839128 | 0.063573158 | 0.949310106 | 0.975117538 | 0.019125293 |
| ENSMUSG00000041716 | Gm20604 | 100859931 | protein_coding | 60.71678525 | -0.019129216 | 0.186135997 | -0.102770108 | 0.91814543 | 0.959342719 | 0.019129216 |
| ENSMUSG00000024317 | Rnf138 | 56515 | protein_coding | 346.2850109 | 0.019190442 | 0.111516664 | 0.172085869 | 0.863370022 | 0.929084741 | 0.019190442 |
| ENSMUSG00000079164 | Tlr5 | 53791 | protein_coding | 50.02563357 | -0.019195429 | 0.286610148 | -0.066974002 | 0.9466024 | 0.973812302 | 0.019195429 |
| ENSMUSG00000032212 | Sltm | 66660 | protein_coding | 1067.282779 | 0.01922212 | 0.065774879 | 0.292241056 | 0.770102323 | 0.874968957 | 0.01922212 |
| ENSMUSG00000020196 | Cabin1 | 104248 | protein_coding | 866.4179341 | -0.019226147 | 0.090619747 | -0.212162886 | 0.83197996 | 0.910655176 | 0.019226147 |
| ENSMUSG00000021715 | Cwc27 | 67285 | protein_coding | 173.3315152 | 0.019233903 | 0.106321027 | 0.180904039 | 0.856442899 | 0.924779506 | 0.019233903 |
| ENSMUSG00000022517 | Mgrn1 | 17237 | protein_coding | 1187.148982 | 0.019246489 | 0.069731061 | 0.276010265 | 0.782540182 | 0.88192882 | 0.019246489 |
| ENSMUSG00000020611 | Gna13 | 14674 | protein_coding | 2636.848981 | -0.019256839 | 0.079314526 | -0.24279082 | 0.808167447 | 0.897538077 | 0.019256839 |
| ENSMUSG00000031878 | Nae1 | 234664 | protein_coding | 463.2783865 | -0.019436005 | 0.086834964 | -0.223826947 | 0.822891961 | 0.906288656 | 0.019436005 |
| ENSMUSG00000034544 | Rsrc1 | 66880 | protein_coding | 308.2774277 | -0.019453004 | 0.088243267 | -0.220447458 | 0.825522688 | 0.907949549 | 0.019453004 |
| ENSMUSG00000046873 | Mbtps2 | 270669 | protein_coding | 231.6247158 | -0.019461927 | 0.101375038 | -0.19197948 | 0.847758279 | 0.919641345 | 0.019461927 |
| ENSMUSG00000020634 | Ubxn2a | 217379 | protein_coding | 1335.337591 | 0.0195289 | 0.069945441 | 0.279201906 | 0.780089881 | 0.880499284 | 0.0195289 |
| ENSMUSG00000031979 | Cog2 | 76332 | protein_coding | 858.0342922 | -0.019530007 | 0.08856016 | -0.220528132 | 0.825459865 | 0.907943149 | 0.019530007 |
| ENSMUSG00000040620 | Dhx33 | 216877 | protein_coding | 413.7561974 | -0.019549791 | 0.136713294 | -0.142998461 | 0.886291397 | 0.94220607 | 0.019549791 |
| ENSMUSG00000045103 | Dmd | 13405 | protein_coding | 367.3707468 | 0.019572982 | 0.104585792 | 0.18714762 | 0.851544893 | 0.921798204 | 0.019572982 |
| ENSMUSG00000030254 | Rad18 | 58186 | protein_coding | 144.058862 | 0.019628216 | 0.150611129 | 0.130323807 | 0.896310246 | 0.948192606 | 0.019628216 |
| ENSMUSG00000086316 | Nbdy | 70123 | protein_coding | 83.63631086 | -0.019659441 | 0.144942459 | -0.135636178 | 0.892108904 | 0.945409888 | 0.019659441 |
| ENSMUSG00000012519 | Mlkl | 74568 | protein_coding | 292.6466262 | 0.019678977 | 0.14000467 | 0.140559435 | 0.888217996 | 0.94326533 | 0.019678977 |
| ENSMUSG00000001729 | Akt1 | 11651 | protein_coding | 2037.283375 | -0.019680037 | 0.091586285 | -0.214879736 | 0.829861084 | 0.909506443 | 0.019680037 |
| ENSMUSG00000046731 | Kctd11 | 216858 | protein_coding | 107.8183302 | -0.019688274 | 0.174756076 | -0.112661454 | 0.910298962 | 0.955759393 | 0.019688274 |
| ENSMUSG00000078923 | Ube2v1 | 66589 | protein_coding | 12.31982715 | -0.019692051 | 0.302065806 | -0.065191263 | 0.948021718 | 0.974453154 | 0.019692051 |
| ENSMUSG00000041781 | Cpsf2 | 51786 | protein_coding | 972.8813831 | -0.019714621 | 0.099769006 | -0.19760266 | 0.843355953 | 0.917115541 | 0.019714621 |
| ENSMUSG00000036450 | Hif1an | 319594 | protein_coding | 834.7191431 | -0.019723271 | 0.078910681 | -0.249944251 | 0.802630462 | 0.894034894 | 0.019723271 |
| ENSMUSG00000035150 | Eif2s3x | 26905 | protein_coding | 682.9009662 | 0.019739069 | 0.081451712 | 0.242340748 | 0.808516142 | 0.897800133 | 0.019739069 |
| ENSMUSG00000070394 | Tmem256 | 69186 | protein_coding | 447.1794529 | 0.019782884 | 0.091817603 | 0.215458511 | 0.829409856 | 0.909405742 | 0.019782884 |

| ensembl_gene_id | external_gene_name | entrezgene_id | gene_biotype | baseMean | log2FoldChange | lfcSE | stat | pvalue | padj | absLog2FC |
|---|---|---|---|---|---|---|---|---|---|---|
| ENSMUSG00000027479 | Mapre1 | 13589 | protein_coding | 1428.74189 | 0.019838383 | 0.075654709 | 0.262222706 | 0.793149749 | 0.888104689 | 0.019838383 |
| ENSMUSG00000022299 | Slc25a32 | 69906 | protein_coding | 441.9973303 | -0.019845539 | 0.085647632 | -0.231711478 | 0.816762116 | 0.902675156 | 0.019845539 |
| ENSMUSG00000073491 | Ifi213 | 623121 | protein_coding | 7.132859646 | -0.019852264 | 0.723880145 | -0.027424794 | 0.978120923 | 0.990307021 | 0.019852264 |
| ENSMUSG00000027323 | Rad51 | 19361 | protein_coding | 110.4402635 | -0.019873152 | 0.154209537 | -0.128871098 | 0.897459647 | 0.948710356 | 0.019873152 |
| ENSMUSG00000053291 | Rab4b | 19342 | protein_coding | 33.60583134 | 0.01993053 | 0.214666513 | 0.092844151 | 0.926027375 | 0.963686666 | 0.01993053 |
| ENSMUSG00000074466 | Gm15417 | 545539 | lncRNA | 61.67867691 | 0.020010795 | 0.165666619 | 0.120789542 | 0.903857733 | 0.952525195 | 0.020010795 |
| ENSMUSG00000027439 | Gzf1 | 74533 | protein_coding | 488.5703061 | -0.020049164 | 0.110355063 | -0.181678698 | 0.855834884 | 0.924583171 | 0.020049164 |
| ENSMUSG00000058594 | Fbh1 | 50755 | protein_coding | 1478.219405 | 0.02006614 | 0.096904575 | 0.207071136 | 0.835954304 | 0.913007852 | 0.02006614 |
| ENSMUSG00000029482 | Aacs | 78894 | protein_coding | 1310.967993 | 0.020066199 | 0.11046421 | 0.181653401 | 0.855854738 | 0.924583171 | 0.020066199 |
| ENSMUSG00000021891 | Mettl6 | 67011 | protein_coding | 282.1998638 | 0.020072309 | 0.099858653 | 0.201007212 | 0.840692934 | 0.915931326 | 0.020072309 |
| ENSMUSG00000055436 | Srsf11 | 69207 | protein_coding | 1266.405355 | -0.02011865 | 0.094903755 | -0.21199003 | 0.832114813 | 0.910655176 | 0.02011865 |
| ENSMUSG00000031482 | Slc25a15 | 18408 | protein_coding | 575.2737915 | -0.020121577 | 0.082612962 | -0.243564404 | 0.8075682 | 0.897060205 | 0.020121577 |
| ENSMUSG00000038416 | Cdc16 | 69957 | protein_coding | 651.6252425 | 0.02015217 | 0.088349489 | 0.228096062 | 0.819571562 | 0.904339164 | 0.02015217 |
| ENSMUSG00000033671 | Cep350 | 74081 | protein_coding | 1830.620894 | -0.020189232 | 0.120417432 | -0.167660377 | 0.866850469 | 0.931031306 | 0.020189232 |
| ENSMUSG00000027562 | Car2 | 12349 | protein_coding | 14587.07894 | -0.020218058 | 0.08317978 | -0.243064574 | 0.807955374 | 0.897427699 | 0.020218058 |
| ENSMUSG00000048410 | Zfp407 | 240476 | protein_coding | 500.9842967 | 0.020286346 | 0.105977368 | 0.191421491 | 0.848195384 | 0.919955452 | 0.020286346 |
| ENSMUSG00000025474 | Tubgcp2 | 74237 | protein_coding | 239.4469756 | -0.020308545 | 0.12285508 | -0.165304883 | 0.868704016 | 0.932204916 | 0.020308545 |
| ENSMUSG00000024665 | Fads2 | 56473 | protein_coding | 2941.364637 | 0.020394422 | 0.217111203 | 0.093935374 | 0.925160494 | 0.963197355 | 0.020394422 |
| ENSMUSG00000018566 | Slc2a4 | 20528 | protein_coding | 8.381831776 | -0.020428528 | 0.425316159 | -0.048031393 | 0.961691223 | 0.981345237 | 0.020428528 |
| ENSMUSG00000002010 | Idh3g | 15929 | protein_coding | 1759.160293 | 0.020453826 | 0.097327797 | 0.210153997 | 0.833547483 | 0.911278944 | 0.020453826 |
| ENSMUSG00000001482 | Def8 | 23854 | protein_coding | 237.449685 | 0.020462257 | 0.096389853 | 0.212286417 | 0.831883591 | 0.910655176 | 0.020462257 |
| ENSMUSG00000022914 | Brwd1 | 93871 | protein_coding | 1406.953164 | 0.020483484 | 0.110430016 | 0.185488376 | 0.852845996 | 0.922175438 | 0.020483484 |
| ENSMUSG00000028139 | Riiad1 | 66353 | protein_coding | 6.974552799 | -0.02049857 | 0.439391836 | -0.046652141 | 0.962790475 | 0.982152343 | 0.02049857 |
| ENSMUSG00000037563 | Rps16 | 20055 | protein_coding | 3137.975418 | -0.02050526 | 0.082492584 | -0.248570954 | 0.803692679 | 0.894376929 | 0.02050526 |
| ENSMUSG00000041741 | Pde3a | 54611 | protein_coding | 1596.745 | 0.020524774 | 0.084143341 | 0.243926302 | 0.807287899 | 0.896811386 | 0.020524774 |
| ENSMUSG00000006058 | Snf8 | 27681 | protein_coding | 627.9476651 | 0.020539523 | 0.07143446 | 0.287529622 | 0.773706824 | 0.877099132 | 0.020539523 |
| ENSMUSG00000000958 | Slc7a7 | 20540 | protein_coding | 516.65978 | 0.020567555 | 0.073671918 | 0.279177679 | 0.780108473 | 0.880499284 | 0.020567555 |
| ENSMUSG00000033032 | Afap1l1 | 106877 | protein_coding | 250.0456585 | 0.020584718 | 0.139818176 | 0.147224909 | 0.8829545 | 0.940966992 | 0.020584718 |
| ENSMUSG00000023156 | Rpp14 | 67053 | protein_coding | 335.1155109 | 0.020669754 | 0.101092197 | 0.204464385 | 0.837990621 | 0.914526523 | 0.020669754 |
| ENSMUSG00000033589 | Reep4 | 72549 | protein_coding | 482.7450676 | 0.020707294 | 0.092423815 | 0.224047173 | 0.822720598 | 0.906181708 | 0.020707294 |
| ENSMUSG00000028291 | Akirin2 | 433693 | protein_coding | 478.6108475 | 0.020736683 | 0.073796798 | 0.280997061 | 0.778712654 | 0.879764214 | 0.020736683 |
| ENSMUSG00000014349 | Ube2z | 268470 | protein_coding | 1533.992066 | 0.020784886 | 0.108394191 | 0.191752764 | 0.847935873 | 0.919771355 | 0.020784886 |
| ENSMUSG00000022210 | Dhrs4 | 28200 | protein_coding | 519.8734903 | -0.020798495 | 0.083040833 | -0.25046106 | 0.802230815 | 0.893981767 | 0.020798495 |
| ENSMUSG00000030722 | Nfatc2ip | 18020 | protein_coding | 67.55302252 | 0.020807013 | 0.185018107 | 0.112459332 | 0.910459213 | 0.955759393 | 0.020807013 |
| ENSMUSG00000038371 | Sbf2 | 319934 | protein_coding | 1064.286331 | 0.020879264 | 0.056385884 | 0.37029239 | 0.711164644 | 0.837163904 | 0.020879264 |
| ENSMUSG00000111356 | Rpl32l | 621697 | processed_pseudogene | 16.56917505 | -0.020931177 | 0.297242572 | -0.07041783 | 0.9438611 | 0.972685016 | 0.020931177 |
| ENSMUSG00000040429 | Mterf1a | 545725 | protein_coding | 41.04556824 | 0.020947558 | 0.229304395 | 0.091352625 | 0.927212404 | 0.963947469 | 0.020947558 |
| ENSMUSG00000048905 | Bnip5 | 207819 | protein_coding | 769.0245534 | -0.020980879 | 0.185413953 | -0.113156957 | 0.90990612 | 0.955560406 | 0.020980879 |

| ensembl_gene_id | external_gene_name | entrezgene_id | gene_biotype | baseMean | log2FoldChange | lfcSE | stat | pvalue | padj | absLog2FC |
|---|---|---|---|---|---|---|---|---|---|---|
| ENSMUSG00000074748 | Atxn7l3b | 382423 | protein_coding | 1571.731246 | -0.020996892 | 0.077679076 | -0.270303065 | 0.786927108 | 0.884494741 | 0.020996892 |
| ENSMUSG00000039046 | Usp6nl | 98910 | protein_coding | 792.6270808 | -0.021022306 | 0.177865434 | -0.118192197 | 0.905915372 | 0.953443904 | 0.021022306 |
| ENSMUSG00000021133 | Susd6 | 217684 | protein_coding | 1134.908735 | -0.02102839 | 0.084998615 | -0.24739686 | 0.804601104 | 0.894700251 | 0.02102839 |
| ENSMUSG00000043088 | Il17re | 57890 | protein_coding | 230.9308592 | -0.021080275 | 0.140268448 | -0.150285226 | 0.880539589 | 0.939399502 | 0.021080275 |
| ENSMUSG00000022426 | Josd1 | 74158 | protein_coding | 668.0393217 | -0.021107429 | 0.08936057 | -0.23620517 | 0.813273468 | 0.900690228 | 0.021107429 |
| ENSMUSG00000026584 | Scyl3 | 240880 | protein_coding | 408.3244856 | -0.021125375 | 0.083720639 | -0.252331749 | 0.80078465 | 0.893137589 | 0.021125375 |
| ENSMUSG00000006191 | Cdkal1 | 68916 | protein_coding | 272.5082783 | -0.021142785 | 0.084768258 | -0.249418657 | 0.803036955 | 0.894084484 | 0.021142785 |
| ENSMUSG00000041426 | Hibch | 227095 | protein_coding | 552.5136343 | -0.021172476 | 0.103976772 | -0.203626975 | 0.838645012 | 0.915052593 | 0.021172476 |
| ENSMUSG00000037339 | Fam53a | 74504 | protein_coding | 344.9337283 | -0.021214266 | 0.084718511 | -0.250408857 | 0.802271181 | 0.893981767 | 0.021214266 |
| ENSMUSG00000051977 | Prdm9 | 213389 | protein_coding | 94.29268542 | -0.021227268 | 0.201222908 | -0.105491309 | 0.915985966 | 0.958406652 | 0.021227268 |
| ENSMUSG00000020224 | Llph | 66225 | protein_coding | 202.5103321 | 0.021234803 | 0.16915946 | 0.125531275 | 0.900102967 | 0.950214744 | 0.021234803 |
| ENSMUSG00000074166 | AW146154 | 101835 | protein_coding | 195.0032719 | 0.021267464 | 0.158911972 | 0.133831727 | 0.893535638 | 0.94649459 | 0.021267464 |
| ENSMUSG00000038766 | Gabpb2 | 213054 | protein_coding | 1289.141513 | -0.021269264 | 0.076018907 | -0.279789126 | 0.779639295 | 0.880278918 | 0.021269264 |
| ENSMUSG00000074754 | Smim26 | 228715 | protein_coding | 93.10054721 | -0.02131574 | 0.124633993 | -0.171026697 | 0.864202774 | 0.9295038 | 0.02131574 |
| ENSMUSG00000028771 | Ptpn12 | 19248 | protein_coding | 503.7341206 | 0.021316547 | 0.205468308 | 0.103746153 | 0.917370799 | 0.959078485 | 0.021316547 |
| ENSMUSG00000069727 | Zfp975 | 434179 | protein_coding | 135.7991798 | 0.02137414 | 0.194514401 | 0.109884616 | 0.912500884 | 0.956960368 | 0.02137414 |
| ENSMUSG00000090266 | Mettl23 | 74319 | protein_coding | 241.8007129 | -0.021420004 | 0.101626019 | -0.210772835 | 0.833064537 | 0.911190103 | 0.021420004 |
| ENSMUSG00000021481 | Zfp346 | 26919 | protein_coding | 118.5059957 | 0.021474513 | 0.126838246 | 0.169306291 | 0.865555728 | 0.930241476 | 0.021474513 |
| ENSMUSG00000042742 | Bmt2 | 101148 | protein_coding | 432.9909489 | -0.021534678 | 0.11870679 | -0.181410669 | 0.856045245 | 0.924704659 | 0.021534678 |
| ENSMUSG00000037103 | Dcaf15 | 212123 | protein_coding | 470.9597916 | -0.02156607 | 0.102441862 | -0.210520089 | 0.833261774 | 0.911238829 | 0.02156607 |
| ENSMUSG00000040446 | Rprd1a | 225283 | protein_coding | 154.9784031 | 0.021587888 | 0.106385099 | 0.202922103 | 0.839195918 | 0.91521483 | 0.021587888 |
| ENSMUSG00000027792 | Bche | 12038 | protein_coding | 1014.55988 | -0.021590763 | 0.100829189 | -0.214132069 | 0.830444068 | 0.909656359 | 0.021590763 |
| ENSMUSG00000050213 | Snip1 | 76793 | protein_coding | 354.6339022 | -0.021623471 | 0.099277047 | -0.217809373 | 0.827577641 | 0.908766402 | 0.021623471 |
| ENSMUSG00000043542 | Zc2hc1a | 67306 | protein_coding | 132.2884556 | 0.021639568 | 0.167754773 | 0.128995244 | 0.897361412 | 0.948703875 | 0.021639568 |
| ENSMUSG00000014791 | Elmo3 | 234683 | protein_coding | 916.1782374 | 0.0217036 | 0.078036407 | 0.278121464 | 0.78091912 | 0.880896827 | 0.0217036 |
| ENSMUSG00000117679 | Apbb3 | 225372 | protein_coding | 151.6263439 | 0.021729337 | 0.115064393 | 0.188845011 | 0.850214293 | 0.920905487 | 0.021729337 |
| ENSMUSG00000027968 | Larp7 | 28036 | protein_coding | 315.5186275 | -0.02180409 | 0.122436176 | -0.178085353 | 0.858655944 | 0.926384278 | 0.02180409 |
| ENSMUSG00000058655 | Eif4b | 75705 | protein_coding | 6411.940809 | -0.021817337 | 0.087464457 | -0.249442319 | 0.803018654 | 0.894084484 | 0.021817337 |
| ENSMUSG00000005069 | Pex5 | 19305 | protein_coding | 555.715909 | 0.021883383 | 0.09610727 | 0.227697474 | 0.819881436 | 0.90449311 | 0.021883383 |
| ENSMUSG00000042211 | Fbxo38 | 107035 | protein_coding | 1116.537455 | -0.021909201 | 0.063257757 | -0.346348049 | 0.729081163 | 0.847668919 | 0.021909201 |
| ENSMUSG00000040842 | Szrd1 | 213491 | protein_coding | 1412.708076 | 0.021926914 | 0.084995113 | 0.257978525 | 0.796423481 | 0.89039329 | 0.021926914 |
| ENSMUSG00000016554 | Eif3d | 55944 | protein_coding | 1188.537524 | 0.021948509 | 0.119250734 | 0.184053452 | 0.853971523 | 0.923049566 | 0.021948509 |
| ENSMUSG00000024070 | Prkd3 | 75292 | protein_coding | 259.2431528 | -0.022029548 | 0.099169214 | -0.222140998 | 0.824204123 | 0.90712571 | 0.022029548 |
| ENSMUSG00000019297 | Nop9 | 67842 | protein_coding | 276.4631507 | -0.022050058 | 0.115391477 | -0.191089137 | 0.848455758 | 0.919988003 | 0.022050058 |
| ENSMUSG00000025893 | Kbtbd3 | 69149 | protein_coding | 86.41124372 | 0.022060668 | 0.166015576 | 0.132883122 | 0.894285815 | 0.947086733 | 0.022060668 |
| ENSMUSG00000003923 | Tfam | 21780 | protein_coding | 741.0999408 | -0.022061775 | 0.078566503 | -0.280803829 | 0.778860867 | 0.879817143 | 0.022061775 |
| ENSMUSG00000018974 | Sart3 | 53890 | protein_coding | 501.364096 | 0.02207779 | 0.092567949 | 0.238503614 | 0.811490512 | 0.899694484 | 0.02207779 |
| ENSMUSG00000026305 | Lrrfip1 | 16978 | protein_coding | 1555.746613 | -0.022097185 | 0.167003847 | -0.132315422 | 0.894734809 | 0.947436248 | 0.022097185 |

| ensembl_gene_id | external_gene_name | entrezgene_id | gene_biotype | baseMean | log2FoldChange | lfcSE | stat | pvalue | padj | absLog2FC |
|---|---|---|---|---|---|---|---|---|---|---|
| ENSMUSG00000078440 | Dohh | 102115 | protein_coding | 81.64136787 | -0.022122812 | 0.1717117 | -0.12883695 | 0.897486668 | 0.948710356 | 0.022122812 |
| ENSMUSG00000032849 | Abcc4 | 239273 | protein_coding | 253.3294541 | 0.022170016 | 0.11112502 | 0.199505173 | 0.841867597 | 0.916434328 | 0.022170016 |
| ENSMUSG00000072762 | 4930522L14Rik | 100041734 | protein_coding | 92.19410365 | 0.022271911 | 0.201934549 | 0.11029272 | 0.912177231 | 0.956746934 | 0.022271911 |
| ENSMUSG00000039621 | Prex1 | 277360 | protein_coding | 130.7033792 | 0.022316352 | 0.181521862 | 0.122940299 | 0.902154374 | 0.951686526 | 0.022316352 |
| ENSMUSG00000028678 | Kif2c | 73804 | protein_coding | 276.0351939 | -0.022318855 | 0.156831871 | -0.142310712 | 0.886834585 | 0.942483462 | 0.022318855 |
| ENSMUSG00000002458 | Rgs19 | 56470 | protein_coding | 30.12057109 | 0.022366331 | 0.247820433 | 0.090252167 | 0.92808683 | 0.964521871 | 0.022366331 |
| ENSMUSG00000059263 | Usp47 | 74996 | protein_coding | 2861.354118 | 0.022384695 | 0.054588716 | 0.41006084 | 0.681761318 | 0.818010868 | 0.022384695 |
| ENSMUSG00000035125 | Gcfc2 | 330361 | protein_coding | 102.2447372 | -0.022409842 | 0.128957461 | -0.173777012 | 0.862040711 | 0.928193452 | 0.022409842 |
| ENSMUSG00000035596 | Mboat7 | 77582 | protein_coding | 550.7254578 | 0.022421179 | 0.083452587 | 0.268669673 | 0.788183892 | 0.884950162 | 0.022421179 |
| ENSMUSG00000062198 | 2700097O09Rik | 72658 | protein_coding | 85.05838382 | -0.022472441 | 0.12085426 | -0.185946619 | 0.852486623 | 0.921945066 | 0.022472441 |
| ENSMUSG00000012535 | Tnpo3 | 320938 | protein_coding | 1286.72085 | -0.022498502 | 0.071886308 | -0.31297339 | 0.754300879 | 0.865108984 | 0.022498502 |
| ENSMUSG00000063801 | Ap3s2 | 11778 | protein_coding | 317.5021547 | -0.022505245 | 0.079784371 | -0.282075862 | 0.777885342 | 0.879295843 | 0.022505245 |
| ENSMUSG00000000085 | Scmh1 | 29871 | protein_coding | 341.7717038 | -0.0225167 | 0.198001398 | -0.113719901 | 0.909459836 | 0.955343683 | 0.0225167 |
| ENSMUSG00000054099 | Slc25a40 | 319653 | protein_coding | 424.5682906 | 0.022534765 | 0.114046043 | 0.197593571 | 0.843363065 | 0.917115541 | 0.022534765 |
| ENSMUSG00000022849 | Hspbap1 | 66667 | protein_coding | 111.5444971 | -0.022551683 | 0.136652523 | -0.165029393 | 0.868920847 | 0.932311974 | 0.022551683 |
| ENSMUSG00000041958 | Pigs | 276846 | protein_coding | 555.6602117 | -0.022637568 | 0.086896916 | -0.260510607 | 0.794469934 | 0.889039236 | 0.022637568 |
| ENSMUSG00000029110 | Rnf4 | 19822 | protein_coding | 1851.06977 | -0.02266559 | 0.052696606 | -0.430114801 | 0.667112134 | 0.807122071 | 0.02266559 |
| ENSMUSG00000072582 | Ptrh2 | 217057 | protein_coding | 226.5734102 | 0.022742709 | 0.096827426 | 0.234878797 | 0.814302807 | 0.900950823 | 0.022742709 |
| ENSMUSG00000021339 | Mrs2 | 380836 | protein_coding | 1201.787052 | 0.022776783 | 0.073312413 | 0.310681125 | 0.756043048 | 0.865975382 | 0.022776783 |
| ENSMUSG00000026878 | Rab14 | 68365 | protein_coding | 1744.97634 | -0.022795365 | 0.062654497 | -0.363826473 | 0.715987593 | 0.84023174 | 0.022795365 |
| ENSMUSG00000021738 | Atxn7 | 246103 | protein_coding | 577.3563268 | 0.022806755 | 0.08751042 | 0.260617593 | 0.794387421 | 0.889039236 | 0.022806755 |
| ENSMUSG00000018932 | Map2k3 | 26397 | protein_coding | 1017.873551 | -0.022858112 | 0.268951105 | -0.084989842 | 0.932269467 | 0.96605218 | 0.022858112 |
| ENSMUSG00000032398 | Snapc5 | 330959 | protein_coding | 439.1300558 | 0.022907013 | 0.071638982 | 0.319756268 | 0.749153102 | 0.861055752 | 0.022907013 |
| ENSMUSG00000048007 | Timm8a1 | 30058 | protein_coding | 88.35415237 | 0.022919582 | 0.163157285 | 0.140475384 | 0.8882844 | 0.94326533 | 0.022919582 |
| ENSMUSG00000051483 | Cbr1 | 12408 | protein_coding | 2884.325175 | 0.02294208 | 0.104930868 | 0.218639955 | 0.826930526 | 0.908412958 | 0.02294208 |
| ENSMUSG00000051378 | Kif18b | 70218 | protein_coding | 165.4428947 | 0.022944806 | 0.113220934 | 0.202655156 | 0.839404576 | 0.915379285 | 0.022944806 |
| ENSMUSG00000005506 | Celf1 | 13046 | protein_coding | 1539.504156 | -0.023003714 | 0.08848399 | -0.259976007 | 0.794882281 | 0.889189108 | 0.023003714 |
| ENSMUSG00000035382 | Pcsk7 | 18554 | protein_coding | 2057.455496 | 0.023034259 | 0.092477382 | 0.249079921 | 0.803298961 | 0.894249532 | 0.023034259 |
| ENSMUSG00000008435 | Rdh13 | 108841 | protein_coding | 652.7403565 | 0.023046193 | 0.080796873 | 0.285236204 | 0.775463185 | 0.877982787 | 0.023046193 |
| ENSMUSG00000058440 | Nrf1 | 18181 | protein_coding | 417.3548538 | 0.023079315 | 0.109964691 | 0.20987932 | 0.833761863 | 0.911278944 | 0.023079315 |
| ENSMUSG00000024360 | Etf1 | 225363 | protein_coding | 2513.511914 | 0.023079395 | 0.122580662 | 0.188279247 | 0.850657753 | 0.921218159 | 0.023079395 |
| ENSMUSG00000028057 | Rit1 | 19769 | protein_coding | 462.3943376 | 0.023102887 | 0.099219403 | 0.232846467 | 0.815880629 | 0.902141385 | 0.023102887 |
| ENSMUSG00000062604 | Srpk2 | 20817 | protein_coding | 296.4037511 | -0.023115543 | 0.122869016 | -0.18813159 | 0.850773498 | 0.921280867 | 0.023115543 |
| ENSMUSG00000021983 | Atp8a2 | 50769 | protein_coding | 577.2336901 | 0.023153517 | 0.082880454 | 0.2793604 | 0.779968258 | 0.880499284 | 0.023153517 |
| ENSMUSG00000056665 | Them6 | 223626 | protein_coding | 15.33555758 | -0.023161346 | 0.282939315 | -0.081859766 | 0.934758229 | 0.967565507 | 0.023161346 |
| ENSMUSG00000028629 | Exo5 | 73172 | protein_coding | 145.0113751 | 0.023209338 | 0.114228684 | 0.203183103 | 0.838991919 | 0.915199917 | 0.023209338 |
| ENSMUSG00000060938 | Rpl26 | 19941 | protein_coding | 5794.371019 | 0.023216971 | 0.105691421 | 0.219667507 | 0.826130112 | 0.908178648 | 0.023216971 |
| ENSMUSG00000033166 | Dis3 | 72662 | protein_coding | 380.8460442 | 0.02323878 | 0.158855002 | 0.146289253 | 0.883693049 | 0.941565516 | 0.02323878 |

| ensembl_gene_id | external_gene_name | entrezgene_id | gene_biotype | baseMean | log2FoldChange | lfcSE | stat | pvalue | padj | absLog2FC |
|---|---|---|---|---|---|---|---|---|---|---|
| ENSMUSG00000051469 | Zfp24 | 59057 | protein_coding | 787.6482292 | 0.023269459 | 0.074314026 | 0.313123382 | 0.754186925 | 0.865108984 | 0.023269459 |
| ENSMUSG00000026435 | Slc45a3 | 212980 | protein_coding | 344.7007047 | 0.023276337 | 0.132630978 | 0.175496988 | 0.860689138 | 0.927608253 | 0.023276337 |
| ENSMUSG00000079037 | Prnp | 19122 | protein_coding | 1792.082108 | 0.023290554 | 0.162834218 | 0.143032306 | 0.886264667 | 0.94220607 | 0.023290554 |
| ENSMUSG00000025555 | Farp1 | 223254 | protein_coding | 580.9272067 | 0.023293047 | 0.113383349 | 0.205436224 | 0.837231321 | 0.91407365 | 0.023293047 |
| ENSMUSG00000059897 | Zfp930 | 234358 | protein_coding | 250.8416136 | -0.023393554 | 0.114504706 | -0.204302119 | 0.838117414 | 0.914602232 | 0.023393554 |
| ENSMUSG00000028952 | Zbtb48 | 100090 | protein_coding | 111.1918592 | -0.023406621 | 0.129261834 | -0.181079137 | 0.85630546 | 0.924704659 | 0.023406621 |
| ENSMUSG00000022993 | Tex49 | 73863 | protein_coding | 39.57340765 | 0.023495921 | 0.180936456 | 0.129857306 | 0.896679323 | 0.948360442 | 0.023495921 |
| ENSMUSG00000037999 | Arap2 | 212285 | protein_coding | 1775.92199 | -0.023604914 | 0.100338039 | -0.235253885 | 0.814011685 | 0.900950463 | 0.023604914 |
| ENSMUSG00000014355 | Anapc1 | 17222 | protein_coding | 1749.418787 | 0.023607677 | 0.066044847 | 0.357449189 | 0.720755558 | 0.843140021 | 0.023607677 |
| ENSMUSG00000047434 | Xxylt1 | 268880 | protein_coding | 138.2077579 | -0.023621193 | 0.112665346 | -0.209658018 | 0.833934594 | 0.911286376 | 0.023621193 |
| ENSMUSG00000005078 | Jkamp | 104771 | protein_coding | 196.2349062 | 0.023687379 | 0.128460043 | 0.18439492 | 0.853703655 | 0.922822654 | 0.023687379 |
| ENSMUSG00000029346 | Srrd | 70118 | protein_coding | 53.13497631 | -0.023712116 | 0.173412744 | -0.136738022 | 0.891237875 | 0.945137528 | 0.023712116 |
| ENSMUSG00000107709 | 9430018G01Rik | 319812 | lncRNA | 7.09997196 | -0.02374437 | 0.706834251 | -0.033592557 | 0.973202058 | 0.988062333 | 0.02374437 |
| ENSMUSG00000028532 | Cachd1 | 320508 | protein_coding | 853.5384029 | -0.023821823 | 0.108679913 | -0.219192511 | 0.826500089 | 0.908397282 | 0.023821823 |
| ENSMUSG00000030059 | Tmf1 | 232286 | protein_coding | 2343.785501 | -0.02382832 | 0.061248494 | -0.389043365 | 0.697244068 | 0.827548697 | 0.02382832 |
| ENSMUSG00000024571 | Naa12 | 27366 | protein_coding | 675.5253338 | -0.023846578 | 0.081925065 | -0.291077921 | 0.770991725 | 0.87556148 | 0.023846578 |
| ENSMUSG00000028224 | Nbn | 27354 | protein_coding | 461.5090905 | 0.023864063 | 0.09539645 | 0.250156721 | 0.802466153 | 0.894011393 | 0.023864063 |
| ENSMUSG00000022434 | Fam118a | 73225 | protein_coding | 224.3071524 | 0.023879067 | 0.170024686 | 0.140444708 | 0.888308636 | 0.94326533 | 0.023879067 |
| ENSMUSG00000021635 | Rad17 | 19356 | protein_coding | 429.2894899 | -0.02390136 | 0.062936834 | -0.37976744 | 0.704118054 | 0.8327907 | 0.02390136 |
| ENSMUSG00000057469 | E2f6 | 50496 | protein_coding | 216.4555869 | -0.023968169 | 0.104128352 | -0.230179091 | 0.817952608 | 0.903241067 | 0.023968169 |
| ENSMUSG00000022295 | Atp6v1c1 | 66335 | protein_coding | 557.2117918 | 0.023978227 | 0.108282209 | 0.221441986 | 0.824748304 | 0.90753651 | 0.023978227 |
| ENSMUSG00000020973 | Dnaaf2 | 109065 | protein_coding | 27.84778213 | -0.024029664 | 0.220734102 | -0.108862492 | 0.913311557 | 0.957282345 | 0.024029664 |
| ENSMUSG00000028407 | Smim27 | 100504309 | protein_coding | 63.08237453 | 0.024041138 | 0.159099374 | 0.151107687 | 0.879890769 | 0.939210573 | 0.024041138 |
| ENSMUSG00000026854 | Usp20 | 74270 | protein_coding | 230.8032012 | 0.024052093 | 0.126912537 | 0.189517076 | 0.849687573 | 0.920572697 | 0.024052093 |
| ENSMUSG00000026922 | Agpat2 | 67512 | protein_coding | 1209.615999 | -0.024099733 | 0.190754734 | -0.126338844 | 0.899463709 | 0.950188262 | 0.024099733 |
| ENSMUSG00000032376 | Usp3 | 235441 | protein_coding | 1364.026656 | -0.024103173 | 0.120061064 | -0.200757618 | 0.840888104 | 0.916056749 | 0.024103173 |
| ENSMUSG00000004054 | Map3k11 | 26403 | protein_coding | 890.3582032 | 0.024113436 | 0.133140764 | 0.181112342 | 0.856279398 | 0.924704659 | 0.024113436 |
| ENSMUSG00000021619 | Atg10 | 66795 | protein_coding | 79.35979632 | -0.024140778 | 0.145869647 | -0.165495551 | 0.868553951 | 0.932161313 | 0.024140778 |
| ENSMUSG00000038291 | Snx25 | 102141 | protein_coding | 442.6296164 | 0.024184604 | 0.114112335 | 0.211936808 | 0.832156334 | 0.910655176 | 0.024184604 |
| ENSMUSG00000017950 | Hnf4a | 15378 | protein_coding | 9793.074718 | 0.024188818 | 0.108857032 | 0.222207214 | 0.824152579 | 0.90712571 | 0.024188818 |
| ENSMUSG00000025785 | Exosc7 | 66446 | protein_coding | 296.4196442 | 0.024246159 | 0.087559084 | 0.276911982 | 0.781847692 | 0.881460224 | 0.024246159 |
| ENSMUSG00000029017 | Pmpcb | 73078 | protein_coding | 1159.851187 | 0.024249602 | 0.09883679 | 0.245349957 | 0.806185473 | 0.895899129 | 0.024249602 |
| ENSMUSG00000055817 | Mta3 | 116871 | protein_coding | 454.6401192 | 0.024284844 | 0.096915529 | 0.250577432 | 0.802140832 | 0.893981767 | 0.024284844 |
| ENSMUSG00000036764 | Dnajc12 | 30045 | protein_coding | 71.91843086 | 0.024296297 | 0.208372327 | 0.116600403 | 0.907176718 | 0.954095762 | 0.024296297 |
| ENSMUSG00000030061 | Uba3 | 22200 | protein_coding | 875.6785384 | -0.024301449 | 0.077732861 | -0.312627748 | 0.754563494 | 0.865216002 | 0.024301449 |
| ENSMUSG00000039414 | Heatr5b | 320473 | protein_coding | 411.3390902 | 0.024318353 | 0.09261684 | 0.262569458 | 0.792882443 | 0.887992656 | 0.024318353 |
| ENSMUSG00000037949 | Ano10 | 102566 | protein_coding | 1058.711395 | -0.024347095 | 0.100844916 | -0.241431056 | 0.809221046 | 0.898394983 | 0.024347095 |
| ENSMUSG00000029556 | Hnf1a | 21405 | protein_coding | 297.6345795 | -0.024407282 | 0.126686192 | -0.192659371 | 0.847225745 | 0.919400582 | 0.024407282 |

| ensembl_gene_id | external_gene_name | entrezgene_id | gene_biotype | baseMean | log2FoldChange | lfcSE | stat | pvalue | padj | absLog2FC |
|---|---|---|---|---|---|---|---|---|---|---|
| ENSMUSG00000039615 | Stub1 | 56424 | protein_coding | 387.3941584 | -0.024452788 | 0.077963179 | -0.313645336 | 0.753790423 | 0.864825603 | 0.024452788 |
| ENSMUSG00000118376 | Gm9926 | 100503565 | lncRNA | 159.7684798 | -0.024463009 | 0.13992862 | -0.174824912 | 0.861217213 | 0.927921787 | 0.024463009 |
| ENSMUSG00000069270 | H2ac6 | 319164 | protein_coding | 281.066182 | -0.02446906 | 0.184690331 | -0.132486959 | 0.894599138 | 0.947355566 | 0.02446906 |
| ENSMUSG00000032528 | Vipr1 | 22354 | protein_coding | 3885.461202 | -0.024472189 | 0.099335259 | -0.246359543 | 0.805403921 | 0.895280456 | 0.024472189 |
| ENSMUSG00000041028 | Ghitm | 66092 | protein_coding | 4915.122678 | 0.024490914 | 0.079997911 | 0.306144415 | 0.759494692 | 0.868410263 | 0.024490914 |
| ENSMUSG00000028975 | Pex14 | 56273 | protein_coding | 244.3110971 | 0.024526898 | 0.129339631 | 0.189631732 | 0.84959772 | 0.920572697 | 0.024526898 |
| ENSMUSG00000056310 | Tyw1 | 100929 | protein_coding | 318.6773658 | -0.024527048 | 0.103328713 | -0.237369144 | 0.812370425 | 0.900072074 | 0.024527048 |
| ENSMUSG00000074579 | Lekr1 | 624866 | protein_coding | 40.41837793 | 0.02457025 | 0.263896383 | 0.093105671 | 0.925819612 | 0.963686666 | 0.02457025 |
| ENSMUSG00000026012 | Cd28 | 12487 | protein_coding | 7.101346509 | -0.024594647 | 0.710631339 | -0.034609573 | 0.972391068 | 0.987518805 | 0.024594647 |
| ENSMUSG00000020288 | Ahsa2 | 268390 | protein_coding | 245.5848753 | -0.02462106 | 0.172379161 | -0.14283084 | 0.88642378 | 0.94220607 | 0.02462106 |
| ENSMUSG00000027423 | Snx5 | 69178 | protein_coding | 1955.716473 | -0.024634076 | 0.09567451 | -0.257477941 | 0.796809842 | 0.890700201 | 0.024634076 |
| ENSMUSG00000033684 | Qsox1 | 104009 | protein_coding | 2110.694085 | -0.024704724 | 0.090424459 | -0.27320842 | 0.784693008 | 0.883167766 | 0.024704724 |
| ENSMUSG00000042271 | Nxt2 | 237082 | protein_coding | 398.8486555 | 0.02477267 | 0.125599547 | 0.197235344 | 0.843643374 | 0.917176415 | 0.02477267 |
| ENSMUSG00000042410 | Agps | 228061 | protein_coding | 1603.861717 | 0.024816247 | 0.072210994 | 0.343663007 | 0.731099734 | 0.848819635 | 0.024816247 |
| ENSMUSG00000027652 | Ralgapb | 228850 | protein_coding | 1904.373059 | -0.024816452 | 0.056832862 | -0.436656741 | 0.662360306 | 0.804505636 | 0.024816452 |
| ENSMUSG00000092558 | Med20 | 56771 | protein_coding | 300.9163738 | 0.024827953 | 0.087216578 | 0.284670109 | 0.775896893 | 0.877982787 | 0.024827953 |
| ENSMUSG00000021037 | Ahsa1 | 217737 | protein_coding | 871.9086969 | 0.024866222 | 0.153485012 | 0.162010752 | 0.871297386 | 0.933729716 | 0.024866222 |
| ENSMUSG00000004994 | Ccdc130 | 67736 | protein_coding | 99.94889267 | 0.024886219 | 0.128253981 | 0.194038569 | 0.846145682 | 0.918830681 | 0.024886219 |
| ENSMUSG00000039361 | Picalm | 233489 | protein_coding | 3947.786423 | 0.024892115 | 0.066163225 | 0.376222823 | 0.70675125 | 0.834197547 | 0.024892115 |
| ENSMUSG00000016528 | Mapkapk2 | 17164 | protein_coding | 1190.921321 | 0.024911753 | 0.080780096 | 0.308389746 | 0.757785784 | 0.867408546 | 0.024911753 |
| ENSMUSG00000039853 | Trim14 | 74735 | protein_coding | 150.6803066 | -0.024927343 | 0.118727539 | -0.209954179 | 0.833703436 | 0.911278944 | 0.024927343 |
| ENSMUSG00000078681 | Tm2d3 | 68634 | protein_coding | 340.1458085 | 0.025007613 | 0.129909707 | 0.192499953 | 0.847350604 | 0.919400582 | 0.025007613 |
| ENSMUSG00000029501 | Ankle2 | 71782 | protein_coding | 779.4871673 | 0.025076011 | 0.067713685 | 0.37032412 | 0.711141004 | 0.837163904 | 0.025076011 |
| ENSMUSG00000090019 | Gimap1 | 16205 | protein_coding | 27.44371644 | 0.025135311 | 0.257445907 | 0.097633367 | 0.922223428 | 0.961522207 | 0.025135311 |
| ENSMUSG00000047992 | Dipk1c | 240479 | protein_coding | 8.670413863 | 0.025169451 | 0.598543221 | 0.042051184 | 0.966457895 | 0.984521816 | 0.025169451 |
| ENSMUSG00000087165 | 2010001A14Rik | 76516 | lncRNA | 53.95867355 | -0.025204147 | 0.168289498 | -0.149766604 | 0.880948758 | 0.93955283 | 0.025204147 |
| ENSMUSG00000030272 | Camk1 | 52163 | protein_coding | 291.7539308 | 0.025206649 | 0.099258333 | 0.253949956 | 0.79953422 | 0.892430347 | 0.025206649 |
| ENSMUSG00000021952 | Xpo4 | 57258 | protein_coding | 368.5808539 | -0.025271856 | 0.116293002 | -0.217311922 | 0.827965267 | 0.90891431 | 0.025271856 |
| ENSMUSG00000002058 | Unc119 | 22248 | protein_coding | 237.3335278 | 0.025272429 | 0.250367486 | 0.100941339 | 0.919597027 | 0.960107027 | 0.025272429 |
| ENSMUSG00000034292 | Traf3ip1 | 74019 | protein_coding | 114.9425552 | -0.02529374 | 0.183807632 | -0.137609844 | 0.890548775 | 0.944864393 | 0.02529374 |
| ENSMUSG00000036955 | Kifbp | 72320 | protein_coding | 306.0963486 | 0.025297741 | 0.075996654 | 0.332879667 | 0.739225118 | 0.854595635 | 0.025297741 |
| ENSMUSG00000028411 | Aptx | 66408 | protein_coding | 142.2923602 | 0.025312554 | 0.117028895 | 0.216293196 | 0.828759213 | 0.909295436 | 0.025312554 |
| ENSMUSG00000022223 | Sdr39u1 | 654795 | protein_coding | 73.00619898 | 0.025353777 | 0.141017946 | 0.179791137 | 0.857316542 | 0.9254719 | 0.025353777 |
| ENSMUSG00000005103 | Wdr1 | 22388 | protein_coding | 5756.185212 | -0.025356143 | 0.147450507 | -0.171963759 | 0.863466021 | 0.929087805 | 0.025356143 |
| ENSMUSG00000010936 | Vac14 | 234729 | protein_coding | 311.3685244 | 0.025356806 | 0.110708063 | 0.229042093 | 0.8188362 | 0.903777566 | 0.025356806 |
| ENSMUSG00000097615 | Gm2061 | 100039138 | lncRNA | 83.21187794 | 0.025380406 | 0.440225659 | 0.057653174 | 0.954024894 | 0.976611016 | 0.025380406 |
| ENSMUSG00000020536 | Llgl1 | 16897 | protein_coding | 378.0945846 | 0.02544852 | 0.083682107 | 0.304109454 | 0.761044504 | 0.869186728 | 0.02544852 |
| ENSMUSG00000005871 | Apc | 11789 | protein_coding | 1988.262116 | -0.025454338 | 0.091321943 | -0.278731895 | 0.780450584 | 0.880641803 | 0.025454338 |

| ensembl_gene_id | external_gene_name | entrezgene_id | gene_biotype | baseMean | log2FoldChange | lfcSE | stat | pvalue | padj | absLog2FC |
|---|---|---|---|---|---|---|---|---|---|---|
| ENSMUSG00000040479 | Dgkz | 104418 | protein_coding | 1733.174722 | -0.025465189 | 0.083774557 | -0.303972828 | 0.761148593 | 0.869186728 | 0.025465189 |
| ENSMUSG00000073678 | Pgap1 | 241062 | protein_coding | 1010.524949 | 0.025476465 | 0.250742993 | 0.101603895 | 0.919071088 | 0.959932084 | 0.025476465 |
| ENSMUSG00000028811 | Yars | 107271 | protein_coding | 907.4617888 | 0.025482594 | 0.090144973 | 0.282684585 | 0.777418634 | 0.879080221 | 0.025482594 |
| ENSMUSG00000055553 | Kxd1 | 75620 | protein_coding | 607.3627822 | -0.02550341 | 0.116563502 | -0.218794128 | 0.826810422 | 0.908412958 | 0.02550341 |
| ENSMUSG00000041297 | Cdk13 | 69562 | protein_coding | 1288.72283 | -0.02550425 | 0.053178356 | -0.479598314 | 0.631513045 | 0.786637975 | 0.02550425 |
| ENSMUSG00000006611 | Hfe | 15216 | protein_coding | 141.6702527 | -0.025523283 | 0.170462932 | -0.149729228 | 0.880978247 | 0.93955283 | 0.025523283 |
| ENSMUSG00000054150 | Syne3 | 212073 | protein_coding | 79.13645608 | -0.025550135 | 0.181317845 | -0.140913518 | 0.887938263 | 0.943170468 | 0.025550135 |
| ENSMUSG00000041147 | Brca2 | 12190 | protein_coding | 133.5419456 | -0.025580888 | 0.167419077 | -0.152795536 | 0.878559518 | 0.938355529 | 0.025580888 |
| ENSMUSG00000013997 | Nit1 | 27045 | protein_coding | 624.3732005 | -0.02559901 | 0.082367895 | -0.310788689 | 0.755961269 | 0.865943984 | 0.02559901 |
| ENSMUSG00000054920 | Klhl5 | 71778 | protein_coding | 272.8476973 | -0.025627625 | 0.177685577 | -0.144230193 | 0.8853187 | 0.94220607 | 0.025627625 |
| ENSMUSG00000069539 | Scyl2 | 213326 | protein_coding | 1136.046705 | 0.025657836 | 0.089495472 | 0.286694235 | 0.774346452 | 0.877412698 | 0.025657836 |
| ENSMUSG00000045394 | Epcam | 17075 | protein_coding | 22668.17527 | -0.025673382 | 0.107672116 | -0.238440398 | 0.811539537 | 0.899694484 | 0.025673382 |
| ENSMUSG00000054484 | Tmem62 | 96957 | protein_coding | 282.0730489 | -0.025695747 | 0.143881614 | -0.178589512 | 0.85826003 | 0.926176542 | 0.025695747 |
| ENSMUSG00000019726 | Lyst | 17101 | protein_coding | 374.6682214 | -0.025710981 | 0.135446657 | -0.189823665 | 0.849447311 | 0.920572697 | 0.025710981 |
| ENSMUSG00000018040 | Rrp7a | 74778 | protein_coding | 608.8576824 | 0.025855257 | 0.094441779 | 0.273769267 | 0.784261943 | 0.882887587 | 0.025855257 |
| ENSMUSG00000019826 | Zbtb24 | 268294 | protein_coding | 217.2482778 | 0.025877816 | 0.108993371 | 0.237425595 | 0.812326635 | 0.900072074 | 0.025877816 |
| ENSMUSG00000041945 | Mfsd9 | 211798 | protein_coding | 126.4587251 | -0.025898538 | 0.158611734 | -0.163282612 | 0.870295923 | 0.933211685 | 0.025898538 |
| ENSMUSG00000036168 | Ccdc38 | 237465 | protein_coding | 7.886588826 | -0.0259529 | 0.527570965 | -0.049193193 | 0.960765336 | 0.980957331 | 0.0259529 |
| ENSMUSG00000028587 | Orc1 | 18392 | protein_coding | 179.7434782 | 0.025960522 | 0.188581976 | 0.137661734 | 0.890507763 | 0.944864393 | 0.025960522 |
| ENSMUSG00000003360 | Ddx23 | 74351 | protein_coding | 1005.873155 | -0.025982245 | 0.083006826 | -0.313013351 | 0.754270519 | 0.865108984 | 0.025982245 |
| ENSMUSG00000026765 | Lypd6b | 71897 | protein_coding | 88.23249082 | -0.025989208 | 0.1901939 | -0.136645854 | 0.89131073 | 0.945137528 | 0.025989208 |
| ENSMUSG00000052406 | Rexo4 | 227656 | protein_coding | 465.7650344 | -0.02601362 | 0.10096609 | -0.257647099 | 0.796679277 | 0.890616755 | 0.02601362 |
| ENSMUSG00000061787 | Rps17 | 20068 | protein_coding | 5243.63402 | -0.026078307 | 0.091758739 | -0.284205157 | 0.776253164 | 0.878261145 | 0.026078307 |
| ENSMUSG00000068452 | Duox2 | 214593 | protein_coding | 1621.58177 | 0.026094296 | 0.252006311 | 0.1035462 | 0.917529484 | 0.95913082 | 0.026094296 |
| ENSMUSG00000053453 | Thoc7 | 66231 | protein_coding | 580.8917605 | -0.02611095 | 0.073429705 | -0.355591102 | 0.722146808 | 0.844044564 | 0.02611095 |
| ENSMUSG00000030830 | Itgal | 16408 | protein_coding | 12.41889114 | 0.026135072 | 0.516685162 | 0.050582199 | 0.959658447 | 0.980212886 | 0.026135072 |
| ENSMUSG00000036452 | Arhgap26 | 71302 | protein_coding | 427.4177889 | -0.026152643 | 0.127760197 | -0.204701023 | 0.837805721 | 0.914450046 | 0.026152643 |
| ENSMUSG00000026389 | Steap3 | 68428 | protein_coding | 307.0120195 | 0.026267463 | 0.171149275 | 0.153476918 | 0.878022192 | 0.938098339 | 0.026267463 |
| ENSMUSG00000029577 | Ube3b | 117146 | protein_coding | 1464.640502 | -0.026336786 | 0.062897494 | -0.418725516 | 0.675416746 | 0.81340072 | 0.026336786 |
| ENSMUSG00000029569 | Tmem168 | 101118 | protein_coding | 1368.840193 | 0.026383583 | 0.096019227 | 0.274773959 | 0.783489906 | 0.882499627 | 0.026383583 |
| ENSMUSG00000022185 | Acin1 | 56215 | protein_coding | 1672.042064 | -0.026389833 | 0.063368653 | -0.41644933 | 0.677081238 | 0.814727654 | 0.026389833 |
| ENSMUSG00000033429 | Mcee | 73724 | protein_coding | 467.1652803 | 0.026422007 | 0.097567676 | 0.270806969 | 0.7865395 | 0.884371111 | 0.026422007 |
| ENSMUSG00000060002 | Chpt1 | 212862 | protein_coding | 2356.843585 | 0.02643371 | 0.119743834 | 0.220752164 | 0.825285412 | 0.907901841 | 0.02643371 |
| ENSMUSG00000029111 | Nelfa | 24116 | protein_coding | 287.3776922 | -0.026466922 | 0.087322977 | -0.303092306 | 0.761819518 | 0.869340971 | 0.026466922 |
| ENSMUSG00000002413 | Braf | 109880 | protein_coding | 536.1224451 | 0.02647582 | 0.117946801 | 0.224472558 | 0.822389618 | 0.906005106 | 0.02647582 |
| ENSMUSG00000018995 | Nars2 | 244141 | protein_coding | 260.4666134 | -0.026480648 | 0.100307785 | -0.263993943 | 0.791784586 | 0.887137372 | 0.026480648 |
| ENSMUSG00000097170 | Gm16982 | 100036523 | lncRNA | 9.62744936 | 0.026496917 | 0.445323355 | 0.059500399 | 0.952553548 | 0.976127816 | 0.026496917 |
| ENSMUSG00000066839 | Ecsit | 26940 | protein_coding | 327.4990595 | -0.0265086 | 0.105695405 | -0.250801824 | 0.801967332 | 0.893955886 | 0.0265086 |

| ensembl_gene_id | external_gene_name | entrezgene_id | gene_biotype | baseMean | log2FoldChange | lfcSE | stat | pvalue | padj | absLog2FC |
|---|---|---|---|---|---|---|---|---|---|---|
| ENSMUSG00000045980 | Tmem104 | 320534 | protein_coding | 85.17154187 | -0.026525631 | 0.130521429 | -0.203228167 | 0.838956698 | 0.915199917 | 0.026525631 |
| ENSMUSG00000020430 | Pes1 | 64934 | protein_coding | 748.924129 | -0.02653057 | 0.106202545 | -0.249811053 | 0.802733472 | 0.894034894 | 0.02653057 |
| ENSMUSG00000033983 | Coil | 12812 | protein_coding | 134.9254055 | -0.02653914 | 0.134072104 | -0.197946773 | 0.843086708 | 0.917115541 | 0.02653914 |
| ENSMUSG00000036241 | Ube2r2 | 67615 | protein_coding | 1100.38718 | 0.026551554 | 0.097018014 | 0.273676539 | 0.784333209 | 0.882887587 | 0.026551554 |
| ENSMUSG00000024121 | Atp6v0c | 11984 | protein_coding | 688.8890003 | -0.026555468 | 0.110858292 | -0.239544266 | 0.810683576 | 0.899310849 | 0.026555468 |
| ENSMUSG00000036180 | Gatad2a | 234366 | protein_coding | 1081.490663 | -0.026584474 | 0.098305288 | -0.270427706 | 0.786831227 | 0.884458731 | 0.026584474 |
| ENSMUSG00000004069 | Dnaja3 | 83945 | protein_coding | 1191.138624 | 0.026591192 | 0.090814 | 0.292809391 | 0.76966785 | 0.874537702 | 0.026591192 |
| ENSMUSG00000041241 | Mul1 | 68350 | protein_coding | 793.1315146 | 0.026610779 | 0.069168689 | 0.384722908 | 0.700442721 | 0.829926899 | 0.026610779 |
| ENSMUSG00000063856 | Gpx1 | 14775 | protein_coding | 2354.277013 | -0.02661178 | 0.12100654 | -0.219920176 | 0.825933323 | 0.908150349 | 0.02661178 |
| ENSMUSG00000052566 | Hook2 | 170833 | protein_coding | 1179.008575 | -0.026652239 | 0.072017396 | -0.370080563 | 0.711322464 | 0.837163904 | 0.026652239 |
| ENSMUSG00000026490 | Cdc42bpa | 226751 | protein_coding | 934.846477 | -0.026669867 | 0.104410092 | -0.255433805 | 0.798388064 | 0.891775953 | 0.026669867 |
| ENSMUSG00000032359 | Ctsh | 13036 | protein_coding | 1015.654136 | -0.026701745 | 0.132483283 | -0.201548035 | 0.840270074 | 0.915836464 | 0.026701745 |
| ENSMUSG00000088252 | Snord13 | 100217422 | snoRNA | 20.97308258 | -0.026702474 | 0.336091672 | -0.079449971 | 0.936674723 | 0.968704702 | 0.026702474 |
| ENSMUSG00000053929 | Cyhr1 | 54151 | protein_coding | 925.0496906 | 0.026726279 | 0.079564347 | 0.335907725 | 0.736940451 | 0.852840632 | 0.026726279 |
| ENSMUSG00000032547 | Ryk | 20187 | protein_coding | 848.1461293 | -0.026762262 | 0.070201999 | -0.381217955 | 0.703041527 | 0.83188808 | 0.026762262 |
| ENSMUSG00000079610 | Ankrd39 | 109346 | protein_coding | 44.07966016 | -0.02676566 | 0.186370663 | -0.143615201 | 0.885804336 | 0.94220607 | 0.02676566 |
| ENSMUSG00000062627 | Mysm1 | 320713 | protein_coding | 574.1540055 | -0.02688028 | 0.072543244 | -0.37054146 | 0.710979091 | 0.837163904 | 0.02688028 |
| ENSMUSG00000021470 | Ercc6l2 | 76251 | protein_coding | 510.714834 | 0.026935067 | 0.083765182 | 0.321554453 | 0.747790254 | 0.859851228 | 0.026935067 |
| ENSMUSG00000026566 | Mpzl1 | 68481 | protein_coding | 414.7682707 | -0.026962153 | 0.160923368 | -0.16754654 | 0.866940032 | 0.931064717 | 0.026962153 |
| ENSMUSG00000046826 | Fam187b | 76415 | protein_coding | 45.56504075 | 0.026967414 | 0.257681299 | 0.104654139 | 0.916650254 | 0.958597942 | 0.026967414 |
| ENSMUSG00000090841 | Myl6 | 17904 | protein_coding | 6189.622906 | 0.026980472 | 0.078485734 | 0.343762755 | 0.731024713 | 0.848794403 | 0.026980472 |
| ENSMUSG00000029101 | Rgs12 | 71729 | protein_coding | 75.25542693 | -0.027006118 | 0.221303189 | -0.122032215 | 0.902873505 | 0.952003786 | 0.027006118 |
| ENSMUSG00000003032 | Klf4 | 16600 | protein_coding | 7591.943022 | 0.027055194 | 0.133177642 | 0.203151173 | 0.839016875 | 0.915199917 | 0.027055194 |
| ENSMUSG00000031388 | Naa10 | 56292 | protein_coding | 238.8028988 | -0.027056739 | 0.102973558 | -0.262754237 | 0.79274001 | 0.887895568 | 0.027056739 |
| ENSMUSG00000041408 | Wapl | 218914 | protein_coding | 3148.788588 | 0.027103639 | 0.062041154 | 0.436865482 | 0.662208906 | 0.804505636 | 0.027103639 |
| ENSMUSG00000018326 | Ywhab | 54401 | protein_coding | 2696.498295 | 0.027131218 | 0.091625842 | 0.296108795 | 0.767146997 | 0.872918801 | 0.027131218 |
| ENSMUSG00000064177 | Ghrl | 58991 | protein_coding | 11.5387716 | -0.027131924 | 0.334005018 | -0.081232083 | 0.935257385 | 0.967926267 | 0.027131924 |
| ENSMUSG00000046591 | Ticrr | 77011 | protein_coding | 110.8295755 | 0.027241557 | 0.148984749 | 0.182847957 | 0.854917319 | 0.923815207 | 0.027241557 |
| ENSMUSG00000062470 | Fbxl12os | 66662 | lncRNA | 18.54950106 | 0.027259907 | 0.309085571 | 0.08819534 | 0.929721421 | 0.965328339 | 0.027259907 |
| ENSMUSG00000032422 | Snx14 | 244962 | protein_coding | 811.7712465 | 0.027372566 | 0.095518917 | 0.286566963 | 0.774443913 | 0.877460709 | 0.027372566 |
| ENSMUSG00000078515 | Ddi2 | 68817 | protein_coding | 1711.074023 | 0.027401994 | 0.086985776 | 0.315016949 | 0.752748785 | 0.863941681 | 0.027401994 |
| ENSMUSG00000020739 | Nup85 | 445007 | protein_coding | 429.8134142 | 0.027409374 | 0.113747462 | 0.240966908 | 0.809580764 | 0.898543824 | 0.027409374 |
| ENSMUSG00000032508 | Myd88 | 17874 | protein_coding | 405.188665 | -0.027444493 | 0.112969053 | -0.242938149 | 0.808053312 | 0.897473897 | 0.027444493 |
| ENSMUSG00000015980 | Lrrc27 | 76612 | protein_coding | 8.000817292 | -0.027445141 | 0.476940445 | -0.057544167 | 0.954111724 | 0.976611016 | 0.027445141 |
| ENSMUSG00000034297 | Med13 | 327987 | protein_coding | 1693.934649 | -0.027497483 | 0.118533424 | -0.231980841 | 0.816552895 | 0.90264989 | 0.027497483 |
| ENSMUSG00000092819 | Gm23639 | 115489002 | snoRNA | 14.69818548 | 0.02751451 | 0.390163314 | 0.070520494 | 0.943779389 | 0.972663741 | 0.02751451 |
| ENSMUSG00000024947 | Men1 | 17283 | protein_coding | 373.0626274 | 0.027525656 | 0.099784914 | 0.27584987 | 0.782663378 | 0.882005256 | 0.027525656 |
| ENSMUSG00000050621 | Rps27rt | 100043813 | protein_coding | 43.46232312 | 0.027535655 | 0.21101775 | 0.130489758 | 0.896178957 | 0.948192606 | 0.027535655 |

| ensembl_gene_id | external_gene_name | entrezgene_id | gene_biotype | baseMean | log2FoldChange | lfcSE | stat | pvalue | padj | absLog2FC |
|---|---|---|---|---|---|---|---|---|---|---|
| ENSMUSG00000020652 | Cenpo | 52504 | protein_coding | 141.9425507 | -0.027549434 | 0.09635674 | -0.285910815 | 0.77494643 | 0.877593098 | 0.027549434 |
| ENSMUSG00000072501 | Phf20l1 | 239510 | protein_coding | 901.5374608 | -0.027557626 | 0.063069035 | -0.436943836 | 0.66215208 | 0.804505636 | 0.027557626 |
| ENSMUSG00000034602 | Mon2 | 67074 | protein_coding | 1904.154628 | -0.027559207 | 0.066845991 | -0.412279131 | 0.680134843 | 0.817167367 | 0.027559207 |
| ENSMUSG00000026608 | Kctd3 | 226823 | protein_coding | 790.7630928 | 0.027585259 | 0.097844734 | 0.281928909 | 0.777998022 | 0.879310181 | 0.027585259 |
| ENSMUSG00000118559 | A930007A09Rik | 432999 | protein_coding | 9.100177758 | -0.027615229 | 0.448049442 | -0.061634335 | 0.950854034 | 0.975614007 | 0.027615229 |
| ENSMUSG00000027305 | Ndufaf1 | 69702 | protein_coding | 230.3504569 | -0.027640695 | 0.089781189 | -0.307867336 | 0.758183281 | 0.867544258 | 0.027640695 |
| ENSMUSG00000023027 | Atf1 | 11908 | protein_coding | 298.194156 | -0.027661485 | 0.079208732 | -0.349222671 | 0.726922147 | 0.846375436 | 0.027661485 |
| ENSMUSG00000027966 | Col11a1 | 12814 | protein_coding | 6.782618846 | 0.027667973 | 1.437497013 | 0.019247326 | 0.984643804 | 0.993212987 | 0.027667973 |
| ENSMUSG00000029106 | Add1 | 11518 | protein_coding | 3645.835642 | 0.027702473 | 0.072021782 | 0.38464021 | 0.700503999 | 0.829926899 | 0.027702473 |
| ENSMUSG00000029775 | Klhdc10 | 76788 | protein_coding | 838.0288358 | 0.027732275 | 0.0671209 | 0.413169006 | 0.679482794 | 0.816753599 | 0.027732275 |
| ENSMUSG00000003199 | Mpnd | 68047 | protein_coding | 812.1303672 | -0.027759833 | 0.09112988 | -0.304618339 | 0.760656851 | 0.869090323 | 0.027759833 |
| ENSMUSG00000032497 | Lrrfip2 | 71268 | protein_coding | 1342.651719 | 0.027796569 | 0.070522322 | 0.394152782 | 0.693468244 | 0.825651233 | 0.027796569 |
| ENSMUSG00000017781 | Pitpna | 18738 | protein_coding | 1740.145323 | 0.027815963 | 0.101782834 | 0.273287371 | 0.784632322 | 0.883161871 | 0.027815963 |
| ENSMUSG00000031600 | Vps37a | 52348 | protein_coding | 745.1697576 | 0.027821751 | 0.057781766 | 0.481497059 | 0.630163269 | 0.785935003 | 0.027821751 |
| ENSMUSG00000030051 | Aplf | 72103 | protein_coding | 166.6205769 | -0.02786115 | 0.125388254 | -0.222199041 | 0.824158941 | 0.90712571 | 0.02786115 |
| ENSMUSG00000021981 | Cab39l | 69008 | protein_coding | 747.2973645 | 0.027911616 | 0.071119272 | 0.392462052 | 0.694716845 | 0.826520011 | 0.027911616 |
| ENSMUSG00000071654 | Uqcc3 | 107197 | protein_coding | 242.5631904 | -0.027928962 | 0.127583541 | -0.218907252 | 0.826722297 | 0.908412958 | 0.027928962 |
| ENSMUSG00000046573 | Lyrm4 | 380840 | protein_coding | 118.0482577 | -0.027953664 | 0.12822563 | -0.218003714 | 0.827426217 | 0.908703155 | 0.027953664 |
| ENSMUSG00000020745 | Pafah1b1 | 18472 | protein_coding | 2612.010607 | 0.027972457 | 0.063507325 | 0.44046033 | 0.659603738 | 0.802969027 | 0.027972457 |
| ENSMUSG00000006288 | Ttc5 | 219022 | protein_coding | 414.0055063 | 0.027973873 | 0.092855648 | 0.301261946 | 0.763214756 | 0.870185014 | 0.027973873 |
| ENSMUSG00000048047 | Zbtb33 | 56805 | protein_coding | 462.6308468 | 0.027979462 | 0.141551369 | 0.197662953 | 0.843308777 | 0.917115541 | 0.027979462 |
| ENSMUSG00000058070 | Eml1 | 68519 | protein_coding | 57.74777881 | -0.02799048 | 0.251377087 | -0.111348574 | 0.911339938 | 0.956372548 | 0.02799048 |
| ENSMUSG00000061904 | Slc25a3 | 18674 | protein_coding | 7486.571895 | 0.028023535 | 0.088115718 | 0.318031061 | 0.750461376 | 0.862088305 | 0.028023535 |
| ENSMUSG00000029147 | Ppm1g | 14208 | protein_coding | 1114.558693 | 0.028044504 | 0.1271045 | 0.220641314 | 0.82537173 | 0.907908903 | 0.028044504 |
| ENSMUSG00000057236 | Rbbp4 | 19646 | protein_coding | 1621.847429 | 0.028046448 | 0.075061167 | 0.373647912 | 0.708666283 | 0.83562604 | 0.028046448 |
| ENSMUSG00000025243 | Slc6a20b | 22599 | protein_coding | 15.06409273 | -0.028060253 | 0.44100141 | -0.063628488 | 0.949266048 | 0.975117538 | 0.028060253 |
| ENSMUSG00000041841 | Rpl37 | 67281 | protein_coding | 4058.279134 | -0.028109981 | 0.108108793 | -0.260015678 | 0.79485168 | 0.889189108 | 0.028109981 |
| ENSMUSG00000070390 | Nlrp1b | 637515 | protein_coding | 245.750037 | -0.028128543 | 0.194326984 | -0.144748518 | 0.884909431 | 0.942104511 | 0.028128543 |
| ENSMUSG00000003198 | Zfp959 | 224893 | protein_coding | 121.0151706 | 0.028131657 | 0.110373788 | 0.254876247 | 0.798818683 | 0.892069269 | 0.028131657 |
| ENSMUSG00000034994 | Eef2 | 13629 | protein_coding | 37971.35299 | 0.028134732 | 0.072904579 | 0.385911727 | 0.699562045 | 0.829246975 | 0.028134732 |
| ENSMUSG00000074925 | Ptar1 | 72351 | protein_coding | 627.7657757 | 0.028156031 | 0.077909901 | 0.361392213 | 0.717806268 | 0.841234511 | 0.028156031 |
| ENSMUSG00000034252 | Senp6 | 215351 | protein_coding | 1929.07544 | 0.028156707 | 0.057643831 | 0.488460023 | 0.625224036 | 0.782844779 | 0.028156707 |
| ENSMUSG00000071078 | Nr2c2ap | 75692 | protein_coding | 33.28356666 | 0.028159928 | 0.241841039 | 0.116439822 | 0.907303976 | 0.954095762 | 0.028159928 |
| ENSMUSG00000024603 | Dctn4 | 67665 | protein_coding | 1322.835716 | 0.028200318 | 0.083987054 | 0.335769813 | 0.737044456 | 0.852861903 | 0.028200318 |
| ENSMUSG00000020211 | Sf3a2 | 20222 | protein_coding | 176.1023084 | -0.028228537 | 0.112330275 | -0.251299455 | 0.801582597 | 0.893676478 | 0.028228537 |
| ENSMUSG00000027805 | Pfn2 | 18645 | protein_coding | 19.8917377 | -0.028270471 | 0.282718285 | -0.0999952 | 0.920348136 | 0.960510098 | 0.028270471 |
| ENSMUSG00000032410 | Xrn1 | 24127 | protein_coding | 737.6823023 | -0.028273459 | 0.131714564 | -0.214657046 | 0.830034714 | 0.909583277 | 0.028273459 |
| ENSMUSG00000053985 | Zfp14 | 243906 | protein_coding | 36.67123871 | -0.028293266 | 0.256358879 | -0.11036585 | 0.912119235 | 0.956746934 | 0.028293266 |

| ensembl_gene_id | external_gene_name | entrezgene_id | gene_biotype | baseMean | log2FoldChange | lfcSE | stat | pvalue | padj | absLog2FC |
|---|---|---|---|---|---|---|---|---|---|---|
| ENSMUSG00000027601 | Mtfr1 | 67472 | protein_coding | 733.6662762 | -0.028345059 | 0.068340767 | -0.414760619 | 0.678317151 | 0.815942199 | 0.028345059 |
| ENSMUSG00000109901 | Chmp1b | 67064 | protein_coding | 1043.753574 | 0.028363447 | 0.103294878 | 0.274587158 | 0.783633433 | 0.882598877 | 0.028363447 |
| ENSMUSG00000029528 | Pxn | 19303 | protein_coding | 1562.214399 | 0.028408076 | 0.095230721 | 0.298307899 | 0.765468176 | 0.87181817 | 0.028408076 |
| ENSMUSG00000068854 | H2bc21 | 319190 | protein_coding | 185.3846942 | 0.028417419 | 0.143664854 | 0.197803556 | 0.843198764 | 0.917115541 | 0.028417419 |
| ENSMUSG00000106583 | 4930447N08Rik | 71647 | lncRNA | 12.20631717 | -0.028488934 | 0.375017261 | -0.075966995 | 0.939445357 | 0.970446184 | 0.028488934 |
| ENSMUSG00000009640 | Dmap1 | 66233 | protein_coding | 204.9465862 | 0.028529871 | 0.122993499 | 0.231962431 | 0.816567193 | 0.90264989 | 0.028529871 |
| ENSMUSG00000091764 | Zfp964 | 636741 | protein_coding | 49.89870619 | -0.028531904 | 0.201673612 | -0.141475645 | 0.887494199 | 0.942901703 | 0.028531904 |
| ENSMUSG00000030650 | Tmc5 | 74424 | protein_coding | 1849.938128 | 0.028537703 | 0.123342635 | 0.231369331 | 0.817027889 | 0.902675156 | 0.028537703 |
| ENSMUSG00000022500 | Litaf | 56722 | protein_coding | 1151.592622 | 0.028538383 | 0.127275049 | 0.224226064 | 0.822581404 | 0.906153717 | 0.028538383 |
| ENSMUSG00000072612 | Gm10382 | 639281 | lncRNA | 205.3601644 | -0.028541625 | 0.24663386 | -0.115724681 | 0.907870745 | 0.954429756 | 0.028541625 |
| ENSMUSG00000027374 | Mrps5 | 77721 | protein_coding | 527.0462157 | -0.028586369 | 0.105127483 | -0.271920983 | 0.785682777 | 0.883803077 | 0.028586369 |
| ENSMUSG00000041837 | Pdcd7 | 50996 | protein_coding | 177.3698136 | -0.028600257 | 0.107129074 | -0.266970075 | 0.789492202 | 0.885815145 | 0.028600257 |
| ENSMUSG00000020743 | Mif4gd | 69674 | protein_coding | 464.0626331 | -0.028600591 | 0.071558424 | -0.399681681 | 0.689390987 | 0.823011515 | 0.028600591 |
| ENSMUSG00000079434 | Neu2 | 23956 | protein_coding | 17.9512965 | 0.028610867 | 0.284675293 | 0.100503514 | 0.919944593 | 0.96034064 | 0.028610867 |
| ENSMUSG00000035234 | Abraxas1 | 70681 | protein_coding | 146.4820891 | 0.02861234 | 0.133069525 | 0.215017975 | 0.829753304 | 0.909462609 | 0.02861234 |
| ENSMUSG00000028953 | Abcf2 | 27407 | protein_coding | 592.2945022 | 0.028627829 | 0.139426939 | 0.205324946 | 0.837318255 | 0.914085243 | 0.028627829 |
| ENSMUSG00000011837 | Snapc2 | 102209 | protein_coding | 142.135533 | 0.028666198 | 0.138529682 | 0.206931809 | 0.836063114 | 0.913048562 | 0.028666198 |
| ENSMUSG00000039148 | Sart1 | 20227 | protein_coding | 697.0599597 | -0.028681378 | 0.084105007 | -0.341018675 | 0.733089523 | 0.850197424 | 0.028681378 |
| ENSMUSG00000032583 | Mon1a | 72825 | protein_coding | 496.7395944 | -0.028685457 | 0.100975519 | -0.284083287 | 0.776346555 | 0.878304419 | 0.028685457 |
| ENSMUSG00000009995 | Tafazzin | 66826 | protein_coding | 622.7852326 | 0.028705462 | 0.082510463 | 0.347900877 | 0.727914627 | 0.846662573 | 0.028705462 |
| ENSMUSG00000062270 | Morf4l1 | 21761 | protein_coding | 1232.745845 | -0.028740862 | 0.057885399 | -0.496513149 | 0.619532413 | 0.778489692 | 0.028740862 |
| ENSMUSG00000036291 | Ap5m1 | 74385 | protein_coding | 802.4468636 | -0.028750023 | 0.071681499 | -0.4010801 | 0.688361152 | 0.822275118 | 0.028750023 |
| ENSMUSG00000033272 | Slc35a4 | 67843 | protein_coding | 683.1309033 | 0.028762315 | 0.08463415 | 0.339842901 | 0.733974838 | 0.850749611 | 0.028762315 |
| ENSMUSG00000037958 | Nsrp1 | 237859 | protein_coding | 291.5622375 | 0.028770886 | 0.104204265 | 0.276100847 | 0.78247061 | 0.881912813 | 0.028770886 |
| ENSMUSG00000008226 | Scrn3 | 74616 | protein_coding | 211.8321587 | 0.028783837 | 0.137113461 | 0.209927143 | 0.833724537 | 0.911278944 | 0.028783837 |
| ENSMUSG00000039197 | Adk | 11534 | protein_coding | 2415.185802 | -0.028850739 | 0.102338312 | -0.281915327 | 0.778008437 | 0.879310181 | 0.028850739 |
| ENSMUSG00000039294 | Cybc1 | 217370 | protein_coding | 283.5393099 | 0.028887264 | 0.105919375 | 0.272728798 | 0.785061696 | 0.883520292 | 0.028887264 |
| ENSMUSG00000033444 | Specc1l | 74392 | protein_coding | 2083.048088 | -0.028893751 | 0.163044317 | -0.177214096 | 0.859340224 | 0.926766895 | 0.028893751 |
| ENSMUSG00000058586 | Serhl | 68607 | protein_coding | 614.4404966 | -0.028900116 | 0.124730629 | -0.231700238 | 0.816770846 | 0.902675156 | 0.028900116 |
| ENSMUSG00000027465 | Tbc1d20 | 67231 | protein_coding | 1089.8746 | 0.029021035 | 0.066512304 | 0.43632581 | 0.662600358 | 0.804658319 | 0.029021035 |
| ENSMUSG00000028822 | Tmem50a | 71817 | protein_coding | 655.9061678 | -0.029031929 | 0.12589039 | -0.230612753 | 0.81761566 | 0.903136927 | 0.029031929 |
| ENSMUSG00000021939 | Ctsb | 13030 | protein_coding | 4221.053464 | 0.029041177 | 0.081694987 | 0.355482976 | 0.722227797 | 0.844044564 | 0.029041177 |
| ENSMUSG00000036323 | Srp72 | 66661 | protein_coding | 1585.868158 | 0.029041524 | 0.050409249 | 0.576114982 | 0.564537465 | 0.741125944 | 0.029041524 |
| ENSMUSG00000047557 | Lxn | 17035 | protein_coding | 165.1983388 | 0.029062198 | 0.096360059 | 0.30160004 | 0.762956978 | 0.870097644 | 0.029062198 |
| ENSMUSG00000040351 | Ankib1 | 70797 | protein_coding | 1604.774768 | -0.029094826 | 0.073212347 | -0.397403261 | 0.691070111 | 0.824089596 | 0.029094826 |
| ENSMUSG00000007216 | Zfp775 | 243372 | protein_coding | 79.94347218 | -0.029114768 | 0.199823622 | -0.145702331 | 0.884156381 | 0.941681233 | 0.029114768 |
| ENSMUSG00000020832 | Eral1 | 57837 | protein_coding | 193.9917504 | -0.029119316 | 0.10226607 | -0.284740733 | 0.775842781 | 0.877982787 | 0.029119316 |
| ENSMUSG00000029012 | Orc5 | 26429 | protein_coding | 255.0085341 | -0.029124251 | 0.097674763 | -0.298175801 | 0.765568991 | 0.871822248 | 0.029124251 |

| ensembl_gene_id | external_gene_name | entrezgene_id | gene_biotype | baseMean | log2FoldChange | lfcSE | stat | pvalue | padj | absLog2FC |
|---|---|---|---|---|---|---|---|---|---|---|
| ENSMUSG00000029655 | N4bp2l2 | 381695 | protein_coding | 1107.385891 | -0.029129641 | 0.063385446 | -0.459563549 | 0.645829528 | 0.795082139 | 0.029129641 |
| ENSMUSG00000077714 | Snord17 | 100313519 | snoRNA | 547.8654204 | 0.029188097 | 0.347377939 | 0.084024039 | 0.933037319 | 0.966437945 | 0.029188097 |
| ENSMUSG00000057440 | Mpp7 | 75739 | protein_coding | 431.7634042 | 0.029201697 | 0.074844979 | 0.390162409 | 0.696416457 | 0.827182798 | 0.029201697 |
| ENSMUSG00000019951 | Uhrf1bp1l | 75089 | protein_coding | 740.4101057 | -0.029291886 | 0.094185998 | -0.311000426 | 0.755800298 | 0.86582186 | 0.029291886 |
| ENSMUSG00000041471 | Shld2 | 75698 | protein_coding | 100.8930851 | 0.029296154 | 0.146323931 | 0.200214372 | 0.841312928 | 0.916315892 | 0.029296154 |
| ENSMUSG00000037536 | Fbxo34 | 78938 | protein_coding | 514.4855976 | -0.02932783 | 0.141564518 | -0.207169358 | 0.835877597 | 0.913007852 | 0.02932783 |
| ENSMUSG00000037600 | Kdf1 | 69073 | protein_coding | 330.7819444 | -0.029334742 | 0.08128345 | -0.360894403 | 0.718178388 | 0.841546779 | 0.029334742 |
| ENSMUSG00000021282 | Eif5 | 217869 | protein_coding | 4180.474264 | 0.029365458 | 0.090977779 | 0.322776155 | 0.746864773 | 0.859293709 | 0.029365458 |
| ENSMUSG00000014313 | Cox6c | 12864 | protein_coding | 5016.061488 | 0.029435829 | 0.09146296 | 0.321833329 | 0.747578964 | 0.859689126 | 0.029435829 |
| ENSMUSG00000024963 | Dnajc4 | 57431 | protein_coding | 222.7919165 | -0.02951265 | 0.08925433 | -0.330657906 | 0.740902902 | 0.855524244 | 0.02951265 |
| ENSMUSG00000018733 | Pex12 | 103737 | protein_coding | 158.9714497 | 0.029514888 | 0.118206192 | 0.249689861 | 0.802827199 | 0.894034894 | 0.029514888 |
| ENSMUSG00000021716 | Srek1ip1 | 67288 | protein_coding | 449.4639467 | 0.029542568 | 0.102837429 | 0.28727447 | 0.773902169 | 0.877099132 | 0.029542568 |
| ENSMUSG00000028894 | Inpp5b | 16330 | protein_coding | 376.050629 | 0.029544879 | 0.075088019 | 0.393469946 | 0.693972418 | 0.826066263 | 0.029544879 |
| ENSMUSG00000024219 | Anks1 | 224650 | protein_coding | 573.4523429 | -0.029627018 | 0.116398961 | -0.254529915 | 0.799086196 | 0.89211142 | 0.029627018 |
| ENSMUSG00000021901 | Bap1 | 104416 | protein_coding | 575.564098 | 0.029696121 | 0.070289976 | 0.422480162 | 0.672674578 | 0.81114094 | 0.029696121 |
| ENSMUSG00000029458 | Brap | 72399 | protein_coding | 730.4302669 | -0.029725823 | 0.059924913 | -0.496051161 | 0.619858316 | 0.778697315 | 0.029725823 |
| ENSMUSG00000039193 | Nlrc4 | 268973 | protein_coding | 1062.149658 | 0.02982695 | 0.170143092 | 0.17530509 | 0.860839912 | 0.927703527 | 0.02982695 |
| ENSMUSG00000024817 | Uhrf2 | 109113 | protein_coding | 1878.446215 | 0.029997495 | 0.081745671 | 0.366961271 | 0.713647907 | 0.838829892 | 0.029997495 |
| ENSMUSG00000037373 | Ctbp1 | 13016 | protein_coding | 2169.820255 | 0.030011569 | 0.080375127 | 0.373393741 | 0.708855417 | 0.835691442 | 0.030011569 |
| ENSMUSG00000020231 | Dip2a | 64451 | protein_coding | 996.0971263 | -0.030035189 | 0.099569621 | -0.301650125 | 0.762918792 | 0.870097644 | 0.030035189 |
| ENSMUSG00000002332 | Dhrs1 | 52585 | protein_coding | 409.7730585 | -0.030055136 | 0.113951812 | -0.263753032 | 0.791970228 | 0.887220551 | 0.030055136 |
| ENSMUSG00000045098 | Kmt5b | 225888 | protein_coding | 1041.046995 | 0.030061673 | 0.087365018 | 0.344092788 | 0.730776507 | 0.848568069 | 0.030061673 |
| ENSMUSG00000028729 | Ebna1bp2 | 69072 | protein_coding | 566.3704196 | -0.030063872 | 0.138270596 | -0.217427802 | 0.827874967 | 0.90891431 | 0.030063872 |
| ENSMUSG00000057842 | Zfp595 | 218314 | protein_coding | 153.0784168 | 0.030126202 | 0.140000262 | 0.215186757 | 0.829621715 | 0.909414769 | 0.030126202 |
| ENSMUSG00000045482 | Trrap | 100683 | protein_coding | 1088.427522 | -0.030139235 | 0.133745116 | -0.225348303 | 0.821708325 | 0.905442422 | 0.030139235 |
| ENSMUSG00000070639 | Lrrc8b | 433926 | protein_coding | 446.9138183 | 0.0301615 | 0.153170921 | 0.19691401 | 0.84389483 | 0.917324565 | 0.0301615 |
| ENSMUSG00000026836 | Acvr1 | 11477 | protein_coding | 155.8976875 | -0.030193077 | 0.183949362 | -0.164137981 | 0.869622523 | 0.932939149 | 0.030193077 |
| ENSMUSG00000054843 | Atrnl1 | 226255 | protein_coding | 2192.172658 | -0.03020848 | 0.074014509 | -0.408142679 | 0.683168928 | 0.818964084 | 0.03020848 |
| ENSMUSG00000014496 | Ankrd28 | 105522 | protein_coding | 818.7671666 | -0.03025295 | 0.08112335 | -0.372925304 | 0.709204039 | 0.835765162 | 0.03025295 |
| ENSMUSG00000022833 | Ccdc14 | 239839 | protein_coding | 92.88425889 | -0.03029468 | 0.153302411 | -0.197613854 | 0.843347195 | 0.917115541 | 0.03029468 |
| ENSMUSG00000035199 | Arl6ip5 | 65106 | protein_coding | 505.1321444 | -0.030300435 | 0.079324633 | -0.381980151 | 0.702476087 | 0.831280767 | 0.030300435 |
| ENSMUSG00000027620 | Rbm39 | 170791 | protein_coding | 4144.502958 | -0.030328178 | 0.055190449 | -0.549518602 | 0.582649602 | 0.75358854 | 0.030328178 |
| ENSMUSG00000030263 | Irag2 | 16970 | protein_coding | 286.6540501 | -0.030328843 | 0.30027575 | -0.101003305 | 0.919547837 | 0.960107027 | 0.030328843 |
| ENSMUSG00000000538 | Tom1l2 | 216810 | protein_coding | 693.1851666 | -0.030357105 | 0.096081445 | -0.315951794 | 0.752039098 | 0.863289099 | 0.030357105 |
| ENSMUSG00000038384 | Setd1b | 208043 | protein_coding | 174.8117618 | 0.030371692 | 0.286326078 | 0.106073788 | 0.915523808 | 0.95821748 | 0.030371692 |
| ENSMUSG00000035967 | Ints6l | 236790 | protein_coding | 418.7079686 | -0.030387475 | 0.179919156 | -0.168895162 | 0.865879104 | 0.930427199 | 0.030387475 |
| ENSMUSG00000111761 | 9230112J17Rik | 100504561 | lncRNA | 9.298339836 | -0.030406874 | 0.375241946 | -0.081032716 | 0.935415934 | 0.96799019 | 0.030406874 |
| ENSMUSG00000052926 | Rnaseh2a | 69724 | protein_coding | 325.7527305 | -0.03046165 | 0.108173048 | -0.28160111 | 0.778249389 | 0.879520098 | 0.03046165 |

| ensembl_gene_id | external_gene_name | entrezgene_id | gene_biotype | baseMean | log2FoldChange | lfcSE | stat | pvalue | padj | absLog2FC |
|---|---|---|---|---|---|---|---|---|---|---|
| ENSMUSG00000035877 | Zhx3 | 320799 | protein_coding | 237.7933023 | -0.030462083 | 0.130530249 | -0.233371832 | 0.815472684 | 0.901878136 | 0.030462083 |
| ENSMUSG00000031467 | Agpat5 | 52123 | protein_coding | 898.8249295 | 0.030473918 | 0.100535635 | 0.303115583 | 0.761801779 | 0.869340971 | 0.030473918 |
| ENSMUSG00000021453 | Gadd45g | 23882 | protein_coding | 173.3913983 | 0.030500686 | 0.316468384 | 0.096378302 | 0.923220124 | 0.962246443 | 0.030500686 |
| ENSMUSG00000063659 | Zbtb18 | 30928 | protein_coding | 516.4959108 | 0.030529568 | 0.09189667 | 0.33221626 | 0.739725967 | 0.854833615 | 0.030529568 |
| ENSMUSG00000015837 | Sqstm1 | 18412 | protein_coding | 2711.94821 | -0.030535547 | 0.12298634 | -0.248284057 | 0.803914634 | 0.894394399 | 0.030535547 |
| ENSMUSG00000037845 | Fdxacb1 | 382137 | protein_coding | 75.75680852 | -0.030554143 | 0.163646054 | -0.186708707 | 0.851889029 | 0.921803579 | 0.030554143 |
| ENSMUSG00000039578 | Ccser1 | 232035 | protein_coding | 375.6629045 | 0.030573368 | 0.096336375 | 0.317360581 | 0.750970014 | 0.862427144 | 0.030573368 |
| ENSMUSG00000067847 | Romo1 | 67067 | protein_coding | 267.6094833 | -0.030579608 | 0.145141105 | -0.210688822 | 0.833130098 | 0.911190103 | 0.030579608 |
| ENSMUSG00000019303 | Psmc3ip | 19183 | protein_coding | 25.65653193 | -0.030581041 | 0.23282184 | -0.131349539 | 0.895498805 | 0.94788821 | 0.030581041 |
| ENSMUSG00000066568 | Lsm14a | 67070 | protein_coding | 828.2941176 | 0.030584176 | 0.079924442 | 0.382663615 | 0.701969196 | 0.83080438 | 0.030584176 |
| ENSMUSG00000025162 | Csnk1d | 104318 | protein_coding | 1507.987965 | -0.030772598 | 0.069373152 | -0.4435808 | 0.65734569 | 0.801674955 | 0.030772598 |
| ENSMUSG00000018501 | Ncor1 | 20185 | protein_coding | 3248.842619 | -0.030820341 | 0.061906502 | -0.497853048 | 0.618587623 | 0.777789999 | 0.030820341 |
| ENSMUSG00000087440 | 4930577N17Rik | 67746 | TEC | 13.52441548 | 0.030823359 | 0.351929472 | 0.087583909 | 0.930207392 | 0.965369999 | 0.030823359 |
| ENSMUSG00000042567 | Nek10 | 674895 | protein_coding | 14.3710347 | 0.030890418 | 0.457233234 | 0.067559433 | 0.946136349 | 0.973769693 | 0.030890418 |
| ENSMUSG00000029642 | Polr1d | 20018 | protein_coding | 1135.581942 | 0.03089773 | 0.098833068 | 0.312625422 | 0.754565262 | 0.865216002 | 0.03089773 |
| ENSMUSG00000017550 | Atad5 | 237877 | protein_coding | 174.3313022 | -0.030928932 | 0.115597835 | -0.267556323 | 0.789040855 | 0.885558253 | 0.030928932 |
| ENSMUSG00000030846 | Tial1 | 21843 | protein_coding | 618.0956858 | -0.030956115 | 0.07212953 | -0.429173952 | 0.667796638 | 0.807344837 | 0.030956115 |
| ENSMUSG00000034863 | Ano8 | 382014 | protein_coding | 178.4441517 | -0.030958988 | 0.097606858 | -0.317180455 | 0.75110668 | 0.862429873 | 0.030958988 |
| ENSMUSG00000046909 | Tefm | 68550 | protein_coding | 127.7551962 | 0.030966614 | 0.136644812 | 0.226621222 | 0.820718289 | 0.904852281 | 0.030966614 |
| ENSMUSG00000066415 | Msl2 | 77853 | protein_coding | 1139.753808 | -0.030987583 | 0.091482354 | -0.338727432 | 0.734815073 | 0.851367123 | 0.030987583 |
| ENSMUSG00000025068 | Gsto1 | 14873 | protein_coding | 4803.993694 | 0.03102811 | 0.087521152 | 0.354521275 | 0.722948263 | 0.844115801 | 0.03102811 |
| ENSMUSG00000025869 | Nop16 | 28126 | protein_coding | 320.7214859 | -0.031043226 | 0.115637919 | -0.268451963 | 0.788351446 | 0.884955829 | 0.031043226 |
| ENSMUSG00000004285 | Atp6v1f | 66144 | protein_coding | 598.1082139 | 0.031062766 | 0.101731764 | 0.30533989 | 0.760107298 | 0.868689179 | 0.031062766 |
| ENSMUSG00000031095 | Cul4b | 72584 | protein_coding | 557.9969568 | -0.031122321 | 0.067369891 | -0.461961872 | 0.644108663 | 0.794001573 | 0.031122321 |
| ENSMUSG00000005609 | Ctr9 | 22083 | protein_coding | 986.3176559 | 0.031138154 | 0.068842956 | 0.452307043 | 0.651047803 | 0.798250711 | 0.031138154 |
| ENSMUSG00000049755 | Zfp672 | 319475 | protein_coding | 498.6285179 | -0.031155037 | 0.087887188 | -0.354488954 | 0.722972481 | 0.844115801 | 0.031155037 |
| ENSMUSG00000036890 | Gtdc1 | 227835 | protein_coding | 75.04217579 | -0.031213995 | 0.135562507 | -0.230255366 | 0.817893342 | 0.903241067 | 0.031213995 |
| ENSMUSG00000025326 | Ube3a | 22215 | protein_coding | 1336.807916 | -0.031229355 | 0.070625059 | -0.442185189 | 0.658355203 | 0.802316884 | 0.031229355 |
| ENSMUSG00000002107 | Celf2 | 14007 | protein_coding | 85.79629898 | -0.03122947 | 0.249915039 | -0.124960348 | 0.900554942 | 0.95050289 | 0.03122947 |
| ENSMUSG00000032324 | Tspan3 | 56434 | protein_coding | 4317.852141 | 0.031288413 | 0.079538336 | 0.39337525 | 0.694042347 | 0.826087767 | 0.031288413 |
| ENSMUSG00000026047 | Poglut2 | 72050 | protein_coding | 101.3621994 | -0.031308757 | 0.134634382 | -0.232546522 | 0.816113558 | 0.902273669 | 0.031308757 |
| ENSMUSG00000022414 | Tab1 | 66513 | protein_coding | 305.1342475 | -0.031311593 | 0.13598976 | -0.230249636 | 0.817897794 | 0.903241067 | 0.031311593 |
| ENSMUSG00000049516 | Spty2d1 | 101685 | protein_coding | 901.3549405 | -0.03136808 | 0.076838049 | -0.408236292 | 0.683100206 | 0.818964084 | 0.03136808 |
| ENSMUSG00000022867 | Usp25 | 30940 | protein_coding | 1105.518628 | 0.031374361 | 0.071252339 | 0.440327448 | 0.659699964 | 0.802969027 | 0.031374361 |
| ENSMUSG00000030243 | Recql | 19691 | protein_coding | 197.0771884 | -0.031382448 | 0.080781809 | -0.388484092 | 0.697657824 | 0.827854714 | 0.031382448 |
| ENSMUSG00000030811 | Fbxl19 | 233902 | protein_coding | 506.256499 | 0.031394629 | 0.096686756 | 0.324704548 | 0.745404692 | 0.858606157 | 0.031394629 |
| ENSMUSG00000030509 | Asb7 | 117589 | protein_coding | 513.8131688 | -0.031395686 | 0.109940376 | -0.285570115 | 0.775207395 | 0.877826204 | 0.031395686 |
| ENSMUSG00000058546 | Rpl23a | 268449 | protein_coding | 1829.452029 | -0.03139578 | 0.115468919 | -0.271898102 | 0.78570037 | 0.883803077 | 0.03139578 |

Supplementary Table 1

| ensembl_gene_id | external_gene_name | entrezgene_id | gene_biotype | baseMean | log2FoldChange | lfcSE | stat | pvalue | padj | absLog2FC |
|---|---|---|---|---|---|---|---|---|---|---|
| ENSMUSG00000020923 | Ubtf | 21429 | protein_coding | 1683.816247 | 0.031414695 | 0.077191737 | 0.406969658 | 0.684030274 | 0.819436988 | 0.031414695 |
| ENSMUSG00000038446 | Cdc40 | 71713 | protein_coding | 360.1603808 | -0.031450207 | 0.068250172 | -0.460807732 | 0.644936554 | 0.794481181 | 0.031450207 |
| ENSMUSG00000042590 | Ipo11 | 76582 | protein_coding | 408.5401786 | 0.031546826 | 0.080562654 | 0.391581258 | 0.695367638 | 0.826592861 | 0.031546826 |
| ENSMUSG00000035236 | Scai | 320271 | protein_coding | 150.9698623 | 0.031591685 | 0.177260154 | 0.178222145 | 0.858548519 | 0.926384278 | 0.031591685 |
| ENSMUSG00000097119 | B230354K17Rik | 105246278 | lncRNA | 208.8347219 | -0.031611205 | 0.135664272 | -0.233010537 | 0.815753224 | 0.902063132 | 0.031611205 |
| ENSMUSG00000004561 | Mettl17 | 52535 | protein_coding | 220.0575659 | 0.031624047 | 0.107195514 | 0.295012788 | 0.767984111 | 0.873434554 | 0.031624047 |
| ENSMUSG00000020024 | Cep83 | 77048 | protein_coding | 403.4092443 | -0.031625211 | 0.077314333 | -0.409047201 | 0.682505023 | 0.81859488 | 0.031625211 |
| ENSMUSG00000044795 | Cyb5d1 | 327951 | protein_coding | 101.1356337 | 0.031627288 | 0.112982205 | 0.279931588 | 0.779529992 | 0.880217901 | 0.031627288 |
| ENSMUSG00000096188 | Cmtm4 | 97487 | protein_coding | 3668.375181 | -0.031633796 | 0.070744856 | -0.447153299 | 0.654764378 | 0.80020617 | 0.031633796 |
| ENSMUSG00000020530 | Ggnbp2 | 217039 | protein_coding | 1647.831502 | -0.031656851 | 0.064001019 | -0.494630416 | 0.620861028 | 0.779403724 | 0.031656851 |
| ENSMUSG00000027067 | Ssrp1 | 20833 | protein_coding | 1440.618301 | 0.031664243 | 0.085364266 | 0.370930889 | 0.710689008 | 0.836895942 | 0.031664243 |
| ENSMUSG00000032299 | Commd4 | 66199 | protein_coding | 284.6396058 | -0.031690837 | 0.091739056 | -0.345445425 | 0.729759532 | 0.848005386 | 0.031690837 |
| ENSMUSG00000005198 | Polr2a | 20020 | protein_coding | 461.3448934 | -0.031695299 | 0.261421458 | -0.121242147 | 0.903499242 | 0.952386474 | 0.031695299 |
| ENSMUSG00000068921 | Dap3 | 65111 | protein_coding | 783.4777537 | -0.031704441 | 0.083327283 | -0.380480916 | 0.70358846 | 0.832349746 | 0.031704441 |
| ENSMUSG00000090659 | Zfp493 | 72958 | protein_coding | 67.28594449 | -0.031733623 | 0.241101371 | -0.13161942 | 0.895285325 | 0.947857168 | 0.031733623 |
| ENSMUSG00000038859 | Baiap2l1 | 66898 | protein_coding | 1622.938985 | 0.031737893 | 0.097516315 | 0.32546239 | 0.744831143 | 0.858257543 | 0.031737893 |
| ENSMUSG00000031629 | Cenpu | 71876 | protein_coding | 74.47015628 | -0.031787907 | 0.18659069 | -0.170361699 | 0.864725692 | 0.929689559 | 0.031787907 |
| ENSMUSG00000026171 | Rnf25 | 57751 | protein_coding | 249.519953 | -0.031811168 | 0.102575177 | -0.310125403 | 0.756465595 | 0.866334771 | 0.031811168 |
| ENSMUSG00000033295 | Ptprf | 19268 | protein_coding | 4910.434178 | -0.031889123 | 0.076364943 | -0.417588509 | 0.676248 | 0.814155566 | 0.031889123 |
| ENSMUSG00000039850 | Endov | 338371 | protein_coding | 207.3414237 | 0.031959296 | 0.122713969 | 0.260437309 | 0.794526467 | 0.889039236 | 0.031959296 |
| ENSMUSG00000063889 | Crem | 12916 | protein_coding | 80.71036094 | 0.03196944 | 0.180345169 | 0.177268068 | 0.859297831 | 0.926766895 | 0.03196944 |
| ENSMUSG00000034159 | Mab21l4 | 71874 | protein_coding | 110.8421842 | 0.032003354 | 0.150354217 | 0.212853052 | 0.831441582 | 0.910435964 | 0.032003354 |
| ENSMUSG00000035770 | Dync1li2 | 234663 | protein_coding | 1820.918802 | 0.032025223 | 0.061077818 | 0.524334757 | 0.600045727 | 0.766564672 | 0.032025223 |
| ENSMUSG00000020390 | Ube2b | 22210 | protein_coding | 936.611875 | -0.03206797 | 0.097940447 | -0.327423152 | 0.743347858 | 0.857217273 | 0.03206797 |
| ENSMUSG00000050668 | Gpatch11 | 53951 | protein_coding | 203.7001615 | 0.032070136 | 0.098608293 | 0.325227578 | 0.745008838 | 0.858274321 | 0.032070136 |
| ENSMUSG00000030595 | Nfkbib | 18036 | protein_coding | 188.8422549 | -0.032077721 | 0.095719471 | -0.335122218 | 0.737532893 | 0.853241133 | 0.032077721 |
| ENSMUSG00000026932 | Nacc2 | 67991 | protein_coding | 376.2374052 | -0.032098397 | 0.0868436 | -0.369611546 | 0.711671947 | 0.837372444 | 0.032098397 |
| ENSMUSG00000032434 | Cmtm6 | 67213 | protein_coding | 2239.157251 | 0.032117473 | 0.051807248 | 0.61994169 | 0.535296177 | 0.720057217 | 0.032117473 |
| ENSMUSG00000004637 | Wwox | 80707 | protein_coding | 143.7733632 | 0.03215196 | 0.115167944 | 0.279174558 | 0.780110868 | 0.880499284 | 0.03215196 |
| ENSMUSG00000032456 | Nmnat3 | 74080 | protein_coding | 153.3383113 | -0.032233946 | 0.211091538 | -0.15270127 | 0.878633859 | 0.938372007 | 0.032233946 |
| ENSMUSG00000047759 | Hs3st3a1 | 15478 | protein_coding | 9.675429188 | 0.032285132 | 0.47783541 | 0.067565382 | 0.946131613 | 0.973769693 | 0.032285132 |
| ENSMUSG00000051278 | Zgrf1 | 71643 | protein_coding | 72.4092669 | 0.032287082 | 0.198652933 | 0.162530104 | 0.870888424 | 0.933600906 | 0.032287082 |
| ENSMUSG00000015971 | Actr8 | 56249 | protein_coding | 416.5185641 | -0.032292949 | 0.073254602 | -0.44083168 | 0.659334857 | 0.802969027 | 0.032292949 |
| ENSMUSG00000027456 | Sdcbp2 | 228765 | protein_coding | 2766.228745 | 0.03230575 | 0.301711678 | 0.107074906 | 0.914729555 | 0.957833724 | 0.03230575 |
| ENSMUSG00000025423 | Pias2 | 17344 | protein_coding | 581.8313086 | -0.032331428 | 0.085559523 | -0.377882278 | 0.705518048 | 0.833694265 | 0.032331428 |
| ENSMUSG00000005893 | Nr2c2 | 22026 | protein_coding | 722.5999234 | 0.03233481 | 0.097501487 | 0.331634016 | 0.740165631 | 0.855106852 | 0.03233481 |
| ENSMUSG00000067713 | Prkag1 | 19082 | protein_coding | 380.2327984 | 0.032342979 | 0.115116215 | 0.280959367 | 0.778741565 | 0.879764214 | 0.032342979 |
| ENSMUSG00000032113 | Chek1 | 12649 | protein_coding | 97.39557275 | -0.032386764 | 0.183628588 | -0.176371034 | 0.86000246 | 0.92724018 | 0.032386764 |

| ensembl_gene_id | external_gene_name | entrezgene_id | gene_biotype | baseMean | log2FoldChange | lfcSE | stat | pvalue | padj | absLog2FC |
|---|---|---|---|---|---|---|---|---|---|---|
| ENSMUSG00000079435 | Rpl36a | 19982 | protein_coding | 2406.699999 | -0.032398246 | 0.090258494 | -0.358949549 | 0.719632832 | 0.842364374 | 0.032398246 |
| ENSMUSG00000063480 | Snu13 | 20826 | protein_coding | 495.491062 | -0.032442449 | 0.107548027 | -0.301655458 | 0.762914727 | 0.870097644 | 0.032442449 |
| ENSMUSG00000054199 | Gon4l | 76022 | protein_coding | 349.6556808 | -0.032475556 | 0.093950273 | -0.345667501 | 0.729592611 | 0.847873275 | 0.032475556 |
| ENSMUSG00000057858 | Fam204a | 76539 | protein_coding | 293.7646319 | 0.032476806 | 0.12453384 | 0.260786995 | 0.794256774 | 0.889039236 | 0.032476806 |
| ENSMUSG00000050335 | Lgals3 | 16854 | protein_coding | 14687.36431 | -0.032564165 | 0.173572361 | -0.187611464 | 0.851181239 | 0.921597088 | 0.032564165 |
| ENSMUSG00000031309 | Rps6ka3 | 110651 | protein_coding | 2070.807595 | 0.032581389 | 0.150788541 | 0.216073374 | 0.828930554 | 0.909374328 | 0.032581389 |
| ENSMUSG00000020397 | Med7 | 66213 | protein_coding | 236.9085056 | 0.032601704 | 0.107132018 | 0.30431336 | 0.760889168 | 0.86915108 | 0.032601704 |
| ENSMUSG00000000826 | Dnajc5 | 13002 | protein_coding | 1590.542185 | 0.032658549 | 0.089480045 | 0.364981371 | 0.715125312 | 0.839800265 | 0.032658549 |
| ENSMUSG00000031433 | Rbm41 | 237073 | protein_coding | 123.8157481 | 0.032672516 | 0.122686998 | 0.266307891 | 0.790002096 | 0.886199971 | 0.032672516 |
| ENSMUSG00000024383 | Map3k2 | 26405 | protein_coding | 1210.491229 | -0.032711386 | 0.081841326 | -0.399692767 | 0.68938282 | 0.823011515 | 0.032711386 |
| ENSMUSG00000001285 | Myg1 | 60315 | protein_coding | 227.1576708 | 0.032809414 | 0.102052598 | 0.321495133 | 0.7478352 | 0.859851228 | 0.032809414 |
| ENSMUSG00000041959 | S100a10 | 20194 | protein_coding | 3672.302676 | -0.032828816 | 0.17015421 | -0.192935669 | 0.84700935 | 0.919400582 | 0.032828816 |
| ENSMUSG00000065990 | Aurkaip1 | 66077 | protein_coding | 680.4078693 | -0.032863339 | 0.099248106 | -0.331123089 | 0.740551512 | 0.855242494 | 0.032863339 |
| ENSMUSG00000034022 | Cpsf1 | 94230 | protein_coding | 821.4995069 | 0.032867667 | 0.083054379 | 0.39573671 | 0.692299271 | 0.824876032 | 0.032867667 |
| ENSMUSG00000024853 | Sf3b2 | 319322 | protein_coding | 2400.760287 | -0.032909933 | 0.057497568 | -0.57237088 | 0.56707074 | 0.743049553 | 0.032909933 |
| ENSMUSG00000031583 | Wrn | 22427 | protein_coding | 497.4166155 | -0.03295064 | 0.076320849 | -0.431738386 | 0.665931565 | 0.80636749 | 0.03295064 |
| ENSMUSG00000021519 | Mterf3 | 66410 | protein_coding | 305.9731412 | -0.03299352 | 0.092633246 | -0.35617364 | 0.721710532 | 0.843885285 | 0.03299352 |
| ENSMUSG00000051224 | Tceanc | 245695 | protein_coding | 95.21981403 | 0.032993776 | 0.167365412 | 0.197136171 | 0.843720979 | 0.917198181 | 0.032993776 |
| ENSMUSG00000050721 | Plekho2 | 102595 | protein_coding | 13.75559562 | 0.03299385 | 0.376764128 | 0.087571633 | 0.930217149 | 0.965369999 | 0.03299385 |
| ENSMUSG00000061374 | Fiz1 | 23877 | protein_coding | 291.2427638 | 0.03299407 | 0.084752239 | 0.389300271 | 0.697054036 | 0.827451522 | 0.03299407 |
| ENSMUSG00000042903 | Foxo4 | 54601 | protein_coding | 365.6851764 | 0.033030748 | 0.092351159 | 0.357664685 | 0.720594265 | 0.843075176 | 0.033030748 |
| ENSMUSG00000005699 | Pard6a | 56513 | protein_coding | 47.99508326 | 0.033035507 | 0.213222674 | 0.154934306 | 0.876873108 | 0.937560238 | 0.033035507 |
| ENSMUSG00000036292 | Gramd1c | 207798 | protein_coding | 193.5912555 | -0.033041062 | 0.123588257 | -0.267347905 | 0.789201306 | 0.885675924 | 0.033041062 |
| ENSMUSG00000008167 | Fbxw9 | 68628 | protein_coding | 262.2353656 | 0.033139647 | 0.084608923 | 0.391680278 | 0.695294464 | 0.826592861 | 0.033139647 |
| ENSMUSG00000018559 | Ctdnep1 | 67181 | protein_coding | 276.9006503 | -0.033181465 | 0.12826197 | -0.258700729 | 0.795866157 | 0.890020091 | 0.033181465 |
| ENSMUSG00000000708 | Kat2b | 18519 | protein_coding | 906.0524924 | 0.03320436 | 0.096865166 | 0.342789482 | 0.731756839 | 0.849260935 | 0.03320436 |
| ENSMUSG00000031198 | Fundc2 | 67391 | protein_coding | 291.5677836 | -0.033232397 | 0.095237546 | -0.348942182 | 0.727132716 | 0.846434985 | 0.033232397 |
| ENSMUSG00000021666 | Gfm2 | 320806 | protein_coding | 734.4524201 | -0.033286933 | 0.073766986 | -0.451244315 | 0.651813474 | 0.798492458 | 0.033286933 |
| ENSMUSG00000040990 | Sh3kbp1 | 58194 | protein_coding | 177.8031788 | -0.03330378 | 0.235524332 | -0.141402714 | 0.88755181 | 0.942901703 | 0.03330378 |
| ENSMUSG00000047180 | Neurl3 | 214854 | protein_coding | 1596.876987 | -0.033320466 | 0.093912265 | -0.354804201 | 0.722736281 | 0.844093231 | 0.033320466 |
| ENSMUSG00000019814 | Ltv1 | 353258 | protein_coding | 380.8493986 | -0.033351088 | 0.158105126 | -0.210942484 | 0.832932153 | 0.911190103 | 0.033351088 |
| ENSMUSG00000057572 | Zbtb8os | 67106 | protein_coding | 259.4697232 | -0.033359231 | 0.089175786 | -0.374083963 | 0.70834185 | 0.835305322 | 0.033359231 |
| ENSMUSG00000021707 | Dhfr | 13361 | protein_coding | 283.1487848 | 0.033366339 | 0.154755173 | 0.215607258 | 0.829293898 | 0.909405742 | 0.033366339 |
| ENSMUSG00000043279 | Trim56 | 384309 | protein_coding | 802.1282543 | -0.033432162 | 0.109815872 | -0.304438343 | 0.76079396 | 0.86915108 | 0.033432162 |
| ENSMUSG00000025525 | Apool | 68117 | protein_coding | 482.2638364 | 0.033455857 | 0.086437559 | 0.387052315 | 0.698717479 | 0.828766651 | 0.033455857 |
| ENSMUSG00000054808 | Actn4 | 60595 | protein_coding | 3783.127494 | -0.033458998 | 0.152028722 | -0.220083399 | 0.825806204 | 0.908116569 | 0.033458998 |
| ENSMUSG00000028530 | Jak1 | 16451 | protein_coding | 1859.24472 | -0.03352994 | 0.065617679 | -0.510989428 | 0.60935846 | 0.772513652 | 0.03352994 |
| ENSMUSG00000014503 | Pkd2l2 | 53871 | protein_coding | 84.61278292 | 0.033531822 | 0.157621967 | 0.212735719 | 0.831533104 | 0.910473588 | 0.033531822 |

| ensembl_gene_id | external_gene_name | entrezgene_id | gene_biotype | baseMean | log2FoldChange | lfcSE | stat | pvalue | padj | absLog2FC |
|---|---|---|---|---|---|---|---|---|---|---|
| ENSMUSG00000027774 | Gfm1 | 28030 | protein_coding | 933.1365495 | -0.033575164 | 0.105830229 | -0.317254949 | 0.751050158 | 0.862427144 | 0.033575164 |
| ENSMUSG00000029461 | Fam168a | 319604 | protein_coding | 189.5367264 | 0.033584806 | 0.140163772 | 0.239611178 | 0.810631699 | 0.899310849 | 0.033584806 |
| ENSMUSG00000021549 | Rasa1 | 218397 | protein_coding | 983.1216704 | -0.03361857 | 0.062741735 | -0.535824675 | 0.592079727 | 0.761026944 | 0.03361857 |
| ENSMUSG00000004264 | Phb2 | 12034 | protein_coding | 2357.956423 | -0.033644747 | 0.092409337 | -0.364083846 | 0.715795399 | 0.840114149 | 0.033644747 |
| ENSMUSG00000043592 | Unc5cl | 76589 | protein_coding | 118.770624 | 0.033655336 | 0.174540396 | 0.192822618 | 0.847097889 | 0.919400582 | 0.033655336 |
| ENSMUSG00000022696 | Sidt1 | 320007 | protein_coding | 640.1991402 | -0.033697466 | 0.09627418 | -0.350015614 | 0.72632698 | 0.846191239 | 0.033697466 |
| ENSMUSG00000049760 | Micos13 | 224904 | protein_coding | 1032.044252 | -0.033725412 | 0.096919744 | -0.347972564 | 0.727860788 | 0.846662573 | 0.033725412 |
| ENSMUSG00000068479 | Mfap1a | 67532 | protein_coding | 293.9753165 | -0.033813979 | 0.085505051 | -0.395461771 | 0.692502129 | 0.824994309 | 0.033813979 |
| ENSMUSG00000023892 | Zfp51 | 22709 | protein_coding | 343.6788465 | 0.033877184 | 0.094233914 | 0.359500982 | 0.719220344 | 0.842333944 | 0.033877184 |
| ENSMUSG00000033285 | Wdr3 | 269470 | protein_coding | 467.9090595 | 0.033895826 | 0.133146851 | 0.254574749 | 0.799051564 | 0.89211142 | 0.033895826 |
| ENSMUSG00000036078 | Sigmar1 | 18391 | protein_coding | 653.4657832 | 0.033926742 | 0.074439994 | 0.455759603 | 0.648562857 | 0.797091758 | 0.033926742 |
| ENSMUSG00000029676 | Pot1a | 101185 | protein_coding | 207.7279963 | 0.033934162 | 0.108855451 | 0.311735989 | 0.755241175 | 0.865722025 | 0.033934162 |
| ENSMUSG00000026709 | Dars2 | 226539 | protein_coding | 421.616633 | -0.033971549 | 0.119412813 | -0.284488306 | 0.776036195 | 0.878078036 | 0.033971549 |
| ENSMUSG00000007029 | Vars | 22321 | protein_coding | 1339.047895 | 0.033992026 | 0.113811614 | 0.298669225 | 0.765192441 | 0.871705624 | 0.033992026 |
| ENSMUSG00000038671 | Arfrp1 | 76688 | protein_coding | 310.7100248 | -0.034019444 | 0.095567798 | -0.355971835 | 0.721861658 | 0.843984213 | 0.034019444 |
| ENSMUSG00000017615 | Tnfaip1 | 21927 | protein_coding | 1380.602236 | 0.034043689 | 0.11272958 | 0.301994284 | 0.76265642 | 0.87004666 | 0.034043689 |
| ENSMUSG00000030403 | Vasp | 22323 | protein_coding | 4140.60236 | 0.034126499 | 0.131740123 | 0.259044077 | 0.795601232 | 0.889848777 | 0.034126499 |
| ENSMUSG00000020775 | Mrpl38 | 60441 | protein_coding | 410.0276622 | -0.034159382 | 0.105563948 | -0.323589468 | 0.746248863 | 0.858895287 | 0.034159382 |
| ENSMUSG00000037533 | Rapgef6 | 192786 | protein_coding | 1296.844265 | 0.034189845 | 0.062754209 | 0.544821546 | 0.585876259 | 0.756020937 | 0.034189845 |
| ENSMUSG00000008668 | Rps18 | 20084 | protein_coding | 7629.473297 | 0.03422895 | 0.113150485 | 0.302508201 | 0.762264682 | 0.869662049 | 0.03422895 |
| ENSMUSG00000041849 | Card6 | 239319 | protein_coding | 59.2060714 | -0.034279333 | 0.203388022 | -0.168541551 | 0.866157258 | 0.930600527 | 0.034279333 |
| ENSMUSG00000041777 | Cir1 | 66935 | protein_coding | 125.4401846 | 0.034298466 | 0.144821855 | 0.236832113 | 0.812787038 | 0.900345778 | 0.034298466 |
| ENSMUSG00000038546 | Ranbp9 | 56705 | protein_coding | 601.938681 | -0.034299205 | 0.081120784 | -0.422816488 | 0.672429157 | 0.810978071 | 0.034299205 |
| ENSMUSG00000032279 | Idh3a | 67834 | protein_coding | 2593.164993 | 0.034380941 | 0.072680302 | 0.473043459 | 0.636182174 | 0.789347063 | 0.034380941 |
| ENSMUSG00000003617 | Cp | 12870 | protein_coding | 9.273719289 | -0.034437759 | 0.6623872 | -0.051990375 | 0.958536363 | 0.97950646 | 0.034437759 |
| ENSMUSG00000024442 | Dele1 | 66839 | protein_coding | 571.3287497 | 0.034480343 | 0.097945313 | 0.352036683 | 0.724810752 | 0.844960894 | 0.034480343 |
| ENSMUSG00000027367 | Stard7 | 99138 | protein_coding | 2426.881282 | 0.034482459 | 0.095050187 | 0.362781607 | 0.716768033 | 0.840627772 | 0.034482459 |
| ENSMUSG00000057342 | Sphk2 | 56632 | protein_coding | 743.1694156 | 0.034485371 | 0.161986325 | 0.212890634 | 0.831412268 | 0.910435964 | 0.034485371 |
| ENSMUSG00000036975 | Tmem177 | 66343 | protein_coding | 127.3084247 | 0.034506601 | 0.150573825 | 0.229167326 | 0.818738867 | 0.90373338 | 0.034506601 |
| ENSMUSG00000020114 | Cand1 | 71902 | protein_coding | 1698.842021 | -0.034516965 | 0.071696251 | -0.481433328 | 0.630208555 | 0.785935003 | 0.034516965 |
| ENSMUSG00000050608 | Micos10 | 433771 | protein_coding | 883.7950481 | -0.034523053 | 0.083532964 | -0.413286574 | 0.679396665 | 0.816753599 | 0.034523053 |
| ENSMUSG00000035476 | Tab3 | 66724 | protein_coding | 210.8321778 | 0.034532486 | 0.113119689 | 0.305273877 | 0.76015757 | 0.868689179 | 0.034532486 |
| ENSMUSG00000025420 | Katnal2 | 71206 | protein_coding | 9.112049809 | -0.034535928 | 0.748810647 | -0.046121043 | 0.963213774 | 0.98246466 | 0.034535928 |
| ENSMUSG00000115667 | Gm10389 | 100038608 | lncRNA | 11.53509184 | 0.034567249 | 0.407729507 | 0.084779857 | 0.932436408 | 0.966103649 | 0.034567249 |
| ENSMUSG00000031818 | Cox4i1 | 12857 | protein_coding | 9904.140921 | 0.034577888 | 0.123494394 | 0.279995607 | 0.779480876 | 0.880217901 | 0.034577888 |
| ENSMUSG00000033623 | Pcgf3 | 69587 | protein_coding | 1183.671985 | -0.034586922 | 0.083995396 | -0.411771643 | 0.680506808 | 0.817182783 | 0.034586922 |
| ENSMUSG00000025337 | Sbds | 66711 | protein_coding | 1150.927306 | -0.034601901 | 0.103314509 | -0.33491812 | 0.737686853 | 0.853357266 | 0.034601901 |
| ENSMUSG00000086914 | Gm16124 | 102636154 | lncRNA | 14.28851373 | 0.034627761 | 0.330126913 | 0.104892268 | 0.916461294 | 0.958597942 | 0.034627761 |

| ensembl_gene_id | external_gene_name | entrezgene_id | gene_biotype | baseMean | log2FoldChange | lfcSE | stat | pvalue | padj | absLog2FC |
|---|---|---|---|---|---|---|---|---|---|---|
| ENSMUSG00000072974 | Gm4787 | 214321 | protein_coding | 12.53920677 | 0.034629175 | 0.44240289 | 0.078275201 | 0.937609144 | 0.969210087 | 0.034629175 |
| ENSMUSG00000031950 | Gabarapl2 | 93739 | protein_coding | 675.6677356 | 0.034633187 | 0.106712085 | 0.324547938 | 0.745523235 | 0.85860846 | 0.034633187 |
| ENSMUSG00000021460 | Auh | 11992 | protein_coding | 300.0269262 | 0.034648983 | 0.10691549 | 0.324078234 | 0.745878805 | 0.858717572 | 0.034648983 |
| ENSMUSG00000058558 | Rpl5 | 100503670 | protein_coding | 6343.967328 | 0.034649021 | 0.082172678 | 0.421661084 | 0.673272413 | 0.811554639 | 0.034649021 |
| ENSMUSG00000028189 | Ctbs | 74245 | protein_coding | 151.0979948 | 0.03465505 | 0.118112801 | 0.293406386 | 0.769211545 | 0.874268694 | 0.03465505 |
| ENSMUSG00000063800 | Prpf38a | 230596 | protein_coding | 640.7274031 | 0.034655091 | 0.079794208 | 0.434305849 | 0.664066357 | 0.805456056 | 0.034655091 |
| ENSMUSG00000025498 | Irf7 | 54123 | protein_coding | 797.9335168 | -0.034690961 | 0.138488087 | -0.250497801 | 0.802202405 | 0.893981767 | 0.034690961 |
| ENSMUSG00000037822 | Smim14 | 68552 | protein_coding | 2320.45102 | -0.034697778 | 0.098330313 | -0.352869601 | 0.724186201 | 0.844771344 | 0.034697778 |
| ENSMUSG00000000957 | Mmp14 | 17387 | protein_coding | 698.4553425 | -0.034729141 | 0.079355155 | -0.437641901 | 0.661645892 | 0.804235025 | 0.034729141 |
| ENSMUSG00000006740 | Kif5b | 16573 | protein_coding | 4607.319303 | 0.034746043 | 0.09341711 | 0.371945168 | 0.709933675 | 0.836191937 | 0.034746043 |
| ENSMUSG00000030990 | Pgap2 | 233575 | protein_coding | 658.9754309 | 0.034777564 | 0.138088288 | 0.251850208 | 0.801156847 | 0.893427589 | 0.034777564 |
| ENSMUSG00000052712 | BC004004 | 80748 | protein_coding | 672.1599728 | -0.034791653 | 0.080138269 | -0.434145304 | 0.664182929 | 0.805536107 | 0.034791653 |
| ENSMUSG00000034057 | Myrfl | 237558 | protein_coding | 14.53758537 | -0.034792458 | 0.320295785 | -0.108626024 | 0.913499119 | 0.957282345 | 0.034792458 |
| ENSMUSG00000031217 | Efnb1 | 13641 | protein_coding | 1578.068433 | -0.034796453 | 0.122083717 | -0.285021243 | 0.775627867 | 0.877982787 | 0.034796453 |
| ENSMUSG00000033400 | Agl | 77559 | protein_coding | 447.6988691 | 0.034801616 | 0.092725283 | 0.375319599 | 0.707422792 | 0.834654091 | 0.034801616 |
| ENSMUSG00000022540 | Rogdi | 66049 | protein_coding | 73.56280713 | -0.034809411 | 0.156472273 | -0.222463766 | 0.823952877 | 0.907037213 | 0.034809411 |
| ENSMUSG00000031311 | Nono | 53610 | protein_coding | 2514.66407 | -0.034833905 | 0.061363121 | -0.567668402 | 0.570260151 | 0.744744605 | 0.034833905 |
| ENSMUSG00000034194 | R3hcc1 | 71843 | protein_coding | 178.2018391 | -0.03484492 | 0.11370574 | -0.306448201 | 0.759263413 | 0.86831717 | 0.03484492 |
| ENSMUSG00000031403 | Dkc1 | 245474 | protein_coding | 426.2440922 | -0.034847113 | 0.151644654 | -0.229794534 | 0.818251432 | 0.903445779 | 0.034847113 |
| ENSMUSG00000059447 | Hadhb | 231086 | protein_coding | 2320.608747 | -0.034855768 | 0.09699598 | -0.359352703 | 0.719331252 | 0.842364374 | 0.034855768 |
| ENSMUSG00000095990 | Zfp97 | 22759 | protein_coding | 145.7387769 | -0.034874148 | 0.180912281 | -0.192768272 | 0.847140452 | 0.919400582 | 0.034874148 |
| ENSMUSG00000067424 | Zfp563 | 240068 | protein_coding | 166.9889472 | 0.034883366 | 0.164713676 | 0.211781844 | 0.832277233 | 0.910724909 | 0.034883366 |
| ENSMUSG00000097779 | 4833407H14Rik | 73899 | lncRNA | 8.711096498 | -0.034962541 | 0.360534575 | -0.09697417 | 0.922746907 | 0.961877857 | 0.034962541 |
| ENSMUSG00000005611 | Irag1 | 17540 | protein_coding | 7.113688268 | 0.035048125 | 0.518008058 | 0.06765942 | 0.946056754 | 0.973769693 | 0.035048125 |
| ENSMUSG00000003970 | Rpl8 | 26961 | protein_coding | 10547.92188 | 0.035065776 | 0.089451715 | 0.392007867 | 0.695052401 | 0.826592861 | 0.035065776 |
| ENSMUSG00000028309 | Rnf20 | 109331 | protein_coding | 1059.287726 | -0.035082854 | 0.0810901 | -0.432640412 | 0.665276026 | 0.806025281 | 0.035082854 |
| ENSMUSG00000024477 | Pggt1b | 225467 | protein_coding | 839.8332773 | 0.035110823 | 0.0990416 | 0.354505817 | 0.722959845 | 0.844115801 | 0.035110823 |
| ENSMUSG00000029686 | Cul1 | 26965 | protein_coding | 1830.996917 | 0.035113463 | 0.095802405 | 0.366519638 | 0.713977362 | 0.839031387 | 0.035113463 |
| ENSMUSG00000008140 | Emc10 | 69683 | protein_coding | 645.0820259 | -0.03515972 | 0.061684124 | -0.569996266 | 0.568680231 | 0.743917437 | 0.03515972 |
| ENSMUSG00000019852 | Arfgef3 | 215821 | protein_coding | 1418.320717 | -0.035184153 | 0.116262144 | -0.302627768 | 0.76217355 | 0.869662049 | 0.035184153 |
| ENSMUSG00000028434 | Epb41l4b | 54357 | protein_coding | 3302.006956 | 0.035248661 | 0.102749995 | 0.343052676 | 0.731558832 | 0.849105085 | 0.035248661 |
| ENSMUSG00000045867 | Cradd | 12905 | protein_coding | 80.06425916 | -0.035272608 | 0.181519586 | -0.19431847 | 0.845926524 | 0.918717965 | 0.035272608 |
| ENSMUSG00000030850 | Ate1 | 11907 | protein_coding | 582.567504 | 0.035337986 | 0.066523958 | 0.531206902 | 0.595275411 | 0.763471032 | 0.035337986 |
| ENSMUSG00000040451 | Sgms1 | 208449 | protein_coding | 682.6622479 | 0.035367122 | 0.081681972 | 0.432985657 | 0.66502519 | 0.806025281 | 0.035367122 |
| ENSMUSG00000032842 | Abcc10 | 224814 | protein_coding | 308.8668369 | -0.035393201 | 0.153692422 | -0.230285918 | 0.817869602 | 0.903241067 | 0.035393201 |
| ENSMUSG00000070780 | Rbm47 | 245945 | protein_coding | 3642.338751 | 0.035407705 | 0.077870714 | 0.454698598 | 0.64932609 | 0.797420867 | 0.035407705 |
| ENSMUSG00000030727 | Rabep2 | 70314 | protein_coding | 298.952631 | -0.035410502 | 0.09502541 | -0.372642454 | 0.709414572 | 0.835827385 | 0.035410502 |
| ENSMUSG00000046861 | Hectd3 | 76608 | protein_coding | 1396.201898 | -0.035442097 | 0.093262345 | -0.380025797 | 0.703926266 | 0.832625775 | 0.035442097 |

| ensembl_gene_id | external_gene_name | entrezgene_id | gene_biotype | baseMean | log2FoldChange | lfcSE | stat | pvalue | padj | absLog2FC |
|---|---|---|---|---|---|---|---|---|---|---|
| ENSMUSG00000025184 | R3hcc1l | 52013 | protein_coding | 357.1728824 | -0.035516094 | 0.072365008 | -0.490790991 | 0.623574282 | 0.781823927 | 0.035516094 |
| ENSMUSG00000047675 | Rps8 | 20116 | protein_coding | 8841.372406 | 0.035545282 | 0.092220749 | 0.385436927 | 0.699913728 | 0.829481091 | 0.035545282 |
| ENSMUSG00000009894 | Snap47 | 67826 | protein_coding | 212.3008962 | -0.035571297 | 0.104191381 | -0.341403448 | 0.73279988 | 0.850024456 | 0.035571297 |
| ENSMUSG00000047044 | D030056L22Rik | 225995 | protein_coding | 289.8932892 | -0.035612025 | 0.113452298 | -0.313894256 | 0.753601353 | 0.864733257 | 0.035612025 |
| ENSMUSG00000020074 | Ccar1 | 67500 | protein_coding | 1285.04882 | 0.03561999 | 0.071363311 | 0.499135895 | 0.617683652 | 0.777436997 | 0.03561999 |
| ENSMUSG00000033009 | Ogfod1 | 270086 | protein_coding | 361.2509724 | 0.035639477 | 0.089374405 | 0.398766033 | 0.690065607 | 0.823384897 | 0.035639477 |
| ENSMUSG00000096370 | Gm21992 | 102902673 | protein_coding | 48.29920653 | 0.035640988 | 0.195506862 | 0.182300446 | 0.855346949 | 0.924159965 | 0.035640988 |
| ENSMUSG00000037257 | Aagab | 66939 | protein_coding | 956.7202145 | -0.035704428 | 0.106872643 | -0.334083887 | 0.738316258 | 0.853775328 | 0.035704428 |
| ENSMUSG00000048000 | Gigyf2 | 227331 | protein_coding | 1248.609306 | -0.035715411 | 0.062631791 | -0.57024412 | 0.568512136 | 0.743758628 | 0.035715411 |
| ENSMUSG00000030327 | Necap1 | 67602 | protein_coding | 552.3773484 | 0.035717328 | 0.073941613 | 0.483047724 | 0.629061851 | 0.785427385 | 0.035717328 |
| ENSMUSG00000036606 | Plxnb2 | 140570 | protein_coding | 4985.936642 | -0.035744774 | 0.087735746 | -0.407414036 | 0.68370392 | 0.819173276 | 0.035744774 |
| ENSMUSG00000030652 | Coq7 | 12850 | protein_coding | 308.9906208 | 0.035746422 | 0.132186029 | 0.270425117 | 0.786833219 | 0.884458731 | 0.035746422 |
| ENSMUSG00000090137 | Uba52 | 22186 | protein_coding | 65.39992794 | 0.035754472 | 0.21123569 | 0.169263404 | 0.86558946 | 0.930241476 | 0.035754472 |
| ENSMUSG00000039318 | Rab3gap2 | 98732 | protein_coding | 807.9791365 | 0.035797992 | 0.08264431 | 0.433157367 | 0.664900449 | 0.806025281 | 0.035797992 |
| ENSMUSG00000056234 | Ncoa4 | 27057 | protein_coding | 892.0452321 | -0.0358621 | 0.07465232 | -0.480388289 | 0.63095132 | 0.786307224 | 0.0358621 |
| ENSMUSG00000042650 | Alkbh5 | 268420 | protein_coding | 2025.4221 | 0.035900149 | 0.066919815 | 0.536465158 | 0.59163711 | 0.760882221 | 0.035900149 |
| ENSMUSG00000033083 | Tbc1d4 | 210789 | protein_coding | 539.2257499 | -0.035902282 | 0.103092127 | -0.348254358 | 0.727649168 | 0.846662573 | 0.035902282 |
| ENSMUSG00000022070 | Bora | 77744 | protein_coding | 272.152661 | -0.035912897 | 0.099320379 | -0.361586386 | 0.71766114 | 0.841126316 | 0.035912897 |
| ENSMUSG00000011008 | Mcoln2 | 68279 | protein_coding | 587.4693844 | 0.035928518 | 0.103142048 | 0.348340166 | 0.727584732 | 0.846662573 | 0.035928518 |
| ENSMUSG00000057788 | Ddx49 | 234374 | protein_coding | 312.3591946 | 0.035967131 | 0.118430047 | 0.303699377 | 0.761356933 | 0.869186728 | 0.035967131 |
| ENSMUSG00000037601 | Nme1 | 18102 | protein_coding | 332.7355185 | 0.035971001 | 0.168393209 | 0.213613133 | 0.830848757 | 0.909974489 | 0.035971001 |
| ENSMUSG00000078622 | Ccdc47 | 67163 | protein_coding | 1407.623315 | 0.036025217 | 0.062900547 | 0.57273297 | 0.56682551 | 0.742894489 | 0.036025217 |
| ENSMUSG00000027713 | 1810062G17Rik | 72282 | protein_coding | 15.83474655 | -0.036053059 | 0.350165974 | -0.102959916 | 0.917994784 | 0.95924824 | 0.036053059 |
| ENSMUSG00000056536 | Pign | 27392 | protein_coding | 793.2673624 | -0.036075568 | 0.063804935 | -0.565404038 | 0.571798979 | 0.74577977 | 0.036075568 |
| ENSMUSG00000033396 | Spg11 | 214585 | protein_coding | 709.2430157 | -0.036092473 | 0.065190383 | -0.553647206 | 0.579820313 | 0.751210179 | 0.036092473 |
| ENSMUSG00000022682 | Rrn3 | 106298 | protein_coding | 677.96705 | -0.036094729 | 0.12262982 | -0.294338921 | 0.768498934 | 0.873738638 | 0.036094729 |
| ENSMUSG00000028228 | Cpne3 | 70568 | protein_coding | 2409.975114 | 0.036130369 | 0.072936748 | 0.495365779 | 0.620341946 | 0.778936265 | 0.036130369 |
| ENSMUSG00000054428 | Atpif1 | 11983 | protein_coding | 1972.781454 | 0.036157715 | 0.109625218 | 0.329830266 | 0.741528217 | 0.856004052 | 0.036157715 |
| ENSMUSG00000029491 | Pde6b | 18587 | protein_coding | 8.830333433 | -0.036176416 | 0.509956609 | -0.070940185 | 0.943445361 | 0.97253954 | 0.036176416 |
| ENSMUSG00000073542 | Cep76 | 225659 | protein_coding | 153.9718835 | -0.036184041 | 0.154068504 | -0.234856836 | 0.814319852 | 0.900950823 | 0.036184041 |
| ENSMUSG00000021395 | Spin1 | 20729 | protein_coding | 921.2675282 | 0.036247523 | 0.070474784 | 0.514333221 | 0.607019039 | 0.771131144 | 0.036247523 |
| ENSMUSG00000025793 | Hgs | 15239 | protein_coding | 907.2205156 | 0.036262079 | 0.089994577 | 0.402936272 | 0.686995107 | 0.821304393 | 0.036262079 |
| ENSMUSG00000057738 | Sptan1 | 20740 | protein_coding | 6258.718206 | -0.036272712 | 0.069502493 | -0.521890803 | 0.601746362 | 0.767564496 | 0.036272712 |
| ENSMUSG00000046862 | Pramel12 | 242736 | protein_coding | 316.9626824 | -0.036299511 | 0.11646718 | -0.311671586 | 0.755290124 | 0.865722025 | 0.036299511 |
| ENSMUSG00000040423 | Rc3h1 | 381305 | protein_coding | 1500.377095 | -0.03639357 | 0.13963049 | -0.260642001 | 0.794368597 | 0.889039236 | 0.03639357 |
| ENSMUSG00000052595 | A1cf | 69865 | protein_coding | 362.2594251 | -0.036414472 | 0.143982325 | -0.252909323 | 0.800338285 | 0.892827301 | 0.036414472 |
| ENSMUSG00000029192 | Tbc1d14 | 100855 | protein_coding | 374.3804102 | -0.036424262 | 0.153308927 | -0.237587354 | 0.812201159 | 0.900009744 | 0.036424262 |
| ENSMUSG00000040128 | Pnrc1 | 108767 | protein_coding | 963.7979401 | -0.036439956 | 0.121982956 | -0.298729894 | 0.765146146 | 0.871705624 | 0.036439956 |

Supplementary Table 1

| ensembl_gene_id | external_gene_name | entrezgene_id | gene_biotype | baseMean | log2FoldChange | lfcSE | stat | pvalue | padj | absLog2FC |
|---|---|---|---|---|---|---|---|---|---|---|
| ENSMUSG00000021510 | Zfp729a | 212281 | protein_coding | 341.6274385 | 0.0364444 | 0.104236798 | 0.34963085 | 0.726615756 | 0.846266142 | 0.0364444 |
| ENSMUSG00000026712 | Mrc1 | 17533 | protein_coding | 20.86796287 | -0.036459612 | 0.443685385 | -0.082174471 | 0.934507974 | 0.967553497 | 0.036459612 |
| ENSMUSG00000031527 | Eri1 | 67276 | protein_coding | 587.3352345 | 0.036477721 | 0.071714087 | 0.508654895 | 0.610994144 | 0.773174617 | 0.036477721 |
| ENSMUSG00000037197 | Rbm17 | 76938 | protein_coding | 829.7390438 | 0.036481056 | 0.073171496 | 0.498569227 | 0.618082889 | 0.777697562 | 0.036481056 |
| ENSMUSG00000020372 | Rack1 | 14694 | protein_coding | 14918.3566 | 0.036481854 | 0.104512179 | 0.349067966 | 0.727038285 | 0.846434985 | 0.036481854 |
| ENSMUSG00000058997 | Vwa8 | 219189 | protein_coding | 3146.328658 | 0.03649344 | 0.082277747 | 0.443539609 | 0.657375476 | 0.801674955 | 0.03649344 |
| ENSMUSG00000027763 | Mbnl1 | 56758 | protein_coding | 2104.567584 | -0.03656602 | 0.089933228 | -0.40659076 | 0.684308585 | 0.819708742 | 0.03656602 |
| ENSMUSG00000034744 | Nagk | 56174 | protein_coding | 474.3974222 | -0.036591188 | 0.096698555 | -0.378404708 | 0.705129971 | 0.833382076 | 0.036591188 |
| ENSMUSG00000027852 | Nras | 18176 | protein_coding | 901.2462527 | -0.036599105 | 0.138502073 | -0.264249513 | 0.791587661 | 0.887042894 | 0.036599105 |
| ENSMUSG00000038235 | F11r | 16456 | protein_coding | 2672.4934 | 0.036666877 | 0.09362054 | 0.391654187 | 0.695313745 | 0.826592861 | 0.036666877 |
| ENSMUSG00000048175 | Asb8 | 78541 | protein_coding | 377.9125365 | 0.036720634 | 0.080648241 | 0.455318472 | 0.648880139 | 0.797283028 | 0.036720634 |
| ENSMUSG00000027676 | Ccdc39 | 51938 | protein_coding | 10.99122287 | 0.036728669 | 0.460730652 | 0.07971831 | 0.936461296 | 0.968704702 | 0.036728669 |
| ENSMUSG00000047747 | Rnf150 | 330812 | protein_coding | 231.7423893 | -0.036744855 | 0.169286958 | -0.217056622 | 0.828164219 | 0.908971863 | 0.036744855 |
| ENSMUSG00000052139 | Babam2 | 107976 | protein_coding | 431.0286915 | -0.03680486 | 0.064780636 | -0.568146015 | 0.569935825 | 0.744741762 | 0.03680486 |
| ENSMUSG00000048550 | Thnsl1 | 208967 | protein_coding | 134.2901476 | -0.036837691 | 0.142718496 | -0.258114347 | 0.79631866 | 0.890338594 | 0.036837691 |
| ENSMUSG00000084883 | Ccdc85c | 668158 | protein_coding | 1022.029319 | -0.036880346 | 0.098048975 | -0.376142086 | 0.706811268 | 0.834197547 | 0.036880346 |
| ENSMUSG00000031938 | 4931406C07Rik | 70984 | protein_coding | 1468.816061 | -0.036881288 | 0.136909775 | -0.26938389 | 0.787634282 | 0.884788751 | 0.036881288 |
| ENSMUSG00000052296 | Ppp6r1 | 243819 | protein_coding | 1787.974617 | -0.036897263 | 0.076433256 | -0.482738338 | 0.629281538 | 0.785542493 | 0.036897263 |
| ENSMUSG00000015120 | Ube2i | 22196 | protein_coding | 850.7284123 | 0.036923378 | 0.072646165 | 0.508263282 | 0.611268717 | 0.773335037 | 0.036923378 |
| ENSMUSG00000019808 | Adat2 | 66757 | protein_coding | 56.11111875 | -0.03692634 | 0.162331617 | -0.227474722 | 0.820054623 | 0.904579103 | 0.03692634 |
| ENSMUSG00000021929 | Kpna3 | 16648 | protein_coding | 678.5225256 | -0.036959129 | 0.077526483 | -0.47672908 | 0.633555053 | 0.788005719 | 0.036959129 |
| ENSMUSG00000066842 | Hmcn1 | 545370 | protein_coding | 402.823791 | 0.037045146 | 0.451712636 | 0.082010427 | 0.934638421 | 0.96756277 | 0.037045146 |
| ENSMUSG00000041712 | Ubr7 | 66622 | protein_coding | 417.5177388 | -0.037097547 | 0.090042961 | -0.411998302 | 0.680340668 | 0.817182783 | 0.037097547 |
| ENSMUSG00000026126 | Ptpn18 | 19253 | protein_coding | 635.9323328 | 0.03717132 | 0.085331201 | 0.435612292 | 0.663118051 | 0.804794609 | 0.03717132 |
| ENSMUSG00000002949 | Timm44 | 21856 | protein_coding | 824.4716661 | -0.037171394 | 0.084655089 | -0.439092254 | 0.660594692 | 0.803570509 | 0.037171394 |
| ENSMUSG00000024789 | Jak2 | 16452 | protein_coding | 1299.593364 | -0.037179563 | 0.071505684 | -0.519952554 | 0.603096645 | 0.76799575 | 0.037179563 |
| ENSMUSG00000020537 | Drg2 | 13495 | protein_coding | 430.8837463 | 0.037180244 | 0.097159989 | 0.382670321 | 0.701964223 | 0.83080438 | 0.037180244 |
| ENSMUSG00000040746 | Rnf167 | 70510 | protein_coding | 743.3806606 | -0.037189788 | 0.124535602 | -0.298627762 | 0.76522408 | 0.871705624 | 0.037189788 |
| ENSMUSG00000102428 | Pcdhga12 | 93724 | protein_coding | 8.199880908 | 0.037195391 | 0.424068747 | 0.087710756 | 0.93010657 | 0.965369999 | 0.037195391 |
| ENSMUSG00000020483 | Dynll2 | 68097 | protein_coding | 1068.579192 | -0.037198456 | 0.130557755 | -0.284919543 | 0.775705783 | 0.877982787 | 0.037198456 |
| ENSMUSG00000038980 | Rbbp8nl | 271887 | protein_coding | 44.91235324 | 0.03722945 | 0.185999223 | 0.200159172 | 0.841356097 | 0.916315892 | 0.03722945 |
| ENSMUSG00000097290 | 1300002E11Rik | 100043489 | lncRNA | 154.2207708 | 0.03728061 | 0.113460951 | 0.328576572 | 0.742475758 | 0.85659524 | 0.03728061 |
| ENSMUSG00000026096 | Osgepl1 | 72085 | protein_coding | 119.9966744 | -0.037294065 | 0.122436968 | -0.304598074 | 0.760672287 | 0.869090323 | 0.037294065 |
| ENSMUSG00000026333 | Gin1 | 252876 | protein_coding | 88.48008075 | 0.037335441 | 0.151983022 | 0.24565534 | 0.805949046 | 0.895698884 | 0.037335441 |
| ENSMUSG00000030982 | Vps35l | 71517 | protein_coding | 680.5333744 | -0.037362923 | 0.057900514 | -0.645295191 | 0.518735908 | 0.705828255 | 0.037362923 |
| ENSMUSG00000033107 | Rnf125 | 67664 | protein_coding | 94.26460876 | -0.037400624 | 0.171081514 | -0.218612892 | 0.82695161 | 0.908412958 | 0.037400624 |
| ENSMUSG00000054237 | Fra10ac1 | 70567 | protein_coding | 147.9472907 | -0.037422563 | 0.120603175 | -0.310295005 | 0.756336629 | 0.866249358 | 0.037422563 |
| ENSMUSG00000041815 | Poldip3 | 73826 | protein_coding | 1179.12025 | -0.037438423 | 0.065930223 | -0.567849178 | 0.570137384 | 0.744744605 | 0.037438423 |

| ensembl_gene_id | external_gene_name | entrezgene_id | gene_biotype | baseMean | log2FoldChange | lfcSE | stat | pvalue | padj | absLog2FC |
|---|---|---|---|---|---|---|---|---|---|---|
| ENSMUSG00000037075 | Rnf139 | 75841 | protein_coding | 502.810046 | 0.03745168 | 0.08785722 | 0.426278904 | 0.669904628 | 0.808780483 | 0.03745168 |
| ENSMUSG00000029759 | Pon3 | 269823 | protein_coding | 886.3846643 | 0.037457029 | 0.067048239 | 0.558657906 | 0.576395213 | 0.748660438 | 0.037457029 |
| ENSMUSG00000029763 | Exoc4 | 20336 | protein_coding | 779.4528073 | 0.037463016 | 0.065174955 | 0.574806925 | 0.565421883 | 0.741604064 | 0.037463016 |
| ENSMUSG00000098449 | Gm7467 | 665055 | lncRNA | 13.0347168 | -0.037475817 | 0.415474478 | -0.090200047 | 0.928128247 | 0.964521871 | 0.037475817 |
| ENSMUSG00000018882 | Mrpl45 | 67036 | protein_coding | 533.8256332 | 0.037527353 | 0.114076359 | 0.328966958 | 0.742180663 | 0.85644089 | 0.037527353 |
| ENSMUSG00000052423 | B4galt3 | 57370 | protein_coding | 242.0029012 | -0.037543994 | 0.115702165 | -0.324488261 | 0.745568408 | 0.85860846 | 0.037543994 |
| ENSMUSG00000026849 | Tor1a | 30931 | protein_coding | 385.235593 | 0.037548256 | 0.09626344 | 0.390057289 | 0.696494185 | 0.827213435 | 0.037548256 |
| ENSMUSG00000024410 | Rmc1 | 76482 | protein_coding | 517.7092669 | -0.037562845 | 0.095580045 | -0.392998822 | 0.694320351 | 0.826233437 | 0.037562845 |
| ENSMUSG00000061132 | Blnk | 17060 | protein_coding | 1167.022452 | -0.037572474 | 0.12904831 | -0.291150456 | 0.770936252 | 0.87556148 | 0.037572474 |
| ENSMUSG00000052299 | Ltn1 | 78913 | protein_coding | 1002.224971 | -0.03763746 | 0.055206116 | -0.681762506 | 0.495389136 | 0.686912002 | 0.03763746 |
| ENSMUSG00000014601 | Strip1 | 229707 | protein_coding | 531.4102902 | 0.037644098 | 0.104934985 | 0.358737341 | 0.719791591 | 0.842383576 | 0.037644098 |
| ENSMUSG00000024240 | Epc1 | 13831 | protein_coding | 444.0431322 | 0.037655852 | 0.08152411 | 0.461898354 | 0.644154215 | 0.794001573 | 0.037655852 |
| ENSMUSG00000031386 | Hcfc1 | 15161 | protein_coding | 1101.082675 | -0.037672301 | 0.103372843 | -0.364431316 | 0.715535954 | 0.840012084 | 0.037672301 |
| ENSMUSG00000022760 | Thap7 | 69009 | protein_coding | 150.987745 | 0.037677027 | 0.125699062 | 0.299739921 | 0.764375545 | 0.87125903 | 0.037677027 |
| ENSMUSG00000069114 | Zbtb10 | 229055 | protein_coding | 631.4297824 | -0.037689676 | 0.131753354 | -0.286062368 | 0.774830353 | 0.877593098 | 0.037689676 |
| ENSMUSG00000027997 | Casp6 | 12368 | protein_coding | 296.8226479 | -0.037716246 | 0.08407958 | -0.448577954 | 0.653736141 | 0.799501631 | 0.037716246 |
| ENSMUSG00000035245 | Eogt | 101351 | protein_coding | 236.5820043 | -0.037742727 | 0.09330623 | -0.404503827 | 0.685842265 | 0.820745689 | 0.037742727 |
| ENSMUSG00000020623 | Map2k6 | 26399 | protein_coding | 308.285205 | -0.037757462 | 0.193715582 | -0.194911849 | 0.845461958 | 0.918463925 | 0.037757462 |
| ENSMUSG00000024422 | Dhx16 | 69192 | protein_coding | 669.2608851 | -0.037758934 | 0.099973214 | -0.377690508 | 0.70566052 | 0.833694265 | 0.037758934 |
| ENSMUSG00000022125 | Cln5 | 211286 | protein_coding | 467.788082 | -0.037759173 | 0.108111284 | -0.349262086 | 0.726892559 | 0.846375436 | 0.037759173 |
| ENSMUSG00000034781 | Gna11 | 14672 | protein_coding | 3238.974497 | 0.037804313 | 0.120407499 | 0.313969755 | 0.75354401 | 0.864733257 | 0.037804313 |
| ENSMUSG00000022575 | Gsdmd | 69146 | protein_coding | 1122.923599 | 0.037810666 | 0.083528629 | 0.452667141 | 0.650788444 | 0.798250711 | 0.037810666 |
| ENSMUSG00000020109 | Dnajb12 | 56709 | protein_coding | 667.4980426 | 0.037813584 | 0.083922124 | 0.450579445 | 0.652292684 | 0.798541656 | 0.037813584 |
| ENSMUSG00000115529 | 9630013A20Rik | 319903 | lncRNA | 9.477110113 | -0.037816422 | 0.560534323 | -0.067464954 | 0.946211561 | 0.973784179 | 0.037816422 |
| ENSMUSG00000048581 | E130311K13Rik | 329659 | protein_coding | 50.81464434 | -0.037850562 | 0.215682878 | -0.175491731 | 0.860693267 | 0.927608253 | 0.037850562 |
| ENSMUSG00000024844 | Banf1 | 23825 | protein_coding | 425.0670441 | 0.037881079 | 0.114052254 | 0.332137925 | 0.739785114 | 0.854833615 | 0.037881079 |
| ENSMUSG00000064120 | Mocs1 | 56738 | protein_coding | 293.1958987 | 0.037940923 | 0.104719418 | 0.362310297 | 0.717120165 | 0.840677852 | 0.037940923 |
| ENSMUSG00000044308 | Ubr3 | 68795 | protein_coding | 1373.378785 | -0.037948569 | 0.075368861 | -0.50350461 | 0.61460954 | 0.775467423 | 0.037948569 |
| ENSMUSG00000010406 | Mrpl52 | 68836 | protein_coding | 547.6557983 | 0.037952203 | 0.093598374 | 0.4054793 | 0.685125231 | 0.820193512 | 0.037952203 |
| ENSMUSG00000031862 | Atp13a1 | 170759 | protein_coding | 1068.728172 | 0.037962886 | 0.08794517 | 0.431665387 | 0.665984628 | 0.806370441 | 0.037962886 |
| ENSMUSG00000003849 | Nqo1 | 18104 | protein_coding | 1159.945693 | 0.038003472 | 0.136792748 | 0.277817888 | 0.781152159 | 0.880987863 | 0.038003472 |
| ENSMUSG00000034868 | Myl12b | 67938 | protein_coding | 642.4252927 | 0.038105962 | 0.108106162 | 0.352486497 | 0.724473443 | 0.844883072 | 0.038105962 |
| ENSMUSG00000094622 | Gm3055 | 100040944 | protein_coding | 12.87990483 | 0.038108788 | 0.314634051 | 0.121120991 | 0.903595203 | 0.952386474 | 0.038108788 |
| ENSMUSG00000040093 | Bmf | 171543 | protein_coding | 142.4386557 | -0.038108845 | 0.292906536 | -0.130105821 | 0.896482705 | 0.948278425 | 0.038108845 |
| ENSMUSG00000024269 | Tpgs2 | 66648 | protein_coding | 107.4289386 | 0.038123455 | 0.120301437 | 0.316899416 | 0.751319925 | 0.862550365 | 0.038123455 |
| ENSMUSG00000028557 | Rnf11 | 29864 | protein_coding | 1400.465173 | -0.038125524 | 0.078498628 | -0.485683956 | 0.62719126 | 0.784304814 | 0.038125524 |
| ENSMUSG00000021488 | Nsd1 | 18193 | protein_coding | 3066.915304 | -0.038130277 | 0.103737711 | -0.367564282 | 0.713198152 | 0.838486874 | 0.038130277 |
| ENSMUSG00000006673 | Qrich1 | 69232 | protein_coding | 1214.427508 | 0.038156258 | 0.064808974 | 0.588749598 | 0.556029261 | 0.735434864 | 0.038156258 |

Supplementary Table 1

| ensembl_gene_id | external_gene_name | entrezgene_id | gene_biotype | baseMean | log2FoldChange | lfcSE | stat | pvalue | padj | absLog2FC |
|---|---|---|---|---|---|---|---|---|---|---|
| ENSMUSG00000055809 | Dnaaf3 | 436022 | protein_coding | 6.819918473 | -0.038158793 | 0.515743425 | -0.073987939 | 0.941019982 | 0.971391204 | 0.038158793 |
| ENSMUSG00000054000 | Tusc1 | 69136 | protein_coding | 37.67474486 | 0.038175219 | 0.19849785 | 0.192320567 | 0.847491109 | 0.919476764 | 0.038175219 |
| ENSMUSG00000026577 | Blzf1 | 66352 | protein_coding | 307.0780857 | 0.03818963 | 0.084008976 | 0.454589872 | 0.649404322 | 0.797420867 | 0.03818963 |
| ENSMUSG00000041685 | Fcho2 | 218503 | protein_coding | 953.0644901 | 0.038209809 | 0.141460953 | 0.270108526 | 0.787076763 | 0.884581166 | 0.038209809 |
| ENSMUSG00000073102 | Drc1 | 381738 | protein_coding | 15.89876454 | 0.038295566 | 0.280876553 | 0.136343052 | 0.891550091 | 0.945195012 | 0.038295566 |
| ENSMUSG00000006599 | Gtf2h1 | 14884 | protein_coding | 867.7564766 | 0.038313289 | 0.091435145 | 0.419021477 | 0.675200437 | 0.813201705 | 0.038313289 |
| ENSMUSG00000044617 | Zbtb39 | 320080 | protein_coding | 237.9004909 | -0.038327872 | 0.12320122 | -0.311099772 | 0.755724774 | 0.865797613 | 0.038327872 |
| ENSMUSG00000040701 | Ap1g2 | 11766 | protein_coding | 461.9249092 | 0.038375216 | 0.073352861 | 0.523159092 | 0.600863546 | 0.766932813 | 0.038375216 |
| ENSMUSG00000029221 | Slc30a9 | 109108 | protein_coding | 1004.854888 | 0.038396048 | 0.070375094 | 0.545591432 | 0.585346816 | 0.75584954 | 0.038396048 |
| ENSMUSG00000052763 | Zfp212 | 232784 | protein_coding | 164.4862102 | -0.038404885 | 0.104382457 | -0.367924711 | 0.712929374 | 0.838411012 | 0.038404885 |
| ENSMUSG00000069678 | Pcgf1 | 69837 | protein_coding | 66.9861094 | -0.038439648 | 0.183030984 | -0.210017166 | 0.833654275 | 0.911278944 | 0.038439648 |
| ENSMUSG00000013787 | Ehmt2 | 110147 | protein_coding | 1733.184735 | -0.038473201 | 0.082976239 | -0.463665282 | 0.642887575 | 0.793238306 | 0.038473201 |
| ENSMUSG00000019210 | Atp6v1e1 | 11973 | protein_coding | 1432.09017 | 0.038487453 | 0.107158949 | 0.359162289 | 0.719473686 | 0.842364374 | 0.038487453 |
| ENSMUSG00000024049 | Myom1 | 17929 | protein_coding | 38.72857794 | -0.038526902 | 0.203923481 | -0.188928228 | 0.850149069 | 0.920905487 | 0.038526902 |
| ENSMUSG00000015755 | Tab2 | 68652 | protein_coding | 1200.385497 | 0.038526965 | 0.066544237 | 0.578967719 | 0.562610955 | 0.739622589 | 0.038526965 |
| ENSMUSG00000026977 | Marchf7 | 57438 | protein_coding | 1754.475955 | -0.038540376 | 0.08774021 | -0.439255566 | 0.660476367 | 0.803487938 | 0.038540376 |
| ENSMUSG00000059689 | Zfp637 | 232337 | protein_coding | 163.7294888 | 0.03857153 | 0.102550636 | 0.376121798 | 0.70682635 | 0.834197547 | 0.03857153 |
| ENSMUSG00000031537 | Ikbkb | 16150 | protein_coding | 664.3499681 | 0.03858882 | 0.062508203 | 0.617340103 | 0.537010421 | 0.721275497 | 0.03858882 |
| ENSMUSG00000025283 | Sat1 | 20229 | protein_coding | 2816.833056 | 0.038637848 | 0.089553706 | 0.431448906 | 0.666141996 | 0.806377087 | 0.038637848 |
| ENSMUSG00000040383 | Aqr | 11834 | protein_coding | 902.3986645 | -0.038675671 | 0.088506102 | -0.436983103 | 0.662123602 | 0.804505636 | 0.038675671 |
| ENSMUSG00000036138 | Acaa1a | 113868 | protein_coding | 1172.80214 | -0.038709362 | 0.131985354 | -0.293285281 | 0.769304104 | 0.874311506 | 0.038709362 |
| ENSMUSG00000051506 | Wdfy4 | 545030 | protein_coding | 12.60345705 | 0.038717571 | 0.630304215 | 0.0614268 | 0.95101931 | 0.975614007 | 0.038717571 |
| ENSMUSG00000028191 | Bcl10 | 12042 | protein_coding | 984.5593626 | 0.038793057 | 0.147270568 | 0.263413505 | 0.792231882 | 0.887388848 | 0.038793057 |
| ENSMUSG00000024981 | Acsl5 | 433256 | protein_coding | 1797.53991 | 0.038809395 | 0.088274785 | 0.439643045 | 0.66019566 | 0.803330517 | 0.038809395 |
| ENSMUSG00000025133 | Ints4 | 101861 | protein_coding | 542.4689851 | 0.038849326 | 0.065457152 | 0.593507742 | 0.552841403 | 0.732884453 | 0.038849326 |
| ENSMUSG00000016984 | Etaa1 | 68145 | protein_coding | 229.1625731 | 0.03886702 | 0.15240105 | 0.255031181 | 0.798699016 | 0.892006327 | 0.03886702 |
| ENSMUSG00000025616 | Usp16 | 74112 | protein_coding | 806.8797187 | 0.038961212 | 0.080307285 | 0.485151657 | 0.627568772 | 0.784362476 | 0.038961212 |
| ENSMUSG00000049482 | Ctu2 | 66965 | protein_coding | 177.2617874 | -0.039001285 | 0.157797013 | -0.247161113 | 0.804783539 | 0.894840643 | 0.039001285 |
| ENSMUSG00000014856 | Tmem208 | 66320 | protein_coding | 497.3865865 | -0.039003471 | 0.127087518 | -0.306902455 | 0.75891762 | 0.868201089 | 0.039003471 |
| ENSMUSG00000008958 | Vps72 | 21427 | protein_coding | 355.8807255 | -0.039008757 | 0.082040867 | -0.475479582 | 0.634445183 | 0.788440577 | 0.039008757 |
| ENSMUSG00000041697 | Cox6a1 | 12861 | protein_coding | 4711.651514 | 0.039048273 | 0.106055683 | 0.368186521 | 0.71273416 | 0.838283978 | 0.039048273 |
| ENSMUSG00000024505 | Dtwd2 | 68857 | protein_coding | 55.01793998 | -0.039092786 | 0.156381196 | -0.249983929 | 0.802599777 | 0.894034894 | 0.039092786 |
| ENSMUSG00000043243 | Niban3 | 100037278 | protein_coding | 10.3239414 | 0.039152831 | 0.390730689 | 0.100204135 | 0.920182263 | 0.96046285 | 0.039152831 |
| ENSMUSG00000046679 | C87436 | 232196 | protein_coding | 180.3536452 | -0.039163606 | 0.142428987 | -0.274969353 | 0.783339784 | 0.882455341 | 0.039163606 |
| ENSMUSG00000023939 | Mrpl14 | 68463 | protein_coding | 260.6338122 | 0.039175174 | 0.101549567 | 0.38577391 | 0.699664119 | 0.829246975 | 0.039175174 |
| ENSMUSG00000024867 | Pip5k1b | 18719 | protein_coding | 988.7469931 | -0.039221827 | 0.081056149 | -0.483884662 | 0.628467726 | 0.784993252 | 0.039221827 |
| ENSMUSG00000024208 | Uqcc2 | 67267 | protein_coding | 321.8115398 | 0.039260925 | 0.106778614 | 0.367685285 | 0.713107914 | 0.838486874 | 0.039260925 |
| ENSMUSG00000029469 | Ift81 | 12589 | protein_coding | 74.67052781 | -0.039278637 | 0.129476008 | -0.303366142 | 0.761610846 | 0.869254376 | 0.039278637 |

| ensembl_gene_id | external_gene_name | entrezgene_id | gene_biotype | baseMean | log2FoldChange | lfcSE | stat | pvalue | padj | absLog2FC |
|---|---|---|---|---|---|---|---|---|---|---|
| ENSMUSG00000034640 | Tiparp | 99929 | protein_coding | 453.3415456 | -0.039281392 | 0.121927398 | -0.322170349 | 0.747323646 | 0.859635374 | 0.039281392 |
| ENSMUSG00000047371 | Zfp768 | 233890 | protein_coding | 424.4746361 | 0.039290489 | 0.11398822 | 0.344689028 | 0.730328168 | 0.848418573 | 0.039290489 |
| ENSMUSG00000039501 | Znfx1 | 98999 | protein_coding | 726.1382373 | 0.039290599 | 0.104966469 | 0.374315714 | 0.708169442 | 0.835163844 | 0.039290599 |
| ENSMUSG00000029310 | Nudt9 | 74167 | protein_coding | 468.8159668 | -0.039359415 | 0.086320413 | -0.4559688 | 0.648412415 | 0.797061488 | 0.039359415 |
| ENSMUSG00000031577 | Tti2 | 234138 | protein_coding | 171.2627975 | -0.039447588 | 0.104270689 | -0.378319047 | 0.705193597 | 0.833382076 | 0.039447588 |
| ENSMUSG00000022422 | Dscc1 | 72107 | protein_coding | 29.71351355 | -0.039467161 | 0.271673761 | -0.145274099 | 0.884494464 | 0.941852375 | 0.039467161 |
| ENSMUSG00000043090 | Zfp866 | 330788 | protein_coding | 285.4944457 | -0.039533512 | 0.101551682 | -0.389294513 | 0.697058295 | 0.827451522 | 0.039533512 |
| ENSMUSG00000031353 | Rbbp7 | 245688 | protein_coding | 2256.688333 | 0.039669645 | 0.090165186 | 0.439966324 | 0.659961498 | 0.803168303 | 0.039669645 |
| ENSMUSG00000020841 | Cpd | 12874 | protein_coding | 6767.779707 | -0.039671314 | 0.071705081 | -0.553256658 | 0.580087675 | 0.751318253 | 0.039671314 |
| ENSMUSG00000025967 | Eef1b2 | 55949 | protein_coding | 6981.153172 | -0.039672082 | 0.090568336 | -0.438034786 | 0.661361067 | 0.804053137 | 0.039672082 |
| ENSMUSG00000022407 | Adsl | 11564 | protein_coding | 621.9163028 | -0.039708321 | 0.114655782 | -0.346326374 | 0.72909745 | 0.847668919 | 0.039708321 |
| ENSMUSG00000056501 | Cebpb | 12608 | protein_coding | 74.72980134 | -0.039802585 | 0.279938771 | -0.142183183 | 0.886935315 | 0.942483462 | 0.039802585 |
| ENSMUSG00000040471 | Ggt6 | 71522 | protein_coding | 598.705544 | 0.039872354 | 0.137836174 | 0.289273512 | 0.772372081 | 0.876172816 | 0.039872354 |
| ENSMUSG00000025894 | Aasdhppt | 67618 | protein_coding | 371.9171425 | 0.039882041 | 0.10928481 | 0.364936724 | 0.71515864 | 0.839800265 | 0.039882041 |
| ENSMUSG00000039480 | Nt5dc1 | 319638 | protein_coding | 401.8220843 | 0.039882534 | 0.087796892 | 0.45425906 | 0.649642379 | 0.797543288 | 0.039882534 |
| ENSMUSG00000024926 | Kat5 | 81601 | protein_coding | 337.4344423 | 0.03989979 | 0.082920803 | 0.481179492 | 0.630388935 | 0.78600125 | 0.03989979 |
| ENSMUSG00000032540 | Abhd5 | 67469 | protein_coding | 398.0332228 | -0.03990816 | 0.069001082 | -0.578370061 | 0.563014303 | 0.739770497 | 0.03990816 |
| ENSMUSG00000015804 | Med28 | 66999 | protein_coding | 606.5686072 | 0.039912425 | 0.104808999 | 0.38081105 | 0.70334346 | 0.832121713 | 0.039912425 |
| ENSMUSG00000006728 | Cdk4 | 12567 | protein_coding | 968.1527077 | 0.039918758 | 0.092689038 | 0.430673988 | 0.666705435 | 0.806813871 | 0.039918758 |
| ENSMUSG00000031594 | Fgl1 | 234199 | protein_coding | 23.82381933 | -0.039964753 | 0.276880626 | -0.144339291 | 0.885232554 | 0.94220607 | 0.039964753 |
| ENSMUSG00000036282 | Naa30 | 70646 | protein_coding | 706.3260449 | -0.03997947 | 0.101413484 | -0.394222426 | 0.69341683 | 0.825651233 | 0.03997947 |
| ENSMUSG00000052415 | Tchh | 99681 | protein_coding | 13.05356485 | 0.040038133 | 0.359437961 | 0.111390943 | 0.911306342 | 0.956372548 | 0.040038133 |
| ENSMUSG00000005698 | Ctcf | 13018 | protein_coding | 1185.76829 | -0.040060909 | 0.063354987 | -0.632324483 | 0.527174868 | 0.712562908 | 0.040060909 |
| ENSMUSG00000025103 | Btbd1 | 83962 | protein_coding | 1103.102763 | 0.040082871 | 0.07241514 | 0.553515066 | 0.579910767 | 0.751266262 | 0.040082871 |
| ENSMUSG00000027472 | Pdrg1 | 68559 | protein_coding | 319.0896084 | 0.040100301 | 0.12170946 | 0.329475629 | 0.741796211 | 0.856183337 | 0.040100301 |
| ENSMUSG00000025208 | Mrpl43 | 94067 | protein_coding | 400.8440413 | 0.040139042 | 0.106890887 | 0.375514163 | 0.707278115 | 0.83460704 | 0.040139042 |
| ENSMUSG00000022992 | Kansl2 | 69612 | protein_coding | 255.2268316 | 0.040227963 | 0.098556749 | 0.408170559 | 0.683148461 | 0.818964084 | 0.040227963 |
| ENSMUSG00000027198 | Ext2 | 14043 | protein_coding | 600.8869221 | 0.040288754 | 0.064253659 | 0.6270266 | 0.530641799 | 0.715912466 | 0.040288754 |
| ENSMUSG00000033577 | Myo6 | 17920 | protein_coding | 5040.680707 | 0.040329668 | 0.045410635 | 0.888110649 | 0.374481235 | 0.58289232 | 0.040329668 |
| ENSMUSG00000031776 | Arl2bp | 107566 | protein_coding | 885.5108205 | -0.040332035 | 0.07138847 | -0.564965672 | 0.572097114 | 0.745911176 | 0.040332035 |
| ENSMUSG00000031119 | Gpc4 | 14735 | protein_coding | 296.5896825 | 0.04034993 | 0.146616956 | 0.275206436 | 0.783157643 | 0.882391137 | 0.04034993 |
| ENSMUSG00000038324 | Trpc4ap | 56407 | protein_coding | 1254.361115 | -0.040351428 | 0.072704121 | -0.555008806 | 0.578888637 | 0.750430391 | 0.040351428 |
| ENSMUSG00000064030 | Pym1 | 78428 | protein_coding | 160.184309 | -0.040354936 | 0.152505904 | -0.26461229 | 0.791308152 | 0.887042894 | 0.040354936 |
| ENSMUSG00000068134 | Zfp120 | 104348 | protein_coding | 409.8662968 | 0.040375088 | 0.140730318 | 0.286896871 | 0.774191287 | 0.87729929 | 0.040375088 |
| ENSMUSG00000025730 | Rab40c | 224624 | protein_coding | 296.7260362 | -0.040408962 | 0.11951678 | -0.338102833 | 0.735285696 | 0.851693009 | 0.040408962 |
| ENSMUSG00000034021 | Pds5b | 100710 | protein_coding | 369.5258445 | 0.040545667 | 0.117147873 | 0.346106729 | 0.729262507 | 0.847675196 | 0.040545667 |
| ENSMUSG00000026342 | Slc35f5 | 74150 | protein_coding | 1664.354453 | 0.040565894 | 0.112490733 | 0.360615432 | 0.718386951 | 0.841729245 | 0.040565894 |
| ENSMUSG00000064853 | Gm23442 | 115488838 | snoRNA | 81.83398677 | 0.040615244 | 0.442229232 | 0.091842062 | 0.926823525 | 0.963794322 | 0.040615244 |

| ensembl_gene_id | external_gene_name | entrezgene_id | gene_biotype | baseMean | log2FoldChange | lfcSE | stat | pvalue | padj | absLog2FC |
|---|---|---|---|---|---|---|---|---|---|---|
| ENSMUSG00000025340 | Rabgef1 | 56715 | protein_coding | 401.1879648 | -0.040633766 | 0.100605489 | -0.403892134 | 0.686292041 | 0.821158325 | 0.040633766 |
| ENSMUSG00000034620 | Rxylt1 | 216395 | protein_coding | 372.8585395 | -0.040643793 | 0.09864238 | -0.412031757 | 0.680316147 | 0.817182783 | 0.040643793 |
| ENSMUSG00000002307 | Daxx | 13163 | protein_coding | 305.0802586 | -0.040706788 | 0.08379899 | -0.485767049 | 0.627132339 | 0.784304814 | 0.040706788 |
| ENSMUSG00000032216 | Nedd4 | 17999 | protein_coding | 5926.752883 | -0.040749394 | 0.054777727 | -0.743904427 | 0.456934297 | 0.655739209 | 0.040749394 |
| ENSMUSG00000048992 | Prss32 | 69814 | protein_coding | 5149.887043 | 0.040753716 | 0.092126034 | 0.442369156 | 0.658222095 | 0.802216048 | 0.040753716 |
| ENSMUSG00000068130 | Zfp442 | 668923 | protein_coding | 77.72699879 | 0.040785589 | 0.24654159 | 0.165430868 | 0.868604859 | 0.932161313 | 0.040785589 |
| ENSMUSG00000070699 | Sars2 | 71984 | protein_coding | 140.527609 | 0.040829897 | 0.119613819 | 0.34134766 | 0.732841873 | 0.850024456 | 0.040829897 |
| ENSMUSG00000027551 | Zfp64 | 22722 | protein_coding | 244.7377872 | -0.040852587 | 0.102120406 | -0.40004333 | 0.689124603 | 0.823011515 | 0.040852587 |
| ENSMUSG00000021636 | Marveld2 | 218518 | protein_coding | 578.5955308 | 0.040887421 | 0.078426165 | 0.52134923 | 0.602123512 | 0.767564496 | 0.040887421 |
| ENSMUSG00000036270 | Edc4 | 234699 | protein_coding | 569.2834033 | -0.040906027 | 0.111326698 | -0.367441303 | 0.713289867 | 0.838515386 | 0.040906027 |
| ENSMUSG00000004364 | Cul3 | 26554 | protein_coding | 1910.630116 | -0.040912413 | 0.062588185 | -0.653676295 | 0.513320379 | 0.700974595 | 0.040912413 |
| ENSMUSG00000028273 | Pdlim5 | 56376 | protein_coding | 1526.636511 | -0.040936019 | 0.121882293 | -0.335865187 | 0.73697253 | 0.852840632 | 0.040936019 |
| ENSMUSG00000028672 | Hmgcl | 15356 | protein_coding | 1226.886638 | -0.041025267 | 0.123284232 | -0.332769786 | 0.739308067 | 0.85461201 | 0.041025267 |
| ENSMUSG00000029392 | Rilpl1 | 75695 | protein_coding | 14.96060231 | -0.041050391 | 0.285016397 | -0.144028172 | 0.885478224 | 0.94220607 | 0.041050391 |
| ENSMUSG00000062101 | Zfp119b | 240120 | protein_coding | 76.88694984 | 0.041054373 | 0.203857387 | 0.201387712 | 0.840395423 | 0.915836464 | 0.041054373 |
| ENSMUSG00000028397 | Kdm4c | 76804 | protein_coding | 552.1210502 | -0.04107325 | 0.08089076 | -0.507761948 | 0.611620299 | 0.773595499 | 0.04107325 |
| ENSMUSG00000031796 | Cfap20 | 14894 | protein_coding | 327.687362 | -0.041083841 | 0.077459146 | -0.530393672 | 0.595839012 | 0.763909664 | 0.041083841 |
| ENSMUSG00000041930 | Fam222a | 433940 | protein_coding | 55.35734595 | -0.041150338 | 0.184448983 | -0.223098754 | 0.823458648 | 0.906681139 | 0.041150338 |
| ENSMUSG00000035105 | Egln3 | 112407 | protein_coding | 235.8339472 | 0.041167616 | 0.220141254 | 0.18700546 | 0.851656352 | 0.921798204 | 0.041167616 |
| ENSMUSG00000031163 | Glod5 | 69824 | protein_coding | 350.069814 | 0.041171738 | 0.113616263 | 0.36237539 | 0.717071528 | 0.840677852 | 0.041171738 |
| ENSMUSG00000002345 | Borcs8 | 72368 | protein_coding | 178.3367071 | -0.041190752 | 0.089095414 | -0.462321801 | 0.643850568 | 0.793811583 | 0.041190752 |
| ENSMUSG00000042549 | Map2k3os | 24082 | lncRNA | 25.53299862 | 0.041191223 | 0.210818372 | 0.195387253 | 0.845089795 | 0.918184874 | 0.041191223 |
| ENSMUSG00000020273 | Papolg | 216578 | protein_coding | 309.1966216 | -0.041221118 | 0.100238793 | -0.411229195 | 0.680904482 | 0.817413822 | 0.041221118 |
| ENSMUSG00000024073 | Birc6 | 12211 | protein_coding | 3710.623753 | 0.041223681 | 0.103885921 | 0.396816823 | 0.691502544 | 0.824338032 | 0.041223681 |
| ENSMUSG00000040432 | Ltb4r2 | 57260 | protein_coding | 10.75759275 | 0.041249976 | 0.48225779 | 0.085535116 | 0.93183598 | 0.966021094 | 0.041249976 |
| ENSMUSG00000005982 | Naa60 | 74763 | protein_coding | 377.7394211 | 0.041287122 | 0.071381271 | 0.578402735 | 0.562992248 | 0.739770497 | 0.041287122 |
| ENSMUSG00000008976 | Gabpa | 14390 | protein_coding | 846.1431932 | 0.041289328 | 0.070034908 | 0.589553545 | 0.555490004 | 0.734992922 | 0.041289328 |
| ENSMUSG00000028337 | Coro2a | 107684 | protein_coding | 2929.169794 | -0.041357345 | 0.153565943 | -0.26931326 | 0.787688629 | 0.884788751 | 0.041357345 |
| ENSMUSG00000025156 | Gps1 | 209318 | protein_coding | 717.8407875 | 0.041388806 | 0.091894098 | 0.450396776 | 0.65242437 | 0.79857184 | 0.041388806 |
| ENSMUSG00000030742 | Lat | 16797 | protein_coding | 9.052462012 | 0.041432154 | 0.555658863 | 0.074564012 | 0.940561609 | 0.971067928 | 0.041432154 |
| ENSMUSG00000052812 | Atad2b | 320817 | protein_coding | 717.9083633 | -0.041445349 | 0.069421235 | -0.597012556 | 0.550498998 | 0.731488115 | 0.041445349 |
| ENSMUSG00000022706 | Mrpl40 | 18100 | protein_coding | 490.4478161 | 0.041450699 | 0.086829563 | 0.477380027 | 0.633091534 | 0.787741735 | 0.041450699 |
| ENSMUSG00000021039 | Snw1 | 66354 | protein_coding | 728.4360701 | 0.04148235 | 0.105964648 | 0.391473487 | 0.695447282 | 0.826592861 | 0.04148235 |
| ENSMUSG00000038047 | Haus6 | 230376 | protein_coding | 222.9920573 | -0.041497725 | 0.112249031 | -0.36969339 | 0.711610958 | 0.837372444 | 0.041497725 |
| ENSMUSG00000028070 | Naxe | 246703 | protein_coding | 643.7969181 | 0.041511255 | 0.118355861 | 0.350732569 | 0.72578899 | 0.845790446 | 0.041511255 |
| ENSMUSG00000073563 | Csnk1g3 | 70425 | protein_coding | 891.4499516 | 0.041570271 | 0.075021809 | 0.554109151 | 0.579504148 | 0.750963201 | 0.041570271 |
| ENSMUSG00000037492 | Zmat4 | 320158 | protein_coding | 22.77167807 | -0.041626601 | 0.270691318 | -0.153778856 | 0.877784106 | 0.938093481 | 0.041626601 |
| ENSMUSG00000020235 | Fzr1 | 56371 | protein_coding | 508.4243118 | -0.041674344 | 0.103407109 | -0.403012366 | 0.686939128 | 0.821304393 | 0.041674344 |

| ensembl_gene_id | external_gene_name | entrezgene_id | gene_biotype | baseMean | log2FoldChange | lfcSE | stat | pvalue | padj | absLog2FC |
|---|---|---|---|---|---|---|---|---|---|---|
| ENSMUSG00000032718 | Mansc1 | 67729 | protein_coding | 708.5054104 | -0.041702094 | 0.095212962 | -0.437987574 | 0.661395291 | 0.804053137 | 0.041702094 |
| ENSMUSG00000038344 | Txlng | 353170 | protein_coding | 363.2651968 | 0.041735363 | 0.087852746 | 0.475060429 | 0.634743902 | 0.788741674 | 0.041735363 |
| ENSMUSG00000030725 | Lipt2 | 67164 | protein_coding | 182.6048816 | 0.041742687 | 0.157879527 | 0.264395818 | 0.791474934 | 0.887042894 | 0.041742687 |
| ENSMUSG00000039233 | Tbce | 70430 | protein_coding | 348.1795911 | -0.041805849 | 0.097307567 | -0.429625879 | 0.667467809 | 0.80727602 | 0.041805849 |
| ENSMUSG00000048756 | Foxo3 | 56484 | protein_coding | 1857.760159 | 0.041828172 | 0.072937631 | 0.573478623 | 0.566320669 | 0.7424161 | 0.041828172 |
| ENSMUSG00000031328 | Flna | 192176 | protein_coding | 436.531162 | 0.041847403 | 0.185686473 | 0.22536592 | 0.821694622 | 0.905442422 | 0.041847403 |
| ENSMUSG00000031023 | Akip1 | 57373 | protein_coding | 121.1070977 | -0.041885366 | 0.12272629 | -0.341290901 | 0.732884597 | 0.850024456 | 0.041885366 |
| ENSMUSG00000002205 | Vrk3 | 101568 | protein_coding | 418.2192294 | 0.041902475 | 0.089158602 | 0.469976801 | 0.638371592 | 0.790506101 | 0.041902475 |
| ENSMUSG00000037730 | Mynn | 80732 | protein_coding | 324.1802547 | -0.041949099 | 0.068580664 | -0.611675316 | 0.540752597 | 0.723851362 | 0.041949099 |
| ENSMUSG00000036769 | Wdr44 | 72404 | protein_coding | 514.6611306 | 0.041967666 | 0.097196693 | 0.431780799 | 0.665900736 | 0.80636749 | 0.041967666 |
| ENSMUSG00000030271 | Ogg1 | 18294 | protein_coding | 79.23396453 | 0.042001253 | 0.137529462 | 0.305398223 | 0.760062875 | 0.868689179 | 0.042001253 |
| ENSMUSG00000029622 | Arpc1b | 11867 | protein_coding | 3651.952456 | -0.042038143 | 0.155091601 | -0.271053639 | 0.786349778 | 0.884220209 | 0.042038143 |
| ENSMUSG00000022024 | Sugt1 | 67955 | protein_coding | 1023.858944 | -0.042082837 | 0.086035329 | -0.489134373 | 0.624746568 | 0.782739191 | 0.042082837 |
| ENSMUSG00000094559 | Cyp2d34 | 223706 | protein_coding | 804.4812829 | -0.042085012 | 0.194171052 | -0.216741947 | 0.828409456 | 0.909178379 | 0.042085012 |
| ENSMUSG00000038497 | Tmco3 | 234076 | protein_coding | 1310.956853 | -0.042101734 | 0.086997919 | -0.483939547 | 0.628428773 | 0.784993252 | 0.042101734 |
| ENSMUSG00000087370 | Tmem170b | 621976 | protein_coding | 388.1810093 | -0.042120095 | 0.151407136 | -0.278190948 | 0.780865783 | 0.880896827 | 0.042120095 |
| ENSMUSG00000024747 | Aldh1a7 | 26358 | protein_coding | 283.6338562 | 0.042123098 | 0.118585296 | 0.355213496 | 0.722429655 | 0.844044564 | 0.042123098 |
| ENSMUSG00000025236 | Adpgk | 72141 | protein_coding | 277.7603451 | -0.042128093 | 0.138883005 | -0.303335119 | 0.761634485 | 0.869254376 | 0.042128093 |
| ENSMUSG00000073702 | Rpl31 | 114641 | protein_coding | 4712.910136 | 0.042144578 | 0.093412601 | 0.451165875 | 0.651870003 | 0.798492458 | 0.042144578 |
| ENSMUSG00000025915 | Sgk3 | 170755 | protein_coding | 99.19190169 | -0.042154409 | 0.176828763 | -0.23839113 | 0.811577746 | 0.899694484 | 0.042154409 |
| ENSMUSG00000045822 | Zswim3 | 67538 | protein_coding | 86.70091072 | -0.04216345 | 0.164420159 | -0.256437228 | 0.797613245 | 0.891285161 | 0.04216345 |
| ENSMUSG00000074170 | Plekhf1 | 72287 | protein_coding | 34.34091131 | -0.042165569 | 0.195484691 | -0.215697552 | 0.829223509 | 0.909405742 | 0.042165569 |
| ENSMUSG00000063760 | Rnf217 | 268291 | protein_coding | 194.6372315 | -0.042166735 | 0.126199558 | -0.334127438 | 0.738283396 | 0.853775328 | 0.042166735 |
| ENSMUSG00000037005 | Xpnpep2 | 170745 | protein_coding | 97.69817578 | 0.042205655 | 0.174112259 | 0.242404847 | 0.808466479 | 0.897800133 | 0.042205655 |
| ENSMUSG00000037236 | Matr3 | 17184 | protein_coding | 1415.763194 | -0.042253957 | 0.066816601 | -0.63238711 | 0.527133954 | 0.712562908 | 0.042253957 |
| ENSMUSG00000025246 | Tbl1x | 21372 | protein_coding | 1439.984641 | -0.042285257 | 0.072870125 | -0.580282486 | 0.561724136 | 0.739199176 | 0.042285257 |
| ENSMUSG00000033769 | Exoc6b | 75914 | protein_coding | 693.958829 | -0.042320028 | 0.090891554 | -0.465610127 | 0.641494593 | 0.792501813 | 0.042320028 |
| ENSMUSG00000029545 | Acads | 11409 | protein_coding | 3223.359912 | -0.042328886 | 0.119930565 | -0.352944939 | 0.72412972 | 0.844771344 | 0.042328886 |
| ENSMUSG00000020628 | Trappc12 | 217449 | protein_coding | 518.8963599 | 0.042330042 | 0.08411663 | 0.503230361 | 0.614802321 | 0.775503166 | 0.042330042 |
| ENSMUSG00000056267 | Cep70 | 68121 | protein_coding | 186.6696114 | 0.042422882 | 0.123269485 | 0.344147476 | 0.73073538 | 0.848568069 | 0.042422882 |
| ENSMUSG00000031378 | Abcd1 | 11666 | protein_coding | 280.0400098 | 0.042432638 | 0.101178566 | 0.41938367 | 0.674935757 | 0.812944399 | 0.042432638 |
| ENSMUSG00000026064 | Ptp4a1 | 19243 | protein_coding | 184.2915579 | 0.042522966 | 0.169983333 | 0.250159619 | 0.802463912 | 0.894011393 | 0.042522966 |
| ENSMUSG00000040441 | Slc26a10 | 216441 | protein_coding | 32.90603323 | -0.042546984 | 0.258406905 | -0.164651111 | 0.869218599 | 0.932568628 | 0.042546984 |
| ENSMUSG00000043284 | Tmem11 | 216821 | protein_coding | 397.1329869 | 0.042578821 | 0.095362376 | 0.44649497 | 0.655239745 | 0.800343347 | 0.042578821 |
| ENSMUSG00000032743 | Katnip | 233865 | protein_coding | 614.0492083 | -0.04259429 | 0.08551881 | -0.498069257 | 0.618435228 | 0.777768363 | 0.04259429 |
| ENSMUSG00000002393 | Nr2f6 | 13864 | protein_coding | 504.0028078 | 0.042599619 | 0.146859944 | 0.290069697 | 0.771762918 | 0.87594952 | 0.042599619 |
| ENSMUSG00000074916 | Chst14 | 72136 | protein_coding | 9.212915861 | 0.042600662 | 0.434430801 | 0.098060869 | 0.92188396 | 0.961231193 | 0.042600662 |
| ENSMUSG00000000194 | Gpr107 | 277463 | protein_coding | 1155.469354 | 0.042607248 | 0.055267438 | 0.770928595 | 0.440749257 | 0.642527127 | 0.042607248 |

| ensembl_gene_id | external_gene_name | entrezgene_id | gene_biotype | baseMean | log2FoldChange | lfcSE | stat | pvalue | padj | absLog2FC |
|---|---|---|---|---|---|---|---|---|---|---|
| ENSMUSG00000024812 | Tjp2 | 21873 | protein_coding | 2392.126359 | 0.042688827 | 0.182329727 | 0.234129823 | 0.814884194 | 0.901289872 | 0.042688827 |
| ENSMUSG00000073643 | Wdfy1 | 69368 | protein_coding | 1158.86416 | -0.042689496 | 0.107177584 | -0.398306201 | 0.69040449 | 0.823665847 | 0.042689496 |
| ENSMUSG00000029640 | Usp12 | 22217 | protein_coding | 1392.142831 | -0.042706041 | 0.091551279 | -0.46647126 | 0.640878215 | 0.792047502 | 0.042706041 |
| ENSMUSG00000035064 | Eef2k | 13631 | protein_coding | 1032.9222 | -0.042761039 | 0.115513391 | -0.370182527 | 0.711246495 | 0.837163904 | 0.042761039 |
| ENSMUSG00000030301 | Ccdc91 | 67015 | protein_coding | 546.0091514 | 0.042768421 | 0.083320072 | 0.513302743 | 0.607739565 | 0.771533271 | 0.042768421 |
| ENSMUSG00000060510 | Zfp266 | 77519 | protein_coding | 1087.446215 | -0.042772707 | 0.086967468 | -0.491824219 | 0.622843614 | 0.780969305 | 0.042772707 |
| ENSMUSG00000066613 | Zfp932 | 69504 | protein_coding | 166.8874442 | -0.042782 | 0.099152088 | -0.431478561 | 0.666120438 | 0.806377087 | 0.042782 |
| ENSMUSG00000041301 | Cftr | 12638 | protein_coding | 4254.21799 | -0.042785057 | 0.06940164 | -0.616484812 | 0.537574593 | 0.721598526 | 0.042785057 |
| ENSMUSG00000039865 | Slc44a3 | 213603 | protein_coding | 1384.204517 | -0.042809353 | 0.102341821 | -0.418297747 | 0.675729437 | 0.813715768 | 0.042809353 |
| ENSMUSG00000024104 | Washc2 | 28006 | protein_coding | 762.2884077 | 0.042822566 | 0.082965374 | 0.516149856 | 0.605749749 | 0.770268402 | 0.042822566 |
| ENSMUSG00000029729 | Zkscan1 | 74570 | protein_coding | 1915.235108 | 0.042835099 | 0.14729431 | 0.290812989 | 0.771194351 | 0.875604783 | 0.042835099 |
| ENSMUSG00000063019 | Manbal | 69161 | protein_coding | 170.7762339 | -0.042845358 | 0.106334009 | -0.402931844 | 0.686998365 | 0.821304393 | 0.042845358 |
| ENSMUSG00000028901 | Gmeb1 | 56809 | protein_coding | 235.1792891 | -0.042847111 | 0.136540309 | -0.313805582 | 0.753668705 | 0.864748244 | 0.042847111 |
| ENSMUSG00000032314 | Etfa | 110842 | protein_coding | 1577.091154 | 0.042864735 | 0.085773057 | 0.499745915 | 0.617253998 | 0.77714186 | 0.042864735 |
| ENSMUSG00000028933 | Xrcc2 | 57434 | protein_coding | 55.83816429 | -0.042883311 | 0.146451057 | -0.292816668 | 0.769662287 | 0.874537702 | 0.042883311 |
| ENSMUSG00000018167 | Stard3 | 59045 | protein_coding | 751.1862135 | -0.042897152 | 0.093092847 | -0.460799661 | 0.644942345 | 0.794481181 | 0.042897152 |
| ENSMUSG00000079641 | Rpl39 | 67248 | protein_coding | 4656.053074 | -0.042918713 | 0.095177104 | -0.450935268 | 0.652036204 | 0.798492458 | 0.042918713 |
| ENSMUSG00000057551 | Zfp317 | 244713 | protein_coding | 573.8366568 | -0.0429284 | 0.075509917 | -0.568513402 | 0.569686408 | 0.744683226 | 0.0429284 |
| ENSMUSG00000003282 | Plag1 | 56711 | protein_coding | 23.91018679 | -0.042940368 | 0.289164875 | -0.148497871 | 0.881949866 | 0.940337202 | 0.042940368 |
| ENSMUSG00000024816 | Frmd8 | 67457 | protein_coding | 509.1743299 | 0.042957454 | 0.122848793 | 0.34967746 | 0.726580771 | 0.846266142 | 0.042957454 |
| ENSMUSG00000027300 | Ubox5 | 140629 | protein_coding | 147.9393859 | -0.042972612 | 0.143853963 | -0.298723867 | 0.765150745 | 0.871705624 | 0.042972612 |
| ENSMUSG00000010057 | Nprl2 | 56032 | protein_coding | 203.1149667 | 0.042987113 | 0.098657608 | 0.435720201 | 0.663039747 | 0.804794609 | 0.042987113 |
| ENSMUSG00000062234 | Gak | 231580 | protein_coding | 2596.729086 | -0.042999729 | 0.063781343 | -0.674174094 | 0.500200637 | 0.690757391 | 0.042999729 |
| ENSMUSG00000068039 | Tcp1 | 21454 | protein_coding | 3218.552242 | 0.043018065 | 0.110876721 | 0.387981032 | 0.698030071 | 0.828227686 | 0.043018065 |
| ENSMUSG00000025813 | Homer2 | 26557 | protein_coding | 453.0721779 | -0.04307598 | 0.151220652 | -0.284855138 | 0.775755127 | 0.877982787 | 0.04307598 |
| ENSMUSG00000042225 | Ammecr1 | 56068 | protein_coding | 401.0029191 | -0.043079551 | 0.091169558 | -0.472521217 | 0.636554801 | 0.789583745 | 0.043079551 |
| ENSMUSG00000008690 | Ncaph2 | 52683 | protein_coding | 1250.009477 | -0.043088183 | 0.081579293 | -0.528175492 | 0.597377538 | 0.764750797 | 0.043088183 |
| ENSMUSG00000039202 | Abhd2 | 54608 | protein_coding | 6991.900954 | -0.043105243 | 0.088631528 | -0.48634209 | 0.626724642 | 0.784047572 | 0.043105243 |
| ENSMUSG00000069792 | Wfdc17 | 100034251 | protein_coding | 33.27850848 | 0.043105405 | 0.263334363 | 0.16369077 | 0.869974584 | 0.933188205 | 0.043105405 |
| ENSMUSG00000017686 | Rhot1 | 59040 | protein_coding | 673.191524 | 0.043117623 | 0.082113492 | 0.525097905 | 0.599515133 | 0.766132625 | 0.043117623 |
| ENSMUSG00000039670 | Oxld1 | 66431 | protein_coding | 93.80498207 | 0.043179135 | 0.183351952 | 0.23549864 | 0.813821735 | 0.900884818 | 0.043179135 |
| ENSMUSG00000041498 | Kif14 | 381293 | protein_coding | 184.9664698 | 0.043232396 | 0.148742897 | 0.290651837 | 0.771317611 | 0.875604783 | 0.043232396 |
| ENSMUSG00000043782 | Bicdl2 | 212733 | protein_coding | 243.865535 | 0.043251827 | 0.100388933 | 0.43084258 | 0.666582837 | 0.806726801 | 0.043251827 |
| ENSMUSG00000057265 | Bbof1 | 72873 | protein_coding | 22.54792341 | -0.043258539 | 0.247569888 | -0.174732637 | 0.861289722 | 0.927926916 | 0.043258539 |
| ENSMUSG00000042729 | Wdr74 | 107071 | protein_coding | 287.2928687 | -0.043300417 | 0.116065824 | -0.373067757 | 0.709098015 | 0.835765162 | 0.043300417 |
| ENSMUSG00000023079 | Gtf2ird1 | 57080 | protein_coding | 852.1034643 | 0.04341089 | 0.069762183 | 0.622269659 | 0.533764568 | 0.718743421 | 0.04341089 |
| ENSMUSG00000024604 | Rbm22 | 66810 | protein_coding | 543.5379249 | -0.043414081 | 0.069885932 | -0.621213461 | 0.534459183 | 0.719249909 | 0.043414081 |
| ENSMUSG00000111080 | Gm20300 | 100504586 | lncRNA | 278.1679231 | 0.043416945 | 0.108090449 | 0.401672354 | 0.687925173 | 0.821906591 | 0.043416945 |

| ensembl_gene_id | external_gene_name | entrezgene_id | gene_biotype | baseMean | log2FoldChange | lfcSE | stat | pvalue | padj | absLog2FC |
|---|---|---|---|---|---|---|---|---|---|---|
| ENSMUSG00000022889 | Mrpl39 | 27393 | protein_coding | 511.6443141 | 0.043453526 | 0.091859085 | 0.473045487 | 0.636180727 | 0.789347063 | 0.043453526 |
| ENSMUSG00000026828 | Galnt5 | 241391 | protein_coding | 1046.238895 | -0.043459283 | 0.091310956 | -0.475948178 | 0.634111298 | 0.788253962 | 0.043459283 |
| ENSMUSG00000054452 | Tle5 | 14797 | protein_coding | 4351.796463 | -0.043499929 | 0.071569682 | -0.607798276 | 0.543321273 | 0.726069405 | 0.043499929 |
| ENSMUSG00000036023 | Parp2 | 11546 | protein_coding | 342.9218444 | -0.043514059 | 0.07681784 | -0.566457725 | 0.571082663 | 0.745230585 | 0.043514059 |
| ENSMUSG00000014074 | Rnf168 | 70238 | protein_coding | 459.8950777 | -0.043649173 | 0.073574157 | -0.593267727 | 0.553001993 | 0.732888318 | 0.043649173 |
| ENSMUSG00000066235 | Pomgnt2 | 215494 | protein_coding | 142.4273594 | -0.04369084 | 0.124225454 | -0.351706016 | 0.725058749 | 0.84513251 | 0.04369084 |
| ENSMUSG00000020225 | Tmbim4 | 68212 | protein_coding | 1025.775716 | 0.043695349 | 0.098977059 | 0.441469468 | 0.658873161 | 0.80270244 | 0.043695349 |
| ENSMUSG00000032309 | Fbxo22 | 71999 | protein_coding | 502.135636 | -0.043706774 | 0.093248379 | -0.468713499 | 0.639274437 | 0.790801742 | 0.043706774 |
| ENSMUSG00000004849 | Ap1s1 | 11769 | protein_coding | 765.5388537 | 0.043714861 | 0.091496967 | 0.477773885 | 0.632811151 | 0.787638978 | 0.043714861 |
| ENSMUSG00000036639 | Nudt1 | 17766 | protein_coding | 205.3871609 | -0.043718461 | 0.108748431 | -0.402014637 | 0.687673254 | 0.821761542 | 0.043718461 |
| ENSMUSG00000029924 | Slc37a3 | 72144 | protein_coding | 755.7928194 | 0.043724976 | 0.068491744 | 0.638397767 | 0.523214784 | 0.709452933 | 0.043724976 |
| ENSMUSG00000071267 | Zfp942 | 73233 | protein_coding | 506.3528997 | 0.043729023 | 0.079858811 | 0.547579189 | 0.583980882 | 0.754575119 | 0.043729023 |
| ENSMUSG00000021009 | Ptpn21 | 24000 | protein_coding | 158.0852667 | 0.043758001 | 0.160622917 | 0.272426886 | 0.785293803 | 0.883563366 | 0.043758001 |
| ENSMUSG00000047067 | Dusp28 | 67446 | protein_coding | 102.1446217 | 0.043767958 | 0.123202336 | 0.355252662 | 0.722400316 | 0.844044564 | 0.043767958 |
| ENSMUSG00000070336 | Fbxo47 | 72973 | protein_coding | 12.19264242 | -0.043787824 | 0.379887755 | -0.115265162 | 0.908234951 | 0.954575761 | 0.043787824 |
| ENSMUSG00000031842 | Pde4c | 110385 | protein_coding | 8.520400232 | -0.043837716 | 0.532814071 | -0.082275823 | 0.93442738 | 0.967553497 | 0.043837716 |
| ENSMUSG00000026960 | Arl6ip6 | 65103 | protein_coding | 262.3027523 | 0.043838495 | 0.113608807 | 0.38587233 | 0.699591224 | 0.829246975 | 0.043838495 |
| ENSMUSG00000008892 | Vdac3 | 22335 | protein_coding | 2834.184757 | -0.043928938 | 0.107020884 | -0.41047071 | 0.681460686 | 0.817711752 | 0.043928938 |
| ENSMUSG00000065954 | Tacc1 | 320165 | protein_coding | 174.2088948 | 0.043948835 | 0.139118777 | 0.315908719 | 0.752071794 | 0.863289099 | 0.043948835 |
| ENSMUSG00000097842 | 9330104G04Rik | 320950 | lncRNA | 7.500342277 | -0.043960393 | 0.391147871 | -0.11238817 | 0.910515635 | 0.955759393 | 0.043960393 |
| ENSMUSG00000040767 | Snrnp25 | 78372 | protein_coding | 139.0071766 | -0.043963508 | 0.177579141 | -0.247571352 | 0.804466079 | 0.894700251 | 0.043963508 |
| ENSMUSG00000020877 | Scrn2 | 217140 | protein_coding | 209.7300805 | 0.044045817 | 0.116726215 | 0.377342975 | 0.705918739 | 0.833805983 | 0.044045817 |
| ENSMUSG00000002968 | Med25 | 75613 | protein_coding | 692.9916992 | -0.044062447 | 0.08086249 | -0.544905885 | 0.585818248 | 0.756020937 | 0.044062447 |
| ENSMUSG00000017132 | Cyth1 | 19157 | protein_coding | 703.6789742 | 0.044103754 | 0.1365861 | 0.322900747 | 0.746770411 | 0.859293709 | 0.044103754 |
| ENSMUSG00000019779 | Frk | 14302 | protein_coding | 2209.780181 | -0.04414703 | 0.172562164 | -0.255832615 | 0.798080089 | 0.891619531 | 0.04414703 |
| ENSMUSG00000049517 | Rps23 | 66475 | protein_coding | 5448.103508 | 0.044155911 | 0.09034538 | 0.488745642 | 0.625021786 | 0.782837698 | 0.044155911 |
| ENSMUSG00000019960 | Dusp6 | 67603 | protein_coding | 1379.399203 | -0.044162501 | 0.338947193 | -0.130293161 | 0.896334491 | 0.948192606 | 0.044162501 |
| ENSMUSG00000012296 | Tjap1 | 74094 | protein_coding | 157.0979078 | -0.044189197 | 0.113540456 | -0.389193407 | 0.69713308 | 0.827478627 | 0.044189197 |
| ENSMUSG00000057637 | Prdm2 | 110593 | protein_coding | 493.5723719 | -0.04420322 | 0.103365625 | -0.427639465 | 0.668913628 | 0.808227326 | 0.04420322 |
| ENSMUSG00000029547 | Ints1 | 68510 | protein_coding | 812.456031 | 0.044245523 | 0.110032105 | 0.402114665 | 0.687599641 | 0.821735225 | 0.044245523 |
| ENSMUSG00000026275 | Ppp1r7 | 66385 | protein_coding | 534.8182187 | 0.044284018 | 0.063195038 | 0.700751508 | 0.483458105 | 0.676897141 | 0.044284018 |
| ENSMUSG00000042558 | Adprs | 100206 | protein_coding | 277.0522953 | 0.044295376 | 0.095158289 | 0.465491518 | 0.64157951 | 0.792545248 | 0.044295376 |
| ENSMUSG00000061118 | Dnajc30 | 66114 | protein_coding | 135.6920729 | -0.044322358 | 0.121004627 | -0.366286472 | 0.714151323 | 0.83917017 | 0.044322358 |
| ENSMUSG00000030417 | Pdcd5 | 56330 | protein_coding | 361.6444557 | -0.044418769 | 0.088625627 | -0.501195541 | 0.616233512 | 0.77653223 | 0.044418769 |
| ENSMUSG00000022587 | Ly6e | 17069 | protein_coding | 1941.300775 | 0.044420402 | 0.128848643 | 0.344748696 | 0.730283305 | 0.848418573 | 0.044420402 |
| ENSMUSG00000031220 | Awat2 | 245532 | protein_coding | 10.30659208 | -0.04444813 | 0.359380986 | -0.123679694 | 0.90156889 | 0.951194891 | 0.04444813 |
| ENSMUSG00000036461 | Elf1 | 13709 | protein_coding | 1895.41258 | -0.044475268 | 0.046580811 | -0.954798068 | 0.339679825 | 0.550230881 | 0.044475268 |
| ENSMUSG00000059900 | Tmem40 | 94346 | protein_coding | 16.40223086 | 0.04447737 | 0.510443337 | 0.087134783 | 0.930564377 | 0.965468682 | 0.04447737 |

| ensembl_gene_id | external_gene_name | entrezgene_id | gene_biotype | baseMean | log2FoldChange | lfcSE | stat | pvalue | padj | absLog2FC |
|---|---|---|---|---|---|---|---|---|---|---|
| ENSMUSG00000035873 | Pawr | 114774 | protein_coding | 657.8287987 | -0.044491114 | 0.066285433 | -0.671204989 | 0.502089951 | 0.692106338 | 0.044491114 |
| ENSMUSG00000036503 | Rnf13 | 24017 | protein_coding | 811.9182764 | -0.044505372 | 0.088864812 | -0.500821085 | 0.616497045 | 0.776741412 | 0.044505372 |
| ENSMUSG00000026134 | Prim2 | 19076 | protein_coding | 136.1783222 | 0.044512473 | 0.133500671 | 0.333425087 | 0.738813429 | 0.854226212 | 0.044512473 |
| ENSMUSG00000018761 | Mpdu1 | 24070 | protein_coding | 558.3811437 | -0.044539482 | 0.090819192 | -0.490419272 | 0.623837242 | 0.781968962 | 0.044539482 |
| ENSMUSG00000033931 | Rbm34 | 52202 | protein_coding | 269.723306 | -0.044539774 | 0.104000801 | -0.428263754 | 0.668459105 | 0.807892603 | 0.044539774 |
| ENSMUSG00000020454 | Eif4enif1 | 74203 | protein_coding | 506.761719 | -0.044598558 | 0.077191422 | -0.577765727 | 0.563422298 | 0.740191918 | 0.044598558 |
| ENSMUSG00000039738 | Slx4 | 52864 | protein_coding | 166.6354958 | 0.044628271 | 0.134409523 | 0.332032057 | 0.739865053 | 0.854833615 | 0.044628271 |
| ENSMUSG00000059495 | Arhgef12 | 69632 | protein_coding | 1197.644442 | -0.044639078 | 0.075084507 | -0.594517821 | 0.552165826 | 0.732480826 | 0.044639078 |
| ENSMUSG00000027959 | Sass6 | 72776 | protein_coding | 110.9326199 | -0.044639263 | 0.135342919 | -0.329823413 | 0.741533395 | 0.856004052 | 0.044639263 |
| ENSMUSG00000028749 | Pla2g2f | 26971 | protein_coding | 429.3984974 | -0.044687306 | 0.27833739 | -0.160550857 | 0.872447163 | 0.934575311 | 0.044687306 |
| ENSMUSG00000029474 | Rnf34 | 80751 | protein_coding | 359.735888 | 0.044703564 | 0.096671169 | 0.46242913 | 0.643773614 | 0.793808373 | 0.044703564 |
| ENSMUSG00000021314 | Amph | 218038 | protein_coding | 50.51949373 | -0.044714854 | 0.257558366 | -0.173610568 | 0.862171526 | 0.928259299 | 0.044714854 |
| ENSMUSG00000003166 | Dgcr2 | 13356 | protein_coding | 896.8856212 | -0.044719099 | 0.07462007 | -0.599290493 | 0.548979186 | 0.730633661 | 0.044719099 |
| ENSMUSG00000060470 | Adgrg3 | 54672 | protein_coding | 6.799336518 | -0.044759682 | 0.587374617 | -0.076202956 | 0.939257631 | 0.970326602 | 0.044759682 |
| ENSMUSG00000032567 | Aste1 | 66595 | protein_coding | 219.7729145 | -0.0448168 | 0.099374701 | -0.450988028 | 0.651998177 | 0.798492458 | 0.0448168 |
| ENSMUSG00000005045 | Chd5 | 269610 | protein_coding | 14.51558226 | -0.044823001 | 0.474644814 | -0.094434827 | 0.924763751 | 0.963036092 | 0.044823001 |
| ENSMUSG00000022948 | Setd4 | 224440 | protein_coding | 100.2180152 | -0.044838328 | 0.140749884 | -0.318567427 | 0.750054557 | 0.861843095 | 0.044838328 |
| ENSMUSG00000022191 | Drosha | 14000 | protein_coding | 451.6512988 | 0.044872024 | 0.099953453 | 0.448929199 | 0.653482731 | 0.799253087 | 0.044872024 |
| ENSMUSG00000040618 | Pck2 | 74551 | protein_coding | 523.9030119 | -0.044941288 | 0.10302816 | -0.436203927 | 0.662688779 | 0.804672955 | 0.044941288 |
| ENSMUSG00000031783 | Polr2c | 20021 | protein_coding | 510.8368005 | 0.044990717 | 0.076145803 | 0.590849601 | 0.554621198 | 0.734331374 | 0.044990717 |
| ENSMUSG00000040139 | 9430038I01Rik | 77252 | protein_coding | 132.6110324 | 0.045028282 | 0.164176444 | 0.274267618 | 0.783878968 | 0.882812996 | 0.045028282 |
| ENSMUSG00000029249 | Rest | 19712 | protein_coding | 876.3308155 | 0.045088443 | 0.107080511 | 0.421070483 | 0.673703614 | 0.811873015 | 0.045088443 |
| ENSMUSG00000034435 | Tmem30b | 238257 | protein_coding | 3524.155056 | 0.04521039 | 0.079076463 | 0.571730051 | 0.567504872 | 0.743173362 | 0.04521039 |
| ENSMUSG00000028062 | Lamtor2 | 83409 | protein_coding | 610.3119492 | -0.045268349 | 0.068879545 | -0.657210333 | 0.511045683 | 0.699008761 | 0.045268349 |
| ENSMUSG00000056342 | Usp34 | 17847 | protein_coding | 2059.183974 | 0.045285248 | 0.135038387 | 0.335350922 | 0.737360385 | 0.853165496 | 0.045285248 |
| ENSMUSG00000028830 | AU040320 | 100317 | protein_coding | 1475.153745 | -0.045356724 | 0.065421579 | -0.693299141 | 0.488121842 | 0.680942953 | 0.045356724 |
| ENSMUSG00000024667 | Tmem216 | 68642 | protein_coding | 64.26548242 | -0.045360182 | 0.170935126 | -0.265364895 | 0.790728377 | 0.88651521 | 0.045360182 |
| ENSMUSG00000011752 | Pgam1 | 18648 | protein_coding | 575.4850991 | 0.045386024 | 0.111003575 | 0.408869928 | 0.682635119 | 0.818689268 | 0.045386024 |
| ENSMUSG00000010154 | Spire2 | 234857 | protein_coding | 599.7575721 | -0.045396935 | 0.098977791 | -0.458657798 | 0.646479924 | 0.795575387 | 0.045396935 |
| ENSMUSG00000006373 | Pgrmc1 | 53328 | protein_coding | 2116.825122 | -0.045402327 | 0.103818842 | -0.437322607 | 0.661877402 | 0.804454419 | 0.045402327 |
| ENSMUSG00000022940 | Pigp | 56176 | protein_coding | 356.318833 | -0.045421497 | 0.094307678 | -0.481630951 | 0.630068135 | 0.785882948 | 0.045421497 |
| ENSMUSG00000057835 | Zfp119a | 104349 | protein_coding | 116.8325401 | -0.045424434 | 0.152349306 | -0.298159768 | 0.765581227 | 0.871822248 | 0.045424434 |
| ENSMUSG00000032047 | Acat1 | 110446 | protein_coding | 2573.537474 | -0.045432397 | 0.085785404 | -0.529605215 | 0.596385678 | 0.764096331 | 0.045432397 |
| ENSMUSG00000054737 | Zfp182 | 319535 | protein_coding | 195.6719598 | -0.045434564 | 0.158146667 | -0.287293844 | 0.773887336 | 0.877099132 | 0.045434564 |
| ENSMUSG00000021235 | Coq6 | 217707 | protein_coding | 328.8816389 | -0.045440839 | 0.104550888 | -0.434628911 | 0.663831806 | 0.805232885 | 0.045440839 |
| ENSMUSG00000022663 | Atg3 | 67841 | protein_coding | 485.357054 | 0.045497568 | 0.091494741 | 0.497269768 | 0.618998828 | 0.778169955 | 0.045497568 |
| ENSMUSG00000058799 | Nap1l1 | 53605 | protein_coding | 978.6281319 | -0.04553622 | 0.112081371 | -0.406278221 | 0.684538186 | 0.819798814 | 0.04553622 |
| ENSMUSG00000020640 | Itsn2 | 20403 | protein_coding | 1587.521621 | -0.045559303 | 0.072315385 | -0.630008446 | 0.528689058 | 0.714245914 | 0.045559303 |

| ensembl_gene_id | external_gene_name | entrezgene_id | gene_biotype | baseMean | log2FoldChange | lfcSE | stat | pvalue | padj | absLog2FC |
|---|---|---|---|---|---|---|---|---|---|---|
| ENSMUSG00000029229 | Chic2 | 74277 | protein_coding | 321.7674706 | 0.045562209 | 0.118873407 | 0.383283443 | 0.701509614 | 0.830569028 | 0.045562209 |
| ENSMUSG00000051518 | Rps19bp1 | 66538 | protein_coding | 171.6162253 | 0.045595125 | 0.128671106 | 0.354354031 | 0.723073581 | 0.844115801 | 0.045595125 |
| ENSMUSG00000005732 | Ranbp1 | 19385 | protein_coding | 457.5907724 | 0.045625095 | 0.140346785 | 0.325088281 | 0.745114258 | 0.858333688 | 0.045625095 |
| ENSMUSG00000062373 | Tmem65 | 74868 | protein_coding | 704.5300796 | -0.045637203 | 0.191383062 | -0.238459991 | 0.811524343 | 0.899694484 | 0.045637203 |
| ENSMUSG00000020553 | Pctp | 18559 | protein_coding | 333.505591 | -0.045646223 | 0.117941621 | -0.387023875 | 0.698738533 | 0.828766651 | 0.045646223 |
| ENSMUSG00000026632 | Tatdn3 | 68972 | protein_coding | 68.78198184 | -0.045649975 | 0.150484326 | -0.303353688 | 0.761620335 | 0.869254376 | 0.045649975 |
| ENSMUSG00000000385 | Tmprss2 | 50528 | protein_coding | 6519.475955 | 0.04565192 | 0.125833748 | 0.362795518 | 0.71675764 | 0.840627772 | 0.04565192 |
| ENSMUSG00000075595 | Zfp652 | 268469 | protein_coding | 1102.190482 | 0.045653856 | 0.110268573 | 0.414024182 | 0.678856397 | 0.816247052 | 0.045653856 |
| ENSMUSG00000033294 | Noc4l | 100608 | protein_coding | 219.5099374 | -0.045656391 | 0.171922681 | -0.26556351 | 0.790575392 | 0.88651521 | 0.045656391 |
| ENSMUSG00000028848 | Gpn2 | 100210 | protein_coding | 161.6536858 | -0.045687113 | 0.127469645 | -0.358415625 | 0.720032301 | 0.842603371 | 0.045687113 |
| ENSMUSG00000030499 | Kctd15 | 233107 | protein_coding | 16.16279649 | -0.045691884 | 0.310511056 | -0.147150588 | 0.883013161 | 0.940966992 | 0.045691884 |
| ENSMUSG00000026172 | Bcs1l | 66821 | protein_coding | 167.6752639 | -0.045707856 | 0.114708789 | -0.398468643 | 0.690284768 | 0.823584703 | 0.045707856 |
| ENSMUSG00000028960 | Ube4b | 63958 | protein_coding | 665.2120056 | -0.045739921 | 0.101872566 | -0.448991542 | 0.653437758 | 0.799253087 | 0.045739921 |
| ENSMUSG00000025825 | Iscu | 66383 | protein_coding | 355.5009941 | 0.04575683 | 0.088486368 | 0.517105976 | 0.605082182 | 0.769603849 | 0.04575683 |
| ENSMUSG00000038299 | Wdr36 | 225348 | protein_coding | 828.3426909 | -0.045762744 | 0.127387895 | -0.359239343 | 0.719416047 | 0.842364374 | 0.045762744 |
| ENSMUSG00000037993 | Dhx38 | 64340 | protein_coding | 627.9119332 | 0.045764151 | 0.102386391 | 0.446974942 | 0.654893152 | 0.800302143 | 0.045764151 |
| ENSMUSG00000030965 | Abraxas2 | 109359 | protein_coding | 564.1735891 | 0.045769297 | 0.06779632 | 0.675100016 | 0.499612224 | 0.690064472 | 0.045769297 |
| ENSMUSG00000040112 | Mrps35 | 232536 | protein_coding | 764.9789909 | 0.04578094 | 0.08689201 | 0.52687169 | 0.598282695 | 0.765321556 | 0.04578094 |
| ENSMUSG00000066152 | Slc31a2 | 20530 | protein_coding | 198.8954577 | -0.045853964 | 0.125050069 | -0.366684834 | 0.713854121 | 0.838968951 | 0.045853964 |
| ENSMUSG00000020680 | Taf15 | 70439 | protein_coding | 490.9137325 | 0.045865341 | 0.150866403 | 0.304012954 | 0.761118022 | 0.869186728 | 0.045865341 |
| ENSMUSG00000046516 | Cox17 | 12856 | protein_coding | 244.3132548 | -0.04588481 | 0.109968039 | -0.417255874 | 0.67649126 | 0.814269937 | 0.04588481 |
| ENSMUSG00000038506 | Dcun1d2 | 102323 | protein_coding | 318.5415602 | -0.045931155 | 0.09016431 | -0.509416138 | 0.610460567 | 0.773061998 | 0.045931155 |
| ENSMUSG00000033624 | Pdpr | 319518 | protein_coding | 721.8343508 | -0.045959323 | 0.09697898 | -0.473910146 | 0.635563984 | 0.789030642 | 0.045959323 |
| ENSMUSG00000029822 | Osbpl3 | 71720 | protein_coding | 296.4274213 | -0.045992419 | 0.147485743 | -0.31184315 | 0.755159729 | 0.865722025 | 0.045992419 |
| ENSMUSG00000040354 | Mars1 | 216443 | protein_coding | 583.1260264 | -0.046005624 | 0.151471005 | -0.303725613 | 0.761336943 | 0.869186728 | 0.046005624 |
| ENSMUSG00000033597 | Caskin1 | 268932 | protein_coding | 12.26411521 | -0.046103853 | 0.434169239 | -0.106188668 | 0.915432662 | 0.95821748 | 0.046103853 |
| ENSMUSG00000028030 | Tbck | 271981 | protein_coding | 564.6756936 | -0.046107814 | 0.073748294 | -0.625205157 | 0.531836417 | 0.71689162 | 0.046107814 |
| ENSMUSG00000021590 | Spata9 | 75571 | protein_coding | 7.612789819 | -0.046185275 | 0.523915702 | -0.08815402 | 0.929754262 | 0.965328339 | 0.046185275 |
| ENSMUSG00000039069 | Mtg2 | 52856 | protein_coding | 160.4263037 | -0.046191119 | 0.12069572 | -0.382707182 | 0.701936889 | 0.83080438 | 0.046191119 |
| ENSMUSG00000026708 | Cenpl | 70454 | protein_coding | 124.0420788 | 0.046243474 | 0.154877578 | 0.298580818 | 0.765259903 | 0.871705624 | 0.046243474 |
| ENSMUSG00000039048 | Foxred1 | 235169 | protein_coding | 416.2468011 | 0.046280159 | 0.076826032 | 0.602402047 | 0.546906548 | 0.728832101 | 0.046280159 |
| ENSMUSG00000000743 | Chmp1a | 234852 | protein_coding | 2435.250373 | 0.046284192 | 0.097630891 | 0.474073236 | 0.635447684 | 0.788947751 | 0.046284192 |
| ENSMUSG00000028318 | Polr1e | 64424 | protein_coding | 133.048428 | -0.046317146 | 0.186238043 | -0.248698629 | 0.80359391 | 0.894329499 | 0.046317146 |
| ENSMUSG00000029669 | Tspan12 | 269831 | protein_coding | 893.4496744 | 0.04631858 | 0.103671124 | 0.446783811 | 0.655031161 | 0.800302454 | 0.04631858 |
| ENSMUSG00000028776 | Tinagl1 | 94242 | protein_coding | 603.00123 | 0.046366475 | 0.189230358 | 0.245026622 | 0.806435818 | 0.896077905 | 0.046366475 |
| ENSMUSG00000020250 | Txnrd1 | 50493 | protein_coding | 3787.977261 | 0.046407748 | 0.105644313 | 0.439282978 | 0.660456507 | 0.803487938 | 0.046407748 |
| ENSMUSG00000039725 | Trp53rka | 381406 | protein_coding | 137.8647199 | -0.046416129 | 0.111363263 | -0.416799291 | 0.676825221 | 0.814542661 | 0.046416129 |
| ENSMUSG00000048039 | Isg20l2 | 229504 | protein_coding | 500.6775836 | -0.046431769 | 0.101499757 | -0.45745695 | 0.647342638 | 0.796267939 | 0.046431769 |

| ensembl_gene_id | external_gene_name | entrezgene_id | gene_biotype | baseMean | log2FoldChange | lfcSE | stat | pvalue | padj | absLog2FC |
|---|---|---|---|---|---|---|---|---|---|---|
| ENSMUSG00000033222 | Ttf2 | 74044 | protein_coding | 245.0906388 | 0.046463139 | 0.087620122 | 0.530279321 | 0.595918282 | 0.76392647 | 0.046463139 |
| ENSMUSG00000024787 | Snx15 | 69024 | protein_coding | 322.9224595 | 0.046464171 | 0.075687632 | 0.61389384 | 0.539285481 | 0.722433906 | 0.046464171 |
| ENSMUSG00000020949 | Fkbp3 | 30795 | protein_coding | 329.5263338 | -0.046547283 | 0.079898722 | -0.582578572 | 0.560177034 | 0.738067647 | 0.046547283 |
| ENSMUSG00000046794 | Ppp1r3b | 244416 | protein_coding | 492.3669617 | -0.046549708 | 0.145321529 | -0.320322169 | 0.748724119 | 0.860624794 | 0.046549708 |
| ENSMUSG00000041886 | Macc1 | 238455 | protein_coding | 156.0684565 | 0.046653094 | 0.158657565 | 0.294048975 | 0.76872048 | 0.87383527 | 0.046653094 |
| ENSMUSG00000014782 | Plekhg4 | 102075 | protein_coding | 7.010456309 | -0.046667062 | 0.596384485 | -0.078249961 | 0.937629221 | 0.969210087 | 0.046667062 |
| ENSMUSG00000039529 | Atp8b1 | 54670 | protein_coding | 9872.522284 | 0.046673736 | 0.061017908 | 0.764918657 | 0.444319983 | 0.645252434 | 0.046673736 |
| ENSMUSG00000046562 | Unc119b | 106840 | protein_coding | 637.2850371 | 0.046692311 | 0.087110381 | 0.536013168 | 0.59194945 | 0.761020102 | 0.046692311 |
| ENSMUSG00000028042 | Zbtb7b | 22724 | protein_coding | 2475.409528 | -0.046730663 | 0.128235724 | -0.364412205 | 0.715550223 | 0.840012084 | 0.046730663 |
| ENSMUSG00000068335 | Dok1 | 13448 | protein_coding | 147.6415851 | -0.046759996 | 0.132462241 | -0.353006223 | 0.724083775 | 0.844771344 | 0.046759996 |
| ENSMUSG00000049106 | Dcaf5 | 320808 | protein_coding | 644.3569167 | -0.046795275 | 0.068567803 | -0.682467176 | 0.494943592 | 0.686593061 | 0.046795275 |
| ENSMUSG00000086549 | Gm13648 | 100415901 | lncRNA | 95.585966 | -0.046796673 | 0.198585055 | -0.235650526 | 0.813703864 | 0.900884818 | 0.046796673 |
| ENSMUSG00000092416 | Zfp141 | 434178 | protein_coding | 238.9241729 | -0.046809077 | 0.100731481 | -0.46469164 | 0.642152296 | 0.79292616 | 0.046809077 |
| ENSMUSG00000028937 | Acot7 | 70025 | protein_coding | 1059.144749 | 0.046905951 | 0.101808558 | 0.460727001 | 0.64499448 | 0.79448396 | 0.046905951 |
| ENSMUSG00000027164 | Traf6 | 22034 | protein_coding | 494.8595728 | 0.046943667 | 0.077704468 | 0.604130866 | 0.54575664 | 0.728169923 | 0.046943667 |
| ENSMUSG00000050855 | Zfp940 | 233057 | protein_coding | 34.70100335 | -0.04694879 | 0.247774227 | -0.189482137 | 0.849714954 | 0.920572697 | 0.04694879 |
| ENSMUSG00000050002 | Idnk | 75731 | protein_coding | 303.3161491 | -0.046957619 | 0.09451516 | -0.496826317 | 0.619311535 | 0.778317353 | 0.046957619 |
| ENSMUSG00000028923 | Necap2 | 66147 | protein_coding | 713.8606007 | 0.047005565 | 0.096048028 | 0.489396461 | 0.624561041 | 0.782629869 | 0.047005565 |
| ENSMUSG00000029614 | Rpl6 | 19988 | protein_coding | 5797.476161 | -0.047037025 | 0.072726774 | -0.646763532 | 0.517784998 | 0.704835436 | 0.047037025 |
| ENSMUSG00000031691 | Tnpo2 | 212999 | protein_coding | 1072.988722 | -0.047076678 | 0.083761316 | -0.562033652 | 0.574093091 | 0.747419535 | 0.047076678 |
| ENSMUSG00000070427 | Il18bp | 16068 | protein_coding | 62.31307792 | 0.047095659 | 0.234844787 | 0.200539513 | 0.841058658 | 0.916179887 | 0.047095659 |
| ENSMUSG00000028708 | Mknk1 | 17346 | protein_coding | 175.3337155 | 0.047124501 | 0.106971322 | 0.440533966 | 0.659550418 | 0.802969027 | 0.047124501 |
| ENSMUSG00000040322 | Slc25a24 | 229731 | protein_coding | 5720.891785 | 0.047153784 | 0.184275436 | 0.255887521 | 0.798037691 | 0.891619531 | 0.047153784 |
| ENSMUSG00000034274 | Thoc5 | 107829 | protein_coding | 462.1918957 | -0.047211629 | 0.086948292 | -0.542985119 | 0.587140048 | 0.757262387 | 0.047211629 |
| ENSMUSG00000024151 | Msh2 | 17685 | protein_coding | 559.8500571 | -0.047237932 | 0.103473666 | -0.456521298 | 0.648015159 | 0.796849027 | 0.047237932 |
| ENSMUSG00000022159 | Rab2b | 76338 | protein_coding | 11.98083651 | -0.047252424 | 0.37427938 | -0.126249071 | 0.899534769 | 0.950188262 | 0.047252424 |
| ENSMUSG00000024044 | Epb41l3 | 13823 | protein_coding | 1318.104114 | 0.047283337 | 0.097917713 | 0.482888494 | 0.629174912 | 0.785506976 | 0.047283337 |
| ENSMUSG00000035566 | Pcdh17 | 219228 | protein_coding | 601.8410745 | 0.047283693 | 0.130555211 | 0.362173926 | 0.717222063 | 0.84073543 | 0.047283693 |
| ENSMUSG00000003420 | Fcgrt | 14132 | protein_coding | 772.2515905 | -0.047297645 | 0.137833853 | -0.3431497 | 0.731485844 | 0.849085297 | 0.047297645 |
| ENSMUSG00000029208 | Guf1 | 231279 | protein_coding | 477.0583469 | 0.047350773 | 0.077032474 | 0.614685872 | 0.538762192 | 0.722272825 | 0.047350773 |
| ENSMUSG00000032423 | Syncrip | 56403 | protein_coding | 1691.886725 | -0.047399281 | 0.125273383 | -0.378366735 | 0.705158176 | 0.833382076 | 0.047399281 |
| ENSMUSG00000097709 | 2810429I04Rik | 76937 | lncRNA | 10.89625598 | -0.047414138 | 0.545444197 | -0.086927568 | 0.930729086 | 0.96552383 | 0.047414138 |
| ENSMUSG00000022515 | Anks3 | 72615 | protein_coding | 311.1885986 | 0.047438793 | 0.077098217 | 0.615303372 | 0.538354391 | 0.721975982 | 0.047438793 |
| ENSMUSG00000028495 | Rps6 | 20104 | protein_coding | 3648.206163 | 0.047499929 | 0.09826623 | 0.483379986 | 0.628825956 | 0.785255964 | 0.047499929 |
| ENSMUSG00000048222 | Mfap1b | 100034361 | protein_coding | 182.7388253 | -0.047539528 | 0.099809818 | -0.476301123 | 0.633859866 | 0.788205164 | 0.047539528 |
| ENSMUSG00000053094 | Tmem248 | 71667 | protein_coding | 839.3830124 | -0.047632848 | 0.085404181 | -0.557734384 | 0.577025775 | 0.749051878 | 0.047632848 |
| ENSMUSG00000042737 | Dpm3 | 68563 | protein_coding | 226.5849332 | 0.047638489 | 0.141573021 | 0.336494119 | 0.736498286 | 0.852539586 | 0.047638489 |
| ENSMUSG00000056121 | Fez2 | 225020 | protein_coding | 302.5441842 | 0.047651757 | 0.095249523 | 0.500283415 | 0.61687553 | 0.776978453 | 0.047651757 |

| ensembl_gene_id | external_gene_name | entrezgene_id | gene_biotype | baseMean | log2FoldChange | lfcSE | stat | pvalue | padj | absLog2FC |
|---|---|---|---|---|---|---|---|---|---|---|
| ENSMUSG00000041199 | Rpusd1 | 106707 | protein_coding | 43.41829373 | 0.047661234 | 0.17929043 | 0.26583256 | 0.790368166 | 0.886363083 | 0.047661234 |
| ENSMUSG00000039357 | Fut11 | 73068 | protein_coding | 149.5290514 | -0.047676852 | 0.088968972 | -0.535881797 | 0.592040246 | 0.761026944 | 0.047676852 |
| ENSMUSG00000055991 | Zkscan5 | 22757 | protein_coding | 207.1752599 | -0.047723592 | 0.133414888 | -0.357708143 | 0.720561738 | 0.843075176 | 0.047723592 |
| ENSMUSG00000031585 | Gtf2e2 | 68153 | protein_coding | 354.0076803 | 0.047732208 | 0.092844216 | 0.514110733 | 0.607174574 | 0.771131144 | 0.047732208 |
| ENSMUSG00000047866 | Lonp2 | 66887 | protein_coding | 1385.02165 | -0.047799549 | 0.081037978 | -0.589841331 | 0.55529703 | 0.734920739 | 0.047799549 |
| ENSMUSG00000031575 | Ash2l | 23808 | protein_coding | 711.6194131 | -0.047807352 | 0.06288446 | -0.760241121 | 0.447110469 | 0.647828214 | 0.047807352 |
| ENSMUSG00000078517 | Emc1 | 230866 | protein_coding | 849.1877065 | 0.047868656 | 0.068271157 | 0.701154896 | 0.483206354 | 0.676728138 | 0.047868656 |
| ENSMUSG00000031974 | Abcb10 | 56199 | protein_coding | 469.2740596 | 0.047878432 | 0.084253993 | 0.568263057 | 0.56985636 | 0.744741762 | 0.047878432 |
| ENSMUSG00000020935 | Dcakd | 68087 | protein_coding | 142.1079753 | -0.047883709 | 0.121773647 | -0.393218976 | 0.694157756 | 0.826101669 | 0.047883709 |
| ENSMUSG00000040312 | Cchcr1 | 240084 | protein_coding | 161.4116598 | -0.047903804 | 0.155629728 | -0.307806257 | 0.75822976 | 0.867544258 | 0.047903804 |
| ENSMUSG00000017478 | Zc3h18 | 76014 | protein_coding | 1197.243325 | -0.047922639 | 0.062005751 | -0.772874106 | 0.439596891 | 0.642022673 | 0.047922639 |
| ENSMUSG00000021376 | Tpmt | 22017 | protein_coding | 220.9940824 | -0.047931464 | 0.100570479 | -0.47659576 | 0.633650003 | 0.788005719 | 0.047931464 |
| ENSMUSG00000020644 | Id2 | 15902 | protein_coding | 283.3129537 | 0.047941249 | 0.180632013 | 0.265408376 | 0.790694884 | 0.88651521 | 0.047941249 |
| ENSMUSG00000033430 | Terf2ip | 57321 | protein_coding | 360.1711449 | 0.047945582 | 0.089893207 | 0.533361568 | 0.593783316 | 0.76209791 | 0.047945582 |
| ENSMUSG00000024911 | Fibp | 58249 | protein_coding | 536.9573102 | -0.047950151 | 0.08909345 | -0.538200633 | 0.590438543 | 0.759898603 | 0.047950151 |
| ENSMUSG00000024182 | Axin1 | 12005 | protein_coding | 776.5203253 | -0.047964858 | 0.086023414 | -0.557579108 | 0.577131826 | 0.749071904 | 0.047964858 |
| ENSMUSG00000022858 | Tra2b | 20462 | protein_coding | 1765.541992 | -0.048015584 | 0.082980737 | -0.578635305 | 0.562835278 | 0.739770497 | 0.048015584 |
| ENSMUSG00000026843 | Fubp3 | 320267 | protein_coding | 613.9996489 | 0.048041093 | 0.057404164 | 0.836892139 | 0.402653199 | 0.609934395 | 0.048041093 |
| ENSMUSG00000050945 | Zfp438 | 240186 | protein_coding | 43.75864589 | -0.048080922 | 0.220018474 | -0.218531297 | 0.827015176 | 0.908412958 | 0.048080922 |
| ENSMUSG00000027327 | 1700037H04Rik | 67326 | protein_coding | 198.6407994 | 0.048080991 | 0.106420048 | 0.451803889 | 0.651410268 | 0.798250711 | 0.048080991 |
| ENSMUSG00000021371 | Mcur1 | 76137 | protein_coding | 767.1637197 | -0.048099583 | 0.068078058 | -0.706535775 | 0.479855013 | 0.673583934 | 0.048099583 |
| ENSMUSG00000026433 | Rab29 | 226422 | protein_coding | 151.7918013 | -0.048150209 | 0.143198322 | -0.336248418 | 0.736683544 | 0.852630104 | 0.048150209 |
| ENSMUSG00000034748 | Sirt6 | 50721 | protein_coding | 133.6526353 | 0.048216861 | 0.13107303 | 0.367862567 | 0.712975713 | 0.838411012 | 0.048216861 |
| ENSMUSG00000035529 | Prdm4 | 72843 | protein_coding | 473.7286081 | 0.04823518 | 0.100393895 | 0.480459294 | 0.630900841 | 0.786307224 | 0.04823518 |
| ENSMUSG00000020547 | Bzw2 | 66912 | protein_coding | 675.6814955 | -0.048287357 | 0.090374022 | -0.534305718 | 0.593130037 | 0.761494544 | 0.048287357 |
| ENSMUSG00000040818 | Dennd6a | 211922 | protein_coding | 879.9095476 | -0.048306457 | 0.066019469 | -0.731700172 | 0.464351591 | 0.66131904 | 0.048306457 |
| ENSMUSG00000032415 | Ube2cbp | 70348 | protein_coding | 37.34958248 | -0.048308594 | 0.200433446 | -0.241020625 | 0.809539131 | 0.898543824 | 0.048308594 |
| ENSMUSG00000037816 | Fbxw17 | 109082 | protein_coding | 81.94495139 | 0.048316081 | 0.134395462 | 0.359506788 | 0.719216001 | 0.842333944 | 0.048316081 |
| ENSMUSG00000071757 | Zhx2 | 387609 | protein_coding | 196.6209196 | -0.048329324 | 0.10476358 | -0.461317989 | 0.644570481 | 0.794268808 | 0.048329324 |
| ENSMUSG00000018583 | G3bp1 | 27041 | protein_coding | 2505.586551 | 0.048340374 | 0.108009963 | 0.447554765 | 0.654474556 | 0.800101761 | 0.048340374 |
| ENSMUSG00000020664 | Dld | 13382 | protein_coding | 2651.74266 | 0.048340925 | 0.076272349 | 0.633793586 | 0.526215539 | 0.711870317 | 0.048340925 |
| ENSMUSG00000039841 | Zfp800 | 627049 | protein_coding | 571.5326262 | 0.048341839 | 0.138961998 | 0.347878122 | 0.727931717 | 0.846662573 | 0.048341839 |
| ENSMUSG00000071252 | 2210408I21Rik | 72371 | protein_coding | 64.15761503 | 0.048390569 | 0.224918432 | 0.215147191 | 0.829652562 | 0.909414769 | 0.048390569 |
| ENSMUSG00000024507 | Hsd17b4 | 15488 | protein_coding | 1599.383023 | -0.048396016 | 0.10886437 | -0.444553309 | 0.656642596 | 0.801149235 | 0.048396016 |
| ENSMUSG00000091387 | Gcnt4 | 218476 | protein_coding | 137.5405084 | -0.048442436 | 0.220506609 | -0.219687004 | 0.826114927 | 0.908178648 | 0.048442436 |
| ENSMUSG00000032786 | Alas1 | 11655 | protein_coding | 2021.999876 | -0.048469403 | 0.129470153 | -0.374367386 | 0.708131004 | 0.835163844 | 0.048469403 |
| ENSMUSG00000020485 | Supt4a | 20922 | protein_coding | 316.1620227 | 0.048514541 | 0.12866792 | 0.377052342 | 0.706134707 | 0.833937393 | 0.048514541 |
| ENSMUSG00000032312 | Csk | 12988 | protein_coding | 863.5420533 | -0.048626502 | 0.075166765 | -0.646914924 | 0.517687006 | 0.704835436 | 0.048626502 |

| ensembl_gene_id | external_gene_name | entrezgene_id | gene_biotype | baseMean | log2FoldChange | lfcSE | stat | pvalue | padj | absLog2FC |
|---|---|---|---|---|---|---|---|---|---|---|
| ENSMUSG00000014551 | Mrps25 | 64658 | protein_coding | 370.7174327 | -0.048628948 | 0.092330946 | -0.526680927 | 0.598415183 | 0.765321556 | 0.048628948 |
| ENSMUSG00000002820 | Atg4d | 235040 | protein_coding | 1049.343227 | 0.048638009 | 0.096454952 | 0.504256218 | 0.61408134 | 0.775272711 | 0.048638009 |
| ENSMUSG00000075376 | Rc3h2 | 319817 | protein_coding | 1216.031579 | 0.048695245 | 0.107444512 | 0.453212954 | 0.650395405 | 0.798050845 | 0.048695245 |
| ENSMUSG00000028292 | Rars2 | 109093 | protein_coding | 649.7965204 | -0.048702154 | 0.090064382 | -0.540748214 | 0.588681141 | 0.758433488 | 0.048702154 |
| ENSMUSG00000038781 | Stap2 | 106766 | protein_coding | 890.9686409 | -0.048767697 | 0.086092554 | -0.566456623 | 0.571083412 | 0.745230585 | 0.048767697 |
| ENSMUSG00000033213 | AA467197 | 433470 | protein_coding | 1970.31568 | -0.048791881 | 0.125821054 | -0.387787889 | 0.698173009 | 0.828258604 | 0.048791881 |
| ENSMUSG00000006678 | Pola1 | 18968 | protein_coding | 401.1642725 | -0.048869215 | 0.144997242 | -0.337035477 | 0.736090156 | 0.852237062 | 0.048869215 |
| ENSMUSG00000037474 | Dtl | 76843 | protein_coding | 204.4152053 | 0.048897582 | 0.121544575 | 0.402301641 | 0.687462047 | 0.821632437 | 0.048897582 |
| ENSMUSG00000051000 | Fhip1a | 229488 | protein_coding | 386.835452 | -0.048911624 | 0.183068674 | -0.26717637 | 0.789333369 | 0.885761723 | 0.048911624 |
| ENSMUSG00000000631 | Myo18a | 360013 | protein_coding | 2803.036003 | -0.04897004 | 0.14557674 | -0.336386428 | 0.736579483 | 0.852571616 | 0.04897004 |
| ENSMUSG00000059714 | Flot1 | 14251 | protein_coding | 414.1170789 | 0.048973488 | 0.092338794 | 0.530367418 | 0.595857212 | 0.763909664 | 0.048973488 |
| ENSMUSG00000064994 | Gm22422 | 115486004 | snoRNA | 57.5642211 | 0.048996326 | 0.305972907 | 0.160132891 | 0.872776392 | 0.934685832 | 0.048996326 |
| ENSMUSG00000024548 | Setbp1 | 240427 | protein_coding | 500.1463992 | 0.049023347 | 0.126930226 | 0.386222801 | 0.699331669 | 0.829099726 | 0.049023347 |
| ENSMUSG00000048310 | Pskh1 | 244631 | protein_coding | 318.5608125 | -0.049043407 | 0.108851636 | -0.450552774 | 0.65231191 | 0.798541656 | 0.049043407 |
| ENSMUSG00000030967 | Zranb1 | 360216 | protein_coding | 955.236742 | -0.049048396 | 0.073018666 | -0.67172408 | 0.501759369 | 0.692014516 | 0.049048396 |
| ENSMUSG00000029003 | Mad2l2 | 71890 | protein_coding | 53.09780133 | -0.049092117 | 0.149897266 | -0.327505083 | 0.743285899 | 0.857217273 | 0.049092117 |
| ENSMUSG00000021501 | Caml | 12328 | protein_coding | 354.6674251 | 0.049114383 | 0.101100633 | 0.485796985 | 0.627111112 | 0.784304814 | 0.049114383 |
| ENSMUSG00000053559 | Smagp | 207818 | protein_coding | 1242.604372 | -0.049148547 | 0.112728023 | -0.435992268 | 0.662842339 | 0.804706767 | 0.049148547 |
| ENSMUSG00000061286 | Exosc5 | 27998 | protein_coding | 202.551379 | -0.049186107 | 0.10764529 | -0.456927633 | 0.647723061 | 0.796612842 | 0.049186107 |
| ENSMUSG00000030016 | Zfp638 | 18139 | protein_coding | 1680.732273 | 0.049189405 | 0.067620103 | 0.72743759 | 0.466957934 | 0.663707223 | 0.049189405 |
| ENSMUSG00000026112 | Coa5 | 76178 | protein_coding | 1708.333251 | 0.049191387 | 0.085643701 | 0.574372499 | 0.565715759 | 0.741806257 | 0.049191387 |
| ENSMUSG00000030604 | Zfp626 | 71163 | protein_coding | 226.135863 | -0.049194497 | 0.125573107 | -0.391759812 | 0.695235692 | 0.826592861 | 0.049194497 |
| ENSMUSG00000069045 | Ddx3y | 26900 | protein_coding | 828.4464454 | -0.049195248 | 0.104580408 | -0.470405965 | 0.638065003 | 0.790438741 | 0.049195248 |
| ENSMUSG00000028657 | Ppt1 | 19063 | protein_coding | 555.7190027 | 0.049211847 | 0.083592479 | 0.588711419 | 0.556054876 | 0.735434864 | 0.049211847 |
| ENSMUSG00000064145 | Arih2 | 23807 | protein_coding | 928.7227857 | -0.04923792 | 0.084088948 | -0.585545677 | 0.558180863 | 0.736676494 | 0.04923792 |
| ENSMUSG00000000600 | Krit1 | 79264 | protein_coding | 487.4986153 | -0.049243258 | 0.088034996 | -0.559360029 | 0.575916035 | 0.748585685 | 0.049243258 |
| ENSMUSG00000041264 | Uspl1 | 231915 | protein_coding | 454.422277 | -0.049252061 | 0.134960674 | -0.36493639 | 0.71515889 | 0.839800265 | 0.049252061 |
| ENSMUSG00000050552 | Lamtor4 | 66096 | protein_coding | 458.3232121 | 0.049400887 | 0.097788505 | 0.505180918 | 0.613431771 | 0.774840702 | 0.049400887 |
| ENSMUSG00000015961 | Adss | 11566 | protein_coding | 990.25657 | -0.049405261 | 0.143476303 | -0.344344396 | 0.7305873 | 0.848536435 | 0.049405261 |
| ENSMUSG00000028102 | Pex11b | 18632 | protein_coding | 163.2804113 | -0.049414752 | 0.119640107 | -0.413028318 | 0.679585867 | 0.816754221 | 0.049414752 |
| ENSMUSG00000043313 | Pcdhb19 | 93890 | protein_coding | 7.200033668 | 0.049423186 | 0.529163853 | 0.093398643 | 0.925586868 | 0.963531518 | 0.049423186 |
| ENSMUSG00000034042 | Gpbp1l1 | 77110 | protein_coding | 2481.86009 | 0.049423728 | 0.068339038 | 0.723213695 | 0.469548606 | 0.665743569 | 0.049423728 |
| ENSMUSG00000024095 | Hnrnpll | 72692 | protein_coding | 1024.251806 | 0.049437489 | 0.082847773 | 0.596726833 | 0.550689775 | 0.731597532 | 0.049437489 |
| ENSMUSG00000044026 | Slc35g1 | 240660 | protein_coding | 2977.159848 | 0.049461137 | 0.09704552 | 0.509669456 | 0.610283055 | 0.773061998 | 0.049461137 |
| ENSMUSG00000096914 | Galntl6 | 270049 | protein_coding | 168.8705964 | 0.049478482 | 0.187813306 | 0.263445029 | 0.792207588 | 0.887388848 | 0.049478482 |
| ENSMUSG00000060032 | H2aj | 232440 | protein_coding | 722.9684862 | -0.049505042 | 0.127617026 | -0.387918786 | 0.698076136 | 0.828227686 | 0.049505042 |
| ENSMUSG00000036561 | Ppp6r2 | 71474 | protein_coding | 517.5802204 | 0.049565393 | 0.084289416 | 0.588038168 | 0.556506674 | 0.735604839 | 0.049565393 |
| ENSMUSG00000048216 | Gpr85 | 64450 | protein_coding | 6.796072282 | -0.049598211 | 0.718231987 | -0.069055976 | 0.944945063 | 0.973212655 | 0.049598211 |

| ensembl_gene_id | external_gene_name | entrezgene_id | gene_biotype | baseMean | log2FoldChange | lfcSE | stat | pvalue | padj | absLog2FC |
|---|---|---|---|---|---|---|---|---|---|---|
| ENSMUSG00000028879 | Stx12 | 100226 | protein_coding | 1167.803919 | -0.04959839 | 0.087632253 | -0.565983285 | 0.571405143 | 0.745461637 | 0.04959839 |
| ENSMUSG00000058704 | Memo1 | 76890 | protein_coding | 521.6522622 | -0.049621048 | 0.099964172 | -0.49638833 | 0.619620457 | 0.778521309 | 0.049621048 |
| ENSMUSG00000040006 | Ginm1 | 215751 | protein_coding | 1051.670236 | 0.049674559 | 0.066207341 | 0.750287788 | 0.453081396 | 0.652992964 | 0.049674559 |
| ENSMUSG00000038264 | Sema7a | 20361 | protein_coding | 85.45717469 | -0.049715528 | 0.577354562 | -0.08610918 | 0.931379626 | 0.965868908 | 0.049715528 |
| ENSMUSG00000031756 | Cenpn | 72155 | protein_coding | 53.52604576 | -0.049752713 | 0.162793573 | -0.30561841 | 0.759895202 | 0.868638637 | 0.049752713 |
| ENSMUSG00000029062 | Cdk11b | 12537 | protein_coding | 593.058324 | -0.049772283 | 0.092863825 | -0.535970634 | 0.591978846 | 0.761020102 | 0.049772283 |
| ENSMUSG00000020409 | Slu7 | 193116 | protein_coding | 889.6692174 | 0.049777286 | 0.082523263 | 0.603190963 | 0.546381659 | 0.728630866 | 0.049777286 |
| ENSMUSG00000002731 | Prkra | 23992 | protein_coding | 510.9924104 | 0.049777806 | 0.117323364 | 0.42427871 | 0.671362564 | 0.810172204 | 0.049777806 |
| ENSMUSG00000039130 | Zc3hc1 | 232679 | protein_coding | 142.7471924 | -0.049791004 | 0.122141779 | -0.40764924 | 0.683531209 | 0.819173276 | 0.049791004 |
| ENSMUSG00000026235 | Epha4 | 13838 | protein_coding | 362.4206448 | 0.049810955 | 0.082516554 | 0.603648028 | 0.546077674 | 0.728347446 | 0.049810955 |
| ENSMUSG00000024350 | Dnajc18 | 76594 | protein_coding | 121.8213273 | -0.049848596 | 0.143201281 | -0.348101604 | 0.727763879 | 0.846662573 | 0.049848596 |
| ENSMUSG00000000740 | Rpl13 | 270106 | protein_coding | 5169.265387 | 0.049858982 | 0.116452861 | 0.428147337 | 0.668543855 | 0.807933727 | 0.049858982 |
| ENSMUSG00000009112 | Bcl2l13 | 94044 | protein_coding | 722.1469069 | 0.049864403 | 0.084144701 | 0.592603013 | 0.553446862 | 0.733153326 | 0.049864403 |
| ENSMUSG00000019877 | Serinc1 | 56442 | protein_coding | 1670.555007 | 0.049911628 | 0.066249236 | 0.753391741 | 0.451214544 | 0.651522551 | 0.049911628 |
| ENSMUSG00000039910 | Cited2 | 17684 | protein_coding | 417.6004155 | 0.049936068 | 0.174623154 | 0.285964757 | 0.774905115 | 0.877593098 | 0.049936068 |
| ENSMUSG00000032712 | Resf1 | 67246 | protein_coding | 938.6930802 | -0.050021291 | 0.098903023 | -0.50576099 | 0.613024447 | 0.774633351 | 0.050021291 |
| ENSMUSG00000041817 | Fam169a | 320557 | protein_coding | 10.55975608 | -0.050076505 | 0.475702335 | -0.105268571 | 0.916162701 | 0.958528606 | 0.050076505 |
| ENSMUSG00000042659 | Arrdc4 | 66412 | protein_coding | 1183.314579 | 0.05009468 | 0.076169192 | 0.657676403 | 0.510746088 | 0.699008761 | 0.05009468 |
| ENSMUSG00000038515 | Grtp1 | 66790 | protein_coding | 999.6664368 | -0.050098078 | 0.099820582 | -0.501881245 | 0.615751059 | 0.77604707 | 0.050098078 |
| ENSMUSG00000028890 | Mtf1 | 17764 | protein_coding | 360.5960965 | -0.050098219 | 0.100169191 | -0.500136001 | 0.616979319 | 0.777041716 | 0.050098219 |
| ENSMUSG00000002996 | Hbp1 | 73389 | protein_coding | 751.6352144 | 0.050166909 | 0.184507175 | 0.27189679 | 0.785701379 | 0.883803077 | 0.050166909 |
| ENSMUSG00000027170 | Eif3m | 98221 | protein_coding | 1528.370381 | 0.050198366 | 0.098443214 | 0.509922056 | 0.610106069 | 0.772969255 | 0.050198366 |
| ENSMUSG00000034789 | Rab24 | 19336 | protein_coding | 187.2185845 | 0.050209416 | 0.118102716 | 0.425133456 | 0.67073939 | 0.809542851 | 0.050209416 |
| ENSMUSG00000056962 | Jmjd6 | 107817 | protein_coding | 529.4611451 | -0.050216515 | 0.16424513 | -0.305741274 | 0.759801645 | 0.868594029 | 0.050216515 |
| ENSMUSG00000021796 | Bmpr1a | 12166 | protein_coding | 1990.745903 | 0.050216615 | 0.054235594 | 0.925897762 | 0.354499115 | 0.56404539 | 0.050216615 |
| ENSMUSG00000015605 | Srf | 20807 | protein_coding | 449.1361699 | -0.05022304 | 0.220153072 | -0.22812782 | 0.819546873 | 0.904339164 | 0.05022304 |
| ENSMUSG00000023723 | Mrps23 | 64656 | protein_coding | 718.0656648 | 0.050226267 | 0.096282395 | 0.521655775 | 0.601910022 | 0.767564496 | 0.050226267 |
| ENSMUSG00000032044 | Rpusd4 | 71989 | protein_coding | 206.0622071 | -0.050228081 | 0.114030779 | -0.440478274 | 0.659590744 | 0.802969027 | 0.050228081 |
| ENSMUSG00000031886 | Ces2e | 234673 | protein_coding | 3581.668714 | -0.050237968 | 0.204287637 | -0.245917807 | 0.805745859 | 0.895535555 | 0.050237968 |
| ENSMUSG00000005442 | Cic | 71722 | protein_coding | 557.1957181 | 0.050308184 | 0.162277242 | 0.310013798 | 0.756550463 | 0.866369673 | 0.050308184 |
| ENSMUSG00000021947 | Cryl1 | 68631 | protein_coding | 1482.791079 | 0.050326444 | 0.106104039 | 0.47431224 | 0.635277264 | 0.788920139 | 0.050326444 |
| ENSMUSG00000034226 | Rhov | 228543 | protein_coding | 20.92711945 | 0.050381253 | 0.290768979 | 0.17326901 | 0.862439981 | 0.928297339 | 0.050381253 |
| ENSMUSG00000040174 | Alkbh3 | 69113 | protein_coding | 178.194855 | 0.050385745 | 0.086941863 | 0.579533762 | 0.562229073 | 0.739448679 | 0.050385745 |
| ENSMUSG00000063663 | Brwd3 | 382236 | protein_coding | 558.7638029 | -0.050387092 | 0.128492701 | -0.392139722 | 0.694954979 | 0.826592861 | 0.050387092 |
| ENSMUSG00000028690 | Mmachc | 67096 | protein_coding | 211.8250047 | -0.05039599 | 0.091844871 | -0.548707717 | 0.58320605 | 0.754056602 | 0.05039599 |
| ENSMUSG00000026563 | Tada1 | 27878 | protein_coding | 129.2775668 | 0.050399448 | 0.126252388 | 0.399195997 | 0.689748792 | 0.823191875 | 0.050399448 |
| ENSMUSG00000060475 | Wtap | 60532 | protein_coding | 1022.824801 | 0.050399608 | 0.063783009 | 0.790172944 | 0.429426775 | 0.633442274 | 0.050399608 |
| ENSMUSG00000021890 | Eaf1 | 74427 | protein_coding | 608.1274542 | 0.050406157 | 0.129359169 | 0.389660486 | 0.696787619 | 0.827409428 | 0.050406157 |

| ensembl_gene_id | external_gene_name | entrezgene_id | gene_biotype | baseMean | log2FoldChange | lfcSE | stat | pvalue | padj | absLog2FC |
|---|---|---|---|---|---|---|---|---|---|---|
| ENSMUSG00000007603 | Dus3l | 224907 | protein_coding | 371.3779457 | 0.050483241 | 0.097329389 | 0.518684457 | 0.603980802 | 0.768632647 | 0.050483241 |
| ENSMUSG00000033808 | Tmem87a | 211499 | protein_coding | 1375.073907 | -0.050532246 | 0.055663613 | -0.907814686 | 0.363976138 | 0.572855539 | 0.050532246 |
| ENSMUSG00000033319 | Fem1c | 240263 | protein_coding | 583.5293141 | -0.050578263 | 0.109560248 | -0.461647945 | 0.644333806 | 0.794161484 | 0.050578263 |
| ENSMUSG00000014550 | Rbsn | 78287 | protein_coding | 486.2070512 | 0.050642568 | 0.092300487 | 0.548670653 | 0.58323149 | 0.754056602 | 0.050642568 |
| ENSMUSG00000030042 | Pole4 | 66979 | protein_coding | 344.8331589 | 0.050644865 | 0.126776209 | 0.399482403 | 0.689537787 | 0.823125075 | 0.050644865 |
| ENSMUSG00000024581 | Napg | 108123 | protein_coding | 530.3646504 | -0.050661707 | 0.062708817 | -0.807888093 | 0.419155006 | 0.625238897 | 0.050661707 |
| ENSMUSG00000028161 | Ppp3ca | 19055 | protein_coding | 1204.361054 | -0.050700096 | 0.058900605 | -0.860773777 | 0.38936265 | 0.59764559 | 0.050700096 |
| ENSMUSG00000030166 | Rad52 | 19365 | protein_coding | 138.41921 | -0.050728107 | 0.143135101 | -0.354407175 | 0.723033758 | 0.844115801 | 0.050728107 |
| ENSMUSG00000097527 | 1700112J16Rik | 76628 | lncRNA | 10.92325082 | 0.050776871 | 0.329446275 | 0.154127926 | 0.877508869 | 0.937882888 | 0.050776871 |
| ENSMUSG00000096199 | Ptrhd1 | 69709 | protein_coding | 82.76621622 | 0.050806658 | 0.13138616 | 0.386697181 | 0.698980404 | 0.828868364 | 0.050806658 |
| ENSMUSG00000059316 | Slc27a4 | 26569 | protein_coding | 1534.625829 | 0.050815705 | 0.175623898 | 0.289343909 | 0.772318214 | 0.876172816 | 0.050815705 |
| ENSMUSG00000023827 | Agpat4 | 68262 | protein_coding | 852.3832119 | 0.05081594 | 0.100743952 | 0.504406852 | 0.613975504 | 0.775245093 | 0.05081594 |
| ENSMUSG00000062078 | Qki | 19317 | protein_coding | 17.70876603 | -0.05084684 | 0.402918977 | -0.126196191 | 0.899576627 | 0.950188262 | 0.05084684 |
| ENSMUSG00000017664 | Slc35c2 | 228875 | protein_coding | 260.2029542 | -0.050849644 | 0.094135369 | -0.540175762 | 0.589075826 | 0.758819229 | 0.050849644 |
| ENSMUSG00000030281 | Il17rc | 171095 | protein_coding | 1149.754418 | 0.050902512 | 0.11504005 | 0.442476439 | 0.658144476 | 0.802182829 | 0.050902512 |
| ENSMUSG00000005204 | Senp3 | 80886 | protein_coding | 599.6548375 | -0.050919105 | 0.076959284 | -0.661636941 | 0.508203927 | 0.697515911 | 0.050919105 |
| ENSMUSG00000032020 | Ubash3b | 72828 | protein_coding | 34.47147124 | -0.050927682 | 0.194127359 | -0.262341598 | 0.793058094 | 0.888064491 | 0.050927682 |
| ENSMUSG00000058355 | Abce1 | 24015 | protein_coding | 1951.154349 | 0.050931797 | 0.157097515 | 0.32420498 | 0.745782852 | 0.858669167 | 0.050931797 |
| ENSMUSG00000007610 | Gtpbp3 | 70359 | protein_coding | 227.8303556 | 0.050943946 | 0.145275851 | 0.350670432 | 0.72583561 | 0.845790446 | 0.050943946 |
| ENSMUSG00000021814 | Anxa7 | 11750 | protein_coding | 2386.279677 | -0.050959711 | 0.137865556 | -0.369633378 | 0.711655678 | 0.837372444 | 0.050959711 |
| ENSMUSG00000036371 | Serbp1 | 66870 | protein_coding | 4501.976418 | 0.050975736 | 0.08384755 | 0.607957371 | 0.543215747 | 0.725990523 | 0.050975736 |
| ENSMUSG00000107002 | 0610012G03Rik | 106264 | protein_coding | 501.6385358 | 0.050987981 | 0.119491165 | 0.426709211 | 0.669591141 | 0.8087085 | 0.050987981 |
| ENSMUSG00000016946 | Kctd5 | 69259 | protein_coding | 929.0862038 | 0.051000374 | 0.109592176 | 0.465365098 | 0.641670024 | 0.79259559 | 0.051000374 |
| ENSMUSG00000001441 | Npepps | 19155 | protein_coding | 882.685385 | -0.051054808 | 0.097940627 | -0.521283246 | 0.602169471 | 0.767564496 | 0.051054808 |
| ENSMUSG00000030494 | Rhpn2 | 52428 | protein_coding | 1663.178132 | -0.051110121 | 0.090568418 | -0.564326083 | 0.572532233 | 0.746198626 | 0.051110121 |
| ENSMUSG00000024012 | Mtch1 | 56462 | protein_coding | 1817.169028 | 0.051171523 | 0.079529705 | 0.643426536 | 0.519947369 | 0.706571256 | 0.051171523 |
| ENSMUSG00000082361 | Btc | 12223 | protein_coding | 141.5329014 | -0.051207987 | 0.341668299 | -0.14987632 | 0.880862195 | 0.93955283 | 0.051207987 |
| ENSMUSG00000036006 | Ripor2 | 193385 | protein_coding | 44.26943059 | 0.051320202 | 0.253861737 | 0.202158083 | 0.839793142 | 0.915678064 | 0.051320202 |
| ENSMUSG00000020171 | Yeats4 | 64050 | protein_coding | 361.8901635 | 0.051333536 | 0.138611022 | 0.370342379 | 0.711127401 | 0.837163904 | 0.051333536 |
| ENSMUSG00000024873 | Cnih2 | 12794 | protein_coding | 12.41823217 | 0.051342001 | 0.314061565 | 0.163477506 | 0.870142482 | 0.933211685 | 0.051342001 |
| ENSMUSG00000045268 | Zfp691 | 195522 | protein_coding | 76.04476745 | -0.051369887 | 0.165099525 | -0.31114497 | 0.755690415 | 0.865797613 | 0.051369887 |
| ENSMUSG00000097471 | 5830432E09Rik | 67765 | lncRNA | 10.99772479 | 0.051427412 | 0.443626381 | 0.115925054 | 0.90771194 | 0.954325794 | 0.051427412 |
| ENSMUSG00000029922 | Mkrn1 | 54484 | protein_coding | 2634.174311 | -0.051437191 | 0.093078659 | -0.552620675 | 0.580523181 | 0.751673912 | 0.051437191 |
| ENSMUSG00000019846 | Lama4 | 16775 | protein_coding | 17.01629038 | -0.051458591 | 0.391843359 | -0.131324394 | 0.895518696 | 0.94788821 | 0.051458591 |
| ENSMUSG00000024889 | Rce1 | 19671 | protein_coding | 339.9072928 | -0.051463676 | 0.10797675 | -0.476618124 | 0.633634075 | 0.788005719 | 0.051463676 |
| ENSMUSG00000039630 | Hnrnpu | 51810 | protein_coding | 6255.8274 | -0.051496298 | 0.10942762 | -0.470596897 | 0.637928624 | 0.790438741 | 0.051496298 |
| ENSMUSG00000021770 | Samd8 | 67630 | protein_coding | 506.8834951 | -0.051507522 | 0.121823407 | -0.422804805 | 0.672437681 | 0.810978071 | 0.051507522 |
| ENSMUSG00000020441 | 2310033P09Rik | 67862 | protein_coding | 94.53406392 | -0.051521821 | 0.144776821 | -0.355870651 | 0.721937436 | 0.843984213 | 0.051521821 |

| ensembl_gene_id | external_gene_name | entrezgene_id | gene_biotype | baseMean | log2FoldChange | lfcSE | stat | pvalue | padj | absLog2FC |
|---|---|---|---|---|---|---|---|---|---|---|
| ENSMUSG00000041977 | Arhgef11 | 213498 | protein_coding | 1165.747209 | -0.05153278 | 0.071699961 | -0.718728152 | 0.472308434 | 0.66775508 | 0.05153278 |
| ENSMUSG00000037119 | Fam91a1 | 210998 | protein_coding | 2193.897179 | 0.051607719 | 0.069522209 | 0.742319896 | 0.45789354 | 0.656524287 | 0.051607719 |
| ENSMUSG00000028780 | Sema3c | 20348 | protein_coding | 2251.365324 | -0.051625653 | 0.098006552 | -0.526757155 | 0.59836224 | 0.765321556 | 0.051625653 |
| ENSMUSG00000067722 | BC003965 | 214489 | protein_coding | 340.6772901 | -0.051646082 | 0.108902069 | -0.474243351 | 0.635326383 | 0.788920139 | 0.051646082 |
| ENSMUSG00000021482 | Prxl2c | 66129 | protein_coding | 786.9864242 | -0.05166033 | 0.10821636 | -0.477380041 | 0.633091524 | 0.787741735 | 0.05166033 |
| ENSMUSG00000037475 | Thoc2 | 331401 | protein_coding | 1293.686237 | 0.051677003 | 0.069743588 | 0.740957042 | 0.458719488 | 0.65713215 | 0.051677003 |
| ENSMUSG00000029869 | Ephb6 | 13848 | protein_coding | 30.4939863 | -0.051678533 | 0.506090957 | -0.102113133 | 0.918666877 | 0.959729751 | 0.051678533 |
| ENSMUSG00000029513 | Prkab1 | 19079 | protein_coding | 1323.986387 | -0.051706586 | 0.159878029 | -0.323412707 | 0.746382707 | 0.858987266 | 0.051706586 |
| ENSMUSG00000012848 | Rps5 | 20103 | protein_coding | 5529.65768 | -0.051733923 | 0.112721832 | -0.458952117 | 0.646268552 | 0.795376718 | 0.051733923 |
| ENSMUSG00000037815 | Ctnna1 | 12385 | protein_coding | 7219.595286 | 0.05175767 | 0.126524306 | 0.409072943 | 0.682486132 | 0.81859488 | 0.05175767 |
| ENSMUSG00000015533 | Itga2 | 16398 | protein_coding | 1405.236055 | -0.051766251 | 0.146422262 | -0.353540852 | 0.723683008 | 0.844689486 | 0.051766251 |
| ENSMUSG00000042331 | Specc1 | 432572 | protein_coding | 697.770948 | -0.051920417 | 0.088817488 | -0.584574262 | 0.558834018 | 0.736907711 | 0.051920417 |
| ENSMUSG00000047139 | Cd24a | 12484 | protein_coding | 7729.368974 | -0.05194127 | 0.102236905 | -0.50804815 | 0.611419576 | 0.773464459 | 0.05194127 |
| ENSMUSG00000041444 | Arhgap32 | 330914 | protein_coding | 1638.787563 | -0.051959881 | 0.148160229 | -0.350700598 | 0.725812977 | 0.845790446 | 0.051959881 |
| ENSMUSG00000040731 | Eif4h | 22384 | protein_coding | 3455.183861 | -0.052033324 | 0.094132476 | -0.552766979 | 0.580422982 | 0.751624168 | 0.052033324 |
| ENSMUSG00000025982 | Sf3b1 | 81898 | protein_coding | 5533.077319 | 0.05203998 | 0.066839706 | 0.778578817 | 0.436227862 | 0.639074944 | 0.05203998 |
| ENSMUSG00000030341 | Tnfrsf1a | 21937 | protein_coding | 412.6895262 | 0.052066779 | 0.206923478 | 0.251623353 | 0.801332206 | 0.893560584 | 0.052066779 |
| ENSMUSG00000000384 | Tbrg4 | 21379 | protein_coding | 900.7555342 | -0.052069369 | 0.079049181 | -0.658695876 | 0.510091082 | 0.698626897 | 0.052069369 |
| ENSMUSG00000030681 | Mvp | 78388 | protein_coding | 5328.372832 | -0.052074047 | 0.131164075 | -0.397014556 | 0.691356727 | 0.824307953 | 0.052074047 |
| ENSMUSG00000029366 | Dck | 13178 | protein_coding | 198.0824685 | 0.052083163 | 0.135488975 | 0.384408866 | 0.700675431 | 0.829926899 | 0.052083163 |
| ENSMUSG00000052056 | Zfp217 | 228913 | protein_coding | 793.1632888 | -0.052117934 | 0.082958939 | -0.628237711 | 0.52984823 | 0.715171253 | 0.052117934 |
| ENSMUSG00000053644 | Aldh7a1 | 110695 | protein_coding | 286.0768288 | -0.0521289 | 0.100033052 | -0.521116764 | 0.602285434 | 0.767564496 | 0.0521289 |
| ENSMUSG00000078671 | Chd2 | 244059 | protein_coding | 1152.152357 | -0.052139872 | 0.099003068 | -0.526649051 | 0.598437323 | 0.765321556 | 0.052139872 |
| ENSMUSG00000055866 | Per2 | 18627 | protein_coding | 270.4676149 | 0.052149506 | 0.225401619 | 0.231362604 | 0.817033114 | 0.902675156 | 0.052149506 |
| ENSMUSG00000062797 | Hikeshi | 67669 | protein_coding | 250.5059388 | 0.052212203 | 0.102351368 | 0.510127069 | 0.609962442 | 0.772848762 | 0.052212203 |
| ENSMUSG00000021276 | Cinp | 67236 | protein_coding | 218.7559936 | 0.052287927 | 0.097721477 | 0.535070989 | 0.59260077 | 0.761247936 | 0.052287927 |
| ENSMUSG00000029815 | Malsu1 | 75593 | protein_coding | 236.0914094 | 0.052302789 | 0.098724474 | 0.529785447 | 0.596260696 | 0.764096331 | 0.052302789 |
| ENSMUSG00000020647 | Ncoa1 | 17977 | protein_coding | 727.7559702 | -0.052336274 | 0.097504634 | -0.536756782 | 0.591435628 | 0.76075157 | 0.052336274 |
| ENSMUSG00000024301 | Kifc5b | 16580 | protein_coding | 70.09130109 | 0.052340267 | 0.182955952 | 0.286081247 | 0.774815894 | 0.877593098 | 0.052340267 |
| ENSMUSG00000028582 | Cc2d1b | 319965 | protein_coding | 516.6598714 | 0.052358321 | 0.092242177 | 0.567618015 | 0.570294372 | 0.744744605 | 0.052358321 |
| ENSMUSG00000026154 | Sdhaf4 | 68002 | protein_coding | 132.3620442 | 0.052358764 | 0.132584274 | 0.394909312 | 0.692909817 | 0.825356534 | 0.052358764 |
| ENSMUSG00000019838 | Slc16a10 | 72472 | protein_coding | 130.2718184 | -0.052401831 | 0.166905573 | -0.313960944 | 0.753550702 | 0.864733257 | 0.052401831 |
| ENSMUSG00000020361 | Hspa4 | 15525 | protein_coding | 2861.652745 | -0.052402975 | 0.129549583 | -0.404501305 | 0.685844118 | 0.820745689 | 0.052402975 |
| ENSMUSG00000025239 | Limd1 | 29806 | protein_coding | 1295.338249 | -0.052420943 | 0.093739689 | -0.559218234 | 0.57601279 | 0.748585685 | 0.052420943 |
| ENSMUSG00000026003 | Acadl | 11363 | protein_coding | 1476.238061 | -0.052444171 | 0.090460309 | -0.579747872 | 0.562084655 | 0.739443779 | 0.052444171 |
| ENSMUSG00000036167 | Pphln1 | 223828 | protein_coding | 288.7545111 | 0.052497717 | 0.104937376 | 0.500276632 | 0.616880306 | 0.776978453 | 0.052497717 |
| ENSMUSG00000000275 | Trim25 | 217069 | protein_coding | 1193.420626 | 0.052511179 | 0.112597553 | 0.466361635 | 0.640956668 | 0.792083001 | 0.052511179 |
| ENSMUSG00000059323 | Tonsl | 72749 | protein_coding | 258.7475189 | 0.052543936 | 0.111955241 | 0.469329849 | 0.638833883 | 0.790625182 | 0.052543936 |

Supplementary Table 1

| ensembl_gene_id | external_gene_name | entrezgene_id | gene_biotype | baseMean | log2FoldChange | lfcSE | stat | pvalue | padj | absLog2FC |
|---|---|---|---|---|---|---|---|---|---|---|
| ENSMUSG00000027360 | Hdc | 15186 | protein_coding | 7.080431684 | 0.052586336 | 0.588744685 | 0.089319423 | 0.928828058 | 0.964808485 | 0.052586336 |
| ENSMUSG00000032468 | Armc8 | 74125 | protein_coding | 655.8322077 | -0.052614782 | 0.062917874 | -0.836245376 | 0.403016876 | 0.610253142 | 0.052614782 |
| ENSMUSG00000026980 | Ly75 | 17076 | protein_coding | 540.5890205 | -0.05270679 | 0.088545294 | -0.595252298 | 0.551674837 | 0.732245284 | 0.05270679 |
| ENSMUSG00000038374 | Rbm8a | 60365 | protein_coding | 613.2217359 | 0.052713109 | 0.09973863 | 0.528512466 | 0.597143697 | 0.764635807 | 0.052713109 |
| ENSMUSG00000043091 | Tuba1c | 22146 | protein_coding | 2087.178106 | 0.052714304 | 0.212937319 | 0.247557845 | 0.804476531 | 0.894700251 | 0.052714304 |
| ENSMUSG00000026926 | Pmpca | 66865 | protein_coding | 880.320604 | -0.052722031 | 0.104312702 | -0.505422929 | 0.613261817 | 0.774748908 | 0.052722031 |
| ENSMUSG00000039063 | Echdc3 | 67856 | protein_coding | 110.8978495 | -0.052736377 | 0.17185919 | -0.306858057 | 0.758951416 | 0.868201089 | 0.052736377 |
| ENSMUSG00000023070 | Rgn | 19733 | protein_coding | 35.53701808 | 0.052750083 | 0.19604338 | 0.269073522 | 0.787873106 | 0.884871162 | 0.052750083 |
| ENSMUSG00000034361 | Cpne2 | 234577 | protein_coding | 322.8118684 | -0.052771503 | 0.092056483 | -0.573251352 | 0.566474519 | 0.742495568 | 0.052771503 |
| ENSMUSG00000032290 | Ptpn9 | 56294 | protein_coding | 738.2859463 | 0.052787401 | 0.071651478 | 0.736724524 | 0.46128989 | 0.659378689 | 0.052787401 |
| ENSMUSG00000071451 | Psmg4 | 69666 | protein_coding | 91.58578846 | -0.052874148 | 0.15615491 | -0.338600612 | 0.734910621 | 0.851382382 | 0.052874148 |
| ENSMUSG00000063108 | Zfp26 | 22688 | protein_coding | 556.5539064 | -0.052877237 | 0.064969723 | -0.813875053 | 0.415716533 | 0.622176792 | 0.052877237 |
| ENSMUSG00000028998 | Tomm7 | 66169 | protein_coding | 1488.0073 | 0.052911457 | 0.087093192 | 0.607526894 | 0.5435013 | 0.726249053 | 0.052911457 |
| ENSMUSG00000041488 | Stx3 | 20908 | protein_coding | 953.7728657 | 0.05292326 | 0.119043937 | 0.444569138 | 0.656631154 | 0.801149235 | 0.05292326 |
| ENSMUSG00000001383 | Zmat2 | 66492 | protein_coding | 682.0869407 | 0.052929653 | 0.085179753 | 0.621387726 | 0.534344546 | 0.719249909 | 0.052929653 |
| ENSMUSG00000019577 | Pdk4 | 27273 | protein_coding | 259.4537452 | 0.052932647 | 0.233512449 | 0.226680194 | 0.820672429 | 0.904852281 | 0.052932647 |
| ENSMUSG00000044279 | Crb3 | 224912 | protein_coding | 841.4076673 | -0.05294288 | 0.125362273 | -0.42231908 | 0.672792133 | 0.811159874 | 0.05294288 |
| ENSMUSG00000061666 | Gdpd1 | 66569 | protein_coding | 1664.291346 | 0.052993495 | 0.104054941 | 0.509283789 | 0.610553319 | 0.773102882 | 0.052993495 |
| ENSMUSG00000044636 | Csrnp2 | 207785 | protein_coding | 89.76644143 | 0.053016837 | 0.129313713 | 0.409986192 | 0.681816077 | 0.81801495 | 0.053016837 |
| ENSMUSG00000030336 | Cd27 | 21940 | protein_coding | 7.494270642 | 0.05302193 | 0.444414748 | 0.119307314 | 0.905031889 | 0.953132467 | 0.05302193 |
| ENSMUSG00000007659 | Bcl2l1 | 12048 | protein_coding | 477.3049379 | -0.053025249 | 0.098737871 | -0.537030507 | 0.591246541 | 0.760569771 | 0.053025249 |
| ENSMUSG00000069844 | Sco1 | 52892 | protein_coding | 936.8599038 | -0.053027838 | 0.10299008 | -0.514882965 | 0.606634805 | 0.771091876 | 0.053027838 |
| ENSMUSG00000042323 | Pbrm1 | 66923 | protein_coding | 1774.766519 | 0.053085626 | 0.075214569 | 0.705789145 | 0.480319274 | 0.673965179 | 0.053085626 |
| ENSMUSG00000042133 | Ppig | 228005 | protein_coding | 744.8357121 | -0.053094946 | 0.093098688 | -0.570308203 | 0.568468678 | 0.743758628 | 0.053094946 |
| ENSMUSG00000051451 | Crebzf | 233490 | protein_coding | 599.7530722 | -0.053095995 | 0.124473453 | -0.426564813 | 0.669696331 | 0.808774218 | 0.053095995 |
| ENSMUSG00000037643 | Prkci | 18759 | protein_coding | 1914.491894 | -0.053131425 | 0.067287063 | -0.789623189 | 0.429747862 | 0.633798513 | 0.053131425 |
| ENSMUSG00000059878 | Zfp422 | 67255 | protein_coding | 404.3890565 | -0.053139468 | 0.066181944 | -0.802929992 | 0.4220152 | 0.627572772 | 0.053139468 |
| ENSMUSG00000005354 | Txn2 | 56551 | protein_coding | 1609.708863 | 0.053160599 | 0.084707391 | 0.62757923 | 0.530279619 | 0.715484436 | 0.053160599 |
| ENSMUSG00000015501 | Hivep2 | 15273 | protein_coding | 399.7007036 | -0.053203401 | 0.110108868 | -0.483189067 | 0.628961497 | 0.785363651 | 0.053203401 |
| ENSMUSG00000013736 | Trnt1 | 70047 | protein_coding | 564.8569754 | -0.05324374 | 0.096874963 | -0.549613009 | 0.582584834 | 0.75356596 | 0.05324374 |
| ENSMUSG00000020307 | Cdc34 | 216150 | protein_coding | 603.5958768 | 0.053283178 | 0.109139771 | 0.488210466 | 0.625400772 | 0.782942977 | 0.053283178 |
| ENSMUSG00000027366 | Sppl2a | 66552 | protein_coding | 5864.849068 | 0.053346261 | 0.1323224 | 0.40315367 | 0.68683518 | 0.821304393 | 0.053346261 |
| ENSMUSG00000061455 | Stx17 | 67727 | protein_coding | 795.8768573 | -0.053377089 | 0.106176118 | -0.502722172 | 0.615159619 | 0.775608518 | 0.053377089 |
| ENSMUSG00000026196 | Bard1 | 12021 | protein_coding | 80.78049667 | -0.053384673 | 0.164102597 | -0.325312784 | 0.744944357 | 0.858274321 | 0.053384673 |
| ENSMUSG00000031226 | Pbdc1 | 67683 | protein_coding | 247.6175085 | 0.053397059 | 0.122291259 | 0.436638395 | 0.662373613 | 0.804505636 | 0.053397059 |
| ENSMUSG00000046792 | Zfp787 | 67109 | protein_coding | 314.2968183 | -0.053440906 | 0.088988128 | -0.600539721 | 0.548146597 | 0.730067647 | 0.053440906 |
| ENSMUSG00000040521 | Tsfm | 66399 | protein_coding | 266.3367743 | 0.053448182 | 0.118128819 | 0.452456755 | 0.650939968 | 0.798250711 | 0.053448182 |
| ENSMUSG00000031453 | Rasa3 | 19414 | protein_coding | 485.4163252 | 0.053455941 | 0.086063489 | 0.621122174 | 0.534519241 | 0.719249909 | 0.053455941 |

| ensembl_gene_id | external_gene_name | entrezgene_id | gene_biotype | baseMean | log2FoldChange | lfcSE | stat | pvalue | padj | absLog2FC |
|---|---|---|---|---|---|---|---|---|---|---|
| ENSMUSG00000030733 | Sh2b1 | 20399 | protein_coding | 471.6143875 | -0.053507024 | 0.095365967 | -0.561070428 | 0.574749527 | 0.747748252 | 0.053507024 |
| ENSMUSG00000027667 | Zfp639 | 67778 | protein_coding | 244.3982308 | 0.053624032 | 0.09520496 | 0.563248301 | 0.573265817 | 0.746727014 | 0.053624032 |
| ENSMUSG00000118516 | Cyp2d13 | 68444 | protein_coding | 77.18906248 | -0.053679485 | 0.268485134 | -0.199934663 | 0.84153168 | 0.916381801 | 0.053679485 |
| ENSMUSG00000070319 | Eif3g | 53356 | protein_coding | 1023.163564 | 0.0537057 | 0.095620529 | 0.561654492 | 0.574351446 | 0.747540635 | 0.0537057 |
| ENSMUSG00000028229 | Rmdn1 | 66302 | protein_coding | 173.6354723 | -0.053713567 | 0.104248991 | -0.515243036 | 0.606383199 | 0.770948583 | 0.053713567 |
| ENSMUSG00000062761 | Zfp512 | 269639 | protein_coding | 240.512616 | -0.053721219 | 0.117228066 | -0.458262439 | 0.646763906 | 0.795863372 | 0.053721219 |
| ENSMUSG00000039253 | Fn3krp | 238024 | protein_coding | 187.8512803 | 0.053742514 | 0.122028333 | 0.440410131 | 0.659640089 | 0.802969027 | 0.053742514 |
| ENSMUSG00000027708 | Dcun1d1 | 114893 | protein_coding | 738.5778356 | 0.053761489 | 0.067170374 | 0.800375022 | 0.423493548 | 0.628356333 | 0.053761489 |
| ENSMUSG00000030447 | Cyfip1 | 20430 | protein_coding | 1937.729681 | 0.05379961 | 0.085524878 | 0.629052175 | 0.529314899 | 0.714606519 | 0.05379961 |
| ENSMUSG00000041459 | Tardbp | 230908 | protein_coding | 1487.959795 | 0.053847309 | 0.129053208 | 0.417248896 | 0.676496364 | 0.814269937 | 0.053847309 |
| ENSMUSG00000051355 | Commd1 | 17846 | protein_coding | 406.0586772 | -0.053867194 | 0.119711829 | -0.449973862 | 0.652729287 | 0.798741727 | 0.053867194 |
| ENSMUSG00000029363 | Rfc5 | 72151 | protein_coding | 358.7254024 | 0.053883256 | 0.129257607 | 0.416867192 | 0.676775552 | 0.814542661 | 0.053883256 |
| ENSMUSG00000070426 | Rnf121 | 75212 | protein_coding | 523.8349311 | -0.053883989 | 0.079920893 | -0.674216546 | 0.500173652 | 0.690757391 | 0.053883989 |
| ENSMUSG00000031392 | Irak1 | 16179 | protein_coding | 675.4090559 | -0.053898283 | 0.0680827 | -0.791659015 | 0.428559526 | 0.632720402 | 0.053898283 |
| ENSMUSG00000038034 | Igsf8 | 140559 | protein_coding | 333.9527298 | -0.053922931 | 0.127238453 | -0.423794298 | 0.671715837 | 0.810432975 | 0.053922931 |
| ENSMUSG00000050796 | B3galt6 | 117592 | protein_coding | 120.1733663 | -0.053929428 | 0.152232975 | -0.354255893 | 0.72314712 | 0.844139782 | 0.053929428 |
| ENSMUSG00000027613 | Eif6 | 16418 | protein_coding | 2395.907147 | 0.054002756 | 0.148028567 | 0.364813068 | 0.715250949 | 0.839846452 | 0.054002756 |
| ENSMUSG00000026749 | Nek6 | 59126 | protein_coding | 937.8807587 | -0.054004794 | 0.207748044 | -0.25995332 | 0.794899781 | 0.889189108 | 0.054004794 |
| ENSMUSG00000045259 | Klhdc9 | 68874 | protein_coding | 9.09041445 | -0.054016741 | 0.374072147 | -0.144401931 | 0.885183092 | 0.94220607 | 0.054016741 |
| ENSMUSG00000027714 | Exosc9 | 50911 | protein_coding | 405.7605773 | 0.054019872 | 0.114904561 | 0.470128179 | 0.638263442 | 0.790506101 | 0.054019872 |
| ENSMUSG00000030339 | Ltbr | 17000 | protein_coding | 1390.926882 | 0.054117261 | 0.09468482 | 0.571551611 | 0.567625786 | 0.743180341 | 0.054117261 |
| ENSMUSG00000009145 | Dqx1 | 93838 | protein_coding | 548.9783551 | -0.054134907 | 0.094656356 | -0.571909896 | 0.567383019 | 0.743135986 | 0.054134907 |
| ENSMUSG00000032594 | Ip6k1 | 27399 | protein_coding | 954.7630533 | -0.054282031 | 0.079548376 | -0.68237761 | 0.49500021 | 0.686611804 | 0.054282031 |
| ENSMUSG00000024754 | Cemip2 | 83921 | protein_coding | 794.5992495 | 0.054284974 | 0.147758207 | 0.367390587 | 0.713327692 | 0.838515386 | 0.054284974 |
| ENSMUSG00000054065 | Pkp3 | 56460 | protein_coding | 999.0260952 | -0.054300807 | 0.156330889 | -0.347345348 | 0.728331886 | 0.84702616 | 0.054300807 |
| ENSMUSG00000001507 | Itga3 | 16400 | protein_coding | 1591.033882 | -0.054322444 | 0.153762421 | -0.353288171 | 0.723872413 | 0.844771344 | 0.054322444 |
| ENSMUSG00000009927 | Rps25 | 75617 | protein_coding | 5546.661635 | 0.054399119 | 0.097305061 | 0.559057446 | 0.576122515 | 0.748585685 | 0.054399119 |
| ENSMUSG00000025794 | Rpl14 | 67115 | protein_coding | 6047.459117 | 0.054417658 | 0.09795708 | 0.555525518 | 0.578535261 | 0.750033342 | 0.054417658 |
| ENSMUSG00000102976 | Zc3h11a | 70579 | protein_coding | 176.0585213 | -0.054429259 | 0.220822592 | -0.246484103 | 0.805307509 | 0.895235762 | 0.054429259 |
| ENSMUSG00000017404 | Rpl19 | 19921 | protein_coding | 6235.062032 | 0.054458669 | 0.097840342 | 0.55660751 | 0.577795622 | 0.74942996 | 0.054458669 |
| ENSMUSG00000036737 | Oxsr1 | 108737 | protein_coding | 1134.873515 | -0.054475499 | 0.097289243 | -0.559933428 | 0.575524847 | 0.74838086 | 0.054475499 |
| ENSMUSG00000051671 | Coa6 | 67892 | protein_coding | 206.9394581 | 0.054484896 | 0.124732515 | 0.436813896 | 0.662246321 | 0.804505636 | 0.054484896 |
| ENSMUSG00000038886 | Man2a2 | 140481 | protein_coding | 108.1806123 | -0.054509352 | 0.174856557 | -0.311737537 | 0.755239998 | 0.865722025 | 0.054509352 |
| ENSMUSG00000097383 | 1500026H17Rik | 69002 | lncRNA | 18.34927144 | 0.054512407 | 0.339667886 | 0.160487374 | 0.872497167 | 0.934575311 | 0.054512407 |
| ENSMUSG00000040725 | Hnrnpul1 | 232989 | protein_coding | 2215.851202 | -0.054513499 | 0.050776174 | -1.07360391 | 0.283000244 | 0.492152843 | 0.054513499 |
| ENSMUSG00000033102 | Cdc14b | 218294 | protein_coding | 272.8128366 | 0.054514895 | 0.157892112 | 0.345266744 | 0.729893846 | 0.848099589 | 0.054514895 |
| ENSMUSG00000020876 | Snx11 | 74479 | protein_coding | 234.6649566 | 0.054540398 | 0.118291475 | 0.461067866 | 0.644749915 | 0.794428456 | 0.054540398 |
| ENSMUSG00000046404 | Yod1 | 226418 | protein_coding | 231.8336027 | 0.054552051 | 0.098458121 | 0.5540635 | 0.579535388 | 0.750963201 | 0.054552051 |

| ensembl_gene_id | external_gene_name | entrezgene_id | gene_biotype | baseMean | log2FoldChange | lfcSE | stat | pvalue | padj | absLog2FC |
|---|---|---|---|---|---|---|---|---|---|---|
| ENSMUSG00000096145 | Vkorc1 | 27973 | protein_coding | 126.0756162 | -0.054578602 | 0.140439938 | -0.38862593 | 0.697552882 | 0.827791857 | 0.054578602 |
| ENSMUSG00000028573 | Fggy | 75578 | protein_coding | 137.8001516 | -0.054579472 | 0.126303476 | -0.432129613 | 0.665647213 | 0.806207042 | 0.054579472 |
| ENSMUSG00000037443 | Cep85 | 70012 | protein_coding | 806.7752316 | -0.054593954 | 0.080639396 | -0.677013433 | 0.498397431 | 0.68900582 | 0.054593954 |
| ENSMUSG00000071454 | Dtnb | 13528 | protein_coding | 82.46612581 | 0.054687926 | 0.152376225 | 0.358900651 | 0.719669413 | 0.842364374 | 0.054687926 |
| ENSMUSG00000039004 | Bmp6 | 12161 | protein_coding | 8.333559326 | 0.054690173 | 0.459508489 | 0.119018853 | 0.905260419 | 0.953132467 | 0.054690173 |
| ENSMUSG00000025981 | Coq10b | 67876 | protein_coding | 621.2382008 | 0.054722352 | 0.228220819 | 0.239778089 | 0.810502294 | 0.899253312 | 0.054722352 |
| ENSMUSG00000024145 | Pigf | 18701 | protein_coding | 289.7335543 | -0.054726051 | 0.131474824 | -0.41624738 | 0.677228993 | 0.814843888 | 0.054726051 |
| ENSMUSG00000007850 | Hnrnph1 | 59013 | protein_coding | 2838.643837 | 0.054767803 | 0.11565865 | 0.473529681 | 0.63583533 | 0.78927929 | 0.054767803 |
| ENSMUSG00000068877 | Selenbp2 | 20342 | protein_coding | 90.54368109 | -0.054771025 | 0.160196955 | -0.341898042 | 0.732427623 | 0.849803845 | 0.054771025 |
| ENSMUSG00000037466 | Tedc1 | 104732 | protein_coding | 201.4411908 | 0.054781886 | 0.115119695 | 0.475868933 | 0.634167756 | 0.788253962 | 0.054781886 |
| ENSMUSG00000039831 | Arhgap29 | 214137 | protein_coding | 219.7115483 | 0.054784248 | 0.100904057 | 0.542934049 | 0.587175212 | 0.757262387 | 0.054784248 |
| ENSMUSG00000026740 | Dnajc1 | 13418 | protein_coding | 508.845071 | 0.054787556 | 0.094680886 | 0.578654872 | 0.562822072 | 0.739770497 | 0.054787556 |
| ENSMUSG00000024925 | Rnaseh2c | 68209 | protein_coding | 226.8371315 | -0.054827998 | 0.147089656 | -0.372752236 | 0.709332855 | 0.835827385 | 0.054827998 |
| ENSMUSG00000000326 | Comt | 12846 | protein_coding | 778.2545167 | 0.054834776 | 0.084379389 | 0.649859839 | 0.515782762 | 0.703131447 | 0.054834776 |
| ENSMUSG00000021759 | Plpp1 | 19012 | protein_coding | 315.2137779 | -0.054867503 | 0.129503601 | -0.423675506 | 0.671802481 | 0.810457465 | 0.054867503 |
| ENSMUSG00000022246 | Rai14 | 75646 | protein_coding | 515.1694084 | 0.054968867 | 0.106894223 | 0.514236092 | 0.607086937 | 0.771131144 | 0.054968867 |
| ENSMUSG00000054517 | Trim65 | 338364 | protein_coding | 179.238251 | -0.054983544 | 0.168030846 | -0.327222921 | 0.743499286 | 0.857217273 | 0.054983544 |
| ENSMUSG00000028104 | Polr3gl | 69870 | protein_coding | 151.0442828 | -0.055011598 | 0.104117807 | -0.528359171 | 0.59725007 | 0.764693012 | 0.055011598 |
| ENSMUSG00000035152 | Ap2b1 | 71770 | protein_coding | 2289.81245 | 0.055014977 | 0.068656526 | 0.801307325 | 0.422953752 | 0.627964891 | 0.055014977 |
| ENSMUSG00000015461 | Atf6b | 12915 | protein_coding | 385.6819598 | 0.055093109 | 0.073613335 | 0.748412074 | 0.454211642 | 0.653598611 | 0.055093109 |
| ENSMUSG00000063535 | Zfp773 | 76373 | protein_coding | 22.56008503 | -0.055100222 | 0.354045985 | -0.155630128 | 0.876324575 | 0.937099553 | 0.055100222 |
| ENSMUSG00000029530 | Ccr9 | 12769 | protein_coding | 14.50265818 | -0.055140615 | 0.473831648 | -0.116371743 | 0.907357929 | 0.954095762 | 0.055140615 |
| ENSMUSG00000058254 | Tspan7 | 21912 | protein_coding | 764.0708 | 0.055163225 | 0.069112252 | 0.798168531 | 0.424772697 | 0.629384766 | 0.055163225 |
| ENSMUSG00000035673 | Sbno2 | 216161 | protein_coding | 862.5038771 | -0.055189929 | 0.151942632 | -0.363228727 | 0.71643403 | 0.840429184 | 0.055189929 |
| ENSMUSG00000057193 | Slc44a2 | 68682 | protein_coding | 376.929358 | -0.055232554 | 0.127668857 | -0.432623552 | 0.665288277 | 0.806025281 | 0.055232554 |
| ENSMUSG00000024666 | Tmem138 | 72982 | protein_coding | 29.46584383 | -0.055275633 | 0.240267684 | -0.230058544 | 0.818046278 | 0.90328188 | 0.055275633 |
| ENSMUSG00000067148 | Polr1c | 20016 | protein_coding | 624.3978661 | 0.055313016 | 0.092846466 | 0.595747131 | 0.551344167 | 0.731939481 | 0.055313016 |
| ENSMUSG00000019087 | Atp6ap1 | 54411 | protein_coding | 1411.035351 | 0.055317448 | 0.077011568 | 0.718300502 | 0.472572021 | 0.667968635 | 0.055317448 |
| ENSMUSG00000005034 | Prkacb | 18749 | protein_coding | 1324.641862 | 0.055322395 | 0.205119314 | 0.269708368 | 0.787384623 | 0.884634427 | 0.055322395 |
| ENSMUSG00000044317 | Gpr4 | 319197 | protein_coding | 7.828575672 | 0.055334963 | 0.527821939 | 0.104836421 | 0.916505609 | 0.958597942 | 0.055334963 |
| ENSMUSG00000041997 | Tlk1 | 228012 | protein_coding | 1606.448786 | -0.0553441 | 0.092872982 | -0.595911734 | 0.551234194 | 0.731854484 | 0.0553441 |
| ENSMUSG00000019362 | D8Ertd738e | 101966 | protein_coding | 696.511732 | 0.055379409 | 0.100419568 | 0.551480251 | 0.581304501 | 0.752303496 | 0.055379409 |
| ENSMUSG00000050323 | Ndufaf6 | 76947 | protein_coding | 106.2695566 | 0.055401591 | 0.134664166 | 0.411405594 | 0.680775153 | 0.81740269 | 0.055401591 |
| ENSMUSG00000037085 | Trmt12 | 68260 | protein_coding | 95.72472599 | 0.055406662 | 0.137874835 | 0.401862034 | 0.687785565 | 0.821834094 | 0.055406662 |
| ENSMUSG00000046865 | Fbl | 14113 | protein_coding | 364.7422471 | -0.055427901 | 0.162813368 | -0.340438268 | 0.733526504 | 0.850397421 | 0.055427901 |
| ENSMUSG00000031753 | Cog4 | 102339 | protein_coding | 890.0070861 | 0.055433885 | 0.08331786 | 0.665330158 | 0.505839345 | 0.695769351 | 0.055433885 |
| ENSMUSG00000022876 | Samsn1 | 67742 | protein_coding | 7.064529055 | 0.055434733 | 0.69692255 | 0.079542172 | 0.93660139 | 0.968704702 | 0.055434733 |
| ENSMUSG00000027597 | Ahcy | 269378 | protein_coding | 1414.079865 | 0.055439535 | 0.09702609 | 0.571387917 | 0.567736718 | 0.743232173 | 0.055439535 |

| ensembl_gene_id | external_gene_name | entrezgene_id | gene_biotype | baseMean | log2FoldChange | lfcSE | stat | pvalue | padj | absLog2FC |
|---|---|---|---|---|---|---|---|---|---|---|
| ENSMUSG00000019806 | Aig1 | 66253 | protein_coding | 870.5674013 | 0.055455051 | 0.073866727 | 0.750744664 | 0.452806338 | 0.652874025 | 0.055455051 |
| ENSMUSG00000019471 | Cdc37 | 12539 | protein_coding | 1127.259379 | 0.055486601 | 0.071025089 | 0.781225363 | 0.434669959 | 0.637867413 | 0.055486601 |
| ENSMUSG00000028639 | Ybx1 | 22608 | protein_coding | 6164.489172 | 0.055513645 | 0.064997188 | 0.854093031 | 0.393053445 | 0.600878709 | 0.055513645 |
| ENSMUSG00000062202 | Btbd9 | 224671 | protein_coding | 401.7914949 | -0.055543336 | 0.10201394 | -0.544468097 | 0.586119397 | 0.756234083 | 0.055543336 |
| ENSMUSG00000067825 | Pex26 | 74043 | protein_coding | 370.2345215 | -0.055548311 | 0.095785764 | -0.579922415 | 0.561966939 | 0.739386206 | 0.055548311 |
| ENSMUSG00000060336 | Zfp937 | 245174 | protein_coding | 56.91527236 | -0.055588201 | 0.191257016 | -0.2906466 | 0.771321616 | 0.875604783 | 0.055588201 |
| ENSMUSG00000018012 | Rac3 | 170758 | protein_coding | 63.11023242 | 0.055593565 | 0.14921548 | 0.37257237 | 0.70946674 | 0.835827385 | 0.055593565 |
| ENSMUSG00000087593 | Gm16174 | 102638408 | lncRNA | 13.32488505 | -0.055603344 | 0.31015221 | -0.1792776 | 0.857719735 | 0.925781659 | 0.055603344 |
| ENSMUSG00000027303 | Ptpra | 19262 | protein_coding | 640.6000762 | 0.055618539 | 0.060396647 | 0.920887849 | 0.357108983 | 0.566762842 | 0.055618539 |
| ENSMUSG00000066829 | Zfp810 | 235050 | protein_coding | 219.3383944 | 0.055630773 | 0.187264712 | 0.29707024 | 0.766412884 | 0.872270412 | 0.055630773 |
| ENSMUSG00000028954 | Nub1 | 53312 | protein_coding | 507.2766937 | 0.055658254 | 0.089429892 | 0.622367459 | 0.533700273 | 0.718743421 | 0.055658254 |
| ENSMUSG00000020124 | Usp15 | 14479 | protein_coding | 1530.295312 | 0.055664941 | 0.059683792 | 0.932664288 | 0.350993329 | 0.560913188 | 0.055664941 |
| ENSMUSG00000026069 | Il1rl1 | 17082 | protein_coding | 7.346110471 | -0.055673823 | 0.847807853 | -0.065667973 | 0.947642171 | 0.974312194 | 0.055673823 |
| ENSMUSG00000021595 | Nsun2 | 28114 | protein_coding | 1271.497898 | -0.055692638 | 0.125027054 | -0.445444697 | 0.655998417 | 0.800853376 | 0.055692638 |
| ENSMUSG00000031681 | Smad1 | 17125 | protein_coding | 847.9212173 | 0.055735594 | 0.080843436 | 0.689426341 | 0.490555012 | 0.683003516 | 0.055735594 |
| ENSMUSG00000036707 | Cab39 | 12283 | protein_coding | 4573.017622 | 0.055737627 | 0.12326654 | 0.452171587 | 0.651145375 | 0.798250711 | 0.055737627 |
| ENSMUSG00000039183 | Nubp2 | 26426 | protein_coding | 265.8532135 | 0.055769908 | 0.122829311 | 0.454043972 | 0.649797179 | 0.797624034 | 0.055769908 |
| ENSMUSG00000004455 | Ppp1cc | 19047 | protein_coding | 890.7511591 | -0.055786771 | 0.080324405 | -0.694518319 | 0.487357217 | 0.680216066 | 0.055786771 |
| ENSMUSG00000021621 | Zcchc9 | 69085 | protein_coding | 362.6088775 | 0.05583087 | 0.087669188 | 0.636835717 | 0.524231858 | 0.70998769 | 0.05583087 |
| ENSMUSG00000021302 | Ggps1 | 14593 | protein_coding | 589.3480818 | 0.055832496 | 0.087834544 | 0.635655335 | 0.525001094 | 0.710649905 | 0.055832496 |
| ENSMUSG00000085696 | Hoxaas3 | 72628 | lncRNA | 167.6957759 | 0.055833749 | 0.141734649 | 0.393931541 | 0.693631583 | 0.825783979 | 0.055833749 |
| ENSMUSG00000036112 | Metap2 | 56307 | protein_coding | 2057.128297 | 0.05584902 | 0.084257869 | 0.66283447 | 0.507436574 | 0.696822981 | 0.05584902 |
| ENSMUSG00000044066 | Cep68 | 216543 | protein_coding | 377.4720588 | -0.055909099 | 0.086414092 | -0.646990532 | 0.517638071 | 0.704835436 | 0.055909099 |
| ENSMUSG00000040969 | Arhgef38 | 77669 | protein_coding | 1485.413215 | -0.055958851 | 0.097271121 | -0.575287403 | 0.56509694 | 0.74142208 | 0.055958851 |
| ENSMUSG00000040424 | Hipk4 | 233020 | protein_coding | 6.675983539 | -0.056003168 | 0.464749516 | -0.120501831 | 0.904085629 | 0.952525195 | 0.056003168 |
| ENSMUSG00000062382 | Ftl1-ps1 | 434624 | protein_coding | 9.404984982 | -0.056065698 | 0.390994988 | -0.14339237 | 0.885980308 | 0.94220607 | 0.056065698 |
| ENSMUSG00000020978 | Klhdc2 | 69554 | protein_coding | 504.7082931 | 0.056076273 | 0.083395398 | 0.67241448 | 0.501319867 | 0.691772602 | 0.056076273 |
| ENSMUSG00000024098 | Twsg1 | 65960 | protein_coding | 688.361027 | 0.056096801 | 0.070428481 | 0.796507313 | 0.425737227 | 0.630311228 | 0.056096801 |
| ENSMUSG00000025766 | D3Ertd751e | 73852 | protein_coding | 181.8790984 | -0.056101267 | 0.123738029 | -0.453387428 | 0.650269788 | 0.797971325 | 0.056101267 |
| ENSMUSG00000115338 | Pnp | 18950 | protein_coding | 330.0327189 | 0.056136927 | 0.103045654 | 0.544777241 | 0.585906734 | 0.756020937 | 0.056136927 |
| ENSMUSG00000040565 | Btaf1 | 107182 | protein_coding | 1213.243367 | 0.056147077 | 0.107245749 | 0.523536623 | 0.600600872 | 0.766753883 | 0.056147077 |
| ENSMUSG00000040785 | Ttc3 | 22129 | protein_coding | 1487.005866 | -0.056150717 | 0.060287331 | -0.93138501 | 0.351654442 | 0.561399852 | 0.056150717 |
| ENSMUSG00000017802 | Retreg3 | 67998 | protein_coding | 1357.221087 | 0.056176 | 0.062758567 | 0.89511286 | 0.37072676 | 0.579256017 | 0.056176 |
| ENSMUSG00000025403 | Shmt2 | 108037 | protein_coding | 261.7536048 | 0.05618547 | 0.129072359 | 0.435302112 | 0.663343152 | 0.804794609 | 0.05618547 |
| ENSMUSG00000028649 | Macf1 | 11426 | protein_coding | 2635.839755 | -0.056189522 | 0.127598705 | -0.440361221 | 0.659675507 | 0.802969027 | 0.056189522 |
| ENSMUSG00000019854 | Reps1 | 19707 | protein_coding | 255.0985584 | -0.056198534 | 0.102204671 | -0.549862677 | 0.582413565 | 0.753466061 | 0.056198534 |
| ENSMUSG00000089911 | Mfsd14a | 15247 | protein_coding | 746.2215251 | 0.05621943 | 0.050612227 | 1.110787525 | 0.266659817 | 0.474459067 | 0.05621943 |
| ENSMUSG00000041769 | Ppp2r2d | 52432 | protein_coding | 509.7065641 | 0.056289982 | 0.082386401 | 0.683243606 | 0.494452923 | 0.686031895 | 0.056289982 |

| ensembl_gene_id | external_gene_name | entrezgene_id | gene_biotype | baseMean | log2FoldChange | lfcSE | stat | pvalue | padj | absLog2FC |
|---|---|---|---|---|---|---|---|---|---|---|
| ENSMUSG00000044145 | 1810024B03Rik | 329509 | protein_coding | 12.70739702 | -0.056293884 | 0.35632042 | -0.15798669 | 0.874467287 | 0.935867429 | 0.056293884 |
| ENSMUSG00000048701 | Ccdc6 | 76551 | protein_coding | 1157.025377 | 0.056301321 | 0.057164736 | 0.984896026 | 0.324675146 | 0.535676996 | 0.056301321 |
| ENSMUSG00000025271 | Pfkfb1 | 18639 | protein_coding | 26.90008538 | 0.056351582 | 0.281779898 | 0.199984392 | 0.841492788 | 0.916381801 | 0.056351582 |
| ENSMUSG00000021973 | Micu2 | 68514 | protein_coding | 1042.821841 | 0.056353966 | 0.08312813 | 0.677916928 | 0.497824366 | 0.688791086 | 0.056353966 |
| ENSMUSG00000022033 | Pbk | 52033 | protein_coding | 413.6945906 | 0.056435363 | 0.165523393 | 0.340950982 | 0.733140483 | 0.850197424 | 0.056435363 |
| ENSMUSG00000028850 | Gpatch3 | 242691 | protein_coding | 65.80229797 | -0.056442165 | 0.165373331 | -0.341301494 | 0.732876623 | 0.850024456 | 0.056442165 |
| ENSMUSG00000006390 | Elovl1 | 54325 | protein_coding | 731.4077893 | -0.056447017 | 0.152046645 | -0.371248026 | 0.710452806 | 0.836679652 | 0.056447017 |
| ENSMUSG00000028114 | Mettl14 | 210529 | protein_coding | 243.5083028 | -0.056566208 | 0.07724081 | -0.732335772 | 0.46396365 | 0.661197506 | 0.056566208 |
| ENSMUSG00000038844 | Kif16b | 16558 | protein_coding | 1150.213982 | 0.05658855 | 0.065430827 | 0.864860681 | 0.387115261 | 0.595269092 | 0.05658855 |
| ENSMUSG00000097537 | 2610020C07Rik | 69918 | lncRNA | 21.8431557 | -0.056761815 | 0.377630084 | -0.150310627 | 0.880519549 | 0.939399502 | 0.056761815 |
| ENSMUSG00000020657 | Dnajc27 | 217378 | protein_coding | 21.4781843 | -0.056786439 | 0.294073204 | -0.193103073 | 0.846878246 | 0.919400582 | 0.056786439 |
| ENSMUSG00000085502 | Gm12320 | 102635935 | lncRNA | 49.55654183 | -0.056795929 | 0.278374052 | -0.204027382 | 0.838332101 | 0.914773837 | 0.056795929 |
| ENSMUSG00000026384 | Ptpn4 | 19258 | protein_coding | 492.1399628 | -0.056819037 | 0.127244788 | -0.446533316 | 0.655212052 | 0.800343347 | 0.056819037 |
| ENSMUSG00000022538 | Lsg1 | 224092 | protein_coding | 389.7772905 | -0.056838313 | 0.11383696 | -0.499295776 | 0.617571031 | 0.777356676 | 0.056838313 |
| ENSMUSG00000048728 | Zfp454 | 237758 | protein_coding | 10.14097439 | 0.056860167 | 0.435218518 | 0.130647399 | 0.896054245 | 0.948165086 | 0.056860167 |
| ENSMUSG00000028622 | Mrpl37 | 56280 | protein_coding | 475.2726214 | 0.056878171 | 0.096579862 | 0.588923709 | 0.555912453 | 0.735368619 | 0.056878171 |
| ENSMUSG00000063730 | Hsd3b2 | 15493 | protein_coding | 484.5988465 | -0.056881433 | 0.177329792 | -0.320766367 | 0.748387449 | 0.860361984 | 0.056881433 |
| ENSMUSG00000079104 | Prps1l3 | 328099 | protein_coding | 426.4108597 | -0.056936417 | 0.081311801 | -0.700223295 | 0.483787866 | 0.677192471 | 0.056936417 |
| ENSMUSG00000061536 | Sec22c | 215474 | protein_coding | 94.19432066 | 0.056945079 | 0.191746264 | 0.296981424 | 0.766480691 | 0.872285256 | 0.056945079 |
| ENSMUSG00000005687 | Bcas2 | 68183 | protein_coding | 533.9279297 | 0.056975016 | 0.061123034 | 0.932136589 | 0.351265941 | 0.561179931 | 0.056975016 |
| ENSMUSG00000039828 | Wdr70 | 545085 | protein_coding | 192.9249679 | 0.056977558 | 0.108812187 | 0.523632136 | 0.600534426 | 0.766753883 | 0.056977558 |
| ENSMUSG00000028284 | Map3k7 | 26409 | protein_coding | 878.7032164 | 0.056983776 | 0.056170156 | 1.01448492 | 0.310351421 | 0.520454563 | 0.056983776 |
| ENSMUSG00000040616 | Tmem51 | 214359 | protein_coding | 352.9118425 | 0.056985608 | 0.09748165 | 0.584577793 | 0.558831643 | 0.736907711 | 0.056985608 |
| ENSMUSG00000029207 | Apbb2 | 11787 | protein_coding | 298.7727841 | 0.057048982 | 0.111984394 | 0.509436893 | 0.610446022 | 0.773061998 | 0.057048982 |
| ENSMUSG00000039756 | Dnttip2 | 99480 | protein_coding | 782.2495247 | -0.057063528 | 0.08748952 | -0.652232717 | 0.514251057 | 0.70188439 | 0.057063528 |
| ENSMUSG00000002015 | Bcap31 | 27061 | protein_coding | 1894.976735 | 0.057074485 | 0.087117154 | 0.655146351 | 0.512373532 | 0.699981612 | 0.057074485 |
| ENSMUSG00000033373 | Fntb | 110606 | protein_coding | 181.3697795 | 0.057094709 | 0.097533958 | 0.585382877 | 0.558290299 | 0.736678642 | 0.057094709 |
| ENSMUSG00000030884 | Uqcrc2 | 67003 | protein_coding | 5487.693863 | 0.057135197 | 0.094747886 | 0.603023445 | 0.546493092 | 0.728657457 | 0.057135197 |
| ENSMUSG00000031562 | Dctd | 320685 | protein_coding | 66.13397909 | -0.0571396 | 0.289273928 | -0.197527653 | 0.843414644 | 0.917115541 | 0.0571396 |
| ENSMUSG00000037458 | Azin1 | 54375 | protein_coding | 3287.275616 | -0.057140799 | 0.153423619 | -0.372438086 | 0.709566701 | 0.83582918 | 0.057140799 |
| ENSMUSG00000039599 | Fam149b | 105428 | protein_coding | 459.4373316 | 0.057141641 | 0.080108935 | 0.713299226 | 0.475660616 | 0.670351322 | 0.057141641 |
| ENSMUSG00000035473 | Galm | 319625 | protein_coding | 860.2261032 | -0.057161218 | 0.088367854 | -0.646855339 | 0.517725573 | 0.704835436 | 0.057161218 |
| ENSMUSG00000020346 | Mgat1 | 17308 | protein_coding | 1039.486894 | -0.057162813 | 0.079774994 | -0.716550517 | 0.473651486 | 0.668798783 | 0.057162813 |
| ENSMUSG00000046138 | 9930021J03Rik | 240613 | protein_coding | 813.5120785 | -0.057171182 | 0.089556585 | -0.638380544 | 0.523225993 | 0.709452933 | 0.057171182 |
| ENSMUSG00000041870 | Ankrd13a | 68420 | protein_coding | 3113.231611 | -0.057229252 | 0.059721266 | -0.958272599 | 0.337925317 | 0.548511589 | 0.057229252 |
| ENSMUSG00000062944 | 9130023H24Rik | 100043133 | protein_coding | 118.7335248 | 0.057256465 | 0.177912524 | 0.321823687 | 0.747586268 | 0.859689126 | 0.057256465 |
| ENSMUSG00000097006 | 9530082P21Rik | 638247 | lncRNA | 83.02856953 | 0.057269903 | 0.185766074 | 0.30829043 | 0.757861347 | 0.867408546 | 0.057269903 |
| ENSMUSG00000002803 | Btbd6 | 399566 | protein_coding | 271.7038711 | -0.057281488 | 0.122734975 | -0.466708756 | 0.640708264 | 0.791898908 | 0.057281488 |

| ensembl_gene_id | external_gene_name | entrezgene_id | gene_biotype | baseMean | log2FoldChange | lfcSE | stat | pvalue | padj | absLog2FC |
|---|---|---|---|---|---|---|---|---|---|---|
| ENSMUSG00000036188 | Ankmy2 | 217473 | protein_coding | 486.4549356 | 0.057291944 | 0.119722443 | 0.478539717 | 0.632266113 | 0.787249165 | 0.057291944 |
| ENSMUSG00000043154 | Ppp2r3a | 235542 | protein_coding | 390.0640489 | -0.057303199 | 0.131631561 | -0.435330238 | 0.663322739 | 0.804794609 | 0.057303199 |
| ENSMUSG00000027893 | Ahcyl1 | 229709 | protein_coding | 1788.844287 | -0.057312454 | 0.073948828 | -0.775028562 | 0.438322782 | 0.640830788 | 0.057312454 |
| ENSMUSG00000031299 | Pdha1 | 18597 | protein_coding | 3989.060477 | 0.057325807 | 0.072502174 | 0.790677072 | 0.429132459 | 0.633370351 | 0.057325807 |
| ENSMUSG00000041161 | Otud3 | 73162 | protein_coding | 216.3901762 | -0.057337342 | 0.114944169 | -0.498827756 | 0.617900732 | 0.777648772 | 0.057337342 |
| ENSMUSG00000030714 | Sgf29 | 75565 | protein_coding | 337.9967805 | -0.057340387 | 0.091762245 | -0.624879946 | 0.532049855 | 0.717022599 | 0.057340387 |
| ENSMUSG00000019432 | Ddx39b | 53817 | protein_coding | 1576.84061 | -0.057367161 | 0.093321708 | -0.614724726 | 0.538736528 | 0.722272825 | 0.057367161 |
| ENSMUSG00000044792 | Isca1 | 69046 | protein_coding | 821.3849181 | 0.057381968 | 0.067700724 | 0.847582792 | 0.396670361 | 0.604651202 | 0.057381968 |
| ENSMUSG00000022247 | Brix1 | 67832 | protein_coding | 444.5982015 | -0.057399807 | 0.124135889 | -0.462394941 | 0.643798127 | 0.793808373 | 0.057399807 |
| ENSMUSG00000040985 | Sun3 | 194974 | protein_coding | 9.280313588 | -0.057410418 | 0.366621618 | -0.156593107 | 0.875565535 | 0.936728093 | 0.057410418 |
| ENSMUSG00000047242 | Taf9b | 407786 | protein_coding | 96.2902304 | -0.0574243 | 0.13179736 | -0.435701443 | 0.663053359 | 0.804794609 | 0.0574243 |
| ENSMUSG00000073402 | Gm8909 | 667977 | protein_coding | 89.66486963 | -0.057466389 | 0.285549609 | -0.201248354 | 0.840504383 | 0.915836464 | 0.057466389 |
| ENSMUSG00000003031 | Cdkn1b | 12576 | protein_coding | 1196.655376 | 0.057482735 | 0.144505883 | 0.397788199 | 0.690786317 | 0.823979937 | 0.057482735 |
| ENSMUSG00000018415 | Gid4 | 66771 | protein_coding | 508.2308056 | -0.057508477 | 0.083690791 | -0.687154186 | 0.491985571 | 0.683859515 | 0.057508477 |
| ENSMUSG00000054385 | Ceacam2 | 26367 | protein_coding | 80.6064998 | 0.05751828 | 0.173593294 | 0.331339295 | 0.740388213 | 0.855177916 | 0.05751828 |
| ENSMUSG00000048429 | Timm29 | 69773 | protein_coding | 415.8616751 | -0.057519114 | 0.110462432 | -0.520712002 | 0.602567413 | 0.767709311 | 0.057519114 |
| ENSMUSG00000026755 | Arpc5l | 74192 | protein_coding | 590.1062165 | 0.057546335 | 0.109344344 | 0.526285428 | 0.598689906 | 0.765385089 | 0.057546335 |
| ENSMUSG00000050471 | Fam118b | 109229 | protein_coding | 460.9326948 | 0.057565744 | 0.095452957 | 0.603079736 | 0.546455647 | 0.728657457 | 0.057565744 |
| ENSMUSG00000027550 | Lrrcc1 | 71710 | protein_coding | 362.6489943 | -0.057576674 | 0.11442395 | -0.503187261 | 0.61483262 | 0.775503166 | 0.057576674 |
| ENSMUSG00000019952 | Poc1b | 382406 | protein_coding | 318.0977197 | 0.057599347 | 0.089143149 | 0.646144407 | 0.518185839 | 0.705260535 | 0.057599347 |
| ENSMUSG00000005312 | Ubqln1 | 56085 | protein_coding | 1995.234244 | 0.057610383 | 0.07566602 | 0.761377206 | 0.446431798 | 0.647239525 | 0.057610383 |
| ENSMUSG00000015087 | Rabl6 | 227624 | protein_coding | 848.494283 | 0.057622312 | 0.065031368 | 0.886069499 | 0.375580078 | 0.583911261 | 0.057622312 |
| ENSMUSG00000052131 | Akr1b7 | 11997 | protein_coding | 28.24457114 | -0.057627674 | 0.244629377 | -0.235571354 | 0.813765304 | 0.900884818 | 0.057627674 |
| ENSMUSG00000041453 | Rpl21 | 19933 | protein_coding | 7109.84206 | 0.057629368 | 0.088479808 | 0.651327905 | 0.514834839 | 0.702380241 | 0.057629368 |
| ENSMUSG00000042750 | Bex2 | 12069 | protein_coding | 35.79023311 | 0.057672017 | 0.205054704 | 0.281251859 | 0.778517231 | 0.879697961 | 0.057672017 |
| ENSMUSG00000048003 | Catsper4 | 329954 | protein_coding | 14.66612811 | -0.057726884 | 0.295112694 | -0.195609628 | 0.844915724 | 0.918121002 | 0.057726884 |
| ENSMUSG00000033392 | Clasp2 | 76499 | protein_coding | 439.173187 | -0.057740273 | 0.078940897 | -0.731436748 | 0.464512426 | 0.661429816 | 0.057740273 |
| ENSMUSG00000022912 | Pros1 | 19128 | protein_coding | 128.5751629 | 0.057743429 | 0.282356232 | 0.204505594 | 0.837958421 | 0.914526523 | 0.057743429 |
| ENSMUSG00000054161 | Fam83e | 73813 | protein_coding | 3979.902992 | -0.057822796 | 0.099356625 | -0.581972225 | 0.56058539 | 0.738310882 | 0.057822796 |
| ENSMUSG00000027406 | Idh3b | 170718 | protein_coding | 3033.027518 | -0.057826567 | 0.074904638 | -0.77200249 | 0.440112953 | 0.642313808 | 0.057826567 |
| ENSMUSG00000021211 | Akr1c12 | 622402 | protein_coding | 2275.196269 | 0.057834086 | 0.125626624 | 0.460364882 | 0.645254338 | 0.794681133 | 0.057834086 |
| ENSMUSG00000028403 | Zdhhc21 | 68268 | protein_coding | 1156.604413 | 0.057836281 | 0.123681281 | 0.467623556 | 0.640053816 | 0.791371033 | 0.057836281 |
| ENSMUSG00000030278 | Cidec | 14311 | protein_coding | 62.50887628 | -0.057913021 | 0.356696945 | -0.162359172 | 0.87102302 | 0.933600906 | 0.057913021 |
| ENSMUSG00000058638 | Zfp110 | 65020 | protein_coding | 884.5465904 | 0.057936162 | 0.080821134 | 0.716844218 | 0.473470224 | 0.668798783 | 0.057936162 |
| ENSMUSG00000033967 | Rnf225 | 381845 | protein_coding | 9.738669715 | -0.057948001 | 0.379686375 | -0.152620702 | 0.878697399 | 0.938376947 | 0.057948001 |
| ENSMUSG00000041881 | Ndufa7 | 66416 | protein_coding | 1319.50268 | 0.058088916 | 0.116079629 | 0.500422999 | 0.616777262 | 0.776971547 | 0.058088916 |
| ENSMUSG00000028986 | Klhl7 | 52323 | protein_coding | 360.4704608 | 0.058124996 | 0.069414329 | 0.837363078 | 0.402388513 | 0.609921868 | 0.058124996 |
| ENSMUSG00000020153 | Ndufs7 | 75406 | protein_coding | 957.5115904 | 0.058150697 | 0.101739035 | 0.571567214 | 0.567615212 | 0.743180341 | 0.058150697 |

| ensembl_gene_id | external_gene_name | entrezgene_id | gene_biotype | baseMean | log2FoldChange | lfcSE | stat | pvalue | padj | absLog2FC |
|---|---|---|---|---|---|---|---|---|---|---|
| ENSMUSG00000024247 | Pkdcc | 106522 | protein_coding | 1114.07389 | 0.05818742 | 0.093501982 | 0.622312157 | 0.533736629 | 0.718743421 | 0.05818742 |
| ENSMUSG00000048897 | Zfp710 | 209225 | protein_coding | 533.209307 | 0.058190593 | 0.086942025 | 0.669303401 | 0.503301957 | 0.693356995 | 0.058190593 |
| ENSMUSG00000034892 | Rps29 | 20090 | protein_coding | 4402.131108 | 0.058223182 | 0.075179711 | 0.774453387 | 0.438662723 | 0.641246315 | 0.058223182 |
| ENSMUSG00000018160 | Med1 | 19014 | protein_coding | 954.8917428 | -0.058231686 | 0.069078316 | -0.842980691 | 0.399239243 | 0.606939679 | 0.058231686 |
| ENSMUSG00000038836 | Agbl3 | 76223 | protein_coding | 24.371184 | 0.058241921 | 0.407592215 | 0.142892624 | 0.886374984 | 0.94220607 | 0.058241921 |
| ENSMUSG00000029098 | Acox3 | 80911 | protein_coding | 494.3282235 | -0.058259714 | 0.093666553 | -0.621990582 | 0.533948061 | 0.718850876 | 0.058259714 |
| ENSMUSG00000056952 | Tatdn2 | 381801 | protein_coding | 936.7741081 | 0.058284546 | 0.147005344 | 0.396479093 | 0.691751628 | 0.824470213 | 0.058284546 |
| ENSMUSG00000025352 | Gdf11 | 14561 | protein_coding | 22.36251907 | 0.058293179 | 0.334416159 | 0.174313283 | 0.86161926 | 0.927978334 | 0.058293179 |
| ENSMUSG00000051184 | Zfp524 | 66056 | protein_coding | 70.48523096 | 0.058358166 | 0.200339219 | 0.291296761 | 0.770824364 | 0.87556148 | 0.058358166 |
| ENSMUSG00000021779 | Thrb | 21834 | protein_coding | 525.1794987 | 0.058360877 | 0.160447672 | 0.363737758 | 0.716053845 | 0.84023174 | 0.058360877 |
| ENSMUSG00000055334 | Snupn | 66069 | protein_coding | 118.2296944 | 0.058365504 | 0.124038042 | 0.470545188 | 0.637965557 | 0.790438741 | 0.058365504 |
| ENSMUSG00000005804 | Bloc1s6 | 18457 | protein_coding | 398.8218972 | 0.058379682 | 0.102756907 | 0.568133896 | 0.569944053 | 0.744741762 | 0.058379682 |
| ENSMUSG00000059866 | Tnip2 | 231130 | protein_coding | 170.6931624 | -0.058384438 | 0.148809045 | -0.392344688 | 0.694803548 | 0.826561425 | 0.058384438 |
| ENSMUSG00000034158 | Lrrc58 | 320184 | protein_coding | 1205.445697 | 0.058388156 | 0.098173001 | 0.594747593 | 0.552012203 | 0.732399032 | 0.058388156 |
| ENSMUSG00000056167 | Cnot10 | 78893 | protein_coding | 593.3653197 | 0.058401315 | 0.080597478 | 0.724604752 | 0.468694543 | 0.66509336 | 0.058401315 |
| ENSMUSG00000033720 | Sfxn5 | 94282 | protein_coding | 296.0366885 | -0.058407382 | 0.179583876 | -0.32523734 | 0.74500145 | 0.858274321 | 0.058407382 |
| ENSMUSG00000021391 | Cenpp | 66336 | protein_coding | 92.65111395 | 0.058443862 | 0.196718759 | 0.297093485 | 0.766395138 | 0.872270412 | 0.058443862 |
| ENSMUSG00000018293 | Pfn1 | 18643 | protein_coding | 5278.882731 | 0.058456732 | 0.131652037 | 0.444024518 | 0.657024858 | 0.801431454 | 0.058456732 |
| ENSMUSG00000025872 | Thoc3 | 73666 | protein_coding | 358.4256379 | -0.058458067 | 0.106431712 | -0.549254222 | 0.582830999 | 0.753761948 | 0.058458067 |
| ENSMUSG00000096696 | Zfp960 | 449000 | protein_coding | 80.38796819 | 0.058473363 | 0.208410021 | 0.280568866 | 0.779041098 | 0.879915375 | 0.058473363 |
| ENSMUSG00000003190 | Bcl2l12 | 75736 | protein_coding | 80.31754891 | 0.058477519 | 0.133366706 | 0.438471647 | 0.661044419 | 0.803875058 | 0.058477519 |
| ENSMUSG00000032253 | Phip | 83946 | protein_coding | 1335.203711 | 0.058484103 | 0.100029323 | 0.584669586 | 0.558769908 | 0.736907711 | 0.058484103 |
| ENSMUSG00000055692 | Tmem191 | 224019 | protein_coding | 52.03208362 | -0.058497202 | 0.235697371 | -0.248187756 | 0.803989139 | 0.894394399 | 0.058497202 |
| ENSMUSG00000024142 | Mlst8 | 56716 | protein_coding | 310.1603439 | 0.058508557 | 0.082406525 | 0.709999081 | 0.477704706 | 0.672281371 | 0.058508557 |
| ENSMUSG00000027706 | Sec62 | 69276 | protein_coding | 1459.493543 | 0.058520448 | 0.078588067 | 0.744648016 | 0.456484533 | 0.655389006 | 0.058520448 |
| ENSMUSG00000038319 | Kcnh2 | 16511 | protein_coding | 47.22967734 | 0.0585274 | 0.188907101 | 0.30982107 | 0.756697028 | 0.866475216 | 0.0585274 |
| ENSMUSG00000001755 | Coasy | 71743 | protein_coding | 436.3468151 | -0.058541396 | 0.110527972 | -0.529652311 | 0.596353018 | 0.764096331 | 0.058541396 |
| ENSMUSG00000002372 | Ranbp3 | 71810 | protein_coding | 637.7482439 | 0.058556675 | 0.091566823 | 0.639496633 | 0.522499903 | 0.708770095 | 0.058556675 |
| ENSMUSG00000067219 | Nipal1 | 70701 | protein_coding | 1153.413694 | -0.058612165 | 0.248286209 | -0.236066938 | 0.813380729 | 0.900690228 | 0.058612165 |
| ENSMUSG00000006412 | Pfdn2 | 18637 | protein_coding | 152.1502998 | 0.058657333 | 0.123055571 | 0.476673524 | 0.633594619 | 0.788005719 | 0.058657333 |
| ENSMUSG00000022848 | Slc49a4 | 224132 | protein_coding | 593.6924972 | 0.058768163 | 0.089109722 | 0.659503378 | 0.509572578 | 0.698579013 | 0.058768163 |
| ENSMUSG00000066900 | Suds3 | 71954 | protein_coding | 1394.097314 | -0.058782325 | 0.073810183 | -0.796398581 | 0.425800403 | 0.630311228 | 0.058782325 |
| ENSMUSG00000024566 | Atp9b | 50771 | protein_coding | 728.659258 | -0.058784402 | 0.065803369 | -0.893334235 | 0.37167821 | 0.579898015 | 0.058784402 |
| ENSMUSG00000056201 | Cfl1 | 12631 | protein_coding | 2986.557015 | -0.058791493 | 0.115080065 | -0.510874695 | 0.609438801 | 0.772513652 | 0.058791493 |
| ENSMUSG00000004668 | Abca13 | 268379 | protein_coding | 22.59259631 | -0.058810535 | 0.636368556 | -0.09241584 | 0.926367655 | 0.963740176 | 0.058810535 |
| ENSMUSG00000010051 | Hyal1 | 15586 | protein_coding | 27.70429344 | 0.058812638 | 0.250546242 | 0.234737657 | 0.814412358 | 0.900950823 | 0.058812638 |
| ENSMUSG00000025001 | Hells | 15201 | protein_coding | 373.9319598 | -0.058824984 | 0.151044439 | -0.389454819 | 0.696939727 | 0.827434106 | 0.058824984 |
| ENSMUSG00000035133 | Arhgap5 | 11855 | protein_coding | 5576.000686 | 0.058827809 | 0.061456804 | 0.957222066 | 0.338455182 | 0.549140688 | 0.058827809 |

| ensembl_gene_id | external_gene_name | entrezgene_id | gene_biotype | baseMean | log2FoldChange | lfcSE | stat | pvalue | padj | absLog2FC |
|---|---|---|---|---|---|---|---|---|---|---|
| ENSMUSG00000028898 | Trnau1ap | 71787 | protein_coding | 286.4757303 | 0.058842995 | 0.11313792 | 0.520099675 | 0.602994105 | 0.76799575 | 0.058842995 |
| ENSMUSG00000041974 | Spidr | 224008 | protein_coding | 209.2615381 | -0.058850729 | 0.090920509 | -0.647276722 | 0.517452864 | 0.704744696 | 0.058850729 |
| ENSMUSG00000003581 | Rnf215 | 71673 | protein_coding | 181.8258069 | 0.058859827 | 0.121120954 | 0.485959072 | 0.626996185 | 0.784262229 | 0.058859827 |
| ENSMUSG00000069135 | Cep43 | 75296 | protein_coding | 291.2413709 | -0.058904004 | 0.09497708 | -0.620191776 | 0.535131535 | 0.719896545 | 0.058904004 |
| ENSMUSG00000021340 | Gpld1 | 14756 | protein_coding | 170.0891548 | -0.058918645 | 0.10934514 | -0.538831854 | 0.590002882 | 0.759583406 | 0.058918645 |
| ENSMUSG00000042428 | Mgat3 | 17309 | protein_coding | 1999.522512 | 0.058952266 | 0.07593059 | 0.776396785 | 0.437514742 | 0.640155687 | 0.058952266 |
| ENSMUSG00000089669 | Tnfsf13 | 69583 | protein_coding | 172.2720525 | 0.058956587 | 0.127336957 | 0.462996667 | 0.643366756 | 0.793445869 | 0.058956587 |
| ENSMUSG00000118100 | Gm50340 | 115489003 | lncRNA | 11.05961916 | 0.058961226 | 0.330983631 | 0.178139403 | 0.858613498 | 0.926384278 | 0.058961226 |
| ENSMUSG00000097392 | Thoc2l | 100042165 | protein_coding | 122.3700459 | -0.058984087 | 0.169576196 | -0.347832349 | 0.727966094 | 0.846662573 | 0.058984087 |
| ENSMUSG00000000148 | Brat1 | 231841 | protein_coding | 211.2306109 | -0.059042478 | 0.131227067 | -0.449926064 | 0.652763753 | 0.798741727 | 0.059042478 |
| ENSMUSG00000028256 | Odf2l | 52184 | protein_coding | 253.3704803 | 0.059063311 | 0.093334889 | 0.63281064 | 0.526857307 | 0.712375483 | 0.059063311 |
| ENSMUSG00000030881 | Arfip2 | 76932 | protein_coding | 870.9675273 | 0.059068357 | 0.066229999 | 0.891867107 | 0.372464167 | 0.580792824 | 0.059068357 |
| ENSMUSG00000037851 | Iars | 105148 | protein_coding | 816.3684016 | 0.059099129 | 0.123083575 | 0.480154474 | 0.631117556 | 0.786391335 | 0.059099129 |
| ENSMUSG00000110062 | B430319F04Rik | 619306 | TEC | 13.06711949 | -0.059175815 | 0.392807472 | -0.150648394 | 0.880253084 | 0.939320286 | 0.059175815 |
| ENSMUSG00000060636 | Rpl35a | 57808 | protein_coding | 3808.812982 | 0.059190807 | 0.085486183 | 0.692402035 | 0.488684888 | 0.681333909 | 0.059190807 |
| ENSMUSG00000037519 | Ppfia1 | 233977 | protein_coding | 963.6264461 | -0.059197841 | 0.054311046 | -1.089977936 | 0.275722863 | 0.484028727 | 0.059197841 |
| ENSMUSG00000025613 | Cct8 | 12469 | protein_coding | 2201.938955 | 0.05921184 | 0.134915252 | 0.438881736 | 0.660747231 | 0.803653446 | 0.05921184 |
| ENSMUSG00000030697 | Ppp4c | 56420 | protein_coding | 1436.754567 | -0.059213339 | 0.099853274 | -0.593003484 | 0.553178821 | 0.732969991 | 0.059213339 |
| ENSMUSG00000018347 | Zkscan6 | 52712 | protein_coding | 277.4129422 | -0.059219795 | 0.104020384 | -0.569309521 | 0.569146107 | 0.74446573 | 0.059219795 |
| ENSMUSG00000037818 | Abhd18 | 269423 | protein_coding | 151.7836961 | -0.059246497 | 0.124064676 | -0.477545252 | 0.632973906 | 0.787741735 | 0.059246497 |
| ENSMUSG00000027177 | Hipk3 | 15259 | protein_coding | 1526.45588 | -0.05948017 | 0.08379387 | -0.709839153 | 0.477803887 | 0.672321665 | 0.05948017 |
| ENSMUSG00000010067 | Rassf1 | 56289 | protein_coding | 233.5242732 | 0.059484983 | 0.108958651 | 0.545940892 | 0.585106568 | 0.755622136 | 0.059484983 |
| ENSMUSG00000026554 | Dcaf8 | 98193 | protein_coding | 1529.876971 | -0.059492442 | 0.088512509 | -0.672135983 | 0.501497131 | 0.691924848 | 0.059492442 |
| ENSMUSG00000030127 | Cops7a | 26894 | protein_coding | 900.9412896 | 0.059505423 | 0.067746056 | 0.878359961 | 0.379748405 | 0.587657295 | 0.059505423 |
| ENSMUSG00000064901 | Snora21 | 100302498 | snoRNA | 13.68534825 | 0.05956848 | 0.370627897 | 0.16072314 | 0.872311463 | 0.934502084 | 0.05956848 |
| ENSMUSG00000020132 | Rab21 | 216344 | protein_coding | 732.604931 | 0.059586919 | 0.104261226 | 0.571515616 | 0.567650178 | 0.743180341 | 0.059586919 |
| ENSMUSG00000056919 | Cep162 | 382090 | protein_coding | 263.4782851 | -0.059597163 | 0.114288196 | -0.521463851 | 0.602043681 | 0.767564496 | 0.059597163 |
| ENSMUSG00000049739 | Zfp646 | 233905 | protein_coding | 839.9912161 | -0.059610721 | 0.059616249 | -0.999907279 | 0.317355381 | 0.528637996 | 0.059610721 |
| ENSMUSG00000019889 | Ptprk | 19272 | protein_coding | 2512.986491 | 0.05964126 | 0.063966581 | 0.93238155 | 0.351139376 | 0.561034004 | 0.05964126 |
| ENSMUSG00000020170 | Frs2 | 327826 | protein_coding | 413.6073623 | -0.059648134 | 0.141031545 | -0.422941792 | 0.67233773 | 0.810978071 | 0.059648134 |
| ENSMUSG00000033933 | Vhl | 22346 | protein_coding | 286.2445433 | -0.05973748 | 0.077957811 | -0.766279594 | 0.443509958 | 0.644487419 | 0.05973748 |
| ENSMUSG00000015027 | Galns | 50917 | protein_coding | 334.8909765 | -0.059780086 | 0.080999674 | -0.738028729 | 0.460496991 | 0.658781284 | 0.059780086 |
| ENSMUSG00000000440 | Pparg | 19016 | protein_coding | 1387.429793 | -0.059790153 | 0.199741727 | -0.299337321 | 0.764682682 | 0.871422033 | 0.059790153 |
| ENSMUSG00000034850 | Tmem127 | 69470 | protein_coding | 1347.457832 | -0.059795534 | 0.081263301 | -0.735824578 | 0.461837462 | 0.659744134 | 0.059795534 |
| ENSMUSG00000060301 | 2610008E11Rik | 72128 | protein_coding | 292.6574228 | 0.059812116 | 0.156991674 | 0.380989098 | 0.703211338 | 0.832027202 | 0.059812116 |
| ENSMUSG00000061983 | Rps12 | 20042 | protein_coding | 8420.373046 | 0.059820968 | 0.08050371 | 0.743083372 | 0.457431206 | 0.656097636 | 0.059820968 |
| ENSMUSG00000024287 | Thoc1 | 225160 | protein_coding | 412.7640503 | 0.059823102 | 0.101333413 | 0.590359098 | 0.554949926 | 0.734641854 | 0.059823102 |
| ENSMUSG00000028861 | Mrps15 | 66407 | protein_coding | 421.8219335 | 0.059847707 | 0.098396382 | 0.608230771 | 0.543034429 | 0.7259297 | 0.059847707 |

| ensembl_gene_id | external_gene_name | entrezgene_id | gene_biotype | baseMean | log2FoldChange | lfcSE | stat | pvalue | padj | absLog2FC |
|---|---|---|---|---|---|---|---|---|---|---|
| ENSMUSG00000031985 | Gnpat | 14712 | protein_coding | 632.1276709 | 0.059860232 | 0.061894149 | 0.967138798 | 0.333474652 | 0.544296824 | 0.059860232 |
| ENSMUSG00000071652 | Ints5 | 109077 | protein_coding | 318.2861183 | -0.059946664 | 0.095262094 | -0.629281396 | 0.529164849 | 0.714548497 | 0.059946664 |
| ENSMUSG00000040848 | Sft2d2 | 108735 | protein_coding | 3016.864169 | 0.059973786 | 0.058621017 | 1.02307652 | 0.306271668 | 0.516330983 | 0.059973786 |
| ENSMUSG00000021065 | Fut8 | 53618 | protein_coding | 1947.78976 | -0.059987137 | 0.078147789 | -0.767611439 | 0.442718064 | 0.643949429 | 0.059987137 |
| ENSMUSG00000032198 | Dock6 | 319899 | protein_coding | 263.0857312 | 0.059990226 | 0.202741254 | 0.295895508 | 0.767309882 | 0.872979414 | 0.059990226 |
| ENSMUSG00000063172 | Hspb11 | 72938 | protein_coding | 194.401904 | -0.05999583 | 0.149377358 | -0.401639386 | 0.687949439 | 0.821906591 | 0.05999583 |
| ENSMUSG00000021993 | Mipep | 70478 | protein_coding | 361.6686826 | 0.060002008 | 0.104213523 | 0.575760289 | 0.564777218 | 0.741139569 | 0.060002008 |
| ENSMUSG00000026254 | Eif4e2 | 26987 | protein_coding | 978.9770376 | -0.060010924 | 0.093679358 | -0.640599217 | 0.521783108 | 0.708038963 | 0.060010924 |
| ENSMUSG00000022386 | Trmu | 72026 | protein_coding | 147.0784274 | -0.060019407 | 0.128398967 | -0.467444623 | 0.640181803 | 0.791371033 | 0.060019407 |
| ENSMUSG00000024766 | Lipo3 | 381236 | protein_coding | 312.2187037 | 0.060085769 | 0.112733098 | 0.532991377 | 0.594039549 | 0.762315657 | 0.060085769 |
| ENSMUSG00000025078 | Nhlrc2 | 66866 | protein_coding | 1208.537503 | -0.060101352 | 0.070102263 | -0.85733826 | 0.391257969 | 0.59924777 | 0.060101352 |
| ENSMUSG00000040249 | Lrp1 | 16971 | protein_coding | 2985.158454 | -0.060109866 | 0.156543746 | -0.383981271 | 0.700992329 | 0.830141697 | 0.060109866 |
| ENSMUSG00000024533 | Spire1 | 68166 | protein_coding | 59.99413795 | 0.060150638 | 0.243059453 | 0.247472943 | 0.804542229 | 0.894700251 | 0.060150638 |
| ENSMUSG00000024958 | Gpr137 | 107173 | protein_coding | 244.222522 | 0.060152112 | 0.112483609 | 0.534763357 | 0.592813504 | 0.761355708 | 0.060152112 |
| ENSMUSG00000037570 | Mcrs1 | 51812 | protein_coding | 516.3219693 | 0.060177744 | 0.123166763 | 0.488587529 | 0.625133744 | 0.782844779 | 0.060177744 |
| ENSMUSG00000007867 | Ift43 | 76411 | protein_coding | 172.0135477 | -0.060193782 | 0.1116172 | -0.539287686 | 0.589688364 | 0.759394898 | 0.060193782 |
| ENSMUSG00000023266 | Frs3 | 107971 | protein_coding | 22.07030941 | 0.060201652 | 0.334284512 | 0.180091061 | 0.85708108 | 0.925280429 | 0.060201652 |
| ENSMUSG00000028255 | Clca1 | 23844 | protein_coding | 34107.38699 | -0.060211859 | 0.2420225 | -0.248786202 | 0.803526165 | 0.89432847 | 0.060211859 |
| ENSMUSG00000022023 | Wbp4 | 22380 | protein_coding | 427.1313201 | -0.060241102 | 0.060639851 | -0.993424316 | 0.320503217 | 0.531989916 | 0.060241102 |
| ENSMUSG00000028790 | Khdrbs1 | 20218 | protein_coding | 1244.542469 | 0.06028659 | 0.102514681 | 0.588077623 | 0.556480192 | 0.735604839 | 0.06028659 |
| ENSMUSG00000058900 | Rsl1 | 380855 | protein_coding | 119.0282806 | 0.060326656 | 0.122964993 | 0.49060025 | 0.62370921 | 0.781870006 | 0.060326656 |
| ENSMUSG00000022801 | Lrch3 | 70144 | protein_coding | 371.6621033 | -0.060334665 | 0.08813885 | -0.684541092 | 0.493633551 | 0.685372663 | 0.060334665 |
| ENSMUSG00000024908 | Ppp6r3 | 52036 | protein_coding | 2818.982627 | 0.06035664 | 0.053508883 | 1.127974202 | 0.259330814 | 0.4664368 | 0.06035664 |
| ENSMUSG00000004667 | Polr2e | 66420 | protein_coding | 552.2632113 | -0.06041051 | 0.073281823 | -0.824358724 | 0.409735761 | 0.616674969 | 0.06041051 |
| ENSMUSG00000001054 | Rmnd5b | 66089 | protein_coding | 243.4376325 | -0.060424842 | 0.130081726 | -0.464514453 | 0.642279207 | 0.792956415 | 0.060424842 |
| ENSMUSG00000031828 | Klhl36 | 234796 | protein_coding | 46.80441201 | 0.060438524 | 0.20838432 | 0.290033931 | 0.771790279 | 0.87594952 | 0.060438524 |
| ENSMUSG00000074580 | 4931440P22Rik | 71004 | transcribed_processed_pseudogene | 22.43481824 | -0.060441891 | 0.273433739 | -0.22104767 | 0.825055316 | 0.907748917 | 0.060441891 |
| ENSMUSG00000025439 | Clns1a | 12729 | protein_coding | 791.4811499 | -0.060442995 | 0.088757145 | -0.680993004 | 0.495875917 | 0.687162611 | 0.060442995 |
| ENSMUSG00000013663 | Pten | 19211 | protein_coding | 2290.178484 | -0.060481191 | 0.051769789 | -1.168271922 | 0.242697098 | 0.447623827 | 0.060481191 |
| ENSMUSG00000027427 | Polr3f | 70408 | protein_coding | 352.1619057 | -0.060481681 | 0.079167128 | -0.763974672 | 0.444882336 | 0.645657025 | 0.060481681 |
| ENSMUSG00000031731 | Ap1g1 | 11765 | protein_coding | 1599.002302 | -0.060514067 | 0.075817249 | -0.798156983 | 0.424779397 | 0.629384766 | 0.060514067 |
| ENSMUSG00000024077 | Strn | 268980 | protein_coding | 995.4988391 | -0.060521602 | 0.0691533 | -0.875180255 | 0.381475835 | 0.589230712 | 0.060521602 |
| ENSMUSG00000022529 | Zfp263 | 74120 | protein_coding | 527.2726362 | 0.060540239 | 0.078206141 | 0.774111064 | 0.438865115 | 0.641483303 | 0.060540239 |
| ENSMUSG00000048677 | Tpcn2 | 233979 | protein_coding | 149.7112141 | -0.060568498 | 0.113523828 | -0.533531141 | 0.59366596 | 0.762081835 | 0.060568498 |
| ENSMUSG00000085457 | 1110046J04Rik | 68808 | lncRNA | 8.227006133 | 0.060606588 | 0.402155197 | 0.150704476 | 0.880208843 | 0.939320286 | 0.060606588 |
| ENSMUSG00000086714 | 0610009E02Rik | 100125929 | lncRNA | 9.0360709 | -0.060614091 | 0.389874859 | -0.155470633 | 0.876450304 | 0.937171082 | 0.060614091 |
| ENSMUSG00000045767 | B230219D22Rik | 78521 | protein_coding | 885.5684098 | 0.060690845 | 0.06782679 | 0.894791642 | 0.370898479 | 0.579410648 | 0.060690845 |
| ENSMUSG00000018507 | Trpv2 | 22368 | protein_coding | 6.987944572 | -0.060710301 | 0.57992881 | -0.104685782 | 0.916625144 | 0.958597942 | 0.060710301 |

| ensembl_gene_id | external_gene_name | entrezgene_id | gene_biotype | baseMean | log2FoldChange | lfcSE | stat | pvalue | padj | absLog2FC |
|---|---|---|---|---|---|---|---|---|---|---|
| ENSMUSG00000026455 | Klhl12 | 240756 | protein_coding | 363.6291325 | -0.060711284 | 0.081488285 | -0.745030821 | 0.456253088 | 0.655296022 | 0.060711284 |
| ENSMUSG00000024759 | Atl3 | 109168 | protein_coding | 1146.57538 | 0.060711598 | 0.083469321 | 0.727352248 | 0.467010199 | 0.663707223 | 0.060711598 |
| ENSMUSG00000022552 | Sharpin | 106025 | protein_coding | 372.0908299 | 0.060718876 | 0.088142698 | 0.68887017 | 0.490904973 | 0.683251871 | 0.060718876 |
| ENSMUSG00000038286 | Bphl | 68021 | protein_coding | 407.2213564 | 0.060761199 | 0.098507837 | 0.616815892 | 0.537356169 | 0.721426926 | 0.060761199 |
| ENSMUSG00000030703 | Gdpd3 | 68616 | protein_coding | 30.09789446 | 0.060791759 | 0.246426478 | 0.246693291 | 0.805145599 | 0.895118246 | 0.060791759 |
| ENSMUSG00000029201 | Ugdh | 22235 | protein_coding | 8287.765284 | -0.060865655 | 0.165329035 | -0.368148611 | 0.712762426 | 0.838283978 | 0.060865655 |
| ENSMUSG00000045128 | Rpl18a | 76808 | protein_coding | 3688.605192 | -0.060901039 | 0.128356585 | -0.474467582 | 0.63516651 | 0.788885209 | 0.060901039 |
| ENSMUSG00000050891 | Tatdn1 | 69694 | protein_coding | 159.5954882 | 0.060904323 | 0.11501485 | 0.529534434 | 0.596434764 | 0.764096331 | 0.060904323 |
| ENSMUSG00000039748 | Exo1 | 26909 | protein_coding | 119.5059416 | 0.060908009 | 0.182089698 | 0.334494537 | 0.738006413 | 0.853608159 | 0.060908009 |
| ENSMUSG00000061755 | Bod1l | 665775 | protein_coding | 984.0008317 | -0.060911333 | 0.091351667 | -0.666778565 | 0.504913586 | 0.694916064 | 0.060911333 |
| ENSMUSG00000063065 | Mapk3 | 26417 | protein_coding | 2646.54395 | 0.060929629 | 0.076282789 | 0.798733628 | 0.424444883 | 0.62900615 | 0.060929629 |
| ENSMUSG00000021416 | Eci3 | 69123 | protein_coding | 140.6129339 | -0.060935433 | 0.119701586 | -0.509061202 | 0.610709326 | 0.77314515 | 0.060935433 |
| ENSMUSG00000049091 | Sephs2 | 20768 | protein_coding | 1201.398036 | 0.060937447 | 0.113124477 | 0.538676053 | 0.5901104 | 0.759660426 | 0.060937447 |
| ENSMUSG00000030064 | Frmd4b | 232288 | protein_coding | 998.1179122 | -0.060956075 | 0.130302383 | -0.467804763 | 0.639924214 | 0.791371033 | 0.060956075 |
| ENSMUSG00000071037 | Camkmt | 73582 | protein_coding | 100.7734247 | -0.060973275 | 0.112520966 | -0.541883681 | 0.587898639 | 0.757854445 | 0.060973275 |
| ENSMUSG00000030865 | Chp2 | 70261 | protein_coding | 1385.918655 | -0.060984577 | 0.119090875 | -0.512084382 | 0.608591954 | 0.772321229 | 0.060984577 |
| ENSMUSG00000004591 | Pkn2 | 109333 | protein_coding | 2376.898552 | 0.060993257 | 0.054833928 | 1.112326977 | 0.265997589 | 0.473810064 | 0.060993257 |
| ENSMUSG00000005615 | Pcyt1a | 13026 | protein_coding | 927.3046667 | -0.061026493 | 0.104075409 | -0.58636803 | 0.557628224 | 0.736454853 | 0.061026493 |
| ENSMUSG00000006705 | Pknox1 | 18771 | protein_coding | 294.146462 | -0.061073884 | 0.135155647 | -0.451878151 | 0.651356765 | 0.798250711 | 0.061073884 |
| ENSMUSG00000087598 | Zfp111 | 56707 | protein_coding | 214.8390107 | -0.061138822 | 0.137523175 | -0.444571044 | 0.656629777 | 0.801149235 | 0.061138822 |
| ENSMUSG00000030094 | Xpc | 22591 | protein_coding | 309.8256695 | -0.061153583 | 0.116087156 | -0.526790261 | 0.598339247 | 0.765321556 | 0.061153583 |
| ENSMUSG00000071291 | Zfp58 | 238693 | protein_coding | 65.84534119 | -0.061162375 | 0.167037569 | -0.366159393 | 0.714246141 | 0.83917017 | 0.061162375 |
| ENSMUSG00000021928 | Ebpl | 68177 | protein_coding | 227.0726682 | 0.061169718 | 0.168236428 | 0.363593775 | 0.716161376 | 0.840296015 | 0.061169718 |
| ENSMUSG00000031697 | Orc6 | 56452 | protein_coding | 169.0267363 | 0.061172405 | 0.119719936 | 0.510962564 | 0.609377271 | 0.772513652 | 0.061172405 |
| ENSMUSG00000035834 | Polr3g | 67486 | protein_coding | 221.7631418 | 0.061181378 | 0.124116387 | 0.492935537 | 0.622058137 | 0.780291542 | 0.061181378 |
| ENSMUSG00000032551 | 1110059G10Rik | 66202 | protein_coding | 185.315292 | 0.061213953 | 0.12709409 | 0.481642798 | 0.630059718 | 0.785882948 | 0.061213953 |
| ENSMUSG00000030276 | Ttll3 | 101100 | protein_coding | 86.84342426 | -0.061232192 | 0.224641077 | -0.272577896 | 0.785177705 | 0.883563366 | 0.061232192 |
| ENSMUSG00000068250 | Amn1 | 232566 | protein_coding | 230.9823625 | 0.06126599 | 0.130049938 | 0.471095877 | 0.637572269 | 0.790351042 | 0.06126599 |
| ENSMUSG00000024400 | Wdr33 | 74320 | protein_coding | 865.8603718 | -0.061300601 | 0.067999435 | -0.901486914 | 0.367329488 | 0.576038603 | 0.061300601 |
| ENSMUSG00000074457 | S100a16 | 67860 | protein_coding | 1553.319995 | 0.061331759 | 0.201207251 | 0.304818832 | 0.760504138 | 0.869022879 | 0.061331759 |
| ENSMUSG00000118522 | Gm31160 | 102633301 | protein_coding | 23.49585273 | 0.061372703 | 0.280859519 | 0.218517439 | 0.827025972 | 0.908412958 | 0.061372703 |
| ENSMUSG00000001415 | Smg5 | 229512 | protein_coding | 1012.924607 | -0.061399771 | 0.101914835 | -0.602461567 | 0.546866939 | 0.728832101 | 0.061399771 |
| ENSMUSG00000001472 | Tcf25 | 66855 | protein_coding | 2834.708267 | 0.06141564 | 0.076330469 | 0.804601892 | 0.42104945 | 0.626854726 | 0.06141564 |
| ENSMUSG00000018707 | Dync1h1 | 13424 | protein_coding | 6172.633462 | -0.061417459 | 0.120008431 | -0.511776199 | 0.60880765 | 0.772360968 | 0.061417459 |
| ENSMUSG00000028651 | Ppie | 56031 | protein_coding | 258.5820663 | -0.061458328 | 0.146281208 | -0.42013823 | 0.674384476 | 0.812440509 | 0.061458328 |
| ENSMUSG00000022641 | Bbx | 70508 | protein_coding | 623.6333541 | -0.061460749 | 0.099187793 | -0.619640255 | 0.535494658 | 0.720263377 | 0.061460749 |
| ENSMUSG00000031446 | Cul4a | 99375 | protein_coding | 1354.942176 | 0.061464178 | 0.049900563 | 1.231733162 | 0.218048778 | 0.417839381 | 0.061464178 |
| ENSMUSG00000045252 | Zfp574 | 232976 | protein_coding | 224.2834763 | -0.061472714 | 0.096867247 | -0.634607834 | 0.525684218 | 0.711354843 | 0.061472714 |

| ensembl_gene_id | external_gene_name | entrezgene_id | gene_biotype | baseMean | log2FoldChange | lfcSE | stat | pvalue | padj | absLog2FC |
|---|---|---|---|---|---|---|---|---|---|---|
| ENSMUSG00000032952 | Ap4b1 | 67489 | protein_coding | 72.83311466 | 0.061521787 | 0.193622392 | 0.317741076 | 0.75068135 | 0.862252293 | 0.061521787 |
| ENSMUSG00000078546 | Zfp995 | 70081 | protein_coding | 243.2980717 | 0.061537442 | 0.12651607 | 0.486400206 | 0.626683444 | 0.784047572 | 0.061537442 |
| ENSMUSG00000058444 | Map2k5 | 23938 | protein_coding | 183.4492845 | 0.061554433 | 0.116188987 | 0.529778554 | 0.596265476 | 0.764096331 | 0.061554433 |
| ENSMUSG00000020189 | Osbpl8 | 237542 | protein_coding | 405.1018463 | -0.061589416 | 0.086415352 | -0.712713831 | 0.476022856 | 0.670570635 | 0.061589416 |
| ENSMUSG00000031754 | Nudt21 | 68219 | protein_coding | 454.7880331 | -0.061629651 | 0.088364835 | -0.697445437 | 0.485524075 | 0.678688781 | 0.061629651 |
| ENSMUSG00000027677 | Ttc14 | 67120 | protein_coding | 608.5485452 | -0.06184591 | 0.105625091 | -0.585522904 | 0.55819617 | 0.736676494 | 0.06184591 |
| ENSMUSG00000031360 | Ctps2 | 55936 | protein_coding | 449.6650678 | -0.06188917 | 0.080848012 | -0.765500206 | 0.443973744 | 0.644867207 | 0.06188917 |
| ENSMUSG00000028851 | Nudc | 18221 | protein_coding | 674.7345155 | 0.061940514 | 0.114383266 | 0.541517271 | 0.588151097 | 0.757944002 | 0.061940514 |
| ENSMUSG00000034343 | Ube2f | 67921 | protein_coding | 629.4316629 | 0.062010939 | 0.13182855 | 0.470390813 | 0.638075827 | 0.790438741 | 0.062010939 |
| ENSMUSG00000055932 | Fto | 26383 | protein_coding | 452.4107909 | -0.062059844 | 0.082618912 | -0.751157846 | 0.452557666 | 0.652717363 | 0.062059844 |
| ENSMUSG00000018861 | Fdxr | 14149 | protein_coding | 177.9914773 | -0.062115952 | 0.14428137 | -0.430519563 | 0.666817739 | 0.806888471 | 0.062115952 |
| ENSMUSG00000041135 | Ripk2 | 192656 | protein_coding | 251.347977 | 0.062253172 | 0.12847352 | 0.484560332 | 0.627988258 | 0.78470262 | 0.062253172 |
| ENSMUSG00000024712 | Rfk | 54391 | protein_coding | 6205.435849 | -0.062257287 | 0.129624373 | -0.480289979 | 0.631021213 | 0.786332803 | 0.062257287 |
| ENSMUSG00000020630 | Rnaseh1 | 19819 | protein_coding | 131.9147363 | -0.062302424 | 0.10100291 | -0.616837914 | 0.537341642 | 0.721426926 | 0.062302424 |
| ENSMUSG00000035278 | Plekhj1 | 78670 | protein_coding | 202.2386488 | -0.062307982 | 0.107701476 | -0.578524871 | 0.562909811 | 0.739770497 | 0.062307982 |
| ENSMUSG00000054716 | Zfp771 | 244216 | protein_coding | 85.49165248 | 0.062330306 | 0.184468873 | 0.337890643 | 0.735445599 | 0.851816281 | 0.062330306 |
| ENSMUSG00000085704 | 4921531C22Rik | 66730 | lncRNA | 52.58372485 | -0.062347043 | 0.186019551 | -0.335163925 | 0.737501433 | 0.853241133 | 0.062347043 |
| ENSMUSG00000025290 | Rps24 | 20088 | protein_coding | 10417.13289 | 0.062370973 | 0.088230946 | 0.706905869 | 0.479624977 | 0.673436762 | 0.062370973 |
| ENSMUSG00000036644 | Tbc1d9b | 76795 | protein_coding | 922.4384603 | -0.062377326 | 0.067626625 | -0.922378218 | 0.356331327 | 0.565952498 | 0.062377326 |
| ENSMUSG00000003992 | Ssbp2 | 66970 | protein_coding | 54.99400262 | -0.062383861 | 0.163066472 | -0.38256706 | 0.702040798 | 0.830827389 | 0.062383861 |
| ENSMUSG00000005674 | Tomm40l | 641376 | protein_coding | 131.312365 | 0.062417755 | 0.140906079 | 0.44297418 | 0.657784411 | 0.801989444 | 0.062417755 |
| ENSMUSG00000049940 | Pgrmc2 | 70804 | protein_coding | 478.6216266 | 0.062437747 | 0.092265564 | 0.676717776 | 0.498585036 | 0.689179521 | 0.062437747 |
| ENSMUSG00000035530 | Eif1 | 20918 | protein_coding | 2342.030645 | -0.062441621 | 0.1013815 | -0.615907452 | 0.537955603 | 0.721865534 | 0.062441621 |
| ENSMUSG00000002221 | Paxip1 | 55982 | protein_coding | 364.6663676 | -0.062473497 | 0.091849391 | -0.680173237 | 0.496394776 | 0.687408798 | 0.062473497 |
| ENSMUSG00000005718 | Tfap4 | 83383 | protein_coding | 214.8053261 | 0.06250507 | 0.185484497 | 0.336982717 | 0.736129929 | 0.852237062 | 0.06250507 |
| ENSMUSG00000035021 | Baz1a | 217578 | protein_coding | 623.3259602 | -0.062524403 | 0.212089667 | -0.294801739 | 0.768145338 | 0.873555543 | 0.062524403 |
| ENSMUSG00000022466 | Rpap3 | 71919 | protein_coding | 170.2629571 | -0.062533942 | 0.094712673 | -0.660248942 | 0.50909409 | 0.698075987 | 0.062533942 |
| ENSMUSG00000071648 | Rom1 | 19881 | protein_coding | 29.18010095 | -0.062542409 | 0.200818731 | -0.311437131 | 0.75546833 | 0.8657529 | 0.062542409 |
| ENSMUSG00000021022 | Ppp2r3c | 59032 | protein_coding | 208.1166665 | 0.062550343 | 0.106578449 | 0.586894848 | 0.55727433 | 0.736238102 | 0.062550343 |
| ENSMUSG00000024163 | Mapk8ip3 | 30957 | protein_coding | 452.6079122 | 0.06257677 | 0.177776718 | 0.351996427 | 0.724840942 | 0.844960894 | 0.06257677 |
| ENSMUSG00000040482 | Dxo | 112403 | protein_coding | 305.36106 | 0.062614708 | 0.084185793 | 0.743768107 | 0.457016778 | 0.655781639 | 0.062614708 |
| ENSMUSG00000041303 | Gtf3c3 | 98488 | protein_coding | 470.8556003 | 0.062626475 | 0.073786073 | 0.848757385 | 0.396016307 | 0.604293902 | 0.062626475 |
| ENSMUSG00000025269 | Apex2 | 77622 | protein_coding | 101.594362 | 0.0626895 | 0.139025865 | 0.450919691 | 0.652047431 | 0.798492458 | 0.0626895 |
| ENSMUSG00000064672 | Gm22806 | 115488214 | snoRNA | 7.960382008 | -0.062695491 | 0.455266703 | -0.137711567 | 0.890468377 | 0.944864393 | 0.062695491 |
| ENSMUSG00000022793 | B4galt4 | 56375 | protein_coding | 948.8184979 | 0.062732894 | 0.082776702 | 0.757856886 | 0.448536663 | 0.64942205 | 0.062732894 |
| ENSMUSG00000032463 | Faim | 23873 | protein_coding | 133.1807198 | 0.062747405 | 0.100999161 | 0.621266593 | 0.53442423 | 0.719249909 | 0.062747405 |
| ENSMUSG00000030265 | Kras | 16653 | protein_coding | 1583.825935 | -0.062782794 | 0.061788683 | -1.016088895 | 0.309587057 | 0.51950091 | 0.062782794 |
| ENSMUSG00000030316 | Tamm41 | 68971 | protein_coding | 201.4268496 | 0.062833788 | 0.098757172 | 0.636245312 | 0.524616544 | 0.710274892 | 0.062833788 |

| ensembl_gene_id | external_gene_name | entrezgene_id | gene_biotype | baseMean | log2FoldChange | lfcSE | stat | pvalue | padj | absLog2FC |
|---|---|---|---|---|---|---|---|---|---|---|
| ENSMUSG00000024054 | Smchd1 | 74355 | protein_coding | 1171.861751 | 0.062836176 | 0.079851014 | 0.786917695 | 0.431330048 | 0.635308397 | 0.062836176 |
| ENSMUSG00000025264 | Tsr2 | 69499 | protein_coding | 152.6227823 | -0.062843061 | 0.143346874 | -0.438398547 | 0.6610974 | 0.803875058 | 0.062843061 |
| ENSMUSG00000042148 | Cox10 | 70383 | protein_coding | 524.1004531 | -0.062872525 | 0.105145958 | -0.597954754 | 0.549870124 | 0.731093174 | 0.062872525 |
| ENSMUSG00000020184 | Mdm2 | 17246 | protein_coding | 712.0719358 | -0.062903316 | 0.112919739 | -0.557062185 | 0.577484943 | 0.749220507 | 0.062903316 |
| ENSMUSG00000056204 | Pgpep1 | 66522 | protein_coding | 607.288456 | -0.062913459 | 0.118505934 | -0.530888679 | 0.595495924 | 0.763692357 | 0.062913459 |
| ENSMUSG00000022744 | Cldnd1 | 224250 | protein_coding | 440.6166437 | 0.062926765 | 0.159521285 | 0.394472532 | 0.693232203 | 0.825617057 | 0.062926765 |
| ENSMUSG00000034353 | Ramp1 | 51801 | protein_coding | 894.4966079 | -0.062930489 | 0.147197147 | -0.427525192 | 0.66899684 | 0.808235867 | 0.062930489 |
| ENSMUSG00000027291 | Vps39 | 269338 | protein_coding | 425.2722979 | -0.062948604 | 0.081814801 | -0.769403617 | 0.441653742 | 0.643011318 | 0.062948604 |
| ENSMUSG00000080717 | B230307C23Rik | 245305 | protein_coding | 104.9127377 | 0.062954498 | 0.117179708 | 0.537247437 | 0.591096708 | 0.760438443 | 0.062954498 |
| ENSMUSG00000032120 | C2cd2l | 71764 | protein_coding | 593.8501207 | 0.0630229 | 0.123210061 | 0.511507744 | 0.608995567 | 0.772360968 | 0.0630229 |
| ENSMUSG00000029047 | Pex10 | 668173 | protein_coding | 98.08989331 | -0.063035848 | 0.125019279 | -0.50420902 | 0.614114502 | 0.775272711 | 0.063035848 |
| ENSMUSG00000018740 | Slc25a35 | 71998 | protein_coding | 207.4436736 | 0.063061803 | 0.125028516 | 0.504379363 | 0.613994817 | 0.775245093 | 0.063061803 |
| ENSMUSG00000031605 | Klhl2 | 77113 | protein_coding | 646.7431265 | 0.063062942 | 0.118562201 | 0.531897532 | 0.594796967 | 0.762965747 | 0.063062942 |
| ENSMUSG00000059326 | Csf2ra | 12982 | protein_coding | 141.6478063 | -0.06308543 | 0.131842258 | -0.478491726 | 0.632300262 | 0.787249165 | 0.06308543 |
| ENSMUSG00000001819 | Hoxd13 | 15433 | protein_coding | 121.2785814 | -0.063121233 | 0.794979139 | -0.079399861 | 0.936714579 | 0.968704702 | 0.063121233 |
| ENSMUSG00000024831 | Ighmbp2 | 20589 | protein_coding | 155.5187334 | 0.063134758 | 0.118572328 | 0.532457774 | 0.59440898 | 0.762640729 | 0.063134758 |
| ENSMUSG00000073043 | Atoh1 | 11921 | protein_coding | 1080.292111 | 0.063138198 | 0.246269628 | 0.256378339 | 0.797658712 | 0.891285161 | 0.063138198 |
| ENSMUSG00000020903 | Stx8 | 55943 | protein_coding | 356.0974894 | 0.063301029 | 0.099417736 | 0.636717665 | 0.524308765 | 0.710014454 | 0.063301029 |
| ENSMUSG00000081534 | Slc48a1 | 67739 | protein_coding | 323.4893691 | -0.063321324 | 0.09603991 | -0.659323026 | 0.509688359 | 0.698579013 | 0.063321324 |
| ENSMUSG00000029388 | Eif2b1 | 209354 | protein_coding | 1119.877485 | 0.063321829 | 0.093013798 | 0.680778877 | 0.496011418 | 0.687222389 | 0.063321829 |
| ENSMUSG00000039656 | Rxrb | 20182 | protein_coding | 534.8798546 | -0.063352221 | 0.108125518 | -0.585913684 | 0.557933521 | 0.736573984 | 0.063352221 |
| ENSMUSG00000037058 | Paip2 | 67869 | protein_coding | 1400.754323 | 0.063396673 | 0.092235376 | 0.687335773 | 0.491871161 | 0.683859515 | 0.063396673 |
| ENSMUSG00000026502 | Desi2 | 78825 | protein_coding | 715.9441564 | 0.063406353 | 0.118680378 | 0.534261466 | 0.593160648 | 0.761494544 | 0.063406353 |
| ENSMUSG00000029250 | Polr2b | 231329 | protein_coding | 1125.428238 | 0.063445741 | 0.079334632 | 0.799723142 | 0.423871222 | 0.628565387 | 0.063445741 |
| ENSMUSG00000037337 | Map4k1 | 26411 | protein_coding | 18.59595856 | -0.063478907 | 0.389369429 | -0.163030023 | 0.870494796 | 0.933277015 | 0.063478907 |
| ENSMUSG00000067336 | Bmpr2 | 12168 | protein_coding | 730.6289723 | -0.063514088 | 0.175425585 | -0.362057152 | 0.717309323 | 0.84077584 | 0.063514088 |
| ENSMUSG00000007656 | Arpp19 | 59046 | protein_coding | 1194.815502 | 0.063552127 | 0.066827781 | 0.950983643 | 0.341612679 | 0.551941783 | 0.063552127 |
| ENSMUSG00000031256 | Cstf2 | 108062 | protein_coding | 1096.61857 | 0.063553566 | 0.077472077 | 0.820341582 | 0.412021408 | 0.618889382 | 0.063553566 |
| ENSMUSG00000000915 | Hip1r | 29816 | protein_coding | 1259.617489 | -0.063564045 | 0.103486008 | -0.6142284 | 0.539064409 | 0.722272825 | 0.063564045 |
| ENSMUSG00000057234 | Mettl15 | 76894 | protein_coding | 77.97956514 | -0.063568311 | 0.191729071 | -0.331552805 | 0.740226962 | 0.855115684 | 0.063568311 |
| ENSMUSG00000050812 | Ecpas | 230249 | protein_coding | 2206.317605 | 0.063568933 | 0.072275053 | 0.879541843 | 0.379107556 | 0.587178874 | 0.063568933 |
| ENSMUSG00000031775 | Pllp | 67801 | protein_coding | 199.4399747 | 0.06357115 | 0.146702337 | 0.433334272 | 0.664771944 | 0.805943638 | 0.06357115 |
| ENSMUSG00000029063 | Nadk | 192185 | protein_coding | 2549.316831 | -0.063584382 | 0.110159498 | -0.577202903 | 0.563802396 | 0.740333435 | 0.063584382 |
| ENSMUSG00000020962 | Gtf2a1 | 83602 | protein_coding | 619.2599389 | -0.063607892 | 0.087260557 | -0.728942078 | 0.466037094 | 0.662771324 | 0.063607892 |
| ENSMUSG00000031792 | Usb1 | 101985 | protein_coding | 195.776276 | -0.063624356 | 0.108686238 | -0.585394774 | 0.558282302 | 0.736678642 | 0.063624356 |
| ENSMUSG00000041560 | Nop53 | 68077 | protein_coding | 1607.593425 | -0.063631258 | 0.092990452 | -0.684277324 | 0.493800063 | 0.685544095 | 0.063631258 |
| ENSMUSG00000000804 | Usp32 | 237898 | protein_coding | 709.917692 | -0.063688087 | 0.098063146 | -0.649459955 | 0.516041122 | 0.703363254 | 0.063688087 |
| ENSMUSG00000024695 | Zfp91 | 109910 | protein_coding | 1477.902261 | 0.063697459 | 0.066052609 | 0.964344322 | 0.334873327 | 0.545413471 | 0.063697459 |

| ensembl_gene_id | external_gene_name | entrezgene_id | gene_biotype | baseMean | log2FoldChange | lfcSE | stat | pvalue | padj | absLog2FC |
|---|---|---|---|---|---|---|---|---|---|---|
| ENSMUSG00000029097 | Trmt44 | 78890 | protein_coding | 144.9883823 | -0.063737008 | 0.114490272 | -0.55670239 | 0.577730784 | 0.749417373 | 0.063737008 |
| ENSMUSG00000033161 | Atp1a1 | 11928 | protein_coding | 29139.22692 | -0.063748404 | 0.107994466 | -0.590293252 | 0.554994063 | 0.734641854 | 0.063748404 |
| ENSMUSG00000029726 | Mepce | 231803 | protein_coding | 444.9003515 | -0.06376073 | 0.100794682 | -0.632580294 | 0.527007759 | 0.712457952 | 0.06376073 |
| ENSMUSG00000051329 | Nup160 | 59015 | protein_coding | 516.3013651 | -0.063762728 | 0.102646086 | -0.621190064 | 0.534474576 | 0.719249909 | 0.063762728 |
| ENSMUSG00000043162 | Pyurf | 66459 | protein_coding | 206.2898101 | -0.063816333 | 0.09858083 | -0.647350337 | 0.51740523 | 0.704740081 | 0.063816333 |
| ENSMUSG00000067928 | Zfp760 | 240034 | protein_coding | 127.6599619 | -0.063841032 | 0.20806163 | -0.306837123 | 0.758967351 | 0.868201089 | 0.063841032 |
| ENSMUSG00000039163 | Cmc1 | 67899 | protein_coding | 262.0044739 | 0.063841112 | 0.094175891 | 0.677892313 | 0.497839974 | 0.688791086 | 0.063841112 |
| ENSMUSG00000033542 | Arhgef5 | 54324 | protein_coding | 2198.068683 | -0.063846056 | 0.110529475 | -0.57763828 | 0.563508357 | 0.740191918 | 0.063846056 |
| ENSMUSG00000021911 | Parg | 26430 | protein_coding | 873.9205789 | 0.063862336 | 0.092669521 | 0.689140667 | 0.49073475 | 0.68307464 | 0.063862336 |
| ENSMUSG00000025226 | Fbxl15 | 68431 | protein_coding | 60.45507602 | 0.063879085 | 0.151672806 | 0.42116373 | 0.673635527 | 0.811873015 | 0.063879085 |
| ENSMUSG00000062190 | Lancl2 | 71835 | protein_coding | 243.4332521 | 0.063945245 | 0.09625747 | 0.664314624 | 0.506488963 | 0.696345153 | 0.063945245 |
| ENSMUSG00000086877 | A230072C01Rik | 320742 | lncRNA | 11.90011222 | -0.063959839 | 0.43346331 | -0.147555371 | 0.882693678 | 0.940754985 | 0.063959839 |
| ENSMUSG00000021687 | Scamp1 | 107767 | protein_coding | 2245.851959 | -0.063988124 | 0.127477184 | -0.501957465 | 0.615697441 | 0.776040899 | 0.063988124 |
| ENSMUSG00000024811 | Tnks2 | 74493 | protein_coding | 2710.067906 | 0.064072681 | 0.070983576 | 0.902640929 | 0.366716497 | 0.575473927 | 0.064072681 |
| ENSMUSG00000020669 | Sh3yl1 | 24057 | protein_coding | 544.1247493 | 0.064157865 | 0.083136704 | 0.771715281 | 0.440283078 | 0.642313808 | 0.064157865 |
| ENSMUSG00000027193 | Api5 | 11800 | protein_coding | 2117.090169 | 0.06417312 | 0.073853095 | 0.868929326 | 0.384885788 | 0.592830561 | 0.06417312 |
| ENSMUSG00000036499 | Eea1 | 216238 | protein_coding | 1947.621614 | -0.064195032 | 0.070075638 | -0.91608202 | 0.359623877 | 0.569481508 | 0.064195032 |
| ENSMUSG00000017688 | Hnf4g | 30942 | protein_coding | 534.5918859 | 0.064202518 | 0.094751529 | 0.677588198 | 0.498032831 | 0.688846377 | 0.064202518 |
| ENSMUSG00000022555 | Dgat1 | 13350 | protein_coding | 408.3504687 | -0.064240505 | 0.103417399 | -0.62117696 | 0.534483197 | 0.719249909 | 0.064240505 |
| ENSMUSG00000031832 | Taf1c | 21341 | protein_coding | 178.1761707 | -0.064262583 | 0.139744866 | -0.459856486 | 0.645619235 | 0.795045584 | 0.064262583 |
| ENSMUSG00000024036 | Slc37a1 | 224674 | protein_coding | 2099.65013 | -0.064265504 | 0.10844843 | -0.592590448 | 0.553455273 | 0.733153326 | 0.064265504 |
| ENSMUSG00000037375 | Hhat | 226861 | protein_coding | 50.17037791 | -0.064268873 | 0.238357305 | -0.269632485 | 0.787443006 | 0.884637622 | 0.064268873 |
| ENSMUSG00000025521 | Tmem192 | 73067 | protein_coding | 156.9056063 | -0.064280803 | 0.114870913 | -0.559591638 | 0.575758009 | 0.74850421 | 0.064280803 |
| ENSMUSG00000071646 | Mta2 | 23942 | protein_coding | 1727.302421 | 0.064307252 | 0.110773861 | 0.580527316 | 0.561559071 | 0.739199176 | 0.064307252 |
| ENSMUSG00000002981 | Clptm1 | 56457 | protein_coding | 2294.808733 | -0.064329415 | 0.086351389 | -0.744972561 | 0.456288308 | 0.655296022 | 0.064329415 |
| ENSMUSG00000022811 | Zfp148 | 22661 | protein_coding | 1360.26687 | -0.064332343 | 0.051975118 | -1.237752724 | 0.215807763 | 0.414792572 | 0.064332343 |
| ENSMUSG00000025260 | Hsd17b10 | 15108 | protein_coding | 694.5002104 | -0.064399928 | 0.080146637 | -0.803526263 | 0.421670624 | 0.627406765 | 0.064399928 |
| ENSMUSG00000005148 | Klf5 | 12224 | protein_coding | 4819.012883 | -0.064419056 | 0.107700505 | -0.598131426 | 0.549752243 | 0.731093174 | 0.064419056 |
| ENSMUSG00000039263 | Npepl1 | 228961 | protein_coding | 539.5923804 | -0.064423459 | 0.090321001 | -0.713272201 | 0.475677336 | 0.670351322 | 0.064423459 |
| ENSMUSG00000029625 | Cpsf4 | 54188 | protein_coding | 216.829328 | -0.064473518 | 0.108539108 | -0.594011861 | 0.55250418 | 0.732685587 | 0.064473518 |
| ENSMUSG00000050714 | Zbtb26 | 320633 | protein_coding | 154.5207433 | -0.064480706 | 0.120553958 | -0.534870089 | 0.592739693 | 0.761322298 | 0.064480706 |
| ENSMUSG00000019917 | Septin10 | 103080 | protein_coding | 315.1158825 | -0.064496308 | 0.13015984 | -0.495516187 | 0.620235799 | 0.778936265 | 0.064496308 |
| ENSMUSG00000042599 | Kdm7a | 338523 | protein_coding | 1567.757311 | 0.064568033 | 0.112227287 | 0.575332746 | 0.565066279 | 0.74142208 | 0.064568033 |
| ENSMUSG00000021413 | Prpf4b | 19134 | protein_coding | 1839.735788 | -0.06457975 | 0.065489868 | -0.986102929 | 0.324082607 | 0.534921011 | 0.06457975 |
| ENSMUSG00000038780 | Smurf1 | 75788 | protein_coding | 415.2875556 | -0.064591858 | 0.108499163 | -0.595321253 | 0.551628752 | 0.732245284 | 0.064591858 |
| ENSMUSG00000094806 | Cyp2d10 | 13101 | protein_coding | 149.942139 | -0.064666793 | 0.211041542 | -0.306417366 | 0.759286888 | 0.86831717 | 0.064666793 |
| ENSMUSG00000019558 | Slc6a8 | 102857 | protein_coding | 1864.437248 | 0.064671708 | 0.139523222 | 0.463519313 | 0.642992175 | 0.793244471 | 0.064671708 |
| ENSMUSG00000028081 | Rps3a1 | 20091 | protein_coding | 8505.11513 | 0.06467277 | 0.090675594 | 0.713232383 | 0.475701971 | 0.670351322 | 0.06467277 |

| ensembl_gene_id | external_gene_name | entrezgene_id | gene_biotype | baseMean | log2FoldChange | lfcSE | stat | pvalue | padj | absLog2FC |
|---|---|---|---|---|---|---|---|---|---|---|
| ENSMUSG00000039952 | Dag1 | 13138 | protein_coding | 2410.553632 | -0.064732849 | 0.083685849 | -0.773522044 | 0.439213488 | 0.641698077 | 0.064732849 |
| ENSMUSG00000029500 | Pgam5 | 72542 | protein_coding | 458.6050228 | 0.064756553 | 0.1487747 | 0.435265893 | 0.663369439 | 0.804794609 | 0.064756553 |
| ENSMUSG00000027424 | Mgme1 | 74528 | protein_coding | 155.7703289 | -0.064773165 | 0.112638452 | -0.575053758 | 0.565254941 | 0.741523966 | 0.064773165 |
| ENSMUSG00000073987 | Ggh | 14590 | protein_coding | 2283.72753 | -0.064846705 | 0.255165046 | -0.254136315 | 0.799390249 | 0.892332181 | 0.064846705 |
| ENSMUSG00000020257 | Wdr82 | 77305 | protein_coding | 1295.55829 | -0.064847523 | 0.056725711 | -1.143176916 | 0.252965145 | 0.459205682 | 0.064847523 |
| ENSMUSG00000039959 | Hip1 | 215114 | protein_coding | 9.593962728 | 0.064875334 | 0.758968829 | 0.085478259 | 0.931881179 | 0.966021094 | 0.064875334 |
| ENSMUSG00000035024 | Ncapd3 | 78658 | protein_coding | 528.0315659 | 0.064940268 | 0.08238043 | 0.788297274 | 0.430522842 | 0.634530196 | 0.064940268 |
| ENSMUSG00000005897 | Nr2c1 | 22025 | protein_coding | 165.6429768 | -0.064948262 | 0.103919826 | -0.624984322 | 0.531981348 | 0.716990939 | 0.064948262 |
| ENSMUSG00000033382 | Trappc8 | 75964 | protein_coding | 1439.00677 | 0.064985574 | 0.065275268 | 0.995561956 | 0.319463027 | 0.530650443 | 0.064985574 |
| ENSMUSG00000026857 | Ntmt1 | 66617 | protein_coding | 200.0454831 | -0.065036314 | 0.140452798 | -0.463047481 | 0.643330334 | 0.793445869 | 0.065036314 |
| ENSMUSG00000057522 | Spop | 20747 | protein_coding | 935.3987981 | -0.065040184 | 0.069569025 | -0.934901481 | 0.349839074 | 0.560080085 | 0.065040184 |
| ENSMUSG00000022124 | Fbxl3 | 50789 | protein_coding | 1578.89081 | -0.065057009 | 0.07674052 | -0.847753039 | 0.396575521 | 0.604651202 | 0.065057009 |
| ENSMUSG00000002326 | Gmpr2 | 105446 | protein_coding | 78.16511782 | 0.065059804 | 0.130718829 | 0.497707979 | 0.618689883 | 0.77784296 | 0.065059804 |
| ENSMUSG00000026484 | Rnf2 | 19821 | protein_coding | 455.9626426 | -0.065102114 | 0.096950152 | -0.67150089 | 0.501901493 | 0.692086138 | 0.065102114 |
| ENSMUSG00000018844 | Fndc8 | 78919 | protein_coding | 7.63855384 | 0.065104959 | 0.455327394 | 0.142984938 | 0.886302076 | 0.94220607 | 0.065104959 |
| ENSMUSG00000032745 | Gpbp1 | 73274 | protein_coding | 1330.797834 | 0.065123429 | 0.071247554 | 0.914044418 | 0.360693506 | 0.570122672 | 0.065123429 |
| ENSMUSG00000066877 | Nck2 | 17974 | protein_coding | 1026.408302 | 0.065134295 | 0.076894113 | 0.847064787 | 0.396959011 | 0.604875455 | 0.065134295 |
| ENSMUSG00000028522 | Mier1 | 71148 | protein_coding | 2867.770609 | 0.065204385 | 0.138464441 | 0.470910687 | 0.637704516 | 0.790351042 | 0.065204385 |
| ENSMUSG00000046694 | Tent5b | 100342 | protein_coding | 13.94409345 | 0.065216233 | 0.326507751 | 0.199738699 | 0.841684946 | 0.916390622 | 0.065216233 |
| ENSMUSG00000036813 | Entpd8 | 72090 | protein_coding | 1390.719895 | 0.06523274 | 0.123204879 | 0.529465555 | 0.596482533 | 0.764096331 | 0.06523274 |
| ENSMUSG00000022800 | Fyttd1 | 69823 | protein_coding | 897.8597347 | 0.065284601 | 0.066128662 | 0.987236082 | 0.323526917 | 0.534446871 | 0.065284601 |
| ENSMUSG00000049134 | Nrap | 18175 | protein_coding | 156.8485702 | 0.065339984 | 0.292083599 | 0.223703024 | 0.822988393 | 0.906288656 | 0.065339984 |
| ENSMUSG00000029227 | Fip1l1 | 66899 | protein_coding | 778.4803493 | -0.065364254 | 0.049618765 | -1.317329331 | 0.187728258 | 0.378477508 | 0.065364254 |
| ENSMUSG00000041323 | Ak7 | 78801 | protein_coding | 13.83901487 | -0.065372699 | 0.34223288 | -0.191018172 | 0.848511356 | 0.919988003 | 0.065372699 |
| ENSMUSG00000051726 | Kcnf1 | 382571 | protein_coding | 274.2595066 | -0.065393187 | 0.342180716 | -0.191107166 | 0.848441633 | 0.919988003 | 0.065393187 |
| ENSMUSG00000045210 | Vcpip1 | 70675 | protein_coding | 1022.503741 | 0.065393952 | 0.108070398 | 0.605105126 | 0.545109148 | 0.727725967 | 0.065393952 |
| ENSMUSG00000037824 | Tspan14 | 52588 | protein_coding | 543.2863248 | -0.065427708 | 0.074324109 | -0.880302617 | 0.378695395 | 0.58680059 | 0.065427708 |
| ENSMUSG00000000441 | Raf1 | 110157 | protein_coding | 821.8439699 | 0.065486632 | 0.086979232 | 0.752899633 | 0.451510229 | 0.651785414 | 0.065486632 |
| ENSMUSG00000052446 | Zfp961 | 234413 | protein_coding | 273.8810222 | 0.065494424 | 0.099371925 | 0.659083781 | 0.50984197 | 0.698582542 | 0.065494424 |
| ENSMUSG00000021112 | Pals1 | 56217 | protein_coding | 1242.967689 | 0.065503935 | 0.054823753 | 1.194809382 | 0.232161514 | 0.435607091 | 0.065503935 |
| ENSMUSG00000033423 | Eri3 | 140546 | protein_coding | 405.3469242 | 0.06554095 | 0.104166421 | 0.629194602 | 0.529221662 | 0.714548497 | 0.06554095 |
| ENSMUSG00000020738 | Sumo2 | 170930 | protein_coding | 239.6717201 | 0.065566057 | 0.101071811 | 0.648707654 | 0.516527356 | 0.703829408 | 0.065566057 |
| ENSMUSG00000039230 | Tbcd | 108903 | protein_coding | 973.9978361 | 0.065579389 | 0.072908254 | 0.899478261 | 0.36839797 | 0.577202721 | 0.065579389 |
| ENSMUSG00000055239 | Kcmf1 | 74287 | protein_coding | 939.7609682 | 0.065708618 | 0.087566645 | 0.750384101 | 0.453023404 | 0.652992964 | 0.065708618 |
| ENSMUSG00000038544 | Inip | 66209 | protein_coding | 301.3349796 | 0.065722043 | 0.103491172 | 0.635049748 | 0.52539597 | 0.711101939 | 0.065722043 |
| ENSMUSG00000032370 | Lactb | 80907 | protein_coding | 191.7723042 | -0.06577492 | 0.09659554 | -0.680931231 | 0.495915006 | 0.687162611 | 0.06577492 |
| ENSMUSG00000031812 | Map1lc3b | 67443 | protein_coding | 1239.117583 | -0.065807645 | 0.100389051 | -0.655526117 | 0.512129077 | 0.699887768 | 0.065807645 |
| ENSMUSG00000024096 | Ralbp1 | 19765 | protein_coding | 1286.983915 | 0.065838506 | 0.060071713 | 1.095998464 | 0.273079447 | 0.481045059 | 0.065838506 |

| ensembl_gene_id | external_gene_name | entrezgene_id | gene_biotype | baseMean | log2FoldChange | lfcSE | stat | pvalue | padj | absLog2FC |
|---|---|---|---|---|---|---|---|---|---|---|
| ENSMUSG00000079334 | Naa80 | 56441 | protein_coding | 123.9527649 | -0.065860981 | 0.135729768 | -0.48523608 | 0.627508891 | 0.784362476 | 0.065860981 |
| ENSMUSG00000024218 | Taf11 | 68776 | protein_coding | 230.5645293 | -0.065863465 | 0.088306838 | -0.745847848 | 0.455759332 | 0.655174606 | 0.065863465 |
| ENSMUSG00000036748 | Cuedc2 | 67116 | protein_coding | 251.8539357 | -0.065864832 | 0.100219119 | -0.657208244 | 0.511047025 | 0.699008761 | 0.065864832 |
| ENSMUSG00000062296 | Trank1 | 320429 | protein_coding | 8.249767117 | 0.065887445 | 0.748800506 | 0.087990652 | 0.929884106 | 0.965328339 | 0.065887445 |
| ENSMUSG00000057181 | 5730455P16Rik | 70591 | protein_coding | 186.5459505 | 0.065919494 | 0.107056657 | 0.615743999 | 0.538063493 | 0.721865534 | 0.065919494 |
| ENSMUSG00000068856 | Sf3b4 | 107701 | protein_coding | 178.0828241 | -0.065932045 | 0.146424012 | -0.450281644 | 0.652507373 | 0.798612067 | 0.065932045 |
| ENSMUSG00000035429 | Ptprh | 545902 | protein_coding | 851.3400006 | -0.065939557 | 0.165259722 | -0.399005613 | 0.689889069 | 0.823235922 | 0.065939557 |
| ENSMUSG00000003068 | Stk11 | 20869 | protein_coding | 676.4592222 | 0.065943107 | 0.075076491 | 0.878345618 | 0.379756187 | 0.587657295 | 0.065943107 |
| ENSMUSG00000014980 | Tsen15 | 66637 | protein_coding | 97.18303293 | 0.06594846 | 0.117710069 | 0.560261845 | 0.575300849 | 0.748337437 | 0.06594846 |
| ENSMUSG00000025132 | Arhgdia | 192662 | protein_coding | 2496.103525 | 0.065964311 | 0.126455606 | 0.521640069 | 0.601920959 | 0.767564496 | 0.065964311 |
| ENSMUSG00000025138 | Sirt7 | 209011 | protein_coding | 717.0849698 | -0.065965125 | 0.084594352 | -0.779781667 | 0.435519399 | 0.638409073 | 0.065965125 |
| ENSMUSG00000072623 | Zfp9 | 22750 | protein_coding | 142.9071355 | 0.065978458 | 0.123316131 | 0.535035097 | 0.592625588 | 0.761247936 | 0.065978458 |
| ENSMUSG00000086629 | 2810403D21Rik | 69964 | lncRNA | 45.12481134 | 0.065990249 | 0.236104561 | 0.279495868 | 0.779864308 | 0.880470563 | 0.065990249 |
| ENSMUSG00000022312 | Eif3h | 68135 | protein_coding | 2465.37148 | 0.065998436 | 0.091884126 | 0.718278972 | 0.472585293 | 0.667968635 | 0.065998436 |
| ENSMUSG00000006818 | Sod2 | 20656 | protein_coding | 1433.630173 | -0.06609084 | 0.078424891 | -0.842727843 | 0.399380671 | 0.606939679 | 0.06609084 |
| ENSMUSG00000027993 | Trim2 | 80890 | protein_coding | 5499.51002 | -0.066105542 | 0.086040039 | -0.768311393 | 0.442302208 | 0.643610382 | 0.066105542 |
| ENSMUSG00000021134 | Srsf5 | 20384 | protein_coding | 3520.37841 | -0.066133094 | 0.095339047 | -0.693662211 | 0.48789407 | 0.680726764 | 0.066133094 |
| ENSMUSG00000042369 | Rbm45 | 241490 | protein_coding | 579.0532335 | 0.066138099 | 0.107159952 | 0.617190453 | 0.537109112 | 0.721277627 | 0.066138099 |
| ENSMUSG00000023032 | Slc4a8 | 59033 | protein_coding | 28.62582015 | -0.066164743 | 0.313135617 | -0.211297404 | 0.832655208 | 0.911075919 | 0.066164743 |
| ENSMUSG00000015745 | Plekho1 | 67220 | protein_coding | 16.51204918 | 0.066195352 | 0.304335615 | 0.217507739 | 0.827812676 | 0.90891431 | 0.066195352 |
| ENSMUSG00000052819 | Best2 | 212989 | protein_coding | 403.7316001 | -0.066239448 | 0.436435479 | -0.151773746 | 0.87936539 | 0.938838523 | 0.066239448 |
| ENSMUSG00000004980 | Hnrnpa2b1 | 53379 | protein_coding | 6001.268098 | -0.066244237 | 0.131442375 | -0.503979309 | 0.614275917 | 0.775275463 | 0.066244237 |
| ENSMUSG00000000838 | Fmr1 | 14265 | protein_coding | 1086.179694 | 0.066292784 | 0.099607493 | 0.665540128 | 0.505705086 | 0.695644755 | 0.066292784 |
| ENSMUSG00000040843 | Tiprl | 226591 | protein_coding | 455.1076786 | 0.066337396 | 0.080151589 | 0.82764917 | 0.407869214 | 0.615028007 | 0.066337396 |
| ENSMUSG00000056069 | Otulinl | 223433 | protein_coding | 22.02717377 | 0.066378323 | 0.278845692 | 0.238046795 | 0.8118448 | 0.899802657 | 0.066378323 |
| ENSMUSG00000038084 | Opa1 | 74143 | protein_coding | 1854.646925 | 0.06642467 | 0.071829658 | 0.924752703 | 0.355094558 | 0.564634733 | 0.06642467 |
| ENSMUSG00000089542 | Gm25835 | 115487914 | snoRNA | 139.5984653 | 0.066428019 | 0.336044288 | 0.197676381 | 0.843298271 | 0.917115541 | 0.066428019 |
| ENSMUSG00000006304 | Arpc2 | 76709 | protein_coding | 3318.167787 | 0.066432237 | 0.093239735 | 0.712488479 | 0.476162343 | 0.670644071 | 0.066432237 |
| ENSMUSG00000002455 | Prpf6 | 68879 | protein_coding | 818.7555152 | 0.066433154 | 0.057336346 | 1.158656933 | 0.246596052 | 0.452144805 | 0.066433154 |
| ENSMUSG00000024833 | Pola2 | 18969 | protein_coding | 277.951119 | 0.066449575 | 0.098378207 | 0.675450156 | 0.499389809 | 0.689876919 | 0.066449575 |
| ENSMUSG00000022200 | Golph3 | 66629 | protein_coding | 3026.938318 | 0.066470636 | 0.083577888 | 0.795313663 | 0.426431068 | 0.631010224 | 0.066470636 |
| ENSMUSG00000037104 | Socs5 | 56468 | protein_coding | 161.1521426 | -0.06650259 | 0.139761462 | -0.475829235 | 0.63419604 | 0.788253962 | 0.06650259 |
| ENSMUSG00000021028 | Mbip | 217588 | protein_coding | 138.4538189 | 0.066511416 | 0.117581643 | 0.565661569 | 0.571623865 | 0.745680614 | 0.066511416 |
| ENSMUSG00000034889 | Cactin | 70312 | protein_coding | 340.0177085 | -0.066515467 | 0.107374754 | -0.619470264 | 0.535606605 | 0.720292301 | 0.066515467 |
| ENSMUSG00000056515 | Rab31 | 106572 | protein_coding | 98.18809257 | 0.066571004 | 0.305432504 | 0.217956515 | 0.827462993 | 0.908703155 | 0.066571004 |
| ENSMUSG00000021171 | Esyt2 | 52635 | protein_coding | 2951.640452 | -0.066585035 | 0.083105808 | -0.801207956 | 0.423011266 | 0.627991746 | 0.066585035 |
| ENSMUSG00000032344 | Cgas | 214763 | protein_coding | 71.21888202 | -0.066597605 | 0.178562433 | -0.372965377 | 0.709174213 | 0.835765162 | 0.066597605 |
| ENSMUSG00000039364 | Sectm1b | 58210 | protein_coding | 1414.203714 | -0.066609207 | 0.167878718 | -0.396769809 | 0.691537216 | 0.824338032 | 0.066609207 |

| ensembl_gene_id | external_gene_name | entrezgene_id | gene_biotype | baseMean | log2FoldChange | lfcSE | stat | pvalue | padj | absLog2FC |
|---|---|---|---|---|---|---|---|---|---|---|
| ENSMUSG00000030279 | C2cd5 | 74741 | protein_coding | 456.4965658 | 0.066625256 | 0.072767146 | 0.915595285 | 0.359879205 | 0.569481508 | 0.066625256 |
| ENSMUSG00000043140 | Tmem186 | 66690 | protein_coding | 342.74643 | -0.066713177 | 0.09943435 | -0.670926865 | 0.502267121 | 0.692239286 | 0.066713177 |
| ENSMUSG00000026626 | Ppp2r5a | 226849 | protein_coding | 1852.733922 | 0.066715089 | 0.072858327 | 0.915682405 | 0.359833496 | 0.569481508 | 0.066715089 |
| ENSMUSG00000043962 | Thrap3 | 230753 | protein_coding | 1777.03906 | 0.066728483 | 0.061230686 | 1.089788265 | 0.275806424 | 0.484028727 | 0.066728483 |
| ENSMUSG00000034216 | Vps18 | 228545 | protein_coding | 274.0094231 | -0.066730125 | 0.081504439 | -0.818729946 | 0.412940507 | 0.619566809 | 0.066730125 |
| ENSMUSG00000021337 | Scgn | 214189 | protein_coding | 21.89260001 | -0.066745376 | 0.258442739 | -0.258259824 | 0.796206391 | 0.890338064 | 0.066745376 |
| ENSMUSG00000071359 | Tbpl1 | 237336 | protein_coding | 364.4550793 | 0.06676608 | 0.092232458 | 0.723889201 | 0.46913376 | 0.665392923 | 0.06676608 |
| ENSMUSG00000016018 | Mtrex | 72198 | protein_coding | 826.2997047 | 0.066849302 | 0.07824566 | 0.854351564 | 0.392910225 | 0.600832763 | 0.066849302 |
| ENSMUSG00000004771 | Rab11a | 53869 | protein_coding | 1551.515881 | 0.066859208 | 0.122471469 | 0.545916598 | 0.585123269 | 0.755622136 | 0.066859208 |
| ENSMUSG00000032024 | Clmp | 71566 | protein_coding | 52.45719667 | 0.06686884 | 0.351811867 | 0.190069882 | 0.849254371 | 0.92046263 | 0.06686884 |
| ENSMUSG00000036402 | Gng12 | 14701 | protein_coding | 2908.381298 | -0.066905604 | 0.058148862 | -1.150591799 | 0.249900208 | 0.45620751 | 0.066905604 |
| ENSMUSG00000102440 | Pcdhga9 | 93717 | protein_coding | 14.02229028 | -0.066929471 | 0.43767834 | -0.152919313 | 0.878461906 | 0.93834699 | 0.066929471 |
| ENSMUSG00000028878 | Fam76a | 230789 | protein_coding | 339.4218302 | -0.066931722 | 0.102783706 | -0.651190001 | 0.514923844 | 0.702381347 | 0.066931722 |
| ENSMUSG00000087679 | Tmem250-ps | 109299 | protein_coding | 661.29981 | -0.066939695 | 0.081890928 | -0.817425047 | 0.413685567 | 0.62021684 | 0.066939695 |
| ENSMUSG00000028249 | Sdcbp | 53378 | protein_coding | 4017.880606 | 0.0669438 | 0.116418764 | 0.575025866 | 0.565273803 | 0.741523966 | 0.0669438 |
| ENSMUSG00000031078 | Cttn | 13043 | protein_coding | 3658.904281 | 0.067010394 | 0.088101743 | 0.760602366 | 0.446894606 | 0.647692201 | 0.067010394 |
| ENSMUSG00000027981 | Rnpc3 | 67225 | protein_coding | 201.215712 | -0.067057902 | 0.092760835 | -0.722911797 | 0.469734075 | 0.665888604 | 0.067057902 |
| ENSMUSG00000048826 | Dact2 | 240025 | protein_coding | 264.7551995 | -0.067066531 | 0.134131939 | -0.50000419 | 0.617072127 | 0.777097157 | 0.067066531 |
| ENSMUSG00000029072 | Tas1r3 | 83771 | protein_coding | 40.2814954 | 0.06708137 | 0.217722244 | 0.308105269 | 0.758002232 | 0.867408546 | 0.06708137 |
| ENSMUSG00000041671 | Pyroxd1 | 232491 | protein_coding | 274.7401867 | -0.067198431 | 0.087984514 | -0.763752937 | 0.445014488 | 0.645771004 | 0.067198431 |
| ENSMUSG00000015542 | Nat9 | 66176 | protein_coding | 226.8960885 | 0.067199806 | 0.126352302 | 0.531844728 | 0.594833541 | 0.762965747 | 0.067199806 |
| ENSMUSG00000061613 | U2af1 | 108121 | protein_coding | 581.280555 | -0.06728568 | 0.129113399 | -0.521136311 | 0.602271818 | 0.767564496 | 0.06728568 |
| ENSMUSG00000022404 | Slc25a17 | 20524 | protein_coding | 752.5551664 | 0.06730564 | 0.097850364 | 0.687842507 | 0.491551965 | 0.683614747 | 0.06730564 |
| ENSMUSG00000027339 | Rassf2 | 215653 | protein_coding | 21.02738063 | 0.067313266 | 0.367038378 | 0.183395716 | 0.854487537 | 0.923481985 | 0.067313266 |
| ENSMUSG00000024056 | Ndc80 | 67052 | protein_coding | 235.4811153 | 0.067344534 | 0.172518342 | 0.390361591 | 0.696269186 | 0.827131234 | 0.067344534 |
| ENSMUSG00000061718 | Ppp1r1b | 19049 | protein_coding | 2185.665412 | -0.067357275 | 0.08612871 | -0.782053686 | 0.434183023 | 0.637531118 | 0.067357275 |
| ENSMUSG00000047423 | AI837181 | 107242 | protein_coding | 226.7965739 | 0.067367082 | 0.137286796 | 0.490703285 | 0.623636323 | 0.781840169 | 0.067367082 |
| ENSMUSG00000029233 | Srd5a3 | 57357 | protein_coding | 204.5338435 | 0.067399772 | 0.109116052 | 0.61768888 | 0.536780444 | 0.721206302 | 0.067399772 |
| ENSMUSG00000040859 | Bsdc1 | 100383 | protein_coding | 1197.413705 | 0.067428821 | 0.109579571 | 0.615341164 | 0.538329438 | 0.721975982 | 0.067428821 |
| ENSMUSG00000021054 | Sgpp1 | 81535 | protein_coding | 614.1185007 | 0.067464779 | 0.092384541 | 0.730260471 | 0.465230985 | 0.662210133 | 0.067464779 |
| ENSMUSG00000033917 | Gde1 | 56209 | protein_coding | 653.0727969 | 0.067515202 | 0.113599475 | 0.594326711 | 0.552293617 | 0.732528331 | 0.067515202 |
| ENSMUSG00000066440 | Zfyve26 | 211978 | protein_coding | 332.2256738 | -0.06752724 | 0.134898676 | -0.500577484 | 0.616668512 | 0.776895995 | 0.06752724 |
| ENSMUSG00000018362 | Kpna2 | 16647 | protein_coding | 2111.237231 | 0.067544414 | 0.111100828 | 0.607955986 | 0.543216666 | 0.725990523 | 0.067544414 |
| ENSMUSG00000096956 | Snhg18 | 100616095 | lncRNA | 532.6228671 | 0.067583006 | 0.10277405 | 0.657588236 | 0.510802756 | 0.699008761 | 0.067583006 |
| ENSMUSG00000051034 | Zfp11 | 22648 | protein_coding | 39.74177512 | 0.067642588 | 0.211087656 | 0.320447863 | 0.748628847 | 0.860577387 | 0.067642588 |
| ENSMUSG00000040010 | Slc7a5 | 20539 | protein_coding | 178.3880244 | -0.06764487 | 0.328730907 | -0.205775814 | 0.836966035 | 0.913846655 | 0.06764487 |
| ENSMUSG00000021097 | Clmn | 94040 | protein_coding | 2247.721558 | -0.067715671 | 0.146726982 | -0.461507968 | 0.644434206 | 0.794206246 | 0.067715671 |
| ENSMUSG00000019897 | Ccdc59 | 52713 | protein_coding | 304.6557651 | 0.067716234 | 0.097416716 | 0.695119241 | 0.486980578 | 0.679869107 | 0.067716234 |

| ensembl_gene_id | external_gene_name | entrezgene_id | gene_biotype | baseMean | log2FoldChange | lfcSE | stat | pvalue | padj | absLog2FC |
|---|---|---|---|---|---|---|---|---|---|---|
| ENSMUSG00000059975 | Zfp74 | 72723 | protein_coding | 67.68566766 | 0.067721833 | 0.209050998 | 0.323948864 | 0.745976749 | 0.858768261 | 0.067721833 |
| ENSMUSG00000013646 | Sh3bp5l | 79566 | protein_coding | 416.0565133 | 0.067729751 | 0.125757385 | 0.538574743 | 0.590180319 | 0.759689035 | 0.067729751 |
| ENSMUSG00000020921 | Tmem101 | 76547 | protein_coding | 128.0214554 | 0.067734428 | 0.134569089 | 0.503343144 | 0.614723038 | 0.775487767 | 0.067734428 |
| ENSMUSG00000048186 | Bend7 | 209645 | protein_coding | 128.6333707 | 0.067740855 | 0.10329233 | 0.655816889 | 0.511941949 | 0.699812166 | 0.067740855 |
| ENSMUSG00000041775 | Mapk1ip1 | 69546 | protein_coding | 89.42510665 | 0.067791731 | 0.179982887 | 0.376656534 | 0.706428869 | 0.834011211 | 0.067791731 |
| ENSMUSG00000072945 | Ripply1 | 622473 | protein_coding | 14.022409 | -0.067802997 | 0.29544713 | -0.229492828 | 0.818485895 | 0.903579383 | 0.067802997 |
| ENSMUSG00000036565 | Ttyh3 | 78339 | protein_coding | 329.1055241 | -0.067828853 | 0.149609527 | -0.453372549 | 0.6502805 | 0.797971325 | 0.067828853 |
| ENSMUSG00000029557 | Mrm2 | 68017 | protein_coding | 87.51424155 | 0.067905171 | 0.167646046 | 0.405050835 | 0.685440145 | 0.82050884 | 0.067905171 |
| ENSMUSG00000030431 | Tmem238 | 664968 | protein_coding | 177.7322481 | -0.067985884 | 0.197437356 | -0.344341544 | 0.730589444 | 0.848536435 | 0.067985884 |
| ENSMUSG00000101856 | 1700096K18Rik | 73571 | lncRNA | 35.91922401 | 0.068007027 | 0.220418637 | 0.308535737 | 0.757674711 | 0.867407662 | 0.068007027 |
| ENSMUSG00000052407 | Ccdc171 | 320226 | protein_coding | 79.87174718 | -0.068028858 | 0.181654764 | -0.374495312 | 0.708035844 | 0.835135786 | 0.068028858 |
| ENSMUSG00000004768 | Rab23 | 19335 | protein_coding | 88.94866088 | -0.068053161 | 0.141428317 | -0.481184834 | 0.630385139 | 0.78600125 | 0.068053161 |
| ENSMUSG00000033126 | Ybey | 216119 | protein_coding | 83.77703704 | -0.068068335 | 0.20176405 | -0.337366021 | 0.735840996 | 0.852088359 | 0.068068335 |
| ENSMUSG00000049305 | Ccdc71 | 72454 | protein_coding | 422.7105897 | 0.06807312 | 0.097535873 | 0.697929056 | 0.485221562 | 0.678589433 | 0.06807312 |
| ENSMUSG00000035443 | Thyn1 | 77862 | protein_coding | 335.93736 | 0.068097882 | 0.104693016 | 0.650452957 | 0.51539968 | 0.702849837 | 0.068097882 |
| ENSMUSG00000052544 | St6galnac3 | 20447 | protein_coding | 6.851462354 | 0.06815079 | 0.832586329 | 0.081854323 | 0.934762558 | 0.967565507 | 0.06815079 |
| ENSMUSG00000032555 | Topbp1 | 235559 | protein_coding | 945.0751371 | 0.068169531 | 0.084678703 | 0.8050375 | 0.42079804 | 0.626692023 | 0.068169531 |
| ENSMUSG00000028234 | Rps20 | 67427 | protein_coding | 5266.48999 | 0.068174925 | 0.122109308 | 0.558310631 | 0.576632287 | 0.748785184 | 0.068174925 |
| ENSMUSG00000018900 | Slc22a5 | 20520 | protein_coding | 440.1940823 | -0.068181311 | 0.09078213 | -0.75104331 | 0.452626591 | 0.652733032 | 0.068181311 |
| ENSMUSG00000030386 | Zfp606 | 67370 | protein_coding | 273.9486198 | 0.068214959 | 0.162013364 | 0.421045259 | 0.673722033 | 0.811873015 | 0.068214959 |
| ENSMUSG00000053483 | Usp21 | 30941 | protein_coding | 211.413379 | -0.068215101 | 0.091020155 | -0.749450506 | 0.45358572 | 0.653148289 | 0.068215101 |
| ENSMUSG00000039703 | Nploc4 | 217365 | protein_coding | 878.9192828 | 0.068272101 | 0.114978391 | 0.593782016 | 0.552657919 | 0.73282845 | 0.068272101 |
| ENSMUSG00000040123 | Zmym5 | 219105 | protein_coding | 592.6519091 | -0.068302536 | 0.069583021 | -0.98159774 | 0.326298062 | 0.537352722 | 0.068302536 |
| ENSMUSG00000007892 | Rplp1 | 56040 | protein_coding | 8660.982034 | 0.068319153 | 0.098045771 | 0.696808765 | 0.48592248 | 0.678962412 | 0.068319153 |
| ENSMUSG00000032187 | Smarca4 | 20586 | protein_coding | 1773.155274 | -0.068352904 | 0.086599544 | -0.789298656 | 0.429937472 | 0.633960753 | 0.068352904 |
| ENSMUSG00000040356 | Skiv2l | 108077 | protein_coding | 990.7054794 | 0.068353423 | 0.085144812 | 0.802790233 | 0.422095988 | 0.627572772 | 0.068353423 |
| ENSMUSG00000001962 | Fam50a | 108160 | protein_coding | 289.9121733 | 0.068379147 | 0.104621422 | 0.653586481 | 0.513378256 | 0.700993473 | 0.068379147 |
| ENSMUSG00000047227 | Gm527 | 217648 | protein_coding | 30.70777567 | -0.068404107 | 0.215347138 | -0.317645767 | 0.750753653 | 0.862273161 | 0.068404107 |
| ENSMUSG00000024805 | Pcgf5 | 76073 | protein_coding | 520.315493 | 0.068473252 | 0.138257207 | 0.495259911 | 0.620416665 | 0.778968677 | 0.068473252 |
| ENSMUSG00000066151 | Fkbp15 | 338355 | protein_coding | 740.5983769 | -0.068487359 | 0.084161775 | -0.813758495 | 0.415783316 | 0.622176792 | 0.068487359 |
| ENSMUSG00000039958 | Etfbkmt | 320204 | protein_coding | 240.8367524 | -0.068542048 | 0.215298098 | -0.318358817 | 0.750212774 | 0.861962711 | 0.068542048 |
| ENSMUSG00000028656 | Cap1 | 12331 | protein_coding | 3434.281081 | 0.068558612 | 0.121288104 | 0.565254211 | 0.571900868 | 0.745802933 | 0.068558612 |
| ENSMUSG00000032905 | Atg12 | 67526 | protein_coding | 849.5937244 | 0.068561332 | 0.082678341 | 0.82925384 | 0.406960785 | 0.614065118 | 0.068561332 |
| ENSMUSG00000096910 | Zfp955b | 100043468 | protein_coding | 276.1375633 | -0.068570399 | 0.117958782 | -0.581308126 | 0.561032805 | 0.738767404 | 0.068570399 |
| ENSMUSG00000054733 | Msra | 110265 | protein_coding | 496.4163216 | -0.068577139 | 0.095952263 | -0.714700584 | 0.474794077 | 0.669646611 | 0.068577139 |
| ENSMUSG00000074698 | Csnk2a1 | 12995 | protein_coding | 1943.47588 | -0.068605738 | 0.070697915 | -0.970406809 | 0.331843756 | 0.542683099 | 0.068605738 |
| ENSMUSG00000039254 | Pomt1 | 99011 | protein_coding | 349.8729794 | -0.068646835 | 0.079099463 | -0.867854622 | 0.385473923 | 0.593449512 | 0.068646835 |
| ENSMUSG00000022635 | Zcrb1 | 67197 | protein_coding | 400.5095794 | 0.068672779 | 0.088179977 | 0.778779741 | 0.436109474 | 0.638980011 | 0.068672779 |

| ensembl_gene_id | external_gene_name | entrezgene_id | gene_biotype | baseMean | log2FoldChange | lfcSE | stat | pvalue | padj | absLog2FC |
|---|---|---|---|---|---|---|---|---|---|---|
| ENSMUSG00000049323 | Smcr8 | 237782 | protein_coding | 563.0864397 | -0.068676744 | 0.113577436 | -0.604668907 | 0.545399011 | 0.727868911 | 0.068676744 |
| ENSMUSG00000010392 | Gosr1 | 53334 | protein_coding | 825.9351221 | 0.068681527 | 0.074557722 | 0.921185967 | 0.356953344 | 0.566572283 | 0.068681527 |
| ENSMUSG00000050730 | Arhgap42 | 71544 | protein_coding | 522.8981794 | 0.068689922 | 0.107127969 | 0.641195042 | 0.521395971 | 0.707815138 | 0.068689922 |
| ENSMUSG00000027207 | Galk2 | 69976 | protein_coding | 824.2226158 | 0.068782562 | 0.082471598 | 0.834015155 | 0.404272444 | 0.611350704 | 0.068782562 |
| ENSMUSG00000037957 | Wdr20 | 69641 | protein_coding | 418.1972766 | 0.068810975 | 0.073257006 | 0.939309127 | 0.347572054 | 0.557627687 | 0.068810975 |
| ENSMUSG00000020777 | Acox1 | 11430 | protein_coding | 3826.559322 | -0.06881442 | 0.137694568 | -0.49976133 | 0.617243143 | 0.77714186 | 0.06881442 |
| ENSMUSG00000062981 | Mrpl42 | 67270 | protein_coding | 901.6836061 | 0.068842033 | 0.094593 | 0.727770897 | 0.466753843 | 0.66367213 | 0.068842033 |
| ENSMUSG00000033751 | Gadd45gip1 | 102060 | protein_coding | 187.0580651 | 0.068865414 | 0.133673501 | 0.515176256 | 0.606429859 | 0.770948583 | 0.068865414 |
| ENSMUSG00000036537 | Rnf113a1 | 69942 | protein_coding | 29.36434853 | 0.068868308 | 0.238455356 | 0.288810068 | 0.772726727 | 0.876325476 | 0.068868308 |
| ENSMUSG00000052117 | D630039A03Rik | 242484 | protein_coding | 550.4382528 | -0.068878493 | 0.238416182 | -0.288900244 | 0.772657717 | 0.876325476 | 0.068878493 |
| ENSMUSG00000029195 | Klb | 83379 | protein_coding | 91.69076303 | 0.068964553 | 0.362761497 | 0.190109903 | 0.84922301 | 0.92046263 | 0.068964553 |
| ENSMUSG00000028927 | Padi2 | 18600 | protein_coding | 134.4546716 | 0.068987356 | 0.192215485 | 0.358906337 | 0.719665159 | 0.842364374 | 0.068987356 |
| ENSMUSG00000047921 | Trappc9 | 76510 | protein_coding | 516.0980072 | -0.068994102 | 0.097838107 | -0.705186397 | 0.480694246 | 0.674079929 | 0.068994102 |
| ENSMUSG00000022387 | Brd1 | 223770 | protein_coding | 1371.61911 | -0.06900092 | 0.062855361 | -1.097773035 | 0.272303609 | 0.48047545 | 0.06900092 |
| ENSMUSG00000028991 | Mtor | 56717 | protein_coding | 945.4567483 | 0.069002632 | 0.077663133 | 0.888486333 | 0.374279204 | 0.582634789 | 0.069002632 |
| ENSMUSG00000021948 | Prkcd | 18753 | protein_coding | 2512.602289 | -0.069098376 | 0.09473896 | -0.729355441 | 0.465784267 | 0.662531353 | 0.069098376 |
| ENSMUSG00000032598 | Nckipsd | 80987 | protein_coding | 84.40848727 | 0.069170756 | 0.136091536 | 0.508266407 | 0.611266525 | 0.773335037 | 0.069170756 |
| ENSMUSG00000027351 | Spred1 | 114715 | protein_coding | 821.1686338 | -0.069188863 | 0.09396905 | -0.736294164 | 0.461551697 | 0.65958881 | 0.069188863 |
| ENSMUSG00000013465 | Nelfb | 58202 | protein_coding | 886.1011189 | -0.069205496 | 0.095344428 | -0.725847306 | 0.467932383 | 0.664576638 | 0.069205496 |
| ENSMUSG00000070003 | Ssbp4 | 76900 | protein_coding | 374.51107 | 0.069245738 | 0.10478563 | 0.660832391 | 0.508719808 | 0.697877891 | 0.069245738 |
| ENSMUSG00000031029 | Eif3f | 66085 | protein_coding | 2434.110715 | 0.06926313 | 0.096685716 | 0.716373969 | 0.473760464 | 0.668798783 | 0.06926313 |
| ENSMUSG00000025968 | Ndufs1 | 227197 | protein_coding | 2743.592392 | 0.069281979 | 0.087284911 | 0.793745195 | 0.427343783 | 0.631814032 | 0.069281979 |
| ENSMUSG00000027475 | Kif3b | 16569 | protein_coding | 782.1578733 | 0.069293408 | 0.078156069 | 0.886603045 | 0.375292654 | 0.583813008 | 0.069293408 |
| ENSMUSG00000024299 | Adamts10 | 224697 | protein_coding | 46.91224615 | -0.069296836 | 0.272107031 | -0.254667571 | 0.798979865 | 0.89211142 | 0.069296836 |
| ENSMUSG00000052917 | Senp7 | 66315 | protein_coding | 260.6689041 | 0.069307291 | 0.144202052 | 0.480626247 | 0.630782158 | 0.786280968 | 0.069307291 |
| ENSMUSG00000091803 | Cox16 | 66272 | protein_coding | 141.572545 | -0.06931263 | 0.118628687 | -0.584282196 | 0.559030469 | 0.737105737 | 0.06931263 |
| ENSMUSG00000010097 | Nxf1 | 53319 | protein_coding | 896.2338628 | -0.069339912 | 0.086418234 | -0.802375945 | 0.422335524 | 0.627648847 | 0.069339912 |
| ENSMUSG00000030188 | Magohb | 66441 | protein_coding | 163.4019737 | -0.069369707 | 0.132031301 | -0.5254035 | 0.599302721 | 0.76592263 | 0.069369707 |
| ENSMUSG00000020691 | Mettl2 | 52686 | protein_coding | 235.5505834 | -0.069372443 | 0.115655565 | -0.599819327 | 0.548626652 | 0.730340602 | 0.069372443 |
| ENSMUSG00000043923 | Ccdc84 | 382073 | protein_coding | 159.0012609 | -0.069400461 | 0.140765489 | -0.493021847 | 0.621997151 | 0.780276492 | 0.069400461 |
| ENSMUSG00000038538 | Ubn2 | 320538 | protein_coding | 789.830266 | 0.069431857 | 0.155760081 | 0.44576156 | 0.655769491 | 0.800758698 | 0.069431857 |
| ENSMUSG00000004530 | Coro1c | 23790 | protein_coding | 1879.358514 | -0.069439954 | 0.135728727 | -0.511608381 | 0.608925119 | 0.772360968 | 0.069439954 |
| ENSMUSG00000059070 | Rpl18 | 19899 | protein_coding | 6145.099951 | 0.069455913 | 0.090748088 | 0.765370538 | 0.444050932 | 0.644920511 | 0.069455913 |
| ENSMUSG00000031174 | Rpgr | 19893 | protein_coding | 118.9624421 | 0.069484385 | 0.108956195 | 0.637727712 | 0.523650943 | 0.709654498 | 0.069484385 |
| ENSMUSG00000044533 | Rps2 | 16898 | protein_coding | 11168.60623 | 0.069530146 | 0.116437466 | 0.597145821 | 0.550410028 | 0.731458975 | 0.069530146 |
| ENSMUSG00000005846 | Rsl1d1 | 66409 | protein_coding | 999.9068081 | -0.069530832 | 0.134354182 | -0.517518927 | 0.60479396 | 0.769421581 | 0.069530832 |
| ENSMUSG00000021518 | Ptdss1 | 19210 | protein_coding | 1464.831638 | 0.069531312 | 0.111508089 | 0.623553971 | 0.532920547 | 0.718013744 | 0.069531312 |
| ENSMUSG00000085196 | Gm14963 | 105246847 | lncRNA | 25.66566449 | 0.069547134 | 0.284301658 | 0.24462444 | 0.806747238 | 0.896335799 | 0.069547134 |

| ensembl_gene_id | external_gene_name | entrezgene_id | gene_biotype | baseMean | log2FoldChange | lfcSE | stat | pvalue | padj | absLog2FC |
|---|---|---|---|---|---|---|---|---|---|---|
| ENSMUSG00000027487 | Cdk5rap1 | 66971 | protein_coding | 208.4971823 | -0.069578801 | 0.247828125 | -0.280754256 | 0.778898891 | 0.879817143 | 0.069578801 |
| ENSMUSG00000008035 | Mid1ip1 | 68041 | protein_coding | 1087.235156 | -0.069589736 | 0.206401408 | -0.337157273 | 0.735998344 | 0.852208609 | 0.069589736 |
| ENSMUSG00000037966 | Ninj1 | 18081 | protein_coding | 184.2116479 | -0.069601762 | 0.115253735 | -0.603900273 | 0.545909947 | 0.728245698 | 0.069601762 |
| ENSMUSG00000005501 | Usp40 | 227334 | protein_coding | 480.3226629 | -0.06961977 | 0.072401858 | -0.961574359 | 0.336263457 | 0.546990445 | 0.06961977 |
| ENSMUSG00000063260 | Syt10 | 54526 | protein_coding | 12.43147212 | -0.069653702 | 0.386657055 | -0.180143363 | 0.857040021 | 0.925280429 | 0.069653702 |
| ENSMUSG00000080059 | Rps19-ps3 | 277692 | processed_pseudogene | 8.586371161 | 0.069669575 | 0.365306091 | 0.190715614 | 0.848748405 | 0.920151416 | 0.069669575 |
| ENSMUSG00000026083 | Eif5b | 226982 | protein_coding | 666.3401732 | -0.069714785 | 0.148368133 | -0.469877083 | 0.638442838 | 0.790509758 | 0.069714785 |
| ENSMUSG00000020226 | Slc5a4b | 64454 | protein_coding | 14.39870447 | 0.069755909 | 0.356582091 | 0.195623702 | 0.844904707 | 0.918121002 | 0.069755909 |
| ENSMUSG00000024999 | Noc3l | 57753 | protein_coding | 259.1046554 | -0.069765236 | 0.178808996 | -0.390166256 | 0.696413612 | 0.827182798 | 0.069765236 |
| ENSMUSG00000038848 | Ythdf1 | 228994 | protein_coding | 530.6561988 | -0.069787411 | 0.096783426 | -0.721067789 | 0.470867808 | 0.666534157 | 0.069787411 |
| ENSMUSG00000021640 | Naip1 | 17940 | protein_coding | 3098.146394 | 0.069852824 | 0.130551166 | 0.5350609 | 0.592607746 | 0.761247936 | 0.069852824 |
| ENSMUSG00000074733 | Zfp950 | 414758 | protein_coding | 358.5285784 | -0.069922033 | 0.161260103 | -0.433597843 | 0.664580501 | 0.80582704 | 0.069922033 |
| ENSMUSG00000037379 | Spon2 | 100689 | protein_coding | 22.62363776 | 0.069938881 | 0.280734386 | 0.249128304 | 0.803261536 | 0.894249532 | 0.069938881 |
| ENSMUSG00000024847 | Aip | 11632 | protein_coding | 286.6510285 | 0.069944429 | 0.089386757 | 0.782492079 | 0.433925439 | 0.637453749 | 0.069944429 |
| ENSMUSG00000018509 | Cenpv | 73139 | protein_coding | 389.3358538 | 0.069945514 | 0.097371989 | 0.71833301 | 0.472551981 | 0.667968635 | 0.069945514 |
| ENSMUSG00000058216 | Gstp3 | 225884 | protein_coding | 43.2171476 | -0.069989809 | 0.313005237 | -0.22360587 | 0.823063995 | 0.906309251 | 0.069989809 |
| ENSMUSG00000021124 | Vti1b | 53612 | protein_coding | 795.5938791 | 0.070107027 | 0.075739185 | 0.925637461 | 0.354634418 | 0.564128335 | 0.070107027 |
| ENSMUSG00000021987 | Mtmr6 | 219135 | protein_coding | 2072.737354 | -0.070117187 | 0.086872017 | -0.807132021 | 0.419590425 | 0.625680983 | 0.070117187 |
| ENSMUSG00000024974 | Smc3 | 13006 | protein_coding | 1223.327229 | -0.070195624 | 0.062945494 | -1.115181085 | 0.26477283 | 0.472220655 | 0.070195624 |
| ENSMUSG00000030469 | Zfp719 | 210105 | protein_coding | 224.0927918 | -0.070224358 | 0.100998807 | -0.69529889 | 0.48686801 | 0.679771534 | 0.070224358 |
| ENSMUSG00000027722 | Spata5 | 57815 | protein_coding | 296.6372381 | -0.070267513 | 0.127359172 | -0.551727151 | 0.581135305 | 0.75217968 | 0.070267513 |
| ENSMUSG00000039232 | Stx11 | 74732 | protein_coding | 8.296484602 | -0.070268409 | 0.619486172 | -0.113430149 | 0.909689538 | 0.955395949 | 0.070268409 |
| ENSMUSG00000071076 | Jund | 16478 | protein_coding | 1295.145998 | -0.070306135 | 0.144329306 | -0.487123068 | 0.62617112 | 0.783599434 | 0.070306135 |
| ENSMUSG00000049600 | Zbtb45 | 232879 | protein_coding | 111.9124015 | 0.070318812 | 0.138010058 | 0.509519471 | 0.610388154 | 0.773061998 | 0.070318812 |
| ENSMUSG00000028247 | Coq3 | 230027 | protein_coding | 172.3993685 | -0.070347681 | 0.16831578 | -0.4179506 | 0.675983236 | 0.813959856 | 0.070347681 |
| ENSMUSG00000028643 | Svbp | 69216 | protein_coding | 141.0175015 | 0.070382448 | 0.124188108 | 0.566740643 | 0.570890403 | 0.74521737 | 0.070382448 |
| ENSMUSG00000026317 | Cln8 | 26889 | protein_coding | 232.0193208 | 0.070442234 | 0.089536807 | 0.786740517 | 0.431433781 | 0.635343681 | 0.070442234 |
| ENSMUSG00000037487 | Ubr5 | 70790 | protein_coding | 3377.072882 | -0.070452297 | 0.071399727 | -0.986730626 | 0.323774712 | 0.534726009 | 0.070452297 |
| ENSMUSG00000053205 | Styx | 56291 | protein_coding | 295.0713073 | -0.07046804 | 0.174908604 | -0.402884925 | 0.687032882 | 0.821304393 | 0.07046804 |
| ENSMUSG00000015342 | Xk | 22439 | protein_coding | 382.6550229 | 0.070555044 | 0.08959349 | 0.7875019 | 0.430988117 | 0.635002887 | 0.070555044 |
| ENSMUSG00000030200 | Bcl2l14 | 66813 | protein_coding | 1253.394392 | 0.070740967 | 0.109169794 | 0.647990295 | 0.516991227 | 0.704236404 | 0.070740967 |
| ENSMUSG00000026153 | Fam135a | 68187 | protein_coding | 1395.597428 | -0.07075332 | 0.091623574 | -0.772217418 | 0.439985667 | 0.642313808 | 0.07075332 |
| ENSMUSG00000025024 | Smndc1 | 76479 | protein_coding | 401.8777639 | -0.070755333 | 0.084838183 | -0.834003403 | 0.404279066 | 0.611350704 | 0.070755333 |
| ENSMUSG00000010095 | Slc3a2 | 17254 | protein_coding | 529.1217269 | 0.070770503 | 0.08055125 | 0.878577343 | 0.379630485 | 0.587634049 | 0.070770503 |
| ENSMUSG00000021226 | Acot2 | 171210 | protein_coding | 94.68359008 | 0.070787801 | 0.145216742 | 0.487463084 | 0.625930198 | 0.783359487 | 0.070787801 |
| ENSMUSG00000008301 | Phax | 56698 | protein_coding | 425.9699038 | -0.070888811 | 0.095674209 | -0.740939608 | 0.458730059 | 0.65713215 | 0.070888811 |
| ENSMUSG00000030649 | Anapc15 | 75430 | protein_coding | 77.13769321 | 0.070890437 | 0.132637058 | 0.53446931 | 0.593016877 | 0.7614941 | 0.070890437 |
| ENSMUSG00000007411 | Mark3 | 17169 | protein_coding | 954.0442114 | -0.070893213 | 0.082520338 | -0.859099883 | 0.390285412 | 0.59854285 | 0.070893213 |

Supplementary Table 1

| ensembl_gene_id | external_gene_name | entrezgene_id | gene_biotype | baseMean | log2FoldChange | lfcSE | stat | pvalue | padj | absLog2FC |
|---|---|---|---|---|---|---|---|---|---|---|
| ENSMUSG00000025764 | Jade1 | 269424 | protein_coding | 688.514274 | 0.070895222 | 0.08433023 | 0.840685733 | 0.400524015 | 0.608097085 | 0.070895222 |
| ENSMUSG00000066306 | Numa1 | 101706 | protein_coding | 2497.609671 | -0.0708992 | 0.08469265 | -0.837135213 | 0.402516569 | 0.609921868 | 0.0708992 |
| ENSMUSG00000022673 | Mcm4 | 17217 | protein_coding | 435.4296557 | 0.070908502 | 0.149941781 | 0.472906891 | 0.636279608 | 0.789347063 | 0.070908502 |
| ENSMUSG00000040407 | Akap9 | 100986 | protein_coding | 3793.830279 | -0.070953966 | 0.106907816 | -0.663692975 | 0.506886836 | 0.696603457 | 0.070953966 |
| ENSMUSG00000021400 | Wrnip1 | 78903 | protein_coding | 407.1359672 | -0.070972212 | 0.088461017 | -0.8022993 | 0.422379847 | 0.627648847 | 0.070972212 |
| ENSMUSG00000035273 | Hpse | 15442 | protein_coding | 222.2692399 | 0.071006774 | 0.09211361 | 0.770860837 | 0.440789422 | 0.642527127 | 0.071006774 |
| ENSMUSG00000033114 | Slc35d2 | 70484 | protein_coding | 108.2914956 | 0.071021265 | 0.171468499 | 0.414194241 | 0.678731859 | 0.81615892 | 0.071021265 |
| ENSMUSG00000022300 | Dcaf13 | 223499 | protein_coding | 624.1888802 | 0.071054946 | 0.092279313 | 0.769998646 | 0.441300696 | 0.642981907 | 0.071054946 |
| ENSMUSG00000062661 | Ncs1 | 14299 | protein_coding | 10.64099385 | 0.071102939 | 0.523837421 | 0.135734745 | 0.89203098 | 0.945409888 | 0.071102939 |
| ENSMUSG00000071379 | Hpcal1 | 53602 | protein_coding | 787.0374557 | -0.071119355 | 0.087619234 | -0.811686555 | 0.416971511 | 0.623101485 | 0.071119355 |
| ENSMUSG00000029536 | Gatc | 384281 | protein_coding | 167.972687 | -0.07111974 | 0.092278944 | -0.770703879 | 0.440882472 | 0.64259116 | 0.07111974 |
| ENSMUSG00000030766 | Arhgap17 | 70497 | protein_coding | 387.5103035 | -0.071145311 | 0.151445704 | -0.469774373 | 0.638516226 | 0.790539172 | 0.071145311 |
| ENSMUSG00000057649 | Brd9 | 105246 | protein_coding | 535.5594741 | -0.071145608 | 0.070936525 | -1.002947478 | 0.315886203 | 0.526825078 | 0.071145608 |
| ENSMUSG00000079487 | Med12 | 59024 | protein_coding | 634.6354726 | -0.071145813 | 0.104401294 | -0.681464861 | 0.495577394 | 0.68695983 | 0.071145813 |
| ENSMUSG00000057177 | Gsk3a | 606496 | protein_coding | 386.8888146 | 0.071175339 | 0.099647263 | 0.714272889 | 0.475058453 | 0.669859335 | 0.071175339 |
| ENSMUSG00000020522 | Mfap3 | 216760 | protein_coding | 391.1400333 | 0.071210304 | 0.063419006 | 1.122854309 | 0.261499375 | 0.469208297 | 0.071210304 |
| ENSMUSG00000026615 | Eprs | 107508 | protein_coding | 2597.210912 | 0.071254181 | 0.067460047 | 1.056242677 | 0.290857357 | 0.499573396 | 0.071254181 |
| ENSMUSG00000052151 | Plpp2 | 50784 | protein_coding | 2494.465617 | 0.071259285 | 0.163007273 | 0.437154019 | 0.661999653 | 0.804480848 | 0.071259285 |
| ENSMUSG00000025583 | Rptor | 74370 | protein_coding | 594.3507076 | -0.071285572 | 0.094103692 | -0.757521525 | 0.448737475 | 0.649476647 | 0.071285572 |
| ENSMUSG00000031706 | Rfx1 | 19724 | protein_coding | 285.9243662 | 0.071374292 | 0.1279395 | 0.557875337 | 0.576929514 | 0.749014624 | 0.071374292 |
| ENSMUSG00000023262 | Acy1 | 109652 | protein_coding | 508.9006071 | 0.071437181 | 0.091184633 | 0.783434432 | 0.433372043 | 0.637255346 | 0.071437181 |
| ENSMUSG00000024101 | Washc1 | 68767 | protein_coding | 301.813284 | -0.071467754 | 0.089582791 | -0.797784416 | 0.424995606 | 0.629529435 | 0.071467754 |
| ENSMUSG00000028760 | Eif4g3 | 230861 | protein_coding | 767.3712737 | 0.071504678 | 0.108495201 | 0.659058443 | 0.50985824 | 0.698582542 | 0.071504678 |
| ENSMUSG00000020577 | Tspan13 | 66109 | protein_coding | 2063.351553 | 0.071515841 | 0.103502042 | 0.690960676 | 0.489590254 | 0.682008993 | 0.071515841 |
| ENSMUSG00000037808 | Fam76b | 72826 | protein_coding | 447.5222674 | -0.071551338 | 0.12021474 | -0.595196044 | 0.551712434 | 0.732245284 | 0.071551338 |
| ENSMUSG00000022228 | Zscan26 | 432731 | protein_coding | 545.2887832 | -0.071578628 | 0.10982527 | -0.651750075 | 0.514562413 | 0.702183862 | 0.071578628 |
| ENSMUSG00000038910 | Plcl2 | 224860 | protein_coding | 1323.412588 | 0.071634435 | 0.11563766 | 0.619473229 | 0.535604653 | 0.720292301 | 0.071634435 |
| ENSMUSG00000037740 | Mrps26 | 99045 | protein_coding | 410.4675838 | 0.071651013 | 0.109873874 | 0.652120569 | 0.514323396 | 0.701922976 | 0.071651013 |
| ENSMUSG00000025580 | Eif4a3 | 192170 | protein_coding | 1197.71236 | 0.071656363 | 0.129729682 | 0.552351332 | 0.580707667 | 0.751809455 | 0.071656363 |
| ENSMUSG00000036985 | Zdhhc9 | 208884 | protein_coding | 332.9352197 | 0.07170025 | 0.108878059 | 0.658537184 | 0.510193012 | 0.698638812 | 0.07170025 |
| ENSMUSG00000020649 | Rrm2 | 20135 | protein_coding | 306.4890225 | 0.07170512 | 0.162860303 | 0.44028605 | 0.659729943 | 0.802969027 | 0.07170512 |
| ENSMUSG00000030744 | Rps3 | 27050 | protein_coding | 12062.23946 | 0.071719297 | 0.094775658 | 0.75672698 | 0.449213448 | 0.649765039 | 0.071719297 |
| ENSMUSG00000028860 | Sytl1 | 269589 | protein_coding | 634.075788 | -0.071739986 | 0.210589969 | -0.340661933 | 0.7333581 | 0.850292692 | 0.071739986 |
| ENSMUSG00000001289 | Pfdn5 | 56612 | protein_coding | 1420.122233 | 0.071766127 | 0.085968956 | 0.834791188 | 0.403835288 | 0.611027688 | 0.071766127 |
| ENSMUSG00000039238 | Zfp750 | 319530 | protein_coding | 120.7598709 | -0.071788122 | 0.356486841 | -0.201376639 | 0.840404081 | 0.915836464 | 0.071788122 |
| ENSMUSG00000035953 | Pip4p1 | 219024 | protein_coding | 341.4762073 | -0.07181537 | 0.11070807 | -0.648691372 | 0.516537882 | 0.703829408 | 0.07181537 |
| ENSMUSG00000014786 | Slc9a5 | 277973 | protein_coding | 73.25380032 | 0.071835028 | 0.190199963 | 0.377681609 | 0.705667132 | 0.833694265 | 0.071835028 |
| ENSMUSG00000034394 | Lif | 16878 | protein_coding | 39.24346669 | -0.07183785 | 0.250922428 | -0.286295055 | 0.774652145 | 0.877593098 | 0.07183785 |

| ensembl_gene_id | external_gene_name | entrezgene_id | gene_biotype | baseMean | log2FoldChange | lfcSE | stat | pvalue | padj | absLog2FC |
|---|---|---|---|---|---|---|---|---|---|---|
| ENSMUSG00000073910 | Mob3b | 214944 | protein_coding | 347.0101403 | 0.07184596 | 0.145494141 | 0.493806548 | 0.621442808 | 0.779949653 | 0.07184596 |
| ENSMUSG00000040234 | Tm7sf3 | 67623 | protein_coding | 816.7600615 | 0.071882352 | 0.0786961 | 0.913416958 | 0.36102329 | 0.570474103 | 0.071882352 |
| ENSMUSG00000038056 | Kmt2c | 231051 | protein_coding | 1851.256396 | -0.071887818 | 0.166112667 | -0.432765422 | 0.665185196 | 0.806025281 | 0.071887818 |
| ENSMUSG00000016503 | Gtf3a | 66596 | protein_coding | 201.211533 | 0.071903134 | 0.126582398 | 0.568034224 | 0.570011729 | 0.744741762 | 0.071903134 |
| ENSMUSG00000043635 | Adamts3 | 330119 | protein_coding | 28.11035132 | 0.071907016 | 0.258199454 | 0.278494065 | 0.78063312 | 0.880776676 | 0.071907016 |
| ENSMUSG00000042063 | Zfp386 | 56220 | protein_coding | 658.6180821 | 0.071965032 | 0.099473935 | 0.723456163 | 0.469399677 | 0.665598871 | 0.071965032 |
| ENSMUSG00000038876 | Rnf146 | 68031 | protein_coding | 195.4287861 | 0.071976494 | 0.107152699 | 0.671718912 | 0.50176266 | 0.692014516 | 0.071976494 |
| ENSMUSG00000003528 | Slc25a1 | 13358 | protein_coding | 1130.242992 | 0.071994192 | 0.102097921 | 0.705148463 | 0.48071785 | 0.674079929 | 0.071994192 |
| ENSMUSG00000063235 | Ptpmt1 | 66461 | protein_coding | 409.5617708 | 0.072046972 | 0.101342423 | 0.710926087 | 0.477130039 | 0.671650664 | 0.072046972 |
| ENSMUSG00000030353 | Tead4 | 21679 | protein_coding | 48.03907278 | -0.072091317 | 0.329871581 | -0.218543583 | 0.827005604 | 0.908412958 | 0.072091317 |
| ENSMUSG00000020668 | Kif3c | 16570 | protein_coding | 19.2737421 | -0.072125473 | 0.33495002 | -0.215332046 | 0.829508447 | 0.909406905 | 0.072125473 |
| ENSMUSG00000024290 | Rock1 | 19877 | protein_coding | 2265.474219 | -0.072138434 | 0.056836036 | -1.269237585 | 0.20435634 | 0.399772183 | 0.072138434 |
| ENSMUSG00000004070 | Hmox2 | 15369 | protein_coding | 347.4759668 | 0.072141454 | 0.086309664 | 0.835844451 | 0.403242416 | 0.610478586 | 0.072141454 |
| ENSMUSG00000074158 | Zfp976 | 208111 | protein_coding | 144.8559437 | 0.072164295 | 0.128076433 | 0.563447105 | 0.573130469 | 0.74661177 | 0.072164295 |
| ENSMUSG00000052144 | Ppp4r2 | 232314 | protein_coding | 979.4380603 | -0.072217169 | 0.081046462 | -0.891058868 | 0.37289759 | 0.581051928 | 0.072217169 |
| ENSMUSG00000026696 | Vamp4 | 53330 | protein_coding | 308.8945858 | -0.072249918 | 0.096628704 | -0.747706579 | 0.454637162 | 0.653915542 | 0.072249918 |
| ENSMUSG00000036718 | Micall2 | 231830 | protein_coding | 323.2399485 | -0.072276246 | 0.13756028 | -0.525415085 | 0.599294669 | 0.76592263 | 0.072276246 |
| ENSMUSG00000016510 | Mtif3 | 76366 | protein_coding | 388.8431208 | -0.072295851 | 0.096029531 | -0.752850191 | 0.451539942 | 0.651785414 | 0.072295851 |
| ENSMUSG00000027646 | Src | 20779 | protein_coding | 1784.573132 | -0.072332784 | 0.089380531 | -0.809267789 | 0.41836113 | 0.624549748 | 0.072332784 |
| ENSMUSG00000002781 | Tmem143 | 70209 | protein_coding | 229.5002876 | 0.072377815 | 0.124056642 | 0.583425554 | 0.559606859 | 0.737621487 | 0.072377815 |
| ENSMUSG00000031109 | Enox2 | 209224 | protein_coding | 389.37536 | 0.072378125 | 0.11723969 | 0.617351726 | 0.537002756 | 0.721275497 | 0.072378125 |
| ENSMUSG00000003200 | Sh3gl1 | 20405 | protein_coding | 376.9434116 | -0.072486908 | 0.105434159 | -0.68750876 | 0.491762182 | 0.683847393 | 0.072486908 |
| ENSMUSG00000035078 | Mtmr9 | 210376 | protein_coding | 376.5627517 | 0.072506778 | 0.109757192 | 0.660610719 | 0.508861993 | 0.697877891 | 0.072506778 |
| ENSMUSG00000003345 | Csnk1g2 | 103236 | protein_coding | 1165.728972 | -0.072515637 | 0.076581254 | -0.946911066 | 0.343684099 | 0.55376472 | 0.072515637 |
| ENSMUSG00000022905 | Kpna1 | 16646 | protein_coding | 1073.171173 | 0.072556711 | 0.094931979 | 0.764302101 | 0.444687234 | 0.645550333 | 0.072556711 |
| ENSMUSG00000020707 | Rnf135 | 71956 | protein_coding | 110.5583784 | 0.07255742 | 0.158021044 | 0.459163022 | 0.646117103 | 0.795304816 | 0.07255742 |
| ENSMUSG00000024513 | Mbd2 | 17191 | protein_coding | 1671.35424 | 0.072559192 | 0.05044254 | 1.438452394 | 0.150305736 | 0.328041762 | 0.072559192 |
| ENSMUSG00000020609 | Apob | 238055 | protein_coding | 2625.968026 | -0.07261938 | 0.170626983 | -0.425603139 | 0.670397053 | 0.809252312 | 0.07261938 |
| ENSMUSG00000045624 | Esf1 | 66580 | protein_coding | 441.8516484 | -0.072662651 | 0.10494994 | -0.692355335 | 0.488714208 | 0.681333909 | 0.072662651 |
| ENSMUSG00000022096 | Hr | 15460 | protein_coding | 368.1513833 | -0.072664237 | 0.1531379 | -0.474501978 | 0.635141988 | 0.788885209 | 0.072664237 |
| ENSMUSG00000028414 | Fktn | 246179 | protein_coding | 383.686044 | 0.07266593 | 0.09035901 | 0.804191305 | 0.4212865 | 0.627009098 | 0.07266593 |
| ENSMUSG00000038957 | Edc3 | 353190 | protein_coding | 209.1298424 | -0.072713011 | 0.120517552 | -0.603339593 | 0.546282799 | 0.728560029 | 0.072713011 |
| ENSMUSG00000040472 | Rabggta | 56187 | protein_coding | 242.6001065 | -0.072716375 | 0.106710621 | -0.681435217 | 0.495596146 | 0.68695983 | 0.072716375 |
| ENSMUSG00000027540 | Ptpn1 | 19246 | protein_coding | 386.8049141 | -0.072726172 | 0.140323907 | -0.518273569 | 0.604267411 | 0.768935953 | 0.072726172 |
| ENSMUSG00000050777 | Tmem37 | 170706 | protein_coding | 112.3939581 | 0.07276464 | 0.18577432 | 0.391682982 | 0.695292466 | 0.826592861 | 0.07276464 |
| ENSMUSG00000055128 | Cgrrf1 | 68755 | protein_coding | 246.8313971 | 0.072842485 | 0.112012064 | 0.650309282 | 0.515492463 | 0.702916184 | 0.072842485 |
| ENSMUSG00000038827 | Abitram | 230234 | protein_coding | 133.1265059 | -0.072891761 | 0.153175733 | -0.475870159 | 0.634166883 | 0.788253962 | 0.072891761 |
| ENSMUSG00000053347 | Zfp943 | 74670 | protein_coding | 502.7215121 | 0.072892725 | 0.101771768 | 0.716237184 | 0.473844907 | 0.66884592 | 0.072892725 |

| ensembl_gene_id | external_gene_name | entrezgene_id | gene_biotype | baseMean | log2FoldChange | lfcSE | stat | pvalue | padj | absLog2FC |
|---|---|---|---|---|---|---|---|---|---|---|
| ENSMUSG00000034341 | Wbp2 | 22378 | protein_coding | 632.3007411 | 0.072921416 | 0.061328563 | 1.189028606 | 0.234428411 | 0.438316627 | 0.072921416 |
| ENSMUSG00000028437 | Ubap1 | 67123 | protein_coding | 885.7980149 | -0.07298694 | 0.069896704 | -1.044211463 | 0.296387565 | 0.505716243 | 0.07298694 |
| ENSMUSG00000041215 | Yeats2 | 208146 | protein_coding | 482.3094292 | -0.072991134 | 0.06685392 | -1.091800362 | 0.27492086 | 0.483060061 | 0.072991134 |
| ENSMUSG00000028820 | Sfpq | 71514 | protein_coding | 2688.343837 | 0.072998626 | 0.132375853 | 0.551449715 | 0.581325429 | 0.752303496 | 0.072998626 |
| ENSMUSG00000000303 | Cdh1 | 12550 | protein_coding | 13345.08879 | -0.073002637 | 0.079027298 | -0.923764808 | 0.355608782 | 0.565274111 | 0.073002637 |
| ENSMUSG00000002233 | Rhoc | 11853 | protein_coding | 887.4590482 | -0.073055678 | 0.116713696 | -0.625939201 | 0.531354821 | 0.716560792 | 0.073055678 |
| ENSMUSG00000022255 | Mtdh | 67154 | protein_coding | 2162.926795 | 0.073073725 | 0.064984942 | 1.12447164 | 0.260812995 | 0.468187692 | 0.073073725 |
| ENSMUSG00000004626 | Stxbp2 | 20911 | protein_coding | 1248.479196 | -0.073108937 | 0.063947944 | -1.143257033 | 0.252931889 | 0.459205682 | 0.073108937 |
| ENSMUSG00000061315 | Naca | 17938 | protein_coding | 4473.665931 | 0.073109972 | 0.083462596 | 0.875960914 | 0.381051283 | 0.588917604 | 0.073109972 |
| ENSMUSG00000004631 | Sgce | 20392 | protein_coding | 10.45842709 | 0.073156289 | 0.744507801 | 0.09826128 | 0.921724824 | 0.961128183 | 0.073156289 |
| ENSMUSG00000024384 | Iws1 | 73473 | protein_coding | 1068.349128 | -0.073173442 | 0.093393878 | -0.783492925 | 0.433337707 | 0.637255346 | 0.073173442 |
| ENSMUSG00000032412 | Atp1b3 | 11933 | protein_coding | 676.4639677 | -0.073213474 | 0.124544152 | -0.587851559 | 0.556631933 | 0.735709355 | 0.073213474 |
| ENSMUSG00000059890 | Ube4a | 140630 | protein_coding | 787.3183052 | 0.073253915 | 0.078475085 | 0.93346716 | 0.350578818 | 0.560627608 | 0.073253915 |
| ENSMUSG00000033192 | Lpcat2 | 270084 | protein_coding | 397.6548498 | 0.073268625 | 0.105439599 | 0.694887168 | 0.487126015 | 0.680012547 | 0.073268625 |
| ENSMUSG00000030745 | Il21r | 60504 | protein_coding | 10.60555644 | -0.073276832 | 0.50997908 | -0.143685957 | 0.88574846 | 0.94220607 | 0.073276832 |
| ENSMUSG00000027809 | Etfdh | 66841 | protein_coding | 1776.038831 | -0.073282733 | 0.089676352 | -0.817191281 | 0.413819124 | 0.620277776 | 0.073282733 |
| ENSMUSG00000029650 | Slc46a3 | 71706 | protein_coding | 69.92805475 | 0.073375024 | 0.179892827 | 0.407881881 | 0.683360396 | 0.819127171 | 0.073375024 |
| ENSMUSG00000001783 | Rtcb | 28088 | protein_coding | 2245.683457 | 0.073415324 | 0.073164093 | 1.003433793 | 0.315651606 | 0.526626038 | 0.073415324 |
| ENSMUSG00000109511 | Nup62 | 18226 | protein_coding | 355.3469682 | -0.073443196 | 0.094159434 | -0.779987652 | 0.435398143 | 0.638336365 | 0.073443196 |
| ENSMUSG00000035764 | Fbxo45 | 268882 | protein_coding | 353.1814249 | -0.073504924 | 0.07730114 | -0.950890555 | 0.341659937 | 0.551941783 | 0.073504924 |
| ENSMUSG00000031644 | Nek1 | 18004 | protein_coding | 402.7047867 | 0.073519186 | 0.101899547 | 0.721486879 | 0.47061001 | 0.666501035 | 0.073519186 |
| ENSMUSG00000031858 | Mau2 | 74549 | protein_coding | 1356.001213 | -0.073536743 | 0.073732661 | -0.997342863 | 0.318598115 | 0.529987548 | 0.073536743 |
| ENSMUSG00000036636 | Clcn7 | 26373 | protein_coding | 365.6274365 | -0.073558068 | 0.08004812 | -0.918923122 | 0.358135785 | 0.567770158 | 0.073558068 |
| ENSMUSG00000029104 | Htt | 15194 | protein_coding | 1078.018392 | 0.073607514 | 0.09735745 | 0.756054256 | 0.449616667 | 0.650217384 | 0.073607514 |
| ENSMUSG00000019797 | Mtres1 | 67851 | protein_coding | 387.7231753 | -0.073617811 | 0.087577262 | -0.840604167 | 0.400569724 | 0.608101981 | 0.073617811 |
| ENSMUSG00000038528 | Mfsd4b5 | 215928 | protein_coding | 13.70218971 | 0.073627175 | 0.374399124 | 0.19665424 | 0.844098122 | 0.917482932 | 0.073627175 |
| ENSMUSG00000038520 | Tbc1d17 | 233204 | protein_coding | 392.9015912 | -0.073649193 | 0.158506778 | -0.464643806 | 0.642186556 | 0.79292616 | 0.073649193 |
| ENSMUSG00000064210 | Ano6 | 105722 | protein_coding | 478.4714597 | 0.073730361 | 0.166612846 | 0.442525067 | 0.658109295 | 0.802182829 | 0.073730361 |
| ENSMUSG00000116564 | Riok2 | 67045 | protein_coding | 357.431163 | -0.073766133 | 0.116687594 | -0.632167742 | 0.527277274 | 0.71264085 | 0.073766133 |
| ENSMUSG00000050394 | Armcx6 | 278097 | protein_coding | 22.44381115 | -0.073784085 | 0.327315433 | -0.225421954 | 0.821651034 | 0.905442422 | 0.073784085 |
| ENSMUSG00000000544 | Gpa33 | 59290 | protein_coding | 5909.828685 | 0.073818462 | 0.067680069 | 1.090697201 | 0.275406143 | 0.48364598 | 0.073818462 |
| ENSMUSG00000024242 | Map4k3 | 225028 | protein_coding | 1416.907712 | 0.073822656 | 0.061865127 | 1.193283832 | 0.232758232 | 0.436366893 | 0.073822656 |
| ENSMUSG00000022297 | Fzd6 | 14368 | protein_coding | 148.3334091 | -0.073844275 | 0.132299847 | -0.558158427 | 0.576736206 | 0.748859077 | 0.073844275 |
| ENSMUSG00000040029 | Ipo8 | 320727 | protein_coding | 1171.605195 | 0.073846954 | 0.064476124 | 1.145337992 | 0.252069174 | 0.458393532 | 0.073846954 |
| ENSMUSG00000032263 | Bckdhb | 12040 | protein_coding | 113.9309221 | -0.073908261 | 0.120337348 | -0.614175583 | 0.539099306 | 0.722272825 | 0.073908261 |
| ENSMUSG00000038095 | Sbno1 | 243272 | protein_coding | 2020.540818 | -0.073922564 | 0.066411636 | -1.113096564 | 0.265666959 | 0.473326994 | 0.073922564 |
| ENSMUSG00000028962 | Slc4a2 | 20535 | protein_coding | 2429.437187 | -0.073923104 | 0.121317101 | -0.609337873 | 0.542300509 | 0.725173014 | 0.073923104 |
| ENSMUSG00000037408 | Cnnm4 | 94220 | protein_coding | 5714.458371 | 0.073940937 | 0.1029703 | 0.718080228 | 0.472707821 | 0.668082535 | 0.073940937 |

| ensembl_gene_id | external_gene_name | entrezgene_id | gene_biotype | baseMean | log2FoldChange | lfcSE | stat | pvalue | padj | absLog2FC |
|---|---|---|---|---|---|---|---|---|---|---|
| ENSMUSG00000022855 | Senp2 | 75826 | protein_coding | 651.4764266 | -0.073961496 | 0.089246338 | -0.828734236 | 0.407254809 | 0.614342654 | 0.073961496 |
| ENSMUSG00000036526 | Card11 | 108723 | protein_coding | 33.68491405 | -0.073965205 | 0.340407018 | -0.217284608 | 0.827986552 | 0.90891431 | 0.073965205 |
| ENSMUSG00000022911 | Arl13b | 68146 | protein_coding | 254.634808 | -0.074011637 | 0.115127938 | -0.642864264 | 0.520312179 | 0.706946378 | 0.074011637 |
| ENSMUSG00000078495 | Zfp984 | 100041677 | protein_coding | 240.0760662 | -0.074069028 | 0.142553101 | -0.519589031 | 0.603350045 | 0.768032832 | 0.074069028 |
| ENSMUSG00000033790 | Tubgcp5 | 233276 | protein_coding | 218.3586504 | -0.074095174 | 0.089301437 | -0.829719839 | 0.406697202 | 0.613841849 | 0.074095174 |
| ENSMUSG00000026792 | Lrsam1 | 227738 | protein_coding | 226.682647 | -0.07415546 | 0.111407084 | -0.665626077 | 0.505650134 | 0.69562924 | 0.07415546 |
| ENSMUSG00000014748 | Tex261 | 21766 | protein_coding | 832.4442983 | -0.074164224 | 0.068259979 | -1.086496443 | 0.277259414 | 0.485615564 | 0.074164224 |
| ENSMUSG00000027247 | Arhgap1 | 228359 | protein_coding | 1460.549431 | -0.074171853 | 0.087803732 | -0.844746017 | 0.398252661 | 0.606149922 | 0.074171853 |
| ENSMUSG00000027878 | Notch2 | 18129 | protein_coding | 207.9072949 | -0.074181051 | 0.130387935 | -0.568925725 | 0.569406547 | 0.744530893 | 0.074181051 |
| ENSMUSG00000055137 | Sugct | 192136 | protein_coding | 31.15859437 | -0.074192776 | 0.257000018 | -0.288687823 | 0.772820282 | 0.876369176 | 0.074192776 |
| ENSMUSG00000043866 | Taf10 | 24075 | protein_coding | 679.7915377 | 0.074231002 | 0.116408966 | 0.63767427 | 0.523685738 | 0.709654498 | 0.074231002 |
| ENSMUSG00000008136 | Fhl2 | 14200 | protein_coding | 290.3822254 | 0.074261059 | 0.470525468 | 0.157825801 | 0.874594067 | 0.935920993 | 0.074261059 |
| ENSMUSG00000044134 | Pheta1 | 231717 | protein_coding | 756.9470114 | 0.074268871 | 0.158631218 | 0.46818572 | 0.639651783 | 0.791145644 | 0.074268871 |
| ENSMUSG00000083307 | AA414768 | 245350 | processed_pseudogene | 14.14285412 | -0.074336536 | 0.399708765 | -0.185976747 | 0.852462996 | 0.921945066 | 0.074336536 |
| ENSMUSG00000085427 | 6430710C18Rik | 76218 | lncRNA | 6.917484388 | 0.074357625 | 0.492870058 | 0.150866591 | 0.880080955 | 0.939320286 | 0.074357625 |
| ENSMUSG00000051413 | Plagl2 | 54711 | protein_coding | 407.86694 | -0.07436834 | 0.166978825 | -0.445375873 | 0.656048145 | 0.800853376 | 0.07436834 |
| ENSMUSG00000028121 | Bcar3 | 29815 | protein_coding | 940.9336395 | 0.074371035 | 0.145544598 | 0.51098451 | 0.609361903 | 0.772513652 | 0.074371035 |
| ENSMUSG00000043881 | Kbtbd7 | 211255 | protein_coding | 287.0118846 | -0.074372057 | 0.093891826 | -0.792103632 | 0.428300253 | 0.632483444 | 0.074372057 |
| ENSMUSG00000027296 | Itpka | 228550 | protein_coding | 112.8105514 | -0.074395094 | 0.193506018 | -0.38445881 | 0.700638419 | 0.829926899 | 0.074395094 |
| ENSMUSG00000038175 | Mylip | 218203 | protein_coding | 301.2656839 | -0.074414677 | 0.250032665 | -0.297619821 | 0.765993345 | 0.872166845 | 0.074414677 |
| ENSMUSG00000020875 | Hoxb9 | 15417 | protein_coding | 601.3603267 | 0.074447513 | 0.130340401 | 0.571177564 | 0.567879285 | 0.743297005 | 0.074447513 |
| ENSMUSG00000087366 | Junos | 230451 | lncRNA | 27.62398234 | 0.07450122 | 0.258305403 | 0.288423004 | 0.773022961 | 0.876474212 | 0.07450122 |
| ENSMUSG00000067194 | Eif1ax | 66235 | protein_coding | 1376.184052 | -0.074583997 | 0.083551597 | -0.892669906 | 0.372033971 | 0.580273902 | 0.074583997 |
| ENSMUSG00000043587 | Pxylp1 | 235534 | protein_coding | 645.9070428 | -0.074619139 | 0.165487855 | -0.450904019 | 0.652058727 | 0.798492458 | 0.074619139 |
| ENSMUSG00000011096 | Akt1s1 | 67605 | protein_coding | 481.2304126 | 0.074619552 | 0.102675303 | 0.726752686 | 0.467377472 | 0.664025366 | 0.074619552 |
| ENSMUSG00000032570 | Atp2c1 | 235574 | protein_coding | 1446.194129 | 0.074626248 | 0.064416047 | 1.158504002 | 0.24665842 | 0.452168679 | 0.074626248 |
| ENSMUSG00000029190 | D5Ertd579e | 320661 | protein_coding | 627.6453577 | -0.074628953 | 0.115702353 | -0.64500808 | 0.51892195 | 0.70596078 | 0.074628953 |
| ENSMUSG00000061306 | Slc38a10 | 72055 | protein_coding | 1524.185374 | 0.074638577 | 0.073149637 | 1.020354711 | 0.307560265 | 0.517462754 | 0.074638577 |
| ENSMUSG00000043716 | Rpl7 | 19989 | protein_coding | 15131.22271 | 0.074640682 | 0.085628045 | 0.871684986 | 0.383380253 | 0.590968806 | 0.074640682 |
| ENSMUSG00000020917 | Acly | 104112 | protein_coding | 5992.363627 | -0.074668921 | 0.112838745 | -0.661731224 | 0.50814349 | 0.697493064 | 0.074668921 |
| ENSMUSG00000097571 | Jpx | 70252 | lncRNA | 30.4391574 | -0.074683425 | 0.227399736 | -0.328423536 | 0.74259145 | 0.856666663 | 0.074683425 |
| ENSMUSG00000021044 | Adck1 | 72113 | protein_coding | 174.0088815 | 0.074732622 | 0.131317666 | 0.569098005 | 0.569289632 | 0.744489545 | 0.074732622 |
| ENSMUSG00000037890 | Wdr19 | 213081 | protein_coding | 139.1403213 | -0.074766609 | 0.193799958 | -0.385792699 | 0.699650203 | 0.829246975 | 0.074766609 |
| ENSMUSG00000030924 | Rexo5 | 434234 | protein_coding | 101.2660748 | 0.074787154 | 0.120067238 | 0.62287728 | 0.533365169 | 0.718433955 | 0.074787154 |
| ENSMUSG00000063334 | Krr1 | 52705 | protein_coding | 522.9284914 | 0.074805782 | 0.101767342 | 0.735066681 | 0.462298885 | 0.660166455 | 0.074805782 |
| ENSMUSG00000029674 | Limk1 | 16885 | protein_coding | 34.94375313 | 0.074806641 | 0.218265414 | 0.342732455 | 0.731799745 | 0.849260935 | 0.074806641 |
| ENSMUSG00000033352 | Map2k4 | 26398 | protein_coding | 847.3690334 | 0.074824308 | 0.124181885 | 0.602538028 | 0.546816058 | 0.728832101 | 0.074824308 |
| ENSMUSG00000020840 | Blmh | 104184 | protein_coding | 950.1411274 | 0.074888922 | 0.152174926 | 0.492123923 | 0.622631741 | 0.780806269 | 0.074888922 |

| ensembl_gene_id | external_gene_name | entrezgene_id | gene_biotype | baseMean | log2FoldChange | lfcSE | stat | pvalue | padj | absLog2FC |
|---|---|---|---|---|---|---|---|---|---|---|
| ENSMUSG00000024642 | Tle4 | 21888 | protein_coding | 912.4478673 | 0.074912424 | 0.191847559 | 0.390478901 | 0.696182454 | 0.827131234 | 0.074912424 |
| ENSMUSG00000036398 | Ppp1r11 | 76497 | protein_coding | 397.4314137 | 0.07495745 | 0.134122863 | 0.558871537 | 0.576249396 | 0.748585685 | 0.07495745 |
| ENSMUSG00000032599 | Ip6k2 | 76500 | protein_coding | 216.9884653 | -0.075006029 | 0.115226813 | -0.650942487 | 0.515083613 | 0.702478962 | 0.075006029 |
| ENSMUSG00000038215 | Cep44 | 382010 | protein_coding | 146.5230081 | 0.075026177 | 0.14921091 | 0.50281965 | 0.615091077 | 0.775590195 | 0.075026177 |
| ENSMUSG00000005299 | Letm1 | 56384 | protein_coding | 1761.914758 | -0.075038468 | 0.094555134 | -0.793594863 | 0.427431324 | 0.631844652 | 0.075038468 |
| ENSMUSG00000041571 | Selenow | 20364 | protein_coding | 616.2258602 | 0.075077027 | 0.083527907 | 0.898825671 | 0.368745524 | 0.577633626 | 0.075077027 |
| ENSMUSG00000058318 | Phf21a | 192285 | protein_coding | 223.947714 | -0.075081917 | 0.130342807 | -0.57603422 | 0.564592052 | 0.741125944 | 0.075081917 |
| ENSMUSG00000033706 | Smyd5 | 232187 | protein_coding | 157.3827598 | 0.075154767 | 0.150962595 | 0.497837011 | 0.618598927 | 0.777789999 | 0.075154767 |
| ENSMUSG00000009470 | Tnpo1 | 238799 | protein_coding | 2636.446273 | 0.075168156 | 0.067350352 | 1.116076652 | 0.264389325 | 0.472048706 | 0.075168156 |
| ENSMUSG00000059810 | Rgs3 | 50780 | protein_coding | 273.1129366 | -0.075212474 | 0.132550681 | -0.567424274 | 0.570425962 | 0.744855281 | 0.075212474 |
| ENSMUSG00000014470 | Rnf166 | 68718 | protein_coding | 238.0057361 | 0.075250475 | 0.085406852 | 0.881082414 | 0.378273215 | 0.586330845 | 0.075250475 |
| ENSMUSG00000034751 | Mast4 | 328329 | protein_coding | 325.7762152 | -0.075353071 | 0.210852253 | -0.357373799 | 0.720811989 | 0.843144111 | 0.075353071 |
| ENSMUSG00000039483 | Asb6 | 72323 | protein_coding | 214.1749927 | 0.07535343 | 0.102099335 | 0.738040363 | 0.460489922 | 0.658781284 | 0.07535343 |
| ENSMUSG00000000416 | Cttnbp2 | 30785 | protein_coding | 167.2051816 | 0.075365818 | 0.280640895 | 0.268548949 | 0.788276802 | 0.884950162 | 0.075365818 |
| ENSMUSG00000019861 | Gopc | 94221 | protein_coding | 722.1574644 | 0.075384358 | 0.068071907 | 1.107422441 | 0.268111328 | 0.476031349 | 0.075384358 |
| ENSMUSG00000024235 | Map3k8 | 26410 | protein_coding | 22.11625066 | -0.075413079 | 0.259017784 | -0.291150199 | 0.770936449 | 0.87556148 | 0.075413079 |
| ENSMUSG00000022824 | Muc13 | 17063 | protein_coding | 14859.25981 | -0.075426033 | 0.091652248 | -0.822958895 | 0.410531369 | 0.617522301 | 0.075426033 |
| ENSMUSG00000115987 | Vps28 | 66914 | protein_coding | 1081.602754 | 0.075464123 | 0.096648313 | 0.780811588 | 0.434913318 | 0.638013252 | 0.075464123 |
| ENSMUSG00000040028 | Elavl1 | 15568 | protein_coding | 1095.355265 | 0.075482254 | 0.115531102 | 0.653350075 | 0.513530617 | 0.701141408 | 0.075482254 |
| ENSMUSG00000032577 | Mapkapk3 | 102626 | protein_coding | 1069.252278 | -0.075487458 | 0.15005651 | -0.503060203 | 0.614921946 | 0.775554419 | 0.075487458 |
| ENSMUSG00000005131 | 4930550C14Rik | 75311 | protein_coding | 7.592536177 | -0.075528179 | 0.498456234 | -0.151524193 | 0.879562228 | 0.938985735 | 0.075528179 |
| ENSMUSG00000010122 | Slc47a1 | 67473 | protein_coding | 27.39428938 | 0.075648731 | 0.244317215 | 0.30963324 | 0.756839876 | 0.866576489 | 0.075648731 |
| ENSMUSG00000049307 | Fut4 | 14345 | protein_coding | 1481.390956 | -0.075669659 | 0.170726584 | -0.443221301 | 0.657605673 | 0.801868244 | 0.075669659 |
| ENSMUSG00000058690 | Ccser2 | 72972 | protein_coding | 881.1032946 | 0.075703428 | 0.09538058 | 0.793698553 | 0.427370943 | 0.631814032 | 0.075703428 |
| ENSMUSG00000003346 | Abhd17a | 216169 | protein_coding | 753.1799569 | 0.075729445 | 0.141727212 | 0.534332424 | 0.593111563 | 0.761494544 | 0.075729445 |
| ENSMUSG00000038861 | Pi4kb | 107650 | protein_coding | 669.7730515 | -0.075742169 | 0.064160845 | -1.180504547 | 0.237799603 | 0.442071905 | 0.075742169 |
| ENSMUSG00000016619 | Nup50 | 18141 | protein_coding | 1056.963765 | 0.075838817 | 0.08565943 | 0.885352805 | 0.375966378 | 0.584199414 | 0.075838817 |
| ENSMUSG00000074217 | Misp3 | 70134 | protein_coding | 37.38837151 | 0.075872844 | 0.207218206 | 0.36614951 | 0.714253515 | 0.83917017 | 0.075872844 |
| ENSMUSG00000020948 | Klhl28 | 66689 | protein_coding | 229.9425859 | -0.075887376 | 0.132154485 | -0.574232315 | 0.565810604 | 0.741869551 | 0.075887376 |
| ENSMUSG00000000881 | Dlg3 | 53310 | protein_coding | 951.5761318 | -0.07593688 | 0.073928774 | -1.027162707 | 0.304343855 | 0.513961388 | 0.07593688 |
| ENSMUSG00000028619 | Tceanc2 | 66526 | protein_coding | 249.6105996 | 0.075948085 | 0.115433763 | 0.65793649 | 0.510578942 | 0.698968748 | 0.075948085 |
| ENSMUSG00000051495 | Irf2bp2 | 270110 | protein_coding | 1957.273299 | 0.075954595 | 0.123090674 | 0.617062141 | 0.537193739 | 0.721330456 | 0.075954595 |
| ENSMUSG00000070661 | Rnf186 | 66825 | protein_coding | 1630.886436 | 0.075983471 | 0.243893162 | 0.311544083 | 0.755387036 | 0.865722025 | 0.075983471 |
| ENSMUSG00000020839 | Tmigd1 | 66601 | protein_coding | 1102.972889 | -0.075986821 | 0.334796247 | -0.226964376 | 0.820451443 | 0.904808599 | 0.075986821 |
| ENSMUSG00000028673 | Fuca1 | 71665 | protein_coding | 3519.34729 | 0.075996876 | 0.076143215 | 0.998078101 | 0.318241488 | 0.52956022 | 0.075996876 |
| ENSMUSG00000055313 | Pgbd1 | 319207 | protein_coding | 17.50072964 | 0.075999379 | 0.319151103 | 0.238129771 | 0.811780445 | 0.899793941 | 0.075999379 |
| ENSMUSG00000031703 | Itfg1 | 71927 | protein_coding | 1747.364583 | 0.076021372 | 0.083044467 | 0.915429711 | 0.359966087 | 0.569481508 | 0.076021372 |
| ENSMUSG00000086993 | Rsf1os2 | 102641859 | lncRNA | 6.725184143 | -0.076021401 | 0.565383919 | -0.134459787 | 0.893039008 | 0.946206699 | 0.076021401 |

| ensembl_gene_id | external_gene_name | entrezgene_id | gene_biotype | baseMean | log2FoldChange | lfcSE | stat | pvalue | padj | absLog2FC |
|---|---|---|---|---|---|---|---|---|---|---|
| ENSMUSG00000035824 | Tk2 | 57813 | protein_coding | 182.1308898 | 0.076035206 | 0.102510229 | 0.741732871 | 0.4582492 | 0.656797737 | 0.076035206 |
| ENSMUSG00000007080 | Pole | 18973 | protein_coding | 257.7632154 | -0.076078616 | 0.171412762 | -0.443832857 | 0.657163431 | 0.801539102 | 0.076078616 |
| ENSMUSG00000008393 | Carhsp1 | 52502 | protein_coding | 1673.530834 | 0.076084147 | 0.083534816 | 0.910807618 | 0.362396748 | 0.571554287 | 0.076084147 |
| ENSMUSG00000038342 | Mlxip | 208104 | protein_coding | 837.9889985 | -0.076086249 | 0.159881655 | -0.47589105 | 0.634151999 | 0.788253962 | 0.076086249 |
| ENSMUSG00000036315 | Polr1h | 66136 | protein_coding | 228.1625823 | 0.076097414 | 0.128448616 | 0.592434677 | 0.553559551 | 0.73323048 | 0.076097414 |
| ENSMUSG00000003382 | Etv3 | 27049 | protein_coding | 854.5742623 | -0.076132732 | 0.078160697 | -0.974053908 | 0.330029774 | 0.540678415 | 0.076132732 |
| ENSMUSG00000017734 | Dbndd2 | 52840 | protein_coding | 36.34767981 | -0.076148773 | 0.207681421 | -0.366661458 | 0.713871559 | 0.838968951 | 0.076148773 |
| ENSMUSG00000030942 | Thumpd1 | 233802 | protein_coding | 515.9331426 | -0.076180785 | 0.088597599 | -0.859851568 | 0.389870869 | 0.598195087 | 0.076180785 |
| ENSMUSG00000047459 | Dynlrb1 | 67068 | protein_coding | 559.396839 | 0.076182508 | 0.092173194 | 0.826514787 | 0.408512133 | 0.61573664 | 0.076182508 |
| ENSMUSG00000027956 | Tmem144 | 70652 | protein_coding | 150.092479 | -0.076187342 | 0.12033141 | -0.633145923 | 0.526638356 | 0.712260825 | 0.076187342 |
| ENSMUSG00000002660 | Clpp | 53895 | protein_coding | 363.3775158 | -0.076190787 | 0.069654446 | -1.093839534 | 0.274025362 | 0.481911887 | 0.076190787 |
| ENSMUSG00000030761 | Myo7a | 17921 | protein_coding | 201.7500057 | 0.076197544 | 0.11836686 | 0.643740521 | 0.519743709 | 0.706451576 | 0.076197544 |
| ENSMUSG00000021243 | Fcf1 | 73736 | protein_coding | 378.9589161 | 0.076310151 | 0.091375781 | 0.83512448 | 0.403647625 | 0.610859797 | 0.076310151 |
| ENSMUSG00000028826 | Maco1 | 66146 | protein_coding | 802.2334436 | 0.076345122 | 0.083672598 | 0.91242682 | 0.361544076 | 0.570843978 | 0.076345122 |
| ENSMUSG00000035704 | Alg8 | 381903 | protein_coding | 161.6281204 | 0.076389657 | 0.149336051 | 0.511528569 | 0.608980989 | 0.772360968 | 0.076389657 |
| ENSMUSG00000034371 | Tkfc | 225913 | protein_coding | 196.7480292 | -0.076601918 | 0.127822683 | -0.599282661 | 0.548984408 | 0.730633661 | 0.076601918 |
| ENSMUSG00000038540 | Tmc3 | 233424 | protein_coding | 7.203800532 | 0.076628404 | 0.593355793 | 0.129144107 | 0.897243622 | 0.948703875 | 0.076628404 |
| ENSMUSG00000060739 | Nsa2 | 59050 | protein_coding | 902.2249062 | -0.076706858 | 0.103390861 | -0.741911396 | 0.458141021 | 0.65670178 | 0.076706858 |
| ENSMUSG00000051435 | Fhad1 | 329977 | protein_coding | 51.36347331 | 0.076745289 | 0.202303415 | 0.379357359 | 0.704422511 | 0.833088933 | 0.076745289 |
| ENSMUSG00000094441 | Zfp955a | 77652 | protein_coding | 246.788182 | -0.076802805 | 0.124303932 | -0.617863038 | 0.536665625 | 0.721189041 | 0.076802805 |
| ENSMUSG00000039201 | Tbc1d25 | 209815 | protein_coding | 102.6110593 | -0.076812158 | 0.110728126 | -0.693700512 | 0.487870045 | 0.680726764 | 0.076812158 |
| ENSMUSG00000021147 | Wdr37 | 207615 | protein_coding | 323.6950648 | -0.076815481 | 0.074678103 | -1.02862121 | 0.303657707 | 0.51328239 | 0.076815481 |
| ENSMUSG00000037343 | Taf2 | 319944 | protein_coding | 629.4949143 | 0.076840609 | 0.06825646 | 1.125763169 | 0.260265778 | 0.467610721 | 0.076840609 |
| ENSMUSG00000072774 | Zfp951 | 626391 | protein_coding | 55.86396555 | 0.076890768 | 0.246575539 | 0.311834533 | 0.755166278 | 0.865722025 | 0.076890768 |
| ENSMUSG00000031667 | Aktip | 14339 | protein_coding | 700.0638848 | -0.076902934 | 0.084077011 | -0.914672544 | 0.360363562 | 0.569883918 | 0.076902934 |
| ENSMUSG00000103793 | Pcdhga6 | 93714 | protein_coding | 10.49167812 | 0.076935319 | 0.365875825 | 0.210277133 | 0.833451382 | 0.911278944 | 0.076935319 |
| ENSMUSG00000036019 | Tmtc2 | 278279 | protein_coding | 648.8851361 | 0.076938435 | 0.069516553 | 1.106764235 | 0.268395875 | 0.476324178 | 0.076938435 |
| ENSMUSG00000028556 | Dock7 | 67299 | protein_coding | 1057.091656 | -0.07693983 | 0.067552173 | -1.138968993 | 0.254716087 | 0.461298219 | 0.07693983 |
| ENSMUSG00000090093 | Gm14399 | 100043761 | protein_coding | 60.65816414 | -0.076991012 | 0.177398572 | -0.434000181 | 0.66428831 | 0.805541241 | 0.076991012 |
| ENSMUSG00000002881 | Nab1 | 17936 | protein_coding | 1254.236709 | -0.076993177 | 0.211815375 | -0.36349192 | 0.716237447 | 0.840316478 | 0.076993177 |
| ENSMUSG00000031004 | Mki67 | 17345 | protein_coding | 3762.316293 | 0.076997289 | 0.117026927 | 0.657945064 | 0.510573432 | 0.698968748 | 0.076997289 |
| ENSMUSG00000072568 | Lratd2 | 399603 | protein_coding | 938.7700861 | 0.077018723 | 0.120943368 | 0.636816425 | 0.524244426 | 0.70998769 | 0.077018723 |
| ENSMUSG00000055485 | Soga1 | 320706 | protein_coding | 68.3304836 | -0.077034378 | 0.230304971 | -0.33448856 | 0.738010922 | 0.853608159 | 0.077034378 |
| ENSMUSG00000029420 | Rimbp2 | 231760 | protein_coding | 65.68127415 | 0.077057083 | 0.235779412 | 0.326818539 | 0.743805137 | 0.857445771 | 0.077057083 |
| ENSMUSG00000022702 | Hira | 15260 | protein_coding | 93.05620658 | -0.077072724 | 0.136327648 | -0.56534918 | 0.571836284 | 0.74577977 | 0.077072724 |
| ENSMUSG00000022771 | Ppil2 | 66053 | protein_coding | 978.522801 | 0.077090593 | 0.068168772 | 1.130878416 | 0.258106269 | 0.465311631 | 0.077090593 |
| ENSMUSG00000075600 | Zc3h3 | 223642 | protein_coding | 65.80019372 | -0.077099826 | 0.21260119 | -0.362650023 | 0.716866338 | 0.840627772 | 0.077099826 |
| ENSMUSG00000075227 | Znhit2 | 29805 | protein_coding | 136.5166418 | 0.077145583 | 0.143435799 | 0.537840509 | 0.590687162 | 0.760015009 | 0.077145583 |

| ensembl_gene_id | external_gene_name | entrezgene_id | gene_biotype | baseMean | log2FoldChange | lfcSE | stat | pvalue | padj | absLog2FC |
|---|---|---|---|---|---|---|---|---|---|---|
| ENSMUSG00000038417 | Fig4 | 103199 | protein_coding | 410.8650269 | 0.077162052 | 0.088363744 | 0.873232037 | 0.382536616 | 0.590239583 | 0.077162052 |
| ENSMUSG00000042111 | Ccdc115 | 69668 | protein_coding | 263.7031695 | 0.077180197 | 0.11081423 | 0.696482727 | 0.486126571 | 0.678983247 | 0.077180197 |
| ENSMUSG00000020456 | Ogdh | 18293 | protein_coding | 5215.335042 | -0.077191475 | 0.068313176 | -1.129964676 | 0.258491109 | 0.465593847 | 0.077191475 |
| ENSMUSG00000015937 | Macroh2a1 | 26914 | protein_coding | 1125.031382 | 0.077205922 | 0.071137742 | 1.085301833 | 0.277787994 | 0.486220568 | 0.077205922 |
| ENSMUSG00000017868 | Sgk2 | 27219 | protein_coding | 447.7355626 | -0.077259724 | 0.292375892 | -0.264247929 | 0.791588881 | 0.887042894 | 0.077259724 |
| ENSMUSG00000041966 | Dcaf17 | 75763 | protein_coding | 319.6447911 | -0.07728612 | 0.113565373 | -0.680543 | 0.496160705 | 0.687323683 | 0.07728612 |
| ENSMUSG00000032030 | Cul5 | 75717 | protein_coding | 707.569846 | -0.077355905 | 0.066639115 | -1.160818323 | 0.24571579 | 0.451102017 | 0.077355905 |
| ENSMUSG00000040782 | Cop1 | 26374 | protein_coding | 1586.123832 | 0.077377193 | 0.057443051 | 1.347024426 | 0.177972366 | 0.365160872 | 0.077377193 |
| ENSMUSG00000021458 | Aopep | 72061 | protein_coding | 870.7552545 | -0.077438272 | 0.117815408 | -0.657284758 | 0.510997835 | 0.699008761 | 0.077438272 |
| ENSMUSG00000025077 | Dclre1a | 55947 | protein_coding | 261.0061261 | -0.07747998 | 0.094035169 | -0.823946837 | 0.409969766 | 0.616911485 | 0.07747998 |
| ENSMUSG00000021207 | Akr1c21 | 77337 | protein_coding | 11.48913058 | -0.0775089 | 0.404981677 | -0.191388658 | 0.848221106 | 0.919955452 | 0.0775089 |
| ENSMUSG00000030685 | Kctd13 | 233877 | protein_coding | 48.88226538 | 0.077529896 | 0.197533341 | 0.392490177 | 0.694696068 | 0.826520011 | 0.077529896 |
| ENSMUSG00000037525 | Bcdin3d | 75284 | protein_coding | 69.78481014 | -0.077532734 | 0.147261966 | -0.526495306 | 0.598544113 | 0.765321556 | 0.077532734 |
| ENSMUSG00000026491 | Ahctf1 | 226747 | protein_coding | 1308.431314 | -0.077576807 | 0.090256524 | -0.859514676 | 0.390056627 | 0.598422457 | 0.077576807 |
| ENSMUSG00000053158 | Fes | 14159 | protein_coding | 105.6081114 | -0.077590977 | 0.14325714 | -0.541620312 | 0.588080096 | 0.757944002 | 0.077590977 |
| ENSMUSG00000061288 | Taok3 | 330177 | protein_coding | 789.1624385 | -0.07763112 | 0.08490311 | -0.914349546 | 0.360533204 | 0.57009559 | 0.07763112 |
| ENSMUSG00000028016 | Ints12 | 71793 | protein_coding | 278.0822062 | 0.077662531 | 0.073329597 | 1.059088467 | 0.289559492 | 0.498385347 | 0.077662531 |
| ENSMUSG00000035851 | Ythdc1 | 231386 | protein_coding | 1051.404492 | -0.07773443 | 0.069029981 | -1.126096656 | 0.26012461 | 0.467426081 | 0.07773443 |
| ENSMUSG00000026198 | Abcb6 | 74104 | protein_coding | 91.47318093 | -0.077735482 | 0.157389111 | -0.493906356 | 0.621372316 | 0.779922636 | 0.077735482 |
| ENSMUSG00000037722 | Gnpnat1 | 54342 | protein_coding | 53.70481954 | -0.077752936 | 0.148815415 | -0.522479044 | 0.601336833 | 0.767290962 | 0.077752936 |
| ENSMUSG00000074406 | Zfp628 | 232816 | protein_coding | 84.21829957 | -0.077790112 | 0.154523699 | -0.503418649 | 0.614669963 | 0.775482231 | 0.077790112 |
| ENSMUSG00000073775 | Kti12 | 100087 | protein_coding | 242.3381027 | -0.077803574 | 0.181290572 | -0.429165032 | 0.667803128 | 0.807344837 | 0.077803574 |
| ENSMUSG00000016253 | Nelfcd | 57314 | protein_coding | 505.0085967 | 0.077851279 | 0.083951465 | 0.92733675 | 0.353751719 | 0.563568452 | 0.077851279 |
| ENSMUSG00000040690 | Col16a1 | 107581 | protein_coding | 265.2181787 | -0.077947375 | 0.161738758 | -0.481933803 | 0.629852972 | 0.785799181 | 0.077947375 |
| ENSMUSG00000026149 | Tm4sf20 | 66261 | protein_coding | 2816.848833 | 0.077963245 | 0.097151063 | 0.802495034 | 0.42226666 | 0.627648847 | 0.077963245 |
| ENSMUSG00000035248 | Tut7 | 214290 | protein_coding | 1738.109508 | 0.078025793 | 0.056062978 | 1.391752554 | 0.163997342 | 0.347129523 | 0.078025793 |
| ENSMUSG00000014859 | E2f4 | 104394 | protein_coding | 437.7044325 | -0.078026482 | 0.131490889 | -0.593398388 | 0.552914568 | 0.732884453 | 0.078026482 |
| ENSMUSG00000024560 | Cxxc1 | 74322 | protein_coding | 510.6321095 | -0.078082045 | 0.081343972 | -0.959899582 | 0.337105757 | 0.547955964 | 0.078082045 |
| ENSMUSG00000035933 | Cog5 | 238123 | protein_coding | 757.2878382 | 0.078104432 | 0.062157675 | 1.256553302 | 0.208915433 | 0.405789022 | 0.078104432 |
| ENSMUSG00000001517 | Foxm1 | 14235 | protein_coding | 204.2228646 | -0.078111222 | 0.134069985 | -0.582615283 | 0.560152315 | 0.738067647 | 0.078111222 |
| ENSMUSG00000022533 | Atp13a3 | 224088 | protein_coding | 3574.122263 | -0.078126966 | 0.085034235 | -0.918770731 | 0.358215505 | 0.567840024 | 0.078126966 |
| ENSMUSG00000004897 | Hdgf | 15191 | protein_coding | 2013.272973 | -0.07814719 | 0.097413651 | -0.802220114 | 0.422425643 | 0.627648847 | 0.07814719 |
| ENSMUSG00000032382 | Snx1 | 56440 | protein_coding | 1813.603371 | 0.078202625 | 0.086795468 | 0.900998939 | 0.367588884 | 0.576275172 | 0.078202625 |
| ENSMUSG00000027860 | Vangl1 | 229658 | protein_coding | 205.9922327 | -0.078252739 | 0.095406762 | -0.820201175 | 0.412101432 | 0.618910638 | 0.078252739 |
| ENSMUSG00000032477 | Cdc25a | 12530 | protein_coding | 796.2069492 | -0.07826683 | 0.100712566 | -0.777130731 | 0.437081645 | 0.639698285 | 0.07826683 |
| ENSMUSG00000071253 | Slc25a16 | 73132 | protein_coding | 558.8423454 | -0.078273978 | 0.094041722 | -0.832332462 | 0.405221312 | 0.612206925 | 0.078273978 |
| ENSMUSG00000033799 | Tasor2 | 105203 | protein_coding | 669.7119071 | -0.078282155 | 0.101705021 | -0.769698037 | 0.441479035 | 0.642981907 | 0.078282155 |
| ENSMUSG00000026439 | Rbbp5 | 213464 | protein_coding | 497.5851976 | 0.078311895 | 0.077274525 | 1.013424475 | 0.310857454 | 0.520864457 | 0.078311895 |

| ensembl_gene_id | external_gene_name | entrezgene_id | gene_biotype | baseMean | log2FoldChange | lfcSE | stat | pvalue | padj | absLog2FC |
|---|---|---|---|---|---|---|---|---|---|---|
| ENSMUSG00000033047 | Eif3l | 223691 | protein_coding | 2624.850702 | -0.07839499 | 0.088570212 | -0.885116879 | 0.376093597 | 0.58431709 | 0.07839499 |
| ENSMUSG00000024570 | Rbfa | 68731 | protein_coding | 116.8046954 | -0.078402385 | 0.110565007 | -0.709106675 | 0.478258281 | 0.672644406 | 0.078402385 |
| ENSMUSG00000040652 | Oaz2 | 18247 | protein_coding | 348.8673356 | -0.078452441 | 0.098740338 | -0.794532842 | 0.426885297 | 0.631389081 | 0.078452441 |
| ENSMUSG00000021786 | Oxsm | 71147 | protein_coding | 373.712982 | 0.078488692 | 0.127425725 | 0.615956408 | 0.537923291 | 0.721865534 | 0.078488692 |
| ENSMUSG00000040331 | Nsmce4a | 67872 | protein_coding | 756.1128329 | 0.078508077 | 0.084801626 | 0.925785045 | 0.354557701 | 0.564062631 | 0.078508077 |
| ENSMUSG00000027030 | Stk39 | 53416 | protein_coding | 732.7575571 | -0.078568333 | 0.076586429 | -1.025878003 | 0.304949092 | 0.514591497 | 0.078568333 |
| ENSMUSG00000038331 | Satb2 | 212712 | protein_coding | 2291.14827 | -0.078634158 | 0.131918929 | -0.596079411 | 0.551122178 | 0.73176676 | 0.078634158 |
| ENSMUSG00000020922 | Lsm12 | 268490 | protein_coding | 542.4031673 | 0.078637065 | 0.102553943 | 0.766787341 | 0.443207964 | 0.644271437 | 0.078637065 |
| ENSMUSG00000009079 | Ewsr1 | 14030 | protein_coding | 2314.998416 | -0.078809421 | 0.083293443 | -0.946165976 | 0.344063937 | 0.554008329 | 0.078809421 |
| ENSMUSG00000049225 | Pdp1 | 381511 | protein_coding | 294.2027863 | 0.078821023 | 0.205115289 | 0.384276682 | 0.700773389 | 0.829944125 | 0.078821023 |
| ENSMUSG00000079316 | Rab9 | 56382 | protein_coding | 464.8545372 | 0.078830567 | 0.095114125 | 0.828799799 | 0.407217702 | 0.614342654 | 0.078830567 |
| ENSMUSG00000014077 | Chp1 | 56398 | protein_coding | 4323.362469 | -0.078859195 | 0.158161982 | -0.498597668 | 0.618062848 | 0.777697562 | 0.078859195 |
| ENSMUSG00000024588 | Fech | 14151 | protein_coding | 740.6247932 | -0.078889467 | 0.074253267 | -1.062437651 | 0.288037049 | 0.497257878 | 0.078889467 |
| ENSMUSG00000041144 | Dnah7b | 227058 | protein_coding | 36.59801803 | -0.078909275 | 0.330588076 | -0.238693651 | 0.81134314 | 0.899694484 | 0.078909275 |
| ENSMUSG00000044030 | Irf2bp1 | 272359 | protein_coding | 498.4305019 | -0.078959129 | 0.09687607 | -0.815052968 | 0.415041992 | 0.621431551 | 0.078959129 |
| ENSMUSG00000025094 | Slc18a2 | 214084 | protein_coding | 60.73273948 | -0.078975752 | 0.179161201 | -0.440808343 | 0.659351753 | 0.802969027 | 0.078975752 |
| ENSMUSG00000036591 | Arhgap21 | 71435 | protein_coding | 1912.72439 | 0.078976308 | 0.109492907 | 0.72129154 | 0.470730161 | 0.666501035 | 0.078976308 |
| ENSMUSG00000037640 | Zfp60 | 22718 | protein_coding | 210.5444891 | 0.079030538 | 0.184289811 | 0.428838346 | 0.668040871 | 0.807570962 | 0.079030538 |
| ENSMUSG00000001150 | Mcm3ap | 54387 | protein_coding | 543.208474 | 0.079033603 | 0.06753752 | 1.170217724 | 0.241913362 | 0.446609087 | 0.079033603 |
| ENSMUSG00000046971 | Pla2g4f | 271844 | protein_coding | 71.06173966 | 0.079050347 | 0.332480577 | 0.237759292 | 0.812067793 | 0.899987184 | 0.079050347 |
| ENSMUSG00000038155 | Gstp2 | 14869 | protein_coding | 321.104728 | -0.079088945 | 0.165119398 | -0.47898034 | 0.631952615 | 0.787062417 | 0.079088945 |
| ENSMUSG00000028223 | Decr1 | 67460 | protein_coding | 1441.419939 | 0.079102259 | 0.099522767 | 0.794815712 | 0.42672071 | 0.631223973 | 0.079102259 |
| ENSMUSG00000117284 | Gm7072 | 631624 | protein_coding | 656.5945613 | 0.079141296 | 0.109922095 | 0.71997623 | 0.471539631 | 0.667069775 | 0.079141296 |
| ENSMUSG00000028188 | Spata1 | 70951 | protein_coding | 30.37661977 | -0.079174916 | 0.259236054 | -0.305416299 | 0.760049109 | 0.868689179 | 0.079174916 |
| ENSMUSG00000047669 | Msl3l2 | 73390 | protein_coding | 13.61942001 | 0.079182162 | 0.362364396 | 0.218515294 | 0.827027643 | 0.908412958 | 0.079182162 |
| ENSMUSG00000025402 | Nab2 | 17937 | protein_coding | 27.50258709 | 0.079200114 | 0.629289314 | 0.125856442 | 0.899845562 | 0.950214744 | 0.079200114 |
| ENSMUSG00000056612 | Ppp1r14b | 18938 | protein_coding | 482.7174346 | 0.07921132 | 0.114631464 | 0.691008535 | 0.489560178 | 0.682008993 | 0.07921132 |
| ENSMUSG00000062376 | Borcs7 | 66439 | protein_coding | 80.86907088 | 0.079219996 | 0.139208103 | 0.569076041 | 0.569304537 | 0.744489545 | 0.079219996 |
| ENSMUSG00000021585 | Cast | 12380 | protein_coding | 2692.612785 | 0.079259305 | 0.12424319 | 0.637936817 | 0.52351481 | 0.709654498 | 0.079259305 |
| ENSMUSG00000038365 | Fbxo25 | 66822 | protein_coding | 1071.769388 | 0.079328399 | 0.148008704 | 0.535971173 | 0.591978473 | 0.761020102 | 0.079328399 |
| ENSMUSG00000050965 | Prkca | 18750 | protein_coding | 1776.124833 | 0.079332926 | 0.074197236 | 1.069216726 | 0.284972024 | 0.49392051 | 0.079332926 |
| ENSMUSG00000009741 | Ubp1 | 22221 | protein_coding | 858.2440147 | -0.079339274 | 0.079812456 | -0.994071336 | 0.320188139 | 0.531665726 | 0.079339274 |
| ENSMUSG00000026219 | Trip12 | 14897 | protein_coding | 3149.820062 | -0.07936242 | 0.07252816 | -1.094229065 | 0.273854527 | 0.481829591 | 0.07936242 |
| ENSMUSG00000026404 | Ddx59 | 67997 | protein_coding | 61.2717031 | 0.07937303 | 0.181821916 | 0.436542696 | 0.662443029 | 0.804528598 | 0.07937303 |
| ENSMUSG00000028519 | Dab1 | 13131 | protein_coding | 37.42312715 | 0.079414503 | 0.298065199 | 0.266433328 | 0.789905501 | 0.886154023 | 0.079414503 |
| ENSMUSG00000014402 | Tsg101 | 22088 | protein_coding | 758.0926577 | 0.079468671 | 0.082929838 | 0.958263909 | 0.337929698 | 0.548511589 | 0.079468671 |
| ENSMUSG00000037366 | Pafah2 | 100163 | protein_coding | 891.5030506 | 0.079489579 | 0.103984969 | 0.764433361 | 0.444609034 | 0.645495643 | 0.079489579 |
| ENSMUSG00000022550 | Adck5 | 268822 | protein_coding | 803.9536924 | -0.079491719 | 0.133290996 | -0.59637726 | 0.550923228 | 0.731624577 | 0.079491719 |

| ensembl_gene_id | external_gene_name | entrezgene_id | gene_biotype | baseMean | log2FoldChange | lfcSE | stat | pvalue | padj | absLog2FC |
|---|---|---|---|---|---|---|---|---|---|---|
| ENSMUSG00000044452 | Zfp507 | 668501 | protein_coding | 521.0040188 | -0.079508542 | 0.107242442 | -0.741390629 | 0.458456625 | 0.657035911 | 0.079508542 |
| ENSMUSG00000041577 | Prelp | 116847 | protein_coding | 91.22301189 | 0.079513116 | 0.255190977 | 0.311582786 | 0.755357618 | 0.865722025 | 0.079513116 |
| ENSMUSG00000032026 | Rexo2 | 104444 | protein_coding | 299.4616213 | 0.079514975 | 0.110888677 | 0.717070279 | 0.473330734 | 0.668725557 | 0.079514975 |
| ENSMUSG00000025934 | Gsta3 | 14859 | protein_coding | 300.456237 | -0.079521675 | 0.207524435 | -0.383191864 | 0.70157751 | 0.830587675 | 0.079521675 |
| ENSMUSG00000020120 | Plek | 56193 | protein_coding | 62.45247885 | -0.079590084 | 0.142807662 | -0.557323628 | 0.577306335 | 0.749071904 | 0.079590084 |
| ENSMUSG00000078963 | Hsbp1l1 | 66255 | protein_coding | 57.19964512 | 0.079624333 | 0.196214177 | 0.405803163 | 0.684887235 | 0.820024075 | 0.079624333 |
| ENSMUSG00000074576 | Mocs3 | 69372 | protein_coding | 109.8399735 | -0.079681697 | 0.169103801 | -0.471199919 | 0.637497976 | 0.790322834 | 0.079681697 |
| ENSMUSG00000018433 | Nol11 | 68979 | protein_coding | 388.8649558 | 0.079705584 | 0.155439565 | 0.512775395 | 0.608108443 | 0.77191196 | 0.079705584 |
| ENSMUSG00000059183 | Mtfmt | 69606 | protein_coding | 215.0361545 | -0.079741223 | 0.097043835 | -0.821703129 | 0.411245878 | 0.618130069 | 0.079741223 |
| ENSMUSG00000076431 | Sox4 | 20677 | protein_coding | 578.4578038 | 0.079766 | 0.114966958 | 0.693816742 | 0.487797142 | 0.680726764 | 0.079766 |
| ENSMUSG00000002768 | Mea1 | 17256 | protein_coding | 696.1501634 | 0.079774484 | 0.081145805 | 0.98310054 | 0.325557959 | 0.536577728 | 0.079774484 |
| ENSMUSG00000030539 | Sema4b | 20352 | protein_coding | 1148.270681 | 0.079840407 | 0.197755151 | 0.403733639 | 0.6864086 | 0.821236098 | 0.079840407 |
| ENSMUSG00000017631 | Abr | 109934 | protein_coding | 2674.208569 | -0.08003375 | 0.071531556 | -1.118859347 | 0.263200148 | 0.470933719 | 0.08003375 |
| ENSMUSG00000043940 | Wdfy3 | 72145 | protein_coding | 679.6850071 | -0.080042946 | 0.144298297 | -0.554704713 | 0.579096652 | 0.750577873 | 0.080042946 |
| ENSMUSG00000014837 | 4931428F04Rik | 74356 | protein_coding | 26.12887652 | -0.08006238 | 0.293802595 | -0.272503994 | 0.785234521 | 0.883563366 | 0.08006238 |
| ENSMUSG00000027639 | Samhd1 | 56045 | protein_coding | 612.0601955 | 0.08008 | 0.109229921 | 0.733132452 | 0.463477649 | 0.66105374 | 0.08008 |
| ENSMUSG00000027655 | Dhx35 | 71715 | protein_coding | 224.5436778 | -0.080083498 | 0.108900643 | -0.73538132 | 0.462107295 | 0.659976112 | 0.080083498 |
| ENSMUSG00000020911 | Krt19 | 16669 | protein_coding | 34649.02189 | 0.08009814 | 0.142996623 | 0.56014008 | 0.575383894 | 0.74838086 | 0.08009814 |
| ENSMUSG00000037349 | Nudt22 | 68323 | protein_coding | 360.1313065 | -0.080101969 | 0.13941903 | -0.574541145 | 0.565601666 | 0.741778785 | 0.080101969 |
| ENSMUSG00000020256 | Aldh1l2 | 216188 | protein_coding | 37.05083586 | 0.080105504 | 0.177732116 | 0.450709221 | 0.652199135 | 0.798541612 | 0.080105504 |
| ENSMUSG00000051076 | Vtcn1 | 242122 | protein_coding | 18.26685354 | -0.080120444 | 0.372965119 | -0.214820206 | 0.829907499 | 0.909506443 | 0.080120444 |
| ENSMUSG00000033124 | Atg9a | 245860 | protein_coding | 421.3906821 | -0.08018109 | 0.09948317 | -0.80597643 | 0.42025644 | 0.626236745 | 0.08018109 |
| ENSMUSG00000031931 | Ankrd49 | 56503 | protein_coding | 334.6577045 | 0.080181632 | 0.113781031 | 0.704701228 | 0.480996187 | 0.674232796 | 0.080181632 |
| ENSMUSG00000113450 | Zfp935 | 71508 | protein_coding | 47.25731547 | -0.080265019 | 0.183769937 | -0.436769041 | 0.662278853 | 0.804505636 | 0.080265019 |
| ENSMUSG00000062743 | Zfp677 | 210503 | protein_coding | 85.1926908 | 0.080266844 | 0.167220715 | 0.480005391 | 0.63122356 | 0.786413345 | 0.080266844 |
| ENSMUSG00000025737 | Wdr24 | 268933 | protein_coding | 402.6985064 | 0.08031372 | 0.089621675 | 0.896141697 | 0.370177089 | 0.578860317 | 0.08031372 |
| ENSMUSG00000046541 | Zfp526 | 210172 | protein_coding | 150.6283291 | 0.080317327 | 0.106009336 | 0.757643902 | 0.44866419 | 0.649476647 | 0.080317327 |
| ENSMUSG00000018427 | Ypel2 | 77864 | protein_coding | 280.441558 | -0.080334031 | 0.244290523 | -0.328846286 | 0.742271876 | 0.856484095 | 0.080334031 |
| ENSMUSG00000044876 | Zfp444 | 72667 | protein_coding | 240.1292347 | 0.080335405 | 0.166565337 | 0.482305662 | 0.629588826 | 0.785654248 | 0.080335405 |
| ENSMUSG00000010609 | Psen2 | 19165 | protein_coding | 172.5732021 | -0.080358411 | 0.098875418 | -0.812723855 | 0.416376401 | 0.622668606 | 0.080358411 |
| ENSMUSG00000022109 | Med4 | 67381 | protein_coding | 196.7136067 | 0.080363915 | 0.096981014 | 0.828656165 | 0.407298997 | 0.614342654 | 0.080363915 |
| ENSMUSG00000079418 | Atg4a | 666468 | protein_coding | 320.8717568 | 0.08037909 | 0.09109828 | 0.882333786 | 0.377596331 | 0.585680627 | 0.08037909 |
| ENSMUSG00000022000 | Zc3h13 | 67302 | protein_coding | 376.5766969 | -0.080411667 | 0.095375614 | -0.843105103 | 0.399169665 | 0.606939679 | 0.080411667 |
| ENSMUSG00000022913 | Psmg1 | 56088 | protein_coding | 196.0344682 | 0.080419667 | 0.112196127 | 0.71677757 | 0.473511354 | 0.668798783 | 0.080419667 |
| ENSMUSG00000009293 | Ube2g2 | 22213 | protein_coding | 417.2927516 | 0.080464125 | 0.154532682 | 0.520693254 | 0.602580475 | 0.767709311 | 0.080464125 |
| ENSMUSG00000053838 | Nudcd3 | 209586 | protein_coding | 576.1982703 | -0.080467613 | 0.063669317 | -1.263836595 | 0.206288683 | 0.402470131 | 0.080467613 |
| ENSMUSG00000090215 | Trim34b | 434218 | protein_coding | 6.788809771 | 0.080480179 | 0.551416467 | 0.145951716 | 0.883959506 | 0.941599221 | 0.080480179 |
| ENSMUSG00000074354 | Arhgap20os | 408028 | lncRNA | 11.3725097 | 0.080483727 | 0.457098933 | 0.176075071 | 0.860234966 | 0.927365315 | 0.080483727 |

| ensembl_gene_id | external_gene_name | entrezgene_id | gene_biotype | baseMean | log2FoldChange | lfcSE | stat | pvalue | padj | absLog2FC |
|---|---|---|---|---|---|---|---|---|---|---|
| ENSMUSG00000021645 | Smn1 | 20595 | protein_coding | 110.8694532 | -0.080483911 | 0.170175774 | -0.472945763 | 0.636251875 | 0.789347063 | 0.080483911 |
| ENSMUSG00000037172 | Dennd11 | 243780 | protein_coding | 994.1182393 | -0.080497105 | 0.061169679 | -1.315964169 | 0.188186069 | 0.3792772 | 0.080497105 |
| ENSMUSG00000014245 | Pigl | 327942 | protein_coding | 91.14416605 | -0.080498412 | 0.121678547 | -0.661566188 | 0.508249283 | 0.697518058 | 0.080498412 |
| ENSMUSG00000063884 | Ptcd3 | 69956 | protein_coding | 795.7695115 | -0.080529167 | 0.079075901 | -1.018378122 | 0.308498295 | 0.518563181 | 0.080529167 |
| ENSMUSG00000024074 | Crim1 | 50766 | protein_coding | 804.2618235 | -0.080555872 | 0.064709987 | -1.244875415 | 0.213177549 | 0.411275679 | 0.080555872 |
| ENSMUSG00000028792 | Ak2 | 11637 | protein_coding | 2476.925376 | 0.080556689 | 0.085019625 | 0.947506988 | 0.343380497 | 0.553579409 | 0.080556689 |
| ENSMUSG00000001305 | Rrp15 | 67223 | protein_coding | 170.2919178 | -0.080567393 | 0.235448554 | -0.342186826 | 0.732210299 | 0.849675481 | 0.080567393 |
| ENSMUSG00000026753 | Ppp6c | 67857 | protein_coding | 1153.385228 | 0.080589155 | 0.075210793 | 1.071510514 | 0.283939946 | 0.493087163 | 0.080589155 |
| ENSMUSG00000055531 | Cpsf6 | 432508 | protein_coding | 1248.005451 | 0.080612616 | 0.067961578 | 1.186149864 | 0.235563122 | 0.439707037 | 0.080612616 |
| ENSMUSG00000034303 | Ccdc15 | 245902 | protein_coding | 49.24246873 | -0.080653978 | 0.21070698 | -0.38277791 | 0.701884441 | 0.83080438 | 0.080653978 |
| ENSMUSG00000074781 | Ube2n | 93765 | protein_coding | 756.225611 | 0.080699018 | 0.087742302 | 0.91972761 | 0.357715121 | 0.567385625 | 0.080699018 |
| ENSMUSG00000096935 | 1700113A16Rik | 76642 | lncRNA | 10.31970807 | 0.08070296 | 0.373044949 | 0.216335753 | 0.828726042 | 0.909295436 | 0.08070296 |
| ENSMUSG00000040134 | Rdh7 | 54150 | protein_coding | 7.287193923 | -0.080729101 | 0.535393843 | -0.150784515 | 0.880145702 | 0.939320286 | 0.080729101 |
| ENSMUSG00000005899 | Smpd4 | 77626 | protein_coding | 341.2490771 | -0.080738868 | 0.091073523 | -0.886524041 | 0.375335205 | 0.583822183 | 0.080738868 |
| ENSMUSG00000038301 | Snx10 | 71982 | protein_coding | 237.4967181 | -0.080753879 | 0.156570866 | -0.515765683 | 0.606018073 | 0.770548086 | 0.080753879 |
| ENSMUSG00000028988 | Ctnnbip1 | 67087 | protein_coding | 255.4537551 | -0.080757 | 0.093844764 | -0.860538149 | 0.389492464 | 0.597729643 | 0.080757 |
| ENSMUSG00000042506 | Usp22 | 216825 | protein_coding | 1071.881941 | 0.080776451 | 0.071186867 | 1.134710014 | 0.25649684 | 0.463387217 | 0.080776451 |
| ENSMUSG00000042419 | Nfkbil1 | 18038 | protein_coding | 135.1526614 | 0.080779509 | 0.194734645 | 0.414818374 | 0.678274868 | 0.815942199 | 0.080779509 |
| ENSMUSG00000030532 | Hddc3 | 68695 | protein_coding | 24.62364986 | -0.0808052 | 0.26343511 | -0.306736637 | 0.759043841 | 0.868226233 | 0.0808052 |
| ENSMUSG00000048603 | Gm9828 | 105246056 | lncRNA | 12.90086045 | -0.080813153 | 0.308290612 | -0.262133031 | 0.793218881 | 0.888119664 | 0.080813153 |
| ENSMUSG00000029804 | Herc3 | 73998 | protein_coding | 329.3817874 | -0.080817068 | 0.113588001 | -0.711493003 | 0.476778785 | 0.671334201 | 0.080817068 |
| ENSMUSG00000009621 | Vav2 | 22325 | protein_coding | 255.9347218 | -0.080842849 | 0.18266315 | -0.442578863 | 0.658070376 | 0.802182829 | 0.080842849 |
| ENSMUSG00000026532 | Spta1 | 20739 | protein_coding | 14.89236859 | -0.080927943 | 0.563239838 | -0.143682918 | 0.885750861 | 0.94220607 | 0.080927943 |
| ENSMUSG00000038811 | Gngt2 | 14710 | protein_coding | 26.19148776 | -0.080943229 | 0.216795661 | -0.373361851 | 0.708879149 | 0.835691442 | 0.080943229 |
| ENSMUSG00000056076 | Eif3b | 27979 | protein_coding | 1975.025248 | 0.080985865 | 0.14467711 | 0.559769714 | 0.575636524 | 0.748407359 | 0.080985865 |
| ENSMUSG00000028696 | Ipp | 16351 | protein_coding | 194.9835243 | 0.080997761 | 0.10324683 | 0.78450603 | 0.432743245 | 0.636683393 | 0.080997761 |
| ENSMUSG00000087177 | E130307A14Rik | 327744 | lncRNA | 43.12070056 | 0.081081117 | 0.233604939 | 0.347086486 | 0.728526346 | 0.847190453 | 0.081081117 |
| ENSMUSG00000021175 | Cdca7l | 217946 | protein_coding | 142.8521054 | -0.081106082 | 0.121633139 | -0.666809085 | 0.504894089 | 0.694916064 | 0.081106082 |
| ENSMUSG00000036309 | Skp1 | 21402 | protein_coding | 2997.954661 | 0.081110039 | 0.084267154 | 0.96253446 | 0.335781202 | 0.54646736 | 0.081110039 |
| ENSMUSG00000067924 | Rtl8b | 553127 | protein_coding | 9.036967617 | -0.081134307 | 0.408284781 | -0.19871989 | 0.842481865 | 0.91678972 | 0.081134307 |
| ENSMUSG00000029833 | Trim24 | 21848 | protein_coding | 908.0560888 | 0.081136118 | 0.086631883 | 0.936561859 | 0.348983979 | 0.55916073 | 0.081136118 |
| ENSMUSG00000028709 | Mob3c | 100465 | protein_coding | 300.3132817 | 0.081143869 | 0.11112139 | 0.730227264 | 0.46525128 | 0.662210133 | 0.081143869 |
| ENSMUSG00000056153 | Socs6 | 54607 | protein_coding | 983.6151712 | 0.081170145 | 0.080878228 | 1.003609347 | 0.315566948 | 0.526539948 | 0.081170145 |
| ENSMUSG00000042851 | Zc3h6 | 78751 | protein_coding | 98.3525506 | 0.081191799 | 0.293506446 | 0.27662697 | 0.782066553 | 0.881644565 | 0.081191799 |
| ENSMUSG00000021428 | Riok1 | 71340 | protein_coding | 330.2024342 | -0.081204223 | 0.131964784 | -0.615347673 | 0.538325141 | 0.721975982 | 0.081204223 |
| ENSMUSG00000030172 | Erc1 | 111173 | protein_coding | 409.4988266 | -0.081211237 | 0.106736206 | -0.760859319 | 0.446741099 | 0.64752864 | 0.081211237 |
| ENSMUSG00000027363 | Usp8 | 84092 | protein_coding | 1013.528747 | 0.081211767 | 0.04611192 | 1.761188168 | 0.07820656 | 0.210346857 | 0.081211767 |
| ENSMUSG00000027162 | Lin7c | 22343 | protein_coding | 4401.835641 | 0.081255231 | 0.079432277 | 1.022949789 | 0.306331587 | 0.516331653 | 0.081255231 |

| ensembl_gene_id | external_gene_name | entrezgene_id | gene_biotype | baseMean | log2FoldChange | lfcSE | stat | pvalue | padj | absLog2FC |
|---|---|---|---|---|---|---|---|---|---|---|
| ENSMUSG00000026036 | Nif3l1 | 65102 | protein_coding | 306.7977661 | 0.081277314 | 0.105644412 | 0.76934797 | 0.441686767 | 0.643011318 | 0.081277314 |
| ENSMUSG00000075576 | Gm12359 | 100037262 | lncRNA | 13.82329697 | -0.081325528 | 0.303003468 | -0.268398012 | 0.78839297 | 0.884955829 | 0.081325528 |
| ENSMUSG00000045838 | Ccdc9b | 214239 | protein_coding | 231.5953765 | -0.081345508 | 0.206862947 | -0.393233827 | 0.694146788 | 0.826101669 | 0.081345508 |
| ENSMUSG00000021932 | Rnaseh2b | 67153 | protein_coding | 239.1193328 | 0.081396332 | 0.09524848 | 0.8545683 | 0.392790183 | 0.600764551 | 0.081396332 |
| ENSMUSG00000006850 | Tmco6 | 71983 | protein_coding | 186.9696275 | 0.081439205 | 0.141862119 | 0.574072947 | 0.565918439 | 0.741949864 | 0.081439205 |
| ENSMUSG00000020850 | Prpf8 | 192159 | protein_coding | 3051.946306 | -0.081506989 | 0.069016512 | -1.18097809 | 0.237611427 | 0.441993884 | 0.081506989 |
| ENSMUSG00000019986 | Ahi1 | 52906 | protein_coding | 31.05234748 | 0.081518676 | 0.236715558 | 0.344373968 | 0.730565063 | 0.848536435 | 0.081518676 |
| ENSMUSG00000023307 | Marchf5 | 69104 | protein_coding | 604.5988504 | 0.081614923 | 0.072556712 | 1.124843183 | 0.260655492 | 0.468010447 | 0.081614923 |
| ENSMUSG00000022105 | Rb1 | 19645 | protein_coding | 792.5040806 | -0.081652368 | 0.079709488 | -1.024374517 | 0.305658413 | 0.515569968 | 0.081652368 |
| ENSMUSG00000022752 | Tomm70a | 28185 | protein_coding | 2173.328182 | 0.08165356 | 0.102962508 | 0.793041675 | 0.427753543 | 0.632086318 | 0.08165356 |
| ENSMUSG00000055065 | Ddx17 | 67040 | protein_coding | 3387.203206 | -0.081676211 | 0.107875802 | -0.757131895 | 0.448970847 | 0.649580486 | 0.081676211 |
| ENSMUSG00000044475 | Ascc1 | 69090 | protein_coding | 308.7549164 | 0.081703695 | 0.100955127 | 0.809307039 | 0.418338559 | 0.624549748 | 0.081703695 |
| ENSMUSG00000033216 | Eefsec | 65967 | protein_coding | 189.5232972 | 0.081707056 | 0.105941177 | 0.771249273 | 0.440559193 | 0.642426699 | 0.081707056 |
| ENSMUSG00000087281 | Gm16015 | 115490183 | lncRNA | 10.46507977 | -0.081708989 | 0.374430358 | -0.218222128 | 0.827256045 | 0.908601178 | 0.081708989 |
| ENSMUSG00000051341 | Zfp52 | 22710 | protein_coding | 129.0089943 | -0.08172385 | 0.15040373 | -0.543363187 | 0.586879767 | 0.757031173 | 0.08172385 |
| ENSMUSG00000035575 | Utp6 | 216987 | protein_coding | 713.5745267 | -0.081729365 | 0.111594542 | -0.732377801 | 0.463938004 | 0.661197506 | 0.081729365 |
| ENSMUSG00000058446 | Znrf2 | 387524 | protein_coding | 1346.588795 | -0.081750642 | 0.075445202 | -1.083576413 | 0.278552654 | 0.487064321 | 0.081750642 |
| ENSMUSG00000025200 | Cwf19l1 | 72502 | protein_coding | 148.6711268 | -0.08178501 | 0.091511604 | -0.893711904 | 0.371476055 | 0.579753122 | 0.08178501 |
| ENSMUSG00000084957 | Bbip1 | 100503572 | protein_coding | 331.8679499 | 0.081825512 | 0.095703711 | 0.854987865 | 0.392557866 | 0.600621715 | 0.081825512 |
| ENSMUSG00000037062 | Sh3glb1 | 54673 | protein_coding | 2105.538939 | 0.081837751 | 0.061954616 | 1.320930631 | 0.186524495 | 0.376929987 | 0.081837751 |
| ENSMUSG00000036087 | Slain2 | 75991 | protein_coding | 1310.080736 | 0.081844831 | 0.060170717 | 1.360210336 | 0.173763373 | 0.359707067 | 0.081844831 |
| ENSMUSG00000020220 | Vps13d | 230895 | protein_coding | 3584.976947 | -0.081858193 | 0.125995826 | -0.649689717 | 0.515892667 | 0.703221087 | 0.081858193 |
| ENSMUSG00000026415 | Fcamr | 64435 | protein_coding | 51.58230329 | -0.081937735 | 0.30499747 | -0.268650539 | 0.788198617 | 0.884950162 | 0.081937735 |
| ENSMUSG00000020698 | Cct6b | 12467 | protein_coding | 9.747463129 | 0.081941764 | 0.413125415 | 0.198345978 | 0.842774381 | 0.917045385 | 0.081941764 |
| ENSMUSG00000027680 | Fxr1 | 14359 | protein_coding | 842.246598 | 0.081946462 | 0.073271706 | 1.118391627 | 0.263399769 | 0.471131469 | 0.081946462 |
| ENSMUSG00000045538 | Ddx28 | 71986 | protein_coding | 129.8152347 | 0.081966504 | 0.146403143 | 0.559868469 | 0.575569157 | 0.74838086 | 0.081966504 |
| ENSMUSG00000027641 | Rbl1 | 19650 | protein_coding | 564.1540666 | -0.082031203 | 0.079348685 | -1.033806703 | 0.301226537 | 0.510581706 | 0.082031203 |
| ENSMUSG00000033313 | Fbxl8 | 50788 | protein_coding | 141.0807134 | -0.082065855 | 0.16345549 | -0.502068517 | 0.615619325 | 0.776003847 | 0.082065855 |
| ENSMUSG00000097145 | 9230114K14Rik | 414108 | lncRNA | 14.8085852 | 0.082072059 | 0.362999951 | 0.226093857 | 0.821128422 | 0.905179148 | 0.082072059 |
| ENSMUSG00000002625 | Akap8l | 54194 | protein_coding | 409.0635143 | -0.082072712 | 0.115100001 | -0.713055703 | 0.475811289 | 0.67038675 | 0.082072712 |
| ENSMUSG00000047731 | Wbp1l | 226178 | protein_coding | 844.097706 | -0.082086173 | 0.073839886 | -1.111677945 | 0.266276646 | 0.474095061 | 0.082086173 |
| ENSMUSG00000020770 | Unk | 217331 | protein_coding | 188.381199 | -0.082138598 | 0.100470424 | -0.817540076 | 0.413619857 | 0.62021684 | 0.082138598 |
| ENSMUSG00000057322 | Rpl38 | 67671 | protein_coding | 4105.685082 | 0.08214255 | 0.093215248 | 0.881213667 | 0.378202184 | 0.586330845 | 0.08214255 |
| ENSMUSG00000001228 | Uhrf1 | 18140 | protein_coding | 528.5475835 | -0.082145301 | 0.162418767 | -0.505762375 | 0.613023475 | 0.774633351 | 0.082145301 |
| ENSMUSG00000021794 | Glud1 | 14661 | protein_coding | 3415.081047 | -0.082161907 | 0.092462525 | -0.888596835 | 0.374219793 | 0.582599243 | 0.082161907 |
| ENSMUSG00000113902 | Ndufb1 | 102631912 | protein_coding | 1792.959815 | 0.082199149 | 0.105967838 | 0.775699027 | 0.437926715 | 0.640519699 | 0.082199149 |
| ENSMUSG00000028567 | Txndc12 | 66073 | protein_coding | 520.9143219 | 0.082257392 | 0.100623323 | 0.817478395 | 0.413655091 | 0.62021684 | 0.082257392 |
| ENSMUSG00000023072 | Cep89 | 72140 | protein_coding | 251.4950503 | 0.082282433 | 0.12454339 | 0.660672822 | 0.508822157 | 0.697877891 | 0.082282433 |

| ensembl_gene_id | external_gene_name | entrezgene_id | gene_biotype | baseMean | log2FoldChange | lfcSE | stat | pvalue | padj | absLog2FC |
|---|---|---|---|---|---|---|---|---|---|---|
| ENSMUSG00000023232 | Serinc2 | 230779 | protein_coding | 1486.283775 | -0.082291789 | 0.083931018 | -0.980469329 | 0.326854502 | 0.53793537 | 0.082291789 |
| ENSMUSG00000090534 | Gm4675 | 105245869 | lncRNA | 10.39827657 | -0.082293973 | 0.36095464 | -0.227989791 | 0.819654177 | 0.904367673 | 0.082293973 |
| ENSMUSG00000002870 | Mcm2 | 17216 | protein_coding | 444.3249119 | 0.0822987 | 0.147365195 | 0.558467686 | 0.576525064 | 0.748706989 | 0.0822987 |
| ENSMUSG00000096916 | Zfp850 | 100043772 | protein_coding | 122.3823732 | -0.082319092 | 0.118783717 | -0.693016634 | 0.488299113 | 0.680942953 | 0.082319092 |
| ENSMUSG00000003184 | Irf3 | 54131 | protein_coding | 766.7598238 | -0.082384256 | 0.107362278 | -0.767348252 | 0.442874487 | 0.644034039 | 0.082384256 |
| ENSMUSG00000024293 | Esco1 | 77805 | protein_coding | 991.005535 | 0.082409441 | 0.072370423 | 1.138717145 | 0.254821149 | 0.461436024 | 0.082409441 |
| ENSMUSG00000036667 | Tcaf1 | 77574 | protein_coding | 296.9147274 | -0.082467084 | 0.108920727 | -0.757129388 | 0.448972349 | 0.649580486 | 0.082467084 |
| ENSMUSG00000032307 | Ube2q2 | 109161 | protein_coding | 538.8985205 | -0.082489689 | 0.070137418 | -1.176115282 | 0.239548812 | 0.443718279 | 0.082489689 |
| ENSMUSG00000021103 | Mnat1 | 17420 | protein_coding | 186.4883916 | 0.082538565 | 0.093704554 | 0.88083835 | 0.37840532 | 0.58646116 | 0.082538565 |
| ENSMUSG00000041203 | Trir | 68544 | protein_coding | 768.3859097 | 0.082553038 | 0.099799274 | 0.827190767 | 0.408128944 | 0.615303154 | 0.082553038 |
| ENSMUSG00000045374 | Wdr81 | 192652 | protein_coding | 376.36015 | -0.082587868 | 0.112653784 | -0.733112238 | 0.463489977 | 0.66105374 | 0.082587868 |
| ENSMUSG00000033379 | Atp6v0b | 114143 | protein_coding | 1063.336781 | 0.082661051 | 0.106117216 | 0.778959853 | 0.436003365 | 0.638923911 | 0.082661051 |
| ENSMUSG00000021111 | Papola | 18789 | protein_coding | 2385.488786 | 0.082671828 | 0.060322591 | 1.370495304 | 0.17053234 | 0.355901352 | 0.082671828 |
| ENSMUSG00000021840 | Mapk1ip1l | 218975 | protein_coding | 1061.505136 | -0.082692577 | 0.124917625 | -0.661976856 | 0.507986054 | 0.697397162 | 0.082692577 |
| ENSMUSG00000042203 | Tbc1d22b | 381085 | protein_coding | 313.2098268 | 0.082704827 | 0.106368407 | 0.777531876 | 0.436845035 | 0.639587069 | 0.082704827 |
| ENSMUSG00000022176 | Rem2 | 140743 | protein_coding | 13.18291654 | 0.082727382 | 0.325886136 | 0.253853642 | 0.79960863 | 0.892450863 | 0.082727382 |
| ENSMUSG00000046311 | Zfp62 | 22720 | protein_coding | 642.500041 | -0.08274235 | 0.068568163 | -1.206716735 | 0.227541252 | 0.430167439 | 0.08274235 |
| ENSMUSG00000033579 | Fa2h | 338521 | protein_coding | 815.3114059 | 0.082783316 | 0.196079528 | 0.422192548 | 0.67288448 | 0.811209809 | 0.082783316 |
| ENSMUSG00000053113 | Socs3 | 12702 | protein_coding | 243.4983497 | -0.08278891 | 0.466549505 | -0.177449358 | 0.85915544 | 0.926766074 | 0.08278891 |
| ENSMUSG00000039089 | L3mbtl3 | 237339 | protein_coding | 340.1223578 | 0.082806638 | 0.135166675 | 0.612626138 | 0.540123572 | 0.723156983 | 0.082806638 |
| ENSMUSG00000037991 | Rmi2 | 223970 | protein_coding | 29.78638243 | -0.082827843 | 0.244554247 | -0.338689039 | 0.734843999 | 0.851367123 | 0.082827843 |
| ENSMUSG00000042772 | Smg7 | 226517 | protein_coding | 973.5139172 | -0.082830308 | 0.10736045 | -0.771516032 | 0.440401123 | 0.642313808 | 0.082830308 |
| ENSMUSG00000024137 | E4f1 | 13560 | protein_coding | 228.8218857 | -0.082841592 | 0.104068072 | -0.796032733 | 0.42601301 | 0.630567347 | 0.082841592 |
| ENSMUSG00000024431 | Nr3c1 | 14815 | protein_coding | 1961.128481 | -0.082982056 | 0.10325215 | -0.803683567 | 0.421579747 | 0.627386921 | 0.082982056 |
| ENSMUSG00000035696 | Rnf38 | 73469 | protein_coding | 521.7536938 | -0.083068058 | 0.122674792 | -0.6771404 | 0.498316878 | 0.688992295 | 0.083068058 |
| ENSMUSG00000005371 | Fbxo11 | 225055 | protein_coding | 607.0202474 | -0.083076497 | 0.104468743 | -0.795228259 | 0.426480736 | 0.631025097 | 0.083076497 |
| ENSMUSG00000024493 | Lars | 107045 | protein_coding | 887.1639363 | -0.083115953 | 0.089425797 | -0.92944045 | 0.352660873 | 0.56239311 | 0.083115953 |
| ENSMUSG00000063808 | Gpatch1 | 67471 | protein_coding | 265.49349 | -0.083141477 | 0.092650656 | -0.897365229 | 0.36952406 | 0.578227258 | 0.083141477 |
| ENSMUSG00000087651 | 1500009L16Rik | 69784 | protein_coding | 74.17563374 | -0.083165034 | 0.207448025 | -0.400895762 | 0.68849687 | 0.822375564 | 0.083165034 |
| ENSMUSG00000019731 | Slc35e1 | 270066 | protein_coding | 1840.320267 | -0.083173122 | 0.096596588 | -0.86103582 | 0.389218315 | 0.597539212 | 0.083173122 |
| ENSMUSG00000038268 | Ovca2 | 246257 | protein_coding | 138.1363417 | 0.083176441 | 0.120076935 | 0.692692908 | 0.488502291 | 0.681157707 | 0.083176441 |
| ENSMUSG00000068299 | Nat8f4 | 75541 | protein_coding | 182.345047 | 0.083209686 | 0.114021637 | 0.729771015 | 0.465530164 | 0.662405258 | 0.083209686 |
| ENSMUSG00000027522 | Stx16 | 228960 | protein_coding | 620.6215133 | -0.083239721 | 0.06550501 | -1.270738238 | 0.203821787 | 0.399074792 | 0.083239721 |
| ENSMUSG00000020783 | Ncbp3 | 66874 | protein_coding | 601.7227452 | -0.083280291 | 0.072135839 | -1.154492579 | 0.248298279 | 0.454115856 | 0.083280291 |
| ENSMUSG00000037020 | Wdr62 | 233064 | protein_coding | 86.1233649 | -0.083342216 | 0.177182815 | -0.470374152 | 0.638087728 | 0.790438741 | 0.083342216 |
| ENSMUSG00000025645 | Ccdc51 | 66658 | protein_coding | 207.7339464 | 0.083348626 | 0.1110185 | 0.750763396 | 0.452795062 | 0.652874025 | 0.083348626 |
| ENSMUSG00000039994 | Timeless | 21853 | protein_coding | 98.48897909 | -0.083357757 | 0.149130805 | -0.558957333 | 0.57619084 | 0.748585685 | 0.083357757 |
| ENSMUSG00000071655 | Ubxn1 | 225896 | protein_coding | 1056.661524 | -0.08337783 | 0.071677196 | -1.163240678 | 0.244731867 | 0.450073799 | 0.08337783 |

| ensembl_gene_id | external_gene_name | entrezgene_id | gene_biotype | baseMean | log2FoldChange | lfcSE | stat | pvalue | padj | absLog2FC |
|---|---|---|---|---|---|---|---|---|---|---|
| ENSMUSG00000049164 | Zfp518a | 72672 | protein_coding | 791.6659722 | -0.083454802 | 0.081036007 | -1.029848404 | 0.303081174 | 0.512579841 | 0.083454802 |
| ENSMUSG00000109293 | Dcst2 | 329702 | protein_coding | 23.75648989 | -0.0834707 | 0.221466549 | -0.376899808 | 0.706248064 | 0.833943348 | 0.0834707 |
| ENSMUSG00000039531 | Zup1 | 72580 | protein_coding | 272.2277645 | 0.083487584 | 0.115900704 | 0.72033716 | 0.471317431 | 0.66688674 | 0.083487584 |
| ENSMUSG00000031879 | Ciao2b | 68523 | protein_coding | 236.359746 | 0.083649165 | 0.115285076 | 0.725585375 | 0.468092989 | 0.664686203 | 0.083649165 |
| ENSMUSG00000025193 | Cutc | 66388 | protein_coding | 156.1716754 | -0.083686891 | 0.161904276 | -0.516891172 | 0.60523213 | 0.769671647 | 0.083686891 |
| ENSMUSG00000022748 | Cmss1 | 66497 | protein_coding | 80.94420354 | 0.083791767 | 0.141599085 | 0.591753595 | 0.554015602 | 0.733651522 | 0.083791767 |
| ENSMUSG00000031622 | Sin3b | 20467 | protein_coding | 932.1676805 | 0.083793432 | 0.085916083 | 0.975293902 | 0.329414494 | 0.540139393 | 0.083793432 |
| ENSMUSG00000026781 | Acbd5 | 74159 | protein_coding | 1338.833265 | 0.083794805 | 0.085882802 | 0.975687829 | 0.329219185 | 0.539905737 | 0.083794805 |
| ENSMUSG00000025036 | Sfxn2 | 94279 | protein_coding | 211.4022453 | -0.083861615 | 0.10269763 | -0.816587636 | 0.414164123 | 0.620546222 | 0.083861615 |
| ENSMUSG00000004865 | Srpk1 | 20815 | protein_coding | 1533.748093 | -0.083898454 | 0.088244387 | -0.950751163 | 0.341730709 | 0.551941783 | 0.083898454 |
| ENSMUSG00000029735 | Tpk1 | 29807 | protein_coding | 530.0511546 | 0.083905835 | 0.071480281 | 1.173831918 | 0.240462357 | 0.444953398 | 0.083905835 |
| ENSMUSG00000098905 | Zfp953 | 629016 | protein_coding | 200.6168361 | -0.083909569 | 0.163979636 | -0.511707252 | 0.60885591 | 0.772360968 | 0.083909569 |
| ENSMUSG00000036099 | Vezt | 215008 | protein_coding | 279.7540451 | -0.083931647 | 0.098956122 | -0.848170331 | 0.396343118 | 0.604537352 | 0.083931647 |
| ENSMUSG00000034211 | Mrps17 | 66258 | protein_coding | 633.4518303 | 0.083947939 | 0.084579855 | 0.992528756 | 0.320939659 | 0.532256186 | 0.083947939 |
| ENSMUSG00000058587 | Tmod3 | 50875 | protein_coding | 1742.578427 | 0.083958134 | 0.078838018 | 1.064944749 | 0.286900933 | 0.495744752 | 0.083958134 |
| ENSMUSG00000044150 | Bclaf3 | 382252 | protein_coding | 195.6043193 | 0.083960905 | 0.131968932 | 0.636217201 | 0.524634863 | 0.710274892 | 0.083960905 |
| ENSMUSG00000020513 | Tubd1 | 56427 | protein_coding | 80.1200667 | 0.083975478 | 0.161442578 | 0.52015694 | 0.602954194 | 0.76799575 | 0.083975478 |
| ENSMUSG00000026192 | Atic | 108147 | protein_coding | 1067.590533 | -0.083980536 | 0.12194833 | -0.688656708 | 0.491039326 | 0.683319446 | 0.083980536 |
| ENSMUSG00000020297 | Nsg2 | 18197 | protein_coding | 12.39909481 | 0.083987967 | 0.413682685 | 0.203025096 | 0.839115417 | 0.915199917 | 0.083987967 |
| ENSMUSG00000024608 | Rps14 | 20044 | protein_coding | 8190.499903 | 0.084029035 | 0.111751569 | 0.751927123 | 0.452094886 | 0.652202459 | 0.084029035 |
| ENSMUSG00000068184 | Ndufaf2 | 75597 | protein_coding | 202.6035682 | -0.084046492 | 0.124839817 | -0.673234661 | 0.500798014 | 0.691282673 | 0.084046492 |
| ENSMUSG00000036211 | H1f6 | 107970 | protein_coding | 11.66379683 | 0.084060222 | 0.449926109 | 0.186831172 | 0.851793006 | 0.921803579 | 0.084060222 |
| ENSMUSG00000036964 | Trim17 | 56631 | protein_coding | 9.198660526 | 0.084109415 | 0.407810411 | 0.206246366 | 0.836598471 | 0.913507949 | 0.084109415 |
| ENSMUSG00000034024 | Cct2 | 12461 | protein_coding | 2966.195826 | -0.084115642 | 0.088938091 | -0.945777459 | 0.344262105 | 0.55406481 | 0.084115642 |
| ENSMUSG00000040964 | Arhgef10l | 72754 | protein_coding | 632.0867104 | 0.084153713 | 0.123046896 | 0.683915771 | 0.494028355 | 0.685741493 | 0.084153713 |
| ENSMUSG00000078572 | Ndufaf8 | 208501 | protein_coding | 223.588298 | 0.084201952 | 0.102890931 | 0.818361254 | 0.413150939 | 0.619765319 | 0.084201952 |
| ENSMUSG00000005378 | Bud23 | 66138 | protein_coding | 351.5066027 | 0.084224625 | 0.116251208 | 0.724505375 | 0.468755528 | 0.66509336 | 0.084224625 |
| ENSMUSG00000032475 | Nck1 | 17973 | protein_coding | 654.5865408 | 0.084228884 | 0.104197385 | 0.808358899 | 0.418884005 | 0.625068806 | 0.084228884 |
| ENSMUSG00000029836 | Cbx3 | 12417 | protein_coding | 414.1216569 | 0.084236962 | 0.079486117 | 1.059769499 | 0.289249476 | 0.498106697 | 0.084236962 |
| ENSMUSG00000043872 | Zmym1 | 68310 | protein_coding | 92.47576316 | 0.084273187 | 0.139496682 | 0.604123237 | 0.545761712 | 0.728169923 | 0.084273187 |
| ENSMUSG00000019790 | Stxbp5 | 78808 | protein_coding | 623.4292676 | 0.084341487 | 0.066918472 | 1.260361824 | 0.207538867 | 0.404082965 | 0.084341487 |
| ENSMUSG00000022674 | Ube2v2 | 70620 | protein_coding | 159.680444 | 0.084363331 | 0.100768747 | 0.837197385 | 0.402481627 | 0.609921868 | 0.084363331 |
| ENSMUSG00000074794 | Arrdc3 | 105171 | protein_coding | 1144.124226 | 0.084404804 | 0.195183823 | 0.432437496 | 0.665423471 | 0.806118498 | 0.084404804 |
| ENSMUSG00000002748 | Baz1b | 22385 | protein_coding | 1790.541812 | -0.084435414 | 0.074708557 | -1.130197367 | 0.258393069 | 0.465469894 | 0.084435414 |
| ENSMUSG00000049878 | Rlf | 109263 | protein_coding | 629.6118668 | -0.084488261 | 0.098109406 | -0.861163718 | 0.38914788 | 0.597488667 | 0.084488261 |
| ENSMUSG00000034940 | Synrg | 217030 | protein_coding | 627.6694047 | -0.08454624 | 0.077185401 | -1.095365686 | 0.273356461 | 0.481320109 | 0.08454624 |
| ENSMUSG00000113722 | Snhg10 | 69434 | lncRNA | 7.68820376 | 0.084575662 | 0.391920734 | 0.215797877 | 0.829145303 | 0.909405742 | 0.084575662 |
| ENSMUSG00000029919 | Hpgds | 54486 | protein_coding | 149.2072805 | -0.08460263 | 0.177474013 | -0.476704325 | 0.633572682 | 0.788005719 | 0.08460263 |

| ensembl_gene_id | external_gene_name | entrezgene_id | gene_biotype | baseMean | log2FoldChange | lfcSE | stat | pvalue | padj | absLog2FC |
|---|---|---|---|---|---|---|---|---|---|---|
| ENSMUSG00000042444 | Mindy2 | 235461 | protein_coding | 492.9691984 | -0.084612995 | 0.087332455 | -0.968860835 | 0.332614629 | 0.543595917 | 0.084612995 |
| ENSMUSG00000020178 | Adora2a | 11540 | protein_coding | 8.131819185 | 0.084627419 | 0.475306411 | 0.178048132 | 0.858685175 | 0.926384278 | 0.084627419 |
| ENSMUSG00000022817 | Itgb5 | 16419 | protein_coding | 383.2884334 | -0.084667096 | 0.118375969 | -0.715238883 | 0.474461447 | 0.669373067 | 0.084667096 |
| ENSMUSG00000073676 | Hspe1 | 15528 | protein_coding | 1216.283499 | 0.084667146 | 0.11363454 | 0.745082844 | 0.45622164 | 0.655296022 | 0.084667146 |
| ENSMUSG00000018965 | Ywhah | 22629 | protein_coding | 2097.974378 | 0.084701592 | 0.100448461 | 0.843234341 | 0.399097396 | 0.606939679 | 0.084701592 |
| ENSMUSG00000044469 | Tnfaip8l1 | 66443 | protein_coding | 244.7213371 | -0.084705996 | 0.1764997 | -0.479921476 | 0.63128323 | 0.786413345 | 0.084705996 |
| ENSMUSG00000024273 | 2700062C07Rik | 68046 | protein_coding | 123.0422172 | 0.084713454 | 0.119036398 | 0.711660089 | 0.476675288 | 0.67124781 | 0.084713454 |
| ENSMUSG00000086682 | Gm16023 | 100502734 | lncRNA | 24.26440589 | -0.084721937 | 0.259739291 | -0.326180673 | 0.744287663 | 0.857877842 | 0.084721937 |
| ENSMUSG00000042251 | Pm20d1 | 212933 | protein_coding | 268.2835022 | -0.084738977 | 0.158325636 | -0.535219561 | 0.592498042 | 0.761247936 | 0.084738977 |
| ENSMUSG00000015653 | Steap2 | 74051 | protein_coding | 375.8381991 | 0.084771467 | 0.126639048 | 0.669394379 | 0.503243936 | 0.693337031 | 0.084771467 |
| ENSMUSG00000032357 | Tinag | 26944 | protein_coding | 483.4869662 | -0.084791156 | 0.109597238 | -0.773661431 | 0.439131034 | 0.641642073 | 0.084791156 |
| ENSMUSG00000033228 | Scaf11 | 72193 | protein_coding | 2022.137954 | -0.084826295 | 0.06238579 | -1.359705399 | 0.173923169 | 0.359819228 | 0.084826295 |
| ENSMUSG00000027361 | Gabpb1 | 14391 | protein_coding | 250.7634459 | 0.084862955 | 0.117494088 | 0.722274253 | 0.470125879 | 0.66616046 | 0.084862955 |
| ENSMUSG00000028089 | Chd1l | 68058 | protein_coding | 265.5957407 | -0.084891362 | 0.103536999 | -0.819913296 | 0.412265536 | 0.619020123 | 0.084891362 |
| ENSMUSG00000001576 | Ergic1 | 67458 | protein_coding | 1539.748255 | 0.084932322 | 0.084195076 | 1.008756404 | 0.313091474 | 0.523451348 | 0.084932322 |
| ENSMUSG00000038495 | Otud7b | 229603 | protein_coding | 758.6097708 | 0.084973568 | 0.162261409 | 0.523683161 | 0.60049893 | 0.766753883 | 0.084973568 |
| ENSMUSG00000004508 | Gab2 | 14389 | protein_coding | 260.3009516 | -0.084999499 | 0.096930915 | -0.87690804 | 0.380536589 | 0.588407609 | 0.084999499 |
| ENSMUSG00000028959 | Fastk | 66587 | protein_coding | 699.3684341 | -0.08500559 | 0.092819632 | -0.915814772 | 0.359764054 | 0.569481508 | 0.08500559 |
| ENSMUSG00000028243 | Ubxn2b | 68053 | protein_coding | 161.2965504 | 0.085018344 | 0.096969866 | 0.876750147 | 0.380622362 | 0.588425991 | 0.085018344 |
| ENSMUSG00000044950 | Pwwp2a | 70802 | protein_coding | 322.9426361 | -0.085022298 | 0.083219969 | -1.021657411 | 0.306943075 | 0.516893761 | 0.085022298 |
| ENSMUSG00000041765 | Ubac2 | 68889 | protein_coding | 455.0582202 | 0.085076927 | 0.09703596 | 0.876756691 | 0.380618807 | 0.588425991 | 0.085076927 |
| ENSMUSG00000030545 | Pex11a | 18631 | protein_coding | 214.0176114 | -0.085088781 | 0.101782965 | -0.835982536 | 0.403164728 | 0.610418991 | 0.085088781 |
| ENSMUSG00000023931 | Efhb | 211482 | protein_coding | 34.85180358 | 0.085144067 | 0.31336821 | 0.271706142 | 0.785847978 | 0.883905557 | 0.085144067 |
| ENSMUSG00000042680 | Garem1 | 381126 | protein_coding | 319.0829159 | -0.085150018 | 0.114111918 | -0.746197412 | 0.455548171 | 0.654989283 | 0.085150018 |
| ENSMUSG00000019494 | Cops6 | 26893 | protein_coding | 588.2140702 | 0.085242433 | 0.074156392 | 1.149495421 | 0.250351754 | 0.456634994 | 0.085242433 |
| ENSMUSG00000021234 | Fam161b | 217705 | protein_coding | 38.62473455 | -0.085251607 | 0.266897169 | -0.319417426 | 0.749409998 | 0.86128887 | 0.085251607 |
| ENSMUSG00000028882 | Ppp1r8 | 100336 | protein_coding | 440.8068931 | 0.085259859 | 0.090001299 | 0.947318095 | 0.343476713 | 0.553678483 | 0.085259859 |
| ENSMUSG00000038982 | Bloc1s5 | 17828 | protein_coding | 209.3660443 | 0.085272018 | 0.11392455 | 0.748495546 | 0.454161311 | 0.653598611 | 0.085272018 |
| ENSMUSG00000026917 | Wdr5 | 140858 | protein_coding | 539.9604248 | -0.085298537 | 0.103109263 | -0.827263573 | 0.408087685 | 0.615299191 | 0.085298537 |
| ENSMUSG00000028251 | Tstd3 | 77032 | protein_coding | 93.18488982 | 0.085345756 | 0.153088707 | 0.557492177 | 0.577191203 | 0.749071904 | 0.085345756 |
| ENSMUSG00000024325 | Ring1 | 19763 | protein_coding | 224.3528739 | -0.085370672 | 0.095286002 | -0.895941377 | 0.370284074 | 0.578905057 | 0.085370672 |
| ENSMUSG00000022119 | Rbm26 | 74213 | protein_coding | 688.7885659 | -0.085414902 | 0.104955211 | -0.8138224 | 0.415746701 | 0.622176792 | 0.085414902 |
| ENSMUSG00000065725 | Gm26165 | 115486130 | snoRNA | 17.205092 | 0.085417906 | 0.407113818 | 0.209813331 | 0.833813368 | 0.911278944 | 0.085417906 |
| ENSMUSG00000041119 | Pde9a | 18585 | protein_coding | 1592.797019 | 0.085425086 | 0.207934832 | 0.410826242 | 0.681199949 | 0.817460468 | 0.085425086 |
| ENSMUSG00000038094 | Atp13a4 | 224079 | protein_coding | 13.29831718 | -0.085436084 | 0.495337374 | -0.172480594 | 0.863059717 | 0.928838863 | 0.085436084 |
| ENSMUSG00000001095 | Slc13a2 | 20500 | protein_coding | 515.6146722 | -0.085451585 | 0.193397393 | -0.441844556 | 0.658601694 | 0.802544735 | 0.085451585 |
| ENSMUSG00000008575 | Nfib | 18028 | protein_coding | 817.9598374 | -0.085538824 | 0.164373234 | -0.52039387 | 0.602789081 | 0.76791366 | 0.085538824 |
| ENSMUSG00000028832 | Stmn1 | 16765 | protein_coding | 762.9023117 | -0.085546475 | 0.116891053 | -0.731847927 | 0.464261392 | 0.661249706 | 0.085546475 |

| ensembl_gene_id | external_gene_name | entrezgene_id | gene_biotype | baseMean | log2FoldChange | lfcSE | stat | pvalue | padj | absLog2FC |
|---|---|---|---|---|---|---|---|---|---|---|
| ENSMUSG00000001225 | Slc26a3 | 13487 | protein_coding | 5991.123857 | -0.08555628 | 0.180719274 | -0.47342089 | 0.635912929 | 0.78927929 | 0.08555628 |
| ENSMUSG00000038212 | Mfsd14b | 66631 | protein_coding | 1530.76257 | 0.08557973 | 0.056836307 | 1.505722916 | 0.132138319 | 0.300071163 | 0.08557973 |
| ENSMUSG00000015714 | Cers2 | 76893 | protein_coding | 1006.405709 | 0.085586711 | 0.117790112 | 0.72660353 | 0.467468865 | 0.664086135 | 0.085586711 |
| ENSMUSG00000034290 | Nek9 | 217718 | protein_coding | 920.118152 | -0.08559339 | 0.062467095 | -1.370215639 | 0.170619598 | 0.355943543 | 0.08559339 |
| ENSMUSG00000060038 | Dhps | 330817 | protein_coding | 236.7354971 | 0.085612459 | 0.117118037 | 0.730992946 | 0.464783462 | 0.661697442 | 0.085612459 |
| ENSMUSG00000022180 | Slc7a8 | 50934 | protein_coding | 38.78843532 | -0.085614845 | 0.258728733 | -0.330905824 | 0.740715623 | 0.855370002 | 0.085614845 |
| ENSMUSG00000020228 | Helb | 117599 | protein_coding | 455.0096815 | 0.08566053 | 0.082481071 | 1.038547736 | 0.299015123 | 0.507887459 | 0.08566053 |
| ENSMUSG00000040653 | Ppp1r14c | 76142 | protein_coding | 26.84299351 | -0.085666178 | 0.265700202 | -0.322416685 | 0.747137046 | 0.859482801 | 0.085666178 |
| ENSMUSG00000024191 | Bnip1 | 224630 | protein_coding | 245.7527819 | 0.085692326 | 0.117354692 | 0.730199403 | 0.465268307 | 0.662210133 | 0.085692326 |
| ENSMUSG00000026121 | Sema4c | 20353 | protein_coding | 63.50580094 | 0.085764425 | 0.198995702 | 0.43098632 | 0.666478318 | 0.806685297 | 0.085764425 |
| ENSMUSG00000063785 | Utp14a | 72554 | protein_coding | 397.987326 | -0.085784545 | 0.086140365 | -0.995869299 | 0.319313654 | 0.530636656 | 0.085784545 |
| ENSMUSG00000023039 | Krt7 | 110310 | protein_coding | 2304.138205 | 0.085790089 | 0.206902729 | 0.414639719 | 0.678405667 | 0.815951481 | 0.085790089 |
| ENSMUSG00000037049 | Smpd1 | 20597 | protein_coding | 355.6641161 | 0.085816936 | 0.115265642 | 0.744514444 | 0.456565306 | 0.655398985 | 0.085816936 |
| ENSMUSG00000031887 | Tradd | 71609 | protein_coding | 715.1816319 | 0.08581898 | 0.108979746 | 0.78747642 | 0.431003027 | 0.635002887 | 0.08581898 |
| ENSMUSG00000020910 | Adprm | 66358 | protein_coding | 191.0356854 | -0.085864147 | 0.131849488 | -0.65122852 | 0.514898983 | 0.702381347 | 0.085864147 |
| ENSMUSG00000074093 | Svip | 75744 | protein_coding | 348.1145159 | 0.085871862 | 0.10307971 | 0.833062707 | 0.404809364 | 0.611862171 | 0.085871862 |
| ENSMUSG00000024472 | Dcp2 | 70640 | protein_coding | 954.5147214 | -0.085875716 | 0.132471964 | -0.648255777 | 0.516819532 | 0.704062735 | 0.085875716 |
| ENSMUSG00000031963 | Bmper | 73230 | protein_coding | 13.49254247 | 0.08590785 | 0.350648779 | 0.24499686 | 0.806458863 | 0.896077905 | 0.08590785 |
| ENSMUSG00000034032 | Rpap1 | 68925 | protein_coding | 349.3514557 | 0.085912607 | 0.078459336 | 1.094995331 | 0.273518682 | 0.481499289 | 0.085912607 |
| ENSMUSG00000020280 | Pus10 | 74467 | protein_coding | 501.7482623 | 0.085951484 | 0.084914524 | 1.012211802 | 0.311436794 | 0.521451199 | 0.085951484 |
| ENSMUSG00000068876 | Cgn | 70737 | protein_coding | 2168.041686 | -0.085960013 | 0.077485769 | -1.109365153 | 0.267272689 | 0.475177965 | 0.085960013 |
| ENSMUSG00000055675 | Kbtbd11 | 74901 | protein_coding | 906.755524 | -0.08598117 | 0.210514079 | -0.4084343 | 0.682954856 | 0.818949405 | 0.08598117 |
| ENSMUSG00000021500 | Ddx46 | 212880 | protein_coding | 1073.526949 | -0.086011822 | 0.063363203 | -1.357441197 | 0.174641066 | 0.360929401 | 0.086011822 |
| ENSMUSG00000001134 | Uxt | 22294 | protein_coding | 184.8665758 | -0.086076603 | 0.123423664 | -0.697407612 | 0.485547739 | 0.678688781 | 0.086076603 |
| ENSMUSG00000029270 | Dipk1a | 67266 | protein_coding | 156.2231756 | -0.086078822 | 0.177752207 | -0.484263029 | 0.628199211 | 0.784780943 | 0.086078822 |
| ENSMUSG00000039285 | Azi2 | 27215 | protein_coding | 2077.947945 | 0.086118616 | 0.087899584 | 0.979738599 | 0.327215168 | 0.538362069 | 0.086118616 |
| ENSMUSG00000041012 | Cmtm8 | 70031 | protein_coding | 81.10136742 | -0.086181718 | 0.174087005 | -0.495049691 | 0.620565045 | 0.779093563 | 0.086181718 |
| ENSMUSG00000041702 | Btbd7 | 238386 | protein_coding | 1144.330349 | -0.086207141 | 0.078585359 | -1.096987303 | 0.272646942 | 0.48070849 | 0.086207141 |
| ENSMUSG00000032612 | Usp4 | 22258 | protein_coding | 1532.687834 | -0.086251749 | 0.082741015 | -1.04243039 | 0.297212183 | 0.506180721 | 0.086251749 |
| ENSMUSG00000021975 | Ints9 | 210925 | protein_coding | 234.1036738 | -0.086315735 | 0.076232059 | -1.132276051 | 0.257518395 | 0.464524689 | 0.086315735 |
| ENSMUSG00000036819 | Jmjd4 | 194952 | protein_coding | 198.0982211 | -0.086332424 | 0.080696855 | -1.069836287 | 0.284693005 | 0.493695176 | 0.086332424 |
| ENSMUSG00000028936 | Rpl22 | 19934 | protein_coding | 2679.143001 | -0.086471915 | 0.081680207 | -1.058664258 | 0.289752711 | 0.498400322 | 0.086471915 |
| ENSMUSG00000030079 | Ruvbl1 | 56505 | protein_coding | 233.6080077 | 0.086500295 | 0.156540913 | 0.55257308 | 0.58055578 | 0.751673912 | 0.086500295 |
| ENSMUSG00000048200 | Cracr2b | 213573 | protein_coding | 267.3330447 | 0.086505675 | 0.116798931 | 0.740637552 | 0.458913233 | 0.657276321 | 0.086505675 |
| ENSMUSG00000043421 | Hilpda | 69573 | protein_coding | 142.7318726 | -0.086537977 | 0.193249202 | -0.447805093 | 0.654293867 | 0.799999441 | 0.086537977 |
| ENSMUSG00000030987 | Stim1 | 20866 | protein_coding | 1500.450656 | -0.086673341 | 0.108057631 | -0.80210292 | 0.422493426 | 0.627691007 | 0.086673341 |
| ENSMUSG00000020246 | Hcfc2 | 67933 | protein_coding | 289.2422689 | 0.086697268 | 0.087222558 | 0.993977587 | 0.320233779 | 0.531665726 | 0.086697268 |
| ENSMUSG00000043929 | Klhl15 | 236904 | protein_coding | 189.0397276 | -0.086707573 | 0.13602644 | -0.637431764 | 0.523843644 | 0.709654498 | 0.086707573 |

| ensembl_gene_id | external_gene_name | entrezgene_id | gene_biotype | baseMean | log2FoldChange | lfcSE | stat | pvalue | padj | absLog2FC |
|---|---|---|---|---|---|---|---|---|---|---|
| ENSMUSG00000045427 | Hnrnph2 | 56258 | protein_coding | 831.706438 | 0.086765654 | 0.080441154 | 1.078622693 | 0.280755961 | 0.489264873 | 0.086765654 |
| ENSMUSG00000039678 | Tbc1d13 | 70296 | protein_coding | 1174.227876 | -0.08678305 | 0.095796872 | -0.905906929 | 0.364985117 | 0.573814056 | 0.08678305 |
| ENSMUSG00000023967 | Mrps18a | 68565 | protein_coding | 498.5297575 | 0.086786522 | 0.083450058 | 1.039981568 | 0.298348464 | 0.507160548 | 0.086786522 |
| ENSMUSG00000023755 | Rhebl1 | 69159 | protein_coding | 17.49824238 | 0.086824482 | 0.286987713 | 0.302537278 | 0.762242521 | 0.869662049 | 0.086824482 |
| ENSMUSG00000027219 | Slc28a2 | 269346 | protein_coding | 47.24335123 | -0.086826292 | 0.278858503 | -0.311363256 | 0.755524484 | 0.865754963 | 0.086826292 |
| ENSMUSG00000042284 | Itga1 | 109700 | protein_coding | 318.8886917 | -0.086886545 | 0.17384372 | -0.499796856 | 0.617218124 | 0.77714186 | 0.086886545 |
| ENSMUSG00000019478 | Rab4a | 19341 | protein_coding | 372.5666959 | 0.086935199 | 0.09281238 | 0.936676761 | 0.348924854 | 0.559122241 | 0.086935199 |
| ENSMUSG00000032469 | Dbr1 | 83703 | protein_coding | 237.7120535 | -0.086948351 | 0.092390735 | -0.941093837 | 0.346656774 | 0.556720013 | 0.086948351 |
| ENSMUSG00000090946 | Ccdc71l | 72123 | protein_coding | 1669.992506 | 0.08695324 | 0.3482849 | 0.249661241 | 0.802849334 | 0.894034894 | 0.08695324 |
| ENSMUSG00000039166 | Akap7 | 432442 | protein_coding | 711.2986529 | 0.08696491 | 0.109727721 | 0.792551872 | 0.428038958 | 0.632390628 | 0.08696491 |
| ENSMUSG00000025409 | Mbd6 | 110962 | protein_coding | 133.4677547 | -0.087046978 | 0.229735042 | -0.378901613 | 0.704760926 | 0.833286574 | 0.087046978 |
| ENSMUSG00000026970 | Rbms1 | 56878 | protein_coding | 232.9479198 | -0.0871606 | 0.118884587 | -0.73315307 | 0.463465075 | 0.66105374 | 0.0871606 |
| ENSMUSG00000037787 | Coa8 | 68020 | protein_coding | 215.8529504 | 0.08717276 | 0.09662637 | 0.902163253 | 0.366970152 | 0.57565559 | 0.08717276 |
| ENSMUSG00000039509 | Nup133 | 234865 | protein_coding | 333.0735163 | 0.087182103 | 0.097139009 | 0.89749838 | 0.369453037 | 0.578177536 | 0.087182103 |
| ENSMUSG00000035776 | Cd99l2 | 171486 | protein_coding | 377.8367789 | -0.087189696 | 0.119093466 | -0.7321115 | 0.464100515 | 0.661249706 | 0.087189696 |
| ENSMUSG00000028907 | Utp11 | 67205 | protein_coding | 484.1403401 | 0.087230646 | 0.089892273 | 0.970390928 | 0.331851669 | 0.542683099 | 0.087230646 |
| ENSMUSG00000028459 | Cd72 | 12517 | protein_coding | 10.34961669 | 0.087321149 | 0.415148937 | 0.210336921 | 0.833404721 | 0.911278944 | 0.087321149 |
| ENSMUSG00000039901 | Armh3 | 71617 | protein_coding | 275.8740192 | -0.087328423 | 0.080064225 | -1.090729631 | 0.275391868 | 0.48364598 | 0.087328423 |
| ENSMUSG00000025234 | Arih1 | 23806 | protein_coding | 1720.601435 | 0.087363967 | 0.062070095 | 1.407504972 | 0.159277704 | 0.340451304 | 0.087363967 |
| ENSMUSG00000032560 | Dnajc13 | 235567 | protein_coding | 2038.332808 | 0.087431143 | 0.069176035 | 1.263893524 | 0.206268246 | 0.402470131 | 0.087431143 |
| ENSMUSG00000007739 | Cct4 | 12464 | protein_coding | 2651.409229 | -0.087456929 | 0.101921765 | -0.858079027 | 0.390848827 | 0.599060828 | 0.087456929 |
| ENSMUSG00000021494 | Ddx41 | 72935 | protein_coding | 470.6263168 | 0.087486467 | 0.089191512 | 0.980883327 | 0.326650281 | 0.537710383 | 0.087486467 |
| ENSMUSG00000038260 | Trpm4 | 68667 | protein_coding | 3236.597373 | -0.087536779 | 0.119362554 | -0.733368853 | 0.463333491 | 0.661051222 | 0.087536779 |
| ENSMUSG00000002957 | Ap2a2 | 11772 | protein_coding | 1576.445989 | 0.087538959 | 0.114623771 | 0.763706849 | 0.445041958 | 0.645771004 | 0.087538959 |
| ENSMUSG00000045665 | Mfsd5 | 106073 | protein_coding | 242.1811773 | 0.08755007 | 0.116699309 | 0.750219263 | 0.453122659 | 0.652992964 | 0.08755007 |
| ENSMUSG00000048921 | Zfp689 | 71131 | protein_coding | 37.08889186 | 0.087561454 | 0.212255842 | 0.412527886 | 0.679952546 | 0.817009969 | 0.087561454 |
| ENSMUSG00000072692 | Rpl37rt | 100502825 | transcribed_processed_pseudogene | 113.675311 | 0.087574286 | 0.142951745 | 0.612614322 | 0.540131386 | 0.723156983 | 0.087574286 |
| ENSMUSG00000026739 | Bmi1 | 12151 | protein_coding | 283.1541738 | 0.087603247 | 0.106972479 | 0.818932569 | 0.412824887 | 0.619566809 | 0.087603247 |
| ENSMUSG00000067369 | Trmt2b | 215201 | protein_coding | 736.6463347 | 0.087607746 | 0.084727783 | 1.033990767 | 0.30114048 | 0.510544498 | 0.087607746 |
| ENSMUSG00000028580 | Pum1 | 80912 | protein_coding | 1133.116031 | -0.087644119 | 0.068656283 | -1.276563711 | 0.20175632 | 0.396639678 | 0.087644119 |
| ENSMUSG00000001829 | Clpb | 20480 | protein_coding | 446.6437885 | -0.087727192 | 0.084590952 | -1.037075366 | 0.299700734 | 0.508644997 | 0.087727192 |
| ENSMUSG00000025217 | Btrc | 12234 | protein_coding | 256.0945029 | -0.087840669 | 0.085160142 | -1.031476308 | 0.302317513 | 0.51166014 | 0.087840669 |
| ENSMUSG00000020475 | Pgam2 | 56012 | protein_coding | 22.14820102 | -0.087855627 | 0.246151683 | -0.356916623 | 0.721154225 | 0.843482486 | 0.087855627 |
| ENSMUSG00000022151 | Ttc33 | 67515 | protein_coding | 329.7416522 | -0.087895295 | 0.121665291 | -0.722435251 | 0.47002692 | 0.66616046 | 0.087895295 |
| ENSMUSG00000034401 | Spata6 | 67946 | protein_coding | 103.9956483 | -0.088004192 | 0.19006118 | -0.463030863 | 0.643342245 | 0.793445869 | 0.088004192 |
| ENSMUSG00000034543 | Morc2a | 74522 | protein_coding | 528.5720056 | -0.088008647 | 0.084300639 | -1.04398553 | 0.296492084 | 0.505840394 | 0.088008647 |
| ENSMUSG00000027883 | Gpsm2 | 76123 | protein_coding | 1066.848438 | 0.088038104 | 0.104482092 | 0.84261429 | 0.399444196 | 0.606939679 | 0.088038104 |
| ENSMUSG00000001761 | Smo | 319757 | protein_coding | 366.8522768 | -0.08804983 | 0.090594332 | -0.971913233 | 0.331093716 | 0.541777348 | 0.08804983 |

| ensembl_gene_id | external_gene_name | entrezgene_id | gene_biotype | baseMean | log2FoldChange | lfcSE | stat | pvalue | padj | absLog2FC |
|---|---|---|---|---|---|---|---|---|---|---|
| ENSMUSG00000025607 | Copg2 | 54160 | protein_coding | 912.7459869 | 0.088063819 | 0.081009298 | 1.087082852 | 0.277000195 | 0.485411397 | 0.088063819 |
| ENSMUSG00000007034 | Slc44a4 | 70129 | protein_coding | 5047.407727 | -0.088064884 | 0.113566877 | -0.775445147 | 0.438076666 | 0.640572992 | 0.088064884 |
| ENSMUSG00000005936 | Kctd20 | 66989 | protein_coding | 587.437289 | -0.088088545 | 0.076369361 | -1.153454007 | 0.248724085 | 0.454842391 | 0.088088545 |
| ENSMUSG00000025609 | Mkln1 | 27418 | protein_coding | 1254.717055 | -0.088120308 | 0.042285075 | -2.083957723 | 0.037164013 | 0.126268857 | 0.088120308 |
| ENSMUSG00000030086 | Chchd6 | 66098 | protein_coding | 54.42698589 | 0.088131475 | 0.186544888 | 0.472441119 | 0.63661196 | 0.789593156 | 0.088131475 |
| ENSMUSG00000078521 | Aunip | 69885 | protein_coding | 35.42607758 | -0.088141755 | 0.246514694 | -0.357551727 | 0.720678809 | 0.843112161 | 0.088141755 |
| ENSMUSG00000074064 | Mlycd | 56690 | protein_coding | 472.8447206 | 0.088155038 | 0.145433151 | 0.606155046 | 0.544411801 | 0.727160685 | 0.088155038 |
| ENSMUSG00000019945 | Cabcoco1 | 73287 | protein_coding | 59.40961421 | -0.088221309 | 0.193496205 | -0.455933019 | 0.648438145 | 0.797061488 | 0.088221309 |
| ENSMUSG00000037149 | Ddx1 | 104721 | protein_coding | 1987.390646 | 0.088223262 | 0.090319317 | 0.976792831 | 0.328671726 | 0.539310112 | 0.088223262 |
| ENSMUSG00000033628 | Pik3c3 | 225326 | protein_coding | 797.0333977 | -0.088236327 | 0.175683289 | -0.502246558 | 0.615494096 | 0.775907398 | 0.088236327 |
| ENSMUSG00000063802 | Hspbp1 | 66245 | protein_coding | 226.7002194 | 0.088243477 | 0.152034623 | 0.580416984 | 0.561633454 | 0.739199176 | 0.088243477 |
| ENSMUSG00000086003 | B230206L02Rik | 100039440 | lncRNA | 9.228926844 | 0.088297887 | 0.369202562 | 0.23915838 | 0.810982774 | 0.899410502 | 0.088297887 |
| ENSMUSG00000032254 | Kif23 | 71819 | protein_coding | 611.732773 | 0.088361802 | 0.119394108 | 0.740085116 | 0.459248349 | 0.657578902 | 0.088361802 |
| ENSMUSG00000063457 | Rps15 | 20054 | protein_coding | 6599.953491 | 0.088370141 | 0.097059365 | 0.910475164 | 0.362571974 | 0.571673569 | 0.088370141 |
| ENSMUSG00000010045 | Tmem115 | 56395 | protein_coding | 565.5128527 | -0.088416849 | 0.100050681 | -0.883720612 | 0.376847051 | 0.58486016 | 0.088416849 |
| ENSMUSG00000038241 | Cep250 | 16328 | protein_coding | 263.7409227 | -0.088501889 | 0.168380561 | -0.52560633 | 0.599161758 | 0.76592263 | 0.088501889 |
| ENSMUSG00000069089 | Cdk7 | 12572 | protein_coding | 463.5156393 | 0.088505064 | 0.074115808 | 1.194145576 | 0.232421028 | 0.435959788 | 0.088505064 |
| ENSMUSG00000010554 | Mettl16 | 67493 | protein_coding | 372.0740949 | 0.088508051 | 0.120850257 | 0.732377849 | 0.463937975 | 0.661197506 | 0.088508051 |
| ENSMUSG00000022471 | Xrcc6 | 14375 | protein_coding | 368.5452016 | -0.088510506 | 0.082519754 | -1.07259779 | 0.283451617 | 0.492615095 | 0.088510506 |
| ENSMUSG00000026274 | Pask | 269224 | protein_coding | 96.19714331 | 0.088564766 | 0.146222962 | 0.605683024 | 0.544725259 | 0.727457359 | 0.088564766 |
| ENSMUSG00000038888 | Ctu1 | 233189 | protein_coding | 178.3242474 | 0.088569706 | 0.15994648 | 0.553745892 | 0.579752763 | 0.751183764 | 0.088569706 |
| ENSMUSG00000039047 | Pigk | 329777 | protein_coding | 632.3935545 | 0.088629808 | 0.077916989 | 1.137490147 | 0.255333441 | 0.462094773 | 0.088629808 |
| ENSMUSG00000026202 | Tuba4a | 22145 | protein_coding | 1601.61472 | -0.088635474 | 0.190245601 | -0.465900253 | 0.6412869 | 0.792368144 | 0.088635474 |
| ENSMUSG00000033712 | Ccar2 | 219158 | protein_coding | 821.7751986 | -0.088668327 | 0.064513496 | -1.374415152 | 0.169312835 | 0.354093686 | 0.088668327 |
| ENSMUSG00000002897 | Il17ra | 16172 | protein_coding | 273.3265668 | -0.088731684 | 0.083775701 | -1.059157758 | 0.289527939 | 0.498385347 | 0.088731684 |
| ENSMUSG00000006273 | Atp6v1b2 | 11966 | protein_coding | 952.758559 | -0.088742093 | 0.064983853 | -1.365602211 | 0.172063852 | 0.357719532 | 0.088742093 |
| ENSMUSG00000053091 | Lins1 | 72635 | protein_coding | 118.0718935 | -0.088799952 | 0.151353632 | -0.58670513 | 0.557401762 | 0.736238102 | 0.088799952 |
| ENSMUSG00000007646 | Rad51c | 114714 | protein_coding | 26.63131485 | 0.08880126 | 0.272507056 | 0.325867743 | 0.744524422 | 0.858026553 | 0.08880126 |
| ENSMUSG00000020733 | Slc9a3r1 | 26941 | protein_coding | 1748.474094 | 0.088814614 | 0.143520354 | 0.618829393 | 0.536028756 | 0.720677474 | 0.088814614 |
| ENSMUSG00000034333 | Zbed4 | 223773 | protein_coding | 241.3143462 | 0.088825893 | 0.097880554 | 0.907492741 | 0.364146286 | 0.573019054 | 0.088825893 |
| ENSMUSG00000047878 | A4galt | 239559 | protein_coding | 157.3917264 | 0.088831494 | 0.305204313 | 0.29105583 | 0.771008621 | 0.87556148 | 0.088831494 |
| ENSMUSG00000054951 | 9130008F23Rik | 71583 | protein_coding | 299.8806531 | 0.088831634 | 0.195010353 | 0.455522655 | 0.648733273 | 0.79717818 | 0.088831634 |
| ENSMUSG00000023959 | Clic5 | 224796 | protein_coding | 1745.703376 | 0.088929832 | 0.09933848 | 0.895220384 | 0.37066929 | 0.57922304 | 0.088929832 |
| ENSMUSG00000029634 | Rnf6 | 74132 | protein_coding | 958.5329243 | 0.088934595 | 0.062356832 | 1.426220555 | 0.153804688 | 0.332898637 | 0.088934595 |
| ENSMUSG00000021388 | Aspn | 66695 | protein_coding | 7.241881397 | 0.088949737 | 0.640571039 | 0.138860066 | 0.889560728 | 0.944280125 | 0.088949737 |
| ENSMUSG00000046947 | Adck2 | 57869 | protein_coding | 240.6506291 | -0.088967957 | 0.118058576 | -0.753591646 | 0.451094462 | 0.651506507 | 0.088967957 |
| ENSMUSG00000059920 | 4930453N24Rik | 67609 | protein_coding | 567.8754571 | 0.089072216 | 0.094967318 | 0.937924937 | 0.348282987 | 0.558374576 | 0.089072216 |
| ENSMUSG00000054519 | Zfp867 | 237775 | protein_coding | 93.03704777 | -0.089103677 | 0.184225036 | -0.483667577 | 0.628621807 | 0.78512408 | 0.089103677 |

| ensembl_gene_id | external_gene_name | entrezgene_id | gene_biotype | baseMean | log2FoldChange | lfcSE | stat | pvalue | padj | absLog2FC |
|---|---|---|---|---|---|---|---|---|---|---|
| ENSMUSG00000061371 | Zfp873 | 408062 | protein_coding | 113.4814935 | 0.089108906 | 0.18920324 | 0.470969238 | 0.637662702 | 0.790351042 | 0.089108906 |
| ENSMUSG00000016496 | Cd274 | 60533 | protein_coding | 30.45531718 | 0.089161397 | 0.258563766 | 0.344833303 | 0.730219695 | 0.848416318 | 0.089161397 |
| ENSMUSG00000027236 | Eif3j1 | 78655 | protein_coding | 443.9721644 | -0.089190657 | 0.127040144 | -0.702066717 | 0.482637558 | 0.676057604 | 0.089190657 |
| ENSMUSG00000038797 | Zscan2 | 22691 | protein_coding | 46.05521434 | -0.08920329 | 0.212196612 | -0.420380369 | 0.674207607 | 0.812374519 | 0.08920329 |
| ENSMUSG00000048833 | Slc39a9 | 328133 | protein_coding | 1642.536146 | 0.089234015 | 0.095131095 | 0.938011017 | 0.348238748 | 0.558359853 | 0.089234015 |
| ENSMUSG00000028059 | Arhgef2 | 16800 | protein_coding | 800.1685177 | 0.089237095 | 0.084138021 | 1.060603685 | 0.288870047 | 0.497873797 | 0.089237095 |
| ENSMUSG00000048787 | Dcun1d3 | 233805 | protein_coding | 199.4955648 | -0.089269305 | 0.127612886 | -0.699532058 | 0.484219586 | 0.67767767 | 0.089269305 |
| ENSMUSG00000001157 | Gmcl1 | 23885 | protein_coding | 625.4432473 | 0.08928098 | 0.07050608 | 1.266287678 | 0.205410107 | 0.401099937 | 0.08928098 |
| ENSMUSG00000002257 | Def6 | 23853 | protein_coding | 12.22099931 | 0.089315612 | 0.37305007 | 0.239419905 | 0.810779997 | 0.899310849 | 0.089315612 |
| ENSMUSG00000009566 | Fpgs | 14287 | protein_coding | 222.555599 | -0.089382524 | 0.138603606 | -0.644878773 | 0.519005749 | 0.706014482 | 0.089382524 |
| ENSMUSG00000051359 | Ncald | 52589 | protein_coding | 436.2026343 | 0.089387349 | 0.097256284 | 0.91909072 | 0.358048123 | 0.567687686 | 0.089387349 |
| ENSMUSG00000000296 | Tpd52l1 | 21987 | protein_coding | 104.0556716 | -0.089409459 | 0.179358589 | -0.498495553 | 0.618134803 | 0.777697562 | 0.089409459 |
| ENSMUSG00000022636 | Alcam | 11658 | protein_coding | 838.4631311 | 0.08945185 | 0.10767352 | 0.830769256 | 0.406103994 | 0.613120797 | 0.08945185 |
| ENSMUSG00000026078 | Pdcl3 | 68833 | protein_coding | 556.5737949 | 0.089452437 | 0.07166208 | 1.248253424 | 0.211938265 | 0.409772837 | 0.089452437 |
| ENSMUSG00000034254 | Agpat1 | 55979 | protein_coding | 1209.325676 | 0.089472683 | 0.070336157 | 1.272072382 | 0.203347403 | 0.398390828 | 0.089472683 |
| ENSMUSG00000035356 | Nfkbiz | 80859 | protein_coding | 1144.590828 | -0.089482077 | 0.195495044 | -0.457720437 | 0.647153304 | 0.796158015 | 0.089482077 |
| ENSMUSG00000017421 | Zfp207 | 22680 | protein_coding | 2016.953282 | 0.089496542 | 0.0528258 | 1.694182415 | 0.090230625 | 0.232179871 | 0.089496542 |
| ENSMUSG00000052336 | Cx3cr1 | 13051 | protein_coding | 7.056943036 | 0.089519833 | 0.553999562 | 0.161588274 | 0.871630092 | 0.933960586 | 0.089519833 |
| ENSMUSG00000062157 | Ifnlr1 | 242700 | protein_coding | 162.986753 | -0.08953009 | 0.134803185 | -0.664154117 | 0.506591677 | 0.696383208 | 0.08953009 |
| ENSMUSG00000028463 | Car9 | 230099 | protein_coding | 713.9821823 | 0.089561897 | 0.11168811 | 0.801892846 | 0.422614945 | 0.627750586 | 0.089561897 |
| ENSMUSG00000010290 | AI597479 | 98404 | protein_coding | 325.0301306 | 0.089656813 | 0.099268264 | 0.903176999 | 0.366431963 | 0.575230748 | 0.089656813 |
| ENSMUSG00000022370 | Mrpl13 | 68537 | protein_coding | 469.1426528 | 0.089661573 | 0.092096671 | 0.973559331 | 0.330275387 | 0.54082722 | 0.089661573 |
| ENSMUSG00000021464 | Ror2 | 26564 | protein_coding | 100.7364781 | -0.089715273 | 0.184817997 | -0.485424983 | 0.627374914 | 0.784304814 | 0.089715273 |
| ENSMUSG00000022952 | Runx1 | 12394 | protein_coding | 225.4857684 | -0.08972439 | 0.226784053 | -0.395638003 | 0.692372098 | 0.824901098 | 0.08972439 |
| ENSMUSG00000033715 | Akr1c14 | 105387 | protein_coding | 3900.362847 | -0.089746334 | 0.19332059 | -0.464235776 | 0.642478832 | 0.793041137 | 0.089746334 |
| ENSMUSG00000039671 | Zmynd8 | 228880 | protein_coding | 1005.34176 | -0.08974694 | 0.10990811 | -0.816563395 | 0.414177981 | 0.620546222 | 0.08974694 |
| ENSMUSG00000047822 | Angptl8 | 624219 | protein_coding | 13.36779916 | -0.089752413 | 0.354517788 | -0.253167586 | 0.800138713 | 0.892792254 | 0.089752413 |
| ENSMUSG00000028455 | Stoml2 | 66592 | protein_coding | 681.636061 | 0.089761751 | 0.076103843 | 1.179464097 | 0.238213425 | 0.442395134 | 0.089761751 |
| ENSMUSG00000010608 | Rbm25 | 67039 | protein_coding | 1616.100907 | -0.089842007 | 0.09339582 | -0.961948901 | 0.336075273 | 0.54678314 | 0.089842007 |
| ENSMUSG00000015733 | Capza2 | 12343 | protein_coding | 2688.239638 | 0.089870351 | 0.099233057 | 0.905649325 | 0.365121492 | 0.573932215 | 0.089870351 |
| ENSMUSG00000025508 | Rplp2 | 67186 | protein_coding | 4916.895875 | 0.089919085 | 0.088264926 | 1.018740846 | 0.308326015 | 0.518367825 | 0.089919085 |
| ENSMUSG00000070934 | Rraga | 68441 | protein_coding | 305.5595625 | 0.0899548 | 0.106732628 | 0.842805067 | 0.399337474 | 0.606939679 | 0.0899548 |
| ENSMUSG00000086915 | Gm16364 | 100504626 | lncRNA | 7.239988798 | -0.089970564 | 0.481033646 | -0.187035906 | 0.851632481 | 0.921798204 | 0.089970564 |
| ENSMUSG00000058173 | Smco4 | 170748 | protein_coding | 176.1399298 | 0.089973237 | 0.119958541 | 0.750036105 | 0.45323296 | 0.65307532 | 0.089973237 |
| ENSMUSG00000032407 | U2surp | 67958 | protein_coding | 1711.363564 | -0.089995931 | 0.062035076 | -1.450726539 | 0.146856022 | 0.323254455 | 0.089995931 |
| ENSMUSG00000038535 | Zfp280d | 235469 | protein_coding | 397.6254615 | -0.090004534 | 0.104570747 | -0.860704708 | 0.389400699 | 0.5976464 | 0.090004534 |
| ENSMUSG00000028538 | St3gal3 | 20441 | protein_coding | 937.2444376 | -0.090032133 | 0.095372397 | -0.944006188 | 0.345166486 | 0.555110237 | 0.090032133 |
| ENSMUSG00000090231 | Cfb | 14962 | protein_coding | 94.46295091 | 0.090045598 | 0.149153183 | 0.603712208 | 0.546034996 | 0.728347446 | 0.090045598 |

| ensembl_gene_id | external_gene_name | entrezgene_id | gene_biotype | baseMean | log2FoldChange | lfcSE | stat | pvalue | padj | absLog2FC |
|---|---|---|---|---|---|---|---|---|---|---|
| ENSMUSG00000035868 | Zfp983 | 73229 | protein_coding | 154.3161403 | 0.090098681 | 0.163876998 | 0.549794551 | 0.582460297 | 0.753466061 | 0.090098681 |
| ENSMUSG00000027411 | Vps16 | 80743 | protein_coding | 547.7156723 | -0.090127465 | 0.076346451 | -1.180506285 | 0.237798912 | 0.442071905 | 0.090127465 |
| ENSMUSG00000046567 | 4930430F08Rik | 68281 | protein_coding | 57.76169293 | 0.090228243 | 0.170075531 | 0.53051866 | 0.595752375 | 0.763909664 | 0.090228243 |
| ENSMUSG00000030929 | Eri2 | 71151 | protein_coding | 153.1564112 | 0.090251019 | 0.114183151 | 0.79040575 | 0.429290845 | 0.633417748 | 0.090251019 |
| ENSMUSG00000038527 | C1rl | 232371 | protein_coding | 10.90459731 | 0.090272365 | 0.452889786 | 0.19932524 | 0.842008337 | 0.916524895 | 0.090272365 |
| ENSMUSG00000015165 | Hnrnpl | 15388 | protein_coding | 2732.839205 | 0.090338854 | 0.060461162 | 1.49416338 | 0.135132921 | 0.304621103 | 0.090338854 |
| ENSMUSG00000026174 | Cnot9 | 58184 | protein_coding | 528.8884852 | 0.090379938 | 0.070425785 | 1.283335892 | 0.199374423 | 0.393706399 | 0.090379938 |
| ENSMUSG00000026526 | Fh1 | 14194 | protein_coding | 1980.697323 | 0.090417876 | 0.095491835 | 0.946865 | 0.343707575 | 0.55376472 | 0.090417876 |
| ENSMUSG00000026313 | Hdac4 | 208727 | protein_coding | 609.6860761 | -0.090444957 | 0.126072122 | -0.717406478 | 0.473123324 | 0.668535437 | 0.090444957 |
| ENSMUSG00000021806 | Nid2 | 18074 | protein_coding | 98.34852293 | 0.090456775 | 0.179264255 | 0.504600179 | 0.613839684 | 0.775232992 | 0.090456775 |
| ENSMUSG00000003235 | Eif2b5 | 224045 | protein_coding | 883.9291527 | 0.090477432 | 0.119415187 | 0.757671066 | 0.448647924 | 0.649476647 | 0.090477432 |
| ENSMUSG00000017291 | Taok1 | 216965 | protein_coding | 2399.406083 | -0.090478941 | 0.095979298 | -0.942692263 | 0.345838332 | 0.555701235 | 0.090478941 |
| ENSMUSG00000025958 | Creb1 | 12912 | protein_coding | 997.6107018 | 0.090499275 | 0.069894488 | 1.294798456 | 0.195389765 | 0.388583865 | 0.090499275 |
| ENSMUSG00000038668 | Lpar1 | 14745 | protein_coding | 502.8157986 | -0.090533879 | 0.061750423 | -1.466125644 | 0.142614061 | 0.316152646 | 0.090533879 |
| ENSMUSG00000028391 | Wdr31 | 71354 | protein_coding | 52.10085906 | -0.090596358 | 0.184880375 | -0.490026903 | 0.624114861 | 0.782255391 | 0.090596358 |
| ENSMUSG00000034312 | Iqsec1 | 232227 | protein_coding | 274.5761219 | -0.09059707 | 0.16036991 | -0.564925613 | 0.572124362 | 0.745911176 | 0.09059707 |
| ENSMUSG00000015341 | Golga7 | 57437 | protein_coding | 597.3912718 | -0.090705245 | 0.084368617 | -1.075106451 | 0.282327072 | 0.491304009 | 0.090705245 |
| ENSMUSG00000029345 | Tfip11 | 54723 | protein_coding | 458.2967466 | -0.090725091 | 0.082774261 | -1.096054383 | 0.273054976 | 0.481045059 | 0.090725091 |
| ENSMUSG00000002550 | Uck1 | 22245 | protein_coding | 414.9102286 | -0.090725217 | 0.100468955 | -0.903017418 | 0.366516651 | 0.575230748 | 0.090725217 |
| ENSMUSG00000061991 | H2ac10 | 319173 | protein_coding | 97.91090455 | 0.090726966 | 0.300919649 | 0.301498974 | 0.763034032 | 0.870097644 | 0.090726966 |
| ENSMUSG00000030688 | Stard10 | 56018 | protein_coding | 6173.787291 | 0.090742721 | 0.087260817 | 1.039902259 | 0.298385313 | 0.50716906 | 0.090742721 |
| ENSMUSG00000015748 | Prpf3 | 70767 | protein_coding | 326.6106073 | 0.090764914 | 0.136896407 | 0.663018962 | 0.50731841 | 0.696822981 | 0.090764914 |
| ENSMUSG00000018569 | Cldn7 | 53624 | protein_coding | 7591.607334 | 0.090766982 | 0.111978797 | 0.810572934 | 0.417610965 | 0.623870517 | 0.090766982 |
| ENSMUSG00000030779 | Rbbp6 | 19647 | protein_coding | 845.9963513 | -0.090844606 | 0.10639179 | -0.85386857 | 0.393177816 | 0.601011157 | 0.090844606 |
| ENSMUSG00000042155 | Klhl23 | 277396 | protein_coding | 74.24918285 | -0.090886223 | 0.161526065 | -0.56267218 | 0.573658132 | 0.747054756 | 0.090886223 |
| ENSMUSG00000003934 | Efnb3 | 13643 | protein_coding | 8.966001019 | 0.090899102 | 0.516740661 | 0.175908552 | 0.860365786 | 0.927443574 | 0.090899102 |
| ENSMUSG00000026355 | Mcm6 | 17219 | protein_coding | 877.054822 | -0.090941718 | 0.121548595 | -0.748192263 | 0.454344197 | 0.653612209 | 0.090941718 |
| ENSMUSG00000085396 | Firre | 103012 | lncRNA | 794.4417976 | -0.09100745 | 0.183638004 | -0.495580696 | 0.620190275 | 0.778936265 | 0.09100745 |
| ENSMUSG00000025868 | Higd2a | 67044 | protein_coding | 1523.876976 | -0.09101346 | 0.076734374 | -1.186084601 | 0.235588891 | 0.439707037 | 0.09101346 |
| ENSMUSG00000038181 | Chpf2 | 100910 | protein_coding | 242.3083203 | 0.091018173 | 0.101141252 | 0.899911471 | 0.368167365 | 0.577011688 | 0.091018173 |
| ENSMUSG00000028156 | Eif4e | 13684 | protein_coding | 1018.500595 | 0.091025494 | 0.15513574 | 0.586747412 | 0.55737336 | 0.736238102 | 0.091025494 |
| ENSMUSG00000023025 | Larp4 | 207214 | protein_coding | 2307.782616 | -0.09102815 | 0.078493336 | -1.159692703 | 0.246173942 | 0.451682819 | 0.09102815 |
| ENSMUSG00000032215 | Rsl24d1 | 225215 | protein_coding | 268.3788562 | -0.091072884 | 0.124747265 | -0.730059167 | 0.465354019 | 0.6622384 | 0.091072884 |
| ENSMUSG00000044857 | Lemd2 | 224640 | protein_coding | 440.2299064 | -0.091092912 | 0.099865173 | -0.912158956 | 0.361685046 | 0.570896781 | 0.091092912 |
| ENSMUSG00000039813 | Tbc1d2 | 381605 | protein_coding | 616.1020756 | 0.091181326 | 0.193105644 | 0.47218364 | 0.636795716 | 0.789636603 | 0.091181326 |
| ENSMUSG00000070565 | Rasal2 | 226525 | protein_coding | 172.5472892 | -0.091284794 | 0.122473736 | -0.745341791 | 0.456065123 | 0.655296022 | 0.091284794 |
| ENSMUSG00000029580 | Actb | 11461 | protein_coding | 49830.20546 | 0.091324748 | 0.194549492 | 0.469416535 | 0.638771932 | 0.790625182 | 0.091324748 |
| ENSMUSG00000020929 | Eftud2 | 20624 | protein_coding | 785.1667489 | 0.091377946 | 0.091341067 | 1.000403751 | 0.317115156 | 0.528403619 | 0.091377946 |

| ensembl_gene_id | external_gene_name | entrezgene_id | gene_biotype | baseMean | log2FoldChange | lfcSE | stat | pvalue | padj | absLog2FC |
|---|---|---|---|---|---|---|---|---|---|---|
| ENSMUSG00000048668 | Rhno1 | 72440 | protein_coding | 146.5259742 | 0.091452715 | 0.117155845 | 0.780607357 | 0.435033463 | 0.63804908 | 0.091452715 |
| ENSMUSG00000001687 | Ubl3 | 24109 | protein_coding | 1630.703086 | 0.091489019 | 0.059212816 | 1.54508813 | 0.122324948 | 0.284935912 | 0.091489019 |
| ENSMUSG00000032590 | Apeh | 235606 | protein_coding | 961.1401093 | -0.091523892 | 0.076229281 | -1.200639575 | 0.229891043 | 0.433125054 | 0.091523892 |
| ENSMUSG00000067367 | Lyar | 17089 | protein_coding | 214.1489105 | -0.091582769 | 0.215623177 | -0.424735273 | 0.671029667 | 0.809831834 | 0.091582769 |
| ENSMUSG00000003662 | Ciao1 | 26371 | protein_coding | 574.9777914 | 0.091600985 | 0.106282363 | 0.861864398 | 0.388762143 | 0.597126655 | 0.091600985 |
| ENSMUSG00000000216 | Scnn1g | 20278 | protein_coding | 25.71219802 | 0.0916066 | 0.370207132 | 0.24744688 | 0.804562397 | 0.894700251 | 0.0916066 |
| ENSMUSG00000031536 | Polb | 18970 | protein_coding | 284.1521382 | -0.09162731 | 0.117363032 | -0.780716967 | 0.434968979 | 0.638013252 | 0.09162731 |
| ENSMUSG00000022707 | Gbe1 | 74185 | protein_coding | 369.0820547 | -0.091656645 | 0.108103514 | -0.847859991 | 0.396515948 | 0.604651202 | 0.091656645 |
| ENSMUSG00000031351 | Zfp185 | 22673 | protein_coding | 15.95802291 | 0.091661935 | 0.306010009 | 0.299539011 | 0.764528811 | 0.871371371 | 0.091661935 |
| ENSMUSG00000074129 | Rpl13a | 22121 | protein_coding | 387.007174 | 0.091733074 | 0.123786725 | 0.741057444 | 0.458658611 | 0.65713215 | 0.091733074 |
| ENSMUSG00000032350 | Gclc | 14629 | protein_coding | 1561.050535 | -0.091738753 | 0.094835289 | -0.967348277 | 0.333369957 | 0.544260768 | 0.091738753 |
| ENSMUSG00000029030 | Tprgl | 67808 | protein_coding | 891.0421786 | -0.091778814 | 0.083999545 | -1.092610841 | 0.274564701 | 0.48259397 | 0.091778814 |
| ENSMUSG00000028383 | Hsdl2 | 72479 | protein_coding | 629.6674668 | 0.091778981 | 0.139674318 | 0.657092745 | 0.511121283 | 0.699050301 | 0.091778981 |
| ENSMUSG00000039840 | Epg5 | 100502841 | protein_coding | 638.7880159 | -0.091787561 | 0.092971685 | -0.987263613 | 0.323513424 | 0.534446871 | 0.091787561 |
| ENSMUSG00000044350 | Lacc1 | 210808 | protein_coding | 136.4055267 | 0.091882579 | 0.120164668 | 0.764638893 | 0.444486603 | 0.645376719 | 0.091882579 |
| ENSMUSG00000026986 | Hnmt | 140483 | protein_coding | 32.02705619 | -0.091932158 | 0.256954963 | -0.357775376 | 0.72051142 | 0.843075176 | 0.091932158 |
| ENSMUSG00000024997 | Prdx3 | 11757 | protein_coding | 1850.198742 | 0.09193791 | 0.082461827 | 1.114914785 | 0.26488694 | 0.472307085 | 0.09193791 |
| ENSMUSG00000021408 | Ripk1 | 19766 | protein_coding | 864.997138 | -0.091964232 | 0.061931595 | -1.484932394 | 0.137561734 | 0.308388369 | 0.091964232 |
| ENSMUSG00000043639 | Rbm20 | 73713 | protein_coding | 54.43522286 | 0.091992442 | 0.223295546 | 0.411976162 | 0.680356896 | 0.817182783 | 0.091992442 |
| ENSMUSG00000024740 | Ddb1 | 13194 | protein_coding | 5977.777413 | 0.09202681 | 0.067106556 | 1.371353484 | 0.170264791 | 0.355529311 | 0.09202681 |
| ENSMUSG00000029454 | Mapkapk5 | 17165 | protein_coding | 20.90735851 | -0.092031249 | 0.332781397 | -0.276551665 | 0.782124384 | 0.881647359 | 0.092031249 |
| ENSMUSG00000041629 | Fam104a | 28081 | protein_coding | 222.0125599 | -0.092065271 | 0.132067411 | -0.697108164 | 0.485735106 | 0.678784766 | 0.092065271 |
| ENSMUSG00000038375 | Trp53inp2 | 68728 | protein_coding | 1158.061273 | 0.092075985 | 0.106828275 | 0.861906506 | 0.388738969 | 0.597126655 | 0.092075985 |
| ENSMUSG00000040940 | Arhgef1 | 16801 | protein_coding | 558.1445791 | -0.092078194 | 0.11804601 | -0.78001953 | 0.435379379 | 0.638336365 | 0.092078194 |
| ENSMUSG00000001435 | Col18a1 | 12822 | protein_coding | 18.36626104 | -0.092091348 | 0.331183873 | -0.278067127 | 0.78096083 | 0.880896827 | 0.092091348 |
| ENSMUSG00000028677 | Rnf220 | 66743 | protein_coding | 476.7048473 | -0.092094276 | 0.066266134 | -1.389763828 | 0.164600605 | 0.347990237 | 0.092094276 |
| ENSMUSG00000054967 | Zfp647 | 239546 | protein_coding | 11.13014401 | 0.092100109 | 0.382507017 | 0.240780181 | 0.809725491 | 0.898579225 | 0.092100109 |
| ENSMUSG00000034518 | Hmgxb4 | 70823 | protein_coding | 296.025382 | -0.092152144 | 0.103981868 | -0.886232822 | 0.375492081 | 0.583911261 | 0.092152144 |
| ENSMUSG00000029681 | Bcl7b | 12054 | protein_coding | 195.4355056 | 0.092182731 | 0.12028374 | 0.766377327 | 0.443451819 | 0.644466916 | 0.092182731 |
| ENSMUSG00000020594 | Pum2 | 80913 | protein_coding | 2435.973258 | 0.092188688 | 0.055278505 | 1.667713137 | 0.095372686 | 0.240851081 | 0.092188688 |
| ENSMUSG00000025421 | Hdhd2 | 76987 | protein_coding | 135.1723647 | -0.092203768 | 0.124936512 | -0.738004986 | 0.460511419 | 0.658781284 | 0.092203768 |
| ENSMUSG00000055401 | Fbxo6 | 50762 | protein_coding | 206.819178 | 0.092241256 | 0.113258666 | 0.814430003 | 0.415398656 | 0.621841803 | 0.092241256 |
| ENSMUSG00000033972 | Zfp944 | 319615 | protein_coding | 312.0181722 | 0.092249095 | 0.079384169 | 1.16205909 | 0.245211464 | 0.450668923 | 0.092249095 |
| ENSMUSG00000027810 | Eif2a | 229317 | protein_coding | 1121.149322 | 0.092249704 | 0.082120617 | 1.12334402 | 0.261291415 | 0.468993651 | 0.092249704 |
| ENSMUSG00000028187 | Rpf1 | 70285 | protein_coding | 523.4780691 | -0.092309789 | 0.086271912 | -1.069986599 | 0.284625341 | 0.493695176 | 0.092309789 |
| ENSMUSG00000060121 | Gemin2 | 66603 | protein_coding | 102.6286122 | -0.092389116 | 0.119732117 | -0.771631853 | 0.440332503 | 0.642313808 | 0.092389116 |
| ENSMUSG00000032783 | Troap | 78733 | protein_coding | 63.83118284 | 0.092432152 | 0.182428572 | 0.506675852 | 0.612382277 | 0.774101099 | 0.092432152 |
| ENSMUSG00000022562 | Oplah | 75475 | protein_coding | 437.8699614 | -0.092544858 | 0.10726344 | -0.862781001 | 0.388257887 | 0.596697378 | 0.092544858 |

| ensembl_gene_id | external_gene_name | entrezgene_id | gene_biotype | baseMean | log2FoldChange | lfcSE | stat | pvalue | padj | absLog2FC |
|---|---|---|---|---|---|---|---|---|---|---|
| ENSMUSG00000039205 | Ciz1 | 68379 | protein_coding | 394.651458 | -0.092559465 | 0.085362312 | -1.084313004 | 0.278226042 | 0.48677333 | 0.092559465 |
| ENSMUSG00000028750 | Pla2g2c | 18781 | protein_coding | 42.67598731 | 0.092613713 | 0.311689937 | 0.297134113 | 0.766364122 | 0.872270412 | 0.092613713 |
| ENSMUSG00000070697 | Utp3 | 65961 | protein_coding | 409.1438693 | -0.09263069 | 0.109801171 | -0.843622059 | 0.398880634 | 0.606758737 | 0.09263069 |
| ENSMUSG00000059486 | Kbtbd2 | 210973 | protein_coding | 715.1638505 | -0.092651091 | 0.096408917 | -0.961022013 | 0.3365411 | 0.547261525 | 0.092651091 |
| ENSMUSG00000068417 | Pnp2 | 667034 | protein_coding | 231.917284 | -0.092670102 | 0.141399694 | -0.655376962 | 0.51222508 | 0.699958912 | 0.092670102 |
| ENSMUSG00000051502 | Ufsp1 | 70240 | protein_coding | 59.3371556 | -0.092672549 | 0.157904538 | -0.586889715 | 0.557277778 | 0.736238102 | 0.092672549 |
| ENSMUSG00000029472 | Anapc5 | 59008 | protein_coding | 2264.946079 | 0.092679168 | 0.083618485 | 1.10835741 | 0.267707491 | 0.47563245 | 0.092679168 |
| ENSMUSG00000054770 | Kctd18 | 51960 | protein_coding | 111.6289149 | 0.092723434 | 0.128391766 | 0.722191435 | 0.470176789 | 0.66616046 | 0.092723434 |
| ENSMUSG00000042055 | Wdr11 | 207425 | protein_coding | 730.0979423 | -0.092820551 | 0.077387239 | -1.199429664 | 0.230360919 | 0.433458782 | 0.092820551 |
| ENSMUSG00000019295 | Tmem129 | 68366 | protein_coding | 335.7374678 | 0.092825833 | 0.10601402 | 0.875599591 | 0.381247748 | 0.589049774 | 0.092825833 |
| ENSMUSG00000026289 | Atg16l1 | 77040 | protein_coding | 484.715146 | -0.092848545 | 0.093628182 | -0.991673047 | 0.321357043 | 0.532354566 | 0.092848545 |
| ENSMUSG00000022369 | Mtbp | 105837 | protein_coding | 113.9118283 | 0.092862569 | 0.129625833 | 0.716389371 | 0.473750956 | 0.668798783 | 0.092862569 |
| ENSMUSG00000026321 | Tnfrsf11a | 21934 | protein_coding | 1881.553346 | -0.092872203 | 0.093566932 | -0.992575061 | 0.320917083 | 0.532256186 | 0.092872203 |
| ENSMUSG00000031246 | Sh3bgrl | 56726 | protein_coding | 2625.395031 | 0.092873528 | 0.079976221 | 1.161264266 | 0.245534447 | 0.450976122 | 0.092873528 |
| ENSMUSG00000031357 | Syap1 | 67043 | protein_coding | 1302.249502 | 0.092930359 | 0.070485051 | 1.31844068 | 0.18735617 | 0.378024934 | 0.092930359 |
| ENSMUSG00000022787 | Wdr53 | 68980 | protein_coding | 124.3373339 | 0.092944123 | 0.102928052 | 0.903000898 | 0.366525418 | 0.575230748 | 0.092944123 |
| ENSMUSG00000087497 | 2810001G20Rik | 66456 | lncRNA | 58.92163911 | 0.092975401 | 0.181611275 | 0.511947293 | 0.608687898 | 0.772339747 | 0.092975401 |
| ENSMUSG00000041143 | Tmco4 | 77056 | protein_coding | 809.7937485 | 0.093014288 | 0.102828988 | 0.904553177 | 0.36570215 | 0.574448111 | 0.093014288 |
| ENSMUSG00000057531 | Dtnbp1 | 94245 | protein_coding | 178.797751 | -0.093017647 | 0.09802026 | -0.948963475 | 0.342639189 | 0.552958586 | 0.093017647 |
| ENSMUSG00000014763 | Fam120b | 67544 | protein_coding | 816.6300498 | 0.093090179 | 0.067093642 | 1.387466474 | 0.165299567 | 0.348987948 | 0.093090179 |
| ENSMUSG00000072763 | 5430403G16Rik | 77200 | protein_coding | 42.70860976 | -0.093124989 | 0.264002224 | -0.352743199 | 0.72428097 | 0.844771344 | 0.093124989 |
| ENSMUSG00000021643 | Serf1 | 20365 | protein_coding | 103.7175076 | 0.093139312 | 0.140558964 | 0.66263516 | 0.507564246 | 0.696938216 | 0.093139312 |
| ENSMUSG00000020238 | Ncln | 103425 | protein_coding | 1042.877781 | 0.093181659 | 0.102650087 | 0.907760155 | 0.364004954 | 0.572855539 | 0.093181659 |
| ENSMUSG00000035914 | Cd276 | 102657 | protein_coding | 69.51427485 | -0.093194307 | 0.211649621 | -0.440323524 | 0.659702806 | 0.802969027 | 0.093194307 |
| ENSMUSG00000034300 | Fam53c | 66306 | protein_coding | 265.9235074 | -0.093209985 | 0.090740759 | -1.027211867 | 0.304320711 | 0.513961388 | 0.093209985 |
| ENSMUSG00000020079 | Supv3l1 | 338359 | protein_coding | 349.8575628 | -0.093249122 | 0.113573819 | -0.821044167 | 0.411621111 | 0.618518967 | 0.093249122 |
| ENSMUSG00000026175 | Vil1 | 22349 | protein_coding | 9568.283981 | 0.093267727 | 0.15171548 | 0.61475419 | 0.538717066 | 0.722272825 | 0.093267727 |
| ENSMUSG00000094410 | Zbed6 | 667118 | protein_coding | 76.36279737 | -0.093336722 | 0.192270662 | -0.485444433 | 0.627361121 | 0.784304814 | 0.093336722 |
| ENSMUSG00000015290 | Ubl4a | 27643 | protein_coding | 565.717977 | 0.093347736 | 0.085631638 | 1.090108036 | 0.275665557 | 0.483994818 | 0.093347736 |
| ENSMUSG00000038010 | Ccdc138 | 76138 | protein_coding | 79.07012124 | 0.093348072 | 0.193525359 | 0.482355762 | 0.629553241 | 0.785654248 | 0.093348072 |
| ENSMUSG00000066037 | Hnrnpr | 74326 | protein_coding | 825.3430574 | 0.093357265 | 0.090491266 | 1.031671558 | 0.302226006 | 0.511596086 | 0.093357265 |
| ENSMUSG00000030034 | Ino80b | 70020 | protein_coding | 169.5486743 | 0.093381342 | 0.107512982 | 0.868558752 | 0.385088524 | 0.592939799 | 0.093381342 |
| ENSMUSG00000033499 | Larp4b | 217980 | protein_coding | 2433.843935 | -0.093396627 | 0.066050493 | -1.414018633 | 0.157356432 | 0.33818806 | 0.093396627 |
| ENSMUSG00000026434 | Nucks1 | 98415 | protein_coding | 2434.927274 | 0.093468988 | 0.078769586 | 1.186612663 | 0.235380439 | 0.439523989 | 0.093468988 |
| ENSMUSG00000029434 | Vps33a | 77573 | protein_coding | 461.3112942 | 0.093477767 | 0.100140042 | 0.93347042 | 0.350577136 | 0.560627608 | 0.093477767 |
| ENSMUSG00000059273 | Zc3h4 | 330474 | protein_coding | 682.00969 | -0.093482339 | 0.089336349 | -1.046408777 | 0.295372342 | 0.504599164 | 0.093482339 |
| ENSMUSG00000029064 | Gnb1 | 14688 | protein_coding | 3558.668953 | 0.093484877 | 0.084439611 | 1.107121119 | 0.268241566 | 0.476156432 | 0.093484877 |
| ENSMUSG00000029319 | Coq2 | 71883 | protein_coding | 608.6066005 | -0.093490466 | 0.088156057 | -1.060510984 | 0.288912196 | 0.497873797 | 0.093490466 |

| ensembl_gene_id | external_gene_name | entrezgene_id | gene_biotype | baseMean | log2FoldChange | lfcSE | stat | pvalue | padj | absLog2FC |
|---|---|---|---|---|---|---|---|---|---|---|
| ENSMUSG00000005802 | Slc30a4 | 22785 | protein_coding | 2767.914813 | -0.093538784 | 0.132115169 | -0.708009415 | 0.478939411 | 0.672927641 | 0.093538784 |
| ENSMUSG00000036845 | Lin37 | 75660 | protein_coding | 122.1593098 | 0.093542075 | 0.111740966 | 0.837133222 | 0.402517688 | 0.609921868 | 0.093542075 |
| ENSMUSG00000020865 | Abcc3 | 76408 | protein_coding | 4541.61556 | -0.093572119 | 0.147596949 | -0.633970551 | 0.526100041 | 0.711774523 | 0.093572119 |
| ENSMUSG00000020605 | Hs1bp3 | 58240 | protein_coding | 210.7757084 | -0.093579881 | 0.16275723 | -0.574966049 | 0.565314259 | 0.741523966 | 0.093579881 |
| ENSMUSG00000054200 | Ffar4 | 107221 | protein_coding | 433.5927054 | 0.093582087 | 0.253342263 | 0.369389954 | 0.711837085 | 0.837504882 | 0.093582087 |
| ENSMUSG00000014426 | Map3k4 | 26407 | protein_coding | 794.9399657 | -0.093592542 | 0.075458634 | -1.240315876 | 0.214858582 | 0.41356707 | 0.093592542 |
| ENSMUSG00000018999 | Slc35b4 | 58246 | protein_coding | 299.7993932 | 0.093663061 | 0.080980896 | 1.156606877 | 0.247433011 | 0.453157756 | 0.093663061 |
| ENSMUSG00000049047 | Armcx3 | 71703 | protein_coding | 216.3083771 | 0.093683351 | 0.104114319 | 0.899812364 | 0.368220114 | 0.577037581 | 0.093683351 |
| ENSMUSG00000014418 | Hps5 | 246694 | protein_coding | 951.2359502 | 0.093693983 | 0.093819979 | 0.998657046 | 0.317960855 | 0.529209253 | 0.093693983 |
| ENSMUSG00000071019 | Sdr16c6 | 242286 | protein_coding | 8.016456635 | 0.093721213 | 0.615480112 | 0.152273342 | 0.87897135 | 0.938606574 | 0.093721213 |
| ENSMUSG00000031295 | Phka2 | 110094 | protein_coding | 236.579674 | -0.093726514 | 0.084840828 | -1.104733609 | 0.269275036 | 0.47724633 | 0.093726514 |
| ENSMUSG00000030019 | Fbxl14 | 101358 | protein_coding | 562.257299 | -0.093734733 | 0.112968315 | -0.829743571 | 0.406683781 | 0.613841849 | 0.093734733 |
| ENSMUSG00000067199 | Frat1 | 14296 | protein_coding | 256.0216596 | -0.09387615 | 0.117487074 | -0.799033853 | 0.424270783 | 0.628865162 | 0.09387615 |
| ENSMUSG00000023022 | Lima1 | 65970 | protein_coding | 5073.294806 | -0.093884008 | 0.101270797 | -0.927059043 | 0.35389588 | 0.563659971 | 0.093884008 |
| ENSMUSG00000038400 | Pmepa1 | 65112 | protein_coding | 138.3529918 | 0.093887544 | 0.311875937 | 0.301041321 | 0.763382986 | 0.870314523 | 0.093887544 |
| ENSMUSG00000087385 | Frg2f1 | 433752 | protein_coding | 46.30233648 | -0.09392214 | 0.18156725 | -0.517285686 | 0.604956745 | 0.769522541 | 0.09392214 |
| ENSMUSG00000024498 | Tcerg1 | 56070 | protein_coding | 711.0161593 | -0.093926311 | 0.123526223 | -0.760375482 | 0.447030175 | 0.647770799 | 0.093926311 |
| ENSMUSG00000019984 | Med23 | 70208 | protein_coding | 539.7082423 | 0.094002152 | 0.087917926 | 1.069203473 | 0.284977994 | 0.49392051 | 0.094002152 |
| ENSMUSG00000035161 | Ints6 | 18130 | protein_coding | 927.1054068 | 0.094070247 | 0.160456479 | 0.586266428 | 0.557696488 | 0.736454853 | 0.094070247 |
| ENSMUSG00000025155 | Dus1l | 68730 | protein_coding | 603.6896922 | 0.094076423 | 0.090260636 | 1.042275212 | 0.297284102 | 0.506180721 | 0.094076423 |
| ENSMUSG00000052539 | Magi3 | 99470 | protein_coding | 1712.282809 | 0.094094722 | 0.088220056 | 1.066591051 | 0.286156543 | 0.495047401 | 0.094094722 |
| ENSMUSG00000028896 | Rcc1 | 100088 | protein_coding | 488.3321439 | 0.094108642 | 0.116601947 | 0.807093231 | 0.419612772 | 0.625680983 | 0.094108642 |
| ENSMUSG00000024382 | Ercc3 | 13872 | protein_coding | 474.4540375 | 0.094117799 | 0.073024217 | 1.288857352 | 0.197447683 | 0.391319924 | 0.094117799 |
| ENSMUSG00000034616 | Ssh3 | 245857 | protein_coding | 1053.530558 | -0.094122102 | 0.092110301 | -1.021841219 | 0.306856057 | 0.516878519 | 0.094122102 |
| ENSMUSG00000029322 | Plac8 | 231507 | protein_coding | 7196.765391 | 0.094143139 | 0.1829756 | 0.514511986 | 0.606894082 | 0.771131144 | 0.094143139 |
| ENSMUSG00000019338 | Zfp687 | 78266 | protein_coding | 403.1512778 | -0.094149428 | 0.107982683 | -0.871893768 | 0.383266334 | 0.590907575 | 0.094149428 |
| ENSMUSG00000003135 | Cnot11 | 52846 | protein_coding | 387.0643461 | 0.094152915 | 0.085592612 | 1.100012172 | 0.271326819 | 0.479295236 | 0.094152915 |
| ENSMUSG00000049411 | Tmem241 | 338363 | protein_coding | 130.6356262 | 0.094158276 | 0.112247899 | 0.838842212 | 0.40155786 | 0.60912358 | 0.094158276 |
| ENSMUSG00000055660 | Mettl4 | 76781 | protein_coding | 401.8762768 | 0.094191065 | 0.105630408 | 0.891704074 | 0.372551569 | 0.580796848 | 0.094191065 |
| ENSMUSG00000022391 | Rangap1 | 19387 | protein_coding | 1778.951478 | 0.094205849 | 0.150029244 | 0.627916579 | 0.530058589 | 0.715246799 | 0.094205849 |
| ENSMUSG00000014850 | Msh3 | 17686 | protein_coding | 466.672069 | -0.094224849 | 0.102200081 | -0.92196453 | 0.356547077 | 0.566040252 | 0.094224849 |
| ENSMUSG00000025937 | Lactb2 | 212442 | protein_coding | 467.613766 | 0.094232549 | 0.0823089 | 1.144864637 | 0.252265235 | 0.458547106 | 0.094232549 |
| ENSMUSG00000047539 | Fbxo28 | 67948 | protein_coding | 560.7011572 | -0.094317716 | 0.067491883 | -1.397467535 | 0.162273023 | 0.344670584 | 0.094317716 |
| ENSMUSG00000031701 | Dnaja2 | 56445 | protein_coding | 2775.042553 | 0.094324278 | 0.102291765 | 0.922110174 | 0.356471111 | 0.565983821 | 0.094324278 |
| ENSMUSG00000024597 | Slc12a2 | 20496 | protein_coding | 9067.343519 | -0.094329832 | 0.098385345 | -0.958779298 | 0.33766994 | 0.548479169 | 0.094329832 |
| ENSMUSG00000020097 | Sgpl1 | 20397 | protein_coding | 1953.027371 | 0.094386072 | 0.074725719 | 1.263100211 | 0.206553168 | 0.402789314 | 0.094386072 |
| ENSMUSG00000034981 | Parm1 | 231440 | protein_coding | 1968.558149 | -0.094412912 | 0.108882204 | -0.867110594 | 0.385881416 | 0.593825534 | 0.094412912 |
| ENSMUSG00000042498 | Radx | 102871 | protein_coding | 80.10732918 | 0.094413161 | 0.189961733 | 0.497011473 | 0.619180961 | 0.778276069 | 0.094413161 |

| ensembl_gene_id | external_gene_name | entrezgene_id | gene_biotype | baseMean | log2FoldChange | lfcSE | stat | pvalue | padj | absLog2FC |
|---|---|---|---|---|---|---|---|---|---|---|
| ENSMUSG00000056692 | Ilrun | 224647 | protein_coding | 1691.999667 | -0.094496644 | 0.066708889 | -1.416552499 | 0.156613805 | 0.337237351 | 0.094496644 |
| ENSMUSG00000035354 | Uvrag | 78610 | protein_coding | 249.4534306 | 0.094531818 | 0.089738085 | 1.053419152 | 0.29214893 | 0.501006478 | 0.094531818 |
| ENSMUSG00000009905 | Kdsr | 70750 | protein_coding | 571.8355398 | -0.09457985 | 0.113749903 | -0.831471915 | 0.405707088 | 0.612753887 | 0.09457985 |
| ENSMUSG00000000399 | Ndufa9 | 66108 | protein_coding | 2252.09155 | 0.094598749 | 0.112586652 | 0.840230592 | 0.400779109 | 0.608194516 | 0.094598749 |
| ENSMUSG00000030513 | Pcsk6 | 18553 | protein_coding | 7.890953818 | -0.094643991 | 0.406815283 | -0.232646105 | 0.816036223 | 0.902250795 | 0.094643991 |
| ENSMUSG00000027739 | Rab33b | 19338 | protein_coding | 285.2682475 | 0.094646225 | 0.087073692 | 1.086966942 | 0.27705142 | 0.485411397 | 0.094646225 |
| ENSMUSG00000030647 | Ndufc2 | 68197 | protein_coding | 1127.690344 | 0.094662393 | 0.115854316 | 0.817081277 | 0.413881982 | 0.620277776 | 0.094662393 |
| ENSMUSG00000021592 | Arsk | 77041 | protein_coding | 179.3775946 | 0.094815988 | 0.126543805 | 0.74927404 | 0.453692051 | 0.653172937 | 0.094815988 |
| ENSMUSG00000019518 | Ap4m1 | 11781 | protein_coding | 317.6655175 | -0.094860058 | 0.116691576 | -0.812912652 | 0.41626814 | 0.622565158 | 0.094860058 |
| ENSMUSG00000014301 | Pam16 | 66449 | protein_coding | 132.7351134 | -0.094910839 | 0.104326451 | -0.909748562 | 0.362955127 | 0.571937996 | 0.094910839 |
| ENSMUSG00000029089 | 5730480H06Rik | 66768 | protein_coding | 104.8705077 | 0.094940636 | 0.121484594 | 0.781503503 | 0.434506417 | 0.637804646 | 0.094940636 |
| ENSMUSG00000023118 | Sympk | 68188 | protein_coding | 1258.616791 | 0.094973714 | 0.080434828 | 1.180753614 | 0.237700616 | 0.442042664 | 0.094973714 |
| ENSMUSG00000024601 | Isoc1 | 66307 | protein_coding | 772.6752079 | -0.0949764 | 0.073940304 | -1.284501081 | 0.198966686 | 0.393175365 | 0.0949764 |
| ENSMUSG00000053040 | Aph1c | 68318 | protein_coding | 8.215958913 | 0.095001128 | 0.471649512 | 0.201423144 | 0.84036772 | 0.915836464 | 0.095001128 |
| ENSMUSG00000031652 | N4bp1 | 80750 | protein_coding | 714.895448 | -0.09501414 | 0.095826691 | -0.991520615 | 0.321431431 | 0.532354566 | 0.09501414 |
| ENSMUSG00000019066 | Rab3d | 19340 | protein_coding | 2562.589865 | 0.09502435 | 0.074591095 | 1.273936928 | 0.20268577 | 0.397632583 | 0.09502435 |
| ENSMUSG00000068744 | Psrc1 | 56742 | protein_coding | 58.01682531 | 0.095033006 | 0.212758074 | 0.446671681 | 0.655112132 | 0.800324132 | 0.095033006 |
| ENSMUSG00000020859 | Spag9 | 70834 | protein_coding | 1590.378998 | -0.095034201 | 0.064918308 | -1.463904465 | 0.143220053 | 0.31713229 | 0.095034201 |
| ENSMUSG00000102252 | Snrpn | 20646 | protein_coding | 111.903406 | -0.095040373 | 0.139400921 | -0.681777224 | 0.495379828 | 0.686912002 | 0.095040373 |
| ENSMUSG00000022503 | Nubp1 | 26425 | protein_coding | 280.3973157 | 0.095081137 | 0.11893933 | 0.799408712 | 0.42405346 | 0.628617629 | 0.095081137 |
| ENSMUSG00000033629 | Hacd3 | 57874 | protein_coding | 1113.18495 | -0.095135064 | 0.083607199 | -1.137881245 | 0.255170074 | 0.461857833 | 0.095135064 |
| ENSMUSG00000055900 | Tmem69 | 230657 | protein_coding | 138.5014828 | -0.09515891 | 0.117914456 | -0.807016484 | 0.419656986 | 0.625680983 | 0.09515891 |
| ENSMUSG00000096472 | Cdkn2d | 12581 | protein_coding | 89.49425228 | -0.095176901 | 0.169569401 | -0.561285825 | 0.574602703 | 0.747743363 | 0.095176901 |
| ENSMUSG00000048720 | Tbc1d12 | 209478 | protein_coding | 218.3367459 | -0.095205054 | 0.110795865 | -0.859283462 | 0.390184146 | 0.598507455 | 0.095205054 |
| ENSMUSG00000023572 | Ccndbp1 | 17151 | protein_coding | 418.2290947 | -0.095249255 | 0.105314129 | -0.904429983 | 0.365767446 | 0.574494022 | 0.095249255 |
| ENSMUSG00000020541 | Tom1l1 | 71943 | protein_coding | 1214.25762 | -0.095269374 | 0.097490113 | -0.977220879 | 0.328459813 | 0.539240147 | 0.095269374 |
| ENSMUSG00000028164 | Manba | 110173 | protein_coding | 472.0604513 | 0.095310818 | 0.101804293 | 0.936216099 | 0.349161936 | 0.559301241 | 0.095310818 |
| ENSMUSG00000035642 | Aamdc | 66273 | protein_coding | 426.7167274 | 0.095317725 | 0.09716778 | 0.980960202 | 0.326612368 | 0.537703544 | 0.095317725 |
| ENSMUSG00000037361 | Sf3b6 | 66055 | protein_coding | 707.7374856 | 0.095321195 | 0.082232189 | 1.159171321 | 0.246386359 | 0.451916389 | 0.095321195 |
| ENSMUSG00000110231 | Gm6329 | 622523 | lncRNA | 8.820599516 | -0.095354914 | 0.514171482 | -0.185453526 | 0.852873329 | 0.922175438 | 0.095354914 |
| ENSMUSG00000036932 | Aifm1 | 26926 | protein_coding | 1111.320438 | -0.095364495 | 0.078079617 | -1.221375029 | 0.221944057 | 0.423092298 | 0.095364495 |
| ENSMUSG00000069565 | Dazap1 | 70248 | protein_coding | 855.3177722 | 0.095414569 | 0.081516517 | 1.170493699 | 0.241802348 | 0.446591536 | 0.095414569 |
| ENSMUSG00000011179 | Odc1 | 18263 | protein_coding | 3086.190095 | 0.095437367 | 0.127539958 | 0.748293856 | 0.45428293 | 0.653612209 | 0.095437367 |
| ENSMUSG00000042572 | Ube2q1 | 70093 | protein_coding | 943.6763492 | -0.095547271 | 0.087898403 | -1.087019426 | 0.277028225 | 0.485411397 | 0.095547271 |
| ENSMUSG00000045594 | Glb1 | 12091 | protein_coding | 1325.80297 | -0.095578676 | 0.081984223 | -1.165817919 | 0.243688071 | 0.448775996 | 0.095578676 |
| ENSMUSG00000026618 | Iars2 | 381314 | protein_coding | 1846.426311 | -0.095610694 | 0.068932098 | -1.387027198 | 0.16543347 | 0.349003763 | 0.095610694 |
| ENSMUSG00000033862 | Cdk10 | 234854 | protein_coding | 252.1095011 | -0.095689868 | 0.091454253 | -1.046314025 | 0.295416072 | 0.504599164 | 0.095689868 |
| ENSMUSG00000032966 | Fkbp1a | 14225 | protein_coding | 484.2068073 | -0.095708325 | 0.127099814 | -0.753017031 | 0.45143968 | 0.651785414 | 0.095708325 |

| ensembl_gene_id | external_gene_name | entrezgene_id | gene_biotype | baseMean | log2FoldChange | lfcSE | stat | pvalue | padj | absLog2FC |
|---|---|---|---|---|---|---|---|---|---|---|
| ENSMUSG00000049109 | Themis | 210757 | protein_coding | 9.473177626 | 0.095733524 | 0.913868156 | 0.104756385 | 0.91656912 | 0.958597942 | 0.095733524 |
| ENSMUSG00000022551 | Cyc1 | 66445 | protein_coding | 3238.678368 | 0.095829976 | 0.095165735 | 1.006979832 | 0.313944466 | 0.524547095 | 0.095829976 |
| ENSMUSG00000015149 | Sirt2 | 64383 | protein_coding | 413.6347434 | -0.095837539 | 0.070111643 | -1.366927584 | 0.171648005 | 0.357153419 | 0.095837539 |
| ENSMUSG00000032637 | Atxn2l | 233871 | protein_coding | 482.4746101 | -0.095843166 | 0.167757563 | -0.571319492 | 0.567783091 | 0.743232173 | 0.095843166 |
| ENSMUSG00000019590 | Cyb561 | 13056 | protein_coding | 1448.549972 | 0.095852114 | 0.070196237 | 1.36548792 | 0.172099747 | 0.357719532 | 0.095852114 |
| ENSMUSG00000030207 | Fam234b | 74525 | protein_coding | 7.751941657 | -0.095899246 | 0.497896492 | -0.1926088 | 0.847265352 | 0.919400582 | 0.095899246 |
| ENSMUSG00000026197 | Zfand2b | 68818 | protein_coding | 329.977043 | -0.095923298 | 0.089846843 | -1.067631261 | 0.285686877 | 0.494484454 | 0.095923298 |
| ENSMUSG00000041396 | Mettl18 | 69962 | protein_coding | 68.72769427 | -0.095948082 | 0.17342604 | -0.553250722 | 0.580091739 | 0.751318253 | 0.095948082 |
| ENSMUSG00000024359 | Hspa9 | 15526 | protein_coding | 5699.720448 | -0.095953295 | 0.090752772 | -1.057304287 | 0.290372737 | 0.499035062 | 0.095953295 |
| ENSMUSG00000019813 | Cep57l1 | 103268 | protein_coding | 64.25967454 | -0.095967451 | 0.253312199 | -0.378850492 | 0.70479889 | 0.833286574 | 0.095967451 |
| ENSMUSG00000035203 | Epn1 | 13854 | protein_coding | 2178.595586 | 0.095968223 | 0.093420322 | 1.027273517 | 0.304291689 | 0.513961388 | 0.095968223 |
| ENSMUSG00000065400 | Mir200a | 387242 | miRNA | 26.57977055 | 0.09600427 | 0.360836906 | 0.266060007 | 0.790192995 | 0.886289277 | 0.09600427 |
| ENSMUSG00000022286 | Grhl2 | 252973 | protein_coding | 1036.970259 | -0.09602106 | 0.075226227 | -1.276430632 | 0.201803334 | 0.396683142 | 0.09602106 |
| ENSMUSG00000062590 | Armc9 | 78795 | protein_coding | 81.44890676 | 0.096112194 | 0.153727685 | 0.625210703 | 0.531832778 | 0.71689162 | 0.096112194 |
| ENSMUSG00000043311 | D17H6S53E | 114585 | protein_coding | 423.4715608 | 0.09614392 | 0.101691211 | 0.945449651 | 0.344429364 | 0.554204557 | 0.09614392 |
| ENSMUSG00000038037 | Socs1 | 12703 | protein_coding | 33.09846239 | 0.096173371 | 0.297181044 | 0.323618792 | 0.746226659 | 0.858895287 | 0.096173371 |
| ENSMUSG00000028274 | Rngtt | 24018 | protein_coding | 317.9675711 | 0.096210127 | 0.076395794 | 1.259364183 | 0.20789882 | 0.404310463 | 0.096210127 |
| ENSMUSG00000000759 | Tubgcp3 | 259279 | protein_coding | 539.5129084 | -0.096229261 | 0.063316545 | -1.519812252 | 0.128558169 | 0.294756659 | 0.096229261 |
| ENSMUSG00000019845 | Tube1 | 71924 | protein_coding | 71.13777135 | 0.096239027 | 0.172616163 | 0.557531956 | 0.577164032 | 0.749071904 | 0.096239027 |
| ENSMUSG00000021240 | Abcd4 | 19300 | protein_coding | 162.7109213 | 0.09626592 | 0.135079326 | 0.712662125 | 0.476054859 | 0.670570635 | 0.09626592 |
| ENSMUSG00000039989 | Cbx4 | 12418 | protein_coding | 215.3464547 | 0.096297562 | 0.145757322 | 0.660670491 | 0.508823652 | 0.697877891 | 0.096297562 |
| ENSMUSG00000110444 | Gm10033 | 378466 | protein_coding | 148.82482 | 0.096313631 | 0.156471249 | 0.615535644 | 0.538201037 | 0.72195268 | 0.096313631 |
| ENSMUSG00000022234 | Cct5 | 12465 | protein_coding | 3836.893461 | 0.096363013 | 0.112719574 | 0.854891561 | 0.392611183 | 0.600621715 | 0.096363013 |
| ENSMUSG00000054321 | Taf4b | 72504 | protein_coding | 161.1725976 | 0.096408064 | 0.185021566 | 0.521063926 | 0.60232224 | 0.767564496 | 0.096408064 |
| ENSMUSG00000030138 | Bms1 | 213895 | protein_coding | 704.0912436 | 0.09640996 | 0.098901942 | 0.974803515 | 0.329657733 | 0.540426963 | 0.09640996 |
| ENSMUSG00000004568 | Arhgef18 | 102098 | protein_coding | 748.2340936 | -0.096410344 | 0.092224285 | -1.045389986 | 0.295842763 | 0.505111323 | 0.096410344 |
| ENSMUSG00000034218 | Atm | 11920 | protein_coding | 910.0313427 | -0.096435971 | 0.125153154 | -0.77054367 | 0.440977461 | 0.642624793 | 0.096435971 |
| ENSMUSG00000032114 | Slc37a4 | 14385 | protein_coding | 446.7381543 | 0.096446733 | 0.15608314 | 0.617918971 | 0.536628753 | 0.721189041 | 0.096446733 |
| ENSMUSG00000050106 | Tmc8 | 217356 | protein_coding | 115.2607653 | 0.096492963 | 0.165900595 | 0.581631203 | 0.560815121 | 0.738541814 | 0.096492963 |
| ENSMUSG00000047260 | Emc6 | 66048 | protein_coding | 594.1784311 | -0.096512919 | 0.096844888 | -0.996572159 | 0.318972225 | 0.530313463 | 0.096512919 |
| ENSMUSG00000006763 | Saal1 | 78935 | protein_coding | 191.8375575 | 0.096553185 | 0.093212291 | 1.035841776 | 0.300275964 | 0.509295662 | 0.096553185 |
| ENSMUSG00000028330 | Ncbp1 | 433702 | protein_coding | 848.2329978 | 0.096563846 | 0.114113762 | 0.846206843 | 0.397437365 | 0.605546427 | 0.096563846 |
| ENSMUSG00000042303 | Sgsm3 | 105835 | protein_coding | 1588.122609 | -0.096666465 | 0.11710435 | -0.825472883 | 0.409103172 | 0.616072168 | 0.096666465 |
| ENSMUSG00000033405 | Nudt15 | 214254 | protein_coding | 33.68747116 | 0.096710294 | 0.239685407 | 0.40348845 | 0.68658893 | 0.821304393 | 0.096710294 |
| ENSMUSG00000041415 | Dicer1 | 192119 | protein_coding | 1005.52103 | -0.096716062 | 0.077702472 | -1.244697354 | 0.213243019 | 0.411305377 | 0.096716062 |
| ENSMUSG00000033632 | AW554918 | 225289 | protein_coding | 125.0510759 | -0.096740548 | 0.153247814 | -0.631268696 | 0.527864851 | 0.713284795 | 0.096740548 |
| ENSMUSG00000021994 | Wnt5a | 22418 | protein_coding | 65.47179848 | 0.096746595 | 0.202413162 | 0.477965928 | 0.632674456 | 0.787591929 | 0.096746595 |
| ENSMUSG00000054894 | Dmac2l | 68055 | protein_coding | 143.1583928 | 0.096752935 | 0.135041051 | 0.716470544 | 0.473700849 | 0.668798783 | 0.096752935 |

Supplementary Table 1

| ensembl_gene_id | external_gene_name | entrezgene_id | gene_biotype | baseMean | log2FoldChange | lfcSE | stat | pvalue | padj | absLog2FC |
|---|---|---|---|---|---|---|---|---|---|---|
| ENSMUSG00000041740 | Rnf10 | 50849 | protein_coding | 1538.790476 | -0.096796513 | 0.101352023 | -0.955052605 | 0.339551095 | 0.550134253 | 0.096796513 |
| ENSMUSG00000020482 | Ccdc117 | 104479 | protein_coding | 581.2382228 | -0.096835104 | 0.099234231 | -0.975823593 | 0.329151891 | 0.539888897 | 0.096835104 |
| ENSMUSG00000043424 | Eif3j2 | 100042807 | protein_coding | 9.870407201 | -0.09685499 | 0.472499618 | -0.204984271 | 0.837584414 | 0.914271145 | 0.09685499 |
| ENSMUSG00000039990 | Edrf1 | 214764 | protein_coding | 550.0568248 | 0.096877554 | 0.081341186 | 1.191002473 | 0.233652614 | 0.437425815 | 0.096877554 |
| ENSMUSG00000038534 | Osbpl7 | 71240 | protein_coding | 583.2122572 | -0.096958034 | 0.152059953 | -0.637630303 | 0.523714365 | 0.709654498 | 0.096958034 |
| ENSMUSG00000029387 | Gtf2h3 | 209357 | protein_coding | 489.6684991 | 0.096991986 | 0.092974829 | 1.043206939 | 0.29685246 | 0.506180721 | 0.096991986 |
| ENSMUSG00000017316 | Ppy | 19064 | protein_coding | 19.39025421 | 0.097043967 | 0.354032581 | 0.27411027 | 0.783999881 | 0.882817412 | 0.097043967 |
| ENSMUSG00000045795 | Whamm | 434204 | protein_coding | 240.7124307 | 0.097103807 | 0.088149669 | 1.101578802 | 0.270644828 | 0.478449592 | 0.097103807 |
| ENSMUSG00000038533 | Cbfa2t2 | 12396 | protein_coding | 373.8563458 | -0.097165604 | 0.100816997 | -0.963781968 | 0.335155249 | 0.545732244 | 0.097165604 |
| ENSMUSG00000022330 | Osr2 | 107587 | protein_coding | 447.4328049 | -0.097183265 | 0.24010048 | -0.404760811 | 0.685653337 | 0.820640694 | 0.097183265 |
| ENSMUSG00000071644 | Eef1g | 67160 | protein_coding | 6655.771592 | 0.097208319 | 0.062397727 | 1.557882375 | 0.119261134 | 0.279886746 | 0.097208319 |
| ENSMUSG00000063714 | Sp3os | 100503849 | lncRNA | 39.95526066 | 0.097218568 | 0.241097642 | 0.403233177 | 0.686776695 | 0.821304393 | 0.097218568 |
| ENSMUSG00000043099 | Hic1 | 15248 | protein_coding | 12.0717582 | 0.097254737 | 0.487139968 | 0.199644339 | 0.841758749 | 0.916390622 | 0.097254737 |
| ENSMUSG00000026798 | Coq4 | 227683 | protein_coding | 174.7305962 | -0.097271552 | 0.112687111 | -0.863200338 | 0.388027328 | 0.596497533 | 0.097271552 |
| ENSMUSG00000062040 | Zfp27 | 22689 | protein_coding | 175.7592974 | 0.097309349 | 0.11666554 | 0.834088193 | 0.404231288 | 0.611350704 | 0.097309349 |
| ENSMUSG00000024370 | Cdc23 | 52563 | protein_coding | 367.1413657 | -0.097337774 | 0.088715021 | -1.097196091 | 0.272555682 | 0.480653997 | 0.097337774 |
| ENSMUSG00000029647 | Pan3 | 72587 | protein_coding | 1325.643824 | -0.097364328 | 0.05730731 | -1.698986185 | 0.089321787 | 0.23066106 | 0.097364328 |
| ENSMUSG00000018500 | Adora2b | 11541 | protein_coding | 25.41616228 | -0.097374697 | 0.29634694 | -0.328583441 | 0.742470566 | 0.85659524 | 0.097374697 |
| ENSMUSG00000024339 | Tap2 | 21355 | protein_coding | 220.9962402 | 0.097381364 | 0.097413984 | 0.999665143 | 0.317472586 | 0.528771321 | 0.097381364 |
| ENSMUSG00000028405 | Aco1 | 11428 | protein_coding | 1800.023442 | 0.097389722 | 0.088182131 | 1.10441561 | 0.269412893 | 0.477331318 | 0.097389722 |
| ENSMUSG00000038605 | Samd10 | 229011 | protein_coding | 154.7240789 | -0.09740666 | 0.137865666 | -0.706533124 | 0.479856661 | 0.673583934 | 0.09740666 |
| ENSMUSG00000027573 | Gid8 | 76425 | protein_coding | 907.2244997 | 0.097419071 | 0.068946904 | 1.412957869 | 0.157668112 | 0.338500834 | 0.097419071 |
| ENSMUSG00000037296 | Lsm1 | 67207 | protein_coding | 246.8523298 | 0.097425986 | 0.109514798 | 0.889614806 | 0.373672753 | 0.582145894 | 0.097425986 |
| ENSMUSG00000024749 | Tmc1 | 13409 | protein_coding | 25.34955346 | -0.097428288 | 0.267493947 | -0.364226141 | 0.715689148 | 0.840074597 | 0.097428288 |
| ENSMUSG00000003099 | Ppp5c | 19060 | protein_coding | 1018.441473 | 0.097441078 | 0.085288597 | 1.142486589 | 0.253251817 | 0.459429949 | 0.097441078 |
| ENSMUSG00000051306 | Usp42 | 76800 | protein_coding | 263.3360509 | -0.097445595 | 0.126727993 | -0.768935045 | 0.441931872 | 0.643250558 | 0.097445595 |
| ENSMUSG00000031916 | Cog8 | 97484 | protein_coding | 433.0190023 | -0.097449857 | 0.120930188 | -0.805835651 | 0.420337618 | 0.626299119 | 0.097449857 |
| ENSMUSG00000058240 | Cryzl1 | 66609 | protein_coding | 420.8474688 | 0.097464579 | 0.086494876 | 1.12682489 | 0.259816525 | 0.467083251 | 0.097464579 |
| ENSMUSG00000020400 | Tnip1 | 57783 | protein_coding | 715.5909004 | -0.097498437 | 0.127858368 | -0.762550302 | 0.445731633 | 0.646536149 | 0.097498437 |
| ENSMUSG00000026353 | Ubxn4 | 67812 | protein_coding | 1230.00745 | 0.097507107 | 0.118198131 | 0.824946268 | 0.409402097 | 0.616397006 | 0.097507107 |
| ENSMUSG00000012405 | Rpl15 | 66480 | protein_coding | 3689.202899 | 0.097544035 | 0.101663233 | 0.959481927 | 0.33731602 | 0.548018112 | 0.097544035 |
| ENSMUSG00000025485 | Ric8a | 101489 | protein_coding | 745.66288 | -0.097611076 | 0.101899023 | -0.957919641 | 0.338103282 | 0.548737423 | 0.097611076 |
| ENSMUSG00000000976 | Heatr6 | 217026 | protein_coding | 323.3387257 | 0.097689979 | 0.093162712 | 1.048595259 | 0.294364439 | 0.503612973 | 0.097689979 |
| ENSMUSG00000039068 | Zzz3 | 108946 | protein_coding | 1134.716274 | -0.097701979 | 0.071298066 | -1.370331397 | 0.170583477 | 0.35593569 | 0.097701979 |
| ENSMUSG00000027951 | Adar | 56417 | protein_coding | 718.1465664 | 0.09773529 | 0.093282503 | 1.047734427 | 0.294760981 | 0.50397643 | 0.09773529 |
| ENSMUSG00000028454 | Pigo | 56703 | protein_coding | 398.3037053 | 0.097765168 | 0.100437675 | 0.97339139 | 0.330358816 | 0.540908226 | 0.097765168 |
| ENSMUSG00000026576 | Atp1b1 | 11931 | protein_coding | 8365.496717 | -0.097769103 | 0.095952727 | -1.01892991 | 0.308236243 | 0.518271601 | 0.097769103 |
| ENSMUSG00000067071 | Hes6 | 55927 | protein_coding | 732.3925204 | 0.097794149 | 0.140122929 | 0.697916817 | 0.485229217 | 0.678589433 | 0.097794149 |

| ensembl_gene_id | external_gene_name | entrezgene_id | gene_biotype | baseMean | log2FoldChange | lfcSE | stat | pvalue | padj | absLog2FC |
|---|---|---|---|---|---|---|---|---|---|---|
| ENSMUSG00000029538 | Srsf9 | 108014 | protein_coding | 341.8052189 | 0.097802362 | 0.112049739 | 0.872847745 | 0.382746071 | 0.59039119 | 0.097802362 |
| ENSMUSG00000032534 | Cep63 | 28135 | protein_coding | 394.5343397 | -0.097880915 | 0.088053672 | -1.111605149 | 0.266307958 | 0.474097815 | 0.097880915 |
| ENSMUSG00000066150 | Slc31a1 | 20529 | protein_coding | 1824.753053 | 0.097901721 | 0.07974029 | 1.227757282 | 0.219538096 | 0.419876136 | 0.097901721 |
| ENSMUSG00000027673 | Ndufb5 | 66046 | protein_coding | 1656.499088 | 0.097906274 | 0.125973765 | 0.777195739 | 0.437043296 | 0.639698285 | 0.097906274 |
| ENSMUSG00000030536 | Iqgap1 | 29875 | protein_coding | 6691.300047 | -0.097926451 | 0.083683891 | -1.170194758 | 0.241922602 | 0.446609087 | 0.097926451 |
| ENSMUSG00000001767 | Crnkl1 | 66877 | protein_coding | 519.377446 | 0.098073095 | 0.082079475 | 1.194855295 | 0.232143572 | 0.435607091 | 0.098073095 |
| ENSMUSG00000026303 | Mlph | 171531 | protein_coding | 693.3783957 | 0.098092453 | 0.152339902 | 0.643905188 | 0.519636917 | 0.706433139 | 0.098092453 |
| ENSMUSG00000027109 | Sp3 | 20687 | protein_coding | 2608.079599 | 0.098093395 | 0.076257501 | 1.286344208 | 0.19832296 | 0.392554755 | 0.098093395 |
| ENSMUSG00000042814 | Mcts2 | 66405 | protein_coding | 109.5726379 | 0.098100128 | 0.154278308 | 0.635864692 | 0.524864618 | 0.710525551 | 0.098100128 |
| ENSMUSG00000039680 | Mrps6 | 121022 | protein_coding | 372.9601227 | 0.098108106 | 0.12241293 | 0.801452147 | 0.422869937 | 0.627898979 | 0.098108106 |
| ENSMUSG00000033099 | Nol12 | 97961 | protein_coding | 162.3036184 | -0.098153545 | 0.160550829 | -0.611354957 | 0.540964615 | 0.724074319 | 0.098153545 |
| ENSMUSG00000034427 | Myo15b | 217328 | protein_coding | 5864.106817 | -0.09816672 | 0.153167527 | -0.64091079 | 0.521580645 | 0.707881371 | 0.09816672 |
| ENSMUSG00000025812 | Pard3 | 93742 | protein_coding | 934.2517638 | -0.098192886 | 0.066684536 | -1.472498604 | 0.140886284 | 0.313281688 | 0.098192886 |
| ENSMUSG00000078861 | Zfp931 | 353208 | protein_coding | 110.899989 | -0.098193444 | 0.151958853 | -0.646184426 | 0.518159925 | 0.705260535 | 0.098193444 |
| ENSMUSG00000024556 | Me2 | 107029 | protein_coding | 2566.054116 | 0.098222481 | 0.131033854 | 0.749596216 | 0.453497931 | 0.653148289 | 0.098222481 |
| ENSMUSG00000049606 | Zfp644 | 52397 | protein_coding | 1536.83993 | -0.098246948 | 0.056116527 | -1.750766706 | 0.079986104 | 0.213797392 | 0.098246948 |
| ENSMUSG00000085995 | Gm2788 | 108167326 | lncRNA | 14.44327167 | -0.098295594 | 0.432300214 | -0.227378084 | 0.820129761 | 0.904579103 | 0.098295594 |
| ENSMUSG00000013973 | Dedd | 21945 | protein_coding | 126.5649769 | -0.098309386 | 0.114680593 | -0.857245185 | 0.391309396 | 0.59924777 | 0.098309386 |
| ENSMUSG00000038637 | Lrrc56 | 70552 | protein_coding | 101.0079696 | -0.098333639 | 0.153443398 | -0.640846329 | 0.521622529 | 0.707881371 | 0.098333639 |
| ENSMUSG00000020610 | Amz2 | 13929 | protein_coding | 476.4994224 | 0.098334392 | 0.08301078 | 1.184597855 | 0.236176485 | 0.440339349 | 0.098334392 |
| ENSMUSG00000006906 | Stambp | 70527 | protein_coding | 179.1670602 | -0.098365552 | 0.194040752 | -0.506932441 | 0.612202223 | 0.774085655 | 0.098365552 |
| ENSMUSG00000029202 | Pds5a | 71521 | protein_coding | 2237.896552 | 0.098373959 | 0.05319362 | 1.849356346 | 0.064406375 | 0.184607654 | 0.098373959 |
| ENSMUSG00000061882 | Ccdc62 | 208908 | protein_coding | 28.24074409 | -0.098418317 | 0.267731737 | -0.367600487 | 0.713171151 | 0.838486874 | 0.098418317 |
| ENSMUSG00000047603 | Zfp235 | 56525 | protein_coding | 173.2535703 | -0.098435023 | 0.15644812 | -0.629186356 | 0.52922706 | 0.714548497 | 0.098435023 |
| ENSMUSG00000040865 | Ino80d | 227195 | protein_coding | 726.026296 | -0.09843935 | 0.169825333 | -0.579650564 | 0.562150287 | 0.739444197 | 0.09843935 |
| ENSMUSG00000017667 | Zfp334 | 228876 | protein_coding | 48.24803876 | 0.098441234 | 0.182506972 | 0.539383419 | 0.58962232 | 0.759394898 | 0.098441234 |
| ENSMUSG00000025779 | Ly96 | 17087 | protein_coding | 67.13955869 | 0.098485933 | 0.175892412 | 0.559921442 | 0.575533022 | 0.74838086 | 0.098485933 |
| ENSMUSG00000037617 | Spag1 | 26942 | protein_coding | 49.05005994 | 0.098504272 | 0.207774615 | 0.474091947 | 0.635434341 | 0.788947751 | 0.098504272 |
| ENSMUSG00000025027 | Xpnpep1 | 170750 | protein_coding | 1001.581033 | 0.098508934 | 0.097128883 | 1.014208445 | 0.3104833 | 0.520484087 | 0.098508934 |
| ENSMUSG00000048874 | Phf3 | 213109 | protein_coding | 1279.158295 | -0.098628661 | 0.061762682 | -1.59689732 | 0.110288597 | 0.264838953 | 0.098628661 |
| ENSMUSG00000077167 | Gm24119 | 115487534 | snoRNA | 30.99043607 | -0.098633706 | 0.344934572 | -0.28594903 | 0.77491716 | 0.877593098 | 0.098633706 |
| ENSMUSG00000022809 | Nr1i2 | 18171 | protein_coding | 1027.653066 | -0.098680236 | 0.100715095 | -0.979795893 | 0.32718688 | 0.538362069 | 0.098680236 |
| ENSMUSG00000086541 | Has2os | 594843 | lncRNA | 18.81383331 | -0.098682718 | 0.328927657 | -0.30001344 | 0.764166904 | 0.871145895 | 0.098682718 |
| ENSMUSG00000029920 | Smarcad1 | 13990 | protein_coding | 590.7460455 | -0.098685926 | 0.052464683 | -1.880997278 | 0.059972289 | 0.175886323 | 0.098685926 |
| ENSMUSG00000020451 | Limk2 | 16886 | protein_coding | 2013.930497 | -0.098710118 | 0.118606387 | -0.832249604 | 0.40526807 | 0.612206925 | 0.098710118 |
| ENSMUSG00000019763 | Rmnd1 | 66084 | protein_coding | 307.6194161 | -0.098730653 | 0.117212055 | -0.842325071 | 0.399606021 | 0.607108423 | 0.098730653 |
| ENSMUSG00000037410 | Tbc1d2b | 67016 | protein_coding | 358.4378012 | -0.098888366 | 0.145008174 | -0.681950287 | 0.495270386 | 0.686866932 | 0.098888366 |
| ENSMUSG00000020064 | Herc4 | 67345 | protein_coding | 1041.31108 | 0.098923502 | 0.074976349 | 1.319396091 | 0.187036728 | 0.377677612 | 0.098923502 |

| ensembl_gene_id | external_gene_name | entrezgene_id | gene_biotype | baseMean | log2FoldChange | lfcSE | stat | pvalue | padj | absLog2FC |
|---|---|---|---|---|---|---|---|---|---|---|
| ENSMUSG00000032977 | Fam207a | 108707 | protein_coding | 380.1588319 | 0.09895461 | 0.135181801 | 0.732011331 | 0.464161652 | 0.661249706 | 0.09895461 |
| ENSMUSG00000030057 | Cnbp | 12785 | protein_coding | 4100.282479 | 0.098994728 | 0.094946063 | 1.042641737 | 0.297114251 | 0.506180721 | 0.098994728 |
| ENSMUSG00000001847 | Rac1 | 19353 | protein_coding | 3796.38398 | 0.099060399 | 0.080162433 | 1.235745915 | 0.216553025 | 0.415723343 | 0.099060399 |
| ENSMUSG00000010517 | Faf1 | 14084 | protein_coding | 1034.959912 | 0.099084037 | 0.085534058 | 1.158416183 | 0.246694238 | 0.452168679 | 0.099084037 |
| ENSMUSG00000020780 | Srp68 | 217337 | protein_coding | 706.2510581 | 0.099102246 | 0.079601652 | 1.24497725 | 0.213140113 | 0.411253267 | 0.099102246 |
| ENSMUSG00000058979 | Hdhd5 | 214932 | protein_coding | 221.0599095 | 0.099156242 | 0.09289389 | 1.067414034 | 0.285784914 | 0.494565635 | 0.099156242 |
| ENSMUSG00000030917 | Tmem159 | 233806 | protein_coding | 340.1777387 | 0.099162671 | 0.114462636 | 0.866332232 | 0.386307995 | 0.594388886 | 0.099162671 |
| ENSMUSG00000047641 | Krt87 | 406219 | protein_coding | 69.25852059 | -0.099172662 | 0.189321535 | -0.523831915 | 0.600395453 | 0.766753883 | 0.099172662 |
| ENSMUSG00000030986 | Dhx32 | 101437 | protein_coding | 1862.345684 | 0.09918037 | 0.079329319 | 1.250235996 | 0.211213351 | 0.40867423 | 0.09918037 |
| ENSMUSG00000029730 | Mcm7 | 17220 | protein_coding | 612.3385149 | 0.09918673 | 0.118538102 | 0.836749772 | 0.402733236 | 0.609997621 | 0.09918673 |
| ENSMUSG00000000278 | Scpep1 | 74617 | protein_coding | 455.5028913 | 0.099218045 | 0.109122478 | 0.909235631 | 0.36322576 | 0.57208147 | 0.099218045 |
| ENSMUSG00000039676 | Capsl | 75568 | protein_coding | 70.05057747 | 0.099218394 | 0.194467254 | 0.510206179 | 0.609907024 | 0.772840022 | 0.099218394 |
| ENSMUSG00000019143 | Hars2 | 70791 | protein_coding | 336.0211598 | -0.099275242 | 0.091677666 | -1.082872706 | 0.278864928 | 0.487423953 | 0.099275242 |
| ENSMUSG00000008822 | Acyp1 | 66204 | protein_coding | 90.53016034 | -0.099347066 | 0.170995287 | -0.580993005 | 0.561245168 | 0.738863793 | 0.099347066 |
| ENSMUSG00000097789 | Ctxnd1 | 100039239 | protein_coding | 190.6450612 | 0.099357758 | 0.15967518 | 0.622249235 | 0.533777996 | 0.718743421 | 0.099357758 |
| ENSMUSG00000027665 | Pik3ca | 18706 | protein_coding | 1204.376759 | -0.099362227 | 0.074144578 | -1.340114545 | 0.180208108 | 0.368125204 | 0.099362227 |
| ENSMUSG00000036864 | Proser3 | 333193 | protein_coding | 34.52569124 | -0.099383928 | 0.229548108 | -0.432954681 | 0.665047694 | 0.806025281 | 0.099383928 |
| ENSMUSG00000037608 | Bclaf1 | 72567 | protein_coding | 2460.526353 | 0.099388736 | 0.058228873 | 1.706863476 | 0.087847431 | 0.228409056 | 0.099388736 |
| ENSMUSG00000071528 | Atp5md | 66477 | protein_coding | 2166.007727 | 0.099393524 | 0.093268945 | 1.065665793 | 0.286574746 | 0.495534183 | 0.099393524 |
| ENSMUSG00000063077 | Kif1b | 16561 | protein_coding | 1303.052932 | -0.099430397 | 0.096306931 | -1.032432406 | 0.301869597 | 0.511236464 | 0.099430397 |
| ENSMUSG00000014030 | Pax5 | 18507 | protein_coding | 7.702472103 | 0.099441329 | 1.079322351 | 0.092133114 | 0.926592279 | 0.963740176 | 0.099441329 |
| ENSMUSG00000048440 | Cyp4f16 | 70101 | protein_coding | 518.721601 | 0.099444989 | 0.080864734 | 1.229769571 | 0.218783406 | 0.41894471 | 0.099444989 |
| ENSMUSG00000028559 | Osbpl9 | 100273 | protein_coding | 980.9615794 | 0.099475197 | 0.056455012 | 1.762025972 | 0.07806491 | 0.210143295 | 0.099475197 |
| ENSMUSG00000055839 | Elob | 67673 | protein_coding | 1181.681011 | 0.099548339 | 0.124530381 | 0.799389985 | 0.424064316 | 0.628617629 | 0.099548339 |
| ENSMUSG00000031370 | Zrsr2 | 22184 | protein_coding | 287.2445579 | -0.099561678 | 0.102353786 | -0.972721009 | 0.330691981 | 0.541342432 | 0.099561678 |
| ENSMUSG00000037355 | Uvssa | 71101 | protein_coding | 295.8267942 | -0.099586912 | 0.14546475 | -0.684611994 | 0.493588797 | 0.685372663 | 0.099586912 |
| ENSMUSG00000106918 | Mrpl33 | 66845 | protein_coding | 330.1081383 | 0.099764561 | 0.101621635 | 0.981725607 | 0.326235047 | 0.537352722 | 0.099764561 |
| ENSMUSG00000074227 | Spint2 | 20733 | protein_coding | 9168.675697 | -0.099799571 | 0.082571255 | -1.208647866 | 0.226798159 | 0.429234918 | 0.099799571 |
| ENSMUSG00000000884 | Gnb1l | 13972 | protein_coding | 38.40666231 | -0.099830704 | 0.205581663 | -0.485601208 | 0.62724994 | 0.784304814 | 0.099830704 |
| ENSMUSG00000039849 | Pcif1 | 228866 | protein_coding | 487.0672216 | -0.099847238 | 0.073718034 | -1.354447921 | 0.175593518 | 0.361771253 | 0.099847238 |
| ENSMUSG00000003948 | Mmd | 67468 | protein_coding | 177.7851845 | -0.099859105 | 0.16166724 | -0.617682994 | 0.536784324 | 0.721206302 | 0.099859105 |
| ENSMUSG00000026860 | Sh3glb2 | 227700 | protein_coding | 680.6415188 | -0.099899493 | 0.093421252 | -1.069344398 | 0.284914512 | 0.49392051 | 0.099899493 |
| ENSMUSG00000027007 | Itprid2 | 70599 | protein_coding | 2332.244501 | -0.099955145 | 0.21237962 | -0.470643769 | 0.637895146 | 0.790438741 | 0.099955145 |
| ENSMUSG00000060733 | Ipmk | 69718 | protein_coding | 3131.762664 | -0.099955999 | 0.111551904 | -0.896049235 | 0.370226468 | 0.578871815 | 0.099955999 |
| ENSMUSG00000051978 | Erich1 | 234086 | protein_coding | 111.3492917 | -0.099989932 | 0.133636765 | -0.748221735 | 0.454326423 | 0.653612209 | 0.099989932 |
| ENSMUSG00000042350 | Arel1 | 68497 | protein_coding | 803.7068236 | -0.100019231 | 0.059944863 | -1.668520479 | 0.09521245 | 0.24060668 | 0.100019231 |
| ENSMUSG00000028163 | Nfkb1 | 18033 | protein_coding | 1232.399926 | -0.100044617 | 0.11518877 | -0.868527518 | 0.385105614 | 0.592939799 | 0.100044617 |
| ENSMUSG00000037857 | Nufip2 | 68564 | protein_coding | 1568.849051 | -0.10004532 | 0.111549975 | -0.896865457 | 0.369790714 | 0.578418334 | 0.10004532 |

| ensembl_gene_id | external_gene_name | entrezgene_id | gene_biotype | baseMean | log2FoldChange | lfcSE | stat | pvalue | padj | absLog2FC |
|---|---|---|---|---|---|---|---|---|---|---|
| ENSMUSG00000032435 | Dync1li1 | 235661 | protein_coding | 877.5462191 | 0.100064794 | 0.120066207 | 0.833413467 | 0.404611582 | 0.611621266 | 0.100064794 |
| ENSMUSG00000019261 | Map1s | 270058 | protein_coding | 127.3971533 | 0.1000899 | 0.208750228 | 0.479472053 | 0.631602845 | 0.786688311 | 0.1000899 |
| ENSMUSG00000022019 | Tdrd3 | 219249 | protein_coding | 569.0408843 | 0.100118348 | 0.079954439 | 1.252192486 | 0.210499733 | 0.407739238 | 0.100118348 |
| ENSMUSG00000027845 | Dclre1b | 140917 | protein_coding | 145.2006836 | -0.100139365 | 0.141933528 | -0.705537065 | 0.480476074 | 0.67397824 | 0.100139365 |
| ENSMUSG00000030155 | Clec2e | 232409 | protein_coding | 446.3100453 | -0.100170203 | 0.195630628 | -0.512037427 | 0.608624816 | 0.772321229 | 0.100170203 |
| ENSMUSG00000042826 | Fgf11 | 14166 | protein_coding | 29.18328644 | 0.100193103 | 0.301247972 | 0.332593452 | 0.739441187 | 0.854703862 | 0.100193103 |
| ENSMUSG00000037725 | Ckap2 | 80986 | protein_coding | 319.9575649 | 0.100213034 | 0.137399723 | 0.729353975 | 0.465785164 | 0.662531353 | 0.100213034 |
| ENSMUSG00000042118 | Bhmt2 | 64918 | protein_coding | 6.869469556 | -0.100225855 | 0.464446563 | -0.215796312 | 0.829146523 | 0.909405742 | 0.100225855 |
| ENSMUSG00000015133 | Lrrk1 | 233328 | protein_coding | 638.8111021 | -0.100253086 | 0.141950837 | -0.706252167 | 0.480031334 | 0.673742347 | 0.100253086 |
| ENSMUSG00000036501 | Fam13b | 225358 | protein_coding | 742.6748763 | -0.100258696 | 0.076003964 | -1.319124565 | 0.187127472 | 0.377782297 | 0.100258696 |
| ENSMUSG00000033781 | Asb13 | 142688 | protein_coding | 245.0523207 | -0.100288756 | 0.109355606 | -0.91708838 | 0.35909633 | 0.568896592 | 0.100288756 |
| ENSMUSG00000044294 | Krt84 | 16680 | protein_coding | 29.82328623 | -0.100338547 | 0.3328706 | -0.301434093 | 0.7630835 | 0.870097644 | 0.100338547 |
| ENSMUSG00000030657 | Xylt1 | 233781 | protein_coding | 121.7669708 | -0.100372441 | 0.189528249 | -0.529590929 | 0.596395585 | 0.764096331 | 0.100372441 |
| ENSMUSG00000038224 | Serpinf2 | 18816 | protein_coding | 15.35898279 | -0.100375227 | 0.423210071 | -0.237175895 | 0.812520336 | 0.900175552 | 0.100375227 |
| ENSMUSG00000002108 | Nr1h3 | 22259 | protein_coding | 174.299412 | -0.100375385 | 0.138536382 | -0.724541696 | 0.468733239 | 0.66509336 | 0.100375385 |
| ENSMUSG00000040549 | Ckap5 | 75786 | protein_coding | 1544.665159 | 0.100419645 | 0.076535136 | 1.312072465 | 0.189495685 | 0.381096879 | 0.100419645 |
| ENSMUSG00000040464 | Gtpbp10 | 207704 | protein_coding | 271.0502204 | 0.100426179 | 0.081813484 | 1.227501563 | 0.219634135 | 0.419876136 | 0.100426179 |
| ENSMUSG00000021902 | Phf7 | 71838 | protein_coding | 106.5548341 | -0.100430482 | 0.172543689 | -0.582058276 | 0.560527428 | 0.738310882 | 0.100430482 |
| ENSMUSG00000037940 | Inpp4b | 234515 | protein_coding | 373.2776932 | 0.100451264 | 0.166073529 | 0.604860174 | 0.545271907 | 0.727864934 | 0.100451264 |
| ENSMUSG00000057948 | Unc13d | 70450 | protein_coding | 245.2624355 | -0.100455518 | 0.102821975 | -0.976984907 | 0.328576624 | 0.53926517 | 0.100455518 |
| ENSMUSG00000045205 | Dpy19l4 | 381510 | protein_coding | 500.3157468 | 0.100457395 | 0.077460616 | 1.296883496 | 0.194671276 | 0.387784402 | 0.100457395 |
| ENSMUSG00000074272 | Ceacam1 | 26365 | protein_coding | 22265.84574 | -0.100488209 | 0.191442851 | -0.52489925 | 0.599653231 | 0.766220177 | 0.100488209 |
| ENSMUSG00000046675 | Tmem251 | 320351 | protein_coding | 187.2668707 | 0.100660026 | 0.163280382 | 0.616485733 | 0.537573985 | 0.721598526 | 0.100660026 |
| ENSMUSG00000058290 | Espl1 | 105988 | protein_coding | 219.7977385 | 0.10069314 | 0.102410723 | 0.983228488 | 0.325494997 | 0.536573088 | 0.10069314 |
| ENSMUSG00000002767 | Mrpl2 | 27398 | protein_coding | 524.4434059 | 0.100702318 | 0.109556288 | 0.919183372 | 0.357999667 | 0.567667366 | 0.100702318 |
| ENSMUSG00000036275 | 9530068E07Rik | 213673 | protein_coding | 1279.654394 | 0.100710396 | 0.051646047 | 1.950011694 | 0.051174725 | 0.157967253 | 0.100710396 |
| ENSMUSG00000073016 | Uprt | 331487 | protein_coding | 303.1329447 | 0.10075009 | 0.131268053 | 0.767514163 | 0.442775876 | 0.643949429 | 0.10075009 |
| ENSMUSG00000028085 | Gatb | 229487 | protein_coding | 210.0719355 | -0.10081145 | 0.101099428 | -0.997151544 | 0.318690957 | 0.53008663 | 0.10081145 |
| ENSMUSG00000097804 | Gm16685 | 102634900 | lncRNA | 9.435813105 | -0.100817226 | 0.536414028 | -0.187946661 | 0.850918464 | 0.921375207 | 0.100817226 |
| ENSMUSG00000033902 | Mapkbp1 | 26390 | protein_coding | 114.4789689 | -0.100822107 | 0.135366467 | -0.744808588 | 0.456387442 | 0.655308679 | 0.100822107 |
| ENSMUSG00000021809 | Nudt13 | 67725 | protein_coding | 163.9848042 | 0.10084732 | 0.135468741 | 0.744432397 | 0.456614925 | 0.655398985 | 0.10084732 |
| ENSMUSG00000024795 | Kif20b | 240641 | protein_coding | 607.5818662 | 0.100913625 | 0.127766145 | 0.789830712 | 0.42962664 | 0.633678408 | 0.100913625 |
| ENSMUSG00000028076 | Cd1d1 | 12479 | protein_coding | 104.2358405 | 0.100946108 | 0.170527414 | 0.591964104 | 0.553874626 | 0.733578739 | 0.100946108 |
| ENSMUSG00000028633 | Ctps | 51797 | protein_coding | 245.7707536 | -0.100969723 | 0.25172404 | -0.401112755 | 0.688337111 | 0.822275118 | 0.100969723 |
| ENSMUSG00000057113 | Npm1 | 18148 | protein_coding | 4496.634868 | -0.100969969 | 0.078047272 | -1.293702727 | 0.195768123 | 0.389044874 | 0.100969969 |
| ENSMUSG00000041439 | Mfsd6 | 98682 | protein_coding | 3357.785163 | 0.10097825 | 0.117621031 | 0.858505055 | 0.39061364 | 0.598873238 | 0.10097825 |
| ENSMUSG00000035051 | Dhx57 | 106794 | protein_coding | 333.0145992 | 0.100985455 | 0.073091084 | 1.381638484 | 0.167082731 | 0.35127842 | 0.100985455 |
| ENSMUSG00000024835 | Coro1b | 23789 | protein_coding | 2861.651118 | 0.101010852 | 0.101111495 | 0.999004634 | 0.317792446 | 0.529144688 | 0.101010852 |

| ensembl_gene_id | external_gene_name | entrezgene_id | gene_biotype | baseMean | log2FoldChange | lfcSE | stat | pvalue | padj | absLog2FC |
|---|---|---|---|---|---|---|---|---|---|---|
| ENSMUSG00000020180 | Snrpd3 | 67332 | protein_coding | 472.0029786 | 0.101049407 | 0.125629252 | 0.804346169 | 0.421197081 | 0.626934595 | 0.101049407 |
| ENSMUSG00000039515 | Ptpa | 110854 | protein_coding | 1069.144685 | 0.101082073 | 0.086200212 | 1.172642978 | 0.240939007 | 0.445516195 | 0.101082073 |
| ENSMUSG00000021357 | Exoc2 | 66482 | protein_coding | 415.4844272 | -0.101094988 | 0.061623968 | -1.64051409 | 0.100898321 | 0.2497061 | 0.101094988 |
| ENSMUSG00000048373 | Fgfbp1 | 14181 | protein_coding | 1475.899329 | 0.101105104 | 0.138025162 | 0.732512117 | 0.46385605 | 0.661197506 | 0.101105104 |
| ENSMUSG00000022403 | St13 | 70356 | protein_coding | 2153.817846 | 0.101108153 | 0.084563212 | 1.195651751 | 0.231832487 | 0.435246093 | 0.101108153 |
| ENSMUSG00000020186 | Csrp2 | 13008 | protein_coding | 2205.388376 | -0.101113796 | 0.127623737 | -0.792280482 | 0.42819715 | 0.632390628 | 0.101113796 |
| ENSMUSG00000033278 | Ptprm | 19274 | protein_coding | 65.83955777 | 0.101115898 | 0.374851052 | 0.269749538 | 0.787352947 | 0.884634427 | 0.101115898 |
| ENSMUSG00000047879 | Usp14 | 59025 | protein_coding | 1000.301651 | 0.101127261 | 0.111888216 | 0.903824054 | 0.366088705 | 0.57488523 | 0.101127261 |
| ENSMUSG00000032481 | Smarcc1 | 20588 | protein_coding | 900.2016132 | -0.101134743 | 0.086649142 | -1.167175356 | 0.243139561 | 0.44812855 | 0.101134743 |
| ENSMUSG00000053470 | Kdm3a | 104263 | protein_coding | 990.9487365 | -0.101161165 | 0.072061562 | -1.403815881 | 0.160373684 | 0.341958773 | 0.101161165 |
| ENSMUSG00000006315 | Tmem147 | 69804 | protein_coding | 486.5903432 | -0.101167025 | 0.104518266 | -0.967936316 | 0.333076176 | 0.543959534 | 0.101167025 |
| ENSMUSG00000056531 | Ccdc18 | 73254 | protein_coding | 148.5384238 | 0.101188515 | 0.157617294 | 0.641988659 | 0.520880547 | 0.707356582 | 0.101188515 |
| ENSMUSG00000086382 | Chrna1os | 102633437 | lncRNA | 9.571422144 | 0.101225886 | 0.390931338 | 0.258935206 | 0.795685234 | 0.889880242 | 0.101225886 |
| ENSMUSG00000037703 | Lzts3 | 241638 | protein_coding | 994.1516197 | 0.10128515 | 0.149772071 | 0.676261935 | 0.498874356 | 0.689463838 | 0.10128515 |
| ENSMUSG00000005823 | Gpr108 | 78308 | protein_coding | 1255.094606 | 0.101294283 | 0.069006742 | 1.467889657 | 0.142134198 | 0.315396142 | 0.101294283 |
| ENSMUSG00000062867 | Impdh2 | 23918 | protein_coding | 1460.610957 | -0.101316201 | 0.085642461 | -1.183013664 | 0.236803732 | 0.440992616 | 0.101316201 |
| ENSMUSG00000038119 | Cdon | 57810 | protein_coding | 314.1209782 | -0.101321175 | 0.154863598 | -0.65426076 | 0.512943822 | 0.700640528 | 0.101321175 |
| ENSMUSG00000013858 | Tmem259 | 216157 | protein_coding | 929.7548015 | -0.101331718 | 0.09483519 | -1.068503341 | 0.285293524 | 0.494306021 | 0.101331718 |
| ENSMUSG00000042705 | Commd10 | 69456 | protein_coding | 309.3890532 | 0.101366794 | 0.087233617 | 1.162015259 | 0.245229267 | 0.450668923 | 0.101366794 |
| ENSMUSG00000041258 | Zfp236 | 329002 | protein_coding | 620.851642 | 0.101444965 | 0.100185205 | 1.012574308 | 0.311263537 | 0.521380336 | 0.101444965 |
| ENSMUSG00000026883 | Dab2ip | 69601 | protein_coding | 1491.428151 | -0.10145072 | 0.079356824 | -1.278412045 | 0.201104179 | 0.395694119 | 0.10145072 |
| ENSMUSG00000016534 | Lamp2 | 16784 | protein_coding | 2117.843103 | 0.101479112 | 0.069497128 | 1.460191462 | 0.144237461 | 0.318731171 | 0.101479112 |
| ENSMUSG00000025059 | Gk | 14933 | protein_coding | 777.6639355 | 0.101492528 | 0.200036214 | 0.50737077 | 0.611894691 | 0.773819661 | 0.101492528 |
| ENSMUSG00000075420 | Smim6 | 68528 | protein_coding | 1408.109649 | 0.101502679 | 0.125993125 | 0.805620774 | 0.420461543 | 0.626390006 | 0.101502679 |
| ENSMUSG00000026675 | Hsd17b7 | 15490 | protein_coding | 374.0229252 | 0.101541977 | 0.173893874 | 0.583930733 | 0.559266915 | 0.737295433 | 0.101541977 |
| ENSMUSG00000006715 | Gmnn | 57441 | protein_coding | 244.392058 | 0.101547503 | 0.131903611 | 0.769861431 | 0.441382095 | 0.642981907 | 0.101547503 |
| ENSMUSG00000037916 | Ndufv1 | 17995 | protein_coding | 2789.548092 | 0.101560826 | 0.118390498 | 0.857846086 | 0.390977458 | 0.599142673 | 0.101560826 |
| ENSMUSG00000034957 | Cebpa | 12606 | protein_coding | 621.3005193 | -0.10162366 | 0.140844551 | -0.721530646 | 0.470583093 | 0.666501035 | 0.10162366 |
| ENSMUSG00000022048 | Dpysl2 | 12934 | protein_coding | 1276.330112 | 0.101686944 | 0.081349544 | 1.250000163 | 0.211299488 | 0.408728456 | 0.101686944 |
| ENSMUSG00000031934 | Panx1 | 55991 | protein_coding | 387.7895737 | 0.101735262 | 0.12619345 | 0.806184962 | 0.420136209 | 0.626116161 | 0.101735262 |
| ENSMUSG00000055493 | Epm2a | 13853 | protein_coding | 19.59149093 | -0.101777407 | 0.257432782 | -0.39535527 | 0.692580715 | 0.825026224 | 0.101777407 |
| ENSMUSG00000028990 | Lzic | 69151 | protein_coding | 301.0224923 | 0.101800136 | 0.096362079 | 1.056433581 | 0.290770171 | 0.49950251 | 0.101800136 |
| ENSMUSG00000049232 | Tigd2 | 68140 | protein_coding | 280.3313286 | -0.101833296 | 0.124760701 | -0.816228949 | 0.414369203 | 0.620774329 | 0.101833296 |
| ENSMUSG00000039126 | Prune2 | 353211 | protein_coding | 45.79486399 | 0.101833322 | 0.349616421 | 0.291271565 | 0.770843633 | 0.87556148 | 0.101833322 |
| ENSMUSG00000022594 | Lynx1 | 23936 | protein_coding | 92.72871362 | 0.10186614 | 0.144491052 | 0.704999647 | 0.480810456 | 0.674150435 | 0.10186614 |
| ENSMUSG00000053226 | Dand5 | 23863 | protein_coding | 31.5517706 | -0.102002603 | 0.224240598 | -0.454880179 | 0.649195443 | 0.797377007 | 0.102002603 |
| ENSMUSG00000045106 | Ccdc73 | 211936 | protein_coding | 24.33347357 | -0.102033292 | 0.312909098 | -0.326079658 | 0.744364087 | 0.857903848 | 0.102033292 |
| ENSMUSG00000028179 | Cth | 107869 | protein_coding | 1539.817969 | 0.102034086 | 0.115120614 | 0.88632333 | 0.375443321 | 0.583911261 | 0.102034086 |

| ensembl_gene_id | external_gene_name | entrezgene_id | gene_biotype | baseMean | log2FoldChange | lfcSE | stat | pvalue | padj | absLog2FC |
|---|---|---|---|---|---|---|---|---|---|---|
| ENSMUSG00000021895 | Arhgef3 | 71704 | protein_coding | 576.2129098 | -0.102035123 | 0.122999621 | -0.829556402 | 0.406789635 | 0.613912577 | 0.102035123 |
| ENSMUSG00000032741 | Tpcn1 | 252972 | protein_coding | 963.5628656 | -0.102066083 | 0.10528011 | -0.969471653 | 0.332309917 | 0.543320915 | 0.102066083 |
| ENSMUSG00000031245 | Hmgn5 | 50887 | protein_coding | 240.3226579 | -0.102083835 | 0.193670713 | -0.527100014 | 0.598124137 | 0.765321556 | 0.102083835 |
| ENSMUSG00000037070 | Rbmxl1 | 19656 | protein_coding | 201.0596676 | -0.102087071 | 0.158844824 | -0.642684278 | 0.520428985 | 0.706984466 | 0.102087071 |
| ENSMUSG00000020982 | Nemf | 66244 | protein_coding | 962.0056618 | 0.102158593 | 0.079902231 | 1.278544934 | 0.201057352 | 0.395684678 | 0.102158593 |
| ENSMUSG00000042719 | Naa25 | 231713 | protein_coding | 452.5416545 | -0.102176332 | 0.121569659 | -0.840475603 | 0.400641775 | 0.608101981 | 0.102176332 |
| ENSMUSG00000027684 | Mecom | 14013 | protein_coding | 2352.65171 | -0.10222902 | 0.129650472 | -0.788497096 | 0.430405997 | 0.634416687 | 0.10222902 |
| ENSMUSG00000063358 | Mapk1 | 26413 | protein_coding | 2143.864382 | 0.102279767 | 0.079974918 | 1.278898043 | 0.200932962 | 0.395606949 | 0.102279767 |
| ENSMUSG00000036968 | Cnpy4 | 66455 | protein_coding | 160.8256771 | 0.10232266 | 0.096032818 | 1.065496798 | 0.286651174 | 0.495534183 | 0.10232266 |
| ENSMUSG00000036390 | Gadd45a | 13197 | protein_coding | 358.6044464 | 0.102347819 | 0.204942683 | 0.499397284 | 0.617499532 | 0.777328108 | 0.102347819 |
| ENSMUSG00000042050 | Dync2i1 | 217935 | protein_coding | 124.5950629 | -0.102354019 | 0.133482 | -0.766800161 | 0.443200341 | 0.644271437 | 0.102354019 |
| ENSMUSG00000048920 | Fkrp | 243853 | protein_coding | 228.7555776 | 0.102386797 | 0.088934013 | 1.151267034 | 0.249622395 | 0.456013936 | 0.102386797 |
| ENSMUSG00000073591 | Pcdhb22 | 93893 | protein_coding | 11.14529615 | -0.102442887 | 0.429271596 | -0.238643525 | 0.811382012 | 0.899694484 | 0.102442887 |
| ENSMUSG00000021241 | Isca2 | 74316 | protein_coding | 234.5252375 | 0.102503939 | 0.097414767 | 1.052242305 | 0.292688395 | 0.501823547 | 0.102503939 |
| ENSMUSG00000021094 | Dhrs7 | 66375 | protein_coding | 472.6153433 | 0.102555966 | 0.096099039 | 1.067190343 | 0.285885892 | 0.494632902 | 0.102555966 |
| ENSMUSG00000037001 | Zfp39 | 22698 | protein_coding | 70.94168863 | 0.102562201 | 0.223396928 | 0.459103007 | 0.646160198 | 0.795304816 | 0.102562201 |
| ENSMUSG00000026667 | Uhmk1 | 16589 | protein_coding | 1944.663344 | -0.10265115 | 0.098956218 | -1.03733906 | 0.299577868 | 0.508544846 | 0.10265115 |
| ENSMUSG00000024672 | Ms4a7 | 109225 | protein_coding | 24.48392028 | -0.102688041 | 0.282298736 | -0.363756643 | 0.716039742 | 0.84023174 | 0.102688041 |
| ENSMUSG00000038080 | Kdm1b | 218214 | protein_coding | 565.0614641 | -0.10281658 | 0.076575727 | -1.342678457 | 0.179376113 | 0.367085742 | 0.10281658 |
| ENSMUSG00000086287 | Gm15972 | 100502834 | lncRNA | 18.54583983 | -0.102835058 | 0.370152635 | -0.277817982 | 0.781152087 | 0.880987863 | 0.102835058 |
| ENSMUSG00000022194 | Pabpn1 | 12050 | protein_coding | 258.4399798 | -0.102857667 | 0.145956485 | -0.704714604 | 0.480987861 | 0.674232796 | 0.102857667 |
| ENSMUSG00000001930 | Vwf | 22371 | protein_coding | 54.81274584 | -0.102870853 | 0.219185236 | -0.46933295 | 0.638831667 | 0.790625182 | 0.102870853 |
| ENSMUSG00000075706 | Gpx4 | 625249 | protein_coding | 1037.095016 | 0.102873737 | 0.081721545 | 1.25883249 | 0.208090842 | 0.404598503 | 0.102873737 |
| ENSMUSG00000099881 | 2810013P06Rik | 100503178 | lncRNA | 229.6812667 | 0.102878144 | 0.161394365 | 0.637433304 | 0.523842641 | 0.709654498 | 0.102878144 |
| ENSMUSG00000021604 | Irx4 | 50916 | protein_coding | 16.13077513 | 0.103082876 | 0.334465777 | 0.308201565 | 0.757928961 | 0.867408546 | 0.103082876 |
| ENSMUSG00000036430 | Tbcc | 72726 | protein_coding | 147.1783688 | 0.103083536 | 0.160621513 | 0.641779139 | 0.521016596 | 0.707440424 | 0.103083536 |
| ENSMUSG00000034613 | Ppm1h | 319468 | protein_coding | 777.5672059 | -0.103110913 | 0.062494828 | -1.649911149 | 0.09896111 | 0.246636295 | 0.103110913 |
| ENSMUSG00000057969 | Sema3b | 20347 | protein_coding | 962.8137072 | -0.103148286 | 0.095295902 | -1.08240002 | 0.279074819 | 0.487504149 | 0.103148286 |
| ENSMUSG00000029718 | Pcolce | 18542 | protein_coding | 39.14562424 | -0.103160416 | 0.268348147 | -0.384427532 | 0.700661598 | 0.829926899 | 0.103160416 |
| ENSMUSG00000031012 | Cask | 12361 | protein_coding | 1359.596001 | 0.103161603 | 0.042844061 | 2.407839065 | 0.016047253 | 0.067998336 | 0.103161603 |
| ENSMUSG00000025417 | Pip4k2c | 117150 | protein_coding | 1007.36172 | -0.103171509 | 0.080166837 | -1.28695995 | 0.198108248 | 0.392178495 | 0.103171509 |
| ENSMUSG00000029687 | Ezh2 | 14056 | protein_coding | 487.451823 | -0.103247833 | 0.084826786 | -1.217160737 | 0.223543066 | 0.425169501 | 0.103247833 |
| ENSMUSG00000085436 | Zfp335os | 629777 | lncRNA | 13.46722295 | -0.103305996 | 0.35072515 | -0.294549723 | 0.768337874 | 0.873665822 | 0.103305996 |
| ENSMUSG00000006642 | Tcf23 | 69852 | protein_coding | 82.60878765 | -0.103321026 | 0.186236113 | -0.554785133 | 0.579041638 | 0.750567643 | 0.103321026 |
| ENSMUSG00000052656 | Rnf103 | 22644 | protein_coding | 1688.523482 | 0.103336046 | 0.092744194 | 1.114205014 | 0.265191245 | 0.472743833 | 0.103336046 |
| ENSMUSG00000039634 | Zfp189 | 230162 | protein_coding | 41.48064049 | -0.103354673 | 0.220954067 | -0.467765426 | 0.639952348 | 0.791371033 | 0.103354673 |
| ENSMUSG00000000339 | Rtca | 66368 | protein_coding | 466.9006733 | 0.103359677 | 0.104676298 | 0.987421979 | 0.323435814 | 0.534407224 | 0.103359677 |
| ENSMUSG00000024381 | Bin1 | 30948 | protein_coding | 182.9728307 | -0.103368918 | 0.117922981 | -0.87657993 | 0.380714844 | 0.588499925 | 0.103368918 |

| ensembl_gene_id | external_gene_name | entrezgene_id | gene_biotype | baseMean | log2FoldChange | lfcSE | stat | pvalue | padj | absLog2FC |
|---|---|---|---|---|---|---|---|---|---|---|
| ENSMUSG00000006456 | Rbm14 | 56275 | protein_coding | 489.0812511 | 0.103375649 | 0.251478159 | 0.411072078 | 0.681019684 | 0.817455242 | 0.103375649 |
| ENSMUSG00000015787 | Abo | 80908 | protein_coding | 170.4532151 | -0.103402392 | 0.152505347 | -0.678024702 | 0.497756031 | 0.688791086 | 0.103402392 |
| ENSMUSG00000039050 | Osbpl2 | 228983 | protein_coding | 1266.418227 | 0.103476753 | 0.101445042 | 1.020027702 | 0.307715323 | 0.517614285 | 0.103476753 |
| ENSMUSG00000071855 | Ccdc112 | 240261 | protein_coding | 49.2631519 | 0.103500514 | 0.211477573 | 0.489416031 | 0.624547189 | 0.782629869 | 0.103500514 |
| ENSMUSG00000079499 | 6530402F18Rik | 76220 | lncRNA | 7.793840991 | 0.103571625 | 0.571162982 | 0.181334625 | 0.85610493 | 0.924704659 | 0.103571625 |
| ENSMUSG00000015289 | Lage3 | 66192 | protein_coding | 161.5151111 | -0.103593803 | 0.115840452 | -0.894280029 | 0.371172082 | 0.579501943 | 0.103593803 |
| ENSMUSG00000025873 | Faf2 | 76577 | protein_coding | 1041.397326 | 0.103598553 | 0.050424726 | 2.054518901 | 0.039925496 | 0.132945456 | 0.103598553 |
| ENSMUSG00000026585 | Kifap3 | 16579 | protein_coding | 337.5221198 | 0.103606114 | 0.092068217 | 1.125319008 | 0.260453878 | 0.467786902 | 0.103606114 |
| ENSMUSG00000031320 | Rps4x | 20102 | protein_coding | 12400.60319 | 0.10362409 | 0.103771264 | 0.998581743 | 0.317997348 | 0.529209253 | 0.10362409 |
| ENSMUSG00000042548 | Asxl1 | 228790 | protein_coding | 760.4956042 | -0.103666809 | 0.114180067 | -0.907923868 | 0.363918446 | 0.572855539 | 0.103666809 |
| ENSMUSG00000029445 | Hpd | 15445 | protein_coding | 873.3190435 | -0.103743448 | 0.135474486 | -0.765778493 | 0.443808114 | 0.644803031 | 0.103743448 |
| ENSMUSG00000037280 | Galnt6 | 207839 | protein_coding | 1976.030185 | -0.103750244 | 0.11699563 | -0.886787337 | 0.375193407 | 0.583715626 | 0.103750244 |
| ENSMUSG00000102869 | Norad | 347740 | lncRNA | 1801.66644 | 0.103767765 | 0.067415871 | 1.539218638 | 0.123750929 | 0.286966176 | 0.103767765 |
| ENSMUSG00000024212 | Mllt1 | 64144 | protein_coding | 529.4242277 | -0.103771556 | 0.096962202 | -1.0702269 | 0.284517189 | 0.493695176 | 0.103771556 |
| ENSMUSG00000016921 | Srsf6 | 67996 | protein_coding | 2135.259961 | -0.103804563 | 0.101416801 | -1.023544048 | 0.306050685 | 0.51606768 | 0.103804563 |
| ENSMUSG00000020156 | Pwwp3a | 68114 | protein_coding | 331.6672879 | -0.103805414 | 0.077171877 | -1.345119728 | 0.178586573 | 0.36594969 | 0.103805414 |
| ENSMUSG00000024423 | Impact | 16210 | protein_coding | 110.1585993 | -0.103886489 | 0.15384108 | -0.675284447 | 0.499495063 | 0.689962481 | 0.103886489 |
| ENSMUSG00000001056 | Nhp2 | 52530 | protein_coding | 364.7156362 | -0.103911869 | 0.130238514 | -0.79785822 | 0.424952771 | 0.629529435 | 0.103911869 |
| ENSMUSG00000032869 | Psmf1 | 228769 | protein_coding | 338.1432117 | 0.103919647 | 0.084648286 | 1.227663918 | 0.219573157 | 0.419876136 | 0.103919647 |
| ENSMUSG00000038880 | Mrps34 | 79044 | protein_coding | 126.9406216 | 0.104003995 | 0.198934356 | 0.522805596 | 0.601109545 | 0.767123855 | 0.104003995 |
| ENSMUSG00000023147 | Get1 | 71446 | protein_coding | 143.1765418 | -0.104007085 | 0.135240717 | -0.769051565 | 0.4418627 | 0.643208653 | 0.104007085 |
| ENSMUSG00000026672 | Optn | 71648 | protein_coding | 546.3698514 | 0.104015098 | 0.098959375 | 1.051088871 | 0.293217775 | 0.502250159 | 0.104015098 |
| ENSMUSG00000054892 | Txk | 22165 | protein_coding | 25.22034242 | 0.104029489 | 0.456630337 | 0.227819924 | 0.819786237 | 0.904450729 | 0.104029489 |
| ENSMUSG00000063145 | Bbs5 | 72569 | protein_coding | 73.92505542 | 0.104061815 | 0.165343114 | 0.629368906 | 0.52910757 | 0.714548497 | 0.104061815 |
| ENSMUSG00000024664 | Fads3 | 60527 | protein_coding | 325.2708108 | 0.104106137 | 0.087486547 | 1.189967375 | 0.234059215 | 0.437878222 | 0.104106137 |
| ENSMUSG00000079562 | Maea | 59003 | protein_coding | 1329.827284 | 0.104124848 | 0.091574585 | 1.137049627 | 0.25551754 | 0.46222412 | 0.104124848 |
| ENSMUSG00000030216 | Wbp11 | 60321 | protein_coding | 568.5121155 | -0.104139643 | 0.074866079 | -1.391012374 | 0.164221675 | 0.347512001 | 0.104139643 |
| ENSMUSG00000027428 | Rbbp9 | 26450 | protein_coding | 164.0413697 | 0.104228758 | 0.135682985 | 0.768178544 | 0.442381119 | 0.643610382 | 0.104228758 |
| ENSMUSG00000005881 | Ergic3 | 66366 | protein_coding | 1371.272447 | -0.104237625 | 0.084675153 | -1.231029669 | 0.218311768 | 0.418242704 | 0.104237625 |
| ENSMUSG00000029312 | Klhl8 | 246293 | protein_coding | 67.95341694 | -0.104257145 | 0.155480601 | -0.670547607 | 0.502508769 | 0.692503865 | 0.104257145 |
| ENSMUSG00000021496 | Pcbd2 | 72562 | protein_coding | 117.6199141 | -0.104262318 | 0.14444106 | -0.721832959 | 0.470397183 | 0.666413442 | 0.104262318 |
| ENSMUSG00000054051 | Ercc6 | 319955 | protein_coding | 393.125933 | -0.104271204 | 0.118088195 | -0.882994309 | 0.377239347 | 0.585247665 | 0.104271204 |
| ENSMUSG00000005150 | Wdr83 | 67836 | protein_coding | 175.8400158 | 0.104281771 | 0.110257248 | 0.94580422 | 0.344248453 | 0.55406481 | 0.104281771 |
| ENSMUSG00000106847 | Peg13 | 353342 | lncRNA | 152.5869317 | -0.104283848 | 0.140154051 | -0.744065888 | 0.456836615 | 0.655658101 | 0.104283848 |
| ENSMUSG00000022885 | St6gal1 | 20440 | protein_coding | 1689.505578 | -0.104291391 | 0.134005704 | -0.778260836 | 0.436415258 | 0.639251631 | 0.104291391 |
| ENSMUSG00000022335 | Zfat | 380993 | protein_coding | 83.98252291 | 0.104297671 | 0.126884248 | 0.821990697 | 0.411082191 | 0.618117355 | 0.104297671 |
| ENSMUSG00000046314 | Stxbp6 | 217517 | protein_coding | 352.4335471 | -0.104315663 | 0.119269847 | -0.874618909 | 0.381781296 | 0.589473873 | 0.104315663 |
| ENSMUSG00000032580 | Rbm5 | 83486 | protein_coding | 1565.037226 | -0.104392282 | 0.051561985 | -2.024597824 | 0.042908678 | 0.139507945 | 0.104392282 |

| ensembl_gene_id | external_gene_name | entrezgene_id | gene_biotype | baseMean | log2FoldChange | lfcSE | stat | pvalue | padj | absLog2FC |
|---|---|---|---|---|---|---|---|---|---|---|
| ENSMUSG00000074922 | Fam122a | 68034 | protein_coding | 65.14179712 | -0.104448582 | 0.166684873 | -0.626623038 | 0.530906363 | 0.716208737 | 0.104448582 |
| ENSMUSG00000035258 | Abi3bp | 320712 | protein_coding | 18.12577068 | -0.104459886 | 0.608447282 | -0.171682722 | 0.86368697 | 0.929200004 | 0.104459886 |
| ENSMUSG00000016427 | Ndufa1 | 54405 | protein_coding | 1269.786357 | 0.10448583 | 0.087888338 | 1.18884748 | 0.23449969 | 0.438393318 | 0.10448583 |
| ENSMUSG00000020434 | 4921536K21Rik | 67430 | protein_coding | 37.60670467 | 0.104569187 | 0.232117791 | 0.450500527 | 0.652349575 | 0.798541656 | 0.104569187 |
| ENSMUSG00000020807 | 4933427D14Rik | 74477 | protein_coding | 7.682781001 | -0.104589356 | 0.395698643 | -0.264315679 | 0.79153668 | 0.887042894 | 0.104589356 |
| ENSMUSG00000001785 | Pwp1 | 103136 | protein_coding | 436.5277608 | -0.104626386 | 0.119833536 | -0.87309771 | 0.382609822 | 0.590295355 | 0.104626386 |
| ENSMUSG00000036817 | Sun1 | 77053 | protein_coding | 1259.33075 | -0.104627856 | 0.108059791 | -0.968240406 | 0.33292432 | 0.543823051 | 0.104627856 |
| ENSMUSG00000038690 | Atp5j2 | 57423 | protein_coding | 3218.089497 | 0.104674432 | 0.12319378 | 0.849673026 | 0.395506899 | 0.603774085 | 0.104674432 |
| ENSMUSG00000004562 | Arhgef40 | 268739 | protein_coding | 282.7327733 | 0.104718072 | 0.197819661 | 0.529361292 | 0.596554844 | 0.764127508 | 0.104718072 |
| ENSMUSG00000020523 | Fam114a2 | 67726 | protein_coding | 1109.94877 | 0.104769767 | 0.077220445 | 1.356762026 | 0.174856837 | 0.361108705 | 0.104769767 |
| ENSMUSG00000057134 | Ado | 211488 | protein_coding | 489.9806371 | -0.10479269 | 0.115982766 | -0.903519496 | 0.366250246 | 0.575082208 | 0.10479269 |
| ENSMUSG00000029203 | Ube2k | 53323 | protein_coding | 956.9636445 | 0.104794712 | 0.07293661 | 1.436791642 | 0.150777206 | 0.328803305 | 0.104794712 |
| ENSMUSG00000020604 | Arsg | 74008 | protein_coding | 107.9915526 | -0.104824605 | 0.159018829 | -0.659196179 | 0.5097698 | 0.698581552 | 0.104824605 |
| ENSMUSG00000057092 | Fxyd3 | 17178 | protein_coding | 488.849908 | -0.104828906 | 0.113081409 | -0.92702158 | 0.35391533 | 0.563659971 | 0.104828906 |
| ENSMUSG00000027978 | Prss12 | 19142 | protein_coding | 16.73552225 | -0.104905787 | 0.324165997 | -0.323617493 | 0.746227643 | 0.858895287 | 0.104905787 |
| ENSMUSG00000005813 | Metap1 | 75624 | protein_coding | 1019.020733 | 0.104940056 | 0.077282234 | 1.35788072 | 0.174501536 | 0.360734648 | 0.104940056 |
| ENSMUSG00000040410 | Fbxl4 | 269514 | protein_coding | 242.9590489 | 0.105037019 | 0.117496131 | 0.893961514 | 0.371342483 | 0.579649458 | 0.105037019 |
| ENSMUSG00000026646 | Suv39h2 | 64707 | protein_coding | 76.34138772 | 0.105068762 | 0.151980061 | 0.69133254 | 0.489356587 | 0.681871378 | 0.105068762 |
| ENSMUSG00000042293 | Gm5617 | 434402 | protein_coding | 59.88132021 | -0.10509469 | 0.162093945 | -0.648356672 | 0.516754288 | 0.704034068 | 0.10509469 |
| ENSMUSG00000038398 | Upf3a | 67031 | protein_coding | 377.9738966 | -0.105122517 | 0.090741845 | -1.158478946 | 0.246668639 | 0.452168679 | 0.105122517 |
| ENSMUSG00000031835 | Mbtps1 | 56453 | protein_coding | 1884.144648 | 0.105253205 | 0.074884381 | 1.405542848 | 0.159859917 | 0.341274462 | 0.105253205 |
| ENSMUSG00000029486 | Mrpl1 | 94061 | protein_coding | 440.0401382 | 0.105280203 | 0.09283598 | 1.134045262 | 0.256775562 | 0.463695525 | 0.105280203 |
| ENSMUSG00000056268 | Dennd1b | 329260 | protein_coding | 1321.12273 | -0.105305093 | 0.073154679 | -1.439485408 | 0.150013041 | 0.327540469 | 0.105305093 |
| ENSMUSG00000042079 | Hnrnpf | 98758 | protein_coding | 1701.152533 | 0.105319553 | 0.101939096 | 1.033161539 | 0.301528308 | 0.510865285 | 0.105319553 |
| ENSMUSG00000089715 | Cbx6 | 494448 | protein_coding | 101.467948 | 0.105321759 | 0.21889578 | 0.48115025 | 0.630409717 | 0.78600125 | 0.105321759 |
| ENSMUSG00000026787 | Gad2 | 14417 | protein_coding | 43.88146959 | 0.105339531 | 0.290462459 | 0.36266143 | 0.716857815 | 0.840627772 | 0.105339531 |
| ENSMUSG00000033306 | Lpp | 210126 | protein_coding | 2181.147587 | 0.105373609 | 0.184524348 | 0.571055311 | 0.56796215 | 0.743305197 | 0.105373609 |
| ENSMUSG00000027394 | Ttl | 69737 | protein_coding | 113.2296799 | -0.105382591 | 0.138169771 | -0.76270367 | 0.445640141 | 0.646462311 | 0.105382591 |
| ENSMUSG00000070803 | Cited4 | 56222 | protein_coding | 98.53338327 | -0.105410952 | 0.149306455 | -0.706003985 | 0.480185659 | 0.673867593 | 0.105410952 |
| ENSMUSG00000095432 | Zfp748 | 212276 | protein_coding | 376.4775507 | -0.105449091 | 0.082681764 | -1.275360918 | 0.202181525 | 0.397230459 | 0.105449091 |
| ENSMUSG00000022723 | Crybg3 | 224273 | protein_coding | 411.8320571 | -0.105463583 | 0.141048848 | -0.747709636 | 0.454635317 | 0.653915542 | 0.105463583 |
| ENSMUSG00000036918 | Ttc7 | 225049 | protein_coding | 1099.705236 | 0.105494207 | 0.087449166 | 1.206348918 | 0.227682983 | 0.430280937 | 0.105494207 |
| ENSMUSG00000004931 | Apba3 | 57267 | protein_coding | 363.4631154 | 0.10549944 | 0.098742496 | 1.06842995 | 0.285326613 | 0.494309581 | 0.10549944 |
| ENSMUSG00000024259 | Slc25a46 | 67453 | protein_coding | 1092.73313 | 0.105506846 | 0.078081924 | 1.351232661 | 0.176620915 | 0.363136432 | 0.105506846 |
| ENSMUSG00000034259 | Exosc4 | 109075 | protein_coding | 200.8072701 | 0.105508149 | 0.11006579 | 0.958591662 | 0.337764494 | 0.548479169 | 0.105508149 |
| ENSMUSG00000007836 | Hnrnpa0 | 77134 | protein_coding | 1378.759161 | 0.105521391 | 0.142381139 | 0.741119165 | 0.45862119 | 0.65713215 | 0.105521391 |
| ENSMUSG00000036376 | Abt1 | 30946 | protein_coding | 188.3058202 | -0.105525196 | 0.101116307 | -1.043602158 | 0.296669494 | 0.506034665 | 0.105525196 |
| ENSMUSG00000035329 | Fbxo33 | 70611 | protein_coding | 500.9019899 | 0.105557559 | 0.111085549 | 0.950236647 | 0.341992021 | 0.552123345 | 0.105557559 |

| ensembl_gene_id | external_gene_name | entrezgene_id | gene_biotype | baseMean | log2FoldChange | lfcSE | stat | pvalue | padj | absLog2FC |
|---|---|---|---|---|---|---|---|---|---|---|
| ENSMUSG00000024862 | Klc2 | 16594 | protein_coding | 138.4844762 | -0.105678864 | 0.120911788 | -0.874016217 | 0.382109422 | 0.589866144 | 0.105678864 |
| ENSMUSG00000036805 | Noxa1 | 241275 | protein_coding | 941.5317939 | 0.1057217 | 0.103995457 | 1.016599219 | 0.309344126 | 0.519147954 | 0.1057217 |
| ENSMUSG00000068457 | Uty | 22290 | protein_coding | 773.3929453 | 0.105733636 | 0.093982461 | 1.125035838 | 0.260573848 | 0.467916601 | 0.105733636 |
| ENSMUSG00000029022 | Miip | 28010 | protein_coding | 241.3975956 | -0.105741956 | 0.119589115 | -0.884210538 | 0.376582571 | 0.58462681 | 0.105741956 |
| ENSMUSG00000042389 | Tsen2 | 381802 | protein_coding | 74.73905429 | 0.105786862 | 0.146701074 | 0.721104893 | 0.470844981 | 0.666534157 | 0.105786862 |
| ENSMUSG00000043747 | 1520401A03Rik | 320309 | protein_coding | 7.798305365 | 0.105789935 | 0.651767991 | 0.16231226 | 0.871059961 | 0.933600906 | 0.105789935 |
| ENSMUSG00000033773 | Rpap2 | 231571 | protein_coding | 213.6063801 | -0.105810813 | 0.123427717 | -0.857269461 | 0.391295982 | 0.59924777 | 0.105810813 |
| ENSMUSG00000022570 | Gfus | 22122 | protein_coding | 1491.580939 | 0.105811415 | 0.093290583 | 1.134213249 | 0.256705108 | 0.463635215 | 0.105811415 |
| ENSMUSG00000117042 | 2700054A10Rik | 72578 | lncRNA | 12.72262681 | 0.105811505 | 0.449925409 | 0.235175659 | 0.814072397 | 0.900950823 | 0.105811505 |
| ENSMUSG00000001552 | Jup | 16480 | protein_coding | 3299.350659 | 0.105817368 | 0.128520237 | 0.823351794 | 0.410307968 | 0.617244552 | 0.105817368 |
| ENSMUSG00000026827 | Gpd2 | 14571 | protein_coding | 897.2603374 | -0.10581964 | 0.132371754 | -0.799412539 | 0.424051242 | 0.628617629 | 0.10581964 |
| ENSMUSG00000040365 | Trim41 | 211007 | protein_coding | 586.6974268 | -0.105881681 | 0.069887038 | -1.515040325 | 0.129762192 | 0.296492928 | 0.105881681 |
| ENSMUSG00000028410 | Dnaja1 | 15502 | protein_coding | 2321.286002 | 0.105901355 | 0.231609478 | 0.457241026 | 0.647497813 | 0.79639731 | 0.105901355 |
| ENSMUSG00000021326 | Trim27 | 19720 | protein_coding | 581.0157439 | 0.105919361 | 0.080030334 | 1.32349018 | 0.18567242 | 0.376066926 | 0.105919361 |
| ENSMUSG00000029699 | Ssc4d | 109267 | protein_coding | 42.76816378 | 0.105964625 | 0.184840136 | 0.573277144 | 0.566457058 | 0.742495568 | 0.105964625 |
| ENSMUSG00000059288 | Cdyl | 12593 | protein_coding | 170.9459349 | -0.10597161 | 0.139667027 | -0.758744655 | 0.448005319 | 0.648829671 | 0.10597161 |
| ENSMUSG00000056716 | Gm5420 | 432436 | lncRNA | 7.239804821 | 0.106012537 | 0.454639793 | 0.233179187 | 0.815622267 | 0.901980939 | 0.106012537 |
| ENSMUSG00000071632 | 2510002D24Rik | 72307 | protein_coding | 208.6931244 | -0.106059573 | 0.099911641 | -1.061533697 | 0.288447427 | 0.497443766 | 0.106059573 |
| ENSMUSG00000027508 | Pag1 | 94212 | protein_coding | 338.8412984 | 0.106069649 | 0.139951017 | 0.757905531 | 0.448507539 | 0.64942205 | 0.106069649 |
| ENSMUSG00000025505 | Tmem80 | 71448 | protein_coding | 92.82831116 | -0.106094939 | 0.125233099 | -0.847179698 | 0.396894968 | 0.60483573 | 0.106094939 |
| ENSMUSG00000035754 | Wdr18 | 216156 | protein_coding | 419.1469781 | -0.106116924 | 0.125738529 | -0.84394915 | 0.398697821 | 0.606538577 | 0.106116924 |
| ENSMUSG00000028018 | Gstcd | 67553 | protein_coding | 154.2022318 | 0.106141044 | 0.116500995 | 0.911074142 | 0.36225631 | 0.571538089 | 0.106141044 |
| ENSMUSG00000024726 | Carnmt1 | 67383 | protein_coding | 148.8308006 | -0.106144041 | 0.116919868 | -0.90783579 | 0.363964986 | 0.572855539 | 0.106144041 |
| ENSMUSG00000036850 | Mrpl41 | 107733 | protein_coding | 289.5033635 | 0.106185572 | 0.098088237 | 1.082551546 | 0.279007524 | 0.487498009 | 0.106185572 |
| ENSMUSG00000037351 | Actr1b | 226977 | protein_coding | 2405.753751 | 0.106198049 | 0.079648357 | 1.333336331 | 0.182421456 | 0.371513739 | 0.106198049 |
| ENSMUSG00000027610 | Gss | 14854 | protein_coding | 2310.107087 | 0.106243172 | 0.108423224 | 0.979893121 | 0.327138879 | 0.53834777 | 0.106243172 |
| ENSMUSG00000005102 | Eif2ak4 | 27103 | protein_coding | 440.1288947 | -0.106281457 | 0.099060307 | -1.072896501 | 0.283317556 | 0.492489581 | 0.106281457 |
| ENSMUSG00000047215 | Rpl9 | 20005 | protein_coding | 9732.301307 | 0.1062915 | 0.119971659 | 0.885971744 | 0.375632754 | 0.583911261 | 0.1062915 |
| ENSMUSG00000036478 | Btg1 | 12226 | protein_coding | 3054.004596 | 0.106342874 | 0.085214035 | 1.24795022 | 0.212049288 | 0.409893332 | 0.106342874 |
| ENSMUSG00000044221 | Grsf1 | 231413 | protein_coding | 1102.193029 | -0.106395749 | 0.074304594 | -1.431886554 | 0.152176291 | 0.330632105 | 0.106395749 |
| ENSMUSG00000025437 | Usp33 | 170822 | protein_coding | 1312.679193 | -0.106401237 | 0.097737825 | -1.088639299 | 0.276312979 | 0.48470409 | 0.106401237 |
| ENSMUSG00000028565 | Nfia | 18027 | protein_coding | 422.9801955 | -0.106427546 | 0.163066956 | -0.652661634 | 0.513974441 | 0.701627091 | 0.106427546 |
| ENSMUSG00000034579 | Pla2g3 | 237625 | protein_coding | 318.633913 | 0.106449383 | 0.197398478 | 0.539261418 | 0.589706487 | 0.759394898 | 0.106449383 |
| ENSMUSG00000053329 | Gatd3a | 28295 | protein_coding | 586.3498136 | -0.106456791 | 0.103700256 | -1.026581753 | 0.30461745 | 0.514140801 | 0.106456791 |
| ENSMUSG00000057411 | Antkmt | 214917 | protein_coding | 190.3648004 | -0.106458595 | 0.114565519 | -0.929237661 | 0.352765934 | 0.562504334 | 0.106458595 |
| ENSMUSG00000013846 | St3gal1 | 20442 | protein_coding | 162.4769166 | -0.106464077 | 0.201439818 | -0.528515552 | 0.597141556 | 0.764635807 | 0.106464077 |
| ENSMUSG00000066324 | Bpnt2 | 242291 | protein_coding | 3073.255266 | 0.10649082 | 0.065286171 | 1.631139009 | 0.102860991 | 0.253030096 | 0.10649082 |
| ENSMUSG00000048307 | Ankrd46 | 68839 | protein_coding | 273.312396 | 0.106511064 | 0.106117437 | 1.003709355 | 0.315518727 | 0.526539948 | 0.106511064 |

| ensembl_gene_id | external_gene_name | entrezgene_id | gene_biotype | baseMean | log2FoldChange | lfcSE | stat | pvalue | padj | absLog2FC |
|---|---|---|---|---|---|---|---|---|---|---|
| ENSMUSG00000052137 | Rbm12b2 | 77604 | protein_coding | 192.7575675 | -0.10652815 | 0.13696073 | -0.777800685 | 0.436686524 | 0.639413767 | 0.10652815 |
| ENSMUSG00000010755 | Cars | 27267 | protein_coding | 473.7288975 | 0.1065367 | 0.220826783 | 0.482444648 | 0.62949011 | 0.785654167 | 0.1065367 |
| ENSMUSG00000019857 | Asf1a | 66403 | protein_coding | 186.5682747 | 0.106536859 | 0.096708926 | 1.101623844 | 0.270625238 | 0.478449592 | 0.106536859 |
| ENSMUSG00000019942 | Cdk1 | 12534 | protein_coding | 608.4556986 | 0.106622389 | 0.140277724 | 0.760080688 | 0.447206356 | 0.647908207 | 0.106622389 |
| ENSMUSG00000032380 | Dapk2 | 13143 | protein_coding | 567.8747974 | -0.10662371 | 0.156562051 | -0.681031637 | 0.495851472 | 0.687162611 | 0.10662371 |
| ENSMUSG00000117924 | Tmem223 | 66836 | protein_coding | 214.5568453 | 0.106631289 | 0.135853544 | 0.784898839 | 0.432512883 | 0.636403252 | 0.106631289 |
| ENSMUSG00000113204 | Gm46430 | 108168101 | lncRNA | 174.3837796 | -0.106692453 | 0.172828828 | -0.617330188 | 0.537016959 | 0.721275497 | 0.106692453 |
| ENSMUSG00000041757 | Plekha6 | 240753 | protein_coding | 4306.939714 | -0.106699553 | 0.11195753 | -0.953035969 | 0.340571847 | 0.550892928 | 0.106699553 |
| ENSMUSG00000036086 | Zranb3 | 226409 | protein_coding | 96.31810106 | -0.106712949 | 0.166982919 | -0.63906506 | 0.522780609 | 0.709030103 | 0.106712949 |
| ENSMUSG00000019302 | Atp6v0a1 | 11975 | protein_coding | 713.9695021 | -0.106724679 | 0.071262455 | -1.497628442 | 0.134229813 | 0.303178171 | 0.106724679 |
| ENSMUSG00000025921 | Rdh10 | 98711 | protein_coding | 396.8138313 | -0.106771847 | 0.286295667 | -0.372942587 | 0.709191175 | 0.835765162 | 0.106771847 |
| ENSMUSG00000022895 | Ets2 | 23872 | protein_coding | 3510.546869 | -0.10678554 | 0.12475979 | -0.855929144 | 0.392036974 | 0.600073507 | 0.10678554 |
| ENSMUSG00000005566 | Trim28 | 21849 | protein_coding | 2111.649084 | 0.1067878 | 0.090486753 | 1.180148442 | 0.237941181 | 0.442159071 | 0.1067878 |
| ENSMUSG00000025742 | Prps2 | 110639 | protein_coding | 777.1257291 | -0.106790387 | 0.069308466 | -1.540798594 | 0.123365812 | 0.286480631 | 0.106790387 |
| ENSMUSG00000028416 | Bag1 | 12017 | protein_coding | 2902.205191 | 0.106825441 | 0.109242165 | 0.977877365 | 0.328134981 | 0.538929056 | 0.106825441 |
| ENSMUSG00000020459 | Mtif2 | 76784 | protein_coding | 548.4683298 | 0.106828435 | 0.089358231 | 1.195507503 | 0.231888806 | 0.43530052 | 0.106828435 |
| ENSMUSG00000021639 | Gtf2h2 | 23894 | protein_coding | 256.9237097 | 0.10684417 | 0.109342979 | 0.977147055 | 0.328496355 | 0.539244558 | 0.10684417 |
| ENSMUSG00000023988 | Bysl | 53414 | protein_coding | 253.0265242 | -0.10685061 | 0.226220879 | -0.472328686 | 0.636692198 | 0.789631187 | 0.10685061 |
| ENSMUSG00000040034 | Nup43 | 69912 | protein_coding | 113.433599 | 0.106867107 | 0.129405693 | 0.825830034 | 0.408900514 | 0.615890871 | 0.106867107 |
| ENSMUSG00000031807 | Pgls | 66171 | protein_coding | 731.8997584 | 0.10688444 | 0.130147416 | 0.821256722 | 0.411500053 | 0.6183954 | 0.10688444 |
| ENSMUSG00000031715 | Smarca5 | 93762 | protein_coding | 2353.279198 | 0.106887525 | 0.081030706 | 1.319099031 | 0.187136007 | 0.377782297 | 0.106887525 |
| ENSMUSG00000024436 | Mrps18b | 66973 | protein_coding | 222.4406401 | 0.10689937 | 0.168892787 | 0.63294219 | 0.526771395 | 0.712369266 | 0.10689937 |
| ENSMUSG00000023904 | Hcfc1r1 | 353502 | protein_coding | 481.5215081 | 0.106915553 | 0.091708471 | 1.165819814 | 0.243687304 | 0.448775996 | 0.106915553 |
| ENSMUSG00000032038 | St3gal4 | 20443 | protein_coding | 2295.060261 | 0.10696172 | 0.230732619 | 0.463574332 | 0.642952748 | 0.793244471 | 0.10696172 |
| ENSMUSG00000043572 | Pars2 | 230577 | protein_coding | 97.85545878 | -0.106968187 | 0.179784688 | -0.59497941 | 0.551857234 | 0.732326245 | 0.106968187 |
| ENSMUSG00000038250 | Usp38 | 74841 | protein_coding | 830.1491959 | -0.106975793 | 0.092798871 | -1.152770415 | 0.249004631 | 0.455198641 | 0.106975793 |
| ENSMUSG00000042978 | Sbk1 | 104175 | protein_coding | 199.2656943 | -0.1069778 | 0.150733738 | -0.709713708 | 0.47788169 | 0.672352226 | 0.1069778 |
| ENSMUSG00000039804 | Ncoa5 | 228869 | protein_coding | 409.0659526 | -0.107003687 | 0.096853085 | -1.104804119 | 0.269244475 | 0.477245271 | 0.107003687 |
| ENSMUSG00000029780 | Nt5c3 | 107569 | protein_coding | 1105.284376 | 0.107005606 | 0.09081418 | 1.178291827 | 0.238680285 | 0.442830469 | 0.107005606 |
| ENSMUSG00000042099 | Kank3 | 80880 | protein_coding | 31.72852136 | -0.107012972 | 0.204418696 | -0.523498946 | 0.600627084 | 0.766753883 | 0.107012972 |
| ENSMUSG00000029629 | Phf14 | 75725 | protein_coding | 611.7990294 | -0.107091762 | 0.071321618 | -1.501532989 | 0.133217761 | 0.301797813 | 0.107091762 |
| ENSMUSG00000026730 | Pter | 19212 | protein_coding | 163.0420833 | -0.107098484 | 0.166901079 | -0.641688387 | 0.521075531 | 0.707440424 | 0.107098484 |
| ENSMUSG00000038280 | Ostm1 | 14628 | protein_coding | 241.886668 | -0.107109365 | 0.132649919 | -0.807458952 | 0.419402114 | 0.625548916 | 0.107109365 |
| ENSMUSG00000024914 | Drap1 | 66556 | protein_coding | 826.1350381 | 0.107123428 | 0.088840957 | 1.205788768 | 0.227898947 | 0.430550876 | 0.107123428 |
| ENSMUSG00000044719 | E230025N22Rik | 240216 | protein_coding | 57.73594664 | 0.107209201 | 0.170570082 | 0.62853462 | 0.529653775 | 0.715003418 | 0.107209201 |
| ENSMUSG00000026721 | Rabgap1l | 29809 | protein_coding | 1378.215261 | -0.107219684 | 0.123357743 | -0.869176762 | 0.384750456 | 0.592679426 | 0.107219684 |
| ENSMUSG00000060862 | Zbtb40 | 230848 | protein_coding | 268.0879014 | 0.10723431 | 0.139924741 | 0.76637133 | 0.443455387 | 0.644466916 | 0.10723431 |
| ENSMUSG00000024983 | Vti1a | 53611 | protein_coding | 420.048711 | 0.107289114 | 0.082754471 | 1.296475145 | 0.194811838 | 0.387870369 | 0.107289114 |

| ensembl_gene_id | external_gene_name | entrezgene_id | gene_biotype | baseMean | log2FoldChange | lfcSE | stat | pvalue | padj | absLog2FC |
|---|---|---|---|---|---|---|---|---|---|---|
| ENSMUSG00000095026 | Gm3336 | 100502950 | protein_coding | 629.2944983 | -0.107329528 | 0.103815876 | -1.033845036 | 0.301208614 | 0.510581706 | 0.107329528 |
| ENSMUSG00000021461 | Fancc | 14088 | protein_coding | 159.2644346 | -0.107342724 | 0.113773053 | -0.943481094 | 0.345434881 | 0.555317601 | 0.107342724 |
| ENSMUSG00000024764 | Naa40 | 70999 | protein_coding | 460.1023665 | -0.107345303 | 0.083502948 | -1.285527106 | 0.198608151 | 0.392728818 | 0.107345303 |
| ENSMUSG00000031393 | Mecp2 | 17257 | protein_coding | 352.8251091 | -0.107353019 | 0.127603668 | -0.841300411 | 0.40017966 | 0.607690116 | 0.107353019 |
| ENSMUSG00000033159 | Cnppd1 | 69171 | protein_coding | 855.1308101 | -0.107428732 | 0.105902083 | -1.014415663 | 0.310384453 | 0.520455161 | 0.107428732 |
| ENSMUSG00000026972 | Arrdc1 | 215705 | protein_coding | 690.5055962 | -0.107447399 | 0.112348151 | -0.956378886 | 0.338880848 | 0.549715714 | 0.107447399 |
| ENSMUSG00000025142 | Aspscr1 | 68938 | protein_coding | 471.5891998 | 0.107568452 | 0.079415815 | 1.354496599 | 0.175577998 | 0.361771253 | 0.107568452 |
| ENSMUSG00000021591 | Glrx | 93692 | protein_coding | 878.6252834 | -0.107602996 | 0.101545874 | -1.059649116 | 0.28930426 | 0.498112447 | 0.107602996 |
| ENSMUSG00000027010 | Slc25a12 | 78830 | protein_coding | 336.2143028 | 0.107604325 | 0.160311639 | 0.671219669 | 0.5020806 | 0.692106338 | 0.107604325 |
| ENSMUSG00000037089 | Slc35b2 | 73836 | protein_coding | 721.1386506 | 0.107623266 | 0.106572195 | 1.009862526 | 0.312561159 | 0.522784221 | 0.107623266 |
| ENSMUSG00000033952 | Aspm | 12316 | protein_coding | 544.5009854 | 0.107626088 | 0.107525519 | 1.000935309 | 0.316858085 | 0.528193536 | 0.107626088 |
| ENSMUSG00000022010 | Tsc22d1 | 21807 | protein_coding | 5942.383947 | -0.10764107 | 0.148602619 | -0.724355135 | 0.468847736 | 0.665105695 | 0.10764107 |
| ENSMUSG00000022544 | Eef2kmt | 70511 | protein_coding | 181.7777023 | -0.107658262 | 0.127884877 | -0.84183732 | 0.399879019 | 0.607349386 | 0.107658262 |
| ENSMUSG00000030259 | Rassf8 | 71323 | protein_coding | 12.00976221 | 0.107673444 | 0.37789808 | 0.284927207 | 0.775699912 | 0.877982787 | 0.107673444 |
| ENSMUSG00000000568 | Hnrnpd | 11991 | protein_coding | 679.7842367 | -0.107673788 | 0.12787613 | -0.84201632 | 0.399778818 | 0.607255104 | 0.107673788 |
| ENSMUSG00000003778 | Brd8 | 78656 | protein_coding | 903.402729 | -0.10767646 | 0.089661288 | -1.200924759 | 0.229780389 | 0.433077983 | 0.10767646 |
| ENSMUSG00000020166 | Cnot2 | 72068 | protein_coding | 911.6433428 | 0.107713555 | 0.076213419 | 1.413314814 | 0.15756318 | 0.33836677 | 0.107713555 |
| ENSMUSG00000002017 | Fam98a | 72722 | protein_coding | 545.2101944 | 0.107749542 | 0.090887582 | 1.185525449 | 0.235809759 | 0.440009526 | 0.107749542 |
| ENSMUSG00000030741 | Spns1 | 73658 | protein_coding | 406.5299823 | 0.107762789 | 0.110417603 | 0.975956604 | 0.32908597 | 0.539888897 | 0.107762789 |
| ENSMUSG00000028420 | Tmem38b | 52076 | protein_coding | 163.9335887 | -0.107801079 | 0.130893017 | -0.823581589 | 0.410177341 | 0.617106327 | 0.107801079 |
| ENSMUSG00000017390 | Aldoc | 11676 | protein_coding | 171.7087436 | -0.107809957 | 0.156437081 | -0.689158588 | 0.490723474 | 0.68307464 | 0.107809957 |
| ENSMUSG00000115518 | Gm10791 | 100038614 | lncRNA | 14.76113271 | -0.107822485 | 0.404119304 | -0.266808548 | 0.789616572 | 0.885892285 | 0.107822485 |
| ENSMUSG00000030780 | Rusf1 | 233913 | protein_coding | 338.114454 | -0.107828678 | 0.087853897 | -1.227363629 | 0.21968595 | 0.419876136 | 0.107828678 |
| ENSMUSG00000024078 | Ttc27 | 74196 | protein_coding | 190.9734848 | -0.107835308 | 0.148246836 | -0.727403775 | 0.466978642 | 0.663707223 | 0.107835308 |
| ENSMUSG00000022221 | Ripk3 | 56532 | protein_coding | 241.7301541 | -0.107835733 | 0.228577938 | -0.471767897 | 0.637092468 | 0.789943081 | 0.107835733 |
| ENSMUSG00000019810 | Fuca2 | 66848 | protein_coding | 1529.191765 | 0.107837551 | 0.076700503 | 1.40595624 | 0.15973712 | 0.341103746 | 0.107837551 |
| ENSMUSG00000035007 | Rundc1 | 217201 | protein_coding | 254.6227513 | -0.107868012 | 0.121275577 | -0.889445462 | 0.373763721 | 0.582173729 | 0.107868012 |
| ENSMUSG00000025857 | Dnaaf5 | 433956 | protein_coding | 118.9919085 | -0.107876375 | 0.14282104 | -0.755325515 | 0.450053693 | 0.650554063 | 0.107876375 |
| ENSMUSG00000031511 | Arhgef7 | 54126 | protein_coding | 500.2147115 | -0.107939273 | 0.129687259 | -0.832304371 | 0.405237164 | 0.612206925 | 0.107939273 |
| ENSMUSG00000041362 | Shtn1 | 71653 | protein_coding | 812.6905097 | 0.107959535 | 0.075672032 | 1.426676828 | 0.153673068 | 0.332794647 | 0.107959535 |
| ENSMUSG00000039000 | Ube3c | 100763 | protein_coding | 1475.014262 | 0.108009655 | 0.079718706 | 1.35488469 | 0.175454298 | 0.361733769 | 0.108009655 |
| ENSMUSG00000040913 | Fbxw4 | 30838 | protein_coding | 91.89659293 | 0.108042469 | 0.140433223 | 0.769351203 | 0.441684848 | 0.643011318 | 0.108042469 |
| ENSMUSG00000034485 | Uaca | 72565 | protein_coding | 1146.165845 | 0.108083859 | 0.085699921 | 1.261189715 | 0.207240503 | 0.403733701 | 0.108083859 |
| ENSMUSG00000028069 | Gpatch4 | 66614 | protein_coding | 127.0489464 | -0.108092549 | 0.281104436 | -0.38452808 | 0.700587088 | 0.829926899 | 0.108092549 |
| ENSMUSG00000035215 | Lsm7 | 66094 | protein_coding | 120.7153655 | 0.108134699 | 0.138957651 | 0.778184567 | 0.436460214 | 0.639258693 | 0.108134699 |
| ENSMUSG00000036834 | Plch1 | 269437 | protein_coding | 782.6684904 | 0.10815457 | 0.133156908 | 0.81223402 | 0.416657362 | 0.622854854 | 0.10815457 |
| ENSMUSG00000090069 | E430024P14Rik | 100040822 | lncRNA | 13.12409865 | -0.108201741 | 0.47848626 | -0.226133434 | 0.821097641 | 0.905179148 | 0.108201741 |
| ENSMUSG00000020140 | Lgr5 | 14160 | protein_coding | 184.740842 | 0.108244101 | 0.205634835 | 0.526389906 | 0.598617328 | 0.765353732 | 0.108244101 |

| ensembl_gene_id | external_gene_name | entrezgene_id | gene_biotype | baseMean | log2FoldChange | lfcSE | stat | pvalue | padj | absLog2FC |
|---|---|---|---|---|---|---|---|---|---|---|
| ENSMUSG00000078995 | Zfp456 | 408065 | protein_coding | 80.28449492 | 0.108263768 | 0.163501619 | 0.662157162 | 0.507870505 | 0.697298629 | 0.108263768 |
| ENSMUSG00000057541 | Pus7 | 78697 | protein_coding | 334.2344216 | -0.108299695 | 0.124855765 | -0.867398429 | 0.385723743 | 0.593661974 | 0.108299695 |
| ENSMUSG00000041846 | Ppp4r3a | 68734 | protein_coding | 1781.478033 | -0.108304097 | 0.078666708 | -1.37674628 | 0.168590705 | 0.353377122 | 0.108304097 |
| ENSMUSG00000004798 | Ulk2 | 29869 | protein_coding | 301.8351156 | 0.108332469 | 0.092785313 | 1.167560522 | 0.242984082 | 0.447893815 | 0.108332469 |
| ENSMUSG00000024613 | Tcof1 | 21453 | protein_coding | 424.7155635 | -0.108368609 | 0.112776285 | -0.960916642 | 0.336594083 | 0.547291808 | 0.108368609 |
| ENSMUSG00000001127 | Araf | 11836 | protein_coding | 1046.466206 | 0.108398385 | 0.089545378 | 1.210541373 | 0.226071225 | 0.428372488 | 0.108398385 |
| ENSMUSG00000032436 | Cmtm7 | 102545 | protein_coding | 69.0659559 | 0.108515117 | 0.158800925 | 0.683340586 | 0.494391654 | 0.686031895 | 0.108515117 |
| ENSMUSG00000026568 | Mpc2 | 70456 | protein_coding | 1088.49606 | 0.108515585 | 0.098994851 | 1.096174035 | 0.27300262 | 0.481016121 | 0.108515585 |
| ENSMUSG00000037098 | Rab11fip3 | 215445 | protein_coding | 214.0102942 | -0.108537841 | 0.114733139 | -0.94600254 | 0.344147291 | 0.55406481 | 0.108537841 |
| ENSMUSG00000029196 | Tada2b | 231151 | protein_coding | 255.991413 | -0.108620865 | 0.102855941 | -1.056048527 | 0.290946045 | 0.499573396 | 0.108620865 |
| ENSMUSG00000029998 | Pcyox1 | 66881 | protein_coding | 1335.090024 | 0.108639472 | 0.093756133 | 1.158745231 | 0.246560048 | 0.452130836 | 0.108639472 |
| ENSMUSG00000005054 | Cstb | 13014 | protein_coding | 578.3191187 | 0.108687634 | 0.094438796 | 1.150879068 | 0.24978199 | 0.456096244 | 0.108687634 |
| ENSMUSG00000034714 | Ttyh2 | 117160 | protein_coding | 315.4286844 | -0.108696823 | 0.166471756 | -0.652944532 | 0.513792038 | 0.701438209 | 0.108696823 |
| ENSMUSG00000018363 | Smurf2 | 66313 | protein_coding | 1040.317793 | -0.108700863 | 0.062603601 | -1.736335611 | 0.08250451 | 0.21870309 | 0.108700863 |
| ENSMUSG00000020801 | Med31 | 67279 | protein_coding | 80.17665332 | 0.10871081 | 0.13013773 | 0.835351977 | 0.40351956 | 0.610808226 | 0.10871081 |
| ENSMUSG00000026280 | Atg4b | 66615 | protein_coding | 677.5024423 | 0.108732856 | 0.079765571 | 1.363155252 | 0.172833587 | 0.358543028 | 0.108732856 |
| ENSMUSG00000005886 | Ncoa2 | 17978 | protein_coding | 853.6000411 | -0.10876507 | 0.110009038 | -0.988692128 | 0.322813796 | 0.533749157 | 0.10876507 |
| ENSMUSG00000059291 | Rpl11 | 67025 | protein_coding | 4817.827825 | 0.108778287 | 0.071612637 | 1.51898174 | 0.128767092 | 0.294980907 | 0.108778287 |
| ENSMUSG00000042613 | Pbxip1 | 229534 | protein_coding | 969.7594988 | -0.108789073 | 0.108780736 | -1.000076638 | 0.317273421 | 0.528556746 | 0.108789073 |
| ENSMUSG00000063047 | Zfp780b | 338354 | protein_coding | 231.5635237 | 0.108817471 | 0.111274041 | 0.977923242 | 0.328112289 | 0.538929056 | 0.108817471 |
| ENSMUSG00000048707 | Tprn | 97031 | protein_coding | 1269.830959 | 0.108853766 | 0.111545482 | 0.975868886 | 0.329129442 | 0.539888897 | 0.108853766 |
| ENSMUSG00000022361 | Zhx1 | 22770 | protein_coding | 567.5319635 | -0.108923183 | 0.0974228 | -1.118046112 | 0.2635473 | 0.471131469 | 0.108923183 |
| ENSMUSG00000004996 | Mri1 | 67873 | protein_coding | 199.6601716 | -0.108979068 | 0.11573752 | -0.941605349 | 0.34639473 | 0.556466586 | 0.108979068 |
| ENSMUSG00000056592 | Zfp658 | 210104 | protein_coding | 48.40994658 | -0.108991149 | 0.253749934 | -0.429521881 | 0.667543474 | 0.80727602 | 0.108991149 |
| ENSMUSG00000091183 | Gm5141 | 380850 | protein_coding | 39.37114613 | -0.108993666 | 0.232135522 | -0.469526015 | 0.638693694 | 0.790625182 | 0.108993666 |
| ENSMUSG00000020544 | Cox11 | 69802 | protein_coding | 78.70199264 | -0.108994125 | 0.180782331 | -0.602902531 | 0.546573532 | 0.72870371 | 0.108994125 |
| ENSMUSG00000031257 | Nox1 | 237038 | protein_coding | 562.2807776 | 0.109013891 | 0.169761575 | 0.642158811 | 0.520770074 | 0.70726686 | 0.109013891 |
| ENSMUSG00000032563 | Mrpl3 | 94062 | protein_coding | 533.2090457 | 0.109015038 | 0.097749168 | 1.115252844 | 0.264742087 | 0.472220655 | 0.109015038 |
| ENSMUSG00000040675 | Mthfd1l | 270685 | protein_coding | 241.1233745 | -0.109024507 | 0.130893021 | -0.832928342 | 0.404885144 | 0.611918642 | 0.109024507 |
| ENSMUSG00000056313 | Tcim | 69068 | protein_coding | 1569.395819 | -0.109029364 | 0.137450716 | -0.793225143 | 0.427646661 | 0.631987012 | 0.109029364 |
| ENSMUSG00000024309 | Pfdn6 | 14976 | protein_coding | 408.6529473 | 0.109056393 | 0.117124024 | 0.931118905 | 0.35179206 | 0.561401056 | 0.109056393 |
| ENSMUSG00000022881 | Rfc4 | 106344 | protein_coding | 169.7672965 | 0.109064418 | 0.111103625 | 0.981645896 | 0.326274328 | 0.537352722 | 0.109064418 |
| ENSMUSG00000047656 | Trpt1 | 107328 | protein_coding | 96.68692741 | -0.109124767 | 0.168231761 | -0.648657346 | 0.51655988 | 0.703829408 | 0.109124767 |
| ENSMUSG00000052459 | Atp6v1a | 11964 | protein_coding | 1311.815507 | 0.109127618 | 0.066494252 | 1.641158657 | 0.100764485 | 0.249491174 | 0.109127618 |
| ENSMUSG00000024769 | Cdc42bpg | 240505 | protein_coding | 1460.793889 | -0.109169864 | 0.115373589 | -0.946229249 | 0.344031671 | 0.554008329 | 0.109169864 |
| ENSMUSG00000033417 | Cacul1 | 78832 | protein_coding | 1030.786618 | 0.109188832 | 0.071362477 | 1.530059449 | 0.126002014 | 0.290484887 | 0.109188832 |
| ENSMUSG00000041225 | Arhgap12 | 75415 | protein_coding | 1668.782168 | -0.109202976 | 0.085942083 | -1.27065778 | 0.203850422 | 0.399081799 | 0.109202976 |
| ENSMUSG00000029782 | Tmem209 | 72649 | protein_coding | 346.6307029 | 0.109217078 | 0.098565887 | 1.108061636 | 0.267835198 | 0.475806271 | 0.109217078 |

| ensembl_gene_id | external_gene_name | entrezgene_id | gene_biotype | baseMean | log2FoldChange | lfcSE | stat | pvalue | padj | absLog2FC |
|---|---|---|---|---|---|---|---|---|---|---|
| ENSMUSG00000039765 | Cc2d2a | 231214 | protein_coding | 393.0421183 | -0.109232364 | 0.124415737 | -0.877962607 | 0.379964012 | 0.58786467 | 0.109232364 |
| ENSMUSG00000049562 | Ap5b1 | 381201 | protein_coding | 156.7258116 | -0.109252727 | 0.122555076 | -0.891458196 | 0.372683409 | 0.580888671 | 0.109252727 |
| ENSMUSG00000028333 | Anp32b | 67628 | protein_coding | 499.1117861 | -0.109298938 | 0.147789376 | -0.739558831 | 0.45956773 | 0.657917922 | 0.109298938 |
| ENSMUSG00000030400 | Ercc2 | 13871 | protein_coding | 236.9784753 | -0.109322078 | 0.098807998 | -1.106409207 | 0.268549442 | 0.476490533 | 0.109322078 |
| ENSMUSG00000040021 | Lats1 | 16798 | protein_coding | 1379.552113 | 0.109345049 | 0.086322542 | 1.266703301 | 0.205261399 | 0.401062215 | 0.109345049 |
| ENSMUSG00000059586 | Nsmce2 | 68501 | protein_coding | 221.7066574 | 0.109419335 | 0.092339812 | 1.184963805 | 0.236031758 | 0.440122237 | 0.109419335 |
| ENSMUSG00000020570 | Sypl | 19027 | protein_coding | 3252.463897 | 0.10942751 | 0.098061294 | 1.115909298 | 0.264460961 | 0.472048706 | 0.10942751 |
| ENSMUSG00000031166 | Wdr13 | 73447 | protein_coding | 1001.147339 | -0.109429234 | 0.064813503 | -1.68837091 | 0.091340051 | 0.234126864 | 0.109429234 |
| ENSMUSG00000042073 | Abhd14b | 76491 | protein_coding | 389.693836 | 0.109443679 | 0.120986209 | 0.904596321 | 0.365679285 | 0.574448111 | 0.109443679 |
| ENSMUSG00000025287 | Acot9 | 56360 | protein_coding | 305.4083387 | -0.109477395 | 0.193269182 | -0.566450348 | 0.571087677 | 0.745230585 | 0.109477395 |
| ENSMUSG00000063273 | Naa15 | 74838 | protein_coding | 944.1310165 | -0.109486809 | 0.125705913 | -0.870975811 | 0.383767361 | 0.591451042 | 0.109486809 |
| ENSMUSG00000061028 | Clasrp | 53609 | protein_coding | 344.5124849 | 0.10954041 | 0.133281601 | 0.821871955 | 0.411149776 | 0.618130069 | 0.10954041 |
| ENSMUSG00000028944 | Prkag2 | 108099 | protein_coding | 516.649801 | -0.109603251 | 0.119117717 | -0.920125518 | 0.357507171 | 0.567168746 | 0.109603251 |
| ENSMUSG00000074519 | Zfp971 | 626848 | protein_coding | 186.3029077 | -0.109607186 | 0.129146593 | -0.848703658 | 0.39604621 | 0.604293902 | 0.109607186 |
| ENSMUSG00000020656 | Grhl1 | 195733 | protein_coding | 68.06948148 | -0.109768586 | 0.433377175 | -0.253286495 | 0.800046831 | 0.892772335 | 0.109768586 |
| ENSMUSG00000047126 | Cltc | 67300 | protein_coding | 8431.326598 | 0.109820881 | 0.065426639 | 1.678534638 | 0.093242774 | 0.23759853 | 0.109820881 |
| ENSMUSG00000038126 | Mphosph9 | 269702 | protein_coding | 194.2750698 | 0.109838031 | 0.110990005 | 0.989620917 | 0.322359442 | 0.533292605 | 0.109838031 |
| ENSMUSG00000003423 | Pih1d1 | 68845 | protein_coding | 369.7727886 | 0.109848456 | 0.109251448 | 1.005464528 | 0.314673222 | 0.525434016 | 0.109848456 |
| ENSMUSG00000022761 | Lztr1 | 66863 | protein_coding | 758.5291982 | 0.109891624 | 0.092497288 | 1.188052388 | 0.234812771 | 0.438721151 | 0.109891624 |
| ENSMUSG00000040455 | Usp45 | 77593 | protein_coding | 543.4296465 | -0.109900913 | 0.091682826 | -1.198707736 | 0.230641609 | 0.433777252 | 0.109900913 |
| ENSMUSG00000038150 | Ormdl3 | 66612 | protein_coding | 304.0523892 | 0.109963877 | 0.098102738 | 1.120905281 | 0.262328183 | 0.470066494 | 0.109963877 |
| ENSMUSG00000025646 | Atrip | 235610 | protein_coding | 152.4918149 | -0.10997651 | 0.107624569 | -1.021853192 | 0.30685039 | 0.516878519 | 0.10997651 |
| ENSMUSG00000024312 | Wdr46 | 57315 | protein_coding | 246.2860461 | -0.109997949 | 0.141491177 | -0.777419143 | 0.436911522 | 0.639621225 | 0.109997949 |
| ENSMUSG00000017679 | Ttpal | 76080 | protein_coding | 407.3935313 | 0.109998742 | 0.119278743 | 0.922199034 | 0.356424767 | 0.565983821 | 0.109998742 |
| ENSMUSG00000018604 | Tbx3 | 21386 | protein_coding | 199.3742931 | -0.110009414 | 0.237226082 | -0.463732374 | 0.6428395 | 0.793238306 | 0.110009414 |
| ENSMUSG00000034621 | Gpatch8 | 237943 | protein_coding | 615.1850367 | -0.110029719 | 0.110976821 | -0.991465767 | 0.3214582 | 0.532354566 | 0.110029719 |
| ENSMUSG00000113679 | Gm10432 | 100038713 | lncRNA | 7.768430457 | -0.110032694 | 0.506432142 | -0.217270361 | 0.827997654 | 0.90891431 | 0.110032694 |
| ENSMUSG00000044864 | Ankrd50 | 99696 | protein_coding | 647.7083357 | 0.110046326 | 0.126051278 | 0.873028241 | 0.382647685 | 0.590296593 | 0.110046326 |
| ENSMUSG00000033998 | Kcnk1 | 16525 | protein_coding | 1497.586936 | 0.110056409 | 0.142623359 | 0.771657668 | 0.440317209 | 0.642313808 | 0.110056409 |
| ENSMUSG00000098973 | Mir6236 | 102465959 | miRNA | 9774.261101 | 0.11009296 | 0.416387786 | 0.264400071 | 0.791471657 | 0.887042894 | 0.11009296 |
| ENSMUSG00000079215 | Zfp664 | 269704 | protein_coding | 926.550114 | -0.11012844 | 0.089904671 | -1.224946813 | 0.220595261 | 0.421299918 | 0.11012844 |
| ENSMUSG00000064215 | Ifi27 | 52668 | protein_coding | 309.0310448 | -0.110230672 | 0.137485686 | -0.801761082 | 0.422691175 | 0.627750586 | 0.110230672 |
| ENSMUSG00000002546 | Golga2 | 99412 | protein_coding | 1275.963758 | 0.110292494 | 0.054672434 | 2.017332809 | 0.043660794 | 0.141347383 | 0.110292494 |
| ENSMUSG00000075232 | Amd1 | 11702 | protein_coding | 901.9079461 | -0.110322435 | 0.194285235 | -0.567837465 | 0.570145338 | 0.744744605 | 0.110322435 |
| ENSMUSG00000002055 | Spag5 | 54141 | protein_coding | 399.5533076 | 0.110421537 | 0.116462313 | 0.948131064 | 0.343062736 | 0.553235117 | 0.110421537 |
| ENSMUSG00000014195 | Dnajc7 | 56354 | protein_coding | 1253.950011 | -0.110436693 | 0.05381766 | -2.052053034 | 0.040164509 | 0.133446035 | 0.110436693 |
| ENSMUSG00000020283 | Pex13 | 72129 | protein_coding | 602.6858453 | -0.110447395 | 0.118591701 | -0.931324819 | 0.351685567 | 0.561399852 | 0.110447395 |
| ENSMUSG00000031373 | Car5b | 56078 | protein_coding | 9.543251545 | -0.110458797 | 0.398120383 | -0.277450744 | 0.781434022 | 0.881180963 | 0.110458797 |

| ensembl_gene_id | external_gene_name | entrezgene_id | gene_biotype | baseMean | log2FoldChange | lfcSE | stat | pvalue | padj | absLog2FC |
|---|---|---|---|---|---|---|---|---|---|---|
| ENSMUSG00000075000 | Nrbf2 | 641340 | protein_coding | 148.2494971 | -0.110492548 | 0.104057754 | -1.061838686 | 0.288308924 | 0.497392119 | 0.110492548 |
| ENSMUSG00000006019 | Dhx34 | 71723 | protein_coding | 148.6771684 | -0.1105298 | 0.161645999 | -0.683776901 | 0.494116055 | 0.685803461 | 0.1105298 |
| ENSMUSG00000043991 | Pura | 19290 | protein_coding | 1024.449764 | -0.110568183 | 0.110220595 | -1.003153571 | 0.315786771 | 0.526796364 | 0.110568183 |
| ENSMUSG00000039747 | Orai2 | 269717 | protein_coding | 380.790093 | -0.110568434 | 0.099819943 | -1.107678796 | 0.26800056 | 0.475940787 | 0.110568434 |
| ENSMUSG00000001034 | Mapk7 | 23939 | protein_coding | 156.7255494 | -0.11058384 | 0.138104314 | -0.800726903 | 0.423289764 | 0.628228988 | 0.11058384 |
| ENSMUSG00000027635 | Dsn1 | 66934 | protein_coding | 133.7182215 | 0.110595888 | 0.14903242 | 0.742092814 | 0.458031104 | 0.656603314 | 0.110595888 |
| ENSMUSG00000026228 | Htr2b | 15559 | protein_coding | 9.685074496 | 0.11063396 | 0.470164961 | 0.235308815 | 0.813969054 | 0.900950463 | 0.11063396 |
| ENSMUSG00000020736 | Nt5c | 50773 | protein_coding | 1155.913332 | 0.110692482 | 0.126123138 | 0.877654041 | 0.380131493 | 0.588066668 | 0.110692482 |
| ENSMUSG00000035847 | Ids | 15931 | protein_coding | 420.6148769 | 0.110709033 | 0.125596393 | 0.881466657 | 0.378065293 | 0.586205601 | 0.110709033 |
| ENSMUSG00000053646 | Plxnb1 | 235611 | protein_coding | 196.6810131 | 0.110720428 | 0.158360016 | 0.699169092 | 0.484446364 | 0.677861368 | 0.110720428 |
| ENSMUSG00000038697 | Taf5l | 102162 | protein_coding | 484.6313524 | -0.110728762 | 0.103113064 | -1.073857747 | 0.282886444 | 0.492008656 | 0.110728762 |
| ENSMUSG00000046032 | Snx12 | 55988 | protein_coding | 358.9753715 | 0.110750882 | 0.103727302 | 1.06771197 | 0.285650457 | 0.494484454 | 0.110750882 |
| ENSMUSG00000022261 | Sdc2 | 15529 | protein_coding | 645.5564691 | 0.110773008 | 0.097170021 | 1.139991605 | 0.254289799 | 0.460735743 | 0.110773008 |
| ENSMUSG00000026260 | Ndufa10 | 67273 | protein_coding | 2308.454122 | 0.110784254 | 0.099249719 | 1.116217311 | 0.264329126 | 0.472048706 | 0.110784254 |
| ENSMUSG00000030365 | Clec2i | 93675 | protein_coding | 7.978345231 | 0.110792089 | 0.611638508 | 0.18113982 | 0.85625783 | 0.924704659 | 0.110792089 |
| ENSMUSG00000057497 | Fam136a | 66488 | protein_coding | 622.358249 | 0.110808004 | 0.090339738 | 1.226569901 | 0.219984288 | 0.420334661 | 0.110808004 |
| ENSMUSG00000004207 | Psap | 19156 | protein_coding | 8174.497932 | -0.110811133 | 0.101802936 | -1.08848661 | 0.276380344 | 0.48476362 | 0.110811133 |
| ENSMUSG00000073158 | 9030624G23Rik | 66808 | protein_coding | 7.699589022 | 0.110870651 | 0.460368707 | 0.240830119 | 0.809686785 | 0.898579225 | 0.110870651 |
| ENSMUSG00000022710 | Usp7 | 252870 | protein_coding | 3207.296688 | 0.110876739 | 0.074785161 | 1.482603474 | 0.138179794 | 0.309205922 | 0.110876739 |
| ENSMUSG00000024294 | Mib1 | 225164 | protein_coding | 1998.502576 | 0.110950756 | 0.0518464 | 2.13998958 | 0.032355609 | 0.114052042 | 0.110950756 |
| ENSMUSG00000024782 | Ak3 | 56248 | protein_coding | 3989.412006 | 0.110960053 | 0.092602909 | 1.198235074 | 0.230825515 | 0.434020635 | 0.110960053 |
| ENSMUSG00000053964 | Lgals4 | 16855 | protein_coding | 51448.11438 | 0.111011601 | 0.089054174 | 1.246562582 | 0.212557929 | 0.410577576 | 0.111011601 |
| ENSMUSG00000097099 | Gm9917 | 100038755 | lncRNA | 94.14026402 | -0.111016106 | 0.119968549 | -0.925376757 | 0.354769965 | 0.564287598 | 0.111016106 |
| ENSMUSG00000046410 | Kcnk6 | 52150 | protein_coding | 836.5934176 | 0.111018463 | 0.13968227 | 0.794792806 | 0.426734037 | 0.631223973 | 0.111018463 |
| ENSMUSG00000041718 | Alg13 | 67574 | protein_coding | 97.19011909 | 0.111060352 | 0.132425222 | 0.838664647 | 0.401657522 | 0.6091793 | 0.111060352 |
| ENSMUSG00000042638 | Gucy2c | 14917 | protein_coding | 5947.893451 | 0.11107777 | 0.094382456 | 1.176890016 | 0.239239408 | 0.443424073 | 0.11107777 |
| ENSMUSG00000029447 | Cct6a | 12466 | protein_coding | 2359.018008 | 0.1111215 | 0.114470466 | 0.970743838 | 0.331675856 | 0.542506986 | 0.1111215 |
| ENSMUSG00000053846 | Lipg | 16891 | protein_coding | 650.6678799 | -0.111204209 | 0.205264214 | -0.541761309 | 0.587982948 | 0.757901789 | 0.111204209 |
| ENSMUSG00000021962 | Dcp1a | 75901 | protein_coding | 413.1948365 | -0.111236062 | 0.092359564 | -1.204380546 | 0.228442528 | 0.431373157 | 0.111236062 |
| ENSMUSG00000032504 | Pdcd6ip | 18571 | protein_coding | 2795.525255 | 0.111249976 | 0.090642525 | 1.227348605 | 0.219691595 | 0.419876136 | 0.111249976 |
| ENSMUSG00000028478 | Clta | 12757 | protein_coding | 1910.167246 | 0.111337098 | 0.089396497 | 1.245430218 | 0.212973652 | 0.411031664 | 0.111337098 |
| ENSMUSG00000027011 | Ube2e3 | 22193 | protein_coding | 537.7146663 | 0.1113518 | 0.072122747 | 1.543920671 | 0.122607552 | 0.285427227 | 0.1113518 |
| ENSMUSG00000048027 | Rgmb | 68799 | protein_coding | 1082.242678 | -0.111375108 | 0.104511135 | -1.065676956 | 0.286569698 | 0.495534183 | 0.111375108 |
| ENSMUSG00000030472 | Ceacam18 | 72431 | protein_coding | 86.15650372 | -0.111384443 | 0.144692923 | -0.769798833 | 0.441419232 | 0.642981907 | 0.111384443 |
| ENSMUSG00000002835 | Chaf1a | 27221 | protein_coding | 154.8579065 | 0.111419982 | 0.167858238 | 0.663774286 | 0.506834786 | 0.696603457 | 0.111419982 |
| ENSMUSG00000025019 | Lcor | 212391 | protein_coding | 1136.982603 | -0.111423678 | 0.098823252 | -1.127504669 | 0.259529168 | 0.46672469 | 0.111423678 |
| ENSMUSG00000000374 | Trappc10 | 216131 | protein_coding | 2030.74383 | 0.111469166 | 0.068782249 | 1.62060949 | 0.105101416 | 0.256707373 | 0.111469166 |
| ENSMUSG00000059423 | Zfp933 | 242747 | protein_coding | 202.3099418 | 0.111481151 | 0.101119077 | 1.102473979 | 0.270255665 | 0.478375791 | 0.111481151 |

| ensembl_gene_id | external_gene_name | entrezgene_id | gene_biotype | baseMean | log2FoldChange | lfcSE | stat | pvalue | padj | absLog2FC |
|---|---|---|---|---|---|---|---|---|---|---|
| ENSMUSG00000048277 | Syngr2 | 20973 | protein_coding | 2314.893331 | 0.111495751 | 0.071324005 | 1.563228979 | 0.117998746 | 0.277742579 | 0.111495751 |
| ENSMUSG00000040532 | Abhd11 | 68758 | protein_coding | 568.5473357 | 0.111582619 | 0.089197569 | 1.250960321 | 0.210948954 | 0.408311453 | 0.111582619 |
| ENSMUSG00000041238 | Rbbp8 | 225182 | protein_coding | 613.2142595 | 0.111635941 | 0.082353701 | 1.355566776 | 0.175237049 | 0.361473948 | 0.111635941 |
| ENSMUSG00000057530 | Ece1 | 230857 | protein_coding | 2089.523565 | -0.111660078 | 0.068000843 | -1.642039608 | 0.100581796 | 0.249193785 | 0.111660078 |
| ENSMUSG00000001173 | Ocrl | 320634 | protein_coding | 379.7778941 | -0.111688002 | 0.080312761 | -1.390663211 | 0.164327579 | 0.347643734 | 0.111688002 |
| ENSMUSG00000027937 | Jtb | 23922 | protein_coding | 724.1737921 | 0.111731218 | 0.090036583 | 1.240953558 | 0.214622906 | 0.41316336 | 0.111731218 |
| ENSMUSG00000015536 | Mocs2 | 17434 | protein_coding | 539.6307732 | 0.111779225 | 0.091175248 | 1.22598213 | 0.2202054 | 0.420656225 | 0.111779225 |
| ENSMUSG00000046836 | Brox | 71678 | protein_coding | 1004.223616 | 0.11177978 | 0.078621165 | 1.421751768 | 0.155098326 | 0.335015749 | 0.11177978 |
| ENSMUSG00000041747 | Utp15 | 105372 | protein_coding | 312.0319104 | -0.111792554 | 0.164726608 | -0.678655113 | 0.497356414 | 0.68832157 | 0.111792554 |
| ENSMUSG00000022217 | Emc9 | 85308 | protein_coding | 39.86543704 | -0.111808214 | 0.190288814 | -0.587571131 | 0.556820193 | 0.735858521 | 0.111808214 |
| ENSMUSG00000039159 | Ube2h | 22214 | protein_coding | 936.0084459 | 0.111850847 | 0.063707773 | 1.755686025 | 0.079142034 | 0.212360903 | 0.111850847 |
| ENSMUSG00000039660 | Spout1 | 227695 | protein_coding | 216.1754368 | -0.111898489 | 0.102935646 | -1.087072292 | 0.277004862 | 0.485411397 | 0.111898489 |
| ENSMUSG00000026135 | Zfp142 | 77264 | protein_coding | 146.1107915 | 0.111969726 | 0.170720788 | 0.655864624 | 0.511911231 | 0.699812166 | 0.111969726 |
| ENSMUSG00000026398 | Nr5a2 | 26424 | protein_coding | 1443.050755 | -0.112062147 | 0.199524184 | -0.56164694 | 0.574356592 | 0.747540635 | 0.112062147 |
| ENSMUSG00000030739 | Myh14 | 71960 | protein_coding | 7798.97205 | 0.112087086 | 0.127120852 | 0.881736426 | 0.377919358 | 0.586067523 | 0.112087086 |
| ENSMUSG00000021866 | Anxa11 | 11744 | protein_coding | 1027.069044 | -0.112158743 | 0.072348612 | -1.55025424 | 0.121080506 | 0.282823038 | 0.112158743 |
| ENSMUSG00000039041 | Adrm1 | 56436 | protein_coding | 512.1642543 | 0.112177597 | 0.152658768 | 0.734825772 | 0.462445609 | 0.660257589 | 0.112177597 |
| ENSMUSG00000064264 | Zfp428 | 232969 | protein_coding | 19.00798721 | -0.112180867 | 0.250220008 | -0.448328924 | 0.653915829 | 0.799659987 | 0.112180867 |
| ENSMUSG00000024654 | Asrgl1 | 66514 | protein_coding | 106.7747757 | 0.112187943 | 0.197482409 | 0.568090816 | 0.569973303 | 0.744741762 | 0.112187943 |
| ENSMUSG00000047368 | Abhd17b | 226016 | protein_coding | 1065.918108 | 0.11234214 | 0.122697767 | 0.915600524 | 0.359876456 | 0.569481508 | 0.11234214 |
| ENSMUSG00000023460 | Rab12 | 19328 | protein_coding | 109.1295358 | -0.112342653 | 0.152026498 | -0.738967578 | 0.459926685 | 0.658313465 | 0.112342653 |
| ENSMUSG00000006527 | Sfmbt1 | 54650 | protein_coding | 162.9820158 | -0.112356812 | 0.117745225 | -0.954236675 | 0.339963854 | 0.550365001 | 0.112356812 |
| ENSMUSG00000007036 | Abhd16a | 193742 | protein_coding | 737.1645202 | 0.112384856 | 0.080192184 | 1.401443997 | 0.161081343 | 0.343100245 | 0.112384856 |
| ENSMUSG00000045962 | Wnk1 | 232341 | protein_coding | 1273.733779 | -0.112420488 | 0.146837756 | -0.765610226 | 0.443908258 | 0.644830892 | 0.112420488 |
| ENSMUSG00000000792 | Slc5a5 | 114479 | protein_coding | 6.680762145 | 0.112431598 | 0.533514848 | 0.210737524 | 0.833092092 | 0.911190103 | 0.112431598 |
| ENSMUSG00000065176 | Rnu12 | 104307 | snRNA | 21.32391789 | -0.112442628 | 0.313125732 | -0.359097372 | 0.719522248 | 0.842364374 | 0.112442628 |
| ENSMUSG00000018217 | Pmp22 | 18858 | protein_coding | 2954.729327 | -0.112480337 | 0.248748261 | -0.452185422 | 0.651135409 | 0.798250711 | 0.112480337 |
| ENSMUSG00000042364 | Snx18 | 170625 | protein_coding | 260.2101815 | -0.112492015 | 0.079751714 | -1.410527874 | 0.158383872 | 0.339625513 | 0.112492015 |
| ENSMUSG00000045827 | Serpinb9 | 20723 | protein_coding | 68.68945488 | 0.112526086 | 0.213706549 | 0.526544866 | 0.598509688 | 0.765321556 | 0.112526086 |
| ENSMUSG00000020464 | Pnpt1 | 71701 | protein_coding | 759.7595746 | 0.112567305 | 0.086584503 | 1.300086061 | 0.193571475 | 0.387069668 | 0.112567305 |
| ENSMUSG00000059811 | Atl2 | 56298 | protein_coding | 1427.060288 | 0.112588567 | 0.076692753 | 1.468047009 | 0.142091454 | 0.315345225 | 0.112588567 |
| ENSMUSG00000039449 | Prpf18 | 67229 | protein_coding | 640.0948198 | 0.112599052 | 0.082588122 | 1.36338071 | 0.172762557 | 0.358489058 | 0.112599052 |
| ENSMUSG00000024544 | Ldlrad4 | 52662 | protein_coding | 47.73033942 | 0.112629229 | 0.265602369 | 0.424052048 | 0.671527855 | 0.810310279 | 0.112629229 |
| ENSMUSG00000034101 | Ctnnd1 | 12388 | protein_coding | 784.7313187 | -0.11269454 | 0.10079135 | -1.11809734 | 0.263525422 | 0.471131469 | 0.11269454 |
| ENSMUSG00000024387 | Csnk2b | 13001 | protein_coding | 932.9943741 | 0.112725644 | 0.080495059 | 1.400404521 | 0.161392217 | 0.343403485 | 0.112725644 |
| ENSMUSG00000035401 | Emsy | 233545 | protein_coding | 362.6122802 | -0.112732395 | 0.124174229 | -0.907856612 | 0.363953983 | 0.572855539 | 0.112732395 |
| ENSMUSG00000058291 | Zfp68 | 24135 | protein_coding | 666.3729318 | 0.112733998 | 0.100023972 | 1.127069791 | 0.259712975 | 0.466949798 | 0.112733998 |
| ENSMUSG00000048758 | Rpl29 | 19944 | protein_coding | 3118.919926 | 0.112753303 | 0.090762289 | 1.242292419 | 0.214128691 | 0.412514121 | 0.112753303 |

| ensembl_gene_id | external_gene_name | entrezgene_id | gene_biotype | baseMean | log2FoldChange | lfcSE | stat | pvalue | padj | absLog2FC |
|---|---|---|---|---|---|---|---|---|---|---|
| ENSMUSG00000058835 | Abi1 | 11308 | protein_coding | 2170.28938 | 0.112824852 | 0.070292335 | 1.605080437 | 0.108476104 | 0.262119878 | 0.112824852 |
| ENSMUSG00000006732 | Mettl1 | 17299 | protein_coding | 179.7713698 | -0.112825993 | 0.15651007 | -0.720886476 | 0.470979364 | 0.666543809 | 0.112825993 |
| ENSMUSG00000030317 | Timp4 | 110595 | protein_coding | 44.57391971 | 0.112838033 | 0.407227684 | 0.277088316 | 0.781712293 | 0.881369958 | 0.112838033 |
| ENSMUSG00000024277 | Mapre2 | 212307 | protein_coding | 966.9695943 | -0.112857659 | 0.070554926 | -1.599571641 | 0.109693644 | 0.263887685 | 0.112857659 |
| ENSMUSG00000020677 | Ddx52 | 78394 | protein_coding | 543.9118184 | 0.112933998 | 0.119237705 | 0.947133273 | 0.343570872 | 0.55376472 | 0.112933998 |
| ENSMUSG00000003348 | Mob3a | 208228 | protein_coding | 471.290795 | 0.112951156 | 0.233220983 | 0.484309578 | 0.62816618 | 0.784780943 | 0.112951156 |
| ENSMUSG00000097316 | Gm10516 | 100038712 | lncRNA | 145.8825232 | 0.112968695 | 0.1464095 | 0.771594019 | 0.440354918 | 0.642313808 | 0.112968695 |
| ENSMUSG00000062691 | Cebpzos | 68554 | protein_coding | 198.0469699 | 0.112995406 | 0.128912764 | 0.876526123 | 0.380744081 | 0.588499925 | 0.112995406 |
| ENSMUSG00000073838 | Tufm | 233870 | protein_coding | 1408.491714 | -0.113015681 | 0.088407512 | -1.278349299 | 0.201126293 | 0.395694119 | 0.113015681 |
| ENSMUSG00000074892 | B3galt5 | 93961 | protein_coding | 6186.009123 | 0.113017785 | 0.106080222 | 1.065399213 | 0.286695313 | 0.495548881 | 0.113017785 |
| ENSMUSG00000038943 | Prc1 | 233406 | protein_coding | 631.03901 | 0.113026069 | 0.118856695 | 0.950944071 | 0.341632768 | 0.551941783 | 0.113026069 |
| ENSMUSG00000052783 | Grk4 | 14772 | protein_coding | 13.8212882 | -0.113045944 | 0.481113125 | -0.234967492 | 0.814233964 | 0.900950823 | 0.113045944 |
| ENSMUSG00000028614 | Ndc1 | 72787 | protein_coding | 381.1433778 | 0.113092508 | 0.107108547 | 1.055868191 | 0.291028439 | 0.499573396 | 0.113092508 |
| ENSMUSG00000001380 | Hars | 15115 | protein_coding | 521.9040293 | 0.11310945 | 0.102585514 | 1.102586962 | 0.270206575 | 0.478375791 | 0.11310945 |
| ENSMUSG00000028221 | Pip4p2 | 72519 | protein_coding | 342.9306888 | 0.113117098 | 0.078046304 | 1.449358814 | 0.147237404 | 0.323788123 | 0.113117098 |
| ENSMUSG00000026483 | Niban1 | 63913 | protein_coding | 1252.719855 | 0.113145256 | 0.087269251 | 1.2965077 | 0.194800629 | 0.387870369 | 0.113145256 |
| ENSMUSG00000026020 | Nop58 | 55989 | protein_coding | 1204.01814 | -0.113179893 | 0.146675591 | -0.771634141 | 0.440331147 | 0.642313808 | 0.113179893 |
| ENSMUSG00000002477 | Snrpd1 | 20641 | protein_coding | 543.3574974 | 0.113195341 | 0.125834587 | 0.899556663 | 0.368356229 | 0.577194099 | 0.113195341 |
| ENSMUSG00000025257 | Ribc1 | 66611 | protein_coding | 29.2894197 | 0.113221499 | 0.254612007 | 0.444682482 | 0.656549231 | 0.801149235 | 0.113221499 |
| ENSMUSG00000057329 | Bcl2 | 12043 | protein_coding | 264.7214843 | -0.11325004 | 0.157323417 | -0.719854945 | 0.471614311 | 0.667069775 | 0.11325004 |
| ENSMUSG00000049076 | Acap2 | 78618 | protein_coding | 1310.55195 | -0.113259787 | 0.091897394 | -1.232459187 | 0.217777603 | 0.417470414 | 0.113259787 |
| ENSMUSG00000024135 | Srbd1 | 78586 | protein_coding | 300.9984695 | 0.113345248 | 0.105656442 | 1.072771769 | 0.28337353 | 0.492533129 | 0.113345248 |
| ENSMUSG00000055302 | Mrfap1 | 67568 | protein_coding | 2255.886416 | 0.113346181 | 0.09980403 | 1.135687419 | 0.256087408 | 0.462992081 | 0.113346181 |
| ENSMUSG00000030822 | Prr14 | 233895 | protein_coding | 397.0236702 | -0.113352954 | 0.070488296 | -1.608110297 | 0.107811024 | 0.261133515 | 0.113352954 |
| ENSMUSG00000024160 | Spsb3 | 79043 | protein_coding | 63.72766806 | 0.113398957 | 0.155927522 | 0.727254274 | 0.467070204 | 0.663707223 | 0.113398957 |
| ENSMUSG00000025199 | Chuk | 12675 | protein_coding | 1537.314995 | 0.113410706 | 0.127266981 | 0.891124348 | 0.372862464 | 0.581051928 | 0.113410706 |
| ENSMUSG00000036617 | Etl4 | 208618 | protein_coding | 1574.599962 | -0.113412108 | 0.127817871 | -0.887294612 | 0.374920306 | 0.583397242 | 0.113412108 |
| ENSMUSG00000020493 | Prr11 | 270906 | protein_coding | 118.3496648 | 0.113413002 | 0.166969238 | 0.679244894 | 0.496982706 | 0.687983708 | 0.113413002 |
| ENSMUSG00000051351 | Zfp46 | 22704 | protein_coding | 273.8493886 | 0.11343707 | 0.118744322 | 0.955305214 | 0.339423372 | 0.550085499 | 0.11343707 |
| ENSMUSG00000020022 | Ndufa12 | 66414 | protein_coding | 804.2932017 | 0.113455217 | 0.080516523 | 1.409092363 | 0.158807859 | 0.340076845 | 0.113455217 |
| ENSMUSG00000074211 | Sdhaf1 | 68332 | protein_coding | 98.28244902 | 0.113467831 | 0.131581019 | 0.86234194 | 0.38849938 | 0.596973148 | 0.113467831 |
| ENSMUSG00000026966 | Ssna1 | 68475 | protein_coding | 341.3901006 | -0.113500584 | 0.094663114 | -1.198994829 | 0.230529956 | 0.433618463 | 0.113500584 |
| ENSMUSG00000030802 | Bckdk | 12041 | protein_coding | 667.4402573 | 0.113522324 | 0.076733747 | 1.479431517 | 0.139025022 | 0.310420576 | 0.113522324 |
| ENSMUSG00000021432 | Slc35b3 | 108652 | protein_coding | 824.3574598 | 0.113526845 | 0.071118747 | 1.596299857 | 0.110421862 | 0.265081713 | 0.113526845 |
| ENSMUSG00000044348 | Slc25a53 | 67062 | protein_coding | 34.10496345 | -0.113549951 | 0.234346928 | -0.484537824 | 0.628004228 | 0.78470262 | 0.113549951 |
| ENSMUSG00000032777 | Gtf3c1 | 233863 | protein_coding | 1457.495338 | -0.113579576 | 0.069775641 | -1.627782625 | 0.103570986 | 0.254232438 | 0.113579576 |
| ENSMUSG00000005681 | Apoa2 | 11807 | protein_coding | 9.270903823 | 0.113611784 | 0.482476994 | 0.235476064 | 0.813839254 | 0.900884818 | 0.113611784 |
| ENSMUSG00000028107 | Tars2 | 71807 | protein_coding | 424.3464842 | -0.113655541 | 0.085490226 | -1.329456555 | 0.18369739 | 0.373110435 | 0.113655541 |

Supplementary Table 1

| ensembl_gene_id | external_gene_name | entrezgene_id | gene_biotype | baseMean | log2FoldChange | lfcSE | stat | pvalue | padj | absLog2FC |
|---|---|---|---|---|---|---|---|---|---|---|
| ENSMUSG00000017210 | Med24 | 23989 | protein_coding | 464.5971875 | 0.113673914 | 0.078611226 | 1.446026474 | 0.148169777 | 0.324966971 | 0.113673914 |
| ENSMUSG00000036352 | Ubac1 | 98766 | protein_coding | 495.3105782 | 0.113700005 | 0.13888529 | 0.818661252 | 0.41297971 | 0.619566809 | 0.113700005 |
| ENSMUSG00000050854 | Tmem125 | 230678 | protein_coding | 251.7191219 | 0.113703139 | 0.189403943 | 0.600320862 | 0.548292418 | 0.73018501 | 0.113703139 |
| ENSMUSG00000031781 | Ciapin1 | 109006 | protein_coding | 267.9413021 | -0.113713868 | 0.125597648 | -0.905382142 | 0.365262973 | 0.574097951 | 0.113713868 |
| ENSMUSG00000038141 | Tmem181a | 77106 | protein_coding | 632.2800267 | -0.113883954 | 0.06566556 | -1.734302633 | 0.082864401 | 0.21942879 | 0.113883954 |
| ENSMUSG00000002812 | Flii | 14248 | protein_coding | 2584.298914 | 0.113910873 | 0.081583483 | 1.396249208 | 0.162639463 | 0.345172734 | 0.113910873 |
| ENSMUSG00000062116 | Zfp954 | 232853 | protein_coding | 285.2403042 | 0.113926971 | 0.09891718 | 1.151740998 | 0.24942752 | 0.45576248 | 0.113926971 |
| ENSMUSG00000027133 | Nop10 | 66181 | protein_coding | 464.7897119 | 0.113953357 | 0.114886426 | 0.991878337 | 0.321256878 | 0.532354566 | 0.113953357 |
| ENSMUSG00000020463 | Ppp4r3b | 104570 | protein_coding | 2087.658368 | 0.113968918 | 0.050213498 | 2.26968691 | 0.023226586 | 0.089447066 | 0.113968918 |
| ENSMUSG00000033885 | Pxk | 218699 | protein_coding | 683.636508 | -0.114045875 | 0.055326396 | -2.061328473 | 0.039271714 | 0.131466972 | 0.114045875 |
| ENSMUSG00000016256 | Ctsz | 64138 | protein_coding | 1124.00011 | -0.114094087 | 0.09168611 | -1.244398817 | 0.213352818 | 0.41141449 | 0.114094087 |
| ENSMUSG00000003344 | Btbd2 | 208198 | protein_coding | 347.4598448 | -0.114099737 | 0.10789582 | -1.057499148 | 0.290283843 | 0.498965961 | 0.114099737 |
| ENSMUSG00000029610 | Aimp2 | 231872 | protein_coding | 157.1631798 | 0.114102231 | 0.150256111 | 0.75938496 | 0.447622308 | 0.648451853 | 0.114102231 |
| ENSMUSG00000020888 | Dvl2 | 13543 | protein_coding | 124.268916 | -0.11412331 | 0.174072015 | -0.655609753 | 0.512075249 | 0.699874255 | 0.11412331 |
| ENSMUSG00000069769 | Msi2 | 76626 | protein_coding | 1318.181283 | 0.114171162 | 0.085997312 | 1.327613151 | 0.184305938 | 0.373994868 | 0.114171162 |
| ENSMUSG00000026743 | Mllt10 | 17354 | protein_coding | 594.707128 | -0.114173123 | 0.084260336 | -1.355004363 | 0.175416167 | 0.36173339 | 0.114173123 |
| ENSMUSG00000034707 | Gns | 75612 | protein_coding | 894.4799087 | -0.114205809 | 0.065259753 | -1.750019013 | 0.080115033 | 0.213992665 | 0.114205809 |
| ENSMUSG00000032228 | Tcf12 | 21406 | protein_coding | 1158.350216 | -0.11421929 | 0.071563858 | -1.596047134 | 0.11047827 | 0.265134532 | 0.11421929 |
| ENSMUSG00000025364 | Pa2g4 | 18813 | protein_coding | 1204.405128 | -0.114226054 | 0.108080743 | -1.056858521 | 0.290576161 | 0.499223071 | 0.114226054 |
| ENSMUSG00000024589 | Nedd4l | 83814 | protein_coding | 1261.302871 | -0.114228072 | 0.081658246 | -1.398855318 | 0.161856374 | 0.344106824 | 0.114228072 |
| ENSMUSG00000048120 | Entpd1 | 12495 | protein_coding | 7.330821291 | -0.114336988 | 0.616251239 | -0.185536322 | 0.852808394 | 0.922175438 | 0.114336988 |
| ENSMUSG00000061477 | Rps7 | 20115 | protein_coding | 6646.505184 | 0.114382671 | 0.103496483 | 1.105184133 | 0.269079811 | 0.477165801 | 0.114382671 |
| ENSMUSG00000018927 | Ccl6 | 20305 | protein_coding | 4890.11694 | -0.114403328 | 0.115966293 | -0.986522244 | 0.323876905 | 0.53475937 | 0.114403328 |
| ENSMUSG00000029290 | Zfp326 | 54367 | protein_coding | 352.4979212 | 0.114413995 | 0.103167019 | 1.109017163 | 0.267422778 | 0.475338691 | 0.114413995 |
| ENSMUSG00000022353 | Mtss1 | 211401 | protein_coding | 647.7548051 | -0.114419034 | 0.107914847 | -1.06027148 | 0.28902111 | 0.497873797 | 0.114419034 |
| ENSMUSG00000003161 | Sri | 109552 | protein_coding | 5237.959064 | 0.114507936 | 0.09375628 | 1.221336162 | 0.221958767 | 0.423092298 | 0.114507936 |
| ENSMUSG00000024228 | Nudt12 | 67993 | protein_coding | 276.3301593 | 0.114524798 | 0.115093885 | 0.99505546 | 0.319709291 | 0.531004128 | 0.114524798 |
| ENSMUSG00000022412 | Mief1 | 239555 | protein_coding | 703.9823816 | -0.114568807 | 0.088724772 | -1.291283203 | 0.196605493 | 0.390214607 | 0.114568807 |
| ENSMUSG00000060600 | Eno3 | 13808 | protein_coding | 639.6110733 | 0.114575208 | 0.247911342 | 0.462162025 | 0.643965133 | 0.793891381 | 0.114575208 |
| ENSMUSG00000021577 | Sdha | 66945 | protein_coding | 6299.326415 | -0.114581091 | 0.068450311 | -1.6739309 | 0.094144181 | 0.23893061 | 0.114581091 |
| ENSMUSG00000026335 | Pam | 18484 | protein_coding | 2777.393562 | -0.114596537 | 0.165864917 | -0.690902807 | 0.489626622 | 0.682008993 | 0.114596537 |
| ENSMUSG00000031669 | Gins3 | 78833 | protein_coding | 60.47670473 | 0.114644348 | 0.156802074 | 0.731140513 | 0.464693331 | 0.661628263 | 0.114644348 |
| ENSMUSG00000038909 | Kat7 | 217127 | protein_coding | 988.922539 | 0.114660444 | 0.081858946 | 1.400707557 | 0.161301542 | 0.343340043 | 0.114660444 |
| ENSMUSG00000015095 | Fbxw5 | 30839 | protein_coding | 471.2692069 | 0.114660693 | 0.085068645 | 1.347860817 | 0.177703153 | 0.364750868 | 0.114660693 |
| ENSMUSG00000026858 | Miga2 | 108958 | protein_coding | 588.8691265 | 0.114663622 | 0.089510646 | 1.281005413 | 0.200191765 | 0.394795779 | 0.114663622 |
| ENSMUSG00000022197 | Pdzd2 | 68070 | protein_coding | 45.10717337 | -0.114664573 | 0.255110576 | -0.449470085 | 0.653092583 | 0.799021328 | 0.114664573 |
| ENSMUSG00000003644 | Rps6ka1 | 20111 | protein_coding | 3598.211826 | 0.114672619 | 0.080250851 | 1.42892714 | 0.153025176 | 0.331907598 | 0.114672619 |
| ENSMUSG00000037697 | Ddhd1 | 114874 | protein_coding | 272.8940974 | -0.114696077 | 0.077877549 | -1.472774609 | 0.140811821 | 0.313223305 | 0.114696077 |

| ensembl_gene_id | external_gene_name | entrezgene_id | gene_biotype | baseMean | log2FoldChange | lfcSE | stat | pvalue | padj | absLog2FC |
|---|---|---|---|---|---|---|---|---|---|---|
| ENSMUSG00000035772 | Mrps2 | 118451 | protein_coding | 412.7554274 | -0.114723399 | 0.0939672 | -1.220887698 | 0.222128543 | 0.423234241 | 0.114723399 |
| ENSMUSG00000023175 | Bsg | 12215 | protein_coding | 7736.540598 | -0.114733332 | 0.090421533 | -1.268871782 | 0.204486798 | 0.399885294 | 0.114733332 |
| ENSMUSG00000039771 | Polr2j | 20022 | protein_coding | 310.5122491 | 0.114749686 | 0.095360054 | 1.203330761 | 0.22884835 | 0.431883474 | 0.114749686 |
| ENSMUSG00000032264 | Zw10 | 26951 | protein_coding | 466.1057238 | 0.114775593 | 0.078570231 | 1.460802544 | 0.144069637 | 0.318448679 | 0.114775593 |
| ENSMUSG00000061759 | Armt1 | 73419 | protein_coding | 242.1530782 | 0.114849889 | 0.104199906 | 1.102207214 | 0.270371596 | 0.478375791 | 0.114849889 |
| ENSMUSG00000035399 | Oser1 | 66680 | protein_coding | 189.5531892 | -0.114885503 | 0.100296843 | -1.145454835 | 0.252020795 | 0.458393532 | 0.114885503 |
| ENSMUSG00000051111 | Sv2c | 75209 | protein_coding | 127.6046541 | -0.114973883 | 0.170901212 | -0.672750544 | 0.501106006 | 0.691618675 | 0.114973883 |
| ENSMUSG00000038642 | Ctss | 13040 | protein_coding | 2309.656725 | -0.11498436 | 0.106417918 | -1.080498117 | 0.279920425 | 0.488182692 | 0.11498436 |
| ENSMUSG00000024644 | Cndp2 | 66054 | protein_coding | 770.5675871 | -0.115003701 | 0.080395858 | -1.430467994 | 0.152582745 | 0.33119896 | 0.115003701 |
| ENSMUSG00000022781 | Pak2 | 224105 | protein_coding | 2199.933138 | -0.115003867 | 0.072044251 | -1.596294864 | 0.110422976 | 0.265081713 | 0.115003867 |
| ENSMUSG00000074282 | Zfp94 | 22756 | protein_coding | 32.13007822 | -0.115015038 | 0.267891207 | -0.429334876 | 0.66767954 | 0.807317977 | 0.115015038 |
| ENSMUSG00000027329 | Spef1 | 70997 | protein_coding | 130.1515705 | -0.115080231 | 0.16601914 | -0.69317448 | 0.488200062 | 0.680942953 | 0.115080231 |
| ENSMUSG00000111375 | Btbd8 | 100503185 | protein_coding | 129.990799 | -0.11510222 | 0.149187529 | -0.77152709 | 0.440394572 | 0.642313808 | 0.11510222 |
| ENSMUSG00000020340 | Cyfip2 | 76884 | protein_coding | 23.62416845 | 0.115143845 | 0.379069968 | 0.30375354 | 0.761315665 | 0.869186728 | 0.115143845 |
| ENSMUSG00000042699 | Dhx9 | 13211 | protein_coding | 1515.290859 | -0.115280645 | 0.118304413 | -0.974440779 | 0.32983773 | 0.54061076 | 0.115280645 |
| ENSMUSG00000034848 | Ttc21b | 73668 | protein_coding | 212.9885124 | 0.11529752 | 0.131815834 | 0.874686419 | 0.381744552 | 0.589473873 | 0.11529752 |
| ENSMUSG00000026234 | Ncl | 17975 | protein_coding | 3179.164209 | 0.115350852 | 0.13809336 | 0.835310632 | 0.403542833 | 0.610808226 | 0.115350852 |
| ENSMUSG00000024039 | Cbs | 12411 | protein_coding | 1301.961957 | -0.115367627 | 0.109038479 | -1.058045081 | 0.29003489 | 0.49866966 | 0.115367627 |
| ENSMUSG00000028597 | Gpx7 | 67305 | protein_coding | 25.6217012 | 0.115451077 | 0.246304926 | 0.468732311 | 0.639260988 | 0.790801742 | 0.115451077 |
| ENSMUSG00000050428 | Fbxo46 | 243867 | protein_coding | 252.4779779 | -0.115490218 | 0.093405415 | -1.236440274 | 0.216294953 | 0.415528407 | 0.115490218 |
| ENSMUSG00000044042 | Fmn1 | 14260 | protein_coding | 781.4800773 | -0.115495515 | 0.140368648 | -0.822801367 | 0.410620959 | 0.617598738 | 0.115495515 |
| ENSMUSG00000018809 | Smyd4 | 319822 | protein_coding | 94.84866247 | -0.115536316 | 0.175371525 | -0.658808871 | 0.51001851 | 0.698626897 | 0.115536316 |
| ENSMUSG00000025862 | Stag2 | 20843 | protein_coding | 2551.085102 | -0.115560753 | 0.052150925 | -2.21589077 | 0.026698984 | 0.099013135 | 0.115560753 |
| ENSMUSG00000029176 | Anapc4 | 52206 | protein_coding | 602.2136498 | 0.115646296 | 0.092063327 | 1.256160287 | 0.209057862 | 0.405828413 | 0.115646296 |
| ENSMUSG00000039164 | Naif1 | 71254 | protein_coding | 113.3522908 | -0.115651534 | 0.127603284 | -0.906336654 | 0.36475769 | 0.57358678 | 0.115651534 |
| ENSMUSG00000085175 | Gm11423 | 100503272 | lncRNA | 27.33223071 | 0.115654801 | 0.295274697 | 0.391685445 | 0.695290646 | 0.826592861 | 0.115654801 |
| ENSMUSG00000033644 | Piwil2 | 57746 | protein_coding | 11.30568601 | -0.115660262 | 0.435090991 | -0.265830055 | 0.790370095 | 0.886363083 | 0.115660262 |
| ENSMUSG00000038058 | Nod1 | 107607 | protein_coding | 212.7597097 | 0.115681648 | 0.147741155 | 0.783002187 | 0.433625828 | 0.637334334 | 0.115681648 |
| ENSMUSG00000095315 | Gm10130 | 102633750 | protein_coding | 63.88318444 | -0.115712207 | 0.188694294 | -0.61322579 | 0.539727054 | 0.722964639 | 0.115712207 |
| ENSMUSG00000017776 | Crk | 12928 | protein_coding | 3107.744541 | 0.115761978 | 0.098710966 | 1.172736752 | 0.240901388 | 0.445498353 | 0.115761978 |
| ENSMUSG00000025510 | Cd151 | 12476 | protein_coding | 1432.338351 | 0.115947585 | 0.100315945 | 1.15582408 | 0.247753122 | 0.453535423 | 0.115947585 |
| ENSMUSG00000044156 | Hepacam2 | 101202 | protein_coding | 3150.92828 | 0.116008856 | 0.127176427 | 0.912188356 | 0.361669573 | 0.570896781 | 0.116008856 |
| ENSMUSG00000038745 | Nlrp6 | 101613 | protein_coding | 2782.139554 | 0.116028106 | 0.170055021 | 0.682297443 | 0.495050889 | 0.686622306 | 0.116028106 |
| ENSMUSG00000045328 | Cenpe | 229841 | protein_coding | 807.5634281 | 0.116044816 | 0.098797625 | 1.174570915 | 0.240166425 | 0.444603768 | 0.116044816 |
| ENSMUSG00000035048 | Anapc13 | 69010 | protein_coding | 457.5965188 | 0.116167223 | 0.085169997 | 1.363945366 | 0.172584761 | 0.35830684 | 0.116167223 |
| ENSMUSG00000031729 | Ist1 | 71955 | protein_coding | 1911.313047 | -0.11624927 | 0.132185034 | -0.879443511 | 0.379160849 | 0.587178874 | 0.11624927 |
| ENSMUSG00000024750 | Zfand5 | 22682 | protein_coding | 2112.745912 | 0.116249373 | 0.089019533 | 1.305886127 | 0.191591285 | 0.384285374 | 0.116249373 |
| ENSMUSG00000042508 | Dmtf1 | 23857 | protein_coding | 508.9265505 | -0.116330181 | 0.097788159 | -1.189614182 | 0.234198069 | 0.438035087 | 0.116330181 |

| ensembl_gene_id | external_gene_name | entrezgene_id | gene_biotype | baseMean | log2FoldChange | lfcSE | stat | pvalue | padj | absLog2FC |
|---|---|---|---|---|---|---|---|---|---|---|
| ENSMUSG00000009090 | Ap1b1 | 11764 | protein_coding | 1506.89698 | -0.116334065 | 0.079420382 | -1.464788535 | 0.142978621 | 0.316696353 | 0.116334065 |
| ENSMUSG00000078619 | Smarcd2 | 83796 | protein_coding | 1184.195935 | -0.116347702 | 0.064097328 | -1.815172429 | 0.069497415 | 0.194817816 | 0.116347702 |
| ENSMUSG00000057778 | Cyb5d2 | 192986 | protein_coding | 117.295059 | 0.116392969 | 0.123929724 | 0.939185256 | 0.347635638 | 0.557673526 | 0.116392969 |
| ENSMUSG00000030682 | Cdipt | 52858 | protein_coding | 859.4724453 | 0.116412834 | 0.089119311 | 1.306258248 | 0.191464751 | 0.384086971 | 0.116412834 |
| ENSMUSG00000118401 | Gpr52 | 620246 | protein_coding | 8.655998735 | -0.116489484 | 0.394533201 | -0.295259014 | 0.767796024 | 0.873407732 | 0.116489484 |
| ENSMUSG00000027420 | Bfsp1 | 12075 | protein_coding | 70.92050841 | -0.116493248 | 0.188177976 | -0.619058887 | 0.535877566 | 0.720595853 | 0.116493248 |
| ENSMUSG00000051124 | Gimap9 | 317758 | protein_coding | 104.6609722 | -0.116495979 | 0.131504093 | -0.885873405 | 0.375685749 | 0.583911261 | 0.116495979 |
| ENSMUSG00000029038 | Ssu72 | 68991 | protein_coding | 821.0065808 | 0.116501443 | 0.07275174 | 1.601356112 | 0.109298068 | 0.263373619 | 0.116501443 |
| ENSMUSG00000021076 | Actr10 | 56444 | protein_coding | 859.3308463 | 0.116518573 | 0.073208912 | 1.591590016 | 0.111476855 | 0.266607109 | 0.116518573 |
| ENSMUSG00000023845 | Lnpep | 240028 | protein_coding | 2925.001376 | 0.116522197 | 0.124034141 | 0.939436477 | 0.347506692 | 0.557627687 | 0.116522197 |
| ENSMUSG00000090626 | Tex9 | 21778 | protein_coding | 168.3364492 | 0.116538354 | 0.205306777 | 0.567630334 | 0.570286006 | 0.744744605 | 0.116538354 |
| ENSMUSG00000013418 | B4galnt2 | 14422 | protein_coding | 6916.537001 | 0.116548692 | 0.189681868 | 0.614442979 | 0.538922642 | 0.722272825 | 0.116548692 |
| ENSMUSG00000038122 | Tbc1d32 | 544696 | protein_coding | 429.2512004 | 0.116669425 | 0.117926931 | 0.989336566 | 0.322498499 | 0.533463272 | 0.116669425 |
| ENSMUSG00000031834 | Pik3r2 | 18709 | protein_coding | 584.9018596 | 0.11667173 | 0.105168281 | 1.109381349 | 0.267265705 | 0.475177965 | 0.11667173 |
| ENSMUSG00000021497 | Txndc15 | 69672 | protein_coding | 540.7648336 | 0.116673545 | 0.10957233 | 1.064808477 | 0.286962608 | 0.495795685 | 0.116673545 |
| ENSMUSG00000028886 | Eya3 | 14050 | protein_coding | 574.1924927 | -0.116687234 | 0.067475993 | -1.729314795 | 0.08375277 | 0.220874619 | 0.116687234 |
| ENSMUSG00000029847 | Slc23a4 | 243753 | protein_coding | 72.51959254 | -0.116702798 | 0.159529622 | -0.731543125 | 0.464447473 | 0.661396455 | 0.116702798 |
| ENSMUSG00000040044 | Orc3 | 50793 | protein_coding | 478.2671895 | -0.116709789 | 0.074585433 | -1.564779941 | 0.117634517 | 0.277049029 | 0.116709789 |
| ENSMUSG00000024404 | Riok3 | 66878 | protein_coding | 2199.005395 | 0.116711269 | 0.113083487 | 1.032080566 | 0.302034377 | 0.511352419 | 0.116711269 |
| ENSMUSG00000021018 | Polr2h | 245841 | protein_coding | 281.0547753 | -0.116747976 | 0.114265366 | -1.021726705 | 0.306910268 | 0.516893761 | 0.116747976 |
| ENSMUSG00000026959 | Grin1 | 14810 | protein_coding | 34.86610994 | -0.116765649 | 0.328571356 | -0.355373792 | 0.72230958 | 0.844044564 | 0.116765649 |
| ENSMUSG00000038903 | Ccdc68 | 381175 | protein_coding | 418.7802055 | -0.116823605 | 0.13184331 | -0.886079125 | 0.375574891 | 0.583911261 | 0.116823605 |
| ENSMUSG00000003531 | Dgcr6 | 13353 | protein_coding | 220.1823156 | 0.116856093 | 0.124388245 | 0.939446431 | 0.347501584 | 0.557627687 | 0.116856093 |
| ENSMUSG00000019979 | Apaf1 | 11783 | protein_coding | 660.7763846 | 0.116939358 | 0.103410914 | 1.130822202 | 0.258129933 | 0.465311631 | 0.116939358 |
| ENSMUSG00000046330 | Rpl37a | 19981 | protein_coding | 4252.915668 | 0.11696102 | 0.112775811 | 1.037110879 | 0.299684186 | 0.508644997 | 0.11696102 |
| ENSMUSG00000033883 | Zfp267 | 241944 | protein_coding | 261.4038618 | 0.11696796 | 0.10298323 | 1.13579619 | 0.256041872 | 0.462962304 | 0.11696796 |
| ENSMUSG00000051396 | Gm45902 | 119065974 | protein_coding | 60.16877215 | 0.116979451 | 0.193224104 | 0.60540817 | 0.544907824 | 0.727614171 | 0.116979451 |
| ENSMUSG00000024645 | Timm21 | 67105 | protein_coding | 306.6872525 | 0.117021219 | 0.109862976 | 1.065156099 | 0.286805296 | 0.495685196 | 0.117021219 |
| ENSMUSG00000048329 | Mfsd6l | 215723 | protein_coding | 215.0590269 | 0.117027373 | 0.172275229 | 0.6793047 | 0.496944819 | 0.687983708 | 0.117027373 |
| ENSMUSG00000064179 | Tnnt1 | 21955 | protein_coding | 20.52690083 | 0.117034794 | 0.254899198 | 0.459141476 | 0.646132574 | 0.795304816 | 0.117034794 |
| ENSMUSG00000031995 | St14 | 19143 | protein_coding | 4372.24759 | 0.117036151 | 0.092874975 | 1.260147328 | 0.20761622 | 0.404082965 | 0.117036151 |
| ENSMUSG00000034610 | Tut4 | 230594 | protein_coding | 364.5804576 | -0.117074045 | 0.125054151 | -0.9361868 | 0.349177018 | 0.559301241 | 0.117074045 |
| ENSMUSG00000006498 | Ptbp1 | 19205 | protein_coding | 2644.703072 | 0.117087248 | 0.116472325 | 1.005279556 | 0.314762256 | 0.525527591 | 0.117087248 |
| ENSMUSG00000009681 | Bcr | 110279 | protein_coding | 465.627333 | -0.117098447 | 0.108183143 | -1.082409362 | 0.27907067 | 0.487504149 | 0.117098447 |
| ENSMUSG00000033020 | Polr2f | 69833 | protein_coding | 383.9746571 | -0.117107464 | 0.090098929 | -1.299765328 | 0.193681412 | 0.387103913 | 0.117107464 |
| ENSMUSG00000026975 | Dph7 | 67228 | protein_coding | 96.84717015 | 0.117124995 | 0.139674283 | 0.838558056 | 0.401717356 | 0.609212039 | 0.117124995 |
| ENSMUSG00000031105 | Slc25a14 | 20523 | protein_coding | 77.05887361 | 0.117145096 | 0.130992812 | 0.894286444 | 0.371168651 | 0.579501943 | 0.117145096 |
| ENSMUSG00000017747 | Ghdc | 80860 | protein_coding | 158.4989012 | -0.117149746 | 0.141811788 | -0.826093147 | 0.408751255 | 0.615833332 | 0.117149746 |

| ensembl_gene_id | external_gene_name | entrezgene_id | gene_biotype | baseMean | log2FoldChange | lfcSE | stat | pvalue | padj | absLog2FC |
|---|---|---|---|---|---|---|---|---|---|---|
| ENSMUSG00000020098 | Pcbd1 | 13180 | protein_coding | 503.2500311 | 0.117150721 | 0.093135863 | 1.257847589 | 0.208446881 | 0.405142395 | 0.117150721 |
| ENSMUSG00000021278 | Amn | 93835 | protein_coding | 89.10075842 | -0.117153565 | 0.292945452 | -0.399915971 | 0.689218409 | 0.823011515 | 0.117153565 |
| ENSMUSG00000037053 | Azgp1 | 12007 | protein_coding | 77.79448949 | -0.117153858 | 0.191485297 | -0.611816466 | 0.540659194 | 0.723787161 | 0.117153858 |
| ENSMUSG00000022721 | Trmt2a | 15547 | protein_coding | 363.2685903 | -0.117180107 | 0.111853127 | -1.04762477 | 0.294811521 | 0.503979247 | 0.117180107 |
| ENSMUSG00000035004 | Igsf6 | 80719 | protein_coding | 12.77673931 | -0.117190368 | 0.368267072 | -0.318221141 | 0.750317198 | 0.862020508 | 0.117190368 |
| ENSMUSG00000006333 | Rps9 | 76846 | protein_coding | 10610.87302 | 0.117225463 | 0.093299982 | 1.256436078 | 0.208957908 | 0.405789022 | 0.117225463 |
| ENSMUSG00000036777 | Anln | 68743 | protein_coding | 539.691365 | 0.117266868 | 0.105865973 | 1.107691783 | 0.267994949 | 0.475940787 | 0.117266868 |
| ENSMUSG00000041684 | Bivm | 246229 | protein_coding | 295.3148691 | 0.117268345 | 0.144195123 | 0.813261524 | 0.416068132 | 0.622441365 | 0.117268345 |
| ENSMUSG00000057841 | Rpl32 | 19951 | protein_coding | 8404.194645 | 0.11731087 | 0.097718237 | 1.200501292 | 0.229944711 | 0.433131429 | 0.11731087 |
| ENSMUSG00000025979 | Mob4 | 19070 | protein_coding | 1305.866061 | 0.117371732 | 0.083803189 | 1.400564029 | 0.161344484 | 0.343385214 | 0.117371732 |
| ENSMUSG00000039156 | Stim2 | 116873 | protein_coding | 823.9410899 | -0.117438929 | 0.11998861 | -0.978750636 | 0.327703206 | 0.538721067 | 0.117438929 |
| ENSMUSG00000024099 | Ndufv2 | 72900 | protein_coding | 2137.485451 | 0.117484408 | 0.086182431 | 1.363206015 | 0.172817592 | 0.358543028 | 0.117484408 |
| ENSMUSG00000022421 | Nptxr | 73340 | protein_coding | 8.479367399 | 0.117498545 | 0.463938436 | 0.253263227 | 0.800064811 | 0.892772335 | 0.117498545 |
| ENSMUSG00000021900 | Btd | 26363 | protein_coding | 152.9665653 | 0.11750429 | 0.136415053 | 0.861373344 | 0.389032452 | 0.597369026 | 0.11750429 |
| ENSMUSG00000023036 | Pcdhgc4 | 93707 | protein_coding | 121.0583184 | 0.117526465 | 0.153670908 | 0.764793194 | 0.444394701 | 0.645302106 | 0.117526465 |
| ENSMUSG00000028152 | Tspan5 | 56224 | protein_coding | 158.898061 | 0.117538602 | 0.11063602 | 1.062390011 | 0.288058667 | 0.497257878 | 0.117538602 |
| ENSMUSG00000031557 | Plekha2 | 83436 | protein_coding | 1005.747604 | 0.117586267 | 0.08738066 | 1.345678398 | 0.178406256 | 0.365695904 | 0.117586267 |
| ENSMUSG00000001100 | Poldip2 | 67811 | protein_coding | 747.6227128 | 0.117605926 | 0.082746097 | 1.421286677 | 0.155233435 | 0.335187361 | 0.117605926 |
| ENSMUSG00000027775 | Mfsd1 | 66868 | protein_coding | 920.073155 | -0.11760919 | 0.099318095 | -1.1841668 | 0.236347042 | 0.440601745 | 0.11760919 |
| ENSMUSG00000028807 | Zbtb8a | 73680 | protein_coding | 218.7721165 | -0.117615141 | 0.119732131 | -0.982318945 | 0.325942744 | 0.53715634 | 0.117615141 |
| ENSMUSG00000019055 | Plod1 | 18822 | protein_coding | 166.9818778 | 0.117694419 | 0.104278415 | 1.128655618 | 0.259043138 | 0.466219108 | 0.117694419 |
| ENSMUSG00000089847 | Timm10b | 14356 | protein_coding | 119.8515622 | -0.117694483 | 0.115419953 | -1.01970656 | 0.30786765 | 0.517815831 | 0.117694483 |
| ENSMUSG00000015846 | Rxra | 20181 | protein_coding | 1380.296247 | -0.117750795 | 0.113604933 | -1.036493677 | 0.299971888 | 0.508888297 | 0.117750795 |
| ENSMUSG00000028607 | Cpt2 | 12896 | protein_coding | 1125.265495 | 0.117837962 | 0.092124774 | 1.279112634 | 0.200857395 | 0.395507058 | 0.117837962 |
| ENSMUSG00000040167 | Ikzf5 | 67143 | protein_coding | 49.42239201 | 0.117848466 | 0.265051857 | 0.444624183 | 0.656591368 | 0.801149235 | 0.117848466 |
| ENSMUSG00000023905 | Tnfrsf12a | 27279 | protein_coding | 263.9482816 | 0.117891341 | 0.85953793 | 0.137156647 | 0.890906978 | 0.945079343 | 0.117891341 |
| ENSMUSG00000024942 | Capn1 | 12333 | protein_coding | 1705.785906 | -0.117968478 | 0.091251874 | -1.292778685 | 0.196087615 | 0.389582578 | 0.117968478 |
| ENSMUSG00000033054 | Npat | 244879 | protein_coding | 474.7937512 | -0.117971344 | 0.063370423 | -1.861615214 | 0.062657347 | 0.180967229 | 0.117971344 |
| ENSMUSG00000038541 | Srd5a2 | 94224 | protein_coding | 75.76736864 | 0.118064406 | 0.264456809 | 0.446441165 | 0.655278603 | 0.800343347 | 0.118064406 |
| ENSMUSG00000026803 | Ttf1 | 22130 | protein_coding | 424.3630111 | 0.118071912 | 0.058013845 | 2.035236814 | 0.041827051 | 0.137167235 | 0.118071912 |
| ENSMUSG00000020018 | Snrpf | 69878 | protein_coding | 192.2278422 | 0.118078057 | 0.126442964 | 0.933844424 | 0.35038415 | 0.560502034 | 0.118078057 |
| ENSMUSG00000019978 | Epb41l2 | 13822 | protein_coding | 617.6785303 | -0.118097025 | 0.116558028 | -1.013203699 | 0.310962874 | 0.520986289 | 0.118097025 |
| ENSMUSG00000036022 | Fam122b | 78755 | protein_coding | 54.45564841 | 0.118155442 | 0.201578404 | 0.586151291 | 0.557773851 | 0.736485277 | 0.118155442 |
| ENSMUSG00000024948 | Map4k2 | 26412 | protein_coding | 131.4308193 | -0.118197352 | 0.149992367 | -0.788022449 | 0.430683575 | 0.634649642 | 0.118197352 |
| ENSMUSG00000022614 | Lmf2 | 105847 | protein_coding | 236.5568659 | -0.118202942 | 0.112970795 | -1.04631416 | 0.29541601 | 0.504599164 | 0.118202942 |
| ENSMUSG00000047090 | Tmem198b | 73827 | protein_coding | 29.67396235 | 0.118211554 | 0.225487138 | 0.524249656 | 0.600104908 | 0.766578793 | 0.118211554 |
| ENSMUSG00000050271 | Prag1 | 244418 | protein_coding | 419.2861948 | -0.118226773 | 0.185754549 | -0.636467716 | 0.524471617 | 0.710174608 | 0.118226773 |
| ENSMUSG00000109032 | Gm7972 | 666190 | transcribed_unprocessed_pseudogene | 7.80347043 | 0.118260026 | 0.521266236 | 0.226870682 | 0.8205243 | 0.904826298 | 0.118260026 |

| ensembl_gene_id | external_gene_name | entrezgene_id | gene_biotype | baseMean | log2FoldChange | lfcSE | stat | pvalue | padj | absLog2FC |
|---|---|---|---|---|---|---|---|---|---|---|
| ENSMUSG00000096255 | Dynlt1b | 21648 | protein_coding | 56.58594367 | 0.118273879 | 0.158662165 | 0.745444758 | 0.456002895 | 0.655296022 | 0.118273879 |
| ENSMUSG00000042015 | Wdr41 | 218460 | protein_coding | 394.4588723 | -0.118280582 | 0.082853781 | -1.427582189 | 0.153412153 | 0.33241039 | 0.118280582 |
| ENSMUSG00000040498 | Igsf23 | 70080 | protein_coding | 114.2839237 | 0.11835064 | 0.151127689 | 0.783116852 | 0.433558496 | 0.637295787 | 0.11835064 |
| ENSMUSG00000027628 | Aar2 | 68295 | protein_coding | 423.7790473 | 0.118415708 | 0.120594557 | 0.981932445 | 0.326133131 | 0.537335907 | 0.118415708 |
| ENSMUSG00000039270 | Megf9 | 230316 | protein_coding | 289.5227061 | -0.118464405 | 0.093740504 | -1.263748324 | 0.206320374 | 0.40248266 | 0.118464405 |
| ENSMUSG00000021540 | Smad5 | 17129 | protein_coding | 694.9148556 | -0.118474852 | 0.137418598 | -0.862145692 | 0.38860735 | 0.597061626 | 0.118474852 |
| ENSMUSG00000025224 | Gbf1 | 107338 | protein_coding | 1279.338698 | 0.118504284 | 0.083115884 | 1.425771803 | 0.153934222 | 0.333043235 | 0.118504284 |
| ENSMUSG00000040025 | Ythdf2 | 213541 | protein_coding | 1262.010224 | -0.118548972 | 0.062576313 | -1.894470374 | 0.058162603 | 0.172324022 | 0.118548972 |
| ENSMUSG00000024736 | Tmem132a | 98170 | protein_coding | 51.52426066 | -0.118560764 | 0.181947909 | -0.651619274 | 0.514646811 | 0.702183862 | 0.118560764 |
| ENSMUSG00000020814 | Mxra7 | 67622 | protein_coding | 27.10586061 | -0.118573535 | 0.292037721 | -0.406021301 | 0.68472695 | 0.819963225 | 0.118573535 |
| ENSMUSG00000025006 | Sorbs1 | 20411 | protein_coding | 197.5320275 | 0.118585926 | 0.173992032 | 0.681559519 | 0.49551752 | 0.68695983 | 0.118585926 |
| ENSMUSG00000018858 | Mrpl58 | 68572 | protein_coding | 391.9259304 | 0.118587436 | 0.111001583 | 1.068340038 | 0.285367154 | 0.49432605 | 0.118587436 |
| ENSMUSG00000025451 | Paip1 | 218693 | protein_coding | 995.3458287 | 0.118626684 | 0.076109924 | 1.558623085 | 0.119085616 | 0.279680874 | 0.118626684 |
| ENSMUSG00000047786 | Lix1 | 66643 | protein_coding | 13.41583266 | 0.118629648 | 0.589502688 | 0.201236823 | 0.8405134 | 0.915836464 | 0.118629648 |
| ENSMUSG00000044296 | Zfp879 | 214779 | protein_coding | 12.15643289 | -0.118637833 | 0.420484815 | -0.282145344 | 0.777832065 | 0.879295843 | 0.118637833 |
| ENSMUSG00000029535 | Triap1 | 69076 | protein_coding | 368.7191043 | 0.118658423 | 0.077228603 | 1.53645694 | 0.124426351 | 0.288103346 | 0.118658423 |
| ENSMUSG00000020520 | Galnt10 | 171212 | protein_coding | 2308.582467 | -0.118674084 | 0.084165194 | -1.410013777 | 0.158535615 | 0.339813656 | 0.118674084 |
| ENSMUSG00000096010 | H4c16 | 320332 | protein_coding | 687.3076982 | 0.118811097 | 0.132290106 | 0.89811023 | 0.369126786 | 0.577871395 | 0.118811097 |
| ENSMUSG00000010607 | Pigyl | 66268 | protein_coding | 139.3506301 | 0.118897941 | 0.116773797 | 1.018190239 | 0.308587557 | 0.518563181 | 0.118897941 |
| ENSMUSG00000021474 | Sfxn1 | 14057 | protein_coding | 840.5473277 | -0.118942116 | 0.064991029 | -1.830131309 | 0.067230306 | 0.190000766 | 0.118942116 |
| ENSMUSG00000030695 | Aldoa | 11674 | protein_coding | 11884.50044 | 0.118948468 | 0.09609769 | 1.237786971 | 0.215795061 | 0.414792572 | 0.118948468 |
| ENSMUSG00000114362 | 4933433G19Rik | 71226 | lncRNA | 16.48094427 | -0.118959301 | 0.349087726 | -0.340771938 | 0.733275278 | 0.850291833 | 0.118959301 |
| ENSMUSG00000048379 | Socs4 | 67296 | protein_coding | 643.273885 | -0.118985929 | 0.116210661 | -1.023881355 | 0.305891318 | 0.515908186 | 0.118985929 |
| ENSMUSG00000012640 | Zfp715 | 69930 | protein_coding | 441.2928779 | -0.118990325 | 0.073349895 | -1.62222898 | 0.104754329 | 0.256243913 | 0.118990325 |
| ENSMUSG00000032180 | Tmed1 | 17083 | protein_coding | 102.1068581 | 0.11900938 | 0.145643898 | 0.817125756 | 0.413856566 | 0.620277776 | 0.11900938 |
| ENSMUSG00000090862 | Rps13 | 68052 | protein_coding | 3535.756243 | 0.119012169 | 0.101150712 | 1.176582613 | 0.239362141 | 0.443424073 | 0.119012169 |
| ENSMUSG00000043262 | Uevld | 54122 | protein_coding | 400.3890606 | 0.119017845 | 0.098361435 | 1.210005178 | 0.226276906 | 0.428594317 | 0.119017845 |
| ENSMUSG00000087138 | Gm15545 | 100502630 | lncRNA | 23.69972956 | -0.119072313 | 0.234093943 | -0.508651833 | 0.61099629 | 0.773174617 | 0.119072313 |
| ENSMUSG00000019080 | Mfsd3 | 69572 | protein_coding | 48.99692611 | -0.11914728 | 0.18550856 | -0.64227376 | 0.520695449 | 0.707225813 | 0.11914728 |
| ENSMUSG00000045078 | Rnf216 | 108086 | protein_coding | 553.2863053 | -0.119164965 | 0.072686568 | -1.639435844 | 0.101122522 | 0.250100293 | 0.119164965 |
| ENSMUSG00000030330 | Ing4 | 28019 | protein_coding | 25.2359177 | -0.11920728 | 0.344532904 | -0.345996794 | 0.729345124 | 0.847709365 | 0.11920728 |
| ENSMUSG00000048012 | Zfp473 | 243963 | protein_coding | 37.93135381 | 0.119227763 | 0.267384631 | 0.445903575 | 0.6556669 | 0.800694808 | 0.119227763 |
| ENSMUSG00000029131 | Dnajb6 | 23950 | protein_coding | 669.1179493 | -0.119264581 | 0.092541579 | -1.288767514 | 0.197478923 | 0.391321758 | 0.119264581 |
| ENSMUSG00000034509 | Mad2l1bp | 66591 | protein_coding | 220.7463705 | 0.119264989 | 0.105491366 | 1.13056635 | 0.258237657 | 0.465369065 | 0.119264989 |
| ENSMUSG00000006676 | Usp19 | 71472 | protein_coding | 1363.821229 | -0.119271602 | 0.082787152 | -1.440701829 | 0.149668938 | 0.327058149 | 0.119271602 |
| ENSMUSG00000024006 | Stk38 | 106504 | protein_coding | 1129.98395 | -0.119272025 | 0.112188817 | -1.063136489 | 0.287720059 | 0.496834898 | 0.119272025 |
| ENSMUSG00000026634 | Angel2 | 52477 | protein_coding | 757.4359517 | -0.11928354 | 0.087176826 | -1.368294134 | 0.171220026 | 0.356588987 | 0.11928354 |
| ENSMUSG00000027823 | Gmps | 229363 | protein_coding | 1141.341781 | 0.119290039 | 0.079465855 | 1.501148379 | 0.133317188 | 0.301829505 | 0.119290039 |

| ensembl_gene_id | external_gene_name | entrezgene_id | gene_biotype | baseMean | log2FoldChange | lfcSE | stat | pvalue | padj | absLog2FC |
|---|---|---|---|---|---|---|---|---|---|---|
| ENSMUSG00000021810 | Ecd | 70601 | protein_coding | 587.2068947 | 0.11933474 | 0.121959967 | 0.97847468 | 0.327839608 | 0.538806428 | 0.11933474 |
| ENSMUSG00000022556 | Hsf1 | 15499 | protein_coding | 259.7163882 | -0.119353596 | 0.139167913 | -0.857622949 | 0.391100699 | 0.599220382 | 0.119353596 |
| ENSMUSG00000031379 | Pir | 69656 | protein_coding | 126.0553507 | 0.119377248 | 0.141621461 | 0.842931907 | 0.399266528 | 0.606939679 | 0.119377248 |
| ENSMUSG00000001910 | Nacc1 | 66830 | protein_coding | 1117.114625 | -0.119383424 | 0.085878193 | -1.390148305 | 0.164483847 | 0.34783573 | 0.119383424 |
| ENSMUSG00000063229 | Ldha | 16828 | protein_coding | 10228.13545 | 0.119397724 | 0.122357909 | 0.975807156 | 0.329160038 | 0.539888897 | 0.119397724 |
| ENSMUSG00000041354 | Rgl2 | 19732 | protein_coding | 392.5534891 | -0.119422001 | 0.109958661 | -1.086062713 | 0.277451248 | 0.485791302 | 0.119422001 |
| ENSMUSG00000017999 | Ddx27 | 228889 | protein_coding | 508.8741489 | -0.119422309 | 0.108112661 | -1.104609836 | 0.269328687 | 0.47728831 | 0.119422309 |
| ENSMUSG00000008373 | Prpf31 | 68988 | protein_coding | 434.6814164 | 0.119476622 | 0.103956231 | 1.149297366 | 0.250433384 | 0.456709748 | 0.119476622 |
| ENSMUSG00000064280 | Ccdc146 | 75172 | protein_coding | 6.99962812 | 0.119556648 | 0.74562321 | 0.16034459 | 0.872609635 | 0.934632928 | 0.119556648 |
| ENSMUSG00000021576 | Pdcd6 | 18570 | protein_coding | 1056.707937 | 0.11958528 | 0.0955809 | 1.251142011 | 0.21088267 | 0.408273388 | 0.11958528 |
| ENSMUSG00000052997 | Uba2 | 50995 | protein_coding | 1221.947133 | 0.119597718 | 0.089250464 | 1.340023482 | 0.180237711 | 0.368125204 | 0.119597718 |
| ENSMUSG00000027864 | Ptgfrn | 19221 | protein_coding | 2283.432462 | 0.119681127 | 0.11770865 | 1.016757281 | 0.309268909 | 0.519076415 | 0.119681127 |
| ENSMUSG00000071172 | Srsf3 | 20383 | protein_coding | 2879.038347 | 0.119724059 | 0.112556815 | 1.063676682 | 0.287475191 | 0.496519719 | 0.119724059 |
| ENSMUSG00000019927 | Ube2d1 | 216080 | protein_coding | 153.2729945 | -0.119729661 | 0.114513118 | -1.045554114 | 0.295766944 | 0.505090156 | 0.119729661 |
| ENSMUSG00000023345 | Poc1a | 70235 | protein_coding | 180.4384141 | 0.119746431 | 0.111982898 | 1.069327846 | 0.284921968 | 0.49392051 | 0.119746431 |
| ENSMUSG00000038759 | Nup205 | 70699 | protein_coding | 912.4825264 | 0.119759224 | 0.091010711 | 1.315880542 | 0.188214141 | 0.379285783 | 0.119759224 |
| ENSMUSG00000022971 | Ifnar2 | 15976 | protein_coding | 46.86776469 | 0.119834014 | 0.18698398 | 0.640878507 | 0.521601621 | 0.707881371 | 0.119834014 |
| ENSMUSG00000034126 | Pomt2 | 217734 | protein_coding | 257.3400875 | 0.119867607 | 0.089701743 | 1.336290724 | 0.181454267 | 0.370064477 | 0.119867607 |
| ENSMUSG00000002804 | Nudt14 | 66174 | protein_coding | 284.9139083 | 0.119868372 | 0.113208361 | 1.058829682 | 0.289677353 | 0.498400322 | 0.119868372 |
| ENSMUSG00000038542 | Pcid2 | 234069 | protein_coding | 206.6916827 | -0.119878517 | 0.109227897 | -1.097508244 | 0.272419279 | 0.480519853 | 0.119878517 |
| ENSMUSG00000034893 | Cog3 | 338337 | protein_coding | 1483.48322 | -0.1198867 | 0.060189155 | -1.991832246 | 0.046389467 | 0.147807847 | 0.1198867 |
| ENSMUSG00000018102 | H2bc4 | 68024 | protein_coding | 2718.407897 | 0.119889808 | 0.113073174 | 1.060285159 | 0.289014889 | 0.497873797 | 0.119889808 |
| ENSMUSG00000036533 | Cdc42ep3 | 260409 | protein_coding | 234.3182546 | 0.119900677 | 0.198963376 | 0.602626871 | 0.54675694 | 0.728826223 | 0.119900677 |
| ENSMUSG00000062931 | Zfp938 | 237411 | protein_coding | 126.6923409 | -0.119916739 | 0.109076458 | -1.09938241 | 0.2716013 | 0.479714516 | 0.119916739 |
| ENSMUSG00000029125 | Stx18 | 71116 | protein_coding | 297.2189609 | 0.119923125 | 0.108090184 | 1.10947285 | 0.26722625 | 0.475177965 | 0.119923125 |
| ENSMUSG00000022831 | Hcls1 | 15163 | protein_coding | 21.86086783 | 0.119940821 | 0.30793687 | 0.389498083 | 0.696907729 | 0.827434106 | 0.119940821 |
| ENSMUSG00000037106 | Fer1l6 | 631797 | protein_coding | 1771.497929 | -0.119975805 | 0.158091335 | -0.758901841 | 0.447911278 | 0.648752463 | 0.119975805 |
| ENSMUSG00000026810 | Dpm2 | 13481 | protein_coding | 264.2306477 | 0.119983036 | 0.094265802 | 1.272816155 | 0.203083287 | 0.398020253 | 0.119983036 |
| ENSMUSG00000074886 | Grk6 | 26385 | protein_coding | 1076.206842 | 0.120042757 | 0.093528508 | 1.283488427 | 0.199321011 | 0.393698545 | 0.120042757 |
| ENSMUSG00000037225 | Fgf2 | 14173 | protein_coding | 12.20270721 | -0.120050295 | 0.420571076 | -0.285445913 | 0.775302536 | 0.877871529 | 0.120050295 |
| ENSMUSG00000026029 | Casp8 | 12370 | protein_coding | 1784.531384 | 0.120106821 | 0.21993389 | 0.546104201 | 0.584994312 | 0.755578122 | 0.120106821 |
| ENSMUSG00000017119 | Nbr1 | 17966 | protein_coding | 3558.98821 | 0.120132917 | 0.074847224 | 1.605041725 | 0.108484623 | 0.262119878 | 0.120132917 |
| ENSMUSG00000000037 | Scml2 | 107815 | protein_coding | 29.17932451 | 0.12016354 | 0.265855152 | 0.45198876 | 0.65127708 | 0.798250711 | 0.12016354 |
| ENSMUSG00000029550 | Sppl3 | 74585 | protein_coding | 555.3535442 | -0.120190087 | 0.08423414 | -1.426857171 | 0.153621068 | 0.332727274 | 0.120190087 |
| ENSMUSG00000038721 | Hoxb7 | 15415 | protein_coding | 146.5098937 | 0.120209857 | 0.242554227 | 0.495599928 | 0.620176704 | 0.778936265 | 0.120209857 |
| ENSMUSG00000021614 | Vcan | 13003 | protein_coding | 96.80480683 | -0.120249062 | 0.159746712 | -0.752748275 | 0.451601195 | 0.651785414 | 0.120249062 |
| ENSMUSG00000029273 | Sult1d1 | 53315 | protein_coding | 7551.467526 | -0.120279654 | 0.179511904 | -0.670037203 | 0.502834075 | 0.692892218 | 0.120279654 |
| ENSMUSG00000039210 | Gpatch2 | 67769 | protein_coding | 773.9099168 | 0.120303844 | 0.108232862 | 1.11152789 | 0.266341192 | 0.474103991 | 0.120303844 |

| ensembl_gene_id | external_gene_name | entrezgene_id | gene_biotype | baseMean | log2FoldChange | lfcSE | stat | pvalue | padj | absLog2FC |
|---|---|---|---|---|---|---|---|---|---|---|
| ENSMUSG00000030557 | Mef2a | 17258 | protein_coding | 616.509909 | 0.120305084 | 0.102806454 | 1.170209448 | 0.241916692 | 0.446609087 | 0.120305084 |
| ENSMUSG00000038022 | Mindy4 | 330323 | protein_coding | 11.80246426 | -0.120312677 | 0.47062736 | -0.255643184 | 0.79822637 | 0.891657874 | 0.120312677 |
| ENSMUSG00000002635 | Pdcd2l | 68079 | protein_coding | 159.1679377 | -0.120314064 | 0.11461815 | -1.049694689 | 0.293858506 | 0.502963495 | 0.120314064 |
| ENSMUSG00000021027 | Ralgapa1 | 56784 | protein_coding | 1221.305884 | -0.120373728 | 0.073446859 | -1.638922735 | 0.101229352 | 0.250175504 | 0.120373728 |
| ENSMUSG00000038069 | Cdkn2aip | 70925 | protein_coding | 330.8250976 | -0.120392252 | 0.073103352 | -1.646877303 | 0.099583268 | 0.247567082 | 0.120392252 |
| ENSMUSG00000052369 | Tmem106c | 380967 | protein_coding | 194.7032517 | -0.120405429 | 0.109795454 | -1.096634015 | 0.272801412 | 0.480874379 | 0.120405429 |
| ENSMUSG00000023084 | Lrrc71 | 74485 | protein_coding | 16.20957702 | 0.120480656 | 0.342224032 | 0.352052004 | 0.724799263 | 0.844960894 | 0.120480656 |
| ENSMUSG00000029173 | Sepsecs | 211006 | protein_coding | 804.5943013 | -0.120570745 | 0.077389995 | -1.557962963 | 0.119242028 | 0.279883145 | 0.120570745 |
| ENSMUSG00000052516 | Robo2 | 268902 | protein_coding | 336.9750662 | -0.120616639 | 0.509471608 | -0.2367485 | 0.812851906 | 0.900355019 | 0.120616639 |
| ENSMUSG00000045983 | Eif4g1 | 208643 | protein_coding | 6264.980673 | 0.120634227 | 0.103821158 | 1.161942608 | 0.245258778 | 0.450668923 | 0.120634227 |
| ENSMUSG00000029466 | Anapc7 | 56317 | protein_coding | 430.51755 | -0.120679964 | 0.070994426 | -1.699851251 | 0.089158909 | 0.230503772 | 0.120679964 |
| ENSMUSG00000043065 | Spice1 | 212514 | protein_coding | 181.746817 | -0.12068851 | 0.132393691 | -0.911588079 | 0.3619856 | 0.571259786 | 0.12068851 |
| ENSMUSG00000026127 | Imp4 | 27993 | protein_coding | 632.3742451 | -0.120734727 | 0.093879082 | -1.286066344 | 0.198419909 | 0.392646374 | 0.120734727 |
| ENSMUSG00000040250 | Ints13 | 71177 | protein_coding | 477.7665825 | 0.120879318 | 0.092615822 | 1.305169192 | 0.191835242 | 0.384442846 | 0.120879318 |
| ENSMUSG00000078584 | AU022252 | 230696 | protein_coding | 276.442421 | 0.120882943 | 0.099841202 | 1.210752076 | 0.225990437 | 0.428372488 | 0.120882943 |
| ENSMUSG00000034453 | Polr3b | 70428 | protein_coding | 450.1797229 | -0.120898498 | 0.102650326 | -1.177770227 | 0.238888221 | 0.443009848 | 0.120898498 |
| ENSMUSG00000061615 | H2ac4 | 319172 | protein_coding | 388.8963428 | 0.120928166 | 0.277303239 | 0.436086379 | 0.662774059 | 0.804685206 | 0.120928166 |
| ENSMUSG00000007415 | Gatad1 | 67210 | protein_coding | 1042.276877 | 0.120928504 | 0.079019103 | 1.530370509 | 0.125925044 | 0.290349465 | 0.120928504 |
| ENSMUSG00000043964 | Orai3 | 269999 | protein_coding | 204.7346314 | -0.120987789 | 0.162059557 | -0.746563737 | 0.455326945 | 0.654730304 | 0.120987789 |
| ENSMUSG00000074829 | 2010315B03Rik | 630836 | protein_coding | 764.1989404 | 0.121016002 | 0.139221121 | 0.869235943 | 0.384718092 | 0.592679426 | 0.121016002 |
| ENSMUSG00000066036 | Ubr4 | 69116 | protein_coding | 2781.776899 | -0.121053353 | 0.15946541 | -0.759119819 | 0.447780884 | 0.648622583 | 0.121053353 |
| ENSMUSG00000020078 | Vps26a | 30930 | protein_coding | 907.1942256 | -0.121109582 | 0.088038957 | -1.375636271 | 0.168934271 | 0.353910966 | 0.121109582 |
| ENSMUSG00000024446 | Rpp21 | 67676 | protein_coding | 226.9284155 | -0.121127369 | 0.100983625 | -1.199475351 | 0.230343164 | 0.433458782 | 0.121127369 |
| ENSMUSG00000017176 | Nt5c3b | 68106 | protein_coding | 115.8266061 | -0.121163829 | 0.136857967 | -0.885325357 | 0.375981177 | 0.584199414 | 0.121163829 |
| ENSMUSG00000047881 | Rell1 | 100532 | protein_coding | 735.2565578 | 0.121169331 | 0.095073753 | 1.274477206 | 0.202494346 | 0.39748185 | 0.121169331 |
| ENSMUSG00000020549 | Elac2 | 68626 | protein_coding | 373.2812105 | -0.121222351 | 0.122640416 | -0.988437219 | 0.322938567 | 0.533862532 | 0.121222351 |
| ENSMUSG00000005983 | 1700037C18Rik | 73261 | protein_coding | 36.46787513 | 0.121304478 | 0.195413903 | 0.620756639 | 0.534759757 | 0.719457168 | 0.121304478 |
| ENSMUSG00000056209 | Npm3 | 18150 | protein_coding | 251.3181771 | -0.121367913 | 0.104314534 | -1.163480373 | 0.244634657 | 0.449946976 | 0.121367913 |
| ENSMUSG00000003429 | Rps11 | 27207 | protein_coding | 6309.401644 | 0.121419856 | 0.110757004 | 1.096272483 | 0.272959548 | 0.480993437 | 0.121419856 |
| ENSMUSG00000040390 | Map3k10 | 269881 | protein_coding | 113.2925471 | -0.121448387 | 0.145403091 | -0.835253134 | 0.403575199 | 0.610808226 | 0.121448387 |
| ENSMUSG00000002416 | Ndufb2 | 68198 | protein_coding | 593.3489567 | 0.121501321 | 0.096729103 | 1.25609892 | 0.209080108 | 0.405828413 | 0.121501321 |
| ENSMUSG00000021660 | Btf3 | 218490 | protein_coding | 2859.942098 | 0.121521204 | 0.07662329 | 1.58595648 | 0.112749182 | 0.268480935 | 0.121521204 |
| ENSMUSG00000030774 | Pak1 | 18479 | protein_coding | 1287.889688 | 0.121564985 | 0.051533138 | 2.35896723 | 0.018325874 | 0.074921592 | 0.121564985 |
| ENSMUSG00000022961 | Son | 20658 | protein_coding | 4476.346075 | -0.1216567 | 0.073638308 | -1.652084397 | 0.098517347 | 0.245876576 | 0.1216567 |
| ENSMUSG00000030201 | Lrp6 | 16974 | protein_coding | 2510.575398 | -0.121678818 | 0.095592437 | -1.272891685 | 0.20305648 | 0.398020253 | 0.121678818 |
| ENSMUSG00000025278 | Flnb | 286940 | protein_coding | 8719.568491 | -0.121694874 | 0.222636694 | -0.546607444 | 0.584648454 | 0.75519265 | 0.121694874 |
| ENSMUSG00000051934 | Spats2 | 72572 | protein_coding | 186.2934207 | 0.121727448 | 0.122253474 | 0.995697256 | 0.319397264 | 0.530636656 | 0.121727448 |
| ENSMUSG00000006215 | Zbtb17 | 22642 | protein_coding | 304.3286886 | -0.121814392 | 0.125756314 | -0.968654277 | 0.332717713 | 0.543652824 | 0.121814392 |

| ensembl_gene_id | external_gene_name | entrezgene_id | gene_biotype | baseMean | log2FoldChange | lfcSE | stat | pvalue | padj | absLog2FC |
|---|---|---|---|---|---|---|---|---|---|---|
| ENSMUSG00000087354 | 4930404I05Rik | 67394 | lncRNA | 8.786777999 | -0.121815902 | 0.481491899 | -0.252996784 | 0.800270698 | 0.892814435 | 0.121815902 |
| ENSMUSG00000029505 | Ep400 | 75560 | protein_coding | 1028.450184 | -0.121835275 | 0.119748859 | -1.017423268 | 0.308952118 | 0.518872768 | 0.121835275 |
| ENSMUSG00000018196 | Glrx2 | 69367 | protein_coding | 263.9321791 | 0.121835418 | 0.07984934 | 1.525816208 | 0.127055643 | 0.292313212 | 0.121835418 |
| ENSMUSG00000031483 | Erlin2 | 244373 | protein_coding | 1998.193924 | 0.121886449 | 0.045561496 | 2.675207345 | 0.007468305 | 0.038397403 | 0.121886449 |
| ENSMUSG00000022408 | Fam83f | 213956 | protein_coding | 453.3369456 | 0.12189376 | 0.088863037 | 1.371703735 | 0.170155685 | 0.355358419 | 0.12189376 |
| ENSMUSG00000027018 | Hat1 | 107435 | protein_coding | 332.2179456 | 0.121916657 | 0.103789081 | 1.174657839 | 0.240131633 | 0.444591033 | 0.121916657 |
| ENSMUSG00000038121 | Fam210a | 108654 | protein_coding | 834.3490586 | -0.12195484 | 0.072059594 | -1.69241641 | 0.090566606 | 0.23274361 | 0.12195484 |
| ENSMUSG00000028015 | Ctso | 229445 | protein_coding | 402.3254244 | 0.121955149 | 0.102765581 | 1.186731468 | 0.235333559 | 0.439487973 | 0.121955149 |
| ENSMUSG00000021597 | Slf1 | 105377 | protein_coding | 291.6292437 | 0.121979944 | 0.102182047 | 1.193751223 | 0.232575297 | 0.43617795 | 0.121979944 |
| ENSMUSG00000084910 | C630043F03Rik | 68285 | lncRNA | 47.46580967 | 0.122018696 | 0.196468644 | 0.621059388 | 0.534560549 | 0.719249909 | 0.122018696 |
| ENSMUSG00000029781 | Fkbp9 | 27055 | protein_coding | 1059.545255 | 0.122022925 | 0.128587893 | 0.94894567 | 0.342648245 | 0.552958586 | 0.122022925 |
| ENSMUSG00000022175 | Lrp10 | 65107 | protein_coding | 2976.273551 | 0.122034912 | 0.066487654 | 1.835452215 | 0.066438713 | 0.188432305 | 0.122034912 |
| ENSMUSG00000040302 | Rbm48 | 269623 | protein_coding | 91.81200327 | 0.122045695 | 0.162241365 | 0.752247709 | 0.451902108 | 0.652077527 | 0.122045695 |
| ENSMUSG00000018189 | Uchl5 | 56207 | protein_coding | 359.85369 | 0.122076746 | 0.098636685 | 1.237640397 | 0.215849429 | 0.414822599 | 0.122076746 |
| ENSMUSG00000039473 | Ubn1 | 170644 | protein_coding | 1083.397761 | -0.122142151 | 0.062884749 | -1.942317554 | 0.052098669 | 0.159956041 | 0.122142151 |
| ENSMUSG00000037826 | Ppm1k | 243382 | protein_coding | 258.3716262 | 0.12217226 | 0.177910427 | 0.686706572 | 0.492267655 | 0.684071633 | 0.12217226 |
| ENSMUSG00000000690 | Hoxb6 | 15414 | protein_coding | 284.1818671 | -0.122182545 | 0.246091029 | -0.496493294 | 0.619546418 | 0.778489692 | 0.122182545 |
| ENSMUSG00000116056 | Gm4544 | 100043600 | lncRNA | 17.83577026 | -0.122202503 | 0.360803614 | -0.33869534 | 0.734839252 | 0.851367123 | 0.122202503 |
| ENSMUSG00000038563 | Efl1 | 101592 | protein_coding | 448.4134403 | -0.122205491 | 0.147980544 | -0.825821339 | 0.408905447 | 0.615890871 | 0.122205491 |
| ENSMUSG00000042709 | Atpaf2 | 246782 | protein_coding | 355.6417647 | 0.122207977 | 0.089672622 | 1.362823723 | 0.172938073 | 0.358619662 | 0.122207977 |
| ENSMUSG00000066233 | Tmem42 | 66079 | protein_coding | 101.6126161 | 0.122229024 | 0.149027519 | 0.820177535 | 0.412114906 | 0.618910638 | 0.122229024 |
| ENSMUSG00000031153 | Gripap1 | 54645 | protein_coding | 382.431045 | 0.122267132 | 0.092496669 | 1.321854428 | 0.18621663 | 0.376785884 | 0.122267132 |
| ENSMUSG00000023965 | Fbxl17 | 50758 | protein_coding | 294.6357828 | -0.122269024 | 0.089283128 | -1.36945274 | 0.170857798 | 0.356297977 | 0.122269024 |
| ENSMUSG00000037022 | Mmaa | 109136 | protein_coding | 301.7304293 | 0.122359654 | 0.085437522 | 1.432153596 | 0.152099869 | 0.330511147 | 0.122359654 |
| ENSMUSG00000028078 | Dclk2 | 70762 | protein_coding | 33.19571748 | -0.122385786 | 0.229485263 | -0.533305644 | 0.593822022 | 0.76209791 | 0.122385786 |
| ENSMUSG00000059796 | Eif4a1 | 13681 | protein_coding | 5473.509132 | 0.122391166 | 0.15923107 | 0.768638725 | 0.44210781 | 0.643447843 | 0.122391166 |
| ENSMUSG00000021709 | Erbin | 59079 | protein_coding | 8545.949992 | -0.122410898 | 0.092891022 | -1.317790416 | 0.187573817 | 0.378355839 | 0.122410898 |
| ENSMUSG00000040287 | Stac3 | 237611 | protein_coding | 8.884559034 | -0.122424906 | 0.543057432 | -0.225436389 | 0.821639806 | 0.905442422 | 0.122424906 |
| ENSMUSG00000029430 | Ran | 19384 | protein_coding | 1992.260746 | 0.122430107 | 0.140109518 | 0.873817204 | 0.38221781 | 0.589976284 | 0.122430107 |
| ENSMUSG00000034243 | Golgb1 | 224139 | protein_coding | 3726.159993 | 0.122466059 | 0.090743821 | 1.349580142 | 0.177150697 | 0.364084685 | 0.122466059 |
| ENSMUSG00000033554 | Dph5 | 69740 | protein_coding | 138.8464694 | -0.122525835 | 0.144053508 | -0.8505578 | 0.39501504 | 0.603182778 | 0.122525835 |
| ENSMUSG00000048280 | Zfp738 | 408068 | protein_coding | 263.8087506 | -0.122537945 | 0.114278114 | -1.07227833 | 0.283595037 | 0.492751979 | 0.122537945 |
| ENSMUSG00000028551 | Cdkn2c | 12580 | protein_coding | 156.9647643 | 0.12255189 | 0.150236826 | 0.815724705 | 0.414657606 | 0.621031156 | 0.12255189 |
| ENSMUSG00000033732 | Sf3b3 | 101943 | protein_coding | 1630.01405 | -0.122555864 | 0.121851065 | -1.005784098 | 0.314519438 | 0.525232293 | 0.122555864 |
| ENSMUSG00000090176 | Cd200r2 | 271375 | protein_coding | 92.79740795 | 0.122556302 | 0.157060489 | 0.780312752 | 0.435206809 | 0.638244049 | 0.122556302 |
| ENSMUSG00000029165 | Agbl5 | 231093 | protein_coding | 138.9226992 | -0.122634987 | 0.119077677 | -1.02987386 | 0.303069223 | 0.512579841 | 0.122634987 |
| ENSMUSG00000022789 | Dnm1l | 74006 | protein_coding | 1991.669063 | 0.12263842 | 0.081091003 | 1.512355444 | 0.130443463 | 0.297324482 | 0.12263842 |
| ENSMUSG00000034120 | Srsf2 | 20382 | protein_coding | 2003.393928 | 0.122677998 | 0.137333461 | 0.893285566 | 0.371704265 | 0.579898015 | 0.122677998 |

| ensembl_gene_id | external_gene_name | entrezgene_id | gene_biotype | baseMean | log2FoldChange | lfcSE | stat | pvalue | padj | absLog2FC |
|---|---|---|---|---|---|---|---|---|---|---|
| ENSMUSG00000038510 | Rpf2 | 67239 | protein_coding | 200.1124068 | -0.122680131 | 0.180570189 | -0.67940412 | 0.49688184 | 0.68796366 | 0.122680131 |
| ENSMUSG00000034190 | Chmp7 | 105513 | protein_coding | 568.0474484 | 0.122731988 | 0.090754828 | 1.352346655 | 0.176264446 | 0.362590739 | 0.122731988 |
| ENSMUSG00000040888 | Gfer | 11692 | protein_coding | 481.5013463 | 0.122738196 | 0.088708439 | 1.383613528 | 0.166476824 | 0.350415963 | 0.122738196 |
| ENSMUSG00000020457 | Drg1 | 13494 | protein_coding | 697.844689 | 0.122782106 | 0.086389558 | 1.421260967 | 0.155240906 | 0.335187361 | 0.122782106 |
| ENSMUSG00000034432 | Cops8 | 108679 | protein_coding | 404.7395694 | 0.12280092 | 0.104237838 | 1.178083917 | 0.238763153 | 0.442885358 | 0.12280092 |
| ENSMUSG00000030161 | Gabarapl1 | 57436 | protein_coding | 355.7864991 | -0.122853209 | 0.102802887 | -1.195036573 | 0.232072741 | 0.435543139 | 0.122853209 |
| ENSMUSG00000008348 | Ubc | 22190 | protein_coding | 1110.151635 | -0.122883424 | 0.194671997 | -0.631233181 | 0.527888069 | 0.713284795 | 0.122883424 |
| ENSMUSG00000015766 | Eps8 | 13860 | protein_coding | 2038.277913 | -0.122942626 | 0.203110259 | -0.605299934 | 0.544979725 | 0.727614171 | 0.122942626 |
| ENSMUSG00000035597 | Prpf39 | 328110 | protein_coding | 624.5144701 | -0.122977643 | 0.139303901 | -0.882801145 | 0.377343722 | 0.585345812 | 0.122977643 |
| ENSMUSG00000064262 | Gimap8 | 243374 | protein_coding | 44.02162786 | 0.123001357 | 0.242334662 | 0.507568153 | 0.61175623 | 0.773705989 | 0.123001357 |
| ENSMUSG00000021385 | Ippk | 75678 | protein_coding | 585.1465699 | -0.123038889 | 0.164691032 | -0.747089187 | 0.455009727 | 0.654392319 | 0.123038889 |
| ENSMUSG00000027342 | Pcna | 18538 | protein_coding | 1199.549411 | 0.123050407 | 0.096702299 | 1.272466194 | 0.203207528 | 0.398214752 | 0.123050407 |
| ENSMUSG00000028431 | Elp1 | 230233 | protein_coding | 385.6565433 | -0.123117291 | 0.130855721 | -0.940862886 | 0.346775129 | 0.556853943 | 0.123117291 |
| ENSMUSG00000032231 | Anxa2 | 12306 | protein_coding | 8617.07665 | 0.123129954 | 0.305742913 | 0.402723821 | 0.687151408 | 0.821322799 | 0.123129954 |
| ENSMUSG00000035376 | Hacd2 | 70757 | protein_coding | 279.7678247 | -0.123168598 | 0.102104493 | -1.206299487 | 0.227702035 | 0.430280937 | 0.123168598 |
| ENSMUSG00000040396 | Abhd13 | 68904 | protein_coding | 1501.371754 | 0.123233825 | 0.081619412 | 1.509859263 | 0.131079339 | 0.298432206 | 0.123233825 |
| ENSMUSG00000027938 | Creb3l4 | 78284 | protein_coding | 737.6233927 | 0.123270174 | 0.139435314 | 0.884067104 | 0.37665999 | 0.58462681 | 0.123270174 |
| ENSMUSG00000031851 | Ntpcr | 66566 | protein_coding | 193.7854396 | 0.123317164 | 0.137610635 | 0.896131056 | 0.370182772 | 0.578860317 | 0.123317164 |
| ENSMUSG00000104585 | 4921511C10Rik | 70924 | lncRNA | 7.612959077 | 0.123318072 | 0.470060949 | 0.262344857 | 0.793055582 | 0.888064491 | 0.123318072 |
| ENSMUSG00000025786 | Zdhhc3 | 69035 | protein_coding | 1588.628348 | 0.12335838 | 0.073550941 | 1.677182894 | 0.093506723 | 0.237985125 | 0.12335838 |
| ENSMUSG00000030619 | Eed | 13626 | protein_coding | 555.0438035 | -0.12336931 | 0.066890247 | -1.844354239 | 0.065131532 | 0.185783495 | 0.12336931 |
| ENSMUSG00000006127 | Inpp5k | 19062 | protein_coding | 353.7572829 | -0.123378257 | 0.124392367 | -0.991847491 | 0.321271927 | 0.532354566 | 0.123378257 |
| ENSMUSG00000032384 | Csnk1g1 | 214897 | protein_coding | 492.6978819 | -0.123443038 | 0.113613391 | -1.086518386 | 0.277249711 | 0.485615564 | 0.123443038 |
| ENSMUSG00000060261 | Gtf2i | 14886 | protein_coding | 823.758807 | -0.123454852 | 0.092815689 | -1.330107587 | 0.183482826 | 0.372769701 | 0.123454852 |
| ENSMUSG00000041995 | Zbed3 | 72114 | protein_coding | 104.5139565 | 0.123464905 | 0.146892827 | 0.840510104 | 0.400622439 | 0.608101981 | 0.123464905 |
| ENSMUSG00000049969 | Plekhf2 | 71801 | protein_coding | 1207.528766 | 0.123468189 | 0.090816888 | 1.359528962 | 0.173979032 | 0.359888054 | 0.123468189 |
| ENSMUSG00000029405 | G3bp2 | 23881 | protein_coding | 2288.447209 | 0.123472489 | 0.072701325 | 1.698352665 | 0.08944122 | 0.230894612 | 0.123472489 |
| ENSMUSG00000025980 | Hspd1 | 15510 | protein_coding | 4678.668315 | -0.123561553 | 0.142876684 | -0.864812571 | 0.38714167 | 0.595269092 | 0.123561553 |
| ENSMUSG00000044674 | Fzd1 | 14362 | protein_coding | 419.2460497 | 0.123653273 | 0.105893461 | 1.167713966 | 0.242922161 | 0.447865081 | 0.123653273 |
| ENSMUSG00000017286 | Glod4 | 67201 | protein_coding | 868.6639136 | 0.123693339 | 0.083079742 | 1.488850776 | 0.136526668 | 0.30684306 | 0.123693339 |
| ENSMUSG00000035305 | Ror1 | 26563 | protein_coding | 112.1158777 | -0.123702095 | 0.34263835 | -0.361028166 | 0.718078391 | 0.841491511 | 0.123702095 |
| ENSMUSG00000041737 | Tmem45b | 235135 | protein_coding | 3983.47491 | -0.123709048 | 0.091787399 | -1.347778112 | 0.17772976 | 0.364757069 | 0.123709048 |
| ENSMUSG00000091512 | Lamtor3 | 56692 | protein_coding | 403.7361652 | 0.123750418 | 0.118620017 | 1.04325072 | 0.296832188 | 0.506180721 | 0.123750418 |
| ENSMUSG00000031604 | Msmo1 | 66234 | protein_coding | 1922.068301 | 0.123794829 | 0.177072959 | 0.699117638 | 0.484478517 | 0.677861368 | 0.123794829 |
| ENSMUSG00000026087 | Mrpl30 | 107734 | protein_coding | 798.4732809 | 0.123833479 | 0.095354355 | 1.298666213 | 0.194058504 | 0.387387373 | 0.123833479 |
| ENSMUSG00000025495 | Ptdss2 | 27388 | protein_coding | 342.7136533 | -0.123835272 | 0.093929536 | -1.318384803 | 0.187374865 | 0.378024934 | 0.123835272 |
| ENSMUSG00000057880 | Abat | 268860 | protein_coding | 2924.311445 | -0.12386366 | 0.143501275 | -0.863153724 | 0.388052953 | 0.596497533 | 0.12386366 |
| ENSMUSG00000074398 | Gm15441 | 100038464 | lncRNA | 10.6145458 | 0.123871843 | 0.373801341 | 0.331384158 | 0.74035433 | 0.855177916 | 0.123871843 |

| ensembl_gene_id | external_gene_name | entrezgene_id | gene_biotype | baseMean | log2FoldChange | lfcSE | stat | pvalue | padj | absLog2FC |
|---|---|---|---|---|---|---|---|---|---|---|
| ENSMUSG00000023170 | Gps2 | 56310 | protein_coding | 261.4503005 | -0.12387966 | 0.096715935 | -1.280860908 | 0.200242526 | 0.394795779 | 0.12387966 |
| ENSMUSG00000056820 | Tsnax | 53424 | protein_coding | 825.5682189 | -0.123910401 | 0.069445105 | -1.784292804 | 0.074376099 | 0.203305733 | 0.123910401 |
| ENSMUSG00000026679 | Enkur | 71233 | protein_coding | 10.00640478 | -0.123922285 | 0.388399286 | -0.319058993 | 0.749681779 | 0.86153906 | 0.123922285 |
| ENSMUSG00000075271 | Ttc30a1 | 78802 | protein_coding | 77.53861354 | 0.123931906 | 0.197288617 | 0.628175652 | 0.529888879 | 0.715171253 | 0.123931906 |
| ENSMUSG00000032913 | Lrig2 | 269473 | protein_coding | 468.7247179 | -0.123968893 | 0.098385984 | -1.260025957 | 0.207659999 | 0.404082965 | 0.123968893 |
| ENSMUSG00000008333 | Snrpb2 | 20639 | protein_coding | 464.9482402 | 0.124063379 | 0.094322875 | 1.315305319 | 0.188407311 | 0.379627028 | 0.124063379 |
| ENSMUSG00000025962 | Fastkd2 | 75619 | protein_coding | 441.1657548 | 0.124082408 | 0.116117012 | 1.068598007 | 0.285250847 | 0.494285846 | 0.124082408 |
| ENSMUSG00000027663 | Zmat3 | 22401 | protein_coding | 212.5719197 | -0.124117267 | 0.163125678 | -0.760868975 | 0.446735331 | 0.64752864 | 0.124117267 |
| ENSMUSG00000038764 | Ptpn3 | 545622 | protein_coding | 1217.840859 | -0.124181821 | 0.077047901 | -1.611748276 | 0.107016728 | 0.260000372 | 0.124181821 |
| ENSMUSG00000020755 | Sap30bp | 57230 | protein_coding | 437.6338272 | -0.124237441 | 0.125936594 | -0.986507869 | 0.323883956 | 0.53475937 | 0.124237441 |
| ENSMUSG00000033728 | Lrrc14 | 223664 | protein_coding | 186.0869024 | -0.124237948 | 0.108763776 | -1.142273215 | 0.253340471 | 0.459486111 | 0.124237948 |
| ENSMUSG00000026049 | Tex30 | 75623 | protein_coding | 126.5831101 | 0.12424699 | 0.175188908 | 0.709217219 | 0.47818969 | 0.672607328 | 0.12424699 |
| ENSMUSG00000021417 | Eci2 | 23986 | protein_coding | 633.1215119 | 0.124276462 | 0.101785992 | 1.2209584 | 0.222101771 | 0.423234241 | 0.124276462 |
| ENSMUSG00000022668 | Gtpbp8 | 66067 | protein_coding | 357.8608417 | -0.124358235 | 0.104588706 | -1.189021646 | 0.234431149 | 0.438316627 | 0.124358235 |
| ENSMUSG00000042672 | Dcst1 | 77772 | protein_coding | 21.49122558 | -0.124364549 | 0.291381152 | -0.426810549 | 0.669517323 | 0.808680664 | 0.124364549 |
| ENSMUSG00000035798 | Zdhhc17 | 320150 | protein_coding | 401.2903399 | -0.124386142 | 0.097826857 | -1.271492778 | 0.203553395 | 0.398696321 | 0.124386142 |
| ENSMUSG00000005873 | Reep5 | 13476 | protein_coding | 620.2464289 | -0.124480579 | 0.101060512 | -1.231743003 | 0.218045101 | 0.417839381 | 0.124480579 |
| ENSMUSG00000016495 | Plgrkt | 67759 | protein_coding | 1293.326593 | 0.124483341 | 0.086861274 | 1.433128197 | 0.151821205 | 0.330085742 | 0.124483341 |
| ENSMUSG00000034379 | Wdr5b | 69544 | protein_coding | 66.77230133 | -0.124533211 | 0.176707085 | -0.704743735 | 0.480969729 | 0.674232796 | 0.124533211 |
| ENSMUSG00000033022 | Cdo1 | 12583 | protein_coding | 538.1894398 | 0.124627204 | 0.118046574 | 1.055746053 | 0.291084251 | 0.499573396 | 0.124627204 |
| ENSMUSG00000055415 | Atp10b | 319767 | protein_coding | 3273.820879 | 0.124641723 | 0.110095233 | 1.132126425 | 0.257581287 | 0.464532918 | 0.124641723 |
| ENSMUSG00000029276 | Glmn | 170823 | protein_coding | 97.79190937 | -0.124656124 | 0.142045311 | -0.877579994 | 0.38017169 | 0.588071737 | 0.124656124 |
| ENSMUSG00000035547 | Capn5 | 12337 | protein_coding | 1364.938595 | 0.124665124 | 0.109865495 | 1.134706805 | 0.256498185 | 0.463387217 | 0.124665124 |
| ENSMUSG00000014668 | Chfr | 231600 | protein_coding | 847.3855295 | 0.124679378 | 0.053420353 | 2.333930267 | 0.019599374 | 0.078614662 | 0.124679378 |
| ENSMUSG00000029427 | Zcchc8 | 70650 | protein_coding | 486.7828125 | -0.124687508 | 0.086685803 | -1.438384409 | 0.150325014 | 0.328041762 | 0.124687508 |
| ENSMUSG00000026094 | Stk17b | 98267 | protein_coding | 1096.722215 | -0.12469745 | 0.134037692 | -0.930316303 | 0.352207339 | 0.561894882 | 0.12469745 |
| ENSMUSG00000011306 | Sugp1 | 70616 | protein_coding | 355.150058 | -0.124755198 | 0.085823515 | -1.453624892 | 0.146050332 | 0.322066965 | 0.124755198 |
| ENSMUSG00000030788 | Rnf141 | 67150 | protein_coding | 1336.581421 | 0.124850237 | 0.079660666 | 1.567275844 | 0.117050229 | 0.27604028 | 0.124850237 |
| ENSMUSG00000024969 | Mark2 | 13728 | protein_coding | 1122.417211 | -0.124887292 | 0.117666127 | -1.061369954 | 0.288521806 | 0.497464313 | 0.124887292 |
| ENSMUSG00000078429 | Ctdsp2 | 52468 | protein_coding | 1677.966957 | -0.124905627 | 0.097535991 | -1.280610632 | 0.200330464 | 0.394859996 | 0.124905627 |
| ENSMUSG00000028068 | Iqgap3 | 404710 | protein_coding | 400.4957991 | -0.124907242 | 0.135843868 | -0.91949121 | 0.357838701 | 0.567468621 | 0.124907242 |
| ENSMUSG00000018574 | Acadvl | 11370 | protein_coding | 1671.562119 | -0.124937839 | 0.109633398 | -1.13959652 | 0.254454436 | 0.46097006 | 0.124937839 |
| ENSMUSG00000027359 | Slc27a2 | 26458 | protein_coding | 119.1018586 | -0.124995504 | 0.17013217 | -0.734696466 | 0.462524373 | 0.660310855 | 0.124995504 |
| ENSMUSG00000001521 | Tulp3 | 22158 | protein_coding | 29.05873991 | 0.124998883 | 0.283665854 | 0.440655373 | 0.659462509 | 0.802969027 | 0.124998883 |
| ENSMUSG00000029559 | 2210016L21Rik | 72357 | protein_coding | 319.8419186 | -0.125007275 | 0.074506484 | -1.677803973 | 0.093385374 | 0.237876576 | 0.125007275 |
| ENSMUSG00000026784 | Pdss1 | 56075 | protein_coding | 524.0582555 | 0.125019975 | 0.069073689 | 1.809950738 | 0.070303427 | 0.196109617 | 0.125019975 |
| ENSMUSG00000020849 | Ywhae | 22627 | protein_coding | 2431.264395 | 0.125086435 | 0.115994469 | 1.078382753 | 0.28086298 | 0.489344296 | 0.125086435 |
| ENSMUSG00000039191 | Rbpj | 19664 | protein_coding | 330.5547475 | -0.125115463 | 0.082358989 | -1.519147622 | 0.128725342 | 0.294927682 | 0.125115463 |

| ensembl_gene_id | external_gene_name | entrezgene_id | gene_biotype | baseMean | log2FoldChange | lfcSE | stat | pvalue | padj | absLog2FC |
|---|---|---|---|---|---|---|---|---|---|---|
| ENSMUSG00000006276 | Eps15l1 | 13859 | protein_coding | 667.5630039 | -0.125122484 | 0.093013133 | -1.345213091 | 0.17855643 | 0.365935 | 0.125122484 |
| ENSMUSG00000032305 | Fam219b | 78323 | protein_coding | 169.3820176 | -0.125170055 | 0.109381972 | -1.144338994 | 0.252483078 | 0.458766879 | 0.125170055 |
| ENSMUSG00000026571 | Dcaf6 | 74106 | protein_coding | 440.7828858 | -0.125172154 | 0.071533918 | -1.749829413 | 0.080147754 | 0.213992665 | 0.125172154 |
| ENSMUSG00000037461 | Ints7 | 77065 | protein_coding | 455.6823745 | -0.125249244 | 0.08542319 | -1.466220643 | 0.142588186 | 0.316152646 | 0.125249244 |
| ENSMUSG00000047388 | Atmin | 234776 | protein_coding | 434.5999345 | -0.125372337 | 0.083025844 | -1.510039899 | 0.131033244 | 0.298386829 | 0.125372337 |
| ENSMUSG00000024683 | Mrpl16 | 94063 | protein_coding | 351.2679563 | 0.125419096 | 0.104131763 | 1.204426893 | 0.228424623 | 0.431373157 | 0.125419096 |
| ENSMUSG00000029068 | Ccnl2 | 56036 | protein_coding | 1506.455092 | -0.125439701 | 0.134454199 | -0.932954882 | 0.350843264 | 0.560729632 | 0.125439701 |
| ENSMUSG00000024251 | Thada | 240174 | protein_coding | 273.0736856 | -0.125466828 | 0.10819744 | -1.15960995 | 0.246207648 | 0.451692631 | 0.125466828 |
| ENSMUSG00000059142 | Zfp945 | 240041 | protein_coding | 226.37776 | -0.125526531 | 0.158435696 | -0.792286931 | 0.42819339 | 0.632390628 | 0.125526531 |
| ENSMUSG00000022141 | Nipbl | 71175 | protein_coding | 1803.963878 | -0.125601093 | 0.103299047 | -1.21589788 | 0.224023826 | 0.425704068 | 0.125601093 |
| ENSMUSG00000046822 | Slc39a3 | 106947 | protein_coding | 361.340042 | 0.125605086 | 0.131491914 | 0.955230496 | 0.339461147 | 0.550085499 | 0.125605086 |
| ENSMUSG00000045934 | Mtmr11 | 194126 | protein_coding | 629.1304842 | 0.125674269 | 0.180194757 | 0.697435768 | 0.485530124 | 0.678688781 | 0.125674269 |
| ENSMUSG00000018666 | Cbx1 | 12412 | protein_coding | 775.1324597 | 0.125702982 | 0.078137277 | 1.608745369 | 0.107672031 | 0.260995299 | 0.125702982 |
| ENSMUSG00000004085 | Map3k20 | 65964 | protein_coding | 447.3898133 | -0.125720012 | 0.072829041 | -1.726234619 | 0.084305212 | 0.222026028 | 0.125720012 |
| ENSMUSG00000037997 | Parp11 | 101187 | protein_coding | 144.4007911 | 0.125770764 | 0.102317604 | 1.229219203 | 0.218989631 | 0.41918842 | 0.125770764 |
| ENSMUSG00000031068 | Glrx3 | 30926 | protein_coding | 1005.157133 | 0.125782123 | 0.088631078 | 1.419164986 | 0.155850921 | 0.336179961 | 0.125782123 |
| ENSMUSG00000022898 | Vps26c | 13185 | protein_coding | 591.366379 | 0.125786758 | 0.080822165 | 1.556339873 | 0.119627295 | 0.280498094 | 0.125786758 |
| ENSMUSG00000053536 | Cstf2t | 83410 | protein_coding | 869.0233074 | 0.125786913 | 0.087598546 | 1.435947495 | 0.151017282 | 0.329101556 | 0.125786913 |
| ENSMUSG00000058773 | H1f5 | 56702 | protein_coding | 1542.696653 | 0.125788083 | 0.145393792 | 0.86515443 | 0.386954034 | 0.595095476 | 0.125788083 |
| ENSMUSG00000027593 | Raly | 19383 | protein_coding | 1519.308123 | 0.125888724 | 0.099666516 | 1.263099468 | 0.206553435 | 0.402789314 | 0.125888724 |
| ENSMUSG00000063015 | Ccni | 12453 | protein_coding | 2714.168782 | -0.125892463 | 0.097405383 | -1.292458984 | 0.196198241 | 0.389705149 | 0.125892463 |
| ENSMUSG00000029076 | Sdf4 | 20318 | protein_coding | 2498.363139 | 0.12589626 | 0.07519387 | 1.674288879 | 0.094073839 | 0.238828197 | 0.12589626 |
| ENSMUSG00000036686 | Cc2d1a | 212139 | protein_coding | 617.2072604 | -0.125897157 | 0.096981517 | -1.298156196 | 0.194233667 | 0.387494219 | 0.125897157 |
| ENSMUSG00000026705 | Klhl20 | 226541 | protein_coding | 174.2401528 | -0.125914479 | 0.116502036 | -1.080792097 | 0.279789605 | 0.488114877 | 0.125914479 |
| ENSMUSG00000025235 | Bbs4 | 102774 | protein_coding | 85.93071718 | -0.125982257 | 0.131157507 | -0.960541713 | 0.336782648 | 0.547486631 | 0.125982257 |
| ENSMUSG00000035949 | Fbxw2 | 30050 | protein_coding | 654.3752263 | 0.126042191 | 0.065454221 | 1.925654129 | 0.054147575 | 0.163841675 | 0.126042191 |
| ENSMUSG00000039967 | Zfp292 | 30046 | protein_coding | 1261.628129 | -0.126075051 | 0.116983893 | -1.077712899 | 0.2811619 | 0.489758011 | 0.126075051 |
| ENSMUSG00000016831 | Tox4 | 268741 | protein_coding | 1069.714763 | 0.12615761 | 0.090940482 | 1.387254681 | 0.165364117 | 0.348987948 | 0.12615761 |
| ENSMUSG00000029627 | Zkscan14 | 67235 | protein_coding | 141.9764893 | 0.126236375 | 0.23332797 | 0.541025474 | 0.588490025 | 0.758309935 | 0.126236375 |
| ENSMUSG00000017843 | Ppp2r5c | 26931 | protein_coding | 2004.723901 | 0.126248177 | 0.069160152 | 1.82544679 | 0.067933634 | 0.191580545 | 0.126248177 |
| ENSMUSG00000028394 | Pole3 | 59001 | protein_coding | 441.7576871 | 0.126456178 | 0.089230919 | 1.417178923 | 0.156430622 | 0.337070746 | 0.126456178 |
| ENSMUSG00000021456 | Fbp2 | 14120 | protein_coding | 2068.011246 | 0.126495534 | 0.095120418 | 1.329846272 | 0.183568927 | 0.372897062 | 0.126495534 |
| ENSMUSG00000060244 | Alyref2 | 56009 | protein_coding | 32.78194281 | 0.126506968 | 0.242228961 | 0.522261946 | 0.601487961 | 0.767360852 | 0.126506968 |
| ENSMUSG00000030610 | Det1 | 76375 | protein_coding | 170.4823358 | 0.126526406 | 0.104556443 | 1.210125389 | 0.226230782 | 0.428568494 | 0.126526406 |
| ENSMUSG00000020432 | Tcn2 | 21452 | protein_coding | 1013.177769 | -0.126528336 | 0.08515085 | -1.48593158 | 0.137297219 | 0.308157609 | 0.126528336 |
| ENSMUSG00000024941 | Scyl1 | 78891 | protein_coding | 726.6060165 | -0.126573436 | 0.089725838 | -1.410668753 | 0.158342309 | 0.339625513 | 0.126573436 |
| ENSMUSG00000034575 | Tent4a | 210106 | protein_coding | 376.2803981 | -0.1266313 | 0.120718387 | -1.048981042 | 0.294186844 | 0.503417281 | 0.1266313 |
| ENSMUSG00000039308 | Ndst2 | 17423 | protein_coding | 252.5864669 | -0.12665649 | 0.100310374 | -1.26264597 | 0.206716439 | 0.403050253 | 0.12665649 |

| ensembl_gene_id | external_gene_name | entrezgene_id | gene_biotype | baseMean | log2FoldChange | lfcSE | stat | pvalue | padj | absLog2FC |
|---|---|---|---|---|---|---|---|---|---|---|
| ENSMUSG00000058748 | Zfp958 | 233987 | protein_coding | 271.0205831 | -0.126664748 | 0.13127576 | -0.964875373 | 0.334607238 | 0.545293316 | 0.126664748 |
| ENSMUSG00000021711 | Trappc13 | 66975 | protein_coding | 776.9060385 | 0.126773764 | 0.076200329 | 1.663690503 | 0.096174295 | 0.242153679 | 0.126773764 |
| ENSMUSG00000016344 | Ppdpf | 66496 | protein_coding | 509.3699232 | -0.126805372 | 0.10748248 | -1.179777137 | 0.238088865 | 0.442256933 | 0.126805372 |
| ENSMUSG00000034636 | Zyg11b | 414872 | protein_coding | 1231.248324 | 0.126830473 | 0.075029629 | 1.690405178 | 0.090950465 | 0.233315994 | 0.126830473 |
| ENSMUSG00000034681 | Rnps1 | 19826 | protein_coding | 523.7977422 | -0.126862709 | 0.115704379 | -1.09643827 | 0.272887024 | 0.480918845 | 0.126862709 |
| ENSMUSG00000029402 | Snrnp35 | 76167 | protein_coding | 92.46773832 | 0.126867112 | 0.123254297 | 1.029311876 | 0.303333145 | 0.51284262 | 0.126867112 |
| ENSMUSG00000049339 | Retreg2 | 227298 | protein_coding | 915.18643 | -0.126868035 | 0.091455869 | -1.387204968 | 0.165379271 | 0.348987948 | 0.126868035 |
| ENSMUSG00000062901 | Klhl24 | 75785 | protein_coding | 1308.205098 | -0.126904709 | 0.328529969 | -0.386280463 | 0.699288968 | 0.829099726 | 0.126904709 |
| ENSMUSG00000030413 | Pglyrp1 | 21946 | protein_coding | 924.3855726 | 0.126930394 | 0.104467212 | 1.215026142 | 0.224356121 | 0.425999491 | 0.126930394 |
| ENSMUSG00000030157 | Clec2d | 93694 | protein_coding | 1806.116454 | 0.126930564 | 0.145765218 | 0.870787733 | 0.383870064 | 0.591552088 | 0.126930564 |
| ENSMUSG00000022219 | Cideb | 12684 | protein_coding | 67.61759584 | 0.126944024 | 0.168205893 | 0.754694274 | 0.450432443 | 0.650983391 | 0.126944024 |
| ENSMUSG00000044147 | Arf6 | 11845 | protein_coding | 2400.638775 | 0.127041698 | 0.149814879 | 0.84799119 | 0.396442877 | 0.604609551 | 0.127041698 |
| ENSMUSG00000032261 | Sh3bgrl2 | 212531 | protein_coding | 3822.109462 | 0.12704177 | 0.062272027 | 2.040109754 | 0.041339395 | 0.136100432 | 0.12704177 |
| ENSMUSG00000021254 | Gpatch2l | 70373 | protein_coding | 243.8321479 | -0.127082495 | 0.088137591 | -1.441864851 | 0.149340504 | 0.326699017 | 0.127082495 |
| ENSMUSG00000042423 | Fbrs | 14123 | protein_coding | 306.2136319 | -0.127104185 | 0.157128544 | -0.808918494 | 0.41856203 | 0.624763941 | 0.127104185 |
| ENSMUSG00000018446 | C1qbp | 12261 | protein_coding | 688.8289849 | -0.127144292 | 0.1077396 | -1.180107327 | 0.237957531 | 0.442159071 | 0.127144292 |
| ENSMUSG00000032561 | Acpp | 56318 | protein_coding | 527.4965868 | -0.127149084 | 0.287805941 | -0.44178756 | 0.658642941 | 0.802544735 | 0.127149084 |
| ENSMUSG00000026116 | Tmem131 | 56030 | protein_coding | 1740.700538 | -0.12716711 | 0.161363664 | -0.788077725 | 0.430651244 | 0.634649642 | 0.12716711 |
| ENSMUSG00000055371 | Stam2 | 56324 | protein_coding | 1268.960784 | 0.127178326 | 0.059784609 | 2.12727537 | 0.03339721 | 0.116747735 | 0.127178326 |
| ENSMUSG00000052798 | Nup107 | 103468 | protein_coding | 277.921536 | 0.127232155 | 0.115421315 | 1.102328066 | 0.270319072 | 0.478375791 | 0.127232155 |
| ENSMUSG00000019990 | Pde7b | 29863 | protein_coding | 22.63329216 | 0.127241495 | 0.418914652 | 0.303740855 | 0.76132533 | 0.869186728 | 0.127241495 |
| ENSMUSG00000027935 | Rab13 | 68328 | protein_coding | 162.99773 | -0.127247958 | 0.110855315 | -1.147874221 | 0.251020492 | 0.457413842 | 0.127247958 |
| ENSMUSG00000030421 | Uri1 | 19777 | protein_coding | 570.5550158 | 0.127257944 | 0.10252778 | 1.241204528 | 0.214530202 | 0.41303482 | 0.127257944 |
| ENSMUSG00000022946 | Dop1b | 70028 | protein_coding | 1160.445826 | -0.127319571 | 0.118755794 | -1.072112504 | 0.283669503 | 0.492778694 | 0.127319571 |
| ENSMUSG00000097767 | Miat | 330166 | lncRNA | 8.66397306 | 0.12733475 | 0.554691505 | 0.229559582 | 0.818434017 | 0.903579383 | 0.12733475 |
| ENSMUSG00000030611 | Mrps11 | 67994 | protein_coding | 198.5401318 | 0.127339486 | 0.104622953 | 1.217127626 | 0.223555661 | 0.425169501 | 0.127339486 |
| ENSMUSG00000029534 | St7 | 64213 | protein_coding | 157.4135779 | 0.127346018 | 0.111327633 | 1.143885084 | 0.252671298 | 0.459036748 | 0.127346018 |
| ENSMUSG00000046230 | Vps13a | 271564 | protein_coding | 1602.787068 | 0.127445492 | 0.083533328 | 1.525684368 | 0.127088489 | 0.292313212 | 0.127445492 |
| ENSMUSG00000094840 | Muc3a | 619309 | protein_coding | 307.8347859 | -0.127461993 | 0.23040163 | -0.553216542 | 0.580115141 | 0.751318253 | 0.127461993 |
| ENSMUSG00000036327 | Qsox2 | 227638 | protein_coding | 431.5874861 | 0.127476378 | 0.103111829 | 1.236292472 | 0.216349869 | 0.41558378 | 0.127476378 |
| ENSMUSG00000026971 | Itgb6 | 16420 | protein_coding | 630.3962989 | 0.127496926 | 0.185812362 | 0.686159548 | 0.492612504 | 0.684312922 | 0.127496926 |
| ENSMUSG00000043336 | Filip1l | 78749 | protein_coding | 505.9659232 | -0.12751202 | 0.109123807 | -1.168507802 | 0.242601994 | 0.447621897 | 0.12751202 |
| ENSMUSG00000023206 | Il15ra | 16169 | protein_coding | 108.6630195 | -0.127558212 | 0.19179557 | -0.66507382 | 0.506003278 | 0.695934739 | 0.127558212 |
| ENSMUSG00000003458 | Ncstn | 59287 | protein_coding | 1663.70758 | 0.127601066 | 0.062556492 | 2.039773367 | 0.041372904 | 0.136154465 | 0.127601066 |
| ENSMUSG00000050379 | Septin6 | 56526 | protein_coding | 90.5981792 | -0.127667206 | 0.144575511 | -0.883048624 | 0.377210001 | 0.585247665 | 0.127667206 |
| ENSMUSG00000038884 | Shfl | 319278 | protein_coding | 25.10619837 | -0.127667243 | 0.327668316 | -0.389623399 | 0.696815047 | 0.827409428 | 0.127667243 |
| ENSMUSG00000027107 | Chrna1 | 11435 | protein_coding | 36.73803185 | -0.12767603 | 0.235310634 | -0.542585043 | 0.587415539 | 0.757354275 | 0.12767603 |
| ENSMUSG00000031516 | Dctn6 | 22428 | protein_coding | 548.0865549 | 0.127693567 | 0.076103454 | 1.677894506 | 0.093367695 | 0.237869586 | 0.127693567 |

| ensembl_gene_id | external_gene_name | entrezgene_id | gene_biotype | baseMean | log2FoldChange | lfcSE | stat | pvalue | padj | absLog2FC |
|---|---|---|---|---|---|---|---|---|---|---|
| ENSMUSG00000046519 | Golph3l | 229593 | protein_coding | 5930.253833 | 0.127750528 | 0.128904527 | 0.991047644 | 0.321662317 | 0.532526492 | 0.127750528 |
| ENSMUSG00000038276 | Asic3 | 171209 | protein_coding | 8.55468655 | -0.127754874 | 0.48004175 | -0.266132839 | 0.790136905 | 0.886288776 | 0.127754874 |
| ENSMUSG00000053199 | Arhgap20 | 244867 | protein_coding | 548.2674205 | -0.127814716 | 0.252253444 | -0.506691658 | 0.612371185 | 0.774101099 | 0.127814716 |
| ENSMUSG00000022201 | Zfr | 22763 | protein_coding | 1299.635463 | 0.12782183 | 0.077029194 | 1.659394618 | 0.0970363 | 0.243592464 | 0.12782183 |
| ENSMUSG00000020128 | Vps54 | 245944 | protein_coding | 1972.753472 | 0.127836812 | 0.063914006 | 2.000137669 | 0.0454854 | 0.145743604 | 0.127836812 |
| ENSMUSG00000038949 | Cnst | 226744 | protein_coding | 652.8594666 | -0.127859312 | 0.095585896 | -1.337637846 | 0.181014521 | 0.369358835 | 0.127859312 |
| ENSMUSG00000030347 | D6Wsu163e | 28040 | protein_coding | 239.912695 | 0.127862978 | 0.131478241 | 0.972502952 | 0.330800397 | 0.541464261 | 0.127862978 |
| ENSMUSG00000031948 | Kars | 85305 | protein_coding | 1548.134071 | 0.127889197 | 0.085738385 | 1.491621246 | 0.135798465 | 0.30556627 | 0.127889197 |
| ENSMUSG00000024730 | Ms4a8a | 64381 | protein_coding | 4183.646406 | 0.127900304 | 0.127551463 | 1.0027349 | 0.315988786 | 0.526912624 | 0.127900304 |
| ENSMUSG00000043333 | Rhbdl2 | 230726 | protein_coding | 546.9097377 | 0.127928284 | 0.129324402 | 0.989204531 | 0.322563082 | 0.533463272 | 0.127928284 |
| ENSMUSG00000022214 | Dcaf11 | 28199 | protein_coding | 1350.611101 | 0.127953334 | 0.070671587 | 1.810534339 | 0.070212964 | 0.19599581 | 0.127953334 |
| ENSMUSG00000026103 | Gls | 14660 | protein_coding | 2106.745094 | -0.128043366 | 0.090567656 | -1.413786911 | 0.157424478 | 0.33818806 | 0.128043366 |
| ENSMUSG00000032850 | Rnft2 | 269695 | protein_coding | 9.523356498 | -0.128082575 | 0.453397314 | -0.282495223 | 0.777563809 | 0.879181966 | 0.128082575 |
| ENSMUSG00000029330 | Cds1 | 74596 | protein_coding | 1912.571249 | 0.128099515 | 0.107764822 | 1.188695092 | 0.234559673 | 0.438420238 | 0.128099515 |
| ENSMUSG00000090150 | Acad11 | 102632 | protein_coding | 1636.794124 | 0.128105135 | 0.109277109 | 1.17229616 | 0.241078173 | 0.445618329 | 0.128105135 |
| ENSMUSG00000030655 | Smg1 | 233789 | protein_coding | 2906.655678 | 0.12811342 | 0.129301882 | 0.990808628 | 0.321779036 | 0.532664361 | 0.12811342 |
| ENSMUSG00000061981 | Flot2 | 14252 | protein_coding | 169.5208131 | -0.128163033 | 0.133716535 | -0.9584681 | 0.33782677 | 0.548511589 | 0.128163033 |
| ENSMUSG00000020277 | Pfkl | 18641 | protein_coding | 3128.540136 | -0.1281828 | 0.131562424 | -0.974311633 | 0.329901831 | 0.540660187 | 0.1281828 |
| ENSMUSG00000029635 | Cdk8 | 264064 | protein_coding | 630.0749897 | -0.128227348 | 0.072764891 | -1.762214537 | 0.078033057 | 0.210143295 | 0.128227348 |
| ENSMUSG00000039953 | Clstn1 | 65945 | protein_coding | 2983.104298 | -0.128270766 | 0.074578814 | -1.71993571 | 0.085444129 | 0.223877953 | 0.128270766 |
| ENSMUSG00000027288 | Zfp106 | 20402 | protein_coding | 2431.375391 | 0.128279892 | 0.083616791 | 1.534140343 | 0.12499513 | 0.28916811 | 0.128279892 |
| ENSMUSG00000033596 | Rfwd3 | 234736 | protein_coding | 851.6339856 | 0.128338139 | 0.101709149 | 1.261815089 | 0.207015331 | 0.403443069 | 0.128338139 |
| ENSMUSG00000058709 | Egln2 | 112406 | protein_coding | 883.4577306 | 0.128348233 | 0.074617839 | 1.720074371 | 0.085418924 | 0.223848736 | 0.128348233 |
| ENSMUSG00000072872 | Rybp | 56353 | protein_coding | 306.4971841 | 0.128420665 | 0.138631242 | 0.92634721 | 0.354265569 | 0.563829056 | 0.128420665 |
| ENSMUSG00000030213 | Atf7ip | 54343 | protein_coding | 866.8266454 | -0.128425504 | 0.16353863 | -0.785291553 | 0.432282648 | 0.636123246 | 0.128425504 |
| ENSMUSG00000019792 | Trmt11 | 73681 | protein_coding | 252.6047809 | 0.128505318 | 0.11299271 | 1.137288574 | 0.25541767 | 0.462147863 | 0.128505318 |
| ENSMUSG00000030980 | Knop1 | 66356 | protein_coding | 531.7641493 | -0.128546695 | 0.096606039 | -1.330627935 | 0.183311466 | 0.372564123 | 0.128546695 |
| ENSMUSG00000003226 | Ranbp2 | 19386 | protein_coding | 3460.30557 | -0.128590785 | 0.102851954 | -1.250251252 | 0.21120778 | 0.40867423 | 0.128590785 |
| ENSMUSG00000037363 | Letm2 | 270035 | protein_coding | 24.80778027 | -0.128603444 | 0.239855329 | -0.536170887 | 0.591840452 | 0.761020102 | 0.128603444 |
| ENSMUSG00000026887 | Mrrf | 67871 | protein_coding | 217.3731217 | -0.128627276 | 0.132169693 | -0.973197956 | 0.330454927 | 0.541009977 | 0.128627276 |
| ENSMUSG00000023977 | Ubr2 | 224826 | protein_coding | 1073.552443 | -0.128676707 | 0.098139955 | -1.311155147 | 0.189805353 | 0.381382573 | 0.128676707 |
| ENSMUSG00000034593 | Myo5a | 17918 | protein_coding | 35.70228993 | 0.128682501 | 0.229354467 | 0.561063853 | 0.574754009 | 0.747748252 | 0.128682501 |
| ENSMUSG00000054619 | Mettl7a1 | 70152 | protein_coding | 2316.17067 | -0.128686256 | 0.160972198 | -0.799431559 | 0.424040217 | 0.628617629 | 0.128686256 |
| ENSMUSG00000098332 | Pigbos1 | 66364 | protein_coding | 160.7261643 | 0.128689347 | 0.142181497 | 0.905106145 | 0.365409157 | 0.574271046 | 0.128689347 |
| ENSMUSG00000024330 | Col11a2 | 12815 | protein_coding | 46.59426077 | 0.128734307 | 0.348901488 | 0.368970359 | 0.712149818 | 0.837810939 | 0.128734307 |
| ENSMUSG00000024826 | Dpf2 | 19708 | protein_coding | 468.8361079 | -0.128811866 | 0.057607651 | -2.236020132 | 0.025350452 | 0.095524487 | 0.128811866 |
| ENSMUSG00000018565 | Elp5 | 54351 | protein_coding | 468.3422836 | 0.128834384 | 0.066773286 | 1.929430045 | 0.053677497 | 0.162790398 | 0.128834384 |
| ENSMUSG00000017724 | Etv4 | 18612 | protein_coding | 153.8676746 | 0.128867734 | 0.24881952 | 0.517916496 | 0.604516532 | 0.769130068 | 0.128867734 |

| ensembl_gene_id | external_gene_name | entrezgene_id | gene_biotype | baseMean | log2FoldChange | lfcSE | stat | pvalue | padj | absLog2FC |
|---|---|---|---|---|---|---|---|---|---|---|
| ENSMUSG00000089682 | Bcl2l2 | 12050 | protein_coding | 668.7754085 | 0.128872669 | 0.077304246 | 1.667083952 | 0.095497713 | 0.241098045 | 0.128872669 |
| ENSMUSG00000032562 | Gnai2 | 14678 | protein_coding | 1503.331482 | 0.128926419 | 0.089870343 | 1.434582481 | 0.15140611 | 0.329588153 | 0.128926419 |
| ENSMUSG00000031634 | Ufsp2 | 192169 | protein_coding | 698.4219225 | 0.128938806 | 0.070279703 | 1.834652129 | 0.06655725 | 0.188667907 | 0.128938806 |
| ENSMUSG00000097073 | 9430037G07Rik | 320692 | lncRNA | 15.72361596 | 0.128995537 | 0.312267531 | 0.413093019 | 0.679538464 | 0.816754221 | 0.128995537 |
| ENSMUSG00000100164 | 2610306M01Rik | 67170 | lncRNA | 35.0746445 | 0.129007973 | 0.200279893 | 0.644138417 | 0.51948568 | 0.706398276 | 0.129007973 |
| ENSMUSG00000018372 | Cep95 | 320162 | protein_coding | 208.8249017 | -0.129010979 | 0.123706353 | -1.042880789 | 0.297003508 | 0.506180721 | 0.129010979 |
| ENSMUSG00000060093 | H4c1 | 326619 | protein_coding | 680.1939302 | 0.129012471 | 0.139782058 | 0.922954439 | 0.356030949 | 0.565615496 | 0.129012471 |
| ENSMUSG00000096433 | Zfp994 | 240038 | protein_coding | 375.7070648 | 0.129022574 | 0.144042266 | 0.89572719 | 0.370398485 | 0.578970272 | 0.129022574 |
| ENSMUSG00000029209 | Gnpda2 | 67980 | protein_coding | 120.1067827 | 0.129032602 | 0.117050431 | 1.102367594 | 0.270301894 | 0.478375791 | 0.129032602 |
| ENSMUSG00000053604 | Rpia | 19895 | protein_coding | 417.1735048 | -0.12904923 | 0.102807967 | -1.255245424 | 0.20938968 | 0.406330266 | 0.12904923 |
| ENSMUSG00000001995 | Sipa1l2 | 244668 | protein_coding | 311.1404186 | 0.129091801 | 0.112282027 | 1.14971028 | 0.250263219 | 0.456634994 | 0.129091801 |
| ENSMUSG00000039585 | Myo9a | 270163 | protein_coding | 352.1793852 | -0.129124004 | 0.166456408 | -0.775722634 | 0.437912773 | 0.640519699 | 0.129124004 |
| ENSMUSG00000026638 | Irf6 | 54139 | protein_coding | 2990.581978 | -0.129129093 | 0.068130537 | -1.89531887 | 0.05805017 | 0.172151015 | 0.129129093 |
| ENSMUSG00000078695 | Cisd3 | 217149 | protein_coding | 1102.017621 | -0.129132278 | 0.07960984 | -1.622064287 | 0.104789584 | 0.256270305 | 0.129132278 |
| ENSMUSG00000064090 | Vrk2 | 69922 | protein_coding | 134.6337316 | 0.129139747 | 0.126599989 | 1.020061285 | 0.307699397 | 0.517614285 | 0.129139747 |
| ENSMUSG00000032058 | Ppp2r1b | 73699 | protein_coding | 492.558153 | -0.12915472 | 0.108655434 | -1.188663237 | 0.234572213 | 0.438420238 | 0.12915472 |
| ENSMUSG00000061451 | Tmem151a | 381199 | protein_coding | 11.21620206 | 0.129161969 | 0.428359068 | 0.301527336 | 0.763012408 | 0.870097644 | 0.129161969 |
| ENSMUSG00000057672 | Pkn1 | 320795 | protein_coding | 808.3982858 | -0.129203571 | 0.073192181 | -1.765264653 | 0.077519297 | 0.209204908 | 0.129203571 |
| ENSMUSG00000029638 | Glcci1 | 170772 | protein_coding | 29.07660286 | -0.129224955 | 0.278448824 | -0.464088707 | 0.642584192 | 0.793052742 | 0.129224955 |
| ENSMUSG00000078941 | Ak6 | 108143 | protein_coding | 201.1905812 | 0.129261067 | 0.138171834 | 0.935509526 | 0.349525776 | 0.559747292 | 0.129261067 |
| ENSMUSG00000041272 | Tox | 252838 | protein_coding | 262.0434931 | -0.12928384 | 0.099636209 | -1.297558799 | 0.194438988 | 0.387709589 | 0.12928384 |
| ENSMUSG00000044783 | Hjurp | 212427 | protein_coding | 2125.07767 | -0.129292336 | 0.065031583 | -1.98814685 | 0.046795451 | 0.148655354 | 0.129292336 |
| ENSMUSG00000015016 | Acsf3 | 257633 | protein_coding | 339.8256639 | -0.129328289 | 0.072228482 | -1.790544189 | 0.07336647 | 0.201653345 | 0.129328289 |
| ENSMUSG00000020697 | Lig3 | 16882 | protein_coding | 490.1137982 | -0.129333446 | 0.082621057 | -1.565381164 | 0.117493562 | 0.276798914 | 0.129333446 |
| ENSMUSG00000015023 | Ddx19a | 13680 | protein_coding | 359.9151678 | 0.129348447 | 0.078260486 | 1.652793803 | 0.098372836 | 0.245631373 | 0.129348447 |
| ENSMUSG00000005510 | Ndufs3 | 68349 | protein_coding | 1215.882561 | 0.129356637 | 0.094118938 | 1.374395413 | 0.169318959 | 0.354093686 | 0.129356637 |
| ENSMUSG00000022545 | Ercc4 | 50505 | protein_coding | 202.5999029 | -0.129359455 | 0.113439779 | -1.140335921 | 0.254146378 | 0.46058067 | 0.129359455 |
| ENSMUSG00000038954 | Supt3 | 109115 | protein_coding | 46.83554539 | 0.129365522 | 0.185164846 | 0.698650551 | 0.484770444 | 0.678210244 | 0.129365522 |
| ENSMUSG00000035898 | Uba6 | 231380 | protein_coding | 562.5764116 | 0.129366856 | 0.134211185 | 0.963905181 | 0.335093466 | 0.545687427 | 0.129366856 |
| ENSMUSG00000021033 | Gstz1 | 14874 | protein_coding | 462.4678445 | -0.12937143 | 0.081636244 | -1.584730312 | 0.113027622 | 0.268862599 | 0.12937143 |
| ENSMUSG00000029761 | Cald1 | 109624 | protein_coding | 27.01692585 | -0.129397737 | 0.371569328 | -0.34824655 | 0.727655031 | 0.846662573 | 0.129397737 |
| ENSMUSG00000037896 | Rcor1 | 217864 | protein_coding | 1156.807631 | -0.129429639 | 0.10251151 | -1.262586406 | 0.206737856 | 0.403050253 | 0.129429639 |
| ENSMUSG00000072946 | Ptgr2 | 77219 | protein_coding | 641.7495433 | 0.129551767 | 0.113898776 | 1.13742897 | 0.255359002 | 0.462094773 | 0.129551767 |
| ENSMUSG00000032329 | Hmg20a | 66867 | protein_coding | 334.8101231 | -0.129565537 | 0.104960565 | -1.234421112 | 0.217046026 | 0.416418827 | 0.129565537 |
| ENSMUSG00000051517 | Arhgef39 | 230098 | protein_coding | 252.7037789 | -0.129581008 | 0.152136751 | -0.851740341 | 0.394358225 | 0.602295312 | 0.129581008 |
| ENSMUSG00000035142 | Nubpl | 76826 | protein_coding | 126.1610485 | -0.129672148 | 0.144520723 | -0.897256431 | 0.3695821 | 0.578227258 | 0.129672148 |
| ENSMUSG00000029994 | Anxa4 | 11746 | protein_coding | 5691.308973 | 0.129700113 | 0.078248677 | 1.65753745 | 0.097410864 | 0.244032752 | 0.129700113 |
| ENSMUSG00000031826 | Usp10 | 22224 | protein_coding | 634.676186 | -0.129732915 | 0.10877711 | -1.192649029 | 0.233006856 | 0.436545136 | 0.129732915 |

| ensembl_gene_id | external_gene_name | entrezgene_id | gene_biotype | baseMean | log2FoldChange | lfcSE | stat | pvalue | padj | absLog2FC |
|---|---|---|---|---|---|---|---|---|---|---|
| ENSMUSG00000057506 | Bloc1s2 | 73689 | protein_coding | 279.5893021 | 0.129784774 | 0.092219498 | 1.407346347 | 0.159324713 | 0.340451304 | 0.129784774 |
| ENSMUSG00000028029 | Aimp1 | 13722 | protein_coding | 686.7108185 | 0.129790481 | 0.116393884 | 1.115097086 | 0.26480882 | 0.472220655 | 0.129790481 |
| ENSMUSG00000018736 | Ndel1 | 83431 | protein_coding | 808.2130839 | 0.12986511 | 0.113435237 | 1.144839236 | 0.252275759 | 0.458547106 | 0.12986511 |
| ENSMUSG00000074746 | Pdzd8 | 107368 | protein_coding | 1517.211541 | -0.130029136 | 0.067204195 | -1.934836612 | 0.053010349 | 0.161728103 | 0.130029136 |
| ENSMUSG00000021772 | Nkiras1 | 69721 | protein_coding | 116.6898653 | -0.130088025 | 0.110138381 | -1.18113254 | 0.237550075 | 0.441993884 | 0.130088025 |
| ENSMUSG00000026304 | Rab17 | 19329 | protein_coding | 228.5984573 | -0.130123375 | 0.192004443 | -0.677710232 | 0.497955438 | 0.688846377 | 0.130123375 |
| ENSMUSG00000033940 | Brk1 | 101314 | protein_coding | 981.2434138 | 0.130142008 | 0.076102358 | 1.710091654 | 0.087248926 | 0.227372528 | 0.130142008 |
| ENSMUSG00000005907 | Pex1 | 71382 | protein_coding | 236.9242255 | -0.130204402 | 0.088181272 | -1.47655391 | 0.139795254 | 0.311725998 | 0.130204402 |
| ENSMUSG00000062580 | Timm17a | 21854 | protein_coding | 867.9621388 | 0.13020773 | 0.106652579 | 1.220858713 | 0.222139519 | 0.423234241 | 0.13020773 |
| ENSMUSG00000025995 | Wdr75 | 73674 | protein_coding | 490.3511592 | -0.130214989 | 0.153302085 | -0.849401288 | 0.395658037 | 0.603875164 | 0.130214989 |
| ENSMUSG00000022698 | Naa50 | 72117 | protein_coding | 1389.151517 | -0.130271516 | 0.106964932 | -1.217889951 | 0.223265796 | 0.424719645 | 0.130271516 |
| ENSMUSG00000054823 | Nsd3 | 234135 | protein_coding | 729.9493036 | -0.130312637 | 0.131932101 | -0.987725017 | 0.323287339 | 0.534217319 | 0.130312637 |
| ENSMUSG00000001751 | Naglu | 27419 | protein_coding | 371.8990016 | 0.130367405 | 0.103194334 | 1.263319415 | 0.206474412 | 0.402733826 | 0.130367405 |
| ENSMUSG00000026279 | Thap4 | 67026 | protein_coding | 1069.749598 | 0.130379018 | 0.094519504 | 1.379387452 | 0.167775326 | 0.35225587 | 0.130379018 |
| ENSMUSG00000024124 | Prss30 | 30943 | protein_coding | 1279.144819 | -0.130414259 | 0.200130043 | -0.651647582 | 0.514628545 | 0.702183862 | 0.130414259 |
| ENSMUSG00000057335 | Cep170 | 545389 | protein_coding | 32.72551395 | -0.130460513 | 0.426013006 | -0.306235987 | 0.759424974 | 0.868410263 | 0.130460513 |
| ENSMUSG00000093904 | Tomm20 | 67952 | protein_coding | 745.8735929 | -0.130516396 | 0.117385209 | -1.111864064 | 0.266196602 | 0.474058528 | 0.130516396 |
| ENSMUSG00000009035 | Tmem184b | 223693 | protein_coding | 266.659389 | -0.130542415 | 0.120457869 | -1.083718453 | 0.278489651 | 0.487034766 | 0.130542415 |
| ENSMUSG00000020315 | Sptbn1 | 20742 | protein_coding | 9761.463792 | -0.13057997 | 0.064102641 | -2.037045075 | 0.041645525 | 0.136769057 | 0.13057997 |
| ENSMUSG00000036249 | Rbm43 | 71684 | protein_coding | 113.5861567 | 0.13059925 | 0.144171663 | 0.905859359 | 0.365010298 | 0.573814056 | 0.13059925 |
| ENSMUSG00000025158 | Rfng | 19719 | protein_coding | 161.8829606 | 0.13059935 | 0.116283088 | 1.123115596 | 0.261388403 | 0.469062019 | 0.13059935 |
| ENSMUSG00000018765 | Fxr2 | 23879 | protein_coding | 708.7857423 | -0.130620104 | 0.07927153 | -1.647755552 | 0.099402843 | 0.247335126 | 0.130620104 |
| ENSMUSG00000012350 | Ehf | 13661 | protein_coding | 8857.042558 | -0.130633801 | 0.081487543 | -1.60311375 | 0.108909543 | 0.26286402 | 0.130633801 |
| ENSMUSG00000022177 | Haus4 | 219072 | protein_coding | 187.4199987 | 0.130644119 | 0.131638942 | 0.99244279 | 0.320981574 | 0.532256186 | 0.130644119 |
| ENSMUSG00000058503 | Fam133b | 68152 | protein_coding | 330.2975667 | 0.130672045 | 0.088105383 | 1.483133496 | 0.138038946 | 0.309064427 | 0.130672045 |
| ENSMUSG00000002728 | Naa20 | 67877 | protein_coding | 724.0194266 | 0.130733758 | 0.098791087 | 1.32333556 | 0.185723811 | 0.37607537 | 0.130733758 |
| ENSMUSG00000002524 | Puf60 | 67959 | protein_coding | 1283.312434 | 0.13076772 | 0.09511835 | 1.374789626 | 0.169196675 | 0.354093686 | 0.13076772 |
| ENSMUSG00000026504 | Sdccag8 | 76816 | protein_coding | 121.3860858 | 0.130768055 | 0.170750407 | 0.765843299 | 0.443769547 | 0.644803031 | 0.130768055 |
| ENSMUSG00000045691 | Thtpa | 105663 | protein_coding | 101.6646244 | -0.13077635 | 0.154743764 | -0.84511548 | 0.398046366 | 0.606068495 | 0.13077635 |
| ENSMUSG00000025856 | Pdgfa | 18590 | protein_coding | 409.0399122 | -0.130787149 | 0.078795166 | -1.65983721 | 0.096947206 | 0.243413537 | 0.130787149 |
| ENSMUSG00000042505 | Sdhaf3 | 71238 | protein_coding | 92.42257015 | 0.130813803 | 0.142967335 | 0.914990847 | 0.360196435 | 0.569732753 | 0.130813803 |
| ENSMUSG00000028496 | Mllt3 | 70122 | protein_coding | 839.5944233 | 0.130817897 | 0.073264612 | 1.785553657 | 0.074171556 | 0.2029556 | 0.130817897 |
| ENSMUSG00000032673 | Prorsd1 | 67939 | protein_coding | 169.549653 | 0.130841958 | 0.112650459 | 1.161486245 | 0.245444214 | 0.450915275 | 0.130841958 |
| ENSMUSG00000034308 | Sdr42e1 | 74032 | protein_coding | 614.8810771 | 0.130884252 | 0.136239391 | 0.960693173 | 0.336706465 | 0.547418656 | 0.130884252 |
| ENSMUSG00000002250 | Ppard | 19015 | protein_coding | 944.3695514 | -0.130899604 | 0.098005452 | -1.335635938 | 0.181668298 | 0.370302264 | 0.130899604 |
| ENSMUSG00000046179 | E2f8 | 108961 | protein_coding | 194.2704788 | -0.130909087 | 0.171748049 | -0.762215857 | 0.445931184 | 0.646707812 | 0.130909087 |
| ENSMUSG00000106464 | C130083M11Rik | 414111 | lncRNA | 15.29060076 | 0.130925594 | 0.345962165 | 0.378439052 | 0.705104462 | 0.833382076 | 0.130925594 |
| ENSMUSG00000031782 | Coq9 | 67914 | protein_coding | 1082.397936 | 0.130930745 | 0.122871965 | 1.065586806 | 0.286610466 | 0.495534183 | 0.130930745 |

| ensembl_gene_id | external_gene_name | entrezgene_id | gene_biotype | baseMean | log2FoldChange | lfcSE | stat | pvalue | padj | absLog2FC |
|---|---|---|---|---|---|---|---|---|---|---|
| ENSMUSG00000020085 | Aifm2 | 71361 | protein_coding | 148.3198454 | -0.130939571 | 0.120309566 | -1.088355451 | 0.27643822 | 0.48476362 | 0.130939571 |
| ENSMUSG00000039662 | Icmt | 57295 | protein_coding | 781.3033191 | 0.130970508 | 0.083208393 | 1.574005969 | 0.115486069 | 0.272915913 | 0.130970508 |
| ENSMUSG00000086158 | Pierce2 | 546143 | protein_coding | 84.46418899 | -0.130973643 | 0.161936106 | -0.808798273 | 0.41863119 | 0.62480862 | 0.130973643 |
| ENSMUSG00000059981 | Taok2 | 381921 | protein_coding | 758.7234665 | -0.130995825 | 0.087293338 | -1.500639426 | 0.133448848 | 0.301885137 | 0.130995825 |
| ENSMUSG00000040729 | Cep126 | 234915 | protein_coding | 55.82617746 | 0.131029319 | 0.200449877 | 0.653676228 | 0.513320422 | 0.700974595 | 0.131029319 |
| ENSMUSG00000000204 | Slfn4 | 20558 | protein_coding | 584.3622114 | -0.131038625 | 0.254832606 | -0.514214516 | 0.60710202 | 0.771131144 | 0.131038625 |
| ENSMUSG00000015994 | Fnta | 14272 | protein_coding | 787.6955918 | 0.131090856 | 0.085934732 | 1.525470003 | 0.12714191 | 0.292351139 | 0.131090856 |
| ENSMUSG00000025196 | Cpn1 | 93721 | protein_coding | 231.4114606 | 0.131151239 | 0.262601608 | 0.49943045 | 0.617476172 | 0.777328108 | 0.131151239 |
| ENSMUSG00000020260 | Pofut2 | 80294 | protein_coding | 487.3664996 | 0.13118643 | 0.105893892 | 1.238847932 | 0.215401819 | 0.414262276 | 0.13118643 |
| ENSMUSG00000074067 | Gm10619 | 100579147 | lncRNA | 15.05624002 | 0.131194285 | 0.374589465 | 0.35023485 | 0.726162454 | 0.846109405 | 0.131194285 |
| ENSMUSG00000026492 | Tfb2m | 15278 | protein_coding | 243.077798 | -0.131202659 | 0.077366196 | -1.695865459 | 0.089911362 | 0.231845261 | 0.131202659 |
| ENSMUSG00000001700 | Gramd3 | 107022 | protein_coding | 1086.252171 | -0.131211062 | 0.137440217 | -0.954677346 | 0.339740889 | 0.550260138 | 0.131211062 |
| ENSMUSG00000027377 | Mall | 228576 | protein_coding | 969.243932 | 0.131212506 | 0.197780299 | 0.663425559 | 0.507058041 | 0.696603457 | 0.131212506 |
| ENSMUSG00000049728 | Zfp668 | 244219 | protein_coding | 101.0905112 | -0.131267058 | 0.116520408 | -1.126558524 | 0.259929184 | 0.467180325 | 0.131267058 |
| ENSMUSG00000025102 | 3110040N11Rik | 67290 | protein_coding | 177.2944813 | 0.131277559 | 0.108433992 | 1.210667947 | 0.226022692 | 0.428372488 | 0.131277559 |
| ENSMUSG00000037029 | Zfp146 | 26465 | protein_coding | 552.7207091 | -0.131331188 | 0.072488517 | -1.811751625 | 0.070024583 | 0.195681974 | 0.131331188 |
| ENSMUSG00000027984 | Hadh | 15107 | protein_coding | 4627.70894 | -0.131332608 | 0.086577876 | -1.516930347 | 0.129284271 | 0.295859555 | 0.131332608 |
| ENSMUSG00000058022 | Adtrp | 109254 | protein_coding | 515.6473901 | 0.131382434 | 0.144412185 | 0.909773883 | 0.362941771 | 0.571937996 | 0.131382434 |
| ENSMUSG00000096795 | Zfp433 | 73610 | protein_coding | 19.01635182 | 0.131393331 | 0.255193931 | 0.514876396 | 0.606639396 | 0.771091876 | 0.131393331 |
| ENSMUSG00000039536 | Stau1 | 20853 | protein_coding | 598.3191241 | -0.131401988 | 0.086495749 | -1.519172792 | 0.128719008 | 0.294927682 | 0.131401988 |
| ENSMUSG00000025262 | Fam120c | 207375 | protein_coding | 139.3374475 | -0.131418461 | 0.173670328 | -0.756712226 | 0.449222288 | 0.649765039 | 0.131418461 |
| ENSMUSG00000029713 | Gnb2 | 14693 | protein_coding | 3554.030337 | 0.131469642 | 0.09733328 | 1.350716247 | 0.176786346 | 0.363429648 | 0.131469642 |
| ENSMUSG00000040254 | Sema3d | 108151 | protein_coding | 8.580214813 | -0.131480904 | 0.622844952 | -0.211097326 | 0.832811326 | 0.911184147 | 0.131480904 |
| ENSMUSG00000028385 | Snx30 | 209131 | protein_coding | 223.2686443 | -0.1315119 | 0.12930429 | -1.017072977 | 0.309118714 | 0.518988393 | 0.1315119 |
| ENSMUSG00000060803 | Gstp1 | 14870 | protein_coding | 1909.555462 | -0.131536371 | 0.123985936 | -1.060897512 | 0.28873648 | 0.497726694 | 0.131536371 |
| ENSMUSG00000079659 | Tmem243 | 652925 | protein_coding | 268.9615849 | -0.131553154 | 0.115579829 | -1.138201669 | 0.255036282 | 0.461720607 | 0.131553154 |
| ENSMUSG00000066042 | Med18 | 67219 | protein_coding | 99.07836195 | 0.131579744 | 0.121417116 | 1.083700125 | 0.27849778 | 0.487034766 | 0.131579744 |
| ENSMUSG00000031865 | Dctn1 | 13191 | protein_coding | 1259.250375 | 0.131580353 | 0.066849463 | 1.96830832 | 0.049032573 | 0.153468144 | 0.131580353 |
| ENSMUSG00000014767 | Tbp | 21374 | protein_coding | 237.4847708 | -0.131624278 | 0.091776466 | -1.434183336 | 0.151519951 | 0.329655755 | 0.131624278 |
| ENSMUSG00000071659 | Hnrnpul2 | 68693 | protein_coding | 2263.315975 | 0.131669603 | 0.068058297 | 1.934659098 | 0.053032143 | 0.161757175 | 0.131669603 |
| ENSMUSG00000041625 | Ggact | 223267 | protein_coding | 554.8498349 | 0.131697469 | 0.08667506 | 1.519439032 | 0.128652023 | 0.294886952 | 0.131697469 |
| ENSMUSG00000000346 | Dazap2 | 23994 | protein_coding | 2856.252812 | 0.131730503 | 0.062077337 | 2.122038582 | 0.033834498 | 0.117950512 | 0.131730503 |
| ENSMUSG00000024213 | Nudt3 | 56409 | protein_coding | 558.9094661 | -0.13175519 | 0.060752032 | -2.168737168 | 0.030102644 | 0.10830696 | 0.13175519 |
| ENSMUSG00000027167 | Elp4 | 77766 | protein_coding | 229.6266368 | 0.131758339 | 0.10010972 | 1.316139321 | 0.188127286 | 0.379206708 | 0.131758339 |
| ENSMUSG00000092274 | Neat1 | 66961 | lncRNA | 11055.72729 | -0.131799088 | 0.304552195 | -0.432763546 | 0.665186559 | 0.806025281 | 0.131799088 |
| ENSMUSG00000098557 | Kctd12 | 239217 | protein_coding | 746.2370005 | 0.131901995 | 0.128863819 | 1.023576639 | 0.306035284 | 0.51606768 | 0.131901995 |
| ENSMUSG00000059248 | Septin9 | 53860 | protein_coding | 617.4896938 | 0.131917955 | 0.099299019 | 1.328492032 | 0.184015614 | 0.373591738 | 0.131917955 |
| ENSMUSG00000045751 | Mms22l | 212377 | protein_coding | 170.0136594 | 0.13194549 | 0.116168884 | 1.135807503 | 0.256037136 | 0.462962304 | 0.13194549 |

| ensembl_gene_id | external_gene_name | entrezgene_id | gene_biotype | baseMean | log2FoldChange | lfcSE | stat | pvalue | padj | absLog2FC |
|---|---|---|---|---|---|---|---|---|---|---|
| ENSMUSG00000026068 | Il18rap | 16174 | protein_coding | 7.763255806 | -0.131980747 | 0.74521253 | -0.177104841 | 0.859426039 | 0.926766895 | 0.131980747 |
| ENSMUSG00000049421 | Zfp260 | 26466 | protein_coding | 875.7061638 | 0.132003922 | 0.091572988 | 1.441515944 | 0.149438976 | 0.326824662 | 0.132003922 |
| ENSMUSG00000051346 | Spryd4 | 66701 | protein_coding | 234.2199411 | -0.13203392 | 0.125529437 | -1.051816397 | 0.292883794 | 0.501996457 | 0.13203392 |
| ENSMUSG00000027630 | Tbl1xr1 | 81004 | protein_coding | 2082.481517 | 0.132037161 | 0.069411802 | 1.902229253 | 0.05714119 | 0.170375304 | 0.132037161 |
| ENSMUSG00000027203 | Dut | 110074 | protein_coding | 302.3714633 | 0.132055058 | 0.112929098 | 1.169362553 | 0.24225759 | 0.447072054 | 0.132055058 |
| ENSMUSG00000037415 | Ranbp10 | 74334 | protein_coding | 313.4558648 | -0.132067244 | 0.112245491 | -1.176592868 | 0.239358046 | 0.443424073 | 0.132067244 |
| ENSMUSG00000037260 | Hgsnat | 52120 | protein_coding | 439.2715442 | -0.132110154 | 0.096474079 | -1.369384968 | 0.170878971 | 0.356297977 | 0.132110154 |
| ENSMUSG00000055408 | Hottip | 791364 | lncRNA | 166.930264 | 0.13214238 | 0.262260327 | 0.503859587 | 0.614360051 | 0.775275463 | 0.13214238 |
| ENSMUSG00000053565 | Eif3k | 73830 | protein_coding | 1508.933331 | 0.132144381 | 0.097467795 | 1.355774803 | 0.175170831 | 0.361418538 | 0.132144381 |
| ENSMUSG00000021708 | Rasgrf2 | 19418 | protein_coding | 195.033638 | 0.132184609 | 0.176362088 | 0.749506938 | 0.453551719 | 0.653148289 | 0.132184609 |
| ENSMUSG00000028158 | Mttp | 17777 | protein_coding | 106.7928556 | 0.132261019 | 0.196410123 | 0.673392068 | 0.500697894 | 0.691223994 | 0.132261019 |
| ENSMUSG00000020740 | Gga3 | 260302 | protein_coding | 168.2022396 | -0.132282083 | 0.135820568 | -0.973947356 | 0.330082679 | 0.540678415 | 0.132282083 |
| ENSMUSG00000000942 | Hoxa4 | 15401 | protein_coding | 7.826032458 | 0.132298687 | 0.460424181 | 0.287340874 | 0.773851328 | 0.877099132 | 0.132298687 |
| ENSMUSG00000021681 | Aggf1 | 66549 | protein_coding | 787.8956324 | -0.132338898 | 0.098093931 | -1.349103824 | 0.17730362 | 0.364187739 | 0.132338898 |
| ENSMUSG00000033557 | Fam20b | 215015 | protein_coding | 2888.748684 | 0.132351573 | 0.077812236 | 1.700909518 | 0.088959979 | 0.230174553 | 0.132351573 |
| ENSMUSG00000039523 | Cep104 | 230967 | protein_coding | 270.2747682 | -0.132366664 | 0.130005303 | -1.018163572 | 0.308600228 | 0.518563181 | 0.132366664 |
| ENSMUSG00000024393 | Prrc2a | 53761 | protein_coding | 1718.027119 | -0.132385877 | 0.120609768 | -1.0976381 | 0.272362549 | 0.480491985 | 0.132385877 |
| ENSMUSG00000068882 | Ssb | 20823 | protein_coding | 1636.800416 | 0.132399481 | 0.082313279 | 1.608482634 | 0.107729516 | 0.261043697 | 0.132399481 |
| ENSMUSG00000033590 | Myo5c | 208943 | protein_coding | 2840.593705 | 0.132407629 | 0.144896519 | 0.913808211 | 0.360817631 | 0.570262277 | 0.132407629 |
| ENSMUSG00000037134 | Prmt9 | 102182 | protein_coding | 291.1283557 | 0.132463484 | 0.068845471 | 1.92406969 | 0.054345849 | 0.164254398 | 0.132463484 |
| ENSMUSG00000032363 | Adamts7 | 108153 | protein_coding | 7.789566017 | -0.132551825 | 0.612954458 | -0.216250691 | 0.828792342 | 0.909295436 | 0.132551825 |
| ENSMUSG00000032518 | Rpsa | 16785 | protein_coding | 13212.04243 | 0.132641519 | 0.081921205 | 1.61913535 | 0.105418144 | 0.257136325 | 0.132641519 |
| ENSMUSG00000031465 | Angpt2 | 11601 | protein_coding | 46.29849213 | 0.132647883 | 0.201576004 | 0.658053939 | 0.510503472 | 0.698968748 | 0.132647883 |
| ENSMUSG00000001348 | Acp5 | 11433 | protein_coding | 1887.410308 | -0.132657961 | 0.100094376 | -1.325328826 | 0.185062112 | 0.375027 | 0.132657961 |
| ENSMUSG00000000827 | Tpd52l2 | 66314 | protein_coding | 776.0538056 | 0.132691085 | 0.071845575 | 1.846892937 | 0.064762659 | 0.185249604 | 0.132691085 |
| ENSMUSG00000037315 | Jade3 | 382207 | protein_coding | 214.4636458 | -0.132701269 | 0.084970901 | -1.561726033 | 0.118352543 | 0.278369656 | 0.132701269 |
| ENSMUSG00000042380 | Smim12 | 80284 | protein_coding | 177.6621633 | 0.132706118 | 0.115248758 | 1.151475477 | 0.249536678 | 0.455909637 | 0.132706118 |
| ENSMUSG00000050174 | Nudt6 | 229228 | protein_coding | 26.85907305 | -0.132732089 | 0.25525253 | -0.520003031 | 0.603061463 | 0.76799575 | 0.132732089 |
| ENSMUSG00000091994 | E130317F20Rik | 414101 | lncRNA | 24.17639763 | -0.132738853 | 0.269479718 | -0.492574558 | 0.622313229 | 0.780488591 | 0.132738853 |
| ENSMUSG00000032582 | Rbm6 | 19654 | protein_coding | 1316.890752 | -0.132798435 | 0.063104071 | -2.104435309 | 0.035340492 | 0.121866932 | 0.132798435 |
| ENSMUSG00000013495 | Tmem175 | 72392 | protein_coding | 185.0272479 | 0.132910099 | 0.11410811 | 1.164773467 | 0.244110702 | 0.449310115 | 0.132910099 |
| ENSMUSG00000037664 | Cdkn1c | 12577 | protein_coding | 71.61624356 | 0.132935054 | 0.210786802 | 0.630661183 | 0.528262085 | 0.71372962 | 0.132935054 |
| ENSMUSG00000030199 | Etv6 | 14011 | protein_coding | 554.7806992 | -0.132946983 | 0.098223575 | -1.353513996 | 0.175891482 | 0.362010534 | 0.132946983 |
| ENSMUSG00000026374 | Tsn | 22099 | protein_coding | 849.6444139 | 0.132974547 | 0.07714647 | 1.723663404 | 0.084768625 | 0.22276764 | 0.132974547 |
| ENSMUSG00000039307 | Hexdc | 238023 | protein_coding | 154.6572007 | -0.132985552 | 0.142846351 | -0.930969193 | 0.3518695 | 0.561468383 | 0.132985552 |
| ENSMUSG00000037531 | Mrpl47 | 74600 | protein_coding | 365.8765657 | 0.13308365 | 0.104509442 | 1.273412701 | 0.202871633 | 0.397850205 | 0.13308365 |
| ENSMUSG00000021963 | Sap18 | 20220 | protein_coding | 656.1054485 | 0.133089492 | 0.081760413 | 1.627798667 | 0.103567583 | 0.254232438 | 0.133089492 |
| ENSMUSG00000040040 | Ift88 | 21821 | protein_coding | 192.4236118 | 0.133094384 | 0.120451654 | 1.104961034 | 0.269176474 | 0.477206176 | 0.133094384 |

| ensembl_gene_id | external_gene_name | entrezgene_id | gene_biotype | baseMean | log2FoldChange | lfcSE | stat | pvalue | padj | absLog2FC |
|---|---|---|---|---|---|---|---|---|---|---|
| ENSMUSG00000025757 | Hspa4l | 18415 | protein_coding | 1226.871438 | 0.133094781 | 0.143112846 | 0.929998841 | 0.352371684 | 0.562044481 | 0.133094781 |
| ENSMUSG00000105402 | Gm3716 | 100042198 | lncRNA | 10.37858844 | -0.133158335 | 0.451337241 | -0.295030684 | 0.76797044 | 0.873434554 | 0.133158335 |
| ENSMUSG00000028869 | Gnl2 | 230737 | protein_coding | 567.3490091 | -0.133193045 | 0.141463233 | -0.94153825 | 0.346429097 | 0.556466586 | 0.133193045 |
| ENSMUSG00000061410 | Zcchc14 | 142682 | protein_coding | 477.4086213 | -0.13320747 | 0.091004349 | -1.463748389 | 0.143262709 | 0.317149191 | 0.13320747 |
| ENSMUSG00000021944 | Gata4 | 14463 | protein_coding | 9.704740351 | -0.133225136 | 0.491486201 | -0.271065873 | 0.786340369 | 0.884220209 | 0.133225136 |
| ENSMUSG00000000938 | Hoxa10 | 15395 | protein_coding | 368.7738758 | 0.133250683 | 0.218684729 | 0.609327789 | 0.542307191 | 0.725173014 | 0.133250683 |
| ENSMUSG00000027650 | Tti1 | 75425 | protein_coding | 251.2755729 | -0.133270225 | 0.099539082 | -1.338873353 | 0.180611906 | 0.368771496 | 0.133270225 |
| ENSMUSG00000031402 | Mpp1 | 17524 | protein_coding | 337.7242992 | -0.133301096 | 0.18488002 | -0.721014069 | 0.470900858 | 0.666534157 | 0.133301096 |
| ENSMUSG00000038393 | Txnip | 56338 | protein_coding | 11795.03423 | -0.133302936 | 0.123313913 | -1.081004832 | 0.279694964 | 0.488003219 | 0.133302936 |
| ENSMUSG00000087687 | Pet100 | 100503890 | protein_coding | 406.6856017 | 0.133375647 | 0.110967413 | 1.201935264 | 0.229388611 | 0.432740253 | 0.133375647 |
| ENSMUSG00000001098 | Kctd10 | 330171 | protein_coding | 979.0945837 | -0.133388662 | 0.121391337 | -1.098831802 | 0.271841438 | 0.480062946 | 0.133388662 |
| ENSMUSG00000023944 | Hsp90ab1 | 15516 | protein_coding | 15395.44047 | -0.133395966 | 0.119264549 | -1.118487992 | 0.263358632 | 0.471131469 | 0.133395966 |
| ENSMUSG00000024542 | Cep192 | 70799 | protein_coding | 550.3060421 | 0.133408263 | 0.080593086 | 1.65533137 | 0.097857299 | 0.244766184 | 0.133408263 |
| ENSMUSG00000031631 | Cfap97 | 66756 | protein_coding | 299.7965481 | -0.13344746 | 0.14121311 | -0.945007581 | 0.344655006 | 0.554399609 | 0.13344746 |
| ENSMUSG00000034617 | Mtrr | 210009 | protein_coding | 210.2474946 | 0.133525509 | 0.123119266 | 1.08452165 | 0.278133573 | 0.486665006 | 0.133525509 |
| ENSMUSG00000034354 | Mtmr3 | 74302 | protein_coding | 1381.416499 | -0.133535638 | 0.075038315 | -1.779566042 | 0.075147008 | 0.204535465 | 0.133535638 |
| ENSMUSG00000023110 | Prmt5 | 27374 | protein_coding | 595.6808666 | 0.133538186 | 0.106650283 | 1.252112816 | 0.210528758 | 0.407745873 | 0.133538186 |
| ENSMUSG00000034639 | Setmar | 74729 | protein_coding | 55.77486097 | 0.133561092 | 0.16177846 | 0.82558019 | 0.409042277 | 0.616038708 | 0.133561092 |
| ENSMUSG00000075229 | Ccdc58 | 381045 | protein_coding | 349.1101329 | 0.133561376 | 0.147598663 | 0.904895569 | 0.365520715 | 0.574333032 | 0.133561376 |
| ENSMUSG00000092118 | Fancf | 100040608 | protein_coding | 32.36520683 | 0.133567384 | 0.221745578 | 0.602345197 | 0.546944381 | 0.728832101 | 0.133567384 |
| ENSMUSG00000008450 | Nutf2 | 68051 | protein_coding | 154.0293812 | 0.133573108 | 0.127633754 | 1.046534349 | 0.295314395 | 0.504587823 | 0.133573108 |
| ENSMUSG00000042595 | Fam199x | 245622 | protein_coding | 646.5066639 | 0.133653 | 0.097945591 | 1.364563719 | 0.172390214 | 0.358089637 | 0.133653 |
| ENSMUSG00000050296 | Abca12 | 74591 | protein_coding | 919.9758139 | -0.133712748 | 0.356375755 | -0.375201584 | 0.707510553 | 0.834695806 | 0.133712748 |
| ENSMUSG00000024936 | Kcnk7 | 16530 | protein_coding | 7.199724937 | -0.133814176 | 0.405808665 | -0.329746966 | 0.741591163 | 0.8560087 | 0.133814176 |
| ENSMUSG00000039367 | Sec24c | 218811 | protein_coding | 1926.513582 | 0.13383795 | 0.084325097 | 1.587166284 | 0.112474989 | 0.267988275 | 0.13383795 |
| ENSMUSG00000003234 | Abcf3 | 27406 | protein_coding | 714.7102194 | 0.133962941 | 0.073061465 | 1.833564944 | 0.0667186 | 0.188923353 | 0.133962941 |
| ENSMUSG00000031021 | Tmem9b | 56786 | protein_coding | 797.8536001 | 0.13396324 | 0.108200598 | 1.23810073 | 0.215678713 | 0.414744724 | 0.13396324 |
| ENSMUSG00000022607 | Ptk2 | 14083 | protein_coding | 1210.946894 | 0.133966666 | 0.085932286 | 1.55897943 | 0.119001249 | 0.279586936 | 0.133966666 |
| ENSMUSG00000015757 | Ppil4 | 67418 | protein_coding | 611.0260085 | -0.133990196 | 0.064076806 | -2.091087303 | 0.036520239 | 0.124693325 | 0.133990196 |
| ENSMUSG00000074656 | Eif2s2 | 67204 | protein_coding | 3312.467874 | 0.134022221 | 0.128757806 | 1.040886181 | 0.297928376 | 0.506735399 | 0.134022221 |
| ENSMUSG00000035268 | Pkig | 18769 | protein_coding | 309.9328153 | 0.134056548 | 0.138495791 | 0.967946726 | 0.333070977 | 0.543959534 | 0.134056548 |
| ENSMUSG00000034820 | Cpsf7 | 269061 | protein_coding | 565.8040937 | -0.134077107 | 0.083753981 | -1.600844581 | 0.109411347 | 0.263566838 | 0.134077107 |
| ENSMUSG00000066979 | Bub3 | 12237 | protein_coding | 753.1955475 | 0.13408694 | 0.096667557 | 1.387093504 | 0.165413253 | 0.349003763 | 0.13408694 |
| ENSMUSG00000042535 | Gtpbp1 | 14904 | protein_coding | 564.9818162 | -0.134087367 | 0.086666729 | -1.547160805 | 0.121824475 | 0.284157212 | 0.134087367 |
| ENSMUSG00000037190 | Cyb561d2 | 56368 | protein_coding | 132.0360857 | 0.134113521 | 0.105865119 | 1.266833896 | 0.205214689 | 0.401062215 | 0.134113521 |
| ENSMUSG00000025139 | Tollip | 54473 | protein_coding | 863.3834002 | -0.134126717 | 0.087130684 | -1.539374088 | 0.123712996 | 0.286952957 | 0.134126717 |
| ENSMUSG00000046185 | Zfp84 | 74352 | protein_coding | 451.4837589 | -0.134140314 | 0.077173062 | -1.738175353 | 0.082179921 | 0.218196662 | 0.134140314 |
| ENSMUSG00000040345 | Arhgap9 | 216445 | protein_coding | 45.87207142 | 0.13417353 | 0.234598617 | 0.571928051 | 0.567370719 | 0.743135986 | 0.13417353 |

Supplementary Table 1

| ensembl_gene_id | external_gene_name | entrezgene_id | gene_biotype | baseMean | log2FoldChange | lfcSE | stat | pvalue | padj | absLog2FC |
|---|---|---|---|---|---|---|---|---|---|---|
| ENSMUSG00000056666 | Retsat | 67442 | protein_coding | 1481.485023 | 0.134240932 | 0.104045778 | 1.290210275 | 0.19697766 | 0.390571413 | 0.134240932 |
| ENSMUSG00000116802 | Gm5165 | 102640673 | lncRNA | 203.3897525 | 0.134273495 | 0.120354017 | 1.115654459 | 0.264570071 | 0.472124579 | 0.134273495 |
| ENSMUSG00000006299 | Aamp | 227290 | protein_coding | 1489.325134 | 0.134351319 | 0.089919752 | 1.494124667 | 0.135143038 | 0.304621103 | 0.134351319 |
| ENSMUSG00000023952 | Gtpbp2 | 56055 | protein_coding | 1403.072199 | 0.134376111 | 0.140826364 | 0.954197116 | 0.339983874 | 0.550365001 | 0.134376111 |
| ENSMUSG00000053877 | Srcap | 100043597 | protein_coding | 110.2632567 | -0.134387489 | 0.203517729 | -0.660323254 | 0.50904641 | 0.698070704 | 0.134387489 |
| ENSMUSG00000054702 | Ap1s3 | 252903 | protein_coding | 171.6517175 | -0.134393974 | 0.145071792 | -0.926396324 | 0.354240054 | 0.563829056 | 0.134393974 |
| ENSMUSG00000049001 | Ndnf | 68169 | protein_coding | 39.92395538 | 0.134408025 | 0.284221154 | 0.472899442 | 0.636284923 | 0.789347063 | 0.134408025 |
| ENSMUSG00000056941 | Commd7 | 99311 | protein_coding | 440.880235 | 0.134478184 | 0.095859599 | 1.402866115 | 0.160656767 | 0.342287451 | 0.134478184 |
| ENSMUSG00000020431 | Adcy1 | 432530 | protein_coding | 11.48526623 | 0.134529712 | 0.436615281 | 0.308119569 | 0.757991351 | 0.867408546 | 0.134529712 |
| ENSMUSG00000033478 | Fhip2a | 226252 | protein_coding | 349.1314005 | -0.134614648 | 0.070396189 | -1.912243408 | 0.055844981 | 0.167482368 | 0.134614648 |
| ENSMUSG00000025481 | Urah | 76974 | protein_coding | 165.3658573 | 0.134638446 | 0.101191617 | 1.330529651 | 0.183343824 | 0.372582345 | 0.134638446 |
| ENSMUSG00000022938 | Fam3b | 52793 | protein_coding | 1651.36275 | 0.134659876 | 0.13500395 | 0.997451378 | 0.318545464 | 0.529987548 | 0.134659876 |
| ENSMUSG00000029141 | Slc4a1ap | 20534 | protein_coding | 289.6173787 | -0.134684004 | 0.090994557 | -1.480132533 | 0.138837881 | 0.31014926 | 0.134684004 |
| ENSMUSG00000038784 | Cnot4 | 53621 | protein_coding | 296.7529469 | -0.134708321 | 0.084603335 | -1.59223417 | 0.111332098 | 0.266434468 | 0.134708321 |
| ENSMUSG00000015002 | Efr3a | 76740 | protein_coding | 1536.863128 | 0.134729607 | 0.057969032 | 2.324165196 | 0.020116642 | 0.08014454 | 0.134729607 |
| ENSMUSG00000066278 | Vps37b | 330192 | protein_coding | 226.4819042 | -0.134773967 | 0.233353632 | -0.577552473 | 0.563566302 | 0.740206486 | 0.134773967 |
| ENSMUSG00000028484 | Psip1 | 101739 | protein_coding | 363.5811724 | 0.134792089 | 0.095345026 | 1.41372964 | 0.157441299 | 0.33818806 | 0.134792089 |
| ENSMUSG00000039158 | Akna | 100182 | protein_coding | 86.83457967 | -0.134917738 | 0.160551227 | -0.840340743 | 0.400717364 | 0.608158758 | 0.134917738 |
| ENSMUSG00000097583 | 6430590A07Rik | 619719 | lncRNA | 24.82910057 | -0.134920852 | 0.319427376 | -0.422383494 | 0.672745123 | 0.811159874 | 0.134920852 |
| ENSMUSG00000026617 | Bpnt1 | 23827 | protein_coding | 2189.976738 | 0.13494512 | 0.083371706 | 1.618596121 | 0.10553419 | 0.257261797 | 0.13494512 |
| ENSMUSG00000029328 | Hnrnpdl | 50926 | protein_coding | 1799.657484 | -0.135014257 | 0.062611735 | -2.156373038 | 0.031054548 | 0.110830572 | 0.135014257 |
| ENSMUSG00000087088 | Gm16638 | 102631889 | lncRNA | 12.18286647 | -0.135049434 | 0.408900707 | -0.330274396 | 0.741192638 | 0.855772195 | 0.135049434 |
| ENSMUSG00000042041 | 2010003K11Rik | 69861 | protein_coding | 155.3522534 | 0.13505948 | 0.300480493 | 0.449478361 | 0.653086615 | 0.799021328 | 0.13505948 |
| ENSMUSG00000021113 | Snapc1 | 75627 | protein_coding | 205.6597713 | -0.135086459 | 0.113228352 | -1.193044463 | 0.23285196 | 0.436411363 | 0.135086459 |
| ENSMUSG00000059995 | Atxn7l3 | 217218 | protein_coding | 697.4175915 | -0.135100648 | 0.086031396 | -1.570364474 | 0.116330343 | 0.274545814 | 0.135100648 |
| ENSMUSG00000027285 | Haus2 | 66296 | protein_coding | 202.8675303 | -0.135127109 | 0.095151289 | -1.42012905 | 0.155570114 | 0.335658113 | 0.135127109 |
| ENSMUSG00000044986 | Tst | 22117 | protein_coding | 2147.563125 | 0.135133021 | 0.118482049 | 1.140535819 | 0.254063139 | 0.460534616 | 0.135133021 |
| ENSMUSG00000034105 | Meak7 | 74347 | protein_coding | 214.4631698 | 0.135195348 | 0.15794599 | 0.855959358 | 0.39202026 | 0.600073507 | 0.135195348 |
| ENSMUSG00000029684 | Wasl | 73178 | protein_coding | 1051.313164 | -0.135219037 | 0.117694851 | -1.148895093 | 0.250599241 | 0.456959899 | 0.135219037 |
| ENSMUSG00000047466 | 8030462N17Rik | 212163 | protein_coding | 442.4241697 | -0.135247905 | 0.085900677 | -1.574468445 | 0.115379191 | 0.272752947 | 0.135247905 |
| ENSMUSG00000026142 | Rhbdd1 | 76867 | protein_coding | 421.22185 | 0.135309568 | 0.082169947 | 1.646703847 | 0.099618933 | 0.247613326 | 0.135309568 |
| ENSMUSG00000027939 | Nup210l | 77595 | protein_coding | 44.15468733 | 0.135311931 | 0.210074173 | 0.644115023 | 0.519500849 | 0.706398276 | 0.135311931 |
| ENSMUSG00000073609 | D2hgdh | 98314 | protein_coding | 236.1450621 | -0.135330236 | 0.108735814 | -1.24457831 | 0.213286797 | 0.411336979 | 0.135330236 |
| ENSMUSG00000024451 | Arap3 | 106952 | protein_coding | 13.9502637 | 0.135371566 | 0.587060046 | 0.230592368 | 0.817631498 | 0.903136927 | 0.135371566 |
| ENSMUSG00000037503 | Fam168b | 214469 | protein_coding | 1782.974413 | -0.135394066 | 0.094987492 | -1.425388364 | 0.15404497 | 0.333192324 | 0.135394066 |
| ENSMUSG00000022972 | Cfap298 | 68001 | protein_coding | 71.14710057 | -0.135423118 | 0.174987882 | -0.77389998 | 0.438989941 | 0.641548012 | 0.135423118 |
| ENSMUSG00000034403 | Pja1 | 18744 | protein_coding | 901.690267 | 0.135495555 | 0.083646356 | 1.619862032 | 0.105261917 | 0.256873268 | 0.135495555 |
| ENSMUSG00000021546 | Hnrnpk | 15387 | protein_coding | 6172.837035 | 0.135533556 | 0.089824064 | 1.508878024 | 0.131329957 | 0.298831936 | 0.135533556 |

| ensembl_gene_id | external_gene_name | entrezgene_id | gene_biotype | baseMean | log2FoldChange | lfcSE | stat | pvalue | padj | absLog2FC |
|---|---|---|---|---|---|---|---|---|---|---|
| ENSMUSG00000042029 | Ncapg2 | 76044 | protein_coding | 449.514396 | 0.135658432 | 0.138652123 | 0.978408617 | 0.327872268 | 0.538806428 | 0.135658432 |
| ENSMUSG00000036073 | Galt | 14430 | protein_coding | 252.0534246 | -0.135674533 | 0.082397485 | -1.646585847 | 0.099643201 | 0.247613326 | 0.135674533 |
| ENSMUSG00000027012 | Dync1i2 | 13427 | protein_coding | 1454.987398 | 0.135714646 | 0.06517865 | 2.082194785 | 0.037324681 | 0.126652647 | 0.135714646 |
| ENSMUSG00000086782 | E130102H24Rik | 77866 | lncRNA | 11.48145509 | -0.135726738 | 0.474717071 | -0.2859108 | 0.774946441 | 0.877593098 | 0.135726738 |
| ENSMUSG00000044229 | Nxpe4 | 244853 | protein_coding | 897.4347879 | -0.135765078 | 0.385725766 | -0.35197306 | 0.724858466 | 0.844960894 | 0.135765078 |
| ENSMUSG00000018428 | Akap1 | 11640 | protein_coding | 1415.720732 | -0.135773089 | 0.094690942 | -1.433855091 | 0.15161362 | 0.329769457 | 0.135773089 |
| ENSMUSG00000046897 | Zfp740 | 68744 | protein_coding | 1011.849432 | -0.135779469 | 0.078741422 | -1.724371558 | 0.084640789 | 0.222541843 | 0.135779469 |
| ENSMUSG00000033187 | BC016579 | 212998 | protein_coding | 613.1421813 | -0.135837906 | 0.208235573 | -0.652328052 | 0.514189567 | 0.701860607 | 0.135837906 |
| ENSMUSG00000050786 | Ccdc126 | 57895 | protein_coding | 37.8323515 | -0.135862665 | 0.210619381 | -0.645062504 | 0.518886681 | 0.70596078 | 0.135862665 |
| ENSMUSG00000022477 | Aco2 | 11429 | protein_coding | 7091.226484 | 0.1358898 | 0.094717021 | 1.434692511 | 0.151374739 | 0.329588153 | 0.1358898 |
| ENSMUSG00000021548 | Ccnh | 66671 | protein_coding | 243.5205866 | 0.13589461 | 0.092252584 | 1.473071037 | 0.140731882 | 0.313156946 | 0.13589461 |
| ENSMUSG00000032112 | Trappc4 | 60409 | protein_coding | 364.8462282 | 0.135895696 | 0.088529553 | 1.535031991 | 0.12477597 | 0.28878693 | 0.135895696 |
| ENSMUSG00000033618 | Map3k13 | 71751 | protein_coding | 302.3446316 | -0.135935063 | 0.236035898 | -0.575908426 | 0.56467708 | 0.741139569 | 0.135935063 |
| ENSMUSG00000056537 | Rlim | 19820 | protein_coding | 2083.223935 | -0.135936152 | 0.088902833 | -1.529041846 | 0.12625407 | 0.290939646 | 0.135936152 |
| ENSMUSG00000000959 | Oxa1l | 69089 | protein_coding | 704.4712986 | -0.135994231 | 0.102126049 | -1.331631178 | 0.182981415 | 0.372178262 | 0.135994231 |
| ENSMUSG00000026203 | Dnajb2 | 56812 | protein_coding | 328.3562722 | 0.136045492 | 0.11962989 | 1.137219899 | 0.25544637 | 0.462147863 | 0.136045492 |
| ENSMUSG00000026510 | Trp53bp2 | 209456 | protein_coding | 577.1459604 | -0.136137128 | 0.073246724 | -1.858610466 | 0.063082373 | 0.181816733 | 0.136137128 |
| ENSMUSG00000037514 | Pank2 | 74450 | protein_coding | 633.2715238 | 0.136153244 | 0.054294824 | 2.50766527 | 0.012153173 | 0.055418192 | 0.136153244 |
| ENSMUSG00000027546 | Atp9a | 11981 | protein_coding | 3642.483192 | -0.136156601 | 0.089626784 | -1.519150808 | 0.12872454 | 0.294927682 | 0.136156601 |
| ENSMUSG00000019820 | Utrn | 22288 | protein_coding | 2587.11359 | 0.136164003 | 0.099007005 | 1.375296648 | 0.169039495 | 0.354084834 | 0.136164003 |
| ENSMUSG00000030796 | Tead2 | 21677 | protein_coding | 44.4085876 | -0.136195251 | 0.202549795 | -0.672403796 | 0.501326667 | 0.691772602 | 0.136195251 |
| ENSMUSG00000005949 | Ctns | 83429 | protein_coding | 212.2459821 | -0.136300252 | 0.190617761 | -0.715044868 | 0.474581319 | 0.669482885 | 0.136300252 |
| ENSMUSG00000000088 | Cox5a | 12858 | protein_coding | 2646.29851 | 0.136346709 | 0.09983419 | 1.365731608 | 0.172023219 | 0.357719532 | 0.136346709 |
| ENSMUSG00000020062 | Slc5a8 | 216225 | protein_coding | 6745.395502 | -0.136350613 | 0.174549261 | -0.781158353 | 0.434709365 | 0.637867413 | 0.136350613 |
| ENSMUSG00000025762 | Larp1b | 214048 | protein_coding | 988.6873381 | 0.136355177 | 0.09640673 | 1.414374052 | 0.157252105 | 0.33818806 | 0.136355177 |
| ENSMUSG00000038618 | Rassf7 | 66985 | protein_coding | 874.5522073 | 0.136376061 | 0.112645278 | 1.21066825 | 0.226022575 | 0.428372488 | 0.136376061 |
| ENSMUSG00000047003 | Zfp41 | 22701 | protein_coding | 94.14872132 | 0.136458982 | 0.222112724 | 0.614368144 | 0.538972081 | 0.722272825 | 0.136458982 |
| ENSMUSG00000018678 | Sp2 | 78912 | protein_coding | 80.11186458 | 0.13645966 | 0.184933534 | 0.737884887 | 0.460584404 | 0.658781284 | 0.13645966 |
| ENSMUSG00000022828 | Gtf2e1 | 74197 | protein_coding | 178.6377662 | 0.136459954 | 0.09545893 | 1.429514819 | 0.152856319 | 0.331702377 | 0.136459954 |
| ENSMUSG00000005882 | Uqcc1 | 56046 | protein_coding | 778.6982999 | 0.136494293 | 0.101244259 | 1.348168232 | 0.17760428 | 0.364683959 | 0.136494293 |
| ENSMUSG00000030528 | Blm | 12144 | protein_coding | 109.8796701 | 0.136525771 | 0.134421336 | 1.015655512 | 0.30979346 | 0.519699986 | 0.136525771 |
| ENSMUSG00000028676 | Srsf10 | 14105 | protein_coding | 1339.404955 | 0.136544009 | 0.09831079 | 1.38890155 | 0.164862689 | 0.34831316 | 0.136544009 |
| ENSMUSG00000045671 | Spred2 | 114716 | protein_coding | 335.0213072 | -0.136544347 | 0.155024677 | -0.880791047 | 0.378430927 | 0.58646116 | 0.136544347 |
| ENSMUSG00000032397 | Tipin | 66131 | protein_coding | 167.3848359 | 0.136564925 | 0.139807737 | 0.976805204 | 0.328665599 | 0.539310112 | 0.136564925 |
| ENSMUSG00000090035 | Galnt4 | 14426 | protein_coding | 5189.819478 | 0.136571627 | 0.106579389 | 1.28140749 | 0.200050575 | 0.394660109 | 0.136571627 |
| ENSMUSG00000026589 | Sec16b | 89867 | protein_coding | 473.2170583 | -0.136606418 | 0.091500696 | -1.492954965 | 0.135448976 | 0.305073452 | 0.136606418 |
| ENSMUSG00000030232 | Aebp2 | 11569 | protein_coding | 694.9326441 | -0.13663145 | 0.120965376 | -1.129508746 | 0.258683282 | 0.465729322 | 0.13663145 |
| ENSMUSG00000026851 | BC005624 | 227707 | protein_coding | 364.9437737 | 0.136646673 | 0.071077966 | 1.922489922 | 0.054544141 | 0.164666238 | 0.136646673 |

| ensembl_gene_id | external_gene_name | entrezgene_id | gene_biotype | baseMean | log2FoldChange | lfcSE | stat | pvalue | padj | absLog2FC |
|---|---|---|---|---|---|---|---|---|---|---|
| ENSMUSG00000038346 | Zfp384 | 269800 | protein_coding | 350.6692045 | -0.13668207 | 0.101498195 | -1.346645326 | 0.178094489 | 0.365317323 | 0.13668207 |
| ENSMUSG00000060519 | Tor3a | 30935 | protein_coding | 454.5145759 | -0.136739423 | 0.123135505 | -1.110479249 | 0.266792565 | 0.474540288 | 0.136739423 |
| ENSMUSG00000024957 | Kcnk4 | 16528 | protein_coding | 9.956148937 | -0.136756952 | 0.490394806 | -0.278871127 | 0.780343727 | 0.880637302 | 0.136756952 |
| ENSMUSG00000038742 | Angptl6 | 70726 | protein_coding | 12.51482762 | 0.136758704 | 0.330966979 | 0.413209513 | 0.679453119 | 0.816753599 | 0.136758704 |
| ENSMUSG00000028955 | Vamp3 | 22319 | protein_coding | 1574.828561 | 0.136767077 | 0.082129411 | 1.665263085 | 0.09586028 | 0.241630406 | 0.136767077 |
| ENSMUSG00000022788 | Fgd4 | 224014 | protein_coding | 1377.203628 | 0.136777292 | 0.091877813 | 1.488686857 | 0.136569847 | 0.306896802 | 0.136777292 |
| ENSMUSG00000024845 | Tmem134 | 66990 | protein_coding | 736.8819297 | 0.136802816 | 0.082205169 | 1.664163183 | 0.096079823 | 0.242030591 | 0.136802816 |
| ENSMUSG00000020868 | Xylt2 | 217119 | protein_coding | 185.6045934 | -0.136848129 | 0.143099432 | -0.956314971 | 0.338913128 | 0.549715714 | 0.136848129 |
| ENSMUSG00000042997 | Nhlrc3 | 212114 | protein_coding | 973.7005593 | 0.1368603 | 0.078154647 | 1.751147318 | 0.079920537 | 0.213693859 | 0.1368603 |
| ENSMUSG00000027455 | Nsfl1c | 386649 | protein_coding | 681.324792 | 0.136878513 | 0.097091019 | 1.409795823 | 0.15859998 | 0.339860149 | 0.136878513 |
| ENSMUSG00000003072 | Atp5d | 66043 | protein_coding | 3972.393421 | 0.136903924 | 0.074962874 | 1.8262897 | 0.067806636 | 0.191357919 | 0.136903924 |
| ENSMUSG00000071072 | Ptges3 | 56351 | protein_coding | 1794.129937 | 0.136941517 | 0.126530104 | 1.08228408 | 0.279126318 | 0.487504149 | 0.136941517 |
| ENSMUSG00000022263 | Trio | 223435 | protein_coding | 1242.371565 | -0.136941877 | 0.147261857 | -0.92992089 | 0.352412046 | 0.562052575 | 0.136941877 |
| ENSMUSG00000061517 | Sox21 | 223227 | protein_coding | 63.23429232 | 0.136963671 | 0.176726768 | 0.775002408 | 0.438338237 | 0.640830788 | 0.136963671 |
| ENSMUSG00000034673 | Pbx2 | 18515 | protein_coding | 455.8663338 | -0.136964477 | 0.095662896 | -1.431740865 | 0.152217997 | 0.330657805 | 0.136964477 |
| ENSMUSG00000018820 | Zfyve27 | 319740 | protein_coding | 371.8966538 | -0.136968852 | 0.087436438 | -1.56649626 | 0.117232483 | 0.276388247 | 0.136968852 |
| ENSMUSG00000022865 | Cxadr | 13052 | protein_coding | 3529.875202 | -0.136982015 | 0.069515276 | -1.970531138 | 0.04877753 | 0.153089359 | 0.136982015 |
| ENSMUSG00000027324 | Rpusd2 | 271842 | protein_coding | 61.26504589 | -0.136991829 | 0.180586274 | -0.758594912 | 0.448094918 | 0.648900432 | 0.136991829 |
| ENSMUSG00000017548 | Suz12 | 52615 | protein_coding | 905.9515352 | 0.136996759 | 0.080781685 | 1.695888846 | 0.089906932 | 0.231845261 | 0.136996759 |
| ENSMUSG00000055198 | Gm12830 | 433746 | lncRNA | 7.555810242 | -0.137007089 | 0.474901343 | -0.288495898 | 0.77296717 | 0.876473346 | 0.137007089 |
| ENSMUSG00000006005 | Tpr | 108989 | protein_coding | 2788.103226 | -0.137023961 | 0.068640524 | -1.996254585 | 0.045906219 | 0.146708317 | 0.137023961 |
| ENSMUSG00000037638 | Zbtb42 | 382639 | protein_coding | 263.0554038 | -0.137075652 | 0.096743976 | -1.416890822 | 0.15651485 | 0.337161014 | 0.137075652 |
| ENSMUSG00000058301 | Upf1 | 19704 | protein_coding | 1250.877884 | -0.137106486 | 0.108808997 | -1.260065711 | 0.207645658 | 0.404082965 | 0.137106486 |
| ENSMUSG00000071350 | Setdb2 | 239122 | protein_coding | 174.2376071 | 0.137128179 | 0.109448608 | 1.252900167 | 0.210242041 | 0.407438281 | 0.137128179 |
| ENSMUSG00000024991 | Eif3a | 13669 | protein_coding | 4098.740652 | -0.137186297 | 0.091918574 | -1.492476343 | 0.135574315 | 0.305305766 | 0.137186297 |
| ENSMUSG00000042745 | Id1 | 15901 | protein_coding | 3264.692332 | 0.137205444 | 0.210767223 | 0.650980938 | 0.515058792 | 0.702478962 | 0.137205444 |
| ENSMUSG00000039100 | Marchf6 | 223455 | protein_coding | 1760.001409 | -0.13724137 | 0.080462686 | -1.705652351 | 0.088072826 | 0.228545168 | 0.13724137 |
| ENSMUSG00000034152 | Exoc3 | 211446 | protein_coding | 986.4144058 | 0.13726932 | 0.086543507 | 1.586130782 | 0.112709645 | 0.268426918 | 0.13726932 |
| ENSMUSG00000029171 | Pgm2 | 66681 | protein_coding | 1218.461462 | -0.13728485 | 0.11141818 | -1.232158433 | 0.217889907 | 0.417635432 | 0.13728485 |
| ENSMUSG00000026519 | Tmem63a | 208795 | protein_coding | 1688.279356 | -0.137319975 | 0.08377155 | -1.639219686 | 0.101167515 | 0.250100293 | 0.137319975 |
| ENSMUSG00000027552 | E2f5 | 13559 | protein_coding | 482.5580022 | 0.137320975 | 0.112157376 | 1.224359734 | 0.220816552 | 0.421469867 | 0.137320975 |
| ENSMUSG00000050359 | Sprr1a | 20753 | protein_coding | 174.3597323 | -0.137328478 | 0.820864219 | -0.167297434 | 0.867136023 | 0.931212417 | 0.137328478 |
| ENSMUSG00000025049 | Taf5 | 226182 | protein_coding | 154.2893097 | -0.137370103 | 0.10026278 | -1.370100683 | 0.170655475 | 0.3559461 | 0.137370103 |
| ENSMUSG00000032294 | Pkm | 18746 | protein_coding | 7712.952853 | 0.13738801 | 0.10824988 | 1.269174708 | 0.20437876 | 0.399772183 | 0.13738801 |
| ENSMUSG00000040891 | Foxa3 | 15377 | protein_coding | 932.2189364 | -0.137437514 | 0.140067911 | -0.981220563 | 0.326483986 | 0.537603321 | 0.137437514 |
| ENSMUSG00000032485 | Scap | 235623 | protein_coding | 806.8089123 | -0.13750689 | 0.080134598 | -1.715949075 | 0.086171363 | 0.225301621 | 0.13750689 |
| ENSMUSG00000038717 | Atp5l | 27425 | protein_coding | 452.2438912 | 0.137523495 | 0.092959199 | 1.479396297 | 0.139034429 | 0.310420576 | 0.137523495 |
| ENSMUSG00000026426 | Arl8a | 68724 | protein_coding | 137.1767381 | -0.137541233 | 0.13244052 | -1.038513234 | 0.299031177 | 0.507887459 | 0.137541233 |

| ensembl_gene_id | external_gene_name | entrezgene_id | gene_biotype | baseMean | log2FoldChange | lfcSE | stat | pvalue | padj | absLog2FC |
|---|---|---|---|---|---|---|---|---|---|---|
| ENSMUSG00000047888 | Tnrc6b | 213988 | protein_coding | 1172.551268 | 0.137547208 | 0.170855367 | 0.805050557 | 0.420790506 | 0.626692023 | 0.137547208 |
| ENSMUSG00000003731 | Kpna6 | 16650 | protein_coding | 891.1201868 | -0.137588039 | 0.091592029 | -1.502183547 | 0.133049713 | 0.301581873 | 0.137588039 |
| ENSMUSG00000041957 | Pkp2 | 67451 | protein_coding | 921.7114976 | 0.137618564 | 0.122838069 | 1.120325037 | 0.262575276 | 0.470344017 | 0.137618564 |
| ENSMUSG00000050270 | Tmem220 | 338369 | protein_coding | 94.99008921 | 0.137621277 | 0.160771724 | 0.856004241 | 0.391995434 | 0.600073507 | 0.137621277 |
| ENSMUSG00000019841 | Rev3l | 19714 | protein_coding | 869.5707328 | 0.137624255 | 0.132417475 | 1.03932095 | 0.298655495 | 0.507472252 | 0.137624255 |
| ENSMUSG00000022400 | Rbx1 | 56438 | protein_coding | 800.6884747 | 0.137628127 | 0.091948322 | 1.496798682 | 0.134445649 | 0.303493524 | 0.137628127 |
| ENSMUSG00000027463 | Slc52a3 | 69698 | protein_coding | 461.9603108 | 0.137636485 | 0.094488371 | 1.456649994 | 0.14521302 | 0.320620041 | 0.137636485 |
| ENSMUSG00000033544 | Angptl1 | 72713 | protein_coding | 10.23172613 | -0.137811204 | 0.38986322 | -0.353486035 | 0.723724097 | 0.844689486 | 0.137811204 |
| ENSMUSG00000079614 | Seh1l | 72124 | protein_coding | 712.8156526 | 0.137836295 | 0.080876073 | 1.704290155 | 0.088326893 | 0.22904457 | 0.137836295 |
| ENSMUSG00000038582 | Pptc7 | 320717 | protein_coding | 635.9254003 | -0.13786684 | 0.104955507 | -1.313574139 | 0.188989556 | 0.380319097 | 0.13786684 |
| ENSMUSG00000062098 | Btbd3 | 228662 | protein_coding | 445.6225515 | -0.137878316 | 0.083114318 | -1.658899677 | 0.09713601 | 0.243675627 | 0.137878316 |
| ENSMUSG00000030970 | Ctbp2 | 13017 | protein_coding | 910.8318091 | 0.137892603 | 0.080831676 | 1.705922852 | 0.088022444 | 0.228545168 | 0.137892603 |
| ENSMUSG00000033658 | Ddx19b | 234733 | protein_coding | 553.394292 | -0.137924406 | 0.070526347 | -1.955643679 | 0.050507144 | 0.156513188 | 0.137924406 |
| ENSMUSG00000047547 | Cltb | 74325 | protein_coding | 1813.497375 | 0.137945218 | 0.092343225 | 1.493831496 | 0.135219667 | 0.304722532 | 0.137945218 |
| ENSMUSG00000026820 | Ptges2 | 96979 | protein_coding | 455.8597865 | -0.137974603 | 0.101917957 | -1.353781095 | 0.175806227 | 0.361907388 | 0.137974603 |
| ENSMUSG00000035293 | G2e3 | 217558 | protein_coding | 641.037573 | 0.137980356 | 0.114890222 | 1.200975628 | 0.229760656 | 0.433077983 | 0.137980356 |
| ENSMUSG00000020137 | Thap2 | 66816 | protein_coding | 339.715235 | 0.138028979 | 0.131329448 | 1.05101317 | 0.293252541 | 0.502250159 | 0.138028979 |
| ENSMUSG00000031993 | Snx19 | 102607 | protein_coding | 598.3417653 | 0.138036846 | 0.077651652 | 1.777642106 | 0.075462654 | 0.205144798 | 0.138036846 |
| ENSMUSG00000060227 | Golm2 | 319996 | protein_coding | 391.4589696 | -0.138040827 | 0.164471576 | -0.839298989 | 0.401301551 | 0.608871318 | 0.138040827 |
| ENSMUSG00000026048 | Ercc5 | 22592 | protein_coding | 272.8264808 | -0.138059239 | 0.096167287 | -1.435615407 | 0.151111808 | 0.329262501 | 0.138059239 |
| ENSMUSG00000030879 | Mrpl17 | 27397 | protein_coding | 454.803894 | 0.138087245 | 0.100509495 | 1.373872642 | 0.169481224 | 0.354357698 | 0.138087245 |
| ENSMUSG00000026596 | Abl2 | 11352 | protein_coding | 447.4994521 | -0.138104303 | 0.241880114 | -0.570961791 | 0.568025544 | 0.743305197 | 0.138104303 |
| ENSMUSG00000000605 | Clcn4 | 12727 | protein_coding | 1652.840972 | 0.138141506 | 0.059535957 | 2.320303781 | 0.02032445 | 0.08083092 | 0.138141506 |
| ENSMUSG00000078713 | Tomm5 | 68512 | protein_coding | 515.8485946 | 0.138164767 | 0.108881397 | 1.268947411 | 0.204459821 | 0.399881635 | 0.138164767 |
| ENSMUSG00000015474 | Ppt2 | 54397 | protein_coding | 146.591075 | -0.138168604 | 0.105327675 | -1.311797726 | 0.189588392 | 0.381182159 | 0.138168604 |
| ENSMUSG00000045802 | Hsf3 | 245525 | protein_coding | 14.204466 | -0.138185855 | 0.348719373 | -0.396266642 | 0.691908333 | 0.824533588 | 0.138185855 |
| ENSMUSG00000042215 | Bag2 | 213539 | protein_coding | 45.93580573 | -0.138194848 | 0.238184009 | -0.580202038 | 0.56177838 | 0.739199176 | 0.138194848 |
| ENSMUSG00000041491 | Cep78 | 208518 | protein_coding | 126.5759558 | 0.13826009 | 0.157304551 | 0.878932552 | 0.379437848 | 0.587392948 | 0.13826009 |
| ENSMUSG00000036779 | Tent4b | 214627 | protein_coding | 574.2229878 | -0.138328364 | 0.089409223 | -1.547137531 | 0.121830085 | 0.284157212 | 0.138328364 |
| ENSMUSG00000034109 | Golim4 | 73124 | protein_coding | 1430.747553 | 0.138387845 | 0.071206714 | 1.943466263 | 0.051959848 | 0.159679038 | 0.138387845 |
| ENSMUSG00000008475 | Arpc5 | 67771 | protein_coding | 3420.124963 | 0.138394336 | 0.08104257 | 1.70767457 | 0.087696743 | 0.228091726 | 0.138394336 |
| ENSMUSG00000000693 | Loxl3 | 16950 | protein_coding | 10.8453789 | 0.138414733 | 0.524499284 | 0.263898803 | 0.791857898 | 0.887157108 | 0.138414733 |
| ENSMUSG00000020687 | Cdc27 | 217232 | protein_coding | 1175.726399 | 0.138440496 | 0.06116048 | 2.263561291 | 0.023601111 | 0.09049947 | 0.138440496 |
| ENSMUSG00000059534 | Uqcr10 | 66152 | protein_coding | 2836.759661 | 0.13844923 | 0.126545905 | 1.094063297 | 0.273927218 | 0.481845691 | 0.13844923 |
| ENSMUSG00000027556 | Car1 | 12346 | protein_coding | 96852.12922 | -0.13851197 | 0.397572311 | -0.348394409 | 0.727544001 | 0.846662573 | 0.13851197 |
| ENSMUSG00000049721 | Gal3st1 | 53897 | protein_coding | 532.1035643 | 0.138517359 | 0.144916364 | 0.95584346 | 0.339151328 | 0.549942578 | 0.138517359 |
| ENSMUSG00000025885 | Myo5b | 17919 | protein_coding | 3511.293596 | 0.138579694 | 0.077998709 | 1.77669215 | 0.075618905 | 0.205239445 | 0.138579694 |
| ENSMUSG00000022671 | Mzt2 | 72083 | protein_coding | 153.1882119 | 0.138686587 | 0.127252605 | 1.089852641 | 0.275778061 | 0.484028727 | 0.138686587 |

| ensembl_gene_id | external_gene_name | entrezgene_id | gene_biotype | baseMean | log2FoldChange | lfcSE | stat | pvalue | padj | absLog2FC |
|---|---|---|---|---|---|---|---|---|---|---|
| ENSMUSG00000020083 | Fam241b | 69894 | protein_coding | 193.0768561 | 0.138699525 | 0.101458788 | 1.367052843 | 0.171608743 | 0.357153419 | 0.138699525 |
| ENSMUSG00000035674 | Ndufa3 | 66091 | protein_coding | 1346.494263 | 0.138765845 | 0.099036817 | 1.401154136 | 0.161167986 | 0.343238893 | 0.138765845 |
| ENSMUSG00000050312 | Nsun3 | 106338 | protein_coding | 254.9108665 | 0.138810268 | 0.15689472 | 0.884735116 | 0.376299512 | 0.584500916 | 0.138810268 |
| ENSMUSG00000056383 | AI987944 | 233168 | protein_coding | 254.9626298 | 0.138823255 | 0.119743605 | 1.159337526 | 0.246318631 | 0.451844197 | 0.138823255 |
| ENSMUSG00000036552 | Ermard | 381062 | protein_coding | 74.2383764 | -0.138847286 | 0.146074116 | -0.950526279 | 0.341844908 | 0.551941783 | 0.138847286 |
| ENSMUSG00000018334 | Ksr1 | 16706 | protein_coding | 264.7938319 | 0.138883925 | 0.177233193 | 0.783622542 | 0.433261625 | 0.637254688 | 0.138883925 |
| ENSMUSG00000098678 | Mroh6 | 223645 | protein_coding | 85.42458095 | -0.138898225 | 0.614842238 | -0.225908723 | 0.821272413 | 0.905275225 | 0.138898225 |
| ENSMUSG00000039105 | Atp6v1g1 | 66290 | protein_coding | 1125.363414 | 0.138942909 | 0.10063711 | 1.380632945 | 0.167391848 | 0.351650931 | 0.138942909 |
| ENSMUSG00000042179 | Pnliprp1 | 18946 | protein_coding | 7.439001819 | -0.138973723 | 0.661761357 | -0.210005799 | 0.833663147 | 0.911278944 | 0.138973723 |
| ENSMUSG00000024790 | Sac3d1 | 66406 | protein_coding | 142.1122854 | -0.13898153 | 0.129740269 | -1.071228931 | 0.284066506 | 0.493199402 | 0.13898153 |
| ENSMUSG00000015839 | Nfe2l2 | 18024 | protein_coding | 3621.583716 | -0.139024163 | 0.092748463 | -1.498937644 | 0.133889809 | 0.302624788 | 0.139024163 |
| ENSMUSG00000028430 | Nol6 | 230082 | protein_coding | 312.5007816 | -0.139032205 | 0.147438327 | -0.942985505 | 0.345688317 | 0.555637797 | 0.139032205 |
| ENSMUSG00000030583 | Sipa1l3 | 74206 | protein_coding | 856.3799012 | -0.139036804 | 0.146989998 | -0.945892958 | 0.344203185 | 0.55406481 | 0.139036804 |
| ENSMUSG00000022325 | Pop1 | 67724 | protein_coding | 122.6091608 | -0.139115474 | 0.201344645 | -0.690932078 | 0.489608226 | 0.682008993 | 0.139115474 |
| ENSMUSG00000021719 | Rgs7bp | 52882 | protein_coding | 9.043889988 | 0.139203583 | 0.648749275 | 0.214572237 | 0.830100841 | 0.909593162 | 0.139203583 |
| ENSMUSG00000038683 | Pak1ip1 | 68083 | protein_coding | 644.7689353 | 0.139299048 | 0.117102872 | 1.189544255 | 0.234225567 | 0.438035087 | 0.139299048 |
| ENSMUSG00000060376 | Bckdha | 12039 | protein_coding | 187.8014194 | -0.139358234 | 0.125358447 | -1.111678049 | 0.266276601 | 0.474095061 | 0.139358234 |
| ENSMUSG00000024238 | Zeb1 | 21417 | protein_coding | 9.52257714 | 0.139368513 | 0.473187575 | 0.294531219 | 0.768352012 | 0.873665822 | 0.139368513 |
| ENSMUSG00000031533 | Mrps31 | 57312 | protein_coding | 541.6614052 | -0.139380993 | 0.097175187 | -1.43432698 | 0.151478975 | 0.329655755 | 0.139380993 |
| ENSMUSG00000029370 | Rassf6 | 73246 | protein_coding | 1043.909192 | -0.139425804 | 0.121122007 | -1.151118678 | 0.249683415 | 0.456020804 | 0.139425804 |
| ENSMUSG00000054277 | Arfgap3 | 66251 | protein_coding | 1980.621915 | -0.139450921 | 0.097273124 | -1.433601751 | 0.151685944 | 0.329836684 | 0.139450921 |
| ENSMUSG00000034785 | Dio1 | 13370 | protein_coding | 718.9146824 | 0.139501503 | 0.112924112 | 1.235356216 | 0.21669796 | 0.415901327 | 0.139501503 |
| ENSMUSG00000032199 | Polr2m | 28015 | protein_coding | 1494.382915 | 0.139568767 | 0.081784166 | 1.706549983 | 0.087905729 | 0.228448759 | 0.139568767 |
| ENSMUSG00000029623 | Pdap1 | 231887 | protein_coding | 763.7644341 | 0.139575082 | 0.104135427 | 1.340322756 | 0.180140436 | 0.368125204 | 0.139575082 |
| ENSMUSG00000019920 | Lims1 | 110829 | protein_coding | 1783.581173 | 0.139577732 | 0.08270795 | 1.687597536 | 0.091488513 | 0.234352176 | 0.139577732 |
| ENSMUSG00000024976 | Shoc2 | 56392 | protein_coding | 1469.235758 | 0.139656591 | 0.066337432 | 2.105245668 | 0.035269929 | 0.121702649 | 0.139656591 |
| ENSMUSG00000099689 | Zfp383 | 73729 | protein_coding | 39.7318099 | 0.139677634 | 0.262232055 | 0.532648972 | 0.594276595 | 0.762558415 | 0.139677634 |
| ENSMUSG00000073982 | Rhog | 56212 | protein_coding | 201.0819272 | 0.139703533 | 0.118862317 | 1.175339134 | 0.239859064 | 0.444189649 | 0.139703533 |
| ENSMUSG00000079480 | Pin4 | 69713 | protein_coding | 158.2449413 | 0.139757166 | 0.146994679 | 0.950763435 | 0.341724478 | 0.551941783 | 0.139757166 |
| ENSMUSG00000068284 | Usf3 | 207806 | protein_coding | 2367.718362 | 0.139804157 | 0.193246843 | 0.723448592 | 0.469404327 | 0.665598871 | 0.139804157 |
| ENSMUSG00000016833 | Mrps18c | 68735 | protein_coding | 341.4949063 | 0.139883255 | 0.105368008 | 1.327568564 | 0.184320675 | 0.373994868 | 0.139883255 |
| ENSMUSG00000059363 | Fxn | 14297 | protein_coding | 85.32241916 | 0.139916383 | 0.134253266 | 1.04218234 | 0.297327149 | 0.506180721 | 0.139916383 |
| ENSMUSG00000057766 | Ankrd29 | 225187 | protein_coding | 10.80525406 | -0.139951567 | 0.41122361 | -0.3403296 | 0.733608329 | 0.850430383 | 0.139951567 |
| ENSMUSG00000003746 | Man1a | 17155 | protein_coding | 1483.555557 | -0.139964541 | 0.08114265 | -1.724919531 | 0.084541975 | 0.222427947 | 0.139964541 |
| ENSMUSG00000026339 | Ccdc93 | 70829 | protein_coding | 1045.256174 | -0.139972918 | 0.091778479 | -1.525116995 | 0.127229919 | 0.292469065 | 0.139972918 |
| ENSMUSG00000035086 | Becn1 | 56208 | protein_coding | 1330.106945 | 0.139989255 | 0.0759133 | 1.844067589 | 0.065173291 | 0.185802787 | 0.139989255 |
| ENSMUSG00000059713 | Rcan3 | 53902 | protein_coding | 529.7902105 | 0.140081465 | 0.090178229 | 1.553384507 | 0.120331302 | 0.2816099 | 0.140081465 |
| ENSMUSG00000020034 | Tcp11l2 | 216198 | protein_coding | 119.4746867 | -0.140097227 | 0.182026216 | -0.769654119 | 0.441505093 | 0.642981907 | 0.140097227 |

| ensembl_gene_id | external_gene_name | entrezgene_id | gene_biotype | baseMean | log2FoldChange | lfcSE | stat | pvalue | padj | absLog2FC |
|---|---|---|---|---|---|---|---|---|---|---|
| ENSMUSG00000040209 | Zfp704 | 170753 | protein_coding | 923.5468896 | 0.140103062 | 0.190171052 | 0.736721287 | 0.461291859 | 0.659378689 | 0.140103062 |
| ENSMUSG00000000751 | Rpa1 | 68275 | protein_coding | 628.5275168 | 0.140108536 | 0.096439239 | 1.452816691 | 0.146274656 | 0.322293917 | 0.140108536 |
| ENSMUSG00000059991 | Nptx2 | 53324 | protein_coding | 12.81772744 | 0.140115825 | 0.518147446 | 0.270416898 | 0.786839541 | 0.884458731 | 0.140115825 |
| ENSMUSG00000032702 | Kank1 | 107351 | protein_coding | 549.9924672 | -0.140132419 | 0.088752372 | -1.578914636 | 0.114355638 | 0.270969445 | 0.140132419 |
| ENSMUSG00000035367 | Rmi1 | 74386 | protein_coding | 180.106128 | -0.140155114 | 0.093324972 | -1.501796474 | 0.133149679 | 0.301722591 | 0.140155114 |
| ENSMUSG00000020776 | Fbf1 | 217335 | protein_coding | 197.2343817 | -0.140160033 | 0.125238556 | -1.119144434 | 0.263078526 | 0.47092771 | 0.140160033 |
| ENSMUSG00000003955 | Fam162a | 70186 | protein_coding | 1483.378852 | -0.140200065 | 0.193501534 | -0.724542393 | 0.468732811 | 0.66509336 | 0.140200065 |
| ENSMUSG00000020898 | Ctc1 | 68964 | protein_coding | 294.2922486 | -0.140297101 | 0.111749593 | -1.255459611 | 0.209311961 | 0.406228941 | 0.140297101 |
| ENSMUSG00000036873 | 2410004B18Rik | 66421 | protein_coding | 208.9914662 | -0.140301092 | 0.102185659 | -1.373001789 | 0.169751791 | 0.354737145 | 0.140301092 |
| ENSMUSG00000022960 | Donson | 60364 | protein_coding | 132.7857014 | 0.140308418 | 0.130891858 | 1.07194152 | 0.2837463 | 0.492858349 | 0.140308418 |
| ENSMUSG00000015942 | Gtf2ird2 | 114674 | protein_coding | 135.1067613 | 0.140364062 | 0.147963902 | 0.948637205 | 0.342805162 | 0.552987707 | 0.140364062 |
| ENSMUSG00000032235 | Ice2 | 93697 | protein_coding | 285.662013 | -0.140400481 | 0.132469019 | -1.059874088 | 0.289201885 | 0.498097711 | 0.140400481 |
| ENSMUSG00000049090 | Zadh2 | 225791 | protein_coding | 539.8764114 | -0.140430808 | 0.112507493 | -1.248190708 | 0.211961226 | 0.409772837 | 0.140430808 |
| ENSMUSG00000001166 | Oas1c | 114643 | protein_coding | 35.16146143 | -0.140488147 | 0.26602562 | -0.528100064 | 0.597429886 | 0.764756347 | 0.140488147 |
| ENSMUSG00000032867 | Fbxw8 | 231672 | protein_coding | 620.3128721 | 0.140523587 | 0.119243316 | 1.178460919 | 0.238612904 | 0.442830469 | 0.140523587 |
| ENSMUSG00000024590 | Lmnb1 | 16906 | protein_coding | 579.0449969 | 0.140528739 | 0.096151007 | 1.461542049 | 0.143866745 | 0.318044345 | 0.140528739 |
| ENSMUSG00000032244 | Fem1b | 14155 | protein_coding | 1021.396614 | -0.140529468 | 0.092197637 | -1.52421985 | 0.127453801 | 0.292913287 | 0.140529468 |
| ENSMUSG00000033029 | 1700088E04Rik | 27660 | protein_coding | 16.01579108 | -0.140530715 | 0.328646418 | -0.427604584 | 0.668939028 | 0.808227326 | 0.140530715 |
| ENSMUSG00000026077 | Npas2 | 18143 | protein_coding | 238.8810315 | 0.140560921 | 0.157342495 | 0.893343664 | 0.371673162 | 0.579898015 | 0.140560921 |
| ENSMUSG00000001524 | Gtf2h4 | 14885 | protein_coding | 184.3501988 | 0.140567732 | 0.117080953 | 1.200602896 | 0.229905277 | 0.433125054 | 0.140567732 |
| ENSMUSG00000045576 | St7l | 229681 | protein_coding | 440.1545575 | -0.140573055 | 0.0681386 | -2.063045825 | 0.039108275 | 0.131057563 | 0.140573055 |
| ENSMUSG00000072770 | Acrbp | 54137 | protein_coding | 34.08926113 | 0.140616273 | 0.313789285 | 0.448123244 | 0.654064254 | 0.799780088 | 0.140616273 |
| ENSMUSG00000091896 | Ube2d2a | 56550 | protein_coding | 687.813856 | 0.140783764 | 0.056180377 | 2.505924173 | 0.012213181 | 0.055580441 | 0.140783764 |
| ENSMUSG00000054091 | 1810037I17Rik | 67704 | protein_coding | 1161.657179 | 0.140837141 | 0.095642396 | 1.472538811 | 0.140875434 | 0.313281688 | 0.140837141 |
| ENSMUSG00000018171 | Vmp1 | 75909 | protein_coding | 1596.643463 | -0.140855639 | 0.139455696 | -1.010038618 | 0.312476789 | 0.522697993 | 0.140855639 |
| ENSMUSG00000070372 | Capza1 | 12340 | protein_coding | 1425.480259 | 0.140886946 | 0.088984438 | 1.58327624 | 0.113358516 | 0.269408302 | 0.140886946 |
| ENSMUSG00000022749 | Tbc1d23 | 67581 | protein_coding | 573.8273901 | 0.140966416 | 0.062627598 | 2.25086735 | 0.02439394 | 0.092670933 | 0.140966416 |
| ENSMUSG00000031918 | Mtmr2 | 77116 | protein_coding | 935.3247179 | 0.140976709 | 0.077938046 | 1.808830433 | 0.070477351 | 0.196218011 | 0.140976709 |
| ENSMUSG00000064068 | Mtx1 | 17827 | protein_coding | 333.1718777 | 0.14100922 | 0.101021724 | 1.395830671 | 0.162765492 | 0.345262415 | 0.14100922 |
| ENSMUSG00000017264 | Exosc10 | 50912 | protein_coding | 509.8727471 | -0.141053421 | 0.13117053 | -1.075343836 | 0.282220817 | 0.491172768 | 0.141053421 |
| ENSMUSG00000019774 | Mtrf1l | 108853 | protein_coding | 168.6868667 | -0.141098836 | 0.098786164 | -1.428325901 | 0.153198076 | 0.332082057 | 0.141098836 |
| ENSMUSG00000027273 | Snap25 | 20614 | protein_coding | 37.67891528 | -0.141189977 | 0.299551574 | -0.471337789 | 0.637399534 | 0.790262301 | 0.141189977 |
| ENSMUSG00000026648 | Dclre1c | 227525 | protein_coding | 129.122178 | -0.141262456 | 0.126435565 | -1.117268358 | 0.263879601 | 0.471566731 | 0.141262456 |
| ENSMUSG00000031820 | Babam1 | 68251 | protein_coding | 656.8513378 | 0.141284402 | 0.088826864 | 1.590559388 | 0.11170877 | 0.266961334 | 0.141284402 |
| ENSMUSG00000024254 | Abcg8 | 67470 | protein_coding | 24.34946156 | 0.141288794 | 0.316247983 | 0.446765835 | 0.655044142 | 0.800302454 | 0.141288794 |
| ENSMUSG00000036442 | Thap11 | 59016 | protein_coding | 329.5972142 | -0.141321582 | 0.103244467 | -1.368805382 | 0.171060118 | 0.356489017 | 0.141321582 |
| ENSMUSG00000027255 | Arfgap2 | 77038 | protein_coding | 464.7837748 | 0.141325002 | 0.073236565 | 1.9297055 | 0.053643338 | 0.162748779 | 0.141325002 |
| ENSMUSG00000059669 | Taf1b | 21340 | protein_coding | 96.67219067 | -0.141330493 | 0.122953134 | -1.149466375 | 0.250363724 | 0.456634994 | 0.141330493 |

| ensembl_gene_id | external_gene_name | entrezgene_id | gene_biotype | baseMean | log2FoldChange | lfcSE | stat | pvalue | padj | absLog2FC |
|---|---|---|---|---|---|---|---|---|---|---|
| ENSMUSG00000032366 | Tpm1 | 22003 | protein_coding | 3589.959372 | -0.141341158 | 0.087487132 | -1.615565103 | 0.106188374 | 0.258500447 | 0.141341158 |
| ENSMUSG00000024052 | Lpin2 | 64898 | protein_coding | 928.0824435 | -0.141346528 | 0.19761835 | -0.715250014 | 0.47445457 | 0.669373067 | 0.141346528 |
| ENSMUSG00000027315 | Spint1 | 20732 | protein_coding | 2848.18652 | -0.141463996 | 0.127767614 | -1.107197606 | 0.268208502 | 0.476150805 | 0.141463996 |
| ENSMUSG00000032624 | Eml4 | 78798 | protein_coding | 1554.98091 | -0.141468846 | 0.127062809 | -1.113377286 | 0.265546426 | 0.473165172 | 0.141468846 |
| ENSMUSG00000002103 | Acp2 | 11432 | protein_coding | 351.5019742 | 0.141474629 | 0.100076113 | 1.413670298 | 0.15745873 | 0.33818806 | 0.141474629 |
| ENSMUSG00000038267 | Slc22a23 | 73102 | protein_coding | 1445.808102 | -0.141516632 | 0.084918256 | -1.666504208 | 0.095613031 | 0.241194783 | 0.141516632 |
| ENSMUSG00000043153 | Crppa | 75847 | protein_coding | 44.6758015 | -0.141549596 | 0.188161367 | -0.752277674 | 0.451884091 | 0.652077527 | 0.141549596 |
| ENSMUSG00000040505 | Abcg5 | 27409 | protein_coding | 40.22632736 | -0.14163893 | 0.287018084 | -0.49348434 | 0.621670402 | 0.780112359 | 0.14163893 |
| ENSMUSG00000036377 | Cracd | 320827 | protein_coding | 723.7978839 | -0.141738098 | 0.157592234 | -0.899397737 | 0.368440844 | 0.577213117 | 0.141738098 |
| ENSMUSG00000007987 | Ift22 | 67286 | protein_coding | 84.53489084 | 0.141758331 | 0.142129556 | 0.997388126 | 0.318576153 | 0.529987548 | 0.141758331 |
| ENSMUSG00000024067 | Dpy30 | 66310 | protein_coding | 575.8192396 | 0.141819436 | 0.088800047 | 1.597064874 | 0.110251247 | 0.264838953 | 0.141819436 |
| ENSMUSG00000059555 | Tor4a | 227612 | protein_coding | 672.9615326 | 0.141897363 | 0.079459494 | 1.785782369 | 0.074134502 | 0.2029556 | 0.141897363 |
| ENSMUSG00000000131 | Xpo6 | 74204 | protein_coding | 763.3315193 | -0.141958456 | 0.096035353 | -1.478189559 | 0.139357048 | 0.311010096 | 0.141958456 |
| ENSMUSG00000018166 | Erbb3 | 13867 | protein_coding | 5868.620275 | -0.141990169 | 0.095901264 | -1.480587035 | 0.138716652 | 0.309971779 | 0.141990169 |
| ENSMUSG00000067851 | Arfgef1 | 211673 | protein_coding | 3550.225714 | 0.142043097 | 0.086696664 | 1.638391724 | 0.101340005 | 0.25035457 | 0.142043097 |
| ENSMUSG00000026939 | Tmem141 | 51875 | protein_coding | 99.70156301 | -0.142059406 | 0.129448792 | -1.097417776 | 0.272458806 | 0.48053636 | 0.142059406 |
| ENSMUSG00000038718 | Pbx3 | 18516 | protein_coding | 71.33782958 | 0.142108445 | 0.145156529 | 0.979001398 | 0.327579289 | 0.538682935 | 0.142108445 |
| ENSMUSG00000103711 | Tstd1 | 226654 | protein_coding | 1921.284401 | 0.142147038 | 0.119421889 | 1.190292995 | 0.233931253 | 0.437742522 | 0.142147038 |
| ENSMUSG00000028645 | Slc2a1 | 20525 | protein_coding | 1207.954071 | -0.142188482 | 0.195924176 | -0.725732193 | 0.468002962 | 0.664617616 | 0.142188482 |
| ENSMUSG00000042359 | Osbpl6 | 99031 | protein_coding | 225.0468032 | -0.142206304 | 0.158414957 | -0.897682311 | 0.369354942 | 0.578133404 | 0.142206304 |
| ENSMUSG00000042167 | Tent2 | 100715 | protein_coding | 405.2219981 | 0.142213744 | 0.069052845 | 2.05949144 | 0.039447184 | 0.131832721 | 0.142213744 |
| ENSMUSG00000059482 | Cfap418 | 67157 | protein_coding | 60.83802719 | 0.142252627 | 0.196459466 | 0.724081309 | 0.469015819 | 0.665284884 | 0.142252627 |
| ENSMUSG00000032615 | Nt5m | 103850 | protein_coding | 140.8302031 | 0.142252816 | 0.147796826 | 0.962488981 | 0.335804036 | 0.54646736 | 0.142252816 |
| ENSMUSG00000029054 | Gabrd | 14403 | protein_coding | 9.310340186 | 0.142286016 | 0.431976662 | 0.329383573 | 0.741865783 | 0.856201593 | 0.142286016 |
| ENSMUSG00000000171 | Sdhd | 66925 | protein_coding | 4478.036414 | 0.142290679 | 0.102727909 | 1.385121926 | 0.166015189 | 0.349853777 | 0.142290679 |
| ENSMUSG00000069303 | H2bc24 | 665622 | protein_coding | 54.93229381 | -0.142308002 | 2.033614648 | -0.06997786 | 0.944211281 | 0.972731213 | 0.142308002 |
| ENSMUSG00000100199 | Gm20324 | 100504637 | lncRNA | 12.3164724 | -0.14241345 | 0.362442593 | -0.392926915 | 0.694373462 | 0.826234909 | 0.14241345 |
| ENSMUSG00000026037 | Orc2 | 18393 | protein_coding | 303.958198 | 0.142418759 | 0.117078436 | 1.216438854 | 0.22381779 | 0.425414053 | 0.142418759 |
| ENSMUSG00000053600 | Zfp472 | 224691 | protein_coding | 218.4051398 | -0.142563817 | 0.098123785 | -1.452897656 | 0.146252172 | 0.322288959 | 0.142563817 |
| ENSMUSG00000043909 | Trp53bp1 | 27223 | protein_coding | 483.1399214 | -0.142581249 | 0.08656017 | -1.647192329 | 0.09951852 | 0.247522016 | 0.142581249 |
| ENSMUSG00000038014 | Fam120a | 218236 | protein_coding | 7466.295478 | 0.142598744 | 0.052097377 | 2.737157826 | 0.006197255 | 0.032980248 | 0.142598744 |
| ENSMUSG00000027811 | 4930579G24Rik | 75939 | protein_coding | 38.44073017 | -0.142604299 | 0.202086394 | -0.705660067 | 0.48039956 | 0.673965179 | 0.142604299 |
| ENSMUSG00000066894 | Vsig10 | 231668 | protein_coding | 2006.769962 | -0.142615063 | 0.091069389 | -1.566004394 | 0.117347588 | 0.276577743 | 0.142615063 |
| ENSMUSG00000058056 | Palld | 72333 | protein_coding | 232.7075668 | -0.142698469 | 0.261599848 | -0.545483762 | 0.585420846 | 0.755883855 | 0.142698469 |
| ENSMUSG00000025237 | Parp6 | 67287 | protein_coding | 350.836169 | -0.142727066 | 0.083421479 | -1.710915074 | 0.087096791 | 0.227050359 | 0.142727066 |
| ENSMUSG00000034612 | Chst11 | 58250 | protein_coding | 99.03621221 | -0.142732672 | 0.307803363 | -0.463713818 | 0.642852796 | 0.793238306 | 0.142732672 |
| ENSMUSG00000054021 | Sirt5 | 68346 | protein_coding | 153.1466921 | 0.142771132 | 0.109530521 | 1.303482638 | 0.19241004 | 0.385434609 | 0.142771132 |
| ENSMUSG00000040771 | Oard1 | 106821 | protein_coding | 368.6947489 | 0.142774964 | 0.11579926 | 1.232952302 | 0.21759356 | 0.417218037 | 0.142774964 |

| ensembl_gene_id | external_gene_name | entrezgene_id | gene_biotype | baseMean | log2FoldChange | lfcSE | stat | pvalue | padj | absLog2FC |
|---|---|---|---|---|---|---|---|---|---|---|
| ENSMUSG00000003808 | Farsa | 66590 | protein_coding | 403.1753993 | 0.142794571 | 0.118925247 | 1.20070863 | 0.229864245 | 0.433125054 | 0.142794571 |
| ENSMUSG00000073406 | H2-Bl | 14963 | protein_coding | 15.49818297 | -0.142798152 | 0.429972384 | -0.332110055 | 0.739806158 | 0.854833615 | 0.142798152 |
| ENSMUSG00000064302 | Clasp1 | 76707 | protein_coding | 620.8830207 | -0.142799239 | 0.09974102 | -1.431700202 | 0.152229639 | 0.330657805 | 0.142799239 |
| ENSMUSG00000002007 | Srpk3 | 56504 | protein_coding | 8.841153217 | -0.142852985 | 0.521273485 | -0.274046137 | 0.784049166 | 0.882817412 | 0.142852985 |
| ENSMUSG00000026201 | Stk16 | 20872 | protein_coding | 707.4207748 | 0.142861832 | 0.080038338 | 1.784917516 | 0.074274697 | 0.203098262 | 0.142861832 |
| ENSMUSG00000020100 | Slc29a3 | 71279 | protein_coding | 107.9635059 | -0.142872369 | 0.190507548 | -0.749956474 | 0.45328092 | 0.653082762 | 0.142872369 |
| ENSMUSG00000050107 | Haspin | 14841 | protein_coding | 148.293247 | 0.142888919 | 0.128881057 | 1.108688296 | 0.267564673 | 0.47543558 | 0.142888919 |
| ENSMUSG00000064380 | Gm26448 | 115487256 | snoRNA | 359.6010873 | 0.14292699 | 0.222985193 | 0.640970766 | 0.521541677 | 0.707881371 | 0.14292699 |
| ENSMUSG00000023830 | Igf2r | 16004 | protein_coding | 3130.067898 | -0.142984371 | 0.066437248 | -2.152171785 | 0.031383827 | 0.111737326 | 0.142984371 |
| ENSMUSG00000034055 | Phka1 | 18679 | protein_coding | 269.497321 | -0.143006459 | 0.094160655 | -1.518749614 | 0.128825533 | 0.29502992 | 0.143006459 |
| ENSMUSG00000025494 | Sigirr | 24058 | protein_coding | 356.8526884 | -0.143066625 | 0.126645583 | -1.129661386 | 0.258618934 | 0.465714909 | 0.143066625 |
| ENSMUSG00000031660 | Brd7 | 26992 | protein_coding | 597.4905347 | -0.143128859 | 0.075735943 | -1.889840581 | 0.058779284 | 0.173602159 | 0.143128859 |
| ENSMUSG00000079737 | Bfar | 66598 | transcribed_processed_pseudogene | 102.4528528 | -0.143167509 | 0.15236494 | -0.939635513 | 0.347404554 | 0.557627687 | 0.143167509 |
| ENSMUSG00000090038 | Gm16573 | 102635191 | lncRNA | 6.735802356 | 0.143232949 | 0.541583261 | 0.264470783 | 0.791417176 | 0.887042894 | 0.143232949 |
| ENSMUSG00000041842 | Fhdc1 | 229474 | protein_coding | 269.8201558 | -0.14332734 | 0.241643109 | -0.593136467 | 0.553089827 | 0.732913042 | 0.14332734 |
| ENSMUSG00000047719 | Ubiad1 | 71707 | protein_coding | 265.688066 | 0.143332991 | 0.121428999 | 1.180385187 | 0.237847051 | 0.44210851 | 0.143332991 |
| ENSMUSG00000024525 | Impa2 | 114663 | protein_coding | 284.8831472 | -0.143342146 | 0.084581946 | -1.69471326 | 0.090129829 | 0.232127689 | 0.143342146 |
| ENSMUSG00000058793 | Cds2 | 110911 | protein_coding | 1037.64731 | -0.143344464 | 0.077043552 | -1.860564068 | 0.062805763 | 0.181128045 | 0.143344464 |
| ENSMUSG00000020864 | Ankrd40 | 71452 | protein_coding | 634.088915 | -0.14340477 | 0.072945826 | -1.965907821 | 0.049309259 | 0.154029787 | 0.14340477 |
| ENSMUSG00000062683 | Atp5g2 | 67942 | protein_coding | 608.3951156 | 0.14340981 | 0.095038767 | 1.508961184 | 0.131308702 | 0.298826263 | 0.14340981 |
| ENSMUSG00000039007 | Cpq | 54381 | protein_coding | 781.1799068 | -0.143443752 | 0.118843578 | -1.206996238 | 0.227433593 | 0.430079813 | 0.143443752 |
| ENSMUSG00000027346 | Gpcpd1 | 74182 | protein_coding | 594.2834841 | -0.143546089 | 0.148186872 | -0.9686829 | 0.332703427 | 0.543652824 | 0.143546089 |
| ENSMUSG00000059325 | Hopx | 74318 | protein_coding | 318.0627753 | -0.143549309 | 0.106029574 | -1.353861033 | 0.175780717 | 0.361907388 | 0.143549309 |
| ENSMUSG00000032066 | Bco2 | 170752 | protein_coding | 240.6498436 | -0.143586851 | 0.207869028 | -0.690756349 | 0.489718671 | 0.682077562 | 0.143586851 |
| ENSMUSG00000031358 | Msl3 | 17692 | protein_coding | 679.693046 | 0.143746661 | 0.078969219 | 1.82028724 | 0.068715274 | 0.193101072 | 0.143746661 |
| ENSMUSG00000078870 | Gm14410 | 100043403 | protein_coding | 62.07351442 | -0.143779613 | 0.168883339 | -0.851354629 | 0.394572387 | 0.60256462 | 0.143779613 |
| ENSMUSG00000037646 | Vps13b | 666173 | protein_coding | 2358.464225 | -0.143822177 | 0.092337324 | -1.557573586 | 0.119334364 | 0.280017347 | 0.143822177 |
| ENSMUSG00000030814 | Bcl7c | 12055 | protein_coding | 91.86855628 | -0.143893384 | 0.146914404 | -0.979436869 | 0.327364168 | 0.538495968 | 0.143893384 |
| ENSMUSG00000054226 | Tprkb | 69786 | protein_coding | 250.5725182 | -0.143937578 | 0.10179755 | -1.413959147 | 0.157373898 | 0.33818806 | 0.143937578 |
| ENSMUSG00000018412 | Kansl1 | 76719 | protein_coding | 794.9938246 | -0.143960172 | 0.087292277 | -1.649174218 | 0.099111948 | 0.246819123 | 0.143960172 |
| ENSMUSG00000073705 | Cenps | 69928 | protein_coding | 24.93972969 | 0.144055858 | 0.282159011 | 0.510548492 | 0.609667251 | 0.772659132 | 0.144055858 |
| ENSMUSG00000028322 | Exosc3 | 66362 | protein_coding | 285.8110162 | 0.144140156 | 0.134494073 | 1.07172125 | 0.283845253 | 0.492976469 | 0.144140156 |
| ENSMUSG00000027709 | Mccc1 | 72039 | protein_coding | 374.3319 | -0.144182643 | 0.112304581 | -1.283853618 | 0.199193179 | 0.393494847 | 0.144182643 |
| ENSMUSG00000001924 | Uba1 | 22201 | protein_coding | 4372.072843 | 0.144192971 | 0.079344556 | 1.817301378 | 0.069170979 | 0.194107709 | 0.144192971 |
| ENSMUSG00000021377 | Dek | 110052 | protein_coding | 903.4421406 | -0.14419396 | 0.088094674 | -1.636806794 | 0.101670846 | 0.250948811 | 0.14419396 |
| ENSMUSG00000064181 | Rab3ip | 216363 | protein_coding | 590.0100336 | -0.14420163 | 0.094055786 | -1.53315001 | 0.125238898 | 0.289329844 | 0.14420163 |
| ENSMUSG00000019373 | Cops3 | 26572 | protein_coding | 849.2014462 | 0.144212997 | 0.067058209 | 2.150564404 | 0.031510597 | 0.11205644 | 0.144212997 |
| ENSMUSG00000069808 | Tlcd3a | 116972 | protein_coding | 702.9563119 | 0.144213899 | 0.150024442 | 0.961269356 | 0.336416751 | 0.547115174 | 0.144213899 |

| ensembl_gene_id | external_gene_name | entrezgene_id | gene_biotype | baseMean | log2FoldChange | lfcSE | stat | pvalue | padj | absLog2FC |
|---|---|---|---|---|---|---|---|---|---|---|
| ENSMUSG00000055044 | Pdlim1 | 54132 | protein_coding | 2676.364444 | -0.144226743 | 0.091383191 | -1.578263372 | 0.114505117 | 0.271202602 | 0.144226743 |
| ENSMUSG00000057110 | Cntrl | 26920 | protein_coding | 1047.54093 | -0.144243424 | 0.093975234 | -1.534908911 | 0.124806204 | 0.288814939 | 0.144243424 |
| ENSMUSG00000039637 | Coro7 | 78885 | protein_coding | 299.7933815 | 0.144295098 | 0.117371967 | 1.229382976 | 0.21892825 | 0.419121294 | 0.144295098 |
| ENSMUSG00000097039 | Pvt1 | 19296 | lncRNA | 26.36421673 | 0.144329745 | 0.254515667 | 0.567076072 | 0.570662499 | 0.745091029 | 0.144329745 |
| ENSMUSG00000036893 | Ehmt1 | 77683 | protein_coding | 823.7713025 | -0.144373794 | 0.075199632 | -1.919873679 | 0.054873857 | 0.165223214 | 0.144373794 |
| ENSMUSG00000028576 | Ift74 | 67694 | protein_coding | 157.6417549 | 0.144427662 | 0.1422044 | 1.015634275 | 0.309803577 | 0.519699986 | 0.144427662 |
| ENSMUSG00000021038 | Vipas39 | 104799 | protein_coding | 867.0680094 | 0.144525394 | 0.127318726 | 1.135146404 | 0.256313981 | 0.463348653 | 0.144525394 |
| ENSMUSG00000050410 | Tcf19 | 106795 | protein_coding | 145.3735189 | 0.144555017 | 0.160057088 | 0.90314661 | 0.366448089 | 0.575230748 | 0.144555017 |
| ENSMUSG00000063316 | Rpl27 | 19942 | protein_coding | 1724.872817 | 0.144592214 | 0.140654306 | 1.027997061 | 0.30395121 | 0.513560504 | 0.144592214 |
| ENSMUSG00000025008 | Tctn3 | 67590 | protein_coding | 70.39016919 | 0.14460756 | 0.156081349 | 0.926488408 | 0.354192218 | 0.563829056 | 0.14460756 |
| ENSMUSG00000034928 | Rnf44 | 105239 | protein_coding | 789.0431886 | -0.144625019 | 0.081531221 | -1.773860581 | 0.076086214 | 0.206077178 | 0.144625019 |
| ENSMUSG00000004221 | Ikbkg | 16151 | protein_coding | 830.9218966 | 0.144643231 | 0.089351268 | 1.618815656 | 0.105486933 | 0.257231468 | 0.144643231 |
| ENSMUSG00000037204 | Atg101 | 68118 | protein_coding | 461.6960013 | 0.144655453 | 0.111297603 | 1.299717597 | 0.193697777 | 0.387103913 | 0.144655453 |
| ENSMUSG00000039043 | Arpin | 70420 | protein_coding | 885.0424941 | 0.144682177 | 0.067778662 | 2.134627225 | 0.032791472 | 0.115120688 | 0.144682177 |
| ENSMUSG00000052593 | Adam17 | 11491 | protein_coding | 705.779067 | 0.144697458 | 0.074822187 | 1.933884379 | 0.053127345 | 0.161919549 | 0.144697458 |
| ENSMUSG00000066148 | Prpf4 | 70052 | protein_coding | 344.734843 | -0.144841959 | 0.150052956 | -0.965272279 | 0.334408453 | 0.545293316 | 0.144841959 |
| ENSMUSG00000118346 | Tmem179b | 67706 | protein_coding | 216.1241682 | -0.14490138 | 0.096403646 | -1.503069503 | 0.132821122 | 0.301192317 | 0.14490138 |
| ENSMUSG00000028943 | Espn | 56226 | protein_coding | 370.0656069 | -0.144919997 | 0.124802376 | -1.161195819 | 0.245562275 | 0.450976122 | 0.144919997 |
| ENSMUSG00000007891 | Ctsd | 13033 | protein_coding | 1987.224554 | -0.144922866 | 0.095744928 | -1.513634924 | 0.130118458 | 0.296838557 | 0.144922866 |
| ENSMUSG00000031845 | Bco1 | 63857 | protein_coding | 30.01247761 | 0.14496259 | 0.207807274 | 0.697581885 | 0.485438713 | 0.678688781 | 0.14496259 |
| ENSMUSG00000047213 | Ythdf3 | 229096 | protein_coding | 2116.993747 | 0.145041929 | 0.087131078 | 1.66464059 | 0.095984482 | 0.241905195 | 0.145041929 |
| ENSMUSG00000034902 | Pip5k1c | 18717 | protein_coding | 434.7094508 | -0.145066601 | 0.076824736 | -1.888279851 | 0.058988391 | 0.173993906 | 0.145066601 |
| ENSMUSG00000023971 | Rrp36 | 224823 | protein_coding | 320.3257998 | 0.14509198 | 0.131823358 | 1.100654556 | 0.271047031 | 0.479054274 | 0.14509198 |
| ENSMUSG00000040009 | Gnaz | 14687 | protein_coding | 10.27430968 | 0.145109535 | 0.44343917 | 0.32723662 | 0.743488926 | 0.857217273 | 0.145109535 |
| ENSMUSG00000041278 | Ttc1 | 66827 | protein_coding | 430.9685923 | 0.145169545 | 0.081709386 | 1.776656916 | 0.075624705 | 0.205239445 | 0.145169545 |
| ENSMUSG00000032512 | Wdr48 | 67561 | protein_coding | 1032.909988 | -0.145183127 | 0.073944627 | -1.963403322 | 0.049599328 | 0.154534306 | 0.145183127 |
| ENSMUSG00000032119 | Hinfp | 102423 | protein_coding | 410.8394131 | 0.145202045 | 0.092331718 | 1.572612834 | 0.115808494 | 0.273516859 | 0.145202045 |
| ENSMUSG00000032409 | Atr | 245000 | protein_coding | 478.8972656 | -0.145226675 | 0.07413632 | -1.958913994 | 0.05012286 | 0.15580736 | 0.145226675 |
| ENSMUSG00000037111 | Setd7 | 73251 | protein_coding | 530.4357068 | -0.145266393 | 0.121613655 | -1.194490807 | 0.232286035 | 0.435789395 | 0.145266393 |
| ENSMUSG00000030615 | Tmem126a | 66271 | protein_coding | 455.1745448 | 0.145288756 | 0.079129454 | 1.836089464 | 0.066344426 | 0.188299006 | 0.145288756 |
| ENSMUSG00000078653 | Cntd1 | 68107 | protein_coding | 25.27688256 | 0.145295284 | 0.250902098 | 0.579091549 | 0.562527402 | 0.739622589 | 0.145295284 |
| ENSMUSG00000025912 | Mybl1 | 17864 | protein_coding | 52.87436263 | 0.145322316 | 0.257446217 | 0.564476409 | 0.572429951 | 0.746187434 | 0.145322316 |
| ENSMUSG00000039697 | Ncoa7 | 211329 | protein_coding | 907.5196524 | 0.145346201 | 0.384116499 | 0.378390935 | 0.705140201 | 0.833382076 | 0.145346201 |
| ENSMUSG00000028536 | 2610528J11Rik | 66451 | protein_coding | 458.1251753 | 0.145359181 | 0.102804775 | 1.413934141 | 0.157381241 | 0.33818806 | 0.145359181 |
| ENSMUSG00000020134 | Peli1 | 67245 | protein_coding | 855.079435 | 0.145388526 | 0.092288776 | 1.575365211 | 0.11517217 | 0.272458387 | 0.145388526 |
| ENSMUSG00000068663 | Clec16a | 74374 | protein_coding | 437.8156162 | 0.145426421 | 0.075253345 | 1.932491122 | 0.053298917 | 0.162074293 | 0.145426421 |
| ENSMUSG00000023284 | Zfp605 | 675812 | protein_coding | 66.25432507 | -0.145578471 | 0.195373423 | -0.745129346 | 0.45619353 | 0.655296022 | 0.145578471 |
| ENSMUSG00000040813 | Tex264 | 21767 | protein_coding | 601.4804142 | -0.145592521 | 0.094431993 | -1.541771145 | 0.123129216 | 0.286098053 | 0.145592521 |

| ensembl_gene_id | external_gene_name | entrezgene_id | gene_biotype | baseMean | log2FoldChange | lfcSE | stat | pvalue | padj | absLog2FC |
|---|---|---|---|---|---|---|---|---|---|---|
| ENSMUSG00000021285 | Ppp1r13b | 21981 | protein_coding | 658.4367243 | -0.145619751 | 0.096777475 | -1.504686401 | 0.132404721 | 0.300504759 | 0.145619751 |
| ENSMUSG00000078607 | 1810010H24Rik | 69066 | protein_coding | 15.21398501 | -0.145645682 | 0.334403364 | -0.435538925 | 0.663171291 | 0.804794609 | 0.145645682 |
| ENSMUSG00000031696 | Vps35 | 65114 | protein_coding | 2290.910924 | 0.145653608 | 0.07530241 | 1.93424896 | 0.053082525 | 0.16183945 | 0.145653608 |
| ENSMUSG00000054612 | Mgmt | 17314 | protein_coding | 48.56685315 | 0.145670733 | 0.173888044 | 0.837727137 | 0.402183969 | 0.609745503 | 0.145670733 |
| ENSMUSG00000029191 | Rfc1 | 19687 | protein_coding | 691.5027529 | -0.145677097 | 0.074814726 | -1.947171428 | 0.051514186 | 0.158593398 | 0.145677097 |
| ENSMUSG00000079436 | Kcnj13 | 100040591 | protein_coding | 10.71439935 | -0.145692018 | 0.364841834 | -0.399329254 | 0.689650614 | 0.823191875 | 0.145692018 |
| ENSMUSG00000026728 | Vim | 22352 | protein_coding | 120.5556946 | 0.145704932 | 0.20212458 | 0.720866961 | 0.470991372 | 0.666543809 | 0.145704932 |
| ENSMUSG00000006307 | Kmt2b | 75410 | protein_coding | 582.2668791 | -0.145730778 | 0.161527464 | -0.902204337 | 0.366948332 | 0.57565559 | 0.145730778 |
| ENSMUSG00000019158 | Tmem160 | 69094 | protein_coding | 110.5262508 | -0.145750262 | 0.133526102 | -1.091548843 | 0.275031452 | 0.483201078 | 0.145750262 |
| ENSMUSG00000058298 | Mcm9 | 71567 | protein_coding | 240.3412262 | -0.145751709 | 0.09725253 | -1.498693226 | 0.133953235 | 0.302702223 | 0.145751709 |
| ENSMUSG00000047945 | Marcksl1 | 17357 | protein_coding | 187.4891091 | 0.145782741 | 0.204174049 | 0.714012101 | 0.475219697 | 0.670024255 | 0.145782741 |
| ENSMUSG00000025351 | Cd63 | 12512 | protein_coding | 3061.275955 | -0.145783033 | 0.057558598 | -2.532775938 | 0.011316326 | 0.052488772 | 0.145783033 |
| ENSMUSG00000018572 | Phf23 | 78246 | protein_coding | 473.4903089 | -0.145784305 | 0.112884813 | -1.291443026 | 0.196550099 | 0.390160765 | 0.145784305 |
| ENSMUSG00000004642 | Slbp | 20492 | protein_coding | 622.6171838 | -0.145784732 | 0.081850786 | -1.781103636 | 0.074895522 | 0.204201575 | 0.145784732 |
| ENSMUSG00000074578 | Zfas1 | 68949 | lncRNA | 524.6218319 | -0.145873235 | 0.136283671 | -1.070364729 | 0.284455169 | 0.493658959 | 0.145873235 |
| ENSMUSG00000017801 | Mlx | 21428 | protein_coding | 1457.904834 | 0.145892531 | 0.090394696 | 1.61395013 | 0.106538244 | 0.259074984 | 0.145892531 |
| ENSMUSG00000045005 | Fzd5 | 14367 | protein_coding | 2231.047924 | -0.145894567 | 0.122570418 | -1.190291821 | 0.233931714 | 0.437742522 | 0.145894567 |
| ENSMUSG00000047407 | Tgif1 | 21815 | protein_coding | 603.8252952 | -0.145938952 | 0.092212926 | -1.582630097 | 0.113505799 | 0.269547674 | 0.145938952 |
| ENSMUSG00000033174 | Mgll | 23945 | protein_coding | 445.8891585 | 0.145950566 | 0.266081126 | 0.548519048 | 0.583335554 | 0.754056602 | 0.145950566 |
| ENSMUSG00000003051 | Elf3 | 13710 | protein_coding | 5398.717419 | -0.145988951 | 0.13077334 | -1.116351015 | 0.264271913 | 0.472003031 | 0.145988951 |
| ENSMUSG00000069729 | Arid1b | 239985 | protein_coding | 901.9669019 | -0.146006246 | 0.091636178 | -1.593325367 | 0.111087219 | 0.266114788 | 0.146006246 |
| ENSMUSG00000013593 | Ndufs2 | 226646 | protein_coding | 3944.984156 | 0.146118805 | 0.102583336 | 1.424391247 | 0.154333247 | 0.333675067 | 0.146118805 |
| ENSMUSG00000021668 | Polk | 27015 | protein_coding | 236.237683 | -0.14617275 | 0.09093216 | -1.607492334 | 0.10794641 | 0.261342212 | 0.14617275 |
| ENSMUSG00000019254 | Ppp1r12c | 232807 | protein_coding | 400.2116027 | -0.146202101 | 0.091626035 | -1.595639277 | 0.110569353 | 0.265233231 | 0.146202101 |
| ENSMUSG00000015889 | Lta4h | 16993 | protein_coding | 1618.152439 | 0.14620609 | 0.06746045 | 2.167286027 | 0.030213051 | 0.108557068 | 0.14620609 |
| ENSMUSG00000033004 | Mycbp2 | 105689 | protein_coding | 1049.02494 | -0.146263312 | 0.117000914 | -1.25010401 | 0.211261555 | 0.408717849 | 0.146263312 |
| ENSMUSG00000035226 | Rims4 | 241770 | protein_coding | 121.6133663 | -0.14626409 | 0.406339451 | -0.359955426 | 0.718880467 | 0.842183589 | 0.14626409 |
| ENSMUSG00000085227 | 6330418K02Rik | 69004 | lncRNA | 9.962059924 | -0.146301635 | 0.340380753 | -0.429817591 | 0.667328336 | 0.807272755 | 0.146301635 |
| ENSMUSG00000040147 | Maob | 109731 | protein_coding | 414.7684205 | 0.146357471 | 0.139928735 | 1.045942931 | 0.295587382 | 0.504837638 | 0.146357471 |
| ENSMUSG00000028618 | Tmem59 | 56374 | protein_coding | 4906.222514 | -0.146423489 | 0.103031477 | -1.42115297 | 0.155272293 | 0.33520969 | 0.146423489 |
| ENSMUSG00000020151 | Ptprr | 19279 | protein_coding | 315.4270735 | -0.146424561 | 0.196881441 | -0.743719472 | 0.457046207 | 0.655781639 | 0.146424561 |
| ENSMUSG00000030867 | Plk1 | 18817 | protein_coding | 840.8867484 | -0.146436951 | 0.070248145 | -2.084566807 | 0.03710864 | 0.126134533 | 0.146436951 |
| ENSMUSG00000024331 | Dsc2 | 13506 | protein_coding | 4960.833953 | 0.146440389 | 0.119771417 | 1.222665583 | 0.221456031 | 0.42238674 | 0.146440389 |
| ENSMUSG00000028689 | Ccdc163 | 68394 | protein_coding | 93.76077894 | -0.146537246 | 0.180247684 | -0.812977139 | 0.416231165 | 0.622565158 | 0.146537246 |
| ENSMUSG00000021893 | Capn7 | 12339 | protein_coding | 1329.413567 | 0.14658981 | 0.061951384 | 2.366207195 | 0.017971384 | 0.073824443 | 0.14658981 |
| ENSMUSG00000021749 | Fam3d | 18300 | protein_coding | 12313.16246 | -0.146593394 | 0.113536834 | -1.29115274 | 0.196650719 | 0.390214607 | 0.146593394 |
| ENSMUSG00000040282 | Cdin1 | 399568 | protein_coding | 62.33807511 | 0.146598676 | 0.207110149 | 0.707829511 | 0.479051137 | 0.672927641 | 0.146598676 |
| ENSMUSG00000037935 | Smarce1 | 57376 | protein_coding | 1048.582404 | 0.146655249 | 0.085679288 | 1.711676795 | 0.086956247 | 0.2269068 | 0.146655249 |

| ensembl_gene_id | external_gene_name | entrezgene_id | gene_biotype | baseMean | log2FoldChange | lfcSE | stat | pvalue | padj | absLog2FC |
|---|---|---|---|---|---|---|---|---|---|---|
| ENSMUSG00000029594 | Rbm19 | 74111 | protein_coding | 200.7433087 | -0.146716029 | 0.189135168 | -0.775720511 | 0.437914027 | 0.640519699 | 0.146716029 |
| ENSMUSG00000022108 | Itm2b | 16432 | protein_coding | 7776.421086 | -0.146734158 | 0.074291598 | -1.975111077 | 0.048255544 | 0.152035744 | 0.146734158 |
| ENSMUSG00000022617 | Chkb | 12651 | protein_coding | 89.70105195 | -0.146767383 | 0.213554662 | -0.687259092 | 0.491919473 | 0.683859515 | 0.146767383 |
| ENSMUSG00000055204 | Ankrd17 | 81702 | protein_coding | 2100.435714 | -0.146792882 | 0.10627042 | -1.381314597 | 0.167182252 | 0.351303286 | 0.146792882 |
| ENSMUSG00000029432 | Nipsnap2 | 14467 | protein_coding | 1293.414622 | 0.14679708 | 0.081245143 | 1.806841298 | 0.070787027 | 0.196873716 | 0.14679708 |
| ENSMUSG00000007833 | Aldh16a1 | 69748 | protein_coding | 1027.872418 | 0.14685619 | 0.108612024 | 1.352117237 | 0.176337814 | 0.362694822 | 0.14685619 |
| ENSMUSG00000051444 | Bbs12 | 241950 | protein_coding | 15.58318829 | -0.146909598 | 0.351658199 | -0.41776247 | 0.676120793 | 0.81406395 | 0.146909598 |
| ENSMUSG00000024619 | Cdx1 | 12590 | protein_coding | 2936.62786 | -0.14699289 | 0.105126414 | -1.398248875 | 0.162038344 | 0.344355804 | 0.14699289 |
| ENSMUSG00000025949 | Pikfyve | 18711 | protein_coding | 692.8286379 | -0.147 | 0.067047204 | -2.19248515 | 0.028344494 | 0.103323822 | 0.147 |
| ENSMUSG00000040822 | 1700123O20Rik | 58248 | protein_coding | 442.4007449 | 0.147017355 | 0.094899139 | 1.549195886 | 0.121334638 | 0.283292014 | 0.147017355 |
| ENSMUSG00000000282 | Mnt | 17428 | protein_coding | 149.6468903 | -0.147143785 | 0.139946632 | -1.05142784 | 0.293062135 | 0.502178856 | 0.147143785 |
| ENSMUSG00000030612 | Mrpl46 | 67308 | protein_coding | 435.0974433 | 0.147155694 | 0.099952776 | 1.472252197 | 0.140952787 | 0.313342078 | 0.147155694 |
| ENSMUSG00000027014 | Cwc22 | 80744 | protein_coding | 460.4342041 | 0.147208193 | 0.110331388 | 1.334236754 | 0.182126278 | 0.370960019 | 0.147208193 |
| ENSMUSG00000020781 | Tsen54 | 76265 | protein_coding | 224.2646455 | -0.147236463 | 0.109139403 | -1.349067884 | 0.177315163 | 0.364187739 | 0.147236463 |
| ENSMUSG00000079109 | Pms2 | 18861 | protein_coding | 131.812669 | -0.147296206 | 0.108868901 | -1.35296861 | 0.176065659 | 0.362228599 | 0.147296206 |
| ENSMUSG00000041361 | Myzap | 102371 | protein_coding | 188.3552295 | -0.147345721 | 0.195384233 | -0.754133119 | 0.450769292 | 0.651233858 | 0.147345721 |
| ENSMUSG00000115219 | Eef1akmt4 | 110599566 | protein_coding | 74.66725716 | -0.14743777 | 0.152466918 | -0.967014824 | 0.333536623 | 0.544302048 | 0.14743777 |
| ENSMUSG00000025135 | Anapc11 | 66156 | protein_coding | 238.8066434 | 0.147443366 | 0.119313691 | 1.235762344 | 0.216546916 | 0.415723343 | 0.147443366 |
| ENSMUSG00000026655 | Fam107b | 66540 | protein_coding | 3318.891242 | -0.147447547 | 0.142998538 | -1.031112267 | 0.302488176 | 0.511794313 | 0.147447547 |
| ENSMUSG00000044702 | Palb2 | 233826 | protein_coding | 76.56171473 | 0.147473459 | 0.165396794 | 0.891634326 | 0.372588965 | 0.580798301 | 0.147473459 |
| ENSMUSG00000028133 | Rwdd3 | 66568 | protein_coding | 37.84548819 | -0.147513943 | 0.181574009 | -0.812417725 | 0.416551979 | 0.622755765 | 0.147513943 |
| ENSMUSG00000019873 | Reep3 | 28193 | protein_coding | 1225.33481 | -0.147567137 | 0.079856185 | -1.847911169 | 0.064615195 | 0.185006439 | 0.147567137 |
| ENSMUSG00000047714 | Ppp1r2 | 66849 | protein_coding | 989.2400896 | 0.147582944 | 0.104004682 | 1.419002881 | 0.155898176 | 0.336191915 | 0.147582944 |
| ENSMUSG00000025702 | Marchf8 | 71779 | protein_coding | 1808.587768 | 0.147667486 | 0.080180652 | 1.841684771 | 0.065521273 | 0.186523628 | 0.147667486 |
| ENSMUSG00000032265 | Tent5a | 212943 | protein_coding | 1709.853626 | -0.147669036 | 0.100846274 | -1.464298384 | 0.143112439 | 0.316948682 | 0.147669036 |
| ENSMUSG00000046556 | Zfp319 | 79233 | protein_coding | 183.8672131 | -0.147711608 | 0.120491446 | -1.225909491 | 0.220232737 | 0.420657997 | 0.147711608 |
| ENSMUSG00000034574 | Daam1 | 208846 | protein_coding | 1063.691046 | -0.147737161 | 0.102325529 | -1.443795715 | 0.148796448 | 0.325956515 | 0.147737161 |
| ENSMUSG00000033088 | Triobp | 110253 | protein_coding | 648.2693299 | 0.14775091 | 0.105869632 | 1.395592936 | 0.162837111 | 0.345362118 | 0.14775091 |
| ENSMUSG00000029617 | Ccz1 | 231874 | protein_coding | 1130.592135 | 0.147764334 | 0.082750122 | 1.78566908 | 0.074152855 | 0.2029556 | 0.147764334 |
| ENSMUSG00000029440 | Psmd9 | 67151 | protein_coding | 262.466614 | 0.147774296 | 0.092337607 | 1.600369562 | 0.109516623 | 0.263740668 | 0.147774296 |
| ENSMUSG00000042306 | S100a14 | 66166 | protein_coding | 1377.115966 | 0.147780451 | 0.247178262 | 0.597869931 | 0.549926724 | 0.731093637 | 0.147780451 |
| ENSMUSG00000055333 | Fat2 | 245827 | protein_coding | 46.77878397 | -0.147815048 | 0.281702635 | -0.524720146 | 0.59977775 | 0.76628379 | 0.147815048 |
| ENSMUSG00000036009 | Mettl25 | 216292 | protein_coding | 87.85970766 | 0.147835584 | 0.129906328 | 1.138016799 | 0.255113468 | 0.461807855 | 0.147835584 |
| ENSMUSG00000036880 | Acaa2 | 52538 | protein_coding | 1852.271782 | -0.147875783 | 0.130146862 | -1.136222421 | 0.255863489 | 0.462744823 | 0.147875783 |
| ENSMUSG00000057897 | Camk2b | 12323 | protein_coding | 348.4226591 | 0.147907854 | 0.105821655 | 1.397708752 | 0.162200545 | 0.344608548 | 0.147907854 |
| ENSMUSG00000029060 | Mib2 | 76580 | protein_coding | 348.5819455 | -0.147952808 | 0.138986878 | -1.064509181 | 0.287098098 | 0.495975976 | 0.147952808 |
| ENSMUSG00000034382 | AI661453 | 224833 | protein_coding | 379.0367721 | -0.147956917 | 0.149104485 | -0.992303597 | 0.321049449 | 0.532286419 | 0.147956917 |
| ENSMUSG00000040525 | Cblc | 80794 | protein_coding | 869.8472705 | 0.147971649 | 0.095684702 | 1.546450445 | 0.121995819 | 0.284449738 | 0.147971649 |

| ensembl_gene_id | external_gene_name | entrezgene_id | gene_biotype | baseMean | log2FoldChange | lfcSE | stat | pvalue | padj | absLog2FC |
|---|---|---|---|---|---|---|---|---|---|---|
| ENSMUSG00000054715 | Zscan22 | 232878 | protein_coding | 87.49741913 | 0.147980049 | 0.146794184 | 1.008078419 | 0.313416817 | 0.523835181 | 0.147980049 |
| ENSMUSG00000117226 | Gm31235 | 102633401 | lncRNA | 66.38550937 | 0.148013369 | 0.21751816 | 0.680464422 | 0.496210442 | 0.687332804 | 0.148013369 |
| ENSMUSG00000024335 | Brd2 | 14312 | protein_coding | 2738.363545 | -0.148022307 | 0.075728256 | -1.954650945 | 0.050624284 | 0.156693275 | 0.148022307 |
| ENSMUSG00000038170 | Pde4dip | 83679 | protein_coding | 2020.673792 | 0.148027869 | 0.077639602 | 1.906602618 | 0.056572066 | 0.169058136 | 0.148027869 |
| ENSMUSG00000051989 | Smim11 | 68936 | protein_coding | 231.3006885 | 0.148066937 | 0.115042557 | 1.28706229 | 0.198072578 | 0.392156622 | 0.148066937 |
| ENSMUSG00000046785 | Epm2aip1 | 77781 | protein_coding | 385.5373017 | -0.148088253 | 0.135259826 | -1.094842846 | 0.273585492 | 0.481510466 | 0.148088253 |
| ENSMUSG00000030373 | Psg28 | 114871 | protein_coding | 7.233225838 | 0.148106537 | 0.551362392 | 0.268619222 | 0.788222719 | 0.884950162 | 0.148106537 |
| ENSMUSG00000022884 | Eif4a2 | 13682 | protein_coding | 5578.857941 | 0.148146236 | 0.116939532 | 1.266861884 | 0.20520468 | 0.401062215 | 0.148146236 |
| ENSMUSG00000025025 | Mxi1 | 17859 | protein_coding | 640.8261731 | -0.148164075 | 0.120100723 | -1.233665136 | 0.21732771 | 0.416808642 | 0.148164075 |
| ENSMUSG00000029502 | Golga3 | 269682 | protein_coding | 1433.469219 | -0.148164583 | 0.106638349 | -1.389411821 | 0.164707558 | 0.348123936 | 0.148164583 |
| ENSMUSG00000021583 | Erap1 | 80898 | protein_coding | 1891.245521 | 0.148169195 | 0.083841087 | 1.767262325 | 0.077184306 | 0.208477468 | 0.148169195 |
| ENSMUSG00000057103 | Nat8f1 | 66116 | protein_coding | 17.47968998 | -0.148193673 | 0.27860585 | -0.531911564 | 0.594787248 | 0.762965747 | 0.148193673 |
| ENSMUSG00000013701 | Timm23 | 53600 | protein_coding | 525.6783518 | 0.148246593 | 0.104477181 | 1.418937521 | 0.155917232 | 0.336191915 | 0.148246593 |
| ENSMUSG00000094724 | Rnaset2b | 68195 | protein_coding | 11.75175192 | -0.148258113 | 0.33141331 | -0.447351113 | 0.654621568 | 0.800150191 | 0.148258113 |
| ENSMUSG00000040188 | Scamp2 | 24044 | protein_coding | 2313.29837 | 0.14826389 | 0.111444917 | 1.330378216 | 0.183393688 | 0.372636135 | 0.14826389 |
| ENSMUSG00000114613 | Gm10732 | 100038690 | lncRNA | 28.8363417 | 0.148290465 | 0.265479086 | 0.558576825 | 0.57645056 | 0.748671276 | 0.148290465 |
| ENSMUSG00000032939 | Nup93 | 71805 | protein_coding | 354.3510953 | 0.148291461 | 0.107113073 | 1.384438495 | 0.166224229 | 0.350047605 | 0.148291461 |
| ENSMUSG00000040987 | Mill2 | 243864 | protein_coding | 87.16723666 | 0.148306138 | 0.262590366 | 0.564781337 | 0.572222502 | 0.745978067 | 0.148306138 |
| ENSMUSG00000004565 | Pnpla6 | 50767 | protein_coding | 679.383593 | -0.148310885 | 0.112306194 | -1.320594 | 0.186636774 | 0.377061205 | 0.148310885 |
| ENSMUSG00000037960 | Card19 | 68480 | protein_coding | 305.7301349 | 0.148372861 | 0.164115427 | 0.904076258 | 0.365954967 | 0.574731878 | 0.148372861 |
| ENSMUSG00000038271 | Iffo1 | 320678 | protein_coding | 14.87052343 | 0.148439512 | 0.453523766 | 0.327302608 | 0.74343902 | 0.857217273 | 0.148439512 |
| ENSMUSG00000025428 | Atp5a1 | 11946 | protein_coding | 25337.09721 | 0.148495524 | 0.102498611 | 1.44875645 | 0.14740561 | 0.323979102 | 0.148495524 |
| ENSMUSG00000026150 | Mff | 75734 | protein_coding | 972.560701 | 0.14852163 | 0.066465639 | 2.234562563 | 0.025446081 | 0.095839484 | 0.14852163 |
| ENSMUSG00000066357 | Wdr6 | 83669 | protein_coding | 311.7101596 | 0.148554943 | 0.131964941 | 1.125715223 | 0.260286078 | 0.467610721 | 0.148554943 |
| ENSMUSG00000072772 | Grcc10 | 14790 | protein_coding | 210.5806658 | -0.148568247 | 0.143807334 | -1.03310619 | 0.301554207 | 0.510865285 | 0.148568247 |
| ENSMUSG00000026469 | Xpr1 | 19775 | protein_coding | 1433.395918 | -0.148602469 | 0.138358014 | -1.074043087 | 0.282803371 | 0.491917887 | 0.148602469 |
| ENSMUSG00000020715 | Ern1 | 78943 | protein_coding | 609.4328211 | -0.148605842 | 0.11543604 | -1.287343552 | 0.19797457 | 0.392060046 | 0.148605842 |
| ENSMUSG00000030752 | Kdm8 | 77035 | protein_coding | 74.91915382 | -0.1486254 | 0.175235971 | -0.848144355 | 0.396357582 | 0.604537352 | 0.1486254 |
| ENSMUSG00000030869 | Ndufab1 | 70316 | protein_coding | 931.7218087 | 0.148683972 | 0.091591178 | 1.623343822 | 0.104515927 | 0.255797431 | 0.148683972 |
| ENSMUSG00000017858 | Ift52 | 245866 | protein_coding | 416.0836574 | 0.148774621 | 0.089632736 | 1.65982461 | 0.096949741 | 0.243413537 | 0.148774621 |
| ENSMUSG00000018841 | Rad51d | 19364 | protein_coding | 307.3802289 | -0.148833825 | 0.109923876 | -1.353971771 | 0.175745384 | 0.361907388 | 0.148833825 |
| ENSMUSG00000101970 | Chaserr | 69170 | lncRNA | 104.834939 | -0.148834541 | 0.201403503 | -0.73898685 | 0.459914982 | 0.658313465 | 0.148834541 |
| ENSMUSG00000061898 | Rbak | 57782 | protein_coding | 167.2398534 | -0.148866243 | 0.101726966 | -1.463390175 | 0.143360645 | 0.317321896 | 0.148866243 |
| ENSMUSG00000024270 | Slc39a6 | 106957 | protein_coding | 57.841948 | 0.148879712 | 0.16272526 | 0.914914575 | 0.360236478 | 0.569739511 | 0.148879712 |
| ENSMUSG00000024777 | Ppp2r5b | 225849 | protein_coding | 389.8115364 | -0.148931724 | 0.110906953 | -1.342852901 | 0.17931961 | 0.367026824 | 0.148931724 |
| ENSMUSG00000067212 | H2-T23 | 15040 | protein_coding | 2174.614673 | 0.149022609 | 0.11127116 | 1.339274338 | 0.18048138 | 0.368552233 | 0.149022609 |
| ENSMUSG00000034083 | Ccdc174 | 232236 | protein_coding | 337.0274359 | -0.149195867 | 0.071279359 | -2.093114602 | 0.036338926 | 0.124287397 | 0.149195867 |
| ENSMUSG00000075419 | Dolk | 227697 | protein_coding | 206.6975734 | 0.149268361 | 0.104384946 | 1.429979772 | 0.152722825 | 0.331457849 | 0.149268361 |

| ensembl_gene_id | external_gene_name | entrezgene_id | gene_biotype | baseMean | log2FoldChange | lfcSE | stat | pvalue | padj | absLog2FC |
|---|---|---|---|---|---|---|---|---|---|---|
| ENSMUSG00000021420 | Fars2 | 69955 | protein_coding | 570.8059136 | -0.149293107 | 0.088243451 | -1.691832149 | 0.090677982 | 0.232877234 | 0.149293107 |
| ENSMUSG00000029578 | Wipi2 | 74781 | protein_coding | 1092.117727 | -0.149310849 | 0.075022317 | -1.990219112 | 0.046566804 | 0.148195015 | 0.149310849 |
| ENSMUSG00000028786 | Tmem54 | 66260 | protein_coding | 1387.801661 | 0.149316818 | 0.142553536 | 1.047443802 | 0.294894939 | 0.503979247 | 0.149316818 |
| ENSMUSG00000017009 | Sdc4 | 20971 | protein_coding | 4985.690625 | -0.149429012 | 0.098567058 | -1.516013725 | 0.129515882 | 0.29611967 | 0.149429012 |
| ENSMUSG00000013643 | Lypd8 | 70163 | protein_coding | 42998.74807 | 0.149527361 | 0.128000644 | 1.168176628 | 0.242735526 | 0.447642869 | 0.149527361 |
| ENSMUSG00000034064 | Poglut1 | 224143 | protein_coding | 255.3374826 | -0.149544593 | 0.085438677 | -1.750314944 | 0.080063984 | 0.213969653 | 0.149544593 |
| ENSMUSG00000020271 | Fbxw11 | 103583 | protein_coding | 1040.973625 | -0.149596257 | 0.057229251 | -2.613982418 | 0.008949364 | 0.044036291 | 0.149596257 |
| ENSMUSG00000039220 | Ppp1r10 | 52040 | protein_coding | 1246.142216 | 0.149632289 | 0.096461886 | 1.551206338 | 0.120852244 | 0.282455545 | 0.149632289 |
| ENSMUSG00000027998 | Plrg1 | 53317 | protein_coding | 570.1619816 | 0.14964626 | 0.108505027 | 1.379164303 | 0.167844102 | 0.352275774 | 0.14964626 |
| ENSMUSG00000047638 | Nr1h4 | 20186 | protein_coding | 1453.409264 | 0.149659863 | 0.191427151 | 0.781811055 | 0.434325623 | 0.637656791 | 0.149659863 |
| ENSMUSG00000047417 | Rexo1 | 66932 | protein_coding | 372.6001536 | -0.149666796 | 0.120696487 | -1.240026115 | 0.214965735 | 0.413723324 | 0.149666796 |
| ENSMUSG00000025809 | Itgb1 | 16412 | protein_coding | 4263.776897 | 0.149710455 | 0.096817158 | 1.546321519 | 0.122026938 | 0.284449738 | 0.149710455 |
| ENSMUSG00000031823 | Zdhhc7 | 102193 | protein_coding | 790.6449307 | 0.14976378 | 0.107554584 | 1.392444418 | 0.163787862 | 0.34682439 | 0.14976378 |
| ENSMUSG00000047454 | Gphn | 268566 | protein_coding | 324.0537158 | -0.149766714 | 0.088749077 | -1.687529834 | 0.091501518 | 0.234352176 | 0.149766714 |
| ENSMUSG00000032333 | Stoml1 | 69106 | protein_coding | 129.671991 | -0.149790678 | 0.133723012 | -1.120156322 | 0.262647153 | 0.470416684 | 0.149790678 |
| ENSMUSG00000078945 | Naip2 | 17948 | protein_coding | 2279.302053 | 0.149793332 | 0.100583467 | 1.489244075 | 0.136423108 | 0.306740157 | 0.149793332 |
| ENSMUSG00000040720 | Virma | 66185 | protein_coding | 940.2109931 | 0.149855234 | 0.062084316 | 2.413737391 | 0.015789838 | 0.06726412 | 0.149855234 |
| ENSMUSG00000094936 | Rbm4 | 19653 | protein_coding | 86.45446151 | -0.149864141 | 0.211130737 | -0.709816787 | 0.477817758 | 0.672321665 | 0.149864141 |
| ENSMUSG00000063364 | 3300002I08Rik | 69277 | protein_coding | 24.65689 | 0.149898595 | 0.26813107 | 0.559049702 | 0.576127801 | 0.748585685 | 0.149898595 |
| ENSMUSG00000045930 | Clec14a | 66864 | protein_coding | 330.5657517 | 0.150037826 | 0.21153169 | 0.709292429 | 0.478143026 | 0.672607328 | 0.150037826 |
| ENSMUSG00000039735 | Fnbp1l | 214459 | protein_coding | 1134.352158 | -0.150044309 | 0.095237706 | -1.575471691 | 0.115147608 | 0.272440745 | 0.150044309 |
| ENSMUSG00000037254 | Itih2 | 16425 | protein_coding | 145.7572365 | 0.150046458 | 0.489447362 | 0.306563012 | 0.759176011 | 0.868315058 | 0.150046458 |
| ENSMUSG00000001247 | Lsr | 54135 | protein_coding | 754.5337685 | -0.150064881 | 0.135918587 | -1.104079177 | 0.269558793 | 0.477483592 | 0.150064881 |
| ENSMUSG00000005374 | Tbl2 | 27368 | protein_coding | 438.1723028 | 0.150084282 | 0.083118583 | 1.805664584 | 0.070970748 | 0.197315774 | 0.150084282 |
| ENSMUSG00000047443 | Erfe | 227358 | protein_coding | 98.67178965 | -0.150091348 | 0.176955019 | -0.848189269 | 0.396332572 | 0.604537352 | 0.150091348 |
| ENSMUSG00000042608 | Stk40 | 74178 | protein_coding | 579.9731293 | -0.150116843 | 0.370850507 | -0.404790719 | 0.685631352 | 0.820640694 | 0.150116843 |
| ENSMUSG00000059278 | Naa38 | 78304 | protein_coding | 238.8873486 | 0.150135377 | 0.100062873 | 1.500410424 | 0.133508121 | 0.301976335 | 0.150135377 |
| ENSMUSG00000023883 | Phf10 | 72057 | protein_coding | 500.2239569 | 0.150157478 | 0.07429692 | 2.021045791 | 0.043275025 | 0.14037911 | 0.150157478 |
| ENSMUSG00000029911 | Ssbp1 | 381760 | protein_coding | 384.6606705 | 0.150236127 | 0.078643356 | 1.910347358 | 0.056088504 | 0.16786503 | 0.150236127 |
| ENSMUSG00000027326 | Knl1 | 76464 | protein_coding | 579.1783873 | 0.150287026 | 0.112114198 | 1.340481662 | 0.180088802 | 0.368080898 | 0.150287026 |
| ENSMUSG00000036572 | Upf3b | 68134 | protein_coding | 361.1489256 | -0.150319303 | 0.090184558 | -1.666796474 | 0.095554882 | 0.241146755 | 0.150319303 |
| ENSMUSG00000007035 | Msh5 | 17687 | protein_coding | 28.49667563 | -0.150334152 | 0.274147074 | -0.548370443 | 0.583437567 | 0.754056602 | 0.150334152 |
| ENSMUSG00000048772 | Tmem53 | 68777 | protein_coding | 42.60436438 | -0.150355537 | 0.224732403 | -0.669042537 | 0.503468344 | 0.693526228 | 0.150355537 |
| ENSMUSG00000014547 | Wdfy2 | 268752 | protein_coding | 273.5747528 | 0.150359357 | 0.1536821 | 0.978379118 | 0.327886852 | 0.538806428 | 0.150359357 |
| ENSMUSG00000029146 | Snx17 | 266781 | protein_coding | 821.937144 | 0.150376758 | 0.085617773 | 1.75637316 | 0.079024713 | 0.212095289 | 0.150376758 |
| ENSMUSG00000031278 | Acsl4 | 50790 | protein_coding | 291.8735818 | 0.150381198 | 0.084600928 | 1.777536024 | 0.075480089 | 0.205144798 | 0.150381198 |
| ENSMUSG00000024457 | Trim26 | 22670 | protein_coding | 850.5551476 | 0.150396853 | 0.067009635 | 2.244406378 | 0.024806267 | 0.093873655 | 0.150396853 |
| ENSMUSG00000049717 | Lig4 | 319583 | protein_coding | 240.3146655 | 0.1504058 | 0.137862309 | 1.090985645 | 0.275279199 | 0.483529674 | 0.1504058 |

| ensembl_gene_id | external_gene_name | entrezgene_id | gene_biotype | baseMean | log2FoldChange | lfcSE | stat | pvalue | padj | absLog2FC |
|---|---|---|---|---|---|---|---|---|---|---|
| ENSMUSG00000015943 | Bola1 | 69168 | protein_coding | 154.3601239 | 0.150440808 | 0.102866381 | 1.46248761 | 0.143607637 | 0.317603783 | 0.150440808 |
| ENSMUSG00000034947 | Tmem106a | 217203 | protein_coding | 168.1178212 | -0.15055651 | 0.130329789 | -1.155196455 | 0.248009986 | 0.453849156 | 0.15055651 |
| ENSMUSG00000036528 | Ppfibp2 | 19024 | protein_coding | 441.7395088 | -0.150581627 | 0.148543688 | -1.013719462 | 0.310716635 | 0.520683278 | 0.150581627 |
| ENSMUSG00000035666 | Gtf3c4 | 269252 | protein_coding | 619.0463778 | -0.150588192 | 0.086060337 | -1.749797836 | 0.080153204 | 0.213992665 | 0.150588192 |
| ENSMUSG00000055240 | Zfp101 | 22643 | protein_coding | 212.4053543 | 0.15065361 | 0.131474341 | 1.145878421 | 0.251845461 | 0.458216818 | 0.15065361 |
| ENSMUSG00000046096 | Mosmo | 233812 | protein_coding | 126.3869653 | -0.150672718 | 0.117728581 | -1.279831255 | 0.200604489 | 0.395157223 | 0.150672718 |
| ENSMUSG00000079283 | 2310009B15Rik | 69549 | protein_coding | 82.38940942 | 0.150741094 | 0.1280905 | 1.176832737 | 0.239262273 | 0.443424073 | 0.150741094 |
| ENSMUSG00000028312 | Smc2 | 14211 | protein_coding | 794.5556791 | 0.150746013 | 0.116332452 | 1.295820819 | 0.195037224 | 0.388054296 | 0.150746013 |
| ENSMUSG00000068329 | Htra2 | 64704 | protein_coding | 198.7893802 | -0.150755101 | 0.110501044 | -1.36428667 | 0.17247736 | 0.358210338 | 0.150755101 |
| ENSMUSG00000047694 | Yipf6 | 77929 | protein_coding | 2532.972489 | 0.150784334 | 0.059135945 | 2.549791567 | 0.010778734 | 0.050629217 | 0.150784334 |
| ENSMUSG00000021067 | Sav1 | 64010 | protein_coding | 332.5257119 | 0.150802755 | 0.081429582 | 1.851940683 | 0.064034341 | 0.183905334 | 0.150802755 |
| ENSMUSG00000035378 | Shq1 | 72171 | protein_coding | 113.9850988 | -0.150813096 | 0.155624611 | -0.96908256 | 0.332503998 | 0.543470876 | 0.150813096 |
| ENSMUSG00000034801 | Sos2 | 20663 | protein_coding | 663.016748 | -0.15089614 | 0.072654856 | -2.076889949 | 0.037811718 | 0.128000666 | 0.15089614 |
| ENSMUSG00000050890 | Pdik1l | 230809 | protein_coding | 453.2386313 | 0.150984772 | 0.161755284 | 0.933414776 | 0.350605854 | 0.560627608 | 0.150984772 |
| ENSMUSG00000097354 | 2310001H17Rik | 76432 | lncRNA | 126.2650023 | 0.150986493 | 0.154378233 | 0.978029675 | 0.328059648 | 0.538929056 | 0.150986493 |
| ENSMUSG00000020629 | Adi1 | 104923 | protein_coding | 282.2748447 | -0.150996843 | 0.125980118 | -1.198576772 | 0.230692555 | 0.433821843 | 0.150996843 |
| ENSMUSG00000023186 | Vwa5a | 67776 | protein_coding | 1153.199262 | 0.151019438 | 0.116466612 | 1.2966758 | 0.194742759 | 0.387829791 | 0.151019438 |
| ENSMUSG00000003444 | Med29 | 67224 | protein_coding | 95.36543213 | 0.151028552 | 0.146319667 | 1.032182176 | 0.301986783 | 0.511326192 | 0.151028552 |
| ENSMUSG00000071708 | Sms | 20603 | protein_coding | 173.3305204 | -0.151042384 | 0.115057364 | -1.312757212 | 0.189264772 | 0.380811412 | 0.151042384 |
| ENSMUSG00000096950 | Gm9530 | 671336 | lncRNA | 8.903641388 | 0.151050336 | 0.432874505 | 0.348947177 | 0.727128967 | 0.846434985 | 0.151050336 |
| ENSMUSG00000038368 | Focad | 230393 | protein_coding | 265.1802309 | 0.151088059 | 0.082486804 | 1.831663396 | 0.067001585 | 0.189556173 | 0.151088059 |
| ENSMUSG00000030854 | Ptpn5 | 19259 | protein_coding | 7.222605817 | -0.151121517 | 0.684666676 | -0.220722759 | 0.82530831 | 0.907901841 | 0.151121517 |
| ENSMUSG00000044496 | 2510039O18Rik | 77034 | protein_coding | 1237.063928 | 0.15114746 | 0.111106459 | 1.360384101 | 0.173708407 | 0.359707067 | 0.15114746 |
| ENSMUSG00000059475 | Zfp426 | 235028 | protein_coding | 349.808489 | 0.15117387 | 0.104282108 | 1.449662582 | 0.147152635 | 0.323691089 | 0.15117387 |
| ENSMUSG00000041126 | H2az2 | 77605 | protein_coding | 580.9765288 | 0.151188823 | 0.107063367 | 1.412143358 | 0.157907754 | 0.338832643 | 0.151188823 |
| ENSMUSG00000028958 | Tmub1 | 64295 | protein_coding | 226.885288 | 0.151221013 | 0.112695783 | 1.341851573 | 0.179644126 | 0.367454944 | 0.151221013 |
| ENSMUSG00000037326 | Capn15 | 50817 | protein_coding | 259.1893802 | -0.151244195 | 0.138853499 | -1.089235751 | 0.276049937 | 0.48434935 | 0.151244195 |
| ENSMUSG00000080440 | Gm25848 | 115488713 | snoRNA | 27.65887122 | -0.151253303 | 0.254819884 | -0.593569469 | 0.552800106 | 0.732884453 | 0.151253303 |
| ENSMUSG00000026245 | Farsb | 23874 | protein_coding | 929.07735 | 0.15129751 | 0.106878814 | 1.415598696 | 0.156893033 | 0.337701653 | 0.15129751 |
| ENSMUSG00000026489 | Coq8a | 67426 | protein_coding | 490.4798297 | -0.151312739 | 0.242417329 | -0.624182848 | 0.53250751 | 0.717578648 | 0.151312739 |
| ENSMUSG00000030849 | Fgfr2 | 14183 | protein_coding | 1086.573402 | -0.151359638 | 0.179934249 | -0.841194153 | 0.400239176 | 0.607722554 | 0.151359638 |
| ENSMUSG00000096687 | Mfsd4b4 | 213332 | protein_coding | 38.99033947 | 0.151367102 | 0.222940146 | 0.678958478 | 0.497164171 | 0.688115304 | 0.151367102 |
| ENSMUSG00000027086 | Fastkd1 | 320720 | protein_coding | 450.492351 | -0.15137434 | 0.131817705 | -1.148361212 | 0.250819479 | 0.457152153 | 0.15137434 |
| ENSMUSG00000035561 | Aldh1b1 | 72535 | protein_coding | 4703.159383 | 0.151383359 | 0.136074152 | 1.11250635 | 0.265920502 | 0.47372573 | 0.151383359 |
| ENSMUSG00000063445 | Nmral1 | 67824 | protein_coding | 84.75000051 | 0.151390678 | 0.147589274 | 1.025756643 | 0.305006307 | 0.514633524 | 0.151390678 |
| ENSMUSG00000025372 | Baiap2 | 108100 | protein_coding | 282.478837 | -0.151425551 | 0.221108939 | -0.684845901 | 0.493441167 | 0.68528476 | 0.151425551 |
| ENSMUSG00000033970 | Rfc3 | 69263 | protein_coding | 206.4847078 | 0.151461244 | 0.123169839 | 1.229694257 | 0.218811618 | 0.418948365 | 0.151461244 |
| ENSMUSG00000034275 | Igsf9b | 235086 | protein_coding | 359.2045192 | 0.151468838 | 0.150864648 | 1.004004853 | 0.315376275 | 0.526387227 | 0.151468838 |

| ensembl_gene_id | external_gene_name | entrezgene_id | gene_biotype | baseMean | log2FoldChange | lfcSE | stat | pvalue | padj | absLog2FC |
|---|---|---|---|---|---|---|---|---|---|---|
| ENSMUSG00000020142 | Slc1a4 | 55963 | protein_coding | 926.8554466 | -0.151471504 | 0.269145229 | -0.562787251 | 0.573579763 | 0.747013774 | 0.151471504 |
| ENSMUSG00000050846 | Zfp623 | 78834 | protein_coding | 273.6335999 | -0.151534693 | 0.07697335 | -1.968664397 | 0.048991642 | 0.153468144 | 0.151534693 |
| ENSMUSG00000048170 | Mcmbp | 210711 | protein_coding | 1117.203556 | -0.151540825 | 0.069076913 | -2.193798461 | 0.028249902 | 0.103131889 | 0.151540825 |
| ENSMUSG00000023043 | Krt18 | 16668 | protein_coding | 9178.378796 | -0.151550986 | 0.212429851 | -0.713416618 | 0.475587993 | 0.670351322 | 0.151550986 |
| ENSMUSG00000045180 | Shroom2 | 110380 | protein_coding | 675.3455022 | -0.151709438 | 0.108684906 | -1.395864832 | 0.162755203 | 0.345262415 | 0.151709438 |
| ENSMUSG00000026707 | Nsun6 | 74455 | protein_coding | 175.2437249 | 0.151717652 | 0.092297808 | 1.643783915 | 0.100220845 | 0.248454104 | 0.151717652 |
| ENSMUSG00000090015 | Gm15446 | 100642166 | protein_coding | 117.9553983 | 0.151721165 | 0.612406351 | 0.247745904 | 0.804331012 | 0.894700251 | 0.151721165 |
| ENSMUSG00000046949 | Nqo2 | 18105 | protein_coding | 439.1813108 | 0.15172264 | 0.116692901 | 1.300187407 | 0.193536746 | 0.387069668 | 0.15172264 |
| ENSMUSG00000030600 | Lrfn1 | 80749 | protein_coding | 7.362182298 | -0.15174385 | 0.528597375 | -0.287068868 | 0.774059589 | 0.877212461 | 0.15174385 |
| ENSMUSG00000089917 | Uckl1 | 68556 | protein_coding | 480.9087087 | -0.15179984 | 0.091417129 | -1.660518564 | 0.096810176 | 0.243178125 | 0.15179984 |
| ENSMUSG00000054008 | Ndst1 | 15531 | protein_coding | 686.0152108 | -0.151804598 | 0.071941489 | -2.110111977 | 0.034848712 | 0.120617479 | 0.151804598 |
| ENSMUSG00000074006 | Omp | 18378 | protein_coding | 80.18736124 | -0.151810841 | 0.267799206 | -0.566883089 | 0.570793615 | 0.745152094 | 0.151810841 |
| ENSMUSG00000050619 | Zscan29 | 99334 | protein_coding | 443.6844148 | -0.151819868 | 0.108874815 | -1.394444325 | 0.163183471 | 0.345682448 | 0.151819868 |
| ENSMUSG00000025371 | Chmp6 | 208092 | protein_coding | 791.4462175 | 0.151867898 | 0.122876876 | 1.235935533 | 0.216482528 | 0.415692112 | 0.151867898 |
| ENSMUSG00000023150 | Ivns1abp | 117198 | protein_coding | 7397.748726 | -0.151872819 | 0.069531764 | -2.184222137 | 0.028945929 | 0.10509477 | 0.151872819 |
| ENSMUSG00000049658 | Bdp1 | 544971 | protein_coding | 1095.574739 | -0.151895676 | 0.107769859 | -1.409444879 | 0.158703662 | 0.339990844 | 0.151895676 |
| ENSMUSG00000036941 | Elac1 | 114615 | protein_coding | 157.896967 | 0.151925407 | 0.139449704 | 1.089463813 | 0.275949404 | 0.484226299 | 0.151925407 |
| ENSMUSG00000050229 | Pigm | 67556 | protein_coding | 483.6398801 | -0.151930973 | 0.094775639 | -1.603059334 | 0.108921555 | 0.26286402 | 0.151930973 |
| ENSMUSG00000078942 | Naip6 | 17952 | protein_coding | 1650.219073 | -0.151965729 | 0.130865864 | -1.161232762 | 0.245547255 | 0.450976122 | 0.151965729 |
| ENSMUSG00000036615 | Rfxap | 170767 | protein_coding | 152.6561327 | 0.15196689 | 0.135579915 | 1.120865806 | 0.262344988 | 0.470066494 | 0.15196689 |
| ENSMUSG00000047141 | Zfp654 | 72020 | protein_coding | 540.1276496 | 0.151998307 | 0.084799424 | 1.792445029 | 0.073061707 | 0.201128046 | 0.151998307 |
| ENSMUSG00000056220 | Pla2g4a | 18783 | protein_coding | 1149.70559 | 0.152040103 | 0.092377964 | 1.645848164 | 0.099795023 | 0.247822752 | 0.152040103 |
| ENSMUSG00000066643 | Wdr35 | 74682 | protein_coding | 46.81411939 | 0.152062269 | 0.231672065 | 0.656368597 | 0.511586991 | 0.699447003 | 0.152062269 |
| ENSMUSG00000042700 | Sipa1l1 | 217692 | protein_coding | 1383.48126 | -0.152068567 | 0.113792653 | -1.336365423 | 0.181429862 | 0.370062088 | 0.152068567 |
| ENSMUSG00000070709 | Zfp974 | 73430 | protein_coding | 152.867178 | -0.152093413 | 0.158665861 | -0.958576801 | 0.337771984 | 0.548479169 | 0.152093413 |
| ENSMUSG00000030423 | Pop4 | 66161 | protein_coding | 240.928057 | 0.152145272 | 0.129812626 | 1.172037546 | 0.241181982 | 0.445706765 | 0.152145272 |
| ENSMUSG00000038406 | Scaf1 | 233208 | protein_coding | 578.7046219 | -0.152187233 | 0.089277103 | -1.704661416 | 0.08825759 | 0.228952262 | 0.152187233 |
| ENSMUSG00000025651 | Uqcrc1 | 22273 | protein_coding | 5259.900319 | 0.152234736 | 0.11150494 | 1.365273471 | 0.172167114 | 0.357766182 | 0.152234736 |
| ENSMUSG00000090925 | 1810064F22Rik | 69862 | lncRNA | 8.393135145 | -0.152254017 | 0.480350854 | -0.316964186 | 0.751270777 | 0.862550365 | 0.152254017 |
| ENSMUSG00000022313 | Utp23 | 78581 | protein_coding | 103.9285752 | 0.15228146 | 0.115830576 | 1.314691381 | 0.188613645 | 0.379779427 | 0.15228146 |
| ENSMUSG00000036257 | Pnpla8 | 67452 | protein_coding | 1398.877107 | 0.152321795 | 0.070587671 | 2.157909353 | 0.03093488 | 0.110502528 | 0.152321795 |
| ENSMUSG00000024276 | Zfp397 | 69256 | protein_coding | 653.3170691 | -0.152363221 | 0.070978161 | -2.146621134 | 0.031823456 | 0.112791279 | 0.152363221 |
| ENSMUSG00000027358 | Bmp2 | 12156 | protein_coding | 220.1588972 | -0.152371869 | 0.213124475 | -0.714943083 | 0.474644214 | 0.669512313 | 0.152371869 |
| ENSMUSG00000028382 | Ptbp3 | 230257 | protein_coding | 5198.772289 | 0.15238815 | 0.082618387 | 1.844482277 | 0.065112886 | 0.185783495 | 0.15238815 |
| ENSMUSG00000037017 | Zscan21 | 22697 | protein_coding | 288.1461359 | -0.152423164 | 0.102887602 | -1.481453179 | 0.138485853 | 0.309629796 | 0.152423164 |
| ENSMUSG00000047037 | Nipa1 | 233280 | protein_coding | 80.50018634 | -0.152433206 | 0.16364129 | -0.931508218 | 0.351590736 | 0.561399852 | 0.152433206 |
| ENSMUSG00000031529 | Tnks | 21951 | protein_coding | 671.6612414 | -0.152438039 | 0.141834784 | -1.074757788 | 0.282483184 | 0.491521974 | 0.152438039 |
| ENSMUSG00000050071 | Bex1 | 19716 | protein_coding | 30.08637096 | 0.152448385 | 0.248124107 | 0.614403763 | 0.538948549 | 0.722272825 | 0.152448385 |

| ensembl_gene_id | external_gene_name | entrezgene_id | gene_biotype | baseMean | log2FoldChange | lfcSE | stat | pvalue | padj | absLog2FC |
|---|---|---|---|---|---|---|---|---|---|---|
| ENSMUSG00000029326 | Enoph1 | 67870 | protein_coding | 395.4669061 | -0.152470806 | 0.088562065 | -1.721626596 | 0.085137182 | 0.223382746 | 0.152470806 |
| ENSMUSG00000029470 | P2rx4 | 18438 | protein_coding | 1960.276243 | 0.152485857 | 0.086353532 | 1.765832314 | 0.077423986 | 0.209018515 | 0.152485857 |
| ENSMUSG00000038696 | Mapkap1 | 227743 | protein_coding | 1011.21243 | 0.152527507 | 0.080423139 | 1.896562469 | 0.057885708 | 0.171823249 | 0.152527507 |
| ENSMUSG00000046807 | Lrrc75b | 192734 | protein_coding | 8.969932191 | 0.152528437 | 0.554334003 | 0.275156198 | 0.783196238 | 0.882391137 | 0.152528437 |
| ENSMUSG00000023074 | Mospd1 | 70380 | protein_coding | 352.6429188 | 0.152599424 | 0.123521419 | 1.235408605 | 0.216678472 | 0.415901327 | 0.152599424 |
| ENSMUSG00000070605 | Zfp992 | 433791 | protein_coding | 62.31338925 | -0.152878316 | 0.22626517 | -0.675659961 | 0.499256563 | 0.689758658 | 0.152878316 |
| ENSMUSG00000039611 | Pgap4 | 67063 | protein_coding | 742.8504471 | 0.152943013 | 0.151304016 | 1.010832474 | 0.312096618 | 0.522338315 | 0.152943013 |
| ENSMUSG00000045659 | Plekha7 | 233765 | protein_coding | 836.5122324 | -0.152971239 | 0.116090926 | -1.317684719 | 0.187609212 | 0.378355839 | 0.152971239 |
| ENSMUSG00000039410 | Prdm16 | 70673 | protein_coding | 118.6562859 | -0.153035708 | 0.173939249 | -0.87982275 | 0.378955338 | 0.587102483 | 0.153035708 |
| ENSMUSG00000030083 | Abtb1 | 80283 | protein_coding | 299.3036801 | -0.153097753 | 0.146769857 | -1.043114412 | 0.296895306 | 0.506180721 | 0.153097753 |
| ENSMUSG00000059436 | Max | 17187 | protein_coding | 1080.229877 | 0.153124061 | 0.070135984 | 2.183245336 | 0.029017748 | 0.105259551 | 0.153124061 |
| ENSMUSG00000087305 | A430035B10Rik | 320312 | lncRNA | 42.37774868 | 0.153130653 | 0.233544789 | 0.655680024 | 0.512030025 | 0.699872499 | 0.153130653 |
| ENSMUSG00000045215 | Asxl3 | 211961 | protein_coding | 47.7293337 | 0.153187759 | 0.298453875 | 0.513271132 | 0.607761674 | 0.771533271 | 0.153187759 |
| ENSMUSG00000112160 | BC024063 | 666584 | protein_coding | 52.6459454 | 0.153192376 | 0.189891769 | 0.806735211 | 0.419819053 | 0.625792035 | 0.153192376 |
| ENSMUSG00000005803 | Sqor | 59010 | protein_coding | 7648.183911 | -0.15332827 | 0.147109307 | -1.042274441 | 0.297284459 | 0.506180721 | 0.15332827 |
| ENSMUSG00000040929 | Rfx3 | 19726 | protein_coding | 144.4245565 | -0.153333228 | 0.201020778 | -0.762773033 | 0.445598766 | 0.646461167 | 0.153333228 |
| ENSMUSG00000033326 | Kdm4a | 230674 | protein_coding | 980.4939836 | 0.153341968 | 0.088878058 | 1.725307378 | 0.084472093 | 0.222281761 | 0.153341968 |
| ENSMUSG00000079020 | Slc45a4 | 106068 | protein_coding | 293.6349341 | -0.153345927 | 0.078437992 | -1.954995568 | 0.050583593 | 0.156635171 | 0.153345927 |
| ENSMUSG00000062328 | Rpl17 | 319195 | protein_coding | 10397.72271 | 0.153471441 | 0.09026195 | 1.70028945 | 0.089076494 | 0.230363759 | 0.153471441 |
| ENSMUSG00000047036 | Zfp445 | 235682 | protein_coding | 1345.70891 | -0.153505234 | 0.060816774 | -2.524060786 | 0.011600787 | 0.053495515 | 0.153505234 |
| ENSMUSG00000001445 | Mrpl10 | 107732 | protein_coding | 636.6942525 | 0.153509196 | 0.092157014 | 1.665735362 | 0.095766135 | 0.241464688 | 0.153509196 |
| ENSMUSG00000028096 | Gpr89 | 67549 | protein_coding | 605.271337 | 0.153586528 | 0.118374344 | 1.297464658 | 0.194471358 | 0.387725596 | 0.153586528 |
| ENSMUSG00000029670 | Ing3 | 71777 | protein_coding | 223.5568255 | 0.153590317 | 0.086199978 | 1.781790667 | 0.074783375 | 0.204173739 | 0.153590317 |
| ENSMUSG00000037938 | Chchd5 | 66170 | protein_coding | 164.9872367 | 0.153595511 | 0.174690326 | 0.879244517 | 0.379268712 | 0.587245265 | 0.153595511 |
| ENSMUSG00000025925 | Terf1 | 21749 | protein_coding | 217.0685794 | 0.153609295 | 0.099031061 | 1.551122372 | 0.120872361 | 0.282461115 | 0.153609295 |
| ENSMUSG00000026516 | Nvl | 67459 | protein_coding | 507.4169923 | -0.153612774 | 0.076137849 | -2.017561257 | 0.043636975 | 0.141298992 | 0.153612774 |
| ENSMUSG00000029102 | Hgfac | 54426 | protein_coding | 2418.614655 | -0.153622261 | 0.120031712 | -1.279847287 | 0.20059885 | 0.395157223 | 0.153622261 |
| ENSMUSG00000004789 | Dlst | 78920 | protein_coding | 2405.951977 | -0.153634476 | 0.066249958 | -2.319012423 | 0.020394363 | 0.081068483 | 0.153634476 |
| ENSMUSG00000025041 | Nt5c2 | 76952 | protein_coding | 412.4166613 | -0.153723785 | 0.149687803 | -1.026962663 | 0.304438045 | 0.513961388 | 0.153723785 |
| ENSMUSG00000078862 | Gm14326 | 628147 | protein_coding | 117.3677797 | -0.153724885 | 0.14663197 | -1.048372225 | 0.294467145 | 0.503630558 | 0.153724885 |
| ENSMUSG00000037440 | Vnn1 | 22361 | protein_coding | 128.7646157 | -0.153734275 | 0.285858025 | -0.537799402 | 0.590715545 | 0.760015009 | 0.153734275 |
| ENSMUSG00000022634 | Yaf2 | 67057 | protein_coding | 139.8369077 | 0.153740007 | 0.10759359 | 1.428895602 | 0.153034242 | 0.331907598 | 0.153740007 |
| ENSMUSG00000004788 | Eif2b2 | 217715 | protein_coding | 386.8809788 | 0.153750235 | 0.094295181 | 1.630520603 | 0.102991514 | 0.253233844 | 0.153750235 |
| ENSMUSG00000073481 | Mtarc2 | 67247 | protein_coding | 1133.809924 | -0.153755128 | 0.086519668 | -1.777111855 | 0.075549838 | 0.205186208 | 0.153755128 |
| ENSMUSG00000026017 | Carf | 241066 | protein_coding | 129.6945766 | -0.153821401 | 0.16455717 | -0.934759642 | 0.349912183 | 0.560140829 | 0.153821401 |
| ENSMUSG00000001998 | Ap4e1 | 108011 | protein_coding | 202.0750664 | -0.153841692 | 0.088901123 | -1.730480857 | 0.083544398 | 0.220642543 | 0.153841692 |
| ENSMUSG00000054717 | Hmgb2 | 97165 | protein_coding | 1657.319414 | 0.153909087 | 0.107466519 | 1.432158481 | 0.152098471 | 0.330511147 | 0.153909087 |
| ENSMUSG00000038467 | Chmp4b | 75608 | protein_coding | 2763.379907 | 0.153909488 | 0.108848561 | 1.413978162 | 0.157368315 | 0.33818806 | 0.153909488 |

| ensembl_gene_id | external_gene_name | entrezgene_id | gene_biotype | baseMean | log2FoldChange | lfcSE | stat | pvalue | padj | absLog2FC |
|---|---|---|---|---|---|---|---|---|---|---|
| ENSMUSG00000028423 | Nfx1 | 74164 | protein_coding | 1188.248459 | -0.153933156 | 0.054418394 | -2.828697158 | 0.00467379 | 0.026653824 | 0.153933156 |
| ENSMUSG00000078784 | Rbis | 69126 | protein_coding | 530.8087057 | 0.154022034 | 0.086090574 | 1.789069649 | 0.073603599 | 0.202052993 | 0.154022034 |
| ENSMUSG00000026641 | Usf1 | 22278 | protein_coding | 502.0781138 | 0.154177055 | 0.091121777 | 1.691989114 | 0.090648049 | 0.232840208 | 0.154177055 |
| ENSMUSG00000027091 | Zc3h15 | 69082 | protein_coding | 1379.703018 | 0.154198017 | 0.089571382 | 1.721509853 | 0.085158346 | 0.223386386 | 0.154198017 |
| ENSMUSG00000109408 | A930037H05Rik | 78439 | lncRNA | 39.59950388 | -0.154211609 | 0.257728099 | -0.598350004 | 0.549606418 | 0.731093174 | 0.154211609 |
| ENSMUSG00000027784 | Ppm1l | 242083 | protein_coding | 186.8507395 | 0.154216096 | 0.106915514 | 1.442410842 | 0.149186507 | 0.326586405 | 0.154216096 |
| ENSMUSG00000084803 | 5830444B04Rik | 641454 | lncRNA | 72.0503745 | 0.154250377 | 0.312431109 | 0.493710044 | 0.621510971 | 0.779973743 | 0.154250377 |
| ENSMUSG00000042302 | Ehbp1 | 216565 | protein_coding | 180.1750368 | -0.154292374 | 0.159509359 | -0.96729355 | 0.333397307 | 0.544260768 | 0.154292374 |
| ENSMUSG00000008601 | Rab25 | 53868 | protein_coding | 1514.884386 | -0.154350468 | 0.081957159 | -1.883306726 | 0.059658813 | 0.175360554 | 0.154350468 |
| ENSMUSG00000024319 | Vps52 | 224705 | protein_coding | 521.537613 | 0.154402236 | 0.072152004 | 2.139957697 | 0.032358186 | 0.114052042 | 0.154402236 |
| ENSMUSG00000072969 | Armcx5 | 494468 | protein_coding | 75.24752633 | -0.154445472 | 0.14538675 | -1.062307755 | 0.288095995 | 0.497268423 | 0.154445472 |
| ENSMUSG00000026162 | Nhej1 | 75570 | protein_coding | 52.73894153 | 0.15451606 | 0.21312839 | 0.724990508 | 0.468457854 | 0.665026443 | 0.15451606 |
| ENSMUSG00000023353 | Agap3 | 213990 | protein_coding | 399.1370174 | -0.154556827 | 0.089972755 | -1.717818101 | 0.085829799 | 0.224666727 | 0.154556827 |
| ENSMUSG00000025207 | Sema4g | 26456 | protein_coding | 2811.921974 | 0.154564104 | 0.175354925 | 0.881435773 | 0.378082003 | 0.586205601 | 0.154564104 |
| ENSMUSG00000052155 | Acvr2a | 11480 | protein_coding | 920.6820672 | 0.154566511 | 0.078417868 | 1.971062408 | 0.048716738 | 0.152928696 | 0.154566511 |
| ENSMUSG00000037685 | Atp8a1 | 11980 | protein_coding | 1878.71969 | 0.154800378 | 0.085517477 | 1.810160727 | 0.070270866 | 0.196106105 | 0.154800378 |
| ENSMUSG00000035930 | Chst4 | 26887 | protein_coding | 1012.009181 | -0.154853967 | 0.106781342 | -1.450196865 | 0.147003629 | 0.323461056 | 0.154853967 |
| ENSMUSG00000032737 | Inppl1 | 16332 | protein_coding | 1265.630222 | -0.154863675 | 0.089945943 | -1.721741639 | 0.085116331 | 0.223382746 | 0.154863675 |
| ENSMUSG00000034973 | Dop1a | 320615 | protein_coding | 949.4367146 | -0.154908211 | 0.059565276 | -2.600646257 | 0.009304835 | 0.045378875 | 0.154908211 |
| ENSMUSG00000002052 | Supt6 | 20926 | protein_coding | 1282.293138 | -0.154947856 | 0.122943809 | -1.260314424 | 0.207555959 | 0.404082965 | 0.154947856 |
| ENSMUSG00000037032 | Apbb1 | 11785 | protein_coding | 11.32479894 | -0.154976437 | 0.472070874 | -0.328290614 | 0.74269194 | 0.856720541 | 0.154976437 |
| ENSMUSG00000020590 | Snx13 | 217463 | protein_coding | 2433.876325 | 0.155029646 | 0.085482393 | 1.813585712 | 0.069741532 | 0.195227236 | 0.155029646 |
| ENSMUSG00000030522 | Mtmr10 | 233315 | protein_coding | 271.7997164 | -0.155044134 | 0.102157884 | -1.517691312 | 0.129092235 | 0.295598207 | 0.155044134 |
| ENSMUSG00000029149 | Krtcap3 | 69815 | protein_coding | 610.0595394 | -0.155054669 | 0.076423885 | -2.02887709 | 0.042470811 | 0.138536775 | 0.155054669 |
| ENSMUSG00000039701 | Usp53 | 99526 | protein_coding | 1826.458608 | -0.15506152 | 0.135699102 | -1.142686412 | 0.253168814 | 0.459383441 | 0.15506152 |
| ENSMUSG00000117698 | BC037704 | 100502982 | lncRNA | 9.469053596 | -0.155093626 | 0.352997216 | -0.439362181 | 0.660399125 | 0.803487938 | 0.155093626 |
| ENSMUSG00000025950 | Idh1 | 15926 | protein_coding | 3957.086396 | 0.155123182 | 0.091611292 | 1.693275777 | 0.090402987 | 0.232435637 | 0.155123182 |
| ENSMUSG00000029265 | Dr1 | 13486 | protein_coding | 786.863523 | 0.155230207 | 0.096090247 | 1.615462665 | 0.106210539 | 0.25850587 | 0.155230207 |
| ENSMUSG00000006932 | Ctnnb1 | 12387 | protein_coding | 11367.04594 | 0.155238431 | 0.130781216 | 1.187008622 | 0.23522422 | 0.439335292 | 0.155238431 |
| ENSMUSG00000015522 | Arnt | 11863 | protein_coding | 509.9230726 | -0.15535379 | 0.072033858 | -2.156677338 | 0.031030814 | 0.110795518 | 0.15535379 |
| ENSMUSG00000038695 | Josd2 | 66124 | protein_coding | 190.7653845 | 0.155380138 | 0.108704297 | 1.429383586 | 0.152894013 | 0.331738979 | 0.155380138 |
| ENSMUSG00000018846 | Pank3 | 211347 | protein_coding | 5781.420442 | 0.155407198 | 0.096756613 | 1.606166161 | 0.108237406 | 0.261882891 | 0.155407198 |
| ENSMUSG00000062526 | Mppe1 | 225651 | protein_coding | 214.1105721 | 0.155410648 | 0.146376205 | 1.061720708 | 0.288362496 | 0.497392119 | 0.155410648 |
| ENSMUSG00000026000 | Lancl1 | 14768 | protein_coding | 288.7160822 | 0.155438841 | 0.132750601 | 1.170908753 | 0.241635456 | 0.446389417 | 0.155438841 |
| ENSMUSG00000025484 | Bet1l | 54399 | protein_coding | 297.5198255 | -0.155460756 | 0.074993167 | -2.072998933 | 0.038172379 | 0.12887455 | 0.155460756 |
| ENSMUSG00000027882 | Stxbp3 | 20912 | protein_coding | 1031.815013 | 0.155539089 | 0.080691097 | 1.927586755 | 0.053906548 | 0.163143358 | 0.155539089 |
| ENSMUSG00000018042 | Cyb5r3 | 109754 | protein_coding | 1325.102822 | 0.155547125 | 0.063556041 | 2.447401107 | 0.014389061 | 0.062551571 | 0.155547125 |
| ENSMUSG00000019832 | Rab32 | 67844 | protein_coding | 291.3614636 | 0.155547308 | 0.191430683 | 0.8125516 | 0.416475191 | 0.622741858 | 0.155547308 |

| ensembl_gene_id | external_gene_name | entrezgene_id | gene_biotype | baseMean | log2FoldChange | lfcSE | stat | pvalue | padj | absLog2FC |
|---|---|---|---|---|---|---|---|---|---|---|
| ENSMUSG00000024767 | Otub1 | 107260 | protein_coding | 726.8270104 | 0.155571937 | 0.079897326 | 1.947148227 | 0.051516967 | 0.158593398 | 0.155571937 |
| ENSMUSG00000019715 | Gle1 | 74412 | protein_coding | 378.4366792 | 0.155609247 | 0.102326114 | 1.520718808 | 0.128330417 | 0.29436159 | 0.155609247 |
| ENSMUSG00000048878 | Hexim1 | 192231 | protein_coding | 1020.789594 | -0.15568986 | 0.073633009 | -2.114403062 | 0.034480856 | 0.119575676 | 0.15568986 |
| ENSMUSG00000025722 | Wdr73 | 71968 | protein_coding | 178.0225768 | -0.155740715 | 0.163543408 | -0.952289774 | 0.340950043 | 0.551279289 | 0.155740715 |
| ENSMUSG00000087022 | 9130024F11Rik | 329160 | lncRNA | 197.4594264 | 0.155790817 | 0.152985365 | 1.018338036 | 0.308517339 | 0.518563181 | 0.155790817 |
| ENSMUSG00000025157 | Zdhhc16 | 74168 | protein_coding | 191.9354186 | 0.155928809 | 0.085975259 | 1.813647436 | 0.069732022 | 0.195227236 | 0.155928809 |
| ENSMUSG00000018209 | Stk4 | 58231 | protein_coding | 896.9416421 | -0.155928895 | 0.057338574 | -2.719441455 | 0.006539227 | 0.034546205 | 0.155928895 |
| ENSMUSG00000026643 | Nmt2 | 18108 | protein_coding | 633.4314086 | -0.15593243 | 0.086476386 | -1.803179303 | 0.07136006 | 0.198062246 | 0.15593243 |
| ENSMUSG00000024188 | Luc7l | 66978 | protein_coding | 757.1332601 | -0.155935726 | 0.085479589 | -1.824245151 | 0.068115018 | 0.191922168 | 0.155935726 |
| ENSMUSG00000041231 | Ublcp1 | 79560 | protein_coding | 443.0775594 | 0.155954227 | 0.103973452 | 1.499942771 | 0.133629228 | 0.302207346 | 0.155954227 |
| ENSMUSG00000030706 | Mrpl48 | 52443 | protein_coding | 442.8126847 | 0.155962304 | 0.078195224 | 1.994524671 | 0.046094747 | 0.147074744 | 0.155962304 |
| ENSMUSG00000074030 | Exoc8 | 102058 | protein_coding | 523.1690472 | -0.155998544 | 0.082409605 | -1.892965558 | 0.05836245 | 0.172734546 | 0.155998544 |
| ENSMUSG00000021615 | Xrcc4 | 108138 | protein_coding | 151.2369766 | 0.156021738 | 0.106174341 | 1.469486291 | 0.141700938 | 0.314741672 | 0.156021738 |
| ENSMUSG00000033653 | Vps8 | 209018 | protein_coding | 360.7025157 | -0.156022628 | 0.082596462 | -1.888974701 | 0.058895219 | 0.173815646 | 0.156022628 |
| ENSMUSG00000060147 | Serpinb6a | 20719 | protein_coding | 5951.127374 | -0.156113562 | 0.106086682 | -1.471566074 | 0.141138093 | 0.313710234 | 0.156113562 |
| ENSMUSG00000021066 | Atl1 | 73991 | protein_coding | 40.95345288 | -0.156182358 | 0.223191697 | -0.699767777 | 0.484072341 | 0.677531129 | 0.156182358 |
| ENSMUSG00000025933 | Tmem14a | 75712 | protein_coding | 50.52618878 | 0.156198423 | 0.208624254 | 0.748706921 | 0.454033871 | 0.653519926 | 0.156198423 |
| ENSMUSG00000027165 | Iftap | 68170 | protein_coding | 75.06855359 | -0.156211537 | 0.151808797 | -1.029001876 | 0.303478793 | 0.513034411 | 0.156211537 |
| ENSMUSG00000045160 | Bola3 | 78653 | protein_coding | 310.8108467 | 0.15624748 | 0.082760723 | 1.887942422 | 0.059033681 | 0.174063026 | 0.15624748 |
| ENSMUSG00000002320 | Tm9sf1 | 74140 | protein_coding | 1261.147039 | 0.156281519 | 0.078984035 | 1.978646943 | 0.04785577 | 0.151211258 | 0.156281519 |
| ENSMUSG00000022427 | Tomm22 | 223696 | protein_coding | 1466.878559 | 0.156292251 | 0.109531393 | 1.42691741 | 0.153603702 | 0.332727274 | 0.156292251 |
| ENSMUSG00000036980 | Taf6 | 21343 | protein_coding | 230.7179458 | 0.156330498 | 0.096645419 | 1.6175676 | 0.105755816 | 0.257683745 | 0.156330498 |
| ENSMUSG00000040850 | Psme4 | 103554 | protein_coding | 3075.407255 | 0.156369288 | 0.05804082 | 2.694126097 | 0.007057346 | 0.036675173 | 0.156369288 |
| ENSMUSG00000021177 | Tdp1 | 104884 | protein_coding | 69.23448379 | -0.156370073 | 0.171663482 | -0.910910528 | 0.362342518 | 0.571538089 | 0.156370073 |
| ENSMUSG00000011958 | Bnip2 | 12175 | protein_coding | 249.8774188 | -0.156388488 | 0.118373536 | -1.321144014 | 0.18645335 | 0.376888975 | 0.156388488 |
| ENSMUSG00000015575 | Atp6v0e | 11974 | protein_coding | 589.5092824 | 0.156389438 | 0.108451409 | 1.442023104 | 0.149295856 | 0.326699017 | 0.156389438 |
| ENSMUSG00000015291 | Gdi1 | 14567 | protein_coding | 548.3856028 | -0.156551781 | 0.09662674 | -1.620170368 | 0.105195685 | 0.256772746 | 0.156551781 |
| ENSMUSG00000057777 | Mab21l2 | 23937 | protein_coding | 7.759412983 | 0.156575153 | 0.447357991 | 0.349999678 | 0.726338939 | 0.846191239 | 0.156575153 |
| ENSMUSG00000036698 | Ago2 | 239528 | protein_coding | 1031.916488 | -0.156580555 | 0.160200485 | -0.977403753 | 0.328369306 | 0.539240147 | 0.156580555 |
| ENSMUSG00000047648 | Fbxo30 | 71865 | protein_coding | 296.0550867 | -0.156606522 | 0.114679883 | -1.365597156 | 0.17206544 | 0.357719532 | 0.156606522 |
| ENSMUSG00000005505 | Kbtbd4 | 67136 | protein_coding | 102.0168759 | -0.156623518 | 0.124656805 | -1.256437774 | 0.208957293 | 0.405789022 | 0.156623518 |
| ENSMUSG00000031673 | Cdh11 | 12552 | protein_coding | 9.199844711 | 0.156652791 | 0.557226269 | 0.281129587 | 0.778611009 | 0.879741516 | 0.156652791 |
| ENSMUSG00000029029 | Wrap73 | 59002 | protein_coding | 147.3941503 | 0.156657003 | 0.130092247 | 1.204199386 | 0.228512523 | 0.431454181 | 0.156657003 |
| ENSMUSG00000057982 | Zfp809 | 235047 | protein_coding | 329.9683744 | 0.15665728 | 0.10172916 | 1.539944692 | 0.123573835 | 0.286838247 | 0.15665728 |
| ENSMUSG00000047193 | Dync2h1 | 110350 | protein_coding | 167.9793088 | -0.156661242 | 0.200449782 | -0.781548579 | 0.434479917 | 0.637804646 | 0.156661242 |
| ENSMUSG00000029823 | Luc7l2 | 192196 | protein_coding | 1937.1665 | 0.156666638 | 0.117721234 | 1.330827354 | 0.183245825 | 0.372478242 | 0.156666638 |
| ENSMUSG00000030888 | Rrp8 | 101867 | protein_coding | 192.5199468 | -0.156731274 | 0.161961163 | -0.967708996 | 0.333189724 | 0.544089186 | 0.156731274 |
| ENSMUSG00000023078 | Cxcl13 | 55985 | protein_coding | 7.368627534 | -0.156754634 | 0.478769966 | -0.327411168 | 0.743356921 | 0.857217273 | 0.156754634 |

| ensembl_gene_id | external_gene_name | entrezgene_id | gene_biotype | baseMean | log2FoldChange | lfcSE | stat | pvalue | padj | absLog2FC |
|---|---|---|---|---|---|---|---|---|---|---|
| ENSMUSG00000063354 | Slc39a4 | 72027 | protein_coding | 1346.5592 | 0.156763945 | 0.151184076 | 1.036907781 | 0.299778837 | 0.508687887 | 0.156763945 |
| ENSMUSG00000040918 | Slc19a2 | 116914 | protein_coding | 168.3004323 | 0.156788285 | 0.102960643 | 1.522798242 | 0.12780919 | 0.293439621 | 0.156788285 |
| ENSMUSG00000094870 | Zfp131 | 72465 | protein_coding | 522.0855208 | -0.156796833 | 0.115094674 | -1.362329177 | 0.173094024 | 0.358895202 | 0.156796833 |
| ENSMUSG00000062397 | Zfp706 | 68036 | protein_coding | 1969.086577 | 0.156905077 | 0.069829542 | 2.246972713 | 0.024641771 | 0.093406504 | 0.156905077 |
| ENSMUSG00000038236 | Hoxa7 | 15404 | protein_coding | 259.9235691 | 0.156953279 | 0.164089669 | 0.956509203 | 0.338815037 | 0.549668559 | 0.156953279 |
| ENSMUSG00000089832 | Shkbp1 | 192192 | protein_coding | 203.1472994 | 0.156961021 | 0.093925963 | 1.671114305 | 0.094699102 | 0.239804023 | 0.156961021 |
| ENSMUSG00000029703 | Lrwd1 | 71735 | protein_coding | 213.7298229 | 0.156963293 | 0.099401746 | 1.579079831 | 0.114317747 | 0.270919964 | 0.156963293 |
| ENSMUSG00000038459 | Abhd17c | 70178 | protein_coding | 1315.891133 | 0.156965916 | 0.079911733 | 1.964241182 | 0.049502129 | 0.15435981 | 0.156965916 |
| ENSMUSG00000020267 | Hint1 | 15254 | protein_coding | 4445.714712 | 0.157038326 | 0.10629856 | 1.477332579 | 0.139586509 | 0.311347698 | 0.157038326 |
| ENSMUSG00000040242 | Fgfr1op2 | 67529 | protein_coding | 1416.153132 | 0.157042352 | 0.054443643 | 2.884493795 | 0.003920435 | 0.023110543 | 0.157042352 |
| ENSMUSG00000031286 | Glt28d2 | 320302 | protein_coding | 152.8883058 | 0.157075698 | 0.195799935 | 0.802225482 | 0.422422539 | 0.627648847 | 0.157075698 |
| ENSMUSG00000021288 | Klc1 | 16593 | protein_coding | 891.682549 | -0.157167081 | 0.13126002 | -1.197372057 | 0.23116157 | 0.434447373 | 0.157167081 |
| ENSMUSG00000035437 | Rabgap1 | 227800 | protein_coding | 770.4382118 | -0.157173853 | 0.087805273 | -1.790027488 | 0.073449493 | 0.201846708 | 0.157173853 |
| ENSMUSG00000038838 | Vars2 | 68915 | protein_coding | 380.1223167 | -0.157197135 | 0.083469368 | -1.883291302 | 0.059660902 | 0.175360554 | 0.157197135 |
| ENSMUSG00000024177 | Nme4 | 56520 | protein_coding | 51.47341066 | -0.1572071 | 0.175556892 | -0.895476661 | 0.370532337 | 0.579122663 | 0.1572071 |
| ENSMUSG00000075273 | Ttc30b | 72421 | protein_coding | 189.2180917 | 0.157210137 | 0.231504796 | 0.679079394 | 0.497087558 | 0.688069056 | 0.157210137 |
| ENSMUSG00000067916 | Zfp991 | 666532 | protein_coding | 78.23228709 | 0.157229771 | 0.198279972 | 0.792968499 | 0.427796176 | 0.632090677 | 0.157229771 |
| ENSMUSG00000047221 | Fam185a | 330050 | protein_coding | 194.5439817 | 0.157235301 | 0.15308349 | 1.027121218 | 0.304363389 | 0.513961388 | 0.157235301 |
| ENSMUSG00000028577 | Plaa | 18786 | protein_coding | 1108.023677 | 0.157242303 | 0.096617566 | 1.627471164 | 0.103637068 | 0.254232438 | 0.157242303 |
| ENSMUSG00000055480 | Zfp458 | 238690 | protein_coding | 65.74180368 | -0.157277196 | 0.186097488 | -0.845133364 | 0.398036382 | 0.606068495 | 0.157277196 |
| ENSMUSG00000029066 | Mrpl20 | 66448 | protein_coding | 536.3218613 | 0.15735628 | 0.094860637 | 1.658815336 | 0.097153009 | 0.243675627 | 0.15735628 |
| ENSMUSG00000050373 | Snx21 | 101113 | protein_coding | 161.5262498 | 0.157367394 | 0.14419444 | 1.091355497 | 0.275116487 | 0.483297166 | 0.157367394 |
| ENSMUSG00000035842 | Ddx11 | 320209 | protein_coding | 165.6974532 | -0.157408662 | 0.139448759 | -1.128792132 | 0.258985532 | 0.466168105 | 0.157408662 |
| ENSMUSG00000071573 | Rnls | 67795 | protein_coding | 35.01452959 | -0.157585007 | 0.229716236 | -0.685998559 | 0.492714017 | 0.68439425 | 0.157585007 |
| ENSMUSG00000010066 | Cacna2d2 | 56808 | protein_coding | 109.4421069 | -0.157628985 | 0.17500955 | -0.900687909 | 0.367754279 | 0.576477165 | 0.157628985 |
| ENSMUSG00000091476 | Catspere2 | 545391 | protein_coding | 34.18313582 | 0.157670792 | 0.220324384 | 0.71563024 | 0.474219696 | 0.669150542 | 0.157670792 |
| ENSMUSG00000039356 | Exosc2 | 227715 | protein_coding | 280.6531946 | -0.157798683 | 0.138540128 | -1.139010662 | 0.254698707 | 0.461298219 | 0.157798683 |
| ENSMUSG00000029364 | Wsb2 | 59043 | protein_coding | 842.1503555 | -0.157821173 | 0.191013681 | -0.826229683 | 0.408673813 | 0.61583316 | 0.157821173 |
| ENSMUSG00000036403 | Cep135 | 381644 | protein_coding | 154.8947712 | -0.15790478 | 0.123268249 | -1.280985023 | 0.200198927 | 0.394795779 | 0.15790478 |
| ENSMUSG00000090247 | Bloc1s1 | 14533 | protein_coding | 109.0362133 | 0.158022184 | 0.128731872 | 1.22752961 | 0.2196236 | 0.419876136 | 0.158022184 |
| ENSMUSG00000026987 | Baz2b | 407823 | protein_coding | 893.0558641 | -0.158038457 | 0.101735825 | -1.553419909 | 0.12032285 | 0.2816099 | 0.158038457 |
| ENSMUSG00000029404 | Arl6ip4 | 65105 | protein_coding | 268.0287219 | 0.158054516 | 0.077446081 | 2.040832979 | 0.041267431 | 0.136004065 | 0.158054516 |
| ENSMUSG00000015305 | Sash1 | 70097 | protein_coding | 1142.830608 | -0.158055508 | 0.151859635 | -1.040800001 | 0.297968379 | 0.506735399 | 0.158055508 |
| ENSMUSG00000020803 | Txndc17 | 52700 | protein_coding | 1585.904924 | 0.158072629 | 0.081951208 | 1.928862703 | 0.053747909 | 0.162897492 | 0.158072629 |
| ENSMUSG00000043895 | S1pr2 | 14739 | protein_coding | 32.54512047 | 0.158099127 | 0.283688319 | 0.557298685 | 0.577323374 | 0.749071904 | 0.158099127 |
| ENSMUSG00000004038 | Gstm3 | 14864 | protein_coding | 709.7065085 | 0.15811925 | 0.101704611 | 1.554691069 | 0.120019663 | 0.28112841 | 0.15811925 |
| ENSMUSG00000028017 | Egf | 13645 | protein_coding | 153.9285582 | -0.158238499 | 0.116356284 | -1.359948027 | 0.173846371 | 0.359707067 | 0.158238499 |
| ENSMUSG00000029055 | Plch2 | 269615 | protein_coding | 695.2851803 | -0.15824017 | 0.18959736 | -0.834611672 | 0.403936388 | 0.611122605 | 0.15824017 |

Supplementary Table 1

| ensembl_gene_id | external_gene_name | entrezgene_id | gene_biotype | baseMean | log2FoldChange | lfcSE | stat | pvalue | padj | absLog2FC |
|---|---|---|---|---|---|---|---|---|---|---|
| ENSMUSG00000038128 | Camk4 | 12326 | protein_coding | 7.42569706 | 0.158266969 | 0.625486604 | 0.253030151 | 0.800244914 | 0.892814435 | 0.158266969 |
| ENSMUSG00000026623 | Lpgat1 | 226856 | protein_coding | 1182.38984 | 0.158297005 | 0.121954874 | 1.297996545 | 0.194288522 | 0.387555114 | 0.158297005 |
| ENSMUSG00000030074 | Gxylt2 | 232313 | protein_coding | 7.72322292 | -0.158334064 | 0.627057129 | -0.252503411 | 0.800651978 | 0.89305215 | 0.158334064 |
| ENSMUSG00000044709 | Gemin7 | 69731 | protein_coding | 248.2013988 | 0.158359612 | 0.124716404 | 1.269757685 | 0.204170958 | 0.399572245 | 0.158359612 |
| ENSMUSG00000034601 | 2700049A03Rik | 76967 | protein_coding | 193.4320573 | -0.158405698 | 0.153766838 | -1.030168145 | 0.30293108 | 0.512434818 | 0.158405698 |
| ENSMUSG00000046605 | B3gntl1 | 210004 | protein_coding | 99.36696147 | -0.158514365 | 0.142281771 | -1.114087661 | 0.265241582 | 0.472780653 | 0.158514365 |
| ENSMUSG00000000579 | Dynlt1c | 100040563 | protein_coding | 32.59509108 | -0.158520296 | 0.205351843 | -0.771944839 | 0.440147098 | 0.642313808 | 0.158520296 |
| ENSMUSG00000020592 | Sdc1 | 20969 | protein_coding | 3427.75529 | -0.158535554 | 0.129168102 | -1.22735839 | 0.219687918 | 0.419876136 | 0.158535554 |
| ENSMUSG00000037573 | Tob1 | 22057 | protein_coding | 1860.559513 | -0.15859257 | 0.152742613 | -1.038299443 | 0.299130668 | 0.508002269 | 0.15859257 |
| ENSMUSG00000044471 | Lncpint | 232685 | lncRNA | 143.8823778 | -0.158604995 | 0.239255232 | -0.662911292 | 0.507387369 | 0.696822981 | 0.158604995 |
| ENSMUSG00000040631 | Dok4 | 114255 | protein_coding | 1019.354273 | -0.158653372 | 0.113724738 | -1.395064742 | 0.162996317 | 0.345469773 | 0.158653372 |
| ENSMUSG00000028327 | Stra6l | 74152 | protein_coding | 388.7142885 | -0.158678195 | 0.139149624 | -1.140342248 | 0.254143743 | 0.46058067 | 0.158678195 |
| ENSMUSG00000032249 | Anp32a | 11737 | protein_coding | 1321.533329 | 0.158686888 | 0.083008557 | 1.911693125 | 0.055915567 | 0.167567854 | 0.158686888 |
| ENSMUSG00000024962 | Vegfb | 22340 | protein_coding | 114.6554027 | -0.158697801 | 0.128482419 | -1.235171337 | 0.216766744 | 0.415983218 | 0.158697801 |
| ENSMUSG00000036192 | Rorb | 225998 | protein_coding | 15.83458872 | 0.158744596 | 0.585109863 | 0.271307332 | 0.786154668 | 0.884188077 | 0.158744596 |
| ENSMUSG00000051452 | Gm11437 | 628813 | protein_coding | 76.50085474 | 0.158757544 | 0.175990021 | 0.902082648 | 0.367012966 | 0.57565559 | 0.158757544 |
| ENSMUSG00000020986 | Sec23a | 20334 | protein_coding | 1609.438368 | 0.158796551 | 0.088200577 | 1.800402627 | 0.071797086 | 0.198953373 | 0.158796551 |
| ENSMUSG00000022536 | Glyr1 | 74022 | protein_coding | 2144.780331 | 0.158799301 | 0.058986936 | 2.692109694 | 0.007100159 | 0.03684957 | 0.158799301 |
| ENSMUSG00000044177 | Wfikkn2 | 278507 | protein_coding | 22.8309379 | -0.158807522 | 0.312107267 | -0.508823532 | 0.610875924 | 0.77314515 | 0.158807522 |
| ENSMUSG00000006526 | Stimate | 69179 | protein_coding | 163.1960631 | -0.15881878 | 0.115150659 | -1.379225971 | 0.167825093 | 0.352275774 | 0.15881878 |
| ENSMUSG00000098274 | Rpl24 | 68193 | protein_coding | 183.6884294 | 0.158915957 | 0.205472963 | 0.773415416 | 0.43927657 | 0.641731377 | 0.158915957 |
| ENSMUSG00000020087 | Tysnd1 | 71767 | protein_coding | 326.0252078 | 0.158980819 | 0.110773935 | 1.435182556 | 0.151235083 | 0.329440974 | 0.158980819 |
| ENSMUSG00000022314 | Rad21 | 19357 | protein_coding | 2559.253004 | 0.158981357 | 0.078996763 | 2.01250471 | 0.044166761 | 0.142435344 | 0.158981357 |
| ENSMUSG00000069743 | Zfp820 | 75424 | protein_coding | 183.5869563 | 0.158988339 | 0.145002209 | 1.096454598 | 0.272879882 | 0.480918845 | 0.158988339 |
| ENSMUSG00000056492 | Adgrf5 | 224792 | protein_coding | 26.98879366 | -0.158996537 | 0.316214625 | -0.50281209 | 0.615096393 | 0.775590195 | 0.158996537 |
| ENSMUSG00000028182 | Lrriq3 | 74435 | protein_coding | 38.99936855 | -0.1590162 | 0.220890943 | -0.719885559 | 0.47159546 | 0.667069775 | 0.1590162 |
| ENSMUSG00000056131 | Pgm3 | 109785 | protein_coding | 743.3452215 | 0.159151501 | 0.071704832 | 2.219536622 | 0.026450239 | 0.098431865 | 0.159151501 |
| ENSMUSG00000041720 | Pi4ka | 224020 | protein_coding | 1059.817952 | -0.159151953 | 0.059207481 | -2.688037873 | 0.007187324 | 0.037192886 | 0.159151953 |
| ENSMUSG00000085492 | Trmt61b | 68789 | lncRNA | 68.96540949 | 0.159219562 | 0.166310377 | 0.957363961 | 0.338383582 | 0.549136393 | 0.159219562 |
| ENSMUSG00000006471 | Ndor1 | 78797 | protein_coding | 67.82688512 | -0.159254457 | 0.160895407 | -0.989801141 | 0.322271327 | 0.533229858 | 0.159254457 |
| ENSMUSG00000016559 | H3f3b | 15081 | protein_coding | 10559.53061 | 0.159255193 | 0.098245275 | 1.620995952 | 0.105018507 | 0.256593769 | 0.159255193 |
| ENSMUSG00000074826 | Gm10767 | 100038538 | protein_coding | 38.31395803 | 0.159281229 | 0.214628442 | 0.74212545 | 0.458011332 | 0.656603314 | 0.159281229 |
| ENSMUSG00000013150 | Gfod2 | 70575 | protein_coding | 241.4188705 | -0.159345421 | 0.131156182 | -1.214928788 | 0.224393253 | 0.425999491 | 0.159345421 |
| ENSMUSG00000054720 | Lrrc8c | 100604 | protein_coding | 12.87207449 | 0.159415167 | 0.449716904 | 0.35447893 | 0.722979992 | 0.844115801 | 0.159415167 |
| ENSMUSG00000041153 | Osgin2 | 209212 | protein_coding | 139.2997601 | 0.159439077 | 0.125595416 | 1.269465734 | 0.204275004 | 0.399716495 | 0.159439077 |
| ENSMUSG00000026830 | Ermn | 77767 | protein_coding | 17.23540446 | 0.159443953 | 0.319919509 | 0.498387713 | 0.618210795 | 0.777731739 | 0.159443953 |
| ENSMUSG00000046352 | Gjb2 | 14619 | protein_coding | 173.622354 | 0.159456308 | 0.127495661 | 1.25068027 | 0.211051152 | 0.408459628 | 0.159456308 |
| ENSMUSG00000041064 | Pif1 | 208084 | protein_coding | 111.7040047 | 0.159573211 | 0.122224697 | 1.30557256 | 0.191697957 | 0.38440955 | 0.159573211 |

| ensembl_gene_id | external_gene_name | entrezgene_id | gene_biotype | baseMean | log2FoldChange | lfcSE | stat | pvalue | padj | absLog2FC |
|---|---|---|---|---|---|---|---|---|---|---|
| ENSMUSG00000061207 | Stk19 | 54402 | protein_coding | 203.2675289 | 0.15971796 | 0.094857188 | 1.683772874 | 0.092225571 | 0.23575171 | 0.15971796 |
| ENSMUSG00000021958 | Pinx1 | 72400 | protein_coding | 93.57082517 | -0.159806598 | 0.203387719 | -0.78572393 | 0.43202924 | 0.6359266 | 0.159806598 |
| ENSMUSG00000039246 | Lyplal1 | 226791 | protein_coding | 115.1458793 | 0.159829036 | 0.151591944 | 1.054337272 | 0.29172853 | 0.5005493 | 0.159829036 |
| ENSMUSG00000028289 | Epha7 | 13841 | protein_coding | 29.82476561 | 0.159839393 | 0.281084363 | 0.56865274 | 0.569591826 | 0.7446207 | 0.159839393 |
| ENSMUSG00000050697 | Prkaa1 | 105787 | protein_coding | 1374.49769 | -0.159856305 | 0.094010606 | -1.700407128 | 0.089054372 | 0.230363759 | 0.159856305 |
| ENSMUSG00000026027 | Stradb | 227154 | protein_coding | 234.5639121 | -0.159904015 | 0.118520664 | -1.349165699 | 0.17728375 | 0.364187739 | 0.159904015 |
| ENSMUSG00000018923 | Med11 | 66172 | protein_coding | 176.4423744 | 0.159925096 | 0.115281937 | 1.387251981 | 0.16536494 | 0.348987948 | 0.159925096 |
| ENSMUSG00000026021 | Sumo1 | 22218 | protein_coding | 1062.957868 | 0.159941075 | 0.07923663 | 2.018524445 | 0.043536671 | 0.141060231 | 0.159941075 |
| ENSMUSG00000038253 | Hoxa5 | 15402 | protein_coding | 88.63781919 | -0.159984172 | 0.238086747 | -0.671957485 | 0.501610762 | 0.691924848 | 0.159984172 |
| ENSMUSG00000039182 | AW209491 | 105351 | protein_coding | 168.729794 | -0.160022143 | 0.122203883 | -1.309468553 | 0.190375683 | 0.382287428 | 0.160022143 |
| ENSMUSG00000023707 | Ogfod2 | 66627 | protein_coding | 293.2502703 | 0.160039692 | 0.079854743 | 2.004135076 | 0.045055593 | 0.144715766 | 0.160039692 |
| ENSMUSG00000028127 | Abcd3 | 19299 | protein_coding | 4488.609441 | 0.160053768 | 0.122793998 | 1.30343316 | 0.192426922 | 0.385434609 | 0.160053768 |
| ENSMUSG00000003464 | Pex19 | 19298 | protein_coding | 927.8152924 | -0.16019195 | 0.101466977 | -1.578759463 | 0.11439124 | 0.271013487 | 0.16019195 |
| ENSMUSG00000032002 | Dcun1d5 | 76863 | protein_coding | 635.8339401 | 0.160193796 | 0.121948379 | 1.313619725 | 0.188974207 | 0.380319097 | 0.160193796 |
| ENSMUSG00000031708 | Tecr | 106529 | protein_coding | 998.2794374 | 0.160227941 | 0.074331319 | 2.155591258 | 0.031115596 | 0.111023568 | 0.160227941 |
| ENSMUSG00000025971 | Maip1 | 68115 | protein_coding | 212.1299769 | 0.160238587 | 0.094476196 | 1.696073658 | 0.089871932 | 0.231845261 | 0.160238587 |
| ENSMUSG00000033819 | Ppp1r16a | 73062 | protein_coding | 575.1615086 | 0.160238921 | 0.116882382 | 1.370941608 | 0.170393159 | 0.355657481 | 0.160238921 |
| ENSMUSG00000056999 | Ide | 15925 | protein_coding | 952.8616094 | 0.160309285 | 0.073592685 | 2.178331785 | 0.029381344 | 0.10623974 | 0.160309285 |
| ENSMUSG00000028307 | Aldob | 230163 | protein_coding | 2753.385981 | 0.160458649 | 0.084811916 | 1.891935187 | 0.058499617 | 0.172968609 | 0.160458649 |
| ENSMUSG00000020376 | Rnf130 | 59044 | protein_coding | 1014.253928 | -0.160479235 | 0.106538234 | -1.506306512 | 0.131988507 | 0.299859213 | 0.160479235 |
| ENSMUSG00000058454 | Dhcr7 | 13360 | protein_coding | 772.8239368 | 0.160487991 | 0.076583549 | 2.095593533 | 0.036118263 | 0.123851819 | 0.160487991 |
| ENSMUSG00000020514 | Mrpl22 | 216767 | protein_coding | 210.8619293 | 0.16049748 | 0.094418741 | 1.699847698 | 0.089159577 | 0.230503772 | 0.16049748 |
| ENSMUSG00000031438 | Rnf128 | 66889 | protein_coding | 3520.57322 | 0.160530879 | 0.108450572 | 1.4802216 | 0.138814118 | 0.310146061 | 0.160530879 |
| ENSMUSG00000032839 | Trpc1 | 22063 | protein_coding | 19.98367131 | -0.160542843 | 0.272440861 | -0.589275935 | 0.555676185 | 0.735117133 | 0.160542843 |
| ENSMUSG00000033364 | Usp37 | 319651 | protein_coding | 654.9329717 | -0.160546412 | 0.079825334 | -2.011221284 | 0.044302089 | 0.142756153 | 0.160546412 |
| ENSMUSG00000113836 | Gm3325 | 100041420 | transcribed_processed_pseudogene | 105.090329 | 0.160562523 | 0.144544809 | 1.110814871 | 0.266648044 | 0.474459067 | 0.160562523 |
| ENSMUSG00000031148 | Gpkow | 209416 | protein_coding | 442.2300223 | -0.160591499 | 0.105960955 | -1.515572392 | 0.129627512 | 0.296227693 | 0.160591499 |
| ENSMUSG00000022560 | Slc52a2 | 52710 | protein_coding | 234.1198079 | -0.160596379 | 0.143527671 | -1.118922773 | 0.263173087 | 0.470933719 | 0.160596379 |
| ENSMUSG00000020116 | Pno1 | 66249 | protein_coding | 467.4016873 | 0.160610006 | 0.131967071 | 1.217046075 | 0.223586686 | 0.425177738 | 0.160610006 |
| ENSMUSG00000038256 | Bcl9 | 77578 | protein_coding | 138.7982855 | -0.160626493 | 0.19200741 | -0.836564033 | 0.40283767 | 0.61003978 | 0.160626493 |
| ENSMUSG00000073733 | Cplane2 | 76166 | protein_coding | 8.580931179 | -0.160629107 | 0.497110504 | -0.323125553 | 0.746600159 | 0.859175444 | 0.160629107 |
| ENSMUSG00000006134 | Crkl | 12929 | protein_coding | 711.3194038 | -0.160633325 | 0.071303298 | -2.252817609 | 0.024270651 | 0.092377156 | 0.160633325 |
| ENSMUSG00000030342 | Cd9 | 12527 | protein_coding | 3235.217586 | -0.160667568 | 0.104755493 | -1.533738829 | 0.125093917 | 0.289178051 | 0.160667568 |
| ENSMUSG00000032806 | Slc10a3 | 214601 | protein_coding | 242.7022424 | 0.160751211 | 0.10457122 | 1.537241423 | 0.1242342 | 0.287742088 | 0.160751211 |
| ENSMUSG00000002658 | Gtf2f1 | 98053 | protein_coding | 673.8919501 | 0.160821611 | 0.108234881 | 1.485857518 | 0.137316812 | 0.308157609 | 0.160821611 |
| ENSMUSG00000040852 | Plekhh2 | 213556 | protein_coding | 56.95832927 | -0.160841098 | 0.269364053 | -0.597114189 | 0.550431145 | 0.731458975 | 0.160841098 |
| ENSMUSG00000046139 | Patl1 | 225929 | protein_coding | 188.7009524 | -0.160910174 | 0.15919635 | -1.010765473 | 0.312128692 | 0.522338315 | 0.160910174 |
| ENSMUSG00000027076 | Timm10 | 30059 | protein_coding | 190.0598978 | -0.160927308 | 0.128432124 | -1.253014457 | 0.210200446 | 0.40740724 | 0.160927308 |

| ensembl_gene_id | external_gene_name | entrezgene_id | gene_biotype | baseMean | log2FoldChange | lfcSE | stat | pvalue | padj | absLog2FC |
|---|---|---|---|---|---|---|---|---|---|---|
| ENSMUSG00000032688 | Malt1 | 240354 | protein_coding | 539.2489085 | 0.160941293 | 0.077877067 | 2.066607001 | 0.038771201 | 0.130292153 | 0.160941293 |
| ENSMUSG00000019988 | Nedd1 | 17997 | protein_coding | 400.6895577 | 0.16094792 | 0.081874377 | 1.965791076 | 0.049322749 | 0.154041715 | 0.16094792 |
| ENSMUSG00000025159 | Mms19 | 72199 | protein_coding | 372.9096901 | -0.161193593 | 0.093572343 | -1.722662783 | 0.084949524 | 0.223059021 | 0.161193593 |
| ENSMUSG00000001983 | Taco1 | 70207 | protein_coding | 177.486207 | -0.161197114 | 0.119881846 | -1.344633233 | 0.178743706 | 0.366177459 | 0.161197114 |
| ENSMUSG00000020477 | Mrps24 | 64660 | protein_coding | 534.8056567 | 0.161203018 | 0.125502848 | 1.284457048 | 0.198982083 | 0.393175365 | 0.161203018 |
| ENSMUSG00000021279 | Cdc42bpb | 217866 | protein_coding | 1406.159867 | -0.161250597 | 0.096021664 | -1.679314756 | 0.093090717 | 0.237277794 | 0.161250597 |
| ENSMUSG00000020462 | Cfap36 | 216618 | protein_coding | 220.4235429 | 0.161303692 | 0.107450948 | 1.50118445 | 0.133307861 | 0.301829505 | 0.161303692 |
| ENSMUSG00000025781 | Atp5c1 | 11949 | protein_coding | 6845.558391 | 0.161337421 | 0.098671575 | 1.635095228 | 0.102029086 | 0.251605402 | 0.161337421 |
| ENSMUSG00000035392 | Dennd1a | 227801 | protein_coding | 166.1934711 | -0.161348754 | 0.140675757 | -1.146954943 | 0.251400243 | 0.457843937 | 0.161348754 |
| ENSMUSG00000074115 | Saa1 | 20208 | protein_coding | 1480.553323 | 0.161382124 | 0.213514466 | 0.755836956 | 0.449746956 | 0.65028772 | 0.161382124 |
| ENSMUSG00000024829 | Mrpl21 | 353242 | protein_coding | 577.5022811 | 0.161412778 | 0.093735945 | 1.721994466 | 0.085070521 | 0.223303108 | 0.161412778 |
| ENSMUSG00000028969 | Cdk5 | 12568 | protein_coding | 307.6308982 | 0.161511483 | 0.087680575 | 1.842044057 | 0.065468706 | 0.186411431 | 0.161511483 |
| ENSMUSG00000035885 | Cox8a | 12868 | protein_coding | 5876.891224 | 0.161520612 | 0.1090068 | 1.481748039 | 0.13840735 | 0.309584646 | 0.161520612 |
| ENSMUSG00000006777 | Krt23 | 94179 | protein_coding | 123.6141541 | -0.161534256 | 0.15686425 | -1.029771002 | 0.303117516 | 0.512586877 | 0.161534256 |
| ENSMUSG00000024734 | Zp1 | 22786 | protein_coding | 7.557587693 | -0.161618265 | 0.41047382 | -0.393735867 | 0.693776058 | 0.825894249 | 0.161618265 |
| ENSMUSG00000070574 | 2310016G11Rik | 69578 | lncRNA | 52.07549527 | -0.161700473 | 0.192776363 | -0.838798236 | 0.401582541 | 0.60912358 | 0.161700473 |
| ENSMUSG00000028755 | Cda | 72269 | protein_coding | 187.2276671 | -0.161711412 | 0.144242823 | -1.121105428 | 0.262242988 | 0.470034424 | 0.161711412 |
| ENSMUSG00000021945 | Zmym2 | 76007 | protein_coding | 1292.794459 | -0.161738528 | 0.081584747 | -1.98246038 | 0.047427738 | 0.150154841 | 0.161738528 |
| ENSMUSG00000047514 | Tspyl1 | 22110 | protein_coding | 658.5609757 | 0.161785506 | 0.06356377 | 2.545247191 | 0.01092004 | 0.051142136 | 0.161785506 |
| ENSMUSG00000021916 | Glt8d1 | 76485 | protein_coding | 132.2726587 | 0.161786942 | 0.11342777 | 1.426343311 | 0.153769268 | 0.332888202 | 0.161786942 |
| ENSMUSG00000032489 | Kif9 | 16578 | protein_coding | 27.57944632 | 0.161794107 | 0.261981216 | 0.617579035 | 0.536852868 | 0.721237557 | 0.161794107 |
| ENSMUSG00000022774 | Ncbp2 | 68092 | protein_coding | 364.0241119 | 0.1618508 | 0.114986792 | 1.407559929 | 0.15926142 | 0.340451304 | 0.1618508 |
| ENSMUSG00000028893 | Sesn2 | 230784 | protein_coding | 245.5201809 | -0.161893059 | 0.124847361 | -1.296727926 | 0.194724817 | 0.387829791 | 0.161893059 |
| ENSMUSG00000046240 | Hepacam | 72927 | protein_coding | 6.979225855 | -0.161923133 | 0.627234927 | -0.258153885 | 0.796288146 | 0.890338594 | 0.161923133 |
| ENSMUSG00000031833 | Mast3 | 546071 | protein_coding | 150.1914682 | -0.161933199 | 0.116663227 | -1.388039768 | 0.165124936 | 0.348728451 | 0.161933199 |
| ENSMUSG00000033530 | Ttc7b | 104718 | protein_coding | 51.98268924 | 0.161949727 | 0.16251402 | 0.996527726 | 0.318993803 | 0.530313463 | 0.161949727 |
| ENSMUSG00000057363 | Uxs1 | 67883 | protein_coding | 268.2879915 | 0.161988584 | 0.083502589 | 1.939922898 | 0.052389061 | 0.160348378 | 0.161988584 |
| ENSMUSG00000102047 | 2010010A06Rik | 100503117 | transcribed_processed_pseudogene | 14.22528877 | -0.162004331 | 0.320458639 | -0.505538973 | 0.613180332 | 0.774707411 | 0.162004331 |
| ENSMUSG00000020689 | Itgb3 | 16416 | protein_coding | 6.711659503 | 0.162009311 | 0.949550508 | 0.170616844 | 0.864525052 | 0.929652738 | 0.162009311 |
| ENSMUSG00000021996 | Esd | 13885 | protein_coding | 2717.441395 | 0.162072401 | 0.100118609 | 1.618803963 | 0.105489449 | 0.257231468 | 0.162072401 |
| ENSMUSG00000033987 | Dnah17 | 69926 | protein_coding | 29.75915528 | 0.162126946 | 0.346727559 | 0.467591749 | 0.640076566 | 0.791371033 | 0.162126946 |
| ENSMUSG00000041733 | Coq5 | 52064 | protein_coding | 602.7544721 | -0.162137849 | 0.09696944 | -1.672050992 | 0.094514267 | 0.239458186 | 0.162137849 |
| ENSMUSG00000025647 | Shisa5 | 66940 | protein_coding | 1656.798359 | -0.162139631 | 0.096901727 | -1.673237789 | 0.094280494 | 0.239110819 | 0.162139631 |
| ENSMUSG00000029135 | Fosl2 | 14284 | protein_coding | 2658.03928 | 0.162144482 | 0.097799261 | 1.657931566 | 0.09733128 | 0.243900172 | 0.162144482 |
| ENSMUSG00000035958 | Tdp2 | 56196 | protein_coding | 447.0234503 | -0.162148573 | 0.096677798 | -1.677205908 | 0.093502224 | 0.237985125 | 0.162148573 |
| ENSMUSG00000073607 | Gm10548 | 633395 | transcribed_processed_pseudogene | 11.01171156 | -0.16218293 | 0.451530627 | -0.359184782 | 0.71945686 | 0.842364374 | 0.16218293 |
| ENSMUSG00000047084 | Ngrn | 83485 | protein_coding | 284.5450413 | 0.162226646 | 0.119256407 | 1.360318079 | 0.17372929 | 0.359707067 | 0.162226646 |
| ENSMUSG00000027131 | Emc4 | 68032 | protein_coding | 362.471045 | 0.162259237 | 0.096191868 | 1.686829045 | 0.091636229 | 0.234584033 | 0.162259237 |

| ensembl_gene_id | external_gene_name | entrezgene_id | gene_biotype | baseMean | log2FoldChange | lfcSE | stat | pvalue | padj | absLog2FC |
|---|---|---|---|---|---|---|---|---|---|---|
| ENSMUSG00000026070 | Il18r1 | 16182 | protein_coding | 17.21437047 | 0.162266695 | 0.433302018 | 0.374488667 | 0.708040787 | 0.835135786 | 0.162266695 |
| ENSMUSG00000030095 | Tmem43 | 74122 | protein_coding | 549.7599315 | -0.162279427 | 0.177776155 | -0.912830112 | 0.361331899 | 0.570811871 | 0.162279427 |
| ENSMUSG00000007041 | Clic1 | 114584 | protein_coding | 2706.564531 | 0.162295276 | 0.121758237 | 1.332930571 | 0.182554589 | 0.371642318 | 0.162295276 |
| ENSMUSG00000036940 | Kdm1a | 99982 | protein_coding | 1044.451456 | -0.162341469 | 0.082898724 | -1.958310821 | 0.050193552 | 0.155875003 | 0.162341469 |
| ENSMUSG00000028779 | Pef1 | 67898 | protein_coding | 786.1743359 | 0.162464783 | 0.098094188 | 1.656212114 | 0.097678871 | 0.244435044 | 0.162464783 |
| ENSMUSG00000073155 | 1810058I24Rik | 67705 | protein_coding | 160.6344887 | -0.162475097 | 0.107838136 | -1.506657131 | 0.131898565 | 0.299697624 | 0.162475097 |
| ENSMUSG00000055670 | Zzef1 | 195018 | protein_coding | 2832.072985 | 0.162478768 | 0.116855658 | 1.390422776 | 0.164400534 | 0.347730593 | 0.162478768 |
| ENSMUSG00000021120 | Pigh | 110417 | protein_coding | 198.9592203 | 0.162560071 | 0.127805071 | 1.271937567 | 0.203395303 | 0.398435664 | 0.162560071 |
| ENSMUSG00000092335 | Zfp977 | 637776 | protein_coding | 10.09100225 | 0.162595973 | 0.363516529 | 0.447286327 | 0.654668338 | 0.800150191 | 0.162595973 |
| ENSMUSG00000054836 | Elp6 | 72341 | protein_coding | 82.92418523 | 0.162605159 | 0.147140643 | 1.105100234 | 0.269116159 | 0.477177133 | 0.162605159 |
| ENSMUSG00000028599 | Tnfrsf1b | 21938 | protein_coding | 332.4116543 | -0.162639512 | 0.319612674 | -0.508864402 | 0.610847274 | 0.77314515 | 0.162639512 |
| ENSMUSG00000022338 | Eny2 | 223527 | protein_coding | 834.6218503 | 0.162649287 | 0.081345293 | 1.999492318 | 0.045555112 | 0.14590827 | 0.162649287 |
| ENSMUSG00000035498 | Cdcp1 | 109332 | protein_coding | 1872.616114 | 0.162697547 | 0.108399615 | 1.500905212 | 0.13338008 | 0.301865663 | 0.162697547 |
| ENSMUSG00000055717 | Slain1 | 105439 | protein_coding | 82.75359361 | -0.162728646 | 0.203383293 | -0.800108229 | 0.423648094 | 0.628365637 | 0.162728646 |
| ENSMUSG00000030979 | Uros | 22276 | protein_coding | 251.9170055 | 0.162731451 | 0.094034666 | 1.730547445 | 0.083532512 | 0.220642543 | 0.162731451 |
| ENSMUSG00000020954 | Strn3 | 94186 | protein_coding | 1028.209312 | 0.162786153 | 0.07095448 | 2.294233633 | 0.021777082 | 0.085048665 | 0.162786153 |
| ENSMUSG00000048497 | Mmgt2 | 216829 | protein_coding | 254.6602906 | 0.162875923 | 0.087934932 | 1.85223231 | 0.063992471 | 0.183863228 | 0.162875923 |
| ENSMUSG00000028847 | Trappc3 | 27096 | protein_coding | 537.4098728 | 0.162898757 | 0.085136348 | 1.913386701 | 0.055698566 | 0.167169164 | 0.162898757 |
| ENSMUSG00000040146 | Rgl3 | 71746 | protein_coding | 61.21746324 | 0.162955644 | 0.164202016 | 0.992409518 | 0.320997798 | 0.532256186 | 0.162955644 |
| ENSMUSG00000055629 | B4galnt4 | 330671 | protein_coding | 69.67338901 | 0.163048748 | 0.210959348 | 0.772891789 | 0.439586425 | 0.642022673 | 0.163048748 |
| ENSMUSG00000100153 | Ppp1ccb | 434233 | transcribed_processed_pseudogene | 12.3216217 | 0.163049596 | 0.36648119 | 0.444905769 | 0.656387853 | 0.801149235 | 0.163049596 |
| ENSMUSG00000026270 | Capn10 | 23830 | protein_coding | 301.8816038 | 0.163088313 | 0.103357644 | 1.577902769 | 0.114587949 | 0.271358437 | 0.163088313 |
| ENSMUSG00000024386 | Proc | 19123 | protein_coding | 50.11450471 | 0.163107831 | 0.192013624 | 0.849459677 | 0.395625559 | 0.603875164 | 0.163107831 |
| ENSMUSG00000026771 | Spopl | 76857 | protein_coding | 386.221697 | -0.163182584 | 0.101918327 | -1.601111295 | 0.109352271 | 0.263464373 | 0.163182584 |
| ENSMUSG00000016239 | Lonrf3 | 74365 | protein_coding | 128.8259851 | -0.163212808 | 0.173714394 | -0.939546829 | 0.347450061 | 0.557627687 | 0.163212808 |
| ENSMUSG00000030613 | Ccdc90b | 66365 | protein_coding | 293.6654893 | 0.163243087 | 0.129628688 | 1.259312955 | 0.207917316 | 0.404310463 | 0.163243087 |
| ENSMUSG00000005683 | Cs | 12974 | protein_coding | 7431.023489 | 0.163264417 | 0.078758123 | 2.07298512 | 0.038173664 | 0.12887455 | 0.163264417 |
| ENSMUSG00000015656 | Hspa8 | 15481 | protein_coding | 28467.66348 | 0.163272915 | 0.20867791 | 0.782415903 | 0.433970191 | 0.637453749 | 0.163272915 |
| ENSMUSG00000031843 | Mphosph6 | 68533 | protein_coding | 238.2169371 | 0.163278424 | 0.124520365 | 1.3112588 | 0.189770343 | 0.381360335 | 0.163278424 |
| ENSMUSG00000047821 | Trim16 | 94092 | protein_coding | 1659.624167 | 0.163355275 | 0.145568459 | 1.122188669 | 0.261782229 | 0.469610018 | 0.163355275 |
| ENSMUSG00000053870 | Fpgt | 75540 | protein_coding | 981.2489153 | -0.163398487 | 0.097688361 | -1.672650508 | 0.094396118 | 0.239303097 | 0.163398487 |
| ENSMUSG00000047617 | Paxx | 227622 | protein_coding | 113.7451062 | -0.163409489 | 0.145993247 | -1.119294844 | 0.263014375 | 0.470865793 | 0.163409489 |
| ENSMUSG00000001739 | Cldn15 | 60363 | protein_coding | 2781.033237 | 0.163411186 | 0.111543627 | 1.464997971 | 0.142921471 | 0.316613797 | 0.163411186 |
| ENSMUSG00000028329 | Xpa | 22590 | protein_coding | 146.2447459 | 0.163415195 | 0.132525388 | 1.233085953 | 0.217543698 | 0.417172648 | 0.163415195 |
| ENSMUSG00000037151 | Lrrc20 | 216011 | protein_coding | 109.9974361 | 0.16341944 | 0.144338492 | 1.132195844 | 0.257552107 | 0.464532891 | 0.16341944 |
| ENSMUSG00000026676 | Ccdc3 | 74186 | protein_coding | 26.57334206 | 0.16344758 | 0.248858612 | 0.656788925 | 0.511316646 | 0.699220189 | 0.16344758 |
| ENSMUSG00000018395 | Kif3a | 16568 | protein_coding | 110.4585221 | 0.163454676 | 0.118907171 | 1.374641032 | 0.169242761 | 0.354093686 | 0.163454676 |
| ENSMUSG00000001630 | Stk38l | 232533 | protein_coding | 1184.819674 | -0.163462114 | 0.11761545 | -1.389801369 | 0.164589202 | 0.347990237 | 0.163462114 |

| ensembl_gene_id | external_gene_name | entrezgene_id | gene_biotype | baseMean | log2FoldChange | lfcSE | stat | pvalue | padj | absLog2FC |
|---|---|---|---|---|---|---|---|---|---|---|
| ENSMUSG00000047832 | Cdca4 | 71963 | protein_coding | 347.794554 | -0.163464572 | 0.200094598 | -0.816936458 | 0.413964742 | 0.620343438 | 0.163464572 |
| ENSMUSG00000043885 | Slc36a4 | 234967 | protein_coding | 301.3587521 | 0.163484796 | 0.131873866 | 1.239705797 | 0.215084232 | 0.413901371 | 0.163484796 |
| ENSMUSG00000032440 | Tgfbr2 | 21813 | protein_coding | 1176.444207 | -0.163504874 | 0.205044866 | -0.797410232 | 0.425212819 | 0.629734057 | 0.163504874 |
| ENSMUSG00000018651 | Tada2a | 217031 | protein_coding | 140.8424559 | -0.163506384 | 0.137955114 | -1.18521437 | 0.2359327 | 0.440061526 | 0.163506384 |
| ENSMUSG00000036555 | Iqce | 74239 | protein_coding | 138.3756378 | -0.163506888 | 0.154622131 | -1.057461092 | 0.290301203 | 0.498965961 | 0.163506888 |
| ENSMUSG00000097814 | Panct2 | 329217 | lncRNA | 32.76306312 | -0.16354078 | 0.289146325 | -0.565598683 | 0.571666624 | 0.745680614 | 0.16354078 |
| ENSMUSG00000071669 | Snx29 | 74478 | protein_coding | 86.42780661 | -0.163755901 | 0.208238385 | -0.786386723 | 0.431640959 | 0.635531262 | 0.163755901 |
| ENSMUSG00000030868 | Dctn5 | 59288 | protein_coding | 572.8273948 | 0.163768954 | 0.071060789 | 2.304631807 | 0.021187203 | 0.083408248 | 0.163768954 |
| ENSMUSG00000019822 | Smpd2 | 20598 | protein_coding | 252.0392713 | -0.16381034 | 0.094319612 | -1.736758 | 0.082429895 | 0.218593229 | 0.16381034 |
| ENSMUSG00000026123 | Plekhb2 | 226971 | protein_coding | 4636.391189 | 0.163822728 | 0.091416518 | 1.792047345 | 0.073125382 | 0.201199014 | 0.163822728 |
| ENSMUSG00000028066 | Pmf1 | 67037 | protein_coding | 220.8463832 | 0.163837204 | 0.114690776 | 1.428512471 | 0.153144408 | 0.332056102 | 0.163837204 |
| ENSMUSG00000072066 | 6720489N17Rik | 211378 | protein_coding | 43.59332049 | 0.163852505 | 0.222751617 | 0.735583908 | 0.461983958 | 0.659894224 | 0.163852505 |
| ENSMUSG00000049321 | Zfp2 | 22678 | protein_coding | 73.81448482 | -0.163894372 | 0.160501272 | -1.021140641 | 0.307187811 | 0.517163878 | 0.163894372 |
| ENSMUSG00000048647 | Exd1 | 241624 | protein_coding | 35.82830594 | 0.163904966 | 0.217055818 | 0.755128185 | 0.450172073 | 0.650666133 | 0.163904966 |
| ENSMUSG00000037152 | Ndufc1 | 66377 | protein_coding | 816.9341332 | 0.163910104 | 0.113747596 | 1.440998398 | 0.149585135 | 0.327026097 | 0.163910104 |
| ENSMUSG00000025404 | R3hdm2 | 71750 | protein_coding | 328.775337 | -0.164120133 | 0.17495135 | -0.938090123 | 0.348198097 | 0.55835088 | 0.164120133 |
| ENSMUSG00000029020 | Mfn2 | 170731 | protein_coding | 1381.354685 | 0.164208709 | 0.060557454 | 2.711618418 | 0.006695562 | 0.035152464 | 0.164208709 |
| ENSMUSG00000054400 | Cklf | 75458 | protein_coding | 90.45019714 | 0.164272515 | 0.130595676 | 1.257871011 | 0.208438409 | 0.405142395 | 0.164272515 |
| ENSMUSG00000097254 | C430042M11Rik | 320021 | lncRNA | 43.72735988 | 0.164275836 | 0.217310293 | 0.755950556 | 0.449678841 | 0.65024826 | 0.164275836 |
| ENSMUSG00000067873 | Htatsf1 | 72459 | protein_coding | 611.9912593 | 0.164313714 | 0.079542407 | 2.065737272 | 0.038853294 | 0.130465137 | 0.164313714 |
| ENSMUSG00000026032 | Ndufb3 | 66495 | protein_coding | 706.1892404 | 0.16432642 | 0.102392959 | 1.604860546 | 0.108524498 | 0.262144101 | 0.16432642 |
| ENSMUSG00000092074 | Dynlt1a | 100310872 | protein_coding | 43.50180035 | 0.164408305 | 0.174583306 | 0.941718361 | 0.346336852 | 0.556430642 | 0.164408305 |
| ENSMUSG00000028293 | Slc35a1 | 24060 | protein_coding | 3259.483666 | 0.164422399 | 0.06305107 | 2.607765426 | 0.009113538 | 0.044637497 | 0.164422399 |
| ENSMUSG00000027589 | Pcmtd2 | 245867 | protein_coding | 790.9634562 | 0.164582924 | 0.148368162 | 1.109287342 | 0.267306244 | 0.475184581 | 0.164582924 |
| ENSMUSG00000032883 | Acsl3 | 74205 | protein_coding | 3292.042322 | -0.164620536 | 0.110343835 | -1.491887031 | 0.135728763 | 0.305532863 | 0.164620536 |
| ENSMUSG00000028064 | Sema4a | 20351 | protein_coding | 1751.672477 | 0.164635435 | 0.120438377 | 1.36696823 | 0.171635264 | 0.357153419 | 0.164635435 |
| ENSMUSG00000020863 | Luc7l3 | 67684 | protein_coding | 1093.781603 | -0.164639644 | 0.099492723 | -1.654790818 | 0.097966937 | 0.244886594 | 0.164639644 |
| ENSMUSG00000069206 | Zfp874a | 238692 | protein_coding | 268.5570224 | 0.164643822 | 0.176329785 | 0.933726667 | 0.350444905 | 0.560542926 | 0.164643822 |
| ENSMUSG00000039428 | Tmem135 | 72759 | protein_coding | 772.6192695 | 0.164695127 | 0.072033418 | 2.286371127 | 0.022232556 | 0.086455115 | 0.164695127 |
| ENSMUSG00000032898 | Fbxo21 | 231670 | protein_coding | 322.0792225 | -0.16470722 | 0.108723581 | -1.514917176 | 0.12979338 | 0.296521652 | 0.16470722 |
| ENSMUSG00000078877 | Gm14295 | 628147 | protein_coding | 82.42663005 | 0.164776636 | 0.170379988 | 0.967112617 | 0.333487739 | 0.544296824 | 0.164776636 |
| ENSMUSG00000052146 | Rps10 | 67097 | protein_coding | 118.8407703 | -0.164816749 | 0.139771451 | -1.179187506 | 0.23832352 | 0.442477798 | 0.164816749 |
| ENSMUSG00000020782 | Llgl2 | 217325 | protein_coding | 2223.628055 | -0.164821015 | 0.113241891 | -1.455477412 | 0.145537139 | 0.321157599 | 0.164821015 |
| ENSMUSG00000041308 | Sntb2 | 20650 | protein_coding | 224.6120347 | -0.164865747 | 0.134762649 | -1.223378638 | 0.221186718 | 0.422024682 | 0.164865747 |
| ENSMUSG00000071506 | Tmem139 | 109218 | protein_coding | 440.8995198 | -0.164910309 | 0.141331368 | -1.166834449 | 0.243277232 | 0.448330412 | 0.164910309 |
| ENSMUSG00000034853 | Acot11 | 329910 | protein_coding | 691.2467945 | -0.164932153 | 0.128394822 | -1.284570121 | 0.198942545 | 0.393175365 | 0.164932153 |
| ENSMUSG00000025466 | Fuom | 69064 | protein_coding | 345.6076353 | -0.164975997 | 0.104630119 | -1.576754364 | 0.114852056 | 0.271903025 | 0.164975997 |
| ENSMUSG00000009073 | Nf2 | 18016 | protein_coding | 1474.58811 | -0.165034138 | 0.088250116 | -1.870072769 | 0.061473713 | 0.178514732 | 0.165034138 |

| ensembl_gene_id | external_gene_name | entrezgene_id | gene_biotype | baseMean | log2FoldChange | lfcSE | stat | pvalue | padj | absLog2FC |
|---|---|---|---|---|---|---|---|---|---|---|
| ENSMUSG00000031891 | Hsd11b2 | 15484 | protein_coding | 2310.168207 | 0.165046839 | 0.138674468 | 1.190174676 | 0.233977744 | 0.437777224 | 0.165046839 |
| ENSMUSG00000002985 | Apoe | 11816 | protein_coding | 47.17052051 | 0.165074258 | 0.246834347 | 0.668765347 | 0.503645175 | 0.693709819 | 0.165074258 |
| ENSMUSG00000023852 | Chd1 | 12648 | protein_coding | 1302.937594 | -0.165207889 | 0.100489407 | -1.644032881 | 0.100169411 | 0.248398364 | 0.165207889 |
| ENSMUSG00000070002 | Ell | 13716 | protein_coding | 295.8538063 | -0.165280848 | 0.100797783 | -1.639727023 | 0.101061938 | 0.250010507 | 0.165280848 |
| ENSMUSG00000039834 | Zfp335 | 329559 | protein_coding | 291.4501358 | -0.165317167 | 0.152169901 | -1.086398598 | 0.277302682 | 0.485637945 | 0.165317167 |
| ENSMUSG00000031007 | Atp6ap2 | 70495 | protein_coding | 2837.16984 | 0.165331484 | 0.054612987 | 3.02732907 | 0.002467252 | 0.016143473 | 0.165331484 |
| ENSMUSG00000038607 | Gng10 | 14700 | protein_coding | 10.57603292 | 0.165365094 | 0.363339399 | 0.455125688 | 0.649018818 | 0.797283028 | 0.165365094 |
| ENSMUSG00000021646 | Mccc2 | 78038 | protein_coding | 427.1848803 | 0.165367768 | 0.093117072 | 1.775912458 | 0.075747347 | 0.205338463 | 0.165367768 |
| ENSMUSG00000035572 | Dcaf10 | 242418 | protein_coding | 549.7354682 | 0.165376883 | 0.068057088 | 2.429972939 | 0.01509995 | 0.064985681 | 0.165376883 |
| ENSMUSG00000002771 | Grin2d | 14814 | protein_coding | 29.07107527 | -0.165394435 | 0.209483216 | -0.789535494 | 0.429799093 | 0.633815383 | 0.165394435 |
| ENSMUSG00000053436 | Mapk14 | 26416 | protein_coding | 820.140646 | -0.165422923 | 0.092159949 | -1.794954587 | 0.072660936 | 0.200493734 | 0.165422923 |
| ENSMUSG00000031896 | Ctrl | 109660 | protein_coding | 10.16444288 | 0.165430333 | 0.399665657 | 0.413921813 | 0.678931368 | 0.816275576 | 0.165430333 |
| ENSMUSG00000034709 | Ppp1r21 | 73825 | protein_coding | 375.35015 | 0.165447558 | 0.084216014 | 1.964561732 | 0.049464985 | 0.154314062 | 0.165447558 |
| ENSMUSG00000032606 | Nicn1 | 66257 | protein_coding | 114.5520122 | -0.165455218 | 0.177435906 | -0.932478784 | 0.351089146 | 0.561010024 | 0.165455218 |
| ENSMUSG00000035580 | Kcnh8 | 211468 | protein_coding | 36.58705595 | -0.165471967 | 0.22214246 | -0.744891216 | 0.456337486 | 0.655296022 | 0.165471967 |
| ENSMUSG00000029238 | Clock | 12753 | protein_coding | 2269.247058 | -0.165488419 | 0.092952631 | -1.780352186 | 0.075018341 | 0.204289989 | 0.165488419 |
| ENSMUSG00000042828 | Trim72 | 434246 | protein_coding | 77.21733559 | -0.165488949 | 0.27453723 | -0.602792375 | 0.54664682 | 0.728740421 | 0.165488949 |
| ENSMUSG00000004934 | Pias4 | 59004 | protein_coding | 319.4078089 | -0.165509645 | 0.085684321 | -1.931621123 | 0.053406287 | 0.16224392 | 0.165509645 |
| ENSMUSG00000046798 | Cldn12 | 64945 | protein_coding | 1448.619062 | 0.165557356 | 0.093526442 | 1.770166306 | 0.07669944 | 0.207484492 | 0.165557356 |
| ENSMUSG00000001016 | Ilf2 | 67781 | protein_coding | 774.9423788 | 0.165565628 | 0.093679017 | 1.767371531 | 0.077166028 | 0.208463448 | 0.165565628 |
| ENSMUSG00000005087 | Cd44 | 12505 | protein_coding | 933.9068613 | -0.165590922 | 0.236250764 | -0.700911687 | 0.483358131 | 0.676828862 | 0.165590922 |
| ENSMUSG00000063888 | Rpl7l1 | 66229 | protein_coding | 1019.20622 | 0.165597992 | 0.129981624 | 1.274010792 | 0.202659591 | 0.397630201 | 0.165597992 |
| ENSMUSG00000043207 | Zmpste24 | 230709 | protein_coding | 839.226731 | 0.165602989 | 0.074139891 | 2.233655693 | 0.025505736 | 0.09599607 | 0.165602989 |
| ENSMUSG00000026594 | Ralgps2 | 78255 | protein_coding | 997.2374724 | 0.165603949 | 0.069419897 | 2.385540126 | 0.017054062 | 0.071165076 | 0.165603949 |
| ENSMUSG00000027433 | Xrn2 | 24128 | protein_coding | 1441.823704 | -0.165639598 | 0.045733564 | -3.62183882 | 0.000292516 | 0.002779953 | 0.165639598 |
| ENSMUSG00000037720 | Tmem33 | 67878 | protein_coding | 1127.101007 | 0.165686142 | 0.083738549 | 1.978612518 | 0.047859649 | 0.151211258 | 0.165686142 |
| ENSMUSG00000026271 | Gpr35 | 64095 | protein_coding | 370.0745707 | 0.16576544 | 0.11741345 | 1.41180963 | 0.158006022 | 0.338952178 | 0.16576544 |
| ENSMUSG00000031337 | Mtm1 | 17772 | protein_coding | 1131.917898 | 0.16590213 | 0.099458205 | 1.668058756 | 0.095304063 | 0.240775631 | 0.16590213 |
| ENSMUSG00000028465 | Tln1 | 21894 | protein_coding | 1790.448368 | 0.165916249 | 0.098448852 | 1.685304054 | 0.091929924 | 0.235146913 | 0.165916249 |
| ENSMUSG00000028136 | Snx27 | 76742 | protein_coding | 615.1116716 | -0.165972087 | 0.074601898 | -2.224770299 | 0.026096662 | 0.09755964 | 0.165972087 |
| ENSMUSG00000038332 | Sesn1 | 140742 | protein_coding | 202.1004309 | -0.165974233 | 0.160929116 | -1.031349931 | 0.302376751 | 0.51166014 | 0.165974233 |
| ENSMUSG00000010080 | Epn3 | 71889 | protein_coding | 530.1983318 | -0.165989317 | 0.115441839 | -1.43786099 | 0.1504735 | 0.32827584 | 0.165989317 |
| ENSMUSG00000104063 | Pcdhgb7 | 93704 | protein_coding | 11.56180101 | 0.16602689 | 0.462781586 | 0.358758634 | 0.719775661 | 0.842383576 | 0.16602689 |
| ENSMUSG00000037400 | Atp11b | 76295 | protein_coding | 2971.675332 | 0.166073556 | 0.085903816 | 1.933250055 | 0.053205401 | 0.161983995 | 0.166073556 |
| ENSMUSG00000033943 | Mga | 29808 | protein_coding | 1136.858176 | -0.166090002 | 0.09724726 | -1.707914473 | 0.087652213 | 0.22801314 | 0.166090002 |
| ENSMUSG00000021338 | Carmil1 | 68732 | protein_coding | 612.6156565 | -0.166138464 | 0.111778906 | -1.486313199 | 0.137196297 | 0.308087216 | 0.166138464 |
| ENSMUSG00000026457 | Adipor1 | 72674 | protein_coding | 2962.883408 | 0.166222552 | 0.08432967 | 1.971104017 | 0.048711979 | 0.152928696 | 0.166222552 |
| ENSMUSG00000060923 | Acyp2 | 75572 | protein_coding | 48.4424107 | 0.166243818 | 0.174739159 | 0.951382732 | 0.341410123 | 0.551911138 | 0.166243818 |

Supplementary Table 1

| ensembl_gene_id | external_gene_name | entrezgene_id | gene_biotype | baseMean | log2FoldChange | lfcSE | stat | pvalue | padj | absLog2FC |
|---|---|---|---|---|---|---|---|---|---|---|
| ENSMUSG00000028743 | Akr7a5 | 110198 | protein_coding | 580.5284208 | -0.166291953 | 0.124577952 | -1.334842569 | 0.181927878 | 0.370603304 | 0.166291953 |
| ENSMUSG00000019837 | Gtf3c6 | 67371 | protein_coding | 504.6013613 | 0.16633545 | 0.099373311 | 1.6738443 | 0.094161203 | 0.238935741 | 0.16633545 |
| ENSMUSG00000031161 | Hdac6 | 15185 | protein_coding | 240.6261643 | 0.16635035 | 0.126813423 | 1.311772416 | 0.189596935 | 0.381182159 | 0.16635035 |
| ENSMUSG00000026425 | Srgap2 | 14270 | protein_coding | 176.0978985 | 0.166466731 | 0.129335976 | 1.287087601 | 0.198063757 | 0.392156622 | 0.166466731 |
| ENSMUSG00000046079 | Lrrc8d | 231549 | protein_coding | 358.3621311 | -0.166539088 | 0.089914029 | -1.852203593 | 0.063996593 | 0.183863228 | 0.166539088 |
| ENSMUSG00000021189 | Atxn3 | 110616 | protein_coding | 783.5752539 | -0.166621403 | 0.057135981 | -2.916225472 | 0.003542944 | 0.021432589 | 0.166621403 |
| ENSMUSG00000069835 | Sat2 | 69215 | protein_coding | 17.61601604 | -0.166665922 | 0.316151723 | -0.527170691 | 0.59807506 | 0.765321556 | 0.166665922 |
| ENSMUSG00000031513 | Leprotl1 | 68192 | protein_coding | 599.8740481 | 0.166703077 | 0.088680842 | 1.87980936 | 0.060134064 | 0.176101375 | 0.166703077 |
| ENSMUSG00000104960 | Snhg8 | 69895 | lncRNA | 180.9098674 | -0.16675394 | 0.112170867 | -1.486606492 | 0.137118772 | 0.307999972 | 0.16675394 |
| ENSMUSG00000051853 | Arf3 | 11842 | protein_coding | 1350.38439 | -0.166961265 | 0.078443323 | -2.128431839 | 0.033301296 | 0.116550876 | 0.166961265 |
| ENSMUSG00000029587 | Zfp12 | 231866 | protein_coding | 272.2643764 | -0.166974677 | 0.178956979 | -0.933043672 | 0.35079742 | 0.560712626 | 0.166974677 |
| ENSMUSG00000028926 | Cdk14 | 18647 | protein_coding | 34.81649307 | -0.167037379 | 0.320054894 | -0.52190228 | 0.601738371 | 0.767564496 | 0.167037379 |
| ENSMUSG00000031112 | Stk26 | 70415 | protein_coding | 100.4772739 | -0.167067488 | 0.27650297 | -0.60421589 | 0.545700118 | 0.728169923 | 0.167067488 |
| ENSMUSG00000020671 | Rab10 | 19325 | protein_coding | 2447.213196 | 0.167086286 | 0.056974743 | 2.932637822 | 0.003360957 | 0.020573914 | 0.167086286 |
| ENSMUSG00000026999 | Nup35 | 69482 | protein_coding | 133.6037301 | 0.167115619 | 0.111178507 | 1.503128822 | 0.132805828 | 0.301192317 | 0.167115619 |
| ENSMUSG00000097162 | 2310010J17Rik | 78329 | lncRNA | 32.58858774 | -0.167165096 | 0.19982015 | -0.836577771 | 0.402829945 | 0.61003978 | 0.167165096 |
| ENSMUSG00000041037 | Irgq | 210146 | protein_coding | 470.2394241 | -0.167214242 | 0.090009975 | -1.857730125 | 0.063207348 | 0.182100725 | 0.167214242 |
| ENSMUSG00000006711 | D130043K22Rik | 210108 | protein_coding | 39.59722696 | -0.167253538 | 0.214957608 | -0.778076846 | 0.436523711 | 0.639292909 | 0.167253538 |
| ENSMUSG00000039208 | Metrnl | 210029 | protein_coding | 727.9256828 | 0.167270967 | 0.233176924 | 0.717356436 | 0.473154193 | 0.668535437 | 0.167270967 |
| ENSMUSG00000021242 | Npc2 | 67963 | protein_coding | 1922.320571 | -0.167276291 | 0.134739685 | -1.241477527 | 0.214429395 | 0.412890644 | 0.167276291 |
| ENSMUSG00000019966 | Kitl | 17311 | protein_coding | 2446.075754 | 0.167306625 | 0.123649711 | 1.353069282 | 0.176033498 | 0.362209218 | 0.167306625 |
| ENSMUSG00000021706 | Zfyve16 | 218441 | protein_coding | 545.6472773 | 0.167383267 | 0.06525798 | 2.564947119 | 0.010319153 | 0.048910622 | 0.167383267 |
| ENSMUSG00000032298 | Neil1 | 72774 | protein_coding | 98.99042611 | -0.167405103 | 0.184592115 | -0.906891949 | 0.364463939 | 0.573238039 | 0.167405103 |
| ENSMUSG00000076437 | Selenoh | 72657 | protein_coding | 226.9630863 | 0.167444591 | 0.159880507 | 1.04731086 | 0.294956229 | 0.504029913 | 0.167444591 |
| ENSMUSG00000061411 | Nol4l | 329540 | protein_coding | 700.0443123 | -0.167476725 | 0.147605727 | -1.134622135 | 0.256533675 | 0.463387217 | 0.167476725 |
| ENSMUSG00000012443 | Kif11 | 16551 | protein_coding | 1177.499341 | 0.16748569 | 0.117927937 | 1.420237599 | 0.155538521 | 0.335647956 | 0.16748569 |
| ENSMUSG00000047804 | Akap10 | 56697 | protein_coding | 557.105021 | -0.167576461 | 0.084845933 | -1.975067688 | 0.048260467 | 0.152035744 | 0.167576461 |
| ENSMUSG00000022853 | Ehhadh | 74147 | protein_coding | 965.5708488 | 0.167657742 | 0.112393714 | 1.491700356 | 0.135777716 | 0.30556627 | 0.167657742 |
| ENSMUSG00000090935 | Synj2bp | 24071 | protein_coding | 1167.223949 | 0.167671309 | 0.064434824 | 2.602184648 | 0.009263196 | 0.045217343 | 0.167671309 |
| ENSMUSG00000026102 | Inpp1 | 16329 | protein_coding | 429.2764051 | 0.167808809 | 0.131765501 | 1.273541308 | 0.202826024 | 0.397809742 | 0.167808809 |
| ENSMUSG00000032313 | Tmem266 | 244886 | protein_coding | 15.05116047 | 0.16785017 | 0.390655914 | 0.429662431 | 0.667441216 | 0.80727602 | 0.16785017 |
| ENSMUSG00000023932 | Cdc5l | 71702 | protein_coding | 1259.871283 | 0.167921822 | 0.082190016 | 2.0430927 | 0.041043262 | 0.135573846 | 0.167921822 |
| ENSMUSG00000025410 | Dctn2 | 69654 | protein_coding | 1675.113634 | 0.167981511 | 0.077719398 | 2.161384604 | 0.030665642 | 0.10981168 | 0.167981511 |
| ENSMUSG00000085399 | Foxd2os | 100040736 | lncRNA | 476.8873922 | -0.168112778 | 0.06595132 | -2.549043415 | 0.010801885 | 0.050708054 | 0.168112778 |
| ENSMUSG00000028399 | Ptprd | 19266 | protein_coding | 1621.529356 | -0.168120418 | 0.067038417 | -2.507822017 | 0.012147784 | 0.055409479 | 0.168120418 |
| ENSMUSG00000034813 | Grip1 | 74053 | protein_coding | 44.91425347 | -0.168152277 | 0.220674235 | -0.76199325 | 0.446064033 | 0.64684158 | 0.168152277 |
| ENSMUSG00000017188 | Coa3 | 52469 | protein_coding | 1088.1841 | 0.168233697 | 0.105032527 | 1.601729494 | 0.10921544 | 0.263294011 | 0.168233697 |
| ENSMUSG00000022149 | C9 | 12279 | protein_coding | 10.11711108 | 0.168274241 | 0.372813958 | 0.451362503 | 0.651728304 | 0.798492458 | 0.168274241 |

| ensembl_gene_id | external_gene_name | entrezgene_id | gene_biotype | baseMean | log2FoldChange | lfcSE | stat | pvalue | padj | absLog2FC |
|---|---|---|---|---|---|---|---|---|---|---|
| ENSMUSG00000036775 | Decr2 | 26378 | protein_coding | 825.7867367 | 0.168284709 | 0.089120609 | 1.888280517 | 0.058988301 | 0.173993906 | 0.168284709 |
| ENSMUSG00000031913 | Vps4a | 116733 | protein_coding | 740.3792223 | 0.168285492 | 0.100577741 | 1.673188229 | 0.094290247 | 0.239110819 | 0.168285492 |
| ENSMUSG00000019804 | Snx3 | 54198 | protein_coding | 983.3102424 | 0.168305892 | 0.105973586 | 1.588187191 | 0.112244017 | 0.267798487 | 0.168305892 |
| ENSMUSG00000040048 | Ndufb10 | 68342 | protein_coding | 1563.458389 | 0.168334473 | 0.098692594 | 1.705644419 | 0.088074304 | 0.228545168 | 0.168334473 |
| ENSMUSG00000027330 | Cdc25b | 12531 | protein_coding | 256.6338502 | -0.168379677 | 0.098681461 | -1.706294926 | 0.087953182 | 0.228497519 | 0.168379677 |
| ENSMUSG00000003039 | Fam32a | 67922 | protein_coding | 1367.750656 | 0.168416404 | 0.075219415 | 2.239001786 | 0.0251558 | 0.094903264 | 0.168416404 |
| ENSMUSG00000036948 | Trappc14 | 231807 | protein_coding | 224.0282724 | -0.16842841 | 0.1198428 | -1.405411177 | 0.159899045 | 0.341312247 | 0.16842841 |
| ENSMUSG00000034127 | Tspan8 | 216350 | protein_coding | 9018.147708 | -0.168484414 | 0.14575627 | -1.155932532 | 0.247708755 | 0.453506327 | 0.168484414 |
| ENSMUSG00000031984 | 2810004N23Rik | 66523 | protein_coding | 337.9331954 | -0.168484491 | 0.10724292 | -1.571054682 | 0.116169948 | 0.274248545 | 0.168484491 |
| ENSMUSG00000005836 | Gata6 | 14465 | protein_coding | 658.8018795 | -0.168587881 | 0.101932996 | -1.653908824 | 0.09814604 | 0.245257314 | 0.168587881 |
| ENSMUSG00000046603 | Tcaim | 382117 | protein_coding | 150.3373931 | 0.168599401 | 0.12765099 | 1.320784117 | 0.186573357 | 0.376980899 | 0.168599401 |
| ENSMUSG00000062312 | Erbb2 | 13866 | protein_coding | 2042.205584 | -0.168624289 | 0.097672642 | -1.726422936 | 0.084271352 | 0.222024104 | 0.168624289 |
| ENSMUSG00000038991 | Txndc5 | 105245 | protein_coding | 5574.457181 | 0.168656799 | 0.094599959 | 1.782842192 | 0.074611995 | 0.203845602 | 0.168656799 |
| ENSMUSG00000030515 | Tarsl2 | 272396 | protein_coding | 147.5630888 | -0.168658616 | 0.132046793 | -1.277264013 | 0.201509054 | 0.39630031 | 0.168658616 |
| ENSMUSG00000069895 | Atxn1l | 52335 | protein_coding | 530.3655041 | -0.168676018 | 0.084810164 | -1.988865617 | 0.046716038 | 0.14856351 | 0.168676018 |
| ENSMUSG00000046707 | Csnk2a2 | 13000 | protein_coding | 382.5370534 | -0.16874768 | 0.083456631 | -2.021980504 | 0.043178366 | 0.140241642 | 0.16874768 |
| ENSMUSG00000005625 | Psmd4 | 19185 | protein_coding | 1220.208634 | 0.168780441 | 0.100852545 | 1.673536761 | 0.094221676 | 0.239013036 | 0.168780441 |
| ENSMUSG00000030084 | Plxna1 | 18844 | protein_coding | 409.4295913 | -0.16878046 | 0.121259606 | -1.391893523 | 0.163954644 | 0.347108299 | 0.16878046 |
| ENSMUSG00000071723 | Gspt2 | 14853 | protein_coding | 38.84222351 | 0.168882408 | 0.234882396 | 0.719008364 | 0.472135766 | 0.667570216 | 0.168882408 |
| ENSMUSG00000071180 | Smim15 | 75616 | protein_coding | 1375.430252 | 0.168885817 | 0.131867015 | 1.280728294 | 0.200289118 | 0.39482736 | 0.168885817 |
| ENSMUSG00000007038 | Neu1 | 18010 | protein_coding | 1588.915872 | -0.168924698 | 0.167011768 | -1.011453864 | 0.311799252 | 0.521948343 | 0.168924698 |
| ENSMUSG00000040651 | Tasor | 218850 | protein_coding | 882.3004198 | -0.168933885 | 0.058929184 | -2.866727027 | 0.004147406 | 0.024162355 | 0.168933885 |
| ENSMUSG00000043372 | Hexim2 | 71059 | protein_coding | 44.60878263 | 0.168947187 | 0.22136918 | 0.763192 | 0.445348899 | 0.646157521 | 0.168947187 |
| ENSMUSG00000026991 | Pkp4 | 227937 | protein_coding | 1024.681522 | 0.169000157 | 0.08047764 | 2.099964126 | 0.035731997 | 0.122871168 | 0.169000157 |
| ENSMUSG00000051950 | B3glct | 381694 | protein_coding | 247.0740233 | -0.169017153 | 0.102203961 | -1.653724089 | 0.098183586 | 0.245274177 | 0.169017153 |
| ENSMUSG00000028572 | Hook1 | 77963 | protein_coding | 3945.228431 | 0.169077084 | 0.057703868 | 2.930082348 | 0.003388722 | 0.020708342 | 0.169077084 |
| ENSMUSG00000048755 | Mcat | 223722 | protein_coding | 209.7449497 | 0.169092565 | 0.102841415 | 1.644206913 | 0.100133469 | 0.248353457 | 0.169092565 |
| ENSMUSG00000071456 | 1110002L01Rik | 100043040 | lncRNA | 22.08045021 | 0.169174015 | 0.292872177 | 0.577637714 | 0.563508739 | 0.740191918 | 0.169174015 |
| ENSMUSG00000073062 | Zxdb | 668166 | protein_coding | 85.92877684 | 0.169179078 | 0.187348058 | 0.903020181 | 0.366515184 | 0.575230748 | 0.169179078 |
| ENSMUSG00000031077 | Fadd | 14082 | protein_coding | 190.3702855 | -0.169181748 | 0.097094316 | -1.742447498 | 0.081430171 | 0.216711739 | 0.169181748 |
| ENSMUSG00000029254 | Stap1 | 56792 | protein_coding | 9.021811471 | 0.169221498 | 0.609743186 | 0.277529133 | 0.781373839 | 0.881175482 | 0.169221498 |
| ENSMUSG00000062075 | Lmnb2 | 16907 | protein_coding | 861.1473658 | -0.169252848 | 0.117475956 | -1.440744586 | 0.149656854 | 0.327058149 | 0.169252848 |
| ENSMUSG00000018752 | Tnfsfm13 | 619441 | protein_coding | 14.00005093 | 0.169344709 | 0.336035218 | 0.503949288 | 0.614297014 | 0.775275463 | 0.169344709 |
| ENSMUSG00000030319 | Cand2 | 67088 | protein_coding | 101.9364088 | 0.169427856 | 0.143217139 | 1.183013829 | 0.236803667 | 0.440992616 | 0.169427856 |
| ENSMUSG00000061232 | H2-K1 | 14972 | protein_coding | 7271.66419 | 0.169432366 | 0.111252856 | 1.522948462 | 0.1277716 | 0.293439621 | 0.169432366 |
| ENSMUSG00000031266 | Gla | 11605 | protein_coding | 317.5214367 | 0.169442528 | 0.110629229 | 1.531625313 | 0.125614922 | 0.289844196 | 0.169442528 |
| ENSMUSG00000030801 | Kat8 | 67773 | protein_coding | 228.7183261 | 0.169485886 | 0.105804852 | 1.601872522 | 0.109183802 | 0.263257583 | 0.169485886 |
| ENSMUSG00000027834 | Serpini1 | 20713 | protein_coding | 10.85141768 | 0.169537911 | 0.440519562 | 0.384858983 | 0.700341896 | 0.82992677 | 0.169537911 |

| ensembl_gene_id | external_gene_name | entrezgene_id | gene_biotype | baseMean | log2FoldChange | lfcSE | stat | pvalue | padj | absLog2FC |
|---|---|---|---|---|---|---|---|---|---|---|
| ENSMUSG00000079426 | Arpc4 | 68089 | protein_coding | 1397.494785 | 0.169556846 | 0.141758117 | 1.196099737 | 0.23165764 | 0.435066931 | 0.169556846 |
| ENSMUSG00000031803 | B3gnt3 | 72297 | protein_coding | 3189.616901 | 0.169577454 | 0.112965126 | 1.501148716 | 0.133317101 | 0.301829505 | 0.169577454 |
| ENSMUSG00000051037 | Zfp455 | 218311 | protein_coding | 35.08736332 | 0.169603267 | 0.250388524 | 0.677360385 | 0.498177327 | 0.688918949 | 0.169603267 |
| ENSMUSG00000031969 | Acad8 | 66948 | protein_coding | 637.254598 | -0.169603582 | 0.090320611 | -1.877794894 | 0.060409229 | 0.176550129 | 0.169603582 |
| ENSMUSG00000024537 | Psmg2 | 107047 | protein_coding | 165.5335327 | 0.169626972 | 0.139066646 | 1.219753103 | 0.222558484 | 0.423917664 | 0.169626972 |
| ENSMUSG00000032425 | Zfp949 | 71640 | protein_coding | 183.9218469 | -0.169630032 | 0.161671439 | -1.049226959 | 0.294073674 | 0.503277692 | 0.169630032 |
| ENSMUSG00000031902 | Nfatc3 | 18021 | protein_coding | 970.252983 | -0.169643351 | 0.080388376 | -2.110297038 | 0.034832779 | 0.120612282 | 0.169643351 |
| ENSMUSG00000021007 | Spata7 | 104871 | protein_coding | 84.61507034 | -0.169655895 | 0.21319069 | -0.795794106 | 0.426151718 | 0.630714046 | 0.169655895 |
| ENSMUSG00000051314 | Ffar2 | 233079 | protein_coding | 322.3578583 | -0.169676025 | 0.147871624 | -1.147454935 | 0.251193648 | 0.457572327 | 0.169676025 |
| ENSMUSG00000058831 | Opn1sw | 12057 | protein_coding | 9.077652249 | -0.169696271 | 0.480604665 | -0.353089106 | 0.72402164 | 0.844771344 | 0.169696271 |
| ENSMUSG00000028049 | Scamp3 | 24045 | protein_coding | 169.3996118 | -0.169744302 | 0.095315092 | -1.780875382 | 0.074932811 | 0.204201575 | 0.169744302 |
| ENSMUSG00000025534 | Gusb | 110006 | protein_coding | 582.2951567 | 0.169754684 | 0.089296529 | 1.901022203 | 0.057299104 | 0.170750259 | 0.169754684 |
| ENSMUSG00000032724 | Abtb2 | 99382 | protein_coding | 58.69137723 | -0.169771524 | 0.216586794 | -0.783849841 | 0.433128225 | 0.637190945 | 0.169771524 |
| ENSMUSG00000031967 | Afg3l1 | 114896 | protein_coding | 485.6694713 | -0.169779847 | 0.08832786 | -1.922155089 | 0.054586246 | 0.164730907 | 0.169779847 |
| ENSMUSG00000061825 | Ces2c | 234671 | protein_coding | 1536.102801 | -0.169781482 | 0.259145644 | -0.65515854 | 0.512365685 | 0.699981612 | 0.169781482 |
| ENSMUSG00000019842 | Traf3ip2 | 103213 | protein_coding | 376.432417 | -0.169826198 | 0.097388969 | -1.743792953 | 0.0811952 | 0.216267081 | 0.169826198 |
| ENSMUSG00000049612 | Omg | 18377 | protein_coding | 7.461799582 | 0.169859066 | 0.596097949 | 0.284951603 | 0.775681221 | 0.877982787 | 0.169859066 |
| ENSMUSG00000037971 | 1110032A03Rik | 68721 | protein_coding | 320.7217148 | 0.169918088 | 0.131329836 | 1.293827005 | 0.195725182 | 0.389008074 | 0.169918088 |
| ENSMUSG00000055612 | Cdca7 | 66953 | protein_coding | 683.9033724 | -0.169949707 | 0.143641609 | -1.183150949 | 0.236749329 | 0.440992616 | 0.169949707 |
| ENSMUSG00000013622 | Atraid | 381629 | protein_coding | 650.4694081 | 0.169961574 | 0.114845735 | 1.479911934 | 0.13889675 | 0.310200146 | 0.169961574 |
| ENSMUSG00000052534 | Pbx1 | 18514 | protein_coding | 601.7061205 | -0.16996705 | 0.104477053 | -1.626836189 | 0.103771893 | 0.254408349 | 0.16996705 |
| ENSMUSG00000042489 | Clspn | 269582 | protein_coding | 182.7631774 | 0.169986236 | 0.163830518 | 1.037573693 | 0.299468571 | 0.50846769 | 0.169986236 |
| ENSMUSG00000036285 | Noa1 | 56412 | protein_coding | 195.1709677 | -0.170045667 | 0.132386609 | -1.284462747 | 0.19898009 | 0.393175365 | 0.170045667 |
| ENSMUSG00000024236 | Svil | 225115 | protein_coding | 152.447502 | 0.170171681 | 0.207859644 | 0.818685518 | 0.412965861 | 0.619566809 | 0.170171681 |
| ENSMUSG00000025069 | Gsto2 | 68214 | protein_coding | 13.26861056 | -0.170191874 | 0.313534838 | -0.542816468 | 0.587256173 | 0.7572714 | 0.170191874 |
| ENSMUSG00000034269 | Setd5 | 72895 | protein_coding | 1875.342208 | -0.170205993 | 0.113420469 | -1.500663806 | 0.133442539 | 0.301885137 | 0.170205993 |
| ENSMUSG00000038146 | Notch3 | 18131 | protein_coding | 11.93232147 | -0.170286457 | 0.535490939 | -0.318000632 | 0.750484458 | 0.862088305 | 0.170286457 |
| ENSMUSG00000033096 | Apmap | 71881 | protein_coding | 592.507787 | 0.170315372 | 0.072719237 | 2.342095163 | 0.019175825 | 0.07759973 | 0.170315372 |
| ENSMUSG00000079469 | Pigb | 55981 | protein_coding | 128.643727 | -0.170324433 | 0.117580207 | -1.448580823 | 0.14745468 | 0.324042238 | 0.170324433 |
| ENSMUSG00000020300 | Cpeb4 | 67579 | protein_coding | 1252.141134 | -0.170343502 | 0.116459105 | -1.462689435 | 0.143552378 | 0.317569761 | 0.170343502 |
| ENSMUSG00000039055 | Eme1 | 268465 | protein_coding | 60.07329328 | 0.170361042 | 0.187086215 | 0.910601791 | 0.362505227 | 0.571624913 | 0.170361042 |
| ENSMUSG00000035284 | Vps13c | 320528 | protein_coding | 1207.0948 | -0.170373896 | 0.241601989 | -0.70518416 | 0.480695638 | 0.674079929 | 0.170373896 |
| ENSMUSG00000026347 | Tmem163 | 72160 | protein_coding | 7.389017207 | -0.170386494 | 0.552052566 | -0.308641792 | 0.757594026 | 0.867377634 | 0.170386494 |
| ENSMUSG00000028468 | Rgp1 | 242406 | protein_coding | 753.150986 | -0.170443603 | 0.087188192 | -1.954893198 | 0.050595678 | 0.156635171 | 0.170443603 |
| ENSMUSG00000001569 | Nom1 | 433864 | protein_coding | 491.0133418 | -0.170492466 | 0.094125161 | -1.811337843 | 0.070088571 | 0.195751493 | 0.170492466 |
| ENSMUSG00000041975 | Mettl8 | 228019 | protein_coding | 157.9345124 | 0.170631954 | 0.138444354 | 1.232494856 | 0.217764287 | 0.417470414 | 0.170631954 |
| ENSMUSG00000049086 | Bmyc | 107771 | protein_coding | 20.15734584 | -0.170644714 | 0.270991306 | -0.629705495 | 0.528887286 | 0.714392529 | 0.170644714 |
| ENSMUSG00000037110 | Ralgapa2 | 241694 | protein_coding | 2305.879502 | -0.170656469 | 0.095013144 | -1.796135367 | 0.072472991 | 0.200112777 | 0.170656469 |

| ensembl_gene_id | external_gene_name | entrezgene_id | gene_biotype | baseMean | log2FoldChange | lfcSE | stat | pvalue | padj | absLog2FC |
|---|---|---|---|---|---|---|---|---|---|---|
| ENSMUSG00000079509 | Zfx | 22764 | protein_coding | 830.1124605 | 0.170708463 | 0.077843162 | 2.192979544 | 0.028308853 | 0.103239441 | 0.170708463 |
| ENSMUSG00000028480 | Glipr2 | 384009 | protein_coding | 134.9267606 | -0.170729158 | 0.247721666 | -0.689197523 | 0.490698975 | 0.68307464 | 0.170729158 |
| ENSMUSG00000001418 | Glmp | 56700 | protein_coding | 717.3968271 | -0.170782509 | 0.068275713 | -2.50136542 | 0.012371545 | 0.0561219 | 0.170782509 |
| ENSMUSG00000022462 | Slc38a2 | 67760 | protein_coding | 3205.242832 | -0.170851758 | 0.109391142 | -1.561842719 | 0.118325045 | 0.278369656 | 0.170851758 |
| ENSMUSG00000026870 | Cutal | 77996 | protein_coding | 1283.306289 | 0.17086922 | 0.107321756 | 1.592120996 | 0.11135752 | 0.266437314 | 0.17086922 |
| ENSMUSG00000048899 | Rimkla | 194237 | protein_coding | 18.28886932 | -0.170893979 | 0.32259247 | -0.529751915 | 0.596283948 | 0.764096331 | 0.170893979 |
| ENSMUSG00000029426 | Scarb2 | 12492 | protein_coding | 2937.19246 | -0.170967722 | 0.116027802 | -1.473506513 | 0.140614509 | 0.312983217 | 0.170967722 |
| ENSMUSG00000021431 | Snrnp48 | 67797 | protein_coding | 468.2305254 | 0.171005287 | 0.102664119 | 1.665677245 | 0.095777717 | 0.241464688 | 0.171005287 |
| ENSMUSG00000023914 | Mep1a | 17287 | protein_coding | 4459.839276 | -0.171079666 | 0.160171138 | -1.068105452 | 0.285472947 | 0.494401772 | 0.171079666 |
| ENSMUSG00000054792 | Klhl18 | 270201 | protein_coding | 215.7606617 | -0.171080351 | 0.103904851 | -1.646509763 | 0.099658852 | 0.247613326 | 0.171080351 |
| ENSMUSG00000041058 | Wwp1 | 107568 | protein_coding | 1516.691428 | 0.171083027 | 0.06815406 | 2.510239711 | 0.012064923 | 0.055126246 | 0.171083027 |
| ENSMUSG00000060743 | H3f3a | 15078 | protein_coding | 2254.562602 | 0.171150271 | 0.088739488 | 1.928682203 | 0.053770327 | 0.162897492 | 0.171150271 |
| ENSMUSG00000042770 | Hebp1 | 15199 | protein_coding | 6.747468289 | -0.171209929 | 0.484905063 | -0.353079277 | 0.724029008 | 0.844771344 | 0.171209929 |
| ENSMUSG00000021692 | Dimt1 | 66254 | protein_coding | 86.14894701 | -0.171223064 | 0.183749407 | -0.931829204 | 0.3514248 | 0.561349008 | 0.171223064 |
| ENSMUSG00000020070 | Rufy2 | 70432 | protein_coding | 142.2894536 | -0.17123042 | 0.148526132 | -1.152863933 | 0.248966238 | 0.455180698 | 0.17123042 |
| ENSMUSG00000006281 | Tep1 | 21745 | protein_coding | 2385.269989 | -0.171248037 | 0.086693934 | -1.975317403 | 0.04823214 | 0.152006632 | 0.171248037 |
| ENSMUSG00000086330 | 1700007J10Rik | 69320 | lncRNA | 7.624166322 | -0.171250688 | 0.459613767 | -0.372596951 | 0.709448442 | 0.835827385 | 0.171250688 |
| ENSMUSG00000052033 | Pfdn4 | 109054 | protein_coding | 173.0723865 | 0.171307504 | 0.142798102 | 1.199648328 | 0.230275949 | 0.433458782 | 0.171307504 |
| ENSMUSG00000022982 | Sod1 | 20655 | protein_coding | 3396.254652 | 0.17132076 | 0.092617634 | 1.849763944 | 0.06434758 | 0.184489941 | 0.17132076 |
| ENSMUSG00000059149 | Mfsd4a | 213006 | protein_coding | 1764.5798 | -0.171320832 | 0.169570056 | -1.010324797 | 0.312339705 | 0.52254097 | 0.171320832 |
| ENSMUSG00000041638 | Gcn1 | 231659 | protein_coding | 1238.220106 | -0.171342273 | 0.113668007 | -1.507392248 | 0.131710145 | 0.29948311 | 0.171342273 |
| ENSMUSG00000021373 | Cap2 | 67252 | protein_coding | 12.58252538 | 0.171355203 | 0.451956012 | 0.379141328 | 0.704582918 | 0.833154916 | 0.171355203 |
| ENSMUSG00000060152 | Pop5 | 117109 | protein_coding | 287.5779731 | -0.17139541 | 0.106464048 | -1.609890033 | 0.107421865 | 0.260626881 | 0.17139541 |
| ENSMUSG00000031601 | Cnot7 | 18983 | protein_coding | 722.9180672 | 0.171408781 | 0.06507785 | 2.633903556 | 0.008440946 | 0.042182001 | 0.171408781 |
| ENSMUSG00000020069 | Hnrnph3 | 432467 | protein_coding | 215.3548379 | -0.171414246 | 0.107462683 | -1.595104844 | 0.110688792 | 0.265435629 | 0.171414246 |
| ENSMUSG00000036057 | Ptpn23 | 104831 | protein_coding | 211.6899393 | -0.171481565 | 0.16463664 | -1.041575949 | 0.297608324 | 0.506551121 | 0.171481565 |
| ENSMUSG00000020290 | Xpo1 | 103573 | protein_coding | 1915.883325 | 0.171611004 | 0.093487017 | 1.8356667 | 0.066406966 | 0.188409361 | 0.171611004 |
| ENSMUSG00000009563 | Tor2a | 30933 | protein_coding | 436.7947415 | 0.171628767 | 0.10862307 | 1.580039743 | 0.114097765 | 0.270508479 | 0.171628767 |
| ENSMUSG00000074749 | Kiz | 228730 | protein_coding | 333.4791903 | -0.171642054 | 0.138543396 | -1.238904626 | 0.21538082 | 0.414262276 | 0.171642054 |
| ENSMUSG00000030446 | Zfp273 | 212569 | protein_coding | 104.695309 | 0.171662398 | 0.23058419 | 0.744467337 | 0.456593794 | 0.655398985 | 0.171662398 |
| ENSMUSG00000058624 | Gda | 14544 | protein_coding | 1089.169875 | 0.171699504 | 0.399483032 | 0.429804246 | 0.667338044 | 0.807272755 | 0.171699504 |
| ENSMUSG00000021638 | Ocln | 18260 | protein_coding | 1931.932065 | -0.171721389 | 0.070459357 | -2.437169388 | 0.014802743 | 0.063889164 | 0.171721389 |
| ENSMUSG00000028415 | Spink4 | 20731 | protein_coding | 9096.433817 | 0.171752459 | 0.148956266 | 1.153039506 | 0.24889417 | 0.455101175 | 0.171752459 |
| ENSMUSG00000004096 | Cwc15 | 66070 | protein_coding | 919.1528885 | 0.171785445 | 0.075538411 | 2.274146915 | 0.022957155 | 0.088689858 | 0.171785445 |
| ENSMUSG00000009828 | Cilk1 | 56542 | protein_coding | 1401.000847 | 0.17180107 | 0.069169589 | 2.483765936 | 0.013000123 | 0.058377625 | 0.17180107 |
| ENSMUSG00000045302 | Preb | 50907 | protein_coding | 1059.724779 | -0.171857432 | 0.10406728 | -1.65140698 | 0.098655501 | 0.246144238 | 0.171857432 |
| ENSMUSG00000029992 | Gfpt1 | 14583 | protein_coding | 14141.83172 | -0.171859196 | 0.098619869 | -1.742642709 | 0.081396045 | 0.216657119 | 0.171859196 |
| ENSMUSG00000040220 | Gas8 | 104346 | protein_coding | 107.2264029 | 0.171875338 | 0.151638806 | 1.133452201 | 0.257024403 | 0.463812519 | 0.171875338 |

| ensembl_gene_id | external_gene_name | entrezgene_id | gene_biotype | baseMean | log2FoldChange | lfcSE | stat | pvalue | padj | absLog2FC |
|---|---|---|---|---|---|---|---|---|---|---|
| ENSMUSG00000046417 | Lrrc75a | 192976 | protein_coding | 704.9158702 | 0.171885492 | 0.104510899 | 1.644665711 | 0.100038768 | 0.248195871 | 0.171885492 |
| ENSMUSG00000048930 | Tada3 | 101206 | protein_coding | 263.8219152 | 0.171890176 | 0.094594956 | 1.81711777 | 0.069199082 | 0.194128339 | 0.171890176 |
| ENSMUSG00000050908 | Tvp23a | 383103 | protein_coding | 7.276618784 | -0.171896705 | 0.456778621 | -0.376323885 | 0.706676125 | 0.834197547 | 0.171896705 |
| ENSMUSG00000019794 | Katna1 | 23924 | protein_coding | 614.6863298 | 0.171906094 | 0.09056435 | 1.898165155 | 0.057674329 | 0.17155684 | 0.171906094 |
| ENSMUSG00000029715 | Pop7 | 74097 | protein_coding | 186.794723 | 0.171923986 | 0.160605691 | 1.070472564 | 0.284406652 | 0.493628544 | 0.171923986 |
| ENSMUSG00000000355 | Mcts1 | 68995 | protein_coding | 298.4089796 | 0.172016191 | 0.098936356 | 1.738655007 | 0.082095465 | 0.218045116 | 0.172016191 |
| ENSMUSG00000032872 | Cyb5r4 | 266690 | protein_coding | 1211.155925 | 0.172054776 | 0.086607767 | 1.986597522 | 0.046967016 | 0.14896269 | 0.172054776 |
| ENSMUSG00000037465 | Klf10 | 21847 | protein_coding | 1633.371598 | -0.17213731 | 0.157632006 | -1.092020045 | 0.274824291 | 0.482943656 | 0.17213731 |
| ENSMUSG00000051499 | Zfp786 | 330301 | protein_coding | 27.04201766 | -0.172148843 | 0.24554281 | -0.70109503 | 0.483243712 | 0.676728138 | 0.172148843 |
| ENSMUSG00000042229 | Rabif | 98710 | protein_coding | 486.5469224 | 0.17221814 | 0.086035189 | 2.001717472 | 0.045315126 | 0.145409528 | 0.17221814 |
| ENSMUSG00000041360 | Pum3 | 52874 | protein_coding | 563.1787041 | -0.172253513 | 0.158416035 | -1.087348974 | 0.276882613 | 0.48532918 | 0.172253513 |
| ENSMUSG00000022009 | Nufip1 | 27275 | protein_coding | 221.4217625 | -0.172254613 | 0.146190099 | -1.178291924 | 0.238680246 | 0.442830469 | 0.172254613 |
| ENSMUSG00000047963 | Stbd1 | 52331 | protein_coding | 7.305648275 | 0.172294285 | 0.472798784 | 0.364413553 | 0.715549217 | 0.840012084 | 0.172294285 |
| ENSMUSG00000004864 | Mapk13 | 26415 | protein_coding | 2295.244794 | 0.172352752 | 0.079520854 | 2.167390604 | 0.030205083 | 0.108557068 | 0.172352752 |
| ENSMUSG00000056211 | R3hdm1 | 226412 | protein_coding | 421.2330691 | -0.172366205 | 0.146602693 | -1.17573696 | 0.239700004 | 0.443946706 | 0.172366205 |
| ENSMUSG00000071550 | Cfap44 | 212517 | protein_coding | 16.28321924 | -0.172385488 | 0.433702574 | -0.397473979 | 0.691017971 | 0.824089117 | 0.172385488 |
| ENSMUSG00000057594 | Arl16 | 70317 | protein_coding | 90.99429402 | 0.17247718 | 0.127217246 | 1.35576886 | 0.175172722 | 0.361418538 | 0.17247718 |
| ENSMUSG00000036687 | Tmem184a | 231832 | protein_coding | 431.3557081 | -0.172500568 | 0.10842039 | -1.591034374 | 0.11160184 | 0.266785848 | 0.172500568 |
| ENSMUSG00000046962 | Zbtb21 | 114565 | protein_coding | 300.1198914 | -0.172503949 | 0.101729997 | -1.695703867 | 0.089941976 | 0.231849133 | 0.172503949 |
| ENSMUSG00000078994 | Zfp429 | 72807 | protein_coding | 151.4942082 | 0.172519226 | 0.115873798 | 1.488854506 | 0.136525686 | 0.30684306 | 0.172519226 |
| ENSMUSG00000087881 | Gm22442 | 115487906 | scaRNA | 549.5553587 | -0.172540943 | 0.108331835 | -1.592707659 | 0.111225788 | 0.266326572 | 0.172540943 |
| ENSMUSG00000118537 | Shld3 | 113002583 | protein_coding | 52.37137516 | 0.17257734 | 0.185945666 | 0.928106279 | 0.353352443 | 0.563101322 | 0.17257734 |
| ENSMUSG00000074500 | Zfp558 | 72230 | protein_coding | 34.61848126 | 0.172600644 | 0.213006802 | 0.810305783 | 0.417764452 | 0.624041283 | 0.172600644 |
| ENSMUSG00000030224 | Strap | 20901 | protein_coding | 1452.450435 | 0.172678098 | 0.102281647 | 1.688260823 | 0.091361172 | 0.234143323 | 0.172678098 |
| ENSMUSG00000003929 | Zfp81 | 224694 | protein_coding | 193.0154551 | -0.172719099 | 0.117445619 | -1.470630413 | 0.141391095 | 0.31409726 | 0.172719099 |
| ENSMUSG00000022685 | Parn | 74108 | protein_coding | 362.4123956 | 0.172726684 | 0.078793115 | 2.192154532 | 0.02836835 | 0.103374762 | 0.172726684 |
| ENSMUSG00000003812 | Dnase2a | 13423 | protein_coding | 119.2408846 | -0.172742509 | 0.163890981 | -1.054008633 | 0.291878964 | 0.500705239 | 0.172742509 |
| ENSMUSG00000043079 | Synpo | 104027 | protein_coding | 381.6414304 | -0.172774758 | 0.108165207 | -1.597322857 | 0.110193758 | 0.264770883 | 0.172774758 |
| ENSMUSG00000046991 | Wdr27 | 71682 | protein_coding | 19.70620286 | 0.172796404 | 0.347011172 | 0.497956313 | 0.618514835 | 0.777789999 | 0.172796404 |
| ENSMUSG00000015672 | Mrpl32 | 75398 | protein_coding | 400.998197 | 0.172811511 | 0.100947687 | 1.711891734 | 0.086916622 | 0.226840564 | 0.172811511 |
| ENSMUSG00000045973 | Slc25a51 | 230125 | protein_coding | 1051.473935 | -0.172819077 | 0.074791318 | -2.310683664 | 0.020850334 | 0.082489847 | 0.172819077 |
| ENSMUSG00000028126 | Pip5k1a | 18720 | protein_coding | 265.6971294 | -0.172872391 | 0.091222628 | -1.895060412 | 0.058084399 | 0.172192406 | 0.172872391 |
| ENSMUSG00000051154 | Commd3 | 12238 | protein_coding | 410.6861307 | 0.172880536 | 0.083456611 | 2.071501984 | 0.038311909 | 0.129224785 | 0.172880536 |
| ENSMUSG00000037016 | Frem2 | 242022 | protein_coding | 15.45031191 | 0.172920332 | 0.421937592 | 0.409824427 | 0.681934746 | 0.818095702 | 0.172920332 |
| ENSMUSG00000045038 | Prkce | 18754 | protein_coding | 273.0451602 | -0.172927549 | 0.130853668 | -1.321533833 | 0.186323429 | 0.376888975 | 0.172927549 |
| ENSMUSG00000035311 | Gnptab | 432486 | protein_coding | 672.8112107 | -0.17294986 | 0.103783924 | -1.666441716 | 0.095625468 | 0.241194783 | 0.17294986 |
| ENSMUSG00000029090 | Adgra3 | 70693 | protein_coding | 1528.646145 | 0.173018392 | 0.12128665 | 1.426524617 | 0.153716966 | 0.332844459 | 0.173018392 |
| ENSMUSG00000068391 | Chrac1 | 93696 | protein_coding | 184.2634182 | 0.1730194 | 0.116059466 | 1.490782324 | 0.136018654 | 0.306003548 | 0.1730194 |

| ensembl_gene_id | external_gene_name | entrezgene_id | gene_biotype | baseMean | log2FoldChange | lfcSE | stat | pvalue | padj | absLog2FC |
|---|---|---|---|---|---|---|---|---|---|---|
| ENSMUSG00000034757 | Tmub2 | 72053 | protein_coding | 513.6663178 | 0.173021766 | 0.106176166 | 1.62957255 | 0.10319187 | 0.253530789 | 0.173021766 |
| ENSMUSG00000025357 | Dgka | 13139 | protein_coding | 1352.047208 | -0.173037855 | 0.105602846 | -1.638571897 | 0.101302449 | 0.250317315 | 0.173037855 |
| ENSMUSG00000034007 | Scaper | 244891 | protein_coding | 447.0280632 | -0.173058884 | 0.0939948 | -1.841153806 | 0.065599023 | 0.186649023 | 0.173058884 |
| ENSMUSG00000031935 | Med17 | 234959 | protein_coding | 531.7267847 | -0.173068797 | 0.078378184 | -2.208124609 | 0.027235588 | 0.100441862 | 0.173068797 |
| ENSMUSG00000022179 | 4931414P19Rik | 74359 | protein_coding | 96.34289613 | 0.173093788 | 0.239989337 | 0.721256161 | 0.470751924 | 0.666501035 | 0.173093788 |
| ENSMUSG00000031540 | Kat6a | 244349 | protein_coding | 1024.555133 | -0.173121881 | 0.14522765 | -1.192072458 | 0.233232835 | 0.436793957 | 0.173121881 |
| ENSMUSG00000087406 | E130215H24Rik | 109344 | lncRNA | 11.3378326 | -0.173149811 | 0.380891754 | -0.454590599 | 0.649403799 | 0.797420867 | 0.173149811 |
| ENSMUSG00000080797 | Gm15760 | 100042948 | transcribed_processed_pseudogene | 12.86376659 | -0.173171383 | 0.353959122 | -0.489241194 | 0.624670949 | 0.782706016 | 0.173171383 |
| ENSMUSG00000024972 | Lgals12 | 56072 | protein_coding | 299.9800958 | -0.173187782 | 0.143349178 | -1.208153291 | 0.226988305 | 0.429514754 | 0.173187782 |
| ENSMUSG00000031959 | Wdr59 | 319481 | protein_coding | 224.1754301 | 0.173281383 | 0.117187593 | 1.478666624 | 0.139229437 | 0.310786074 | 0.173281383 |
| ENSMUSG00000065637 | Gm26397 | 115487043 | snoRNA | 66.7984054 | -0.173398196 | 0.280353952 | -0.618497422 | 0.536247497 | 0.720849871 | 0.173398196 |
| ENSMUSG00000002343 | Armc6 | 76813 | protein_coding | 107.9291272 | 0.173399896 | 0.211271915 | 0.820742766 | 0.411792805 | 0.618718593 | 0.173399896 |
| ENSMUSG00000027942 | 4933434E20Rik | 99650 | protein_coding | 174.5281076 | 0.173444472 | 0.095770631 | 1.811040307 | 0.070134613 | 0.195811412 | 0.173444472 |
| ENSMUSG00000037216 | Lipt1 | 623661 | protein_coding | 41.98241206 | -0.173461666 | 0.180068125 | -0.96331134 | 0.335391304 | 0.546004977 | 0.173461666 |
| ENSMUSG00000069170 | Adgrv1 | 110789 | protein_coding | 30.51145004 | -0.173491361 | 0.525362935 | -0.330231445 | 0.741225088 | 0.855772195 | 0.173491361 |
| ENSMUSG00000053119 | Chmp3 | 66700 | protein_coding | 2403.433135 | 0.173512097 | 0.065339655 | 2.655540445 | 0.007918144 | 0.0401401 | 0.173512097 |
| ENSMUSG00000025204 | Ndufb8 | 67264 | protein_coding | 2200.479754 | 0.173559616 | 0.106990939 | 1.622189847 | 0.104762705 | 0.256243913 | 0.173559616 |
| ENSMUSG00000029605 | Oas1b | 23961 | protein_coding | 17.15167072 | 0.173673847 | 0.329717939 | 0.526734599 | 0.598377905 | 0.765321556 | 0.173673847 |
| ENSMUSG00000009863 | Sdhb | 67680 | protein_coding | 2972.739263 | 0.17367515 | 0.108827281 | 1.5958788 | 0.110515856 | 0.265184777 | 0.17367515 |
| ENSMUSG00000047507 | Baiap3 | 545192 | protein_coding | 41.87412686 | -0.173802298 | 0.210455459 | -0.825838865 | 0.408895504 | 0.615890871 | 0.173802298 |
| ENSMUSG00000025082 | Vwa2 | 240675 | protein_coding | 14.89588405 | 0.173804952 | 0.322503336 | 0.53892451 | 0.589938944 | 0.759562485 | 0.173804952 |
| ENSMUSG00000031400 | G6pdx | 14381 | protein_coding | 849.6265915 | 0.173846803 | 0.096703422 | 1.797731659 | 0.072219542 | 0.19967979 | 0.173846803 |
| ENSMUSG00000027282 | Mtch2 | 56428 | protein_coding | 3590.258639 | 0.173878131 | 0.116190783 | 1.496488164 | 0.13452649 | 0.303632979 | 0.173878131 |
| ENSMUSG00000042712 | Tceal9 | 22381 | protein_coding | 1497.490065 | 0.173918119 | 0.106797771 | 1.62848079 | 0.103422979 | 0.254020233 | 0.173918119 |
| ENSMUSG00000058407 | Txndc9 | 98258 | protein_coding | 1066.069595 | 0.17395119 | 0.093798233 | 1.854525231 | 0.063664054 | 0.183139074 | 0.17395119 |
| ENSMUSG00000031816 | Mthfsd | 234814 | protein_coding | 248.6039684 | -0.173984773 | 0.0860589 | -2.021694118 | 0.043207962 | 0.140280558 | 0.173984773 |
| ENSMUSG00000004393 | Ddx56 | 52513 | protein_coding | 357.4703601 | -0.174045733 | 0.127653436 | -1.363423803 | 0.172748983 | 0.358489058 | 0.174045733 |
| ENSMUSG00000026155 | Smap1 | 98366 | protein_coding | 349.9820869 | 0.174088634 | 0.09703817 | 1.794022239 | 0.072809619 | 0.200676868 | 0.174088634 |
| ENSMUSG00000019370 | Calm3 | 12315 | protein_coding | 5869.783989 | 0.17415624 | 0.070440887 | 2.472374319 | 0.01342189 | 0.059732847 | 0.17415624 |
| ENSMUSG00000037157 | Il22ra1 | 230828 | protein_coding | 347.6696338 | -0.174294742 | 0.114235565 | -1.525748509 | 0.127072509 | 0.292313212 | 0.174294742 |
| ENSMUSG00000023938 | Aars2 | 224805 | protein_coding | 241.2582599 | -0.174298578 | 0.115669784 | -1.506863522 | 0.131845643 | 0.299620118 | 0.174298578 |
| ENSMUSG00000110080 | Gm6145 | 620376 | lncRNA | 13.7380245 | -0.174337585 | 0.385244359 | -0.452537671 | 0.65088169 | 0.798250711 | 0.174337585 |
| ENSMUSG00000006262 | Mob1b | 68473 | protein_coding | 1560.198295 | 0.17437286 | 0.111376984 | 1.565609471 | 0.117440071 | 0.276713825 | 0.17437286 |
| ENSMUSG00000027425 | Kat14 | 228714 | protein_coding | 383.0419522 | -0.174428551 | 0.085747385 | -2.034214228 | 0.041930001 | 0.137306959 | 0.174428551 |
| ENSMUSG00000032846 | Zswim6 | 67263 | protein_coding | 312.3915723 | -0.174430016 | 0.140410268 | -1.242288171 | 0.214130258 | 0.412514121 | 0.174430016 |
| ENSMUSG00000026821 | Ralgds | 19730 | protein_coding | 1071.15459 | -0.174454022 | 0.103348029 | -1.688024665 | 0.091406494 | 0.234221789 | 0.174454022 |
| ENSMUSG00000060601 | Nr1h2 | 22260 | protein_coding | 598.6266955 | -0.174494493 | 0.081120931 | -2.151041553 | 0.03147292 | 0.111947448 | 0.174494493 |
| ENSMUSG00000015143 | Actn1 | 109711 | protein_coding | 1428.862931 | -0.174544951 | 0.094514269 | -1.846757672 | 0.064782269 | 0.185251702 | 0.174544951 |

| ensembl_gene_id | external_gene_name | entrezgene_id | gene_biotype | baseMean | log2FoldChange | lfcSE | stat | pvalue | padj | absLog2FC |
|---|---|---|---|---|---|---|---|---|---|---|
| ENSMUSG00000033845 | Mrpl15 | 27395 | protein_coding | 806.4623008 | 0.174569686 | 0.084881374 | 2.056631259 | 0.039721709 | 0.132481904 | 0.174569686 |
| ENSMUSG00000035139 | Secisbp2 | 75420 | protein_coding | 281.0660831 | -0.174608681 | 0.123284095 | -1.416311504 | 0.156684321 | 0.337343589 | 0.174608681 |
| ENSMUSG00000006050 | Sra1 | 24068 | protein_coding | 1058.188107 | 0.174620485 | 0.095777809 | 1.823183124 | 0.06827566 | 0.192272757 | 0.174620485 |
| ENSMUSG00000024313 | Mep1b | 17288 | protein_coding | 1283.479268 | 0.174660269 | 0.135241148 | 1.291472841 | 0.196539767 | 0.390160765 | 0.174660269 |
| ENSMUSG00000030588 | Yif1b | 77254 | protein_coding | 258.4753655 | 0.174675049 | 0.140196035 | 1.245934302 | 0.212788516 | 0.410823694 | 0.174675049 |
| ENSMUSG00000055943 | Emc7 | 73024 | protein_coding | 811.0058487 | 0.174848061 | 0.07863406 | 2.223566484 | 0.026177626 | 0.097739621 | 0.174848061 |
| ENSMUSG00000011832 | Evi5l | 213027 | protein_coding | 161.1703923 | 0.174850021 | 0.12180736 | 1.43546352 | 0.151155057 | 0.329311687 | 0.174850021 |
| ENSMUSG00000087330 | Bloodlinc | 105463053 | lncRNA | 9.355238384 | 0.174857135 | 0.450961367 | 0.387743048 | 0.698206195 | 0.828258604 | 0.174857135 |
| ENSMUSG00000027651 | Rprd1b | 70470 | protein_coding | 461.6132629 | -0.174940394 | 0.070890783 | -2.467745256 | 0.013596704 | 0.060191302 | 0.174940394 |
| ENSMUSG00000009356 | Lpo | 76113 | protein_coding | 333.7357141 | -0.174944885 | 0.171442754 | -1.020427406 | 0.307525802 | 0.51745943 | 0.174944885 |
| ENSMUSG00000020646 | Mboat2 | 67216 | protein_coding | 216.3933248 | -0.174969661 | 0.192231242 | -0.910204081 | 0.362714893 | 0.57178571 | 0.174969661 |
| ENSMUSG00000026383 | Epb41l5 | 226352 | protein_coding | 339.3190528 | -0.175000229 | 0.101691864 | -1.720887225 | 0.08527129 | 0.223632151 | 0.175000229 |
| ENSMUSG00000034867 | Ankrd27 | 245886 | protein_coding | 453.7103822 | -0.175068765 | 0.085854303 | -2.039137921 | 0.041436264 | 0.136250376 | 0.175068765 |
| ENSMUSG00000029484 | Anxa3 | 11745 | protein_coding | 1333.740092 | -0.175084608 | 0.188685912 | -0.927915638 | 0.353451332 | 0.563202563 | 0.175084608 |
| ENSMUSG00000048087 | Ahcyl | 11615 | protein_coding | 13.37095037 | 0.175107192 | 0.342901028 | 0.510663947 | 0.60958639 | 0.772618129 | 0.175107192 |
| ENSMUSG00000045968 | Teddm2 | 240817 | protein_coding | 15.4752188 | -0.175161942 | 0.30038864 | -0.58311773 | 0.559814049 | 0.73771144 | 0.175161942 |
| ENSMUSG00000029507 | Pus1 | 56361 | protein_coding | 309.325692 | -0.175261366 | 0.171435786 | -1.022314941 | 0.306631864 | 0.516664797 | 0.175261366 |
| ENSMUSG00000030978 | Rrm1 | 20133 | protein_coding | 927.6471361 | 0.175297653 | 0.122790146 | 1.427619874 | 0.1534013 | 0.33241039 | 0.175297653 |
| ENSMUSG00000058921 | Slc10a5 | 241877 | protein_coding | 990.0454763 | 0.175419102 | 0.093126377 | 1.883667196 | 0.059610006 | 0.175340384 | 0.175419102 |
| ENSMUSG00000048234 | Rnf149 | 67702 | protein_coding | 680.4253402 | -0.175428992 | 0.097802409 | -1.79370829 | 0.072859741 | 0.200728816 | 0.175428992 |
| ENSMUSG00000029708 | Gcc1 | 74375 | protein_coding | 434.7050769 | -0.175509699 | 0.085986567 | -2.04112927 | 0.04123798 | 0.135935128 | 0.175509699 |
| ENSMUSG00000021598 | Med10 | 28077 | protein_coding | 314.8569495 | 0.175527675 | 0.11387228 | 1.541443407 | 0.123208906 | 0.286241462 | 0.175527675 |
| ENSMUSG00000043003 | Rasef | 242505 | protein_coding | 2409.23384 | 0.175648045 | 0.093535649 | 1.877872747 | 0.060398575 | 0.176550129 | 0.175648045 |
| ENSMUSG00000025747 | Tyms | 22171 | protein_coding | 483.849816 | 0.175689917 | 0.089048129 | 1.97297707 | 0.048498175 | 0.152534832 | 0.175689917 |
| ENSMUSG00000056917 | Sipa1 | 20469 | protein_coding | 58.79516515 | -0.175713724 | 0.179611889 | -0.978296731 | 0.327927586 | 0.538810666 | 0.175713724 |
| ENSMUSG00000054648 | Zfp869 | 66869 | protein_coding | 610.405456 | 0.175720429 | 0.090060234 | 1.95114338 | 0.051039992 | 0.157643008 | 0.175720429 |
| ENSMUSG00000044730 | 9930104L06Rik | 194268 | protein_coding | 113.7646959 | -0.175725578 | 0.126898976 | -1.384767503 | 0.166123571 | 0.349909579 | 0.175725578 |
| ENSMUSG00000034160 | Ogt | 108155 | protein_coding | 1470.979576 | -0.175754135 | 0.108860686 | -1.614486741 | 0.106421891 | 0.258831558 | 0.175754135 |
| ENSMUSG00000034773 | Hrob | 217216 | protein_coding | 30.38604351 | -0.175778815 | 0.250887044 | -0.700629307 | 0.483534384 | 0.676897141 | 0.175778815 |
| ENSMUSG00000027160 | Ccdc34 | 68201 | protein_coding | 326.7997182 | 0.175883291 | 0.153643325 | 1.14475062 | 0.252312476 | 0.458561515 | 0.175883291 |
| ENSMUSG00000025358 | Cdk2 | 12566 | protein_coding | 285.8113926 | -0.17591651 | 0.12994831 | -1.353742186 | 0.175818644 | 0.361907388 | 0.17591651 |
| ENSMUSG00000038705 | Gmeb2 | 229004 | protein_coding | 381.6234916 | -0.176012401 | 0.104728326 | -1.680657059 | 0.092829546 | 0.236839501 | 0.176012401 |
| ENSMUSG00000096215 | Smim22 | 432995 | protein_coding | 260.761336 | -0.176022263 | 0.101861282 | -1.728058585 | 0.083977721 | 0.221383176 | 0.176022263 |
| ENSMUSG00000056216 | Cebpg | 12611 | protein_coding | 1601.028468 | 0.17605787 | 0.065851362 | 2.673564577 | 0.007504983 | 0.038548649 | 0.17605787 |
| ENSMUSG00000028331 | Trmo | 74753 | protein_coding | 76.86930035 | -0.176076185 | 0.166073795 | -1.060228583 | 0.28904062 | 0.497873797 | 0.176076185 |
| ENSMUSG00000025470 | Zfp511 | 69752 | protein_coding | 139.3322728 | -0.17609001 | 0.129871528 | -1.35587848 | 0.175137835 | 0.361418538 | 0.17609001 |
| ENSMUSG00000055210 | Foxd2 | 17301 | protein_coding | 696.086822 | -0.176212994 | 0.154053896 | -1.143839906 | 0.252690037 | 0.459036748 | 0.176212994 |
| ENSMUSG00000031750 | Il34 | 76527 | protein_coding | 28.29973503 | -0.176213797 | 0.286925663 | -0.61414443 | 0.539119891 | 0.722272825 | 0.176213797 |

| ensembl_gene_id | external_gene_name | entrezgene_id | gene_biotype | baseMean | log2FoldChange | lfcSE | stat | pvalue | padj | absLog2FC |
|---|---|---|---|---|---|---|---|---|---|---|
| ENSMUSG00000041219 | Arhgap11a | 228482 | protein_coding | 589.4905553 | 0.176280503 | 0.109448398 | 1.61062662 | 0.107261128 | 0.260276546 | 0.176280503 |
| ENSMUSG00000027907 | S100a11 | 20195 | protein_coding | 389.4379251 | 0.176406879 | 0.214955873 | 0.820665547 | 0.4118368 | 0.61872633 | 0.176406879 |
| ENSMUSG00000035890 | Rnf126 | 70294 | protein_coding | 293.0213508 | 0.17642926 | 0.111555267 | 1.581541274 | 0.11375433 | 0.269866479 | 0.17642926 |
| ENSMUSG00000032041 | Tirap | 117149 | protein_coding | 291.8966738 | 0.176491941 | 0.171902735 | 1.026696527 | 0.304563385 | 0.514104027 | 0.176491941 |
| ENSMUSG00000039242 | B3galnt2 | 97884 | protein_coding | 480.0057799 | 0.176517574 | 0.070908527 | 2.489370212 | 0.012796963 | 0.057627942 | 0.176517574 |
| ENSMUSG00000069208 | Zfp825 | 235956 | protein_coding | 110.1521266 | 0.17652404 | 0.163527458 | 1.079476453 | 0.280375385 | 0.488708594 | 0.17652404 |
| ENSMUSG00000042447 | Mios | 252875 | protein_coding | 297.0917259 | -0.176542519 | 0.086522624 | -2.040420293 | 0.041308482 | 0.136047522 | 0.176542519 |
| ENSMUSG00000042275 | Pelo | 105083 | protein_coding | 40.10084027 | 0.176545444 | 0.19807282 | 0.89131585 | 0.372759748 | 0.580950809 | 0.176545444 |
| ENSMUSG00000022401 | Xpnpep3 | 321003 | protein_coding | 230.227778 | -0.176562246 | 0.091916615 | -1.920895868 | 0.054744837 | 0.165037012 | 0.176562246 |
| ENSMUSG00000043391 | 2510009E07Rik | 72190 | protein_coding | 468.0150804 | -0.176690116 | 0.137402703 | -1.285928969 | 0.198467852 | 0.392646374 | 0.176690116 |
| ENSMUSG00000051319 | Mtln | 67885 | protein_coding | 150.7467836 | 0.17676336 | 0.099529556 | 1.775988624 | 0.075734792 | 0.205338463 | 0.17676336 |
| ENSMUSG00000021678 | F2rl1 | 14063 | protein_coding | 1176.031744 | 0.176788402 | 0.103624213 | 1.706053023 | 0.087998208 | 0.22853994 | 0.176788402 |
| ENSMUSG00000073684 | Faap20 | 67513 | protein_coding | 156.4551448 | -0.176825735 | 0.106568077 | -1.659274888 | 0.097060413 | 0.243614601 | 0.176825735 |
| ENSMUSG00000032666 | 1700025G04Rik | 69399 | protein_coding | 260.7181336 | -0.176840278 | 0.251068595 | -0.704350451 | 0.481214555 | 0.67440173 | 0.176840278 |
| ENSMUSG00000024225 | Clps | 109791 | protein_coding | 111.6748217 | -0.176889797 | 0.177964265 | -0.993962455 | 0.320241146 | 0.531665726 | 0.176889797 |
| ENSMUSG00000019470 | Xab2 | 67439 | protein_coding | 532.1427305 | -0.176900752 | 0.103965731 | -1.701529437 | 0.088843615 | 0.229985552 | 0.176900752 |
| ENSMUSG00000071537 | Klrg2 | 74253 | protein_coding | 7.511539182 | 0.1769242 | 0.541881182 | 0.326499988 | 0.744046098 | 0.857661474 | 0.1769242 |
| ENSMUSG00000022346 | Myc | 17869 | protein_coding | 276.6383576 | -0.176949049 | 0.22928893 | -0.771729575 | 0.44027461 | 0.642313808 | 0.176949049 |
| ENSMUSG00000022283 | Pabpc1 | 18458 | protein_coding | 4647.671685 | -0.176950422 | 0.116438004 | -1.519696457 | 0.128587283 | 0.294780978 | 0.176950422 |
| ENSMUSG00000030521 | Mphosph10 | 67973 | protein_coding | 390.2014142 | -0.177025088 | 0.181757695 | -0.973961998 | 0.330075408 | 0.540678415 | 0.177025088 |
| ENSMUSG00000017740 | Slc12a5 | 57138 | protein_coding | 12.81920097 | -0.177090883 | 0.599945545 | -0.295178261 | 0.767857708 | 0.873415523 | 0.177090883 |
| ENSMUSG00000035047 | Kri1 | 215194 | protein_coding | 178.2588806 | -0.177147692 | 0.11464572 | -1.54517493 | 0.122303957 | 0.284928685 | 0.177147692 |
| ENSMUSG00000018126 | Baiap2l2 | 207495 | protein_coding | 655.7881139 | -0.177183318 | 0.13728902 | -1.290586222 | 0.196847196 | 0.390507179 | 0.177183318 |
| ENSMUSG00000027230 | Creb3l1 | 26427 | protein_coding | 1036.642314 | -0.177225218 | 0.113861971 | -1.556491747 | 0.119591204 | 0.280496052 | 0.177225218 |
| ENSMUSG00000019173 | Rab5c | 19345 | protein_coding | 1719.55985 | 0.177276397 | 0.071179092 | 2.49056841 | 0.012753894 | 0.057466485 | 0.177276397 |
| ENSMUSG00000042626 | Shc1 | 20416 | protein_coding | 1067.3173 | 0.17746392 | 0.10842999 | 1.636668232 | 0.10169981 | 0.250948811 | 0.17746392 |
| ENSMUSG00000049299 | Trappc1 | 245828 | protein_coding | 380.2839962 | 0.177500304 | 0.081329492 | 2.18248386 | 0.029073841 | 0.105343072 | 0.177500304 |
| ENSMUSG00000095677 | Dynlt1f | 100040531 | protein_coding | 98.65833299 | 0.177534869 | 0.140056474 | 1.267594877 | 0.204942662 | 0.400678375 | 0.177534869 |
| ENSMUSG00000025858 | Get4 | 67604 | protein_coding | 270.4280228 | 0.17754638 | 0.100479375 | 1.766993266 | 0.077229356 | 0.208563781 | 0.17754638 |
| ENSMUSG00000032085 | Tagln | 21345 | protein_coding | 28.49601045 | -0.177569032 | 0.220591447 | -0.804967892 | 0.420838209 | 0.626693249 | 0.177569032 |
| ENSMUSG00000036661 | Dennd3 | 105841 | protein_coding | 163.1343196 | -0.177591331 | 0.135304282 | -1.312532961 | 0.189340372 | 0.380880708 | 0.177591331 |
| ENSMUSG00000042520 | Ubap2l | 74383 | protein_coding | 910.1868272 | -0.177619493 | 0.072408851 | -2.453008047 | 0.014166717 | 0.061903744 | 0.177619493 |
| ENSMUSG00000067150 | Xpo5 | 72322 | protein_coding | 595.8972064 | -0.17763773 | 0.099135866 | -1.791861385 | 0.073155173 | 0.201211465 | 0.17763773 |
| ENSMUSG00000041623 | D11Wsu47e | 276852 | protein_coding | 115.4019536 | -0.177681852 | 0.14122647 | -1.258134202 | 0.208343226 | 0.405039778 | 0.177681852 |
| ENSMUSG00000049401 | Ogfr | 72075 | protein_coding | 679.9730234 | 0.177687599 | 0.088435299 | 2.009238412 | 0.044511856 | 0.143287155 | 0.177687599 |
| ENSMUSG00000002222 | Rmnd5a | 68477 | protein_coding | 3146.012487 | 0.177724892 | 0.069662119 | 2.551241562 | 0.01073399 | 0.050463693 | 0.177724892 |
| ENSMUSG00000074865 | Zfp934 | 77117 | protein_coding | 75.22253629 | -0.177799076 | 0.197920029 | -0.898337966 | 0.369005398 | 0.577870217 | 0.177799076 |
| ENSMUSG00000024369 | Nelfe | 27632 | protein_coding | 311.2439933 | 0.177837898 | 0.109106431 | 1.629948814 | 0.103112316 | 0.253408395 | 0.177837898 |

| ensembl_gene_id | external_gene_name | entrezgene_id | gene_biotype | baseMean | log2FoldChange | lfcSE | stat | pvalue | padj | absLog2FC |
|---|---|---|---|---|---|---|---|---|---|---|
| ENSMUSG00000027088 | Phospho2 | 73373 | protein_coding | 446.3376638 | 0.177870965 | 0.094717685 | 1.877906587 | 0.060393945 | 0.176550129 | 0.177870965 |
| ENSMUSG00000056394 | Lig1 | 16881 | protein_coding | 358.8786089 | -0.17790743 | 0.127850449 | -1.391527607 | 0.164065494 | 0.347227636 | 0.17790743 |
| ENSMUSG00000035726 | Supt16 | 114741 | protein_coding | 1477.516852 | -0.177926146 | 0.084541177 | -2.104609295 | 0.035325332 | 0.121841032 | 0.177926146 |
| ENSMUSG00000032621 | Srek1 | 218543 | protein_coding | 684.175281 | -0.177927036 | 0.113191812 | -1.571907302 | 0.115972051 | 0.273862539 | 0.177927036 |
| ENSMUSG00000020905 | Usp43 | 216835 | protein_coding | 425.9675101 | -0.17801012 | 0.119981036 | -1.483652125 | 0.137901233 | 0.308886351 | 0.17801012 |
| ENSMUSG00000028668 | Eloa | 27224 | protein_coding | 1071.185918 | -0.178052264 | 0.077518167 | -2.296910144 | 0.021623896 | 0.08462541 | 0.178052264 |
| ENSMUSG00000038323 | 1700066M21Rik | 73467 | protein_coding | 177.3884565 | 0.178063977 | 0.105994543 | 1.679935322 | 0.0929699 | 0.237045714 | 0.178063977 |
| ENSMUSG00000021012 | Zc3h14 | 75553 | protein_coding | 1095.252704 | 0.178090661 | 0.056184952 | 3.169721706 | 0.00152585 | 0.010806465 | 0.178090661 |
| ENSMUSG00000027502 | Rtf2 | 66404 | protein_coding | 963.9097431 | 0.178112514 | 0.092388921 | 1.927855772 | 0.053873068 | 0.16307302 | 0.178112514 |
| ENSMUSG00000026181 | Ppm1f | 68606 | protein_coding | 113.0541764 | -0.178131398 | 0.125745157 | -1.416606432 | 0.156598027 | 0.337237351 | 0.178131398 |
| ENSMUSG00000032462 | Pik3cb | 74769 | protein_coding | 213.3122629 | -0.178226018 | 0.103774003 | -1.7174438 | 0.085898115 | 0.224799607 | 0.178226018 |
| ENSMUSG00000049287 | Iba57 | 216792 | protein_coding | 247.5565146 | -0.178246989 | 0.111043399 | -1.605201122 | 0.108449551 | 0.262119878 | 0.178246989 |
| ENSMUSG00000104011 | Gm32391 | 102634922 | lncRNA | 14.21675597 | -0.17827616 | 0.320679111 | -0.555933186 | 0.578256531 | 0.749855086 | 0.17827616 |
| ENSMUSG00000024614 | Tmx3 | 67988 | protein_coding | 908.0386235 | 0.178306361 | 0.091940863 | 1.939359231 | 0.052457611 | 0.160434875 | 0.178306361 |
| ENSMUSG00000024561 | Mbd1 | 17190 | protein_coding | 837.5438528 | -0.178306986 | 0.084718824 | -2.10469148 | 0.035318173 | 0.121841032 | 0.178306986 |
| ENSMUSG00000002565 | Scin | 20259 | protein_coding | 2744.685063 | -0.178429624 | 0.128772089 | -1.385623431 | 0.16586192 | 0.349682152 | 0.178429624 |
| ENSMUSG00000013653 | 1810065E05Rik | 69864 | protein_coding | 14398.44956 | 0.178576153 | 0.331283457 | 0.539043376 | 0.589856925 | 0.759518278 | 0.178576153 |
| ENSMUSG00000028470 | Hint2 | 68917 | protein_coding | 334.3536963 | 0.178633515 | 0.127202589 | 1.404322951 | 0.160222703 | 0.341809566 | 0.178633515 |
| ENSMUSG00000026273 | Mterf4 | 69821 | protein_coding | 214.8541867 | 0.178664672 | 0.111688374 | 1.599671174 | 0.10967155 | 0.263874388 | 0.178664672 |
| ENSMUSG00000049288 | Lix1l | 280411 | protein_coding | 8.990063211 | 0.178679568 | 0.545741939 | 0.327406702 | 0.743360298 | 0.857217273 | 0.178679568 |
| ENSMUSG00000034566 | Atp5h | 71679 | protein_coding | 3054.709906 | 0.178740823 | 0.093411035 | 1.913487238 | 0.055685706 | 0.167162066 | 0.178740823 |
| ENSMUSG00000035517 | Tdrd7 | 100121 | protein_coding | 839.0596377 | 0.178835655 | 0.095838219 | 1.866016041 | 0.062039121 | 0.179697968 | 0.178835655 |
| ENSMUSG00000032278 | Paqr5 | 74090 | protein_coding | 858.9204124 | -0.178878484 | 0.129487566 | -1.381433677 | 0.167145657 | 0.351303286 | 0.178878484 |
| ENSMUSG00000020021 | Fgd6 | 13998 | protein_coding | 382.6124781 | -0.178890935 | 0.171554101 | -1.042766884 | 0.297056272 | 0.506180721 | 0.178890935 |
| ENSMUSG00000052428 | Tmco1 | 68944 | protein_coding | 890.3086515 | 0.178896099 | 0.081098249 | 2.205918149 | 0.027389731 | 0.100800286 | 0.178896099 |
| ENSMUSG00000060681 | Slc9a6 | 236794 | protein_coding | 358.7817239 | 0.179030389 | 0.089792527 | 1.993822813 | 0.046171422 | 0.147230912 | 0.179030389 |
| ENSMUSG00000036568 | Bicral | 210982 | protein_coding | 332.2030066 | -0.179086073 | 0.100324293 | -1.785071871 | 0.07424966 | 0.203069209 | 0.179086073 |
| ENSMUSG00000021978 | Extl3 | 54616 | protein_coding | 558.4141477 | -0.179116053 | 0.08365556 | -2.14111354 | 0.032264883 | 0.113823933 | 0.179116053 |
| ENSMUSG00000032348 | Gsta4 | 14860 | protein_coding | 686.8491699 | -0.179119308 | 0.100781038 | -1.777311607 | 0.075516985 | 0.205186208 | 0.179119308 |
| ENSMUSG00000015597 | Zfp318 | 57908 | protein_coding | 294.1219874 | -0.179157302 | 0.116540798 | -1.53729256 | 0.124221683 | 0.287742088 | 0.179157302 |
| ENSMUSG00000031785 | Adgrg1 | 14766 | protein_coding | 623.0973854 | 0.179167612 | 0.098226529 | 1.824024672 | 0.068148342 | 0.191982101 | 0.179167612 |
| ENSMUSG00000093803 | Ppp2r3d | 19054 | protein_coding | 262.8160795 | -0.179193125 | 0.121070687 | -1.480070274 | 0.138854493 | 0.31014926 | 0.179193125 |
| ENSMUSG00000027654 | Fam83d | 71878 | protein_coding | 202.2570593 | 0.17919935 | 0.120805252 | 1.483373833 | 0.137975116 | 0.309008386 | 0.17919935 |
| ENSMUSG00000026848 | Tor1b | 30934 | protein_coding | 527.5977727 | -0.179243407 | 0.083276567 | -2.152387067 | 0.031366881 | 0.111737326 | 0.179243407 |
| ENSMUSG00000001017 | Chtop | 66511 | protein_coding | 1361.664116 | 0.179250088 | 0.092764412 | 1.932315263 | 0.053320606 | 0.162109297 | 0.179250088 |
| ENSMUSG00000061079 | Zfp143 | 20841 | protein_coding | 268.1458795 | 0.179289445 | 0.130437943 | 1.374519109 | 0.169280582 | 0.354093686 | 0.179289445 |
| ENSMUSG00000037553 | Zdhhc18 | 503610 | protein_coding | 298.0429976 | -0.179359772 | 0.142214577 | -1.261191197 | 0.207239969 | 0.403733701 | 0.179359772 |
| ENSMUSG00000008683 | Rps15a | 267019 | protein_coding | 5915.482544 | 0.1793926 | 0.089906885 | 1.99531548 | 0.046008483 | 0.146917224 | 0.1793926 |

| ensembl_gene_id | external_gene_name | entrezgene_id | gene_biotype | baseMean | log2FoldChange | lfcSE | stat | pvalue | padj | absLog2FC |
|---|---|---|---|---|---|---|---|---|---|---|
| ENSMUSG00000038225 | Primpol | 408022 | protein_coding | 162.0587698 | -0.179396096 | 0.134114405 | -1.33763481 | 0.181015511 | 0.369358835 | 0.179396096 |
| ENSMUSG00000053289 | Ddx10 | 77591 | protein_coding | 635.1408152 | 0.179415378 | 0.113276654 | 1.583868984 | 0.113223537 | 0.269195376 | 0.179415378 |
| ENSMUSG00000073468 | Sft2d1 | 106489 | protein_coding | 311.815146 | 0.179465678 | 0.109998489 | 1.631528568 | 0.102778837 | 0.252945197 | 0.179465678 |
| ENSMUSG00000030660 | Pik3c2a | 18704 | protein_coding | 2339.586118 | 0.179479511 | 0.06769672 | 2.65122907 | 0.008019943 | 0.040591563 | 0.179479511 |
| ENSMUSG00000029923 | Rab19 | 19331 | protein_coding | 130.47887 | 0.179492405 | 0.22280766 | 0.805593509 | 0.420477269 | 0.626390006 | 0.179492405 |
| ENSMUSG00000021830 | Txndc16 | 70561 | protein_coding | 458.139906 | -0.179586339 | 0.155827645 | -1.152467773 | 0.249128905 | 0.45537356 | 0.179586339 |
| ENSMUSG00000027905 | Ddx20 | 53975 | protein_coding | 290.1528378 | -0.17963214 | 0.099805317 | -1.799825362 | 0.071888218 | 0.199136615 | 0.17963214 |
| ENSMUSG00000021811 | Dnajc9 | 108671 | protein_coding | 292.2534925 | 0.179665914 | 0.103621078 | 1.733874211 | 0.082940405 | 0.219554789 | 0.179665914 |
| ENSMUSG00000110754 | 4930405J17Rik | 73811 | lncRNA | 15.1324435 | 0.179722238 | 0.318435382 | 0.564391547 | 0.57248769 | 0.746198626 | 0.179722238 |
| ENSMUSG00000032988 | Slc16a8 | 57274 | protein_coding | 18.32224909 | -0.179740329 | 0.32575173 | -0.551770912 | 0.581105319 | 0.75217968 | 0.179740329 |
| ENSMUSG00000055447 | Cd47 | 16423 | protein_coding | 484.9939445 | 0.179781328 | 0.134079519 | 1.340855999 | 0.17996721 | 0.368002072 | 0.179781328 |
| ENSMUSG00000006720 | Zfp184 | 193452 | protein_coding | 19.65414828 | -0.179824253 | 0.254301109 | -0.707131218 | 0.479484937 | 0.673299487 | 0.179824253 |
| ENSMUSG00000029551 | Psmg3 | 66506 | protein_coding | 101.8546164 | 0.17985245 | 0.140968723 | 1.275832298 | 0.202014808 | 0.397018359 | 0.17985245 |
| ENSMUSG00000102543 | Pcdhgc5 | 93708 | protein_coding | 18.16570949 | -0.179869542 | 0.327797519 | -0.548721486 | 0.583196599 | 0.754056602 | 0.179869542 |
| ENSMUSG00000027582 | Zgpat | 229007 | protein_coding | 368.0963693 | -0.179894278 | 0.100065804 | -1.797759768 | 0.072215085 | 0.19967979 | 0.179894278 |
| ENSMUSG00000033918 | Parl | 381038 | protein_coding | 511.1492909 | -0.179913649 | 0.087627984 | -2.053152904 | 0.04005775 | 0.133272807 | 0.179913649 |
| ENSMUSG00000023328 | Ache | 11423 | protein_coding | 46.14302373 | -0.17993572 | 0.214498069 | -0.838868717 | 0.401542984 | 0.60912358 | 0.17993572 |
| ENSMUSG00000018377 | Vezf1 | 22344 | protein_coding | 640.4169601 | -0.179991976 | 0.088158967 | -2.041675199 | 0.04118376 | 0.135812616 | 0.179991976 |
| ENSMUSG00000030678 | Maz | 17188 | protein_coding | 628.4747165 | -0.179994672 | 0.071636162 | -2.512623068 | 0.01198373 | 0.054849669 | 0.179994672 |
| ENSMUSG00000020817 | Rabep1 | 54189 | protein_coding | 684.9360641 | -0.179994965 | 0.109055044 | -1.650496469 | 0.098841435 | 0.246453722 | 0.179994965 |
| ENSMUSG00000025940 | Tmem70 | 70397 | protein_coding | 558.3072824 | 0.180002501 | 0.098973592 | 1.818692208 | 0.068958401 | 0.193613505 | 0.180002501 |
| ENSMUSG00000060373 | Hnrnpc | 15381 | protein_coding | 2622.454814 | 0.180005861 | 0.070665996 | 2.547276926 | 0.010856724 | 0.050860559 | 0.180005861 |
| ENSMUSG00000029263 | Pigg | 433931 | protein_coding | 416.3407361 | 0.180037201 | 0.073983932 | 2.433463532 | 0.014955142 | 0.064519366 | 0.180037201 |
| ENSMUSG00000079043 | Fastkd5 | 380601 | protein_coding | 109.3992943 | -0.180038687 | 0.12378454 | -1.454452126 | 0.145820997 | 0.321650304 | 0.180038687 |
| ENSMUSG00000037286 | Stag1 | 20842 | protein_coding | 1045.895628 | 0.180057261 | 0.05226624 | 3.445001243 | 0.000571057 | 0.00484592 | 0.180057261 |
| ENSMUSG00000002718 | Cse1l | 110750 | protein_coding | 991.5828676 | 0.18006435 | 0.105573678 | 1.705579963 | 0.088086313 | 0.228545168 | 0.18006435 |
| ENSMUSG00000004947 | Dtx2 | 74198 | protein_coding | 98.06794852 | -0.180068171 | 0.146826457 | -1.226401394 | 0.220047662 | 0.420405321 | 0.180068171 |
| ENSMUSG00000030075 | Cntn3 | 18488 | protein_coding | 28.44379158 | -0.180082187 | 0.349979002 | -0.514551404 | 0.606866531 | 0.771131144 | 0.180082187 |
| ENSMUSG00000037885 | Stk35 | 67333 | protein_coding | 402.7100702 | -0.180110295 | 0.097844403 | -1.840782822 | 0.065653391 | 0.186770354 | 0.180110295 |
| ENSMUSG00000034724 | Cnot6l | 231464 | protein_coding | 782.0035463 | -0.180117968 | 0.148292335 | -1.214614142 | 0.224513293 | 0.426176585 | 0.180117968 |
| ENSMUSG00000039981 | Zc3h12d | 237256 | protein_coding | 248.1485678 | 0.180121067 | 0.117544942 | 1.532359144 | 0.125433834 | 0.289636141 | 0.180121067 |
| ENSMUSG00000038812 | Trmt112 | 67674 | protein_coding | 118.3654888 | -0.180133111 | 0.129659838 | -1.389274539 | 0.164749283 | 0.348165926 | 0.180133111 |
| ENSMUSG00000042185 | Nfrkb | 235134 | protein_coding | 369.7491379 | -0.180180773 | 0.079323081 | -2.271479765 | 0.023117951 | 0.08913646 | 0.180180773 |
| ENSMUSG00000028973 | Abcb8 | 74610 | protein_coding | 831.6256633 | 0.180301432 | 0.092798952 | 1.942925304 | 0.052025184 | 0.159787338 | 0.180301432 |
| ENSMUSG00000022489 | Pde1b | 18574 | protein_coding | 86.10512889 | -0.180309188 | 0.224887832 | -0.801773873 | 0.422683774 | 0.627750586 | 0.180309188 |
| ENSMUSG00000034906 | Ncaph | 215387 | protein_coding | 361.2987487 | 0.180318074 | 0.140703908 | 1.281542756 | 0.200003092 | 0.394660109 | 0.180318074 |
| ENSMUSG00000016308 | Ube2a | 22209 | protein_coding | 502.7019939 | 0.180320322 | 0.081038144 | 2.225128989 | 0.02607258 | 0.097509572 | 0.180320322 |
| ENSMUSG00000048351 | Coa7 | 69893 | protein_coding | 220.0629268 | -0.180340202 | 0.107673326 | -1.674882797 | 0.09395723 | 0.23876049 | 0.180340202 |

| ensembl_gene_id | external_gene_name | entrezgene_id | gene_biotype | baseMean | log2FoldChange | lfcSE | stat | pvalue | padj | absLog2FC |
|---|---|---|---|---|---|---|---|---|---|---|
| ENSMUSG00000002395 | Use1 | 67023 | protein_coding | 822.0199753 | 0.180524241 | 0.107149318 | 1.684791315 | 0.092028841 | 0.235362136 | 0.180524241 |
| ENSMUSG00000015337 | Endog | 13804 | protein_coding | 105.5550151 | 0.180559783 | 0.191820434 | 0.941295873 | 0.346553257 | 0.55660989 | 0.180559783 |
| ENSMUSG00000001018 | Snapin | 20615 | protein_coding | 270.6664405 | -0.180561607 | 0.108995893 | -1.656590927 | 0.097602208 | 0.24431997 | 0.180561607 |
| ENSMUSG00000031608 | Galnt7 | 108150 | protein_coding | 4913.446739 | 0.180570335 | 0.062204374 | 2.902855909 | 0.003697767 | 0.022100577 | 0.180570335 |
| ENSMUSG00000031171 | Ftsj1 | 54632 | protein_coding | 217.9730732 | 0.180586452 | 0.112660293 | 1.602929011 | 0.108950328 | 0.262893625 | 0.180586452 |
| ENSMUSG00000039988 | Ankrd13c | 433667 | protein_coding | 1355.711678 | 0.180599992 | 0.067517796 | 2.674850224 | 0.007476264 | 0.038418309 | 0.180599992 |
| ENSMUSG00000042426 | Dhx29 | 218629 | protein_coding | 288.9067647 | 0.180624018 | 0.157706002 | 1.14532114 | 0.252076152 | 0.458393532 | 0.180624018 |
| ENSMUSG00000051579 | Tceal8 | 66684 | protein_coding | 321.6141792 | 0.18062558 | 0.119593841 | 1.5103251 | 0.13096049 | 0.29828953 | 0.18062558 |
| ENSMUSG00000058192 | Zfp846 | 244721 | protein_coding | 159.775592 | 0.180686923 | 0.137644618 | 1.312706057 | 0.189282016 | 0.380811412 | 0.180686923 |
| ENSMUSG00000044573 | Acp1 | 11431 | protein_coding | 800.0839368 | 0.180754097 | 0.076351864 | 2.367382894 | 0.017914388 | 0.073712317 | 0.180754097 |
| ENSMUSG00000024217 | Snrpc | 20630 | protein_coding | 290.8594695 | 0.180778451 | 0.128447811 | 1.407407792 | 0.159306502 | 0.340451304 | 0.180778451 |
| ENSMUSG00000054469 | Lclat1 | 225010 | protein_coding | 1307.206574 | 0.180832998 | 0.098324623 | 1.839142544 | 0.065894223 | 0.187321646 | 0.180832998 |
| ENSMUSG00000026785 | Pkn3 | 263803 | protein_coding | 94.81641489 | -0.180898536 | 0.168345923 | -1.074564398 | 0.282569798 | 0.491565285 | 0.180898536 |
| ENSMUSG00000035451 | Foxa1 | 15375 | protein_coding | 625.3842078 | -0.180934131 | 0.115788478 | -1.562626387 | 0.118140498 | 0.277994069 | 0.180934131 |
| ENSMUSG00000027881 | Prpf38b | 66921 | protein_coding | 1140.909806 | 0.180949393 | 0.115860112 | 1.561791977 | 0.118337002 | 0.278369656 | 0.180949393 |
| ENSMUSG00000022235 | Cmbl | 69574 | protein_coding | 676.6484321 | 0.180972579 | 0.145265737 | 1.245803603 | 0.212836507 | 0.410824034 | 0.180972579 |
| ENSMUSG00000038975 | Rabggtb | 19352 | protein_coding | 679.5101721 | -0.181051815 | 0.084528059 | -2.1419138 | 0.032200419 | 0.113672047 | 0.181051815 |
| ENSMUSG00000010914 | Pdhx | 27402 | protein_coding | 449.8880656 | 0.181054638 | 0.090404612 | 2.00271462 | 0.045207929 | 0.14514652 | 0.181054638 |
| ENSMUSG00000005981 | Trap1 | 68015 | protein_coding | 1445.923609 | -0.181123696 | 0.067211188 | -2.694844442 | 0.00704215 | 0.036625014 | 0.181123696 |
| ENSMUSG00000029413 | Naaa | 67111 | protein_coding | 222.3531765 | 0.181135486 | 0.146753879 | 1.234280744 | 0.217098309 | 0.41646897 | 0.181135486 |
| ENSMUSG00000016487 | Ppfibp1 | 67533 | protein_coding | 478.1615945 | 0.181316731 | 0.124053798 | 1.461597579 | 0.143851518 | 0.318044345 | 0.181316731 |
| ENSMUSG00000036469 | Marchf1 | 72925 | protein_coding | 7.413037037 | -0.181341158 | 0.562188296 | -0.322563025 | 0.7470262 | 0.859417357 | 0.181341158 |
| ENSMUSG00000022571 | Pycrl | 66194 | protein_coding | 606.9139031 | 0.181351182 | 0.115026009 | 1.576610233 | 0.114885237 | 0.271936836 | 0.181351182 |
| ENSMUSG00000032083 | Apoa1 | 11806 | protein_coding | 66.12362347 | -0.181368548 | 0.712745055 | -0.254464828 | 0.799136473 | 0.89211142 | 0.181368548 |
| ENSMUSG00000046070 | Igfals | 16005 | protein_coding | 17.90282218 | 0.18144039 | 0.281722458 | 0.644039498 | 0.519549822 | 0.706398276 | 0.18144039 |
| ENSMUSG00000073427 | Gm4924 | 237412 | protein_coding | 77.84176819 | -0.1814447 | 0.148942526 | -1.218219572 | 0.223140544 | 0.424532081 | 0.1814447 |
| ENSMUSG00000000787 | Ddx3x | 13205 | protein_coding | 6297.616183 | -0.18148206 | 0.125773933 | -1.442922678 | 0.149042254 | 0.326360327 | 0.18148206 |
| ENSMUSG00000019054 | Fis1 | 66437 | protein_coding | 868.6356811 | 0.181504835 | 0.109544032 | 1.65691213 | 0.097537242 | 0.244195722 | 0.181504835 |
| ENSMUSG00000019578 | Ubxn6 | 66530 | protein_coding | 189.7987236 | 0.181527048 | 0.139813744 | 1.298349098 | 0.194167402 | 0.387494219 | 0.181527048 |
| ENSMUSG00000045664 | Cdc42ep2 | 104252 | protein_coding | 148.759519 | 0.181536293 | 0.181749769 | 0.998825443 | 0.317879259 | 0.529178599 | 0.181536293 |
| ENSMUSG00000049038 | Mterf2 | 74238 | protein_coding | 87.5193086 | -0.181585995 | 0.184213085 | -0.985738856 | 0.324261277 | 0.535105016 | 0.181585995 |
| ENSMUSG00000051235 | Gen1 | 209334 | protein_coding | 135.5227341 | 0.18162983 | 0.135085304 | 1.344556548 | 0.178768483 | 0.366181122 | 0.18162983 |
| ENSMUSG00000042726 | Trafd1 | 231712 | protein_coding | 425.0344514 | 0.181678178 | 0.099668796 | 1.822819032 | 0.068330804 | 0.192394034 | 0.181678178 |
| ENSMUSG00000030435 | U2af2 | 22185 | protein_coding | 1511.578179 | 0.181707491 | 0.12315254 | 1.475466856 | 0.140087073 | 0.312201888 | 0.181707491 |
| ENSMUSG00000073599 | Ecscr | 68545 | protein_coding | 47.42982046 | -0.18172888 | 0.217102708 | -0.837064086 | 0.402556547 | 0.609921868 | 0.18172888 |
| ENSMUSG00000003153 | Slc2a3 | 20527 | protein_coding | 7.380548462 | -0.181759077 | 0.623706201 | -0.291417781 | 0.770731817 | 0.87556148 | 0.181759077 |
| ENSMUSG00000065582 | Mir194-2 | 723957 | miRNA | 9.320124105 | 0.181776399 | 0.496917555 | 0.365807965 | 0.714508376 | 0.839407687 | 0.181776399 |
| ENSMUSG00000028060 | Khdc4 | 74200 | protein_coding | 987.5202233 | -0.18182525 | 0.106893832 | -1.700989166 | 0.088945022 | 0.230173242 | 0.18182525 |

| ensembl_gene_id | external_gene_name | entrezgene_id | gene_biotype | baseMean | log2FoldChange | lfcSE | stat | pvalue | padj | absLog2FC |
|---|---|---|---|---|---|---|---|---|---|---|
| ENSMUSG00000020515 | Cnot8 | 69125 | protein_coding | 546.5111824 | 0.181826133 | 0.099919587 | 1.819724626 | 0.068800951 | 0.193307735 | 0.181826133 |
| ENSMUSG00000066640 | Fbxl18 | 231863 | protein_coding | 287.7298381 | -0.181828502 | 0.10449597 | -1.740052775 | 0.081849752 | 0.217573906 | 0.181828502 |
| ENSMUSG00000071054 | Safb | 224903 | protein_coding | 736.116918 | -0.181865355 | 0.10601339 | -1.715494192 | 0.086254659 | 0.225482392 | 0.181865355 |
| ENSMUSG00000042202 | Slc35e2 | 320541 | protein_coding | 484.3575983 | 0.18187212 | 0.083763011 | 2.171270098 | 0.029910761 | 0.107713906 | 0.18187212 |
| ENSMUSG00000071302 | 2610044O15Rik8 | 72139 | protein_coding | 200.2890979 | 0.181882756 | 0.157316456 | 1.156158483 | 0.247616338 | 0.453389244 | 0.181882756 |
| ENSMUSG00000054874 | Pcnx3 | 104401 | protein_coding | 809.4777082 | -0.181889823 | 0.114497537 | -1.588591579 | 0.112152631 | 0.267660638 | 0.181889823 |
| ENSMUSG00000024943 | Smc5 | 226026 | protein_coding | 479.7977189 | -0.182045458 | 0.081796245 | -2.225596765 | 0.026041203 | 0.097481618 | 0.182045458 |
| ENSMUSG00000036109 | Mbnl3 | 171170 | protein_coding | 463.0326277 | 0.182076535 | 0.107916226 | 1.687202579 | 0.091564406 | 0.234452231 | 0.182076535 |
| ENSMUSG00000071014 | Ndufb6 | 230075 | protein_coding | 654.4393627 | 0.182122094 | 0.132313567 | 1.376443084 | 0.168684497 | 0.353480683 | 0.182122094 |
| ENSMUSG00000059842 | Zfp341 | 228807 | protein_coding | 40.41583034 | -0.182143147 | 0.18023504 | -1.010586768 | 0.312214252 | 0.522423427 | 0.182143147 |
| ENSMUSG00000029571 | Tmem106b | 71900 | protein_coding | 1321.601652 | 0.182197884 | 0.113655328 | 1.603073859 | 0.108918349 | 0.26286402 | 0.182197884 |
| ENSMUSG00000036529 | Sbf1 | 77980 | protein_coding | 1313.580109 | -0.182238304 | 0.10760161 | -1.693639193 | 0.090333866 | 0.232295417 | 0.182238304 |
| ENSMUSG00000042305 | Tmem183a | 57439 | protein_coding | 2428.630223 | 0.182354165 | 0.067664066 | 2.69499269 | 0.007039018 | 0.036625014 | 0.182354165 |
| ENSMUSG00000030805 | Stx4a | 20909 | protein_coding | 306.4172249 | -0.182375995 | 0.085553985 | -2.131706596 | 0.033030974 | 0.115782869 | 0.182375995 |
| ENSMUSG00000039607 | Rbms3 | 207181 | protein_coding | 73.68222201 | -0.182418252 | 0.180755486 | -1.009198975 | 0.312879218 | 0.523151394 | 0.182418252 |
| ENSMUSG00000024594 | Prrc1 | 73137 | protein_coding | 834.9295087 | 0.182430928 | 0.061454922 | 2.968532419 | 0.002992255 | 0.018741897 | 0.182430928 |
| ENSMUSG00000074825 | Itpripl1 | 73338 | protein_coding | 60.1248396 | -0.18245745 | 0.203595043 | -0.896178254 | 0.370157568 | 0.578860317 | 0.18245745 |
| ENSMUSG00000033701 | Acbd6 | 72482 | protein_coding | 206.7555254 | -0.182477417 | 0.110998141 | -1.643968227 | 0.100182766 | 0.248398364 | 0.182477417 |
| ENSMUSG00000025993 | Slc40a1 | 53945 | protein_coding | 2132.452878 | 0.182480851 | 0.149653174 | 1.219358373 | 0.222708202 | 0.424013417 | 0.182480851 |
| ENSMUSG00000036693 | Nop14 | 75416 | protein_coding | 268.1968728 | -0.182558733 | 0.156031902 | -1.170009022 | 0.241997338 | 0.446695284 | 0.182558733 |
| ENSMUSG00000031885 | Cbfb | 12400 | protein_coding | 857.3019548 | 0.182571727 | 0.113550971 | 1.60783942 | 0.107870353 | 0.261197777 | 0.182571727 |
| ENSMUSG00000028902 | Sf3a3 | 75062 | protein_coding | 569.2204648 | 0.182619522 | 0.099550872 | 1.834434173 | 0.066589571 | 0.188667907 | 0.182619522 |
| ENSMUSG00000032349 | Elovl5 | 68801 | protein_coding | 921.7098782 | 0.182628094 | 0.071663745 | 2.548402874 | 0.010821742 | 0.050786299 | 0.182628094 |
| ENSMUSG00000024248 | Cox7a2l | 20463 | protein_coding | 2949.532544 | 0.182666181 | 0.107657767 | 1.696730164 | 0.089747687 | 0.231610687 | 0.182666181 |
| ENSMUSG00000004268 | Emg1 | 14791 | protein_coding | 804.5709755 | 0.182690146 | 0.124859186 | 1.463169445 | 0.143421019 | 0.317411421 | 0.182690146 |
| ENSMUSG00000053768 | Chchd3 | 66075 | protein_coding | 2477.970105 | 0.182772709 | 0.095431944 | 1.91521518 | 0.055465067 | 0.166688225 | 0.182772709 |
| ENSMUSG00000002846 | Timmdc1 | 76916 | protein_coding | 343.4868262 | 0.182780623 | 0.086491999 | 2.113266267 | 0.034577984 | 0.119886403 | 0.182780623 |
| ENSMUSG00000036860 | Mrpl55 | 67212 | protein_coding | 264.5212614 | 0.182790128 | 0.08700826 | 2.100836499 | 0.035655321 | 0.12266387 | 0.182790128 |
| ENSMUSG00000000581 | C1d | 57316 | protein_coding | 596.8827708 | 0.182851278 | 0.078449095 | 2.330827107 | 0.019762478 | 0.0791294 | 0.182851278 |
| ENSMUSG00000023960 | Enpp5 | 83965 | protein_coding | 840.4431396 | 0.182900019 | 0.096732468 | 1.890782107 | 0.058653437 | 0.173294739 | 0.182900019 |
| ENSMUSG00000031546 | Gins4 | 109145 | protein_coding | 247.9332033 | 0.182906815 | 0.117727925 | 1.553640009 | 0.12027031 | 0.281549897 | 0.182906815 |
| ENSMUSG00000033075 | Senp1 | 223870 | protein_coding | 425.3318932 | -0.182974811 | 0.100935465 | -1.812790096 | 0.069864202 | 0.195399091 | 0.182974811 |
| ENSMUSG00000050029 | Rap2c | 72065 | protein_coding | 736.1527009 | 0.182984972 | 0.104103753 | 1.757717333 | 0.078795617 | 0.211670171 | 0.182984972 |
| ENSMUSG00000022792 | Yars2 | 70120 | protein_coding | 227.8126175 | -0.183018204 | 0.153520274 | -1.192143544 | 0.233204966 | 0.43679312 | 0.183018204 |
| ENSMUSG00000031799 | Tpm4 | 326618 | protein_coding | 2299.999687 | -0.183029348 | 0.145863925 | -1.254795168 | 0.209553127 | 0.406597906 | 0.183029348 |
| ENSMUSG00000028458 | Tesk1 | 21754 | protein_coding | 227.9695627 | -0.183054963 | 0.112105065 | -1.632887533 | 0.102492653 | 0.252435901 | 0.183054963 |
| ENSMUSG00000052833 | Sae1 | 56459 | protein_coding | 1073.688235 | 0.183096557 | 0.103920105 | 1.761897347 | 0.078086643 | 0.210164354 | 0.183096557 |
| ENSMUSG00000031169 | Porcn | 53627 | protein_coding | 28.82502108 | 0.183099745 | 0.221639423 | 0.82611542 | 0.408738621 | 0.615833332 | 0.183099745 |

Supplementary Table 1

| ensembl_gene_id | external_gene_name | entrezgene_id | gene_biotype | baseMean | log2FoldChange | lfcSE | stat | pvalue | padj | absLog2FC |
|---|---|---|---|---|---|---|---|---|---|---|
| ENSMUSG00000056673 | Kdm5d | 20592 | protein_coding | 856.2591503 | -0.183145887 | 0.109567905 | -1.671528597 | 0.094617315 | 0.239635012 | 0.183145887 |
| ENSMUSG00000029422 | Rsrc2 | 208606 | protein_coding | 1299.344858 | -0.183164501 | 0.060766118 | -3.014253781 | 0.002576122 | 0.016659551 | 0.183164501 |
| ENSMUSG00000059518 | Znhit1 | 70103 | protein_coding | 238.3131957 | 0.183191367 | 0.081561856 | 2.246042146 | 0.024701308 | 0.093565384 | 0.183191367 |
| ENSMUSG00000090264 | Eif4ebp3 | 108112 | protein_coding | 269.3910153 | 0.183207597 | 0.14181745 | 1.291855104 | 0.196407328 | 0.389983519 | 0.183207597 |
| ENSMUSG00000029644 | Pdx1 | 18609 | protein_coding | 8.1108244 | -0.183207686 | 0.432943631 | -0.423167527 | 0.672173037 | 0.810843088 | 0.183207686 |
| ENSMUSG00000041390 | Mdfic | 16543 | protein_coding | 77.42086512 | -0.183210314 | 0.141238805 | -1.297166975 | 0.194573741 | 0.387730172 | 0.183210314 |
| ENSMUSG00000037486 | Asxl2 | 75302 | protein_coding | 815.5325779 | -0.183277993 | 0.096592475 | -1.897435522 | 0.057770481 | 0.171719678 | 0.183277993 |
| ENSMUSG00000049957 | Ccdc137 | 67291 | protein_coding | 245.776492 | -0.183286306 | 0.149868665 | -1.222979506 | 0.221337437 | 0.422211102 | 0.183286306 |
| ENSMUSG00000071369 | Map3k5 | 26408 | protein_coding | 874.8164603 | 0.18329428 | 0.090882598 | 2.016824824 | 0.043713797 | 0.141432737 | 0.18329428 |
| ENSMUSG00000092607 | Scnm1 | 69269 | protein_coding | 224.2462617 | 0.183312028 | 0.102260777 | 1.792593738 | 0.073037908 | 0.201097287 | 0.183312028 |
| ENSMUSG00000093805 | Gal3st2b | 100041596 | protein_coding | 156.39359 | 0.183353195 | 0.359938119 | 0.509401992 | 0.61047048 | 0.773061998 | 0.183353195 |
| ENSMUSG00000014956 | Ppp1cb | 19046 | protein_coding | 6145.61811 | 0.183363951 | 0.081344806 | 2.254156842 | 0.024186303 | 0.092184227 | 0.183363951 |
| ENSMUSG00000052684 | Jun | 16476 | protein_coding | 8622.487307 | -0.18338547 | 0.266363541 | -0.688478122 | 0.491151743 | 0.683356478 | 0.18338547 |
| ENSMUSG00000021608 | Lpcat1 | 210992 | protein_coding | 195.4830085 | -0.183411044 | 0.134857889 | -1.360031999 | 0.173819798 | 0.359707067 | 0.183411044 |
| ENSMUSG00000039316 | Rftn1 | 76438 | protein_coding | 8.398836952 | 0.183415155 | 0.608468224 | 0.301437525 | 0.763080884 | 0.870097644 | 0.183415155 |
| ENSMUSG00000019907 | Ppp1r12a | 17931 | protein_coding | 1562.074663 | -0.183471077 | 0.064612416 | -2.839563777 | 0.004517526 | 0.025873844 | 0.183471077 |
| ENSMUSG00000022637 | Cblb | 208650 | protein_coding | 314.6147632 | -0.183513957 | 0.142726869 | -1.285770222 | 0.198523266 | 0.39266606 | 0.183513957 |
| ENSMUSG00000074212 | Dnajb14 | 70604 | protein_coding | 403.4019069 | 0.183543095 | 0.107367918 | 1.709478021 | 0.08736244 | 0.22744507 | 0.183543095 |
| ENSMUSG00000030842 | Lamtor1 | 66508 | protein_coding | 750.6343795 | 0.183577361 | 0.084005887 | 2.185291628 | 0.028867471 | 0.104882711 | 0.183577361 |
| ENSMUSG00000021607 | Mrpl36 | 94066 | protein_coding | 513.9356835 | 0.183652012 | 0.109837424 | 1.672034955 | 0.094517429 | 0.239458186 | 0.183652012 |
| ENSMUSG00000021710 | Nln | 75805 | protein_coding | 736.1595278 | -0.1837536 | 0.063654922 | -2.8867147 | 0.00389287 | 0.023007659 | 0.1837536 |
| ENSMUSG00000023094 | Msrb2 | 76467 | protein_coding | 82.85544903 | 0.183816002 | 0.143511574 | 1.280844441 | 0.200248311 | 0.394795779 | 0.183816002 |
| ENSMUSG00000031497 | Tnfsf13b | 24099 | protein_coding | 59.80102645 | 0.183852965 | 0.205823404 | 0.893255876 | 0.371720162 | 0.579898015 | 0.183852965 |
| ENSMUSG00000046432 | Bex3 | 12070 | protein_coding | 211.2299762 | 0.184023496 | 0.104558367 | 1.760007367 | 0.078406557 | 0.210778 | 0.184023496 |
| ENSMUSG00000045100 | Slc25a26 | 67582 | protein_coding | 87.25174017 | -0.184024668 | 0.125513636 | -1.46617271 | 0.142601241 | 0.316152646 | 0.184024668 |
| ENSMUSG00000025465 | Echs1 | 93747 | protein_coding | 1680.544255 | -0.184109965 | 0.103371981 | -1.781043202 | 0.074905393 | 0.204201575 | 0.184109965 |
| ENSMUSG00000045211 | Nudt18 | 213484 | protein_coding | 67.99827231 | 0.184206351 | 0.153989532 | 1.196226449 | 0.231608202 | 0.435030123 | 0.184206351 |
| ENSMUSG00000001052 | Sec24b | 99683 | protein_coding | 347.2184656 | -0.184210558 | 0.136065742 | -1.35383496 | 0.175789038 | 0.361907388 | 0.184210558 |
| ENSMUSG00000062300 | Nectin2 | 19294 | protein_coding | 1396.025869 | -0.18421391 | 0.126418966 | -1.457169878 | 0.145069493 | 0.320347551 | 0.18421391 |
| ENSMUSG00000028706 | Nsun4 | 72181 | protein_coding | 180.8126752 | -0.184281912 | 0.093165614 | -1.978003516 | 0.04792831 | 0.151318557 | 0.184281912 |
| ENSMUSG00000112489 | 9230116L04Rik | 321017 | lncRNA | 67.31161037 | 0.18428594 | 0.229530202 | 0.802883187 | 0.422042255 | 0.627572772 | 0.18428594 |
| ENSMUSG00000025787 | Tgm4 | 331046 | protein_coding | 10.26932351 | -0.184368352 | 0.382701326 | -0.481755195 | 0.629979862 | 0.785882948 | 0.184368352 |
| ENSMUSG00000066798 | Zbtb6 | 241322 | protein_coding | 594.9918872 | 0.18439052 | 0.115769156 | 1.592743065 | 0.111217842 | 0.266326572 | 0.18439052 |
| ENSMUSG00000049775 | Tmsb4x | 19241 | protein_coding | 6690.479238 | 0.184484552 | 0.15726652 | 1.173069461 | 0.240767952 | 0.445303291 | 0.184484552 |
| ENSMUSG00000026495 | Efcab2 | 68226 | protein_coding | 53.46302153 | 0.184539459 | 0.211270648 | 0.873474192 | 0.382404667 | 0.590207513 | 0.184539459 |
| ENSMUSG00000037331 | Larp1 | 73158 | protein_coding | 1931.670809 | -0.184543089 | 0.087266664 | -2.114703148 | 0.034455256 | 0.119538949 | 0.184543089 |
| ENSMUSG00000021213 | Akr1c13 | 27384 | protein_coding | 3377.188323 | 0.184558274 | 0.110216929 | 1.674500246 | 0.094032326 | 0.238820528 | 0.184558274 |
| ENSMUSG00000020821 | Kif1c | 16562 | protein_coding | 4043.516648 | -0.184575241 | 0.05493249 | -3.360037771 | 0.000779318 | 0.006259697 | 0.184575241 |

| ensembl_gene_id | external_gene_name | entrezgene_id | gene_biotype | baseMean | log2FoldChange | lfcSE | stat | pvalue | padj | absLog2FC |
|---|---|---|---|---|---|---|---|---|---|---|
| ENSMUSG00000022718 | Dgcr8 | 94223 | protein_coding | 351.2527712 | 0.184628723 | 0.103794209 | 1.778795988 | 0.075273215 | 0.204774 | 0.184628723 |
| ENSMUSG00000027637 | Rab5if | 67388 | protein_coding | 354.647155 | 0.184646012 | 0.123439771 | 1.495838903 | 0.134695641 | 0.303885576 | 0.184646012 |
| ENSMUSG00000060098 | Prmt7 | 214572 | protein_coding | 331.4303001 | -0.184648374 | 0.105410575 | -1.751706353 | 0.079824314 | 0.213474537 | 0.184648374 |
| ENSMUSG00000025373 | Rnf41 | 67588 | protein_coding | 376.4929436 | -0.184682424 | 0.07789521 | -2.370908611 | 0.017744418 | 0.07314521 | 0.184682424 |
| ENSMUSG00000056648 | Hoxb8 | 15416 | protein_coding | 57.85658953 | -0.184737077 | 0.261371653 | -0.706798443 | 0.479691742 | 0.673471139 | 0.184737077 |
| ENSMUSG00000020659 | Cbll1 | 104836 | protein_coding | 150.5126248 | -0.18478725 | 0.15916252 | -1.160997263 | 0.245643013 | 0.451020393 | 0.18478725 |
| ENSMUSG00000033446 | Lpar6 | 67168 | protein_coding | 277.456375 | 0.184802583 | 0.094036461 | 1.965222646 | 0.049388474 | 0.1541261 | 0.184802583 |
| ENSMUSG00000004535 | Tax1bp1 | 52440 | protein_coding | 8419.135222 | 0.184889305 | 0.097030356 | 1.905479002 | 0.056717834 | 0.169428931 | 0.184889305 |
| ENSMUSG00000017485 | Top2b | 21974 | protein_coding | 2652.236232 | -0.184977305 | 0.047966389 | -3.856394199 | 0.000115072 | 0.001270177 | 0.184977305 |
| ENSMUSG00000048799 | Cep120 | 225523 | protein_coding | 241.9090517 | -0.185025103 | 0.117998578 | -1.568028244 | 0.116874541 | 0.275666769 | 0.185025103 |
| ENSMUSG00000022160 | Mettl3 | 56335 | protein_coding | 256.2908477 | -0.185050875 | 0.108908171 | -1.699145932 | 0.089291691 | 0.230651255 | 0.185050875 |
| ENSMUSG00000096959 | 4930509G22Rik | 105246071 | lncRNA | 8.216123282 | -0.185057237 | 0.406132749 | -0.455657018 | 0.648636635 | 0.797120926 | 0.185057237 |
| ENSMUSG00000045039 | Megf8 | 269878 | protein_coding | 448.9574179 | -0.185105186 | 0.092675321 | -1.997351437 | 0.045787019 | 0.146360187 | 0.185105186 |
| ENSMUSG00000054381 | Zfp747 | 269997 | protein_coding | 232.562293 | -0.185122772 | 0.092985112 | -1.99088615 | 0.046493405 | 0.148079776 | 0.185122772 |
| ENSMUSG00000030816 | Rnf40 | 233900 | protein_coding | 860.0733781 | -0.18516408 | 0.062590644 | -2.958334804 | 0.00309306 | 0.019247562 | 0.18516408 |
| ENSMUSG00000074604 | Mgst2 | 211666 | protein_coding | 1520.713613 | 0.185168201 | 0.097832951 | 1.892697699 | 0.058398083 | 0.172773032 | 0.185168201 |
| ENSMUSG00000041261 | Car8 | 12319 | protein_coding | 1445.235149 | 0.185254043 | 0.261560574 | 0.708264399 | 0.478781081 | 0.672927641 | 0.185254043 |
| ENSMUSG00000029433 | Diablo | 66593 | protein_coding | 95.67768323 | -0.185262153 | 0.115357197 | -1.60598695 | 0.108276777 | 0.261882891 | 0.185262153 |
| ENSMUSG00000024780 | Cdc37l1 | 67072 | protein_coding | 669.7543859 | -0.185265292 | 0.072849426 | -2.543126328 | 0.01098655 | 0.051378088 | 0.185265292 |
| ENSMUSG00000042312 | S100a13 | 20196 | protein_coding | 512.5025785 | 0.185272068 | 0.102271939 | 1.81156307 | 0.070053736 | 0.195688513 | 0.185272068 |
| ENSMUSG00000049751 | Rpl36al | 66483 | protein_coding | 264.5323986 | 0.185284401 | 0.142965021 | 1.296012132 | 0.194971305 | 0.388054296 | 0.185284401 |
| ENSMUSG00000024085 | Man2a1 | 17158 | protein_coding | 2137.485575 | 0.185366181 | 0.055367903 | 3.347899597 | 0.000814265 | 0.006471863 | 0.185366181 |
| ENSMUSG00000033502 | Cdc14a | 229776 | protein_coding | 701.6661848 | -0.185379649 | 0.089500765 | -2.071263286 | 0.038334198 | 0.129224785 | 0.185379649 |
| ENSMUSG00000019088 | Dnase1l1 | 69537 | protein_coding | 191.4357158 | 0.185395977 | 0.100678598 | 1.841463639 | 0.065553644 | 0.186553233 | 0.185395977 |
| ENSMUSG00000029152 | Ociad1 | 68095 | protein_coding | 1531.739068 | 0.18541651 | 0.095378099 | 1.944015577 | 0.051893573 | 0.159506142 | 0.18541651 |
| ENSMUSG00000045854 | Lyrm2 | 108755 | protein_coding | 157.9434692 | 0.185440343 | 0.128762671 | 1.440171611 | 0.149818853 | 0.327251055 | 0.185440343 |
| ENSMUSG00000060675 | Plaat3 | 225845 | protein_coding | 667.4313909 | -0.18546333 | 0.114761524 | -1.616075873 | 0.10607791 | 0.258271011 | 0.18546333 |
| ENSMUSG00000100594 | 2810414N06Rik | 102640772 | lncRNA | 14.89621553 | 0.185497074 | 0.304037143 | 0.6101132 | 0.541786823 | 0.724809384 | 0.185497074 |
| ENSMUSG00000052681 | Rap1b | 215449 | protein_coding | 1633.752191 | 0.185514793 | 0.094166967 | 1.970062327 | 0.048831228 | 0.153137192 | 0.185514793 |
| ENSMUSG00000024846 | Cst6 | 73720 | protein_coding | 17.30816399 | -0.18566555 | 0.417821467 | -0.44436575 | 0.656778171 | 0.801253272 | 0.18566555 |
| ENSMUSG00000061559 | Wdr61 | 66317 | protein_coding | 722.3107823 | 0.18582716 | 0.100733744 | 1.844735954 | 0.065075958 | 0.185724755 | 0.18582716 |
| ENSMUSG00000040681 | Hmgn1 | 15312 | protein_coding | 1228.00947 | 0.185919388 | 0.115389029 | 1.611239725 | 0.107127483 | 0.26012589 | 0.185919388 |
| ENSMUSG00000009555 | Cdk9 | 107951 | protein_coding | 439.6822441 | -0.185924335 | 0.078899609 | -2.356467127 | 0.018449701 | 0.075256593 | 0.185924335 |
| ENSMUSG00000021192 | Golga5 | 27277 | protein_coding | 932.8262287 | 0.185940937 | 0.071402876 | 2.604109898 | 0.009211322 | 0.045060789 | 0.185940937 |
| ENSMUSG00000020362 | Cnot6 | 104625 | protein_coding | 1099.358321 | 0.185955179 | 0.095008028 | 1.957257546 | 0.050317196 | 0.156046009 | 0.185955179 |
| ENSMUSG00000022564 | Grina | 66168 | protein_coding | 418.3374378 | -0.186009466 | 0.126301487 | -1.472741697 | 0.140820699 | 0.313223305 | 0.186009466 |
| ENSMUSG00000075704 | Txnrd2 | 26462 | protein_coding | 138.4509976 | -0.186041419 | 0.107802961 | -1.725754254 | 0.084391633 | 0.222216884 | 0.186041419 |
| ENSMUSG00000019773 | Fbxo5 | 67141 | protein_coding | 151.7673651 | 0.186067898 | 0.149412381 | 1.24533119 | 0.213010035 | 0.411052077 | 0.186067898 |

| ensembl_gene_id | external_gene_name | entrezgene_id | gene_biotype | baseMean | log2FoldChange | lfcSE | stat | pvalue | padj | absLog2FC |
|---|---|---|---|---|---|---|---|---|---|---|
| ENSMUSG00000027304 | Rtf1 | 76246 | protein_coding | 400.1106897 | -0.186107206 | 0.073188613 | -2.542843737 | 0.010995439 | 0.051404565 | 0.186107206 |
| ENSMUSG00000038518 | Jarid2 | 16468 | protein_coding | 748.2926654 | -0.18614646 | 0.063491017 | -2.931855059 | 0.00336944 | 0.020609998 | 0.18614646 |
| ENSMUSG00000020774 | Aspa | 11484 | protein_coding | 58.76330836 | -0.186183029 | 0.232457042 | -0.800935207 | 0.423169157 | 0.628167598 | 0.186183029 |
| ENSMUSG00000003437 | Paf1 | 54624 | protein_coding | 488.6043149 | 0.186219247 | 0.087623672 | 2.125216183 | 0.033568578 | 0.117228747 | 0.186219247 |
| ENSMUSG00000056632 | Dsg3 | 13512 | protein_coding | 22.14973336 | 0.186221261 | 0.388845975 | 0.478907518 | 0.632004423 | 0.787065398 | 0.186221261 |
| ENSMUSG00000056019 | Zfp709 | 236193 | protein_coding | 165.1349043 | 0.18622155 | 0.120002244 | 1.551817223 | 0.120705964 | 0.282237902 | 0.18622155 |
| ENSMUSG00000024580 | Grpel2 | 17714 | protein_coding | 1130.83114 | -0.186263484 | 0.078431035 | -2.374869651 | 0.017555148 | 0.072634077 | 0.186263484 |
| ENSMUSG00000070493 | Chchd2 | 14004 | protein_coding | 3959.987772 | 0.1864451 | 0.121608218 | 1.533162004 | 0.125235943 | 0.289329844 | 0.1864451 |
| ENSMUSG00000033486 | Catsper2 | 212670 | protein_coding | 25.75578469 | 0.18652015 | 0.255496254 | 0.73003086 | 0.465371322 | 0.6622384 | 0.18652015 |
| ENSMUSG00000010376 | Nedd8 | 18002 | protein_coding | 797.4588858 | 0.186530944 | 0.103574893 | 1.800928175 | 0.071714202 | 0.198792269 | 0.186530944 |
| ENSMUSG00000028525 | Pde4b | 18578 | protein_coding | 52.79400096 | 0.186571545 | 0.254892619 | 0.731961349 | 0.464192159 | 0.661249706 | 0.186571545 |
| ENSMUSG00000004110 | Cacna1e | 12290 | protein_coding | 39.72562109 | -0.186668591 | 0.575096843 | -0.324586361 | 0.745494151 | 0.85860846 | 0.186668591 |
| ENSMUSG00000074165 | Zfp788 | 67607 | protein_coding | 158.4854801 | -0.186687097 | 0.115166847 | -1.621014226 | 0.105014588 | 0.256593769 | 0.186687097 |
| ENSMUSG00000021594 | Srd5a1 | 78925 | protein_coding | 138.5300833 | 0.186716012 | 0.120485548 | 1.549696335 | 0.121214418 | 0.283052815 | 0.186716012 |
| ENSMUSG00000003316 | Glg1 | 20340 | protein_coding | 1649.723106 | -0.186732348 | 0.065163243 | -2.865608582 | 0.004162085 | 0.02423015 | 0.186732348 |
| ENSMUSG00000038664 | Herc1 | 235439 | protein_coding | 1734.101118 | -0.186780626 | 0.116751464 | -1.599813996 | 0.109639853 | 0.263874388 | 0.186780626 |
| ENSMUSG00000002329 | Mdp1 | 67881 | protein_coding | 1164.282935 | 0.186782863 | 0.10757423 | 1.736316059 | 0.082507965 | 0.21870309 | 0.186782863 |
| ENSMUSG00000045382 | Cxcr4 | 12767 | protein_coding | 13.74806263 | 0.18682891 | 0.332647513 | 0.561642288 | 0.574359762 | 0.747540635 | 0.18682891 |
| ENSMUSG00000009535 | Rnmt | 67897 | protein_coding | 389.541776 | -0.186864824 | 0.0966546 | -1.933325724 | 0.053196085 | 0.161983995 | 0.186864824 |
| ENSMUSG00000087259 | 2610035D17Rik | 72386 | lncRNA | 299.5703528 | 0.186927359 | 0.22453023 | 0.832526464 | 0.405111847 | 0.6121451 | 0.186927359 |
| ENSMUSG00000063882 | Uqcrh | 66576 | protein_coding | 5767.845407 | 0.186946069 | 0.09441517 | 1.980042714 | 0.047698729 | 0.150832908 | 0.186946069 |
| ENSMUSG00000034563 | Ccpg1 | 72278 | protein_coding | 12.62640425 | -0.187028674 | 0.338096796 | -0.553180852 | 0.580139577 | 0.751318253 | 0.187028674 |
| ENSMUSG00000060591 | Ifitm2 | 80876 | protein_coding | 266.1066545 | -0.187055439 | 0.132790959 | -1.408645888 | 0.158939903 | 0.340262253 | 0.187055439 |
| ENSMUSG00000017760 | Ctsa | 19025 | protein_coding | 1754.302443 | 0.187065363 | 0.057535987 | 3.251275794 | 0.001148884 | 0.008575172 | 0.187065363 |
| ENSMUSG00000030314 | Atg7 | 74244 | protein_coding | 428.9638848 | -0.187076663 | 0.078714519 | -2.376647482 | 0.017470776 | 0.072417303 | 0.187076663 |
| ENSMUSG00000003271 | Sult2b1 | 54200 | protein_coding | 183.5673207 | -0.187208091 | 0.165729969 | -1.12959709 | 0.258646038 | 0.465714909 | 0.187208091 |
| ENSMUSG00000032584 | Mst1r | 19882 | protein_coding | 872.3328954 | -0.187268635 | 0.163042839 | -1.148585462 | 0.250726955 | 0.457061914 | 0.187268635 |
| ENSMUSG00000078773 | Rad54b | 623474 | protein_coding | 78.82570674 | 0.187268871 | 0.203970614 | 0.918116912 | 0.358557662 | 0.568269297 | 0.187268871 |
| ENSMUSG00000023087 | Noct | 12457 | protein_coding | 394.366337 | -0.18728236 | 0.172810601 | -1.083743467 | 0.278478557 | 0.487034766 | 0.18728236 |
| ENSMUSG00000060508 | Nlrp9b | 243874 | protein_coding | 195.8797778 | 0.187324592 | 0.202896249 | 0.923253109 | 0.355875318 | 0.565481052 | 0.187324592 |
| ENSMUSG00000027649 | Ctnnbl1 | 66642 | protein_coding | 373.9437937 | 0.187408747 | 0.106376746 | 1.761745442 | 0.078112316 | 0.210164354 | 0.187408747 |
| ENSMUSG00000049823 | Zbtb12 | 193736 | protein_coding | 34.57095487 | -0.187456377 | 0.222390879 | -0.842913961 | 0.399276565 | 0.606939679 | 0.187456377 |
| ENSMUSG00000060450 | Rnf14 | 56736 | protein_coding | 913.7108383 | 0.187474689 | 0.078150388 | 2.398896473 | 0.016444563 | 0.069220139 | 0.187474689 |
| ENSMUSG00000027605 | Acss2 | 60525 | protein_coding | 1810.870346 | -0.187551964 | 0.14367317 | -1.305407009 | 0.191754293 | 0.384424123 | 0.187551964 |
| ENSMUSG00000037710 | Cisd1 | 52637 | protein_coding | 1034.866241 | 0.187608196 | 0.133043126 | 1.410130692 | 0.158501096 | 0.339785391 | 0.187608196 |
| ENSMUSG00000070738 | Dgkd | 227333 | protein_coding | 1857.79939 | -0.187630827 | 0.096048044 | -1.953510137 | 0.050759177 | 0.157049762 | 0.187630827 |
| ENSMUSG00000030882 | Dnhd1 | 77505 | protein_coding | 14.5065263 | -0.187675495 | 0.438714789 | -0.427784748 | 0.668807842 | 0.808191436 | 0.187675495 |
| ENSMUSG00000039844 | Rapgef1 | 107746 | protein_coding | 534.0134937 | -0.187701574 | 0.086337769 | -2.174037814 | 0.029702295 | 0.107081034 | 0.187701574 |

| ensembl_gene_id | external_gene_name | entrezgene_id | gene_biotype | baseMean | log2FoldChange | lfcSE | stat | pvalue | padj | absLog2FC |
|---|---|---|---|---|---|---|---|---|---|---|
| ENSMUSG00000040812 | Agbl2 | 271813 | protein_coding | 39.5723421 | -0.187721952 | 0.358930202 | -0.523004058 | 0.600971429 | 0.767009048 | 0.187721952 |
| ENSMUSG00000028252 | Ccnc | 51813 | protein_coding | 438.7651331 | 0.187725335 | 0.115362373 | 1.627266587 | 0.103680491 | 0.254260641 | 0.187725335 |
| ENSMUSG00000006494 | Pdk1 | 228026 | protein_coding | 987.2988274 | -0.187740487 | 0.097742103 | -1.920773973 | 0.054760209 | 0.165037012 | 0.187740487 |
| ENSMUSG00000026567 | Adcy10 | 271639 | protein_coding | 7.183870839 | -0.187758347 | 0.820267562 | -0.228898906 | 0.818947491 | 0.903777566 | 0.187758347 |
| ENSMUSG00000029840 | Mtpn | 14489 | protein_coding | 2847.914165 | 0.187931192 | 0.077515557 | 2.424431946 | 0.015332357 | 0.065666517 | 0.187931192 |
| ENSMUSG00000110605 | Gm32856 | 102635552 | lncRNA | 37.52197791 | 0.188044481 | 0.212265657 | 0.88589216 | 0.375675641 | 0.583911261 | 0.188044481 |
| ENSMUSG00000009376 | Met | 17295 | protein_coding | 1099.169332 | -0.188060312 | 0.196753873 | -0.955815047 | 0.339165685 | 0.549942578 | 0.188060312 |
| ENSMUSG00000034285 | Nipsnap1 | 18082 | protein_coding | 869.4650333 | -0.188147165 | 0.087062949 | -2.161047465 | 0.030691673 | 0.109880191 | 0.188147165 |
| ENSMUSG00000027695 | Pld1 | 18805 | protein_coding | 2170.801889 | -0.188249429 | 0.116575788 | -1.614824408 | 0.106348726 | 0.258732626 | 0.188249429 |
| ENSMUSG00000024982 | Zdhhc6 | 66980 | protein_coding | 679.985354 | 0.188257617 | 0.073878186 | 2.54821655 | 0.010827524 | 0.050790108 | 0.188257617 |
| ENSMUSG00000029050 | Ski | 20481 | protein_coding | 744.0799358 | -0.188285717 | 0.09306559 | -2.023150737 | 0.043057609 | 0.139901465 | 0.188285717 |
| ENSMUSG00000042082 | Arsb | 11881 | protein_coding | 222.8547092 | -0.188306188 | 0.110148299 | -1.70956964 | 0.087345484 | 0.22744507 | 0.188306188 |
| ENSMUSG00000022390 | Zc3h7b | 20286 | protein_coding | 495.9146518 | -0.188420557 | 0.078542998 | -2.39894786 | 0.016442256 | 0.069220139 | 0.188420557 |
| ENSMUSG00000027259 | Adal | 75894 | protein_coding | 154.3642944 | 0.188425667 | 0.116677663 | 1.61492493 | 0.106326953 | 0.258719173 | 0.188425667 |
| ENSMUSG00000037341 | Slc9a7 | 236727 | protein_coding | 13.32326483 | -0.188446851 | 0.352100585 | -0.535207434 | 0.592506426 | 0.761247936 | 0.188446851 |
| ENSMUSG00000078768 | Zfp566 | 72556 | protein_coding | 24.28311103 | -0.188550645 | 0.271991991 | -0.693221313 | 0.488170675 | 0.680942953 | 0.188550645 |
| ENSMUSG00000091754 | Gm3636 | 100041151 | protein_coding | 8.050077223 | -0.188588182 | 0.457883777 | -0.411869106 | 0.680435366 | 0.817182783 | 0.188588182 |
| ENSMUSG00000020412 | Ascc2 | 75452 | protein_coding | 736.0158191 | -0.188655885 | 0.076742124 | -2.458309396 | 0.013959285 | 0.061268531 | 0.188655885 |
| ENSMUSG00000020741 | Cluh | 74148 | protein_coding | 2969.037282 | -0.188798246 | 0.082511214 | -2.288152557 | 0.022128638 | 0.086126011 | 0.188798246 |
| ENSMUSG00000028964 | Park7 | 57320 | protein_coding | 816.2725468 | 0.188918274 | 0.098802758 | 1.912074905 | 0.055866587 | 0.167515625 | 0.188918274 |
| ENSMUSG00000022008 | Gpalpp1 | 67467 | protein_coding | 752.193519 | 0.188921278 | 0.067752114 | 2.788418938 | 0.0052966 | 0.029272813 | 0.188921278 |
| ENSMUSG00000019699 | Akt3 | 23797 | protein_coding | 20.59934571 | 0.188925088 | 0.4044302 | 0.467138921 | 0.640400489 | 0.791579931 | 0.188925088 |
| ENSMUSG00000031303 | Map3k15 | 270672 | protein_coding | 58.22949053 | -0.188935281 | 0.172439755 | -1.095659644 | 0.273227749 | 0.481253079 | 0.188935281 |
| ENSMUSG00000033760 | Rbm4b | 66704 | protein_coding | 159.2143373 | 0.188972854 | 0.172492694 | 1.095541204 | 0.273279604 | 0.481257732 | 0.188972854 |
| ENSMUSG00000027983 | Cyp2u1 | 71519 | protein_coding | 21.40188171 | 0.189011772 | 0.273122923 | 0.692039212 | 0.488912704 | 0.681431708 | 0.189011772 |
| ENSMUSG00000073448 | Gm10509 | 100310809 | protein_coding | 53.76449385 | 0.189037911 | 0.164978465 | 1.145833859 | 0.251863903 | 0.458216818 | 0.189037911 |
| ENSMUSG00000040506 | Ambra1 | 228361 | protein_coding | 614.9843569 | -0.189054806 | 0.084241451 | -2.244201671 | 0.024819429 | 0.093882597 | 0.189054806 |
| ENSMUSG00000007050 | Lsm2 | 27756 | protein_coding | 116.6882681 | 0.189112452 | 0.166776037 | 1.133930603 | 0.256823659 | 0.463695525 | 0.189112452 |
| ENSMUSG00000034345 | Gtf2h5 | 66467 | protein_coding | 385.0129094 | 0.189175772 | 0.093236891 | 2.028979847 | 0.042460343 | 0.138531001 | 0.189175772 |
| ENSMUSG00000022890 | Atp5j | 11957 | protein_coding | 2334.670696 | 0.189176633 | 0.072364015 | 2.614236289 | 0.008942717 | 0.044019314 | 0.189176633 |
| ENSMUSG00000019734 | Tmc4 | 353499 | protein_coding | 2732.221403 | -0.189261015 | 0.056422919 | -3.354328694 | 0.000795578 | 0.006345501 | 0.189261015 |
| ENSMUSG00000036898 | Zfp157 | 72154 | protein_coding | 201.9129095 | 0.18927208 | 0.164842419 | 1.148200087 | 0.250885973 | 0.457221028 | 0.18927208 |
| ENSMUSG00000042487 | Leo1 | 235497 | protein_coding | 481.4718836 | 0.189276238 | 0.078966552 | 2.39691659 | 0.016533687 | 0.069540737 | 0.189276238 |
| ENSMUSG00000042992 | Borcs5 | 67774 | protein_coding | 99.82628944 | -0.189358599 | 0.150321093 | -1.259694131 | 0.207779723 | 0.404190942 | 0.189358599 |
| ENSMUSG00000021771 | Vdac2 | 22334 | protein_coding | 6428.158985 | 0.18947834 | 0.115843275 | 1.635643851 | 0.101914147 | 0.251399804 | 0.18947834 |
| ENSMUSG00000024937 | Ehbp1l1 | 114601 | protein_coding | 602.6214174 | 0.189552893 | 0.160746824 | 1.179201483 | 0.238317956 | 0.442477798 | 0.189552893 |
| ENSMUSG00000034525 | Ice1 | 218333 | protein_coding | 707.5923629 | -0.189579849 | 0.100780521 | -1.881115991 | 0.059956142 | 0.175886323 | 0.189579849 |
| ENSMUSG00000028497 | Hacd4 | 66775 | protein_coding | 65.36338462 | -0.189607163 | 0.143608266 | -1.320308143 | 0.186732158 | 0.377206064 | 0.189607163 |

Supplementary Table 1

| ensembl_gene_id | external_gene_name | entrezgene_id | gene_biotype | baseMean | log2FoldChange | lfcSE | stat | pvalue | padj | absLog2FC |
|---|---|---|---|---|---|---|---|---|---|---|
| ENSMUSG00000100738 | 2010106C02Rik | 72104 | lncRNA | 7.123140384 | -0.18964162 | 0.461570912 | -0.410861291 | 0.681174248 | 0.817460468 | 0.18964162 |
| ENSMUSG00000041729 | Coro2b | 235431 | protein_coding | 11.29097356 | -0.18965849 | 0.500996344 | -0.378562622 | 0.705012683 | 0.833382076 | 0.18965849 |
| ENSMUSG00000020133 | 2310011J03Rik | 66374 | protein_coding | 417.7571434 | 0.189661268 | 0.091977109 | 2.06204859 | 0.03920311 | 0.131310866 | 0.189661268 |
| ENSMUSG00000049800 | Sertad2 | 58172 | protein_coding | 675.106323 | -0.189677241 | 0.093175081 | -2.035707827 | 0.041779703 | 0.137068405 | 0.189677241 |
| ENSMUSG00000026853 | Crat | 12908 | protein_coding | 594.9797615 | -0.189702596 | 0.076929398 | -2.465931119 | 0.013665761 | 0.060312619 | 0.189702596 |
| ENSMUSG00000039835 | Nhsl1 | 215819 | protein_coding | 880.1232788 | 0.189739145 | 0.116582368 | 1.627511507 | 0.103628507 | 0.254232438 | 0.189739145 |
| ENSMUSG00000023942 | Slc29a1 | 63959 | protein_coding | 237.227759 | -0.189748572 | 0.111916237 | -1.695451673 | 0.08998977 | 0.2319348 | 0.189748572 |
| ENSMUSG00000036792 | Mbd5 | 109241 | protein_coding | 191.8533476 | -0.189787314 | 0.165093012 | -1.149578117 | 0.250317676 | 0.456634994 | 0.189787314 |
| ENSMUSG00000033938 | Ndufb7 | 66916 | protein_coding | 950.7852546 | 0.189790865 | 0.105897422 | 1.792214213 | 0.073098659 | 0.201174597 | 0.189790865 |
| ENSMUSG00000046667 | Rbm12b1 | 72397 | protein_coding | 23.61931831 | -0.189861237 | 0.261911346 | -0.724906499 | 0.468509393 | 0.665040337 | 0.189861237 |
| ENSMUSG00000089804 | Gm16136 | 105245924 | lncRNA | 83.57441976 | -0.189892731 | 0.160385839 | -1.18397442 | 0.236423189 | 0.440601745 | 0.189892731 |
| ENSMUSG00000005370 | Msh6 | 17688 | protein_coding | 555.6825351 | -0.189935397 | 0.077503015 | -2.450683967 | 0.014258508 | 0.062136666 | 0.189935397 |
| ENSMUSG00000019320 | Noxo1 | 71893 | protein_coding | 44.13995719 | -0.189960921 | 0.189777393 | -1.000967068 | 0.31684273 | 0.528193536 | 0.189960921 |
| ENSMUSG00000028086 | Fbxw7 | 50754 | protein_coding | 305.9682266 | -0.190011069 | 0.090004184 | -2.111135962 | 0.034760627 | 0.120414804 | 0.190011069 |
| ENSMUSG00000030498 | Gas2 | 14453 | protein_coding | 56.45365111 | 0.190065254 | 0.163170624 | 1.164825191 | 0.24408976 | 0.449310115 | 0.190065254 |
| ENSMUSG00000022765 | Snap29 | 67474 | protein_coding | 652.290311 | 0.190107341 | 0.081791617 | 2.324288827 | 0.020110019 | 0.0801382 | 0.190107341 |
| ENSMUSG00000091931 | Gon7 | 100233175 | protein_coding | 71.10794777 | 0.190142558 | 0.141467537 | 1.344072017 | 0.1789251 | 0.366360585 | 0.190142558 |
| ENSMUSG00000026565 | Pou2f1 | 18986 | protein_coding | 295.9849744 | -0.190162377 | 0.207043081 | -0.918467671 | 0.358374077 | 0.568034859 | 0.190162377 |
| ENSMUSG00000070473 | Cldn3 | 12739 | protein_coding | 5326.442826 | 0.190188936 | 0.114449676 | 1.661769103 | 0.09655908 | 0.242795263 | 0.190188936 |
| ENSMUSG00000057604 | Lmcd1 | 30937 | protein_coding | 10.19701664 | 0.19019505 | 0.353356203 | 0.53825304 | 0.590402367 | 0.759898603 | 0.19019505 |
| ENSMUSG00000022704 | Qtrt2 | 106248 | protein_coding | 225.0693585 | -0.190215258 | 0.179674907 | -1.058663458 | 0.289753075 | 0.498400322 | 0.190215258 |
| ENSMUSG00000034108 | Ccs | 12460 | protein_coding | 533.4756452 | -0.190229835 | 0.089621369 | -2.122594609 | 0.033787837 | 0.117813632 | 0.190229835 |
| ENSMUSG00000069301 | H2ac11 | 319167 | protein_coding | 43.02635361 | 0.190300761 | 0.401964005 | 0.473427368 | 0.635908308 | 0.78927929 | 0.190300761 |
| ENSMUSG00000008730 | Hipk1 | 15257 | protein_coding | 2240.933558 | -0.190302699 | 0.106023388 | -1.794912451 | 0.07266765 | 0.200493734 | 0.190302699 |
| ENSMUSG00000029352 | Crybb3 | 12962 | protein_coding | 10.27322661 | -0.190321554 | 0.4030336 | -0.472222551 | 0.636767945 | 0.789636603 | 0.190321554 |
| ENSMUSG00000022378 | Cyrib | 223601 | protein_coding | 570.6292168 | 0.190379504 | 0.081824819 | 2.326671855 | 0.019982737 | 0.079690795 | 0.190379504 |
| ENSMUSG00000030304 | Ergic2 | 67456 | protein_coding | 572.0823577 | 0.190414132 | 0.084122101 | 2.263544664 | 0.023602135 | 0.09049947 | 0.190414132 |
| ENSMUSG00000020904 | Cfap52 | 71860 | protein_coding | 27.565013 | -0.190447941 | 0.252735513 | -0.753546422 | 0.451121627 | 0.651506507 | 0.190447941 |
| ENSMUSG00000027035 | Cers6 | 241447 | protein_coding | 2938.24287 | -0.190462152 | 0.079793193 | -2.386947358 | 0.016988924 | 0.070949024 | 0.190462152 |
| ENSMUSG00000090330 | 9130221H12Rik | 77124 | lncRNA | 41.89502561 | -0.190539116 | 0.19174056 | -0.993734012 | 0.32035238 | 0.531794966 | 0.190539116 |
| ENSMUSG00000035711 | Dok3 | 27261 | protein_coding | 8.780899217 | 0.1905484 | 0.523197706 | 0.36419961 | 0.715708958 | 0.840074597 | 0.1905484 |
| ENSMUSG00000002668 | Dennd1c | 70785 | protein_coding | 310.0787716 | -0.190586502 | 0.117625755 | -1.620278668 | 0.105172429 | 0.256772746 | 0.190586502 |
| ENSMUSG00000034263 | Ints14 | 69882 | protein_coding | 439.8595921 | 0.190622326 | 0.091259374 | 2.088797204 | 0.036725982 | 0.125245747 | 0.190622326 |
| ENSMUSG00000022711 | Pmm2 | 54128 | protein_coding | 2780.571498 | 0.190623209 | 0.078065641 | 2.441832361 | 0.01461293 | 0.063197056 | 0.190623209 |
| ENSMUSG00000033014 | Trim33 | 94093 | protein_coding | 928.3331737 | -0.190661209 | 0.062511702 | -3.050008306 | 0.00228835 | 0.015132223 | 0.190661209 |
| ENSMUSG00000044062 | Plekhd1os | 68354 | lncRNA | 15.24705658 | -0.190843494 | 0.321623752 | -0.593375001 | 0.552930216 | 0.732884453 | 0.190843494 |
| ENSMUSG00000020455 | Trim11 | 94091 | protein_coding | 302.9252306 | -0.190843662 | 0.110967464 | -1.71981638 | 0.085465824 | 0.223897968 | 0.190843662 |
| ENSMUSG00000020248 | Nfyb | 18045 | protein_coding | 470.7590514 | 0.190876672 | 0.098224686 | 1.943265797 | 0.051984052 | 0.159722604 | 0.190876672 |

| ensembl_gene_id | external_gene_name | entrezgene_id | gene_biotype | baseMean | log2FoldChange | lfcSE | stat | pvalue | padj | absLog2FC |
|---|---|---|---|---|---|---|---|---|---|---|
| ENSMUSG00000060314 | Zfp941 | 407812 | protein_coding | 14.89900528 | 0.190911412 | 0.378750763 | 0.504055519 | 0.614222364 | 0.775275463 | 0.190911412 |
| ENSMUSG00000038023 | Atp6v0a2 | 21871 | protein_coding | 2399.938428 | 0.190944867 | 0.110370224 | 1.730039674 | 0.083623187 | 0.220715026 | 0.190944867 |
| ENSMUSG00000072875 | Gpr27 | 14761 | protein_coding | 14.19686988 | 0.190967341 | 0.415495828 | 0.459613138 | 0.645793927 | 0.795082139 | 0.190967341 |
| ENSMUSG00000020150 | Gamt | 14431 | protein_coding | 18.37257953 | 0.191121073 | 0.319445963 | 0.598289211 | 0.549646975 | 0.731093174 | 0.191121073 |
| ENSMUSG00000021972 | Hmbox1 | 219150 | protein_coding | 288.9405498 | -0.191239726 | 0.174232308 | -1.097613459 | 0.272373313 | 0.480491985 | 0.191239726 |
| ENSMUSG00000020843 | Timm22 | 56322 | protein_coding | 302.4562004 | 0.191265956 | 0.10202127 | 1.874765493 | 0.060824992 | 0.177344744 | 0.191265956 |
| ENSMUSG00000062352 | Itgb1bp1 | 16413 | protein_coding | 105.4441944 | 0.191268937 | 0.126100509 | 1.516797497 | 0.129317819 | 0.295859555 | 0.191268937 |
| ENSMUSG00000028646 | Rragc | 54170 | protein_coding | 779.4855417 | 0.191288424 | 0.120713251 | 1.584651417 | 0.113045556 | 0.268865106 | 0.191288424 |
| ENSMUSG00000021520 | Uqcrb | 67530 | protein_coding | 3870.456211 | 0.191429682 | 0.096821321 | 1.977143883 | 0.048025369 | 0.151504867 | 0.191429682 |
| ENSMUSG00000078348 | Sf3b5 | 66125 | protein_coding | 345.7614114 | 0.1915047 | 0.12735254 | 1.503736797 | 0.132649151 | 0.300930875 | 0.1915047 |
| ENSMUSG00000025507 | Pidd1 | 57913 | protein_coding | 50.55667439 | 0.191590652 | 0.17904997 | 1.070040123 | 0.284601249 | 0.493695176 | 0.191590652 |
| ENSMUSG00000023806 | Rsph3b | 100037282 | protein_coding | 162.628868 | 0.191698937 | 0.114013046 | 1.681377202 | 0.092689671 | 0.236617233 | 0.191698937 |
| ENSMUSG00000039842 | Mcph1 | 244329 | protein_coding | 221.3346918 | 0.19176695 | 0.088084615 | 2.177076565 | 0.029474855 | 0.106502223 | 0.19176695 |
| ENSMUSG00000037174 | Elf2 | 69257 | protein_coding | 968.6898576 | -0.191796805 | 0.057653667 | -3.326706075 | 0.00087879 | 0.006895256 | 0.191796805 |
| ENSMUSG00000052310 | Slc39a1 | 30791 | protein_coding | 1822.922105 | 0.191831369 | 0.077361594 | 2.479671895 | 0.013150333 | 0.058853189 | 0.191831369 |
| ENSMUSG00000057137 | Tmem140 | 68487 | protein_coding | 43.30571043 | 0.191861513 | 0.184062296 | 1.042372701 | 0.297238918 | 0.506180721 | 0.191861513 |
| ENSMUSG00000095253 | Zfp799 | 240064 | protein_coding | 232.2350614 | -0.19189083 | 0.177621985 | -1.080332651 | 0.279994075 | 0.488257677 | 0.19189083 |
| ENSMUSG00000055862 | Izumo4 | 71564 | protein_coding | 118.1650192 | -0.191918826 | 0.165345761 | -1.1607121 | 0.245758999 | 0.451129347 | 0.191918826 |
| ENSMUSG00000023092 | Fhl1 | 14199 | protein_coding | 983.1019661 | -0.191971614 | 0.235973702 | -0.813529697 | 0.415914427 | 0.622269866 | 0.191971614 |
| ENSMUSG00000034041 | Lyl1 | 17095 | protein_coding | 7.91407151 | 0.192152601 | 0.404952143 | 0.474506937 | 0.635138452 | 0.788885209 | 0.192152601 |
| ENSMUSG00000054499 | Dedd2 | 67379 | protein_coding | 579.058604 | -0.192174422 | 0.147962624 | -1.298803822 | 0.194011263 | 0.387387373 | 0.192174422 |
| ENSMUSG00000047412 | Zbtb44 | 235132 | protein_coding | 745.1319341 | -0.192224572 | 0.121273355 | -1.585051983 | 0.112954524 | 0.268809153 | 0.192224572 |
| ENSMUSG00000024900 | Cpt1a | 12894 | protein_coding | 4387.450243 | -0.192274815 | 0.117695214 | -1.633667236 | 0.102328741 | 0.252149164 | 0.192274815 |
| ENSMUSG00000021224 | Numb | 18222 | protein_coding | 786.9252468 | 0.192282759 | 0.097684973 | 1.968396506 | 0.049022434 | 0.153468144 | 0.192282759 |
| ENSMUSG00000063200 | Nol7 | 70078 | protein_coding | 699.9770108 | 0.192318071 | 0.097035203 | 1.98194124 | 0.047485818 | 0.150249126 | 0.192318071 |
| ENSMUSG00000026436 | Elk4 | 13714 | protein_coding | 621.2016978 | -0.192333047 | 0.132557919 | -1.450935927 | 0.146797703 | 0.323178135 | 0.192333047 |
| ENSMUSG00000027575 | Arfgap1 | 228998 | protein_coding | 713.3590694 | -0.192348579 | 0.085502802 | -2.249617276 | 0.024473251 | 0.092900367 | 0.192348579 |
| ENSMUSG00000025780 | Itih5 | 209378 | protein_coding | 2873.038485 | -0.192360843 | 0.26745376 | -0.719230279 | 0.471999046 | 0.667439386 | 0.192360843 |
| ENSMUSG00000024349 | Sting1 | 72512 | protein_coding | 76.15818162 | 0.192387794 | 0.235347567 | 0.817462429 | 0.413664212 | 0.62021684 | 0.192387794 |
| ENSMUSG00000027263 | Tubgcp4 | 51885 | protein_coding | 332.5158027 | -0.192398799 | 0.090490419 | -2.126178664 | 0.033488386 | 0.117000002 | 0.192398799 |
| ENSMUSG00000020694 | Tlk2 | 24086 | protein_coding | 580.7382001 | -0.192445244 | 0.06779717 | -2.838543902 | 0.004531988 | 0.025928704 | 0.192445244 |
| ENSMUSG00000064655 | Gm25788 | 115490049 | snoRNA | 43.30755724 | 0.192448143 | 0.311485488 | 0.617839837 | 0.536680921 | 0.721189041 | 0.192448143 |
| ENSMUSG00000027957 | Slc35a3 | 229782 | protein_coding | 6192.95048 | 0.192484203 | 0.059596099 | 3.229812136 | 0.001238716 | 0.009106042 | 0.192484203 |
| ENSMUSG00000009549 | Srp14 | 20813 | protein_coding | 469.1387695 | 0.192597006 | 0.09793838 | 1.966512068 | 0.04923949 | 0.153872198 | 0.192597006 |
| ENSMUSG00000063275 | Hacd1 | 30963 | protein_coding | 343.8263531 | -0.192634732 | 0.09544742 | -2.018228802 | 0.043567438 | 0.141131208 | 0.192634732 |
| ENSMUSG00000028082 | Sh3d19 | 27059 | protein_coding | 1580.680483 | -0.192690363 | 0.070211967 | -2.744409119 | 0.006061994 | 0.032477442 | 0.192690363 |
| ENSMUSG00000089235 | Gm23119 | 115486069 | scaRNA | 48.54305169 | -0.192708659 | 0.261515296 | -0.736892495 | 0.461187728 | 0.659378689 | 0.192708659 |
| ENSMUSG00000020882 | Cacnb1 | 12295 | protein_coding | 28.99071965 | -0.192729305 | 0.343445354 | -0.561164395 | 0.574685473 | 0.747748252 | 0.192729305 |

| ensembl_gene_id | external_gene_name | entrezgene_id | gene_biotype | baseMean | log2FoldChange | lfcSE | stat | pvalue | padj | absLog2FC |
|---|---|---|---|---|---|---|---|---|---|---|
| ENSMUSG00000021062 | Rab15 | 104886 | protein_coding | 1482.667452 | -0.192807615 | 0.094809087 | -2.033640673 | 0.041987838 | 0.137439844 | 0.192807615 |
| ENSMUSG00000048486 | Fitm2 | 228859 | protein_coding | 314.6043479 | -0.192877341 | 0.141808626 | -1.360124173 | 0.173790633 | 0.359707067 | 0.192877341 |
| ENSMUSG00000020526 | Znhit3 | 448850 | protein_coding | 90.5161024 | 0.193024776 | 0.128201453 | 1.505636423 | 0.132160533 | 0.300078827 | 0.193024776 |
| ENSMUSG00000026377 | Nifk | 67949 | protein_coding | 571.42564 | -0.193028776 | 0.138240967 | -1.396321084 | 0.162617828 | 0.345172734 | 0.193028776 |
| ENSMUSG00000036782 | Klhl13 | 67455 | protein_coding | 44.40996491 | -0.193067068 | 0.210890085 | -0.915486701 | 0.359936181 | 0.569481508 | 0.193067068 |
| ENSMUSG00000028568 | Btf3l4 | 70533 | protein_coding | 230.8199859 | -0.193084232 | 0.10226734 | -1.888034168 | 0.059021364 | 0.17405893 | 0.193084232 |
| ENSMUSG00000038871 | Bpgm | 12183 | protein_coding | 683.2114128 | 0.193104718 | 0.078835976 | 2.449449185 | 0.01430749 | 0.062247936 | 0.193104718 |
| ENSMUSG00000020023 | Tmcc3 | 319880 | protein_coding | 1820.313683 | 0.193139191 | 0.141061853 | 1.36918088 | 0.170942741 | 0.356337649 | 0.193139191 |
| ENSMUSG00000027459 | Fam110a | 73847 | protein_coding | 141.2172552 | 0.193201885 | 0.116461578 | 1.658932391 | 0.097129417 | 0.243675627 | 0.193201885 |
| ENSMUSG00000039653 | Baat | 12012 | protein_coding | 15.02210008 | 0.193286624 | 0.377211766 | 0.512408789 | 0.608364941 | 0.772114485 | 0.193286624 |
| ENSMUSG00000078954 | Arhgap8 | 73167 | protein_coding | 57.30813609 | -0.193289841 | 0.228144517 | -0.847225453 | 0.396869469 | 0.60483573 | 0.193289841 |
| ENSMUSG00000045996 | Polr2k | 17749 | protein_coding | 142.8586852 | 0.19330481 | 0.137300429 | 1.407896616 | 0.159161685 | 0.340376923 | 0.19330481 |
| ENSMUSG00000022280 | Rnf19a | 30945 | protein_coding | 547.1560688 | -0.193326025 | 0.092437911 | -2.091414914 | 0.036490886 | 0.124625545 | 0.193326025 |
| ENSMUSG00000069049 | Eif2s3y | 26908 | protein_coding | 660.3238567 | 0.193475726 | 0.21123945 | 0.915907167 | 0.359715588 | 0.569481508 | 0.193475726 |
| ENSMUSG00000038545 | Cul7 | 66515 | protein_coding | 158.4732118 | 0.193500497 | 0.121462972 | 1.593082189 | 0.111141754 | 0.266205393 | 0.193500497 |
| ENSMUSG00000023106 | Denr | 68184 | protein_coding | 517.6652611 | 0.193502765 | 0.101798939 | 1.900832833 | 0.057323912 | 0.170792232 | 0.193502765 |
| ENSMUSG00000017978 | Cadps2 | 320405 | protein_coding | 1142.955695 | -0.193510247 | 0.09425553 | -2.053038651 | 0.040068829 | 0.133272807 | 0.193510247 |
| ENSMUSG00000019179 | Mdh2 | 17448 | protein_coding | 3534.055985 | 0.193514709 | 0.083165234 | 2.326870235 | 0.019972173 | 0.079668612 | 0.193514709 |
| ENSMUSG00000033543 | Gtf2a2 | 235459 | protein_coding | 537.7273554 | 0.193578307 | 0.088591656 | 2.185062512 | 0.028884263 | 0.10489479 | 0.193578307 |
| ENSMUSG00000026979 | Psd4 | 215632 | protein_coding | 1051.075475 | -0.193579628 | 0.100370468 | -1.928651251 | 0.053774172 | 0.162897492 | 0.193579628 |
| ENSMUSG00000026925 | Inpp5e | 64436 | protein_coding | 214.2883349 | -0.193580996 | 0.105979107 | -1.82659585 | 0.067760558 | 0.191295669 | 0.193580996 |
| ENSMUSG00000037434 | Slc30a1 | 22782 | protein_coding | 1019.421911 | 0.193617923 | 0.147781508 | 1.310163403 | 0.190140563 | 0.381863434 | 0.193617923 |
| ENSMUSG00000013698 | Pea15a | 18611 | protein_coding | 169.3903007 | -0.193663312 | 0.137116362 | -1.412401187 | 0.157831867 | 0.338737828 | 0.193663312 |
| ENSMUSG00000002496 | Tsc2 | 22084 | protein_coding | 765.0030879 | -0.193691711 | 0.078143037 | -2.478681649 | 0.013186895 | 0.058934022 | 0.193691711 |
| ENSMUSG00000031262 | Cenpi | 102920 | protein_coding | 137.7754687 | 0.193717626 | 0.127896819 | 1.514639905 | 0.129863621 | 0.296554516 | 0.193717626 |
| ENSMUSG00000017778 | Cox7c | 12867 | protein_coding | 2103.687955 | 0.193756681 | 0.089983127 | 2.153255708 | 0.031298588 | 0.111526601 | 0.193756681 |
| ENSMUSG00000021575 | Ahrr | 11624 | protein_coding | 8.941409252 | 0.19377631 | 0.470556216 | 0.411802679 | 0.680484058 | 0.817182783 | 0.19377631 |
| ENSMUSG00000054455 | Vapb | 56491 | protein_coding | 1016.674297 | 0.193807658 | 0.092211001 | 2.101784544 | 0.035572154 | 0.122506651 | 0.193807658 |
| ENSMUSG00000043190 | Rfesd | 218341 | protein_coding | 138.7061044 | 0.193868159 | 0.179765589 | 1.078449775 | 0.280833084 | 0.489344296 | 0.193868159 |
| ENSMUSG00000004105 | Angptl2 | 26360 | protein_coding | 15.7022149 | -0.193939108 | 0.42513785 | -0.45617935 | 0.648261014 | 0.796966766 | 0.193939108 |
| ENSMUSG00000050866 | Clrn3 | 212070 | protein_coding | 1783.872199 | 0.193988152 | 0.127478627 | 1.521730796 | 0.128076548 | 0.293948596 | 0.193988152 |
| ENSMUSG00000078812 | Eif5a | 276770 | protein_coding | 3373.845469 | 0.194117801 | 0.088130813 | 2.202609901 | 0.027622255 | 0.101360933 | 0.194117801 |
| ENSMUSG00000022195 | 6030458C11Rik | 77877 | protein_coding | 497.5677419 | 0.194172904 | 0.081897239 | 2.370933445 | 0.017743226 | 0.07314521 | 0.194172904 |
| ENSMUSG00000035212 | Leprot | 230514 | protein_coding | 1067.845323 | 0.19422797 | 0.090365485 | 2.14936013 | 0.031605863 | 0.112244859 | 0.19422797 |
| ENSMUSG00000022892 | App | 11820 | protein_coding | 4257.584294 | -0.194229155 | 0.056148563 | -3.459200818 | 0.000541781 | 0.004659549 | 0.194229155 |
| ENSMUSG00000025192 | Entpd7 | 93685 | protein_coding | 302.5940903 | 0.194257162 | 0.122531907 | 1.585359824 | 0.112884604 | 0.268723056 | 0.194257162 |
| ENSMUSG00000022420 | Dnal4 | 54152 | protein_coding | 162.1243018 | 0.194292307 | 0.116400174 | 1.669175407 | 0.095082622 | 0.240431181 | 0.194292307 |
| ENSMUSG00000078578 | Ube2d3 | 66105 | protein_coding | 5414.24844 | 0.194314228 | 0.083718851 | 2.32103315 | 0.020285055 | 0.080714552 | 0.194314228 |

| ensembl_gene_id | external_gene_name | entrezgene_id | gene_biotype | baseMean | log2FoldChange | lfcSE | stat | pvalue | padj | absLog2FC |
|---|---|---|---|---|---|---|---|---|---|---|
| ENSMUSG00000038024 | Dennd4c | 329877 | protein_coding | 2034.271458 | -0.194322768 | 0.105015792 | -1.850414732 | 0.064253797 | 0.184274927 | 0.194322768 |
| ENSMUSG00000074340 | Ovgp1 | 12659 | protein_coding | 8.511079235 | 0.1943789 | 0.532327636 | 0.365148992 | 0.715000192 | 0.83979965 | 0.1943789 |
| ENSMUSG00000031441 | Atp11a | 50770 | protein_coding | 816.7663106 | -0.194380328 | 0.159781913 | -1.216535239 | 0.223781096 | 0.425414053 | 0.194380328 |
| ENSMUSG00000038178 | Slc43a2 | 215113 | protein_coding | 123.4329262 | -0.194425715 | 0.139170122 | -1.39703632 | 0.16240265 | 0.34485394 | 0.194425715 |
| ENSMUSG00000026826 | Nr4a2 | 18227 | protein_coding | 116.2362932 | 0.194428875 | 0.198014462 | 0.981892299 | 0.32615291 | 0.537335907 | 0.194428875 |
| ENSMUSG00000024528 | Srfbp1 | 67222 | protein_coding | 171.852947 | -0.194442249 | 0.116171103 | -1.673757442 | 0.094178279 | 0.238941006 | 0.194442249 |
| ENSMUSG00000028744 | Slc66a1 | 212555 | protein_coding | 79.10353877 | -0.194443543 | 0.134335084 | -1.447451682 | 0.14777046 | 0.324601832 | 0.194443543 |
| ENSMUSG00000098789 | Jmjd7 | 433466 | protein_coding | 20.70346991 | -0.194519656 | 0.281988142 | -0.68981502 | 0.490310522 | 0.68272279 | 0.194519656 |
| ENSMUSG00000025369 | Smarcc2 | 68094 | protein_coding | 1062.510227 | -0.194522523 | 0.0735378 | -2.645204548 | 0.008164155 | 0.041190579 | 0.194522523 |
| ENSMUSG00000057143 | Trim12c | 319236 | protein_coding | 289.2528786 | 0.19456792 | 0.108197505 | 1.798266232 | 0.072134828 | 0.199564716 | 0.19456792 |
| ENSMUSG00000059439 | Bcas3 | 192197 | protein_coding | 266.1039596 | -0.1945775 | 0.088054424 | -2.209741329 | 0.02712312 | 0.100096629 | 0.1945775 |
| ENSMUSG00000078786 | BC024978 | 414069 | protein_coding | 92.40981143 | 0.19460882 | 0.208269339 | 0.93440936 | 0.350092771 | 0.560204708 | 0.19460882 |
| ENSMUSG00000068115 | Ninl | 78177 | protein_coding | 38.60716585 | 0.194612123 | 0.192540103 | 1.010761502 | 0.312130593 | 0.522338315 | 0.194612123 |
| ENSMUSG00000044442 | N6amt1 | 67768 | protein_coding | 117.6841011 | -0.194616667 | 0.132044365 | -1.473873326 | 0.140515701 | 0.312850724 | 0.194616667 |
| ENSMUSG00000019998 | Stx7 | 53331 | protein_coding | 1491.135411 | 0.194656312 | 0.059796499 | 3.255312854 | 0.001132675 | 0.008497998 | 0.194656312 |
| ENSMUSG00000020790 | Ankfy1 | 11736 | protein_coding | 1829.514225 | 0.194670097 | 0.063838405 | 3.049419827 | 0.002292838 | 0.015153663 | 0.194670097 |
| ENSMUSG00000004748 | Mtfp1 | 67900 | protein_coding | 166.8064676 | 0.194678972 | 0.125474771 | 1.551538766 | 0.120772625 | 0.282352321 | 0.194678972 |
| ENSMUSG00000018411 | Mapt | 17762 | protein_coding | 84.57367675 | 0.194717388 | 0.187005984 | 1.041236136 | 0.297765968 | 0.506657018 | 0.194717388 |
| ENSMUSG00000021906 | Oxnad1 | 218885 | protein_coding | 404.0998296 | 0.194726105 | 0.103218232 | 1.886547578 | 0.059221203 | 0.174551319 | 0.194726105 |
| ENSMUSG00000031683 | Lsm6 | 78651 | protein_coding | 405.8838 | 0.19480141 | 0.097179388 | 2.004554802 | 0.045010663 | 0.144600612 | 0.19480141 |
| ENSMUSG00000085154 | C130046K22Rik | 399609 | lncRNA | 7.220655594 | 0.194841854 | 0.395105157 | 0.493139232 | 0.621914212 | 0.780233898 | 0.194841854 |
| ENSMUSG00000017493 | Igfbp4 | 16010 | protein_coding | 28.49497326 | -0.194884665 | 0.335773584 | -0.580404995 | 0.561641538 | 0.739199176 | 0.194884665 |
| ENSMUSG00000047446 | Arl4a | 11861 | protein_coding | 1748.078224 | -0.194942053 | 0.115085129 | -1.693894369 | 0.090285357 | 0.232259644 | 0.194942053 |
| ENSMUSG00000038074 | Fkbp14 | 231997 | protein_coding | 60.97331284 | 0.194951195 | 0.155185986 | 1.256242265 | 0.209028147 | 0.405828413 | 0.194951195 |
| ENSMUSG00000032413 | Rasa2 | 114713 | protein_coding | 1110.750771 | -0.19497235 | 0.154494512 | -1.262001788 | 0.206948143 | 0.403361481 | 0.19497235 |
| ENSMUSG00000003604 | Aven | 74268 | protein_coding | 326.2639368 | 0.195035357 | 0.140028602 | 1.392825139 | 0.163672675 | 0.34662656 | 0.195035357 |
| ENSMUSG00000024830 | Rps6kb2 | 58988 | protein_coding | 182.7162535 | -0.195108966 | 0.095554198 | -2.041867032 | 0.041164723 | 0.1357952 | 0.195108966 |
| ENSMUSG00000061119 | Prcp | 72461 | protein_coding | 868.3044766 | -0.195132276 | 0.096405497 | -2.02407832 | 0.042962094 | 0.139653109 | 0.195132276 |
| ENSMUSG00000015176 | Nolc1 | 70769 | protein_coding | 638.3473501 | -0.195138763 | 0.209454618 | -0.931651758 | 0.351516526 | 0.56137402 | 0.195138763 |
| ENSMUSG00000023051 | Tarbp2 | 21357 | protein_coding | 243.2846393 | 0.195143853 | 0.112034651 | 1.74181694 | 0.081540482 | 0.216896591 | 0.195143853 |
| ENSMUSG00000057147 | Dph6 | 66632 | protein_coding | 89.08303524 | -0.195234422 | 0.163195654 | -1.196321207 | 0.231571236 | 0.435011988 | 0.195234422 |
| ENSMUSG00000016409 | Nkap | 67050 | protein_coding | 275.3601463 | -0.195268024 | 0.079851107 | -2.445401579 | 0.014469093 | 0.062813769 | 0.195268024 |
| ENSMUSG00000071203 | Naip5 | 17951 | protein_coding | 921.8877431 | 0.195277981 | 0.088143382 | 2.215458229 | 0.026728628 | 0.099076936 | 0.195277981 |
| ENSMUSG00000068328 | Aup1 | 11993 | protein_coding | 1132.769813 | 0.195337882 | 0.08744241 | 2.233903232 | 0.025489441 | 0.095957413 | 0.195337882 |
| ENSMUSG00000033411 | Ctdspl2 | 329506 | protein_coding | 552.178369 | -0.195353376 | 0.092119365 | -2.120654825 | 0.033950861 | 0.118226784 | 0.195353376 |
| ENSMUSG00000070923 | Klhl9 | 242521 | protein_coding | 2105.548183 | 0.195378001 | 0.067304651 | 2.902890042 | 0.003697364 | 0.022100577 | 0.195378001 |
| ENSMUSG00000027349 | Fam98b | 68215 | protein_coding | 140.1586699 | 0.195382375 | 0.165716453 | 1.179016151 | 0.238391745 | 0.44255288 | 0.195382375 |
| ENSMUSG00000096957 | E230013L22Rik | 102132 | lncRNA | 27.61224368 | 0.195396411 | 0.241400113 | 0.809429658 | 0.418268048 | 0.624549748 | 0.195396411 |

| ensembl_gene_id | external_gene_name | entrezgene_id | gene_biotype | baseMean | log2FoldChange | lfcSE | stat | pvalue | padj | absLog2FC |
|---|---|---|---|---|---|---|---|---|---|---|
| ENSMUSG00000038619 | Ensa | 56205 | protein_coding | 1481.710015 | 0.195500501 | 0.088562113 | 2.207495892 | 0.027279434 | 0.100464004 | 0.195500501 |
| ENSMUSG00000041926 | Rnpep | 215615 | protein_coding | 1047.971915 | 0.195530327 | 0.105763744 | 1.848746275 | 0.064494458 | 0.184760383 | 0.195530327 |
| ENSMUSG00000020111 | Micu1 | 216001 | protein_coding | 1481.009816 | -0.195579984 | 0.06015394 | -3.251324562 | 0.001148686 | 0.008575172 | 0.195579984 |
| ENSMUSG00000036672 | Cenpt | 320394 | protein_coding | 95.27231373 | 0.195598994 | 0.146768801 | 1.332701449 | 0.182629798 | 0.371747913 | 0.195598994 |
| ENSMUSG00000032352 | Lrrc1 | 214345 | protein_coding | 1014.542378 | 0.195719534 | 0.128058226 | 1.528363623 | 0.126422281 | 0.291200882 | 0.195719534 |
| ENSMUSG00000052396 | Mogat2 | 233549 | protein_coding | 71.32456541 | 0.195735463 | 0.275593307 | 0.710233005 | 0.477559657 | 0.672136628 | 0.195735463 |
| ENSMUSG00000026812 | Tsc1 | 64930 | protein_coding | 456.6522933 | -0.195747612 | 0.106730649 | -1.834033744 | 0.066648985 | 0.188759786 | 0.195747612 |
| ENSMUSG00000037674 | Rfx7 | 319758 | protein_coding | 620.1728437 | 0.195786067 | 0.115771764 | 1.691138327 | 0.090810387 | 0.233144397 | 0.195786067 |
| ENSMUSG00000050357 | Carmil2 | 234695 | protein_coding | 12.15875567 | -0.19586311 | 0.391011681 | -0.500913706 | 0.616431856 | 0.776720719 | 0.19586311 |
| ENSMUSG00000056050 | Mia3 | 338366 | protein_coding | 4952.848837 | 0.195939815 | 0.083690129 | 2.341253596 | 0.019219108 | 0.077695925 | 0.195939815 |
| ENSMUSG00000057789 | Bak1 | 12018 | protein_coding | 766.0080231 | 0.195940584 | 0.097338879 | 2.012973494 | 0.044117418 | 0.142305029 | 0.195940584 |
| ENSMUSG00000028300 | C9orf72 | 73205 | protein_coding | 255.5100457 | 0.196032409 | 0.177927227 | 1.101756106 | 0.270567718 | 0.478449592 | 0.196032409 |
| ENSMUSG00000024975 | Pdcd4 | 18569 | protein_coding | 2710.655725 | 0.196076137 | 0.166462447 | 1.177900119 | 0.238836428 | 0.442965373 | 0.196076137 |
| ENSMUSG00000040836 | Gpr161 | 240888 | protein_coding | 8.202575371 | -0.19608221 | 0.577608041 | -0.339472785 | 0.734253596 | 0.850992598 | 0.19608221 |
| ENSMUSG00000061838 | Suclg2 | 20917 | protein_coding | 3887.220685 | -0.196126182 | 0.092655287 | -2.116729528 | 0.034282809 | 0.11920564 | 0.196126182 |
| ENSMUSG00000067787 | Blcap | 53619 | protein_coding | 378.9242366 | -0.196182298 | 0.10263415 | -1.911471939 | 0.05594396 | 0.167596009 | 0.196182298 |
| ENSMUSG00000040697 | Dnajc16 | 214063 | protein_coding | 361.210999 | -0.196232044 | 0.084305781 | -2.327622638 | 0.01993215 | 0.079588691 | 0.196232044 |
| ENSMUSG00000029344 | Tpst2 | 22022 | protein_coding | 156.7268768 | 0.196246137 | 0.112274492 | 1.747913827 | 0.08047895 | 0.214718379 | 0.196246137 |
| ENSMUSG00000044791 | Setd2 | 235626 | protein_coding | 1532.584062 | -0.196249131 | 0.083983907 | -2.336746856 | 0.019452351 | 0.078439447 | 0.196249131 |
| ENSMUSG00000028540 | Dph2 | 67728 | protein_coding | 109.4303145 | -0.196271765 | 0.170877372 | -1.148611793 | 0.250716092 | 0.457061914 | 0.196271765 |
| ENSMUSG00000049799 | Lrrc19 | 100061 | protein_coding | 1082.402367 | 0.196309726 | 0.13801359 | 1.422394168 | 0.154911855 | 0.334658352 | 0.196309726 |
| ENSMUSG00000078234 | Klhdc7a | 242721 | protein_coding | 175.3807955 | 0.196389373 | 0.243905372 | 0.805186746 | 0.420711924 | 0.626680962 | 0.196389373 |
| ENSMUSG00000035093 | Secisbp2l | 70354 | protein_coding | 1464.609436 | 0.196490585 | 0.113815627 | 1.72639373 | 0.084276603 | 0.222024104 | 0.196490585 |
| ENSMUSG00000026269 | Rnpepl1 | 108657 | protein_coding | 872.7219997 | -0.196552922 | 0.07709252 | -2.549571898 | 0.010785527 | 0.050646189 | 0.196552922 |
| ENSMUSG00000071256 | Zfp213 | 449521 | protein_coding | 150.1263246 | 0.196552983 | 0.136084517 | 1.444344943 | 0.14864197 | 0.325707703 | 0.196552983 |
| ENSMUSG00000038481 | Cdk19 | 78334 | protein_coding | 533.0765749 | -0.196608441 | 0.059873501 | -3.283730492 | 0.001024428 | 0.007814701 | 0.196608441 |
| ENSMUSG00000031445 | Proz | 66901 | protein_coding | 19.98219162 | 0.19668003 | 0.240695919 | 0.817130725 | 0.413853726 | 0.620277776 | 0.19668003 |
| ENSMUSG00000039427 | Alg1 | 208211 | protein_coding | 147.1444017 | -0.196705119 | 0.107241658 | -1.834223024 | 0.066620895 | 0.188713786 | 0.196705119 |
| ENSMUSG00000024246 | Thumpd2 | 72167 | protein_coding | 41.45544459 | -0.196740356 | 0.201650614 | -0.975649676 | 0.329238098 | 0.539905737 | 0.196740356 |
| ENSMUSG00000006021 | Kptn | 70394 | protein_coding | 204.036008 | -0.196792576 | 0.095383346 | -2.063175441 | 0.039095963 | 0.131043875 | 0.196792576 |
| ENSMUSG00000020413 | Hus1 | 15574 | protein_coding | 128.9540767 | -0.196883964 | 0.120295455 | -1.636670014 | 0.101699438 | 0.250948811 | 0.196883964 |
| ENSMUSG00000036840 | Siah1a | 20437 | protein_coding | 233.561148 | -0.196926542 | 0.108521421 | -1.814632901 | 0.069580343 | 0.194981651 | 0.196926542 |
| ENSMUSG00000063787 | Chchd1 | 66121 | protein_coding | 403.0005509 | 0.196948161 | 0.099149196 | 1.986381817 | 0.046990944 | 0.149008911 | 0.196948161 |
| ENSMUSG00000057278 | Snrpg | 68011 | protein_coding | 317.8867797 | 0.196958465 | 0.130963281 | 1.503921283 | 0.132601636 | 0.300865935 | 0.196958465 |
| ENSMUSG00000022394 | L3mbtl2 | 214669 | protein_coding | 227.8837989 | -0.196963911 | 0.104263079 | -1.889105073 | 0.058877751 | 0.173815646 | 0.196963911 |
| ENSMUSG00000027134 | Lpcat4 | 99010 | protein_coding | 218.6540816 | 0.196975659 | 0.327071618 | 0.602240146 | 0.547014296 | 0.728864289 | 0.196975659 |
| ENSMUSG00000020793 | Galr2 | 14428 | protein_coding | 7.174468628 | -0.197016692 | 0.44744858 | -0.440311357 | 0.659711616 | 0.802969027 | 0.197016692 |
| ENSMUSG00000056486 | Chn1 | 108699 | protein_coding | 7.438402961 | -0.197083505 | 0.46009268 | -0.428356098 | 0.668391883 | 0.807885474 | 0.197083505 |

| ensembl_gene_id | external_gene_name | entrezgene_id | gene_biotype | baseMean | log2FoldChange | lfcSE | stat | pvalue | padj | absLog2FC |
|---|---|---|---|---|---|---|---|---|---|---|
| ENSMUSG00000038203 | Hoxa13 | 15398 | protein_coding | 456.3348078 | 0.197116759 | 0.24276945 | 0.811950429 | 0.416820076 | 0.622981156 | 0.197116759 |
| ENSMUSG00000040824 | Snrpd2 | 107686 | protein_coding | 578.19811 | 0.197252954 | 0.112717326 | 1.749979008 | 0.080121936 | 0.213992665 | 0.197252954 |
| ENSMUSG00000023913 | Pla2g7 | 27226 | protein_coding | 8.380639724 | -0.19729832 | 0.504326563 | -0.391211438 | 0.695640955 | 0.826693212 | 0.19729832 |
| ENSMUSG00000073147 | 5031425E22Rik | 269630 | lncRNA | 201.1602892 | -0.19730746 | 0.137043744 | -1.43974073 | 0.149940765 | 0.327427544 | 0.19730746 |
| ENSMUSG00000050017 | Pitpnb | 56305 | protein_coding | 403.9741325 | 0.197334374 | 0.105299679 | 1.874026364 | 0.060926791 | 0.177509042 | 0.197334374 |
| ENSMUSG00000039485 | Tspyl4 | 72480 | protein_coding | 13.12553587 | -0.197376299 | 0.332936476 | -0.592834711 | 0.553291776 | 0.733058676 | 0.197376299 |
| ENSMUSG00000033106 | Slc7a6os | 66432 | protein_coding | 280.9241885 | -0.197402753 | 0.122116545 | -1.616511116 | 0.105983852 | 0.258081455 | 0.197402753 |
| ENSMUSG00000035569 | Ankrd11 | 77087 | protein_coding | 2478.43221 | -0.197414012 | 0.087638674 | -2.252590128 | 0.024285004 | 0.092383937 | 0.197414012 |
| ENSMUSG00000071533 | Pcnp | 76302 | protein_coding | 797.5860283 | 0.197415435 | 0.062915326 | 3.137795637 | 0.001702235 | 0.011814031 | 0.197415435 |
| ENSMUSG00000041483 | Zfp281 | 226442 | protein_coding | 174.2255978 | 0.197417697 | 0.179424612 | 1.100282144 | 0.271209209 | 0.479281529 | 0.197417697 |
| ENSMUSG00000038172 | Ttc39b | 69863 | protein_coding | 1166.889188 | -0.19742285 | 0.090346448 | -2.185175564 | 0.028875977 | 0.10488861 | 0.19742285 |
| ENSMUSG00000000001 | Gnai3 | 14679 | protein_coding | 3442.738063 | 0.19742465 | 0.101079551 | 1.953161123 | 0.050800506 | 0.157122878 | 0.19742465 |
| ENSMUSG00000036752 | Tubb4b | 227613 | protein_coding | 8086.446297 | 0.197451701 | 0.162495696 | 1.215119571 | 0.22432049 | 0.425999491 | 0.197451701 |
| ENSMUSG00000030246 | Ldhb | 16832 | protein_coding | 87.38919429 | 0.197512246 | 0.174052662 | 1.1347844 | 0.256465664 | 0.463387217 | 0.197512246 |
| ENSMUSG00000021290 | Atp5mpl | 70257 | protein_coding | 2314.793654 | 0.197547649 | 0.123563896 | 1.59874895 | 0.109876396 | 0.264167734 | 0.197547649 |
| ENSMUSG00000028741 | Mrto4 | 69902 | protein_coding | 238.1443198 | -0.197589263 | 0.230393081 | -0.857618043 | 0.391103409 | 0.599220382 | 0.197589263 |
| ENSMUSG00000024127 | Prepl | 213760 | protein_coding | 710.3772515 | 0.197600951 | 0.063050239 | 3.134023798 | 0.001724269 | 0.011956531 | 0.197600951 |
| ENSMUSG00000023791 | Pigx | 72084 | protein_coding | 314.269553 | 0.197638955 | 0.101889122 | 1.939745397 | 0.05241064 | 0.160383605 | 0.197638955 |
| ENSMUSG00000049811 | Fam161a | 73873 | protein_coding | 99.15155708 | -0.197683668 | 0.222248366 | -0.889471861 | 0.37374954 | 0.582173729 | 0.197683668 |
| ENSMUSG00000057230 | Aak1 | 269774 | protein_coding | 519.2023113 | -0.197698822 | 0.141347664 | -1.398670594 | 0.161911786 | 0.344178691 | 0.197698822 |
| ENSMUSG00000039899 | Fgl2 | 14190 | protein_coding | 928.8924671 | 0.197725706 | 0.216045422 | 0.915204331 | 0.360084371 | 0.569612063 | 0.197725706 |
| ENSMUSG00000022419 | Deptor | 97998 | protein_coding | 1731.053777 | -0.197744468 | 0.169913271 | -1.163796489 | 0.244506495 | 0.449763189 | 0.197744468 |
| ENSMUSG00000042460 | C1galt1 | 94192 | protein_coding | 5965.097566 | 0.197745329 | 0.068584855 | 2.883221493 | 0.003936306 | 0.023169799 | 0.197745329 |
| ENSMUSG00000081984 | Dnajb3 | 15504 | protein_coding | 14.33112033 | 0.19778175 | 0.462864835 | 0.427299149 | 0.669161453 | 0.808373426 | 0.19778175 |
| ENSMUSG00000030505 | Prmt3 | 71974 | protein_coding | 215.7073974 | -0.197791562 | 0.15780852 | -1.25336428 | 0.210073165 | 0.407309235 | 0.197791562 |
| ENSMUSG00000058761 | Rnf169 | 108937 | protein_coding | 393.0528955 | -0.197812364 | 0.094721503 | -2.088357529 | 0.036765596 | 0.125262547 | 0.197812364 |
| ENSMUSG00000021265 | Slc25a29 | 214663 | protein_coding | 62.66164169 | -0.197842908 | 0.17280627 | -1.144882692 | 0.252257754 | 0.458547106 | 0.197842908 |
| ENSMUSG00000007815 | Rhoa | 11848 | protein_coding | 3359.779152 | 0.197981887 | 0.070416783 | 2.811572459 | 0.004929999 | 0.02773854 | 0.197981887 |
| ENSMUSG00000020380 | Rad50 | 19360 | protein_coding | 573.9438843 | -0.198001263 | 0.093329441 | -2.121530591 | 0.033877176 | 0.118021799 | 0.198001263 |
| ENSMUSG00000034189 | Hsdl1 | 72552 | protein_coding | 383.9465972 | -0.198030284 | 0.093428188 | -2.119598894 | 0.034039886 | 0.1184591 | 0.198030284 |
| ENSMUSG00000021114 | Atp6v1d | 73834 | protein_coding | 594.3189623 | 0.198071331 | 0.092113028 | 2.150307461 | 0.031530903 | 0.112057438 | 0.198071331 |
| ENSMUSG00000035614 | Togaram1 | 328108 | protein_coding | 709.4229875 | -0.198126259 | 0.072185219 | -2.744692911 | 0.006056754 | 0.032468905 | 0.198126259 |
| ENSMUSG00000029338 | Antxr2 | 71914 | protein_coding | 757.455158 | -0.198159617 | 0.11972741 | -1.655089811 | 0.097906281 | 0.244811813 | 0.198159617 |
| ENSMUSG00000051403 | Ppp1r37 | 232947 | protein_coding | 1298.001598 | -0.198213693 | 0.090407883 | -2.192438151 | 0.028347884 | 0.103323822 | 0.198213693 |
| ENSMUSG00000041936 | Agrn | 11603 | protein_coding | 692.7418523 | -0.198223076 | 0.117719579 | -1.683858188 | 0.092209079 | 0.235747384 | 0.198223076 |
| ENSMUSG00000043257 | Pigv | 230801 | protein_coding | 287.2314919 | 0.198234496 | 0.141704813 | 1.398925637 | 0.161835284 | 0.344106824 | 0.198234496 |
| ENSMUSG00000053192 | Mllt11 | 56772 | protein_coding | 59.64722045 | -0.198310372 | 0.150381538 | -1.318714876 | 0.187264451 | 0.377945791 | 0.198310372 |
| ENSMUSG00000022820 | Ndufb4 | 68194 | protein_coding | 476.339601 | 0.198326967 | 0.085221302 | 2.327199448 | 0.019954652 | 0.079644384 | 0.198326967 |

| ensembl_gene_id | external_gene_name | entrezgene_id | gene_biotype | baseMean | log2FoldChange | lfcSE | stat | pvalue | padj | absLog2FC |
|---|---|---|---|---|---|---|---|---|---|---|
| ENSMUSG00000013833 | Med16 | 216154 | protein_coding | 283.2917084 | -0.198352103 | 0.098629333 | -2.011086316 | 0.04431634 | 0.142773193 | 0.198352103 |
| ENSMUSG00000020063 | Sirt1 | 93759 | protein_coding | 407.3916173 | -0.198352437 | 0.091701193 | -2.163030069 | 0.030538867 | 0.109480773 | 0.198352437 |
| ENSMUSG00000027863 | Cd2 | 12481 | protein_coding | 8.902775857 | 0.198399772 | 0.461863981 | 0.429563205 | 0.667513408 | 0.80727602 | 0.198399772 |
| ENSMUSG00000036202 | Rif1 | 51869 | protein_coding | 971.1667078 | -0.198441243 | 0.083778242 | -2.368648917 | 0.017853192 | 0.073498484 | 0.198441243 |
| ENSMUSG00000071722 | Spin4 | 270624 | protein_coding | 15.49094364 | 0.198483385 | 0.330885759 | 0.599854722 | 0.54860306 | 0.730340602 | 0.198483385 |
| ENSMUSG00000030909 | Anks4b | 72074 | protein_coding | 1991.405513 | 0.198486998 | 0.128574389 | 1.543752211 | 0.122648372 | 0.285438819 | 0.198486998 |
| ENSMUSG00000042396 | Rbm7 | 67010 | protein_coding | 929.3428269 | 0.198521827 | 0.079175745 | 2.507356614 | 0.012163792 | 0.055448884 | 0.198521827 |
| ENSMUSG00000042258 | Isl1 | 16392 | protein_coding | 14.83816376 | 0.198585645 | 0.331238445 | 0.599524749 | 0.548823011 | 0.730540943 | 0.198585645 |
| ENSMUSG00000039087 | Rreb1 | 68750 | protein_coding | 935.1308035 | -0.198707918 | 0.151887123 | -1.308260467 | 0.190784979 | 0.382916223 | 0.198707918 |
| ENSMUSG00000046743 | Fat4 | 329628 | protein_coding | 14.29592189 | -0.198735987 | 0.493092016 | -0.403040367 | 0.686918529 | 0.821304393 | 0.198735987 |
| ENSMUSG00000031788 | Kifc3 | 16582 | protein_coding | 217.6693189 | 0.198801716 | 0.139102454 | 1.429174755 | 0.152954012 | 0.331823958 | 0.198801716 |
| ENSMUSG00000035629 | Rubcn | 100502698 | protein_coding | 432.0186122 | -0.198878206 | 0.084678749 | -2.34862003 | 0.018843125 | 0.076545088 | 0.198878206 |
| ENSMUSG00000031365 | Zfp275 | 27081 | protein_coding | 269.3681091 | -0.198888025 | 0.146776531 | -1.35503969 | 0.175404912 | 0.36173339 | 0.198888025 |
| ENSMUSG00000021365 | Nedd9 | 18003 | protein_coding | 1807.979032 | -0.198908243 | 0.119259391 | -1.667862306 | 0.095343064 | 0.240822125 | 0.198908243 |
| ENSMUSG00000023908 | Pkmyt1 | 268930 | protein_coding | 146.3876464 | -0.198921527 | 0.155566219 | -1.278693589 | 0.201004978 | 0.395650926 | 0.198921527 |
| ENSMUSG00000054079 | Utp18 | 217109 | protein_coding | 332.6290528 | -0.198971133 | 0.162510165 | -1.224361155 | 0.220816016 | 0.421469867 | 0.198971133 |
| ENSMUSG00000010660 | Plcd1 | 18799 | protein_coding | 444.591232 | 0.198993048 | 0.125444816 | 1.586299497 | 0.112671386 | 0.26837593 | 0.198993048 |
| ENSMUSG00000016541 | Atxn10 | 54138 | protein_coding | 325.1356918 | 0.199013617 | 0.091069761 | 2.185287576 | 0.028867768 | 0.104882711 | 0.199013617 |
| ENSMUSG00000097360 | 9430065F17Rik | 77857 | lncRNA | 7.762132681 | 0.199096003 | 0.591112688 | 0.336815648 | 0.736255877 | 0.852320925 | 0.199096003 |
| ENSMUSG00000050711 | Scg2 | 20254 | protein_coding | 26.15069414 | 0.199179544 | 0.423716524 | 0.470077357 | 0.638299751 | 0.790506101 | 0.199179544 |
| ENSMUSG00000024366 | Gfra3 | 14587 | protein_coding | 26.35480827 | 0.199224584 | 0.242198869 | 0.82256612 | 0.410754772 | 0.617741668 | 0.199224584 |
| ENSMUSG00000032388 | Spg21 | 27965 | protein_coding | 524.7444949 | 0.199314908 | 0.1067537 | 1.867053872 | 0.061894065 | 0.179473634 | 0.199314908 |
| ENSMUSG00000048154 | Kmt2d | 381022 | protein_coding | 1051.546637 | -0.199323482 | 0.27594748 | -0.722323981 | 0.470095312 | 0.66616046 | 0.199323482 |
| ENSMUSG00000001674 | Ddx18 | 66942 | protein_coding | 269.1079657 | -0.199356225 | 0.15459164 | -1.289566665 | 0.197201156 | 0.390965896 | 0.199356225 |
| ENSMUSG00000019732 | Calr3 | 73316 | protein_coding | 10.73906281 | -0.199372671 | 0.356776222 | -0.558817148 | 0.576286518 | 0.748585685 | 0.199372671 |
| ENSMUSG00000003779 | Kif20a | 19348 | protein_coding | 396.7811304 | 0.199438361 | 0.148376542 | 1.344136742 | 0.178904173 | 0.366360585 | 0.199438361 |
| ENSMUSG00000037025 | Foxa2 | 15376 | protein_coding | 375.8360906 | -0.199471179 | 0.229629429 | -0.868665572 | 0.385030077 | 0.59293813 | 0.199471179 |
| ENSMUSG00000086728 | Man2c1os | 76794 | lncRNA | 19.28928626 | -0.199471659 | 0.333550491 | -0.598025381 | 0.549822998 | 0.731093174 | 0.199471659 |
| ENSMUSG00000035595 | Fam174c | 69770 | protein_coding | 110.7296811 | 0.199481461 | 0.161518981 | 1.235034172 | 0.216817787 | 0.416031045 | 0.199481461 |
| ENSMUSG00000029600 | Rita1 | 100764 | protein_coding | 159.536246 | 0.199502911 | 0.14163377 | 1.408582937 | 0.158958527 | 0.340262253 | 0.199502911 |
| ENSMUSG00000055737 | Ghr | 14600 | protein_coding | 308.3756469 | -0.199537542 | 0.147036623 | -1.35706015 | 0.174762099 | 0.361071142 | 0.199537542 |
| ENSMUSG00000001604 | Tcea3 | 21401 | protein_coding | 640.5992999 | -0.199590294 | 0.109613616 | -1.82085311 | 0.068629189 | 0.192961291 | 0.199590294 |
| ENSMUSG00000017831 | Rab5a | 271457 | protein_coding | 1412.536212 | 0.199592201 | 0.0627216 | 3.182192421 | 0.001461647 | 0.01043528 | 0.199592201 |
| ENSMUSG00000029185 | Fam114a1 | 68303 | protein_coding | 1445.1347 | 0.199624542 | 0.08173934 | 2.442208879 | 0.014597698 | 0.06316548 | 0.199624542 |
| ENSMUSG00000028233 | Tgs1 | 116940 | protein_coding | 202.1276185 | -0.199676479 | 0.113688723 | -1.756343756 | 0.07902973 | 0.212095289 | 0.199676479 |
| ENSMUSG00000045917 | Tmem268 | 230279 | protein_coding | 345.3350675 | 0.199696554 | 0.074644653 | 2.675296154 | 0.007466326 | 0.038397403 | 0.199696554 |
| ENSMUSG00000028467 | Gba2 | 230101 | protein_coding | 184.3314455 | -0.199704825 | 0.11522257 | -1.733209258 | 0.083058482 | 0.219701126 | 0.199704825 |
| ENSMUSG00000022440 | C1qtnf6 | 72709 | protein_coding | 183.1619975 | -0.199844866 | 0.146108146 | -1.367787301 | 0.171378664 | 0.35687271 | 0.199844866 |

| ensembl_gene_id | external_gene_name | entrezgene_id | gene_biotype | baseMean | log2FoldChange | lfcSE | stat | pvalue | padj | absLog2FC |
|---|---|---|---|---|---|---|---|---|---|---|
| ENSMUSG00000022701 | Ccdc191 | 212153 | protein_coding | 115.2928278 | -0.199926445 | 0.173830528 | -1.150122752 | 0.250093317 | 0.45640311 | 0.199926445 |
| ENSMUSG00000026869 | Psmd5 | 66998 | protein_coding | 808.0031788 | 0.199934308 | 0.099244782 | 2.014557388 | 0.043951046 | 0.141883312 | 0.199934308 |
| ENSMUSG00000079507 | H2-Q1 | 15006 | protein_coding | 211.0623537 | 0.20001932 | 0.281279391 | 0.711105494 | 0.477018866 | 0.671553517 | 0.20001932 |
| ENSMUSG00000063362 | Alg11 | 207958 | protein_coding | 548.6557896 | 0.200024555 | 0.071561303 | 2.795149694 | 0.005187569 | 0.028840348 | 0.200024555 |
| ENSMUSG00000040723 | Rcsd1 | 226594 | protein_coding | 15.66725488 | 0.200058198 | 0.444425422 | 0.450150212 | 0.652602133 | 0.798666675 | 0.200058198 |
| ENSMUSG00000046111 | Cep295 | 319675 | protein_coding | 312.6084373 | -0.200211599 | 0.12500609 | -1.601614763 | 0.109240824 | 0.263315352 | 0.200211599 |
| ENSMUSG00000094338 | H2bc13 | 319185 | protein_coding | 485.9692653 | 0.200212438 | 0.169233446 | 1.183054789 | 0.236787434 | 0.440992616 | 0.200212438 |
| ENSMUSG00000058402 | Zfp420 | 233058 | protein_coding | 64.27177189 | 0.200259143 | 0.189670653 | 1.055825661 | 0.291047872 | 0.499573396 | 0.200259143 |
| ENSMUSG00000021115 | Vrk1 | 22367 | protein_coding | 293.2402508 | 0.200355994 | 0.111307292 | 1.800025769 | 0.071856569 | 0.199083569 | 0.200355994 |
| ENSMUSG00000024220 | Zfp523 | 224656 | protein_coding | 225.9856706 | 0.200486129 | 0.117474847 | 1.706630263 | 0.087890797 | 0.228448759 | 0.200486129 |
| ENSMUSG00000103567 | Pcdhga5 | 93713 | protein_coding | 9.770646819 | -0.200501976 | 0.359755113 | -0.557329051 | 0.57730263 | 0.749071904 | 0.200501976 |
| ENSMUSG00000020152 | Actr2 | 66713 | protein_coding | 4095.169367 | 0.200527239 | 0.094038314 | 2.132399338 | 0.032974032 | 0.11561651 | 0.200527239 |
| ENSMUSG00000057130 | Txnl4a | 27366 | protein_coding | 332.232776 | 0.200556684 | 0.103280171 | 1.941870175 | 0.052152819 | 0.159965985 | 0.200556684 |
| ENSMUSG00000021536 | Adcy2 | 210044 | protein_coding | 37.77725449 | 0.200593229 | 0.175831988 | 1.140823302 | 0.253943461 | 0.460370072 | 0.200593229 |
| ENSMUSG00000029088 | Kcnip4 | 80334 | protein_coding | 9.252183932 | -0.200597017 | 0.455630979 | -0.440262025 | 0.659747342 | 0.802969027 | 0.200597017 |
| ENSMUSG00000071266 | Zfp946 | 74149 | protein_coding | 482.4261657 | 0.200600664 | 0.093225191 | 2.151786037 | 0.03141421 | 0.111788547 | 0.200600664 |
| ENSMUSG00000091337 | Eid1 | 58521 | protein_coding | 290.6542022 | -0.20062722 | 0.143232823 | -1.400707015 | 0.161301704 | 0.343340043 | 0.20062722 |
| ENSMUSG00000042298 | Ttc19 | 72795 | protein_coding | 346.8258407 | -0.200634697 | 0.083106637 | -2.414183792 | 0.015770505 | 0.067199733 | 0.200634697 |
| ENSMUSG00000031921 | Terf2 | 21750 | protein_coding | 219.3210011 | -0.200647841 | 0.103447508 | -1.939610199 | 0.05242708 | 0.1604031 | 0.200647841 |
| ENSMUSG00000043061 | Tmem18 | 211986 | protein_coding | 387.805706 | 0.200658028 | 0.097773754 | 2.052268835 | 0.040143543 | 0.133429475 | 0.200658028 |
| ENSMUSG00000058351 | Smim4 | 66487 | protein_coding | 84.35622374 | -0.200673286 | 0.163846291 | -1.224765507 | 0.220663584 | 0.421379879 | 0.200673286 |
| ENSMUSG00000024079 | Eif2ak2 | 19106 | protein_coding | 826.8703105 | 0.200742748 | 0.074127302 | 2.708081144 | 0.006767347 | 0.035469006 | 0.200742748 |
| ENSMUSG00000030465 | Psd3 | 234353 | protein_coding | 1057.328544 | -0.200756873 | 0.097467285 | -2.059735974 | 0.039423788 | 0.131832721 | 0.200756873 |
| ENSMUSG00000090110 | Cmc4 | 105886298 | protein_coding | 36.63323385 | 0.200779165 | 0.189215489 | 1.061113792 | 0.288638191 | 0.497611116 | 0.200779165 |
| ENSMUSG00000043683 | Fem1a | 14154 | protein_coding | 777.6677808 | -0.200900865 | 0.102748985 | -1.955258873 | 0.050552523 | 0.156623339 | 0.200900865 |
| ENSMUSG00000031355 | Arhgap6 | 11856 | protein_coding | 35.70883931 | 0.200914205 | 0.251490337 | 0.798894331 | 0.424351687 | 0.628926553 | 0.200914205 |
| ENSMUSG00000029048 | Rer1 | 67830 | protein_coding | 2329.066965 | 0.20092888 | 0.076034987 | 2.642584535 | 0.008227592 | 0.041435132 | 0.20092888 |
| ENSMUSG00000014498 | Ankrd52 | 237615 | protein_coding | 404.6392178 | -0.200961831 | 0.14921204 | -1.346820472 | 0.17803806 | 0.365248611 | 0.200961831 |
| ENSMUSG00000021236 | Entpd5 | 12499 | protein_coding | 4542.003867 | -0.201062522 | 0.11950294 | -1.682490168 | 0.09247383 | 0.236203505 | 0.201062522 |
| ENSMUSG00000025198 | Erlin1 | 226144 | protein_coding | 768.8364578 | 0.20107078 | 0.09395074 | 2.140172402 | 0.032340837 | 0.114041365 | 0.20107078 |
| ENSMUSG00000091144 | Phf11c | 628705 | protein_coding | 26.9684438 | 0.201100476 | 0.268290813 | 0.749561542 | 0.453518821 | 0.653148289 | 0.201100476 |
| ENSMUSG00000002486 | Tchp | 77832 | protein_coding | 84.7979297 | -0.201121751 | 0.152431514 | -1.319423687 | 0.187027507 | 0.377677612 | 0.201121751 |
| ENSMUSG00000015112 | Slc25a13 | 50799 | protein_coding | 760.8820445 | 0.201129038 | 0.06983658 | 2.879995543 | 0.003976808 | 0.023373652 | 0.201129038 |
| ENSMUSG00000037344 | Slc12a9 | 83704 | protein_coding | 196.638443 | -0.201131375 | 0.105082627 | -1.914030707 | 0.055616232 | 0.167016468 | 0.201131375 |
| ENSMUSG00000038072 | Galnt11 | 231050 | protein_coding | 285.3152997 | 0.20115089 | 0.088553466 | 2.27151911 | 0.023115571 | 0.08913646 | 0.20115089 |
| ENSMUSG00000034664 | Itga2b | 16399 | protein_coding | 16.6104274 | 0.201151001 | 0.316981465 | 0.634582847 | 0.525700518 | 0.711354843 | 0.201151001 |
| ENSMUSG00000031333 | Abcb7 | 11306 | protein_coding | 1442.097199 | 0.201171951 | 0.059648517 | 3.372622816 | 0.000744559 | 0.006038356 | 0.201171951 |
| ENSMUSG00000087260 | Lamtor5 | 68576 | protein_coding | 594.4346627 | 0.201204561 | 0.081515164 | 2.468308356 | 0.013575332 | 0.06014684 | 0.201204561 |

| ensembl_gene_id | external_gene_name | entrezgene_id | gene_biotype | baseMean | log2FoldChange | lfcSE | stat | pvalue | padj | absLog2FC |
|---|---|---|---|---|---|---|---|---|---|---|
| ENSMUSG00000013155 | Enkd1 | 102124 | protein_coding | 23.92441734 | -0.201304727 | 0.27909391 | -0.721279542 | 0.470737541 | 0.666501035 | 0.201304727 |
| ENSMUSG00000021550 | 2210016F16Rik | 70153 | protein_coding | 970.1465676 | 0.201309114 | 0.092073618 | 2.186393018 | 0.028786864 | 0.104636506 | 0.201309114 |
| ENSMUSG00000025503 | Taldo1 | 21351 | protein_coding | 3883.265793 | 0.201353151 | 0.11222516 | 1.794189026 | 0.072783003 | 0.200676868 | 0.201353151 |
| ENSMUSG00000026098 | Pms1 | 227099 | protein_coding | 90.80329552 | -0.20137478 | 0.122785525 | -1.640053096 | 0.100994128 | 0.249904375 | 0.20137478 |
| ENSMUSG00000032479 | Map4 | 17758 | protein_coding | 917.29644 | 0.201390964 | 0.09746092 | 2.066376599 | 0.038792934 | 0.130312171 | 0.201390964 |
| ENSMUSG00000028132 | Tlcd4 | 99887 | protein_coding | 762.1435048 | 0.201419926 | 0.084326718 | 2.388565921 | 0.016914274 | 0.070743298 | 0.201419926 |
| ENSMUSG00000062937 | Mtap | 66902 | protein_coding | 84.87881406 | 0.201452238 | 0.19532845 | 1.031351234 | 0.30237614 | 0.51166014 | 0.201452238 |
| ENSMUSG00000025225 | Nfkb2 | 18034 | protein_coding | 530.6587489 | -0.201508838 | 0.125518771 | -1.605407993 | 0.108404047 | 0.262051852 | 0.201508838 |
| ENSMUSG00000051007 | Gatd1 | 213350 | protein_coding | 378.0151251 | 0.201515844 | 0.109848596 | 1.834487211 | 0.066581705 | 0.188667907 | 0.201515844 |
| ENSMUSG00000021868 | Ppif | 105675 | protein_coding | 1055.573634 | -0.201530616 | 0.112612849 | -1.789588116 | 0.07352015 | 0.201905024 | 0.201530616 |
| ENSMUSG00000034826 | Nup54 | 269113 | protein_coding | 293.5981606 | -0.201548827 | 0.12092263 | -1.666758549 | 0.095562426 | 0.241146755 | 0.201548827 |
| ENSMUSG00000042333 | Tnfrsf14 | 230979 | protein_coding | 135.0409498 | -0.201560621 | 0.12039868 | -1.674109897 | 0.094109003 | 0.238879395 | 0.201560621 |
| ENSMUSG00000041237 | Pklr | 18770 | protein_coding | 621.0190157 | 0.201632762 | 0.06678489 | 3.019137422 | 0.002534955 | 0.016480313 | 0.201632762 |
| ENSMUSG00000024866 | Acy3 | 71670 | protein_coding | 176.5779626 | 0.201652458 | 0.121607512 | 1.658223692 | 0.097272324 | 0.243839091 | 0.201652458 |
| ENSMUSG00000024424 | Ttc39c | 72747 | protein_coding | 21.11739112 | -0.201719865 | 0.306247992 | -0.658681428 | 0.510100361 | 0.698626897 | 0.201719865 |
| ENSMUSG00000028186 | Uox | 22262 | protein_coding | 32.32833645 | 0.201774484 | 0.277440234 | 0.727271895 | 0.467059412 | 0.663707223 | 0.201774484 |
| ENSMUSG00000069601 | Ank3 | 11735 | protein_coding | 1183.803119 | -0.201837111 | 0.122658902 | -1.645515385 | 0.099863573 | 0.247877043 | 0.201837111 |
| ENSMUSG00000049313 | Sorl1 | 20660 | protein_coding | 2618.11374 | -0.201854958 | 0.092829326 | -2.17447403 | 0.029669554 | 0.10701464 | 0.201854958 |
| ENSMUSG00000044433 | Camsap3 | 69697 | protein_coding | 857.7450366 | -0.201931914 | 0.100651989 | -2.006238673 | 0.04483079 | 0.144139043 | 0.201931914 |
| ENSMUSG00000035638 | Muc20 | 224116 | protein_coding | 21.87173491 | 0.201935392 | 0.292466664 | 0.690456099 | 0.489907409 | 0.682280773 | 0.201935392 |
| ENSMUSG00000034738 | Nostrin | 329416 | protein_coding | 911.7404264 | 0.201971306 | 0.183197832 | 1.102476506 | 0.270254567 | 0.478375791 | 0.201971306 |
| ENSMUSG00000033538 | Casp4 | 12363 | protein_coding | 340.9206423 | -0.201995445 | 0.094282339 | -2.142452629 | 0.032157077 | 0.113544207 | 0.201995445 |
| ENSMUSG00000025650 | Col7a1 | 12836 | protein_coding | 58.71179903 | 0.202014466 | 0.344065671 | 0.587139266 | 0.557110178 | 0.736158198 | 0.202014466 |
| ENSMUSG00000090641 | Zfp712 | 78251 | protein_coding | 82.25873887 | 0.202076937 | 0.200800517 | 1.006356656 | 0.314244036 | 0.524882446 | 0.202076937 |
| ENSMUSG00000078490 | Cfap74 | 544678 | protein_coding | 32.99078497 | -0.20214062 | 0.361508046 | -0.559159393 | 0.576052944 | 0.748585685 | 0.20214062 |
| ENSMUSG00000097276 | 4930525G20Rik | 320163 | lncRNA | 11.44718157 | 0.202259414 | 0.367469845 | 0.550410916 | 0.582037563 | 0.753102705 | 0.202259414 |
| ENSMUSG00000038007 | Acer2 | 230379 | protein_coding | 128.3866237 | -0.202316672 | 0.111817082 | -1.809353885 | 0.070396043 | 0.19615605 | 0.202316672 |
| ENSMUSG00000004018 | Fancl | 67030 | protein_coding | 149.8986033 | 0.202334963 | 0.106852159 | 1.893597332 | 0.058278478 | 0.172571035 | 0.202334963 |
| ENSMUSG00000031024 | Denn2b | 76954 | protein_coding | 578.6970202 | 0.202490441 | 0.129066325 | 1.568886711 | 0.116674339 | 0.275235318 | 0.202490441 |
| ENSMUSG00000034560 | Washc4 | 319277 | protein_coding | 722.1307298 | 0.20249935 | 0.062664664 | 3.231475862 | 0.001231527 | 0.009081373 | 0.20249935 |
| ENSMUSG00000026283 | Ing5 | 66262 | protein_coding | 380.9204602 | -0.202594326 | 0.085073741 | -2.381396701 | 0.017247127 | 0.071703093 | 0.202594326 |
| ENSMUSG00000029621 | Arpc1a | 56443 | protein_coding | 1118.670777 | 0.202607738 | 0.085114664 | 2.380409306 | 0.017293417 | 0.071813535 | 0.202607738 |
| ENSMUSG00000054387 | Mdm4 | 17248 | protein_coding | 1152.103591 | -0.202633699 | 0.08904717 | -2.275577082 | 0.022871334 | 0.088529438 | 0.202633699 |
| ENSMUSG00000005320 | Fgfr4 | 14186 | protein_coding | 265.7612321 | 0.202711378 | 0.211249045 | 0.959584828 | 0.337264208 | 0.548018112 | 0.202711378 |
| ENSMUSG00000031506 | Ptpn7 | 320139 | protein_coding | 10.64487205 | 0.202746729 | 0.689304214 | 0.294132438 | 0.768656705 | 0.873825137 | 0.202746729 |
| ENSMUSG00000062619 | 2310039H08Rik | 67101 | protein_coding | 257.3412484 | 0.202785822 | 0.094921663 | 2.136349238 | 0.032650958 | 0.11477918 | 0.202785822 |
| ENSMUSG00000029862 | Clcn1 | 12723 | protein_coding | 12.0159631 | -0.202804941 | 0.414925791 | -0.488774006 | 0.625001703 | 0.782837698 | 0.202804941 |
| ENSMUSG00000059013 | Sh2d3c | 27387 | protein_coding | 36.74596087 | 0.202810292 | 0.247646421 | 0.818951034 | 0.412814351 | 0.619566809 | 0.202810292 |

| ensembl_gene_id | external_gene_name | entrezgene_id | gene_biotype | baseMean | log2FoldChange | lfcSE | stat | pvalue | padj | absLog2FC |
|---|---|---|---|---|---|---|---|---|---|---|
| ENSMUSG00000023048 | Prr13 | 66151 | protein_coding | 1527.091298 | 0.202832607 | 0.127115953 | 1.595650286 | 0.110566894 | 0.265233231 | 0.202832607 |
| ENSMUSG00000024998 | Plce1 | 74055 | protein_coding | 1459.89111 | -0.202924259 | 0.11914086 | -1.703229774 | 0.088525077 | 0.229361781 | 0.202924259 |
| ENSMUSG00000032802 | Srxn1 | 76650 | protein_coding | 1626.315398 | 0.203007916 | 0.218688546 | 0.928296975 | 0.353253543 | 0.563000043 | 0.203007916 |
| ENSMUSG00000039031 | Arhgap18 | 73910 | protein_coding | 845.5567159 | -0.203025439 | 0.095297487 | -2.130438532 | 0.033135426 | 0.116072368 | 0.203025439 |
| ENSMUSG00000006920 | Ezh1 | 14055 | protein_coding | 597.8726978 | -0.203207004 | 0.100805636 | -2.015829783 | 0.043817778 | 0.141596585 | 0.203207004 |
| ENSMUSG00000031732 | Phlpp2 | 244650 | protein_coding | 433.5530901 | -0.203216265 | 0.146520138 | -1.386951094 | 0.165456676 | 0.349003763 | 0.203216265 |
| ENSMUSG00000039826 | Trub2 | 227682 | protein_coding | 163.5986679 | -0.203306888 | 0.105350227 | -1.929819184 | 0.053629246 | 0.162748779 | 0.203306888 |
| ENSMUSG00000035285 | Nat14 | 269854 | protein_coding | 15.69852451 | -0.203321619 | 0.335162966 | -0.606635101 | 0.544093101 | 0.726820743 | 0.203321619 |
| ENSMUSG00000062510 | Nsl1 | 381318 | protein_coding | 100.7632348 | 0.203329125 | 0.131442583 | 1.546904511 | 0.121886273 | 0.284246641 | 0.203329125 |
| ENSMUSG00000062006 | Rpl34 | 68436 | protein_coding | 3705.71778 | 0.203362834 | 0.114931353 | 1.769428703 | 0.076822359 | 0.207781718 | 0.203362834 |
| ENSMUSG00000014361 | Mertk | 17289 | protein_coding | 606.270904 | -0.203373484 | 0.165464549 | -1.229106088 | 0.219032033 | 0.419207437 | 0.203373484 |
| ENSMUSG00000007812 | Zfp655 | 72611 | protein_coding | 825.6481895 | -0.203432307 | 0.063486665 | -3.20433129 | 0.001353766 | 0.009819119 | 0.203432307 |
| ENSMUSG00000038855 | Itpkb | 320404 | protein_coding | 150.4868702 | -0.203583162 | 0.137253819 | -1.483260457 | 0.138005224 | 0.309032365 | 0.203583162 |
| ENSMUSG00000025347 | Mettl7b | 71664 | protein_coding | 3390.056184 | -0.203584666 | 0.206880913 | -0.984066938 | 0.325082602 | 0.53623816 | 0.203584666 |
| ENSMUSG00000001416 | Cct3 | 12462 | protein_coding | 2433.719966 | 0.20368539 | 0.132313178 | 1.539418776 | 0.123702093 | 0.286952957 | 0.20368539 |
| ENSMUSG00000023923 | Tbc1d5 | 72238 | protein_coding | 279.6310142 | -0.203740772 | 0.10841631 | -1.879244673 | 0.060211092 | 0.176229746 | 0.203740772 |
| ENSMUSG00000074785 | Plxnc1 | 54712 | protein_coding | 13.82628797 | 0.203761147 | 0.600424475 | 0.339361827 | 0.734337171 | 0.851027537 | 0.203761147 |
| ENSMUSG00000028438 | Kif24 | 109242 | protein_coding | 113.4356786 | 0.203806645 | 0.17253976 | 1.181215538 | 0.23751711 | 0.441993884 | 0.203806645 |
| ENSMUSG00000062908 | Acadm | 11364 | protein_coding | 2111.533272 | -0.203807537 | 0.110124792 | -1.85069623 | 0.064213267 | 0.184274927 | 0.203807537 |
| ENSMUSG00000020538 | Srebf1 | 20787 | protein_coding | 1322.23382 | 0.203834662 | 0.117458781 | 1.735371857 | 0.082674962 | 0.219036392 | 0.203834662 |
| ENSMUSG00000110277 | Gm45871 | 108168395 | protein_coding | 506.7196469 | 0.203907842 | 0.259156179 | 0.786814508 | 0.431390459 | 0.635338625 | 0.203907842 |
| ENSMUSG00000006024 | Napa | 108124 | protein_coding | 1098.570746 | 0.203953083 | 0.080750743 | 2.525711537 | 0.011546424 | 0.053359893 | 0.203953083 |
| ENSMUSG00000040997 | Abhd4 | 105501 | protein_coding | 601.0989271 | -0.203957798 | 0.089685687 | -2.27413989 | 0.022957577 | 0.088689858 | 0.203957798 |
| ENSMUSG00000020709 | Adap2 | 216991 | protein_coding | 32.85531399 | -0.203963284 | 0.225481763 | -0.904566654 | 0.365695008 | 0.574448111 | 0.203963284 |
| ENSMUSG00000058297 | Spock2 | 94214 | protein_coding | 18.90060648 | -0.204003395 | 0.302958967 | -0.673369721 | 0.500712107 | 0.691223994 | 0.204003395 |
| ENSMUSG00000024193 | Phf1 | 21652 | protein_coding | 128.4954547 | -0.204028419 | 0.108562691 | -1.879360375 | 0.060195303 | 0.176215914 | 0.204028419 |
| ENSMUSG00000046010 | Zfp830 | 66983 | protein_coding | 203.5484048 | -0.204050498 | 0.095403584 | -2.138813755 | 0.032450755 | 0.114226658 | 0.204050498 |
| ENSMUSG00000007827 | Ankrd26 | 232339 | protein_coding | 160.0932186 | 0.204125827 | 0.130362462 | 1.565832858 | 0.117387752 | 0.27663147 | 0.204125827 |
| ENSMUSG00000022521 | Crebbp | 12914 | protein_coding | 1019.657325 | -0.204157717 | 0.147402328 | -1.385037262 | 0.166041074 | 0.349853777 | 0.204157717 |
| ENSMUSG00000075463 | 4930594M22Rik | 105245866 | lncRNA | 12.35953452 | -0.204317981 | 0.390234776 | -0.523577072 | 0.600572732 | 0.766753883 | 0.204317981 |
| ENSMUSG00000049285 | Mblac1 | 330216 | protein_coding | 35.2639098 | 0.204318448 | 0.226763946 | 0.901018224 | 0.36757863 | 0.576275172 | 0.204318448 |
| ENSMUSG00000047583 | Tyw3 | 209584 | protein_coding | 36.56578872 | -0.204388443 | 0.191826351 | -1.065486794 | 0.286655699 | 0.495534183 | 0.204388443 |
| ENSMUSG00000020679 | Hnf1b | 21410 | protein_coding | 239.3404976 | -0.204401673 | 0.181331712 | -1.127225187 | 0.259647284 | 0.466884391 | 0.204401673 |
| ENSMUSG00000042510 | AA986860 | 212439 | protein_coding | 512.6668608 | 0.204413781 | 0.120917937 | 1.690516619 | 0.090929162 | 0.233298918 | 0.204413781 |
| ENSMUSG00000078570 | 1110065P20Rik | 68920 | protein_coding | 68.91532114 | 0.20444851 | 0.191298341 | 1.068741679 | 0.285186085 | 0.494227393 | 0.20444851 |
| ENSMUSG00000026427 | Eif2d | 16865 | protein_coding | 522.3945157 | 0.204552772 | 0.077234779 | 2.648454168 | 0.008086081 | 0.04083548 | 0.204552772 |
| ENSMUSG00000043702 | Pde12 | 211948 | protein_coding | 496.8251276 | -0.204577753 | 0.124511797 | -1.643039112 | 0.100374841 | 0.248719737 | 0.204577753 |
| ENSMUSG00000020925 | Ccdc43 | 52715 | protein_coding | 402.2949787 | 0.204645126 | 0.095255578 | 2.148379455 | 0.031683624 | 0.112470862 | 0.204645126 |

| ensembl_gene_id | external_gene_name | entrezgene_id | gene_biotype | baseMean | log2FoldChange | lfcSE | stat | pvalue | padj | absLog2FC |
|---|---|---|---|---|---|---|---|---|---|---|
| ENSMUSG00000021116 | Eif2s1 | 13665 | protein_coding | 1312.533794 | 0.204663138 | 0.163334285 | 1.253032315 | 0.210193947 | 0.40740724 | 0.204663138 |
| ENSMUSG00000015880 | Ncapg | 54392 | protein_coding | 390.7663693 | 0.20467417 | 0.128367784 | 1.594435648 | 0.110838494 | 0.265598845 | 0.20467417 |
| ENSMUSG00000040084 | Bub1b | 12236 | protein_coding | 644.8844678 | 0.204723511 | 0.109235528 | 1.874147675 | 0.060910074 | 0.177492802 | 0.204723511 |
| ENSMUSG00000053898 | Ech1 | 51798 | protein_coding | 3106.53894 | 0.204752589 | 0.119389056 | 1.715003002 | 0.086344677 | 0.225677181 | 0.204752589 |
| ENSMUSG00000054611 | Kdm2a | 225876 | protein_coding | 1356.738872 | -0.204845434 | 0.060957458 | -3.360465515 | 0.000778112 | 0.006253166 | 0.204845434 |
| ENSMUSG00000025323 | Sp4 | 20688 | protein_coding | 164.9619908 | -0.204866595 | 0.188266431 | -1.088173785 | 0.276518395 | 0.484850836 | 0.204866595 |
| ENSMUSG00000019775 | Rgs17 | 56533 | protein_coding | 458.4362875 | -0.20503368 | 0.152879157 | -1.34114868 | 0.179872185 | 0.367874186 | 0.20503368 |
| ENSMUSG00000099843 | Gm7160 | 635504 | lncRNA | 22.06853145 | -0.205058429 | 0.271158125 | -0.75623192 | 0.449510158 | 0.650122382 | 0.205058429 |
| ENSMUSG00000049092 | Gpr137c | 70713 | protein_coding | 13.06599408 | -0.205061291 | 0.453224137 | -0.452450066 | 0.650944786 | 0.798250711 | 0.205061291 |
| ENSMUSG00000027016 | Zfp385b | 241494 | protein_coding | 34.74978126 | -0.205082047 | 0.303387383 | -0.675974211 | 0.499057019 | 0.689656446 | 0.205082047 |
| ENSMUSG00000004383 | Large1 | 16795 | protein_coding | 216.702799 | -0.205257155 | 0.102244689 | -2.007509221 | 0.04469547 | 0.143762004 | 0.205257155 |
| ENSMUSG00000027534 | Snx16 | 74718 | protein_coding | 138.5606319 | -0.205281247 | 0.10817285 | -1.897715065 | 0.057733627 | 0.171691787 | 0.205281247 |
| ENSMUSG00000035504 | Reep6 | 70335 | protein_coding | 173.9584412 | -0.205283185 | 0.170986667 | -1.200580074 | 0.229914134 | 0.433125054 | 0.205283185 |
| ENSMUSG00000013419 | Zfp651 | 270210 | protein_coding | 8.608931919 | 0.205342554 | 0.508210408 | 0.404050273 | 0.686175751 | 0.821080862 | 0.205342554 |
| ENSMUSG00000021143 | Pacs2 | 217893 | protein_coding | 464.7493312 | -0.205367104 | 0.083411026 | -2.462109786 | 0.013812238 | 0.060807443 | 0.205367104 |
| ENSMUSG00000040820 | Hlcs | 110948 | protein_coding | 192.0147534 | -0.205456387 | 0.110149016 | -1.865258489 | 0.062145181 | 0.17987433 | 0.205456387 |
| ENSMUSG00000018239 | Zcchc10 | 67966 | protein_coding | 36.05857184 | -0.20547771 | 0.24515118 | -0.838167329 | 0.401936733 | 0.609486699 | 0.20547771 |
| ENSMUSG00000037253 | Mex3c | 240396 | protein_coding | 1075.2777 | 0.20557471 | 0.181971687 | 1.129707118 | 0.258599657 | 0.465714909 | 0.20557471 |
| ENSMUSG00000047281 | Sfn | 55948 | protein_coding | 3124.129317 | 0.205604916 | 0.229977141 | 0.894023271 | 0.37130944 | 0.579649458 | 0.205604916 |
| ENSMUSG00000032652 | Crebl2 | 232430 | protein_coding | 263.7546813 | -0.205607357 | 0.135760761 | -1.514482943 | 0.129903397 | 0.296561471 | 0.205607357 |
| ENSMUSG00000052632 | Asap2 | 211914 | protein_coding | 439.2896174 | -0.205617964 | 0.099718168 | -2.061990993 | 0.039208593 | 0.131310866 | 0.205617964 |
| ENSMUSG00000065523 | Mir192 | 387187 | miRNA | 9.067830026 | -0.205686416 | 0.447412409 | -0.459724432 | 0.645714031 | 0.795062854 | 0.205686416 |
| ENSMUSG00000004843 | Chmp2b | 68942 | protein_coding | 2302.7857 | 0.205738155 | 0.111716178 | 1.84161469 | 0.065531531 | 0.186523628 | 0.205738155 |
| ENSMUSG00000038987 | Cfap157 | 227736 | protein_coding | 52.96643314 | 0.205777279 | 0.225124906 | 0.914058258 | 0.360686235 | 0.570122672 | 0.205777279 |
| ENSMUSG00000039745 | Htatip2 | 53415 | protein_coding | 332.1096863 | 0.205813375 | 0.106464482 | 1.933164669 | 0.053215916 | 0.161983995 | 0.205813375 |
| ENSMUSG00000021696 | Elovl7 | 74559 | protein_coding | 1470.094737 | 0.205894196 | 0.09727079 | 2.116711451 | 0.034284344 | 0.11920564 | 0.205894196 |
| ENSMUSG00000034342 | Cbl | 12402 | protein_coding | 791.5434649 | -0.205917408 | 0.229503584 | -0.897229596 | 0.369596416 | 0.578227258 | 0.205917408 |
| ENSMUSG00000036632 | Alg5 | 66248 | protein_coding | 654.8362069 | 0.20595281 | 0.092766017 | 2.220132088 | 0.026409803 | 0.098329906 | 0.20595281 |
| ENSMUSG00000022020 | Naa16 | 66897 | protein_coding | 211.5042614 | -0.206170889 | 0.134088781 | -1.537570016 | 0.124153783 | 0.287706527 | 0.206170889 |
| ENSMUSG00000024146 | Cript | 56724 | protein_coding | 646.1804485 | 0.20619909 | 0.1108161 | 1.860732238 | 0.062781999 | 0.181128045 | 0.20619909 |
| ENSMUSG00000035852 | Misp | 78906 | protein_coding | 5626.258126 | 0.206255857 | 0.16478592 | 1.251659466 | 0.210693976 | 0.408016249 | 0.206255857 |
| ENSMUSG00000037210 | Fam193a | 231128 | protein_coding | 412.347485 | -0.206273711 | 0.102427634 | -2.013848248 | 0.044025468 | 0.142065976 | 0.206273711 |
| ENSMUSG00000040481 | Bptf | 207165 | protein_coding | 1171.895493 | -0.20628053 | 0.102389657 | -2.014661806 | 0.043940097 | 0.14187672 | 0.20628053 |
| ENSMUSG00000024425 | Ndfip1 | 65113 | protein_coding | 1521.469557 | 0.206324867 | 0.089479919 | 2.305823133 | 0.021120517 | 0.083248601 | 0.206324867 |
| ENSMUSG00000039131 | Gipc2 | 54120 | protein_coding | 2541.272351 | -0.206397925 | 0.07767517 | -2.657193087 | 0.00787943 | 0.03998202 | 0.206397925 |
| ENSMUSG00000028088 | Fmo5 | 14263 | protein_coding | 383.9932571 | -0.206442241 | 0.193423365 | -1.067307671 | 0.285832925 | 0.494594985 | 0.206442241 |
| ENSMUSG00000049680 | Urgcp | 72046 | protein_coding | 1144.582241 | 0.206477551 | 0.171114144 | 1.206665601 | 0.227560952 | 0.430167439 | 0.206477551 |
| ENSMUSG00000066880 | Zfp617 | 170938 | protein_coding | 310.8248537 | 0.206487057 | 0.110814504 | 1.863357686 | 0.062411959 | 0.180449752 | 0.206487057 |

Supplementary Table 1

| ensembl_gene_id | external_gene_name | entrezgene_id | gene_biotype | baseMean | log2FoldChange | lfcSE | stat | pvalue | padj | absLog2FC |
|---|---|---|---|---|---|---|---|---|---|---|
| ENSMUSG00000018541 | Cwc25 | 67480 | protein_coding | 251.1332958 | -0.206501656 | 0.076465167 | -2.700597723 | 0.0069215 | 0.036134268 | 0.206501656 |
| ENSMUSG00000039231 | Suv39h1 | 20937 | protein_coding | 179.2014182 | 0.206507752 | 0.119735842 | 1.724694531 | 0.084582537 | 0.222462129 | 0.206507752 |
| ENSMUSG00000079508 | Apoo | 68316 | protein_coding | 259.3813432 | 0.20650782 | 0.095139158 | 2.170587003 | 0.029962405 | 0.107875496 | 0.20650782 |
| ENSMUSG00000021108 | Prkch | 18755 | protein_coding | 117.8478477 | -0.20659326 | 0.161225506 | -1.281393153 | 0.200055608 | 0.394660109 | 0.20659326 |
| ENSMUSG00000032217 | Rnf111 | 93836 | protein_coding | 608.2555945 | -0.206679968 | 0.083260515 | -2.482328706 | 0.013052681 | 0.058531278 | 0.206679968 |
| ENSMUSG00000002489 | Tiam1 | 21844 | protein_coding | 254.4406417 | 0.206818944 | 0.141618476 | 1.460395208 | 0.144181489 | 0.318651694 | 0.206818944 |
| ENSMUSG00000028248 | Pnisr | 66625 | protein_coding | 940.7625 | -0.206837051 | 0.146140742 | -1.41532777 | 0.156972416 | 0.337781228 | 0.206837051 |
| ENSMUSG00000029291 | Rufy3 | 52822 | protein_coding | 420.3778695 | -0.206869197 | 0.080668261 | -2.564443502 | 0.01033414 | 0.048938762 | 0.206869197 |
| ENSMUSG00000049327 | Kmt5a | 67956 | protein_coding | 1977.364452 | 0.20687255 | 0.065419581 | 3.162242035 | 0.001565594 | 0.011024217 | 0.20687255 |
| ENSMUSG00000027102 | Hoxd8 | 15437 | protein_coding | 44.5309803 | 0.206911416 | 0.275727028 | 0.75042123 | 0.453001049 | 0.652992964 | 0.206911416 |
| ENSMUSG00000020956 | Dtd2 | 328092 | protein_coding | 128.2706713 | 0.207031317 | 0.111509284 | 1.856628526 | 0.063364023 | 0.182407763 | 0.207031317 |
| ENSMUSG00000059182 | Skap2 | 54353 | protein_coding | 841.0437364 | 0.207086094 | 0.124485143 | 1.663540644 | 0.096204262 | 0.242190846 | 0.207086094 |
| ENSMUSG00000009248 | Ascl2 | 17173 | protein_coding | 72.11481403 | -0.207098377 | 0.209345074 | -0.989267973 | 0.322532049 | 0.533463272 | 0.207098377 |
| ENSMUSG00000042632 | Pla2g6 | 53357 | protein_coding | 144.2627452 | 0.207136636 | 0.102252783 | 2.025731039 | 0.042792354 | 0.139212399 | 0.207136636 |
| ENSMUSG00000026797 | Stxbp1 | 20910 | protein_coding | 102.4522461 | -0.207142907 | 0.190383129 | -1.088031842 | 0.276581051 | 0.484907317 | 0.207142907 |
| ENSMUSG00000057894 | Zfp329 | 67230 | protein_coding | 216.0061231 | -0.207190892 | 0.130906629 | -1.582737974 | 0.113481199 | 0.269539001 | 0.207190892 |
| ENSMUSG00000002109 | Ddb2 | 107986 | protein_coding | 106.0827761 | -0.207236616 | 0.152767187 | -1.356551889 | 0.174923637 | 0.361185338 | 0.207236616 |
| ENSMUSG00000040195 | Nemp1 | 72243 | protein_coding | 169.3457897 | -0.207257658 | 0.174232528 | -1.189546294 | 0.234224765 | 0.438035087 | 0.207257658 |
| ENSMUSG00000032059 | Alg9 | 102580 | protein_coding | 594.9016547 | 0.207314152 | 0.105312314 | 1.968565148 | 0.049003048 | 0.153468144 | 0.207314152 |
| ENSMUSG00000019969 | Psen1 | 19164 | protein_coding | 2429.011462 | 0.207365152 | 0.06978809 | 2.971354437 | 0.002964894 | 0.018599775 | 0.207365152 |
| ENSMUSG00000041954 | Tnfrsf18 | 21936 | protein_coding | 22.09919489 | 0.207370442 | 0.292910601 | 0.707964961 | 0.478967017 | 0.672927641 | 0.207370442 |
| ENSMUSG00000025531 | Chm | 12662 | protein_coding | 488.5083566 | 0.207376579 | 0.070951522 | 2.922792546 | 0.003469076 | 0.02104969 | 0.207376579 |
| ENSMUSG00000024792 | Zfpl1 | 81909 | protein_coding | 265.8614511 | 0.207506322 | 0.091669338 | 2.263639368 | 0.023596304 | 0.09049947 | 0.207506322 |
| ENSMUSG00000059820 | Nkapd1 | 270156 | protein_coding | 331.4057721 | -0.207511224 | 0.089956053 | -2.306806665 | 0.021065601 | 0.08309383 | 0.207511224 |
| ENSMUSG00000003868 | Ruvbl2 | 20174 | protein_coding | 560.3109232 | 0.207555321 | 0.099080177 | 2.094821862 | 0.036186831 | 0.123980177 | 0.207555321 |
| ENSMUSG00000034006 | Slc66a2 | 66943 | protein_coding | 745.1758362 | -0.20759838 | 0.176942548 | -1.173253024 | 0.240694354 | 0.445218869 | 0.20759838 |
| ENSMUSG00000022472 | Desi1 | 28075 | protein_coding | 275.393821 | -0.207598854 | 0.112393993 | -1.847063604 | 0.064737923 | 0.185224651 | 0.207598854 |
| ENSMUSG00000004677 | Myo9b | 17925 | protein_coding | 588.5866912 | -0.207611327 | 0.147453035 | -1.407982727 | 0.159136185 | 0.340376923 | 0.207611327 |
| ENSMUSG00000031570 | Plpp5 | 71910 | protein_coding | 655.5051669 | 0.207654242 | 0.069740915 | 2.977509558 | 0.002906006 | 0.018295202 | 0.207654242 |
| ENSMUSG00000001518 | Itfg2 | 101142 | protein_coding | 157.487271 | -0.207746955 | 0.101037836 | -2.056130296 | 0.039769959 | 0.132498476 | 0.207746955 |
| ENSMUSG00000012076 | Brms1l | 52592 | protein_coding | 110.187378 | 0.207778056 | 0.116643294 | 1.781311628 | 0.074861556 | 0.204201575 | 0.207778056 |
| ENSMUSG00000024479 | Mal2 | 105853 | protein_coding | 5572.379805 | 0.20783563 | 0.110512192 | 1.880657922 | 0.060018467 | 0.175956956 | 0.20783563 |
| ENSMUSG00000038070 | Cntln | 338349 | protein_coding | 62.66251569 | -0.207874425 | 0.219527283 | -0.946918405 | 0.343680359 | 0.55376472 | 0.207874425 |
| ENSMUSG00000029253 | Cenpc1 | 12617 | protein_coding | 445.0100778 | 0.207910225 | 0.078010984 | 2.665140402 | 0.007695619 | 0.039261952 | 0.207910225 |
| ENSMUSG00000021712 | Trim23 | 81003 | protein_coding | 242.7236716 | 0.207963886 | 0.16502102 | 1.260226636 | 0.207587617 | 0.404082965 | 0.207963886 |
| ENSMUSG00000095193 | Gm20939 | 100044193 | lncRNA | 61.9132833 | 0.20796585 | 0.190742402 | 1.090296899 | 0.275582381 | 0.483902123 | 0.20796585 |
| ENSMUSG00000060678 | H4c3 | 319155 | protein_coding | 3020.776388 | 0.207975704 | 0.130704359 | 1.591191798 | 0.111566418 | 0.266741205 | 0.207975704 |
| ENSMUSG00000024231 | Cul2 | 71745 | protein_coding | 707.4046534 | 0.208122504 | 0.080139353 | 2.597007549 | 0.009403986 | 0.045736392 | 0.208122504 |

| ensembl_gene_id | external_gene_name | entrezgene_id | gene_biotype | baseMean | log2FoldChange | lfcSE | stat | pvalue | padj | absLog2FC |
|---|---|---|---|---|---|---|---|---|---|---|
| ENSMUSG00000047415 | Gpr68 | 238377 | protein_coding | 18.94518912 | -0.208208252 | 0.488391312 | -0.426314407 | 0.669878762 | 0.808780483 | 0.208208252 |
| ENSMUSG00000048327 | Ckap2l | 70466 | protein_coding | 311.0570198 | 0.208234305 | 0.130160066 | 1.599832502 | 0.109635746 | 0.263874388 | 0.208234305 |
| ENSMUSG00000021392 | Nol8 | 70930 | protein_coding | 360.2459051 | -0.208254992 | 0.161014463 | -1.293393081 | 0.195875142 | 0.389208989 | 0.208254992 |
| ENSMUSG00000037946 | Fgd3 | 30938 | protein_coding | 164.0219446 | 0.20840124 | 0.191937775 | 1.085775013 | 0.277578544 | 0.485907357 | 0.20840124 |
| ENSMUSG00000017428 | Psmd11 | 69077 | protein_coding | 713.3021586 | 0.208488127 | 0.096148082 | 2.1684065 | 0.030127772 | 0.108372887 | 0.208488127 |
| ENSMUSG00000027765 | P2ry1 | 18441 | protein_coding | 455.6390185 | -0.208519705 | 0.091425404 | -2.280763293 | 0.022562458 | 0.087545633 | 0.208519705 |
| ENSMUSG00000025401 | Myo1a | 432516 | protein_coding | 6675.332796 | -0.208527477 | 0.100658303 | -2.071637131 | 0.038299294 | 0.129216514 | 0.208527477 |
| ENSMUSG00000007950 | Abhd8 | 64296 | protein_coding | 105.5087823 | 0.208538317 | 0.178735465 | 1.166742802 | 0.243314252 | 0.448346762 | 0.208538317 |
| ENSMUSG00000049184 | Purg | 75029 | protein_coding | 21.61907139 | 0.208542762 | 0.293608441 | 0.710275092 | 0.477533562 | 0.672136628 | 0.208542762 |
| ENSMUSG00000050520 | Cldn8 | 54420 | protein_coding | 1441.12065 | 0.208545518 | 0.25061206 | 0.832144784 | 0.405327226 | 0.612238223 | 0.208545518 |
| ENSMUSG00000029490 | Mfsd7a | 243197 | protein_coding | 182.7135222 | 0.208557804 | 0.143577458 | 1.452580416 | 0.146340287 | 0.322393927 | 0.208557804 |
| ENSMUSG00000000561 | Wdr77 | 70465 | protein_coding | 250.8178839 | -0.208560816 | 0.167560034 | -1.244693086 | 0.213244589 | 0.411305377 | 0.208560816 |
| ENSMUSG00000026761 | Orc4 | 26428 | protein_coding | 599.3313835 | 0.208574933 | 0.053969055 | 3.864713481 | 0.00011122 | 0.0012345 | 0.208574933 |
| ENSMUSG00000022040 | Ephx2 | 13850 | protein_coding | 429.9189105 | -0.208582094 | 0.105545396 | -1.976231112 | 0.048128608 | 0.151740393 | 0.208582094 |
| ENSMUSG00000037221 | Mospd3 | 68929 | protein_coding | 35.19035956 | 0.208587917 | 0.26961363 | 0.773654943 | 0.439134872 | 0.641642073 | 0.208587917 |
| ENSMUSG00000032899 | Styk1 | 243659 | protein_coding | 1642.868348 | -0.208613138 | 0.101745509 | -2.050342471 | 0.040331023 | 0.133776037 | 0.208613138 |
| ENSMUSG00000034321 | Exosc1 | 66583 | protein_coding | 150.3637638 | -0.208672675 | 0.144905663 | -1.440058797 | 0.149850765 | 0.327275879 | 0.208672675 |
| ENSMUSG00000044080 | S100a1 | 20193 | protein_coding | 1002.195836 | -0.208681844 | 0.075484206 | -2.764576264 | 0.005699678 | 0.030921142 | 0.208681844 |
| ENSMUSG00000025140 | Pycr1 | 209027 | protein_coding | 111.5024978 | 0.208752849 | 0.139385158 | 1.49766913 | 0.134219236 | 0.303178171 | 0.208752849 |
| ENSMUSG00000030905 | Crym | 12971 | protein_coding | 54.97163732 | -0.208905281 | 0.158134295 | -1.321062462 | 0.186480538 | 0.376888975 | 0.208905281 |
| ENSMUSG00000109336 | Samd4b | 233033 | protein_coding | 309.5796363 | 0.208908554 | 0.153284782 | 1.362878629 | 0.172920765 | 0.358619662 | 0.208908554 |
| ENSMUSG00000056851 | Pcbp2 | 18521 | protein_coding | 752.2633857 | -0.208980798 | 0.180582985 | -1.157256307 | 0.247167659 | 0.452801103 | 0.208980798 |
| ENSMUSG00000063870 | Chd4 | 107932 | protein_coding | 3602.779869 | -0.20905983 | 0.083092264 | -2.515996332 | 0.011869642 | 0.054436987 | 0.20905983 |
| ENSMUSG00000021263 | Degs2 | 70059 | protein_coding | 2237.591313 | 0.209100107 | 0.13155262 | 1.589478846 | 0.111952326 | 0.267382914 | 0.209100107 |
| ENSMUSG00000023805 | Synj2 | 20975 | protein_coding | 696.7554991 | -0.209123953 | 0.095964181 | -2.179187593 | 0.029317735 | 0.106081983 | 0.209123953 |
| ENSMUSG00000022956 | Atp5o | 28080 | protein_coding | 3896.579696 | 0.20912588 | 0.102018806 | 2.049875782 | 0.040376554 | 0.133871306 | 0.20912588 |
| ENSMUSG00000021785 | Ngly1 | 59007 | protein_coding | 1083.62331 | 0.209175894 | 0.081524025 | 2.565819025 | 0.010293252 | 0.04883852 | 0.209175894 |
| ENSMUSG00000085421 | 4732490B19Rik | 319871 | lncRNA | 24.2719024 | -0.209177691 | 0.24257591 | -0.862318485 | 0.388512284 | 0.596973148 | 0.209177691 |
| ENSMUSG00000021792 | Prxl2a | 70564 | protein_coding | 131.5821241 | -0.209183035 | 0.123458602 | -1.694357714 | 0.09019733 | 0.232146031 | 0.209183035 |
| ENSMUSG00000093674 | Rpl41 | 67945 | protein_coding | 7212.608683 | 0.209210313 | 0.105472461 | 1.983553913 | 0.047305591 | 0.149827691 | 0.209210313 |
| ENSMUSG00000019188 | H13 | 14950 | protein_coding | 1471.95174 | 0.209224373 | 0.079482649 | 2.632327639 | 0.008480204 | 0.042327446 | 0.209224373 |
| ENSMUSG00000039062 | Anpep | 16790 | protein_coding | 111.3136977 | -0.209229446 | 0.357721416 | -0.584894941 | 0.558618361 | 0.736907711 | 0.209229446 |
| ENSMUSG00000028699 | Tspan1 | 66805 | protein_coding | 16786.67873 | 0.209279854 | 0.091966247 | 2.275615907 | 0.022869008 | 0.088529438 | 0.209279854 |
| ENSMUSG00000028024 | Enpep | 13809 | protein_coding | 209.7041036 | 0.209342093 | 0.271365915 | 0.77143842 | 0.44044711 | 0.642322062 | 0.209342093 |
| ENSMUSG00000020993 | Trappc6b | 78232 | protein_coding | 467.2982954 | 0.209428657 | 0.076572556 | 2.735035473 | 0.006237355 | 0.033149348 | 0.209428657 |
| ENSMUSG00000001794 | Capns1 | 12336 | protein_coding | 2117.213624 | 0.209494935 | 0.122718148 | 1.707122691 | 0.08779925 | 0.228321055 | 0.209494935 |
| ENSMUSG00000040688 | Tbl3 | 213773 | protein_coding | 402.8722239 | -0.209500762 | 0.131146096 | -1.597460908 | 0.110163005 | 0.264736926 | 0.209500762 |
| ENSMUSG00000021967 | Mrpl57 | 67840 | protein_coding | 430.342156 | 0.209549379 | 0.095113897 | 2.203141552 | 0.027584773 | 0.101309196 | 0.209549379 |

| ensembl_gene_id | external_gene_name | entrezgene_id | gene_biotype | baseMean | log2FoldChange | lfcSE | stat | pvalue | padj | absLog2FC |
|---|---|---|---|---|---|---|---|---|---|---|
| ENSMUSG00000022755 | Adgrg7 | 239853 | protein_coding | 4686.311724 | 0.20959547 | 0.126215664 | 1.660613773 | 0.096791041 | 0.243168407 | 0.20959547 |
| ENSMUSG00000021773 | Comtd1 | 69156 | protein_coding | 72.72153288 | -0.209612428 | 0.161912085 | -1.294606439 | 0.19545603 | 0.388667122 | 0.209612428 |
| ENSMUSG00000039065 | Atpsckmt | 68073 | protein_coding | 178.1547719 | 0.2096353 | 0.11891808 | 1.762854738 | 0.077924993 | 0.209908554 | 0.2096353 |
| ENSMUSG00000090191 | 9230105E05Rik | 320626 | lncRNA | 113.2536685 | 0.209692738 | 0.193954514 | 1.08114389 | 0.279633113 | 0.488002215 | 0.209692738 |
| ENSMUSG00000040102 | Klhl42 | 232539 | protein_coding | 412.0036814 | -0.209708325 | 0.094379123 | -2.221977889 | 0.0262848 | 0.098047844 | 0.209708325 |
| ENSMUSG00000031367 | Ap1s2 | 108012 | protein_coding | 46.48216865 | 0.209723319 | 0.219954438 | 0.953485278 | 0.340344252 | 0.550691106 | 0.209723319 |
| ENSMUSG00000028232 | Tmem68 | 72098 | protein_coding | 260.9097283 | 0.209725883 | 0.094528479 | 2.21865288 | 0.026510349 | 0.098566022 | 0.209725883 |
| ENSMUSG00000062210 | Tnfaip8 | 106869 | protein_coding | 812.8838948 | -0.209769025 | 0.132528687 | -1.582819769 | 0.113462549 | 0.269534897 | 0.209769025 |
| ENSMUSG00000071424 | Grid2 | 14804 | protein_coding | 68.25644008 | -0.209779683 | 0.225297091 | -0.931124686 | 0.351789071 | 0.561401056 | 0.209779683 |
| ENSMUSG00000026189 | Pecr | 111175 | protein_coding | 91.91427359 | 0.209827405 | 0.127506105 | 1.645626334 | 0.099840714 | 0.24785893 | 0.209827405 |
| ENSMUSG00000020974 | Pole2 | 18974 | protein_coding | 66.88602137 | -0.209877006 | 0.137696929 | -1.52419525 | 0.127459944 | 0.292913287 | 0.209877006 |
| ENSMUSG00000025137 | Pcyt2 | 68671 | protein_coding | 818.4746895 | -0.209945423 | 0.089731198 | -2.339714916 | 0.019298465 | 0.077937613 | 0.209945423 |
| ENSMUSG00000019977 | Hbs1l | 56422 | protein_coding | 754.8873844 | 0.209990297 | 0.084439541 | 2.486871605 | 0.01288719 | 0.057937359 | 0.209990297 |
| ENSMUSG00000028800 | Hdac1 | 433759 | protein_coding | 1033.218364 | 0.210038009 | 0.096968423 | 2.166045425 | 0.030307716 | 0.108823557 | 0.210038009 |
| ENSMUSG00000057093 | Zfp607b | 112415 | protein_coding | 20.52539298 | -0.21006873 | 0.298264456 | -0.704303599 | 0.481243725 | 0.67440173 | 0.21006873 |
| ENSMUSG00000020289 | Nprl3 | 17168 | protein_coding | 133.0486021 | -0.210127758 | 0.101457744 | -2.071086444 | 0.038350718 | 0.12925312 | 0.210127758 |
| ENSMUSG00000004040 | Stat3 | 20848 | protein_coding | 1624.941551 | 0.210231828 | 0.142933278 | 1.470838914 | 0.141334686 | 0.314015745 | 0.210231828 |
| ENSMUSG00000025324 | Atp10a | 11982 | protein_coding | 25.9256998 | -0.21024462 | 0.405228708 | -0.518829529 | 0.603879624 | 0.768565294 | 0.21024462 |
| ENSMUSG00000028889 | Yrdc | 230734 | protein_coding | 382.1257986 | 0.210294551 | 0.148234502 | 1.418661296 | 0.155997786 | 0.336320078 | 0.210294551 |
| ENSMUSG00000022557 | Bop1 | 12181 | protein_coding | 558.7044685 | -0.210300007 | 0.162139243 | -1.297033359 | 0.194619709 | 0.387730172 | 0.210300007 |
| ENSMUSG00000067370 | B3galt4 | 54218 | protein_coding | 117.9097399 | 0.210307257 | 0.149884365 | 1.403130053 | 0.160578061 | 0.342211313 | 0.210307257 |
| ENSMUSG00000022836 | Mylk | 107589 | protein_coding | 722.4995107 | -0.210322421 | 0.174296086 | -1.206696177 | 0.227549172 | 0.430167439 | 0.210322421 |
| ENSMUSG00000056429 | Tgoln1 | 22134 | protein_coding | 17482.30631 | -0.210324295 | 0.064788945 | -3.246299127 | 0.001169159 | 0.00870611 | 0.210324295 |
| ENSMUSG00000025091 | Pnliprp2 | 18947 | protein_coding | 26.03518313 | -0.210326846 | 0.226499273 | -0.928598328 | 0.353097289 | 0.56288012 | 0.210326846 |
| ENSMUSG00000027985 | Lef1 | 16842 | protein_coding | 8.871710872 | -0.210350915 | 0.847102019 | -0.248318278 | 0.803888158 | 0.894394399 | 0.210350915 |
| ENSMUSG00000034203 | Chchd4 | 72170 | protein_coding | 382.8648947 | 0.210387673 | 0.129779214 | 1.621119954 | 0.104991915 | 0.256593769 | 0.210387673 |
| ENSMUSG00000048490 | Nrip1 | 268903 | protein_coding | 1945.697082 | -0.210448282 | 0.165049148 | -1.275064335 | 0.202286472 | 0.397338627 | 0.210448282 |
| ENSMUSG00000008859 | Rala | 56044 | protein_coding | 1121.52501 | 0.21047624 | 0.062311985 | 3.377781008 | 0.000730732 | 0.005941351 | 0.21047624 |
| ENSMUSG00000021068 | Nin | 18080 | protein_coding | 143.823293 | -0.210498647 | 0.194272225 | -1.083524146 | 0.278575839 | 0.487064321 | 0.210498647 |
| ENSMUSG00000038188 | Scarf1 | 380713 | protein_coding | 45.53908762 | -0.210531737 | 0.240415451 | -0.875699697 | 0.38119331 | 0.589022802 | 0.210531737 |
| ENSMUSG00000034795 | Ccdc122 | 108811 | protein_coding | 217.5605399 | 0.210574756 | 0.197733474 | 1.064942382 | 0.286902004 | 0.495744752 | 0.210574756 |
| ENSMUSG00000003308 | Keap1 | 50868 | protein_coding | 686.1190821 | 0.210611913 | 0.076646833 | 2.747822783 | 0.005999243 | 0.032217106 | 0.210611913 |
| ENSMUSG00000029004 | Kmt2e | 69188 | protein_coding | 674.9198018 | -0.210705781 | 0.191926395 | -1.097846813 | 0.272271386 | 0.48047545 | 0.210705781 |
| ENSMUSG00000021555 | Naa35 | 78689 | protein_coding | 799.566196 | 0.210843604 | 0.068565445 | 3.075070905 | 0.002104525 | 0.014112378 | 0.210843604 |
| ENSMUSG00000028381 | Ugcg | 22234 | protein_coding | 1399.607281 | -0.210911304 | 0.155503407 | -1.356313075 | 0.174999577 | 0.361280983 | 0.210911304 |
| ENSMUSG00000021693 | Kif2a | 16563 | protein_coding | 397.8156973 | -0.210992252 | 0.114742444 | -1.838833517 | 0.065939677 | 0.187383974 | 0.210992252 |
| ENSMUSG00000089984 | Fbxo24 | 71176 | protein_coding | 13.67346673 | -0.211005122 | 0.37173796 | -0.567617905 | 0.570294447 | 0.744744605 | 0.211005122 |
| ENSMUSG00000001376 | Vps50 | 73288 | protein_coding | 778.6706011 | 0.211071244 | 0.086330115 | 2.444931821 | 0.014487953 | 0.062846872 | 0.211071244 |

Supplementary Table 1

| ensembl_gene_id | external_gene_name | entrezgene_id | gene_biotype | baseMean | log2FoldChange | lfcSE | stat | pvalue | padj | absLog2FC |
|---|---|---|---|---|---|---|---|---|---|---|
| ENSMUSG00000091086 | Rpl6l | 432502 | processed_pseudogene | 9.489958162 | 0.211274206 | 0.398323124 | 0.530409091 | 0.595828325 | 0.763909664 | 0.211274206 |
| ENSMUSG00000044337 | Ackr3 | 12778 | protein_coding | 20.34037294 | 0.211288364 | 0.335730678 | 0.629338866 | 0.529127232 | 0.714548497 | 0.211288364 |
| ENSMUSG00000060716 | Plekhh1 | 211945 | protein_coding | 1454.052364 | -0.211293042 | 0.094958028 | -2.225120373 | 0.026073158 | 0.097509572 | 0.211293042 |
| ENSMUSG00000005656 | Snx6 | 72183 | protein_coding | 840.5921635 | 0.21131221 | 0.086136334 | 2.453229666 | 0.014157991 | 0.061884875 | 0.21131221 |
| ENSMUSG00000021843 | Ktn1 | 16709 | protein_coding | 1259.071156 | -0.211363716 | 0.092903372 | -2.275091961 | 0.022900414 | 0.088538846 | 0.211363716 |
| ENSMUSG00000049792 | Bag5 | 70369 | protein_coding | 245.2568355 | -0.211389591 | 0.113871164 | -1.856392653 | 0.063397611 | 0.182471477 | 0.211389591 |
| ENSMUSG00000044715 | Gskip | 66787 | protein_coding | 239.3317178 | 0.211473861 | 0.11689243 | 1.809132215 | 0.070430465 | 0.19615605 | 0.211473861 |
| ENSMUSG00000031520 | Vegfc | 22341 | protein_coding | 54.81568077 | 0.211549115 | 0.214497308 | 0.986255339 | 0.32400783 | 0.534853012 | 0.211549115 |
| ENSMUSG00000018548 | Trim37 | 68729 | protein_coding | 515.0468014 | 0.211566792 | 0.115997963 | 1.823883689 | 0.068169658 | 0.19200819 | 0.211566792 |
| ENSMUSG00000029465 | Arpc3 | 56378 | protein_coding | 2444.389071 | 0.211615635 | 0.098915279 | 2.13936246 | 0.032406325 | 0.114129198 | 0.211615635 |
| ENSMUSG00000031555 | Adam9 | 11502 | protein_coding | 1551.913027 | 0.211649301 | 0.058163371 | 3.63887609 | 0.000273831 | 0.002639364 | 0.211649301 |
| ENSMUSG00000071647 | Eml3 | 225898 | protein_coding | 304.3304816 | -0.211674165 | 0.075204738 | -2.814638675 | 0.00488321 | 0.027561788 | 0.211674165 |
| ENSMUSG00000026343 | Gpr39 | 71111 | protein_coding | 119.2443381 | -0.211710199 | 0.235578837 | -0.898680891 | 0.368822659 | 0.577640836 | 0.211710199 |
| ENSMUSG00000049488 | Tmem67 | 329795 | protein_coding | 94.08618965 | 0.211715148 | 0.181218111 | 1.168289121 | 0.242690162 | 0.447623827 | 0.211715148 |
| ENSMUSG00000053110 | Yap1 | 22601 | protein_coding | 1038.140137 | -0.21172575 | 0.104340191 | -2.029186919 | 0.042439255 | 0.138490568 | 0.21172575 |
| ENSMUSG00000026187 | Xrcc5 | 22596 | protein_coding | 407.2272876 | -0.211733864 | 0.087964125 | -2.407047919 | 0.01608206 | 0.068096417 | 0.211733864 |
| ENSMUSG00000022523 | Fgf12 | 14167 | protein_coding | 37.41473439 | -0.211901604 | 0.262591061 | -0.806964271 | 0.419687068 | 0.625680983 | 0.211901604 |
| ENSMUSG00000037995 | Igsf9 | 93842 | protein_coding | 1009.082121 | 0.211972868 | 0.114328371 | 1.854070576 | 0.063729064 | 0.183259889 | 0.211972868 |
| ENSMUSG00000020708 | Psmc5 | 19184 | protein_coding | 1022.099427 | 0.212019623 | 0.108087054 | 1.961563526 | 0.049813322 | 0.154996405 | 0.212019623 |
| ENSMUSG00000040385 | Ppp1ca | 19045 | protein_coding | 2834.961432 | 0.212102915 | 0.083340257 | 2.545023538 | 0.010927037 | 0.051159861 | 0.212102915 |
| ENSMUSG00000001105 | Ift20 | 55978 | protein_coding | 468.071669 | 0.21213487 | 0.07883896 | 2.690736535 | 0.007129447 | 0.036965442 | 0.21213487 |
| ENSMUSG00000020154 | Ptprb | 19263 | protein_coding | 11.07691028 | -0.212191499 | 0.649216293 | -0.326842535 | 0.743786986 | 0.857445771 | 0.212191499 |
| ENSMUSG00000032902 | Slc16a1 | 20501 | protein_coding | 6167.447021 | -0.212273695 | 0.129926169 | -1.63380247 | 0.102300334 | 0.252118167 | 0.212273695 |
| ENSMUSG00000054426 | A930005H10Rik | 68161 | lncRNA | 22.68193169 | -0.212291731 | 0.271450421 | -0.782064475 | 0.434176683 | 0.637531118 | 0.212291731 |
| ENSMUSG00000027341 | Tmem230 | 70612 | protein_coding | 239.8930134 | 0.212294042 | 0.092501769 | 2.295026841 | 0.021731586 | 0.084900835 | 0.212294042 |
| ENSMUSG00000034278 | Dnajc17 | 69408 | protein_coding | 88.24091219 | -0.21229803 | 0.158913813 | -1.335931888 | 0.181571537 | 0.370208838 | 0.21229803 |
| ENSMUSG00000054128 | H2-T3 | 15043 | protein_coding | 655.6881548 | 0.212298634 | 0.220046906 | 0.964788088 | 0.334650963 | 0.545293316 | 0.212298634 |
| ENSMUSG00000041476 | Smpx | 66106 | protein_coding | 101.3734374 | 0.212444744 | 0.241657882 | 0.879113656 | 0.379339654 | 0.587298018 | 0.212444744 |
| ENSMUSG00000029723 | Spacdr | 78829 | protein_coding | 741.1486844 | -0.21259044 | 0.11165236 | -1.90403893 | 0.056905114 | 0.169853087 | 0.21259044 |
| ENSMUSG00000022553 | Maf1 | 68877 | protein_coding | 789.208745 | 0.212613053 | 0.127867859 | 1.662756019 | 0.096361284 | 0.242547808 | 0.212613053 |
| ENSMUSG00000032267 | Usp28 | 235323 | protein_coding | 285.5285337 | -0.212632597 | 0.148211908 | -1.434652584 | 0.151386122 | 0.329588153 | 0.212632597 |
| ENSMUSG00000024727 | Trpm6 | 225997 | protein_coding | 1408.494119 | 0.212643051 | 0.200547077 | 1.060314884 | 0.28900137 | 0.497873797 | 0.212643051 |
| ENSMUSG00000027243 | Harbi1 | 241547 | protein_coding | 223.3384616 | 0.212659279 | 0.163057624 | 1.304197098 | 0.192166389 | 0.385058026 | 0.212659279 |
| ENSMUSG00000008763 | Man1a2 | 17156 | protein_coding | 2463.516734 | 0.212743321 | 0.054423528 | 3.909032171 | 9.27E-05 | 0.001058819 | 0.212743321 |
| ENSMUSG00000007721 | Ccdc124 | 234388 | protein_coding | 377.6810537 | 0.212743693 | 0.113116759 | 1.880744241 | 0.060006718 | 0.175954899 | 0.212743693 |
| ENSMUSG00000078908 | Mon1b | 270096 | protein_coding | 475.6981184 | 0.212773143 | 0.092652802 | 2.296456654 | 0.021649785 | 0.084685111 | 0.212773143 |
| ENSMUSG00000062906 | Hdac10 | 170787 | protein_coding | 225.3368754 | -0.212851601 | 0.122882975 | -1.732148826 | 0.083247068 | 0.220076231 | 0.212851601 |
| ENSMUSG00000041057 | Wdr43 | 72515 | protein_coding | 851.5295265 | -0.212854832 | 0.180249581 | -1.180889467 | 0.237646636 | 0.441993884 | 0.212854832 |

| ensembl_gene_id | external_gene_name | entrezgene_id | gene_biotype | baseMean | log2FoldChange | lfcSE | stat | pvalue | padj | absLog2FC |
|---|---|---|---|---|---|---|---|---|---|---|
| ENSMUSG00000022805 | Cfap91 | 320214 | protein_coding | 14.06224903 | -0.212915308 | 0.356166994 | -0.597796292 | 0.549975865 | 0.731097946 | 0.212915308 |
| ENSMUSG00000021140 | Pcnx | 54604 | protein_coding | 728.7966391 | -0.21312123 | 0.113779056 | -1.873114764 | 0.061052539 | 0.17777785 | 0.21312123 |
| ENSMUSG00000032336 | Nptn | 20320 | protein_coding | 1387.724706 | 0.213137095 | 0.07988329 | 2.668106118 | 0.007628016 | 0.039042108 | 0.213137095 |
| ENSMUSG00000035765 | Dym | 69190 | protein_coding | 661.6981042 | -0.213151777 | 0.071412778 | -2.984784844 | 0.002837779 | 0.017943682 | 0.213151777 |
| ENSMUSG00000028211 | Trp53inp1 | 60599 | protein_coding | 1516.601081 | -0.213166739 | 0.213416471 | -0.998829835 | 0.317877131 | 0.529178599 | 0.213166739 |
| ENSMUSG00000051169 | Rpusd3 | 101122 | protein_coding | 36.23466316 | -0.21317228 | 0.214885842 | -0.99202571 | 0.321184984 | 0.532354566 | 0.21317228 |
| ENSMUSG00000021457 | Syk | 20963 | protein_coding | 893.6042114 | 0.21320662 | 0.146862122 | 1.451746827 | 0.146572014 | 0.322770501 | 0.21320662 |
| ENSMUSG00000019132 | BC005537 | 79555 | protein_coding | 1919.687619 | -0.213214431 | 0.096552786 | -2.208268033 | 0.027225594 | 0.100428269 | 0.213214431 |
| ENSMUSG00000028483 | Snapc3 | 77634 | protein_coding | 215.3758915 | -0.213228829 | 0.144585759 | -1.474756785 | 0.140277943 | 0.312550067 | 0.213228829 |
| ENSMUSG00000035585 | Tsen34 | 66078 | protein_coding | 383.1393 | 0.213237208 | 0.107574376 | 1.982230488 | 0.04745345 | 0.150206389 | 0.213237208 |
| ENSMUSG00000042197 | Zfp451 | 98403 | protein_coding | 461.7395384 | -0.213280875 | 0.11205024 | -1.903439697 | 0.056983195 | 0.169999688 | 0.213280875 |
| ENSMUSG00000027108 | Ola1 | 67059 | protein_coding | 1346.547593 | 0.213282344 | 0.083884632 | 2.542567556 | 0.011004133 | 0.051423605 | 0.213282344 |
| ENSMUSG00000027166 | Dnajc24 | 99349 | protein_coding | 102.9378991 | 0.213325437 | 0.125940234 | 1.693862474 | 0.090291419 | 0.232259644 | 0.213325437 |
| ENSMUSG00000025228 | Actr1a | 54130 | protein_coding | 2257.848472 | 0.213360959 | 0.092480103 | 2.307101219 | 0.021049178 | 0.08304975 | 0.213360959 |
| ENSMUSG00000028854 | Slc9a1 | 20544 | protein_coding | 506.7898426 | -0.213376764 | 0.15488974 | -1.377604252 | 0.168325507 | 0.352960595 | 0.213376764 |
| ENSMUSG00000070814 | Zswim9 | 321008 | protein_coding | 70.91963288 | -0.213407435 | 0.213250668 | -1.000735127 | 0.31695488 | 0.528247079 | 0.213407435 |
| ENSMUSG00000028563 | Tm2d1 | 94043 | protein_coding | 273.555149 | 0.213410717 | 0.117621996 | 1.814377617 | 0.069619609 | 0.195057368 | 0.213410717 |
| ENSMUSG00000041477 | Dcp1b | 319618 | protein_coding | 141.6114298 | -0.21357252 | 0.194507289 | -1.098018081 | 0.272196594 | 0.4804721 | 0.21357252 |
| ENSMUSG00000039382 | Wdr45 | 54636 | protein_coding | 154.0381425 | -0.213620166 | 0.126210451 | -1.692571135 | 0.090537129 | 0.232705405 | 0.213620166 |
| ENSMUSG00000032625 | Thsd7a | 330267 | protein_coding | 9.03568615 | -0.21367015 | 0.990957651 | -0.21561986 | 0.829284074 | 0.909405742 | 0.21367015 |
| ENSMUSG00000028688 | Toe1 | 68276 | protein_coding | 188.9247988 | 0.213685302 | 0.115889654 | 1.843868658 | 0.065202284 | 0.185852179 | 0.213685302 |
| ENSMUSG00000040370 | Etfrf1 | 67636 | protein_coding | 386.6080663 | 0.213709349 | 0.110454415 | 1.934819431 | 0.053012458 | 0.161728103 | 0.213709349 |
| ENSMUSG00000050144 | Slc25a44 | 229517 | protein_coding | 602.2686969 | 0.213746642 | 0.089855577 | 2.378779928 | 0.017370043 | 0.072087035 | 0.213746642 |
| ENSMUSG00000022089 | Bin3 | 57784 | protein_coding | 366.1076831 | 0.213815605 | 0.082411267 | 2.594494825 | 0.009473004 | 0.045973799 | 0.213815605 |
| ENSMUSG00000060397 | Zfp128 | 243833 | protein_coding | 32.14107015 | 0.213816889 | 0.28435261 | 0.751942768 | 0.452085477 | 0.652202459 | 0.213816889 |
| ENSMUSG00000040236 | Trappc5 | 66682 | protein_coding | 369.9787974 | 0.213841655 | 0.111850143 | 1.911858579 | 0.055894336 | 0.167567284 | 0.213841655 |
| ENSMUSG00000077222 | Gm22270 | 115490266 | snoRNA | 10.41902176 | 0.213856317 | 0.350494528 | 0.610155935 | 0.541758517 | 0.724809384 | 0.213856317 |
| ENSMUSG00000025173 | Wdr45b | 66840 | protein_coding | 695.7223209 | 0.213859348 | 0.093425245 | 2.289095928 | 0.022073779 | 0.086026709 | 0.213859348 |
| ENSMUSG00000027944 | Hax1 | 23897 | protein_coding | 337.6346969 | 0.213867484 | 0.086649304 | 2.468196207 | 0.013579586 | 0.060148958 | 0.213867484 |
| ENSMUSG00000067942 | Zfp160 | 224585 | protein_coding | 389.5138432 | 0.213904339 | 0.109322565 | 1.956634848 | 0.050390415 | 0.156204476 | 0.213904339 |
| ENSMUSG00000023915 | Tnfrsf21 | 94185 | protein_coding | 984.6259431 | -0.213912634 | 0.159142973 | -1.344153807 | 0.178898656 | 0.366360585 | 0.213912634 |
| ENSMUSG00000047409 | Ctdspl | 69274 | protein_coding | 1292.265116 | 0.213918848 | 0.100821306 | 2.121762308 | 0.033857703 | 0.11800558 | 0.213918848 |
| ENSMUSG00000037270 | Bltp1 | 229227 | protein_coding | 1956.923223 | 0.214027335 | 0.17069345 | 1.253869643 | 0.209889391 | 0.407185609 | 0.214027335 |
| ENSMUSG00000042557 | Sin3a | 20466 | protein_coding | 844.8550374 | -0.214049793 | 0.091044999 | -2.351032957 | 0.018721376 | 0.076205998 | 0.214049793 |
| ENSMUSG00000033948 | Zswim5 | 74464 | protein_coding | 185.5907564 | -0.214117094 | 0.099883292 | -2.143672779 | 0.032059115 | 0.113349074 | 0.214117094 |
| ENSMUSG00000042046 | Dstyk | 213452 | protein_coding | 199.9878036 | -0.214184268 | 0.114456132 | -1.871321909 | 0.061300476 | 0.178191396 | 0.214184268 |
| ENSMUSG00000010054 | Tusc2 | 80385 | protein_coding | 517.2875911 | 0.214215832 | 0.093113541 | 2.300587339 | 0.021414967 | 0.084108607 | 0.214215832 |
| ENSMUSG00000027447 | Cst3 | 13010 | protein_coding | 841.3097793 | -0.214229247 | 0.101058095 | -2.119862316 | 0.034017658 | 0.118433499 | 0.214229247 |

| ensembl_gene_id | external_gene_name | entrezgene_id | gene_biotype | baseMean | log2FoldChange | lfcSE | stat | pvalue | padj | absLog2FC |
|---|---|---|---|---|---|---|---|---|---|---|
| ENSMUSG00000048271 | Rbm33 | 381626 | protein_coding | 827.5294697 | -0.214275072 | 0.118956224 | -1.801293497 | 0.071656632 | 0.198667857 | 0.214275072 |
| ENSMUSG00000023169 | Slc38a1 | 105727 | protein_coding | 798.3833172 | -0.214321164 | 0.108775573 | -1.970305995 | 0.048803312 | 0.153109937 | 0.214321164 |
| ENSMUSG00000033991 | Ttc37 | 218343 | protein_coding | 688.9137744 | 0.214341221 | 0.066612252 | 3.217744691 | 0.001292028 | 0.009414189 | 0.214341221 |
| ENSMUSG00000048807 | Slc35e4 | 103710 | protein_coding | 122.8428286 | -0.214378838 | 0.157566144 | -1.360564099 | 0.173651484 | 0.359677611 | 0.214378838 |
| ENSMUSG00000033282 | Rpgrip1l | 244585 | protein_coding | 175.0427269 | 0.214414362 | 0.114239202 | 1.876889523 | 0.060533237 | 0.176782802 | 0.214414362 |
| ENSMUSG00000044024 | Rell2 | 225392 | protein_coding | 7.463728021 | -0.214520771 | 0.414719795 | -0.517266776 | 0.604969943 | 0.769522541 | 0.214520771 |
| ENSMUSG00000039081 | Zfp503 | 218820 | protein_coding | 42.15493576 | -0.214555745 | 0.231341212 | -0.927442812 | 0.353696671 | 0.563537119 | 0.214555745 |
| ENSMUSG00000049939 | Lrrc4 | 192198 | protein_coding | 14.74143337 | -0.214629165 | 0.410626621 | -0.522686923 | 0.60119214 | 0.767167794 | 0.214629165 |
| ENSMUSG00000020230 | Prmt2 | 15468 | protein_coding | 180.4123839 | 0.214629951 | 0.0997283 | 2.152146906 | 0.031385786 | 0.111737326 | 0.214629951 |
| ENSMUSG00000058239 | Usf2 | 22282 | protein_coding | 228.289174 | -0.214663103 | 0.083432683 | -2.572889852 | 0.01008533 | 0.048182104 | 0.214663103 |
| ENSMUSG00000026072 | Il1r1 | 16177 | protein_coding | 453.6561831 | -0.214688983 | 0.158207232 | -1.357011187 | 0.174777656 | 0.361071142 | 0.214688983 |
| ENSMUSG00000036120 | Rfxank | 19727 | protein_coding | 194.0524499 | -0.214707254 | 0.109128668 | -1.967468839 | 0.049129185 | 0.153635373 | 0.214707254 |
| ENSMUSG00000040363 | Bcor | 71458 | protein_coding | 557.1034701 | -0.214745387 | 0.102499924 | -2.095078503 | 0.036164015 | 0.123928663 | 0.214745387 |
| ENSMUSG00000049686 | Orai1 | 109305 | protein_coding | 243.0579298 | 0.214763653 | 0.128834239 | 1.666976534 | 0.095519071 | 0.24111375 | 0.214763653 |
| ENSMUSG00000066406 | Akap13 | 75547 | protein_coding | 2563.221901 | -0.214792755 | 0.138105396 | -1.55528141 | 0.119879063 | 0.280881688 | 0.214792755 |
| ENSMUSG00000036550 | Cnot1 | 234594 | protein_coding | 4414.691487 | 0.214986309 | 0.072424799 | 2.968407374 | 0.002993473 | 0.018742154 | 0.214986309 |
| ENSMUSG00000029397 | Rchy1 | 68098 | protein_coding | 634.9797566 | 0.215001378 | 0.069588404 | 3.089615019 | 0.002004161 | 0.013526388 | 0.215001378 |
| ENSMUSG00000029162 | Khk | 16548 | protein_coding | 188.7369164 | -0.215015328 | 0.123806403 | -1.73670604 | 0.082439071 | 0.218593229 | 0.215015328 |
| ENSMUSG00000060012 | Kif13b | 16554 | protein_coding | 2440.187841 | -0.215056373 | 0.091800061 | -2.342660449 | 0.019146799 | 0.077541373 | 0.215056373 |
| ENSMUSG00000015127 | Unkl | 74154 | protein_coding | 60.01646532 | -0.215065936 | 0.153287833 | -1.403020264 | 0.160610796 | 0.342235286 | 0.215065936 |
| ENSMUSG00000059355 | Wdr83os | 414077 | protein_coding | 165.4712213 | -0.215082439 | 0.104529602 | -2.05762228 | 0.039626406 | 0.132320628 | 0.215082439 |
| ENSMUSG00000019948 | Actr6 | 67019 | protein_coding | 88.55945484 | 0.21509393 | 0.14592164 | 1.474037229 | 0.140471568 | 0.312850724 | 0.21509393 |
| ENSMUSG00000029826 | Zc3hav1 | 78781 | protein_coding | 1544.569883 | 0.215127853 | 0.10590917 | 2.031248605 | 0.042229782 | 0.138118269 | 0.215127853 |
| ENSMUSG00000025377 | Tepsin | 78777 | protein_coding | 327.8709889 | 0.215219789 | 0.097029255 | 2.218091748 | 0.026548577 | 0.098616078 | 0.215219789 |
| ENSMUSG00000026380 | Tfcp2l1 | 81879 | protein_coding | 6873.697965 | -0.215244756 | 0.117462553 | -1.832454266 | 0.06688377 | 0.189323739 | 0.215244756 |
| ENSMUSG00000019782 | Rwdd1 | 66521 | protein_coding | 302.3430324 | 0.215311643 | 0.090674637 | 2.374552027 | 0.01757026 | 0.072634077 | 0.215311643 |
| ENSMUSG00000085287 | 4833418N02Rik | 74597 | lncRNA | 53.01924732 | 0.215333643 | 0.187057372 | 1.151163627 | 0.249664926 | 0.456020804 | 0.215333643 |
| ENSMUSG00000022751 | Nit2 | 52633 | protein_coding | 237.8789378 | -0.215638874 | 0.092381889 | -2.334211573 | 0.019584646 | 0.078608426 | 0.215638874 |
| ENSMUSG00000032316 | Clk3 | 102414 | protein_coding | 593.0138436 | -0.215653955 | 0.083336038 | -2.587763482 | 0.009660128 | 0.046678138 | 0.215653955 |
| ENSMUSG00000024037 | Wdr4 | 57773 | protein_coding | 228.1123443 | -0.215761737 | 0.147122205 | -1.466547738 | 0.142499126 | 0.316117838 | 0.215761737 |
| ENSMUSG00000030029 | Lrig1 | 16206 | protein_coding | 2269.274491 | -0.215773326 | 0.118947298 | -1.814024615 | 0.069673936 | 0.195142335 | 0.215773326 |
| ENSMUSG00000061533 | Cep128 | 75216 | protein_coding | 149.6827964 | 0.215988191 | 0.14324216 | 1.507853484 | 0.13159203 | 0.299299993 | 0.215988191 |
| ENSMUSG00000039058 | Ak5 | 229949 | protein_coding | 6.792064315 | -0.216011085 | 0.608704417 | -0.354870245 | 0.722686801 | 0.844093231 | 0.216011085 |
| ENSMUSG00000064373 | Selenop | 20363 | protein_coding | 4350.929377 | -0.216017872 | 0.163517358 | -1.321069972 | 0.186478034 | 0.376888975 | 0.216017872 |
| ENSMUSG00000097281 | Gm26685 | 102635083 | lncRNA | 12.54504383 | -0.216078332 | 0.45023699 | -0.479921322 | 0.631283339 | 0.786413345 | 0.216078332 |
| ENSMUSG00000004043 | Stat5a | 20850 | protein_coding | 154.5218085 | -0.216086227 | 0.114783275 | -1.882558469 | 0.059760229 | 0.175536148 | 0.216086227 |
| ENSMUSG00000037926 | Ssh2 | 237860 | protein_coding | 726.5214249 | 0.216207939 | 0.136031628 | 1.589394632 | 0.111971326 | 0.267388198 | 0.216207939 |
| ENSMUSG00000031730 | Dhodh | 56749 | protein_coding | 179.060342 | 0.216213754 | 0.103368238 | 2.091684616 | 0.036466738 | 0.124617506 | 0.216213754 |

| ensembl_gene_id | external_gene_name | entrezgene_id | gene_biotype | baseMean | log2FoldChange | lfcSE | stat | pvalue | padj | absLog2FC |
|---|---|---|---|---|---|---|---|---|---|---|
| ENSMUSG00000049624 | Slc17a5 | 235504 | protein_coding | 423.8649227 | -0.21621703 | 0.093107851 | -2.32222126 | 0.020221025 | 0.080499996 | 0.21621703 |
| ENSMUSG00000043020 | Dnai3 | 242253 | protein_coding | 14.99492348 | -0.216233348 | 0.320915299 | -0.673801931 | 0.500437247 | 0.690964327 | 0.216233348 |
| ENSMUSG00000022681 | Ntan1 | 18203 | protein_coding | 408.4945237 | -0.216241749 | 0.095245046 | -2.270372671 | 0.023184981 | 0.089343223 | 0.216241749 |
| ENSMUSG00000038112 | AW551984 | 244810 | protein_coding | 28.2642348 | -0.216292038 | 0.341163304 | -0.633983888 | 0.526091337 | 0.711774523 | 0.216292038 |
| ENSMUSG00000084771 | A230072E10Rik | 331547 | lncRNA | 9.243586935 | 0.216292724 | 0.38020018 | 0.568891694 | 0.569429642 | 0.744530893 | 0.216292724 |
| ENSMUSG00000005107 | Slc2a9 | 117591 | protein_coding | 101.952285 | 0.216337424 | 0.115488441 | 1.873238754 | 0.061035423 | 0.177760508 | 0.216337424 |
| ENSMUSG00000003380 | Rabac1 | 14470 | protein_coding | 479.9040109 | 0.216348332 | 0.154023143 | 1.40464821 | 0.160125913 | 0.341659149 | 0.216348332 |
| ENSMUSG00000028106 | Rprd2 | 75137 | protein_coding | 617.6923764 | -0.216356759 | 0.110457053 | -1.95874101 | 0.050143125 | 0.155811367 | 0.216356759 |
| ENSMUSG00000049382 | Krt8 | 16691 | protein_coding | 67970.2979 | 0.216439711 | 0.151276269 | 1.430757864 | 0.152499623 | 0.331153919 | 0.216439711 |
| ENSMUSG00000029462 | Vps29 | 56433 | protein_coding | 547.9442395 | 0.216494425 | 0.096469008 | 2.244186289 | 0.024820418 | 0.093882597 | 0.216494425 |
| ENSMUSG00000005503 | Evx1 | 14028 | protein_coding | 15.33107606 | 0.216558431 | 0.346401937 | 0.625165185 | 0.531862649 | 0.71689162 | 0.216558431 |
| ENSMUSG00000043705 | Capn13 | 381122 | protein_coding | 494.6479826 | -0.216673527 | 0.24678652 | -0.877979586 | 0.379954797 | 0.58786467 | 0.216673527 |
| ENSMUSG00000072571 | Tmem253 | 619301 | protein_coding | 110.1131861 | 0.21692603 | 0.163510612 | 1.326678602 | 0.184615022 | 0.374457819 | 0.21692603 |
| ENSMUSG00000027583 | Zbtb46 | 72147 | protein_coding | 11.58883368 | 0.216963909 | 0.487096879 | 0.445422499 | 0.656014456 | 0.800853376 | 0.216963909 |
| ENSMUSG00000039983 | Ccdc32 | 269336 | protein_coding | 502.5202137 | 0.216973618 | 0.160152201 | 1.354796352 | 0.175482449 | 0.361733769 | 0.216973618 |
| ENSMUSG00000020472 | Zkscan17 | 268417 | protein_coding | 243.4110568 | -0.217062968 | 0.12865481 | -1.687173356 | 0.091570023 | 0.234452231 | 0.217062968 |
| ENSMUSG00000028793 | Rnf19b | 75234 | protein_coding | 562.1380224 | 0.217192647 | 0.105147802 | 2.065593793 | 0.038866851 | 0.130468115 | 0.217192647 |
| ENSMUSG00000024140 | Epas1 | 13819 | protein_coding | 956.2192433 | -0.217196194 | 0.128792504 | -1.686404008 | 0.09171801 | 0.234730223 | 0.217196194 |
| ENSMUSG00000019891 | Dcbld1 | 66686 | protein_coding | 577.2714557 | 0.21720066 | 0.063576173 | 3.416384621 | 0.000634585 | 0.005311444 | 0.21720066 |
| ENSMUSG00000024772 | Ehd1 | 13660 | protein_coding | 803.5686918 | 0.217202816 | 0.090338715 | 2.404315973 | 0.016202762 | 0.068510111 | 0.217202816 |
| ENSMUSG00000004099 | Dnmt1 | 13433 | protein_coding | 1001.395251 | -0.217341968 | 0.108331137 | -2.006274233 | 0.044826998 | 0.144139043 | 0.217341968 |
| ENSMUSG00000034918 | Cdhr2 | 268663 | protein_coding | 7122.44632 | -0.217375475 | 0.124584576 | -1.744802453 | 0.081019262 | 0.215979046 | 0.217375475 |
| ENSMUSG00000026309 | Ilkap | 67444 | protein_coding | 538.909252 | -0.217388327 | 0.103043648 | -2.109672285 | 0.034886593 | 0.120719891 | 0.217388327 |
| ENSMUSG00000014353 | Tmem87b | 72477 | protein_coding | 1609.132779 | -0.21738928 | 0.07348483 | -2.958287873 | 0.003093531 | 0.019247562 | 0.21738928 |
| ENSMUSG00000062007 | Hsh2d | 209488 | protein_coding | 56.63030929 | 0.21740621 | 0.24713107 | 0.879720262 | 0.37901087 | 0.587131408 | 0.21740621 |
| ENSMUSG00000022999 | Lmbr1l | 74775 | protein_coding | 204.1601319 | -0.217434136 | 0.109902135 | -1.978434141 | 0.047879751 | 0.151225199 | 0.217434136 |
| ENSMUSG00000052253 | Zfp622 | 52521 | protein_coding | 789.8161073 | -0.217450011 | 0.093146143 | -2.334503663 | 0.019569364 | 0.078608426 | 0.217450011 |
| ENSMUSG00000042228 | Lyn | 17096 | protein_coding | 378.7102451 | 0.217457002 | 0.086505059 | 2.513806775 | 0.011943585 | 0.054713093 | 0.217457002 |
| ENSMUSG00000031216 | Stard8 | 236920 | protein_coding | 57.23124276 | -0.217502154 | 0.204416471 | -1.064014815 | 0.287321988 | 0.496308929 | 0.217502154 |
| ENSMUSG00000031430 | Vsig1 | 78789 | protein_coding | 110.8531017 | 0.217546355 | 0.307177156 | 0.708211374 | 0.478814004 | 0.672927641 | 0.217546355 |
| ENSMUSG00000028857 | Tmem222 | 52174 | protein_coding | 344.3486034 | 0.217611351 | 0.115195921 | 1.88905431 | 0.058884552 | 0.173815646 | 0.217611351 |
| ENSMUSG00000056459 | Zbtb25 | 109929 | protein_coding | 32.45508649 | 0.217966306 | 0.207520311 | 1.050337217 | 0.2935631 | 0.502565892 | 0.217966306 |
| ENSMUSG00000027995 | Tlr2 | 24088 | protein_coding | 385.6989783 | -0.217983958 | 0.163671305 | -1.331839797 | 0.182912837 | 0.372086293 | 0.217983958 |
| ENSMUSG00000043219 | Hoxa6 | 15403 | protein_coding | 19.82867319 | -0.21800542 | 0.391950085 | -0.556207099 | 0.578069287 | 0.749673311 | 0.21800542 |
| ENSMUSG00000031447 | Lamp1 | 16783 | protein_coding | 5289.548274 | -0.218012401 | 0.075304579 | -2.895074945 | 0.00379068 | 0.022520684 | 0.218012401 |
| ENSMUSG00000031075 | Ano1 | 101772 | protein_coding | 1749.586784 | -0.218039991 | 0.278618338 | -0.78257588 | 0.43387621 | 0.637453749 | 0.218039991 |
| ENSMUSG00000037012 | Hk1 | 15275 | protein_coding | 2544.331972 | -0.218171168 | 0.082540938 | -2.643187415 | 0.008212956 | 0.041397457 | 0.218171168 |
| ENSMUSG00000017453 | Pipox | 19193 | protein_coding | 51.23994525 | -0.218197961 | 0.182552116 | -1.195263932 | 0.231983927 | 0.435427762 | 0.218197961 |

| ensembl_gene_id | external_gene_name | entrezgene_id | gene_biotype | baseMean | log2FoldChange | lfcSE | stat | pvalue | padj | absLog2FC |
|---|---|---|---|---|---|---|---|---|---|---|
| ENSMUSG00000053390 | Zfp952 | 240067 | protein_coding | 215.3491474 | 0.218207125 | 0.090335613 | 2.415516057 | 0.01571293 | 0.066990242 | 0.218207125 |
| ENSMUSG00000063296 | Tmem117 | 320709 | protein_coding | 494.994653 | -0.218274047 | 0.091343008 | -2.389608706 | 0.016866332 | 0.07056657 | 0.218274047 |
| ENSMUSG00000050088 | 1600012H06Rik | 67912 | protein_coding | 579.9115741 | 0.218342804 | 0.080422524 | 2.714945915 | 0.00662866 | 0.034879844 | 0.218342804 |
| ENSMUSG00000005583 | Mef2c | 17260 | protein_coding | 9.583936158 | 0.218420171 | 0.995437969 | 0.219421177 | 0.826321975 | 0.908326875 | 0.218420171 |
| ENSMUSG00000108659 | Gm34121 | 102637256 | lncRNA | 13.84894621 | 0.218471322 | 0.375221904 | 0.582245651 | 0.560401228 | 0.738240903 | 0.218471322 |
| ENSMUSG00000025878 | Uimc1 | 20184 | protein_coding | 373.8207429 | -0.218478405 | 0.067146625 | -3.253751107 | 0.00113892 | 0.00853681 | 0.218478405 |
| ENSMUSG00000050390 | C77080 | 97130 | protein_coding | 1740.476013 | -0.218490723 | 0.114792682 | -1.903350624 | 0.056994809 | 0.170002495 | 0.218490723 |
| ENSMUSG00000051065 | Mb21d2 | 239796 | protein_coding | 189.6715962 | 0.218491462 | 0.132237157 | 1.652269799 | 0.098479563 | 0.245820793 | 0.218491462 |
| ENSMUSG00000027618 | Nfs1 | 18041 | protein_coding | 551.4140371 | -0.218534962 | 0.091633976 | -2.384868281 | 0.017085237 | 0.071239179 | 0.218534962 |
| ENSMUSG00000019996 | Map7 | 17761 | protein_coding | 2227.503949 | 0.218541983 | 0.076517579 | 2.856101662 | 0.004288778 | 0.024831572 | 0.218541983 |
| ENSMUSG00000047528 | Flacc1 | 108812 | protein_coding | 22.78525827 | -0.218549873 | 0.339169036 | -0.64436859 | 0.519336448 | 0.706343689 | 0.218549873 |
| ENSMUSG00000030493 | Faap24 | 101831 | protein_coding | 94.67012737 | 0.21867484 | 0.166307041 | 1.314886241 | 0.188548138 | 0.379766659 | 0.21867484 |
| ENSMUSG00000028842 | Ago3 | 214150 | protein_coding | 289.5168009 | -0.218750374 | 0.146384815 | -1.494351542 | 0.13508376 | 0.304588637 | 0.218750374 |
| ENSMUSG00000026158 | Ogfrl1 | 70155 | protein_coding | 55.00770004 | 0.218975621 | 0.246806655 | 0.887235483 | 0.374952132 | 0.583397242 | 0.218975621 |
| ENSMUSG00000109865 | Hspa14 | 50497 | protein_coding | 471.2842231 | 0.218993634 | 0.098567165 | 2.22177064 | 0.02629881 | 0.098054177 | 0.218993634 |
| ENSMUSG00000033916 | Chmp2a | 68953 | protein_coding | 1933.18261 | 0.219033639 | 0.092860368 | 2.358741883 | 0.018337005 | 0.074947862 | 0.219033639 |
| ENSMUSG00000001089 | Luzp1 | 269593 | protein_coding | 2068.285213 | -0.219055417 | 0.078729374 | -2.782384842 | 0.005396101 | 0.029627404 | 0.219055417 |
| ENSMUSG00000037860 | Aim2 | 383619 | protein_coding | 880.064912 | 0.219110454 | 0.055857844 | 3.922644286 | 8.76E-05 | 0.001006503 | 0.219110454 |
| ENSMUSG00000028621 | Cyb5rl | 230582 | protein_coding | 65.56203643 | -0.219112813 | 0.141932687 | -1.54377979 | 0.122641689 | 0.285438819 | 0.219112813 |
| ENSMUSG00000028636 | Ppcs | 106564 | protein_coding | 406.7656135 | 0.219155088 | 0.152289333 | 1.439070507 | 0.150130547 | 0.327752105 | 0.219155088 |
| ENSMUSG00000021764 | Ndufs4 | 17993 | protein_coding | 1482.938553 | 0.219215555 | 0.097216598 | 2.254919022 | 0.024138412 | 0.092046116 | 0.219215555 |
| ENSMUSG00000044018 | Mrpl50 | 28028 | protein_coding | 774.6228846 | 0.21925107 | 0.095193363 | 2.30321804 | 0.021266578 | 0.083700039 | 0.21925107 |
| ENSMUSG00000020130 | Tbc1d15 | 66687 | protein_coding | 1143.897296 | -0.219257617 | 0.122644363 | -1.787751284 | 0.073816144 | 0.202470043 | 0.219257617 |
| ENSMUSG00000020262 | Adarb1 | 110532 | protein_coding | 7.29962067 | -0.219280779 | 0.55620257 | -0.394246253 | 0.69339924 | 0.825651233 | 0.219280779 |
| ENSMUSG00000025558 | Dock9 | 105445 | protein_coding | 662.5305342 | -0.2192904 | 0.092707098 | -2.365411107 | 0.018010066 | 0.073921367 | 0.2192904 |
| ENSMUSG00000037316 | Bag4 | 67384 | protein_coding | 407.0762002 | -0.219310979 | 0.10861353 | -2.019186552 | 0.043467834 | 0.140894516 | 0.219310979 |
| ENSMUSG00000048922 | Cdca2 | 108912 | protein_coding | 296.6210371 | 0.219323878 | 0.118360535 | 1.853015262 | 0.063880171 | 0.183628111 | 0.219323878 |
| ENSMUSG00000079022 | Col22a1 | 69700 | protein_coding | 6.7609165 | 0.219352282 | 1.418987568 | 0.154583653 | 0.877149558 | 0.937792869 | 0.219352282 |
| ENSMUSG00000014232 | Cluap1 | 76779 | protein_coding | 113.0018225 | 0.219353236 | 0.127128065 | 1.72545092 | 0.084446242 | 0.222281761 | 0.219353236 |
| ENSMUSG00000070315 | 4930581F22Rik | 78934 | lncRNA | 47.53768701 | -0.219360544 | 0.217123505 | -1.010303068 | 0.312350112 | 0.52254097 | 0.219360544 |
| ENSMUSG00000022437 | Samm50 | 68653 | protein_coding | 1697.532494 | 0.219364556 | 0.108253082 | 2.026404726 | 0.042723327 | 0.139104079 | 0.219364556 |
| ENSMUSG00000019961 | Tmpo | 21917 | protein_coding | 2495.564519 | 0.219413593 | 0.086460065 | 2.537744933 | 0.011156927 | 0.051915725 | 0.219413593 |
| ENSMUSG00000022962 | Gart | 14450 | protein_coding | 591.1095415 | -0.219465891 | 0.10703736 | -2.050367196 | 0.040328612 | 0.133776037 | 0.219465891 |
| ENSMUSG00000030397 | Mark4 | 232944 | protein_coding | 214.6120464 | -0.219493628 | 0.111405884 | -1.97021576 | 0.048813648 | 0.153112207 | 0.219493628 |
| ENSMUSG00000030180 | Kdm5a | 214899 | protein_coding | 1152.16586 | -0.219586733 | 0.05654802 | -3.883190512 | 0.000103095 | 0.001157218 | 0.219586733 |
| ENSMUSG00000040699 | Limd2 | 67803 | protein_coding | 73.18843922 | 0.219590686 | 0.137915745 | 1.59220897 | 0.111337758 | 0.266434468 | 0.219590686 |
| ENSMUSG00000033377 | Palmd | 114301 | protein_coding | 69.86214906 | -0.219682268 | 0.217826927 | -1.008517504 | 0.313206089 | 0.523588013 | 0.219682268 |
| ENSMUSG00000060216 | Arrb2 | 216869 | protein_coding | 48.09972885 | 0.219756795 | 0.170297828 | 1.290426287 | 0.19690269 | 0.390519991 | 0.219756795 |

| ensembl_gene_id | external_gene_name | entrezgene_id | gene_biotype | baseMean | log2FoldChange | lfcSE | stat | pvalue | padj | absLog2FC |
|---|---|---|---|---|---|---|---|---|---|---|
| ENSMUSG00000043384 | Gprasp1 | 67298 | protein_coding | 154.8926649 | -0.219805407 | 0.121320396 | -1.811776201 | 0.070020784 | 0.195681974 | 0.219805407 |
| ENSMUSG00000025577 | Cbx2 | 12416 | protein_coding | 74.25878728 | -0.219812614 | 0.213514306 | -1.029498297 | 0.303245579 | 0.512749001 | 0.219812614 |
| ENSMUSG00000024013 | Fgd2 | 26382 | protein_coding | 30.59709084 | -0.21981445 | 0.294790621 | -0.74566297 | 0.455871034 | 0.655276042 | 0.21981445 |
| ENSMUSG00000038816 | Ctnnal1 | 54366 | protein_coding | 326.0559687 | -0.219848334 | 0.253836865 | -0.866100887 | 0.386434839 | 0.594526621 | 0.219848334 |
| ENSMUSG00000025171 | Ubtd1 | 226122 | protein_coding | 54.61988874 | -0.219985679 | 0.168023332 | -1.309256731 | 0.190447401 | 0.382383235 | 0.219985679 |
| ENSMUSG00000029999 | Tgfa | 21802 | protein_coding | 480.7299676 | -0.220009481 | 0.132412861 | -1.661541637 | 0.096604714 | 0.242853517 | 0.220009481 |
| ENSMUSG00000070643 | Sox13 | 20668 | protein_coding | 426.2006262 | -0.22003625 | 0.112943303 | -1.948200953 | 0.051390923 | 0.158327779 | 0.22003625 |
| ENSMUSG00000024824 | Rad9a | 19367 | protein_coding | 90.13724247 | -0.220088409 | 0.125169989 | -1.758316118 | 0.078693737 | 0.21147863 | 0.220088409 |
| ENSMUSG00000020241 | Col6a2 | 12834 | protein_coding | 16.26724935 | -0.220108418 | 0.491808481 | -0.44754905 | 0.654478681 | 0.800101761 | 0.220108418 |
| ENSMUSG00000040121 | Rep15 | 66532 | protein_coding | 1857.757054 | 0.220144608 | 0.173818106 | 1.266522879 | 0.205325944 | 0.401099937 | 0.220144608 |
| ENSMUSG00000024456 | Diaph1 | 13367 | protein_coding | 1401.032095 | -0.22023424 | 0.072180298 | -3.051168352 | 0.002279527 | 0.015084466 | 0.22023424 |
| ENSMUSG00000050244 | Heatr1 | 217995 | protein_coding | 596.9103153 | -0.22027531 | 0.222067018 | -0.991931681 | 0.321230853 | 0.532354566 | 0.22027531 |
| ENSMUSG00000030876 | Mettl9 | 59052 | protein_coding | 1062.360439 | 0.220309896 | 0.070567998 | 3.121951909 | 0.001796563 | 0.012371661 | 0.220309896 |
| ENSMUSG00000026961 | Lrrc26 | 227618 | protein_coding | 383.8716577 | 0.220336821 | 0.128632593 | 1.712915953 | 0.086728003 | 0.226496742 | 0.220336821 |
| ENSMUSG00000037364 | Srrt | 83701 | protein_coding | 1172.771037 | 0.220344822 | 0.09213847 | 2.391453028 | 0.016781831 | 0.07034237 | 0.220344822 |
| ENSMUSG00000017837 | Nkiras2 | 71966 | protein_coding | 726.196536 | 0.220462032 | 0.121856707 | 1.809190791 | 0.070421368 | 0.19615605 | 0.220462032 |
| ENSMUSG00000041417 | Pik3r1 | 18708 | protein_coding | 2182.993517 | -0.220504976 | 0.132821012 | -1.660166361 | 0.09688099 | 0.24331763 | 0.220504976 |
| ENSMUSG00000009646 | Pla2g12b | 69836 | protein_coding | 56.54063241 | 0.220570499 | 0.222163307 | 0.992830466 | 0.32079258 | 0.532256186 | 0.220570499 |
| ENSMUSG00000042628 | Zfyve1 | 217695 | protein_coding | 654.8774253 | -0.220611293 | 0.147342222 | -1.497271382 | 0.134322658 | 0.303287236 | 0.220611293 |
| ENSMUSG00000063021 | H2ac15 | 319169 | protein_coding | 378.6515003 | 0.220694873 | 0.19747474 | 1.117585334 | 0.263744135 | 0.471430433 | 0.220694873 |
| ENSMUSG00000049960 | Mrps16 | 66242 | protein_coding | 554.2572902 | 0.220809071 | 0.108026442 | 2.044027997 | 0.040950781 | 0.135324491 | 0.220809071 |
| ENSMUSG00000034349 | Smc4 | 70099 | protein_coding | 1749.870618 | 0.220835717 | 0.090503362 | 2.440083011 | 0.014683887 | 0.06343503 | 0.220835717 |
| ENSMUSG00000046020 | Pofut1 | 140484 | protein_coding | 761.2859895 | 0.220837807 | 0.101482368 | 2.176119967 | 0.029546291 | 0.106666665 | 0.220837807 |
| ENSMUSG00000027222 | Pex16 | 18633 | protein_coding | 296.9080345 | -0.220889838 | 0.104462758 | -2.114531943 | 0.034469859 | 0.119563572 | 0.220889838 |
| ENSMUSG00000084834 | 4930565N06Rik | 68306 | lncRNA | 56.43262949 | -0.220925718 | 0.27691943 | -0.797797823 | 0.424987825 | 0.629529435 | 0.220925718 |
| ENSMUSG00000044320 | 1700001O22Rik | 73598 | protein_coding | 28.67409676 | -0.220932021 | 0.262183676 | -0.842661238 | 0.399417931 | 0.606939679 | 0.220932021 |
| ENSMUSG00000026767 | Mindy3 | 66960 | protein_coding | 547.2545453 | 0.220963629 | 0.086915832 | 2.542271331 | 0.011013464 | 0.051443535 | 0.220963629 |
| ENSMUSG00000085169 | Gm10785 | 100038392 | lncRNA | 13.76339756 | -0.221009437 | 0.30479149 | -0.725116825 | 0.468380364 | 0.664975705 | 0.221009437 |
| ENSMUSG00000038046 | Mrm3 | 67390 | protein_coding | 114.8560996 | -0.221069401 | 0.135550856 | -1.630896382 | 0.102912185 | 0.253116937 | 0.221069401 |
| ENSMUSG00000030291 | Med21 | 108098 | protein_coding | 316.6589485 | 0.221093353 | 0.09348824 | 2.364932256 | 0.018033369 | 0.073991629 | 0.221093353 |
| ENSMUSG00000023089 | Ndufa5 | 68202 | protein_coding | 910.1993077 | 0.221181646 | 0.108479651 | 2.038922906 | 0.041457722 | 0.136292797 | 0.221181646 |
| ENSMUSG00000042918 | Mamstr | 74490 | protein_coding | 19.22865604 | 0.22120716 | 0.367737744 | 0.601535098 | 0.547483642 | 0.729306636 | 0.22120716 |
| ENSMUSG00000022455 | Wbp2nl | 74716 | protein_coding | 7.895322219 | 0.221217898 | 0.54436732 | 0.406376154 | 0.684466238 | 0.819774287 | 0.221217898 |
| ENSMUSG00000036678 | Aaas | 223921 | protein_coding | 174.4692983 | 0.221245761 | 0.125874676 | 1.757666975 | 0.07880419 | 0.211670171 | 0.221245761 |
| ENSMUSG00000017561 | Crlf3 | 54394 | protein_coding | 179.5449682 | -0.22129477 | 0.138391234 | -1.599051937 | 0.109809063 | 0.264125454 | 0.22129477 |
| ENSMUSG00000016477 | E2f3 | 13557 | protein_coding | 256.4048824 | -0.221302413 | 0.178534391 | -1.23955061 | 0.215141658 | 0.413911865 | 0.221302413 |
| ENSMUSG00000021218 | Gdi2 | 14569 | protein_coding | 7189.81804 | 0.221408282 | 0.077292878 | 2.86453663 | 0.004176199 | 0.024303431 | 0.221408282 |
| ENSMUSG00000031521 | Aga | 11593 | protein_coding | 326.0431984 | 0.221415853 | 0.108433712 | 2.041946641 | 0.041156825 | 0.1357952 | 0.221415853 |

| ensembl_gene_id | external_gene_name | entrezgene_id | gene_biotype | baseMean | log2FoldChange | lfcSE | stat | pvalue | padj | absLog2FC |
|---|---|---|---|---|---|---|---|---|---|---|
| ENSMUSG00000058076 | Sdhc | 66052 | protein_coding | 2397.281864 | 0.221439409 | 0.089049861 | 2.486690109 | 0.012893766 | 0.057937359 | 0.221439409 |
| ENSMUSG00000057101 | Zfp180 | 210135 | protein_coding | 465.1615251 | -0.221444163 | 0.068830143 | -3.217255621 | 0.001294232 | 0.009421633 | 0.221444163 |
| ENSMUSG00000052419 | Smim30 | 66520 | protein_coding | 337.6486458 | 0.221534521 | 0.089975259 | 2.462171535 | 0.01380986 | 0.060807443 | 0.221534521 |
| ENSMUSG00000028700 | Pomgnt1 | 68273 | protein_coding | 602.4825381 | -0.221541903 | 0.08637468 | -2.564894069 | 0.010320731 | 0.048910622 | 0.221541903 |
| ENSMUSG00000026004 | Kansl1l | 68691 | protein_coding | 228.5100685 | -0.221545146 | 0.193596624 | -1.144364715 | 0.252472415 | 0.458766879 | 0.221545146 |
| ENSMUSG00000067365 | Tmem128 | 66309 | protein_coding | 508.6321266 | 0.221548965 | 0.093849319 | 2.360688055 | 0.018241067 | 0.074632345 | 0.221548965 |
| ENSMUSG00000035247 | Hectd1 | 207304 | protein_coding | 4248.470612 | 0.22158148 | 0.071580292 | 3.095565467 | 0.00196438 | 0.013325658 | 0.22158148 |
| ENSMUSG00000026605 | Cenpf | 108000 | protein_coding | 994.4616755 | 0.221615889 | 0.112562927 | 1.96881776 | 0.048974022 | 0.15346414 | 0.221615889 |
| ENSMUSG00000036459 | Wtip | 101543 | protein_coding | 163.0122809 | 0.221618121 | 0.143896216 | 1.540124731 | 0.123529953 | 0.286797469 | 0.221618121 |
| ENSMUSG00000037742 | Eef1a1 | 13627 | protein_coding | 104262.2081 | 0.221667696 | 0.092252039 | 2.402848751 | 0.016267914 | 0.068749095 | 0.221667696 |
| ENSMUSG00000006301 | Tmbim1 | 69660 | protein_coding | 2123.858464 | -0.221730605 | 0.094702733 | -2.341332697 | 0.019215036 | 0.077695925 | 0.221730605 |
| ENSMUSG00000031310 | Zmym3 | 56364 | protein_coding | 294.7288436 | -0.221741262 | 0.163025735 | -1.360161098 | 0.17377895 | 0.359707067 | 0.221741262 |
| ENSMUSG00000041515 | Irf8 | 15900 | protein_coding | 935.5644857 | 0.221786611 | 0.132809422 | 1.669961418 | 0.094926997 | 0.240190185 | 0.221786611 |
| ENSMUSG00000000409 | Lck | 16818 | protein_coding | 84.25267617 | 0.221805537 | 0.153927753 | 1.440971707 | 0.149592676 | 0.327026097 | 0.221805537 |
| ENSMUSG00000040712 | Camta2 | 216874 | protein_coding | 494.3112333 | -0.221858541 | 0.136902911 | -1.620553867 | 0.105113353 | 0.256707373 | 0.221858541 |
| ENSMUSG00000044934 | Zfp367 | 238673 | protein_coding | 275.72176 | 0.221891146 | 0.15239283 | 1.45604715 | 0.145379586 | 0.320854378 | 0.221891146 |
| ENSMUSG00000022453 | Naga | 17939 | protein_coding | 1617.995676 | 0.221895995 | 0.091472314 | 2.425826856 | 0.015273555 | 0.065520384 | 0.221895995 |
| ENSMUSG00000011256 | Adam19 | 11492 | protein_coding | 13.04897215 | 0.221948208 | 0.518157104 | 0.428341533 | 0.668402485 | 0.807885474 | 0.221948208 |
| ENSMUSG00000022856 | Tmem41a | 66664 | protein_coding | 208.9034894 | -0.221989953 | 0.102100647 | -2.174226709 | 0.029688113 | 0.10705734 | 0.221989953 |
| ENSMUSG00000062031 | Pgghg | 212974 | protein_coding | 400.7087226 | -0.222012517 | 0.085636113 | -2.59251043 | 0.009527829 | 0.046169536 | 0.222012517 |
| ENSMUSG00000061859 | Patj | 12695 | protein_coding | 1660.597225 | -0.222037342 | 0.073000273 | -3.041596027 | 0.002353275 | 0.015501639 | 0.222037342 |
| ENSMUSG00000026662 | Sephs1 | 109079 | protein_coding | 539.9547988 | 0.222088022 | 0.073902424 | 3.005152063 | 0.002654481 | 0.01706289 | 0.222088022 |
| ENSMUSG00000021697 | Depdc1b | 218581 | protein_coding | 71.91608016 | 0.222088532 | 0.184635999 | 1.202845237 | 0.229036215 | 0.432186807 | 0.222088532 |
| ENSMUSG00000035168 | Tanc1 | 66860 | protein_coding | 570.9615855 | -0.222202168 | 0.080882288 | -2.747229004 | 0.006010115 | 0.032253746 | 0.222202168 |
| ENSMUSG00000003299 | Mrpl4 | 66163 | protein_coding | 426.3264169 | 0.222281722 | 0.083561249 | 2.660105323 | 0.007811622 | 0.039739225 | 0.222281722 |
| ENSMUSG00000021650 | Ptcd2 | 68927 | protein_coding | 703.1116095 | 0.222297552 | 0.115667192 | 1.921872131 | 0.054621849 | 0.164775912 | 0.222297552 |
| ENSMUSG00000027901 | Dennd2d | 72121 | protein_coding | 1177.316103 | 0.222364921 | 0.080183954 | 2.773184785 | 0.005551059 | 0.030311028 | 0.222364921 |
| ENSMUSG00000048796 | Cyb561d1 | 72023 | protein_coding | 259.8222056 | -0.222395123 | 0.130886835 | -1.699140509 | 0.089292712 | 0.230651255 | 0.222395123 |
| ENSMUSG00000091537 | Tma7 | 66167 | protein_coding | 897.2370474 | 0.222453369 | 0.097205823 | 2.288477799 | 0.022109712 | 0.086124599 | 0.222453369 |
| ENSMUSG00000108621 | Gm33989 | 102637087 | lncRNA | 378.9251409 | -0.222464199 | 0.165020253 | -1.348102401 | 0.177625449 | 0.364683959 | 0.222464199 |
| ENSMUSG00000079427 | Mthfsl | 100039707 | protein_coding | 81.22352679 | 0.222500037 | 0.125204764 | 1.777089227 | 0.07555356 | 0.205186208 | 0.222500037 |
| ENSMUSG00000040957 | Cables1 | 63955 | protein_coding | 493.2829661 | -0.222558131 | 0.124981304 | -1.780731392 | 0.074956342 | 0.204225901 | 0.222558131 |
| ENSMUSG00000040263 | Klhdc4 | 234825 | protein_coding | 342.0000409 | 0.222602606 | 0.118405207 | 1.880006894 | 0.060107138 | 0.176054892 | 0.222602606 |
| ENSMUSG00000020476 | Dbnl | 13169 | protein_coding | 1116.669162 | 0.222646393 | 0.075318144 | 2.956079109 | 0.003115772 | 0.019348152 | 0.222646393 |
| ENSMUSG00000019312 | Grb7 | 14786 | protein_coding | 727.8865187 | -0.222665717 | 0.15673227 | -1.420675632 | 0.15541108 | 0.335418385 | 0.222665717 |
| ENSMUSG00000047230 | Cldn2 | 12738 | protein_coding | 2614.607544 | 0.222678169 | 0.127110832 | 1.751842591 | 0.079800878 | 0.213474537 | 0.222678169 |
| ENSMUSG00000079554 | Aox2 | 213043 | protein_coding | 10.91625441 | -0.222688389 | 0.456417362 | -0.487905166 | 0.625617015 | 0.783090594 | 0.222688389 |
| ENSMUSG00000018697 | Aatf | 56321 | protein_coding | 347.8990702 | -0.222746408 | 0.121149728 | -1.838604274 | 0.065973412 | 0.187446398 | 0.222746408 |

| ensembl_gene_id | external_gene_name | entrezgene_id | gene_biotype | baseMean | log2FoldChange | lfcSE | stat | pvalue | padj | absLog2FC |
|---|---|---|---|---|---|---|---|---|---|---|
| ENSMUSG00000051223 | Bzw1 | 66882 | protein_coding | 4858.38995 | 0.222775358 | 0.128822341 | 1.729322385 | 0.083751413 | 0.220874619 | 0.222775358 |
| ENSMUSG00000062127 | Cttnbp2nl | 80281 | protein_coding | 1072.260465 | -0.222801844 | 0.115191208 | -1.934191407 | 0.053089599 | 0.16183945 | 0.222801844 |
| ENSMUSG00000022272 | Myo10 | 17909 | protein_coding | 1494.276153 | -0.222807212 | 0.104367213 | -2.134839159 | 0.03277415 | 0.115085244 | 0.222807212 |
| ENSMUSG00000024527 | Afg3l2 | 69597 | protein_coding | 1948.270883 | -0.222842126 | 0.061100261 | -3.647155033 | 0.00026516 | 0.002572148 | 0.222842126 |
| ENSMUSG00000040728 | Esrp1 | 207920 | protein_coding | 4361.787807 | 0.222843104 | 0.048570803 | 4.588005383 | 4.48E-06 | 7.30E-05 | 0.222843104 |
| ENSMUSG00000036160 | Surf6 | 20935 | protein_coding | 521.4849422 | -0.222855921 | 0.19661111 | -1.133485898 | 0.25701026 | 0.463812519 | 0.222855921 |
| ENSMUSG00000007570 | Fance | 72775 | protein_coding | 145.176768 | -0.22291155 | 0.111687796 | -1.995845189 | 0.045950776 | 0.146821257 | 0.22291155 |
| ENSMUSG00000091996 | Smim35 | 408059 | protein_coding | 9.467926502 | -0.222974934 | 0.466600573 | -0.477871111 | 0.632741944 | 0.787614387 | 0.222974934 |
| ENSMUSG00000037661 | Gpr160 | 71862 | protein_coding | 473.9076177 | 0.223013229 | 0.166410515 | 1.34013905 | 0.180200143 | 0.368125204 | 0.223013229 |
| ENSMUSG00000047098 | Rnf31 | 268749 | protein_coding | 379.2738939 | -0.223027574 | 0.102890284 | -2.167625213 | 0.030187214 | 0.108537684 | 0.223027574 |
| ENSMUSG00000057554 | Lgals8 | 56048 | protein_coding | 821.1830752 | 0.223033411 | 0.08608518 | 2.590845628 | 0.009574043 | 0.046351171 | 0.223033411 |
| ENSMUSG00000030067 | Foxp1 | 108655 | protein_coding | 1590.458057 | -0.223134479 | 0.11650524 | -1.915231276 | 0.055463016 | 0.166688225 | 0.223134479 |
| ENSMUSG00000024376 | Epb41l4a | 13824 | protein_coding | 108.0786881 | -0.223137046 | 0.161007774 | -1.385877463 | 0.165784323 | 0.349604218 | 0.223137046 |
| ENSMUSG00000037419 | Endod1 | 71946 | protein_coding | 4044.415594 | 0.223173687 | 0.111496918 | 2.001613063 | 0.045326363 | 0.145409528 | 0.223173687 |
| ENSMUSG00000037773 | Pced1a | 319513 | protein_coding | 106.058916 | -0.223176382 | 0.12225318 | -1.825526181 | 0.067921664 | 0.191580545 | 0.223176382 |
| ENSMUSG00000067786 | Nnat | 18111 | protein_coding | 25.33192693 | -0.223176996 | 0.384973865 | -0.579719864 | 0.562103546 | 0.739443779 | 0.223176996 |
| ENSMUSG00000034663 | Bmp2k | 140780 | protein_coding | 500.7896643 | -0.223246505 | 0.100877334 | -2.213049224 | 0.026894252 | 0.099482501 | 0.223246505 |
| ENSMUSG00000048495 | Tyw5 | 68736 | protein_coding | 91.11483702 | 0.223260197 | 0.121919092 | 1.831216045 | 0.067068303 | 0.189677544 | 0.223260197 |
| ENSMUSG00000041079 | Rwdd2b | 53858 | protein_coding | 244.1358082 | 0.223278473 | 0.132897641 | 1.680078522 | 0.092942039 | 0.237012616 | 0.223278473 |
| ENSMUSG00000058230 | Arhgap35 | 232906 | protein_coding | 954.5856058 | -0.223400033 | 0.112380735 | -1.987885494 | 0.046824355 | 0.148717514 | 0.223400033 |
| ENSMUSG00000036825 | Ssx2ip | 99167 | protein_coding | 1067.088329 | 0.223480239 | 0.106839221 | 2.091743434 | 0.036461473 | 0.124617506 | 0.223480239 |
| ENSMUSG00000023882 | Zfp54 | 22712 | protein_coding | 285.9281131 | 0.223503531 | 0.11326794 | 1.973228539 | 0.04846953 | 0.152483247 | 0.223503531 |
| ENSMUSG00000067276 | Capn6 | 12338 | protein_coding | 21.15233419 | 0.223507412 | 0.254440427 | 0.878427279 | 0.379711886 | 0.587657295 | 0.223507412 |
| ENSMUSG00000021486 | Prelid1 | 66494 | protein_coding | 2643.396982 | 0.223533756 | 0.09856236 | 2.2679424 | 0.023332718 | 0.089698043 | 0.223533756 |
| ENSMUSG00000021951 | Eef1akmt1 | 68043 | protein_coding | 177.119474 | 0.223554124 | 0.13366647 | 1.672477208 | 0.094430259 | 0.239351553 | 0.223554124 |
| ENSMUSG00000051890 | Klhdc1 | 271005 | protein_coding | 56.23527577 | -0.223579585 | 0.22389056 | -0.998611038 | 0.317983151 | 0.529209253 | 0.223579585 |
| ENSMUSG00000103421 | Golt1a | 68338 | protein_coding | 51.171872 | 0.223584966 | 0.195236685 | 1.145199563 | 0.252126499 | 0.458432749 | 0.223584966 |
| ENSMUSG00000027566 | Psma7 | 26444 | protein_coding | 1838.404806 | 0.223594778 | 0.091575156 | 2.441653255 | 0.014620181 | 0.063210492 | 0.223594778 |
| ENSMUSG00000035266 | Helq | 191578 | protein_coding | 125.0401626 | -0.223763547 | 0.109309231 | -2.047069082 | 0.040651302 | 0.134577456 | 0.223763547 |
| ENSMUSG00000041920 | Slc16a6 | 104681 | protein_coding | 91.90224311 | 0.22384042 | 0.246363653 | 0.908577288 | 0.3635733 | 0.572459027 | 0.22384042 |
| ENSMUSG00000031928 | Mre11a | 17535 | protein_coding | 385.4313462 | 0.223852053 | 0.074172013 | 3.01801238 | 0.002544385 | 0.016523768 | 0.223852053 |
| ENSMUSG00000027404 | Snrpb | 20638 | protein_coding | 1278.420917 | 0.223962113 | 0.151018434 | 1.483011757 | 0.138071287 | 0.309078789 | 0.223962113 |
| ENSMUSG00000055319 | Sec23ip | 207352 | protein_coding | 844.4715158 | 0.223985359 | 0.090360355 | 2.478801218 | 0.013182475 | 0.058930806 | 0.223985359 |
| ENSMUSG00000023026 | Dip2b | 239667 | protein_coding | 649.7812118 | -0.224058303 | 0.089205398 | -2.511712388 | 0.012014696 | 0.054959818 | 0.224058303 |
| ENSMUSG00000020758 | Itgb4 | 192897 | protein_coding | 2036.498199 | -0.224072416 | 0.141173025 | -1.58721835 | 0.1124632 | 0.267988275 | 0.224072416 |
| ENSMUSG00000021476 | Habp4 | 56541 | protein_coding | 172.7803564 | 0.224074886 | 0.099055217 | 2.262120998 | 0.023689929 | 0.090758727 | 0.224074886 |
| ENSMUSG00000050350 | Gpr18 | 110168 | protein_coding | 12.8156328 | 0.224079443 | 0.381006326 | 0.588125256 | 0.556448222 | 0.735604839 | 0.224079443 |
| ENSMUSG00000004127 | Trmt10a | 108943 | protein_coding | 79.28970236 | 0.224082959 | 0.150843855 | 1.485529252 | 0.13740368 | 0.308196452 | 0.224082959 |

| ensembl_gene_id | external_gene_name | entrezgene_id | gene_biotype | baseMean | log2FoldChange | lfcSE | stat | pvalue | padj | absLog2FC |
|---|---|---|---|---|---|---|---|---|---|---|
| ENSMUSG00000097277 | 2900076A07Rik | 100504421 | lncRNA | 46.96613559 | -0.224183429 | 0.204148548 | -1.098138738 | 0.272143912 | 0.480459791 | 0.224183429 |
| ENSMUSG00000018909 | Arrb1 | 109689 | protein_coding | 1423.098167 | -0.224206017 | 0.089045849 | -2.517871644 | 0.011806634 | 0.054210454 | 0.224206017 |
| ENSMUSG00000028439 | Fam219a | 71901 | protein_coding | 24.12270458 | 0.224258808 | 0.300289945 | 0.746807583 | 0.455179718 | 0.654577694 | 0.224258808 |
| ENSMUSG00000086427 | Hoxa11os | 15397 | lncRNA | 270.2873957 | -0.224304516 | 0.254566953 | -0.881121896 | 0.378251847 | 0.586330845 | 0.224304516 |
| ENSMUSG00000045757 | Zfp764 | 233893 | protein_coding | 140.8037883 | -0.224357227 | 0.121454836 | -1.8472482 | 0.064711177 | 0.185181391 | 0.224357227 |
| ENSMUSG00000033356 | Pus7l | 78895 | protein_coding | 64.76256099 | 0.224359357 | 0.228050674 | 0.983813611 | 0.325207166 | 0.53627702 | 0.224359357 |
| ENSMUSG00000019916 | P4ha1 | 18451 | protein_coding | 267.203075 | 0.224367341 | 0.177737881 | 1.262349591 | 0.206823019 | 0.403166938 | 0.224367341 |
| ENSMUSG00000024187 | Fam234a | 106581 | protein_coding | 2734.905863 | -0.224390639 | 0.085052985 | -2.638245308 | 0.008333627 | 0.04180725 | 0.224390639 |
| ENSMUSG00000025134 | Alyref | 21681 | protein_coding | 724.2221156 | 0.224418556 | 0.133384208 | 1.682497201 | 0.092472467 | 0.236203505 | 0.224418556 |
| ENSMUSG00000024169 | Ift140 | 106633 | protein_coding | 312.6958973 | -0.224449792 | 0.107863732 | -2.080864327 | 0.037446326 | 0.127011304 | 0.224449792 |
| ENSMUSG00000037656 | Slc20a2 | 20516 | protein_coding | 234.8720841 | -0.224513605 | 0.093243061 | -2.407831787 | 0.016047573 | 0.067998336 | 0.224513605 |
| ENSMUSG00000025733 | Rhot2 | 214952 | protein_coding | 513.5878375 | -0.224519807 | 0.079551156 | -2.822332431 | 0.004767573 | 0.027101321 | 0.224519807 |
| ENSMUSG00000014778 | Fhod1 | 234686 | protein_coding | 298.583571 | -0.224536711 | 0.116211348 | -1.932141004 | 0.053342104 | 0.162114111 | 0.224536711 |
| ENSMUSG00000035441 | Myo1d | 338367 | protein_coding | 3979.954379 | 0.224543887 | 0.074702279 | 3.005850564 | 0.002648391 | 0.017043869 | 0.224543887 |
| ENSMUSG00000029014 | Dnajc2 | 22791 | protein_coding | 567.9789202 | -0.224566511 | 0.12503038 | -1.796095562 | 0.07247932 | 0.200112777 | 0.224566511 |
| ENSMUSG00000021258 | Ccnk | 12454 | protein_coding | 139.3952624 | -0.224667076 | 0.163864534 | -1.371053701 | 0.170358216 | 0.35563115 | 0.224667076 |
| ENSMUSG00000028863 | Meaf6 | 70088 | protein_coding | 176.2492252 | -0.224695799 | 0.107582547 | -2.088589701 | 0.036744673 | 0.125245747 | 0.224695799 |
| ENSMUSG00000018425 | Dhx40 | 67487 | protein_coding | 651.8055216 | 0.224777596 | 0.087969206 | 2.555185002 | 0.010613137 | 0.050028426 | 0.224777596 |
| ENSMUSG00000041779 | Tram2 | 170829 | protein_coding | 22.17700546 | -0.224790987 | 0.43718208 | -0.514181614 | 0.607125021 | 0.771131144 | 0.224790987 |
| ENSMUSG00000039463 | Slc9a8 | 77031 | protein_coding | 539.6537276 | -0.224822451 | 0.072698877 | -3.092516126 | 0.001984675 | 0.01342331 | 0.224822451 |
| ENSMUSG00000028948 | Nol9 | 74035 | protein_coding | 415.5038438 | -0.224833559 | 0.175952308 | -1.27780966 | 0.201316548 | 0.395970606 | 0.224833559 |
| ENSMUSG00000042345 | Ubash3a | 328795 | protein_coding | 7.347606234 | 0.224968203 | 0.94943791 | 0.236948831 | 0.812696487 | 0.900308085 | 0.224968203 |
| ENSMUSG00000042759 | Apobr | 171504 | protein_coding | 8.348095973 | 0.22499476 | 0.452859923 | 0.496830804 | 0.619308371 | 0.778317353 | 0.22499476 |
| ENSMUSG00000013878 | Rnf170 | 77733 | protein_coding | 318.8401301 | 0.225036873 | 0.101524531 | 2.216576333 | 0.026652056 | 0.09893124 | 0.225036873 |
| ENSMUSG00000050973 | Gdpgp1 | 269952 | protein_coding | 182.7531758 | -0.225081794 | 0.099638619 | -2.258981468 | 0.023884538 | 0.091340436 | 0.225081794 |
| ENSMUSG00000026429 | Ube2t | 67196 | protein_coding | 66.79620944 | 0.225091078 | 0.201062211 | 1.119509612 | 0.262922793 | 0.470754749 | 0.225091078 |
| ENSMUSG00000021936 | Mapk8 | 26419 | protein_coding | 473.2731217 | 0.22512523 | 0.078226731 | 2.877855531 | 0.004003885 | 0.023489456 | 0.22512523 |
| ENSMUSG00000055760 | Gemin6 | 67242 | protein_coding | 84.81225662 | 0.225142764 | 0.173381835 | 1.298537207 | 0.194102799 | 0.387427242 | 0.225142764 |
| ENSMUSG00000001665 | Gstt3 | 103140 | protein_coding | 649.3656018 | 0.225235851 | 0.117759399 | 1.912678341 | 0.055789244 | 0.167378239 | 0.225235851 |
| ENSMUSG00000024899 | Papss2 | 23972 | protein_coding | 9637.783021 | -0.22532083 | 0.204450388 | -1.102080716 | 0.270426582 | 0.478375791 | 0.22532083 |
| ENSMUSG00000086550 | Pla2g10os | 76684 | lncRNA | 82.86191549 | -0.225354547 | 0.130077113 | -1.732468857 | 0.083190117 | 0.219962183 | 0.225354547 |
| ENSMUSG00000024174 | Pot1b | 72836 | protein_coding | 98.45256653 | -0.225385279 | 0.146429984 | -1.539201694 | 0.123755064 | 0.286966176 | 0.225385279 |
| ENSMUSG00000030313 | Dennd5b | 320560 | protein_coding | 320.5355439 | -0.225427723 | 0.139272899 | -1.618604378 | 0.105532413 | 0.257261797 | 0.225427723 |
| ENSMUSG00000022241 | Tars | 110960 | protein_coding | 1650.515484 | 0.225466945 | 0.13008109 | 1.73327994 | 0.083045924 | 0.219701126 | 0.225466945 |
| ENSMUSG00000022174 | Dad1 | 13135 | protein_coding | 930.8383695 | 0.225476741 | 0.077189275 | 2.921088995 | 0.003488102 | 0.021157077 | 0.225476741 |
| ENSMUSG00000034075 | Zdhhc5 | 228136 | protein_coding | 2368.466564 | 0.225659256 | 0.09441423 | 2.390097936 | 0.016843881 | 0.070491153 | 0.225659256 |
| ENSMUSG00000026965 | Anapc2 | 99152 | protein_coding | 651.1859799 | -0.225686271 | 0.096633883 | -2.335477617 | 0.019518484 | 0.078530312 | 0.225686271 |
| ENSMUSG00000092183 | 4930515G01Rik | 67642 | lncRNA | 13.97084607 | -0.225697386 | 0.328043834 | -0.688009841 | 0.491446584 | 0.683557312 | 0.225697386 |

| ensembl_gene_id | external_gene_name | entrezgene_id | gene_biotype | baseMean | log2FoldChange | lfcSE | stat | pvalue | padj | absLog2FC |
|---|---|---|---|---|---|---|---|---|---|---|
| ENSMUSG00000074405 | Zfp865 | 319748 | protein_coding | 135.751349 | -0.225700055 | 0.119733752 | -1.88501614 | 0.059427659 | 0.17500132 | 0.225700055 |
| ENSMUSG00000046442 | Ppm1e | 320472 | protein_coding | 13.80426998 | 0.225814806 | 0.475481308 | 0.474918367 | 0.634845159 | 0.788753018 | 0.225814806 |
| ENSMUSG00000063952 | Brpf3 | 268936 | protein_coding | 569.2695679 | -0.225820536 | 0.17687677 | -1.276711101 | 0.201704261 | 0.396586282 | 0.225820536 |
| ENSMUSG00000027454 | Gins1 | 69270 | protein_coding | 45.64063627 | 0.225872585 | 0.227187834 | 0.994210742 | 0.320120279 | 0.531631301 | 0.225872585 |
| ENSMUSG00000031264 | Btk | 12229 | protein_coding | 9.262875106 | 0.225901617 | 0.397199952 | 0.568735256 | 0.569535818 | 0.744608594 | 0.225901617 |
| ENSMUSG00000047854 | Stx19 | 68159 | protein_coding | 38.78489884 | -0.226009749 | 0.201314464 | -1.122670195 | 0.261577591 | 0.469295773 | 0.226009749 |
| ENSMUSG00000074622 | Mafb | 16658 | protein_coding | 103.9772942 | 0.226016813 | 0.240742534 | 0.938832078 | 0.347816966 | 0.557852043 | 0.226016813 |
| ENSMUSG00000031903 | Pla2g15 | 192654 | protein_coding | 79.62164024 | 0.226026968 | 0.139296379 | 1.622633487 | 0.104667778 | 0.256090378 | 0.226026968 |
| ENSMUSG00000035683 | Melk | 17279 | protein_coding | 448.1878063 | 0.226036871 | 0.128871089 | 1.753976581 | 0.07943452 | 0.212894672 | 0.226036871 |
| ENSMUSG00000022034 | Esco2 | 71988 | protein_coding | 262.2436356 | 0.226055993 | 0.164432294 | 1.3747664 | 0.169203878 | 0.354093686 | 0.226055993 |
| ENSMUSG00000031563 | Wwc2 | 52357 | protein_coding | 255.1882955 | -0.226083729 | 0.201567797 | -1.121626238 | 0.262021391 | 0.469933196 | 0.226083729 |
| ENSMUSG00000030527 | Crtc3 | 70461 | protein_coding | 132.8330074 | -0.226104556 | 0.124718777 | -1.812915108 | 0.069844916 | 0.195399091 | 0.226104556 |
| ENSMUSG00000041915 | Ammecr1l | 225339 | protein_coding | 357.8148352 | -0.22615891 | 0.087820041 | -2.575253963 | 0.010016651 | 0.047983523 | 0.22615891 |
| ENSMUSG00000026417 | Pigr | 18703 | protein_coding | 59938.28686 | 0.226209645 | 0.110914044 | 2.039504075 | 0.041399745 | 0.136190758 | 0.226209645 |
| ENSMUSG00000056763 | Cspp1 | 211660 | protein_coding | 463.5484381 | -0.226237596 | 0.148829231 | -1.520115328 | 0.128481993 | 0.294624414 | 0.226237596 |
| ENSMUSG00000032411 | Tfdp2 | 211586 | protein_coding | 1087.043312 | 0.226253099 | 0.100380263 | 2.253960002 | 0.024198684 | 0.092187669 | 0.226253099 |
| ENSMUSG00000029521 | Chek2 | 50883 | protein_coding | 269.8112209 | 0.226283323 | 0.147796308 | 1.531048547 | 0.125757395 | 0.29008889 | 0.226283323 |
| ENSMUSG00000056687 | Gm11696 | 100036768 | lncRNA | 28.67451745 | 0.226324939 | 0.26272831 | 0.861441002 | 0.388995202 | 0.597369026 | 0.226324939 |
| ENSMUSG00000049470 | Aff4 | 93736 | protein_coding | 2163.291667 | -0.226378212 | 0.098708787 | -2.293394733 | 0.02182529 | 0.085183341 | 0.226378212 |
| ENSMUSG00000026039 | Sgo2a | 68549 | protein_coding | 120.109869 | 0.226392538 | 0.158024417 | 1.432642767 | 0.151959953 | 0.330297235 | 0.226392538 |
| ENSMUSG00000025538 | Sumf2 | 67902 | protein_coding | 75.93678808 | -0.226505708 | 0.157623227 | -1.437007174 | 0.150715955 | 0.328759755 | 0.226505708 |
| ENSMUSG00000074358 | Ccdc61 | 232933 | protein_coding | 206.6157406 | 0.226518977 | 0.124742662 | 1.815890199 | 0.069387217 | 0.194577393 | 0.226518977 |
| ENSMUSG00000024346 | Pfdn1 | 67199 | protein_coding | 626.9310788 | 0.226601291 | 0.098024484 | 2.311680519 | 0.020795296 | 0.08233345 | 0.226601291 |
| ENSMUSG00000007837 | Prrg2 | 65116 | protein_coding | 439.799261 | 0.226697049 | 0.073150639 | 3.09904402 | 0.001941462 | 0.013209569 | 0.226697049 |
| ENSMUSG00000030890 | Ilk | 16202 | protein_coding | 11.35166415 | 0.226778091 | 0.393648529 | 0.57609282 | 0.564552445 | 0.741125944 | 0.226778091 |
| ENSMUSG00000029153 | Ociad2 | 433904 | protein_coding | 3177.261243 | 0.226815381 | 0.109715669 | 2.067301621 | 0.038705742 | 0.130201701 | 0.226815381 |
| ENSMUSG00000060802 | B2m | 12010 | protein_coding | 3381.007778 | 0.226932035 | 0.108464536 | 2.092223355 | 0.03641854 | 0.124532956 | 0.226932035 |
| ENSMUSG00000043831 | Lysmd4 | 75099 | protein_coding | 219.7609379 | 0.226974321 | 0.137130534 | 1.655169823 | 0.097890055 | 0.244809671 | 0.226974321 |
| ENSMUSG00000034659 | Tmem109 | 68539 | protein_coding | 355.0829596 | -0.226985739 | 0.099465165 | -2.282062651 | 0.022485642 | 0.087332675 | 0.226985739 |
| ENSMUSG00000032870 | Smap2 | 69780 | protein_coding | 341.416308 | 0.227010037 | 0.155163971 | 1.463033176 | 0.143458301 | 0.317449823 | 0.227010037 |
| ENSMUSG00000028920 | Fbxo42 | 213499 | protein_coding | 390.2621314 | -0.227016792 | 0.092508073 | -2.454021416 | 0.014126857 | 0.061849978 | 0.227016792 |
| ENSMUSG00000045975 | C2cd2 | 207781 | protein_coding | 248.8015615 | -0.227056099 | 0.095151856 | -2.386249813 | 0.017021184 | 0.071065118 | 0.227056099 |
| ENSMUSG00000037376 | Trmt6 | 66926 | protein_coding | 339.9801647 | 0.227063252 | 0.139991368 | 1.62198038 | 0.104807549 | 0.256274892 | 0.227063252 |
| ENSMUSG00000038116 | Phf20 | 228829 | protein_coding | 273.837545 | 0.227066918 | 0.115535713 | 1.965339653 | 0.049374939 | 0.15411851 | 0.227066918 |
| ENSMUSG00000029394 | Cdk2ap1 | 13445 | protein_coding | 211.2223371 | 0.227172163 | 0.153716767 | 1.477861959 | 0.13944473 | 0.311118597 | 0.227172163 |
| ENSMUSG00000038598 | Shoc1 | 100155 | protein_coding | 24.76753448 | 0.22720849 | 0.308716963 | 0.735976695 | 0.46174488 | 0.659671043 | 0.22720849 |
| ENSMUSG00000016028 | Celsr1 | 12614 | protein_coding | 1461.419588 | -0.227295191 | 0.09095468 | -2.498993919 | 0.012454645 | 0.056405341 | 0.227295191 |
| ENSMUSG00000051730 | Mettl5 | 75422 | protein_coding | 123.6013491 | -0.227321968 | 0.136626154 | -1.663824687 | 0.096147469 | 0.242153679 | 0.227321968 |

| ensembl_gene_id | external_gene_name | entrezgene_id | gene_biotype | baseMean | log2FoldChange | lfcSE | stat | pvalue | padj | absLog2FC |
|---|---|---|---|---|---|---|---|---|---|---|
| ENSMUSG00000002763 | Pex6 | 224824 | protein_coding | 558.4648025 | -0.227352453 | 0.098200988 | -2.315174793 | 0.020603368 | 0.081756464 | 0.227352453 |
| ENSMUSG00000044312 | Neurog3 | 11925 | protein_coding | 11.18771437 | -0.227391318 | 0.333795059 | -0.681230329 | 0.49572576 | 0.687079699 | 0.227391318 |
| ENSMUSG00000028549 | Itgb3bp | 67733 | protein_coding | 115.615704 | 0.227416244 | 0.109087583 | 2.084712459 | 0.037095409 | 0.126116473 | 0.227416244 |
| ENSMUSG00000001833 | Septin7 | 235072 | protein_coding | 1907.993476 | 0.227466196 | 0.083872688 | 2.712041334 | 0.006687025 | 0.035128938 | 0.227466196 |
| ENSMUSG00000032782 | Cntrob | 216846 | protein_coding | 168.1047712 | -0.227501092 | 0.166668627 | -1.364990496 | 0.172256037 | 0.357883248 | 0.227501092 |
| ENSMUSG00000024735 | Prpf19 | 28000 | protein_coding | 946.4321473 | 0.227521859 | 0.087494353 | 2.600417647 | 0.009311036 | 0.045381331 | 0.227521859 |
| ENSMUSG00000035235 | Trim13 | 66597 | protein_coding | 59.03466842 | 0.227570347 | 0.156363354 | 1.455394381 | 0.145560111 | 0.321163797 | 0.227570347 |
| ENSMUSG00000038914 | Dido1 | 23856 | protein_coding | 1288.51398 | -0.227580755 | 0.117474199 | -1.937282885 | 0.052710773 | 0.161085418 | 0.227580755 |
| ENSMUSG00000059208 | Hnrnpm | 76936 | protein_coding | 2308.667788 | 0.227610763 | 0.085021841 | 2.67708581 | 0.007426561 | 0.038232148 | 0.227610763 |
| ENSMUSG00000054556 | Gm4876 | 232599 | lncRNA | 56.46271013 | -0.227689644 | 0.299068515 | -0.761329369 | 0.446460363 | 0.647239525 | 0.227689644 |
| ENSMUSG00000075284 | Wipf1 | 215280 | protein_coding | 229.4769367 | -0.227811347 | 0.126722736 | -1.797714867 | 0.072222204 | 0.19967979 | 0.227811347 |
| ENSMUSG00000058325 | Dock1 | 330662 | protein_coding | 1733.584872 | -0.22785671 | 0.062099901 | -3.669195995 | 0.000243314 | 0.002395759 | 0.22785671 |
| ENSMUSG00000032295 | Man2c1 | 73744 | protein_coding | 454.540469 | -0.227872525 | 0.098750683 | -2.307553921 | 0.02102396 | 0.083011808 | 0.227872525 |
| ENSMUSG00000057219 | Armc7 | 276905 | protein_coding | 86.18508553 | -0.227928796 | 0.149757596 | -1.521984878 | 0.12801287 | 0.293844789 | 0.227928796 |
| ENSMUSG00000031684 | Slc10a7 | 76775 | protein_coding | 493.8067132 | 0.228031346 | 0.074830159 | 3.047318726 | 0.002308927 | 0.015228404 | 0.228031346 |
| ENSMUSG00000018547 | Pip4k2b | 108083 | protein_coding | 150.3185582 | -0.228031467 | 0.124242821 | -1.835369362 | 0.06645098 | 0.188433543 | 0.228031467 |
| ENSMUSG00000031848 | Lsm4 | 50783 | protein_coding | 524.3241877 | 0.228054564 | 0.114454798 | 1.992529513 | 0.04631299 | 0.147593699 | 0.228054564 |
| ENSMUSG00000049892 | Rasd1 | 19416 | protein_coding | 351.5917711 | 0.228136061 | 0.407359027 | 0.560036836 | 0.575454313 | 0.74838086 | 0.228136061 |
| ENSMUSG00000020719 | Ddx5 | 13207 | protein_coding | 11833.53836 | -0.228215415 | 0.064974415 | -3.512388913 | 0.000444098 | 0.003954864 | 0.228215415 |
| ENSMUSG00000031864 | Ints10 | 70885 | protein_coding | 585.69499 | 0.228280261 | 0.115377115 | 1.978557542 | 0.047865844 | 0.151211258 | 0.228280261 |
| ENSMUSG00000032330 | Cox7a2 | 12866 | protein_coding | 3691.059316 | 0.228290062 | 0.135015959 | 1.690837622 | 0.090867819 | 0.233216666 | 0.228290062 |
| ENSMUSG00000031490 | Eif4ebp1 | 13685 | protein_coding | 172.5879858 | 0.2282933 | 0.164542101 | 1.387446118 | 0.165305771 | 0.348987948 | 0.2282933 |
| ENSMUSG00000021185 | Dglucy | 217830 | protein_coding | 444.6994512 | 0.228339503 | 0.105776063 | 2.158706762 | 0.030872924 | 0.110425237 | 0.228339503 |
| ENSMUSG00000032182 | Yipf2 | 74766 | protein_coding | 232.4494464 | 0.228344061 | 0.100147878 | 2.280068884 | 0.022603603 | 0.087662575 | 0.228344061 |
| ENSMUSG00000000532 | Acvr1b | 11479 | protein_coding | 1389.027346 | -0.228353221 | 0.131593492 | -1.735292654 | 0.082688982 | 0.219037105 | 0.228353221 |
| ENSMUSG00000001065 | Zfp276 | 57247 | protein_coding | 278.4916518 | -0.22855297 | 0.115823633 | -1.973284422 | 0.048463167 | 0.152483247 | 0.22855297 |
| ENSMUSG00000056973 | Ces1d | 104158 | protein_coding | 1131.456943 | -0.228606336 | 0.163724676 | -1.396285163 | 0.16262864 | 0.345172734 | 0.228606336 |
| ENSMUSG00000047298 | Kcnv2 | 240595 | protein_coding | 7.390953592 | 0.228648366 | 0.470914395 | 0.485541254 | 0.627292456 | 0.784304814 | 0.228648366 |
| ENSMUSG00000020167 | Tcf3 | 21423 | protein_coding | 419.2005115 | 0.228693566 | 0.131602434 | 1.73776091 | 0.082252951 | 0.218281407 | 0.228693566 |
| ENSMUSG00000058135 | Gstm1 | 14862 | protein_coding | 2843.011147 | 0.228825943 | 0.109554993 | 2.088685657 | 0.036736029 | 0.125245747 | 0.228825943 |
| ENSMUSG00000068036 | Afdn | 17356 | protein_coding | 2574.346203 | -0.228863865 | 0.080233903 | -2.852458347 | 0.00433825 | 0.02508022 | 0.228863865 |
| ENSMUSG00000038039 | Gcc2 | 70297 | protein_coding | 2680.003103 | -0.228942057 | 0.090697956 | -2.524225107 | 0.011595365 | 0.0534901 | 0.228942057 |
| ENSMUSG00000026095 | Asnsd1 | 70396 | protein_coding | 128.5603783 | 0.2289613 | 0.131737825 | 1.738007286 | 0.08220953 | 0.218221429 | 0.2289613 |
| ENSMUSG00000033900 | Map9 | 213582 | protein_coding | 38.94403788 | 0.228999821 | 0.237251734 | 0.965218744 | 0.334435261 | 0.545293316 | 0.228999821 |
| ENSMUSG00000032042 | Srpr | 67398 | protein_coding | 1950.1991 | 0.229015403 | 0.084317764 | 2.716099086 | 0.006605615 | 0.034792262 | 0.229015403 |
| ENSMUSG00000073096 | Lrrc61 | 243371 | protein_coding | 40.68537125 | -0.229042595 | 0.202951346 | -1.128559134 | 0.259083858 | 0.466239712 | 0.229042595 |
| ENSMUSG00000032897 | Nfyc | 18046 | protein_coding | 434.1623623 | 0.229238625 | 0.078010804 | 2.938549715 | 0.003297518 | 0.02027127 | 0.229238625 |
| ENSMUSG00000021366 | Hivep1 | 110521 | protein_coding | 680.7356096 | -0.229281214 | 0.124861431 | -1.836285328 | 0.066315469 | 0.188259104 | 0.229281214 |

| ensembl_gene_id | external_gene_name | entrezgene_id | gene_biotype | baseMean | log2FoldChange | lfcSE | stat | pvalue | padj | absLog2FC |
|---|---|---|---|---|---|---|---|---|---|---|
| ENSMUSG00000041130 | Zfp598 | 213753 | protein_coding | 796.2123468 | -0.229368796 | 0.110496392 | -2.075803485 | 0.03791213 | 0.128181789 | 0.229368796 |
| ENSMUSG00000041440 | Gk5 | 235533 | protein_coding | 330.6682223 | -0.229383689 | 0.088236064 | -2.599659123 | 0.00933164 | 0.04546784 | 0.229383689 |
| ENSMUSG00000066090 | Insl5 | 23919 | protein_coding | 298.5929011 | -0.229452504 | 0.272174013 | -0.843036048 | 0.399208284 | 0.606939679 | 0.229452504 |
| ENSMUSG00000026895 | Ndufa8 | 68375 | protein_coding | 1065.657578 | 0.229489092 | 0.103182318 | 2.224112591 | 0.02614087 | 0.097654439 | 0.229489092 |
| ENSMUSG00000022292 | Rrm2b | 382985 | protein_coding | 110.8382034 | -0.229504344 | 0.111179672 | -2.064265348 | 0.038992565 | 0.13075233 | 0.229504344 |
| ENSMUSG00000036339 | Tmem260 | 218989 | protein_coding | 500.2474604 | 0.229571339 | 0.109564605 | 2.095305678 | 0.036143828 | 0.123892113 | 0.229571339 |
| ENSMUSG00000024878 | Cbwd1 | 226043 | protein_coding | 244.3190348 | 0.229578282 | 0.103170906 | 2.225223078 | 0.026066266 | 0.097509572 | 0.229578282 |
| ENSMUSG00000040412 | Elapor1 | 229722 | protein_coding | 5160.940678 | -0.229605797 | 0.097781386 | -2.348154448 | 0.018866696 | 0.076601768 | 0.229605797 |
| ENSMUSG00000036136 | Fam110c | 104943 | protein_coding | 539.0993258 | -0.229645793 | 0.104007263 | -2.207978417 | 0.027245778 | 0.100441862 | 0.229645793 |
| ENSMUSG00000095217 | H2bc15 | 319187 | protein_coding | 1235.685269 | 0.229648775 | 0.148730369 | 1.544061086 | 0.122573535 | 0.285427227 | 0.229648775 |
| ENSMUSG00000002948 | Map2k7 | 26400 | protein_coding | 238.712152 | -0.229699679 | 0.10505644 | -2.18644073 | 0.028783376 | 0.104636506 | 0.229699679 |
| ENSMUSG00000016257 | Prelid3b | 66390 | protein_coding | 2148.291913 | 0.229725386 | 0.08350448 | 2.751054636 | 0.005940373 | 0.031944047 | 0.229725386 |
| ENSMUSG00000042414 | Prdm14 | 383491 | protein_coding | 22.81758884 | 0.22979616 | 0.360521475 | 0.637399367 | 0.52386474 | 0.709654498 | 0.22979616 |
| ENSMUSG00000054115 | Skp2 | 27401 | protein_coding | 110.3029643 | 0.229880116 | 0.140965877 | 1.630750082 | 0.102943064 | 0.253153795 | 0.229880116 |
| ENSMUSG00000033703 | Fcsk | 234730 | protein_coding | 676.4371507 | 0.229888518 | 0.11180708 | 2.056117715 | 0.039771171 | 0.132498476 | 0.229888518 |
| ENSMUSG00000036751 | Cox6b1 | 110323 | protein_coding | 5795.344618 | 0.229933718 | 0.093063877 | 2.470708551 | 0.013484567 | 0.059911347 | 0.229933718 |
| ENSMUSG00000090386 | Mir99ahg | 77994 | lncRNA | 14.74754795 | -0.229943134 | 0.733890586 | -0.31332073 | 0.754037003 | 0.865046195 | 0.229943134 |
| ENSMUSG00000031917 | Nip7 | 66164 | protein_coding | 342.6872913 | -0.229946319 | 0.172482318 | -1.333158792 | 0.182479699 | 0.371537345 | 0.229946319 |
| ENSMUSG00000036368 | Rmdn2 | 381110 | protein_coding | 105.4380054 | 0.230012129 | 0.128054992 | 1.796198065 | 0.072463022 | 0.200112777 | 0.230012129 |
| ENSMUSG00000030340 | Scnn1a | 20276 | protein_coding | 558.5513722 | -0.230020308 | 0.122709273 | -1.874514475 | 0.060859549 | 0.177378023 | 0.230020308 |
| ENSMUSG00000020496 | Rnf187 | 108660 | protein_coding | 717.9475283 | 0.230029626 | 0.112044455 | 2.053021067 | 0.040070534 | 0.133272807 | 0.230029626 |
| ENSMUSG00000028550 | Atg4c | 242557 | protein_coding | 197.0400017 | 0.230035872 | 0.122610408 | 1.876152891 | 0.060634289 | 0.177004261 | 0.230035872 |
| ENSMUSG00000035936 | Aldh5a1 | 214579 | protein_coding | 290.7256981 | -0.230146775 | 0.091822348 | -2.5064353 | 0.012195537 | 0.055531881 | 0.230146775 |
| ENSMUSG00000060708 | Bloc1s4 | 117197 | protein_coding | 96.86619389 | -0.230170677 | 0.128281385 | -1.794264049 | 0.072771033 | 0.200676868 | 0.230170677 |
| ENSMUSG00000024327 | Slc39a7 | 14977 | protein_coding | 196.3658145 | 0.230187475 | 0.124318641 | 1.851592599 | 0.064084347 | 0.184015769 | 0.230187475 |
| ENSMUSG00000040321 | Zfp770 | 228491 | protein_coding | 143.087608 | -0.230233107 | 0.111229253 | -2.069897086 | 0.038461983 | 0.129548715 | 0.230233107 |
| ENSMUSG00000001419 | Mef2d | 17261 | protein_coding | 540.9638521 | -0.230269514 | 0.110528464 | -2.083350361 | 0.037219299 | 0.126374067 | 0.230269514 |
| ENSMUSG00000022329 | Stk3 | 56274 | protein_coding | 468.3043111 | 0.230324524 | 0.089214022 | 2.581707673 | 0.009831282 | 0.047323258 | 0.230324524 |
| ENSMUSG00000036273 | Lrrk2 | 66725 | protein_coding | 226.9242848 | -0.230325561 | 0.322995537 | -0.713092087 | 0.475788776 | 0.67038675 | 0.230325561 |
| ENSMUSG00000002028 | Kmt2a | 214162 | protein_coding | 622.2136541 | -0.230373072 | 0.245218257 | -0.939461341 | 0.347493932 | 0.557627687 | 0.230373072 |
| ENSMUSG00000048330 | Ric3 | 320360 | protein_coding | 17.5677388 | -0.230434378 | 0.40112936 | -0.574464002 | 0.565653853 | 0.74178615 | 0.230434378 |
| ENSMUSG00000039943 | Plcb4 | 18798 | protein_coding | 537.8538453 | 0.230454916 | 0.101277491 | 2.275480099 | 0.022877145 | 0.088529438 | 0.230454916 |
| ENSMUSG00000025154 | Arhgap19 | 71085 | protein_coding | 216.1369302 | 0.230495644 | 0.131699107 | 1.750168617 | 0.080089222 | 0.213992665 | 0.230495644 |
| ENSMUSG00000091491 | Vmn2r97 | 627367 | protein_coding | 8.226162902 | -0.230513136 | 0.98090929 | -0.234999442 | 0.814209166 | 0.900950823 | 0.230513136 |
| ENSMUSG00000089844 | A530032D15Rik | 381287 | protein_coding | 8.405196763 | -0.230536306 | 0.786388596 | -0.293158251 | 0.769401193 | 0.874359461 | 0.230536306 |
| ENSMUSG00000060703 | Cd302 | 66205 | protein_coding | 60.90657671 | 0.230638197 | 0.207125978 | 1.113516515 | 0.265486659 | 0.473111604 | 0.230638197 |
| ENSMUSG00000028945 | Rheb | 19744 | protein_coding | 707.6525252 | 0.230660927 | 0.072849779 | 3.166254335 | 0.001544157 | 0.010906091 | 0.230660927 |
| ENSMUSG00000036093 | Arl5a | 75423 | protein_coding | 984.4639443 | 0.23066115 | 0.061070542 | 3.776962567 | 0.000158753 | 0.001673469 | 0.23066115 |

| ensembl_gene_id | external_gene_name | entrezgene_id | gene_biotype | baseMean | log2FoldChange | lfcSE | stat | pvalue | padj | absLog2FC |
|---|---|---|---|---|---|---|---|---|---|---|
| ENSMUSG00000045327 | 6330549D23Rik | 229613 | transcribed_processed_pseudogene | 16.58049863 | -0.230738306 | 0.29887185 | -0.77203091 | 0.44009612 | 0.642313808 | 0.230738306 |
| ENSMUSG00000031156 | Slc35a2 | 22232 | protein_coding | 1202.394244 | 0.230784977 | 0.093153336 | 2.477474104 | 0.013231601 | 0.059084088 | 0.230784977 |
| ENSMUSG00000033157 | Abhd10 | 213012 | protein_coding | 159.4425218 | 0.230863532 | 0.102548845 | 2.251254329 | 0.024369434 | 0.09261664 | 0.230863532 |
| ENSMUSG00000038803 | Ost4 | 67695 | protein_coding | 429.7401816 | 0.230890472 | 0.102551385 | 2.251461278 | 0.024356337 | 0.092588958 | 0.230890472 |
| ENSMUSG00000025147 | Mob2 | 101513 | protein_coding | 755.7863085 | 0.230891209 | 0.084671158 | 2.726916877 | 0.006392914 | 0.033896916 | 0.230891209 |
| ENSMUSG00000039914 | Coq10a | 210582 | protein_coding | 164.3393979 | -0.230897498 | 0.106802354 | -2.16191395 | 0.03062481 | 0.109714792 | 0.230897498 |
| ENSMUSG00000037526 | Atg14 | 100504663 | protein_coding | 203.628396 | -0.230982889 | 0.173352799 | -1.332443951 | 0.182714348 | 0.371824982 | 0.230982889 |
| ENSMUSG00000031545 | Gpat4 | 102247 | protein_coding | 1347.920375 | 0.231013736 | 0.108139846 | 2.136249913 | 0.032659049 | 0.114782289 | 0.231013736 |
| ENSMUSG00000026942 | Traf2 | 22030 | protein_coding | 341.0361387 | 0.231092067 | 0.115102723 | 2.007702878 | 0.044674875 | 0.143724784 | 0.231092067 |
| ENSMUSG00000009076 | Zmat5 | 67178 | protein_coding | 104.2155233 | 0.231147268 | 0.145310444 | 1.590713385 | 0.111674093 | 0.266918511 | 0.231147268 |
| ENSMUSG00000027963 | Extl2 | 58193 | protein_coding | 150.0932123 | 0.231224889 | 0.117526937 | 1.967420366 | 0.049134768 | 0.153635373 | 0.231224889 |
| ENSMUSG00000030282 | Cmas | 12764 | protein_coding | 2961.180431 | 0.23125822 | 0.100053106 | 2.311354726 | 0.020813269 | 0.082384134 | 0.23125822 |
| ENSMUSG00000020219 | Timm13 | 30055 | protein_coding | 677.9716762 | 0.231267066 | 0.121926955 | 1.896767335 | 0.057858652 | 0.171774951 | 0.231267066 |
| ENSMUSG00000029725 | Ppp1r35 | 69871 | protein_coding | 60.85718357 | -0.231444088 | 0.155745982 | -1.486035691 | 0.137269681 | 0.308157609 | 0.231444088 |
| ENSMUSG00000025612 | Bach1 | 12013 | protein_coding | 1054.911184 | 0.231447214 | 0.232279594 | 0.996416475 | 0.319047832 | 0.530340919 | 0.231447214 |
| ENSMUSG00000021374 | Nup153 | 218210 | protein_coding | 1040.90795 | -0.231458628 | 0.12215045 | -1.894865128 | 0.058110272 | 0.172233051 | 0.231458628 |
| ENSMUSG00000051627 | H1f4 | 50709 | protein_coding | 3290.063632 | 0.231470647 | 0.13350471 | 1.733801356 | 0.082953335 | 0.219554789 | 0.231470647 |
| ENSMUSG00000041112 | Elmo1 | 140580 | protein_coding | 227.4560993 | -0.231502568 | 0.133241716 | -1.737463126 | 0.082305457 | 0.218347988 | 0.231502568 |
| ENSMUSG00000002871 | Tpra1 | 24100 | protein_coding | 305.7091983 | -0.231523859 | 0.094670398 | -2.445578161 | 0.01446201 | 0.0628091 | 0.231523859 |
| ENSMUSG00000037455 | Slc18b1 | 76306 | protein_coding | 401.3324259 | 0.231557891 | 0.086920271 | 2.66402633 | 0.007721152 | 0.039379605 | 0.231557891 |
| ENSMUSG00000055723 | Rras2 | 66922 | protein_coding | 917.0818223 | -0.231595203 | 0.100899399 | -2.295308047 | 0.021715477 | 0.084858713 | 0.231595203 |
| ENSMUSG00000025393 | Atp5b | 11947 | protein_coding | 28371.85426 | 0.231599642 | 0.110218766 | 2.101272327 | 0.035617068 | 0.12260831 | 0.231599642 |
| ENSMUSG00000063317 | Usp31 | 76179 | protein_coding | 264.6312189 | -0.231633071 | 0.156942365 | -1.475911693 | 0.1399676 | 0.312022944 | 0.231633071 |
| ENSMUSG00000023921 | Mmut | 17850 | protein_coding | 1426.821003 | 0.231649702 | 0.082485639 | 2.80836404 | 0.00497939 | 0.027964962 | 0.231649702 |
| ENSMUSG00000002845 | Tmem39a | 67846 | protein_coding | 367.1392844 | 0.231707488 | 0.104398612 | 2.219449881 | 0.026456133 | 0.098431865 | 0.231707488 |
| ENSMUSG00000017715 | Pgs1 | 74451 | protein_coding | 583.1745223 | 0.231727767 | 0.156407771 | 1.481561719 | 0.138456951 | 0.309608637 | 0.231727767 |
| ENSMUSG00000027531 | Impa1 | 55980 | protein_coding | 1276.064876 | 0.231737623 | 0.078264536 | 2.960953123 | 0.003066886 | 0.019119124 | 0.231737623 |
| ENSMUSG00000002778 | Kdelr1 | 68137 | protein_coding | 1419.935132 | 0.231768936 | 0.069012237 | 3.358374502 | 0.000784023 | 0.00628165 | 0.231768936 |
| ENSMUSG00000028836 | Slc30a2 | 230810 | protein_coding | 9.383380546 | 0.231794258 | 0.48219783 | 0.480703654 | 0.630727134 | 0.786273913 | 0.231794258 |
| ENSMUSG00000070044 | Fam149a | 212326 | protein_coding | 98.0449065 | 0.23183599 | 0.217031246 | 1.068214805 | 0.285423628 | 0.494370111 | 0.23183599 |
| ENSMUSG00000032403 | 2300009A05Rik | 69478 | protein_coding | 67.70971688 | 0.231922522 | 0.136925268 | 1.693789065 | 0.090305373 | 0.232259644 | 0.231922522 |
| ENSMUSG00000074182 | Znhit6 | 229937 | protein_coding | 93.69400614 | -0.231924719 | 0.175283901 | -1.323137599 | 0.185789623 | 0.376160813 | 0.231924719 |
| ENSMUSG00000054434 | Tmem120b | 330189 | protein_coding | 39.58832808 | -0.231949863 | 0.190216473 | -1.219399453 | 0.222692617 | 0.424013417 | 0.231949863 |
| ENSMUSG00000034154 | Ino80 | 68142 | protein_coding | 672.6916268 | -0.231968226 | 0.081906124 | -2.832123104 | 0.004624004 | 0.026407723 | 0.231968226 |
| ENSMUSG00000033488 | Cryzl2 | 226527 | protein_coding | 124.0906617 | -0.231971531 | 0.159277962 | -1.456394394 | 0.145283624 | 0.320687024 | 0.231971531 |
| ENSMUSG00000028115 | Bnipl | 171388 | protein_coding | 90.7393697 | 0.232000489 | 0.220671725 | 1.051337631 | 0.29310355 | 0.502178856 | 0.232000489 |
| ENSMUSG00000060771 | Tsga10 | 211484 | protein_coding | 81.11325275 | 0.232023237 | 0.167757628 | 1.38308606 | 0.16663848 | 0.350562195 | 0.232023237 |
| ENSMUSG00000021957 | Tkt | 21881 | protein_coding | 6855.877569 | 0.232049975 | 0.105884887 | 2.191530655 | 0.028413414 | 0.10349162 | 0.232049975 |

| ensembl_gene_id | external_gene_name | entrezgene_id | gene_biotype | baseMean | log2FoldChange | lfcSE | stat | pvalue | padj | absLog2FC |
|---|---|---|---|---|---|---|---|---|---|---|
| ENSMUSG00000027961 | Lrrc39 | 109245 | protein_coding | 10.21702004 | -0.232086953 | 0.445308444 | -0.521182466 | 0.602239668 | 0.767564496 | 0.232086953 |
| ENSMUSG00000042208 | Sanbr | 71675 | protein_coding | 447.4882781 | 0.232108057 | 0.082295791 | 2.820412268 | 0.004796199 | 0.02721548 | 0.232108057 |
| ENSMUSG00000028409 | Smu1 | 74255 | protein_coding | 1377.674928 | 0.232111535 | 0.083412159 | 2.78270624 | 0.005390759 | 0.02960828 | 0.232111535 |
| ENSMUSG00000035992 | Fnip1 | 216742 | protein_coding | 722.0903723 | 0.232179278 | 0.058627693 | 3.960232187 | 7.49E-05 | 0.000873729 | 0.232179278 |
| ENSMUSG00000031996 | Aplp2 | 11804 | protein_coding | 2472.755313 | -0.232458961 | 0.068515678 | -3.392784966 | 0.000691859 | 0.005700949 | 0.232458961 |
| ENSMUSG00000001248 | Gramd1a | 52857 | protein_coding | 146.6426144 | 0.232506223 | 0.178595185 | 1.301861655 | 0.192963683 | 0.386218338 | 0.232506223 |
| ENSMUSG00000022354 | Ndufb9 | 66218 | protein_coding | 2448.27035 | 0.232578605 | 0.090481055 | 2.57046742 | 0.010156138 | 0.048368275 | 0.232578605 |
| ENSMUSG00000051736 | Fam229b | 66337 | protein_coding | 10.35512532 | -0.232672407 | 0.442754935 | -0.52551059 | 0.599228294 | 0.76592263 | 0.232672407 |
| ENSMUSG00000026839 | Upp2 | 76654 | protein_coding | 46.21972895 | 0.232824836 | 0.21934559 | 1.061452095 | 0.288484493 | 0.497453827 | 0.232824836 |
| ENSMUSG00000009575 | Cbx5 | 12419 | protein_coding | 1966.879556 | 0.232872499 | 0.082924032 | 2.808263091 | 0.004980952 | 0.027964962 | 0.232872499 |
| ENSMUSG00000054939 | Zfp174 | 385674 | protein_coding | 33.86691549 | -0.232917114 | 0.285524392 | -0.815752071 | 0.414641951 | 0.621031156 | 0.232917114 |
| ENSMUSG00000022199 | Slc22a17 | 59049 | protein_coding | 14.01775619 | -0.232936132 | 0.455523088 | -0.511359661 | 0.609099236 | 0.772403867 | 0.232936132 |
| ENSMUSG00000061244 | Exoc5 | 105504 | protein_coding | 1546.984904 | 0.232949618 | 0.119301039 | 1.952620192 | 0.050864617 | 0.157192785 | 0.232949618 |
| ENSMUSG00000085347 | Hk1os | 70162 | lncRNA | 8.613482876 | -0.232983863 | 0.402297541 | -0.579133201 | 0.562499299 | 0.739622589 | 0.232983863 |
| ENSMUSG00000093979 | Gm2237 | 100039441 | protein_coding | 7.567381015 | -0.233095311 | 0.411374773 | -0.566625195 | 0.570968854 | 0.745230585 | 0.233095311 |
| ENSMUSG00000018076 | Med13l | 76199 | protein_coding | 811.5825278 | -0.233140515 | 0.115880383 | -2.011906664 | 0.044229777 | 0.142609693 | 0.233140515 |
| ENSMUSG00000035725 | Prkx | 19108 | protein_coding | 487.1736018 | -0.23315552 | 0.216800035 | -1.07544042 | 0.282177593 | 0.491172768 | 0.23315552 |
| ENSMUSG00000047284 | Neurl4 | 216860 | protein_coding | 616.7760603 | -0.233338136 | 0.089570449 | -2.605079452 | 0.009185296 | 0.044947278 | 0.233338136 |
| ENSMUSG00000040648 | Ppip5k2 | 227399 | protein_coding | 1846.847646 | 0.233350628 | 0.062344218 | 3.74293936 | 0.00018188 | 0.001880252 | 0.233350628 |
| ENSMUSG00000037395 | Rcor3 | 214742 | protein_coding | 256.705569 | -0.233412559 | 0.158944663 | -1.468514609 | 0.141964491 | 0.315151277 | 0.233412559 |
| ENSMUSG00000034708 | Grn | 14824 | protein_coding | 963.9482918 | -0.233496589 | 0.0835734 | -2.793910368 | 0.005207491 | 0.028930909 | 0.233496589 |
| ENSMUSG00000041035 | Dpcd | 226162 | protein_coding | 201.8337758 | 0.23349746 | 0.097770148 | 2.38822856 | 0.016929809 | 0.07076185 | 0.23349746 |
| ENSMUSG00000028261 | Ndufaf4 | 68493 | protein_coding | 564.4469526 | 0.233507284 | 0.155726203 | 1.499473305 | 0.133750889 | 0.302403053 | 0.233507284 |
| ENSMUSG00000051246 | Msantd1 | 403174 | protein_coding | 26.29794472 | 0.233573662 | 0.339219684 | 0.688561642 | 0.491099167 | 0.683343018 | 0.233573662 |
| ENSMUSG00000070520 | Nsmce3 | 66647 | protein_coding | 247.7800365 | 0.23367853 | 0.083616603 | 2.794642708 | 0.00519571 | 0.028875532 | 0.23367853 |
| ENSMUSG00000024132 | Eci1 | 13177 | protein_coding | 757.3785746 | 0.233776894 | 0.087632856 | 2.667685439 | 0.007637573 | 0.039065915 | 0.233776894 |
| ENSMUSG00000024298 | Zfp871 | 208292 | protein_coding | 894.6612491 | -0.233830447 | 0.162518306 | -1.438794508 | 0.150208753 | 0.327877897 | 0.233830447 |
| ENSMUSG00000024164 | C3 | 12266 | protein_coding | 78.19682772 | -0.23383582 | 0.369636441 | -0.6326103 | 0.526988159 | 0.712457952 | 0.23383582 |
| ENSMUSG00000042272 | Sestd1 | 228071 | protein_coding | 628.925971 | 0.233873113 | 0.075570462 | 3.094768847 | 0.001969663 | 0.01335581 | 0.233873113 |
| ENSMUSG00000051910 | Sox6 | 20679 | protein_coding | 222.2726265 | 0.233996153 | 0.171131723 | 1.367345275 | 0.171517107 | 0.357067635 | 0.233996153 |
| ENSMUSG00000031196 | F8 | 14069 | protein_coding | 20.67707194 | -0.233999823 | 0.285781684 | -0.81880623 | 0.412896976 | 0.619566809 | 0.233999823 |
| ENSMUSG00000020721 | Helz | 78455 | protein_coding | 450.260922 | -0.23405832 | 0.176426027 | -1.326665478 | 0.184619365 | 0.374457819 | 0.23405832 |
| ENSMUSG00000031766 | Slc12a3 | 20497 | protein_coding | 8.839987862 | -0.234066088 | 0.549298097 | -0.426118512 | 0.670021491 | 0.808860263 | 0.234066088 |
| ENSMUSG00000002409 | Dyrk1b | 13549 | protein_coding | 129.9248256 | -0.234267901 | 0.191738927 | -1.221806679 | 0.221780742 | 0.422854161 | 0.234267901 |
| ENSMUSG00000052485 | Tmem171 | 380863 | protein_coding | 1303.438166 | -0.234314386 | 0.132853532 | -1.763704608 | 0.077781725 | 0.209628987 | 0.234314386 |
| ENSMUSG00000028184 | Adgrl2 | 99633 | protein_coding | 331.7762989 | -0.234340798 | 0.142778662 | -1.641287252 | 0.100737801 | 0.249487959 | 0.234340798 |
| ENSMUSG00000053046 | Brsk2 | 75770 | protein_coding | 18.88006285 | 0.234380288 | 0.416183211 | 0.563166129 | 0.573321765 | 0.746738823 | 0.234380288 |
| ENSMUSG00000042032 | Mat2b | 108645 | protein_coding | 1450.417951 | 0.234388246 | 0.121410144 | 1.930549119 | 0.053538835 | 0.162524598 | 0.234388246 |

| ensembl_gene_id | external_gene_name | entrezgene_id | gene_biotype | baseMean | log2FoldChange | lfcSE | stat | pvalue | padj | absLog2FC |
|---|---|---|---|---|---|---|---|---|---|---|
| ENSMUSG00000028357 | Kif12 | 16552 | protein_coding | 288.0444602 | -0.234390089 | 0.137402501 | -1.705864795 | 0.088033256 | 0.228545168 | 0.234390089 |
| ENSMUSG00000035671 | Zswim4 | 212168 | protein_coding | 213.2740847 | -0.234431669 | 0.178718902 | -1.311734047 | 0.189609885 | 0.381182159 | 0.234431669 |
| ENSMUSG00000011158 | Brf1 | 72308 | protein_coding | 364.5420611 | -0.234434744 | 0.07566912 | -3.098156088 | 0.001947288 | 0.013232256 | 0.234434744 |
| ENSMUSG00000040586 | Ofd1 | 237222 | protein_coding | 186.078977 | -0.234481744 | 0.093815672 | -2.499387776 | 0.01244081 | 0.056374741 | 0.234481744 |
| ENSMUSG00000004891 | Nes | 18008 | protein_coding | 17.34997524 | 0.234497993 | 0.793314464 | 0.295592736 | 0.767541122 | 0.873180129 | 0.234497993 |
| ENSMUSG00000010025 | Aldh3a2 | 11671 | protein_coding | 2312.016236 | 0.234551186 | 0.089371286 | 2.624457989 | 0.0086787 | 0.04311473 | 0.234551186 |
| ENSMUSG00000034936 | Arl4d | 80981 | protein_coding | 30.20805888 | 0.234556831 | 0.272245098 | 0.861564939 | 0.388926972 | 0.597322226 | 0.234556831 |
| ENSMUSG00000029267 | Mtf2 | 17765 | protein_coding | 622.7078019 | 0.234569543 | 0.088483625 | 2.650993839 | 0.008025531 | 0.040606941 | 0.234569543 |
| ENSMUSG00000070939 | Tgfbrap1 | 73122 | protein_coding | 369.7409836 | -0.234596619 | 0.091010387 | -2.577690604 | 0.009946301 | 0.047759252 | 0.234596619 |
| ENSMUSG00000042831 | Alkbh6 | 233065 | protein_coding | 237.0847619 | 0.234612759 | 0.107684245 | 2.178710174 | 0.029353205 | 0.106162093 | 0.234612759 |
| ENSMUSG00000061143 | Maml3 | 433586 | protein_coding | 293.8340778 | -0.234843143 | 0.146050285 | -1.607960866 | 0.10784375 | 0.261173065 | 0.234843143 |
| ENSMUSG00000020788 | Atp2a3 | 53313 | protein_coding | 13190.27198 | -0.234882187 | 0.080005245 | -2.935834869 | 0.003326514 | 0.020410894 | 0.234882187 |
| ENSMUSG00000026277 | Stk25 | 59041 | protein_coding | 2713.023271 | -0.234939287 | 0.16684562 | -1.40812379 | 0.159094418 | 0.340376923 | 0.234939287 |
| ENSMUSG00000005973 | Rcn1 | 19672 | protein_coding | 84.17906976 | 0.235041967 | 0.229879002 | 1.022459488 | 0.306563477 | 0.516604218 | 0.235041967 |
| ENSMUSG00000005534 | Insr | 16337 | protein_coding | 1565.2276 | -0.235069615 | 0.136399174 | -1.723394715 | 0.08481717 | 0.222821686 | 0.235069615 |
| ENSMUSG00000029228 | Lnx1 | 16924 | protein_coding | 340.4463512 | -0.235091948 | 0.118911496 | -1.977032972 | 0.048037903 | 0.151514401 | 0.235091948 |
| ENSMUSG00000038451 | Spsb2 | 14794 | protein_coding | 166.1605326 | -0.235115964 | 0.111799476 | -2.103014892 | 0.035464468 | 0.122241515 | 0.235115964 |
| ENSMUSG00000041782 | Lad1 | 16763 | protein_coding | 4077.601971 | -0.235117751 | 0.095272875 | -2.467835157 | 0.01359329 | 0.060191302 | 0.235117751 |
| ENSMUSG00000021190 | Lgmn | 19141 | protein_coding | 474.8187237 | -0.235122127 | 0.241168994 | -0.974926846 | 0.329596549 | 0.540382277 | 0.235122127 |
| ENSMUSG00000046269 | Usp27x | 54651 | protein_coding | 28.83207264 | -0.235139046 | 0.252454818 | -0.93141041 | 0.351641308 | 0.561399852 | 0.235139046 |
| ENSMUSG00000022568 | Scrib | 105782 | protein_coding | 996.3214492 | -0.235160156 | 0.081780575 | -2.875501381 | 0.004033864 | 0.023647914 | 0.235160156 |
| ENSMUSG00000002064 | Sdf2 | 20316 | protein_coding | 763.5679108 | 0.235165222 | 0.115828292 | 2.030291724 | 0.042326895 | 0.138233724 | 0.235165222 |
| ENSMUSG00000023015 | Racgap1 | 26934 | protein_coding | 730.9589043 | 0.235217204 | 0.125421768 | 1.875409728 | 0.060736378 | 0.17718114 | 0.235217204 |
| ENSMUSG00000012114 | Med15 | 94112 | protein_coding | 592.6072956 | -0.235260096 | 0.093891435 | -2.505660877 | 0.012222278 | 0.055603864 | 0.235260096 |
| ENSMUSG00000020227 | Irak3 | 73914 | protein_coding | 84.86918045 | -0.235313583 | 0.16517522 | -1.424630061 | 0.154264166 | 0.333575839 | 0.235313583 |
| ENSMUSG00000066621 | Tecpr1 | 70381 | protein_coding | 785.2754677 | 0.235340608 | 0.091960723 | 2.559142648 | 0.010493067 | 0.049594534 | 0.235340608 |
| ENSMUSG00000029439 | Sfswap | 231769 | protein_coding | 521.9716441 | -0.235384972 | 0.115584106 | -2.03648217 | 0.041701962 | 0.136897949 | 0.235384972 |
| ENSMUSG00000022957 | Itsn1 | 16443 | protein_coding | 469.6898543 | -0.235416771 | 0.147848986 | -1.592278559 | 0.111322128 | 0.266434468 | 0.235416771 |
| ENSMUSG00000042854 | Trp53rkb | 76367 | protein_coding | 41.36133778 | -0.235426675 | 0.177537668 | -1.326066054 | 0.184817819 | 0.374764891 | 0.235426675 |
| ENSMUSG00000118047 | 4930554I06Rik | 75354 | lncRNA | 9.453845938 | -0.23555011 | 0.454646 | -0.518095638 | 0.604391543 | 0.769032473 | 0.23555011 |
| ENSMUSG00000024944 | Arl2 | 56327 | protein_coding | 166.0084996 | -0.235583911 | 0.120355636 | -1.957398245 | 0.050300665 | 0.156025119 | 0.235583911 |
| ENSMUSG00000027281 | Slx4ip | 74243 | protein_coding | 226.3228759 | 0.235658894 | 0.099349799 | 2.372011795 | 0.017691526 | 0.073002753 | 0.235658894 |
| ENSMUSG00000068735 | Trp53i11 | 277414 | protein_coding | 1654.632832 | 0.235662388 | 0.160735788 | 1.466147587 | 0.142608084 | 0.316152646 | 0.235662388 |
| ENSMUSG00000000308 | Ckmt1 | 12716 | protein_coding | 16185.907 | 0.235662911 | 0.177000946 | 1.33142176 | 0.183050273 | 0.372217515 | 0.235662911 |
| ENSMUSG00000036298 | Slc2a13 | 239606 | protein_coding | 858.3918643 | 0.235706896 | 0.142220855 | 1.657330047 | 0.097452766 | 0.244099332 | 0.235706896 |
| ENSMUSG00000036944 | Tmem71 | 213068 | protein_coding | 10.33891072 | 0.235719067 | 0.394148648 | 0.598046114 | 0.549809164 | 0.731093174 | 0.235719067 |
| ENSMUSG00000026796 | Niban2 | 227737 | protein_coding | 1558.978453 | -0.235945794 | 0.141369887 | -1.668996125 | 0.095118148 | 0.240482835 | 0.235945794 |
| ENSMUSG00000020946 | Gosr2 | 56494 | protein_coding | 956.4519649 | 0.235990806 | 0.098037902 | 2.40713847 | 0.016078072 | 0.068096417 | 0.235990806 |

| ensembl_gene_id | external_gene_name | entrezgene_id | gene_biotype | baseMean | log2FoldChange | lfcSE | stat | pvalue | padj | absLog2FC |
|---|---|---|---|---|---|---|---|---|---|---|
| ENSMUSG00000040187 | Arntl2 | 272322 | protein_coding | 28.76166479 | -0.236006499 | 0.24218245 | -0.974498769 | 0.32980895 | 0.54061076 | 0.236006499 |
| ENSMUSG00000031755 | Bbs2 | 67378 | protein_coding | 149.6583527 | -0.236029662 | 0.112647942 | -2.095286064 | 0.036145571 | 0.123892113 | 0.236029662 |
| ENSMUSG00000061650 | Med9 | 192191 | protein_coding | 231.0711879 | 0.236054886 | 0.118095804 | 1.998842275 | 0.045625422 | 0.146061871 | 0.236054886 |
| ENSMUSG00000018924 | Alox15 | 11687 | protein_coding | 8.126529653 | -0.236076487 | 0.5152711 | -0.458159767 | 0.646837662 | 0.795892645 | 0.236076487 |
| ENSMUSG00000057963 | Itpk1 | 217837 | protein_coding | 2254.414896 | 0.236096065 | 0.113227594 | 2.08514601 | 0.037056049 | 0.126036461 | 0.236096065 |
| ENSMUSG00000021109 | Hif1a | 15251 | protein_coding | 1400.782348 | -0.23615489 | 0.198180293 | -1.191616411 | 0.233411687 | 0.437077516 | 0.23615489 |
| ENSMUSG00000002475 | Abhd3 | 106861 | protein_coding | 112.0969011 | 0.236168747 | 0.23222082 | 1.017000746 | 0.309153075 | 0.518991376 | 0.236168747 |
| ENSMUSG00000027679 | Dnajc19 | 67713 | protein_coding | 356.9051431 | 0.236213225 | 0.069387387 | 3.404267479 | 0.000663418 | 0.005499701 | 0.236213225 |
| ENSMUSG00000024870 | Rab1b | 76308 | protein_coding | 1231.822448 | 0.236215265 | 0.090336787 | 2.614829172 | 0.008927209 | 0.043995232 | 0.236215265 |
| ENSMUSG00000022514 | Il1rap | 16180 | protein_coding | 184.3934604 | 0.236232733 | 0.09009536 | 2.622029956 | 0.008740775 | 0.043317692 | 0.236232733 |
| ENSMUSG00000074649 | BC029722 | 613262 | protein_coding | 54.82332567 | -0.236248143 | 0.152125652 | -1.552980315 | 0.120427837 | 0.281762672 | 0.236248143 |
| ENSMUSG00000026479 | Lamc2 | 16782 | protein_coding | 855.4654295 | -0.236288977 | 0.263102193 | -0.898088208 | 0.369138525 | 0.577871395 | 0.236288977 |
| ENSMUSG00000034793 | G6pc3 | 68401 | protein_coding | 376.3158206 | -0.236319684 | 0.084148052 | -2.808379718 | 0.004979148 | 0.027964962 | 0.236319684 |
| ENSMUSG00000030750 | Nsmce1 | 67711 | protein_coding | 266.9489589 | 0.236327197 | 0.126223897 | 1.872285696 | 0.061167089 | 0.177981255 | 0.236327197 |
| ENSMUSG00000048249 | Crebrf | 77128 | protein_coding | 729.2472811 | -0.23637547 | 0.178068479 | -1.32744139 | 0.184362716 | 0.374032523 | 0.23637547 |
| ENSMUSG00000025880 | Smad7 | 17131 | protein_coding | 706.9631427 | -0.236406985 | 0.139526574 | -1.694350964 | 0.090198612 | 0.232146031 | 0.236406985 |
| ENSMUSG00000006641 | Slc5a6 | 330064 | protein_coding | 305.7258811 | -0.236456915 | 0.200714129 | -1.178078079 | 0.23876548 | 0.442885358 | 0.236456915 |
| ENSMUSG00000015092 | Edf1 | 59022 | protein_coding | 1944.285694 | 0.236499276 | 0.121465508 | 1.947048835 | 0.05152888 | 0.158599421 | 0.236499276 |
| ENSMUSG00000024181 | Mrpl28 | 68611 | protein_coding | 719.335647 | 0.236529664 | 0.121788954 | 1.942127401 | 0.052121679 | 0.159960328 | 0.236529664 |
| ENSMUSG00000038736 | Nudcd1 | 67429 | protein_coding | 824.7519622 | 0.236542824 | 0.096178795 | 2.45940722 | 0.013916666 | 0.061148872 | 0.236542824 |
| ENSMUSG00000061518 | Cox5b | 12859 | protein_coding | 4236.026211 | 0.23659303 | 0.091437788 | 2.587475436 | 0.009668209 | 0.046679733 | 0.23659303 |
| ENSMUSG00000055963 | Triqk | 208820 | protein_coding | 181.3099572 | 0.236706575 | 0.17826115 | 1.327864065 | 0.184223018 | 0.373891976 | 0.236706575 |
| ENSMUSG00000021477 | Ctsl | 13039 | protein_coding | 1087.287033 | -0.236773131 | 0.103480241 | -2.288099923 | 0.022131703 | 0.086126011 | 0.236773131 |
| ENSMUSG00000042644 | Itpr3 | 16440 | protein_coding | 2770.939593 | -0.23680602 | 0.089671633 | -2.640813057 | 0.008270734 | 0.041570292 | 0.23680602 |
| ENSMUSG00000028788 | Ptp4a2 | 19244 | protein_coding | 7259.461198 | 0.23701782 | 0.084157453 | 2.816361612 | 0.004857096 | 0.027482711 | 0.23701782 |
| ENSMUSG00000025280 | Polr3a | 218832 | protein_coding | 332.7157653 | -0.237080539 | 0.094568829 | -2.506962828 | 0.012177352 | 0.055480795 | 0.237080539 |
| ENSMUSG00000043460 | Elfn2 | 207393 | protein_coding | 24.41293802 | -0.23712746 | 0.432367767 | -0.548439264 | 0.583390323 | 0.754056602 | 0.23712746 |
| ENSMUSG00000051375 | Pcdh1 | 75599 | protein_coding | 1225.030985 | -0.237150586 | 0.116341694 | -2.038397226 | 0.041510223 | 0.136380949 | 0.237150586 |
| ENSMUSG00000053178 | Mterf1b | 208595 | protein_coding | 16.24816862 | 0.237222336 | 0.28695919 | 0.82667621 | 0.408420609 | 0.615684597 | 0.237222336 |
| ENSMUSG00000024081 | Cebpz | 12607 | protein_coding | 668.3294823 | -0.23724254 | 0.119316316 | -1.988349523 | 0.046773047 | 0.148643475 | 0.23724254 |
| ENSMUSG00000038009 | Dnajc22 | 72778 | protein_coding | 966.7768862 | 0.237244831 | 0.113463164 | 2.090941441 | 0.036533314 | 0.124711234 | 0.237244831 |
| ENSMUSG00000063146 | Clip2 | 269713 | protein_coding | 1333.340256 | 0.237246661 | 0.135623822 | 1.749299337 | 0.080239291 | 0.214186596 | 0.237246661 |
| ENSMUSG00000021752 | Kctd6 | 71393 | protein_coding | 116.9587861 | -0.237312323 | 0.137933665 | -1.720481525 | 0.085344949 | 0.223728497 | 0.237312323 |
| ENSMUSG00000079560 | Hoxa3 | 15400 | protein_coding | 73.33892926 | -0.237314549 | 0.189332449 | -1.253427768 | 0.210050071 | 0.407309235 | 0.237314549 |
| ENSMUSG00000032221 | Mns1 | 17427 | protein_coding | 39.83943372 | 0.237362954 | 0.206479657 | 1.149570653 | 0.250320751 | 0.456634994 | 0.237362954 |
| ENSMUSG00000020692 | Nle1 | 217011 | protein_coding | 102.0347374 | -0.237369997 | 0.220291485 | -1.077526878 | 0.28124495 | 0.489849049 | 0.237369997 |
| ENSMUSG00000040359 | Ufl1 | 67490 | protein_coding | 507.6642823 | 0.237406032 | 0.081812637 | 2.901825934 | 0.003709946 | 0.02216505 | 0.237406032 |
| ENSMUSG00000030691 | Fchsd2 | 207278 | protein_coding | 880.8650099 | -0.23741538 | 0.156982214 | -1.512371207 | 0.130439455 | 0.297324482 | 0.23741538 |

| ensembl_gene_id | external_gene_name | entrezgene_id | gene_biotype | baseMean | log2FoldChange | lfcSE | stat | pvalue | padj | absLog2FC |
|---|---|---|---|---|---|---|---|---|---|---|
| ENSMUSG00000049493 | Pls1 | 102502 | protein_coding | 4075.680254 | 0.237424111 | 0.130136323 | 1.824426155 | 0.068087671 | 0.191900312 | 0.237424111 |
| ENSMUSG00000031839 | Hsbp1 | 68196 | protein_coding | 1057.333556 | 0.237439844 | 0.077529176 | 3.062586949 | 0.002194327 | 0.014614647 | 0.237439844 |
| ENSMUSG00000021572 | Cep72 | 74470 | protein_coding | 99.12411082 | -0.237448667 | 0.181837977 | -1.305825501 | 0.191611906 | 0.384285374 | 0.237448667 |
| ENSMUSG00000021918 | Nek4 | 23955 | protein_coding | 289.2676509 | 0.23745175 | 0.093562247 | 2.537901305 | 0.011151943 | 0.051907699 | 0.23745175 |
| ENSMUSG00000018661 | Cog1 | 16834 | protein_coding | 903.1137762 | 0.237549235 | 0.09289509 | 2.5571775 | 0.010552536 | 0.049786963 | 0.237549235 |
| ENSMUSG00000041852 | Tcf20 | 21411 | protein_coding | 2863.983482 | -0.237565838 | 0.06979464 | -3.403783428 | 0.000664594 | 0.005504762 | 0.237565838 |
| ENSMUSG00000030352 | Tspan9 | 109246 | protein_coding | 133.8947378 | 0.237640365 | 0.135400774 | 1.755088679 | 0.07924414 | 0.212599068 | 0.237640365 |
| ENSMUSG00000025374 | Nabp2 | 69917 | protein_coding | 291.1938541 | 0.237663492 | 0.079296001 | 2.997168714 | 0.002724999 | 0.017388158 | 0.237663492 |
| ENSMUSG00000026281 | Dtymk | 21915 | protein_coding | 315.8851041 | 0.237685038 | 0.103870357 | 2.288285561 | 0.022120897 | 0.086126011 | 0.237685038 |
| ENSMUSG00000044676 | Zfp612 | 234725 | protein_coding | 200.7947906 | -0.237749633 | 0.114560293 | -2.07532319 | 0.037956592 | 0.128304879 | 0.237749633 |
| ENSMUSG00000094112 | 9330182O14Rik | 328531 | protein_coding | 9.793103474 | -0.237790035 | 0.436179119 | -0.545166021 | 0.585639339 | 0.75599124 | 0.237790035 |
| ENSMUSG00000038866 | Zcchc2 | 227449 | protein_coding | 509.5577827 | -0.2378114 | 0.140219938 | -1.695988485 | 0.089888061 | 0.231845261 | 0.2378114 |
| ENSMUSG00000041845 | Rhod | 11854 | protein_coding | 95.77293251 | 0.237845658 | 0.212205421 | 1.12082744 | 0.262361322 | 0.470066494 | 0.237845658 |
| ENSMUSG00000029186 | Pi4k2b | 67073 | protein_coding | 2767.149081 | 0.23794257 | 0.113827891 | 2.090371423 | 0.036584448 | 0.124832281 | 0.23794257 |
| ENSMUSG00000021838 | Samd4 | 74480 | protein_coding | 7.30420592 | -0.238023122 | 0.610411686 | -0.389938671 | 0.696581897 | 0.827255924 | 0.238023122 |
| ENSMUSG00000022474 | Pmm1 | 29858 | protein_coding | 402.0398943 | -0.238115783 | 0.12881151 | -1.848559831 | 0.064521397 | 0.184804319 | 0.238115783 |
| ENSMUSG00000030726 | Pold3 | 67967 | protein_coding | 477.8758135 | -0.238243422 | 0.075123411 | -3.171360527 | 0.001517267 | 0.010760031 | 0.238243422 |
| ENSMUSG00000055373 | Fut9 | 14348 | protein_coding | 244.521768 | -0.238313318 | 0.522383658 | -0.456203625 | 0.648243559 | 0.796966766 | 0.238313318 |
| ENSMUSG00000025817 | Nudt5 | 53893 | protein_coding | 428.1575913 | 0.238361919 | 0.0834452 | 2.856508441 | 0.004283286 | 0.024808793 | 0.238361919 |
| ENSMUSG00000029769 | Ccdc136 | 232664 | protein_coding | 8.102613068 | -0.238369844 | 0.545152315 | -0.437253658 | 0.6619274 | 0.804454419 | 0.238369844 |
| ENSMUSG00000080465 | Snord94 | 115490493 | snoRNA | 51.38476184 | 0.238420884 | 0.317416499 | 0.751129462 | 0.452574746 | 0.652717363 | 0.238420884 |
| ENSMUSG00000066512 | Klk1b5 | 16622 | protein_coding | 8.06030354 | 0.238451128 | 0.379615148 | 0.628139127 | 0.529912803 | 0.715171253 | 0.238451128 |
| ENSMUSG00000030031 | Kbtbd8 | 243574 | protein_coding | 248.1848192 | 0.238531781 | 0.158800712 | 1.502082567 | 0.133075786 | 0.301598054 | 0.238531781 |
| ENSMUSG00000073758 | Sh3d21 | 66938 | protein_coding | 440.1800166 | 0.238546417 | 0.229279823 | 1.04041609 | 0.29814663 | 0.506925653 | 0.238546417 |
| ENSMUSG00000035919 | Bbs9 | 319845 | protein_coding | 78.78545308 | -0.238668342 | 0.140746437 | -1.695732753 | 0.089936503 | 0.231849133 | 0.238668342 |
| ENSMUSG00000022683 | Pla2g10 | 26565 | protein_coding | 786.1383618 | 0.23882137 | 0.13626586 | 1.752613383 | 0.079668393 | 0.213270278 | 0.23882137 |
| ENSMUSG00000029705 | Cux1 | 13047 | protein_coding | 1485.181482 | -0.238827461 | 0.067971255 | -3.51365384 | 0.000441988 | 0.003946183 | 0.238827461 |
| ENSMUSG00000004356 | Utp20 | 70683 | protein_coding | 1888.860475 | 0.238831075 | 0.087893359 | 2.717282361 | 0.006582043 | 0.034714827 | 0.238831075 |
| ENSMUSG00000037752 | Xkr8 | 381560 | protein_coding | 20.8497733 | -0.238877486 | 0.3296801 | -0.724573567 | 0.46871368 | 0.66509336 | 0.238877486 |
| ENSMUSG00000018169 | Mfng | 17305 | protein_coding | 8.191095312 | -0.238906967 | 0.539559928 | -0.442781153 | 0.657924037 | 0.802098282 | 0.238906967 |
| ENSMUSG00000039262 | Prrc2b | 227723 | protein_coding | 909.8368918 | -0.238960218 | 0.122234178 | -1.954937822 | 0.05059041 | 0.156635171 | 0.238960218 |
| ENSMUSG00000073079 | Srp54a | 24067 | protein_coding | 56.91942391 | 0.238990702 | 0.163088665 | 1.465403514 | 0.142810859 | 0.316412765 | 0.238990702 |
| ENSMUSG00000022052 | Ppp2r2a | 71978 | protein_coding | 342.985167 | -0.239008775 | 0.187558908 | -1.274313107 | 0.202552473 | 0.39748185 | 0.239008775 |
| ENSMUSG00000071074 | Yipf3 | 28064 | protein_coding | 615.7688918 | -0.239132205 | 0.120132669 | -1.99056766 | 0.046528439 | 0.148132115 | 0.239132205 |
| ENSMUSG00000085112 | A530072M11Rik | 100415915 | lncRNA | 27.48589776 | 0.239197679 | 0.231862416 | 1.031636276 | 0.30224254 | 0.511596086 | 0.239197679 |
| ENSMUSG00000028684 | Urod | 22275 | protein_coding | 381.4743134 | -0.239216649 | 0.100414176 | -2.382299564 | 0.017204895 | 0.071663068 | 0.239216649 |
| ENSMUSG00000031847 | 1700030J22Rik | 69528 | protein_coding | 18.17755414 | -0.239278391 | 0.306793125 | -0.779934004 | 0.435429722 | 0.638336365 | 0.239278391 |
| ENSMUSG00000038187 | Btbd10 | 68815 | protein_coding | 237.6271909 | -0.239305637 | 0.133993411 | -1.785950774 | 0.074107229 | 0.20295391 | 0.239305637 |

| ensembl_gene_id | external_gene_name | entrezgene_id | gene_biotype | baseMean | log2FoldChange | lfcSE | stat | pvalue | padj | absLog2FC |
|---|---|---|---|---|---|---|---|---|---|---|
| ENSMUSG00000040296 | Rigi | 230073 | protein_coding | 342.7307078 | 0.239334983 | 0.086672569 | 2.761369425 | 0.005755953 | 0.031131007 | 0.239334983 |
| ENSMUSG00000028013 | Ppa2 | 74776 | protein_coding | 1309.066473 | -0.239380218 | 0.081713809 | -2.929495284 | 0.00339513 | 0.020719397 | 0.239380218 |
| ENSMUSG00000075470 | Alg10b | 380959 | protein_coding | 1055.733915 | 0.239435241 | 0.068149102 | 3.513402752 | 0.000442406 | 0.003947702 | 0.239435241 |
| ENSMUSG00000029166 | Mapre3 | 100732 | protein_coding | 84.81557662 | -0.239502427 | 0.262814194 | -0.91129944 | 0.362137621 | 0.571441255 | 0.239502427 |
| ENSMUSG00000002280 | Ciao3 | 67563 | protein_coding | 360.5091414 | 0.239513704 | 0.111945372 | 2.139558795 | 0.03239044 | 0.114115223 | 0.239513704 |
| ENSMUSG00000049300 | Prmt6 | 99890 | protein_coding | 133.4725799 | -0.239549329 | 0.153586756 | -1.559700423 | 0.118830692 | 0.279370519 | 0.239549329 |
| ENSMUSG00000029776 | Hibadh | 58875 | protein_coding | 956.2545788 | -0.239573927 | 0.107899666 | -2.220339828 | 0.026395708 | 0.098300407 | 0.239573927 |
| ENSMUSG00000029646 | Cdx2 | 12591 | protein_coding | 3305.943864 | 0.239619245 | 0.186364012 | 1.285759215 | 0.198527108 | 0.39266606 | 0.239619245 |
| ENSMUSG00000057229 | Dmac2 | 66349 | protein_coding | 505.6644691 | 0.239630718 | 0.095106959 | 2.519591839 | 0.011749099 | 0.053982933 | 0.239630718 |
| ENSMUSG00000042198 | Chchd7 | 66433 | protein_coding | 356.5326653 | 0.239671778 | 0.086035516 | 2.785730687 | 0.005340722 | 0.029445145 | 0.239671778 |
| ENSMUSG00000038930 | Rccd1 | 269955 | protein_coding | 76.950984 | -0.239672812 | 0.165978939 | -1.443995327 | 0.14874029 | 0.325878313 | 0.239672812 |
| ENSMUSG00000078317 | F8a | 14070 | protein_coding | 233.4351879 | 0.239709722 | 0.173785991 | 1.379338578 | 0.167790387 | 0.35225587 | 0.239709722 |
| ENSMUSG00000026806 | Ddx31 | 227674 | protein_coding | 144.2491368 | -0.239749747 | 0.198148539 | -1.209949608 | 0.22629823 | 0.428594317 | 0.239749747 |
| ENSMUSG00000063383 | Zfp947 | 210853 | protein_coding | 115.9869475 | 0.239773933 | 0.186828395 | 1.283391278 | 0.199355028 | 0.393706399 | 0.239773933 |
| ENSMUSG00000036934 | 4921524J17Rik | 66714 | protein_coding | 104.092692 | -0.239800076 | 0.170718587 | -1.404651247 | 0.16012501 | 0.341659149 | 0.239800076 |
| ENSMUSG00000038482 | Tfdp1 | 21781 | protein_coding | 816.4291346 | 0.239869518 | 0.083031709 | 2.88889052 | 0.003866036 | 0.022883028 | 0.239869518 |
| ENSMUSG00000034951 | Cog7 | 233824 | protein_coding | 555.2705174 | 0.2399068 | 0.061422678 | 3.905834244 | 9.39E-05 | 0.001068324 | 0.2399068 |
| ENSMUSG00000036768 | Kif15 | 209737 | protein_coding | 486.779195 | 0.239909708 | 0.128327976 | 1.869504335 | 0.061552681 | 0.178676602 | 0.239909708 |
| ENSMUSG00000028040 | Efna4 | 13639 | protein_coding | 94.86539013 | 0.239955241 | 0.226431576 | 1.059725172 | 0.289269648 | 0.498106697 | 0.239955241 |
| ENSMUSG00000006717 | Acot13 | 66834 | protein_coding | 463.5698275 | 0.240006051 | 0.097804912 | 2.453926349 | 0.014130592 | 0.061849978 | 0.240006051 |
| ENSMUSG00000025016 | Tm9sf3 | 107358 | protein_coding | 11395.61095 | 0.240025323 | 0.056707972 | 4.232655746 | 2.31E-05 | 0.000312917 | 0.240025323 |
| ENSMUSG00000038965 | Ube2l3 | 22195 | protein_coding | 1050.140615 | 0.2400578 | 0.070456387 | 3.407182954 | 0.000656371 | 0.005462215 | 0.2400578 |
| ENSMUSG00000003721 | Insig2 | 72999 | protein_coding | 491.9698655 | 0.240063368 | 0.108320844 | 2.216225058 | 0.026676093 | 0.098951281 | 0.240063368 |
| ENSMUSG00000041992 | Rapgef5 | 217944 | protein_coding | 569.075756 | -0.240088126 | 0.111920724 | -2.145162386 | 0.031939866 | 0.11305293 | 0.240088126 |
| ENSMUSG00000063904 | Dpp3 | 75221 | protein_coding | 1773.752221 | 0.24010094 | 0.101478331 | 2.366031629 | 0.017979908 | 0.073829332 | 0.24010094 |
| ENSMUSG00000048439 | Nupl2 | 231042 | protein_coding | 136.0762222 | 0.240105994 | 0.134824407 | 1.780879295 | 0.074932172 | 0.204201575 | 0.240105994 |
| ENSMUSG00000050931 | Sgms2 | 74442 | protein_coding | 1293.955631 | -0.240125672 | 0.130768564 | -1.836264496 | 0.066318548 | 0.188259104 | 0.240125672 |
| ENSMUSG00000047205 | Dusp18 | 75219 | protein_coding | 186.1648903 | -0.24013653 | 0.108996238 | -2.203163473 | 0.027583229 | 0.101309196 | 0.24013653 |
| ENSMUSG00000005410 | Mcm5 | 17218 | protein_coding | 561.2262332 | 0.240242624 | 0.145864332 | 1.6470279 | 0.099552311 | 0.247566483 | 0.240242624 |
| ENSMUSG00000038722 | Bud31 | 231889 | protein_coding | 539.1710433 | 0.240248342 | 0.12093136 | 1.986650466 | 0.046961145 | 0.14896269 | 0.240248342 |
| ENSMUSG00000051705 | Senp8 | 71599 | protein_coding | 111.9993034 | 0.240275985 | 0.228087759 | 1.053436561 | 0.292140955 | 0.501006478 | 0.240275985 |
| ENSMUSG00000018001 | Cyth3 | 19159 | protein_coding | 233.4587605 | -0.240305712 | 0.08243834 | -2.914975137 | 0.003557169 | 0.021477856 | 0.240305712 |
| ENSMUSG00000021182 | Ccdc88c | 68339 | protein_coding | 1224.505511 | 0.240365728 | 0.099805734 | 2.408335854 | 0.016025431 | 0.067940661 | 0.240365728 |
| ENSMUSG00000066149 | Cdc26 | 66440 | protein_coding | 315.7801348 | 0.240370025 | 0.097013749 | 2.477690306 | 0.013223587 | 0.05906486 | 0.240370025 |
| ENSMUSG00000057421 | Las1l | 76130 | protein_coding | 380.2685339 | 0.240400689 | 0.111371655 | 2.158544637 | 0.030885512 | 0.110425237 | 0.240400689 |
| ENSMUSG00000036912 | Piwil4 | 330890 | protein_coding | 128.6679349 | 0.240409501 | 0.118026392 | 2.036913067 | 0.041658754 | 0.1367843 | 0.240409501 |
| ENSMUSG00000042541 | Sem1 | 20422 | protein_coding | 1691.724628 | 0.240474214 | 0.112613893 | 2.135386753 | 0.032729432 | 0.114953559 | 0.240474214 |
| ENSMUSG00000018919 | Tm4sf5 | 75604 | protein_coding | 26.22914333 | 0.240545421 | 0.264521152 | 0.909361762 | 0.363159199 | 0.572033201 | 0.240545421 |

| ensembl_gene_id | external_gene_name | entrezgene_id | gene_biotype | baseMean | log2FoldChange | lfcSE | stat | pvalue | padj | absLog2FC |
|---|---|---|---|---|---|---|---|---|---|---|
| ENSMUSG00000027099 | Mtx2 | 53375 | protein_coding | 772.5911669 | 0.24060411 | 0.087011658 | 2.765193948 | 0.005688896 | 0.030883687 | 0.24060411 |
| ENSMUSG00000029777 | Gars | 353172 | protein_coding | 2292.39917 | 0.240625425 | 0.139179349 | 1.728887414 | 0.083829247 | 0.221028347 | 0.240625425 |
| ENSMUSG00000087963 | Gm25394 | 115488237 | snoRNA | 11.74125693 | 0.240629844 | 0.443234049 | 0.542895665 | 0.587201641 | 0.757262387 | 0.240629844 |
| ENSMUSG00000063450 | Syne2 | 319565 | protein_coding | 2763.266089 | -0.240705778 | 0.148422688 | -1.621758656 | 0.104855034 | 0.256351647 | 0.240705778 |
| ENSMUSG00000001964 | Emd | 13726 | protein_coding | 266.5072378 | 0.240811089 | 0.088238484 | 2.729093675 | 0.006350866 | 0.033718865 | 0.240811089 |
| ENSMUSG00000003452 | Bicd1 | 12121 | protein_coding | 301.369532 | -0.241057169 | 0.167173228 | -1.441960369 | 0.149313554 | 0.326699017 | 0.241057169 |
| ENSMUSG00000020650 | Bcap29 | 12033 | protein_coding | 135.0832893 | -0.241075291 | 0.13560826 | -1.777733089 | 0.075447702 | 0.205144798 | 0.241075291 |
| ENSMUSG00000037251 | Pomk | 74653 | protein_coding | 181.9464566 | -0.24109281 | 0.119295313 | -2.020974703 | 0.043282384 | 0.14037911 | 0.24109281 |
| ENSMUSG00000040649 | Rimklb | 108653 | protein_coding | 8.987255133 | 0.241096967 | 0.500261517 | 0.481941863 | 0.629847246 | 0.785799181 | 0.241096967 |
| ENSMUSG00000026088 | Mitd1 | 69028 | protein_coding | 287.4608008 | 0.241199859 | 0.082839118 | 2.911666181 | 0.003595066 | 0.02161654 | 0.241199859 |
| ENSMUSG00000031930 | Wwp2 | 66894 | protein_coding | 1484.731017 | -0.241239109 | 0.078771258 | -3.062527074 | 0.002194766 | 0.014614647 | 0.241239109 |
| ENSMUSG00000041696 | Rasl12 | 70784 | protein_coding | 11.24687039 | -0.241308545 | 0.400867055 | -0.601966517 | 0.547196426 | 0.729045978 | 0.241308545 |
| ENSMUSG00000043987 | Cep164 | 214552 | protein_coding | 454.141936 | -0.24131746 | 0.097651299 | -2.471216085 | 0.013465443 | 0.059843072 | 0.24131746 |
| ENSMUSG00000035357 | Pdzrn3 | 55983 | protein_coding | 94.40226546 | -0.241317469 | 0.220030459 | -1.09674574 | 0.272752555 | 0.480841472 | 0.241317469 |
| ENSMUSG00000091264 | Smim13 | 108934 | protein_coding | 218.3940256 | -0.241379784 | 0.099667176 | -2.421858362 | 0.015441368 | 0.066062343 | 0.241379784 |
| ENSMUSG00000027524 | Edn3 | 13616 | protein_coding | 116.5163284 | 0.241395094 | 0.17131045 | 1.409108982 | 0.158802945 | 0.340076845 | 0.241395094 |
| ENSMUSG00000036231 | Agr3 | 403205 | protein_coding | 28.32219124 | -0.241462204 | 0.242225105 | -0.996850445 | 0.318837108 | 0.530163635 | 0.241462204 |
| ENSMUSG00000032420 | Nt5e | 23959 | protein_coding | 445.9064651 | -0.241592029 | 0.291249945 | -0.829500687 | 0.406821148 | 0.613912577 | 0.241592029 |
| ENSMUSG00000061186 | Sfmbt2 | 353282 | protein_coding | 9.598636612 | -0.241695181 | 0.694423596 | -0.34805151 | 0.727801499 | 0.846662573 | 0.241695181 |
| ENSMUSG00000069305 | H4c18 | 319161 | protein_coding | 1171.851718 | 0.241784154 | 0.140539652 | 1.720398125 | 0.085360098 | 0.223731386 | 0.241784154 |
| ENSMUSG00000086468 | Etaa1os | 102633673 | lncRNA | 11.48154302 | -0.241875164 | 0.357457553 | -0.676654228 | 0.498625364 | 0.689179521 | 0.241875164 |
| ENSMUSG00000024082 | Ndufaf7 | 73694 | protein_coding | 283.9005142 | -0.241892648 | 0.090560844 | -2.67105116 | 0.007561412 | 0.038813462 | 0.241892648 |
| ENSMUSG00000021131 | Erh | 13877 | protein_coding | 276.6957299 | 0.241947255 | 0.134489206 | 1.79900872 | 0.072017301 | 0.199424822 | 0.241947255 |
| ENSMUSG00000047635 | Mtrfr | 72650 | protein_coding | 99.59747183 | 0.242014865 | 0.13873948 | 1.744383538 | 0.081092234 | 0.216066691 | 0.242014865 |
| ENSMUSG00000004317 | Clcn5 | 12728 | protein_coding | 563.2064137 | -0.24212833 | 0.127432665 | -1.900049169 | 0.057426667 | 0.171066384 | 0.24212833 |
| ENSMUSG00000030380 | Mzf1 | 109889 | protein_coding | 16.67444438 | -0.24213529 | 0.350154979 | -0.691508916 | 0.489245779 | 0.681776621 | 0.24213529 |
| ENSMUSG00000058656 | Samd12 | 320679 | protein_coding | 311.3441853 | 0.242225096 | 0.197983621 | 1.223460276 | 0.221155899 | 0.422016433 | 0.242225096 |
| ENSMUSG00000025473 | Adam8 | 11501 | protein_coding | 16.70297222 | -0.242253585 | 0.452426288 | -0.535454265 | 0.592335775 | 0.761247936 | 0.242253585 |
| ENSMUSG00000032050 | Rdx | 19684 | protein_coding | 179.8287947 | 0.242479301 | 0.504699983 | 0.480442459 | 0.630912809 | 0.786307224 | 0.242479301 |
| ENSMUSG00000038462 | Uqcrfs1 | 66694 | protein_coding | 3899.542692 | 0.242521644 | 0.122834315 | 1.974380236 | 0.048338524 | 0.152161266 | 0.242521644 |
| ENSMUSG00000037143 | Cfap61 | 78774 | protein_coding | 7.144372495 | -0.242529089 | 0.736692752 | -0.329213351 | 0.741994432 | 0.856288025 | 0.242529089 |
| ENSMUSG00000003873 | Bax | 12028 | protein_coding | 234.0198023 | 0.242664653 | 0.10055096 | 2.413349938 | 0.015806635 | 0.067304582 | 0.242664653 |
| ENSMUSG00000021832 | Psmc6 | 67089 | protein_coding | 1584.378305 | 0.242675614 | 0.084764048 | 2.862954501 | 0.004197108 | 0.02440728 | 0.242675614 |
| ENSMUSG00000025366 | Esyt1 | 23943 | protein_coding | 622.7692117 | 0.242772238 | 0.119543606 | 2.030825786 | 0.04227267 | 0.138201784 | 0.242772238 |
| ENSMUSG00000028032 | Papss1 | 23971 | protein_coding | 2376.189866 | 0.242839732 | 0.123977776 | 1.958735989 | 0.050143713 | 0.155811367 | 0.242839732 |
| ENSMUSG00000031488 | Rab11fip1 | 75767 | protein_coding | 2350.935952 | -0.242880666 | 0.061186171 | -3.969535297 | 7.20E-05 | 0.000845887 | 0.242880666 |
| ENSMUSG00000034723 | Tmx4 | 52837 | protein_coding | 276.6998869 | 0.242965485 | 0.204843392 | 1.186103602 | 0.235581388 | 0.439707037 | 0.242965485 |
| ENSMUSG00000045435 | Tmem60 | 212090 | protein_coding | 163.3130758 | 0.242971194 | 0.127239173 | 1.909562825 | 0.056189525 | 0.168072631 | 0.242971194 |

| ensembl_gene_id | external_gene_name | entrezgene_id | gene_biotype | baseMean | log2FoldChange | lfcSE | stat | pvalue | padj | absLog2FC |
|---|---|---|---|---|---|---|---|---|---|---|
| ENSMUSG00000061808 | Ttr | 22139 | protein_coding | 253.1433267 | -0.243038315 | 0.301275231 | -0.806698626 | 0.419840136 | 0.625792035 | 0.243038315 |
| ENSMUSG00000016128 | Stard13 | 243362 | protein_coding | 353.4145366 | -0.243044156 | 0.104309273 | -2.330034041 | 0.019804352 | 0.079157771 | 0.243044156 |
| ENSMUSG00000021807 | Rtraf | 68045 | protein_coding | 2033.303226 | 0.243152267 | 0.078625198 | 3.092548854 | 0.001984456 | 0.01342331 | 0.243152267 |
| ENSMUSG00000045503 | Sys1 | 66460 | protein_coding | 323.5763567 | 0.243191843 | 0.104116866 | 2.335758385 | 0.019503838 | 0.078508245 | 0.243191843 |
| ENSMUSG00000024091 | Vapa | 30960 | protein_coding | 2242.583684 | 0.243393762 | 0.089268748 | 2.72652824 | 0.006400448 | 0.033925567 | 0.243393762 |
| ENSMUSG00000098176 | Ccdc166 | 223648 | protein_coding | 83.80994907 | -0.243650841 | 0.272631159 | -0.893701374 | 0.37148169 | 0.579753122 | 0.243650841 |
| ENSMUSG00000110803 | Gm20275 | 100504539 | lncRNA | 57.50525024 | -0.243659109 | 0.190368653 | -1.27993294 | 0.200568722 | 0.395157223 | 0.243659109 |
| ENSMUSG00000002679 | Med6 | 69792 | protein_coding | 253.805161 | 0.243675636 | 0.096304524 | 2.53026158 | 0.011397752 | 0.052804942 | 0.243675636 |
| ENSMUSG00000033068 | Entpd6 | 12497 | protein_coding | 818.4832155 | 0.244054852 | 0.093922009 | 2.598484161 | 0.009363636 | 0.045581908 | 0.244054852 |
| ENSMUSG00000032078 | Zpr1 | 22687 | protein_coding | 394.019991 | -0.24410382 | 0.103708106 | -2.353758354 | 0.018584688 | 0.075746395 | 0.24410382 |
| ENSMUSG00000033906 | Zdhhc15 | 108672 | protein_coding | 130.3271154 | -0.244158698 | 0.152053914 | -1.605737675 | 0.10833156 | 0.261956138 | 0.244158698 |
| ENSMUSG00000021671 | Poc5 | 67463 | protein_coding | 228.3125942 | 0.244220479 | 0.091077446 | 2.681459453 | 0.00733018 | 0.037833799 | 0.244220479 |
| ENSMUSG00000035623 | Rsf1 | 233532 | protein_coding | 751.5793046 | -0.244254275 | 0.127161775 | -1.920815227 | 0.054755006 | 0.165037012 | 0.244254275 |
| ENSMUSG00000021578 | Ccdc127 | 67433 | protein_coding | 564.5448033 | 0.24425697 | 0.069113581 | 3.534138537 | 0.000409107 | 0.003700312 | 0.24425697 |
| ENSMUSG00000054509 | Parp4 | 328417 | protein_coding | 385.1369676 | -0.244326666 | 0.221243881 | -1.104331854 | 0.26944921 | 0.477342567 | 0.244326666 |
| ENSMUSG00000030452 | Nipa2 | 93790 | protein_coding | 759.187383 | 0.244342284 | 0.085806212 | 2.847606004 | 0.004404942 | 0.025356675 | 0.244342284 |
| ENSMUSG00000036990 | Otud4 | 73945 | protein_coding | 1580.900653 | -0.244369213 | 0.080746361 | -3.026380514 | 0.002475006 | 0.016183045 | 0.244369213 |
| ENSMUSG00000067931 | Zfp948 | 381066 | protein_coding | 343.0733125 | -0.244430197 | 0.142926173 | -1.710184994 | 0.08723167 | 0.227364758 | 0.244430197 |
| ENSMUSG00000032096 | Arcn1 | 213827 | protein_coding | 4679.444796 | 0.244459588 | 0.071938692 | 3.398165583 | 0.000678393 | 0.005613221 | 0.244459588 |
| ENSMUSG00000061578 | Ksr2 | 333050 | protein_coding | 22.11677559 | -0.244627356 | 0.649540583 | -0.376615969 | 0.706459019 | 0.834011211 | 0.244627356 |
| ENSMUSG00000028980 | H6pd | 100198 | protein_coding | 413.9415148 | 0.2446785 | 0.096174453 | 2.544111154 | 0.010955622 | 0.051278619 | 0.2446785 |
| ENSMUSG00000074912 | Gm14207 | 100043609 | lncRNA | 54.82561404 | -0.244745613 | 0.182572381 | -1.340540182 | 0.18006979 | 0.368080898 | 0.244745613 |
| ENSMUSG00000037390 | Muc3 | 666339 | protein_coding | 5677.212182 | -0.244841057 | 0.194901533 | -1.256229506 | 0.209032772 | 0.405828413 | 0.244841057 |
| ENSMUSG00000025500 | Lmntd2 | 72000 | protein_coding | 62.95319931 | -0.244862665 | 0.232838873 | -1.051639969 | 0.292964763 | 0.502081208 | 0.244862665 |
| ENSMUSG00000020561 | Polr1f | 28071 | protein_coding | 370.6216959 | 0.24497759 | 0.090667877 | 2.701922631 | 0.00689398 | 0.036026023 | 0.24497759 |
| ENSMUSG00000064138 | Fam172a | 68675 | protein_coding | 226.9339251 | -0.245054977 | 0.088476404 | -2.769721266 | 0.005610428 | 0.030520116 | 0.245054977 |
| ENSMUSG00000033904 | Ccp110 | 101565 | protein_coding | 91.01093282 | -0.245069621 | 0.176896252 | -1.385386167 | 0.165934419 | 0.349742414 | 0.245069621 |
| ENSMUSG00000046312 | Myorg | 329828 | protein_coding | 239.6746408 | -0.245073994 | 0.189136532 | -1.295751756 | 0.195061024 | 0.388054296 | 0.245073994 |
| ENSMUSG00000037762 | Slc16a9 | 66859 | protein_coding | 286.3020037 | -0.245127211 | 0.219530441 | -1.116597819 | 0.264166325 | 0.471920281 | 0.245127211 |
| ENSMUSG00000037348 | Paqr7 | 71904 | protein_coding | 32.81566359 | -0.245211759 | 0.252173313 | -0.972393772 | 0.330854689 | 0.541497481 | 0.245211759 |
| ENSMUSG00000026278 | Bok | 51800 | protein_coding | 205.128463 | -0.245382308 | 0.201716516 | -1.21647108 | 0.223805521 | 0.425414053 | 0.245382308 |
| ENSMUSG00000057388 | Mrpl18 | 67681 | protein_coding | 744.3285945 | 0.245413551 | 0.092781996 | 2.645055732 | 0.008167746 | 0.04119565 | 0.245413551 |
| ENSMUSG00000026035 | Ppil3 | 70225 | protein_coding | 194.1455948 | 0.245506371 | 0.136873967 | 1.793667384 | 0.072866274 | 0.200728816 | 0.245506371 |
| ENSMUSG00000061979 | Rcc1l | 94254 | protein_coding | 227.6616754 | -0.24573263 | 0.170604307 | -1.440365921 | 0.1497639 | 0.327175888 | 0.24573263 |
| ENSMUSG00000038793 | Lefty1 | 13590 | protein_coding | 53.01968035 | -0.245743687 | 0.219784278 | -1.118113134 | 0.263518678 | 0.471131469 | 0.245743687 |
| ENSMUSG00000021816 | Ppp3cb | 19056 | protein_coding | 769.1674305 | 0.245791071 | 0.065184103 | 3.770721101 | 0.000162777 | 0.00170735 | 0.245791071 |
| ENSMUSG00000055125 | M5C1000I18Rik | 102633151 | lncRNA | 10.44786584 | -0.2458479 | 0.403427814 | -0.609397498 | 0.542260996 | 0.725173014 | 0.2458479 |
| ENSMUSG00000025504 | Eps8l2 | 98845 | protein_coding | 1155.121249 | -0.245923583 | 0.153288534 | -1.60431819 | 0.108643934 | 0.262353028 | 0.245923583 |

| ensembl_gene_id | external_gene_name | entrezgene_id | gene_biotype | baseMean | log2FoldChange | lfcSE | stat | pvalue | padj | absLog2FC |
|---|---|---|---|---|---|---|---|---|---|---|
| ENSMUSG00000063439 | B9d2 | 232987 | protein_coding | 17.50105445 | -0.245942309 | 0.29496829 | -0.833792369 | 0.404397996 | 0.611414455 | 0.245942309 |
| ENSMUSG00000070420 | Zscan25 | 666311 | protein_coding | 138.4026103 | -0.246020035 | 0.125662066 | -1.957790783 | 0.050254567 | 0.155973256 | 0.246020035 |
| ENSMUSG00000000563 | Atp5pb | 11950 | protein_coding | 5781.31848 | 0.246064379 | 0.092618945 | 2.656739162 | 0.007890046 | 0.04002314 | 0.246064379 |
| ENSMUSG00000073664 | Nbeal1 | 269198 | protein_coding | 2777.599585 | 0.246091754 | 0.138581672 | 1.775788607 | 0.075767766 | 0.205347793 | 0.246091754 |
| ENSMUSG00000074909 | Ranbp6 | 240614 | protein_coding | 320.4844436 | -0.24614339 | 0.103886433 | -2.369350673 | 0.017819349 | 0.073411269 | 0.24614339 |
| ENSMUSG00000060279 | Ap2a1 | 11771 | protein_coding | 688.9720339 | 0.24620345 | 0.086653701 | 2.841234086 | 0.004493931 | 0.025776282 | 0.24620345 |
| ENSMUSG00000042292 | Mrtfa | 223701 | protein_coding | 278.6868951 | -0.246242497 | 0.126130473 | -1.95228394 | 0.050904503 | 0.157285534 | 0.246242497 |
| ENSMUSG00000037214 | Thap1 | 73754 | protein_coding | 126.3684467 | 0.24624978 | 0.128925325 | 1.910018683 | 0.056130808 | 0.16796008 | 0.24624978 |
| ENSMUSG00000030407 | Qpctl | 67369 | protein_coding | 162.8444952 | 0.246273605 | 0.117581655 | 2.094490035 | 0.03621635 | 0.124027955 | 0.246273605 |
| ENSMUSG00000025277 | Abhd6 | 66082 | protein_coding | 636.668267 | 0.246352031 | 0.167133144 | 1.473986692 | 0.140485174 | 0.312850724 | 0.246352031 |
| ENSMUSG00000026888 | Grb14 | 50915 | protein_coding | 28.13710119 | -0.246389111 | 0.255460241 | -0.964491029 | 0.334799804 | 0.545376485 | 0.246389111 |
| ENSMUSG00000024511 | Rab27b | 80718 | protein_coding | 1625.350439 | 0.246400058 | 0.101570991 | 2.425890058 | 0.015270896 | 0.065520384 | 0.246400058 |
| ENSMUSG00000032033 | Barx2 | 12023 | protein_coding | 24.75039921 | -0.246454777 | 0.243290848 | -1.013004718 | 0.311057907 | 0.521090697 | 0.246454777 |
| ENSMUSG00000022983 | Scaf4 | 224432 | protein_coding | 263.3346897 | -0.246487145 | 0.170478727 | -1.445852798 | 0.148218494 | 0.325003327 | 0.246487145 |
| ENSMUSG00000001763 | Tspan33 | 232670 | protein_coding | 215.996615 | -0.246487422 | 0.181341811 | -1.359242088 | 0.174069888 | 0.360029241 | 0.246487422 |
| ENSMUSG00000031448 | Adprhl1 | 234072 | protein_coding | 7.26431673 | 0.246832143 | 0.539235526 | 0.457744586 | 0.647135952 | 0.796158015 | 0.246832143 |
| ENSMUSG00000060961 | Slc4a4 | 54403 | protein_coding | 1471.595806 | -0.246856093 | 0.080998845 | -3.047649545 | 0.002306387 | 0.015217953 | 0.246856093 |
| ENSMUSG00000045281 | Gpr20 | 239530 | protein_coding | 220.4445929 | -0.246863671 | 0.161721223 | -1.526476648 | 0.126891202 | 0.292110039 | 0.246863671 |
| ENSMUSG00000018381 | Abi3 | 66610 | protein_coding | 9.875423333 | -0.246863809 | 0.352939419 | -0.699450944 | 0.48427026 | 0.677689045 | 0.246863809 |
| ENSMUSG00000049233 | Apoo-ps | 621156 | processed_pseudogene | 9.568390497 | -0.246903909 | 0.356691465 | -0.692205824 | 0.488808082 | 0.681360016 | 0.246903909 |
| ENSMUSG00000000134 | Tfe3 | 209446 | protein_coding | 292.6476846 | -0.24690443 | 0.166104877 | -1.486436972 | 0.137163577 | 0.30805717 | 0.24690443 |
| ENSMUSG00000030748 | Il4ra | 16190 | protein_coding | 679.027481 | 0.246975712 | 0.1006628 | 2.453495355 | 0.014147537 | 0.061872603 | 0.246975712 |
| ENSMUSG00000040621 | Gemin8 | 237221 | protein_coding | 24.9179055 | 0.247029103 | 0.28090909 | 0.87939163 | 0.379188969 | 0.587178874 | 0.247029103 |
| ENSMUSG00000029156 | Sgcb | 24051 | protein_coding | 39.89766757 | -0.247035572 | 0.206231514 | -1.197855592 | 0.23097324 | 0.43419589 | 0.247035572 |
| ENSMUSG00000028849 | Map7d1 | 245877 | protein_coding | 115.3162142 | 0.247208733 | 0.271767463 | 0.909633295 | 0.363015933 | 0.571977227 | 0.247208733 |
| ENSMUSG00000079658 | Eloc | 67923 | protein_coding | 576.8591534 | 0.247315722 | 0.090007922 | 2.747710607 | 0.006001295 | 0.032217267 | 0.247315722 |
| ENSMUSG00000029385 | Ccng2 | 12452 | protein_coding | 1287.631583 | -0.247339615 | 0.204832521 | -1.207521222 | 0.227231477 | 0.429748631 | 0.247339615 |
| ENSMUSG00000032803 | Cdv3 | 321022 | protein_coding | 2449.814376 | 0.247366853 | 0.147604307 | 1.675878285 | 0.093762036 | 0.238454687 | 0.247366853 |
| ENSMUSG00000002608 | Ccdc97 | 52132 | protein_coding | 196.4417115 | -0.247511653 | 0.119395132 | -2.073046432 | 0.038167959 | 0.12887455 | 0.247511653 |
| ENSMUSG00000033416 | Gucd1 | 68778 | protein_coding | 1955.292292 | 0.247520562 | 0.121201401 | 2.042225254 | 0.041129192 | 0.135745085 | 0.247520562 |
| ENSMUSG00000007880 | Arid1a | 93760 | protein_coding | 613.4401241 | -0.247532223 | 0.185754368 | -1.332578206 | 0.182670261 | 0.371782765 | 0.247532223 |
| ENSMUSG00000033762 | Recql4 | 79456 | protein_coding | 58.77579111 | 0.247579852 | 0.180838883 | 1.369063159 | 0.170979533 | 0.356367705 | 0.247579852 |
| ENSMUSG00000039176 | Polg | 18975 | protein_coding | 849.6907511 | -0.247635624 | 0.096829369 | -2.557443315 | 0.010544474 | 0.049778419 | 0.247635624 |
| ENSMUSG00000039886 | Tmem120a | 215210 | protein_coding | 464.1469036 | 0.247643336 | 0.106923535 | 2.316078829 | 0.020553965 | 0.081601085 | 0.247643336 |
| ENSMUSG00000001467 | Cyp51 | 13121 | protein_coding | 2387.022281 | 0.24776879 | 0.146164804 | 1.695133052 | 0.090050184 | 0.23201542 | 0.24776879 |
| ENSMUSG00000017697 | Ada | 11486 | protein_coding | 106.1147134 | 0.247812239 | 0.130160419 | 1.9038986 | 0.056923391 | 0.169853087 | 0.247812239 |
| ENSMUSG00000027071 | P2rx3 | 228139 | protein_coding | 8.467255304 | -0.247866204 | 0.433231586 | -0.572133271 | 0.567231691 | 0.743121103 | 0.247866204 |
| ENSMUSG00000032010 | Usp2 | 53376 | protein_coding | 162.2885671 | -0.247894053 | 0.15670938 | -1.581871187 | 0.11367898 | 0.269807598 | 0.247894053 |

| ensembl_gene_id | external_gene_name | entrezgene_id | gene_biotype | baseMean | log2FoldChange | lfcSE | stat | pvalue | padj | absLog2FC |
|---|---|---|---|---|---|---|---|---|---|---|
| ENSMUSG00000024241 | Sos1 | 20662 | protein_coding | 540.9851988 | -0.247924207 | 0.134207387 | -1.84732161 | 0.064700543 | 0.185181391 | 0.247924207 |
| ENSMUSG00000039221 | Rpl22l1 | 68028 | protein_coding | 1069.55553 | 0.247998447 | 0.152643551 | 1.624689976 | 0.104228634 | 0.255423713 | 0.247998447 |
| ENSMUSG00000034210 | Efcab14 | 230648 | protein_coding | 968.736967 | -0.248110099 | 0.086827242 | -2.857514467 | 0.004269732 | 0.024757294 | 0.248110099 |
| ENSMUSG00000020961 | Ston2 | 108800 | protein_coding | 512.3611543 | -0.248215346 | 0.121924555 | -2.035810958 | 0.041769342 | 0.137062644 | 0.248215346 |
| ENSMUSG00000000916 | Nsun5 | 100609 | protein_coding | 100.5101487 | -0.248328814 | 0.191407719 | -1.297381396 | 0.19449999 | 0.387730172 | 0.248328814 |
| ENSMUSG00000003573 | Homer3 | 26558 | protein_coding | 19.72505582 | -0.248373479 | 0.290415782 | -0.855234096 | 0.392421564 | 0.600604465 | 0.248373479 |
| ENSMUSG00000028974 | Dffa | 13347 | protein_coding | 254.6187592 | -0.248400206 | 0.089336785 | -2.78049189 | 0.005427662 | 0.029759653 | 0.248400206 |
| ENSMUSG00000032135 | Mcam | 84004 | protein_coding | 8.320593248 | -0.248438765 | 0.485876424 | -0.51132089 | 0.60912638 | 0.772403867 | 0.248438765 |
| ENSMUSG00000020234 | Tektip1 | 432479 | protein_coding | 10.9766867 | -0.248470944 | 0.409381773 | -0.606941883 | 0.543889482 | 0.726645837 | 0.248470944 |
| ENSMUSG00000023473 | Celsr3 | 107934 | protein_coding | 19.08329308 | -0.248507393 | 0.430846971 | -0.57678807 | 0.56408263 | 0.740640354 | 0.248507393 |
| ENSMUSG00000002105 | Slc39a13 | 68427 | protein_coding | 85.43033139 | -0.248529963 | 0.139177929 | -1.785699538 | 0.07414792 | 0.2029556 | 0.248529963 |
| ENSMUSG00000030762 | Aqp8 | 11833 | protein_coding | 1544.933448 | 0.248642229 | 0.314665601 | 0.790179252 | 0.429423091 | 0.633442274 | 0.248642229 |
| ENSMUSG00000038967 | Pdk2 | 18604 | protein_coding | 443.6539591 | 0.248699852 | 0.130625286 | 1.903918141 | 0.056920846 | 0.169853087 | 0.248699852 |
| ENSMUSG00000021850 | ccdc198 | 67082 | protein_coding | 116.1211079 | 0.248720962 | 0.206617208 | 1.203776607 | 0.228675934 | 0.431660378 | 0.248720962 |
| ENSMUSG00000045314 | Sowahb | 78088 | protein_coding | 468.4013153 | -0.248743945 | 0.210094429 | -1.183962595 | 0.23642787 | 0.440601745 | 0.248743945 |
| ENSMUSG00000020385 | Clk4 | 12750 | protein_coding | 583.9795123 | -0.248747415 | 0.120309885 | -2.067555917 | 0.038681801 | 0.130176152 | 0.248747415 |
| ENSMUSG00000055707 | Klhl26 | 234378 | protein_coding | 168.392642 | -0.248897655 | 0.140126024 | -1.776241465 | 0.075693126 | 0.205338463 | 0.248897655 |
| ENSMUSG00000026754 | Golga1 | 76899 | protein_coding | 681.4256984 | 0.248916249 | 0.086447657 | 2.87938686 | 0.003984492 | 0.023401547 | 0.248916249 |
| ENSMUSG00000051695 | Pcbp1 | 23983 | protein_coding | 1156.926985 | 0.248998852 | 0.115729626 | 2.151556696 | 0.031432286 | 0.111827887 | 0.248998852 |
| ENSMUSG00000034487 | Poglut3 | 68304 | protein_coding | 288.4674668 | 0.249167067 | 0.124710767 | 1.997959537 | 0.045721047 | 0.146233903 | 0.249167067 |
| ENSMUSG00000037979 | Ccdc92 | 215707 | protein_coding | 26.14166319 | 0.249285181 | 0.251468148 | 0.991319111 | 0.321529784 | 0.532417756 | 0.249285181 |
| ENSMUSG00000020828 | Pld2 | 18806 | protein_coding | 45.33898433 | -0.249302018 | 0.254682886 | -0.978872285 | 0.327643088 | 0.538721067 | 0.249302018 |
| ENSMUSG00000089706 | B230216N24Rik | 78603 | lncRNA | 16.63833337 | -0.249333612 | 0.427329896 | -0.583468685 | 0.559577832 | 0.737621487 | 0.249333612 |
| ENSMUSG00000018287 | Spag7 | 216873 | protein_coding | 415.1401406 | 0.249370854 | 0.079411846 | 3.140222355 | 0.001688196 | 0.01174218 | 0.249370854 |
| ENSMUSG00000073471 | Rsph3a | 66832 | protein_coding | 139.8102345 | 0.249375193 | 0.107469385 | 2.320430073 | 0.020317624 | 0.080823953 | 0.249375193 |
| ENSMUSG00000050064 | Zfp697 | 242109 | protein_coding | 71.43753747 | -0.249471938 | 0.179388218 | -1.390681846 | 0.164321925 | 0.347643734 | 0.249471938 |
| ENSMUSG00000025497 | Cdhr5 | 72040 | protein_coding | 5454.930251 | -0.249518099 | 0.115402643 | -2.162152378 | 0.030606433 | 0.109673625 | 0.249518099 |
| ENSMUSG00000035011 | Zbtb7a | 16969 | protein_coding | 1951.02172 | -0.249665501 | 0.134394954 | -1.85769996 | 0.063211634 | 0.182100725 | 0.249665501 |
| ENSMUSG00000032602 | Slc25a20 | 57279 | protein_coding | 2063.089011 | 0.249739752 | 0.170106426 | 1.468138257 | 0.142066672 | 0.315334162 | 0.249739752 |
| ENSMUSG00000041650 | Pcca | 110821 | protein_coding | 628.5694541 | -0.249745748 | 0.110123228 | -2.267875292 | 0.023336809 | 0.089698043 | 0.249745748 |
| ENSMUSG00000015711 | Prune1 | 229589 | protein_coding | 272.0056843 | -0.249747105 | 0.078864905 | -3.166771162 | 0.001541416 | 0.010892488 | 0.249747105 |
| ENSMUSG00000025571 | Tnrc6c | 217351 | protein_coding | 541.8743834 | -0.24977132 | 0.097027757 | -2.574225446 | 0.010046479 | 0.048054148 | 0.24977132 |
| ENSMUSG00000031827 | Cotl1 | 72042 | protein_coding | 688.811716 | -0.24982338 | 0.13537549 | -1.845410719 | 0.064977815 | 0.185602458 | 0.24982338 |
| ENSMUSG00000016637 | Ift27 | 67042 | protein_coding | 107.3537026 | 0.249843558 | 0.118091308 | 2.115681185 | 0.034371932 | 0.119325161 | 0.249843558 |
| ENSMUSG00000058385 | H2bc8 | 319181 | protein_coding | 1276.176515 | 0.2499068 | 0.140698653 | 1.776184733 | 0.075702474 | 0.205338463 | 0.2499068 |
| ENSMUSG00000033060 | Lmo7 | 380928 | protein_coding | 3975.551409 | -0.249910515 | 0.226273971 | -1.104459845 | 0.269393713 | 0.477331318 | 0.249910515 |
| ENSMUSG00000024421 | Lama3 | 16774 | protein_coding | 795.857437 | 0.249955101 | 0.254232637 | 0.983174718 | 0.325521456 | 0.536573088 | 0.249955101 |
| ENSMUSG00000028010 | Gar1 | 68147 | protein_coding | 142.2972022 | -0.250066174 | 0.223640472 | -1.118161538 | 0.263498008 | 0.471131469 | 0.250066174 |

| ensembl_gene_id | external_gene_name | entrezgene_id | gene_biotype | baseMean | log2FoldChange | lfcSE | stat | pvalue | padj | absLog2FC |
|---|---|---|---|---|---|---|---|---|---|---|
| ENSMUSG00000020720 | Psmd12 | 66997 | protein_coding | 923.3001128 | 0.250227863 | 0.122411702 | 2.044149853 | 0.040938745 | 0.135312791 | 0.250227863 |
| ENSMUSG00000021087 | Rtn1 | 104001 | protein_coding | 8.329381525 | -0.250252782 | 0.538079055 | -0.465085529 | 0.641870208 | 0.792729067 | 0.250252782 |
| ENSMUSG00000103088 | Pcdhgb6 | 93703 | protein_coding | 33.5606092 | 0.250377629 | 0.191343806 | 1.308522258 | 0.19069623 | 0.382834579 | 0.250377629 |
| ENSMUSG00000034557 | Zfyve9 | 230597 | protein_coding | 612.1848919 | -0.250414858 | 0.084718063 | -2.95586146 | 0.003117971 | 0.019354265 | 0.250414858 |
| ENSMUSG00000023961 | Enpp4 | 224794 | protein_coding | 249.9577294 | -0.250414943 | 0.110916894 | -2.257680817 | 0.023965566 | 0.091562374 | 0.250414943 |
| ENSMUSG00000053675 | Tgm5 | 74176 | protein_coding | 8.923767833 | 0.250451134 | 0.428240944 | 0.584836964 | 0.558657347 | 0.736907711 | 0.250451134 |
| ENSMUSG00000021950 | Anxa8 | 11752 | protein_coding | 194.9712576 | -0.250464448 | 0.292918229 | -0.855066101 | 0.392514555 | 0.600621715 | 0.250464448 |
| ENSMUSG00000068101 | Cenpm | 66570 | protein_coding | 46.2439595 | 0.25051085 | 0.223872944 | 1.118986714 | 0.263145807 | 0.470933719 | 0.25051085 |
| ENSMUSG00000011960 | Ccnt1 | 12455 | protein_coding | 757.198169 | -0.250522512 | 0.11999009 | -2.087860012 | 0.036810464 | 0.125334988 | 0.250522512 |
| ENSMUSG00000036986 | Pml | 18854 | protein_coding | 357.7949267 | -0.250530365 | 0.102508701 | -2.443991225 | 0.01452578 | 0.062956904 | 0.250530365 |
| ENSMUSG00000017307 | Acot8 | 170789 | protein_coding | 391.9351194 | 0.250535515 | 0.106929617 | 2.342994588 | 0.019129661 | 0.077491667 | 0.250535515 |
| ENSMUSG00000047473 | Zfp30 | 22693 | protein_coding | 17.50318448 | -0.250539491 | 0.364698088 | -0.686977803 | 0.492096716 | 0.683894642 | 0.250539491 |
| ENSMUSG00000017221 | Psmd3 | 22123 | protein_coding | 1248.876281 | 0.250599073 | 0.105233826 | 2.381354754 | 0.017249092 | 0.071703093 | 0.250599073 |
| ENSMUSG00000050605 | Zfp61 | 22719 | protein_coding | 64.32921157 | 0.250669211 | 0.239251653 | 1.047721961 | 0.294766726 | 0.50397643 | 0.250669211 |
| ENSMUSG00000043671 | Dpy19l3 | 233115 | protein_coding | 86.58868199 | -0.250691055 | 0.183415466 | -1.366793436 | 0.171690061 | 0.357194265 | 0.250691055 |
| ENSMUSG00000023456 | Tpi1 | 21991 | protein_coding | 5939.661388 | 0.250699267 | 0.102103992 | 2.455332664 | 0.014075427 | 0.061710266 | 0.250699267 |
| ENSMUSG00000023829 | Slc22a1 | 20517 | protein_coding | 453.0434308 | -0.250702249 | 0.116590454 | -2.150281089 | 0.031532987 | 0.112057438 | 0.250702249 |
| ENSMUSG00000030007 | Cct7 | 12468 | protein_coding | 2589.82406 | 0.250704427 | 0.114232988 | 2.194676264 | 0.028186829 | 0.102996055 | 0.250704427 |
| ENSMUSG00000010476 | Ebf3 | 13593 | protein_coding | 12.57782746 | 0.25082415 | 0.560525465 | 0.447480383 | 0.654528249 | 0.800101761 | 0.25082415 |
| ENSMUSG00000036835 | Psenen | 66340 | protein_coding | 274.5499848 | 0.251015954 | 0.126340291 | 1.986824251 | 0.046941876 | 0.14896269 | 0.251015954 |
| ENSMUSG00000027761 | Aadac | 67758 | protein_coding | 834.8266265 | -0.251035297 | 0.147707812 | -1.699539738 | 0.089217534 | 0.230541367 | 0.251035297 |
| ENSMUSG00000097080 | 1700086O06Rik | 73516 | lncRNA | 23.71691427 | -0.251118866 | 0.254141883 | -0.988105 | 0.323101227 | 0.534020582 | 0.251118866 |
| ENSMUSG00000000738 | Spg7 | 234847 | protein_coding | 1102.13957 | -0.251198965 | 0.100964083 | -2.488003238 | 0.012846256 | 0.057808688 | 0.251198965 |
| ENSMUSG00000005610 | Eif4g2 | 13690 | protein_coding | 9676.214488 | 0.251201343 | 0.083268824 | 3.016751419 | 0.002554992 | 0.016549784 | 0.251201343 |
| ENSMUSG00000030798 | Cd37 | 12493 | protein_coding | 10.73529944 | -0.25121971 | 0.456419761 | -0.550413744 | 0.582035624 | 0.753102705 | 0.25121971 |
| ENSMUSG00000021504 | B4galt7 | 218271 | protein_coding | 253.3917818 | 0.251299707 | 0.079225292 | 3.171963147 | 0.001514122 | 0.010749598 | 0.251299707 |
| ENSMUSG00000027122 | Arl14ep | 212772 | protein_coding | 254.2493274 | 0.251401231 | 0.084986171 | 2.958142796 | 0.003094987 | 0.019249104 | 0.251401231 |
| ENSMUSG00000030516 | Tjp1 | 21872 | protein_coding | 1767.523733 | -0.251413759 | 0.107895432 | -2.330161281 | 0.019797629 | 0.07915076 | 0.251413759 |
| ENSMUSG00000029649 | Pomp | 66537 | protein_coding | 1016.027428 | 0.251446747 | 0.107729609 | 2.334054209 | 0.019592884 | 0.078608426 | 0.251446747 |
| ENSMUSG00000025195 | Dnmbp | 71972 | protein_coding | 2142.063641 | -0.251451619 | 0.081441687 | -3.087505019 | 0.002018444 | 0.013605489 | 0.251451619 |
| ENSMUSG00000040124 | Gorab | 98376 | protein_coding | 322.1745432 | 0.251452142 | 0.114535143 | 2.19541474 | 0.028133861 | 0.102873309 | 0.251452142 |
| ENSMUSG00000010453 | Kansl3 | 226976 | protein_coding | 955.1350907 | -0.251507814 | 0.071819354 | -3.501950378 | 0.000461866 | 0.004075676 | 0.251507814 |
| ENSMUSG00000056602 | Fry | 320365 | protein_coding | 272.4619716 | -0.251584814 | 0.147029582 | -1.711116977 | 0.087059521 | 0.227050359 | 0.251584814 |
| ENSMUSG00000057396 | Zfp759 | 268670 | protein_coding | 155.3211874 | 0.251715456 | 0.197697862 | 1.27323307 | 0.202935349 | 0.397926165 | 0.251715456 |
| ENSMUSG00000068394 | Cep152 | 99100 | protein_coding | 197.8190665 | -0.251719425 | 0.144835571 | -1.73796688 | 0.08221665 | 0.218221429 | 0.251719425 |
| ENSMUSG00000021798 | Ldb3 | 24131 | protein_coding | 180.0052321 | 0.251926747 | 0.115736699 | 2.176723099 | 0.029501234 | 0.106528146 | 0.251926747 |
| ENSMUSG00000032328 | Tmem30a | 69981 | protein_coding | 2452.989232 | 0.252206566 | 0.078133808 | 3.227880138 | 0.001247112 | 0.009150884 | 0.252206566 |
| ENSMUSG00000086213 | A330040F15Rik | 74333 | lncRNA | 7.909847037 | -0.252217576 | 0.456894121 | -0.552026312 | 0.580930326 | 0.752036593 | 0.252217576 |

| ensembl_gene_id | external_gene_name | entrezgene_id | gene_biotype | baseMean | log2FoldChange | lfcSE | stat | pvalue | padj | absLog2FC |
|---|---|---|---|---|---|---|---|---|---|---|
| ENSMUSG00000060798 | Intu | 380614 | protein_coding | 145.71292 | 0.252254235 | 0.136732098 | 1.844879434 | 0.065055079 | 0.185724755 | 0.252254235 |
| ENSMUSG00000044465 | Fhip1b | 74349 | protein_coding | 836.0445392 | -0.25237847 | 0.138494487 | -1.822299753 | 0.068409516 | 0.192513563 | 0.25237847 |
| ENSMUSG00000026341 | Actr3 | 74117 | protein_coding | 5058.032622 | 0.252427972 | 0.088585874 | 2.84952849 | 0.004378408 | 0.025238355 | 0.252427972 |
| ENSMUSG00000038611 | Phrf1 | 101471 | protein_coding | 846.4767892 | -0.252452514 | 0.09810292 | -2.573343523 | 0.010072118 | 0.048133422 | 0.252452514 |
| ENSMUSG00000022016 | Akap11 | 219181 | protein_coding | 989.9315417 | -0.252530657 | 0.090982438 | -2.775597832 | 0.005510032 | 0.030128317 | 0.252530657 |
| ENSMUSG00000014633 | Cmc2 | 66531 | protein_coding | 173.7898795 | 0.252536135 | 0.120327033 | 2.098748121 | 0.03583911 | 0.12315973 | 0.252536135 |
| ENSMUSG00000047907 | Tshz2 | 228911 | protein_coding | 205.5553191 | -0.25254252 | 0.108604238 | -2.325346829 | 0.020053422 | 0.079938547 | 0.25254252 |
| ENSMUSG00000003154 | Foxj2 | 60611 | protein_coding | 203.4691908 | -0.252560795 | 0.128644586 | -1.963244653 | 0.049617753 | 0.154534306 | 0.252560795 |
| ENSMUSG00000022013 | Dnajc15 | 66148 | protein_coding | 450.4570297 | 0.252590448 | 0.0783097 | 3.225532049 | 0.001257388 | 0.009203892 | 0.252590448 |
| ENSMUSG00000074671 | Tspyl3 | 241732 | protein_coding | 8.602047514 | -0.252607042 | 0.396653756 | -0.636845202 | 0.524225679 | 0.70998769 | 0.252607042 |
| ENSMUSG00000039879 | Heca | 380629 | protein_coding | 1076.352012 | -0.252610107 | 0.112281569 | -2.249791386 | 0.024462191 | 0.09288052 | 0.252610107 |
| ENSMUSG00000022003 | Slc25a30 | 67554 | protein_coding | 69.98531325 | -0.252619228 | 0.135742923 | -1.861012144 | 0.062742462 | 0.181128045 | 0.252619228 |
| ENSMUSG00000031907 | Zfp90 | 22751 | protein_coding | 88.88395116 | -0.252633553 | 0.162982929 | -1.550061442 | 0.12112677 | 0.282889617 | 0.252633553 |
| ENSMUSG00000000876 | Pxmp4 | 59038 | protein_coding | 452.9279541 | 0.252652785 | 0.139459452 | 1.811657663 | 0.070039109 | 0.195681974 | 0.252652785 |
| ENSMUSG00000040740 | Slc25a34 | 384071 | protein_coding | 458.4491803 | -0.252662765 | 0.168754256 | -1.497223071 | 0.134335224 | 0.303287236 | 0.252662765 |
| ENSMUSG00000038564 | Ift172 | 67661 | protein_coding | 430.0017368 | 0.252671171 | 0.073066923 | 3.458078731 | 0.000544042 | 0.004671432 | 0.252671171 |
| ENSMUSG00000035891 | Cerk | 223753 | protein_coding | 372.5538628 | -0.252865546 | 0.084389942 | -2.996394364 | 0.002731929 | 0.017412629 | 0.252865546 |
| ENSMUSG00000035164 | Zc3h12c | 244871 | protein_coding | 49.0076728 | 0.252865861 | 0.199710622 | 1.266161303 | 0.205455339 | 0.401139082 | 0.252865861 |
| ENSMUSG00000036766 | Dner | 227325 | protein_coding | 8.181372177 | -0.252959332 | 0.544761073 | -0.464349134 | 0.642397627 | 0.793002356 | 0.252959332 |
| ENSMUSG00000036304 | Zdhhc23 | 332175 | protein_coding | 276.1226293 | -0.253084973 | 0.108418773 | -2.334327951 | 0.019578556 | 0.078608426 | 0.253084973 |
| ENSMUSG00000031906 | Smpd3 | 58994 | protein_coding | 2089.232172 | 0.253209833 | 0.086096198 | 2.941010633 | 0.003271433 | 0.020126452 | 0.253209833 |
| ENSMUSG00000044906 | 4930503L19Rik | 269033 | protein_coding | 8.751516049 | 0.253216472 | 0.473751433 | 0.534492255 | 0.593001007 | 0.7614941 | 0.253216472 |
| ENSMUSG00000041353 | Tmem29 | 382245 | protein_coding | 133.4914573 | 0.253262006 | 0.105502013 | 2.400541924 | 0.016370815 | 0.069048936 | 0.253262006 |
| ENSMUSG00000038020 | Rapgefl1 | 268480 | protein_coding | 740.4364686 | -0.253268777 | 0.205816875 | -1.230553991 | 0.218489722 | 0.41848296 | 0.253268777 |
| ENSMUSG00000026601 | Axdnd1 | 77352 | protein_coding | 9.959212318 | -0.253361226 | 0.556531489 | -0.455250478 | 0.648929049 | 0.797283028 | 0.253361226 |
| ENSMUSG00000097431 | Gm26782 | 102640338 | lncRNA | 36.34640892 | -0.253404664 | 0.227247063 | -1.115106447 | 0.264804809 | 0.472220655 | 0.253404664 |
| ENSMUSG00000010721 | Lmbr1 | 56873 | protein_coding | 322.5069454 | -0.253507949 | 0.087847416 | -2.885775835 | 0.003904501 | 0.023033667 | 0.253507949 |
| ENSMUSG00000049119 | Fam110b | 242297 | protein_coding | 20.8140918 | -0.253563482 | 0.313737541 | -0.808202553 | 0.418973988 | 0.625144514 | 0.253563482 |
| ENSMUSG00000034317 | Trim59 | 66949 | protein_coding | 257.2379055 | 0.25358892 | 0.120352095 | 2.107058633 | 0.035112497 | 0.121317106 | 0.25358892 |
| ENSMUSG00000067714 | Lpar5 | 381810 | protein_coding | 83.65275062 | -0.253619939 | 0.162976443 | -1.556175449 | 0.119666378 | 0.280537081 | 0.253619939 |
| ENSMUSG00000055013 | Agap1 | 347722 | protein_coding | 823.7445125 | -0.253654153 | 0.083272991 | -3.046055503 | 0.00231865 | 0.015286201 | 0.253654153 |
| ENSMUSG00000030663 | 1110004F10Rik | 56372 | protein_coding | 685.050051 | 0.253762939 | 0.081110452 | 3.128609597 | 0.001756355 | 0.012147296 | 0.253762939 |
| ENSMUSG00000053769 | Lysmd1 | 217779 | protein_coding | 23.22632575 | -0.253895079 | 0.242920971 | -1.045175628 | 0.295941806 | 0.505172123 | 0.253895079 |
| ENSMUSG00000040213 | Kyat3 | 229905 | protein_coding | 128.6186918 | 0.253959896 | 0.128749784 | 1.972507357 | 0.048551717 | 0.152651353 | 0.253959896 |
| ENSMUSG00000056770 | Setd3 | 52690 | protein_coding | 2166.048204 | 0.254006973 | 0.06943393 | 3.658254312 | 0.000253939 | 0.00248302 | 0.254006973 |
| ENSMUSG00000022372 | Sla | 20491 | protein_coding | 19.59553395 | 0.254026971 | 0.393573188 | 0.645437696 | 0.518643582 | 0.705762919 | 0.254026971 |
| ENSMUSG00000024097 | Srsf7 | 225027 | protein_coding | 1666.639411 | 0.254032708 | 0.102281331 | 2.483666434 | 0.013003755 | 0.058377625 | 0.254032708 |
| ENSMUSG00000035601 | Trmt10b | 69934 | protein_coding | 166.0778335 | 0.254050513 | 0.12043819 | 2.109385011 | 0.034911362 | 0.120753133 | 0.254050513 |

| ensembl_gene_id | external_gene_name | entrezgene_id | gene_biotype | baseMean | log2FoldChange | lfcSE | stat | pvalue | padj | absLog2FC |
|---|---|---|---|---|---|---|---|---|---|---|
| ENSMUSG00000032657 | Entrep3 | 68521 | protein_coding | 21.13723204 | -0.254051505 | 0.296047311 | -0.858144953 | 0.390812428 | 0.599060828 | 0.254051505 |
| ENSMUSG00000024483 | Ankhd1 | 108857 | protein_coding | 1334.3891 | 0.25406604 | 0.105276418 | 2.413323373 | 0.015807788 | 0.067304582 | 0.25406604 |
| ENSMUSG00000017718 | Afmid | 71562 | protein_coding | 57.62490396 | 0.254084201 | 0.197700246 | 1.285199214 | 0.198722678 | 0.392857741 | 0.254084201 |
| ENSMUSG00000044968 | Napepld | 242864 | protein_coding | 196.8194784 | -0.254117447 | 0.217884829 | -1.166292523 | 0.243496195 | 0.448578232 | 0.254117447 |
| ENSMUSG00000019992 | Mtfr2 | 71804 | protein_coding | 33.36879025 | 0.254140557 | 0.244052637 | 1.041335015 | 0.297720091 | 0.506633078 | 0.254140557 |
| ENSMUSG00000097727 | F630040K05Rik | 433111 | lncRNA | 9.351497944 | -0.254147111 | 0.695699864 | -0.365311428 | 0.714878949 | 0.839719166 | 0.254147111 |
| ENSMUSG00000030056 | Isy1 | 57905 | protein_coding | 215.9538233 | 0.254187175 | 0.101613999 | 2.501497593 | 0.012366928 | 0.056119782 | 0.254187175 |
| ENSMUSG00000020072 | Pbld2 | 67307 | protein_coding | 336.8593441 | -0.25421533 | 0.122003175 | -2.083677986 | 0.037189468 | 0.12630146 | 0.25421533 |
| ENSMUSG00000039987 | Phtf2 | 68770 | protein_coding | 84.99203906 | -0.254281893 | 0.1762407 | -1.44281028 | 0.149073922 | 0.326384802 | 0.254281893 |
| ENSMUSG00000039477 | Tnrc18 | 231861 | protein_coding | 1044.130912 | -0.254301568 | 0.142562251 | -1.783793159 | 0.074457282 | 0.203492722 | 0.254301568 |
| ENSMUSG00000047228 | A2ml1 | 232400 | protein_coding | 14.79013842 | -0.254389349 | 0.494185668 | -0.514764725 | 0.606717438 | 0.771129536 | 0.254389349 |
| ENSMUSG00000077394 | Gm24339 | 115487283 | snoRNA | 258.5438452 | -0.254403205 | 0.112689662 | -2.257555838 | 0.023973365 | 0.091570204 | 0.254403205 |
| ENSMUSG00000118506 | Cfap141 | 73545 | protein_coding | 7.687275203 | 0.254506801 | 0.465277643 | 0.546999851 | 0.584378834 | 0.754966832 | 0.254506801 |
| ENSMUSG00000046982 | Tshz1 | 110796 | protein_coding | 812.1454697 | -0.254523357 | 0.07007851 | -3.631974453 | 0.000281261 | 0.002692262 | 0.254523357 |
| ENSMUSG00000044447 | Dock5 | 68813 | protein_coding | 1331.122591 | -0.254537517 | 0.179502237 | -1.418018634 | 0.156185325 | 0.336633268 | 0.254537517 |
| ENSMUSG00000004394 | Tmed4 | 103694 | protein_coding | 881.5727324 | 0.25459513 | 0.091169014 | 2.792562074 | 0.005229243 | 0.029031505 | 0.25459513 |
| ENSMUSG00000061313 | Ddhd2 | 72108 | protein_coding | 295.4995456 | -0.254620839 | 0.096383374 | -2.641750637 | 0.008247876 | 0.041468486 | 0.254620839 |
| ENSMUSG00000064127 | Med14 | 26896 | protein_coding | 808.1895336 | 0.254659559 | 0.077144487 | 3.301072691 | 0.000963159 | 0.007454422 | 0.254659559 |
| ENSMUSG00000052102 | Gnpda1 | 26384 | protein_coding | 171.4016451 | -0.254676371 | 0.150280542 | -1.694672957 | 0.090137479 | 0.232127689 | 0.254676371 |
| ENSMUSG00000023018 | Smarcd1 | 83797 | protein_coding | 138.9172362 | -0.254745795 | 0.112529092 | -2.263821653 | 0.023585086 | 0.09049947 | 0.254745795 |
| ENSMUSG00000025034 | Trim8 | 93679 | protein_coding | 430.5025543 | -0.254806729 | 0.137663226 | -1.850942589 | 0.064177813 | 0.184242305 | 0.254806729 |
| ENSMUSG00000075703 | Selenoi | 28042 | protein_coding | 1304.250391 | 0.254832473 | 0.091522136 | 2.78438073 | 0.005363004 | 0.02952711 | 0.254832473 |
| ENSMUSG00000040749 | Siah1b | 20438 | protein_coding | 60.51790834 | 0.254861571 | 0.189866764 | 1.342317982 | 0.179492915 | 0.367192802 | 0.254861571 |
| ENSMUSG00000029082 | Bst1 | 12182 | protein_coding | 7.161956898 | -0.254862622 | 0.731102424 | -0.348600433 | 0.727389303 | 0.846662573 | 0.254862622 |
| ENSMUSG00000034247 | Plekhm1 | 353047 | protein_coding | 417.6461548 | -0.255061927 | 0.129885579 | -1.96374323 | 0.049559877 | 0.154449171 | 0.255061927 |
| ENSMUSG00000038990 | Cables2 | 252966 | protein_coding | 356.5997094 | 0.255073624 | 0.116247484 | 2.194229196 | 0.028218937 | 0.103042463 | 0.255073624 |
| ENSMUSG00000033713 | Foxn3 | 71375 | protein_coding | 1074.187397 | -0.255073974 | 0.140644721 | -1.813605042 | 0.069738554 | 0.195227236 | 0.255073974 |
| ENSMUSG00000030873 | Scnn1b | 20277 | protein_coding | 177.7268172 | 0.255197441 | 0.190447515 | 1.339988296 | 0.18024915 | 0.368125204 | 0.255197441 |
| ENSMUSG00000048732 | Klhl11 | 217194 | protein_coding | 179.3034785 | -0.255215701 | 0.146390408 | -1.743390873 | 0.081265361 | 0.216345425 | 0.255215701 |
| ENSMUSG00000019082 | Slc25a22 | 68267 | protein_coding | 201.9254186 | -0.255224522 | 0.169458292 | -1.506119992 | 0.132036373 | 0.299882412 | 0.255224522 |
| ENSMUSG00000011884 | Gltp | 56356 | protein_coding | 628.3407256 | 0.255261207 | 0.085808115 | 2.974790988 | 0.002931882 | 0.018424685 | 0.255261207 |
| ENSMUSG00000042625 | Safb2 | 224902 | protein_coding | 716.8942775 | -0.255268896 | 0.125418361 | -2.03533911 | 0.041816764 | 0.13716174 | 0.255268896 |
| ENSMUSG00000034112 | Atp2c2 | 69047 | protein_coding | 993.1200419 | -0.255359705 | 0.156480195 | -1.631897921 | 0.102700992 | 0.252792675 | 0.255359705 |
| ENSMUSG00000041328 | Pcf11 | 74737 | protein_coding | 1740.296549 | -0.255380762 | 0.10916074 | -2.339492784 | 0.019309945 | 0.077944449 | 0.255380762 |
| ENSMUSG00000003546 | Klc4 | 74764 | protein_coding | 1873.90429 | 0.255397741 | 0.105610167 | 2.418306375 | 0.015592942 | 0.066589104 | 0.255397741 |
| ENSMUSG00000021733 | Slc4a7 | 218756 | protein_coding | 4874.807828 | 0.255416587 | 0.11533914 | 2.214483199 | 0.026795557 | 0.099278817 | 0.255416587 |
| ENSMUSG00000022656 | Nectin3 | 58998 | protein_coding | 2189.454709 | 0.25542388 | 0.0672574 | 3.797706734 | 0.000146041 | 0.001561168 | 0.25542388 |
| ENSMUSG00000064289 | Tank | 21353 | protein_coding | 545.6961984 | 0.255445776 | 0.084369421 | 3.027705684 | 0.00246418 | 0.016132122 | 0.255445776 |

| ensembl_gene_id | external_gene_name | entrezgene_id | gene_biotype | baseMean | log2FoldChange | lfcSE | stat | pvalue | padj | absLog2FC |
|---|---|---|---|---|---|---|---|---|---|---|
| ENSMUSG00000039461 | Tcta | 102791 | protein_coding | 469.1837885 | 0.255450191 | 0.152976392 | 1.669866752 | 0.09494573 | 0.240199426 | 0.255450191 |
| ENSMUSG00000032726 | Bmp8a | 12163 | protein_coding | 73.56737672 | -0.255497819 | 0.133321856 | -1.916398603 | 0.055314379 | 0.166360918 | 0.255497819 |
| ENSMUSG00000022351 | Sqle | 20775 | protein_coding | 1688.991851 | 0.255540875 | 0.138198197 | 1.849089784 | 0.064444849 | 0.184669522 | 0.255540875 |
| ENSMUSG00000040433 | Zbtb38 | 245007 | protein_coding | 936.1677075 | -0.255618895 | 0.099563692 | -2.567390693 | 0.010246709 | 0.048675687 | 0.255618895 |
| ENSMUSG00000024019 | Cmtr1 | 74157 | protein_coding | 52.81033242 | -0.255679828 | 0.174357018 | -1.466415468 | 0.142535135 | 0.316152646 | 0.255679828 |
| ENSMUSG00000025572 | Tmc6 | 217353 | protein_coding | 915.7145603 | 0.255705677 | 0.096223514 | 2.657413623 | 0.007874277 | 0.03998202 | 0.255705677 |
| ENSMUSG00000024851 | Pitpnm1 | 18739 | protein_coding | 318.7415159 | -0.255812337 | 0.182733696 | -1.399918798 | 0.161537636 | 0.343612643 | 0.255812337 |
| ENSMUSG00000028007 | Snx7 | 76561 | protein_coding | 1030.171189 | 0.25588746 | 0.12859858 | 1.989815593 | 0.046611253 | 0.148306839 | 0.25588746 |
| ENSMUSG00000028648 | Ndufs5 | 595136 | protein_coding | 349.3268875 | 0.255926037 | 0.134411551 | 1.904047948 | 0.05690394 | 0.169853087 | 0.255926037 |
| ENSMUSG00000037426 | Depdc5 | 277854 | protein_coding | 370.6335223 | -0.256001567 | 0.111011883 | -2.306073542 | 0.021106523 | 0.083214036 | 0.256001567 |
| ENSMUSG00000044390 | Tigd3 | 332359 | protein_coding | 6.810591165 | -0.256003936 | 0.524225564 | -0.488346913 | 0.625304138 | 0.782883533 | 0.256003936 |
| ENSMUSG00000040883 | Tmem205 | 235043 | protein_coding | 312.0302031 | -0.256039471 | 0.114928496 | -2.227815382 | 0.025892829 | 0.097063068 | 0.256039471 |
| ENSMUSG00000040703 | Cyp2s1 | 74134 | protein_coding | 956.4480233 | -0.256039479 | 0.115485522 | -2.217069939 | 0.026618313 | 0.09885206 | 0.256039479 |
| ENSMUSG00000047843 | Bri3 | 55950 | protein_coding | 215.1910426 | -0.25607538 | 0.095735968 | -2.674808496 | 0.007477195 | 0.038418309 | 0.25607538 |
| ENSMUSG00000038729 | Pakap | 677884 | protein_coding | 14.89418521 | -0.256124872 | 0.389620088 | -0.657370808 | 0.510942517 | 0.699008761 | 0.256124872 |
| ENSMUSG00000028873 | Cdca8 | 52276 | protein_coding | 324.5869787 | 0.256188129 | 0.140353092 | 1.825311623 | 0.067954017 | 0.191604104 | 0.256188129 |
| ENSMUSG00000027111 | Itga6 | 16403 | protein_coding | 5338.650482 | -0.256202771 | 0.196774388 | -1.302012797 | 0.192912011 | 0.386163443 | 0.256202771 |
| ENSMUSG00000073131 | Vma21 | 67048 | protein_coding | 326.1807077 | 0.256270776 | 0.126782368 | 2.021343972 | 0.04324417 | 0.140369502 | 0.256270776 |
| ENSMUSG00000022018 | Rgcc | 66214 | protein_coding | 109.7057924 | -0.256280699 | 0.154574248 | -1.657977977 | 0.097321912 | 0.243900172 | 0.256280699 |
| ENSMUSG00000030541 | Idh2 | 269951 | protein_coding | 1656.318518 | -0.256364155 | 0.097726281 | -2.623287725 | 0.008708569 | 0.043211867 | 0.256364155 |
| ENSMUSG00000071551 | Akr1c19 | 432720 | protein_coding | 4320.846637 | -0.256405414 | 0.181871411 | -1.409817038 | 0.158593714 | 0.339860149 | 0.256405414 |
| ENSMUSG00000006931 | P3h4 | 66180 | protein_coding | 42.78179521 | -0.256439419 | 0.202582143 | -1.265854015 | 0.205565352 | 0.401304686 | 0.256439419 |
| ENSMUSG00000040659 | Efhd2 | 27984 | protein_coding | 1413.110203 | -0.256493192 | 0.15006038 | -1.709266575 | 0.087401582 | 0.227509788 | 0.256493192 |
| ENSMUSG00000049882 | Vcpkmt | 207965 | protein_coding | 62.30556788 | -0.256523903 | 0.190600205 | -1.345874227 | 0.178343082 | 0.3656389 | 0.256523903 |
| ENSMUSG00000038733 | Wdr26 | 226757 | protein_coding | 1759.422625 | -0.256630805 | 0.079745568 | -3.218119969 | 0.001290338 | 0.009406183 | 0.256630805 |
| ENSMUSG00000020792 | Exoc7 | 53413 | protein_coding | 603.0555174 | 0.25665793 | 0.090329596 | 2.841349246 | 0.004492309 | 0.025776282 | 0.25665793 |
| ENSMUSG00000052031 | Tagap1 | 380608 | protein_coding | 131.8639253 | -0.256669489 | 0.135145846 | -1.89920369 | 0.0575377 | 0.171329048 | 0.256669489 |
| ENSMUSG00000001062 | Vps9d1 | 72325 | protein_coding | 207.4108065 | -0.256747821 | 0.105835831 | -2.425906421 | 0.015270207 | 0.065520384 | 0.256747821 |
| ENSMUSG00000038147 | Cd84 | 12523 | protein_coding | 12.80019182 | -0.256777107 | 0.563323334 | -0.45582544 | 0.648515509 | 0.797091758 | 0.256777107 |
| ENSMUSG00000025220 | Oga | 76055 | protein_coding | 2105.503388 | -0.256782535 | 0.098914261 | -2.596011271 | 0.009431297 | 0.045827243 | 0.256782535 |
| ENSMUSG00000034875 | Nudt19 | 110959 | protein_coding | 1135.851214 | 0.256962478 | 0.120924294 | 2.124986385 | 0.033587749 | 0.117269984 | 0.256962478 |
| ENSMUSG00000019659 | Ccdc12 | 72654 | protein_coding | 601.0241317 | 0.256994867 | 0.114320777 | 2.248015398 | 0.024575208 | 0.093220747 | 0.256994867 |
| ENSMUSG00000062585 | Cnr2 | 12802 | protein_coding | 7.084743815 | -0.257038904 | 0.419510465 | -0.612711542 | 0.540067089 | 0.723156983 | 0.257038904 |
| ENSMUSG00000005161 | Prdx2 | 21672 | protein_coding | 2419.345988 | 0.257039348 | 0.096563221 | 2.661876284 | 0.007770643 | 0.039571347 | 0.257039348 |
| ENSMUSG00000020847 | Rph3al | 380714 | protein_coding | 307.9871002 | 0.257078609 | 0.13551922 | 1.896990032 | 0.057829254 | 0.171719678 | 0.257078609 |
| ENSMUSG00000030956 | Fam53b | 77938 | protein_coding | 308.6606046 | -0.257125735 | 0.144769229 | -1.776107647 | 0.075715176 | 0.205338463 | 0.257125735 |
| ENSMUSG00000034667 | Xpot | 73192 | protein_coding | 846.7545065 | 0.257126617 | 0.108459517 | 2.370715122 | 0.017753709 | 0.073164574 | 0.257126617 |
| ENSMUSG00000011658 | Fuz | 70300 | protein_coding | 72.52138668 | -0.257312405 | 0.149869533 | -1.716909365 | 0.085995733 | 0.224953201 | 0.257312405 |

| ensembl_gene_id | external_gene_name | entrezgene_id | gene_biotype | baseMean | log2FoldChange | lfcSE | stat | pvalue | padj | absLog2FC |
|---|---|---|---|---|---|---|---|---|---|---|
| ENSMUSG00000030512 | Snrpa1 | 68981 | protein_coding | 316.3762546 | 0.257363096 | 0.120881113 | 2.129059611 | 0.033249329 | 0.116423875 | 0.257363096 |
| ENSMUSG00000037108 | Zcwpw1 | 381678 | protein_coding | 16.96755734 | -0.257385682 | 0.279706183 | -0.920200185 | 0.357468157 | 0.567163344 | 0.257385682 |
| ENSMUSG00000006289 | Osgep | 66246 | protein_coding | 378.3542017 | 0.257460724 | 0.108907262 | 2.36403634 | 0.018077039 | 0.074132616 | 0.257460724 |
| ENSMUSG00000044098 | Rsbn1 | 229675 | protein_coding | 364.7461946 | -0.257510515 | 0.101661434 | -2.533020678 | 0.011308428 | 0.052467417 | 0.257510515 |
| ENSMUSG00000048779 | P2ry6 | 233571 | protein_coding | 149.7509263 | -0.257562455 | 0.14524896 | -1.773248186 | 0.076187591 | 0.206310089 | 0.257562455 |
| ENSMUSG00000036854 | Hspb6 | 243912 | protein_coding | 24.81917734 | 0.257583222 | 0.241664079 | 1.065873021 | 0.286481046 | 0.495501152 | 0.257583222 |
| ENSMUSG00000101355 | H3c10 | 319152 | protein_coding | 332.2715672 | 0.257636725 | 0.279179392 | 0.922835754 | 0.356092805 | 0.565657346 | 0.257636725 |
| ENSMUSG00000019647 | Sema6a | 20358 | protein_coding | 276.7086557 | -0.257821157 | 0.136018251 | -1.895489431 | 0.058027591 | 0.172116101 | 0.257821157 |
| ENSMUSG00000027379 | Bub1 | 12235 | protein_coding | 313.8990379 | 0.25793488 | 0.134621901 | 1.915994933 | 0.055365741 | 0.166470253 | 0.25793488 |
| ENSMUSG00000032386 | Trip4 | 56404 | protein_coding | 487.2736187 | 0.257941151 | 0.087376166 | 2.95207677 | 0.003156444 | 0.019539777 | 0.257941151 |
| ENSMUSG00000071866 | Ppia | 268373 | protein_coding | 6607.7015 | 0.258131783 | 0.104527584 | 2.469508755 | 0.013529871 | 0.060028909 | 0.258131783 |
| ENSMUSG00000025314 | Ptprj | 19271 | protein_coding | 1522.322839 | -0.258168362 | 0.077747048 | -3.320619487 | 0.000898179 | 0.007019723 | 0.258168362 |
| ENSMUSG00000036026 | Tmem63b | 224807 | protein_coding | 885.973385 | -0.258173295 | 0.067936049 | -3.800240048 | 0.000144556 | 0.001547371 | 0.258173295 |
| ENSMUSG00000041319 | Thoc6 | 386612 | protein_coding | 167.3523545 | 0.258204181 | 0.142097309 | 1.817094098 | 0.069202706 | 0.194128339 | 0.258204181 |
| ENSMUSG00000030228 | Pik3c2g | 18705 | protein_coding | 32.59802735 | -0.258270022 | 0.31358349 | -0.82360848 | 0.410162056 | 0.617106327 | 0.258270022 |
| ENSMUSG00000028670 | Lypla2 | 26394 | protein_coding | 734.0540798 | -0.25832859 | 0.113543764 | -2.275145558 | 0.022897199 | 0.088538846 | 0.25832859 |
| ENSMUSG00000041115 | Iqsec2 | 245666 | protein_coding | 197.4013061 | -0.258451398 | 0.167137938 | -1.546335923 | 0.12202346 | 0.284449738 | 0.258451398 |
| ENSMUSG00000005417 | Mprip | 26936 | protein_coding | 2013.476245 | -0.258495441 | 0.08704943 | -2.969524797 | 0.002982607 | 0.018696171 | 0.258495441 |
| ENSMUSG00000044167 | Foxo1 | 56458 | protein_coding | 248.4833549 | -0.258526227 | 0.138512254 | -1.866450224 | 0.061978402 | 0.179611609 | 0.258526227 |
| ENSMUSG00000036867 | Smad6 | 17130 | protein_coding | 231.4069526 | -0.25853991 | 0.253990113 | -1.017913282 | 0.308719169 | 0.518602032 | 0.25853991 |
| ENSMUSG00000025630 | Hprt | 15452 | protein_coding | 1423.884067 | 0.258561138 | 0.097771567 | 2.644543246 | 0.008180125 | 0.041245026 | 0.258561138 |
| ENSMUSG00000024592 | C330018D20Rik | 77422 | protein_coding | 260.9971252 | 0.258569031 | 0.133070474 | 1.943098437 | 0.052004266 | 0.159753895 | 0.258569031 |
| ENSMUSG00000028217 | Cdh17 | 12557 | protein_coding | 12119.406 | 0.258600508 | 0.151142234 | 1.710974496 | 0.087085821 | 0.227050359 | 0.258600508 |
| ENSMUSG00000027130 | Slc12a6 | 107723 | protein_coding | 497.5818006 | -0.258619344 | 0.157280834 | -1.644315692 | 0.10011101 | 0.248336421 | 0.258619344 |
| ENSMUSG00000055733 | Nap1l3 | 54561 | protein_coding | 12.89136266 | -0.258629355 | 0.431047138 | -0.600002489 | 0.548504577 | 0.730300167 | 0.258629355 |
| ENSMUSG00000020527 | Myo19 | 66196 | protein_coding | 641.4801763 | -0.258683825 | 0.133716757 | -1.934565502 | 0.053043637 | 0.161761257 | 0.258683825 |
| ENSMUSG00000035829 | Ppp1r26 | 241289 | protein_coding | 79.78925079 | -0.258855764 | 0.161176342 | -1.606040693 | 0.108264969 | 0.261882891 | 0.258855764 |
| ENSMUSG00000029094 | Afap1 | 70292 | protein_coding | 176.3121405 | 0.258880303 | 0.14570064 | 1.776795919 | 0.075601824 | 0.205239445 | 0.258880303 |
| ENSMUSG00000047040 | Prr15l | 217138 | protein_coding | 3555.730784 | 0.258882173 | 0.149704251 | 1.729290726 | 0.083757076 | 0.220874619 | 0.258882173 |
| ENSMUSG00000033257 | Ttll4 | 67534 | protein_coding | 458.3813653 | -0.258899206 | 0.094499248 | -2.739695954 | 0.006149604 | 0.032847006 | 0.258899206 |
| ENSMUSG00000054312 | Mrps21 | 66292 | protein_coding | 585.0987599 | 0.25890093 | 0.100825399 | 2.567814575 | 0.010234188 | 0.048630714 | 0.25890093 |
| ENSMUSG00000039396 | Neil3 | 234258 | protein_coding | 128.9883591 | -0.259001851 | 0.206593241 | -1.25368018 | 0.209958275 | 0.407185609 | 0.259001851 |
| ENSMUSG00000047632 | Fgfbp3 | 72514 | protein_coding | 13.32941712 | -0.259264513 | 0.376850976 | -0.687976229 | 0.491467751 | 0.683557312 | 0.259264513 |
| ENSMUSG00000020778 | Ten1 | 69535 | protein_coding | 228.6169254 | 0.259280184 | 0.094751116 | 2.736434097 | 0.006210903 | 0.033031818 | 0.259280184 |
| ENSMUSG00000032239 | Rp9 | 55934 | protein_coding | 431.9922509 | -0.259304731 | 0.088471141 | -2.930952722 | 0.003379242 | 0.020662022 | 0.259304731 |
| ENSMUSG00000071064 | Zfp827 | 622675 | protein_coding | 323.2911488 | -0.259317698 | 0.08952698 | -2.896531298 | 0.00377313 | 0.022458302 | 0.259317698 |
| ENSMUSG00000015697 | Setdb1 | 84505 | protein_coding | 609.9839351 | -0.259348109 | 0.065853294 | -3.938270839 | 8.21E-05 | 0.000949329 | 0.259348109 |
| ENSMUSG00000084798 | 4930533B01Rik | 75166 | lncRNA | 8.336896123 | -0.259403243 | 0.436046273 | -0.594898431 | 0.551911366 | 0.732326245 | 0.259403243 |

| ensembl_gene_id | external_gene_name | entrezgene_id | gene_biotype | baseMean | log2FoldChange | lfcSE | stat | pvalue | padj | absLog2FC |
|---|---|---|---|---|---|---|---|---|---|---|
| ENSMUSG00000028138 | Adh5 | 11532 | protein_coding | 1707.622723 | -0.259450415 | 0.148668747 | -1.745157746 | 0.080957414 | 0.215886437 | 0.259450415 |
| ENSMUSG00000028012 | Rrh | 20132 | protein_coding | 6.801611611 | -0.259501113 | 0.534851562 | -0.485183425 | 0.627546239 | 0.784362476 | 0.259501113 |
| ENSMUSG00000037652 | Phc3 | 241915 | protein_coding | 596.4753735 | -0.259543447 | 0.184650666 | -1.40559172 | 0.159845396 | 0.341274462 | 0.259543447 |
| ENSMUSG00000025956 | Mettl21a | 67099 | protein_coding | 130.6799012 | 0.259555533 | 0.137211826 | 1.891641123 | 0.058538813 | 0.173020265 | 0.259555533 |
| ENSMUSG00000050912 | Tmem123 | 71929 | protein_coding | 456.6096947 | 0.25965032 | 0.09382104 | 2.767506295 | 0.005648695 | 0.030707311 | 0.25965032 |
| ENSMUSG00000045216 | Hs6st1 | 50785 | protein_coding | 1037.364917 | 0.259667137 | 0.102841766 | 2.524919085 | 0.011572493 | 0.053443511 | 0.259667137 |
| ENSMUSG00000008604 | Ubqln4 | 94232 | protein_coding | 629.7689087 | -0.259735648 | 0.065133954 | -3.987715015 | 6.67E-05 | 0.000791804 | 0.259735648 |
| ENSMUSG00000028271 | Gtf2b | 229906 | protein_coding | 593.8127769 | 0.259761128 | 0.091517387 | 2.838380076 | 0.004534315 | 0.025932703 | 0.259761128 |
| ENSMUSG00000063052 | Lrrc40 | 67144 | protein_coding | 523.2737033 | 0.259767927 | 0.081072571 | 3.204140729 | 0.001354662 | 0.009821149 | 0.259767927 |
| ENSMUSG00000021737 | Psmd6 | 66413 | protein_coding | 1014.524879 | 0.259817972 | 0.138845134 | 1.871278912 | 0.061306432 | 0.178191396 | 0.259817972 |
| ENSMUSG00000047115 | Fam221a | 231946 | protein_coding | 58.94451649 | -0.260008738 | 0.200927073 | -1.29404532 | 0.195649767 | 0.388961796 | 0.260008738 |
| ENSMUSG00000030120 | Mlf2 | 30853 | protein_coding | 1929.901512 | 0.260069723 | 0.101016964 | 2.574515337 | 0.010038064 | 0.048042752 | 0.260069723 |
| ENSMUSG00000030861 | Acadsb | 66885 | protein_coding | 423.8342537 | -0.260123374 | 0.095877247 | -2.71308764 | 0.006665947 | 0.035041324 | 0.260123374 |
| ENSMUSG00000052062 | Pard3b | 72823 | protein_coding | 531.4295498 | -0.260135921 | 0.136249349 | -1.909263588 | 0.056228097 | 0.168156427 | 0.260135921 |
| ENSMUSG00000000056 | Narf | 67608 | protein_coding | 684.7441843 | 0.260184424 | 0.150418453 | 1.729737407 | 0.083677202 | 0.220777788 | 0.260184424 |
| ENSMUSG00000031503 | Col4a2 | 12827 | protein_coding | 9.977803124 | -0.26023823 | 0.464129028 | -0.560702336 | 0.575000475 | 0.748007805 | 0.26023823 |
| ENSMUSG00000009585 | Apobec3 | 80287 | protein_coding | 253.8643697 | -0.26026485 | 0.130989962 | -1.986906831 | 0.046932723 | 0.14896269 | 0.26026485 |
| ENSMUSG00000058833 | Rex1bd | 66462 | protein_coding | 250.4222301 | 0.260284473 | 0.127908503 | 2.034927053 | 0.041858213 | 0.137241174 | 0.260284473 |
| ENSMUSG00000093606 | B130034C11Rik | 414095 | lncRNA | 12.48392398 | 0.260285906 | 0.385235992 | 0.675653136 | 0.499260897 | 0.689758658 | 0.260285906 |
| ENSMUSG00000051339 | 2900026A02Rik | 243219 | protein_coding | 2674.857447 | -0.260323577 | 0.119484611 | -2.178720556 | 0.029352434 | 0.106162093 | 0.260323577 |
| ENSMUSG00000041506 | Rrp9 | 27966 | protein_coding | 233.5453717 | 0.260428638 | 0.245924675 | 1.058977258 | 0.289610137 | 0.498385347 | 0.260428638 |
| ENSMUSG00000005225 | Plekha8 | 231999 | protein_coding | 1537.197238 | -0.260429369 | 0.077617073 | -3.355310355 | 0.00079276 | 0.006329664 | 0.260429369 |
| ENSMUSG00000030533 | Unc45a | 101869 | protein_coding | 607.8941145 | -0.26046864 | 0.115963119 | -2.24613344 | 0.024695462 | 0.093565384 | 0.26046864 |
| ENSMUSG00000069267 | H3c2 | 319150 | protein_coding | 809.1205031 | 0.260560768 | 0.186750736 | 1.395232881 | 0.162945625 | 0.345459967 | 0.260560768 |
| ENSMUSG00000019428 | Fkbp8 | 14232 | protein_coding | 2909.562467 | 0.260608422 | 0.095707367 | 2.722971401 | 0.006469766 | 0.034236023 | 0.260608422 |
| ENSMUSG00000022974 | Paxbp1 | 67367 | protein_coding | 517.8125383 | -0.260610285 | 0.128105562 | -2.034340125 | 0.041917314 | 0.137293642 | 0.260610285 |
| ENSMUSG00000000439 | Mkrn2 | 67027 | protein_coding | 226.1167157 | -0.260698837 | 0.101128232 | -2.577903652 | 0.009940171 | 0.047759252 | 0.260698837 |
| ENSMUSG00000018848 | Rars | 104458 | protein_coding | 1318.009424 | 0.260771469 | 0.118754564 | 2.195885875 | 0.028100113 | 0.102820721 | 0.260771469 |
| ENSMUSG00000028603 | Scp2 | 20280 | protein_coding | 2438.569823 | 0.260773682 | 0.079841976 | 3.266122586 | 0.00109031 | 0.008238638 | 0.260773682 |
| ENSMUSG00000027080 | Med19 | 381379 | protein_coding | 360.0118739 | 0.260809857 | 0.100871653 | 2.585561437 | 0.009722054 | 0.046896694 | 0.260809857 |
| ENSMUSG00000089661 | Mia | 12587 | protein_coding | 8.917494646 | 0.260931045 | 0.371268327 | 0.70280987 | 0.482174246 | 0.67549148 | 0.260931045 |
| ENSMUSG00000062929 | Cfl2 | 12632 | protein_coding | 327.8978029 | -0.260934226 | 0.11067254 | -2.357714263 | 0.01838784 | 0.075059334 | 0.260934226 |
| ENSMUSG00000023963 | Cyp39a1 | 56050 | protein_coding | 338.0944741 | -0.260969771 | 0.126397647 | -2.064672699 | 0.038953979 | 0.130677966 | 0.260969771 |
| ENSMUSG00000039200 | Atf7ip2 | 75329 | protein_coding | 21.19989969 | -0.261045798 | 0.311012957 | -0.839340587 | 0.401278213 | 0.608871318 | 0.261045798 |
| ENSMUSG00000037887 | Dusp8 | 18218 | protein_coding | 146.9929995 | -0.261083269 | 0.228495727 | -1.142617733 | 0.25319734 | 0.459383441 | 0.261083269 |
| ENSMUSG00000051379 | Flrt3 | 71436 | protein_coding | 1117.435298 | 0.26116274 | 0.332531554 | 0.785377318 | 0.432232376 | 0.636108037 | 0.26116274 |
| ENSMUSG00000097047 | 1110020A21Rik | 68531 | lncRNA | 27.91849998 | 0.261198774 | 0.249071476 | 1.048690033 | 0.294320803 | 0.50359241 | 0.261198774 |
| ENSMUSG00000021675 | F2rl2 | 14064 | protein_coding | 19.3832473 | 0.261202162 | 0.379435728 | 0.688396327 | 0.491203236 | 0.683368429 | 0.261202162 |

| ensembl_gene_id | external_gene_name | entrezgene_id | gene_biotype | baseMean | log2FoldChange | lfcSE | stat | pvalue | padj | absLog2FC |
|---|---|---|---|---|---|---|---|---|---|---|
| ENSMUSG00000023994 | Nfya | 18044 | protein_coding | 356.0472085 | -0.261251426 | 0.124765623 | -2.093937575 | 0.036265542 | 0.124139798 | 0.261251426 |
| ENSMUSG00000025528 | 2010106E10Rik | 67715 | protein_coding | 30.41667136 | 0.261345575 | 0.403983815 | 0.646920905 | 0.517683135 | 0.704835436 | 0.261345575 |
| ENSMUSG00000020766 | Galk1 | 14635 | protein_coding | 320.8377685 | 0.261345831 | 0.154481605 | 1.691760198 | 0.090691705 | 0.232877234 | 0.261345831 |
| ENSMUSG00000031626 | Sorbs2 | 234214 | protein_coding | 727.4715904 | -0.261346092 | 0.216353283 | -1.20795991 | 0.227062683 | 0.429544279 | 0.261346092 |
| ENSMUSG00000068040 | Tm9sf4 | 99237 | protein_coding | 2640.6424 | 0.26151199 | 0.068780065 | 3.802148073 | 0.000143447 | 0.001536532 | 0.26151199 |
| ENSMUSG00000027569 | Mrgbp | 73247 | protein_coding | 256.0286259 | -0.26152051 | 0.111508306 | -2.345300714 | 0.019011739 | 0.077112039 | 0.26152051 |
| ENSMUSG00000031889 | Phaf1 | 234678 | protein_coding | 218.186387 | -0.261583759 | 0.080163985 | -3.263108206 | 0.001101974 | 0.00830679 | 0.261583759 |
| ENSMUSG00000037148 | Arhgap10 | 78514 | protein_coding | 42.47672873 | 0.261606993 | 0.211467559 | 1.237102246 | 0.216049126 | 0.41510621 | 0.261606993 |
| ENSMUSG00000030315 | Vgll4 | 232334 | protein_coding | 675.3435404 | -0.261611498 | 0.088612873 | -2.95229677 | 0.003154196 | 0.019533452 | 0.261611498 |
| ENSMUSG00000028716 | Pdzk1ip1 | 67182 | protein_coding | 460.0257633 | -0.261709578 | 0.21781149 | -1.201541654 | 0.229541159 | 0.432847922 | 0.261709578 |
| ENSMUSG00000032579 | Hemk1 | 69536 | protein_coding | 94.54758309 | -0.261722032 | 0.135031429 | -1.938230487 | 0.052595109 | 0.160824514 | 0.261722032 |
| ENSMUSG00000027132 | Katnbl1 | 72425 | protein_coding | 262.2772686 | 0.261741191 | 0.128940802 | 2.029933018 | 0.042363348 | 0.138299531 | 0.261741191 |
| ENSMUSG00000041341 | Atg2b | 76559 | protein_coding | 686.3666172 | -0.261796102 | 0.074023955 | -3.536640288 | 0.000405251 | 0.003673786 | 0.261796102 |
| ENSMUSG00000031399 | Fam3a | 66294 | protein_coding | 409.1578447 | 0.26194078 | 0.087299164 | 3.000495858 | 0.002695404 | 0.017241292 | 0.26194078 |
| ENSMUSG00000068587 | Mgam | 232714 | protein_coding | 7590.093872 | 0.262020298 | 0.105825655 | 2.475961978 | 0.013287773 | 0.05925186 | 0.262020298 |
| ENSMUSG00000038712 | Mindy1 | 75007 | protein_coding | 1203.221374 | -0.262059126 | 0.070582095 | -3.712827247 | 0.000204957 | 0.002087789 | 0.262059126 |
| ENSMUSG00000028977 | Casz1 | 69743 | protein_coding | 940.0424564 | -0.262106655 | 0.134958292 | -1.942130801 | 0.052121268 | 0.159960328 | 0.262106655 |
| ENSMUSG00000036863 | Syde2 | 214804 | protein_coding | 242.4081548 | -0.262224593 | 0.142830099 | -1.8359197 | 0.066369534 | 0.188336706 | 0.262224593 |
| ENSMUSG00000047187 | Rab2a | 59021 | protein_coding | 2841.591299 | 0.262265591 | 0.074284225 | 3.530569128 | 0.000414667 | 0.003742102 | 0.262265591 |
| ENSMUSG00000025574 | Tk1 | 21877 | protein_coding | 423.5459379 | 0.262413697 | 0.121105289 | 2.166822755 | 0.030248372 | 0.108659464 | 0.262413697 |
| ENSMUSG00000029260 | Ugt2b34 | 100727 | protein_coding | 9877.973047 | 0.262533206 | 0.101685205 | 2.581823058 | 0.009827996 | 0.047321741 | 0.262533206 |
| ENSMUSG00000060402 | Chst8 | 68947 | protein_coding | 13.24449947 | -0.262546494 | 0.34201898 | -0.767637206 | 0.442702752 | 0.643949429 | 0.262546494 |
| ENSMUSG00000005686 | Ampd3 | 11717 | protein_coding | 696.8274873 | -0.262547758 | 0.125955938 | -2.084441287 | 0.037120046 | 0.126146381 | 0.262547758 |
| ENSMUSG00000028992 | Nmnat1 | 66454 | protein_coding | 148.2686418 | 0.262550256 | 0.101989705 | 2.574281941 | 0.010044838 | 0.048054148 | 0.262550256 |
| ENSMUSG00000097357 | Gm16793 | 100504714 | lncRNA | 7.88104489 | -0.262601218 | 0.419348388 | -0.626212537 | 0.531175545 | 0.716450469 | 0.262601218 |
| ENSMUSG00000047181 | Samd14 | 217125 | protein_coding | 24.82449528 | -0.262630899 | 0.252201173 | -1.041354788 | 0.297710918 | 0.506633078 | 0.262630899 |
| ENSMUSG00000015671 | Psma2 | 19166 | protein_coding | 1670.156393 | 0.262637934 | 0.103910633 | 2.527536673 | 0.011486583 | 0.053139207 | 0.262637934 |
| ENSMUSG00000049643 | 2310022A10Rik | 66367 | protein_coding | 63.77520797 | -0.262642588 | 0.170313256 | -1.542114773 | 0.123045706 | 0.285966245 | 0.262642588 |
| ENSMUSG00000024414 | Mrpl27 | 94064 | protein_coding | 389.2029599 | 0.262662189 | 0.105267884 | 2.495178772 | 0.012589369 | 0.056902231 | 0.262662189 |
| ENSMUSG00000055305 | Zfp93 | 22755 | protein_coding | 101.9616969 | 0.2627082 | 0.173764177 | 1.511866285 | 0.130567883 | 0.297522923 | 0.2627082 |
| ENSMUSG00000087819 | Gm25117 | 115487522 | snoRNA | 45.8890102 | -0.262882354 | 0.283786016 | -0.926340055 | 0.354269286 | 0.563829056 | 0.262882354 |
| ENSMUSG00000002297 | Dbf4 | 27214 | protein_coding | 481.026683 | 0.262897234 | 0.111927266 | 2.348822093 | 0.018832903 | 0.07652308 | 0.262897234 |
| ENSMUSG00000032405 | Pias1 | 56469 | protein_coding | 333.6173947 | -0.262977721 | 0.084050392 | -3.128810151 | 0.001755157 | 0.012144281 | 0.262977721 |
| ENSMUSG00000022582 | Ly6g | 546644 | protein_coding | 100.7731926 | 0.263023842 | 0.34525242 | 0.761830551 | 0.446161144 | 0.646923504 | 0.263023842 |
| ENSMUSG00000033237 | Arid2 | 77044 | protein_coding | 712.6399087 | -0.26303444 | 0.09954184 | -2.642451053 | 0.008230836 | 0.041435132 | 0.26303444 |
| ENSMUSG00000052560 | Cpne8 | 66871 | protein_coding | 292.5037175 | 0.263037792 | 0.121016549 | 2.173568778 | 0.029737535 | 0.107162775 | 0.263037792 |
| ENSMUSG00000030591 | Psmd8 | 57296 | protein_coding | 1848.746818 | 0.263154354 | 0.094371976 | 2.788479853 | 0.005295604 | 0.029272813 | 0.263154354 |
| ENSMUSG00000031666 | Rbl2 | 19651 | protein_coding | 606.2692639 | -0.263159226 | 0.122778772 | -2.143360946 | 0.032084127 | 0.113374214 | 0.263159226 |

| ensembl_gene_id | external_gene_name | entrezgene_id | gene_biotype | baseMean | log2FoldChange | lfcSE | stat | pvalue | padj | absLog2FC |
|---|---|---|---|---|---|---|---|---|---|---|
| ENSMUSG00000003119 | Cdk12 | 69131 | protein_coding | 944.294804 | -0.263192456 | 0.084263814 | -3.123433919 | 0.00178754 | 0.012335011 | 0.263192456 |
| ENSMUSG00000058006 | Mdn1 | 100019 | protein_coding | 603.1354935 | -0.263216513 | 0.197083422 | -1.335558877 | 0.181693499 | 0.370302264 | 0.263216513 |
| ENSMUSG00000022686 | B3gnt5 | 108105 | protein_coding | 1538.904868 | 0.263375485 | 0.266891462 | 0.986826194 | 0.323727851 | 0.534723345 | 0.263375485 |
| ENSMUSG00000050856 | Atp5k | 11958 | protein_coding | 2615.057392 | 0.263410746 | 0.109355833 | 2.408748925 | 0.016007306 | 0.067918053 | 0.263410746 |
| ENSMUSG00000001909 | Trmt1 | 212528 | protein_coding | 517.702603 | -0.263477026 | 0.103301758 | -2.55055703 | 0.010755093 | 0.050533072 | 0.263477026 |
| ENSMUSG00000061482 | H4c4 | 319156 | protein_coding | 3240.196857 | 0.263667068 | 0.109457748 | 2.408847906 | 0.016002966 | 0.067917728 | 0.263667068 |
| ENSMUSG00000027499 | Pkia | 18767 | protein_coding | 138.1985643 | 0.263686575 | 0.137802567 | 1.913509891 | 0.055682809 | 0.167162066 | 0.263686575 |
| ENSMUSG00000028051 | Hcn3 | 15168 | protein_coding | 14.15504286 | -0.263716164 | 0.336697227 | -0.783244243 | 0.433483699 | 0.637295787 | 0.263716164 |
| ENSMUSG00000003500 | Impdh1 | 23917 | protein_coding | 287.2042119 | -0.26372719 | 0.141590621 | -1.862603522 | 0.062518068 | 0.180690944 | 0.26372719 |
| ENSMUSG00000029918 | Mrps33 | 14548 | protein_coding | 136.05891 | 0.263736939 | 0.126878136 | 2.078663413 | 0.037648298 | 0.127587679 | 0.263736939 |
| ENSMUSG00000026259 | Ngef | 53972 | protein_coding | 633.8112631 | -0.263772867 | 0.130887613 | -2.015262252 | 0.043877178 | 0.141731026 | 0.263772867 |
| ENSMUSG00000041598 | Cdc42ep4 | 56699 | protein_coding | 1005.294891 | -0.263850584 | 0.140157787 | -1.882525327 | 0.059764724 | 0.175536148 | 0.263850584 |
| ENSMUSG00000031955 | Bcar1 | 12927 | protein_coding | 787.6124045 | -0.263873778 | 0.104653403 | -2.521406565 | 0.011688671 | 0.053777342 | 0.263873778 |
| ENSMUSG00000038894 | Irs2 | 384783 | protein_coding | 592.1807751 | -0.263934457 | 0.180463928 | -1.462533039 | 0.143595198 | 0.317603783 | 0.263934457 |
| ENSMUSG00000048534 | Jaml | 270152 | protein_coding | 35.13560723 | 0.264041607 | 0.257325555 | 1.02609944 | 0.304844714 | 0.514469868 | 0.264041607 |
| ENSMUSG00000024007 | Ppil1 | 68816 | protein_coding | 377.9847106 | 0.264045865 | 0.123215696 | 2.142956408 | 0.032116599 | 0.113451583 | 0.264045865 |
| ENSMUSG00000022742 | Cpox | 12892 | protein_coding | 391.5047532 | 0.264113044 | 0.095373382 | 2.769253215 | 0.005618495 | 0.030553564 | 0.264113044 |
| ENSMUSG00000027272 | Ubr1 | 22222 | protein_coding | 538.923082 | -0.264153127 | 0.089433431 | -2.953628466 | 0.003140619 | 0.01947208 | 0.264153127 |
| ENSMUSG00000033467 | Crlf2 | 57914 | protein_coding | 35.80505784 | 0.264159284 | 0.210703521 | 1.253701327 | 0.209950585 | 0.407185609 | 0.264159284 |
| ENSMUSG00000026437 | Cdk18 | 18557 | protein_coding | 300.4768769 | 0.264164628 | 0.133536688 | 1.97821761 | 0.047904163 | 0.151272305 | 0.264164628 |
| ENSMUSG00000022337 | Emc2 | 66736 | protein_coding | 746.3269585 | 0.264177264 | 0.080884299 | 3.26611304 | 0.001090347 | 0.008238638 | 0.264177264 |
| ENSMUSG00000062248 | Cks2 | 66197 | protein_coding | 494.2244663 | 0.264186776 | 0.116430617 | 2.269049006 | 0.023265346 | 0.08954201 | 0.264186776 |
| ENSMUSG00000029056 | Pank4 | 269614 | protein_coding | 282.4314025 | -0.26420998 | 0.093337393 | -2.830698105 | 0.004644654 | 0.026506645 | 0.26420998 |
| ENSMUSG00000006731 | B4galnt1 | 14421 | protein_coding | 1120.114741 | 0.264255909 | 0.144655392 | 1.826796116 | 0.06773043 | 0.191244512 | 0.264255909 |
| ENSMUSG00000019883 | Echdc1 | 52665 | protein_coding | 213.1104183 | 0.264257395 | 0.10599201 | 2.493182234 | 0.012660385 | 0.05714214 | 0.264257395 |
| ENSMUSG00000020381 | Mrnip | 68067 | protein_coding | 20.85650752 | -0.264302291 | 0.269252531 | -0.98161488 | 0.326289614 | 0.537352722 | 0.264302291 |
| ENSMUSG00000037013 | Ss18 | 268996 | protein_coding | 1002.055157 | 0.26436821 | 0.078224334 | 3.37961598 | 0.000725872 | 0.005904844 | 0.26436821 |
| ENSMUSG00000022095 | Fhip2b | 239170 | protein_coding | 225.8118221 | -0.264424298 | 0.109925812 | -2.405479584 | 0.016151254 | 0.0683286 | 0.264424298 |
| ENSMUSG00000025348 | Itga7 | 16404 | protein_coding | 379.4679803 | -0.264467614 | 0.141509001 | -1.868910194 | 0.06163531 | 0.178853564 | 0.264467614 |
| ENSMUSG00000044149 | Nkrf | 77286 | protein_coding | 102.0307222 | -0.264696165 | 0.148733676 | -1.779665318 | 0.07513075 | 0.204526163 | 0.264696165 |
| ENSMUSG00000046324 | Ermp1 | 226090 | protein_coding | 2028.369065 | -0.26477638 | 0.166043648 | -1.594619151 | 0.110797427 | 0.265540389 | 0.26477638 |
| ENSMUSG00000044991 | Shld1 | 73747 | protein_coding | 61.77736387 | 0.264831648 | 0.174629958 | 1.516530446 | 0.129385278 | 0.295971378 | 0.264831648 |
| ENSMUSG00000005262 | Ufd1 | 22230 | protein_coding | 536.7933521 | 0.264847185 | 0.06368093 | 4.15897167 | 3.20E-05 | 0.000418056 | 0.264847185 |
| ENSMUSG00000042406 | Atf4 | 11911 | protein_coding | 3680.968507 | 0.264855273 | 0.148715321 | 1.780954853 | 0.074919826 | 0.204201575 | 0.264855273 |
| ENSMUSG00000035242 | Oaz1 | 18245 | protein_coding | 2430.304415 | 0.264864969 | 0.101310145 | 2.614397295 | 0.008938503 | 0.044019314 | 0.264864969 |
| ENSMUSG00000022216 | Psme1 | 19186 | protein_coding | 1366.2724 | 0.264923114 | 0.091954004 | 2.881039465 | 0.00396366 | 0.023313581 | 0.264923114 |
| ENSMUSG00000002910 | Arrdc2 | 70807 | protein_coding | 221.1554422 | 0.264936754 | 0.142219913 | 1.862866804 | 0.062481008 | 0.180616606 | 0.264936754 |
| ENSMUSG00000098923 | Tmem185b | 226351 | protein_coding | 191.2759416 | -0.265036044 | 0.088923773 | -2.980485811 | 0.002877916 | 0.018161427 | 0.265036044 |

| ensembl_gene_id | external_gene_name | entrezgene_id | gene_biotype | baseMean | log2FoldChange | lfcSE | stat | pvalue | padj | absLog2FC |
|---|---|---|---|---|---|---|---|---|---|---|
| ENSMUSG00000054280 | Prr14l | 215476 | protein_coding | 1184.072224 | -0.265038933 | 0.066019404 | -4.014561139 | 5.96E-05 | 0.000719193 | 0.265038933 |
| ENSMUSG00000007817 | Zmiz1 | 328365 | protein_coding | 596.3114926 | -0.265065708 | 0.191558522 | -1.383732268 | 0.166440449 | 0.350391683 | 0.265065708 |
| ENSMUSG00000117975 | Itprip | 414801 | protein_coding | 35.99907996 | 0.265142964 | 0.34023125 | 0.779302207 | 0.435801715 | 0.638764122 | 0.265142964 |
| ENSMUSG00000012609 | Ttll5 | 320244 | protein_coding | 299.903071 | 0.265143224 | 0.108190589 | 2.450705051 | 0.014257673 | 0.062136666 | 0.265143224 |
| ENSMUSG00000022554 | Hgh1 | 59053 | protein_coding | 91.25788344 | -0.265158475 | 0.21039882 | -1.260265982 | 0.207573427 | 0.404082965 | 0.265158475 |
| ENSMUSG00000029415 | Sdad1 | 231452 | protein_coding | 362.0491444 | -0.265301449 | 0.171112319 | -1.550452068 | 0.12103305 | 0.282753654 | 0.265301449 |
| ENSMUSG00000026199 | Ankzf1 | 52231 | protein_coding | 322.4665168 | -0.265319788 | 0.153882825 | -1.724167648 | 0.084677583 | 0.222601838 | 0.265319788 |
| ENSMUSG00000038351 | Sgsm2 | 97761 | protein_coding | 384.3427213 | -0.265477759 | 0.130755269 | -2.030340815 | 0.042321908 | 0.138233724 | 0.265477759 |
| ENSMUSG00000020329 | Polrmt | 216151 | protein_coding | 326.4428435 | -0.265512159 | 0.083718685 | -3.171480271 | 0.001516642 | 0.010760031 | 0.265512159 |
| ENSMUSG00000037126 | Psd | 73728 | protein_coding | 67.00757367 | -0.265521491 | 0.168631859 | -1.574563036 | 0.115357341 | 0.272752947 | 0.265521491 |
| ENSMUSG00000042129 | Rassf4 | 213391 | protein_coding | 348.4727801 | 0.265568224 | 0.196058105 | 1.35453836 | 0.175564684 | 0.361771253 | 0.265568224 |
| ENSMUSG00000025575 | Cant1 | 76025 | protein_coding | 3577.326221 | 0.265596867 | 0.106384745 | 2.496569104 | 0.012540123 | 0.056727942 | 0.265596867 |
| ENSMUSG00000021819 | Zswim8 | 268721 | protein_coding | 539.9911953 | -0.265705264 | 0.11539075 | -2.302656524 | 0.021298176 | 0.083782996 | 0.265705264 |
| ENSMUSG00000027806 | Tsc22d2 | 72033 | protein_coding | 444.8813329 | -0.265751878 | 0.151585627 | -1.753146938 | 0.079576788 | 0.213132518 | 0.265751878 |
| ENSMUSG00000024232 | Bambi | 68010 | protein_coding | 113.2979731 | -0.265761891 | 0.162467761 | -1.635782323 | 0.101885153 | 0.251367211 | 0.265761891 |
| ENSMUSG00000073436 | Eme2 | 193838 | protein_coding | 112.8074003 | -0.265794148 | 0.153662341 | -1.729728615 | 0.083678774 | 0.220777788 | 0.265794148 |
| ENSMUSG00000029863 | Casp2 | 12366 | protein_coding | 717.6849508 | -0.265970248 | 0.073336864 | -3.626692411 | 0.000287075 | 0.002734766 | 0.265970248 |
| ENSMUSG00000025076 | Casp7 | 12369 | protein_coding | 1997.034566 | 0.266168662 | 0.096592978 | 2.755569493 | 0.005859006 | 0.031581131 | 0.266168662 |
| ENSMUSG00000022708 | Zbtb20 | 56490 | protein_coding | 320.1043077 | -0.266210009 | 0.25753766 | -1.033674099 | 0.301288546 | 0.510586416 | 0.266210009 |
| ENSMUSG00000033985 | Tesk2 | 230661 | protein_coding | 348.6582312 | 0.266355237 | 0.129433404 | 2.05785546 | 0.03960401 | 0.132273575 | 0.266355237 |
| ENSMUSG00000024105 | Themis3 | 74556 | protein_coding | 595.1109792 | 0.266358663 | 0.105294836 | 2.529646011 | 0.011417766 | 0.052851546 | 0.266358663 |
| ENSMUSG00000042473 | Tbc1d8b | 245638 | protein_coding | 820.7788704 | 0.266380732 | 0.070753907 | 3.764890771 | 0.000166622 | 0.001740297 | 0.266380732 |
| ENSMUSG00000031805 | Jak3 | 16453 | protein_coding | 117.7062121 | 0.266486211 | 0.154718709 | 1.722391647 | 0.084998596 | 0.223151085 | 0.266486211 |
| ENSMUSG00000019189 | Rnf145 | 74315 | protein_coding | 1072.698054 | 0.266549969 | 0.112529178 | 2.368718706 | 0.017849823 | 0.073498484 | 0.266549969 |
| ENSMUSG00000021139 | Gm20498 | 105940408 | protein_coding | 9.423590659 | -0.266597601 | 0.382815299 | -0.696413133 | 0.486170141 | 0.678983247 | 0.266597601 |
| ENSMUSG00000003134 | Tbc1d8 | 54610 | protein_coding | 239.0582169 | -0.26661322 | 0.124323829 | -2.144506171 | 0.031992352 | 0.113163263 | 0.26661322 |
| ENSMUSG00000025035 | Arl3 | 56350 | protein_coding | 59.72184885 | -0.266643579 | 0.193288234 | -1.379512726 | 0.167736725 | 0.35225587 | 0.266643579 |
| ENSMUSG00000043015 | Nemp2 | 227094 | protein_coding | 71.18674785 | -0.266800545 | 0.180969915 | -1.474281212 | 0.140405892 | 0.312781125 | 0.266800545 |
| ENSMUSG00000031451 | Gas6 | 14456 | protein_coding | 4185.873555 | -0.266800799 | 0.10909748 | -2.445526683 | 0.014464075 | 0.0628091 | 0.266800799 |
| ENSMUSG00000049577 | Zfpm1 | 22761 | protein_coding | 53.2434921 | 0.266843009 | 0.240192873 | 1.110953065 | 0.266588553 | 0.474438264 | 0.266843009 |
| ENSMUSG00000034282 | Evpl | 14027 | protein_coding | 674.7559794 | -0.266847692 | 0.136372337 | -1.956758222 | 0.050375901 | 0.156197655 | 0.266847692 |
| ENSMUSG00000029596 | Sdsl | 257635 | protein_coding | 205.7338673 | 0.266916051 | 0.103206361 | 2.586236424 | 0.009703035 | 0.046819128 | 0.266916051 |
| ENSMUSG00000025086 | Trub1 | 72133 | protein_coding | 130.0999754 | -0.266946939 | 0.105750918 | -2.524299028 | 0.011592927 | 0.0534901 | 0.266946939 |
| ENSMUSG00000019214 | Chtf18 | 214901 | protein_coding | 95.2899956 | 0.266983081 | 0.192985091 | 1.38343889 | 0.166530333 | 0.350441953 | 0.266983081 |
| ENSMUSG00000029790 | Cep41 | 83922 | protein_coding | 68.09731012 | -0.267000832 | 0.139664751 | -1.91172669 | 0.055911259 | 0.167567854 | 0.267000832 |
| ENSMUSG00000042606 | Hirip3 | 233876 | protein_coding | 129.7904541 | -0.267005572 | 0.180048675 | -1.482963269 | 0.13808417 | 0.309078789 | 0.267005572 |
| ENSMUSG00000027333 | Smox | 228608 | protein_coding | 273.2728812 | -0.26705934 | 0.194459233 | -1.373343585 | 0.169645559 | 0.354608197 | 0.26705934 |
| ENSMUSG00000045636 | Mtus1 | 102103 | protein_coding | 3007.715239 | -0.267224803 | 0.102883787 | -2.597346089 | 0.009394721 | 0.04570529 | 0.267224803 |

| ensembl_gene_id | external_gene_name | entrezgene_id | gene_biotype | baseMean | log2FoldChange | lfcSE | stat | pvalue | padj | absLog2FC |
|---|---|---|---|---|---|---|---|---|---|---|
| ENSMUSG00000022812 | Gsk3b | 56637 | protein_coding | 1231.364754 | 0.267275705 | 0.089503602 | 2.986200537 | 0.002824673 | 0.017875009 | 0.267275705 |
| ENSMUSG00000039737 | Prkrip1 | 66801 | protein_coding | 261.8384465 | -0.267286677 | 0.105446613 | -2.534805723 | 0.01125097 | 0.052276971 | 0.267286677 |
| ENSMUSG00000010362 | Rdm1 | 66599 | protein_coding | 142.9799455 | -0.267287366 | 0.114819789 | -2.327885891 | 0.019918164 | 0.079552786 | 0.267287366 |
| ENSMUSG00000085148 | Mir22hg | 100042498 | lncRNA | 191.3867711 | -0.267460899 | 0.154108601 | -1.735535179 | 0.082646056 | 0.218996237 | 0.267460899 |
| ENSMUSG00000040811 | Eml2 | 72205 | protein_coding | 234.9065274 | -0.267480684 | 0.115912041 | -2.307617751 | 0.021020406 | 0.083011808 | 0.267480684 |
| ENSMUSG00000029480 | Dhx37 | 208144 | protein_coding | 175.3077738 | -0.267533407 | 0.137877971 | -1.940363674 | 0.052335508 | 0.160276865 | 0.267533407 |
| ENSMUSG00000036995 | Asap3 | 230837 | protein_coding | 22.9550472 | 0.26755959 | 0.272710948 | 0.981110554 | 0.326538226 | 0.537637055 | 0.26755959 |
| ENSMUSG00000046201 | Scaf8 | 106583 | protein_coding | 940.9286885 | -0.267576661 | 0.0705762 | -3.791315787 | 0.000149851 | 0.00159441 | 0.267576661 |
| ENSMUSG00000028447 | Dctn3 | 53598 | protein_coding | 546.603534 | 0.267590872 | 0.097555773 | 2.74295272 | 0.006088945 | 0.032567065 | 0.267590872 |
| ENSMUSG00000041949 | Tango6 | 272538 | protein_coding | 75.50912716 | -0.267591825 | 0.133042779 | -2.011321668 | 0.044291491 | 0.142750885 | 0.267591825 |
| ENSMUSG00000031958 | Ldhd | 52815 | protein_coding | 83.0041949 | -0.267705228 | 0.135788254 | -1.971490327 | 0.048667818 | 0.152854694 | 0.267705228 |
| ENSMUSG00000000058 | Cav2 | 12390 | protein_coding | 125.5011312 | -0.267707252 | 0.185077581 | -1.446459641 | 0.148048324 | 0.324809051 | 0.267707252 |
| ENSMUSG00000100680 | 1810044D09Rik | 69798 | lncRNA | 19.58795471 | -0.267829927 | 0.243832364 | -1.098418284 | 0.272021882 | 0.480297588 | 0.267829927 |
| ENSMUSG00000042104 | Uggt2 | 66435 | protein_coding | 31.60527572 | 0.267926208 | 0.378468238 | 0.70792257 | 0.478993343 | 0.672927641 | 0.267926208 |
| ENSMUSG00000022475 | Hdac7 | 56233 | protein_coding | 78.19426538 | -0.26794239 | 0.149352085 | -1.794031795 | 0.072808094 | 0.200676868 | 0.26794239 |
| ENSMUSG00000097177 | 9330159M07Rik | 319673 | lncRNA | 24.51429137 | 0.267957427 | 0.27059369 | 0.990257485 | 0.322048282 | 0.532943899 | 0.267957427 |
| ENSMUSG00000079110 | Capn3 | 12335 | protein_coding | 13.36530639 | 0.267974693 | 0.490167592 | 0.546700144 | 0.584584755 | 0.755171612 | 0.267974693 |
| ENSMUSG00000061393 | Acvr2b | 11481 | protein_coding | 55.70266835 | -0.268085922 | 0.215611527 | -1.243374719 | 0.213729782 | 0.412041633 | 0.268085922 |
| ENSMUSG00000030720 | Cln3 | 12752 | protein_coding | 256.2040772 | -0.268106388 | 0.108256684 | -2.476580456 | 0.013264772 | 0.059182436 | 0.268106388 |
| ENSMUSG00000030729 | Pgm2l1 | 70974 | protein_coding | 351.7741412 | -0.268164899 | 0.079073797 | -3.391324431 | 0.000695557 | 0.005722538 | 0.268164899 |
| ENSMUSG00000002396 | Ocel1 | 77090 | protein_coding | 298.5840457 | -0.268229261 | 0.1058033 | -2.535169129 | 0.011239305 | 0.052238006 | 0.268229261 |
| ENSMUSG00000039219 | Arid4b | 94246 | protein_coding | 889.3572411 | -0.268231861 | 0.087095276 | -3.079752138 | 0.002071729 | 0.013929297 | 0.268231861 |
| ENSMUSG00000090112 | Shprh | 268281 | protein_coding | 519.6836903 | -0.268261208 | 0.089211073 | -3.007039368 | 0.002638056 | 0.016991086 | 0.268261208 |
| ENSMUSG00000030058 | Copg1 | 54161 | protein_coding | 2239.253899 | 0.26832563 | 0.071210639 | 3.768055378 | 0.000164524 | 0.001722907 | 0.26832563 |
| ENSMUSG00000021216 | Tubal3 | 238463 | protein_coding | 268.5906004 | -0.268373251 | 0.13986322 | -1.918826496 | 0.055006295 | 0.165528108 | 0.268373251 |
| ENSMUSG00000022807 | Osbpl11 | 106326 | protein_coding | 338.9012655 | -0.268387751 | 0.083268486 | -3.223161182 | 0.001267842 | 0.009267639 | 0.268387751 |
| ENSMUSG00000017146 | Brca1 | 12189 | protein_coding | 240.7516018 | 0.268478306 | 0.13045843 | 2.057960584 | 0.039593917 | 0.132267599 | 0.268478306 |
| ENSMUSG00000030609 | Aen | 68048 | protein_coding | 257.0700159 | -0.268498399 | 0.26861146 | -0.999579092 | 0.317514245 | 0.528771321 | 0.268498399 |
| ENSMUSG00000051166 | Eml5 | 319670 | protein_coding | 54.57968451 | -0.26853629 | 0.306751166 | -0.875420601 | 0.381345094 | 0.589085895 | 0.26853629 |
| ENSMUSG00000006219 | Fblim1 | 74202 | protein_coding | 555.6276476 | -0.268538758 | 0.098032651 | -2.739278744 | 0.006157414 | 0.032847006 | 0.268538758 |
| ENSMUSG00000030103 | Bhlhe40 | 20893 | protein_coding | 1957.012987 | -0.268558448 | 0.221869944 | -1.210431856 | 0.226113224 | 0.428396745 | 0.268558448 |
| ENSMUSG00000031352 | Hccs | 15159 | protein_coding | 458.4538367 | 0.26859372 | 0.087730747 | 3.061568819 | 0.002201804 | 0.014655382 | 0.26859372 |
| ENSMUSG00000022620 | Arsa | 11883 | protein_coding | 279.6520125 | 0.268680506 | 0.126986609 | 2.115817633 | 0.034360321 | 0.119325161 | 0.268680506 |
| ENSMUSG00000036181 | H1f2 | 50708 | protein_coding | 5487.596367 | 0.268695332 | 0.177082512 | 1.517345383 | 0.129179506 | 0.295737287 | 0.268695332 |
| ENSMUSG00000036391 | Sec24a | 77371 | protein_coding | 945.2307941 | 0.268698936 | 0.14893621 | 1.804120941 | 0.07121235 | 0.197814842 | 0.268698936 |
| ENSMUSG00000026547 | Tagln2 | 21346 | protein_coding | 2279.567313 | -0.268721422 | 0.105179832 | -2.554875926 | 0.010622566 | 0.050058054 | 0.268721422 |
| ENSMUSG00000040795 | Iqcc | 230767 | protein_coding | 101.0156354 | -0.268726423 | 0.218329423 | -1.230830091 | 0.218386419 | 0.418335403 | 0.268726423 |
| ENSMUSG00000031458 | Coprs | 66423 | protein_coding | 146.360649 | -0.268835647 | 0.137278268 | -1.958326322 | 0.050191734 | 0.155875003 | 0.268835647 |

| ensembl_gene_id | external_gene_name | entrezgene_id | gene_biotype | baseMean | log2FoldChange | lfcSE | stat | pvalue | padj | absLog2FC |
|---|---|---|---|---|---|---|---|---|---|---|
| ENSMUSG00000031976 | Urb2 | 382038 | protein_coding | 336.033841 | -0.268842838 | 0.13007836 | -2.066776039 | 0.038755262 | 0.130285736 | 0.268842838 |
| ENSMUSG00000028173 | Wls | 68151 | protein_coding | 2072.412891 | -0.26889206 | 0.089041279 | -3.019858456 | 0.002528928 | 0.016461288 | 0.26889206 |
| ENSMUSG00000034235 | Usp54 | 78787 | protein_coding | 151.9148827 | -0.26890265 | 0.144136814 | -1.865607005 | 0.062096369 | 0.179798393 | 0.26890265 |
| ENSMUSG00000026879 | Gsn | 227753 | protein_coding | 1958.598121 | -0.269016193 | 0.186233832 | -1.444507641 | 0.148596232 | 0.325697094 | 0.269016193 |
| ENSMUSG00000021361 | Tmem14c | 66154 | protein_coding | 1438.078451 | 0.26903942 | 0.107420523 | 2.504543949 | 0.012260937 | 0.055718176 | 0.26903942 |
| ENSMUSG00000031813 | Mvb12a | 73711 | protein_coding | 733.5452533 | 0.269137285 | 0.11414208 | 2.357914672 | 0.018377917 | 0.075038056 | 0.269137285 |
| ENSMUSG00000008036 | Ap2s1 | 232910 | protein_coding | 1018.781139 | 0.269196901 | 0.109749783 | 2.452823984 | 0.014173968 | 0.061903744 | 0.269196901 |
| ENSMUSG00000030225 | Dera | 232449 | protein_coding | 677.1895904 | 0.269314515 | 0.108499973 | 2.482162039 | 0.013058788 | 0.058542181 | 0.269314515 |
| ENSMUSG00000046441 | Cmtr2 | 234728 | protein_coding | 205.6059267 | -0.269331285 | 0.14007634 | -1.922746441 | 0.054511901 | 0.164600107 | 0.269331285 |
| ENSMUSG00000018322 | Tomm34 | 67145 | protein_coding | 295.4050893 | -0.269333433 | 0.100627072 | -2.67655042 | 0.007438437 | 0.038268549 | 0.269333433 |
| ENSMUSG00000005621 | Zfp592 | 233410 | protein_coding | 556.5548258 | -0.26934318 | 0.135372107 | -1.989650492 | 0.04662945 | 0.148335107 | 0.26934318 |
| ENSMUSG00000020846 | Rflnb | 76566 | protein_coding | 1280.770933 | -0.26935831 | 0.186834045 | -1.441698224 | 0.149387525 | 0.326757003 | 0.26935831 |
| ENSMUSG00000006998 | Psmd2 | 21762 | protein_coding | 2864.936818 | 0.269391156 | 0.110882914 | 2.429510067 | 0.015119245 | 0.065051144 | 0.269391156 |
| ENSMUSG00000038762 | Abcf1 | 224742 | protein_coding | 1465.383893 | 0.269477442 | 0.107216965 | 2.513384364 | 0.011957897 | 0.054762906 | 0.269477442 |
| ENSMUSG00000045287 | Rtn4rl1 | 237847 | protein_coding | 65.74345636 | -0.269529971 | 0.22788615 | -1.182739589 | 0.236912369 | 0.44114335 | 0.269529971 |
| ENSMUSG00000018379 | Srsf1 | 110809 | protein_coding | 3761.716909 | 0.269709982 | 0.097136456 | 2.776609249 | 0.005492917 | 0.030054686 | 0.269709982 |
| ENSMUSG00000099474 | 1700097N02Rik | 67522 | lncRNA | 8.943160186 | 0.269750869 | 0.486698082 | 0.554246831 | 0.579409933 | 0.750922816 | 0.269750869 |
| ENSMUSG00000038025 | Phf2 | 18676 | protein_coding | 386.9052774 | -0.269831884 | 0.127052063 | -2.123789872 | 0.033687718 | 0.11756748 | 0.269831884 |
| ENSMUSG00000059588 | Calcrl | 54598 | protein_coding | 70.89926534 | -0.269942725 | 0.229070565 | -1.178426068 | 0.23862679 | 0.442830469 | 0.269942725 |
| ENSMUSG00000049922 | Slc35c1 | 228368 | protein_coding | 2270.471776 | 0.270078421 | 0.080022234 | 3.375042242 | 0.000738044 | 0.005994676 | 0.270078421 |
| ENSMUSG00000007777 | 0610009B22Rik | 66050 | protein_coding | 261.9684843 | 0.270094614 | 0.112881565 | 2.392725641 | 0.016723741 | 0.070155143 | 0.270094614 |
| ENSMUSG00000044461 | Shisa2 | 219134 | protein_coding | 798.9254642 | -0.270135651 | 0.150415571 | -1.795928769 | 0.072505846 | 0.200151321 | 0.270135651 |
| ENSMUSG00000024253 | Dync2li1 | 213575 | protein_coding | 27.00534678 | 0.270379622 | 0.26437405 | 1.022716194 | 0.306442052 | 0.516454239 | 0.270379622 |
| ENSMUSG00000054662 | Ano9 | 71345 | protein_coding | 1100.351136 | -0.270397599 | 0.119910116 | -2.255002386 | 0.024133179 | 0.092046116 | 0.270397599 |
| ENSMUSG00000071660 | Ttc9c | 70387 | protein_coding | 748.4154351 | 0.27043276 | 0.071392189 | 3.78798805 | 0.000151872 | 0.00161268 | 0.27043276 |
| ENSMUSG00000041530 | Ago1 | 236511 | protein_coding | 506.9050123 | -0.270504391 | 0.118821754 | -2.276556123 | 0.022812746 | 0.088387597 | 0.270504391 |
| ENSMUSG00000033739 | Fkbpl | 56299 | protein_coding | 85.60388156 | 0.270562336 | 0.15875838 | 1.704239711 | 0.088336313 | 0.22904457 | 0.270562336 |
| ENSMUSG00000041406 | BC055324 | 381306 | protein_coding | 87.66031587 | 0.270638785 | 0.140734987 | 1.923038406 | 0.054475227 | 0.164520556 | 0.270638785 |
| ENSMUSG00000038383 | Pigu | 228812 | protein_coding | 455.9779163 | 0.2707688 | 0.070251702 | 3.854266783 | 0.000116077 | 0.001278613 | 0.2707688 |
| ENSMUSG00000008855 | Hdac5 | 15184 | protein_coding | 477.7318356 | -0.270801855 | 0.176800122 | -1.531683649 | 0.125600519 | 0.289844196 | 0.270801855 |
| ENSMUSG00000059474 | Mbtd1 | 103537 | protein_coding | 545.059988 | -0.270878707 | 0.124139505 | -2.182050805 | 0.029105783 | 0.10541085 | 0.270878707 |
| ENSMUSG00000034187 | Nsf | 18195 | protein_coding | 817.3025383 | 0.271028345 | 0.076235651 | 3.55513914 | 0.000377779 | 0.003464171 | 0.271028345 |
| ENSMUSG00000022526 | Zfp251 | 71591 | protein_coding | 139.3422097 | -0.271056955 | 0.168250349 | -1.611033543 | 0.107172412 | 0.260176472 | 0.271056955 |
| ENSMUSG00000045438 | Cox19 | 68033 | protein_coding | 312.7878554 | 0.271119309 | 0.10372933 | 2.613718887 | 0.008956269 | 0.044053355 | 0.271119309 |
| ENSMUSG00000066800 | Rnasel | 24014 | protein_coding | 5020.395924 | -0.271143861 | 0.113520713 | -2.388496816 | 0.016917455 | 0.070743298 | 0.271143861 |
| ENSMUSG00000002308 | Cd320 | 54219 | protein_coding | 270.080783 | -0.27121868 | 0.146680318 | -1.849046163 | 0.064451147 | 0.184669522 | 0.27121868 |
| ENSMUSG00000047642 | D930020B18Rik | 216393 | protein_coding | 26.90179466 | 0.271260831 | 0.217068625 | 1.249654716 | 0.211425706 | 0.408886431 | 0.271260831 |
| ENSMUSG00000034023 | Fancd2 | 211651 | protein_coding | 173.780566 | 0.271317829 | 0.177790732 | 1.526051592 | 0.126997016 | 0.292271128 | 0.271317829 |

| ensembl_gene_id | external_gene_name | entrezgene_id | gene_biotype | baseMean | log2FoldChange | lfcSE | stat | pvalue | padj | absLog2FC |
|---|---|---|---|---|---|---|---|---|---|---|
| ENSMUSG00000026409 | Pfkfb2 | 18640 | protein_coding | 390.340835 | 0.271328192 | 0.088136567 | 3.078497398 | 0.002080473 | 0.013976289 | 0.271328192 |
| ENSMUSG00000036151 | Tm6sf2 | 107770 | protein_coding | 84.92641732 | -0.271340829 | 0.15341229 | -1.768703332 | 0.076943396 | 0.208003125 | 0.271340829 |
| ENSMUSG00000041605 | Inava | 67313 | protein_coding | 1197.704132 | -0.271445324 | 0.135843037 | -1.998227727 | 0.045691977 | 0.146170277 | 0.271445324 |
| ENSMUSG00000020107 | Anapc16 | 52717 | protein_coding | 509.7275558 | 0.271489334 | 0.104757454 | 2.591599196 | 0.0095531 | 0.046277911 | 0.271489334 |
| ENSMUSG00000027692 | Tnik | 665113 | protein_coding | 471.0840704 | -0.271538128 | 0.091736358 | -2.959983733 | 0.003076553 | 0.019156894 | 0.271538128 |
| ENSMUSG00000009772 | Nuak2 | 74137 | protein_coding | 550.6066223 | 0.271584922 | 0.18023145 | 1.506867548 | 0.131844611 | 0.299620118 | 0.271584922 |
| ENSMUSG00000021198 | Unc79 | 217843 | protein_coding | 39.62608597 | -0.271622756 | 0.307102168 | -0.88447033 | 0.376442372 | 0.58462681 | 0.271622756 |
| ENSMUSG00000024120 | Lrpprc | 72416 | protein_coding | 1332.594788 | -0.27162558 | 0.099196522 | -2.73825708 | 0.006176577 | 0.032925208 | 0.27162558 |
| ENSMUSG00000020163 | Uqcr11 | 66594 | protein_coding | 2540.514936 | 0.271653563 | 0.101779797 | 2.669032279 | 0.007607014 | 0.038984723 | 0.271653563 |
| ENSMUSG00000026090 | Cracdl | 72097 | protein_coding | 669.6064788 | -0.271670933 | 0.117443323 | -2.31320884 | 0.02071116 | 0.082061531 | 0.271670933 |
| ENSMUSG00000061046 | Haghl | 68977 | protein_coding | 87.72528758 | -0.271762552 | 0.127694755 | -2.128220161 | 0.033318834 | 0.116561034 | 0.271762552 |
| ENSMUSG00000026737 | Pip4k2a | 18718 | protein_coding | 26.43757664 | 0.271841563 | 0.228955686 | 1.18731082 | 0.235105042 | 0.439164197 | 0.271841563 |
| ENSMUSG00000025209 | Twnk | 226153 | protein_coding | 297.3978047 | -0.271897209 | 0.116534327 | -2.333194139 | 0.019637958 | 0.078749597 | 0.271897209 |
| ENSMUSG00000034739 | Mfrp | 259172 | protein_coding | 8.133050028 | -0.27192471 | 0.563896546 | -0.482224465 | 0.629646499 | 0.785664664 | 0.27192471 |
| ENSMUSG00000037989 | Wnk2 | 75607 | protein_coding | 192.1904751 | -0.271929296 | 0.172652646 | -1.575007983 | 0.115254602 | 0.272612904 | 0.271929296 |
| ENSMUSG00000005575 | Ube2m | 22192 | protein_coding | 781.3249722 | 0.271937299 | 0.104958784 | 2.59089606 | 0.00957264 | 0.046351171 | 0.271937299 |
| ENSMUSG00000066441 | Rdh11 | 17252 | protein_coding | 622.5761842 | 0.272064429 | 0.122948054 | 2.212840472 | 0.026908646 | 0.099512633 | 0.272064429 |
| ENSMUSG00000026933 | Camsap1 | 227634 | protein_coding | 346.627805 | -0.272108712 | 0.11286366 | -2.410950622 | 0.015911002 | 0.067635561 | 0.272108712 |
| ENSMUSG00000060981 | H4c8 | 69386 | protein_coding | 2865.131269 | 0.272153576 | 0.153425715 | 1.773845912 | 0.076088641 | 0.206077178 | 0.272153576 |
| ENSMUSG00000031765 | Mt1 | 17748 | protein_coding | 3206.543097 | -0.272176689 | 0.191235973 | -1.423250473 | 0.154663559 | 0.334257959 | 0.272176689 |
| ENSMUSG00000078656 | Vps25 | 28084 | protein_coding | 8.746310499 | -0.272202465 | 0.376157393 | -0.723639814 | 0.469286892 | 0.665550852 | 0.272202465 |
| ENSMUSG00000029376 | Mthfd2l | 665563 | protein_coding | 152.9330099 | 0.272211851 | 0.155960575 | 1.745388858 | 0.080917204 | 0.215815342 | 0.272211851 |
| ENSMUSG00000045248 | Med26 | 70625 | protein_coding | 118.971538 | -0.272431162 | 0.113455466 | -2.401216722 | 0.016340655 | 0.068946745 | 0.272431162 |
| ENSMUSG00000053749 | Gm9920 | 100503794 | lncRNA | 33.69654606 | 0.27243935 | 0.252378147 | 1.079488666 | 0.280369944 | 0.488708594 | 0.27243935 |
| ENSMUSG00000039852 | Rere | 68703 | protein_coding | 686.569778 | -0.272559973 | 0.19378283 | -1.406522822 | 0.159568933 | 0.340835988 | 0.272559973 |
| ENSMUSG00000032220 | Myo1e | 71602 | protein_coding | 1493.071043 | -0.272577696 | 0.137184378 | -1.98694414 | 0.046928587 | 0.14896269 | 0.272577696 |
| ENSMUSG00000097415 | AU020206 | 108167440 | lncRNA | 90.13492222 | 0.272584547 | 0.167842142 | 1.624053073 | 0.104364482 | 0.255544576 | 0.272584547 |
| ENSMUSG00000026111 | Unc50 | 67387 | protein_coding | 611.3236901 | 0.272611542 | 0.101071772 | 2.697207522 | 0.006992368 | 0.036444515 | 0.272611542 |
| ENSMUSG00000073616 | Cops9 | 66915 | protein_coding | 614.6043734 | 0.272654946 | 0.11046059 | 2.468345908 | 0.013573908 | 0.06014684 | 0.272654946 |
| ENSMUSG00000059839 | Zfp874b | 408067 | protein_coding | 127.9689614 | -0.272698118 | 0.178653677 | -1.526406415 | 0.126908681 | 0.292110039 | 0.272698118 |
| ENSMUSG00000002057 | Foxn1 | 15218 | protein_coding | 21.05949115 | -0.272727889 | 0.576214641 | -0.473309544 | 0.635992354 | 0.789316364 | 0.272727889 |
| ENSMUSG00000041187 | Prkd2 | 101540 | protein_coding | 162.0648061 | -0.272739052 | 0.110823522 | -2.461021342 | 0.013854212 | 0.060958535 | 0.272739052 |
| ENSMUSG00000002068 | Ccne1 | 12447 | protein_coding | 70.09363424 | 0.272783292 | 0.224370548 | 1.215771384 | 0.224072023 | 0.425744862 | 0.272783292 |
| ENSMUSG00000030510 | Cers3 | 545975 | protein_coding | 16.54342963 | 0.272856509 | 0.361998769 | 0.753749826 | 0.450999457 | 0.651506507 | 0.272856509 |
| ENSMUSG00000029071 | Dvl1 | 13542 | protein_coding | 693.2191723 | -0.272978986 | 0.120341927 | -2.268361428 | 0.023307187 | 0.089627443 | 0.272978986 |
| ENSMUSG00000024902 | Mrpl11 | 66419 | protein_coding | 436.7030718 | 0.27317812 | 0.109756042 | 2.488957467 | 0.012811829 | 0.05767858 | 0.27317812 |
| ENSMUSG00000006423 | Steep1 | 77644 | protein_coding | 616.3902836 | 0.273211538 | 0.064330178 | 4.24701977 | 2.17E-05 | 0.000296947 | 0.273211538 |
| ENSMUSG00000025888 | Casp1 | 12362 | protein_coding | 2667.234854 | 0.273248097 | 0.121568416 | 2.247689868 | 0.024595972 | 0.093277298 | 0.273248097 |

| ensembl_gene_id | external_gene_name | entrezgene_id | gene_biotype | baseMean | log2FoldChange | lfcSE | stat | pvalue | padj | absLog2FC |
|---|---|---|---|---|---|---|---|---|---|---|
| ENSMUSG00000022519 | Srl | 106393 | protein_coding | 128.7241506 | 0.273268868 | 0.158355489 | 1.725667167 | 0.084407308 | 0.222221422 | 0.273268868 |
| ENSMUSG00000053730 | Tmem39b | 230770 | protein_coding | 54.1792452 | -0.273385338 | 0.148102227 | -1.845923208 | 0.064903356 | 0.185504307 | 0.273385338 |
| ENSMUSG00000018899 | Irf1 | 16362 | protein_coding | 1436.46194 | 0.273393592 | 0.151364455 | 1.806194138 | 0.07088802 | 0.197120179 | 0.273393592 |
| ENSMUSG00000061894 | Zscan20 | 269585 | protein_coding | 97.40970142 | -0.273448258 | 0.163873208 | -1.668657506 | 0.095185275 | 0.240576176 | 0.273448258 |
| ENSMUSG00000040661 | Rad54l2 | 81000 | protein_coding | 471.3663001 | -0.273499573 | 0.107697099 | -2.539525913 | 0.011100282 | 0.051720096 | 0.273499573 |
| ENSMUSG00000031787 | Katnb1 | 74187 | protein_coding | 205.4276433 | 0.273598286 | 0.127303347 | 2.149183764 | 0.031619836 | 0.112269448 | 0.273598286 |
| ENSMUSG00000027900 | Dram2 | 67171 | protein_coding | 2791.37031 | 0.273686482 | 0.080803404 | 3.387066259 | 0.000706443 | 0.005782234 | 0.273686482 |
| ENSMUSG00000046782 | Ttc6 | 70846 | protein_coding | 112.1411027 | -0.273689651 | 0.188664861 | -1.450665743 | 0.146872959 | 0.323254455 | 0.273689651 |
| ENSMUSG00000030055 | Rab43 | 69834 | protein_coding | 71.60125203 | -0.273738164 | 0.177238894 | -1.544458768 | 0.122477233 | 0.28524892 | 0.273738164 |
| ENSMUSG00000025083 | Afap1l2 | 226250 | protein_coding | 29.94940659 | -0.273857669 | 0.356999436 | -0.767109527 | 0.443016398 | 0.644181595 | 0.273857669 |
| ENSMUSG00000022257 | Laptm4b | 114128 | protein_coding | 1074.456512 | 0.273875961 | 0.07647445 | 3.581274033 | 0.000341923 | 0.00317005 | 0.273875961 |
| ENSMUSG00000023020 | Cox14 | 66379 | protein_coding | 868.2792606 | 0.273902919 | 0.092362765 | 2.965512333 | 0.003021792 | 0.01888976 | 0.273902919 |
| ENSMUSG00000042523 | Dnal1 | 105000 | protein_coding | 24.48608226 | -0.273956887 | 0.301387138 | -0.908986656 | 0.36335717 | 0.572231857 | 0.273956887 |
| ENSMUSG00000047030 | Spata2 | 263876 | protein_coding | 614.2564698 | -0.273971613 | 0.094065166 | -2.912572472 | 0.00358465 | 0.021562746 | 0.273971613 |
| ENSMUSG00000024691 | Fam111a | 107373 | protein_coding | 231.0981958 | 0.274112234 | 0.145726767 | 1.881001276 | 0.059971745 | 0.175886323 | 0.274112234 |
| ENSMUSG00000040560 | Wdr7 | 104082 | protein_coding | 465.56413 | 0.274279304 | 0.070176489 | 3.908421552 | 9.29E-05 | 0.001059978 | 0.274279304 |
| ENSMUSG00000026432 | Avpr1b | 26361 | protein_coding | 13.541157 | -0.27429264 | 0.527648937 | -0.519839272 | 0.603175605 | 0.76799575 | 0.27429264 |
| ENSMUSG00000024392 | Bag6 | 224727 | protein_coding | 1423.55598 | -0.274341552 | 0.057199304 | -4.796239327 | 1.62E-06 | 2.90E-05 | 0.274341552 |
| ENSMUSG00000067925 | Rtl8a | 66158 | protein_coding | 10.2509028 | -0.274371632 | 0.396637307 | -0.691744391 | 0.489097864 | 0.681630132 | 0.274371632 |
| ENSMUSG00000028719 | Cmpk1 | 66588 | protein_coding | 9947.618535 | 0.274380889 | 0.093735113 | 2.927194293 | 0.003420351 | 0.020809532 | 0.274380889 |
| ENSMUSG00000032320 | Rcn2 | 26611 | protein_coding | 344.5742491 | 0.274478871 | 0.097287378 | 2.821320474 | 0.00478264 | 0.027167577 | 0.274478871 |
| ENSMUSG00000000776 | Polr3d | 67065 | protein_coding | 266.9695417 | -0.274510658 | 0.163300485 | -1.681015567 | 0.092759891 | 0.236699702 | 0.274510658 |
| ENSMUSG00000060538 | Tmem219 | 68742 | protein_coding | 242.4787647 | -0.274535189 | 0.095622092 | -2.871043527 | 0.004091192 | 0.023896041 | 0.274535189 |
| ENSMUSG00000001768 | Rin2 | 74030 | protein_coding | 1392.710213 | -0.274624292 | 0.065584755 | -4.187319001 | 2.82E-05 | 0.000374353 | 0.274624292 |
| ENSMUSG00000044566 | Cage1 | 71213 | protein_coding | 42.35256437 | 0.274643916 | 0.260053648 | 1.056104841 | 0.290920319 | 0.499573396 | 0.274643916 |
| ENSMUSG00000049604 | Hoxb13 | 15408 | protein_coding | 1718.329756 | -0.274667164 | 0.468511906 | -0.586254395 | 0.557704573 | 0.736454853 | 0.274667164 |
| ENSMUSG00000065778 | Gm22154 | 115490050 | snoRNA | 83.71625107 | 0.274764089 | 0.351172224 | 0.782419765 | 0.433967922 | 0.637453749 | 0.274764089 |
| ENSMUSG00000050732 | Vamp8 | 22320 | protein_coding | 1263.122941 | 0.274928822 | 0.086833789 | 3.166150222 | 0.00154471 | 0.010906091 | 0.274928822 |
| ENSMUSG00000045598 | Zfp553 | 233887 | protein_coding | 243.3642839 | -0.274993421 | 0.093917368 | -2.928035862 | 0.003411107 | 0.020774708 | 0.274993421 |
| ENSMUSG00000003233 | Dvl3 | 13544 | protein_coding | 200.8402388 | -0.275063051 | 0.162831587 | -1.689248727 | 0.091171775 | 0.233733145 | 0.275063051 |
| ENSMUSG00000032458 | Copb2 | 50797 | protein_coding | 4736.728789 | 0.275130287 | 0.068506211 | 4.016136366 | 5.92E-05 | 0.000716579 | 0.275130287 |
| ENSMUSG00000040043 | Rbms2 | 56516 | protein_coding | 145.8560185 | -0.27522241 | 0.132446865 | -2.077983573 | 0.037710872 | 0.127745379 | 0.27522241 |
| ENSMUSG00000040774 | Cept1 | 99712 | protein_coding | 2000.94338 | 0.275228641 | 0.076751479 | 3.585971835 | 0.000335825 | 0.003118964 | 0.275228641 |
| ENSMUSG00000034586 | Hid1 | 217310 | protein_coding | 1101.432891 | -0.27524839 | 0.136531192 | -2.016011035 | 0.043798822 | 0.141596585 | 0.27524839 |
| ENSMUSG00000022476 | Polr3h | 78929 | protein_coding | 110.2233072 | -0.275260812 | 0.11213307 | -2.45476926 | 0.014097504 | 0.061772753 | 0.275260812 |
| ENSMUSG00000045237 | Eola1 | 68618 | protein_coding | 257.3295669 | 0.27533831 | 0.133291214 | 2.065689862 | 0.038857773 | 0.130465137 | 0.27533831 |
| ENSMUSG00000052794 | 1700030K09Rik | 72254 | protein_coding | 60.58948739 | 0.275422306 | 0.243100107 | 1.132958393 | 0.257231726 | 0.464112702 | 0.275422306 |
| ENSMUSG00000035198 | Tubg1 | 103733 | protein_coding | 198.3664734 | 0.275677312 | 0.116910369 | 2.358022776 | 0.018372566 | 0.075038056 | 0.275677312 |

| ensembl_gene_id | external_gene_name | entrezgene_id | gene_biotype | baseMean | log2FoldChange | lfcSE | stat | pvalue | padj | absLog2FC |
|---|---|---|---|---|---|---|---|---|---|---|
| ENSMUSG00000072694 | 1500011B03Rik | 66236 | protein_coding | 91.26238351 | 0.275773213 | 0.125120486 | 2.204061235 | 0.027520038 | 0.101162973 | 0.275773213 |
| ENSMUSG00000039108 | Lsm14b | 241846 | protein_coding | 409.0897884 | -0.275783733 | 0.096182202 | -2.867305242 | 0.004139835 | 0.024127074 | 0.275783733 |
| ENSMUSG00000051185 | Fam174a | 67698 | protein_coding | 193.6617975 | 0.275865159 | 0.126390704 | 2.182638041 | 0.029062476 | 0.105343072 | 0.275865159 |
| ENSMUSG00000032207 | Lipc | 15450 | protein_coding | 8.598479443 | 0.275984855 | 0.427563392 | 0.645482894 | 0.5186143 | 0.705762919 | 0.275984855 |
| ENSMUSG00000033767 | Tmem131l | 229473 | protein_coding | 298.2114138 | -0.276018926 | 0.121272887 | -2.276015131 | 0.022845104 | 0.088491444 | 0.276018926 |
| ENSMUSG00000059970 | Hspa2 | 15512 | protein_coding | 28.70375409 | 0.276081888 | 0.239993898 | 1.150370448 | 0.249991326 | 0.45632155 | 0.276081888 |
| ENSMUSG00000038145 | Snrk | 20623 | protein_coding | 683.4108756 | -0.276093971 | 0.097360513 | -2.835790015 | 0.004571249 | 0.026134549 | 0.276093971 |
| ENSMUSG00000033735 | Spr | 20751 | protein_coding | 768.0835424 | 0.276100572 | 0.110974893 | 2.487955297 | 0.012847988 | 0.057808688 | 0.276100572 |
| ENSMUSG00000053819 | Camk2d | 108058 | protein_coding | 1689.894552 | 0.276242386 | 0.066436892 | 4.157966732 | 3.21E-05 | 0.00041921 | 0.276242386 |
| ENSMUSG00000074300 | Smim31 | 407795 | protein_coding | 1660.890058 | 0.276276939 | 0.115145871 | 2.399364694 | 0.016423548 | 0.069186532 | 0.276276939 |
| ENSMUSG00000022965 | Ifngr2 | 15980 | protein_coding | 1703.012119 | -0.276296723 | 0.084023151 | -3.288340401 | 0.001007799 | 0.007713707 | 0.276296723 |
| ENSMUSG00000042766 | Trim46 | 360213 | protein_coding | 131.1616332 | -0.276309187 | 0.250783349 | -1.101784419 | 0.270555406 | 0.478449592 | 0.276309187 |
| ENSMUSG00000056214 | Pard6g | 93737 | protein_coding | 14.95050052 | -0.276339674 | 0.301837215 | -0.915525522 | 0.35991581 | 0.569481508 | 0.276339674 |
| ENSMUSG00000032812 | Arap1 | 69710 | protein_coding | 1110.768642 | -0.276344331 | 0.082282447 | -3.358484579 | 0.000783711 | 0.00628165 | 0.276344331 |
| ENSMUSG00000052298 | Cdc42se2 | 72729 | protein_coding | 1051.682459 | 0.276350584 | 0.095892649 | 2.881874532 | 0.003953171 | 0.023260476 | 0.276350584 |
| ENSMUSG00000042744 | Hectd4 | 269700 | protein_coding | 1127.985518 | -0.27635266 | 0.194493652 | -1.420882668 | 0.155350873 | 0.33533388 | 0.27635266 |
| ENSMUSG00000035944 | Ttc38 | 239570 | protein_coding | 1131.897995 | 0.276410004 | 0.126593624 | 2.18344334 | 0.029003177 | 0.105248302 | 0.276410004 |
| ENSMUSG00000032000 | Birc3 | 11796 | protein_coding | 589.5686976 | -0.276432395 | 0.117437262 | -2.353872949 | 0.01857896 | 0.07574243 | 0.276432395 |
| ENSMUSG00000044501 | Zfp758 | 224598 | protein_coding | 427.5579264 | 0.276433163 | 0.090693926 | 3.047978782 | 0.002303862 | 0.015211282 | 0.276433163 |
| ENSMUSG00000015790 | Surf1 | 20930 | protein_coding | 870.7088392 | -0.276504496 | 0.095707474 | -2.889058552 | 0.003863971 | 0.022879306 | 0.276504496 |
| ENSMUSG00000028799 | Zfp362 | 230761 | protein_coding | 63.49460076 | -0.276513624 | 0.230354426 | -1.200383377 | 0.229990482 | 0.433166418 | 0.276513624 |
| ENSMUSG00000020453 | Patz1 | 56218 | protein_coding | 107.0319009 | -0.276522832 | 0.116504273 | -2.373499483 | 0.017620418 | 0.072765884 | 0.276522832 |
| ENSMUSG00000078350 | Smim1 | 68859 | protein_coding | 8.939773077 | -0.276654189 | 0.372373299 | -0.742948514 | 0.457512852 | 0.656155656 | 0.276654189 |
| ENSMUSG00000021118 | Plek2 | 27260 | protein_coding | 680.8198859 | -0.276931398 | 0.385040557 | -0.719226565 | 0.472001334 | 0.667439386 | 0.276931398 |
| ENSMUSG00000029772 | Ahcyl2 | 74340 | protein_coding | 9622.74832 | -0.276976484 | 0.10932577 | -2.533496758 | 0.011293079 | 0.052426743 | 0.276976484 |
| ENSMUSG00000027870 | Hao2 | 56185 | protein_coding | 7465.673433 | -0.276985938 | 0.143286318 | -1.933094121 | 0.053224604 | 0.161983995 | 0.276985938 |
| ENSMUSG00000050910 | Cdr2l | 237988 | protein_coding | 98.69975872 | -0.277024673 | 0.1751717 | -1.581446506 | 0.113775981 | 0.269876966 | 0.277024673 |
| ENSMUSG00000078486 | Perm1 | 74183 | protein_coding | 17.52706869 | -0.27704826 | 0.342836038 | -0.808107169 | 0.419028891 | 0.62516787 | 0.27704826 |
| ENSMUSG00000085683 | Tmem238l | 71576 | protein_coding | 1107.439338 | 0.277065364 | 0.115769806 | 2.393243754 | 0.016700142 | 0.070092191 | 0.277065364 |
| ENSMUSG00000030060 | Hmces | 232210 | protein_coding | 137.7730996 | -0.277185889 | 0.100992968 | -2.744605825 | 0.006058362 | 0.032468905 | 0.277185889 |
| ENSMUSG00000074283 | Zfp109 | 56869 | protein_coding | 65.07923214 | -0.277251285 | 0.160833644 | -1.723838855 | 0.084736938 | 0.222721114 | 0.277251285 |
| ENSMUSG00000047767 | Atg16l2 | 73683 | protein_coding | 183.4343471 | -0.27725322 | 0.183666117 | -1.509550181 | 0.131158242 | 0.298526504 | 0.27725322 |
| ENSMUSG00000039770 | Ypel5 | 383295 | protein_coding | 619.5935908 | 0.277276185 | 0.092404062 | 3.000692586 | 0.002693664 | 0.017241292 | 0.277276185 |
| ENSMUSG00000051586 | Mical3 | 194401 | protein_coding | 378.8991759 | 0.277286078 | 0.152907244 | 1.813426689 | 0.069766036 | 0.195227236 | 0.277286078 |
| ENSMUSG00000058331 | Zfp85 | 22746 | protein_coding | 71.68187443 | 0.277307719 | 0.246550992 | 1.124747936 | 0.260695862 | 0.468030173 | 0.277307719 |
| ENSMUSG00000027884 | Clcc1 | 229725 | protein_coding | 507.7576476 | 0.277328331 | 0.090394265 | 3.067985886 | 0.002155068 | 0.014398457 | 0.277328331 |
| ENSMUSG00000026229 | Psmd1 | 70247 | protein_coding | 1721.55933 | 0.277350966 | 0.105531301 | 2.628139367 | 0.008585333 | 0.042733497 | 0.277350966 |
| ENSMUSG00000020121 | Srgap1 | 117600 | protein_coding | 221.1769217 | -0.277410374 | 0.22445475 | -1.235930067 | 0.21648456 | 0.415692112 | 0.277410374 |

| ensembl_gene_id | external_gene_name | entrezgene_id | gene_biotype | baseMean | log2FoldChange | lfcSE | stat | pvalue | padj | absLog2FC |
|---|---|---|---|---|---|---|---|---|---|---|
| ENSMUSG00000025436 | Atp23 | 68876 | protein_coding | 62.92151375 | -0.277478386 | 0.166084585 | -1.670705236 | 0.094779914 | 0.239939491 | 0.277478386 |
| ENSMUSG00000026880 | Stom | 13830 | protein_coding | 463.7952813 | -0.277615352 | 0.466847189 | -0.594660006 | 0.552070761 | 0.732415715 | 0.277615352 |
| ENSMUSG00000048644 | Ctxn1 | 330695 | protein_coding | 8.451105195 | -0.277646227 | 0.542532872 | -0.51175927 | 0.608819499 | 0.772360968 | 0.277646227 |
| ENSMUSG00000025557 | Slc15a1 | 56643 | protein_coding | 299.34207 | -0.277696129 | 0.26639901 | -1.042406759 | 0.297223134 | 0.506180721 | 0.277696129 |
| ENSMUSG00000022861 | Dgkg | 110197 | protein_coding | 104.8823173 | -0.277897344 | 0.23345081 | -1.190389289 | 0.233893421 | 0.437742522 | 0.277897344 |
| ENSMUSG00000060639 | H4c9 | 319158 | protein_coding | 128.6003391 | 0.27789943 | 0.150785106 | 1.843016443 | 0.06532661 | 0.186108536 | 0.27789943 |
| ENSMUSG00000028145 | Them4 | 75778 | protein_coding | 157.2150718 | -0.277921992 | 0.142854976 | -1.945483457 | 0.051716819 | 0.159085649 | 0.277921992 |
| ENSMUSG00000020042 | Btbd11 | 74007 | protein_coding | 22.88153348 | -0.277965332 | 0.269287036 | -1.032226937 | 0.301965819 | 0.511326192 | 0.277965332 |
| ENSMUSG00000115902 | D730005E14Rik | 109361 | lncRNA | 45.08962302 | 0.277975088 | 0.287955415 | 0.96534072 | 0.334374183 | 0.545293316 | 0.277975088 |
| ENSMUSG00000023330 | Dtwd1 | 69185 | protein_coding | 73.3268202 | 0.277995114 | 0.129050899 | 2.154150932 | 0.031228337 | 0.111301177 | 0.277995114 |
| ENSMUSG00000030701 | Plekhb1 | 27276 | protein_coding | 22.8381451 | -0.278046179 | 0.282738685 | -0.983403384 | 0.325408946 | 0.536498673 | 0.278046179 |
| ENSMUSG00000106864 | Gtf3c2 | 71752 | protein_coding | 316.9695849 | -0.278081513 | 0.084145816 | -3.304757464 | 0.000950586 | 0.00736906 | 0.278081513 |
| ENSMUSG00000072889 | Nfxl1 | 100978 | protein_coding | 359.5585826 | 0.278118635 | 0.102047107 | 2.725394613 | 0.006422468 | 0.034016631 | 0.278118635 |
| ENSMUSG00000070972 | Dnajc25 | 72429 | protein_coding | 173.4779142 | 0.278121635 | 0.115769631 | 2.402371265 | 0.016289167 | 0.068793316 | 0.278121635 |
| ENSMUSG00000030222 | Rerg | 232441 | protein_coding | 8.37506222 | -0.278142139 | 0.591853892 | -0.469950679 | 0.638390255 | 0.790506101 | 0.278142139 |
| ENSMUSG00000100372 | 1700003I22Rik | 69356 | lncRNA | 16.78980763 | 0.27820599 | 0.32640775 | 0.852326544 | 0.394032877 | 0.602029321 | 0.27820599 |
| ENSMUSG00000061360 | Phf5a | 68479 | protein_coding | 537.0322945 | 0.278223232 | 0.114865264 | 2.422170317 | 0.015428118 | 0.066023391 | 0.278223232 |
| ENSMUSG00000071645 | Tut1 | 70044 | protein_coding | 327.1298294 | -0.278383186 | 0.088091145 | -3.160172168 | 0.001576759 | 0.011083241 | 0.278383186 |
| ENSMUSG00000022090 | Pdlim2 | 213019 | protein_coding | 318.3725375 | -0.278398406 | 0.294977784 | -0.943794486 | 0.345274679 | 0.555172126 | 0.278398406 |
| ENSMUSG00000029179 | Zcchc4 | 78796 | protein_coding | 166.3626736 | 0.278450905 | 0.110801986 | 2.513049775 | 0.011969245 | 0.054799117 | 0.278450905 |
| ENSMUSG00000035790 | Cep19 | 66994 | protein_coding | 390.2069885 | 0.278457174 | 0.102473127 | 2.717367788 | 0.006580344 | 0.034714827 | 0.278457174 |
| ENSMUSG00000117655 | 6720468P15Rik | 78073 | lncRNA | 26.20194186 | 0.278471395 | 0.250374384 | 1.112219992 | 0.266043574 | 0.473838986 | 0.278471395 |
| ENSMUSG00000038214 | Bend3 | 331623 | protein_coding | 124.3776187 | -0.278500002 | 0.181584505 | -1.533721185 | 0.12509826 | 0.289178051 | 0.278500002 |
| ENSMUSG00000089281 | Scarna6 | 100217415 | scaRNA | 342.0543629 | -0.278547922 | 0.187841682 | -1.482886652 | 0.138104528 | 0.309080923 | 0.278547922 |
| ENSMUSG00000028996 | Rbp7 | 63954 | protein_coding | 22.89863819 | -0.278578125 | 0.234994908 | -1.18546452 | 0.235833835 | 0.440009526 | 0.278578125 |
| ENSMUSG00000078676 | Casc3 | 192160 | protein_coding | 494.7974394 | -0.278599582 | 0.072518709 | -3.841761449 | 0.000122155 | 0.001339076 | 0.278599582 |
| ENSMUSG00000034653 | Ythdc2 | 240255 | protein_coding | 643.8755195 | 0.278755798 | 0.157144191 | 1.773885475 | 0.076082096 | 0.206077178 | 0.278755798 |
| ENSMUSG00000025759 | Mfsd8 | 72175 | protein_coding | 224.8334132 | 0.278761984 | 0.126817059 | 2.198142641 | 0.027938943 | 0.102325012 | 0.278761984 |
| ENSMUSG00000030760 | Acer3 | 66190 | protein_coding | 478.3308842 | -0.278888719 | 0.077466415 | -3.600124262 | 0.000318065 | 0.002983593 | 0.278888719 |
| ENSMUSG00000042377 | Fam83g | 69640 | protein_coding | 617.4180096 | -0.279270248 | 0.203170797 | -1.374559001 | 0.169268207 | 0.354093686 | 0.279270248 |
| ENSMUSG00000025241 | Fyco1 | 17281 | protein_coding | 574.1477026 | -0.279273218 | 0.137730257 | -2.0276824 | 0.042592673 | 0.138773685 | 0.279273218 |
| ENSMUSG00000026156 | B3gat2 | 280645 | protein_coding | 29.27348512 | -0.279470413 | 0.257823049 | -1.083962099 | 0.278381604 | 0.486992003 | 0.279470413 |
| ENSMUSG00000071662 | Polr2g | 67710 | protein_coding | 354.1406232 | 0.279510218 | 0.099281263 | 2.815337059 | 0.00487261 | 0.02753137 | 0.279510218 |
| ENSMUSG00000014177 | Tvp23b | 67510 | protein_coding | 917.5489327 | 0.279533487 | 0.07944669 | 3.518503892 | 0.000433987 | 0.003885639 | 0.279533487 |
| ENSMUSG00000028367 | Txn1 | 22166 | protein_coding | 9652.06576 | 0.279626203 | 0.095755637 | 2.92020619 | 0.003497999 | 0.02120903 | 0.279626203 |
| ENSMUSG00000020918 | Kat2a | 14534 | protein_coding | 272.9476763 | -0.279633327 | 0.112207365 | -2.49211206 | 0.012698597 | 0.057282143 | 0.279633327 |
| ENSMUSG00000038347 | Tcte2 | 21646 | protein_coding | 16.73908156 | -0.279640011 | 0.29477203 | -0.948665351 | 0.342790842 | 0.552987707 | 0.279640011 |
| ENSMUSG00000061531 | Tmem236 | 625286 | protein_coding | 1297.832471 | 0.279675748 | 0.157342615 | 1.777495234 | 0.075486794 | 0.205144798 | 0.279675748 |

| ensembl_gene_id | external_gene_name | entrezgene_id | gene_biotype | baseMean | log2FoldChange | lfcSE | stat | pvalue | padj | absLog2FC |
|---|---|---|---|---|---|---|---|---|---|---|
| ENSMUSG00000071650 | Ganab | 14376 | protein_coding | 3989.451321 | 0.279688529 | 0.063056346 | 4.43553342 | 9.18E-06 | 0.00013827 | 0.279688529 |
| ENSMUSG00000039018 | Mtg1 | 212508 | protein_coding | 108.7108358 | -0.27973816 | 0.124080355 | -2.254491942 | 0.024165237 | 0.092126351 | 0.27973816 |
| ENSMUSG00000074024 | 4632427E13Rik | 666737 | lncRNA | 182.6929252 | -0.279759338 | 0.195472402 | -1.431196096 | 0.152374022 | 0.330926292 | 0.279759338 |
| ENSMUSG00000106617 | Gm36266 | 108169076 | lncRNA | 33.56938886 | -0.280006237 | 0.250254159 | -1.118887446 | 0.263188159 | 0.470933719 | 0.280006237 |
| ENSMUSG00000063253 | Scoc | 56367 | protein_coding | 83.60511188 | 0.280050751 | 0.152437103 | 1.837156086 | 0.066186856 | 0.187985779 | 0.280050751 |
| ENSMUSG00000032105 | Pdzd3 | 170761 | protein_coding | 250.857478 | -0.280062223 | 0.132757422 | -2.109578658 | 0.034894664 | 0.120721592 | 0.280062223 |
| ENSMUSG00000033105 | Lss | 16987 | protein_coding | 899.5176474 | 0.280066158 | 0.163077064 | 1.717385332 | 0.08590879 | 0.224799607 | 0.280066158 |
| ENSMUSG00000014907 | Naf1 | 234344 | protein_coding | 125.1773933 | -0.280077368 | 0.144326206 | -1.940585672 | 0.052308553 | 0.160225123 | 0.280077368 |
| ENSMUSG00000089862 | Umad1 | 100036521 | protein_coding | 336.9579352 | 0.280081961 | 0.136198387 | 2.056426416 | 0.039741432 | 0.132481904 | 0.280081961 |
| ENSMUSG00000020440 | Arf5 | 11844 | protein_coding | 1046.999469 | 0.280162908 | 0.105434737 | 2.657216367 | 0.007878886 | 0.03998202 | 0.280162908 |
| ENSMUSG00000022106 | Rcbtb2 | 105670 | protein_coding | 277.1187793 | 0.280221443 | 0.10131563 | 2.765826386 | 0.005677875 | 0.030834368 | 0.280221443 |
| ENSMUSG00000038599 | Capn8 | 170725 | protein_coding | 369.541938 | 0.280274994 | 0.13894055 | 2.017229625 | 0.043671556 | 0.141353494 | 0.280274994 |
| ENSMUSG00000070737 | Tmem35b | 100039968 | protein_coding | 107.0390973 | -0.280277597 | 0.126708999 | -2.211978618 | 0.026968142 | 0.099660429 | 0.280277597 |
| ENSMUSG00000024170 | Telo2 | 71718 | protein_coding | 158.4041047 | -0.280302763 | 0.11439579 | -2.450289161 | 0.014274153 | 0.06215383 | 0.280302763 |
| ENSMUSG00000060073 | Psma3 | 19167 | protein_coding | 1430.301784 | 0.280380965 | 0.111827313 | 2.5072673 | 0.012166866 | 0.055448884 | 0.280380965 |
| ENSMUSG00000029878 | Dbpht2 | 386753 | lncRNA | 12.1946638 | -0.280416757 | 0.381993708 | -0.73408737 | 0.46289549 | 0.660676356 | 0.280416757 |
| ENSMUSG00000037580 | Gch1 | 14528 | protein_coding | 301.9703678 | 0.280481044 | 0.174633772 | 1.606109979 | 0.108249748 | 0.261882891 | 0.280481044 |
| ENSMUSG00000062082 | Cd200r4 | 239849 | protein_coding | 6.936542108 | 0.280490467 | 0.602038182 | 0.465901459 | 0.641286037 | 0.792368144 | 0.280490467 |
| ENSMUSG00000000552 | Zfp385a | 29813 | protein_coding | 7.614331239 | 0.280701893 | 0.598460279 | 0.469040139 | 0.639040945 | 0.790697132 | 0.280701893 |
| ENSMUSG00000039563 | 2210406O10Rik | 76710 | lncRNA | 6.703213131 | 0.280761562 | 0.558209184 | 0.502968367 | 0.614986513 | 0.77557444 | 0.280761562 |
| ENSMUSG00000030872 | Gga2 | 74105 | protein_coding | 165.7784507 | 0.280838658 | 0.127194664 | 2.20794371 | 0.027248197 | 0.100441862 | 0.280838658 |
| ENSMUSG00000045165 | AI467606 | 101602 | protein_coding | 22.76980185 | 0.280841466 | 0.285422002 | 0.983951704 | 0.32513926 | 0.536276083 | 0.280841466 |
| ENSMUSG00000026750 | Psmb7 | 19177 | protein_coding | 832.3662963 | 0.280863834 | 0.088134741 | 3.186755097 | 0.001438785 | 0.010321491 | 0.280863834 |
| ENSMUSG00000034480 | Diaph2 | 54004 | protein_coding | 243.5922349 | -0.280866649 | 0.11815501 | -2.377103173 | 0.017449207 | 0.072377855 | 0.280866649 |
| ENSMUSG00000027787 | Nmd3 | 97112 | protein_coding | 473.1040245 | -0.2808848 | 0.150994506 | -1.860231925 | 0.06285272 | 0.181230653 | 0.2808848 |
| ENSMUSG00000053684 | Kics2 | 270802 | protein_coding | 46.26192003 | -0.280952843 | 0.216607533 | -1.297059433 | 0.194610738 | 0.387730172 | 0.280952843 |
| ENSMUSG00000007617 | Homer1 | 26556 | protein_coding | 519.8992877 | 0.28096088 | 0.126414222 | 2.222541691 | 0.02624672 | 0.097959566 | 0.28096088 |
| ENSMUSG00000029657 | Hsph1 | 15505 | protein_coding | 1201.263813 | 0.280976458 | 0.268951046 | 1.044712272 | 0.296155971 | 0.505483531 | 0.280976458 |
| ENSMUSG00000046062 | Ppp1r15b | 108954 | protein_coding | 2704.359267 | -0.281038804 | 0.113996006 | -2.465339038 | 0.013688366 | 0.060378924 | 0.281038804 |
| ENSMUSG00000028691 | Prdx1 | 18477 | protein_coding | 8326.280845 | 0.281061678 | 0.095833182 | 2.932822148 | 0.003358962 | 0.020573914 | 0.281061678 |
| ENSMUSG00000040687 | Madd | 228355 | protein_coding | 533.4031767 | -0.281242466 | 0.088341741 | -3.183573954 | 0.001454689 | 0.010399593 | 0.281242466 |
| ENSMUSG00000054814 | Usp46 | 69727 | protein_coding | 401.548901 | -0.281287445 | 0.103778023 | -2.710472186 | 0.006718748 | 0.035249085 | 0.281287445 |
| ENSMUSG00000033434 | Gtpbp6 | 107999 | protein_coding | 86.66273422 | -0.281303846 | 0.171445463 | -1.640777426 | 0.100843626 | 0.249609522 | 0.281303846 |
| ENSMUSG00000006699 | Cdc42 | 12540 | protein_coding | 5836.062132 | 0.281398676 | 0.101060186 | 2.784466253 | 0.00536159 | 0.02952711 | 0.281398676 |
| ENSMUSG00000071516 | H2ac13 | 319191 | protein_coding | 150.4634158 | 0.281512807 | 0.310353211 | 0.907072319 | 0.364368555 | 0.573201219 | 0.281512807 |
| ENSMUSG00000049489 | Ccnq | 69109 | protein_coding | 133.0053506 | 0.281533823 | 0.164242252 | 1.714137625 | 0.086503453 | 0.22594736 | 0.281533823 |
| ENSMUSG00000090173 | Fbxw10 | 213980 | protein_coding | 14.42051043 | 0.28154744 | 0.405869509 | 0.693689557 | 0.487876917 | 0.680726764 | 0.28154744 |
| ENSMUSG00000028961 | Pgd | 110208 | protein_coding | 2212.320412 | 0.281550135 | 0.090866392 | 3.098506824 | 0.001944985 | 0.013225891 | 0.281550135 |

| ensembl_gene_id | external_gene_name | entrezgene_id | gene_biotype | baseMean | log2FoldChange | lfcSE | stat | pvalue | padj | absLog2FC |
|---|---|---|---|---|---|---|---|---|---|---|
| ENSMUSG00000027387 | Zc3h8 | 57432 | protein_coding | 52.00199954 | -0.281565913 | 0.176500295 | -1.595271628 | 0.110651507 | 0.265390334 | 0.281565913 |
| ENSMUSG00000028578 | Caap1 | 67770 | protein_coding | 178.4963559 | 0.281588799 | 0.107738215 | 2.613638994 | 0.008958364 | 0.044053355 | 0.281588799 |
| ENSMUSG00000045679 | Pqlc3 | 217430 | protein_coding | 440.2329836 | 0.281610509 | 0.088981638 | 3.164815983 | 0.001551811 | 0.01094178 | 0.281610509 |
| ENSMUSG00000039099 | Wdr93 | 626359 | protein_coding | 6.708011045 | 0.281644173 | 0.557421377 | 0.505262599 | 0.613374408 | 0.774829691 | 0.281644173 |
| ENSMUSG00000039917 | Rhbdd2 | 215160 | protein_coding | 162.5953382 | -0.281657056 | 0.097489876 | -2.889090301 | 0.003863581 | 0.022879306 | 0.281657056 |
| ENSMUSG00000033066 | Gas7 | 14457 | protein_coding | 124.5936541 | -0.281726574 | 0.154067126 | -1.828596287 | 0.067460109 | 0.190515004 | 0.281726574 |
| ENSMUSG00000058145 | Adamts17 | 233332 | protein_coding | 82.95739535 | 0.281730761 | 0.26268893 | 1.072488134 | 0.283500841 | 0.49264689 | 0.281730761 |
| ENSMUSG00000027699 | Ect2 | 13605 | protein_coding | 762.7927503 | 0.281759114 | 0.119043254 | 2.366863349 | 0.017939555 | 0.073758708 | 0.281759114 |
| ENSMUSG00000038279 | Nop2 | 110109 | protein_coding | 485.4772313 | -0.281819193 | 0.178192895 | -1.581540016 | 0.113754617 | 0.269866479 | 0.281819193 |
| ENSMUSG00000027523 | Gnas | 14683 | protein_coding | 116.9813997 | -0.281843811 | 0.236548365 | -1.191484923 | 0.233463272 | 0.437122722 | 0.281843811 |
| ENSMUSG00000074505 | Fat3 | 270120 | protein_coding | 9.507242266 | -0.281879275 | 1.798052432 | -0.156769219 | 0.875426733 | 0.936642507 | 0.281879275 |
| ENSMUSG00000038290 | Smg6 | 103677 | protein_coding | 560.0466944 | -0.282014247 | 0.124790985 | -2.259892793 | 0.023827905 | 0.091167638 | 0.282014247 |
| ENSMUSG00000020802 | Ube2o | 217342 | protein_coding | 245.3496144 | -0.282045927 | 0.118828862 | -2.373547317 | 0.017618136 | 0.072765884 | 0.282045927 |
| ENSMUSG00000070777 | Ceacam20 | 71601 | protein_coding | 843.6244645 | 0.282060223 | 0.136196396 | 2.070981551 | 0.03836052 | 0.129258804 | 0.282060223 |
| ENSMUSG00000069308 | H2bc22 | 319188 | protein_coding | 52.19882208 | 0.282073854 | 0.194682052 | 1.448895014 | 0.147366904 | 0.323938731 | 0.282073854 |
| ENSMUSG00000026842 | Abl1 | 11350 | protein_coding | 1074.16728 | -0.282121821 | 0.08211534 | -3.435677456 | 0.000591074 | 0.00499715 | 0.282121821 |
| ENSMUSG00000039810 | Zc3h10 | 103284 | protein_coding | 196.0744224 | -0.282128112 | 0.089772787 | -3.142690794 | 0.001674026 | 0.011648703 | 0.282128112 |
| ENSMUSG00000038240 | Pdss2 | 71365 | protein_coding | 471.7729479 | 0.282233803 | 0.091935413 | 3.069913903 | 0.002141205 | 0.014317847 | 0.282233803 |
| ENSMUSG00000010048 | Ifrd2 | 15983 | protein_coding | 331.8977992 | -0.282388541 | 0.141507728 | -1.995569748 | 0.045980775 | 0.146858188 | 0.282388541 |
| ENSMUSG00000096878 | Gm21083 | 105242399 | protein_coding | 7.512132338 | -0.282469144 | 0.472740881 | -0.597513682 | 0.550164475 | 0.731287638 | 0.282469144 |
| ENSMUSG00000023452 | Pisd | 320951 | protein_coding | 704.867656 | -0.282492377 | 0.081574255 | -3.463008979 | 0.000534171 | 0.004601552 | 0.282492377 |
| ENSMUSG00000037606 | Osbpl5 | 79196 | protein_coding | 341.4336239 | -0.282505452 | 0.127191022 | -2.221111588 | 0.026343404 | 0.098174484 | 0.282505452 |
| ENSMUSG00000056228 | Cars2 | 71941 | protein_coding | 280.1450018 | 0.282515041 | 0.087478857 | 3.229523691 | 0.001239966 | 0.009109861 | 0.282515041 |
| ENSMUSG00000030946 | Lhpp | 76429 | protein_coding | 155.647136 | -0.282576755 | 0.119569093 | -2.363292624 | 0.018113361 | 0.074243338 | 0.282576755 |
| ENSMUSG00000026361 | Cdc73 | 214498 | protein_coding | 690.0696878 | 0.282584836 | 0.157136714 | 1.798337437 | 0.072123551 | 0.199564716 | 0.282584836 |
| ENSMUSG00000029246 | Ppat | 231327 | protein_coding | 344.477949 | -0.282602319 | 0.116407304 | -2.42770263 | 0.015194796 | 0.065323269 | 0.282602319 |
| ENSMUSG00000060860 | Ube2s | 77891 | protein_coding | 434.6827627 | 0.282622267 | 0.131423279 | 2.150473419 | 0.031517786 | 0.112056987 | 0.282622267 |
| ENSMUSG00000038244 | Mical2 | 320878 | protein_coding | 1205.360611 | -0.282684181 | 0.115663493 | -2.444022518 | 0.01452452 | 0.062956904 | 0.282684181 |
| ENSMUSG00000025969 | Nrp2 | 18187 | protein_coding | 1600.070836 | -0.28272377 | 0.163239488 | -1.731956979 | 0.083281223 | 0.220129989 | 0.28272377 |
| ENSMUSG00000020747 | Tmem94 | 71947 | protein_coding | 1628.369028 | -0.282734002 | 0.077672951 | -3.640057449 | 0.000272577 | 0.002629685 | 0.282734002 |
| ENSMUSG00000029575 | Mmab | 77697 | protein_coding | 256.2441738 | -0.282748461 | 0.116492342 | -2.427184961 | 0.015216496 | 0.065363632 | 0.282748461 |
| ENSMUSG00000054545 | Ugt1a6a | 94284 | protein_coding | 92.98877873 | -0.282760136 | 0.215472389 | -1.312280135 | 0.189425632 | 0.3810041 | 0.282760136 |
| ENSMUSG00000039067 | Psmd7 | 17463 | protein_coding | 919.0229864 | 0.28289976 | 0.102022453 | 2.77291665 | 0.005555635 | 0.030317273 | 0.28289976 |
| ENSMUSG00000047789 | Slc38a9 | 268706 | protein_coding | 182.5605642 | -0.28300454 | 0.121072212 | -2.337485501 | 0.019413954 | 0.07830475 | 0.28300454 |
| ENSMUSG00000038740 | Mvb12b | 72543 | protein_coding | 78.63838173 | -0.283024549 | 0.195164689 | -1.450183178 | 0.147007445 | 0.323461056 | 0.283024549 |
| ENSMUSG00000014769 | Psmb1 | 19170 | protein_coding | 2201.02247 | 0.283118176 | 0.087643959 | 3.230321624 | 0.00123651 | 0.009105472 | 0.283118176 |
| ENSMUSG00000037621 | Atoh8 | 71093 | protein_coding | 27.96522272 | 0.283181017 | 0.261847788 | 1.081471871 | 0.279487266 | 0.487908064 | 0.283181017 |
| ENSMUSG00000085741 | 5430405H02Rik | 74487 | lncRNA | 17.57151452 | 0.283221502 | 0.271165679 | 1.044459252 | 0.296272962 | 0.505629031 | 0.283221502 |

| ensembl_gene_id | external_gene_name | entrezgene_id | gene_biotype | baseMean | log2FoldChange | lfcSE | stat | pvalue | padj | absLog2FC |
|---|---|---|---|---|---|---|---|---|---|---|
| ENSMUSG00000029128 | Rab28 | 100972 | protein_coding | 376.9978028 | 0.283251653 | 0.079079404 | 3.581863796 | 0.000341152 | 0.003166587 | 0.283251653 |
| ENSMUSG00000045441 | Gprin3 | 243385 | protein_coding | 476.2577905 | -0.283319842 | 0.166284465 | -1.703826285 | 0.088413546 | 0.229170213 | 0.283319842 |
| ENSMUSG00000019989 | Enpp3 | 209558 | protein_coding | 29.32805184 | 0.283334018 | 0.274413738 | 1.032506683 | 0.301834818 | 0.511231921 | 0.283334018 |
| ENSMUSG00000052533 | Nup188 | 227699 | protein_coding | 334.5842271 | -0.283346766 | 0.088752945 | -3.192533682 | 0.001410305 | 0.010155216 | 0.283346766 |
| ENSMUSG00000047844 | Bex4 | 406217 | protein_coding | 22.62427082 | -0.283400153 | 0.299709701 | -0.945582181 | 0.344361736 | 0.554151721 | 0.283400153 |
| ENSMUSG00000021713 | Ppwd1 | 238831 | protein_coding | 251.2318868 | 0.283425846 | 0.087005209 | 3.257573322 | 0.001123692 | 0.008435802 | 0.283425846 |
| ENSMUSG00000072640 | Lyrm9 | 66274 | protein_coding | 123.9466419 | -0.28372786 | 0.134654305 | -2.107083467 | 0.035110345 | 0.121317106 | 0.28372786 |
| ENSMUSG00000021209 | Ppp4r4 | 74521 | protein_coding | 11.78753321 | -0.283737288 | 0.39758555 | -0.713650907 | 0.475443071 | 0.670223668 | 0.283737288 |
| ENSMUSG00000022120 | Obi1 | 72486 | protein_coding | 178.6241427 | -0.283858164 | 0.108273486 | -2.62167752 | 0.008749818 | 0.043349021 | 0.283858164 |
| ENSMUSG00000023277 | Twf2 | 23999 | protein_coding | 211.6941823 | 0.283902357 | 0.108220468 | 2.623370253 | 0.00870646 | 0.043211867 | 0.283902357 |
| ENSMUSG00000028745 | Capzb | 12345 | protein_coding | 1985.532077 | 0.2839038 | 0.078071413 | 3.636462939 | 0.000276407 | 0.002656981 | 0.2839038 |
| ENSMUSG00000006567 | Atp7b | 11979 | protein_coding | 19.21856129 | -0.283927435 | 0.317120019 | -0.895331164 | 0.370610086 | 0.579187348 | 0.283927435 |
| ENSMUSG00000052915 | Msl1 | 74026 | protein_coding | 714.3395237 | -0.283963562 | 0.066947647 | -4.24157643 | 2.22E-05 | 0.00030268 | 0.283963562 |
| ENSMUSG00000025529 | Zfp711 | 245595 | protein_coding | 53.24632016 | 0.284010128 | 0.294233232 | 0.965255103 | 0.334417054 | 0.545293316 | 0.284010128 |
| ENSMUSG00000027115 | Kif18a | 228421 | protein_coding | 214.4862343 | 0.284093458 | 0.16885433 | 1.682476594 | 0.09247646 | 0.236203505 | 0.284093458 |
| ENSMUSG00000003863 | Ppfia3 | 76787 | protein_coding | 332.6883291 | -0.284140852 | 0.136731705 | -2.078090465 | 0.037701028 | 0.127739198 | 0.284140852 |
| ENSMUSG00000026074 | Map4k4 | 26921 | protein_coding | 329.0029702 | -0.284143934 | 0.130526319 | -2.176909117 | 0.029487349 | 0.106502223 | 0.284143934 |
| ENSMUSG00000030357 | Fkbp4 | 14228 | protein_coding | 4191.891677 | 0.284268169 | 0.103235914 | 2.753578273 | 0.005894767 | 0.031741668 | 0.284268169 |
| ENSMUSG00000068086 | Cyp2d9 | 13105 | protein_coding | 56.6149181 | 0.284284502 | 0.258914836 | 1.097984598 | 0.272211215 | 0.4804721 | 0.284284502 |
| ENSMUSG00000024388 | Myo7b | 17922 | protein_coding | 3101.843067 | -0.284356262 | 0.121036478 | -2.349343493 | 0.018806548 | 0.076455004 | 0.284356262 |
| ENSMUSG00000097350 | 4732491K20Rik | 224523 | lncRNA | 21.39610762 | 0.284398678 | 0.272807372 | 1.042488977 | 0.297185033 | 0.506180721 | 0.284398678 |
| ENSMUSG00000090171 | Ugt1a2 | 22236 | protein_coding | 8.117072177 | -0.284453443 | 0.575863026 | -0.493960248 | 0.621334254 | 0.779922636 | 0.284453443 |
| ENSMUSG00000018199 | Ro60 | 20822 | protein_coding | 454.202552 | 0.284560678 | 0.109095839 | 2.608355005 | 0.009097855 | 0.044587886 | 0.284560678 |
| ENSMUSG00000040841 | Six5 | 20475 | protein_coding | 46.68807037 | -0.284598923 | 0.201299883 | -1.413805707 | 0.157418957 | 0.33818806 | 0.284598923 |
| ENSMUSG00000013275 | Slc41a1 | 98396 | protein_coding | 212.112915 | -0.284616432 | 0.111155918 | -2.560515316 | 0.010451706 | 0.049435191 | 0.284616432 |
| ENSMUSG00000040016 | Ptger3 | 19218 | protein_coding | 96.08639922 | -0.284661864 | 0.246306829 | -1.155720547 | 0.247795481 | 0.453560839 | 0.284661864 |
| ENSMUSG00000032340 | Neo1 | 18007 | protein_coding | 1584.229945 | -0.28471474 | 0.115394937 | -2.467307036 | 0.013613357 | 0.060214816 | 0.28471474 |
| ENSMUSG00000022750 | Klhl22 | 224023 | protein_coding | 404.9116111 | 0.284742774 | 0.099525821 | 2.860993959 | 0.004223151 | 0.024505045 | 0.284742774 |
| ENSMUSG00000038302 | Afg1l | 215951 | protein_coding | 408.0621998 | 0.28485416 | 0.129877095 | 2.193259401 | 0.028288695 | 0.103239441 | 0.28485416 |
| ENSMUSG00000030878 | Cdr2 | 12585 | protein_coding | 128.9480811 | 0.284904788 | 0.242732071 | 1.173741841 | 0.240498445 | 0.444959837 | 0.284904788 |
| ENSMUSG00000025911 | Adhfe1 | 76187 | protein_coding | 34.76660586 | -0.284913542 | 0.20376064 | -1.398275652 | 0.162030306 | 0.344355804 | 0.284913542 |
| ENSMUSG00000020458 | Rtn4 | 68585 | protein_coding | 1453.529435 | -0.285125399 | 0.179619591 | -1.587384746 | 0.112425532 | 0.267988275 | 0.285125399 |
| ENSMUSG00000048612 | Myof | 226101 | protein_coding | 896.7353898 | 0.285143669 | 0.191779157 | 1.486833468 | 0.1370588 | 0.307908683 | 0.285143669 |
| ENSMUSG00000051391 | Ywhag | 22628 | protein_coding | 907.7247324 | -0.285247386 | 0.173588379 | -1.643240104 | 0.100333265 | 0.248655399 | 0.285247386 |
| ENSMUSG00000031107 | Rbmx2 | 209003 | protein_coding | 67.46115753 | 0.285291616 | 0.157630632 | 1.809874216 | 0.070315295 | 0.196109617 | 0.285291616 |
| ENSMUSG00000032018 | Sc5d | 235293 | protein_coding | 1189.211441 | 0.28534807 | 0.096833397 | 2.946793954 | 0.003210871 | 0.019822771 | 0.28534807 |
| ENSMUSG00000112627 | 4933412E12Rik | 71086 | lncRNA | 9.518902727 | 0.285390988 | 0.475209558 | 0.600558182 | 0.548134298 | 0.730067647 | 0.285390988 |
| ENSMUSG00000030451 | Herc2 | 15204 | protein_coding | 2377.222177 | 0.285410274 | 0.089490023 | 3.189297119 | 0.001426192 | 0.010246454 | 0.285410274 |

Supplementary Table 1

| ensembl_gene_id | external_gene_name | entrezgene_id | gene_biotype | baseMean | log2FoldChange | lfcSE | stat | pvalue | padj | absLog2FC |
|---|---|---|---|---|---|---|---|---|---|---|
| ENSMUSG00000033871 | Ppargc1b | 170826 | protein_coding | 492.92035 | -0.285464441 | 0.152956022 | -1.866317113 | 0.061997012 | 0.179611609 | 0.285464441 |
| ENSMUSG00000026773 | Pfkfb3 | 170768 | protein_coding | 29.90112548 | 0.285510808 | 0.353772532 | 0.807046287 | 0.419639816 | 0.625680983 | 0.285510808 |
| ENSMUSG00000040945 | Rcc2 | 108911 | protein_coding | 908.9681064 | -0.28560561 | 0.086561506 | -3.299452882 | 0.000968735 | 0.007490296 | 0.28560561 |
| ENSMUSG00000074918 | Inafm2 | 100043272 | protein_coding | 311.1875517 | -0.28568817 | 0.133884723 | -2.133836961 | 0.032856129 | 0.115296855 | 0.28568817 |
| ENSMUSG00000065822 | Snord15a | 449630 | snoRNA | 32.51976788 | -0.285730849 | 0.379579642 | -0.752755989 | 0.451596558 | 0.651785414 | 0.285730849 |
| ENSMUSG00000021193 | Pitrm1 | 69617 | protein_coding | 642.9327712 | 0.285775677 | 0.090023589 | 3.174453294 | 0.00150119 | 0.010684076 | 0.285775677 |
| ENSMUSG00000039648 | Kyat1 | 70266 | protein_coding | 214.645331 | -0.285950397 | 0.13672062 | -2.091494296 | 0.036483777 | 0.124625545 | 0.285950397 |
| ENSMUSG00000097855 | A930007I19Rik | 77779 | lncRNA | 11.24832989 | -0.285957927 | 0.472805495 | -0.60481092 | 0.545304637 | 0.727864934 | 0.285957927 |
| ENSMUSG00000043286 | Pnpla1 | 433091 | protein_coding | 27.48361133 | 0.285992945 | 0.22822126 | 1.253138927 | 0.210155151 | 0.40740724 | 0.285992945 |
| ENSMUSG00000026683 | Nuf2 | 66977 | protein_coding | 242.8849797 | 0.286015531 | 0.127179923 | 2.248904731 | 0.024518558 | 0.093028011 | 0.286015531 |
| ENSMUSG00000032185 | Carm1 | 59035 | protein_coding | 281.315995 | -0.286074448 | 0.114616266 | -2.49593238 | 0.012562655 | 0.056813732 | 0.286074448 |
| ENSMUSG00000014039 | Prdm15 | 114604 | protein_coding | 321.8846019 | -0.286079633 | 0.103905321 | -2.753272211 | 0.005900281 | 0.031760623 | 0.286079633 |
| ENSMUSG00000038646 | Ramac | 67148 | protein_coding | 566.5179946 | 0.28621968 | 0.101674477 | 2.815059274 | 0.004876824 | 0.027535643 | 0.28621968 |
| ENSMUSG00000100826 | Snhg14 | 16353 | lncRNA | 59.86146347 | -0.286329707 | 0.333340727 | -0.858970069 | 0.390357029 | 0.598595048 | 0.286329707 |
| ENSMUSG00000038002 | Cramp1 | 57354 | protein_coding | 503.2843493 | -0.286332879 | 0.150770387 | -1.899132079 | 0.057547112 | 0.171329048 | 0.286332879 |
| ENSMUSG00000003848 | Nob1 | 67619 | protein_coding | 219.5091959 | -0.286356891 | 0.110280744 | -2.596617324 | 0.009414675 | 0.045760433 | 0.286356891 |
| ENSMUSG00000067006 | Serpinb5 | 20724 | protein_coding | 1197.352162 | -0.286417458 | 0.093063541 | -3.077654826 | 0.002086364 | 0.01400405 | 0.286417458 |
| ENSMUSG00000022364 | Tbc1d31 | 210544 | protein_coding | 124.554113 | 0.286420065 | 0.137772166 | 2.078939991 | 0.037622866 | 0.127555772 | 0.286420065 |
| ENSMUSG00000047989 | Ino80c | 225280 | protein_coding | 406.8772765 | 0.286553303 | 0.161032544 | 1.779474477 | 0.075162006 | 0.204541334 | 0.286553303 |
| ENSMUSG00000063652 | Slc22a21 | 56517 | protein_coding | 130.1261718 | -0.286555919 | 0.126876131 | -2.258548683 | 0.023911473 | 0.091399554 | 0.286555919 |
| ENSMUSG00000021149 | Gtpbp4 | 69237 | protein_coding | 692.4725418 | -0.286588158 | 0.18808382 | -1.523725744 | 0.127577236 | 0.293118459 | 0.286588158 |
| ENSMUSG00000069456 | Rdh16 | 19683 | protein_coding | 275.8277965 | 0.286611867 | 0.191689971 | 1.495184472 | 0.134866305 | 0.304184439 | 0.286611867 |
| ENSMUSG00000048481 | Mypop | 232934 | protein_coding | 18.65587158 | -0.286631283 | 0.297240056 | -0.964309073 | 0.334890994 | 0.545413471 | 0.286631283 |
| ENSMUSG00000032009 | Sesn3 | 75747 | protein_coding | 595.4465113 | 0.28675563 | 0.151579136 | 1.891788255 | 0.058519199 | 0.172994395 | 0.28675563 |
| ENSMUSG00000073557 | Ppp1r12b | 329251 | protein_coding | 152.1300538 | -0.28677828 | 0.18229719 | -1.573136043 | 0.115687321 | 0.273311724 | 0.28677828 |
| ENSMUSG00000056698 | Elmod3 | 232089 | protein_coding | 141.6563016 | -0.286903905 | 0.156391843 | -1.834519628 | 0.066576897 | 0.188667907 | 0.286903905 |
| ENSMUSG00000036199 | Ndufa13 | 67184 | protein_coding | 1802.165699 | 0.28691405 | 0.098657665 | 2.908177993 | 0.003635414 | 0.021826184 | 0.28691405 |
| ENSMUSG00000032258 | Lca5 | 75782 | protein_coding | 42.69366458 | 0.286919706 | 0.219527612 | 1.306986871 | 0.19121717 | 0.38368696 | 0.286919706 |
| ENSMUSG00000034595 | Ppp1r18 | 76448 | protein_coding | 12.61446044 | 0.286950978 | 0.485779357 | 0.590702289 | 0.554719914 | 0.73440103 | 0.286950978 |
| ENSMUSG00000044352 | Sowaha | 237761 | protein_coding | 48.94826113 | -0.286961779 | 0.181623027 | -1.579985663 | 0.11411015 | 0.270508479 | 0.286961779 |
| ENSMUSG00000024462 | Gabbr1 | 54393 | protein_coding | 95.92247182 | -0.28703252 | 0.247348291 | -1.160438661 | 0.245870254 | 0.451177581 | 0.28703252 |
| ENSMUSG00000073639 | Rab18 | 19330 | protein_coding | 1269.366216 | 0.287088743 | 0.080180405 | 3.580534952 | 0.000342891 | 0.003171647 | 0.287088743 |
| ENSMUSG00000031016 | Wee1 | 22390 | protein_coding | 200.7635808 | -0.28716095 | 0.142466525 | -2.015638063 | 0.043837837 | 0.14163267 | 0.28716095 |
| ENSMUSG00000024713 | Pcsk5 | 18552 | protein_coding | 613.1991042 | -0.287258533 | 0.176502966 | -1.627499752 | 0.103631001 | 0.254232438 | 0.287258533 |
| ENSMUSG00000025429 | Pstpip2 | 19201 | protein_coding | 148.9682083 | -0.287263683 | 0.151586316 | -1.895050224 | 0.058085748 | 0.172192406 | 0.287263683 |
| ENSMUSG00000024913 | Lrp5 | 16973 | protein_coding | 946.6966343 | -0.287330525 | 0.105815777 | -2.715384539 | 0.006619886 | 0.034845187 | 0.287330525 |
| ENSMUSG00000035365 | Parpbp | 75317 | protein_coding | 164.8507331 | 0.287348789 | 0.167013837 | 1.720508872 | 0.085339982 | 0.223728497 | 0.287348789 |
| ENSMUSG00000009378 | Slc16a12 | 240638 | protein_coding | 67.56895666 | 0.287438715 | 0.433350136 | 0.663294391 | 0.507142028 | 0.696658751 | 0.287438715 |

| ensembl_gene_id | external_gene_name | entrezgene_id | gene_biotype | baseMean | log2FoldChange | lfcSE | stat | pvalue | padj | absLog2FC |
|---|---|---|---|---|---|---|---|---|---|---|
| ENSMUSG00000041540 | Sox5 | 20678 | protein_coding | 13.60064617 | -0.287440507 | 0.553229984 | -0.519567839 | 0.603364819 | 0.768032832 | 0.287440507 |
| ENSMUSG00000028199 | Cryz | 12972 | protein_coding | 215.1201523 | -0.287498089 | 0.184494101 | -1.558305044 | 0.119160954 | 0.279775305 | 0.287498089 |
| ENSMUSG00000022322 | Shcbp1 | 20419 | protein_coding | 254.9793746 | 0.287541752 | 0.124131387 | 2.316430664 | 0.020534766 | 0.081545189 | 0.287541752 |
| ENSMUSG00000062963 | Ufc1 | 66155 | protein_coding | 1035.102144 | 0.287566475 | 0.11178156 | 2.572575256 | 0.010094501 | 0.048209507 | 0.287566475 |
| ENSMUSG00000020308 | Tpgs1 | 110012 | protein_coding | 149.2795549 | 0.287593558 | 0.165450452 | 1.738245823 | 0.082167508 | 0.218196662 | 0.287593558 |
| ENSMUSG00000000127 | Fer | 14158 | protein_coding | 279.614768 | 0.287657641 | 0.087470429 | 3.288627322 | 0.001006772 | 0.007709553 | 0.287657641 |
| ENSMUSG00000030894 | Tpp1 | 12751 | protein_coding | 373.7571981 | -0.287701148 | 0.074971119 | -3.837493044 | 0.000124297 | 0.001358819 | 0.287701148 |
| ENSMUSG00000034329 | Brip1 | 237911 | protein_coding | 175.5924959 | 0.28786546 | 0.136830285 | 2.103813934 | 0.035394681 | 0.122027376 | 0.28786546 |
| ENSMUSG00000034430 | Zxdc | 80292 | protein_coding | 313.6292585 | -0.287959395 | 0.093799927 | -3.069931966 | 0.002141075 | 0.014317847 | 0.287959395 |
| ENSMUSG00000033220 | Rac2 | 19354 | protein_coding | 45.15081904 | -0.28798663 | 0.245862644 | -1.171331382 | 0.241465602 | 0.446127376 | 0.28798663 |
| ENSMUSG00000027469 | Tpx2 | 72119 | protein_coding | 638.1303054 | 0.288180544 | 0.110881597 | 2.598993447 | 0.009349756 | 0.045528251 | 0.288180544 |
| ENSMUSG00000026790 | Odf2 | 18286 | protein_coding | 525.2380749 | -0.288276691 | 0.096148734 | -2.998236998 | 0.002715464 | 0.017349343 | 0.288276691 |
| ENSMUSG00000020386 | Sar1b | 66397 | protein_coding | 913.0976923 | 0.288295911 | 0.10072432 | 2.862227416 | 0.004206749 | 0.024454418 | 0.288295911 |
| ENSMUSG00000026430 | Rassf5 | 54354 | protein_coding | 21.72285783 | 0.288436432 | 0.293285614 | 0.983466005 | 0.325378138 | 0.536498673 | 0.288436432 |
| ENSMUSG00000053641 | Dennd4a | 102442 | protein_coding | 211.1755599 | -0.288437624 | 0.125491008 | -2.29847244 | 0.021534915 | 0.084360089 | 0.288437624 |
| ENSMUSG00000025413 | Ttc4 | 72354 | protein_coding | 672.307028 | 0.288444761 | 0.093034309 | 3.100412792 | 0.001932511 | 0.013159913 | 0.288444761 |
| ENSMUSG00000001158 | Snrnp27 | 66618 | protein_coding | 317.3781611 | 0.288445935 | 0.098834164 | 2.918484092 | 0.003517378 | 0.021302205 | 0.288445935 |
| ENSMUSG00000038059 | Smim3 | 106878 | protein_coding | 161.3458311 | -0.28850377 | 0.39573116 | -0.729039811 | 0.46597731 | 0.662745477 | 0.28850377 |
| ENSMUSG00000061136 | Prpf40a | 56194 | protein_coding | 1991.936378 | 0.288518983 | 0.06510365 | 4.431686768 | 9.35E-06 | 0.000140363 | 0.288518983 |
| ENSMUSG00000028553 | Angptl3 | 30924 | protein_coding | 10.95759837 | -0.288564338 | 0.424200984 | -0.680253815 | 0.496343763 | 0.687397918 | 0.288564338 |
| ENSMUSG00000053164 | Gpr21 | 338346 | protein_coding | 15.33409989 | -0.288638518 | 0.368312903 | -0.783677453 | 0.433229396 | 0.637254688 | 0.288638518 |
| ENSMUSG00000061461 | Smim20 | 66278 | protein_coding | 470.3314206 | 0.28866149 | 0.120345674 | 2.398602965 | 0.016457749 | 0.069257337 | 0.28866149 |
| ENSMUSG00000054405 | Dnajc8 | 68598 | protein_coding | 573.0636167 | -0.288722755 | 0.08455166 | -3.414749701 | 0.000638406 | 0.005337815 | 0.288722755 |
| ENSMUSG00000049657 | Zbtb5 | 230119 | protein_coding | 145.6380424 | 0.288754529 | 0.167725018 | 1.721594865 | 0.085142934 | 0.223382746 | 0.288754529 |
| ENSMUSG00000028555 | Ttc39a | 230603 | protein_coding | 2043.135759 | 0.28883275 | 0.084846578 | 3.404176764 | 0.000663638 | 0.005499701 | 0.28883275 |
| ENSMUSG00000085007 | Smim43 | 100503068 | protein_coding | 11.05720759 | -0.288910316 | 0.433398527 | -0.666615823 | 0.50501756 | 0.694999109 | 0.288910316 |
| ENSMUSG00000030754 | Copb1 | 70349 | protein_coding | 3393.795354 | 0.289013899 | 0.071287408 | 4.054206885 | 5.03E-05 | 0.000623058 | 0.289013899 |
| ENSMUSG00000008496 | Pou2f2 | 18987 | protein_coding | 6.781213225 | -0.289039027 | 1.078995928 | -0.267877774 | 0.788793403 | 0.885342916 | 0.289039027 |
| ENSMUSG00000058486 | Wdr91 | 101240 | protein_coding | 57.68170033 | -0.289051975 | 0.174041893 | -1.660818374 | 0.09674993 | 0.243141824 | 0.289051975 |
| ENSMUSG00000035342 | Lzts2 | 226154 | protein_coding | 804.8103281 | -0.289102002 | 0.117183337 | -2.467091402 | 0.013621558 | 0.060217646 | 0.289102002 |
| ENSMUSG00000070167 | Snora57 | 118027464 | snoRNA | 66.37033169 | 0.289102975 | 0.226126001 | 1.278503903 | 0.201071809 | 0.395684678 | 0.289102975 |
| ENSMUSG00000031844 | Hsd17b2 | 15486 | protein_coding | 1912.276739 | -0.289111502 | 0.151046992 | -1.914050042 | 0.055613762 | 0.167016468 | 0.289111502 |
| ENSMUSG00000034171 | Faah | 14073 | protein_coding | 418.6748968 | -0.289123113 | 0.082251806 | -3.515097436 | 0.000439593 | 0.003926995 | 0.289123113 |
| ENSMUSG00000056411 | Gm12500 | 791415 | lncRNA | 9.373367428 | 0.289125353 | 0.400850265 | 0.721280185 | 0.470737146 | 0.666501035 | 0.289125353 |
| ENSMUSG00000026924 | Sec16a | 227648 | protein_coding | 2425.383085 | 0.289169715 | 0.086603318 | 3.339014259 | 0.000840762 | 0.006642691 | 0.289169715 |
| ENSMUSG00000018567 | Gabarap | 56486 | protein_coding | 3148.230138 | 0.289193467 | 0.089945665 | 3.21520182 | 0.001303528 | 0.009476312 | 0.289193467 |
| ENSMUSG00000061758 | Akr1b10 | 67861 | protein_coding | 107.2542254 | 0.289208133 | 0.117765627 | 2.455794101 | 0.014057367 | 0.061665037 | 0.289208133 |
| ENSMUSG00000051234 | Rnf7 | 19823 | protein_coding | 695.3294361 | 0.289278954 | 0.087648118 | 3.300458242 | 0.000965271 | 0.007467136 | 0.289278954 |

| ensembl_gene_id | external_gene_name | entrezgene_id | gene_biotype | baseMean | log2FoldChange | lfcSE | stat | pvalue | padj | absLog2FC |
|---|---|---|---|---|---|---|---|---|---|---|
| ENSMUSG00000041528 | Rnf123 | 84585 | protein_coding | 380.0391304 | -0.289285283 | 0.095799905 | -3.01968237 | 0.002530399 | 0.016464132 | 0.289285283 |
| ENSMUSG00000034462 | Pkd2 | 18764 | protein_coding | 237.4244768 | -0.289419279 | 0.10421083 | -2.777247603 | 0.00548214 | 0.030017025 | 0.289419279 |
| ENSMUSG00000074305 | Peak1 | 244895 | protein_coding | 302.9257356 | -0.289449556 | 0.196433157 | -1.473526977 | 0.140608995 | 0.312983217 | 0.289449556 |
| ENSMUSG00000015750 | Aph1a | 226548 | protein_coding | 921.2505919 | 0.289533571 | 0.112579029 | 2.57182509 | 0.010116399 | 0.048243712 | 0.289533571 |
| ENSMUSG00000003038 | Hmgn2 | 15331 | protein_coding | 797.4829476 | 0.289534171 | 0.083861968 | 3.452508675 | 0.0005554 | 0.004746713 | 0.289534171 |
| ENSMUSG00000028453 | Fancg | 60534 | protein_coding | 113.9386626 | 0.289580045 | 0.188777639 | 1.533974291 | 0.125035978 | 0.289176702 | 0.289580045 |
| ENSMUSG00000032513 | Gorasp1 | 74498 | protein_coding | 524.6652838 | 0.289695036 | 0.115227278 | 2.514118533 | 0.011933032 | 0.054680475 | 0.289695036 |
| ENSMUSG00000002346 | Slc25a42 | 73095 | protein_coding | 57.88327811 | -0.289706395 | 0.255335631 | -1.134610136 | 0.256538704 | 0.463387217 | 0.289706395 |
| ENSMUSG00000027602 | Map1lc3a | 66734 | protein_coding | 235.3328444 | -0.289715485 | 0.144458712 | -2.005524484 | 0.044907006 | 0.144325825 | 0.289715485 |
| ENSMUSG00000003549 | Ercc1 | 13870 | protein_coding | 128.2670668 | 0.289772186 | 0.184581123 | 1.56989069 | 0.116440544 | 0.274745608 | 0.289772186 |
| ENSMUSG00000021277 | Traf3 | 22031 | protein_coding | 380.4330276 | -0.289836046 | 0.124063209 | -2.336196585 | 0.019480998 | 0.078455964 | 0.289836046 |
| ENSMUSG00000097404 | Gm10814 | 100502985 | lncRNA | 8.763593777 | 0.289920656 | 0.371136211 | 0.781170491 | 0.434702227 | 0.637867413 | 0.289920656 |
| ENSMUSG00000037211 | Spry1 | 24063 | protein_coding | 133.9632272 | -0.290123791 | 0.141298097 | -2.053274581 | 0.040045955 | 0.133272807 | 0.290123791 |
| ENSMUSG00000000276 | Dgke | 56077 | protein_coding | 151.1130269 | -0.290135523 | 0.153293789 | -1.892676305 | 0.058400929 | 0.172773032 | 0.290135523 |
| ENSMUSG00000051147 | Nat2 | 17961 | protein_coding | 381.3250917 | 0.290203747 | 0.155092315 | 1.871167802 | 0.061321826 | 0.178203621 | 0.290203747 |
| ENSMUSG00000021969 | Zdhhc20 | 75965 | protein_coding | 1474.868826 | 0.290208597 | 0.090111833 | 3.220538143 | 0.001279502 | 0.009348579 | 0.290208597 |
| ENSMUSG00000024989 | Cep55 | 74107 | protein_coding | 298.9651181 | 0.290223317 | 0.173991543 | 1.668031166 | 0.09530954 | 0.240775631 | 0.290223317 |
| ENSMUSG00000029541 | Cyp2w1 | 545817 | protein_coding | 54.18702161 | -0.290244301 | 0.219656212 | -1.32135713 | 0.186382313 | 0.376888975 | 0.290244301 |
| ENSMUSG00000053560 | Ier2 | 15936 | protein_coding | 2801.739485 | -0.290322916 | 0.270769149 | -1.072215637 | 0.283623188 | 0.492751979 | 0.290322916 |
| ENSMUSG00000031986 | Sprtn | 244666 | protein_coding | 284.9548849 | -0.29034754 | 0.084028402 | -3.455350022 | 0.000549579 | 0.00470882 | 0.29034754 |
| ENSMUSG00000029651 | Mtus2 | 77521 | protein_coding | 40.68767446 | -0.29039612 | 0.340842177 | -0.85199585 | 0.394216396 | 0.602136435 | 0.29039612 |
| ENSMUSG00000027438 | Napb | 17957 | protein_coding | 35.9535187 | -0.290419077 | 0.229273354 | -1.266693541 | 0.205264891 | 0.401062215 | 0.290419077 |
| ENSMUSG00000089943 | Ugt1a5 | 394433 | protein_coding | 11.47208783 | 0.290472108 | 0.441655887 | 0.657688749 | 0.510738154 | 0.699008761 | 0.290472108 |
| ENSMUSG00000039178 | Tbc1d19 | 67249 | protein_coding | 114.012658 | -0.290503277 | 0.195535211 | -1.485682684 | 0.137363073 | 0.308157609 | 0.290503277 |
| ENSMUSG00000030538 | Cib1 | 23991 | protein_coding | 1928.29863 | 0.290537337 | 0.108863285 | 2.668827573 | 0.007611651 | 0.038992079 | 0.290537337 |
| ENSMUSG00000031176 | Dynlt3 | 67117 | protein_coding | 1369.236757 | 0.290620134 | 0.067780504 | 4.287665607 | 1.81E-05 | 0.000252944 | 0.290620134 |
| ENSMUSG00000039496 | Cdnf | 227526 | protein_coding | 17.42520198 | -0.290669619 | 0.29298781 | -0.992087756 | 0.321154719 | 0.532354566 | 0.290669619 |
| ENSMUSG00000022559 | Fbxl6 | 30840 | protein_coding | 188.4860232 | -0.290679715 | 0.119725076 | -2.42789334 | 0.015186809 | 0.065306558 | 0.290679715 |
| ENSMUSG00000033318 | Gstt2 | 14872 | protein_coding | 604.46747 | 0.290694287 | 0.091979872 | 3.160411945 | 0.001575462 | 0.011079012 | 0.290694287 |
| ENSMUSG00000032643 | Fhl3 | 14201 | protein_coding | 11.74051838 | -0.290713953 | 0.574074396 | -0.506404665 | 0.6125726 | 0.774246657 | 0.290713953 |
| ENSMUSG00000079317 | Trappc2 | 66226 | protein_coding | 55.83577302 | 0.290770876 | 0.159630717 | 1.821522085 | 0.068527533 | 0.192777559 | 0.290770876 |
| ENSMUSG00000024340 | Btnl2 | 547431 | protein_coding | 52.35488248 | 0.290860954 | 0.246513666 | 1.17989789 | 0.238040829 | 0.442256933 | 0.290860954 |
| ENSMUSG00000071337 | Tia1 | 21841 | protein_coding | 511.0387261 | 0.290868136 | 0.110300755 | 2.637045747 | 0.008363155 | 0.041902591 | 0.290868136 |
| ENSMUSG00000062825 | Actg1 | 11465 | protein_coding | 11809.63922 | -0.290915666 | 0.230915792 | -1.259834434 | 0.207729095 | 0.404141813 | 0.290915666 |
| ENSMUSG00000053754 | Chd8 | 67772 | protein_coding | 992.8811594 | -0.290930049 | 0.129622764 | -2.244436388 | 0.024804337 | 0.093873655 | 0.290930049 |
| ENSMUSG00000007613 | Tgfbr1 | 21812 | protein_coding | 904.2124197 | 0.291091607 | 0.099575522 | 2.923324929 | 0.003463149 | 0.021025924 | 0.291091607 |
| ENSMUSG00000027506 | Tpd52 | 21985 | protein_coding | 7128.04504 | 0.291143927 | 0.069416463 | 4.194162496 | 2.74E-05 | 0.000364139 | 0.291143927 |
| ENSMUSG00000084786 | Ubl5 | 66177 | protein_coding | 1043.992607 | 0.291225141 | 0.101527585 | 2.868433663 | 0.004125097 | 0.024049979 | 0.291225141 |

| ensembl_gene_id | external_gene_name | entrezgene_id | gene_biotype | baseMean | log2FoldChange | lfcSE | stat | pvalue | padj | absLog2FC |
|---|---|---|---|---|---|---|---|---|---|---|
| ENSMUSG00000030101 | Sumf1 | 58911 | protein_coding | 500.9599013 | 0.291228557 | 0.090102234 | 3.232201293 | 0.001228405 | 0.009066744 | 0.291228557 |
| ENSMUSG00000095440 | Fignl2 | 668225 | protein_coding | 25.62213643 | -0.291254164 | 0.280371973 | -1.038813404 | 0.298891526 | 0.507812718 | 0.291254164 |
| ENSMUSG00000066057 | Gm1976 | 100038746 | lncRNA | 43.53240919 | -0.291273946 | 0.207564018 | -1.403296915 | 0.160528318 | 0.342171494 | 0.291273946 |
| ENSMUSG00000027002 | Nckap1 | 50884 | protein_coding | 4167.985144 | 0.291278032 | 0.091032375 | 3.19971913 | 0.001375616 | 0.009950412 | 0.291278032 |
| ENSMUSG00000026698 | Pigc | 67292 | protein_coding | 356.5821088 | 0.291367932 | 0.118655698 | 2.455574718 | 0.014065951 | 0.061685701 | 0.291367932 |
| ENSMUSG00000037541 | Shank2 | 210274 | protein_coding | 302.6755315 | -0.29141919 | 0.22732264 | -1.281962899 | 0.199855661 | 0.394461087 | 0.29141919 |
| ENSMUSG00000075334 | Rprm | 67874 | protein_coding | 22.37636067 | -0.291447041 | 0.465336169 | -0.62631504 | 0.531108324 | 0.71642051 | 0.291447041 |
| ENSMUSG00000021662 | Arhgef28 | 110596 | protein_coding | 1427.339172 | -0.291464814 | 0.121094663 | -2.406917101 | 0.016087821 | 0.068096417 | 0.291464814 |
| ENSMUSG00000054850 | Smim10l2a | 320237 | protein_coding | 34.0913829 | 0.291550489 | 0.210218454 | 1.38689294 | 0.165474411 | 0.349003763 | 0.291550489 |
| ENSMUSG00000037750 | Fam222b | 216971 | protein_coding | 103.933761 | -0.29167299 | 0.186451629 | -1.564335968 | 0.117738689 | 0.277253378 | 0.29167299 |
| ENSMUSG00000032093 | Cd3e | 12501 | protein_coding | 13.55252944 | 0.291693423 | 0.453821958 | 0.642748589 | 0.520387247 | 0.706984466 | 0.291693423 |
| ENSMUSG00000020303 | Stc2 | 20856 | protein_coding | 435.3241428 | 0.291701737 | 0.209991717 | 1.38911068 | 0.164799096 | 0.348224995 | 0.291701737 |
| ENSMUSG00000032965 | Ift57 | 73916 | protein_coding | 8.025769314 | 0.291738481 | 0.418833702 | 0.696549679 | 0.486084657 | 0.678983247 | 0.291738481 |
| ENSMUSG00000024955 | Esrra | 26379 | protein_coding | 1660.348166 | 0.291782143 | 0.10402534 | 2.804914109 | 0.005032999 | 0.028207362 | 0.291782143 |
| ENSMUSG00000030346 | Rad51ap1 | 19362 | protein_coding | 94.78812862 | 0.29196677 | 0.164316608 | 1.776854897 | 0.075592117 | 0.205239445 | 0.29196677 |
| ENSMUSG00000068798 | Rap1a | 109905 | protein_coding | 951.4125143 | 0.292028095 | 0.09636698 | 3.030375089 | 0.002442502 | 0.016016538 | 0.292028095 |
| ENSMUSG00000059734 | Ndufs8 | 225887 | protein_coding | 965.985988 | 0.292066465 | 0.111230456 | 2.625777825 | 0.008645123 | 0.042963966 | 0.292066465 |
| ENSMUSG00000101225 | 1700008J07Rik | 629159 | lncRNA | 26.20954197 | -0.292081602 | 0.232959916 | -1.253784803 | 0.209920234 | 0.407185609 | 0.292081602 |
| ENSMUSG00000030471 | Zdhhc13 | 243983 | protein_coding | 2212.336068 | 0.292188962 | 0.189449155 | 1.542308076 | 0.122998747 | 0.285966245 | 0.292188962 |
| ENSMUSG00000024587 | Nars | 70223 | protein_coding | 2820.719622 | 0.292253313 | 0.143351219 | 2.038722203 | 0.041477761 | 0.136328617 | 0.292253313 |
| ENSMUSG00000003309 | Ap1m2 | 11768 | protein_coding | 2055.294946 | 0.292262247 | 0.108188824 | 2.701408841 | 0.006904641 | 0.036069896 | 0.292262247 |
| ENSMUSG00000027184 | Caprin1 | 53872 | protein_coding | 6192.338666 | 0.292300351 | 0.068811325 | 4.247852385 | 2.16E-05 | 0.000296101 | 0.292300351 |
| ENSMUSG00000028809 | Srrm1 | 51796 | protein_coding | 853.1779023 | -0.29234747 | 0.113082175 | -2.585265715 | 0.009730398 | 0.046922729 | 0.29234747 |
| ENSMUSG00000061731 | Ext1 | 14042 | protein_coding | 481.4162727 | 0.292561674 | 0.097353663 | 3.005142945 | 0.002654561 | 0.01706289 | 0.292561674 |
| ENSMUSG00000027452 | Acss1 | 68738 | protein_coding | 2024.164067 | 0.292566041 | 0.078799956 | 3.712769064 | 0.000205004 | 0.002087789 | 0.292566041 |
| ENSMUSG00000006342 | Susd2 | 71733 | protein_coding | 7.447666439 | -0.292578332 | 0.641277922 | -0.45624264 | 0.648215506 | 0.796966766 | 0.292578332 |
| ENSMUSG00000005779 | Psmb4 | 19172 | protein_coding | 2031.326304 | 0.292592656 | 0.122105965 | 2.396219185 | 0.016565181 | 0.069635842 | 0.292592656 |
| ENSMUSG00000019868 | Vta1 | 66201 | protein_coding | 441.391497 | 0.292643134 | 0.113797848 | 2.571605165 | 0.010122827 | 0.048249367 | 0.292643134 |
| ENSMUSG00000073889 | Il11ra1 | 16157 | protein_coding | 106.4353879 | -0.292644143 | 0.184970857 | -1.582109464 | 0.113624583 | 0.269718683 | 0.292644143 |
| ENSMUSG00000092618 | Btnl6 | 624681 | protein_coding | 1327.387476 | 0.292648858 | 0.136102784 | 2.150204795 | 0.031539019 | 0.112057438 | 0.292648858 |
| ENSMUSG00000028041 | Adam15 | 11490 | protein_coding | 534.7227715 | -0.292701638 | 0.126817822 | -2.30804814 | 0.020996459 | 0.082944345 | 0.292701638 |
| ENSMUSG00000034424 | Gcsh | 68133 | protein_coding | 646.7870041 | 0.292978386 | 0.126762189 | 2.311244304 | 0.020819364 | 0.082387786 | 0.292978386 |
| ENSMUSG00000038608 | Dock10 | 210293 | protein_coding | 58.00789595 | 0.293064886 | 0.346960982 | 0.84466237 | 0.398299375 | 0.606149922 | 0.293064886 |
| ENSMUSG00000053175 | Bcl3 | 12051 | protein_coding | 330.6525013 | -0.293132566 | 0.426551183 | -0.687215458 | 0.491946965 | 0.683859515 | 0.293132566 |
| ENSMUSG00000020091 | Eif4ebp2 | 13688 | protein_coding | 1050.360463 | -0.293222437 | 0.170508093 | -1.719698056 | 0.085487341 | 0.223917509 | 0.293222437 |
| ENSMUSG00000062753 | Smim29 | 106672 | protein_coding | 77.16138231 | -0.293375045 | 0.142444724 | -2.059571161 | 0.039439556 | 0.131832721 | 0.293375045 |
| ENSMUSG00000030602 | Pak4 | 70584 | protein_coding | 1035.866183 | 0.293394989 | 0.098510041 | 2.978325718 | 0.002898278 | 0.018253765 | 0.293394989 |
| ENSMUSG00000029456 | Acad10 | 71985 | protein_coding | 216.5673077 | -0.293584549 | 0.099653724 | -2.946046933 | 0.003218636 | 0.01986301 | 0.293584549 |

| ensembl_gene_id | external_gene_name | entrezgene_id | gene_biotype | baseMean | log2FoldChange | lfcSE | stat | pvalue | padj | absLog2FC |
|---|---|---|---|---|---|---|---|---|---|---|
| ENSMUSG00000000420 | Galnt1 | 14423 | protein_coding | 1630.137214 | 0.293838535 | 0.082720196 | 3.552198226 | 0.000382027 | 0.003497971 | 0.293838535 |
| ENSMUSG00000096740 | Lbhd1 | 102308570 | protein_coding | 91.77316696 | 0.293857956 | 0.128151008 | 2.293060038 | 0.021844549 | 0.085237622 | 0.293857956 |
| ENSMUSG00000027495 | Fam210b | 67017 | protein_coding | 331.6734268 | -0.293992423 | 0.106033058 | -2.772648715 | 0.005560211 | 0.030329807 | 0.293992423 |
| ENSMUSG00000028194 | Ddah1 | 69219 | protein_coding | 916.3511278 | 0.294104668 | 0.164656146 | 1.78617485 | 0.074070952 | 0.202942265 | 0.294104668 |
| ENSMUSG00000038756 | Ttll6 | 237930 | protein_coding | 324.5446138 | -0.294140253 | 0.258121383 | -1.139542374 | 0.254477005 | 0.46097006 | 0.294140253 |
| ENSMUSG00000024563 | Smad2 | 17126 | protein_coding | 874.1222982 | 0.294170212 | 0.062492329 | 4.707301189 | 2.51E-06 | 4.30E-05 | 0.294170212 |
| ENSMUSG00000027671 | Actl6a | 56456 | protein_coding | 564.0053725 | 0.294321292 | 0.127046773 | 2.316637295 | 0.020523498 | 0.081520767 | 0.294321292 |
| ENSMUSG00000067813 | Xkr9 | 381246 | protein_coding | 47.30898097 | 0.29435243 | 0.197932912 | 1.487132317 | 0.136979868 | 0.307774768 | 0.29435243 |
| ENSMUSG00000035900 | Gramd4 | 223752 | protein_coding | 551.7562402 | -0.294411828 | 0.078126027 | -3.768421846 | 0.000164283 | 0.001721512 | 0.294411828 |
| ENSMUSG00000032860 | P2ry2 | 18442 | protein_coding | 189.9099528 | -0.294491899 | 0.242375296 | -1.215024402 | 0.224356785 | 0.425999491 | 0.294491899 |
| ENSMUSG00000058396 | Gpr182 | 11536 | protein_coding | 8.906101741 | 0.294523732 | 0.441122848 | 0.667668278 | 0.504345363 | 0.69455412 | 0.294523732 |
| ENSMUSG00000027185 | Nat10 | 98956 | protein_coding | 385.5757215 | -0.294551242 | 0.141843863 | -2.07658784 | 0.037839616 | 0.128018141 | 0.294551242 |
| ENSMUSG00000033610 | Pank1 | 75735 | protein_coding | 512.2102989 | -0.294584959 | 0.135584318 | -2.172706724 | 0.029802397 | 0.10737222 | 0.294584959 |
| ENSMUSG00000103472 | Pcdhga7 | 93715 | protein_coding | 14.6153414 | -0.294615268 | 0.280309662 | -1.05103501 | 0.293242511 | 0.502250159 | 0.294615268 |
| ENSMUSG00000002428 | Hltf | 20585 | protein_coding | 885.6322171 | 0.294842456 | 0.057148909 | 5.159196597 | 2.48E-07 | 5.35E-06 | 0.294842456 |
| ENSMUSG00000030275 | Etnk1 | 75320 | protein_coding | 1487.481562 | 0.294848158 | 0.075050004 | 3.928689431 | 8.54E-05 | 0.000984379 | 0.294848158 |
| ENSMUSG00000013523 | Bcas1 | 76960 | protein_coding | 1326.922987 | 0.29485923 | 0.151246068 | 1.94953319 | 0.051231783 | 0.158099111 | 0.29485923 |
| ENSMUSG00000035478 | Mbd3 | 17192 | protein_coding | 335.7249088 | 0.294954373 | 0.09471537 | 3.114113078 | 0.001844988 | 0.012655884 | 0.294954373 |
| ENSMUSG00000028433 | Ubap2 | 68926 | protein_coding | 339.5085401 | -0.294964913 | 0.133131279 | -2.215594378 | 0.026719294 | 0.099065391 | 0.294964913 |
| ENSMUSG00000078496 | Zfp982 | 195531 | protein_coding | 8.390227184 | 0.295010436 | 0.447404903 | 0.65938132 | 0.509650934 | 0.698579013 | 0.295010436 |
| ENSMUSG00000026480 | Ncf2 | 17970 | protein_coding | 392.8516779 | 0.295084732 | 0.115063081 | 2.564547451 | 0.010331045 | 0.048938762 | 0.295084732 |
| ENSMUSG00000038127 | Ccdc50 | 67501 | protein_coding | 814.8856591 | -0.295148652 | 0.090049525 | -3.277625871 | 0.00104684 | 0.007958986 | 0.295148652 |
| ENSMUSG00000069631 | Strada | 72149 | protein_coding | 81.48578854 | -0.295212916 | 0.133467863 | -2.211865158 | 0.026975983 | 0.099660429 | 0.295212916 |
| ENSMUSG00000046727 | Cystm1 | 66060 | protein_coding | 2587.658346 | 0.295346925 | 0.104032761 | 2.838979967 | 0.0045258 | 0.025902601 | 0.295346925 |
| ENSMUSG00000036904 | Fzd8 | 14370 | protein_coding | 295.4343941 | 0.295382108 | 0.156107775 | 1.892167824 | 0.058468624 | 0.172941178 | 0.295382108 |
| ENSMUSG00000046572 | Zfp518b | 100515 | protein_coding | 70.8521631 | 0.29540078 | 0.186522001 | 1.583731559 | 0.11325482 | 0.269202025 | 0.29540078 |
| ENSMUSG00000063550 | Nup98 | 269966 | protein_coding | 1557.545101 | 0.29541266 | 0.092566734 | 3.191347972 | 0.001416106 | 0.010187777 | 0.29541266 |
| ENSMUSG00000048232 | Fbxo10 | 269529 | protein_coding | 52.68835943 | -0.295419987 | 0.202343851 | -1.459989944 | 0.144292838 | 0.31880931 | 0.295419987 |
| ENSMUSG00000024800 | Rpp30 | 54364 | protein_coding | 118.4215547 | 0.295475786 | 0.145866889 | 2.025653586 | 0.042800296 | 0.139212399 | 0.295475786 |
| ENSMUSG00000090258 | Churc1 | 211151 | protein_coding | 259.5389681 | 0.295559911 | 0.109799073 | 2.691825207 | 0.007106218 | 0.036860261 | 0.295559911 |
| ENSMUSG00000090100 | Ttbk2 | 140810 | protein_coding | 124.0890842 | -0.295573906 | 0.181102683 | -1.632079111 | 0.102662822 | 0.252737778 | 0.295573906 |
| ENSMUSG00000039328 | Rnf122 | 68867 | protein_coding | 43.02679763 | -0.295637144 | 0.279704155 | -1.05696372 | 0.290528145 | 0.499223071 | 0.295637144 |
| ENSMUSG00000053253 | Ndfip2 | 76273 | protein_coding | 2175.242301 | 0.295670505 | 0.087195781 | 3.390880885 | 0.000696684 | 0.005728848 | 0.295670505 |
| ENSMUSG00000069495 | Epc2 | 227867 | protein_coding | 621.297948 | 0.295725648 | 0.079502313 | 3.719711238 | 0.000199451 | 0.002039057 | 0.295725648 |
| ENSMUSG00000107283 | Mpv17 | 17527 | protein_coding | 184.4565192 | 0.295778727 | 0.122221809 | 2.420015949 | 0.015519826 | 0.066326749 | 0.295778727 |
| ENSMUSG00000020075 | Ddx21 | 56200 | protein_coding | 1581.735744 | -0.295803771 | 0.183887128 | -1.608615973 | 0.107700339 | 0.261024195 | 0.295803771 |
| ENSMUSG00000040459 | Arglu1 | 234023 | protein_coding | 892.8395652 | -0.295862582 | 0.115264557 | -2.566813147 | 0.01026379 | 0.048727765 | 0.295862582 |
| ENSMUSG00000029279 | Brdt | 114642 | protein_coding | 19.95716142 | -0.295884563 | 0.357438538 | -0.827791441 | 0.407788623 | 0.614964702 | 0.295884563 |

| ensembl_gene_id | external_gene_name | entrezgene_id | gene_biotype | baseMean | log2FoldChange | lfcSE | stat | pvalue | padj | absLog2FC |
|---|---|---|---|---|---|---|---|---|---|---|
| ENSMUSG00000050122 | Vwa3b | 70853 | protein_coding | 28.9113366 | -0.295984694 | 0.273791578 | -1.081058432 | 0.279671122 | 0.488003219 | 0.295984694 |
| ENSMUSG00000050989 | Selenon | 74777 | protein_coding | 30.26288801 | 0.29600093 | 0.22470462 | 1.317289026 | 0.187741762 | 0.378477508 | 0.29600093 |
| ENSMUSG00000032549 | Rab6b | 270192 | protein_coding | 15.01622652 | -0.296090181 | 0.339625625 | -0.871813432 | 0.383310166 | 0.590917956 | 0.296090181 |
| ENSMUSG00000002147 | Stat6 | 20852 | protein_coding | 1193.490614 | -0.296154146 | 0.083136418 | -3.562267341 | 0.000367666 | 0.003379215 | 0.296154146 |
| ENSMUSG00000032648 | Pygm | 19309 | protein_coding | 11.84776804 | 0.296203476 | 0.495373872 | 0.59793924 | 0.549880476 | 0.731093174 | 0.296203476 |
| ENSMUSG00000031904 | Slc7a6 | 330836 | protein_coding | 252.0862823 | -0.296226444 | 0.190133442 | -1.55799233 | 0.119235066 | 0.279883145 | 0.296226444 |
| ENSMUSG00000022827 | Rabl3 | 67657 | protein_coding | 188.0825993 | 0.296298133 | 0.105674932 | 2.803863952 | 0.005049421 | 0.028269656 | 0.296298133 |
| ENSMUSG00000026288 | Inpp5d | 16331 | protein_coding | 71.89041411 | -0.296427669 | 0.159254358 | -1.861347306 | 0.062695147 | 0.181038488 | 0.296427669 |
| ENSMUSG00000002741 | Ykt6 | 56418 | protein_coding | 1408.907037 | 0.29644759 | 0.107301874 | 2.762743822 | 0.005731773 | 0.031042396 | 0.29644759 |
| ENSMUSG00000024072 | Yipf4 | 67864 | protein_coding | 935.2013372 | 0.296456381 | 0.092959884 | 3.189078648 | 0.00142727 | 0.010249578 | 0.296456381 |
| ENSMUSG00000069540 | Gm4925 | 237433 | transcribed_processed_pseudogene | 9.260429049 | 0.296464335 | 0.40447677 | 0.732957631 | 0.463584271 | 0.66105374 | 0.296464335 |
| ENSMUSG00000022463 | Srebf2 | 20788 | protein_coding | 1583.057167 | -0.296511517 | 0.147090446 | -2.015844847 | 0.043816202 | 0.141596585 | 0.296511517 |
| ENSMUSG00000069309 | H2ac22 | 319170 | protein_coding | 55.29067951 | 0.296566669 | 0.273895503 | 1.082773048 | 0.278909171 | 0.487423953 | 0.296566669 |
| ENSMUSG00000035875 | AI182371 | 98870 | protein_coding | 11.26517808 | -0.296643718 | 0.448904786 | -0.660816564 | 0.508729959 | 0.697877891 | 0.296643718 |
| ENSMUSG00000055976 | Cldn23 | 71908 | protein_coding | 765.5420892 | -0.296848087 | 0.166632382 | -1.781454983 | 0.074838153 | 0.204201575 | 0.296848087 |
| ENSMUSG00000044847 | Lsm11 | 72290 | protein_coding | 175.0748298 | -0.29691752 | 0.107465892 | -2.762900052 | 0.005729031 | 0.031038095 | 0.29691752 |
| ENSMUSG00000075031 | H2bc3 | 319178 | protein_coding | 2042.600698 | 0.296979403 | 0.156748708 | 1.894621059 | 0.058142623 | 0.172296874 | 0.296979403 |
| ENSMUSG00000078779 | Zfp59 | 22717 | protein_coding | 129.7427687 | -0.296991754 | 0.147249918 | -2.016923053 | 0.043703544 | 0.141428291 | 0.296991754 |
| ENSMUSG00000059772 | Slx1b | 75764 | protein_coding | 154.0126481 | -0.297048239 | 0.158794753 | -1.870642662 | 0.061394627 | 0.178382638 | 0.297048239 |
| ENSMUSG00000037761 | Actr5 | 109275 | protein_coding | 153.9265771 | -0.297123437 | 0.125818796 | -2.36151868 | 0.018200255 | 0.074522792 | 0.297123437 |
| ENSMUSG00000078776 | 9530053A07Rik | 319482 | protein_coding | 13.48535614 | -0.297142051 | 0.480160212 | -0.618839386 | 0.536022173 | 0.720677474 | 0.297142051 |
| ENSMUSG00000040170 | Fmo2 | 55990 | protein_coding | 21.12944287 | -0.297164353 | 0.360123365 | -0.825173766 | 0.409272945 | 0.616269566 | 0.297164353 |
| ENSMUSG00000048174 | Tmem81 | 74626 | protein_coding | 29.8331136 | -0.297222545 | 0.199475848 | -1.490017698 | 0.136219583 | 0.306325782 | 0.297222545 |
| ENSMUSG00000028282 | Casp8ap2 | 26885 | protein_coding | 541.5547768 | 0.297267074 | 0.091390507 | 3.252712827 | 0.00114309 | 0.008551964 | 0.297267074 |
| ENSMUSG00000030105 | Arl8b | 67166 | protein_coding | 1498.694502 | 0.297319459 | 0.075249526 | 3.951114056 | 7.78E-05 | 0.000905048 | 0.297319459 |
| ENSMUSG00000034175 | Rhbdd3 | 279766 | protein_coding | 92.33745928 | -0.297347873 | 0.124336164 | -2.391483415 | 0.016780442 | 0.07034237 | 0.297347873 |
| ENSMUSG00000022949 | Clic6 | 209195 | protein_coding | 678.1865083 | -0.297395452 | 0.232371252 | -1.279828934 | 0.200605306 | 0.395157223 | 0.297395452 |
| ENSMUSG00000000567 | Sox9 | 20682 | protein_coding | 465.218555 | -0.297526647 | 0.156365235 | -1.902767243 | 0.057070923 | 0.170197653 | 0.297526647 |
| ENSMUSG00000022994 | Adcy6 | 11512 | protein_coding | 2986.677717 | -0.297552257 | 0.093080054 | -3.196734902 | 0.001389926 | 0.010035694 | 0.297552257 |
| ENSMUSG00000030880 | Polr3e | 26939 | protein_coding | 385.7456052 | -0.297572501 | 0.150197689 | -1.981205595 | 0.047568222 | 0.150479968 | 0.297572501 |
| ENSMUSG00000012117 | Dhdds | 67422 | protein_coding | 423.7183925 | 0.297587018 | 0.085922565 | 3.463432652 | 0.00053333 | 0.004596797 | 0.297587018 |
| ENSMUSG00000011257 | Pabpc4 | 230721 | protein_coding | 820.8162491 | -0.297610421 | 0.111454739 | -2.670235684 | 0.007579802 | 0.038882799 | 0.297610421 |
| ENSMUSG00000100632 | 9430060I03Rik | 100037260 | lncRNA | 16.89163962 | -0.297632355 | 0.281858553 | -1.055963536 | 0.290984875 | 0.499573396 | 0.297632355 |
| ENSMUSG00000024524 | Gnal | 14680 | protein_coding | 64.86492483 | -0.297757332 | 0.229046395 | -1.299986983 | 0.193605431 | 0.387069668 | 0.297757332 |
| ENSMUSG00000043953 | Ccrl2 | 54199 | protein_coding | 16.16868373 | -0.297861885 | 0.287475998 | -1.036127841 | 0.300142505 | 0.509123517 | 0.297861885 |
| ENSMUSG00000032092 | Mpzl2 | 14012 | protein_coding | 215.9290824 | -0.297918852 | 0.110069427 | -2.706644878 | 0.006796692 | 0.03557598 | 0.297918852 |
| ENSMUSG00000038485 | Socs7 | 192157 | protein_coding | 369.3522231 | -0.297920779 | 0.119999818 | -2.482676918 | 0.01303993 | 0.058523528 | 0.297920779 |
| ENSMUSG00000002885 | Adgre5 | 26364 | protein_coding | 476.8329156 | -0.29792212 | 0.150714437 | -1.976732457 | 0.04807188 | 0.151591547 | 0.29792212 |

| ensembl_gene_id | external_gene_name | entrezgene_id | gene_biotype | baseMean | log2FoldChange | lfcSE | stat | pvalue | padj | absLog2FC |
|---|---|---|---|---|---|---|---|---|---|---|
| ENSMUSG00000070462 | Tlnrd1 | 80889 | protein_coding | 1042.770379 | 0.297947038 | 0.140806058 | 2.116010079 | 0.03434395 | 0.119325161 | 0.297947038 |
| ENSMUSG00000028693 | Nasp | 50927 | protein_coding | 618.5150644 | -0.297993111 | 0.099456745 | -2.996208173 | 0.002733597 | 0.017416296 | 0.297993111 |
| ENSMUSG00000026558 | Uck2 | 80914 | protein_coding | 990.699347 | -0.297997048 | 0.084557504 | -3.524193976 | 0.000424773 | 0.003818164 | 0.297997048 |
| ENSMUSG00000070544 | Top1 | 21969 | protein_coding | 2133.233715 | 0.298087795 | 0.134668869 | 2.213487027 | 0.026864087 | 0.099440198 | 0.298087795 |
| ENSMUSG00000034761 | Map4k5 | 399510 | protein_coding | 496.4710051 | -0.298094792 | 0.081047386 | -3.678030946 | 0.000235041 | 0.002338963 | 0.298094792 |
| ENSMUSG00000049659 | Aftph | 216549 | protein_coding | 2506.330401 | 0.298114339 | 0.065546064 | 4.548165349 | 5.41E-06 | 8.68E-05 | 0.298114339 |
| ENSMUSG00000026458 | Ppfia4 | 68507 | protein_coding | 20.51601311 | -0.298194943 | 0.325371552 | -0.916475153 | 0.359417734 | 0.569292528 | 0.298194943 |
| ENSMUSG00000025047 | Pdcd11 | 18572 | protein_coding | 560.4086482 | -0.298256261 | 0.161583351 | -1.845835347 | 0.064916117 | 0.185504307 | 0.298256261 |
| ENSMUSG00000038206 | Fbxo8 | 50753 | protein_coding | 587.372564 | 0.298292545 | 0.115669988 | 2.578824032 | 0.009913727 | 0.047662496 | 0.298292545 |
| ENSMUSG00000037234 | Hook3 | 320191 | protein_coding | 497.681981 | -0.298339638 | 0.103377876 | -2.885913798 | 0.00390279 | 0.023032103 | 0.298339638 |
| ENSMUSG00000028426 | Rad23b | 19359 | protein_coding | 2539.970396 | 0.298379673 | 0.102751628 | 2.903892409 | 0.003685548 | 0.02206895 | 0.298379673 |
| ENSMUSG00000036622 | Atp13a2 | 74772 | protein_coding | 405.3083953 | 0.298407777 | 0.124815156 | 2.390797616 | 0.016811817 | 0.070412471 | 0.298407777 |
| ENSMUSG00000026473 | Glul | 14645 | protein_coding | 7944.42879 | -0.298453464 | 0.13322597 | -2.240204849 | 0.025077627 | 0.094653185 | 0.298453464 |
| ENSMUSG00000039114 | Nrn1 | 68404 | protein_coding | 18.49284903 | -0.29855071 | 0.345536126 | -0.864021698 | 0.387575971 | 0.595879351 | 0.29855071 |
| ENSMUSG00000070985 | Acnat1 | 230161 | protein_coding | 358.5453394 | 0.298587258 | 0.127864722 | 2.335180922 | 0.019533971 | 0.078530312 | 0.298587258 |
| ENSMUSG00000039195 | Bbln | 73737 | protein_coding | 129.4780523 | 0.298604503 | 0.153233734 | 1.948686458 | 0.05133288 | 0.158205488 | 0.298604503 |
| ENSMUSG00000074513 | Arfip1 | 99889 | protein_coding | 352.3518775 | 0.298629659 | 0.10992852 | 2.71658038 | 0.006596018 | 0.034754009 | 0.298629659 |
| ENSMUSG00000033453 | Adamts15 | 235130 | protein_coding | 437.393333 | 0.298643302 | 0.315064774 | 0.94787906 | 0.343191026 | 0.553385976 | 0.298643302 |
| ENSMUSG00000039789 | Zfp597 | 71063 | protein_coding | 210.6413136 | -0.29881383 | 0.124940604 | -2.391647077 | 0.016772962 | 0.070342217 | 0.29881383 |
| ENSMUSG00000013160 | Atp6v0d1 | 11972 | protein_coding | 1228.280859 | 0.298817942 | 0.086957879 | 3.436352688 | 0.000589603 | 0.004987358 | 0.298817942 |
| ENSMUSG00000075486 | Commd6 | 66200 | protein_coding | 429.2063832 | 0.298875838 | 0.091265349 | 3.274800795 | 0.001057365 | 0.008027507 | 0.298875838 |
| ENSMUSG00000049539 | H1f1 | 80838 | protein_coding | 1420.246938 | 0.29890016 | 0.16343536 | 1.82885858 | 0.067420797 | 0.190437746 | 0.29890016 |
| ENSMUSG00000022747 | St3gal6 | 54613 | protein_coding | 1559.474547 | -0.298919012 | 0.252070492 | -1.185854835 | 0.235679632 | 0.43982486 | 0.298919012 |
| ENSMUSG00000102748 | Pcdhgb2 | 93700 | protein_coding | 9.659628871 | -0.298953427 | 0.359024597 | -0.832682301 | 0.405023929 | 0.612070317 | 0.298953427 |
| ENSMUSG00000038425 | Poli | 26447 | protein_coding | 85.52366576 | -0.298987673 | 0.240625911 | -1.242541472 | 0.214036849 | 0.412483839 | 0.298987673 |
| ENSMUSG00000045690 | Wdr89 | 72338 | protein_coding | 27.23593128 | -0.299071952 | 0.236146412 | -1.266468332 | 0.20534546 | 0.401099937 | 0.299071952 |
| ENSMUSG00000069020 | Urm1 | 68205 | protein_coding | 308.2182205 | 0.299100819 | 0.08907871 | 3.357713872 | 0.000785899 | 0.006290441 | 0.299100819 |
| ENSMUSG00000042569 | Dhrs7b | 216820 | protein_coding | 629.509536 | 0.299111312 | 0.107729434 | 2.776505005 | 0.005494679 | 0.030054686 | 0.299111312 |
| ENSMUSG00000005483 | Dnajb1 | 81489 | protein_coding | 2271.954652 | -0.299115041 | 0.256393443 | -1.166625158 | 0.243361779 | 0.448382465 | 0.299115041 |
| ENSMUSG00000003824 | Syce2 | 71846 | protein_coding | 53.13618177 | -0.299119156 | 0.205691424 | -1.454213063 | 0.145887244 | 0.321751873 | 0.299119156 |
| ENSMUSG00000084128 | Esrp2 | 77411 | protein_coding | 866.7482899 | -0.299139975 | 0.107846574 | -2.773755015 | 0.005541339 | 0.030268329 | 0.299139975 |
| ENSMUSG00000017929 | B4galt5 | 56336 | protein_coding | 543.4692502 | -0.299149038 | 0.11418687 | -2.619819931 | 0.008797621 | 0.043545215 | 0.299149038 |
| ENSMUSG00000046691 | Chtf8 | 214987 | protein_coding | 43.93382756 | 0.29922112 | 0.188475286 | 1.587588092 | 0.112379513 | 0.267961209 | 0.29922112 |
| ENSMUSG00000021803 | Cdhr1 | 170677 | protein_coding | 103.5048712 | 0.299378064 | 0.166792233 | 1.794916101 | 0.072667068 | 0.200493734 | 0.299378064 |
| ENSMUSG00000040181 | Fmo1 | 14261 | protein_coding | 49.40135684 | -0.299380628 | 0.271760972 | -1.101632166 | 0.270621618 | 0.478449592 | 0.299380628 |
| ENSMUSG00000086962 | Gm12248 | 676527 | protein_coding | 10.22863697 | -0.299381884 | 0.511975553 | -0.584758163 | 0.558710339 | 0.736907711 | 0.299381884 |
| ENSMUSG00000010651 | Acaa1b | 235674 | protein_coding | 141.9359084 | 0.299508513 | 0.306129993 | 0.978370365 | 0.32789118 | 0.538806428 | 0.299508513 |
| ENSMUSG00000024069 | Slc30a6 | 210148 | protein_coding | 282.3064191 | 0.299526245 | 0.092450996 | 3.239837966 | 0.001195976 | 0.008864361 | 0.299526245 |

Supplementary Table 1

| ensembl_gene_id | external_gene_name | entrezgene_id | gene_biotype | baseMean | log2FoldChange | lfcSE | stat | pvalue | padj | absLog2FC |
|---|---|---|---|---|---|---|---|---|---|---|
| ENSMUSG00000031167 | Rbm3 | 19652 | protein_coding | 2078.381834 | 0.299606571 | 0.103814751 | 2.885973035 | 0.003902056 | 0.023032103 | 0.299606571 |
| ENSMUSG00000031231 | Cox7b | 66142 | protein_coding | 6471.413238 | 0.299635913 | 0.098860307 | 3.030902118 | 0.002438243 | 0.015995194 | 0.299635913 |
| ENSMUSG00000036513 | Commd2 | 52245 | protein_coding | 212.3792269 | 0.299709734 | 0.099083576 | 3.024817488 | 0.002487832 | 0.016253564 | 0.299709734 |
| ENSMUSG00000045519 | Zfp560 | 434377 | protein_coding | 88.69265568 | 0.299721517 | 0.142876858 | 2.097761117 | 0.035926253 | 0.123347635 | 0.299721517 |
| ENSMUSG00000039304 | Tnfsf10 | 22035 | protein_coding | 176.6813724 | 0.299790961 | 0.23073655 | 1.299278165 | 0.193848485 | 0.387259334 | 0.299790961 |
| ENSMUSG00000034457 | Eda2r | 245527 | protein_coding | 62.3776321 | 0.299835964 | 0.170158476 | 1.762098316 | 0.078052688 | 0.210143295 | 0.299835964 |
| ENSMUSG00000021982 | Cdadc1 | 71891 | protein_coding | 342.2281892 | -0.299927446 | 0.107257661 | -2.796326551 | 0.005168715 | 0.028745561 | 0.299927446 |
| ENSMUSG00000032252 | Glce | 93683 | protein_coding | 759.8674062 | 0.299932332 | 0.072898825 | 4.114364427 | 3.88E-05 | 0.000494721 | 0.299932332 |
| ENSMUSG00000042328 | Hps4 | 192232 | protein_coding | 291.5934241 | -0.300085971 | 0.159521154 | -1.881167257 | 0.05994917 | 0.175886323 | 0.300085971 |
| ENSMUSG00000030122 | Ptms | 69202 | protein_coding | 1894.493765 | 0.300314965 | 0.087110115 | 3.447532641 | 0.000565732 | 0.004805856 | 0.300314965 |
| ENSMUSG00000035184 | Fam124a | 629059 | protein_coding | 96.96055068 | -0.300354983 | 0.194772832 | -1.542078431 | 0.123054536 | 0.285966245 | 0.300354983 |
| ENSMUSG00000085184 | 4933439K11Rik | 71319 | lncRNA | 34.16732543 | -0.300392119 | 0.246283314 | -1.219701466 | 0.222578065 | 0.423917664 | 0.300392119 |
| ENSMUSG00000023393 | Slc17a9 | 228993 | protein_coding | 963.1045114 | 0.300472646 | 0.122335248 | 2.456141229 | 0.014043795 | 0.061622472 | 0.300472646 |
| ENSMUSG00000041220 | Elovl6 | 170439 | protein_coding | 1414.229541 | 0.300511728 | 0.124686397 | 2.410140437 | 0.015946381 | 0.067728187 | 0.300511728 |
| ENSMUSG00000073633 | Fbxo36 | 66153 | protein_coding | 82.25519459 | 0.300550533 | 0.155198691 | 1.936553275 | 0.052799974 | 0.161248547 | 0.300550533 |
| ENSMUSG00000028718 | Stil | 20460 | protein_coding | 196.5215781 | 0.300598149 | 0.118338846 | 2.540147724 | 0.011080566 | 0.051650936 | 0.300598149 |
| ENSMUSG00000038298 | Pdzk1 | 59020 | protein_coding | 6.789480887 | -0.300646207 | 0.690367915 | -0.435486935 | 0.663209021 | 0.804794609 | 0.300646207 |
| ENSMUSG00000022051 | Bnip3l | 12177 | protein_coding | 1057.69929 | 0.300716731 | 0.07443931 | 4.039757115 | 5.35E-05 | 0.000658619 | 0.300716731 |
| ENSMUSG00000068922 | Msto1 | 229524 | protein_coding | 118.8390988 | -0.300769889 | 0.121517728 | -2.475111197 | 0.01331947 | 0.05933981 | 0.300769889 |
| ENSMUSG00000001774 | Chordc1 | 66917 | protein_coding | 1132.677853 | 0.300780888 | 0.165210253 | 1.820594561 | 0.06866851 | 0.193003711 | 0.300780888 |
| ENSMUSG00000029762 | Akr1b8 | 14187 | protein_coding | 1154.465319 | -0.300786242 | 0.135584311 | -2.218444305 | 0.026524553 | 0.098572814 | 0.300786242 |
| ENSMUSG00000051146 | Camk2n2 | 73047 | protein_coding | 9.566843978 | -0.300786272 | 0.374429147 | -0.803319599 | 0.421790033 | 0.627406765 | 0.300786272 |
| ENSMUSG00000059554 | Ccdc28a | 215814 | protein_coding | 224.5286535 | 0.30081763 | 0.156674265 | 1.920019408 | 0.054855448 | 0.165200538 | 0.30081763 |
| ENSMUSG00000070047 | Fat1 | 14107 | protein_coding | 5147.240759 | -0.30093427 | 0.156387718 | -1.924283269 | 0.054319087 | 0.164204671 | 0.30093427 |
| ENSMUSG00000054942 | Miga1 | 215708 | protein_coding | 283.4624618 | -0.300935907 | 0.156870642 | -1.918369832 | 0.055064134 | 0.165670857 | 0.300935907 |
| ENSMUSG00000001229 | Dpp9 | 224897 | protein_coding | 550.1942887 | 0.301036255 | 0.121490019 | 2.477868208 | 0.013216996 | 0.059051979 | 0.301036255 |
| ENSMUSG00000004610 | Etfb | 110826 | protein_coding | 545.6272571 | 0.301040866 | 0.105977978 | 2.840598322 | 0.004502899 | 0.025808627 | 0.301040866 |
| ENSMUSG00000009569 | Mrtfb | 239719 | protein_coding | 725.1729434 | -0.301052164 | 0.164113709 | -1.83441204 | 0.066592854 | 0.188667907 | 0.301052164 |
| ENSMUSG00000015776 | Med22 | 20933 | protein_coding | 595.0917439 | -0.301224853 | 0.133106903 | -2.263029533 | 0.023633869 | 0.090577541 | 0.301224853 |
| ENSMUSG00000047721 | Bola2 | 66162 | protein_coding | 313.7924902 | -0.301292551 | 0.077504395 | -3.887425354 | 0.000101313 | 0.00113963 | 0.301292551 |
| ENSMUSG00000031971 | Ccsap | 73420 | protein_coding | 40.02738309 | -0.30139999 | 0.193753318 | -1.555586207 | 0.119806521 | 0.280753017 | 0.30139999 |
| ENSMUSG00000022516 | Nudt16l1 | 66911 | protein_coding | 197.8684405 | 0.301402391 | 0.096507587 | 3.123095284 | 0.001789598 | 0.012335011 | 0.301402391 |
| ENSMUSG00000004270 | Lpcat3 | 14792 | protein_coding | 1061.814321 | 0.301416938 | 0.070727778 | 4.261648623 | 2.03E-05 | 0.000281059 | 0.301416938 |
| ENSMUSG00000026019 | Wdr12 | 57750 | protein_coding | 193.837462 | -0.301424781 | 0.147166511 | -2.048188678 | 0.040541515 | 0.134362309 | 0.301424781 |
| ENSMUSG00000042857 | Gm9776 | 328309 | lncRNA | 28.94396606 | -0.301481334 | 0.269248109 | -1.119715696 | 0.262834934 | 0.470650347 | 0.301481334 |
| ENSMUSG00000028101 | Pias3 | 229615 | protein_coding | 174.4419705 | -0.301530907 | 0.122903294 | -2.453399717 | 0.014151299 | 0.061872603 | 0.301530907 |
| ENSMUSG00000068264 | Ap5s1 | 69596 | protein_coding | 196.3663449 | 0.301619822 | 0.160401359 | 1.880406898 | 0.060052644 | 0.17598711 | 0.301619822 |
| ENSMUSG00000032127 | Vps11 | 71732 | protein_coding | 475.4751658 | 0.301636516 | 0.072908605 | 4.137186752 | 3.52E-05 | 0.000454187 | 0.301636516 |

| ensembl_gene_id | external_gene_name | entrezgene_id | gene_biotype | baseMean | log2FoldChange | lfcSE | stat | pvalue | padj | absLog2FC |
|---|---|---|---|---|---|---|---|---|---|---|
| ENSMUSG00000074634 | Tmem267 | 633640 | protein_coding | 56.97901685 | 0.30179511 | 0.266948138 | 1.130538357 | 0.258249445 | 0.465369065 | 0.30179511 |
| ENSMUSG00000020681 | Ace | 11421 | protein_coding | 263.8994258 | -0.30180479 | 0.196292534 | -1.537525574 | 0.124164657 | 0.287706527 | 0.30180479 |
| ENSMUSG00000046791 | Riox1 | 71952 | protein_coding | 181.4021688 | -0.301846238 | 0.149587597 | -2.017856052 | 0.043606255 | 0.14122823 | 0.301846238 |
| ENSMUSG00000040268 | Plekha1 | 101476 | protein_coding | 963.7579119 | -0.301857839 | 0.08390031 | -3.597815518 | 0.000320901 | 0.003003121 | 0.301857839 |
| ENSMUSG00000002844 | Adprh | 11544 | protein_coding | 273.2589722 | 0.301858462 | 0.122197458 | 2.470251564 | 0.013501807 | 0.059966278 | 0.301858462 |
| ENSMUSG00000034570 | Inpp5j | 170835 | protein_coding | 662.1704261 | -0.30198502 | 0.147797866 | -2.043229909 | 0.041029684 | 0.135557107 | 0.30198502 |
| ENSMUSG00000063810 | Alms1 | 236266 | protein_coding | 247.1532848 | -0.302038024 | 0.124559142 | -2.424856335 | 0.015314446 | 0.065607448 | 0.302038024 |
| ENSMUSG00000036882 | Arhgap33 | 233071 | protein_coding | 23.31086156 | -0.302039536 | 0.427687473 | -0.706215533 | 0.480054112 | 0.673742347 | 0.302039536 |
| ENSMUSG00000029213 | Commd8 | 27784 | protein_coding | 490.8924801 | 0.302053188 | 0.071153471 | 4.245094223 | 2.19E-05 | 0.000299251 | 0.302053188 |
| ENSMUSG00000073434 | Wdr90 | 106618 | protein_coding | 182.2907633 | -0.302067479 | 0.167465696 | -1.80375734 | 0.071269356 | 0.197938676 | 0.302067479 |
| ENSMUSG00000048285 | Frmd6 | 319710 | protein_coding | 63.86406976 | 0.302191177 | 0.409099758 | 0.738673566 | 0.460105241 | 0.658509864 | 0.302191177 |
| ENSMUSG00000025821 | Zfp282 | 101095 | protein_coding | 315.0142899 | -0.302198134 | 0.100160749 | -3.017131325 | 0.002551792 | 0.016542619 | 0.302198134 |
| ENSMUSG00000001366 | Fbxo9 | 71538 | protein_coding | 687.6722274 | 0.302234416 | 0.087411491 | 3.457605089 | 0.000545 | 0.004677131 | 0.302234416 |
| ENSMUSG00000041438 | Utp4 | 21771 | protein_coding | 319.0694729 | -0.302303028 | 0.164925459 | -1.83296763 | 0.066807385 | 0.189141136 | 0.302303028 |
| ENSMUSG00000048924 | Ccdc125 | 76041 | protein_coding | 125.1922019 | -0.302315674 | 0.112484403 | -2.687623055 | 0.007196258 | 0.037227022 | 0.302315674 |
| ENSMUSG00000031493 | Ggn | 243897 | protein_coding | 7.766842453 | -0.302323602 | 0.491811712 | -0.614714115 | 0.538743537 | 0.722272825 | 0.302323602 |
| ENSMUSG00000005447 | Pafah1b3 | 18476 | protein_coding | 617.7761883 | 0.302449703 | 0.105140004 | 2.876637718 | 0.004019368 | 0.023571608 | 0.302449703 |
| ENSMUSG00000024271 | Elp2 | 58523 | protein_coding | 407.2545428 | 0.302459348 | 0.115250148 | 2.62437274 | 0.008680873 | 0.04311473 | 0.302459348 |
| ENSMUSG00000041229 | Phf8 | 320595 | protein_coding | 520.9162268 | 0.302508635 | 0.077059989 | 3.92562522 | 8.65E-05 | 0.000995555 | 0.302508635 |
| ENSMUSG00000011877 | Git1 | 216963 | protein_coding | 492.2601636 | -0.30251388 | 0.106352238 | -2.844452412 | 0.004448783 | 0.025562849 | 0.30251388 |
| ENSMUSG00000039115 | Itga9 | 104099 | protein_coding | 7.151269553 | -0.302533195 | 0.735532429 | -0.411311838 | 0.68084389 | 0.81740269 | 0.302533195 |
| ENSMUSG00000027242 | Wdr76 | 241627 | protein_coding | 185.3912707 | -0.302544022 | 0.149016535 | -2.03027149 | 0.042328951 | 0.138233724 | 0.302544022 |
| ENSMUSG00000029217 | Tec | 21682 | protein_coding | 253.5675569 | -0.30257296 | 0.139783891 | -2.164576747 | 0.030420113 | 0.109128731 | 0.30257296 |
| ENSMUSG00000021483 | Cdk20 | 105278 | protein_coding | 37.63654756 | -0.302586914 | 0.195815637 | -1.545264308 | 0.122282345 | 0.28492001 | 0.302586914 |
| ENSMUSG00000047909 | Ankrd16 | 320816 | protein_coding | 224.0852912 | -0.302658644 | 0.157354864 | -1.923414613 | 0.054428001 | 0.164409102 | 0.302658644 |
| ENSMUSG00000068923 | Syt11 | 229521 | protein_coding | 38.89373174 | 0.302671775 | 0.221677709 | 1.36536856 | 0.17213724 | 0.357750778 | 0.302671775 |
| ENSMUSG00000106779 | Gm8579 | 667335 | lncRNA | 7.506278352 | 0.302864462 | 0.427298975 | 0.708788178 | 0.478455935 | 0.672843015 | 0.302864462 |
| ENSMUSG00000050705 | 2310061I04Rik | 69662 | protein_coding | 187.4584632 | -0.302880733 | 0.118581952 | -2.554189139 | 0.010643542 | 0.050142071 | 0.302880733 |
| ENSMUSG00000029499 | Pxmp2 | 19301 | protein_coding | 60.54240014 | -0.302961627 | 0.154076611 | -1.966305108 | 0.049263377 | 0.153916648 | 0.302961627 |
| ENSMUSG00000039686 | Zer1 | 227693 | protein_coding | 251.0296881 | -0.30304273 | 0.108480612 | -2.793519722 | 0.005213785 | 0.028955776 | 0.30304273 |
| ENSMUSG00000021485 | Mxd3 | 17121 | protein_coding | 52.98719022 | 0.303080929 | 0.230444117 | 1.315203588 | 0.188441489 | 0.379647868 | 0.303080929 |
| ENSMUSG00000021411 | Pxdc1 | 66895 | protein_coding | 31.98980077 | 0.303236928 | 0.256783141 | 1.180906683 | 0.237639796 | 0.441993884 | 0.303236928 |
| ENSMUSG00000068011 | Mkrn2os | 70291 | protein_coding | 433.1318918 | 0.303278875 | 0.108193282 | 2.803121118 | 0.005061066 | 0.028305006 | 0.303278875 |
| ENSMUSG00000039168 | Dap | 223453 | protein_coding | 578.9996637 | 0.303293266 | 0.078440519 | 3.866538224 | 0.000110391 | 0.00122787 | 0.303293266 |
| ENSMUSG00000046711 | Hmga1 | 15361 | protein_coding | 352.2636564 | -0.30340129 | 0.139035891 | -2.182179646 | 0.029096277 | 0.105400386 | 0.30340129 |
| ENSMUSG00000035151 | Elmod2 | 244548 | protein_coding | 525.0657218 | 0.30349102 | 0.085589402 | 3.545894865 | 0.000391282 | 0.003569124 | 0.30349102 |
| ENSMUSG00000031987 | Egln1 | 112405 | protein_coding | 580.435789 | -0.30358733 | 0.086544319 | -3.507882806 | 0.000451688 | 0.004001383 | 0.30358733 |
| ENSMUSG00000041939 | Mvk | 17855 | protein_coding | 401.0102713 | 0.303632449 | 0.123879025 | 2.451040029 | 0.014244411 | 0.06210922 | 0.303632449 |

| ensembl_gene_id | external_gene_name | entrezgene_id | gene_biotype | baseMean | log2FoldChange | lfcSE | stat | pvalue | padj | absLog2FC |
|---|---|---|---|---|---|---|---|---|---|---|
| ENSMUSG00000022031 | Elp3 | 74195 | protein_coding | 723.2637627 | 0.303633412 | 0.082663568 | 3.673122501 | 0.000239605 | 0.002370448 | 0.303633412 |
| ENSMUSG00000022791 | Tnk2 | 51789 | protein_coding | 638.2870416 | -0.303666262 | 0.1108603 | -2.739179517 | 0.006159273 | 0.032847006 | 0.303666262 |
| ENSMUSG00000023990 | Tfeb | 21425 | protein_coding | 202.9449702 | -0.30369609 | 0.113755923 | -2.669716731 | 0.007591526 | 0.038918449 | 0.30369609 |
| ENSMUSG00000028899 | Taf12 | 66464 | protein_coding | 332.1633509 | 0.303710951 | 0.108870357 | 2.789656968 | 0.005276391 | 0.029221691 | 0.303710951 |
| ENSMUSG00000057054 | Inca1 | 103844 | protein_coding | 19.24111271 | -0.303729878 | 0.284490969 | -1.06762573 | 0.285689372 | 0.494484454 | 0.303729878 |
| ENSMUSG00000031954 | Cfdp1 | 23837 | protein_coding | 827.1080322 | 0.30389174 | 0.086517869 | 3.512473703 | 0.000443956 | 0.003954864 | 0.30389174 |
| ENSMUSG00000109771 | Gm35315 | 102638847 | protein_coding | 45.50108473 | 0.303898224 | 0.222085382 | 1.368384636 | 0.171191711 | 0.356588987 | 0.303898224 |
| ENSMUSG00000031657 | Heatr3 | 234549 | protein_coding | 313.5547483 | -0.303903622 | 0.142401796 | -2.134127738 | 0.032832326 | 0.115238714 | 0.303903622 |
| ENSMUSG00000090000 | Ier3ip1 | 66191 | protein_coding | 662.7980157 | 0.304061541 | 0.098391585 | 3.090320603 | 0.001999405 | 0.013505738 | 0.304061541 |
| ENSMUSG00000050994 | Adgb | 215772 | protein_coding | 11.9669916 | -0.304099052 | 0.603756127 | -0.503678619 | 0.614487236 | 0.77537453 | 0.304099052 |
| ENSMUSG00000019461 | Plscr3 | 70310 | protein_coding | 105.0590363 | -0.304106341 | 0.132206166 | -2.300243247 | 0.021434443 | 0.084132037 | 0.304106341 |
| ENSMUSG00000053617 | Sh3pxd2a | 14218 | protein_coding | 241.2195065 | -0.304184763 | 0.133703552 | -2.275068667 | 0.022901811 | 0.088538846 | 0.304184763 |
| ENSMUSG00000025321 | Itgb8 | 320910 | protein_coding | 12.43905401 | 0.304257974 | 0.640784051 | 0.474821391 | 0.634914285 | 0.788777375 | 0.304257974 |
| ENSMUSG00000052713 | Zfp608 | 269023 | protein_coding | 289.0407239 | 0.304296768 | 0.203626228 | 1.49438887 | 0.135074009 | 0.304588637 | 0.304296768 |
| ENSMUSG00000024668 | Sdhaf2 | 66072 | protein_coding | 550.5628696 | 0.304386292 | 0.074587507 | 4.080928635 | 4.49E-05 | 0.000560367 | 0.304386292 |
| ENSMUSG00000002319 | Ipo4 | 75751 | protein_coding | 133.0401638 | -0.304596234 | 0.146714951 | -2.076109028 | 0.037883868 | 0.128140636 | 0.304596234 |
| ENSMUSG00000027331 | Knstrn | 51944 | protein_coding | 512.7903823 | 0.304626686 | 0.13725289 | 2.219455524 | 0.02645575 | 0.098431865 | 0.304626686 |
| ENSMUSG00000019850 | Tnfaip3 | 21929 | protein_coding | 489.879433 | -0.304657075 | 0.114718848 | -2.655684577 | 0.007914761 | 0.040135724 | 0.304657075 |
| ENSMUSG00000030203 | Dusp16 | 70686 | protein_coding | 550.4011049 | -0.304724558 | 0.113579502 | -2.682918598 | 0.007298275 | 0.037681338 | 0.304724558 |
| ENSMUSG00000042675 | Ypel3 | 66090 | protein_coding | 244.9400533 | -0.304811612 | 0.170148532 | -1.791444264 | 0.073222031 | 0.201325827 | 0.304811612 |
| ENSMUSG00000028911 | Srsf4 | 57317 | protein_coding | 176.6966095 | -0.304833127 | 0.105438917 | -2.891087415 | 0.003839113 | 0.02276597 | 0.304833127 |
| ENSMUSG00000021335 | Slc17a1 | 20504 | protein_coding | 26.50497982 | -0.304914548 | 0.23859961 | -1.277933973 | 0.201272708 | 0.395933271 | 0.304914548 |
| ENSMUSG00000027378 | Nphp1 | 53885 | protein_coding | 89.53615068 | 0.304927303 | 0.169246774 | 1.801672758 | 0.071596907 | 0.198571396 | 0.304927303 |
| ENSMUSG00000074194 | Zfp791 | 244556 | protein_coding | 76.64885226 | -0.305043704 | 0.262857915 | -1.160488945 | 0.245849792 | 0.451177581 | 0.305043704 |
| ENSMUSG00000022178 | Ajuba | 16475 | protein_coding | 61.54053206 | -0.305091392 | 0.165763728 | -1.840519607 | 0.065691988 | 0.186846784 | 0.305091392 |
| ENSMUSG00000031340 | Gabre | 14404 | protein_coding | 29.67264351 | -0.305099204 | 0.292697479 | -1.042370456 | 0.297239958 | 0.506180721 | 0.305099204 |
| ENSMUSG00000004936 | Map2k1 | 26395 | protein_coding | 2910.165499 | 0.305148791 | 0.100981437 | 3.021830546 | 0.002512512 | 0.016387913 | 0.305148791 |
| ENSMUSG00000026213 | Stk11ip | 71728 | protein_coding | 233.7092956 | -0.305183105 | 0.104781129 | -2.912576978 | 0.003584598 | 0.021562746 | 0.305183105 |
| ENSMUSG00000029127 | Zbtb49 | 75079 | protein_coding | 80.80107251 | -0.305273738 | 0.127479396 | -2.394690803 | 0.016634386 | 0.069882561 | 0.305273738 |
| ENSMUSG00000045009 | Prrt3 | 210673 | protein_coding | 10.22803421 | -0.305276848 | 0.430877181 | -0.708500849 | 0.478634285 | 0.67287616 | 0.305276848 |
| ENSMUSG00000003680 | Taf6l | 225895 | protein_coding | 88.14572346 | 0.305289241 | 0.12335453 | 2.474892829 | 0.013327616 | 0.059346455 | 0.305289241 |
| ENSMUSG00000030614 | Tmem126b | 68472 | protein_coding | 307.1482525 | 0.305327181 | 0.095916575 | 3.183257748 | 0.001456279 | 0.010406287 | 0.305327181 |
| ENSMUSG00000028393 | Alad | 17025 | protein_coding | 436.9103431 | -0.305356418 | 0.078649249 | -3.882509016 | 0.000103384 | 0.001158925 | 0.305356418 |
| ENSMUSG00000036054 | Sugp2 | 234373 | protein_coding | 296.7767774 | -0.305447874 | 0.1378189 | -2.21629889 | 0.026671039 | 0.098951281 | 0.305447874 |
| ENSMUSG00000029167 | Ppargc1a | 19017 | protein_coding | 448.7185502 | -0.305512599 | 0.26283009 | -1.162395823 | 0.245074719 | 0.450555293 | 0.305512599 |
| ENSMUSG00000005514 | Por | 18984 | protein_coding | 1233.410879 | -0.305565144 | 0.137003016 | -2.230353419 | 0.025723989 | 0.096652975 | 0.305565144 |
| ENSMUSG00000028098 | Rnf115 | 67845 | protein_coding | 866.4406225 | 0.305676302 | 0.077550649 | 3.94163436 | 8.09E-05 | 0.000938155 | 0.305676302 |
| ENSMUSG00000047517 | Dmbt1 | 12945 | protein_coding | 81706.07716 | 0.305728714 | 0.164004643 | 1.864146698 | 0.062301106 | 0.18017115 | 0.305728714 |

| ensembl_gene_id | external_gene_name | entrezgene_id | gene_biotype | baseMean | log2FoldChange | lfcSE | stat | pvalue | padj | absLog2FC |
|---|---|---|---|---|---|---|---|---|---|---|
| ENSMUSG00000030204 | Ddx47 | 67755 | protein_coding | 640.6716917 | -0.305830117 | 0.109592584 | -2.79060959 | 0.005260889 | 0.029184127 | 0.305830117 |
| ENSMUSG00000046806 | Cyren | 78412 | protein_coding | 23.86464848 | 0.305841365 | 0.229098106 | 1.334979893 | 0.181882927 | 0.370561793 | 0.305841365 |
| ENSMUSG00000018585 | Atox1 | 11927 | protein_coding | 648.9918254 | 0.306083894 | 0.11874223 | 2.577717247 | 0.009945534 | 0.047759252 | 0.306083894 |
| ENSMUSG00000062861 | Zfp28 | 22690 | protein_coding | 38.91057082 | -0.306100112 | 0.229292613 | -1.334975897 | 0.181884235 | 0.370561793 | 0.306100112 |
| ENSMUSG00000004880 | Lbr | 98386 | protein_coding | 1353.084227 | 0.306118595 | 0.0853275 | 3.587572523 | 0.000333771 | 0.003105317 | 0.306118595 |
| ENSMUSG00000033365 | Ipo13 | 230673 | protein_coding | 205.5102753 | -0.306163005 | 0.137370566 | -2.228738037 | 0.02583134 | 0.09694665 | 0.306163005 |
| ENSMUSG00000020641 | Rsad2 | 58185 | protein_coding | 78.30978954 | 0.306187598 | 0.24736608 | 1.237791369 | 0.21579343 | 0.414792572 | 0.306187598 |
| ENSMUSG00000027634 | Ndrg3 | 29812 | protein_coding | 748.9850737 | -0.306201735 | 0.11604701 | -2.638600814 | 0.008324894 | 0.041789763 | 0.306201735 |
| ENSMUSG00000112640 | Gm32687 | 102635315 | protein_coding | 32.39279295 | 0.306204212 | 0.235035477 | 1.302799967 | 0.192643062 | 0.385770517 | 0.306204212 |
| ENSMUSG00000005580 | Adcy9 | 11515 | protein_coding | 250.1781775 | -0.30624066 | 0.105141846 | -2.912642977 | 0.003583841 | 0.021562746 | 0.30624066 |
| ENSMUSG00000037686 | Aspg | 104816 | protein_coding | 527.9582464 | -0.306347643 | 0.169538703 | -1.806948136 | 0.070770366 | 0.196861752 | 0.306347643 |
| ENSMUSG00000036875 | Dna2 | 327762 | protein_coding | 83.34155291 | -0.306385557 | 0.202057576 | -1.516327982 | 0.12943644 | 0.296015171 | 0.306385557 |
| ENSMUSG00000075543 | Urad | 231903 | protein_coding | 11.48988742 | 0.306389166 | 0.318335668 | 0.962471995 | 0.335812564 | 0.54646736 | 0.306389166 |
| ENSMUSG00000057133 | Chd6 | 71389 | protein_coding | 790.1356148 | -0.306449881 | 0.151102923 | -2.028087043 | 0.042551365 | 0.138714315 | 0.306449881 |
| ENSMUSG00000025578 | Cbx8 | 30951 | protein_coding | 97.81362909 | 0.306670832 | 0.146602813 | 2.091848212 | 0.036452096 | 0.124617506 | 0.306670832 |
| ENSMUSG00000078813 | Leng1 | 69757 | protein_coding | 204.524676 | 0.306915666 | 0.117609702 | 2.609611798 | 0.009064502 | 0.044465473 | 0.306915666 |
| ENSMUSG00000069310 | H3c3 | 319148 | protein_coding | 877.7902547 | 0.30693595 | 0.224780227 | 1.365493549 | 0.172097979 | 0.357719532 | 0.30693595 |
| ENSMUSG00000020415 | Pttg1 | 30939 | protein_coding | 217.7335117 | 0.306944875 | 0.110694746 | 2.772894705 | 0.00555601 | 0.030317273 | 0.306944875 |
| ENSMUSG00000079553 | Kifc1 | 100502766 | protein_coding | 203.6197507 | 0.306995719 | 0.169234898 | 1.814021351 | 0.069674439 | 0.195142335 | 0.306995719 |
| ENSMUSG00000049439 | Cyp20a1 | 77951 | protein_coding | 378.9741296 | 0.307006931 | 0.084868519 | 3.617441846 | 0.000297529 | 0.002819358 | 0.307006931 |
| ENSMUSG00000037681 | Esyt3 | 272636 | protein_coding | 8.879473697 | -0.307071076 | 0.581233068 | -0.528309715 | 0.59728439 | 0.764693012 | 0.307071076 |
| ENSMUSG00000005043 | Sgsh | 27029 | protein_coding | 156.9166544 | -0.307105455 | 0.110052852 | -2.790527013 | 0.005262231 | 0.029184127 | 0.307105455 |
| ENSMUSG00000020265 | Sumo3 | 20610 | protein_coding | 1149.209018 | 0.307113238 | 0.072559479 | 4.23257224 | 2.31E-05 | 0.000312917 | 0.307113238 |
| ENSMUSG00000061751 | Kalrn | 545156 | protein_coding | 482.4145358 | -0.307128319 | 0.114741551 | -2.676696606 | 0.007435193 | 0.038264217 | 0.307128319 |
| ENSMUSG00000020844 | Nxn | 18230 | protein_coding | 33.69752592 | 0.307152775 | 0.210397002 | 1.459872392 | 0.144325148 | 0.318836472 | 0.307152775 |
| ENSMUSG00000028362 | Tnfsf8 | 21949 | protein_coding | 7.867545366 | 0.307172029 | 0.583183334 | 0.526716062 | 0.59839078 | 0.765321556 | 0.307172029 |
| ENSMUSG00000029175 | Slc35f6 | 74919 | protein_coding | 584.8909514 | 0.307257316 | 0.118876321 | 2.584680559 | 0.009746925 | 0.046988205 | 0.307257316 |
| ENSMUSG00000028457 | Atp8b5 | 320571 | protein_coding | 39.21806687 | -0.30743427 | 0.321806273 | -0.955339579 | 0.339405999 | 0.550085499 | 0.30743427 |
| ENSMUSG00000024965 | Fermt3 | 108101 | protein_coding | 62.38676446 | -0.307522067 | 0.17294494 | -1.778150128 | 0.075379201 | 0.205027309 | 0.307522067 |
| ENSMUSG00000070705 | Eid2b | 434156 | protein_coding | 25.76316012 | -0.307683259 | 0.260800354 | -1.179765497 | 0.238093496 | 0.442256933 | 0.307683259 |
| ENSMUSG00000085795 | Zfp703 | 353310 | protein_coding | 147.570174 | -0.307715799 | 0.207261272 | -1.484675819 | 0.13762972 | 0.308453738 | 0.307715799 |
| ENSMUSG00000068739 | Sars | 20226 | protein_coding | 1616.35237 | 0.307733368 | 0.083841576 | 3.670414862 | 0.000242157 | 0.002391405 | 0.307733368 |
| ENSMUSG00000021271 | Zfp839 | 72805 | protein_coding | 138.3870242 | -0.307752512 | 0.120507298 | -2.55380808 | 0.010655197 | 0.050182134 | 0.307752512 |
| ENSMUSG00000018659 | Pnpo | 103711 | protein_coding | 1172.000335 | 0.307764575 | 0.144555102 | 2.129046786 | 0.03325039 | 0.116423875 | 0.307764575 |
| ENSMUSG00000030530 | Furin | 18550 | protein_coding | 736.3077359 | -0.307765499 | 0.096180173 | -3.199885063 | 0.001374824 | 0.009949203 | 0.307765499 |
| ENSMUSG00000045251 | Zfp688 | 69234 | protein_coding | 78.28047322 | 0.307821301 | 0.190401372 | 1.616696868 | 0.10594373 | 0.258023201 | 0.307821301 |
| ENSMUSG00000038354 | Ankrd35 | 213121 | protein_coding | 27.04813589 | -0.30786768 | 0.390381124 | -0.788633621 | 0.430326174 | 0.634416687 | 0.30786768 |
| ENSMUSG00000025608 | Podxl | 27205 | protein_coding | 11.80575815 | 0.308150184 | 0.430145539 | 0.716385866 | 0.47375312 | 0.668798783 | 0.308150184 |

| ensembl_gene_id | external_gene_name | entrezgene_id | gene_biotype | baseMean | log2FoldChange | lfcSE | stat | pvalue | padj | absLog2FC |
|---|---|---|---|---|---|---|---|---|---|---|
| ENSMUSG00000042390 | Gatad2b | 229542 | protein_coding | 381.418498 | -0.308325119 | 0.187148398 | -1.647490022 | 0.099457365 | 0.247408545 | 0.308325119 |
| ENSMUSG00000027356 | Fermt1 | 241639 | protein_coding | 3117.094306 | 0.308571302 | 0.119542903 | 2.581259899 | 0.009844044 | 0.047370373 | 0.308571302 |
| ENSMUSG00000021557 | Agtpbp1 | 67269 | protein_coding | 85.03469505 | -0.308575464 | 0.159248238 | -1.937700961 | 0.052659717 | 0.160991164 | 0.308575464 |
| ENSMUSG00000027879 | Sec22b | 20333 | protein_coding | 850.1745731 | 0.308621071 | 0.077109469 | 4.002375748 | 6.27E-05 | 0.000752137 | 0.308621071 |
| ENSMUSG00000054752 | Fsd1l | 319636 | protein_coding | 282.1423061 | -0.308739046 | 0.124738736 | -2.475085565 | 0.013320426 | 0.05933981 | 0.308739046 |
| ENSMUSG00000034473 | Sec22a | 317717 | protein_coding | 556.5962529 | 0.308800935 | 0.110428142 | 2.796397098 | 0.005167586 | 0.028745561 | 0.308800935 |
| ENSMUSG00000061887 | Ssbp3 | 72475 | protein_coding | 729.2310252 | 0.30881103 | 0.082779589 | 3.730521406 | 0.000191084 | 0.001962337 | 0.30881103 |
| ENSMUSG00000084799 | Ino80dos | 100380944 | lncRNA | 23.50803005 | -0.308835973 | 0.305905857 | -1.009578489 | 0.31269728 | 0.522956979 | 0.308835973 |
| ENSMUSG00000035000 | Dpp4 | 13482 | protein_coding | 1037.527327 | 0.308852563 | 0.091626758 | 3.370768226 | 0.000749589 | 0.006066795 | 0.308852563 |
| ENSMUSG00000027896 | Slc16a4 | 229699 | protein_coding | 20.59121921 | 0.308931847 | 0.44852439 | 0.688773795 | 0.490965629 | 0.683276587 | 0.308931847 |
| ENSMUSG00000049881 | 2810025M15Rik | 69953 | transcribed_processed_pseudogene | 99.71414916 | 0.308979548 | 0.186869198 | 1.653453603 | 0.098238581 | 0.24537047 | 0.308979548 |
| ENSMUSG00000113035 | 5830428M24Rik | 76062 | lncRNA | 24.45418305 | -0.309001871 | 0.311646208 | -0.991514938 | 0.321434202 | 0.532354566 | 0.309001871 |
| ENSMUSG00000030871 | Ears2 | 67417 | protein_coding | 240.8524709 | 0.309002668 | 0.107158503 | 2.883603805 | 0.00393153 | 0.023150247 | 0.309002668 |
| ENSMUSG00000031758 | Cdyl2 | 75796 | protein_coding | 25.02027198 | 0.309031356 | 0.254513918 | 1.214202189 | 0.224670525 | 0.426373421 | 0.309031356 |
| ENSMUSG00000032840 | Spring1 | 76792 | protein_coding | 187.2901123 | 0.309143477 | 0.113792635 | 2.71672658 | 0.006593105 | 0.034750157 | 0.309143477 |
| ENSMUSG00000028053 | Ash1l | 192195 | protein_coding | 1443.08034 | -0.309185546 | 0.134431935 | -2.299941195 | 0.021451552 | 0.084157714 | 0.309185546 |
| ENSMUSG00000014554 | Dguok | 27369 | protein_coding | 262.1878566 | 0.309195858 | 0.101445345 | 3.047905829 | 0.002304421 | 0.015211282 | 0.309195858 |
| ENSMUSG00000028999 | Rint1 | 72772 | protein_coding | 419.7618218 | 0.309216982 | 0.080160844 | 3.857456673 | 0.000114573 | 0.001267289 | 0.309216982 |
| ENSMUSG00000021569 | Trip13 | 69716 | protein_coding | 140.8425278 | 0.309219558 | 0.147111446 | 2.101940847 | 0.035558458 | 0.122485968 | 0.309219558 |
| ENSMUSG00000069237 | Fam8a1 | 97863 | protein_coding | 1146.999875 | 0.309488981 | 0.074801666 | 4.137461086 | 3.51E-05 | 0.000454013 | 0.309488981 |
| ENSMUSG00000026811 | St6galnac6 | 50935 | protein_coding | 7811.338582 | 0.30954929 | 0.104787449 | 2.954068387 | 0.003136145 | 0.019451915 | 0.30954929 |
| ENSMUSG00000004896 | Rrnad1 | 229503 | protein_coding | 142.7714865 | -0.309575684 | 0.123615031 | -2.504353074 | 0.012267555 | 0.055732347 | 0.309575684 |
| ENSMUSG00000053862 | Slc51b | 330962 | protein_coding | 402.3261756 | 0.30959685 | 0.248330381 | 1.246713543 | 0.212502551 | 0.410520393 | 0.30959685 |
| ENSMUSG00000114279 | H2bc14 | 319186 | protein_coding | 690.460872 | 0.309837035 | 0.172453864 | 1.796637245 | 0.072393228 | 0.199979072 | 0.309837035 |
| ENSMUSG00000036046 | 5031439G07Rik | 223739 | protein_coding | 618.4472476 | -0.309965109 | 0.089327826 | -3.469972617 | 0.000520511 | 0.004498484 | 0.309965109 |
| ENSMUSG00000067206 | Lrrc66 | 231296 | protein_coding | 959.3650647 | 0.309966011 | 0.103423109 | 2.997067224 | 0.002725906 | 0.017388158 | 0.309966011 |
| ENSMUSG00000045193 | Cirbp | 12696 | protein_coding | 310.4062544 | -0.310020038 | 0.118840745 | -2.608701577 | 0.009088646 | 0.044556467 | 0.310020038 |
| ENSMUSG00000021368 | Tbc1d7 | 67046 | protein_coding | 90.04068671 | 0.310050473 | 0.139630658 | 2.220504269 | 0.026384556 | 0.098300407 | 0.310050473 |
| ENSMUSG00000024906 | Mus81 | 71711 | protein_coding | 143.5131263 | 0.310064959 | 0.124761141 | 2.4852687 | 0.012945368 | 0.058148287 | 0.310064959 |
| ENSMUSG00000021040 | Slirp | 380773 | protein_coding | 179.9866313 | 0.310084706 | 0.146548225 | 2.115922634 | 0.034351388 | 0.119325161 | 0.310084706 |
| ENSMUSG00000013822 | Elof1 | 66126 | protein_coding | 749.1206396 | 0.310094149 | 0.101597497 | 3.05218295 | 0.002271836 | 0.015046071 | 0.310094149 |
| ENSMUSG00000097494 | 4933406C10Rik | 74076 | lncRNA | 44.64720847 | -0.310148045 | 0.190958051 | -1.624168466 | 0.104339859 | 0.255523566 | 0.310148045 |
| ENSMUSG00000098912 | 1500004A13Rik | 319830 | lncRNA | 32.73322195 | -0.310186431 | 0.258650693 | -1.199248405 | 0.230431371 | 0.433485954 | 0.310186431 |
| ENSMUSG00000055632 | Hmcn2 | 665700 | protein_coding | 11.96969173 | -0.310249164 | 0.794593778 | -0.390450029 | 0.696203799 | 0.827131234 | 0.310249164 |
| ENSMUSG00000015970 | Chdh | 218865 | protein_coding | 1336.999697 | -0.310292567 | 0.102253597 | -3.034539384 | 0.002409033 | 0.015823125 | 0.310292567 |
| ENSMUSG00000040225 | Prrc2c | 226562 | protein_coding | 2270.008694 | -0.310344974 | 0.15997349 | -1.939977515 | 0.052382423 | 0.160348378 | 0.310344974 |
| ENSMUSG00000033200 | Tpsg1 | 26945 | protein_coding | 438.9848866 | -0.310397046 | 0.145811327 | -2.128758109 | 0.033274278 | 0.116481914 | 0.310397046 |
| ENSMUSG00000021767 | Kat6b | 54169 | protein_coding | 702.7712163 | 0.310417483 | 0.112346775 | 2.763029763 | 0.005726754 | 0.03103632 | 0.310417483 |

| ensembl_gene_id | external_gene_name | entrezgene_id | gene_biotype | baseMean | log2FoldChange | lfcSE | stat | pvalue | padj | absLog2FC |
|---|---|---|---|---|---|---|---|---|---|---|
| ENSMUSG00000091625 | Lsm5 | 66373 | protein_coding | 148.1997081 | 0.310479574 | 0.156641627 | 1.982101308 | 0.047467903 | 0.150222285 | 0.310479574 |
| ENSMUSG00000021072 | Tmx1 | 72736 | protein_coding | 1324.540153 | 0.310494783 | 0.083787219 | 3.705753523 | 0.000210763 | 0.002136878 | 0.310494783 |
| ENSMUSG00000028175 | Depdc1a | 76131 | protein_coding | 131.1342701 | 0.31061743 | 0.147288348 | 2.10890701 | 0.034952609 | 0.120869552 | 0.31061743 |
| ENSMUSG00000014599 | Csf1 | 12977 | protein_coding | 9.518876489 | -0.310622083 | 0.542777283 | -0.572282763 | 0.567130425 | 0.743049553 | 0.310622083 |
| ENSMUSG00000033323 | Ctdp1 | 67655 | protein_coding | 387.1018759 | -0.310652999 | 0.092227327 | -3.36834004 | 0.000756223 | 0.006101436 | 0.310652999 |
| ENSMUSG00000030834 | Abcc6 | 27421 | protein_coding | 129.3106145 | 0.310931003 | 0.174215778 | 1.784746522 | 0.074302441 | 0.203139252 | 0.310931003 |
| ENSMUSG00000048264 | Dip2c | 208440 | protein_coding | 279.4861543 | -0.311009016 | 0.081742965 | -3.804719048 | 0.000141965 | 0.001526821 | 0.311009016 |
| ENSMUSG00000042207 | Kdm5b | 75605 | protein_coding | 1174.448553 | -0.31101527 | 0.09313405 | -3.339436745 | 0.000839485 | 0.006635887 | 0.31101527 |
| ENSMUSG00000032046 | Abhd12 | 76192 | protein_coding | 508.1037213 | 0.311024433 | 0.085524697 | 3.636662197 | 0.000276194 | 0.00265653 | 0.311024433 |
| ENSMUSG00000033826 | Dnah8 | 13417 | protein_coding | 461.6728811 | -0.311034466 | 0.201958546 | -1.540090637 | 0.123538262 | 0.286797469 | 0.311034466 |
| ENSMUSG00000102805 | Gm37240 | 99889 | protein_coding | 8.850736871 | -0.311041217 | 0.404284792 | -0.769361657 | 0.441678644 | 0.643011318 | 0.311041217 |
| ENSMUSG00000032816 | Igdcc4 | 56741 | protein_coding | 826.9859613 | 0.311078023 | 0.118268663 | 2.630265824 | 0.008531813 | 0.042533556 | 0.311078023 |
| ENSMUSG00000025532 | Crcp | 12909 | protein_coding | 260.6429345 | 0.31109267 | 0.118322164 | 2.629200303 | 0.008558593 | 0.042642282 | 0.31109267 |
| ENSMUSG00000061111 | Mcrip1 | 192173 | protein_coding | 92.47775243 | -0.311174541 | 0.145480878 | -2.138937738 | 0.032440711 | 0.114216545 | 0.311174541 |
| ENSMUSG00000008200 | Fnbp4 | 55935 | protein_coding | 636.640742 | -0.311181431 | 0.091367147 | -3.405835046 | 0.00065962 | 0.005477805 | 0.311181431 |
| ENSMUSG00000030554 | Synm | 233335 | protein_coding | 8.73501763 | -0.311190329 | 0.467011685 | -0.666343773 | 0.505191392 | 0.695118219 | 0.311190329 |
| ENSMUSG00000003062 | Stard3nl | 76205 | protein_coding | 514.1135842 | 0.311214388 | 0.125394996 | 2.481872473 | 0.013069404 | 0.058573288 | 0.311214388 |
| ENSMUSG00000014303 | Glis2 | 83396 | protein_coding | 12.05015302 | -0.311255749 | 0.33643565 | -0.925156856 | 0.354884322 | 0.564413128 | 0.311255749 |
| ENSMUSG00000026478 | Lamc1 | 226519 | protein_coding | 143.8520909 | -0.311360225 | 0.172018317 | -1.810041113 | 0.070289412 | 0.196106105 | 0.311360225 |
| ENSMUSG00000031700 | Gpt2 | 108682 | protein_coding | 542.4653005 | -0.311362734 | 0.168262824 | -1.850454703 | 0.064248041 | 0.184274927 | 0.311362734 |
| ENSMUSG00000050930 | Map10 | 74393 | protein_coding | 53.23343124 | -0.31142233 | 0.174141752 | -1.78832661 | 0.073723329 | 0.20225029 | 0.31142233 |
| ENSMUSG00000040511 | Pvr | 52118 | protein_coding | 201.9462281 | -0.311475955 | 0.250811452 | -1.24187294 | 0.214283446 | 0.412709398 | 0.311475955 |
| ENSMUSG00000052395 | Rft1 | 328370 | protein_coding | 279.6998034 | 0.311487012 | 0.125851676 | 2.475032686 | 0.013322398 | 0.05933981 | 0.311487012 |
| ENSMUSG00000039458 | Mtmr12 | 268783 | protein_coding | 604.0711577 | -0.311623582 | 0.080295168 | -3.880975531 | 0.000104038 | 0.001164527 | 0.311623582 |
| ENSMUSG00000026322 | Htr4 | 15562 | protein_coding | 420.7780815 | -0.311635795 | 0.17617684 | -1.768880601 | 0.076913802 | 0.207958418 | 0.311635795 |
| ENSMUSG00000021699 | Pde4d | 238871 | protein_coding | 319.5757143 | -0.311656186 | 0.166530795 | -1.871462788 | 0.061280963 | 0.178191396 | 0.311656186 |
| ENSMUSG00000074628 | Tldc2 | 383766 | protein_coding | 58.07940952 | 0.31173207 | 0.203227909 | 1.533903844 | 0.12505331 | 0.289176702 | 0.31173207 |
| ENSMUSG00000013091 | Tmem190 | 78052 | protein_coding | 11.90751836 | -0.311767967 | 0.342801474 | -0.909470908 | 0.363101607 | 0.571999051 | 0.311767967 |
| ENSMUSG00000037169 | Mycn | 18109 | protein_coding | 32.82473662 | -0.31181972 | 0.322962698 | -0.965497631 | 0.334295622 | 0.545293316 | 0.31181972 |
| ENSMUSG00000020899 | Pfas | 237823 | protein_coding | 422.9480034 | -0.311941925 | 0.100184327 | -3.113679909 | 0.001847698 | 0.012669022 | 0.311941925 |
| ENSMUSG00000099583 | H3c4 | 319149 | protein_coding | 402.7666043 | 0.311973025 | 0.190603973 | 1.636760344 | 0.101680555 | 0.250948811 | 0.311973025 |
| ENSMUSG00000037568 | Vash2 | 226841 | protein_coding | 7.044471765 | 0.312007148 | 0.418809099 | 0.744986555 | 0.456279848 | 0.655296022 | 0.312007148 |
| ENSMUSG00000050565 | Tor1aip2 | 240832 | protein_coding | 3462.058005 | 0.312011403 | 0.080964738 | 3.853670264 | 0.00011636 | 0.001279963 | 0.312011403 |
| ENSMUSG00000068749 | Psma5 | 26442 | protein_coding | 1002.121776 | 0.312073866 | 0.097931149 | 3.186666042 | 0.001439229 | 0.010321491 | 0.312073866 |
| ENSMUSG00000037275 | Gemin5 | 216766 | protein_coding | 353.901713 | -0.312077725 | 0.119525036 | -2.610982073 | 0.009028263 | 0.044314998 | 0.312077725 |
| ENSMUSG00000079003 | Samd1 | 666704 | protein_coding | 198.6804291 | -0.312118126 | 0.102771479 | -3.037011133 | 0.002389366 | 0.015719879 | 0.312118126 |
| ENSMUSG00000027384 | Ndufaf5 | 69487 | protein_coding | 171.6856645 | 0.312157893 | 0.126370439 | 2.470181274 | 0.01350446 | 0.059966278 | 0.312157893 |
| ENSMUSG00000028833 | Ncdn | 26562 | protein_coding | 325.6616188 | -0.312459284 | 0.140366784 | -2.226020119 | 0.026012834 | 0.097421213 | 0.312459284 |

Supplementary Table 1

| ensembl_gene_id | external_gene_name | entrezgene_id | gene_biotype | baseMean | log2FoldChange | lfcSE | stat | pvalue | padj | absLog2FC |
|---|---|---|---|---|---|---|---|---|---|---|
| ENSMUSG00000038369 | Ncoa6 | 56406 | protein_coding | 351.262375 | -0.312501507 | 0.19698756 | -1.586402244 | 0.112648092 | 0.268360575 | 0.312501507 |
| ENSMUSG00000036333 | Kidins220 | 77480 | protein_coding | 1707.92821 | -0.312541318 | 0.079209827 | -3.945739177 | 7.96E-05 | 0.000923571 | 0.312541318 |
| ENSMUSG00000024993 | Dennd10 | 67894 | protein_coding | 396.9186465 | 0.312544932 | 0.075699963 | 4.128732951 | 3.65E-05 | 0.000468173 | 0.312544932 |
| ENSMUSG00000001855 | Nup214 | 227720 | protein_coding | 298.7072254 | -0.312731597 | 0.130156393 | -2.402737127 | 0.01627288 | 0.068751841 | 0.312731597 |
| ENSMUSG00000032322 | Pstpip1 | 19200 | protein_coding | 24.65373628 | -0.312816452 | 0.281954072 | -1.109458889 | 0.26723227 | 0.475177965 | 0.312816452 |
| ENSMUSG00000027286 | Lrrc57 | 66606 | protein_coding | 107.575881 | -0.312822286 | 0.115697397 | -2.703797101 | 0.006855213 | 0.03584696 | 0.312822286 |
| ENSMUSG00000067235 | H2-Q10 | 15007 | protein_coding | 80.50345541 | -0.312832425 | 0.200749232 | -1.558324389 | 0.11915637 | 0.279775305 | 0.312832425 |
| ENSMUSG00000019982 | Myb | 17863 | protein_coding | 1615.721872 | -0.312850187 | 0.165609966 | -1.889078271 | 0.058881342 | 0.173815646 | 0.312850187 |
| ENSMUSG00000041431 | Ccnb1 | 268697 | protein_coding | 588.1141463 | 0.31287552 | 0.135292418 | 2.312587241 | 0.020745344 | 0.082176532 | 0.31287552 |
| ENSMUSG00000039117 | Taf4 | 228980 | protein_coding | 425.9574742 | -0.312909987 | 0.093849178 | -3.334179291 | 0.000855515 | 0.006735859 | 0.312909987 |
| ENSMUSG00000094483 | Purb | 19291 | protein_coding | 4145.748088 | 0.31298188 | 0.081143926 | 3.857120233 | 0.000114731 | 0.001267289 | 0.31298188 |
| ENSMUSG00000029601 | Iqcd | 75732 | protein_coding | 15.43548576 | -0.313152939 | 0.387608998 | -0.807909363 | 0.419142761 | 0.625238897 | 0.313152939 |
| ENSMUSG00000016520 | Lnx2 | 140887 | protein_coding | 459.6827055 | -0.313276818 | 0.073581708 | -4.257536659 | 2.07E-05 | 0.000285532 | 0.313276818 |
| ENSMUSG00000030245 | Golt1b | 66964 | protein_coding | 593.0571127 | 0.313483153 | 0.111448 | 2.812819902 | 0.004910915 | 0.027659495 | 0.313483153 |
| ENSMUSG00000022433 | Csnk1e | 27373 | protein_coding | 163.980517 | -0.313494193 | 0.173308985 | -1.808874437 | 0.070470513 | 0.196218011 | 0.313494193 |
| ENSMUSG00000049858 | Suox | 211389 | protein_coding | 370.2401593 | 0.313597576 | 0.130719194 | 2.399017063 | 0.016439149 | 0.069220139 | 0.313597576 |
| ENSMUSG00000028245 | Nsmaf | 18201 | protein_coding | 284.6521139 | -0.313742823 | 0.110126387 | -2.848934146 | 0.004386596 | 0.025269331 | 0.313742823 |
| ENSMUSG00000036810 | Cnep1r1 | 382030 | protein_coding | 298.4668861 | 0.313823802 | 0.093860003 | 3.343530716 | 0.000827195 | 0.006571355 | 0.313823802 |
| ENSMUSG00000032402 | Smad3 | 17127 | protein_coding | 838.1021629 | 0.313849678 | 0.127074816 | 2.469802343 | 0.013518773 | 0.060013102 | 0.313849678 |
| ENSMUSG00000021748 | Pdhb | 68263 | protein_coding | 1528.810174 | 0.313921549 | 0.066614972 | 4.712477377 | 2.45E-06 | 4.21E-05 | 0.313921549 |
| ENSMUSG00000036781 | Rps27l | 67941 | protein_coding | 1642.492804 | 0.313996466 | 0.097046532 | 3.235524844 | 0.001214193 | 0.008978491 | 0.313996466 |
| ENSMUSG00000029084 | Cd38 | 12494 | protein_coding | 2665.635444 | 0.314077803 | 0.187238643 | 1.677419783 | 0.093460423 | 0.237985125 | 0.314077803 |
| ENSMUSG00000032609 | Klhdc8b | 78267 | protein_coding | 40.08769811 | -0.314120791 | 0.235613964 | -1.333201077 | 0.182465826 | 0.371537345 | 0.314120791 |
| ENSMUSG00000053553 | 3110082I17Rik | 73212 | protein_coding | 50.78269925 | -0.314162593 | 0.182363008 | -1.722732025 | 0.084936996 | 0.223059021 | 0.314162593 |
| ENSMUSG00000075520 | Malrd1 | 102635496 | protein_coding | 499.8531851 | -0.314165502 | 0.211640618 | -1.484429143 | 0.137695108 | 0.308495056 | 0.314165502 |
| ENSMUSG00000044827 | Tlr1 | 21897 | protein_coding | 1077.968991 | 0.314293364 | 0.108456695 | 2.897869642 | 0.003757067 | 0.02239617 | 0.314293364 |
| ENSMUSG00000036372 | Tmem258 | 69038 | protein_coding | 699.3489933 | 0.31432546 | 0.111630146 | 2.815775766 | 0.004865962 | 0.027513325 | 0.31432546 |
| ENSMUSG00000085628 | Appbp2os | 654810 | lncRNA | 10.81437796 | -0.314594905 | 0.360740372 | -0.872081223 | 0.383164069 | 0.590807095 | 0.314594905 |
| ENSMUSG00000026005 | Rpe | 66646 | protein_coding | 806.5148527 | 0.314618583 | 0.071377299 | 4.40782417 | 1.04E-05 | 0.000154584 | 0.314618583 |
| ENSMUSG00000022021 | Diaph3 | 56419 | protein_coding | 195.3059625 | 0.314675215 | 0.124690573 | 2.523648805 | 0.011614389 | 0.053528355 | 0.314675215 |
| ENSMUSG00000047735 | Samd9l | 209086 | protein_coding | 2362.819138 | 0.314708543 | 0.146599262 | 2.146726651 | 0.03181505 | 0.112791279 | 0.314708543 |
| ENSMUSG00000028837 | Psmb2 | 26445 | protein_coding | 1081.096892 | 0.314798532 | 0.089108456 | 3.532757134 | 0.00041125 | 0.003717589 | 0.314798532 |
| ENSMUSG00000029036 | Atad3a | 108888 | protein_coding | 755.3842087 | -0.314873558 | 0.123573426 | -2.54806852 | 0.010832119 | 0.050790108 | 0.314873558 |
| ENSMUSG00000069266 | H4c2 | 326620 | protein_coding | 478.7822148 | 0.31488884 | 0.132205322 | 2.381816666 | 0.017227472 | 0.071685549 | 0.31488884 |
| ENSMUSG00000049932 | H2ax | 15270 | protein_coding | 789.6394248 | 0.31497527 | 0.19060243 | 1.652524943 | 0.098427585 | 0.245729556 | 0.31497527 |
| ENSMUSG00000024909 | Efemp2 | 58859 | protein_coding | 22.44018732 | 0.314979843 | 0.268318249 | 1.173903915 | 0.240433514 | 0.444953398 | 0.314979843 |
| ENSMUSG00000107476 | Zfp862-ps | 58894 | transcribed_unitary_pseudogene | 81.66270013 | -0.315007041 | 0.247829066 | -1.271065766 | 0.203705253 | 0.398895656 | 0.315007041 |
| ENSMUSG00000019878 | Hsf2 | 15500 | protein_coding | 158.5685929 | -0.315152989 | 0.168864239 | -1.866309829 | 0.06199803 | 0.179611609 | 0.315152989 |

| ensembl_gene_id | external_gene_name | entrezgene_id | gene_biotype | baseMean | log2FoldChange | lfcSE | stat | pvalue | padj | absLog2FC |
|---|---|---|---|---|---|---|---|---|---|---|
| ENSMUSG00000087368 | BC065397 | 436230 | lncRNA | 14.59608817 | -0.315278051 | 0.326655412 | -0.965170144 | 0.334459598 | 0.545293316 | 0.315278051 |
| ENSMUSG00000031103 | Elf4 | 56501 | protein_coding | 617.3708235 | -0.315453833 | 0.141687408 | -2.226406973 | 0.025986934 | 0.097347103 | 0.315453833 |
| ENSMUSG00000027276 | Jag1 | 16449 | protein_coding | 496.6197297 | -0.315518036 | 0.111279674 | -2.835360903 | 0.004577394 | 0.026160293 | 0.315518036 |
| ENSMUSG00000017311 | Pyy | 217212 | protein_coding | 329.9434676 | -0.315613424 | 0.162135142 | -1.946607136 | 0.051581852 | 0.158701129 | 0.315613424 |
| ENSMUSG00000023008 | Fmnl3 | 22379 | protein_coding | 26.03112194 | -0.315637832 | 0.276349558 | -1.14216876 | 0.253383879 | 0.459512515 | 0.315637832 |
| ENSMUSG00000025232 | Hexa | 15211 | protein_coding | 1963.332886 | -0.315687083 | 0.087351483 | -3.613986537 | 0.000301525 | 0.002848568 | 0.315687083 |
| ENSMUSG00000035773 | Kiss1r | 114229 | protein_coding | 17.3548511 | -0.315710234 | 0.305280703 | -1.034163739 | 0.301059623 | 0.510461749 | 0.315710234 |
| ENSMUSG00000053293 | Pom121 | 107939 | protein_coding | 167.4888147 | -0.315801084 | 0.211368701 | -1.494076853 | 0.135155533 | 0.304621103 | 0.315801084 |
| ENSMUSG00000090236 | Car15 | 80733 | protein_coding | 9.996815047 | -0.315811891 | 0.339260495 | -0.93088319 | 0.351913991 | 0.561483126 | 0.315811891 |
| ENSMUSG00000029851 | Tcaf2 | 232748 | protein_coding | 395.4626026 | -0.315850849 | 0.089823958 | -3.516331902 | 0.000437554 | 0.003910973 | 0.315850849 |
| ENSMUSG00000043183 | Simc1 | 319719 | protein_coding | 87.60409186 | -0.315868051 | 0.182839121 | -1.727573664 | 0.084064688 | 0.221575765 | 0.315868051 |
| ENSMUSG00000024130 | Abca3 | 27410 | protein_coding | 850.1996994 | -0.315870882 | 0.08694715 | -3.632906677 | 0.000280246 | 0.002685779 | 0.315870882 |
| ENSMUSG00000017721 | Pigt | 78928 | protein_coding | 888.5822471 | 0.316041172 | 0.068866857 | 4.58916216 | 4.45E-06 | 7.26E-05 | 0.316041172 |
| ENSMUSG00000013033 | Adgrl1 | 330814 | protein_coding | 48.35232649 | -0.316047626 | 0.200744259 | -1.574379399 | 0.115399764 | 0.272753737 | 0.316047626 |
| ENSMUSG00000021427 | Ssr1 | 107513 | protein_coding | 3073.192682 | 0.316071472 | 0.079024188 | 3.999680079 | 6.34E-05 | 0.000757902 | 0.316071472 |
| ENSMUSG00000032358 | Fam83b | 208994 | protein_coding | 705.5815644 | -0.316114099 | 0.056885791 | -5.556995766 | 2.74E-08 | 6.87E-07 | 0.316114099 |
| ENSMUSG00000054893 | Zfp667 | 384763 | protein_coding | 84.42062359 | -0.316187878 | 0.179558942 | -1.76091413 | 0.078252938 | 0.210400539 | 0.316187878 |
| ENSMUSG00000029560 | Snx8 | 231834 | protein_coding | 74.19548437 | 0.316347187 | 0.172867015 | 1.830003177 | 0.067249464 | 0.190021192 | 0.316347187 |
| ENSMUSG00000026659 | Dusp12 | 80915 | protein_coding | 95.63609275 | 0.316435158 | 0.120390778 | 2.628400303 | 0.00857875 | 0.042714075 | 0.316435158 |
| ENSMUSG00000091575 | 2010016I18Rik | 69206 | lncRNA | 100.5713401 | -0.316487271 | 0.254887239 | -1.24167562 | 0.214356269 | 0.412799739 | 0.316487271 |
| ENSMUSG00000010358 | Ifi35 | 70110 | protein_coding | 278.8964392 | 0.316529192 | 0.111197992 | 2.846536937 | 0.00441976 | 0.025405247 | 0.316529192 |
| ENSMUSG00000048895 | Cdk5r1 | 12569 | protein_coding | 14.39945537 | 0.316563789 | 0.38811075 | 0.815653236 | 0.414698492 | 0.621033996 | 0.316563789 |
| ENSMUSG00000060548 | Tnfrsf19 | 29820 | protein_coding | 52.24745235 | -0.316608542 | 0.180135066 | -1.757617492 | 0.078812615 | 0.211670171 | 0.316608542 |
| ENSMUSG00000101972 | H3c11 | 319153 | protein_coding | 494.8191387 | 0.316658452 | 0.210480897 | 1.504452213 | 0.132464969 | 0.300598664 | 0.316658452 |
| ENSMUSG00000027478 | Dnmt3b | 13436 | protein_coding | 45.62523645 | -0.316660451 | 0.226926326 | -1.395432856 | 0.162885349 | 0.345418432 | 0.316660451 |
| ENSMUSG00000030486 | Zfp108 | 54678 | protein_coding | 50.39480628 | 0.316851608 | 0.262363212 | 1.207683067 | 0.227169194 | 0.429681857 | 0.316851608 |
| ENSMUSG00000001227 | Sema6b | 20359 | protein_coding | 42.11915291 | 0.316961416 | 0.187028887 | 1.694719039 | 0.090128732 | 0.232127689 | 0.316961416 |
| ENSMUSG00000002617 | Zfp40 | 22700 | protein_coding | 118.9771227 | 0.317018 | 0.142496745 | 2.22473853 | 0.026098796 | 0.09755964 | 0.317018 |
| ENSMUSG00000058571 | Gpc6 | 23888 | protein_coding | 14.20782277 | 0.317158378 | 0.363408414 | 0.872732622 | 0.382808832 | 0.59043082 | 0.317158378 |
| ENSMUSG00000036822 | Topors | 106021 | protein_coding | 698.3539443 | -0.317213059 | 0.076717293 | -4.134831252 | 3.55E-05 | 0.000458127 | 0.317213059 |
| ENSMUSG00000029925 | Tbxas1 | 21391 | protein_coding | 14.89279994 | -0.317276209 | 0.425303731 | -0.745999119 | 0.455667947 | 0.655102361 | 0.317276209 |
| ENSMUSG00000007908 | Hmgcll1 | 208982 | protein_coding | 10.91095095 | -0.317283656 | 0.411431024 | -0.771170957 | 0.440605607 | 0.642435564 | 0.317283656 |
| ENSMUSG00000052748 | Swt1 | 66875 | protein_coding | 426.2633538 | -0.317298988 | 0.088097264 | -3.601689475 | 0.000316156 | 0.002967431 | 0.317298988 |
| ENSMUSG00000024799 | Tm7sf2 | 73166 | protein_coding | 384.3871374 | 0.317315144 | 0.103779683 | 3.05758445 | 0.002231288 | 0.014820663 | 0.317315144 |
| ENSMUSG00000036646 | Man1b1 | 227619 | protein_coding | 1737.033349 | 0.317480909 | 0.077399014 | 4.101872766 | 4.10E-05 | 0.000517653 | 0.317480909 |
| ENSMUSG00000031712 | Il15 | 16168 | protein_coding | 176.8742992 | -0.317497437 | 0.164457897 | -1.930569731 | 0.053536284 | 0.162524598 | 0.317497437 |
| ENSMUSG00000030677 | Kif22 | 110033 | protein_coding | 311.6517262 | 0.317622961 | 0.126446687 | 2.511912077 | 0.0120079 | 0.054944508 | 0.317622961 |
| ENSMUSG00000056476 | Med12l | 329650 | protein_coding | 26.588346 | -0.317678527 | 0.399580154 | -0.795030793 | 0.42659559 | 0.631136407 | 0.317678527 |

| ensembl_gene_id | external_gene_name | entrezgene_id | gene_biotype | baseMean | log2FoldChange | lfcSE | stat | pvalue | padj | absLog2FC |
|---|---|---|---|---|---|---|---|---|---|---|
| ENSMUSG00000015668 | Pdzd11 | 72621 | protein_coding | 479.402668 | 0.317770908 | 0.083650019 | 3.798814525 | 0.00014539 | 0.001555251 | 0.317770908 |
| ENSMUSG00000019864 | Rtn4ip1 | 170728 | protein_coding | 302.8218908 | -0.317824787 | 0.095069143 | -3.343090902 | 0.000828508 | 0.006575073 | 0.317824787 |
| ENSMUSG00000071478 | H2ac7 | 319165 | protein_coding | 91.87106587 | 0.317848823 | 0.302572718 | 1.050487383 | 0.293494089 | 0.502507368 | 0.317848823 |
| ENSMUSG00000033177 | Tmprss7 | 208171 | protein_coding | 31.91650079 | 0.317883333 | 0.258791362 | 1.228338269 | 0.219320011 | 0.419619115 | 0.317883333 |
| ENSMUSG00000000983 | Wfdc18 | 14038 | protein_coding | 74.74149459 | 0.317933609 | 0.237170938 | 1.340525159 | 0.18007467 | 0.368080898 | 0.317933609 |
| ENSMUSG00000027284 | Cdan1 | 68968 | protein_coding | 132.9649416 | -0.317935557 | 0.147721144 | -2.152268449 | 0.031376217 | 0.111737326 | 0.317935557 |
| ENSMUSG00000047797 | Gjb1 | 14618 | protein_coding | 572.7658669 | 0.317953653 | 0.105601485 | 3.010882406 | 0.002604897 | 0.016804699 | 0.317953653 |
| ENSMUSG00000042784 | Muc1 | 17829 | protein_coding | 32.10396368 | -0.317954923 | 0.288463432 | -1.102236494 | 0.27035887 | 0.478375791 | 0.317954923 |
| ENSMUSG00000041346 | Wrap53 | 216853 | protein_coding | 197.1308473 | 0.3179652 | 0.09599283 | 3.31238489 | 0.000925042 | 0.007190857 | 0.3179652 |
| ENSMUSG00000019872 | Smpdl3a | 57319 | protein_coding | 2213.863083 | -0.317982867 | 0.153110623 | -2.0768178 | 0.037818379 | 0.128000666 | 0.317982867 |
| ENSMUSG00000034245 | Hdac11 | 232232 | protein_coding | 415.8400308 | -0.318137724 | 0.201557455 | -1.578397207 | 0.114474386 | 0.271170141 | 0.318137724 |
| ENSMUSG00000036214 | Polr1has | 76416 | protein_coding | 78.4131943 | 0.318273051 | 0.200452003 | 1.587776851 | 0.112336808 | 0.267939604 | 0.318273051 |
| ENSMUSG00000114456 | H2bc9 | 319182 | protein_coding | 748.6476836 | 0.318283209 | 0.159251525 | 1.99861954 | 0.045649535 | 0.146063838 | 0.318283209 |
| ENSMUSG00000078485 | Plekhn1 | 231002 | protein_coding | 188.169699 | -0.318284629 | 0.123043041 | -2.58677474 | 0.00968789 | 0.046760217 | 0.318284629 |
| ENSMUSG00000085355 | 3010003L21Rik | 109163 | lncRNA | 37.10976044 | -0.318289606 | 0.232141272 | -1.37110305 | 0.170342834 | 0.35563115 | 0.318289606 |
| ENSMUSG00000025986 | Slc39a10 | 227059 | protein_coding | 185.1988558 | 0.31832222 | 0.166973681 | 1.906421525 | 0.056595538 | 0.169096555 | 0.31832222 |
| ENSMUSG00000064141 | Zfp69 | 381549 | protein_coding | 19.92593362 | -0.318346145 | 0.299834776 | -1.061738567 | 0.288354386 | 0.497392119 | 0.318346145 |
| ENSMUSG00000070371 | Prss36 | 77613 | protein_coding | 29.57671935 | -0.318347668 | 0.248945445 | -1.278784868 | 0.200972824 | 0.395636527 | 0.318347668 |
| ENSMUSG00000071477 | Zfp777 | 72306 | protein_coding | 288.4426095 | -0.318365754 | 0.092881703 | -3.427647691 | 0.000608835 | 0.005125541 | 0.318365754 |
| ENSMUSG00000040247 | Tbc1d10c | 108995 | protein_coding | 8.485928522 | 0.318473302 | 0.54795099 | 0.581207641 | 0.561100519 | 0.738795491 | 0.318473302 |
| ENSMUSG00000069272 | H2ac8 | 319166 | protein_coding | 240.2564432 | 0.31857801 | 0.254409816 | 1.252223735 | 0.21048835 | 0.407739238 | 0.31857801 |
| ENSMUSG00000020585 | Laptm4a | 17775 | protein_coding | 3192.758231 | 0.318631919 | 0.086369019 | 3.689192273 | 0.000224967 | 0.002250512 | 0.318631919 |
| ENSMUSG00000059552 | Trp53 | 22059 | protein_coding | 322.15419 | -0.318786815 | 0.127011197 | -2.509911121 | 0.012076155 | 0.055145929 | 0.318786815 |
| ENSMUSG00000007379 | Dennd2c | 329727 | protein_coding | 43.54932057 | -0.318791177 | 0.234251945 | -1.3608902 | 0.173548391 | 0.35951083 | 0.318791177 |
| ENSMUSG00000030629 | Zfand6 | 65098 | protein_coding | 1353.876475 | 0.318828337 | 0.093201218 | 3.420860188 | 0.000624234 | 0.005233053 | 0.318828337 |
| ENSMUSG00000028683 | Eif2b3 | 108067 | protein_coding | 143.306927 | -0.318887795 | 0.148585818 | -2.146152303 | 0.03186083 | 0.112843642 | 0.318887795 |
| ENSMUSG00000026413 | Pkp1 | 18772 | protein_coding | 555.8352862 | -0.319028518 | 0.091305746 | -3.494068346 | 0.000475719 | 0.004172841 | 0.319028518 |
| ENSMUSG00000029313 | Aff1 | 17355 | protein_coding | 1295.191918 | -0.319122822 | 0.108703347 | -2.935722125 | 0.003327723 | 0.020410894 | 0.319122822 |
| ENSMUSG00000044600 | Smim7 | 66818 | protein_coding | 1733.492605 | 0.319146114 | 0.114417789 | 2.789305031 | 0.005282129 | 0.029233409 | 0.319146114 |
| ENSMUSG00000061458 | Nol10 | 217431 | protein_coding | 104.5661372 | -0.319205331 | 0.201700156 | -1.582573547 | 0.113518697 | 0.269547674 | 0.319205331 |
| ENSMUSG00000076432 | Ywhaq | 22630 | protein_coding | 1969.405001 | 0.319211975 | 0.083364032 | 3.829133124 | 0.000128595 | 0.001401963 | 0.319211975 |
| ENSMUSG00000029598 | Plbd2 | 71772 | protein_coding | 245.2003381 | -0.319230857 | 0.092576583 | -3.448289486 | 0.000564149 | 0.004802654 | 0.319230857 |
| ENSMUSG00000027953 | Slc50a1 | 19729 | protein_coding | 480.5678859 | -0.319329152 | 0.093290437 | -3.422956984 | 0.000619439 | 0.005195588 | 0.319329152 |
| ENSMUSG00000116220 | A430088P11Rik | 106059 | lncRNA | 24.98570449 | -0.319386751 | 0.330296616 | -0.966969492 | 0.333559285 | 0.544302048 | 0.319386751 |
| ENSMUSG00000023348 | Trip6 | 22051 | protein_coding | 74.93141011 | -0.319395094 | 0.151522454 | -2.107906024 | 0.035039119 | 0.121116122 | 0.319395094 |
| ENSMUSG00000023966 | Rsph9 | 75564 | protein_coding | 15.99131042 | -0.319416651 | 0.373571523 | -0.855034794 | 0.392531886 | 0.600621715 | 0.319416651 |
| ENSMUSG00000034320 | Slc26a2 | 13521 | protein_coding | 14146.2236 | 0.319421521 | 0.123108071 | 2.594643228 | 0.009468915 | 0.045973799 | 0.319421521 |
| ENSMUSG00000044881 | Coa4 | 68185 | protein_coding | 266.3159858 | 0.319499365 | 0.152768455 | 2.091396195 | 0.036492563 | 0.124625545 | 0.319499365 |

| ensembl_gene_id | external_gene_name | entrezgene_id | gene_biotype | baseMean | log2FoldChange | lfcSE | stat | pvalue | padj | absLog2FC |
|---|---|---|---|---|---|---|---|---|---|---|
| ENSMUSG00000020599 | Rgs9 | 19739 | protein_coding | 8.563244154 | -0.319531798 | 0.805896816 | -0.396492195 | 0.691741965 | 0.824470213 | 0.319531798 |
| ENSMUSG00000078566 | Bnip3 | 12176 | protein_coding | 59.31428811 | -0.319566826 | 0.186753638 | -1.711167872 | 0.087050128 | 0.227050359 | 0.319566826 |
| ENSMUSG00000011427 | Zfp790 | 233056 | protein_coding | 339.7799989 | 0.319681799 | 0.088601112 | 3.608101406 | 0.000308446 | 0.002903621 | 0.319681799 |
| ENSMUSG00000021556 | Golm1 | 105348 | protein_coding | 9511.219049 | 0.319792897 | 0.070435303 | 4.540236009 | 5.62E-06 | 8.96E-05 | 0.319792897 |
| ENSMUSG00000000823 | Zfp512b | 269401 | protein_coding | 211.0494893 | -0.319963845 | 0.156949475 | -2.03864234 | 0.041485736 | 0.136328617 | 0.319963845 |
| ENSMUSG00000068396 | Rpl34-ps1 | 619547 | processed_pseudogene | 10.16456596 | 0.320076138 | 0.389521639 | 0.821715935 | 0.411238588 | 0.618130069 | 0.320076138 |
| ENSMUSG00000021606 | Ndufs6 | 407785 | protein_coding | 1044.42863 | 0.320149853 | 0.08384774 | 3.818228777 | 0.000134413 | 0.001456418 | 0.320149853 |
| ENSMUSG00000021823 | Vcl | 22330 | protein_coding | 840.4377604 | -0.320234412 | 0.21565644 | -1.484928586 | 0.137562743 | 0.308388369 | 0.320234412 |
| ENSMUSG00000097519 | 4930558J18Rik | 75323 | lncRNA | 8.704143828 | 0.320261183 | 0.459030579 | 0.697690302 | 0.485370894 | 0.678688781 | 0.320261183 |
| ENSMUSG00000040350 | Trim7 | 94089 | protein_coding | 41.69772214 | -0.320284463 | 0.228147858 | -1.403846021 | 0.160364706 | 0.341958773 | 0.320284463 |
| ENSMUSG00000017716 | Birc5 | 11799 | protein_coding | 405.4286133 | 0.320326594 | 0.138757235 | 2.308539761 | 0.020969134 | 0.08285695 | 0.320326594 |
| ENSMUSG00000026603 | Smyd2 | 226830 | protein_coding | 249.3652827 | -0.320427293 | 0.085640539 | -3.741537571 | 0.000182898 | 0.001888008 | 0.320427293 |
| ENSMUSG00000029027 | Dffb | 13368 | protein_coding | 122.2054139 | -0.320454586 | 0.114953682 | -2.787684402 | 0.005308623 | 0.029308752 | 0.320454586 |
| ENSMUSG00000039187 | Fanci | 208836 | protein_coding | 102.9298877 | 0.320791697 | 0.1544022 | 2.077636824 | 0.037742822 | 0.127826423 | 0.320791697 |
| ENSMUSG00000044700 | Tmem201 | 230917 | protein_coding | 150.5400817 | -0.320799337 | 0.127015381 | -2.525673135 | 0.011547687 | 0.053359893 | 0.320799337 |
| ENSMUSG00000031618 | Nr3c2 | 110784 | protein_coding | 2555.027504 | -0.320832615 | 0.079890333 | -4.015912851 | 5.92E-05 | 0.000716713 | 0.320832615 |
| ENSMUSG00000046753 | Ccdc66 | 320234 | protein_coding | 236.5882535 | 0.32093866 | 0.088822443 | 3.613260914 | 0.00030237 | 0.002854862 | 0.32093866 |
| ENSMUSG00000064288 | H4c12 | 319160 | protein_coding | 898.4375468 | 0.320980761 | 0.127823263 | 2.511129451 | 0.012034556 | 0.055016701 | 0.320980761 |
| ENSMUSG00000032232 | Cgnl1 | 68178 | protein_coding | 626.7458541 | -0.321290868 | 0.100951868 | -3.182614377 | 0.001459519 | 0.010424759 | 0.321290868 |
| ENSMUSG00000036208 | Nepro | 212547 | protein_coding | 247.5946362 | -0.321402753 | 0.09293887 | -3.458216692 | 0.000543764 | 0.004671432 | 0.321402753 |
| ENSMUSG00000092032 | Vmn2r59 | 628444 | protein_coding | 9.581709492 | -0.321513732 | 1.649112583 | -0.194961663 | 0.84542296 | 0.918463925 | 0.321513732 |
| ENSMUSG00000041075 | Fzd7 | 14369 | protein_coding | 263.2011071 | 0.321534646 | 0.141469063 | 2.272826569 | 0.023036633 | 0.088930561 | 0.321534646 |
| ENSMUSG00000020420 | Zfp607a | 545938 | protein_coding | 99.85157521 | -0.321534668 | 0.169860194 | -1.892937128 | 0.058366231 | 0.172734546 | 0.321534668 |
| ENSMUSG00000019971 | Cep290 | 216274 | protein_coding | 347.9213482 | 0.321573521 | 0.108024079 | 2.976868912 | 0.002912085 | 0.018318992 | 0.321573521 |
| ENSMUSG00000045409 | Trim39 | 79263 | protein_coding | 292.3660427 | -0.321616049 | 0.13694483 | -2.348508139 | 0.018848787 | 0.076548568 | 0.321616049 |
| ENSMUSG00000070034 | Sp110 | 109032 | protein_coding | 77.25130034 | -0.321623282 | 0.464050254 | -0.69307856 | 0.488260252 | 0.680942953 | 0.321623282 |
| ENSMUSG00000090622 | A930033H14Rik | 320700 | lncRNA | 6.75564743 | -0.321810267 | 0.516257579 | -0.62335214 | 0.533053142 | 0.718131645 | 0.321810267 |
| ENSMUSG00000099759 | 1700030C10Rik | 69513 | lncRNA | 17.86276265 | 0.32183894 | 0.321631678 | 1.000644407 | 0.316998753 | 0.528264924 | 0.32183894 |
| ENSMUSG00000010911 | Apip | 56369 | protein_coding | 119.6976914 | 0.321966676 | 0.119218961 | 2.700633142 | 0.006920763 | 0.036134268 | 0.321966676 |
| ENSMUSG00000044770 | Scml4 | 268297 | protein_coding | 775.7303769 | -0.322074528 | 0.197590307 | -1.630011777 | 0.103099008 | 0.253408395 | 0.322074528 |
| ENSMUSG00000044813 | Shb | 230126 | protein_coding | 324.3889971 | 0.322128041 | 0.292793845 | 1.1001872 | 0.271250567 | 0.479281529 | 0.322128041 |
| ENSMUSG00000001082 | Mfsd10 | 68294 | protein_coding | 221.1517315 | 0.322128608 | 0.107470076 | 2.997379555 | 0.002723114 | 0.017388158 | 0.322128608 |
| ENSMUSG00000040562 | Gstm2 | 14863 | protein_coding | 1153.641476 | 0.322159232 | 0.11546404 | 2.790126108 | 0.005268752 | 0.0291898 | 0.322159232 |
| ENSMUSG00000029198 | Grpel1 | 17713 | protein_coding | 1122.507195 | 0.322169445 | 0.113160347 | 2.847017116 | 0.004413099 | 0.025386934 | 0.322169445 |
| ENSMUSG00000021820 | Camk2g | 12325 | protein_coding | 150.4201522 | -0.322176872 | 0.116949496 | -2.754837639 | 0.005872127 | 0.031630448 | 0.322176872 |
| ENSMUSG00000057914 | Cacnb2 | 12296 | protein_coding | 10.69063111 | -0.322222971 | 0.61987755 | -0.519817133 | 0.603191036 | 0.76799575 | 0.322222971 |
| ENSMUSG00000032908 | Sgpp2 | 433323 | protein_coding | 1837.793077 | 0.322331497 | 0.089709667 | 3.593051982 | 0.000326827 | 0.003047837 | 0.322331497 |
| ENSMUSG00000038806 | Sde2 | 208768 | protein_coding | 347.5871507 | -0.322332609 | 0.116322739 | -2.771019758 | 0.005588104 | 0.030440258 | 0.322332609 |

Supplementary Table 1

| ensembl_gene_id | external_gene_name | entrezgene_id | gene_biotype | baseMean | log2FoldChange | lfcSE | stat | pvalue | padj | absLog2FC |
|---|---|---|---|---|---|---|---|---|---|---|
| ENSMUSG00000036123 | Slc9a3 | 105243 | protein_coding | 2292.186002 | -0.322378219 | 0.346968811 | -0.92912737 | 0.352823082 | 0.562539144 | 0.322378219 |
| ENSMUSG00000035835 | Plppr3 | 216152 | protein_coding | 44.68060914 | -0.322420303 | 0.227801658 | -1.415355384 | 0.156964324 | 0.337781228 | 0.322420303 |
| ENSMUSG00000033721 | Vav3 | 57257 | protein_coding | 403.4108788 | -0.322428228 | 0.17762376 | -1.815231403 | 0.069488356 | 0.194817816 | 0.322428228 |
| ENSMUSG00000033717 | Adra2a | 11551 | protein_coding | 513.1535369 | -0.32247062 | 0.275736658 | -1.169487662 | 0.242207209 | 0.447030873 | 0.32247062 |
| ENSMUSG00000044345 | Marveld1 | 277010 | protein_coding | 51.27935738 | -0.322628695 | 0.156978688 | -2.055238827 | 0.039855943 | 0.132753127 | 0.322628695 |
| ENSMUSG00000021704 | Mtx3 | 382793 | protein_coding | 295.1518716 | -0.322690923 | 0.112776064 | -2.861342292 | 0.004218513 | 0.024495982 | 0.322690923 |
| ENSMUSG00000040283 | Btnl9 | 237754 | protein_coding | 10.72584528 | -0.322704273 | 0.46239737 | -0.697893835 | 0.48524359 | 0.678589433 | 0.322704273 |
| ENSMUSG00000043753 | Dmrta1 | 242523 | protein_coding | 14.08719206 | -0.322706885 | 0.390459889 | -0.826478963 | 0.408532447 | 0.61573664 | 0.322706885 |
| ENSMUSG00000030555 | Ttc23 | 67009 | protein_coding | 186.8734831 | 0.32290481 | 0.14333475 | 2.252801997 | 0.024271636 | 0.092377156 | 0.32290481 |
| ENSMUSG00000030838 | Ush1c | 72088 | protein_coding | 2549.769256 | 0.322945234 | 0.065891732 | 4.901149568 | 9.53E-07 | 1.81E-05 | 0.322945234 |
| ENSMUSG00000024827 | Gldc | 104174 | protein_coding | 7.42380447 | -0.322972808 | 0.491764597 | -0.656763033 | 0.511333298 | 0.699220189 | 0.322972808 |
| ENSMUSG00000041429 | Nthl1 | 18207 | protein_coding | 31.94209186 | -0.322982168 | 0.207829099 | -1.55407577 | 0.120166344 | 0.281430603 | 0.322982168 |
| ENSMUSG00000008540 | Mgst1 | 56615 | protein_coding | 2381.361056 | 0.323070698 | 0.114839105 | 2.813246389 | 0.004904406 | 0.027652167 | 0.323070698 |
| ENSMUSG00000038567 | Cyp24a1 | 13081 | protein_coding | 12.14180546 | 0.323184903 | 0.544585767 | 0.593450881 | 0.552879446 | 0.732884453 | 0.323184903 |
| ENSMUSG00000025145 | Lrrc45 | 217366 | protein_coding | 515.4266706 | 0.323191663 | 0.09935208 | 3.252993417 | 0.001141961 | 0.008548225 | 0.323191663 |
| ENSMUSG00000015944 | Castor2 | 80909 | protein_coding | 44.94506028 | -0.323508434 | 0.277054955 | -1.167668827 | 0.242940376 | 0.447865081 | 0.323508434 |
| ENSMUSG00000097325 | Gm16897 | 320400 | lncRNA | 7.726041219 | 0.323539964 | 0.430120748 | 0.752207294 | 0.451926408 | 0.652077527 | 0.323539964 |
| ENSMUSG00000054693 | Adam10 | 11487 | protein_coding | 1824.032212 | 0.323567355 | 0.070359994 | 4.598740536 | 4.25E-06 | 6.97E-05 | 0.323567355 |
| ENSMUSG00000026764 | Kif5c | 16574 | protein_coding | 46.22499276 | 0.323748195 | 0.383114782 | 0.845042297 | 0.398087224 | 0.606072768 | 0.323748195 |
| ENSMUSG00000033581 | Igf2bp2 | 319765 | protein_coding | 509.6440361 | -0.323749839 | 0.132614807 | -2.441279712 | 0.014635314 | 0.063242345 | 0.323749839 |
| ENSMUSG00000003518 | Dusp3 | 72349 | protein_coding | 338.6805026 | -0.323777644 | 0.101341507 | -3.194916415 | 0.001398713 | 0.010085427 | 0.323777644 |
| ENSMUSG00000066571 | Garre1 | 233103 | protein_coding | 265.426637 | -0.323840393 | 0.121690066 | -2.661190042 | 0.0077865 | 0.039636742 | 0.323840393 |
| ENSMUSG00000060429 | Sntb1 | 20649 | protein_coding | 9.147279421 | -0.323890377 | 0.600629694 | -0.539251356 | 0.589713429 | 0.759394898 | 0.323890377 |
| ENSMUSG00000027087 | Itgav | 16410 | protein_coding | 787.8079897 | 0.32389741 | 0.219755295 | 1.473900371 | 0.140508418 | 0.312850724 | 0.32389741 |
| ENSMUSG00000034192 | Lsm3 | 67678 | protein_coding | 259.1665411 | 0.323902565 | 0.101972191 | 3.176381337 | 0.001491248 | 0.010618058 | 0.323902565 |
| ENSMUSG00000039616 | Mocos | 68591 | protein_coding | 134.7917162 | 0.323976349 | 0.141565069 | 2.288533122 | 0.022106494 | 0.086124599 | 0.323976349 |
| ENSMUSG00000040936 | Ulk4 | 209012 | protein_coding | 33.67841838 | -0.323995469 | 0.250664117 | -1.292548262 | 0.196167344 | 0.389692373 | 0.323995469 |
| ENSMUSG00000068917 | Clk2 | 12748 | protein_coding | 322.3549836 | -0.324020026 | 0.107388066 | -3.017281507 | 0.002550528 | 0.016542619 | 0.324020026 |
| ENSMUSG00000020261 | Slc36a1 | 215335 | protein_coding | 434.3314751 | -0.324026164 | 0.193677399 | -1.673020005 | 0.094323358 | 0.239156709 | 0.324026164 |
| ENSMUSG00000033276 | Stk36 | 269209 | protein_coding | 12.55435255 | 0.324098535 | 0.387170814 | 0.83709444 | 0.402539486 | 0.609921868 | 0.324098535 |
| ENSMUSG00000036908 | Unc93b1 | 54445 | protein_coding | 402.0188086 | -0.324156517 | 0.121735224 | -2.662799683 | 0.007749353 | 0.039498143 | 0.324156517 |
| ENSMUSG00000047117 | Ankdd1b | 271144 | protein_coding | 23.39192465 | 0.324279608 | 0.433270199 | 0.748446602 | 0.454190822 | 0.653598611 | 0.324279608 |
| ENSMUSG00000035337 | Uchl4 | 93841 | protein_coding | 13.65082823 | -0.324348457 | 0.340512599 | -0.952529973 | 0.340828272 | 0.551168324 | 0.324348457 |
| ENSMUSG00000022508 | Bcl6 | 12053 | protein_coding | 70.25662127 | 0.324385619 | 0.220815394 | 1.469035348 | 0.141823202 | 0.314924867 | 0.324385619 |
| ENSMUSG00000002083 | Bbc3 | 170770 | protein_coding | 78.0996906 | -0.324529734 | 0.175383177 | -1.850404007 | 0.064255342 | 0.184274927 | 0.324529734 |
| ENSMUSG00000030922 | Lyrm1 | 73919 | protein_coding | 31.67167279 | 0.324561044 | 0.232706385 | 1.394723415 | 0.163099261 | 0.345641968 | 0.324561044 |
| ENSMUSG00000027875 | Hmgcs2 | 15360 | protein_coding | 18284.91087 | -0.324638088 | 0.212227147 | -1.529672776 | 0.126097746 | 0.290663516 | 0.324638088 |
| ENSMUSG00000021559 | Dapk1 | 69635 | protein_coding | 239.2111992 | -0.324638707 | 0.167940129 | -1.933062145 | 0.053228543 | 0.161983995 | 0.324638707 |

| ensembl_gene_id | external_gene_name | entrezgene_id | gene_biotype | baseMean | log2FoldChange | lfcSE | stat | pvalue | padj | absLog2FC |
|---|---|---|---|---|---|---|---|---|---|---|
| ENSMUSG00000037266 | Rsrp1 | 27981 | protein_coding | 1937.730535 | -0.324698863 | 0.148746821 | -2.182896157 | 0.029043458 | 0.105304849 | 0.324698863 |
| ENSMUSG00000037112 | Sik2 | 235344 | protein_coding | 247.6275214 | -0.324702197 | 0.114065713 | -2.846624008 | 0.004418552 | 0.025405247 | 0.324702197 |
| ENSMUSG00000040260 | Daam2 | 76441 | protein_coding | 16.57414709 | -0.324708655 | 0.499509343 | -0.650055218 | 0.515656554 | 0.703079747 | 0.324708655 |
| ENSMUSG00000009207 | Lnpk | 69605 | protein_coding | 311.8684798 | 0.324793786 | 0.098610044 | 3.293719086 | 0.000988713 | 0.00761151 | 0.324793786 |
| ENSMUSG00000000295 | Hddc2 | 69692 | protein_coding | 198.2377607 | 0.324824261 | 0.111512868 | 2.912885902 | 0.003581054 | 0.021562746 | 0.324824261 |
| ENSMUSG00000038984 | Tspyl5 | 239364 | protein_coding | 8.887231312 | 0.324922166 | 0.378927883 | 0.857477586 | 0.391180997 | 0.59924777 | 0.324922166 |
| ENSMUSG00000027931 | Npr1 | 18160 | protein_coding | 8.257921314 | 0.324925979 | 0.518931714 | 0.626144 | 0.531220495 | 0.716450469 | 0.324925979 |
| ENSMUSG00000033450 | Tagap | 72536 | protein_coding | 39.20179122 | -0.324936597 | 0.458160897 | -0.709219401 | 0.478188336 | 0.672607328 | 0.324936597 |
| ENSMUSG00000066191 | Anks6 | 75691 | protein_coding | 140.0882893 | -0.32499679 | 0.181928296 | -1.786400445 | 0.074034444 | 0.202894119 | 0.32499679 |
| ENSMUSG00000038650 | Rnh1 | 107702 | protein_coding | 1436.049692 | 0.325011741 | 0.126218021 | 2.575002672 | 0.010023931 | 0.048003962 | 0.325011741 |
| ENSMUSG00000024927 | Rela | 19697 | protein_coding | 348.6190297 | -0.325107908 | 0.127336213 | -2.553145736 | 0.010675482 | 0.050247953 | 0.325107908 |
| ENSMUSG00000027835 | Pdcd10 | 56426 | protein_coding | 922.0679741 | 0.325126027 | 0.086334152 | 3.765902822 | 0.000165948 | 0.0017344 | 0.325126027 |
| ENSMUSG00000021913 | Ogdhl | 239017 | protein_coding | 24.54631184 | -0.325545969 | 0.239577636 | -1.358832878 | 0.174199552 | 0.360225341 | 0.325545969 |
| ENSMUSG00000020330 | Hmmr | 15366 | protein_coding | 630.5293537 | 0.325563443 | 0.136064102 | 2.392721059 | 0.01672395 | 0.070155143 | 0.325563443 |
| ENSMUSG00000004071 | Cdip1 | 66626 | protein_coding | 400.7748364 | -0.325572162 | 0.085574499 | -3.804546493 | 0.000142064 | 0.001526855 | 0.325572162 |
| ENSMUSG00000060621 | Nkpd1 | 69547 | protein_coding | 9.37899689 | -0.32566043 | 0.396070052 | -0.822229372 | 0.410946365 | 0.617971459 | 0.32566043 |
| ENSMUSG00000027615 | Hps3 | 12807 | protein_coding | 208.6722163 | -0.325668798 | 0.117728574 | -2.766268083 | 0.00567019 | 0.030803133 | 0.325668798 |
| ENSMUSG00000097101 | 1810034E14Rik | 66503 | lncRNA | 82.07229485 | 0.32567664 | 0.165693288 | 1.965539122 | 0.049351872 | 0.154102455 | 0.32567664 |
| ENSMUSG00000043668 | Tox3 | 244579 | protein_coding | 257.6527502 | -0.325805554 | 0.112111697 | -2.90607994 | 0.003659879 | 0.021948252 | 0.325805554 |
| ENSMUSG00000071637 | Cebpd | 12609 | protein_coding | 313.7893134 | 0.325841436 | 0.21122658 | 1.54261569 | 0.122924049 | 0.28587155 | 0.325841436 |
| ENSMUSG00000027599 | Armc1 | 74252 | protein_coding | 571.0136655 | 0.325893801 | 0.091910544 | 3.545771638 | 0.000391465 | 0.003569124 | 0.325893801 |
| ENSMUSG00000035274 | Tpbg | 21983 | protein_coding | 89.19761911 | -0.325916841 | 0.204089624 | -1.596929992 | 0.110281313 | 0.264838953 | 0.325916841 |
| ENSMUSG00000024222 | Fkbp5 | 14229 | protein_coding | 2050.186899 | -0.325947979 | 0.254618663 | -1.280141745 | 0.20049529 | 0.395135978 | 0.325947979 |
| ENSMUSG00000038208 | Pgap3 | 320655 | protein_coding | 391.8294632 | 0.325964998 | 0.082374045 | 3.957132363 | 7.59E-05 | 0.000883846 | 0.325964998 |
| ENSMUSG00000021468 | Sptlc1 | 268656 | protein_coding | 999.4140821 | 0.325973823 | 0.083018126 | 3.926537946 | 8.62E-05 | 0.000992503 | 0.325973823 |
| ENSMUSG00000117322 | 6330415G19Rik | 320020 | lncRNA | 8.390495413 | -0.325979474 | 0.410869792 | -0.793388759 | 0.427551358 | 0.63196344 | 0.325979474 |
| ENSMUSG00000023495 | Pcbp4 | 59092 | protein_coding | 36.96094341 | -0.325984741 | 0.233769496 | -1.39447082 | 0.163175475 | 0.345682448 | 0.325984741 |
| ENSMUSG00000042010 | Acacb | 100705 | protein_coding | 372.5576781 | 0.326054397 | 0.146570989 | 2.224549346 | 0.026111506 | 0.09758425 | 0.326054397 |
| ENSMUSG00000021782 | Dlg5 | 71228 | protein_coding | 338.4219784 | -0.326099686 | 0.144084718 | -2.263249634 | 0.023620305 | 0.090547345 | 0.326099686 |
| ENSMUSG00000073437 | D330041H03Rik | 654822 | lncRNA | 22.63293755 | -0.32615922 | 0.268091434 | -1.216596947 | 0.223757605 | 0.425414053 | 0.32615922 |
| ENSMUSG00000085208 | Brip1os | 74038 | lncRNA | 944.2501067 | 0.32640862 | 0.07545732 | 4.325738289 | 1.52E-05 | 0.000216584 | 0.32640862 |
| ENSMUSG00000041609 | Bicdl1 | 75665 | protein_coding | 235.134986 | 0.326488568 | 0.198092906 | 1.648158809 | 0.099320087 | 0.247182867 | 0.326488568 |
| ENSMUSG00000063894 | Zkscan8 | 93681 | protein_coding | 226.3033844 | -0.326568512 | 0.205080238 | -1.592393865 | 0.111296234 | 0.266434468 | 0.326568512 |
| ENSMUSG00000070871 | Ccnyl1 | 227210 | protein_coding | 191.4029333 | 0.326611072 | 0.101010744 | 3.233429021 | 0.001223137 | 0.009036239 | 0.326611072 |
| ENSMUSG00000037847 | Nmrk1 | 225994 | protein_coding | 68.85410394 | 0.326681393 | 0.182523176 | 1.789807743 | 0.073484824 | 0.201894694 | 0.326681393 |
| ENSMUSG00000044254 | Pcsk9 | 100102 | protein_coding | 332.4265854 | -0.326771965 | 0.1975859 | -1.653822289 | 0.098163626 | 0.245262782 | 0.326771965 |
| ENSMUSG00000113634 | Gm36756 | 102640764 | lncRNA | 10.39091384 | -0.326790427 | 0.461512962 | -0.708085045 | 0.478892446 | 0.672927641 | 0.326790427 |
| ENSMUSG00000020284 | Cfap410 | 67884 | protein_coding | 122.8993831 | -0.326874957 | 0.131244315 | -2.490583731 | 0.012753344 | 0.057466485 | 0.326874957 |

| ensembl_gene_id | external_gene_name | entrezgene_id | gene_biotype | baseMean | log2FoldChange | lfcSE | stat | pvalue | padj | absLog2FC |
|---|---|---|---|---|---|---|---|---|---|---|
| ENSMUSG00000023017 | Asic1 | 11419 | protein_coding | 50.73843694 | -0.326902838 | 0.19522917 | -1.674456939 | 0.094040831 | 0.238820528 | 0.326902838 |
| ENSMUSG00000033961 | Zfp446 | 269870 | protein_coding | 61.57150417 | 0.327122198 | 0.19051946 | 1.717001498 | 0.085978898 | 0.224946106 | 0.327122198 |
| ENSMUSG00000026627 | Pacc1 | 66950 | protein_coding | 72.97988667 | -0.327193359 | 0.182845847 | -1.789449226 | 0.073542498 | 0.201928099 | 0.327193359 |
| ENSMUSG00000015094 | Npdc1 | 18146 | protein_coding | 411.3381412 | -0.327246234 | 0.086285885 | -3.792581287 | 0.000149089 | 0.001589489 | 0.327246234 |
| ENSMUSG00000022945 | Chaf1b | 110749 | protein_coding | 119.0516103 | -0.327495926 | 0.204820343 | -1.598942375 | 0.109833407 | 0.264138909 | 0.327495926 |
| ENSMUSG00000112980 | D430020J02Rik | 319545 | lncRNA | 37.53658184 | -0.327510009 | 0.258017485 | -1.269332615 | 0.204322459 | 0.399760241 | 0.327510009 |
| ENSMUSG00000034949 | Zfr2 | 103406 | protein_coding | 61.97777794 | -0.327539377 | 0.237645952 | -1.378266174 | 0.168121123 | 0.352676319 | 0.327539377 |
| ENSMUSG00000038776 | Ephx1 | 13849 | protein_coding | 273.7231636 | 0.32756508 | 0.117755615 | 2.781736401 | 0.005406894 | 0.029676431 | 0.32756508 |
| ENSMUSG00000041566 | Tssk1 | 22114 | protein_coding | 7.288503244 | -0.327668345 | 0.495356001 | -0.661480519 | 0.508304204 | 0.69753333 | 0.327668345 |
| ENSMUSG00000020088 | Sar1a | 20224 | protein_coding | 1739.140498 | 0.327806793 | 0.093152666 | 3.519027475 | 0.000433132 | 0.003880311 | 0.327806793 |
| ENSMUSG00000037692 | Ahdc1 | 230793 | protein_coding | 262.3639081 | -0.3278103 | 0.150163883 | -2.183016941 | 0.029034562 | 0.105296565 | 0.3278103 |
| ENSMUSG00000022841 | Ap2m1 | 11773 | protein_coding | 1531.311579 | 0.32786126 | 0.093745728 | 3.497346128 | 0.000469912 | 0.004139797 | 0.32786126 |
| ENSMUSG00000020744 | Slc25a19 | 67283 | protein_coding | 222.5658346 | 0.327873728 | 0.088284032 | 3.713850881 | 0.000204129 | 0.002082875 | 0.327873728 |
| ENSMUSG00000040669 | Phc1 | 13619 | protein_coding | 117.3178334 | -0.328158517 | 0.118994949 | -2.757751653 | 0.00582004 | 0.031424268 | 0.328158517 |
| ENSMUSG00000034173 | Chchd2l | 71970 | protein_coding | 14.98110356 | -0.328492897 | 0.322950138 | -1.017162893 | 0.309075945 | 0.51897129 | 0.328492897 |
| ENSMUSG00000038422 | Hdhd3 | 72748 | protein_coding | 200.0718662 | 0.328585518 | 0.144400818 | 2.275510087 | 0.022875348 | 0.088529438 | 0.328585518 |
| ENSMUSG00000036585 | Fgf1 | 14164 | protein_coding | 537.2278532 | 0.328624684 | 0.157423912 | 2.087514405 | 0.03684166 | 0.125387599 | 0.328624684 |
| ENSMUSG00000038526 | Car14 | 23831 | protein_coding | 8.36698339 | -0.328757841 | 0.603216971 | -0.545007611 | 0.585748283 | 0.756000214 | 0.328757841 |
| ENSMUSG00000025188 | Hps1 | 192236 | protein_coding | 219.1503893 | -0.328912611 | 0.09755877 | -3.37143049 | 0.000747789 | 0.006061468 | 0.328912611 |
| ENSMUSG00000025545 | Clybl | 69634 | protein_coding | 491.4739474 | -0.3291036 | 0.089830497 | -3.663606573 | 0.000248689 | 0.002437608 | 0.3291036 |
| ENSMUSG00000078453 | Abracl | 73112 | protein_coding | 619.3246614 | 0.329292026 | 0.091955262 | 3.58100255 | 0.000342278 | 0.003171464 | 0.329292026 |
| ENSMUSG00000030091 | Nup210 | 54563 | protein_coding | 424.0438976 | -0.329433995 | 0.109387035 | -3.011636571 | 0.002598435 | 0.016776599 | 0.329433995 |
| ENSMUSG00000024935 | Slc1a1 | 20510 | protein_coding | 358.8900151 | -0.329438007 | 0.258035106 | -1.276717776 | 0.201701903 | 0.396586282 | 0.329438007 |
| ENSMUSG00000034390 | Cmip | 74440 | protein_coding | 607.0487489 | -0.329440952 | 0.127727903 | -2.579240276 | 0.009901789 | 0.047619472 | 0.329440952 |
| ENSMUSG00000025758 | Plk4 | 20873 | protein_coding | 321.0912207 | 0.329455049 | 0.114743817 | 2.87122268 | 0.004088874 | 0.023891262 | 0.329455049 |
| ENSMUSG00000079685 | Ulbp1 | 77777 | protein_coding | 77.01393815 | -0.329526585 | 0.253305111 | -1.300907763 | 0.193290029 | 0.386677164 | 0.329526585 |
| ENSMUSG00000058706 | 0610030E20Rik | 68364 | protein_coding | 247.8148041 | -0.329681835 | 0.102144944 | -3.227588377 | 0.001248385 | 0.009154194 | 0.329681835 |
| ENSMUSG00000038102 | Trappc11 | 320714 | protein_coding | 1111.48803 | 0.32979092 | 0.064598085 | 5.105273952 | 3.30E-07 | 6.93E-06 | 0.32979092 |
| ENSMUSG00000020056 | Washc3 | 67282 | protein_coding | 361.8872603 | 0.329811108 | 0.101324147 | 3.255009975 | 0.001133884 | 0.008503061 | 0.329811108 |
| ENSMUSG00000033186 | Mzt1 | 76789 | protein_coding | 807.2915938 | 0.329868253 | 0.100440359 | 3.284220159 | 0.00102265 | 0.007806918 | 0.329868253 |
| ENSMUSG00000085929 | Gm13421 | 105244102 | lncRNA | 18.57697152 | 0.329878235 | 0.33267944 | 0.991579867 | 0.321402515 | 0.532354566 | 0.329878235 |
| ENSMUSG00000028785 | Hpca | 15444 | protein_coding | 28.79588899 | -0.329904162 | 0.260302416 | -1.267388012 | 0.205016584 | 0.400773705 | 0.329904162 |
| ENSMUSG00000097180 | 2700038G22Rik | 67194 | lncRNA | 9.69026119 | -0.329948969 | 0.35962216 | -0.91748787 | 0.358887046 | 0.56873474 | 0.329948969 |
| ENSMUSG00000078789 | Dph1 | 116905 | protein_coding | 126.9399448 | -0.330187302 | 0.14978264 | -2.204443063 | 0.0274932 | 0.101087648 | 0.330187302 |
| ENSMUSG00000023055 | Calcoco1 | 67488 | protein_coding | 200.8077101 | -0.330217527 | 0.20824322 | -1.585730025 | 0.112800565 | 0.268563139 | 0.330217527 |
| ENSMUSG00000110206 | Flt3l | 14256 | protein_coding | 15.28452423 | -0.33022259 | 0.312035324 | -1.058285917 | 0.289925111 | 0.498588553 | 0.33022259 |
| ENSMUSG00000019866 | Crybg1 | 11630 | protein_coding | 1857.347536 | 0.33032914 | 0.09903858 | 3.335358186 | 0.000851896 | 0.006717326 | 0.33032914 |
| ENSMUSG00000031431 | Tsc22d3 | 14605 | protein_coding | 754.579337 | 0.330361085 | 0.287074553 | 1.150784984 | 0.249820704 | 0.456114646 | 0.330361085 |

| ensembl_gene_id | external_gene_name | entrezgene_id | gene_biotype | baseMean | log2FoldChange | lfcSE | stat | pvalue | padj | absLog2FC |
|---|---|---|---|---|---|---|---|---|---|---|
| ENSMUSG00000002249 | Tead3 | 21678 | protein_coding | 199.9679674 | -0.330455142 | 0.134858188 | -2.450389896 | 0.01427016 | 0.06215383 | 0.330455142 |
| ENSMUSG00000059456 | Ptk2b | 19229 | protein_coding | 672.3015102 | -0.33046047 | 0.120451526 | -2.743514177 | 0.006078542 | 0.032536352 | 0.33046047 |
| ENSMUSG00000024818 | Slc25a45 | 107375 | protein_coding | 30.35770775 | -0.33047171 | 0.245182758 | -1.347858686 | 0.177703838 | 0.364750868 | 0.33047171 |
| ENSMUSG00000031628 | Casp3 | 12367 | protein_coding | 1076.072663 | -0.330487342 | 0.094145542 | -3.510387599 | 0.000447454 | 0.003977146 | 0.330487342 |
| ENSMUSG00000027488 | Snta1 | 20648 | protein_coding | 51.0585814 | 0.33053825 | 0.191396913 | 1.726977958 | 0.084171622 | 0.221820913 | 0.33053825 |
| ENSMUSG00000035757 | Selenoo | 223776 | protein_coding | 319.5231347 | -0.330549981 | 0.107010687 | -3.088943633 | 0.002008695 | 0.013545513 | 0.330549981 |
| ENSMUSG00000045639 | Zfp629 | 320683 | protein_coding | 344.2376107 | -0.330550605 | 0.134806187 | -2.452043275 | 0.014204758 | 0.062004215 | 0.330550605 |
| ENSMUSG00000020122 | Egfr | 13649 | protein_coding | 899.5507285 | -0.330801192 | 0.136289627 | -2.427192731 | 0.01521617 | 0.065363632 | 0.330801192 |
| ENSMUSG00000036894 | Rap2b | 74012 | protein_coding | 67.37368807 | 0.33080873 | 0.259721197 | 1.27370709 | 0.202767243 | 0.397743429 | 0.33080873 |
| ENSMUSG00000022773 | Ypel1 | 106369 | protein_coding | 18.9230675 | -0.330814514 | 0.311476919 | -1.062083559 | 0.288197753 | 0.497390163 | 0.330814514 |
| ENSMUSG00000064307 | Lrrc51 | 69358 | protein_coding | 56.90276648 | -0.33086654 | 0.204222364 | -1.620128836 | 0.105204604 | 0.256772746 | 0.33086654 |
| ENSMUSG00000039470 | Zdhhc2 | 70546 | protein_coding | 332.6746524 | 0.330889746 | 0.110069445 | 3.006190741 | 0.00264543 | 0.017031693 | 0.330889746 |
| ENSMUSG00000032998 | Foxj3 | 230700 | protein_coding | 559.1047037 | -0.331008794 | 0.073889428 | -4.479785599 | 7.47E-06 | 0.000115769 | 0.331008794 |
| ENSMUSG00000014606 | Slc25a11 | 67863 | protein_coding | 1355.934655 | 0.331034425 | 0.099399188 | 3.330353424 | 0.000867358 | 0.006822361 | 0.331034425 |
| ENSMUSG00000032586 | Traip | 22036 | protein_coding | 92.54093128 | 0.331083305 | 0.164797427 | 2.009032007 | 0.044533739 | 0.143328632 | 0.331083305 |
| ENSMUSG00000021223 | Papln | 170721 | protein_coding | 23.46442013 | -0.331091538 | 0.282827005 | -1.170650367 | 0.241739343 | 0.446529544 | 0.331091538 |
| ENSMUSG00000037933 | Bicd2 | 76895 | protein_coding | 274.639929 | -0.331109587 | 0.10191804 | -3.248782918 | 0.001158999 | 0.008644173 | 0.331109587 |
| ENSMUSG00000049550 | Clip1 | 56430 | protein_coding | 810.2248613 | -0.331260217 | 0.055350267 | -5.984798904 | 2.17E-09 | 6.33E-08 | 0.331260217 |
| ENSMUSG00000004937 | Sgta | 52551 | protein_coding | 931.8908758 | 0.331325403 | 0.063810573 | 5.192327622 | 2.08E-07 | 4.54E-06 | 0.331325403 |
| ENSMUSG00000062410 | Hsd3b3 | 15494 | protein_coding | 2220.838822 | -0.331341011 | 0.146558139 | -2.260816169 | 0.023770643 | 0.090992261 | 0.331341011 |
| ENSMUSG00000031173 | Otc | 18416 | protein_coding | 471.7475699 | -0.331543025 | 0.127459829 | -2.601156989 | 0.009290993 | 0.045339133 | 0.331543025 |
| ENSMUSG00000048997 | Atxn7l2 | 72522 | protein_coding | 44.01843294 | -0.331570007 | 0.16626357 | -1.99424328 | 0.046125474 | 0.147143312 | 0.331570007 |
| ENSMUSG00000050310 | Rictor | 78757 | protein_coding | 877.4068166 | -0.331624943 | 0.109917842 | -3.017025606 | 0.002552682 | 0.016542619 | 0.331624943 |
| ENSMUSG00000032855 | Pkd1 | 18763 | protein_coding | 621.074054 | -0.331669044 | 0.142526457 | -2.32707001 | 0.019961539 | 0.079646142 | 0.331669044 |
| ENSMUSG00000074647 | Fam83c | 71405 | protein_coding | 151.3765637 | -0.331680496 | 0.214346659 | -1.54740222 | 0.121766286 | 0.284133227 | 0.331680496 |
| ENSMUSG00000051316 | Taf7 | 24074 | protein_coding | 227.0343792 | -0.3317999 | 0.085791796 | -3.867501512 | 0.000109956 | 0.001224742 | 0.3317999 |
| ENSMUSG00000033427 | Upb1 | 103149 | protein_coding | 99.69183743 | -0.331853078 | 0.17377009 | -1.909724956 | 0.056168636 | 0.168041705 | 0.331853078 |
| ENSMUSG00000026031 | Cflar | 12633 | protein_coding | 1203.82282 | 0.331853163 | 0.08141473 | 4.076082594 | 4.58E-05 | 0.000571259 | 0.331853163 |
| ENSMUSG00000026782 | Abi2 | 329165 | protein_coding | 96.67053206 | 0.331967875 | 0.186495274 | 1.780033716 | 0.075070443 | 0.204396926 | 0.331967875 |
| ENSMUSG00000056718 | Gm13199 | 791411 | protein_coding | 6.927610529 | -0.331975027 | 0.463766265 | -0.715824009 | 0.474100026 | 0.669040951 | 0.331975027 |
| ENSMUSG00000043621 | Ubxn10 | 212190 | protein_coding | 106.4673254 | 0.332015683 | 0.193689827 | 1.714161695 | 0.086499034 | 0.22594736 | 0.332015683 |
| ENSMUSG00000065738 | Gm24494 | 115489817 | snoRNA | 9.253396684 | -0.332127539 | 0.423863726 | -0.783571509 | 0.433291579 | 0.637254688 | 0.332127539 |
| ENSMUSG00000002814 | Top3a | 21975 | protein_coding | 145.3092364 | -0.332238074 | 0.100009846 | -3.322053652 | 0.000893575 | 0.006990601 | 0.332238074 |
| ENSMUSG00000017376 | Nlk | 18099 | protein_coding | 243.2378949 | 0.332324396 | 0.081539412 | 4.075629063 | 4.59E-05 | 0.000571492 | 0.332324396 |
| ENSMUSG00000040430 | Pitpnc1 | 71795 | protein_coding | 43.7481005 | -0.332336947 | 0.206296736 | -1.610965611 | 0.107187218 | 0.260176472 | 0.332336947 |
| ENSMUSG00000027827 | Kcnab1 | 16497 | protein_coding | 20.9735309 | 0.332360248 | 0.389825372 | 0.852587522 | 0.393888085 | 0.60186583 | 0.332360248 |
| ENSMUSG00000113800 | 2210039B01Rik | 73644 | lncRNA | 16.06031106 | -0.332402145 | 0.325719384 | -1.020516926 | 0.307483366 | 0.51745943 | 0.332402145 |
| ENSMUSG00000059791 | Nrm | 106582 | protein_coding | 98.77546535 | -0.332557733 | 0.157150665 | -2.116171332 | 0.034330238 | 0.119325161 | 0.332557733 |

| ensembl_gene_id | external_gene_name | entrezgene_id | gene_biotype | baseMean | log2FoldChange | lfcSE | stat | pvalue | padj | absLog2FC |
|---|---|---|---|---|---|---|---|---|---|---|
| ENSMUSG00000019943 | Atp2b1 | 67972 | protein_coding | 5608.731204 | 0.33258783 | 0.234365103 | 1.419101332 | 0.155869476 | 0.336179961 | 0.33258783 |
| ENSMUSG00000109324 | Prmt1 | 15469 | protein_coding | 350.2995974 | 0.332638729 | 0.122208013 | 2.721906038 | 0.006490659 | 0.034323779 | 0.332638729 |
| ENSMUSG00000097267 | 1700109K24Rik | 74303 | lncRNA | 12.11011487 | -0.332748214 | 0.403441953 | -0.824773456 | 0.409500219 | 0.616436962 | 0.332748214 |
| ENSMUSG00000028932 | Psmc2 | 19181 | protein_coding | 983.0523205 | 0.332767446 | 0.105168719 | 3.164129494 | 0.001555476 | 0.010962662 | 0.332767446 |
| ENSMUSG00000048489 | Depp1 | 213393 | protein_coding | 21.61481604 | -0.332798093 | 0.484224909 | -0.687279996 | 0.491906302 | 0.683859515 | 0.332798093 |
| ENSMUSG00000061778 | Mospd2 | 76763 | protein_coding | 401.8505414 | 0.332874708 | 0.08046769 | 4.1367499 | 3.52E-05 | 0.000454683 | 0.332874708 |
| ENSMUSG00000024771 | Lipk | 240633 | protein_coding | 7.204934214 | 0.332885506 | 0.457625108 | 0.727419672 | 0.466968907 | 0.663707223 | 0.332885506 |
| ENSMUSG00000078435 | AU041133 | 216177 | protein_coding | 112.8143805 | 0.332890629 | 0.155358531 | 2.142725126 | 0.032135177 | 0.113492038 | 0.332890629 |
| ENSMUSG00000058317 | Ube2e2 | 218793 | protein_coding | 110.386887 | 0.332964135 | 0.127948066 | 2.602338162 | 0.00925905 | 0.045210961 | 0.332964135 |
| ENSMUSG00000028759 | Hp1bp3 | 15441 | protein_coding | 2071.787629 | 0.33298085 | 0.064756508 | 5.142044555 | 2.72E-07 | 5.80E-06 | 0.33298085 |
| ENSMUSG00000035621 | Midn | 59090 | protein_coding | 1483.427001 | -0.33298299 | 0.203243045 | -1.638348757 | 0.101348962 | 0.25035457 | 0.33298299 |
| ENSMUSG00000025227 | Mfsd13a | 75146 | protein_coding | 126.7204101 | -0.332985384 | 0.137282042 | -2.42555675 | 0.015284926 | 0.065534172 | 0.332985384 |
| ENSMUSG00000035521 | Gnptg | 214505 | protein_coding | 204.8016893 | -0.333097711 | 0.091458196 | -3.642076115 | 0.000270448 | 0.002615481 | 0.333097711 |
| ENSMUSG00000033909 | Usp36 | 72344 | protein_coding | 357.1300574 | -0.333115158 | 0.160772695 | -2.071963513 | 0.038268844 | 0.129141133 | 0.333115158 |
| ENSMUSG00000042408 | Zmym6 | 100177 | protein_coding | 218.2656934 | -0.333158354 | 0.161945891 | -2.057220173 | 0.039665052 | 0.132421913 | 0.333158354 |
| ENSMUSG00000000486 | Septin1 | 54204 | protein_coding | 21.66169318 | 0.333205534 | 0.324423923 | 1.027068322 | 0.304388294 | 0.513961388 | 0.333205534 |
| ENSMUSG00000018931 | Natd1 | 24083 | protein_coding | 109.6094683 | -0.333215831 | 0.139367381 | -2.39091694 | 0.016806355 | 0.070408105 | 0.333215831 |
| ENSMUSG00000029275 | Gfi1 | 14581 | protein_coding | 152.2868274 | -0.333271152 | 0.300600862 | -1.108683286 | 0.267566835 | 0.47543558 | 0.333271152 |
| ENSMUSG00000071553 | Cpa2 | 232680 | protein_coding | 38.13472589 | 0.333388447 | 0.249071529 | 1.338524915 | 0.180725384 | 0.368908615 | 0.333388447 |
| ENSMUSG00000041355 | Ssr2 | 66256 | protein_coding | 1724.997306 | 0.333420596 | 0.083984982 | 3.970002594 | 7.19E-05 | 0.000845476 | 0.333420596 |
| ENSMUSG00000023940 | Sgo1 | 72415 | protein_coding | 244.0422167 | 0.33342069 | 0.134069394 | 2.486926218 | 0.012885212 | 0.057937359 | 0.33342069 |
| ENSMUSG00000020919 | Stat5b | 20851 | protein_coding | 319.0791165 | -0.333499843 | 0.091366749 | -3.650122697 | 0.000262115 | 0.002551937 | 0.333499843 |
| ENSMUSG00000040414 | Slc25a28 | 246696 | protein_coding | 270.4354487 | 0.333659283 | 0.109982966 | 3.033736003 | 0.002415457 | 0.01585878 | 0.333659283 |
| ENSMUSG00000015405 | Ace2 | 70008 | protein_coding | 105.5360626 | -0.333704238 | 0.210983159 | -1.581662915 | 0.113726543 | 0.269866479 | 0.333704238 |
| ENSMUSG00000002984 | Tomm40 | 53333 | protein_coding | 728.9341896 | 0.333706786 | 0.115092832 | 2.899457586 | 0.003738089 | 0.022316448 | 0.333706786 |
| ENSMUSG00000023122 | Sult1c2 | 69083 | protein_coding | 884.8068904 | -0.333729534 | 0.134723848 | -2.477137775 | 0.013244077 | 0.059123221 | 0.333729534 |
| ENSMUSG00000036206 | Sh3bp4 | 98402 | protein_coding | 236.7128777 | -0.333912582 | 0.208372702 | -1.602477574 | 0.109050043 | 0.2630147 | 0.333912582 |
| ENSMUSG00000089824 | Rbm12 | 75710 | protein_coding | 231.337641 | -0.333931873 | 0.164727115 | -2.027182188 | 0.042643784 | 0.13890188 | 0.333931873 |
| ENSMUSG00000032946 | Rasgrp2 | 19395 | protein_coding | 11.19046864 | 0.334001269 | 0.738197414 | 0.452455214 | 0.650941078 | 0.798250711 | 0.334001269 |
| ENSMUSG00000035781 | R3hdm4 | 109284 | protein_coding | 1254.789555 | -0.33406903 | 0.069925772 | -4.77748073 | 1.78E-06 | 3.13E-05 | 0.33406903 |
| ENSMUSG00000097184 | 4632428C04Rik | 100043102 | lncRNA | 29.9887629 | -0.334190986 | 0.270122656 | -1.23718236 | 0.216019389 | 0.415099146 | 0.334190986 |
| ENSMUSG00000030335 | Mrpl51 | 66493 | protein_coding | 1335.13168 | 0.33419356 | 0.072494452 | 4.609919143 | 4.03E-06 | 6.64E-05 | 0.33419356 |
| ENSMUSG00000021375 | Kif13a | 16553 | protein_coding | 1346.109963 | -0.334226384 | 0.086445587 | -3.866320944 | 0.00011049 | 0.001228107 | 0.334226384 |
| ENSMUSG00000045107 | Saysd1 | 67509 | protein_coding | 132.324588 | -0.334276358 | 0.142132334 | -2.35186708 | 0.018679449 | 0.076074215 | 0.334276358 |
| ENSMUSG00000038612 | Mcl1 | 17210 | protein_coding | 4569.078075 | -0.334474221 | 0.068910221 | -4.853767928 | 1.21E-06 | 2.23E-05 | 0.334474221 |
| ENSMUSG00000038412 | Higd1a | 56295 | protein_coding | 3478.800676 | 0.3345359 | 0.214614899 | 1.558772952 | 0.119050128 | 0.279638762 | 0.3345359 |
| ENSMUSG00000039715 | Dync2i2 | 71820 | protein_coding | 96.99076155 | 0.334554304 | 0.135601151 | 2.467193699 | 0.013617667 | 0.060217158 | 0.334554304 |
| ENSMUSG00000027375 | Mal | 17153 | protein_coding | 1254.780029 | -0.334585022 | 0.536123294 | -0.624082232 | 0.532573582 | 0.717606972 | 0.334585022 |

Supplementary Table 1

| ensembl_gene_id | external_gene_name | entrezgene_id | gene_biotype | baseMean | log2FoldChange | lfcSE | stat | pvalue | padj | absLog2FC |
|---|---|---|---|---|---|---|---|---|---|---|
| ENSMUSG00000021221 | Dpf3 | 70127 | protein_coding | 47.22186774 | -0.334618115 | 0.168286639 | -1.988381948 | 0.046769464 | 0.148643475 | 0.334618115 |
| ENSMUSG00000023057 | Fabp2 | 14079 | protein_coding | 6447.811352 | 0.334627075 | 0.186863712 | 1.790754729 | 0.073332663 | 0.201595211 | 0.334627075 |
| ENSMUSG00000027297 | Ltk | 17005 | protein_coding | 13.92758109 | -0.334744082 | 0.31127988 | -1.075379759 | 0.28220474 | 0.491172768 | 0.334744082 |
| ENSMUSG00000014704 | Hoxa2 | 15399 | protein_coding | 24.76858243 | -0.334814799 | 0.252509808 | -1.325947697 | 0.184857023 | 0.374796672 | 0.334814799 |
| ENSMUSG00000026028 | Trak2 | 70827 | protein_coding | 1258.943058 | -0.334845826 | 0.113437231 | -2.951815933 | 0.003159112 | 0.019548692 | 0.334845826 |
| ENSMUSG00000026556 | Vangl2 | 93840 | protein_coding | 34.04655975 | -0.334910752 | 0.243686466 | -1.374351057 | 0.169332723 | 0.354093686 | 0.334910752 |
| ENSMUSG00000032565 | Nudt16 | 75686 | protein_coding | 256.7061962 | 0.334916442 | 0.148398162 | 2.256877289 | 0.024015744 | 0.091687329 | 0.334916442 |
| ENSMUSG00000030763 | Lcmt1 | 30949 | protein_coding | 274.4807758 | 0.334916932 | 0.116027899 | 2.886520687 | 0.003895271 | 0.02301331 | 0.334916932 |
| ENSMUSG00000053460 | Ggcx | 56316 | protein_coding | 381.8674233 | 0.334949912 | 0.096020945 | 3.488300533 | 0.000486101 | 0.004244859 | 0.334949912 |
| ENSMUSG00000054766 | Set | 56086 | protein_coding | 2730.260197 | 0.334985927 | 0.127802535 | 2.621121146 | 0.008764111 | 0.043392839 | 0.334985927 |
| ENSMUSG00000052914 | Cyp2j6 | 13110 | protein_coding | 1546.670829 | 0.335047464 | 0.111024965 | 3.017766896 | 0.002546447 | 0.016528039 | 0.335047464 |
| ENSMUSG00000037239 | Spred3 | 101809 | protein_coding | 13.70419134 | -0.3352261 | 0.50274364 | -0.666793318 | 0.504904161 | 0.694916064 | 0.3352261 |
| ENSMUSG00000024050 | Wiz | 22404 | protein_coding | 225.9259949 | -0.33548655 | 0.140476295 | -2.388207564 | 0.016930777 | 0.07076185 | 0.33548655 |
| ENSMUSG00000015932 | Dstn | 56431 | protein_coding | 19689.70082 | 0.335523819 | 0.142565928 | 2.353464274 | 0.018599395 | 0.075767561 | 0.335523819 |
| ENSMUSG00000040624 | Plekhg1 | 213783 | protein_coding | 243.001488 | -0.335557349 | 0.192951944 | -1.739072138 | 0.082022075 | 0.217959227 | 0.335557349 |
| ENSMUSG00000001583 | Tnk1 | 83813 | protein_coding | 313.7000209 | -0.335671287 | 0.102067212 | -3.288727892 | 0.001006413 | 0.007709553 | 0.335671287 |
| ENSMUSG00000059409 | Ppp2r5d | 21770 | protein_coding | 895.7162327 | 0.335733265 | 0.08899254 | 3.772600106 | 0.000161555 | 0.001699637 | 0.335733265 |
| ENSMUSG00000035811 | Ugt2b35 | 243085 | protein_coding | 846.4218478 | 0.335740291 | 0.129335822 | 2.595880134 | 0.009434897 | 0.045830755 | 0.335740291 |
| ENSMUSG00000023021 | Cers5 | 71949 | protein_coding | 423.7450625 | -0.335779242 | 0.128255254 | -2.618054478 | 0.008843269 | 0.043730389 | 0.335779242 |
| ENSMUSG00000008682 | Rpl10 | 110954 | protein_coding | 9223.892735 | 0.336015759 | 0.105869385 | 3.173870891 | 0.001504206 | 0.010700755 | 0.336015759 |
| ENSMUSG00000020994 | Pnn | 18949 | protein_coding | 779.4319909 | 0.336026764 | 0.124443632 | 2.700232697 | 0.006929099 | 0.036162088 | 0.336026764 |
| ENSMUSG00000051256 | Jagn1 | 67767 | protein_coding | 422.5416716 | 0.336299673 | 0.088757898 | 3.788954897 | 0.000151282 | 0.001607489 | 0.336299673 |
| ENSMUSG00000041642 | Kif21b | 16565 | protein_coding | 512.4334179 | -0.336353339 | 0.14388405 | -2.337669383 | 0.019404406 | 0.078286062 | 0.336353339 |
| ENSMUSG00000085333 | 1700030A11Rik | 74179 | lncRNA | 14.76842945 | -0.336417404 | 0.45708043 | -0.736013581 | 0.461722433 | 0.659671043 | 0.336417404 |
| ENSMUSG00000051220 | Ercc6l | 236930 | protein_coding | 198.2472799 | 0.33642691 | 0.148878756 | 2.259737512 | 0.023837547 | 0.091182623 | 0.33642691 |
| ENSMUSG00000019689 | Fmc1 | 66117 | protein_coding | 388.0668521 | 0.336554501 | 0.141302642 | 2.38179907 | 0.017228295 | 0.071685549 | 0.336554501 |
| ENSMUSG00000046329 | Slc25a23 | 66972 | protein_coding | 127.1585879 | -0.336624596 | 0.169311014 | -1.988202583 | 0.046789289 | 0.148655354 | 0.336624596 |
| ENSMUSG00000051786 | Tubgcp6 | 328580 | protein_coding | 272.17138 | -0.336828797 | 0.088135343 | -3.821722219 | 0.000132523 | 0.001438872 | 0.336828797 |
| ENSMUSG00000020572 | Nampt | 59027 | protein_coding | 1464.230193 | 0.336836099 | 0.074217025 | 4.538528709 | 5.66E-06 | 9.00E-05 | 0.336836099 |
| ENSMUSG00000024026 | Glo1 | 109801 | protein_coding | 1126.225845 | 0.336854596 | 0.091975717 | 3.662429683 | 0.000249834 | 0.002447332 | 0.336854596 |
| ENSMUSG00000051703 | Tmem198 | 319998 | protein_coding | 12.50874952 | -0.336859838 | 0.312129167 | -1.079232169 | 0.280484242 | 0.488844842 | 0.336859838 |
| ENSMUSG00000038152 | 5033430I15Rik | 76000 | lncRNA | 21.23892733 | -0.336932543 | 0.236333474 | -1.425665766 | 0.153964843 | 0.333064243 | 0.336932543 |
| ENSMUSG00000004151 | Etv1 | 14009 | protein_coding | 24.25559227 | -0.336970276 | 0.45779749 | -0.736068421 | 0.46168906 | 0.659671043 | 0.336970276 |
| ENSMUSG00000023066 | Rttn | 246102 | protein_coding | 227.5161902 | 0.337071581 | 0.119548359 | 2.819541691 | 0.004809228 | 0.027279697 | 0.337071581 |
| ENSMUSG00000007564 | Ppp2r1a | 51792 | protein_coding | 3196.321697 | 0.337153972 | 0.104413724 | 3.229019715 | 0.001242153 | 0.009121723 | 0.337153972 |
| ENSMUSG00000027890 | Gstm4 | 14865 | protein_coding | 181.1702357 | 0.337253785 | 0.122722646 | 2.748097409 | 0.00599422 | 0.032200989 | 0.337253785 |
| ENSMUSG00000031158 | Timm17b | 21855 | protein_coding | 370.7830839 | 0.337267826 | 0.093357956 | 3.612630766 | 0.000303106 | 0.002860116 | 0.337267826 |
| ENSMUSG00000000706 | Btn1a1 | 12231 | protein_coding | 68.80112485 | 0.337330279 | 0.305169863 | 1.105385296 | 0.268992672 | 0.477123143 | 0.337330279 |

| ensembl_gene_id | external_gene_name | entrezgene_id | gene_biotype | baseMean | log2FoldChange | lfcSE | stat | pvalue | padj | absLog2FC |
|---|---|---|---|---|---|---|---|---|---|---|
| ENSMUSG00000034607 | Pof1b | 69693 | protein_coding | 3952.143202 | 0.337364101 | 0.155085509 | 2.175342508 | 0.029604459 | 0.106828235 | 0.337364101 |
| ENSMUSG00000029468 | P2rx7 | 18439 | protein_coding | 118.9645766 | -0.337376484 | 0.209649833 | -1.609238026 | 0.107564305 | 0.260853265 | 0.337376484 |
| ENSMUSG00000055296 | Tmem245 | 242474 | protein_coding | 589.6595742 | -0.337415471 | 0.165167225 | -2.042871829 | 0.041065128 | 0.135583278 | 0.337415471 |
| ENSMUSG00000045071 | E130308A19Rik | 230259 | protein_coding | 118.0506018 | -0.337492648 | 0.127917288 | -2.638366191 | 0.008330657 | 0.041805515 | 0.337492648 |
| ENSMUSG00000026775 | Yme1l1 | 27377 | protein_coding | 3395.295841 | 0.337688098 | 0.076742323 | 4.400285057 | 1.08E-05 | 0.000159293 | 0.337688098 |
| ENSMUSG00000027405 | Nop56 | 67134 | protein_coding | 713.6972603 | -0.337806713 | 0.189211758 | -1.785336788 | 0.074206705 | 0.203016901 | 0.337806713 |
| ENSMUSG00000024002 | Brd4 | 57261 | protein_coding | 379.4048106 | -0.33780853 | 0.191530491 | -1.763732383 | 0.077777046 | 0.209628987 | 0.33780853 |
| ENSMUSG00000047515 | BC049715 | 320135 | protein_coding | 27.9431515 | -0.33785457 | 0.294712209 | -1.1463881 | 0.251634604 | 0.45811361 | 0.33785457 |
| ENSMUSG00000004446 | Bid | 12122 | protein_coding | 158.7249768 | -0.337905114 | 0.109354016 | -3.090011031 | 0.002001491 | 0.013514092 | 0.337905114 |
| ENSMUSG00000062519 | Zfp398 | 272347 | protein_coding | 277.7356611 | -0.338045033 | 0.172907898 | -1.955058364 | 0.050576182 | 0.156635171 | 0.338045033 |
| ENSMUSG00000063511 | Snrnp70 | 20637 | protein_coding | 1939.688499 | 0.338148911 | 0.116194935 | 2.910186335 | 0.003612133 | 0.021710966 | 0.338148911 |
| ENSMUSG00000044562 | Rasip1 | 69903 | protein_coding | 24.37122371 | 0.338177396 | 0.24578098 | 1.375929889 | 0.168843341 | 0.353766999 | 0.338177396 |
| ENSMUSG00000062949 | Atp11c | 320940 | protein_coding | 483.2471191 | 0.33818195 | 0.078720114 | 4.296004294 | 1.74E-05 | 0.000244263 | 0.33818195 |
| ENSMUSG00000028970 | Abcb1b | 18669 | protein_coding | 8.178339981 | -0.338187293 | 0.760623412 | -0.444618568 | 0.656595426 | 0.801149235 | 0.338187293 |
| ENSMUSG00000023800 | Tiam2 | 24001 | protein_coding | 439.6862211 | 0.338727624 | 0.13064984 | 2.592637113 | 0.009524321 | 0.04616658 | 0.338727624 |
| ENSMUSG00000036620 | Mgat4b | 103534 | protein_coding | 3349.151792 | 0.338742686 | 0.094287132 | 3.592671432 | 0.000327305 | 0.003050507 | 0.338742686 |
| ENSMUSG00000086844 | B230206H07Rik | 320871 | lncRNA | 16.54033567 | -0.338889489 | 0.37137349 | -0.912530105 | 0.361489729 | 0.570814753 | 0.338889489 |
| ENSMUSG00000030102 | Itpr1 | 16438 | protein_coding | 571.7488504 | -0.338918174 | 0.14732397 | -2.300495795 | 0.021420147 | 0.084108607 | 0.338918174 |
| ENSMUSG00000024292 | Cyp4f14 | 64385 | protein_coding | 1412.305461 | 0.339073915 | 0.182233661 | 1.860654684 | 0.062792957 | 0.181128045 | 0.339073915 |
| ENSMUSG00000038976 | Ppp1r9b | 217124 | protein_coding | 603.9166987 | -0.339139014 | 0.127626282 | -2.657281937 | 0.007877353 | 0.03998202 | 0.339139014 |
| ENSMUSG00000026315 | Serpinb8 | 20725 | protein_coding | 9.354170615 | -0.339189003 | 0.55056021 | -0.616079761 | 0.537841879 | 0.721865534 | 0.339189003 |
| ENSMUSG00000116270 | Gm1600 | 381033 | lncRNA | 7.963669758 | 0.339214891 | 0.659611618 | 0.514264579 | 0.607067023 | 0.771131144 | 0.339214891 |
| ENSMUSG00000038429 | Usp5 | 22225 | protein_coding | 1285.482953 | 0.339264106 | 0.08145618 | 4.164989138 | 3.11E-05 | 0.000408522 | 0.339264106 |
| ENSMUSG00000027466 | Rbck1 | 24105 | protein_coding | 988.7923627 | 0.339307822 | 0.074614421 | 4.547483142 | 5.43E-06 | 8.70E-05 | 0.339307822 |
| ENSMUSG00000027274 | Mkks | 59030 | protein_coding | 385.2066641 | 0.339353806 | 0.143230815 | 2.369279301 | 0.017822789 | 0.073411269 | 0.339353806 |
| ENSMUSG00000031838 | Ifi30 | 65972 | protein_coding | 1062.262226 | -0.33946663 | 0.081346681 | -4.173085187 | 3.01E-05 | 0.000396554 | 0.33946663 |
| ENSMUSG00000079184 | Mphosph8 | 75339 | protein_coding | 267.1115022 | 0.339646875 | 0.129188552 | 2.629078734 | 0.008561654 | 0.042642282 | 0.339646875 |
| ENSMUSG00000007594 | Hapln4 | 330790 | protein_coding | 24.14107976 | 0.339906793 | 0.282014247 | 1.205282341 | 0.228094324 | 0.43086888 | 0.339906793 |
| ENSMUSG00000026393 | Nek7 | 59125 | protein_coding | 1062.97248 | 0.33993502 | 0.106960854 | 3.178125518 | 0.001482306 | 0.010560955 | 0.33993502 |
| ENSMUSG00000054889 | Dsp | 109620 | protein_coding | 9882.125794 | -0.339999412 | 0.093256676 | -3.645845261 | 0.000266514 | 0.002583713 | 0.339999412 |
| ENSMUSG00000024429 | Gnl1 | 14670 | protein_coding | 382.7952186 | -0.340003483 | 0.097378051 | -3.491582318 | 0.000480168 | 0.004199334 | 0.340003483 |
| ENSMUSG00000031099 | Smarca1 | 93761 | protein_coding | 6.864493572 | -0.340044517 | 0.48776129 | -0.697153553 | 0.485706702 | 0.678784766 | 0.340044517 |
| ENSMUSG00000016933 | Plcg1 | 18803 | protein_coding | 303.4690334 | -0.340121285 | 0.100796316 | -3.374342425 | 0.000739923 | 0.006006875 | 0.340121285 |
| ENSMUSG00000053004 | Hrh1 | 15465 | protein_coding | 137.7995774 | -0.340184063 | 0.222044746 | -1.532051845 | 0.125509641 | 0.289727184 | 0.340184063 |
| ENSMUSG00000026553 | Copa | 12847 | protein_coding | 5964.177769 | 0.340243207 | 0.070023261 | 4.859002579 | 1.18E-06 | 2.18E-05 | 0.340243207 |
| ENSMUSG00000033147 | Slc22a15 | 242126 | protein_coding | 229.8578118 | -0.340304307 | 0.147659822 | -2.304650672 | 0.021186146 | 0.083408248 | 0.340304307 |
| ENSMUSG00000001151 | Pcnt | 18541 | protein_coding | 882.3240539 | 0.340325359 | 0.108036566 | 3.150094181 | 0.001632178 | 0.011417364 | 0.340325359 |
| ENSMUSG00000025509 | Pnpla2 | 66853 | protein_coding | 750.0335745 | -0.340785719 | 0.089549703 | -3.805548296 | 0.00014149 | 0.001523772 | 0.340785719 |

| ensembl_gene_id | external_gene_name | entrezgene_id | gene_biotype | baseMean | log2FoldChange | lfcSE | stat | pvalue | padj | absLog2FC |
|---|---|---|---|---|---|---|---|---|---|---|
| ENSMUSG00000032827 | Ppp1r9a | 243725 | protein_coding | 683.7620415 | -0.340822061 | 0.115469314 | -2.95162455 | 0.00316107 | 0.019553215 | 0.340822061 |
| ENSMUSG00000029283 | Cdc7 | 12545 | protein_coding | 111.6965793 | 0.341028453 | 0.16230923 | 2.101103269 | 0.035631902 | 0.122632873 | 0.341028453 |
| ENSMUSG00000051550 | Zfp579 | 68490 | protein_coding | 60.92638456 | -0.341050112 | 0.206841568 | -1.648847069 | 0.099178969 | 0.246947416 | 0.341050112 |
| ENSMUSG00000023972 | Ptk7 | 71461 | protein_coding | 254.0663472 | -0.341159696 | 0.117422832 | -2.905394885 | 0.0036679 | 0.021979803 | 0.341159696 |
| ENSMUSG00000052707 | Tnrc6a | 233833 | protein_coding | 788.1627221 | -0.341200284 | 0.113704118 | -3.000773324 | 0.00269295 | 0.017241292 | 0.341200284 |
| ENSMUSG00000029103 | Lrpap1 | 16976 | protein_coding | 598.3647398 | -0.34155702 | 0.076186288 | -4.48318232 | 7.35E-06 | 0.000114163 | 0.34155702 |
| ENSMUSG00000027778 | Ift80 | 68259 | protein_coding | 152.3784854 | 0.341651201 | 0.115213305 | 2.965379754 | 0.003023095 | 0.018890489 | 0.341651201 |
| ENSMUSG00000026131 | Dst | 13518 | protein_coding | 4493.510038 | -0.341913282 | 0.171325933 | -1.995689015 | 0.045967783 | 0.14684614 | 0.341913282 |
| ENSMUSG00000025223 | Ldb1 | 16825 | protein_coding | 382.9961177 | -0.341929672 | 0.113422182 | -3.014663142 | 0.002572648 | 0.016643842 | 0.341929672 |
| ENSMUSG00000053965 | Pde5a | 242202 | protein_coding | 1874.630112 | -0.342026148 | 0.064385498 | -5.312161273 | 1.08E-07 | 2.47E-06 | 0.342026148 |
| ENSMUSG00000058886 | Deaf1 | 54006 | protein_coding | 234.6663253 | -0.342048271 | 0.11304922 | -3.025657954 | 0.002480928 | 0.016215108 | 0.342048271 |
| ENSMUSG00000070324 | Fbxw22 | 382156 | protein_coding | 14.15680132 | 0.342087162 | 0.514518716 | 0.664868256 | 0.506134761 | 0.696024236 | 0.342087162 |
| ENSMUSG00000030960 | Eef1akmt2 | 72096 | protein_coding | 207.9498207 | 0.34219702 | 0.082475656 | 4.149066958 | 3.34E-05 | 0.000434419 | 0.34219702 |
| ENSMUSG00000017639 | Rab11fip4 | 268451 | protein_coding | 433.7622725 | -0.342202098 | 0.117388693 | -2.915119768 | 0.003555521 | 0.021476046 | 0.342202098 |
| ENSMUSG00000057286 | St6galnac2 | 20446 | protein_coding | 694.0958119 | -0.342293996 | 0.102480885 | -3.340076499 | 0.000837553 | 0.006623906 | 0.342293996 |
| ENSMUSG00000027496 | Aurka | 20878 | protein_coding | 401.4271108 | 0.34232735 | 0.128287666 | 2.668435415 | 0.007620543 | 0.039016395 | 0.34232735 |
| ENSMUSG00000016382 | Pls3 | 102866 | protein_coding | 291.3777576 | -0.342405707 | 0.125239453 | -2.734008319 | 0.006256846 | 0.033241844 | 0.342405707 |
| ENSMUSG00000024215 | Spdef | 30051 | protein_coding | 1781.103677 | -0.34249716 | 0.225786525 | -1.516906995 | 0.129290167 | 0.295859555 | 0.34249716 |
| ENSMUSG00000032372 | Plscr2 | 18828 | protein_coding | 32.93195206 | -0.34256368 | 0.304977778 | -1.123241445 | 0.261334965 | 0.469018966 | 0.34256368 |
| ENSMUSG00000020198 | Ap3d1 | 11776 | protein_coding | 2144.809701 | 0.342565832 | 0.083150157 | 4.11984587 | 3.79E-05 | 0.00048426 | 0.342565832 |
| ENSMUSG00000020424 | Castor1 | 71962 | protein_coding | 21.66350898 | 0.342623082 | 0.292317122 | 1.172093787 | 0.241159404 | 0.445706765 | 0.342623082 |
| ENSMUSG00000087396 | 4933407K13Rik | 74396 | lncRNA | 54.64097168 | -0.342623965 | 0.191201405 | -1.79195318 | 0.073140466 | 0.201205759 | 0.342623965 |
| ENSMUSG00000051427 | Ccdc157 | 216516 | protein_coding | 89.18686643 | -0.342666279 | 0.12848735 | -2.666926191 | 0.007654848 | 0.039124542 | 0.342666279 |
| ENSMUSG00000018068 | Ints2 | 70422 | protein_coding | 438.16438 | 0.34266737 | 0.131078259 | 2.614219714 | 0.00894315 | 0.044019314 | 0.34266737 |
| ENSMUSG00000026788 | Zbtb43 | 71834 | protein_coding | 531.7180816 | -0.342696151 | 0.075867897 | -4.517011362 | 6.27E-06 | 9.88E-05 | 0.342696151 |
| ENSMUSG00000074063 | Osgin1 | 71839 | protein_coding | 182.8102439 | -0.342758355 | 0.17459285 | -1.96318666 | 0.049624489 | 0.154534306 | 0.342758355 |
| ENSMUSG00000032076 | Cadm1 | 54725 | protein_coding | 30.85176396 | 0.342791112 | 0.25864767 | 1.325320703 | 0.185064805 | 0.375027 | 0.342791112 |
| ENSMUSG00000069274 | H4c6 | 319157 | protein_coding | 1450.225332 | 0.342836988 | 0.179446212 | 1.910527864 | 0.056065282 | 0.167827066 | 0.342836988 |
| ENSMUSG00000068966 | Zbtb34 | 241311 | protein_coding | 60.46183548 | -0.342916273 | 0.29450358 | -1.164387453 | 0.24426703 | 0.449478426 | 0.342916273 |
| ENSMUSG00000031633 | Slc25a4 | 11739 | protein_coding | 339.8830434 | 0.343029465 | 0.143249318 | 2.394632445 | 0.016637034 | 0.069882561 | 0.343029465 |
| ENSMUSG00000044367 | Slc16a13 | 69309 | protein_coding | 92.75185117 | -0.343034363 | 0.114555393 | -2.994484621 | 0.002749089 | 0.017500996 | 0.343034363 |
| ENSMUSG00000090213 | Peds1 | 407243 | protein_coding | 526.686181 | 0.343078579 | 0.102356393 | 3.35180411 | 0.000802868 | 0.006394041 | 0.343078579 |
| ENSMUSG00000028466 | Creb3 | 12913 | protein_coding | 676.0908937 | 0.343082511 | 0.101936408 | 3.36565234 | 0.000763629 | 0.006146074 | 0.343082511 |
| ENSMUSG00000102018 | Iqank1 | 432964 | protein_coding | 37.33023205 | -0.343320382 | 0.237543871 | -1.445292527 | 0.148375736 | 0.325258564 | 0.343320382 |
| ENSMUSG00000039103 | Nexn | 68810 | protein_coding | 28.19679139 | -0.34336092 | 0.383282519 | -0.895842892 | 0.370336678 | 0.578930475 | 0.34336092 |
| ENSMUSG00000060288 | Ppih | 66101 | protein_coding | 306.9112462 | 0.343423159 | 0.097412755 | 3.525443437 | 0.000422774 | 0.003804492 | 0.343423159 |
| ENSMUSG00000079523 | Tmsb10 | 19240 | protein_coding | 2392.619169 | 0.343636389 | 0.141610492 | 2.426630843 | 0.015239754 | 0.065433283 | 0.343636389 |
| ENSMUSG00000022360 | Atad2 | 70472 | protein_coding | 591.9228113 | 0.343767048 | 0.109327384 | 3.14438192 | 0.001664381 | 0.011591716 | 0.343767048 |

| ensembl_gene_id | external_gene_name | entrezgene_id | gene_biotype | baseMean | log2FoldChange | lfcSE | stat | pvalue | padj | absLog2FC |
|---|---|---|---|---|---|---|---|---|---|---|
| ENSMUSG00000043602 | Zfp3 | 193043 | protein_coding | 46.06169335 | 0.343809652 | 0.24977476 | 1.376478762 | 0.168673458 | 0.353480683 | 0.343809652 |
| ENSMUSG00000050272 | Dscam | 13508 | protein_coding | 6.96491196 | -0.343971935 | 0.806866458 | -0.426305905 | 0.669884956 | 0.808780483 | 0.343971935 |
| ENSMUSG00000023909 | Paqr4 | 76498 | protein_coding | 2136.390473 | 0.344043264 | 0.106925224 | 3.217606207 | 0.001292652 | 0.009414428 | 0.344043264 |
| ENSMUSG00000090115 | Usp49 | 224836 | protein_coding | 134.4702359 | -0.344178372 | 0.134653459 | -2.556030685 | 0.010587378 | 0.049921776 | 0.344178372 |
| ENSMUSG00000052676 | Zmat1 | 215693 | protein_coding | 17.00022055 | 0.34422089 | 0.360367243 | 0.955194725 | 0.339479233 | 0.550085499 | 0.34422089 |
| ENSMUSG00000025231 | Sufu | 24069 | protein_coding | 96.50023109 | -0.344233174 | 0.156267166 | -2.202850313 | 0.027605301 | 0.101359435 | 0.344233174 |
| ENSMUSG00000039741 | Bahcc1 | 268515 | protein_coding | 180.7507431 | -0.344473154 | 0.173049593 | -1.990603663 | 0.046524477 | 0.148132115 | 0.344473154 |
| ENSMUSG00000026799 | Med27 | 68975 | protein_coding | 115.09061 | 0.344574848 | 0.149762446 | 2.300809429 | 0.021402405 | 0.084105999 | 0.344574848 |
| ENSMUSG00000038065 | Mturn | 68235 | protein_coding | 24.11721649 | 0.344741388 | 0.234140001 | 1.472372882 | 0.140920212 | 0.313313392 | 0.344741388 |
| ENSMUSG00000039960 | Rhou | 69581 | protein_coding | 517.4531543 | -0.34478709 | 0.127949276 | -2.694717014 | 0.007044843 | 0.036625014 | 0.34478709 |
| ENSMUSG00000032226 | Gcnt3 | 72077 | protein_coding | 6813.890661 | -0.344935796 | 0.216775777 | -1.59121006 | 0.11156231 | 0.266741205 | 0.344935796 |
| ENSMUSG00000006958 | Chrd | 12667 | protein_coding | 12.09088962 | 0.34502238 | 0.419854433 | 0.821766671 | 0.411209706 | 0.618130069 | 0.34502238 |
| ENSMUSG00000021024 | Psma6 | 26443 | protein_coding | 1623.523642 | 0.345312281 | 0.081495386 | 4.237200373 | 2.26E-05 | 0.000307847 | 0.345312281 |
| ENSMUSG00000044155 | Lsm8 | 76522 | protein_coding | 380.8363729 | 0.345352316 | 0.093527794 | 3.692509995 | 0.000222052 | 0.002228631 | 0.345352316 |
| ENSMUSG00000025202 | Scd3 | 30049 | protein_coding | 6.820895395 | 0.345359525 | 0.748408992 | 0.461458279 | 0.644469848 | 0.794206246 | 0.345359525 |
| ENSMUSG00000095567 | Noc2l | 57741 | protein_coding | 562.1878555 | -0.345399584 | 0.130711602 | -2.642455435 | 0.00823073 | 0.041435132 | 0.345399584 |
| ENSMUSG00000031010 | Usp9x | 22284 | protein_coding | 4334.772333 | 0.345431845 | 0.060108476 | 5.746807585 | 9.09E-09 | 2.47E-07 | 0.345431845 |
| ENSMUSG00000050079 | Rspry1 | 67610 | protein_coding | 728.4830965 | 0.34544059 | 0.098182882 | 3.518338251 | 0.000434258 | 0.003885881 | 0.34544059 |
| ENSMUSG00000029130 | Rnf32 | 56874 | protein_coding | 202.7661689 | -0.345505172 | 0.17842357 | -1.936432347 | 0.052814771 | 0.161248547 | 0.345505172 |
| ENSMUSG00000050530 | Fam171a1 | 269233 | protein_coding | 260.5401745 | -0.3457045 | 0.097612746 | -3.541591803 | 0.00039772 | 0.00361787 | 0.3457045 |
| ENSMUSG00000026463 | Atp2b4 | 381290 | protein_coding | 23.8601368 | -0.345718475 | 0.496350419 | -0.69652097 | 0.486102629 | 0.678983247 | 0.345718475 |
| ENSMUSG00000038658 | Ric1 | 226089 | protein_coding | 495.3181849 | -0.345851987 | 0.102682671 | -3.368163137 | 0.000756708 | 0.006101436 | 0.345851987 |
| ENSMUSG00000097321 | 1700028E10Rik | 625175 | lncRNA | 9.448430517 | -0.345875228 | 0.472303207 | -0.73231607 | 0.463975673 | 0.661197506 | 0.345875228 |
| ENSMUSG00000020287 | Mpg | 268395 | protein_coding | 73.52266221 | -0.346001874 | 0.139560586 | -2.479223415 | 0.01316688 | 0.058894151 | 0.346001874 |
| ENSMUSG00000035578 | Iqcg | 69707 | protein_coding | 16.33987414 | -0.346072803 | 0.306137837 | -1.130447664 | 0.25828764 | 0.46538524 | 0.346072803 |
| ENSMUSG00000055370 | Gm9968 | 791329 | lncRNA | 11.63908462 | -0.346087596 | 0.427619752 | -0.809334916 | 0.418322528 | 0.624549748 | 0.346087596 |
| ENSMUSG00000005268 | Prlr | 19116 | protein_coding | 403.3696899 | -0.346096787 | 0.20446484 | -1.692695852 | 0.090513375 | 0.232681898 | 0.346096787 |
| ENSMUSG00000073423 | Zfp414 | 328801 | protein_coding | 67.14208873 | -0.346102377 | 0.193398636 | -1.78958024 | 0.073521417 | 0.201905024 | 0.346102377 |
| ENSMUSG00000034998 | Foxn2 | 14236 | protein_coding | 441.0648586 | -0.346253391 | 0.140054919 | -2.472268688 | 0.013425857 | 0.059733812 | 0.346253391 |
| ENSMUSG00000058812 | 0610039K10Rik | 68386 | lncRNA | 13.58453812 | -0.346275251 | 0.340206112 | -1.017839595 | 0.308754192 | 0.518602032 | 0.346275251 |
| ENSMUSG00000042507 | Mideas | 238317 | protein_coding | 783.5173973 | -0.346365792 | 0.16926385 | -2.046306949 | 0.04072618 | 0.134778018 | 0.346365792 |
| ENSMUSG00000023030 | Slc11a2 | 18174 | protein_coding | 467.3231821 | 0.346385805 | 0.117116059 | 2.957628592 | 0.003100154 | 0.01925868 | 0.346385805 |
| ENSMUSG00000028737 | Aldh4a1 | 212647 | protein_coding | 212.5382315 | 0.346401378 | 0.131867332 | 2.626893044 | 0.008616841 | 0.042854354 | 0.346401378 |
| ENSMUSG00000066551 | Hmgb1 | 15289 | protein_coding | 2521.27789 | 0.346483633 | 0.092515254 | 3.745151409 | 0.000180285 | 0.001867087 | 0.346483633 |
| ENSMUSG00000021245 | Mlh3 | 217716 | protein_coding | 248.8964092 | 0.346523054 | 0.112263057 | 3.086706015 | 0.002023876 | 0.013630573 | 0.346523054 |
| ENSMUSG00000061665 | Cd2ap | 12488 | protein_coding | 6703.463554 | 0.346536704 | 0.060515903 | 5.726374158 | 1.03E-08 | 2.75E-07 | 0.346536704 |
| ENSMUSG00000024959 | Bad | 12015 | protein_coding | 396.7191211 | 0.346593197 | 0.085555851 | 4.051075344 | 5.10E-05 | 0.000630966 | 0.346593197 |
| ENSMUSG00000086247 | Gm15787 | 100504007 | lncRNA | 36.42582828 | -0.346613475 | 0.247524615 | -1.400319219 | 0.161417748 | 0.343403485 | 0.346613475 |

Supplementary Table 1

| ensembl_gene_id | external_gene_name | entrezgene_id | gene_biotype | baseMean | log2FoldChange | lfcSE | stat | pvalue | padj | absLog2FC |
|---|---|---|---|---|---|---|---|---|---|---|
| ENSMUSG00000021495 | Fam193b | 212483 | protein_coding | 351.332144 | -0.346788817 | 0.165556084 | -2.094690869 | 0.036198481 | 0.123993422 | 0.346788817 |
| ENSMUSG00000021910 | Nisch | 64652 | protein_coding | 1643.242566 | -0.34681165 | 0.089392083 | -3.879668507 | 0.000104599 | 0.00116998 | 0.34681165 |
| ENSMUSG00000030587 | 2200002D01Rik | 72275 | protein_coding | 2630.017963 | -0.346877694 | 0.151061468 | -2.296268529 | 0.021660532 | 0.084706349 | 0.346877694 |
| ENSMUSG00000039206 | Daglb | 231871 | protein_coding | 592.2366621 | 0.347029438 | 0.176769611 | 1.963173625 | 0.049626003 | 0.154534306 | 0.347029438 |
| ENSMUSG00000041180 | Hectd2 | 226098 | protein_coding | 14.84847873 | -0.347042426 | 0.593628083 | -0.584612548 | 0.558808269 | 0.736907711 | 0.347042426 |
| ENSMUSG00000026200 | Glb1l | 74577 | protein_coding | 33.31984992 | -0.347103528 | 0.229927442 | -1.509622011 | 0.131139901 | 0.298526504 | 0.347103528 |
| ENSMUSG00000044627 | Swi5 | 72931 | protein_coding | 1093.344905 | 0.347109833 | 0.083970089 | 4.133731885 | 3.57E-05 | 0.000459584 | 0.347109833 |
| ENSMUSG00000035171 | 1110059E24Rik | 66206 | protein_coding | 936.4457851 | 0.347124388 | 0.118745524 | 2.923262925 | 0.003463839 | 0.021025924 | 0.347124388 |
| ENSMUSG00000038379 | Ttk | 22137 | protein_coding | 222.7932321 | 0.347214647 | 0.137554037 | 2.524205428 | 0.011596014 | 0.0534901 | 0.347214647 |
| ENSMUSG00000024659 | Anxa1 | 16952 | protein_coding | 2079.806133 | 0.347371728 | 0.185296678 | 1.874678664 | 0.060836944 | 0.177344744 | 0.347371728 |
| ENSMUSG00000026637 | Traf5 | 22033 | protein_coding | 34.1963008 | 0.347484018 | 0.227837197 | 1.525141736 | 0.127223749 | 0.292469065 | 0.347484018 |
| ENSMUSG00000056116 | H2-T22 | 15039 | protein_coding | 546.6300633 | 0.347523668 | 0.095471429 | 3.640080293 | 0.000272553 | 0.002629685 | 0.347523668 |
| ENSMUSG00000038388 | Pals2 | 56524 | protein_coding | 1553.086374 | -0.347538265 | 0.229514384 | -1.514233051 | 0.129966742 | 0.296662405 | 0.347538265 |
| ENSMUSG00000042216 | Sgsm1 | 52850 | protein_coding | 397.0476719 | 0.347560203 | 0.148540882 | 2.339828596 | 0.019292592 | 0.077933655 | 0.347560203 |
| ENSMUSG00000003810 | Mast2 | 17776 | protein_coding | 1214.025919 | -0.347569357 | 0.105662589 | -3.289426858 | 0.001003916 | 0.007698787 | 0.347569357 |
| ENSMUSG00000017677 | Wsb1 | 78889 | protein_coding | 1793.141605 | -0.347590338 | 0.134281146 | -2.588526744 | 0.009638746 | 0.046607476 | 0.347590338 |
| ENSMUSG00000040557 | Mettl27 | 79565 | protein_coding | 154.4260171 | -0.347591779 | 0.129188412 | -2.690580167 | 0.00713279 | 0.036970736 | 0.347591779 |
| ENSMUSG00000029865 | Sval1 | 71578 | protein_coding | 3923.44205 | 0.347626009 | 0.346838655 | 1.002270087 | 0.316213166 | 0.527231582 | 0.347626009 |
| ENSMUSG00000069094 | Pde7a | 18583 | protein_coding | 1040.272749 | -0.347653883 | 0.223563603 | -1.555055822 | 0.119932775 | 0.280966207 | 0.347653883 |
| ENSMUSG00000027962 | Vcam1 | 22329 | protein_coding | 9.155650124 | -0.347731977 | 0.555950262 | -0.625473177 | 0.531660548 | 0.716861865 | 0.347731977 |
| ENSMUSG00000023067 | Cdkn1a | 12575 | protein_coding | 1452.496234 | 0.347966437 | 0.188000224 | 1.850883096 | 0.064186373 | 0.184242305 | 0.347966437 |
| ENSMUSG00000053841 | Txlna | 109658 | protein_coding | 1153.073583 | 0.347977481 | 0.09246027 | 3.763535218 | 0.000167528 | 0.001746327 | 0.347977481 |
| ENSMUSG00000020092 | Pald1 | 27355 | protein_coding | 16.97539043 | -0.348113839 | 0.301092878 | -1.156167631 | 0.247612597 | 0.453389244 | 0.348113839 |
| ENSMUSG00000056260 | Lrif1 | 321000 | protein_coding | 293.9142955 | 0.348174577 | 0.113362423 | 3.071340298 | 0.002131001 | 0.014267587 | 0.348174577 |
| ENSMUSG00000029204 | Rhoh | 74734 | protein_coding | 18.11823453 | 0.348284593 | 0.59792764 | 0.582486189 | 0.560239242 | 0.738088556 | 0.348284593 |
| ENSMUSG00000036820 | Amdhd2 | 245847 | protein_coding | 300.3450091 | -0.348344742 | 0.111547667 | -3.122833058 | 0.001791193 | 0.012340019 | 0.348344742 |
| ENSMUSG00000039128 | Cdc123 | 98828 | protein_coding | 958.7516507 | 0.348352363 | 0.075570268 | 4.609648366 | 4.03E-06 | 6.64E-05 | 0.348352363 |
| ENSMUSG00000041654 | Slc39a11 | 69806 | protein_coding | 2022.694151 | 0.348493066 | 0.082584743 | 4.219823819 | 2.44E-05 | 0.000328632 | 0.348493066 |
| ENSMUSG00000026082 | Rev1 | 56210 | protein_coding | 143.1476488 | -0.348544044 | 0.138970986 | -2.508034611 | 0.012140477 | 0.055392015 | 0.348544044 |
| ENSMUSG00000021892 | Sh3bp5 | 24056 | protein_coding | 85.20584121 | -0.348630979 | 0.176657514 | -1.973485141 | 0.048440316 | 0.152451561 | 0.348630979 |
| ENSMUSG00000085747 | Slc13a2os | 790911 | lncRNA | 49.39453194 | -0.348837563 | 0.289458184 | -1.205139746 | 0.228149358 | 0.430921733 | 0.348837563 |
| ENSMUSG00000027253 | Lrp4 | 228357 | protein_coding | 737.5114094 | -0.348857896 | 0.105988068 | -3.291482747 | 0.000996607 | 0.007657482 | 0.348857896 |
| ENSMUSG00000020253 | Ppm1m | 67905 | protein_coding | 51.87139917 | -0.34896705 | 0.17042067 | -2.047680309 | 0.040591335 | 0.134477991 | 0.34896705 |
| ENSMUSG00000045948 | Mrps12 | 24030 | protein_coding | 396.0139636 | 0.349043318 | 0.093992465 | 3.71352445 | 0.000204393 | 0.00208423 | 0.349043318 |
| ENSMUSG00000029269 | Sult1b1 | 56362 | protein_coding | 2640.960337 | 0.349140396 | 0.121942524 | 2.86315539 | 0.004194448 | 0.024400719 | 0.349140396 |
| ENSMUSG00000050043 | Tmx2 | 66958 | protein_coding | 639.3966391 | 0.349219306 | 0.09685305 | 3.605661428 | 0.000311359 | 0.002927672 | 0.349219306 |
| ENSMUSG00000059883 | Irak4 | 266632 | protein_coding | 535.968013 | 0.34926718 | 0.12203909 | 2.861928762 | 0.004210715 | 0.024468542 | 0.34926718 |
| ENSMUSG00000020003 | Pex7 | 18634 | protein_coding | 743.7493592 | 0.349272013 | 0.095829058 | 3.644740121 | 0.000267662 | 0.002591685 | 0.349272013 |

| ensembl_gene_id | external_gene_name | entrezgene_id | gene_biotype | baseMean | log2FoldChange | lfcSE | stat | pvalue | padj | absLog2FC |
|---|---|---|---|---|---|---|---|---|---|---|
| ENSMUSG00000029119 | Man2b2 | 17160 | protein_coding | 419.3558195 | -0.349279812 | 0.094538962 | -3.694559422 | 0.000220268 | 0.002212955 | 0.349279812 |
| ENSMUSG00000020869 | Lrrc59 | 98238 | protein_coding | 2698.595974 | 0.349293534 | 0.154800824 | 2.256406165 | 0.024045206 | 0.091756598 | 0.349293534 |
| ENSMUSG00000062939 | Stat4 | 20849 | protein_coding | 12.54927113 | 0.349307617 | 0.486688272 | 0.717723514 | 0.472927782 | 0.668334106 | 0.349307617 |
| ENSMUSG00000022305 | Lrp12 | 239393 | protein_coding | 219.3113846 | -0.349345707 | 0.151557854 | -2.305032022 | 0.02116478 | 0.083382897 | 0.349345707 |
| ENSMUSG00000037353 | Letmd1 | 68614 | protein_coding | 277.8073726 | -0.349495729 | 0.093950446 | -3.72000074 | 0.000199222 | 0.002038029 | 0.349495729 |
| ENSMUSG00000020486 | Septin4 | 18952 | protein_coding | 15.64666031 | 0.349507473 | 0.490853002 | 0.712041022 | 0.476439374 | 0.67097492 | 0.349507473 |
| ENSMUSG00000027742 | Cog6 | 67542 | protein_coding | 1083.990351 | 0.349575463 | 0.073785169 | 4.737747 | 2.16E-06 | 3.75E-05 | 0.349575463 |
| ENSMUSG00000037791 | Phf12 | 268448 | protein_coding | 351.9961021 | -0.349690829 | 0.116702446 | -2.996431023 | 0.0027316 | 0.017412629 | 0.349690829 |
| ENSMUSG00000025487 | Psmd13 | 23997 | protein_coding | 1072.164923 | 0.34987348 | 0.091992048 | 3.803301353 | 0.00014278 | 0.001533518 | 0.34987348 |
| ENSMUSG00000026946 | Nmi | 64685 | protein_coding | 405.5055416 | 0.349920214 | 0.101414216 | 3.45040594 | 0.000559744 | 0.004772808 | 0.349920214 |
| ENSMUSG00000026766 | Mmadhc | 109129 | protein_coding | 936.1338821 | 0.350047133 | 0.117394096 | 2.981812065 | 0.002865478 | 0.01809728 | 0.350047133 |
| ENSMUSG00000028266 | Lmo4 | 16911 | protein_coding | 391.6589763 | -0.350297722 | 0.103114593 | -3.397169233 | 0.000680868 | 0.005630775 | 0.350297722 |
| ENSMUSG00000074247 | Dda1 | 66498 | protein_coding | 481.9272364 | 0.350334438 | 0.113473296 | 3.087373409 | 0.002019338 | 0.013605757 | 0.350334438 |
| ENSMUSG00000034858 | Atosa | 235493 | protein_coding | 205.612527 | -0.35041608 | 0.142414466 | -2.460537114 | 0.013872922 | 0.061024001 | 0.35041608 |
| ENSMUSG00000020521 | Rnft1 | 76892 | protein_coding | 966.9979411 | 0.35062102 | 0.101569247 | 3.452039197 | 0.000556367 | 0.004746713 | 0.35062102 |
| ENSMUSG00000042213 | Zfand4 | 67492 | protein_coding | 14.98877789 | 0.350701082 | 0.412724105 | 0.849722801 | 0.395479219 | 0.603774085 | 0.350701082 |
| ENSMUSG00000037706 | Cd81 | 12520 | protein_coding | 1044.548294 | -0.350712339 | 0.096230293 | -3.644510787 | 0.000267901 | 0.002592421 | 0.350712339 |
| ENSMUSG00000078938 | Syce3 | 75459 | protein_coding | 7.528488777 | -0.350715767 | 0.503609266 | -0.696404515 | 0.486175536 | 0.678983247 | 0.350715767 |
| ENSMUSG00000048652 | Samd13 | 75015 | protein_coding | 13.60636389 | -0.350815406 | 0.32769909 | -1.07054129 | 0.284375734 | 0.493628544 | 0.350815406 |
| ENSMUSG00000002379 | Ndufa11 | 69875 | protein_coding | 877.692417 | 0.350838305 | 0.09572843 | 3.664933255 | 0.000247403 | 0.002427995 | 0.350838305 |
| ENSMUSG00000085451 | Gm40614 | 105245120 | lncRNA | 14.12623164 | -0.350881023 | 0.310194089 | -1.131166053 | 0.257985207 | 0.465156031 | 0.350881023 |
| ENSMUSG00000061130 | Ppm1b | 19043 | protein_coding | 1392.931078 | 0.350913059 | 0.07383028 | 4.752969399 | 2.00E-06 | 3.51E-05 | 0.350913059 |
| ENSMUSG00000020268 | Lyrm7 | 75530 | protein_coding | 69.20994606 | 0.350951625 | 0.191584258 | 1.831839568 | 0.066975326 | 0.189515544 | 0.350951625 |
| ENSMUSG00000031214 | Ophn1 | 94190 | protein_coding | 330.5753013 | -0.351022483 | 0.111803998 | -3.1396237 | 0.00169165 | 0.011755933 | 0.351022483 |
| ENSMUSG00000102758 | Naaladl2 | 635702 | protein_coding | 255.4070744 | 0.351132664 | 0.153899254 | 2.281574832 | 0.022514454 | 0.087423263 | 0.351132664 |
| ENSMUSG00000033863 | Klf9 | 16601 | protein_coding | 1112.072313 | -0.351145651 | 0.10452721 | -3.359370731 | 0.000781202 | 0.006265356 | 0.351145651 |
| ENSMUSG00000040434 | Large2 | 228366 | protein_coding | 101.5794966 | -0.351470543 | 0.149986309 | -2.343350846 | 0.019111402 | 0.077457102 | 0.351470543 |
| ENSMUSG00000003559 | As3mt | 57344 | protein_coding | 708.6133441 | 0.351582922 | 0.105599019 | 3.329414671 | 0.000870287 | 0.006839397 | 0.351582922 |
| ENSMUSG00000021493 | Pdlim7 | 67399 | protein_coding | 69.82079259 | 0.35164978 | 0.257763158 | 1.364236002 | 0.172493301 | 0.358210338 | 0.35164978 |
| ENSMUSG00000027075 | Slc43a1 | 72401 | protein_coding | 104.2040649 | 0.351686956 | 0.170197542 | 2.066345682 | 0.038795851 | 0.130312171 | 0.351686956 |
| ENSMUSG00000037499 | Nenf | 66208 | protein_coding | 199.3233091 | -0.351710414 | 0.151132092 | -2.327172264 | 0.019956099 | 0.079644384 | 0.351710414 |
| ENSMUSG00000052605 | Isoc2b | 67441 | protein_coding | 194.2443933 | 0.351749572 | 0.145523268 | 2.417136289 | 0.015643159 | 0.066728504 | 0.351749572 |
| ENSMUSG00000024480 | Ap3s1 | 11777 | protein_coding | 699.2313786 | 0.351790844 | 0.10847876 | 3.242946776 | 0.001183003 | 0.00879686 | 0.351790844 |
| ENSMUSG00000044952 | Kctd21 | 622320 | protein_coding | 135.4218725 | -0.351821149 | 0.142573747 | -2.467643279 | 0.013600578 | 0.060191721 | 0.351821149 |
| ENSMUSG00000015759 | Cnih1 | 12793 | protein_coding | 917.6164217 | 0.351874264 | 0.09778967 | 3.598276412 | 0.000320333 | 0.002999568 | 0.351874264 |
| ENSMUSG00000017756 | Slc12a7 | 20499 | protein_coding | 1145.171263 | -0.351965302 | 0.111736692 | -3.149952757 | 0.001632969 | 0.011417877 | 0.351965302 |
| ENSMUSG00000036002 | Atosb | 230088 | protein_coding | 528.9436573 | 0.35209842 | 0.155431437 | 2.26529734 | 0.023494439 | 0.090238589 | 0.35209842 |
| ENSMUSG00000042570 | Mier2 | 70427 | protein_coding | 359.7385798 | 0.352130896 | 0.148029667 | 2.378785979 | 0.017369758 | 0.072087035 | 0.352130896 |

| ensembl_gene_id | external_gene_name | entrezgene_id | gene_biotype | baseMean | log2FoldChange | lfcSE | stat | pvalue | padj | absLog2FC |
|---|---|---|---|---|---|---|---|---|---|---|
| ENSMUSG00000055024 | Ep300 | 328572 | protein_coding | 773.0181896 | -0.352141243 | 0.191651107 | -1.837407826 | 0.066149712 | 0.18791379 | 0.352141243 |
| ENSMUSG00000018920 | Cxcl16 | 66102 | protein_coding | 36.70346375 | -0.352156776 | 0.283399502 | -1.242616069 | 0.214009345 | 0.412480742 | 0.352156776 |
| ENSMUSG00000047996 | Prrg1 | 546336 | protein_coding | 72.530616 | 0.352191881 | 0.232186075 | 1.516851866 | 0.129304089 | 0.295859555 | 0.352191881 |
| ENSMUSG00000075389 | Reno1 | 100042332 | lncRNA | 24.13530962 | -0.35228299 | 0.251032342 | -1.403337066 | 0.16051635 | 0.342171494 | 0.35228299 |
| ENSMUSG00000016252 | Atp5e | 67126 | protein_coding | 2012.476518 | 0.352285231 | 0.164775804 | 2.137967003 | 0.032519421 | 0.114367265 | 0.352285231 |
| ENSMUSG00000037145 | Lypd8l | 78354 | protein_coding | 457.4330964 | 0.352329247 | 0.273678542 | 1.287383526 | 0.197960643 | 0.392060046 | 0.352329247 |
| ENSMUSG00000026611 | Spata17 | 74717 | protein_coding | 18.13452502 | 0.352336096 | 0.322270422 | 1.093293311 | 0.274265039 | 0.482226906 | 0.352336096 |
| ENSMUSG00000021102 | Glrx5 | 73046 | protein_coding | 580.3944966 | 0.352339962 | 0.10501006 | 3.3552972 | 0.000792798 | 0.006329664 | 0.352339962 |
| ENSMUSG00000030737 | Slco2b1 | 101488 | protein_coding | 207.6793741 | -0.352348269 | 0.179415938 | -1.963862706 | 0.049546016 | 0.154436192 | 0.352348269 |
| ENSMUSG00000022014 | Epsti1 | 108670 | protein_coding | 611.2449018 | 0.352363738 | 0.09244261 | 3.811702606 | 0.000138013 | 0.001491363 | 0.352363738 |
| ENSMUSG00000030303 | Far2 | 330450 | protein_coding | 282.4307725 | 0.352413539 | 0.145229174 | 2.426602923 | 0.015240927 | 0.065433283 | 0.352413539 |
| ENSMUSG00000037953 | A4gnt | 333424 | protein_coding | 406.7225444 | -0.352533678 | 0.230129295 | -1.531893967 | 0.125548603 | 0.289775125 | 0.352533678 |
| ENSMUSG00000079038 | D130040H23Rik | 211135 | protein_coding | 45.94015244 | -0.352835904 | 0.230387562 | -1.531488509 | 0.125648704 | 0.289880151 | 0.352835904 |
| ENSMUSG00000040007 | Bahd1 | 228536 | protein_coding | 385.3020313 | -0.352912403 | 0.120491987 | -2.928928408 | 0.003401328 | 0.020741587 | 0.352912403 |
| ENSMUSG00000073888 | Ccl27a | 20301 | protein_coding | 20.24423233 | -0.352916006 | 0.294887747 | -1.196780844 | 0.231391986 | 0.434726535 | 0.352916006 |
| ENSMUSG00000085990 | Gm16731 | 105247126 | lncRNA | 13.05209703 | 0.352943034 | 0.417487831 | 0.845397178 | 0.397889119 | 0.606002858 | 0.352943034 |
| ENSMUSG00000029389 | Ddx55 | 67848 | protein_coding | 212.6868955 | -0.35297351 | 0.099672235 | -3.541342373 | 0.000398097 | 0.003619226 | 0.35297351 |
| ENSMUSG00000022450 | Ndufa6 | 67130 | protein_coding | 2335.20189 | 0.352996908 | 0.097743736 | 3.611452989 | 0.000304486 | 0.002868041 | 0.352996908 |
| ENSMUSG00000055762 | Eef1d | 66656 | protein_coding | 1560.248383 | 0.353021309 | 0.098386587 | 3.588104034 | 0.000333091 | 0.003100806 | 0.353021309 |
| ENSMUSG00000040269 | Mrps28 | 66230 | protein_coding | 285.2531291 | 0.353108374 | 0.131608022 | 2.683030792 | 0.007295827 | 0.037680913 | 0.353108374 |
| ENSMUSG00000032301 | Psma4 | 26441 | protein_coding | 1792.743882 | 0.353247023 | 0.101545678 | 3.478700718 | 0.000503851 | 0.004380643 | 0.353247023 |
| ENSMUSG00000087231 | E230016M11Rik | 320172 | lncRNA | 20.4700998 | -0.353414839 | 0.347170095 | -1.017987562 | 0.308683867 | 0.518602032 | 0.353414839 |
| ENSMUSG00000073000 | Gm10451 | 100041694 | lncRNA | 27.74763493 | -0.353565099 | 0.301557592 | -1.17246293 | 0.241011247 | 0.445587011 | 0.353565099 |
| ENSMUSG00000072618 | Gm10384 | 100038735 | lncRNA | 43.16951941 | -0.35366334 | 0.218272651 | -1.620282426 | 0.105171622 | 0.256772746 | 0.35366334 |
| ENSMUSG00000049866 | Arl4c | 320982 | protein_coding | 18.45368476 | 0.353716126 | 0.350390842 | 1.009490215 | 0.312739592 | 0.522972832 | 0.353716126 |
| ENSMUSG00000021514 | Zfp369 | 170936 | protein_coding | 244.2471666 | -0.353736588 | 0.137942082 | -2.564384877 | 0.010335886 | 0.048938762 | 0.353736588 |
| ENSMUSG00000001901 | Kcnh6 | 192775 | protein_coding | 42.17301529 | -0.353748372 | 0.219462014 | -1.61188884 | 0.106986131 | 0.25996569 | 0.353748372 |
| ENSMUSG00000019278 | Dpep1 | 13479 | protein_coding | 3510.86576 | 0.35378959 | 0.156861959 | 2.255419946 | 0.024106982 | 0.091970301 | 0.35378959 |
| ENSMUSG00000022479 | Vdr | 22337 | protein_coding | 3079.774001 | -0.353837373 | 0.120807953 | -2.928924499 | 0.00340137 | 0.020741587 | 0.353837373 |
| ENSMUSG00000029524 | Sirt4 | 75387 | protein_coding | 61.66530809 | -0.353930421 | 0.184389438 | -1.919472317 | 0.054924586 | 0.165344701 | 0.353930421 |
| ENSMUSG00000101337 | Dnah7c | 100101919 | protein_coding | 8.387184132 | -0.353948661 | 1.182338662 | -0.29936318 | 0.764662953 | 0.871422033 | 0.353948661 |
| ENSMUSG00000027176 | Cstf3 | 228410 | protein_coding | 363.6756041 | 0.354016205 | 0.087621043 | 4.040310311 | 5.34E-05 | 0.000657575 | 0.354016205 |
| ENSMUSG00000033256 | Shf | 435684 | protein_coding | 153.4727422 | -0.354132402 | 0.259331468 | -1.36555893 | 0.172077445 | 0.357719532 | 0.354132402 |
| ENSMUSG00000041460 | Cacna2d4 | 319734 | protein_coding | 16.05448702 | -0.354250773 | 0.502211888 | -0.705381097 | 0.480573104 | 0.674054985 | 0.354250773 |
| ENSMUSG00000041762 | Gpr155 | 68526 | protein_coding | 14.89511104 | 0.354352131 | 0.365864925 | 0.968532665 | 0.332778414 | 0.543696233 | 0.354352131 |
| ENSMUSG00000038622 | Med30 | 69790 | protein_coding | 202.4193594 | 0.354515055 | 0.121007121 | 2.929704058 | 0.00339285 | 0.020713419 | 0.354515055 |
| ENSMUSG00000006464 | Bbs1 | 52028 | protein_coding | 35.98592898 | -0.354596899 | 0.235607289 | -1.505033657 | 0.132315424 | 0.300387695 | 0.354596899 |
| ENSMUSG00000026669 | Mcm10 | 70024 | protein_coding | 271.9767295 | 0.354622911 | 0.120863896 | 2.934068163 | 0.003345507 | 0.02051087 | 0.354622911 |

| ensembl_gene_id | external_gene_name | entrezgene_id | gene_biotype | baseMean | log2FoldChange | lfcSE | stat | pvalue | padj | absLog2FC |
|---|---|---|---|---|---|---|---|---|---|---|
| ENSMUSG00000093661 | Eif4e3 | 66892 | protein_coding | 129.3451575 | 0.354639797 | 0.231280361 | 1.533376183 | 0.125183193 | 0.28930904 | 0.354639797 |
| ENSMUSG00000026239 | Pde6d | 18582 | protein_coding | 82.14692942 | 0.354651678 | 0.154587488 | 2.294180994 | 0.021780105 | 0.085048665 | 0.354651678 |
| ENSMUSG00000002102 | Psmc3 | 19182 | protein_coding | 1816.229456 | 0.354847652 | 0.095801539 | 3.703986974 | 0.000212237 | 0.002150454 | 0.354847652 |
| ENSMUSG00000027189 | Trim44 | 80985 | protein_coding | 703.7001268 | -0.354871834 | 0.0894424 | -3.967601882 | 7.26E-05 | 0.000850199 | 0.354871834 |
| ENSMUSG00000096210 | H1f0 | 14958 | protein_coding | 3543.496535 | 0.355136147 | 0.124528003 | 2.851857713 | 0.004346455 | 0.02511075 | 0.355136147 |
| ENSMUSG00000031983 | 2310022B05Rik | 69551 | protein_coding | 205.8091363 | -0.35530458 | 0.124692609 | -2.849443784 | 0.004379574 | 0.025238355 | 0.35530458 |
| ENSMUSG00000067274 | Rplp0 | 11837 | protein_coding | 22800.94805 | 0.35536802 | 0.106828621 | 3.326524437 | 0.000879363 | 0.006896354 | 0.35536802 |
| ENSMUSG00000043467 | Zbtb37 | 240869 | protein_coding | 145.315222 | -0.355383207 | 0.202208719 | -1.757506836 | 0.078831458 | 0.211670171 | 0.355383207 |
| ENSMUSG00000027954 | Efna1 | 13636 | protein_coding | 841.5291074 | 0.355461986 | 0.190932059 | 1.861719756 | 0.062642602 | 0.180967229 | 0.355461986 |
| ENSMUSG00000026994 | Galnt3 | 14425 | protein_coding | 1490.392489 | 0.355681262 | 0.180867219 | 1.966532489 | 0.049237133 | 0.153872198 | 0.355681262 |
| ENSMUSG00000039033 | Tasp1 | 75812 | protein_coding | 231.5670097 | 0.355702723 | 0.086844002 | 4.095881306 | 4.21E-05 | 0.000529128 | 0.355702723 |
| ENSMUSG00000004952 | Rasa4 | 54153 | protein_coding | 423.8951335 | -0.355727064 | 0.140793478 | -2.526587667 | 0.011517664 | 0.053267523 | 0.355727064 |
| ENSMUSG00000031558 | Slit2 | 20563 | protein_coding | 108.5620024 | -0.355761174 | 0.169560479 | -2.098137351 | 0.035893014 | 0.123265184 | 0.355761174 |
| ENSMUSG00000052609 | Plekhg3 | 263406 | protein_coding | 731.6642717 | -0.355807119 | 0.189233154 | -1.880257826 | 0.060072948 | 0.17598711 | 0.355807119 |
| ENSMUSG00000020275 | Rel | 19696 | protein_coding | 99.38310026 | -0.355945385 | 0.22561558 | -1.577663147 | 0.114643017 | 0.271448487 | 0.355945385 |
| ENSMUSG00000050511 | Oprd1 | 18386 | protein_coding | 14.6271382 | -0.356103132 | 0.298397014 | -1.193387051 | 0.232717824 | 0.436353122 | 0.356103132 |
| ENSMUSG00000037313 | Tacc3 | 21335 | protein_coding | 497.1139251 | 0.356150109 | 0.127157506 | 2.800857933 | 0.005096695 | 0.028464292 | 0.356150109 |
| ENSMUSG00000035929 | H2-Q4 | 15015 | protein_coding | 226.9618349 | 0.356303178 | 0.106616766 | 3.341905695 | 0.000832053 | 0.006596787 | 0.356303178 |
| ENSMUSG00000010592 | Dazl | 13164 | protein_coding | 9.290865167 | 0.356402392 | 0.622762977 | 0.572292197 | 0.567124035 | 0.743049553 | 0.356402392 |
| ENSMUSG00000020894 | Vamp2 | 22318 | protein_coding | 163.3034298 | -0.356436605 | 0.143061088 | -2.491499338 | 0.01272052 | 0.057364793 | 0.356436605 |
| ENSMUSG00000028982 | Slc25a33 | 70556 | protein_coding | 85.90739443 | -0.356455444 | 0.180537388 | -1.974413439 | 0.048334751 | 0.152161266 | 0.356455444 |
| ENSMUSG00000030603 | Psmc4 | 23996 | protein_coding | 1075.462621 | 0.356497612 | 0.111479435 | 3.197877821 | 0.001384429 | 0.010005077 | 0.356497612 |
| ENSMUSG00000037242 | Clic4 | 29876 | protein_coding | 802.706119 | -0.356595374 | 0.134187455 | -2.657441966 | 0.007873615 | 0.03998202 | 0.356595374 |
| ENSMUSG00000020272 | Stk10 | 20868 | protein_coding | 370.2315768 | -0.356610973 | 0.177522305 | -2.008823469 | 0.044555858 | 0.143370851 | 0.356610973 |
| ENSMUSG00000006010 | Odr4 | 226499 | protein_coding | 722.2828767 | 0.356661774 | 0.100450844 | 3.55061002 | 0.000384339 | 0.003516231 | 0.356661774 |
| ENSMUSG00000046994 | Mars2 | 212679 | protein_coding | 370.4866576 | -0.356695754 | 0.120094618 | -2.970122723 | 0.002976808 | 0.018667165 | 0.356695754 |
| ENSMUSG00000038349 | Plcl1 | 227120 | protein_coding | 15.09777124 | 0.356740924 | 0.359904305 | 0.991210492 | 0.321582808 | 0.532450205 | 0.356740924 |
| ENSMUSG00000089704 | Galnt2 | 108148 | protein_coding | 11.3700093 | 0.356875158 | 0.381876815 | 0.934529525 | 0.350030813 | 0.56016184 | 0.356875158 |
| ENSMUSG00000045414 | Dipk2a | 68861 | protein_coding | 574.1325935 | 0.357249957 | 0.098437321 | 3.629212496 | 0.000284287 | 0.002711453 | 0.357249957 |
| ENSMUSG00000052214 | Opa3 | 403187 | protein_coding | 988.2034333 | 0.357263403 | 0.084210977 | 4.242480213 | 2.21E-05 | 0.00030198 | 0.357263403 |
| ENSMUSG00000034647 | Ankrd12 | 106585 | protein_coding | 356.7855993 | -0.357366859 | 0.101836563 | -3.50921957 | 0.000449424 | 0.003990201 | 0.357366859 |
| ENSMUSG00000027332 | Ivd | 56357 | protein_coding | 1133.006143 | 0.357429169 | 0.084203743 | 4.244813322 | 2.19E-05 | 0.000299369 | 0.357429169 |
| ENSMUSG00000026600 | Soat1 | 20652 | protein_coding | 1170.75399 | -0.357484331 | 0.095510047 | -3.742897646 | 0.00018191 | 0.001880252 | 0.357484331 |
| ENSMUSG00000039699 | Batf2 | 74481 | protein_coding | 137.1610553 | 0.3574955 | 0.16319698 | 2.190576686 | 0.02848244 | 0.103719319 | 0.3574955 |
| ENSMUSG00000058258 | Idi1 | 319554 | protein_coding | 398.1970025 | 0.357654285 | 0.181333882 | 1.972352215 | 0.048569413 | 0.152676852 | 0.357654285 |
| ENSMUSG00000024118 | Tedc2 | 72016 | protein_coding | 55.42873706 | -0.357711267 | 0.190101275 | -1.881687888 | 0.059878405 | 0.175740416 | 0.357711267 |
| ENSMUSG00000022629 | Kif21a | 16564 | protein_coding | 1394.3025 | 0.357823931 | 0.077300527 | 4.628997283 | 3.67E-06 | 6.10E-05 | 0.357823931 |
| ENSMUSG00000069910 | Spdl1 | 70385 | protein_coding | 143.9107566 | 0.35786907 | 0.153303224 | 2.3343871 | 0.019575462 | 0.078608426 | 0.35786907 |

| ensembl_gene_id | external_gene_name | entrezgene_id | gene_biotype | baseMean | log2FoldChange | lfcSE | stat | pvalue | padj | absLog2FC |
|---|---|---|---|---|---|---|---|---|---|---|
| ENSMUSG00000038013 | Wipf2 | 68524 | protein_coding | 461.1436948 | -0.35790446 | 0.116206378 | -3.079903758 | 0.002070675 | 0.013928087 | 0.35790446 |
| ENSMUSG00000072964 | Bhlhb9 | 70237 | protein_coding | 46.89164349 | 0.357933359 | 0.199006943 | 1.798597345 | 0.072082398 | 0.19950103 | 0.357933359 |
| ENSMUSG00000004815 | Dgkq | 110524 | protein_coding | 420.7313555 | -0.357935265 | 0.134619284 | -2.658870668 | 0.007840305 | 0.039859682 | 0.357935265 |
| ENSMUSG00000026944 | Abca2 | 11305 | protein_coding | 107.9353118 | -0.357956019 | 0.127075225 | -2.816882825 | 0.004849221 | 0.02745766 | 0.357956019 |
| ENSMUSG00000004567 | Mcoln1 | 94178 | protein_coding | 347.2475119 | -0.357989221 | 0.092340755 | -3.87682798 | 0.000105827 | 0.001181229 | 0.357989221 |
| ENSMUSG00000005142 | Man2b1 | 17159 | protein_coding | 1633.852527 | -0.358020223 | 0.085890292 | -4.168343292 | 3.07E-05 | 0.000403555 | 0.358020223 |
| ENSMUSG00000023764 | Sfi1 | 78887 | protein_coding | 558.6215118 | -0.358066295 | 0.22856842 | -1.566560659 | 0.117217419 | 0.276388247 | 0.358066295 |
| ENSMUSG00000032202 | Rab27a | 11891 | protein_coding | 744.952215 | 0.358088861 | 0.147909695 | 2.420996556 | 0.015478024 | 0.066183615 | 0.358088861 |
| ENSMUSG00000038831 | Ralgps1 | 241308 | protein_coding | 169.668841 | 0.358102993 | 0.141145527 | 2.537118966 | 0.011176897 | 0.051980373 | 0.358102993 |
| ENSMUSG00000038000 | Acd | 497652 | protein_coding | 129.2475116 | -0.358133921 | 0.115186041 | -3.109178149 | 0.001876086 | 0.012819516 | 0.358133921 |
| ENSMUSG00000035297 | Cops4 | 26891 | protein_coding | 1185.226569 | 0.358203445 | 0.070757442 | 5.062413685 | 4.14E-07 | 8.60E-06 | 0.358203445 |
| ENSMUSG00000036155 | Mgat5 | 107895 | protein_coding | 2216.600302 | -0.358214039 | 0.112925035 | -3.172140167 | 0.0015132 | 0.010749598 | 0.358214039 |
| ENSMUSG00000002274 | Metrn | 70083 | protein_coding | 19.51021035 | 0.35830925 | 0.251214201 | 1.426309696 | 0.153778967 | 0.332888202 | 0.35830925 |
| ENSMUSG00000004945 | Tmem242 | 70544 | protein_coding | 495.0770084 | 0.358310308 | 0.112492297 | 3.185198618 | 0.001446547 | 0.010350673 | 0.358310308 |
| ENSMUSG00000072704 | Smim10l1 | 381820 | protein_coding | 2121.669453 | 0.358377674 | 0.07983012 | 4.489253844 | 7.15E-06 | 0.000111066 | 0.358377674 |
| ENSMUSG00000016179 | Camk1g | 215303 | protein_coding | 7.714863957 | -0.358447406 | 0.547103382 | -0.655173075 | 0.512356327 | 0.699981612 | 0.358447406 |
| ENSMUSG00000020471 | Pold2 | 18972 | protein_coding | 471.9411338 | 0.35850834 | 0.114208804 | 3.139060468 | 0.001694905 | 0.011773416 | 0.35850834 |
| ENSMUSG00000028914 | Casp9 | 12371 | protein_coding | 282.6873701 | -0.358528029 | 0.08041589 | -4.458422672 | 8.26E-06 | 0.000125846 | 0.358528029 |
| ENSMUSG00000014226 | Cacybp | 12301 | protein_coding | 812.418429 | 0.358585296 | 0.148707571 | 2.411345262 | 0.015893795 | 0.067616549 | 0.358585296 |
| ENSMUSG00000050944 | Efcab5 | 319634 | protein_coding | 73.81769942 | 0.358622437 | 0.182556026 | 1.964451377 | 0.04947777 | 0.154314062 | 0.358622437 |
| ENSMUSG00000038060 | Dlec1 | 320256 | protein_coding | 9.894196414 | -0.358632361 | 0.489286819 | -0.732969593 | 0.463576975 | 0.66105374 | 0.358632361 |
| ENSMUSG00000072494 | Ppp1r3e | 105651 | protein_coding | 37.58536897 | -0.358708162 | 0.259962951 | -1.379843399 | 0.167634867 | 0.352115015 | 0.358708162 |
| ENSMUSG00000003228 | Grk5 | 14773 | protein_coding | 179.3591266 | 0.358724815 | 0.105131806 | 3.412143544 | 0.000644542 | 0.005379931 | 0.358724815 |
| ENSMUSG00000069804 | Gm10277 | 791303 | protein_coding | 27.14468999 | -0.358761296 | 0.404884213 | -0.886083688 | 0.375572432 | 0.583911261 | 0.358761296 |
| ENSMUSG00000006575 | Rundc3a | 51799 | protein_coding | 105.5452101 | -0.358785249 | 0.133860618 | -2.680289788 | 0.007355845 | 0.037941677 | 0.358785249 |
| ENSMUSG00000003269 | Cyth2 | 19158 | protein_coding | 824.5658402 | 0.358910914 | 0.119544631 | 3.002317301 | 0.002679327 | 0.017187405 | 0.358910914 |
| ENSMUSG00000009614 | Sardh | 192166 | protein_coding | 73.36000135 | 0.359136385 | 0.191572276 | 1.874678282 | 0.060836996 | 0.177344744 | 0.359136385 |
| ENSMUSG00000048118 | Arid4a | 238247 | protein_coding | 884.2125909 | 0.359316229 | 0.178799361 | 2.009605786 | 0.044472928 | 0.143190783 | 0.359316229 |
| ENSMUSG00000029189 | Sel1l3 | 231238 | protein_coding | 3346.585661 | 0.35932755 | 0.108415803 | 3.314346624 | 0.000918576 | 0.007154558 | 0.35932755 |
| ENSMUSG00000087141 | Plcxd2 | 433022 | protein_coding | 271.4656508 | 0.359409401 | 0.153213925 | 2.345801145 | 0.018986234 | 0.077047831 | 0.359409401 |
| ENSMUSG00000035455 | Fignl1 | 60530 | protein_coding | 149.9946789 | 0.359556851 | 0.163845922 | 2.194481546 | 0.028200809 | 0.103023507 | 0.359556851 |
| ENSMUSG00000074529 | Zfp972 | 78703 | protein_coding | 14.07035678 | 0.3596242 | 0.29933183 | 1.20142318 | 0.229587089 | 0.432847922 | 0.3596242 |
| ENSMUSG00000020827 | Mink1 | 50932 | protein_coding | 788.3553744 | -0.359637608 | 0.110906987 | -3.242695665 | 0.001184046 | 0.008800507 | 0.359637608 |
| ENSMUSG00000049904 | Tmem17 | 103765 | protein_coding | 13.92482783 | 0.359696445 | 0.353625917 | 1.017166523 | 0.309074219 | 0.51897129 | 0.359696445 |
| ENSMUSG00000078439 | Smim24 | 72273 | protein_coding | 1327.535919 | -0.359733559 | 0.10449715 | -3.442520289 | 0.000576321 | 0.004885383 | 0.359733559 |
| ENSMUSG00000089857 | Zfp882 | 382019 | protein_coding | 42.06990392 | 0.359820061 | 0.267395839 | 1.345645702 | 0.178416805 | 0.365695904 | 0.359820061 |
| ENSMUSG00000020857 | Nme2 | 18103 | protein_coding | 146.2106015 | 0.35999205 | 0.145638081 | 2.471826387 | 0.013442478 | 0.059757687 | 0.35999205 |
| ENSMUSG00000034066 | Farp2 | 227377 | protein_coding | 473.4132092 | -0.360067298 | 0.123625609 | -2.912562378 | 0.003584766 | 0.021562746 | 0.360067298 |

| ensembl_gene_id | external_gene_name | entrezgene_id | gene_biotype | baseMean | log2FoldChange | lfcSE | stat | pvalue | padj | absLog2FC |
|---|---|---|---|---|---|---|---|---|---|---|
| ENSMUSG00000020473 | Aebp1 | 11568 | protein_coding | 7.549391821 | -0.360204486 | 0.793043959 | -0.454204943 | 0.649681326 | 0.797543288 | 0.360204486 |
| ENSMUSG00000016493 | Cd46 | 17221 | protein_coding | 39.2162975 | -0.360210448 | 0.283687687 | -1.269742976 | 0.204176199 | 0.399572245 | 0.360210448 |
| ENSMUSG00000020728 | Cep112 | 76380 | protein_coding | 27.35305814 | 0.360308854 | 0.34465424 | 1.045421214 | 0.295828336 | 0.505111323 | 0.360308854 |
| ENSMUSG00000048832 | Vps37c | 107305 | protein_coding | 513.1309 | 0.360318463 | 0.089417081 | 4.029637936 | 5.59E-05 | 0.000681826 | 0.360318463 |
| ENSMUSG00000019893 | Ros1 | 19886 | protein_coding | 16.63660598 | -0.360544608 | 0.59518696 | -0.60576698 | 0.544669499 | 0.727443886 | 0.360544608 |
| ENSMUSG00000089736 | Tgfbr3l | 100044509 | protein_coding | 11.93381343 | -0.360561692 | 0.365426995 | -0.986685976 | 0.323796607 | 0.534726009 | 0.360561692 |
| ENSMUSG00000089774 | Slc5a3 | 53881 | protein_coding | 1176.689579 | -0.360568357 | 0.178339807 | -2.021805245 | 0.043196476 | 0.140271858 | 0.360568357 |
| ENSMUSG00000052212 | Cd177 | 68891 | protein_coding | 3618.385294 | 0.360582837 | 0.138300825 | 2.607235617 | 0.009127653 | 0.044683106 | 0.360582837 |
| ENSMUSG00000028345 | Tex10 | 269536 | protein_coding | 312.2538716 | -0.360615352 | 0.074145184 | -4.863638247 | 1.15E-06 | 2.14E-05 | 0.360615352 |
| ENSMUSG00000046387 | Pcdhb17 | 93888 | protein_coding | 8.284301213 | 0.360649075 | 0.425487023 | 0.847614745 | 0.39665256 | 0.604651202 | 0.360649075 |
| ENSMUSG00000026034 | Clk1 | 12747 | protein_coding | 2179.027599 | -0.360865731 | 0.114638238 | -3.147865296 | 0.001644675 | 0.01147957 | 0.360865731 |
| ENSMUSG00000074221 | Zfp568 | 243905 | protein_coding | 316.2631214 | -0.361003416 | 0.126340113 | -2.857393484 | 0.00427136 | 0.024757721 | 0.361003416 |
| ENSMUSG00000008090 | Fgfrl1 | 116701 | protein_coding | 228.1475238 | -0.361020585 | 0.152116747 | -2.373312557 | 0.017629339 | 0.072783853 | 0.361020585 |
| ENSMUSG00000034429 | Zfp707 | 69020 | protein_coding | 112.7184969 | -0.361051442 | 0.225863869 | -1.598535626 | 0.109923823 | 0.264241874 | 0.361051442 |
| ENSMUSG00000026065 | Slc9a4 | 110895 | protein_coding | 8.054418566 | 0.361059343 | 0.442493808 | 0.815964734 | 0.414520306 | 0.620942296 | 0.361059343 |
| ENSMUSG00000008734 | Gprc5b | 64297 | protein_coding | 8.186722218 | 0.361142368 | 0.685592853 | 0.526759238 | 0.598360793 | 0.765321556 | 0.361142368 |
| ENSMUSG00000041073 | Nacad | 192950 | protein_coding | 66.08736194 | -0.361162553 | 0.235977007 | -1.530498916 | 0.125893282 | 0.290318255 | 0.361162553 |
| ENSMUSG00000001627 | Ifrd1 | 15982 | protein_coding | 982.8872847 | -0.361176291 | 0.156408364 | -2.309187836 | 0.02093316 | 0.082735329 | 0.361176291 |
| ENSMUSG00000034163 | Zfc3h1 | 216345 | protein_coding | 984.0318287 | -0.361206321 | 0.126246557 | -2.861118185 | 0.004221497 | 0.024504372 | 0.361206321 |
| ENSMUSG00000038045 | Sult6b1 | 73671 | protein_coding | 9.611535303 | 0.361378459 | 0.378429397 | 0.954942882 | 0.339606582 | 0.55016819 | 0.361378459 |
| ENSMUSG00000056487 | Mettl7a2 | 393082 | protein_coding | 24.65918151 | -0.361521644 | 0.587160495 | -0.615711799 | 0.538084749 | 0.721865534 | 0.361521644 |
| ENSMUSG00000020696 | Rffl | 67338 | protein_coding | 372.4332131 | -0.361728038 | 0.08503851 | -4.253696815 | 2.10E-05 | 0.000289721 | 0.361728038 |
| ENSMUSG00000022664 | Slc35a5 | 74102 | protein_coding | 437.0879482 | 0.361927549 | 0.103751952 | 3.488392675 | 0.000485934 | 0.004244859 | 0.361927549 |
| ENSMUSG00000015968 | Cacna1d | 12289 | protein_coding | 33.54389996 | -0.361934211 | 0.320842796 | -1.128073359 | 0.259288939 | 0.4664368 | 0.361934211 |
| ENSMUSG00000015452 | Ager | 11596 | protein_coding | 10.81876491 | 0.361995039 | 0.445531202 | 0.812502105 | 0.416503579 | 0.622741858 | 0.361995039 |
| ENSMUSG00000004709 | Cd244a | 18106 | protein_coding | 20.84341887 | 0.362012952 | 0.343375927 | 1.054275863 | 0.291756636 | 0.5005493 | 0.362012952 |
| ENSMUSG00000078580 | E430018J23Rik | 101604 | protein_coding | 129.9783842 | -0.362106879 | 0.164481829 | -2.201500801 | 0.02770059 | 0.101592217 | 0.362106879 |
| ENSMUSG00000020834 | Dhrs13 | 70451 | protein_coding | 68.49241198 | -0.362157008 | 0.139087104 | -2.603814423 | 0.009219266 | 0.045080317 | 0.362157008 |
| ENSMUSG00000037243 | Zfp692 | 103836 | protein_coding | 199.8100372 | -0.362168999 | 0.140944555 | -2.569584899 | 0.010182044 | 0.048469694 | 0.362168999 |
| ENSMUSG00000074797 | Itpa | 16434 | protein_coding | 503.6769458 | 0.362180236 | 0.105427319 | 3.435354692 | 0.000591779 | 0.005000452 | 0.362180236 |
| ENSMUSG00000030866 | Ern2 | 26918 | protein_coding | 1632.206493 | -0.362255131 | 0.105478605 | -3.434394407 | 0.000593879 | 0.005015541 | 0.362255131 |
| ENSMUSG00000029212 | Gabrb1 | 14400 | protein_coding | 11.69718379 | -0.362407259 | 0.849761013 | -0.426481391 | 0.669757105 | 0.808780483 | 0.362407259 |
| ENSMUSG00000027793 | Ccna1 | 12427 | protein_coding | 14.14591406 | 0.362454126 | 0.301573285 | 1.201877435 | 0.229411018 | 0.432740253 | 0.362454126 |
| ENSMUSG00000052331 | Ankrd44 | 329154 | protein_coding | 208.2331456 | -0.362468531 | 0.170373416 | -2.127494657 | 0.033379005 | 0.116720261 | 0.362468531 |
| ENSMUSG00000027796 | Smad9 | 55994 | protein_coding | 12.87542662 | 0.362481499 | 0.379924557 | 0.9540881 | 0.340039049 | 0.550365001 | 0.362481499 |
| ENSMUSG00000028344 | Invs | 16348 | protein_coding | 100.9401124 | 0.36252366 | 0.153982725 | 2.354313835 | 0.018556936 | 0.075672013 | 0.36252366 |
| ENSMUSG00000081769 | Gm12216 | 622459 | lncRNA | 18.02740923 | -0.362541428 | 0.330234998 | -1.097828608 | 0.272279337 | 0.48047545 | 0.362541428 |
| ENSMUSG00000094091 | Ankrd40cl | 102639132 | protein_coding | 352.6012489 | 0.362576939 | 0.218581198 | 1.658774603 | 0.09716122 | 0.243675627 | 0.362576939 |

| ensembl_gene_id | external_gene_name | entrezgene_id | gene_biotype | baseMean | log2FoldChange | lfcSE | stat | pvalue | padj | absLog2FC |
|---|---|---|---|---|---|---|---|---|---|---|
| ENSMUSG00000079108 | Srp54c | 100101806 | protein_coding | 137.5271837 | 0.362580176 | 0.113194401 | 3.203163534 | 0.001359268 | 0.009850053 | 0.362580176 |
| ENSMUSG00000031711 | Zfp330 | 30932 | protein_coding | 667.0597712 | 0.362662983 | 0.10568658 | 3.431495107 | 0.000600264 | 0.005066775 | 0.362662983 |
| ENSMUSG00000041736 | Tspo | 12257 | protein_coding | 675.957411 | -0.362674589 | 0.133414429 | -2.71840603 | 0.006559729 | 0.034620065 | 0.362674589 |
| ENSMUSG00000037447 | Arid5a | 214855 | protein_coding | 118.7504038 | -0.362784457 | 0.365291877 | -0.993135846 | 0.320643757 | 0.532167736 | 0.362784457 |
| ENSMUSG00000000711 | Rab5b | 19344 | protein_coding | 1067.865308 | 0.36278889 | 0.077025951 | 4.709956635 | 2.48E-06 | 4.25E-05 | 0.36278889 |
| ENSMUSG00000070423 | Or51e1 | 259097 | protein_coding | 11.26553402 | -0.362791943 | 0.390753595 | -0.928441727 | 0.353178482 | 0.562936742 | 0.362791943 |
| ENSMUSG00000026276 | Septin2 | 18000 | protein_coding | 57.13005361 | -0.362858679 | 0.177493428 | -2.044349936 | 0.040918989 | 0.135275563 | 0.362858679 |
| ENSMUSG00000025104 | Hdgfl3 | 29877 | protein_coding | 6.823307425 | -0.362923095 | 0.927140575 | -0.391443438 | 0.695469489 | 0.826592861 | 0.362923095 |
| ENSMUSG00000042105 | Inpp5f | 101490 | protein_coding | 234.5242458 | -0.362992646 | 0.118512511 | -3.062905704 | 0.002191991 | 0.014608383 | 0.362992646 |
| ENSMUSG00000035045 | Zc3h12b | 547176 | protein_coding | 7.584147721 | -0.363023944 | 0.564959803 | -0.642565969 | 0.520505771 | 0.707028476 | 0.363023944 |
| ENSMUSG00000078502 | Zfp268 | 433801 | protein_coding | 35.66065139 | -0.363055961 | 0.303322309 | -1.196931284 | 0.231333339 | 0.434667622 | 0.363055961 |
| ENSMUSG00000024487 | Yipf5 | 67180 | protein_coding | 709.6519567 | 0.363366178 | 0.066440693 | 5.469030519 | 4.53E-08 | 1.09E-06 | 0.363366178 |
| ENSMUSG00000021186 | Fbln5 | 23876 | protein_coding | 6.874883663 | -0.363408801 | 0.688635331 | -0.527723143 | 0.597691508 | 0.765029761 | 0.363408801 |
| ENSMUSG00000042363 | Lgalsl | 216551 | protein_coding | 724.7278282 | 0.363541506 | 0.064462029 | 5.639622428 | 1.70E-08 | 4.45E-07 | 0.363541506 |
| ENSMUSG00000057596 | Trim30d | 209387 | protein_coding | 337.1908514 | 0.363555317 | 0.141145502 | 2.575748516 | 0.010002337 | 0.047943791 | 0.363555317 |
| ENSMUSG00000033498 | Strc | 140476 | protein_coding | 25.00068156 | -0.363680549 | 0.28740538 | -1.265392279 | 0.205730742 | 0.40157834 | 0.363680549 |
| ENSMUSG00000032997 | Chpf | 74241 | protein_coding | 201.9756271 | 0.363707313 | 0.14746195 | 2.466448553 | 0.013646033 | 0.060242243 | 0.363707313 |
| ENSMUSG00000030409 | Dmpk | 13400 | protein_coding | 54.30869166 | -0.363778353 | 0.222727755 | -1.633287032 | 0.102408643 | 0.252307017 | 0.363778353 |
| ENSMUSG00000022037 | Clu | 12759 | protein_coding | 41.57578124 | -0.363941814 | 0.247053695 | -1.473128395 | 0.140716418 | 0.313156946 | 0.363941814 |
| ENSMUSG00000034656 | Cacna1a | 12286 | protein_coding | 109.6823944 | -0.364042837 | 0.220552637 | -1.650593901 | 0.098821525 | 0.246442657 | 0.364042837 |
| ENSMUSG00000026395 | Ptprc | 19264 | protein_coding | 226.9983176 | 0.364078129 | 0.190471624 | 1.911456001 | 0.055946006 | 0.167596009 | 0.364078129 |
| ENSMUSG00000040177 | 2310057M21Rik | 68277 | protein_coding | 108.9413138 | -0.364113216 | 0.191454717 | -1.901824207 | 0.057194141 | 0.170469362 | 0.364113216 |
| ENSMUSG00000026974 | Zmynd19 | 67187 | protein_coding | 279.1657576 | 0.364124743 | 0.173460842 | 2.099175462 | 0.035801436 | 0.123083375 | 0.364124743 |
| ENSMUSG00000041632 | Mrps27 | 218506 | protein_coding | 402.2914486 | 0.36417005 | 0.086099864 | 4.229623959 | 2.34E-05 | 0.000316681 | 0.36417005 |
| ENSMUSG00000077391 | Gm24336 | 115489055 | snoRNA | 470.983737 | -0.364236127 | 0.117522926 | -3.099277207 | 0.001939934 | 0.013204817 | 0.364236127 |
| ENSMUSG00000028048 | Gba | 14466 | protein_coding | 736.192578 | 0.364290076 | 0.073169825 | 4.978692732 | 6.40E-07 | 1.27E-05 | 0.364290076 |
| ENSMUSG00000014592 | Camta1 | 100072 | protein_coding | 170.58557 | 0.364315146 | 0.196471554 | 1.854289529 | 0.063697749 | 0.183202917 | 0.364315146 |
| ENSMUSG00000027660 | Skil | 20482 | protein_coding | 1644.214093 | 0.364336494 | 0.097608369 | 3.732635823 | 0.000189486 | 0.001948444 | 0.364336494 |
| ENSMUSG00000051721 | Wdcp | 238037 | protein_coding | 47.37195576 | -0.36439796 | 0.225907786 | -1.613038516 | 0.10673614 | 0.259516599 | 0.36439796 |
| ENSMUSG00000028389 | Zfp37 | 22696 | protein_coding | 153.0432188 | 0.364513417 | 0.23959972 | 1.521343249 | 0.128173722 | 0.294121538 | 0.364513417 |
| ENSMUSG00000072825 | Cep170b | 217882 | protein_coding | 1787.526214 | -0.364549305 | 0.130957923 | -2.78371325 | 0.005374052 | 0.029547084 | 0.364549305 |
| ENSMUSG00000100210 | H3c7 | 260423 | protein_coding | 1257.146353 | 0.364607547 | 0.188660345 | 1.932613586 | 0.053283817 | 0.162074293 | 0.364607547 |
| ENSMUSG00000051177 | Plcb1 | 18795 | protein_coding | 137.4355019 | -0.364671806 | 0.137607512 | -2.650086481 | 0.008047117 | 0.040690311 | 0.364671806 |
| ENSMUSG00000028295 | Smim8 | 66291 | protein_coding | 216.5392269 | 0.364921081 | 0.120823135 | 3.02029146 | 0.002525315 | 0.016444491 | 0.364921081 |
| ENSMUSG00000063410 | Stk24 | 223255 | protein_coding | 1058.457907 | 0.364939646 | 0.072922163 | 5.004509379 | 5.60E-07 | 1.13E-05 | 0.364939646 |
| ENSMUSG00000047502 | Mroh7 | 381538 | protein_coding | 8.695231223 | 0.364971694 | 0.464396348 | 0.785905608 | 0.431922789 | 0.635828667 | 0.364971694 |
| ENSMUSG00000040272 | Accs | 329470 | protein_coding | 56.31258401 | -0.364971836 | 0.216446308 | -1.686200328 | 0.091757221 | 0.234780564 | 0.364971836 |
| ENSMUSG00000040570 | Rundc3b | 242819 | protein_coding | 379.9002097 | -0.364977369 | 0.087003151 | -4.194990213 | 2.73E-05 | 0.000363419 | 0.364977369 |

Supplementary Table 1

| ensembl_gene_id | external_gene_name | entrezgene_id | gene_biotype | baseMean | log2FoldChange | lfcSE | stat | pvalue | padj | absLog2FC |
|---|---|---|---|---|---|---|---|---|---|---|
| ENSMUSG00000025959 | Klf7 | 93691 | protein_coding | 243.4809328 | 0.365182023 | 0.126450943 | 2.887934369 | 0.003877808 | 0.022944176 | 0.365182023 |
| ENSMUSG00000026525 | Opn3 | 13603 | protein_coding | 19.07915213 | 0.365199768 | 0.275295295 | 1.326574682 | 0.184649416 | 0.374471084 | 0.365199768 |
| ENSMUSG00000031960 | Aars | 234734 | protein_coding | 1487.753121 | 0.365401829 | 0.180272232 | 2.026944608 | 0.042668078 | 0.138952595 | 0.365401829 |
| ENSMUSG00000014504 | Srp19 | 66384 | protein_coding | 402.8186206 | 0.365442251 | 0.104589982 | 3.4940464 | 0.000475758 | 0.004172841 | 0.365442251 |
| ENSMUSG00000047153 | Khnyn | 219094 | protein_coding | 1158.295621 | -0.365488433 | 0.106692806 | -3.425614575 | 0.00061341 | 0.005159802 | 0.365488433 |
| ENSMUSG00000022385 | Gtse1 | 29870 | protein_coding | 154.7182663 | 0.365609515 | 0.175255677 | 2.086149346 | 0.036965097 | 0.125753965 | 0.365609515 |
| ENSMUSG00000030306 | Tmtc1 | 387314 | protein_coding | 8.113900768 | 0.365710826 | 0.53267914 | 0.686549929 | 0.49236639 | 0.684090358 | 0.365710826 |
| ENSMUSG00000067567 | Hdac8 | 70315 | protein_coding | 111.1215205 | -0.365785409 | 0.143826456 | -2.543241475 | 0.01098293 | 0.051376242 | 0.365785409 |
| ENSMUSG00000041351 | Rap1gap | 110351 | protein_coding | 1857.659868 | -0.365820217 | 0.126322703 | -2.895918216 | 0.003780509 | 0.022477025 | 0.365820217 |
| ENSMUSG00000026896 | Ifih1 | 71586 | protein_coding | 1292.533381 | 0.365872181 | 0.087537934 | 4.17958436 | 2.92E-05 | 0.00038635 | 0.365872181 |
| ENSMUSG00000013539 | Tango2 | 27883 | protein_coding | 242.2668735 | 0.366067076 | 0.093497891 | 3.91524421 | 9.03E-05 | 0.001034893 | 0.366067076 |
| ENSMUSG00000007338 | Mrpl49 | 18120 | protein_coding | 478.581453 | -0.366067604 | 0.098928519 | -3.700324293 | 0.000215324 | 0.002172323 | 0.366067604 |
| ENSMUSG00000040717 | Il17rd | 171463 | protein_coding | 73.67757504 | -0.366219975 | 0.194288759 | -1.884926213 | 0.0594398 | 0.17500132 | 0.366219975 |
| ENSMUSG00000060260 | Pwwp2b | 101631 | protein_coding | 217.2185173 | -0.366341145 | 0.106978938 | -3.424423089 | 0.000616106 | 0.005175812 | 0.366341145 |
| ENSMUSG00000116895 | Gm3435 | 100041621 | protein_coding | 8.276869686 | -0.36644571 | 0.369174721 | -0.992607807 | 0.320901119 | 0.532256186 | 0.36644571 |
| ENSMUSG00000032553 | Srprb | 20818 | protein_coding | 645.8313018 | 0.366455417 | 0.119444394 | 3.068000141 | 0.002154965 | 0.014398457 | 0.366455417 |
| ENSMUSG00000061039 | Or8b53 | 258783 | protein_coding | 7.257603366 | -0.36660884 | 0.60135915 | -0.60963376 | 0.542104443 | 0.725112493 | 0.36660884 |
| ENSMUSG00000022012 | Enox1 | 239188 | protein_coding | 9.51960813 | -0.366694636 | 0.555795827 | -0.659765004 | 0.509404645 | 0.698441696 | 0.366694636 |
| ENSMUSG00000045775 | Slc16a5 | 217316 | protein_coding | 1282.495018 | 0.366704876 | 0.200849858 | 1.825766177 | 0.06788549 | 0.191512591 | 0.366704876 |
| ENSMUSG00000032184 | Lysmd2 | 70082 | protein_coding | 39.32773571 | 0.366785172 | 0.389017313 | 0.942850509 | 0.345757372 | 0.555667786 | 0.366785172 |
| ENSMUSG00000028063 | Lmna | 16905 | protein_coding | 4941.091401 | -0.366796155 | 0.154984335 | -2.366665996 | 0.017949123 | 0.073779002 | 0.366796155 |
| ENSMUSG00000055884 | Fancm | 104806 | protein_coding | 170.1671935 | 0.366838173 | 0.103617905 | 3.540297149 | 0.000399677 | 0.003631518 | 0.366838173 |
| ENSMUSG00000045316 | Fahd1 | 68636 | protein_coding | 675.3218688 | 0.366895665 | 0.144304555 | 2.542509251 | 0.011005969 | 0.051423605 | 0.366895665 |
| ENSMUSG00000041168 | Lonp1 | 74142 | protein_coding | 1036.221577 | -0.366959042 | 0.094295776 | -3.891574531 | 9.96E-05 | 0.001124282 | 0.366959042 |
| ENSMUSG00000039633 | Lonrf1 | 244421 | protein_coding | 69.26701768 | -0.366966568 | 0.198390191 | -1.849721335 | 0.064353724 | 0.184489941 | 0.366966568 |
| ENSMUSG00000057378 | Ryr3 | 20192 | protein_coding | 27.90351404 | -0.367017831 | 0.544470318 | -0.674082349 | 0.500258961 | 0.690778043 | 0.367017831 |
| ENSMUSG00000025289 | Prdx4 | 53381 | protein_coding | 362.7868915 | 0.36703909 | 0.131990512 | 2.780799034 | 0.005422529 | 0.029741752 | 0.36703909 |
| ENSMUSG00000021990 | Spata13 | 219140 | protein_coding | 692.9923149 | -0.367068703 | 0.106565289 | -3.444542848 | 0.000572026 | 0.00485156 | 0.367068703 |
| ENSMUSG00000046434 | Hnrnpa1 | 15382 | protein_coding | 2427.422271 | -0.367151662 | 0.074442993 | -4.931984139 | 8.14E-07 | 1.57E-05 | 0.367151662 |
| ENSMUSG00000027854 | Sike1 | 66641 | protein_coding | 863.9555125 | -0.367175099 | 0.093663857 | -3.92013643 | 8.85E-05 | 0.001016234 | 0.367175099 |
| ENSMUSG00000022240 | Ctnnd2 | 18163 | protein_coding | 12.16072088 | -0.367244789 | 0.580012189 | -0.633167365 | 0.526624355 | 0.712260825 | 0.367244789 |
| ENSMUSG00000031429 | Psmd10 | 53380 | protein_coding | 331.7451825 | 0.367246936 | 0.08587479 | 4.276539556 | 1.90E-05 | 0.000264984 | 0.367246936 |
| ENSMUSG00000022838 | Eaf2 | 106389 | protein_coding | 9.179989594 | -0.367351005 | 0.423484668 | -0.867448181 | 0.385696492 | 0.593661974 | 0.367351005 |
| ENSMUSG00000059834 | Sclt1 | 67161 | protein_coding | 237.2902487 | 0.367391145 | 0.135831178 | 2.704763004 | 0.006835314 | 0.035754639 | 0.367391145 |
| ENSMUSG00000021196 | Pfkp | 56421 | protein_coding | 2376.451393 | 0.367528646 | 0.113771102 | 3.230421785 | 0.001236077 | 0.009105472 | 0.367528646 |
| ENSMUSG00000070436 | Serpinh1 | 12406 | protein_coding | 12.36994107 | 0.367631002 | 0.381014851 | 0.964873156 | 0.334608348 | 0.545293316 | 0.367631002 |
| ENSMUSG00000027163 | Commd9 | 76501 | protein_coding | 299.6875768 | 0.367672135 | 0.140531845 | 2.616290526 | 0.008889089 | 0.043888843 | 0.367672135 |
| ENSMUSG00000039153 | Runx2 | 12393 | protein_coding | 8.122714466 | -0.36780441 | 1.031266584 | -0.356653086 | 0.72135153 | 0.843589367 | 0.36780441 |

Supplementary Table 1

| ensembl_gene_id | external_gene_name | entrezgene_id | gene_biotype | baseMean | log2FoldChange | lfcSE | stat | pvalue | padj | absLog2FC |
|---|---|---|---|---|---|---|---|---|---|---|
| ENSMUSG00000032526 | Ss18l2 | 26901 | protein_coding | 235.4074273 | 0.367807701 | 0.099216813 | 3.707110607 | 0.000209637 | 0.002128174 | 0.367807701 |
| ENSMUSG00000030800 | Prss8 | 76560 | protein_coding | 716.8063351 | 0.368022459 | 0.085656082 | 4.296512895 | 1.74E-05 | 0.000243919 | 0.368022459 |
| ENSMUSG00000057246 | BC051142 | 407788 | protein_coding | 29.69581636 | -0.368112371 | 0.318628171 | -1.155303909 | 0.247965996 | 0.453820795 | 0.368112371 |
| ENSMUSG00000030823 | 9130019O22Rik | 78921 | protein_coding | 101.7596944 | -0.368123579 | 0.132267613 | -2.783172477 | 0.005383018 | 0.029586167 | 0.368123579 |
| ENSMUSG00000054676 | 1600014C10Rik | 72244 | protein_coding | 144.0621373 | 0.368142865 | 0.112131114 | 3.283146412 | 0.001026553 | 0.007823417 | 0.368142865 |
| ENSMUSG00000026531 | Mptx1 | 66289 | protein_coding | 19903.62136 | -0.368164358 | 0.487400137 | -0.755363673 | 0.450030803 | 0.650554063 | 0.368164358 |
| ENSMUSG00000107722 | 2900060B14Rik | 68204 | lncRNA | 11.1685244 | 0.368206754 | 0.379918345 | 0.969173399 | 0.332458681 | 0.543456188 | 0.368206754 |
| ENSMUSG00000103906 | Tigd5 | 105734 | protein_coding | 53.49498026 | 0.368224816 | 0.186720015 | 1.972069333 | 0.048601692 | 0.152718042 | 0.368224816 |
| ENSMUSG00000001211 | Agpat3 | 28169 | protein_coding | 785.9583504 | 0.368230184 | 0.059127377 | 6.227744305 | 4.73E-10 | 1.48E-08 | 0.368230184 |
| ENSMUSG00000020706 | Ftsj3 | 56095 | protein_coding | 602.3851011 | -0.368376895 | 0.165129149 | -2.230841109 | 0.025691656 | 0.096581707 | 0.368376895 |
| ENSMUSG00000022285 | Ywhaz | 22631 | protein_coding | 4011.977791 | 0.368479308 | 0.115655207 | 3.186015713 | 0.001442468 | 0.010332564 | 0.368479308 |
| ENSMUSG00000053279 | Aldh1a1 | 11668 | protein_coding | 2912.384379 | -0.368528959 | 0.299906243 | -1.228813896 | 0.21914159 | 0.419328117 | 0.368528959 |
| ENSMUSG00000028637 | Ccdc30 | 73332 | protein_coding | 12.48060856 | 0.368588736 | 0.615961405 | 0.59839583 | 0.549575847 | 0.731093174 | 0.368588736 |
| ENSMUSG00000027804 | Ppid | 67738 | protein_coding | 562.7220047 | -0.368681827 | 0.150305653 | -2.452880645 | 0.014171735 | 0.061903744 | 0.368681827 |
| ENSMUSG00000033508 | Asprv1 | 67855 | protein_coding | 7.421503491 | -0.369036779 | 0.406776205 | -0.907223125 | 0.364288817 | 0.573132387 | 0.369036779 |
| ENSMUSG00000037628 | Cdkn3 | 72391 | protein_coding | 181.121624 | 0.369070203 | 0.156585239 | 2.3569923 | 0.018423629 | 0.075186155 | 0.369070203 |
| ENSMUSG00000049791 | Fzd4 | 14366 | protein_coding | 99.31841382 | -0.369205017 | 0.197933225 | -1.86530087 | 0.062139243 | 0.17987433 | 0.369205017 |
| ENSMUSG00000038843 | Gcnt1 | 14537 | protein_coding | 410.5746881 | -0.369267988 | 0.16859066 | -2.190322936 | 0.028500825 | 0.103762545 | 0.369267988 |
| ENSMUSG00000050022 | Amz1 | 231842 | protein_coding | 7.692881991 | 0.369412566 | 0.474896693 | 0.777879845 | 0.436639851 | 0.639404206 | 0.369412566 |
| ENSMUSG00000001036 | Epn2 | 13855 | protein_coding | 216.3163205 | -0.369414674 | 0.092694394 | -3.985296828 | 6.74E-05 | 0.000798721 | 0.369414674 |
| ENSMUSG00000036613 | Eipr1 | 380752 | protein_coding | 277.9312102 | 0.369494274 | 0.112097804 | 3.296177637 | 0.0009801 | 0.007556165 | 0.369494274 |
| ENSMUSG00000050592 | Fam78a | 241303 | protein_coding | 42.49192244 | -0.369574242 | 0.279730785 | -1.321178294 | 0.186441922 | 0.376888975 | 0.369574242 |
| ENSMUSG00000022995 | Enah | 13800 | protein_coding | 12.25870913 | -0.369604614 | 0.827045136 | -0.446897754 | 0.654948886 | 0.800302454 | 0.369604614 |
| ENSMUSG00000038296 | Galnt18 | 233733 | protein_coding | 12.35222703 | 0.369635868 | 0.42384925 | 0.872092773 | 0.383157769 | 0.590807095 | 0.369635868 |
| ENSMUSG00000033039 | Micall1 | 27008 | protein_coding | 135.2767342 | -0.369731204 | 0.152737574 | -2.420695802 | 0.015490834 | 0.066220614 | 0.369731204 |
| ENSMUSG00000025265 | Fgd1 | 14163 | protein_coding | 34.85591933 | -0.369792031 | 0.271567774 | -1.361693346 | 0.173294682 | 0.359125383 | 0.369792031 |
| ENSMUSG00000006398 | Cdc20 | 107995 | protein_coding | 725.5315957 | 0.369882218 | 0.146681838 | 2.52166337 | 0.011680142 | 0.053769164 | 0.369882218 |
| ENSMUSG00000041679 | Fbxl9 | 234684 | protein_coding | 37.94437191 | -0.369936365 | 0.237690053 | -1.556381345 | 0.119617439 | 0.280498094 | 0.369936365 |
| ENSMUSG00000061607 | Mdc1 | 240087 | protein_coding | 290.1483535 | -0.369949814 | 0.165036444 | -2.241624973 | 0.024985621 | 0.094350632 | 0.369949814 |
| ENSMUSG00000049536 | Tceal1 | 237052 | protein_coding | 15.19135011 | -0.370228424 | 0.397328012 | -0.931795425 | 0.351442259 | 0.561349008 | 0.370228424 |
| ENSMUSG00000020363 | Gfpt2 | 14584 | protein_coding | 11.7454114 | 0.370276744 | 0.482927566 | 0.766733503 | 0.44323998 | 0.644271437 | 0.370276744 |
| ENSMUSG00000028772 | Zcchc17 | 619605 | protein_coding | 386.3439034 | 0.370406235 | 0.108558694 | 3.412036573 | 0.000644795 | 0.005379931 | 0.370406235 |
| ENSMUSG00000052565 | H1f3 | 14957 | protein_coding | 2762.860646 | 0.370421635 | 0.158540351 | 2.336450194 | 0.019467791 | 0.07844244 | 0.370421635 |
| ENSMUSG00000104900 | 4930596I21Rik | 78229 | lncRNA | 8.057999101 | -0.370500187 | 0.474346268 | -0.781075371 | 0.434758167 | 0.637880258 | 0.370500187 |
| ENSMUSG00000023980 | Taf8 | 63856 | protein_coding | 271.3857258 | 0.370538225 | 0.103093338 | 3.594201446 | 0.000325388 | 0.003037972 | 0.370538225 |
| ENSMUSG00000047246 | H2bc6 | 319179 | protein_coding | 934.1567342 | 0.37056111 | 0.152786044 | 2.42535967 | 0.015293227 | 0.065534172 | 0.37056111 |
| ENSMUSG00000021824 | Ap3m1 | 55946 | protein_coding | 1253.433141 | 0.370571196 | 0.076736905 | 4.829113151 | 1.37E-06 | 2.51E-05 | 0.370571196 |
| ENSMUSG00000053801 | Grwd1 | 101612 | protein_coding | 211.1084862 | -0.370616358 | 0.265641301 | -1.395175961 | 0.162962784 | 0.345459967 | 0.370616358 |

Supplementary Table 1

| ensembl_gene_id | external_gene_name | entrezgene_id | gene_biotype | baseMean | log2FoldChange | lfcSE | stat | pvalue | padj | absLog2FC |
|---|---|---|---|---|---|---|---|---|---|---|
| ENSMUSG00000001300 | Efnb2 | 13642 | protein_coding | 1789.445387 | 0.370739511 | 0.191803067 | 1.932917533 | 0.053246358 | 0.162007256 | 0.370739511 |
| ENSMUSG00000040761 | Spen | 56381 | protein_coding | 620.8251737 | -0.370742052 | 0.212467073 | -1.744938864 | 0.080995511 | 0.215951876 | 0.370742052 |
| ENSMUSG00000061589 | Dot1l | 208266 | protein_coding | 357.553693 | -0.370879305 | 0.193111209 | -1.920547792 | 0.054788742 | 0.165091769 | 0.370879305 |
| ENSMUSG00000004558 | Ndrg2 | 29811 | protein_coding | 1365.362889 | -0.370912579 | 0.152137767 | -2.438004629 | 0.014768585 | 0.063783628 | 0.370912579 |
| ENSMUSG00000046338 | Gpat2 | 215456 | protein_coding | 11.16680798 | -0.371051115 | 0.479378755 | -0.774024946 | 0.438916039 | 0.641498869 | 0.371051115 |
| ENSMUSG00000001632 | Brpf1 | 78783 | protein_coding | 462.5826917 | -0.371063403 | 0.116117265 | -3.195592007 | 0.001395443 | 0.010070962 | 0.371063403 |
| ENSMUSG00000032530 | Lyzl4 | 69032 | protein_coding | 20.9842319 | -0.371144265 | 0.228565052 | -1.623801457 | 0.104418189 | 0.255636784 | 0.371144265 |
| ENSMUSG00000028840 | Zfp593 | 68040 | protein_coding | 10.52850658 | -0.371241686 | 0.358035859 | -1.036884091 | 0.299789879 | 0.508687887 | 0.371241686 |
| ENSMUSG00000022863 | Btg3 | 12228 | protein_coding | 93.54050456 | -0.371255577 | 0.14999935 | -2.475047905 | 0.013321831 | 0.05933981 | 0.371255577 |
| ENSMUSG00000005628 | Tmod4 | 50874 | protein_coding | 7.710163765 | -0.371361363 | 0.446975318 | -0.830831924 | 0.406068586 | 0.613120797 | 0.371361363 |
| ENSMUSG00000086358 | Gm13270 | 105244084 | lncRNA | 7.615662211 | 0.371370623 | 0.606222 | 0.612598393 | 0.540141921 | 0.723156983 | 0.371370623 |
| ENSMUSG00000034731 | Dgkh | 380921 | protein_coding | 1116.522162 | -0.371396466 | 0.142004149 | -2.615391652 | 0.008912519 | 0.043963644 | 0.371396466 |
| ENSMUSG00000038774 | Ascc3 | 77987 | protein_coding | 1418.940306 | 0.371430431 | 0.05954587 | 6.23771939 | 4.44E-10 | 1.40E-08 | 0.371430431 |
| ENSMUSG00000039887 | Alg14 | 66789 | protein_coding | 293.0422434 | 0.371488126 | 0.119047055 | 3.12051504 | 0.001805351 | 0.012421437 | 0.371488126 |
| ENSMUSG00000021676 | Iqgap2 | 544963 | protein_coding | 6451.718286 | 0.371580417 | 0.124730084 | 2.979076137 | 0.002891189 | 0.018216322 | 0.371580417 |
| ENSMUSG00000044361 | BC024139 | 271278 | protein_coding | 25.81371654 | -0.371610127 | 0.332773023 | -1.116707491 | 0.264119414 | 0.471889403 | 0.371610127 |
| ENSMUSG00000052397 | Ezr | 22350 | protein_coding | 6354.331959 | -0.37163082 | 0.148817568 | -2.49722412 | 0.012516982 | 0.056655438 | 0.37163082 |
| ENSMUSG00000009545 | Kcnq1 | 16535 | protein_coding | 991.9985723 | -0.371803793 | 0.089134108 | -4.171285304 | 3.03E-05 | 0.000399369 | 0.371803793 |
| ENSMUSG00000004113 | Cacna1b | 12287 | protein_coding | 130.2351576 | -0.371884237 | 0.151841525 | -2.449160318 | 0.01431897 | 0.062280872 | 0.371884237 |
| ENSMUSG00000039086 | Ss18l1 | 269397 | protein_coding | 116.7330587 | -0.371988646 | 0.142646604 | -2.607763771 | 0.009113582 | 0.044637497 | 0.371988646 |
| ENSMUSG00000049013 | Prss48 | 368202 | protein_coding | 12.12668366 | -0.372056353 | 0.435871527 | -0.853591779 | 0.393331215 | 0.601187947 | 0.372056353 |
| ENSMUSG00000013076 | Amotl1 | 75723 | protein_coding | 174.9215581 | 0.372170605 | 0.138089821 | 2.695134234 | 0.007036028 | 0.036625014 | 0.372170605 |
| ENSMUSG00000040605 | Bace2 | 56175 | protein_coding | 990.584094 | -0.372172619 | 0.128594437 | -2.894158013 | 0.003801768 | 0.022561311 | 0.372172619 |
| ENSMUSG00000021703 | Serinc5 | 218442 | protein_coding | 1370.984894 | -0.372274253 | 0.13272747 | -2.804801857 | 0.005034752 | 0.028207362 | 0.372274253 |
| ENSMUSG00000033016 | Nfatc1 | 18018 | protein_coding | 146.9198203 | -0.372307358 | 0.131649348 | -2.828022792 | 0.004683647 | 0.026691863 | 0.372307358 |
| ENSMUSG00000039345 | Mettl22 | 239706 | protein_coding | 58.4401634 | -0.372332329 | 0.169584106 | -2.195561469 | 0.028123347 | 0.102873309 | 0.372332329 |
| ENSMUSG00000024968 | Rcor2 | 104383 | protein_coding | 16.53482632 | -0.37255819 | 0.319829019 | -1.164866749 | 0.244072935 | 0.449310115 | 0.37255819 |
| ENSMUSG00000002006 | Pdzd4 | 245469 | protein_coding | 28.07431208 | -0.372597206 | 0.303696093 | -1.226875204 | 0.2198695 | 0.420165734 | 0.372597206 |
| ENSMUSG00000027082 | Tfpi | 21788 | protein_coding | 57.0102442 | -0.372629369 | 0.214185634 | -1.73974959 | 0.081902998 | 0.217679116 | 0.372629369 |
| ENSMUSG00000090489 | Gm17415 | 108167670 | processed_pseudogene | 22.65362703 | 0.37278556 | 0.346384936 | 1.076217586 | 0.281829956 | 0.490653355 | 0.37278556 |
| ENSMUSG00000032245 | Cln6 | 76524 | protein_coding | 297.4877942 | -0.372809275 | 0.098050963 | -3.802199034 | 0.000143417 | 0.001536532 | 0.372809275 |
| ENSMUSG00000019890 | Nts | 67405 | protein_coding | 154.6113149 | -0.372820507 | 0.350762406 | -1.062886161 | 0.28783358 | 0.496977047 | 0.372820507 |
| ENSMUSG00000031229 | Atrx | 22589 | protein_coding | 2515.762154 | 0.372981022 | 0.07308546 | 5.103354624 | 3.34E-07 | 6.99E-06 | 0.372981022 |
| ENSMUSG00000031749 | St3gal2 | 20444 | protein_coding | 12.47856351 | -0.373045472 | 0.326320776 | -1.143186396 | 0.252961209 | 0.459205682 | 0.373045472 |
| ENSMUSG00000027999 | Pla2g12a | 66350 | protein_coding | 126.9666126 | -0.373078076 | 0.167863368 | -2.22251037 | 0.026248834 | 0.097959566 | 0.373078076 |
| ENSMUSG00000014444 | Piezo1 | 234839 | protein_coding | 198.098169 | -0.373286037 | 0.165705301 | -2.252710289 | 0.024277422 | 0.092377156 | 0.373286037 |
| ENSMUSG00000079547 | H2-DMb1 | 14999 | protein_coding | 51.84097805 | 0.373422005 | 0.439497529 | 0.849656667 | 0.395515997 | 0.603774085 | 0.373422005 |
| ENSMUSG00000055653 | Gpc3 | 14734 | protein_coding | 8.835545597 | 0.373573494 | 0.466990351 | 0.79995977 | 0.423734106 | 0.628420563 | 0.373573494 |

| ensembl_gene_id | external_gene_name | entrezgene_id | gene_biotype | baseMean | log2FoldChange | lfcSE | stat | pvalue | padj | absLog2FC |
|---|---|---|---|---|---|---|---|---|---|---|
| ENSMUSG00000096002 | Vmn2r53 | 637908 | protein_coding | 10.72816955 | -0.373600449 | 1.734041497 | -0.215450697 | 0.829415948 | 0.909405742 | 0.373600449 |
| ENSMUSG00000040860 | Crocc | 230872 | protein_coding | 105.3629808 | -0.373609253 | 0.180066711 | -2.074837991 | 0.038001552 | 0.128429604 | 0.373609253 |
| ENSMUSG00000025245 | Lztfl1 | 93730 | protein_coding | 407.7622306 | 0.373675976 | 0.080663588 | 4.632523606 | 3.61E-06 | 6.00E-05 | 0.373675976 |
| ENSMUSG00000021250 | Fos | 14281 | protein_coding | 16880.71975 | -0.373800852 | 0.31766457 | -1.17671559 | 0.239309043 | 0.443424073 | 0.373800852 |
| ENSMUSG00000032596 | Uba7 | 74153 | protein_coding | 462.8409367 | 0.373812582 | 0.093996169 | 3.976891678 | 6.98E-05 | 0.00082563 | 0.373812582 |
| ENSMUSG00000030286 | Emc3 | 66087 | protein_coding | 951.0182069 | 0.373818395 | 0.085431981 | 4.37562599 | 1.21E-05 | 0.000176939 | 0.373818395 |
| ENSMUSG00000085645 | Hoxb5os | 75395 | lncRNA | 26.46338998 | -0.37392705 | 0.3039543 | -1.230208126 | 0.218619178 | 0.418680566 | 0.37392705 |
| ENSMUSG00000051343 | Rab11fip5 | 52055 | protein_coding | 30.89682189 | -0.373963645 | 0.253105118 | -1.477503293 | 0.139540776 | 0.311289283 | 0.373963645 |
| ENSMUSG00000079197 | Psme2 | 19188 | protein_coding | 1721.297898 | 0.373983284 | 0.124635031 | 3.000627359 | 0.002694241 | 0.017241292 | 0.373983284 |
| ENSMUSG00000039929 | Urb1 | 207932 | protein_coding | 204.5316555 | -0.373997071 | 0.250509877 | -1.492943413 | 0.135452 | 0.305073452 | 0.373997071 |
| ENSMUSG00000036534 | Slc38a7 | 234595 | protein_coding | 138.4369248 | -0.374292558 | 0.117145921 | -3.195096794 | 0.00139784 | 0.01008369 | 0.374292558 |
| ENSMUSG00000014959 | Gorasp2 | 70231 | protein_coding | 1806.023571 | 0.374359321 | 0.087642416 | 4.271439986 | 1.94E-05 | 0.000270335 | 0.374359321 |
| ENSMUSG00000029464 | Gpn3 | 68080 | protein_coding | 254.195909 | 0.374527921 | 0.103282862 | 3.626234925 | 0.000287584 | 0.002737976 | 0.374527921 |
| ENSMUSG00000020015 | Cdk17 | 237459 | protein_coding | 266.4308017 | -0.374588172 | 0.0958367 | -3.908608823 | 9.28E-05 | 0.001059916 | 0.374588172 |
| ENSMUSG00000025060 | Slk | 20874 | protein_coding | 2826.870265 | -0.37461129 | 0.083906885 | -4.464607272 | 8.02E-06 | 0.000122854 | 0.37461129 |
| ENSMUSG00000042121 | Ssh1 | 231637 | protein_coding | 119.4373755 | -0.374689793 | 0.191854115 | -1.952993256 | 0.050820394 | 0.157122878 | 0.374689793 |
| ENSMUSG00000049690 | Nckap5 | 210356 | protein_coding | 66.0815021 | 0.374691739 | 0.275238793 | 1.361333317 | 0.173408379 | 0.359267515 | 0.374691739 |
| ENSMUSG00000020190 | Mknk2 | 17347 | protein_coding | 2224.767956 | -0.375181892 | 0.121372868 | -3.091151248 | 0.00199382 | 0.013479445 | 0.375181892 |
| ENSMUSG00000001948 | Spa17 | 20686 | protein_coding | 16.31140891 | -0.375636211 | 0.309177087 | -1.214954883 | 0.224383299 | 0.425999491 | 0.375636211 |
| ENSMUSG00000044786 | Zfp36 | 22695 | protein_coding | 2321.509253 | -0.375931177 | 0.260491993 | -1.443158285 | 0.148975888 | 0.326304722 | 0.375931177 |
| ENSMUSG00000036580 | Spg20 | 229285 | protein_coding | 203.2497379 | 0.375960192 | 0.134844342 | 2.788105053 | 0.005301735 | 0.029280871 | 0.375960192 |
| ENSMUSG00000027314 | Dll4 | 54485 | protein_coding | 150.3661542 | -0.376028528 | 0.30931672 | -1.215674756 | 0.224108844 | 0.425764035 | 0.376028528 |
| ENSMUSG00000095098 | Ccdc85b | 240514 | protein_coding | 139.5498985 | -0.376104816 | 0.122258187 | -3.076315983 | 0.002095756 | 0.014061165 | 0.376104816 |
| ENSMUSG00000044340 | Phlpp1 | 98432 | protein_coding | 449.7618596 | -0.376112666 | 0.153493817 | -2.450344079 | 0.014271976 | 0.06215383 | 0.376112666 |
| ENSMUSG00000017144 | Rnd3 | 74194 | protein_coding | 949.1783442 | -0.37617346 | 0.148687977 | -2.529952112 | 0.01140781 | 0.052836171 | 0.37617346 |
| ENSMUSG00000020752 | Recql5 | 170472 | protein_coding | 525.2784562 | -0.376190007 | 0.146247803 | -2.572278004 | 0.010103173 | 0.048209507 | 0.376190007 |
| ENSMUSG00000033953 | Ppp3r1 | 19058 | protein_coding | 1194.451721 | 0.376194079 | 0.083316204 | 4.515257051 | 6.32E-06 | 9.93E-05 | 0.376194079 |
| ENSMUSG00000050954 | Zfp169 | 67911 | protein_coding | 122.3397106 | -0.37620388 | 0.176823548 | -2.127566624 | 0.033373032 | 0.116720261 | 0.37620388 |
| ENSMUSG00000001918 | Slc1a5 | 20514 | protein_coding | 1977.048733 | -0.37622802 | 0.091051133 | -4.132052032 | 3.60E-05 | 0.000461834 | 0.37622802 |
| ENSMUSG00000042688 | Mapk6 | 50772 | protein_coding | 1685.611561 | -0.376443138 | 0.078995097 | -4.765398757 | 1.88E-06 | 3.31E-05 | 0.376443138 |
| ENSMUSG00000054364 | Rhob | 11852 | protein_coding | 3871.936731 | 0.37662223 | 0.298880569 | 1.260109454 | 0.20762988 | 0.404082965 | 0.37662223 |
| ENSMUSG00000064128 | Cenpj | 219103 | protein_coding | 99.00227645 | 0.37674197 | 0.126971826 | 2.967130446 | 0.003005934 | 0.018798004 | 0.37674197 |
| ENSMUSG00000075028 | Prdm11 | 100042784 | protein_coding | 23.59550896 | -0.376760702 | 0.285521688 | -1.319551956 | 0.186984652 | 0.377668215 | 0.376760702 |
| ENSMUSG00000042670 | Immp1l | 66541 | protein_coding | 345.4075079 | 0.37677003 | 0.120633213 | 3.123269475 | 0.001788539 | 0.012335011 | 0.37677003 |
| ENSMUSG00000064254 | Ethe1 | 66071 | protein_coding | 5067.352287 | -0.376926563 | 0.15426718 | -2.443336049 | 0.01455218 | 0.063044022 | 0.376926563 |
| ENSMUSG00000114828 | AI463229 | 100042802 | lncRNA | 38.52618302 | -0.376930447 | 0.235581576 | -1.599999686 | 0.109598653 | 0.263874388 | 0.376930447 |
| ENSMUSG00000047749 | Zc3hav1l | 209032 | protein_coding | 26.6371058 | -0.377062848 | 0.265522889 | -1.420076625 | 0.155585375 | 0.335658113 | 0.377062848 |
| ENSMUSG00000031723 | Txnl4b | 234723 | protein_coding | 117.3338076 | 0.377089609 | 0.157745799 | 2.390489072 | 0.01682595 | 0.07043462 | 0.377089609 |

| ensembl_gene_id | external_gene_name | entrezgene_id | gene_biotype | baseMean | log2FoldChange | lfcSE | stat | pvalue | padj | absLog2FC |
|---|---|---|---|---|---|---|---|---|---|---|
| ENSMUSG00000073791 | Efcab7 | 230500 | protein_coding | 53.69272979 | 0.377090437 | 0.194617417 | 1.937598614 | 0.052672212 | 0.160998463 | 0.377090437 |
| ENSMUSG00000036611 | Eepd1 | 67484 | protein_coding | 112.5404032 | -0.377292569 | 0.155248917 | -2.430242838 | 0.015088709 | 0.064954855 | 0.377292569 |
| ENSMUSG00000000394 | Gcg | 14526 | protein_coding | 704.1520501 | -0.377418022 | 0.182372251 | -2.069492589 | 0.038499886 | 0.129646128 | 0.377418022 |
| ENSMUSG00000044734 | Serpinb1a | 66222 | protein_coding | 4997.925037 | 0.377451344 | 0.115835641 | 3.258507832 | 0.001119998 | 0.008418747 | 0.377451344 |
| ENSMUSG00000020086 | Macroh2a2 | 404634 | protein_coding | 14.46089145 | -0.377509649 | 0.355581508 | -1.061668395 | 0.288386252 | 0.497392119 | 0.377509649 |
| ENSMUSG00000050188 | Lsm10 | 116748 | protein_coding | 110.5698122 | 0.377605863 | 0.146838235 | 2.571577242 | 0.010123643 | 0.048249367 | 0.377605863 |
| ENSMUSG00000057706 | Mex3b | 108797 | protein_coding | 30.05920571 | 0.377792281 | 0.222502285 | 1.6979254 | 0.089521843 | 0.231065291 | 0.377792281 |
| ENSMUSG00000004633 | Chn2 | 69993 | protein_coding | 345.1626507 | -0.377839535 | 0.193835237 | -1.949281978 | 0.05126176 | 0.158099111 | 0.377839535 |
| ENSMUSG00000047146 | Tet1 | 52463 | protein_coding | 31.71040532 | -0.378319025 | 0.396384566 | -0.954424207 | 0.339868958 | 0.550313384 | 0.378319025 |
| ENSMUSG00000040435 | Ppp1r15a | 17872 | protein_coding | 414.6580291 | -0.378332914 | 0.229534132 | -1.648264292 | 0.099298449 | 0.247167635 | 0.378332914 |
| ENSMUSG00000026727 | Rsu1 | 20163 | protein_coding | 259.3598847 | 0.378376919 | 0.10899399 | 3.471539299 | 0.000517484 | 0.004482044 | 0.378376919 |
| ENSMUSG00000032498 | Mlh1 | 17350 | protein_coding | 301.6601923 | 0.378435557 | 0.08296316 | 4.56148916 | 5.08E-06 | 8.21E-05 | 0.378435557 |
| ENSMUSG00000012422 | Tmem167 | 66074 | protein_coding | 1045.591064 | 0.378479325 | 0.087460864 | 4.3274135 | 1.51E-05 | 0.000215136 | 0.378479325 |
| ENSMUSG00000089762 | Ier5l | 72500 | protein_coding | 17.84907122 | -0.378534553 | 0.340586653 | -1.111419222 | 0.266387942 | 0.474134221 | 0.378534553 |
| ENSMUSG00000037185 | Krt80 | 74127 | protein_coding | 6.972112141 | -0.378536422 | 0.709652405 | -0.533411033 | 0.593749082 | 0.76209791 | 0.378536422 |
| ENSMUSG00000055923 | Aasdh | 231326 | protein_coding | 130.5108467 | -0.378575121 | 0.159356066 | -2.375655537 | 0.017517808 | 0.07253019 | 0.378575121 |
| ENSMUSG00000032218 | Ccnb2 | 12442 | protein_coding | 623.505495 | 0.378585214 | 0.147225205 | 2.571470114 | 0.010126776 | 0.04824986 | 0.378585214 |
| ENSMUSG00000028347 | Tmeff1 | 230157 | protein_coding | 173.9628062 | -0.378627307 | 0.153482121 | -2.466914737 | 0.01362828 | 0.060230441 | 0.378627307 |
| ENSMUSG00000044252 | Osbpl1a | 64291 | protein_coding | 193.4383493 | -0.378714852 | 0.153419417 | -2.46849362 | 0.013568307 | 0.06014684 | 0.378714852 |
| ENSMUSG00000002833 | Hdgfl2 | 15193 | protein_coding | 462.5338705 | 0.378926327 | 0.076137254 | 4.976884595 | 6.46E-07 | 1.28E-05 | 0.378926327 |
| ENSMUSG00000042439 | Zfp532 | 328977 | protein_coding | 132.88927 | -0.37903761 | 0.131256604 | -2.887760309 | 0.003879954 | 0.022948351 | 0.37903761 |
| ENSMUSG00000033685 | Ucp2 | 22228 | protein_coding | 2697.993324 | -0.379080314 | 0.064985895 | -5.83327066 | 5.44E-09 | 1.50E-07 | 0.379080314 |
| ENSMUSG00000031327 | Chic1 | 12212 | protein_coding | 349.0323752 | 0.379085281 | 0.144534116 | 2.622808305 | 0.008720832 | 0.043232312 | 0.379085281 |
| ENSMUSG00000056025 | Clca3a1 | 12722 | protein_coding | 45.11095713 | 0.379132708 | 0.26793122 | 1.415037443 | 0.157057518 | 0.337873062 | 0.379132708 |
| ENSMUSG00000031059 | Ndufb11 | 104130 | protein_coding | 1342.171242 | 0.379185478 | 0.104747683 | 3.619989174 | 0.000294615 | 0.002798231 | 0.379185478 |
| ENSMUSG00000024875 | Yif1a | 68090 | protein_coding | 803.9143044 | 0.379324857 | 0.111881319 | 3.390421755 | 0.000697852 | 0.005735491 | 0.379324857 |
| ENSMUSG00000062866 | Phactr2 | 215789 | protein_coding | 300.8280763 | -0.379383038 | 0.155995182 | -2.43201767 | 0.015014975 | 0.064742425 | 0.379383038 |
| ENSMUSG00000032021 | Crtam | 54698 | protein_coding | 6.913429709 | 0.379656038 | 0.693473354 | 0.547470261 | 0.584055697 | 0.754610572 | 0.379656038 |
| ENSMUSG00000079470 | Utp14b | 195434 | protein_coding | 1698.842926 | -0.379656665 | 0.168135081 | -2.258045519 | 0.023942822 | 0.091497424 | 0.379656665 |
| ENSMUSG00000022659 | Gcsam | 14525 | protein_coding | 7.433080659 | -0.379657559 | 0.515244877 | -0.73684878 | 0.461214315 | 0.659378689 | 0.379657559 |
| ENSMUSG00000020823 | Sec14l1 | 74136 | protein_coding | 401.7576705 | -0.379698228 | 0.102364185 | -3.709287865 | 0.000207843 | 0.002115351 | 0.379698228 |
| ENSMUSG00000068732 | Tmem167b | 67495 | protein_coding | 1173.000684 | 0.379815151 | 0.063094778 | 6.019755757 | 1.75E-09 | 5.15E-08 | 0.379815151 |
| ENSMUSG00000028218 | Cibar1 | 68099 | protein_coding | 118.587347 | 0.379825312 | 0.151414751 | 2.508509308 | 0.012124177 | 0.055333495 | 0.379825312 |
| ENSMUSG00000021876 | Rnase4 | 58809 | protein_coding | 2363.724129 | -0.379982557 | 0.129691334 | -2.929899366 | 0.003390718 | 0.020708342 | 0.379982557 |
| ENSMUSG00000029320 | 1700016H13Rik | 74218 | protein_coding | 13.40522778 | -0.3801011 | 0.335356489 | -1.133424019 | 0.257036232 | 0.463812519 | 0.3801011 |
| ENSMUSG00000038593 | Tctn1 | 654470 | protein_coding | 64.25174089 | -0.380211467 | 0.173039255 | -2.197255567 | 0.028002199 | 0.10248599 | 0.380211467 |
| ENSMUSG00000032392 | Parp16 | 214424 | protein_coding | 214.2743721 | 0.380391319 | 0.133404848 | 2.851405522 | 0.004352642 | 0.025133906 | 0.380391319 |
| ENSMUSG00000047613 | A430005L14Rik | 97159 | protein_coding | 284.0518334 | 0.380423183 | 0.105764124 | 3.596901936 | 0.00032203 | 0.003011915 | 0.380423183 |

| ensembl_gene_id | external_gene_name | entrezgene_id | gene_biotype | baseMean | log2FoldChange | lfcSE | stat | pvalue | padj | absLog2FC |
|---|---|---|---|---|---|---|---|---|---|---|
| ENSMUSG00000039824 | Myl6b | 216459 | protein_coding | 15.72638389 | -0.380450376 | 0.284869296 | -1.335526088 | 0.181704223 | 0.370302264 | 0.380450376 |
| ENSMUSG00000043323 | Fbrsl1 | 381668 | protein_coding | 292.1457852 | -0.380459408 | 0.132962096 | -2.861412543 | 0.004217578 | 0.024495982 | 0.380459408 |
| ENSMUSG00000052392 | Acot4 | 171282 | protein_coding | 117.6065219 | 0.380502375 | 0.14769718 | 2.576233171 | 0.009988326 | 0.047891049 | 0.380502375 |
| ENSMUSG00000089789 | Rdh1 | 107605 | protein_coding | 49.44240765 | -0.380522141 | 0.342652928 | -1.110517698 | 0.266776006 | 0.474540288 | 0.380522141 |
| ENSMUSG00000057409 | Zfp53 | 24132 | protein_coding | 304.8640073 | 0.38089765 | 0.10938515 | 3.482169653 | 0.000497368 | 0.004331375 | 0.38089765 |
| ENSMUSG00000020474 | Polm | 54125 | protein_coding | 106.5275559 | -0.380914096 | 0.133518258 | -2.852898943 | 0.00433224 | 0.025055887 | 0.380914096 |
| ENSMUSG00000067995 | Gtf2f2 | 68705 | protein_coding | 362.5697902 | 0.380947811 | 0.165942528 | 2.295661132 | 0.021695264 | 0.084821348 | 0.380947811 |
| ENSMUSG00000020897 | Aurkb | 20877 | protein_coding | 380.4611183 | 0.38096244 | 0.151229659 | 2.519098716 | 0.011765567 | 0.05403747 | 0.38096244 |
| ENSMUSG00000091844 | Gm8251 | 102636082 | protein_coding | 21.2851822 | -0.381268706 | 0.489488292 | -0.77891282 | 0.436031072 | 0.638923911 | 0.381268706 |
| ENSMUSG00000089417 | Gm22009 | 115489854 | scaRNA | 1391.807156 | -0.381339862 | 0.09864348 | -3.865839504 | 0.000110708 | 0.001229674 | 0.381339862 |
| ENSMUSG00000028687 | Mutyh | 70603 | protein_coding | 24.19930007 | 0.381368034 | 0.248059851 | 1.537403305 | 0.124194578 | 0.287733997 | 0.381368034 |
| ENSMUSG00000029821 | Gsdme | 54722 | protein_coding | 501.386165 | -0.381602109 | 0.148345256 | -2.572391724 | 0.010099854 | 0.048209507 | 0.381602109 |
| ENSMUSG00000067653 | Ankrd23 | 78321 | protein_coding | 7.325082947 | -0.381698601 | 0.488090148 | -0.782024802 | 0.434199998 | 0.637531118 | 0.381698601 |
| ENSMUSG00000035172 | Plekhh3 | 217198 | protein_coding | 253.0373813 | -0.381702863 | 0.214294655 | -1.78120571 | 0.074878851 | 0.204201575 | 0.381702863 |
| ENSMUSG00000020811 | Wscd1 | 216881 | protein_coding | 11.88962176 | 0.381712967 | 0.341904527 | 1.116431449 | 0.264237498 | 0.47199449 | 0.381712967 |
| ENSMUSG00000031642 | Sh3rf1 | 59009 | protein_coding | 546.5352846 | -0.381785475 | 0.106723423 | -3.57733535 | 0.000347115 | 0.003206986 | 0.381785475 |
| ENSMUSG00000034833 | Tespa1 | 67596 | protein_coding | 8.569294112 | 0.381995587 | 0.717473057 | 0.532418023 | 0.594436505 | 0.762640729 | 0.381995587 |
| ENSMUSG00000034472 | Rasd2 | 75141 | protein_coding | 244.4642093 | -0.382114283 | 0.271589561 | -1.406954969 | 0.159440742 | 0.340607851 | 0.382114283 |
| ENSMUSG00000022184 | Fbxo4 | 106052 | protein_coding | 197.8219172 | 0.382199274 | 0.158829294 | 2.406352535 | 0.016112708 | 0.068183636 | 0.382199274 |
| ENSMUSG00000021974 | Fgf9 | 14180 | protein_coding | 25.6575822 | -0.382215494 | 0.374345843 | -1.021022409 | 0.307243822 | 0.517203508 | 0.382215494 |
| ENSMUSG00000042797 | Aqp11 | 66333 | protein_coding | 287.4151988 | -0.38228619 | 0.131232914 | -2.913035901 | 0.003579334 | 0.021562746 | 0.38228619 |
| ENSMUSG00000031347 | Cetn2 | 26370 | protein_coding | 296.177027 | 0.38251361 | 0.109291225 | 3.499948043 | 0.000465349 | 0.004101869 | 0.38251361 |
| ENSMUSG00000024055 | Cyp4f13 | 170716 | protein_coding | 435.2956509 | -0.382530605 | 0.090516146 | -4.226103546 | 2.38E-05 | 0.000320805 | 0.382530605 |
| ENSMUSG00000029009 | Mthfr | 17769 | protein_coding | 244.8820782 | -0.382554358 | 0.121343578 | -3.152654349 | 0.001617933 | 0.011332643 | 0.382554358 |
| ENSMUSG00000036641 | Ccdc148 | 227933 | protein_coding | 16.44578347 | -0.382568204 | 0.418491968 | -0.914159012 | 0.360633298 | 0.570122672 | 0.382568204 |
| ENSMUSG00000031090 | Nadsyn1 | 78914 | protein_coding | 586.492046 | -0.382854617 | 0.068281817 | -5.606977608 | 2.06E-08 | 5.32E-07 | 0.382854617 |
| ENSMUSG00000060935 | Tmem263 | 103266 | protein_coding | 619.8392377 | 0.38291687 | 0.096327633 | 3.975150813 | 7.03E-05 | 0.000829231 | 0.38291687 |
| ENSMUSG00000026107 | Nabp1 | 109019 | protein_coding | 310.765911 | 0.38305757 | 0.118672918 | 3.227843192 | 0.001247273 | 0.009150884 | 0.38305757 |
| ENSMUSG00000071862 | Lrrtm2 | 107065 | protein_coding | 59.47098689 | -0.383087736 | 0.312104082 | -1.227435839 | 0.219658823 | 0.419876136 | 0.383087736 |
| ENSMUSG00000024646 | Cyb5a | 109672 | protein_coding | 1131.446432 | 0.38316419 | 0.098665891 | 3.883451379 | 0.000102984 | 0.001156792 | 0.38316419 |
| ENSMUSG00000056832 | Ttc26 | 264134 | protein_coding | 46.54720043 | 0.383217039 | 0.191605893 | 2.000027416 | 0.045497304 | 0.145752424 | 0.383217039 |
| ENSMUSG00000054640 | Slc8a1 | 20541 | protein_coding | 1617.106751 | -0.383231254 | 0.177502701 | -2.159016467 | 0.030848889 | 0.110377388 | 0.383231254 |
| ENSMUSG00000020027 | Socs2 | 216233 | protein_coding | 506.071891 | -0.383304894 | 0.128623061 | -2.980063539 | 0.002881886 | 0.018172082 | 0.383304894 |
| ENSMUSG00000039199 | Zdhhc1 | 70796 | protein_coding | 133.8635545 | -0.383331121 | 0.136683886 | -2.804508512 | 0.005039336 | 0.028223115 | 0.383331121 |
| ENSMUSG00000015709 | Arnt2 | 11864 | protein_coding | 48.41491692 | 0.383382639 | 0.241902946 | 1.584861388 | 0.112997831 | 0.268850022 | 0.383382639 |
| ENSMUSG00000030410 | Dmwd | 13401 | protein_coding | 17.41130662 | 0.383486251 | 0.29566288 | 1.297038881 | 0.194617809 | 0.387730172 | 0.383486251 |
| ENSMUSG00000052760 | A630001G21Rik | 319997 | protein_coding | 30.35154375 | -0.383576917 | 0.403772143 | -0.949983607 | 0.342120582 | 0.552274919 | 0.383576917 |
| ENSMUSG00000079465 | Col4a3 | 12828 | protein_coding | 13.07561767 | -0.383795334 | 0.546982863 | -0.70165879 | 0.482891979 | 0.676354506 | 0.383795334 |

| ensembl_gene_id | external_gene_name | entrezgene_id | gene_biotype | baseMean | log2FoldChange | lfcSE | stat | pvalue | padj | absLog2FC |
|---|---|---|---|---|---|---|---|---|---|---|
| ENSMUSG00000001750 | Tcirg1 | 27060 | protein_coding | 329.1000131 | -0.383810333 | 0.127143252 | -3.018723577 | 0.00253842 | 0.016496107 | 0.383810333 |
| ENSMUSG00000031568 | Rwdd4a | 192174 | protein_coding | 381.5735265 | 0.383820509 | 0.120054657 | 3.197048068 | 0.001388418 | 0.010029351 | 0.383820509 |
| ENSMUSG00000065653 | Gm23105 | 115487521 | snoRNA | 32.42798913 | 0.38383517 | 0.436426022 | 0.879496527 | 0.379132115 | 0.587178874 | 0.38383517 |
| ENSMUSG00000027800 | Tm4sf1 | 17112 | protein_coding | 19.98408995 | -0.384009931 | 0.338669808 | -1.133877075 | 0.256846115 | 0.463695525 | 0.384009931 |
| ENSMUSG00000042605 | Atxn2 | 20239 | protein_coding | 468.8210918 | -0.38404972 | 0.130415486 | -2.944816859 | 0.003231459 | 0.019918992 | 0.38404972 |
| ENSMUSG00000031200 | Mtcp1 | 17763 | protein_coding | 57.20348253 | 0.38440899 | 0.200313406 | 1.919037759 | 0.054979555 | 0.165478904 | 0.38440899 |
| ENSMUSG00000028630 | Dyrk2 | 69181 | protein_coding | 1458.688228 | -0.384501 | 0.095158663 | -4.040630531 | 5.33E-05 | 0.000657186 | 0.384501 |
| ENSMUSG00000040054 | Baz2a | 116848 | protein_coding | 1022.180322 | -0.384580252 | 0.147943008 | -2.599516242 | 0.009335526 | 0.045472862 | 0.384580252 |
| ENSMUSG00000040606 | Kazn | 71529 | protein_coding | 143.4690914 | -0.384940703 | 0.137306832 | -2.803507278 | 0.005055009 | 0.028281063 | 0.384940703 |
| ENSMUSG00000066258 | Trim12a | 76681 | protein_coding | 194.518945 | 0.384984957 | 0.142034595 | 2.710501316 | 0.006718158 | 0.035249085 | 0.384984957 |
| ENSMUSG00000006154 | Eps8l1 | 67425 | protein_coding | 69.25661448 | -0.385043278 | 0.192777888 | -1.997341507 | 0.045788097 | 0.146360187 | 0.385043278 |
| ENSMUSG00000020532 | Acaca | 107476 | protein_coding | 1228.740816 | 0.385134828 | 0.080150416 | 4.805150725 | 1.55E-06 | 2.79E-05 | 0.385134828 |
| ENSMUSG00000074738 | Fndc10 | 230991 | protein_coding | 23.82211028 | -0.385217327 | 0.24830765 | -1.551371157 | 0.120812763 | 0.28240471 | 0.385217327 |
| ENSMUSG00000014075 | Dynlt2b | 66061 | protein_coding | 38.24426446 | -0.385252301 | 0.23789459 | -1.619424392 | 0.105355982 | 0.257024059 | 0.385252301 |
| ENSMUSG00000092203 | 1110038B12Rik | 68763 | lncRNA | 833.9744564 | -0.385290924 | 0.105884074 | -3.638799593 | 0.000273912 | 0.002639364 | 0.385290924 |
| ENSMUSG00000001436 | Slc19a1 | 20509 | protein_coding | 113.4934859 | -0.38533126 | 0.159150626 | -2.421173388 | 0.015470496 | 0.06616919 | 0.38533126 |
| ENSMUSG00000002808 | Epdr1 | 105298 | protein_coding | 26.35295877 | 0.385404952 | 0.33615928 | 1.146495054 | 0.251590372 | 0.458085443 | 0.385404952 |
| ENSMUSG00000056305 | Usp39 | 28035 | protein_coding | 390.7870044 | 0.385581116 | 0.16056615 | 2.401384827 | 0.01633315 | 0.068933336 | 0.385581116 |
| ENSMUSG00000097011 | Gm4651 | 100043793 | lncRNA | 16.81631238 | 0.385670851 | 0.302999996 | 1.272841109 | 0.20307443 | 0.398020253 | 0.385670851 |
| ENSMUSG00000006585 | Cdt1 | 67177 | protein_coding | 174.1237405 | -0.38574967 | 0.133486876 | -2.889794732 | 0.003854935 | 0.022842782 | 0.38574967 |
| ENSMUSG00000030654 | Arl6ip1 | 54208 | protein_coding | 2622.875406 | 0.385945638 | 0.113856417 | 3.389757452 | 0.000699545 | 0.005740521 | 0.385945638 |
| ENSMUSG00000038217 | Tlcd2 | 380712 | protein_coding | 60.55724311 | -0.386225131 | 0.143304016 | -2.695145203 | 0.007035796 | 0.036625014 | 0.386225131 |
| ENSMUSG00000031391 | L1cam | 16728 | protein_coding | 1469.2878 | -0.386266562 | 0.087234513 | -4.42790987 | 9.52E-06 | 0.000142173 | 0.386266562 |
| ENSMUSG00000021938 | Pspc1 | 66645 | protein_coding | 451.8342164 | 0.386285764 | 0.142628622 | 2.708332725 | 0.006762219 | 0.035459758 | 0.386285764 |
| ENSMUSG00000052456 | Get3 | 56495 | protein_coding | 731.5721526 | 0.386327926 | 0.10013845 | 3.857937958 | 0.000114348 | 0.001265691 | 0.386327926 |
| ENSMUSG00000001025 | S100a6 | 20200 | protein_coding | 12044.01792 | -0.386447097 | 0.139932999 | -2.761658077 | 0.005750867 | 0.031114066 | 0.386447097 |
| ENSMUSG00000032834 | Pwp2 | 110816 | protein_coding | 250.062007 | -0.386524707 | 0.226173051 | -1.708977731 | 0.087455075 | 0.227574649 | 0.386524707 |
| ENSMUSG00000056185 | Snx32 | 225861 | protein_coding | 44.19338914 | -0.386580648 | 0.245614227 | -1.573934264 | 0.115502648 | 0.272915913 | 0.386580648 |
| ENSMUSG00000029757 | Dync1i1 | 13426 | protein_coding | 51.49010155 | 0.3866888 | 0.187301514 | 2.064525755 | 0.038967895 | 0.13069712 | 0.3866888 |
| ENSMUSG00000042446 | Zmym4 | 67785 | protein_coding | 382.1875652 | -0.386690682 | 0.110247325 | -3.507483585 | 0.000452366 | 0.004002939 | 0.386690682 |
| ENSMUSG00000066735 | Vkorc1l1 | 69568 | protein_coding | 735.446637 | 0.386696361 | 0.065734432 | 5.882706373 | 4.04E-09 | 1.14E-07 | 0.386696361 |
| ENSMUSG00000073125 | Xlr3b | 574437 | protein_coding | 27.47921435 | 0.386756435 | 0.76963077 | 0.502522053 | 0.615300344 | 0.775724543 | 0.386756435 |
| ENSMUSG00000055681 | Cope | 59042 | protein_coding | 1392.669368 | 0.386892404 | 0.09937508 | 3.893253782 | 9.89E-05 | 0.001118109 | 0.386892404 |
| ENSMUSG00000007097 | Atp1a2 | 98660 | protein_coding | 6.904720274 | -0.387139369 | 0.784801631 | -0.493295826 | 0.621803577 | 0.780188222 | 0.387139369 |
| ENSMUSG00000022973 | Synj1 | 104015 | protein_coding | 384.9124959 | -0.387395703 | 0.147922682 | -2.618906706 | 0.008821207 | 0.043648395 | 0.387395703 |
| ENSMUSG00000079429 | Mroh2a | 100040766 | protein_coding | 163.9391983 | -0.387396707 | 0.204213236 | -1.897020557 | 0.057825225 | 0.171719678 | 0.387396707 |
| ENSMUSG00000032815 | Fanca | 14087 | protein_coding | 144.3657754 | 0.387405734 | 0.124342998 | 3.115621638 | 0.001835576 | 0.012596751 | 0.387405734 |
| ENSMUSG00000052675 | Zfp112 | 57745 | protein_coding | 32.93102113 | 0.387564003 | 0.263835039 | 1.468963353 | 0.141842729 | 0.314924867 | 0.387564003 |

| ensembl_gene_id | external_gene_name | entrezgene_id | gene_biotype | baseMean | log2FoldChange | lfcSE | stat | pvalue | padj | absLog2FC |
|---|---|---|---|---|---|---|---|---|---|---|
| ENSMUSG00000028698 | Pik3r3 | 18710 | protein_coding | 412.3390367 | 0.387865231 | 0.108290993 | 3.581694278 | 0.000341373 | 0.003166798 | 0.387865231 |
| ENSMUSG00000034168 | Irf2bpl | 238330 | protein_coding | 482.126746 | -0.387903764 | 0.165032418 | -2.3504701 | 0.018749714 | 0.076282361 | 0.387903764 |
| ENSMUSG00000054474 | Thnsl2 | 232078 | protein_coding | 284.8133122 | 0.38810882 | 0.144027211 | 2.694690948 | 0.007045395 | 0.036625014 | 0.38810882 |
| ENSMUSG00000040596 | Pogk | 71592 | protein_coding | 345.2214717 | -0.388197226 | 0.144310343 | -2.690016657 | 0.007144845 | 0.036997105 | 0.388197226 |
| ENSMUSG00000021336 | Slc17a4 | 319848 | protein_coding | 1049.138485 | -0.388216531 | 0.150357484 | -2.581956821 | 0.009824188 | 0.047317708 | 0.388216531 |
| ENSMUSG00000030704 | Rab6a | 19346 | protein_coding | 2207.869786 | 0.388279923 | 0.08685551 | 4.470412102 | 7.81E-06 | 0.000119912 | 0.388279923 |
| ENSMUSG00000042499 | Hoxd11 | 15431 | protein_coding | 10.191434 | -0.388465215 | 0.712666795 | -0.545086734 | 0.585693866 | 0.75599124 | 0.388465215 |
| ENSMUSG00000038160 | Atg5 | 11793 | protein_coding | 762.4696281 | 0.388541565 | 0.089931195 | 4.320431475 | 1.56E-05 | 0.00022166 | 0.388541565 |
| ENSMUSG00000027001 | Dusp19 | 68082 | protein_coding | 123.8144506 | 0.388553856 | 0.134051058 | 2.89855121 | 0.003748911 | 0.0223574 | 0.388553856 |
| ENSMUSG00000032591 | Mst1 | 15235 | protein_coding | 15.24599039 | -0.388672266 | 0.367746256 | -1.056903394 | 0.290555679 | 0.499223071 | 0.388672266 |
| ENSMUSG00000030030 | 1700003E16Rik | 71837 | protein_coding | 39.21178698 | -0.388678538 | 0.261611489 | -1.485708977 | 0.137356115 | 0.308157609 | 0.388678538 |
| ENSMUSG00000053332 | Gas5 | 14455 | lncRNA | 1354.147469 | -0.388692378 | 0.078101709 | -4.976746132 | 6.47E-07 | 1.28E-05 | 0.388692378 |
| ENSMUSG00000051149 | Adnp | 11538 | protein_coding | 16.09495892 | -0.38874724 | 0.286330503 | -1.357687136 | 0.17456298 | 0.360814839 | 0.38874724 |
| ENSMUSG00000047014 | Catsperb | 271036 | protein_coding | 6.692614476 | -0.388836341 | 0.768016062 | -0.50628673 | 0.612655378 | 0.77428984 | 0.388836341 |
| ENSMUSG00000030852 | Tacc2 | 57752 | protein_coding | 658.0420913 | -0.388849902 | 0.125602053 | -3.095888106 | 0.001962244 | 0.013316839 | 0.388849902 |
| ENSMUSG00000029916 | Agk | 69923 | protein_coding | 120.1989552 | -0.388935573 | 0.113539548 | -3.425551521 | 0.000613553 | 0.005159802 | 0.388935573 |
| ENSMUSG00000027994 | Mcub | 66815 | protein_coding | 61.03983781 | -0.389032101 | 0.186688791 | -2.083853559 | 0.03717349 | 0.126274119 | 0.389032101 |
| ENSMUSG00000063727 | Tnfrsf11b | 18383 | protein_coding | 21.20859749 | 0.389041006 | 0.414165297 | 0.939337527 | 0.347557477 | 0.557627687 | 0.389041006 |
| ENSMUSG00000041298 | Katnal1 | 231912 | protein_coding | 61.3911785 | 0.389058734 | 0.200912089 | 1.936462542 | 0.052811076 | 0.161248547 | 0.389058734 |
| ENSMUSG00000032308 | Ulk3 | 71742 | protein_coding | 95.46796255 | -0.389105791 | 0.16904937 | -2.301728724 | 0.021350475 | 0.083926547 | 0.389105791 |
| ENSMUSG00000049422 | Chchd10 | 103172 | protein_coding | 2128.827283 | 0.38912044 | 0.095245927 | 4.085428662 | 4.40E-05 | 0.000550912 | 0.38912044 |
| ENSMUSG00000096054 | Syne1 | 64009 | protein_coding | 263.9097862 | -0.389157034 | 0.228082665 | -1.706210478 | 0.087968899 | 0.228501079 | 0.389157034 |
| ENSMUSG00000034687 | Fras1 | 231470 | protein_coding | 13.58248998 | -0.389314161 | 0.753016389 | -0.517006225 | 0.605151813 | 0.769630955 | 0.389314161 |
| ENSMUSG00000020528 | Prpsap2 | 212627 | protein_coding | 319.2199799 | 0.389492638 | 0.105426008 | 3.694464442 | 0.000220351 | 0.002212955 | 0.389492638 |
| ENSMUSG00000041794 | Myrip | 245049 | protein_coding | 92.73666103 | -0.389596062 | 0.301086748 | -1.293966154 | 0.195677112 | 0.388961796 | 0.389596062 |
| ENSMUSG00000054604 | Cggbp1 | 106143 | protein_coding | 2195.116728 | 0.389713547 | 0.069445369 | 5.61180033 | 2.00E-08 | 5.19E-07 | 0.389713547 |
| ENSMUSG00000107456 | Gm10400 | 100093700 | TEC | 20.89287293 | 0.389838521 | 0.456367737 | 0.854220158 | 0.392983016 | 0.600878709 | 0.389838521 |
| ENSMUSG00000022270 | Retreg1 | 66270 | protein_coding | 1795.651081 | -0.389916338 | 0.106080478 | -3.675665356 | 0.00023723 | 0.002354271 | 0.389916338 |
| ENSMUSG00000035314 | Gdpd5 | 233552 | protein_coding | 77.64375903 | -0.389957895 | 0.192535488 | -2.025381915 | 0.042828163 | 0.139274598 | 0.389957895 |
| ENSMUSG00000024038 | Ndufv3 | 78330 | protein_coding | 1377.749094 | 0.390333791 | 0.101780342 | 3.83506071 | 0.000125533 | 0.001370455 | 0.390333791 |
| ENSMUSG00000002227 | Mov10 | 17454 | protein_coding | 349.0239536 | 0.390400704 | 0.106367035 | 3.670316693 | 0.00024225 | 0.002391405 | 0.390400704 |
| ENSMUSG00000053054 | Adh6a | 69117 | protein_coding | 11.20183843 | -0.390404105 | 0.445162892 | -0.876991573 | 0.380491215 | 0.588394571 | 0.390404105 |
| ENSMUSG00000032381 | Ciao2a | 68250 | protein_coding | 1344.493637 | 0.390524747 | 0.086423538 | 4.518731302 | 6.22E-06 | 9.81E-05 | 0.390524747 |
| ENSMUSG00000069255 | Dusp22 | 105352 | protein_coding | 44.64505576 | 0.39084517 | 0.204128629 | 1.914700417 | 0.05553072 | 0.166854049 | 0.39084517 |
| ENSMUSG00000000682 | Cd52 | 23833 | protein_coding | 27.79185776 | 0.390874237 | 0.320708411 | 1.218783868 | 0.222926236 | 0.424276387 | 0.390874237 |
| ENSMUSG00000049764 | Zfp280b | 64453 | protein_coding | 246.5367399 | 0.391054803 | 0.120046238 | 3.257534829 | 0.001123845 | 0.008435802 | 0.391054803 |
| ENSMUSG00000032067 | Pts | 19286 | protein_coding | 766.3234734 | 0.391083062 | 0.120427143 | 3.24746608 | 0.001164375 | 0.008674542 | 0.391083062 |
| ENSMUSG00000031093 | Dock11 | 75974 | protein_coding | 238.9462177 | -0.391235976 | 0.136318441 | -2.870015035 | 0.004104523 | 0.023956335 | 0.391235976 |

| ensembl_gene_id | external_gene_name | entrezgene_id | gene_biotype | baseMean | log2FoldChange | lfcSE | stat | pvalue | padj | absLog2FC |
|---|---|---|---|---|---|---|---|---|---|---|
| ENSMUSG00000046761 | Fam83h | 105732 | protein_coding | 1318.353931 | -0.39124482 | 0.171565565 | -2.28043909 | 0.022581659 | 0.0875988 | 0.39124482 |
| ENSMUSG00000022075 | Rhobtb2 | 246710 | protein_coding | 212.5279435 | -0.391628411 | 0.103479484 | -3.784599561 | 0.000153956 | 0.001629379 | 0.391628411 |
| ENSMUSG00000022205 | Sub1 | 20024 | protein_coding | 1134.275502 | 0.391636894 | 0.099619414 | 3.931331027 | 8.45E-05 | 0.000974328 | 0.391636894 |
| ENSMUSG00000037139 | Myom3 | 242702 | protein_coding | 37.88652868 | -0.39198109 | 0.322768493 | -1.214434178 | 0.224581971 | 0.426256152 | 0.39198109 |
| ENSMUSG00000039942 | Ptger4 | 19219 | protein_coding | 531.2039326 | -0.392034909 | 0.111052256 | -3.5301841 | 0.000415271 | 0.00374543 | 0.392034909 |
| ENSMUSG00000072949 | Acot1 | 26897 | protein_coding | 101.9699828 | 0.392287879 | 0.171009944 | 2.293947757 | 0.0217935 | 0.085080114 | 0.392287879 |
| ENSMUSG00000036957 | Lrfn3 | 233067 | protein_coding | 134.4262676 | -0.392354968 | 0.203485888 | -1.928167949 | 0.053834239 | 0.163014057 | 0.392354968 |
| ENSMUSG00000042747 | Krtcap2 | 66059 | protein_coding | 685.1624328 | 0.392561628 | 0.098941386 | 3.967618027 | 7.26E-05 | 0.000850199 | 0.392561628 |
| ENSMUSG00000053182 | Cd200l1 | 208166 | protein_coding | 249.6468328 | -0.392566413 | 0.141656126 | -2.771263227 | 0.005583927 | 0.030427911 | 0.392566413 |
| ENSMUSG00000046295 | Ankle1 | 234396 | protein_coding | 62.27954099 | -0.392653616 | 0.236841459 | -1.657875348 | 0.097342629 | 0.243900172 | 0.392653616 |
| ENSMUSG00000038615 | Nfe2l1 | 18023 | protein_coding | 2577.697696 | 0.392667593 | 0.074629009 | 5.261594616 | 1.43E-07 | 3.20E-06 | 0.392667593 |
| ENSMUSG00000039512 | Uhrf1bp1 | 224648 | protein_coding | 210.8402557 | -0.392699966 | 0.123545379 | -3.178588851 | 0.001479939 | 0.010551683 | 0.392699966 |
| ENSMUSG00000065219 | Snord32a | 27209 | snoRNA | 6.770534444 | -0.392775527 | 0.440450036 | -0.891759553 | 0.372521825 | 0.580796848 | 0.392775527 |
| ENSMUSG00000010110 | Stx5a | 56389 | protein_coding | 1178.286321 | 0.392815016 | 0.082650663 | 4.752714662 | 2.01E-06 | 3.51E-05 | 0.392815016 |
| ENSMUSG00000020593 | Lpin1 | 14245 | protein_coding | 168.6562456 | 0.392842329 | 0.163663932 | 2.400298727 | 0.016381697 | 0.069048936 | 0.392842329 |
| ENSMUSG00000028692 | Akr1a1 | 58810 | protein_coding | 6592.823591 | 0.393091851 | 0.090726445 | 4.332715219 | 1.47E-05 | 0.000210395 | 0.393091851 |
| ENSMUSG00000026110 | Mgat4a | 269181 | protein_coding | 4591.0907 | -0.393158082 | 0.108771755 | -3.614523673 | 0.0003009 | 0.002844356 | 0.393158082 |
| ENSMUSG00000006538 | Ihh | 16147 | protein_coding | 967.5477506 | -0.39323635 | 0.154161387 | -2.550809626 | 0.010747301 | 0.050511365 | 0.39323635 |
| ENSMUSG00000097577 | 6230400D17Rik | 76133 | lncRNA | 26.31900004 | -0.39328032 | 0.291395488 | -1.349644506 | 0.177130041 | 0.364084685 | 0.39328032 |
| ENSMUSG00000038335 | Tsr1 | 104662 | protein_coding | 711.1526608 | -0.393326559 | 0.121393317 | -3.240100602 | 0.001194875 | 0.008860323 | 0.393326559 |
| ENSMUSG00000040097 | Flywch1 | 224613 | protein_coding | 301.8760185 | 0.393351768 | 0.082962849 | 4.741300158 | 2.12E-06 | 3.69E-05 | 0.393351768 |
| ENSMUSG00000018286 | Psmb6 | 19175 | protein_coding | 1533.576608 | 0.393395149 | 0.093840801 | 4.19215463 | 2.76E-05 | 0.000367071 | 0.393395149 |
| ENSMUSG00000040389 | Wdr47 | 99512 | protein_coding | 89.78249383 | -0.393547088 | 0.122045967 | -3.224580838 | 0.001261572 | 0.009230282 | 0.393547088 |
| ENSMUSG00000024725 | Ostf1 | 20409 | protein_coding | 1074.941101 | 0.393561692 | 0.088097474 | 4.467343637 | 7.92E-06 | 0.000121527 | 0.393561692 |
| ENSMUSG00000028803 | Nipal3 | 74552 | protein_coding | 1298.184122 | 0.393604048 | 0.079954072 | 4.922876809 | 8.53E-07 | 1.63E-05 | 0.393604048 |
| ENSMUSG00000103585 | Pcdhgb4 | 93701 | protein_coding | 12.8408057 | -0.393634555 | 0.348886419 | -1.128259898 | 0.259210174 | 0.466414331 | 0.393634555 |
| ENSMUSG00000042787 | Exog | 208194 | protein_coding | 135.2099627 | 0.393747868 | 0.138919959 | 2.834350601 | 0.004591892 | 0.026233738 | 0.393747868 |
| ENSMUSG00000022601 | Zbtb11 | 271377 | protein_coding | 486.2638234 | -0.393955399 | 0.110312944 | -3.571252701 | 0.000355278 | 0.003272913 | 0.393955399 |
| ENSMUSG00000022621 | Rabl2 | 68708 | protein_coding | 65.06928267 | 0.394005956 | 0.157995689 | 2.493776623 | 0.012639206 | 0.057062718 | 0.394005956 |
| ENSMUSG00000021884 | Hacl1 | 56794 | protein_coding | 20.64790433 | -0.394054065 | 0.274615047 | -1.434932533 | 0.151306324 | 0.329506034 | 0.394054065 |
| ENSMUSG00000052337 | Immt | 76614 | protein_coding | 4382.202774 | 0.394233014 | 0.069864654 | 5.642810675 | 1.67E-08 | 4.38E-07 | 0.394233014 |
| ENSMUSG00000026649 | Cfap126 | 75472 | protein_coding | 11.25462896 | 0.394422573 | 0.373460408 | 1.05612955 | 0.290909032 | 0.499573396 | 0.394422573 |
| ENSMUSG00000031145 | Prickle3 | 54630 | protein_coding | 121.7968979 | -0.39445008 | 0.164334539 | -2.400287141 | 0.016382216 | 0.069048936 | 0.39445008 |
| ENSMUSG00000022978 | Mis18a | 66578 | protein_coding | 168.8592889 | 0.39446078 | 0.136438177 | 2.891132009 | 0.003838568 | 0.02276597 | 0.39446078 |
| ENSMUSG00000072258 | Taf1a | 21339 | protein_coding | 135.4907641 | -0.394869247 | 0.115926442 | -3.406205172 | 0.000658727 | 0.005473238 | 0.394869247 |
| ENSMUSG00000032536 | Trak1 | 67095 | protein_coding | 2570.589188 | -0.39493077 | 0.124992925 | -3.159624984 | 0.001579723 | 0.011099176 | 0.39493077 |
| ENSMUSG00000044148 | 1810030O07Rik | 69155 | protein_coding | 507.9228944 | 0.395026535 | 0.08016205 | 4.927849677 | 8.31E-07 | 1.59E-05 | 0.395026535 |
| ENSMUSG00000034201 | Gas2l1 | 78926 | protein_coding | 393.7127632 | -0.395238507 | 0.10289221 | -3.841287004 | 0.000122391 | 0.001340744 | 0.395238507 |

| ensembl_gene_id | external_gene_name | entrezgene_id | gene_biotype | baseMean | log2FoldChange | lfcSE | stat | pvalue | padj | absLog2FC |
|---|---|---|---|---|---|---|---|---|---|---|
| ENSMUSG00000052302 | Tbc1d30 | 74694 | protein_coding | 306.1378045 | -0.395381425 | 0.131643587 | -3.003423373 | 0.002669608 | 0.017145771 | 0.395381425 |
| ENSMUSG00000043822 | Adamtsl5 | 66548 | protein_coding | 53.59821783 | -0.395435463 | 0.195653623 | -2.02109962 | 0.043269454 | 0.14037911 | 0.395435463 |
| ENSMUSG00000037579 | Kcnh3 | 16512 | protein_coding | 306.5429044 | -0.395563012 | 0.221151536 | -1.78865144 | 0.073670968 | 0.20214146 | 0.395563012 |
| ENSMUSG00000026447 | Pik3c2b | 240752 | protein_coding | 812.6172416 | -0.395574668 | 0.125299805 | -3.1570254 | 0.001593875 | 0.011183804 | 0.395574668 |
| ENSMUSG00000047810 | Ccdc88b | 78317 | protein_coding | 29.70098266 | 0.395733971 | 0.291237808 | 1.35880013 | 0.174209932 | 0.360225341 | 0.395733971 |
| ENSMUSG00000066838 | Zfp772 | 232855 | protein_coding | 90.54728721 | -0.395868703 | 0.147854482 | -2.677421055 | 0.007419133 | 0.038218615 | 0.395868703 |
| ENSMUSG00000075040 | Zfp408 | 381410 | protein_coding | 189.7739483 | -0.396022704 | 0.107739203 | -3.675753069 | 0.000237149 | 0.002354271 | 0.396022704 |
| ENSMUSG00000041293 | Adgrf1 | 77596 | protein_coding | 28.26842147 | -0.396098952 | 0.411346008 | -0.962933744 | 0.335580774 | 0.546257595 | 0.396098952 |
| ENSMUSG00000028378 | Ptgr1 | 67103 | protein_coding | 408.0061456 | 0.396251466 | 0.121177911 | 3.269997504 | 0.001075484 | 0.008141784 | 0.396251466 |
| ENSMUSG00000029174 | Tbc1d1 | 57915 | protein_coding | 686.3572092 | 0.396477502 | 0.138209191 | 2.86867681 | 0.004121927 | 0.0240403 | 0.396477502 |
| ENSMUSG00000031822 | Gse1 | 382034 | protein_coding | 473.3378168 | -0.396683139 | 0.163644595 | -2.424052797 | 0.015348374 | 0.065717446 | 0.396683139 |
| ENSMUSG00000054723 | Vmac | 106639 | protein_coding | 289.7321481 | 0.396716464 | 0.093389934 | 4.247957423 | 2.16E-05 | 0.000296101 | 0.396716464 |
| ENSMUSG00000024661 | Fth1 | 14319 | protein_coding | 9589.571294 | -0.396856906 | 0.105663763 | -3.755846805 | 0.000172756 | 0.001797298 | 0.396856906 |
| ENSMUSG00000087590 | Epb41l4aos | 69749 | lncRNA | 141.3579816 | -0.396892345 | 0.134819752 | -2.943873869 | 0.003241321 | 0.019962569 | 0.396892345 |
| ENSMUSG00000060176 | Kif27 | 75050 | protein_coding | 47.70442384 | -0.397027673 | 0.305830587 | -1.298194785 | 0.19422041 | 0.387494219 | 0.397027673 |
| ENSMUSG00000021360 | Gcnt2 | 14538 | protein_coding | 113.0798852 | -0.397193562 | 0.244077543 | -1.627325303 | 0.103668027 | 0.254260641 | 0.397193562 |
| ENSMUSG00000020564 | Atxn7l1 | 380753 | protein_coding | 144.0024059 | -0.397240896 | 0.150347671 | -2.642148659 | 0.008238189 | 0.041445948 | 0.397240896 |
| ENSMUSG00000032311 | Nrg4 | 83961 | protein_coding | 20.94212597 | -0.397299912 | 0.270754378 | -1.467381303 | 0.142272358 | 0.315658743 | 0.397299912 |
| ENSMUSG00000031621 | Isx | 71597 | protein_coding | 1355.672998 | -0.397342337 | 0.236336131 | -1.68125938 | 0.092712544 | 0.236617233 | 0.397342337 |
| ENSMUSG00000051098 | Mblac2 | 72852 | protein_coding | 75.24140026 | -0.397466533 | 0.187199569 | -2.123223549 | 0.033735124 | 0.117681351 | 0.397466533 |
| ENSMUSG00000031949 | Adat1 | 30947 | protein_coding | 46.62221779 | -0.397518922 | 0.19865284 | -2.00107344 | 0.045384476 | 0.14556664 | 0.397518922 |
| ENSMUSG00000041890 | Git2 | 26431 | protein_coding | 287.2749833 | -0.397715629 | 0.116157552 | -3.423932593 | 0.000617219 | 0.005179699 | 0.397715629 |
| ENSMUSG00000029163 | Emilin1 | 100952 | protein_coding | 33.59051929 | 0.397803474 | 0.238107151 | 1.670691001 | 0.094782727 | 0.239939491 | 0.397803474 |
| ENSMUSG00000021763 | Cspg4b | 408066 | protein_coding | 7.338471292 | -0.397907539 | 0.582115061 | -0.683554792 | 0.49425634 | 0.685938397 | 0.397907539 |
| ENSMUSG00000037336 | Mfsd2b | 432628 | protein_coding | 14.62983193 | 0.397915805 | 0.339400427 | 1.172408085 | 0.241033255 | 0.445587011 | 0.397915805 |
| ENSMUSG00000061273 | Mmgt1 | 236792 | protein_coding | 823.915178 | 0.397967713 | 0.076352619 | 5.212233971 | 1.87E-07 | 4.10E-06 | 0.397967713 |
| ENSMUSG00000099481 | Xndc1 | 102443350 | protein_coding | 42.42874189 | -0.398043702 | 0.199152641 | -1.998686529 | 0.045642281 | 0.146063838 | 0.398043702 |
| ENSMUSG00000042564 | Fam227a | 75729 | protein_coding | 21.41298291 | -0.398090285 | 0.353368134 | -1.126559662 | 0.259928702 | 0.467180325 | 0.398090285 |
| ENSMUSG00000022131 | Gpr180 | 58245 | protein_coding | 397.3925573 | 0.398283107 | 0.100327807 | 3.969817717 | 7.19E-05 | 0.000845508 | 0.398283107 |
| ENSMUSG00000038879 | Nipal2 | 223473 | protein_coding | 692.5427905 | 0.398287756 | 0.093168633 | 4.274912517 | 1.91E-05 | 0.000266693 | 0.398287756 |
| ENSMUSG00000024773 | Atg2a | 329015 | protein_coding | 632.5716257 | -0.398438997 | 0.093788176 | -4.248286018 | 2.15E-05 | 0.000296039 | 0.398438997 |
| ENSMUSG00000044067 | Gpr22 | 73010 | protein_coding | 19.1835405 | -0.39856107 | 0.451378576 | -0.882986236 | 0.377243708 | 0.585247665 | 0.39856107 |
| ENSMUSG00000028093 | Acp6 | 66659 | protein_coding | 475.3532704 | -0.398734007 | 0.07826561 | -5.094625916 | 3.49E-07 | 7.31E-06 | 0.398734007 |
| ENSMUSG00000026692 | Fmo4 | 226564 | protein_coding | 69.63276644 | -0.39893172 | 0.294553889 | -1.354359034 | 0.17562186 | 0.36178285 | 0.39893172 |
| ENSMUSG00000026411 | Tmem9 | 66241 | protein_coding | 127.0748056 | -0.39894888 | 0.148038617 | -2.694897371 | 0.007041031 | 0.036625014 | 0.39894888 |
| ENSMUSG00000056144 | Trim34a | 94094 | protein_coding | 83.69327494 | 0.398957801 | 0.160038067 | 2.492893156 | 0.012670697 | 0.057172481 | 0.398957801 |
| ENSMUSG00000040111 | Gramd1b | 235283 | protein_coding | 157.723681 | -0.399248269 | 0.14105789 | -2.830385946 | 0.004649188 | 0.02652302 | 0.399248269 |
| ENSMUSG00000032607 | Amt | 434437 | protein_coding | 276.1183302 | -0.399314012 | 0.109329075 | -3.652404562 | 0.000259796 | 0.002533217 | 0.399314012 |

| ensembl_gene_id | external_gene_name | entrezgene_id | gene_biotype | baseMean | log2FoldChange | lfcSE | stat | pvalue | padj | absLog2FC |
|---|---|---|---|---|---|---|---|---|---|---|
| ENSMUSG00000045331 | 2310079G19Rik | 69699 | protein_coding | 9.866840876 | -0.399394867 | 0.49745556 | -0.802875471 | 0.422046715 | 0.627572772 | 0.399394867 |
| ENSMUSG00000032640 | Chsy1 | 269941 | protein_coding | 125.1639195 | 0.399413676 | 0.232104089 | 1.720838601 | 0.085280116 | 0.223632151 | 0.399413676 |
| ENSMUSG00000020229 | Slc5a4a | 64452 | protein_coding | 7.157988581 | 0.399419339 | 0.522776475 | 0.764034645 | 0.444846597 | 0.645657025 | 0.399419339 |
| ENSMUSG00000018398 | Septin8 | 20362 | protein_coding | 510.5353384 | -0.399442209 | 0.070356594 | -5.677395447 | 1.37E-08 | 3.61E-07 | 0.399442209 |
| ENSMUSG00000032285 | Dnaja4 | 58233 | protein_coding | 59.72159754 | -0.399496619 | 0.182164584 | -2.193053169 | 0.028303549 | 0.103239441 | 0.399496619 |
| ENSMUSG00000022244 | Amacr | 17117 | protein_coding | 149.2096458 | 0.399670972 | 0.137072853 | 2.915755835 | 0.003548281 | 0.021456725 | 0.399670972 |
| ENSMUSG00000044860 | Gm1123 | 382097 | protein_coding | 2771.468507 | -0.399678156 | 0.120525174 | -3.316138379 | 0.000912706 | 0.007122776 | 0.399678156 |
| ENSMUSG00000046959 | Slc26a1 | 231583 | protein_coding | 31.36360008 | -0.39982069 | 0.401487004 | -0.995849645 | 0.319323205 | 0.530636656 | 0.39982069 |
| ENSMUSG00000028517 | Plpp3 | 67916 | protein_coding | 76.87480292 | -0.399972913 | 0.195568801 | -2.045177503 | 0.040837362 | 0.135061772 | 0.399972913 |
| ENSMUSG00000021178 | Psmc1 | 19179 | protein_coding | 1298.003318 | 0.400023997 | 0.113079328 | 3.537551957 | 0.000403855 | 0.003665298 | 0.400023997 |
| ENSMUSG00000035325 | Sec31a | 69162 | protein_coding | 2690.2182 | 0.400143194 | 0.057067647 | 7.011734561 | 2.35E-12 | 9.33E-11 | 0.400143194 |
| ENSMUSG00000036053 | Fmnl2 | 71409 | protein_coding | 77.57922647 | 0.400164008 | 0.200040267 | 2.000417282 | 0.045455224 | 0.14569496 | 0.400164008 |
| ENSMUSG00000025736 | Jmjd8 | 72106 | protein_coding | 244.6051624 | 0.400200509 | 0.102806118 | 3.8927694 | 9.91E-05 | 0.00111955 | 0.400200509 |
| ENSMUSG00000020392 | Cdkn2aipnl | 52626 | protein_coding | 487.496538 | 0.400257267 | 0.099097482 | 4.039025601 | 5.37E-05 | 0.000659181 | 0.400257267 |
| ENSMUSG00000073413 | Ly6g6d | 114654 | protein_coding | 26.22429393 | -0.400274526 | 0.275795908 | -1.451343238 | 0.146684307 | 0.322973133 | 0.400274526 |
| ENSMUSG00000046380 | Jrk | 16469 | protein_coding | 60.83057039 | -0.400276138 | 0.228440115 | -1.752214744 | 0.079736889 | 0.213417774 | 0.400276138 |
| ENSMUSG00000062833 | Sval3 | 387564 | protein_coding | 11.15832811 | -0.400360391 | 0.466456322 | -0.858301994 | 0.390725729 | 0.598987431 | 0.400360391 |
| ENSMUSG00000002825 | Qtrt1 | 60507 | protein_coding | 171.1916089 | -0.400388811 | 0.115205833 | -3.475421351 | 0.000510052 | 0.004427303 | 0.400388811 |
| ENSMUSG00000022130 | Tgds | 76355 | protein_coding | 255.3301575 | 0.400501195 | 0.108645177 | 3.686322819 | 0.000227518 | 0.002270615 | 0.400501195 |
| ENSMUSG00000022604 | Cep97 | 74201 | protein_coding | 85.17663197 | -0.400555612 | 0.266161331 | -1.504935412 | 0.132340683 | 0.300402223 | 0.400555612 |
| ENSMUSG00000027353 | Mcm8 | 66634 | protein_coding | 110.0514893 | 0.400663726 | 0.153210951 | 2.615111548 | 0.008919832 | 0.043986094 | 0.400663726 |
| ENSMUSG00000027395 | Polr1b | 20017 | protein_coding | 399.6670676 | -0.400762186 | 0.173772728 | -2.306243286 | 0.021097042 | 0.083197249 | 0.400762186 |
| ENSMUSG00000035469 | Rcbtb1 | 71330 | protein_coding | 854.7276824 | 0.400936345 | 0.064279308 | 6.237409195 | 4.45E-10 | 1.40E-08 | 0.400936345 |
| ENSMUSG00000064326 | Siva1 | 30954 | protein_coding | 96.24242509 | -0.400982009 | 0.13839229 | -2.89743027 | 0.003762334 | 0.022419173 | 0.400982009 |
| ENSMUSG00000032246 | Calml4 | 75600 | protein_coding | 1884.008652 | 0.401086929 | 0.083943724 | 4.778045443 | 1.77E-06 | 3.13E-05 | 0.401086929 |
| ENSMUSG00000079057 | Cyp4v3 | 102294 | protein_coding | 151.6418639 | -0.401213842 | 0.138626637 | -2.894204539 | 0.003801205 | 0.022561311 | 0.401213842 |
| ENSMUSG00000030551 | Nr2f2 | 11819 | protein_coding | 149.4736407 | -0.401363402 | 0.230569633 | -1.740747022 | 0.081727932 | 0.217322621 | 0.401363402 |
| ENSMUSG00000039997 | Ifi203 | 15950 | protein_coding | 14.11679994 | 0.401413498 | 0.420873889 | 0.953761944 | 0.340204157 | 0.550544115 | 0.401413498 |
| ENSMUSG00000070327 | Rnf213 | 672511 | protein_coding | 2242.437546 | 0.401439476 | 0.117081981 | 3.428704179 | 0.00060647 | 0.005111035 | 0.401439476 |
| ENSMUSG00000074569 | Gcnt7 | 654821 | protein_coding | 15.36459013 | -0.401622979 | 0.416153711 | -0.965083257 | 0.334503112 | 0.545293316 | 0.401622979 |
| ENSMUSG00000021877 | Arf4 | 11843 | protein_coding | 4142.372861 | 0.401793398 | 0.07737117 | 5.19306346 | 2.07E-07 | 4.53E-06 | 0.401793398 |
| ENSMUSG00000021144 | Mta1 | 116870 | protein_coding | 220.2213593 | -0.401828424 | 0.09954839 | -4.036513562 | 5.43E-05 | 0.000665729 | 0.401828424 |
| ENSMUSG00000027195 | Hsd17b12 | 56348 | protein_coding | 1684.275679 | 0.402129229 | 0.102593722 | 3.919628021 | 8.87E-05 | 0.001016981 | 0.402129229 |
| ENSMUSG00000028128 | F3 | 14066 | protein_coding | 532.3516351 | -0.402143893 | 0.713034306 | -0.563989543 | 0.572761248 | 0.746436031 | 0.402143893 |
| ENSMUSG00000022186 | Oxct1 | 67041 | protein_coding | 2508.973233 | -0.402155334 | 0.105605203 | -3.808101504 | 0.000140038 | 0.001509565 | 0.402155334 |
| ENSMUSG00000037432 | Fer1l5 | 100534273 | protein_coding | 44.83908539 | 0.402260838 | 0.186687472 | 2.154728618 | 0.031183077 | 0.111164738 | 0.402260838 |
| ENSMUSG00000005667 | Mthfd2 | 17768 | protein_coding | 158.2460734 | 0.402404757 | 0.224318313 | 1.793900606 | 0.072829035 | 0.200695651 | 0.402404757 |
| ENSMUSG00000001119 | Col6a1 | 12833 | protein_coding | 16.43490487 | -0.402559721 | 0.531510899 | -0.75738752 | 0.448817731 | 0.649533783 | 0.402559721 |

| ensembl_gene_id | external_gene_name | entrezgene_id | gene_biotype | baseMean | log2FoldChange | lfcSE | stat | pvalue | padj | absLog2FC |
|---|---|---|---|---|---|---|---|---|---|---|
| ENSMUSG00000029859 | Epha1 | 13835 | protein_coding | 458.9775271 | -0.402806077 | 0.105777983 | -3.808033242 | 0.000140076 | 0.001509565 | 0.402806077 |
| ENSMUSG00000045257 | Morn2 | 378462 | protein_coding | 44.3459448 | 0.402893919 | 0.222250769 | 1.812789765 | 0.069864253 | 0.195399091 | 0.402893919 |
| ENSMUSG00000026687 | Aldh9a1 | 56752 | protein_coding | 901.684438 | 0.403172059 | 0.070779213 | 5.696193056 | 1.23E-08 | 3.26E-07 | 0.403172059 |
| ENSMUSG00000022114 | Spry2 | 24064 | protein_coding | 481.8650112 | -0.403239528 | 0.218487093 | -1.845598855 | 0.064950473 | 0.185566122 | 0.403239528 |
| ENSMUSG00000020029 | Nudt4 | 71207 | protein_coding | 6233.828565 | -0.40343241 | 0.118172176 | -3.413937397 | 0.000640313 | 0.005350946 | 0.40343241 |
| ENSMUSG00000033308 | Dpyd | 99586 | protein_coding | 617.2239451 | -0.403649145 | 0.110965412 | -3.637612264 | 0.000275177 | 0.002648353 | 0.403649145 |
| ENSMUSG00000024885 | Aldh3b1 | 67689 | protein_coding | 94.50767213 | -0.403885996 | 0.163389061 | -2.471928005 | 0.013438658 | 0.059757381 | 0.403885996 |
| ENSMUSG00000004661 | Arid3b | 56380 | protein_coding | 67.44574873 | -0.403904758 | 0.264736928 | -1.525683478 | 0.127088711 | 0.292313212 | 0.403904758 |
| ENSMUSG00000025169 | Ogfod3 | 66179 | protein_coding | 117.3750552 | -0.403939015 | 0.124646437 | -3.240678389 | 0.001192456 | 0.008850627 | 0.403939015 |
| ENSMUSG00000012126 | Ubxn11 | 67586 | protein_coding | 19.37502888 | -0.404395707 | 0.280636509 | -1.440994646 | 0.149586195 | 0.327026097 | 0.404395707 |
| ENSMUSG00000029630 | Cyp3a25 | 56388 | protein_coding | 9.74809022 | -0.404412473 | 0.548037997 | -0.7379278 | 0.460558325 | 0.658781284 | 0.404412473 |
| ENSMUSG00000037876 | Jmjd1c | 108829 | protein_coding | 1826.762518 | 0.404459902 | 0.098058265 | 4.124689579 | 3.71E-05 | 0.000475707 | 0.404459902 |
| ENSMUSG00000031919 | Tmed6 | 66269 | protein_coding | 55.91218184 | -0.404673176 | 0.198302023 | -2.040691113 | 0.041281539 | 0.136022416 | 0.404673176 |
| ENSMUSG00000022139 | Mbnl2 | 105559 | protein_coding | 3440.319119 | 0.404817054 | 0.099223443 | 4.079852925 | 4.51E-05 | 0.000562526 | 0.404817054 |
| ENSMUSG00000031504 | Rab20 | 19332 | protein_coding | 255.2091483 | 0.404854467 | 0.16738134 | 2.418755085 | 0.015573722 | 0.066539228 | 0.404854467 |
| ENSMUSG00000023984 | Gm20517 | 112694759 | protein_coding | 7.026818699 | -0.404913109 | 0.454017396 | -0.89184492 | 0.372476061 | 0.580792824 | 0.404913109 |
| ENSMUSG00000022814 | Umps | 22247 | protein_coding | 409.2855901 | -0.404938598 | 0.135264336 | -2.9936834 | 0.002756318 | 0.017540005 | 0.404938598 |
| ENSMUSG00000068762 | Gstm6 | 14867 | protein_coding | 200.6270074 | 0.405035918 | 0.126071979 | 3.21273545 | 0.001314773 | 0.009549343 | 0.405035918 |
| ENSMUSG00000052085 | Dock8 | 76088 | protein_coding | 376.1356988 | -0.405117485 | 0.120575482 | -3.359866186 | 0.000779802 | 0.006260429 | 0.405117485 |
| ENSMUSG00000024922 | Ovol1 | 18426 | protein_coding | 312.2352474 | -0.405134105 | 0.229952408 | -1.761817185 | 0.07810019 | 0.210164354 | 0.405134105 |
| ENSMUSG00000041921 | Metap1d | 66559 | protein_coding | 170.1451549 | -0.405134891 | 0.107702835 | -3.761599146 | 0.00016883 | 0.001758752 | 0.405134891 |
| ENSMUSG00000018405 | Mrm1 | 217038 | protein_coding | 139.1427349 | -0.405296796 | 0.1104573 | -3.669262188 | 0.000243252 | 0.002395759 | 0.405296796 |
| ENSMUSG00000070691 | Runx3 | 12399 | protein_coding | 7.74572483 | 0.405327043 | 0.593187698 | 0.683303184 | 0.494415282 | 0.686031895 | 0.405327043 |
| ENSMUSG00000053774 | Ubxn7 | 224111 | protein_coding | 652.6524944 | -0.405348731 | 0.138179165 | -2.933501085 | 0.003351625 | 0.02054047 | 0.405348731 |
| ENSMUSG00000033287 | Kctd17 | 72844 | protein_coding | 237.2516222 | -0.405364666 | 0.110731355 | -3.660793884 | 0.000251435 | 0.002461497 | 0.405364666 |
| ENSMUSG00000059493 | Nhs | 195727 | protein_coding | 26.7348829 | -0.405373755 | 0.257198961 | -1.576109612 | 0.115000544 | 0.272133215 | 0.405373755 |
| ENSMUSG00000021327 | Zkscan3 | 72739 | protein_coding | 823.8633442 | 0.405416159 | 0.117367784 | 3.454237145 | 0.000551852 | 0.004720675 | 0.405416159 |
| ENSMUSG00000001542 | Ell2 | 192657 | protein_coding | 1016.516595 | 0.40545935 | 0.076183561 | 5.322137011 | 1.03E-07 | 2.35E-06 | 0.40545935 |
| ENSMUSG00000033059 | Pygb | 110078 | protein_coding | 3170.386232 | 0.405514599 | 0.116308637 | 3.486539004 | 0.000489314 | 0.004270572 | 0.405514599 |
| ENSMUSG00000032091 | Tmprss4 | 214523 | protein_coding | 2285.017072 | 0.40558751 | 0.099508553 | 4.075906001 | 4.58E-05 | 0.000571259 | 0.40558751 |
| ENSMUSG00000074653 | Lrrc31 | 320352 | protein_coding | 216.3787133 | -0.405601801 | 0.297478867 | -1.363464253 | 0.172736243 | 0.358489058 | 0.405601801 |
| ENSMUSG00000049502 | Dtx3l | 209200 | protein_coding | 806.7552323 | 0.405725639 | 0.120870274 | 3.356703236 | 0.000788777 | 0.00630705 | 0.405725639 |
| ENSMUSG00000039878 | Slc39a5 | 72002 | protein_coding | 1312.124208 | 0.405728059 | 0.114453236 | 3.544924307 | 0.000392726 | 0.003578568 | 0.405728059 |
| ENSMUSG00000085004 | 5430427M07Rik | 71349 | lncRNA | 44.92979771 | -0.405742111 | 0.395857475 | -1.024970189 | 0.305377254 | 0.515150276 | 0.405742111 |
| ENSMUSG00000039934 | Gsap | 212167 | protein_coding | 81.74151369 | 0.405778483 | 0.220882945 | 1.837074759 | 0.06619886 | 0.18798635 | 0.405778483 |
| ENSMUSG00000046668 | Cxxc5 | 67393 | protein_coding | 155.0078557 | -0.406051199 | 0.16564714 | -2.451302195 | 0.01423404 | 0.062097996 | 0.406051199 |
| ENSMUSG00000065952 | Rps23rg1 | 546049 | protein_coding | 26.98147119 | 0.406247472 | 0.347689296 | 1.168420992 | 0.242636991 | 0.447621897 | 0.406247472 |
| ENSMUSG00000073198 | Bnip3l-ps | 100043324 | transcribed_processed_pseudogene | 7.085460499 | 0.406561765 | 0.4173973 | 0.974040237 | 0.330036561 | 0.540678415 | 0.406561765 |

| ensembl_gene_id | external_gene_name | entrezgene_id | gene_biotype | baseMean | log2FoldChange | lfcSE | stat | pvalue | padj | absLog2FC |
|---|---|---|---|---|---|---|---|---|---|---|
| ENSMUSG00000033576 | Apol6 | 71939 | protein_coding | 144.7216207 | 0.406678875 | 0.182022861 | 2.234218669 | 0.025468689 | 0.095901956 | 0.406678875 |
| ENSMUSG00000044949 | Ubtd2 | 327900 | protein_coding | 31.94383969 | -0.406729014 | 0.313844199 | -1.295958361 | 0.194989831 | 0.388054296 | 0.406729014 |
| ENSMUSG00000075415 | Fnbp1 | 14269 | protein_coding | 196.4214724 | -0.406757256 | 0.158398467 | -2.567936814 | 0.01023058 | 0.048628076 | 0.406757256 |
| ENSMUSG00000001473 | Tubb6 | 67951 | protein_coding | 59.88469614 | -0.406779391 | 0.146841487 | -2.770193898 | 0.005602293 | 0.030486274 | 0.406779391 |
| ENSMUSG00000045019 | Acer1 | 171168 | protein_coding | 715.1944627 | 0.406839955 | 0.173673021 | 2.342562778 | 0.019151812 | 0.077541957 | 0.406839955 |
| ENSMUSG00000031774 | Psme3ip1 | 102122 | protein_coding | 599.8372851 | 0.40691743 | 0.078594321 | 5.177440638 | 2.25E-07 | 4.90E-06 | 0.40691743 |
| ENSMUSG00000026941 | Mamdc4 | 381352 | protein_coding | 52.42294404 | -0.406921641 | 0.224976951 | -1.808725916 | 0.070493595 | 0.196228937 | 0.406921641 |
| ENSMUSG00000028398 | Dmac1 | 66928 | protein_coding | 273.1757467 | 0.407001709 | 0.107806825 | 3.775287039 | 0.000159824 | 0.001682531 | 0.407001709 |
| ENSMUSG00000019923 | Zwint | 52696 | protein_coding | 523.395368 | 0.407048968 | 0.119482644 | 3.406762309 | 0.000657383 | 0.005467783 | 0.407048968 |
| ENSMUSG00000085037 | 4933421O10Rik | 100504166 | lncRNA | 94.63986818 | 0.407324289 | 0.163801774 | 2.486690336 | 0.012893758 | 0.057937359 | 0.407324289 |
| ENSMUSG00000022837 | Iqcb1 | 320299 | protein_coding | 43.15810336 | 0.407372911 | 0.208617976 | 1.952721998 | 0.050852546 | 0.157185979 | 0.407372911 |
| ENSMUSG00000048445 | Ccdc57 | 71276 | protein_coding | 41.36253486 | -0.407561383 | 0.250055547 | -1.629883395 | 0.103126144 | 0.253408395 | 0.407561383 |
| ENSMUSG00000032641 | Gpr19 | 14760 | protein_coding | 47.40075896 | -0.407621694 | 0.207718507 | -1.962375428 | 0.049718791 | 0.154762734 | 0.407621694 |
| ENSMUSG00000031591 | Asah1 | 11886 | protein_coding | 2318.971352 | 0.40762493 | 0.10452683 | 3.899715779 | 9.63E-05 | 0.001093341 | 0.40762493 |
| ENSMUSG00000028601 | Echdc2 | 52430 | protein_coding | 178.32902 | -0.407657239 | 0.122255688 | -3.334464392 | 0.000854638 | 0.006732285 | 0.407657239 |
| ENSMUSG00000092341 | Malat1 | 72289 | lncRNA | 48086.88132 | -0.407839119 | 0.199236529 | -2.047009753 | 0.040657127 | 0.134577456 | 0.407839119 |
| ENSMUSG00000029436 | Mmp17 | 23948 | protein_coding | 7.46190741 | -0.407844477 | 0.538361333 | -0.757566437 | 0.448710579 | 0.649476647 | 0.407844477 |
| ENSMUSG00000069744 | Psmb3 | 26446 | protein_coding | 1058.663124 | 0.408303107 | 0.094060094 | 4.340874986 | 1.42E-05 | 0.000203827 | 0.408303107 |
| ENSMUSG00000005251 | Ripk4 | 72388 | protein_coding | 747.5600201 | -0.408419509 | 0.132675118 | -3.078342922 | 0.002081552 | 0.013977641 | 0.408419509 |
| ENSMUSG00000030047 | Arhgap25 | 232201 | protein_coding | 11.04341021 | 0.408429481 | 0.515938659 | 0.791624108 | 0.428579885 | 0.632720402 | 0.408429481 |
| ENSMUSG00000018339 | Gpx3 | 14778 | protein_coding | 17.93998185 | -0.408590912 | 0.30142256 | -1.355541906 | 0.175244967 | 0.361473948 | 0.408590912 |
| ENSMUSG00000056724 | Nbeal2 | 235627 | protein_coding | 1239.629226 | 0.408649289 | 0.15886281 | 2.572340802 | 0.01010134 | 0.048209507 | 0.408649289 |
| ENSMUSG00000033813 | Tcea1 | 21399 | protein_coding | 985.0459632 | 0.408681608 | 0.069990034 | 5.839140004 | 5.25E-09 | 1.45E-07 | 0.408681608 |
| ENSMUSG00000015335 | Zdhhc12 | 66220 | protein_coding | 189.266229 | 0.408727091 | 0.104196433 | 3.922659154 | 8.76E-05 | 0.001006503 | 0.408727091 |
| ENSMUSG00000046733 | Gprc5a | 232431 | protein_coding | 1159.348359 | -0.408820048 | 0.410314989 | -0.996356603 | 0.319076911 | 0.530340919 | 0.408820048 |
| ENSMUSG00000020700 | Map3k3 | 26406 | protein_coding | 445.8596893 | 0.40882229 | 0.121784074 | 3.356943796 | 0.000788091 | 0.00630473 | 0.40882229 |
| ENSMUSG00000054582 | Pabpc1l | 381404 | protein_coding | 8.639238468 | -0.409096185 | 0.560691462 | -0.72962799 | 0.465617608 | 0.662470501 | 0.409096185 |
| ENSMUSG00000028664 | Ephb2 | 13844 | protein_coding | 1268.911303 | -0.409312035 | 0.165572161 | -2.472106609 | 0.013431945 | 0.059744213 | 0.409312035 |
| ENSMUSG00000026418 | Tnni1 | 21952 | protein_coding | 258.2295161 | -0.409319897 | 0.282973287 | -1.446496598 | 0.148037966 | 0.324809051 | 0.409319897 |
| ENSMUSG00000019505 | Ubb | 22187 | protein_coding | 4887.737053 | 0.409381798 | 0.12309845 | 3.325645429 | 0.000882141 | 0.006911331 | 0.409381798 |
| ENSMUSG00000025544 | Tm9sf2 | 68059 | protein_coding | 11198.43874 | 0.409415469 | 0.083151069 | 4.923754754 | 8.49E-07 | 1.63E-05 | 0.409415469 |
| ENSMUSG00000101360 | 4933417O13Rik | 71153 | lncRNA | 11.65513889 | -0.409445151 | 0.460642718 | -0.888856232 | 0.37408035 | 0.582496023 | 0.409445151 |
| ENSMUSG00000044646 | Zbtb7c | 207259 | protein_coding | 349.6200565 | 0.409707863 | 0.17208583 | 2.380834398 | 0.017273475 | 0.071761063 | 0.409707863 |
| ENSMUSG00000027200 | Sema6d | 214968 | protein_coding | 653.9895194 | 0.409788387 | 0.156713244 | 2.61489314 | 0.008925538 | 0.043995232 | 0.409788387 |
| ENSMUSG00000024854 | Pold4 | 69745 | protein_coding | 147.4295334 | 0.409970944 | 0.149835427 | 2.736141589 | 0.006216427 | 0.03304915 | 0.409970944 |
| ENSMUSG00000024892 | Pcx | 18563 | protein_coding | 627.6194615 | -0.410018319 | 0.240882083 | -1.702153659 | 0.088726567 | 0.22975291 | 0.410018319 |
| ENSMUSG00000071711 | Mpst | 246221 | protein_coding | 3025.233762 | 0.41006306 | 0.237028367 | 1.730016812 | 0.083627272 | 0.220715026 | 0.41006306 |
| ENSMUSG00000039801 | Cplane1 | 73692 | protein_coding | 127.6281854 | 0.410087958 | 0.145675838 | 2.815071888 | 0.004876632 | 0.027535643 | 0.410087958 |

| ensembl_gene_id | external_gene_name | entrezgene_id | gene_biotype | baseMean | log2FoldChange | lfcSE | stat | pvalue | padj | absLog2FC |
|---|---|---|---|---|---|---|---|---|---|---|
| ENSMUSG00000060579 | Fhit | 14198 | protein_coding | 92.99597745 | 0.410178574 | 0.143803242 | 2.852359725 | 0.004339596 | 0.02508022 | 0.410178574 |
| ENSMUSG00000055320 | Tead1 | 21676 | protein_coding | 211.2241315 | -0.410293615 | 0.222350724 | -1.845254238 | 0.065000564 | 0.185609355 | 0.410293615 |
| ENSMUSG00000032101 | Ddx25 | 30959 | protein_coding | 11.09901436 | -0.410300024 | 0.422293591 | -0.971598984 | 0.331250088 | 0.541977546 | 0.410300024 |
| ENSMUSG00000063430 | Wscd2 | 320916 | protein_coding | 7.6144742 | -0.410330376 | 0.839359638 | -0.48886122 | 0.624939953 | 0.782837698 | 0.410330376 |
| ENSMUSG00000022265 | Ank | 11732 | protein_coding | 1133.322657 | -0.410474528 | 0.134583148 | -3.049969735 | 0.002288644 | 0.015132223 | 0.410474528 |
| ENSMUSG00000021754 | Map3k1 | 26401 | protein_coding | 716.81698 | -0.410487364 | 0.082210224 | -4.993142515 | 5.94E-07 | 1.20E-05 | 0.410487364 |
| ENSMUSG00000032400 | Zwilch | 68014 | protein_coding | 153.5436226 | 0.410550374 | 0.141402198 | 2.903422859 | 0.003691079 | 0.022085183 | 0.410550374 |
| ENSMUSG00000031972 | Acta1 | 11459 | protein_coding | 167.5104837 | 0.410692766 | 0.302705588 | 1.356739955 | 0.174863852 | 0.361108705 | 0.410692766 |
| ENSMUSG00000036437 | Npy1r | 18166 | protein_coding | 60.98578476 | -0.41072061 | 0.422940043 | -0.971108358 | 0.331494322 | 0.542265745 | 0.41072061 |
| ENSMUSG00000034832 | Tet3 | 194388 | protein_coding | 1101.142612 | -0.410737998 | 0.172956659 | -2.374803031 | 0.017558317 | 0.072634077 | 0.410737998 |
| ENSMUSG00000104445 | Rhbg | 58176 | protein_coding | 19.38501942 | 0.410947406 | 0.291797314 | 1.408331693 | 0.159032875 | 0.340375657 | 0.410947406 |
| ENSMUSG00000034880 | Mrpl34 | 94065 | protein_coding | 433.3084961 | 0.411015889 | 0.109004048 | 3.770647949 | 0.000162824 | 0.00170735 | 0.411015889 |
| ENSMUSG00000044982 | Sft2d3 | 67158 | protein_coding | 73.19495044 | 0.411049728 | 0.152652994 | 2.692706629 | 0.00708746 | 0.036795654 | 0.411049728 |
| ENSMUSG00000033033 | Calhm2 | 72691 | protein_coding | 24.34072906 | -0.411272779 | 0.317310918 | -1.296119218 | 0.194934415 | 0.388054296 | 0.411272779 |
| ENSMUSG00000042210 | Abhd14a | 68644 | protein_coding | 127.4093324 | -0.411391118 | 0.121722555 | -3.379744343 | 0.000725533 | 0.005904844 | 0.411391118 |
| ENSMUSG00000054414 | Slc30a7 | 66500 | protein_coding | 1239.589543 | 0.411415829 | 0.058996525 | 6.973560323 | 3.09E-12 | 1.20E-10 | 0.411415829 |
| ENSMUSG00000023044 | Csad | 246277 | protein_coding | 378.8971 | -0.411425379 | 0.139352361 | -2.952410541 | 0.003153034 | 0.019533452 | 0.411425379 |
| ENSMUSG00000037594 | Clba1 | 217887 | protein_coding | 25.32391904 | 0.411526981 | 0.248685049 | 1.654811905 | 0.097962658 | 0.244886594 | 0.411526981 |
| ENSMUSG00000021998 | Lcp1 | 18826 | protein_coding | 239.1280247 | 0.411720077 | 0.112278397 | 3.666957202 | 0.000245454 | 0.002411838 | 0.411720077 |
| ENSMUSG00000029554 | Mad1l1 | 17120 | protein_coding | 224.8098461 | 0.411858798 | 0.136854412 | 3.00946673 | 0.002617067 | 0.016862722 | 0.411858798 |
| ENSMUSG00000010601 | Apol7a | 75761 | protein_coding | 115.8604691 | 0.41210064 | 0.175448229 | 2.348844682 | 0.01883176 | 0.07652308 | 0.41210064 |
| ENSMUSG00000025512 | Chid1 | 68038 | protein_coding | 753.8300841 | 0.412134491 | 0.090299372 | 4.564090356 | 5.02E-06 | 8.11E-05 | 0.412134491 |
| ENSMUSG00000063698 | Sfxn4 | 94281 | protein_coding | 8.131968287 | -0.412226261 | 0.59729451 | -0.690155784 | 0.490096226 | 0.68248406 | 0.412226261 |
| ENSMUSG00000029475 | Kdm2b | 30841 | protein_coding | 190.0344702 | -0.412236826 | 0.099082658 | -4.160534613 | 3.18E-05 | 0.00041623 | 0.412236826 |
| ENSMUSG00000030089 | Slc41a3 | 71699 | protein_coding | 30.2306853 | 0.412248612 | 0.285025612 | 1.446356379 | 0.14807727 | 0.324827814 | 0.412248612 |
| ENSMUSG00000032740 | Ccdc88a | 108686 | protein_coding | 25.69330163 | 0.412283883 | 0.37713965 | 1.093186259 | 0.274312029 | 0.482256291 | 0.412283883 |
| ENSMUSG00000000782 | Tcf7 | 21414 | protein_coding | 38.76279087 | -0.412285993 | 0.254574635 | -1.619509317 | 0.105337723 | 0.257018881 | 0.412285993 |
| ENSMUSG00000031201 | Brcc3 | 210766 | protein_coding | 555.2192669 | 0.412310336 | 0.07029352 | 5.86555254 | 4.48E-09 | 1.26E-07 | 0.412310336 |
| ENSMUSG00000028756 | Pink1 | 68943 | protein_coding | 1483.743133 | 0.412426456 | 0.128790905 | 3.202294874 | 0.001363374 | 0.009875314 | 0.412426456 |
| ENSMUSG00000026914 | Psmd14 | 59029 | protein_coding | 1194.857497 | 0.412490294 | 0.113084249 | 3.647637021 | 0.000264663 | 0.002568894 | 0.412490294 |
| ENSMUSG00000087968 | Gm25395 | 115485773 | scaRNA | 584.3227892 | -0.412549384 | 0.211929892 | -1.946631408 | 0.05157894 | 0.158701129 | 0.412549384 |
| ENSMUSG00000052488 | Cherp | 27967 | protein_coding | 165.771125 | -0.412624337 | 0.178280779 | -2.314463393 | 0.020642317 | 0.081849845 | 0.412624337 |
| ENSMUSG00000028917 | Plekhm2 | 69582 | protein_coding | 197.1283852 | -0.412679247 | 0.1153903 | -3.576377282 | 0.000348389 | 0.003213165 | 0.412679247 |
| ENSMUSG00000028609 | Magoh | 17149 | protein_coding | 679.5624354 | 0.412836422 | 0.109082281 | 3.784633207 | 0.000153935 | 0.001629379 | 0.412836422 |
| ENSMUSG00000044894 | Uqcrq | 22272 | protein_coding | 1213.255908 | 0.412883606 | 0.121002856 | 3.412180651 | 0.000644454 | 0.005379931 | 0.412883606 |
| ENSMUSG00000026494 | Kif26b | 269152 | protein_coding | 23.05294547 | -0.413034484 | 0.430416378 | -0.959616095 | 0.337248466 | 0.548018112 | 0.413034484 |
| ENSMUSG00000062729 | Ppox | 19044 | protein_coding | 214.1020542 | 0.413458201 | 0.112651564 | 3.670239339 | 0.000242323 | 0.002391405 | 0.413458201 |
| ENSMUSG00000047710 | Champ1 | 101994 | protein_coding | 282.3289871 | -0.41346709 | 0.095776341 | -4.31700652 | 1.58E-05 | 0.000224725 | 0.41346709 |

| ensembl_gene_id | external_gene_name | entrezgene_id | gene_biotype | baseMean | log2FoldChange | lfcSE | stat | pvalue | padj | absLog2FC |
|---|---|---|---|---|---|---|---|---|---|---|
| ENSMUSG00000056413 | Adap1 | 231821 | protein_coding | 501.9816734 | -0.413690073 | 0.181953984 | -2.273597231 | 0.022990214 | 0.088794406 | 0.413690073 |
| ENSMUSG00000066026 | Dhrs3 | 20148 | protein_coding | 418.1740465 | -0.413706382 | 0.15015955 | -2.755112019 | 0.005867204 | 0.031614625 | 0.413706382 |
| ENSMUSG00000024583 | Txnl1 | 53382 | protein_coding | 1446.7776 | 0.413955259 | 0.090446186 | 4.576812784 | 4.72E-06 | 7.66E-05 | 0.413955259 |
| ENSMUSG00000026640 | Plxna2 | 18845 | protein_coding | 911.0350414 | -0.414110597 | 0.223547336 | -1.852451493 | 0.063961017 | 0.183827332 | 0.414110597 |
| ENSMUSG00000037544 | Dlgap5 | 218977 | protein_coding | 383.4540242 | 0.414262597 | 0.138453119 | 2.992078471 | 0.00277085 | 0.017611374 | 0.414262597 |
| ENSMUSG00000015659 | Serac1 | 321007 | protein_coding | 95.80812565 | -0.414319379 | 0.179584443 | -2.307100616 | 0.021049212 | 0.08304975 | 0.414319379 |
| ENSMUSG00000041235 | Chd7 | 320790 | protein_coding | 1302.647226 | -0.414476943 | 0.132672617 | -3.124057944 | 0.001783753 | 0.012315397 | 0.414476943 |
| ENSMUSG00000026778 | Prkcq | 18761 | protein_coding | 11.87238725 | 0.414487306 | 0.59487063 | 0.696768819 | 0.485947482 | 0.678962412 | 0.414487306 |
| ENSMUSG00000018849 | Wwc1 | 211652 | protein_coding | 576.3813634 | -0.41449692 | 0.144356239 | -2.871347466 | 0.00408726 | 0.023890596 | 0.41449692 |
| ENSMUSG00000058794 | Nfe2 | 18022 | protein_coding | 7.734686313 | -0.41463812 | 0.527168135 | -0.786538663 | 0.431551977 | 0.63545899 | 0.41463812 |
| ENSMUSG00000073002 | Vamp5 | 53620 | protein_coding | 10.85366579 | -0.414654672 | 0.370855387 | -1.118103408 | 0.263522831 | 0.471131469 | 0.414654672 |
| ENSMUSG00000003762 | Coq8b | 76889 | protein_coding | 265.5701107 | 0.414845638 | 0.119834572 | 3.461819319 | 0.000536537 | 0.00461944 | 0.414845638 |
| ENSMUSG00000042660 | Wdr55 | 67936 | protein_coding | 248.8597751 | -0.414926204 | 0.119068617 | -3.484765472 | 0.000492569 | 0.004294496 | 0.414926204 |
| ENSMUSG00000049553 | Polr1a | 20019 | protein_coding | 411.8554892 | -0.415013451 | 0.143340122 | -2.89530556 | 0.003787896 | 0.022512541 | 0.415013451 |
| ENSMUSG00000047757 | Fancb | 237211 | protein_coding | 63.07657723 | 0.415089947 | 0.188028201 | 2.207594096 | 0.027272581 | 0.100461996 | 0.415089947 |
| ENSMUSG00000032593 | Amigo3 | 320844 | protein_coding | 158.7602584 | 0.415225214 | 0.190999462 | 2.173960128 | 0.02970813 | 0.107081034 | 0.415225214 |
| ENSMUSG00000032754 | Slc8b1 | 170756 | protein_coding | 480.0340331 | -0.415250153 | 0.220576029 | -1.882571529 | 0.059758458 | 0.175536148 | 0.415250153 |
| ENSMUSG00000030839 | Sergef | 27414 | protein_coding | 60.06827226 | -0.415376342 | 0.172300055 | -2.410773123 | 0.015918747 | 0.067638334 | 0.415376342 |
| ENSMUSG00000025579 | Gaa | 14387 | protein_coding | 564.3306685 | -0.415527884 | 0.084117917 | -4.9398261 | 7.82E-07 | 1.51E-05 | 0.415527884 |
| ENSMUSG00000038872 | Zfhx3 | 11906 | protein_coding | 255.9161368 | -0.415566705 | 0.274541151 | -1.513677286 | 0.130107709 | 0.296838557 | 0.415566705 |
| ENSMUSG00000050212 | Eva1b | 230752 | protein_coding | 16.32999248 | -0.415626513 | 0.318048253 | -1.30680332 | 0.191279517 | 0.383763718 | 0.415626513 |
| ENSMUSG00000040680 | Kremen2 | 73016 | protein_coding | 101.4936379 | -0.415657047 | 0.151837041 | -2.737520728 | 0.006190422 | 0.032965911 | 0.415657047 |
| ENSMUSG00000028358 | Zfp618 | 72701 | protein_coding | 328.9109987 | -0.415789862 | 0.272017759 | -1.5285394 | 0.126378669 | 0.291142528 | 0.415789862 |
| ENSMUSG00000010830 | Kdelr3 | 105785 | protein_coding | 344.8012881 | 0.415797111 | 0.151797916 | 2.739149013 | 0.006159845 | 0.032847006 | 0.415797111 |
| ENSMUSG00000031808 | Slc27a1 | 26457 | protein_coding | 359.8535926 | -0.415840528 | 0.134059971 | -3.10189928 | 0.001922834 | 0.013099613 | 0.415840528 |
| ENSMUSG00000021965 | Ska3 | 219114 | protein_coding | 113.719652 | 0.415959069 | 0.138582612 | 3.001524243 | 0.002686316 | 0.01721837 | 0.415959069 |
| ENSMUSG00000038702 | Dsel | 319901 | protein_coding | 143.0029445 | 0.416122097 | 0.123445673 | 3.370892519 | 0.000749251 | 0.006066795 | 0.416122097 |
| ENSMUSG00000062115 | Rai1 | 19377 | protein_coding | 205.4088833 | -0.416160069 | 0.155813256 | -2.670890012 | 0.007565043 | 0.038819589 | 0.416160069 |
| ENSMUSG00000037325 | Bbs7 | 71492 | protein_coding | 124.1304125 | 0.416223145 | 0.129643743 | 3.210514716 | 0.001324975 | 0.009614669 | 0.416223145 |
| ENSMUSG00000027698 | Nceh1 | 320024 | protein_coding | 992.9312587 | -0.416382835 | 0.238770538 | -1.743861863 | 0.08118318 | 0.216267081 | 0.416382835 |
| ENSMUSG00000057058 | Skap1 | 78473 | protein_coding | 15.81123437 | 0.416385706 | 0.359698286 | 1.157597137 | 0.247028478 | 0.452677129 | 0.416385706 |
| ENSMUSG00000094651 | Gal3st2 | 381334 | protein_coding | 247.1847364 | 0.416407576 | 0.196851402 | 2.115339652 | 0.034401009 | 0.119376748 | 0.416407576 |
| ENSMUSG00000027210 | Meis2 | 17536 | protein_coding | 8.854218769 | -0.416409923 | 0.920008504 | -0.452615297 | 0.650825782 | 0.798250711 | 0.416409923 |
| ENSMUSG00000075502 | Kbtbd6 | 432879 | protein_coding | 13.06790991 | 0.416449207 | 0.316046838 | 1.317681929 | 0.187610147 | 0.378355839 | 0.416449207 |
| ENSMUSG00000032773 | Chrm1 | 12669 | protein_coding | 102.1115485 | -0.416476758 | 0.139022266 | -2.9957558 | 0.002737656 | 0.017435178 | 0.416476758 |
| ENSMUSG00000041912 | Tdrkh | 72634 | protein_coding | 7.758963328 | 0.417059088 | 0.525758356 | 0.793252421 | 0.427630772 | 0.631987012 | 0.417059088 |
| ENSMUSG00000030616 | Sytl2 | 83671 | protein_coding | 2025.593239 | -0.417367656 | 0.188578746 | -2.213227442 | 0.026881969 | 0.099482501 | 0.417367656 |
| ENSMUSG00000070305 | Mpzl3 | 319742 | protein_coding | 444.5905238 | -0.417453795 | 0.183956054 | -2.26931262 | 0.023249322 | 0.089512979 | 0.417453795 |

| ensembl_gene_id | external_gene_name | entrezgene_id | gene_biotype | baseMean | log2FoldChange | lfcSE | stat | pvalue | padj | absLog2FC |
|---|---|---|---|---|---|---|---|---|---|---|
| ENSMUSG00000029993 | Nfu1 | 56748 | protein_coding | 331.5887915 | -0.41747733 | 0.103515787 | -4.032982241 | 5.51E-05 | 0.00067374 | 0.41747733 |
| ENSMUSG00000034800 | Zfp661 | 72180 | protein_coding | 74.88601706 | -0.417595016 | 0.189486774 | -2.203821444 | 0.027536904 | 0.101201616 | 0.417595016 |
| ENSMUSG00000030189 | Ybx3 | 56449 | protein_coding | 1998.812618 | -0.417666428 | 0.08594957 | -4.859435943 | 1.18E-06 | 2.18E-05 | 0.417666428 |
| ENSMUSG00000031150 | Ccdc120 | 54648 | protein_coding | 134.727371 | -0.41775332 | 0.201143795 | -2.076888921 | 0.037811813 | 0.128000666 | 0.41775332 |
| ENSMUSG00000030518 | Entrep2 | 70638 | protein_coding | 51.8450006 | -0.417908505 | 0.223865309 | -1.86678547 | 0.061931552 | 0.179549648 | 0.417908505 |
| ENSMUSG00000027429 | Sec23b | 27054 | protein_coding | 1188.024738 | 0.417957851 | 0.131170561 | 3.186369303 | 0.001440706 | 0.010327435 | 0.417957851 |
| ENSMUSG00000030795 | Fus | 233908 | protein_coding | 614.6036466 | -0.418067612 | 0.179950103 | -2.323241866 | 0.020166163 | 0.08030166 | 0.418067612 |
| ENSMUSG00000043998 | Mgat2 | 217664 | protein_coding | 935.452981 | 0.418193152 | 0.09904984 | 4.222047744 | 2.42E-05 | 0.000326231 | 0.418193152 |
| ENSMUSG00000087424 | 5730405O15Rik | 70489 | lncRNA | 8.120751127 | 0.418649491 | 0.416197797 | 1.005890696 | 0.314468152 | 0.525201712 | 0.418649491 |
| ENSMUSG00000036186 | Dipk1b | 56279 | protein_coding | 37.63059115 | -0.418770055 | 0.195813013 | -2.138622193 | 0.032466279 | 0.114255412 | 0.418770055 |
| ENSMUSG00000022947 | Cbr3 | 109857 | protein_coding | 524.4739532 | -0.418927674 | 0.14404381 | -2.908335134 | 0.003633587 | 0.021823445 | 0.418927674 |
| ENSMUSG00000041607 | Mbp | 17196 | protein_coding | 1171.74933 | -0.418931205 | 0.082900262 | -5.053436428 | 4.34E-07 | 9.00E-06 | 0.418931205 |
| ENSMUSG00000074627 | Mroh8 | 629499 | protein_coding | 13.39549832 | -0.419572384 | 0.418297271 | -1.003048342 | 0.315837537 | 0.526825078 | 0.419572384 |
| ENSMUSG00000044201 | Cdc25c | 12532 | protein_coding | 125.145837 | 0.419761457 | 0.127483868 | 3.292663326 | 0.000992432 | 0.007636455 | 0.419761457 |
| ENSMUSG00000043252 | Tmem64 | 100201 | protein_coding | 186.116897 | 0.41987469 | 0.157680466 | 2.662819946 | 0.007748886 | 0.039498143 | 0.41987469 |
| ENSMUSG00000039005 | Tlr4 | 21898 | protein_coding | 593.1006677 | 0.419924483 | 0.181365575 | 2.315348345 | 0.020593876 | 0.081739162 | 0.419924483 |
| ENSMUSG00000026179 | Pnkd | 56695 | protein_coding | 277.0629551 | 0.420014688 | 0.090813705 | 4.625014357 | 3.75E-06 | 6.19E-05 | 0.420014688 |
| ENSMUSG00000029722 | Agfg2 | 231801 | protein_coding | 308.6278861 | -0.420107105 | 0.098502034 | -4.26495869 | 2.00E-05 | 0.000277649 | 0.420107105 |
| ENSMUSG00000073926 | Or52d3 | 57250 | protein_coding | 6.773992699 | -0.420140399 | 0.590676219 | -0.711287141 | 0.476906318 | 0.671454418 | 0.420140399 |
| ENSMUSG00000027936 | Crtc2 | 74343 | protein_coding | 136.8341775 | -0.420319525 | 0.142448402 | -2.950679129 | 0.003170761 | 0.019590336 | 0.420319525 |
| ENSMUSG00000031939 | Taf1d | 75316 | protein_coding | 1906.349021 | -0.42060877 | 0.089815952 | -4.683007417 | 2.83E-06 | 4.80E-05 | 0.42060877 |
| ENSMUSG00000063931 | Pepd | 18624 | protein_coding | 575.1418967 | 0.420698381 | 0.111779641 | 3.763640485 | 0.000167458 | 0.001746327 | 0.420698381 |
| ENSMUSG00000046634 | Pkd1l1 | 171395 | protein_coding | 8.737099757 | -0.420783069 | 0.68926289 | -0.610482699 | 0.541542101 | 0.724720445 | 0.420783069 |
| ENSMUSG00000030751 | Psma1 | 26440 | protein_coding | 1676.101371 | 0.420785297 | 0.10040323 | 4.190953809 | 2.78E-05 | 0.000368712 | 0.420785297 |
| ENSMUSG00000036504 | Phpt1 | 75454 | protein_coding | 15.46532449 | 0.420855286 | 0.305210079 | 1.378903633 | 0.167924468 | 0.352398014 | 0.420855286 |
| ENSMUSG00000022528 | Hes1 | 15205 | protein_coding | 981.3252457 | 0.421382145 | 0.12662426 | 3.32781527 | 0.000875299 | 0.006871247 | 0.421382145 |
| ENSMUSG00000038936 | Sccpdh | 109232 | protein_coding | 305.3369187 | 0.421407625 | 0.151832923 | 2.775469363 | 0.005512209 | 0.03012988 | 0.421407625 |
| ENSMUSG00000032431 | Crtap | 56693 | protein_coding | 84.60114142 | -0.421520344 | 0.133352012 | -3.160959756 | 0.001572502 | 0.01106797 | 0.421520344 |
| ENSMUSG00000026657 | Frmd4a | 209630 | protein_coding | 161.3194365 | -0.42158314 | 0.199059705 | -2.117872826 | 0.03418584 | 0.118915058 | 0.42158314 |
| ENSMUSG00000021214 | Akr1c18 | 105349 | protein_coding | 64.89577908 | -0.421648416 | 0.299488163 | -1.407896766 | 0.159161641 | 0.340376923 | 0.421648416 |
| ENSMUSG00000092837 | Rpph1 | 85029 | ribozyme | 135.1988302 | 0.421712167 | 0.174760201 | 2.413090413 | 0.015817895 | 0.067329619 | 0.421712167 |
| ENSMUSG00000045467 | Ttll13 | 269954 | protein_coding | 9.502360942 | -0.421789706 | 0.514669062 | -0.81953577 | 0.412480802 | 0.619284967 | 0.421789706 |
| ENSMUSG00000026768 | Itga8 | 241226 | protein_coding | 11.74725094 | -0.422036146 | 0.515458335 | -0.81875899 | 0.412923933 | 0.619566809 | 0.422036146 |
| ENSMUSG00000002324 | Rec8 | 56739 | protein_coding | 51.5497063 | 0.422089982 | 0.265470029 | 1.589972262 | 0.111841058 | 0.267157224 | 0.422089982 |
| ENSMUSG00000028608 | Czib | 74098 | protein_coding | 314.3682331 | 0.422135448 | 0.119109589 | 3.544092894 | 0.000393966 | 0.00358577 | 0.422135448 |
| ENSMUSG00000032604 | Qars | 97541 | protein_coding | 2667.751566 | 0.422450748 | 0.08854555 | 4.770999194 | 1.83E-06 | 3.23E-05 | 0.422450748 |
| ENSMUSG00000072082 | Ccnf | 12449 | protein_coding | 291.8733195 | 0.422492222 | 0.131245692 | 3.219094028 | 0.001285963 | 0.009382877 | 0.422492222 |
| ENSMUSG00000033055 | Ankrd54 | 223690 | protein_coding | 234.8301247 | 0.422677603 | 0.107358608 | 3.937063004 | 8.25E-05 | 0.000953426 | 0.422677603 |

| ensembl_gene_id | external_gene_name | entrezgene_id | gene_biotype | baseMean | log2FoldChange | lfcSE | stat | pvalue | padj | absLog2FC |
|---|---|---|---|---|---|---|---|---|---|---|
| ENSMUSG00000028461 | Ccdc107 | 622404 | protein_coding | 468.5847 | -0.422681994 | 0.088072393 | -4.799256393 | 1.59E-06 | 2.86E-05 | 0.422681994 |
| ENSMUSG00000026311 | Asb1 | 65247 | protein_coding | 70.83634029 | -0.422738571 | 0.186880575 | -2.262078711 | 0.023692541 | 0.090758727 | 0.422738571 |
| ENSMUSG00000027201 | Myef2 | 17876 | protein_coding | 237.8174195 | 0.422802325 | 0.12665112 | 3.338322838 | 0.000842858 | 0.006655942 | 0.422802325 |
| ENSMUSG00000079017 | Ifi27l2a | 76933 | protein_coding | 10.06306949 | 0.42283246 | 0.392342421 | 1.077712828 | 0.281161932 | 0.489758011 | 0.42283246 |
| ENSMUSG00000038502 | Ptov1 | 84113 | protein_coding | 684.7736707 | 0.42293876 | 0.094530615 | 4.474092955 | 7.67E-06 | 0.000118092 | 0.42293876 |
| ENSMUSG00000039753 | Fbxl5 | 242960 | protein_coding | 2156.003563 | 0.422942142 | 0.174156923 | 2.428511798 | 0.015160932 | 0.065212887 | 0.422942142 |
| ENSMUSG00000068742 | Cry2 | 12953 | protein_coding | 190.0772259 | -0.423137691 | 0.207236127 | -2.041814317 | 0.041169953 | 0.1357952 | 0.423137691 |
| ENSMUSG00000029673 | Auts2 | 319974 | protein_coding | 399.7031628 | -0.42315291 | 0.181582014 | -2.330367971 | 0.019786711 | 0.07915076 | 0.42315291 |
| ENSMUSG00000029446 | Psph | 100678 | protein_coding | 99.02358092 | 0.423283949 | 0.120686338 | 3.507306256 | 0.000452668 | 0.004003383 | 0.423283949 |
| ENSMUSG00000024454 | Hdac3 | 15183 | protein_coding | 1045.689799 | 0.423557319 | 0.073679319 | 5.748659507 | 9.00E-09 | 2.45E-07 | 0.423557319 |
| ENSMUSG00000091405 | H4c14 | 97122 | protein_coding | 535.5509893 | 0.423630628 | 0.151310063 | 2.79975184 | 0.005114191 | 0.028531989 | 0.423630628 |
| ENSMUSG00000024921 | Smarca2 | 67155 | protein_coding | 498.6285039 | -0.423693118 | 0.115847366 | -3.657339257 | 0.000254847 | 0.002490308 | 0.423693118 |
| ENSMUSG00000037992 | Rara | 19401 | protein_coding | 232.3872073 | -0.423698015 | 0.110542782 | -3.832887219 | 0.000126648 | 0.001381678 | 0.423698015 |
| ENSMUSG00000062822 | 4833420G17Rik | 67392 | protein_coding | 395.7069581 | -0.423720627 | 0.132455153 | -3.198974277 | 0.001379175 | 0.009971627 | 0.423720627 |
| ENSMUSG00000024622 | Hmgxb3 | 106894 | protein_coding | 550.5450867 | -0.423763112 | 0.099890565 | -4.242273664 | 2.21E-05 | 0.000302 | 0.423763112 |
| ENSMUSG00000040714 | Klc3 | 232943 | protein_coding | 15.48379909 | -0.423784072 | 0.29759432 | -1.424032798 | 0.154436978 | 0.333813568 | 0.423784072 |
| ENSMUSG00000029471 | Camkk2 | 207565 | protein_coding | 52.34226246 | -0.423911534 | 0.18030187 | -2.351121121 | 0.01871694 | 0.076205998 | 0.423911534 |
| ENSMUSG00000058267 | Mrps14 | 64659 | protein_coding | 526.5694194 | 0.424339768 | 0.090884796 | 4.668985223 | 3.03E-06 | 5.10E-05 | 0.424339768 |
| ENSMUSG00000020546 | Stxbp4 | 20913 | protein_coding | 42.23958404 | 0.424382337 | 0.212015133 | 2.001660595 | 0.045321247 | 0.145409528 | 0.424382337 |
| ENSMUSG00000079260 | Tmppe | 100504715 | protein_coding | 115.041885 | -0.42485701 | 0.187578326 | -2.264957892 | 0.023515264 | 0.090275035 | 0.42485701 |
| ENSMUSG00000004263 | Atn1 | 13498 | protein_coding | 8.474121292 | -0.424881789 | 0.471053896 | -0.901981265 | 0.36706682 | 0.57568337 | 0.424881789 |
| ENSMUSG00000035299 | Mid1 | 17318 | protein_coding | 91.26531095 | -0.424898423 | 0.408826306 | -1.039312825 | 0.298659272 | 0.507472252 | 0.424898423 |
| ENSMUSG00000034324 | Tmem132c | 208213 | protein_coding | 26.90266818 | -0.424942264 | 0.305305847 | -1.391857603 | 0.163965523 | 0.347108299 | 0.424942264 |
| ENSMUSG00000038807 | Rap1gap2 | 380711 | protein_coding | 1539.76971 | -0.425005666 | 0.079178304 | -5.367703619 | 7.97E-08 | 1.85E-06 | 0.425005666 |
| ENSMUSG00000032292 | Nr2e3 | 23958 | protein_coding | 53.82537256 | 0.425084866 | 0.27299335 | 1.557125348 | 0.119440728 | 0.280225646 | 0.425084866 |
| ENSMUSG00000031613 | Hpgd | 15446 | protein_coding | 2828.672844 | -0.425133428 | 0.16655195 | -2.552557492 | 0.010693526 | 0.05030905 | 0.425133428 |
| ENSMUSG00000073856 | Iqck | 434232 | protein_coding | 12.21835073 | -0.425308084 | 0.428471527 | -0.99261691 | 0.320896681 | 0.532256186 | 0.425308084 |
| ENSMUSG00000056493 | Foxk1 | 17425 | protein_coding | 408.0513069 | -0.425581067 | 0.154183581 | -2.760223001 | 0.005776192 | 0.031229865 | 0.425581067 |
| ENSMUSG00000030935 | Acsm3 | 20216 | protein_coding | 2520.510294 | -0.425594657 | 0.178467865 | -2.384713106 | 0.017092445 | 0.07125058 | 0.425594657 |
| ENSMUSG00000026385 | Dbi | 13167 | protein_coding | 3661.541066 | 0.425610431 | 0.120852276 | 3.521741131 | 0.000428723 | 0.003844985 | 0.425610431 |
| ENSMUSG00000021670 | Hmgcr | 15357 | protein_coding | 3132.394898 | 0.425685099 | 0.225965701 | 1.883848285 | 0.0595855 | 0.175321947 | 0.425685099 |
| ENSMUSG00000025991 | Cps1 | 227231 | protein_coding | 444.5756461 | -0.425800792 | 0.258835728 | -1.645061891 | 0.099957048 | 0.248070406 | 0.425800792 |
| ENSMUSG00000031502 | Col4a1 | 12826 | protein_coding | 18.87852412 | -0.42585229 | 0.410022674 | -1.038606684 | 0.298987696 | 0.507887459 | 0.42585229 |
| ENSMUSG00000054191 | Klf1 | 16596 | protein_coding | 7.180413728 | -0.425923237 | 0.455693521 | -0.934670381 | 0.349958196 | 0.560158189 | 0.425923237 |
| ENSMUSG00000017897 | Eya2 | 14049 | protein_coding | 37.59935076 | -0.425930446 | 0.361942259 | -1.176791146 | 0.239278877 | 0.443424073 | 0.425930446 |
| ENSMUSG00000017607 | Tns4 | 217169 | protein_coding | 1536.010504 | -0.425933451 | 0.250242084 | -1.702085611 | 0.088739321 | 0.22975291 | 0.425933451 |
| ENSMUSG00000028675 | Pnrc2 | 52830 | protein_coding | 1963.328281 | 0.426134096 | 0.086298281 | 4.937921014 | 7.90E-07 | 1.53E-05 | 0.426134096 |
| ENSMUSG00000019899 | Lama2 | 16773 | protein_coding | 10.35159549 | -0.426198292 | 0.783284291 | -0.544116992 | 0.586360969 | 0.75648449 | 0.426198292 |

| ensembl_gene_id | external_gene_name | entrezgene_id | gene_biotype | baseMean | log2FoldChange | lfcSE | stat | pvalue | padj | absLog2FC |
|---|---|---|---|---|---|---|---|---|---|---|
| ENSMUSG00000041935 | Rimoc1 | 106064 | protein_coding | 490.4117501 | -0.426288774 | 0.129485267 | -3.292179751 | 0.00099414 | 0.007642215 | 0.426288774 |
| ENSMUSG00000041375 | Ccdc9 | 243846 | protein_coding | 799.0986486 | -0.426359109 | 0.1067823 | -3.992788212 | 6.53E-05 | 0.000775625 | 0.426359109 |
| ENSMUSG00000029247 | Paics | 67054 | protein_coding | 1238.926992 | -0.426586134 | 0.18651223 | -2.287175134 | 0.022185603 | 0.086314675 | 0.426586134 |
| ENSMUSG00000063049 | Ing2 | 69260 | protein_coding | 124.9359118 | -0.426789809 | 0.131839875 | -3.237183045 | 0.00120716 | 0.008930626 | 0.426789809 |
| ENSMUSG00000018770 | Atp5g3 | 228033 | protein_coding | 5911.756623 | 0.427184746 | 0.123895234 | 3.447951392 | 0.000564856 | 0.004803535 | 0.427184746 |
| ENSMUSG00000041674 | BC006965 | 217294 | lncRNA | 9.331205585 | 0.427443812 | 0.553701222 | 0.771975562 | 0.440128902 | 0.642313808 | 0.427443812 |
| ENSMUSG00000015747 | Vps45 | 22365 | protein_coding | 303.4435939 | 0.427884437 | 0.093850882 | 4.559194634 | 5.14E-06 | 8.27E-05 | 0.427884437 |
| ENSMUSG00000007476 | Lrrc8a | 241296 | protein_coding | 174.3357961 | -0.42806335 | 0.13469266 | -3.178074803 | 0.001482565 | 0.010560955 | 0.42806335 |
| ENSMUSG00000071547 | Nt5dc2 | 70021 | protein_coding | 43.28195091 | 0.428073676 | 0.245527464 | 1.743485916 | 0.081248772 | 0.21633742 | 0.428073676 |
| ENSMUSG00000070808 | Bicra | 243842 | protein_coding | 84.84145744 | -0.428117217 | 0.265650965 | -1.611577875 | 0.107053829 | 0.260022383 | 0.428117217 |
| ENSMUSG00000019437 | Tlcd1 | 68385 | protein_coding | 150.8074311 | -0.428217589 | 0.109852182 | -3.898125488 | 9.69E-05 | 0.001099468 | 0.428217589 |
| ENSMUSG00000026405 | C4bp | 12269 | protein_coding | 163.7074791 | 0.428315567 | 0.225766574 | 1.897161119 | 0.057806677 | 0.171719678 | 0.428315567 |
| ENSMUSG00000072919 | Noxred1 | 71275 | protein_coding | 7.348008081 | 0.428377807 | 0.484518343 | 0.884131248 | 0.376625367 | 0.58462681 | 0.428377807 |
| ENSMUSG00000032109 | Nlrx1 | 270151 | protein_coding | 258.9862956 | 0.428749347 | 0.162686947 | 2.635425619 | 0.008403184 | 0.042037034 | 0.428749347 |
| ENSMUSG00000003526 | Prodh | 19125 | protein_coding | 217.1653028 | -0.428762624 | 0.1424669 | -3.009559587 | 0.002616268 | 0.016862722 | 0.428762624 |
| ENSMUSG00000020519 | Sap30l | 50724 | protein_coding | 278.1840062 | 0.428843363 | 0.095805229 | 4.476199952 | 7.60E-06 | 0.00011716 | 0.428843363 |
| ENSMUSG00000073728 | Tmem51os1 | 100038693 | lncRNA | 16.70115314 | -0.428845498 | 0.280195896 | -1.530520266 | 0.125888001 | 0.290318255 | 0.428845498 |
| ENSMUSG00000079657 | Rab26 | 328778 | protein_coding | 10.60448389 | 0.428889863 | 0.430764736 | 0.995647571 | 0.319421412 | 0.530636656 | 0.428889863 |
| ENSMUSG00000075700 | Selenot | 69227 | protein_coding | 2164.399438 | 0.428994647 | 0.09192411 | 4.666834928 | 3.06E-06 | 5.14E-05 | 0.428994647 |
| ENSMUSG00000028277 | Ube2j1 | 56228 | protein_coding | 1062.582955 | 0.429010563 | 0.094574882 | 4.536199798 | 5.73E-06 | 9.09E-05 | 0.429010563 |
| ENSMUSG00000008999 | Bmp7 | 12162 | protein_coding | 47.09581648 | -0.429083759 | 0.299425769 | -1.433022149 | 0.151851508 | 0.330106567 | 0.429083759 |
| ENSMUSG00000032557 | Uba5 | 66663 | protein_coding | 633.0601867 | 0.429442273 | 0.117390147 | 3.658248023 | 0.000253945 | 0.00248302 | 0.429442273 |
| ENSMUSG00000078532 | Nkain1 | 67149 | protein_coding | 89.2232898 | 0.429748633 | 0.207755066 | 2.068535037 | 0.038589739 | 0.129921237 | 0.429748633 |
| ENSMUSG00000041836 | Ptpre | 19267 | protein_coding | 157.3928798 | -0.429861291 | 0.19278041 | -2.229797572 | 0.025760885 | 0.096750618 | 0.429861291 |
| ENSMUSG00000035901 | Dennd5a | 19347 | protein_coding | 53.85095572 | 0.430009083 | 0.211486332 | 2.033271276 | 0.042025123 | 0.137533628 | 0.430009083 |
| ENSMUSG00000105504 | Gbp5 | 229898 | protein_coding | 8.353227382 | 0.430027395 | 0.59286679 | 0.725335611 | 0.468246164 | 0.664844437 | 0.430027395 |
| ENSMUSG00000031595 | Pdgfrl | 68797 | protein_coding | 11.20464538 | 0.430113405 | 0.44154158 | 0.974117557 | 0.329998173 | 0.540678415 | 0.430113405 |
| ENSMUSG00000038633 | Degs1 | 13244 | protein_coding | 426.8344131 | 0.430177086 | 0.121318336 | 3.545853815 | 0.000391343 | 0.003569124 | 0.430177086 |
| ENSMUSG00000037493 | Cib2 | 56506 | protein_coding | 35.13977601 | -0.430217808 | 0.22301227 | -1.929121693 | 0.053715757 | 0.162875418 | 0.430217808 |
| ENSMUSG00000025584 | Pde8a | 18584 | protein_coding | 131.5069284 | -0.430225862 | 0.104986305 | -4.097923665 | 4.17E-05 | 0.000524897 | 0.430225862 |
| ENSMUSG00000031538 | Plat | 18791 | protein_coding | 14.66159929 | 0.430538636 | 0.427332355 | 1.007503015 | 0.31369311 | 0.524237105 | 0.430538636 |
| ENSMUSG00000086370 | Ftx | 78878 | lncRNA | 89.58246814 | -0.430939678 | 0.165188715 | -2.608771902 | 0.009086779 | 0.044556467 | 0.430939678 |
| ENSMUSG00000049538 | Adamts16 | 271127 | protein_coding | 14.80411607 | -0.431232448 | 0.39690028 | -1.086500741 | 0.277257513 | 0.485615564 | 0.431232448 |
| ENSMUSG00000032028 | Nxpe2 | 78252 | protein_coding | 1658.012453 | -0.431272167 | 0.313084247 | -1.377495582 | 0.16835908 | 0.352984523 | 0.431272167 |
| ENSMUSG00000017774 | Myo1c | 17913 | protein_coding | 1151.417486 | -0.431658078 | 0.119752638 | -3.604580945 | 0.000312657 | 0.002938053 | 0.431658078 |
| ENSMUSG00000028518 | Prkaa2 | 108079 | protein_coding | 2337.135981 | -0.431727016 | 0.118682488 | -3.637664025 | 0.000275122 | 0.002648353 | 0.431727016 |
| ENSMUSG00000028795 | Ccdc28b | 66264 | protein_coding | 130.43142 | -0.431792665 | 0.186777836 | -2.311798203 | 0.020788807 | 0.082328222 | 0.431792665 |
| ENSMUSG00000003123 | Lipe | 16890 | protein_coding | 76.89905279 | -0.431873254 | 0.180884092 | -2.38756902 | 0.016960218 | 0.070866303 | 0.431873254 |

| ensembl_gene_id | external_gene_name | entrezgene_id | gene_biotype | baseMean | log2FoldChange | lfcSE | stat | pvalue | padj | absLog2FC |
|---|---|---|---|---|---|---|---|---|---|---|
| ENSMUSG00000021125 | Arg2 | 11847 | protein_coding | 146.3361352 | -0.432088718 | 0.240287617 | -1.79821467 | 0.072142996 | 0.199564716 | 0.432088718 |
| ENSMUSG00000056735 | A930024E05Rik | 109202 | lncRNA | 9.451633542 | 0.432388981 | 0.461980933 | 0.935945512 | 0.349301243 | 0.559443961 | 0.432388981 |
| ENSMUSG00000043631 | Ecm2 | 407800 | protein_coding | 11.19793023 | -0.432527951 | 0.54631772 | -0.791715032 | 0.428526856 | 0.632720402 | 0.432527951 |
| ENSMUSG00000005682 | Pan2 | 103135 | protein_coding | 574.9037856 | 0.432621706 | 0.09703001 | 4.45863818 | 8.25E-06 | 0.00012584 | 0.432621706 |
| ENSMUSG00000031349 | Nsdhl | 18194 | protein_coding | 553.0360924 | 0.432706786 | 0.099402319 | 4.353085436 | 1.34E-05 | 0.000193668 | 0.432706786 |
| ENSMUSG00000032531 | Amotl2 | 56332 | protein_coding | 84.88278301 | -0.432838643 | 0.210101437 | -2.060141277 | 0.039385037 | 0.131735587 | 0.432838643 |
| ENSMUSG00000005481 | Ddx39a | 68278 | protein_coding | 912.4437559 | 0.433048563 | 0.200652102 | 2.158205956 | 0.030911823 | 0.110460495 | 0.433048563 |
| ENSMUSG00000021301 | Hecw1 | 94253 | protein_coding | 9.59403201 | -0.433189855 | 1.871581083 | -0.231456633 | 0.816960073 | 0.902675156 | 0.433189855 |
| ENSMUSG00000063972 | Nr6a1 | 14536 | protein_coding | 158.3441829 | -0.433221613 | 0.140628314 | -3.08061443 | 0.00206574 | 0.013906636 | 0.433221613 |
| ENSMUSG00000051510 | Mafg | 17134 | protein_coding | 341.6224366 | -0.433274888 | 0.134140281 | -3.230013272 | 0.001237845 | 0.009106042 | 0.433274888 |
| ENSMUSG00000045629 | Sh3tc2 | 225608 | protein_coding | 17.32865966 | -0.433321061 | 0.306632726 | -1.413159861 | 0.157608726 | 0.338418951 | 0.433321061 |
| ENSMUSG00000018143 | Mafk | 17135 | protein_coding | 960.2451816 | -0.433396707 | 0.200740963 | -2.15898489 | 0.030851339 | 0.110377388 | 0.433396707 |
| ENSMUSG00000052837 | Junb | 16477 | protein_coding | 3484.114165 | -0.433429899 | 0.24511428 | -1.768276819 | 0.077014637 | 0.208125067 | 0.433429899 |
| ENSMUSG00000079516 | Reg3a | 19694 | protein_coding | 20.53874444 | 0.433480543 | 0.37626725 | 1.152054937 | 0.249298499 | 0.455578992 | 0.433480543 |
| ENSMUSG00000035762 | Tmem161b | 72745 | protein_coding | 384.9864585 | 0.433596718 | 0.074107154 | 5.850942767 | 4.89E-09 | 1.37E-07 | 0.433596718 |
| ENSMUSG00000031627 | Irf2 | 16363 | protein_coding | 384.1506204 | -0.433804545 | 0.155074261 | -2.797398761 | 0.005151591 | 0.028700434 | 0.433804545 |
| ENSMUSG00000022687 | Boc | 117606 | protein_coding | 861.3485718 | -0.433871837 | 0.265830536 | -1.632136936 | 0.102650642 | 0.252737778 | 0.433871837 |
| ENSMUSG00000036885 | Arhgef26 | 622434 | protein_coding | 184.1881991 | -0.434057514 | 0.206142045 | -2.105623402 | 0.035237078 | 0.121615641 | 0.434057514 |
| ENSMUSG00000029381 | Shroom3 | 27428 | protein_coding | 2421.800836 | -0.434078292 | 0.117341514 | -3.699272976 | 0.000216218 | 0.002178317 | 0.434078292 |
| ENSMUSG00000036564 | Ndrg4 | 234593 | protein_coding | 7.109628667 | -0.434342623 | 0.606122222 | -0.716592475 | 0.473625589 | 0.668798783 | 0.434342623 |
| ENSMUSG00000041594 | Tmtc4 | 70551 | protein_coding | 477.1601184 | 0.434375081 | 0.0855993 | 5.074516746 | 3.88E-07 | 8.10E-06 | 0.434375081 |
| ENSMUSG00000019256 | Ahr | 11622 | protein_coding | 237.6863569 | -0.434470384 | 0.114030452 | -3.81012596 | 0.000138896 | 0.001499229 | 0.434470384 |
| ENSMUSG00000060441 | Trim5 | 667823 | protein_coding | 22.99017487 | -0.434484722 | 0.332852161 | -1.305338442 | 0.191777629 | 0.384424123 | 0.434484722 |
| ENSMUSG00000040722 | Scamp5 | 56807 | protein_coding | 568.9182512 | -0.434624946 | 0.111236741 | -3.907206744 | 9.34E-05 | 0.001064557 | 0.434624946 |
| ENSMUSG00000089960 | Ugt1a1 | 394436 | protein_coding | 280.4054453 | -0.434790016 | 0.218127119 | -1.9932873 | 0.046229996 | 0.147358691 | 0.434790016 |
| ENSMUSG00000028212 | Ccne2 | 12448 | protein_coding | 183.5396176 | 0.434818185 | 0.157663013 | 2.75789595 | 0.005817471 | 0.031421052 | 0.434818185 |
| ENSMUSG00000071192 | Wfikkn1 | 215001 | protein_coding | 18.28706717 | -0.434943056 | 0.356516603 | -1.219979805 | 0.222472531 | 0.42381802 | 0.434943056 |
| ENSMUSG00000006418 | Rnf114 | 81018 | protein_coding | 1376.249311 | 0.435112165 | 0.09343308 | 4.656939105 | 3.21E-06 | 5.39E-05 | 0.435112165 |
| ENSMUSG00000074136 | 4930513N10Rik | 319960 | lncRNA | 24.39161598 | 0.43515767 | 0.291739643 | 1.491595953 | 0.1358051 | 0.30556627 | 0.43515767 |
| ENSMUSG00000031609 | Sap30 | 60406 | protein_coding | 144.3780838 | -0.435282497 | 0.1433036 | -3.037484734 | 0.002385615 | 0.015701683 | 0.435282497 |
| ENSMUSG00000091586 | Cyp4f17 | 208285 | protein_coding | 43.17821834 | -0.435407901 | 0.214082814 | -2.033829309 | 0.041968808 | 0.137405792 | 0.435407901 |
| ENSMUSG00000040463 | Mybbp1a | 18432 | protein_coding | 871.3467066 | -0.435450238 | 0.204112517 | -2.133383315 | 0.032893295 | 0.11540185 | 0.435450238 |
| ENSMUSG00000029603 | Dtx1 | 14357 | protein_coding | 13.30190084 | 0.435546708 | 0.427418128 | 1.019017864 | 0.308194486 | 0.518256099 | 0.435546708 |
| ENSMUSG00000039286 | Fndc3b | 72007 | protein_coding | 832.23321 | 0.435607332 | 0.092946774 | 4.686632097 | 2.78E-06 | 4.73E-05 | 0.435607332 |
| ENSMUSG00000055978 | Fut2 | 14344 | protein_coding | 3228.029404 | 0.435660123 | 0.164303397 | 2.651558831 | 0.008012115 | 0.040564836 | 0.435660123 |
| ENSMUSG00000024810 | Il33 | 77125 | protein_coding | 17.54085698 | 0.43566323 | 0.349250033 | 1.247425023 | 0.212241696 | 0.410115945 | 0.43566323 |
| ENSMUSG00000005267 | Zfp287 | 170740 | protein_coding | 40.37746242 | 0.435702302 | 0.235463551 | 1.850402326 | 0.064255584 | 0.184274927 | 0.435702302 |
| ENSMUSG00000051212 | Gpr183 | 321019 | protein_coding | 18.65373088 | 0.435721321 | 0.332413825 | 1.310779782 | 0.189932175 | 0.381541138 | 0.435721321 |

| ensembl_gene_id | external_gene_name | entrezgene_id | gene_biotype | baseMean | log2FoldChange | lfcSE | stat | pvalue | padj | absLog2FC |
|---|---|---|---|---|---|---|---|---|---|---|
| ENSMUSG00000033439 | Trmt13 | 229780 | protein_coding | 87.90638792 | -0.435986546 | 0.176930725 | -2.464165255 | 0.013733278 | 0.060536076 | 0.435986546 |
| ENSMUSG00000020883 | Fbxl20 | 72194 | protein_coding | 460.6688184 | -0.436250953 | 0.181141443 | -2.408344248 | 0.016025062 | 0.067940661 | 0.436250953 |
| ENSMUSG00000021451 | Sema4d | 20354 | protein_coding | 115.0915286 | -0.43628785 | 0.177141517 | -2.462933917 | 0.013780531 | 0.060718202 | 0.43628785 |
| ENSMUSG00000031508 | Ankrd10 | 102334 | protein_coding | 800.5920985 | -0.436295445 | 0.145376625 | -3.00113891 | 0.002689718 | 0.01723324 | 0.436295445 |
| ENSMUSG00000032717 | Mdfi | 17240 | protein_coding | 7.252011298 | 0.436482269 | 0.78109094 | 0.558811076 | 0.576290663 | 0.748585685 | 0.436482269 |
| ENSMUSG00000022055 | Nefl | 18039 | protein_coding | 8.170330627 | 0.43665646 | 0.489016066 | 0.892928659 | 0.371895378 | 0.580114542 | 0.43665646 |
| ENSMUSG00000054659 | Pm20d2 | 242377 | protein_coding | 26.6964491 | -0.436839751 | 0.217247171 | -2.010796039 | 0.044347005 | 0.142814212 | 0.436839751 |
| ENSMUSG00000043432 | Leng9 | 243813 | protein_coding | 604.4729907 | 0.436865186 | 0.192983836 | 2.263739777 | 0.023590124 | 0.09049947 | 0.436865186 |
| ENSMUSG00000021203 | Otub2 | 68149 | protein_coding | 27.06395041 | -0.436907214 | 0.237004906 | -1.843452193 | 0.065263016 | 0.185992005 | 0.436907214 |
| ENSMUSG00000032733 | Snx33 | 235406 | protein_coding | 347.7208285 | -0.436990983 | 0.091236097 | -4.789672044 | 1.67E-06 | 2.98E-05 | 0.436990983 |
| ENSMUSG00000086938 | 4930481A15Rik | 74931 | lncRNA | 93.46756271 | 0.437012052 | 0.193624686 | 2.257005865 | 0.024007708 | 0.0916794 | 0.437012052 |
| ENSMUSG00000021079 | Timm9 | 30056 | protein_coding | 177.7689412 | -0.437015999 | 0.15151583 | -2.884292679 | 0.00392294 | 0.023116753 | 0.437015999 |
| ENSMUSG00000035200 | Chrnb4 | 108015 | protein_coding | 77.84229829 | -0.437037499 | 0.16387731 | -2.666857902 | 0.007656404 | 0.039124542 | 0.437037499 |
| ENSMUSG00000033458 | Fan1 | 330554 | protein_coding | 122.3070955 | -0.43708821 | 0.190281934 | -2.29705574 | 0.02161559 | 0.084615534 | 0.43708821 |
| ENSMUSG00000070287 | Slc35g2 | 245020 | protein_coding | 107.9499369 | -0.437110071 | 0.167372775 | -2.611596012 | 0.009012068 | 0.044262786 | 0.437110071 |
| ENSMUSG00000039481 | Nrtn | 18188 | protein_coding | 11.56368618 | -0.437275978 | 0.482132913 | -0.906961476 | 0.36442717 | 0.573236812 | 0.437275978 |
| ENSMUSG00000048076 | Arf1 | 11840 | protein_coding | 5992.0105 | 0.437419593 | 0.090923121 | 4.810873036 | 1.50E-06 | 2.72E-05 | 0.437419593 |
| ENSMUSG00000073802 | Cdkn2b | 12579 | protein_coding | 510.8231986 | 0.437554446 | 0.129241025 | 3.385569287 | 0.000710308 | 0.005804916 | 0.437554446 |
| ENSMUSG00000032959 | Pebp1 | 23980 | protein_coding | 1242.30777 | 0.437604853 | 0.112028622 | 3.906187945 | 9.38E-05 | 0.001067727 | 0.437604853 |
| ENSMUSG00000029817 | Tra2a | 101214 | protein_coding | 1426.469812 | 0.437673111 | 0.074736809 | 5.856192083 | 4.74E-09 | 1.33E-07 | 0.437673111 |
| ENSMUSG00000055022 | Cntn1 | 12805 | protein_coding | 8.889263162 | -0.437732741 | 0.836558785 | -0.523254013 | 0.600797499 | 0.766909966 | 0.437732741 |
| ENSMUSG00000029727 | Cyp3a13 | 13113 | protein_coding | 1361.896083 | -0.437748596 | 0.212860032 | -2.056509117 | 0.039733468 | 0.132481904 | 0.437748596 |
| ENSMUSG00000026482 | Rgl1 | 19731 | protein_coding | 10.73406939 | -0.438040402 | 0.416990611 | -1.050480252 | 0.293497366 | 0.502507368 | 0.438040402 |
| ENSMUSG00000117286 | Gm1043 | 381634 | protein_coding | 14.15958636 | -0.438160668 | 0.506332891 | -0.865360864 | 0.386840755 | 0.59503617 | 0.438160668 |
| ENSMUSG00000000340 | Dbt | 13171 | protein_coding | 562.239757 | -0.438232882 | 0.107745806 | -4.067284847 | 4.76E-05 | 0.000590952 | 0.438232882 |
| ENSMUSG00000026109 | Tmeff2 | 56363 | protein_coding | 7.959955533 | -0.438427836 | 0.704815796 | -0.622045985 | 0.533911631 | 0.718850876 | 0.438427836 |
| ENSMUSG00000035840 | Lysmd3 | 80289 | protein_coding | 729.41474 | 0.438519438 | 0.119363821 | 3.673805289 | 0.000238965 | 0.002365589 | 0.438519438 |
| ENSMUSG00000034248 | Slc25a37 | 67712 | protein_coding | 344.227218 | -0.438706346 | 0.167657194 | -2.616686685 | 0.00887878 | 0.043853148 | 0.438706346 |
| ENSMUSG00000106943 | Dancr | 70036 | lncRNA | 34.73487046 | -0.438798052 | 0.215059451 | -2.040356983 | 0.041314783 | 0.136047522 | 0.438798052 |
| ENSMUSG00000034327 | Kctd9 | 105440 | protein_coding | 296.6267493 | 0.438805873 | 0.130686054 | 3.357710022 | 0.00078591 | 0.006290441 | 0.438805873 |
| ENSMUSG00000048371 | Pdp2 | 382051 | protein_coding | 391.2799226 | -0.438826217 | 0.154375966 | -2.84258119 | 0.004474983 | 0.025694856 | 0.438826217 |
| ENSMUSG00000022677 | Cep20 | 66086 | protein_coding | 618.7149036 | 0.438998972 | 0.09722009 | 4.51551702 | 6.32E-06 | 9.93E-05 | 0.438998972 |
| ENSMUSG00000061436 | Hipk2 | 15258 | protein_coding | 391.6904929 | -0.439005026 | 0.196731685 | -2.231491209 | 0.025648609 | 0.096442646 | 0.439005026 |
| ENSMUSG00000090083 | Rnf8 | 58230 | protein_coding | 108.0072119 | -0.439081851 | 0.127083338 | -3.455070182 | 0.00055015 | 0.004711174 | 0.439081851 |
| ENSMUSG00000025626 | Phf6 | 70998 | protein_coding | 461.9532194 | 0.439410568 | 0.081592152 | 5.385451359 | 7.23E-08 | 1.69E-06 | 0.439410568 |
| ENSMUSG00000025871 | 4833439L19Rik | 97820 | protein_coding | 2864.724507 | 0.439426815 | 0.081258715 | 5.407749959 | 6.38E-08 | 1.51E-06 | 0.439426815 |
| ENSMUSG00000032116 | Stt3a | 16430 | protein_coding | 3320.586963 | 0.439459246 | 0.071098472 | 6.180994215 | 6.37E-10 | 1.95E-08 | 0.439459246 |
| ENSMUSG00000041649 | Klf8 | 245671 | protein_coding | 24.55917431 | -0.439614889 | 0.264438114 | -1.662449042 | 0.096422773 | 0.242664232 | 0.439614889 |

| ensembl_gene_id | external_gene_name | entrezgene_id | gene_biotype | baseMean | log2FoldChange | lfcSE | stat | pvalue | padj | absLog2FC |
|---|---|---|---|---|---|---|---|---|---|---|
| ENSMUSG00000029001 | Fbxo44 | 230903 | protein_coding | 31.29994058 | -0.439780943 | 0.224590196 | -1.958148443 | 0.050212597 | 0.155903751 | 0.439780943 |
| ENSMUSG00000017405 | Nek8 | 140859 | protein_coding | 128.1646398 | -0.439789601 | 0.156417147 | -2.81164571 | 0.004928876 | 0.02773854 | 0.439789601 |
| ENSMUSG00000079442 | St6galnac4 | 20448 | protein_coding | 441.9439282 | 0.439794914 | 0.12948366 | 3.396528287 | 0.000682465 | 0.005638124 | 0.439794914 |
| ENSMUSG00000002797 | Ggct | 110175 | protein_coding | 69.47313462 | 0.439872325 | 0.178850192 | 2.459445633 | 0.013915177 | 0.061148872 | 0.439872325 |
| ENSMUSG00000099083 | Atf7 | 223922 | protein_coding | 162.4765588 | -0.439994634 | 0.199712999 | -2.203134679 | 0.027585258 | 0.101309196 | 0.439994634 |
| ENSMUSG00000022802 | Lmln | 239833 | protein_coding | 91.26782887 | -0.440036532 | 0.2162732 | -2.034632731 | 0.041887841 | 0.137293642 | 0.440036532 |
| ENSMUSG00000074476 | Spc24 | 67629 | protein_coding | 225.8348998 | 0.44006181 | 0.143200747 | 3.073041305 | 0.002118892 | 0.014198445 | 0.44006181 |
| ENSMUSG00000043155 | Hpdl | 242642 | protein_coding | 93.26277496 | -0.440281227 | 0.207520977 | -2.121622756 | 0.03386943 | 0.118020625 | 0.440281227 |
| ENSMUSG00000048763 | Hoxb3 | 15410 | protein_coding | 50.72628699 | -0.44034586 | 0.276606057 | -1.591960296 | 0.111393626 | 0.266448067 | 0.44034586 |
| ENSMUSG00000049303 | Syt12 | 171180 | protein_coding | 14.85822916 | -0.440349488 | 0.502786484 | -0.875818069 | 0.381128945 | 0.588980483 | 0.440349488 |
| ENSMUSG00000085282 | Gm15663 | 327812 | lncRNA | 28.79534076 | -0.440507998 | 0.205681242 | -2.141702349 | 0.032217442 | 0.113706938 | 0.440507998 |
| ENSMUSG00000028919 | Arhgef19 | 213649 | protein_coding | 210.5238068 | -0.440585175 | 0.179717377 | -2.451544651 | 0.014224454 | 0.062073178 | 0.440585175 |
| ENSMUSG00000031227 | Magee1 | 107528 | protein_coding | 10.10941711 | 0.440694407 | 0.38532246 | 1.14370288 | 0.252746879 | 0.459036748 | 0.440694407 |
| ENSMUSG00000069633 | Pex11g | 69129 | protein_coding | 137.4444367 | 0.440714816 | 0.119143958 | 3.699011042 | 0.000216441 | 0.002179188 | 0.440714816 |
| ENSMUSG00000050947 | Amigo1 | 229715 | protein_coding | 170.7753635 | -0.44075682 | 0.121028685 | -3.641755006 | 0.000270786 | 0.002617157 | 0.44075682 |
| ENSMUSG00000000378 | Ccm2 | 216527 | protein_coding | 282.820983 | 0.440843919 | 0.125649125 | 3.508531555 | 0.000450588 | 0.003993858 | 0.440843919 |
| ENSMUSG00000085551 | 1810063I02Rik | 69854 | lncRNA | 13.32447793 | 0.440849106 | 0.492968621 | 0.894274174 | 0.371175215 | 0.579501943 | 0.440849106 |
| ENSMUSG00000040857 | Erf | 13875 | protein_coding | 339.9409591 | -0.440851359 | 0.135912372 | -3.243644081 | 0.001180111 | 0.008779453 | 0.440851359 |
| ENSMUSG00000056598 | Drc3 | 74665 | protein_coding | 29.80047284 | 0.441002433 | 0.205417557 | 2.146858525 | 0.031804546 | 0.112791279 | 0.441002433 |
| ENSMUSG00000034584 | Exph5 | 320051 | protein_coding | 1123.269603 | -0.44101488 | 0.207618246 | -2.124162436 | 0.033656563 | 0.117484492 | 0.44101488 |
| ENSMUSG00000037278 | Tmem97 | 69071 | protein_coding | 845.848121 | 0.441234898 | 0.108796991 | 4.055579971 | 5.00E-05 | 0.000619891 | 0.441234898 |
| ENSMUSG00000020143 | Dock2 | 94176 | protein_coding | 39.52487709 | 0.441356068 | 0.470611364 | 0.937835552 | 0.348328927 | 0.558392025 | 0.441356068 |
| ENSMUSG00000025499 | Hras | 15461 | protein_coding | 689.8837047 | 0.441361227 | 0.117038071 | 3.771091109 | 0.000162535 | 0.00170735 | 0.441361227 |
| ENSMUSG00000035722 | Abca7 | 27403 | protein_coding | 1060.399115 | 0.441563655 | 0.107629525 | 4.102625707 | 4.08E-05 | 0.000516791 | 0.441563655 |
| ENSMUSG00000052934 | Fbxo31 | 76454 | protein_coding | 229.7001034 | -0.441590833 | 0.15626317 | -2.82594314 | 0.004714163 | 0.026845617 | 0.441590833 |
| ENSMUSG00000078919 | Dpm1 | 13480 | protein_coding | 60.80048762 | -0.4417617 | 0.14763503 | -2.992255283 | 0.002769246 | 0.017611374 | 0.4417617 |
| ENSMUSG00000034591 | Slc41a2 | 338365 | protein_coding | 463.3853691 | 0.441864623 | 0.131531974 | 3.35937042 | 0.000781203 | 0.006265356 | 0.441864623 |
| ENSMUSG00000064451 | Snora23 | 100379145 | snoRNA | 29.78279671 | 0.442193843 | 0.365530618 | 1.209731336 | 0.226382001 | 0.428651036 | 0.442193843 |
| ENSMUSG00000026388 | 3110009E18Rik | 73103 | protein_coding | 15.63269976 | 0.442262516 | 0.305604888 | 1.447170948 | 0.147849051 | 0.324686593 | 0.442262516 |
| ENSMUSG00000036553 | Sh3tc1 | 231147 | protein_coding | 389.6456409 | 0.442386398 | 0.140553634 | 3.147456137 | 0.001646978 | 0.011490612 | 0.442386398 |
| ENSMUSG00000023088 | Abcc1 | 17250 | protein_coding | 668.9409645 | -0.442544326 | 0.115851425 | -3.81992994 | 0.00013349 | 0.001447394 | 0.442544326 |
| ENSMUSG00000068551 | Zfp467 | 68910 | protein_coding | 294.6281612 | -0.442827714 | 0.189818447 | -2.332901363 | 0.019653323 | 0.078791373 | 0.442827714 |
| ENSMUSG00000031292 | Cdkl5 | 382253 | protein_coding | 67.99878413 | 0.442893836 | 0.172023999 | 2.574604933 | 0.010035464 | 0.048042752 | 0.442893836 |
| ENSMUSG00000020405 | Fabp6 | 16204 | protein_coding | 101.4628892 | 0.442905841 | 0.32248817 | 1.373401822 | 0.169627464 | 0.354608197 | 0.442905841 |
| ENSMUSG00000025176 | Hoga1 | 67432 | protein_coding | 58.96307583 | 0.442996574 | 0.233306639 | 1.898773976 | 0.0575942 | 0.171405161 | 0.442996574 |
| ENSMUSG00000036943 | Rab8b | 235442 | protein_coding | 203.6537738 | 0.443262406 | 0.129732376 | 3.416744688 | 0.000633747 | 0.005307213 | 0.443262406 |
| ENSMUSG00000099632 | 2900093K20Rik | 108168734 | lncRNA | 10.73244401 | 0.443285952 | 0.379545605 | 1.167938572 | 0.242831544 | 0.447768098 | 0.443285952 |
| ENSMUSG00000032178 | Ilf3 | 16201 | protein_coding | 509.5105129 | -0.443492273 | 0.07256621 | -6.111553503 | 9.87E-10 | 2.98E-08 | 0.443492273 |

Supplementary Table 1

| ensembl_gene_id | external_gene_name | entrezgene_id | gene_biotype | baseMean | log2FoldChange | lfcSE | stat | pvalue | padj | absLog2FC |
|---|---|---|---|---|---|---|---|---|---|---|
| ENSMUSG00000075702 | Selenom | 114679 | protein_coding | 1187.14533 | 0.443547199 | 0.101955062 | 4.350418606 | 1.36E-05 | 0.000195684 | 0.443547199 |
| ENSMUSG00000055725 | Paqr3 | 231474 | protein_coding | 10.58661661 | -0.44361935 | 0.495735167 | -0.894871656 | 0.3708557 | 0.579400646 | 0.44361935 |
| ENSMUSG00000050288 | Fzd2 | 57265 | protein_coding | 32.73645368 | 0.443656686 | 0.208678454 | 2.12603015 | 0.033500749 | 0.117017529 | 0.443656686 |
| ENSMUSG00000031242 | Chmp1b2 | 67028 | protein_coding | 571.7578453 | 0.443853879 | 0.087498013 | 5.072730955 | 3.92E-07 | 8.16E-06 | 0.443853879 |
| ENSMUSG00000051951 | Xkr4 | 497097 | protein_coding | 47.89391006 | -0.444474365 | 0.264198158 | -1.682352249 | 0.092500555 | 0.236227167 | 0.444474365 |
| ENSMUSG00000061024 | Rrs1 | 59014 | protein_coding | 268.685681 | -0.444507257 | 0.212696349 | -2.089867831 | 0.036629674 | 0.12495983 | 0.444507257 |
| ENSMUSG00000035351 | Nup37 | 69736 | protein_coding | 188.3487913 | 0.444590266 | 0.137142415 | 3.241814475 | 0.001187713 | 0.008819531 | 0.444590266 |
| ENSMUSG00000002043 | Trappc6a | 67091 | protein_coding | 773.0666035 | 0.444926964 | 0.128085086 | 3.473682832 | 0.000513367 | 0.004451234 | 0.444926964 |
| ENSMUSG00000035049 | Rrp12 | 107094 | protein_coding | 208.1253944 | -0.444957311 | 0.247081778 | -1.800850369 | 0.071726467 | 0.198792269 | 0.444957311 |
| ENSMUSG00000039814 | Xkr5 | 319581 | protein_coding | 26.57925415 | 0.445014468 | 0.28347539 | 1.56985221 | 0.116449498 | 0.274745608 | 0.445014468 |
| ENSMUSG00000052496 | Pkdrej | 18766 | protein_coding | 47.52695949 | 0.445060528 | 0.221055898 | 2.013339308 | 0.044078946 | 0.142209732 | 0.445060528 |
| ENSMUSG00000053647 | Gper1 | 76854 | protein_coding | 10.57744334 | -0.445186132 | 0.384393368 | -1.15815248 | 0.246801817 | 0.452313811 | 0.445186132 |
| ENSMUSG00000054978 | Kbtbd13 | 74492 | protein_coding | 37.21639379 | 0.445311421 | 0.190279344 | 2.340303533 | 0.019268073 | 0.077854355 | 0.445311421 |
| ENSMUSG00000029455 | Aldh2 | 11669 | protein_coding | 5661.805787 | -0.445344816 | 0.133555404 | -3.334532355 | 0.00085443 | 0.006732285 | 0.445344816 |
| ENSMUSG00000022574 | Naprt | 223646 | protein_coding | 630.8833572 | -0.445467904 | 0.136064165 | -3.273954644 | 0.001060536 | 0.008043916 | 0.445467904 |
| ENSMUSG00000024143 | Rhoq | 104215 | protein_coding | 660.8011356 | 0.445543877 | 0.105991473 | 4.203582272 | 2.63E-05 | 0.000351062 | 0.445543877 |
| ENSMUSG00000047766 | Lrrc49 | 102747 | protein_coding | 116.8405385 | 0.445728885 | 0.124010982 | 3.594269447 | 0.000325303 | 0.003037972 | 0.445728885 |
| ENSMUSG00000033805 | Ephx4 | 384214 | protein_coding | 85.13698113 | -0.445814096 | 0.124272536 | -3.587390334 | 0.000334004 | 0.003105673 | 0.445814096 |
| ENSMUSG00000075590 | Nrbp2 | 223649 | protein_coding | 325.51512 | -0.445862576 | 0.168257341 | -2.649884841 | 0.008051921 | 0.040701682 | 0.445862576 |
| ENSMUSG00000044244 | Il20rb | 213208 | protein_coding | 17.65399328 | -0.446046379 | 0.281920582 | -1.582170326 | 0.113610693 | 0.269718683 | 0.446046379 |
| ENSMUSG00000021553 | Slc28a3 | 114304 | protein_coding | 71.08444307 | -0.446571514 | 0.393439686 | -1.135044403 | 0.256356714 | 0.463348653 | 0.446571514 |
| ENSMUSG00000050555 | Hyls1 | 76832 | protein_coding | 138.4157111 | 0.446732468 | 0.135217719 | 3.303801251 | 0.000953834 | 0.00738584 | 0.446732468 |
| ENSMUSG00000062124 | Defb45 | 433490 | protein_coding | 13.46625534 | -0.446882101 | 0.592198984 | -0.754614773 | 0.450480157 | 0.650993281 | 0.446882101 |
| ENSMUSG00000005672 | Kit | 16590 | protein_coding | 1124.861726 | -0.446895237 | 0.152630618 | -2.927952754 | 0.003412019 | 0.020774708 | 0.446895237 |
| ENSMUSG00000114469 | C730002L08Rik | 320155 | lncRNA | 11.62172516 | -0.446900259 | 0.515415277 | -0.86706832 | 0.385904577 | 0.593825534 | 0.446900259 |
| ENSMUSG00000048078 | Tenm4 | 23966 | protein_coding | 259.6610327 | -0.446967658 | 0.194520579 | -2.29779111 | 0.021573681 | 0.084491171 | 0.446967658 |
| ENSMUSG00000025808 | Ccdc7a | 74703 | protein_coding | 11.02097648 | -0.447073785 | 0.845730638 | -0.528624322 | 0.597066085 | 0.764635807 | 0.447073785 |
| ENSMUSG00000069805 | Fbp1 | 14121 | protein_coding | 28.0058297 | -0.447104863 | 0.248811236 | -1.796964118 | 0.072341316 | 0.199870336 | 0.447104863 |
| ENSMUSG00000039218 | Srrm2 | 75956 | protein_coding | 3502.059574 | -0.447138887 | 0.205918035 | -2.17144112 | 0.029897843 | 0.107691735 | 0.447138887 |
| ENSMUSG00000104713 | Gbp6 | 100702 | protein_coding | 18.90263318 | 0.447229979 | 0.308441723 | 1.449965895 | 0.14706803 | 0.323549666 | 0.447229979 |
| ENSMUSG00000020105 | Lrig3 | 320398 | protein_coding | 1515.160825 | -0.447403637 | 0.143528522 | -3.11717582 | 0.001825926 | 0.012535929 | 0.447403637 |
| ENSMUSG00000025010 | Ccnj | 240665 | protein_coding | 80.85540001 | -0.447469503 | 0.158430863 | -2.824383433 | 0.004737168 | 0.026966978 | 0.447469503 |
| ENSMUSG00000024403 | Atp6v1g2 | 66237 | protein_coding | 16.2717259 | -0.447517647 | 0.325487482 | -1.374915079 | 0.169157774 | 0.354093686 | 0.447517647 |
| ENSMUSG00000000154 | Slc22a18 | 18400 | protein_coding | 917.5168369 | 0.447710431 | 0.125164531 | 3.576975274 | 0.000347593 | 0.003207684 | 0.447710431 |
| ENSMUSG00000026819 | Slc25a25 | 227731 | protein_coding | 714.6444266 | -0.447906405 | 0.177931571 | -2.517295851 | 0.011825948 | 0.054267848 | 0.447906405 |
| ENSMUSG00000027347 | Rasgrp1 | 19419 | protein_coding | 23.64867778 | 0.447992863 | 0.318175165 | 1.408007013 | 0.159128993 | 0.340376923 | 0.447992863 |
| ENSMUSG00000045991 | Onecut2 | 225631 | protein_coding | 7.106549784 | -0.448219173 | 0.938705753 | -0.477486338 | 0.633015847 | 0.787741735 | 0.448219173 |
| ENSMUSG00000069833 | Ahnak | 66395 | protein_coding | 16078.954 | -0.448341238 | 0.170257807 | -2.633307965 | 0.008455764 | 0.042220503 | 0.448341238 |

| ensembl_gene_id | external_gene_name | entrezgene_id | gene_biotype | baseMean | log2FoldChange | lfcSE | stat | pvalue | padj | absLog2FC |
|---|---|---|---|---|---|---|---|---|---|---|
| ENSMUSG00000027636 | Sla2 | 77799 | protein_coding | 8.833750877 | 0.448398365 | 0.598337199 | 0.749407469 | 0.453611651 | 0.653148289 | 0.448398365 |
| ENSMUSG00000069302 | H2ac12 | 319168 | protein_coding | 51.21733569 | 0.448926799 | 0.317459185 | 1.414124462 | 0.157325362 | 0.33818806 | 0.448926799 |
| ENSMUSG00000027422 | Rrbp1 | 81910 | protein_coding | 7366.047734 | 0.448943858 | 0.103428897 | 4.340603747 | 1.42E-05 | 0.000203895 | 0.448943858 |
| ENSMUSG00000030717 | Nupr1 | 56312 | protein_coding | 3198.755418 | 0.449192668 | 0.170554175 | 2.633724256 | 0.008445404 | 0.042182001 | 0.449192668 |
| ENSMUSG00000056752 | Dnah9 | 237806 | protein_coding | 27.8449737 | -0.449246558 | 0.459407183 | -0.977883182 | 0.328132104 | 0.538929056 | 0.449246558 |
| ENSMUSG00000048126 | Col6a3 | 12835 | protein_coding | 14.89124831 | -0.449347694 | 0.492588444 | -0.912217286 | 0.361654346 | 0.570896781 | 0.449347694 |
| ENSMUSG00000027244 | Atg13 | 51897 | protein_coding | 414.0325096 | 0.449505144 | 0.09215274 | 4.877827204 | 1.07E-06 | 2.01E-05 | 0.449505144 |
| ENSMUSG00000009733 | Tfcp2 | 21422 | protein_coding | 184.0699914 | 0.44966798 | 0.096877886 | 4.641595725 | 3.46E-06 | 5.76E-05 | 0.44966798 |
| ENSMUSG00000020621 | Rdh14 | 105014 | protein_coding | 443.0355459 | 0.449751201 | 0.103491236 | 4.345790202 | 1.39E-05 | 0.000199676 | 0.449751201 |
| ENSMUSG00000025422 | Agap2 | 216439 | protein_coding | 14.62757823 | 0.449781834 | 0.526410201 | 0.854432215 | 0.392865553 | 0.600822134 | 0.449781834 |
| ENSMUSG00000024413 | Npc1 | 18145 | protein_coding | 610.2091961 | -0.44998268 | 0.148528731 | -3.02960025 | 0.002448776 | 0.01604447 | 0.44998268 |
| ENSMUSG00000055799 | Tcf7l1 | 21415 | protein_coding | 108.6148111 | -0.45005854 | 0.259025255 | -1.737508336 | 0.082297484 | 0.218347988 | 0.45005854 |
| ENSMUSG00000025602 | Zfp202 | 80902 | protein_coding | 40.85417244 | -0.450163869 | 0.198328401 | -2.269790247 | 0.023220313 | 0.089444532 | 0.450163869 |
| ENSMUSG00000029178 | Klf3 | 16599 | protein_coding | 3242.145902 | -0.450310704 | 0.204152125 | -2.205760558 | 0.027400769 | 0.100817615 | 0.450310704 |
| ENSMUSG00000027940 | Tpm3 | 59069 | protein_coding | 3998.653323 | 0.450442766 | 0.121666541 | 3.702273128 | 0.000213676 | 0.002160913 | 0.450442766 |
| ENSMUSG00000032776 | Mctp2 | 244049 | protein_coding | 1183.341598 | 0.450644628 | 0.108886083 | 4.138679751 | 3.49E-05 | 0.000452481 | 0.450644628 |
| ENSMUSG00000069268 | H2bc7 | 319180 | protein_coding | 291.366854 | 0.4511745 | 0.150458714 | 2.998659825 | 0.002711699 | 0.017332239 | 0.4511745 |
| ENSMUSG00000038415 | Foxq1 | 15220 | protein_coding | 46.73859427 | -0.451565423 | 0.382486419 | -1.180605115 | 0.237759631 | 0.442071905 | 0.451565423 |
| ENSMUSG00000040943 | Tet2 | 214133 | protein_coding | 510.7122032 | -0.45183613 | 0.152589205 | -2.961127747 | 0.003065148 | 0.019115769 | 0.45183613 |
| ENSMUSG00000033955 | Tnks1bp1 | 228140 | protein_coding | 2032.30968 | -0.452018253 | 0.20856125 | -2.167316568 | 0.030210724 | 0.108557068 | 0.452018253 |
| ENSMUSG00000022129 | Dct | 13190 | protein_coding | 19.85004998 | -0.452043337 | 0.299364312 | -1.510010775 | 0.131040675 | 0.298386829 | 0.452043337 |
| ENSMUSG00000005980 | Dnase1 | 13419 | protein_coding | 15.77404185 | -0.452076887 | 0.325584726 | -1.388507665 | 0.164982512 | 0.348473871 | 0.452076887 |
| ENSMUSG00000036246 | Gmip | 78816 | protein_coding | 167.0374493 | -0.452145952 | 0.13593958 | -3.326080248 | 0.000880766 | 0.006903955 | 0.452145952 |
| ENSMUSG00000036242 | Armh4 | 67419 | protein_coding | 145.9503975 | -0.452228597 | 0.130662709 | -3.461037967 | 0.000538097 | 0.004630366 | 0.452228597 |
| ENSMUSG00000033065 | Pfkm | 18642 | protein_coding | 161.6106819 | -0.452297169 | 0.18703202 | -2.41828736 | 0.015593757 | 0.066589104 | 0.452297169 |
| ENSMUSG00000051650 | B3gnt2 | 53625 | protein_coding | 41.45621861 | -0.452339386 | 0.243226089 | -1.859748632 | 0.062921098 | 0.181394977 | 0.452339386 |
| ENSMUSG00000035165 | Kcne3 | 57442 | protein_coding | 2662.017867 | -0.452360461 | 0.116108359 | -3.896019747 | 9.78E-05 | 0.001106995 | 0.452360461 |
| ENSMUSG00000022546 | Gpt | 76282 | protein_coding | 619.5928578 | -0.452586468 | 0.233203126 | -1.940739287 | 0.052289908 | 0.160198819 | 0.452586468 |
| ENSMUSG00000068547 | Clca4a | 99663 | protein_coding | 1730.145518 | -0.452596967 | 0.50361052 | -0.898704354 | 0.368810157 | 0.577640836 | 0.452596967 |
| ENSMUSG00000022452 | Smdt1 | 69029 | protein_coding | 1397.87934 | 0.45260484 | 0.120363803 | 3.760306922 | 0.000169705 | 0.001766707 | 0.45260484 |
| ENSMUSG00000022900 | Ildr1 | 106347 | protein_coding | 3794.769597 | 0.452777352 | 0.162503384 | 2.786264135 | 0.005331941 | 0.029406907 | 0.452777352 |
| ENSMUSG00000060780 | Lrrtm1 | 74342 | protein_coding | 22.33273518 | 0.452871483 | 0.47184187 | 0.959795034 | 0.337158382 | 0.547985583 | 0.452871483 |
| ENSMUSG00000020718 | Polg2 | 50776 | protein_coding | 148.1395797 | -0.453042456 | 0.225995464 | -2.004652874 | 0.04500017 | 0.144596067 | 0.453042456 |
| ENSMUSG00000032633 | Flcn | 216805 | protein_coding | 307.6465078 | -0.453082981 | 0.138335781 | -3.275240709 | 0.00105572 | 0.008018839 | 0.453082981 |
| ENSMUSG00000031304 | Il2rg | 16186 | protein_coding | 8.473056849 | 0.453145224 | 0.451829391 | 1.002912234 | 0.315903209 | 0.526825078 | 0.453145224 |
| ENSMUSG00000027317 | Ppp1r14d | 72112 | protein_coding | 1757.161714 | 0.453158865 | 0.096417603 | 4.69995987 | 2.60E-06 | 4.46E-05 | 0.453158865 |
| ENSMUSG00000032589 | Bsn | 12217 | protein_coding | 161.9614097 | -0.453167132 | 0.150790205 | -3.005282291 | 0.002653345 | 0.01706289 | 0.453167132 |
| ENSMUSG00000091971 | Hspa1a | 193740 | protein_coding | 1030.3004 | 0.453169287 | 0.542852267 | 0.834793025 | 0.403834253 | 0.611027688 | 0.453169287 |

| ensembl_gene_id | external_gene_name | entrezgene_id | gene_biotype | baseMean | log2FoldChange | lfcSE | stat | pvalue | padj | absLog2FC |
|---|---|---|---|---|---|---|---|---|---|---|
| ENSMUSG00000035126 | Dnai4 | 242584 | protein_coding | 42.53232867 | 0.453171968 | 0.27474998 | 1.64939764 | 0.099066198 | 0.246812223 | 0.453171968 |
| ENSMUSG00000029093 | Sorcs2 | 81840 | protein_coding | 286.6653311 | -0.453200337 | 0.117834009 | -3.846091119 | 0.000120017 | 0.001318368 | 0.453200337 |
| ENSMUSG00000050315 | Synpo2 | 118449 | protein_coding | 28.55312724 | -0.453440932 | 0.330510205 | -1.371942302 | 0.1700814 | 0.355286104 | 0.453440932 |
| ENSMUSG00000024539 | Ptpn2 | 19255 | protein_coding | 637.8922527 | 0.453693315 | 0.093809651 | 4.836318124 | 1.32E-06 | 2.42E-05 | 0.453693315 |
| ENSMUSG00000021048 | Mthfd1 | 108156 | protein_coding | 928.3509341 | -0.453721986 | 0.094410813 | -4.805826469 | 1.54E-06 | 2.78E-05 | 0.453721986 |
| ENSMUSG00000024871 | Doc2g | 60425 | protein_coding | 10.55378799 | -0.453889525 | 0.364337737 | -1.245793336 | 0.212840277 | 0.410824034 | 0.453889525 |
| ENSMUSG00000029161 | Cgref1 | 68567 | protein_coding | 1730.104491 | 0.453934275 | 0.129282827 | 3.51117226 | 0.000446135 | 0.003969856 | 0.453934275 |
| ENSMUSG00000051811 | Cox6b2 | 333182 | protein_coding | 18.90554231 | -0.454143652 | 0.268402169 | -1.692026758 | 0.090640872 | 0.232840208 | 0.454143652 |
| ENSMUSG00000021281 | Tnfaip2 | 21928 | protein_coding | 7.081554588 | 0.454490309 | 0.575078996 | 0.790309352 | 0.429347126 | 0.633442111 | 0.454490309 |
| ENSMUSG00000046804 | Phgr1 | 53906 | protein_coding | 2996.695533 | -0.454553524 | 0.148007032 | -3.07116167 | 0.002132277 | 0.014270127 | 0.454553524 |
| ENSMUSG00000086013 | Gm15706 | 100039495 | lncRNA | 40.82595988 | -0.454585916 | 0.180734467 | -2.515214306 | 0.011896005 | 0.054542189 | 0.454585916 |
| ENSMUSG00000051615 | Rap2a | 76108 | protein_coding | 1166.307612 | 0.454624951 | 0.058960599 | 7.71065696 | 1.25E-14 | 6.15E-13 | 0.454624951 |
| ENSMUSG00000028885 | Smpdl3b | 100340 | protein_coding | 76.52880078 | 0.454690726 | 0.169002724 | 2.69043431 | 0.007135908 | 0.036974869 | 0.454690726 |
| ENSMUSG00000062727 | H2bc12 | 319184 | protein_coding | 532.8766342 | 0.454748371 | 0.149367112 | 3.044501321 | 0.002330664 | 0.015359047 | 0.454748371 |
| ENSMUSG00000027104 | Atf2 | 11909 | protein_coding | 931.8052962 | 0.455001559 | 0.07318243 | 6.217360651 | 5.06E-10 | 1.58E-08 | 0.455001559 |
| ENSMUSG00000000149 | Gna12 | 14673 | protein_coding | 361.8748863 | 0.455020266 | 0.161573685 | 2.816178054 | 0.004859872 | 0.027488653 | 0.455020266 |
| ENSMUSG00000039763 | Dnajc28 | 246738 | protein_coding | 93.50995396 | -0.455133496 | 0.287118418 | -1.585176946 | 0.112926137 | 0.268781756 | 0.455133496 |
| ENSMUSG00000099517 | H3c8 | 97908 | protein_coding | 256.6042944 | 0.455260471 | 0.188814208 | 2.411155795 | 0.015902054 | 0.067621779 | 0.455260471 |
| ENSMUSG00000020422 | Tns3 | 319939 | protein_coding | 1753.752383 | -0.455301055 | 0.107893768 | -4.219901315 | 2.44E-05 | 0.000328632 | 0.455301055 |
| ENSMUSG00000074224 | Wdr87-ps | 114675 | transcribed_unitary_pseudogene | 11.52677181 | -0.455322873 | 1.775319779 | -0.256473723 | 0.797585068 | 0.891285161 | 0.455322873 |
| ENSMUSG00000021587 | Pcsk1 | 18548 | protein_coding | 451.9501125 | -0.455584211 | 0.186141116 | -2.447520579 | 0.014384291 | 0.062547908 | 0.455584211 |
| ENSMUSG00000021275 | Tecpr2 | 104859 | protein_coding | 275.9345304 | -0.45558491 | 0.105075409 | -4.335790028 | 1.45E-05 | 0.000207848 | 0.45558491 |
| ENSMUSG00000048442 | Smim5 | 66528 | protein_coding | 71.79667029 | -0.455631687 | 0.251783397 | -1.80961768 | 0.070355096 | 0.196138387 | 0.455631687 |
| ENSMUSG00000032014 | Oaf | 102644 | protein_coding | 149.5398216 | -0.455759361 | 0.166204956 | -2.742152651 | 0.006103797 | 0.032635541 | 0.455759361 |
| ENSMUSG00000038902 | Pogz | 229584 | protein_coding | 216.1075909 | -0.456097459 | 0.152826847 | -2.984406661 | 0.002841289 | 0.017958749 | 0.456097459 |
| ENSMUSG00000000530 | Acvrl1 | 11482 | protein_coding | 68.73823089 | -0.456166081 | 0.15395493 | -2.96298457 | 0.003046718 | 0.019023179 | 0.456166081 |
| ENSMUSG00000020589 | Cyria | 76820 | protein_coding | 303.7285835 | -0.456385939 | 0.139822586 | -3.264035885 | 0.001098373 | 0.008290853 | 0.456385939 |
| ENSMUSG00000029314 | Gpat3 | 231510 | protein_coding | 294.8873937 | 0.456520436 | 0.232913361 | 1.960043999 | 0.049990648 | 0.155487412 | 0.456520436 |
| ENSMUSG00000015377 | Dennd6b | 69440 | protein_coding | 88.03730433 | -0.456567171 | 0.143139687 | -3.189661657 | 0.001424394 | 0.010238157 | 0.456567171 |
| ENSMUSG00000003037 | Rab8a | 17274 | protein_coding | 1767.872236 | 0.456627921 | 0.079017091 | 5.778850085 | 7.52E-09 | 2.06E-07 | 0.456627921 |
| ENSMUSG00000048251 | Bcl11b | 58208 | protein_coding | 309.4625795 | -0.456725424 | 0.199190428 | -2.292908496 | 0.021853274 | 0.085250782 | 0.456725424 |
| ENSMUSG00000032727 | Mier3 | 218613 | protein_coding | 1412.500643 | -0.456843532 | 0.147443787 | -3.098425112 | 0.001945521 | 0.013225891 | 0.456843532 |
| ENSMUSG00000027715 | Ccna2 | 12428 | protein_coding | 1377.196155 | 0.457205429 | 0.118985079 | 3.842544228 | 0.000121765 | 0.001336651 | 0.457205429 |
| ENSMUSG00000021774 | Ube2e1 | 22194 | protein_coding | 242.3523012 | 0.457308037 | 0.105392752 | 4.339084298 | 1.43E-05 | 0.000205125 | 0.457308037 |
| ENSMUSG00000018796 | Acsl1 | 14081 | protein_coding | 440.8568058 | -0.457337696 | 0.100882649 | -4.533363275 | 5.81E-06 | 9.19E-05 | 0.457337696 |
| ENSMUSG00000034362 | Csta1 | 209294 | protein_coding | 13.33003533 | 0.457405299 | 0.495352271 | 0.923393967 | 0.355801934 | 0.565477271 | 0.457405299 |
| ENSMUSG00000044551 | 9930012K11Rik | 268759 | protein_coding | 33.11205337 | -0.457478536 | 0.210289458 | -2.175470608 | 0.029594868 | 0.106817825 | 0.457478536 |
| ENSMUSG00000028931 | Kcnab2 | 16498 | protein_coding | 12.42877968 | -0.457593938 | 0.546098088 | -0.837933601 | 0.402067996 | 0.609627705 | 0.457593938 |

| ensembl_gene_id | external_gene_name | entrezgene_id | gene_biotype | baseMean | log2FoldChange | lfcSE | stat | pvalue | padj | absLog2FC |
|---|---|---|---|---|---|---|---|---|---|---|
| ENSMUSG00000028634 | Hivep3 | 16656 | protein_coding | 93.4740183 | -0.45817302 | 0.242037273 | -1.892985384 | 0.058359813 | 0.172734546 | 0.45817302 |
| ENSMUSG00000020952 | Scfd1 | 76983 | protein_coding | 1056.914044 | 0.458236829 | 0.076073754 | 6.023586377 | 1.71E-09 | 5.04E-08 | 0.458236829 |
| ENSMUSG00000035112 | Wnk4 | 69847 | protein_coding | 281.2883876 | -0.45834142 | 0.168294372 | -2.723450666 | 0.006460386 | 0.034197751 | 0.45834142 |
| ENSMUSG00000035104 | Eva1a | 232146 | protein_coding | 31.54592407 | 0.458463272 | 0.321270202 | 1.427033287 | 0.1535703 | 0.332707799 | 0.458463272 |
| ENSMUSG00000040543 | Pitpnm3 | 327958 | protein_coding | 187.7238254 | -0.458532031 | 0.170344293 | -2.691795675 | 0.007106847 | 0.036860261 | 0.458532031 |
| ENSMUSG00000021208 | Ifi27l2b | 217845 | protein_coding | 1402.875177 | 0.45868501 | 0.1028876 | 4.458117502 | 8.27E-06 | 0.000125857 | 0.45868501 |
| ENSMUSG00000032548 | Slco2a1 | 24059 | protein_coding | 342.0291106 | -0.458694914 | 0.207761984 | -2.207790398 | 0.027258888 | 0.10045802 | 0.458694914 |
| ENSMUSG00000050332 | Amer1 | 72345 | protein_coding | 196.1170479 | -0.458812956 | 0.15989275 | -2.869504447 | 0.004111156 | 0.023986258 | 0.458812956 |
| ENSMUSG00000067377 | Tspan6 | 56496 | protein_coding | 186.462293 | -0.459139991 | 0.139542165 | -3.290331568 | 0.001000694 | 0.007681471 | 0.459139991 |
| ENSMUSG00000038079 | Tmem237 | 381259 | protein_coding | 24.8840952 | 0.4592786 | 0.224984847 | 2.041375707 | 0.041213497 | 0.135882547 | 0.4592786 |
| ENSMUSG00000069662 | Marcks | 17118 | protein_coding | 1192.555205 | 0.459377745 | 0.09895403 | 4.642334872 | 3.44E-06 | 5.75E-05 | 0.459377745 |
| ENSMUSG00000098134 | Rnf113a2 | 66381 | protein_coding | 167.0277353 | 0.459587942 | 0.100947752 | 4.552730821 | 5.30E-06 | 8.51E-05 | 0.459587942 |
| ENSMUSG00000045319 | Proser2 | 227545 | protein_coding | 156.1154451 | -0.459610413 | 0.143928674 | -3.193320699 | 0.001406467 | 0.010132157 | 0.459610413 |
| ENSMUSG00000051439 | Cd14 | 12475 | protein_coding | 361.0555374 | 0.459920048 | 0.1389986 | 3.308810655 | 0.000936932 | 0.007272636 | 0.459920048 |
| ENSMUSG00000040022 | Rab11fip2 | 74998 | protein_coding | 86.22408372 | -0.459991064 | 0.172299898 | -2.669711759 | 0.007591639 | 0.038918449 | 0.459991064 |
| ENSMUSG00000070604 | Vsig10l | 75690 | protein_coding | 23.44493408 | -0.460054175 | 0.273636789 | -1.681258492 | 0.092712716 | 0.236617233 | 0.460054175 |
| ENSMUSG00000027746 | Ufm1 | 67890 | protein_coding | 1209.42898 | 0.460099363 | 0.089765129 | 5.125591287 | 2.97E-07 | 6.27E-06 | 0.460099363 |
| ENSMUSG00000064427 | Gm22748 | 115488064 | snoRNA | 62.02602984 | -0.460099486 | 0.163201642 | -2.819208681 | 0.004814221 | 0.02728858 | 0.460099486 |
| ENSMUSG00000044359 | P2ry4 | 57385 | protein_coding | 55.08738435 | 0.460219995 | 0.220960613 | 2.082814615 | 0.037268125 | 0.126487682 | 0.460219995 |
| ENSMUSG00000032012 | Nectin1 | 58235 | protein_coding | 167.6416258 | -0.460395852 | 0.191863341 | -2.399603025 | 0.01641286 | 0.069159799 | 0.460395852 |
| ENSMUSG00000034334 | Fam151b | 73942 | protein_coding | 51.47044467 | 0.460648928 | 0.18002099 | 2.55886231 | 0.010501533 | 0.049605104 | 0.460648928 |
| ENSMUSG00000035559 | Mpv17l2 | 234384 | protein_coding | 323.4058022 | 0.46075008 | 0.098472501 | 4.678972062 | 2.88E-06 | 4.89E-05 | 0.46075008 |
| ENSMUSG00000068959 | Zfp619 | 70227 | protein_coding | 42.47546891 | -0.460831086 | 0.192764954 | -2.39063728 | 0.01681916 | 0.070424706 | 0.460831086 |
| ENSMUSG00000020444 | Guk1 | 14923 | protein_coding | 911.5504808 | 0.460867721 | 0.120370978 | 3.828727884 | 0.000128807 | 0.001403313 | 0.460867721 |
| ENSMUSG00000022193 | Psmb5 | 19173 | protein_coding | 704.6501581 | 0.460957048 | 0.103686061 | 4.445699286 | 8.76E-06 | 0.00013239 | 0.460957048 |
| ENSMUSG00000066637 | Ttc32 | 75516 | protein_coding | 138.1338229 | 0.46110394 | 0.127294705 | 3.622334013 | 0.000291957 | 0.00277795 | 0.46110394 |
| ENSMUSG00000042757 | Tmem108 | 81907 | protein_coding | 9.023687723 | -0.461347034 | 0.687636385 | -0.670917136 | 0.502273319 | 0.692239286 | 0.461347034 |
| ENSMUSG00000034311 | Kif4 | 16571 | protein_coding | 595.4055516 | 0.461353233 | 0.126405864 | 3.649777136 | 0.000262468 | 0.002553257 | 0.461353233 |
| ENSMUSG00000030287 | Itpr2 | 16439 | protein_coding | 875.445578 | -0.461514487 | 0.106927379 | -4.316148866 | 1.59E-05 | 0.000225398 | 0.461514487 |
| ENSMUSG00000031478 | Nek3 | 23954 | protein_coding | 126.6164533 | -0.461608608 | 0.115823046 | -3.985464224 | 6.73E-05 | 0.000798721 | 0.461608608 |
| ENSMUSG00000028159 | Dapp1 | 26377 | protein_coding | 121.0183447 | 0.462022528 | 0.19519559 | 2.36697217 | 0.017934281 | 0.073758708 | 0.462022528 |
| ENSMUSG00000051735 | Rinl | 320435 | protein_coding | 24.29410196 | 0.462511405 | 0.252713851 | 1.830178292 | 0.067223283 | 0.190000766 | 0.462511405 |
| ENSMUSG00000002580 | Mien1 | 103742 | protein_coding | 662.7229418 | 0.462597976 | 0.075381883 | 6.136726165 | 8.42E-10 | 2.55E-08 | 0.462597976 |
| ENSMUSG00000020328 | Nudcd2 | 52653 | protein_coding | 522.5132378 | 0.462623513 | 0.109146969 | 4.238537413 | 2.25E-05 | 0.000306281 | 0.462623513 |
| ENSMUSG00000049804 | Armcx4 | 100503043 | protein_coding | 9.653261587 | -0.462658368 | 0.54293006 | -0.852150953 | 0.394130315 | 0.602062688 | 0.462658368 |
| ENSMUSG00000097061 | 9330151L19Rik | 414085 | lncRNA | 32.83699683 | -0.462738897 | 0.23829633 | -1.9418633 | 0.052153651 | 0.159965985 | 0.462738897 |
| ENSMUSG00000021702 | Thbs4 | 21828 | protein_coding | 2135.277522 | -0.462807753 | 0.324515961 | -1.426147891 | 0.153825657 | 0.332898787 | 0.462807753 |
| ENSMUSG00000021714 | Cenpk | 60411 | protein_coding | 111.2464636 | 0.462813508 | 0.14805191 | 3.126021864 | 0.001771884 | 0.012238753 | 0.462813508 |

| ensembl_gene_id | external_gene_name | entrezgene_id | gene_biotype | baseMean | log2FoldChange | lfcSE | stat | pvalue | padj | absLog2FC |
|---|---|---|---|---|---|---|---|---|---|---|
| ENSMUSG00000052384 | Nrros | 224109 | protein_coding | 21.5872042 | 0.463016865 | 0.305157739 | 1.517303368 | 0.129190108 | 0.295737287 | 0.463016865 |
| ENSMUSG00000006782 | Cnp | 12799 | protein_coding | 473.3088976 | 0.463189865 | 0.093749767 | 4.940704153 | 7.78E-07 | 1.51E-05 | 0.463189865 |
| ENSMUSG00000078153 | Psme2b | 621823 | protein_coding | 44.88864722 | 0.463365336 | 0.185722138 | 2.494938627 | 0.012597892 | 0.056924601 | 0.463365336 |
| ENSMUSG00000026918 | Brd3 | 67382 | protein_coding | 517.4606828 | -0.463406051 | 0.09198519 | -5.037833288 | 4.71E-07 | 9.65E-06 | 0.463406051 |
| ENSMUSG00000030000 | Add2 | 11519 | protein_coding | 7.027797055 | -0.463494533 | 0.884285783 | -0.524145634 | 0.600177251 | 0.766609723 | 0.463494533 |
| ENSMUSG00000069814 | Ccdc92b | 432582 | protein_coding | 18.88720664 | -0.463504828 | 0.391082494 | -1.185184289 | 0.23594459 | 0.440061526 | 0.463504828 |
| ENSMUSG00000038828 | Tmem214 | 68796 | protein_coding | 865.9731208 | 0.463643052 | 0.101069163 | 4.587383912 | 4.49E-06 | 7.31E-05 | 0.463643052 |
| ENSMUSG00000021917 | Spcs1 | 69019 | protein_coding | 1149.176721 | 0.463789649 | 0.118506579 | 3.913619417 | 9.09E-05 | 0.001040388 | 0.463789649 |
| ENSMUSG00000028334 | Nans | 94181 | protein_coding | 6214.961988 | 0.463880758 | 0.107605079 | 4.310955956 | 1.63E-05 | 0.000230142 | 0.463880758 |
| ENSMUSG00000038085 | Cnbd2 | 70873 | protein_coding | 40.77654203 | -0.464113532 | 0.195701789 | -2.37153444 | 0.017714396 | 0.073059268 | 0.464113532 |
| ENSMUSG00000020658 | Efr3b | 668212 | protein_coding | 16.10212517 | -0.464214575 | 0.369435316 | -1.256551701 | 0.208916014 | 0.405789022 | 0.464214575 |
| ENSMUSG00000089900 | Tbc1d22bos | 74092 | lncRNA | 7.081020454 | -0.464397018 | 0.438541763 | -1.058957338 | 0.289619209 | 0.498385347 | 0.464397018 |
| ENSMUSG00000021244 | Ylpm1 | 56531 | protein_coding | 397.6659548 | -0.464898007 | 0.14907035 | -3.118648381 | 0.001816826 | 0.012484215 | 0.464898007 |
| ENSMUSG00000050240 | Hic2 | 58180 | protein_coding | 122.6580502 | -0.465289571 | 0.137194677 | -3.391454974 | 0.000695226 | 0.005722538 | 0.465289571 |
| ENSMUSG00000017386 | Traf4 | 22032 | protein_coding | 747.6575554 | -0.465492221 | 0.115627696 | -4.025784782 | 5.68E-05 | 0.000690444 | 0.465492221 |
| ENSMUSG00000022092 | Ppp3cc | 19057 | protein_coding | 73.15045056 | -0.465498558 | 0.157244193 | -2.960354524 | 0.003072852 | 0.019141333 | 0.465498558 |
| ENSMUSG00000058470 | Gm8369 | 666926 | protein_coding | 6.877876782 | -0.465596667 | 1.050578509 | -0.443181222 | 0.65763466 | 0.801868244 | 0.465596667 |
| ENSMUSG00000042138 | Msantd2 | 235184 | protein_coding | 103.2643617 | -0.465627236 | 0.145802895 | -3.193539041 | 0.001405403 | 0.010129079 | 0.465627236 |
| ENSMUSG00000033184 | Tmed7 | 66676 | protein_coding | 3336.003335 | 0.465795956 | 0.075890099 | 6.137769819 | 8.37E-10 | 2.53E-08 | 0.465795956 |
| ENSMUSG00000025464 | Paox | 212503 | protein_coding | 108.3078584 | -0.465835994 | 0.148020013 | -3.147114941 | 0.001648901 | 0.011498993 | 0.465835994 |
| ENSMUSG00000046876 | Atxn1 | 20238 | protein_coding | 714.9525633 | -0.465842066 | 0.149783855 | -3.110095327 | 0.00187027 | 0.012792498 | 0.465842066 |
| ENSMUSG00000032491 | Nradd | 67169 | protein_coding | 20.99993815 | -0.466106835 | 0.280078336 | -1.664201675 | 0.096072133 | 0.242030591 | 0.466106835 |
| ENSMUSG00000033792 | Atp7a | 11977 | protein_coding | 573.0237395 | -0.466170033 | 0.107101968 | -4.352581384 | 1.35E-05 | 0.000193938 | 0.466170033 |
| ENSMUSG00000000861 | Bcl11a | 14025 | protein_coding | 75.37174304 | -0.466302056 | 0.201580886 | -2.313225547 | 0.020710242 | 0.082061531 | 0.466302056 |
| ENSMUSG00000025665 | Rps6ka6 | 67071 | protein_coding | 83.58697888 | 0.466306717 | 0.174063429 | 2.678947085 | 0.007385406 | 0.038069499 | 0.466306717 |
| ENSMUSG00000042763 | Maneal | 215090 | protein_coding | 8.123358773 | -0.46641254 | 0.48505888 | -0.961558605 | 0.336271374 | 0.546990445 | 0.46641254 |
| ENSMUSG00000055748 | Gsdmc4 | 74548 | protein_coding | 8238.760836 | -0.466709514 | 0.132624961 | -3.519017162 | 0.000433149 | 0.003880311 | 0.466709514 |
| ENSMUSG00000040520 | Manea | 242362 | protein_coding | 424.4451654 | 0.466717766 | 0.140768461 | 3.315499537 | 0.000914795 | 0.007130953 | 0.466717766 |
| ENSMUSG00000032375 | Aph1b | 208117 | protein_coding | 245.9069364 | 0.466780609 | 0.134669561 | 3.466118135 | 0.000528031 | 0.004558526 | 0.466780609 |
| ENSMUSG00000078866 | Zfp970 | 628308 | protein_coding | 81.68406613 | -0.466875364 | 0.147416887 | -3.167041274 | 0.001539985 | 0.010887206 | 0.466875364 |
| ENSMUSG00000043461 | Sptssb | 66183 | protein_coding | 1391.00396 | 0.466917851 | 0.259148291 | 1.80174004 | 0.071586315 | 0.198571396 | 0.466917851 |
| ENSMUSG00000019487 | Trip10 | 106628 | protein_coding | 546.1747021 | -0.467036817 | 0.095582036 | -4.886240502 | 1.03E-06 | 1.94E-05 | 0.467036817 |
| ENSMUSG00000020359 | Phykpl | 72947 | protein_coding | 336.8294169 | -0.467057703 | 0.094714806 | -4.931200547 | 8.17E-07 | 1.57E-05 | 0.467057703 |
| ENSMUSG00000045817 | Zfp36l2 | 12193 | protein_coding | 1904.646712 | -0.467094653 | 0.16252642 | -2.87396137 | 0.004053585 | 0.023737319 | 0.467094653 |
| ENSMUSG00000022229 | Atp12a | 192113 | protein_coding | 442.5119776 | 0.467357518 | 0.359687244 | 1.299344154 | 0.193825848 | 0.387259334 | 0.467357518 |
| ENSMUSG00000030088 | Aldh1l1 | 107747 | protein_coding | 1109.591368 | -0.467377095 | 0.337600286 | -1.384409656 | 0.166233054 | 0.350047605 | 0.467377095 |
| ENSMUSG00000002997 | Prkar2b | 19088 | protein_coding | 78.07640753 | -0.467423717 | 0.211321478 | -2.211908231 | 0.026973006 | 0.099660429 | 0.467423717 |
| ENSMUSG00000022652 | Morc1 | 17450 | protein_coding | 7.56842711 | -0.467432699 | 0.726166499 | -0.643699069 | 0.519770594 | 0.706451576 | 0.467432699 |

| ensembl_gene_id | external_gene_name | entrezgene_id | gene_biotype | baseMean | log2FoldChange | lfcSE | stat | pvalue | padj | absLog2FC |
|---|---|---|---|---|---|---|---|---|---|---|
| ENSMUSG00000075270 | Pde11a | 241489 | protein_coding | 55.10831552 | 0.467443335 | 0.250046006 | 1.86942932 | 0.061563109 | 0.178676602 | 0.467443335 |
| ENSMUSG00000006587 | Snai3 | 30927 | protein_coding | 12.08224589 | 0.46774341 | 0.389334012 | 1.201393651 | 0.229598538 | 0.432847922 | 0.46774341 |
| ENSMUSG00000034255 | Arhgap27 | 544817 | protein_coding | 1170.938334 | -0.467900391 | 0.167086239 | -2.800352641 | 0.005104681 | 0.028488912 | 0.467900391 |
| ENSMUSG00000085028 | Slc2a4rg-ps | 329584 | transcribed_unitary_pseudogene | 55.69991949 | -0.46808129 | 0.211149829 | -2.216820601 | 0.026635353 | 0.098892286 | 0.46808129 |
| ENSMUSG00000028862 | Map3k6 | 53608 | protein_coding | 109.2585942 | -0.468183114 | 0.288254954 | -1.624197984 | 0.104333561 | 0.255523566 | 0.468183114 |
| ENSMUSG00000097331 | F420014N23Rik | 100038591 | lncRNA | 9.697922899 | -0.4682096 | 0.414254986 | -1.130244936 | 0.25837303 | 0.465469894 | 0.4682096 |
| ENSMUSG00000027306 | Nusap1 | 108907 | protein_coding | 734.5836442 | 0.468545453 | 0.143560284 | 3.263754009 | 0.001099466 | 0.008291805 | 0.468545453 |
| ENSMUSG00000034993 | Vat1 | 26949 | protein_coding | 596.5584315 | -0.468678646 | 0.163834229 | -2.860688193 | 0.004227226 | 0.024519757 | 0.468678646 |
| ENSMUSG00000109186 | Gm34821 | 102638204 | lncRNA | 19.01533357 | -0.468699248 | 0.280257447 | -1.672388201 | 0.094447797 | 0.239357918 | 0.468699248 |
| ENSMUSG00000031853 | Map3k21 | 234878 | protein_coding | 1265.632301 | -0.468906942 | 0.098394045 | -4.765602854 | 1.88E-06 | 3.31E-05 | 0.468906942 |
| ENSMUSG00000001143 | Lman2l | 214895 | protein_coding | 150.3390469 | 0.469008922 | 0.143915356 | 3.258922023 | 0.001118364 | 0.008410435 | 0.469008922 |
| ENSMUSG00000037300 | Ttc13 | 234875 | protein_coding | 782.0641616 | 0.469051895 | 0.084698591 | 5.537894899 | 3.06E-08 | 7.61E-07 | 0.469051895 |
| ENSMUSG00000023249 | Parp3 | 235587 | protein_coding | 150.4312099 | 0.469120323 | 0.158457982 | 2.960534494 | 0.003071057 | 0.019137637 | 0.469120323 |
| ENSMUSG00000034908 | Sidt2 | 214597 | protein_coding | 733.8396206 | -0.469414658 | 0.07951406 | -5.90354279 | 3.56E-09 | 1.02E-07 | 0.469414658 |
| ENSMUSG00000044365 | Cxxc4 | 319478 | protein_coding | 31.3864122 | 0.469433346 | 0.33289475 | 1.410155451 | 0.158493787 | 0.339785391 | 0.469433346 |
| ENSMUSG00000042808 | Gpx2 | 14776 | protein_coding | 4583.260449 | 0.46948852 | 0.13849142 | 3.390018822 | 0.000698878 | 0.005739537 | 0.46948852 |
| ENSMUSG00000054520 | Sh3bp2 | 24055 | protein_coding | 90.90546388 | -0.469681839 | 0.126810656 | -3.703804198 | 0.00021239 | 0.002150637 | 0.469681839 |
| ENSMUSG00000044052 | Ccr10 | 12777 | protein_coding | 9.799197031 | -0.470296512 | 0.521334853 | -0.902100656 | 0.3670034 | 0.57565559 | 0.470296512 |
| ENSMUSG00000019975 | Ikbip | 67454 | protein_coding | 255.0006872 | 0.470918441 | 0.142334748 | 3.308527599 | 0.000937879 | 0.007276446 | 0.470918441 |
| ENSMUSG00000043019 | Edem3 | 66967 | protein_coding | 3026.154066 | 0.471040044 | 0.072469617 | 6.499827984 | 8.04E-11 | 2.80E-09 | 0.471040044 |
| ENSMUSG00000054931 | Zkscan4 | 544922 | protein_coding | 9.680250468 | 0.471205832 | 0.493698324 | 0.954440816 | 0.339860554 | 0.550313384 | 0.471205832 |
| ENSMUSG00000047342 | Zfp286 | 192651 | protein_coding | 14.4299702 | 0.471389928 | 0.474876363 | 0.992658226 | 0.320876539 | 0.532256186 | 0.471389928 |
| ENSMUSG00000037369 | Kdm6a | 22289 | protein_coding | 762.1985378 | 0.471445245 | 0.065226107 | 7.227861183 | 4.91E-13 | 2.09E-11 | 0.471445245 |
| ENSMUSG00000031425 | Plp1 | 18823 | protein_coding | 7.869253176 | -0.471541799 | 0.514577828 | -0.916366335 | 0.359474786 | 0.56932628 | 0.471541799 |
| ENSMUSG00000007591 | Tssk4 | 71099 | protein_coding | 27.83091168 | 0.47157589 | 0.259269517 | 1.818863614 | 0.06893224 | 0.19360831 | 0.47157589 |
| ENSMUSG00000016181 | Utp25 | 215193 | protein_coding | 138.6426421 | -0.471594797 | 0.155709246 | -3.028688465 | 0.002456178 | 0.016086351 | 0.471594797 |
| ENSMUSG00000060615 | Ang4 | 219033 | protein_coding | 375.7702355 | -0.471598715 | 0.555829578 | -0.848459193 | 0.396182289 | 0.60444363 | 0.471598715 |
| ENSMUSG00000047654 | Tssk6 | 83984 | protein_coding | 14.1050709 | -0.471620606 | 0.340876923 | -1.383550996 | 0.166495982 | 0.350415963 | 0.471620606 |
| ENSMUSG00000059005 | Hnrnpa3 | 229279 | protein_coding | 1980.265479 | 0.471771053 | 0.127613767 | 3.696866439 | 0.000218277 | 0.002194898 | 0.471771053 |
| ENSMUSG00000024201 | Kdm4b | 193796 | protein_coding | 487.4633965 | 0.471946481 | 0.076067095 | 6.204344723 | 5.49E-10 | 1.71E-08 | 0.471946481 |
| ENSMUSG00000029134 | Plb1 | 665270 | protein_coding | 18.68343324 | 0.47200598 | 0.641072557 | 0.736275441 | 0.461563089 | 0.65958881 | 0.47200598 |
| ENSMUSG00000044072 | Eml6 | 237711 | protein_coding | 74.54712154 | 0.472131947 | 0.192055504 | 2.458309907 | 0.013959266 | 0.061268531 | 0.472131947 |
| ENSMUSG00000086894 | Gm15708 | 100504231 | lncRNA | 29.5202672 | -0.472144904 | 0.272747312 | -1.731070788 | 0.08343914 | 0.220437655 | 0.472144904 |
| ENSMUSG00000036745 | Ttll7 | 70892 | protein_coding | 180.4906568 | 0.472357985 | 0.11803296 | 4.001915937 | 6.28E-05 | 0.000752468 | 0.472357985 |
| ENSMUSG00000038508 | Gdf15 | 23886 | protein_coding | 173.9137829 | 0.472453788 | 0.199866655 | 2.363844979 | 0.018086379 | 0.074151826 | 0.472453788 |
| ENSMUSG00000027661 | Slc2a10 | 170441 | protein_coding | 196.1889078 | -0.472459277 | 0.240856587 | -1.961579226 | 0.049811493 | 0.154996405 | 0.472459277 |
| ENSMUSG00000033307 | Mif | 17319 | protein_coding | 694.884389 | 0.472494539 | 0.131571877 | 3.591151466 | 0.00032922 | 0.003066561 | 0.472494539 |
| ENSMUSG00000051839 | Gypa | 14934 | protein_coding | 12.10282849 | 0.472805011 | 0.437578331 | 1.080503711 | 0.279917935 | 0.488182692 | 0.472805011 |

| ensembl_gene_id | external_gene_name | entrezgene_id | gene_biotype | baseMean | log2FoldChange | lfcSE | stat | pvalue | padj | absLog2FC |
|---|---|---|---|---|---|---|---|---|---|---|
| ENSMUSG00000047022 | Mipol1 | 73490 | protein_coding | 61.32474686 | -0.47301787 | 0.245919272 | -1.923468074 | 0.054421292 | 0.164409102 | 0.47301787 |
| ENSMUSG00000029864 | Gstk1 | 76263 | protein_coding | 673.87073 | -0.473273065 | 0.120096694 | -3.940766821 | 8.12E-05 | 0.00094087 | 0.473273065 |
| ENSMUSG00000021287 | Xrcc3 | 74335 | protein_coding | 115.4777981 | 0.473344878 | 0.131047779 | 3.612002304 | 0.000303842 | 0.002863665 | 0.473344878 |
| ENSMUSG00000046402 | Rbp1 | 19659 | protein_coding | 9.26875898 | -0.473427322 | 0.413161673 | -1.145864568 | 0.251851194 | 0.458216818 | 0.473427322 |
| ENSMUSG00000084925 | 1810062O18Rik | 75602 | lncRNA | 9.475226891 | -0.473431931 | 0.36371848 | -1.301643873 | 0.193038155 | 0.38631885 | 0.473431931 |
| ENSMUSG00000078202 | Nrarp | 67122 | protein_coding | 65.27685717 | -0.473598719 | 0.249769974 | -1.89613952 | 0.057941598 | 0.171957103 | 0.473598719 |
| ENSMUSG00000045294 | Insig1 | 231070 | protein_coding | 1841.191766 | 0.474157398 | 0.216577985 | 2.189314841 | 0.028573964 | 0.104005051 | 0.474157398 |
| ENSMUSG00000057367 | Birc2 | 11797 | protein_coding | 700.7094567 | 0.474258502 | 0.117497819 | 4.036317505 | 5.43E-05 | 0.000665772 | 0.474258502 |
| ENSMUSG00000001525 | Tubb5 | 22154 | protein_coding | 3793.408316 | 0.474586257 | 0.130558985 | 3.635033289 | 0.000277945 | 0.002668538 | 0.474586257 |
| ENSMUSG00000059060 | Rad51b | 19363 | protein_coding | 38.50072734 | -0.474622534 | 0.213884568 | -2.21905927 | 0.026482693 | 0.098486184 | 0.474622534 |
| ENSMUSG00000035863 | Palm | 18483 | protein_coding | 23.97375009 | -0.474725349 | 0.250093021 | -1.89819511 | 0.057670385 | 0.17155684 | 0.474725349 |
| ENSMUSG00000047787 | Flrt1 | 396184 | protein_coding | 7.834131872 | -0.474868747 | 0.552644491 | -0.859266226 | 0.390193653 | 0.598507455 | 0.474868747 |
| ENSMUSG00000024158 | Hagh | 14651 | protein_coding | 887.1643165 | 0.475067249 | 0.136874933 | 3.470812646 | 0.000518886 | 0.004491696 | 0.475067249 |
| ENSMUSG00000027401 | Tgm3 | 21818 | protein_coding | 4576.675737 | -0.475249662 | 0.429001845 | -1.107803305 | 0.267946772 | 0.475940787 | 0.475249662 |
| ENSMUSG00000050860 | Phospho1 | 237928 | protein_coding | 19.87220957 | -0.475415047 | 0.385240006 | -1.234074963 | 0.217174972 | 0.416565866 | 0.475415047 |
| ENSMUSG00000007458 | M6pr | 17113 | protein_coding | 3700.516042 | 0.475673955 | 0.092246454 | 5.156555447 | 2.52E-07 | 5.41E-06 | 0.475673955 |
| ENSMUSG00000040600 | Eps8l3 | 99662 | protein_coding | 1962.576732 | -0.475896714 | 0.150035172 | -3.171901011 | 0.001514446 | 0.010749598 | 0.475896714 |
| ENSMUSG00000024197 | Plin3 | 66905 | protein_coding | 1360.232833 | 0.476114452 | 0.096214953 | 4.948445518 | 7.48E-07 | 1.46E-05 | 0.476114452 |
| ENSMUSG00000071713 | Csf2rb | 12983 | protein_coding | 7.868414661 | 0.476471868 | 0.621292076 | 0.766904788 | 0.443138127 | 0.644271437 | 0.476471868 |
| ENSMUSG00000086712 | Mexis | 381524 | lncRNA | 34.24603333 | 0.476517914 | 0.216337178 | 2.202663081 | 0.027618504 | 0.101360933 | 0.476517914 |
| ENSMUSG00000109644 | 0610005C13Rik | 71661 | lncRNA | 239.5642685 | 0.47653885 | 0.150278662 | 3.17103469 | 0.00151897 | 0.010767314 | 0.47653885 |
| ENSMUSG00000029710 | Ephb4 | 13846 | protein_coding | 1417.994369 | -0.476593299 | 0.127112545 | -3.749380504 | 0.000177272 | 0.00184067 | 0.476593299 |
| ENSMUSG00000021509 | Slc25a48 | 328258 | protein_coding | 13.06497155 | 0.476616835 | 0.415321843 | 1.147584321 | 0.251140205 | 0.457527301 | 0.476616835 |
| ENSMUSG00000056215 | Lrguk | 74354 | protein_coding | 8.878701456 | -0.476687646 | 0.672587018 | -0.708737507 | 0.478487385 | 0.672843015 | 0.476687646 |
| ENSMUSG00000049504 | Proser1 | 212127 | protein_coding | 86.04487238 | -0.476856951 | 0.214259597 | -2.225603694 | 0.026040739 | 0.097481618 | 0.476856951 |
| ENSMUSG00000030789 | Itgax | 16411 | protein_coding | 14.02817838 | 0.477151303 | 0.505381996 | 0.944139893 | 0.345098166 | 0.555056406 | 0.477151303 |
| ENSMUSG00000039985 | Sinhcaf | 56306 | protein_coding | 252.3276688 | -0.477240994 | 0.091320727 | -5.225987691 | 1.73E-07 | 3.83E-06 | 0.477240994 |
| ENSMUSG00000038569 | Rad9b | 231724 | protein_coding | 37.77213208 | -0.47773576 | 0.2021888 | -2.362820097 | 0.018136471 | 0.074299824 | 0.47773576 |
| ENSMUSG00000047238 | Mageh1 | 75625 | protein_coding | 72.63542459 | 0.477748772 | 0.185597304 | 2.574114819 | 0.010049692 | 0.048055085 | 0.477748772 |
| ENSMUSG00000040563 | Plppr2 | 235044 | protein_coding | 12.38927625 | -0.477811705 | 0.402316318 | -1.187651812 | 0.234970616 | 0.438964575 | 0.477811705 |
| ENSMUSG00000070883 | Ccdc173 | 75051 | protein_coding | 13.09461947 | 0.477899344 | 0.321904598 | 1.484599311 | 0.137649998 | 0.308453738 | 0.477899344 |
| ENSMUSG00000114744 | Gm38409 | 328436 | lncRNA | 7.708723523 | 0.477904599 | 0.431784101 | 1.106813793 | 0.268374444 | 0.476324178 | 0.477904599 |
| ENSMUSG00000026701 | Prdx6 | 11758 | protein_coding | 11583.14712 | -0.478373962 | 0.148557076 | -3.220135817 | 0.001281299 | 0.009353129 | 0.478373962 |
| ENSMUSG00000059674 | Cdh24 | 239096 | protein_coding | 35.69633805 | 0.478398415 | 0.254396594 | 1.880522095 | 0.060036958 | 0.175978776 | 0.478398415 |
| ENSMUSG00000013236 | Ptprs | 19280 | protein_coding | 431.7257053 | -0.478630673 | 0.120361595 | -3.976606266 | 6.99E-05 | 0.000826008 | 0.478630673 |
| ENSMUSG00000069520 | Tmem19 | 67226 | protein_coding | 637.0686585 | 0.478641392 | 0.105615838 | 4.531909253 | 5.85E-06 | 9.24E-05 | 0.478641392 |
| ENSMUSG00000036281 | Snapc4 | 227644 | protein_coding | 177.4920466 | -0.479064695 | 0.142104963 | -3.371203115 | 0.000748407 | 0.006063387 | 0.479064695 |
| ENSMUSG00000001911 | Nfix | 18032 | protein_coding | 245.3878158 | -0.479077156 | 0.124900038 | -3.835684631 | 0.000125215 | 0.001367918 | 0.479077156 |

| ensembl_gene_id | external_gene_name | entrezgene_id | gene_biotype | baseMean | log2FoldChange | lfcSE | stat | pvalue | padj | absLog2FC |
|---|---|---|---|---|---|---|---|---|---|---|
| ENSMUSG00000003623 | Crot | 74114 | protein_coding | 1107.409334 | 0.479457861 | 0.124125514 | 3.862685807 | 0.000112147 | 0.001243062 | 0.479457861 |
| ENSMUSG00000073077 | Cfap47 | 636104 | protein_coding | 55.60105986 | -0.479544559 | 0.291553099 | -1.644793218 | 0.100012461 | 0.248169261 | 0.479544559 |
| ENSMUSG00000073530 | Pappa2 | 23850 | protein_coding | 92.57474388 | -0.479576628 | 0.332028108 | -1.444385631 | 0.14863053 | 0.325707703 | 0.479576628 |
| ENSMUSG00000022311 | Csmd3 | 239420 | protein_coding | 9.158964367 | -0.479880249 | 1.70574102 | -0.28133242 | 0.778455446 | 0.879690553 | 0.479880249 |
| ENSMUSG00000036815 | Dpp10 | 269109 | protein_coding | 11.11979999 | -0.48008527 | 1.779910848 | -0.269724335 | 0.787372338 | 0.884634427 | 0.48008527 |
| ENSMUSG00000008206 | Cers4 | 67260 | protein_coding | 539.8998527 | -0.480087191 | 0.177049361 | -2.71160081 | 0.006695918 | 0.035152464 | 0.480087191 |
| ENSMUSG00000029034 | Ints11 | 71957 | protein_coding | 416.188117 | 0.480453456 | 0.100437475 | 4.783607469 | 1.72E-06 | 3.05E-05 | 0.480453456 |
| ENSMUSG00000011034 | Slc5a1 | 20537 | protein_coding | 2888.230109 | 0.480549192 | 0.11402225 | 4.214521234 | 2.50E-05 | 0.000335599 | 0.480549192 |
| ENSMUSG00000031762 | Mt2 | 17750 | protein_coding | 605.3728393 | -0.48056567 | 0.228750904 | -2.100825231 | 0.035656311 | 0.12266387 | 0.48056567 |
| ENSMUSG00000052373 | Mpp3 | 13384 | protein_coding | 121.5202525 | -0.480660971 | 0.248353235 | -1.935392431 | 0.052942157 | 0.161575529 | 0.480660971 |
| ENSMUSG00000029675 | Eln | 13717 | protein_coding | 11.40633388 | -0.48067001 | 0.789650066 | -0.608712682 | 0.542714899 | 0.725563448 | 0.48067001 |
| ENSMUSG00000055980 | Irs1 | 16367 | protein_coding | 77.21200774 | -0.480766084 | 0.351354051 | -1.368323725 | 0.171210768 | 0.356588987 | 0.480766084 |
| ENSMUSG00000039462 | Col10a1 | 12813 | protein_coding | 7.746438832 | -0.481070761 | 0.439127262 | -1.095515588 | 0.27329082 | 0.481257732 | 0.481070761 |
| ENSMUSG00000051331 | Cacna1c | 12288 | protein_coding | 46.19724524 | -0.481127731 | 0.373302677 | -1.288840826 | 0.19745343 | 0.391319924 | 0.481127731 |
| ENSMUSG00000021248 | Tmed10 | 68581 | protein_coding | 2243.94577 | 0.481163204 | 0.052014397 | 9.250577338 | 2.23E-20 | 1.62E-18 | 0.481163204 |
| ENSMUSG00000056124 | B4galt6 | 56386 | protein_coding | 493.7118696 | -0.481328055 | 0.244378966 | -1.969596905 | 0.048884587 | 0.153244184 | 0.481328055 |
| ENSMUSG00000078970 | Dnaaf10 | 103784 | protein_coding | 347.5981559 | 0.481443057 | 0.090889607 | 5.297008906 | 1.18E-07 | 2.67E-06 | 0.481443057 |
| ENSMUSG00000015652 | Steap1 | 70358 | protein_coding | 81.47785731 | 0.481569624 | 0.146414309 | 3.289088528 | 0.001005124 | 0.007704339 | 0.481569624 |
| ENSMUSG00000023991 | Foxp4 | 74123 | protein_coding | 418.2584494 | -0.48166646 | 0.149000704 | -3.232645524 | 0.001226496 | 0.009056855 | 0.48166646 |
| ENSMUSG00000034177 | Rnf43 | 207742 | protein_coding | 517.381755 | -0.48169005 | 0.109542606 | -4.397284943 | 1.10E-05 | 0.000161212 | 0.48169005 |
| ENSMUSG00000041164 | Zmiz2 | 52915 | protein_coding | 325.2540893 | -0.481695805 | 0.11483225 | -4.194778066 | 2.73E-05 | 0.000363455 | 0.481695805 |
| ENSMUSG00000029516 | Cit | 12704 | protein_coding | 341.689419 | 0.481793254 | 0.175883405 | 2.739276365 | 0.006157459 | 0.032847006 | 0.481793254 |
| ENSMUSG00000043008 | Klhl6 | 239743 | protein_coding | 6.988425945 | 0.481916641 | 0.550491796 | 0.875429287 | 0.38134037 | 0.589085895 | 0.481916641 |
| ENSMUSG00000038500 | Prr3 | 75210 | protein_coding | 46.79468728 | -0.48194589 | 0.213811757 | -2.254066366 | 0.024191993 | 0.092184227 | 0.48194589 |
| ENSMUSG00000037894 | H2az1 | 51788 | protein_coding | 1866.642235 | 0.482062759 | 0.10154097 | 4.747470492 | 2.06E-06 | 3.59E-05 | 0.482062759 |
| ENSMUSG00000043505 | Gimap5 | 317757 | protein_coding | 14.01080598 | 0.482083374 | 0.394028974 | 1.223471892 | 0.221151514 | 0.422016433 | 0.482083374 |
| ENSMUSG00000018830 | Myh11 | 17880 | protein_coding | 12.19074174 | -0.482155403 | 0.729710879 | -0.660748547 | 0.508773585 | 0.697877891 | 0.482155403 |
| ENSMUSG00000030966 | Trim21 | 20821 | protein_coding | 207.6444243 | 0.482242819 | 0.11694033 | 4.123836669 | 3.73E-05 | 0.000477089 | 0.482242819 |
| ENSMUSG00000022844 | Pdia5 | 72599 | protein_coding | 957.1490425 | 0.482318834 | 0.085121312 | 5.666252344 | 1.46E-08 | 3.84E-07 | 0.482318834 |
| ENSMUSG00000036295 | Lrrn3 | 16981 | protein_coding | 18.83617837 | -0.482371699 | 0.275769691 | -1.749183161 | 0.080259364 | 0.214204281 | 0.482371699 |
| ENSMUSG00000036833 | Pnpla7 | 241274 | protein_coding | 163.9715251 | -0.482558516 | 0.112813322 | -4.27749584 | 1.89E-05 | 0.00026408 | 0.482558516 |
| ENSMUSG00000038507 | Parp12 | 243771 | protein_coding | 965.2406999 | 0.482701696 | 0.152784978 | 3.159353114 | 0.001581198 | 0.011099744 | 0.482701696 |
| ENSMUSG00000019843 | Fyn | 14360 | protein_coding | 39.06250168 | 0.483221874 | 0.262719833 | 1.839304887 | 0.065870355 | 0.18728722 | 0.483221874 |
| ENSMUSG00000116903 | Gm19522 | 100503040 | lncRNA | 18.1642203 | 0.4832658 | 0.32246545 | 1.498659156 | 0.133962078 | 0.302702223 | 0.4832658 |
| ENSMUSG00000025002 | Cyp2c55 | 72082 | protein_coding | 12385.13685 | -0.483300845 | 0.351498172 | -1.374973994 | 0.169139507 | 0.354093686 | 0.483300845 |
| ENSMUSG00000063524 | Eno1 | 13806 | protein_coding | 3290.381357 | 0.483330537 | 0.098076282 | 4.928108259 | 8.30E-07 | 1.59E-05 | 0.483330537 |
| ENSMUSG00000022906 | Parp9 | 80285 | protein_coding | 746.4533956 | 0.483789208 | 0.12576947 | 3.846634694 | 0.000119751 | 0.001316355 | 0.483789208 |
| ENSMUSG00000035067 | Xkr6 | 219149 | protein_coding | 53.43348198 | -0.483917165 | 0.179574857 | -2.69479354 | 0.007043226 | 0.036625014 | 0.483917165 |

| ensembl_gene_id | external_gene_name | entrezgene_id | gene_biotype | baseMean | log2FoldChange | lfcSE | stat | pvalue | padj | absLog2FC |
|---|---|---|---|---|---|---|---|---|---|---|
| ENSMUSG00000027357 | Crls1 | 66586 | protein_coding | 753.2550204 | 0.483991784 | 0.092045794 | 5.258162964 | 1.46E-07 | 3.25E-06 | 0.483991784 |
| ENSMUSG00000074088 | Snrnp40 | 66585 | protein_coding | 527.8279248 | 0.483997127 | 0.131227127 | 3.688239917 | 0.000225811 | 0.002256407 | 0.483997127 |
| ENSMUSG00000052291 | 5330438D12Rik | 102633868 | lncRNA | 20.9448208 | -0.484076372 | 0.292378553 | -1.655649388 | 0.097792842 | 0.244643379 | 0.484076372 |
| ENSMUSG00000035769 | Xylb | 102448 | protein_coding | 84.75053478 | -0.484181543 | 0.193249616 | -2.505472215 | 0.012228801 | 0.055603864 | 0.484181543 |
| ENSMUSG00000045667 | Smtnl2 | 276829 | protein_coding | 12.20997049 | -0.484239005 | 0.344052435 | -1.407456992 | 0.159291922 | 0.340451304 | 0.484239005 |
| ENSMUSG00000065434 | Mir7-1 | 723902 | miRNA | 7.116043266 | -0.484312318 | 0.474064714 | -1.021616466 | 0.306962461 | 0.516893761 | 0.484312318 |
| ENSMUSG00000031790 | Mmp15 | 17388 | protein_coding | 1417.06456 | -0.484364902 | 0.112308568 | -4.31280455 | 1.61E-05 | 0.000228429 | 0.484364902 |
| ENSMUSG00000050957 | Insl6 | 27356 | protein_coding | 70.37172052 | 0.484430135 | 0.15332824 | 3.159431904 | 0.00158077 | 0.011099744 | 0.484430135 |
| ENSMUSG00000021690 | Jmy | 57748 | protein_coding | 399.9202556 | -0.484517846 | 0.194050415 | -2.496865804 | 0.012529636 | 0.056696604 | 0.484517846 |
| ENSMUSG00000032503 | Arpp21 | 74100 | protein_coding | 11.0266359 | -0.484562376 | 1.902927967 | -0.25464042 | 0.799000838 | 0.89211142 | 0.484562376 |
| ENSMUSG00000039577 | Nphp4 | 260305 | protein_coding | 7.128674733 | -0.484702206 | 0.641644099 | -0.755406629 | 0.450005037 | 0.650554063 | 0.484702206 |
| ENSMUSG00000005470 | Asf1b | 66929 | protein_coding | 175.3675057 | 0.484901501 | 0.149731151 | 3.238481095 | 0.00120168 | 0.008898353 | 0.484901501 |
| ENSMUSG00000100094 | 1810008I18Rik | 100503969 | lncRNA | 33.52135017 | 0.48491019 | 0.246221546 | 1.969405998 | 0.048906487 | 0.153282671 | 0.48491019 |
| ENSMUSG00000049709 | Nlrp10 | 244202 | protein_coding | 34.61222234 | -0.484923565 | 0.338106402 | -1.434233611 | 0.151505608 | 0.329655755 | 0.484923565 |
| ENSMUSG00000067455 | H4c11 | 319159 | protein_coding | 615.2448992 | 0.485096626 | 0.175637182 | 2.761924441 | 0.005746177 | 0.031099258 | 0.485096626 |
| ENSMUSG00000034220 | Gpc1 | 14733 | protein_coding | 28.74824124 | -0.485201191 | 0.258416369 | -1.877594643 | 0.060436639 | 0.176597833 | 0.485201191 |
| ENSMUSG00000044017 | Adgrd1 | 243277 | protein_coding | 13.35854851 | -0.485271553 | 0.453579753 | -1.069870402 | 0.284677647 | 0.493695176 | 0.485271553 |
| ENSMUSG00000000142 | Axin2 | 12006 | protein_coding | 387.5101276 | -0.485320362 | 0.152384294 | -3.184845034 | 0.001448315 | 0.010358674 | 0.485320362 |
| ENSMUSG00000057315 | Arhgap24 | 231532 | protein_coding | 269.8946945 | -0.48539109 | 0.15595182 | -3.11244262 | 0.001855461 | 0.012711303 | 0.48539109 |
| ENSMUSG00000053070 | Cfap300 | 234912 | protein_coding | 21.58987554 | -0.485440989 | 0.337984743 | -1.436280778 | 0.150922462 | 0.328975834 | 0.485440989 |
| ENSMUSG00000078974 | Sec61g | 20335 | protein_coding | 686.6908019 | 0.485550293 | 0.10713404 | 4.532175708 | 5.84E-06 | 9.23E-05 | 0.485550293 |
| ENSMUSG00000044453 | Ffar1 | 233081 | protein_coding | 8.623675007 | -0.485711496 | 0.506648821 | -0.958674877 | 0.337722559 | 0.548479169 | 0.485711496 |
| ENSMUSG00000038725 | Pkhd1l1 | 192190 | protein_coding | 11.67950629 | -0.485712592 | 0.823707761 | -0.58966616 | 0.555414487 | 0.734954054 | 0.485712592 |
| ENSMUSG00000118215 | Vmn1r55 | 384522 | protein_coding | 10.39984913 | -0.48574908 | 1.620076455 | -0.299830961 | 0.764306098 | 0.871242219 | 0.48574908 |
| ENSMUSG00000025876 | Unc5a | 107448 | protein_coding | 69.91994529 | 0.485854209 | 0.147328724 | 3.297756181 | 0.000974607 | 0.007524743 | 0.485854209 |
| ENSMUSG00000037295 | Ldlrap1 | 100017 | protein_coding | 209.3559732 | -0.486014016 | 0.106474126 | -4.56462086 | 5.00E-06 | 8.10E-05 | 0.486014016 |
| ENSMUSG00000087230 | Mroh3 | 76422 | protein_coding | 99.90777822 | -0.486048913 | 0.21594064 | -2.250845014 | 0.024395355 | 0.092670933 | 0.486048913 |
| ENSMUSG00000042579 | 4632404H12Rik | 74034 | lncRNA | 66.22251049 | -0.48616964 | 0.218773492 | -2.222251127 | 0.026266339 | 0.098001932 | 0.48616964 |
| ENSMUSG00000059854 | Hydin | 244653 | protein_coding | 14.04680449 | -0.48678645 | 0.728077857 | -0.668591202 | 0.503756286 | 0.693802865 | 0.48678645 |
| ENSMUSG00000002228 | Ppm1j | 71887 | protein_coding | 84.01602586 | -0.487069933 | 0.204314584 | -2.383921515 | 0.017129255 | 0.071366669 | 0.487069933 |
| ENSMUSG00000023243 | Kcnk5 | 16529 | protein_coding | 894.6607383 | -0.487092805 | 0.28098673 | -1.733508214 | 0.083005379 | 0.219656035 | 0.487092805 |
| ENSMUSG00000041774 | Ydjc | 69101 | protein_coding | 291.2749911 | 0.487793631 | 0.147189235 | 3.314057783 | 0.000919525 | 0.007158453 | 0.487793631 |
| ENSMUSG00000043415 | Otud1 | 71198 | protein_coding | 24.69790496 | -0.488029998 | 0.279738417 | -1.744594123 | 0.081055545 | 0.216039611 | 0.488029998 |
| ENSMUSG00000042594 | Sh2b3 | 16923 | protein_coding | 163.6478635 | -0.488072326 | 0.11616925 | -4.201390018 | 2.65E-05 | 0.000353886 | 0.488072326 |
| ENSMUSG00000002983 | Relb | 19698 | protein_coding | 434.0375307 | -0.488442409 | 0.083506196 | -5.849175692 | 4.94E-09 | 1.38E-07 | 0.488442409 |
| ENSMUSG00000058392 | Rrp1b | 72462 | protein_coding | 174.154646 | -0.488470322 | 0.28545901 | -1.711175003 | 0.087048812 | 0.227050359 | 0.488470322 |
| ENSMUSG00000042870 | Tom1 | 21968 | protein_coding | 164.5201001 | -0.488602187 | 0.108198937 | -4.515776221 | 6.31E-06 | 9.93E-05 | 0.488602187 |
| ENSMUSG00000031831 | Dnaaf1 | 68270 | protein_coding | 20.16412943 | 0.488658882 | 0.286788132 | 1.703902037 | 0.08839939 | 0.229170213 | 0.488658882 |

| ensembl_gene_id | external_gene_name | entrezgene_id | gene_biotype | baseMean | log2FoldChange | lfcSE | stat | pvalue | padj | absLog2FC |
|---|---|---|---|---|---|---|---|---|---|---|
| ENSMUSG00000090210 | Itga10 | 213119 | protein_coding | 14.45267261 | -0.488754104 | 0.393205289 | -1.242999819 | 0.2138679 | 0.412258007 | 0.488754104 |
| ENSMUSG00000018821 | Avpi1 | 69534 | protein_coding | 105.1852757 | 0.48885857 | 0.161350576 | 3.029791281 | 0.002447228 | 0.016040925 | 0.48885857 |
| ENSMUSG00000070283 | Ndufaf3 | 66706 | protein_coding | 250.0364274 | 0.4891927 | 0.108356035 | 4.514678867 | 6.34E-06 | 9.95E-05 | 0.4891927 |
| ENSMUSG00000033021 | Gmppa | 69080 | protein_coding | 903.7488549 | 0.489387628 | 0.07361372 | 6.648049123 | 2.97E-11 | 1.07E-09 | 0.489387628 |
| ENSMUSG00000066043 | Phactr4 | 100169 | protein_coding | 321.0898611 | -0.489630313 | 0.175798553 | -2.785178286 | 0.00534983 | 0.029474953 | 0.489630313 |
| ENSMUSG00000030082 | Sec61a1 | 53421 | protein_coding | 3750.187631 | 0.48977614 | 0.107405562 | 4.560063097 | 5.11E-06 | 8.24E-05 | 0.48977614 |
| ENSMUSG00000038524 | Fchsd1 | 319262 | protein_coding | 92.87148914 | -0.489865932 | 0.142889361 | -3.428288363 | 0.0006074 | 0.005116164 | 0.489865932 |
| ENSMUSG00000004032 | Gstm5 | 14866 | protein_coding | 482.7255881 | 0.489885748 | 0.112249594 | 4.364254059 | 1.28E-05 | 0.000184703 | 0.489885748 |
| ENSMUSG00000032306 | Mpi | 110119 | protein_coding | 1156.591686 | 0.489955618 | 0.071118133 | 6.889320623 | 5.61E-12 | 2.16E-10 | 0.489955618 |
| ENSMUSG00000096299 | Gm21814 | 436100 | lncRNA | 8.477950736 | -0.489961773 | 0.457836718 | -1.070167056 | 0.28454412 | 0.493695176 | 0.489961773 |
| ENSMUSG00000005233 | Spc25 | 66442 | protein_coding | 281.012697 | 0.49027318 | 0.11215684 | 4.371317717 | 1.23E-05 | 0.000179972 | 0.49027318 |
| ENSMUSG00000021831 | Ero1a | 50527 | protein_coding | 779.4826781 | 0.490519058 | 0.082602452 | 5.938311092 | 2.88E-09 | 8.29E-08 | 0.490519058 |
| ENSMUSG00000030077 | Chl1 | 12661 | protein_coding | 8.04597894 | -0.490701961 | 1.086075137 | -0.451812166 | 0.651404304 | 0.798250711 | 0.490701961 |
| ENSMUSG00000097852 | 4933405D12Rik | 74057 | lncRNA | 12.72977805 | 0.490773279 | 0.371119609 | 1.322412686 | 0.186030767 | 0.376457636 | 0.490773279 |
| ENSMUSG00000053399 | Adamts18 | 208936 | protein_coding | 51.94076746 | 0.490775683 | 0.30912304 | 1.587638639 | 0.112368076 | 0.267961209 | 0.490775683 |
| ENSMUSG00000022822 | Abcc5 | 27416 | protein_coding | 221.3791435 | -0.490785328 | 0.189177677 | -2.594308878 | 0.009478129 | 0.045984662 | 0.490785328 |
| ENSMUSG00000087030 | Gm16143 | 102632251 | lncRNA | 7.18163764 | -0.490795567 | 0.452418739 | -1.084825902 | 0.277998771 | 0.486536032 | 0.490795567 |
| ENSMUSG00000097207 | 6030443J06Rik | 320719 | lncRNA | 19.42764546 | -0.49114486 | 0.370628281 | -1.32516833 | 0.185115327 | 0.37508166 | 0.49114486 |
| ENSMUSG00000029449 | Rhof | 23912 | protein_coding | 321.5606368 | 0.491243095 | 0.268538259 | 1.829322559 | 0.067351301 | 0.190275191 | 0.491243095 |
| ENSMUSG00000029478 | Ncor2 | 20602 | protein_coding | 256.6777718 | -0.49127131 | 0.237656987 | -2.067144403 | 0.038720549 | 0.130224009 | 0.49127131 |
| ENSMUSG00000030401 | Rtn2 | 20167 | protein_coding | 28.49486629 | -0.491598311 | 0.28363961 | -1.733179337 | 0.083063798 | 0.219701126 | 0.491598311 |
| ENSMUSG00000025161 | Slc16a3 | 80879 | protein_coding | 40.69979359 | 0.491905219 | 0.301991991 | 1.628868426 | 0.103340875 | 0.253857721 | 0.491905219 |
| ENSMUSG00000020534 | Shmt1 | 20425 | protein_coding | 1177.286441 | 0.492086703 | 0.104744895 | 4.697954056 | 2.63E-06 | 4.49E-05 | 0.492086703 |
| ENSMUSG00000042367 | Gjb3 | 14620 | protein_coding | 521.024121 | -0.492354601 | 0.122237842 | -4.027841083 | 5.63E-05 | 0.000686005 | 0.492354601 |
| ENSMUSG00000073411 | H2-D1 | 14964 | protein_coding | 6520.537666 | 0.492690003 | 0.121978028 | 4.039170097 | 5.36E-05 | 0.000659181 | 0.492690003 |
| ENSMUSG00000046658 | Zfp316 | 54201 | protein_coding | 123.4471495 | -0.492848007 | 0.12894634 | -3.822117058 | 0.000132311 | 0.001437549 | 0.492848007 |
| ENSMUSG00000003752 | Itpkc | 233011 | protein_coding | 383.4547907 | -0.492897144 | 0.111704024 | -4.412528101 | 1.02E-05 | 0.000151666 | 0.492897144 |
| ENSMUSG00000012819 | Cdh23 | 22295 | protein_coding | 14.11841656 | -0.493068416 | 0.629624541 | -0.783114989 | 0.43355959 | 0.637295787 | 0.493068416 |
| ENSMUSG00000029512 | Ulk1 | 22241 | protein_coding | 228.9333563 | -0.4932965 | 0.154637909 | -3.190010158 | 0.001422678 | 0.010230435 | 0.4932965 |
| ENSMUSG00000001156 | Mxd1 | 17119 | protein_coding | 5761.299687 | 0.493321467 | 0.195669102 | 2.521202694 | 0.011695446 | 0.053792973 | 0.493321467 |
| ENSMUSG00000086494 | 2210417A02Rik | 70138 | lncRNA | 9.859974899 | -0.493366464 | 0.390003361 | -1.265031313 | 0.205860103 | 0.401732385 | 0.493366464 |
| ENSMUSG00000006651 | Aplp1 | 11803 | protein_coding | 222.9675013 | 0.49338594 | 0.22302264 | 2.21226841 | 0.026948125 | 0.099635498 | 0.49338594 |
| ENSMUSG00000050627 | Gpd1l | 333433 | protein_coding | 2665.939132 | 0.493489354 | 0.098295972 | 5.020443311 | 5.16E-07 | 1.05E-05 | 0.493489354 |
| ENSMUSG00000078157 | Fem1al | 216622 | protein_coding | 35.01514126 | -0.493609025 | 0.317937568 | -1.552534444 | 0.120534398 | 0.281919513 | 0.493609025 |
| ENSMUSG00000039155 | Cdh26 | 381409 | protein_coding | 39.65681893 | 0.49374667 | 0.230588252 | 2.141248157 | 0.032254031 | 0.113810858 | 0.49374667 |
| ENSMUSG00000025477 | Inpp5a | 212111 | protein_coding | 348.7864685 | -0.494371813 | 0.150426478 | -3.286468042 | 0.001014523 | 0.007761441 | 0.494371813 |
| ENSMUSG00000019464 | Ptger1 | 19216 | protein_coding | 13.2900381 | -0.494381548 | 0.322126907 | -1.534741546 | 0.124847326 | 0.288868131 | 0.494381548 |
| ENSMUSG00000022967 | Ifnar1 | 15975 | protein_coding | 776.0088312 | 0.494401905 | 0.089969891 | 5.495192866 | 3.90E-08 | 9.49E-07 | 0.494401905 |

Supplementary Table 1

| ensembl_gene_id | external_gene_name | entrezgene_id | gene_biotype | baseMean | log2FoldChange | lfcSE | stat | pvalue | padj | absLog2FC |
|---|---|---|---|---|---|---|---|---|---|---|
| ENSMUSG00000030323 | Ift122 | 81896 | protein_coding | 107.9078501 | 0.494411725 | 0.13978368 | 3.536977462 | 0.000404734 | 0.003671188 | 0.494411725 |
| ENSMUSG00000020525 | Ppm1d | 53892 | protein_coding | 197.3595908 | -0.49461282 | 0.093132112 | -5.31087302 | 1.09E-07 | 2.48E-06 | 0.49461282 |
| ENSMUSG00000028841 | Cnksr1 | 194231 | protein_coding | 511.2783008 | 0.49506819 | 0.136440991 | 3.62844177 | 0.000285137 | 0.002717931 | 0.49506819 |
| ENSMUSG00000068205 | Macrod2 | 72899 | protein_coding | 128.7276104 | 0.495311 | 0.124713924 | 3.971577368 | 7.14E-05 | 0.000840526 | 0.495311 |
| ENSMUSG00000074259 | Gramd2 | 546134 | protein_coding | 40.11929338 | 0.495378691 | 0.247609834 | 2.00064223 | 0.045430959 | 0.145686394 | 0.495378691 |
| ENSMUSG00000091956 | C2cd4b | 75697 | protein_coding | 59.74819246 | -0.495481723 | 0.398789379 | -1.242464693 | 0.214065159 | 0.41248849 | 0.495481723 |
| ENSMUSG00000018570 | 2810408A11Rik | 70419 | protein_coding | 78.16634245 | -0.495735748 | 0.151415867 | -3.274001311 | 0.001060361 | 0.008043916 | 0.495735748 |
| ENSMUSG00000001440 | Kpnb1 | 16211 | protein_coding | 4253.295004 | 0.495736998 | 0.117209346 | 4.229500583 | 2.34E-05 | 0.000316681 | 0.495736998 |
| ENSMUSG00000118087 | 4833438C02Rik | 100503392 | lncRNA | 84.63914434 | -0.495926373 | 0.163376604 | -3.035479746 | 0.002401534 | 0.015780374 | 0.495926373 |
| ENSMUSG00000022451 | Twf1 | 19230 | protein_coding | 2685.836207 | 0.496025285 | 0.098190681 | 5.051653371 | 4.38E-07 | 9.06E-06 | 0.496025285 |
| ENSMUSG00000016319 | Slc25a5 | 11740 | protein_coding | 18131.71651 | 0.496102354 | 0.097164101 | 5.105819395 | 3.29E-07 | 6.92E-06 | 0.496102354 |
| ENSMUSG00000078716 | Tmem8b | 242409 | protein_coding | 26.01299025 | -0.49616752 | 0.353580824 | -1.403264789 | 0.160537894 | 0.342171494 | 0.49616752 |
| ENSMUSG00000001313 | Rnd2 | 11858 | protein_coding | 24.95462531 | -0.496338214 | 0.362272552 | -1.370068507 | 0.170665518 | 0.3559461 | 0.496338214 |
| ENSMUSG00000020601 | Trib2 | 217410 | protein_coding | 67.64633467 | 0.496463033 | 0.223594439 | 2.2203729 | 0.026393465 | 0.098300407 | 0.496463033 |
| ENSMUSG00000032788 | Pdxk | 216134 | protein_coding | 393.0145212 | 0.496467756 | 0.146281509 | 3.393920117 | 0.000688998 | 0.00568031 | 0.496467756 |
| ENSMUSG00000023868 | Pde10a | 23984 | protein_coding | 10.08559287 | -0.496523597 | 0.642629342 | -0.772643832 | 0.439733196 | 0.642162863 | 0.496523597 |
| ENSMUSG00000029073 | Cptp | 79554 | protein_coding | 431.4862216 | 0.496708324 | 0.091496145 | 5.428734967 | 5.68E-08 | 1.35E-06 | 0.496708324 |
| ENSMUSG00000021466 | Ptch1 | 19206 | protein_coding | 252.2418368 | -0.496725955 | 0.243158663 | -2.042805917 | 0.041071655 | 0.135583278 | 0.496725955 |
| ENSMUSG00000016624 | Phf21b | 271305 | protein_coding | 89.9095649 | -0.496796563 | 0.191821932 | -2.589884053 | 0.009600826 | 0.046452601 | 0.496796563 |
| ENSMUSG00000063903 | Klk1 | 16612 | protein_coding | 19151.11649 | 0.4969811 | 0.116650187 | 4.2604398 | 2.04E-05 | 0.000282338 | 0.4969811 |
| ENSMUSG00000039713 | Plekhg5 | 269608 | protein_coding | 149.3809074 | -0.497029236 | 0.157170657 | -3.162353879 | 0.001564993 | 0.011024217 | 0.497029236 |
| ENSMUSG00000034206 | Polq | 77782 | protein_coding | 148.3156663 | 0.497038832 | 0.171948245 | 2.890630446 | 0.003844699 | 0.022790611 | 0.497038832 |
| ENSMUSG00000022445 | Cyp2d26 | 76279 | protein_coding | 365.1926545 | -0.497190663 | 0.223312831 | -2.226431244 | 0.02598531 | 0.097347103 | 0.497190663 |
| ENSMUSG00000065208 | Gm24616 | 115488691 | snoRNA | 8.761122408 | 0.497564635 | 0.472265922 | 1.053568788 | 0.292080385 | 0.50099681 | 0.497564635 |
| ENSMUSG00000022679 | Mpv17l | 93734 | protein_coding | 140.6212659 | 0.497608602 | 0.12394061 | 4.014895533 | 5.95E-05 | 0.000718719 | 0.497608602 |
| ENSMUSG00000002504 | Slc9a3r2 | 65962 | protein_coding | 134.7649442 | -0.497888547 | 0.190421542 | -2.614665036 | 0.0089315 | 0.044002763 | 0.497888547 |
| ENSMUSG00000003575 | Crtc1 | 382056 | protein_coding | 105.2326291 | -0.497898295 | 0.146622898 | -3.395774476 | 0.000684347 | 0.005648709 | 0.497898295 |
| ENSMUSG00000098188 | Sowahc | 268301 | protein_coding | 1178.334804 | -0.497939425 | 0.089703793 | -5.550929465 | 2.84E-08 | 7.11E-07 | 0.497939425 |
| ENSMUSG00000055639 | Dach1 | 13134 | protein_coding | 55.54651146 | -0.497988415 | 0.25932139 | -1.920352252 | 0.05481342 | 0.16513489 | 0.497988415 |
| ENSMUSG00000026956 | Uap1l1 | 227620 | protein_coding | 209.7426151 | -0.498000368 | 0.106504854 | -4.675846685 | 2.93E-06 | 4.94E-05 | 0.498000368 |
| ENSMUSG00000035778 | Ggta1 | 14594 | protein_coding | 10.31665461 | 0.498067304 | 0.541646921 | 0.91954239 | 0.357811944 | 0.567468621 | 0.498067304 |
| ENSMUSG00000025903 | Lypla1 | 18777 | protein_coding | 2002.225217 | 0.498388273 | 0.165595087 | 3.009680313 | 0.002615228 | 0.016862722 | 0.498388273 |
| ENSMUSG00000046598 | Bdh1 | 71911 | protein_coding | 1655.541635 | -0.498667858 | 0.091940501 | -5.423810526 | 5.83E-08 | 1.38E-06 | 0.498667858 |
| ENSMUSG00000027381 | Bcl2l11 | 12125 | protein_coding | 116.0297475 | -0.498698264 | 0.113372946 | -4.39874132 | 1.09E-05 | 0.000160282 | 0.498698264 |
| ENSMUSG00000037370 | Enpp1 | 18605 | protein_coding | 441.065447 | 0.498782277 | 0.07596638 | 6.565829201 | 5.17E-11 | 1.83E-09 | 0.498782277 |
| ENSMUSG00000015957 | Wnt11 | 22411 | protein_coding | 14.93720236 | -0.49880088 | 0.36065619 | -1.38303707 | 0.1666535 | 0.350562195 | 0.49880088 |
| ENSMUSG00000035545 | Leng8 | 232798 | protein_coding | 715.3382305 | -0.498811444 | 0.198120753 | -2.517714257 | 0.01181191 | 0.054219051 | 0.498811444 |
| ENSMUSG00000063838 | Cdc42ep5 | 58804 | protein_coding | 1320.190468 | 0.498853699 | 0.202673081 | 2.461371274 | 0.013840705 | 0.060915932 | 0.498853699 |

| ensembl_gene_id | external_gene_name | entrezgene_id | gene_biotype | baseMean | log2FoldChange | lfcSE | stat | pvalue | padj | absLog2FC |
|---|---|---|---|---|---|---|---|---|---|---|
| ENSMUSG00000030036 | Mogs | 57377 | protein_coding | 1357.032106 | 0.498898212 | 0.124407325 | 4.010199653 | 6.07E-05 | 0.000730394 | 0.498898212 |
| ENSMUSG00000066952 | Myo1h | 231646 | protein_coding | 9.922971272 | -0.499357594 | 0.439233022 | -1.136885364 | 0.255586211 | 0.462295841 | 0.499357594 |
| ENSMUSG00000041000 | Trim62 | 67525 | protein_coding | 54.7723872 | -0.499414971 | 0.191805969 | -2.603750932 | 0.009220974 | 0.045080317 | 0.499414971 |
| ENSMUSG00000029810 | Tmem176b | 65963 | protein_coding | 689.0361527 | 0.499878247 | 0.09148828 | 5.463850092 | 4.66E-08 | 1.12E-06 | 0.499878247 |
| ENSMUSG00000034111 | Tmed8 | 382620 | protein_coding | 92.68304192 | -0.500051103 | 0.14251546 | -3.508749874 | 0.000450218 | 0.003993858 | 0.500051103 |
| ENSMUSG00000027194 | Ttc17 | 74569 | protein_coding | 1181.703701 | 0.500503614 | 0.06736847 | 7.429345122 | 1.09E-13 | 4.92E-12 | 0.500503614 |
| ENSMUSG00000053012 | Krcc1 | 57896 | protein_coding | 1844.716232 | 0.500528984 | 0.089332393 | 5.602995378 | 2.11E-08 | 5.41E-07 | 0.500528984 |
| ENSMUSG00000031662 | Snx20 | 71607 | protein_coding | 237.5759358 | 0.500539581 | 0.177891375 | 2.813737203 | 0.004896924 | 0.027619763 | 0.500539581 |
| ENSMUSG00000019853 | Hebp2 | 56016 | protein_coding | 127.9637761 | 0.500783179 | 0.141812406 | 3.531307262 | 0.000413511 | 0.003734964 | 0.500783179 |
| ENSMUSG00000060206 | Zfp462 | 242466 | protein_coding | 292.456942 | -0.500956192 | 0.135096698 | -3.708130533 | 0.000208795 | 0.002120974 | 0.500956192 |
| ENSMUSG00000033233 | Trim45 | 229644 | protein_coding | 44.56100956 | 0.501020126 | 0.213935967 | 2.341916293 | 0.019185017 | 0.077616869 | 0.501020126 |
| ENSMUSG00000048521 | Cxcr6 | 80901 | protein_coding | 18.94143337 | 0.501123074 | 0.366156815 | 1.368602341 | 0.171123612 | 0.356574684 | 0.501123074 |
| ENSMUSG00000019817 | Plagl1 | 22634 | protein_coding | 25.53416893 | -0.501289479 | 0.362840864 | -1.381568421 | 0.167104256 | 0.35127842 | 0.501289479 |
| ENSMUSG00000024371 | C2 | 12263 | protein_coding | 17.49613693 | 0.501298972 | 0.276673806 | 1.811877238 | 0.070005167 | 0.195681974 | 0.501298972 |
| ENSMUSG00000056458 | Mok | 26448 | protein_coding | 7.48079864 | -0.501322016 | 0.458061802 | -1.094441872 | 0.273761228 | 0.481766528 | 0.501322016 |
| ENSMUSG00000031530 | Dusp4 | 319520 | protein_coding | 85.66215468 | -0.501377661 | 0.353755495 | -1.417299993 | 0.156395237 | 0.337040095 | 0.501377661 |
| ENSMUSG00000034587 | 8430429K09Rik | 71523 | lncRNA | 39.6074227 | -0.501576186 | 0.181954186 | -2.75660701 | 0.00584045 | 0.031523784 | 0.501576186 |
| ENSMUSG00000083674 | Zfp133-ps | 668917 | transcribed_unitary_pseudogene | 6.739143504 | 0.50169481 | 0.450443903 | 1.113778666 | 0.265374151 | 0.472964024 | 0.50169481 |
| ENSMUSG00000074207 | Adh1 | 11522 | protein_coding | 6345.843332 | -0.501796859 | 0.340061347 | -1.475606867 | 0.14004946 | 0.312161742 | 0.501796859 |
| ENSMUSG00000042724 | Map3k9 | 338372 | protein_coding | 204.4983371 | -0.50182432 | 0.133947724 | -3.746419162 | 0.000179377 | 0.00185889 | 0.50182432 |
| ENSMUSG00000054342 | Kcnn4 | 16534 | protein_coding | 699.0322174 | -0.501957415 | 0.179893137 | -2.790308871 | 0.005265778 | 0.0291898 | 0.501957415 |
| ENSMUSG00000063694 | Cycs | 13063 | protein_coding | 3035.408589 | 0.501957827 | 0.149798818 | 3.350879755 | 0.000805553 | 0.006409013 | 0.501957827 |
| ENSMUSG00000067629 | Syngap1 | 240057 | protein_coding | 16.77752936 | 0.501989335 | 0.430983651 | 1.164752615 | 0.244119145 | 0.449310115 | 0.501989335 |
| ENSMUSG00000068855 | H2ac20 | 319176 | protein_coding | 328.8439219 | 0.502488841 | 0.204812004 | 2.453414988 | 0.014150698 | 0.061872603 | 0.502488841 |
| ENSMUSG00000040380 | Cbln3 | 56410 | protein_coding | 59.66515998 | 0.502580361 | 0.211135132 | 2.380372969 | 0.017295123 | 0.071813535 | 0.502580361 |
| ENSMUSG00000097534 | Gm16675 | 100503498 | lncRNA | 25.24115824 | -0.502598698 | 0.237567672 | -2.115602234 | 0.034378652 | 0.119325161 | 0.502598698 |
| ENSMUSG00000021725 | Parp8 | 52552 | protein_coding | 11.63574955 | 0.502709808 | 0.632909147 | 0.794284314 | 0.427029934 | 0.631485729 | 0.502709808 |
| ENSMUSG00000040189 | Odad1 | 211535 | protein_coding | 80.22389155 | 0.502750303 | 0.259544248 | 1.937050455 | 0.052739176 | 0.1611413 | 0.502750303 |
| ENSMUSG00000091618 | H60c | 670558 | protein_coding | 54.85299969 | 0.502873746 | 0.360203649 | 1.396081766 | 0.162689874 | 0.345233721 | 0.502873746 |
| ENSMUSG00000014776 | Nol3 | 78688 | protein_coding | 25.94811316 | 0.503143012 | 0.251723216 | 1.998794627 | 0.045630579 | 0.146061871 | 0.503143012 |
| ENSMUSG00000064109 | Hcst | 23900 | protein_coding | 14.86200824 | -0.503256564 | 0.311263557 | -1.616818138 | 0.105917543 | 0.257998872 | 0.503256564 |
| ENSMUSG00000062794 | Zfp599 | 235048 | protein_coding | 108.2817319 | -0.503275192 | 0.138480189 | -3.634275751 | 0.000278763 | 0.002674777 | 0.503275192 |
| ENSMUSG00000007030 | Vwa7 | 27762 | protein_coding | 7.617872102 | -0.503277299 | 0.478405389 | -1.051989193 | 0.292804508 | 0.501914572 | 0.503277299 |
| ENSMUSG00000063179 | Pstk | 214580 | protein_coding | 258.936771 | 0.50362641 | 0.125896542 | 4.000319638 | 6.33E-05 | 0.000756424 | 0.50362641 |
| ENSMUSG00000040952 | Rps19 | 20085 | protein_coding | 6213.497698 | 0.503654265 | 0.114648919 | 4.393013628 | 1.12E-05 | 0.000163808 | 0.503654265 |
| ENSMUSG00000032527 | Pccb | 66904 | protein_coding | 2781.664268 | 0.503661815 | 0.077949864 | 6.461355902 | 1.04E-10 | 3.54E-09 | 0.503661815 |
| ENSMUSG00000023912 | Slc25a27 | 74011 | protein_coding | 19.38009117 | -0.503895096 | 0.287422037 | -1.753154011 | 0.079575575 | 0.213132518 | 0.503895096 |
| ENSMUSG00000022790 | Igsf11 | 207683 | protein_coding | 21.45918254 | 0.504053379 | 0.333161219 | 1.512941335 | 0.130294561 | 0.297112637 | 0.504053379 |

| ensembl_gene_id | external_gene_name | entrezgene_id | gene_biotype | baseMean | log2FoldChange | lfcSE | stat | pvalue | padj | absLog2FC |
|---|---|---|---|---|---|---|---|---|---|---|
| ENSMUSG00000039236 | Isg20 | 57444 | protein_coding | 284.5008784 | 0.50424431 | 0.155212737 | 3.248730221 | 0.001159214 | 0.008644173 | 0.50424431 |
| ENSMUSG00000048349 | Pou4f1 | 18996 | protein_coding | 7.531747719 | -0.504276 | 0.458369373 | -1.10015204 | 0.271265883 | 0.479281529 | 0.504276 |
| ENSMUSG00000024033 | Rsph1 | 22092 | protein_coding | 16.37902503 | -0.504344929 | 0.288227126 | -1.749817706 | 0.080149774 | 0.213992665 | 0.504344929 |
| ENSMUSG00000025735 | Rhbdl1 | 214951 | protein_coding | 11.19196206 | -0.504601063 | 0.44211781 | -1.141327156 | 0.253733806 | 0.460042351 | 0.504601063 |
| ENSMUSG00000113585 | Gm19605 | 102638940 | lncRNA | 59.58380232 | 0.504892778 | 0.15574224 | 3.241848691 | 0.001187571 | 0.008819531 | 0.504892778 |
| ENSMUSG00000052040 | Klf13 | 50794 | protein_coding | 875.9506756 | -0.505028429 | 0.136569742 | -3.697952585 | 0.000217345 | 0.00218691 | 0.505028429 |
| ENSMUSG00000004328 | Hif3a | 53417 | protein_coding | 54.16306125 | -0.505064866 | 0.218659263 | -2.309826068 | 0.020897785 | 0.082621898 | 0.505064866 |
| ENSMUSG00000018554 | Ybx2 | 53422 | protein_coding | 101.348161 | -0.505255506 | 0.22894248 | -2.206910251 | 0.02732033 | 0.100568111 | 0.505255506 |
| ENSMUSG00000116953 | 9030025P20Rik | 100041574 | protein_coding | 10.75776514 | -0.505731891 | 0.341480216 | -1.480999098 | 0.138606814 | 0.309813265 | 0.505731891 |
| ENSMUSG00000028571 | Cyp2j13 | 230459 | protein_coding | 9.265245677 | -0.505872456 | 0.379682936 | -1.332354993 | 0.182743564 | 0.37183693 | 0.505872456 |
| ENSMUSG00000039157 | Fam102a | 98952 | protein_coding | 1388.766883 | -0.505910757 | 0.123062733 | -4.110998871 | 3.94E-05 | 0.00050112 | 0.505910757 |
| ENSMUSG00000004837 | Grap | 71520 | protein_coding | 7.648297254 | 0.50596523 | 0.487678299 | 1.037497939 | 0.299503856 | 0.508473397 | 0.50596523 |
| ENSMUSG00000038055 | Dexi | 58239 | protein_coding | 154.3234977 | 0.505977633 | 0.129533857 | 3.906141953 | 9.38E-05 | 0.001067727 | 0.505977633 |
| ENSMUSG00000027555 | Car13 | 71934 | protein_coding | 755.7203115 | 0.506029146 | 0.078389508 | 6.455317315 | 1.08E-10 | 3.68E-09 | 0.506029146 |
| ENSMUSG00000040003 | Magi2 | 50791 | protein_coding | 12.75902042 | -0.506367519 | 0.555986626 | -0.910754855 | 0.362424554 | 0.571554287 | 0.506367519 |
| ENSMUSG00000006442 | Srm | 20810 | protein_coding | 218.2236891 | 0.506620204 | 0.281010004 | 1.802854691 | 0.071411038 | 0.198159411 | 0.506620204 |
| ENSMUSG00000039109 | F13a1 | 74145 | protein_coding | 14.02910249 | 0.506632034 | 0.530712714 | 0.954625771 | 0.33976698 | 0.550260138 | 0.506632034 |
| ENSMUSG00000040183 | Ankrd6 | 140577 | protein_coding | 7.312733113 | -0.506790596 | 0.569936776 | -0.889204938 | 0.373892949 | 0.58231807 | 0.506790596 |
| ENSMUSG00000005958 | Ephb3 | 13845 | protein_coding | 448.383931 | -0.506977741 | 0.159891184 | -3.170767315 | 0.001520369 | 0.010772435 | 0.506977741 |
| ENSMUSG00000020364 | Zfp354a | 21408 | protein_coding | 27.02706006 | -0.507034711 | 0.236260823 | -2.146080353 | 0.031866568 | 0.112843642 | 0.507034711 |
| ENSMUSG00000034107 | Ano7 | 404545 | protein_coding | 599.2454928 | -0.50704256 | 0.163362311 | -3.103791545 | 0.001910579 | 0.013032848 | 0.50704256 |
| ENSMUSG00000018819 | Lsp1 | 16985 | protein_coding | 28.46460533 | 0.507050216 | 0.29288626 | 1.731218858 | 0.083412737 | 0.220404459 | 0.507050216 |
| ENSMUSG00000027865 | Gdap2 | 14547 | protein_coding | 384.8972548 | 0.507115609 | 0.113264364 | 4.477274142 | 7.56E-06 | 0.000116702 | 0.507115609 |
| ENSMUSG00000048827 | Pkd1l3 | 244646 | protein_coding | 23.88322134 | -0.507151958 | 0.282122503 | -1.797630297 | 0.072235614 | 0.199682204 | 0.507151958 |
| ENSMUSG00000075225 | Ccdc162 | 75973 | protein_coding | 17.44374455 | -0.507238197 | 0.425045965 | -1.193372573 | 0.232723492 | 0.436353122 | 0.507238197 |
| ENSMUSG00000030124 | Lag3 | 16768 | protein_coding | 7.240918211 | -0.507569204 | 0.623049489 | -0.81465311 | 0.4152709 | 0.621715848 | 0.507569204 |
| ENSMUSG00000046027 | Stard5 | 170460 | protein_coding | 713.5710189 | -0.507867201 | 0.178234074 | -2.849439449 | 0.004379634 | 0.025238355 | 0.507867201 |
| ENSMUSG00000092260 | Zfp963 | 620419 | protein_coding | 51.62877456 | 0.508040659 | 0.217661147 | 2.334089786 | 0.019591021 | 0.078608426 | 0.508040659 |
| ENSMUSG00000027223 | Mapk8ip1 | 19099 | protein_coding | 24.89428339 | -0.508118769 | 0.263444187 | -1.928753011 | 0.053761532 | 0.162897492 | 0.508118769 |
| ENSMUSG00000051747 | Ttn | 22138 | protein_coding | 66.17802065 | -0.50842918 | 0.697617351 | -0.728808106 | 0.466119052 | 0.6628287 | 0.50842918 |
| ENSMUSG00000073700 | Klhl21 | 242785 | protein_coding | 383.3328278 | 0.508699767 | 0.1705084 | 2.983429361 | 0.002850378 | 0.018009054 | 0.508699767 |
| ENSMUSG00000033386 | Frrs1 | 20321 | protein_coding | 445.7801841 | -0.509252881 | 0.127397381 | -3.997357547 | 6.41E-05 | 0.000763656 | 0.509252881 |
| ENSMUSG00000005553 | Atp4a | 11944 | protein_coding | 8.696563895 | -0.509273823 | 0.446665208 | -1.140168998 | 0.254215901 | 0.460654251 | 0.509273823 |
| ENSMUSG00000060166 | Zdhhc8 | 27801 | protein_coding | 61.93502316 | -0.509476421 | 0.180567515 | -2.821528675 | 0.004779536 | 0.027159635 | 0.509476421 |
| ENSMUSG00000031659 | Adcy7 | 11513 | protein_coding | 35.32808073 | 0.509663548 | 0.300406997 | 1.696576821 | 0.089776695 | 0.231648015 | 0.509663548 |
| ENSMUSG00000036983 | Tfb1m | 224481 | protein_coding | 135.3474985 | 0.509904538 | 0.181660012 | 2.806916797 | 0.005001816 | 0.028062319 | 0.509904538 |
| ENSMUSG00000044712 | Slc38a6 | 625098 | protein_coding | 74.96472624 | 0.510074603 | 0.161768239 | 3.153119589 | 0.001615356 | 0.011319575 | 0.510074603 |
| ENSMUSG00000032449 | Slc25a36 | 192287 | protein_coding | 968.4197712 | -0.51008614 | 0.141698571 | -3.599797336 | 0.000318465 | 0.002985589 | 0.51008614 |

| ensembl_gene_id | external_gene_name | entrezgene_id | gene_biotype | baseMean | log2FoldChange | lfcSE | stat | pvalue | padj | absLog2FC |
|---|---|---|---|---|---|---|---|---|---|---|
| ENSMUSG00000027309 | Dnaaf9 | 228602 | protein_coding | 207.2244329 | -0.510177332 | 0.11349765 | -4.495047534 | 6.96E-06 | 0.000108508 | 0.510177332 |
| ENSMUSG00000031389 | Arhgap4 | 171207 | protein_coding | 33.70512433 | -0.510220442 | 0.228227301 | -2.235580232 | 0.025379281 | 0.095610497 | 0.510220442 |
| ENSMUSG00000034614 | Pik3ip1 | 216505 | protein_coding | 41.58555399 | -0.510690631 | 0.197353607 | -2.587693424 | 0.009662093 | 0.046678138 | 0.510690631 |
| ENSMUSG00000047730 | Fcgbp | 215384 | protein_coding | 78165.01314 | 0.510831677 | 0.162052931 | 3.15225201 | 0.001620164 | 0.011338299 | 0.510831677 |
| ENSMUSG00000026097 | Ormdl1 | 227102 | protein_coding | 349.3652584 | 0.510846424 | 0.102469066 | 4.985372126 | 6.18E-07 | 1.24E-05 | 0.510846424 |
| ENSMUSG00000037902 | Sirpa | 19261 | protein_coding | 21.28853637 | -0.510974149 | 0.274480769 | -1.861602733 | 0.062659108 | 0.180967229 | 0.510974149 |
| ENSMUSG00000039693 | Msantd3 | 66665 | protein_coding | 32.21501677 | 0.511015888 | 0.22199612 | 2.301913605 | 0.021340045 | 0.083906254 | 0.511015888 |
| ENSMUSG00000018750 | Zbtb4 | 75580 | protein_coding | 285.1607515 | -0.511431343 | 0.144643302 | -3.535810743 | 0.000406526 | 0.00368115 | 0.511431343 |
| ENSMUSG00000042647 | Acad12 | 338350 | protein_coding | 145.0903977 | -0.511488142 | 0.146435091 | -3.492934241 | 0.000477744 | 0.004187952 | 0.511488142 |
| ENSMUSG00000049122 | Frmd3 | 242506 | protein_coding | 25.37484744 | 0.511608508 | 0.285161382 | 1.794101659 | 0.072796944 | 0.200676868 | 0.511608508 |
| ENSMUSG00000005533 | Igf1r | 16001 | protein_coding | 275.0875352 | -0.511709029 | 0.184683656 | -2.770732613 | 0.005593033 | 0.030456697 | 0.511709029 |
| ENSMUSG00000031089 | Slc6a14 | 56774 | protein_coding | 11509.68426 | -0.511729919 | 0.151935158 | -3.368080988 | 0.000756934 | 0.006101436 | 0.511729919 |
| ENSMUSG00000014773 | Dll1 | 13388 | protein_coding | 201.7413722 | -0.511785108 | 0.228638365 | -2.238404332 | 0.0251947 | 0.095027511 | 0.511785108 |
| ENSMUSG00000007655 | Cav1 | 12389 | protein_coding | 7.359783798 | 0.511938439 | 0.8013837 | 0.638818133 | 0.522941251 | 0.70918759 | 0.511938439 |
| ENSMUSG00000029368 | Alb | 11657 | protein_coding | 19.10560239 | 0.512122001 | 0.330973662 | 1.547319499 | 0.121786222 | 0.28413812 | 0.512122001 |
| ENSMUSG00000047496 | Rnf152 | 320311 | protein_coding | 210.5249022 | -0.512439306 | 0.284158298 | -1.803358583 | 0.071331918 | 0.198043365 | 0.512439306 |
| ENSMUSG00000072980 | Oip5 | 70645 | protein_coding | 67.89077923 | 0.513032136 | 0.15948318 | 3.216841643 | 0.001296101 | 0.009430927 | 0.513032136 |
| ENSMUSG00000020258 | Glyctk | 235582 | protein_coding | 14.60900254 | -0.513056773 | 0.28373032 | -1.808255012 | 0.07056682 | 0.196398447 | 0.513056773 |
| ENSMUSG00000024696 | Lpxn | 107321 | protein_coding | 22.77040991 | 0.513382551 | 0.2564761 | 2.001677939 | 0.045319381 | 0.145409528 | 0.513382551 |
| ENSMUSG00000035632 | Cnot3 | 232791 | protein_coding | 393.8876717 | 0.513676735 | 0.119452454 | 4.300261033 | 1.71E-05 | 0.000240254 | 0.513676735 |
| ENSMUSG00000032332 | Col12a1 | 12816 | protein_coding | 9.039552108 | -0.513774705 | 0.924391215 | -0.55579791 | 0.578349015 | 0.749869458 | 0.513774705 |
| ENSMUSG00000031821 | Gins2 | 272551 | protein_coding | 111.8602083 | 0.513893117 | 0.140805132 | 3.649676051 | 0.000262571 | 0.002553257 | 0.513893117 |
| ENSMUSG00000020818 | Mfsd11 | 69900 | protein_coding | 331.152646 | 0.514167563 | 0.064001169 | 8.033721469 | 9.46E-16 | 5.12E-14 | 0.514167563 |
| ENSMUSG00000002688 | Prkd1 | 18760 | protein_coding | 12.34439739 | 0.514340709 | 0.494181023 | 1.040794132 | 0.297971104 | 0.506735399 | 0.514340709 |
| ENSMUSG00000112280 | A830082N09Rik | 414093 | lncRNA | 7.966425408 | -0.514958779 | 0.69732819 | -0.738474059 | 0.460226425 | 0.658624124 | 0.514958779 |
| ENSMUSG00000026670 | Uap1 | 107652 | protein_coding | 4235.907321 | 0.515002911 | 0.103301317 | 4.9854438 | 6.18E-07 | 1.24E-05 | 0.515002911 |
| ENSMUSG00000090124 | Ugt1a7c | 394432 | protein_coding | 2533.461065 | -0.515004651 | 0.11346833 | -4.538752374 | 5.66E-06 | 9.00E-05 | 0.515004651 |
| ENSMUSG00000038248 | Sobp | 109205 | protein_coding | 23.60078533 | -0.515154249 | 0.334861046 | -1.538411993 | 0.123947912 | 0.287329695 | 0.515154249 |
| ENSMUSG00000073295 | Nudt11 | 58242 | protein_coding | 27.00072189 | 0.51518421 | 0.276878629 | 1.860686077 | 0.062788521 | 0.181128045 | 0.51518421 |
| ENSMUSG00000021418 | Rpp40 | 208366 | protein_coding | 56.60006098 | -0.515250942 | 0.161004547 | -3.200226018 | 0.001373199 | 0.009941958 | 0.515250942 |
| ENSMUSG00000038895 | Zfp653 | 319601 | protein_coding | 52.42149386 | -0.515335561 | 0.199092385 | -2.588424275 | 0.009641614 | 0.046607476 | 0.515335561 |
| ENSMUSG00000038252 | Ncapd2 | 68298 | protein_coding | 968.1094951 | 0.515369708 | 0.105213125 | 4.898340442 | 9.66E-07 | 1.83E-05 | 0.515369708 |
| ENSMUSG00000025453 | Nnt | 18115 | protein_coding | 8.292290428 | -0.515476265 | 0.404518909 | -1.274294608 | 0.202559027 | 0.39748185 | 0.515476265 |
| ENSMUSG00000041189 | Chrnb1 | 11443 | protein_coding | 22.95923466 | -0.515530661 | 0.26658774 | -1.933812336 | 0.053136205 | 0.161919549 | 0.515530661 |
| ENSMUSG00000031910 | Has3 | 15118 | protein_coding | 14.42175723 | -0.515680163 | 0.360959548 | -1.428636995 | 0.153108595 | 0.33202365 | 0.515680163 |
| ENSMUSG00000061479 | Snrpa | 53607 | protein_coding | 468.0920508 | 0.515769196 | 0.100444363 | 5.134874504 | 2.82E-07 | 6.00E-06 | 0.515769196 |
| ENSMUSG00000085078 | C030013C21Rik | 77417 | lncRNA | 9.70066542 | -0.515779682 | 0.46940616 | -1.098791889 | 0.271858851 | 0.480062946 | 0.515779682 |
| ENSMUSG00000024986 | Hhex | 15242 | protein_coding | 12.63776134 | -0.515859256 | 0.430064025 | -1.199494089 | 0.230335882 | 0.433458782 | 0.515859256 |

| ensembl_gene_id | external_gene_name | entrezgene_id | gene_biotype | baseMean | log2FoldChange | lfcSE | stat | pvalue | padj | absLog2FC |
|---|---|---|---|---|---|---|---|---|---|---|
| ENSMUSG00000042524 | Sun2 | 223697 | protein_coding | 513.6019209 | -0.516139108 | 0.215334889 | -2.396913522 | 0.016533825 | 0.069540737 | 0.516139108 |
| ENSMUSG00000019970 | Sgk1 | 20393 | protein_coding | 2291.982639 | -0.516190082 | 0.211803923 | -2.437112946 | 0.014805054 | 0.063889164 | 0.516190082 |
| ENSMUSG00000055184 | Fam72a | 108900 | protein_coding | 26.81312796 | 0.516404855 | 0.260978643 | 1.978724578 | 0.047847024 | 0.151211258 | 0.516404855 |
| ENSMUSG00000020598 | Nrcam | 319504 | protein_coding | 8.060987755 | -0.516424908 | 1.304534528 | -0.395869099 | 0.692201598 | 0.824821355 | 0.516424908 |
| ENSMUSG00000022383 | Ppara | 19013 | protein_coding | 314.874471 | -0.51645751 | 0.197232375 | -2.618523005 | 0.008831134 | 0.043683943 | 0.51645751 |
| ENSMUSG00000026614 | Slc30a10 | 226781 | protein_coding | 479.7501372 | -0.516555755 | 0.308014402 | -1.677050656 | 0.093532577 | 0.237985125 | 0.516555755 |
| ENSMUSG00000038437 | Mllt6 | 246198 | protein_coding | 782.8595781 | -0.516568953 | 0.084075633 | -6.144098264 | 8.04E-10 | 2.44E-08 | 0.516568953 |
| ENSMUSG00000058908 | Pla2g2a | 18780 | protein_coding | 33.05902476 | 0.516814417 | 0.294747321 | 1.75341515 | 0.079530772 | 0.213080932 | 0.516814417 |
| ENSMUSG00000000301 | Pemt | 18618 | protein_coding | 45.75916357 | 0.516973342 | 0.179852561 | 2.874428575 | 0.004047593 | 0.023710946 | 0.516973342 |
| ENSMUSG00000042035 | Igsf3 | 78908 | protein_coding | 380.0604892 | -0.517106022 | 0.15349526 | -3.368872914 | 0.000754762 | 0.006094139 | 0.517106022 |
| ENSMUSG00000028108 | Ecm1 | 13601 | protein_coding | 12.62095663 | -0.517993209 | 0.373353152 | -1.387408159 | 0.165317339 | 0.348987948 | 0.517993209 |
| ENSMUSG00000029298 | Gbp9 | 236573 | protein_coding | 18.64657627 | 0.51809149 | 0.447869536 | 1.156791091 | 0.247357722 | 0.453071965 | 0.51809149 |
| ENSMUSG00000038692 | Hoxb4 | 15412 | protein_coding | 47.48363456 | -0.518155335 | 0.298679788 | -1.734818877 | 0.082772891 | 0.219222916 | 0.518155335 |
| ENSMUSG00000036523 | Greb1 | 268527 | protein_coding | 146.5514506 | -0.518188318 | 0.137061055 | -3.780711573 | 0.000156381 | 0.001652842 | 0.518188318 |
| ENSMUSG00000027559 | Car3 | 12350 | protein_coding | 16.77785283 | 0.518386422 | 0.358236641 | 1.447050253 | 0.147882849 | 0.324714368 | 0.518386422 |
| ENSMUSG00000057561 | Eif1a | 13664 | protein_coding | 1941.015279 | 0.518479781 | 0.151427777 | 3.423941048 | 0.0006172 | 0.005179699 | 0.518479781 |
| ENSMUSG00000015850 | Adamtsl4 | 229595 | protein_coding | 51.55957464 | -0.518688637 | 0.185322377 | -2.798845158 | 0.005128572 | 0.028602207 | 0.518688637 |
| ENSMUSG00000034135 | Sik3 | 70661 | protein_coding | 276.8110352 | -0.518768184 | 0.165894316 | -3.127100405 | 0.001765396 | 0.012199373 | 0.518768184 |
| ENSMUSG00000028635 | Edn2 | 13615 | protein_coding | 170.4184426 | -0.51878018 | 0.329410819 | -1.574872925 | 0.115285779 | 0.272646161 | 0.51878018 |
| ENSMUSG00000031015 | Swap70 | 20947 | protein_coding | 603.4499872 | -0.518856846 | 0.094304427 | -5.501935174 | 3.76E-08 | 9.18E-07 | 0.518856846 |
| ENSMUSG00000052861 | Dnah6 | 330355 | protein_coding | 9.61593539 | -0.519141505 | 1.092927986 | -0.475000651 | 0.634786509 | 0.788741674 | 0.519141505 |
| ENSMUSG00000020108 | Ddit4 | 74747 | protein_coding | 825.105367 | 0.519354852 | 0.167396887 | 3.102535895 | 0.001918703 | 0.013077064 | 0.519354852 |
| ENSMUSG00000041124 | Msantd4 | 78100 | protein_coding | 729.7432981 | 0.519360558 | 0.074064638 | 7.012260845 | 2.34E-12 | 9.31E-11 | 0.519360558 |
| ENSMUSG00000043518 | Rai2 | 24004 | protein_coding | 20.71723252 | -0.519396715 | 0.34990733 | -1.484383638 | 0.137707173 | 0.308495056 | 0.519396715 |
| ENSMUSG00000021025 | Nfkbia | 18035 | protein_coding | 977.7840722 | -0.519574382 | 0.129498711 | -4.01219733 | 6.02E-05 | 0.000724786 | 0.519574382 |
| ENSMUSG00000032714 | Syde1 | 71709 | protein_coding | 8.324880664 | -0.519579653 | 0.425537016 | -1.220997547 | 0.222086949 | 0.423234241 | 0.519579653 |
| ENSMUSG00000110344 | Smim36 | 121811722 | protein_coding | 20.94914709 | -0.519753033 | 0.304598644 | -1.706353732 | 0.08794224 | 0.228497519 | 0.519753033 |
| ENSMUSG00000047819 | Tigd4 | 403175 | protein_coding | 6.999208879 | 0.519900036 | 0.508241706 | 1.022938556 | 0.306336898 | 0.516331653 | 0.519900036 |
| ENSMUSG00000024029 | Tff3 | 21786 | protein_coding | 8188.669153 | 0.519953214 | 0.172061073 | 3.02191079 | 0.002511846 | 0.016387913 | 0.519953214 |
| ENSMUSG00000052942 | Glis3 | 226075 | protein_coding | 145.7108349 | -0.520040078 | 0.179369096 | -2.899273554 | 0.003740284 | 0.022321186 | 0.520040078 |
| ENSMUSG00000012483 | Rpa3 | 68240 | protein_coding | 283.3056793 | 0.520060549 | 0.104184821 | 4.991711305 | 5.98E-07 | 1.21E-05 | 0.520060549 |
| ENSMUSG00000079440 | Alpi | 76768 | protein_coding | 189.5330817 | -0.520438934 | 0.177473448 | -2.932488994 | 0.003362568 | 0.02057587 | 0.520438934 |
| ENSMUSG00000097587 | 4930578M01Rik | 75951 | lncRNA | 9.313422687 | 0.520562323 | 0.56398629 | 0.923005278 | 0.356004455 | 0.565615496 | 0.520562323 |
| ENSMUSG00000024742 | Fen1 | 14156 | protein_coding | 32.06690831 | 0.520580497 | 0.242861924 | 2.143524551 | 0.032071002 | 0.113365938 | 0.520580497 |
| ENSMUSG00000000934 | Top1mt | 72960 | protein_coding | 84.24375564 | -0.520601366 | 0.149682614 | -3.478034965 | 0.000505104 | 0.004386749 | 0.520601366 |
| ENSMUSG00000033031 | Cip2a | 224171 | protein_coding | 630.3625367 | 0.520678159 | 0.106331562 | 4.896741392 | 9.74E-07 | 1.84E-05 | 0.520678159 |
| ENSMUSG00000057666 | Gapdh | 14433 | protein_coding | 8029.539766 | 0.520700069 | 0.093506591 | 5.568592171 | 2.57E-08 | 6.53E-07 | 0.520700069 |
| ENSMUSG00000049598 | Vsig8 | 240916 | protein_coding | 17.97429685 | -0.520804913 | 0.387777377 | -1.343051305 | 0.179255362 | 0.366942475 | 0.520804913 |

| ensembl_gene_id | external_gene_name | entrezgene_id | gene_biotype | baseMean | log2FoldChange | lfcSE | stat | pvalue | padj | absLog2FC |
|---|---|---|---|---|---|---|---|---|---|---|
| ENSMUSG00000041141 | Pnma8a | 71691 | protein_coding | 12.48186515 | -0.521199595 | 0.339213593 | -1.536493836 | 0.124417308 | 0.288103346 | 0.521199595 |
| ENSMUSG00000050824 | Sstr5 | 20609 | protein_coding | 8.323036237 | -0.521399423 | 0.431341239 | -1.2087864 | 0.226744918 | 0.429204895 | 0.521399423 |
| ENSMUSG00000078612 | Fyb2 | 242594 | protein_coding | 124.0356563 | -0.521442659 | 0.279609911 | -1.864893336 | 0.062196357 | 0.179957052 | 0.521442659 |
| ENSMUSG00000021036 | Sptlc2 | 20773 | protein_coding | 4042.603703 | 0.521472846 | 0.093160234 | 5.597590546 | 2.17E-08 | 5.56E-07 | 0.521472846 |
| ENSMUSG00000104667 | Gm4961 | 242914 | lncRNA | 9.079810125 | 0.521651671 | 0.551570967 | 0.945756216 | 0.344272942 | 0.55406481 | 0.521651671 |
| ENSMUSG00000037843 | Vstm2l | 277432 | protein_coding | 12.60379779 | -0.521782004 | 0.413547052 | -1.261723428 | 0.207048324 | 0.403458001 | 0.521782004 |
| ENSMUSG00000022074 | Tnfrsf10b | 21933 | protein_coding | 116.0829301 | -0.521867646 | 0.244040725 | -2.138444906 | 0.032480651 | 0.114256142 | 0.521867646 |
| ENSMUSG00000020263 | Appl2 | 216190 | protein_coding | 2236.814186 | -0.521919022 | 0.109710772 | -4.757226778 | 1.96E-06 | 3.45E-05 | 0.521919022 |
| ENSMUSG00000022371 | Col14a1 | 12818 | protein_coding | 17.17836261 | -0.522046675 | 0.458547214 | -1.138479657 | 0.254920249 | 0.461563001 | 0.522046675 |
| ENSMUSG00000028581 | Laptm5 | 16792 | protein_coding | 37.11041119 | 0.522122272 | 0.346238959 | 1.507982446 | 0.13155902 | 0.299267647 | 0.522122272 |
| ENSMUSG00000028195 | Ccn1 | 16007 | protein_coding | 293.7665729 | -0.522123571 | 0.435312208 | -1.199423223 | 0.230363422 | 0.433458782 | 0.522123571 |
| ENSMUSG00000039431 | Mtmr7 | 54384 | protein_coding | 309.6217226 | 0.522345522 | 0.093220923 | 5.603307776 | 2.10E-08 | 5.41E-07 | 0.522345522 |
| ENSMUSG00000063576 | Klhdc3 | 71765 | protein_coding | 338.0767124 | -0.522444771 | 0.128332002 | -4.07104046 | 4.68E-05 | 0.000581958 | 0.522444771 |
| ENSMUSG00000026694 | Eef1aknmt | 71449 | protein_coding | 157.5561277 | -0.522478757 | 0.140197981 | -3.726720985 | 0.000193987 | 0.001988304 | 0.522478757 |
| ENSMUSG00000057614 | Gnai1 | 14677 | protein_coding | 194.7763407 | -0.522635826 | 0.119559741 | -4.371336227 | 1.23E-05 | 0.000179972 | 0.522635826 |
| ENSMUSG00000032062 | 2310030G06Rik | 66952 | protein_coding | 287.0547421 | 0.522669172 | 0.095555466 | 5.469798783 | 4.51E-08 | 1.09E-06 | 0.522669172 |
| ENSMUSG00000031728 | Zfp821 | 75871 | protein_coding | 183.4038812 | -0.522709308 | 0.101920889 | -5.128578757 | 2.92E-07 | 6.19E-06 | 0.522709308 |
| ENSMUSG00000069300 | H2bc11 | 319183 | protein_coding | 442.9101977 | 0.522710725 | 0.187767734 | 2.783815481 | 0.005372359 | 0.029547084 | 0.522710725 |
| ENSMUSG00000026248 | Mrpl44 | 69163 | protein_coding | 482.2974819 | 0.522763988 | 0.112289191 | 4.655514794 | 3.23E-06 | 5.41E-05 | 0.522763988 |
| ENSMUSG00000037669 | Ldah | 68832 | protein_coding | 889.3591437 | 0.522859076 | 0.08027692 | 6.51319305 | 7.36E-11 | 2.58E-09 | 0.522859076 |
| ENSMUSG00000029832 | Nfe2l3 | 18025 | protein_coding | 59.51371024 | -0.523096627 | 0.292776676 | -1.786674519 | 0.07399011 | 0.202842422 | 0.523096627 |
| ENSMUSG00000030491 | Tdrd12 | 71981 | protein_coding | 21.94525509 | -0.523169964 | 0.254983174 | -2.051782299 | 0.040190824 | 0.13350562 | 0.523169964 |
| ENSMUSG00000024018 | Ccdc167 | 68597 | protein_coding | 80.98964742 | 0.523238599 | 0.178749918 | 2.927210295 | 0.003420175 | 0.020809532 | 0.523238599 |
| ENSMUSG00000031980 | Agt | 11606 | protein_coding | 99.75755208 | -0.523604145 | 0.144087612 | -3.633928955 | 0.000279138 | 0.002676765 | 0.523604145 |
| ENSMUSG00000036902 | Neto2 | 74513 | protein_coding | 65.19759208 | 0.523818686 | 0.298977881 | 1.752031567 | 0.07976838 | 0.213466189 | 0.523818686 |
| ENSMUSG00000071636 | Rimbp3 | 239731 | protein_coding | 9.164504992 | -0.523855468 | 0.358337775 | -1.461904117 | 0.143767487 | 0.317913165 | 0.523855468 |
| ENSMUSG00000042354 | Gnl3 | 30877 | protein_coding | 482.0843875 | -0.524007645 | 0.212219055 | -2.469182822 | 0.013542201 | 0.060066884 | 0.524007645 |
| ENSMUSG00000018537 | Pcgf2 | 22658 | protein_coding | 91.45288862 | -0.524012385 | 0.231000757 | -2.268444451 | 0.023302131 | 0.089627443 | 0.524012385 |
| ENSMUSG00000111329 | A830035O19Rik | 100101457 | lncRNA | 10.9890384 | -0.524036164 | 0.395380017 | -1.325398711 | 0.185038944 | 0.375027 | 0.524036164 |
| ENSMUSG00000021815 | Mss51 | 74843 | protein_coding | 8.353222048 | 0.524414064 | 0.4334489 | 1.20986364 | 0.226331221 | 0.428605837 | 0.524414064 |
| ENSMUSG00000068893 | Sprr2a2 | 100303744 | protein_coding | 259.6755142 | 0.524466129 | 0.198942138 | 2.636274718 | 0.008382184 | 0.041971526 | 0.524466129 |
| ENSMUSG00000029334 | Prkg2 | 19092 | protein_coding | 705.3146379 | -0.524508492 | 0.203491543 | -2.577544431 | 0.009950509 | 0.047759252 | 0.524508492 |
| ENSMUSG00000052512 | Nav2 | 78286 | protein_coding | 244.7598224 | -0.524530131 | 0.255875699 | -2.049941179 | 0.040370171 | 0.133871306 | 0.524530131 |
| ENSMUSG00000018800 | Abca5 | 217265 | protein_coding | 45.47667751 | -0.524715939 | 0.287933596 | -1.822350524 | 0.068401817 | 0.192513563 | 0.524715939 |
| ENSMUSG00000036989 | Trim3 | 55992 | protein_coding | 97.91683608 | -0.525135933 | 0.155090803 | -3.385990155 | 0.000709219 | 0.005798996 | 0.525135933 |
| ENSMUSG00000036411 | 9530077C05Rik | 68283 | protein_coding | 10.37057771 | -0.525973027 | 0.463879909 | -1.133856019 | 0.256854948 | 0.463695525 | 0.525973027 |
| ENSMUSG00000005968 | Tuft1 | 22156 | protein_coding | 539.8071756 | -0.526055524 | 0.180984823 | -2.906627837 | 0.003653476 | 0.021918103 | 0.526055524 |
| ENSMUSG00000007207 | Stx1a | 20907 | protein_coding | 19.39772775 | -0.526123354 | 0.316698567 | -1.661274819 | 0.096658265 | 0.242949794 | 0.526123354 |

| ensembl_gene_id | external_gene_name | entrezgene_id | gene_biotype | baseMean | log2FoldChange | lfcSE | stat | pvalue | padj | absLog2FC |
|---|---|---|---|---|---|---|---|---|---|---|
| ENSMUSG00000006930 | Hap1 | 15114 | protein_coding | 35.06701561 | -0.526198743 | 0.303152639 | -1.735755113 | 0.082607144 | 0.218929549 | 0.526198743 |
| ENSMUSG00000040550 | Otud6b | 72201 | protein_coding | 748.8655591 | 0.526318591 | 0.094907743 | 5.545581164 | 2.93E-08 | 7.29E-07 | 0.526318591 |
| ENSMUSG00000028294 | Cfap206 | 69329 | protein_coding | 11.41449875 | -0.526467107 | 0.408735969 | -1.288037134 | 0.197733037 | 0.391727838 | 0.526467107 |
| ENSMUSG00000111709 | Gm3776 | 100042295 | protein_coding | 25.26393353 | 0.526610541 | 0.224872038 | 2.341823139 | 0.019189806 | 0.077616869 | 0.526610541 |
| ENSMUSG00000024620 | Pdgfrb | 18596 | protein_coding | 116.565403 | -0.526633165 | 0.173448598 | -3.036249184 | 0.002395413 | 0.015746653 | 0.526633165 |
| ENSMUSG00000041482 | Piezo2 | 667742 | protein_coding | 45.43438944 | -0.526819528 | 0.415421254 | -1.268157376 | 0.204741755 | 0.400334726 | 0.526819528 |
| ENSMUSG00000025050 | Pcgf6 | 71041 | protein_coding | 179.3226177 | -0.52723939 | 0.116083782 | -4.541886746 | 5.58E-06 | 8.90E-05 | 0.52723939 |
| ENSMUSG00000037736 | Limch1 | 77569 | protein_coding | 42.36189788 | 0.527440564 | 0.243906238 | 2.162472625 | 0.030581765 | 0.10960989 | 0.527440564 |
| ENSMUSG00000045007 | Tubg2 | 103768 | protein_coding | 12.42483955 | -0.527555583 | 0.433051017 | -1.218229637 | 0.22313672 | 0.424532081 | 0.527555583 |
| ENSMUSG00000011114 | Tbrg1 | 21376 | protein_coding | 1766.732183 | 0.52762607 | 0.090948906 | 5.801345971 | 6.58E-09 | 1.81E-07 | 0.52762607 |
| ENSMUSG00000101174 | Hoxd4 | 15436 | protein_coding | 27.99483992 | -0.527718937 | 0.297157834 | -1.775887679 | 0.075751432 | 0.205338463 | 0.527718937 |
| ENSMUSG00000090565 | 4930405O22Rik | 414079 | lncRNA | 9.31976174 | -0.527867968 | 0.419790622 | -1.257455362 | 0.208588793 | 0.405368753 | 0.527867968 |
| ENSMUSG00000031432 | Prps1 | 19139 | protein_coding | 216.047972 | 0.527990715 | 0.158448185 | 3.33226106 | 0.000861434 | 0.006779112 | 0.527990715 |
| ENSMUSG00000002032 | Tmem25 | 71687 | protein_coding | 56.71250388 | -0.528116971 | 0.185064228 | -2.853695589 | 0.004321392 | 0.025011311 | 0.528116971 |
| ENSMUSG00000044468 | Tent5c | 74645 | protein_coding | 164.6469514 | -0.528121085 | 0.266921094 | -1.978566314 | 0.047864855 | 0.151211258 | 0.528121085 |
| ENSMUSG00000029177 | Cenpa | 12615 | protein_coding | 447.6241525 | 0.528295136 | 0.128763604 | 4.102829683 | 4.08E-05 | 0.000516745 | 0.528295136 |
| ENSMUSG00000020838 | Slc6a4 | 15567 | protein_coding | 142.1026313 | -0.528336398 | 0.33881123 | -1.559382783 | 0.118905809 | 0.279464619 | 0.528336398 |
| ENSMUSG00000033487 | Fndc3a | 319448 | protein_coding | 2761.542932 | 0.528804323 | 0.051659845 | 10.23627394 | 1.36E-24 | 1.21E-22 | 0.528804323 |
| ENSMUSG00000044854 | 1700056E22Rik | 73363 | protein_coding | 11.47302551 | 0.528831131 | 0.436442812 | 1.211684821 | 0.225633052 | 0.427894183 | 0.528831131 |
| ENSMUSG00000070867 | Trabd2b | 666048 | protein_coding | 16.07341192 | -0.528885396 | 0.362818011 | -1.457715386 | 0.144919009 | 0.32005962 | 0.528885396 |
| ENSMUSG00000060950 | Trmt61a | 328162 | protein_coding | 96.75855775 | -0.528937068 | 0.365267618 | -1.448080919 | 0.147594421 | 0.324304586 | 0.528937068 |
| ENSMUSG00000022840 | Adcy5 | 224129 | protein_coding | 51.301074 | -0.529058424 | 0.173985324 | -3.040822136 | 0.002359332 | 0.015535111 | 0.529058424 |
| ENSMUSG00000095789 | Nupr1l | 69034 | protein_coding | 7.519702152 | -0.529145811 | 0.403504621 | -1.311374848 | 0.189731152 | 0.38132969 | 0.529145811 |
| ENSMUSG00000015451 | C4a | 625018 | lncRNA | 8.312464062 | -0.529163489 | 0.559236197 | -0.946225391 | 0.344033638 | 0.554008329 | 0.529163489 |
| ENSMUSG00000059742 | Kcnh7 | 170738 | protein_coding | 32.19342622 | -0.529254697 | 0.338984211 | -1.561296013 | 0.118453924 | 0.278566972 | 0.529254697 |
| ENSMUSG00000033460 | Armcx1 | 78248 | protein_coding | 50.93634018 | -0.529569959 | 0.265100229 | -1.997621657 | 0.045757693 | 0.146321731 | 0.529569959 |
| ENSMUSG00000001891 | Ugp2 | 216558 | protein_coding | 4586.25154 | 0.52960101 | 0.105866185 | 5.002551184 | 5.66E-07 | 1.14E-05 | 0.52960101 |
| ENSMUSG00000028995 | Hycc1 | 84652 | protein_coding | 428.3604872 | 0.52976986 | 0.130863427 | 4.048265204 | 5.16E-05 | 0.000637103 | 0.52976986 |
| ENSMUSG00000024924 | Vldlr | 22359 | protein_coding | 357.951729 | -0.530167705 | 0.235432445 | -2.251888876 | 0.024329295 | 0.092530328 | 0.530167705 |
| ENSMUSG00000002799 | Jag2 | 16450 | protein_coding | 51.72116425 | -0.530209807 | 0.23791961 | -2.22852503 | 0.025845524 | 0.096959775 | 0.530209807 |
| ENSMUSG00000019987 | Arg1 | 11846 | protein_coding | 24.87308146 | -0.530238136 | 0.285775048 | -1.855438882 | 0.063533579 | 0.182796758 | 0.530238136 |
| ENSMUSG00000024732 | Ccdc86 | 108673 | protein_coding | 170.0027461 | -0.530516918 | 0.216857271 | -2.446387501 | 0.014429582 | 0.06271061 | 0.530516918 |
| ENSMUSG00000005677 | Nr1i3 | 12355 | protein_coding | 7.416297255 | 0.530518016 | 0.455510056 | 1.164668066 | 0.24415338 | 0.449321206 | 0.530518016 |
| ENSMUSG00000061762 | Tac1 | 21333 | protein_coding | 72.33829424 | -0.530577088 | 0.274447346 | -1.933256398 | 0.05320462 | 0.161983995 | 0.530577088 |
| ENSMUSG00000065546 | Mir196a-1 | 387191 | miRNA | 33.66453723 | -0.530579319 | 0.262676059 | -2.019899798 | 0.043393782 | 0.14071176 | 0.530579319 |
| ENSMUSG00000086968 | 4933431E20Rik | 329735 | lncRNA | 341.8817373 | -0.531144962 | 0.140306212 | -3.785612585 | 0.00015333 | 0.001624915 | 0.531144962 |
| ENSMUSG00000022661 | Cd200 | 17470 | protein_coding | 48.51888493 | -0.531257904 | 0.220336358 | -2.411122292 | 0.015903515 | 0.067621779 | 0.531257904 |
| ENSMUSG00000031840 | Rab3a | 19339 | protein_coding | 103.1186491 | 0.531286038 | 0.151395731 | 3.509253759 | 0.000449366 | 0.003990201 | 0.531286038 |

| ensembl_gene_id | external_gene_name | entrezgene_id | gene_biotype | baseMean | log2FoldChange | lfcSE | stat | pvalue | padj | absLog2FC |
|---|---|---|---|---|---|---|---|---|---|---|
| ENSMUSG00000020389 | Cdkl3 | 213084 | protein_coding | 43.02407612 | 0.53128724 | 0.203445524 | 2.611447178 | 0.009015991 | 0.044268407 | 0.53128724 |
| ENSMUSG00000025026 | Add3 | 27360 | protein_coding | 1676.208475 | -0.531341832 | 0.119815192 | -4.434678302 | 9.22E-06 | 0.000138583 | 0.531341832 |
| ENSMUSG00000025350 | Rdh5 | 19682 | protein_coding | 44.29418685 | -0.531422823 | 0.209986624 | -2.530746063 | 0.011382022 | 0.052747409 | 0.531422823 |
| ENSMUSG00000042340 | Ctf1 | 13019 | protein_coding | 27.64931256 | -0.531443926 | 0.244098068 | -2.177173828 | 0.0294676 | 0.106502223 | 0.531443926 |
| ENSMUSG00000014294 | Ndufa2 | 17991 | protein_coding | 949.4420539 | 0.531459881 | 0.121832771 | 4.362207956 | 1.29E-05 | 0.00018627 | 0.531459881 |
| ENSMUSG00000026921 | Egfl7 | 353156 | protein_coding | 16.79856357 | -0.531784583 | 0.283453096 | -1.876093758 | 0.060642407 | 0.177004261 | 0.531784583 |
| ENSMUSG00000032086 | Bace1 | 23821 | protein_coding | 285.0177077 | -0.531860227 | 0.093500879 | -5.68829123 | 1.28E-08 | 3.40E-07 | 0.531860227 |
| ENSMUSG00000025648 | Pfkfb4 | 270198 | protein_coding | 308.3309767 | -0.532491528 | 0.117143692 | -4.545627004 | 5.48E-06 | 8.77E-05 | 0.532491528 |
| ENSMUSG00000034573 | Ptpn13 | 19249 | protein_coding | 321.7070165 | -0.532584419 | 0.194627883 | -2.736424039 | 0.006211093 | 0.033031818 | 0.532584419 |
| ENSMUSG00000027690 | Slc2a2 | 20526 | protein_coding | 8.709309496 | -0.53304827 | 0.563017322 | -0.946770639 | 0.343755666 | 0.55376472 | 0.53304827 |
| ENSMUSG00000041773 | Enc1 | 13803 | protein_coding | 227.2642054 | -0.533096041 | 0.153252566 | -3.478545624 | 0.000504143 | 0.004380788 | 0.533096041 |
| ENSMUSG00000068874 | Selenbp1 | 20341 | protein_coding | 13741.01848 | -0.53334721 | 0.174477075 | -3.056832597 | 0.002236892 | 0.014845504 | 0.53334721 |
| ENSMUSG00000053007 | Creb5 | 231991 | protein_coding | 13.97577619 | -0.533448494 | 0.994493219 | -0.536402344 | 0.591680512 | 0.760882221 | 0.533448494 |
| ENSMUSG00000038352 | Arl5c | 217151 | protein_coding | 33.35550483 | 0.533449376 | 0.369680979 | 1.44299925 | 0.149020683 | 0.326357959 | 0.533449376 |
| ENSMUSG00000032051 | Fdx1 | 14148 | protein_coding | 530.0476329 | 0.533569433 | 0.107439862 | 4.966214783 | 6.83E-07 | 1.34E-05 | 0.533569433 |
| ENSMUSG00000027257 | Pacsin3 | 80708 | protein_coding | 65.98478689 | -0.533709428 | 0.182364778 | -2.926603675 | 0.003426853 | 0.020841126 | 0.533709428 |
| ENSMUSG00000056895 | H2bc27 | 78303 | protein_coding | 21.28997238 | 0.534000305 | 0.308372642 | 1.731672117 | 0.083331958 | 0.220227547 | 0.534000305 |
| ENSMUSG00000070304 | Scn2b | 72821 | protein_coding | 61.1123263 | -0.534046927 | 0.256157838 | -2.084835395 | 0.037084245 | 0.126105433 | 0.534046927 |
| ENSMUSG00000031157 | Pqbp1 | 54633 | protein_coding | 393.6938393 | 0.534067116 | 0.113212869 | 4.717371086 | 2.39E-06 | 4.11E-05 | 0.534067116 |
| ENSMUSG00000023467 | Tulp2 | 56734 | protein_coding | 26.16928256 | -0.534092307 | 0.277412285 | -1.925265518 | 0.054196149 | 0.16392637 | 0.534092307 |
| ENSMUSG00000028003 | Lrat | 79235 | protein_coding | 196.9755076 | -0.534344926 | 0.196739862 | -2.715997264 | 0.006607646 | 0.034792262 | 0.534344926 |
| ENSMUSG00000026104 | Stat1 | 20846 | protein_coding | 282.0064423 | 0.534388992 | 0.14586005 | 3.663710471 | 0.000248588 | 0.002437608 | 0.534388992 |
| ENSMUSG00000028542 | Slc6a9 | 14664 | protein_coding | 164.2742483 | 0.534431396 | 0.117879373 | 4.533714272 | 5.80E-06 | 9.19E-05 | 0.534431396 |
| ENSMUSG00000037846 | Rtkn2 | 170799 | protein_coding | 10.97440444 | -0.534471604 | 0.631895655 | -0.845822566 | 0.397651735 | 0.605699229 | 0.534471604 |
| ENSMUSG00000030793 | Pycard | 66824 | protein_coding | 2284.559353 | 0.53466508 | 0.114836047 | 4.655899386 | 3.23E-06 | 5.41E-05 | 0.53466508 |
| ENSMUSG00000023886 | Smoc2 | 64074 | protein_coding | 514.9137765 | 0.53470809 | 0.13085712 | 4.0861979 | 4.38E-05 | 0.000549535 | 0.53470809 |
| ENSMUSG00000040339 | Fam102b | 329739 | protein_coding | 1097.94717 | 0.534716932 | 0.161143702 | 3.318261436 | 0.000905797 | 0.007072318 | 0.534716932 |
| ENSMUSG00000018378 | Cuedc1 | 70393 | protein_coding | 88.71339811 | -0.534984678 | 0.19409575 | -2.756292597 | 0.005846067 | 0.031543416 | 0.534984678 |
| ENSMUSG00000024966 | Stip1 | 20867 | protein_coding | 1385.694257 | 0.535111694 | 0.187956577 | 2.84699637 | 0.004413387 | 0.025386934 | 0.535111694 |
| ENSMUSG00000058046 | 4933430I17Rik | 214106 | protein_coding | 8.664759714 | -0.535301578 | 0.565528308 | -0.94655134 | 0.343867449 | 0.553859918 | 0.535301578 |
| ENSMUSG00000029910 | Mad2l1 | 56150 | protein_coding | 591.5747562 | 0.536010541 | 0.116203072 | 4.612705442 | 3.97E-06 | 6.55E-05 | 0.536010541 |
| ENSMUSG00000024521 | Pmaip1 | 58801 | protein_coding | 268.7376018 | -0.536085895 | 0.236128293 | -2.270316229 | 0.023188403 | 0.089343223 | 0.536085895 |
| ENSMUSG00000027624 | Epb41l1 | 13821 | protein_coding | 845.8966935 | -0.536172888 | 0.097140454 | -5.519563309 | 3.40E-08 | 8.34E-07 | 0.536172888 |
| ENSMUSG00000021262 | Evl | 14026 | protein_coding | 81.69225904 | -0.536217496 | 0.235688218 | -2.275113708 | 0.02289911 | 0.088538846 | 0.536217496 |
| ENSMUSG00000085887 | Arhgap27os3 | 653030 | lncRNA | 24.58635588 | -0.536477954 | 0.30826306 | -1.740325145 | 0.081801941 | 0.217483111 | 0.536477954 |
| ENSMUSG00000005355 | Casp14 | 12365 | protein_coding | 42.37000531 | -0.536716725 | 0.564630841 | -0.950562183 | 0.341826673 | 0.551941783 | 0.536716725 |
| ENSMUSG00000058248 | Kcnh1 | 16510 | protein_coding | 23.76149706 | -0.536809955 | 0.320116513 | -1.676920538 | 0.093558021 | 0.238011846 | 0.536809955 |
| ENSMUSG00000028246 | Faxc | 76132 | protein_coding | 9.425107732 | -0.536853647 | 0.48167553 | -1.114554536 | 0.265041362 | 0.472529529 | 0.536853647 |

Supplementary Table 1

| ensembl_gene_id | external_gene_name | entrezgene_id | gene_biotype | baseMean | log2FoldChange | lfcSE | stat | pvalue | padj | absLog2FC |
|---|---|---|---|---|---|---|---|---|---|---|
| ENSMUSG00000027371 | Fahd2a | 68126 | protein_coding | 130.1698408 | -0.536914925 | 0.115357768 | -4.654345631 | 3.25E-06 | 5.44E-05 | 0.536914925 |
| ENSMUSG00000029234 | Tmem165 | 21982 | protein_coding | 990.2476033 | 0.536977284 | 0.090795173 | 5.914161164 | 3.34E-09 | 9.57E-08 | 0.536977284 |
| ENSMUSG00000037593 | Rskr | 216976 | protein_coding | 10.20324982 | -0.537188151 | 0.481378775 | -1.115936512 | 0.264449312 | 0.472048706 | 0.537188151 |
| ENSMUSG00000020635 | Fkbp1b | 14226 | protein_coding | 15.14201279 | -0.537499541 | 0.380464031 | -1.412747322 | 0.157730032 | 0.338588134 | 0.537499541 |
| ENSMUSG00000000628 | Hk2 | 15277 | protein_coding | 3293.549651 | 0.537753673 | 0.242770723 | 2.21506805 | 0.026755394 | 0.099153075 | 0.537753673 |
| ENSMUSG00000022324 | Matn2 | 17181 | protein_coding | 139.4909369 | -0.538254044 | 0.277537451 | -1.939392476 | 0.052453566 | 0.160434875 | 0.538254044 |
| ENSMUSG00000020387 | Jade2 | 76901 | protein_coding | 179.2815666 | -0.538280558 | 0.192181746 | -2.800893258 | 0.005096137 | 0.028464292 | 0.538280558 |
| ENSMUSG00000018476 | Kdm6b | 216850 | protein_coding | 273.2613914 | -0.538491447 | 0.225191984 | -2.39125495 | 0.016790889 | 0.070361819 | 0.538491447 |
| ENSMUSG00000116632 | Magef1 | 76222 | transcribed_processed_pseudogene | 34.43052691 | -0.538538023 | 0.234219767 | -2.299285114 | 0.021488755 | 0.084241418 | 0.538538023 |
| ENSMUSG00000028028 | Alpk1 | 71481 | protein_coding | 450.2601382 | 0.53868357 | 0.104181329 | 5.170634449 | 2.33E-07 | 5.04E-06 | 0.53868357 |
| ENSMUSG00000031982 | Arv1 | 68865 | protein_coding | 31.27797109 | -0.538940603 | 0.193236208 | -2.789024943 | 0.005286699 | 0.02924854 | 0.538940603 |
| ENSMUSG00000006360 | Crip1 | 12925 | protein_coding | 4409.80203 | -0.538986266 | 0.129024081 | -4.17740829 | 2.95E-05 | 0.00038974 | 0.538986266 |
| ENSMUSG00000045102 | Poln | 272158 | protein_coding | 6.956395542 | -0.539297585 | 0.851227321 | -0.633552955 | 0.526372611 | 0.712022332 | 0.539297585 |
| ENSMUSG00000026575 | Nme7 | 171567 | protein_coding | 196.2278359 | 0.539525252 | 0.127466562 | 4.232680664 | 2.31E-05 | 0.000312917 | 0.539525252 |
| ENSMUSG00000026604 | Ptpn14 | 19250 | protein_coding | 63.09671112 | -0.539857272 | 0.218037203 | -2.475986968 | 0.013286843 | 0.05925186 | 0.539857272 |
| ENSMUSG00000051844 | B230319C09Rik | 320775 | lncRNA | 33.91510752 | -0.539906391 | 0.386646419 | -1.396382753 | 0.162599266 | 0.345172734 | 0.539906391 |
| ENSMUSG00000039936 | Pik3cd | 18707 | protein_coding | 52.29659291 | 0.539973944 | 0.316560068 | 1.705755079 | 0.08805369 | 0.228545168 | 0.539973944 |
| ENSMUSG00000020149 | Rab1a | 19324 | protein_coding | 4600.892942 | 0.540070464 | 0.101279407 | 5.332480523 | 9.69E-08 | 2.22E-06 | 0.540070464 |
| ENSMUSG00000020895 | Tmem107 | 66910 | protein_coding | 102.6118038 | 0.540140668 | 0.172951283 | 3.123079861 | 0.001789692 | 0.012335011 | 0.540140668 |
| ENSMUSG00000028360 | Slc44a5 | 242259 | protein_coding | 7.009519004 | -0.540407957 | 0.648025908 | -0.833929555 | 0.404320682 | 0.611355593 | 0.540407957 |
| ENSMUSG00000029700 | Slc13a1 | 55961 | protein_coding | 261.4866583 | -0.540409627 | 0.189528691 | -2.851334146 | 0.004353619 | 0.025133906 | 0.540409627 |
| ENSMUSG00000028669 | Pithd1 | 66193 | protein_coding | 838.3203069 | 0.540445089 | 0.107404232 | 5.031878881 | 4.86E-07 | 9.92E-06 | 0.540445089 |
| ENSMUSG00000037622 | Wdtc1 | 230796 | protein_coding | 374.6417221 | -0.541132777 | 0.075944967 | -7.1253277 | 1.04E-12 | 4.30E-11 | 0.541132777 |
| ENSMUSG00000025178 | Pi4k2a | 84095 | protein_coding | 379.1292954 | 0.541779539 | 0.092642709 | 5.848053689 | 4.97E-09 | 1.38E-07 | 0.541779539 |
| ENSMUSG00000032192 | Gnb5 | 14697 | protein_coding | 104.8156414 | -0.542082647 | 0.13692856 | -3.958872035 | 7.53E-05 | 0.000878076 | 0.542082647 |
| ENSMUSG00000026587 | Astn1 | 11899 | protein_coding | 8.003791025 | -0.542145467 | 1.106784144 | -0.489838484 | 0.624248196 | 0.782360946 | 0.542145467 |
| ENSMUSG00000039652 | Cpeb3 | 208922 | protein_coding | 72.89437267 | -0.542348034 | 0.222483358 | -2.437701582 | 0.01478097 | 0.063819815 | 0.542348034 |
| ENSMUSG00000024426 | Atat1 | 73242 | protein_coding | 58.01707828 | -0.542561618 | 0.166309703 | -3.262356975 | 0.001104899 | 0.008324897 | 0.542561618 |
| ENSMUSG00000023832 | Acat2 | 110460 | protein_coding | 449.7816332 | 0.542773144 | 0.126190077 | 4.301234746 | 1.70E-05 | 0.000239412 | 0.542773144 |
| ENSMUSG00000019947 | Arid5b | 71371 | protein_coding | 454.6529566 | -0.542879699 | 0.196522786 | -2.762426233 | 0.005737352 | 0.03106205 | 0.542879699 |
| ENSMUSG00000025467 | Prap1 | 22264 | protein_coding | 196.2886508 | -0.542942154 | 0.248670792 | -2.183377263 | 0.029008039 | 0.105248302 | 0.542942154 |
| ENSMUSG00000032501 | Trib1 | 211770 | protein_coding | 1524.311434 | -0.543115376 | 0.235384542 | -2.307353623 | 0.021035114 | 0.083035267 | 0.543115376 |
| ENSMUSG00000062017 | Abca14 | 67928 | protein_coding | 10.92048577 | 0.543137568 | 0.549200851 | 0.988959806 | 0.322682808 | 0.533605872 | 0.543137568 |
| ENSMUSG00000040428 | Plekha4 | 69217 | protein_coding | 13.63144983 | -0.543198931 | 0.473306996 | -1.147667233 | 0.251105963 | 0.457517247 | 0.543198931 |
| ENSMUSG00000006476 | Nsmf | 56876 | protein_coding | 517.6661456 | -0.543261353 | 0.129806985 | -4.185147297 | 2.85E-05 | 0.000377636 | 0.543261353 |
| ENSMUSG00000056148 | Rdh9 | 103142 | protein_coding | 28.15846939 | 0.543393011 | 0.277296319 | 1.959611344 | 0.050041234 | 0.155611958 | 0.543393011 |
| ENSMUSG00000044456 | Rin3 | 217835 | protein_coding | 36.92358221 | -0.543580654 | 0.209513001 | -2.594496054 | 0.00947297 | 0.045973799 | 0.543580654 |
| ENSMUSG00000052698 | Tln2 | 70549 | protein_coding | 736.3911245 | -0.543624522 | 0.139841723 | -3.887427237 | 0.000101312 | 0.00113963 | 0.543624522 |

| ensembl_gene_id | external_gene_name | entrezgene_id | gene_biotype | baseMean | log2FoldChange | lfcSE | stat | pvalue | padj | absLog2FC |
|---|---|---|---|---|---|---|---|---|---|---|
| ENSMUSG00000001566 | Mnx1 | 15285 | protein_coding | 14.15956512 | -0.544036743 | 0.318791676 | -1.706558808 | 0.087904087 | 0.228448759 | 0.544036743 |
| ENSMUSG00000045348 | Nyap1 | 243300 | protein_coding | 78.44963519 | -0.544087503 | 0.21539475 | -2.526001694 | 0.011536893 | 0.053340965 | 0.544087503 |
| ENSMUSG00000040721 | Zfhx2 | 239102 | protein_coding | 50.98749403 | -0.544270635 | 0.305523872 | -1.78143407 | 0.074841567 | 0.204201575 | 0.544270635 |
| ENSMUSG00000044763 | Trmt10c | 52575 | protein_coding | 398.9538627 | -0.544572985 | 0.112319615 | -4.848422816 | 1.24E-06 | 2.29E-05 | 0.544572985 |
| ENSMUSG00000022098 | Bmp1 | 12153 | protein_coding | 62.03649727 | -0.544659832 | 0.1674346 | -3.252970598 | 0.001142053 | 0.008548225 | 0.544659832 |
| ENSMUSG00000098207 | Arl14 | 71619 | protein_coding | 283.6618588 | -0.544757424 | 0.226687578 | -2.403119877 | 0.016255858 | 0.068716375 | 0.544757424 |
| ENSMUSG00000028710 | Atpaf1 | 230649 | protein_coding | 148.3993854 | -0.544839878 | 0.100896357 | -5.399995559 | 6.66E-08 | 1.57E-06 | 0.544839878 |
| ENSMUSG00000024063 | Lbh | 77889 | protein_coding | 285.8392556 | -0.544900829 | 0.141267113 | -3.857237672 | 0.000114676 | 0.001267289 | 0.544900829 |
| ENSMUSG00000053469 | Tg | 21819 | protein_coding | 6.818261909 | -0.54496631 | 1.781545675 | -0.305895222 | 0.759684424 | 0.868522359 | 0.54496631 |
| ENSMUSG00000052752 | Traf7 | 224619 | protein_coding | 1181.180736 | 0.5452745 | 0.081916313 | 6.65648246 | 2.80E-11 | 1.01E-09 | 0.5452745 |
| ENSMUSG00000031647 | Mfap3l | 71306 | protein_coding | 1027.121684 | -0.54540134 | 0.193323972 | -2.821178017 | 0.004784764 | 0.027169956 | 0.54540134 |
| ENSMUSG00000078762 | Haus5 | 71909 | protein_coding | 15.65898624 | -0.545549391 | 0.3871046 | -1.409307434 | 0.158744282 | 0.34003213 | 0.545549391 |
| ENSMUSG00000033170 | Card10 | 105844 | protein_coding | 319.4794402 | -0.545561142 | 0.13644134 | -3.99850325 | 6.37E-05 | 0.000760539 | 0.545561142 |
| ENSMUSG00000031981 | Capn9 | 73647 | protein_coding | 547.850359 | 0.54563229 | 0.153372717 | 3.5575577 | 0.000374319 | 0.003436398 | 0.54563229 |
| ENSMUSG00000021127 | Zfp36l1 | 12192 | protein_coding | 316.8240758 | -0.545812475 | 0.153877027 | -3.547069269 | 0.000389542 | 0.00355974 | 0.545812475 |
| ENSMUSG00000041827 | Oasl1 | 231655 | protein_coding | 192.6272273 | 0.546092917 | 0.182007627 | 3.000384796 | 0.002696387 | 0.017241292 | 0.546092917 |
| ENSMUSG00000047109 | Cldn14 | 56173 | protein_coding | 24.37558543 | -0.546261258 | 0.37544848 | -1.454956636 | 0.145681267 | 0.321386596 | 0.546261258 |
| ENSMUSG00000021013 | Ttc8 | 76260 | protein_coding | 23.40894275 | 0.546323535 | 0.280145955 | 1.950138938 | 0.051159561 | 0.157951055 | 0.546323535 |
| ENSMUSG00000002980 | Bcam | 57278 | protein_coding | 47.88270593 | -0.546606872 | 0.193283113 | -2.828011533 | 0.004683812 | 0.026691863 | 0.546606872 |
| ENSMUSG00000046159 | Chrm3 | 12671 | protein_coding | 34.71823499 | -0.546897536 | 0.215423887 | -2.538704239 | 0.011126384 | 0.051803871 | 0.546897536 |
| ENSMUSG00000040794 | C1qtnf4 | 67445 | protein_coding | 17.28417712 | -0.546949149 | 0.301392391 | -1.814741066 | 0.069563711 | 0.194969345 | 0.546949149 |
| ENSMUSG00000052557 | Gan | 209239 | protein_coding | 478.9079941 | -0.547452503 | 0.146413761 | -3.739078205 | 0.000184696 | 0.001902873 | 0.547452503 |
| ENSMUSG00000001424 | Snd1 | 56463 | protein_coding | 2131.194816 | 0.547499957 | 0.087356078 | 6.267451215 | 3.67E-10 | 1.17E-08 | 0.547499957 |
| ENSMUSG00000025810 | Nrp1 | 18186 | protein_coding | 110.6486933 | 0.547564791 | 0.114225571 | 4.79371463 | 1.64E-06 | 2.93E-05 | 0.547564791 |
| ENSMUSG00000030786 | Itgam | 16409 | protein_coding | 18.44198841 | 0.547567609 | 0.320223471 | 1.709954637 | 0.087274262 | 0.227399911 | 0.547567609 |
| ENSMUSG00000049526 | Tmem202 | 73893 | protein_coding | 6.730564045 | -0.547895204 | 0.402230586 | -1.362142071 | 0.173153053 | 0.358895202 | 0.547895204 |
| ENSMUSG00000035372 | 1810055G02Rik | 72056 | protein_coding | 200.0679591 | -0.547916797 | 0.174983142 | -3.13125477 | 0.001740611 | 0.012059352 | 0.547916797 |
| ENSMUSG00000018387 | Shroom1 | 71774 | protein_coding | 12.27514125 | -0.548111182 | 0.32867234 | -1.667652297 | 0.09538477 | 0.240851081 | 0.548111182 |
| ENSMUSG00000017466 | Timp2 | 21858 | protein_coding | 91.66569687 | -0.54812917 | 0.19123864 | -2.866205132 | 0.00415425 | 0.024193378 | 0.54812917 |
| ENSMUSG00000051498 | Tlr6 | 21899 | protein_coding | 20.00935821 | 0.548136649 | 0.289123757 | 1.895854755 | 0.057979253 | 0.172019679 | 0.548136649 |
| ENSMUSG00000055172 | C1ra | 50909 | protein_coding | 24.1785499 | -0.548267042 | 0.340886932 | -1.608354534 | 0.107757553 | 0.261043697 | 0.548267042 |
| ENSMUSG00000051675 | Trim32 | 69807 | protein_coding | 328.102314 | 0.548355359 | 0.089162937 | 6.15003698 | 7.75E-10 | 2.36E-08 | 0.548355359 |
| ENSMUSG00000021665 | Hexb | 15212 | protein_coding | 1199.396584 | -0.54858383 | 0.281249519 | -1.950523618 | 0.051113741 | 0.157840184 | 0.54858383 |
| ENSMUSG00000045176 | Borcs6 | 71923 | protein_coding | 162.6623447 | 0.549113785 | 0.119278063 | 4.603644377 | 4.15E-06 | 6.82E-05 | 0.549113785 |
| ENSMUSG00000015224 | Cyp2j9 | 74519 | protein_coding | 54.49471672 | 0.549157226 | 0.189745944 | 2.894171091 | 0.00380161 | 0.022561311 | 0.549157226 |
| ENSMUSG00000040613 | Apobec1 | 11810 | protein_coding | 402.8687186 | 0.549520071 | 0.232115986 | 2.367437418 | 0.017911749 | 0.073712317 | 0.549520071 |
| ENSMUSG00000043644 | 0610009L18Rik | 66838 | lncRNA | 27.90503934 | -0.549537896 | 0.240525153 | -2.284741899 | 0.022327966 | 0.086762588 | 0.549537896 |
| ENSMUSG00000085826 | Gm15638 | 100503378 | lncRNA | 20.87592997 | -0.54960546 | 0.359329019 | -1.529532634 | 0.126132455 | 0.290701454 | 0.54960546 |

Supplementary Table 1

| ensembl_gene_id | external_gene_name | entrezgene_id | gene_biotype | baseMean | log2FoldChange | lfcSE | stat | pvalue | padj | absLog2FC |
|---|---|---|---|---|---|---|---|---|---|---|
| ENSMUSG00000038570 | Saxo2 | 330577 | protein_coding | 12.79725609 | 0.549827372 | 0.330389949 | 1.664177053 | 0.096077052 | 0.242030591 | 0.549827372 |
| ENSMUSG00000034471 | Caskin2 | 140721 | protein_coding | 834.9370864 | -0.549854656 | 0.137127401 | -4.009808782 | 6.08E-05 | 0.000731051 | 0.549854656 |
| ENSMUSG00000078651 | Aoc2 | 237940 | protein_coding | 22.25485749 | -0.549972479 | 0.394356268 | -1.394608185 | 0.163134026 | 0.34566965 | 0.549972479 |
| ENSMUSG00000061171 | Slc38a11 | 320106 | protein_coding | 29.771306 | -0.550322357 | 0.282628816 | -1.947155868 | 0.051516051 | 0.158593398 | 0.550322357 |
| ENSMUSG00000025037 | Maoa | 17161 | protein_coding | 12535.14412 | -0.550380461 | 0.154162957 | -3.570121319 | 0.000356816 | 0.003285182 | 0.550380461 |
| ENSMUSG00000031574 | Star | 20845 | protein_coding | 37.29140046 | -0.550394792 | 0.185154406 | -2.972625957 | 0.00295264 | 0.018530203 | 0.550394792 |
| ENSMUSG00000026958 | Dpp7 | 83768 | protein_coding | 174.7360158 | -0.550443202 | 0.113016245 | -4.87047859 | 1.11E-06 | 2.08E-05 | 0.550443202 |
| ENSMUSG00000039568 | Ubald1 | 207740 | protein_coding | 283.0583429 | -0.550626662 | 0.164123569 | -3.354951797 | 0.000793788 | 0.006334398 | 0.550626662 |
| ENSMUSG00000038775 | Vill | 22351 | protein_coding | 84.17067884 | -0.55064385 | 0.195849866 | -2.811561024 | 0.004930174 | 0.02773854 | 0.55064385 |
| ENSMUSG00000047728 | Ly6g2 | 223631 | protein_coding | 117.3625485 | 0.550707229 | 0.197477122 | 2.788714073 | 0.005291776 | 0.029266462 | 0.550707229 |
| ENSMUSG00000028150 | Rorc | 19885 | protein_coding | 530.9139389 | 0.550970356 | 0.165906457 | 3.320969927 | 0.000897052 | 0.007014357 | 0.550970356 |
| ENSMUSG00000000811 | Txnrd3 | 232223 | protein_coding | 153.4923112 | -0.551013501 | 0.118860175 | -4.635812648 | 3.56E-06 | 5.91E-05 | 0.551013501 |
| ENSMUSG00000092060 | Bend4 | 666938 | protein_coding | 7.154373769 | -0.551043307 | 0.685909515 | -0.803376094 | 0.421757388 | 0.627406765 | 0.551043307 |
| ENSMUSG00000039264 | Gimap3 | 83408 | protein_coding | 49.43330796 | 0.551074732 | 0.248028924 | 2.221816403 | 0.026295716 | 0.098054177 | 0.551074732 |
| ENSMUSG00000078451 | Ppil6 | 73075 | protein_coding | 16.6543665 | -0.551302557 | 0.302055814 | -1.825167841 | 0.067975705 | 0.191631333 | 0.551302557 |
| ENSMUSG00000035818 | Plekhs1 | 226245 | protein_coding | 16.3251583 | -0.551530994 | 0.334227943 | -1.650164224 | 0.098909352 | 0.246545878 | 0.551530994 |
| ENSMUSG00000024990 | Rbp4 | 19662 | protein_coding | 97.05144102 | -0.551572298 | 0.13278505 | -4.153873481 | 3.27E-05 | 0.000426086 | 0.551572298 |
| ENSMUSG00000062110 | Scfd2 | 212986 | protein_coding | 255.1165129 | 0.552120953 | 0.107360416 | 5.142686413 | 2.71E-07 | 5.79E-06 | 0.552120953 |
| ENSMUSG00000021257 | Angel1 | 68737 | protein_coding | 68.26545482 | -0.55221238 | 0.140871275 | -3.919978572 | 8.86E-05 | 0.001016234 | 0.55221238 |
| ENSMUSG00000034265 | Zdhhc14 | 224454 | protein_coding | 87.69700588 | -0.552454452 | 0.175449696 | -3.148791152 | 0.001639473 | 0.011453302 | 0.552454452 |
| ENSMUSG00000037552 | Plekhg2 | 101497 | protein_coding | 163.5348172 | -0.552794644 | 0.192552918 | -2.870871292 | 0.004093422 | 0.0239003 | 0.552794644 |
| ENSMUSG00000061959 | Ces1e | 13897 | protein_coding | 424.3723511 | 0.552922057 | 0.101148803 | 5.46642216 | 4.59E-08 | 1.11E-06 | 0.552922057 |
| ENSMUSG00000035960 | Apex1 | 11792 | protein_coding | 183.1796008 | 0.553031709 | 0.128220858 | 4.313118129 | 1.61E-05 | 0.000228308 | 0.553031709 |
| ENSMUSG00000052572 | Dlg2 | 23859 | protein_coding | 16.33673078 | -0.553034482 | 0.571611596 | -0.967500461 | 0.333293911 | 0.544203528 | 0.553034482 |
| ENSMUSG00000115149 | 9330188P03Rik | 380930 | lncRNA | 21.13565436 | -0.553044527 | 0.345322059 | -1.601532576 | 0.109259011 | 0.263319341 | 0.553044527 |
| ENSMUSG00000022469 | Rapgef3 | 223864 | protein_coding | 331.8785199 | -0.553328041 | 0.232357091 | -2.381369294 | 0.017248411 | 0.071703093 | 0.553328041 |
| ENSMUSG00000079173 | Zan | 22635 | protein_coding | 836.7134517 | -0.553365911 | 0.25640754 | -2.158149913 | 0.030916178 | 0.110460495 | 0.553365911 |
| ENSMUSG00000084781 | D930015M05Rik | 320507 | lncRNA | 8.260531652 | -0.553511914 | 0.583677981 | -0.948317278 | 0.342967957 | 0.553138277 | 0.553511914 |
| ENSMUSG00000063873 | Slc24a3 | 94249 | protein_coding | 49.45918814 | -0.553520175 | 0.190632057 | -2.903604896 | 0.003688934 | 0.022080923 | 0.553520175 |
| ENSMUSG00000032656 | Marchf3 | 320253 | protein_coding | 91.12154451 | 0.553666107 | 0.222361168 | 2.489940634 | 0.012776443 | 0.057551807 | 0.553666107 |
| ENSMUSG00000110218 | Tincr | 100504425 | protein_coding | 37.42455962 | -0.553738119 | 0.195580274 | -2.831257512 | 0.004636538 | 0.02646981 | 0.553738119 |
| ENSMUSG00000096981 | Gm16845 | 100503652 | lncRNA | 13.38041321 | -0.553850175 | 0.295809941 | -1.872317655 | 0.06116267 | 0.177981255 | 0.553850175 |
| ENSMUSG00000037972 | Snn | 20621 | protein_coding | 22.3560004 | 0.554341089 | 0.226046428 | 2.452332883 | 0.014193329 | 0.061971312 | 0.554341089 |
| ENSMUSG00000085057 | Gm13415 | 102638285 | lncRNA | 12.70208142 | -0.554375911 | 0.339847694 | -1.631248117 | 0.102837976 | 0.253012555 | 0.554375911 |
| ENSMUSG00000029860 | Zyx | 22793 | protein_coding | 471.2944451 | -0.554551568 | 0.242654678 | -2.285352887 | 0.022292144 | 0.086648725 | 0.554551568 |
| ENSMUSG00000048279 | Sacs | 50720 | protein_coding | 20.47176118 | 0.554970085 | 0.385891217 | 1.438151638 | 0.150391034 | 0.328140875 | 0.554970085 |
| ENSMUSG00000029428 | Stx2 | 13852 | protein_coding | 13.02290592 | 0.555255296 | 0.399353913 | 1.390384012 | 0.164412299 | 0.347730593 | 0.555255296 |
| ENSMUSG00000034758 | Tle6 | 114606 | protein_coding | 45.70257956 | -0.555339134 | 0.223877915 | -2.480544509 | 0.013118188 | 0.058742341 | 0.555339134 |

| ensembl_gene_id | external_gene_name | entrezgene_id | gene_biotype | baseMean | log2FoldChange | lfcSE | stat | pvalue | padj | absLog2FC |
|---|---|---|---|---|---|---|---|---|---|---|
| ENSMUSG00000004044 | Cavin1 | 19285 | protein_coding | 58.78810934 | -0.555617038 | 0.457168038 | -1.215345325 | 0.224234412 | 0.425951784 | 0.555617038 |
| ENSMUSG00000031371 | Haus7 | 73738 | protein_coding | 146.0582469 | 0.555713676 | 0.133152826 | 4.173502678 | 3.00E-05 | 0.000396156 | 0.555713676 |
| ENSMUSG00000026238 | Ptma | 19231 | protein_coding | 7272.790017 | 0.555759516 | 0.106245151 | 5.230916529 | 1.69E-07 | 3.73E-06 | 0.555759516 |
| ENSMUSG00000042115 | Klhdc8a | 213417 | protein_coding | 126.6584274 | -0.556772126 | 0.226460474 | -2.458584123 | 0.013948609 | 0.061268531 | 0.556772126 |
| ENSMUSG00000026511 | Srp9 | 27058 | protein_coding | 879.1714334 | 0.556784842 | 0.082062814 | 6.784861683 | 1.16E-11 | 4.31E-10 | 0.556784842 |
| ENSMUSG00000035637 | Grhpr | 76238 | protein_coding | 284.0088218 | 0.556831826 | 0.103709714 | 5.369138591 | 7.91E-08 | 1.84E-06 | 0.556831826 |
| ENSMUSG00000015247 | Nipsnap3b | 66536 | protein_coding | 901.1823084 | 0.557055029 | 0.088604927 | 6.286953169 | 3.24E-10 | 1.04E-08 | 0.557055029 |
| ENSMUSG00000035275 | Raver2 | 242570 | protein_coding | 9.868577516 | -0.557274954 | 0.415796422 | -1.340259138 | 0.180161111 | 0.368125204 | 0.557274954 |
| ENSMUSG00000055805 | Fmnl1 | 57778 | protein_coding | 14.99084099 | 0.557311161 | 0.463892633 | 1.201379632 | 0.229603973 | 0.432847922 | 0.557311161 |
| ENSMUSG00000005373 | Mlxipl | 58805 | protein_coding | 166.6698865 | -0.557575339 | 0.177320143 | -3.144455726 | 0.001663961 | 0.011591716 | 0.557575339 |
| ENSMUSG00000023341 | Mx2 | 17858 | protein_coding | 20.39403316 | 0.557582508 | 0.28852773 | 1.932509256 | 0.053296681 | 0.162074293 | 0.557582508 |
| ENSMUSG00000024194 | Cuta | 67675 | protein_coding | 393.6632825 | 0.557653348 | 0.098796755 | 5.644450016 | 1.66E-08 | 4.35E-07 | 0.557653348 |
| ENSMUSG00000022941 | Ripply3 | 170765 | protein_coding | 7.012319303 | 0.557702941 | 0.451150946 | 1.236178149 | 0.216392351 | 0.415615262 | 0.557702941 |
| ENSMUSG00000046449 | Nexmif | 245555 | protein_coding | 64.92612245 | -0.557823021 | 0.194206864 | -2.872313628 | 0.004074783 | 0.02384392 | 0.557823021 |
| ENSMUSG00000072115 | Ang | 11727 | protein_coding | 278.8304195 | -0.55807707 | 0.122921884 | -4.540095324 | 5.62E-06 | 8.96E-05 | 0.55807707 |
| ENSMUSG00000020988 | L2hgdh | 217666 | protein_coding | 504.7212546 | 0.558131988 | 0.091606554 | 6.092708035 | 1.11E-09 | 3.34E-08 | 0.558131988 |
| ENSMUSG00000022102 | Dok2 | 13449 | protein_coding | 9.125963344 | -0.558195535 | 0.553725654 | -1.008072375 | 0.313419718 | 0.523835181 | 0.558195535 |
| ENSMUSG00000020251 | Glt8d2 | 74782 | protein_coding | 7.149418634 | -0.558258216 | 0.51594094 | -1.082019613 | 0.279243813 | 0.487589943 | 0.558258216 |
| ENSMUSG00000019900 | Rfx6 | 320995 | protein_coding | 16.66169274 | -0.558301576 | 0.33694494 | -1.65695195 | 0.09752919 | 0.244195722 | 0.558301576 |
| ENSMUSG00000027227 | Sord | 20322 | protein_coding | 1363.179503 | -0.558321564 | 0.100335613 | -5.564540337 | 2.63E-08 | 6.65E-07 | 0.558321564 |
| ENSMUSG00000075025 | Gm10804 | 100038525 | lncRNA | 11.15102948 | -0.558495273 | 0.419213001 | -1.332247024 | 0.182779029 | 0.371861588 | 0.558495273 |
| ENSMUSG00000034919 | Ttc22 | 230576 | protein_coding | 728.0697906 | 0.558617524 | 0.149699486 | 3.731592789 | 0.000190273 | 0.001955269 | 0.558617524 |
| ENSMUSG00000004665 | Cnn2 | 12798 | protein_coding | 150.5490315 | 0.55863776 | 0.266891462 | 2.093127129 | 0.036337808 | 0.124287397 | 0.55863776 |
| ENSMUSG00000060904 | Arl1 | 104303 | protein_coding | 4434.290874 | 0.55881878 | 0.083229645 | 6.714179558 | 1.89E-11 | 6.89E-10 | 0.55881878 |
| ENSMUSG00000073293 | Nudt10 | 102954 | protein_coding | 21.12342805 | 0.558821956 | 0.309188185 | 1.807384573 | 0.070702338 | 0.196741236 | 0.558821956 |
| ENSMUSG00000032289 | Thsd4 | 207596 | protein_coding | 322.2539612 | -0.559674429 | 0.161800065 | -3.459049463 | 0.000542085 | 0.004659652 | 0.559674429 |
| ENSMUSG00000057406 | Nsd2 | 107823 | protein_coding | 1283.397301 | 0.559724801 | 0.090405394 | 6.191276602 | 5.97E-10 | 1.84E-08 | 0.559724801 |
| ENSMUSG00000016206 | H2-M3 | 14991 | protein_coding | 81.17200447 | 0.559826343 | 0.141109733 | 3.967312046 | 7.27E-05 | 0.000850199 | 0.559826343 |
| ENSMUSG00000035849 | Krt222 | 268481 | protein_coding | 16.69265145 | -0.559952617 | 0.347535226 | -1.611211112 | 0.107133717 | 0.26012589 | 0.559952617 |
| ENSMUSG00000071856 | Mcc | 328949 | protein_coding | 376.5162468 | -0.560028088 | 0.151791743 | -3.689450283 | 0.000224739 | 0.002249934 | 0.560028088 |
| ENSMUSG00000031725 | Ces1f | 234564 | protein_coding | 1827.111822 | -0.560293259 | 0.171662644 | -3.263920713 | 0.001098819 | 0.008290853 | 0.560293259 |
| ENSMUSG00000024940 | Ltbp3 | 16998 | protein_coding | 9.8354548 | -0.560631742 | 0.709063446 | -0.790665131 | 0.429139429 | 0.633370351 | 0.560631742 |
| ENSMUSG00000028165 | Cisd2 | 67006 | protein_coding | 961.8127539 | 0.560829076 | 0.101063277 | 5.549286448 | 2.87E-08 | 7.16E-07 | 0.560829076 |
| ENSMUSG00000045094 | Arhgef37 | 328967 | protein_coding | 158.4604916 | -0.560899214 | 0.147419896 | -3.804772816 | 0.000141934 | 0.001526821 | 0.560899214 |
| ENSMUSG00000027006 | Dnajc10 | 66861 | protein_coding | 3361.363449 | 0.561614126 | 0.083371756 | 6.736263577 | 1.63E-11 | 5.98E-10 | 0.561614126 |
| ENSMUSG00000029074 | Ttll10 | 330010 | protein_coding | 167.5233582 | -0.561826047 | 0.239038882 | -2.350354224 | 0.018755553 | 0.07628663 | 0.561826047 |
| ENSMUSG00000025855 | Prkar1b | 19085 | protein_coding | 7.667310347 | -0.561841758 | 0.534420734 | -1.051309805 | 0.293116325 | 0.502178856 | 0.561841758 |
| ENSMUSG00000045555 | Mettl24 | 327747 | protein_coding | 9.431720653 | 0.56186573 | 0.432632051 | 1.29871499 | 0.194041758 | 0.387387373 | 0.56186573 |

| ensembl_gene_id | external_gene_name | entrezgene_id | gene_biotype | baseMean | log2FoldChange | lfcSE | stat | pvalue | padj | absLog2FC |
|---|---|---|---|---|---|---|---|---|---|---|
| ENSMUSG00000048546 | Tob2 | 57259 | protein_coding | 450.6044111 | -0.56189907 | 0.113651273 | -4.944063156 | 7.65E-07 | 1.49E-05 | 0.56189907 |
| ENSMUSG00000038679 | Trps1 | 83925 | protein_coding | 8.409349168 | -0.561993277 | 0.864722155 | -0.649911968 | 0.515749087 | 0.703131447 | 0.561993277 |
| ENSMUSG00000035239 | Neu3 | 50877 | protein_coding | 39.66841373 | -0.562229863 | 0.514698354 | -1.092348282 | 0.274680046 | 0.482743437 | 0.562229863 |
| ENSMUSG00000026832 | Cytip | 227929 | protein_coding | 132.5510416 | -0.56276683 | 0.199817441 | -2.816404958 | 0.004856441 | 0.027482711 | 0.56276683 |
| ENSMUSG00000114384 | 4930455J16Rik | 67585 | lncRNA | 7.439079415 | -0.562771772 | 1.215636784 | -0.462944014 | 0.643404498 | 0.793445869 | 0.562771772 |
| ENSMUSG00000038242 | Aox4 | 71872 | protein_coding | 7.375665426 | -0.562832058 | 1.339536386 | -0.420169295 | 0.674361784 | 0.812440509 | 0.562832058 |
| ENSMUSG00000032123 | Dpagt1 | 13478 | protein_coding | 547.8857169 | 0.562917676 | 0.062904567 | 8.948756836 | 3.60E-19 | 2.42E-17 | 0.562917676 |
| ENSMUSG00000037754 | Ppp1r16b | 228852 | protein_coding | 19.02871932 | 0.563470729 | 0.367397811 | 1.53368015 | 0.125108359 | 0.289178051 | 0.563470729 |
| ENSMUSG00000025732 | Mcrip2 | 68241 | protein_coding | 247.5677182 | -0.563568101 | 0.106438424 | -5.294780608 | 1.19E-07 | 2.70E-06 | 0.563568101 |
| ENSMUSG00000025591 | Tma16 | 66282 | protein_coding | 134.6239633 | -0.563594142 | 0.133556319 | -4.21989873 | 2.44E-05 | 0.000328632 | 0.563594142 |
| ENSMUSG00000028874 | Fgr | 14191 | protein_coding | 22.91154153 | -0.563602736 | 0.293421577 | -1.92079513 | 0.054757541 | 0.165037012 | 0.563602736 |
| ENSMUSG00000027224 | Duoxa1 | 213696 | protein_coding | 16.1835188 | 0.564188257 | 0.393420048 | 1.43406077 | 0.151554922 | 0.329686806 | 0.564188257 |
| ENSMUSG00000047671 | Dynlt4 | 242646 | protein_coding | 27.45931109 | -0.564194958 | 0.327099107 | -1.724844076 | 0.084555576 | 0.222427947 | 0.564194958 |
| ENSMUSG00000062093 | Gm10110 | 328451 | transcribed_processed_pseudogene | 8.027780495 | -0.564248169 | 0.412769537 | -1.366981133 | 0.171631219 | 0.357153419 | 0.564248169 |
| ENSMUSG00000002384 | Bmp8b | 12164 | protein_coding | 21.63111616 | -0.564250936 | 0.45951031 | -1.227939664 | 0.219469619 | 0.419854923 | 0.564250936 |
| ENSMUSG00000091705 | H2-Q2 | 15013 | protein_coding | 2012.05617 | 0.564331931 | 0.200864616 | 2.809513904 | 0.004961638 | 0.027886014 | 0.564331931 |
| ENSMUSG00000024621 | Csf1r | 12978 | protein_coding | 8.300184378 | 0.564418397 | 0.515951854 | 1.093936174 | 0.273982972 | 0.481890546 | 0.564418397 |
| ENSMUSG00000023935 | Spats1 | 71020 | protein_coding | 6.963191234 | 0.564602357 | 0.593705228 | 0.950980941 | 0.341614051 | 0.551941783 | 0.564602357 |
| ENSMUSG00000085440 | Sorbs2os | 319940 | lncRNA | 7.589726749 | -0.564698866 | 0.557784641 | -1.01239587 | 0.311348812 | 0.521451199 | 0.564698866 |
| ENSMUSG00000038239 | Hrc | 15464 | protein_coding | 34.68933154 | -0.564744697 | 0.299790977 | -1.883794846 | 0.059592731 | 0.175321947 | 0.564744697 |
| ENSMUSG00000004347 | Pde1c | 18575 | protein_coding | 65.16982357 | -0.56479416 | 0.21333232 | -2.647485197 | 0.008109291 | 0.040939711 | 0.56479416 |
| ENSMUSG00000022565 | Plec | 18810 | protein_coding | 9642.558436 | -0.564834924 | 0.307734126 | -1.835464044 | 0.066436962 | 0.188432305 | 0.564834924 |
| ENSMUSG00000043004 | Gng2 | 14702 | protein_coding | 19.36503784 | 0.564853016 | 0.326013624 | 1.732605554 | 0.083165801 | 0.219934399 | 0.564853016 |
| ENSMUSG00000038530 | Rgs4 | 19736 | protein_coding | 15.22006286 | -0.565302238 | 0.401643544 | -1.407472485 | 0.159287331 | 0.340451304 | 0.565302238 |
| ENSMUSG00000116660 | 9030404E10Rik | 71528 | lncRNA | 12.1213319 | -0.565441853 | 0.383029319 | -1.476236479 | 0.13988042 | 0.311872246 | 0.565441853 |
| ENSMUSG00000020102 | Slc16a7 | 20503 | protein_coding | 119.9419645 | -0.565783588 | 0.196440796 | -2.880173564 | 0.003974563 | 0.023369081 | 0.565783588 |
| ENSMUSG00000074811 | Hps6 | 20170 | protein_coding | 113.8546625 | -0.56578911 | 0.128360998 | -4.407796114 | 1.04E-05 | 0.000154584 | 0.56578911 |
| ENSMUSG00000053950 | Adnp2 | 240442 | protein_coding | 464.9913253 | 0.566069686 | 0.106691245 | 5.305680765 | 1.12E-07 | 2.55E-06 | 0.566069686 |
| ENSMUSG00000017412 | Cacnb4 | 12298 | protein_coding | 7.91842999 | -0.566108043 | 0.608642103 | -0.930116469 | 0.352310785 | 0.562003624 | 0.566108043 |
| ENSMUSG00000016024 | Lbp | 16803 | protein_coding | 11.68543758 | 0.566115665 | 0.50490037 | 1.121242326 | 0.262184728 | 0.470034424 | 0.566115665 |
| ENSMUSG00000026211 | Obsl1 | 98733 | protein_coding | 9.589050084 | 0.566206718 | 0.495337421 | 1.143072771 | 0.253008378 | 0.459205682 | 0.566206718 |
| ENSMUSG00000035158 | Mitf | 17342 | protein_coding | 32.80484517 | 0.566542978 | 0.257633607 | 2.199025908 | 0.02787608 | 0.102165256 | 0.566542978 |
| ENSMUSG00000021596 | Mctp1 | 78771 | protein_coding | 64.82526255 | -0.566743166 | 0.185646826 | -3.052802891 | 0.002267148 | 0.015027521 | 0.566743166 |
| ENSMUSG00000038304 | Cd160 | 54215 | protein_coding | 32.65294363 | 0.566822543 | 0.228852391 | 2.476804111 | 0.013256464 | 0.059161937 | 0.566822543 |
| ENSMUSG00000030223 | Ptpro | 19277 | protein_coding | 160.6232261 | 0.566931608 | 0.223283895 | 2.539061802 | 0.011115019 | 0.051766087 | 0.566931608 |
| ENSMUSG00000008305 | Tle1 | 21885 | protein_coding | 902.3855323 | 0.567023001 | 0.116664036 | 4.860306748 | 1.17E-06 | 2.17E-05 | 0.567023001 |
| ENSMUSG00000021701 | Plk2 | 20620 | protein_coding | 550.8601972 | -0.567023433 | 0.21609367 | -2.623970575 | 0.008691129 | 0.043152216 | 0.567023433 |
| ENSMUSG00000070806 | Zmynd12 | 332934 | protein_coding | 11.26760039 | 0.567051754 | 0.354473683 | 1.599700574 | 0.109665025 | 0.263874388 | 0.567051754 |

| ensembl_gene_id | external_gene_name | entrezgene_id | gene_biotype | baseMean | log2FoldChange | lfcSE | stat | pvalue | padj | absLog2FC |
|---|---|---|---|---|---|---|---|---|---|---|
| ENSMUSG00000000167 | Pih1d2 | 72614 | protein_coding | 22.73045708 | 0.567298995 | 0.240590503 | 2.357944253 | 0.018376452 | 0.075038056 | 0.567298995 |
| ENSMUSG00000042677 | Zc3h12a | 230738 | protein_coding | 251.6147209 | -0.56769202 | 0.155910202 | -3.641147347 | 0.000271426 | 0.002621752 | 0.56769202 |
| ENSMUSG00000110266 | Gm32742 | 102635385 | protein_coding | 20.46335551 | -0.567736114 | 0.316441796 | -1.794124926 | 0.072793231 | 0.200676868 | 0.567736114 |
| ENSMUSG00000027245 | Hypk | 67693 | protein_coding | 21.44364841 | -0.567878875 | 0.295428446 | -1.922221382 | 0.054577907 | 0.164730907 | 0.567878875 |
| ENSMUSG00000037235 | Mxd4 | 17122 | protein_coding | 1674.287894 | 0.568044269 | 0.114139391 | 4.976759245 | 6.47E-07 | 1.28E-05 | 0.568044269 |
| ENSMUSG00000041891 | Lman1 | 70361 | protein_coding | 1697.005307 | 0.568136159 | 0.065129065 | 8.723235321 | 2.70E-18 | 1.74E-16 | 0.568136159 |
| ENSMUSG00000078552 | Dcdc2b | 100504491 | protein_coding | 13.74903509 | -0.56825568 | 0.31800527 | -1.786937935 | 0.07394752 | 0.202773241 | 0.56825568 |
| ENSMUSG00000020366 | Mapk9 | 26420 | protein_coding | 1028.444764 | 0.568321599 | 0.069388559 | 8.190422231 | 2.60E-16 | 1.47E-14 | 0.568321599 |
| ENSMUSG00000034768 | Asb16 | 217217 | protein_coding | 11.76749587 | -0.568325623 | 0.388240561 | -1.46384917 | 0.143235164 | 0.31713229 | 0.568325623 |
| ENSMUSG00000072572 | Slc39a2 | 214922 | protein_coding | 7.485031055 | 0.56855708 | 0.505204347 | 1.125400214 | 0.260419481 | 0.467786902 | 0.56855708 |
| ENSMUSG00000035062 | Zc4h2 | 245522 | protein_coding | 13.84532665 | 0.568603765 | 0.315337942 | 1.803156832 | 0.071363588 | 0.198062246 | 0.568603765 |
| ENSMUSG00000091945 | Vmn2r114 | 666002 | protein_coding | 10.113125 | -0.568652344 | 1.702827229 | -0.333946001 | 0.738420307 | 0.853833658 | 0.568652344 |
| ENSMUSG00000022768 | Ccdc116 | 76872 | protein_coding | 126.9817489 | 0.568809552 | 0.214441777 | 2.652512767 | 0.007989511 | 0.040463251 | 0.568809552 |
| ENSMUSG00000015647 | Lama5 | 16776 | protein_coding | 90.84469038 | -0.569243886 | 0.288074505 | -1.97603008 | 0.048151371 | 0.151782115 | 0.569243886 |
| ENSMUSG00000085667 | Gm12992 | 545681 | lncRNA | 28.35828845 | -0.569523607 | 0.307256052 | -1.853579785 | 0.063799302 | 0.183428751 | 0.569523607 |
| ENSMUSG00000030269 | Mtmr14 | 97287 | protein_coding | 576.4806784 | 0.5695414 | 0.108766347 | 5.236375166 | 1.64E-07 | 3.64E-06 | 0.5695414 |
| ENSMUSG00000030364 | Clec2h | 94071 | protein_coding | 4999.002312 | 0.569594752 | 0.168074561 | 3.38894089 | 0.000701631 | 0.005750958 | 0.569594752 |
| ENSMUSG00000010175 | Prox1 | 19130 | protein_coding | 24.9701284 | -0.569649256 | 0.288033031 | -1.977721977 | 0.04796008 | 0.151388852 | 0.569649256 |
| ENSMUSG00000024474 | Ik | 24010 | protein_coding | 1566.616442 | 0.569735001 | 0.074965972 | 7.599914811 | 2.96E-14 | 1.41E-12 | 0.569735001 |
| ENSMUSG00000020282 | Rhbdf1 | 13650 | protein_coding | 281.7925613 | -0.569842621 | 0.175441661 | -3.248046203 | 0.001162004 | 0.008660926 | 0.569842621 |
| ENSMUSG00000020941 | Map3k14 | 53859 | protein_coding | 101.9265193 | -0.570787319 | 0.189189985 | -3.017005991 | 0.002552848 | 0.016542619 | 0.570787319 |
| ENSMUSG00000022339 | Ebag9 | 55960 | protein_coding | 435.590207 | 0.570850485 | 0.113669548 | 5.022017728 | 5.11E-07 | 1.04E-05 | 0.570850485 |
| ENSMUSG00000031364 | Grpr | 14829 | protein_coding | 36.24968068 | 0.571022752 | 0.30686582 | 1.860822271 | 0.062769279 | 0.181128045 | 0.571022752 |
| ENSMUSG00000040524 | Zfp609 | 214812 | protein_coding | 252.4903093 | -0.571224537 | 0.146879814 | -3.889060862 | 0.000100633 | 0.001134381 | 0.571224537 |
| ENSMUSG00000059602 | Syn3 | 27204 | protein_coding | 34.67887204 | -0.571642901 | 0.309933295 | -1.84440623 | 0.06512396 | 0.185783495 | 0.571642901 |
| ENSMUSG00000086390 | 1810019D21Rik | 69771 | lncRNA | 131.9665339 | 0.571673107 | 0.211083699 | 2.708276908 | 0.006763357 | 0.035459758 | 0.571673107 |
| ENSMUSG00000009030 | Pdcl | 67466 | protein_coding | 878.4881815 | 0.571724112 | 0.073234851 | 7.80672191 | 5.87E-15 | 2.98E-13 | 0.571724112 |
| ENSMUSG00000028419 | Chmp5 | 76959 | protein_coding | 1866.780237 | 0.572088959 | 0.083413868 | 6.858439433 | 6.96E-12 | 2.64E-10 | 0.572088959 |
| ENSMUSG00000036966 | Spryd3 | 223918 | protein_coding | 516.7388346 | 0.572252778 | 0.116375655 | 4.917289415 | 8.78E-07 | 1.67E-05 | 0.572252778 |
| ENSMUSG00000072972 | Adam4 | 11498 | protein_coding | 17.79329889 | 0.572584617 | 0.355210368 | 1.611959189 | 0.106970821 | 0.25996569 | 0.572584617 |
| ENSMUSG00000030041 | M1ap | 110958 | protein_coding | 7.045125674 | -0.572618246 | 0.550159261 | -1.040822697 | 0.297957844 | 0.506735399 | 0.572618246 |
| ENSMUSG00000004100 | Ppan | 235036 | protein_coding | 274.592476 | -0.572660544 | 0.197064847 | -2.90594975 | 0.003661402 | 0.021949122 | 0.572660544 |
| ENSMUSG00000027712 | Anxa5 | 11747 | protein_coding | 793.2745373 | 0.572823357 | 0.115144356 | 4.974827932 | 6.53E-07 | 1.29E-05 | 0.572823357 |
| ENSMUSG00000068231 | Vmn1r43 | 113847 | protein_coding | 6.932962362 | -0.572884686 | 0.989403628 | -0.5790202 | 0.562575544 | 0.739622589 | 0.572884686 |
| ENSMUSG00000020661 | Dnmt3a | 13435 | protein_coding | 603.7931113 | -0.572902639 | 0.133429462 | -4.293674221 | 1.76E-05 | 0.000246408 | 0.572902639 |
| ENSMUSG00000074884 | Serf2 | 378702 | protein_coding | 1449.920708 | 0.573180207 | 0.075907744 | 7.551010939 | 4.32E-14 | 2.03E-12 | 0.573180207 |
| ENSMUSG00000020773 | Trim47 | 217333 | protein_coding | 33.97813576 | 0.573216968 | 0.266024696 | 2.154750953 | 0.031181328 | 0.111164738 | 0.573216968 |
| ENSMUSG00000056553 | Ptprn2 | 19276 | protein_coding | 692.4374866 | -0.573663104 | 0.117283732 | -4.891241904 | 1.00E-06 | 1.89E-05 | 0.573663104 |

| ensembl_gene_id | external_gene_name | entrezgene_id | gene_biotype | baseMean | log2FoldChange | lfcSE | stat | pvalue | padj | absLog2FC |
|---|---|---|---|---|---|---|---|---|---|---|
| ENSMUSG00000073792 | Alg6 | 320438 | protein_coding | 414.7298438 | 0.573742787 | 0.082159961 | 6.983240784 | 2.88E-12 | 1.13E-10 | 0.573742787 |
| ENSMUSG00000064945 | Rny3 | 19874 | misc_RNA | 69.29897272 | 0.573862418 | 0.356538019 | 1.609540605 | 0.107498184 | 0.260760999 | 0.573862418 |
| ENSMUSG00000029504 | Ddx51 | 69663 | protein_coding | 127.0534758 | -0.573913339 | 0.170355394 | -3.368917921 | 0.000754639 | 0.006094139 | 0.573913339 |
| ENSMUSG00000040877 | Wdr25 | 212198 | protein_coding | 60.74991451 | -0.574073249 | 0.185516102 | -3.094465889 | 0.001971676 | 0.013363769 | 0.574073249 |
| ENSMUSG00000039716 | Dock3 | 208869 | protein_coding | 18.39599125 | -0.574176707 | 0.628570604 | -0.913464142 | 0.360998484 | 0.570474103 | 0.574176707 |
| ENSMUSG00000000805 | Car4 | 12351 | protein_coding | 1775.988674 | -0.574273865 | 0.242118468 | -2.371871385 | 0.01769825 | 0.073011585 | 0.574273865 |
| ENSMUSG00000044807 | Zfp354c | 30944 | protein_coding | 7.209587189 | -0.574414511 | 0.717921505 | -0.800107682 | 0.423648411 | 0.628365637 | 0.574414511 |
| ENSMUSG00000022043 | Trim35 | 66854 | protein_coding | 349.2579536 | 0.57476358 | 0.115270576 | 4.986212459 | 6.16E-07 | 1.24E-05 | 0.57476358 |
| ENSMUSG00000023191 | P3h3 | 14789 | protein_coding | 6.694457545 | -0.575020625 | 0.437392924 | -1.314654611 | 0.188626007 | 0.379779427 | 0.575020625 |
| ENSMUSG00000005947 | Itgae | 16407 | protein_coding | 56.14946037 | 0.575171115 | 0.263535339 | 2.182519875 | 0.029071186 | 0.105343072 | 0.575171115 |
| ENSMUSG00000026923 | Notch1 | 18128 | protein_coding | 464.1021154 | -0.575859149 | 0.166657262 | -3.45534988 | 0.000549579 | 0.00470882 | 0.575859149 |
| ENSMUSG00000046447 | Camk2n1 | 66259 | protein_coding | 741.6057709 | 0.576019271 | 0.121499647 | 4.74091311 | 2.13E-06 | 3.70E-05 | 0.576019271 |
| ENSMUSG00000026834 | Acvr1c | 269275 | protein_coding | 299.9033943 | -0.576269962 | 0.218640117 | -2.635700944 | 0.008396369 | 0.04201614 | 0.576269962 |
| ENSMUSG00000020654 | Adcy3 | 104111 | protein_coding | 18.59461259 | 0.576306718 | 0.283499899 | 2.032828655 | 0.042069837 | 0.137651677 | 0.576306718 |
| ENSMUSG00000054027 | Nt5dc3 | 103466 | protein_coding | 98.24079685 | -0.576365785 | 0.258575531 | -2.22900358 | 0.025813667 | 0.096903154 | 0.576365785 |
| ENSMUSG00000039646 | Vasn | 246154 | protein_coding | 74.84002064 | -0.576441747 | 0.16927674 | -3.405321656 | 0.000660862 | 0.005485255 | 0.576441747 |
| ENSMUSG00000028137 | Celf3 | 78784 | protein_coding | 33.51222073 | -0.576688098 | 0.246990735 | -2.334857211 | 0.019550881 | 0.07857846 | 0.576688098 |
| ENSMUSG00000052087 | Rgs14 | 51791 | protein_coding | 28.6651987 | 0.576885028 | 0.26057962 | 2.213853203 | 0.026838879 | 0.099393086 | 0.576885028 |
| ENSMUSG00000096852 | Cyp2d12 | 380997 | protein_coding | 60.31330023 | -0.577054088 | 0.344512012 | -1.674989747 | 0.093936243 | 0.23874525 | 0.577054088 |
| ENSMUSG00000017057 | Il13ra1 | 16164 | protein_coding | 3188.934555 | 0.577104462 | 0.100683704 | 5.731855689 | 9.93E-09 | 2.68E-07 | 0.577104462 |
| ENSMUSG00000027340 | Slc23a2 | 54338 | protein_coding | 924.403514 | -0.577367321 | 0.131401701 | -4.393910556 | 1.11E-05 | 0.000163386 | 0.577367321 |
| ENSMUSG00000050982 | Apol10a | 245282 | protein_coding | 3075.408995 | 0.577624354 | 0.230137294 | 2.509911985 | 0.012076126 | 0.055145929 | 0.577624354 |
| ENSMUSG00000021756 | Il6st | 16195 | protein_coding | 914.3458923 | 0.578127362 | 0.131211676 | 4.406066445 | 1.05E-05 | 0.000155678 | 0.578127362 |
| ENSMUSG00000020936 | Nmt1 | 18107 | protein_coding | 1588.976782 | 0.57831257 | 0.084682361 | 6.829197537 | 8.54E-12 | 3.20E-10 | 0.57831257 |
| ENSMUSG00000039683 | Sdk1 | 330222 | protein_coding | 9.438931773 | 0.578866916 | 0.628799111 | 0.920591182 | 0.357263908 | 0.566895749 | 0.578866916 |
| ENSMUSG00000074824 | Rslcan18 | 432770 | protein_coding | 12.08805739 | -0.578891515 | 0.482712888 | -1.199246031 | 0.230432293 | 0.433485954 | 0.578891515 |
| ENSMUSG00000030641 | Ddias | 74041 | protein_coding | 132.8736004 | 0.578910808 | 0.146024237 | 3.964484397 | 7.36E-05 | 0.00085956 | 0.578910808 |
| ENSMUSG00000019235 | Rps6kl1 | 238323 | protein_coding | 26.72210745 | -0.578969699 | 0.217227814 | -2.665265047 | 0.007692767 | 0.039259976 | 0.578969699 |
| ENSMUSG00000023505 | Cdca3 | 14793 | protein_coding | 682.0467007 | 0.579435175 | 0.129976655 | 4.457994214 | 8.27E-06 | 0.000125857 | 0.579435175 |
| ENSMUSG00000035735 | Dagla | 269060 | protein_coding | 160.7705768 | -0.579751988 | 0.160294722 | -3.61678777 | 0.000298282 | 0.002824632 | 0.579751988 |
| ENSMUSG00000051790 | Nlgn2 | 216856 | protein_coding | 22.04465221 | -0.579954172 | 0.29695897 | -1.952977452 | 0.050822267 | 0.157122878 | 0.579954172 |
| ENSMUSG00000032238 | Rora | 19883 | protein_coding | 29.98790122 | -0.5801263 | 0.253321376 | -2.290080328 | 0.022016661 | 0.085825103 | 0.5801263 |
| ENSMUSG00000020492 | Ska2 | 66140 | protein_coding | 112.8294652 | 0.580277059 | 0.16500107 | 3.516807862 | 0.00043677 | 0.003906159 | 0.580277059 |
| ENSMUSG00000031637 | Lrp2bp | 67620 | protein_coding | 7.437054945 | -0.580353413 | 0.576231204 | -1.007153742 | 0.313860899 | 0.524462484 | 0.580353413 |
| ENSMUSG00000031897 | Psmb10 | 19171 | protein_coding | 1292.263988 | 0.580456149 | 0.126932945 | 4.572935344 | 4.81E-06 | 7.79E-05 | 0.580456149 |
| ENSMUSG00000054708 | Ankrd24 | 70615 | protein_coding | 104.0022539 | -0.580836552 | 0.178601964 | -3.252128579 | 0.001145442 | 0.008557504 | 0.580836552 |
| ENSMUSG00000026620 | Mark1 | 226778 | protein_coding | 85.57083122 | 0.580855854 | 0.206065428 | 2.81879333 | 0.004820454 | 0.027314194 | 0.580855854 |
| ENSMUSG00000036158 | Prickle1 | 106042 | protein_coding | 20.94866161 | -0.581254762 | 0.298400356 | -1.947902375 | 0.051426646 | 0.158407196 | 0.581254762 |

| ensembl_gene_id | external_gene_name | entrezgene_id | gene_biotype | baseMean | log2FoldChange | lfcSE | stat | pvalue | padj | absLog2FC |
|---|---|---|---|---|---|---|---|---|---|---|
| ENSMUSG00000054871 | Tmem158 | 72309 | protein_coding | 11.66853563 | 0.581896702 | 0.447643992 | 1.299909554 | 0.19363197 | 0.387069668 | 0.581896702 |
| ENSMUSG00000063751 | Platr8 | 100038454 | lncRNA | 8.738215364 | -0.581932096 | 0.389169238 | -1.495318848 | 0.134831248 | 0.304148438 | 0.581932096 |
| ENSMUSG00000019944 | Rhobtb1 | 69288 | protein_coding | 50.94113725 | -0.581952428 | 0.232392877 | -2.504174982 | 0.012273732 | 0.05574451 | 0.581952428 |
| ENSMUSG00000073771 | Btbd19 | 78611 | protein_coding | 60.45705365 | -0.582091956 | 0.207063746 | -2.811172734 | 0.004936128 | 0.02776195 | 0.582091956 |
| ENSMUSG00000062012 | Zfp13 | 22654 | protein_coding | 36.95484513 | -0.582239827 | 0.211492666 | -2.753002443 | 0.005905146 | 0.031776067 | 0.582239827 |
| ENSMUSG00000043439 | Epop | 103551 | protein_coding | 53.25688822 | -0.582483738 | 0.380170767 | -1.532163408 | 0.125482116 | 0.28970563 | 0.582483738 |
| ENSMUSG00000053198 | Prx | 19153 | protein_coding | 53.76731539 | 0.582770573 | 0.165289907 | 3.525748075 | 0.000422288 | 0.003802267 | 0.582770573 |
| ENSMUSG00000029632 | Ndufa4 | 17992 | protein_coding | 2345.854043 | 0.583171593 | 0.111486471 | 5.230873193 | 1.69E-07 | 3.73E-06 | 0.583171593 |
| ENSMUSG00000102101 | Zbtb11os1 | 66391 | lncRNA | 12.06326763 | -0.583175845 | 0.356211677 | -1.637160942 | 0.101596846 | 0.250850187 | 0.583175845 |
| ENSMUSG00000044641 | Pard6b | 58220 | protein_coding | 384.8533911 | -0.583317968 | 0.318358623 | -1.832266903 | 0.066911666 | 0.189369049 | 0.583317968 |
| ENSMUSG00000056035 | Cyp3a11 | 13112 | protein_coding | 10.44655972 | -0.583542591 | 0.618357478 | -0.943697799 | 0.345324099 | 0.555195544 | 0.583542591 |
| ENSMUSG00000031922 | Cep57 | 74360 | protein_coding | 427.2213122 | 0.583586932 | 0.07619952 | 7.658669444 | 1.88E-14 | 9.12E-13 | 0.583586932 |
| ENSMUSG00000020886 | Dlg4 | 13385 | protein_coding | 15.70303996 | -0.584014029 | 0.691942618 | -0.844020897 | 0.398657729 | 0.606535515 | 0.584014029 |
| ENSMUSG00000053219 | Raet1e | 379043 | protein_coding | 104.1448377 | -0.584077717 | 0.196368198 | -2.974400758 | 0.002935614 | 0.018437878 | 0.584077717 |
| ENSMUSG00000040711 | Sh3pxd2b | 268396 | protein_coding | 142.4066671 | -0.584174686 | 0.320810373 | -1.820934529 | 0.06861681 | 0.192960548 | 0.584174686 |
| ENSMUSG00000028039 | Efna3 | 13638 | protein_coding | 45.71982817 | -0.584402574 | 0.32819973 | -1.780630877 | 0.074972772 | 0.20423573 | 0.584402574 |
| ENSMUSG00000032482 | Cspg5 | 29873 | protein_coding | 17.77321902 | -0.584420294 | 0.397198268 | -1.471356602 | 0.141194703 | 0.313792275 | 0.584420294 |
| ENSMUSG00000016526 | Dyrk3 | 226419 | protein_coding | 17.29643234 | -0.584617505 | 0.326808508 | -1.788868682 | 0.073635967 | 0.202080234 | 0.584617505 |
| ENSMUSG00000010342 | Tex14 | 83560 | protein_coding | 13.95652217 | -0.584640048 | 0.417131871 | -1.401571273 | 0.16104331 | 0.343065112 | 0.584640048 |
| ENSMUSG00000023010 | Tmbim6 | 110213 | protein_coding | 14133.62838 | 0.584689353 | 0.107395084 | 5.444284171 | 5.20E-08 | 1.24E-06 | 0.584689353 |
| ENSMUSG00000033669 | Zfp7 | 223669 | protein_coding | 375.2287182 | -0.584726928 | 0.162444771 | -3.599542944 | 0.000318777 | 0.002986753 | 0.584726928 |
| ENSMUSG00000095864 | Vmn1r77 | 628946 | protein_coding | 7.880668893 | -0.584898443 | 1.811767641 | -0.322833033 | 0.746821694 | 0.859293709 | 0.584898443 |
| ENSMUSG00000026546 | Cfap45 | 71870 | protein_coding | 37.5462461 | -0.585009192 | 0.232473778 | -2.516452375 | 0.011854292 | 0.054382247 | 0.585009192 |
| ENSMUSG00000023235 | Ccl25 | 20300 | protein_coding | 109.0402128 | -0.585486118 | 0.172294063 | -3.398179305 | 0.000678359 | 0.005613221 | 0.585486118 |
| ENSMUSG00000043633 | Fam221b | 242408 | protein_coding | 21.47335831 | 0.585996927 | 0.31529704 | 1.858555119 | 0.063090224 | 0.181816733 | 0.585996927 |
| ENSMUSG00000049676 | Catsperg1 | 320225 | protein_coding | 11.93097201 | -0.586072295 | 0.554114911 | -1.057672847 | 0.290204618 | 0.498907627 | 0.586072295 |
| ENSMUSG00000017418 | Arl5b | 75869 | protein_coding | 726.8802726 | -0.586153244 | 0.277384301 | -2.113144982 | 0.034588361 | 0.11989628 | 0.586153244 |
| ENSMUSG00000015749 | Anp32e | 66471 | protein_coding | 984.9224404 | 0.586230428 | 0.129004649 | 4.544258141 | 5.51E-06 | 8.81E-05 | 0.586230428 |
| ENSMUSG00000030165 | Klrd1 | 16643 | protein_coding | 27.64955931 | 0.586370132 | 0.324477404 | 1.807121623 | 0.070743318 | 0.196820885 | 0.586370132 |
| ENSMUSG00000020255 | Nopchap1 | 28109 | protein_coding | 901.6980749 | 0.586426151 | 0.137289648 | 4.271452086 | 1.94E-05 | 0.000270335 | 0.586426151 |
| ENSMUSG00000006800 | Sulf2 | 72043 | protein_coding | 347.8215942 | 0.58645368 | 0.213768789 | 2.743401793 | 0.006080623 | 0.032536352 | 0.58645368 |
| ENSMUSG00000070583 | Fv1 | 14349 | protein_coding | 69.01641892 | -0.586678175 | 0.173881848 | -3.374004719 | 0.000740831 | 0.006011185 | 0.586678175 |
| ENSMUSG00000038773 | Kdm3b | 277250 | protein_coding | 1467.784559 | 0.586865577 | 0.0895283 | 6.555084561 | 5.56E-11 | 1.96E-09 | 0.586865577 |
| ENSMUSG00000034522 | Zfp395 | 380912 | protein_coding | 96.34245576 | -0.586898511 | 0.222140132 | -2.642019282 | 0.008241336 | 0.041448691 | 0.586898511 |
| ENSMUSG00000024206 | Rfx2 | 19725 | protein_coding | 22.9280249 | -0.586922392 | 0.311732139 | -1.882777933 | 0.059730469 | 0.17553264 | 0.586922392 |
| ENSMUSG00000022428 | Cby1 | 73739 | protein_coding | 174.5417114 | 0.587209891 | 0.125829386 | 4.666715065 | 3.06E-06 | 5.14E-05 | 0.587209891 |
| ENSMUSG00000079084 | Ccdc82 | 66396 | protein_coding | 256.9781397 | -0.587392592 | 0.086532084 | -6.788147981 | 1.14E-11 | 4.23E-10 | 0.587392592 |
| ENSMUSG00000040713 | Creg1 | 433375 | protein_coding | 794.4903949 | 0.587422615 | 0.111330696 | 5.276376038 | 1.32E-07 | 2.96E-06 | 0.587422615 |

| ensembl_gene_id | external_gene_name | entrezgene_id | gene_biotype | baseMean | log2FoldChange | lfcSE | stat | pvalue | padj | absLog2FC |
|---|---|---|---|---|---|---|---|---|---|---|
| ENSMUSG00000036264 | Fstl4 | 320027 | protein_coding | 42.40288178 | -0.587662911 | 0.199852797 | -2.940478795 | 0.003277054 | 0.020153251 | 0.587662911 |
| ENSMUSG00000030322 | Mbd4 | 17193 | protein_coding | 63.36141735 | 0.587682838 | 0.178343041 | 3.295238407 | 0.000983382 | 0.007577799 | 0.587682838 |
| ENSMUSG00000026024 | Als2 | 74018 | protein_coding | 247.535153 | 0.587684346 | 0.122884084 | 4.782428505 | 1.73E-06 | 3.07E-05 | 0.587684346 |
| ENSMUSG00000021280 | Exoc3l4 | 74190 | protein_coding | 33.86422143 | 0.587735661 | 0.228033215 | 2.577412507 | 0.009954308 | 0.047759252 | 0.587735661 |
| ENSMUSG00000029714 | Gigyf1 | 57330 | protein_coding | 220.6768766 | -0.587760836 | 0.170143793 | -3.454494737 | 0.000551325 | 0.004718702 | 0.587760836 |
| ENSMUSG00000002265 | Peg3 | 18616 | protein_coding | 51.80334921 | -0.58808378 | 0.240720987 | -2.44301001 | 0.014565333 | 0.063059698 | 0.58808378 |
| ENSMUSG00000039851 | 4932438H23Rik | 74387 | protein_coding | 17.39476696 | 0.588213104 | 0.333663633 | 1.762892465 | 0.077918628 | 0.209908554 | 0.588213104 |
| ENSMUSG00000022615 | Tymp | 72962 | protein_coding | 35.00230084 | -0.588446297 | 0.209599116 | -2.807484631 | 0.004993006 | 0.028022764 | 0.588446297 |
| ENSMUSG00000040938 | Slc16a11 | 216867 | protein_coding | 39.02775955 | -0.588616976 | 0.196722375 | -2.99212012 | 0.002770472 | 0.017611374 | 0.588616976 |
| ENSMUSG00000025165 | Sectm1a | 209588 | protein_coding | 86.05041835 | 0.588689311 | 0.162052513 | 3.63270708 | 0.000280463 | 0.002686242 | 0.588689311 |
| ENSMUSG00000025270 | Alas2 | 11656 | protein_coding | 28.11966873 | 0.589084866 | 0.444401243 | 1.3255698 | 0.184982235 | 0.375002805 | 0.589084866 |
| ENSMUSG00000008307 | 1700109H08Rik | 77036 | protein_coding | 24.12434812 | -0.589317414 | 0.249236494 | -2.364490868 | 0.018054872 | 0.07406078 | 0.589317414 |
| ENSMUSG00000042589 | Cux2 | 13048 | protein_coding | 33.24222227 | -0.589607766 | 0.323688978 | -1.821525619 | 0.068526996 | 0.192777559 | 0.589607766 |
| ENSMUSG00000031672 | Got2 | 14719 | protein_coding | 1113.086016 | 0.589621317 | 0.085677121 | 6.881899255 | 5.91E-12 | 2.26E-10 | 0.589621317 |
| ENSMUSG00000037801 | Iqch | 78250 | protein_coding | 13.85203986 | -0.589660851 | 0.508756182 | -1.159024443 | 0.246446222 | 0.451974142 | 0.589660851 |
| ENSMUSG00000037318 | Traf3ip3 | 215243 | protein_coding | 12.02313354 | 0.589879924 | 0.506022138 | 1.1657196 | 0.243727833 | 0.448797332 | 0.589879924 |
| ENSMUSG00000025359 | Pmel | 20431 | protein_coding | 32.39748124 | -0.589915658 | 0.201750188 | -2.923990629 | 0.003455751 | 0.021000843 | 0.589915658 |
| ENSMUSG00000015478 | Rnf5 | 54197 | protein_coding | 264.6007553 | 0.590040982 | 0.128841794 | 4.579577504 | 4.66E-06 | 7.56E-05 | 0.590040982 |
| ENSMUSG00000031953 | Tmem170 | 66817 | protein_coding | 581.5589755 | 0.590125734 | 0.079631732 | 7.41068569 | 1.26E-13 | 5.62E-12 | 0.590125734 |
| ENSMUSG00000042308 | Setd1a | 233904 | protein_coding | 89.99171366 | -0.590484151 | 0.154247413 | -3.828162434 | 0.000129104 | 0.001405348 | 0.590484151 |
| ENSMUSG00000034528 | Hsd17b13 | 243168 | protein_coding | 348.3261956 | 0.590561232 | 0.170490418 | 3.463896914 | 0.00053241 | 0.004591356 | 0.590561232 |
| ENSMUSG00000028949 | Smarcd3 | 66993 | protein_coding | 10.06995287 | -0.590932135 | 0.441478653 | -1.338529351 | 0.180723939 | 0.368908615 | 0.590932135 |
| ENSMUSG00000090700 | Cyp4f40 | 631304 | protein_coding | 1060.575632 | 0.591282748 | 0.126148819 | 4.687184158 | 2.77E-06 | 4.72E-05 | 0.591282748 |
| ENSMUSG00000032300 | 1700017B05Rik | 74211 | protein_coding | 146.2294953 | -0.591680113 | 0.257411677 | -2.298575262 | 0.02152907 | 0.084357939 | 0.591680113 |
| ENSMUSG00000053907 | Mat2a | 232087 | protein_coding | 3341.282756 | -0.592158564 | 0.11346734 | -5.218757787 | 1.80E-07 | 3.97E-06 | 0.592158564 |
| ENSMUSG00000024617 | Camk2a | 12322 | protein_coding | 11.15107925 | -0.592254537 | 0.621710458 | -0.95262116 | 0.340782052 | 0.551168324 | 0.592254537 |
| ENSMUSG00000029607 | Ankrd61 | 66729 | protein_coding | 10.18886712 | -0.592386539 | 0.351277212 | -1.686379073 | 0.091722809 | 0.234730223 | 0.592386539 |
| ENSMUSG00000047777 | Phf13 | 230936 | protein_coding | 266.2554727 | 0.592400744 | 0.140134666 | 4.227367583 | 2.36E-05 | 0.000319426 | 0.592400744 |
| ENSMUSG00000020395 | Itk | 16428 | protein_coding | 20.87918479 | 0.592454664 | 0.380743264 | 1.556047656 | 0.119696761 | 0.280537081 | 0.592454664 |
| ENSMUSG00000049971 | Glt1d1 | 319804 | protein_coding | 298.2945555 | -0.592607614 | 0.160554759 | -3.690999991 | 0.000223374 | 0.002240493 | 0.592607614 |
| ENSMUSG00000015202 | Cnksr3 | 215748 | protein_coding | 352.9742561 | -0.592703661 | 0.21070621 | -2.812938745 | 0.0049091 | 0.027659055 | 0.592703661 |
| ENSMUSG00000035948 | Acss3 | 380660 | protein_coding | 19.58584019 | -0.592866797 | 0.304150354 | -1.949255655 | 0.051264902 | 0.158099111 | 0.592866797 |
| ENSMUSG00000028527 | Ak4 | 11639 | protein_coding | 129.3473352 | 0.592959974 | 0.32124336 | 1.845827955 | 0.06491719 | 0.185504307 | 0.592959974 |
| ENSMUSG00000022253 | Nadk2 | 68646 | protein_coding | 641.3891312 | 0.593101453 | 0.12076483 | 4.911210097 | 9.05E-07 | 1.72E-05 | 0.593101453 |
| ENSMUSG00000028369 | Svep1 | 64817 | protein_coding | 20.33721781 | -0.593266847 | 0.465541989 | -1.274357334 | 0.202536806 | 0.39748185 | 0.593266847 |
| ENSMUSG00000037892 | Pcdh18 | 73173 | protein_coding | 63.39858703 | 0.593416203 | 0.219819479 | 2.699561503 | 0.006943092 | 0.036211385 | 0.593416203 |
| ENSMUSG00000057182 | Scn3a | 20269 | protein_coding | 56.58731137 | -0.593490113 | 0.235500487 | -2.520122661 | 0.011731395 | 0.05394274 | 0.593490113 |
| ENSMUSG00000027829 | Ccnl1 | 56706 | protein_coding | 1518.907936 | -0.59383321 | 0.118626729 | -5.005897207 | 5.56E-07 | 1.13E-05 | 0.59383321 |

Supplementary Table 1

| ensembl_gene_id | external_gene_name | entrezgene_id | gene_biotype | baseMean | log2FoldChange | lfcSE | stat | pvalue | padj | absLog2FC |
|---|---|---|---|---|---|---|---|---|---|---|
| ENSMUSG00000024575 | Pde6a | 225600 | protein_coding | 19.7596927 | -0.593919148 | 0.287546146 | -2.065474204 | 0.038878154 | 0.130478359 | 0.593919148 |
| ENSMUSG00000037965 | Zc3h7a | 106205 | protein_coding | 1632.622032 | 0.594050952 | 0.104035743 | 5.710065941 | 1.13E-08 | 3.02E-07 | 0.594050952 |
| ENSMUSG00000027843 | Ptpn22 | 19260 | protein_coding | 29.8908474 | 0.594148602 | 0.349593835 | 1.699539702 | 0.089217541 | 0.230541367 | 0.594148602 |
| ENSMUSG00000049281 | Scn3b | 235281 | protein_coding | 9.21176606 | 0.594257463 | 0.671710051 | 0.884693421 | 0.376322005 | 0.584500916 | 0.594257463 |
| ENSMUSG00000024579 | Pcyox1l | 240334 | protein_coding | 136.1033499 | 0.594431827 | 0.122644039 | 4.846805699 | 1.25E-06 | 2.30E-05 | 0.594431827 |
| ENSMUSG00000079111 | Kdelr2 | 66913 | protein_coding | 3260.147471 | 0.594434025 | 0.075569004 | 7.866109022 | 3.66E-15 | 1.89E-13 | 0.594434025 |
| ENSMUSG00000051557 | Pusl1 | 433813 | protein_coding | 65.04059448 | -0.594448151 | 0.211836256 | -2.806168138 | 0.005013452 | 0.028117701 | 0.594448151 |
| ENSMUSG00000044505 | Lingo4 | 320747 | protein_coding | 9.468657343 | 0.594597318 | 0.396107538 | 1.50110074 | 0.133329507 | 0.301829505 | 0.594597318 |
| ENSMUSG00000078878 | Gm14305 | 100043387 | protein_coding | 72.57051494 | -0.594780646 | 0.213397909 | -2.787190597 | 0.00531672 | 0.029343282 | 0.594780646 |
| ENSMUSG00000015396 | Cd83 | 12522 | protein_coding | 91.19922678 | -0.595568689 | 0.385190782 | -1.546165476 | 0.122064609 | 0.284495916 | 0.595568689 |
| ENSMUSG00000055850 | Rnf181 | 66510 | protein_coding | 964.4316398 | 0.595769887 | 0.080413497 | 7.408829502 | 1.27E-13 | 5.68E-12 | 0.595769887 |
| ENSMUSG00000045282 | Tmem86b | 68255 | protein_coding | 34.956638 | -0.595805203 | 0.28897891 | -2.06176016 | 0.039230576 | 0.131356866 | 0.595805203 |
| ENSMUSG00000085720 | Gm7854 | 665934 | lncRNA | 8.823579819 | -0.595916793 | 0.442235167 | -1.347511092 | 0.177815683 | 0.364886396 | 0.595916793 |
| ENSMUSG00000086401 | Gm15559 | 100504049 | lncRNA | 7.457223567 | -0.596381343 | 0.477124731 | -1.249948502 | 0.21131836 | 0.408728456 | 0.596381343 |
| ENSMUSG00000059895 | Ptp4a3 | 19245 | protein_coding | 95.70590322 | -0.596514511 | 0.148516887 | -4.016475977 | 5.91E-05 | 0.000716092 | 0.596514511 |
| ENSMUSG00000021200 | Asb2 | 65256 | protein_coding | 11.3901958 | 0.59673122 | 0.418956735 | 1.424326595 | 0.154351952 | 0.333675067 | 0.59673122 |
| ENSMUSG00000019802 | Sec63 | 140740 | protein_coding | 1458.755588 | 0.596841027 | 0.053428857 | 11.17076156 | 5.67E-29 | 6.18E-27 | 0.596841027 |
| ENSMUSG00000101031 | Ms4a12 | 381213 | protein_coding | 353.0922908 | 0.59684205 | 0.122744957 | 4.862456809 | 1.16E-06 | 2.15E-05 | 0.59684205 |
| ENSMUSG00000031556 | Tm2d2 | 69742 | protein_coding | 1341.755233 | 0.596870401 | 0.11689205 | 5.106167606 | 3.29E-07 | 6.92E-06 | 0.596870401 |
| ENSMUSG00000029095 | Ablim2 | 231148 | protein_coding | 78.64650735 | -0.596997383 | 0.240489999 | -2.482420826 | 0.013049306 | 0.058531278 | 0.596997383 |
| ENSMUSG00000050914 | Ankrd37 | 654824 | protein_coding | 109.7496791 | 0.597089477 | 0.17515181 | 3.408982617 | 0.000652056 | 0.005429145 | 0.597089477 |
| ENSMUSG00000007039 | Ddah2 | 51793 | protein_coding | 19.67130753 | -0.597131798 | 0.291098465 | -2.051305215 | 0.040237233 | 0.133614527 | 0.597131798 |
| ENSMUSG00000063568 | Jazf1 | 231986 | protein_coding | 15.22409607 | -0.597187888 | 0.376103539 | -1.58782842 | 0.112325143 | 0.267939604 | 0.597187888 |
| ENSMUSG00000019888 | Mgat4c | 67569 | protein_coding | 11346.31879 | 0.597331204 | 0.217736067 | 2.743372801 | 0.00608116 | 0.032536352 | 0.597331204 |
| ENSMUSG00000074277 | Phldb3 | 232970 | protein_coding | 69.70350686 | -0.597566677 | 0.157525121 | -3.793469088 | 0.000148557 | 0.001585937 | 0.597566677 |
| ENSMUSG00000048537 | Phldb1 | 102693 | protein_coding | 1092.848241 | -0.597636106 | 0.101526512 | -5.886502864 | 3.94E-09 | 1.12E-07 | 0.597636106 |
| ENSMUSG00000026885 | Ttll11 | 74410 | protein_coding | 8.857029251 | -0.597828942 | 0.411312933 | -1.453464976 | 0.146094697 | 0.322075618 | 0.597828942 |
| ENSMUSG00000013083 | 2200002J24Rik | 69147 | protein_coding | 10.00206756 | -0.598025719 | 0.341530059 | -1.751019283 | 0.079942588 | 0.213716943 | 0.598025719 |
| ENSMUSG00000085958 | A930019D19Rik | 77806 | lncRNA | 13.20984826 | -0.598176375 | 0.344010437 | -1.738832053 | 0.082064309 | 0.217998718 | 0.598176375 |
| ENSMUSG00000053931 | Cnn3 | 71994 | protein_coding | 350.7451205 | -0.598218047 | 0.092070525 | -6.497389347 | 8.17E-11 | 2.84E-09 | 0.598218047 |
| ENSMUSG00000079036 | Alkbh1 | 211064 | protein_coding | 288.8400474 | -0.598432294 | 0.127390495 | -4.69762124 | 2.63E-06 | 4.49E-05 | 0.598432294 |
| ENSMUSG00000027902 | Chil6 | 229688 | protein_coding | 14.10385744 | 0.598620002 | 0.288754605 | 2.073109806 | 0.038162062 | 0.12887455 | 0.598620002 |
| ENSMUSG00000073598 | Smim33 | 73449 | protein_coding | 319.4265811 | 0.598735953 | 0.141678577 | 4.226016139 | 2.38E-05 | 0.000320805 | 0.598735953 |
| ENSMUSG00000024553 | Galr1 | 14427 | protein_coding | 7.58683679 | -0.598825089 | 0.53405815 | -1.121273195 | 0.262171592 | 0.470034424 | 0.598825089 |
| ENSMUSG00000062961 | Ccdc177 | 380768 | protein_coding | 7.545683145 | -0.599105301 | 0.540257423 | -1.108925626 | 0.267462268 | 0.475355836 | 0.599105301 |
| ENSMUSG00000075296 | Aldh3b2 | 621603 | protein_coding | 146.1623473 | 0.599291371 | 0.136035122 | 4.405416507 | 1.06E-05 | 0.000155898 | 0.599291371 |
| ENSMUSG00000020469 | Myl7 | 17898 | protein_coding | 43.01849254 | 0.599463647 | 0.318568828 | 1.881739817 | 0.059871351 | 0.175740416 | 0.599463647 |
| ENSMUSG00000039234 | Sec24d | 69608 | protein_coding | 1079.012301 | 0.599748432 | 0.100798044 | 5.950000689 | 2.68E-09 | 7.77E-08 | 0.599748432 |

| ensembl_gene_id | external_gene_name | entrezgene_id | gene_biotype | baseMean | log2FoldChange | lfcSE | stat | pvalue | padj | absLog2FC |
|---|---|---|---|---|---|---|---|---|---|---|
| ENSMUSG00000056758 | Hmga2 | 15364 | protein_coding | 112.351598 | -0.600270088 | 0.199220494 | -3.013094058 | 0.002585988 | 0.016716562 | 0.600270088 |
| ENSMUSG00000000817 | Fasl | 14103 | protein_coding | 7.878609713 | 0.600341703 | 0.494681203 | 1.213593115 | 0.224903137 | 0.426713183 | 0.600341703 |
| ENSMUSG00000041372 | B4galnt3 | 330406 | protein_coding | 573.446178 | -0.600349961 | 0.296310976 | -2.026080742 | 0.042756511 | 0.139148668 | 0.600349961 |
| ENSMUSG00000040658 | Dnph1 | 381101 | protein_coding | 34.40521172 | -0.600511609 | 0.249098348 | -2.410741036 | 0.015920148 | 0.067638334 | 0.600511609 |
| ENSMUSG00000097726 | 9530036O11Rik | 654796 | lncRNA | 20.73481244 | -0.600777902 | 0.333485622 | -1.801510658 | 0.071622429 | 0.198607599 | 0.600777902 |
| ENSMUSG00000047123 | Ticam1 | 106759 | protein_coding | 421.1459232 | 0.601021922 | 0.108655594 | 5.531440206 | 3.18E-08 | 7.87E-07 | 0.601021922 |
| ENSMUSG00000037072 | Selenof | 93684 | protein_coding | 3318.532175 | 0.601313007 | 0.092698298 | 6.486775056 | 8.77E-11 | 3.02E-09 | 0.601313007 |
| ENSMUSG00000031383 | Dusp9 | 75590 | protein_coding | 18.54019095 | -0.601616268 | 0.512528444 | -1.173820253 | 0.24046703 | 0.444953398 | 0.601616268 |
| ENSMUSG00000108084 | Gm7932 | 666105 | lncRNA | 17.27860361 | -0.601711994 | 0.286738587 | -2.098468853 | 0.035863749 | 0.123217814 | 0.601711994 |
| ENSMUSG00000074682 | Zcchc3 | 67917 | protein_coding | 22.69151676 | 0.601726649 | 0.269760031 | 2.230599719 | 0.025707655 | 0.09661905 | 0.601726649 |
| ENSMUSG00000009013 | Dynll1 | 56455 | protein_coding | 1677.487133 | 0.601728087 | 0.174403422 | 3.45020803 | 0.000560155 | 0.004773753 | 0.601728087 |
| ENSMUSG00000021303 | Gng4 | 14706 | protein_coding | 13.67233462 | -0.601919576 | 0.395227683 | -1.522969167 | 0.127766419 | 0.293439621 | 0.601919576 |
| ENSMUSG00000025044 | Msr1 | 20288 | protein_coding | 8.805582126 | -0.602068432 | 0.607052875 | -0.991789113 | 0.32130041 | 0.532354566 | 0.602068432 |
| ENSMUSG00000059654 | Reg1 | 19692 | protein_coding | 36.46972805 | 0.602130618 | 0.345183153 | 1.744380085 | 0.081092835 | 0.216066691 | 0.602130618 |
| ENSMUSG00000024673 | Ms4a1 | 12482 | protein_coding | 10.76847401 | 0.602168481 | 0.471990228 | 1.275807094 | 0.202023719 | 0.397018359 | 0.602168481 |
| ENSMUSG00000043263 | Ifi209 | 236312 | protein_coding | 11.26755986 | 0.60218267 | 0.508069809 | 1.185236083 | 0.235924117 | 0.440061526 | 0.60218267 |
| ENSMUSG00000032194 | Kank2 | 235041 | protein_coding | 263.4563566 | -0.602275543 | 0.28312754 | -2.127223456 | 0.033401521 | 0.116747735 | 0.602275543 |
| ENSMUSG00000096141 | Dnah7a | 627872 | protein_coding | 11.40962989 | -0.602657039 | 0.850401353 | -0.70867366 | 0.478527014 | 0.672843015 | 0.602657039 |
| ENSMUSG00000068373 | D430041D05Rik | 241589 | protein_coding | 7.932991528 | -0.602755734 | 1.858241568 | -0.324368879 | 0.745658778 | 0.858650449 | 0.602755734 |
| ENSMUSG00000017764 | Zswim1 | 71971 | protein_coding | 96.19538992 | 0.602808494 | 0.137131931 | 4.395828829 | 1.10E-05 | 0.000162147 | 0.602808494 |
| ENSMUSG00000034422 | Parp14 | 547253 | protein_coding | 870.929811 | 0.602878863 | 0.10754128 | 5.606022755 | 2.07E-08 | 5.34E-07 | 0.602878863 |
| ENSMUSG00000049044 | Rapgef4 | 56508 | protein_coding | 88.60670491 | -0.603038994 | 0.175250555 | -3.441010468 | 0.000579546 | 0.004910112 | 0.603038994 |
| ENSMUSG00000004187 | Kifc2 | 16581 | protein_coding | 119.3961739 | -0.603115671 | 0.235504976 | -2.560946611 | 0.01043874 | 0.049396389 | 0.603115671 |
| ENSMUSG00000048731 | Ggnbp1 | 70772 | protein_coding | 16.08045071 | -0.603116041 | 0.352724105 | -1.709880424 | 0.087287987 | 0.227399911 | 0.603116041 |
| ENSMUSG00000023019 | Gpd1 | 14555 | protein_coding | 456.271784 | -0.603381449 | 0.14713536 | -4.100859573 | 4.12E-05 | 0.000518906 | 0.603381449 |
| ENSMUSG00000026723 | Trdmt1 | 13434 | protein_coding | 65.99114521 | -0.603424654 | 0.162817812 | -3.706134149 | 0.000210447 | 0.00213503 | 0.603424654 |
| ENSMUSG00000032911 | Cspg4 | 121021 | protein_coding | 33.62230567 | -0.603499946 | 0.194093894 | -3.109319589 | 0.001875188 | 0.012818879 | 0.603499946 |
| ENSMUSG00000044033 | Ccdc141 | 545428 | protein_coding | 38.16789633 | -0.603554038 | 0.230570475 | -2.617655348 | 0.008853618 | 0.043754398 | 0.603554038 |
| ENSMUSG00000037321 | Tap1 | 21354 | protein_coding | 275.2791564 | 0.603836559 | 0.172893792 | 3.492528856 | 0.00047847 | 0.004192007 | 0.603836559 |
| ENSMUSG00000068015 | Lrch1 | 380916 | protein_coding | 378.8210897 | -0.603836886 | 0.134837548 | -4.47825471 | 7.53E-06 | 0.000116489 | 0.603836886 |
| ENSMUSG00000049357 | Brd8dc | 271508 | protein_coding | 11.05515855 | 0.604286475 | 0.473966505 | 1.274956075 | 0.20232479 | 0.397343604 | 0.604286475 |
| ENSMUSG00000042404 | Dennd4b | 229541 | protein_coding | 29.10445367 | -0.604348401 | 0.258034919 | -2.342118665 | 0.019174617 | 0.07759973 | 0.604348401 |
| ENSMUSG00000001249 | Hpn | 15451 | protein_coding | 74.14532299 | -0.604622026 | 0.263922152 | -2.290910488 | 0.021968591 | 0.085658683 | 0.604622026 |
| ENSMUSG00000063142 | Kcnma1 | 16531 | protein_coding | 264.5967855 | -0.604901607 | 0.132073793 | -4.580027512 | 4.65E-06 | 7.56E-05 | 0.604901607 |
| ENSMUSG00000087273 | Gm13203 | 115489366 | lncRNA | 7.736871888 | 0.604933346 | 0.405937371 | 1.490213487 | 0.136168111 | 0.306253272 | 0.604933346 |
| ENSMUSG00000086429 | Gt(ROSA)26Sor | 14910 | lncRNA | 88.01388217 | 0.605132719 | 0.197303033 | 3.067021875 | 0.00216203 | 0.01442682 | 0.605132719 |
| ENSMUSG00000061080 | Lsamp | 268890 | protein_coding | 10.21759831 | -0.605346177 | 1.793654419 | -0.337493204 | 0.735745134 | 0.852039298 | 0.605346177 |
| ENSMUSG00000022505 | Emp2 | 13731 | protein_coding | 522.1505456 | -0.605903964 | 0.138615811 | -4.371102823 | 1.24E-05 | 0.000179984 | 0.605903964 |

| ensembl_gene_id | external_gene_name | entrezgene_id | gene_biotype | baseMean | log2FoldChange | lfcSE | stat | pvalue | padj | absLog2FC |
|---|---|---|---|---|---|---|---|---|---|---|
| ENSMUSG00000021565 | Slc6a19 | 74338 | protein_coding | 33.33776516 | -0.606221766 | 0.295261563 | -2.053168585 | 0.04005623 | 0.133272807 | 0.606221766 |
| ENSMUSG00000034157 | Cipc | 217732 | protein_coding | 1293.852891 | -0.606436401 | 0.164387548 | -3.689065322 | 0.000225079 | 0.002250512 | 0.606436401 |
| ENSMUSG00000042743 | Sgtb | 218544 | protein_coding | 47.00078775 | -0.606556518 | 0.218248854 | -2.779196804 | 0.00544935 | 0.029858013 | 0.606556518 |
| ENSMUSG00000020808 | Pimreg | 109212 | protein_coding | 58.64757272 | 0.606591566 | 0.240229702 | 2.525048154 | 0.011568244 | 0.053439381 | 0.606591566 |
| ENSMUSG00000029049 | Morn1 | 76866 | protein_coding | 39.65575155 | -0.606803854 | 0.194533743 | -3.119273016 | 0.001812979 | 0.012463154 | 0.606803854 |
| ENSMUSG00000001175 | Calm1 | 12313 | protein_coding | 19104.66618 | 0.607211712 | 0.108946832 | 5.573468303 | 2.50E-08 | 6.36E-07 | 0.607211712 |
| ENSMUSG00000052751 | Repin1 | 58887 | protein_coding | 241.3734293 | -0.607275728 | 0.184816804 | -3.28582529 | 0.001016841 | 0.007767979 | 0.607275728 |
| ENSMUSG00000025040 | Fundc1 | 72018 | protein_coding | 353.8948297 | 0.607546098 | 0.111817921 | 5.433351749 | 5.53E-08 | 1.32E-06 | 0.607546098 |
| ENSMUSG00000066438 | Plekhd1 | 217682 | protein_coding | 69.47313279 | -0.607890535 | 0.224708657 | -2.705238609 | 0.006825534 | 0.035715213 | 0.607890535 |
| ENSMUSG00000043487 | Acot6 | 217700 | protein_coding | 24.78347991 | -0.608192754 | 0.260540592 | -2.334349322 | 0.019577438 | 0.078608426 | 0.608192754 |
| ENSMUSG00000031586 | Rbpms | 19663 | protein_coding | 43.53890467 | -0.609940809 | 0.242903344 | -2.511043273 | 0.012037494 | 0.055016701 | 0.609940809 |
| ENSMUSG00000033209 | Ttc28 | 209683 | protein_coding | 6.74356991 | -0.61017473 | 0.722850779 | -0.84412267 | 0.398600861 | 0.606506927 | 0.61017473 |
| ENSMUSG00000031951 | Tmem231 | 234740 | protein_coding | 62.09881818 | 0.610346046 | 0.223921338 | 2.725716324 | 0.006416212 | 0.033997811 | 0.610346046 |
| ENSMUSG00000073755 | 5730409E04Rik | 230757 | protein_coding | 10.01434873 | -0.610424441 | 0.450625888 | -1.354614677 | 0.175540355 | 0.361771253 | 0.610424441 |
| ENSMUSG00000030096 | Slc6a6 | 21366 | protein_coding | 2055.228569 | -0.610425735 | 0.147535453 | -4.13748507 | 3.51E-05 | 0.000454013 | 0.610425735 |
| ENSMUSG00000017830 | Dhx58 | 80861 | protein_coding | 106.4139718 | 0.610622828 | 0.186543751 | 3.273349144 | 0.001062811 | 0.00805635 | 0.610622828 |
| ENSMUSG00000025782 | Taf3 | 209361 | protein_coding | 96.6741198 | -0.610771728 | 0.163358263 | -3.738848092 | 0.000184865 | 0.001903385 | 0.610771728 |
| ENSMUSG00000098488 | Pla2g4b | 211429 | protein_coding | 8.59008127 | -0.611270044 | 0.461626899 | -1.3241647 | 0.185448352 | 0.37566086 | 0.611270044 |
| ENSMUSG00000022377 | Asap1 | 13196 | protein_coding | 291.4529738 | -0.611322211 | 0.203632889 | -3.002079944 | 0.002681417 | 0.017193888 | 0.611322211 |
| ENSMUSG00000053580 | Tanc2 | 77097 | protein_coding | 136.4383296 | -0.611560243 | 0.153808047 | -3.976126445 | 7.00E-05 | 0.00082645 | 0.611560243 |
| ENSMUSG00000028671 | Gale | 74246 | protein_coding | 3387.346788 | 0.611787271 | 0.127657944 | 4.792394834 | 1.65E-06 | 2.95E-05 | 0.611787271 |
| ENSMUSG00000047344 | Lancl3 | 236285 | protein_coding | 79.31776437 | -0.611802672 | 0.226036177 | -2.706658203 | 0.006796419 | 0.03557598 | 0.611802672 |
| ENSMUSG00000026639 | Lamb3 | 16780 | protein_coding | 2801.196145 | 0.612360978 | 0.170712594 | 3.587087292 | 0.000334392 | 0.003107469 | 0.612360978 |
| ENSMUSG00000028023 | Pitx2 | 18741 | protein_coding | 134.1890995 | -0.612497934 | 0.7595009 | -0.806447938 | 0.419984615 | 0.625948812 | 0.612497934 |
| ENSMUSG00000035041 | Creb3l3 | 208677 | protein_coding | 111.8058362 | 0.612640385 | 0.438910444 | 1.395820934 | 0.162768425 | 0.345262415 | 0.612640385 |
| ENSMUSG00000055116 | Arntl | 11865 | protein_coding | 883.4211167 | 0.612644761 | 0.107506618 | 5.69867022 | 1.21E-08 | 3.22E-07 | 0.612644761 |
| ENSMUSG00000005469 | Prkaca | 18747 | protein_coding | 791.7100511 | 0.612789983 | 0.077454371 | 7.911625612 | 2.54E-15 | 1.33E-13 | 0.612789983 |
| ENSMUSG00000023571 | C1qtnf12 | 67389 | protein_coding | 77.79896712 | -0.61281857 | 0.144062751 | -4.253830827 | 2.10E-05 | 0.000289721 | 0.61281857 |
| ENSMUSG00000001988 | Npas1 | 18142 | protein_coding | 7.272494434 | 0.612869004 | 0.48935995 | 1.25238897 | 0.210428164 | 0.407699767 | 0.612869004 |
| ENSMUSG00000025931 | Paqr8 | 74229 | protein_coding | 636.4751456 | 0.613117117 | 0.244704981 | 2.505535912 | 0.012226598 | 0.055603864 | 0.613117117 |
| ENSMUSG00000055818 | A230083G16Rik | 442825 | lncRNA | 11.03693763 | -0.613161518 | 0.412792756 | -1.485397962 | 0.137438435 | 0.308196452 | 0.613161518 |
| ENSMUSG00000029570 | Lfng | 16848 | protein_coding | 211.1099602 | -0.613387413 | 0.19244962 | -3.187262277 | 0.001436265 | 0.01030952 | 0.613387413 |
| ENSMUSG00000015085 | Entpd2 | 12496 | protein_coding | 91.52667741 | -0.613493852 | 0.132653891 | -4.624770891 | 3.75E-06 | 6.19E-05 | 0.613493852 |
| ENSMUSG00000030309 | Caprin2 | 232560 | protein_coding | 120.9258272 | -0.613674693 | 0.162048257 | -3.78698731 | 0.000152485 | 0.00161703 | 0.613674693 |
| ENSMUSG00000044921 | Rassf9 | 237504 | protein_coding | 142.5802324 | -0.613702077 | 0.292141994 | -2.100697915 | 0.035667492 | 0.122675841 | 0.613702077 |
| ENSMUSG00000070563 | Spaca4 | 69363 | protein_coding | 102.6316169 | -0.613762402 | 0.256104922 | -2.396527167 | 0.016551266 | 0.069595716 | 0.613762402 |
| ENSMUSG00000028073 | Pear1 | 73182 | protein_coding | 53.95562103 | -0.61379365 | 0.371235009 | -1.653382993 | 0.098252942 | 0.24537047 | 0.61379365 |
| ENSMUSG00000055833 | 1700034H15Rik | 98736 | lncRNA | 7.84180282 | -0.614079146 | 0.464697638 | -1.321459582 | 0.186348171 | 0.376888975 | 0.614079146 |

| ensembl_gene_id | external_gene_name | entrezgene_id | gene_biotype | baseMean | log2FoldChange | lfcSE | stat | pvalue | padj | absLog2FC |
|---|---|---|---|---|---|---|---|---|---|---|
| ENSMUSG00000041670 | Rims1 | 116837 | protein_coding | 10.13426447 | -0.614251159 | 0.640965729 | -0.958321375 | 0.337900729 | 0.548511589 | 0.614251159 |
| ENSMUSG00000054598 | 9130230L23Rik | 231253 | lncRNA | 59.02391587 | -0.614254857 | 0.21301236 | -2.883658284 | 0.00393085 | 0.023150247 | 0.614254857 |
| ENSMUSG00000034883 | Lrr1 | 69706 | protein_coding | 21.16343967 | 0.614303171 | 0.286206586 | 2.146362807 | 0.031844044 | 0.112839141 | 0.614303171 |
| ENSMUSG00000052727 | Map1b | 17755 | protein_coding | 62.9868901 | -0.614429267 | 0.21968101 | -2.796915704 | 0.005159299 | 0.028723285 | 0.614429267 |
| ENSMUSG00000030298 | Sec13 | 110379 | protein_coding | 1701.337905 | 0.614434184 | 0.11054499 | 5.558227325 | 2.73E-08 | 6.85E-07 | 0.614434184 |
| ENSMUSG00000021720 | Rnf180 | 71816 | protein_coding | 82.85243058 | -0.61446822 | 0.1986659 | -3.092972774 | 0.001981623 | 0.013414064 | 0.61446822 |
| ENSMUSG00000057156 | Homez | 239099 | protein_coding | 135.9864086 | 0.614517827 | 0.126428738 | 4.860586594 | 1.17E-06 | 2.17E-05 | 0.614517827 |
| ENSMUSG00000042320 | Prox2 | 73422 | protein_coding | 12.87085156 | 0.614628973 | 0.387229948 | 1.587245449 | 0.112457065 | 0.267988275 | 0.614628973 |
| ENSMUSG00000031740 | Mmp2 | 17390 | protein_coding | 11.30172582 | -0.614689874 | 0.311964212 | -1.970385863 | 0.048794165 | 0.153109937 | 0.614689874 |
| ENSMUSG00000042622 | Maff | 17133 | protein_coding | 296.6494859 | -0.61471435 | 0.216355175 | -2.841227853 | 0.004494019 | 0.025776282 | 0.61471435 |
| ENSMUSG00000038167 | Plekhg6 | 213522 | protein_coding | 222.5731972 | -0.614828739 | 0.18428595 | -3.336275719 | 0.000849089 | 0.00669851 | 0.614828739 |
| ENSMUSG00000010538 | Tsacc | 76927 | protein_coding | 19.79236497 | -0.614925847 | 0.263887067 | -2.330261402 | 0.019792339 | 0.07915076 | 0.614925847 |
| ENSMUSG00000041570 | Camsap2 | 67886 | protein_coding | 158.9993786 | -0.614996083 | 0.235454601 | -2.611951856 | 0.009002693 | 0.044230381 | 0.614996083 |
| ENSMUSG00000096918 | Gm16863 | 328646 | lncRNA | 10.09327017 | -0.615507944 | 0.457207912 | -1.346232049 | 0.178227692 | 0.365449367 | 0.615507944 |
| ENSMUSG00000037905 | Bri3bp | 76809 | protein_coding | 299.4262625 | -0.615740693 | 0.136920619 | -4.497063309 | 6.89E-06 | 0.00010759 | 0.615740693 |
| ENSMUSG00000087331 | 1810021B22Rik | 69120 | lncRNA | 22.77660429 | 0.615900633 | 0.309936757 | 1.987181642 | 0.046902271 | 0.148935283 | 0.615900633 |
| ENSMUSG00000086554 | 9530034E10Rik | 78615 | lncRNA | 9.411759714 | -0.615969156 | 0.444481103 | -1.385816296 | 0.165803005 | 0.349604218 | 0.615969156 |
| ENSMUSG00000038312 | Edem2 | 108687 | protein_coding | 866.6171973 | 0.616402479 | 0.091431568 | 6.741681182 | 1.57E-11 | 5.77E-10 | 0.616402479 |
| ENSMUSG00000029136 | Rbks | 71336 | protein_coding | 116.4647628 | 0.616590696 | 0.140332828 | 4.393773743 | 1.11E-05 | 0.000163386 | 0.616590696 |
| ENSMUSG00000095105 | Edaradd | 171211 | protein_coding | 23.63236104 | 0.616826046 | 0.306163643 | 2.01469397 | 0.043936724 | 0.14187672 | 0.616826046 |
| ENSMUSG00000070822 | Zscan18 | 232875 | protein_coding | 7.069394784 | 0.617458238 | 0.514695106 | 1.199658265 | 0.230272088 | 0.433458782 | 0.617458238 |
| ENSMUSG00000087026 | A230103J11Rik | 320466 | lncRNA | 13.19928878 | -0.617854404 | 0.350776402 | -1.761391019 | 0.078172244 | 0.21029007 | 0.617854404 |
| ENSMUSG00000058569 | Tmed9 | 67511 | protein_coding | 1746.39783 | 0.617895687 | 0.095956881 | 6.439305671 | 1.20E-10 | 4.06E-09 | 0.617895687 |
| ENSMUSG00000027009 | Itga4 | 16401 | protein_coding | 34.86948452 | 0.618015161 | 0.375075343 | 1.647709377 | 0.099412323 | 0.247335126 | 0.618015161 |
| ENSMUSG00000053128 | Rnf26 | 213211 | protein_coding | 45.14696216 | 0.618295746 | 0.168246365 | 3.674942665 | 0.000237903 | 0.002359475 | 0.618295746 |
| ENSMUSG00000020309 | Chac2 | 68044 | protein_coding | 205.9431697 | 0.618807828 | 0.118515233 | 5.221335783 | 1.78E-07 | 3.92E-06 | 0.618807828 |
| ENSMUSG00000042097 | Zfp239 | 22685 | protein_coding | 236.1218014 | 0.61882797 | 0.134799963 | 4.590713221 | 4.42E-06 | 7.23E-05 | 0.61882797 |
| ENSMUSG00000020423 | Btg2 | 12227 | protein_coding | 5867.240294 | -0.618977288 | 0.26797883 | -2.309799204 | 0.020899273 | 0.082621898 | 0.618977288 |
| ENSMUSG00000040502 | Marchf9 | 216438 | protein_coding | 51.46889711 | -0.619038293 | 0.187675419 | -3.298451646 | 0.000972196 | 0.007513412 | 0.619038293 |
| ENSMUSG00000027168 | Pax6 | 18508 | protein_coding | 20.68842064 | -0.619234615 | 0.482778251 | -1.28264812 | 0.199615384 | 0.39408451 | 0.619234615 |
| ENSMUSG00000039145 | Camk1d | 227541 | protein_coding | 1413.960178 | -0.619280143 | 0.099176267 | -6.244237302 | 4.26E-10 | 1.35E-08 | 0.619280143 |
| ENSMUSG00000028521 | Slc35d1 | 242585 | protein_coding | 479.3481989 | -0.619915098 | 0.221504217 | -2.798660481 | 0.005131506 | 0.028608552 | 0.619915098 |
| ENSMUSG00000036330 | Slc18a1 | 110877 | protein_coding | 69.6245064 | -0.620198713 | 0.155890907 | -3.978414926 | 6.94E-05 | 0.000820969 | 0.620198713 |
| ENSMUSG00000056899 | Immp2l | 93757 | protein_coding | 77.35293319 | -0.620592144 | 0.163715142 | -3.790682628 | 0.000150234 | 0.001597413 | 0.620592144 |
| ENSMUSG00000036832 | Lpar3 | 65086 | protein_coding | 7.455984669 | 0.620707202 | 0.517486921 | 1.199464522 | 0.230347372 | 0.433458782 | 0.620707202 |
| ENSMUSG00000087203 | Gm13986 | 329496 | lncRNA | 15.03187445 | -0.620740278 | 0.374123671 | -1.659184719 | 0.097078576 | 0.243621799 | 0.620740278 |
| ENSMUSG00000031875 | Cmtm3 | 68119 | protein_coding | 12.17494683 | -0.620799027 | 0.406245424 | -1.528137894 | 0.126478305 | 0.291287802 | 0.620799027 |
| ENSMUSG00000001288 | Rarg | 19411 | protein_coding | 123.2859369 | -0.621083421 | 0.12975098 | -4.786733954 | 1.70E-06 | 3.02E-05 | 0.621083421 |

| ensembl_gene_id | external_gene_name | entrezgene_id | gene_biotype | baseMean | log2FoldChange | lfcSE | stat | pvalue | padj | absLog2FC |
|---|---|---|---|---|---|---|---|---|---|---|
| ENSMUSG00000021999 | Cpb2 | 56373 | protein_coding | 7.566892674 | -0.621103055 | 0.416324629 | -1.491871995 | 0.135732706 | 0.305532863 | 0.621103055 |
| ENSMUSG00000032353 | Tmed3 | 66111 | protein_coding | 1840.838726 | 0.621565517 | 0.105899287 | 5.869402259 | 4.37E-09 | 1.23E-07 | 0.621565517 |
| ENSMUSG00000068617 | Efcab1 | 66793 | protein_coding | 8.661126691 | -0.621844115 | 0.455208223 | -1.366065207 | 0.171918498 | 0.357622806 | 0.621844115 |
| ENSMUSG00000019738 | Polr2i | 69920 | protein_coding | 317.5410888 | 0.622190353 | 0.11193984 | 5.558256585 | 2.72E-08 | 6.85E-07 | 0.622190353 |
| ENSMUSG00000020393 | Kremen1 | 84035 | protein_coding | 309.8144399 | -0.622752436 | 0.110751719 | -5.622959541 | 1.88E-08 | 4.88E-07 | 0.622752436 |
| ENSMUSG00000030787 | Lyve1 | 114332 | protein_coding | 33.81867118 | 0.622765956 | 0.221186523 | 2.815569174 | 0.004869091 | 0.027521252 | 0.622765956 |
| ENSMUSG00000037405 | Icam1 | 15894 | protein_coding | 22.56740408 | 0.622856755 | 0.238413771 | 2.612503259 | 0.008988184 | 0.044186354 | 0.622856755 |
| ENSMUSG00000038700 | Hoxb5 | 15413 | protein_coding | 49.44185753 | -0.623505663 | 0.291448673 | -2.139332652 | 0.032408737 | 0.114129198 | 0.623505663 |
| ENSMUSG00000058057 | Mettl7a3 | 668178 | protein_coding | 9.666684385 | 0.623721543 | 0.57688023 | 1.081197639 | 0.279609208 | 0.488002215 | 0.623721543 |
| ENSMUSG00000026786 | Apbb1ip | 54519 | protein_coding | 31.12077672 | 0.624167165 | 0.323295428 | 1.930640244 | 0.053527557 | 0.162524598 | 0.624167165 |
| ENSMUSG00000039084 | Chad | 12643 | protein_coding | 28.11007077 | -0.624180221 | 0.350559444 | -1.780526049 | 0.07498991 | 0.204247485 | 0.624180221 |
| ENSMUSG00000025867 | Cplx2 | 12890 | protein_coding | 71.55096795 | -0.624262001 | 0.207876845 | -3.00303769 | 0.002672993 | 0.017160595 | 0.624262001 |
| ENSMUSG00000029359 | Tesc | 57816 | protein_coding | 33.26057242 | -0.624464263 | 0.266296552 | -2.344995676 | 0.0190273 | 0.077155509 | 0.624464263 |
| ENSMUSG00000107455 | Gm43868 | 115490363 | TEC | 11.44748425 | -0.624668049 | 0.338710811 | -1.844251879 | 0.065146441 | 0.18579275 | 0.624668049 |
| ENSMUSG00000042351 | Grap2 | 17444 | protein_coding | 31.67213183 | 0.624885927 | 0.284934497 | 2.193086245 | 0.028301166 | 0.103239441 | 0.624885927 |
| ENSMUSG00000037679 | Inf2 | 70435 | protein_coding | 203.4144382 | -0.625063249 | 0.22773567 | -2.744687505 | 0.006056854 | 0.032468905 | 0.625063249 |
| ENSMUSG00000027889 | Ampd2 | 109674 | protein_coding | 524.7112406 | 0.625218084 | 0.129483324 | 4.828560647 | 1.38E-06 | 2.51E-05 | 0.625218084 |
| ENSMUSG00000096992 | Gm26788 | 108167617 | lncRNA | 25.2499964 | -0.6252485 | 0.291058835 | -2.148185953 | 0.031698986 | 0.112500323 | 0.6252485 |
| ENSMUSG00000031197 | Vbp1 | 22327 | protein_coding | 746.2692267 | 0.625271124 | 0.114722955 | 5.450270377 | 5.03E-08 | 1.20E-06 | 0.625271124 |
| ENSMUSG00000038587 | Akap12 | 83397 | protein_coding | 8.285282902 | -0.625619581 | 0.530694492 | -1.178869559 | 0.238450121 | 0.442609664 | 0.625619581 |
| ENSMUSG00000020812 | Snhg16 | 66293 | lncRNA | 79.5613364 | -0.625732908 | 0.199670153 | -3.133832968 | 0.00172539 | 0.011959103 | 0.625732908 |
| ENSMUSG00000059921 | Unc5c | 22253 | protein_coding | 7.790792341 | -0.625792817 | 1.8330007 | -0.34140348 | 0.732799856 | 0.850024456 | 0.625792817 |
| ENSMUSG00000045045 | Lrfn4 | 225875 | protein_coding | 257.5498704 | -0.625800109 | 0.242210827 | -2.583699983 | 0.009774678 | 0.047107737 | 0.625800109 |
| ENSMUSG00000016356 | Col20a1 | 73368 | protein_coding | 49.79069145 | -0.626187333 | 0.273809575 | -2.286944617 | 0.022199056 | 0.086345925 | 0.626187333 |
| ENSMUSG00000034974 | Dapk3 | 13144 | protein_coding | 415.2559109 | 0.626213598 | 0.156789227 | 3.993983589 | 6.50E-05 | 0.000772875 | 0.626213598 |
| ENSMUSG00000045752 | Tssc4 | 56844 | protein_coding | 250.9628529 | 0.626417063 | 0.132994159 | 4.710109597 | 2.48E-06 | 4.25E-05 | 0.626417063 |
| ENSMUSG00000055188 | Rbm3os | 70226 | lncRNA | 16.31504426 | -0.626479011 | 0.314056588 | -1.99479659 | 0.046065069 | 0.147009502 | 0.626479011 |
| ENSMUSG00000041797 | Abca9 | 217262 | protein_coding | 8.063867638 | -0.626498074 | 0.89041989 | -0.703598472 | 0.481682863 | 0.674898368 | 0.626498074 |
| ENSMUSG00000042616 | Oscp1 | 230751 | protein_coding | 29.30819385 | 0.626508889 | 0.235300915 | 2.662585863 | 0.007754278 | 0.039510603 | 0.626508889 |
| ENSMUSG00000040274 | Cdk6 | 12571 | protein_coding | 726.9749908 | -0.626582493 | 0.137382178 | -4.56087174 | 5.09E-06 | 8.22E-05 | 0.626582493 |
| ENSMUSG00000024785 | Rcl1 | 59028 | protein_coding | 214.9236246 | 0.626799727 | 0.25842839 | 2.425429061 | 0.015290304 | 0.065534172 | 0.626799727 |
| ENSMUSG00000028838 | Extl1 | 56219 | protein_coding | 17.53467784 | -0.62696994 | 0.322855923 | -1.941949625 | 0.052143199 | 0.159965985 | 0.62696994 |
| ENSMUSG00000043501 | Lgals2 | 107753 | protein_coding | 122.0250035 | 0.627284036 | 0.276457926 | 2.269003628 | 0.023268105 | 0.08954201 | 0.627284036 |
| ENSMUSG00000006411 | Nectin4 | 71740 | protein_coding | 48.43822029 | -0.627503165 | 0.254207583 | -2.46846753 | 0.013569296 | 0.06014684 | 0.627503165 |
| ENSMUSG00000040229 | Gpr34 | 23890 | protein_coding | 10.38163268 | 0.627600108 | 0.417976148 | 1.501521345 | 0.13322077 | 0.301797813 | 0.627600108 |
| ENSMUSG00000056445 | Hoxaas2 | 105242657 | lncRNA | 16.10365298 | -0.627701987 | 0.352267119 | -1.781892076 | 0.074766833 | 0.204163572 | 0.627701987 |
| ENSMUSG00000020953 | Coch | 12810 | protein_coding | 10.84507006 | -0.627839556 | 0.477948604 | -1.313613118 | 0.188976432 | 0.380319097 | 0.627839556 |
| ENSMUSG00000026686 | Lmx1a | 110648 | protein_coding | 27.70458974 | -0.627860388 | 0.266229612 | -2.358341669 | 0.018356789 | 0.075009474 | 0.627860388 |

| ensembl_gene_id | external_gene_name | entrezgene_id | gene_biotype | baseMean | log2FoldChange | lfcSE | stat | pvalue | padj | absLog2FC |
|---|---|---|---|---|---|---|---|---|---|---|
| ENSMUSG00000033249 | Hsf4 | 26386 | protein_coding | 8.070303214 | -0.62796322 | 0.442723672 | -1.418408951 | 0.156071404 | 0.336433255 | 0.62796322 |
| ENSMUSG00000030428 | Ttyh1 | 57776 | protein_coding | 79.28245965 | -0.627975213 | 0.281352442 | -2.231987783 | 0.02561577 | 0.09634191 | 0.627975213 |
| ENSMUSG00000058260 | Serpina9 | 71907 | protein_coding | 31.40630031 | -0.628016137 | 0.526740912 | -1.192267626 | 0.233156324 | 0.436753373 | 0.628016137 |
| ENSMUSG00000040998 | Npnt | 114249 | protein_coding | 7.65793556 | -0.628300362 | 0.722029966 | -0.870185991 | 0.38419877 | 0.591944091 | 0.628300362 |
| ENSMUSG00000003847 | Nfat5 | 54446 | protein_coding | 441.5289123 | -0.628421166 | 0.224218762 | -2.802714456 | 0.005067451 | 0.02832083 | 0.628421166 |
| ENSMUSG00000047604 | Frat2 | 212398 | protein_coding | 361.7829828 | -0.6285239 | 0.169857847 | -3.700293575 | 0.00021535 | 0.002172323 | 0.6285239 |
| ENSMUSG00000086040 | Wipf3 | 330319 | protein_coding | 15.59762034 | -0.628846912 | 0.356076743 | -1.766043202 | 0.077388601 | 0.208958406 | 0.628846912 |
| ENSMUSG00000091811 | Inafm1 | 66300 | protein_coding | 238.6188808 | -0.62910491 | 0.126727032 | -4.964251897 | 6.90E-07 | 1.35E-05 | 0.62910491 |
| ENSMUSG00000029369 | Afm | 280662 | protein_coding | 52.78167217 | 0.629599265 | 0.179605404 | 3.505458362 | 0.000455822 | 0.00402904 | 0.629599265 |
| ENSMUSG00000022225 | Cma1 | 17228 | protein_coding | 8.707957996 | 0.629683883 | 0.863155643 | 0.729513719 | 0.465687479 | 0.662510733 | 0.629683883 |
| ENSMUSG00000090698 | Apold1 | 381823 | protein_coding | 9.616238733 | -0.629773645 | 0.503307607 | -1.251269873 | 0.210836033 | 0.408241713 | 0.629773645 |
| ENSMUSG00000023919 | Cenpq | 83815 | protein_coding | 152.1533487 | 0.630102798 | 0.12771093 | 4.933820439 | 8.06E-07 | 1.55E-05 | 0.630102798 |
| ENSMUSG00000074183 | Gsta1 | 14857 | protein_coding | 129.587945 | 0.630156093 | 0.239446278 | 2.631722233 | 0.008495329 | 0.042364933 | 0.630156093 |
| ENSMUSG00000085315 | A430018G15Rik | 115489452 | lncRNA | 10.99375853 | -0.63024895 | 0.381743634 | -1.650974354 | 0.098743812 | 0.246287415 | 0.63024895 |
| ENSMUSG00000033594 | Spata2l | 78779 | protein_coding | 91.70859017 | -0.630322414 | 0.174944874 | -3.602977324 | 0.000314593 | 0.002954503 | 0.630322414 |
| ENSMUSG00000021589 | Rhobtb3 | 73296 | protein_coding | 210.0250871 | 0.63075459 | 0.181738576 | 3.470669821 | 0.000519162 | 0.004491696 | 0.63075459 |
| ENSMUSG00000097743 | Gm16973 | 100503823 | lncRNA | 39.81044541 | -0.630896999 | 0.214723864 | -2.93817831 | 0.003301471 | 0.020287742 | 0.630896999 |
| ENSMUSG00000045725 | Prr15 | 78004 | protein_coding | 480.6269262 | 0.631480891 | 0.206645209 | 3.055869986 | 0.002244085 | 0.014887044 | 0.631480891 |
| ENSMUSG00000050675 | Gp1ba | 14723 | protein_coding | 12.5295343 | -0.631619837 | 0.483902007 | -1.305263934 | 0.19180299 | 0.38442658 | 0.631619837 |
| ENSMUSG00000033697 | Arhgap39 | 223666 | protein_coding | 587.6090908 | 0.631842604 | 0.145797396 | 4.333702936 | 1.47E-05 | 0.00020964 | 0.631842604 |
| ENSMUSG00000101581 | C430002N11Rik | 319707 | lncRNA | 8.979416443 | -0.632021381 | 0.573074388 | -1.102860981 | 0.270087541 | 0.478260629 | 0.632021381 |
| ENSMUSG00000029408 | Abcb9 | 56325 | protein_coding | 192.7097652 | 0.632449506 | 0.146795619 | 4.308367711 | 1.64E-05 | 0.000232645 | 0.632449506 |
| ENSMUSG00000029417 | Cxcl9 | 17329 | protein_coding | 7.453513191 | 0.632519194 | 0.489868187 | 1.291202838 | 0.196633351 | 0.390214607 | 0.632519194 |
| ENSMUSG00000042156 | Dzip1 | 66573 | protein_coding | 11.21936003 | 0.632572692 | 0.448972424 | 1.408934399 | 0.158854567 | 0.340131138 | 0.632572692 |
| ENSMUSG00000028456 | Unc13b | 22249 | protein_coding | 1011.190723 | 0.632866243 | 0.0694538 | 9.112046303 | 8.08E-20 | 5.65E-18 | 0.632866243 |
| ENSMUSG00000026955 | Sapcd2 | 72080 | protein_coding | 116.5324929 | 0.63300087 | 0.16418177 | 3.855488145 | 0.000115499 | 0.001273663 | 0.63300087 |
| ENSMUSG00000030156 | Cd69 | 12515 | protein_coding | 30.70332318 | 0.633636104 | 0.334228565 | 1.89581673 | 0.057984283 | 0.172019679 | 0.633636104 |
| ENSMUSG00000070369 | Itgad | 381924 | protein_coding | 8.177874058 | 0.633800947 | 0.61717159 | 1.026944464 | 0.304446616 | 0.513961388 | 0.633800947 |
| ENSMUSG00000026579 | F5 | 14067 | protein_coding | 14.16596445 | -0.634064637 | 0.419585836 | -1.511167875 | 0.130745685 | 0.297885463 | 0.634064637 |
| ENSMUSG00000041132 | N4bp2l1 | 100637 | protein_coding | 26.1730758 | -0.634172832 | 0.310065361 | -2.045287582 | 0.040826514 | 0.13505394 | 0.634172832 |
| ENSMUSG00000065701 | Rny1 | 19872 | misc_RNA | 23.08102522 | 0.634291794 | 0.355043843 | 1.786516812 | 0.074015618 | 0.202877432 | 0.634291794 |
| ENSMUSG00000072647 | Adam1a | 280668 | protein_coding | 10.15866291 | -0.634606775 | 0.386091801 | -1.643668095 | 0.10024478 | 0.248474767 | 0.634606775 |
| ENSMUSG00000079355 | Ackr4 | 252837 | protein_coding | 41.86477121 | -0.634648357 | 0.25395724 | -2.499036283 | 0.012453156 | 0.056405341 | 0.634648357 |
| ENSMUSG00000086859 | Snhg20 | 76972 | lncRNA | 31.97481199 | -0.634882462 | 0.251673997 | -2.522638296 | 0.011647814 | 0.053666873 | 0.634882462 |
| ENSMUSG00000050103 | Agmo | 319660 | protein_coding | 122.7530194 | -0.635096176 | 0.117475907 | -5.406182338 | 6.44E-08 | 1.52E-06 | 0.635096176 |
| ENSMUSG00000043760 | Pkhd1 | 241035 | protein_coding | 180.1763031 | 0.635609676 | 0.247611474 | 2.566963745 | 0.010259334 | 0.04872113 | 0.635609676 |
| ENSMUSG00000092008 | Cyp2c69 | 100043108 | protein_coding | 76.88356017 | -0.635754841 | 0.302350487 | -2.102708174 | 0.035491287 | 0.12230749 | 0.635754841 |
| ENSMUSG00000089809 | Rasgef1b | 320292 | protein_coding | 548.1517541 | -0.636196761 | 0.226159203 | -2.813048292 | 0.004907428 | 0.027659055 | 0.636196761 |

| ensembl_gene_id | external_gene_name | entrezgene_id | gene_biotype | baseMean | log2FoldChange | lfcSE | stat | pvalue | padj | absLog2FC |
|---|---|---|---|---|---|---|---|---|---|---|
| ENSMUSG00000049154 | Fam183b | 75429 | protein_coding | 9.853675743 | -0.636218132 | 0.38817638 | -1.638992389 | 0.101214845 | 0.250175504 | 0.636218132 |
| ENSMUSG00000008461 | Fut1 | 14343 | protein_coding | 38.98973827 | 0.636427756 | 0.232345021 | 2.739149538 | 0.006159835 | 0.032847006 | 0.636427756 |
| ENSMUSG00000098702 | 1500015A07Rik | 68982 | lncRNA | 29.74968973 | -0.636642403 | 0.214551422 | -2.967318501 | 0.003004096 | 0.018793888 | 0.636642403 |
| ENSMUSG00000072235 | Tuba1a | 22142 | protein_coding | 91.09146036 | -0.636754327 | 0.148689483 | -4.282443599 | 1.85E-05 | 0.000258728 | 0.636754327 |
| ENSMUSG00000064043 | Trerf1 | 224829 | protein_coding | 10.53199279 | -0.637269823 | 0.691106461 | -0.922100804 | 0.356475998 | 0.565983821 | 0.637269823 |
| ENSMUSG00000028654 | Mycl | 16918 | protein_coding | 201.3566755 | -0.637499017 | 0.102970455 | -6.19108675 | 5.98E-10 | 1.84E-08 | 0.637499017 |
| ENSMUSG00000005413 | Hmox1 | 15368 | protein_coding | 38.04387901 | 0.637992271 | 0.297093446 | 2.147446468 | 0.031757755 | 0.112683786 | 0.637992271 |
| ENSMUSG00000023050 | Map3k12 | 26404 | protein_coding | 44.90366669 | -0.638028207 | 0.302985996 | -2.105800976 | 0.035221644 | 0.121588718 | 0.638028207 |
| ENSMUSG00000040584 | Abcb1a | 18671 | protein_coding | 5559.374545 | -0.638527044 | 0.168980217 | -3.77870887 | 0.000157644 | 0.001663981 | 0.638527044 |
| ENSMUSG00000021286 | Zfyve21 | 68520 | protein_coding | 345.5417535 | 0.639054235 | 0.119515095 | 5.347058736 | 8.94E-08 | 2.06E-06 | 0.639054235 |
| ENSMUSG00000023007 | Prpf40b | 54614 | protein_coding | 112.4930305 | -0.639150346 | 0.149408804 | -4.277862687 | 1.89E-05 | 0.000263877 | 0.639150346 |
| ENSMUSG00000003813 | Rad23a | 19358 | protein_coding | 342.0276949 | -0.639627399 | 0.075633888 | -8.456889033 | 2.75E-17 | 1.65E-15 | 0.639627399 |
| ENSMUSG00000020576 | Nbas | 71169 | protein_coding | 1153.949565 | 0.639680828 | 0.063479441 | 10.07697633 | 6.98E-24 | 5.95E-22 | 0.639680828 |
| ENSMUSG00000033854 | Kcnk10 | 72258 | protein_coding | 23.623118 | -0.639852589 | 0.358233164 | -1.786134432 | 0.074077494 | 0.202942265 | 0.639852589 |
| ENSMUSG00000049295 | Zfp219 | 69890 | protein_coding | 544.9672999 | 0.639880178 | 0.112011024 | 5.712653569 | 1.11E-08 | 2.98E-07 | 0.639880178 |
| ENSMUSG00000092083 | Kcnb2 | 98741 | protein_coding | 19.0306119 | -0.640015347 | 0.420798179 | -1.520955601 | 0.12827098 | 0.294267631 | 0.640015347 |
| ENSMUSG00000035954 | Dock4 | 238130 | protein_coding | 50.54386278 | -0.640385738 | 0.365380796 | -1.752652974 | 0.079661592 | 0.213270278 | 0.640385738 |
| ENSMUSG00000029059 | Prxl2b | 66469 | protein_coding | 112.8275446 | -0.640672943 | 0.120262572 | -5.327284571 | 9.97E-08 | 2.28E-06 | 0.640672943 |
| ENSMUSG00000038473 | Nos1ap | 70729 | protein_coding | 75.43537182 | -0.640678639 | 0.269797081 | -2.374668538 | 0.017564715 | 0.072634077 | 0.640678639 |
| ENSMUSG00000036473 | Tbc1d24 | 224617 | protein_coding | 310.5356574 | -0.640809285 | 0.21951145 | -2.919252205 | 0.003508722 | 0.021265954 | 0.640809285 |
| ENSMUSG00000029442 | Wdr66 | 269701 | protein_coding | 12.95472653 | -0.640932838 | 0.946010544 | -0.677511305 | 0.4980816 | 0.688846377 | 0.640932838 |
| ENSMUSG00000006057 | Atp5g1 | 11951 | protein_coding | 3522.602413 | 0.641174963 | 0.107445639 | 5.967435888 | 2.41E-09 | 6.99E-08 | 0.641174963 |
| ENSMUSG00000090272 | Mndal | 100040462 | protein_coding | 14.46015104 | 0.64161256 | 0.458546682 | 1.399230625 | 0.161743836 | 0.343959388 | 0.64161256 |
| ENSMUSG00000097032 | 4930539J05Rik | 319587 | lncRNA | 14.72101922 | -0.641693017 | 0.345598033 | -1.856761198 | 0.063345136 | 0.182399926 | 0.641693017 |
| ENSMUSG00000015568 | Lpl | 16956 | protein_coding | 345.2612185 | -0.641910847 | 0.181781399 | -3.531224052 | 0.000413641 | 0.003734964 | 0.641910847 |
| ENSMUSG00000085385 | Snhg17 | 68108 | lncRNA | 148.2735592 | -0.641944848 | 0.197378193 | -3.252359537 | 0.001144511 | 0.008557504 | 0.641944848 |
| ENSMUSG00000044927 | H1f10 | 243529 | protein_coding | 6.756201417 | 0.641964036 | 0.572887256 | 1.120576569 | 0.262468143 | 0.470204992 | 0.641964036 |
| ENSMUSG00000095061 | E030018B13Rik | 381994 | protein_coding | 11.8142667 | -0.642689765 | 0.45922092 | -1.399521965 | 0.161656516 | 0.343819601 | 0.642689765 |
| ENSMUSG00000001039 | B9d1 | 27078 | protein_coding | 48.49691179 | 0.642782599 | 0.195400837 | 3.289559088 | 0.001003445 | 0.007698787 | 0.642782599 |
| ENSMUSG00000023826 | Prkn | 50873 | protein_coding | 21.4322388 | -0.64307194 | 0.281022463 | -2.28832931 | 0.022118351 | 0.086126011 | 0.64307194 |
| ENSMUSG00000047649 | Cd3eap | 70333 | protein_coding | 106.6307994 | -0.643919482 | 0.248720461 | -2.588928467 | 0.009627509 | 0.046567557 | 0.643919482 |
| ENSMUSG00000036931 | Nfkbid | 243910 | protein_coding | 43.97393103 | -0.644118433 | 0.207592382 | -3.102803803 | 0.001916967 | 0.013070825 | 0.644118433 |
| ENSMUSG00000037852 | Cpe | 12876 | protein_coding | 104.9372905 | -0.644187435 | 0.143421767 | -4.491559737 | 7.07E-06 | 0.000110085 | 0.644187435 |
| ENSMUSG00000108171 | Gm43915 | 215458 | lncRNA | 10.26459759 | -0.644249345 | 0.532202 | -1.21053537 | 0.226073527 | 0.428372488 | 0.644249345 |
| ENSMUSG00000044966 | Fbxo48 | 319701 | protein_coding | 12.52021179 | -0.644271901 | 0.367251168 | -1.75430865 | 0.079377634 | 0.212813829 | 0.644271901 |
| ENSMUSG00000000197 | Nalcn | 338370 | protein_coding | 8.723947673 | -0.644493357 | 2.059800019 | -0.312891228 | 0.754363302 | 0.865108984 | 0.644493357 |
| ENSMUSG00000037936 | Scarb1 | 20778 | protein_coding | 199.5501096 | -0.64465238 | 0.127579021 | -5.052965425 | 4.35E-07 | 9.01E-06 | 0.64465238 |
| ENSMUSG00000038387 | Rras | 20130 | protein_coding | 48.46576655 | -0.644787919 | 0.186499105 | -3.457324468 | 0.000545568 | 0.004679483 | 0.644787919 |

| ensembl_gene_id | external_gene_name | entrezgene_id | gene_biotype | baseMean | log2FoldChange | lfcSE | stat | pvalue | padj | absLog2FC |
|---|---|---|---|---|---|---|---|---|---|---|
| ENSMUSG00000026205 | Slc23a3 | 22626 | protein_coding | 97.18230733 | 0.644902603 | 0.181793341 | 3.547448988 | 0.000388981 | 0.003556654 | 0.644902603 |
| ENSMUSG00000110622 | Iqcn | 637079 | protein_coding | 19.13872604 | -0.645162761 | 0.42261844 | -1.526584502 | 0.126864363 | 0.292092451 | 0.645162761 |
| ENSMUSG00000015806 | Qdpr | 110391 | protein_coding | 1113.369054 | 0.645185201 | 0.124741891 | 5.172161481 | 2.31E-07 | 5.01E-06 | 0.645185201 |
| ENSMUSG00000002732 | Fkbp7 | 14231 | protein_coding | 40.34786297 | 0.645215297 | 0.177750687 | 3.629889191 | 0.000283543 | 0.002705975 | 0.645215297 |
| ENSMUSG00000024558 | Mapk4 | 225724 | protein_coding | 6.999358459 | -0.645306329 | 0.562976647 | -1.146239959 | 0.251695878 | 0.458172794 | 0.645306329 |
| ENSMUSG00000031639 | Tlr3 | 142980 | protein_coding | 493.607697 | 0.645436199 | 0.119715377 | 5.39142267 | 6.99E-08 | 1.63E-06 | 0.645436199 |
| ENSMUSG00000091076 | Vmn2r115 | 638102 | protein_coding | 10.25168142 | -0.64566732 | 1.602979402 | -0.402792025 | 0.687101228 | 0.821322799 | 0.64566732 |
| ENSMUSG00000019762 | Iyd | 70337 | protein_coding | 74.61935998 | -0.645972644 | 0.191015836 | -3.38177534 | 0.00072019 | 0.005876634 | 0.645972644 |
| ENSMUSG00000043445 | Pgp | 67078 | protein_coding | 1182.177204 | 0.645991135 | 0.191058091 | 3.381124203 | 0.000721899 | 0.005884549 | 0.645991135 |
| ENSMUSG00000025153 | Fasn | 14104 | protein_coding | 4982.931445 | 0.646050658 | 0.1324835 | 4.876461291 | 1.08E-06 | 2.02E-05 | 0.646050658 |
| ENSMUSG00000100017 | 2410022M11Rik | 100503842 | lncRNA | 32.27754371 | 0.646088015 | 0.280324856 | 2.304783183 | 0.021178719 | 0.083408248 | 0.646088015 |
| ENSMUSG00000021179 | Nrde2 | 217827 | protein_coding | 312.29646 | 0.64640724 | 0.134599116 | 4.802462724 | 1.57E-06 | 2.82E-05 | 0.64640724 |
| ENSMUSG00000020600 | Slc7a15 | 328059 | protein_coding | 10.96077335 | 0.646408672 | 0.444811802 | 1.453218349 | 0.146163139 | 0.32213733 | 0.646408672 |
| ENSMUSG00000030270 | Cpne9 | 211232 | protein_coding | 9.498768394 | -0.646865435 | 0.947772054 | -0.682511615 | 0.494915501 | 0.686593061 | 0.646865435 |
| ENSMUSG00000096727 | Psmb9 | 16912 | protein_coding | 178.5123956 | 0.646872681 | 0.152467324 | 4.242697139 | 2.21E-05 | 0.000301948 | 0.646872681 |
| ENSMUSG00000056671 | Prelid2 | 77619 | protein_coding | 503.1482389 | -0.647314861 | 0.12020597 | -5.385047511 | 7.24E-08 | 1.69E-06 | 0.647314861 |
| ENSMUSG00000053297 | AI854703 | 243373 | protein_coding | 236.2700085 | -0.647552809 | 0.580309039 | -1.115875793 | 0.264475305 | 0.472048706 | 0.647552809 |
| ENSMUSG00000023927 | Satb1 | 20230 | protein_coding | 14.55537211 | 0.64757958 | 0.501805559 | 1.290499016 | 0.196877453 | 0.390518565 | 0.64757958 |
| ENSMUSG00000020682 | Mmp28 | 118453 | protein_coding | 206.8459934 | -0.647904605 | 0.279466878 | -2.318359195 | 0.020429808 | 0.081189116 | 0.647904605 |
| ENSMUSG00000031129 | Slc9a9 | 331004 | protein_coding | 34.17321012 | -0.648067448 | 0.211436594 | -3.065067568 | 0.002176208 | 0.014515346 | 0.648067448 |
| ENSMUSG00000060224 | Pyroxd2 | 74580 | protein_coding | 149.4045033 | -0.64828219 | 0.158053128 | -4.101672626 | 4.10E-05 | 0.000517691 | 0.64828219 |
| ENSMUSG00000029335 | Bmp3 | 110075 | protein_coding | 969.4404804 | -0.648371463 | 0.173860269 | -3.729267568 | 0.000192037 | 0.001970856 | 0.648371463 |
| ENSMUSG00000025268 | Maged2 | 80884 | protein_coding | 321.4616211 | 0.648638326 | 0.128919215 | 5.031354927 | 4.87E-07 | 9.93E-06 | 0.648638326 |
| ENSMUSG00000005763 | Cd247 | 12503 | protein_coding | 10.19220977 | 0.648822721 | 0.373107015 | 1.738972187 | 0.082039655 | 0.21796958 | 0.648822721 |
| ENSMUSG00000009418 | Nav1 | 215690 | protein_coding | 79.08126018 | -0.648875194 | 0.222551232 | -2.915621664 | 0.003549807 | 0.021457807 | 0.648875194 |
| ENSMUSG00000017765 | Slc12a4 | 20498 | protein_coding | 8.705340371 | -0.649504487 | 0.551877807 | -1.176899087 | 0.239235786 | 0.443424073 | 0.649504487 |
| ENSMUSG00000052889 | Prkcb | 18751 | protein_coding | 10.7578936 | 0.649848447 | 0.498345405 | 1.304012117 | 0.192229451 | 0.385135936 | 0.649848447 |
| ENSMUSG00000052928 | Ctif | 269037 | protein_coding | 133.964007 | 0.649891054 | 0.133403742 | 4.871610362 | 1.11E-06 | 2.07E-05 | 0.649891054 |
| ENSMUSG00000016664 | Pacsin2 | 23970 | protein_coding | 2543.466619 | 0.651080598 | 0.133848184 | 4.864321483 | 1.15E-06 | 2.14E-05 | 0.651080598 |
| ENSMUSG00000049521 | Cdc42ep1 | 104445 | protein_coding | 233.8835341 | -0.651290422 | 0.158344276 | -4.113128926 | 3.90E-05 | 0.000496979 | 0.651290422 |
| ENSMUSG00000027293 | Ehd4 | 98878 | protein_coding | 1094.09714 | 0.651550032 | 0.237769297 | 2.740261421 | 0.006139033 | 0.032812927 | 0.651550032 |
| ENSMUSG00000090290 | Tarbp1 | 212728 | protein_coding | 224.7397052 | -0.651585243 | 0.118666589 | -5.490890468 | 4.00E-08 | 9.68E-07 | 0.651585243 |
| ENSMUSG00000044948 | Cfap43 | 100048534 | protein_coding | 16.90142763 | -0.651669925 | 0.659394107 | -0.988285939 | 0.32301263 | 0.533929552 | 0.651669925 |
| ENSMUSG00000097787 | 2700046G09Rik | 67188 | lncRNA | 17.67372485 | -0.65170605 | 0.298727316 | -2.181608492 | 0.02913844 | 0.10550513 | 0.65170605 |
| ENSMUSG00000056870 | Gulp1 | 70676 | protein_coding | 125.833768 | 0.651881204 | 0.257720928 | 2.52940733 | 0.011425534 | 0.052872141 | 0.651881204 |
| ENSMUSG00000078606 | Gvin2 | 100042856 | protein_coding | 6.92486126 | 0.652170639 | 0.493463032 | 1.321620053 | 0.186294702 | 0.376888975 | 0.652170639 |
| ENSMUSG00000037509 | Arhgef4 | 226970 | protein_coding | 23.4628109 | 0.652227772 | 0.276327444 | 2.360343816 | 0.018258005 | 0.074663286 | 0.652227772 |
| ENSMUSG00000055067 | Smyd3 | 69726 | protein_coding | 134.3517054 | -0.652552614 | 0.13529053 | -4.823342896 | 1.41E-06 | 2.57E-05 | 0.652552614 |

| ensembl_gene_id | external_gene_name | entrezgene_id | gene_biotype | baseMean | log2FoldChange | lfcSE | stat | pvalue | padj | absLog2FC |
|---|---|---|---|---|---|---|---|---|---|---|
| ENSMUSG00000053141 | Ptprt | 19281 | protein_coding | 137.4270484 | -0.652623882 | 0.158908805 | -4.106908243 | 4.01E-05 | 0.000508922 | 0.652623882 |
| ENSMUSG00000073433 | Arhgdig | 14570 | protein_coding | 14.94614566 | -0.652924689 | 0.346679206 | -1.883368477 | 0.059650449 | 0.175360554 | 0.652924689 |
| ENSMUSG00000024354 | Slc23a1 | 20522 | protein_coding | 14.20881529 | -0.653123419 | 0.433031553 | -1.508258264 | 0.13148844 | 0.299149819 | 0.653123419 |
| ENSMUSG00000054630 | Ugt2b5 | 22238 | protein_coding | 816.8759702 | 0.653269447 | 0.254321808 | 2.56867255 | 0.010208887 | 0.048553944 | 0.653269447 |
| ENSMUSG00000034035 | Ccdc17 | 622665 | protein_coding | 49.02309528 | -0.653380818 | 0.198523072 | -3.291208484 | 0.00099758 | 0.007661257 | 0.653380818 |
| ENSMUSG00000038042 | Ptpdc1 | 218232 | protein_coding | 9.284047093 | -0.653462408 | 0.400911547 | -1.629941599 | 0.103113841 | 0.253408395 | 0.653462408 |
| ENSMUSG00000021055 | Esr2 | 13983 | protein_coding | 18.00763123 | -0.653569837 | 0.485432668 | -1.346365582 | 0.178184645 | 0.36540814 | 0.653569837 |
| ENSMUSG00000079299 | Klrb1 | 100043861 | protein_coding | 7.863071162 | 0.653612007 | 0.463358407 | 1.410597059 | 0.158363459 | 0.339625513 | 0.653612007 |
| ENSMUSG00000040037 | Negr1 | 320840 | protein_coding | 9.367254464 | -0.653911226 | 0.805263194 | -0.812046584 | 0.416764902 | 0.622957146 | 0.653911226 |
| ENSMUSG00000004707 | Ly9 | 17085 | protein_coding | 8.370012993 | 0.653997829 | 0.553201218 | 1.182206054 | 0.237123951 | 0.441445333 | 0.653997829 |
| ENSMUSG00000027544 | Nfatc2 | 18019 | protein_coding | 74.33512555 | -0.654160958 | 0.299540075 | -2.183884603 | 0.028970729 | 0.10516084 | 0.654160958 |
| ENSMUSG00000016087 | Fli1 | 14247 | protein_coding | 12.93359818 | 0.65427911 | 0.458068853 | 1.428342278 | 0.153193365 | 0.332082057 | 0.65427911 |
| ENSMUSG00000038011 | Dnah10 | 56087 | protein_coding | 28.39634735 | 0.654335171 | 0.519010241 | 1.260736533 | 0.207403787 | 0.404002387 | 0.654335171 |
| ENSMUSG00000039349 | C130074G19Rik | 226777 | protein_coding | 286.7144215 | -0.654615937 | 0.195819319 | -3.342958904 | 0.000828902 | 0.006575073 | 0.654615937 |
| ENSMUSG00000032470 | Mras | 17532 | protein_coding | 19.08868386 | -0.654704838 | 0.310767277 | -2.106736736 | 0.035140406 | 0.121360882 | 0.654704838 |
| ENSMUSG00000011171 | Vipr2 | 22355 | protein_coding | 69.83816545 | -0.655163285 | 0.213591318 | -3.067368527 | 0.002159524 | 0.014416137 | 0.655163285 |
| ENSMUSG00000074480 | Mex3a | 72640 | protein_coding | 38.82900285 | -0.655184237 | 0.268346609 | -2.441559596 | 0.014623974 | 0.063210492 | 0.655184237 |
| ENSMUSG00000035759 | Bbs10 | 71769 | protein_coding | 29.19979937 | 0.655449925 | 0.312765976 | 2.095656101 | 0.036112709 | 0.123851819 | 0.655449925 |
| ENSMUSG00000019823 | Mical1 | 171580 | protein_coding | 176.5381252 | -0.65575666 | 0.20691368 | -3.169228156 | 0.001528444 | 0.010815215 | 0.65575666 |
| ENSMUSG00000041798 | Gck | 103988 | protein_coding | 7.170518668 | -0.656074876 | 0.516063882 | -1.27130555 | 0.203619969 | 0.398777679 | 0.656074876 |
| ENSMUSG00000001588 | Acap1 | 216859 | protein_coding | 25.14498062 | 0.656317149 | 0.36191201 | 1.81347159 | 0.069759117 | 0.195227236 | 0.656317149 |
| ENSMUSG00000091239 | Vmn2r76 | 675969 | protein_coding | 18.49303636 | -0.656608247 | 1.89411574 | -0.346656877 | 0.728849111 | 0.847503916 | 0.656608247 |
| ENSMUSG00000021135 | Slc10a1 | 20493 | protein_coding | 22.28163832 | -0.656684225 | 0.318665238 | -2.060733792 | 0.039328444 | 0.131575452 | 0.656684225 |
| ENSMUSG00000050567 | Maml1 | 103806 | protein_coding | 206.8019107 | -0.657138239 | 0.145857631 | -4.505340133 | 6.63E-06 | 0.000103683 | 0.657138239 |
| ENSMUSG00000034818 | Celf5 | 319586 | protein_coding | 39.23303604 | -0.65740982 | 0.306227533 | -2.146801803 | 0.031809064 | 0.112791279 | 0.65740982 |
| ENSMUSG00000000028 | Cdc45 | 12544 | protein_coding | 165.9214416 | 0.657744993 | 0.163049407 | 4.034022605 | 5.48E-05 | 0.000671795 | 0.657744993 |
| ENSMUSG00000053317 | Sec61b | 66212 | protein_coding | 1504.752172 | 0.657795624 | 0.117476811 | 5.599365704 | 2.15E-08 | 5.51E-07 | 0.657795624 |
| ENSMUSG00000034686 | Prr7 | 432763 | protein_coding | 7.358949209 | -0.658028378 | 0.477513371 | -1.378031313 | 0.168193621 | 0.352776928 | 0.658028378 |
| ENSMUSG00000021306 | Gpr137b | 83924 | protein_coding | 77.6993115 | -0.658593627 | 0.293251167 | -2.245834631 | 0.024714602 | 0.093593482 | 0.658593627 |
| ENSMUSG00000018168 | Ikzf3 | 22780 | protein_coding | 22.39633282 | 0.658706488 | 0.384981428 | 1.711008479 | 0.087079548 | 0.227050359 | 0.658706488 |
| ENSMUSG00000049871 | Nlrc3 | 268857 | protein_coding | 17.33524316 | 0.659068536 | 0.326380416 | 2.019326237 | 0.043453323 | 0.14087615 | 0.659068536 |
| ENSMUSG00000031668 | Eif2ak3 | 13666 | protein_coding | 1027.072669 | 0.659167141 | 0.197157071 | 3.343360393 | 0.000827703 | 0.006572113 | 0.659167141 |
| ENSMUSG00000064037 | Gpn1 | 74254 | protein_coding | 297.9636025 | 0.659388268 | 0.14761532 | 4.466936542 | 7.93E-06 | 0.000121641 | 0.659388268 |
| ENSMUSG00000020232 | Hmg20b | 15353 | protein_coding | 680.9810238 | 0.660572702 | 0.090605528 | 7.290644554 | 3.08E-13 | 1.34E-11 | 0.660572702 |
| ENSMUSG00000003809 | Gcdh | 270076 | protein_coding | 759.5651383 | 0.660655826 | 0.09647138 | 6.848205443 | 7.48E-12 | 2.82E-10 | 0.660655826 |
| ENSMUSG00000000305 | Cdh4 | 12561 | protein_coding | 8.027724196 | -0.660678726 | 0.547631502 | -1.206429367 | 0.227651978 | 0.430280937 | 0.660678726 |
| ENSMUSG00000028957 | Per3 | 18628 | protein_coding | 172.6861682 | -0.660806873 | 0.242220075 | -2.728125953 | 0.006369528 | 0.033795418 | 0.660806873 |
| ENSMUSG00000024190 | Dusp1 | 19252 | protein_coding | 2029.241485 | -0.66091811 | 0.414064824 | -1.596170627 | 0.110450704 | 0.26510832 | 0.66091811 |

| ensembl_gene_id | external_gene_name | entrezgene_id | gene_biotype | baseMean | log2FoldChange | lfcSE | stat | pvalue | padj | absLog2FC |
|---|---|---|---|---|---|---|---|---|---|---|
| ENSMUSG00000032766 | Gng11 | 66066 | protein_coding | 326.7501042 | 0.66123909 | 0.131237796 | 5.03848059 | 4.69E-07 | 9.63E-06 | 0.66123909 |
| ENSMUSG00000070348 | Ccnd1 | 12443 | protein_coding | 1192.61509 | 0.66139151 | 0.135566261 | 4.878732386 | 1.07E-06 | 2.01E-05 | 0.66139151 |
| ENSMUSG00000026527 | Rgs7 | 24012 | protein_coding | 8.025234711 | -0.661841364 | 0.55476443 | -1.193013338 | 0.23286415 | 0.436411363 | 0.661841364 |
| ENSMUSG00000045411 | 2410002F23Rik | 668661 | protein_coding | 371.8847545 | -0.662071729 | 0.132015024 | -5.015124087 | 5.30E-07 | 1.08E-05 | 0.662071729 |
| ENSMUSG00000027457 | Snph | 241727 | protein_coding | 71.51415506 | -0.662102507 | 0.192037541 | -3.447776419 | 0.000565222 | 0.004804083 | 0.662102507 |
| ENSMUSG00000028801 | Stpg1 | 78806 | protein_coding | 14.26240212 | 0.662460082 | 0.350246077 | 1.891413281 | 0.058569197 | 0.173077954 | 0.662460082 |
| ENSMUSG00000038372 | Gmds | 218138 | protein_coding | 6163.14184 | 0.662611224 | 0.076707769 | 8.638124075 | 5.71E-18 | 3.60E-16 | 0.662611224 |
| ENSMUSG00000027298 | Tyro3 | 22174 | protein_coding | 140.8960268 | -0.662710747 | 0.104956208 | -6.314164348 | 2.72E-10 | 8.80E-09 | 0.662710747 |
| ENSMUSG00000030930 | Chst15 | 77590 | protein_coding | 112.44809 | -0.66312131 | 0.247051988 | -2.684136705 | 0.007271737 | 0.037568673 | 0.66312131 |
| ENSMUSG00000033149 | Phldb2 | 208177 | protein_coding | 186.4153646 | -0.66314193 | 0.342491246 | -1.936230307 | 0.0528395 | 0.161293131 | 0.66314193 |
| ENSMUSG00000053783 | 1700016K19Rik | 74230 | protein_coding | 14.20476591 | -0.66319067 | 0.331107189 | -2.002948566 | 0.045182809 | 0.145095119 | 0.66319067 |
| ENSMUSG00000054263 | Lifr | 16880 | protein_coding | 67.52706586 | -0.663446363 | 0.387472361 | -1.712241777 | 0.086852121 | 0.226746532 | 0.663446363 |
| ENSMUSG00000020183 | Cpm | 70574 | protein_coding | 673.455926 | -0.663789315 | 0.20021351 | -3.315407207 | 0.000915097 | 0.007130953 | 0.663789315 |
| ENSMUSG00000040163 | 1700034J05Rik | 73344 | protein_coding | 6.981841254 | -0.663789458 | 0.546878057 | -1.213779654 | 0.224831878 | 0.4266288 | 0.663789458 |
| ENSMUSG00000104117 | Gm20743 | 433374 | lncRNA | 7.998236544 | 0.66433653 | 0.554810723 | 1.197411121 | 0.231146351 | 0.434447373 | 0.66433653 |
| ENSMUSG00000048782 | Insc | 233752 | protein_coding | 39.44315392 | 0.664600113 | 0.230317176 | 2.885586409 | 0.003906852 | 0.023039 | 0.664600113 |
| ENSMUSG00000030123 | Plxnd1 | 67784 | protein_coding | 29.5471479 | -0.664790509 | 0.284985585 | -2.332716261 | 0.019663042 | 0.078810502 | 0.664790509 |
| ENSMUSG00000029811 | Aoc1 | 76507 | protein_coding | 3071.681607 | -0.664835704 | 0.158115239 | -4.204754123 | 2.61E-05 | 0.000349835 | 0.664835704 |
| ENSMUSG00000037188 | Grhl3 | 230824 | protein_coding | 13.60484564 | -0.665442813 | 0.459732486 | -1.447456581 | 0.147769089 | 0.324601832 | 0.665442813 |
| ENSMUSG00000032532 | Cck | 12424 | protein_coding | 20.61173419 | -0.666032871 | 0.361527911 | -1.842272344 | 0.065435323 | 0.186349691 | 0.666032871 |
| ENSMUSG00000031398 | Plxna3 | 18846 | protein_coding | 60.36849605 | -0.666043331 | 0.229092529 | -2.907311448 | 0.0036455 | 0.021878497 | 0.666043331 |
| ENSMUSG00000021629 | Slc30a5 | 69048 | protein_coding | 1461.659986 | 0.666141511 | 0.089632838 | 7.431891295 | 1.07E-13 | 4.86E-12 | 0.666141511 |
| ENSMUSG00000036916 | Zfp280c | 208968 | protein_coding | 167.2897846 | -0.666197384 | 0.144007356 | -4.626134403 | 3.73E-06 | 6.16E-05 | 0.666197384 |
| ENSMUSG00000054256 | Msi1 | 17690 | protein_coding | 18.6568671 | -0.666247533 | 0.343019021 | -1.942304924 | 0.052100197 | 0.159956041 | 0.666247533 |
| ENSMUSG00000047793 | Sned1 | 208777 | protein_coding | 193.6886991 | -0.666553692 | 0.188983423 | -3.527048455 | 0.00042022 | 0.003787924 | 0.666553692 |
| ENSMUSG00000025909 | Sntg1 | 71096 | protein_coding | 8.800499579 | -0.667629672 | 0.83406817 | -0.800449767 | 0.423450257 | 0.628350632 | 0.667629672 |
| ENSMUSG00000047875 | Gpr157 | 269604 | protein_coding | 19.54226148 | -0.667656957 | 0.285779261 | -2.336268049 | 0.019477276 | 0.078455964 | 0.667656957 |
| ENSMUSG00000080365 | Gm25776 | 115488078 | snoRNA | 71.82602956 | -0.667741858 | 0.206610583 | -3.231886027 | 0.001229761 | 0.009072549 | 0.667741858 |
| ENSMUSG00000028883 | Sema3a | 20346 | protein_coding | 8.628260096 | -0.668029586 | 1.231140012 | -0.542610572 | 0.587397958 | 0.757354275 | 0.668029586 |
| ENSMUSG00000109564 | Muc16 | 73732 | protein_coding | 53.53473963 | -0.668395415 | 0.530739519 | -1.259366207 | 0.20789809 | 0.404310463 | 0.668395415 |
| ENSMUSG00000035397 | Klf16 | 118445 | protein_coding | 201.4904602 | -0.668425951 | 0.207705242 | -3.218146756 | 0.001290218 | 0.009406183 | 0.668425951 |
| ENSMUSG00000070000 | Fcho1 | 74015 | protein_coding | 74.37728513 | -0.668439669 | 0.206377319 | -3.238920201 | 0.001199831 | 0.008888797 | 0.668439669 |
| ENSMUSG00000024066 | Xdh | 22436 | protein_coding | 927.5628332 | 0.668571354 | 0.100297109 | 6.66590852 | 2.63E-11 | 9.50E-10 | 0.668571354 |
| ENSMUSG00000027876 | Reg4 | 67709 | protein_coding | 3710.026274 | -0.66874071 | 0.20509967 | -3.260564529 | 0.001111907 | 0.008369777 | 0.66874071 |
| ENSMUSG00000003411 | Rab3b | 69908 | protein_coding | 13.65496065 | 0.669145901 | 0.378444352 | 1.76814873 | 0.077036043 | 0.208147598 | 0.669145901 |
| ENSMUSG00000062995 | Ica1 | 15893 | protein_coding | 729.8678004 | 0.669208832 | 0.075337556 | 8.882805168 | 6.52E-19 | 4.27E-17 | 0.669208832 |
| ENSMUSG00000042348 | Arl15 | 218639 | protein_coding | 161.1179342 | 0.669518869 | 0.109050515 | 6.13952962 | 8.28E-10 | 2.51E-08 | 0.669518869 |
| ENSMUSG00000017713 | Tha1 | 71776 | protein_coding | 96.75192469 | -0.669669826 | 0.153295501 | -4.368489764 | 1.25E-05 | 0.000181819 | 0.669669826 |

| ensembl_gene_id | external_gene_name | entrezgene_id | gene_biotype | baseMean | log2FoldChange | lfcSE | stat | pvalue | padj | absLog2FC |
|---|---|---|---|---|---|---|---|---|---|---|
| ENSMUSG00000029188 | Slc34a2 | 20531 | protein_coding | 425.3797292 | 0.670039331 | 0.211715291 | 3.16481313 | 0.001551826 | 0.01094178 | 0.670039331 |
| ENSMUSG00000061013 | Mkx | 210719 | protein_coding | 191.6068652 | -0.670069884 | 0.20276199 | -3.304711527 | 0.000950742 | 0.00736906 | 0.670069884 |
| ENSMUSG00000061175 | Fnip2 | 329679 | protein_coding | 426.7288735 | -0.670570502 | 0.085343536 | -7.857308601 | 3.92E-15 | 2.01E-13 | 0.670570502 |
| ENSMUSG00000071226 | Cecr2 | 330409 | protein_coding | 11.65717401 | -0.670939783 | 0.686488839 | -0.977349878 | 0.328395968 | 0.539240147 | 0.670939783 |
| ENSMUSG00000025384 | Faap100 | 71885 | protein_coding | 257.3809984 | -0.670960321 | 0.182593879 | -3.674604671 | 0.000238218 | 0.00236113 | 0.670960321 |
| ENSMUSG00000050050 | Ccdc158 | 320696 | protein_coding | 6.95377181 | -0.671033668 | 0.764843856 | -0.877347268 | 0.380298046 | 0.588210067 | 0.671033668 |
| ENSMUSG00000030032 | Wdr54 | 75659 | protein_coding | 17.52263134 | -0.671107142 | 0.287946273 | -2.330667923 | 0.019770877 | 0.079143133 | 0.671107142 |
| ENSMUSG00000039395 | Mreg | 381269 | protein_coding | 12.7446407 | -0.671184547 | 0.435941165 | -1.53962186 | 0.123652554 | 0.286937336 | 0.671184547 |
| ENSMUSG00000096917 | 2500002B13Rik | 78353 | lncRNA | 20.64141679 | -0.671268729 | 0.284863888 | -2.35645428 | 0.018450339 | 0.075256593 | 0.671268729 |
| ENSMUSG00000035505 | Cox18 | 231430 | protein_coding | 108.0313825 | 0.671422649 | 0.155931086 | 4.305893501 | 1.66E-05 | 0.000235052 | 0.671422649 |
| ENSMUSG00000029053 | Prkcz | 18762 | protein_coding | 1631.266886 | 0.671727666 | 0.103516616 | 6.489080596 | 8.64E-11 | 2.99E-09 | 0.671727666 |
| ENSMUSG00000043541 | Casc1 | 320662 | protein_coding | 13.60350955 | -0.67179467 | 0.421239457 | -1.594804711 | 0.110755913 | 0.265491014 | 0.67179467 |
| ENSMUSG00000074415 | Mir100hg | 73144 | lncRNA | 14.82817859 | -0.671937856 | 1.631438115 | -0.411868431 | 0.680435861 | 0.817182783 | 0.671937856 |
| ENSMUSG00000006586 | Runx1t1 | 12395 | protein_coding | 15.44432192 | -0.671973244 | 0.449076096 | -1.496346052 | 0.1345635 | 0.303673481 | 0.671973244 |
| ENSMUSG00000039682 | Lap3 | 66988 | protein_coding | 1313.11456 | 0.672258341 | 0.10424478 | 6.448844146 | 1.13E-10 | 3.82E-09 | 0.672258341 |
| ENSMUSG00000015702 | Anxa9 | 71790 | protein_coding | 25.24790833 | 0.67311278 | 0.315550893 | 2.133135402 | 0.03291362 | 0.115447731 | 0.67311278 |
| ENSMUSG00000046574 | Prr12 | 233210 | protein_coding | 92.24890903 | -0.673733372 | 0.299907954 | -2.246467169 | 0.0246741 | 0.093506797 | 0.673733372 |
| ENSMUSG00000022442 | Ttll1 | 319953 | protein_coding | 48.16960152 | -0.674282804 | 0.18480071 | -3.648702454 | 0.000263568 | 0.002559825 | 0.674282804 |
| ENSMUSG00000019122 | Ccl9 | 20308 | protein_coding | 467.1995915 | 0.674293037 | 0.219442123 | 3.07276027 | 0.002120888 | 0.014205847 | 0.674293037 |
| ENSMUSG00000053695 | Defb37 | 353320 | protein_coding | 83.22966325 | 0.674529533 | 0.339767425 | 1.985268403 | 0.047114618 | 0.149341617 | 0.674529533 |
| ENSMUSG00000035181 | Heatr5a | 320487 | protein_coding | 1425.974071 | 0.675020887 | 0.074258262 | 9.090178894 | 9.89E-20 | 6.88E-18 | 0.675020887 |
| ENSMUSG00000021187 | Tc2n | 74413 | protein_coding | 1044.135752 | 0.675403343 | 0.079641195 | 8.480577722 | 2.24E-17 | 1.36E-15 | 0.675403343 |
| ENSMUSG00000086290 | Snhg12 | 100039864 | lncRNA | 173.8745067 | -0.675574022 | 0.130496947 | -5.176933521 | 2.26E-07 | 4.91E-06 | 0.675574022 |
| ENSMUSG00000000489 | Pdgfb | 18591 | protein_coding | 19.50925513 | -0.675579146 | 0.319310935 | -2.115740715 | 0.034366866 | 0.119325161 | 0.675579146 |
| ENSMUSG00000053111 | Fank1 | 66930 | protein_coding | 10.22852046 | -0.675911013 | 0.399811435 | -1.69057449 | 0.0909181 | 0.233298918 | 0.675911013 |
| ENSMUSG00000003541 | Ier3 | 15937 | protein_coding | 200.5825652 | -0.675981464 | 0.276667859 | -2.443295967 | 0.014553796 | 0.063044022 | 0.675981464 |
| ENSMUSG00000010663 | Fads1 | 76267 | protein_coding | 291.7854394 | 0.676079805 | 0.141260882 | 4.786036963 | 1.70E-06 | 3.03E-05 | 0.676079805 |
| ENSMUSG00000030556 | Lrrc28 | 67867 | protein_coding | 122.3131078 | 0.67608492 | 0.121570843 | 5.561242334 | 2.68E-08 | 6.76E-07 | 0.67608492 |
| ENSMUSG00000030718 | Ppme1 | 72590 | protein_coding | 657.7257774 | 0.676174044 | 0.121490877 | 5.565636355 | 2.61E-08 | 6.62E-07 | 0.676174044 |
| ENSMUSG00000007670 | Khsrp | 16549 | protein_coding | 1232.567434 | 0.676308304 | 0.091350915 | 7.403410281 | 1.33E-13 | 5.91E-12 | 0.676308304 |
| ENSMUSG00000040329 | Il7 | 16196 | protein_coding | 147.4824535 | 0.67630881 | 0.139211021 | 4.858155661 | 1.18E-06 | 2.19E-05 | 0.67630881 |
| ENSMUSG00000056737 | Capg | 12332 | protein_coding | 470.4729718 | 0.676309928 | 0.204195774 | 3.312066229 | 0.000926096 | 0.007195542 | 0.676309928 |
| ENSMUSG00000049532 | Sall2 | 50524 | protein_coding | 20.93904086 | -0.676814507 | 0.258554099 | -2.617690104 | 0.008852716 | 0.043754398 | 0.676814507 |
| ENSMUSG00000080115 | Eef1akmt3 | 100504608 | protein_coding | 24.13324638 | 0.677040425 | 0.232401899 | 2.913231034 | 0.003577098 | 0.021562746 | 0.677040425 |
| ENSMUSG00000031539 | Ap3m2 | 64933 | protein_coding | 20.78976851 | 0.677123027 | 0.29453682 | 2.298941865 | 0.021508241 | 0.084297062 | 0.677123027 |
| ENSMUSG00000020101 | Vsir | 74048 | protein_coding | 24.18966432 | 0.67724983 | 0.262708253 | 2.577954148 | 0.009938718 | 0.047759252 | 0.67724983 |
| ENSMUSG00000037709 | Fam13a | 58909 | protein_coding | 1156.759408 | 0.67732575 | 0.113249266 | 5.980840109 | 2.22E-09 | 6.46E-08 | 0.67732575 |
| ENSMUSG00000050921 | P2ry10 | 78826 | protein_coding | 22.65623207 | 0.677328546 | 0.295064448 | 2.295527471 | 0.021702914 | 0.084830433 | 0.677328546 |

| ensembl_gene_id | external_gene_name | entrezgene_id | gene_biotype | baseMean | log2FoldChange | lfcSE | stat | pvalue | padj | absLog2FC |
|---|---|---|---|---|---|---|---|---|---|---|
| ENSMUSG00000069227 | Gprin1 | 26913 | protein_coding | 12.43388476 | -0.677803034 | 0.362140632 | -1.871656959 | 0.061254078 | 0.178191396 | 0.677803034 |
| ENSMUSG00000062232 | Rapgef2 | 76089 | protein_coding | 640.1823431 | 0.678007039 | 0.1228664 | 5.51824617 | 3.42E-08 | 8.39E-07 | 0.678007039 |
| ENSMUSG00000015627 | Gata5 | 14464 | protein_coding | 14.60685788 | -0.678106606 | 0.326212731 | -2.078725145 | 0.03764262 | 0.127587679 | 0.678106606 |
| ENSMUSG00000109864 | Eid3 | 66341 | protein_coding | 26.93302491 | -0.678286056 | 0.394844635 | -1.717855571 | 0.085822962 | 0.224666727 | 0.678286056 |
| ENSMUSG00000032507 | Fbxl2 | 72179 | protein_coding | 11.77722575 | -0.678319139 | 0.349473292 | -1.940975618 | 0.052261235 | 0.16017259 | 0.678319139 |
| ENSMUSG00000020484 | Xbp1 | 22433 | protein_coding | 4693.822232 | 0.678700136 | 0.101145875 | 6.710111852 | 1.94E-11 | 7.07E-10 | 0.678700136 |
| ENSMUSG00000035910 | Dcdc2a | 195208 | protein_coding | 18.15887264 | 0.678888449 | 0.379477366 | 1.78900907 | 0.073613355 | 0.202052993 | 0.678888449 |
| ENSMUSG00000048572 | Tmem252 | 226040 | protein_coding | 255.3572454 | -0.678906977 | 0.239113205 | -2.839270114 | 0.004521686 | 0.025888361 | 0.678906977 |
| ENSMUSG00000074345 | Tnfaip8l3 | 244882 | protein_coding | 7.268697548 | 0.679328469 | 0.660680674 | 1.028225126 | 0.303843942 | 0.513472525 | 0.679328469 |
| ENSMUSG00000001663 | Gstt1 | 14871 | protein_coding | 226.1986245 | -0.679451228 | 0.142204643 | -4.777982036 | 1.77E-06 | 3.13E-05 | 0.679451228 |
| ENSMUSG00000024338 | Psmb8 | 16913 | protein_coding | 199.9185513 | 0.679669362 | 0.22767702 | 2.985234789 | 0.002833607 | 0.017924423 | 0.679669362 |
| ENSMUSG00000067889 | Sptbn2 | 20743 | protein_coding | 241.8771963 | -0.679735696 | 0.131033364 | -5.187500915 | 2.13E-07 | 4.66E-06 | 0.679735696 |
| ENSMUSG00000044951 | Mylk4 | 238564 | protein_coding | 7.950126771 | -0.679975356 | 1.861185761 | -0.365345239 | 0.714853713 | 0.839719166 | 0.679975356 |
| ENSMUSG00000028262 | Clca3a2 | 80797 | protein_coding | 128.9690286 | 0.680287572 | 0.258291093 | 2.633801905 | 0.008443473 | 0.042182001 | 0.680287572 |
| ENSMUSG00000074218 | Cox7a1 | 12865 | protein_coding | 712.7961226 | 0.680535186 | 0.112750452 | 6.035764565 | 1.58E-09 | 4.69E-08 | 0.680535186 |
| ENSMUSG00000070855 | Or10ag54 | 257875 | protein_coding | 6.679981935 | -0.680599563 | 1.495240768 | -0.455177238 | 0.648981735 | 0.797283028 | 0.680599563 |
| ENSMUSG00000018470 | Kcnab3 | 16499 | protein_coding | 7.830016525 | -0.680761563 | 0.446242949 | -1.525540212 | 0.127124412 | 0.292351139 | 0.680761563 |
| ENSMUSG00000048100 | Taf13 | 99730 | protein_coding | 405.1764426 | 0.680796226 | 0.118825682 | 5.729369405 | 1.01E-08 | 2.71E-07 | 0.680796226 |
| ENSMUSG00000037904 | Ankrd9 | 74251 | protein_coding | 321.6553447 | -0.68085846 | 0.173921638 | -3.91474268 | 9.05E-05 | 0.001036301 | 0.68085846 |
| ENSMUSG00000047021 | Cfap65 | 241116 | protein_coding | 10.75930684 | -0.680975659 | 0.726761101 | -0.937000698 | 0.348758199 | 0.558911419 | 0.680975659 |
| ENSMUSG00000028864 | Hgf | 15234 | protein_coding | 8.768199215 | 0.681096743 | 0.640617563 | 1.063187745 | 0.287696819 | 0.496834898 | 0.681096743 |
| ENSMUSG00000034771 | Tle2 | 21886 | protein_coding | 35.87161461 | -0.681179183 | 0.317573614 | -2.144948926 | 0.031956931 | 0.113088201 | 0.681179183 |
| ENSMUSG00000028035 | Dnajb4 | 67035 | protein_coding | 322.9006229 | -0.681551199 | 0.178881677 | -3.810067148 | 0.000138929 | 0.001499229 | 0.681551199 |
| ENSMUSG00000044328 | Trp53i13 | 216964 | protein_coding | 39.38006866 | -0.681726718 | 0.187491313 | -3.636044289 | 0.000276857 | 0.002659695 | 0.681726718 |
| ENSMUSG00000041420 | Meis3 | 17537 | protein_coding | 82.01028322 | -0.68234359 | 0.148210326 | -4.603886968 | 4.15E-06 | 6.82E-05 | 0.68234359 |
| ENSMUSG00000074489 | Bglap3 | 12095 | protein_coding | 31.51217402 | -0.68239039 | 0.2547495 | -2.678672141 | 0.007391473 | 0.038088443 | 0.68239039 |
| ENSMUSG00000018672 | Copz2 | 56358 | protein_coding | 88.84355214 | -0.68318862 | 0.167653246 | -4.075009798 | 4.60E-05 | 0.000572567 | 0.68318862 |
| ENSMUSG00000047904 | Sstr2 | 20606 | protein_coding | 54.54134447 | -0.683390005 | 0.292105603 | -2.339530626 | 0.019307988 | 0.077944449 | 0.683390005 |
| ENSMUSG00000068154 | Insm1 | 53626 | protein_coding | 34.02816914 | -0.683552548 | 0.24908704 | -2.744231691 | 0.006065271 | 0.032484076 | 0.683552548 |
| ENSMUSG00000042834 | Nrep | 27528 | protein_coding | 275.7582799 | -0.683599802 | 0.191101417 | -3.57715717 | 0.000347351 | 0.003207311 | 0.683599802 |
| ENSMUSG00000073600 | Prob1 | 381148 | protein_coding | 15.7041064 | 0.683629867 | 0.387565634 | 1.76390734 | 0.077747581 | 0.209607899 | 0.683629867 |
| ENSMUSG00000063160 | Numbl | 18223 | protein_coding | 13.03135879 | -0.683888164 | 0.447715851 | -1.527504918 | 0.126635505 | 0.291607681 | 0.683888164 |
| ENSMUSG00000055827 | Gsdmc3 | 270328 | protein_coding | 1133.254361 | -0.6840841 | 0.170843926 | -4.004146453 | 6.22E-05 | 0.000747089 | 0.6840841 |
| ENSMUSG00000074923 | Pak6 | 214230 | protein_coding | 28.16427465 | -0.684149809 | 0.269148996 | -2.541899914 | 0.011025174 | 0.051483134 | 0.684149809 |
| ENSMUSG00000056055 | Sag | 20215 | protein_coding | 6.733789125 | -0.684632722 | 0.891236756 | -0.768182772 | 0.442378607 | 0.643610382 | 0.684632722 |
| ENSMUSG00000046879 | Irgm1 | 15944 | protein_coding | 699.8945068 | 0.684748145 | 0.115652604 | 5.920732643 | 3.21E-09 | 9.21E-08 | 0.684748145 |
| ENSMUSG00000020014 | Cfap54 | 380654 | protein_coding | 11.40858284 | -0.685445504 | 1.438108758 | -0.47662981 | 0.633625752 | 0.788005719 | 0.685445504 |
| ENSMUSG00000006014 | Prg4 | 96875 | protein_coding | 12.29286868 | -0.685487139 | 0.63311819 | -1.082715913 | 0.278934538 | 0.487423953 | 0.685487139 |

| ensembl_gene_id | external_gene_name | entrezgene_id | gene_biotype | baseMean | log2FoldChange | lfcSE | stat | pvalue | padj | absLog2FC |
|---|---|---|---|---|---|---|---|---|---|---|
| ENSMUSG00000054169 | Ceacam10 | 26366 | protein_coding | 515.64341 | -0.685888209 | 0.114615261 | -5.98426602 | 2.17E-09 | 6.34E-08 | 0.685888209 |
| ENSMUSG00000015083 | C8g | 69379 | protein_coding | 29.14959512 | -0.68597563 | 0.225030268 | -3.0483705 | 0.002300861 | 0.015200376 | 0.68597563 |
| ENSMUSG00000049744 | Arhgap15 | 76117 | protein_coding | 12.06470987 | 0.687256998 | 0.61299062 | 1.121154184 | 0.262222238 | 0.470034424 | 0.687256998 |
| ENSMUSG00000030257 | Srgap3 | 259302 | protein_coding | 145.8630748 | -0.687531436 | 0.190681508 | -3.605653444 | 0.000311368 | 0.002927672 | 0.687531436 |
| ENSMUSG00000020173 | Cobl | 12808 | protein_coding | 640.1210703 | -0.687927201 | 0.120630972 | -5.702741108 | 1.18E-08 | 3.15E-07 | 0.687927201 |
| ENSMUSG00000068245 | Phf11d | 219132 | protein_coding | 134.6944336 | 0.689254946 | 0.185869878 | 3.70826598 | 0.000208683 | 0.002120974 | 0.689254946 |
| ENSMUSG00000021751 | Acox2 | 93732 | protein_coding | 57.42977028 | 0.689575236 | 0.210327912 | 3.278572151 | 0.001043337 | 0.007936137 | 0.689575236 |
| ENSMUSG00000086999 | Bcas1os2 | 73303 | lncRNA | 18.93058402 | -0.690189618 | 0.374635239 | -1.842297644 | 0.065431624 | 0.186349691 | 0.690189618 |
| ENSMUSG00000097482 | Gm17634 | 100504568 | lncRNA | 8.815105245 | -0.690200177 | 0.37712171 | -1.830178848 | 0.0672232 | 0.190000766 | 0.690200177 |
| ENSMUSG00000033949 | Trim36 | 28105 | protein_coding | 249.7627669 | -0.690677414 | 0.171488374 | -4.027546577 | 5.64E-05 | 0.00068634 | 0.690677414 |
| ENSMUSG00000024743 | Syt7 | 54525 | protein_coding | 747.2375053 | -0.691147215 | 0.114955399 | -6.012307579 | 1.83E-09 | 5.38E-08 | 0.691147215 |
| ENSMUSG00000031714 | Gab1 | 14388 | protein_coding | 388.2916941 | -0.691697697 | 0.180816585 | -3.82541068 | 0.000130554 | 0.001419432 | 0.691697697 |
| ENSMUSG00000022797 | Tfrc | 22042 | protein_coding | 5204.921517 | 0.692185323 | 0.157123468 | 4.405359248 | 1.06E-05 | 0.000155898 | 0.692185323 |
| ENSMUSG00000023367 | Tmem176a | 66058 | protein_coding | 455.6030949 | 0.692450721 | 0.082700658 | 8.372977142 | 5.62E-17 | 3.31E-15 | 0.692450721 |
| ENSMUSG00000009108 | Gnat2 | 14686 | protein_coding | 33.89497774 | 0.692592728 | 0.198749536 | 3.484751429 | 0.000492594 | 0.004294496 | 0.692592728 |
| ENSMUSG00000028047 | Thbs3 | 21827 | protein_coding | 13.98069337 | -0.693308284 | 0.363247585 | -1.908638383 | 0.056308757 | 0.168366037 | 0.693308284 |
| ENSMUSG00000064899 | Snord118 | 100216530 | snoRNA | 68.68809071 | 0.693594996 | 0.281170068 | 2.466816615 | 0.013632015 | 0.060230441 | 0.693594996 |
| ENSMUSG00000040666 | Sh3bgr | 50795 | protein_coding | 72.76485429 | -0.694024428 | 0.417525256 | -1.662233405 | 0.096465985 | 0.242696288 | 0.694024428 |
| ENSMUSG00000051008 | 4930412M03Rik | 100504140 | lncRNA | 11.9869829 | -0.694731817 | 0.376476788 | -1.845351 | 0.064986496 | 0.185602458 | 0.694731817 |
| ENSMUSG00000085980 | Gm12408 | 100040557 | lncRNA | 21.97395229 | 0.69476726 | 0.35241676 | 1.971436489 | 0.048673971 | 0.152854694 | 0.69476726 |
| ENSMUSG00000004655 | Aqp1 | 11826 | protein_coding | 1164.71909 | -0.695014613 | 0.172417251 | -4.031003907 | 5.55E-05 | 0.000678395 | 0.695014613 |
| ENSMUSG00000051344 | Plekhm3 | 241075 | protein_coding | 41.79668175 | -0.69523487 | 0.230753125 | -3.012894711 | 0.002587687 | 0.016720761 | 0.69523487 |
| ENSMUSG00000004319 | Clcn3 | 12725 | protein_coding | 2582.959196 | 0.695439017 | 0.061866202 | 11.24101687 | 2.56E-29 | 2.88E-27 | 0.695439017 |
| ENSMUSG00000040855 | Reps2 | 194590 | protein_coding | 857.4242576 | 0.696351743 | 0.077906839 | 8.938262006 | 3.95E-19 | 2.65E-17 | 0.696351743 |
| ENSMUSG00000035775 | Krt20 | 66809 | protein_coding | 8985.275785 | 0.696572782 | 0.258649694 | 2.693112723 | 0.007078833 | 0.036774838 | 0.696572782 |
| ENSMUSG00000027173 | Depdc7 | 211896 | protein_coding | 13.53382104 | -0.69700556 | 0.323218313 | -2.156454424 | 0.031048199 | 0.110830572 | 0.69700556 |
| ENSMUSG00000034413 | Neurl1b | 240055 | protein_coding | 718.3460698 | -0.697125754 | 0.108481413 | -6.426223007 | 1.31E-10 | 4.39E-09 | 0.697125754 |
| ENSMUSG00000000290 | Itgb2 | 16414 | protein_coding | 23.88870632 | 0.697425688 | 0.398847441 | 1.748602637 | 0.08035973 | 0.214436218 | 0.697425688 |
| ENSMUSG00000025804 | Ccr1 | 12768 | protein_coding | 19.27523169 | 0.69766483 | 0.297460098 | 2.345406437 | 0.019006348 | 0.07710981 | 0.69766483 |
| ENSMUSG00000021176 | Efcab11 | 78767 | protein_coding | 48.07399249 | 0.697690154 | 0.210190292 | 3.319326248 | 0.000902349 | 0.007048858 | 0.697690154 |
| ENSMUSG00000020806 | Rhbdf2 | 217344 | protein_coding | 208.877335 | -0.698027999 | 0.205987265 | -3.388694918 | 0.000702261 | 0.005750958 | 0.698027999 |
| ENSMUSG00000117662 | Gm4013 | 100042757 | lncRNA | 8.171529419 | -0.698451006 | 0.544377911 | -1.283025985 | 0.199482972 | 0.393871921 | 0.698451006 |
| ENSMUSG00000067231 | Cyp2c65 | 72303 | protein_coding | 3093.948682 | -0.698619883 | 0.336803818 | -2.074263547 | 0.038054841 | 0.128582417 | 0.698619883 |
| ENSMUSG00000029706 | Pax4 | 18506 | protein_coding | 8.900643979 | -0.69891002 | 0.485104081 | -1.440742405 | 0.14965747 | 0.327058149 | 0.69891002 |
| ENSMUSG00000014873 | Surf2 | 20931 | protein_coding | 211.2362271 | -0.699333728 | 0.092388122 | -7.569519912 | 3.75E-14 | 1.78E-12 | 0.699333728 |
| ENSMUSG00000097121 | D130020L05Rik | 319760 | lncRNA | 13.50772895 | -0.699400633 | 0.354500886 | -1.972916463 | 0.048505081 | 0.152534832 | 0.699400633 |
| ENSMUSG00000026222 | Sp100 | 20684 | protein_coding | 80.69500277 | 0.699502771 | 0.240442009 | 2.909236929 | 0.003623122 | 0.021768797 | 0.699502771 |
| ENSMUSG00000046589 | Lrrc8e | 72267 | protein_coding | 312.4818947 | -0.699724499 | 0.140496368 | -4.980374286 | 6.35E-07 | 1.26E-05 | 0.699724499 |

| ensembl_gene_id | external_gene_name | entrezgene_id | gene_biotype | baseMean | log2FoldChange | lfcSE | stat | pvalue | padj | absLog2FC |
|---|---|---|---|---|---|---|---|---|---|---|
| ENSMUSG00000048163 | Selplg | 20345 | protein_coding | 25.84970734 | 0.6997968 | 0.300946556 | 2.325319184 | 0.020054899 | 0.079938547 | 0.6997968 |
| ENSMUSG00000090877 | Hspa1b | 15511 | protein_coding | 2889.075229 | 0.700084757 | 0.483130591 | 1.449059053 | 0.147321092 | 0.323882726 | 0.700084757 |
| ENSMUSG00000058063 | Trim31 | 224762 | protein_coding | 1015.351344 | 0.700135044 | 0.141166354 | 4.959645297 | 7.06E-07 | 1.38E-05 | 0.700135044 |
| ENSMUSG00000027533 | Fabp5 | 16592 | protein_coding | 66.19171449 | -0.700435736 | 0.190649673 | -3.67394144 | 0.000238837 | 0.002365589 | 0.700435736 |
| ENSMUSG00000102752 | Gm7694 | 665574 | protein_coding | 78.10590009 | -0.701084508 | 0.162312024 | -4.319362745 | 1.56E-05 | 0.000222537 | 0.701084508 |
| ENSMUSG00000026622 | Nek2 | 18005 | protein_coding | 669.0528428 | 0.701330166 | 0.142801666 | 4.911218363 | 9.05E-07 | 1.72E-05 | 0.701330166 |
| ENSMUSG00000050222 | Il17d | 239114 | protein_coding | 21.64822601 | -0.701604556 | 0.269601292 | -2.602378312 | 0.009257966 | 0.045210961 | 0.701604556 |
| ENSMUSG00000042515 | Pwwp3b | 245631 | protein_coding | 8.997843034 | -0.701702575 | 0.406704155 | -1.725339086 | 0.084466382 | 0.222281761 | 0.701702575 |
| ENSMUSG00000051335 | Gfod1 | 328232 | protein_coding | 70.00302301 | -0.702335821 | 0.138678337 | -5.064495548 | 4.09E-07 | 8.51E-06 | 0.702335821 |
| ENSMUSG00000052142 | Rasal3 | 320484 | protein_coding | 15.19483589 | 0.703447181 | 0.474779527 | 1.481629137 | 0.138439002 | 0.309608637 | 0.703447181 |
| ENSMUSG00000025592 | Dach2 | 93837 | protein_coding | 7.671117836 | -0.70381279 | 0.510495724 | -1.378684986 | 0.167991901 | 0.352493083 | 0.70381279 |
| ENSMUSG00000097768 | 2310043M15Rik | 102633000 | lncRNA | 8.066667901 | -0.704193509 | 0.492279771 | -1.430474196 | 0.152580967 | 0.33119896 | 0.704193509 |
| ENSMUSG00000062991 | Nrg1 | 211323 | protein_coding | 10.43692113 | -0.704249687 | 1.097397039 | -0.641745569 | 0.521038396 | 0.707440424 | 0.704249687 |
| ENSMUSG00000039384 | Dusp10 | 63953 | protein_coding | 77.87690628 | 0.704273338 | 0.315777901 | 2.230280634 | 0.025728818 | 0.096652975 | 0.704273338 |
| ENSMUSG00000092111 | Vmn2r113 | 434701 | protein_coding | 9.054087856 | -0.704324634 | 1.679106049 | -0.419464056 | 0.674877018 | 0.812935124 | 0.704324634 |
| ENSMUSG00000026113 | Inpp4a | 269180 | protein_coding | 176.6400869 | -0.704712151 | 0.177616753 | -3.967599551 | 7.26E-05 | 0.000850199 | 0.704712151 |
| ENSMUSG00000028004 | Npy2r | 18167 | protein_coding | 9.760500518 | -0.705087582 | 0.451187639 | -1.562736923 | 0.118114486 | 0.277973926 | 0.705087582 |
| ENSMUSG00000027364 | Usp50 | 75083 | protein_coding | 8.478939674 | -0.70529257 | 0.567924038 | -1.241878355 | 0.214281447 | 0.412709398 | 0.70529257 |
| ENSMUSG00000037217 | Syn1 | 20964 | protein_coding | 9.298231113 | -0.705512285 | 0.434348178 | -1.62430124 | 0.104311533 | 0.255523566 | 0.705512285 |
| ENSMUSG00000032064 | Dixdc1 | 330938 | protein_coding | 236.1551391 | -0.705714521 | 0.227853424 | -3.09723027 | 0.001953381 | 0.013267995 | 0.705714521 |
| ENSMUSG00000051984 | Sec31b | 240667 | protein_coding | 16.32014479 | -0.705898525 | 0.334475496 | -2.110464093 | 0.034818401 | 0.120588712 | 0.705898525 |
| ENSMUSG00000029752 | Asns | 27053 | protein_coding | 756.7055786 | 0.70611839 | 0.201610998 | 3.502380312 | 0.000461121 | 0.00407136 | 0.70611839 |
| ENSMUSG00000024778 | Fas | 14102 | protein_coding | 131.0301143 | 0.706185833 | 0.172460686 | 4.094764143 | 4.23E-05 | 0.000530848 | 0.706185833 |
| ENSMUSG00000041961 | Znrf3 | 407821 | protein_coding | 177.3442419 | -0.70644057 | 0.149096726 | -4.738136042 | 2.16E-06 | 3.74E-05 | 0.70644057 |
| ENSMUSG00000005089 | Slc1a2 | 20511 | protein_coding | 21.09695504 | -0.706776131 | 0.44596398 | -1.584827838 | 0.113005456 | 0.268850022 | 0.706776131 |
| ENSMUSG00000038213 | Tapbpl | 213233 | protein_coding | 418.6654949 | 0.706830507 | 0.11161974 | 6.332486591 | 2.41E-10 | 7.87E-09 | 0.706830507 |
| ENSMUSG00000030366 | Ceacam12 | 67315 | protein_coding | 62.43096806 | -0.706875039 | 0.390103507 | -1.812019188 | 0.069983232 | 0.195681974 | 0.706875039 |
| ENSMUSG00000033880 | Lgals3bp | 19039 | protein_coding | 1525.843852 | 0.70688127 | 0.144881681 | 4.879024479 | 1.07E-06 | 2.00E-05 | 0.70688127 |
| ENSMUSG00000038453 | Srcin1 | 56013 | protein_coding | 10.87569081 | -0.706970812 | 0.795514098 | -0.888696773 | 0.374166066 | 0.582572541 | 0.706970812 |
| ENSMUSG00000036466 | Megf11 | 214058 | protein_coding | 57.61247811 | -0.707056018 | 0.231102575 | -3.059490004 | 0.002217142 | 0.014745151 | 0.707056018 |
| ENSMUSG00000052353 | Cemip | 80982 | protein_coding | 8.916586016 | -0.707818012 | 0.84127657 | -0.84136185 | 0.400145251 | 0.607690116 | 0.707818012 |
| ENSMUSG00000003402 | Prkcsh | 19089 | protein_coding | 2148.163841 | 0.707851121 | 0.094151397 | 7.518222155 | 5.55E-14 | 2.59E-12 | 0.707851121 |
| ENSMUSG00000021610 | Clptm1l | 218335 | protein_coding | 2379.688273 | 0.707879326 | 0.083881381 | 8.439051858 | 3.20E-17 | 1.92E-15 | 0.707879326 |
| ENSMUSG00000061414 | Cracr2a | 381812 | protein_coding | 891.5482127 | -0.708263395 | 0.147410956 | -4.804686255 | 1.55E-06 | 2.79E-05 | 0.708263395 |
| ENSMUSG00000047343 | Mettl21c | 433294 | protein_coding | 12.5423217 | 0.708536516 | 0.441832739 | 1.603630635 | 0.108795494 | 0.262679192 | 0.708536516 |
| ENSMUSG00000039347 | Atp6v0e2 | 76252 | protein_coding | 212.8579172 | -0.708621126 | 0.110085033 | -6.437034257 | 1.22E-10 | 4.11E-09 | 0.708621126 |
| ENSMUSG00000076435 | Acsf2 | 264895 | protein_coding | 1187.268315 | -0.709054959 | 0.156019342 | -4.544660615 | 5.50E-06 | 8.80E-05 | 0.709054959 |
| ENSMUSG00000030887 | Pdzd9 | 67983 | protein_coding | 6.976089637 | -0.710861343 | 0.443547562 | -1.602672191 | 0.109007046 | 0.262950815 | 0.710861343 |

| ensembl_gene_id | external_gene_name | entrezgene_id | gene_biotype | baseMean | log2FoldChange | lfcSE | stat | pvalue | padj | absLog2FC |
|---|---|---|---|---|---|---|---|---|---|---|
| ENSMUSG00000031133 | Arhgef6 | 73341 | protein_coding | 12.98862699 | 0.711085311 | 0.38217387 | 1.86063299 | 0.062796023 | 0.181128045 | 0.711085311 |
| ENSMUSG00000055994 | Nod2 | 257632 | protein_coding | 53.84961887 | -0.711920828 | 0.208214436 | -3.419171312 | 0.000628122 | 0.005262873 | 0.711920828 |
| ENSMUSG00000030406 | Gipr | 381853 | protein_coding | 9.390386684 | -0.711966192 | 0.434607358 | -1.638182556 | 0.101383617 | 0.250401343 | 0.711966192 |
| ENSMUSG00000097251 | 5033417F24Rik | 75995 | lncRNA | 7.488488303 | -0.712087406 | 0.431612138 | -1.649831744 | 0.098977354 | 0.246638188 | 0.712087406 |
| ENSMUSG00000089371 | Mir1938 | 100316693 | miRNA | 26.68020132 | -0.712420042 | 0.352907373 | -2.018716795 | 0.043516663 | 0.141024092 | 0.712420042 |
| ENSMUSG00000069265 | H3c1 | 360198 | protein_coding | 631.4972244 | 0.712499154 | 0.235814621 | 3.021437563 | 0.002515775 | 0.016402483 | 0.712499154 |
| ENSMUSG00000050503 | Fbxl22 | 74165 | protein_coding | 32.25903677 | -0.712658128 | 0.229341848 | -3.107405531 | 0.001887373 | 0.012880067 | 0.712658128 |
| ENSMUSG00000024883 | Rin1 | 225870 | protein_coding | 93.05899572 | -0.712903962 | 0.276598754 | -2.577393974 | 0.009954842 | 0.047759252 | 0.712903962 |
| ENSMUSG00000024855 | Pacs1 | 107975 | protein_coding | 88.57683875 | -0.712936494 | 0.205560347 | -3.468258863 | 0.000523842 | 0.004524817 | 0.712936494 |
| ENSMUSG00000046688 | Tifa | 211550 | protein_coding | 899.4697874 | 0.713091217 | 0.228036166 | 3.127097027 | 0.001765417 | 0.012199373 | 0.713091217 |
| ENSMUSG00000052273 | Dnah3 | 381917 | protein_coding | 8.947944634 | -0.713180786 | 1.599386238 | -0.445909043 | 0.65566295 | 0.800694808 | 0.713180786 |
| ENSMUSG00000024114 | Prss41 | 71003 | protein_coding | 9.208192172 | -0.713408405 | 0.439990347 | -1.621418312 | 0.104927956 | 0.256490559 | 0.713408405 |
| ENSMUSG00000045092 | S1pr1 | 13609 | protein_coding | 7.299663693 | 0.713509775 | 0.625086294 | 1.141458039 | 0.253679364 | 0.459996006 | 0.713509775 |
| ENSMUSG00000085852 | Gm13807 | 639658 | lncRNA | 17.89107857 | -0.71352032 | 0.300091981 | -2.377672064 | 0.017422313 | 0.072285125 | 0.71352032 |
| ENSMUSG00000053510 | Nrd1 | 230598 | protein_coding | 1718.337763 | 0.713676105 | 0.055862732 | 12.77553172 | 2.25E-37 | 3.69E-35 | 0.713676105 |
| ENSMUSG00000038205 | Prkab2 | 108097 | protein_coding | 56.48078466 | -0.714107001 | 0.255858881 | -2.791019017 | 0.005254239 | 0.029160108 | 0.714107001 |
| ENSMUSG00000074818 | Pdzd7 | 100503041 | protein_coding | 134.6954579 | -0.714108627 | 0.320921623 | -2.22518078 | 0.026069105 | 0.097509572 | 0.714108627 |
| ENSMUSG00000049687 | Pheta2 | 338368 | protein_coding | 8.940861525 | -0.715209956 | 0.418695102 | -1.708188019 | 0.087601461 | 0.227918339 | 0.715209956 |
| ENSMUSG00000072244 | Trim6 | 94088 | protein_coding | 88.8588681 | 0.715413254 | 0.194989325 | 3.668986772 | 0.000243514 | 0.002395759 | 0.715413254 |
| ENSMUSG00000022619 | Mapk8ip2 | 60597 | protein_coding | 10.77613671 | -0.7156726 | 0.404078096 | -1.771124459 | 0.076540008 | 0.207123554 | 0.7156726 |
| ENSMUSG00000095620 | Csta2 | 76770 | protein_coding | 6.979863148 | 0.715696671 | 0.434092732 | 1.648718395 | 0.09920534 | 0.246974469 | 0.715696671 |
| ENSMUSG00000032356 | Rasgrf1 | 19417 | protein_coding | 7.324284894 | -0.716108444 | 1.040475239 | -0.688251308 | 0.491294539 | 0.683435757 | 0.716108444 |
| ENSMUSG00000033855 | Ston1 | 77057 | protein_coding | 62.07619668 | -0.716217452 | 0.211300539 | -3.389567561 | 0.00070003 | 0.00574154 | 0.716217452 |
| ENSMUSG00000024697 | Gna14 | 14675 | protein_coding | 909.2641529 | 0.716473189 | 0.185396806 | 3.864539008 | 0.000111299 | 0.001234522 | 0.716473189 |
| ENSMUSG00000052949 | Rnf157 | 217340 | protein_coding | 28.67696654 | 0.716590264 | 0.257001536 | 2.788272306 | 0.005298998 | 0.029275908 | 0.716590264 |
| ENSMUSG00000048058 | Ldlrad3 | 241576 | protein_coding | 350.1295346 | -0.716654254 | 0.191156024 | -3.749053988 | 0.000177503 | 0.001841867 | 0.716654254 |
| ENSMUSG00000056145 | AI504432 | 229694 | lncRNA | 7.850257836 | 0.71671232 | 0.655664214 | 1.093108797 | 0.274346034 | 0.482262844 | 0.71671232 |
| ENSMUSG00000043029 | Trpv3 | 246788 | protein_coding | 149.0527004 | -0.71748685 | 0.334755215 | -2.143317919 | 0.032087579 | 0.113374214 | 0.71748685 |
| ENSMUSG00000000555 | Itga5 | 16402 | protein_coding | 117.4350304 | -0.717878408 | 0.210838371 | -3.404875529 | 0.000661942 | 0.005491362 | 0.717878408 |
| ENSMUSG00000016494 | Cd34 | 12490 | protein_coding | 13.5902146 | -0.71812242 | 0.495545794 | -1.449154505 | 0.147294439 | 0.323868833 | 0.71812242 |
| ENSMUSG00000056468 | 5730596B20Rik | 77580 | TEC | 23.14345345 | -0.718514689 | 0.365749281 | -1.964500617 | 0.049472065 | 0.154314062 | 0.718514689 |
| ENSMUSG00000044916 | 1700029I15Rik | 75641 | protein_coding | 11.60112515 | -0.718829439 | 0.382581339 | -1.878893102 | 0.060259091 | 0.176337828 | 0.718829439 |
| ENSMUSG00000032177 | Pde4a | 18577 | protein_coding | 20.1247338 | -0.719228025 | 0.404287982 | -1.778999268 | 0.075239882 | 0.204718285 | 0.719228025 |
| ENSMUSG00000090655 | Vmn2r120 | 224916 | protein_coding | 10.16447095 | -0.719584601 | 1.636551682 | -0.439695617 | 0.660157578 | 0.803330517 | 0.719584601 |
| ENSMUSG00000042807 | Hecw2 | 329152 | protein_coding | 9.848301432 | -0.719644757 | 0.767815699 | -0.937262364 | 0.348623617 | 0.558808189 | 0.719644757 |
| ENSMUSG00000030670 | Cyp2r1 | 244209 | protein_coding | 52.88966497 | 0.719957435 | 0.168906421 | 4.262463386 | 2.02E-05 | 0.000280366 | 0.719957435 |
| ENSMUSG00000029798 | Herc6 | 67138 | protein_coding | 67.09952444 | 0.72044566 | 0.21402373 | 3.366195228 | 0.000762127 | 0.006137091 | 0.72044566 |
| ENSMUSG00000027070 | Lrp2 | 14725 | protein_coding | 9.474082994 | -0.72051183 | 1.535950553 | -0.469098324 | 0.638999357 | 0.790697132 | 0.72051183 |

| ensembl_gene_id | external_gene_name | entrezgene_id | gene_biotype | baseMean | log2FoldChange | lfcSE | stat | pvalue | padj | absLog2FC |
|---|---|---|---|---|---|---|---|---|---|---|
| ENSMUSG00000022860 | Chodl | 246048 | protein_coding | 20.38226037 | -0.720661301 | 0.5910953 | -1.219196466 | 0.222769633 | 0.424029005 | 0.720661301 |
| ENSMUSG00000067229 | Cyp2c66 | 69888 | protein_coding | 54.15146694 | -0.721003341 | 0.373230894 | -1.931789017 | 0.053385552 | 0.162213864 | 0.721003341 |
| ENSMUSG00000020319 | Wdpcp | 216560 | protein_coding | 71.82610543 | 0.721213215 | 0.155199717 | 4.647000824 | 3.37E-06 | 5.62E-05 | 0.721213215 |
| ENSMUSG00000018648 | Dusp14 | 56405 | protein_coding | 20.36711346 | 0.721636317 | 0.408960979 | 1.764560325 | 0.077637688 | 0.209453439 | 0.721636317 |
| ENSMUSG00000015222 | Map2 | 17756 | protein_coding | 94.8403795 | 0.721894037 | 0.199641613 | 3.615949731 | 0.000299248 | 0.002830421 | 0.721894037 |
| ENSMUSG00000042607 | Asb4 | 65255 | protein_coding | 35.66068891 | -0.722338692 | 0.228863814 | -3.156194415 | 0.001598423 | 0.011210779 | 0.722338692 |
| ENSMUSG00000078161 | Erich3 | 209601 | protein_coding | 8.424153706 | -0.722855224 | 0.723190641 | -0.999536198 | 0.317535013 | 0.528771321 | 0.722855224 |
| ENSMUSG00000037738 | Nek5 | 330721 | protein_coding | 18.93781842 | -0.722885493 | 0.322337763 | -2.242633587 | 0.024920453 | 0.094151825 | 0.722885493 |
| ENSMUSG00000025791 | Pgm1 | 72157 | protein_coding | 1430.398613 | 0.72288868 | 0.152844808 | 4.729559925 | 2.25E-06 | 3.88E-05 | 0.72288868 |
| ENSMUSG00000056293 | Gsdmc2 | 331063 | protein_coding | 2729.523625 | -0.723250081 | 0.11515236 | -6.280809878 | 3.37E-10 | 1.08E-08 | 0.723250081 |
| ENSMUSG00000023132 | Gzma | 14938 | protein_coding | 28.94709716 | 0.723398144 | 0.379669595 | 1.90533599 | 0.05673641 | 0.169428931 | 0.723398144 |
| ENSMUSG00000114442 | F630042J09Rik | 100038627 | lncRNA | 7.50839214 | -0.72379321 | 0.485536195 | -1.490709069 | 0.136037894 | 0.306003612 | 0.72379321 |
| ENSMUSG00000013089 | Etv5 | 104156 | protein_coding | 459.3444628 | -0.72400548 | 0.1350036 | -5.362860548 | 8.19E-08 | 1.89E-06 | 0.72400548 |
| ENSMUSG00000019929 | Dcn | 13179 | protein_coding | 36.09227333 | -0.724276711 | 0.482592977 | -1.500802428 | 0.13340667 | 0.301875472 | 0.724276711 |
| ENSMUSG00000098112 | Bin2 | 668218 | protein_coding | 16.92276734 | 0.724562136 | 0.35298343 | 2.052680309 | 0.040103593 | 0.133327078 | 0.724562136 |
| ENSMUSG00000032500 | Dclk3 | 245038 | protein_coding | 38.09099099 | -0.724709763 | 0.289799875 | -2.500724899 | 0.012393941 | 0.056178342 | 0.724709763 |
| ENSMUSG00000022498 | Txndc11 | 106200 | protein_coding | 1000.64712 | 0.724919741 | 0.084366993 | 8.592456764 | 8.51E-18 | 5.30E-16 | 0.724919741 |
| ENSMUSG00000064125 | Prr36 | 73072 | protein_coding | 10.98490616 | -0.724950418 | 0.646152189 | -1.121949953 | 0.26188372 | 0.469739177 | 0.724950418 |
| ENSMUSG00000058756 | Thra | 21833 | protein_coding | 454.037352 | -0.725033654 | 0.115490072 | -6.277887272 | 3.43E-10 | 1.10E-08 | 0.725033654 |
| ENSMUSG00000025170 | Rab40b | 217371 | protein_coding | 6.787540918 | -0.725052523 | 0.462964355 | -1.56610874 | 0.117323162 | 0.276561096 | 0.725052523 |
| ENSMUSG00000072915 | Gm12258 | 108167848 | protein_coding | 36.88814226 | 0.725503937 | 0.228077798 | 3.180949415 | 0.001467933 | 0.010475462 | 0.725503937 |
| ENSMUSG00000067158 | Col4a4 | 12829 | protein_coding | 9.319332273 | -0.725963634 | 0.85926046 | -0.84487029 | 0.398183264 | 0.606149922 | 0.725963634 |
| ENSMUSG00000069273 | H3c6 | 319151 | protein_coding | 1576.488817 | 0.726146081 | 0.180913596 | 4.013772854 | 5.98E-05 | 0.000721054 | 0.726146081 |
| ENSMUSG00000060063 | Alox5ap | 11690 | protein_coding | 24.82041386 | -0.72645482 | 0.288102921 | -2.521511473 | 0.011685186 | 0.053776842 | 0.72645482 |
| ENSMUSG00000035699 | Slc51a | 106407 | protein_coding | 614.9626002 | -0.72656568 | 0.424995805 | -1.709583181 | 0.087342978 | 0.22744507 | 0.72656568 |
| ENSMUSG00000054453 | Sytl5 | 236643 | protein_coding | 25.4119865 | 0.726863858 | 0.33957047 | 2.140539068 | 0.032311227 | 0.113962184 | 0.726863858 |
| ENSMUSG00000027434 | Nkx2-2 | 18088 | protein_coding | 12.90769052 | -0.726882958 | 0.370585323 | -1.96144562 | 0.049827063 | 0.155008878 | 0.726882958 |
| ENSMUSG00000014329 | Bicc1 | 83675 | protein_coding | 10.71212095 | -0.726893875 | 0.455721958 | -1.595038074 | 0.110703722 | 0.265435629 | 0.726893875 |
| ENSMUSG00000055254 | Ntrk2 | 18212 | protein_coding | 199.8014756 | 0.72693075 | 0.185806375 | 3.912302532 | 9.14E-05 | 0.001045328 | 0.72693075 |
| ENSMUSG00000102785 | Gm2447 | 329639 | lncRNA | 6.968876296 | 0.727252635 | 0.457846729 | 1.588419419 | 0.112191529 | 0.26771336 | 0.727252635 |
| ENSMUSG00000030747 | Dgat2 | 67800 | protein_coding | 691.1322307 | 0.727782905 | 0.276250609 | 2.634502452 | 0.00842607 | 0.042138284 | 0.727782905 |
| ENSMUSG00000053414 | Hunk | 26559 | protein_coding | 381.3594872 | -0.727955088 | 0.120630098 | -6.034605791 | 1.59E-09 | 4.72E-08 | 0.727955088 |
| ENSMUSG00000022504 | Ciita | 12265 | protein_coding | 21.69593516 | 0.728104516 | 0.407465843 | 1.786909329 | 0.073952144 | 0.202773241 | 0.728104516 |
| ENSMUSG00000001823 | Hoxd12 | 15432 | protein_coding | 6.826515592 | 0.728302339 | 0.832415468 | 0.874926484 | 0.381613908 | 0.589386825 | 0.728302339 |
| ENSMUSG00000048865 | Arhgap30 | 226652 | protein_coding | 28.22051705 | 0.728563935 | 0.386486384 | 1.885095997 | 0.059416878 | 0.17500132 | 0.728563935 |
| ENSMUSG00000020131 | Pcsk4 | 18551 | protein_coding | 27.02973035 | -0.729219199 | 0.275581675 | -2.646109178 | 0.008142353 | 0.0410936 | 0.729219199 |
| ENSMUSG00000097327 | E030030I06Rik | 319887 | transcribed_unprocessed_pseudogene | 65.73132801 | 0.729262963 | 0.164627736 | 4.429769744 | 9.43E-06 | 0.000141411 | 0.729262963 |
| ENSMUSG00000037820 | Tgm2 | 21817 | protein_coding | 390.9426689 | 0.729271172 | 0.266341003 | 2.738110786 | 0.006179326 | 0.032928838 | 0.729271172 |

| ensembl_gene_id | external_gene_name | entrezgene_id | gene_biotype | baseMean | log2FoldChange | lfcSE | stat | pvalue | padj | absLog2FC |
|---|---|---|---|---|---|---|---|---|---|---|
| ENSMUSG00000004612 | Nkg7 | 72310 | protein_coding | 16.14439978 | 0.729350425 | 0.389710222 | 1.871519873 | 0.061273058 | 0.178191396 | 0.729350425 |
| ENSMUSG00000030598 | Fbxo17 | 50760 | protein_coding | 15.22601121 | -0.73014172 | 0.300316502 | -2.431240758 | 0.015047212 | 0.064811248 | 0.73014172 |
| ENSMUSG00000046841 | Ckap4 | 216197 | protein_coding | 580.0373688 | 0.730429637 | 0.13644658 | 5.353227895 | 8.64E-08 | 1.99E-06 | 0.730429637 |
| ENSMUSG00000027254 | Map1a | 17754 | protein_coding | 21.43767568 | -0.730874012 | 0.400986306 | -1.822690705 | 0.068350249 | 0.192414769 | 0.730874012 |
| ENSMUSG00000043535 | Setx | 269254 | protein_coding | 1633.477728 | 0.731048564 | 0.065172452 | 11.21714071 | 3.36E-29 | 3.74E-27 | 0.731048564 |
| ENSMUSG00000024131 | Slc3a1 | 20532 | protein_coding | 522.6602984 | 0.731122274 | 0.245565643 | 2.977298712 | 0.002908005 | 0.018300556 | 0.731122274 |
| ENSMUSG00000034412 | Tbc1d10a | 103724 | protein_coding | 261.4030452 | 0.731391114 | 0.215753046 | 3.389945719 | 0.000699065 | 0.005739537 | 0.731391114 |
| ENSMUSG00000006378 | Gcat | 26912 | protein_coding | 67.49173424 | 0.731881491 | 0.19656669 | 3.72332409 | 0.000196617 | 0.002012669 | 0.731881491 |
| ENSMUSG00000025144 | Cenpx | 20892 | protein_coding | 254.9794691 | 0.732263988 | 0.094217796 | 7.77203482 | 7.72E-15 | 3.86E-13 | 0.732263988 |
| ENSMUSG00000027950 | Chrnb2 | 11444 | protein_coding | 42.96888675 | -0.732406549 | 0.214824331 | -3.40932774 | 0.000651232 | 0.005425116 | 0.732406549 |
| ENSMUSG00000032186 | Tmod2 | 50876 | protein_coding | 16.15938785 | -0.732586685 | 0.475368154 | -1.541093317 | 0.123294076 | 0.286397557 | 0.732586685 |
| ENSMUSG00000021379 | Id4 | 15904 | protein_coding | 121.4114598 | 0.732907099 | 0.163696053 | 4.477243547 | 7.56E-06 | 0.000116702 | 0.732907099 |
| ENSMUSG00000028544 | Slc5a9 | 230612 | protein_coding | 277.5715433 | 0.733543072 | 0.227538147 | 3.223824586 | 0.001264909 | 0.009250442 | 0.733543072 |
| ENSMUSG00000022543 | Dnaaf8 | 74684 | transcribed_unitary_pseudogene | 11.12402713 | 0.733625517 | 0.445677822 | 1.646089352 | 0.099745364 | 0.247738057 | 0.733625517 |
| ENSMUSG00000029121 | Crmp1 | 12933 | protein_coding | 9.101616828 | -0.734462029 | 0.518770674 | -1.415773995 | 0.156841686 | 0.337636758 | 0.734462029 |
| ENSMUSG00000046157 | Tmem229b | 268567 | protein_coding | 34.06020006 | 0.734699799 | 0.219228782 | 3.351292626 | 0.000804353 | 0.006402663 | 0.734699799 |
| ENSMUSG00000032578 | Cish | 12700 | protein_coding | 103.1463384 | -0.735025364 | 0.173606412 | -4.233860694 | 2.30E-05 | 0.000311924 | 0.735025364 |
| ENSMUSG00000064284 | Cdpf1 | 72355 | protein_coding | 315.311136 | 0.735135078 | 0.118574083 | 6.199795605 | 5.65E-10 | 1.75E-08 | 0.735135078 |
| ENSMUSG00000027225 | Duoxa2 | 66811 | protein_coding | 125.7710116 | 0.735734738 | 0.353421994 | 2.081745765 | 0.037365698 | 0.126764822 | 0.735734738 |
| ENSMUSG00000048578 | Mlec | 109154 | protein_coding | 8421.169686 | 0.736334758 | 0.072595922 | 10.1429218 | 3.56E-24 | 3.08E-22 | 0.736334758 |
| ENSMUSG00000026950 | Neb | 17996 | protein_coding | 23.8885372 | -0.736468489 | 0.781021732 | -0.942955181 | 0.345703828 | 0.555637797 | 0.736468489 |
| ENSMUSG00000042195 | Slc35f2 | 72022 | protein_coding | 290.8683505 | 0.736861533 | 0.098896685 | 7.450821389 | 9.28E-14 | 4.22E-12 | 0.736861533 |
| ENSMUSG00000080712 | H2bc26-ps | 382522 | protein_coding | 34.92288957 | 0.737324526 | 0.221517815 | 3.328511184 | 0.000873115 | 0.006857483 | 0.737324526 |
| ENSMUSG00000049807 | Arhgap23 | 58996 | protein_coding | 7.906565051 | -0.737468145 | 0.562262204 | -1.31160896 | 0.189652108 | 0.381218927 | 0.737468145 |
| ENSMUSG00000040928 | S100pbp | 74648 | protein_coding | 216.5071478 | -0.737757537 | 0.184234921 | -4.004439194 | 6.22E-05 | 0.000746727 | 0.737757537 |
| ENSMUSG00000094145 | Vmn2r20 | 667180 | protein_coding | 10.24174017 | -0.738098057 | 1.860132415 | -0.396798664 | 0.691515936 | 0.824338032 | 0.738098057 |
| ENSMUSG00000039908 | Slc26a11 | 268512 | protein_coding | 59.63575466 | -0.738568303 | 0.189199945 | -3.903639106 | 9.48E-05 | 0.001077295 | 0.738568303 |
| ENSMUSG00000021684 | Pde8b | 218461 | protein_coding | 323.0923233 | -0.738652492 | 0.178478434 | -4.138609222 | 3.49E-05 | 0.000452481 | 0.738652492 |
| ENSMUSG00000001670 | Tat | 234724 | protein_coding | 264.3432844 | 0.738660402 | 0.222196176 | 3.324361446 | 0.000886213 | 0.00693982 | 0.738660402 |
| ENSMUSG00000073805 | Insyn2a | 627214 | protein_coding | 27.39977604 | -0.738772808 | 0.400850809 | -1.843011891 | 0.065327274 | 0.186108536 | 0.738772808 |
| ENSMUSG00000031937 | Vstm5 | 69137 | protein_coding | 89.22028173 | -0.738893723 | 0.196322547 | -3.763672247 | 0.000167436 | 0.001746327 | 0.738893723 |
| ENSMUSG00000066113 | Adamtsl1 | 77739 | protein_coding | 7.287786334 | -0.739240896 | 1.047647231 | -0.705620054 | 0.48042445 | 0.673965179 | 0.739240896 |
| ENSMUSG00000006736 | Tspan31 | 67125 | protein_coding | 643.7366411 | 0.739287129 | 0.096654502 | 7.648760456 | 2.03E-14 | 9.76E-13 | 0.739287129 |
| ENSMUSG00000069917 | Hba-a2 | 110257 | protein_coding | 18.85162911 | 0.739784608 | 0.460420017 | 1.606760308 | 0.108106959 | 0.26169113 | 0.739784608 |
| ENSMUSG00000026726 | Cubn | 65969 | protein_coding | 120.0144826 | -0.740412939 | 0.307217357 | -2.410062202 | 0.015949801 | 0.067728187 | 0.740412939 |
| ENSMUSG00000053746 | Ptrh1 | 329384 | protein_coding | 131.3686509 | 0.740488263 | 0.165494934 | 4.474386287 | 7.66E-06 | 0.000118044 | 0.740488263 |
| ENSMUSG00000028370 | Pappa | 18491 | protein_coding | 13.29205473 | 0.740600576 | 0.560530504 | 1.321249372 | 0.186418229 | 0.376888975 | 0.740600576 |
| ENSMUSG00000061740 | Cyp2d22 | 56448 | protein_coding | 119.2300136 | -0.740961678 | 0.164862839 | -4.494412941 | 6.98E-06 | 0.000108725 | 0.740961678 |

| ensembl_gene_id | external_gene_name | entrezgene_id | gene_biotype | baseMean | log2FoldChange | lfcSE | stat | pvalue | padj | absLog2FC |
|---|---|---|---|---|---|---|---|---|---|---|
| ENSMUSG00000032280 | Tle3 | 21887 | protein_coding | 683.5774867 | -0.741540771 | 0.107659929 | -6.887806633 | 5.67E-12 | 2.17E-10 | 0.741540771 |
| ENSMUSG00000028940 | Hes2 | 15206 | protein_coding | 66.56040012 | 0.742038832 | 0.21844644 | 3.396891397 | 0.00068156 | 0.00563357 | 0.742038832 |
| ENSMUSG00000090145 | Ugt1a6b | 394435 | protein_coding | 13.0262551 | -0.742180884 | 0.410202106 | -1.809305396 | 0.070403571 | 0.19615605 | 0.742180884 |
| ENSMUSG00000024598 | Fbn2 | 14119 | protein_coding | 10.64195383 | -0.742469736 | 1.038923465 | -0.714652966 | 0.474823507 | 0.669646611 | 0.742469736 |
| ENSMUSG00000064341 | mt-Nd1 | 17716 | protein_coding | 62755.35166 | -0.742532783 | 0.182971855 | -4.058180337 | 4.95E-05 | 0.000613508 | 0.742532783 |
| ENSMUSG00000021490 | Slc34a1 | 20505 | protein_coding | 17.04054873 | 0.742610184 | 0.374049889 | 1.985323902 | 0.047108446 | 0.149341617 | 0.742610184 |
| ENSMUSG00000002325 | Irf9 | 16391 | protein_coding | 463.0328901 | 0.743068176 | 0.102882308 | 7.222506872 | 5.10E-13 | 2.17E-11 | 0.743068176 |
| ENSMUSG00000071181 | 3830408C21Rik | 100040322 | lncRNA | 8.417889387 | -0.74307431 | 0.716882649 | -1.036535493 | 0.29995239 | 0.508888297 | 0.74307431 |
| ENSMUSG00000028207 | Asph | 65973 | protein_coding | 2394.115415 | 0.743562168 | 0.042526784 | 17.48456139 | 1.88E-68 | 6.80E-66 | 0.743562168 |
| ENSMUSG00000070704 | Ugt2b36 | 231396 | protein_coding | 100.7078435 | 0.744472948 | 0.2268953 | 3.281129882 | 0.001033921 | 0.007872035 | 0.744472948 |
| ENSMUSG00000020135 | Apc2 | 23805 | protein_coding | 15.88001169 | -0.744549603 | 0.416872647 | -1.786036114 | 0.074093411 | 0.202950963 | 0.744549603 |
| ENSMUSG00000054404 | Slfn5 | 327978 | protein_coding | 14.63308492 | 0.745011105 | 0.573567482 | 1.298907503 | 0.193975675 | 0.387387373 | 0.745011105 |
| ENSMUSG00000000253 | Gmpr | 66355 | protein_coding | 151.5955545 | -0.745064083 | 0.250971773 | -2.968716655 | 0.002990462 | 0.018738033 | 0.745064083 |
| ENSMUSG00000029380 | Cxcl1 | 14825 | protein_coding | 23.21398433 | -0.745746642 | 0.399828891 | -1.865164473 | 0.062158354 | 0.179879771 | 0.745746642 |
| ENSMUSG00000032065 | Tex12 | 66654 | protein_coding | 27.27653282 | 0.745961084 | 0.294625502 | 2.531895843 | 0.011344768 | 0.052590068 | 0.745961084 |
| ENSMUSG00000090588 | Muc21 | 672682 | protein_coding | 11.08554562 | -0.74627698 | 1.963753964 | -0.380025703 | 0.703926336 | 0.832625775 | 0.74627698 |
| ENSMUSG00000030731 | Syt3 | 20981 | protein_coding | 7.431135709 | -0.746529461 | 0.513682387 | -1.45328997 | 0.146143261 | 0.32213733 | 0.746529461 |
| ENSMUSG00000022332 | Khdrbs3 | 13992 | protein_coding | 411.7341919 | 0.746718123 | 0.101670884 | 7.34446377 | 2.07E-13 | 9.02E-12 | 0.746718123 |
| ENSMUSG00000000594 | Gm2a | 14667 | protein_coding | 191.612588 | -0.747362042 | 0.188173799 | -3.971658355 | 7.14E-05 | 0.000840526 | 0.747362042 |
| ENSMUSG00000044162 | Tnip3 | 414084 | protein_coding | 320.9709707 | 0.747671196 | 0.285051611 | 2.622932712 | 0.008717649 | 0.043229984 | 0.747671196 |
| ENSMUSG00000072884 | Gm10433 | 100038378 | lncRNA | 25.07191257 | 0.747737537 | 0.24999742 | 2.990981016 | 0.002780828 | 0.01766774 | 0.747737537 |
| ENSMUSG00000031207 | Msn | 17698 | protein_coding | 34.97525772 | 0.748254314 | 0.34121072 | 2.192939053 | 0.02831177 | 0.103239441 | 0.748254314 |
| ENSMUSG00000024027 | Glp1r | 14652 | protein_coding | 20.25990737 | -0.748301374 | 0.397969459 | -1.880298492 | 0.060067409 | 0.17598711 | 0.748301374 |
| ENSMUSG00000020573 | Pik3cg | 30955 | protein_coding | 124.9374222 | -0.748319483 | 0.138450313 | -5.404967791 | 6.48E-08 | 1.53E-06 | 0.748319483 |
| ENSMUSG00000003585 | Sec14l2 | 67815 | protein_coding | 87.54527498 | 0.748442621 | 0.182067923 | 4.110787938 | 3.94E-05 | 0.00050112 | 0.748442621 |
| ENSMUSG00000085024 | C230035I16Rik | 320842 | lncRNA | 7.838599012 | -0.749355324 | 0.410430522 | -1.825778749 | 0.067883596 | 0.191512591 | 0.749355324 |
| ENSMUSG00000026930 | Gpsm1 | 67839 | protein_coding | 84.92434875 | -0.750316962 | 0.172018024 | -4.36185085 | 1.29E-05 | 0.000186405 | 0.750316962 |
| ENSMUSG00000064647 | Gm23301 | 115488997 | snoRNA | 7.295990431 | -0.751162383 | 0.467849063 | -1.605565644 | 0.108369379 | 0.262007813 | 0.751162383 |
| ENSMUSG00000072294 | Klf12 | 16597 | protein_coding | 240.1683382 | 0.751454524 | 0.139001444 | 5.406091503 | 6.44E-08 | 1.52E-06 | 0.751454524 |
| ENSMUSG00000049350 | Zg16 | 69036 | protein_coding | 43505.61411 | 0.751800015 | 0.203208913 | 3.699640945 | 0.000215905 | 0.002176538 | 0.751800015 |
| ENSMUSG00000044125 | 9530080O11Rik | 319247 | lncRNA | 14.56058899 | 0.752088076 | 0.323916045 | 2.321861139 | 0.020240414 | 0.08055705 | 0.752088076 |
| ENSMUSG00000030579 | Tyrobp | 22177 | protein_coding | 13.13042045 | 0.752516173 | 0.380523502 | 1.977581331 | 0.047975957 | 0.151408965 | 0.752516173 |
| ENSMUSG00000023031 | Cela1 | 109901 | protein_coding | 1716.763751 | -0.753498083 | 0.233190021 | -3.231262133 | 0.001232449 | 0.009083961 | 0.753498083 |
| ENSMUSG00000037129 | Tmprss13 | 214531 | protein_coding | 46.57610908 | -0.753843654 | 0.580647056 | -1.298282056 | 0.19419043 | 0.387494219 | 0.753843654 |
| ENSMUSG00000054423 | Cadps | 27062 | protein_coding | 32.92957396 | -0.753998119 | 0.26185846 | -2.879410951 | 0.003984188 | 0.023401547 | 0.753998119 |
| ENSMUSG00000037907 | Ankrd13b | 268445 | protein_coding | 68.53335454 | -0.754028873 | 0.176533625 | -4.271304543 | 1.94E-05 | 0.000270335 | 0.754028873 |
| ENSMUSG00000058589 | Anks1b | 77531 | protein_coding | 11.87370371 | -0.754471579 | 0.827647278 | -0.911585888 | 0.361986754 | 0.571259786 | 0.754471579 |
| ENSMUSG00000063063 | Ctnna2 | 12386 | protein_coding | 12.60801525 | -0.754486162 | 0.520517422 | -1.449492622 | 0.147200059 | 0.323750697 | 0.754486162 |

| ensembl_gene_id | external_gene_name | entrezgene_id | gene_biotype | baseMean | log2FoldChange | lfcSE | stat | pvalue | padj | absLog2FC |
|---|---|---|---|---|---|---|---|---|---|---|
| ENSMUSG00000064202 | Spata6l | 381218 | protein_coding | 24.03440514 | 0.754793884 | 0.277554002 | 2.719448756 | 0.006539083 | 0.034546205 | 0.754793884 |
| ENSMUSG00000045545 | Krt14 | 16664 | protein_coding | 12.51258868 | -0.755350036 | 0.473637316 | -1.594785737 | 0.110760158 | 0.265491014 | 0.755350036 |
| ENSMUSG00000036299 | BC031181 | 407819 | protein_coding | 1268.613514 | 0.755354616 | 0.108179653 | 6.98240929 | 2.90E-12 | 1.14E-10 | 0.755354616 |
| ENSMUSG00000004902 | Slc25a18 | 71803 | protein_coding | 10.62927502 | -0.755640266 | 0.405895404 | -1.861662534 | 0.062650673 | 0.180967229 | 0.755640266 |
| ENSMUSG00000000266 | Mid2 | 23947 | protein_coding | 1126.402006 | 0.755855303 | 0.093544175 | 8.080196385 | 6.47E-16 | 3.53E-14 | 0.755855303 |
| ENSMUSG00000068740 | Celsr2 | 53883 | protein_coding | 52.99672475 | -0.756131167 | 0.286702255 | -2.637339444 | 0.008355917 | 0.041902591 | 0.756131167 |
| ENSMUSG00000030214 | Plbd1 | 66857 | protein_coding | 741.173882 | -0.756153031 | 0.191947347 | -3.939377356 | 8.17E-05 | 0.000945648 | 0.756153031 |
| ENSMUSG00000041763 | Tpp2 | 22019 | protein_coding | 2027.953647 | 0.756335741 | 0.090499744 | 8.357324621 | 6.42E-17 | 3.76E-15 | 0.756335741 |
| ENSMUSG00000039021 | Ttc16 | 338348 | protein_coding | 8.430515752 | -0.756382719 | 0.55616657 | -1.359993139 | 0.173832095 | 0.359707067 | 0.756382719 |
| ENSMUSG00000062859 | Tcp11 | 21463 | protein_coding | 11.243639 | -0.756725064 | 0.431272404 | -1.75463363 | 0.079321995 | 0.212772102 | 0.756725064 |
| ENSMUSG00000029463 | Fam216a | 68948 | protein_coding | 18.64030027 | 0.756880771 | 0.333790847 | 2.267530037 | 0.023357866 | 0.08975732 | 0.756880771 |
| ENSMUSG00000030824 | Nucb1 | 18220 | protein_coding | 5554.511479 | 0.756973563 | 0.071468646 | 10.59168745 | 3.26E-26 | 3.07E-24 | 0.756973563 |
| ENSMUSG00000045672 | Col27a1 | 373864 | protein_coding | 14.87599955 | -0.757374794 | 0.645491404 | -1.173330566 | 0.240663269 | 0.445213072 | 0.757374794 |
| ENSMUSG00000055200 | Sertad3 | 170742 | protein_coding | 232.965567 | 0.757894536 | 0.082939886 | 9.137877667 | 6.37E-20 | 4.49E-18 | 0.757894536 |
| ENSMUSG00000081058 | H3c15 | 97114 | protein_coding | 13.86903453 | 0.758123445 | 0.327539564 | 2.314601132 | 0.020634771 | 0.081840296 | 0.758123445 |
| ENSMUSG00000029797 | Sspo | 243369 | protein_coding | 67.15091977 | -0.759011451 | 0.247931683 | -3.061373368 | 0.002203242 | 0.014658828 | 0.759011451 |
| ENSMUSG00000023176 | Cpn2 | 71756 | protein_coding | 25.61472793 | 0.759052717 | 0.534052114 | 1.421308328 | 0.155227143 | 0.335187361 | 0.759052717 |
| ENSMUSG00000052105 | Mtcl1 | 68617 | protein_coding | 109.6323838 | -0.759340928 | 0.295055055 | -2.573556754 | 0.010065914 | 0.048118209 | 0.759340928 |
| ENSMUSG00000001827 | Folr1 | 14275 | protein_coding | 10.31761248 | -0.760053467 | 0.385386038 | -1.972187347 | 0.048588224 | 0.152705845 | 0.760053467 |
| ENSMUSG00000022123 | Scel | 64929 | protein_coding | 58.0420896 | -0.760777196 | 0.327461698 | -2.323255517 | 0.02016543 | 0.08030166 | 0.760777196 |
| ENSMUSG00000049916 | 2610318N02Rik | 70458 | protein_coding | 37.72817816 | 0.761099094 | 0.273121494 | 2.786668608 | 0.005325291 | 0.029380405 | 0.761099094 |
| ENSMUSG00000031485 | Plpbp | 114863 | protein_coding | 949.6185589 | 0.761149269 | 0.069531684 | 10.94679751 | 6.88E-28 | 7.12E-26 | 0.761149269 |
| ENSMUSG00000022722 | Arl6 | 56297 | protein_coding | 37.83166437 | 0.761434413 | 0.207525486 | 3.669112787 | 0.000243394 | 0.002395759 | 0.761434413 |
| ENSMUSG00000022358 | Fbxo32 | 67731 | protein_coding | 745.2673276 | -0.761443749 | 0.28873837 | -2.637140844 | 0.008360811 | 0.041902591 | 0.761443749 |
| ENSMUSG00000003352 | Cacnb3 | 12297 | protein_coding | 419.6351692 | -0.761527165 | 0.096153538 | -7.919907886 | 2.38E-15 | 1.25E-13 | 0.761527165 |
| ENSMUSG00000085793 | Lin52 | 217708 | protein_coding | 165.0703703 | 0.762420666 | 0.106294065 | 7.172749192 | 7.35E-13 | 3.07E-11 | 0.762420666 |
| ENSMUSG00000032175 | Tyk2 | 54721 | protein_coding | 869.0529649 | 0.762436002 | 0.092404496 | 8.251070371 | 1.57E-16 | 9.03E-15 | 0.762436002 |
| ENSMUSG00000063455 | D630045J12Rik | 330286 | protein_coding | 21.25990434 | -0.762440679 | 0.339302549 | -2.247082086 | 0.024634782 | 0.093402238 | 0.762440679 |
| ENSMUSG00000031596 | Slc7a2 | 11988 | protein_coding | 16.96131739 | -0.762440681 | 0.371691507 | -2.051272807 | 0.040240387 | 0.133614527 | 0.762440681 |
| ENSMUSG00000071716 | Apol7e | 666348 | protein_coding | 9.291675224 | 0.762632239 | 0.418868972 | 1.820694035 | 0.068653379 | 0.19299524 | 0.762632239 |
| ENSMUSG00000029648 | Flt1 | 14254 | protein_coding | 9.08838455 | -0.762911549 | 1.135250048 | -0.672020715 | 0.501570509 | 0.691924848 | 0.762911549 |
| ENSMUSG00000051295 | 9630028B13Rik | 319562 | lncRNA | 10.63156179 | -0.763225653 | 0.435210441 | -1.753693342 | 0.079483067 | 0.212988945 | 0.763225653 |
| ENSMUSG00000058975 | Kcnc1 | 16502 | protein_coding | 8.799585681 | -0.763750731 | 0.568459639 | -1.34354434 | 0.179095778 | 0.366662925 | 0.763750731 |
| ENSMUSG00000000184 | Ccnd2 | 12444 | protein_coding | 1697.534084 | -0.76394176 | 0.151579784 | -5.039865725 | 4.66E-07 | 9.57E-06 | 0.76394176 |
| ENSMUSG00000022899 | Slc15a2 | 57738 | protein_coding | 145.1131729 | -0.764179922 | 0.63220192 | -1.208759257 | 0.226755349 | 0.429204895 | 0.764179922 |
| ENSMUSG00000044197 | Gpr146 | 80290 | protein_coding | 10.97444657 | -0.7642237 | 0.385088055 | -1.984542732 | 0.047195369 | 0.149567816 | 0.7642237 |
| ENSMUSG00000026628 | Atf3 | 11910 | protein_coding | 3925.61582 | -0.76441227 | 0.632727183 | -1.208123012 | 0.22699995 | 0.429514754 | 0.76441227 |
| ENSMUSG00000021700 | Rab3c | 67295 | protein_coding | 44.2634331 | -0.764754596 | 0.260501276 | -2.935703841 | 0.003327919 | 0.020410894 | 0.764754596 |

| ensembl_gene_id | external_gene_name | entrezgene_id | gene_biotype | baseMean | log2FoldChange | lfcSE | stat | pvalue | padj | absLog2FC |
|---|---|---|---|---|---|---|---|---|---|---|
| ENSMUSG00000057098 | Ebf1 | 13591 | protein_coding | 8.017748057 | -0.765642904 | 1.064129099 | -0.719501896 | 0.471831735 | 0.667318049 | 0.765642904 |
| ENSMUSG00000032040 | Dcps | 69305 | protein_coding | 248.6312387 | 0.766445028 | 0.134994246 | 5.677612583 | 1.37E-08 | 3.61E-07 | 0.766445028 |
| ENSMUSG00000080727 | C920021L13Rik | 100042889 | lncRNA | 8.2845169 | -0.766991172 | 0.433687739 | -1.768533216 | 0.076971804 | 0.208044612 | 0.766991172 |
| ENSMUSG00000054435 | Gimap4 | 107526 | protein_coding | 62.89011464 | 0.767108709 | 0.293247941 | 2.615904845 | 0.008899135 | 0.043913482 | 0.767108709 |
| ENSMUSG00000029377 | Ereg | 13874 | protein_coding | 88.67871123 | 0.767621107 | 0.372938596 | 2.058304274 | 0.039560934 | 0.132185139 | 0.767621107 |
| ENSMUSG00000041025 | Iffo2 | 212632 | protein_coding | 217.6482506 | -0.767739203 | 0.11670433 | -6.578498008 | 4.75E-11 | 1.69E-09 | 0.767739203 |
| ENSMUSG00000032514 | Ttc21a | 74052 | protein_coding | 10.83085722 | -0.768053286 | 0.381951361 | -2.010866736 | 0.044339535 | 0.142814212 | 0.768053286 |
| ENSMUSG00000044726 | Erich5 | 239368 | protein_coding | 9.47453691 | -0.768303468 | 0.431421638 | -1.780864474 | 0.074934594 | 0.204201575 | 0.768303468 |
| ENSMUSG00000032932 | Hspa13 | 110920 | protein_coding | 926.993722 | 0.769034052 | 0.099088639 | 7.761071867 | 8.42E-15 | 4.18E-13 | 0.769034052 |
| ENSMUSG00000038156 | Spon1 | 233744 | protein_coding | 21.89523737 | -0.769643451 | 0.394816794 | -1.949368578 | 0.051251424 | 0.158099111 | 0.769643451 |
| ENSMUSG00000023224 | Serping1 | 12258 | protein_coding | 8.275431522 | -0.77068725 | 0.460094165 | -1.675064169 | 0.093921642 | 0.23874525 | 0.77068725 |
| ENSMUSG00000056222 | Spock1 | 20745 | protein_coding | 7.445773127 | -0.770699533 | 1.036833396 | -0.743320514 | 0.457287654 | 0.656009886 | 0.770699533 |
| ENSMUSG00000040734 | Ppp1r13l | 333654 | protein_coding | 41.13141849 | -0.771040597 | 0.24946959 | -3.090719786 | 0.00199672 | 0.013493318 | 0.771040597 |
| ENSMUSG00000040105 | Plpp6 | 74411 | protein_coding | 702.070937 | 0.77135249 | 0.07594854 | 10.15625166 | 3.11E-24 | 2.75E-22 | 0.77135249 |
| ENSMUSG00000031060 | Rbm10 | 236732 | protein_coding | 1117.087514 | 0.771356588 | 0.107826679 | 7.153670996 | 8.45E-13 | 3.52E-11 | 0.771356588 |
| ENSMUSG00000058099 | Nfam1 | 74039 | protein_coding | 10.54929242 | 0.771372331 | 0.622501231 | 1.23914989 | 0.215289994 | 0.414147225 | 0.771372331 |
| ENSMUSG00000020437 | Myo1g | 246177 | protein_coding | 19.1105424 | 0.771829428 | 0.474165236 | 1.627764688 | 0.103574791 | 0.254232438 | 0.771829428 |
| ENSMUSG00000099988 | 1700054A03Rik | 78457 | lncRNA | 9.438179233 | -0.771893025 | 0.526663069 | -1.46562968 | 0.142749201 | 0.316320155 | 0.771893025 |
| ENSMUSG00000002014 | Ssr4 | 20832 | protein_coding | 2445.241705 | 0.771933452 | 0.104895805 | 7.359049802 | 1.85E-13 | 8.13E-12 | 0.771933452 |
| ENSMUSG00000005125 | Ndrg1 | 17988 | protein_coding | 1994.616834 | -0.772034241 | 0.15608725 | -4.946171082 | 7.57E-07 | 1.47E-05 | 0.772034241 |
| ENSMUSG00000103948 | 4930594C11Rik | 77633 | TEC | 15.23083819 | -0.772858323 | 0.39324662 | -1.965327317 | 0.049376366 | 0.15411851 | 0.772858323 |
| ENSMUSG00000001999 | Blvra | 109778 | protein_coding | 301.0143389 | 0.773035204 | 0.110796952 | 6.977043971 | 3.01E-12 | 1.18E-10 | 0.773035204 |
| ENSMUSG00000064370 | mt-Cytb | 17711 | protein_coding | 75438.98575 | -0.773329179 | 0.182545018 | -4.236375155 | 2.27E-05 | 0.000308717 | 0.773329179 |
| ENSMUSG00000032826 | Ank2 | 109676 | protein_coding | 24.74757747 | -0.773504603 | 0.972234088 | -0.795595025 | 0.426267459 | 0.63082673 | 0.773504603 |
| ENSMUSG00000015467 | Egfl8 | 81701 | protein_coding | 11.37001492 | -0.774094851 | 0.420679497 | -1.840105966 | 0.065752682 | 0.186986023 | 0.774094851 |
| ENSMUSG00000030321 | Efcab12 | 212516 | protein_coding | 9.155320908 | 0.774276266 | 0.413311168 | 1.873349493 | 0.06102014 | 0.177748497 | 0.774276266 |
| ENSMUSG00000056822 | Or2l13 | 259071 | protein_coding | 8.069212289 | 0.774684658 | 0.540140586 | 1.434227826 | 0.151507259 | 0.329655755 | 0.774684658 |
| ENSMUSG00000110151 | Gm38416 | 330850 | lncRNA | 37.8523074 | 0.774936268 | 0.235806854 | 3.286317831 | 0.001015064 | 0.007761853 | 0.774936268 |
| ENSMUSG00000026938 | Fcna | 14133 | protein_coding | 60.80305458 | -0.775599062 | 0.168693258 | -4.597688564 | 4.27E-06 | 7.00E-05 | 0.775599062 |
| ENSMUSG00000022518 | 4930562C15Rik | 78809 | protein_coding | 11.95525014 | -0.77562682 | 0.390082704 | -1.988365061 | 0.04677133 | 0.148643475 | 0.77562682 |
| ENSMUSG00000020638 | Cmpk2 | 22169 | protein_coding | 188.1640632 | 0.776286627 | 0.182549225 | 4.252478359 | 2.11E-05 | 0.0002908 | 0.776286627 |
| ENSMUSG00000079227 | Ccr5 | 12774 | protein_coding | 12.26016081 | 0.776889497 | 0.726257425 | 1.069716425 | 0.284746971 | 0.493735004 | 0.776889497 |
| ENSMUSG00000001666 | Ddt | 13202 | protein_coding | 288.9478658 | 0.778267603 | 0.110493472 | 7.043561869 | 1.87E-12 | 7.46E-11 | 0.778267603 |
| ENSMUSG00000029452 | Tmem116 | 77462 | protein_coding | 29.26999826 | 0.778347777 | 0.219001721 | 3.554071511 | 0.000379316 | 0.003476264 | 0.778347777 |
| ENSMUSG00000042429 | Adora1 | 11539 | protein_coding | 12.24327028 | -0.778488677 | 0.481325239 | -1.617385945 | 0.105794998 | 0.257739786 | 0.778488677 |
| ENSMUSG00000028701 | Lurap1 | 68075 | protein_coding | 39.20640765 | -0.778560576 | 0.214435224 | -3.630749476 | 0.000282599 | 0.002700691 | 0.778560576 |
| ENSMUSG00000025213 | Kazald1 | 107250 | protein_coding | 28.54647923 | -0.779133936 | 0.256606788 | -3.036295106 | 0.002395048 | 0.015746653 | 0.779133936 |
| ENSMUSG00000024485 | Slc4a9 | 240215 | protein_coding | 8.941443283 | -0.779353106 | 0.538782903 | -1.44650675 | 0.14803512 | 0.324809051 | 0.779353106 |

| ensembl_gene_id | external_gene_name | entrezgene_id | gene_biotype | baseMean | log2FoldChange | lfcSE | stat | pvalue | padj | absLog2FC |
|---|---|---|---|---|---|---|---|---|---|---|
| ENSMUSG00000074261 | Erich4 | 632778 | protein_coding | 37.52616705 | 0.779551115 | 0.278634702 | 2.797753144 | 0.005145943 | 0.028678997 | 0.779551115 |
| ENSMUSG00000031970 | Dbndd1 | 72185 | protein_coding | 14.70054927 | -0.779665963 | 0.379781931 | -2.05293064 | 0.040079305 | 0.133274148 | 0.779665963 |
| ENSMUSG00000040978 | Gm11992 | 626870 | protein_coding | 28.75049967 | 0.780091163 | 0.358348104 | 2.17690886 | 0.029487368 | 0.106502223 | 0.780091163 |
| ENSMUSG00000021959 | Lats2 | 50523 | protein_coding | 58.45510342 | -0.780138296 | 0.231520583 | -3.369628246 | 0.000752697 | 0.006088853 | 0.780138296 |
| ENSMUSG00000019027 | Dnah1 | 110084 | protein_coding | 9.356638411 | -0.780220347 | 0.637822187 | -1.223256831 | 0.221232707 | 0.42206187 | 0.780220347 |
| ENSMUSG00000032418 | Me1 | 17436 | protein_coding | 286.2502847 | 0.780656015 | 0.212264855 | 3.677745032 | 0.000235305 | 0.002339537 | 0.780656015 |
| ENSMUSG00000020642 | Rnf144a | 108089 | protein_coding | 8.738699128 | -0.780690802 | 0.747922398 | -1.043812572 | 0.296572114 | 0.505922741 | 0.780690802 |
| ENSMUSG00000032757 | Bet1 | 12068 | protein_coding | 719.419528 | 0.781231782 | 0.081505949 | 9.584966419 | 9.25E-22 | 7.05E-20 | 0.781231782 |
| ENSMUSG00000021537 | Cetn3 | 12626 | protein_coding | 756.170726 | 0.781300606 | 0.097752264 | 7.992659966 | 1.32E-15 | 7.08E-14 | 0.781300606 |
| ENSMUSG00000087213 | 2810408I11Rik | 69941 | lncRNA | 21.88700502 | 0.781667798 | 0.266108425 | 2.937403418 | 0.003309732 | 0.020322829 | 0.781667798 |
| ENSMUSG00000025194 | Abcc2 | 12780 | protein_coding | 31.05928175 | -0.782013128 | 0.3819141 | -2.047615229 | 0.040597716 | 0.134477991 | 0.782013128 |
| ENSMUSG00000047501 | Cldn4 | 12740 | protein_coding | 366.5919438 | -0.782097281 | 0.328429556 | -2.381324295 | 0.017250518 | 0.071703093 | 0.782097281 |
| ENSMUSG00000039137 | Whrn | 73750 | protein_coding | 122.0280549 | -0.782792698 | 0.2133541 | -3.668983626 | 0.000243517 | 0.002395759 | 0.782792698 |
| ENSMUSG00000024352 | Spata24 | 71242 | protein_coding | 12.16285831 | 0.783121806 | 0.380021731 | 2.06072901 | 0.0393289 | 0.131575452 | 0.783121806 |
| ENSMUSG00000050556 | Kcnb1 | 16500 | protein_coding | 8.715963952 | -0.783698397 | 0.51029689 | -1.535769495 | 0.124594924 | 0.288451737 | 0.783698397 |
| ENSMUSG00000051980 | Casr | 12374 | protein_coding | 7.410191535 | 0.78376797 | 0.532853336 | 1.470888734 | 0.14132121 | 0.314015745 | 0.78376797 |
| ENSMUSG00000053411 | Cbx7 | 52609 | protein_coding | 189.9002927 | -0.783834698 | 0.123033556 | -6.370901779 | 1.88E-10 | 6.22E-09 | 0.783834698 |
| ENSMUSG00000006241 | Ccdc159 | 67119 | protein_coding | 23.9838306 | -0.783880534 | 0.314241296 | -2.494517887 | 0.012612837 | 0.056975969 | 0.783880534 |
| ENSMUSG00000038209 | Itln1 | 16429 | protein_coding | 37.02461587 | -0.785007463 | 0.467177712 | -1.680318738 | 0.092895317 | 0.236958409 | 0.785007463 |
| ENSMUSG00000038305 | Spats2l | 67198 | protein_coding | 177.6882962 | 0.785060942 | 0.165035567 | 4.756919724 | 1.97E-06 | 3.45E-05 | 0.785060942 |
| ENSMUSG00000034912 | Mdga2 | 320772 | protein_coding | 9.177008519 | -0.785215777 | 1.542004966 | -0.509217411 | 0.61059984 | 0.773102882 | 0.785215777 |
| ENSMUSG00000042042 | Csgalnact2 | 78752 | protein_coding | 225.9710492 | 0.785452767 | 0.10463632 | 7.506502235 | 6.07E-14 | 2.82E-12 | 0.785452767 |
| ENSMUSG00000035640 | Cbarp | 100503659 | protein_coding | 22.78038211 | -0.785490264 | 0.307640567 | -2.553272707 | 0.01067159 | 0.050244485 | 0.785490264 |
| ENSMUSG00000020099 | Unc5b | 107449 | protein_coding | 1436.585938 | 0.785697851 | 0.145629345 | 5.39518909 | 6.85E-08 | 1.60E-06 | 0.785697851 |
| ENSMUSG00000022365 | Derl1 | 67819 | protein_coding | 1677.057673 | 0.786522942 | 0.096008485 | 8.192223259 | 2.56E-16 | 1.45E-14 | 0.786522942 |
| ENSMUSG00000086992 | Gm15941 | 100503538 | lncRNA | 11.25888827 | -0.786603239 | 0.396506474 | -1.983834542 | 0.047274288 | 0.14977207 | 0.786603239 |
| ENSMUSG00000029016 | Clcn6 | 26372 | protein_coding | 43.63980573 | -0.786614833 | 0.208094016 | -3.780093479 | 0.000156769 | 0.001655852 | 0.786614833 |
| ENSMUSG00000054958 | Nt5c1a | 230718 | protein_coding | 20.99903763 | -0.786791682 | 0.711806428 | -1.105345008 | 0.269010122 | 0.477123143 | 0.786791682 |
| ENSMUSG00000020331 | Hcn2 | 15166 | protein_coding | 94.85434823 | -0.787206454 | 0.158577282 | -4.964181794 | 6.90E-07 | 1.35E-05 | 0.787206454 |
| ENSMUSG00000023951 | Vegfa | 22339 | protein_coding | 611.5501878 | -0.787695664 | 0.160481946 | -4.90831327 | 9.19E-07 | 1.74E-05 | 0.787695664 |
| ENSMUSG00000057895 | Zfp105 | 22646 | protein_coding | 15.12014289 | 0.788208011 | 0.363227429 | 2.170012361 | 0.03000591 | 0.107983309 | 0.788208011 |
| ENSMUSG00000052026 | Slc6a7 | 240332 | protein_coding | 59.73638988 | -0.788259753 | 0.266690546 | -2.955709397 | 0.003119509 | 0.019356265 | 0.788259753 |
| ENSMUSG00000024427 | Spry4 | 24066 | protein_coding | 59.0040852 | -0.788380989 | 0.685387268 | -1.150270841 | 0.250032337 | 0.456344112 | 0.788380989 |
| ENSMUSG00000026664 | Phyh | 16922 | protein_coding | 3227.674004 | 0.788404315 | 0.113751856 | 6.930913852 | 4.18E-12 | 1.62E-10 | 0.788404315 |
| ENSMUSG00000046082 | Tmem174 | 68344 | protein_coding | 16.46527351 | 0.78881147 | 0.483124624 | 1.632728763 | 0.102526055 | 0.252479129 | 0.78881147 |
| ENSMUSG00000046523 | Kctd4 | 67516 | protein_coding | 14.24314176 | 0.788823034 | 0.44677277 | 1.765602308 | 0.077462592 | 0.209087303 | 0.788823034 |
| ENSMUSG00000078632 | Lrrc37a | 237954 | protein_coding | 7.494896221 | -0.788882971 | 2.012854537 | -0.391922495 | 0.695115482 | 0.826592861 | 0.788882971 |
| ENSMUSG00000034799 | Unc13a | 382018 | protein_coding | 41.01963284 | -0.789024871 | 0.336039241 | -2.348014086 | 0.018873808 | 0.076611113 | 0.789024871 |

| ensembl_gene_id | external_gene_name | entrezgene_id | gene_biotype | baseMean | log2FoldChange | lfcSE | stat | pvalue | padj | absLog2FC |
|---|---|---|---|---|---|---|---|---|---|---|
| ENSMUSG00000055491 | Pprc1 | 226169 | protein_coding | 220.4443791 | -0.78903373 | 0.254789688 | -3.096804015 | 0.001956192 | 0.013281424 | 0.78903373 |
| ENSMUSG00000018449 | Rpain | 69723 | protein_coding | 151.2226044 | 0.789326766 | 0.106749676 | 7.394184216 | 1.42E-13 | 6.30E-12 | 0.789326766 |
| ENSMUSG00000029672 | Fam3c | 27999 | protein_coding | 338.8527346 | 0.789508172 | 0.20478063 | 3.855385017 | 0.000115548 | 0.001273663 | 0.789508172 |
| ENSMUSG00000099950 | 9130227L01Rik | 329159 | lncRNA | 35.30485129 | 0.789514071 | 0.293954331 | 2.685839225 | 0.007234789 | 0.037389917 | 0.789514071 |
| ENSMUSG00000025731 | Mettl26 | 68347 | protein_coding | 625.7149748 | -0.789735208 | 0.093917689 | -8.40880157 | 4.14E-17 | 2.47E-15 | 0.789735208 |
| ENSMUSG00000064345 | mt-Nd2 | 17717 | protein_coding | 11475.87985 | -0.789753731 | 0.291045645 | -2.713504714 | 0.006657562 | 0.035008798 | 0.789753731 |
| ENSMUSG00000032036 | Kirrel3 | 67703 | protein_coding | 18.72385887 | -0.790209195 | 0.392822962 | -2.011616611 | 0.044260367 | 0.142679442 | 0.790209195 |
| ENSMUSG00000026873 | Phf19 | 74016 | protein_coding | 57.14181263 | -0.790466109 | 0.196580025 | -4.021090684 | 5.79E-05 | 0.000703276 | 0.790466109 |
| ENSMUSG00000046613 | Vwa5b2 | 328643 | protein_coding | 26.70063306 | -0.790544276 | 0.325085208 | -2.43180636 | 0.015023737 | 0.064755004 | 0.790544276 |
| ENSMUSG00000055148 | Klf2 | 16598 | protein_coding | 254.2443136 | -0.7905601 | 0.508053843 | -1.556055744 | 0.119694837 | 0.280537081 | 0.7905601 |
| ENSMUSG00000048970 | C1galt1c1 | 59048 | protein_coding | 1465.089316 | 0.790732501 | 0.10737076 | 7.364504999 | 1.78E-13 | 7.84E-12 | 0.790732501 |
| ENSMUSG00000018401 | Mtmr4 | 170749 | protein_coding | 1470.677853 | 0.790940494 | 0.093024832 | 8.502466258 | 1.86E-17 | 1.13E-15 | 0.790940494 |
| ENSMUSG00000027490 | E2f1 | 13555 | protein_coding | 56.09311407 | 0.791789683 | 0.192417299 | 4.114961017 | 3.87E-05 | 0.000494022 | 0.791789683 |
| ENSMUSG00000029868 | Trpv6 | 64177 | protein_coding | 35.04003225 | -0.792146917 | 0.476547448 | -1.662262424 | 0.096460168 | 0.242696288 | 0.792146917 |
| ENSMUSG00000020032 | Nuak1 | 77976 | protein_coding | 18.50199624 | -0.792323459 | 0.37049808 | -2.13853594 | 0.03247327 | 0.114255412 | 0.792323459 |
| ENSMUSG00000061601 | Pclo | 26875 | protein_coding | 30.17100636 | -0.792445546 | 0.489801527 | -1.617891129 | 0.105686062 | 0.257575797 | 0.792445546 |
| ENSMUSG00000062444 | Ap3b2 | 11775 | protein_coding | 9.168313114 | -0.7926577 | 0.584442227 | -1.356263567 | 0.175015323 | 0.361280983 | 0.7926577 |
| ENSMUSG00000020914 | Top2a | 21973 | protein_coding | 4781.525336 | 0.793441894 | 0.135094774 | 5.873224186 | 4.27E-09 | 1.20E-07 | 0.793441894 |
| ENSMUSG00000059540 | Tcea2 | 21400 | protein_coding | 16.93868273 | -0.793774748 | 0.283820699 | -2.79674721 | 0.00516199 | 0.028728226 | 0.793774748 |
| ENSMUSG00000090254 | Gm1965 | 434065 | lncRNA | 62.99932585 | -0.794713789 | 0.279761394 | -2.840684256 | 0.004501686 | 0.025808627 | 0.794713789 |
| ENSMUSG00000064330 | Pde6h | 78600 | protein_coding | 7.00240935 | -0.795184177 | 1.102491459 | -0.721261077 | 0.4707489 | 0.666501035 | 0.795184177 |
| ENSMUSG00000020407 | Upp1 | 22271 | protein_coding | 282.991221 | 0.79539683 | 0.186801934 | 4.257968927 | 2.06E-05 | 0.000285228 | 0.79539683 |
| ENSMUSG00000032661 | Oas3 | 246727 | protein_coding | 91.78072343 | 0.795842203 | 0.314126648 | 2.533507449 | 0.011292734 | 0.052426743 | 0.795842203 |
| ENSMUSG00000073787 | Gm10575 | 100126795 | lncRNA | 21.83145669 | -0.796691929 | 0.266730821 | -2.986876158 | 0.002818439 | 0.017842644 | 0.796691929 |
| ENSMUSG00000058672 | Tubb2a | 22151 | protein_coding | 214.420049 | -0.796753021 | 0.259108296 | -3.074980746 | 0.002105161 | 0.014112378 | 0.796753021 |
| ENSMUSG00000024687 | Osbp | 76303 | protein_coding | 1844.523571 | 0.796797062 | 0.112118878 | 7.106716342 | 1.19E-12 | 4.89E-11 | 0.796797062 |
| ENSMUSG00000058740 | Kcnt1 | 227632 | protein_coding | 7.363835745 | -0.797065941 | 0.744283043 | -1.070917776 | 0.284206402 | 0.493388509 | 0.797065941 |
| ENSMUSG00000054206 | Gzmm | 16904 | protein_coding | 7.87384229 | -0.797119053 | 0.417076896 | -1.911204048 | 0.055978364 | 0.167629889 | 0.797119053 |
| ENSMUSG00000027510 | Rbm38 | 56190 | protein_coding | 95.98095167 | -0.79716037 | 0.17016646 | -4.684591617 | 2.81E-06 | 4.77E-05 | 0.79716037 |
| ENSMUSG00000100291 | 2310069B03Rik | 69652 | lncRNA | 14.79237766 | 0.797218264 | 0.385511033 | 2.067951878 | 0.038644548 | 0.130078269 | 0.797218264 |
| ENSMUSG00000041377 | Ninj2 | 29862 | protein_coding | 6.736160194 | -0.797318407 | 0.592800682 | -1.345002515 | 0.178624422 | 0.365980165 | 0.797318407 |
| ENSMUSG00000059659 | Gm10069 | 791299 | lncRNA | 10.32519734 | 0.798621257 | 0.396149829 | 2.015957596 | 0.04380441 | 0.141596585 | 0.798621257 |
| ENSMUSG00000106951 | 5930430L01Rik | 319982 | lncRNA | 15.94761711 | -0.798807454 | 0.275002876 | -2.904723996 | 0.003675771 | 0.022018683 | 0.798807454 |
| ENSMUSG00000020205 | Phlda1 | 21664 | protein_coding | 147.4178487 | -0.798940921 | 0.476241158 | -1.677597384 | 0.093425724 | 0.237941306 | 0.798940921 |
| ENSMUSG00000032121 | Tmem218 | 66279 | protein_coding | 143.3243759 | 0.799368435 | 0.163252367 | 4.896519727 | 9.75E-07 | 1.84E-05 | 0.799368435 |
| ENSMUSG00000027199 | Gatm | 67092 | protein_coding | 156.601151 | 0.799690728 | 0.361271438 | 2.213545394 | 0.026860067 | 0.099440198 | 0.799690728 |
| ENSMUSG00000040599 | Mis12 | 67139 | protein_coding | 506.4723079 | 0.800324793 | 0.12272251 | 6.52141806 | 6.96E-11 | 2.45E-09 | 0.800324793 |
| ENSMUSG00000035493 | Tgfbi | 21810 | protein_coding | 72.79577037 | -0.800452811 | 0.25911716 | -3.089154003 | 0.002007274 | 0.013541658 | 0.800452811 |

| ensembl_gene_id | external_gene_name | entrezgene_id | gene_biotype | baseMean | log2FoldChange | lfcSE | stat | pvalue | padj | absLog2FC |
|---|---|---|---|---|---|---|---|---|---|---|
| ENSMUSG00000012123 | Crybg2 | 230806 | protein_coding | 647.8780142 | -0.800613095 | 0.189884772 | -4.216310173 | 2.48E-05 | 0.00033323 | 0.800613095 |
| ENSMUSG00000026817 | Ak1 | 11636 | protein_coding | 14.35314284 | -0.801457381 | 0.358095566 | -2.238110321 | 0.025213862 | 0.095054767 | 0.801457381 |
| ENSMUSG00000031877 | Ces2g | 72361 | protein_coding | 178.9173362 | -0.801784954 | 0.18289307 | -4.383900142 | 1.17E-05 | 0.00017066 | 0.801784954 |
| ENSMUSG00000050241 | Klre1 | 243655 | protein_coding | 7.131062958 | 0.802020817 | 0.707549784 | 1.133518568 | 0.256996548 | 0.463812519 | 0.802020817 |
| ENSMUSG00000020892 | Aloxe3 | 23801 | protein_coding | 6.907516066 | -0.802135705 | 0.881095227 | -0.910384804 | 0.362619609 | 0.571692085 | 0.802135705 |
| ENSMUSG00000022829 | Stxbp5l | 207227 | protein_coding | 9.048310362 | -0.802396269 | 0.643857798 | -1.246232121 | 0.21267919 | 0.410736891 | 0.802396269 |
| ENSMUSG00000041831 | Sytl3 | 83672 | protein_coding | 33.63672013 | -0.802566371 | 0.228800497 | -3.50771254 | 0.000451977 | 0.004001719 | 0.802566371 |
| ENSMUSG00000039110 | Mycbpap | 104601 | protein_coding | 75.80769185 | -0.802802298 | 0.180590303 | -4.445434149 | 8.77E-06 | 0.000132428 | 0.802802298 |
| ENSMUSG00000020439 | Smtn | 29856 | protein_coding | 219.2493857 | 0.803120574 | 0.136443672 | 5.886096173 | 3.95E-09 | 1.12E-07 | 0.803120574 |
| ENSMUSG00000037287 | Tbcel | 272589 | protein_coding | 181.7027403 | -0.803950815 | 0.135220743 | -5.945469578 | 2.76E-09 | 7.97E-08 | 0.803950815 |
| ENSMUSG00000004462 | Tbccd1 | 70573 | protein_coding | 766.7496999 | 0.804752591 | 0.086100536 | 9.346661817 | 9.05E-21 | 6.70E-19 | 0.804752591 |
| ENSMUSG00000078816 | Prkcg | 18752 | protein_coding | 12.56579943 | -0.804932923 | 0.381263634 | -2.111223971 | 0.034753065 | 0.120414798 | 0.804932923 |
| ENSMUSG00000011267 | Zfp296 | 63872 | protein_coding | 7.075012664 | -0.80494073 | 0.43633332 | -1.844784004 | 0.065068965 | 0.185724755 | 0.80494073 |
| ENSMUSG00000078247 | Airn | 104103 | lncRNA | 63.44765319 | -0.805098852 | 0.204780612 | -3.931518932 | 8.44E-05 | 0.000974272 | 0.805098852 |
| ENSMUSG00000093042 | Mir3109 | 100526560 | miRNA | 19.27641376 | -0.805322743 | 0.350224157 | -2.299449445 | 0.021479431 | 0.084234579 | 0.805322743 |
| ENSMUSG00000047031 | Vmn1r68 | 628580 | protein_coding | 6.814969984 | -0.806460483 | 1.777812807 | -0.453625083 | 0.650098697 | 0.797932653 | 0.806460483 |
| ENSMUSG00000006784 | Odad4 | 74407 | protein_coding | 23.21961044 | -0.806641139 | 0.422048194 | -1.91125362 | 0.055971996 | 0.167629889 | 0.806641139 |
| ENSMUSG00000064061 | Dzip3 | 224170 | protein_coding | 152.7413056 | 0.806723365 | 0.170296271 | 4.73717575 | 2.17E-06 | 3.75E-05 | 0.806723365 |
| ENSMUSG00000030683 | Sez6l2 | 233878 | protein_coding | 28.58276416 | -0.807499029 | 0.241681038 | -3.341176607 | 0.000834241 | 0.006607109 | 0.807499029 |
| ENSMUSG00000020866 | Cacna1g | 12291 | protein_coding | 7.252749022 | -0.807824894 | 0.956446327 | -0.844610796 | 0.398328179 | 0.606149922 | 0.807824894 |
| ENSMUSG00000041801 | Phlda3 | 27280 | protein_coding | 31.58137553 | 0.808006416 | 0.275061457 | 2.937548662 | 0.003308182 | 0.020321146 | 0.808006416 |
| ENSMUSG00000085408 | C530005A16Rik | 654318 | lncRNA | 10.49500948 | -0.808103063 | 0.410823684 | -1.967031341 | 0.049179598 | 0.153745364 | 0.808103063 |
| ENSMUSG00000028976 | Slc2a5 | 56485 | protein_coding | 71.52615344 | 0.808977994 | 0.153628426 | 5.265809283 | 1.40E-07 | 3.14E-06 | 0.808977994 |
| ENSMUSG00000032281 | Acsbg1 | 94180 | protein_coding | 7.728938763 | 0.810797408 | 0.537973723 | 1.507131991 | 0.131776828 | 0.299549211 | 0.810797408 |
| ENSMUSG00000092981 | Mir5125 | 100628593 | miRNA | 9.476393084 | -0.811051362 | 0.431012808 | -1.881733785 | 0.05987217 | 0.175740416 | 0.811051362 |
| ENSMUSG00000015950 | Ncf1 | 17969 | protein_coding | 12.53853589 | 0.811695295 | 0.438270891 | 1.852040168 | 0.064020055 | 0.183897464 | 0.811695295 |
| ENSMUSG00000044231 | Nhlrc1 | 105193 | protein_coding | 40.8333426 | -0.811766221 | 0.202551926 | -4.007694405 | 6.13E-05 | 0.000737068 | 0.811766221 |
| ENSMUSG00000021057 | Akap5 | 238276 | protein_coding | 7.452712733 | -0.811888089 | 0.893251323 | -0.908913391 | 0.363395845 | 0.572236185 | 0.811888089 |
| ENSMUSG00000031101 | Sash3 | 74131 | protein_coding | 18.90899158 | 0.812930782 | 0.358064033 | 2.270350295 | 0.023186338 | 0.089343223 | 0.812930782 |
| ENSMUSG00000000901 | Mmp11 | 17385 | protein_coding | 7.067698372 | 0.813041258 | 0.463415184 | 1.75445537 | 0.079352511 | 0.212813829 | 0.813041258 |
| ENSMUSG00000038677 | Scube3 | 268935 | protein_coding | 9.397515762 | -0.813395994 | 0.639330701 | -1.272261746 | 0.203280135 | 0.398308033 | 0.813395994 |
| ENSMUSG00000050747 | Trim15 | 69097 | protein_coding | 229.4644749 | 0.813468639 | 0.365029237 | 2.228502695 | 0.025847012 | 0.096959775 | 0.813468639 |
| ENSMUSG00000057132 | Rpgrip1 | 77945 | protein_coding | 19.51003395 | -0.81358917 | 0.414246157 | -1.964023457 | 0.049527372 | 0.154408295 | 0.81358917 |
| ENSMUSG00000074646 | 6430550D23Rik | 320095 | protein_coding | 14.44470991 | -0.813720709 | 0.368586646 | -2.207678214 | 0.027266713 | 0.100461996 | 0.813720709 |
| ENSMUSG00000025582 | Nptx1 | 18164 | protein_coding | 7.72643935 | -0.813914887 | 0.528856469 | -1.539009041 | 0.123802089 | 0.287033432 | 0.813914887 |
| ENSMUSG00000069874 | Irgm2 | 54396 | protein_coding | 234.8350627 | 0.814277481 | 0.177403366 | 4.58997764 | 4.43E-06 | 7.25E-05 | 0.814277481 |
| ENSMUSG00000052387 | Trpm3 | 226025 | protein_coding | 13.53629606 | -0.814582464 | 1.80135124 | -0.452206347 | 0.651120336 | 0.798250711 | 0.814582464 |
| ENSMUSG00000040760 | Appl1 | 72993 | protein_coding | 1705.066435 | 0.814736783 | 0.163409098 | 4.985871619 | 6.17E-07 | 1.24E-05 | 0.814736783 |

| ensembl_gene_id | external_gene_name | entrezgene_id | gene_biotype | baseMean | log2FoldChange | lfcSE | stat | pvalue | padj | absLog2FC |
|---|---|---|---|---|---|---|---|---|---|---|
| ENSMUSG00000027188 | Pamr1 | 210622 | protein_coding | 12.4288905 | -0.815038366 | 0.451600384 | -1.804777841 | 0.071109453 | 0.197597936 | 0.815038366 |
| ENSMUSG00000100510 | Hand2os1 | 102636514 | lncRNA | 7.091706186 | -0.815119894 | 1.803149004 | -0.452053542 | 0.651230411 | 0.798250711 | 0.815119894 |
| ENSMUSG00000112129 | Pbld1 | 68371 | protein_coding | 153.1938785 | -0.815446622 | 0.239176352 | -3.409394849 | 0.000651072 | 0.005425116 | 0.815446622 |
| ENSMUSG00000060791 | Gmfg | 63986 | protein_coding | 223.6802613 | 0.815802615 | 0.139696985 | 5.839801141 | 5.23E-09 | 1.45E-07 | 0.815802615 |
| ENSMUSG00000040374 | Pex2 | 19302 | protein_coding | 971.0888968 | 0.816205008 | 0.100979967 | 8.082840923 | 6.33E-16 | 3.48E-14 | 0.816205008 |
| ENSMUSG00000066392 | Nrxn3 | 18191 | protein_coding | 8.183113263 | -0.816448203 | 2.04226186 | -0.399776453 | 0.689321175 | 0.823011515 | 0.816448203 |
| ENSMUSG00000030523 | Trpm1 | 17364 | protein_coding | 10.80693829 | -0.816494896 | 1.49787258 | -0.545103039 | 0.585682652 | 0.75599124 | 0.816494896 |
| ENSMUSG00000027287 | Snap23 | 20619 | protein_coding | 1217.556362 | 0.816587229 | 0.075786378 | 10.77485496 | 4.52E-27 | 4.42E-25 | 0.816587229 |
| ENSMUSG00000069515 | Lyz1 | 17110 | protein_coding | 75.06400837 | -0.816651057 | 0.825094277 | -0.989766963 | 0.322288036 | 0.533229858 | 0.816651057 |
| ENSMUSG00000028044 | Cks1b | 54124 | protein_coding | 188.5987613 | 0.816916496 | 0.180960276 | 4.514341569 | 6.35E-06 | 9.96E-05 | 0.816916496 |
| ENSMUSG00000041617 | Ccdc74a | 72315 | protein_coding | 16.80600624 | -0.817525225 | 0.349964124 | -2.33602581 | 0.019489896 | 0.078471959 | 0.817525225 |
| ENSMUSG00000003279 | Dlgap1 | 224997 | protein_coding | 13.64996092 | -0.818313818 | 0.625546086 | -1.30815912 | 0.190819345 | 0.382936943 | 0.818313818 |
| ENSMUSG00000048217 | Nags | 217214 | protein_coding | 77.24374248 | -0.81846179 | 0.285028642 | -2.871507169 | 0.004085195 | 0.023887294 | 0.81846179 |
| ENSMUSG00000027078 | Ube2l6 | 56791 | protein_coding | 86.24052227 | 0.819340147 | 0.16666752 | 4.916015713 | 8.83E-07 | 1.68E-05 | 0.819340147 |
| ENSMUSG00000018581 | Dnah11 | 13411 | protein_coding | 15.53824776 | -0.819590586 | 0.568964236 | -1.44049579 | 0.14972718 | 0.327140539 | 0.819590586 |
| ENSMUSG00000029033 | Acap3 | 140500 | protein_coding | 109.2967059 | -0.819692302 | 0.155937596 | -5.256540574 | 1.47E-07 | 3.27E-06 | 0.819692302 |
| ENSMUSG00000004473 | Clec11a | 20256 | protein_coding | 25.00650481 | 0.819865888 | 0.257356219 | 3.185724015 | 0.001443923 | 0.01033654 | 0.819865888 |
| ENSMUSG00000055730 | Ces2a | 102022 | protein_coding | 1621.235471 | 0.820049639 | 0.159689485 | 5.135276383 | 2.82E-07 | 6.00E-06 | 0.820049639 |
| ENSMUSG00000056629 | Fkbp2 | 14227 | protein_coding | 827.6817801 | 0.820266468 | 0.074630595 | 10.99102137 | 4.22E-28 | 4.42E-26 | 0.820266468 |
| ENSMUSG00000059479 | B3gnt8 | 232984 | protein_coding | 15.27926495 | -0.820610364 | 0.512021983 | -1.602685806 | 0.109004039 | 0.262950815 | 0.820610364 |
| ENSMUSG00000024978 | Gpam | 14732 | protein_coding | 665.2559422 | -0.820709753 | 0.097828092 | -8.389305502 | 4.89E-17 | 2.90E-15 | 0.820709753 |
| ENSMUSG00000036223 | Ska1 | 66468 | protein_coding | 80.42836025 | 0.820886678 | 0.156781784 | 5.23585493 | 1.64E-07 | 3.64E-06 | 0.820886678 |
| ENSMUSG00000079445 | B3gnt7 | 227327 | protein_coding | 3171.369557 | -0.821488047 | 0.388211327 | -2.116084691 | 0.034337605 | 0.119325161 | 0.821488047 |
| ENSMUSG00000055322 | Tns1 | 21961 | protein_coding | 16.89495846 | -0.821838265 | 0.890142709 | -0.923265738 | 0.355868738 | 0.565481052 | 0.821838265 |
| ENSMUSG00000002831 | Plin4 | 57435 | protein_coding | 8.229046664 | -0.822144466 | 0.4513861 | -1.821377455 | 0.0685495 | 0.192805303 | 0.822144466 |
| ENSMUSG00000010307 | Tmem86a | 67893 | protein_coding | 22.53037297 | -0.822398097 | 0.275042483 | -2.990076617 | 0.002789075 | 0.017713071 | 0.822398097 |
| ENSMUSG00000030008 | Pradc1 | 73327 | protein_coding | 139.3215449 | 0.822632839 | 0.130287004 | 6.314005319 | 2.72E-10 | 8.80E-09 | 0.822632839 |
| ENSMUSG00000038210 | Hoxa11 | 15396 | protein_coding | 35.71730293 | 0.822906049 | 0.264595339 | 3.110054966 | 0.001870525 | 0.012792498 | 0.822906049 |
| ENSMUSG00000064363 | mt-Nd4 | 17719 | protein_coding | 14232.88681 | -0.823115301 | 0.296889798 | -2.772460715 | 0.005563423 | 0.030336942 | 0.823115301 |
| ENSMUSG00000060924 | Csmd1 | 94109 | protein_coding | 11.75692179 | -0.823309021 | 1.777842401 | -0.463094491 | 0.643296639 | 0.793445869 | 0.823309021 |
| ENSMUSG00000026586 | Prrx1 | 18933 | protein_coding | 7.807716672 | -0.823487133 | 0.971628104 | -0.847533258 | 0.396697957 | 0.604651202 | 0.823487133 |
| ENSMUSG00000042581 | Thsd7b | 210417 | protein_coding | 7.996095174 | -0.823589193 | 1.377011148 | -0.598099146 | 0.54977378 | 0.731093174 | 0.823589193 |
| ENSMUSG00000097368 | Gm10390 | 100503427 | lncRNA | 11.19270569 | -0.823935931 | 0.335665364 | -2.454634939 | 0.014102772 | 0.061772753 | 0.823935931 |
| ENSMUSG00000024399 | Ltb | 16994 | protein_coding | 10.84202274 | 0.823999951 | 0.477328536 | 1.726274229 | 0.084298089 | 0.222026028 | 0.823999951 |
| ENSMUSG00000013338 | Fer1l4 | 74562 | protein_coding | 75.05232234 | 0.824461125 | 0.340983881 | 2.417888855 | 0.015610844 | 0.066644205 | 0.824461125 |
| ENSMUSG00000027570 | Col9a3 | 12841 | protein_coding | 10.63103222 | -0.824997139 | 0.596735499 | -1.382517279 | 0.166812929 | 0.350851225 | 0.824997139 |
| ENSMUSG00000097397 | Gm16861 | 100503704 | lncRNA | 13.12976631 | -0.825054985 | 0.375289098 | -2.198451779 | 0.027916927 | 0.102291424 | 0.825054985 |
| ENSMUSG00000037613 | Tnfrsf23 | 79201 | protein_coding | 31.17350183 | -0.825160295 | 0.246104534 | -3.352885386 | 0.000799738 | 0.00637549 | 0.825160295 |

| ensembl_gene_id | external_gene_name | entrezgene_id | gene_biotype | baseMean | log2FoldChange | lfcSE | stat | pvalue | padj | absLog2FC |
|---|---|---|---|---|---|---|---|---|---|---|
| ENSMUSG00000020076 | Ddx50 | 94213 | protein_coding | 1731.670994 | 0.825255133 | 0.051899548 | 15.90100812 | 6.24E-57 | 1.74E-54 | 0.825255133 |
| ENSMUSG00000035390 | Brsk1 | 381979 | protein_coding | 18.73914018 | -0.825608394 | 0.404869967 | -2.039193967 | 0.041430673 | 0.136250376 | 0.825608394 |
| ENSMUSG00000051079 | Rgs13 | 246709 | protein_coding | 151.5209839 | -0.825787771 | 0.196489178 | -4.202713758 | 2.64E-05 | 0.000352117 | 0.825787771 |
| ENSMUSG00000019796 | Lrp11 | 237253 | protein_coding | 27.71450671 | -0.825813469 | 0.256440902 | -3.220287651 | 0.00128062 | 0.009352462 | 0.825813469 |
| ENSMUSG00000068270 | Shroom4 | 208431 | protein_coding | 33.96938124 | -0.825984474 | 0.204987714 | -4.02943404 | 5.59E-05 | 0.000681895 | 0.825984474 |
| ENSMUSG00000001403 | Ube2c | 68612 | protein_coding | 713.0179026 | 0.826496478 | 0.174667297 | 4.731832987 | 2.23E-06 | 3.85E-05 | 0.826496478 |
| ENSMUSG00000018599 | Mief2 | 237781 | protein_coding | 211.1267867 | 0.827115208 | 0.138471175 | 5.973194119 | 2.33E-09 | 6.76E-08 | 0.827115208 |
| ENSMUSG00000036103 | Colec12 | 140792 | protein_coding | 6.723509367 | -0.827648534 | 0.893390817 | -0.926412628 | 0.354231584 | 0.563829056 | 0.827648534 |
| ENSMUSG00000076441 | Ass1 | 11898 | protein_coding | 159.215052 | -0.827758465 | 0.167383843 | -4.945271014 | 7.60E-07 | 1.48E-05 | 0.827758465 |
| ENSMUSG00000022883 | Robo1 | 19876 | protein_coding | 17.54711376 | -0.828019382 | 0.664436845 | -1.246197269 | 0.212691981 | 0.410736891 | 0.828019382 |
| ENSMUSG00000025081 | Tdrd1 | 83561 | protein_coding | 48.06102118 | -0.828132482 | 0.266479378 | -3.107679429 | 0.001885625 | 0.012873653 | 0.828132482 |
| ENSMUSG00000020427 | Igfbp3 | 16009 | protein_coding | 10.90411167 | 0.829285297 | 0.486920379 | 1.703123001 | 0.088545052 | 0.229361781 | 0.829285297 |
| ENSMUSG00000086189 | Gm15462 | 108168861 | lncRNA | 24.36952924 | -0.829409695 | 0.44663383 | -1.857023897 | 0.063307754 | 0.182344649 | 0.829409695 |
| ENSMUSG00000074502 | Ubtfl1 | 546118 | protein_coding | 7.74690533 | -0.829442383 | 1.35880678 | -0.610419667 | 0.541583843 | 0.724720445 | 0.829442383 |
| ENSMUSG00000070942 | Il1rl2 | 107527 | protein_coding | 15.76304041 | -0.829569518 | 0.366223108 | -2.265202547 | 0.023500253 | 0.090239158 | 0.829569518 |
| ENSMUSG00000002033 | Cd3g | 12502 | protein_coding | 36.02335648 | 0.829592133 | 0.319018817 | 2.600448899 | 0.009310188 | 0.045381331 | 0.829592133 |
| ENSMUSG00000105814 | Mir703 | 735265 | miRNA | 30.04515073 | 0.829722352 | 0.369978565 | 2.242622761 | 0.024921151 | 0.094151825 | 0.829722352 |
| ENSMUSG00000034765 | Dusp5 | 240672 | protein_coding | 131.6442207 | -0.829792688 | 0.278896945 | -2.975266323 | 0.002927343 | 0.018407705 | 0.829792688 |
| ENSMUSG00000063406 | Tmed5 | 73130 | protein_coding | 954.839766 | 0.829895735 | 0.160387098 | 5.174329772 | 2.29E-07 | 4.97E-06 | 0.829895735 |
| ENSMUSG00000009731 | Kcnd1 | 16506 | protein_coding | 9.234113015 | -0.830980045 | 0.499985289 | -1.662008988 | 0.096510972 | 0.242732792 | 0.830980045 |
| ENSMUSG00000090564 | A430057M04Rik | 319486 | lncRNA | 8.491451811 | -0.83222042 | 0.450279649 | -1.848230141 | 0.064569057 | 0.184907576 | 0.83222042 |
| ENSMUSG00000022032 | Scara5 | 71145 | protein_coding | 23.90473195 | -0.832419385 | 0.356993735 | -2.331747882 | 0.019713958 | 0.078974831 | 0.832419385 |
| ENSMUSG00000039704 | Lmbrd2 | 320506 | protein_coding | 1325.222512 | 0.832716512 | 0.13204575 | 6.306272737 | 2.86E-10 | 9.23E-09 | 0.832716512 |
| ENSMUSG00000044408 | Sptssa | 104725 | protein_coding | 1232.450866 | 0.832831576 | 0.1004931 | 8.287450365 | 1.16E-16 | 6.73E-15 | 0.832831576 |
| ENSMUSG00000026018 | Ica1l | 70375 | protein_coding | 15.74155673 | -0.832946744 | 0.462167546 | -1.802261433 | 0.071504283 | 0.198383596 | 0.832946744 |
| ENSMUSG00000063286 | Gvin-ps7 | 668139 | transcribed_unprocessed_pseudogene | 42.04023083 | 0.833061997 | 0.337326789 | 2.469599283 | 0.013526448 | 0.060028909 | 0.833061997 |
| ENSMUSG00000018669 | Cdk5rap3 | 80280 | protein_coding | 718.8086783 | 0.833500692 | 0.06453906 | 12.91467053 | 3.72E-38 | 6.44E-36 | 0.833500692 |
| ENSMUSG00000032454 | Rbp2 | 19660 | protein_coding | 91.04788325 | -0.833580394 | 0.182971243 | -4.555800021 | 5.22E-06 | 8.40E-05 | 0.833580394 |
| ENSMUSG00000090394 | 4930523C07Rik | 67647 | protein_coding | 125.8183395 | -0.834007089 | 0.141128601 | -5.909553982 | 3.43E-09 | 9.81E-08 | 0.834007089 |
| ENSMUSG00000026969 | Fam166a | 68222 | protein_coding | 28.62361731 | -0.834371115 | 0.251867265 | -3.312741391 | 0.000923864 | 0.007185205 | 0.834371115 |
| ENSMUSG00000037411 | Serpine1 | 18787 | protein_coding | 11.03942699 | -0.83439373 | 0.403988819 | -2.065388175 | 0.038886286 | 0.130478359 | 0.83439373 |
| ENSMUSG00000049436 | Upk1b | 22268 | protein_coding | 49.50316958 | 0.834432305 | 0.236740768 | 3.524666708 | 0.000424016 | 0.00381351 | 0.834432305 |
| ENSMUSG00000024516 | Sec11c | 66286 | protein_coding | 2325.767871 | 0.834663052 | 0.070727851 | 11.80105201 | 3.85E-32 | 4.80E-30 | 0.834663052 |
| ENSMUSG00000078664 | Sprr2a1 | 20755 | protein_coding | 394.9996905 | 0.834863018 | 0.620082009 | 1.346375166 | 0.178181556 | 0.36540814 | 0.834863018 |
| ENSMUSG00000042671 | Rgs8 | 67792 | protein_coding | 15.73780506 | -0.836669379 | 0.468706162 | -1.785061617 | 0.074251323 | 0.203069209 | 0.836669379 |
| ENSMUSG00000051998 | Lax1 | 240754 | protein_coding | 7.058302462 | 0.836845617 | 0.610084598 | 1.371687827 | 0.170160639 | 0.355358419 | 0.836845617 |
| ENSMUSG00000093565 | Rab26os | 75614 | lncRNA | 40.88430314 | 0.837407178 | 0.201058629 | 4.164989996 | 3.11E-05 | 0.000408522 | 0.837407178 |
| ENSMUSG00000048478 | Spata33 | 320869 | protein_coding | 6.917873767 | -0.837769793 | 0.449301989 | -1.864602903 | 0.062237086 | 0.18004219 | 0.837769793 |

| ensembl_gene_id | external_gene_name | entrezgene_id | gene_biotype | baseMean | log2FoldChange | lfcSE | stat | pvalue | padj | absLog2FC |
|---|---|---|---|---|---|---|---|---|---|---|
| ENSMUSG00000073399 | Trim40 | 195359 | protein_coding | 257.1222794 | 0.837967901 | 0.239966679 | 3.49201775 | 0.000479386 | 0.004197728 | 0.837967901 |
| ENSMUSG00000052776 | Oas1a | 246730 | protein_coding | 537.0592309 | 0.838031863 | 0.13165859 | 6.365189403 | 1.95E-10 | 6.45E-09 | 0.838031863 |
| ENSMUSG00000073421 | H2-Ab1 | 14961 | protein_coding | 190.3190643 | 0.838432109 | 0.359026449 | 2.33529343 | 0.019528097 | 0.078530312 | 0.838432109 |
| ENSMUSG00000029384 | 2010109A12Rik | 75610 | protein_coding | 9.097992291 | -0.839167257 | 0.448107686 | -1.872691057 | 0.061111059 | 0.177915728 | 0.839167257 |
| ENSMUSG00000015879 | Fam184b | 58227 | protein_coding | 16.5077578 | -0.83940436 | 0.45837802 | -1.831249151 | 0.067063364 | 0.189677544 | 0.83940436 |
| ENSMUSG00000015437 | Gzmb | 14939 | protein_coding | 20.86942265 | 0.839497679 | 0.334268453 | 2.511447523 | 0.012023716 | 0.054985286 | 0.839497679 |
| ENSMUSG00000028354 | Fmn2 | 54418 | protein_coding | 15.19227466 | 0.840365439 | 0.767557166 | 1.094857134 | 0.273579231 | 0.481510466 | 0.840365439 |
| ENSMUSG00000014867 | Surf4 | 20932 | protein_coding | 5192.552665 | 0.840688275 | 0.099197504 | 8.474893423 | 2.35E-17 | 1.42E-15 | 0.840688275 |
| ENSMUSG00000030662 | Ipo5 | 70572 | protein_coding | 2646.132884 | 0.841209275 | 0.09217545 | 9.126174883 | 7.10E-20 | 4.98E-18 | 0.841209275 |
| ENSMUSG00000103749 | Pcdhgb5 | 93702 | protein_coding | 14.7837062 | -0.841621201 | 0.348177649 | -2.417217776 | 0.015639657 | 0.066728504 | 0.841621201 |
| ENSMUSG00000026496 | Parp1 | 11545 | protein_coding | 1090.628152 | 0.841639049 | 0.09727573 | 8.652096977 | 5.06E-18 | 3.21E-16 | 0.841639049 |
| ENSMUSG00000049551 | Fzd9 | 14371 | protein_coding | 25.79803771 | -0.841833575 | 0.290592374 | -2.896956871 | 0.003768016 | 0.022444634 | 0.841833575 |
| ENSMUSG00000004098 | Col5a3 | 53867 | protein_coding | 8.698404069 | -0.841908538 | 1.162249031 | -0.724378783 | 0.468833222 | 0.665105695 | 0.841908538 |
| ENSMUSG00000024268 | Celf4 | 108013 | protein_coding | 24.92468441 | -0.842163112 | 0.436785151 | -1.928094647 | 0.053843355 | 0.163014057 | 0.842163112 |
| ENSMUSG00000065979 | Cpped1 | 223978 | protein_coding | 1368.147909 | 0.843647515 | 0.059073731 | 14.28126355 | 2.86E-46 | 6.10E-44 | 0.843647515 |
| ENSMUSG00000048458 | Inka2 | 109050 | protein_coding | 16.05531817 | 0.843905364 | 0.407705791 | 2.069888098 | 0.038462825 | 0.129548715 | 0.843905364 |
| ENSMUSG00000028341 | Nr4a3 | 18124 | protein_coding | 6.939230176 | -0.843930017 | 0.548818451 | -1.537721655 | 0.124116687 | 0.287679072 | 0.843930017 |
| ENSMUSG00000021069 | Pygl | 110095 | protein_coding | 98.83690256 | -0.84490481 | 0.127975185 | -6.602098769 | 4.05E-11 | 1.44E-09 | 0.84490481 |
| ENSMUSG00000087543 | Gm16576 | 100504191 | lncRNA | 21.54934795 | -0.845065102 | 0.402976506 | -2.097057991 | 0.035988442 | 0.123494972 | 0.845065102 |
| ENSMUSG00000098132 | Rassf10 | 78748 | protein_coding | 55.61798958 | -0.845137883 | 0.226010276 | -3.739378133 | 0.000184476 | 0.001901835 | 0.845137883 |
| ENSMUSG00000003865 | Gys1 | 14936 | protein_coding | 430.6814984 | 0.845396716 | 0.081433279 | 10.38146475 | 3.01E-25 | 2.76E-23 | 0.845396716 |
| ENSMUSG00000052271 | Bhlha15 | 17341 | protein_coding | 55.45042394 | 0.845881984 | 0.26002003 | 3.253141627 | 0.001141366 | 0.008548225 | 0.845881984 |
| ENSMUSG00000039911 | Spsb1 | 74646 | protein_coding | 41.01273059 | -0.846035629 | 0.322071112 | -2.626859709 | 0.008617686 | 0.042854354 | 0.846035629 |
| ENSMUSG00000087128 | Gm12655 | 115489886 | lncRNA | 8.114831776 | -0.846702805 | 0.434376424 | -1.949237479 | 0.051267071 | 0.158099111 | 0.846702805 |
| ENSMUSG00000035413 | Tmem98 | 103743 | protein_coding | 1624.869644 | 0.846984324 | 0.136981558 | 6.183199668 | 6.28E-10 | 1.93E-08 | 0.846984324 |
| ENSMUSG00000085944 | 1700003D09Rik | 69338 | lncRNA | 23.14616843 | -0.847157904 | 0.391259144 | -2.165209217 | 0.030371667 | 0.109004035 | 0.847157904 |
| ENSMUSG00000025432 | Avil | 11567 | protein_coding | 129.7807395 | -0.847375492 | 0.124459758 | -6.808429521 | 9.87E-12 | 3.69E-10 | 0.847375492 |
| ENSMUSG00000025163 | Cd7 | 12516 | protein_coding | 51.59202945 | 0.847522683 | 0.261752219 | 3.23788156 | 0.001204208 | 0.008912929 | 0.847522683 |
| ENSMUSG00000049580 | Tsku | 244152 | protein_coding | 542.3470706 | -0.847826506 | 0.245580172 | -3.452341038 | 0.000555745 | 0.004746713 | 0.847826506 |
| ENSMUSG00000024300 | Myo1f | 17916 | protein_coding | 21.43934386 | 0.847859862 | 0.334328382 | 2.53600923 | 0.011212378 | 0.052128066 | 0.847859862 |
| ENSMUSG00000039405 | Prss23 | 76453 | protein_coding | 241.1339715 | -0.848210284 | 0.272045618 | -3.117897252 | 0.001821463 | 0.012510678 | 0.848210284 |
| ENSMUSG00000020674 | Pxdn | 69675 | protein_coding | 15.20631511 | -0.848239098 | 0.387816562 | -2.187217309 | 0.028726664 | 0.104513089 | 0.848239098 |
| ENSMUSG00000017499 | Cdc6 | 23834 | protein_coding | 131.8666024 | 0.848366914 | 0.160774199 | 5.276760325 | 1.31E-07 | 2.96E-06 | 0.848366914 |
| ENSMUSG00000041608 | Entpd3 | 215446 | protein_coding | 9.090935519 | -0.848683341 | 0.466697043 | -1.81848879 | 0.068989459 | 0.193666566 | 0.848683341 |
| ENSMUSG00000015363 | Trabd | 67976 | protein_coding | 1531.830985 | 0.848913349 | 0.092631085 | 9.164454314 | 4.98E-20 | 3.54E-18 | 0.848913349 |
| ENSMUSG00000062044 | Lmtk3 | 381983 | protein_coding | 8.91059219 | -0.849354754 | 0.599400454 | -1.417007192 | 0.156480825 | 0.337133312 | 0.849354754 |
| ENSMUSG00000011382 | Dhdh | 71755 | protein_coding | 20.84541589 | -0.849579174 | 0.385253323 | -2.205248138 | 0.027436687 | 0.100903151 | 0.849579174 |
| ENSMUSG00000026872 | Zeb2 | 24136 | protein_coding | 16.36478711 | -0.849660409 | 0.848921725 | -1.000870143 | 0.316889593 | 0.528193536 | 0.849660409 |

Supplementary Table 1

| ensembl_gene_id | external_gene_name | entrezgene_id | gene_biotype | baseMean | log2FoldChange | lfcSE | stat | pvalue | padj | absLog2FC |
|---|---|---|---|---|---|---|---|---|---|---|
| ENSMUSG00000021238 | Aldh6a1 | 104776 | protein_coding | 1073.323628 | 0.849972598 | 0.086834623 | 9.788406608 | 1.26E-22 | 1.00E-20 | 0.849972598 |
| ENSMUSG00000029828 | 4921507P07Rik | 70821 | protein_coding | 14.34300672 | -0.851273755 | 0.397099871 | -2.143727101 | 0.032054759 | 0.113349074 | 0.851273755 |
| ENSMUSG00000000531 | Tamalin | 56149 | protein_coding | 74.10065854 | -0.851312432 | 0.387767172 | -2.195421618 | 0.028133368 | 0.102873309 | 0.851312432 |
| ENSMUSG00000020447 | Npc1l1 | 237636 | protein_coding | 16.24395973 | -0.851434553 | 0.497223989 | -1.712376256 | 0.086827352 | 0.226719026 | 0.851434553 |
| ENSMUSG00000054146 | Krt15 | 16665 | protein_coding | 12.45416498 | -0.8515591 | 0.478107937 | -1.781102203 | 0.074895756 | 0.204201575 | 0.8515591 |
| ENSMUSG00000026399 | Cd55 | 13136 | protein_coding | 277.2927895 | -0.851765402 | 0.124461371 | -6.843612565 | 7.72E-12 | 2.91E-10 | 0.851765402 |
| ENSMUSG00000037418 | Best1 | 24115 | protein_coding | 18.04182148 | -0.851812459 | 0.324960782 | -2.621277726 | 0.008760087 | 0.043386399 | 0.851812459 |
| ENSMUSG00000070564 | Ntn5 | 243967 | protein_coding | 12.19714048 | -0.852026304 | 0.517538058 | -1.646306568 | 0.099700658 | 0.247665639 | 0.852026304 |
| ENSMUSG00000032525 | Nktr | 18087 | protein_coding | 2418.49249 | 0.852235846 | 0.128816149 | 6.615908445 | 3.69E-11 | 1.32E-09 | 0.852235846 |
| ENSMUSG00000057719 | Sh3rf2 | 269016 | protein_coding | 313.9722232 | -0.852478728 | 0.187873758 | -4.537508261 | 5.69E-06 | 9.04E-05 | 0.852478728 |
| ENSMUSG00000042942 | Greb1l | 381157 | protein_coding | 122.7162802 | -0.852777605 | 0.231889119 | -3.67752316 | 0.00023551 | 0.002340111 | 0.852777605 |
| ENSMUSG00000109372 | Gm19410 | 100502846 | protein_coding | 14.84430161 | -0.853177226 | 0.494756152 | -1.724439852 | 0.084628468 | 0.222541843 | 0.853177226 |
| ENSMUSG00000085589 | A430078I02Rik | 320486 | lncRNA | 45.39524686 | 0.853340803 | 0.182301849 | 4.680922364 | 2.86E-06 | 4.84E-05 | 0.853340803 |
| ENSMUSG00000018008 | Cyth4 | 72318 | protein_coding | 17.53163864 | 0.854383667 | 0.381035893 | 2.242265578 | 0.024944213 | 0.094216606 | 0.854383667 |
| ENSMUSG00000026358 | Rgs1 | 50778 | protein_coding | 116.4090795 | 0.854838595 | 0.221818437 | 3.853776118 | 0.00011631 | 0.001279963 | 0.854838595 |
| ENSMUSG00000017167 | Cntnap1 | 53321 | protein_coding | 106.7581827 | -0.855561739 | 0.172396883 | -4.962744815 | 6.95E-07 | 1.36E-05 | 0.855561739 |
| ENSMUSG00000033730 | Egr3 | 13655 | protein_coding | 6.77860391 | -0.85688943 | 0.487000344 | -1.759525308 | 0.078488326 | 0.210962209 | 0.85688943 |
| ENSMUSG00000047369 | Dnah14 | 240960 | protein_coding | 18.09531303 | -0.856917099 | 0.580331651 | -1.476598936 | 0.139783177 | 0.311725998 | 0.856917099 |
| ENSMUSG00000051238 | Swsap1 | 66962 | protein_coding | 175.1097865 | 0.85796512 | 0.122407967 | 7.009062712 | 2.40E-12 | 9.48E-11 | 0.85796512 |
| ENSMUSG00000031671 | Setd6 | 66083 | protein_coding | 101.4407699 | -0.858597745 | 0.17388968 | -4.937600349 | 7.91E-07 | 1.53E-05 | 0.858597745 |
| ENSMUSG00000039096 | Rsad1 | 237926 | protein_coding | 60.76181894 | -0.860835973 | 0.213408559 | -4.033746246 | 5.49E-05 | 0.000672069 | 0.860835973 |
| ENSMUSG00000000392 | Fap | 14089 | protein_coding | 82.93544838 | 0.86086111 | 0.161857454 | 5.318637421 | 1.05E-07 | 2.39E-06 | 0.86086111 |
| ENSMUSG00000031209 | Heph | 15203 | protein_coding | 3547.032797 | 0.860893541 | 0.114558089 | 7.514908342 | 5.70E-14 | 2.65E-12 | 0.860893541 |
| ENSMUSG00000026874 | Hc | 15139 | protein_coding | 20.89541256 | -0.861778123 | 0.29942727 | -2.878088306 | 0.004000931 | 0.023480779 | 0.861778123 |
| ENSMUSG00000024501 | Dpysl3 | 22240 | protein_coding | 191.8765057 | -0.861949872 | 0.185736938 | -4.640702506 | 3.47E-06 | 5.78E-05 | 0.861949872 |
| ENSMUSG00000020653 | Klf11 | 194655 | protein_coding | 243.0513336 | -0.863391949 | 0.240285884 | -3.593186307 | 0.000326659 | 0.003047837 | 0.863391949 |
| ENSMUSG00000045969 | Ing1 | 26356 | protein_coding | 583.3165564 | 0.864254363 | 0.115948562 | 7.453773879 | 9.07E-14 | 4.14E-12 | 0.864254363 |
| ENSMUSG00000068758 | Il3ra | 16188 | protein_coding | 20.36673302 | 0.865241516 | 0.28286755 | 3.058822106 | 0.002222091 | 0.014771894 | 0.865241516 |
| ENSMUSG00000051985 | Igfn1 | 226438 | protein_coding | 8.193382781 | -0.865357334 | 0.774435881 | -1.117403462 | 0.263821856 | 0.47151644 | 0.865357334 |
| ENSMUSG00000001815 | Evx2 | 14029 | protein_coding | 7.581726079 | -0.865544402 | 0.799417678 | -1.082718617 | 0.278933338 | 0.487423953 | 0.865544402 |
| ENSMUSG00000025935 | Tram1 | 72265 | protein_coding | 3946.386188 | 0.866143218 | 0.088529179 | 9.783703221 | 1.32E-22 | 1.04E-20 | 0.866143218 |
| ENSMUSG00000004961 | Syt5 | 53420 | protein_coding | 9.161175302 | -0.866487249 | 0.411334526 | -2.106526911 | 0.035158608 | 0.121397422 | 0.866487249 |
| ENSMUSG00000025875 | Tspan17 | 74257 | protein_coding | 77.06316436 | -0.86716435 | 0.157639152 | -5.500945301 | 3.78E-08 | 9.21E-07 | 0.86716435 |
| ENSMUSG00000066861 | Oas1g | 23960 | protein_coding | 181.6833625 | 0.867746941 | 0.148949576 | 5.825776512 | 5.68E-09 | 1.57E-07 | 0.867746941 |
| ENSMUSG00000029765 | Plxna4 | 243743 | protein_coding | 8.849023065 | -0.868602821 | 1.044783699 | -0.831370955 | 0.405764102 | 0.612781892 | 0.868602821 |
| ENSMUSG00000074102 | Rbm15b | 109095 | protein_coding | 1536.303722 | 0.869065668 | 0.079740991 | 10.89860632 | 1.17E-27 | 1.19E-25 | 0.869065668 |
| ENSMUSG00000023247 | Guca2a | 14915 | protein_coding | 20207.97142 | 0.869112044 | 0.18048696 | 4.815373048 | 1.47E-06 | 2.67E-05 | 0.869112044 |
| ENSMUSG00000032011 | Thy1 | 21838 | protein_coding | 14.83410905 | 0.870176551 | 0.452279933 | 1.923977802 | 0.054357366 | 0.164258039 | 0.870176551 |

| ensembl_gene_id | external_gene_name | entrezgene_id | gene_biotype | baseMean | log2FoldChange | lfcSE | stat | pvalue | padj | absLog2FC |
|---|---|---|---|---|---|---|---|---|---|---|
| ENSMUSG00000056290 | Ms4a4b | 60361 | protein_coding | 15.88945259 | 0.870465384 | 0.544612326 | 1.59832112 | 0.109971529 | 0.264316662 | 0.870465384 |
| ENSMUSG00000061702 | Tmem91 | 320208 | protein_coding | 6.699032575 | -0.870976996 | 0.547077289 | -1.59205475 | 0.111372403 | 0.266437314 | 0.870976996 |
| ENSMUSG00000025189 | Cnnm1 | 83674 | protein_coding | 23.12642298 | -0.871437189 | 0.312877542 | -2.785234066 | 0.00534891 | 0.029474953 | 0.871437189 |
| ENSMUSG00000040612 | Ildr2 | 100039795 | protein_coding | 10.20053744 | -0.871545819 | 0.576799546 | -1.511002956 | 0.130787698 | 0.297938565 | 0.871545819 |
| ENSMUSG00000043366 | Or51e2 | 170639 | protein_coding | 25.51509666 | -0.871715431 | 0.309192563 | -2.819328587 | 0.004812423 | 0.027288098 | 0.871715431 |
| ENSMUSG00000032417 | Rwdd2a | 69519 | protein_coding | 8.829121034 | -0.871757837 | 0.426377988 | -2.044565761 | 0.040897688 | 0.13523321 | 0.871757837 |
| ENSMUSG00000006463 | Zdhhc24 | 70605 | protein_coding | 192.5586217 | 0.872032175 | 0.113153503 | 7.706629927 | 1.29E-14 | 6.33E-13 | 0.872032175 |
| ENSMUSG00000048696 | Mex3d | 237400 | protein_coding | 102.1387484 | -0.872138233 | 0.170522134 | -5.11451631 | 3.15E-07 | 6.64E-06 | 0.872138233 |
| ENSMUSG00000092517 | Art2a | 11871 | protein_coding | 141.1791003 | 0.872326646 | 0.309645102 | 2.817182125 | 0.004844704 | 0.02744184 | 0.872326646 |
| ENSMUSG00000117869 | Snhg4 | 100503380 | lncRNA | 100.5750164 | -0.872633915 | 0.169145014 | -5.159087419 | 2.48E-07 | 5.35E-06 | 0.872633915 |
| ENSMUSG00000032006 | Pdgfd | 71785 | protein_coding | 24.71634232 | -0.873348308 | 0.271599457 | -3.215574576 | 0.001301837 | 0.009468335 | 0.873348308 |
| ENSMUSG00000028443 | Nudt2 | 66401 | protein_coding | 284.2281246 | 0.873828932 | 0.106568086 | 8.199724367 | 2.41E-16 | 1.37E-14 | 0.873828932 |
| ENSMUSG00000070509 | Rgma | 244058 | protein_coding | 14.13080135 | -0.873995678 | 0.405523558 | -2.155227879 | 0.031144006 | 0.111075175 | 0.873995678 |
| ENSMUSG00000044037 | Als2cl | 235633 | protein_coding | 403.3194576 | -0.87423825 | 0.15706456 | -5.566107659 | 2.60E-08 | 6.62E-07 | 0.87423825 |
| ENSMUSG00000029007 | Agtrap | 11610 | protein_coding | 625.9370656 | 0.875139932 | 0.153740551 | 5.692316875 | 1.25E-08 | 3.33E-07 | 0.875139932 |
| ENSMUSG00000087412 | Gm15501 | 100040298 | transcribed_processed_pseudogene | 9.774782257 | -0.875213178 | 0.359792805 | -2.43254775 | 0.014993014 | 0.064665241 | 0.875213178 |
| ENSMUSG00000081683 | Fzd10 | 93897 | protein_coding | 14.80221999 | -0.875223473 | 0.334579964 | -2.615887279 | 0.008899593 | 0.043913482 | 0.875223473 |
| ENSMUSG00000016758 | Bik | 12124 | protein_coding | 40.89143927 | -0.875373881 | 0.239808008 | -3.650311303 | 0.000261923 | 0.002551623 | 0.875373881 |
| ENSMUSG00000002699 | Lcp2 | 16822 | protein_coding | 17.61650861 | 0.875379883 | 0.378363375 | 2.313595713 | 0.020689909 | 0.082018136 | 0.875379883 |
| ENSMUSG00000040473 | Cfap69 | 207686 | protein_coding | 10.94837613 | -0.87573506 | 0.683393615 | -1.281450455 | 0.200035492 | 0.394660109 | 0.87573506 |
| ENSMUSG00000023828 | Slc22a3 | 20519 | protein_coding | 15.34344241 | -0.875747392 | 0.345664408 | -2.533519133 | 0.011292358 | 0.052426743 | 0.875747392 |
| ENSMUSG00000069892 | 9930111J21Rik2 | 245240 | protein_coding | 7.104520793 | 0.87592741 | 0.462390706 | 1.894344756 | 0.058179264 | 0.172341327 | 0.87592741 |
| ENSMUSG00000112188 | Gm47708 | 102641383 | lncRNA | 22.2011689 | -0.875991298 | 0.371034103 | -2.360945504 | 0.018228409 | 0.074602329 | 0.875991298 |
| ENSMUSG00000040537 | Adam22 | 11496 | protein_coding | 54.08939152 | -0.876049452 | 0.231318369 | -3.787202276 | 0.000152353 | 0.001616709 | 0.876049452 |
| ENSMUSG00000061958 | Defa38 | 634825 | protein_coding | 11.5580048 | -0.876608383 | 1.121973026 | -0.781309677 | 0.43462038 | 0.637867413 | 0.876608383 |
| ENSMUSG00000030671 | Pde3b | 18576 | protein_coding | 277.3765612 | 0.876773146 | 0.102207667 | 8.578350021 | 9.62E-18 | 5.94E-16 | 0.876773146 |
| ENSMUSG00000055745 | Rtl6 | 223732 | protein_coding | 15.00082982 | 0.877482381 | 0.357488773 | 2.454573254 | 0.014105192 | 0.061772753 | 0.877482381 |
| ENSMUSG00000010021 | Kif19a | 286942 | protein_coding | 12.23416917 | -0.877512723 | 0.345401899 | -2.540555584 | 0.01106765 | 0.051636066 | 0.877512723 |
| ENSMUSG00000095961 | Vmn2r102 | 224572 | protein_coding | 11.75083338 | -0.878007601 | 1.761267081 | -0.498509062 | 0.618125283 | 0.777697562 | 0.878007601 |
| ENSMUSG00000090799 | Klhl33 | 546611 | protein_coding | 7.575977025 | -0.878904963 | 0.679233904 | -1.293965095 | 0.195677478 | 0.388961796 | 0.878904963 |
| ENSMUSG00000062760 | Shisal1 | 72301 | protein_coding | 7.83753333 | -0.879588509 | 0.562374442 | -1.564062025 | 0.117803003 | 0.277363819 | 0.879588509 |
| ENSMUSG00000026365 | Cfh | 12628 | protein_coding | 7.168850587 | -0.87983144 | 0.745974572 | -1.179438915 | 0.238223447 | 0.442395134 | 0.87983144 |
| ENSMUSG00000030870 | Ubfd1 | 28018 | protein_coding | 1057.134034 | 0.880026715 | 0.118446829 | 7.429719479 | 1.09E-13 | 4.92E-12 | 0.880026715 |
| ENSMUSG00000027867 | Spag17 | 74362 | protein_coding | 44.30906905 | 0.880704572 | 0.298958139 | 2.945912676 | 0.003220033 | 0.019863937 | 0.880704572 |
| ENSMUSG00000024891 | Slc29a2 | 13340 | protein_coding | 17.77247375 | -0.881051253 | 0.340506488 | -2.587472732 | 0.009668285 | 0.046679733 | 0.881051253 |
| ENSMUSG00000033578 | Tmem35a | 67564 | protein_coding | 63.0666813 | -0.881427363 | 0.226560962 | -3.890464419 | 0.000100053 | 0.001128638 | 0.881427363 |
| ENSMUSG00000048960 | Prex2 | 109294 | protein_coding | 10.81493263 | -0.881598625 | 0.870924076 | -1.012256578 | 0.311415391 | 0.521451199 | 0.881598625 |
| ENSMUSG00000010492 | Uckl1os | 100271841 | lncRNA | 24.87351989 | -0.881885698 | 0.269075054 | -3.277471044 | 0.001047415 | 0.007959552 | 0.881885698 |

| ensembl_gene_id | external_gene_name | entrezgene_id | gene_biotype | baseMean | log2FoldChange | lfcSE | stat | pvalue | padj | absLog2FC |
|---|---|---|---|---|---|---|---|---|---|---|
| ENSMUSG00000029343 | Crybb1 | 12960 | protein_coding | 7.313983412 | 0.882898559 | 0.692321128 | 1.275273168 | 0.202212571 | 0.397242456 | 0.882898559 |
| ENSMUSG00000030590 | Fam98c | 73833 | protein_coding | 431.8602971 | 0.883315118 | 0.110009898 | 8.029414919 | 9.79E-16 | 5.29E-14 | 0.883315118 |
| ENSMUSG00000083282 | Ctsf | 56464 | protein_coding | 113.8171781 | -0.883603344 | 0.138840477 | -6.364162407 | 1.96E-10 | 6.48E-09 | 0.883603344 |
| ENSMUSG00000020732 | Rab37 | 58222 | protein_coding | 58.45871939 | -0.884240243 | 0.212583439 | -4.159497304 | 3.19E-05 | 0.000417438 | 0.884240243 |
| ENSMUSG00000030022 | Adamts9 | 101401 | protein_coding | 13.23697618 | -0.884767074 | 0.595499117 | -1.485757154 | 0.137343367 | 0.308157609 | 0.884767074 |
| ENSMUSG00000032437 | Stt3b | 68292 | protein_coding | 6133.498354 | 0.885010211 | 0.077568853 | 11.40935029 | 3.76E-30 | 4.37E-28 | 0.885010211 |
| ENSMUSG00000097908 | 4933404O12Rik | 66752 | lncRNA | 15.13624441 | -0.885655403 | 0.383308672 | -2.310554046 | 0.0208575 | 0.082497706 | 0.885655403 |
| ENSMUSG00000037306 | Man1c1 | 230815 | protein_coding | 18.04318665 | -0.885688455 | 0.321807241 | -2.752232835 | 0.005919042 | 0.031840089 | 0.885688455 |
| ENSMUSG00000039275 | Foxk2 | 68837 | protein_coding | 848.1766429 | 0.886604179 | 0.123392339 | 7.185244914 | 6.71E-13 | 2.83E-11 | 0.886604179 |
| ENSMUSG00000029563 | Foxp2 | 114142 | protein_coding | 44.28859204 | -0.888207616 | 0.530028542 | -1.675773181 | 0.093782629 | 0.238468984 | 0.888207616 |
| ENSMUSG00000038751 | Ptk6 | 20459 | protein_coding | 260.207493 | 0.889119171 | 0.333335181 | 2.667342728 | 0.007645366 | 0.039093224 | 0.889119171 |
| ENSMUSG00000033849 | B3galt2 | 26878 | protein_coding | 191.0073174 | -0.889157468 | 0.206702052 | -4.30163831 | 1.70E-05 | 0.000239188 | 0.889157468 |
| ENSMUSG00000033705 | Stard9 | 668880 | protein_coding | 16.99764331 | -0.891233779 | 0.515068645 | -1.730320388 | 0.083573049 | 0.220676394 | 0.891233779 |
| ENSMUSG00000030208 | Emp1 | 13730 | protein_coding | 9077.474174 | -0.891343039 | 0.403108577 | -2.21117359 | 0.027023819 | 0.099799532 | 0.891343039 |
| ENSMUSG00000031861 | Lpar2 | 53978 | protein_coding | 22.61365362 | -0.891431843 | 0.315700012 | -2.82366743 | 0.004747762 | 0.027017633 | 0.891431843 |
| ENSMUSG00000022204 | Ngdn | 68966 | protein_coding | 583.3044493 | 0.891528508 | 0.119064822 | 7.487757428 | 7.01E-14 | 3.23E-12 | 0.891528508 |
| ENSMUSG00000024910 | Ctsw | 13041 | protein_coding | 45.72537652 | 0.891984868 | 0.244704912 | 3.64514493 | 0.000267241 | 0.002589183 | 0.891984868 |
| ENSMUSG00000034063 | 4930590J08Rik | 381798 | protein_coding | 7.545947752 | -0.892166893 | 0.557622796 | -1.599946952 | 0.109610352 | 0.263874388 | 0.892166893 |
| ENSMUSG00000074802 | Gas2l3 | 237436 | protein_coding | 291.4010869 | 0.892266455 | 0.135502462 | 6.584872625 | 4.55E-11 | 1.62E-09 | 0.892266455 |
| ENSMUSG00000024411 | Aqp4 | 11829 | protein_coding | 13913.30471 | 0.892387062 | 0.252362961 | 3.536125338 | 0.000406042 | 0.003678861 | 0.892387062 |
| ENSMUSG00000096986 | 4930509E16Rik | 75115 | lncRNA | 31.54535951 | 0.892631694 | 0.224995805 | 3.967325934 | 7.27E-05 | 0.000850199 | 0.892631694 |
| ENSMUSG00000032068 | Plet1 | 76509 | protein_coding | 1323.468863 | -0.89314247 | 0.327720934 | -2.725314066 | 0.006424035 | 0.034016631 | 0.89314247 |
| ENSMUSG00000085957 | Syna | 214292 | protein_coding | 13.00168946 | 0.893662304 | 0.471084386 | 1.897032315 | 0.057823673 | 0.171719678 | 0.893662304 |
| ENSMUSG00000026856 | Dolpp1 | 57170 | protein_coding | 772.8566052 | 0.894271058 | 0.112718667 | 7.933655378 | 2.13E-15 | 1.13E-13 | 0.894271058 |
| ENSMUSG00000059401 | Mamld1 | 333639 | protein_coding | 12.4615415 | -0.894865699 | 0.372799398 | -2.400394702 | 0.016377402 | 0.069048936 | 0.894865699 |
| ENSMUSG00000030310 | Slc6a1 | 232333 | protein_coding | 8.395827687 | 0.895262367 | 0.580312212 | 1.542725361 | 0.122897426 | 0.285856281 | 0.895262367 |
| ENSMUSG00000050010 | Shisa3 | 330096 | protein_coding | 7.483553398 | -0.895697416 | 0.551355798 | -1.624536133 | 0.104261435 | 0.255449337 | 0.895697416 |
| ENSMUSG00000039620 | Trmt9b | 319582 | protein_coding | 22.05420908 | -0.896545498 | 0.354015829 | -2.53250116 | 0.0113252 | 0.052514637 | 0.896545498 |
| ENSMUSG00000046697 | Enpp7 | 238011 | protein_coding | 8.616342582 | -0.896888117 | 0.689962046 | -1.299909354 | 0.193632039 | 0.387069668 | 0.896888117 |
| ENSMUSG00000065444 | Mir27a | 387220 | miRNA | 7.056447852 | -0.897077811 | 0.504672838 | -1.777543281 | 0.075478896 | 0.205144798 | 0.897077811 |
| ENSMUSG00000030492 | Slc7a9 | 30962 | protein_coding | 66.28227542 | 0.89722518 | 0.20324145 | 4.414577737 | 1.01E-05 | 0.000150516 | 0.89722518 |
| ENSMUSG00000027801 | Tm4sf4 | 229302 | protein_coding | 19.24746617 | -0.897365386 | 0.307655715 | -2.916784388 | 0.003536602 | 0.021402352 | 0.897365386 |
| ENSMUSG00000063382 | Bcl9l | 80288 | protein_coding | 113.1735161 | -0.897449337 | 0.233599345 | -3.841831564 | 0.00012212 | 0.001339076 | 0.897449337 |
| ENSMUSG00000038580 | Sct | 20287 | protein_coding | 191.1534371 | -0.897803458 | 0.248262503 | -3.6163474 | 0.000298789 | 0.002827758 | 0.897803458 |
| ENSMUSG00000042500 | Ago4 | 76850 | protein_coding | 102.877746 | -0.898221803 | 0.187715737 | -4.785010659 | 1.71E-06 | 3.04E-05 | 0.898221803 |
| ENSMUSG00000040270 | Bach2 | 12014 | protein_coding | 48.10818351 | -0.898373398 | 0.38515472 | -2.332500032 | 0.019674401 | 0.078836192 | 0.898373398 |
| ENSMUSG00000040253 | Gbp7 | 229900 | protein_coding | 102.3618958 | 0.898384122 | 0.243618279 | 3.687671241 | 0.000226316 | 0.002260036 | 0.898384122 |
| ENSMUSG00000021728 | Emb | 13723 | protein_coding | 27.58848674 | 0.898617122 | 0.301380648 | 2.981668298 | 0.002866824 | 0.018098604 | 0.898617122 |

| ensembl_gene_id | external_gene_name | entrezgene_id | gene_biotype | baseMean | log2FoldChange | lfcSE | stat | pvalue | padj | absLog2FC |
|---|---|---|---|---|---|---|---|---|---|---|
| ENSMUSG00000032554 | Trf | 22041 | protein_coding | 257.4507663 | 0.899286158 | 0.554905731 | 1.620610687 | 0.105101159 | 0.256707373 | 0.899286158 |
| ENSMUSG00000096146 | Kcnj11 | 16514 | protein_coding | 9.253961094 | -0.899704676 | 0.442954989 | -2.031142437 | 0.042240548 | 0.138125117 | 0.899704676 |
| ENSMUSG00000030790 | Adm | 11535 | protein_coding | 51.67581191 | 0.89982401 | 0.225019728 | 3.998867198 | 6.36E-05 | 0.00075994 | 0.89982401 |
| ENSMUSG00000059003 | Grin2a | 14811 | protein_coding | 8.514129981 | -0.900562889 | 1.370005487 | -0.657342542 | 0.510960688 | 0.699008761 | 0.900562889 |
| ENSMUSG00000044005 | Gls2 | 216456 | protein_coding | 15.39221386 | 0.901180002 | 0.323846806 | 2.782735491 | 0.005390273 | 0.02960828 | 0.901180002 |
| ENSMUSG00000108622 | Gm36864 | 102640913 | protein_coding | 11.53113338 | -0.901274281 | 0.416305972 | -2.164932386 | 0.030392864 | 0.109055538 | 0.901274281 |
| ENSMUSG00000037416 | Dmxl1 | 240283 | protein_coding | 2601.082219 | 0.901444522 | 0.086978063 | 10.36404457 | 3.61E-25 | 3.27E-23 | 0.901444522 |
| ENSMUSG00000028884 | Rpa2 | 19891 | protein_coding | 238.8972698 | 0.901497484 | 0.139050321 | 6.483246353 | 8.98E-11 | 3.09E-09 | 0.901497484 |
| ENSMUSG00000039474 | Wfs1 | 22393 | protein_coding | 171.2396321 | 0.90177815 | 0.116383977 | 7.748301579 | 9.31E-15 | 4.61E-13 | 0.90177815 |
| ENSMUSG00000043614 | Vps37d | 194309 | protein_coding | 11.57646732 | -0.902007328 | 0.43904658 | -2.054468407 | 0.039930378 | 0.132945456 | 0.902007328 |
| ENSMUSG00000006362 | Cbfa2t3 | 12398 | protein_coding | 241.2839391 | -0.902097919 | 0.194888002 | -4.628801722 | 3.68E-06 | 6.10E-05 | 0.902097919 |
| ENSMUSG00000030149 | Klrk1 | 27007 | protein_coding | 16.73496457 | 0.90227229 | 0.478799538 | 1.88444687 | 0.059504555 | 0.175127227 | 0.90227229 |
| ENSMUSG00000026715 | Serpinc1 | 11905 | protein_coding | 12.16093525 | -0.902342567 | 0.488640918 | -1.846637346 | 0.064799718 | 0.185268337 | 0.902342567 |
| ENSMUSG00000045312 | Lhfpl2 | 218454 | protein_coding | 683.4161641 | -0.902361971 | 0.239526909 | -3.767267625 | 0.000165044 | 0.001726081 | 0.902361971 |
| ENSMUSG00000028541 | B4galt2 | 53418 | protein_coding | 6.940472957 | 0.903474194 | 0.504119774 | 1.792181621 | 0.073103878 | 0.201174597 | 0.903474194 |
| ENSMUSG00000079018 | Ly6c1 | 17067 | protein_coding | 20.63478626 | 0.9035175 | 0.646649609 | 1.397228867 | 0.162344759 | 0.344776976 | 0.9035175 |
| ENSMUSG00000020326 | Ccng1 | 12450 | protein_coding | 3173.564376 | 0.90409176 | 0.083466615 | 10.83177702 | 2.43E-27 | 2.42E-25 | 0.90409176 |
| ENSMUSG00000053835 | H2-T24 | 15042 | protein_coding | 27.37761092 | 0.904432068 | 0.279127445 | 3.240211897 | 0.001194409 | 0.008860323 | 0.904432068 |
| ENSMUSG00000022657 | Cd96 | 84544 | protein_coding | 23.73707666 | 0.904975267 | 0.305960011 | 2.95782205 | 0.003098209 | 0.019254107 | 0.904975267 |
| ENSMUSG00000060090 | Rp2 | 19889 | protein_coding | 1412.589165 | 0.905667797 | 0.085806488 | 10.55477061 | 4.83E-26 | 4.52E-24 | 0.905667797 |
| ENSMUSG00000030877 | Mfsd13b | 74466 | protein_coding | 7.851195862 | 0.905794359 | 0.418146716 | 2.166211822 | 0.030295004 | 0.108802441 | 0.905794359 |
| ENSMUSG00000040026 | Saa3 | 20210 | protein_coding | 115.3651944 | 0.906949268 | 0.368926345 | 2.458347797 | 0.013957793 | 0.061268531 | 0.906949268 |
| ENSMUSG00000020019 | Ntn4 | 57764 | protein_coding | 187.553953 | -0.907741981 | 0.202965379 | -4.472398121 | 7.73E-06 | 0.000118918 | 0.907741981 |
| ENSMUSG00000020900 | Myh10 | 77579 | protein_coding | 24.70567974 | -0.907869623 | 0.303869334 | -2.987697416 | 0.002810877 | 0.017801848 | 0.907869623 |
| ENSMUSG00000034088 | Hdlbp | 110611 | protein_coding | 10023.05754 | 0.90789877 | 0.066493239 | 13.65400133 | 1.91E-42 | 3.62E-40 | 0.90789877 |
| ENSMUSG00000021484 | Lman2 | 66890 | protein_coding | 4384.361367 | 0.908081126 | 0.084095693 | 10.79818825 | 3.51E-27 | 3.45E-25 | 0.908081126 |
| ENSMUSG00000118204 | 4930414N06Rik | 73894 | lncRNA | 11.08340224 | 0.908582875 | 0.433271324 | 2.097029795 | 0.035990938 | 0.123494972 | 0.908582875 |
| ENSMUSG00000024774 | Ankrd22 | 52024 | protein_coding | 99.12291571 | 0.910466188 | 0.204708485 | 4.447623099 | 8.68E-06 | 0.00013139 | 0.910466188 |
| ENSMUSG00000024109 | Nrxn1 | 18189 | protein_coding | 19.24050463 | -0.911388125 | 0.684672138 | -1.331130733 | 0.183145998 | 0.372322841 | 0.911388125 |
| ENSMUSG00000030711 | Sult1a1 | 20887 | protein_coding | 5773.622785 | -0.911400986 | 0.098412152 | -9.261061415 | 2.02E-20 | 1.48E-18 | 0.911400986 |
| ENSMUSG00000070644 | Etnk2 | 214253 | protein_coding | 12.17746482 | -0.911419901 | 0.332033475 | -2.744963891 | 0.006051756 | 0.032466273 | 0.911419901 |
| ENSMUSG00000098766 | Mir6386 | 102465203 | miRNA | 8.525147729 | -0.911577025 | 0.400147941 | -2.278100004 | 0.02272062 | 0.088052077 | 0.911577025 |
| ENSMUSG00000031149 | Praf2 | 54637 | protein_coding | 152.7028414 | 0.911762239 | 0.128495492 | 7.095674897 | 1.29E-12 | 5.26E-11 | 0.911762239 |
| ENSMUSG00000031422 | Morf4l2 | 56397 | protein_coding | 3015.622448 | 0.912193676 | 0.119807002 | 7.613859487 | 2.66E-14 | 1.28E-12 | 0.912193676 |
| ENSMUSG00000050295 | Foxc1 | 17300 | protein_coding | 24.7130344 | -0.912326114 | 0.331812712 | -2.749521282 | 0.005968239 | 0.032072236 | 0.912326114 |
| ENSMUSG00000025007 | Aldh18a1 | 56454 | protein_coding | 1123.700817 | 0.912687461 | 0.164016418 | 5.56461036 | 2.63E-08 | 6.65E-07 | 0.912687461 |
| ENSMUSG00000031098 | Syt8 | 55925 | protein_coding | 15.08701135 | 0.912923709 | 0.4459358 | 2.047208834 | 0.040637584 | 0.134568699 | 0.912923709 |
| ENSMUSG00000022026 | Olfm4 | 380924 | protein_coding | 34.49591559 | -0.913996829 | 0.827981609 | -1.103885424 | 0.269642843 | 0.477579361 | 0.913996829 |

| ensembl_gene_id | external_gene_name | entrezgene_id | gene_biotype | baseMean | log2FoldChange | lfcSE | stat | pvalue | padj | absLog2FC |
|---|---|---|---|---|---|---|---|---|---|---|
| ENSMUSG00000020089 | Ppa1 | 67895 | protein_coding | 4117.250758 | 0.914916264 | 0.10545177 | 8.676158469 | 4.09E-18 | 2.61E-16 | 0.914916264 |
| ENSMUSG00000054793 | Cadm4 | 260299 | protein_coding | 12.53836584 | -0.915082934 | 0.385032492 | -2.376638213 | 0.017471215 | 0.072417303 | 0.915082934 |
| ENSMUSG00000038418 | Egr1 | 13653 | protein_coding | 2257.527043 | -0.91528529 | 0.26867879 | -3.406615345 | 0.000657738 | 0.005467872 | 0.91528529 |
| ENSMUSG00000030844 | Rgs10 | 67865 | protein_coding | 10.20586612 | 0.915285879 | 0.525374451 | 1.742159095 | 0.081480609 | 0.216809747 | 0.915285879 |
| ENSMUSG00000021071 | Trim9 | 94090 | protein_coding | 12.20315183 | -0.915674144 | 0.522734053 | -1.751701728 | 0.07982511 | 0.213474537 | 0.915674144 |
| ENSMUSG00000037649 | H2-DMa | 14998 | protein_coding | 55.51142886 | 0.916168002 | 0.184294543 | 4.971216114 | 6.65E-07 | 1.31E-05 | 0.916168002 |
| ENSMUSG00000027849 | Syt6 | 54524 | protein_coding | 9.408935617 | -0.916213575 | 0.651565027 | -1.406173654 | 0.159672565 | 0.341011615 | 0.916213575 |
| ENSMUSG00000057176 | Ccdc189 | 233899 | protein_coding | 19.69251075 | -0.916562316 | 0.26228522 | -3.494525216 | 0.000474906 | 0.004169955 | 0.916562316 |
| ENSMUSG00000025978 | Rftn2 | 74013 | protein_coding | 71.02667982 | -0.91666065 | 0.205680456 | -4.456722184 | 8.32E-06 | 0.000126485 | 0.91666065 |
| ENSMUSG00000055413 | H2-Q5 | 15016 | protein_coding | 10.57252557 | 0.916676673 | 0.389960146 | 2.350693226 | 0.018738476 | 0.076256117 | 0.916676673 |
| ENSMUSG00000043811 | Rtn4r | 65079 | protein_coding | 45.05899125 | -0.917199631 | 0.337609665 | -2.716745777 | 0.006592723 | 0.034750157 | 0.917199631 |
| ENSMUSG00000025515 | Muc2 | 17831 | protein_coding | 37065.285 | -0.91832721 | 0.303633299 | -3.024461456 | 0.002490762 | 0.016266035 | 0.91832721 |
| ENSMUSG00000073739 | Gm16287 | 100038595 | lncRNA | 11.72559093 | -0.918500578 | 0.376065598 | -2.442394581 | 0.01459019 | 0.063150149 | 0.918500578 |
| ENSMUSG00000023068 | Nus1 | 52014 | protein_coding | 2972.311921 | 0.920226546 | 0.142987035 | 6.435734173 | 1.23E-10 | 4.14E-09 | 0.920226546 |
| ENSMUSG00000020377 | Ltc4s | 17001 | protein_coding | 14.97603752 | -0.920363864 | 0.35534947 | -2.59002459 | 0.009596908 | 0.04644775 | 0.920363864 |
| ENSMUSG00000019039 | Dalrd3 | 67789 | protein_coding | 597.577188 | 0.920495489 | 0.12097273 | 7.609115597 | 2.76E-14 | 1.32E-12 | 0.920495489 |
| ENSMUSG00000014602 | Kif1a | 16560 | protein_coding | 30.94076384 | -0.920855908 | 0.279196044 | -3.298241251 | 0.000972925 | 0.007515398 | 0.920855908 |
| ENSMUSG00000028909 | Ptpru | 19273 | protein_coding | 72.44835931 | -0.920901469 | 0.387578119 | -2.376040914 | 0.017499522 | 0.072492169 | 0.920901469 |
| ENSMUSG00000039579 | Grin3a | 242443 | protein_coding | 98.60381052 | -0.920907574 | 0.245459715 | -3.751766656 | 0.000175593 | 0.001825616 | 0.920907574 |
| ENSMUSG00000022439 | Parvg | 64099 | protein_coding | 7.621434086 | 0.921048109 | 0.58267421 | 1.580725719 | 0.113940765 | 0.270227592 | 0.921048109 |
| ENSMUSG00000006445 | Epha2 | 13836 | protein_coding | 484.4714172 | -0.921305814 | 0.553757543 | -1.663735015 | 0.096165396 | 0.242153679 | 0.921305814 |
| ENSMUSG00000020785 | Camkk1 | 55984 | protein_coding | 26.52523377 | -0.921498296 | 0.477078599 | -1.931543981 | 0.053415816 | 0.16224392 | 0.921498296 |
| ENSMUSG00000074657 | Kif5a | 16572 | protein_coding | 6.780491429 | -0.921955319 | 0.834115596 | -1.105308813 | 0.2690258 | 0.477123143 | 0.921955319 |
| ENSMUSG00000022490 | Ppp1r1a | 58200 | protein_coding | 13.04831706 | 0.922135338 | 0.347912034 | 2.650484173 | 0.008037649 | 0.040655344 | 0.922135338 |
| ENSMUSG00000006179 | Prss16 | 54373 | protein_coding | 24.72615009 | 0.922726509 | 0.326852666 | 2.823065577 | 0.004756684 | 0.027058674 | 0.922726509 |
| ENSMUSG00000002633 | Shh | 20423 | protein_coding | 15.04524057 | -0.923310834 | 0.279410519 | -3.304495608 | 0.000951475 | 0.007371152 | 0.923310834 |
| ENSMUSG00000001985 | Grik3 | 14807 | protein_coding | 6.712072524 | -0.923361282 | 0.647860127 | -1.425247896 | 0.154085556 | 0.333234858 | 0.923361282 |
| ENSMUSG00000030699 | Tbx6 | 21389 | protein_coding | 12.56414186 | -0.923374313 | 0.40059153 | -2.305027049 | 0.021165059 | 0.083382897 | 0.923374313 |
| ENSMUSG00000028100 | Nudt17 | 78373 | protein_coding | 23.84544834 | -0.924355059 | 0.289568939 | -3.192176148 | 0.001412052 | 0.010163201 | 0.924355059 |
| ENSMUSG00000079588 | Tmem182 | 381339 | protein_coding | 56.3633348 | 0.924491455 | 0.207308392 | 4.45949847 | 8.22E-06 | 0.000125577 | 0.924491455 |
| ENSMUSG00000000049 | Apoh | 11818 | protein_coding | 7.5949351 | 0.924529148 | 0.494277228 | 1.870466807 | 0.061419022 | 0.178420971 | 0.924529148 |
| ENSMUSG00000040297 | Suco | 226551 | protein_coding | 1532.876925 | 0.925695417 | 0.129045375 | 7.173410262 | 7.32E-13 | 3.07E-11 | 0.925695417 |
| ENSMUSG00000021879 | Dnah12 | 110083 | protein_coding | 9.025903433 | -0.926430334 | 1.132545816 | -0.818006937 | 0.413353226 | 0.620010377 | 0.926430334 |
| ENSMUSG00000063704 | Mapk15 | 332110 | protein_coding | 34.43332281 | -0.926674185 | 0.256000203 | -3.619818158 | 0.00029481 | 0.002798412 | 0.926674185 |
| ENSMUSG00000046318 | Ccbe1 | 320924 | protein_coding | 8.554013084 | -0.927119259 | 1.244329443 | -0.745075401 | 0.456226139 | 0.655296022 | 0.927119259 |
| ENSMUSG00000045912 | C2cd4c | 237397 | protein_coding | 32.37373235 | -0.927220545 | 0.252101384 | -3.677966895 | 0.0002351 | 0.002338963 | 0.927220545 |
| ENSMUSG00000096971 | 4930556M19Rik | 75259 | lncRNA | 19.44201831 | -0.927665828 | 0.307231925 | -3.019431747 | 0.002532493 | 0.016471032 | 0.927665828 |
| ENSMUSG00000029378 | Areg | 11839 | protein_coding | 926.2907137 | -0.927809427 | 0.360081093 | -2.57666799 | 0.009975772 | 0.047845255 | 0.927809427 |

| ensembl_gene_id | external_gene_name | entrezgene_id | gene_biotype | baseMean | log2FoldChange | lfcSE | stat | pvalue | padj | absLog2FC |
|---|---|---|---|---|---|---|---|---|---|---|
| ENSMUSG00000049555 | Tmie | 20776 | protein_coding | 17.31466684 | -0.928388907 | 0.304136719 | -3.052538052 | 0.00226915 | 0.015034532 | 0.928388907 |
| ENSMUSG00000038665 | Dgki | 320127 | protein_coding | 17.60815953 | -0.928815139 | 0.395254576 | -2.349916219 | 0.018777637 | 0.076356958 | 0.928815139 |
| ENSMUSG00000034330 | Plcg2 | 234779 | protein_coding | 163.9733768 | -0.928946853 | 0.118959024 | -7.808964983 | 5.77E-15 | 2.93E-13 | 0.928946853 |
| ENSMUSG00000022389 | Tef | 21685 | protein_coding | 282.7508964 | -0.929585015 | 0.226434787 | -4.105310079 | 4.04E-05 | 0.000512046 | 0.929585015 |
| ENSMUSG00000020038 | Cry1 | 12952 | protein_coding | 328.0971733 | 0.929627647 | 0.212070332 | 4.3835818 | 1.17E-05 | 0.000170753 | 0.929627647 |
| ENSMUSG00000033847 | Pla2g4c | 232889 | protein_coding | 48.78188013 | 0.930206475 | 0.212965097 | 4.367882297 | 1.25E-05 | 0.000182159 | 0.930206475 |
| ENSMUSG00000046351 | Zfp322a | 218100 | protein_coding | 500.151104 | 0.93048968 | 0.124064483 | 7.500048837 | 6.38E-14 | 2.95E-12 | 0.93048968 |
| ENSMUSG00000058488 | Kl | 16591 | protein_coding | 7.452544531 | -0.930894436 | 0.871934657 | -1.067619492 | 0.285692187 | 0.494484454 | 0.930894436 |
| ENSMUSG00000014846 | Tppp3 | 67971 | protein_coding | 342.9151631 | -0.931524113 | 0.315449535 | -2.95300519 | 0.003146967 | 0.019503847 | 0.931524113 |
| ENSMUSG00000021252 | Erg28 | 58520 | protein_coding | 674.4665291 | 0.931620183 | 0.09773264 | 9.532334176 | 1.54E-21 | 1.16E-19 | 0.931620183 |
| ENSMUSG00000055926 | Gm14137 | 623781 | protein_coding | 34.8883316 | 0.931669066 | 0.279121689 | 3.337859801 | 0.000844263 | 0.00666374 | 0.931669066 |
| ENSMUSG00000110086 | Smim32 | 115488204 | protein_coding | 7.613384029 | -0.932568295 | 0.542443347 | -1.719199434 | 0.085578064 | 0.224118284 | 0.932568295 |
| ENSMUSG00000073418 | C4b | 12268 | protein_coding | 19.27981024 | 0.932577286 | 0.543798085 | 1.714933008 | 0.08635751 | 0.225677181 | 0.932577286 |
| ENSMUSG00000018983 | E2f2 | 242705 | protein_coding | 567.1896833 | 0.933304288 | 0.127261934 | 7.33372705 | 2.24E-13 | 9.74E-12 | 0.933304288 |
| ENSMUSG00000070495 | Ctcfl | 664799 | protein_coding | 12.0942809 | -0.93417314 | 0.425728991 | -2.19429064 | 0.028214522 | 0.103042463 | 0.93417314 |
| ENSMUSG00000015312 | Gadd45b | 17873 | protein_coding | 168.8894845 | 0.934545478 | 0.169408813 | 5.516510382 | 3.46E-08 | 8.46E-07 | 0.934545478 |
| ENSMUSG00000034116 | Vav1 | 22324 | protein_coding | 103.376307 | -0.934926506 | 0.122040912 | -7.660763024 | 1.85E-14 | 9.00E-13 | 0.934926506 |
| ENSMUSG00000026223 | Itm2c | 64294 | protein_coding | 903.2192656 | 0.934967387 | 0.117895518 | 7.930474406 | 2.18E-15 | 1.15E-13 | 0.934967387 |
| ENSMUSG00000070337 | Gpr179 | 217143 | protein_coding | 10.26417565 | -0.935061889 | 0.582703247 | -1.604696547 | 0.108560602 | 0.26219155 | 0.935061889 |
| ENSMUSG00000035697 | Arhgap45 | 70719 | protein_coding | 56.58098141 | 0.935933848 | 0.241305077 | 3.878633049 | 0.000105045 | 0.001173322 | 0.935933848 |
| ENSMUSG00000026080 | Chst10 | 98388 | protein_coding | 9.137624687 | -0.936497109 | 0.603109885 | -1.552780234 | 0.120475646 | 0.281823483 | 0.936497109 |
| ENSMUSG00000041565 | L3mbtl4 | 320858 | protein_coding | 9.995406419 | -0.93678234 | 0.645209041 | -1.451905167 | 0.146527976 | 0.322718141 | 0.93678234 |
| ENSMUSG00000024053 | Emilin2 | 246707 | protein_coding | 7.684497144 | -0.93780807 | 0.555030227 | -1.689652244 | 0.091094505 | 0.233610251 | 0.93780807 |
| ENSMUSG00000065611 | Mir23a | 387216 | miRNA | 10.13144383 | -0.937862645 | 0.5160512 | -1.817382938 | 0.069158498 | 0.194106883 | 0.937862645 |
| ENSMUSG00000009739 | Pou6f1 | 19009 | protein_coding | 32.13178683 | -0.939002302 | 0.380705102 | -2.466482057 | 0.013644756 | 0.060242243 | 0.939002302 |
| ENSMUSG00000025064 | Col17a1 | 12821 | protein_coding | 14.71533892 | -0.939221466 | 0.463512242 | -2.026314262 | 0.042732591 | 0.139105805 | 0.939221466 |
| ENSMUSG00000118339 | 4930592I03Rik | 75853 | lncRNA | 8.270389197 | 0.939315441 | 0.498381016 | 1.884733591 | 0.059465815 | 0.175045555 | 0.939315441 |
| ENSMUSG00000023011 | Faim2 | 72393 | protein_coding | 9.883516844 | -0.939585741 | 0.879932384 | -1.067793115 | 0.285613845 | 0.494484454 | 0.939585741 |
| ENSMUSG00000029407 | Uso1 | 56041 | protein_coding | 1578.500693 | 0.939713059 | 0.120600187 | 7.791970142 | 6.60E-15 | 3.31E-13 | 0.939713059 |
| ENSMUSG00000070953 | Rabepk | 227746 | protein_coding | 489.8371583 | 0.940416326 | 0.103649789 | 9.073017239 | 1.16E-19 | 7.95E-18 | 0.940416326 |
| ENSMUSG00000085719 | 4933424L21Rik | 78762 | lncRNA | 12.19203827 | -0.940516867 | 0.439531866 | -2.139814971 | 0.032369723 | 0.114067466 | 0.940516867 |
| ENSMUSG00000048965 | Mrgpre | 244238 | protein_coding | 12.93943398 | -0.940621138 | 0.360398734 | -2.609945736 | 0.009055659 | 0.044435776 | 0.940621138 |
| ENSMUSG00000040667 | Nup88 | 19069 | protein_coding | 1298.364642 | 0.941283071 | 0.086470701 | 10.88557235 | 1.35E-27 | 1.37E-25 | 0.941283071 |
| ENSMUSG00000042116 | Vwa1 | 246228 | protein_coding | 10.4755941 | -0.942213885 | 0.406655459 | -2.316983247 | 0.020504644 | 0.081466195 | 0.942213885 |
| ENSMUSG00000002602 | Axl | 26362 | protein_coding | 10.99103472 | -0.94278703 | 0.579271873 | -1.627538077 | 0.103622868 | 0.254232438 | 0.94278703 |
| ENSMUSG00000078700 | D030028A08Rik | 319371 | lncRNA | 153.5464962 | 0.942861156 | 0.136123569 | 6.926509216 | 4.31E-12 | 1.67E-10 | 0.942861156 |
| ENSMUSG00000036712 | Cyld | 74256 | protein_coding | 1454.044462 | 0.943133104 | 0.069152346 | 13.63848313 | 2.36E-42 | 4.43E-40 | 0.943133104 |
| ENSMUSG00000028804 | Csmd2 | 329942 | protein_coding | 9.153749561 | -0.943585437 | 1.234969327 | -0.764055767 | 0.44483401 | 0.645657025 | 0.943585437 |

| ensembl_gene_id | external_gene_name | entrezgene_id | gene_biotype | baseMean | log2FoldChange | lfcSE | stat | pvalue | padj | absLog2FC |
|---|---|---|---|---|---|---|---|---|---|---|
| ENSMUSG00000030621 | Me3 | 109264 | protein_coding | 21.57638993 | -0.943657331 | 0.410276362 | -2.30005289 | 0.021445224 | 0.084153615 | 0.943657331 |
| ENSMUSG00000057667 | Bloc1s3 | 232946 | protein_coding | 245.216788 | 0.944358887 | 0.132579226 | 7.122977805 | 1.06E-12 | 4.36E-11 | 0.944358887 |
| ENSMUSG00000050100 | Hmx2 | 15372 | protein_coding | 7.674061957 | -0.94442444 | 0.490676135 | -1.92474093 | 0.054261777 | 0.164093713 | 0.94442444 |
| ENSMUSG00000001506 | Col1a1 | 12842 | protein_coding | 31.05207679 | -0.944756288 | 0.360163649 | -2.623130596 | 0.008712587 | 0.043218338 | 0.944756288 |
| ENSMUSG00000031144 | Syp | 20977 | protein_coding | 13.42390268 | -0.945330165 | 0.340081438 | -2.77971703 | 0.005440628 | 0.029820485 | 0.945330165 |
| ENSMUSG00000054252 | Fgfr3 | 14184 | protein_coding | 543.3154479 | -0.945478249 | 0.194120056 | -4.87058508 | 1.11E-06 | 2.08E-05 | 0.945478249 |
| ENSMUSG00000001520 | Nrip2 | 60345 | protein_coding | 9.381062775 | -0.945565594 | 0.493977069 | -1.914189245 | 0.055595979 | 0.167016468 | 0.945565594 |
| ENSMUSG00000049265 | Kcnk3 | 16527 | protein_coding | 7.583184522 | -0.945728512 | 0.436184587 | -2.168184158 | 0.030144678 | 0.108409218 | 0.945728512 |
| ENSMUSG00000025240 | Sacm1l | 83493 | protein_coding | 1228.778711 | 0.945883431 | 0.101471691 | 9.321648414 | 1.15E-20 | 8.45E-19 | 0.945883431 |
| ENSMUSG00000005237 | Dnah2 | 327954 | protein_coding | 27.93402913 | -0.946308654 | 0.43062913 | -2.19750265 | 0.027984567 | 0.102444999 | 0.946308654 |
| ENSMUSG00000060992 | Copz1 | 56447 | protein_coding | 2209.020014 | 0.946343738 | 0.084593205 | 11.18699476 | 4.72E-29 | 5.22E-27 | 0.946343738 |
| ENSMUSG00000016262 | Sertad4 | 214791 | protein_coding | 7.462001313 | -0.94653149 | 0.710951206 | -1.331359285 | 0.18307082 | 0.372217515 | 0.94653149 |
| ENSMUSG00000030110 | Ret | 19713 | protein_coding | 53.77824868 | 0.946559264 | 0.231557636 | 4.087791192 | 4.35E-05 | 0.000546623 | 0.946559264 |
| ENSMUSG00000039116 | Adgrg6 | 215798 | protein_coding | 9.049587797 | -0.94662809 | 0.68701883 | -1.377877939 | 0.168240977 | 0.352829794 | 0.94662809 |
| ENSMUSG00000019564 | Arid3a | 13496 | protein_coding | 109.1862234 | 0.946642658 | 0.23374414 | 4.049909685 | 5.12E-05 | 0.000633134 | 0.946642658 |
| ENSMUSG00000042249 | Grk3 | 320129 | protein_coding | 107.8014857 | -0.947498048 | 0.244043915 | -3.882489953 | 0.000103392 | 0.001158925 | 0.947498048 |
| ENSMUSG00000020717 | Pecam1 | 18613 | protein_coding | 31.52038184 | -0.947669783 | 0.276347995 | -3.429262379 | 0.000605224 | 0.005103235 | 0.947669783 |
| ENSMUSG00000061474 | Mrps36 | 66128 | protein_coding | 249.3675437 | 0.947762401 | 0.109746251 | 8.635943304 | 5.82E-18 | 3.65E-16 | 0.947762401 |
| ENSMUSG00000117011 | E130008D07Rik | 545207 | lncRNA | 7.935751254 | -0.948133142 | 0.577693032 | -1.641240396 | 0.100747523 | 0.249487959 | 0.948133142 |
| ENSMUSG00000062980 | Cped1 | 214642 | protein_coding | 20.35783901 | -0.94856117 | 0.360371399 | -2.632176618 | 0.008483975 | 0.042327446 | 0.94856117 |
| ENSMUSG00000075289 | Carns1 | 107239 | protein_coding | 7.883147945 | -0.948624407 | 0.449158928 | -2.112001672 | 0.034686306 | 0.120209634 | 0.948624407 |
| ENSMUSG00000091670 | Vmn2r105 | 627743 | protein_coding | 7.960591849 | -0.949149288 | 1.821100709 | -0.521195387 | 0.602230668 | 0.767564496 | 0.949149288 |
| ENSMUSG00000029438 | Bcl7a | 77045 | protein_coding | 105.7327859 | -0.949167362 | 0.170805257 | -5.557014936 | 2.74E-08 | 6.87E-07 | 0.949167362 |
| ENSMUSG00000029552 | Tes | 21753 | protein_coding | 3877.714141 | 0.949393129 | 0.200679386 | 4.73089513 | 2.24E-06 | 3.86E-05 | 0.949393129 |
| ENSMUSG00000027217 | Tspan18 | 241556 | protein_coding | 11.44859185 | -0.949478633 | 0.692898714 | -1.370299315 | 0.170593487 | 0.35593569 | 0.949478633 |
| ENSMUSG00000001281 | Itgb7 | 16421 | protein_coding | 32.42356092 | 0.949794047 | 0.271683641 | 3.495955975 | 0.000472367 | 0.004154531 | 0.949794047 |
| ENSMUSG00000020937 | Plcd3 | 72469 | protein_coding | 187.9609149 | -0.950955263 | 0.232890284 | -4.08327581 | 4.44E-05 | 0.000555607 | 0.950955263 |
| ENSMUSG00000005986 | Ankrd13d | 68423 | protein_coding | 11.19427428 | -0.951070268 | 0.389137176 | -2.44404885 | 0.01452346 | 0.062956904 | 0.951070268 |
| ENSMUSG00000073067 | 9130019P16Rik | 100042056 | lncRNA | 8.117798117 | -0.951712493 | 0.57534861 | -1.654149288 | 0.098097184 | 0.245173692 | 0.951712493 |
| ENSMUSG00000037683 | Armc3 | 70882 | protein_coding | 17.21535657 | 0.951725041 | 0.46935193 | 2.027742893 | 0.042586496 | 0.138773685 | 0.951725041 |
| ENSMUSG00000007989 | Fzd3 | 14365 | protein_coding | 235.3984946 | 0.952502038 | 0.152597055 | 6.241942474 | 4.32E-10 | 1.36E-08 | 0.952502038 |
| ENSMUSG00000075602 | Ly6a | 110454 | protein_coding | 981.7088397 | 0.95262811 | 0.411553572 | 2.314712285 | 0.020628683 | 0.081836529 | 0.95262811 |
| ENSMUSG00000057400 | Ces1c | 13884 | protein_coding | 13.9451701 | 0.954048448 | 0.548061922 | 1.740767621 | 0.08172432 | 0.217322621 | 0.954048448 |
| ENSMUSG00000031576 | Kcnu1 | 16532 | protein_coding | 7.88062286 | -0.954176428 | 0.499290642 | -1.911064114 | 0.055996342 | 0.167652206 | 0.954176428 |
| ENSMUSG00000058163 | Gm5431 | 432555 | protein_coding | 79.50745094 | 0.954318874 | 0.289510489 | 3.296318822 | 0.000979608 | 0.007556025 | 0.954318874 |
| ENSMUSG00000097021 | 4933433G15Rik | 71274 | lncRNA | 9.490441897 | -0.954466268 | 0.380327931 | -2.509587625 | 0.012087222 | 0.055180647 | 0.954466268 |
| ENSMUSG00000064080 | Fbln2 | 14115 | protein_coding | 9.049879541 | -0.955466444 | 0.582854124 | -1.639289154 | 0.101153054 | 0.250100293 | 0.955466444 |
| ENSMUSG00000004366 | Sst | 20604 | protein_coding | 45.91895943 | -0.955966775 | 0.263934681 | -3.621982426 | 0.000292354 | 0.002779953 | 0.955966775 |

| ensembl_gene_id | external_gene_name | entrezgene_id | gene_biotype | baseMean | log2FoldChange | lfcSE | stat | pvalue | padj | absLog2FC |
|---|---|---|---|---|---|---|---|---|---|---|
| ENSMUSG00000031561 | Tenm3 | 23965 | protein_coding | 6.867175811 | -0.956202198 | 0.990796253 | -0.965084592 | 0.334502444 | 0.545293316 | 0.956202198 |
| ENSMUSG00000013483 | Card14 | 170720 | protein_coding | 337.4902574 | 0.95744911 | 0.213206279 | 4.490717218 | 7.10E-06 | 0.000110413 | 0.95744911 |
| ENSMUSG00000062309 | Rpp25 | 102614 | protein_coding | 9.826023871 | -0.957717444 | 0.380103971 | -2.519619673 | 0.01174817 | 0.053982933 | 0.957717444 |
| ENSMUSG00000024970 | Spindoc | 68229 | protein_coding | 244.7770895 | 0.957802109 | 0.190129618 | 5.037627071 | 4.71E-07 | 9.65E-06 | 0.957802109 |
| ENSMUSG00000048022 | Tmem229a | 319832 | protein_coding | 9.08668298 | -0.958460786 | 0.521176388 | -1.839033403 | 0.065910273 | 0.187333838 | 0.958460786 |
| ENSMUSG00000003545 | Fosb | 14282 | protein_coding | 3090.464991 | -0.958475777 | 0.380414594 | -2.519555746 | 0.011750303 | 0.053982933 | 0.958475777 |
| ENSMUSG00000024843 | Chka | 12660 | protein_coding | 1038.174778 | 0.958491397 | 0.225228029 | 4.255648833 | 2.08E-05 | 0.000287703 | 0.958491397 |
| ENSMUSG00000028214 | Gem | 14579 | protein_coding | 13.20500858 | -0.959325767 | 0.438476201 | -2.187862794 | 0.028679598 | 0.104365693 | 0.959325767 |
| ENSMUSG00000021010 | Npas3 | 27386 | protein_coding | 11.10509169 | -0.959357419 | 0.481840478 | -1.991027037 | 0.046477915 | 0.148060046 | 0.959357419 |
| ENSMUSG00000038028 | Tigar | 319801 | protein_coding | 345.6694136 | 0.95954943 | 0.114380496 | 8.38910007 | 4.90E-17 | 2.90E-15 | 0.95954943 |
| ENSMUSG00000001123 | Lgals9 | 16859 | protein_coding | 922.8470126 | 0.959625877 | 0.157305761 | 6.100386085 | 1.06E-09 | 3.19E-08 | 0.959625877 |
| ENSMUSG00000027074 | Slc43a3 | 58207 | protein_coding | 13.02589932 | -0.960140204 | 0.426091179 | -2.253367943 | 0.024235959 | 0.092285526 | 0.960140204 |
| ENSMUSG00000028680 | Plk3 | 12795 | protein_coding | 618.1193253 | -0.960914012 | 0.251020196 | -3.82803467 | 0.000129171 | 0.001405348 | 0.960914012 |
| ENSMUSG00000041734 | Kirrel | 170643 | protein_coding | 9.092928973 | -0.960948087 | 0.545673368 | -1.761031679 | 0.078233041 | 0.210382557 | 0.960948087 |
| ENSMUSG00000025175 | Fn3k | 63828 | protein_coding | 8.603585453 | -0.962291333 | 0.473958643 | -2.030327639 | 0.042323247 | 0.138233724 | 0.962291333 |
| ENSMUSG00000021273 | Fdft1 | 14137 | protein_coding | 1877.204747 | 0.963280901 | 0.159490625 | 6.039733689 | 1.54E-09 | 4.61E-08 | 0.963280901 |
| ENSMUSG00000022758 | P2rx6 | 18440 | protein_coding | 6.913422183 | -0.96334629 | 0.48315528 | -1.993864768 | 0.046166835 | 0.147230912 | 0.96334629 |
| ENSMUSG00000053693 | Mast1 | 56527 | protein_coding | 13.45224968 | -0.965002499 | 0.424527537 | -2.273121094 | 0.023018884 | 0.088883583 | 0.965002499 |
| ENSMUSG00000047108 | Dnajb7 | 57755 | protein_coding | 7.307437088 | -0.965151606 | 0.513813035 | -1.878410122 | 0.060325082 | 0.176466098 | 0.965151606 |
| ENSMUSG00000055053 | Nfic | 18029 | protein_coding | 149.5722014 | -0.965762 | 0.198893216 | -4.855680948 | 1.20E-06 | 2.21E-05 | 0.965762 |
| ENSMUSG00000025425 | St8sia5 | 225742 | protein_coding | 9.611464508 | -0.966227705 | 0.691334856 | -1.397626196 | 0.162225348 | 0.344615276 | 0.966227705 |
| ENSMUSG00000021061 | Sptb | 20741 | protein_coding | 41.1465504 | -0.966643273 | 0.254802446 | -3.793696997 | 0.000148421 | 0.001585544 | 0.966643273 |
| ENSMUSG00000051527 | Usp29 | 57775 | protein_coding | 9.189921534 | -0.967840633 | 1.245041463 | -0.777356146 | 0.436948678 | 0.639621225 | 0.967840633 |
| ENSMUSG00000023045 | Soat2 | 223920 | protein_coding | 28.40364211 | 0.967905518 | 0.216162729 | 4.477670706 | 7.55E-06 | 0.000116695 | 0.967905518 |
| ENSMUSG00000025151 | Maged1 | 94275 | protein_coding | 1808.018225 | 0.968028838 | 0.12926806 | 7.488538476 | 6.96E-14 | 3.22E-12 | 0.968028838 |
| ENSMUSG00000021846 | Peli2 | 93834 | protein_coding | 229.5318875 | -0.96872056 | 0.095458286 | -10.1481034 | 3.38E-24 | 2.94E-22 | 0.96872056 |
| ENSMUSG00000029814 | Igf2bp3 | 140488 | protein_coding | 13.17709793 | -0.968767924 | 0.44454479 | -2.179235807 | 0.029314155 | 0.106081983 | 0.968767924 |
| ENSMUSG00000046814 | Gchfr | 320415 | protein_coding | 42.51773288 | 0.968912143 | 0.236109108 | 4.103662715 | 4.07E-05 | 0.000515297 | 0.968912143 |
| ENSMUSG00000028464 | Tpm2 | 22004 | protein_coding | 43.99260702 | -0.969016322 | 0.255558462 | -3.791759871 | 0.000149584 | 0.00159369 | 0.969016322 |
| ENSMUSG00000039452 | Snx22 | 382083 | protein_coding | 9.930785183 | 0.969428041 | 0.379085687 | 2.557279457 | 0.010549443 | 0.049786963 | 0.969428041 |
| ENSMUSG00000072437 | Nanos1 | 332397 | protein_coding | 57.70078713 | 0.969567585 | 0.320280542 | 3.027244736 | 0.002467941 | 0.016143473 | 0.969567585 |
| ENSMUSG00000028641 | P3h1 | 56401 | protein_coding | 29.50881174 | 0.970212232 | 0.234137219 | 4.14377619 | 3.42E-05 | 0.000443483 | 0.970212232 |
| ENSMUSG00000029406 | Pitpnm2 | 19679 | protein_coding | 37.56269578 | -0.970487415 | 0.299986993 | -3.235098316 | 0.001216009 | 0.008987742 | 0.970487415 |
| ENSMUSG00000033981 | Gria2 | 14800 | protein_coding | 9.275650026 | -0.970836704 | 0.997130279 | -0.973630753 | 0.330239911 | 0.540824728 | 0.970836704 |
| ENSMUSG00000064065 | Ipcef1 | 320495 | protein_coding | 27.06687957 | 0.971024722 | 0.324845556 | 2.989188877 | 0.002797192 | 0.017750465 | 0.971024722 |
| ENSMUSG00000025185 | Loxl4 | 67573 | protein_coding | 8.048355212 | -0.971207833 | 0.484199319 | -2.005801731 | 0.044877406 | 0.144259802 | 0.971207833 |
| ENSMUSG00000028034 | Fubp1 | 51886 | protein_coding | 783.3363799 | -0.9720696 | 0.185455708 | -5.241518901 | 1.59E-07 | 3.54E-06 | 0.9720696 |
| ENSMUSG00000039781 | Cep131 | 12009 | protein_coding | 255.3243226 | 0.972411905 | 0.120201153 | 8.089871695 | 5.97E-16 | 3.29E-14 | 0.972411905 |

| ensembl_gene_id | external_gene_name | entrezgene_id | gene_biotype | baseMean | log2FoldChange | lfcSE | stat | pvalue | padj | absLog2FC |
|---|---|---|---|---|---|---|---|---|---|---|
| ENSMUSG00000030020 | Prickle2 | 243548 | protein_coding | 7.362239941 | -0.973264991 | 0.879446059 | -1.106679575 | 0.268432489 | 0.476336084 | 0.973264991 |
| ENSMUSG00000040710 | St8sia4 | 20452 | protein_coding | 32.97836395 | 0.973301502 | 0.452786758 | 2.149580316 | 0.031588427 | 0.112207953 | 0.973301502 |
| ENSMUSG00000026791 | Slc2a8 | 56017 | protein_coding | 193.7931168 | 0.974190385 | 0.118597854 | 8.214232813 | 2.14E-16 | 1.22E-14 | 0.974190385 |
| ENSMUSG00000045790 | Ccdc149 | 100503884 | protein_coding | 243.7591869 | 0.974885553 | 0.102413051 | 9.519153453 | 1.75E-21 | 1.31E-19 | 0.974885553 |
| ENSMUSG00000030630 | Fah | 14085 | protein_coding | 79.60094983 | -0.975137489 | 0.197259452 | -4.943425924 | 7.68E-07 | 1.49E-05 | 0.975137489 |
| ENSMUSG00000025255 | Zfhx4 | 80892 | protein_coding | 8.058407298 | -0.975564533 | 1.13542058 | -0.859209839 | 0.390224756 | 0.598507455 | 0.975564533 |
| ENSMUSG00000038604 | Ripor1 | 75687 | protein_coding | 64.86673572 | -0.977123261 | 0.20281903 | -4.81770997 | 1.45E-06 | 2.64E-05 | 0.977123261 |
| ENSMUSG00000022488 | Nckap1l | 105855 | protein_coding | 25.86106861 | 0.97794958 | 0.350497672 | 2.790174255 | 0.005267968 | 0.0291898 | 0.97794958 |
| ENSMUSG00000030173 | Klra5 | 16636 | protein_coding | 9.130327271 | 0.97801218 | 0.554437696 | 1.763971296 | 0.077736812 | 0.209607899 | 0.97801218 |
| ENSMUSG00000053581 | Zfand2a | 100494 | protein_coding | 381.8861678 | -0.97825726 | 0.248137955 | -3.94239269 | 8.07E-05 | 0.000935874 | 0.97825726 |
| ENSMUSG00000009487 | Otog | 18419 | protein_coding | 7.635976821 | -0.979339364 | 1.193823168 | -0.820338715 | 0.412023042 | 0.618889382 | 0.979339364 |
| ENSMUSG00000045091 | Aqp12 | 208760 | protein_coding | 9.98558908 | 0.979981324 | 0.454657341 | 2.155428352 | 0.03112833 | 0.111044129 | 0.979981324 |
| ENSMUSG00000023259 | Slc26a6 | 171429 | protein_coding | 56.10399362 | 0.980402137 | 0.155480473 | 6.305628742 | 2.87E-10 | 9.25E-09 | 0.980402137 |
| ENSMUSG00000027840 | Wnt2b | 22414 | protein_coding | 11.38615294 | -0.980697797 | 0.37684581 | -2.602384769 | 0.009257792 | 0.045210961 | 0.980697797 |
| ENSMUSG00000019359 | Gdpd2 | 71584 | protein_coding | 30.68264912 | -0.981134656 | 0.315675493 | -3.108048229 | 0.001883273 | 0.012863112 | 0.981134656 |
| ENSMUSG00000103409 | Lsmem2 | 434436 | protein_coding | 12.16718306 | -0.981283851 | 0.403545962 | -2.431653252 | 0.015030088 | 0.064755004 | 0.981283851 |
| ENSMUSG00000015966 | Il17rb | 50905 | protein_coding | 35.13043757 | -0.981352012 | 0.202719941 | -4.840924898 | 1.29E-06 | 2.37E-05 | 0.981352012 |
| ENSMUSG00000019579 | Mydgf | 28106 | protein_coding | 1516.053696 | 0.98173796 | 0.127153068 | 7.720914437 | 1.15E-14 | 5.70E-13 | 0.98173796 |
| ENSMUSG00000049897 | Stkld1 | 279029 | protein_coding | 8.573333127 | -0.982547245 | 0.412120115 | -2.384128336 | 0.017119631 | 0.071345233 | 0.982547245 |
| ENSMUSG00000033420 | Antxr1 | 69538 | protein_coding | 7.421889722 | -0.983161474 | 0.99271552 | -0.990375847 | 0.321990447 | 0.532903557 | 0.983161474 |
| ENSMUSG00000063434 | Sorcs3 | 66673 | protein_coding | 14.01968231 | 0.983886252 | 0.444576124 | 2.213088376 | 0.026891553 | 0.099482501 | 0.983886252 |
| ENSMUSG00000014725 | Adam28 | 13522 | protein_coding | 7.650391277 | -0.983913481 | 1.365056439 | -0.720785935 | 0.471041231 | 0.666555146 | 0.983913481 |
| ENSMUSG00000016756 | Cmah | 12763 | protein_coding | 857.7355872 | -0.984132667 | 0.218416803 | -4.505755287 | 6.61E-06 | 0.000103583 | 0.984132667 |
| ENSMUSG00000066705 | Fxyd6 | 59095 | protein_coding | 8.619811921 | -0.98463101 | 0.522155246 | -1.885705481 | 0.059334654 | 0.174821008 | 0.98463101 |
| ENSMUSG00000052563 | D930048N14Rik | 97775 | lncRNA | 27.59486372 | -0.984997883 | 0.288801056 | -3.41064502 | 0.000648094 | 0.00540463 | 0.984997883 |
| ENSMUSG00000040212 | Emp3 | 13732 | protein_coding | 12.38665239 | 0.985391904 | 0.439240728 | 2.243398301 | 0.024871141 | 0.094032003 | 0.985391904 |
| ENSMUSG00000022094 | Slc39a14 | 213053 | protein_coding | 1183.687576 | 0.985795616 | 0.139900862 | 7.046387006 | 1.84E-12 | 7.37E-11 | 0.985795616 |
| ENSMUSG00000036800 | Fam135b | 70363 | protein_coding | 10.32727132 | -0.98595081 | 1.919018694 | -0.513778637 | 0.607406765 | 0.771267136 | 0.98595081 |
| ENSMUSG00000026674 | Ddr2 | 18214 | protein_coding | 8.466754274 | -0.986545389 | 0.643749522 | -1.53249883 | 0.125399386 | 0.289598582 | 0.986545389 |
| ENSMUSG00000036676 | Tmtc3 | 237500 | protein_coding | 647.9138853 | 0.986803884 | 0.107351588 | 9.192261603 | 3.85E-20 | 2.77E-18 | 0.986803884 |
| ENSMUSG00000066595 | Flvcr1 | 226844 | protein_coding | 294.9794524 | 0.987870566 | 0.100505526 | 9.829017472 | 8.44E-23 | 6.86E-21 | 0.987870566 |
| ENSMUSG00000010760 | Phlda2 | 22113 | protein_coding | 21.62932777 | -0.988048806 | 0.509018647 | -1.94108568 | 0.052247886 | 0.160162495 | 0.988048806 |
| ENSMUSG00000002289 | Angptl4 | 57875 | protein_coding | 46.83248755 | -0.988918755 | 0.368861208 | -2.681005031 | 0.007340141 | 0.037872941 | 0.988918755 |
| ENSMUSG00000040033 | Stat2 | 20847 | protein_coding | 405.7985175 | 0.989262829 | 0.135331206 | 7.309938759 | 2.67E-13 | 1.16E-11 | 0.989262829 |
| ENSMUSG00000027187 | Cat | 12359 | protein_coding | 2142.01642 | 0.990498313 | 0.096707959 | 10.24215917 | 1.28E-24 | 1.15E-22 | 0.990498313 |
| ENSMUSG00000064351 | mt-Co1 | 17708 | protein_coding | 684969.298 | -0.990946314 | 0.154819397 | -6.400659946 | 1.55E-10 | 5.16E-09 | 0.990946314 |
| ENSMUSG00000031932 | Gpr83 | 14608 | protein_coding | 7.482093513 | -0.9910117 | 0.857949839 | -1.155092822 | 0.248052417 | 0.453861899 | 0.9910117 |
| ENSMUSG00000027220 | Syt13 | 80976 | protein_coding | 24.73438912 | -0.991707413 | 0.262272677 | -3.781207498 | 0.00015607 | 0.001650648 | 0.991707413 |

| ensembl_gene_id | external_gene_name | entrezgene_id | gene_biotype | baseMean | log2FoldChange | lfcSE | stat | pvalue | padj | absLog2FC |
|---|---|---|---|---|---|---|---|---|---|---|
| ENSMUSG00000050150 | Slc9b1 | 74446 | protein_coding | 7.711896619 | -0.992292871 | 0.679977551 | -1.459302398 | 0.144481896 | 0.319138488 | 0.992292871 |
| ENSMUSG00000021775 | Nr1d2 | 353187 | protein_coding | 662.8009445 | -0.992852388 | 0.18766981 | -5.290421436 | 1.22E-07 | 2.76E-06 | 0.992852388 |
| ENSMUSG00000042357 | Gjb5 | 14622 | protein_coding | 8.520412923 | -0.993103499 | 0.396820553 | -2.502651366 | 0.01232669 | 0.055953127 | 0.993103499 |
| ENSMUSG00000027808 | Serp1 | 28146 | protein_coding | 6447.938516 | 0.993550879 | 0.110638098 | 8.980187599 | 2.70E-19 | 1.82E-17 | 0.993550879 |
| ENSMUSG00000035270 | Impg2 | 224224 | protein_coding | 13.01151115 | 0.994611878 | 0.554677283 | 1.793136132 | 0.072951159 | 0.200927904 | 0.994611878 |
| ENSMUSG00000041084 | Ostc | 66357 | protein_coding | 1961.772174 | 0.995412033 | 0.085171614 | 11.68713365 | 1.48E-31 | 1.80E-29 | 0.995412033 |
| ENSMUSG00000030283 | St8sia1 | 20449 | protein_coding | 7.118204151 | -0.995983519 | 0.916863847 | -1.086293808 | 0.277349026 | 0.485665708 | 0.995983519 |
| ENSMUSG00000040415 | Dtx3 | 80904 | protein_coding | 31.18367466 | -0.999075329 | 0.268711399 | -3.718023624 | 0.000200787 | 0.002050503 | 0.999075329 |
| ENSMUSG00000078851 | H2ac25 | 319162 | protein_coding | 105.168149 | 0.99910212 | 0.171887613 | 5.812531251 | 6.15E-09 | 1.70E-07 | 0.99910212 |
| ENSMUSG00000075705 | Msrb1 | 27361 | protein_coding | 777.3145894 | 0.999191713 | 0.112883416 | 8.851536777 | 8.63E-19 | 5.64E-17 | 0.999191713 |
| ENSMUSG00000037795 | N4bp2 | 333789 | protein_coding | 727.6837854 | 0.999250497 | 0.102059274 | 9.790883821 | 1.23E-22 | 9.86E-21 | 0.999250497 |
| ENSMUSG00000040747 | Cd53 | 12508 | protein_coding | 25.43373603 | 1.000409665 | 0.318614362 | 3.139876239 | 0.001690192 | 0.01175093 | 1.000409665 |
| ENSMUSG00000010505 | Myt1 | 17932 | protein_coding | 28.47966847 | -1.001225438 | 0.267180811 | -3.747370305 | 0.000178698 | 0.001853063 | 1.001225438 |
| ENSMUSG00000040875 | Osbpl10 | 74486 | protein_coding | 396.930921 | 1.001365044 | 0.12830503 | 7.804565775 | 5.97E-15 | 3.02E-13 | 1.001365044 |
| ENSMUSG00000021367 | Edn1 | 13614 | protein_coding | 175.7791038 | -1.001821829 | 0.357420114 | -2.802925158 | 0.005064142 | 0.028312269 | 1.001821829 |
| ENSMUSG00000028019 | Pdgfc | 54635 | protein_coding | 48.5943893 | 1.002359606 | 0.209493869 | 4.784672759 | 1.71E-06 | 3.04E-05 | 1.002359606 |
| ENSMUSG00000039903 | Eva1c | 70967 | protein_coding | 9.022687602 | -1.002956286 | 0.424410131 | -2.363177059 | 0.018119011 | 0.074247389 | 1.002956286 |
| ENSMUSG00000038917 | 3930402G23Rik | 665306 | lncRNA | 21.07803508 | 1.003017309 | 0.347520238 | 2.886212655 | 0.003899086 | 0.023027305 | 1.003017309 |
| ENSMUSG00000029771 | Irf5 | 27056 | protein_coding | 10.93228697 | -1.003992548 | 0.417977138 | -2.402027424 | 0.016304486 | 0.0688306 | 1.003992548 |
| ENSMUSG00000079363 | Gbp4 | 17472 | protein_coding | 18.65891673 | 1.005392316 | 0.497426737 | 2.021186723 | 0.04326044 | 0.14037911 | 1.005392316 |
| ENSMUSG00000046805 | Mpeg1 | 17476 | protein_coding | 13.32677646 | 1.005483309 | 0.479504452 | 2.096921737 | 0.036000504 | 0.12350119 | 1.005483309 |
| ENSMUSG00000047150 | 1700001C19Rik | 75462 | protein_coding | 7.067692567 | -1.005929843 | 0.48187206 | -2.087545484 | 0.036838854 | 0.125387599 | 1.005929843 |
| ENSMUSG00000021573 | Tppp | 72948 | protein_coding | 1237.207383 | -1.006468343 | 0.242591455 | -4.14882026 | 3.34E-05 | 0.000434533 | 1.006468343 |
| ENSMUSG00000051457 | Spn | 20737 | protein_coding | 26.20444492 | 1.007053051 | 0.362592818 | 2.777366236 | 0.005480139 | 0.030016388 | 1.007053051 |
| ENSMUSG00000021730 | Hcn1 | 15165 | protein_coding | 9.719716313 | -1.007117973 | 0.919554901 | -1.095223322 | 0.273418811 | 0.481376679 | 1.007117973 |
| ENSMUSG00000024083 | Pja2 | 224938 | protein_coding | 1393.496726 | 1.007871889 | 0.043088999 | 23.39046891 | 5.34E-121 | 4.48E-118 | 1.007871889 |
| ENSMUSG00000026981 | Il1rn | 16181 | protein_coding | 759.5101811 | -1.008519838 | 0.519492488 | -1.941355958 | 0.052215116 | 0.160123676 | 1.008519838 |
| ENSMUSG00000027822 | Slc33a1 | 11416 | protein_coding | 963.5228359 | 1.008789415 | 0.079353622 | 12.71258179 | 5.03E-37 | 8.10E-35 | 1.008789415 |
| ENSMUSG00000032854 | Ugt8a | 22239 | protein_coding | 110.8374592 | -1.009260955 | 0.3426613 | -2.945360199 | 0.003225789 | 0.01989174 | 1.009260955 |
| ENSMUSG00000031760 | Mt3 | 17751 | protein_coding | 35.82413025 | -1.00930195 | 0.251259269 | -4.016974004 | 5.90E-05 | 0.000715125 | 1.00930195 |
| ENSMUSG00000050192 | Eif5a2 | 208691 | protein_coding | 18.65716435 | 1.009871118 | 0.313014642 | 3.226274371 | 0.001254131 | 0.009188498 | 1.009871118 |
| ENSMUSG00000064247 | Plcxd1 | 403178 | protein_coding | 10.65020332 | -1.01051834 | 0.371644837 | -2.719043125 | 0.006547107 | 0.034565524 | 1.01051834 |
| ENSMUSG00000070284 | Gmppb | 331026 | protein_coding | 940.9705583 | 1.010641429 | 0.166821222 | 6.058230587 | 1.38E-09 | 4.12E-08 | 1.010641429 |
| ENSMUSG00000050075 | Gpr171 | 229323 | protein_coding | 12.26549306 | 1.010842015 | 0.413300332 | 2.445780798 | 0.014453885 | 0.062799093 | 1.010842015 |
| ENSMUSG00000004609 | Cd33 | 12489 | protein_coding | 8.979979774 | -1.011241357 | 0.655725065 | -1.542172796 | 0.123031609 | 0.285966245 | 1.011241357 |
| ENSMUSG00000029231 | Pdgfra | 18595 | protein_coding | 7.341921872 | -1.011498342 | 0.704903 | -1.434946853 | 0.151302243 | 0.329506034 | 1.011498342 |
| ENSMUSG00000027955 | Gask1b | 68659 | protein_coding | 23.04513516 | 1.011906128 | 0.339586471 | 2.979818729 | 0.00288419 | 0.018179414 | 1.011906128 |
| ENSMUSG00000042246 | Tmc7 | 209760 | protein_coding | 129.9013671 | -1.012767805 | 0.201709749 | -5.020916486 | 5.14E-07 | 1.05E-05 | 1.012767805 |

| ensembl_gene_id | external_gene_name | entrezgene_id | gene_biotype | baseMean | log2FoldChange | lfcSE | stat | pvalue | padj | absLog2FC |
|---|---|---|---|---|---|---|---|---|---|---|
| ENSMUSG00000099032 | Tcf24 | 100039596 | protein_coding | 15.3028633 | 1.013104792 | 0.339040638 | 2.988151501 | 0.002806704 | 0.01778249 | 1.013104792 |
| ENSMUSG00000021294 | Kif26a | 668303 | protein_coding | 13.72784169 | -1.013853314 | 0.420963733 | -2.40841012 | 0.016022171 | 0.067940661 | 1.013853314 |
| ENSMUSG00000036959 | Bcorl1 | 320376 | protein_coding | 154.503184 | -1.014421497 | 0.245495054 | -4.13214637 | 3.59E-05 | 0.000461834 | 1.014421497 |
| ENSMUSG00000000902 | Smarcb1 | 20587 | protein_coding | 1012.031394 | 1.016363945 | 0.093063817 | 10.92115046 | 9.13E-28 | 9.39E-26 | 1.016363945 |
| ENSMUSG00000086784 | Isoc2a | 664994 | protein_coding | 424.3072792 | 1.017062289 | 0.137491985 | 7.397247842 | 1.39E-13 | 6.17E-12 | 1.017062289 |
| ENSMUSG00000030035 | Wbp1 | 22377 | protein_coding | 1068.382438 | 1.017461785 | 0.089692015 | 11.34395057 | 7.95E-30 | 9.04E-28 | 1.017461785 |
| ENSMUSG00000040364 | Sec1 | 56546 | protein_coding | 15.45966487 | -1.017721248 | 0.603777032 | -1.685591193 | 0.091874567 | 0.235043061 | 1.017721248 |
| ENSMUSG00000022148 | Fyb | 23880 | protein_coding | 62.87332792 | -1.018099537 | 0.196636005 | -5.177584534 | 2.25E-07 | 4.90E-06 | 1.018099537 |
| ENSMUSG00000001802 | Lrp3 | 435965 | protein_coding | 7.283613738 | -1.018599785 | 0.500264254 | -2.036123461 | 0.041737959 | 0.136987887 | 1.018599785 |
| ENSMUSG00000059146 | Ntrk3 | 18213 | protein_coding | 21.20492968 | -1.018778729 | 1.390180123 | -0.732839373 | 0.463656404 | 0.661097413 | 1.018778729 |
| ENSMUSG00000037060 | Cavin3 | 109042 | protein_coding | 10.613021 | -1.018893945 | 0.406715204 | -2.505177913 | 0.012238981 | 0.055634273 | 1.018893945 |
| ENSMUSG00000074445 | Sprr2a3 | 100042514 | protein_coding | 1506.269719 | 1.019095167 | 0.214376023 | 4.753774003 | 2.00E-06 | 3.50E-05 | 1.019095167 |
| ENSMUSG00000022091 | Sorbs3 | 20410 | protein_coding | 31.06627978 | -1.020400455 | 0.238986574 | -4.269697825 | 1.96E-05 | 0.000272052 | 1.020400455 |
| ENSMUSG00000026825 | Dnm1 | 13429 | protein_coding | 30.94278624 | -1.021909615 | 0.305872776 | -3.340962963 | 0.000834884 | 0.006607109 | 1.021909615 |
| ENSMUSG00000027803 | Wwtr1 | 97064 | protein_coding | 16.28636507 | -1.022264488 | 0.279601806 | -3.656144083 | 0.000256037 | 0.002500408 | 1.022264488 |
| ENSMUSG00000057751 | Megf6 | 230971 | protein_coding | 12.69412872 | -1.022475242 | 0.473670747 | -2.1586202 | 0.030879644 | 0.110425237 | 1.022475242 |
| ENSMUSG00000111731 | Gm48727 | 118567821 | lncRNA | 7.558156572 | 1.023799497 | 0.450486447 | 2.272653271 | 0.023047083 | 0.088942434 | 1.023799497 |
| ENSMUSG00000047867 | Gimap6 | 231931 | protein_coding | 46.4634854 | 1.02457258 | 0.263944713 | 3.881769661 | 0.000103699 | 0.001161546 | 1.02457258 |
| ENSMUSG00000072214 | Septin5 | 18951 | protein_coding | 351.28533 | 1.024622318 | 0.13390822 | 7.651676015 | 1.98E-14 | 9.60E-13 | 1.024622318 |
| ENSMUSG00000084174 | Sycn | 68416 | protein_coding | 1594.080059 | -1.024850264 | 0.234373963 | -4.372713808 | 1.23E-05 | 0.000179152 | 1.024850264 |
| ENSMUSG00000069919 | Hba-a1 | 15122 | protein_coding | 54.91259059 | 1.025359692 | 0.401705385 | 2.552516668 | 0.010694779 | 0.05030905 | 1.025359692 |
| ENSMUSG00000047935 | Sox1ot | 434280 | lncRNA | 13.50142037 | 1.025399095 | 0.479637081 | 2.137864512 | 0.032527741 | 0.114371272 | 1.025399095 |
| ENSMUSG00000034028 | Cd226 | 225825 | protein_coding | 18.54417162 | 1.025957182 | 0.418768037 | 2.449941474 | 0.014287944 | 0.062196875 | 1.025957182 |
| ENSMUSG00000051236 | Msrb3 | 320183 | protein_coding | 12.88937871 | 1.026302271 | 0.415039725 | 2.472780819 | 0.013406634 | 0.059681628 | 1.026302271 |
| ENSMUSG00000026209 | Dnpep | 13437 | protein_coding | 1596.423482 | 1.026585904 | 0.110695385 | 9.273972044 | 1.79E-20 | 1.32E-18 | 1.026585904 |
| ENSMUSG00000031565 | Fgfr1 | 14182 | protein_coding | 9.705914893 | -1.027271131 | 0.505903828 | -2.030566038 | 0.042299036 | 0.138233724 | 1.027271131 |
| ENSMUSG00000094921 | Vmn2r112 | 628185 | protein_coding | 6.84645198 | -1.027371276 | 1.99881225 | -0.513990884 | 0.607258364 | 0.771140175 | 1.027371276 |
| ENSMUSG00000033268 | Duox1 | 99439 | protein_coding | 13.55154257 | -1.028842712 | 0.411800113 | -2.49840318 | 0.012475422 | 0.056483376 | 1.028842712 |
| ENSMUSG00000064368 | mt-Nd6 | 17722 | protein_coding | 3956.315271 | -1.02989299 | 0.316670852 | -3.252250667 | 0.00114495 | 0.008557504 | 1.02989299 |
| ENSMUSG00000006356 | Crip2 | 68337 | protein_coding | 19.42850181 | -1.033991217 | 0.328679699 | -3.145893166 | 0.001655805 | 0.011542082 | 1.033991217 |
| ENSMUSG00000116138 | C030006K11Rik | 223665 | protein_coding | 60.90118634 | -1.036537658 | 0.180061593 | -5.75657273 | 8.58E-09 | 2.34E-07 | 1.036537658 |
| ENSMUSG00000030483 | Cyp2b10 | 13088 | protein_coding | 6.799136661 | -1.036673241 | 0.60393871 | -1.716520606 | 0.0860668 | 0.225102132 | 1.036673241 |
| ENSMUSG00000036432 | Siah2 | 20439 | protein_coding | 426.3466698 | 1.036788915 | 0.208094157 | 4.982306723 | 6.28E-07 | 1.25E-05 | 1.036788915 |
| ENSMUSG00000020080 | Hkdc1 | 216019 | protein_coding | 502.3711595 | -1.037784139 | 0.297209534 | -3.491759246 | 0.000479851 | 0.004199334 | 1.037784139 |
| ENSMUSG00000032035 | Ets1 | 23871 | protein_coding | 117.5492661 | 1.038346508 | 0.352725402 | 2.943781491 | 0.003242288 | 0.019962569 | 1.038346508 |
| ENSMUSG00000026837 | Col5a1 | 12831 | protein_coding | 18.79086933 | -1.038965898 | 0.461964204 | -2.249018188 | 0.024511339 | 0.09302278 | 1.038965898 |
| ENSMUSG00000040265 | Dnm3 | 103967 | protein_coding | 13.59723662 | -1.039156523 | 0.889373629 | -1.168413913 | 0.242639845 | 0.447621897 | 1.039156523 |
| ENSMUSG00000064140 | Trim38 | 214158 | protein_coding | 19.54769204 | -1.03957878 | 0.322102685 | -3.227476294 | 0.001248874 | 0.009154194 | 1.03957878 |

| ensembl_gene_id | external_gene_name | entrezgene_id | gene_biotype | baseMean | log2FoldChange | lfcSE | stat | pvalue | padj | absLog2FC |
|---|---|---|---|---|---|---|---|---|---|---|
| ENSMUSG00000078810 | Gp6 | 243816 | protein_coding | 10.93486363 | 1.04071211 | 0.556428363 | 1.870343387 | 0.061436148 | 0.178438178 | 1.04071211 |
| ENSMUSG00000031661 | Nkd1 | 93960 | protein_coding | 15.55971512 | -1.040918635 | 0.315930043 | -3.294775721 | 0.000985003 | 0.007586616 | 1.040918635 |
| ENSMUSG00000036760 | Kcnk9 | 223604 | protein_coding | 7.680375843 | -1.041140134 | 1.82334427 | -0.571005789 | 0.567995719 | 0.743305197 | 1.041140134 |
| ENSMUSG00000038179 | Slamf7 | 75345 | protein_coding | 9.181747044 | 1.042370897 | 0.555328771 | 1.877033843 | 0.060513456 | 0.17675744 | 1.042370897 |
| ENSMUSG00000027513 | Pck1 | 18534 | protein_coding | 2696.392067 | -1.042921479 | 0.235674163 | -4.425268618 | 9.63E-06 | 0.000143788 | 1.042921479 |
| ENSMUSG00000068227 | Il2rb | 16185 | protein_coding | 44.37895496 | 1.043150423 | 0.273605399 | 3.812609058 | 0.000137508 | 0.001486911 | 1.043150423 |
| ENSMUSG00000036278 | Macrod1 | 107227 | protein_coding | 90.49509395 | -1.04338509 | 0.139586567 | -7.474824472 | 7.73E-14 | 3.54E-12 | 1.04338509 |
| ENSMUSG00000013415 | Igf2bp1 | 140486 | protein_coding | 15.66695014 | -1.04360529 | 0.395883618 | -2.636141643 | 0.008385472 | 0.041974795 | 1.04360529 |
| ENSMUSG00000050953 | Gja1 | 14609 | protein_coding | 78.34909383 | -1.04430894 | 0.146226004 | -7.141745738 | 9.22E-13 | 3.83E-11 | 1.04430894 |
| ENSMUSG00000034825 | Nrip3 | 78593 | protein_coding | 8.933253768 | -1.044981891 | 0.559877285 | -1.866448092 | 0.0619787 | 0.179611609 | 1.044981891 |
| ENSMUSG00000078427 | Sarnp | 66118 | protein_coding | 717.2449631 | 1.04552539 | 0.088437565 | 11.82218649 | 3.00E-32 | 3.76E-30 | 1.04552539 |
| ENSMUSG00000071158 | Klrh1 | 232415 | protein_coding | 9.545465666 | 1.045569513 | 0.581712404 | 1.797399378 | 0.072272239 | 0.199726438 | 1.045569513 |
| ENSMUSG00000014164 | Klhl3 | 100503085 | protein_coding | 20.61100877 | -1.045853644 | 0.386368147 | -2.706883713 | 0.006791804 | 0.035573778 | 1.045853644 |
| ENSMUSG00000034438 | Gbp8 | 76074 | protein_coding | 24.14435275 | 1.047146037 | 0.394458719 | 2.654640365 | 0.0079393 | 0.040234544 | 1.047146037 |
| ENSMUSG00000039976 | Tbc1d16 | 207592 | protein_coding | 195.7443053 | -1.049370316 | 0.165265913 | -6.349587143 | 2.16E-10 | 7.08E-09 | 1.049370316 |
| ENSMUSG00000032611 | 1700102P08Rik | 112418 | protein_coding | 8.519430695 | -1.049550036 | 0.456443376 | -2.299409066 | 0.021481722 | 0.084234579 | 1.049550036 |
| ENSMUSG00000043789 | Vwce | 71768 | protein_coding | 7.090219485 | -1.052050802 | 0.560243785 | -1.877844664 | 0.060402418 | 0.176550129 | 1.052050802 |
| ENSMUSG00000031543 | Ank1 | 11733 | protein_coding | 10.7808198 | -1.052113149 | 0.791545428 | -1.329188587 | 0.183785759 | 0.373242328 | 1.052113149 |
| ENSMUSG00000036098 | Myrf | 225908 | protein_coding | 7.968852495 | -1.052313303 | 0.769602962 | -1.367345702 | 0.171516973 | 0.357067635 | 1.052313303 |
| ENSMUSG00000044788 | Fads6 | 328035 | protein_coding | 17.89083494 | -1.052867332 | 0.288236524 | -3.652789444 | 0.000259407 | 0.002531761 | 1.052867332 |
| ENSMUSG00000047990 | C2cd4a | 244911 | protein_coding | 105.0227065 | 1.053379829 | 0.266766367 | 3.948698034 | 7.86E-05 | 0.000913315 | 1.053379829 |
| ENSMUSG00000097990 | Gm19557 | 100503104 | lncRNA | 10.20257483 | -1.053606394 | 0.357871658 | -2.944090066 | 0.003239057 | 0.019958106 | 1.053606394 |
| ENSMUSG00000005465 | Il27ra | 50931 | protein_coding | 7.434971983 | 1.054250183 | 0.516073709 | 2.042828701 | 0.041069398 | 0.135583278 | 1.054250183 |
| ENSMUSG00000035828 | Pim3 | 223775 | protein_coding | 1028.322258 | -1.054346236 | 0.178689307 | -5.900443939 | 3.63E-09 | 1.03E-07 | 1.054346236 |
| ENSMUSG00000040327 | Cul9 | 78309 | protein_coding | 48.83050036 | -1.05514324 | 0.225748076 | -4.673985525 | 2.95E-06 | 4.98E-05 | 1.05514324 |
| ENSMUSG00000073197 | 5730507C01Rik | 236366 | protein_coding | 74.05127922 | 1.05519984 | 0.250127369 | 4.218650051 | 2.46E-05 | 0.000330069 | 1.05519984 |
| ENSMUSG00000026354 | Lct | 226413 | protein_coding | 38.79366138 | -1.055263467 | 0.539055751 | -1.957614707 | 0.05027524 | 0.155976632 | 1.055263467 |
| ENSMUSG00000100084 | 2310068J16Rik | 70281 | lncRNA | 6.850840476 | -1.055523438 | 0.610039629 | -1.730253884 | 0.083584925 | 0.220676394 | 1.055523438 |
| ENSMUSG00000027249 | F2 | 14061 | protein_coding | 12.60436203 | -1.056220659 | 0.349659879 | -3.020708761 | 0.002521838 | 0.016428562 | 1.056220659 |
| ENSMUSG00000030220 | Arhgdib | 11857 | protein_coding | 27.52746408 | 1.056659955 | 0.298548316 | 3.539326462 | 0.000401149 | 0.003642821 | 1.056659955 |
| ENSMUSG00000050822 | Slc29a4 | 243328 | protein_coding | 13.11011554 | -1.056809808 | 0.41342362 | -2.55623955 | 0.010581025 | 0.049906593 | 1.056809808 |
| ENSMUSG00000061356 | Nuggc | 100503545 | protein_coding | 10.78571741 | 1.056850799 | 0.44383632 | 2.381172408 | 0.017257633 | 0.071713951 | 1.056850799 |
| ENSMUSG00000036030 | Prtg | 235472 | protein_coding | 7.010974643 | -1.05712437 | 0.922325712 | -1.146150818 | 0.251732754 | 0.458187556 | 1.05712437 |
| ENSMUSG00000063239 | Grm4 | 268934 | protein_coding | 9.005545957 | -1.057502612 | 0.531720806 | -1.988830604 | 0.046719904 | 0.14856351 | 1.057502612 |
| ENSMUSG00000003882 | Il7r | 16197 | protein_coding | 8.206227679 | 1.05760647 | 0.555081222 | 1.905318409 | 0.056738694 | 0.169428931 | 1.05760647 |
| ENSMUSG00000032519 | Slc25a38 | 208638 | protein_coding | 355.0529251 | 1.057667435 | 0.13457523 | 7.859302437 | 3.86E-15 | 1.98E-13 | 1.057667435 |
| ENSMUSG00000035431 | Sstr1 | 20605 | protein_coding | 165.1338325 | -1.057831291 | 0.210149392 | -5.033710923 | 4.81E-07 | 9.84E-06 | 1.057831291 |
| ENSMUSG00000023073 | Slc10a2 | 20494 | protein_coding | 1084.607398 | -1.059075054 | 0.476277624 | -2.223650664 | 0.026171957 | 0.097739621 | 1.059075054 |

Supplementary Table 1

| ensembl_gene_id | external_gene_name | entrezgene_id | gene_biotype | baseMean | log2FoldChange | lfcSE | stat | pvalue | padj | absLog2FC |
|---|---|---|---|---|---|---|---|---|---|---|
| ENSMUSG00000033857 | Engase | 217364 | protein_coding | 680.4523477 | 1.059091745 | 0.115570225 | 9.164053693 | 5.00E-20 | 3.54E-18 | 1.059091745 |
| ENSMUSG00000029403 | Cdkl2 | 53886 | protein_coding | 230.5212021 | 1.059207001 | 0.089831756 | 11.79100852 | 4.34E-32 | 5.36E-30 | 1.059207001 |
| ENSMUSG00000043557 | Mdga1 | 74762 | protein_coding | 11.73598006 | -1.059318076 | 0.474581488 | -2.232109983 | 0.025607695 | 0.096334285 | 1.059318076 |
| ENSMUSG00000049103 | Ccr2 | 12772 | protein_coding | 10.97933771 | 1.060318615 | 0.541629394 | 1.957645997 | 0.050271566 | 0.155976632 | 1.060318615 |
| ENSMUSG00000009292 | Trpm2 | 28240 | protein_coding | 15.63169226 | -1.060721486 | 0.599511732 | -1.769308971 | 0.076842327 | 0.207800439 | 1.060721486 |
| ENSMUSG00000046840 | Hnf4aos | 68314 | lncRNA | 7.720086907 | -1.061005669 | 0.480103721 | -2.209950941 | 0.027108567 | 0.100066109 | 1.061005669 |
| ENSMUSG00000022763 | Aifm3 | 72168 | protein_coding | 27.55150314 | -1.06129809 | 0.276475341 | -3.838671789 | 0.000123702 | 0.001354172 | 1.06129809 |
| ENSMUSG00000030921 | Trim30a | 20128 | protein_coding | 129.8102576 | 1.061796833 | 0.177142508 | 5.99402621 | 2.05E-09 | 5.99E-08 | 1.061796833 |
| ENSMUSG00000028777 | Gnat3 | 242851 | protein_coding | 41.51691643 | -1.062158176 | 0.240326981 | -4.419637661 | 9.89E-06 | 0.00014731 | 1.062158176 |
| ENSMUSG00000070695 | Cntnap5a | 636808 | protein_coding | 7.38253707 | -1.062830598 | 1.136245043 | -0.935388545 | 0.349588098 | 0.559790814 | 1.062830598 |
| ENSMUSG00000091382 | Vmn1r18 | 171199 | protein_coding | 9.235579283 | -1.063597175 | 2.038585169 | -0.521733009 | 0.601856238 | 0.767564496 | 1.063597175 |
| ENSMUSG00000048856 | Slc25a47 | 104910 | protein_coding | 121.1851892 | 1.064569477 | 0.135009111 | 7.885167644 | 3.14E-15 | 1.64E-13 | 1.064569477 |
| ENSMUSG00000052613 | Pcdh15 | 11994 | protein_coding | 12.33164867 | -1.065050241 | 1.653719766 | -0.644033084 | 0.519553981 | 0.706398276 | 1.065050241 |
| ENSMUSG00000064367 | mt-Nd5 | 17721 | protein_coding | 15480.9135 | -1.068448018 | 0.255444776 | -4.182696687 | 2.88E-05 | 0.000381414 | 1.068448018 |
| ENSMUSG00000031824 | 6430548M08Rik | 234797 | protein_coding | 1558.927635 | 1.068836793 | 0.177000563 | 6.038606756 | 1.55E-09 | 4.63E-08 | 1.068836793 |
| ENSMUSG00000037386 | Rims2 | 116838 | protein_coding | 58.54431497 | -1.06995535 | 0.541068814 | -1.977484788 | 0.047986858 | 0.151413367 | 1.06995535 |
| ENSMUSG00000032446 | Eomes | 13813 | protein_coding | 9.877719598 | -1.070057094 | 0.441193943 | -2.42536669 | 0.015292931 | 0.065534172 | 1.070057094 |
| ENSMUSG00000016995 | Matn4 | 17183 | protein_coding | 7.083083465 | -1.070058924 | 0.493038759 | -2.170334289 | 0.029981531 | 0.10791996 | 1.070058924 |
| ENSMUSG00000007888 | Crlf1 | 12931 | protein_coding | 9.367465543 | -1.071914778 | 0.421985721 | -2.540168361 | 0.011079912 | 0.051650936 | 1.071914778 |
| ENSMUSG00000040488 | Ltbp4 | 108075 | protein_coding | 20.6597107 | -1.071915874 | 0.481128267 | -2.227921216 | 0.025885769 | 0.097059447 | 1.071915874 |
| ENSMUSG00000034701 | Neurod1 | 18012 | protein_coding | 36.38182402 | -1.072210408 | 0.305935268 | -3.504696978 | 0.000457127 | 0.004038338 | 1.072210408 |
| ENSMUSG00000055214 | Pld5 | 319455 | protein_coding | 12.47253154 | -1.072489486 | 0.610207429 | -1.757581823 | 0.078818688 | 0.211670171 | 1.072489486 |
| ENSMUSG00000015829 | Tnr | 21960 | protein_coding | 12.35818896 | -1.072892046 | 0.999439387 | -1.073493861 | 0.283049592 | 0.492184923 | 1.072892046 |
| ENSMUSG00000040828 | Catsperd | 106757 | protein_coding | 9.559587415 | -1.073067143 | 0.432868161 | -2.478969902 | 0.013176242 | 0.05891948 | 1.073067143 |
| ENSMUSG00000051855 | Mest | 17294 | protein_coding | 13.11752133 | -1.073481674 | 0.449981695 | -2.385611874 | 0.017050735 | 0.071165076 | 1.073481674 |
| ENSMUSG00000032690 | Oas2 | 246728 | protein_coding | 35.39223501 | 1.075511744 | 0.380581116 | 2.825972436 | 0.004713732 | 0.026845617 | 1.075511744 |
| ENSMUSG00000086199 | Bcas3os1 | 71489 | lncRNA | 8.274071271 | -1.075629252 | 0.591400987 | -1.818781633 | 0.068944752 | 0.19360931 | 1.075629252 |
| ENSMUSG00000045136 | Tubb2b | 73710 | protein_coding | 23.53546696 | -1.075990672 | 0.247838281 | -4.34150313 | 1.42E-05 | 0.000203429 | 1.075990672 |
| ENSMUSG00000062785 | Kcnc3 | 16504 | protein_coding | 23.76168342 | 1.076095092 | 0.367266259 | 2.930013487 | 0.003389473 | 0.020708342 | 1.076095092 |
| ENSMUSG00000055912 | Tmem150a | 232086 | protein_coding | 7.251947707 | -1.076626192 | 0.45575091 | -2.362312766 | 0.018161313 | 0.074382461 | 1.076626192 |
| ENSMUSG00000016200 | Syt14 | 329324 | protein_coding | 13.14049013 | -1.077750214 | 0.525811016 | -2.049691205 | 0.040394574 | 0.133903178 | 1.077750214 |
| ENSMUSG00000057716 | Tmem178b | 434008 | protein_coding | 6.98947151 | -1.07776261 | 0.855374896 | -1.259988591 | 0.207673478 | 0.404082965 | 1.07776261 |
| ENSMUSG00000109305 | Smim38 | 100040049 | protein_coding | 45.83122743 | -1.078235897 | 0.312348352 | -3.452030054 | 0.000556386 | 0.004746713 | 1.078235897 |
| ENSMUSG00000036594 | H2-Aa | 14960 | protein_coding | 53.23170012 | 1.078322006 | 0.375516423 | 2.87157083 | 0.004084372 | 0.023887294 | 1.078322006 |
| ENSMUSG00000024901 | Peli3 | 240518 | protein_coding | 27.12871838 | -1.078333822 | 0.30848828 | -3.495542268 | 0.0004731 | 0.004158681 | 1.078333822 |
| ENSMUSG00000026610 | Esrrg | 26381 | protein_coding | 79.0532492 | -1.079260745 | 0.306323992 | -3.523265471 | 0.000426264 | 0.003827247 | 1.079260745 |
| ENSMUSG00000029154 | Cwh43 | 231293 | protein_coding | 401.2779002 | -1.079474073 | 0.224970574 | -4.79829008 | 1.60E-06 | 2.87E-05 | 1.079474073 |
| ENSMUSG00000022754 | Tmem45a | 56277 | protein_coding | 11.90147258 | -1.07967613 | 0.326044433 | -3.311438626 | 0.000928176 | 0.007208185 | 1.07967613 |

| ensembl_gene_id | external_gene_name | entrezgene_id | gene_biotype | baseMean | log2FoldChange | lfcSE | stat | pvalue | padj | absLog2FC |
|---|---|---|---|---|---|---|---|---|---|---|
| ENSMUSG00000054074 | Skida1 | 72668 | protein_coding | 7.426638918 | -1.079782971 | 0.672363871 | -1.605950317 | 0.108284827 | 0.261882891 | 1.079782971 |
| ENSMUSG00000040136 | Abcc8 | 20927 | protein_coding | 55.16902274 | -1.079850443 | 0.294376839 | -3.66825884 | 0.000244208 | 0.002401076 | 1.079850443 |
| ENSMUSG00000101304 | Plet1os | 69631 | lncRNA | 39.20605274 | -1.080298489 | 0.318516424 | -3.391657097 | 0.000694713 | 0.005721506 | 1.080298489 |
| ENSMUSG00000064105 | Cnnm2 | 94219 | protein_coding | 133.3522676 | 1.081918029 | 0.111064246 | 9.741371005 | 2.01E-22 | 1.57E-20 | 1.081918029 |
| ENSMUSG00000047619 | Ddi1 | 71829 | protein_coding | 7.291023556 | -1.082434554 | 0.662154696 | -1.634715515 | 0.102108698 | 0.251723781 | 1.082434554 |
| ENSMUSG00000031709 | Tbc1d9 | 71310 | protein_coding | 163.2615555 | 1.082666715 | 0.213236211 | 5.077311719 | 3.83E-07 | 7.99E-06 | 1.082666715 |
| ENSMUSG00000032377 | Plscr4 | 235527 | protein_coding | 37.33107624 | -1.082817697 | 0.254570827 | -4.253502686 | 2.10E-05 | 0.000289722 | 1.082817697 |
| ENSMUSG00000008153 | Clstn3 | 232370 | protein_coding | 6.744445338 | -1.084356209 | 0.79969117 | -1.355968716 | 0.175109122 | 0.361418538 | 1.084356209 |
| ENSMUSG00000020181 | Nav3 | 260315 | protein_coding | 8.300184112 | -1.086131583 | 1.268574081 | -0.856183016 | 0.391896555 | 0.600031561 | 1.086131583 |
| ENSMUSG00000022847 | Thpo | 21832 | protein_coding | 9.09059744 | 1.086600038 | 0.438202004 | 2.479678388 | 0.013150093 | 0.058853189 | 1.086600038 |
| ENSMUSG00000017692 | Rhbdl3 | 246104 | protein_coding | 72.53790738 | -1.088751586 | 0.232825773 | -4.676250275 | 2.92E-06 | 4.94E-05 | 1.088751586 |
| ENSMUSG00000087626 | Gm15050 | 100504154 | lncRNA | 8.610335158 | 1.088774185 | 0.408337409 | 2.666359147 | 0.007667773 | 0.039170073 | 1.088774185 |
| ENSMUSG00000087341 | 0610040F04Rik | 75394 | lncRNA | 44.63019203 | 1.089081587 | 0.369010376 | 2.951357622 | 0.003163804 | 0.019562525 | 1.089081587 |
| ENSMUSG00000063564 | Col23a1 | 237759 | protein_coding | 11.95938081 | -1.090806725 | 0.522870913 | -2.086187425 | 0.036961649 | 0.125753965 | 1.090806725 |
| ENSMUSG00000035934 | Pknox2 | 208076 | protein_coding | 8.04499302 | -1.091342088 | 0.816354405 | -1.336848409 | 0.181272124 | 0.369787704 | 1.091342088 |
| ENSMUSG00000024378 | Stard4 | 170459 | protein_coding | 3474.902169 | 1.09214271 | 0.100868384 | 10.8274037 | 2.55E-27 | 2.53E-25 | 1.09214271 |
| ENSMUSG00000074151 | Nlrc5 | 434341 | protein_coding | 47.90330548 | 1.092508091 | 0.297983733 | 3.666334672 | 0.000246052 | 0.002416223 | 1.092508091 |
| ENSMUSG00000050677 | Ccdc96 | 66717 | protein_coding | 7.830998176 | -1.092609137 | 0.466319459 | -2.343048561 | 0.019126893 | 0.077491667 | 1.092609137 |
| ENSMUSG00000052305 | Hbb-bs | 15129 | protein_coding | 343.2904301 | 1.09330738 | 0.394417104 | 2.771957325 | 0.005572034 | 0.030373497 | 1.09330738 |
| ENSMUSG00000025790 | Slco3a1 | 108116 | protein_coding | 15.88559506 | -1.093412503 | 0.414103098 | -2.64043546 | 0.008279956 | 0.041603515 | 1.093412503 |
| ENSMUSG00000084890 | A830036E02Rik | 353123 | lncRNA | 9.715915859 | -1.09494737 | 0.523750448 | -2.090589847 | 0.036564847 | 0.124792131 | 1.09494737 |
| ENSMUSG00000034449 | Dhrs11 | 192970 | protein_coding | 1628.531484 | 1.095857079 | 0.096142588 | 11.39824816 | 4.27E-30 | 4.89E-28 | 1.095857079 |
| ENSMUSG00000025738 | Fbxl16 | 214931 | protein_coding | 10.07197785 | -1.096359476 | 0.470496526 | -2.330218007 | 0.019794632 | 0.07915076 | 1.096359476 |
| ENSMUSG00000001334 | Fndc5 | 384061 | protein_coding | 151.3760188 | -1.096538996 | 0.247179874 | -4.43619854 | 9.16E-06 | 0.000137974 | 1.096538996 |
| ENSMUSG00000023236 | Scg5 | 20394 | protein_coding | 19.34527558 | -1.097125156 | 0.308863894 | -3.552131467 | 0.000382124 | 0.003497971 | 1.097125156 |
| ENSMUSG00000080316 | Spaca6 | 75202 | protein_coding | 16.03471827 | -1.09755481 | 0.462398837 | -2.373610661 | 0.017615114 | 0.072765884 | 1.09755481 |
| ENSMUSG00000054422 | Fabp1 | 14080 | protein_coding | 28.07288789 | -1.099014345 | 0.534360717 | -2.056690002 | 0.039716054 | 0.132481904 | 1.099014345 |
| ENSMUSG00000054083 | Capn12 | 60594 | protein_coding | 11.10491835 | -1.099150158 | 0.440651472 | -2.494375325 | 0.012617904 | 0.0569827 | 1.099150158 |
| ENSMUSG00000049571 | Cfap46 | 212124 | protein_coding | 8.367655342 | -1.099204703 | 1.224749731 | -0.897493321 | 0.369455736 | 0.578177536 | 1.099204703 |
| ENSMUSG00000040118 | Cacna2d1 | 12293 | protein_coding | 116.7898575 | 1.100277537 | 0.15509656 | 7.094145328 | 1.30E-12 | 5.30E-11 | 1.100277537 |
| ENSMUSG00000025212 | Sfxn3 | 94280 | protein_coding | 142.8478732 | -1.100983378 | 0.221497892 | -4.970626902 | 6.67E-07 | 1.32E-05 | 1.100983378 |
| ENSMUSG00000042286 | Stab1 | 192187 | protein_coding | 7.685705912 | -1.102204712 | 0.731810388 | -1.50613428 | 0.132032706 | 0.299882412 | 1.102204712 |
| ENSMUSG00000031381 | Piga | 18700 | protein_coding | 776.0170877 | 1.102384131 | 0.092128054 | 11.9657811 | 5.37E-33 | 6.95E-31 | 1.102384131 |
| ENSMUSG00000035042 | Ccl5 | 20304 | protein_coding | 29.55996166 | 1.103213957 | 0.40642341 | 2.714444907 | 0.006638694 | 0.03492111 | 1.103213957 |
| ENSMUSG00000018654 | Ikzf1 | 22778 | protein_coding | 18.27243715 | 1.103286555 | 0.388505508 | 2.839822169 | 0.004513869 | 0.025862196 | 1.103286555 |
| ENSMUSG00000039457 | Ppl | 19041 | protein_coding | 182.0001454 | -1.104168287 | 0.21866861 | -5.049505205 | 4.43E-07 | 9.15E-06 | 1.104168287 |
| ENSMUSG00000028763 | Hspg2 | 15530 | protein_coding | 350.032358 | -1.104426553 | 0.202816184 | -5.445455735 | 5.17E-08 | 1.23E-06 | 1.104426553 |
| ENSMUSG00000026117 | Zap70 | 22637 | protein_coding | 12.3792007 | 1.104637813 | 0.434748791 | 2.540864602 | 0.011057873 | 0.051605568 | 1.104637813 |

| ensembl_gene_id | external_gene_name | entrezgene_id | gene_biotype | baseMean | log2FoldChange | lfcSE | stat | pvalue | padj | absLog2FC |
|---|---|---|---|---|---|---|---|---|---|---|
| ENSMUSG00000020620 | Abca8b | 27404 | protein_coding | 9.881253019 | -1.104746692 | 0.714730056 | -1.545683833 | 0.122180944 | 0.284725395 | 1.104746692 |
| ENSMUSG00000069920 | B3gnt9 | 97440 | protein_coding | 63.65168135 | -1.105287249 | 0.20120174 | -5.493427871 | 3.94E-08 | 9.56E-07 | 1.105287249 |
| ENSMUSG00000020077 | Srgn | 19073 | protein_coding | 27.06643523 | 1.105455996 | 0.285193556 | 3.876160495 | 0.000106118 | 0.001183643 | 1.105455996 |
| ENSMUSG00000038503 | Mesd | 67943 | protein_coding | 619.3808994 | 1.10819463 | 0.124222883 | 8.921018452 | 4.62E-19 | 3.07E-17 | 1.10819463 |
| ENSMUSG00000023973 | Cnpy3 | 72029 | protein_coding | 551.7647169 | 1.108997254 | 0.089240911 | 12.42700501 | 1.86E-35 | 2.78E-33 | 1.108997254 |
| ENSMUSG00000027750 | Postn | 50706 | protein_coding | 12.42693142 | -1.109447713 | 0.468882769 | -2.366151597 | 0.017974083 | 0.073824443 | 1.109447713 |
| ENSMUSG00000016918 | Sulf1 | 240725 | protein_coding | 16.21382692 | -1.110027959 | 0.450394807 | -2.464566514 | 0.01371791 | 0.060492489 | 1.110027959 |
| ENSMUSG00000101969 | Gm20125 | 100504235 | lncRNA | 7.206713653 | -1.110587804 | 0.701209193 | -1.583818088 | 0.113235122 | 0.269195376 | 1.110587804 |
| ENSMUSG00000031232 | Magt1 | 67075 | protein_coding | 3441.941297 | 1.111664102 | 0.065972765 | 16.85034885 | 1.04E-63 | 3.32E-61 | 1.111664102 |
| ENSMUSG00000001333 | Sync | 68828 | protein_coding | 13.80657041 | -1.111682024 | 0.353153454 | -3.147872442 | 0.001644634 | 0.01147957 | 1.111682024 |
| ENSMUSG00000021390 | Ogn | 18295 | protein_coding | 11.01749943 | -1.112186453 | 0.457369617 | -2.43170165 | 0.01502808 | 0.064755004 | 1.112186453 |
| ENSMUSG00000029767 | Calu | 12321 | protein_coding | 4060.254624 | 1.112440949 | 0.139050079 | 8.000289957 | 1.24E-15 | 6.68E-14 | 1.112440949 |
| ENSMUSG00000033763 | Mtss2 | 244654 | protein_coding | 8.516577326 | -1.112621404 | 0.572267778 | -1.944232136 | 0.051867464 | 0.159456663 | 1.112621404 |
| ENSMUSG00000027790 | Sis | 69983 | protein_coding | 210.3870921 | -1.112964158 | 0.312151675 | -3.565459513 | 0.000363219 | 0.003340275 | 1.112964158 |
| ENSMUSG00000087476 | Rap1gapos | 69808 | lncRNA | 34.89508034 | 1.113478081 | 0.252468526 | 4.410363936 | 1.03E-05 | 0.000153047 | 1.113478081 |
| ENSMUSG00000017204 | Gsdma | 57911 | protein_coding | 14.01265046 | -1.113624923 | 0.302575425 | -3.680487017 | 0.000232789 | 0.002318863 | 1.113624923 |
| ENSMUSG00000047910 | Pcdhb16 | 93887 | protein_coding | 7.86947302 | 1.114155777 | 0.566048728 | 1.968303648 | 0.049033111 | 0.153468144 | 1.114155777 |
| ENSMUSG00000037095 | Lrg1 | 76905 | protein_coding | 16.52089228 | 1.115221901 | 0.81822721 | 1.362973373 | 0.172890902 | 0.358615222 | 1.115221901 |
| ENSMUSG00000095115 | Itpripl2 | 319622 | protein_coding | 110.5725674 | -1.116321797 | 0.243597736 | -4.582644375 | 4.59E-06 | 7.47E-05 | 1.116321797 |
| ENSMUSG00000043924 | Ncmap | 230822 | protein_coding | 23.6045774 | 1.116674748 | 0.329012849 | 3.394015617 | 0.000688758 | 0.00568031 | 1.116674748 |
| ENSMUSG00000022951 | Rcan1 | 54720 | protein_coding | 441.2405974 | 1.11736602 | 0.125376616 | 8.912076742 | 5.01E-19 | 3.31E-17 | 1.11736602 |
| ENSMUSG00000037071 | Scd1 | 20249 | protein_coding | 1140.236049 | 1.117756732 | 0.113634561 | 9.836415242 | 7.85E-23 | 6.44E-21 | 1.117756732 |
| ENSMUSG00000035845 | Alg12 | 223774 | protein_coding | 317.8034219 | 1.117894093 | 0.098787142 | 11.31619024 | 1.09E-29 | 1.23E-27 | 1.117894093 |
| ENSMUSG00000030098 | Grip2 | 243547 | protein_coding | 7.151219382 | -1.118136677 | 0.812102981 | -1.376840996 | 0.168561413 | 0.353362225 | 1.118136677 |
| ENSMUSG00000045064 | Zc2hc1c | 72350 | protein_coding | 14.99945599 | -1.119197047 | 0.419370814 | -2.66875283 | 0.007613345 | 0.038992079 | 1.119197047 |
| ENSMUSG00000040896 | Kcnd3 | 56543 | protein_coding | 13.24338665 | -1.11940365 | 0.59787404 | -1.872306834 | 0.061164166 | 0.177981255 | 1.11940365 |
| ENSMUSG00000087175 | Gm15133 | 622070 | lncRNA | 17.38394661 | 1.119685275 | 0.308399659 | 3.630630715 | 0.000282729 | 0.002700691 | 1.119685275 |
| ENSMUSG00000004933 | Matk | 17179 | protein_coding | 63.09382551 | -1.120165685 | 0.180050762 | -6.221388205 | 4.93E-10 | 1.54E-08 | 1.120165685 |
| ENSMUSG00000035227 | Spcs2 | 66624 | protein_coding | 1732.999498 | 1.120691966 | 0.079795866 | 14.04448662 | 8.33E-45 | 1.66E-42 | 1.120691966 |
| ENSMUSG00000060586 | H2-Eb1 | 14969 | protein_coding | 41.52546435 | 1.123129879 | 0.406224469 | 2.764801152 | 0.00569575 | 0.030910362 | 1.123129879 |
| ENSMUSG00000031523 | Dlc1 | 50768 | protein_coding | 12.16386384 | -1.124426171 | 0.481212161 | -2.33665369 | 0.019457198 | 0.078439447 | 1.124426171 |
| ENSMUSG00000026166 | Ccl20 | 20297 | protein_coding | 12.06943634 | -1.124562539 | 0.499770331 | -2.250158662 | 0.024438875 | 0.092814117 | 1.124562539 |
| ENSMUSG00000024503 | Spink1 | 20730 | protein_coding | 2508.074254 | 1.124617691 | 0.342662469 | 3.281998446 | 0.001030742 | 0.00785158 | 1.124617691 |
| ENSMUSG00000055385 | Rnf212 | 671564 | protein_coding | 23.43272525 | -1.12548387 | 0.295155949 | -3.813183754 | 0.000137188 | 0.001484464 | 1.12548387 |
| ENSMUSG00000038777 | Sema6c | 20360 | protein_coding | 8.405889309 | -1.125644212 | 0.739198432 | -1.522790314 | 0.127811174 | 0.293439621 | 1.125644212 |
| ENSMUSG00000086564 | Cd101 | 630146 | protein_coding | 9.276910466 | 1.125990982 | 0.496761989 | 2.266660912 | 0.023410948 | 0.089939599 | 1.125990982 |
| ENSMUSG00000075701 | Selenos | 109815 | protein_coding | 1472.807919 | 1.126399936 | 0.073262316 | 15.37488848 | 2.41E-53 | 6.00E-51 | 1.126399936 |
| ENSMUSG00000007872 | Id3 | 15903 | protein_coding | 2314.695741 | 1.127985398 | 0.226078865 | 4.989344752 | 6.06E-07 | 1.22E-05 | 1.127985398 |

| ensembl_gene_id | external_gene_name | entrezgene_id | gene_biotype | baseMean | log2FoldChange | lfcSE | stat | pvalue | padj | absLog2FC |
|---|---|---|---|---|---|---|---|---|---|---|
| ENSMUSG00000026249 | Serpine2 | 20720 | protein_coding | 23.21110188 | 1.129897029 | 0.335397488 | 3.368829731 | 0.00075488 | 0.006094139 | 1.129897029 |
| ENSMUSG00000028757 | Ddost | 13200 | protein_coding | 5228.615336 | 1.129949062 | 0.093351267 | 12.10427127 | 1.00E-33 | 1.35E-31 | 1.129949062 |
| ENSMUSG00000024897 | Apba1 | 319924 | protein_coding | 9.900320771 | -1.130133632 | 0.596769334 | -1.893752858 | 0.058257822 | 0.172541946 | 1.130133632 |
| ENSMUSG00000094002 | Gm9866 | 636791 | lncRNA | 6.788509205 | -1.130720316 | 0.932148774 | -1.21302559 | 0.225120037 | 0.427073839 | 1.130720316 |
| ENSMUSG00000102737 | 1700016A09Rik | 74227 | TEC | 7.94935814 | -1.130726713 | 0.448381485 | -2.52179617 | 0.011675734 | 0.05376441 | 1.130726713 |
| ENSMUSG00000032978 | Guca2b | 14916 | protein_coding | 132.7065857 | -1.132633038 | 0.16047538 | -7.057986312 | 1.69E-12 | 6.81E-11 | 1.132633038 |
| ENSMUSG00000050148 | Ubqln2 | 54609 | protein_coding | 307.6419023 | 1.133942898 | 0.131113981 | 8.64852771 | 5.22E-18 | 3.30E-16 | 1.133942898 |
| ENSMUSG00000036106 | Prr5 | 109270 | protein_coding | 254.8696133 | 1.134671591 | 0.143002181 | 7.934645376 | 2.11E-15 | 1.12E-13 | 1.134671591 |
| ENSMUSG00000078492 | 1700045H11Rik | 73348 | lncRNA | 9.580525578 | -1.134878102 | 0.427629501 | -2.65388169 | 0.007957172 | 0.040312289 | 1.134878102 |
| ENSMUSG00000053977 | Cd8a | 12525 | protein_coding | 30.65043065 | 1.134907891 | 0.277742157 | 4.086192406 | 4.39E-05 | 0.000549535 | 1.134907891 |
| ENSMUSG00000027238 | Frmd5 | 228564 | protein_coding | 16.52593444 | 1.136148227 | 0.524622323 | 2.165649795 | 0.030337958 | 0.108907595 | 1.136148227 |
| ENSMUSG00000022150 | Dab2 | 13132 | protein_coding | 10.82869387 | -1.13650908 | 0.698602375 | -1.626832546 | 0.103772667 | 0.254408349 | 1.13650908 |
| ENSMUSG00000091119 | Ccdc152 | 100039139 | protein_coding | 7.565687423 | -1.136642325 | 0.577630773 | -1.967766225 | 0.049094942 | 0.153601303 | 1.136642325 |
| ENSMUSG00000066372 | Vmn2r65 | 100009609 | protein_coding | 7.971933936 | -1.136789593 | 1.403364941 | -0.810045598 | 0.417913969 | 0.624147558 | 1.136789593 |
| ENSMUSG00000025964 | Adam23 | 23792 | protein_coding | 10.33792974 | -1.137015332 | 1.534345873 | -0.74104239 | 0.458667739 | 0.65713215 | 1.137015332 |
| ENSMUSG00000024486 | Hbegf | 15200 | protein_coding | 2338.798601 | -1.137195826 | 0.348767703 | -3.260611051 | 0.001111724 | 0.008369777 | 1.137195826 |
| ENSMUSG00000020218 | Wif1 | 24117 | protein_coding | 17.39936106 | -1.137608996 | 0.380641748 | -2.988660599 | 0.002802032 | 0.017764119 | 1.137608996 |
| ENSMUSG00000032374 | Plod2 | 26432 | protein_coding | 22.414249 | 1.137725088 | 0.339064085 | 3.355486881 | 0.000792254 | 0.006329664 | 1.137725088 |
| ENSMUSG00000068220 | Lgals1 | 16852 | protein_coding | 16.39414117 | 1.137820506 | 0.341751049 | 3.329384093 | 0.000870383 | 0.006839397 | 1.137820506 |
| ENSMUSG00000032181 | Scg3 | 20255 | protein_coding | 20.5598312 | -1.138641727 | 0.287389751 | -3.962012292 | 7.43E-05 | 0.000867874 | 1.138641727 |
| ENSMUSG00000030643 | Rab30 | 75985 | protein_coding | 14.21528162 | -1.138832459 | 0.455047473 | -2.502667363 | 0.012326133 | 0.055953127 | 1.138832459 |
| ENSMUSG00000022537 | Tmem44 | 224090 | protein_coding | 116.4280215 | -1.138893142 | 0.176257057 | -6.461546358 | 1.04E-10 | 3.54E-09 | 1.138893142 |
| ENSMUSG00000025854 | Fam20c | 80752 | protein_coding | 6.81166096 | -1.13916576 | 0.518994753 | -2.194946581 | 0.02816743 | 0.102948789 | 1.13916576 |
| ENSMUSG00000034855 | Cxcl10 | 15945 | protein_coding | 12.24784254 | 1.139367275 | 0.669782458 | 1.701100501 | 0.088924117 | 0.230156538 | 1.139367275 |
| ENSMUSG00000073968 | Trim68 | 101700 | protein_coding | 237.450656 | 1.139512863 | 0.113011245 | 10.08318121 | 6.56E-24 | 5.61E-22 | 1.139512863 |
| ENSMUSG00000035713 | Usp35 | 244144 | protein_coding | 41.45381491 | -1.139745752 | 0.265325937 | -4.295643938 | 1.74E-05 | 0.000244445 | 1.139745752 |
| ENSMUSG00000028967 | Errfi1 | 74155 | protein_coding | 1882.52123 | -1.139970514 | 0.407056898 | -2.800518844 | 0.005102053 | 0.028484226 | 1.139970514 |
| ENSMUSG00000026043 | Col3a1 | 12825 | protein_coding | 110.1233413 | -1.139986102 | 0.40503655 | -2.814526496 | 0.004884915 | 0.027561788 | 1.139986102 |
| ENSMUSG00000051674 | Dcun1d4 | 100737 | protein_coding | 301.5011488 | 1.140383488 | 0.102564621 | 11.1186828 | 1.02E-28 | 1.10E-26 | 1.140383488 |
| ENSMUSG00000004698 | Hdac9 | 79221 | protein_coding | 37.64911003 | -1.140594055 | 0.243311324 | -4.687796831 | 2.76E-06 | 4.71E-05 | 1.140594055 |
| ENSMUSG00000040061 | Plcb2 | 18796 | protein_coding | 46.1383479 | -1.143209873 | 0.215973281 | -5.293293082 | 1.20E-07 | 2.72E-06 | 1.143209873 |
| ENSMUSG00000029245 | Epha5 | 13839 | protein_coding | 13.51961803 | -1.146007892 | 0.905772898 | -1.26522652 | 0.205790138 | 0.401645058 | 1.146007892 |
| ENSMUSG00000024357 | Sil1 | 81500 | protein_coding | 922.0249509 | 1.146198986 | 0.080647703 | 14.21241946 | 7.67E-46 | 1.59E-43 | 1.146198986 |
| ENSMUSG00000009394 | Syn2 | 20965 | protein_coding | 55.43914165 | -1.146606202 | 0.33567588 | -3.415813496 | 0.000635918 | 0.0053198 | 1.146606202 |
| ENSMUSG00000044716 | Dok7 | 231134 | protein_coding | 55.22036938 | -1.147262626 | 0.180418635 | -6.358892079 | 2.03E-10 | 6.69E-09 | 1.147262626 |
| ENSMUSG00000064036 | Mro | 71263 | protein_coding | 9.562002224 | -1.147933426 | 0.503251549 | -2.28103307 | 0.02254649 | 0.087504994 | 1.147933426 |
| ENSMUSG00000102224 | 4930447F24Rik | 76873 | lncRNA | 23.31084803 | -1.148968175 | 0.259379753 | -4.429675654 | 9.44E-06 | 0.000141411 | 1.148968175 |
| ENSMUSG00000050395 | Tnfsf15 | 326623 | protein_coding | 41.66304346 | -1.150063159 | 0.278292009 | -4.132577007 | 3.59E-05 | 0.000461524 | 1.150063159 |

| ensembl_gene_id | external_gene_name | entrezgene_id | gene_biotype | baseMean | log2FoldChange | lfcSE | stat | pvalue | padj | absLog2FC |
|---|---|---|---|---|---|---|---|---|---|---|
| ENSMUSG00000025190 | Got1 | 14718 | protein_coding | 872.9284798 | 1.150186431 | 0.146035312 | 7.876084307 | 3.38E-15 | 1.75E-13 | 1.150186431 |
| ENSMUSG00000097075 | Cdiptos | 381922 | protein_coding | 9.756003498 | -1.150626129 | 0.576726876 | -1.995097119 | 0.046032288 | 0.146963779 | 1.150626129 |
| ENSMUSG00000062257 | Opcml | 330908 | protein_coding | 9.485837981 | -1.150784943 | 1.090041979 | -1.055725344 | 0.291093715 | 0.499573396 | 1.150784943 |
| ENSMUSG00000073940 | Hbb-bt | 101488143 | protein_coding | 40.48579406 | 1.152541897 | 0.37257371 | 3.093460075 | 0.001978372 | 0.013403447 | 1.152541897 |
| ENSMUSG00000091620 | Vmn2r23 | 435916 | protein_coding | 8.26142222 | -1.152642801 | 1.691450902 | -0.681452119 | 0.495585454 | 0.68695983 | 1.152642801 |
| ENSMUSG00000061947 | Serpina10 | 217847 | protein_coding | 13.89284468 | 1.152889532 | 0.552018106 | 2.08849949 | 0.036752801 | 0.125245747 | 1.152889532 |
| ENSMUSG00000051727 | Kctd14 | 233529 | protein_coding | 356.1974753 | 1.154209085 | 0.130624551 | 8.83608076 | 9.91E-19 | 6.44E-17 | 1.154209085 |
| ENSMUSG00000036961 | Wnt8b | 22423 | protein_coding | 7.026785107 | -1.154444233 | 0.743381645 | -1.552963057 | 0.120431961 | 0.281762672 | 1.154444233 |
| ENSMUSG00000090362 | Vmn2r79 | 621430 | protein_coding | 8.006952846 | -1.154548897 | 1.653229708 | -0.698359636 | 0.484952313 | 0.678405097 | 1.154548897 |
| ENSMUSG00000024401 | Tnf | 21926 | protein_coding | 32.90852777 | -1.15496609 | 0.350807804 | -3.29230444 | 0.0009937 | 0.007642215 | 1.15496609 |
| ENSMUSG00000026193 | Fn1 | 14268 | protein_coding | 35.05032617 | -1.156444056 | 0.409934364 | -2.821046876 | 0.004786721 | 0.027171379 | 1.156444056 |
| ENSMUSG00000025203 | Scd2 | 20250 | protein_coding | 15606.07895 | 1.157129967 | 0.141033455 | 8.204648829 | 2.31E-16 | 1.32E-14 | 1.157129967 |
| ENSMUSG00000028402 | Mpdz | 17475 | protein_coding | 22.43726937 | -1.157892186 | 0.344242366 | -3.36359583 | 0.000769341 | 0.006185798 | 1.157892186 |
| ENSMUSG00000030707 | Coro1a | 12721 | protein_coding | 62.50640425 | 1.158548641 | 0.236751496 | 4.893521947 | 9.90E-07 | 1.87E-05 | 1.158548641 |
| ENSMUSG00000064267 | Hvcn1 | 74096 | protein_coding | 9.432429591 | 1.159248608 | 0.379225528 | 3.056884419 | 0.002236505 | 0.014845504 | 1.159248608 |
| ENSMUSG00000030062 | Rpn1 | 103963 | protein_coding | 7690.565362 | 1.160509116 | 0.091471789 | 12.68707136 | 6.97E-37 | 1.11E-34 | 1.160509116 |
| ENSMUSG00000056749 | Nfil3 | 18030 | protein_coding | 254.0895965 | 1.160581101 | 0.225246357 | 5.152496653 | 2.57E-07 | 5.52E-06 | 1.160581101 |
| ENSMUSG00000020893 | Per1 | 18626 | protein_coding | 447.1139223 | 1.160793168 | 0.206271527 | 5.62750073 | 1.83E-08 | 4.77E-07 | 1.160793168 |
| ENSMUSG00000040483 | Xaf1 | 327959 | protein_coding | 82.19172062 | 1.161641521 | 0.219621509 | 5.289288508 | 1.23E-07 | 2.77E-06 | 1.161641521 |
| ENSMUSG00000034110 | Kctd7 | 212919 | protein_coding | 102.7695114 | 1.161770258 | 0.170120089 | 6.829118556 | 8.54E-12 | 3.20E-10 | 1.161770258 |
| ENSMUSG00000004451 | Ralb | 64143 | protein_coding | 1736.304313 | 1.161900328 | 0.104659586 | 11.10170953 | 1.23E-28 | 1.32E-26 | 1.161900328 |
| ENSMUSG00000020901 | Pik3r5 | 320207 | protein_coding | 95.38829103 | -1.162056461 | 0.160578278 | -7.23669775 | 4.60E-13 | 1.97E-11 | 1.162056461 |
| ENSMUSG00000028343 | Erp44 | 76299 | protein_coding | 1969.777318 | 1.162405837 | 0.085564202 | 13.58518877 | 4.90E-42 | 9.08E-40 | 1.162405837 |
| ENSMUSG00000006345 | Ggt1 | 14598 | protein_coding | 130.5625361 | 1.162946203 | 0.345632157 | 3.364693305 | 0.000766288 | 0.00616436 | 1.162946203 |
| ENSMUSG00000093930 | Hmgcs1 | 208715 | protein_coding | 6346.886757 | 1.165013693 | 0.199293701 | 5.845712566 | 5.04E-09 | 1.40E-07 | 1.165013693 |
| ENSMUSG00000043931 | Gimap7 | 231932 | protein_coding | 14.42378393 | 1.165096205 | 0.455482716 | 2.557937249 | 0.010529509 | 0.049722508 | 1.165096205 |
| ENSMUSG00000016552 | Foxred2 | 239554 | protein_coding | 34.41190254 | -1.16582761 | 0.256077529 | -4.552635344 | 5.30E-06 | 8.51E-05 | 1.16582761 |
| ENSMUSG00000035969 | Rusc2 | 100213 | protein_coding | 63.94674188 | -1.166316943 | 0.254129964 | -4.589450715 | 4.44E-06 | 7.26E-05 | 1.166316943 |
| ENSMUSG00000033526 | Ppip5k1 | 327655 | protein_coding | 2105.98819 | 1.167561069 | 0.06156113 | 18.96588106 | 3.27E-80 | 1.44E-77 | 1.167561069 |
| ENSMUSG00000034459 | Ifit1 | 15957 | protein_coding | 229.0117932 | 1.168839185 | 0.308282064 | 3.791460223 | 0.000149764 | 0.00159441 | 1.168839185 |
| ENSMUSG00000048376 | F2r | 14062 | protein_coding | 7.891734636 | 1.169417524 | 0.45584945 | 2.565359078 | 0.010306908 | 0.048888752 | 1.169417524 |
| ENSMUSG00000036027 | Pou2af2 | 69809 | protein_coding | 11.90784635 | -1.170058302 | 0.318771929 | -3.670518624 | 0.000242059 | 0.002391405 | 1.170058302 |
| ENSMUSG00000036814 | Slc6a20a | 102680 | protein_coding | 36.70118677 | -1.170158623 | 0.281377748 | -4.158675063 | 3.20E-05 | 0.000418256 | 1.170158623 |
| ENSMUSG00000063689 | H2ac21 | 621893 | protein_coding | 29.83144103 | 1.171341794 | 0.268324005 | 4.365400686 | 1.27E-05 | 0.000183904 | 1.171341794 |
| ENSMUSG00000056073 | Grik2 | 14806 | protein_coding | 9.180829413 | -1.172482248 | 1.248608201 | -0.939031353 | 0.347714647 | 0.557744099 | 1.172482248 |
| ENSMUSG00000066108 | Muc5b | 74180 | protein_coding | 11.03782348 | -1.173188689 | 0.760469316 | -1.542716667 | 0.122899536 | 0.285856281 | 1.173188689 |
| ENSMUSG00000024395 | Lims2 | 225341 | protein_coding | 7.338289193 | -1.175491966 | 0.511842144 | -2.296590813 | 0.021642123 | 0.084675937 | 1.175491966 |
| ENSMUSG00000037962 | Rflna | 73121 | protein_coding | 9.569945649 | -1.175682835 | 0.462254385 | -2.543367618 | 0.010978965 | 0.051372784 | 1.175682835 |

| ensembl_gene_id | external_gene_name | entrezgene_id | gene_biotype | baseMean | log2FoldChange | lfcSE | stat | pvalue | padj | absLog2FC |
|---|---|---|---|---|---|---|---|---|---|---|
| ENSMUSG00000026581 | Sell | 20343 | protein_coding | 10.13868376 | 1.176266005 | 0.651724868 | 1.804850579 | 0.071098067 | 0.197597936 | 1.176266005 |
| ENSMUSG00000056043 | Rgs9bp | 243923 | protein_coding | 7.517062126 | -1.176954859 | 0.650232177 | -1.810053241 | 0.070287531 | 0.196106105 | 1.176954859 |
| ENSMUSG00000056004 | Elapor2 | 231014 | protein_coding | 28.85139643 | -1.176963627 | 0.265402281 | -4.434640203 | 9.22E-06 | 0.000138583 | 1.176963627 |
| ENSMUSG00000032679 | Cd59a | 12509 | protein_coding | 16.63818837 | -1.177798235 | 0.294294698 | -4.002104835 | 6.28E-05 | 0.000752433 | 1.177798235 |
| ENSMUSG00000046768 | Rhoj | 80837 | protein_coding | 7.738226138 | 1.177887153 | 0.56035594 | 2.102033847 | 0.035550311 | 0.122484394 | 1.177887153 |
| ENSMUSG00000019232 | Etnppl | 71760 | protein_coding | 75.01223971 | 1.178089124 | 0.233386793 | 5.047796879 | 4.47E-07 | 9.22E-06 | 1.178089124 |
| ENSMUSG00000025887 | Casp12 | 12364 | protein_coding | 43.27581054 | 1.178795338 | 0.203080387 | 5.804574993 | 6.45E-09 | 1.78E-07 | 1.178795338 |
| ENSMUSG00000032094 | Cd3d | 12500 | protein_coding | 15.53231715 | 1.179625217 | 0.443309805 | 2.660949981 | 0.007792053 | 0.03965234 | 1.179625217 |
| ENSMUSG00000046997 | Spsb4 | 211949 | protein_coding | 27.21479928 | -1.179785491 | 0.243961822 | -4.835943111 | 1.33E-06 | 2.42E-05 | 1.179785491 |
| ENSMUSG00000008193 | Spib | 272382 | protein_coding | 77.34992972 | -1.181068356 | 0.200188074 | -5.899793794 | 3.64E-09 | 1.04E-07 | 1.181068356 |
| ENSMUSG00000001946 | Esam | 69524 | protein_coding | 11.8793124 | -1.181288671 | 0.372819942 | -3.168523294 | 0.001532155 | 0.010836661 | 1.181288671 |
| ENSMUSG00000059974 | Ntm | 235106 | protein_coding | 7.760799628 | -1.182218724 | 0.799880156 | -1.477994815 | 0.139409166 | 0.311082824 | 1.182218724 |
| ENSMUSG00000041809 | Efhc1 | 71877 | protein_coding | 15.34094164 | 1.18421369 | 0.350050584 | 3.382978758 | 0.000717042 | 0.005856944 | 1.18421369 |
| ENSMUSG00000050097 | Ces2b | 234669 | protein_coding | 212.93855 | -1.184394796 | 0.281570877 | -4.206382448 | 2.59E-05 | 0.000347617 | 1.184394796 |
| ENSMUSG00000027642 | Rpn2 | 20014 | protein_coding | 8541.612328 | 1.186518607 | 0.0977411 | 12.13940305 | 6.53E-34 | 9.04E-32 | 1.186518607 |
| ENSMUSG00000035279 | Ssc5d | 269855 | protein_coding | 7.371426133 | -1.186673692 | 0.701311606 | -1.692077648 | 0.09063117 | 0.232840208 | 1.186673692 |
| ENSMUSG00000019737 | Syne4 | 233066 | protein_coding | 20.37332952 | 1.187499869 | 0.2674951 | 4.439333165 | 9.02E-06 | 0.000136109 | 1.187499869 |
| ENSMUSG00000036095 | Dgkb | 217480 | protein_coding | 6.83044519 | -1.189663778 | 1.249548422 | -0.952074971 | 0.341058961 | 0.551399425 | 1.189663778 |
| ENSMUSG00000060208 | Defa17 | 23855 | protein_coding | 7.781604249 | -1.19078654 | 1.061761548 | -1.121519744 | 0.262066692 | 0.469961526 | 1.19078654 |
| ENSMUSG00000026173 | Plcd4 | 18802 | protein_coding | 11.60201016 | -1.191764618 | 0.511521696 | -2.329841777 | 0.019814515 | 0.079178524 | 1.191764618 |
| ENSMUSG00000029390 | Tmed2 | 56334 | protein_coding | 2232.803329 | 1.19204135 | 0.098730778 | 12.07365504 | 1.46E-33 | 1.95E-31 | 1.19204135 |
| ENSMUSG00000031377 | Bmx | 12169 | protein_coding | 45.47595128 | -1.192351474 | 0.219451754 | -5.433319405 | 5.53E-08 | 1.32E-06 | 1.192351474 |
| ENSMUSG00000002908 | Kcnn1 | 84036 | protein_coding | 6.990385396 | -1.193440736 | 0.571546461 | -2.088090499 | 0.036789672 | 0.125293737 | 1.193440736 |
| ENSMUSG00000078670 | Fam174b | 100038347 | protein_coding | 2269.029252 | 1.194869287 | 0.086369776 | 13.83434516 | 1.58E-43 | 3.07E-41 | 1.194869287 |
| ENSMUSG00000003378 | Grik5 | 14809 | protein_coding | 31.76857413 | -1.195050534 | 0.258240799 | -4.627659687 | 3.70E-06 | 6.12E-05 | 1.195050534 |
| ENSMUSG00000030826 | Bcat2 | 12036 | protein_coding | 854.7411732 | 1.198266257 | 0.094463526 | 12.68496216 | 7.16E-37 | 1.13E-34 | 1.198266257 |
| ENSMUSG00000029913 | Prdm5 | 70779 | protein_coding | 18.69498665 | 1.198336044 | 0.366756809 | 3.267385946 | 0.001085456 | 0.008209469 | 1.198336044 |
| ENSMUSG00000118384 | 9030625G05Rik | 71545 | lncRNA | 20.21869031 | 1.198590705 | 0.308957524 | 3.879467604 | 0.000104685 | 0.001170125 | 1.198590705 |
| ENSMUSG00000067599 | Klra7 | 16638 | protein_coding | 10.45395006 | 1.198935131 | 0.416441547 | 2.878999799 | 0.003989386 | 0.023421651 | 1.198935131 |
| ENSMUSG00000039629 | Strip2 | 320609 | protein_coding | 113.4680251 | -1.200147264 | 0.1182369 | -10.1503614 | 3.30E-24 | 2.89E-22 | 1.200147264 |
| ENSMUSG00000025724 | Sec11a | 56529 | protein_coding | 1579.836609 | 1.20105839 | 0.093609255 | 12.83055177 | 1.11E-37 | 1.85E-35 | 1.20105839 |
| ENSMUSG00000050158 | Or2m13 | 258458 | protein_coding | 66.76320184 | 1.201233803 | 0.300791483 | 3.993576517 | 6.51E-05 | 0.000773626 | 1.201233803 |
| ENSMUSG00000075592 | Nynrin | 277154 | protein_coding | 16.36814191 | -1.201355931 | 0.381105085 | -3.152295722 | 0.001619921 | 0.011338299 | 1.201355931 |
| ENSMUSG00000074796 | Slc4a11 | 269356 | protein_coding | 13.56565603 | -1.201541573 | 0.749892875 | -1.602284291 | 0.109092758 | 0.263077888 | 1.201541573 |
| ENSMUSG00000030934 | Oat | 18242 | protein_coding | 4804.187354 | 1.201686784 | 0.112262055 | 10.70430062 | 9.72E-27 | 9.38E-25 | 1.201686784 |
| ENSMUSG00000024677 | Ms4a6b | 69774 | protein_coding | 17.72182275 | 1.20228263 | 0.410872571 | 2.92616912 | 0.003431643 | 0.020862296 | 1.20228263 |
| ENSMUSG00000036052 | Dnajb5 | 56323 | protein_coding | 69.94965601 | 1.202508741 | 0.242587469 | 4.957010951 | 7.16E-07 | 1.40E-05 | 1.202508741 |
| ENSMUSG00000026042 | Col5a2 | 12832 | protein_coding | 14.57876244 | -1.202601628 | 0.53869593 | -2.23243125 | 0.025586474 | 0.096277194 | 1.202601628 |

| ensembl_gene_id | external_gene_name | entrezgene_id | gene_biotype | baseMean | log2FoldChange | lfcSE | stat | pvalue | padj | absLog2FC |
|---|---|---|---|---|---|---|---|---|---|---|
| ENSMUSG00000054909 | Wbscr25 | 71304 | lncRNA | 14.08479861 | -1.204274813 | 0.365201596 | -3.297561746 | 0.000975282 | 0.007526306 | 1.204274813 |
| ENSMUSG00000075010 | AW112010 | 107350 | lncRNA | 786.9958654 | 1.204310819 | 0.131136522 | 9.183641586 | 4.17E-20 | 2.99E-18 | 1.204310819 |
| ENSMUSG00000019935 | Slc17a8 | 216227 | protein_coding | 18.47015042 | 1.205268754 | 0.312278817 | 3.859591773 | 0.000113577 | 0.001258031 | 1.205268754 |
| ENSMUSG00000024030 | Abcg1 | 11307 | protein_coding | 93.83969554 | 1.205897593 | 0.179217677 | 6.728675487 | 1.71E-11 | 6.28E-10 | 1.205897593 |
| ENSMUSG00000022054 | Nefm | 18040 | protein_coding | 11.12907321 | -1.205899547 | 0.484029486 | -2.491376212 | 0.01272493 | 0.057368436 | 1.205899547 |
| ENSMUSG00000043833 | 2900005J15Rik | 67261 | lncRNA | 12.24774749 | -1.207455784 | 0.447253123 | -2.699714596 | 0.006939898 | 0.036206583 | 1.207455784 |
| ENSMUSG00000042751 | Nmnat2 | 226518 | protein_coding | 19.15687807 | -1.207568039 | 0.348541683 | -3.464630191 | 0.000530961 | 0.004581337 | 1.207568039 |
| ENSMUSG00000021136 | Smoc1 | 64075 | protein_coding | 8.341580165 | -1.20855475 | 0.555155146 | -2.176967573 | 0.029482987 | 0.106502223 | 1.20855475 |
| ENSMUSG00000027947 | Il6ra | 16194 | protein_coding | 39.74941335 | -1.20895447 | 0.348377967 | -3.470238033 | 0.000519997 | 0.004496481 | 1.20895447 |
| ENSMUSG00000038628 | Polr3k | 67005 | protein_coding | 896.7014994 | 1.209573124 | 0.100525728 | 12.03247319 | 2.40E-33 | 3.16E-31 | 1.209573124 |
| ENSMUSG00000020368 | Canx | 12330 | protein_coding | 18546.93463 | 1.212951661 | 0.085014329 | 14.26761429 | 3.48E-46 | 7.30E-44 | 1.212951661 |
| ENSMUSG00000020916 | Krt36 | 16673 | protein_coding | 27.47606403 | -1.21312346 | 0.546107678 | -2.221399752 | 0.026323898 | 0.098124747 | 1.21312346 |
| ENSMUSG00000030411 | Nova2 | 384569 | protein_coding | 7.3765828 | -1.213787049 | 0.939559727 | -1.291867897 | 0.196402897 | 0.389983519 | 1.213787049 |
| ENSMUSG00000053137 | Mapk11 | 19094 | protein_coding | 28.08361443 | -1.21637098 | 0.598818995 | -2.031283225 | 0.042226272 | 0.138118269 | 1.21637098 |
| ENSMUSG00000022678 | Nde1 | 67203 | protein_coding | 673.7540432 | 1.217244242 | 0.121497405 | 10.01868509 | 1.26E-23 | 1.06E-21 | 1.217244242 |
| ENSMUSG00000005360 | Slc1a3 | 20512 | protein_coding | 24.7205556 | -1.217830376 | 0.391298798 | -3.112277326 | 0.0018565 | 0.012712955 | 1.217830376 |
| ENSMUSG00000035232 | Pdk3 | 236900 | protein_coding | 717.0173681 | 1.218073075 | 0.111055038 | 10.96819286 | 5.43E-28 | 5.66E-26 | 1.218073075 |
| ENSMUSG00000027397 | Slc20a1 | 20515 | protein_coding | 4877.228567 | 1.21848712 | 0.167738875 | 7.264190388 | 3.75E-13 | 1.62E-11 | 1.21848712 |
| ENSMUSG00000097610 | A930012L18Rik | 626275 | lncRNA | 7.542064996 | -1.218817367 | 0.509064481 | -2.394229833 | 0.016655309 | 0.069940879 | 1.218817367 |
| ENSMUSG00000058881 | Zfp516 | 329003 | protein_coding | 352.6526807 | 1.219418189 | 0.192852835 | 6.323050374 | 2.56E-10 | 8.34E-09 | 1.219418189 |
| ENSMUSG00000074652 | Myh7b | 668940 | protein_coding | 7.97048267 | -1.221271292 | 0.654523441 | -1.865893895 | 0.062056212 | 0.17971479 | 1.221271292 |
| ENSMUSG00000039278 | Pcsk1n | 30052 | protein_coding | 18.08299502 | -1.221476261 | 0.405081651 | -3.015382843 | 0.002566551 | 0.016611142 | 1.221476261 |
| ENSMUSG00000036492 | Rnf39 | 386454 | protein_coding | 127.3688345 | -1.221799522 | 0.253853594 | -4.813008556 | 1.49E-06 | 2.69E-05 | 1.221799522 |
| ENSMUSG00000001053 | N4bp3 | 212706 | protein_coding | 180.0552591 | -1.221878591 | 0.634858665 | -1.92464663 | 0.054273582 | 0.164098255 | 1.221878591 |
| ENSMUSG00000028560 | Usp1 | 230484 | protein_coding | 1267.849549 | 1.221901549 | 0.134465593 | 9.087094515 | 1.02E-19 | 7.01E-18 | 1.221901549 |
| ENSMUSG00000037784 | Dzip1l | 72507 | protein_coding | 34.2706044 | -1.221971261 | 0.237317299 | -5.149103192 | 2.62E-07 | 5.60E-06 | 1.221971261 |
| ENSMUSG00000061577 | Adgrg5 | 382045 | protein_coding | 10.8825802 | 1.222309851 | 0.488673247 | 2.501282518 | 0.012374442 | 0.0561219 | 1.222309851 |
| ENSMUSG00000053702 | Nebl | 74103 | protein_coding | 83.37995833 | -1.223877337 | 0.201017484 | -6.088412371 | 1.14E-09 | 3.42E-08 | 1.223877337 |
| ENSMUSG00000029361 | Nos1 | 18125 | protein_coding | 8.675496299 | -1.224959653 | 1.261292229 | -0.971194165 | 0.331451599 | 0.542251548 | 1.224959653 |
| ENSMUSG00000041268 | Dmxl2 | 235380 | protein_coding | 32.50855359 | -1.225562325 | 0.277142116 | -4.422143929 | 9.77E-06 | 0.000145748 | 1.225562325 |
| ENSMUSG00000024165 | Jpt2 | 52009 | protein_coding | 1804.887109 | 1.225587126 | 0.084136661 | 14.56662422 | 4.58E-48 | 1.04E-45 | 1.225587126 |
| ENSMUSG00000029664 | Tfpi2 | 21789 | protein_coding | 263.0940358 | 1.226169779 | 0.157293632 | 7.795419064 | 6.42E-15 | 3.24E-13 | 1.226169779 |
| ENSMUSG00000050463 | Krt78 | 332131 | protein_coding | 10.64419442 | 1.227938954 | 0.553488004 | 2.218546645 | 0.026517583 | 0.098569909 | 1.227938954 |
| ENSMUSG00000020185 | E2f7 | 52679 | protein_coding | 75.77359916 | 1.228235821 | 0.254547821 | 4.825167291 | 1.40E-06 | 2.55E-05 | 1.228235821 |
| ENSMUSG00000022894 | Adamts5 | 23794 | protein_coding | 7.697780505 | -1.228711579 | 0.908835272 | -1.35196291 | 0.176387181 | 0.362749518 | 1.228711579 |
| ENSMUSG00000070802 | Pnma8b | 434128 | protein_coding | 34.26383346 | -1.228948084 | 0.259186599 | -4.74155719 | 2.12E-06 | 3.69E-05 | 1.228948084 |
| ENSMUSG00000062070 | Pgk1 | 18655 | protein_coding | 2792.225627 | 1.230239983 | 0.111453393 | 11.03815635 | 2.50E-28 | 2.64E-26 | 1.230239983 |
| ENSMUSG00000006310 | Zbtb32 | 58206 | protein_coding | 102.9562154 | 1.230786685 | 0.246921952 | 4.984517066 | 6.21E-07 | 1.24E-05 | 1.230786685 |

| ensembl_gene_id | external_gene_name | entrezgene_id | gene_biotype | baseMean | log2FoldChange | lfcSE | stat | pvalue | padj | absLog2FC |
|---|---|---|---|---|---|---|---|---|---|---|
| ENSMUSG00000047798 | Cd300lf | 246746 | protein_coding | 21.46537148 | -1.230789181 | 0.358693637 | -3.431310327 | 0.000600673 | 0.005067542 | 1.230789181 |
| ENSMUSG00000070601 | Vmn2r84 | 625068 | protein_coding | 7.166587582 | -1.230886155 | 1.269097087 | -0.969891246 | 0.332100704 | 0.543034598 | 1.230886155 |
| ENSMUSG00000019433 | Gipc1 | 67903 | protein_coding | 1276.52481 | 1.231967293 | 0.101600117 | 12.12564834 | 7.72E-34 | 1.06E-31 | 1.231967293 |
| ENSMUSG00000025701 | Alox5 | 11689 | protein_coding | 55.61592515 | -1.233019958 | 0.192058613 | -6.420019085 | 1.36E-10 | 4.56E-09 | 1.233019958 |
| ENSMUSG00000020212 | Mdm1 | 17245 | protein_coding | 147.6004117 | 1.233386228 | 0.161217403 | 7.650453399 | 2.00E-14 | 9.67E-13 | 1.233386228 |
| ENSMUSG00000064213 | Defa24 | 503491 | protein_coding | 43.78011084 | -1.233761891 | 1.259788405 | -0.979340568 | 0.327411732 | 0.538518595 | 1.233761891 |
| ENSMUSG00000041895 | Wipi1 | 52639 | protein_coding | 478.333675 | 1.233841126 | 0.108247083 | 11.39837759 | 4.26E-30 | 4.89E-28 | 1.233841126 |
| ENSMUSG00000062488 | Ifit3b | 667370 | protein_coding | 20.23415589 | 1.23594763 | 0.376870226 | 3.279504579 | 0.001039895 | 0.007913737 | 1.23594763 |
| ENSMUSG00000039013 | Siglecf | 233186 | protein_coding | 39.24037236 | -1.236865042 | 0.251388864 | -4.920126632 | 8.65E-07 | 1.65E-05 | 1.236865042 |
| ENSMUSG00000044339 | Alkbh2 | 231642 | protein_coding | 45.64499405 | 1.236964118 | 0.239796424 | 5.158392674 | 2.49E-07 | 5.36E-06 | 1.236964118 |
| ENSMUSG00000003283 | Hck | 15162 | protein_coding | 126.6811012 | -1.238035619 | 0.159303531 | -7.77155167 | 7.75E-15 | 3.86E-13 | 1.238035619 |
| ENSMUSG00000065232 | Gm22973 | 115488085 | snRNA | 105.5181015 | 1.239350718 | 0.30868205 | 4.014975015 | 5.95E-05 | 0.000718719 | 1.239350718 |
| ENSMUSG00000005640 | Insrr | 23920 | protein_coding | 93.20529118 | -1.239448549 | 0.215809386 | -5.743255996 | 9.29E-09 | 2.52E-07 | 1.239448549 |
| ENSMUSG00000031298 | Adgrg2 | 237175 | protein_coding | 15.4183627 | -1.24078233 | 0.356602332 | -3.479456576 | 0.000502432 | 0.00437069 | 1.24078233 |
| ENSMUSG00000020396 | Nefh | 380684 | protein_coding | 7.32475254 | -1.240910388 | 0.454745384 | -2.728802604 | 0.006356474 | 0.033737394 | 1.240910388 |
| ENSMUSG00000041193 | Pla2g5 | 18784 | protein_coding | 350.0500014 | 1.24199194 | 0.308547176 | 4.025290258 | 5.69E-05 | 0.000691369 | 1.24199194 |
| ENSMUSG00000091572 | Vmn2r3 | 637004 | protein_coding | 10.01510084 | -1.244156907 | 1.900333883 | -0.654704375 | 0.512658108 | 0.700310321 | 1.244156907 |
| ENSMUSG00000018593 | Sparc | 20692 | protein_coding | 22.7435835 | -1.245431412 | 0.472259137 | -2.63717801 | 0.008359895 | 0.041902591 | 1.245431412 |
| ENSMUSG00000031748 | Gnao1 | 14681 | protein_coding | 35.49666539 | -1.245467521 | 0.28124343 | -4.428432415 | 9.49E-06 | 0.000142095 | 1.245467521 |
| ENSMUSG00000032323 | Cyp11a1 | 13070 | protein_coding | 6.682522546 | -1.245815406 | 1.145209292 | -1.087849544 | 0.276661533 | 0.484995037 | 1.245815406 |
| ENSMUSG00000057836 | Xlr3a | 22445 | protein_coding | 135.2837193 | 1.24813577 | 0.386442355 | 3.229811 | 0.001238721 | 0.009106042 | 1.24813577 |
| ENSMUSG00000071984 | Fndc1 | 68655 | protein_coding | 9.51389073 | -1.248357059 | 0.788646352 | -1.582911094 | 0.11344173 | 0.269525637 | 1.248357059 |
| ENSMUSG00000060962 | Dmkn | 73712 | protein_coding | 8.364063329 | -1.248493829 | 0.523446426 | -2.385141569 | 0.01707255 | 0.071204917 | 1.248493829 |
| ENSMUSG00000032015 | Pou2f3 | 18988 | protein_coding | 26.30673519 | -1.249198691 | 0.276261887 | -4.521791642 | 6.13E-06 | 9.68E-05 | 1.249198691 |
| ENSMUSG00000070331 | Qrich2 | 217341 | protein_coding | 14.49917033 | 1.249661377 | 0.416159498 | 3.002842378 | 0.002674709 | 0.017164693 | 1.249661377 |
| ENSMUSG00000021266 | Wars1 | 22375 | protein_coding | 2794.548934 | 1.249786332 | 0.158669841 | 7.876647027 | 3.36E-15 | 1.74E-13 | 1.249786332 |
| ENSMUSG00000060181 | Slc35e3 | 215436 | protein_coding | 176.3296477 | -1.251361073 | 0.178600461 | -7.006482888 | 2.44E-12 | 9.63E-11 | 1.251361073 |
| ENSMUSG00000006576 | Slc4a3 | 20536 | protein_coding | 11.77933306 | -1.251944486 | 0.51244121 | -2.443098764 | 0.014561752 | 0.063059698 | 1.251944486 |
| ENSMUSG00000001128 | Cfp | 18636 | protein_coding | 27.27630675 | 1.252101605 | 0.30101581 | 4.159587518 | 3.19E-05 | 0.000417438 | 1.252101605 |
| ENSMUSG00000035941 | Ibtk | 108837 | protein_coding | 3001.905592 | 1.253061211 | 0.106732547 | 11.74019776 | 7.93E-32 | 9.72E-30 | 1.253061211 |
| ENSMUSG00000032515 | Csrnp1 | 215418 | protein_coding | 215.0212874 | -1.253881424 | 0.261959588 | -4.786545258 | 1.70E-06 | 3.02E-05 | 1.253881424 |
| ENSMUSG00000033965 | Slc16a2 | 20502 | protein_coding | 21.96077884 | -1.254766973 | 0.358993087 | -3.495239932 | 0.000473636 | 0.004161098 | 1.254766973 |
| ENSMUSG00000042453 | Reln | 19699 | protein_coding | 13.56303175 | -1.257423483 | 0.869940283 | -1.445413563 | 0.148341756 | 0.325228836 | 1.257423483 |
| ENSMUSG00000026414 | Tnnt2 | 21956 | protein_coding | 39.98228946 | -1.257880794 | 0.234490313 | -5.3643188 | 8.13E-08 | 1.88E-06 | 1.257880794 |
| ENSMUSG00000042641 | Rgsl1 | 240816 | protein_coding | 10.49624242 | -1.25894542 | 0.47360928 | -2.658194157 | 0.007856062 | 0.039927044 | 1.25894542 |
| ENSMUSG00000089712 | Gm15889 | 115486954 | lncRNA | 10.73356447 | -1.259763936 | 0.513945785 | -2.451161142 | 0.014239619 | 0.062105326 | 1.259763936 |
| ENSMUSG00000042682 | Selenok | 80795 | protein_coding | 1359.001463 | 1.262686407 | 0.097905236 | 12.89702632 | 4.68E-38 | 8.01E-36 | 1.262686407 |
| ENSMUSG00000027068 | Dhrs9 | 241452 | protein_coding | 562.8088422 | 1.262960366 | 0.4690263 | 2.692728248 | 0.007087001 | 0.036795654 | 1.262960366 |

| ensembl_gene_id | external_gene_name | entrezgene_id | gene_biotype | baseMean | log2FoldChange | lfcSE | stat | pvalue | padj | absLog2FC |
|---|---|---|---|---|---|---|---|---|---|---|
| ENSMUSG00000030825 | Hsd17b14 | 66065 | protein_coding | 11.10064792 | -1.263093034 | 0.432767733 | -2.918639581 | 0.003515624 | 0.021299683 | 1.263093034 |
| ENSMUSG00000024302 | Dtna | 13527 | protein_coding | 8.629561337 | -1.263761334 | 1.138037796 | -1.110473957 | 0.266794844 | 0.474540288 | 1.263761334 |
| ENSMUSG00000097691 | 9030616G12Rik | 77699 | lncRNA | 16.05009108 | -1.263958684 | 0.324409281 | -3.896185338 | 9.77E-05 | 0.001106995 | 1.263958684 |
| ENSMUSG00000108356 | 9130221F21Rik | 330350 | lncRNA | 21.97542736 | -1.264473947 | 0.617547343 | -2.047574102 | 0.040601749 | 0.134477991 | 1.264473947 |
| ENSMUSG00000071847 | Apcdd1 | 494504 | protein_coding | 8.022090605 | -1.264907023 | 0.759049911 | -1.666434586 | 0.095626887 | 0.241194783 | 1.264907023 |
| ENSMUSG00000034854 | Mfsd12 | 73822 | protein_coding | 312.3451233 | 1.26515379 | 0.133657364 | 9.465649729 | 2.92E-21 | 2.17E-19 | 1.26515379 |
| ENSMUSG00000070291 | Ddx43 | 100048658 | protein_coding | 47.02803651 | 1.265259086 | 0.226347124 | 5.589905732 | 2.27E-08 | 5.80E-07 | 1.265259086 |
| ENSMUSG00000025716 | Myo3a | 667663 | protein_coding | 208.7561973 | -1.265995322 | 0.126312881 | -10.02269377 | 1.21E-23 | 1.03E-21 | 1.265995322 |
| ENSMUSG00000025492 | Ifitm3 | 66141 | protein_coding | 82.58348014 | 1.26652563 | 0.374640033 | 3.380646802 | 0.000723154 | 0.00588973 | 1.26652563 |
| ENSMUSG00000059049 | Frem1 | 329872 | protein_coding | 6.796103817 | -1.267073166 | 1.205675215 | -1.050924121 | 0.293293441 | 0.5022662 | 1.267073166 |
| ENSMUSG00000036097 | Slf2 | 226151 | protein_coding | 1224.007292 | 1.269087959 | 0.102884388 | 12.33508779 | 5.86E-35 | 8.49E-33 | 1.269087959 |
| ENSMUSG00000040152 | Thbs1 | 21825 | protein_coding | 1779.396258 | -1.269793937 | 0.241863543 | -5.250042738 | 1.52E-07 | 3.39E-06 | 1.269793937 |
| ENSMUSG00000031532 | Saraf | 67887 | protein_coding | 2599.244618 | 1.272016276 | 0.097866977 | 12.99740033 | 1.27E-38 | 2.22E-36 | 1.272016276 |
| ENSMUSG00000028906 | Epb41 | 269587 | protein_coding | 2247.972147 | 1.272338513 | 0.069939936 | 18.19187401 | 5.99E-74 | 2.33E-71 | 1.272338513 |
| ENSMUSG00000022064 | Pibf1 | 52023 | protein_coding | 776.5610557 | 1.273026154 | 0.141226373 | 9.014082356 | 1.99E-19 | 1.35E-17 | 1.273026154 |
| ENSMUSG00000049907 | Rasl11b | 68939 | protein_coding | 31.64245311 | 1.274392436 | 0.311223244 | 4.094785526 | 4.23E-05 | 0.000530848 | 1.274392436 |
| ENSMUSG00000027022 | Xirp2 | 241431 | protein_coding | 8.349922166 | -1.274532303 | 1.856866246 | -0.686388859 | 0.492467928 | 0.684171754 | 1.274532303 |
| ENSMUSG00000073409 | H2-Q6 | 110557 | protein_coding | 55.74901954 | 1.276179663 | 0.2636308 | 4.840783647 | 1.29E-06 | 2.37E-05 | 1.276179663 |
| ENSMUSG00000032080 | Apoa4 | 11808 | protein_coding | 26.02436083 | -1.276192772 | 0.697149706 | -1.830586401 | 0.0671623 | 0.189909659 | 1.276192772 |
| ENSMUSG00000038295 | Atg9b | 213948 | protein_coding | 16.71376789 | 1.276799237 | 0.388776601 | 3.284146308 | 0.001022918 | 0.007806918 | 1.276799237 |
| ENSMUSG00000033740 | St18 | 240690 | protein_coding | 48.18358407 | -1.280259678 | 0.239481829 | -5.345957491 | 8.99E-08 | 2.07E-06 | 1.280259678 |
| ENSMUSG00000024912 | Fosl1 | 14283 | protein_coding | 26.3879299 | -1.28135202 | 0.575122174 | -2.227964905 | 0.025882855 | 0.097059447 | 1.28135202 |
| ENSMUSG00000058715 | Fcer1g | 14127 | protein_coding | 11.23539453 | 1.282660572 | 0.37880276 | 3.386090879 | 0.000708959 | 0.005798996 | 1.282660572 |
| ENSMUSG00000110195 | Pde2a | 207728 | protein_coding | 23.5762927 | -1.283037393 | 0.286311883 | -4.481257921 | 7.42E-06 | 0.000115085 | 1.283037393 |
| ENSMUSG00000066513 | Klk1b4 | 18048 | protein_coding | 7.057291275 | -1.285429596 | 0.429993033 | -2.989419588 | 0.00279508 | 0.017744135 | 1.285429596 |
| ENSMUSG00000047250 | Ptgs1 | 19224 | protein_coding | 42.87708326 | -1.286764538 | 0.193632792 | -6.645385441 | 3.02E-11 | 1.08E-09 | 1.286764538 |
| ENSMUSG00000004035 | Gstm7 | 68312 | protein_coding | 223.1289129 | 1.286825662 | 0.132532813 | 9.709487246 | 2.75E-22 | 2.13E-20 | 1.286825662 |
| ENSMUSG00000050663 | Trhde | 237553 | protein_coding | 7.192760553 | -1.287598339 | 1.817084161 | -0.708606881 | 0.478568465 | 0.672843015 | 1.287598339 |
| ENSMUSG00000037362 | Ccn3 | 18133 | protein_coding | 3296.280241 | -1.287818735 | 0.12410573 | -10.37678705 | 3.16E-25 | 2.88E-23 | 1.287818735 |
| ENSMUSG00000111296 | 2010001M07Rik | 115487174 | lncRNA | 9.831139411 | -1.288232079 | 0.441459851 | -2.918118321 | 0.003521507 | 0.021319104 | 1.288232079 |
| ENSMUSG00000035208 | Slfn8 | 276950 | protein_coding | 8.95149991 | 1.28868405 | 0.507460806 | 2.539475041 | 0.011101897 | 0.051720096 | 1.28868405 |
| ENSMUSG00000079339 | Ifit1bl1 | 667373 | protein_coding | 752.9640041 | 1.288800066 | 0.242500336 | 5.31463208 | 1.07E-07 | 2.44E-06 | 1.288800066 |
| ENSMUSG00000030847 | Bag3 | 29810 | protein_coding | 902.2513899 | 1.290093713 | 0.142745294 | 9.037732016 | 1.60E-19 | 1.09E-17 | 1.290093713 |
| ENSMUSG00000046460 | Sh2d7 | 244885 | protein_coding | 145.6643483 | -1.290937693 | 0.150475574 | -8.579051442 | 9.57E-18 | 5.93E-16 | 1.290937693 |
| ENSMUSG00000073405 | H2-T-ps | 667803 | unprocessed_pseudogene | 31.19542152 | 1.291958191 | 0.214949904 | 6.010508335 | 1.85E-09 | 5.43E-08 | 1.291958191 |
| ENSMUSG00000002076 | Hsf2bp | 74377 | protein_coding | 19.3312253 | 1.29520331 | 0.315717513 | 4.102411982 | 4.09E-05 | 0.000516858 | 1.29520331 |
| ENSMUSG00000020175 | Rab36 | 76877 | protein_coding | 13.65974167 | 1.296699517 | 0.500041314 | 2.593184766 | 0.009509167 | 0.046121197 | 1.296699517 |
| ENSMUSG00000059277 | R74862 | 97423 | lncRNA | 6.827146383 | -1.297397399 | 0.543927782 | -2.385238337 | 0.01706806 | 0.071204833 | 1.297397399 |

| ensembl_gene_id | external_gene_name | entrezgene_id | gene_biotype | baseMean | log2FoldChange | lfcSE | stat | pvalue | padj | absLog2FC |
|---|---|---|---|---|---|---|---|---|---|---|
| ENSMUSG00000033327 | Tnxb | 81877 | protein_coding | 21.43786141 | -1.299948808 | 0.443907665 | -2.928421629 | 0.003406877 | 0.020767217 | 1.299948808 |
| ENSMUSG00000051043 | Gprc5c | 70355 | protein_coding | 9.574221658 | -1.300747522 | 0.465050964 | -2.796999948 | 0.005157954 | 0.028723285 | 1.300747522 |
| ENSMUSG00000052477 | C130026I21Rik | 620078 | protein_coding | 26.06423392 | -1.301507039 | 1.911937695 | -0.680726701 | 0.496044438 | 0.687222389 | 1.301507039 |
| ENSMUSG00000053334 | Ficd | 231630 | protein_coding | 166.5029928 | 1.303355799 | 0.098106197 | 13.28515262 | 2.82E-40 | 5.05E-38 | 1.303355799 |
| ENSMUSG00000028278 | Rragd | 52187 | protein_coding | 13.49336171 | -1.303829564 | 0.449083945 | -2.90330923 | 0.003692419 | 0.022085183 | 1.303829564 |
| ENSMUSG00000027514 | Zbp1 | 58203 | protein_coding | 152.2565596 | 1.304516869 | 0.172577559 | 7.559017974 | 4.06E-14 | 1.92E-12 | 1.304516869 |
| ENSMUSG00000078853 | Igtp | 16145 | protein_coding | 134.0650619 | 1.304788626 | 0.159338305 | 8.18879446 | 2.64E-16 | 1.48E-14 | 1.304788626 |
| ENSMUSG00000027674 | Pex5l | 58869 | protein_coding | 47.35801422 | -1.305504405 | 0.200832362 | -6.50046831 | 8.01E-11 | 2.80E-09 | 1.305504405 |
| ENSMUSG00000091006 | Vmn2r69 | 330581 | protein_coding | 8.769241334 | -1.305542187 | 1.683191669 | -0.775634891 | 0.437964593 | 0.640519699 | 1.305542187 |
| ENSMUSG00000075304 | Sp5 | 64406 | protein_coding | 12.20563596 | 1.306394084 | 0.464733218 | 2.811062421 | 0.004937821 | 0.02776195 | 1.306394084 |
| ENSMUSG00000047155 | Cyp4x1 | 81906 | protein_coding | 13.84057469 | -1.306471387 | 0.390063504 | -3.349381248 | 0.000809923 | 0.006440563 | 1.306471387 |
| ENSMUSG00000038550 | Ciart | 229599 | protein_coding | 20.37929934 | -1.307425024 | 0.371209444 | -3.522068319 | 0.000428194 | 0.003842406 | 1.307425024 |
| ENSMUSG00000024538 | Ppic | 19038 | protein_coding | 16.09993975 | -1.311984982 | 0.307804705 | -4.262394173 | 2.02E-05 | 0.000280366 | 1.311984982 |
| ENSMUSG00000044122 | Proca1 | 216974 | protein_coding | 7.734722152 | -1.312127406 | 0.471357018 | -2.783723072 | 0.005373889 | 0.029547084 | 1.312127406 |
| ENSMUSG00000061603 | Akap6 | 238161 | protein_coding | 8.158586739 | -1.312261464 | 1.680765329 | -0.78075234 | 0.43494817 | 0.638013252 | 1.312261464 |
| ENSMUSG00000037849 | Ifi206 | 102639543 | protein_coding | 6.925314976 | 1.314196019 | 0.633057382 | 2.075950863 | 0.037898496 | 0.128162896 | 1.314196019 |
| ENSMUSG00000018442 | Derl2 | 116891 | protein_coding | 1234.341639 | 1.314627357 | 0.092053643 | 14.28110078 | 2.87E-46 | 6.10E-44 | 1.314627357 |
| ENSMUSG00000045275 | Lca5l | 385668 | protein_coding | 6.902863117 | -1.315263839 | 0.666072227 | -1.974656479 | 0.048307145 | 0.152122619 | 1.315263839 |
| ENSMUSG00000038963 | Slco4a1 | 108115 | protein_coding | 83.99139109 | -1.315911593 | 0.202855042 | -6.486955307 | 8.76E-11 | 3.02E-09 | 1.315911593 |
| ENSMUSG00000049608 | Gpr55 | 227326 | protein_coding | 25.92111961 | 1.316971614 | 0.313670147 | 4.19858768 | 2.69E-05 | 0.000357993 | 1.316971614 |
| ENSMUSG00000027820 | Mme | 17380 | protein_coding | 25.34810537 | -1.318230674 | 0.490197605 | -2.689182198 | 0.007162731 | 0.037077668 | 1.318230674 |
| ENSMUSG00000021613 | Hapln1 | 12950 | protein_coding | 24.14548577 | 1.319329688 | 0.306538216 | 4.303964795 | 1.68E-05 | 0.000236898 | 1.319329688 |
| ENSMUSG00000026609 | Ush2a | 22283 | protein_coding | 9.242954184 | -1.321314487 | 1.76396205 | -0.749060609 | 0.453820676 | 0.653272094 | 1.321314487 |
| ENSMUSG00000009246 | Trpm5 | 56843 | protein_coding | 211.6350486 | -1.321776775 | 0.145779415 | -9.066964489 | 1.22E-19 | 8.37E-18 | 1.321776775 |
| ENSMUSG00000086645 | Gm15743 | 100047123 | lncRNA | 9.915599099 | -1.323681076 | 0.440679042 | -3.003730489 | 0.002666915 | 0.017135383 | 1.323681076 |
| ENSMUSG00000001270 | Ckb | 12709 | protein_coding | 315.9162311 | 1.326283984 | 0.25640431 | 5.172627499 | 2.31E-07 | 5.00E-06 | 1.326283984 |
| ENSMUSG00000074364 | Ehd2 | 259300 | protein_coding | 8.57852419 | -1.331439 | 0.594135532 | -2.240968479 | 0.025028117 | 0.094488705 | 1.331439 |
| ENSMUSG00000062151 | Unc13c | 208898 | protein_coding | 7.109717456 | -1.332486631 | 0.920754488 | -1.447168218 | 0.147849816 | 0.324686593 | 1.332486631 |
| ENSMUSG00000043629 | 1700019D03Rik | 67080 | protein_coding | 365.8496632 | 1.333953569 | 0.126765459 | 10.52300507 | 6.77E-26 | 6.30E-24 | 1.333953569 |
| ENSMUSG00000050473 | Slc35d3 | 76157 | protein_coding | 9.061431785 | -1.334593543 | 0.440078601 | -3.032625399 | 0.002424363 | 0.015910697 | 1.334593543 |
| ENSMUSG00000037605 | Adgrl3 | 319387 | protein_coding | 11.21797232 | -1.335213174 | 1.647982397 | -0.810210823 | 0.417819018 | 0.624064265 | 1.335213174 |
| ENSMUSG00000001943 | Vsig2 | 57276 | protein_coding | 168.4842868 | -1.335445517 | 0.187405279 | -7.125975972 | 1.03E-12 | 4.29E-11 | 1.335445517 |
| ENSMUSG00000030302 | Atp2b2 | 11941 | protein_coding | 7.464968277 | -1.336530017 | 1.160016096 | -1.152165062 | 0.249253252 | 0.45554857 | 1.336530017 |
| ENSMUSG00000018620 | Mmp20 | 30800 | protein_coding | 6.922393777 | 1.336709953 | 0.70067976 | 1.907733075 | 0.056425725 | 0.168652456 | 1.336709953 |
| ENSMUSG00000091649 | Phf11b | 236451 | protein_coding | 74.01359135 | 1.339981816 | 0.237466662 | 5.642820785 | 1.67E-08 | 4.38E-07 | 1.339981816 |
| ENSMUSG00000028036 | Ptgfr | 19220 | protein_coding | 13.25261921 | -1.34025772 | 0.413194423 | -3.243649101 | 0.00118009 | 0.008779453 | 1.34025772 |
| ENSMUSG00000060550 | H2-Q7 | 15018 | protein_coding | 77.91823939 | 1.341609132 | 0.21753041 | 6.167455536 | 6.94E-10 | 2.12E-08 | 1.341609132 |
| ENSMUSG00000036110 | Slc17a2 | 218103 | protein_coding | 26.03950281 | -1.342437937 | 0.270258731 | -4.96723245 | 6.79E-07 | 1.34E-05 | 1.342437937 |

| ensembl_gene_id | external_gene_name | entrezgene_id | gene_biotype | baseMean | log2FoldChange | lfcSE | stat | pvalue | padj | absLog2FC |
|---|---|---|---|---|---|---|---|---|---|---|
| ENSMUSG00000050628 | Ubald2 | 319370 | protein_coding | 878.8245133 | 1.348161242 | 0.158028637 | 8.531119875 | 1.45E-17 | 8.88E-16 | 1.348161242 |
| ENSMUSG00000059708 | Akap17b | 338351 | protein_coding | 74.82966834 | 1.348195938 | 0.169697662 | 7.944693659 | 1.95E-15 | 1.04E-13 | 1.348195938 |
| ENSMUSG00000022146 | Osmr | 18414 | protein_coding | 25.82672635 | 1.348954625 | 0.718054063 | 1.878625433 | 0.060295656 | 0.176412419 | 1.348954625 |
| ENSMUSG00000033788 | Dysf | 26903 | protein_coding | 6.90339937 | -1.349818026 | 1.113932064 | -1.211759738 | 0.225604364 | 0.427890726 | 1.349818026 |
| ENSMUSG00000060594 | Layn | 244864 | protein_coding | 6.687054552 | -1.349960541 | 0.50647094 | -2.665425468 | 0.007689097 | 0.039253828 | 1.349960541 |
| ENSMUSG00000094584 | Ms4a18 | 76002 | protein_coding | 16.46779436 | 1.350780232 | 0.379481944 | 3.559537557 | 0.000371508 | 0.003412564 | 1.350780232 |
| ENSMUSG00000074896 | Ifit3 | 15959 | protein_coding | 50.0251221 | 1.351635293 | 0.363540921 | 3.717972898 | 0.000200828 | 0.002050503 | 1.351635293 |
| ENSMUSG00000036111 | Lmo1 | 109594 | protein_coding | 7.416884856 | 1.353400754 | 0.432297472 | 3.130716326 | 0.001743805 | 0.01207623 | 1.353400754 |
| ENSMUSG00000031486 | Adgra2 | 78560 | protein_coding | 11.35898809 | -1.353865741 | 0.388884199 | -3.481411036 | 0.000498779 | 0.004341289 | 1.353865741 |
| ENSMUSG00000066456 | Hmgn3 | 94353 | protein_coding | 8.902620942 | -1.353945848 | 0.412016348 | -3.286145939 | 0.001015684 | 0.007762865 | 1.353945848 |
| ENSMUSG00000054580 | Pla2r1 | 18779 | protein_coding | 7.674463399 | -1.355816132 | 0.876115092 | -1.547531989 | 0.121735017 | 0.284133227 | 1.355816132 |
| ENSMUSG00000030605 | Mfge8 | 17304 | protein_coding | 407.4669461 | 1.356768969 | 0.131580791 | 10.31129965 | 6.26E-25 | 5.64E-23 | 1.356768969 |
| ENSMUSG00000020334 | Slc22a4 | 30805 | protein_coding | 16.7610825 | -1.357708123 | 0.351126719 | -3.866718334 | 0.00011031 | 0.001227822 | 1.357708123 |
| ENSMUSG00000045004 | Spata21 | 329972 | protein_coding | 8.808785443 | -1.358031445 | 0.550392351 | -2.467387932 | 0.013610282 | 0.060214816 | 1.358031445 |
| ENSMUSG00000020044 | Timp3 | 21859 | protein_coding | 41.00949132 | -1.358461226 | 0.247282424 | -5.493561596 | 3.94E-08 | 9.56E-07 | 1.358461226 |
| ENSMUSG00000018906 | P4ha2 | 18452 | protein_coding | 110.6560915 | 1.359561486 | 0.209329907 | 6.494826764 | 8.31E-11 | 2.88E-09 | 1.359561486 |
| ENSMUSG00000025433 | Crisp3 | 11572 | protein_coding | 27.55529231 | -1.361587283 | 0.339259393 | -4.013410718 | 5.98E-05 | 0.000721615 | 1.361587283 |
| ENSMUSG00000001493 | Meox1 | 17285 | protein_coding | 11.49281925 | 1.364254259 | 0.501741665 | 2.719037213 | 0.006547224 | 0.034565524 | 1.364254259 |
| ENSMUSG00000022757 | Tfg | 21787 | protein_coding | 1668.674611 | 1.364857822 | 0.095007969 | 14.36571938 | 8.49E-47 | 1.88E-44 | 1.364857822 |
| ENSMUSG00000039954 | Stk32a | 269019 | protein_coding | 18.01424684 | -1.36513995 | 0.349889995 | -3.901626132 | 9.55E-05 | 0.00108552 | 1.36513995 |
| ENSMUSG00000027797 | Dclk1 | 13175 | protein_coding | 335.2825354 | -1.365441318 | 0.156721168 | -8.71255195 | 2.97E-18 | 1.90E-16 | 1.365441318 |
| ENSMUSG00000079243 | Xirp1 | 22437 | protein_coding | 7.938633334 | -1.365953454 | 0.662253768 | -2.062583136 | 0.039152251 | 0.131177337 | 1.365953454 |
| ENSMUSG00000028339 | Col15a1 | 12819 | protein_coding | 18.23531181 | -1.367121717 | 0.376515182 | -3.630986957 | 0.000282339 | 0.002700691 | 1.367121717 |
| ENSMUSG00000024028 | Tff2 | 21785 | protein_coding | 7.980871792 | 1.368503821 | 0.472431549 | 2.896724033 | 0.003770813 | 0.022452901 | 1.368503821 |
| ENSMUSG00000034675 | Dbn1 | 56320 | protein_coding | 17.07943811 | -1.373681821 | 0.317296975 | -4.3293253 | 1.50E-05 | 0.000213468 | 1.373681821 |
| ENSMUSG00000053093 | Myh7 | 140781 | protein_coding | 8.493843756 | -1.374675196 | 0.775705417 | -1.772161398 | 0.076367769 | 0.206762847 | 1.374675196 |
| ENSMUSG00000026520 | Pycr2 | 69051 | protein_coding | 528.3500027 | 1.375293795 | 0.166822045 | 8.244077046 | 1.66E-16 | 9.54E-15 | 1.375293795 |
| ENSMUSG00000012017 | Scarf2 | 224024 | protein_coding | 34.879826 | 1.375494678 | 0.210404648 | 6.537377826 | 6.26E-11 | 2.21E-09 | 1.375494678 |
| ENSMUSG00000036887 | C1qa | 12259 | protein_coding | 18.66447583 | -1.375934867 | 0.57894753 | -2.376614104 | 0.017472356 | 0.072417303 | 1.375934867 |
| ENSMUSG00000026875 | Traf1 | 22029 | protein_coding | 111.0681562 | -1.37728409 | 0.215288139 | -6.397398827 | 1.58E-10 | 5.26E-09 | 1.37728409 |
| ENSMUSG00000068196 | Col8a1 | 12837 | protein_coding | 557.5024651 | -1.377478505 | 0.200583897 | -6.867343427 | 6.54E-12 | 2.49E-10 | 1.377478505 |
| ENSMUSG00000055945 | Prr18 | 320111 | protein_coding | 7.6003198 | -1.378564449 | 0.447521345 | -3.08044402 | 0.002066922 | 0.013908718 | 1.378564449 |
| ENSMUSG00000026360 | Rgs2 | 19735 | protein_coding | 389.369526 | 1.381195153 | 0.195929417 | 7.049452692 | 1.80E-12 | 7.22E-11 | 1.381195153 |
| ENSMUSG00000031253 | Srpx2 | 68792 | protein_coding | 11.4163372 | -1.382297659 | 0.335255846 | -4.12311277 | 3.74E-05 | 0.000477823 | 1.382297659 |
| ENSMUSG00000049252 | Lrp1b | 94217 | protein_coding | 14.06574045 | -1.382319219 | 1.798800303 | -0.768467304 | 0.442209608 | 0.643537199 | 1.382319219 |
| ENSMUSG00000024793 | Tnfrsf25 | 85030 | protein_coding | 12.1316085 | -1.382577203 | 0.413782429 | -3.341314437 | 0.000833827 | 0.006607109 | 1.382577203 |
| ENSMUSG00000013629 | Cad | 69719 | protein_coding | 1005.345078 | 1.385685457 | 0.109565344 | 12.6471145 | 1.16E-36 | 1.78E-34 | 1.385685457 |
| ENSMUSG00000037627 | Rgs22 | 626596 | protein_coding | 9.504879467 | -1.386783974 | 0.640830949 | -2.16404026 | 0.03046126 | 0.109251733 | 1.386783974 |

| ensembl_gene_id | external_gene_name | entrezgene_id | gene_biotype | baseMean | log2FoldChange | lfcSE | stat | pvalue | padj | absLog2FC |
|---|---|---|---|---|---|---|---|---|---|---|
| ENSMUSG00000037921 | Ddx60 | 234311 | protein_coding | 419.6758284 | 1.388454061 | 0.218400187 | 6.357384954 | 2.05E-10 | 6.74E-09 | 1.388454061 |
| ENSMUSG00000028451 | 1700022I11Rik | 67317 | protein_coding | 8.771509563 | 1.393547871 | 0.464967051 | 2.99708951 | 0.002725707 | 0.017388158 | 1.393547871 |
| ENSMUSG00000022262 | Dnah5 | 110082 | protein_coding | 134.474639 | -1.39384193 | 0.19713076 | -7.070646569 | 1.54E-12 | 6.25E-11 | 1.39384193 |
| ENSMUSG00000009628 | Tex15 | 104271 | protein_coding | 17.33608085 | 1.397105607 | 0.354876994 | 3.936872859 | 8.26E-05 | 0.000953489 | 1.397105607 |
| ENSMUSG00000045896 | Paip2b | 232164 | protein_coding | 1002.450315 | 1.398447489 | 0.125299458 | 11.16084233 | 6.34E-29 | 6.87E-27 | 1.398447489 |
| ENSMUSG00000029661 | Col1a2 | 12843 | protein_coding | 39.93826201 | -1.398581473 | 0.408344282 | -3.425005651 | 0.000614787 | 0.005167452 | 1.398581473 |
| ENSMUSG00000025889 | Snca | 20617 | protein_coding | 8.801434292 | 1.399569335 | 0.576585855 | 2.427339003 | 0.015210036 | 0.065363632 | 1.399569335 |
| ENSMUSG00000099907 | Gm10421 | 100038689 | lncRNA | 6.82006314 | -1.400713363 | 0.614770598 | -2.278432586 | 0.022700817 | 0.087996741 | 1.400713363 |
| ENSMUSG00000019429 | Ffar3 | 233080 | protein_coding | 13.42187105 | -1.403906617 | 0.361137577 | -3.887456484 | 0.0001013 | 0.00113963 | 1.403906617 |
| ENSMUSG00000032355 | Mlip | 69642 | protein_coding | 9.492810205 | -1.407024172 | 0.646871326 | -2.175122185 | 0.029620961 | 0.106863573 | 1.407024172 |
| ENSMUSG00000052631 | Sh2d6 | 71130 | protein_coding | 59.79363914 | -1.407370308 | 0.226719114 | -6.207550308 | 5.38E-10 | 1.68E-08 | 1.407370308 |
| ENSMUSG00000073457 | Smok2b | 236574 | protein_coding | 7.601535885 | -1.407827504 | 1.790533418 | -0.786261507 | 0.431714299 | 0.635580492 | 1.407827504 |
| ENSMUSG00000020053 | Igf1 | 16000 | protein_coding | 6.928893824 | -1.411753292 | 0.932988151 | -1.513152435 | 0.130240943 | 0.297032895 | 1.411753292 |
| ENSMUSG00000030592 | Ryr1 | 20190 | protein_coding | 19.14258636 | -1.412037219 | 1.456956701 | -0.969168966 | 0.332460892 | 0.543456188 | 1.412037219 |
| ENSMUSG00000047394 | Odf3b | 70113 | protein_coding | 37.27073448 | -1.413111792 | 0.234014719 | -6.038559451 | 1.55E-09 | 4.63E-08 | 1.413111792 |
| ENSMUSG00000019768 | Esr1 | 13982 | protein_coding | 149.6947798 | 1.414295429 | 0.245066819 | 5.771060453 | 7.88E-09 | 2.15E-07 | 1.414295429 |
| ENSMUSG00000028452 | Vcp | 269523 | protein_coding | 5404.970674 | 1.414970736 | 0.101943328 | 13.87997391 | 8.38E-44 | 1.65E-41 | 1.414970736 |
| ENSMUSG00000015355 | Cd48 | 12506 | protein_coding | 10.11279609 | 1.415563255 | 0.462935832 | 3.057795826 | 0.002229714 | 0.014816392 | 1.415563255 |
| ENSMUSG00000020312 | Shc2 | 216148 | protein_coding | 11.03552294 | -1.415694559 | 0.451595921 | -3.134870122 | 0.001719302 | 0.011927284 | 1.415694559 |
| ENSMUSG00000100182 | 1810006J02Rik | 69030 | lncRNA | 10.71719957 | -1.416314384 | 0.372500568 | -3.802180474 | 0.000143428 | 0.001536532 | 1.416314384 |
| ENSMUSG00000038600 | Atp6v0a4 | 140494 | protein_coding | 7.258411549 | -1.416777104 | 0.556189677 | -2.54729126 | 0.010856278 | 0.050860559 | 1.416777104 |
| ENSMUSG00000020321 | Mdh1 | 17449 | protein_coding | 9971.704894 | 1.417086593 | 0.111373273 | 12.72375818 | 4.36E-37 | 7.09E-35 | 1.417086593 |
| ENSMUSG00000097115 | 1810019N24Rik | 69086 | lncRNA | 7.942066592 | 1.417367686 | 0.480752177 | 2.948229364 | 0.003195998 | 0.019738605 | 1.417367686 |
| ENSMUSG00000020581 | Agr2 | 23795 | protein_coding | 42897.662 | 1.419456706 | 0.121671908 | 11.66626484 | 1.90E-31 | 2.27E-29 | 1.419456706 |
| ENSMUSG00000024471 | Myot | 58916 | protein_coding | 11.58222071 | 1.420770807 | 0.546298198 | 2.600723949 | 0.009302728 | 0.045378875 | 1.420770807 |
| ENSMUSG00000022602 | Arc | 11838 | protein_coding | 186.754063 | -1.423053307 | 0.491389578 | -2.895977798 | 0.003779791 | 0.022477025 | 1.423053307 |
| ENSMUSG00000026574 | Dpt | 56429 | protein_coding | 7.644035267 | -1.424315374 | 0.539689487 | -2.639138634 | 0.008311698 | 0.041736674 | 1.424315374 |
| ENSMUSG00000073608 | Gal3st2c | 619597 | protein_coding | 1224.34507 | -1.427034838 | 0.32308681 | -4.416877427 | 1.00E-05 | 0.000149064 | 1.427034838 |
| ENSMUSG00000018865 | Sult4a1 | 29859 | protein_coding | 8.434489595 | 1.427310561 | 0.49197194 | 2.901203192 | 0.003717327 | 0.022200822 | 1.427310561 |
| ENSMUSG00000027828 | Ssr3 | 67437 | protein_coding | 7975.810886 | 1.429288745 | 0.088945093 | 16.06933784 | 4.19E-58 | 1.28E-55 | 1.429288745 |
| ENSMUSG00000053310 | Nrgn | 64011 | protein_coding | 6.839542356 | -1.430740643 | 0.478706794 | -2.988761935 | 0.002801103 | 0.017764119 | 1.430740643 |
| ENSMUSG00000048218 | Amigo2 | 105827 | protein_coding | 26.89701435 | -1.431063957 | 0.330039703 | -4.336035761 | 1.45E-05 | 0.000207802 | 1.431063957 |
| ENSMUSG00000023033 | Scn8a | 20273 | protein_coding | 9.52233926 | -1.431312425 | 1.016424138 | -1.408184214 | 0.159076529 | 0.340376923 | 1.431312425 |
| ENSMUSG00000022415 | Syngr1 | 20972 | protein_coding | 26.47279954 | -1.432603862 | 0.301603916 | -4.749951138 | 2.03E-06 | 3.55E-05 | 1.432603862 |
| ENSMUSG00000017723 | Wfdc2 | 67701 | protein_coding | 1081.041912 | -1.432843871 | 0.250068265 | -5.729810911 | 1.01E-08 | 2.71E-07 | 1.432843871 |
| ENSMUSG00000030708 | Dnajb13 | 69387 | protein_coding | 84.91515018 | -1.433759942 | 0.297958689 | -4.811942041 | 1.49E-06 | 2.71E-05 | 1.433759942 |
| ENSMUSG00000060534 | Dcc | 13176 | protein_coding | 7.69481491 | -1.435017915 | 1.986988058 | -0.722207619 | 0.47016684 | 0.66616046 | 1.435017915 |
| ENSMUSG00000072620 | Slfn2 | 20556 | protein_coding | 45.30708085 | 1.43748376 | 0.364050065 | 3.948588114 | 7.86E-05 | 0.000913315 | 1.43748376 |

| ensembl_gene_id | external_gene_name | entrezgene_id | gene_biotype | baseMean | log2FoldChange | lfcSE | stat | pvalue | padj | absLog2FC |
|---|---|---|---|---|---|---|---|---|---|---|
| ENSMUSG00000001642 | Akr1b3 | 11677 | protein_coding | 1069.40729 | 1.43822957 | 0.129614241 | 11.0962311 | 1.31E-28 | 1.39E-26 | 1.43822957 |
| ENSMUSG00000031551 | Ido1 | 15930 | protein_coding | 28.5839718 | 1.441728718 | 0.505257118 | 2.85345553 | 0.004324658 | 0.025021123 | 1.441728718 |
| ENSMUSG00000045730 | Adrb2 | 11555 | protein_coding | 20.77005835 | -1.443644419 | 0.392186598 | -3.681014159 | 0.000232308 | 0.002315522 | 1.443644419 |
| ENSMUSG00000029467 | Atp2a2 | 11938 | protein_coding | 9795.224961 | 1.445344519 | 0.14235373 | 10.15319035 | 3.21E-24 | 2.82E-22 | 1.445344519 |
| ENSMUSG00000030145 | Zfp248 | 72720 | protein_coding | 12.21647264 | 1.44562476 | 0.455578227 | 3.173164726 | 0.001507869 | 0.010717244 | 1.44562476 |
| ENSMUSG00000002603 | Tgfb1 | 21803 | protein_coding | 18.54466677 | -1.450311869 | 0.362925224 | -3.996172696 | 6.44E-05 | 0.000766339 | 1.450311869 |
| ENSMUSG00000014905 | Dnajb9 | 27362 | protein_coding | 1515.002911 | 1.450769275 | 0.113137914 | 12.82301593 | 1.22E-37 | 2.02E-35 | 1.450769275 |
| ENSMUSG00000028125 | Abca4 | 11304 | protein_coding | 8.760846516 | -1.452900614 | 1.23414894 | -1.177249007 | 0.239096134 | 0.443292191 | 1.452900614 |
| ENSMUSG00000015619 | Gata3 | 14462 | protein_coding | 8.002543306 | 1.455150389 | 1.022676124 | 1.422884875 | 0.154769531 | 0.334441608 | 1.455150389 |
| ENSMUSG00000034923 | Ly6g6f | 433099 | protein_coding | 14.15972924 | -1.455384118 | 0.402325929 | -3.617425606 | 0.000297548 | 0.002819358 | 1.455384118 |
| ENSMUSG00000034684 | Sema3f | 20350 | protein_coding | 36.36453244 | -1.45772608 | 0.26694862 | -5.460699072 | 4.74E-08 | 1.14E-06 | 1.45772608 |
| ENSMUSG00000085034 | Gm11240 | 667768 | lncRNA | 15.91217719 | 1.457899709 | 0.383380838 | 3.802745378 | 0.000143101 | 0.001535929 | 1.457899709 |
| ENSMUSG00000002279 | Lmf1 | 76483 | protein_coding | 780.3314608 | 1.463277458 | 0.086362349 | 16.94346529 | 2.15E-64 | 6.99E-62 | 1.463277458 |
| ENSMUSG00000066687 | Zbtb16 | 235320 | protein_coding | 20.10845417 | -1.465421617 | 0.453722437 | -3.229775517 | 0.001238874 | 0.009106042 | 1.465421617 |
| ENSMUSG00000055567 | Unc80 | 329178 | protein_coding | 17.4749582 | -1.46674224 | 0.617675183 | -2.374617404 | 0.017567149 | 0.072634077 | 1.46674224 |
| ENSMUSG00000025150 | Cbr2 | 12409 | protein_coding | 362.2198843 | 1.469978811 | 0.198334807 | 7.411602802 | 1.25E-13 | 5.60E-12 | 1.469978811 |
| ENSMUSG00000028011 | Tdo2 | 56720 | protein_coding | 14.00040146 | -1.470612031 | 0.502230255 | -2.928162961 | 0.003409713 | 0.020774708 | 1.470612031 |
| ENSMUSG00000014852 | Adamts13 | 279028 | protein_coding | 7.86630717 | -1.475148158 | 0.648455611 | -2.274863741 | 0.022914105 | 0.088564879 | 1.475148158 |
| ENSMUSG00000033569 | Adgrb3 | 210933 | protein_coding | 21.44142927 | -1.479514861 | 0.437517036 | -3.381616573 | 0.000720607 | 0.00587702 | 1.479514861 |
| ENSMUSG00000032818 | Loxhd1 | 240411 | protein_coding | 7.770132694 | -1.480142804 | 1.815522522 | -0.815270968 | 0.414917223 | 0.621303143 | 1.480142804 |
| ENSMUSG00000070424 | Art5 | 11875 | protein_coding | 12.38730925 | 1.482314809 | 0.368153048 | 4.026354845 | 5.66E-05 | 0.000689299 | 1.482314809 |
| ENSMUSG00000040447 | Spns2 | 216892 | protein_coding | 44.61512394 | -1.48420304 | 0.210216477 | -7.060355402 | 1.66E-12 | 6.71E-11 | 1.48420304 |
| ENSMUSG00000049336 | Tenm2 | 23964 | protein_coding | 6.709250746 | -1.484411843 | 1.282707788 | -1.157248639 | 0.247170791 | 0.452801103 | 1.484411843 |
| ENSMUSG00000028655 | Mfsd2a | 76574 | protein_coding | 146.8822497 | 1.485876553 | 0.59894792 | 2.480810942 | 0.013108388 | 0.058714961 | 1.485876553 |
| ENSMUSG00000022816 | Fstl1 | 14314 | protein_coding | 16.22236332 | -1.489293239 | 0.580565858 | -2.565244268 | 0.010310319 | 0.048890374 | 1.489293239 |
| ENSMUSG00000026475 | Rgs16 | 19734 | protein_coding | 30.6390495 | -1.493722422 | 0.207606896 | -7.194955702 | 6.25E-13 | 2.65E-11 | 1.493722422 |
| ENSMUSG00000026170 | Cyp27a1 | 104086 | protein_coding | 111.4856779 | -1.496781163 | 0.179449007 | -8.340983265 | 7.37E-17 | 4.30E-15 | 1.496781163 |
| ENSMUSG00000058618 | Defa39 | 382000 | protein_coding | 8.186484483 | -1.496818117 | 0.988738719 | -1.513866189 | 0.130059781 | 0.296789712 | 1.496818117 |
| ENSMUSG00000046718 | Bst2 | 69550 | protein_coding | 172.0633773 | 1.49697556 | 0.121938638 | 12.27646611 | 1.21E-34 | 1.74E-32 | 1.49697556 |
| ENSMUSG00000006369 | Fbln1 | 14114 | protein_coding | 95.03745473 | 1.497472332 | 0.217934118 | 6.871215694 | 6.37E-12 | 2.43E-10 | 1.497472332 |
| ENSMUSG00000041481 | Serpina3g | 20715 | protein_coding | 9.738603229 | 1.497936936 | 0.442035762 | 3.388723416 | 0.000702188 | 0.005750958 | 1.497936936 |
| ENSMUSG00000020000 | Moxd1 | 59012 | protein_coding | 10.14468412 | 1.498721899 | 0.669283686 | 2.23929244 | 0.025136894 | 0.094854408 | 1.498721899 |
| ENSMUSG00000038151 | Prdm1 | 12142 | protein_coding | 42.86575832 | 1.500308085 | 0.279037235 | 5.376730753 | 7.59E-08 | 1.76E-06 | 1.500308085 |
| ENSMUSG00000072663 | Spef2 | 320277 | protein_coding | 7.522357703 | -1.500526028 | 1.30931806 | -1.146036303 | 0.251780132 | 0.458216818 | 1.500526028 |
| ENSMUSG00000000157 | Itgb2l | 16415 | protein_coding | 164.5890566 | 1.502283481 | 0.270865023 | 5.546243902 | 2.92E-08 | 7.28E-07 | 1.502283481 |
| ENSMUSG00000057346 | Apol9a | 223672 | protein_coding | 52.94414573 | 1.506582355 | 0.288793784 | 5.216810195 | 1.82E-07 | 4.00E-06 | 1.506582355 |
| ENSMUSG00000074004 | B3gnt6 | 272411 | protein_coding | 324.0202209 | -1.506926073 | 0.241379099 | -6.242984902 | 4.29E-10 | 1.36E-08 | 1.506926073 |
| ENSMUSG00000024042 | Sik1 | 17691 | protein_coding | 213.4149807 | -1.507760765 | 0.234594045 | -6.427105884 | 1.30E-10 | 4.37E-09 | 1.507760765 |

| ensembl_gene_id | external_gene_name | entrezgene_id | gene_biotype | baseMean | log2FoldChange | lfcSE | stat | pvalue | padj | absLog2FC |
|---|---|---|---|---|---|---|---|---|---|---|
| ENSMUSG00000055150 | Zfp78 | 330463 | protein_coding | 11.83087083 | -1.509525763 | 0.431153231 | -3.501135221 | 0.000463281 | 0.0040859 | 1.509525763 |
| ENSMUSG00000037868 | Egr2 | 13654 | protein_coding | 57.90540136 | -1.510138656 | 0.39348305 | -3.837874736 | 0.000124104 | 0.001357641 | 1.510138656 |
| ENSMUSG00000032171 | Pin1 | 23988 | protein_coding | 497.1910545 | 1.510396397 | 0.157855032 | 9.568249936 | 1.09E-21 | 8.25E-20 | 1.510396397 |
| ENSMUSG00000110125 | Gm33148 | 102635939 | lncRNA | 17.13799066 | -1.511122584 | 1.818276356 | -0.831074209 | 0.405931708 | 0.612976892 | 1.511122584 |
| ENSMUSG00000037003 | Tns2 | 209039 | protein_coding | 287.5975672 | 1.514171382 | 0.143919325 | 10.52097331 | 6.92E-26 | 6.40E-24 | 1.514171382 |
| ENSMUSG00000023034 | Nr4a1 | 15370 | protein_coding | 750.4497555 | -1.516986967 | 0.397516451 | -3.816161475 | 0.000135544 | 0.00146767 | 1.516986967 |
| ENSMUSG00000041096 | Tspyl2 | 52808 | protein_coding | 177.3247717 | 1.517689234 | 0.170327224 | 8.910432522 | 5.08E-19 | 3.35E-17 | 1.517689234 |
| ENSMUSG00000029591 | Ung | 22256 | protein_coding | 273.6068941 | 1.520874966 | 0.146266605 | 10.39796451 | 2.53E-25 | 2.33E-23 | 1.520874966 |
| ENSMUSG00000078920 | Ifi47 | 15953 | protein_coding | 163.6630119 | 1.52714919 | 0.192519071 | 7.932456666 | 2.15E-15 | 1.14E-13 | 1.52714919 |
| ENSMUSG00000071661 | Zbtb3 | 75291 | protein_coding | 40.87188608 | 1.528146417 | 0.241338865 | 6.331953276 | 2.42E-10 | 7.89E-09 | 1.528146417 |
| ENSMUSG00000038539 | Atf5 | 107503 | protein_coding | 170.5192544 | 1.533173259 | 0.18899546 | 8.112222687 | 4.97E-16 | 2.75E-14 | 1.533173259 |
| ENSMUSG00000060487 | Samd5 | 320825 | protein_coding | 88.63394339 | -1.533263719 | 0.258146915 | -5.939500453 | 2.86E-09 | 8.25E-08 | 1.533263719 |
| ENSMUSG00000062028 | Irgc1 | 210145 | protein_coding | 8.798941175 | -1.534166464 | 0.533865909 | -2.873692507 | 0.004057037 | 0.023748803 | 1.534166464 |
| ENSMUSG00000026421 | Csrp1 | 13007 | protein_coding | 977.6166462 | 1.53726416 | 0.145097604 | 10.59469016 | 3.15E-26 | 2.99E-24 | 1.53726416 |
| ENSMUSG00000050737 | Ptges | 64292 | protein_coding | 38.69160918 | 1.539185961 | 0.2481128 | 6.203573376 | 5.52E-10 | 1.71E-08 | 1.539185961 |
| ENSMUSG00000031382 | Asb11 | 68854 | protein_coding | 10.88483544 | -1.540412499 | 0.370972332 | -4.152364919 | 3.29E-05 | 0.000428554 | 1.540412499 |
| ENSMUSG00000039672 | Kcne2 | 246133 | protein_coding | 8.837656548 | -1.541132537 | 0.493862169 | -3.120572165 | 0.001805001 | 0.012421437 | 1.541132537 |
| ENSMUSG00000003273 | Car11 | 12348 | protein_coding | 12.24402506 | -1.541297119 | 0.411966239 | -3.741319005 | 0.000183057 | 0.001888427 | 1.541297119 |
| ENSMUSG00000040328 | Or2v1 | 18356 | protein_coding | 17.8957395 | 1.541512194 | 0.520125111 | 2.963733457 | 0.003039314 | 0.018984391 | 1.541512194 |
| ENSMUSG00000032126 | Hmbs | 15288 | protein_coding | 858.7836213 | 1.542459586 | 0.099495535 | 15.50280211 | 3.32E-54 | 8.40E-52 | 1.542459586 |
| ENSMUSG00000033061 | Resp18 | 19711 | protein_coding | 28.93251924 | -1.543707115 | 0.213325846 | -7.236381061 | 4.61E-13 | 1.97E-11 | 1.543707115 |
| ENSMUSG00000054408 | Spcs3 | 76687 | protein_coding | 3188.686817 | 1.546357513 | 0.076899689 | 20.10876157 | 6.18E-90 | 3.28E-87 | 1.546357513 |
| ENSMUSG00000042349 | Ikbke | 56489 | protein_coding | 1365.507038 | 1.548908596 | 0.096824015 | 15.99715317 | 1.34E-57 | 3.95E-55 | 1.548908596 |
| ENSMUSG00000028613 | Lrp8 | 16975 | protein_coding | 118.3115301 | 1.549995168 | 0.310442015 | 4.992865306 | 5.95E-07 | 1.20E-05 | 1.549995168 |
| ENSMUSG00000040797 | Iqsec3 | 243621 | protein_coding | 8.598835875 | -1.550066435 | 2.080890675 | -0.744905272 | 0.456328988 | 0.655296022 | 1.550066435 |
| ENSMUSG00000025092 | Hspa12a | 73442 | protein_coding | 49.4058032 | 1.550728663 | 0.210766671 | 7.35756113 | 1.87E-13 | 8.20E-12 | 1.550728663 |
| ENSMUSG00000022558 | Mroh1 | 223658 | protein_coding | 1524.330322 | 1.551506883 | 0.091185174 | 17.01490301 | 6.37E-65 | 2.16E-62 | 1.551506883 |
| ENSMUSG00000091017 | Garin4 | 619288 | protein_coding | 16.93499194 | -1.55273308 | 0.563466494 | -2.755679524 | 0.005857035 | 0.031581131 | 1.55273308 |
| ENSMUSG00000004951 | Hspb1 | 15507 | protein_coding | 7.10035731 | -1.554089285 | 0.615586172 | -2.524568216 | 0.011584052 | 0.053481386 | 1.554089285 |
| ENSMUSG00000029616 | Erp29 | 67397 | protein_coding | 1804.259762 | 1.55924356 | 0.109002625 | 14.30464235 | 2.05E-46 | 4.47E-44 | 1.55924356 |
| ENSMUSG00000021313 | Ryr2 | 20191 | protein_coding | 16.24222454 | -1.559913685 | 1.617203057 | -0.964575028 | 0.334757712 | 0.545363694 | 1.559913685 |
| ENSMUSG00000024186 | Rgs11 | 50782 | protein_coding | 32.85222564 | -1.560867598 | 0.272216319 | -5.73392368 | 9.81E-09 | 2.65E-07 | 1.560867598 |
| ENSMUSG00000024008 | Cpne5 | 240058 | protein_coding | 8.939688189 | -1.561095387 | 0.692634318 | -2.253852209 | 0.024205467 | 0.092191458 | 1.561095387 |
| ENSMUSG00000040466 | Blvrb | 233016 | protein_coding | 654.5127382 | 1.561096005 | 0.097957848 | 15.93640572 | 3.54E-57 | 1.03E-54 | 1.561096005 |
| ENSMUSG00000035095 | Fam167a | 219148 | protein_coding | 11.84404129 | -1.561271752 | 0.449301501 | -3.474886572 | 0.000511069 | 0.004433722 | 1.561271752 |
| ENSMUSG00000029373 | Pf4 | 56744 | protein_coding | 7.358908622 | 1.561800201 | 0.601276999 | 2.597472055 | 0.009391276 | 0.045702489 | 1.561800201 |
| ENSMUSG00000026204 | Ptprn | 19275 | protein_coding | 24.74071375 | -1.564609525 | 0.289805119 | -5.398833296 | 6.71E-08 | 1.57E-06 | 1.564609525 |
| ENSMUSG00000074444 | Defa30 | 100038927 | protein_coding | 14.22334218 | -1.565710108 | 1.046579418 | -1.496026084 | 0.134646858 | 0.303818552 | 1.565710108 |

| ensembl_gene_id | external_gene_name | entrezgene_id | gene_biotype | baseMean | log2FoldChange | lfcSE | stat | pvalue | padj | absLog2FC |
|---|---|---|---|---|---|---|---|---|---|---|
| ENSMUSG00000030107 | Usp18 | 24110 | protein_coding | 47.76248648 | 1.568247831 | 0.244763249 | 6.407203038 | 1.48E-10 | 4.95E-09 | 1.568247831 |
| ENSMUSG00000034730 | Adgrb1 | 107831 | protein_coding | 10.77357283 | -1.571586835 | 0.417162832 | -3.767322288 | 0.000165008 | 0.001726081 | 1.571586835 |
| ENSMUSG00000038648 | Creb3l2 | 208647 | protein_coding | 237.0692979 | 1.573424895 | 0.1467958 | 10.71845988 | 8.34E-27 | 8.10E-25 | 1.573424895 |
| ENSMUSG00000040462 | Os9 | 216440 | protein_coding | 4352.603396 | 1.574674616 | 0.08421822 | 18.69755282 | 5.18E-78 | 2.17E-75 | 1.574674616 |
| ENSMUSG00000026307 | Scly | 50880 | protein_coding | 287.978551 | 1.576737083 | 0.116281521 | 13.55965308 | 6.95E-42 | 1.26E-39 | 1.576737083 |
| ENSMUSG00000038738 | Shank1 | 243961 | protein_coding | 38.06611979 | 1.580630026 | 0.542272199 | 2.914827698 | 0.00355885 | 0.021479863 | 1.580630026 |
| ENSMUSG00000032478 | Nme6 | 54369 | protein_coding | 254.1189493 | 1.581688359 | 0.136105385 | 11.62105641 | 3.22E-31 | 3.83E-29 | 1.581688359 |
| ENSMUSG00000024731 | Ms4a10 | 69826 | protein_coding | 44.47462643 | 1.583219112 | 0.231125265 | 6.850047804 | 7.38E-12 | 2.79E-10 | 1.583219112 |
| ENSMUSG00000021919 | Chat | 12647 | protein_coding | 11.80468374 | -1.583724391 | 0.543242275 | -2.915318749 | 0.003553254 | 0.021470499 | 1.583724391 |
| ENSMUSG00000042686 | Jph1 | 57339 | protein_coding | 12.4187578 | 1.591583192 | 0.486620093 | 3.270689423 | 0.001072857 | 0.008125754 | 1.591583192 |
| ENSMUSG00000032383 | Ppib | 19035 | protein_coding | 3703.186292 | 1.591959562 | 0.10139399 | 15.70072901 | 1.50E-55 | 4.04E-53 | 1.591959562 |
| ENSMUSG00000030815 | Phkg2 | 68961 | protein_coding | 270.585691 | 1.59373176 | 0.131644567 | 12.10632382 | 9.78E-34 | 1.33E-31 | 1.59373176 |
| ENSMUSG00000024150 | Mcfd2 | 193813 | protein_coding | 2141.411061 | 1.594903112 | 0.12870559 | 12.39187138 | 2.89E-35 | 4.27E-33 | 1.594903112 |
| ENSMUSG00000036599 | Chst12 | 59031 | protein_coding | 10.36097808 | 1.5950001 | 0.411597627 | 3.875144066 | 0.000106562 | 0.001187763 | 1.5950001 |
| ENSMUSG00000037470 | Uggt1 | 320011 | protein_coding | 5854.008248 | 1.595251255 | 0.062251963 | 25.62571804 | 7.89E-145 | 8.97E-142 | 1.595251255 |
| ENSMUSG00000031924 | Cyb5b | 66427 | protein_coding | 10203.21174 | 1.595505924 | 0.10515729 | 15.17256596 | 5.37E-52 | 1.32E-49 | 1.595505924 |
| ENSMUSG00000026544 | Dusp23 | 68440 | protein_coding | 150.0577562 | 1.60290477 | 0.140149058 | 11.43714264 | 2.73E-30 | 3.19E-28 | 1.60290477 |
| ENSMUSG00000038860 | Garnl3 | 99326 | protein_coding | 12.19474262 | -1.615942598 | 0.436528822 | -3.701800466 | 0.000214075 | 0.00216357 | 1.615942598 |
| ENSMUSG00000027350 | Chgb | 12653 | protein_coding | 2047.425698 | -1.616382649 | 0.184825156 | -8.745468895 | 2.22E-18 | 1.43E-16 | 1.616382649 |
| ENSMUSG00000049999 | Ppp1r3d | 228966 | protein_coding | 95.70922418 | 1.620266538 | 0.218182749 | 7.426189952 | 1.12E-13 | 5.03E-12 | 1.620266538 |
| ENSMUSG00000052135 | Foxo6 | 329934 | protein_coding | 11.53633529 | -1.624044627 | 0.389499105 | -4.169572165 | 3.05E-05 | 0.00040205 | 1.624044627 |
| ENSMUSG00000024610 | Cd74 | 16149 | protein_coding | 159.7623744 | 1.625457317 | 0.408794976 | 3.976216475 | 7.00E-05 | 0.00082645 | 1.625457317 |
| ENSMUSG00000030209 | Grin2b | 14812 | protein_coding | 23.03025393 | -1.626008726 | 0.815684263 | -1.993429074 | 0.046214483 | 0.147338727 | 1.626008726 |
| ENSMUSG00000095562 | Gm55594 | 170942 | lncRNA | 8.59483975 | -1.632106483 | 0.6407121 | -2.547332075 | 0.010855008 | 0.050860559 | 1.632106483 |
| ENSMUSG00000029561 | Oasl2 | 23962 | protein_coding | 294.0054054 | 1.632364973 | 0.218335353 | 7.476411639 | 7.64E-14 | 3.51E-12 | 1.632364973 |
| ENSMUSG00000020873 | Slc35b1 | 110172 | protein_coding | 1691.222346 | 1.633806032 | 0.123242717 | 13.25681611 | 4.12E-40 | 7.29E-38 | 1.633806032 |
| ENSMUSG00000030284 | Creld1 | 171508 | protein_coding | 176.0810413 | 1.638845428 | 0.131797143 | 12.43460507 | 1.70E-35 | 2.55E-33 | 1.638845428 |
| ENSMUSG00000006204 | Cdcp3 | 71395 | protein_coding | 7.741197507 | -1.644426372 | 2.043843574 | -0.804575455 | 0.421064711 | 0.626854726 | 1.644426372 |
| ENSMUSG00000009406 | Elk1 | 13712 | protein_coding | 141.1445575 | 1.645026679 | 0.1674772 | 9.822391826 | 9.02E-23 | 7.29E-21 | 1.645026679 |
| ENSMUSG00000006386 | Tek | 21687 | protein_coding | 7.093306097 | -1.64557041 | 0.439726504 | -3.742258867 | 0.000182374 | 0.001883817 | 1.64557041 |
| ENSMUSG00000028270 | Gbp2 | 14469 | protein_coding | 115.5950857 | 1.647066017 | 0.27943374 | 5.894299012 | 3.76E-09 | 1.07E-07 | 1.647066017 |
| ENSMUSG00000006435 | Neurl1a | 18011 | protein_coding | 34.62253945 | -1.650351289 | 0.193441334 | -8.531533854 | 1.44E-17 | 8.88E-16 | 1.650351289 |
| ENSMUSG00000040046 | Tph1 | 21990 | protein_coding | 348.3052401 | -1.652871882 | 0.255816901 | -6.461151996 | 1.04E-10 | 3.54E-09 | 1.652871882 |
| ENSMUSG00000011751 | Sptbn4 | 80297 | protein_coding | 48.29506733 | 1.657966288 | 0.308113341 | 5.381027257 | 7.41E-08 | 1.72E-06 | 1.657966288 |
| ENSMUSG00000057074 | Ces1g | 12623 | protein_coding | 1322.914261 | -1.661488213 | 0.34762165 | -4.779587849 | 1.76E-06 | 3.11E-05 | 1.661488213 |
| ENSMUSG00000050505 | Pcdh20 | 219257 | protein_coding | 10.4669742 | -1.66340613 | 0.399018413 | -4.168745294 | 3.06E-05 | 0.000403177 | 1.66340613 |
| ENSMUSG00000033389 | Arhgap44 | 216831 | protein_coding | 13.56207156 | -1.664170161 | 0.45837984 | -3.630548327 | 0.00028282 | 0.002700691 | 1.664170161 |
| ENSMUSG00000095385 | D630033O11Rik | 235302 | lncRNA | 23.33988358 | -1.675580588 | 0.303400337 | -5.522672138 | 3.34E-08 | 8.22E-07 | 1.675580588 |

Supplementary Table 1

| ensembl_gene_id | external_gene_name | entrezgene_id | gene_biotype | baseMean | log2FoldChange | lfcSE | stat | pvalue | padj | absLog2FC |
|---|---|---|---|---|---|---|---|---|---|---|
| ENSMUSG00000075266 | Cenpw | 66311 | protein_coding | 139.7089284 | 1.678575746 | 0.170083747 | 9.869113164 | 5.67E-23 | 4.68E-21 | 1.678575746 |
| ENSMUSG00000044937 | Ttc41 | 103220 | protein_coding | 57.13333795 | 1.682755157 | 0.23435381 | 7.180404522 | 6.95E-13 | 2.93E-11 | 1.682755157 |
| ENSMUSG00000056258 | Kcnq3 | 110862 | protein_coding | 7.020388619 | -1.682824073 | 1.774469962 | -0.948353091 | 0.342949732 | 0.553138277 | 1.682824073 |
| ENSMUSG00000022211 | Carmil3 | 268747 | protein_coding | 11.51972722 | -1.684773581 | 0.408439531 | -4.124903325 | 3.71E-05 | 0.000475648 | 1.684773581 |
| ENSMUSG00000024856 | Cdk2ap2 | 52004 | protein_coding | 1551.929999 | 1.685853928 | 0.112295286 | 15.01268651 | 6.06E-51 | 1.46E-48 | 1.685853928 |
| ENSMUSG00000036853 | Mcoln3 | 171166 | protein_coding | 23.08825268 | 1.690170158 | 0.411167489 | 4.110660987 | 3.95E-05 | 0.00050112 | 1.690170158 |
| ENSMUSG00000040204 | Pclaf | 68026 | protein_coding | 1296.32588 | 1.691952642 | 0.175993656 | 9.613713808 | 7.00E-22 | 5.38E-20 | 1.691952642 |
| ENSMUSG00000040694 | Apobec2 | 11811 | protein_coding | 38.75123423 | 1.695266881 | 0.387950985 | 4.369796558 | 1.24E-05 | 0.000180899 | 1.695266881 |
| ENSMUSG00000046546 | Fam43a | 224093 | protein_coding | 85.46003196 | -1.695482993 | 0.30837 | -5.498209927 | 3.84E-08 | 9.34E-07 | 1.695482993 |
| ENSMUSG00000027204 | Fbn1 | 14118 | protein_coding | 81.39476714 | -1.708711749 | 0.342388064 | -4.990570431 | 6.02E-07 | 1.21E-05 | 1.708711749 |
| ENSMUSG00000028356 | Ambp | 11699 | protein_coding | 23.19196332 | -1.712874348 | 0.283690522 | -6.037827191 | 1.56E-09 | 4.64E-08 | 1.712874348 |
| ENSMUSG00000020787 | P2rx1 | 18436 | protein_coding | 24.02857965 | -1.715033835 | 0.368931741 | -4.648648098 | 3.34E-06 | 5.58E-05 | 1.715033835 |
| ENSMUSG00000101009 | 1700108F19Rik | 73272 | lncRNA | 11.75182432 | 1.719183003 | 0.417797114 | 4.114875247 | 3.87E-05 | 0.000494022 | 1.719183003 |
| ENSMUSG00000034040 | Galnt17 | 212996 | protein_coding | 7.08040961 | -1.719676626 | 0.781868207 | -2.199445648 | 0.02784625 | 0.102079417 | 1.719676626 |
| ENSMUSG00000006313 | Upk1a | 109637 | protein_coding | 208.144169 | 1.720275415 | 0.172219235 | 9.98886923 | 1.71E-23 | 1.43E-21 | 1.720275415 |
| ENSMUSG00000015599 | Ttbk1 | 106763 | protein_coding | 8.645465935 | -1.722383667 | 0.977485342 | -1.76205575 | 0.078059879 | 0.210143295 | 1.722383667 |
| ENSMUSG00000052229 | Gpr17 | 574402 | protein_coding | 8.25401229 | -1.730795475 | 0.528769859 | -3.273249119 | 0.001063187 | 0.00805635 | 1.730795475 |
| ENSMUSG00000059824 | Dbp | 13170 | protein_coding | 58.468424 | -1.730841533 | 0.317566749 | -5.450323566 | 5.03E-08 | 1.20E-06 | 1.730841533 |
| ENSMUSG00000059743 | Fdps | 110196 | protein_coding | 639.8077454 | 1.744964764 | 0.172486136 | 10.11655085 | 4.67E-24 | 4.02E-22 | 1.744964764 |
| ENSMUSG00000060371 | Caln1 | 140904 | protein_coding | 7.035174515 | -1.746180907 | 2.035453766 | -0.857882864 | 0.390957147 | 0.599142673 | 1.746180907 |
| ENSMUSG00000066279 | Chrna10 | 504186 | protein_coding | 9.495526741 | 1.75038315 | 0.66386946 | 2.636637555 | 0.008373224 | 0.041939848 | 1.75038315 |
| ENSMUSG00000028602 | Tnfrsf8 | 21941 | protein_coding | 18.12393567 | 1.751957515 | 0.558320024 | 3.137909158 | 0.001701576 | 0.011814031 | 1.751957515 |
| ENSMUSG00000041681 | Iapp | 15874 | protein_coding | 9.62481919 | -1.753216937 | 0.497129134 | -3.526683144 | 0.0004208 | 0.003791008 | 1.753216937 |
| ENSMUSG00000020889 | Nr1d1 | 217166 | protein_coding | 309.5577908 | -1.754728542 | 0.317844402 | -5.520715583 | 3.38E-08 | 8.30E-07 | 1.754728542 |
| ENSMUSG00000033355 | Rtp4 | 67775 | protein_coding | 112.5797958 | 1.759715284 | 0.249778746 | 7.045096162 | 1.85E-12 | 7.41E-11 | 1.759715284 |
| ENSMUSG00000070576 | Mn1 | 433938 | protein_coding | 6.875537651 | -1.767995245 | 0.561311568 | -3.149757365 | 0.001634061 | 0.011420503 | 1.767995245 |
| ENSMUSG00000027971 | Ndst4 | 64580 | protein_coding | 8.198684198 | -1.775213726 | 2.006222332 | -0.884853935 | 0.376235415 | 0.584480415 | 1.775213726 |
| ENSMUSG00000068246 | Apol9b | 71898 | protein_coding | 28.0179342 | 1.780786359 | 0.353224988 | 5.041507317 | 4.62E-07 | 9.51E-06 | 1.780786359 |
| ENSMUSG00000024807 | Syvn1 | 74126 | protein_coding | 1078.464006 | 1.784381558 | 0.144583848 | 12.34149996 | 5.41E-35 | 7.91E-33 | 1.784381558 |
| ENSMUSG00000026573 | Xcl1 | 16963 | protein_coding | 20.76377329 | 1.788860969 | 0.345812524 | 5.172921293 | 2.30E-07 | 5.00E-06 | 1.788860969 |
| ENSMUSG00000045466 | Zfp956 | 101197 | protein_coding | 198.3068836 | 1.802199407 | 0.140281984 | 12.84697689 | 8.94E-38 | 1.52E-35 | 1.802199407 |
| ENSMUSG00000017754 | Pltp | 18830 | protein_coding | 22.28294203 | 1.802968712 | 0.350031358 | 5.150877681 | 2.59E-07 | 5.56E-06 | 1.802968712 |
| ENSMUSG00000017002 | Slpi | 20568 | protein_coding | 28.2881929 | 1.80498269 | 0.633371146 | 2.849802524 | 0.004374638 | 0.025236956 | 1.80498269 |
| ENSMUSG00000043067 | Dpy19l1 | 244745 | protein_coding | 225.8590011 | 1.807513936 | 0.143218552 | 12.62066901 | 1.62E-36 | 2.46E-34 | 1.807513936 |
| ENSMUSG00000012428 | Steap4 | 117167 | protein_coding | 74.26949178 | 1.819370657 | 0.293511005 | 6.198645443 | 5.70E-10 | 1.76E-08 | 1.819370657 |
| ENSMUSG00000041592 | Sdk2 | 237979 | protein_coding | 10.36507076 | -1.82105888 | 0.790748685 | -2.302955307 | 0.021281358 | 0.083737517 | 1.82105888 |
| ENSMUSG00000061100 | Retnla | 57262 | protein_coding | 6.753994897 | -1.823200616 | 0.752530491 | -2.422759793 | 0.015403108 | 0.065934077 | 1.823200616 |
| ENSMUSG00000030109 | Slc6a12 | 14411 | protein_coding | 7.579919837 | -1.823418975 | 0.572328562 | -3.185965365 | 0.001442719 | 0.010332564 | 1.823418975 |

| ensembl_gene_id | external_gene_name | entrezgene_id | gene_biotype | baseMean | log2FoldChange | lfcSE | stat | pvalue | padj | absLog2FC |
|---|---|---|---|---|---|---|---|---|---|---|
| ENSMUSG00000032558 | Nphp3 | 74025 | protein_coding | 231.5053092 | 1.826974623 | 0.113188823 | 16.14094551 | 1.32E-58 | 4.11E-56 | 1.826974623 |
| ENSMUSG00000073988 | Ttpa | 50500 | protein_coding | 77.16077835 | 1.830061058 | 0.235419217 | 7.773626469 | 7.63E-15 | 3.82E-13 | 1.830061058 |
| ENSMUSG00000047497 | Adamts12 | 239337 | protein_coding | 6.838172294 | -1.832070271 | 1.08413369 | -1.689893311 | 0.091048367 | 0.233529533 | 1.832070271 |
| ENSMUSG00000030017 | Reg3g | 19695 | protein_coding | 4398.724292 | 1.833378634 | 0.763169215 | 2.402322576 | 0.016291335 | 0.068793316 | 1.833378634 |
| ENSMUSG00000021194 | Chga | 12652 | protein_coding | 848.4749514 | -1.835113102 | 0.187721455 | -9.775723819 | 1.43E-22 | 1.12E-20 | 1.835113102 |
| ENSMUSG00000031168 | Ebp | 13595 | protein_coding | 2422.085018 | 1.838868117 | 0.108183699 | 16.99764514 | 8.55E-65 | 2.84E-62 | 1.838868117 |
| ENSMUSG00000050821 | Fam131a | 78408 | protein_coding | 7.898404911 | -1.84771926 | 0.45834294 | -4.031302983 | 5.55E-05 | 0.000678052 | 1.84771926 |
| ENSMUSG00000044938 | Klhl31 | 244923 | protein_coding | 9.668579461 | 1.854815453 | 0.692833042 | 2.677146356 | 0.007425219 | 0.038232148 | 1.854815453 |
| ENSMUSG00000004929 | Thop1 | 50492 | protein_coding | 949.9597705 | 1.855080467 | 0.130856541 | 14.17644437 | 1.28E-45 | 2.62E-43 | 1.855080467 |
| ENSMUSG00000005338 | Cadm3 | 94332 | protein_coding | 9.318114736 | -1.859813166 | 1.104793776 | -1.683403009 | 0.092297101 | 0.235858853 | 1.859813166 |
| ENSMUSG00000028268 | Gbp3 | 55932 | protein_coding | 65.25453369 | 1.869107727 | 0.302041865 | 6.188240591 | 6.08E-10 | 1.87E-08 | 1.869107727 |
| ENSMUSG00000020990 | Cdkl1 | 71091 | protein_coding | 15.88080339 | -1.88343955 | 0.422378962 | -4.459122537 | 8.23E-06 | 0.000125677 | 1.88343955 |
| ENSMUSG00000049112 | Oxtr | 18430 | protein_coding | 6.933751816 | -1.889687075 | 0.528013111 | -3.578863925 | 0.000345091 | 0.00319014 | 1.889687075 |
| ENSMUSG00000005397 | Nid1 | 18073 | protein_coding | 17.21995344 | -1.891652396 | 0.454009936 | -4.166544048 | 3.09E-05 | 0.000406417 | 1.891652396 |
| ENSMUSG00000061462 | Obscn | 380698 | protein_coding | 22.73775377 | -1.89189192 | 0.486787818 | -3.886481643 | 0.000101708 | 0.00114326 | 1.89189192 |
| ENSMUSG00000086254 | 3100003L05Rik | 73076 | lncRNA | 22.18553505 | -1.892498226 | 0.332581297 | -5.690332684 | 1.27E-08 | 3.36E-07 | 1.892498226 |
| ENSMUSG00000056091 | St3gal5 | 20454 | protein_coding | 80.14140882 | -1.897493511 | 0.680215548 | -2.789547396 | 0.005278177 | 0.029221691 | 1.897493511 |
| ENSMUSG00000091345 | Col6a5 | 665033 | protein_coding | 14.83153005 | -1.89947404 | 0.629721623 | -3.016370998 | 0.0025582 | 0.016563828 | 1.89947404 |
| ENSMUSG00000030104 | Edem1 | 192193 | protein_coding | 5286.933038 | 1.904141368 | 0.086781522 | 21.94178354 | 1.04E-106 | 6.61E-104 | 1.904141368 |
| ENSMUSG00000020826 | Nos2 | 18126 | protein_coding | 52.93148248 | 1.906394518 | 0.443504581 | 4.298477626 | 1.72E-05 | 0.000241981 | 1.906394518 |
| ENSMUSG00000084984 | Far1os | 69295 | lncRNA | 101.9438508 | 1.907863392 | 0.263163606 | 7.249723542 | 4.18E-13 | 1.80E-11 | 1.907863392 |
| ENSMUSG00000000159 | Igsf5 | 72058 | protein_coding | 721.8907999 | 1.914379132 | 0.158695925 | 12.06319026 | 1.65E-33 | 2.19E-31 | 1.914379132 |
| ENSMUSG00000050762 | Prss27 | 213171 | protein_coding | 19.77060895 | 1.919713249 | 0.399132327 | 4.809716271 | 1.51E-06 | 2.73E-05 | 1.919713249 |
| ENSMUSG00000026207 | Speg | 11790 | protein_coding | 8.771902872 | -1.925797587 | 1.267910122 | -1.518875473 | 0.128793844 | 0.294999762 | 1.925797587 |
| ENSMUSG00000112023 | Lilrb4b | 14727 | protein_coding | 8.037003376 | 1.94417816 | 0.656557742 | 2.961168587 | 0.003064741 | 0.019115769 | 1.94417816 |
| ENSMUSG00000006095 | Tbcb | 66411 | protein_coding | 1369.606495 | 1.946470676 | 0.129810139 | 14.99475072 | 7.95E-51 | 1.89E-48 | 1.946470676 |
| ENSMUSG00000087658 | Hotairm1 | 105242658 | lncRNA | 19.81074743 | 1.949338202 | 0.311655799 | 6.254779177 | 3.98E-10 | 1.26E-08 | 1.949338202 |
| ENSMUSG00000003355 | Fkbp11 | 66120 | protein_coding | 244.9153744 | 1.950341116 | 0.154197631 | 12.64832088 | 1.14E-36 | 1.77E-34 | 1.950341116 |
| ENSMUSG00000027933 | Ints3 | 229543 | protein_coding | 2390.220202 | 1.951163463 | 0.141262334 | 13.81234057 | 2.15E-43 | 4.12E-41 | 1.951163463 |
| ENSMUSG00000006519 | Cyba | 13057 | protein_coding | 345.6165214 | -1.955569863 | 0.095222609 | -20.53682286 | 1.01E-93 | 5.96E-91 | 1.955569863 |
| ENSMUSG00000047104 | Pbp2 | 76400 | protein_coding | 8.645630769 | -1.955578891 | 0.734668105 | -2.661853535 | 0.007771169 | 0.039571347 | 1.955578891 |
| ENSMUSG00000020607 | Lratd1 | 105005 | protein_coding | 198.3064443 | 1.958882171 | 0.275858857 | 7.101030539 | 1.24E-12 | 5.07E-11 | 1.958882171 |
| ENSMUSG00000035459 | Stab2 | 192188 | protein_coding | 7.023739291 | -1.96205518 | 0.84011387 | -2.335463383 | 0.019519227 | 0.078530312 | 1.96205518 |
| ENSMUSG00000024112 | Cacna1h | 58226 | protein_coding | 242.79961 | -1.96609738 | 0.237921493 | -8.263639222 | 1.41E-16 | 8.18E-15 | 1.96609738 |
| ENSMUSG00000025130 | P4hb | 18453 | protein_coding | 30501.87341 | 1.970034733 | 0.12499936 | 15.76035861 | 5.83E-56 | 1.60E-53 | 1.970034733 |
| ENSMUSG00000022208 | Jph4 | 319984 | protein_coding | 8.561301254 | -1.971739312 | 0.786743909 | -2.506202195 | 0.012203581 | 0.055552627 | 1.971739312 |
| ENSMUSG00000030087 | Klf15 | 66277 | protein_coding | 9.611364943 | -1.975707997 | 0.469926748 | -4.204289297 | 2.62E-05 | 0.000350261 | 1.975707997 |
| ENSMUSG00000045932 | Ifit2 | 15958 | protein_coding | 98.92468821 | 1.976133514 | 0.291939133 | 6.768991521 | 1.30E-11 | 4.80E-10 | 1.976133514 |

| ensembl_gene_id | external_gene_name | entrezgene_id | gene_biotype | baseMean | log2FoldChange | lfcSE | stat | pvalue | padj | absLog2FC |
|---|---|---|---|---|---|---|---|---|---|---|
| ENSMUSG00000062421 | Arf2 | 11841 | protein_coding | 411.3163686 | 1.976919623 | 0.249685928 | 7.91762532 | 2.42E-15 | 1.27E-13 | 1.976919623 |
| ENSMUSG00000040276 | Pacsin1 | 23969 | protein_coding | 94.6926578 | -1.981014334 | 0.358631 | -5.523823461 | 3.32E-08 | 8.18E-07 | 1.981014334 |
| ENSMUSG00000029641 | Rasl11a | 68895 | protein_coding | 22.40464891 | 2.001766875 | 0.265576999 | 7.5374256 | 4.79E-14 | 2.25E-12 | 2.001766875 |
| ENSMUSG00000068114 | Ccdc134 | 76457 | protein_coding | 448.5839452 | 2.012777953 | 0.212439849 | 9.474578164 | 2.68E-21 | 2.00E-19 | 2.012777953 |
| ENSMUSG00000070476 | Fam217b | 71532 | protein_coding | 80.16296029 | 2.013389099 | 0.209774728 | 9.597863002 | 8.16E-22 | 6.25E-20 | 2.013389099 |
| ENSMUSG00000022650 | Retnlb | 57263 | protein_coding | 2242.40764 | 2.019901229 | 0.321366696 | 6.285347086 | 3.27E-10 | 1.05E-08 | 2.019901229 |
| ENSMUSG00000041120 | Nbl1 | 17965 | protein_coding | 38.34272916 | -2.023240676 | 0.267915872 | -7.551776082 | 4.29E-14 | 2.02E-12 | 2.023240676 |
| ENSMUSG00000026452 | Syt2 | 20980 | protein_coding | 31.90236878 | -2.032120757 | 0.267187536 | -7.605597127 | 2.84E-14 | 1.35E-12 | 2.032120757 |
| ENSMUSG00000024781 | Lipa | 16889 | protein_coding | 1112.147934 | 2.032655944 | 0.171575323 | 11.84701801 | 2.23E-32 | 2.82E-30 | 2.032655944 |
| ENSMUSG00000053398 | Phgdh | 236539 | protein_coding | 387.8945428 | 2.052249195 | 0.206804666 | 9.923611654 | 3.29E-23 | 2.73E-21 | 2.052249195 |
| ENSMUSG00000037973 | Itprid1 | 232016 | protein_coding | 11.97430385 | -2.057530404 | 0.400429966 | -5.138302774 | 2.77E-07 | 5.91E-06 | 2.057530404 |
| ENSMUSG00000038576 | Susd4 | 96935 | protein_coding | 17.57984529 | 2.070469235 | 0.358680554 | 5.772460229 | 7.81E-09 | 2.14E-07 | 2.070469235 |
| ENSMUSG00000026185 | Igfbp5 | 16011 | protein_coding | 11.00978947 | -2.077151051 | 0.653472002 | -3.178638174 | 0.001479687 | 0.010551683 | 2.077151051 |
| ENSMUSG00000034118 | Tpst1 | 22021 | protein_coding | 78.60893244 | 2.078514555 | 0.224885756 | 9.242535382 | 2.41E-20 | 1.74E-18 | 2.078514555 |
| ENSMUSG00000004500 | Zfp324 | 243834 | protein_coding | 432.7910694 | 2.089111822 | 0.192245794 | 10.86687922 | 1.66E-27 | 1.67E-25 | 2.089111822 |
| ENSMUSG00000047554 | Tmem41b | 233724 | protein_coding | 1368.896243 | 2.092748648 | 0.100234816 | 20.87846057 | 8.41E-97 | 5.15E-94 | 2.092748648 |
| ENSMUSG00000025408 | Ddit3 | 13198 | protein_coding | 615.9903795 | 2.095062685 | 0.154234951 | 13.58357925 | 5.01E-42 | 9.18E-40 | 2.095062685 |
| ENSMUSG00000024640 | Psat1 | 107272 | protein_coding | 297.0869844 | 2.102588132 | 0.25799903 | 8.149597032 | 3.65E-16 | 2.03E-14 | 2.102588132 |
| ENSMUSG00000004892 | Bcan | 12032 | protein_coding | 6.836399285 | -2.113392995 | 0.57909269 | -3.649490024 | 0.000262761 | 0.002553547 | 2.113392995 |
| ENSMUSG00000034532 | Fbxo16 | 50759 | protein_coding | 9.411262643 | 2.121290455 | 0.650752863 | 3.259748171 | 0.001115112 | 0.008389939 | 2.121290455 |
| ENSMUSG00000033768 | Nrxn2 | 18190 | protein_coding | 32.33353407 | 2.135573458 | 0.719643712 | 2.967542721 | 0.003001906 | 0.018787565 | 2.135573458 |
| ENSMUSG00000110404 | Rnf223 | 100045778 | protein_coding | 10.53477055 | 2.142845203 | 0.549726963 | 3.898017285 | 9.70E-05 | 0.001099468 | 2.142845203 |
| ENSMUSG00000022610 | Mapk12 | 29857 | protein_coding | 11.40925355 | -2.14387978 | 0.453742099 | -4.724886192 | 2.30E-06 | 3.97E-05 | 2.14387978 |
| ENSMUSG00000024184 | Pdia2 | 69191 | protein_coding | 13.80442126 | 2.154886693 | 0.387541645 | 5.560400339 | 2.69E-08 | 6.78E-07 | 2.154886693 |
| ENSMUSG00000019139 | Isyna1 | 71780 | protein_coding | 2029.596056 | 2.166313898 | 0.13488938 | 16.05992927 | 4.87E-58 | 1.46E-55 | 2.166313898 |
| ENSMUSG00000035107 | Dcbld2 | 73379 | protein_coding | 272.0262174 | 2.175141118 | 0.124367958 | 17.48956202 | 1.72E-68 | 6.37E-66 | 2.175141118 |
| ENSMUSG00000028037 | Ifi44 | 99899 | protein_coding | 51.40043749 | 2.199036021 | 0.314481535 | 6.992575956 | 2.70E-12 | 1.06E-10 | 2.199036021 |
| ENSMUSG00000051817 | Sox12 | 20667 | protein_coding | 26.98756782 | 2.237742725 | 0.276866275 | 8.082395447 | 6.35E-16 | 3.48E-14 | 2.237742725 |
| ENSMUSG00000002900 | Lamb1 | 16777 | protein_coding | 40.05526536 | -2.240684025 | 0.285538556 | -7.847220544 | 4.25E-15 | 2.17E-13 | 2.240684025 |
| ENSMUSG00000078964 | Ces1b | 382044 | protein_coding | 8.158253626 | -2.241206322 | 0.649888811 | -3.448599643 | 0.000563502 | 0.004799707 | 2.241206322 |
| ENSMUSG00000022665 | Ccdc80 | 67896 | protein_coding | 8.850494692 | -2.251716634 | 0.653535825 | -3.445437188 | 0.000570136 | 0.004840689 | 2.251716634 |
| ENSMUSG00000006517 | Mvd | 192156 | protein_coding | 1451.294583 | 2.253072014 | 0.12503712 | 18.01922506 | 1.38E-72 | 5.22E-70 | 2.253072014 |
| ENSMUSG00000071176 | Arhgef10 | 234094 | protein_coding | 142.9455031 | 2.268630882 | 0.157072557 | 14.44320338 | 2.77E-47 | 6.21E-45 | 2.268630882 |
| ENSMUSG00000048108 | Tmem72 | 319776 | protein_coding | 18.3495073 | 2.27132644 | 0.743692556 | 3.054120174 | 0.002257216 | 0.01496792 | 2.27132644 |
| ENSMUSG00000040808 | S100g | 12309 | protein_coding | 279.8554535 | -2.283257342 | 0.512976117 | -4.451001257 | 8.55E-06 | 0.000129779 | 2.283257342 |
| ENSMUSG00000025353 | Ormdl2 | 66844 | protein_coding | 1455.995967 | 2.294265716 | 0.09136553 | 25.11084566 | 3.79E-139 | 3.35E-136 | 2.294265716 |
| ENSMUSG00000107768 | Gm36582 | 102640545 | lncRNA | 17.10402845 | -2.299952366 | 0.560859512 | -4.100763765 | 4.12E-05 | 0.000518906 | 2.299952366 |
| ENSMUSG00000022548 | Apod | 11815 | protein_coding | 12.31470377 | 2.302934648 | 0.416848516 | 5.524631991 | 3.30E-08 | 8.15E-07 | 2.302934648 |

Supplementary Table 1

| ensembl_gene_id | external_gene_name | entrezgene_id | gene_biotype | baseMean | log2FoldChange | lfcSE | stat | pvalue | padj | absLog2FC |
|---|---|---|---|---|---|---|---|---|---|---|
| ENSMUSG00000020176 | Grb10 | 14783 | protein_coding | 51.43267341 | 2.304170549 | 0.342755938 | 6.722481782 | 1.79E-11 | 6.54E-10 | 2.304170549 |
| ENSMUSG00000022964 | Tmem50b | 77975 | protein_coding | 1721.250315 | 2.309167902 | 0.114633632 | 20.1438954 | 3.04E-90 | 1.67E-87 | 2.309167902 |
| ENSMUSG00000025017 | Pik3ap1 | 83490 | protein_coding | 80.04895426 | 2.315082999 | 0.206969485 | 11.18562475 | 4.80E-29 | 5.27E-27 | 2.315082999 |
| ENSMUSG00000112778 | Gm33677 | 102636674 | lncRNA | 15.83252255 | -2.317090553 | 0.35902238 | -6.453888897 | 1.09E-10 | 3.70E-09 | 2.317090553 |
| ENSMUSG00000044362 | Ccdc89 | 70054 | protein_coding | 29.34497428 | 2.358754267 | 0.34942888 | 6.750312875 | 1.48E-11 | 5.45E-10 | 2.358754267 |
| ENSMUSG00000022864 | D16Ertd472e | 67102 | protein_coding | 54.96748657 | 2.363946623 | 0.321001556 | 7.364284007 | 1.78E-13 | 7.84E-12 | 2.363946623 |
| ENSMUSG00000004460 | Dnajb11 | 67838 | protein_coding | 7018.669833 | 2.379157098 | 0.124491588 | 19.11098688 | 2.05E-81 | 9.31E-79 | 2.379157098 |
| ENSMUSG00000027932 | Slc27a3 | 26568 | protein_coding | 71.50679682 | 2.399120073 | 0.271796617 | 8.826894542 | 1.08E-18 | 6.97E-17 | 2.399120073 |
| ENSMUSG00000099413 | Gm17767 | 100045066 | lncRNA | 19.79974182 | 2.426557898 | 0.353642085 | 6.861620837 | 6.81E-12 | 2.59E-10 | 2.426557898 |
| ENSMUSG00000042448 | Hoxd1 | 15429 | protein_coding | 10.59797159 | 2.443541687 | 0.542392559 | 4.50511654 | 6.63E-06 | 0.000103691 | 2.443541687 |
| ENSMUSG00000031770 | Herpud1 | 64209 | protein_coding | 8120.653899 | 2.453064987 | 0.134681162 | 18.21386864 | 4.01E-74 | 1.60E-71 | 2.453064987 |
| ENSMUSG00000073424 | Cyp4f15 | 106648 | protein_coding | 11.03332111 | -2.459573663 | 0.580215402 | -4.239069929 | 2.24E-05 | 0.000305817 | 2.459573663 |
| ENSMUSG00000035692 | Isg15 | 100038882 | protein_coding | 92.77392563 | 2.466631093 | 0.276150861 | 8.932186873 | 4.18E-19 | 2.78E-17 | 2.466631093 |
| ENSMUSG00000030759 | Far1 | 67420 | protein_coding | 12979.05374 | 2.466728947 | 0.105880574 | 23.29727596 | 4.72E-120 | 3.76E-117 | 2.466728947 |
| ENSMUSG00000032098 | Treh | 58866 | protein_coding | 46.67514744 | 2.467218097 | 0.2317038 | 10.64815554 | 1.78E-26 | 1.70E-24 | 2.467218097 |
| ENSMUSG00000027248 | Pdia3 | 14827 | protein_coding | 22242.81415 | 2.472714668 | 0.080661779 | 30.65534502 | 2.24E-206 | 5.95E-203 | 2.472714668 |
| ENSMUSG00000015243 | Abca1 | 11303 | protein_coding | 608.4833076 | 2.478398055 | 0.208507734 | 11.88636031 | 1.39E-32 | 1.78E-30 | 2.478398055 |
| ENSMUSG00000056367 | Actr3b | 242894 | protein_coding | 10.84108435 | 2.49292929 | 0.566161704 | 4.403210726 | 1.07E-05 | 0.000157304 | 2.49292929 |
| ENSMUSG00000070858 | Gm1673 | 381633 | protein_coding | 11.69837495 | 2.505741822 | 0.443306286 | 5.652394069 | 1.58E-08 | 4.16E-07 | 2.505741822 |
| ENSMUSG00000038143 | Stox2 | 71069 | protein_coding | 436.7512907 | 2.510615345 | 0.125505107 | 20.00408905 | 5.07E-89 | 2.61E-86 | 2.510615345 |
| ENSMUSG00000022136 | Dnajc3 | 100037258 | protein_coding | 8488.77101 | 2.525710306 | 0.099225336 | 25.45428817 | 6.33E-143 | 6.30E-140 | 2.525710306 |
| ENSMUSG00000026663 | Atf6 | 226641 | protein_coding | 5272.646882 | 2.526024891 | 0.081063455 | 31.16108089 | 3.59E-213 | 1.14E-209 | 2.526024891 |
| ENSMUSG00000032122 | Slc37a2 | 56857 | protein_coding | 1452.800784 | -2.5431442 | 0.454073265 | -5.600735384 | 2.13E-08 | 5.47E-07 | 2.5431442 |
| ENSMUSG00000024011 | Pi16 | 74116 | protein_coding | 7.74841779 | -2.545843577 | 0.720278633 | -3.534526028 | 0.000408507 | 0.00369699 | 2.545843577 |
| ENSMUSG00000020571 | Pdia6 | 71853 | protein_coding | 12860.11705 | 2.560376761 | 0.116459852 | 21.98505943 | 4.00E-107 | 2.66E-104 | 2.560376761 |
| ENSMUSG00000058013 | Septin11 | 52398 | protein_coding | 2234.324444 | 2.597369176 | 0.163164087 | 15.91875528 | 4.70E-57 | 1.34E-54 | 2.597369176 |
| ENSMUSG00000031155 | Pim2 | 18715 | protein_coding | 129.0696237 | 2.602636523 | 0.21631805 | 12.03152729 | 2.43E-33 | 3.17E-31 | 2.602636523 |
| ENSMUSG00000040856 | Dlk1 | 13386 | protein_coding | 8.900822318 | -2.61345116 | 0.757947469 | -3.448063708 | 0.000564621 | 0.004803535 | 2.61345116 |
| ENSMUSG00000078921 | Tgtp2 | 100039796 | protein_coding | 9.213500072 | 2.63499817 | 0.530726053 | 4.9648932 | 6.87E-07 | 1.35E-05 | 2.63499817 |
| ENSMUSG00000027313 | Chac1 | 69065 | protein_coding | 38.28549404 | 2.635517041 | 0.327599324 | 8.044940405 | 8.63E-16 | 4.69E-14 | 2.635517041 |
| ENSMUSG00000043410 | Hfm1 | 330149 | protein_coding | 11.12548766 | 2.648039327 | 0.563521908 | 4.699088514 | 2.61E-06 | 4.47E-05 | 2.648039327 |
| ENSMUSG00000046550 | Spin2c | 278240 | protein_coding | 30.61965315 | 2.708089744 | 0.331501865 | 8.169153868 | 3.11E-16 | 1.74E-14 | 2.708089744 |
| ENSMUSG00000007783 | Cpt1c | 78070 | protein_coding | 20.35535319 | 2.736249665 | 0.377610597 | 7.246220542 | 4.29E-13 | 1.84E-11 | 2.736249665 |
| ENSMUSG00000039759 | Thap3 | 69876 | protein_coding | 513.7762188 | 2.738741025 | 0.141911139 | 19.2989856 | 5.48E-83 | 2.57E-80 | 2.738741025 |
| ENSMUSG00000023809 | Rps6ka2 | 20112 | protein_coding | 60.04208604 | 2.753961218 | 0.253816944 | 10.85018667 | 1.99E-27 | 1.99E-25 | 2.753961218 |
| ENSMUSG00000057069 | Ero1b | 67475 | protein_coding | 572.573739 | 2.798471405 | 0.110193024 | 25.39608509 | 2.79E-142 | 2.61E-139 | 2.798471405 |
| ENSMUSG00000078922 | Tgtp1 | 21822 | protein_coding | 10.70520884 | 2.807655279 | 0.600730778 | 4.673733025 | 2.96E-06 | 4.99E-05 | 2.807655279 |
| ENSMUSG00000078349 | AW011738 | 100382 | lncRNA | 85.24903948 | 2.822540307 | 0.181234702 | 15.57395067 | 1.09E-54 | 2.81E-52 | 2.822540307 |

Supplementary Table 1

| ensembl_gene_id | external_gene_name | entrezgene_id | gene_biotype | baseMean | log2FoldChange | lfcSE | stat | pvalue | padj | absLog2FC |
|---|---|---|---|---|---|---|---|---|---|---|
| ENSMUSG00000020964 | Sel1l | 20338 | protein_coding | 7514.70565 | 2.845580188 | 0.100168916 | 28.40781665 | 1.62E-177 | 2.87E-174 | 2.845580188 |
| ENSMUSG00000020388 | Pdlim4 | 30794 | protein_coding | 12.10561455 | -2.869654202 | 0.399817998 | -7.17740125 | 7.10E-13 | 2.99E-11 | 2.869654202 |
| ENSMUSG00000031617 | Tmem184c | 234463 | protein_coding | 214.9476783 | 2.883149591 | 0.157394338 | 18.31800067 | 5.95E-75 | 2.43E-72 | 2.883149591 |
| ENSMUSG00000071356 | Reg3b | 18489 | protein_coding | 28157.10932 | 2.901485631 | 0.783592448 | 3.702799382 | 0.000213233 | 0.002157804 | 2.901485631 |
| ENSMUSG00000000791 | Il12rb1 | 16161 | protein_coding | 17.84130331 | 2.903100613 | 0.491205271 | 5.910157695 | 3.42E-09 | 9.79E-08 | 2.903100613 |
| ENSMUSG00000031734 | Irx3 | 16373 | protein_coding | 13.16798959 | 2.927675027 | 0.593263501 | 4.934864562 | 8.02E-07 | 1.55E-05 | 2.927675027 |
| ENSMUSG00000068341 | Reg3d | 30053 | protein_coding | 9.094218334 | 2.984275002 | 0.817088568 | 3.652327444 | 0.000259874 | 0.002533217 | 2.984275002 |
| ENSMUSG00000053550 | Shisa7 | 232813 | protein_coding | 113.0556444 | 2.999118905 | 0.202446302 | 14.81439212 | 1.18E-49 | 2.77E-47 | 2.999118905 |
| ENSMUSG00000004988 | Fxyd4 | 108017 | protein_coding | 21.67546852 | -3.016056732 | 0.365230636 | -8.257951096 | 1.48E-16 | 8.55E-15 | 3.016056732 |
| ENSMUSG00000019876 | Pkib | 18768 | protein_coding | 98.78953782 | 3.02176161 | 0.311781857 | 9.691909723 | 3.26E-22 | 2.52E-20 | 3.02176161 |
| ENSMUSG00000045349 | Sh2d5 | 230863 | protein_coding | 17.8926631 | 3.046565111 | 0.44126045 | 6.904233345 | 5.05E-12 | 1.95E-10 | 3.046565111 |
| ENSMUSG00000054999 | Naaladl1 | 381204 | protein_coding | 2780.105174 | 3.066419586 | 0.149594673 | 20.49818703 | 2.23E-93 | 1.27E-90 | 3.066419586 |
| ENSMUSG00000097651 | 4930461G14Rik | 74879 | lncRNA | 7.403000604 | 3.103700503 | 0.627198116 | 4.948516943 | 7.48E-07 | 1.46E-05 | 3.103700503 |
| ENSMUSG00000003814 | Calr | 12317 | protein_coding | 45889.9582 | 3.123377789 | 0.118704076 | 26.3123045 | 1.39E-152 | 1.70E-149 | 3.123377789 |
| ENSMUSG00000027737 | Slc7a11 | 26570 | protein_coding | 401.7401999 | 3.169960294 | 0.323854669 | 9.788218607 | 1.27E-22 | 1.00E-20 | 3.169960294 |
| ENSMUSG00000026833 | Olfm1 | 56177 | protein_coding | 48.9215417 | 3.267196342 | 0.274367806 | 11.90808932 | 1.07E-32 | 1.38E-30 | 3.267196342 |
| ENSMUSG00000074195 | Clca4b | 99709 | protein_coding | 629.8847059 | 3.305576326 | 0.518465475 | 6.375692283 | 1.82E-10 | 6.04E-09 | 3.305576326 |
| ENSMUSG00000070407 | Hs3st3b1 | 54710 | protein_coding | 931.2285558 | 3.339771402 | 0.146054925 | 22.86654423 | 1.00E-115 | 7.24E-113 | 3.339771402 |
| ENSMUSG00000094430 | Smbd1 | 381043 | protein_coding | 62.69863534 | 3.354027964 | 0.238073471 | 14.08820544 | 4.49E-45 | 9.05E-43 | 3.354027964 |
| ENSMUSG00000032875 | Arhgef17 | 207212 | protein_coding | 180.0863365 | 3.370570009 | 0.215160689 | 15.66536163 | 2.61E-55 | 6.93E-53 | 3.370570009 |
| ENSMUSG00000022226 | Mcpt2 | 17225 | protein_coding | 27.39951241 | 3.417683595 | 0.396908634 | 8.610756491 | 7.26E-18 | 4.53E-16 | 3.417683595 |
| ENSMUSG00000001120 | Pcbp3 | 59093 | protein_coding | 9.676276815 | 3.428370137 | 0.546812791 | 6.269732884 | 3.62E-10 | 1.15E-08 | 3.428370137 |
| ENSMUSG00000032115 | Hyou1 | 12282 | protein_coding | 15539.2681 | 3.461225354 | 0.156506913 | 22.11547904 | 2.24E-108 | 1.55E-105 | 3.461225354 |
| ENSMUSG00000060445 | Sycp2 | 320558 | protein_coding | 23.99985968 | 3.471217298 | 0.618652886 | 5.610928801 | 2.01E-08 | 5.20E-07 | 3.471217298 |
| ENSMUSG00000053395 | Cacng8 | 81905 | protein_coding | 8.428156654 | 3.480834613 | 0.85260149 | 4.082604424 | 4.45E-05 | 0.000556777 | 3.480834613 |
| ENSMUSG00000097391 | Mirg | 100040724 | lncRNA | 14.55164826 | -3.512464148 | 0.599618469 | -5.857831817 | 4.69E-09 | 1.32E-07 | 3.512464148 |
| ENSMUSG00000028713 | Cyp4b1 | 13120 | protein_coding | 192.6964281 | -3.550591888 | 0.3625465 | -9.793479978 | 1.20E-22 | 9.66E-21 | 3.550591888 |
| ENSMUSG00000027398 | Il1b | 16176 | protein_coding | 8.830385331 | 3.576958264 | 0.741055663 | 4.82684155 | 1.39E-06 | 2.53E-05 | 3.576958264 |
| ENSMUSG00000030559 | Rab38 | 72433 | protein_coding | 59.83603902 | 3.593689191 | 0.293359568 | 12.25011755 | 1.68E-34 | 2.36E-32 | 3.593689191 |
| ENSMUSG00000032575 | Manf | 74840 | protein_coding | 8474.594821 | 3.633618344 | 0.136468573 | 26.62604489 | 3.39E-156 | 4.50E-153 | 3.633618344 |
| ENSMUSG00000021268 | Meg3 | 17263 | lncRNA | 93.0176607 | -3.633855367 | 0.339810869 | -10.69375849 | 1.09E-26 | 1.04E-24 | 3.633855367 |
| ENSMUSG00000019789 | Hey2 | 15214 | protein_coding | 17.44693395 | 3.65231333 | 0.51399653 | 7.105715932 | 1.20E-12 | 4.91E-11 | 3.65231333 |
| ENSMUSG00000085925 | Rtl1 | 353326 | protein_coding | 8.185671988 | -3.669717388 | 0.715310228 | -5.130245934 | 2.89E-07 | 6.15E-06 | 3.669717388 |
| ENSMUSG00000022227 | Mcpt1 | 17224 | protein_coding | 58.3304406 | 3.675920707 | 0.369433554 | 9.950153862 | 2.52E-23 | 2.10E-21 | 3.675920707 |
| ENSMUSG00000026864 | Hspa5 | 14828 | protein_coding | 90500.10627 | 3.733440226 | 0.162664366 | 22.95180139 | 1.41E-116 | 1.07E-113 | 3.733440226 |
| ENSMUSG00000060671 | Atp8b2 | 54667 | protein_coding | 458.0547416 | 3.765782389 | 0.129234413 | 29.13916121 | 1.15E-186 | 2.28E-183 | 3.765782389 |
| ENSMUSG00000042821 | Snai1 | 20613 | protein_coding | 11.86624299 | 3.796247443 | 0.617015006 | 6.152601484 | 7.62E-10 | 2.33E-08 | 3.796247443 |
| ENSMUSG00000030659 | Nucb2 | 53322 | protein_coding | 1075.584718 | 3.921129736 | 0.153961612 | 25.46822995 | 4.44E-143 | 4.71E-140 | 3.921129736 |

Supplementary Table 1

| ensembl_gene_id | external_gene_name | entrezgene_id | gene_biotype | baseMean | log2FoldChange | lfcSE | stat | pvalue | padj | absLog2FC |
|---|---|---|---|---|---|---|---|---|---|---|
| ENSMUSG00000018623 | Mmp7 | 17393 | protein_coding | 189.6443959 | 3.931104673 | 0.322327845 | 12.19598225 | 3.27E-34 | 4.56E-32 | 3.931104673 |
| ENSMUSG00000020048 | Hsp90b1 | 22027 | protein_coding | 78395.5205 | 3.978658323 | 0.100894217 | 39.4339581 | 0 | 0 | 3.978658323 |
| ENSMUSG00000054072 | Iigp1 | 60440 | protein_coding | 89.68914387 | 4.113918773 | 0.491571056 | 8.368919861 | 5.81E-17 | 3.42E-15 | 4.113918773 |
| ENSMUSG00000022769 | Sdf2l1 | 64136 | protein_coding | 4816.345053 | 4.18008538 | 0.148212324 | 28.20335896 | 5.32E-175 | 8.47E-172 | 4.18008538 |
| ENSMUSG00000023272 | Creld2 | 76737 | protein_coding | 4203.092847 | 4.192310133 | 0.213533161 | 19.63306363 | 8.07E-86 | 4.02E-83 | 4.192310133 |
| ENSMUSG00000029844 | Hoxa1 | 15394 | protein_coding | 52.13193115 | 4.205834195 | 0.34316373 | 12.25605688 | 1.56E-34 | 2.22E-32 | 4.205834195 |
| ENSMUSG00000025823 | Pdia4 | 12304 | protein_coding | 32285.43449 | 4.353091684 | 0.123447242 | 35.26276995 | 2.19E-272 | 1.16E-268 | 4.353091684 |
| ENSMUSG00000026442 | Nfasc | 269116 | protein_coding | 223.594189 | 4.526550543 | 0.263890442 | 17.15314321 | 5.95E-66 | 2.06E-63 | 4.526550543 |
| ENSMUSG00000064358 | mt-Co3 | 17710 | protein_coding | 88.88789024 | -4.600550885 | 2.158627693 | -2.131238703 | 0.033069482 | 0.115892345 | 4.600550885 |
| ENSMUSG00000032715 | Trib3 | 228775 | protein_coding | 76.46064037 | 4.602458416 | 0.31199237 | 14.75183006 | 2.99E-49 | 6.91E-47 | 4.602458416 |
| ENSMUSG00000036067 | Slc2a6 | 227659 | protein_coding | 709.9512588 | 4.789414921 | 0.278917092 | 17.17146439 | 4.34E-66 | 1.54E-63 | 4.789414921 |
| ENSMUSG00000021622 | Ckmt2 | 76722 | protein_coding | 610.3971048 | 4.893355557 | 0.16097082 | 30.39902232 | 5.66E-203 | 1.29E-199 | 4.893355557 |
| ENSMUSG00000018340 | Anxa6 | 11749 | protein_coding | 1244.763916 | 5.002379153 | 0.16034743 | 31.19712704 | 1.17E-213 | 4.64E-210 | 5.002379153 |
| ENSMUSG00000035202 | Lars2 | 102436 | protein_coding | 33744.32593 | 5.151473276 | 1.391963825 | 3.700867209 | 0.000214864 | 0.002170166 | 5.151473276 |
| ENSMUSG00000036036 | Zfp57 | 22715 | protein_coding | 82.82103236 | 5.309154424 | 0.539845199 | 9.834586715 | 7.99E-23 | 6.53E-21 | 5.309154424 |
| ENSMUSG00000029151 | Slc30a3 | 22784 | protein_coding | 65.42792787 | 5.543773835 | 0.474649918 | 11.67971094 | 1.62E-31 | 1.95E-29 | 5.543773835 |
| ENSMUSG00000017446 | C1qtnf1 | 56745 | protein_coding | 367.1305856 | 5.869434692 | 0.218849937 | 26.81944884 | 1.92E-158 | 2.78E-155 | 5.869434692 |
| ENSMUSG00000052584 | Serp2 | 72661 | protein_coding | 258.0623384 | 5.887909128 | 0.314422072 | 18.72613168 | 3.03E-78 | 1.31E-75 | 5.887909128 |
| ENSMUSG00000028558 | Calr4 | 108802 | protein_coding | 61.66948757 | 6.042550827 | 0.520276122 | 11.61412291 | 3.49E-31 | 4.12E-29 | 6.042550827 |
| ENSMUSG00000029697 | Fezf1 | 73191 | protein_coding | 28.66334234 | 6.270215158 | 0.738266416 | 8.493160498 | 2.01E-17 | 1.22E-15 | 6.270215158 |
| ENSMUSG00000098682 | Otx2os1 | 606497 | lncRNA | 53.77866784 | 7.647594206 | 0.833977032 | 9.170029761 | 4.73E-20 | 3.38E-18 | 7.647594206 |
| ENSMUSG00000009092 | Derl3 | 70377 | protein_coding | 4279.743615 | 8.577861616 | 0.240042802 | 35.73471706 | 1.14E-279 | 9.10E-276 | 8.577861616 |
