## Supplementary Table 2 for "ATF6 activation alters colonic lipid metabolism causing tumor-associated microbial adaptation"

| pathway | pval | padj | ES | NES | nMoreExtreme | size | Genes_in_Pathway | Percentage_of_Pathway_detected |
|---|---|---|---|---|---|---|---|---|
| KEGG_ABC_TRANSPORTERS | 0.29707113 | 0.650061531 | -0.418667788 | -1.132391375 | 212 | 42 | 43 | 97.67 |
| KEGG_ACUTE_MYELOID_LEUKEMIA | 0.503989362 | 0.765463918 | -0.343677559 | -0.976351633 | 378 | 58 | 59 | 98.31 |
| KEGG_ADHERENS_JUNCTION | 0.211009174 | 0.577172153 | 0.312211673 | 1.133617145 | 45 | 72 | 73 | 98.63 |
| KEGG_ADIPOCYTOKINE_SIGNALING_PATHWAY | 0.47008547 | 0.745841905 | 0.275265614 | 0.982827516 | 109 | 64 | 67 | 95.52 |
| KEGG_ALANINE_ASPARTATE_AND_GLUTAMATE_METABOLISM | 0.459302326 | 0.745841905 | -0.387172273 | -0.997047853 | 315 | 31 | 33 | 93.94 |
| KEGG_ALDOSTERONE_REGULATED_SODIUM_REABSORPTION | 0.03202847 | 0.198576512 | 0.446493565 | 1.470062552 | 8 | 41 | 43 | 95.35 |
| KEGG_ALLOGRAFT_REJECTION | 1 | 1 | -0.143814835 | -0.355888025 | 662 | 25 | 37 | 67.57 |
| KEGG_ALPHA_LINOLENIC_ACID_METABOLISM | 0.701277955 | 0.865410087 | -0.358059943 | -0.825691019 | 438 | 18 | 18 | 100 |
| KEGG_ALZHEIMERS_DISEASE | 0.01459854 | 0.123424021 | 0.323487678 | 1.359399122 | 1 | 154 | 165 | 93.33 |
| KEGG_AMINOACYL_TRNA_BIOSYNTHESIS | 0.055478502 | 0.271552668 | -0.545155271 | -1.467974438 | 39 | 41 | 41 | 100 |
| KEGG_AMINO_SUGAR_AND_NUCLEOTIDE_SUGAR_METABOLISM | 0.147018031 | 0.464587866 | -0.470344699 | -1.288916389 | 105 | 45 | 48 | 93.75 |
| KEGG_AMYOTROPHIC_LATERAL_SCLEROSIS_ALS | 0.502024291 | 0.765463918 | -0.346806404 | -0.969990463 | 371 | 51 | 51 | 100 |
| KEGG_ANTIGEN_PROCESSING_AND_PRESENTATION | 0.066485753 | 0.299917662 | -0.5158562 | -1.447193248 | 48 | 55 | 279 | 19.71 |
| KEGG_APOPTOSIS | 0.301886792 | 0.652917946 | 0.285190608 | 1.077391995 | 63 | 82 | 89 | 92.13 |
| KEGG_ARACHIDONIC_ACID_METABOLISM | 0.669201521 | 0.846520095 | 0.257366726 | 0.890082136 | 175 | 53 | 60 | 88.33 |
| KEGG_ARGININE_AND_PROLINE_METABOLISM | 0.029769959 | 0.198576512 | -0.551106183 | -1.545045298 | 21 | 53 | 59 | 89.83 |
| KEGG_ARRHYTHMOGENIC_RIGHT_VENTRICULAR_CARDIOMYOPATHY_ARVC | 0.702564103 | 0.865410087 | -0.284579355 | -0.839808769 | 547 | 73 | 74 | 98.65 |
| KEGG_ASCORBATE_AND_ALDARATE_METABOLISM | 0.100755668 | 0.395408163 | 0.556511328 | 1.406666921 | 39 | 13 | 31 | 41.94 |
| KEGG_ASTHMA | 0.700767263 | 0.865410087 | 0.316067455 | 0.824869313 | 273 | 14 | 32 | 43.75 |
| KEGG_AUTOIMMUNE_THYROID_DISEASE | 0.993957704 | 1 | 0.152936485 | 0.456489792 | 328 | 26 | 221 | 11.76 |
| KEGG_AXON_GUIDANCE | 0.874109264 | 0.945257692 | -0.241076923 | -0.761048892 | 735 | 129 | 129 | 100 |
| KEGG_BASAL_CELL_CARCINOMA | 0.9797023 | 1 | -0.208426585 | -0.58433116 | 723 | 53 | 55 | 96.36 |
| KEGG_BASAL_TRANSCRIPTION_FACTORS | 0.067723343 | 0.299917662 | -0.55273268 | -1.433133424 | 46 | 32 | 38 | 84.21 |
| KEGG_BASE_EXCISION_REPAIR | 0.031700288 | 0.198576512 | -0.607958905 | -1.576324791 | 21 | 32 | 32 | 100 |
| KEGG_BETA_ALANINE_METABOLISM | 0.019662921 | 0.15901319 | 0.559373911 | 1.625906355 | 6 | 22 | 22 | 100 |
| KEGG_BIOSYNTHESIS_OF_UNSATURATED_FATTY_ACIDS | 0.621417798 | 0.827497627 | -0.353011381 | -0.873571376 | 411 | 25 | 29 | 86.21 |
| KEGG_BLADDER_CANCER | 0.388579387 | 0.720710523 | -0.391276191 | -1.056306395 | 278 | 40 | 42 | 95.24 |
| KEGG_BUTANOATE_METABOLISM | 0.014035088 | 0.123424021 | 0.482078663 | 1.556954296 | 3 | 37 | 38 | 97.37 |
| KEGG_B_CELL_RECEPTOR_SIGNALING_PATHWAY | 0.334841629 | 0.675949367 | 0.279848597 | 1.046349256 | 73 | 78 | 90 | 86.67 |
| KEGG_CALCIUM_SIGNALING_PATHWAY | 0.742009132 | 0.873740801 | -0.261260232 | -0.847666715 | 649 | 159 | 173 | 91.91 |
| KEGG_CARDIAC_MUSCLE_CONTRACTION | 0.154929577 | 0.464587866 | 0.331800163 | 1.198699594 | 32 | 71 | 79 | 89.87 |
| KEGG_CELL_ADHESION_MOLECULES_CAMS | 0.942682927 | 0.985050699 | -0.219687941 | -0.681503417 | 772 | 111 | 129 | 86.05 |
| KEGG_CELL_CYCLE | 0.002430134 | 0.07635468 | -0.550870122 | -1.717995953 | 1 | 122 | 124 | 98.39 |
| KEGG_CHEMOKINE_SIGNALING_PATHWAY | 0.924855491 | 0.982773596 | -0.22697454 | -0.731728399 | 799 | 154 | 178 | 86.52 |
| KEGG_CHRONIC_MYELOID_LEUKEMIA | 0.859693878 | 0.935105621 | -0.25132088 | -0.740693032 | 673 | 72 | 72 | 100 |
| KEGG_CIRCADIAN_RHYTHM_MAMMAL | 0.431404959 | 0.745841905 | -0.472918004 | -1.031372062 | 260 | 13 | 13 | 100 |
| KEGG_CITRATE_CYCLE_TCA_CYCLE | 0.117460317 | 0.414034344 | 0.42920585 | 1.32075881 | 36 | 30 | 31 | 96.77 |
| KEGG_COLORECTAL_CANCER | 0.281818182 | 0.640275387 | -0.392623875 | -1.132416731 | 216 | 63 | 64 | 98.44 |
| KEGG_COMPLEMENT_AND_COAGULATION_CASCADES | 0.585062241 | 0.813579501 | 0.266688861 | 0.93466286 | 140 | 59 | 74 | 79.73 |
| KEGG_CYSTEINE_AND_METHIONINE_METABOLISM | 0.749291785 | 0.873740801 | -0.304076246 | -0.795792556 | 528 | 33 | 38 | 86.84 |

Supplementary Table 2

| pathway | pval | padj | ES | NES | nMoreExtreme | size | Genes_in_Pathway | Percentage_of_Pathway_detected |
|---|---|---|---|---|---|---|---|---|
| KEGG_CYTOKINE_CYTOKINE_RECEPTOR_INTERACTION | 0.994337486 | 1 | -0.188939804 | -0.620446061 | 877 | 190 | 460 | 41.3 |
| KEGG_CYTOSOLIC_DNA_SENSING_PATHWAY | 0.226890756 | 0.61161856 | -0.456301329 | -1.206644983 | 161 | 35 | 221 | 15.84 |
| KEGG_DILATED_CARDIOMYOPATHY | 0.669172932 | 0.846520095 | -0.29075172 | -0.874810565 | 533 | 85 | 90 | 94.44 |
| KEGG_DNA_REPLICATION | 0.029535865 | 0.198576512 | -0.604401057 | -1.593146062 | 20 | 34 | 34 | 100 |
| KEGG_DORSO_VENTRAL_AXIS_FORMATION | 0.60620155 | 0.827497627 | -0.37884654 | -0.899108845 | 390 | 21 | 21 | 100 |
| KEGG_DRUG_METABOLISM_CYTOCHROME_P450 | 0.003773585 | 0.07635468 | 0.526789741 | 1.844185557 | 0 | 55 | 84 | 65.48 |
| KEGG_DRUG_METABOLISM_OTHER_ENZYMES | 0.003571429 | 0.07635468 | 0.634345425 | 2.118781331 | 0 | 43 | 73 | 58.9 |
| KEGG_ECM_RECEPTOR_INTERACTION | 0.720183486 | 0.873740801 | 0.232469782 | 0.880015422 | 156 | 81 | 83 | 97.59 |
| KEGG_ENDOCYTOSIS | 0.751937984 | 0.873740801 | 0.21534177 | 0.918627914 | 96 | 171 | 184 | 92.93 |
| KEGG_ENDOMETRIAL_CANCER | 0.532608696 | 0.776386404 | -0.341703785 | -0.954997295 | 391 | 52 | 52 | 100 |
| KEGG_EPITHELIAL_CELL_SIGNALING_IN_HELICOBACTER_PYLORI_INFECTION | 0.004166667 | 0.07635468 | 0.493085756 | 1.729339218 | 0 | 60 | 64 | 93.75 |
| KEGG_ERBB_SIGNALING_PATHWAY | 0.76875 | 0.882638889 | -0.268364194 | -0.807307249 | 614 | 86 | 86 | 100 |
| KEGG_ETHER_LIPID_METABOLISM | 0.796829971 | 0.907409885 | -0.29143036 | -0.755624923 | 552 | 32 | 32 | 100 |
| KEGG_FATTY_ACID_METABOLISM | 0.003558719 | 0.07635468 | 0.548169768 | 1.86199698 | 0 | 45 | 51 | 88.24 |
| KEGG_FC_EPSILON_RI_SIGNALING_PATHWAY | 0.563063063 | 0.793407043 | 0.259598948 | 0.944866913 | 124 | 73 | 78 | 93.59 |
| KEGG_FC_GAMMA_R_MEDIATED_PHAGOCYTOSIS | 0.319796954 | 0.668339702 | 0.279359481 | 1.078629503 | 62 | 95 | 99 | 95.96 |
| KEGG_FOCAL_ADHESION | 0.756302521 | 0.873740801 | 0.216627406 | 0.929758832 | 89 | 189 | 195 | 96.92 |
| KEGG_FOLATE_BIOSYNTHESIS | 0.740863787 | 0.873740801 | -0.365262332 | -0.773543382 | 445 | 11 | 17 | 64.71 |
| KEGG_FRUCTOSE_AND_MANNOSE_METABOLISM | 0.277777778 | 0.640275387 | -0.431184742 | -1.151770244 | 199 | 38 | 38 | 100 |
| KEGG_GALACTOSE_METABOLISM | 0.254901961 | 0.632156863 | -0.476069178 | -1.178093482 | 168 | 25 | 26 | 96.15 |
| KEGG_GAP_JUNCTION | 0.315533981 | 0.666924095 | 0.28216508 | 1.075265617 | 64 | 83 | 84 | 98.81 |
| KEGG_GLIOMA | 0.814717477 | 0.907409885 | -0.266413252 | -0.759580778 | 619 | 59 | 61 | 96.72 |
| KEGG_GLUTATHIONE_METABOLISM | 0.807692308 | 0.907409885 | -0.269318821 | -0.745072564 | 587 | 47 | 48 | 97.92 |
| KEGG_GLYCEROLIPID_METABOLISM | 0.659340659 | 0.846520095 | -0.315423567 | -0.87366641 | 479 | 48 | 49 | 97.96 |
| KEGG_GLYCEROPHOSPHOLIPID_METABOLISM | 0.290115533 | 0.64239868 | -0.380429921 | -1.128314372 | 225 | 76 | 76 | 100 |
| KEGG_GLYCINE_SERINE_AND_THREONINE_METABOLISM | 0.113372093 | 0.414034344 | -0.522593602 | -1.345785493 | 77 | 31 | 32 | 96.88 |
| KEGG_GLYCOLYSIS_GLUCONEOGENESIS | 0.004 | 0.07635468 | 0.451443563 | 1.583426397 | 0 | 58 | 67 | 86.57 |
| KEGG_GLYCOSAMINOGLYCAN_BIOSYNTHESIS_CHONDROITIN_SULFATE | 0.127131783 | 0.437898363 | -0.572197888 | -1.357985698 | 81 | 21 | 22 | 95.45 |
| KEGG_GLYCOSAMINOGLYCAN_BIOSYNTHESIS_HEPARAN_SULFATE | 0.039694656 | 0.217153121 | -0.643743361 | -1.557359506 | 25 | 23 | 26 | 88.46 |
| KEGG_GLYCOSAMINOGLYCAN_BIOSYNTHESIS_KERATAN_SULFATE | 0.453355155 | 0.745841905 | -0.459789389 | -1.01700232 | 276 | 14 | 14 | 100 |
| KEGG_GLYCOSAMINOGLYCAN_DEGRADATION | 0.285714286 | 0.640275387 | -0.491404143 | -1.163054552 | 183 | 20 | 22 | 90.91 |
| KEGG_GLYCOSPHINGOLIPID_BIOSYNTHESIS_GANGLIO_SERIES | 0.672638436 | 0.846520095 | -0.376241069 | -0.839625867 | 412 | 15 | 15 | 100 |
| KEGG_GLYCOSPHINGOLIPID_BIOSYNTHESIS_GLOBO_SERIES | 0.826446281 | 0.910696183 | -0.314773655 | -0.686480006 | 499 | 13 | 14 | 92.86 |
| KEGG_GLYCOSPHINGOLIPID_BIOSYNTHESIS_LACTO_AND_NEOLACTO_SERIES | 0.743034056 | 0.873740801 | -0.329065664 | -0.783718571 | 479 | 22 | 23 | 95.65 |
| KEGG_GLYCOSYLPHOSPHATIDYLINOSITOL_GPI_ANCHOR_BIOSYNTHESIS | 0.626911315 | 0.827497627 | -0.357263627 | -0.867574028 | 409 | 24 | 24 | 100 |
| KEGG_GLYOXYLATE_AND_DICARBOXYLATE_METABOLISM | 0.155555556 | 0.464587866 | -0.581649065 | -1.320892985 | 97 | 16 | 16 | 100 |
| KEGG_GNRH_SIGNALING_PATHWAY | 0.394052045 | 0.720710523 | -0.346101748 | -1.048129125 | 317 | 90 | 95 | 94.74 |
| KEGG_GRAFT_VERSUS_HOST_DISEASE | 0.983606557 | 1 | -0.203782034 | -0.512919135 | 659 | 28 | 50 | 56 |
| KEGG_HEDGEHOG_SIGNALING_PATHWAY | 0.993178718 | 1 | -0.189822822 | -0.526652826 | 727 | 49 | 52 | 94.23 |
| KEGG_HEMATOPOIETIC_CELL_LINEAGE | 0.538461538 | 0.776386404 | 0.26004374 | 0.969150179 | 118 | 77 | 88 | 87.5 |

Supplementary Table 2

| pathway | pval | padj | ES | NES | nMoreExtreme | size | Genes_in_Pathway | Percentage_of_Pathway_detected |
|---|---|---|---|---|---|---|---|---|
| KEGG_HISTIDINE_METABOLISM | 0.175159236 | 0.50122489 | 0.395600889 | 1.228485587 | 54 | 31 | 33 | 93.94 |
| KEGG_HOMOLOGOUS_RECOMBINATION | 0.008941878 | 0.092399404 | -0.698939802 | -1.740272396 | 5 | 26 | 26 | 100 |
| KEGG_HUNTINGTONS_DISEASE | 0.008064516 | 0.088235294 | 0.374987876 | 1.576761722 | 0 | 164 | 175 | 93.71 |
| KEGG_HYPERTROPHIC_CARDIOMYOPATHY_HCM | 0.399103139 | 0.720710523 | 0.269768146 | 1.017571561 | 88 | 79 | 84 | 94.05 |
| KEGG_INOSITOL_PHOSPHATE_METABOLISM | 0.333333333 | 0.675949367 | 0.311160239 | 1.068097874 | 86 | 51 | 53 | 96.23 |
| KEGG_INSULIN_SIGNALING_PATHWAY | 0.1152019 | 0.414034344 | -0.403471549 | -1.276767036 | 96 | 132 | 137 | 96.35 |
| KEGG_INTESTINAL_IMMUNE_NETWORK_FOR_IGA_PRODUCTION | 0.079861111 | 0.330092593 | 0.430043051 | 1.357213779 | 22 | 35 | 46 | 76.09 |
| KEGG_JAK_STAT_SIGNALING_PATHWAY | 0.82746051 | 0.910696183 | -0.252952626 | -0.788360871 | 680 | 118 | 353 | 33.43 |
| KEGG_LEISHMANIA_INFECTION | 0.200854701 | 0.557596632 | 0.325095917 | 1.1607451 | 46 | 64 | 70 | 91.43 |
| KEGG_LEUKOCYTE_TRANSENDOTHELIAL_MIGRATION | 0.005076142 | 0.07635468 | 0.397588197 | 1.559989831 | 0 | 105 | 112 | 93.75 |
| KEGG_LIMONENE_AND_PINENE_DEGRADATION | 0.278325123 | 0.640275387 | 0.510063865 | 1.1873329 | 112 | 10 | 10 | 100 |
| KEGG_LINOLEIC_ACID_METABOLISM | 0.958715596 | 0.996207268 | -0.241258034 | -0.585867657 | 626 | 24 | 28 | 85.71 |
| KEGG_LONG_TERM_DEPRESSION | 0.738341969 | 0.873740801 | -0.280541127 | -0.813764468 | 569 | 65 | 67 | 97.01 |
| KEGG_LONG_TERM_POTENTIATION | 0.329015544 | 0.675949367 | -0.376356991 | -1.091697141 | 253 | 65 | 68 | 95.59 |
| KEGG_LYSINE_DEGRADATION | 0.619972261 | 0.827497627 | -0.329460568 | -0.887159525 | 446 | 41 | 42 | 97.62 |
| KEGG_LYSOSOME | 0.005347594 | 0.07635468 | 0.426910097 | 1.707510505 | 0 | 114 | 118 | 96.61 |
| KEGG_MAPK_SIGNALING_PATHWAY | 0.244919786 | 0.624042195 | -0.334022427 | -1.123216188 | 228 | 252 | 266 | 94.74 |
| KEGG_MATURITY_ONSET_DIABETES_OF_THE_YOUNG | 0.383285303 | 0.720710523 | 0.364829684 | 1.063966891 | 132 | 23 | 25 | 92 |
| KEGG_MELANOGENESIS | 0.253382534 | 0.632156863 | -0.381332559 | -1.158185666 | 205 | 91 | 97 | 93.81 |
| KEGG_MELANOMA | 0.627296588 | 0.827497627 | -0.309975822 | -0.888530138 | 477 | 61 | 69 | 88.41 |
| KEGG_METABOLISM_OF_XENOBIOTICS_BY_CYTOCHROME_P450 | 0.035714286 | 0.209725159 | 0.431537968 | 1.441382808 | 9 | 43 | 67 | 64.18 |
| KEGG_MISMATCH_REPAIR | 0.037209302 | 0.209725159 | -0.666478199 | -1.581739256 | 23 | 21 | 21 | 100 |
| KEGG_MTOR_SIGNALING_PATHWAY | 0.457489879 | 0.745841905 | -0.357058775 | -0.998665547 | 338 | 51 | 52 | 98.08 |
| KEGG_NATURAL_KILLER_CELL_MEDIATED_CYTOTOXICITY | 0.140625 | 0.458881579 | 0.305388578 | 1.18654796 | 26 | 99 | 316 | 31.33 |
| KEGG_NEUROACTIVE_LIGAND_RECEPTOR_INTERACTION | 0.633663366 | 0.830009762 | 0.220186656 | 0.960284373 | 63 | 212 | 313 | 67.73 |
| KEGG_NEUROTROPHIN_SIGNALING_PATHWAY | 0.413043478 | 0.731677019 | -0.32983333 | -1.030691347 | 341 | 121 | 126 | 96.03 |
| KEGG_NICOTINATE_AND_NICOTINAMIDE_METABOLISM | 0.045977011 | 0.237547893 | 0.498927802 | 1.473028741 | 15 | 24 | 26 | 92.31 |
| KEGG_NITROGEN_METABOLISM | 0.801550388 | 0.907409885 | -0.320088774 | -0.759660226 | 516 | 21 | 23 | 91.3 |
| KEGG_NOD_LIKE_RECEPTOR_SIGNALING_PATHWAY | 0.394270123 | 0.720710523 | -0.377828765 | -1.048264824 | 288 | 49 | 52 | 94.23 |
| KEGG_NON_HOMOLOGOUS_END_JOINING | 0.509001637 | 0.765463918 | -0.459633277 | -0.990481374 | 310 | 12 | 13 | 92.31 |
| KEGG_NON_SMALL_CELL_LUNG_CANCER | 0.47148289 | 0.745841905 | 0.28876469 | 0.998669472 | 123 | 53 | 53 | 100 |
| KEGG_NOTCH_SIGNALING_PATHWAY | 0.995839112 | 1 | -0.177860036 | -0.487401507 | 717 | 45 | 46 | 97.83 |
| KEGG_NUCLEOTIDE_EXCISION_REPAIR | 0.523012552 | 0.776386404 | -0.355674401 | -0.962010059 | 374 | 42 | 42 | 100 |
| KEGG_N_GLYCAN_BIOSYNTHESIS | 0.001390821 | 0.07635468 | -0.666742735 | -1.823520975 | 0 | 44 | 46 | 95.65 |
| KEGG_OLFACTORY_TRANSDUCTION | 0.9375 | 0.985050699 | -0.223717118 | -0.721615942 | 809 | 155 | 650 | 23.85 |
| KEGG_ONE_CARBON_POOL_BY_FOLATE | 0.838658147 | 0.917590678 | -0.308368657 | -0.711102247 | 524 | 18 | 18 | 100 |
| KEGG_OOCYTE_MEIOSIS | 0.062111801 | 0.288819876 | -0.442150083 | -1.35700632 | 49 | 104 | 111 | 93.69 |
| KEGG_OTHER_GLYCAN_DEGRADATION | 0.172043011 | 0.5 | 0.472540533 | 1.262431864 | 63 | 16 | 16 | 100 |
| KEGG_OXIDATIVE_PHOSPHORYLATION | 0.005747126 | 0.07635468 | 0.551242706 | 2.23044642 | 0 | 121 | 132 | 91.67 |
| KEGG_O_GLYCAN_BIOSYNTHESIS | 0.283987915 | 0.640275387 | 0.373079045 | 1.139376819 | 93 | 28 | 29 | 96.55 |

Supplementary Table 2

| pathway | pval | padj | ES | NES | nMoreExtreme | size | Genes_in_Pathway | Percentage_of_Pathway_detected |
|---|---|---|---|---|---|---|---|---|
| KEGG_P53_SIGNALING_PATHWAY | 0.67357513 | 0.846520095 | -0.294014619 | -0.85284697 | 519 | 65 | 66 | 98.48 |
| KEGG_PANCREATIC_CANCER | 0.409207161 | 0.731677019 | -0.354287225 | -1.03750164 | 319 | 70 | 71 | 98.59 |
| KEGG_PANTOTHENATE_AND_COA_BIOSYNTHESIS | 0.510309278 | 0.765463918 | 0.363801257 | 0.962552434 | 197 | 15 | 15 | 100 |
| KEGG_PARKINSONS_DISEASE | 0.005586592 | 0.07635468 | 0.440585334 | 1.77319905 | 0 | 118 | 126 | 93.65 |
| KEGG_PATHOGENIC_ESCHERICHIA_COLI_INFECTION | 0.278195489 | 0.640275387 | 0.320370937 | 1.106431403 | 73 | 52 | 52 | 100 |
| KEGG_PATHWAYS_IN_CANCER | 0.369747899 | 0.720710523 | -0.309363897 | -1.056307246 | 351 | 317 | 340 | 93.24 |
| KEGG_PENTOSE_AND_GLUCURONATE_INTERCONVERSIONS | 0.384408602 | 0.720710523 | 0.395227309 | 1.055883071 | 142 | 16 | 34 | 47.06 |
| KEGG_PENTOSE_PHOSPHATE_PATHWAY | 0.079365079 | 0.330092593 | 0.455783342 | 1.402543474 | 24 | 30 | 31 | 96.77 |
| KEGG_PEROXISOME | 0.342747112 | 0.678201732 | -0.366725693 | -1.089920049 | 266 | 79 | 80 | 98.75 |
| KEGG_PHENYLALANINE_METABOLISM | 0.443213296 | 0.745841905 | 0.362289801 | 1.011955208 | 159 | 19 | 21 | 90.48 |
| KEGG_PHOSPHATIDYLINOSITOL_SIGNALING_SYSTEM | 0.004587156 | 0.07635468 | 0.477175535 | 1.73258854 | 0 | 72 | 75 | 96 |
| KEGG_PORPHYRIN_AND_CHLOROPHYLL_METABOLISM | 0.306451613 | 0.655172414 | -0.449127475 | -1.144200008 | 208 | 29 | 47 | 61.7 |
| KEGG_PPAR_SIGNALING_PATHWAY | 0.113043478 | 0.414034344 | 0.349297369 | 1.246146632 | 25 | 65 | 78 | 83.33 |
| KEGG_PRIMARY_BILE_ACID_BIOSYNTHESIS | 0.284634761 | 0.640275387 | 0.451571015 | 1.141414339 | 112 | 13 | 15 | 86.67 |
| KEGG_PRIMARY_IMMUNODEFICIENCY | 0.46835443 | 0.745841905 | -0.378142793 | -0.996749912 | 332 | 34 | 38 | 89.47 |
| KEGG_PRION_DISEASES | 0.023054755 | 0.176582278 | -0.63884108 | -1.656396549 | 15 | 32 | 34 | 94.12 |
| KEGG_PROGESTERONE_MEDIATED_OOCYTE_MATURATION | 0.036989796 | 0.209725159 | -0.483648711 | -1.447869391 | 28 | 81 | 84 | 96.43 |
| KEGG_PROPANOATE_METABOLISM | 0.088888889 | 0.35942029 | 0.447283736 | 1.376388356 | 27 | 30 | 32 | 93.75 |
| KEGG_PROSTATE_CANCER | 0.070807453 | 0.306283403 | -0.440750284 | -1.340987138 | 56 | 95 | 103 | 92.23 |
| KEGG_PROTEASOME | 0.031900139 | 0.198576512 | -0.57009003 | -1.562255052 | 22 | 45 | 45 | 100 |
| KEGG_PROTEIN_EXPORT | 0.001490313 | 0.07635468 | -0.778129921 | -1.937445856 | 0 | 26 | 26 | 100 |
| KEGG_PROXIMAL_TUBULE_BICARBONATE_RECLAMATION | 0.238764045 | 0.624042195 | 0.406803954 | 1.182438296 | 84 | 22 | 23 | 95.65 |
| KEGG_PURINE_METABOLISM | 0.586127168 | 0.813579501 | -0.290726084 | -0.936838555 | 506 | 153 | 158 | 96.84 |
| KEGG_PYRIMIDINE_METABOLISM | 0.608856089 | 0.827497627 | -0.299420219 | -0.909400986 | 494 | 91 | 96 | 94.79 |
| KEGG_PYRUVATE_METABOLISM | 0.39748954 | 0.720710523 | -0.395863361 | -1.051303404 | 284 | 36 | 39 | 92.31 |
| KEGG_REGULATION_OF_ACTIN_CYTOSKELETON | 0.392857143 | 0.720710523 | 0.238165674 | 1.027112574 | 43 | 196 | 211 | 92.89 |
| KEGG_REGULATION_OF_AUTOPHAGY | 0.117977528 | 0.414034344 | 0.458907476 | 1.333885202 | 41 | 22 | 206 | 10.68 |
| KEGG_RENAL_CELL_CARCINOMA | 0.536363636 | 0.776386404 | 0.26406649 | 0.950530395 | 117 | 70 | 70 | 100 |
| KEGG_RENIN_ANGIOTENSIN_SYSTEM | 0.438291139 | 0.745841905 | -0.442611906 | -1.016012634 | 276 | 17 | 19 | 89.47 |
| KEGG_RETINOL_METABOLISM | 0.00729927 | 0.084854015 | 0.472311524 | 1.607808854 | 1 | 47 | 81 | 58.02 |
| KEGG_RIBOFLAVIN_METABOLISM | 0.440721649 | 0.745841905 | 0.386598289 | 1.022869265 | 170 | 15 | 16 | 93.75 |
| KEGG_RIBOSOME | 0.004273504 | 0.07635468 | 0.718026132 | 2.5636905 | 0 | 64 | 74 | 86.49 |
| KEGG_RIG_I_LIKE_RECEPTOR_SIGNALING_PATHWAY | 0.150815217 | 0.464587866 | -0.455703072 | -1.273603689 | 110 | 52 | 238 | 21.85 |
| KEGG_RNA_DEGRADATION | 0.135752688 | 0.455594185 | -0.461413577 | -1.302080532 | 100 | 56 | 58 | 96.55 |
| KEGG_RNA_POLYMERASE | 0.754147813 | 0.873740801 | -0.314348041 | -0.777894044 | 499 | 25 | 27 | 92.59 |
| KEGG_SELENOAMINO_ACID_METABOLISM | 0.492378049 | 0.763185976 | -0.39684314 | -0.985005153 | 322 | 27 | 27 | 100 |
| KEGG_SMALL_CELL_LUNG_CANCER | 0.529113924 | 0.776386404 | -0.319692361 | -0.958875298 | 417 | 82 | 82 | 100 |
| KEGG_SNARE_INTERACTIONS_IN_VESICULAR_TRANSPORT | 0.460339943 | 0.745841905 | -0.381744968 | -0.999057992 | 324 | 33 | 34 | 97.06 |
| KEGG_SPHINGOLIPID_METABOLISM | 0.912011173 | 0.974908495 | -0.247589468 | -0.664656673 | 652 | 39 | 40 | 97.5 |
| KEGG_SPLICEOSOME | 0.009720535 | 0.095158918 | -0.494667403 | -1.54271681 | 7 | 122 | 154 | 79.22 |

| pathway | pval | padj | ES | NES | nMoreExtreme | size | Genes_in_Pathway | Percentage_of_Pathway_detected |
|---|---|---|---|---|---|---|---|---|
| KEGG_STARCH_AND_SUCROSE_METABOLISM | 0.43697479 | 0.745841905 | -0.380823753 | -1.007051791 | 311 | 35 | 52 | 67.31 |
| KEGG_STEROID_BIOSYNTHESIS | 0.023734177 | 0.176582278 | -0.727694698 | -1.670418255 | 14 | 17 | 17 | 100 |
| KEGG_STEROID_HORMONE_BIOSYNTHESIS | 0.012987013 | 0.120779221 | 0.517322649 | 1.618039579 | 3 | 32 | 69 | 46.38 |
| KEGG_SULFUR_METABOLISM | 0.061749571 | 0.288819876 | -0.755374196 | -1.532879243 | 35 | 9 | 14 | 64.29 |
| KEGG_SYSTEMIC_LUPUS_ERYTHEMATOSUS | 0.102040816 | 0.395408163 | -0.453684063 | -1.337097915 | 79 | 72 | 105 | 68.57 |
| KEGG_TASTE_TRANSDUCTION | 0.474320242 | 0.745841905 | 0.328923713 | 0.981782196 | 156 | 26 | 69 | 37.68 |
| KEGG_TAURINE_AND_HYPOTAURINE_METABOLISM | 0.556650246 | 0.790482378 | 0.408374229 | 0.950618522 | 225 | 10 | 10 | 100 |
| KEGG_TERPENOID_BACKBONE_BIOSYNTHESIS | 0.050488599 | 0.253807553 | -0.676389718 | -1.509442616 | 30 | 15 | 17 | 88.24 |
| KEGG_TGF_BETA_SIGNALING_PATHWAY | 0.907810499 | 0.974908495 | -0.235139092 | -0.699613739 | 708 | 77 | 83 | 92.77 |
| KEGG_THYROID_CANCER | 0.181950509 | 0.512769618 | -0.486755328 | -1.252850847 | 124 | 30 | 31 | 96.77 |
| KEGG_TIGHT_JUNCTION | 0.811664642 | 0.907409885 | -0.256716033 | -0.800090034 | 667 | 118 | 125 | 94.4 |
| KEGG_TOLL_LIKE_RECEPTOR_SIGNALING_PATHWAY | 0.337974684 | 0.675949367 | -0.366372743 | -1.098886977 | 266 | 82 | 267 | 30.71 |
| KEGG_TRYPTOPHAN_METABOLISM | 0.244755245 | 0.624042195 | 0.348364118 | 1.136777462 | 69 | 39 | 45 | 86.67 |
| KEGG_TYPE_II_DIABETES_MELLITUS | 0.744827586 | 0.873740801 | -0.286047747 | -0.786378656 | 539 | 46 | 48 | 95.83 |
| KEGG_TYPE_I_DIABETES_MELLITUS | 0.929936306 | 0.982773596 | 0.220388133 | 0.684385835 | 291 | 31 | 44 | 70.45 |
| KEGG_TYROSINE_METABOLISM | 0.556737589 | 0.790482378 | 0.29835653 | 0.962717442 | 156 | 38 | 44 | 86.36 |
| KEGG_T_CELL_RECEPTOR_SIGNALING_PATHWAY | 0.157360406 | 0.464587866 | 0.291629291 | 1.144246059 | 30 | 105 | 108 | 97.22 |
| KEGG_UBIQUITIN_MEDIATED_PROTEOLYSIS | 0.235849057 | 0.624042195 | -0.361250203 | -1.145221202 | 199 | 133 | 136 | 97.79 |
| KEGG_VALINE_LEUCINE_AND_ISOLEUCINE_BIOSYNTHESIS | 0.156040268 | 0.464587866 | -0.641355559 | -1.336111476 | 92 | 10 | 11 | 90.91 |
| KEGG_VALINE_LEUCINE_AND_ISOLEUCINE_DEGRADATION | 0.00729927 | 0.084854015 | 0.487949572 | 1.66104277 | 1 | 47 | 48 | 97.92 |
| KEGG_VASCULAR_SMOOTH_MUSCLE_CONTRACTION | 0.644688645 | 0.838546069 | -0.288672093 | -0.894614596 | 527 | 110 | 119 | 92.44 |
| KEGG_VASOPRESSIN_REGULATED_WATER_REABSORPTION | 0.045769764 | 0.237547893 | -0.558569963 | -1.504097037 | 32 | 41 | 44 | 93.18 |
| KEGG_VEGF_SIGNALING_PATHWAY | 0.137168142 | 0.455594185 | 0.329636352 | 1.208572136 | 30 | 74 | 74 | 100 |
| KEGG_VIBRIO_CHOLERAE_INFECTION | 0.275579809 | 0.640275387 | -0.416482388 | -1.156863398 | 201 | 50 | 52 | 96.15 |
| KEGG_VIRAL_MYOCARDITIS | 0.477178423 | 0.745841905 | 0.285215957 | 0.999594662 | 114 | 59 | 70 | 84.29 |
| KEGG_WNT_SIGNALING_PATHWAY | 0.618266979 | 0.827497627 | -0.286970985 | -0.91743209 | 527 | 143 | 151 | 94.7 |

Supplementary Table 2

| pathway | pval | padj | ES | NES | nMoreExtreme | size | Genes_in_Pathway | Percentage_of_Pathway_detected |
|---|---|---|---|---|---|---|---|---|
| KEGG_ABC_TRANSPORTERS | 0.368421053 | 0.692185008 | 0.382531378 | 1.069332114 | 251 | 30 | 43 | 69.77 |
| KEGG_ACUTE_MYELOID_LEUKEMIA | 0.318361955 | 0.650717842 | 0.357323963 | 1.108263866 | 240 | 56 | 59 | 94.92 |
| KEGG_ADHERENS_JUNCTION | 0.211453744 | 0.551928783 | -0.280633446 | -1.144286757 | 47 | 65 | 73 | 89.04 |
| KEGG_ADIPOCYTOKINE_SIGNALING_PATHWAY | 0.121338912 | 0.417945142 | -0.324665613 | -1.26817802 | 28 | 55 | 67 | 82.09 |
| KEGG_ALANINE_ASPARTATE_AND_GLUTAMATE_METABOLISM | 0.777443609 | 0.89708642 | 0.283488233 | 0.757648608 | 516 | 25 | 33 | 75.76 |
| KEGG_ALDOSTERONE_REGULATED_SODIUM_REABSORPTION | 0.461538462 | 0.784723751 | -0.278935641 | -0.976772712 | 143 | 31 | 43 | 72.09 |
| KEGG_ALLOGRAFT_REJECTION | 0.614318707 | 0.817927847 | -0.401593943 | -0.858101709 | 265 | 6 | 37 | 16.22 |
| KEGG_ALPHA_LINOLENIC_ACID_METABOLISM | 0.965798046 | 0.989829707 | 0.210850287 | 0.508873573 | 592 | 14 | 18 | 77.78 |
| KEGG_ALZHEIMERS_DISEASE | 0.001123596 | 0.05593985 | 0.473255967 | 1.640148147 | 0 | 140 | 165 | 84.85 |
| KEGG_AMINOACYL_TRNA_BIOSYNTHESIS | 0.099579243 | 0.385869565 | 0.470201024 | 1.392572372 | 70 | 41 | 41 | 100 |
| KEGG_AMINO_SUGAR_AND_NUCLEOTIDE_SUGAR_METABOLISM | 0.033660589 | 0.236095084 | 0.520334968 | 1.541051731 | 23 | 41 | 48 | 85.42 |
| KEGG_AMYOTROPHIC_LATERAL_SCLEROSIS_ALS | 0.737658674 | 0.862921468 | 0.271913813 | 0.80956627 | 522 | 43 | 51 | 84.31 |
| KEGG_ANTIGEN_PROCESSING_AND_PRESENTATION | 0.001390821 | 0.05593985 | 0.709493275 | 2.058442213 | 0 | 36 | 279 | 12.9 |
| KEGG_APOPTOSIS | 0.964197531 | 0.989829707 | 0.191398321 | 0.617201226 | 780 | 74 | 89 | 83.15 |
| KEGG_ARACHIDONIC_ACID_METABOLISM | 0.460361613 | 0.784723751 | 0.343412166 | 0.996336573 | 330 | 36 | 60 | 60 |
| KEGG_ARGININE_AND_PROLINE_METABOLISM | 0.012482663 | 0.135964912 | 0.544849455 | 1.630625805 | 8 | 46 | 59 | 77.97 |
| KEGG_ARRHYTHMOGENIC_RIGHT_VENTRICULAR_CARDIOMYOPATHY_ARVC | 0.216438356 | 0.551928783 | 0.401008435 | 1.204359999 | 157 | 48 | 74 | 64.86 |
| KEGG_ASCORBATE_AND_ALDARATE_METABOLISM | 0.208877285 | 0.551928783 | -0.470256781 | -1.277958347 | 79 | 13 | 31 | 41.94 |
| KEGG_ASTHMA | 0.614349776 | 0.817927847 | -0.44225894 | -0.893148881 | 273 | 5 | 32 | 15.62 |
| KEGG_AUTOIMMUNE_THYROID_DISEASE | 0.656950673 | 0.817927847 | -0.423431503 | -0.855126576 | 292 | 5 | 221 | 2.26 |
| KEGG_AXON_GUIDANCE | 0.072847682 | 0.341449275 | -0.284506507 | -1.222931841 | 10 | 101 | 129 | 78.29 |
| KEGG_BASAL_CELL_CARCINOMA | 0.224832215 | 0.551928783 | -0.329914606 | -1.172844731 | 66 | 35 | 55 | 63.64 |
| KEGG_BASAL_TRANSCRIPTION_FACTORS | 0.894202899 | 0.959337149 | 0.240713246 | 0.678699014 | 616 | 31 | 38 | 81.58 |
| KEGG_BASE_EXCISION_REPAIR | 0.195652174 | 0.551928783 | 0.442018827 | 1.246286802 | 134 | 31 | 32 | 96.88 |
| KEGG_BETA_ALANINE_METABOLISM | 0.508716323 | 0.784794937 | 0.38479367 | 0.974738731 | 320 | 18 | 22 | 81.82 |
| KEGG_BIOSYNTHESIS_OF_UNSATURATED_FATTY_ACIDS | 0.975193798 | 0.991180582 | 0.192117262 | 0.500212505 | 628 | 22 | 29 | 75.86 |
| KEGG_BLADDER_CANCER | 0.367176634 | 0.692185008 | 0.367633131 | 1.073486646 | 263 | 37 | 42 | 88.1 |
| KEGG_BUTANOATE_METABOLISM | 0.907759883 | 0.959337149 | 0.229060658 | 0.632235369 | 619 | 28 | 38 | 73.68 |
| KEGG_B_CELL_RECEPTOR_SIGNALING_PATHWAY | 0.428571429 | 0.763499446 | -0.24957968 | -1.020558458 | 95 | 66 | 90 | 73.33 |
| KEGG_CALCIUM_SIGNALING_PATHWAY | 0.514757969 | 0.784794937 | 0.291092221 | 0.977834847 | 435 | 103 | 173 | 59.54 |
| KEGG_CARDIAC_MUSCLE_CONTRACTION | 0.057746479 | 0.298356808 | 0.492460545 | 1.469587425 | 40 | 45 | 79 | 56.96 |
| KEGG_CELL_ADHESION_MOLECULES_CAMS | 0.699354839 | 0.839225806 | 0.267792211 | 0.840865669 | 541 | 65 | 129 | 50.39 |
| KEGG_CELL_CYCLE | 0.718277066 | 0.856407271 | 0.250866641 | 0.852037079 | 616 | 118 | 124 | 95.16 |
| KEGG_CHEMOKINE_SIGNALING_PATHWAY | 0.884222474 | 0.959337149 | 0.214005104 | 0.738406267 | 778 | 132 | 178 | 74.16 |
| KEGG_CHRONIC_MYELOID_LEUKEMIA | 0.780150754 | 0.89708642 | 0.246155775 | 0.785144284 | 620 | 69 | 72 | 95.83 |
| KEGG_CIRCADIAN_RHYTHM_MAMMAL | 0.539735099 | 0.803125828 | 0.406137831 | 0.943928748 | 325 | 12 | 13 | 92.31 |
| KEGG_CITRATE_CYCLE_TCA_CYCLE | 0.665217391 | 0.817927847 | 0.306535206 | 0.856738096 | 458 | 29 | 31 | 93.55 |
| KEGG_COLORECTAL_CANCER | 0.574967405 | 0.817927847 | 0.292600937 | 0.914616784 | 440 | 58 | 64 | 90.62 |
| KEGG_COMPLEMENT_AND_COAGULATION_CASCADES | 0.230769231 | 0.551928783 | -0.336385107 | -1.150520368 | 71 | 29 | 74 | 39.19 |
| KEGG_CYSTEINE_AND_METHIONINE_METABOLISM | 0.231454006 | 0.551928783 | 0.439757041 | 1.204860888 | 155 | 27 | 38 | 71.05 |
| KEGG_CYTOKINE_CYTOKINE_RECEPTOR_INTERACTION | 0.851351351 | 0.954940642 | -0.194472102 | -0.858144217 | 125 | 112 | 460 | 24.35 |
| KEGG_CYTOSOLIC_DNA_SENSING_PATHWAY | 0.171875 | 0.54184322 | 0.441642794 | 1.268331027 | 120 | 33 | 221 | 14.93 |
| KEGG_DILATED_CARDIOMYOPATHY | 0.268817204 | 0.581395349 | 0.375517931 | 1.151503132 | 199 | 53 | 90 | 58.89 |
| KEGG_DNA_REPLICATION | 0.053824363 | 0.29688747 | 0.511563339 | 1.469072614 | 37 | 34 | 34 | 100 |

| pathway | pval | padj | ES | NES | nMoreExtreme | size | Genes_in_Pathway | Percentage_of_Pathway_detected |
|---|---|---|---|---|---|---|---|---|
| KEGG_DORSO_VENTRAL_AXIS_FORMATION | 0.781333333 | 0.89708642 | -0.251049397 | -0.776806167 | 292 | 19 | 21 | 90.48 |
| KEGG_DRUG_METABOLISM_CYTOCHROME_P450 | 0.003558719 | 0.066192171 | -0.551525336 | -2.103398725 | 0 | 46 | 84 | 54.76 |
| KEGG_DRUG_METABOLISM_OTHER_ENZYMES | 0.028268551 | 0.228606545 | -0.438162103 | -1.547057556 | 7 | 37 | 73 | 50.68 |
| KEGG_ECM_RECEPTOR_INTERACTION | 0.361147327 | 0.692185008 | 0.344174698 | 1.075826887 | 276 | 58 | 83 | 69.88 |
| KEGG_ENDOCYTOSIS | 0.673672566 | 0.818974492 | 0.255456258 | 0.893276545 | 608 | 152 | 184 | 82.61 |
| KEGG_ENDOMETRIAL_CANCER | 0.656207367 | 0.817927847 | 0.285950219 | 0.866429714 | 480 | 51 | 52 | 98.08 |
| KEGG_EPITHELIAL_CELL_SIGNALING_IN_HELICOBACTER_PYLORI_INFECTION | 0.565891473 | 0.817927847 | -0.246142258 | -0.95167333 | 145 | 52 | 64 | 81.25 |
| KEGG_ERBB_SIGNALING_PATHWAY | 0.889585947 | 0.959337149 | 0.220115178 | 0.707983168 | 708 | 78 | 86 | 90.7 |
| KEGG_ETHER_LIPID_METABOLISM | 0.995488722 | 0.995488722 | 0.166862387 | 0.445955213 | 661 | 25 | 32 | 78.12 |
| KEGG_FATTY_ACID_METABOLISM | 0.605006954 | 0.817927847 | 0.309133153 | 0.896883388 | 434 | 36 | 51 | 70.59 |
| KEGG_FC_EPSILON_RI_SIGNALING_PATHWAY | 0.885750963 | 0.959337149 | 0.221079221 | 0.698588149 | 689 | 67 | 78 | 85.9 |
| KEGG_FC_GAMMA_R_MEDIATED_PHAGOCYTOSIS | 0.534567901 | 0.801851852 | 0.292186742 | 0.952459685 | 432 | 85 | 99 | 85.86 |
| KEGG_FOCAL_ADHESION | 0.204646018 | 0.551928783 | 0.336268515 | 1.175859927 | 184 | 152 | 195 | 77.95 |
| KEGG_FOLATE_BIOSYNTHESIS | 0.484193012 | 0.784723751 | 0.439399874 | 0.98359528 | 290 | 10 | 17 | 58.82 |
| KEGG_FRUCTOSE_AND_MANNOSE_METABOLISM | 0.203125 | 0.551928783 | 0.429834819 | 1.236183205 | 142 | 35 | 38 | 92.11 |
| KEGG_GALACTOSE_METABOLISM | 0.265957447 | 0.581395349 | 0.451166412 | 1.184822141 | 174 | 23 | 26 | 88.46 |
| KEGG_GAP_JUNCTION | 0.216450216 | 0.551928783 | -0.27910703 | -1.138827411 | 49 | 64 | 84 | 76.19 |
| KEGG_GLIOMA | 0.668414155 | 0.817927847 | 0.277255288 | 0.860640157 | 509 | 55 | 61 | 90.16 |
| KEGG_GLUTATHIONE_METABOLISM | 0.492977528 | 0.784723751 | 0.32670576 | 0.975689387 | 350 | 44 | 48 | 91.67 |
| KEGG_GLYCEROLIPID_METABOLISM | 0.190409027 | 0.551928783 | 0.413447814 | 1.230954031 | 134 | 43 | 49 | 87.76 |
| KEGG_GLYCEROPHOSPHOLIPID_METABOLISM | 0.665137615 | 0.817927847 | -0.214876772 | -0.895062283 | 144 | 70 | 76 | 92.11 |
| KEGG_GLYCINE_SERINE_AND_THREONINE_METABOLISM | 0.213533835 | 0.551928783 | 0.467503179 | 1.241755383 | 141 | 24 | 32 | 75 |
| KEGG_GLYCOLYSIS_GLUCONEOGENESIS | 0.089041096 | 0.360035736 | 0.473651176 | 1.422530002 | 64 | 48 | 67 | 71.64 |
| KEGG_GLYCOSAMINOGLYCAN_BIOSYNTHESIS_CHONDROITIN_SULFATE | 0.081433225 | 0.341449275 | 0.604403137 | 1.458687999 | 49 | 14 | 22 | 63.64 |
| KEGG_GLYCOSAMINOGLYCAN_BIOSYNTHESIS_HEPARAN_SULFATE | 0.023923445 | 0.211952393 | 0.638675844 | 1.625881701 | 14 | 19 | 26 | 73.08 |
| KEGG_GLYCOSAMINOGLYCAN_BIOSYNTHESIS_KERATAN_SULFATE | 0.299668874 | 0.629759581 | 0.507320063 | 1.15769115 | 180 | 11 | 14 | 78.57 |
| KEGG_GLYCOSAMINOGLYCAN_DEGRADATION | 0.239302694 | 0.556378764 | 0.481238192 | 1.219046834 | 150 | 18 | 22 | 81.82 |
| KEGG_GLYCOSPHINGOLIPID_BIOSYNTHESIS_GANGLIO_SERIES | 0.42218543 | 0.762393107 | 0.447381352 | 1.039785236 | 254 | 12 | 15 | 80 |
| KEGG_GLYCOSPHINGOLIPID_BIOSYNTHESIS_GLOBO_SERIES | 0.32615894 | 0.659408293 | 0.482086384 | 1.120445236 | 196 | 12 | 14 | 85.71 |
| KEGG_GLYCOSPHINGOLIPID_BIOSYNTHESIS_LACTO_AND_NEOLACTO_SERIES | 0.212938005 | 0.551928783 | -0.40402461 | -1.202513313 | 78 | 18 | 23 | 78.26 |
| KEGG_GLYCOSYLPHOSPHATIDYLINOSITOL_GPI_ANCHOR_BIOSYNTHESIS | 0.384962406 | 0.716030075 | 0.401219 | 1.065695113 | 255 | 24 | 24 | 100 |
| KEGG_GLYOXYLATE_AND_DICARBOXYLATE_METABOLISM | 0.055374593 | 0.29688747 | 0.628824977 | 1.517628535 | 33 | 14 | 16 | 87.5 |
| KEGG_GNRH_SIGNALING_PATHWAY | 0.801488834 | 0.91458235 | 0.238425272 | 0.772388112 | 645 | 81 | 95 | 85.26 |
| KEGG_GRAFT_VERSUS_HOST_DISEASE | 0.905982906 | 0.959337149 | 0.282614342 | 0.594995965 | 529 | 8 | 50 | 16 |
| KEGG_HEDGEHOG_SIGNALING_PATHWAY | 0.213414634 | 0.551928783 | -0.346193354 | -1.172808434 | 69 | 27 | 52 | 51.92 |
| KEGG_HEMATOPOIETIC_CELL_LINEAGE | 0.994366197 | 0.995488722 | 0.17283114 | 0.515758009 | 705 | 45 | 88 | 51.14 |
| KEGG_HISTIDINE_METABOLISM | 0.207282913 | 0.551928783 | -0.373060783 | -1.202248921 | 73 | 22 | 33 | 66.67 |
| KEGG_HOMOLOGOUS_RECOMBINATION | 0.367164179 | 0.692185008 | 0.394602913 | 1.071797344 | 245 | 26 | 26 | 100 |
| KEGG_HUNTINGTONS_DISEASE | 0.003351955 | 0.066192171 | 0.442057034 | 1.539389379 | 2 | 150 | 175 | 85.71 |
| KEGG_HYPERTROPHIC_CARDIOMYOPATHY_HCM | 0.454794521 | 0.784723751 | 0.338478485 | 1.016562026 | 331 | 48 | 84 | 57.14 |
| KEGG_INOSITOL_PHOSPHATE_METABOLISM | 0.680763984 | 0.822221435 | 0.277993177 | 0.84231986 | 498 | 51 | 53 | 96.23 |
| KEGG_INSULIN_SIGNALING_PATHWAY | 0.342989571 | 0.678681492 | 0.318285554 | 1.083127308 | 295 | 119 | 137 | 86.86 |
| KEGG_INTESTINAL_IMMUNE_NETWORK_FOR_IGA_PRODUCTION | 0.263736264 | 0.581395349 | -0.365477916 | -1.145970069 | 95 | 20 | 46 | 43.48 |
| KEGG_JAK_STAT_SIGNALING_PATHWAY | 0.852258852 | 0.954940642 | 0.225228133 | 0.740251594 | 697 | 87 | 353 | 24.65 |

| pathway | pval | padj | ES | NES | nMoreExtreme | size | Genes_in_Pathway | Percentage_of_Pathway_detected |
|---|---|---|---|---|---|---|---|---|
| KEGG_LEISHMANIA_INFECTION | 0.003558719 | 0.066192171 | -0.461092151 | -1.758506053 | 0 | 46 | 70 | 65.71 |
| KEGG_LEUKOCYTE_TRANSENDOTHELIAL_MIGRATION | 0.055865922 | 0.29688747 | -0.294097759 | -1.265357495 | 9 | 88 | 112 | 78.57 |
| KEGG_LIMONENE_AND_PINENE_DEGRADATION | 0.477537438 | 0.784723751 | 0.442981056 | 0.991611745 | 286 | 10 | 10 | 100 |
| KEGG_LINOLEIC_ACID_METABOLISM | 0.936 | 0.98359322 | -0.203668295 | -0.630197837 | 350 | 19 | 28 | 67.86 |
| KEGG_LONG_TERM_DEPRESSION | 0.479452055 | 0.784723751 | 0.331634825 | 0.996008268 | 349 | 48 | 67 | 71.64 |
| KEGG_LONG_TERM_POTENTIATION | 0.494132986 | 0.784723751 | 0.314271684 | 0.981284203 | 378 | 57 | 68 | 83.82 |
| KEGG_LYSINE_DEGRADATION | 0.48028169 | 0.784723751 | 0.338859284 | 0.991176776 | 340 | 38 | 42 | 90.48 |
| KEGG_LYSOSOME | 0.012820513 | 0.135964912 | -0.298605444 | -1.316300651 | 1 | 107 | 118 | 90.68 |
| KEGG_MAPK_SIGNALING_PATHWAY | 0.511777302 | 0.784794937 | 0.275581498 | 0.983427124 | 477 | 207 | 266 | 77.82 |
| KEGG_MATURITY_ONSET_DIABETES_OF_THE_YOUNG | 0.967084639 | 0.989829707 | 0.204461969 | 0.527898999 | 616 | 20 | 25 | 80 |
| KEGG_MELANOGENESIS | 0.577777778 | 0.817927847 | 0.28610023 | 0.922586005 | 467 | 74 | 97 | 76.29 |
| KEGG_MELANOMA | 0.230040595 | 0.551928783 | 0.390526755 | 1.185198902 | 169 | 50 | 69 | 72.46 |
| KEGG_METABOLISM_OF_XENOBIOTICS_BY_CYTOCHROME_P450 | 0.081272085 | 0.341449275 | -0.389311156 | -1.374575214 | 22 | 37 | 67 | 55.22 |
| KEGG_MISMATCH_REPAIR | 0.234836703 | 0.552906668 | 0.467069708 | 1.213335096 | 150 | 21 | 21 | 100 |
| KEGG_MTOR_SIGNALING_PATHWAY | 0.120996441 | 0.417945142 | -0.326444943 | -1.244990631 | 33 | 46 | 52 | 88.46 |
| KEGG_NATURAL_KILLER_CELL_MEDIATED_CYTOTOXICITY | 0.64516129 | 0.817927847 | -0.218790882 | -0.928731694 | 119 | 82 | 316 | 25.95 |
| KEGG_NEUROACTIVE_LIGAND_RECEPTOR_INTERACTION | 0.005235602 | 0.088529272 | -0.421030338 | -1.807192653 | 0 | 84 | 313 | 26.84 |
| KEGG_NEUROTROPHIN_SIGNALING_PATHWAY | 0.663924794 | 0.817927847 | 0.260962628 | 0.879253276 | 564 | 109 | 126 | 86.51 |
| KEGG_NICOTINATE_AND_NICOTINAMIDE_METABOLISM | 0.493001555 | 0.784723751 | 0.378035312 | 0.982045087 | 316 | 21 | 26 | 80.77 |
| KEGG_NITROGEN_METABOLISM | 0.509333333 | 0.784794937 | -0.308919652 | -0.955870413 | 190 | 19 | 23 | 82.61 |
| KEGG_NOD_LIKE_RECEPTOR_SIGNALING_PATHWAY | 0.051460362 | 0.29688747 | 0.500092158 | 1.488458197 | 36 | 42 | 52 | 80.77 |
| KEGG_NON_HOMOLOGOUS_END_JOINING | 0.968543046 | 0.989829707 | 0.226394628 | 0.526177031 | 584 | 12 | 13 | 92.31 |
| KEGG_NON_SMALL_CELL_LUNG_CANCER | 0.593451569 | 0.817927847 | 0.30140885 | 0.913269396 | 434 | 51 | 53 | 96.23 |
| KEGG_NOTCH_SIGNALING_PATHWAY | 0.134275618 | 0.451410178 | -0.335942789 | -1.244283428 | 37 | 42 | 46 | 91.3 |
| KEGG_NUCLEOTIDE_EXCISION_REPAIR | 0.862308762 | 0.959337149 | 0.238709946 | 0.710488598 | 619 | 42 | 42 | 100 |
| KEGG_N_GLYCAN_BIOSYNTHESIS | 0.001410437 | 0.05593985 | 0.712867328 | 2.122412747 | 0 | 43 | 46 | 93.48 |
| KEGG_OLFACTORY_TRANSDUCTION | 0.635258359 | 0.817927847 | 0.33139613 | 0.870289679 | 417 | 23 | 650 | 3.54 |
| KEGG_ONE_CARBON_POOL_BY_FOLATE | 0.884210526 | 0.959337149 | -0.232384214 | -0.687148869 | 335 | 17 | 18 | 94.44 |
| KEGG_OOCYTE_MEIOSIS | 0.619804401 | 0.817927847 | 0.273310655 | 0.900223627 | 506 | 92 | 111 | 82.88 |
| KEGG_OTHER_GLYCAN_DEGRADATION | 0.1043257 | 0.39601184 | -0.4809257 | -1.384029727 | 40 | 15 | 16 | 93.75 |
| KEGG_OXIDATIVE_PHOSPHORYLATION | 0.003512881 | 0.066192171 | 0.476189795 | 1.609345334 | 2 | 111 | 132 | 84.09 |
| KEGG_O_GLYCAN_BIOSYNTHESIS | 0.431007752 | 0.763499446 | 0.396778582 | 1.033085763 | 277 | 22 | 29 | 75.86 |
| KEGG_P53_SIGNALING_PATHWAY | 0.604681404 | 0.817927847 | 0.288480346 | 0.900810492 | 464 | 61 | 66 | 92.42 |
| KEGG_PANCREATIC_CANCER | 0.225680934 | 0.551928783 | 0.381349916 | 1.194274319 | 173 | 64 | 71 | 90.14 |
| KEGG_PANTOTHENATE_AND_COA_BIOSYNTHESIS | 0.035541195 | 0.236095084 | 0.662316317 | 1.574567881 | 21 | 13 | 15 | 86.67 |
| KEGG_PARKINSONS_DISEASE | 0.001182033 | 0.05593985 | 0.503222288 | 1.690531972 | 0 | 107 | 126 | 84.92 |
| KEGG_PATHOGENIC_ESCHERICHIA_COLI_INFECTION | 0.051317614 | 0.29688747 | 0.497074252 | 1.487644146 | 36 | 46 | 52 | 88.46 |
| KEGG_PATHWAYS_IN_CANCER | 0.358350951 | 0.692185008 | 0.29087853 | 1.061653659 | 338 | 262 | 340 | 77.06 |
| KEGG_PENTOSE_AND_GLUCURONATE_INTERCONVERSIONS | 0.60904685 | 0.817927847 | 0.355298208 | 0.876039153 | 376 | 16 | 34 | 47.06 |
| KEGG_PENTOSE_PHOSPHATE_PATHWAY | 0.297744361 | 0.629759581 | 0.432972736 | 1.150037581 | 197 | 24 | 31 | 77.42 |
| KEGG_PEROXISOME | 0.07037037 | 0.341449275 | 0.439455145 | 1.417108847 | 56 | 74 | 80 | 92.5 |
| KEGG_PHENYLALANINE_METABOLISM | 0.2247557 | 0.551928783 | 0.512030277 | 1.235752059 | 137 | 14 | 21 | 66.67 |
| KEGG_PHOSPHATIDYLINOSITOL_SIGNALING_SYSTEM | 0.009174312 | 0.121887287 | -0.367719909 | -1.53172545 | 1 | 70 | 75 | 93.33 |
| KEGG_PORPHYRIN_AND_CHLOROPHYLL_METABOLISM | 0.077598829 | 0.341449275 | 0.521440351 | 1.439239005 | 52 | 28 | 47 | 59.57 |

| pathway | pval | padj | ES | NES | nMoreExtreme | size | Genes_in_Pathway | Percentage_of_Pathway_detected |
|---|---|---|---|---|---|---|---|---|
| KEGG_PPAR_SIGNALING_PATHWAY | 0.44486692 | 0.780615539 | -0.260910242 | -1.004148619 | 116 | 50 | 78 | 64.1 |
| KEGG_PRIMARY_BILE_ACID_BIOSYNTHESIS | 0.400690846 | 0.730671543 | 0.524048187 | 1.072932414 | 231 | 7 | 15 | 46.67 |
| KEGG_PRIMARY_IMMUNODEFICIENCY | 0.590111643 | 0.817927847 | 0.355618529 | 0.905300652 | 369 | 19 | 38 | 50 |
| KEGG_PRION_DISEASES | 0.030395137 | 0.23556231 | 0.613404763 | 1.61088132 | 19 | 23 | 34 | 67.65 |
| KEGG_PROGESTERONE_MEDIATED_OOCYTE_MATURATION | 0.609876543 | 0.817927847 | 0.280485227 | 0.904479333 | 493 | 74 | 84 | 88.1 |
| KEGG_PROPANOATE_METABOLISM | 0.034782609 | 0.236095084 | 0.558563476 | 1.561134249 | 23 | 29 | 32 | 90.62 |
| KEGG_PROSTATE_CANCER | 0.051787916 | 0.29688747 | 0.438063155 | 1.426313289 | 41 | 84 | 103 | 81.55 |
| KEGG_PROTEASOME | 0.002781641 | 0.066192171 | 0.598252706 | 1.780620094 | 1 | 42 | 45 | 93.33 |
| KEGG_PROTEIN_EXPORT | 0.001503759 | 0.05593985 | 0.84299277 | 2.239109502 | 0 | 24 | 26 | 92.31 |
| KEGG_PROXIMAL_TUBULE_BICARBONATE_RECLAMATION | 0.099236641 | 0.385869565 | -0.485794395 | -1.398041079 | 38 | 15 | 23 | 65.22 |
| KEGG_PURINE_METABOLISM | 0.472190692 | 0.784723751 | 0.292167251 | 1.006910646 | 415 | 131 | 158 | 82.91 |
| KEGG_PYRIMIDINE_METABOLISM | 0.245421245 | 0.563559897 | 0.358662785 | 1.178807884 | 200 | 87 | 96 | 90.62 |
| KEGG_PYRUVATE_METABOLISM | 0.007082153 | 0.101329266 | 0.607075336 | 1.743357437 | 4 | 34 | 39 | 87.18 |
| KEGG_REGULATION_OF_ACTIN_CYTOSKELETON | 0.524807056 | 0.79361067 | 0.279024933 | 0.978866553 | 475 | 168 | 211 | 79.62 |
| KEGG_REGULATION_OF_AUTOPHAGY | 0.168 | 0.538758621 | -0.411966218 | -1.274720842 | 62 | 19 | 206 | 9.22 |
| KEGG_RENAL_CELL_CARCINOMA | 0.497835498 | 0.784723751 | -0.241987737 | -0.987371289 | 114 | 64 | 70 | 91.43 |
| KEGG_RENIN_ANGIOTENSIN_SYSTEM | 0.582781457 | 0.817927847 | 0.394009056 | 0.915739551 | 351 | 12 | 19 | 63.16 |
| KEGG_RETINOL_METABOLISM | 0.00660066 | 0.101329266 | -0.530072403 | -1.862847126 | 1 | 32 | 81 | 39.51 |
| KEGG_RIBOFLAVIN_METABOLISM | 0.620689655 | 0.817927847 | 0.364389739 | 0.885039554 | 377 | 15 | 16 | 93.75 |
| KEGG_RIBOSOME | 0.013157895 | 0.135964912 | -0.384481907 | -1.552471037 | 2 | 63 | 74 | 85.14 |
| KEGG_RIG_I_LIKE_RECEPTOR_SIGNALING_PATHWAY | 0.11878453 | 0.417945142 | 0.449625478 | 1.347717179 | 85 | 47 | 238 | 19.75 |
| KEGG_RNA_DEGRADATION | 0.96 | 0.989829707 | -0.175429218 | -0.68256045 | 239 | 54 | 58 | 93.1 |
| KEGG_RNA_POLYMERASE | 0.332330827 | 0.664661654 | 0.417095815 | 1.114727268 | 220 | 25 | 27 | 92.59 |
| KEGG_SELENOAMINO_ACID_METABOLISM | 0.203007519 | 0.551928783 | 0.473134269 | 1.256712364 | 134 | 24 | 27 | 88.89 |
| KEGG_SMALL_CELL_LUNG_CANCER | 0.730337079 | 0.859763903 | 0.253273144 | 0.813559956 | 584 | 75 | 82 | 91.46 |
| KEGG_SNARE_INTERACTIONS_IN_VESICULAR_TRANSPORT | 0.082608696 | 0.341449275 | 0.498313084 | 1.405010335 | 56 | 31 | 34 | 91.18 |
| KEGG_SPHINGOLIPID_METABOLISM | 0.310954064 | 0.642638398 | -0.310585684 | -1.096612249 | 87 | 37 | 40 | 92.5 |
| KEGG_SPLICEOSOME | 0.253179191 | 0.574284506 | 0.338488203 | 1.153661286 | 218 | 120 | 154 | 77.92 |
| KEGG_STARCH_AND_SUCROSE_METABOLISM | 0.135908441 | 0.451410178 | 0.464154343 | 1.321867187 | 94 | 32 | 52 | 61.54 |
| KEGG_STEROID_BIOSYNTHESIS | 0.009852217 | 0.122167488 | 0.697100777 | 1.69313703 | 5 | 15 | 17 | 88.24 |
| KEGG_STEROID_HORMONE_BIOSYNTHESIS | 0.025069638 | 0.211952393 | -0.528799919 | -1.681205727 | 8 | 21 | 69 | 30.43 |
| KEGG_SULFUR_METABOLISM | 0.041450777 | 0.265856709 | 0.780190056 | 1.597355396 | 23 | 7 | 14 | 50 |
| KEGG_SYSTEMIC_LUPUS_ERYTHEMATOSUS | 0.113764045 | 0.414904164 | 0.460206398 | 1.374381946 | 80 | 44 | 105 | 41.9 |
| KEGG_TASTE_TRANSDUCTION | 0.650470219 | 0.817927847 | 0.333023505 | 0.859831172 | 414 | 20 | 69 | 28.99 |
| KEGG_TAURINE_AND_HYPOTAURINE_METABOLISM | 0.98108747 | 0.991751465 | -0.228437048 | -0.510768137 | 414 | 7 | 10 | 70 |
| KEGG_TERPENOID_BACKBONE_BIOSYNTHESIS | 0.024429967 | 0.211952393 | 0.677581989 | 1.63530044 | 14 | 14 | 17 | 82.35 |
| KEGG_TGF_BETA_SIGNALING_PATHWAY | 0.081545064 | 0.341449275 | -0.324208208 | -1.299772198 | 18 | 61 | 83 | 73.49 |
| KEGG_THYROID_CANCER | 0.035139092 | 0.236095084 | 0.565133571 | 1.559837624 | 23 | 28 | 31 | 90.32 |
| KEGG_TIGHT_JUNCTION | 0.301336574 | 0.629759581 | 0.338410881 | 1.119057036 | 247 | 94 | 125 | 75.2 |
| KEGG_TOLL_LIKE_RECEPTOR_SIGNALING_PATHWAY | 0.16112532 | 0.525777359 | 0.398600349 | 1.263153065 | 125 | 68 | 267 | 25.47 |
| KEGG_TRYPTOPHAN_METABOLISM | 0.257142857 | 0.576247849 | 0.445074722 | 1.189503494 | 170 | 25 | 45 | 55.56 |
| KEGG_TYPE_II_DIABETES_MELLITUS | 0.881615599 | 0.959337149 | 0.233296534 | 0.693166548 | 632 | 40 | 48 | 83.33 |
| KEGG_TYPE_I_DIABETES_MELLITUS | 0.663907285 | 0.817927847 | 0.366198021 | 0.835654332 | 400 | 11 | 44 | 25 |
| KEGG_TYROSINE_METABOLISM | 0.636498516 | 0.817927847 | 0.319994659 | 0.876731954 | 428 | 27 | 44 | 61.36 |

Supplementary Table 2

| pathway | pval | padj | ES | NES | nMoreExtreme | size | Genes_in_Pathway | Percentage_of_Pathway_detected |
|---|---|---|---|---|---|---|---|---|
| KEGG_T_CELL_RECEPTOR_SIGNALING_PATHWAY | 0.625 | 0.817927847 | -0.217545927 | -0.941814495 | 114 | 92 | 108 | 85.19 |
| KEGG_UBIQUITIN_MEDIATED_PROTEOLYSIS | 0.727066818 | 0.859763903 | 0.245505521 | 0.845857405 | 641 | 129 | 136 | 94.85 |
| KEGG_VALINE_LEUCINE_AND_ISOLEUCINE_BIOSYNTHESIS | 0.017241379 | 0.168784029 | 0.781479954 | 1.676393127 | 9 | 9 | 11 | 81.82 |
| KEGG_VALINE_LEUCINE_AND_ISOLEUCINE_DEGRADATION | 0.109396914 | 0.406956522 | 0.45762671 | 1.355331617 | 77 | 41 | 48 | 85.42 |
| KEGG_VASCULAR_SMOOTH_MUSCLE_CONTRACTION | 0.810473815 | 0.919195913 | 0.237365649 | 0.764181903 | 649 | 77 | 119 | 64.71 |
| KEGG_VASOPRESSIN_REGULATED_WATER_REABSORPTION | 0.06056338 | 0.30445375 | 0.504210621 | 1.474835958 | 42 | 38 | 44 | 86.36 |
| KEGG_VEGF_SIGNALING_PATHWAY | 0.393939394 | 0.725472547 | -0.253771193 | -1.035450776 | 90 | 64 | 74 | 86.49 |
| KEGG_VIBRIO_CHOLERAE_INFECTION | 0.073509015 | 0.341449275 | 0.484123695 | 1.448885708 | 52 | 46 | 52 | 88.46 |
| KEGG_VIRAL_MYOCARDITIS | 0.571014493 | 0.817927847 | 0.326584854 | 0.920816872 | 393 | 31 | 70 | 44.29 |
| KEGG_WNT_SIGNALING_PATHWAY | 0.902522936 | 0.959337149 | 0.210061137 | 0.7181806 | 786 | 121 | 151 | 80.13 |
