## Supplementary Table 3 for "ATF6 activation alters colonic lipid metabolism causing tumor-associated microbial adaptation"

| geneID | Cond1_DF_Detected | Cond1_DF_ValueCol | Cond1_DF_PadjCol | Cond1_DF_Cutoff | Cond1_DF_Cutoff_Specific.x | Cond2_DF_Detected | Cond2_DF_ValueCol | Cond2_DF_PadjCol | Cond2_DF_Cutoff | Cond2_DF_Cutoff_Specific | BG_Method | RG1_All | RG2_Significant | RG3_SignificantChange | Cond1_DF_ensembl_gene_id | Cond1_DF_entrezgene_id | Cond1_DF_gene_biotype |
|---|---|---|---|---|---|---|---|---|---|---|---|---|---|---|---|---|---|
| 0610005C13Rik | TRUE | 0.186539086 | 0.63839585 | No Change | Not Significant | TRUE | 0.472272074 | 5.45E-06 | No Change | Significant Positive | TRUE | Cond1 Not Significant + Cond2 Significant Positive | None | None | ENSMUSG00000109644 | 71661 | lncRNA |
| 0610009B22Rik | TRUE | -0.031420495 | 0.888973789 | No Change | Not Significant | TRUE | 0.169642304 | 0.108948886 | No Change | Not Significant | TRUE | Cond1 Not Significant + Cond2 Not Significant | None | None | ENSMUSG00000007777 | 66050 | protein_coding |
| 0610009E02Rik | TRUE | 0.043932232 | 0.95833356 | No Change | Not Significant | TRUE | 0.144578297 | 0.866155901 | No Change | Not Significant | TRUE | Cond1 Not Significant + Cond2 Not Significant | None | None | ENSMUSG00000086714 | 100125929 | lncRNA |
| 0610009L18Rik | TRUE | -0.356237644 | 0.251451529 | No Change | Not Significant | TRUE | -0.638136414 | 0.000483146 | DOWN | DOWN | TRUE | Cond1 Not Significant + Cond2 DOWN | Cond2_DOWN | Cond2_DOWN | ENSMUSG00000043644 | 66838 | lncRNA |
| 0610010K14Rik | TRUE | 0.127571036 | 0.897695523 | No Change | Not Significant | FALSE | NA | NA | No Change | Not Detected | FALSE | Background = FALSE | Background = FALSE | Background = FALSE | ENSMUSG00000020831 | 104457 | protein_coding |
| 0610012G03Rik | TRUE | -0.118099356 | 0.298143983 | No Change | Not Significant | TRUE | 0.067449961 | 0.523118689 | No Change | Not Significant | TRUE | Cond1 Not Significant + Cond2 Not Significant | None | None | ENSMUSG00000107002 | 106264 | protein_coding |
| 0610030E20Rik | TRUE | -0.356318573 | 0.001548948 | No Change | Significant Negative | TRUE | -0.15781434 | 0.116930286 | No Change | Not Significant | TRUE | Cond1 Significant Negative + Cond2 Not Significant | None | None | ENSMUSG00000058706 | 68364 | protein_coding |
| 0610031O16Rik | TRUE | 0.445215205 | 1 | No Change | Not Significant | FALSE | NA | NA | No Change | Not Detected | FALSE | Background = FALSE | Background = FALSE | Background = FALSE | ENSMUSG00000099146 | 68369 | lncRNA |
| [illegible] | [illegible] | [illegible] | [illegible] | [illegible] | [illegible] | [illegible] | [illegible] | [illegible] | [illegible] | [illegible] | [illegible] | [illegible] | [illegible] | [illegible] | [illegible] | [illegible] | [illegible] |

| geneID | Cond1_DF_baseMean | Cond1_DF_ log2FoldChange | Cond1_DF_lfcSE | Cond1_DF_stat | Cond1_DF_pvalue | Cond1_DF_ padj | Cond1_DF_absLog2FC | Cond1_DF_Cutoff_Specific.y | Cond2_DF_ensembl_gene_id | Cond2_DF_entrezgene_id | Cond2_DF_gene_biotype | Cond2_DF_baseMean | Cond2_DF_ log2FoldChange | Cond2_DF_lfcSE | Cond2_DF_stat | Cond2_DF_pvalue | Cond2_DF_ padj | Cond2_DF_absLog2FC | RG1_Specific_Cond2 | RG1_Specific_Cond1 |
|---|---|---|---|---|---|---|---|---|---|---|---|---|---|---|---|---|---|---|---|---|
| 0610005C13Rik | 209.945329 | 0.186539086 | 0.230519809 | -0.809210657 | 0.418393986 | 0.63839585 | 0.186539086 | Not Significant | ENSMUSG00000109644 | 71661 | lncRNA | 565.9222448 | 0.472272074 | 0.090667207 | 5.20885214 | 1.90E-07 | 5.45E-06 | 0.472272074 | Cond1 No Change + Cond2 Significant Positive | Cond2 No Change + Cond1 Not Significant |
| 0610009B22Rik | 170.8093752 | -0.031420495 | 0.111380025 | 0.282101709 | 0.777865523 | 0.888973789 | 0.031420495 | Not Significant | ENSMUSG00000007777 | 66050 | protein_coding | 308.1654574 | 0.169642304 | 0.074749629 | 2.269473505 | 0.023239547 | 0.108948886 | 0.169642304 | Cond1 No Change + Cond2 Not Significant | Cond2 No Change + Cond1 Not Significant |
| 0610009E02Rik | 7.583474848 | 0.043932232 | 0.412831978 | -0.106416738 | 0.915251714 | 0.95833356 | 0.043932232 | Not Significant | ENSMUSG00000086714 | 100125929 | lncRNA | 12.43099416 | 0.144578297 | 0.412512593 | 0.350482141 | 0.725976891 | 0.866155901 | 0.144578297 | Cond1 No Change + Cond2 Not Significant | Cond2 No Change + Cond1 Not Significant |
| 0610009L18Rik | 27.30046995 | -0.356237644 | 0.217930367 | 1.63463977 | 0.102124586 | 0.251451529 | 0.356237644 | Not Significant | ENSMUSG00000043644 | 66838 | lncRNA | 80.44210233 | -0.638136414 | 0.152245438 | -4.191497761 | 2.77E-05 | 0.000483146 | 0.638136414 | Cond1 No Change + Cond2 DOWN | Cond2 DOWN + Cond1 Not Significant |
| 0610010K14Rik | 5.887441764 | 0.127571036 | 0.489431162 | -0.260651642 | 0.794361161 | 0.897695523 | 0.127571036 | Not Significant | NA | NA | NA | NA | NA | NA | NA | NA | NA | NA | Background = FALSE | Background = FALSE |
| 0610012G03Rik | 416.943223 | -0.118099356 | 0.078043327 | 1.513253727 | 0.130215222 | 0.298143983 | 0.118099356 | Not Significant | ENSMUSG00000107002 | 106264 | protein_coding | 905.3325972 | 0.067449961 | 0.062424678 | 1.08050155 | 0.279918897 | 0.523118689 | 0.067449961 | Cond1 No Change + Cond2 Not Significant | Cond2 No Change + Cond1 Not Significant |
| 0610030E20Rik | 218.4395863 | -0.356318573 | 0.095664175 | 3.724681413 | 0.000195562 | 0.001548948 | 0.356318573 | Significant Negative | ENSMUSG00000058706 | 68364 | protein_coding | 411.6306048 | -0.15781434 | 0.070738116 | -2.230966106 | 0.025683374 | 0.116930286 | 0.15781434 | Cond1 No Change + Cond2 Not Significant | Cond2 No Change + Cond1 Significant Negative |
| 0610031O16Rik | 2.883680877 | 0.445215205 | 0.846881922 | -0.525711074 | 0.599088968 | 1 | 0.445215205 | Not Significant | NA | NA | NA | NA | NA | NA | NA | NA | NA | NA | Background = FALSE | Background = FALSE |
| [illegible] | [illegible] | [illegible] | [illegible] | [illegible] | [illegible] | [illegible] | [illegible] | [illegible] | [illegible] | [illegible] | [illegible] | [illegible] | [illegible] | [illegible] | [illegible] | [illegible] | [illegible] | [illegible] | [illegible] | [illegible] |

Supplementary Table 3

| geneID | Cond1_DF_Detected | Cond1_DF_ValueCol | Cond1_DF_PadjCol | Cond1_DF_Cutoff | Cond1_DF_Cutoff_Specific.x |
|---|---|---|---|---|---|
| 2010109A12Rik | TRUE | 0.022757241 | 0.984122985 | No Change | Not Significant |
| 2010308F09Rik | TRUE | -0.393686544 | 1 | No Change | Not Significant |
| 2010310C07Rik | TRUE | -0.068911277 | 1 | No Change | Not Significant |
| 2010315B03Rik | TRUE | -0.129275634 | 0.297380166 | No Change | Not Significant |
| 2200002D01Rik | TRUE | -0.532823754 | 0.01711111 | DOWN | DOWN |
| 2200002J24Rik | TRUE | -1.314634992 | 0.012282071 | DOWN | DOWN |
| 2210010C04Rik | TRUE | -1.033043676 | 1 | No Change | Not Significant |
| 2210016F16Rik | TRUE | -0.098867361 | 0.548178261 | No Change | Not Significant |
| 2210016L21Rik | TRUE | -0.144208112 | 0.166982469 | No Change | Not Significant |
| 2210039B01Rik | TRUE | -0.162512005 | 0.766833571 | No Change | Not Significant |
| 2210406O10Rik | TRUE | 0.807557196 | 0.358301502 | No Change | Not Significant |
| 2210408F21Rik | TRUE | -0.265341472 | 1 | No Change | Not Significant |
| 2210408I21Rik | TRUE | -0.148524097 | 0.559393767 | No Change | Not Significant |
| 2210417A02Rik | TRUE | -0.627623562 | 1 | No Change | Not Significant |
| 2300009A05Rik | TRUE | -0.107217269 | 0.77652216 | No Change | Not Significant |
| 2310001H17Rik | TRUE | -0.031397545 | 0.932314788 | No Change | Not Significant |
| 2310001K24Rik | TRUE | 0.067146517 | 1 | No Change | Not Significant |
| 2310002F09Rik | TRUE | 1.341503604 | 1 | No Change | Not Significant |
| 2310003N18Rik | TRUE | -0.22892818 | 1 | No Change | Not Significant |
| 2310008N11Rik | TRUE | 1.141135797 | 1 | No Change | Not Significant |
| 2310009B15Rik | TRUE | 0.093005324 | 0.63998633 | No Change | Not Significant |
| 2310010J17Rik | TRUE | 0.091604329 | 0.827323214 | No Change | Not Significant |
| 2310011J03Rik | TRUE | 0.000119126 | 0.999319591 | No Change | Not Significant |
| 2310016G11Rik | TRUE | -0.215311655 | 0.621711048 | No Change | Not Significant |
| 2310022A10Rik | TRUE | 0.075086548 | 0.763289104 | No Change | Not Significant |
| 2310022B05Rik | TRUE | -0.091694937 | 0.549759128 | No Change | Not Significant |
| 2310030G06Rik | TRUE | 0.44338908 | 8.11E-05 | No Change | Significant Positive |
| 2310033P09Rik | TRUE | 0.024078543 | 0.915993052 | No Change | Not Significant |
| 2310034C09Rik | TRUE | 1.771357605 | 1 | No Change | Not Significant |
| 2310039H08Rik | TRUE | -0.013260125 | 0.938630585 | No Change | Not Significant |
| 2310039L15Rik | TRUE | 0.966435978 | 1 | No Change | Not Significant |
| 2310043M15Rik | TRUE | -0.47015899 | 0.559438733 | No Change | Not Significant |
| 2310043O21Rik | TRUE | 0.226070465 | 1 | No Change | Not Significant |
| 2310057M21Rik | TRUE | 0.131277836 | 0.593022866 | No Change | Not Significant |
| 2310061I04Rik | TRUE | -0.146476616 | 0.451758428 | No Change | Not Significant |
| 2310065F04Rik | TRUE | -0.428142042 | 1 | No Change | Not Significant |
| 2310068J16Rik | TRUE | 0.026229054 | 0.97289203 | No Change | Not Significant |
| 2310069B03Rik | TRUE | 1.648332961 | 0.015118795 | UP | UP |
| 2310079G19Rik | TRUE | -2.063323298 | 0.00119646 | DOWN | DOWN |
| 2410002F23Rik | TRUE | -0.232308211 | 0.147350951 | No Change | Not Significant |
| 2410004B18Rik | TRUE | 0.132677766 | 0.316005805 | No Change | Not Significant |
| 2410004P03Rik | TRUE | -0.849415362 | 1 | No Change | Not Significant |
| 2410017I17Rik | TRUE | 2.690516879 | 1 | No Change | Not Significant |
| 2410018L13Rik | TRUE | 0.358068425 | 1 | No Change | Not Significant |
| 2410021H03Rik | TRUE | -0.529249725 | 1 | No Change | Not Significant |
| 2410022M11Rik | TRUE | 0.013225859 | 0.976016991 | No Change | Not Significant |
| 2500002B13Rik | TRUE | -0.145522167 | 0.884482916 | No Change | Not Significant |
| 2510002D24Rik | TRUE | -0.112352711 | 0.374977244 | No Change | Not Significant |
| 2510009E07Rik | TRUE | -0.029074643 | 0.927743965 | No Change | Not Significant |
| 2510039O18Rik | TRUE | 0.105377236 | 0.259060655 | No Change | Not Significant |
| 2610008E11Rik | TRUE | -0.057385417 | 0.807082912 | No Change | Not Significant |
| 2610020C07Rik | TRUE | 0.046748972 | 0.938630585 | No Change | Not Significant |
| 2610028E06Rik | TRUE | -2.112189026 | 1 | No Change | Not Significant |
| 2610028H24Rik | TRUE | -1.060316066 | 1 | No Change | Not Significant |
| 2610035D17Rik | TRUE | 0.04504154 | 0.895392916 | No Change | Not Significant |
| 2610044O15Rik8 | TRUE | -0.094100813 | 0.736796542 | No Change | Not Significant |
| 2610206C17Rik | TRUE | 0.303483748 | 1 | No Change | Not Significant |
| 2610306M01Rik | TRUE | -0.411824729 | 0.165147848 | No Change | Not Significant |
| 2610318N02Rik | TRUE | 0.936618224 | 0.001240778 | UP | UP |
| 2610528A11Rik | TRUE | -0.549188157 | 1.57E-08 | DOWN | DOWN |
| 2610528J11Rik | TRUE | -0.110730221 | 0.387766717 | No Change | Not Significant |
| 2700038G22Rik | TRUE | -0.126147937 | 0.877286805 | No Change | Not Significant |
| 2700046G09Rik | TRUE | -0.378028814 | 0.276057789 | No Change | Not Significant |
| 2700049A03Rik | TRUE | 0.003347456 | 0.985876467 | No Change | Not Significant |
| 2700054A10Rik | TRUE | 0.207315338 | 0.706691156 | No Change | Not Significant |
| 2700062C07Rik | TRUE | -0.059536749 | 0.800207711 | No Change | Not Significant |
| 2700097O09Rik | TRUE | 0.100823035 | 0.714031075 | No Change | Not Significant |
| 2810001G20Rik | TRUE | -0.351042011 | 0.16072315 | No Change | Not Significant |
| 2810004N23Rik | TRUE | -0.156677062 | 0.328001251 | No Change | Not Significant |
| 2810013P06Rik | TRUE | -0.154842553 | 0.439526427 | No Change | Not Significant |
| 2810021J22Rik | TRUE | -0.004513938 | 0.985568894 | No Change | Not Significant |
| 2810025M15Rik | TRUE | 0.327619451 | 0.114584988 | No Change | Not Significant |
| 2810029C07Rik | TRUE | 0.008788029 | 1 | No Change | Not Significant |
| 2810403D21Rik | TRUE | -0.352359755 | 0.313784669 | No Change | Not Significant |
| 2810408A11Rik | TRUE | -0.313259287 | 0.296522686 | No Change | Not Significant |
| 2810408I11Rik | TRUE | 1.608529862 | 0.00043081 | UP | UP |
| 2810414N06Rik | TRUE | 0.034145185 | 0.957266079 | No Change | Not Significant |
| 2810429I04Rik | TRUE | 0.525206505 | 0.654709467 | No Change | Not Significant |
| 2810442N19Rik | TRUE | 0.787779122 | 1 | No Change | Not Significant |
| 2810459M11Rik | TRUE | -0.054083295 | 0.784660848 | No Change | Not Significant |
| 2900005J15Rik | TRUE | -1.179084545 | 0.050703535 | No Change | Not Significant |
| 2900009J06Rik | TRUE | 0.996028858 | 1 | No Change | Not Significant |
| 2900026A02Rik | TRUE | -0.156778446 | 0.595844868 | No Change | Not Significant |
| 2900052N01Rik | TRUE | -0.428229571 | 1 | No Change | Not Significant |
| 2900057B20Rik | TRUE | 0.414138901 | 0.558663295 | No Change | Not Significant |
| 2900060B14Rik | TRUE | 0.880518929 | 0.258037088 | No Change | Not Significant |
| 2900060N12Rik | TRUE | -0.097416817 | 1 | No Change | Not Significant |
| 2900076A07Rik | TRUE | -0.140641719 | 0.727284417 | No Change | Not Significant |
| 2900079G21Rik | TRUE | 0.004301634 | 1 | No Change | Not Significant |
| 2900093K20Rik | TRUE | 0.266920753 | 0.636395319 | No Change | Not Significant |
| 3010003L21Rik | TRUE | 0.078897661 | 0.825971257 | No Change | Not Significant |
| 3100003L05Rik | TRUE | -1.677080597 | 0.037106194 | DOWN | DOWN |
| 3110006O06Rik | TRUE | 0.205472176 | 1 | No Change | Not Significant |
| 3110009E18Rik | TRUE | -0.439583656 | 0.321507788 | No Change | Not Significant |
| 3110015C05Rik | TRUE | 1.062689261 | 1 | No Change | Not Significant |
| 3110040N11Rik | TRUE | -0.035405226 | 0.882132403 | No Change | Not Significant |
| 3110070M22Rik | TRUE | -0.645053728 | 0.475073328 | No Change | Not Significant |
| 3110082I17Rik | TRUE | 0.100549361 | 0.765317902 | No Change | Not Significant |
| 3110099E03Rik | TRUE | 0.967940753 | 1 | No Change | Not Significant |
| 3200001D21Rik | TRUE | -0.783450942 | 1 | No Change | Not Significant |
| 3300002I08Rik | TRUE | 0.027409734 | 0.966412148 | No Change | Not Significant |
| 3300005D01Rik | TRUE | 1.366490021 | 1 | No Change | Not Significant |
| 3425401B19Rik | TRUE | -0.623155065 | 0.344649174 | No Change | Not Significant |
| 3632454L22Rik | TRUE | 1.046795245 | 1 | No Change | Not Significant |
| 3830403N18Rik | TRUE | 0.08345999 | 1 | No Change | Not Significant |
| 3830406C13Rik | TRUE | -0.331464336 | 0.023333491 | No Change | Significant Negative |
| 3830408C21Rik | TRUE | 0.516543546 | 0.666174493 | No Change | Not Significant |
| 3830432H09Rik | TRUE | 0.618772462 | 1 | No Change | Not Significant |
| 3930402G23Rik | TRUE | 2.004322372 | 3.83E-09 | UP | UP |
| 4631405J19Rik | TRUE | 0.957865558 | 1 | No Change | Not Significant |
| 4632404H12Rik | TRUE | -0.396497277 | 0.087507252 | No Change | Not Significant |
| 4632427E13Rik | TRUE | -0.110839329 | 0.728048657 | No Change | Not Significant |
| 4632428C04Rik | TRUE | -0.92722336 | 0.000353491 | DOWN | DOWN |
| 4732414G09Rik | TRUE | -0.676140942 | 1 | No Change | Not Significant |
| 4732471J01Rik | TRUE | 0.324823822 | 0.363275924 | No Change | Not Significant |
| 4732490B19Rik | TRUE | 0.055771267 | 0.91440345 | No Change | Not Significant |
| 4732491K20Rik | TRUE | 0.606616605 | 0.185168116 | No Change | Not Significant |
| 4833407H14Rik | TRUE | 0.23915415 | 0.758030479 | No Change | Not Significant |
| 4833417C18Rik | TRUE | -0.584625111 | 1 | No Change | Not Significant |
| 4833418N02Rik | TRUE | 0.445836499 | 0.14994898 | No Change | Not Significant |
| 4833419F23Rik | TRUE | -1.04309595 | 1 | No Change | Not Significant |
| 4833420G17Rik | TRUE | -0.285964596 | 0.010995173 | No Change | Significant Negative |
| 4833422C13Rik | TRUE | -1.267038999 | 1 | No Change | Not Significant |
| 4833422M21Rik | TRUE | -0.265714533 | 1 | No Change | Not Significant |
| 4833427F10Rik | TRUE | 1.516596284 | 1 | No Change | Not Significant |
| 4833428L15Rik | TRUE | -0.843025252 | 1 | No Change | Not Significant |
| 4833438C02Rik | TRUE | -0.241596158 | 0.346538122 | No Change | Not Significant |
| 4833439L19Rik | TRUE | 0.383177946 | 3.05E-09 | No Change | Significant Positive |
| 4921507G05Rik | TRUE | -1.560512382 | 1 | No Change | Not Significant |
| 4921507P07Rik | TRUE | -0.2399992 | 0.725037586 | No Change | Not Significant |
| 4921511C10Rik | TRUE | 0.196222695 | 0.825190702 | No Change | Not Significant |
| 4921511I17Rik | TRUE | -1.919793723 | 1 | No Change | Not Significant |
| 4921517D22Rik | TRUE | 0.005691297 | 1 | No Change | Not Significant |
| 4921518K17Rik | TRUE | -0.511320371 | 1 | No Change | Not Significant |
| 4921524J17Rik | TRUE | -0.184059112 | 0.37240098 | No Change | Not Significant |
| 4921525O09Rik | TRUE | 1.604988559 | 1 | No Change | Not Significant |
| 4921531C22Rik | TRUE | 0.083504658 | 0.870239999 | No Change | Not Significant |
| 4921536K21Rik | TRUE | 0.24844281 | 0.432209358 | No Change | Not Significant |
| 4921539E11Rik | TRUE | -0.319545924 | 1 | No Change | Not Significant |
| 4930404H11Rik | TRUE | 0 | 1 | No Change | Not Significant |
| 4930404I05Rik | TRUE | -0.197951695 | 1 | No Change | Not Significant |
| 4930405A10Rik | TRUE | 1.333243514 | 1 | No Change | Not Significant |
| 4930405A21Rik | TRUE | 0.477511936 | 1 | No Change | Not Significant |
| 4930405J17Rik | TRUE | -0.298121852 | 0.69793377 | No Change | Not Significant |
| 4930405O22Rik | TRUE | -0.451755639 | 0.568175062 | No Change | Not Significant |
| 4930407I10Rik | TRUE | 0.31932635 | 1 | No Change | Not Significant |
| 4930412M03Rik | TRUE | -0.626437708 | 0.208009073 | No Change | Not Significant |
| 4930413F20Rik | TRUE | -0.790673755 | 0.260075004 | No Change | Not Significant |
| 4930414N06Rik | TRUE | 1.002130212 | 0.097003559 | No Change | Not Significant |
| 4930417O13Rik | TRUE | -0.039134584 | 1 | No Change | Not Significant |
| 4930419G24Rik | TRUE | 0.14983293 | 1 | No Change | Not Significant |
| 4930422M22Rik | TRUE | 0.676126437 | 1 | No Change | Not Significant |
| 4930426D05Rik | TRUE | -1.117854557 | 1 | No Change | Not Significant |
| 4930428E07Rik | TRUE | 2.004454413 | 1 | No Change | Not Significant |
| 4930429F24Rik | TRUE | 1.256206269 | 1 | No Change | Not Significant |
| 4930429H19Rik | TRUE | 0.96373696 | 1 | No Change | Not Significant |
| 4930430F08Rik | TRUE | 0.241135254 | 0.464617944 | No Change | Not Significant |
| 4930432J09Rik | TRUE | -0.428755724 | 1 | No Change | Not Significant |
| 4930432O09Rik | TRUE | 1.623123746 | 1 | No Change | Not Significant |
| 4930438A08Rik | TRUE | 0.231526289 | 1 | No Change | Not Significant |
| 4930442L01Rik | TRUE | 1.111705803 | 1 | No Change | Not Significant |
| 4930443O20Rik | TRUE | -0.930107353 | 1 | No Change | Not Significant |
| 4930445N18Rik | TRUE | 2.20293302 | 1 | No Change | Not Significant |
| 4930447A16Rik | TRUE | -0.284105303 | 1 | No Change | Not Significant |
| 4930447C04Rik | TRUE | 1.681489086 | 1 | No Change | Not Significant |
| 4930447F24Rik | TRUE | -1.539977251 | 9.25E-06 | DOWN | DOWN |
| 4930447N08Rik | TRUE | -0.239353748 | 0.681625181 | No Change | Not Significant |
| 4930448I18Rik | TRUE | -0.816877869 | 1 | No Change | Not Significant |
| 4930449E18Rik | TRUE | 0.198660024 | 1 | No Change | Not Significant |
| 4930451E10Rik | TRUE | 0.027887773 | 1 | No Change | Not Significant |
| 4930451I11Rik | TRUE | 1.862060497 | 1 | No Change | Not Significant |
| 4930452G13Rik | TRUE | 2.211969154 | 1 | No Change | Not Significant |
| 4930453N24Rik | TRUE | -0.059367844 | 0.709429427 | No Change | Not Significant |
| 4930455C13Rik | TRUE | -0.184100068 | 1 | No Change | Not Significant |
| 4930455J16Rik | TRUE | 1.561255414 | 0.698788598 | No Change | Not Significant |
| 4930459C07Rik | TRUE | 1.210937295 | 1 | No Change | Not Significant |
| 4930461G14Rik | TRUE | 0.507917825 | 1 | No Change | Not Significant |
| 4930471M09Rik | TRUE | 0.821056796 | 1 | No Change | Not Significant |
| 4930473A02Rik | TRUE | -0.199750992 | 1 | No Change | Not Significant |
| 4930473H19Rik | TRUE | -0.963910746 | 1 | No Change | Not Significant |
| 4930474M22Rik | TRUE | 0.484557718 | 1 | No Change | Not Significant |
| 4930474N05Rik | TRUE | 0.611342111 | 1 | No Change | Not Significant |
| 4930478L05Rik | TRUE | 1.971979442 | 1 | No Change | Not Significant |
| 4930481A15Rik | TRUE | 0.256211543 | 0.378310591 | No Change | Not Significant |
| 4930483J18Rik | TRUE | 0.46693282 | 1 | No Change | Not Significant |
| 4930483K19Rik | TRUE | -0.423302653 | 1 | No Change | Not Significant |
| 4930483O08Rik | TRUE | -0.97887499 | 1 | No Change | Not Significant |
| 4930486I03Rik | TRUE | -0.894128781 | 1 | No Change | Not Significant |
| 4930500A05Rik | TRUE | 0.234830986 | 1 | No Change | Not Significant |
| 4930500H12Rik | TRUE | -0.427974407 | 1 | No Change | Not Significant |
| 4930502E09Rik | TRUE | 0.644653523 | 1 | No Change | Not Significant |
| 4930503L19Rik | TRUE | 0.182093066 | 1 | No Change | Not Significant |

| geneID | Cond1_DF_Detected | Cond1_DF_ValueCol | Cond1_DF_PadjCol | Cond1_DF_Cutoff | Cond1_DF_Cutoff_Specific.x | Cond2_DF_Detected | Cond2_DF_ValueCol | Cond2_DF_PadjCol | Cond2_DF_Cutoff | Cond2_DF_Cutoff_Specific | BG_Method | RG1_All | RG2_Significant | RG3_SignificantChange | Cond1_DF_ensembl_gene_id | Cond1_DF_entrezgene_id | Cond1_DF_gene_biotype | Cond1_DF_baseMean | Cond1_DF_ log2FoldChange | Cond1_DF_lfcSE | Cond1_DF_stat | Cond1_DF_pvalue | Cond1_DF_ padj | Cond1_DF_absLog2FC | Cond1_DF_Cutoff_Specific.y | Cond2_DF_ensembl_gene_id | Cond2_DF_entrezgene_id | Cond2_DF_gene_biotype | Cond2_DF_baseMean | Cond2_DF_ log2FoldChange | Cond2_DF_lfcSE | Cond2_DF_stat | Cond2_DF_pvalue | Cond2_DF_ padj | Cond2_DF_absLog2FC | RG1_Specific_Cond2 | RG1_Specific_Cond1 |
|---|---|---|---|---|---|---|---|---|---|---|---|---|---|---|---|---|---|---|---|---|---|---|---|---|---|---|---|---|---|---|---|---|---|---|---|---|---|
| 4930505N22Rik | TRUE | -0.012707235 | 1 | No Change | Not Significant | FALSE | NA | NA | No Change | Not Detected | FALSE | Background = FALSE | Background = FALSE | Background = FALSE | ENSMUSG00000097690 | 100174921 | lncRNA | 3.853838291 | -0.012707235 | 0.615118635 | 0.020658185 | 0.983518325 | 1 | 0.012707235 | Not Significant | NA | NA | NA | NA | NA | NA | NA | NA | NA | NA | Background = FALSE | Background = FALSE |
| 4930506C21Rik | TRUE | -1.006204124 | 0.199909758 | No Change | Not Significant | TRUE | -0.415492728 | 0.619709567 | No Change | Not Significant | TRUE | Cond1 Not Significant + Cond2 Not Significant | None | None | ENSMUSG00000087478 | 75060 | lncRNA | 4.825503213 | -1.006204124 | 0.564489036 | 1.782504284 | 0.074667033 | 0.199909758 | 1.006204124 | Not Significant | ENSMUSG00000087478 | 75060 | lncRNA | 9.056892336 | -0.415492728 | 0.471776585 | -0.88069807 | 0.378481261 | 0.619709567 | 0.415492728 | Cond1 No Change + Cond2 Not Significant | Cond2 No Change + Cond1 Not Significant |
| 4930509E16Rik | TRUE | 1.258872688 | 1.87E-05 | UP | UP | TRUE | 0.134056048 | 0.800376096 | No Change | Not Significant | TRUE | Cond1 UP + Cond2 Not Significant | Cond1_UP | Cond1_UP | ENSMUSG00000096986 | 75115 | lncRNA | 20.61589951 | 1.258872688 | 0.260898934 | -4.825135416 | 1.40E-06 | 1.87E-05 | 1.258872688 | UP | ENSMUSG00000096986 | 75115 | lncRNA | 19.36663933 | 0.134056048 | 0.271736378 | 0.493331251 | 0.62177855 | 0.800376096 | 0.134056048 | Cond1 UP + Cond2 Not Significant | Cond2 No Change + Cond1 UP |
| 4930509G22Rik | TRUE | 0.720232678 | 0.466665282 | No Change | Not Significant | TRUE | -0.045518389 | 0.958435249 | No Change | Not Significant | TRUE | Cond1 Not Significant + Cond2 Not Significant | None | None | ENSMUSG00000096959 | 105246071 | lncRNA | 5.380531304 | 0.720232678 | 0.631388722 | -1.140711979 | 0.2539898 | 0.466665282 | 0.720232678 | Not Significant | ENSMUSG00000096959 | 105246071 | lncRNA | 9.334327379 | -0.045518389 | 0.380716381 | -0.119559839 | 0.904831835 | 0.958435249 | 0.045518389 | Cond1 No Change + Cond2 Not Significant | Cond2 No Change + Cond1 Not Significant |
| 4930511M06Rik | TRUE | -0.284180066 | 1 | No Change | Not Significant | FALSE | NA | NA | No Change | Not Detected | FALSE | Background = FALSE | Background = FALSE | Background = FALSE | ENSMUSG00000086607 | 75084 | lncRNA | 2.971304886 | -0.284180066 | 0.876040502 | 0.32439147 | 0.745641677 | 1 | 0.284180066 | Not Significant | NA | NA | NA | NA | NA | NA | NA | NA | NA | NA | Background = FALSE | Background = FALSE |
| 4930513N10Rik | TRUE | 0.462029547 | 0.331941022 | No Change | Not Significant | TRUE | -0.022264259 | 0.960412417 | No Change | Not Significant | TRUE | Cond1 Not Significant + Cond2 Not Significant | None | None | ENSMUSG00000074136 | 319960 | lncRNA | 26.40119587 | 0.462029547 | 0.322264546 | -1.433696484 | 0.151658896 | 0.331941022 | 0.462029547 | Not Significant | ENSMUSG00000074136 | 319960 | lncRNA | 33.93903853 | -0.022264259 | 0.199562633 | -0.111565268 | 0.911168112 | 0.960412417 | 0.022264259 | Cond1 No Change + Cond2 Not Significant | Cond2 No Change + Cond1 Not Significant |
| 4930515G01Rik | TRUE | -0.372178165 | 0.463839794 | No Change | Not Significant | TRUE | -0.030527139 | 0.95873684 | No Change | Not Significant | TRUE | Cond1 Not Significant + Cond2 Not Significant | None | None | ENSMUSG00000092183 | 67642 | lncRNA | 12.02847996 | -0.372178165 | 0.324520184 | 1.146856755 | 0.251440828 | 0.463839794 | 0.372178165 | Not Significant | ENSMUSG00000092183 | 67642 | lncRNA | 25.4682919 | -0.030527139 | 0.261505813 | -0.116735985 | 0.907069274 | 0.95873684 | 0.030527139 | Cond1 No Change + Cond2 Not Significant | Cond2 No Change + Cond1 Not Significant |
| 4930518J21Rik | TRUE | -0.935628363 | 1 | No Change | Not Significant | FALSE | NA | NA | No Change | Not Detected | FALSE | Background = FALSE | Background = FALSE | Background = FALSE | ENSMUSG00000110310 | 75123 | lncRNA | 1.038590796 | -0.935628363 | 1.498501155 | 0.624376138 | 0.532380592 | 1 | 0.935628363 | Not Significant | NA | NA | NA | NA | NA | NA | NA | NA | NA | NA | Background = FALSE | Background = FALSE |
| 4930519G04Rik | TRUE | 1.792641517 | 1 | No Change | Not Significant | FALSE | NA | NA | No Change | Not Detected | FALSE | Background = FALSE | Background = FALSE | Background = FALSE | ENSMUSG00000029564 | 67593 | protein_coding | 1.909338251 | 1.792641517 | 2.381893788 | -0.752611861 | 0.451683188 | 1 | 1.792641517 | Not Significant | NA | NA | NA | NA | NA | NA | NA | NA | NA | NA | Background = FALSE | Background = FALSE |
| 4930519K11Rik | TRUE | -0.298419904 | 1 | No Change | Not Significant | FALSE | NA | NA | No Change | Not Detected | FALSE | Background = FALSE | Background = FALSE | Background = FALSE | ENSMUSG00000114953 | 78019 | lncRNA | 3.677895781 | -0.298419904 | 0.54923255 | 0.5433398 | 0.586895866 | 1 | 0.298419904 | Not Significant | NA | NA | NA | NA | NA | NA | NA | NA | NA | NA | Background = FALSE | Background = FALSE |
| 4930522L14Rik | TRUE | -0.133566613 | 0.728048657 | No Change | Not Significant | TRUE | -0.192713392 | 0.457336743 | No Change | Not Significant | TRUE | Cond1 Not Significant + Cond2 Not Significant | None | None | ENSMUSG00000072762 | 100041734 | protein_coding | 38.25845462 | -0.133566613 | 0.213202854 | 0.626476665 | 0.531002338 | 0.728048657 | 0.133566613 | Not Significant | ENSMUSG00000072762 | 100041734 | protein_coding | 85.62875708 | -0.192713392 | 0.157377792 | -1.224527236 | 0.220753398 | 0.457336743 | 0.192713392 | Cond1 No Change + Cond2 Not Significant | Cond2 No Change + Cond1 Not Significant |
| 4930523C07Rik | TRUE | -0.556252925 | 0.0109583 | DOWN | DOWN | TRUE | -0.236062921 | 0.223773946 | No Change | Not Significant | TRUE | Cond1 DOWN + Cond2 Not Significant | Cond1_DOWN | Cond1_DOWN | ENSMUSG00000090394 | 67647 | protein_coding | 86.8960717 | -0.556252925 | 0.179245608 | 3.103300165 | 0.001913754 | 0.0109583 | 0.556252925 | DOWN | ENSMUSG00000090394 | 67647 | protein_coding | 178.0684326 | -0.236062921 | 0.128939951 | -1.830797349 | 0.067130796 | 0.223773946 | 0.236062921 | Cond1 DOWN + Cond2 Not Significant | Cond2 No Change + Cond1 DOWN |
| 4930523O13Rik | TRUE | 0.611660012 | 1 | No Change | Not Significant | FALSE | NA | NA | No Change | Not Detected | FALSE | Background = FALSE | Background = FALSE | Background = FALSE | ENSMUSG00000115177 | 74726 | lncRNA | 1.260303311 | 0.611660012 | 2.912908344 | -0.209982581 | 0.833681268 | 1 | 0.611660012 | Not Significant | NA | NA | NA | NA | NA | NA | NA | NA | NA | NA | Background = FALSE | Background = FALSE |
| [illegible] | [illegible] | [illegible] | [illegible] | [illegible] | [illegible] | [illegible] | [illegible] | [illegible] | [illegible] | [illegible] | [illegible] | [illegible] | [illegible] | [illegible] | [illegible] | [illegible] | [illegible] | [illegible] | [illegible] | [illegible] | [illegible] | [illegible] | [illegible] | [illegible] | [illegible] | [illegible] | [illegible] | [illegible] | [illegible] | [illegible] | [illegible] | [illegible] | [illegible] | [illegible] | [illegible] | [illegible] | [illegible] |

| geneID | Cond1_DF_Detected | Cond1_DF_ValueCol | Cond1_DF_PadjCol | Cond1_DF_Cutoff | Cond1_DF_Cutoff_Specific.x | Cond2_DF_Detected | Cond2_DF_ValueCol |
|---|---|---|---|---|---|---|---|
| 9030404E10Rik | TRUE | -1.7471369 | 5.62E-05 | DOWN | DOWN | TRUE | -0.52157447 |
| 9030612E09Rik | TRUE | -0.500529123 | 0.493730605 | No Change | Not Significant | TRUE | -0.465336335 |
| 9030616G12Rik | TRUE | -1.629224121 | 5.10E-05 | DOWN | DOWN | TRUE | -0.272280437 |
| 9030624G23Rik | TRUE | -0.395966082 | 0.668637691 | No Change | Not Significant | TRUE | -0.460138348 |
| 9030625G05Rik | TRUE | 2.582208515 | 1.52E-12 | UP | UP | TRUE | 0.30399798 |
| 9130008F23Rik | TRUE | 0.100845033 | 0.687930892 | No Change | Not Significant | TRUE | 0.16013774 |
| 9130019O22Rik | TRUE | -0.380721855 | 0.131176532 | No Change | Not Significant | TRUE | -0.406351881 |
| 9130019P16Rik | TRUE | -1.189320866 | 0.207979566 | No Change | Not Significant | TRUE | -0.548187737 |
| 9130023H24Rik | TRUE | -0.063114579 | 0.826153034 | No Change | Not Significant | TRUE | 0.021786368 |
| 9130024F11Rik | TRUE | 0.386843586 | 0.013502881 | No Change | Significant Positive | TRUE | 0.112191757 |
| 9130209A04Rik | TRUE | -0.313612729 | 0.607019295 | No Change | Not Significant | TRUE | -0.261672266 |
| 9130221F21Rik | TRUE | 1.017750544 | 0.512561987 | No Change | Not Significant | TRUE | 0.358024608 |
| 9130221H12Rik | TRUE | 0.04502611 | 0.907217398 | No Change | Not Significant | TRUE | 0.03481705 |
| 9130227L01Rik | TRUE | 1.460080735 | 1.14E-05 | UP | UP | TRUE | 0.083943842 |
| 9130230L23Rik | TRUE | -0.387092689 | 0.214258446 | No Change | Not Significant | TRUE | -0.249848851 |
| 9130401M01Rik | TRUE | 0.007631306 | 0.981238658 | No Change | Not Significant | TRUE | 0.001952592 |
| 9230020A06Rik | TRUE | 1.886601444 | 1 | No Change | Not Significant | FALSE | NA |
| 9230102K24Rik | TRUE | 0.861089921 | 1 | No Change | Not Significant | FALSE | NA |
| 9230105E05Rik | TRUE | -0.147226743 | 0.535990818 | No Change | Not Significant | TRUE | 0.201960544 |
| 9230112D13Rik | TRUE | 0.544270779 | 1 | No Change | Not Significant | FALSE | NA |
| 9230112J17Rik | TRUE | -0.49250319 | 0.451187917 | No Change | Not Significant | TRUE | -0.324861402 |
| 9230114K14Rik | TRUE | -0.017538295 | 0.980447529 | No Change | Not Significant | TRUE | -0.063248474 |
| 9230116L04Rik | TRUE | -0.020098286 | 0.958030863 | No Change | Not Significant | TRUE | 0.029446066 |
| 9230116N13Rik | TRUE | -0.113767407 | 1 | No Change | Not Significant | TRUE | -0.898376659 |
| 9230117E06Rik | TRUE | 0.042350893 | 1 | No Change | Not Significant | FALSE | NA |
| 9330104G04Rik | TRUE | 0.317761913 | 0.715609679 | No Change | Not Significant | TRUE | 0.171180062 |
| 9330111N05Rik | TRUE | -1.047617789 | 1 | No Change | Not Significant | FALSE | NA |
| 9330136K24Rik | TRUE | -0.111581537 | 1 | No Change | Not Significant | FALSE | NA |
| 9330151L19Rik | TRUE | -0.550331165 | 0.124751997 | No Change | Not Significant | TRUE | -0.177179258 |
| 9330158H04Rik | TRUE | 1.234040885 | 0.833930977 | No Change | Not Significant | FALSE | NA |
| 9330159F19Rik | TRUE | 3.795683596 | 1 | No Change | Not Significant | FALSE | NA |
| 9330159M07Rik | TRUE | 0.060424378 | 0.909608677 | No Change | Not Significant | TRUE | 0.486283565 |
| 9330162B11Rik | TRUE | -0.98507199 | 1 | No Change | Not Significant | FALSE | NA |
| 9330182O14Rik | TRUE | -1.312916722 | 1 | No Change | Not Significant | TRUE | 0.092504803 |
| 9330188P03Rik | TRUE | -0.427361405 | 0.256689693 | No Change | Not Significant | TRUE | -0.294067669 |
| 9330199G10Rik | TRUE | 2.504132081 | 1 | No Change | Not Significant | FALSE | NA |
| 9430015G10Rik | TRUE | 0.374077802 | 0.012123334 | No Change | Significant Positive | TRUE | -0.0575129 |
| 9430018G01Rik | TRUE | -0.664679326 | 0.464375257 | No Change | Not Significant | TRUE | -0.094496208 |
| 9430037G07Rik | TRUE | 0.079780151 | 0.910399889 | No Change | Not Significant | TRUE | 0.253592507 |
| 9430038I01Rik | TRUE | -0.375357306 | 0.052452983 | No Change | Not Significant | TRUE | -0.01686264 |
| 9430041J12Rik | TRUE | -2.142364928 | 1 | No Change | Not Significant | FALSE | NA |
| 9430060I03Rik | TRUE | 0.118676941 | 0.870311987 | No Change | Not Significant | TRUE | 0.298411564 |
| 9430065F17Rik | TRUE | -0.791010115 | 1 | No Change | Not Significant | TRUE | 0.042699878 |
| 9530026P05Rik | TRUE | -2.042783673 | 1 | No Change | Not Significant | TRUE | -1.485320763 |
| 9530034E10Rik | TRUE | -1.016354722 | 0.091355786 | No Change | Not Significant | FALSE | NA |
| 9530036O11Rik | TRUE | -0.224218508 | 0.696363286 | No Change | Not Significant | TRUE | -0.162690993 |
| 9530051G07Rik | TRUE | -1.087849259 | 1 | No Change | Not Significant | FALSE | NA |
| 9530052E02Rik | TRUE | 0.249615533 | 1 | No Change | Not Significant | FALSE | NA |
| 9530053A07Rik | TRUE | 0.345348693 | 0.777473116 | No Change | Not Significant | TRUE | 0.33163034 |
| 9530059O14Rik | TRUE | 0.512642239 | 1 | No Change | Not Significant | FALSE | NA |
| 9530062K07Rik | TRUE | -0.011620401 | 1 | No Change | Not Significant | TRUE | 0.409058265 |
| 9530068E07Rik | TRUE | -0.016948443 | 0.908714429 | No Change | Not Significant | TRUE | 0.00893473 |
| 9530077C05Rik | TRUE | -0.299962366 | 0.696638145 | No Change | Not Significant | TRUE | -0.701574014 |
| 9530080O11Rik | TRUE | 0.616975399 | 0.24588517 | No Change | Not Significant | TRUE | 0.012861131 |
| 9530082P21Rik | TRUE | 0.143875865 | 0.632329199 | No Change | Not Significant | TRUE | -0.523286664 |
| 9630001P10Rik | TRUE | -1.794985629 | 1 | No Change | Not Significant | FALSE | NA |
| 9630013A20Rik | TRUE | 2.396717721 | 1 | No Change | Not Significant | TRUE | -0.119523782 |
| 9630013K17Rik | TRUE | -0.094722778 | 0.938415991 | No Change | Not Significant | FALSE | NA |
| 9630015K15Rik | TRUE | -0.496377971 | 1 | No Change | Not Significant | FALSE | NA |
| 9630028B13Rik | TRUE | 0.104929082 | 0.870990647 | No Change | Not Significant | TRUE | 0.011857861 |
| 9630028H03Rik | TRUE | 0.623723543 | 1 | No Change | Not Significant | FALSE | NA |
| 9630028I04Rik | TRUE | 0.462083447 | 1 | No Change | Not Significant | FALSE | NA |
| 9830107B12Rik | TRUE | -1.237245317 | 1 | No Change | Not Significant | FALSE | NA |
| 9830132P13Rik | TRUE | -0.697927385 | 1 | No Change | Not Significant | FALSE | NA |
| 9830166K06Rik | TRUE | -3.014151465 | 1 | No Change | Not Significant | FALSE | NA |
| 9930012K11Rik | TRUE | -0.25680999 | 0.490619315 | No Change | Not Significant | TRUE | -0.129047999 |
| 9930021J03Rik | TRUE | 0.040233696 | 0.808756278 | No Change | Not Significant | TRUE | -0.17003397 |
| 9930104L06Rik | TRUE | -0.024364311 | 0.909014472 | No Change | Not Significant | TRUE | -0.198553011 |
| 9930111J21Rik1 | TRUE | 1.19189141 | 1 | No Change | Not Significant | FALSE | NA |
| 9930111J21Rik2 | TRUE | -0.243742361 | 1 | No Change | Not Significant | FALSE | NA |
| a | TRUE | 2.660518464 | 1 | No Change | Not Significant | FALSE | NA |
| A130010J15Rik | TRUE | 0.031133076 | 0.895656612 | No Change | Not Significant | TRUE | 0.05167485 |
| A1cf | TRUE | 0.078314089 | 0.794172689 | No Change | Not Significant | TRUE | 0.137602643 |
| A230004M16Rik | TRUE | -0.968538837 | 1 | No Change | Not Significant | FALSE | NA |
| A230020J21Rik | TRUE | 1.417747328 | 1 | No Change | Not Significant | FALSE | NA |
| A230028O05Rik | TRUE | 2.865786344 | 1 | No Change | Not Significant | FALSE | NA |
| A230051N06Rik | TRUE | -2.559196265 | 1 | No Change | Not Significant | FALSE | NA |
| A230057D06Rik | TRUE | 1.202386851 | 1 | No Change | Not Significant | FALSE | NA |
| A230072C01Rik | TRUE | 0.024514094 | 0.977426278 | No Change | Not Significant | TRUE | -0.117818177 |
| A230072E10Rik | TRUE | -0.308074459 | 0.662372535 | No Change | Not Significant | TRUE | 0.530273181 |
| A230072I06Rik | TRUE | 1.758684461 | 1 | No Change | Not Significant | FALSE | NA |
| A230083G16Rik | TRUE | 0.150766756 | 0.801948109 | No Change | Not Significant | TRUE | -0.36000669 |
| A230103J11Rik | TRUE | -0.386234676 | 0.445817123 | No Change | Not Significant | TRUE | -0.233323603 |
| A230103O09Rik | TRUE | 2.953373818 | 1 | No Change | Not Significant | FALSE | NA |
| A230107N01Rik | TRUE | 0.092365976 | 1 | No Change | Not Significant | FALSE | NA |
| A230108P19Rik | TRUE | 1.470027507 | 1 | No Change | Not Significant | FALSE | NA |
| A2m | TRUE | -1.186150004 | 1 | No Change | Not Significant | FALSE | NA |
| A2ml1 | TRUE | -0.057078456 | 0.962730128 | No Change | Not Significant | TRUE | -0.565364531 |
| A330008L17Rik | TRUE | -0.499587708 | 1 | No Change | Not Significant | FALSE | NA |
| A330009N23Rik | TRUE | -0.129040413 | 0.914497068 | No Change | Not Significant | FALSE | NA |
| A330015K06Rik | TRUE | 2.373037377 | 1 | No Change | Not Significant | FALSE | NA |
| A330032B11Rik | TRUE | -0.594587674 | 1 | No Change | Not Significant | FALSE | NA |
| A330033J07Rik | TRUE | -1.646876128 | 1 | No Change | Not Significant | FALSE | NA |
| A330040F15Rik | TRUE | -0.160864149 | 0.846814623 | No Change | Not Significant | TRUE | 0.151473278 |
| A330049N07Rik | TRUE | -1.503002112 | 1 | No Change | Not Significant | FALSE | NA |
| A330074K22Rik | TRUE | 1.124985255 | 1 | No Change | Not Significant | FALSE | NA |
| A330076C08Rik | TRUE | -0.479451038 | 1 | No Change | Not Significant | FALSE | NA |
| A3galt2 | TRUE | 0.864096823 | 1 | No Change | Not Significant | FALSE | NA |
| A430005L14Rik | TRUE | 0.06959224 | 0.668738427 | No Change | Not Significant | TRUE | 0.232654656 |
| A430018G15Rik | TRUE | -0.397192569 | 0.541897747 | No Change | Not Significant | TRUE | -0.222383728 |
| A430033K04Rik | TRUE | 0.106329086 | 0.45166903 | No Change | Not Significant | TRUE | -0.128026492 |
| A430035B10Rik | TRUE | -0.136828443 | 0.76328646 | No Change | Not Significant | TRUE | 0.042742088 |
| A430057M04Rik | TRUE | -0.011216587 | 0.989302036 | No Change | Not Significant | TRUE | -0.444097271 |
| A430078I02Rik | TRUE | 1.543569914 | 6.75E-13 | UP | UP | TRUE | 0.355961441 |
| A430088P11Rik | TRUE | -0.209774966 | 0.62720573 | No Change | Not Significant | TRUE | -0.13918293 |
| A430090L17Rik | TRUE | -0.262835114 | 0.787780401 | No Change | Not Significant | FALSE | NA |
| A430093F15Rik | TRUE | -0.028579537 | 1 | No Change | Not Significant | FALSE | NA |
| A430108G06Rik | TRUE | 1.802917078 | 1 | No Change | Not Significant | FALSE | NA |
| A4galt | TRUE | -0.10606364 | 0.827951097 | No Change | Not Significant | TRUE | 0.325336051 |
| A4gnt | TRUE | -0.187126369 | 0.643957486 | No Change | Not Significant | TRUE | 0.035335478 |
| A530010L16Rik | TRUE | -0.363817815 | 1 | No Change | Not Significant | FALSE | NA |
| A530016L24Rik | TRUE | -0.84048971 | 1 | No Change | Not Significant | FALSE | NA |
| A530032D15Rik | TRUE | -0.229616486 | 0.885199039 | No Change | Not Significant | TRUE | 0.087194178 |
| A530040E14Rik | TRUE | 0.278647845 | 1 | No Change | Not Significant | FALSE | NA |
| A530064D06Rik | TRUE | 2.772972669 | 1 | No Change | Not Significant | FALSE | NA |
| A530065N20Rik | TRUE | -0.090873239 | 1 | No Change | Not Significant | FALSE | NA |
| A530072M11Rik | TRUE | 0.12187632 | 0.779177919 | No Change | Not Significant | TRUE | -0.030008375 |
| A630001G21Rik | TRUE | 0.74812005 | 0.01984355 | UP | UP | TRUE | -0.058566439 |
| A630072M18Rik | TRUE | 0.165749891 | 1 | No Change | Not Significant | FALSE | NA |
| A630095N17Rik | TRUE | 0.051543989 | 1 | No Change | Not Significant | FALSE | NA |
| A730006G06Rik | TRUE | 0.524287384 | 1 | No Change | Not Significant | FALSE | NA |
| A730018C14Rik | TRUE | -0.338774122 | 1 | No Change | Not Significant | FALSE | NA |
| A730020M07Rik | TRUE | -0.766115511 | 1 | No Change | Not Significant | FALSE | NA |
| A730036I17Rik | TRUE | -0.422516346 | 1 | No Change | Not Significant | FALSE | NA |
| A730046J19Rik | TRUE | 0.609377624 | 1 | No Change | Not Significant | FALSE | NA |
| A730049H05Rik | TRUE | -2.831541923 | 1 | No Change | Not Significant | FALSE | NA |
| A730056A06Rik | TRUE | -0.818023029 | 1 | No Change | Not Significant | FALSE | NA |
| A730081D07Rik | TRUE | -0.474390274 | 1 | No Change | Not Significant | FALSE | NA |
| A730082K24Rik | TRUE | 0.753083675 | 1 | No Change | Not Significant | FALSE | NA |
| A830005F24Rik | TRUE | 1.118184821 | 1 | No Change | Not Significant | FALSE | NA |
| A830008E24Rik | TRUE | -0.106679485 | 1 | No Change | Not Significant | FALSE | NA |
| A830009L08Rik | TRUE | -1.532900623 | 1 | No Change | Not Significant | FALSE | NA |
| A830018L16Rik | TRUE | -1.340466696 | 1 | No Change | Not Significant | FALSE | NA |
| A830019L24Rik | TRUE | 1.126460431 | 1 | No Change | Not Significant | FALSE | NA |
| A830035O19Rik | TRUE | -0.290867214 | 0.6397711 | No Change | Not Significant | TRUE | -0.749907575 |
| A830036E02Rik | TRUE | 1.29324271 | 0.201273764 | No Change | Not Significant | FALSE | NA |
| A830082K12Rik | TRUE | 1.7677283 | 1 | No Change | Not Significant | FALSE | NA |
| A830082N09Rik | TRUE | 0.443357974 | 0.663584952 | No Change | Not Significant | FALSE | NA |
| A930002C04Rik | TRUE | -0.428080566 | 1 | No Change | Not Significant | FALSE | NA |
| A930003O13Rik | TRUE | -1.898806956 | 1 | No Change | Not Significant | FALSE | NA |
| A930004D18Rik | TRUE | -0.571531775 | 0.235723563 | No Change | Not Significant | TRUE | -0.084227814 |
| A930005H10Rik | TRUE | -0.068259187 | 0.901709044 | No Change | Not Significant | TRUE | -0.051707834 |
| A930006K02Rik | TRUE | -0.532934316 | 1 | No Change | Not Significant | FALSE | NA |
| A930007A09Rik | TRUE | -0.107052009 | 0.932166328 | No Change | Not Significant | TRUE | 0.111829908 |
| A930007I19Rik | TRUE | 0.574746681 | 0.493321208 | No Change | Not Significant | TRUE | 0.409273327 |
| A930009A15Rik | TRUE | 1.520258197 | 1 | No Change | Not Significant | FALSE | NA |
| A930012L18Rik | TRUE | -0.221377447 | 1 | No Change | Not Significant | TRUE | -0.082051261 |
| A930016O22Rik | TRUE | 0.341357266 | 1 | No Change | Not Significant | FALSE | NA |
| A930019D19Rik | TRUE | -0.058425128 | 0.940676072 | No Change | Not Significant | TRUE | 0.210278603 |
| A930024E05Rik | TRUE | 0.120328276 | 0.882690208 | No Change | Not Significant | TRUE | -0.604778485 |
| A930033H14Rik | TRUE | 0.253347587 | 0.851895956 | No Change | Not Significant | TRUE | -0.724384325 |
| A930037H05Rik | TRUE | -0.221850432 | 0.63998633 | No Change | Not Significant | TRUE | -0.263093054 |
| A930041C12Rik | TRUE | 1.064542774 | 1 | No Change | Not Significant | FALSE | NA |
| AA414768 | TRUE | -0.186502172 | 0.734194205 | No Change | Not Significant | TRUE | -0.080972827 |
| AA467197 | TRUE | -0.48004957 | 0.000479251 | No Change | Significant Negative | TRUE | -0.138310364 |
| AA986860 | TRUE | 0.475896206 | 0.01548252 | No Change | Significant Positive | TRUE | 0.005764742 |
| Aaas | TRUE | 0.425184094 | 0.013385512 | No Change | Significant Positive | TRUE | 0.084180238 |
| Aacs | TRUE | -0.09534827 | 0.692354022 | No Change | Not Significant | TRUE | -0.077484205 |
| Aadac | TRUE | -0.473954241 | 0.019112763 | No Change | Significant Negative | TRUE | 0.2288404 |
| Aagab | TRUE | -0.106290111 | 0.532495959 | No Change | Not Significant | TRUE | 0.162001209 |
| Aak1 | TRUE | -0.0278767 | 0.875092959 | No Change | Not Significant | TRUE | 0.004726204 |
| Aamdc | TRUE | -0.500570725 | 6.88E-07 | DOWN | DOWN | TRUE | 0.034431211 |
| Aamp | TRUE | 0.061430623 | 0.443169915 | No Change | Not Significant | TRUE | 0.128918155 |
| Aanat | TRUE | -0.284077796 | 1 | No Change | Not Significant | FALSE | NA |
| Aar2 | TRUE | -0.031912379 | 0.815684492 | No Change | Not Significant | TRUE | 0.054056359 |
| Aars | TRUE | 0.578521617 | 4.21E-13 | UP | UP | TRUE | 0.269698318 |
| Aars2 | TRUE | -0.27101562 | 0.008643912 | No Change | Significant Negative | TRUE | -0.048592374 |
| Aarsd1 | TRUE | 1.268423171 | 0.049808611 | UP | UP | FALSE | NA |
| Aasdh | TRUE | -0.213207738 | 0.203572456 | No Change | Not Significant | TRUE | -0.104531653 |
| Aasdhppt | TRUE | -0.171651483 | 0.189357722 | No Change | Not Significant | TRUE | -0.038527182 |
| Aass | TRUE | 2.010819434 | 1 | No Change | Not Significant | FALSE | NA |
| Aatf | TRUE | -0.222078377 | 0.088198982 | No Change | Not Significant | TRUE | -0.13947227 |
| Aatk | TRUE | 0.621940972 | 0.764339772 | No Change | Not Significant | FALSE | NA |
| AB124611 | TRUE | 0.100065706 | 1 | No Change | Not Significant | FALSE | NA |
| Abat | TRUE | -0.418034489 | 0.017884376 | No Change | Significant Negative | TRUE | -0.130447697 |
| Abca1 | TRUE | 2.896717358 | 2.45E-97 | UP | UP | TRUE | 1.900838571 |
| Abca12 | TRUE | 0.576938512 | 0.087888475 | No Change | Not Significant | TRUE | 0.037079256 |
| Abca13 | TRUE | -0.265192401 | 0.799093792 | No Change | Not Significant | TRUE | -0.332850218 |
| Abca14 | TRUE | 1.163201987 | 0.292138489 | No Change | Not Significant | TRUE | -0.978544253 |
| Abca15 | TRUE | 1.722440106 | 1 | No Change | Not Significant | FALSE | NA |
| Abca16 | TRUE | -0.731130138 | 1 | No Change | Not Significant | FALSE | NA |
| Abca17 | TRUE | 0.437312016 | 1 | No Change | Not Significant | FALSE | NA |
| Abca2 | TRUE | -0.18788942 | 0.1835581 | No Change | Not Significant | TRUE | 0.128016857 |
| Abca3 | TRUE | -0.261263943 | 0.015558323 | No Change | Significant Negative | TRUE | -0.02243156 |
| Abca4 | TRUE | 0.305404623 | 1 | No Change | Not Significant | FALSE | NA |
| Abca5 | TRUE | -0.056603231 | 0.906297546 | No Change | Not Significant | TRUE | -0.131672822 |
| Abca6 | TRUE | 4.13881597 | 1 | No Change | Not Significant | FALSE | NA |
| Abca7 | TRUE | 0.712226479 | 8.87E-08 | UP | UP | TRUE | 0.153414401 |
| Abca8a | TRUE | 1.789090661 | 1 | No Change | Not Significant | FALSE | NA |
| Abca8b | TRUE | -0.184256913 | 0.904987729 | No Change | Not Significant | FALSE | NA |
| Abca9 | TRUE | -0.143669806 | 1 | No Change | Not Significant | FALSE | NA |
| Abcb10 | TRUE | -0.108349771 | 0.405213026 | No Change | Not Significant | TRUE | -0.115726193 |

| geneID | Cond1_DF_Detected | Cond1_DF_ValueCol | Cond1_DF_PadjCol | Cond1_DF_Cutoff | Cond1_DF_Cutoff_Specific.x | Cond2_DF_Detected | Cond2_DF_ValueCol | Cond2_DF_PadjCol | Cond2_DF_Cutoff | Cond2_DF_Cutoff_Specific | BG_Method | RG1_All | RG2_Significant | RG3_SignificantChange |
|---|---|---|---|---|---|---|---|---|---|---|---|---|---|---|
| Abcb11 | TRUE | 2.828112913 | 1 | No Change | Not Significant | FALSE | NA | NA | No Change | Not Detected | FALSE | Background = FALSE | Background = FALSE | Background = FALSE |
| Abcb1a | TRUE | -0.594954129 | 1.18E-05 | DOWN | DOWN | TRUE | -0.587335577 | 5.40E-14 | DOWN | DOWN | TRUE | Cond1 DOWN + Cond2 DOWN | Core_DOWN | Core_DOWN |
| Abcb1b | TRUE | 0.970374285 | 0.443634979 | No Change | Not Significant | FALSE | NA | NA | No Change | Not Detected | FALSE | Background = FALSE | Background = FALSE | Background = FALSE |
| Abcb4 | TRUE | -0.555492136 | 0.712202101 | No Change | Not Significant | FALSE | NA | NA | No Change | Not Detected | FALSE | Background = FALSE | Background = FALSE | Background = FALSE |
| Abcb5 | TRUE | 2.539944654 | 1 | No Change | Not Significant | FALSE | NA | NA | No Change | Not Detected | FALSE | Background = FALSE | Background = FALSE | Background = FALSE |
| Abcb6 | TRUE | -0.311077401 | 0.121849545 | No Change | Not Significant | TRUE | -0.046912656 | 0.865592838 | No Change | Not Significant | TRUE | Cond1 Not Significant + Cond2 Not Significant | None | None |
| Abcb7 | TRUE | -0.060120583 | 0.717711842 | No Change | Not Significant | TRUE | 0.048332514 | 0.5725114 | No Change | Not Significant | TRUE | Cond1 Not Significant + Cond2 Not Significant | None | None |
| Abcb8 | TRUE | -0.002956157 | 0.987234441 | No Change | Not Significant | TRUE | 0.056137976 | 0.592690624 | No Change | Not Significant | TRUE | Cond1 Not Significant + Cond2 Not Significant | None | None |
| Abcb9 | TRUE | 1.213290249 | 4.53E-36 | UP | UP | TRUE | 0.523330475 | 4.76E-08 | UP | UP | TRUE | Cond1 UP + Cond2 UP | Core_UP | Core_UP |
| Abcc1 | TRUE | -0.264899968 | 0.019641571 | No Change | Significant Negative | TRUE | -0.002246529 | 0.986357608 | No Change | Not Significant | TRUE | Cond1 Significant Negative + Cond2 Not Significant | None | None |
| Abcc10 | TRUE | 0.143115349 | 0.50115173 | No Change | Not Significant | TRUE | 0.023528692 | 0.908143613 | No Change | Not Significant | TRUE | Cond1 Not Significant + Cond2 Not Significant | None | None |
| Abcc12 | TRUE | 2.009529999 | 1 | No Change | Not Significant | FALSE | NA | NA | No Change | Not Detected | FALSE | Background = FALSE | Background = FALSE | Background = FALSE |
| Abcc2 | TRUE | -0.969937068 | 0.127792639 | No Change | Not Significant | TRUE | -1.112744072 | 0.016537282 | DOWN | DOWN | TRUE | Cond1 Not Significant + Cond2 DOWN | Cond2_DOWN | Cond2_DOWN |
| Abcc3 | TRUE | -0.112038452 | 0.636725956 | No Change | Not Significant | TRUE | -0.164247784 | 0.094722877 | No Change | Not Significant | TRUE | Cond1 Not Significant + Cond2 Not Significant | None | None |
| Abcc4 | TRUE | 0.361022037 | 0.124778744 | No Change | Not Significant | TRUE | 0.207692795 | 0.043063317 | No Change | Significant Positive | TRUE | Cond1 Not Significant + Cond2 Significant Positive | None | None |
| Abcc5 | TRUE | -0.131023627 | 0.761753322 | No Change | Not Significant | TRUE | -0.442157325 | 0.051170901 | No Change | Not Significant | TRUE | Cond1 Not Significant + Cond2 Not Significant | None | None |
| Abcc6 | TRUE | 0.399563256 | 0.10016904 | No Change | Not Significant | TRUE | -0.004265959 | 0.986375338 | No Change | Not Significant | TRUE | Cond1 Not Significant + Cond2 Not Significant | None | None |
| Abcc8 | TRUE | -0.97453043 | 0.002313363 | DOWN | DOWN | TRUE | -0.634173859 | 0.012507901 | DOWN | DOWN | TRUE | Cond1 DOWN + Cond2 DOWN | Core_DOWN | Core_DOWN |
| Abcc9 | TRUE | 0.702004972 | 1 | No Change | Not Significant | FALSE | NA | NA | No Change | Not Detected | FALSE | Background = FALSE | Background = FALSE | Background = FALSE |
| Abcd1 | TRUE | -0.121829149 | 0.22515016 | No Change | Not Significant | TRUE | 0.066992129 | 0.639482053 | No Change | Not Significant | TRUE | Cond1 Not Significant + Cond2 Not Significant | None | None |
| Abcd3 | TRUE | 0.105321402 | 0.255483337 | No Change | Not Significant | TRUE | 0.186014673 | 0.002907732 | No Change | Significant Positive | TRUE | Cond1 Not Significant + Cond2 Significant Positive | None | None |
| Abcd4 | TRUE | 0.047068275 | 0.847573958 | No Change | Not Significant | TRUE | -0.066738956 | 0.649970343 | No Change | Not Significant | TRUE | Cond1 Not Significant + Cond2 Not Significant | None | None |
| Abce1 | TRUE | 0.221760713 | 0.160458367 | No Change | Not Significant | TRUE | 0.13224774 | 0.26363254 | No Change | Not Significant | TRUE | Cond1 Not Significant + Cond2 Not Significant | None | None |
| Abcf1 | TRUE | 0.333644807 | 4.48E-07 | No Change | Significant Positive | TRUE | 0.240117473 | 1.21E-07 | No Change | Significant Positive | TRUE | Cond1 Significant Positive + Cond2 Significant Positive | None | None |
| Abcf2 | TRUE | 0.053610205 | 0.753386941 | No Change | Not Significant | TRUE | 0.139206238 | 0.108932668 | No Change | Not Significant | TRUE | Cond1 Not Significant + Cond2 Not Significant | None | None |
| Abcf3 | TRUE | 0.127036762 | 0.161179189 | No Change | Not Significant | TRUE | 0.096058088 | 0.193436559 | No Change | Not Significant | TRUE | Cond1 Not Significant + Cond2 Not Significant | None | None |
| Abcg1 | TRUE | 1.161719463 | 3.12E-09 | UP | UP | TRUE | 0.801900105 | 0.000257257 | UP | UP | TRUE | Cond1 UP + Cond2 UP | Core_UP | Core_UP |
| Abcg2 | TRUE | 0.025885196 | 0.914175515 | No Change | Not Significant | TRUE | 0.049506368 | 0.657014281 | No Change | Not Significant | TRUE | Cond1 Not Significant + Cond2 Not Significant | None | None |
| Abcg3 | TRUE | 1.250368351 | 1 | No Change | Not Significant | FALSE | NA | NA | No Change | Not Detected | FALSE | Background = FALSE | Background = FALSE | Background = FALSE |
| Abcg4 | TRUE | 1.135931174 | 1 | No Change | Not Significant | FALSE | NA | NA | No Change | Not Detected | FALSE | Background = FALSE | Background = FALSE | Background = FALSE |
| Abcg5 | TRUE | -0.216167954 | 0.660955754 | No Change | Not Significant | TRUE | 0.124011128 | 0.725473655 | No Change | Not Significant | TRUE | Cond1 Not Significant + Cond2 Not Significant | None | None |
| Abcg8 | TRUE | -0.490842949 | 0.414416136 | No Change | Not Significant | TRUE | 0.08692652 | 0.892291327 | No Change | Not Significant | TRUE | Cond1 Not Significant + Cond2 Not Significant | None | None |
| Abhd1 | TRUE | -0.059253962 | 0.962738158 | No Change | Not Significant | TRUE | -0.467443365 | 0.517029701 | No Change | Not Significant | TRUE | Cond1 Not Significant + Cond2 Not Significant | None | None |
| Abhd10 | TRUE | 0.171858112 | 0.36728482 | No Change | Not Significant | TRUE | 0.037532099 | 0.852953349 | No Change | Not Significant | TRUE | Cond1 Not Significant + Cond2 Not Significant | None | None |
| Abhd11 | TRUE | 0.110235222 | 0.292957669 | No Change | Not Significant | TRUE | -0.068446349 | 0.449547151 | No Change | Not Significant | TRUE | Cond1 Not Significant + Cond2 Not Significant | None | None |
| Abhd12 | TRUE | -0.011912247 | 0.95200802 | No Change | Not Significant | TRUE | 0.124838885 | 0.317250963 | No Change | Not Significant | TRUE | Cond1 Not Significant + Cond2 Not Significant | None | None |
| Abhd13 | TRUE | 0.089164857 | 0.575712264 | No Change | Not Significant | TRUE | 0.04264926 | 0.485573395 | No Change | Not Significant | TRUE | Cond1 Not Significant + Cond2 Not Significant | None | None |
| Abhd14a | TRUE | -0.297398973 | 0.22515016 | No Change | Not Significant | TRUE | 0.08761883 | 0.542214583 | No Change | Not Significant | TRUE | Cond1 Not Significant + Cond2 Not Significant | None | None |
| Abhd14b | TRUE | -0.295575166 | 0.006384236 | No Change | Significant Negative | TRUE | 0.004743612 | 0.974240941 | No Change | Not Significant | TRUE | Cond1 Significant Negative + Cond2 Not Significant | None | None |
| Abhd15 | TRUE | 1.424560034 | 0.02782961 | UP | UP | FALSE | NA | NA | No Change | Not Detected | FALSE | Background = FALSE | Background = FALSE | Background = FALSE |
| Abhd16a | TRUE | -0.087253526 | 0.494297255 | No Change | Not Significant | TRUE | -0.024923576 | 0.786019643 | No Change | Not Significant | TRUE | Cond1 Not Significant + Cond2 Not Significant | None | None |
| Abhd16b | TRUE | 0.059363746 | 1 | No Change | Not Significant | FALSE | NA | NA | No Change | Not Detected | FALSE | Background = FALSE | Background = FALSE | Background = FALSE |
| Abhd17a | TRUE | -0.022335931 | 0.909231489 | No Change | Not Significant | TRUE | 0.085695051 | 0.604649053 | No Change | Not Significant | TRUE | Cond1 Not Significant + Cond2 Not Significant | None | None |
| Abhd17b | TRUE | 0.152863935 | 0.246352764 | No Change | Not Significant | TRUE | 0.02406725 | 0.816434847 | No Change | Not Significant | TRUE | Cond1 Not Significant + Cond2 Not Significant | None | None |
| Abhd17c | TRUE | 0.013952095 | 0.950335782 | No Change | Not Significant | TRUE | 0.121676437 | 0.249623847 | No Change | Not Significant | TRUE | Cond1 Not Significant + Cond2 Not Significant | None | None |
| Abhd18 | TRUE | -0.171703444 | 0.42248653 | No Change | Not Significant | TRUE | 0.022141297 | 0.927812059 | No Change | Not Significant | TRUE | Cond1 Not Significant + Cond2 Not Significant | None | None |
| Abhd2 | TRUE | -0.131000946 | 0.211290803 | No Change | Not Significant | TRUE | -0.006242627 | 0.957069421 | No Change | Not Significant | TRUE | Cond1 Not Significant + Cond2 Not Significant | None | None |
| Abhd3 | TRUE | -0.195707423 | 0.480830149 | No Change | Not Significant | TRUE | 0.23735145 | 0.126172771 | No Change | Not Significant | TRUE | Cond1 Not Significant + Cond2 Not Significant | None | None |
| Abhd4 | TRUE | -0.407917768 | 3.00E-09 | No Change | Significant Negative | TRUE | -0.034408674 | 0.753179518 | No Change | Not Significant | TRUE | Cond1 Significant Negative + Cond2 Not Significant | None | None |
| Abhd5 | TRUE | -0.069288783 | 0.625062193 | No Change | Not Significant | TRUE | 0.052574476 | 0.566263466 | No Change | Not Significant | TRUE | Cond1 Not Significant + Cond2 Not Significant | None | None |
| Abhd6 | TRUE | 0.177519614 | 0.181743011 | No Change | Not Significant | TRUE | -0.070007649 | 0.508211544 | No Change | Not Significant | TRUE | Cond1 Not Significant + Cond2 Not Significant | None | None |
| Abhd8 | TRUE | -0.006511022 | 0.985415068 | No Change | Not Significant | TRUE | 0.090485515 | 0.749012844 | No Change | Not Significant | TRUE | Cond1 Not Significant + Cond2 Not Significant | None | None |
| Abi1 | TRUE | -0.040773616 | 0.696497169 | No Change | Not Significant | TRUE | -0.008599637 | 0.899716112 | No Change | Not Significant | TRUE | Cond1 Not Significant + Cond2 Not Significant | None | None |
| Abi2 | TRUE | 0.762509824 | 1.87E-05 | UP | UP | TRUE | 0.424737988 | 0.005283881 | No Change | Significant Positive | TRUE | Cond1 UP + Cond2 Significant Positive | Core_UP | Cond1_UP |
| Abi3 | TRUE | -0.75173191 | 0.08204539 | No Change | Not Significant | TRUE | 0.075452918 | 0.838859992 | No Change | Not Significant | TRUE | Cond1 Not Significant + Cond2 Not Significant | None | None |
| Abi3bp | TRUE | 0.170225729 | 0.890384202 | No Change | Not Significant | FALSE | NA | NA | No Change | Not Detected | FALSE | Background = FALSE | Background = FALSE | Background = FALSE |
| Abitram | TRUE | 0.093116051 | 0.667666133 | No Change | Not Significant | TRUE | -0.050660915 | 0.768546546 | No Change | Not Significant | TRUE | Cond1 Not Significant + Cond2 Not Significant | None | None |
| Abl1 | TRUE | -0.064527082 | 0.643978573 | No Change | Not Significant | TRUE | -0.124167307 | 0.133697276 | No Change | Not Significant | TRUE | Cond1 Not Significant + Cond2 Not Significant | None | None |
| Abl2 | TRUE | 0.012493361 | 0.943195324 | No Change | Not Significant | TRUE | -0.079792769 | 0.333247419 | No Change | Not Significant | TRUE | Cond1 Not Significant + Cond2 Not Significant | None | None |
| Ablim1 | TRUE | -0.134864807 | 0.487629303 | No Change | Not Significant | TRUE | -0.093986194 | 0.497115408 | No Change | Not Significant | TRUE | Cond1 Not Significant + Cond2 Not Significant | None | None |
| Ablim2 | TRUE | -0.663828869 | 0.013237523 | DOWN | DOWN | TRUE | -0.759215299 | 2.07E-06 | DOWN | DOWN | TRUE | Cond1 DOWN + Cond2 DOWN | Core_DOWN | Core_DOWN |
| Ablim3 | TRUE | 0.966780389 | 1 | No Change | Not Significant | FALSE | NA | NA | No Change | Not Detected | FALSE | Background = FALSE | Background = FALSE | Background = FALSE |
| Abo | TRUE | -0.118389905 | 0.832872319 | No Change | Not Significant | TRUE | 0.098346782 | 0.60052913 | No Change | Not Significant | TRUE | Cond1 Not Significant + Cond2 Not Significant | None | None |
| Abr | TRUE | -0.061975022 | 0.667636772 | No Change | Not Significant | TRUE | 0.078938958 | 0.301715221 | No Change | Not Significant | TRUE | Cond1 Not Significant + Cond2 Not Significant | None | None |
| Abra | TRUE | 1.101451601 | 1 | No Change | Not Significant | FALSE | NA | NA | No Change | Not Detected | FALSE | Background = FALSE | Background = FALSE | Background = FALSE |
| Abracl | TRUE | 0.079209114 | 0.696186289 | No Change | Not Significant | TRUE | 0.144183666 | 0.223554482 | No Change | Not Significant | TRUE | Cond1 Not Significant + Cond2 Not Significant | None | None |
| Abraxas1 | TRUE | 0.116317973 | 0.503470022 | No Change | Not Significant | TRUE | -0.11035199 | 0.558101668 | No Change | Not Significant | TRUE | Cond1 Not Significant + Cond2 Not Significant | None | None |
| Abraxas2 | TRUE | 0.077407415 | 0.559327232 | No Change | Not Significant | TRUE | 0.064332218 | 0.419779698 | No Change | Not Significant | TRUE | Cond1 Not Significant + Cond2 Not Significant | None | None |
| Abt1 | TRUE | -0.023271642 | 0.910297568 | No Change | Not Significant | TRUE | -0.080490319 | 0.512256092 | No Change | Not Significant | TRUE | Cond1 Not Significant + Cond2 Not Significant | None | None |
| Abtb1 | TRUE | -0.415717512 | 0.001541925 | No Change | Significant Negative | TRUE | -0.118736868 | 0.366672302 | No Change | Not Significant | TRUE | Cond1 Significant Negative + Cond2 Not Significant | None | None |
| Abtb2 | TRUE | -0.255284593 | 0.450500711 | No Change | Not Significant | TRUE | 0.282141586 | 0.154385176 | No Change | Not Significant | TRUE | Cond1 Not Significant + Cond2 Not Significant | None | None |
| Acaa1a | TRUE | -0.234479419 | 0.027207712 | No Change | Significant Negative | TRUE | -0.099826229 | 0.244881547 | No Change | Not Significant | TRUE | Cond1 Significant Negative + Cond2 Not Significant | None | None |
| Acaa1b | TRUE | 1.14002688 | 0.000797335 | UP | UP | TRUE | -0.1641975 | 0.582615186 | No Change | Not Significant | TRUE | Cond1 UP + Cond2 Not Significant | Cond1_UP | Cond1_UP |
| Acaa2 | TRUE | -0.321901941 | 2.73E-05 | No Change | Significant Negative | TRUE | -0.1618255 | 0.003276707 | No Change | Significant Negative | TRUE | Cond1 Significant Negative + Cond2 Significant Negative | None | None |
| Acaca | TRUE | 0.272408031 | 0.027859338 | No Change | Significant Positive | TRUE | 0.052071554 | 0.612036907 | No Change | Not Significant | TRUE | Cond1 Significant Positive + Cond2 Not Significant | None | None |
| Acacb | TRUE | 0.440976235 | 0.110778523 | No Change | Not Significant | TRUE | 0.231119732 | 0.086176324 | No Change | Not Significant | TRUE | Cond1 Not Significant + Cond2 Not Significant | None | None |
| Acad10 | TRUE | -0.320425756 | 0.014624596 | No Change | Significant Negative | TRUE | -0.04872944 | 0.651017488 | No Change | Not Significant | TRUE | Cond1 Significant Negative + Cond2 Not Significant | None | None |
| Acad11 | TRUE | -0.104148292 | 0.447999484 | No Change | Not Significant | TRUE | 0.058327576 | 0.481535591 | No Change | Not Significant | TRUE | Cond1 Not Significant + Cond2 Not Significant | None | None |
| Acad12 | TRUE | -0.796147124 | 3.39E-07 | DOWN | DOWN | TRUE | -0.097207857 | 0.544051223 | No Change | Not Significant | TRUE | Cond1 DOWN + Cond2 Not Significant | Cond1_DOWN | Cond1_DOWN |
| Acad8 | TRUE | -0.336297867 | 0.003770114 | No Change | Significant Negative | TRUE | 0.010485824 | 0.929183351 | No Change | Not Significant | TRUE | Cond1 Significant Negative + Cond2 Not Significant | None | None |
| Acad9 | TRUE | -0.102935359 | 0.545498722 | No Change | Not Significant | TRUE | -0.10564063 | 0.234890673 | No Change | Not Significant | TRUE | Cond1 Not Significant + Cond2 Not Significant | None | None |
| Acadl | TRUE | -0.314209249 | 0.00129073 | No Change | Significant Negative | TRUE | -0.067923973 | 0.543615578 | No Change | Not Significant | TRUE | Cond1 Significant Negative + Cond2 Not Significant | None | None |
| Acadm | TRUE | -0.559956657 | 1.62E-11 | DOWN | DOWN | TRUE | -0.090048028 | 0.138156009 | No Change | Not Significant | TRUE | Cond1 DOWN + Cond2 Not Significant | Cond1_DOWN | Cond1_DOWN |
| Acads | TRUE | -0.288006537 | 0.000261952 | No Change | Significant Negative | TRUE | -0.071243848 | 0.472813526 | No Change | Not Significant | TRUE | Cond1 Significant Negative + Cond2 Not Significant | None | None |
| Acadsb | TRUE | -0.21090754 | 0.26676599 | No Change | Not Significant | TRUE | 0.020946855 | 0.869778854 | No Change | Not Significant | TRUE | Cond1 Not Significant + Cond2 Not Significant | None | None |
| Acadvl | TRUE | -0.351134618 | 8.65E-06 | No Change | Significant Negative | TRUE | 0.046453909 | 0.479283878 | No Change | Not Significant | TRUE | Cond1 Significant Negative + Cond2 Not Significant | None | None |
| Acan | TRUE | 0.94676266 | 1 | No Change | Not Significant | FALSE | NA | NA | No Change | Not Detected | FALSE | Background = FALSE | Background = FALSE | Background = FALSE |
| Acap1 | TRUE | -0.392755286 | 0.559393767 | No Change | Not Significant | TRUE | 0.746416294 | 0.009541901 | UP | UP | TRUE | Cond1 Not Significant + Cond2 UP | Cond2_UP | Cond2_UP |
| Acap2 | TRUE | -0.207100257 | 0.104512461 | No Change | Not Significant | TRUE | -0.135109074 | 0.179634215 | No Change | Not Significant | TRUE | Cond1 Not Significant + Cond2 Not Significant | None | None |
| Acap3 | TRUE | -0.462733605 | 0.00104843 | No Change | Significant Negative | TRUE | -0.122555449 | 0.478679126 | No Change | Not Significant | TRUE | Cond1 Significant Negative + Cond2 Not Significant | None | None |
| Acat1 | TRUE | -0.296849264 | 0.046798532 | No Change | Significant Negative | TRUE | -0.005511861 | 0.961718903 | No Change | Not Significant | TRUE | Cond1 Significant Negative + Cond2 Not Significant | None | None |
| Acat2 | TRUE | 0.401062355 | 0.017740208 | No Change | Significant Positive | TRUE | 0.131872501 | 0.484226665 | No Change | Not Significant | TRUE | Cond1 Significant Positive + Cond2 Not Significant | None | None |
| Acat3 | TRUE | 0.619988018 | 1 | No Change | Not Significant | FALSE | NA | NA | No Change | Not Detected | FALSE | Background = FALSE | Background = FALSE | Background = FALSE |
| Acbd3 | TRUE | 0.091889589 | 0.322785367 | No Change | Not Significant | TRUE | 0.040647375 | 0.547169304 | No Change | Not Significant | TRUE | Cond1 Not Significant + Cond2 Not Significant | None | None |
| Acbd5 | TRUE | -0.043282933 | 0.765922697 | No Change | Not Significant | TRUE | -0.018855958 | 0.86404218 | No Change | Not Significant | TRUE | Cond1 Not Significant + Cond2 Not Significant | None | None |
| Acbd6 | TRUE | -0.15628415 | 0.443634979 | No Change | Not Significant | TRUE | -0.137718198 | 0.405982819 | No Change | Not Significant | TRUE | Cond1 Not Significant + Cond2 Not Significant | None | None |
| Acbd7 | TRUE | 0.747419608 | 1 | No Change | Not Significant | FALSE | NA | NA | No Change | Not Detected | FALSE | Background = FALSE | Background = FALSE | Background = FALSE |
| Accs | TRUE | -0.311444374 | 0.289400593 | No Change | Not Significant | TRUE | -0.119795866 | 0.606826059 | No Change | Not Significant | TRUE | Cond1 Not Significant + Cond2 Not Significant | None | None |
| Accsl | TRUE | 1.805063169 | 1 | No Change | Not Significant | FALSE | NA | NA | No Change | Not Detected | FALSE | Background = FALSE | Background = FALSE | Background = FALSE |
| Acd | TRUE | -0.299636472 | 0.016201536 | No Change | Significant Negative | TRUE | -0.208889281 | 0.089842947 | No Change | Not Significant | TRUE | Cond1 Significant Negative + Cond2 Not Significant | None | None |
| Ace | TRUE | -0.400844034 | 0.18053173 | No Change | Not Significant | TRUE | -0.372785252 | 0.12424678 | No Change | Not Significant | TRUE | Cond1 Not Significant + Cond2 Not Significant | None | None |
| Ace2 | TRUE | -1.052320961 | 2.16E-08 | DOWN | DOWN | TRUE | -0.35154217 | 0.009978907 | No Change | Significant Negative | TRUE | Cond1 DOWN + Cond2 Significant Negative | Core_DOWN | Cond1_DOWN |
| Ace3 | TRUE | 0.89215915 | 0.227592128 | No Change | Not Significant | TRUE | 0.511177679 | 0.499763105 | No Change | Not Significant | TRUE | Cond1 Not Significant + Cond2 Not Significant | None | None |
| Acer1 | TRUE | -0.0901324 | 0.732400135 | No Change | Not Significant | TRUE | 0.16407254 | 0.059203 | No Change | Not Significant | TRUE | Cond1 Not Significant + Cond2 Not Significant | None | None |
| Acer2 | TRUE | 0.052852237 | 0.80530915 | No Change | Not Significant | TRUE | -0.070852783 | 0.698573273 | No Change | Not Significant | TRUE | Cond1 Not Significant + Cond2 Not Significant | None | None |
| Acer3 | TRUE | -0.454192757 | 1.39E-05 | No Change | Significant Negative | TRUE | -0.213366786 | 0.012289435 | No Change | Significant Negative | TRUE | Cond1 Significant Negative + Cond2 Significant Negative | None | None |
| Ache | TRUE | -0.08220532 | 0.888506311 | No Change | Not Significant | TRUE | -0.430782192 | 0.148466815 | No Change | Not Significant | TRUE | Cond1 Not Significant + Cond2 Not Significant | None | None |
| Acin1 | TRUE | 0.140602625 | 0.001599893 | No Change | Significant Positive | TRUE | 0.054053822 | 0.42444695 | No Change | Not Significant | TRUE | Cond1 Significant Positive + Cond2 Not Significant | None | None |
| Ackr2 | TRUE | 1.218547651 | 0.122751949 | No Change | Not Significant | FALSE | NA | NA | No Change | Not Detected | FALSE | Background = FALSE | Background = FALSE | Background = FALSE |
| Ackr3 | TRUE | 0.894161161 | 0.023887859 | UP | UP | TRUE | 0.203431248 | 0.688854258 | No Change | Not Significant | TRUE | Cond1 UP + Cond2 Not Significant | Cond1_UP | Cond1_UP |
| Ackr4 | TRUE | -0.40239054 | 0.114868009 | No Change | Not Significant | TRUE | -0.00991172 | 0.982024691 | No Change | Not Significant | TRUE | Cond1 Not Significant + Cond2 Not Significant | None | None |
| Acly | TRUE | -0.149914517 | 0.62490656 | No Change | Not Significant | TRUE | -0.067272685 | 0.750864769 | No Change | Not Significant | TRUE | Cond1 Not Significant + Cond2 Not Significant | None | None |
| Acmsd | TRUE | 0.184639395 | 1 | No Change | Not Significant | FALSE | NA | NA | No Change | Not Detected | FALSE | Background = FALSE | Background = FALSE | Background = FALSE |
| Acnat1 | TRUE | -0.167437031 | 0.436092673 | No Change | Not Significant | TRUE | 0.166812796 | 0.198761565 | No Change | Not Significant | TRUE | Cond1 Not Significant + Cond2 Not Significant | None | None |
| Acnat2 | TRUE | -0.024195822 | 1 | No Change | Not Significant | FALSE | NA | NA | No Change | Not Detected | FALSE | Background = FALSE | Background = FALSE | Background = FALSE |
| Aco1 | TRUE | -0.045312223 | 0.784161879 | No Change | Not Significant | TRUE | 0.038401257 | 0.751462933 | No Change | Not Significant | TRUE | Cond1 Not Significant + Cond2 Not Significant | None | None |
| Aco2 | TRUE | -0.121270551 | 0.301713259 | No Change | Not Significant | TRUE | 0.122580597 | 0.142262102 | No Change | Not Significant | TRUE | Cond1 Not Significant + Cond2 Not Significant | None | None |
| Acod1 | TRUE | 2.094562872 | 1 | No Change | Not Significant | FALSE | NA | NA | No Change | Not Detected | FALSE | Background = FALSE | Background = FALSE | Background = FALSE |
| Acot1 | TRUE | 0.508825661 | 0.006808522 | UP | UP | TRUE | -0.017849342 | 0.960412417 | No Change | Not Significant | TRUE | Cond1 UP + Cond2 Not Significant | Cond1_UP | Cond1_UP |
| Acot10 | TRUE | 1.835977672 | 1 | No Change | Not Significant | FALSE | NA | NA | No Change | Not Detected | FALSE | Background = FALSE | Background = FALSE | Background = FALSE |
| Acot11 | TRUE | -0.209320032 | 0.055305475 | No Change | Not Significant | TRUE | 0.015284428 | 0.913597373 | No Change | Not Significant | TRUE | Cond1 Not Significant + Cond2 Not Significant | None | None |
| Acot12 | TRUE | -0.249603534 | 1 | No Change | Not Significant | FALSE | NA | NA | No Change | Not Detected | FALSE | Background = FALSE | Background = FALSE | Background = FALSE |
| Acot13 | TRUE | -0.008028876 | 0.969180133 | No Change | Not Significant | TRUE | 0.061062679 | 0.568485113 | No Change | Not Significant | TRUE | Cond1 Not Significant + Cond2 Not Significant | None | None |
| Acot2 | TRUE | -0.316007831 | 0.064388674 | No Change | Not Significant | TRUE | 0.099613176 | 0.692759569 | No Change | Not Significant | TRUE | Cond1 Not Significant + Cond2 Not Significant | None | None |
| Acot3 | TRUE | 0.008011155 | 1 | No Change | Not Significant | FALSE | NA | NA | No Change | Not Detected | FALSE | Background = FALSE | Background = FALSE | Background = FALSE |
| Acot4 | TRUE | 0.340652983 | 0.21796851 | No Change | Not Significant | TRUE | 0.015556872 | 0.949475732 | No Change | Not Significant | TRUE | Cond1 Not Significant + Cond2 Not Significant | None | None |
| Acot5 | TRUE | -0.317817493 | 1 | No Change | Not Significant | FALSE | NA | NA | No Change | Not Detected | FALSE | Background = FALSE | Background = FALSE | Background = FALSE |
| Acot6 | TRUE | -0.04018921 | 0.920280009 | No Change | Not Significant | TRUE | 0.128837341 | 0.676717913 | No Change | Not Significant | TRUE | Cond1 Not Significant + Cond2 Not Significant | None | None |
| Acot7 | TRUE | 0.006145378 | 0.976822768 | No Change | Not Significant | TRUE | 0.269994083 | 0.093396937 | No Change | Not Significant | TRUE | Cond1 Not Significant + Cond2 Not Significant | None | None |
| Acot8 | TRUE | 0.056837418 | 0.675987813 | No Change | Not Significant | TRUE | 0.23563455 | 0.041321095 | No Change | Significant Positive | TRUE | Cond1 Not Significant + Cond2 Significant Positive | None | None |
| Acot9 | TRUE | -0.376979036 | 0.012440868 | No Change | Significant Negative | TRUE | 0.001935648 | 0.992621208 | No Change | Not Significant | TRUE | Cond1 Significant Negative + Cond2 Not Significant | None | None |
| Acox1 | TRUE | -0.190702452 | 0.296522686 | No Change | Not Significant | TRUE | -0.069167977 | 0.4461694 | No Change | Not Significant | TRUE | Cond1 Not Significant + Cond2 Not Significant | None | None |
| Acox2 | TRUE | 1.479468001 | 1.63E-09 | UP | UP | TRUE | 0.33206867 | 0.258509777 | No Change | Not Significant | TRUE | Cond1 UP + Cond2 Not Significant | Cond1_UP | Cond1_UP |
| Acox3 | TRUE | -0.318775112 | 0.00021219 | No Change | Significant Negative | TRUE | -0.042741208 | 0.735121083 | No Change | Not Significant | TRUE | Cond1 Significant Negative + Cond2 Not Significant | None | None |
| Acoxl | TRUE | 0.085509496 | 1 | No Change | Not Significant | FALSE | NA | NA | No Change | Not Detected | FALSE | Background = FALSE | Background = FALSE | Background = FALSE |
| Acp1 | TRUE | 8.73E-05 | 0.999548819 | No Change | Not Significant | TRUE | 0.07407245 | 0.372364958 | No Change | Not Significant | TRUE | Cond1 Not Significant + Cond2 Not Significant | None | None |
| Acp2 | TRUE | -0.011731494 | 0.941973978 | No Change | Not Significant | TRUE | 0.052012085 | 0.588148333 | No Change | Not Significant | TRUE | Cond1 Not Significant + Cond2 Not Significant | None | None |
| Acp4 | TRUE | -0.066789642 | 0.95096289 | No Change | Not Significant | TRUE | -0.386372913 | 0.708970987 | No Change | Not Significant | TRUE | Cond1 Not Significant + Cond2 Not Significant | None | None |
| Acp5 | TRUE | -0.374886724 | 0.002629971 | No Change | Significant Negative | TRUE | -0.021650261 | 0.880449447 | No Change | Not Significant | TRUE | Cond1 Significant Negative + Cond2 Not Significant | None | None |
| Acp6 | TRUE | -0.485462568 | 4.05E-08 | No Change | Significant Negative | TRUE | -0.2868626 | 6.90E-05 | No Change | Significant Negative | TRUE | Cond1 Significant Negative + Cond2 Significant Negative | None | None |
| Acp7 | TRUE | 0.917437094 | 1 | No Change | Not Significant | FALSE | NA | NA | No Change | Not Detected | FALSE | Background = FALSE | Background = FALSE | Background = FALSE |
| Acpp | TRUE | -0.361533542 | 0.382529046 | No Change | Not Significant | TRUE | 0.00880552 | 0.967733155 | No Change | Not Significant | TRUE | Cond1 Not Significant + Cond2 Not Significant | None | None |
| Acrbp | TRUE | -0.051060815 | 0.934688628 | No Change | Not Significant | TRUE | -0.258009068 | 0.464315218 | No Change | Not Significant | TRUE | Cond1 Not Significant + Cond2 Not Significant | None | None |
| Acrv1 | TRUE | 1.090631609 | 1 | No Change | Not Significant | FALSE | NA | NA | No Change | Not Detected | FALSE | Background = FALSE | Background = FALSE | Background = FALSE |
| Acsbg1 | TRUE | 0.036758853 | 0.980697519 | No Change | Not Significant | TRUE | 0.06536155 | 0.95019842 | No Change | Not Significant | TRUE | Cond1 Not Significant + Cond2 Not Significant | None | None |
| Acsbg3 | TRUE | 0.640776847 | 0.659241321 | No Change | Not Significant | FALSE | NA | NA | No Change | Not Detected | FALSE | Background = FALSE | Background = FALSE | Background = FALSE |
| Acsf2 | TRUE | -0.519602816 | 0.004964661 | DOWN | DOWN | TRUE | -0.175077148 | 0.070728973 | No Change | Not Significant | TRUE | Cond1 DOWN + Cond2 Not Significant | Cond1_DOWN | Cond1_DOWN |
| Acsf3 | TRUE | -0.053005182 | 0.749713986 | No Change | Not Significant | TRUE | 0.101925238 | 0.277610848 | No Change | Not Significant | TRUE | Cond1 Not Significant + Cond2 Not Significant | None | None |
| Acsl1 | TRUE | 0.020936564 | 0.95457824 | No Change | Not Significant | TRUE | 0.107364363 | 0.241391445 | No Change | Not Significant | TRUE | Cond1 Not Significant + Cond2 Not Significant | None | None |
| Acsl3 | TRUE | -0.238352532 | 0.055375938 | No Change | Not Significant | TRUE | -0.229603927 | 0.00041284 | No Change | Significant Negative | TRUE | Cond1 Not Significant + Cond2 Significant Negative | None | None |
| Acsl4 | TRUE | 0.130213002 | 0.441179379 | No Change | Not Significant | TRUE | 0.077787081 | 0.569172418 | No Change | Not Significant | TRUE | Cond1 Not Significant + Cond2 Not Significant | None | None |
| Acsl5 | TRUE | -0.036807802 | 0.875699372 | No Change | Not Significant | TRUE | 0.017014877 | 0.914534473 | No Change | Not Significant | TRUE | Cond1 Not Significant + Cond2 Not Significant | None | None |
| Acsl6 | TRUE | 0.790782355 | 1 | No Change | Not Significant | FALSE | NA | NA | No Change | Not Detected | FALSE | Background = FALSE | Background = FALSE | Background = FALSE |
| Acsm1 | TRUE | 0.4970555 | 1 | No Change | Not Significant | FALSE | NA | NA | No Change | Not Detected | FALSE | Background = FALSE | Background = FALSE | Background = FALSE |
| Acsm2 | TRUE | 1.700816114 | 1 | No Change | Not Significant | FALSE | NA | NA | No Change | Not Detected | FALSE | Background = FALSE | Background = FALSE | Background = FALSE |
| Acsm3 | TRUE | -0.57749439 | 0.065583696 | No Change | Not Significant | TRUE | -0.022338417 | 0.900662858 | No Change | Not Significant | TRUE | Cond1 Not Significant + Cond2 Not Significant | None | None |
| Acsm4 | TRUE | -2.242821818 | 1 | No Change | Not Significant | FALSE | NA | NA | No Change | Not Detected | FALSE | Background = FALSE | Background = FALSE | Background = FALSE |
| Acsm5 | TRUE | -0.396527332 | 1 | No Change | Not Significant | FALSE | NA | NA | No Change | Not Detected | FALSE | Background = FALSE | Background = FALSE | Background = FALSE |
| Acss1 | TRUE | 0.302446943 | 0.004334906 | No Change | Significant Positive | TRUE | 0.727079094 | 1.93E-71 | UP | UP | TRUE | Cond1 Significant Positive + Cond2 UP | Core_UP | Cond2_UP |
| Acss2 | TRUE | -0.249188588 | 0.352592954 | No Change | Not Significant | TRUE | -0.068872246 | 0.609913364 | No Change | Not Significant | TRUE | Cond1 Not Significant + Cond2 Not Significant | None | None |
| Acss2os | TRUE | -0.139267811 | 1 | No Change | Not Significant | FALSE | NA | NA | No Change | Not Detected | FALSE | Background = FALSE | Background = FALSE | Background = FALSE |
| Acss3 | TRUE | -0.367394129 | 1 | No Change | Not Significant | TRUE | 0.180128867 | 0.79448284 | No Change | Not Significant | TRUE | Cond1 Not Significant + Cond2 Not Significant | None | None |
| Acta1 | TRUE | 0.567640258 | 0.183701378 | No Change | Not Significant | TRUE | 0.370181756 | 0.021656722 | No Change | Significant Positive | TRUE | Cond1 Not Significant + Cond2 Significant Positive | None | None |
| Acta2 | TRUE | -0.43305208 | 1 | No Change | Not Significant | FALSE | NA | NA | No Change | Not Detected | FALSE | Background = FALSE | Background = FALSE | Background = FALSE |
| Actb | TRUE | 0.104087843 | 0.561491337 | No Change | Not Significant | TRUE | 0.161059554 | 0.14237022 | No Change | Not Significant | TRUE | Cond1 Not Significant + Cond2 Not Significant | None | None |
| Acte1 | TRUE | -0.930937674 | 1 | No Change | Not Significant | FALSE | NA | NA | No Change | Not Detected | FALSE | Background = FALSE | Background = FALSE | Background = FALSE |
| Actg1 | TRUE | -0.596757894 | 4.92E-05 | DOWN | DOWN | TRUE | 0.075553108 | 0.32728228 | No Change | Not Significant | TRUE | Cond1 DOWN + Cond2 Not Significant | Cond1_DOWN | Cond1_DOWN |
| Actg2 | TRUE | 2.909093797 | 1 | No Change | Not Significant | FALSE | NA | NA | No Change | Not Detected | FALSE | Background = FALSE | Background = FALSE | Background = FALSE |
| Actl10 | TRUE | 0.23604865 | 1 | No Change | Not Significant | FALSE | NA | NA | No Change | Not Detected | FALSE | Background = FALSE | Background = FALSE | Background = FALSE |
| Actl11 | TRUE | 0.11015461 | 1 | No Change | Not Significant | FALSE | NA | NA | No Change | Not Detected | FALSE | Background = FALSE | Background = FALSE | Background = FALSE |
| Actl6a | TRUE | 0.376070265 | 5.49E-06 | No Change | Significant Positive | TRUE | 0.084922321 | 0.530775478 | No Change | Not Significant | TRUE | Cond1 Significant Positive + Cond2 Not Significant | None | None |
| Actl6b | TRUE | 1.120034938 | 1 | No Change | Not Significant | FALSE | NA | NA | No Change | Not Detected | FALSE | Background = FALSE | Background = FALSE | Background = FALSE |
| Actl7a | TRUE | 1.401832785 | 1 | No Change | Not Significant | FALSE | NA | NA | No Change | Not Detected | FALSE | Background = FALSE | Background = FALSE | Background = FALSE |
| Actl9 | TRUE | 2.376594709 | 1 | No Change | Not Significant | FALSE | NA | NA | No Change | Not Detected | FALSE | Background = FALSE | Background = FALSE | Background = FALSE |
| Actn1 | TRUE | -0.123598939 | 0.465324602 | No Change | Not Significant | TRUE | 0.064592504 | 0.443170686 | No Change | Not Significant | TRUE | Cond1 Not Significant + Cond2 Not Significant | None | None |
| Actn2 | TRUE | 1.329739681 | 1 | No Change | Not Significant | FALSE | NA | NA | No Change | Not Detected | FALSE | Background = FALSE | Background = FALSE | Background = FALSE |
| Actn3 | TRUE | 1.584770026 | 1 | No Change | Not Significant | FALSE | NA | NA | No Change | Not Detected | FALSE | Background = FALSE | Background = FALSE | Background = FALSE |
| Actn4 | TRUE | 0.082645002 | 0.715485319 | No Change | Not Significant | TRUE | 0.132502188 | 0.281107986 | No Change | Not Significant | TRUE | Cond1 Not Significant + Cond2 Not Significant | None | None |
| Actr10 | TRUE | 0.073540235 | 0.464320478 | No Change | Not Significant | TRUE | 0.061597581 | 0.521098971 | No Change | Not Significant | TRUE | Cond1 Not Significant + Cond2 Not Significant | None | None |
| Actr1a | TRUE | 0.075389501 | 0.253681814 | No Change | Not Significant | TRUE | 0.088969965 | 0.229476957 | No Change | Not Significant | TRUE | Cond1 Not Significant + Cond2 Not Significant | None | None |
| Actr1b | TRUE | -0.024774675 | 0.758644032 | No Change | Not Significant | TRUE | 0.136587522 | 0.020906252 | No Change | Significant Positive | TRUE | Cond1 Not Significant + Cond2 Significant Positive | None | None |
| Actr2 | TRUE | 0.047256835 | 0.733468492 | No Change | Not Significant | TRUE | 0.08811549 | 0.133956365 | No Change | Not Significant | TRUE | Cond1 Not Significant + Cond2 Not Significant | None | None |
| Actr3 | TRUE | -0.086171998 | 0.524360158 | No Change | Not Significant | TRUE | 0.092853447 | 0.233599794 | No Change | Not Significant | TRUE | Cond1 Not Significant + Cond2 Not Significant | None | None |
| Actr3b | TRUE | 3.817230303 | 6.46E-05 | UP | UP | FALSE | NA | NA | No Change | Not Detected | FALSE | Background = FALSE | Background = FALSE | Background = FALSE |
| Actr5 | TRUE | -0.201962493 | 0.192474654 | No Change | Not Significant | TRUE | -0.017833327 | 0.935889087 | No Change | Not Significant | TRUE | Cond1 Not Significant + Cond2 Not Significant | None | None |
| Actr6 | TRUE | 0.272896673 | 0.343330529 | No Change | Not Significant | TRUE | -0.026727529 | 0.935592329 | No Change | Not Significant | TRUE | Cond1 Not Significant + Cond2 Not Significant | None | None |
| Actr8 | TRUE | -0.042635034 | 0.727438372 | No Change | Not Significant | TRUE | -0.002784633 | 0.984186478 | No Change | Not Significant | TRUE | Cond1 Not Significant + Cond2 Not Significant | None | None |
| Acvr1 | TRUE | 0.058528695 | 0.804573016 | No Change | Not Significant | TRUE | 0.194222804 | 0.161612222 | No Change | Not Significant | TRUE | Cond1 Not Significant + Cond2 Not Significant | None | None |
| Acvr1b | TRUE | -0.208322326 | 0.03273648 | No Change | Significant Negative | TRUE | -0.109273679 | 0.127510737 | No Change | Not Significant | TRUE | Cond1 Significant Negative + Cond2 Not Significant | None | None |
| Acvr1c | TRUE | -0.577390317 | 0.001522563 | DOWN | DOWN | TRUE | -0.141428692 | 0.42043505 | No Change | Not Significant | TRUE | Cond1 DOWN + Cond2 Not Significant | Cond1_DOWN | Cond1_DOWN |
| Acvr2a | TRUE | 0.272170651 | 0.002851314 | No Change | Significant Positive | TRUE | 0.231823448 | 5.65E-06 | No Change | Significant Positive | TRUE | Cond1 Significant Positive + Cond2 Significant Positive | None | None |
| Acvr2b | TRUE | 0.077834957 | 0.811872956 | No Change | Not Significant | TRUE | -0.262881069 | 0.254924035 | No Change | Not Significant | TRUE | Cond1 Not Significant + Cond2 Not Significant | None | None |

| geneID | Cond1_DF_Detected | Cond1_DF_ValueCol | Cond1_DF_PadjCol | Cond1_DF_Cutoff | Cond1_DF_Cutoff_Specific.x | Cond2_DF_Detected | Cond2_DF_ValueCol | Cond2_DF_PadjCol | Cond2_DF_Cutoff | Cond2_DF_Cutoff_Specific | BG_Method |
|---|---|---|---|---|---|---|---|---|---|---|---|
| Acvrl1 | TRUE | 0.015958066 | 0.964050667 | No Change | Not Significant | TRUE | 0.112533633 | 0.707337899 | No Change | Not Significant | TRUE |
| Acy1 | TRUE | 0.06468491 | 0.740220295 | No Change | Not Significant | TRUE | 0.233899717 | 0.078738347 | No Change | Not Significant | TRUE |
| Acy3 | TRUE | 0.007117378 | 0.980506535 | No Change | Not Significant | TRUE | 0.027498788 | 0.88389772 | No Change | Not Significant | TRUE |
| Acyp1 | TRUE | -0.326530646 | 0.070752953 | No Change | Not Significant | TRUE | 0.038087012 | 0.841984797 | No Change | Not Significant | TRUE |
| Acyp2 | TRUE | -0.227130817 | 0.380591057 | No Change | Not Significant | TRUE | 0.015055746 | 0.969665987 | No Change | Not Significant | TRUE |
| Ada | TRUE | 0.483890981 | 0.002186165 | No Change | Significant Positive | TRUE | 0.308451967 | 0.056752643 | No Change | Not Significant | TRUE |
| Adad1 | TRUE | 0.683646242 | 1 | No Change | Not Significant | FALSE | NA | NA | No Change | Not Detected | FALSE |
| Adal | TRUE | 0.100546845 | 0.657938595 | No Change | Not Significant | TRUE | -0.19258912 | 0.28619758 | No Change | Not Significant | TRUE |
| Adam10 | TRUE | 0.111941509 | 0.340507018 | No Change | Not Significant | TRUE | -0.004735461 | 0.968095473 | No Change | Not Significant | TRUE |
| Adam11 | TRUE | 0.012480511 | 0.99400894 | No Change | Not Significant | FALSE | NA | NA | No Change | Not Detected | FALSE |
| Adam12 | TRUE | 1.866050382 | 0.375961595 | No Change | Not Significant | FALSE | NA | NA | No Change | Not Detected | FALSE |
| Adam15 | TRUE | -0.361492351 | 0.000708826 | No Change | Significant Negative | TRUE | 0.029365927 | 0.887492981 | No Change | Not Significant | TRUE |
| [illegible] | [illegible] | [illegible] | [illegible] | [illegible] | [illegible] | [illegible] | [illegible] | [illegible] | [illegible] | [illegible] | [illegible] |

| geneID | RG1_All | RG2_Significant | RG3_SignificantChange | Cond1_DF_ensembl_gene_id | Cond1_DF_entrezgene_id | Cond1_DF_gene_biotype | Cond1_DF_baseMean | Cond1_DF_log2FoldChange | Cond1_DF_lfcSE | Cond1_DF_stat | Cond1_DF_pvalue | Cond1_DF_padj | Cond1_DF_absLog2FC | Cond1_DF_Cutoff_Specific.y |
|---|---|---|---|---|---|---|---|---|---|---|---|---|---|---|
| Acvrl1 | Cond1 Not Significant + Cond2 Not Significant | None | None | ENSMUSG00000000530 | 11482 | protein_coding | 74.4878905 | 0.015958066 | 0.169509624 | -0.094142535 | 0.924995932 | 0.964050667 | 0.015958066 | Not Significant |
| Acy1 | Cond1 Not Significant + Cond2 Not Significant | None | None | ENSMUSG00000023262 | 109652 | protein_coding | 470.0688601 | 0.06468491 | 0.107966192 | -0.599121904 | 0.549091596 | 0.740220295 | 0.06468491 | Not Significant |
| Acy3 | Cond1 Not Significant + Cond2 Not Significant | None | None | ENSMUSG00000024866 | 71670 | protein_coding | 184.4799706 | 0.007117378 | 0.141216887 | -0.050400331 | 0.959803373 | 0.980506535 | 0.007117378 | Not Significant |
| Acyp1 | Cond1 Not Significant + Cond2 Not Significant | None | None | ENSMUSG00000008822 | 66204 | protein_coding | 74.6327845 | -0.326530646 | 0.139320234 | 2.343741725 | 0.019091387 | 0.070752953 | 0.326530646 | Not Significant |
| Acyp2 | Cond1 Not Significant + Cond2 Not Significant | None | None | ENSMUSG00000060923 | 75572 | protein_coding | 37.0417996 | -0.227130817 | 0.17143161 | 1.324906284 | 0.185202236 | 0.380591057 | 0.227130817 | Not Significant |
| Ada | Cond1 Significant Positive + Cond2 Not Significant | None | None | ENSMUSG00000017697 | 11486 | protein_coding | 97.23261115 | 0.483890981 | 0.133396241 | -3.62747088 | 0.000286211 | 0.002186165 | 0.483890981 | Significant Positive |
| Adad1 | Background = FALSE | Background = FALSE | Background = FALSE | ENSMUSG00000027719 | 21744 | protein_coding | 1.038165905 | 0.683646242 | 2.233067964 | -0.306146635 | 0.759493002 | 1 | 0.683646242 | Not Significant |
| Adal | Cond1 Not Significant + Cond2 Not Significant | None | None | ENSMUSG00000027259 | 75894 | protein_coding | 93.69636391 | 0.100546845 | 0.130234094 | -0.772047025 | 0.440086576 | 0.657938595 | 0.100546845 | Not Significant |
| Adam10 | Cond1 Not Significant + Cond2 Not Significant | None | None | ENSMUSG00000054693 | 11487 | protein_coding | 1382.500117 | 0.111941509 | 0.079143832 | -1.414405982 | 0.157242735 | 0.340507018 | 0.111941509 | Not Significant |
| Adam11 | Background = FALSE | Background = FALSE | Background = FALSE | ENSMUSG00000020926 | 11488 | protein_coding | 4.708587363 | 0.012480511 | 0.931843851 | -0.013393351 | 0.989313972 | 0.99400894 | 0.012480511 | Not Significant |
| Adam12 | Background = FALSE | Background = FALSE | Background = FALSE | ENSMUSG00000054555 | 11489 | protein_coding | 4.432815474 | 1.866050382 | 1.397363493 | -1.335407996 | 0.181742849 | 0.375961595 | 1.866050382 | Not Significant |
| Adam15 | Cond1 Significant Negative + Cond2 Not Significant | None | None | ENSMUSG00000028041 | 11490 | protein_coding | 595.1730466 | -0.361492351 | 0.091641555 | 3.944633518 | 7.99E-05 | 0.000708826 | 0.361492351 | Significant Negative |
| [illegible] | [illegible] | [illegible] | [illegible] | [illegible] | [illegible] | [illegible] | [illegible] | [illegible] | [illegible] | [illegible] | [illegible] | [illegible] | [illegible] | [illegible] |

| geneID | Cond2_DF_ensembl_gene_id | Cond2_DF_entrezgene_id | Cond2_DF_gene_biotype | Cond2_DF_baseMean | Cond2_DF_log2FoldChange | Cond2_DF_lfcSE | Cond2_DF_stat | Cond2_DF_pvalue | Cond2_DF_padj | Cond2_DF_absLog2FC | RG1_Specific_Cond2 | RG1_Specific_Cond1 |
|---|---|---|---|---|---|---|---|---|---|---|---|---|
| Acvrl1 | ENSMUSG00000000530 | 11482 | protein_coding | 93.34048019 | 0.112533633 | 0.161619953 | 0.696285519 | 0.48625004 | 0.707337899 | 0.112533633 | Cond1 No Change + Cond2 Not Significant | Cond2 No Change + Cond1 Not Significant |
| Acy1 | ENSMUSG00000023262 | 109652 | protein_coding | 965.4495195 | 0.233899717 | 0.095788674 | 2.44183061 | 0.014613001 | 0.078738347 | 0.233899717 | Cond1 No Change + Cond2 Not Significant | Cond2 No Change + Cond1 Not Significant |
| Acy3 | ENSMUSG00000024866 | 71670 | protein_coding | 438.5774404 | 0.027498788 | 0.088219811 | 0.311707632 | 0.755262727 | 0.88389772 | 0.027498788 | Cond1 No Change + Cond2 Not Significant | Cond2 No Change + Cond1 Not Significant |
| Acyp1 | ENSMUSG00000008822 | 66204 | protein_coding | 168.3781209 | 0.038087012 | 0.094237139 | 0.404161375 | 0.686094055 | 0.841984797 | 0.038087012 | Cond1 No Change + Cond2 Not Significant | Cond2 No Change + Cond1 Not Significant |
| Acyp2 | ENSMUSG00000060923 | 75572 | protein_coding | 86.83409667 | 0.015055746 | 0.173583892 | 0.086734696 | 0.930882397 | 0.969665987 | 0.015055746 | Cond1 No Change + Cond2 Not Significant | Cond2 No Change + Cond1 Not Significant |
| Ada | ENSMUSG00000017697 | 11486 | protein_coding | 163.666552 | 0.308451967 | 0.118711658 | 2.598329201 | 0.009367864 | 0.056752643 | 0.308451967 | Cond1 No Change + Cond2 Not Significant | Cond2 No Change + Cond1 Significant Positive |
| Adad1 | NA | NA | NA | NA | NA | NA | NA | NA | NA | NA | Background = FALSE | Background = FALSE |
| Adal | ENSMUSG00000027259 | 75894 | protein_coding | 159.0301054 | -0.19258912 | 0.116743058 | -1.649683702 | 0.099007645 | 0.28619758 | 0.19258912 | Cond1 No Change + Cond2 Not Significant | Cond2 No Change + Cond1 Not Significant |
| Adam10 | ENSMUSG00000054693 | 11487 | protein_coding | 2501.420079 | -0.004735461 | 0.051727058 | -0.091547079 | 0.927057899 | 0.968095473 | 0.004735461 | Cond1 No Change + Cond2 Not Significant | Cond2 No Change + Cond1 Not Significant |
| Adam11 | NA | NA | NA | NA | NA | NA | NA | NA | NA | NA | Background = FALSE | Background = FALSE |
| Adam12 | NA | NA | NA | NA | NA | NA | NA | NA | NA | NA | Background = FALSE | Background = FALSE |
| Adam15 | ENSMUSG00000028041 | 11490 | protein_coding | 797.846257 | 0.029365927 | 0.096718675 | 0.303622094 | 0.761415817 | 0.887492981 | 0.029365927 | Cond1 No Change + Cond2 Not Significant | Cond2 No Change + Cond1 Significant Negative |
| [illegible] | [illegible] | [illegible] | [illegible] | [illegible] | [illegible] | [illegible] | [illegible] | [illegible] | [illegible] | [illegible] | [illegible] | [illegible] |

| geneID | Cond1_DF_Detected | Cond1_DF_ValueCol | Cond1_DF_PadjCol | Cond1_DF_Cutoff | Cond1_DF_Cutoff_Specific.x | Cond2_DF_Detected | Cond2_DF_ValueCol | Cond2_DF_PadjCol | Cond2_DF_Cutoff | Cond2_DF_Cutoff_Specific | BG_Method | RG1_All |
|---|---|---|---|---|---|---|---|---|---|---|---|---|
| Amfr | TRUE | -0.170374538 | 0.100654009 | No Change | Not Significant | TRUE | -0.039848863 | 0.503084198 | No Change | Not Significant | TRUE | Cond1 Not Significant + Cond2 Not Significant |
| Amh | TRUE | 1.046311148 | 1 | No Change | Not Significant | FALSE | NA | NA | No Change | Not Detected | FALSE | Background = FALSE |
| Amhr2 | TRUE | -1.315120014 | 0.104512288 | No Change | Not Significant | TRUE | 0.030015746 | 0.977863038 | No Change | Not Significant | TRUE | Cond1 Not Significant + Cond2 Not Significant |
| Amigo1 | TRUE | -0.29209991 | 0.073420411 | No Change | Not Significant | TRUE | -0.200193925 | 0.120870289 | No Change | Not Significant | TRUE | Cond1 Not Significant + Cond2 Not Significant |
| Amigo2 | TRUE | -1.13580945 | 0.009775631 | DOWN | DOWN | TRUE | -0.306131164 | 0.486948216 | No Change | Not Significant | TRUE | Cond1 DOWN + Cond2 Not Significant |
| Amigo3 | TRUE | 0.871866059 | 4.91E-14 | UP | UP | TRUE | 0.190361236 | 0.376539145 | No Change | Not Significant | TRUE | Cond1 UP + Cond2 Not Significant |
| Ammecr1 | TRUE | 0.088773214 | 0.77323163 | No Change | Not Significant | TRUE | 0.021503841 | 0.910371624 | No Change | Not Significant | TRUE | Cond1 Not Significant + Cond2 Not Significant |
| Ammecr1l | TRUE | 0.031971293 | 0.824897384 | No Change | Not Significant | TRUE | 0.006412054 | 0.959122007 | No Change | Not Significant | TRUE | Cond1 Not Significant + Cond2 Not Significant |
| Amn | TRUE | 0.20743774 | 0.630052795 | No Change | Not Significant | TRUE | -0.341157895 | 0.050864992 | No Change | Not Significant | TRUE | Cond1 Not Significant + Cond2 Not Significant |
| Amn1 | TRUE | 0.074306562 | 0.726359126 | No Change | Not Significant | TRUE | -0.078652825 | 0.669750853 | No Change | Not Significant | TRUE | Cond1 Not Significant + Cond2 Not Significant |
| Amot | TRUE | 0.234256784 | 0.122133027 | No Change | Not Significant | TRUE | 0.087660751 | 0.259435364 | No Change | Not Significant | TRUE | Cond1 Not Significant + Cond2 Not Significant |
| Amotl1 | TRUE | 1.058620504 | 1.77E-08 | UP | UP | TRUE | 0.279106755 | 0.10455009 | No Change | Not Significant | TRUE | Cond1 UP + Cond2 Not Significant |
| [illegible] | [illegible] | [illegible] | [illegible] | [illegible] | [illegible] | [illegible] | [illegible] | [illegible] | [illegible] | [illegible] | [illegible] | [illegible] |

| geneID | RG2_Significant | RG3_SignificantChange | Cond1_DF_ensembl_gene_id | Cond1_DF_entrezgene_id | Cond1_DF_gene_biotype | Cond1_DF_baseMean | Cond1_DF_log2FoldChange | Cond1_DF_lfcSE | Cond1_DF_stat | Cond1_DF_pvalue | Cond1_DF_padj | Cond1_DF_absLog2FC | Cond1_DF_Cutoff_Specific.y |
|---|---|---|---|---|---|---|---|---|---|---|---|---|---|
| Amfr | None | None | ENSMUSG00000031751 | 23802 | protein_coding | 1974.139966 | -0.170374538 | 0.078460778 | 2.171461225 | 0.029896325 | 0.100654009 | 0.170374538 | Not Significant |
| Amh | Background = FALSE | Background = FALSE | ENSMUSG00000035262 | 11705 | protein_coding | 2.337177898 | 1.046311148 | 0.830705295 | -1.259545538 | 0.207833353 | 1 | 1.046311148 | Not Significant |
| Amhr2 | None | None | ENSMUSG00000023047 | 110542 | protein_coding | 4.921331539 | -1.315120014 | 0.611067325 | 2.152168771 | 0.031384064 | 0.104512288 | 1.315120014 | Not Significant |
| Amigo1 | None | None | ENSMUSG00000050947 | 229715 | protein_coding | 141.1691325 | -0.29209991 | 0.125595748 | 2.325714958 | 0.020033762 | 0.073420411 | 0.29209991 | Not Significant |
| Amigo2 | Cond1_DOWN | Cond1_DOWN | ENSMUSG00000048218 | 105827 | protein_coding | 15.85397742 | -1.13580945 | 0.36138999 | 3.142891283 | 0.00167288 | 0.009775631 | 1.13580945 | DOWN |
| Amigo3 | Cond1_UP | Cond1_UP | ENSMUSG00000032593 | 320844 | protein_coding | 184.2765389 | 0.871866059 | 0.109026414 | -7.996833337 | 1.28E-15 | 4.91E-14 | 0.871866059 | UP |
| Ammecr1 | None | None | ENSMUSG00000042225 | 56068 | protein_coding | 276.4674192 | 0.088773214 | 0.166744663 | -0.532390138 | 0.594455814 | 0.77323163 | 0.088773214 | Not Significant |
| Ammecr1l | None | None | ENSMUSG00000041915 | 225339 | protein_coding | 371.2493527 | 0.031971293 | 0.075797799 | -0.421797111 | 0.673173115 | 0.824897384 | 0.031971293 | Not Significant |
| Amn | None | None | ENSMUSG00000021278 | 93835 | protein_coding | 101.6828334 | 0.20743774 | 0.251504523 | -0.824787316 | 0.409492349 | 0.630052795 | 0.20743774 | Not Significant |
| Amn1 | None | None | ENSMUSG00000068250 | 232566 | protein_coding | 178.3719994 | 0.074306562 | 0.117853104 | -0.630501524 | 0.528366506 | 0.726359126 | 0.074306562 | Not Significant |
| Amot | None | None | ENSMUSG00000041688 | 27494 | protein_coding | 546.022296 | 0.234256784 | 0.113153428 | -2.070257954 | 0.038428195 | 0.122133027 | 0.234256784 | Not Significant |
| Amotl1 | Cond1_UP | Cond1_UP | ENSMUSG00000013076 | 75723 | protein_coding | 201.4995652 | 1.058620504 | 0.172215843 | -6.147056422 | 7.89E-10 | 1.77E-08 | 1.058620504 | UP |
| [illegible] | [illegible] | [illegible] | [illegible] | [illegible] | [illegible] | [illegible] | [illegible] | [illegible] | [illegible] | [illegible] | [illegible] | [illegible] | [illegible] |

| geneID | Cond2_DF_ensembl_gene_id | Cond2_DF_entrezgene_id | Cond2_DF_gene_biotype | Cond2_DF_baseMean | Cond2_DF_log2FoldChange | Cond2_DF_lfcSE | Cond2_DF_stat | Cond2_DF_pvalue | Cond2_DF_padj | Cond2_DF_absLog2FC | RG1_Specific_Cond2 | RG1_Specific_Cond1 |
|---|---|---|---|---|---|---|---|---|---|---|---|---|
| Amfr | ENSMUSG00000031751 | 23802 | protein_coding | 4573.508954 | -0.039848863 | 0.035468308 | -1.123506183 | 0.261222576 | 0.503084198 | 0.039848863 | Cond1 No Change + Cond2 Not Significant | Cond2 No Change + Cond1 Not Significant |
| Amh | NA | NA | NA | NA | NA | NA | NA | NA | NA | NA | Background = FALSE | Background = FALSE |
| Amhr2 | ENSMUSG00000023047 | 110542 | protein_coding | 8.126746293 | 0.030015746 | 0.453613966 | 0.066170242 | 0.947242288 | 0.977863038 | 0.030015746 | Cond1 No Change + Cond2 Not Significant | Cond2 No Change + Cond1 Not Significant |
| Amigo1 | ENSMUSG00000050947 | 229715 | protein_coding | 305.6467758 | -0.200193925 | 0.090578936 | -2.210159829 | 0.027094072 | 0.120870289 | 0.200193925 | Cond1 No Change + Cond2 Not Significant | Cond2 No Change + Cond1 Not Significant |
| Amigo2 | ENSMUSG00000048218 | 105827 | protein_coding | 41.34473318 | -0.306131164 | 0.263707635 | -1.160873344 | 0.24569341 | 0.486948216 | 0.306131164 | Cond1 DOWN + Cond2 Not Significant | Cond2 No Change + Cond1 DOWN |
| Amigo3 | ENSMUSG00000032593 | 320844 | protein_coding | 202.3055102 | 0.190361236 | 0.134605317 | 1.414217807 | 0.157297961 | 0.376539145 | 0.190361236 | Cond1 UP + Cond2 Not Significant | Cond2 No Change + Cond1 UP |
| Ammecr1 | ENSMUSG00000042225 | 56068 | protein_coding | 499.2417365 | 0.021503841 | 0.086365642 | 0.248986056 | 0.803371568 | 0.910371624 | 0.021503841 | Cond1 No Change + Cond2 Not Significant | Cond2 No Change + Cond1 Not Significant |
| Ammecr1l | ENSMUSG00000041915 | 225339 | protein_coding | 618.0489396 | 0.006412054 | 0.05543173 | 0.115674804 | 0.907910276 | 0.959122007 | 0.006412054 | Cond1 No Change + Cond2 Not Significant | Cond2 No Change + Cond1 Not Significant |
| Amn | ENSMUSG00000021278 | 93835 | protein_coding | 220.9248607 | -0.341157895 | 0.128732191 | -2.650136626 | 0.008045923 | 0.050864992 | 0.341157895 | Cond1 No Change + Cond2 Not Significant | Cond2 No Change + Cond1 Not Significant |
| Amn1 | ENSMUSG00000068250 | 232566 | protein_coding | 256.7969022 | -0.078652825 | 0.101085279 | -0.778083861 | 0.436519577 | 0.669750853 | 0.078652825 | Cond1 No Change + Cond2 Not Significant | Cond2 No Change + Cond1 Not Significant |
| Amot | ENSMUSG00000041688 | 27494 | protein_coding | 961.5370493 | 0.087660751 | 0.050832578 | 1.724499424 | 0.084617723 | 0.259435364 | 0.087660751 | Cond1 No Change + Cond2 Not Significant | Cond2 No Change + Cond1 Not Significant |
| Amotl1 | ENSMUSG00000013076 | 75723 | protein_coding | 196.7064093 | 0.279106755 | 0.121672855 | 2.293911456 | 0.021795585 | 0.10455009 | 0.279106755 | Cond1 UP + Cond2 Not Significant | Cond2 No Change + Cond1 UP |
| [illegible] | [illegible] | [illegible] | [illegible] | [illegible] | [illegible] | [illegible] | [illegible] | [illegible] | [illegible] | [illegible] | [illegible] | [illegible] |

| geneID | Cond1_DF_Detected | Cond1_DF_ValueCol | Cond1_DF_PadjCol | Cond1_DF_Cutoff | Cond1_DF_Cutoff_Specific.x | Cond2_DF_Detected | Cond2_DF_ValueCol | Cond2_DF_PadjCol | Cond2_DF_Cutoff | Cond2_DF_Cutoff_Specific | BG_Method | RG1_All | RG2_Significant | RG3_SignificantChange | Cond1_DF_ensembl_gene_id | Cond1_DF_entrezgene_id | Cond1_DF_gene_biotype |
|---|---|---|---|---|---|---|---|---|---|---|---|---|---|---|---|---|---|
| Aph1c | TRUE | 0.775495552 | 0.387122324 | No Change | Not Significant | TRUE | 0.638815483 | 0.233254754 | No Change | Not Significant | TRUE | Cond1 Not Significant + Cond2 Not Significant | None | None | ENSMUSG00000053040 | 68318 | protein_coding |
| Api5 | TRUE | -0.057159168 | 0.753307251 | No Change | Not Significant | TRUE | 0.085367157 | 0.237154121 | No Change | Not Significant | TRUE | Cond1 Not Significant + Cond2 Not Significant | None | None | ENSMUSG00000027193 | 11800 | protein_coding |
| Apip | TRUE | 0.35824535 | 0.027285164 | No Change | Significant Positive | TRUE | 0.24846911 | 0.113078271 | No Change | Not Significant | TRUE | Cond1 Significant Positive + Cond2 Not Significant | None | None | ENSMUSG00000010911 | 56369 | protein_coding |
| Aplf | TRUE | 0.001403937 | 0.993753796 | No Change | Not Significant | TRUE | -0.045179839 | 0.777161429 | No Change | Not Significant | TRUE | Cond1 Not Significant + Cond2 Not Significant | None | None | ENSMUSG00000030051 | 72103 | protein_coding |
| Apln | TRUE | -0.840980496 | 1 | No Change | Not Significant | TRUE | 0.089106737 | 0.92580236 | No Change | Not Significant | TRUE | Cond1 Not Significant + Cond2 Not Significant | None | None | ENSMUSG00000037010 | 30878 | protein_coding |
| Aplnr | TRUE | 0.637723598 | 1 | No Change | Not Significant | FALSE | NA | NA | No Change | Not Detected | FALSE | Background = FALSE | Background = FALSE | Background = FALSE | ENSMUSG00000044338 | 23796 | protein_coding |
| Aplp1 | TRUE | 0.943827175 | 0.012608423 | UP | UP | TRUE | 0.174624882 | 0.29617307 | No Change | Not Significant | TRUE | Cond1 UP + Cond2 Not Significant | Cond1_UP | Cond1_UP | ENSMUSG00000006651 | 11803 | protein_coding |
| Aplp2 | TRUE | -0.406533732 | 0.001078713 | No Change | Significant Negative | TRUE | -0.100599568 | 0.142802169 | No Change | Not Significant | TRUE | Cond1 Significant Negative + Cond2 Not Significant | None | None | ENSMUSG00000031996 | 11804 | protein_coding |
| Apmap | TRUE | 0.173820212 | 0.252075246 | No Change | Not Significant | TRUE | 0.094813779 | 0.382615199 | No Change | Not Significant | TRUE | Cond1 Not Significant + Cond2 Not Significant | None | None | ENSMUSG00000033096 | 71881 | protein_coding |
| Apoa1 | TRUE | -0.978576495 | 0.496688848 | No Change | Not Significant | TRUE | 0.21588956 | 0.89645106 | No Change | Not Significant | TRUE | Cond1 Not Significant + Cond2 Not Significant | None | None | ENSMUSG00000032083 | 11806 | protein_coding |
| Apoa2 | TRUE | -0.051807693 | 0.957921881 | No Change | Not Significant | TRUE | -0.188114382 | 0.679535938 | No Change | Not Significant | TRUE | Cond1 Not Significant + Cond2 Not Significant | None | None | ENSMUSG00000005681 | 11807 | protein_coding |
| Apoa4 | TRUE | 0.144449811 | 0.948996349 | No Change | Not Significant | TRUE | 0.244559666 | 0.914838553 | No Change | Not Significant | TRUE | Cond1 Not Significant + Cond2 Not Significant | None | None | ENSMUSG00000032080 | 11808 | protein_coding |
| [illegible] | | | | | | | | | | | | | | | | | |

| geneID | Cond1_DF_baseMean | Cond1_DF_log2FoldChange | Cond1_DF_lfcSE | Cond1_DF_stat | Cond1_DF_pvalue | Cond1_DF_padj | Cond1_DF_absLog2FC | Cond1_DF_Cutoff_Specific.y | Cond2_DF_ensembl_gene_id | Cond2_DF_entrezgene_id | Cond2_DF_gene_biotype | Cond2_DF_baseMean | Cond2_DF_log2FoldChange | Cond2_DF_lfcSE | Cond2_DF_stat | Cond2_DF_pvalue | Cond2_DF_padj | Cond2_DF_absLog2FC | RG1_Specific_Cond2 | RG1_Specific_Cond1 |
|---|---|---|---|---|---|---|---|---|---|---|---|---|---|---|---|---|---|---|---|---|
| Aph1c | 6.957583161 | 0.775495552 | 0.591489862 | -1.311088493 | 0.189827868 | 0.387122324 | 0.775495552 | Not Significant | ENSMUSG00000053040 | 68318 | protein_coding | 11.56822678 | 0.638815483 | 0.35452932 | 1.801869259 | 0.071565978 | 0.233254754 | 0.638815483 | Cond1 No Change + Cond2 Not Significant | Cond2 No Change + Cond1 Not Significant |
| Api5 | 1363.922969 | -0.057159168 | 0.0998409 | 0.572502537 | 0.566981568 | 0.753307251 | 0.057159168 | Not Significant | ENSMUSG00000027193 | 11800 | protein_coding | 2605.590811 | 0.085367157 | 0.047678701 | 1.790467326 | 0.073378815 | 0.237154121 | 0.085367157 | Cond1 No Change + Cond2 Not Significant | Cond2 No Change + Cond1 Not Significant |
| Apip | 83.67163471 | 0.35824535 | 0.129711695 | -2.761858512 | 0.005747338 | 0.027285164 | 0.35824535 | Significant Positive | ENSMUSG00000010911 | 56369 | protein_coding | 131.0491503 | 0.24846911 | 0.110431473 | 2.249984576 | 0.024449924 | 0.113078271 | 0.24846911 | Cond1 No Change + Cond2 Not Significant | Cond2 No Change + Cond1 Significant Positive |
| Aplf | 119.6536241 | 0.001403937 | 0.101829949 | -0.013787071 | 0.988999857 | 0.993753796 | 0.001403937 | Not Significant | ENSMUSG00000030051 | 72103 | protein_coding | 247.2303443 | -0.045179839 | 0.083218887 | -0.542903666 | 0.587196132 | 0.777161429 | 0.045179839 | Cond1 No Change + Cond2 Not Significant | Cond2 No Change + Cond1 Not Significant |
| Apln | 3.604274032 | -0.840980496 | 0.566801467 | 1.483730274 | 0.137880492 | 1 | 0.840980496 | Not Significant | ENSMUSG00000037010 | 30878 | protein_coding | 8.601614382 | 0.089106737 | 0.421131593 | 0.21158882 | 0.832427832 | 0.92580236 | 0.089106737 | Cond1 No Change + Cond2 Not Significant | Cond2 No Change + Cond1 Not Significant |
| Aplnr | 1.79248052 | 0.637723598 | 1.769109495 | -0.360477178 | 0.718490321 | 1 | 0.637723598 | Not Significant | NA | NA | NA | NA | NA | NA | NA | NA | NA | NA | Background = FALSE | Background = FALSE |
| Aplp1 | 252.9501547 | 0.943827175 | 0.309394097 | -3.050566202 | 0.002284103 | 0.012608423 | 0.943827175 | UP | ENSMUSG00000006651 | 11803 | protein_coding | 304.1067939 | 0.174624882 | 0.107602029 | 1.622877223 | 0.104615653 | 0.29617307 | 0.174624882 | Cond1 UP + Cond2 Not Significant | Cond2 No Change + Cond1 UP |
| Aplp2 | 2189.793415 | -0.406533732 | 0.106169768 | 3.829091287 | 0.000128617 | 0.001078713 | 0.406533732 | Significant Negative | ENSMUSG00000031996 | 11804 | protein_coding | 4608.702902 | -0.100599568 | 0.047543163 | -2.11596289 | 0.034347964 | 0.142802169 | 0.100599568 | Cond1 No Change + Cond2 Not Significant | Cond2 No Change + Cond1 Significant Negative |
| Apmap | 436.6915348 | 0.173820212 | 0.106441701 | -1.633008579 | 0.102467192 | 0.252075246 | 0.173820212 | Not Significant | ENSMUSG00000033096 | 71881 | protein_coding | 807.9956262 | 0.094813779 | 0.067794308 | 1.398550733 | 0.161947749 | 0.382615199 | 0.094813779 | Cond1 No Change + Cond2 Not Significant | Cond2 No Change + Cond1 Not Significant |
| Apoa1 | 80.42771587 | -0.978576495 | 0.904465923 | 1.08193849 | 0.27927986 | 0.496688848 | 0.978576495 | Not Significant | ENSMUSG00000032083 | 11806 | protein_coding | 376.3191895 | 0.21588956 | 0.764469436 | 0.282404436 | 0.777633414 | 0.89645106 | 0.21588956 | Cond1 No Change + Cond2 Not Significant | Cond2 No Change + Cond1 Not Significant |
| Apoa2 | 6.008385744 | -0.051807693 | 0.477423468 | 0.108515179 | 0.9135804 | 0.957921881 | 0.051807693 | Not Significant | ENSMUSG00000005681 | 11807 | protein_coding | 24.57446243 | -0.188114382 | 0.249113536 | -0.755135129 | 0.450167907 | 0.679535938 | 0.188114382 | Cond1 No Change + Cond2 Not Significant | Cond2 No Change + Cond1 Not Significant |
| Apoa4 | 19.15411222 | 0.144449811 | 1.106416613 | -0.130556437 | 0.896126206 | 0.948996349 | 0.144449811 | Not Significant | ENSMUSG00000032080 | 11808 | protein_coding | 99.71291572 | 0.244559666 | 1.022469037 | 0.239185401 | 0.810961823 | 0.914838553 | 0.244559666 | Cond1 No Change + Cond2 Not Significant | Cond2 No Change + Cond1 Not Significant |
| [illegible] | | | | | | | | | | | | | | | | | | | | |

Supplementary Table 3

| geneID | Cond1_DF_Detected | Cond1_DF_ValueCol | Cond1_DF_PadjCol | Cond1_DF_Cutoff | Cond1_DF_Cutoff_Specific.x | Cond2_DF_Detected | Cond2_DF_ValueCol | Cond2_DF_PadjCol | Cond2_DF_Cutoff | Cond2_DF_Cutoff_Specific | BG_Method | RG1_All | RG2_Significant | RG3_SignificantChange |
|---|---|---|---|---|---|---|---|---|---|---|---|---|---|---|
| Armc10 | TRUE | -0.140214552 | 0.442685693 | No Change | Not Significant | TRUE | -0.01384908 | 0.930889346 | No Change | Not Significant | TRUE | Cond1 Not Significant + Cond2 Not Significant | None | None |
| Armc12 | TRUE | 0.607046636 | 1 | No Change | Not Significant | FALSE | NA | NA | No Change | Not Detected | FALSE | Background = FALSE | Background = FALSE | Background = FALSE |
| Armc2 | TRUE | 2.42060778 | 1 | No Change | Not Significant | FALSE | NA | NA | No Change | Not Detected | FALSE | Background = FALSE | Background = FALSE | Background = FALSE |
| Armc3 | TRUE | -0.122600446 | 0.892991423 | No Change | Not Significant | TRUE | -0.226435945 | 0.775860848 | No Change | Not Significant | TRUE | Cond1 Not Significant + Cond2 Not Significant | None | None |
| Armc5 | TRUE | 0.067916445 | 0.761994054 | No Change | Not Significant | TRUE | 0.093131826 | 0.661834894 | No Change | Not Significant | TRUE | Cond1 Not Significant + Cond2 Not Significant | None | None |
| Armc6 | TRUE | 0.235861173 | 0.262539089 | No Change | Not Significant | TRUE | 0.175827139 | 0.325080339 | No Change | Not Significant | TRUE | Cond1 Not Significant + Cond2 Not Significant | None | None |
| Armc7 | TRUE | 0.14216663 | 0.533579433 | No Change | Not Significant | TRUE | 0.000824526 | 0.997610035 | No Change | Not Significant | TRUE | Cond1 Not Significant + Cond2 Not Significant | None | None |
| Armc8 | TRUE | -0.086273863 | 0.501193606 | No Change | Not Significant | TRUE | 0.030392345 | 0.792706869 | No Change | Not Significant | TRUE | Cond1 Not Significant + Cond2 Not Significant | None | None |
| Armc9 | TRUE | 0.412267579 | 0.051024976 | No Change | Not Significant | TRUE | 0.109887158 | 0.58688999 | No Change | Not Significant | TRUE | Cond1 Not Significant + Cond2 Not Significant | None | None |
| Armcx1 | TRUE | 0.126768776 | 0.746140914 | No Change | Not Significant | TRUE | 0.132808402 | 0.642103246 | No Change | Not Significant | TRUE | Cond1 Not Significant + Cond2 Not Significant | None | None |
| Armcx2 | TRUE | 0.028809611 | 0.984770701 | No Change | Not Significant | TRUE | 0.476969214 | 0.481155234 | No Change | Not Significant | TRUE | Cond1 Not Significant + Cond2 Not Significant | None | None |
| Armcx3 | TRUE | -0.235595253 | 0.438050261 | No Change | Not Significant | TRUE | 0.352585051 | 0.008027948 | No Change | Significant Positive | TRUE | Cond1 Not Significant + Cond2 Significant Positive | None | None |

| geneID | Cond1_DF_ensembl_gene_id | Cond1_DF_entrezgene_id | Cond1_DF_gene_biotype | Cond1_DF_baseMean | Cond1_DF_log2FoldChange | Cond1_DF_lfcSE | Cond1_DF_stat | Cond1_DF_pvalue | Cond1_DF_padj | Cond1_DF_absLog2FC | Cond1_DF_Cutoff_Specific.y |
|---|---|---|---|---|---|---|---|---|---|---|---|
| Armc10 | ENSMUSG00000038525 | 67211 | protein_coding | 375.7459975 | -0.140214552 | 0.117614965 | 1.19214891 | 0.233202862 | 0.442685693 | 0.140214552 | Not Significant |
| Armc12 | ENSMUSG00000024223 | 67645 | protein_coding | 0.933154481 | 0.607046636 | 2.116458262 | -0.286821926 | 0.774248673 | 1 | 0.607046636 | Not Significant |
| Armc2 | ENSMUSG00000071324 | 213402 | protein_coding | 2.391436592 | 2.42060778 | 1.8842145 | -1.284677397 | 0.19890504 | 1 | 2.42060778 | Not Significant |
| Armc3 | ENSMUSG00000037683 | 70882 | protein_coding | 10.16162001 | -0.122600446 | 0.451201594 | 0.271719887 | 0.785837409 | 0.892991423 | 0.122600446 | Not Significant |
| Armc5 | ENSMUSG00000042178 | 233912 | protein_coding | 309.5404149 | 0.067916445 | 0.122195863 | -0.555799872 | 0.578347673 | 0.761994054 | 0.067916445 | Not Significant |
| Armc6 | ENSMUSG00000002343 | 76813 | protein_coding | 109.92289 | 0.235861173 | 0.146971326 | -1.604810809 | 0.108535446 | 0.262539089 | 0.235861173 | Not Significant |
| Armc7 | ENSMUSG00000057219 | 276905 | protein_coding | 72.83403963 | 0.14216663 | 0.140772545 | -1.009903109 | 0.312541713 | 0.533579433 | 0.14216663 | Not Significant |
| Armc8 | ENSMUSG00000032468 | 74125 | protein_coding | 500.1575359 | -0.086273863 | 0.080381047 | 1.073311015 | 0.283131595 | 0.501193606 | 0.086273863 | Not Significant |
| Armc9 | ENSMUSG00000062590 | 78795 | protein_coding | 82.94303056 | 0.412267579 | 0.165268941 | -2.494525445 | 0.012612568 | 0.051024976 | 0.412267579 | Not Significant |
| Armcx1 | ENSMUSG00000033460 | 78248 | protein_coding | 32.71425468 | 0.126768776 | 0.215754226 | -0.587561033 | 0.556826973 | 0.746140914 | 0.126768776 | Not Significant |
| Armcx2 | ENSMUSG00000033436 | 67416 | protein_coding | 4.787585489 | 0.028809611 | 0.781104876 | -0.036883154 | 0.970578172 | 0.984770701 | 0.028809611 | Not Significant |
| Armcx3 | ENSMUSG00000049047 | 71703 | protein_coding | 124.1914394 | -0.235595253 | 0.19590508 | 1.202599001 | 0.229131534 | 0.438050261 | 0.235595253 | Not Significant |

| geneID | Cond2_DF_ensembl_gene_id | Cond2_DF_entrezgene_id | Cond2_DF_gene_biotype | Cond2_DF_baseMean | Cond2_DF_log2FoldChange | Cond2_DF_lfcSE | Cond2_DF_stat | Cond2_DF_pvalue | Cond2_DF_padj | Cond2_DF_absLog2FC | RG1_Specific_Cond2 | RG1_Specific_Cond1 |
|---|---|---|---|---|---|---|---|---|---|---|---|---|
| Armc10 | ENSMUSG00000038525 | 67211 | protein_coding | 820.3632108 | -0.01384908 | 0.071555727 | -0.19354258 | 0.846534062 | 0.930889346 | 0.01384908 | Cond1 No Change + Cond2 Not Significant | Cond2 No Change + Cond1 Not Significant |
| Armc12 | NA | NA | NA | NA | NA | NA | NA | NA | NA | NA | Background = FALSE | Background = FALSE |
| Armc2 | NA | NA | NA | NA | NA | NA | NA | NA | NA | NA | Background = FALSE | Background = FALSE |
| Armc3 | ENSMUSG00000037683 | 70882 | protein_coding | 11.7366086 | -0.226435945 | 0.414816337 | -0.545870365 | 0.58515505 | 0.775860848 | 0.226435945 | Cond1 No Change + Cond2 Not Significant | Cond2 No Change + Cond1 Not Significant |
| Armc5 | ENSMUSG00000042178 | 233912 | protein_coding | 641.6155004 | 0.093131826 | 0.117101889 | 0.795305927 | 0.426435567 | 0.661834894 | 0.093131826 | Cond1 No Change + Cond2 Not Significant | Cond2 No Change + Cond1 Not Significant |
| Armc6 | ENSMUSG00000002343 | 76813 | protein_coding | 178.5496737 | 0.175827139 | 0.113967456 | 1.542783754 | 0.122883253 | 0.325080339 | 0.175827139 | Cond1 No Change + Cond2 Not Significant | Cond2 No Change + Cond1 Not Significant |
| Armc7 | ENSMUSG00000057219 | 276905 | protein_coding | 135.0818283 | 0.000824526 | 0.127158281 | 0.006484252 | 0.994826352 | 0.997610035 | 0.000824526 | Cond1 No Change + Cond2 Not Significant | Cond2 No Change + Cond1 Not Significant |
| Armc8 | ENSMUSG00000032468 | 74125 | protein_coding | 1000.942996 | 0.030392345 | 0.0597488 | 0.508668708 | 0.610984459 | 0.792706869 | 0.030392345 | Cond1 No Change + Cond2 Not Significant | Cond2 No Change + Cond1 Not Significant |
| Armc9 | ENSMUSG00000062590 | 78795 | protein_coding | 122.160688 | 0.109887158 | 0.116242594 | 0.945326098 | 0.344492419 | 0.58688999 | 0.109887158 | Cond1 No Change + Cond2 Not Significant | Cond2 No Change + Cond1 Not Significant |
| Armcx1 | ENSMUSG00000033460 | 78248 | protein_coding | 73.99092188 | 0.132808402 | 0.15873638 | 0.836660139 | 0.402783631 | 0.642103246 | 0.132808402 | Cond1 No Change + Cond2 Not Significant | Cond2 No Change + Cond1 Not Significant |
| Armcx2 | ENSMUSG00000033436 | 67416 | protein_coding | 10.02264972 | 0.476969214 | 0.406409863 | 1.173616236 | 0.240548775 | 0.481155234 | 0.476969214 | Cond1 No Change + Cond2 Not Significant | Cond2 No Change + Cond1 Not Significant |
| Armcx3 | ENSMUSG00000049047 | 71703 | protein_coding | 239.7910915 | 0.352585051 | 0.104726041 | 3.366737126 | 0.000760632 | 0.008027948 | 0.352585051 | Cond1 No Change + Cond2 Significant Positive | Cond2 No Change + Cond1 Not Significant |

| geneID | Cond1_DF_Detected | Cond1_DF_ValueCol | Cond1_DF_PadjCol | Cond1_DF_Cutoff | Cond1_DF_Cutoff_Specific.x |
|---|---|---|---|---|---|
| Atp2b1 | TRUE | 0.550112654 | 3.50E-08 | UP | UP |
| Atp2b2 | TRUE | 1.11847907 | 0.827306109 | No Change | Not Significant |
| Atp2b3 | TRUE | 0.235750183 | 0.935658236 | No Change | Not Significant |
| Atp2b4 | TRUE | 0.390564948 | 0.736456016 | No Change | Not Significant |
| Atp2c1 | TRUE | -0.075426461 | 0.726648836 | No Change | Not Significant |
| Atp2c2 | TRUE | -0.417013785 | 0.257759428 | No Change | Not Significant |
| Atp4a | TRUE | -0.533476572 | 0.599156327 | No Change | Not Significant |
| Atp4b | TRUE | -0.02344889 | 1 | No Change | Not Significant |
| Atp5a1 | TRUE | -0.083046504 | 0.588321101 | No Change | Not Significant |
| Atp5b | TRUE | 0.013252596 | 0.931099354 | No Change | Not Significant |
| Atp5c1 | TRUE | -0.190761343 | 0.12190868 | No Change | Not Significant |
| Atp5d | TRUE | -0.089569095 | 0.503574403 | No Change | Not Significant |
| Atp5e | TRUE | 0.029424917 | 0.872440598 | No Change | Not Significant |
| Atp5g1 | TRUE | 0.267901041 | 0.048844287 | No Change | Significant Positive |
| Atp5g2 | TRUE | -0.567568289 | 0.009157744 | DOWN | DOWN |
| Atp5g3 | TRUE | 0.181316147 | 0.100633238 | No Change | Not Significant |
| Atp5h | TRUE | -0.381146027 | 0.001114664 | No Change | Significant Negative |
| Atp5j | TRUE | -0.064859737 | 0.522023104 | No Change | Not Significant |
| Atp5j2 | TRUE | -0.294175284 | 0.002263768 | No Change | Significant Negative |
| Atp5k | TRUE | -0.100293981 | 0.569192626 | No Change | Not Significant |
| Atp5l | TRUE | -0.230800596 | 0.112975508 | No Change | Not Significant |
| Atp5md | TRUE | -0.367979712 | 0.006649187 | No Change | Significant Negative |
| Atp5mpl | TRUE | -0.201532603 | 0.020065343 | No Change | Significant Negative |
| Atp5o | TRUE | -0.096354634 | 0.483116267 | No Change | Not Significant |
| Atp5pb | TRUE | -0.096477089 | 0.451804114 | No Change | Not Significant |
| Atp6ap1 | TRUE | -0.186121318 | 0.046543198 | No Change | Significant Negative |
| Atp6ap2 | TRUE | 0.173647109 | 0.441638415 | No Change | Not Significant |
| Atp6v0a1 | TRUE | 0.048549246 | 0.72786082 | No Change | Not Significant |
| Atp6v0a2 | TRUE | 0.10690178 | 0.285476835 | No Change | Not Significant |
| Atp6v0a4 | TRUE | -0.763894332 | 0.560514185 | No Change | Not Significant |
| Atp6v0b | TRUE | -0.218491762 | 0.010159502 | No Change | Significant Negative |
| Atp6v0c | TRUE | -0.329553712 | 0.008943393 | No Change | Significant Negative |
| Atp6v0d1 | TRUE | 0.070551755 | 0.649494967 | No Change | Not Significant |
| Atp6v0d2 | TRUE | -0.343551436 | 1 | No Change | Not Significant |
| Atp6v0e | TRUE | -0.199640393 | 0.03389483 | No Change | Significant Negative |
| Atp6v0e2 | TRUE | -1.35714368 | 1.41E-18 | DOWN | DOWN |
| Atp6v1a | TRUE | -0.103531983 | 0.461760239 | No Change | Not Significant |
| Atp6v1b1 | TRUE | -0.53166271 | 1 | No Change | Not Significant |
| Atp6v1b2 | TRUE | -0.106660774 | 0.429396526 | No Change | Not Significant |
| Atp6v1c1 | TRUE | -0.058885387 | 0.622737045 | No Change | Not Significant |
| Atp6v1c2 | TRUE | 2.103292525 | 0.002305389 | UP | UP |
| Atp6v1d | TRUE | 0.190984481 | 0.013498197 | No Change | Significant Positive |
| Atp6v1e1 | TRUE | -0.278196959 | 0.000111756 | No Change | Significant Negative |
| Atp6v1f | TRUE | -0.404148447 | 1.98E-08 | No Change | Significant Negative |
| Atp6v1fnb | TRUE | -1.340707109 | 1 | No Change | Not Significant |
| Atp6v1g1 | TRUE | -0.356669561 | 0.005018564 | No Change | Significant Negative |
| Atp6v1g2 | TRUE | 0.287888831 | 0.633557127 | No Change | Not Significant |
| Atp6v1h | TRUE | -0.188864273 | 0.256047562 | No Change | Not Significant |
| Atp7a | TRUE | -0.334192883 | 0.000713673 | No Change | Significant Negative |
| Atp7b | TRUE | 0.110256476 | 0.862666522 | No Change | Not Significant |
| Atp8a1 | TRUE | 0.091148908 | 0.644974953 | No Change | Not Significant |
| Atp8a2 | TRUE | 0.020191664 | 0.889454308 | No Change | Not Significant |
| Atp8b1 | TRUE | -0.078110375 | 0.474068167 | No Change | Not Significant |
| Atp8b2 | TRUE | 4.04944735 | 0 | UP | UP |
| Atp8b3 | TRUE | 3.505344774 | 1 | No Change | Not Significant |
| Atp8b4 | TRUE | 0.185715921 | 0.942116626 | No Change | Not Significant |
| Atp8b5 | TRUE | -0.356622989 | 0.404039964 | No Change | Not Significant |
| Atp9a | TRUE | -0.268777541 | 0.022498653 | No Change | Significant Negative |
| Atp9b | TRUE | 0.037262973 | 0.735049182 | No Change | Not Significant |
| Atpaf1 | TRUE | -0.599116347 | 0.000123789 | DOWN | DOWN |
| Atpaf2 | TRUE | -0.079737567 | 0.622742125 | No Change | Not Significant |
| Atpif1 | TRUE | -0.227151539 | 0.030641927 | No Change | Significant Negative |
| Atpsckmt | TRUE | 0.07392467 | 0.697627026 | No Change | Not Significant |
| Atr | TRUE | 0.218553107 | 0.215987092 | No Change | Not Significant |
| Atraid | TRUE | -0.017944344 | 0.919410944 | No Change | Not Significant |
| Atrip | TRUE | 0.038484841 | 0.880259019 | No Change | Not Significant |
| Atrn | TRUE | -0.022446365 | 0.88791795 | No Change | Not Significant |
| Atrnl1 | TRUE | -0.042902399 | 0.756249943 | No Change | Not Significant |
| Atrx | TRUE | 0.502099735 | 1.26E-10 | UP | UP |
| Atxn1 | TRUE | -0.371395024 | 0.023423844 | No Change | Significant Negative |
| Atxn10 | TRUE | 0.382164347 | 0.003940543 | No Change | Significant Positive |
| Atxn1l | TRUE | -0.001134663 | 0.993636457 | No Change | Not Significant |
| Atxn2 | TRUE | 0.057469107 | 0.717206457 | No Change | Not Significant |
| Atxn2l | TRUE | 0.415279316 | 0.042874537 | No Change | Significant Positive |
| Atxn3 | TRUE | -0.262869782 | 0.159023459 | No Change | Not Significant |
| Atxn7 | TRUE | 0.109133206 | 0.172412699 | No Change | Not Significant |
| Atxn7l1 | TRUE | -0.064553385 | 0.833389267 | No Change | Not Significant |
| Atxn7l1os2 | TRUE | -0.048607013 | 0.964583514 | No Change | Not Significant |
| Atxn7l2 | TRUE | -0.467747712 | 0.084474747 | No Change | Not Significant |
| Atxn7l3 | TRUE | 0.057208424 | 0.71375046 | No Change | Not Significant |
| Atxn7l3b | TRUE | -0.157415013 | 0.309078852 | No Change | Not Significant |
| AU018091 | TRUE | 0.413282614 | 1 | No Change | Not Significant |
| AU020206 | TRUE | 0.497592225 | 0.025458165 | No Change | Significant Positive |
| AU021092 | TRUE | -0.181971562 | 1 | No Change | Not Significant |
| AU022252 | TRUE | 0.101110758 | 0.445505084 | No Change | Not Significant |
| AU040320 | TRUE | -0.100353293 | 0.086109128 | No Change | Not Significant |
| AU040972 | TRUE | -0.663966723 | 0.489835334 | No Change | Not Significant |
| AU041133 | TRUE | -0.243200906 | 0.332739097 | No Change | Not Significant |
| Auh | TRUE | -0.249912331 | 0.045254367 | No Change | Significant Negative |
| Aunip | TRUE | 0.346931407 | 0.38964213 | No Change | Not Significant |
| Aup1 | TRUE | 0.071704537 | 0.468471767 | No Change | Not Significant |
| Aurka | TRUE | 0.846052167 | 1.49E-05 | UP | UP |
| Aurkaip1 | TRUE | -0.253221126 | 0.001128742 | No Change | Significant Negative |
| Aurkb | TRUE | 1.100891597 | 7.22E-06 | UP | UP |
| Auts2 | TRUE | -0.152047087 | 0.522337974 | No Change | Not Significant |
| AV039307 | TRUE | -0.652395656 | 1 | No Change | Not Significant |
| Aven | TRUE | 0.032502468 | 0.842293038 | No Change | Not Significant |
| Avil | TRUE | -0.665484301 | 6.46E-05 | DOWN | DOWN |
| Avl9 | TRUE | -0.155790786 | 0.233934706 | No Change | Not Significant |
| Avpi1 | TRUE | 0.107502194 | 0.72936091 | No Change | Not Significant |
| Avpr1a | TRUE | -0.533398855 | 1 | No Change | Not Significant |
| Avpr1b | TRUE | 0.032076551 | 0.972316965 | No Change | Not Significant |
| Avpr2 | TRUE | -0.058586729 | 1 | No Change | Not Significant |
| AW011738 | TRUE | 3.604268243 | 4.79E-32 | UP | UP |
| AW112010 | TRUE | 0.109514569 | 0.669164362 | No Change | Not Significant |
| AW146154 | TRUE | -0.110188149 | 0.545489766 | No Change | Not Significant |
| AW209491 | TRUE | -0.180095485 | 0.225215631 | No Change | Not Significant |
| AW551984 | TRUE | -0.095396961 | 0.853563561 | No Change | Not Significant |
| AW554918 | TRUE | 0.107242396 | 0.560514185 | No Change | Not Significant |
| AW822073 | TRUE | 1.138850891 | 1 | No Change | Not Significant |
| Awat1 | TRUE | -2.580250757 | 1 | No Change | Not Significant |
| Awat2 | TRUE | 0.100445412 | 0.900998399 | No Change | Not Significant |
| Axdnd1 | TRUE | 0.343584538 | 0.800822518 | No Change | Not Significant |
| Axin1 | TRUE | 0.140119047 | 0.101747126 | No Change | Not Significant |
| Axin2 | TRUE | 0.042348738 | 0.880144002 | No Change | Not Significant |
| Axl | TRUE | -0.586506061 | 0.660786267 | No Change | Not Significant |
| AY358078 | TRUE | -0.846770523 | 1 | No Change | Not Significant |
| AY512915 | TRUE | 2.578156424 | 1 | No Change | Not Significant |
| AY512931 | TRUE | 0.924378699 | 1 | No Change | Not Significant |
| AY702103 | TRUE | -0.383808552 | 1 | No Change | Not Significant |
| Azgp1 | TRUE | -0.435776376 | 0.14941403 | No Change | Not Significant |
| Azi2 | TRUE | -0.046336836 | 0.671862287 | No Change | Not Significant |
| Azin1 | TRUE | -0.110290323 | 0.249173195 | No Change | Not Significant |
| Azin2 | TRUE | -0.118968129 | 0.925637253 | No Change | Not Significant |
| B020014A21Rik | TRUE | 0.038419811 | 1 | No Change | Not Significant |
| B130034C11Rik | TRUE | 0.32063232 | 0.567534275 | No Change | Not Significant |
| B230110C06Rik | TRUE | 0.176016293 | 1 | No Change | Not Significant |
| B230206H07Rik | TRUE | -0.005020063 | 0.995401911 | No Change | Not Significant |
| B230206L02Rik | TRUE | -0.263268727 | 0.742102974 | No Change | Not Significant |
| B230208H11Rik | TRUE | 2.031497015 | 1 | No Change | Not Significant |
| B230214G05Rik | TRUE | 1.875157026 | 1 | No Change | Not Significant |
| B230216N24Rik | TRUE | -0.624738192 | 0.187201528 | No Change | Not Significant |
| B230219D22Rik | TRUE | 0.10499056 | 0.482454733 | No Change | Not Significant |
| B230220B15Rik | TRUE | 0.922977537 | 1 | No Change | Not Significant |
| B230307C23Rik | TRUE | -0.004967796 | 0.984403993 | No Change | Not Significant |
| B230319C09Rik | TRUE | -0.129167085 | 0.777353752 | No Change | Not Significant |
| B230354K17Rik | TRUE | 0.030054566 | 0.906704483 | No Change | Not Significant |
| B2m | TRUE | -0.647593839 | 1.74E-06 | DOWN | DOWN |
| B3galnt2 | TRUE | 0.082546348 | 0.639797547 | No Change | Not Significant |
| B3galt1 | TRUE | 1.291168274 | 1 | No Change | Not Significant |
| B3galt2 | TRUE | -0.940432669 | 0.001219773 | DOWN | DOWN |
| B3galt4 | TRUE | 0.086039437 | 0.623781054 | No Change | Not Significant |
| B3galt5 | TRUE | 0.150210324 | 0.50724071 | No Change | Not Significant |
| B3galt6 | TRUE | 0.103858599 | 0.683821668 | No Change | Not Significant |
| B3galt9 | TRUE | -0.158824566 | 1 | No Change | Not Significant |
| B3gat2 | TRUE | -0.247168251 | 0.566362983 | No Change | Not Significant |
| B3gat3 | TRUE | -0.179335563 | 0.190611112 | No Change | Not Significant |
| B3glct | TRUE | -0.120711225 | 0.465324602 | No Change | Not Significant |
| B3gnt2 | TRUE | 0.403989351 | 0.290432101 | No Change | Not Significant |
| B3gnt3 | TRUE | 0.105995821 | 0.281984397 | No Change | Not Significant |
| B3gnt5 | TRUE | 0.361301907 | 0.520943084 | No Change | Not Significant |
| B3gnt6 | TRUE | -1.509727715 | 3.16E-07 | DOWN | DOWN |
| B3gnt7 | TRUE | -0.281607624 | 0.708204999 | No Change | Not Significant |
| B3gnt8 | TRUE | -0.712960945 | 0.053754342 | No Change | Not Significant |
| B3gnt9 | TRUE | -0.838919716 | 2.53E-06 | DOWN | DOWN |
| B3gntl1 | TRUE | 0.051363241 | 0.82296411 | No Change | Not Significant |
| B430212C06Rik | TRUE | 2.361786765 | 1 | No Change | Not Significant |
| B430306N03Rik | TRUE | 2.673643059 | 1 | No Change | Not Significant |
| B430319F04Rik | TRUE | 0.445266052 | 0.287840111 | No Change | Not Significant |
| B430319G15Rik | TRUE | 0.514130511 | 1 | No Change | Not Significant |
| B4galnt1 | TRUE | 0.12084814 | 0.422162995 | No Change | Not Significant |
| B4galnt2 | TRUE | -0.149824261 | 0.387893639 | No Change | Not Significant |
| B4galnt3 | TRUE | -0.5271946 | 0.026283536 | DOWN | DOWN |
| B4galnt4 | TRUE | 0.79146115 | 0.004600301 | UP | UP |
| B4galt1 | TRUE | -0.151459495 | 0.46108514 | No Change | Not Significant |
| B4galt2 | TRUE | 1.851139228 | 0.002930712 | UP | UP |
| B4galt3 | TRUE | 0.162264635 | 0.199366082 | No Change | Not Significant |
| B4galt4 | TRUE | 0.25551756 | 0.093901357 | No Change | Not Significant |
| B4galt5 | TRUE | -0.359573837 | 0.000471385 | No Change | Significant Negative |
| B4galt6 | TRUE | -0.070572776 | 0.660786267 | No Change | Not Significant |
| B4galt7 | TRUE | 0.045793305 | 0.760926598 | No Change | Not Significant |
| B4gat1 | TRUE | 0.018710865 | 1 | No Change | Not Significant |
| B630019A10Rik | TRUE | 0.607287872 | 1 | No Change | Not Significant |
| B930018H19Rik | TRUE | -1.099667559 | 1 | No Change | Not Significant |
| B930025P03Rik | TRUE | -1.193927635 | 1 | No Change | Not Significant |
| B930092H01Rik | TRUE | 0.753082267 | 1 | No Change | Not Significant |
| B9d1 | TRUE | 0.753368555 | 0.000192981 | UP | UP |
| B9d2 | TRUE | -0.285615074 | 0.446089228 | No Change | Not Significant |
| Baalc | TRUE | 2.564408721 | 1 | No Change | Not Significant |
| Baat | TRUE | -0.519744587 | 0.346538122 | No Change | Not Significant |
| Babam1 | TRUE | -0.058616945 | 0.613316042 | No Change | Not Significant |
| Babam2 | TRUE | 0.012021489 | 0.919582966 | No Change | Not Significant |
| Bace1 | TRUE | -0.262601166 | 0.049790441 | No Change | Significant Negative |
| Bace2 | TRUE | -0.64736694 | 0.03055251 | DOWN | DOWN |
| Bach1 | TRUE | 0.03317724 | 0.858992577 | No Change | Not Significant |
| Bach2 | TRUE | -0.455048061 | 0.201270862 | No Change | Not Significant |
| Bach2os | TRUE | 1.089512293 | 1 | No Change | Not Significant |
| Bad | TRUE | 0.231876788 | 0.012122206 | No Change | Significant Positive |
| Bag1 | TRUE | 0.009512225 | 0.953990085 | No Change | Not Significant |
| Bag2 | TRUE | -0.069413042 | 0.850598485 | No Change | Not Significant |
| Bag3 | TRUE | 1.789015341 | 1.71E-54 | UP | UP |
| Bag4 | TRUE | -0.136661497 | 0.32840994 | No Change | Not Significant |

| geneID | Cond1_DF_Detected | Cond1_DF_ValueCol | Cond1_DF_PadjCol | Cond1_DF_Cutoff | Cond1_DF_Cutoff_Specific.x | Cond2_DF_Detected | Cond2_DF_ValueCol | Cond2_DF_PadjCol | Cond2_DF_Cutoff | Cond2_DF_Cutoff_Specific | BG_Method |
|---|---|---|---|---|---|---|---|---|---|---|---|
| Bag5 | TRUE | -0.035940248 | 0.876432521 | No Change | Not Significant | TRUE | -0.030280454 | 0.852953349 | No Change | Not Significant | TRUE |
| Bag6 | TRUE | -0.102197466 | 0.414593589 | No Change | Not Significant | TRUE | -0.016870197 | 0.892047592 | No Change | Not Significant | TRUE |
| Bahcc1 | TRUE | 0.282853021 | 0.230625736 | No Change | Not Significant | TRUE | -0.126090457 | 0.472527059 | No Change | Not Significant | TRUE |
| Bahd1 | TRUE | -0.183361829 | 0.031346224 | No Change | Significant Negative | TRUE | -0.016388895 | 0.943461099 | No Change | Not Significant | TRUE |
| Baiap2 | TRUE | 0.198193977 | 0.156392751 | No Change | Not Significant | TRUE | 0.115050406 | 0.381255129 | No Change | Not Significant | TRUE |
| Baiap2l1 | TRUE | -0.024338443 | 0.905233898 | No Change | Not Significant | TRUE | 0.098017865 | 0.263421446 | No Change | Not Significant | TRUE |
| Baiap2l2 | TRUE | -0.126087469 | 0.668072037 | No Change | Not Significant | TRUE | -0.139955413 | 0.373324674 | No Change | Not Significant | TRUE |
| Baiap3 | TRUE | -0.192223821 | 0.615929821 | No Change | Not Significant | TRUE | -0.114184226 | 0.679535938 | No Change | Not Significant | TRUE |
| Bak1 | TRUE | -0.061415208 | 0.658733455 | No Change | Not Significant | TRUE | 0.103863267 | 0.537983118 | No Change | Not Significant | TRUE |
| Bambi | TRUE | 0.35714635 | 0.225622081 | No Change | Not Significant | TRUE | -0.398275484 | 0.00046847 | No Change | Significant Negative | TRUE |
| Banf1 | TRUE | -0.015141907 | 0.935638301 | No Change | Not Significant | TRUE | -0.03414904 | 0.831111145 | No Change | Not Significant | TRUE |
| Bank1 | TRUE | -2.306229062 | 1 | No Change | Not Significant | FALSE | NA | NA | No Change | Not Detected | FALSE |
| Banp | TRUE | 0.061128306 | 0.888050066 | No Change | Not Significant | TRUE | 0.108289039 | 0.588148333 | No Change | Not Significant | TRUE |
| Bap1 | TRUE | 0.074132354 | 0.375447286 | No Change | Not Significant | TRUE | -0.044397478 | 0.622235448 | No Change | Not Significant | TRUE |
| Bard1 | TRUE | 0.301161035 | 0.181269925 | No Change | Not Significant | TRUE | 0.102057612 | 0.685606255 | No Change | Not Significant | TRUE |
| Barhl1 | TRUE | -0.009268455 | 1 | No Change | Not Significant | FALSE | NA | NA | No Change | Not Detected | FALSE |
| Barx2 | TRUE | -0.162774023 | 0.716025064 | No Change | Not Significant | TRUE | 0.587731332 | 0.076614782 | No Change | Not Significant | TRUE |
| Basp1 | TRUE | -0.261471636 | 1 | No Change | Not Significant | FALSE | NA | NA | No Change | Not Detected | FALSE |
| Batf | TRUE | 0.976447262 | 1 | No Change | Not Significant | FALSE | NA | NA | No Change | Not Detected | FALSE |
| Batf2 | TRUE | -0.37119916 | 0.049536564 | No Change | Significant Negative | TRUE | -0.119844385 | 0.645187136 | No Change | Not Significant | TRUE |
| Bax | TRUE | 0.164710168 | 0.166725366 | No Change | Not Significant | TRUE | 0.085139168 | 0.523431015 | No Change | Not Significant | TRUE |
| Baz1a | TRUE | 0.239177704 | 0.056246986 | No Change | Not Significant | TRUE | -0.089710107 | 0.402088841 | No Change | Not Significant | TRUE |
| Baz1b | TRUE | 0.148559454 | 0.125757237 | No Change | Not Significant | TRUE | 0.009960309 | 0.915984616 | No Change | Not Significant | TRUE |
| Baz2a | TRUE | -0.176648288 | 0.214258446 | No Change | Not Significant | TRUE | -0.133576608 | 0.083135909 | No Change | Not Significant | TRUE |
| Baz2b | TRUE | -0.145466887 | 0.328900667 | No Change | Not Significant | TRUE | -0.1638317 | 0.059914226 | No Change | Not Significant | TRUE |
| BB031773 | TRUE | 3.225384238 | 1 | No Change | Not Significant | FALSE | NA | NA | No Change | Not Detected | FALSE |
| BB557941 | TRUE | 1.017664986 | 1 | No Change | Not Significant | FALSE | NA | NA | No Change | Not Detected | FALSE |
| Bbc3 | TRUE | -0.189342567 | 0.604102573 | No Change | Not Significant | TRUE | -0.265449444 | 0.220501906 | No Change | Not Significant | TRUE |
| Bbip1 | TRUE | 0.069721217 | 0.647057325 | No Change | Not Significant | TRUE | 0.012103677 | 0.938898215 | No Change | Not Significant | TRUE |
| Bbln | TRUE | 0.046804991 | 0.799444652 | No Change | Not Significant | TRUE | 0.160953645 | 0.419784692 | No Change | Not Significant | TRUE |
| Bbof1 | TRUE | 0.512903277 | 0.199909758 | No Change | Not Significant | TRUE | -0.278419739 | 0.551915127 | No Change | Not Significant | TRUE |
| Bbox1 | TRUE | -0.075817678 | 1 | No Change | Not Significant | FALSE | NA | NA | No Change | Not Detected | FALSE |
| Bbs1 | TRUE | -0.179697031 | 0.555300096 | No Change | Not Significant | TRUE | 0.193115547 | 0.500132153 | No Change | Not Significant | TRUE |
| Bbs10 | TRUE | -0.06705613 | 0.892732467 | No Change | Not Significant | TRUE | 0.084329789 | 0.831002845 | No Change | Not Significant | TRUE |
| Bbs12 | TRUE | 0.374953451 | 0.488337435 | No Change | Not Significant | TRUE | 0.002640996 | 0.996976705 | No Change | Not Significant | TRUE |
| Bbs2 | TRUE | -0.075553612 | 0.685755422 | No Change | Not Significant | TRUE | 0.003257395 | 0.989317668 | No Change | Not Significant | TRUE |
| Bbs4 | TRUE | -0.08553112 | 0.78770474 | No Change | Not Significant | TRUE | -0.075758752 | 0.743740778 | No Change | Not Significant | TRUE |
| Bbs5 | TRUE | 0.06559827 | 0.840048595 | No Change | Not Significant | TRUE | 0.244380403 | 0.275350828 | No Change | Not Significant | TRUE |
| Bbs7 | TRUE | 0.42131402 | 0.01096738 | No Change | Significant Positive | TRUE | 0.131972978 | 0.5332246 | No Change | Not Significant | TRUE |
| Bbs9 | TRUE | 0.188041025 | 0.377007156 | No Change | Not Significant | TRUE | 0.023323887 | 0.939297866 | No Change | Not Significant | TRUE |
| Bbx | TRUE | -0.122868724 | 0.445952845 | No Change | Not Significant | TRUE | 0.002700817 | 0.983542327 | No Change | Not Significant | TRUE |
| BC002059 | TRUE | -0.034045883 | 0.90054944 | No Change | Not Significant | TRUE | -0.229618585 | 0.11171095 | No Change | Not Significant | TRUE |
| BC003965 | TRUE | -0.19675496 | 0.071033953 | No Change | Not Significant | TRUE | 0.004921455 | 0.976374805 | No Change | Not Significant | TRUE |
| BC004004 | TRUE | -0.2748639 | 0.000142578 | No Change | Significant Negative | TRUE | -0.129233689 | 0.062318117 | No Change | Not Significant | TRUE |
| BC005537 | TRUE | -0.051463547 | 0.652196761 | No Change | Not Significant | TRUE | -0.078348412 | 0.39047175 | No Change | Not Significant | TRUE |
| BC005624 | TRUE | -0.226963208 | 0.002731683 | No Change | Significant Negative | TRUE | 0.128296483 | 0.104092769 | No Change | Not Significant | TRUE |
| BC006965 | TRUE | -0.003290501 | 0.998042036 | No Change | Not Significant | TRUE | 0.309354285 | 0.711873325 | No Change | Not Significant | TRUE |
| BC016579 | TRUE | 0.208201138 | 0.07883501 | No Change | Not Significant | TRUE | 0.060916398 | 0.605044452 | No Change | Not Significant | TRUE |
| BC024063 | TRUE | -0.718209455 | 0.04055653 | DOWN | DOWN | TRUE | -0.327606186 | 0.337453004 | No Change | Not Significant | TRUE |
| BC024139 | TRUE | 0.166866626 | 0.685318046 | No Change | Not Significant | TRUE | -0.085198274 | 0.849656295 | No Change | Not Significant | TRUE |
| BC024978 | TRUE | 0.550946056 | 0.046464079 | UP | UP | TRUE | 0.095645169 | 0.710842238 | No Change | Not Significant | TRUE |
| BC028528 | TRUE | 0.600784028 | 1 | No Change | Not Significant | FALSE | NA | NA | No Change | Not Detected | FALSE |
| BC029722 | TRUE | -0.196394691 | 0.375025422 | No Change | Not Significant | TRUE | -0.076016341 | 0.738126248 | No Change | Not Significant | TRUE |
| BC031181 | TRUE | 0.55534349 | 1.29E-12 | UP | UP | TRUE | 0.354418495 | 3.01E-06 | No Change | Significant Positive | TRUE |
| BC034090 | TRUE | 0.58616817 | 1 | No Change | Not Significant | FALSE | NA | NA | No Change | Not Detected | FALSE |
| BC035044 | TRUE | -0.108941103 | 1 | No Change | Not Significant | FALSE | NA | NA | No Change | Not Detected | FALSE |
| BC037704 | TRUE | 0.251818163 | 0.668637691 | No Change | Not Significant | TRUE | 0.152824826 | 0.755288004 | No Change | Not Significant | TRUE |
| BC049715 | TRUE | 0.127398317 | 0.761298159 | No Change | Not Significant | TRUE | 0.051164752 | 0.877659754 | No Change | Not Significant | TRUE |
| BC049762 | TRUE | -0.427971386 | 1 | No Change | Not Significant | TRUE | -1.021543748 | 0.132480392 | No Change | Not Significant | TRUE |
| BC050972 | TRUE | 0.007452894 | 1 | No Change | Not Significant | FALSE | NA | NA | No Change | Not Detected | FALSE |
| BC051019 | TRUE | -0.555131892 | 1 | No Change | Not Significant | FALSE | NA | NA | No Change | Not Detected | FALSE |
| BC051142 | TRUE | 0.131840314 | 0.838747423 | No Change | Not Significant | TRUE | -0.339763076 | 0.498869056 | No Change | Not Significant | TRUE |
| BC053393 | TRUE | 3.423462447 | 1 | No Change | Not Significant | FALSE | NA | NA | No Change | Not Detected | FALSE |
| BC055324 | TRUE | 0.828738254 | 0.001407454 | UP | UP | TRUE | 0.286064646 | 0.323066278 | No Change | Not Significant | TRUE |
| BC055402 | TRUE | 2.849042648 | 1 | No Change | Not Significant | FALSE | NA | NA | No Change | Not Detected | FALSE |
| BC065397 | TRUE | -0.031847266 | 0.967478151 | No Change | Not Significant | TRUE | -0.438520896 | 0.352866698 | No Change | Not Significant | TRUE |
| BC107364 | TRUE | 0.744191668 | 1 | No Change | Not Significant | FALSE | NA | NA | No Change | Not Detected | FALSE |
| BC147527 | TRUE | -0.740644823 | 1 | No Change | Not Significant | FALSE | NA | NA | No Change | Not Detected | FALSE |
| Bcam | TRUE | 0.037646522 | 0.908812719 | No Change | Not Significant | TRUE | -0.411510067 | 0.008402281 | No Change | Significant Negative | TRUE |
| Bcan | TRUE | -2.462469808 | 0.077273602 | No Change | Not Significant | FALSE | NA | NA | No Change | Not Detected | FALSE |
| Bcap29 | TRUE | -0.094526413 | 0.714354347 | No Change | Not Significant | TRUE | -0.179141058 | 0.269540799 | No Change | Not Significant | TRUE |
| Bcap31 | TRUE | -0.303757459 | 0.00535797 | No Change | Significant Negative | TRUE | -0.028719561 | 0.769017785 | No Change | Not Significant | TRUE |
| Bcar1 | TRUE | -0.069264578 | 0.560532006 | No Change | Not Significant | TRUE | 0.028330674 | 0.854251145 | No Change | Not Significant | TRUE |
| Bcar3 | TRUE | -0.019111428 | 0.931842966 | No Change | Not Significant | TRUE | 0.048071953 | 0.640442713 | No Change | Not Significant | TRUE |
| Bcas1 | TRUE | 0.167165719 | 0.597748482 | No Change | Not Significant | TRUE | 0.184453903 | 0.04918511 | No Change | Significant Positive | TRUE |
| Bcas1os2 | TRUE | -0.92797127 | 0.134442563 | No Change | Not Significant | TRUE | -0.128467481 | 0.880475742 | No Change | Not Significant | TRUE |
| Bcas2 | TRUE | 0.048004974 | 0.808650118 | No Change | Not Significant | TRUE | -0.093711751 | 0.316511058 | No Change | Not Significant | TRUE |
| Bcas3 | TRUE | -0.294337255 | 0.131848679 | No Change | Not Significant | TRUE | -0.094207862 | 0.572749056 | No Change | Not Significant | TRUE |
| Bcas3os1 | TRUE | -0.721916753 | 0.433049501 | No Change | Not Significant | TRUE | 0.069511015 | 0.950733597 | No Change | Not Significant | TRUE |
| Bcas3os2 | TRUE | -2.153939561 | 1 | No Change | Not Significant | FALSE | NA | NA | No Change | Not Detected | FALSE |
| Bcat1 | TRUE | 2.770558915 | 1 | No Change | Not Significant | FALSE | NA | NA | No Change | Not Detected | FALSE |
| Bcat2 | TRUE | 1.182250361 | 6.04E-21 | UP | UP | TRUE | 1.344101164 | 1.79E-111 | UP | UP | TRUE |
| Bccip | TRUE | -0.036146441 | 0.853791016 | No Change | Not Significant | TRUE | -0.029026379 | 0.857064391 | No Change | Not Significant | TRUE |
| Bcdin3d | TRUE | -0.090088264 | 0.715139121 | No Change | Not Significant | TRUE | -0.007289107 | 0.980585772 | No Change | Not Significant | TRUE |
| Bche | TRUE | -0.116289601 | 0.448589299 | No Change | Not Significant | TRUE | -0.416986176 | 0.007209695 | No Change | Significant Negative | TRUE |
| Bckdha | TRUE | -0.334150474 | 0.037507197 | No Change | Significant Negative | TRUE | -0.095947316 | 0.457199933 | No Change | Not Significant | TRUE |
| Bckdhb | TRUE | -0.024435906 | 0.936580622 | No Change | Not Significant | TRUE | 0.110817775 | 0.561462863 | No Change | Not Significant | TRUE |
| Bckdk | TRUE | 0.138898965 | 0.270127107 | No Change | Not Significant | TRUE | 0.133956734 | 0.183396824 | No Change | Not Significant | TRUE |
| Bcl10 | TRUE | 0.002230816 | 0.985568894 | No Change | Not Significant | TRUE | 0.080915947 | 0.339763458 | No Change | Not Significant | TRUE |
| Bcl11a | TRUE | -0.126154601 | 0.67261003 | No Change | Not Significant | TRUE | -0.105581954 | 0.645300059 | No Change | Not Significant | TRUE |
| Bcl11b | TRUE | -0.042553678 | 0.852123921 | No Change | Not Significant | TRUE | -0.03958566 | 0.853157579 | No Change | Not Significant | TRUE |
| Bcl2 | TRUE | 0.372820247 | 0.054426466 | No Change | Not Significant | TRUE | 0.110403863 | 0.499216262 | No Change | Not Significant | TRUE |
| Bcl2a1b | TRUE | 0.639586945 | 1 | No Change | Not Significant | FALSE | NA | NA | No Change | Not Detected | FALSE |
| Bcl2a1d | TRUE | 1.269723467 | 1 | No Change | Not Significant | FALSE | NA | NA | No Change | Not Detected | FALSE |
| Bcl2l1 | TRUE | 0.011847126 | 0.948680909 | No Change | Not Significant | TRUE | 0.105568158 | 0.142635415 | No Change | Not Significant | TRUE |
| Bcl2l10 | TRUE | -1.071651321 | 1 | No Change | Not Significant | FALSE | NA | NA | No Change | Not Detected | FALSE |
| Bcl2l11 | TRUE | -0.50058175 | 0.008574071 | DOWN | DOWN | TRUE | -0.382742555 | 0.003155171 | No Change | Significant Negative | TRUE |
| Bcl2l12 | TRUE | 0.216683866 | 0.263984696 | No Change | Not Significant | TRUE | 0.137888709 | 0.401524931 | No Change | Not Significant | TRUE |
| Bcl2l13 | TRUE | 0.00458516 | 0.968724317 | No Change | Not Significant | TRUE | 0.057731801 | 0.514726356 | No Change | Not Significant | TRUE |
| Bcl2l14 | TRUE | 0.094493735 | 0.520880615 | No Change | Not Significant | TRUE | 0.106604999 | 0.175287213 | No Change | Not Significant | TRUE |
| Bcl2l15 | TRUE | 1.641477059 | 1 | No Change | Not Significant | FALSE | NA | NA | No Change | Not Detected | FALSE |
| Bcl2l2 | TRUE | 0.11223478 | 0.462014445 | No Change | Not Significant | TRUE | 0.051389118 | 0.50034619 | No Change | Not Significant | TRUE |
| Bcl3 | TRUE | 0.52882196 | 0.007074869 | UP | UP | TRUE | 0.140404896 | 0.445605776 | No Change | Not Significant | TRUE |
| Bcl6 | TRUE | -0.184164335 | 0.527079777 | No Change | Not Significant | TRUE | -0.115197262 | 0.747864433 | No Change | Not Significant | TRUE |
| Bcl6b | TRUE | 0.231535838 | 0.821350533 | No Change | Not Significant | FALSE | NA | NA | No Change | Not Detected | FALSE |
| Bcl7a | TRUE | -0.321494965 | 0.084361596 | No Change | Not Significant | TRUE | -0.309829406 | 0.01731607 | No Change | Significant Negative | TRUE |
| Bcl7b | TRUE | -0.026402473 | 0.885199039 | No Change | Not Significant | TRUE | -0.0596417 | 0.595956629 | No Change | Not Significant | TRUE |
| Bcl7c | TRUE | -0.128611203 | 0.601768559 | No Change | Not Significant | TRUE | 0.076798913 | 0.691625579 | No Change | Not Significant | TRUE |
| Bcl9 | TRUE | 0.243009828 | 0.405918699 | No Change | Not Significant | TRUE | -0.104423431 | 0.517140055 | No Change | Not Significant | TRUE |
| Bcl9l | TRUE | -0.114123842 | 0.727166502 | No Change | Not Significant | TRUE | -0.243059209 | 0.207235951 | No Change | Not Significant | TRUE |
| Bclaf1 | TRUE | 0.099711735 | 0.432138537 | No Change | Not Significant | TRUE | -0.077248736 | 0.5257021 | No Change | Not Significant | TRUE |
| Bclaf3 | TRUE | 0.021652919 | 0.937816815 | No Change | Not Significant | TRUE | -0.044885843 | 0.83235935 | No Change | Not Significant | TRUE |
| Bco1 | TRUE | 0.060465097 | 0.904987729 | No Change | Not Significant | TRUE | -0.115151926 | 0.747912187 | No Change | Not Significant | TRUE |
| Bco2 | TRUE | -0.75998136 | 0.001860579 | DOWN | DOWN | TRUE | -0.196669329 | 0.260220234 | No Change | Not Significant | TRUE |
| Bcor | TRUE | 0.12071086 | 0.208654479 | No Change | Not Significant | TRUE | -0.076452568 | 0.494893811 | No Change | Not Significant | TRUE |
| Bcorl1 | TRUE | -0.5410071 | 0.066168368 | No Change | Not Significant | TRUE | -0.384163717 | 0.022899104 | No Change | Significant Negative | TRUE |
| Bcr | TRUE | -0.048579325 | 0.772538316 | No Change | Not Significant | TRUE | -0.034870618 | 0.797513438 | No Change | Not Significant | TRUE |
| Bcs1l | TRUE | -0.326025129 | 0.021513836 | No Change | Significant Negative | TRUE | 0.073941201 | 0.585130336 | No Change | Not Significant | TRUE |
| Bdh1 | TRUE | -0.646448708 | 1.97E-07 | DOWN | DOWN | TRUE | -0.070085212 | 0.391953183 | No Change | Not Significant | TRUE |
| Bdh2 | TRUE | 0.405653735 | 1 | No Change | Not Significant | FALSE | NA | NA | No Change | Not Detected | FALSE |
| Bdkrb1 | TRUE | 0.118021865 | 1 | No Change | Not Significant | FALSE | NA | NA | No Change | Not Detected | FALSE |
| Bdkrb2 | TRUE | 1.518598082 | 1 | No Change | Not Significant | FALSE | NA | NA | No Change | Not Detected | FALSE |
| Bdnf | TRUE | -0.742673487 | 1 | No Change | Not Significant | FALSE | NA | NA | No Change | Not Detected | FALSE |
| Bdp1 | TRUE | 0.121528755 | 0.239840996 | No Change | Not Significant | TRUE | -0.089732755 | 0.154536441 | No Change | Not Significant | TRUE |
| Bean1 | TRUE | 0.176026823 | 1 | No Change | Not Significant | FALSE | NA | NA | No Change | Not Detected | FALSE |
| Becn1 | TRUE | 0.005277878 | 0.954378365 | No Change | Not Significant | TRUE | 0.038447794 | 0.64854129 | No Change | Not Significant | TRUE |
| Becn2 | TRUE | -0.341282537 | 1 | No Change | Not Significant | FALSE | NA | NA | No Change | Not Detected | FALSE |
| Begain | TRUE | 1.202884266 | 1 | No Change | Not Significant | FALSE | NA | NA | No Change | Not Detected | FALSE |
| Bend3 | TRUE | 0.083839938 | 0.63773884 | No Change | Not Significant | TRUE | -0.093686818 | 0.609985966 | No Change | Not Significant | TRUE |
| Bend4 | TRUE | -0.569086141 | 1 | No Change | Not Significant | FALSE | NA | NA | No Change | Not Detected | FALSE |
| Bend5 | TRUE | -0.569104155 | 1 | No Change | Not Significant | FALSE | NA | NA | No Change | Not Detected | FALSE |
| Bend6 | TRUE | -0.033668619 | 1 | No Change | Not Significant | FALSE | NA | NA | No Change | Not Detected | FALSE |
| Bend7 | TRUE | 0.162444936 | 0.382367026 | No Change | Not Significant | TRUE | 0.182204843 | 0.178465808 | No Change | Not Significant | TRUE |
| Best1 | TRUE | 0.310283159 | 0.804573016 | No Change | Not Significant | TRUE | -1.123919031 | 0.057394659 | No Change | Not Significant | TRUE |
| Best2 | TRUE | 0.190428732 | 0.850157504 | No Change | Not Significant | TRUE | 0.653472359 | 5.52E-05 | UP | UP | TRUE |
| Best3 | TRUE | -0.528102515 | 1 | No Change | Not Significant | FALSE | NA | NA | No Change | Not Detected | FALSE |
| Bet1 | TRUE | 0.597419435 | 4.83E-05 | UP | UP | TRUE | 0.390983686 | 3.86E-06 | No Change | Significant Positive | TRUE |
| Bet1l | TRUE | -0.04093148 | 0.777725809 | No Change | Not Significant | TRUE | 0.031230252 | 0.797543961 | No Change | Not Significant | TRUE |
| Bex1 | TRUE | 0.652908525 | 0.038193919 | UP | UP | TRUE | -0.047227834 | 0.920659938 | No Change | Not Significant | TRUE |
| Bex2 | TRUE | 0.489982418 | 0.098824478 | No Change | Not Significant | TRUE | 0.093403861 | 0.755953522 | No Change | Not Significant | TRUE |
| Bex3 | TRUE | 0.521588167 | 0.003624129 | UP | UP | TRUE | 0.141959517 | 0.215929529 | No Change | Not Significant | TRUE |
| Bex4 | TRUE | 0.309644738 | 0.472341459 | No Change | Not Significant | TRUE | -0.064987512 | 0.889251381 | No Change | Not Significant | TRUE |
| Bfar | TRUE | -0.080216454 | 0.570683233 | No Change | Not Significant | TRUE | -0.039009791 | 0.62056297 | No Change | Not Significant | TRUE |
| Bfsp1 | TRUE | -0.008183653 | 0.982490542 | No Change | Not Significant | TRUE | -0.314730045 | 0.023672009 | No Change | Significant Negative | TRUE |
| Bglap3 | TRUE | -1.27119305 | 0.004304208 | DOWN | DOWN | TRUE | -1.416174944 | 1.24E-06 | DOWN | DOWN | TRUE |
| Bgn | TRUE | 0.60082056 | 0.827372024 | No Change | Not Significant | FALSE | NA | NA | No Change | Not Detected | FALSE |
| Bhlha15 | TRUE | 1.40918012 | 8.22E-09 | UP | UP | TRUE | 1.485474475 | 7.66E-10 | UP | UP | TRUE |
| Bhlha9 | TRUE | 0.201121741 | 1 | No Change | Not Significant | FALSE | NA | NA | No Change | Not Detected | FALSE |
| Bhlhb9 | TRUE | 0.226666937 | 0.606725963 | No Change | Not Significant | TRUE | 0.032150894 | 0.937787345 | No Change | Not Significant | TRUE |
| Bhlhe40 | TRUE | -0.289909527 | 0.048151079 | No Change | Significant Negative | TRUE | -0.15174677 | 0.146771529 | No Change | Not Significant | TRUE |
| Bhlhe41 | TRUE | 1.374163123 | 1 | No Change | Not Significant | FALSE | NA | NA | No Change | Not Detected | FALSE |
| Bhmt | TRUE | 0.632719358 | 1 | No Change | Not Significant | FALSE | NA | NA | No Change | Not Detected | FALSE |
| Bhmt2 | TRUE | -0.006069778 | 0.992998437 | No Change | Not Significant | TRUE | -0.286266689 | 0.576171574 | No Change | Not Significant | TRUE |
| Bicc1 | TRUE | -0.43205503 | 0.570058297 | No Change | Not Significant | TRUE | -0.396780853 | 0.504745134 | No Change | Not Significant | TRUE |
| Bicd1 | TRUE | -0.073389255 | 0.711726983 | No Change | Not Significant | TRUE | 0.056955681 | 0.685606255 | No Change | Not Significant | TRUE |
| Bicd2 | TRUE | -0.077159192 | 0.691115487 | No Change | Not Significant | TRUE | -0.048293974 | 0.790834872 | No Change | Not Significant | TRUE |
| Bicdl1 | TRUE | 0.677614384 | 2.03E-08 | UP | UP | TRUE | 0.263326572 | 0.000248702 | No Change | Significant Positive | TRUE |
| Bicdl2 | TRUE | -0.089499341 | 0.539723961 | No Change | Not Significant | TRUE | 0.106963053 | 0.380615229 | No Change | Not Significant | TRUE |
| Bicra | TRUE | 0.387114297 | 0.370029503 | No Change | Not Significant | TRUE | -0.308502135 | 0.124816482 | No Change | Not Significant | TRUE |
| Bicral | TRUE | -0.121214157 | 0.292089426 | No Change | Not Significant | TRUE | -0.124554382 | 0.397562024 | No Change | Not Significant | TRUE |
| Bid | TRUE | -0.32523774 | 0.018172963 | No Change | Significant Negative | TRUE | -0.312546481 | 0.030837772 | No Change | Significant Negative | TRUE |
| Bik | TRUE | -1.3583055 | 4.43E-11 | DOWN | DOWN | TRUE | -0.180316175 | 0.451769221 | No Change | Not Significant | TRUE |
| Bin1 | TRUE | 0.024769901 | 0.901751383 | No Change | Not Significant | TRUE | 0.117474447 | 0.425519673 | No Change | Not Significant | TRUE |
| Bin2 | TRUE | 0.104052263 | 0.885228942 | No Change | Not Significant | TRUE | 0.373385278 | 0.359247178 | No Change | Not Significant | TRUE |
| Bin3 | TRUE | -0.00340044 | 0.983635967 | No Change | Not Significant | TRUE | 0.07842396 | 0.361886677 | No Change | Not Significant | TRUE |
| Birc2 | TRUE | 0.209677081 | 0.054421559 | No Change | Not Significant | TRUE | 0.00497476 | 0.974315 | No Change | Not Significant | TRUE |
| Birc3 | TRUE | -0.450965353 | 0.000140653 | No Change | Significant Negative | TRUE | -0.115838367 | 0.412210695 | No Change | Not Significant | TRUE |
| Birc5 | TRUE | 0.783701832 | 5.03E-05 | UP | UP | TRUE | 0.049813034 | 0.830998117 | No Change | Not Significant | TRUE |
| Birc6 | TRUE | 0.09535494 | 0.38145272 | No Change | Not Significant | TRUE | 0.035370156 | 0.693047027 | No Change | Not Significant | TRUE |
| Bivm | TRUE | 0.053251406 | 0.788668238 | No Change | Not Significant | TRUE | 0.089236706 | 0.469953022 | No Change | Not Significant | TRUE |
| Blcap | TRUE | -0.21211707 | 0.105941585 | No Change | Not Significant | TRUE | -0.079802807 | 0.371487597 | No Change | Not Significant | TRUE |
| Blk | TRUE | 1.60759819 | 1 | No Change | Not Significant | FALSE | NA | NA | No Change | Not Detected | FALSE |
| Blm | TRUE | 0.618719317 | 0.008334723 | UP | UP | TRUE | 0.058279288 | 0.841819353 | No Change | Not Significant | TRUE |
| Blmh | TRUE | 0.097270548 | 0.297320449 | No Change | Not Significant | TRUE | 0.077508915 | 0.505363174 | No Change | Not Significant | TRUE |
| Blnk | TRUE | -0.127804627 | 0.316544664 | No Change | Not Significant | TRUE | -0.125248018 | 0.088986709 | No Change | Not Significant | TRUE |
| Bloc1s1 | TRUE | -0.14034462 | 0.486449404 | No Change | Not Significant | TRUE | -0.122665264 | 0.555505998 | No Change | Not Significant | TRUE |
| Bloc1s2 | TRUE | -0.090662602 | 0.725856654 | No Change | Not Significant | TRUE | 0.131786239 | 0.25082096 | No Change | Not Significant | TRUE |
| Bloc1s3 | TRUE | 0.680519184 | 3.18E-15 | UP | UP | TRUE | 0.34720506 | 0.010442524 | No Change | Significant Positive | TRUE |
| Bloc1s4 | TRUE | -0.092306725 | 0.72936091 | No Change | Not Significant | TRUE | -0.084397941 | 0.713632533 | No Change | Not Significant | TRUE |
| Bloc1s5 | TRUE | -0.048081644 | 0.802698942 | No Change | Not Significant | TRUE | 0.128957399 | 0.225514314 | No Change | Not Significant | TRUE |
| Bloc1s6 | TRUE | -0.065791159 | 0.64620651 | No Change | Not Significant | TRUE | 0.072165291 | 0.440953549 | No Change | Not Significant | TRUE |
| Bloc1s6os | TRUE | 0.196519823 | 1 | No Change | Not Significant | FALSE | NA | NA | No Change | Not Detected | FALSE |
| Bloodlinc | TRUE | 0.569251038 | 0.433049501 | No Change | Not Significant | TRUE | 0.728405363 | 0.262627924 | No Change | Not Significant | TRUE |
| Bltp1 | TRUE | 0.267672642 | 0.14195729 | No Change | Not Significant | TRUE | 0.096072874 | 0.404658227 | No Change | Not Significant | TRUE |
| Bltp2 | TRUE | -0.060909872 | 0.494858209 | No Change | Not Significant | TRUE | 0.070606581 | 0.29617307 | No Change | Not Significant | TRUE |
| Blvra | TRUE | 0.700368889 | 1.41E-14 | UP | UP | TRUE | 0.428413822 | 9.22E-07 | No Change | Significant Positive | TRUE |
| Blvrb | TRUE | 1.337292591 | 1.08E-47 | UP | UP | TRUE | 0.860889144 | 1.08E-33 | UP | UP | TRUE |
| Blzf1 | TRUE | 0.069626746 | 0.673388737 | No Change | Not Significant | TRUE | 0.114795598 | 0.256862183 | No Change | Not Significant | TRUE |
| Bmerb1 | TRUE | -0.518795814 | 1 | No Change | Not Significant | FALSE | NA | NA | No Change | Not Detected | FALSE |
| Bmf | TRUE | -0.54187613 | 0.003784612 | DOWN | DOWN | TRUE | -0.057387485 | 0.760667465 | No Change | Not Significant | TRUE |
| Bmi1 | TRUE | -0.054832705 | 0.775625491 | No Change | Not Significant | TRUE | -0.166360433 | 0.092350341 | No Change | Not Significant | TRUE |
| Bmp1 | TRUE | 0.402348706 | 0.365696631 | No Change | Not Significant | TRUE | 0.047938675 | 0.895837077 | No Change | Not Significant | TRUE |

| geneID | Cond1_DF_Detected | Cond1_DF_ValueCol | Cond1_DF_PadjCol | Cond1_DF_Cutoff | Cond1_DF_Cutoff_Specific.x | Cond2_DF_Detected | Cond2_DF_ValueCol |
|---|---|---|---|---|---|---|---|
| Bmp10 | TRUE | 2.512713418 | 1 | No Change | Not Significant | FALSE | NA |
| Bmp2 | TRUE | -0.44310701 | 0.179735514 | No Change | Not Significant | TRUE | -0.292454913 |
| Bmp2k | TRUE | -0.017712003 | 0.92744128 | No Change | Not Significant | TRUE | -0.127334379 |
| Bmp3 | TRUE | -0.949790815 | 2.46E-08 | DOWN | DOWN | TRUE | -0.344997762 |
| Bmp4 | TRUE | 0.362360113 | 1 | No Change | Not Significant | FALSE | NA |
| Bmp5 | TRUE | -2.087453219 | 1 | No Change | Not Significant | FALSE | NA |
| Bmp6 | TRUE | -0.027262305 | 0.981095512 | No Change | Not Significant | TRUE | -0.181769995 |
| Bmp7 | TRUE | -0.049705675 | 0.937028043 | No Change | Not Significant | TRUE | -0.182872901 |
| Bmp8a | TRUE | -0.060165004 | 0.840298483 | No Change | Not Significant | TRUE | 0.289379805 |
| Bmp8b | TRUE | -0.060901687 | 0.947328174 | No Change | Not Significant | TRUE | 0.500641778 |
| Bmper | TRUE | 1.43390898 | 0.006304476 | UP | UP | TRUE | 0.13708823 |
| Bmpr1a | TRUE | -0.043149081 | 0.786099952 | No Change | Not Significant | TRUE | -0.072975821 |
| Bmpr1b | TRUE | -0.937098979 | 1 | No Change | Not Significant | FALSE | NA |
| Bmpr2 | TRUE | -0.05291299 | 0.748973617 | No Change | Not Significant | TRUE | -0.035483842 |
| Bms1 | TRUE | 0.284760069 | 0.000413537 | No Change | Significant Positive | TRUE | 0.137258389 |
| Bmt2 | TRUE | -0.003734192 | 0.987811867 | No Change | Not Significant | TRUE | -0.132156546 |
| Bmx | TRUE | -0.667076207 | 0.004486866 | DOWN | DOWN | TRUE | 0.332945595 |
| Bmyc | TRUE | 0.037455388 | 0.943887948 | No Change | Not Significant | TRUE | -0.323767048 |
| Bnc1 | TRUE | -1.645889739 | 1 | No Change | Not Significant | FALSE | NA |
| Bnc2 | TRUE | 1.032553777 | 1 | No Change | Not Significant | FALSE | NA |
| Bnip1 | TRUE | -0.170774652 | 0.202404322 | No Change | Not Significant | TRUE | -0.02287707 |
| Bnip2 | TRUE | 0.032786664 | 0.898299072 | No Change | Not Significant | TRUE | -0.209418054 |
| Bnip3 | TRUE | -0.975338454 | 2.33E-05 | DOWN | DOWN | TRUE | -0.574636342 |
| Bnip3l | TRUE | -0.150125895 | 0.265929452 | No Change | Not Significant | TRUE | 0.099925689 |
| Bnip3l-ps | TRUE | 0.427812265 | 1 | No Change | Not Significant | TRUE | 0.640770694 |
| Bnip5 | TRUE | -0.119815796 | 0.861253255 | No Change | Not Significant | TRUE | 0.117819065 |
| Bnipl | TRUE | -0.109797578 | 0.702914252 | No Change | Not Significant | TRUE | 0.064406072 |
| Boc | TRUE | -0.276228171 | 0.083216624 | No Change | Not Significant | TRUE | -0.207991322 |
| Bod1 | TRUE | -0.056506291 | 0.648871201 | No Change | Not Significant | TRUE | 0.004512988 |
| Bod1l | TRUE | 0.007269901 | 0.956641558 | No Change | Not Significant | TRUE | -0.065274845 |
| Bok | TRUE | -0.316684385 | 0.020161214 | No Change | Significant Negative | TRUE | 0.327132827 |
| Bola1 | TRUE | -0.061509539 | 0.772538316 | No Change | Not Significant | TRUE | 0.161562469 |
| Bola2 | TRUE | -0.285371928 | 0.001809424 | No Change | Significant Negative | TRUE | -0.049257527 |
| Bola3 | TRUE | -0.068540804 | 0.736259883 | No Change | Not Significant | TRUE | 0.071149594 |
| Boll | TRUE | -0.320578684 | 1 | No Change | Not Significant | FALSE | NA |
| Bop1 | TRUE | 0.014929909 | 0.931099354 | No Change | Not Significant | TRUE | 0.018996457 |
| Bora | TRUE | 0.467734146 | 0.000137432 | No Change | Significant Positive | TRUE | -0.257568797 |
| Borcs5 | TRUE | 0.068344066 | 0.762380457 | No Change | Not Significant | TRUE | 0.14573533 |
| Borcs6 | TRUE | 0.532798775 | 2.50E-06 | UP | UP | TRUE | 0.258670202 |
| Borcs7 | TRUE | -0.180392135 | 0.501169049 | No Change | Not Significant | TRUE | -0.024771539 |
| Borcs8 | TRUE | -0.146945391 | 0.204680714 | No Change | Not Significant | TRUE | 0.126456826 |
| Bpgm | TRUE | 0.104217551 | 0.342932541 | No Change | Not Significant | TRUE | 0.257613529 |
| Bphl | TRUE | -0.220533272 | 0.080865411 | No Change | Not Significant | TRUE | -0.00237313 |
| Bpi | TRUE | 1.556172221 | 1 | No Change | Not Significant | FALSE | NA |
| Bpifb1 | TRUE | 2.924583418 | 1 | No Change | Not Significant | FALSE | NA |
| Bpifb4 | TRUE | 3.564681205 | 1 | No Change | Not Significant | FALSE | NA |
| Bpifb5 | TRUE | 1.056793073 | 1 | No Change | Not Significant | FALSE | NA |
| Bpifb9b | TRUE | -0.921339891 | 1 | No Change | Not Significant | FALSE | NA |
| Bpifc | TRUE | 0.470303085 | 1 | No Change | Not Significant | FALSE | NA |
| Bpnt1 | TRUE | -0.071365493 | 0.62019321 | No Change | Not Significant | TRUE | 0.16710602 |
| Bpnt2 | TRUE | 0.069535496 | 0.744656458 | No Change | Not Significant | TRUE | -0.044873146 |
| Bptf | TRUE | 0.020706206 | 0.890758364 | No Change | Not Significant | TRUE | -0.148798862 |
| Braf | TRUE | 0.185841084 | 0.205363124 | No Change | Not Significant | TRUE | -0.000460328 |
| Brap | TRUE | -0.029296812 | 0.774662172 | No Change | Not Significant | TRUE | -0.024679534 |
| Brat1 | TRUE | 0.143462782 | 0.429222486 | No Change | Not Significant | TRUE | 0.017152271 |
| Brca1 | TRUE | 0.566276424 | 0.00592004 | UP | UP | TRUE | 0.25025968 |
| Brca2 | TRUE | 0.26816452 | 0.239722683 | No Change | Not Significant | TRUE | -0.094368698 |
| Brcc3 | TRUE | 0.255694053 | 0.008569101 | No Change | Significant Positive | TRUE | 0.107546216 |
| Brd1 | TRUE | -0.06080027 | 0.514433539 | No Change | Not Significant | TRUE | 0.011175455 |
| Brd2 | TRUE | 0.123490018 | 0.449590928 | No Change | Not Significant | TRUE | -0.043983697 |
| Brd3 | TRUE | -0.086125284 | 0.587183556 | No Change | Not Significant | TRUE | -0.077011969 |
| Brd3os | TRUE | 0.398122754 | 1 | No Change | Not Significant | FALSE | NA |
| Brd4 | TRUE | 0.117094102 | 0.732746487 | No Change | Not Significant | TRUE | -0.242319507 |
| Brd7 | TRUE | -0.175792955 | 0.0473047 | No Change | Significant Negative | TRUE | 0.010095861 |
| Brd8 | TRUE | 0.147545936 | 0.181269925 | No Change | Not Significant | TRUE | -0.091080444 |
| Brd8dc | TRUE | 0.546279411 | 0.612497263 | No Change | Not Significant | FALSE | NA |
| Brd9 | TRUE | 0.178158296 | 0.044908772 | No Change | Significant Positive | TRUE | -0.06560713 |
| Brdt | TRUE | -0.163048896 | 0.847226925 | No Change | Not Significant | TRUE | -0.05637002 |
| Brf1 | TRUE | -0.064024964 | 0.716883239 | No Change | Not Significant | TRUE | -0.189750292 |
| Brf2 | TRUE | -0.036952361 | 0.890541867 | No Change | Not Significant | TRUE | 0.001176739 |
| Bri3 | TRUE | -0.429840708 | 0.000254944 | No Change | Significant Negative | TRUE | -0.11738326 |
| Bri3bp | TRUE | -0.51215617 | 6.95E-05 | DOWN | DOWN | TRUE | -0.282395282 |
| Brinp1 | TRUE | 1.299248433 | 1 | No Change | Not Significant | FALSE | NA |
| Brinp2 | TRUE | 1.467778209 | 1 | No Change | Not Significant | FALSE | NA |
| Brinp3 | TRUE | 2.52104946 | 1 | No Change | Not Significant | FALSE | NA |
| Brip1 | TRUE | 0.625359714 | 0.008063204 | UP | UP | TRUE | 0.017112888 |
| Brip1os | TRUE | 0.209234744 | 0.142788208 | No Change | Not Significant | TRUE | 0.13737204 |
| Brix1 | TRUE | 0.13048472 | 0.431320242 | No Change | Not Significant | TRUE | -0.044740496 |
| Brk1 | TRUE | -0.173205111 | 0.131725022 | No Change | Not Significant | TRUE | 0.045463047 |
| Brme1 | TRUE | -1.045973464 | 0.071918188 | No Change | Not Significant | TRUE | -0.196365661 |
| Brms1 | TRUE | 0.629538821 | 0.47508221 | No Change | Not Significant | TRUE | -0.397076148 |
| Brms1l | TRUE | 0.449066429 | 0.051764384 | No Change | Not Significant | TRUE | 0.148912094 |
| Brox | TRUE | -0.150893215 | 0.256383382 | No Change | Not Significant | TRUE | -0.005321807 |
| Brpf1 | TRUE | -0.070445577 | 0.673388737 | No Change | Not Significant | TRUE | -0.218621976 |
| Brpf3 | TRUE | 0.200108723 | 0.604659935 | No Change | Not Significant | TRUE | -0.197394306 |
| Brsk1 | TRUE | -0.194649664 | 0.732371878 | No Change | Not Significant | TRUE | 0.084695971 |
| Brsk2 | TRUE | 0.959062086 | 0.035388872 | UP | UP | TRUE | -0.042436208 |
| Brwd1 | TRUE | 0.365872768 | 0.00149302 | No Change | Significant Positive | TRUE | -0.005822807 |
| Brwd3 | TRUE | 0.110921861 | 0.42530681 | No Change | Not Significant | TRUE | 0.085857851 |
| Bscl2 | TRUE | -0.240393397 | 0.028908905 | No Change | Significant Negative | TRUE | -0.054911779 |
| Bsdc1 | TRUE | -0.198341418 | 0.054307717 | No Change | Not Significant | TRUE | 0.140760077 |
| Bsg | TRUE | -0.394371247 | 6.93E-08 | No Change | Significant Negative | TRUE | -0.023285095 |
| Bsn | TRUE | -0.172689663 | 0.305611042 | No Change | Not Significant | TRUE | -0.244548315 |
| Bspry | TRUE | -0.125005849 | 0.034960626 | No Change | Significant Negative | TRUE | 0.157028987 |
| Bst1 | TRUE | 1.290982422 | 0.42723932 | No Change | Not Significant | TRUE | 0.551295621 |
| Bst2 | TRUE | 0.911582482 | 0.003827457 | UP | UP | TRUE | 0.461238858 |
| Btaf1 | TRUE | 0.149398896 | 0.124214011 | No Change | Not Significant | TRUE | 0.000223565 |
| Btbd1 | TRUE | -0.213985996 | 0.069759652 | No Change | Not Significant | TRUE | 0.027519191 |
| Btbd10 | TRUE | -0.252325877 | 0.060282943 | No Change | Not Significant | TRUE | -0.135763039 |
| Btbd11 | TRUE | 0.317565868 | 0.587266381 | No Change | Not Significant | TRUE | 0.390588547 |
| Btbd16 | TRUE | -0.947102528 | 1 | No Change | Not Significant | FALSE | NA |
| Btbd17 | TRUE | 0.031969009 | 1 | No Change | Not Significant | FALSE | NA |
| Btbd18 | TRUE | -0.314996763 | 1 | No Change | Not Significant | FALSE | NA |
| Btbd19 | TRUE | -0.124986058 | 0.695380179 | No Change | Not Significant | TRUE | -0.366848866 |
| Btbd2 | TRUE | -0.07768697 | 0.552585717 | No Change | Not Significant | TRUE | -0.016238602 |
| Btbd3 | TRUE | -0.200627796 | 0.108571968 | No Change | Not Significant | TRUE | -0.068634335 |
| Btbd35f12 | TRUE | 3.349923482 | 1 | No Change | Not Significant | FALSE | NA |
| Btbd6 | TRUE | -0.205913948 | 0.071685421 | No Change | Not Significant | TRUE | -0.088149745 |
| Btbd7 | TRUE | -0.042080322 | 0.696186289 | No Change | Not Significant | TRUE | -0.036044747 |
| Btbd8 | TRUE | 0.017168504 | 0.947050968 | No Change | Not Significant | TRUE | -0.022611 |
| Btbd9 | TRUE | -0.101010594 | 0.283211393 | No Change | Not Significant | TRUE | -0.034147701 |
| Btc | TRUE | -0.412942233 | 0.091871348 | No Change | Not Significant | TRUE | 0.151142521 |
| Btd | TRUE | 0.010136004 | 0.971185231 | No Change | Not Significant | TRUE | 0.101608866 |
| Btf3 | TRUE | -0.248312568 | 0.008079258 | No Change | Significant Negative | TRUE | 0.001866035 |
| Btf3l4 | TRUE | -0.275445414 | 0.030671822 | No Change | Significant Negative | TRUE | -0.177396731 |
| Btf3l4b | TRUE | 0.686085123 | 1 | No Change | Not Significant | FALSE | NA |
| Btg1 | TRUE | -0.178855532 | 0.261367195 | No Change | Not Significant | TRUE | -0.123131097 |
| Btg2 | TRUE | -0.418378903 | 0.040813086 | No Change | Significant Negative | TRUE | 0.191052475 |
| Btg3 | TRUE | -0.271385934 | 0.453724307 | No Change | Not Significant | TRUE | -0.107739216 |
| Btk | TRUE | 0.25606685 | 0.777615016 | No Change | Not Significant | TRUE | 0.209956336 |
| Btla | TRUE | -1.964049897 | 1 | No Change | Not Significant | FALSE | NA |
| Btn1a1 | TRUE | 0.439057655 | 0.0821132 | No Change | Not Significant | TRUE | 0.110610058 |
| Btn2a2 | TRUE | 1.994908742 | 1 | No Change | Not Significant | FALSE | NA |
| Btnl1 | TRUE | -0.603370987 | 0.005170946 | DOWN | DOWN | TRUE | -0.174467013 |
| Btnl10 | TRUE | 1.437913241 | 1 | No Change | Not Significant | FALSE | NA |
| Btnl2 | TRUE | -0.158570466 | 0.833786242 | No Change | Not Significant | TRUE | -0.229916187 |
| Btnl4 | TRUE | -0.266506715 | 0.070728647 | No Change | Not Significant | TRUE | -0.235942505 |
| Btnl6 | TRUE | -0.218548376 | 0.387358014 | No Change | Not Significant | TRUE | -0.297800789 |
| Btnl9 | TRUE | 0.328688612 | 0.810260975 | No Change | Not Significant | TRUE | -0.2268161 |
| Btrc | TRUE | -0.011224434 | 0.95132126 | No Change | Not Significant | TRUE | -0.023291725 |
| Bub1 | TRUE | 0.673809074 | 0.008117753 | UP | UP | TRUE | 0.172162672 |
| Bub1b | TRUE | 0.681261323 | 0.000377711 | UP | UP | TRUE | 0.038161129 |
| Bub3 | TRUE | 0.233088273 | 0.089020995 | No Change | Not Significant | TRUE | 0.149746883 |
| Bud13 | TRUE | 0.029551605 | 0.887804402 | No Change | Not Significant | TRUE | -0.052924182 |
| Bud23 | TRUE | 0.144365477 | 0.233822059 | No Change | Not Significant | TRUE | 0.136559977 |
| Bud31 | TRUE | 0.156416099 | 0.148254918 | No Change | Not Significant | TRUE | 0.023190529 |
| Bysl | TRUE | 0.28176839 | 0.074891965 | No Change | Not Significant | TRUE | 0.249155809 |
| Bzw1 | TRUE | 0.263647282 | 0.003569964 | No Change | Significant Positive | TRUE | 0.093427661 |
| Bzw2 | TRUE | -0.030096289 | 0.89588517 | No Change | Not Significant | TRUE | -0.126325987 |
| C030006K11Rik | TRUE | -1.266885626 | 3.46E-16 | DOWN | DOWN | TRUE | -0.193883421 |
| C030013C21Rik | TRUE | 0.110377278 | 0.88791795 | No Change | Not Significant | TRUE | -0.14663865 |
| C030018K13Rik | TRUE | 0.329042794 | 1 | No Change | Not Significant | FALSE | NA |
| C030029H02Rik | TRUE | -1.1877723 | 1 | No Change | Not Significant | FALSE | NA |
| C030034L19Rik | TRUE | 1.448493493 | 1 | No Change | Not Significant | FALSE | NA |
| C130026I21Rik | TRUE | 0.738071763 | 0.888933284 | No Change | Not Significant | FALSE | NA |
| C130046K22Rik | TRUE | -0.160043341 | 0.836797094 | No Change | Not Significant | TRUE | 0.57912283 |
| C130050O18Rik | TRUE | -1.053775367 | 1 | No Change | Not Significant | FALSE | NA |
| C130071C03Rik | TRUE | -0.42850014 | 1 | No Change | Not Significant | FALSE | NA |
| C130074G19Rik | TRUE | -0.887416913 | 4.35E-07 | DOWN | DOWN | TRUE | -0.03761876 |
| C130083M11Rik | TRUE | 0.058831254 | 0.927499565 | No Change | Not Significant | TRUE | -0.348076842 |
| C1d | TRUE | -0.01468401 | 0.930879221 | No Change | Not Significant | TRUE | 0.043231868 |
| C1galt1 | TRUE | 0.158770538 | 0.508958453 | No Change | Not Significant | TRUE | 0.158512737 |
| C1galt1c1 | TRUE | 0.551378957 | 0.00171518 | UP | UP | TRUE | 0.319555097 |
| C1qa | TRUE | 0.414858017 | 0.640805153 | No Change | Not Significant | TRUE | -0.006175594 |
| C1qb | TRUE | -0.186175041 | 1 | No Change | Not Significant | FALSE | NA |
| C1qbp | TRUE | -0.025385037 | 0.902857271 | No Change | Not Significant | TRUE | -0.033264997 |
| C1qc | TRUE | 1.029034737 | 1 | No Change | Not Significant | FALSE | NA |
| C1ql3 | TRUE | -2.290888723 | 1 | No Change | Not Significant | FALSE | NA |
| C1ql4 | TRUE | 1.707575636 | 1 | No Change | Not Significant | FALSE | NA |
| C1qtnf1 | TRUE | 7.710628117 | 1.50E-260 | UP | UP | TRUE | 6.511563321 |
| C1qtnf12 | TRUE | -1.022506899 | 2.89E-08 | DOWN | DOWN | TRUE | -0.872962618 |
| C1qtnf4 | TRUE | -0.605886954 | 0.149272747 | No Change | Not Significant | TRUE | -0.350540421 |
| C1qtnf6 | TRUE | 0.121669318 | 0.577401503 | No Change | Not Significant | TRUE | 0.017205746 |
| C1qtnf9 | TRUE | -0.81580376 | 1 | No Change | Not Significant | FALSE | NA |
| C1ra | TRUE | -0.889107569 | 0.018083738 | DOWN | DOWN | TRUE | -0.554457376 |
| C1rb | TRUE | 0.521055642 | 1 | No Change | Not Significant | FALSE | NA |
| C1rl | TRUE | 1.161072494 | 0.001937524 | UP | UP | TRUE | -0.053495082 |
| C1s1 | TRUE | 0.482696432 | 1 | No Change | Not Significant | FALSE | NA |
| C1s2 | TRUE | -1.52158501 | 1 | No Change | Not Significant | FALSE | NA |
| C2 | TRUE | -0.23678542 | 0.606725963 | No Change | Not Significant | TRUE | -0.326115541 |
| C230024C17Rik | TRUE | -0.210115282 | 1 | No Change | Not Significant | FALSE | NA |
| C230035I16Rik | TRUE | -0.286911267 | 1 | No Change | Not Significant | TRUE | -0.958421491 |
| C2cd2 | TRUE | -0.229414959 | 0.132283734 | No Change | Not Significant | TRUE | -0.049180417 |
| C2cd2l | TRUE | 0.30217044 | 0.050060192 | No Change | Not Significant | TRUE | 0.088694175 |
| C2cd3 | TRUE | 0.093967873 | 0.520523758 | No Change | Not Significant | TRUE | 0.032989153 |
| C2cd4a | TRUE | 1.895176934 | 9.31E-18 | UP | UP | TRUE | 0.853823656 |
| C2cd4b | TRUE | 0.845433632 | 0.000169939 | UP | UP | TRUE | 0.239672365 |
| C2cd4c | TRUE | -0.444098878 | 0.142911457 | No Change | Not Significant | TRUE | -0.024167028 |
| C2cd4d | TRUE | 0.449623403 | 1 | No Change | Not Significant | TRUE | 1.302580274 |
| C2cd5 | TRUE | 0.303018298 | 0.000123434 | No Change | Significant Positive | TRUE | 0.033024057 |
| C2cd6 | TRUE | 2.852027029 | 1 | No Change | Not Significant | FALSE | NA |
| C3 | TRUE | -0.60718107 | 0.057414402 | No Change | Not Significant | TRUE | 0.194286938 |
| C330018D20Rik | TRUE | -0.181911462 | 0.173330403 | No Change | Not Significant | TRUE | 0.093108966 |
| C330022C24Rik | TRUE | 0.663750616 | 1 | No Change | Not Significant | FALSE | NA |
| C330024D21Rik | TRUE | 2.852576153 | 1 | No Change | Not Significant | FALSE | NA |
| C3ar1 | TRUE | 0.963772876 | 1 | No Change | Not Significant | FALSE | NA |
| C430002N11Rik | TRUE | -0.45938012 | 0.653742472 | No Change | Not Significant | TRUE | -0.490290437 |
| C430042M11Rik | TRUE | 0.037882324 | 0.934200477 | No Change | Not Significant | TRUE | 0.150195909 |
| C4a | TRUE | -0.270547007 | 0.83386713 | No Change | Not Significant | TRUE | -0.189382151 |
| C4b | TRUE | 0.590666306 | 0.710248999 | No Change | Not Significant | FALSE | NA |
| C4bp | TRUE | -0.185603197 | 0.726359126 | No Change | Not Significant | TRUE | 0.159294879 |
| C530005A16Rik | TRUE | 0.044569538 | 0.95200802 | No Change | Not Significant | TRUE | -0.392244136 |

Supplementary Table 3

| geneID | Cond1_DF_Detected | Cond1_DF_ValueCol | Cond1_DF_PadjCol | Cond1_DF_Cutoff | Cond1_DF_Cutoff_Specific.x | Cond2_DF_Detected | Cond2_DF_ValueCol | Cond2_DF_PadjCol | Cond2_DF_Cutoff | Cond2_DF_Cutoff_Specific |
|---|---|---|---|---|---|---|---|---|---|---|
| C5ar2 | TRUE | -0.369051302 | 1 | No Change | Not Significant | FALSE | NA | NA | No Change | Not Detected |
| C630043F03Rik | TRUE | 0.069355483 | 0.86640918 | No Change | Not Significant | TRUE | 0.01650121 | 0.968095473 | No Change | Not Significant |
| C7 | TRUE | -1.692868931 | 1 | No Change | Not Significant | FALSE | NA | NA | No Change | Not Detected |
| C730002L08Rik | TRUE | 0.018421344 | 1 | No Change | Not Significant | TRUE | -0.489292671 | 0.442081119 | No Change | Not Significant |
| C730027H18Rik | TRUE | 0.497794706 | 1 | No Change | Not Significant | FALSE | NA | NA | No Change | Not Detected |
| C77080 | TRUE | -0.068366682 | 0.786448489 | No Change | Not Significant | TRUE | -0.043102055 | 0.761525033 | No Change | Not Significant |
| C86187 | TRUE | 1.569448922 | 1 | No Change | Not Significant | FALSE | NA | NA | No Change | Not Detected |
| C87198 | TRUE | 1.457311324 | 1 | No Change | Not Significant | FALSE | NA | NA | No Change | Not Detected |
| C87436 | TRUE | -0.156473941 | 0.383501006 | No Change | Not Significant | TRUE | 0.002928692 | 0.988536579 | No Change | Not Significant |
| C8a | TRUE | 0.510427675 | 1 | No Change | Not Significant | FALSE | NA | NA | No Change | Not Detected |
| C8b | TRUE | -1.736324486 | 1 | No Change | Not Significant | FALSE | NA | NA | No Change | Not Detected |
| C8g | TRUE | -0.15508757 | 0.670174662 | No Change | Not Significant | TRUE | -0.121069043 | 0.725329993 | No Change | Not Significant |
| C9 | TRUE | 0.408835041 | 0.598287036 | No Change | Not Significant | TRUE | -0.029493164 | 0.973498396 | No Change | Not Significant |
| C920021L13Rik | TRUE | -0.453741391 | 0.559860817 | No Change | Not Significant | TRUE | -0.256138951 | 0.616171935 | No Change | Not Significant |
| C9orf72 | TRUE | 0.28771223 | 0.063213005 | No Change | Not Significant | TRUE | -0.027479312 | 0.873419092 | No Change | Not Significant |
| Caap1 | TRUE | 0.216230368 | 0.082564172 | No Change | Not Significant | TRUE | 0.100131022 | 0.504584307 | No Change | Not Significant |
| Cab39 | TRUE | 0.052775941 | 0.60257474 | No Change | Not Significant | TRUE | 0.062074117 | 0.421276292 | No Change | Not Significant |
| Cab39l | TRUE | -0.167341678 | 0.284857762 | No Change | Not Significant | TRUE | 0.006518155 | 0.961349829 | No Change | Not Significant |
| Cabcoco1 | TRUE | 0.37634742 | 0.106546551 | No Change | Not Significant | TRUE | -0.006811754 | [illegible] | [illegible] | [illegible] |
| Cabin1 | TRUE | 0.125938486 | 0.279625468 | No Change | Not Significant | TRUE | 0.032008529 | [illegible] | [illegible] | [illegible] |
| Cables1 | TRUE | -0.304936894 | 0.173816344 | No Change | Not Significant | TRUE | 0.042242083 | [illegible] | [illegible] | [illegible] |
| Cables2 | TRUE | 0.509220928 | 1.25E-07 | UP | UP | TRUE | 0.090248567 | [illegible] | [illegible] | [illegible] |
| Cabp1 | TRUE | 0.752225396 | 0.583394032 | No Change | Not Significant | FALSE | NA | NA | [illegible] | [illegible] |
| Cabp2 | TRUE | 1.189927792 | 1 | No Change | Not Significant | FALSE | NA | NA | [illegible] | [illegible] |
| Cabp4 | TRUE | 0.503197912 | 0.75067522 | No Change | Not Significant | FALSE | NA | NA | [illegible] | [illegible] |
| Cabp7 | TRUE | 0.21637376 | 1 | No Change | Not Significant | FALSE | NA | NA | [illegible] | [illegible] |
| Cabyr | TRUE | -0.045821259 | 1 | No Change | Not Significant | FALSE | NA | NA | [illegible] | [illegible] |
| [illegible] | [illegible] | [illegible] | [illegible] | [illegible] | [illegible] | [illegible] | [illegible] | [illegible] | [illegible] | [illegible] |

| geneID | BG_Method | RG1_All | RG2_Significant | RG3_SignificantChange | Cond1_DF_ensembl_gene_id | Cond1_DF_entrezgene_id | Cond1_DF_gene_biotype |
|---|---|---|---|---|---|---|---|
| C5ar2 | FALSE | Background = FALSE | Background = FALSE | Background = FALSE | ENSMUSG00000074361 | 319430 | protein_coding |
| C630043F03Rik | TRUE | Cond1 Not Significant + Cond2 Not Significant | None | None | ENSMUSG00000084910 | 68285 | lncRNA |
| C7 | FALSE | Background = FALSE | Background = FALSE | Background = FALSE | ENSMUSG00000079105 | 109828 | protein_coding |
| C730002L08Rik | TRUE | Cond1 Not Significant + Cond2 Not Significant | None | None | ENSMUSG00000114469 | 320155 | lncRNA |
| C730027H18Rik | FALSE | Background = FALSE | Background = FALSE | Background = FALSE | ENSMUSG00000112366 | 319572 | lncRNA |
| C77080 | TRUE | Cond1 Not Significant + Cond2 Not Significant | None | None | ENSMUSG00000050390 | 97130 | protein_coding |
| C86187 | FALSE | Background = FALSE | Background = FALSE | Background = FALSE | ENSMUSG00000110001 | 97402 | lncRNA |
| C87198 | FALSE | Background = FALSE | Background = FALSE | Background = FALSE | ENSMUSG00000092515 | 97836 | lncRNA |
| C87436 | TRUE | Cond1 Not Significant + Cond2 Not Significant | None | None | ENSMUSG00000046679 | 232196 | protein_coding |
| C8a | FALSE | Background = FALSE | Background = FALSE | Background = FALSE | ENSMUSG00000035031 | 230558 | protein_coding |
| C8b | FALSE | Background = FALSE | Background = FALSE | Background = FALSE | ENSMUSG00000029656 | 110382 | protein_coding |
| C8g | TRUE | Cond1 Not Significant + Cond2 Not Significant | None | None | ENSMUSG00000015083 | 69379 | protein_coding |
| C9 | TRUE | Cond1 Not Significant + Cond2 Not Significant | None | None | ENSMUSG00000022149 | 12279 | protein_coding |
| C920021L13Rik | TRUE | Cond1 Not Significant + Cond2 Not Significant | None | None | ENSMUSG00000080727 | 100042889 | lncRNA |
| C9orf72 | TRUE | Cond1 Not Significant + Cond2 Not Significant | None | None | ENSMUSG00000028300 | 73205 | protein_coding |
| Caap1 | TRUE | Cond1 Not Significant + Cond2 Not Significant | None | None | ENSMUSG00000028578 | 67770 | protein_coding |
| Cab39 | TRUE | Cond1 Not Significant + Cond2 Not Significant | None | None | ENSMUSG00000036707 | 12283 | protein_coding |
| Cab39l | TRUE | Cond1 Not Significant + Cond2 Not Significant | None | None | ENSMUSG00000021981 | 69008 | protein_coding |

| geneID | Cond1_DF_baseMean | Cond1_DF_log2FoldChange | Cond1_DF_lfcSE | Cond1_DF_stat | Cond1_DF_pvalue | Cond1_DF_padj | Cond1_DF_absLog2FC | Cond1_DF_Cutoff_Specific.y |
|---|---|---|---|---|---|---|---|---|
| C5ar2 | 1.769504278 | -0.369051302 | 1.869579998 | 0.197397973 | 0.843516116 | 1 | 0.369051302 | Not Significant |
| C630043F03Rik | 23.65459368 | 0.069355483 | 0.206373945 | -0.33606705 | 0.736820305 | 0.86640918 | 0.069355483 | Not Significant |
| C7 | 1.342631701 | -1.692868931 | 2.211809329 | 0.765377426 | 0.444046831 | 1 | 1.692868931 | Not Significant |
| C730002L08Rik | 4.338382363 | 0.018421344 | 0.672139094 | -0.027407042 | 0.978135082 | 1 | 0.018421344 | Not Significant |
| C730027H18Rik | 1.115101584 | 0.497794706 | 2.02113682 | -0.246294413 | 0.805454335 | 1 | 0.497794706 | Not Significant |
| C77080 | 2537.273468 | -0.068366682 | 0.135404491 | 0.504907048 | 0.613624124 | 0.786448489 | 0.068366682 | Not Significant |
| C86187 | 1.101541491 | 1.569448922 | 1.84403727 | -0.851093927 | 0.394717179 | 1 | 1.569448922 | Not Significant |
| C87198 | 1.496234869 | 1.457311324 | 2.383530159 | -0.611408804 | 0.540928976 | 1 | 1.457311324 | Not Significant |
| C87436 | 142.7843888 | -0.156473941 | 0.118682469 | 1.318425058 | 0.187361397 | 0.383501006 | 0.156473941 | Not Significant |
| C8a | 1.067705722 | 0.510427675 | 2.277989343 | -0.224069387 | 0.822703313 | 1 | 0.510427675 | Not Significant |
| C8b | 1.777951398 | -1.736324486 | 2.231920874 | 0.777950736 | 0.436598056 | 1 | 1.736324486 | Not Significant |
| C8g | 24.06824754 | -0.15508757 | 0.207457481 | 0.747563157 | 0.454723695 | 0.670174662 | 0.15508757 | Not Significant |
| C9 | 6.463641515 | 0.408835041 | 0.461864913 | -0.885183155 | 0.376057856 | 0.598287036 | 0.408835041 | Not Significant |
| C920021L13Rik | 7.519670238 | -0.453741391 | 0.473741766 | 0.957782116 | 0.338172639 | 0.559860817 | 0.453741391 | Not Significant |
| C9orf72 | 158.0226828 | 0.28771223 | 0.12004144 | -2.396774239 | 0.016540111 | 0.063213005 | 0.28771223 | Not Significant |
| Caap1 | 128.4473468 | 0.216230368 | 0.095287174 | -2.269249456 | 0.02325316 | 0.082564172 | 0.216230368 | Not Significant |
| Cab39 | 3787.699087 | 0.052775941 | 0.060166226 | -0.877168878 | 0.380394918 | 0.60257474 | 0.052775941 | Not Significant |
| Cab39l | 465.6122326 | -0.167341678 | 0.108182978 | 1.546839259 | 0.121902011 | 0.284857762 | 0.167341678 | Not Significant |

| geneID | Cond2_DF_ensembl_gene_id | Cond2_DF_entrezgene_id | Cond2_DF_gene_biotype | Cond2_DF_baseMean | Cond2_DF_log2FoldChange | Cond2_DF_lfcSE | Cond2_DF_stat | Cond2_DF_pvalue | Cond2_DF_padj | Cond2_DF_absLog2FC | RG1_Specific_Cond2 | RG1_Specific_Cond1 |
|---|---|---|---|---|---|---|---|---|---|---|---|---|
| C5ar2 | NA | NA | NA | NA | NA | NA | NA | NA | NA | NA | Background = FALSE | Background = FALSE |
| C630043F03Rik | ENSMUSG00000084910 | 68285 | lncRNA | 51.58932497 | 0.01650121 | 0.181391233 | 0.090970274 | 0.927516211 | 0.968095473 | 0.01650121 | Cond1 No Change + Cond2 Not Significant | Cond2 No Change + Cond1 Not Significant |
| C7 | NA | NA | NA | NA | NA | NA | NA | NA | NA | NA | Background = FALSE | Background = FALSE |
| C730002L08Rik | ENSMUSG00000114469 | 320155 | lncRNA | 11.89754733 | -0.489292671 | 0.387772113 | -1.26180469 | 0.207019074 | 0.442081119 | 0.489292671 | Cond1 No Change + Cond2 Not Significant | Cond2 No Change + Cond1 Not Significant |
| C730027H18Rik | NA | NA | NA | NA | NA | NA | NA | NA | NA | NA | Background = FALSE | Background = FALSE |
| C77080 | ENSMUSG00000050390 | 97130 | protein_coding | 4726.701024 | -0.043102055 | 0.074857261 | -0.575789899 | 0.564757202 | 0.761525033 | 0.043102055 | Cond1 No Change + Cond2 Not Significant | Cond2 No Change + Cond1 Not Significant |
| C86187 | NA | NA | NA | NA | NA | NA | NA | NA | NA | NA | Background = FALSE | Background = FALSE |
| C87198 | NA | NA | NA | NA | NA | NA | NA | NA | NA | NA | Background = FALSE | Background = FALSE |
| C87436 | ENSMUSG00000046679 | 232196 | protein_coding | 297.1354016 | 0.002928692 | 0.081611451 | 0.035885796 | 0.971373422 | 0.988536579 | 0.002928692 | Cond1 No Change + Cond2 Not Significant | Cond2 No Change + Cond1 Not Significant |
| C8a | NA | NA | NA | NA | NA | NA | NA | NA | NA | NA | Background = FALSE | Background = FALSE |
| C8b | NA | NA | NA | NA | NA | NA | NA | NA | NA | NA | Background = FALSE | Background = FALSE |
| C8g | ENSMUSG00000015083 | 69379 | protein_coding | 50.66032756 | -0.121069043 | 0.184283298 | -0.656972414 | 0.511198654 | 0.725329993 | 0.121069043 | Cond1 No Change + Cond2 Not Significant | Cond2 No Change + Cond1 Not Significant |
| C9 | ENSMUSG00000022149 | 12279 | protein_coding | 14.10181921 | -0.029493164 | 0.387896995 | -0.076033495 | 0.93939245 | 0.973498396 | 0.029493164 | Cond1 No Change + Cond2 Not Significant | Cond2 No Change + Cond1 Not Significant |
| C920021L13Rik | ENSMUSG00000080727 | 100042889 | lncRNA | 18.37218659 | -0.256138951 | 0.288310668 | -0.888413021 | 0.374318624 | 0.616171935 | 0.256138951 | Cond1 No Change + Cond2 Not Significant | Cond2 No Change + Cond1 Not Significant |
| C9orf72 | ENSMUSG00000028300 | 73205 | protein_coding | 267.9608266 | -0.027479312 | 0.081631083 | -0.336628049 | 0.736397309 | 0.873419092 | 0.027479312 | Cond1 No Change + Cond2 Not Significant | Cond2 No Change + Cond1 Not Significant |
| Caap1 | ENSMUSG00000028578 | 67770 | protein_coding | 225.424921 | 0.100131022 | 0.089387551 | 1.12018979 | 0.262632894 | 0.504584307 | 0.100131022 | Cond1 No Change + Cond2 Not Significant | Cond2 No Change + Cond1 Not Significant |
| Cab39 | ENSMUSG00000036707 | 12283 | protein_coding | 7020.978477 | 0.062074117 | 0.047424092 | 1.308915236 | 0.190563065 | 0.421276292 | 0.062074117 | Cond1 No Change + Cond2 Not Significant | Cond2 No Change + Cond1 Not Significant |
| Cab39l | ENSMUSG00000021981 | 69008 | protein_coding | 942.9451527 | 0.006518155 | 0.059745597 | 0.109098496 | 0.913124368 | 0.961349829 | 0.006518155 | Cond1 No Change + Cond2 Not Significant | Cond2 No Change + Cond1 Not Significant |

| geneID | Cond1_DF_Detected | Cond1_DF_ValueCol | Cond1_DF_PadjCol | Cond1_DF_Cutoff | Cond1_DF_Cutoff_Specific.x | Cond2_DF_Detected | Cond2_DF_ValueCol | Cond2_DF_PadjCol | Cond2_DF_Cutoff | Cond2_DF_Cutoff_Specific | BG_Method |
|---|---|---|---|---|---|---|---|---|---|---|---|
| Catsper1 | TRUE | 1.901344307 | 1 | No Change | Not Significant | FALSE | NA | NA | No Change | Not Detected | FALSE |
| Catsper2 | TRUE | 0.692327015 | 0.056215521 | No Change | Not Significant | TRUE | 0.805015419 | 0.000714599 | UP | UP | TRUE |
| Catsper3 | TRUE | 2.08287833 | 1 | No Change | Not Significant | FALSE | NA | NA | No Change | Not Detected | FALSE |
| Catsper4 | TRUE | 0.891191736 | 0.165304689 | No Change | Not Significant | TRUE | 0.043326805 | 0.950993787 | No Change | Not Significant | TRUE |
| Catsperb | TRUE | -0.890147185 | 1 | No Change | Not Significant | FALSE | NA | NA | No Change | Not Detected | FALSE |
| Catsperd | TRUE | -1.233214455 | 0.191288267 | No Change | Not Significant | TRUE | -0.634168424 | 0.282646118 | No Change | Not Significant | TRUE |
| Catspere2 | TRUE | 0.053246542 | 0.930374388 | No Change | Not Significant | TRUE | 0.380394884 | 0.23750328 | No Change | Not Significant | TRUE |
| Catsperg1 | TRUE | 0.350574887 | 0.793088519 | No Change | Not Significant | FALSE | NA | NA | No Change | Not Detected | FALSE |
| Catsperg2 | TRUE | 2.142666942 | 1 | No Change | Not Significant | FALSE | NA | NA | No Change | Not Detected | FALSE |
| Catsperz | TRUE | 0.448574867 | 0.529213766 | No Change | Not Significant | TRUE | -0.028207369 | 0.970732425 | No Change | Not Significant | TRUE |
| Cav1 | TRUE | 0.508178023 | 1 | No Change | Not Significant | TRUE | 2.079235365 | 0.00710592 | UP | UP | TRUE |
| Cav2 | TRUE | -0.510403562 | 0.145821162 | No Change | Not Significant | TRUE | 0.355803709 | 0.042967024 | No Change | Significant Positive | TRUE |
| Cavin1 | TRUE | -0.647184841 | 0.02193128 | DOWN | DOWN | TRUE | -0.002701836 | 0.995823024 | No Change | Not Significant | TRUE |
| Cavin2 | TRUE | -0.874371668 | 1 | No Change | Not Significant | FALSE | NA | NA | No Change | Not Detected | FALSE |
| Cavin3 | TRUE | -0.38214037 | 0.490623547 | No Change | Not Significant | TRUE | 0.884107082 | 0.011821691 | UP | UP | TRUE |
| Cavin4 | TRUE | 1.215392238 | 1 | No Change | Not Significant | FALSE | NA | NA | No Change | Not Detected | FALSE |
| Cbarp | TRUE | -0.199842875 | 0.668802273 | No Change | Not Significant | TRUE | -0.564032306 | 0.04505532 | DOWN | DOWN | TRUE |
| Cbfa2t2 | TRUE | -0.018215001 | 0.879422681 | No Change | Not Significant | TRUE | -0.011130241 | 0.927560229 | No Change | Not Significant | TRUE |
| Cbfa2t3 | TRUE | -0.361963822 | 0.003828526 | No Change | Significant Negative | TRUE | -0.208432884 | 0.133784989 | No Change | Not Significant | TRUE |
| Cbfb | TRUE | 0.212476646 | 0.029324606 | No Change | Significant Positive | TRUE | -0.010249337 | 0.940691929 | No Change | Not Significant | TRUE |
| Cbl | TRUE | -0.218044978 | 0.232062395 | No Change | Not Significant | TRUE | -0.226293082 | 0.030989506 | No Change | Significant Negative | TRUE |
| Cblb | TRUE | -0.126296637 | 0.474080016 | No Change | Not Significant | TRUE | -0.137004716 | 0.270928354 | No Change | Not Significant | TRUE |
| Cblc | TRUE | -0.145736187 | 0.184267881 | No Change | Not Significant | TRUE | 0.155615078 | 0.171421356 | No Change | Not Significant | TRUE |
| Cblif | TRUE | -0.315448324 | 1 | No Change | Not Significant | FALSE | NA | NA | No Change | Not Detected | FALSE |
| Cbll1 | TRUE | -0.083722358 | 0.772293296 | No Change | Not Significant | TRUE | -0.238999596 | 0.21752641 | No Change | Not Significant | TRUE |
| Cbln3 | TRUE | 0.990469348 | 4.27E-07 | UP | UP | TRUE | 0.309902693 | 0.168180809 | No Change | Not Significant | TRUE |
| Cbln4 | TRUE | 0.689447912 | 1 | No Change | Not Significant | FALSE | NA | NA | No Change | Not Detected | FALSE |
| Cbr1 | TRUE | -0.409425473 | 0.001225641 | No Change | Significant Negative | TRUE | -0.015836032 | 0.923652088 | No Change | Not Significant | TRUE |
| Cbr2 | TRUE | 1.286626969 | 3.53E-06 | UP | UP | TRUE | 0.646466725 | 0.000170681 | UP | UP | TRUE |
| Cbr3 | TRUE | -0.624685924 | 0.00748498 | DOWN | DOWN | TRUE | 0.269698538 | 0.284673902 | No Change | Not Significant | TRUE |
| Cbr4 | TRUE | -0.241766209 | 0.144615997 | No Change | Not Significant | TRUE | 0.007031667 | 0.968789653 | No Change | Not Significant | TRUE |
| Cbs | TRUE | -0.194737851 | 0.28232699 | No Change | Not Significant | TRUE | 0.060671206 | 0.623211682 | No Change | Not Significant | TRUE |
| Cbwd1 | TRUE | 0.363233685 | 0.007622708 | No Change | Significant Positive | TRUE | 0.101811079 | 0.453747313 | No Change | Not Significant | TRUE |
| Cbx1 | TRUE | 0.131589435 | 0.463479905 | No Change | Not Significant | TRUE | 0.081965743 | 0.394876088 | No Change | Not Significant | TRUE |
| Cbx2 | TRUE | -0.203022728 | 0.365420225 | No Change | Not Significant | TRUE | 0.041084752 | 0.839494578 | No Change | Not Significant | TRUE |
| Cbx3 | TRUE | 0.082198381 | 0.754536534 | No Change | Not Significant | TRUE | -0.005461956 | 0.980775949 | No Change | Not Significant | TRUE |
| Cbx4 | TRUE | 0.739831594 | 9.54E-08 | UP | UP | TRUE | 0.092827094 | 0.605769124 | No Change | Not Significant | TRUE |
| Cbx5 | TRUE | 0.435816628 | 0.017949136 | No Change | Significant Positive | TRUE | 0.15034402 | 0.183993767 | No Change | Not Significant | TRUE |
| Cbx6 | TRUE | 0.412682315 | 0.170033542 | No Change | Not Significant | TRUE | -0.128515031 | 0.548477307 | No Change | Not Significant | TRUE |
| Cbx7 | TRUE | -0.701451449 | 8.41E-09 | DOWN | DOWN | TRUE | -0.666461873 | 9.22E-09 | DOWN | DOWN | TRUE |
| Cbx8 | TRUE | 0.285718204 | 0.061382313 | No Change | Not Significant | TRUE | 0.26062552 | 0.034804603 | No Change | Significant Positive | TRUE |
| Cby1 | TRUE | 0.222237512 | 0.233277344 | No Change | Not Significant | TRUE | 0.294023559 | 0.003775888 | No Change | Significant Positive | TRUE |
| Cby2 | TRUE | 2.217268286 | 1 | No Change | Not Significant | FALSE | NA | NA | No Change | Not Detected | FALSE |
| Cc2d1a | TRUE | -0.086355597 | 0.57545396 | No Change | Not Significant | TRUE | 0.104017345 | 0.219914726 | No Change | Not Significant | TRUE |
| Cc2d1b | TRUE | 0.040816689 | 0.702242166 | No Change | Not Significant | TRUE | 0.098030709 | 0.12411837 | No Change | Not Significant | TRUE |
| Cc2d2a | TRUE | 0.011849771 | 0.956997404 | No Change | Not Significant | TRUE | -0.255922776 | 0.015822676 | No Change | Significant Negative | TRUE |
| Cc2d2b | TRUE | 0.994195709 | 1 | No Change | Not Significant | FALSE | NA | NA | No Change | Not Detected | FALSE |
| Ccar1 | TRUE | 0.182135619 | 0.001108596 | No Change | Significant Positive | TRUE | 0.052403531 | 0.657986673 | No Change | Not Significant | TRUE |
| Ccar2 | TRUE | -3.23E-05 | 0.999654528 | No Change | Not Significant | TRUE | -0.039310425 | 0.655860109 | No Change | Not Significant | TRUE |
| Ccbe1 | TRUE | 0.170767771 | 0.965886893 | No Change | Not Significant | FALSE | NA | NA | No Change | Not Detected | FALSE |
| Ccdc102a | TRUE | -0.676321316 | 0.408576247 | No Change | Not Significant | TRUE | 0.095371823 | 0.926721051 | No Change | Not Significant | TRUE |
| Ccdc103 | TRUE | -0.707948487 | 1 | No Change | Not Significant | FALSE | NA | NA | No Change | Not Detected | FALSE |
| Ccdc106 | TRUE | 0.329518308 | 1 | No Change | Not Significant | FALSE | NA | NA | No Change | Not Detected | FALSE |
| Ccdc107 | TRUE | -0.519251682 | 1.22E-07 | DOWN | DOWN | TRUE | -0.360365713 | 4.67E-06 | No Change | Significant Negative | TRUE |
| Ccdc112 | TRUE | 0.287904226 | 0.347734431 | No Change | Not Significant | TRUE | 0.359751031 | 0.093968897 | No Change | Not Significant | TRUE |
| Ccdc115 | TRUE | 0.011535229 | 0.965151432 | No Change | Not Significant | TRUE | 0.026524308 | 0.87279256 | No Change | Not Significant | TRUE |
| Ccdc116 | TRUE | 0.877013755 | 0.00298846 | UP | UP | TRUE | -0.03922473 | 0.890119548 | No Change | Not Significant | TRUE |
| Ccdc117 | TRUE | -0.041536268 | 0.803690806 | No Change | Not Significant | TRUE | -0.104408871 | 0.21967756 | No Change | Not Significant | TRUE |
| Ccdc12 | TRUE | -0.063366826 | 0.713811425 | No Change | Not Significant | TRUE | 0.180040059 | 0.012866746 | No Change | Significant Positive | TRUE |
| Ccdc120 | TRUE | -0.441077938 | 0.068717608 | No Change | Not Significant | TRUE | -0.143789703 | 0.410507024 | No Change | Not Significant | TRUE |
| Ccdc121rt3 | TRUE | 0.9125183 | 0.360508479 | No Change | Not Significant | FALSE | NA | NA | No Change | Not Detected | FALSE |
| Ccdc122 | TRUE | 0.128658723 | 0.607820554 | No Change | Not Significant | TRUE | -0.030561567 | 0.891239011 | No Change | Not Significant | TRUE |
| Ccdc124 | TRUE | 0.129768613 | 0.383442977 | No Change | Not Significant | TRUE | 0.160066391 | 0.381379516 | No Change | Not Significant | TRUE |
| Ccdc125 | TRUE | -0.079648607 | 0.782404996 | No Change | Not Significant | TRUE | 0.04678464 | 0.846497211 | No Change | Not Significant | TRUE |
| Ccdc126 | TRUE | 0.033173133 | 0.938630585 | No Change | Not Significant | TRUE | 0.358822081 | 0.084790715 | No Change | Not Significant | TRUE |
| Ccdc127 | TRUE | 0.327624074 | 0.007553859 | No Change | Significant Positive | TRUE | 0.169920625 | 0.029032271 | No Change | Significant Positive | TRUE |
| Ccdc13 | TRUE | 0.489531188 | 1 | No Change | Not Significant | FALSE | NA | NA | No Change | Not Detected | FALSE |
| Ccdc130 | TRUE | 0.057945349 | 0.803288086 | No Change | Not Significant | TRUE | 0.015327849 | 0.941223339 | No Change | Not Significant | TRUE |
| Ccdc134 | TRUE | 2.400035664 | 4.94E-77 | UP | UP | TRUE | 1.343611603 | 6.68E-34 | UP | UP | TRUE |
| Ccdc136 | TRUE | -0.356143431 | 0.769577776 | No Change | Not Significant | TRUE | -0.15846539 | 0.865841503 | No Change | Not Significant | TRUE |
| Ccdc137 | TRUE | -0.115779276 | 0.388417088 | No Change | Not Significant | TRUE | -0.023144972 | 0.885925238 | No Change | Not Significant | TRUE |
| Ccdc138 | TRUE | -0.038320542 | 0.911456552 | No Change | Not Significant | TRUE | 0.085405686 | 0.708115319 | No Change | Not Significant | TRUE |
| Ccdc14 | TRUE | -0.089607611 | 0.75067522 | No Change | Not Significant | TRUE | -0.058497246 | 0.815430435 | No Change | Not Significant | TRUE |
| Ccdc141 | TRUE | -0.112855343 | 0.874376659 | No Change | Not Significant | TRUE | -0.493806763 | 0.149514754 | No Change | Not Significant | TRUE |
| Ccdc142 | TRUE | 0.234636888 | 1 | No Change | Not Significant | FALSE | NA | NA | No Change | Not Detected | FALSE |
| Ccdc144b | TRUE | 3.219147383 | 1 | No Change | Not Significant | FALSE | NA | NA | No Change | Not Detected | FALSE |
| Ccdc146 | TRUE | 0.403355433 | 1 | No Change | Not Significant | FALSE | NA | NA | No Change | Not Detected | FALSE |
| Ccdc148 | TRUE | -0.627978064 | 0.575410323 | No Change | Not Significant | TRUE | -0.144980364 | 0.783232176 | No Change | Not Significant | TRUE |
| Ccdc149 | TRUE | 0.905499763 | 5.90E-21 | UP | UP | TRUE | 0.69947706 | 7.38E-15 | UP | UP | TRUE |
| Ccdc15 | TRUE | 0.140493183 | 0.687930892 | No Change | Not Significant | TRUE | -0.180524734 | 0.590760052 | No Change | Not Significant | TRUE |
| Ccdc150 | TRUE | -0.070766411 | 1 | No Change | Not Significant | FALSE | NA | NA | No Change | Not Detected | FALSE |
| Ccdc152 | TRUE | -0.714612007 | 1 | No Change | Not Significant | TRUE | -0.158636229 | 0.892439856 | No Change | Not Significant | TRUE |
| Ccdc153 | TRUE | -0.385949568 | 1 | No Change | Not Significant | FALSE | NA | NA | No Change | Not Detected | FALSE |
| Ccdc154 | TRUE | 0.39518159 | 1 | No Change | Not Significant | FALSE | NA | NA | No Change | Not Detected | FALSE |
| Ccdc157 | TRUE | 0.077995952 | 0.767158627 | No Change | Not Significant | TRUE | 0.084478494 | 0.705097682 | No Change | Not Significant | TRUE |
| Ccdc158 | TRUE | -0.665419524 | 0.462129017 | No Change | Not Significant | TRUE | -0.321981736 | 0.734592691 | No Change | Not Significant | TRUE |
| Ccdc159 | TRUE | -0.2021179 | 0.701011313 | No Change | Not Significant | TRUE | -0.123791702 | 0.792525305 | No Change | Not Significant | TRUE |
| Ccdc160 | TRUE | -0.964315377 | 1 | No Change | Not Significant | TRUE | 0.022894655 | 0.981205596 | No Change | Not Significant | TRUE |
| Ccdc162 | TRUE | -0.335436347 | 0.744442234 | No Change | Not Significant | TRUE | -0.188407091 | 0.827707324 | No Change | Not Significant | TRUE |
| Ccdc163 | TRUE | -0.153792249 | 0.451799148 | No Change | Not Significant | TRUE | -0.143182029 | 0.495031103 | No Change | Not Significant | TRUE |
| Ccdc166 | TRUE | -0.277208705 | 0.231384745 | No Change | Not Significant | TRUE | 0.007612584 | 0.981205596 | No Change | Not Significant | TRUE |
| Ccdc167 | TRUE | 1.034996974 | 6.48E-09 | UP | UP | TRUE | 0.27071475 | 0.306582205 | No Change | Not Significant | TRUE |
| Ccdc169 | TRUE | 1.759312202 | 1 | No Change | Not Significant | FALSE | NA | NA | No Change | Not Detected | FALSE |
| Ccdc17 | TRUE | -0.330149738 | 0.148427209 | No Change | Not Significant | TRUE | -0.00924559 | 0.97697769 | No Change | Not Significant | TRUE |
| Ccdc170 | TRUE | -2.151079925 | 1 | No Change | Not Significant | FALSE | NA | NA | No Change | Not Detected | FALSE |
| Ccdc171 | TRUE | 0.244119833 | 0.360812744 | No Change | Not Significant | TRUE | -0.128357499 | 0.611697618 | No Change | Not Significant | TRUE |
| Ccdc173 | TRUE | 0.425730796 | 0.462725114 | No Change | Not Significant | TRUE | -0.660623368 | 0.286840423 | No Change | Not Significant | TRUE |
| Ccdc174 | TRUE | -0.140756955 | 0.172473724 | No Change | Not Significant | TRUE | -0.055070009 | 0.699497125 | No Change | Not Significant | TRUE |
| Ccdc175 | TRUE | -0.673301145 | 1 | No Change | Not Significant | FALSE | NA | NA | No Change | Not Detected | FALSE |
| Ccdc177 | TRUE | 0.237598968 | 0.727438372 | No Change | Not Significant | TRUE | -0.044092463 | 0.944199804 | No Change | Not Significant | TRUE |
| Ccdc178 | TRUE | -0.428279948 | 1 | No Change | Not Significant | FALSE | NA | NA | No Change | Not Detected | FALSE |
| Ccdc18 | TRUE | 0.491464478 | 0.027715628 | No Change | Significant Positive | TRUE | -0.057506541 | 0.801438232 | No Change | Not Significant | TRUE |
| Ccdc180 | TRUE | 0.936220928 | 1 | No Change | Not Significant | FALSE | NA | NA | No Change | Not Detected | FALSE |
| Ccdc181 | TRUE | -0.060088481 | 0.76182422 | No Change | Not Significant | TRUE | 0.243853982 | 0.052776264 | No Change | Not Significant | TRUE |
| Ccdc183 | TRUE | -0.091623253 | 1 | No Change | Not Significant | FALSE | NA | NA | No Change | Not Detected | FALSE |
| Ccdc184 | TRUE | -0.749186619 | 1 | No Change | Not Significant | FALSE | NA | NA | No Change | Not Detected | FALSE |
| Ccdc186 | TRUE | 0.027722065 | 0.870032583 | No Change | Not Significant | TRUE | -0.079721446 | 0.209031847 | No Change | Not Significant | TRUE |
| Ccdc187 | TRUE | -0.923928782 | 1 | No Change | Not Significant | TRUE | -0.086385478 | 0.929613548 | No Change | Not Significant | TRUE |
| Ccdc188 | TRUE | 1.020863456 | 1 | No Change | Not Significant | FALSE | NA | NA | No Change | Not Detected | FALSE |
| Ccdc189 | TRUE | -1.16506603 | 0.000180539 | DOWN | DOWN | TRUE | -0.658535521 | 0.05184024 | No Change | Not Significant | TRUE |
| Ccdc191 | TRUE | -0.27871388 | 0.114868009 | No Change | Not Significant | TRUE | 0.017483707 | 0.937561241 | No Change | Not Significant | TRUE |
| Ccdc192 | TRUE | 0.346384551 | 1 | No Change | Not Significant | FALSE | NA | NA | No Change | Not Detected | FALSE |
| ccdc198 | TRUE | -0.317645901 | 0.119750527 | No Change | Not Significant | TRUE | -0.115008369 | 0.677580465 | No Change | Not Significant | TRUE |
| Ccdc200 | TRUE | -0.04616548 | 1 | No Change | Not Significant | FALSE | NA | NA | No Change | Not Detected | FALSE |
| Ccdc24 | TRUE | 0.164782867 | 1 | No Change | Not Significant | FALSE | NA | NA | No Change | Not Detected | FALSE |
| Ccdc25 | TRUE | -0.02216458 | 0.913524643 | No Change | Not Significant | TRUE | 0.029355347 | 0.858813602 | No Change | Not Significant | TRUE |
| Ccdc27 | TRUE | -0.29986656 | 1 | No Change | Not Significant | FALSE | NA | NA | No Change | Not Detected | FALSE |
| Ccdc28a | TRUE | -0.169433923 | 0.386024762 | No Change | Not Significant | TRUE | -0.090422405 | 0.470652152 | No Change | Not Significant | TRUE |
| Ccdc28b | TRUE | -0.609058719 | 2.13E-06 | DOWN | DOWN | TRUE | -0.267894979 | 0.005628532 | No Change | Significant Negative | TRUE |
| Ccdc3 | TRUE | 0.991462961 | 0.003668869 | UP | UP | TRUE | 0.692887692 | 0.023394202 | UP | UP | TRUE |
| Ccdc30 | TRUE | 1.062025068 | 0.388417088 | No Change | Not Significant | FALSE | NA | NA | No Change | Not Detected | FALSE |
| Ccdc32 | TRUE | -0.174238427 | 0.407126819 | No Change | Not Significant | TRUE | 0.090480817 | 0.470523296 | No Change | Not Significant | TRUE |
| Ccdc33 | TRUE | 1.375786748 | 1 | No Change | Not Significant | FALSE | NA | NA | No Change | Not Detected | FALSE |
| Ccdc34 | TRUE | 0.322103617 | 0.083737724 | No Change | Not Significant | TRUE | 0.086229607 | 0.586128211 | No Change | Not Significant | TRUE |
| Ccdc34os | TRUE | 0.143220052 | 1 | No Change | Not Significant | FALSE | NA | NA | No Change | Not Detected | FALSE |
| Ccdc38 | TRUE | 0.382238788 | 0.763773846 | No Change | Not Significant | FALSE | NA | NA | No Change | Not Detected | FALSE |
| Ccdc39 | TRUE | 0.220425022 | 0.876249099 | No Change | Not Significant | FALSE | NA | NA | No Change | Not Detected | FALSE |
| Ccdc40 | TRUE | -0.077912531 | 1 | No Change | Not Significant | FALSE | NA | NA | No Change | Not Detected | FALSE |
| Ccdc43 | TRUE | -0.011406851 | 0.94517522 | No Change | Not Significant | TRUE | -0.018634085 | 0.910796813 | No Change | Not Significant | TRUE |
| Ccdc47 | TRUE | -0.034138204 | 0.848979902 | No Change | Not Significant | TRUE | 0.020357747 | 0.822489025 | No Change | Not Significant | TRUE |
| Ccdc50 | TRUE | -0.257350533 | 0.001312529 | No Change | Significant Negative | TRUE | -0.173708458 | 0.037290326 | No Change | Significant Negative | TRUE |
| Ccdc51 | TRUE | -0.164825682 | 0.264619521 | No Change | Not Significant | TRUE | 0.052496355 | 0.685788934 | No Change | Not Significant | TRUE |
| Ccdc57 | TRUE | -0.160201647 | 0.659670632 | No Change | Not Significant | TRUE | -0.264819374 | 0.243800777 | No Change | Not Significant | TRUE |
| Ccdc58 | TRUE | -0.008032453 | 0.962383589 | No Change | Not Significant | TRUE | -0.037792764 | 0.759206347 | No Change | Not Significant | TRUE |
| Ccdc59 | TRUE | 0.087384268 | 0.424883585 | No Change | Not Significant | TRUE | -0.092442734 | 0.463452522 | No Change | Not Significant | TRUE |
| Ccdc6 | TRUE | 0.253030164 | 0.010450353 | No Change | Significant Positive | TRUE | 0.056939524 | 0.580250264 | No Change | Not Significant | TRUE |
| Ccdc60 | TRUE | 0.434549439 | 1 | No Change | Not Significant | FALSE | NA | NA | No Change | Not Detected | FALSE |
| Ccdc61 | TRUE | 0.264992986 | 0.004340608 | No Change | Significant Positive | TRUE | 0.062470442 | 0.668067439 | No Change | Not Significant | TRUE |
| Ccdc62 | TRUE | 0.258752684 | 0.671491432 | No Change | Not Significant | TRUE | -0.137657539 | 0.812013688 | No Change | Not Significant | TRUE |
| Ccdc63 | TRUE | -0.139580788 | 1 | No Change | Not Significant | FALSE | NA | NA | No Change | Not Detected | FALSE |
| Ccdc65 | TRUE | 0.743502144 | 1 | No Change | Not Significant | FALSE | NA | NA | No Change | Not Detected | FALSE |
| Ccdc66 | TRUE | 0.405024683 | 9.65E-05 | No Change | Significant Positive | TRUE | 0.19286477 | 0.245659762 | No Change | Not Significant | TRUE |
| Ccdc68 | TRUE | -0.174607454 | 0.5416756 | No Change | Not Significant | TRUE | -0.001098253 | 0.995814509 | No Change | Not Significant | TRUE |
| Ccdc69 | TRUE | -0.121250271 | 0.903305601 | No Change | Not Significant | TRUE | 0.247977493 | 0.740799392 | No Change | Not Significant | TRUE |
| Ccdc71 | TRUE | -0.006868352 | 0.962436702 | No Change | Not Significant | TRUE | 0.028407375 | 0.778426904 | No Change | Not Significant | TRUE |
| Ccdc71l | TRUE | 0.484978735 | 0.007435174 | No Change | Significant Positive | TRUE | 0.16703593 | 0.201136435 | No Change | Not Significant | TRUE |
| Ccdc73 | TRUE | -0.194918901 | 0.767207271 | No Change | Not Significant | TRUE | -0.100881423 | 0.874869968 | No Change | Not Significant | TRUE |
| Ccdc74a | TRUE | 0.348495088 | 0.597844879 | No Change | Not Significant | TRUE | -0.150992702 | 0.789753 | No Change | Not Significant | TRUE |
| Ccdc77 | TRUE | 0.3603478 | 0.027823083 | No Change | Significant Positive | TRUE | -0.20434264 | 0.248883957 | No Change | Not Significant | TRUE |
| Ccdc78 | TRUE | 0.481294859 | 1 | No Change | Not Significant | FALSE | NA | NA | No Change | Not Detected | FALSE |
| Ccdc7a | TRUE | -0.227145412 | 0.903566201 | No Change | Not Significant | FALSE | NA | NA | No Change | Not Detected | FALSE |
| Ccdc8 | TRUE | 0.132473234 | 1 | No Change | Not Significant | FALSE | NA | NA | No Change | Not Detected | FALSE |
| Ccdc80 | TRUE | -0.654696908 | 1 | No Change | Not Significant | FALSE | NA | NA | No Change | Not Detected | FALSE |
| Ccdc81 | TRUE | 0.913024815 | 1 | No Change | Not Significant | FALSE | NA | NA | No Change | Not Detected | FALSE |
| Ccdc82 | TRUE | -0.441393526 | 0.004881237 | No Change | Significant Negative | TRUE | -0.34235186 | 0.043098347 | No Change | Significant Negative | TRUE |
| Ccdc83 | TRUE | -1.49781879 | 1 | No Change | Not Significant | FALSE | NA | NA | No Change | Not Detected | FALSE |
| Ccdc84 | TRUE | 0.192134521 | 0.405231932 | No Change | Not Significant | TRUE | -0.175740061 | 0.51411852 | No Change | Not Significant | TRUE |
| Ccdc85a | TRUE | -1.091858734 | 1 | No Change | Not Significant | FALSE | NA | NA | No Change | Not Detected | FALSE |
| Ccdc85b | TRUE | -0.290324241 | 0.094754639 | No Change | Not Significant | TRUE | -0.259133636 | 0.054260619 | No Change | Not Significant | TRUE |
| Ccdc85c | TRUE | 0.212380865 | 0.066769512 | No Change | Not Significant | TRUE | 0.137013414 | 0.376633179 | No Change | Not Significant | TRUE |
| Ccdc86 | TRUE | -0.112338445 | 0.673388737 | No Change | Not Significant | TRUE | -0.049744085 | 0.822097164 | No Change | Not Significant | TRUE |
| Ccdc87 | TRUE | 0.559110095 | 1 | No Change | Not Significant | FALSE | NA | NA | No Change | Not Detected | FALSE |
| Ccdc88a | TRUE | 1.07759181 | 0.008911609 | UP | UP | TRUE | 0.686158453 | 0.043714661 | UP | UP | TRUE |
| Ccdc88b | TRUE | 0.361701309 | 0.451933708 | No Change | Not Significant | TRUE | -0.002873605 | 0.995858831 | No Change | Not Significant | TRUE |
| Ccdc88c | TRUE | 0.426123328 | 6.56E-07 | No Change | Significant Positive | TRUE | 0.120954939 | 0.058770253 | No Change | Not Significant | TRUE |
| Ccdc89 | TRUE | 2.214460412 | 1.48E-15 | UP | UP | TRUE | 1.296561065 | 5.38E-05 | UP | UP | TRUE |
| Ccdc9 | TRUE | -0.399862762 | 2.89E-07 | No Change | Significant Negative | TRUE | -0.104069246 | 0.20590328 | No Change | Not Significant | TRUE |
| Ccdc90b | TRUE | -0.254635434 | 0.220716112 | No Change | Not Significant | TRUE | -0.012747305 | 0.935592329 | No Change | Not Significant | TRUE |
| Ccdc91 | TRUE | -0.311548838 | 0.010218283 | No Change | Significant Negative | TRUE | 0.027978482 | 0.752522003 | No Change | Not Significant | TRUE |
| Ccdc92 | TRUE | 0.381332092 | 0.411599641 | No Change | Not Significant | TRUE | 0.183974696 | 0.575843468 | No Change | Not Significant | TRUE |
| Ccdc92b | TRUE | 0.407043986 | 0.379300188 | No Change | Not Significant | TRUE | -0.145637907 | 0.693580708 | No Change | Not Significant | TRUE |
| Ccdc93 | TRUE | -0.27529202 | 0.031346224 | No Change | Significant Negative | TRUE | -0.19936254 | 0.000203809 | No Change | Significant Negative | TRUE |
| Ccdc96 | TRUE | 0.231984455 | 0.786448489 | No Change | Not Significant | TRUE | -0.189601419 | 0.857573161 | No Change | Not Significant | TRUE |
| Ccdc97 | TRUE | -0.044280732 | 0.827306109 | No Change | Not Significant | TRUE | -0.044105912 | 0.81030697 | No Change | Not Significant | TRUE |
| Ccdc9b | TRUE | 0.229829549 | 0.58491775 | No Change | Not Significant | TRUE | -0.322676987 | 0.254853985 | No Change | Not Significant | TRUE |
| Ccer1 | TRUE | 1.106208618 | 1 | No Change | Not Significant | FALSE | NA | NA | No Change | Not Detected | FALSE |
| Ccer2 | TRUE | -0.980973987 | 1 | No Change | Not Significant | FALSE | NA | NA | No Change | Not Detected | FALSE |
| Cchcr1 | TRUE | 0.027749132 | 0.898947765 | No Change | Not Significant | TRUE | 0.053008476 | 0.759431884 | No Change | Not Significant | TRUE |
| Cck | TRUE | -0.467267735 | 0.422731291 | No Change | Not Significant | TRUE | -0.323257287 | 0.448053195 | No Change | Not Significant | TRUE |
| Cckar | TRUE | -0.292593854 | 1 | No Change | Not Significant | FALSE | NA | NA | No Change | Not Detected | FALSE |
| Cckbr | TRUE | 0.00644893 | 1 | No Change | Not Significant | FALSE | NA | NA | No Change | Not Detected | FALSE |
| Ccl1 | TRUE | 0.609544515 | 1 | No Change | Not Significant | FALSE | NA | NA | No Change | Not Detected | FALSE |
| Ccl20 | TRUE | -0.125360955 | 0.87897876 | No Change | Not Significant | TRUE | 0.182181117 | 0.729699066 | No Change | Not Significant | TRUE |
| Ccl22 | TRUE | 0.303356684 | 1 | No Change | Not Significant | FALSE | NA | NA | No Change | Not Detected | FALSE |
| Ccl24 | TRUE | -0.622873649 | 1 | No Change | Not Significant | FALSE | NA | NA | No Change | Not Detected | FALSE |
| Ccl25 | TRUE | -0.205960895 | 0.499165737 | No Change | Not Significant | TRUE | -0.954425004 | 0.000137788 | DOWN | DOWN | TRUE |
| Ccl27a | TRUE | -0.563348988 | 0.189172946 | No Change | Not Significant | TRUE | -0.220663833 | 0.562649386 | No Change | Not Significant | TRUE |
| Ccl28 | TRUE | -0.099565611 | 0.761298159 | No Change | Not Significant | TRUE | 0.0645963 | 0.645753068 | No Change | Not Significant | TRUE |
| Ccl3 | TRUE | 0.826193839 | 1 | No Change | Not Significant | FALSE | NA | NA | No Change | Not Detected | FALSE |
| Ccl4 | TRUE | 2.831592737 | 1 | No Change | Not Significant | FALSE | NA | NA | No Change | Not Detected | FALSE |
| Ccl5 | TRUE | 0.57179397 | 0.286151898 | No Change | Not Significant | TRUE | 0.536224757 | 0.077109679 | No Change | Not Significant | TRUE |
| Ccl6 | TRUE | -0.282361431 | 0.662536791 | No Change | Not Significant | TRUE | 0.196347158 | 0.00175663 | No Change | Significant Positive | TRUE |

Supplementary Table 3

| geneID | Cond1_DF_Detected | Cond1_DF_ValueCol | Cond1_DF_PadjCol | Cond1_DF_Cutoff | Cond1_DF_Cutoff_Specific.x | Cond2_DF_Detected | Cond2_DF_ValueCol | Cond2_DF_PadjCol | Cond2_DF_Cutoff | Cond2_DF_Cutoff_Specific | BG_Method | RG1_All | RG2_Significant | RG3_SignificantChange |
|---|---|---|---|---|---|---|---|---|---|---|---|---|---|---|
| Ccl7 | TRUE | 0.877295989 | 1 | No Change | Not Significant | FALSE | NA | NA | No Change | Not Detected | FALSE | Background = FALSE | Background = FALSE | Background = FALSE |
| Ccl9 | TRUE | 0.698750125 | 3.91E-05 | UP | UP | TRUE | 0.422127024 | 0.001110122 | No Change | Significant Positive | TRUE | Cond1 UP + Cond2 Significant Positive | Core_UP | Cond1_UP |
| Ccm2 | TRUE | 0.66481035 | 7.23E-14 | UP | UP | TRUE | 0.212810957 | 0.024375404 | No Change | Significant Positive | TRUE | Cond1 UP + Cond2 Significant Positive | Core_UP | Cond1_UP |
| Ccm2l | TRUE | -1.440489346 | 1 | No Change | Not Significant | FALSE | NA | NA | No Change | Not Detected | FALSE | Background = FALSE | Background = FALSE | Background = FALSE |
| Ccn1 | TRUE | -0.134371613 | 0.803650332 | No Change | Not Significant | TRUE | 0.012916057 | 0.967127575 | No Change | Not Significant | TRUE | Cond1 Not Significant + Cond2 Not Significant | None | None |
| Ccn2 | TRUE | -0.506338118 | 1 | No Change | Not Significant | FALSE | NA | NA | No Change | Not Detected | FALSE | Background = FALSE | Background = FALSE | Background = FALSE |
| Ccn3 | TRUE | -0.803678623 | 0.181269925 | No Change | Not Significant | TRUE | 0.386570974 | 0.339503983 | No Change | Not Significant | TRUE | Cond1 Not Significant + Cond2 Not Significant | None | None |
| Ccn4 | TRUE | 0.438863489 | 1 | No Change | Not Significant | FALSE | NA | NA | No Change | Not Detected | FALSE | Background = FALSE | Background = FALSE | Background = FALSE |
| Ccn6 | TRUE | 0.805433347 | 1 | No Change | Not Significant | FALSE | NA | NA | No Change | Not Detected | FALSE | Background = FALSE | Background = FALSE | Background = FALSE |
| Ccna1 | TRUE | 0.798676731 | 0.148614618 | No Change | Not Significant | TRUE | -0.176140294 | 0.813324566 | No Change | Not Significant | TRUE | Cond1 Not Significant + Cond2 Not Significant | None | None |
| Ccna2 | TRUE | 0.833112983 | 8.78E-06 | UP | UP | TRUE | 0.170640888 | 0.305832804 | No Change | Not Significant | TRUE | Cond1 UP + Cond2 Not Significant | Cond1_UP | Cond1_UP |
| Ccnb1 | TRUE | 0.66603387 | 0.000635639 | UP | UP | TRUE | 0.042717589 | 0.789753 | No Change | Not Significant | TRUE | Cond1 UP + Cond2 Not Significant | Cond1_UP | Cond1_UP |
| Ccnb1-ps | TRUE | 0.510816437 | 1 | No Change | Not Significant | FALSE | NA | NA | No Change | Not Detected | FALSE | Background = FALSE | Background = FALSE | Background = FALSE |
| Ccnb2 | TRUE | 0.68207059 | 5.78E-07 | UP | UP | TRUE | -0.055861675 | 0.748759819 | No Change | Not Significant | TRUE | Cond1 UP + Cond2 Not Significant | Cond1_UP | Cond1_UP |
| Ccnc | TRUE | 0.081206208 | 0.541378279 | No Change | Not Significant | TRUE | 0.092044916 | 0.44133635 | No Change | Not Significant | TRUE | Cond1 Not Significant + Cond2 Not Significant | None | None |
| Ccnd1 | TRUE | 0.665820005 | 2.16E-06 | UP | UP | TRUE | 0.382577032 | 4.55E-09 | No Change | Significant Positive | TRUE | Cond1 UP + Cond2 Significant Positive | Core_UP | Cond1_UP |
| Ccnd2 | TRUE | -0.313018104 | 0.176840786 | No Change | Not Significant | TRUE | -0.085901987 | 0.680967534 | No Change | Not Significant | TRUE | Cond1 Not Significant + Cond2 Not Significant | None | None |
| Ccnd3 | TRUE | 0.079419905 | 0.716272317 | No Change | Not Significant | TRUE | 0.218805512 | 0.034269516 | No Change | Significant Positive | TRUE | Cond1 Not Significant + Cond2 Significant Positive | None | None |
| Ccndbp1 | TRUE | -0.262711944 | 0.022878813 | No Change | Significant Negative | TRUE | -0.061896535 | 0.447576381 | No Change | Not Significant | TRUE | Cond1 Significant Negative + Cond2 Not Significant | None | None |
| Ccne1 | TRUE | 1.013376708 | 3.70E-06 | UP | UP | TRUE | 0.573412677 | 0.026849159 | UP | UP | TRUE | Cond1 UP + Cond2 UP | Core_UP | Core_UP |
| Ccne2 | TRUE | 0.481776503 | 0.013849316 | No Change | Significant Positive | TRUE | 0.139710296 | 0.589851392 | No Change | Not Significant | TRUE | Cond1 Significant Positive + Cond2 Not Significant | None | None |
| Ccnf | TRUE | 0.944200406 | 4.87E-06 | UP | UP | TRUE | 0.20589312 | 0.258112004 | No Change | Not Significant | TRUE | Cond1 UP + Cond2 Not Significant | Cond1_UP | Cond1_UP |
| Ccng1 | TRUE | 0.743828896 | 1.31E-13 | UP | UP | TRUE | 0.427645846 | 3.58E-18 | No Change | Significant Positive | TRUE | Cond1 UP + Cond2 Significant Positive | Core_UP | Cond1_UP |
| Ccng2 | TRUE | -0.370365461 | 0.16083904 | No Change | Not Significant | TRUE | -0.23679234 | 0.046820968 | No Change | Significant Negative | TRUE | Cond1 Not Significant + Cond2 Significant Negative | None | None |
| Ccnh | TRUE | 0.145652888 | 0.415642124 | No Change | Not Significant | TRUE | 0.000721356 | 0.996976705 | No Change | Not Significant | TRUE | Cond1 Not Significant + Cond2 Not Significant | None | None |
| Ccni | TRUE | -0.406089434 | 6.46E-05 | No Change | Significant Negative | TRUE | -0.129458025 | 0.118482312 | No Change | Not Significant | TRUE | Cond1 Significant Negative + Cond2 Not Significant | None | None |
| Ccnj | TRUE | -0.110564745 | 0.730361684 | No Change | Not Significant | TRUE | 0.018739274 | 0.950656978 | No Change | Not Significant | TRUE | Cond1 Not Significant + Cond2 Not Significant | None | None |
| Ccnjl | TRUE | 0.389902964 | 0.147109787 | No Change | Not Significant | TRUE | 0.49514302 | 0.005779259 | No Change | Significant Positive | TRUE | Cond1 Not Significant + Cond2 Significant Positive | None | None |
| Ccnk | TRUE | 0.162063427 | 0.508154368 | No Change | Not Significant | TRUE | -0.238587753 | 0.093297675 | No Change | Not Significant | TRUE | Cond1 Not Significant + Cond2 Not Significant | None | None |
| Ccnl1 | TRUE | -0.296099288 | 0.098614315 | No Change | Not Significant | TRUE | -0.464197989 | 0.017417983 | No Change | Significant Negative | TRUE | Cond1 Not Significant + Cond2 Significant Negative | None | None |
| Ccnl2 | TRUE | 0.209822827 | 0.212413992 | No Change | Not Significant | TRUE | -0.188440998 | 0.183629264 | No Change | Not Significant | TRUE | Cond1 Not Significant + Cond2 Not Significant | None | None |
| Ccno | TRUE | 0.129887485 | 0.932314788 | No Change | Not Significant | FALSE | NA | NA | No Change | Not Detected | FALSE | Background = FALSE | Background = FALSE | Background = FALSE |
| Ccnq | TRUE | -0.206116429 | 0.346267391 | No Change | Not Significant | TRUE | -0.016632906 | 0.948995857 | No Change | Not Significant | TRUE | Cond1 Not Significant + Cond2 Not Significant | None | None |
| Ccnt1 | TRUE | -0.073157468 | 0.590110273 | No Change | Not Significant | TRUE | -0.044673091 | 0.75524772 | No Change | Not Significant | TRUE | Cond1 Not Significant + Cond2 Not Significant | None | None |
| Ccnt2 | TRUE | -0.15810535 | 0.027715628 | No Change | Significant Negative | TRUE | -0.151131462 | 0.01497706 | No Change | Significant Negative | TRUE | Cond1 Significant Negative + Cond2 Significant Negative | None | None |
| Ccny | TRUE | -0.097604421 | 0.539479894 | No Change | Not Significant | TRUE | 0.025112548 | 0.665528957 | No Change | Not Significant | TRUE | Cond1 Not Significant + Cond2 Not Significant | None | None |
| Ccnyl1 | TRUE | 0.46199229 | 0.000177672 | No Change | Significant Positive | TRUE | 0.128575198 | 0.353306674 | No Change | Not Significant | TRUE | Cond1 Significant Positive + Cond2 Not Significant | None | None |
| Ccp110 | TRUE | 0.240905139 | 0.352338897 | No Change | Not Significant | TRUE | 0.059231465 | 0.888754045 | No Change | Not Significant | TRUE | Cond1 Not Significant + Cond2 Not Significant | None | None |
| Ccpg1 | TRUE | 0.730085713 | 0.128452801 | No Change | Not Significant | TRUE | -0.044686481 | 0.950489771 | No Change | Not Significant | TRUE | Cond1 Not Significant + Cond2 Not Significant | None | None |
| Ccr1 | TRUE | 1.163831839 | 0.039588986 | UP | UP | TRUE | 0.235422377 | 0.629260691 | No Change | Not Significant | TRUE | Cond1 UP + Cond2 Not Significant | Cond1_UP | Cond1_UP |
| Ccr10 | TRUE | -0.2492891 | 0.761475268 | No Change | Not Significant | TRUE | -0.332950509 | 0.54999264 | No Change | Not Significant | TRUE | Cond1 Not Significant + Cond2 Not Significant | None | None |
| Ccr2 | TRUE | 1.025063209 | 0.468613008 | No Change | Not Significant | TRUE | 0.362472963 | 0.749829352 | No Change | Not Significant | TRUE | Cond1 Not Significant + Cond2 Not Significant | None | None |
| Ccr4 | TRUE | 0.78448959 | 1 | No Change | Not Significant | FALSE | NA | NA | No Change | Not Detected | FALSE | Background = FALSE | Background = FALSE | Background = FALSE |
| Ccr5 | TRUE | 1.607539957 | 0.159701725 | No Change | Not Significant | TRUE | 1.230396716 | 0.124739465 | No Change | Not Significant | TRUE | Cond1 Not Significant + Cond2 Not Significant | None | None |
| Ccr6 | TRUE | 1.521196223 | 1 | No Change | Not Significant | FALSE | NA | NA | No Change | Not Detected | FALSE | Background = FALSE | Background = FALSE | Background = FALSE |
| Ccr7 | TRUE | 0.467307482 | 1 | No Change | Not Significant | FALSE | NA | NA | No Change | Not Detected | FALSE | Background = FALSE | Background = FALSE | Background = FALSE |
| Ccr8 | TRUE | -0.416086163 | 1 | No Change | Not Significant | FALSE | NA | NA | No Change | Not Detected | FALSE | Background = FALSE | Background = FALSE | Background = FALSE |
| Ccr9 | TRUE | -0.320306317 | 0.76182422 | No Change | Not Significant | TRUE | 0.060216686 | 0.936097283 | No Change | Not Significant | TRUE | Cond1 Not Significant + Cond2 Not Significant | None | None |
| Ccrl2 | TRUE | -0.807817187 | 0.032763544 | DOWN | DOWN | TRUE | -0.309790897 | 0.514863213 | No Change | Not Significant | TRUE | Cond1 DOWN + Cond2 Not Significant | Cond1_DOWN | Cond1_DOWN |
| Ccs | TRUE | -0.423458229 | 1.09E-05 | No Change | Significant Negative | TRUE | -0.065430163 | 0.486881861 | No Change | Not Significant | TRUE | Cond1 Significant Negative + Cond2 Not Significant | None | None |
| Ccsap | TRUE | 0.539494968 | 0.050176222 | No Change | Not Significant | TRUE | -0.049338858 | 0.895541399 | No Change | Not Significant | TRUE | Cond1 Not Significant + Cond2 Not Significant | None | None |
| Ccser1 | TRUE | 0.009062499 | 0.968634984 | No Change | Not Significant | TRUE | -0.13174636 | 0.234068382 | No Change | Not Significant | TRUE | Cond1 Not Significant + Cond2 Not Significant | None | None |
| Ccser2 | TRUE | 0.062898141 | 0.481401448 | No Change | Not Significant | TRUE | 0.112174486 | 0.118377705 | No Change | Not Significant | TRUE | Cond1 Not Significant + Cond2 Not Significant | None | None |
| Cct2 | TRUE | -0.081920156 | 0.60964446 | No Change | Not Significant | TRUE | -0.033648567 | 0.675687484 | No Change | Not Significant | TRUE | Cond1 Not Significant + Cond2 Not Significant | None | None |
| Cct3 | TRUE | 0.098758567 | 0.604659935 | No Change | Not Significant | TRUE | 0.143880042 | 0.234551147 | No Change | Not Significant | TRUE | Cond1 Not Significant + Cond2 Not Significant | None | None |
| Cct4 | TRUE | -0.187230009 | 0.12191044 | No Change | Not Significant | TRUE | -0.086500533 | 0.175287213 | No Change | Not Significant | TRUE | Cond1 Not Significant + Cond2 Not Significant | None | None |
| Cct5 | TRUE | 0.015347276 | 0.932314788 | No Change | Not Significant | TRUE | 0.035602458 | 0.706927105 | No Change | Not Significant | TRUE | Cond1 Not Significant + Cond2 Not Significant | None | None |
| Cct6a | TRUE | 0.070032574 | 0.636776036 | No Change | Not Significant | TRUE | 0.034121246 | 0.745625688 | No Change | Not Significant | TRUE | Cond1 Not Significant + Cond2 Not Significant | None | None |
| Cct6b | TRUE | -0.514141832 | 0.532389508 | No Change | Not Significant | TRUE | 0.003205206 | 0.996814475 | No Change | Not Significant | TRUE | Cond1 Not Significant + Cond2 Not Significant | None | None |
| Cct7 | TRUE | 0.261070971 | 0.016170677 | No Change | Significant Positive | TRUE | 0.1631668 | 0.064978209 | No Change | Not Significant | TRUE | Cond1 Significant Positive + Cond2 Not Significant | None | None |
| Cct8 | TRUE | 0.109736869 | 0.474068167 | No Change | Not Significant | TRUE | 0.080977882 | 0.400571839 | No Change | Not Significant | TRUE | Cond1 Not Significant + Cond2 Not Significant | None | None |
| Cct8l1 | TRUE | 1.050995534 | 1 | No Change | Not Significant | FALSE | NA | NA | No Change | Not Detected | FALSE | Background = FALSE | Background = FALSE | Background = FALSE |
| Ccz1 | TRUE | 0.015159948 | 0.932932188 | No Change | Not Significant | TRUE | -0.009396705 | 0.935543319 | No Change | Not Significant | TRUE | Cond1 Not Significant + Cond2 Not Significant | None | None |
| Cd101 | TRUE | 1.397339725 | 0.022396188 | UP | UP | TRUE | 1.20342044 | 0.012600696 | UP | UP | TRUE | Cond1 UP + Cond2 UP | Core_UP | Core_UP |
| Cd109 | TRUE | 3.385951792 | 1 | No Change | Not Significant | FALSE | NA | NA | No Change | Not Detected | FALSE | Background = FALSE | Background = FALSE | Background = FALSE |
| Cd14 | TRUE | 0.419793558 | 0.158736855 | No Change | Not Significant | TRUE | 0.360270763 | 0.001579682 | No Change | Significant Positive | TRUE | Cond1 Not Significant + Cond2 Significant Positive | None | None |
| Cd151 | TRUE | -0.205911825 | 0.001242392 | No Change | Significant Negative | TRUE | -0.017213804 | 0.879490473 | No Change | Not Significant | TRUE | Cond1 Significant Negative + Cond2 Not Significant | None | None |
| Cd160 | TRUE | 0.228151048 | 0.708596728 | No Change | Not Significant | TRUE | 0.440045849 | 0.142420193 | No Change | Not Significant | TRUE | Cond1 Not Significant + Cond2 Not Significant | None | None |
| Cd163 | TRUE | 0.552313536 | 1 | No Change | Not Significant | FALSE | NA | NA | No Change | Not Detected | FALSE | Background = FALSE | Background = FALSE | Background = FALSE |
| Cd164 | TRUE | 0.00045761 | 0.99808463 | No Change | Not Significant | TRUE | -0.059024269 | 0.141755849 | No Change | Not Significant | TRUE | Cond1 Not Significant + Cond2 Not Significant | None | None |
| Cd164l2 | TRUE | -0.316493183 | 0.733377497 | No Change | Not Significant | TRUE | 0.000429984 | 0.999454275 | No Change | Not Significant | TRUE | Cond1 Not Significant + Cond2 Not Significant | None | None |
| Cd177 | TRUE | 0.662546294 | 0.038458141 | UP | UP | TRUE | 0.769866233 | 4.56E-12 | UP | UP | TRUE | Cond1 UP + Cond2 UP | Core_UP | Core_UP |
| Cd180 | TRUE | 0.344131581 | 1 | No Change | Not Significant | FALSE | NA | NA | No Change | Not Detected | FALSE | Background = FALSE | Background = FALSE | Background = FALSE |
| Cd19 | TRUE | 0.758548989 | 1 | No Change | Not Significant | FALSE | NA | NA | No Change | Not Detected | FALSE | Background = FALSE | Background = FALSE | Background = FALSE |
| Cd1d1 | TRUE | 0.025306402 | 0.952305972 | No Change | Not Significant | TRUE | 0.202136495 | 0.238421516 | No Change | Not Significant | TRUE | Cond1 Not Significant + Cond2 Not Significant | None | None |
| Cd2 | TRUE | -0.424758546 | 1 | No Change | Not Significant | TRUE | 0.458423949 | 0.472837299 | No Change | Not Significant | TRUE | Cond1 Not Significant + Cond2 Not Significant | None | None |
| Cd200 | TRUE | -0.039034219 | 0.93002838 | No Change | Not Significant | TRUE | -0.769130829 | 0.000114488 | DOWN | DOWN | TRUE | Cond1 Not Significant + Cond2 DOWN | Cond2_DOWN | Cond2_DOWN |
| Cd200l1 | TRUE | -0.279580111 | 0.220499674 | No Change | Not Significant | TRUE | -0.349646307 | 0.03837566 | No Change | Significant Negative | TRUE | Cond1 Not Significant + Cond2 Significant Negative | None | None |
| Cd200r1 | TRUE | -0.114557637 | 0.92788168 | No Change | Not Significant | TRUE | 0.238594759 | 0.783947174 | No Change | Not Significant | TRUE | Cond1 Not Significant + Cond2 Not Significant | None | None |
| Cd200r2 | TRUE | -0.023373302 | 0.95833356 | No Change | Not Significant | TRUE | 0.026644458 | 0.930889346 | No Change | Not Significant | TRUE | Cond1 Not Significant + Cond2 Not Significant | None | None |
| Cd200r3 | TRUE | -0.026798009 | 1 | No Change | Not Significant | FALSE | NA | NA | No Change | Not Detected | FALSE | Background = FALSE | Background = FALSE | Background = FALSE |
| Cd200r4 | TRUE | -0.11850523 | 0.935258087 | No Change | Not Significant | FALSE | NA | NA | No Change | Not Detected | FALSE | Background = FALSE | Background = FALSE | Background = FALSE |
| Cd209f | TRUE | 2.425418171 | 1 | No Change | Not Significant | FALSE | NA | NA | No Change | Not Detected | FALSE | Background = FALSE | Background = FALSE | Background = FALSE |
| Cd209g | TRUE | 1.726335262 | 1 | No Change | Not Significant | FALSE | NA | NA | No Change | Not Detected | FALSE | Background = FALSE | Background = FALSE | Background = FALSE |
| Cd22 | TRUE | 0.494931743 | 1 | No Change | Not Significant | FALSE | NA | NA | No Change | Not Detected | FALSE | Background = FALSE | Background = FALSE | Background = FALSE |
| Cd226 | TRUE | -0.034457624 | 0.974163449 | No Change | Not Significant | TRUE | 0.393786295 | 0.38300506 | No Change | Not Significant | TRUE | Cond1 Not Significant + Cond2 Not Significant | None | None |
| Cd244a | TRUE | -0.200797063 | 0.806110415 | No Change | Not Significant | TRUE | 0.770412165 | 0.007209695 | UP | UP | TRUE | Cond1 Not Significant + Cond2 UP | Cond2_UP | Cond2_UP |
| Cd247 | TRUE | -0.175791099 | 0.839586229 | No Change | Not Significant | TRUE | 0.140662911 | 0.862820371 | No Change | Not Significant | TRUE | Cond1 Not Significant + Cond2 Not Significant | None | None |
| Cd248 | TRUE | -0.054192345 | 1 | No Change | Not Significant | FALSE | NA | NA | No Change | Not Detected | FALSE | Background = FALSE | Background = FALSE | Background = FALSE |
| Cd24a | TRUE | 0.112223957 | 0.67923791 | No Change | Not Significant | TRUE | 0.12779864 | 0.00809297 | No Change | Significant Positive | TRUE | Cond1 Not Significant + Cond2 Significant Positive | None | None |
| Cd27 | TRUE | 0.393661995 | 0.679207466 | No Change | Not Significant | TRUE | 0.222824935 | 0.773321692 | No Change | Not Significant | TRUE | Cond1 Not Significant + Cond2 Not Significant | None | None |
| Cd274 | TRUE | -0.211807789 | 0.662372535 | No Change | Not Significant | TRUE | -0.270761083 | 0.646388536 | No Change | Not Significant | TRUE | Cond1 Not Significant + Cond2 Not Significant | None | None |
| Cd276 | TRUE | 0.20069789 | 0.450375375 | No Change | Not Significant | TRUE | -0.077120008 | 0.789753 | No Change | Not Significant | TRUE | Cond1 Not Significant + Cond2 Not Significant | None | None |
| Cd28 | TRUE | 0.36396969 | 1 | No Change | Not Significant | FALSE | NA | NA | No Change | Not Detected | FALSE | Background = FALSE | Background = FALSE | Background = FALSE |
| Cd2ap | TRUE | 0.197604011 | 0.022575574 | No Change | Significant Positive | TRUE | 0.090419148 | 0.185896496 | No Change | Not Significant | TRUE | Cond1 Significant Positive + Cond2 Not Significant | None | None |
| Cd2bp2 | TRUE | -0.046014757 | 0.685332006 | No Change | Not Significant | TRUE | 0.012351829 | 0.923333171 | No Change | Not Significant | TRUE | Cond1 Not Significant + Cond2 Not Significant | None | None |
| Cd300a | TRUE | 0.077590407 | 0.977047797 | No Change | Not Significant | FALSE | NA | NA | No Change | Not Detected | FALSE | Background = FALSE | Background = FALSE | Background = FALSE |
| Cd300c2 | TRUE | -0.133173146 | 1 | No Change | Not Significant | FALSE | NA | NA | No Change | Not Detected | FALSE | Background = FALSE | Background = FALSE | Background = FALSE |
| Cd300e | TRUE | -0.41753209 | 1 | No Change | Not Significant | FALSE | NA | NA | No Change | Not Detected | FALSE | Background = FALSE | Background = FALSE | Background = FALSE |
| Cd300ld | TRUE | 2.099740825 | 1 | No Change | Not Significant | FALSE | NA | NA | No Change | Not Detected | FALSE | Background = FALSE | Background = FALSE | Background = FALSE |
| Cd300lf | TRUE | -0.435617234 | 0.409799304 | No Change | Not Significant | TRUE | -0.303249082 | 0.419339065 | No Change | Not Significant | TRUE | Cond1 Not Significant + Cond2 Not Significant | None | None |
| Cd300lg | TRUE | -1.29520975 | 1 | No Change | Not Significant | FALSE | NA | NA | No Change | Not Detected | FALSE | Background = FALSE | Background = FALSE | Background = FALSE |
| Cd302 | TRUE | 0.279381909 | 0.298143983 | No Change | Not Significant | TRUE | 0.05576789 | 0.871241669 | No Change | Not Significant | TRUE | Cond1 Not Significant + Cond2 Not Significant | None | None |
| Cd320 | TRUE | -0.610940542 | 0.000354329 | DOWN | DOWN | TRUE | -0.083011808 | 0.561388929 | No Change | Not Significant | TRUE | Cond1 DOWN + Cond2 Not Significant | Cond1_DOWN | Cond1_DOWN |
| Cd33 | TRUE | -0.472446099 | 0.797713998 | No Change | Not Significant | FALSE | NA | NA | No Change | Not Detected | FALSE | Background = FALSE | Background = FALSE | Background = FALSE |
| Cd34 | TRUE | 0.245630213 | 0.874014884 | No Change | Not Significant | TRUE | 1.139000374 | 0.059145001 | No Change | Not Significant | TRUE | Cond1 Not Significant + Cond2 Not Significant | None | None |
| Cd36 | TRUE | -0.288752939 | 1 | No Change | Not Significant | TRUE | 0.16786977 | 0.892332662 | No Change | Not Significant | TRUE | Cond1 Not Significant + Cond2 Not Significant | None | None |
| Cd37 | TRUE | 0.085897477 | 0.905779176 | No Change | Not Significant | TRUE | 0.114688096 | 0.852795671 | No Change | Not Significant | TRUE | Cond1 Not Significant + Cond2 Not Significant | None | None |
| Cd38 | TRUE | -0.051182396 | 0.850681912 | No Change | Not Significant | TRUE | -0.223183571 | 0.001627504 | No Change | Significant Negative | TRUE | Cond1 Not Significant + Cond2 Significant Negative | None | None |
| Cd3d | TRUE | -0.136987282 | 0.890747356 | No Change | Not Significant | TRUE | 0.341067131 | 0.616422248 | No Change | Not Significant | TRUE | Cond1 Not Significant + Cond2 Not Significant | None | None |
| Cd3e | TRUE | -0.560137202 | 0.380604488 | No Change | Not Significant | TRUE | 0.237996767 | 0.685946212 | No Change | Not Significant | TRUE | Cond1 Not Significant + Cond2 Not Significant | None | None |
| Cd3eap | TRUE | -0.371685767 | 0.070451816 | No Change | Not Significant | TRUE | -0.260956676 | 0.043445662 | No Change | Significant Negative | TRUE | Cond1 Not Significant + Cond2 Significant Negative | None | None |
| Cd3g | TRUE | -0.105789813 | 0.888506311 | No Change | Not Significant | TRUE | 0.330958321 | 0.246560224 | No Change | Not Significant | TRUE | Cond1 Not Significant + Cond2 Not Significant | None | None |
| Cd4 | TRUE | -1.30866602 | 1 | No Change | Not Significant | FALSE | NA | NA | No Change | Not Detected | FALSE | Background = FALSE | Background = FALSE | Background = FALSE |
| Cd40 | TRUE | 1.024912637 | 1 | No Change | Not Significant | FALSE | NA | NA | No Change | Not Detected | FALSE | Background = FALSE | Background = FALSE | Background = FALSE |
| Cd40lg | TRUE | 1.686112304 | 1 | No Change | Not Significant | FALSE | NA | NA | No Change | Not Detected | FALSE | Background = FALSE | Background = FALSE | Background = FALSE |
| Cd44 | TRUE | 0.205437056 | 0.518790947 | No Change | Not Significant | TRUE | -0.169931071 | 0.115681275 | No Change | Not Significant | TRUE | Cond1 Not Significant + Cond2 Not Significant | None | None |
| Cd46 | TRUE | -0.42768496 | 0.22586956 | No Change | Not Significant | TRUE | -0.570241943 | 0.086684114 | No Change | Not Significant | TRUE | Cond1 Not Significant + Cond2 Not Significant | None | None |
| Cd47 | TRUE | -0.058189662 | 0.753386941 | No Change | Not Significant | TRUE | -0.127290553 | 0.336509736 | No Change | Not Significant | TRUE | Cond1 Not Significant + Cond2 Not Significant | None | None |
| Cd48 | TRUE | -0.259689033 | 0.80133583 | No Change | Not Significant | TRUE | 0.016253496 | 0.986201549 | No Change | Not Significant | TRUE | Cond1 Not Significant + Cond2 Not Significant | None | None |
| Cd5 | TRUE | 0.434960568 | 0.752646336 | No Change | Not Significant | FALSE | NA | NA | No Change | Not Detected | FALSE | Background = FALSE | Background = FALSE | Background = FALSE |
| Cd52 | TRUE | -0.189347919 | 0.780450414 | No Change | Not Significant | TRUE | 0.164427122 | 0.627104366 | No Change | Not Significant | TRUE | Cond1 Not Significant + Cond2 Not Significant | None | None |
| Cd53 | TRUE | 0.016344329 | 0.982706396 | No Change | Not Significant | TRUE | 0.275734472 | 0.5169296 | No Change | Not Significant | TRUE | Cond1 Not Significant + Cond2 Not Significant | None | None |
| Cd55 | TRUE | -0.92717424 | 1.29E-12 | DOWN | DOWN | TRUE | -0.231150538 | 0.25082096 | No Change | Not Significant | TRUE | Cond1 DOWN + Cond2 Not Significant | Cond1_DOWN | Cond1_DOWN |
| Cd55b | TRUE | 1.520645824 | 1 | No Change | Not Significant | FALSE | NA | NA | No Change | Not Detected | FALSE | Background = FALSE | Background = FALSE | Background = FALSE |
| Cd55os | TRUE | 0.068866993 | 1 | No Change | Not Significant | FALSE | NA | NA | No Change | Not Detected | FALSE | Background = FALSE | Background = FALSE | Background = FALSE |
| Cd59a | TRUE | -0.985896279 | 0.042468102 | DOWN | DOWN | TRUE | 0.334596714 | 0.426250675 | No Change | Not Significant | TRUE | Cond1 DOWN + Cond2 Not Significant | Cond1_DOWN | Cond1_DOWN |
| Cd59b | TRUE | -0.64047218 | 1 | No Change | Not Significant | FALSE | NA | NA | No Change | Not Detected | FALSE | Background = FALSE | Background = FALSE | Background = FALSE |
| Cd5l | TRUE | 0.868511853 | 1 | No Change | Not Significant | FALSE | NA | NA | No Change | Not Detected | FALSE | Background = FALSE | Background = FALSE | Background = FALSE |
| Cd6 | TRUE | -1.162925426 | 0.595844868 | No Change | Not Significant | FALSE | NA | NA | No Change | Not Detected | FALSE | Background = FALSE | Background = FALSE | Background = FALSE |
| Cd63 | TRUE | -0.463254931 | 9.39E-09 | No Change | Significant Negative | TRUE | -0.185194583 | 0.000381741 | No Change | Significant Negative | TRUE | Cond1 Significant Negative + Cond2 Significant Negative | None | None |
| Cd68 | TRUE | 0.94424811 | 0.169705389 | No Change | Not Significant | TRUE | -0.599152433 | 0.503084198 | No Change | Not Significant | TRUE | Cond1 Not Significant + Cond2 Not Significant | None | None |
| Cd69 | TRUE | 0.021606761 | 0.984403993 | No Change | Not Significant | TRUE | 0.149142116 | 0.790773109 | No Change | Not Significant | TRUE | Cond1 Not Significant + Cond2 Not Significant | None | None |
| Cd7 | TRUE | 0.178760942 | 0.761222618 | No Change | Not Significant | TRUE | 0.485530777 | 0.034222527 | No Change | Significant Positive | TRUE | Cond1 Not Significant + Cond2 Significant Positive | None | None |
| Cd72 | TRUE | 0.429537533 | 0.688134988 | No Change | Not Significant | TRUE | -0.23178639 | 0.752144991 | No Change | Not Significant | TRUE | Cond1 Not Significant + Cond2 Not Significant | None | None |
| Cd74 | TRUE | 0.08679968 | 0.894720656 | No Change | Not Significant | FALSE | NA | NA | No Change | Not Detected | FALSE | Background = FALSE | Background = FALSE | Background = FALSE |
| Cd79a | TRUE | 0.010669654 | 1 | No Change | Not Significant | FALSE | NA | NA | No Change | Not Detected | FALSE | Background = FALSE | Background = FALSE | Background = FALSE |
| Cd79b | TRUE | -0.00061115 | 1 | No Change | Not Significant | FALSE | NA | NA | No Change | Not Detected | FALSE | Background = FALSE | Background = FALSE | Background = FALSE |
| Cd80 | TRUE | -0.784101544 | 1 | No Change | Not Significant | FALSE | NA | NA | No Change | Not Detected | FALSE | Background = FALSE | Background = FALSE | Background = FALSE |
| Cd81 | TRUE | -0.238476279 | 0.370029503 | No Change | Not Significant | TRUE | -0.138012796 | 0.203800871 | No Change | Not Significant | TRUE | Cond1 Not Significant + Cond2 Not Significant | None | None |
| Cd82 | TRUE | -0.050829774 | 0.684193826 | No Change | Not Significant | TRUE | 0.002235224 | 0.986873905 | No Change | Not Significant | TRUE | Cond1 Not Significant + Cond2 Not Significant | None | None |
| Cd83 | TRUE | -0.858396212 | 0.003915665 | DOWN | DOWN | TRUE | -0.129701799 | 0.559370353 | No Change | Not Significant | TRUE | Cond1 DOWN + Cond2 Not Significant | Cond1_DOWN | Cond1_DOWN |
| Cd84 | TRUE | 0.78385679 | 0.59267286 | No Change | Not Significant | TRUE | 0.754081669 | 0.238288521 | No Change | Not Significant | TRUE | Cond1 Not Significant + Cond2 Not Significant | None | None |
| Cd86 | TRUE | 2.884195661 | 1 | No Change | Not Significant | FALSE | NA | NA | No Change | Not Detected | FALSE | Background = FALSE | Background = FALSE | Background = FALSE |
| Cd8a | TRUE | -0.224533596 | 0.750868206 | No Change | Not Significant | TRUE | -0.111184616 | 0.82901007 | No Change | Not Significant | TRUE | Cond1 Not Significant + Cond2 Not Significant | None | None |
| Cd8b1 | TRUE | -1.226081399 | 1 | No Change | Not Significant | FALSE | NA | NA | No Change | Not Detected | FALSE | Background = FALSE | Background = FALSE | Background = FALSE |
| Cd9 | TRUE | -0.587787499 | 0.000584065 | DOWN | DOWN | TRUE | -0.049971522 | 0.679801803 | No Change | Not Significant | TRUE | Cond1 DOWN + Cond2 Not Significant | Cond1_DOWN | Cond1_DOWN |
| Cd93 | TRUE | 0.866845278 | 1 | No Change | Not Significant | FALSE | NA | NA | No Change | Not Detected | FALSE | Background = FALSE | Background = FALSE | Background = FALSE |
| Cd96 | TRUE | -0.588853677 | 0.326547965 | No Change | Not Significant | TRUE | 0.428713155 | 0.306028716 | No Change | Not Significant | TRUE | Cond1 Not Significant + Cond2 Not Significant | None | None |
| Cd99l2 | TRUE | -0.309623051 | 0.01737744 | No Change | Significant Negative | TRUE | -0.040622659 | 0.82592067 | No Change | Not Significant | TRUE | Cond1 Significant Negative + Cond2 Not Significant | None | None |
| Cda | TRUE | 0.117318272 | 0.696047947 | No Change | Not Significant | TRUE | -0.088266065 | 0.657926757 | No Change | Not Significant | TRUE | Cond1 Not Significant + Cond2 Not Significant | None | None |
| Cdadc1 | TRUE | -0.324596698 | 0.007882634 | No Change | Significant Negative | TRUE | -0.0340309 | 0.819274771 | No Change | Not Significant | TRUE | Cond1 Significant Negative + Cond2 Not Significant | None | None |
| Cdan1 | TRUE | 0.116305635 | 0.559563944 | No Change | Not Significant | TRUE | -0.122582475 | 0.523452524 | No Change | Not Significant | TRUE | Cond1 Not Significant + Cond2 Not Significant | None | None |
| Cdc123 | TRUE | 0.320629917 | 0.004076571 | No Change | Significant Positive | TRUE | 0.188429921 | 0.013257847 | No Change | Significant Positive | TRUE | Cond1 Significant Positive + Cond2 Significant Positive | None | None |
| Cdc14a | TRUE | -0.172175971 | 0.252796186 | No Change | Not Significant | TRUE | -0.094049274 | 0.311837862 | No Change | Not Significant | TRUE | Cond1 Not Significant + Cond2 Not Significant | None | None |
| Cdc14b | TRUE | -0.393038146 | 0.019029502 | No Change | Significant Negative | TRUE | -0.070934644 | 0.622373434 | No Change | Not Significant | TRUE | Cond1 Significant Negative + Cond2 Not Significant | None | None |
| Cdc16 | TRUE | 0.0530955 | 0.745548913 | No Change | Not Significant | TRUE | 0.000799524 | 0.995858831 | No Change | Not Significant | TRUE | Cond1 Not Significant + Cond2 Not Significant | None | None |
| Cdc20 | TRUE | 0.797609987 | 2.10E-07 | UP | UP | TRUE | 0.046228924 | 0.815430435 | No Change | Not Significant | TRUE | Cond1 UP + Cond2 Not Significant | Cond1_UP | Cond1_UP |
| Cdc20b | TRUE | -1.546126028 | 1 | No Change | Not Significant | FALSE | NA | NA | No Change | Not Detected | FALSE | Background = FALSE | Background = FALSE | Background = FALSE |
| Cdc23 | TRUE | 0.004706802 | 0.984706282 | No Change | Not Significant | TRUE | -0.108654647 | 0.241736064 | No Change | Not Significant | TRUE | Cond1 Not Significant + Cond2 Not Significant | None | None |
| Cdc25a | TRUE | 0.208944955 | 0.124778744 | No Change | Not Significant | TRUE | 0.052886786 | 0.696651125 | No Change | Not Significant | TRUE | Cond1 Not Significant + Cond2 Not Significant | None | None |
| Cdc25b | TRUE | 0.363292342 | 0.028025774 | No Change | Significant Positive | TRUE | -0.213353989 | 0.041292666 | No Change | Significant Negative | TRUE | Cond1 Significant Positive + Cond2 Significant Negative | None | None |
| Cdc25c | TRUE | 0.855671356 | 1.92E-05 | UP | UP | TRUE | 0.009989952 | 0.97221237 | No Change | Not Significant | TRUE | Cond1 UP + Cond2 Not Significant | Cond1_UP | Cond1_UP |
| Cdc26 | TRUE | -0.040610067 | 0.8036518 | No Change | Not Significant | TRUE | 0.218651228 | 0.032609438 | No Change | Significant Positive | TRUE | Cond1 Not Significant + Cond2 Significant Positive | None | None |
| Cdc27 | TRUE | 0.22226015 | 0.048151079 | No Change | Significant Positive | TRUE | 0.093242737 | 0.331476631 | No Change | Not Significant | TRUE | Cond1 Significant Positive + Cond2 Not Significant | None | None |
| Cdc34 | TRUE | 0.012289872 | 0.94399288 | No Change | Not Significant | TRUE | 0.111482117 | 0.424880538 | No Change | Not Significant | TRUE | Cond1 Not Significant + Cond2 Not Significant | None | None |
| Cdc37 | TRUE | 0.01844402 | 0.893375436 | No Change | Not Significant | TRUE | 0.087709231 | 0.352096817 | No Change | Not Significant | TRUE | Cond1 Not Significant + Cond2 Not Significant | None | None |
| Cdc37l1 | TRUE | -0.117622086 | 0.438111017 | No Change | Not Significant | TRUE | -0.117116577 | 0.243202137 | No Change | Not Significant | TRUE | Cond1 Not Significant + Cond2 Not Significant | None | None |
| Cdc40 | TRUE | 0.027435932 | 0.879513202 | No Change | Not Significant | TRUE | -0.01485905 | 0.918128862 | No Change | Not Significant | TRUE | Cond1 Not Significant + Cond2 Not Significant | None | None |
| Cdc42 | TRUE | -0.008225363 | 0.959274295 | No Change | Not Significant | TRUE | 0.109214932 | 0.145506574 | No Change | Not Significant | TRUE | Cond1 Not Significant + Cond2 Not Significant | None | None |
| Cdc42bpa | TRUE | 0.011162061 | 0.968109605 | No Change | Not Significant | TRUE | 0.285777849 | 0.0002563 | No Change | Significant Positive | TRUE | Cond1 Not Significant + Cond2 Significant Positive | None | None |
| Cdc42bpb | TRUE | -0.030960034 | 0.82339418 | No Change | Not Significant | TRUE | 0.021010353 | 0.830562128 | No Change | Not Significant | TRUE | Cond1 Not Significant + Cond2 Not Significant | None | None |
| Cdc42bpg | TRUE | 0.127891327 | 0.40082018 | No Change | Not Significant | TRUE | -0.040856294 | 0.682988492 | No Change | Not Significant | TRUE | Cond1 Not Significant + Cond2 Not Significant | None | None |
| Cdc42ep1 | TRUE | -0.302608899 | 0.069131639 | No Change | Not Significant | TRUE | -0.062866229 | 0.685606255 | No Change | Not Significant | TRUE | Cond1 Not Significant + Cond2 Not Significant | None | None |
| Cdc42ep2 | TRUE | 0.047293877 | 0.922154269 | No Change | Not Significant | TRUE | 0.076376365 | 0.711826106 | No Change | Not Significant | TRUE | Cond1 Not Significant + Cond2 Not Significant | None | None |
| Cdc42ep3 | TRUE | -0.076884892 | 0.807443873 | No Change | Not Significant | TRUE | 0.34937615 | 0.004587415 | No Change | Significant Positive | TRUE | Cond1 Not Significant + Cond2 Significant Positive | None | None |
| Cdc42ep4 | TRUE | -0.26821291 | 1.41E-05 | No Change | Significant Negative | TRUE | 0.067925248 | 0.511644242 | No Change | Not Significant | TRUE | Cond1 Significant Negative + Cond2 Not Significant | None | None |
| Cdc42ep5 | TRUE | 0.188353623 | 0.279268705 | No Change | Not Significant | TRUE | 0.481165856 | 9.88E-08 | No Change | Significant Positive | TRUE | Cond1 Not Significant + Cond2 Significant Positive | None | None |
| Cdc42se1 | TRUE | 0.032125114 | 0.866653357 | No Change | Not Significant | TRUE | 0.019243044 | 0.803431336 | No Change | Not Significant | TRUE | Cond1 Not Significant + Cond2 Not Significant | None | None |
| Cdc42se2 | TRUE | 0.083741491 | 0.603354756 | No Change | Not Significant | TRUE | 0.170573623 | 0.032307417 | No Change | Significant Positive | TRUE | Cond1 Not Significant + Cond2 Significant Positive | None | None |
| Cdc45 | TRUE | 0.928910028 | 0.000136072 | UP | UP | TRUE | 0.442011734 | 0.101895312 | No Change | Not Significant | TRUE | Cond1 UP + Cond2 Not Significant | Cond1_UP | Cond1_UP |
| Cdc5l | TRUE | -0.023929167 | 0.911456552 | No Change | Not Significant | TRUE | 0.003607547 | 0.977252742 | No Change | Not Significant | TRUE | Cond1 Not Significant + Cond2 Not Significant | None | None |
| Cdc6 | TRUE | 0.972588609 | 1.56E-05 | UP | UP | TRUE | 0.748214854 | 0.00088192 | UP | UP | TRUE | Cond1 UP + Cond2 UP | Core_UP | Core_UP |
| Cdc7 | TRUE | 0.526131385 | 0.019692129 | UP | UP | TRUE | -0.209481075 | 0.525566345 | No Change | Not Significant | TRUE | Cond1 UP + Cond2 Not Significant | Cond1_UP | Cond1_UP |
| Cdc73 | TRUE | 0.326925199 | 0.003631345 | No Change | Significant Positive | TRUE | 0.155180779 | 0.170111172 | No Change | Not Significant | TRUE | Cond1 Significant Positive + Cond2 Not Significant | None | None |
| Cdca2 | TRUE | 0.921813687 | 6.78E-07 | UP | UP | TRUE | -0.00384345 | 0.989206294 | No Change | Not Significant | TRUE | Cond1 UP + Cond2 Not Significant | Cond1_UP | Cond1_UP |
| Cdca3 | TRUE | 0.907724988 | 1.19E-08 | UP | UP | TRUE | 0.072986606 | 0.685788934 | No Change | Not Significant | TRUE | Cond1 UP + Cond2 Not Significant | Cond1_UP | Cond1_UP |
| Cdca4 | TRUE | 0.159339116 | 0.326976379 | No Change | Not Significant | TRUE | 0.012255075 | 0.936097283 | No Change | Not Significant | TRUE | Cond1 Not Significant + Cond2 Not Significant | None | None |
| Cdca5 | TRUE | 0.531836714 | 0.090681791 | No Change | Not Significant | TRUE | 0.109547324 | 0.633207999 | No Change | Not Significant | TRUE | Cond1 Not Significant + Cond2 Not Significant | None | None |
| Cdca7 | TRUE | -0.141036577 | 0.464617944 | No Change | Not Significant | TRUE | -0.209535131 | 0.048003141 | No Change | Significant Negative | TRUE | Cond1 Not Significant + Cond2 Significant Negative | None | None |
| Cdca7l | TRUE | 0.159344422 | 0.459757614 | No Change | Not Significant | TRUE | -0.043958602 | 0.88186074 | No Change | Not Significant | TRUE | Cond1 Not Significant + Cond2 Not Significant | None | None |
| Cdca8 | TRUE | 0.774494429 | 8.31E-06 | UP | UP | TRUE | 0.159675374 | 0.226076106 | No Change | Not Significant | TRUE | Cond1 UP + Cond2 Not Significant | Cond1_UP | Cond1_UP |

| geneID | Cond1_DF_Detected | Cond1_DF_ValueCol | Cond1_DF_PadjCol | Cond1_DF_Cutoff | Cond1_DF_Cutoff_Specific.x | Cond2_DF_Detected | Cond2_DF_ValueCol | Cond2_DF_PadjCol | Cond2_DF_Cutoff | Cond2_DF_Cutoff_Specific | BG_Method | RG1_All | RG2_Significant | RG3_SignificantChange |
|---|---|---|---|---|---|---|---|---|---|---|---|---|---|---|
| Cdcp1 | TRUE | 0.148036815 | 0.107027093 | No Change | Not Significant | TRUE | 0.082207527 | 0.143968177 | No Change | Not Significant | TRUE | Cond1 Not Significant + Cond2 Not Significant | None | None |
| Cdcp2 | TRUE | -1.703511717 | 1 | No Change | Not Significant | FALSE | NA | NA | No Change | Not Detected | FALSE | Background = FALSE | Background = FALSE | Background = FALSE |
| Cdcp3 | TRUE | 1.971597298 | 0.559327232 | No Change | Not Significant | FALSE | NA | NA | No Change | Not Detected | FALSE | Background = FALSE | Background = FALSE | Background = FALSE |
| Cdh1 | TRUE | -0.193595812 | 0.060376822 | No Change | Not Significant | TRUE | 0.051015315 | 0.653466789 | No Change | Not Significant | TRUE | Cond1 Not Significant + Cond2 Not Significant | None | None |
| Cdh10 | TRUE | 0.96033746 | 1 | No Change | Not Significant | FALSE | NA | NA | No Change | Not Detected | FALSE | Background = FALSE | Background = FALSE | Background = FALSE |
| Cdh11 | TRUE | 0.62048485 | 0.559613498 | No Change | Not Significant | TRUE | 0.684380385 | 0.299598598 | No Change | Not Significant | TRUE | Cond1 Not Significant + Cond2 Not Significant | None | None |
| Cdh12 | TRUE | 3.93793523 | 1 | No Change | Not Significant | FALSE | NA | NA | No Change | Not Detected | FALSE | Background = FALSE | Background = FALSE | Background = FALSE |
| Cdh13 | TRUE | 1.024353603 | 1 | No Change | Not Significant | FALSE | NA | NA | No Change | Not Detected | FALSE | Background = FALSE | Background = FALSE | Background = FALSE |
| Cdh15 | TRUE | 2.777955963 | 1 | No Change | Not Significant | FALSE | NA | NA | No Change | Not Detected | FALSE | Background = FALSE | Background = FALSE | Background = FALSE |
| Cdh16 | TRUE | 0.329555822 | 1 | No Change | Not Significant | FALSE | NA | NA | No Change | Not Detected | FALSE | Background = FALSE | Background = FALSE | Background = FALSE |
| Cdh17 | TRUE | -0.087428268 | 0.53139398 | No Change | Not Significant | TRUE | 0.093606626 | 0.290200492 | No Change | Not Significant | TRUE | Cond1 Not Significant + Cond2 Not Significant | None | None |
| Cdh2 | TRUE | 0.413066099 | 0.752646336 | No Change | Not Significant | TRUE | 1.167361649 | 0.036914915 | UP | UP | TRUE | Cond1 Not Significant + Cond2 UP | Cond2_UP | Cond2_UP |
| Cdh20 | TRUE | 0.291403192 | 1 | No Change | Not Significant | FALSE | NA | NA | No Change | Not Detected | FALSE | Background = FALSE | Background = FALSE | Background = FALSE |
| Cdh22 | TRUE | 0.607091558 | 1 | No Change | Not Significant | FALSE | NA | NA | No Change | Not Detected | FALSE | Background = FALSE | Background = FALSE | Background = FALSE |
| Cdh23 | TRUE | 0.564561708 | 0.679754841 | No Change | Not Significant | TRUE | 0.118138914 | 0.940691929 | No Change | Not Significant | TRUE | Cond1 Not Significant + Cond2 Not Significant | None | None |
| Cdh24 | TRUE | 0.25944757 | 0.397165509 | No Change | Not Significant | TRUE | 0.066388703 | 0.892439856 | No Change | Not Significant | TRUE | Cond1 Not Significant + Cond2 Not Significant | None | None |
| Cdh26 | TRUE | 0.298036901 | 0.424602824 | No Change | Not Significant | TRUE | 0.099348453 | 0.770206405 | No Change | Not Significant | TRUE | Cond1 Not Significant + Cond2 Not Significant | None | None |
| Cdh3 | TRUE | 0.553759129 | 0.50637921 | No Change | Not Significant | TRUE | 0.381092042 | 0.702150736 | No Change | Not Significant | TRUE | Cond1 Not Significant + Cond2 Not Significant | None | None |
| Cdh4 | TRUE | -0.26063473 | 0.880656876 | No Change | Not Significant | TRUE | 0.929641998 | 0.285746134 | No Change | Not Significant | TRUE | Cond1 Not Significant + Cond2 Not Significant | None | None |
| Cdh6 | TRUE | 0.142284816 | 1 | No Change | Not Significant | FALSE | NA | NA | No Change | Not Detected | FALSE | Background = FALSE | Background = FALSE | Background = FALSE |
| Cdh7 | TRUE | -0.691919105 | 1 | No Change | Not Significant | FALSE | NA | NA | No Change | Not Detected | FALSE | Background = FALSE | Background = FALSE | Background = FALSE |
| Cdh8 | TRUE | 2.655855407 | 1 | No Change | Not Significant | FALSE | NA | NA | No Change | Not Detected | FALSE | Background = FALSE | Background = FALSE | Background = FALSE |
| Cdh9 | TRUE | -1.470249984 | 1 | No Change | Not Significant | FALSE | NA | NA | No Change | Not Detected | FALSE | Background = FALSE | Background = FALSE | Background = FALSE |
| Cdhr1 | TRUE | 0.109234846 | 0.783477444 | No Change | Not Significant | TRUE | -0.095678063 | 0.623117424 | No Change | Not Significant | TRUE | Cond1 Not Significant + Cond2 Not Significant | None | None |
| Cdhr2 | TRUE | -0.412417106 | 0.006903589 | No Change | Significant Negative | TRUE | -0.18504019 | 0.018226234 | No Change | Significant Negative | TRUE | Cond1 Significant Negative + Cond2 Significant Negative | None | None |
| Cdhr3 | TRUE | 3.094690106 | 1 | No Change | Not Significant | FALSE | NA | NA | No Change | Not Detected | FALSE | Background = FALSE | Background = FALSE | Background = FALSE |
| Cdhr4 | TRUE | 1.935370195 | 1 | No Change | Not Significant | FALSE | NA | NA | No Change | Not Detected | FALSE | Background = FALSE | Background = FALSE | Background = FALSE |
| Cdhr5 | TRUE | -0.400697645 | 0.013111071 | No Change | Significant Negative | TRUE | -0.206100007 | 0.026714719 | No Change | Significant Negative | TRUE | Cond1 Significant Negative + Cond2 Significant Negative | None | None |
| Cdin1 | TRUE | 0.232068636 | 0.446932387 | No Change | Not Significant | TRUE | 0.421120581 | 0.040015956 | No Change | Significant Positive | TRUE | Cond1 Not Significant + Cond2 Significant Positive | None | None |
| Cdip1 | TRUE | -0.060657373 | 0.696651931 | No Change | Not Significant | TRUE | -0.225947667 | 0.007789628 | No Change | Significant Negative | TRUE | Cond1 Not Significant + Cond2 Significant Negative | None | None |
| Cdipt | TRUE | 0.068298517 | 0.514433539 | No Change | Not Significant | TRUE | 0.0880111 | 0.336231563 | No Change | Not Significant | TRUE | Cond1 Not Significant + Cond2 Not Significant | None | None |
| Cdiptos | TRUE | -0.617211159 | 0.460735958 | No Change | Not Significant | TRUE | -0.342104633 | 0.642408959 | No Change | Not Significant | TRUE | Cond1 Not Significant + Cond2 Not Significant | None | None |
| Cdk1 | TRUE | 0.630445207 | 0.000938301 | UP | UP | TRUE | 0.077923474 | 0.745625688 | No Change | Not Significant | TRUE | Cond1 UP + Cond2 Not Significant | Cond1_UP | Cond1_UP |
| Cdk10 | TRUE | -0.16052879 | 0.132820779 | No Change | Not Significant | TRUE | 0.007651667 | 0.959446759 | No Change | Not Significant | TRUE | Cond1 Not Significant + Cond2 Not Significant | None | None |
| Cdk11b | TRUE | 0.151512323 | 0.279437874 | No Change | Not Significant | TRUE | 0.090350365 | 0.342017997 | No Change | Not Significant | TRUE | Cond1 Not Significant + Cond2 Not Significant | None | None |
| Cdk12 | TRUE | 0.073470601 | 0.548613253 | No Change | Not Significant | TRUE | -0.168643649 | 0.000448813 | No Change | Significant Negative | TRUE | Cond1 Not Significant + Cond2 Significant Negative | None | None |
| Cdk13 | TRUE | 0.164347696 | 0.015751711 | No Change | Significant Positive | TRUE | -0.071251389 | 0.218011427 | No Change | Not Significant | TRUE | Cond1 Significant Positive + Cond2 Not Significant | None | None |
| Cdk14 | TRUE | 0.83154056 | 0.000752996 | UP | UP | TRUE | 0.790711244 | 8.02E-07 | UP | UP | TRUE | Cond1 UP + Cond2 UP | Core_UP | Core_UP |
| Cdk15 | TRUE | 0.488097818 | 1 | No Change | Not Significant | FALSE | NA | NA | No Change | Not Detected | FALSE | Background = FALSE | Background = FALSE | Background = FALSE |
| Cdk16 | TRUE | 0.032310606 | 0.727020313 | No Change | Not Significant | TRUE | 0.049891034 | 0.571937821 | No Change | Not Significant | TRUE | Cond1 Not Significant + Cond2 Not Significant | None | None |
| Cdk17 | TRUE | -0.187181625 | 0.416065952 | No Change | Not Significant | TRUE | -0.20865078 | 0.033928886 | No Change | Significant Negative | TRUE | Cond1 Not Significant + Cond2 Significant Negative | None | None |
| Cdk18 | TRUE | 0.753736377 | 3.71E-06 | UP | UP | TRUE | 0.189697867 | 0.157771355 | No Change | Not Significant | TRUE | Cond1 UP + Cond2 Not Significant | Cond1_UP | Cond1_UP |
| Cdk19 | TRUE | -0.100414931 | 0.554606509 | No Change | Not Significant | TRUE | -0.120507977 | 0.138361096 | No Change | Not Significant | TRUE | Cond1 Not Significant + Cond2 Not Significant | None | None |
| Cdk2 | TRUE | 0.025774013 | 0.929786937 | No Change | Not Significant | TRUE | -0.127738895 | 0.352173719 | No Change | Not Significant | TRUE | Cond1 Not Significant + Cond2 Not Significant | None | None |
| Cdk20 | TRUE | 0.084040926 | 0.833389267 | No Change | Not Significant | TRUE | 0.034082984 | 0.931678839 | No Change | Not Significant | TRUE | Cond1 Not Significant + Cond2 Not Significant | None | None |
| Cdk2ap1 | TRUE | -0.317892845 | 0.233281352 | No Change | Not Significant | TRUE | -0.004023139 | 0.982572197 | No Change | Not Significant | TRUE | Cond1 Not Significant + Cond2 Not Significant | None | None |
| Cdk2ap2 | TRUE | 1.523140514 | 1.71E-63 | UP | UP | TRUE | 1.329351059 | 3.01E-50 | UP | UP | TRUE | Cond1 UP + Cond2 UP | Core_UP | Core_UP |
| Cdk4 | TRUE | 0.056241416 | 0.800270871 | No Change | Not Significant | TRUE | -0.043786077 | 0.753561 | No Change | Not Significant | TRUE | Cond1 Not Significant + Cond2 Not Significant | None | None |
| Cdk5 | TRUE | -0.223736017 | 0.047698058 | No Change | Significant Negative | TRUE | 0.009287536 | 0.94557102 | No Change | Not Significant | TRUE | Cond1 Significant Negative + Cond2 Not Significant | None | None |
| Cdk5r1 | TRUE | 0.715021065 | 0.160458367 | No Change | Not Significant | TRUE | -0.224810546 | 0.671740242 | No Change | Not Significant | TRUE | Cond1 Not Significant + Cond2 Not Significant | None | None |
| Cdk5r2 | TRUE | -0.547772562 | 1 | No Change | Not Significant | FALSE | NA | NA | No Change | Not Detected | FALSE | Background = FALSE | Background = FALSE | Background = FALSE |
| Cdk5rap1 | TRUE | -0.186539481 | 0.685335967 | No Change | Not Significant | TRUE | -0.078600736 | 0.811381567 | No Change | Not Significant | TRUE | Cond1 Not Significant + Cond2 Not Significant | None | None |
| Cdk5rap2 | TRUE | 0.437215006 | 4.16E-05 | No Change | Significant Positive | TRUE | 0.010988013 | 0.947098467 | No Change | Not Significant | TRUE | Cond1 Significant Positive + Cond2 Not Significant | None | None |
| Cdk5rap3 | TRUE | 0.942495702 | 7.28E-33 | UP | UP | TRUE | 0.620307586 | 6.68E-20 | UP | UP | TRUE | Cond1 UP + Cond2 UP | Core_UP | Core_UP |
| Cdk6 | TRUE | -0.368289224 | 0.008984351 | No Change | Significant Negative | TRUE | -0.376156426 | 0.00012886 | No Change | Significant Negative | TRUE | Cond1 Significant Negative + Cond2 Significant Negative | None | None |
| Cdk7 | TRUE | 0.06350932 | 0.676855263 | No Change | Not Significant | TRUE | 0.067449185 | 0.543683373 | No Change | Not Significant | TRUE | Cond1 Not Significant + Cond2 Not Significant | None | None |
| Cdk8 | TRUE | 0.0226503 | 0.870990647 | No Change | Not Significant | TRUE | -0.137413839 | 0.118917792 | No Change | Not Significant | TRUE | Cond1 Not Significant + Cond2 Not Significant | None | None |
| Cdk9 | TRUE | -0.15434096 | 0.063640522 | No Change | Not Significant | TRUE | -0.198853104 | 0.001710212 | No Change | Significant Negative | TRUE | Cond1 Not Significant + Cond2 Significant Negative | None | None |
| Cdkal1 | TRUE | -0.133461028 | 0.300075862 | No Change | Not Significant | TRUE | -0.010663236 | 0.944776763 | No Change | Not Significant | TRUE | Cond1 Not Significant + Cond2 Not Significant | None | None |
| Cdkl1 | TRUE | -2.319142869 | 7.30E-05 | DOWN | DOWN | TRUE | -1.524893095 | 3.83E-09 | DOWN | DOWN | TRUE | Cond1 DOWN + Cond2 DOWN | Core_DOWN | Core_DOWN |
| Cdkl2 | TRUE | 0.901167182 | 2.25E-11 | UP | UP | TRUE | 0.398121696 | 6.43E-06 | No Change | Significant Positive | TRUE | Cond1 UP + Cond2 Significant Positive | Core_UP | Cond1_UP |
| Cdkl3 | TRUE | 0.502618879 | 0.042188532 | UP | UP | TRUE | 0.123682711 | 0.736439527 | No Change | Not Significant | TRUE | Cond1 UP + Cond2 Not Significant | Cond1_UP | Cond1_UP |
| Cdkl4 | TRUE | -0.226887055 | 1 | No Change | Not Significant | FALSE | NA | NA | No Change | Not Detected | FALSE | Background = FALSE | Background = FALSE | Background = FALSE |
| Cdkl5 | TRUE | 0.663709184 | 0.000150633 | UP | UP | TRUE | 0.178590337 | 0.345432975 | No Change | Not Significant | TRUE | Cond1 UP + Cond2 Not Significant | Cond1_UP | Cond1_UP |
| Cdkn1a | TRUE | 0.364809767 | 0.1468042 | No Change | Not Significant | TRUE | 0.106005576 | 0.326064971 | No Change | Not Significant | TRUE | Cond1 Not Significant + Cond2 Not Significant | None | None |
| Cdkn1b | TRUE | -0.120642609 | 0.249566874 | No Change | Not Significant | TRUE | -0.028892607 | 0.72429229 | No Change | Not Significant | TRUE | Cond1 Not Significant + Cond2 Not Significant | None | None |
| Cdkn1c | TRUE | 0.555515277 | 0.046240775 | UP | UP | TRUE | 0.24894274 | 0.150848528 | No Change | Not Significant | TRUE | Cond1 UP + Cond2 Not Significant | Cond1_UP | Cond1_UP |
| Cdkn2a | TRUE | 0.739565451 | 1 | No Change | Not Significant | FALSE | NA | NA | No Change | Not Detected | FALSE | Background = FALSE | Background = FALSE | Background = FALSE |
| Cdkn2aip | TRUE | -0.209609606 | 0.090782827 | No Change | Not Significant | TRUE | -0.19649353 | 0.04081985 | No Change | Significant Negative | TRUE | Cond1 Not Significant + Cond2 Significant Negative | None | None |
| Cdkn2aipnl | TRUE | 0.29181261 | 0.054759062 | No Change | Not Significant | TRUE | 0.026134247 | 0.829512341 | No Change | Not Significant | TRUE | Cond1 Not Significant + Cond2 Not Significant | None | None |
| Cdkn2b | TRUE | -0.233392585 | 0.167639761 | No Change | Not Significant | TRUE | -0.039909521 | 0.792937111 | No Change | Not Significant | TRUE | Cond1 Not Significant + Cond2 Not Significant | None | None |
| Cdkn2c | TRUE | 0.306931123 | 0.219013029 | No Change | Not Significant | TRUE | 0.1363937 | 0.617324668 | No Change | Not Significant | TRUE | Cond1 Not Significant + Cond2 Not Significant | None | None |
| Cdkn2d | TRUE | 0.29436986 | 0.242436056 | No Change | Not Significant | TRUE | 0.049852824 | 0.840074678 | No Change | Not Significant | TRUE | Cond1 Not Significant + Cond2 Not Significant | None | None |
| Cdkn3 | TRUE | 0.744056177 | 0.00013417 | UP | UP | TRUE | 0.070916221 | 0.786125192 | No Change | Not Significant | TRUE | Cond1 UP + Cond2 Not Significant | Cond1_UP | Cond1_UP |
| Cdnf | TRUE | 0.521651741 | 0.18053173 | No Change | Not Significant | TRUE | -0.002363128 | 0.996814475 | No Change | Not Significant | TRUE | Cond1 Not Significant + Cond2 Not Significant | None | None |
| Cdo1 | TRUE | 0.251526659 | 0.278515327 | No Change | Not Significant | TRUE | -0.02865719 | 0.853157579 | No Change | Not Significant | TRUE | Cond1 Not Significant + Cond2 Not Significant | None | None |
| Cdon | TRUE | 0.17951289 | 0.188479843 | No Change | Not Significant | TRUE | 0.09515671 | 0.469307431 | No Change | Not Significant | TRUE | Cond1 Not Significant + Cond2 Not Significant | None | None |
| Cdpf1 | TRUE | 0.613017293 | 3.14E-10 | UP | UP | TRUE | 0.520923927 | 2.54E-07 | UP | UP | TRUE | Cond1 UP + Cond2 UP | Core_UP | Core_UP |
| Cdr1os | TRUE | -0.305096388 | 0.962580298 | No Change | Not Significant | FALSE | NA | NA | No Change | Not Detected | FALSE | Background = FALSE | Background = FALSE | Background = FALSE |
| Cdr2 | TRUE | 0.772096875 | 4.46E-05 | UP | UP | TRUE | 0.051970463 | 0.819270326 | No Change | Not Significant | TRUE | Cond1 UP + Cond2 Not Significant | Cond1_UP | Cond1_UP |
| Cdr2l | TRUE | 0.445594217 | 0.055305475 | No Change | Not Significant | TRUE | 0.039146199 | 0.891549271 | No Change | Not Significant | TRUE | Cond1 Not Significant + Cond2 Not Significant | None | None |
| Cds1 | TRUE | -0.088647554 | 0.53655334 | No Change | Not Significant | TRUE | -0.177320048 | 0.034530138 | No Change | Significant Negative | TRUE | Cond1 Not Significant + Cond2 Significant Negative | None | None |
| Cds2 | TRUE | -0.263544963 | 0.035388872 | No Change | Significant Negative | TRUE | -0.11306649 | 0.086281495 | No Change | Not Significant | TRUE | Cond1 Significant Negative + Cond2 Not Significant | None | None |
| Cdsn | TRUE | 0.269495136 | 0.675880563 | No Change | Not Significant | TRUE | -0.084063503 | 0.900566916 | No Change | Not Significant | TRUE | Cond1 Not Significant + Cond2 Not Significant | None | None |
| Cdt1 | TRUE | -0.093601819 | 0.709429427 | No Change | Not Significant | TRUE | -0.112436622 | 0.645466424 | No Change | Not Significant | TRUE | Cond1 Not Significant + Cond2 Not Significant | None | None |
| Cdv3 | TRUE | 0.225772383 | 0.049857012 | No Change | Significant Positive | TRUE | 0.068382765 | 0.361756348 | No Change | Not Significant | TRUE | Cond1 Significant Positive + Cond2 Not Significant | None | None |
| Cdx1 | TRUE | -0.047606128 | 0.769577776 | No Change | Not Significant | TRUE | 0.037864239 | 0.810530941 | No Change | Not Significant | TRUE | Cond1 Not Significant + Cond2 Not Significant | None | None |
| Cdx2 | TRUE | 0.277472726 | 0.132476266 | No Change | Not Significant | TRUE | 0.019495427 | 0.891758816 | No Change | Not Significant | TRUE | Cond1 Not Significant + Cond2 Not Significant | None | None |
| Cdyl | TRUE | 0.073575608 | 0.660497294 | No Change | Not Significant | TRUE | -0.025479499 | 0.880695457 | No Change | Not Significant | TRUE | Cond1 Not Significant + Cond2 Not Significant | None | None |
| Cdyl2 | TRUE | 0.160052384 | 0.764064128 | No Change | Not Significant | TRUE | -0.102694144 | 0.761533505 | No Change | Not Significant | TRUE | Cond1 Not Significant + Cond2 Not Significant | None | None |
| Ceacam1 | TRUE | -0.265434745 | 0.109351285 | No Change | Not Significant | TRUE | -0.069517857 | 0.249693708 | No Change | Not Significant | TRUE | Cond1 Not Significant + Cond2 Not Significant | None | None |
| Ceacam10 | TRUE | -0.696602865 | 0.024791549 | DOWN | DOWN | TRUE | 0.331060356 | 0.09547586 | No Change | Not Significant | TRUE | Cond1 DOWN + Cond2 Not Significant | Cond1_DOWN | Cond1_DOWN |
| Ceacam11 | TRUE | 1.686550943 | 1 | No Change | Not Significant | FALSE | NA | NA | No Change | Not Detected | FALSE | Background = FALSE | Background = FALSE | Background = FALSE |
| Ceacam12 | TRUE | -0.402780086 | 0.420139706 | No Change | Not Significant | TRUE | -1.40179365 | 0.001066785 | DOWN | DOWN | TRUE | Cond1 Not Significant + Cond2 DOWN | Cond2_DOWN | Cond2_DOWN |
| Ceacam13 | TRUE | 1.108765806 | 1 | No Change | Not Significant | FALSE | NA | NA | No Change | Not Detected | FALSE | Background = FALSE | Background = FALSE | Background = FALSE |
| Ceacam18 | TRUE | -0.495887862 | 0.012051751 | No Change | Significant Negative | TRUE | -0.405270029 | 0.000167595 | No Change | Significant Negative | TRUE | Cond1 Significant Negative + Cond2 Significant Negative | None | None |
| Ceacam19 | TRUE | 0.380383504 | 1 | No Change | Not Significant | FALSE | NA | NA | No Change | Not Detected | FALSE | Background = FALSE | Background = FALSE | Background = FALSE |
| Ceacam2 | TRUE | -0.344088807 | 0.362441146 | No Change | Not Significant | TRUE | -0.228435017 | 0.226749513 | No Change | Not Significant | TRUE | Cond1 Not Significant + Cond2 Not Significant | None | None |
| Ceacam20 | TRUE | -0.014827299 | 0.975869522 | No Change | Not Significant | TRUE | -0.07774078 | 0.679246742 | No Change | Not Significant | TRUE | Cond1 Not Significant + Cond2 Not Significant | None | None |
| Ceacam3 | TRUE | 0.176011399 | 1 | No Change | Not Significant | FALSE | NA | NA | No Change | Not Detected | FALSE | Background = FALSE | Background = FALSE | Background = FALSE |
| Ceacam9 | TRUE | 1.288357957 | 1 | No Change | Not Significant | FALSE | NA | NA | No Change | Not Detected | FALSE | Background = FALSE | Background = FALSE | Background = FALSE |
| Cebpa | TRUE | 0.041243233 | 0.807915411 | No Change | Not Significant | TRUE | -0.054499887 | 0.746077797 | No Change | Not Significant | TRUE | Cond1 Not Significant + Cond2 Not Significant | None | None |
| Cebpb | TRUE | 0.455458921 | 0.190674731 | No Change | Not Significant | TRUE | -0.373238417 | 0.179737215 | No Change | Not Significant | TRUE | Cond1 Not Significant + Cond2 Not Significant | None | None |
| Cebpd | TRUE | 0.50604147 | 0.041407144 | UP | UP | TRUE | -0.216370954 | 0.365355509 | No Change | Not Significant | TRUE | Cond1 UP + Cond2 Not Significant | Cond1_UP | Cond1_UP |
| Cebpg | TRUE | 0.0349375 | 0.795381772 | No Change | Not Significant | TRUE | 0.056988073 | 0.55760602 | No Change | Not Significant | TRUE | Cond1 Not Significant + Cond2 Not Significant | None | None |
| Cebpz | TRUE | -0.046046933 | 0.852575063 | No Change | Not Significant | TRUE | -0.14065284 | 0.323307444 | No Change | Not Significant | TRUE | Cond1 Not Significant + Cond2 Not Significant | None | None |
| Cebpzos | TRUE | 0.05715031 | 0.797446101 | No Change | Not Significant | TRUE | -0.036397516 | 0.84234163 | No Change | Not Significant | TRUE | Cond1 Not Significant + Cond2 Not Significant | None | None |
| Cecr2 | TRUE | 0.620318356 | 0.707452102 | No Change | Not Significant | TRUE | -0.304349432 | 0.678988684 | No Change | Not Significant | TRUE | Cond1 Not Significant + Cond2 Not Significant | None | None |
| Cel | TRUE | -0.499883509 | 1 | No Change | Not Significant | FALSE | NA | NA | No Change | Not Detected | FALSE | Background = FALSE | Background = FALSE | Background = FALSE |
| Cela1 | TRUE | -1.255656282 | 8.13E-09 | DOWN | DOWN | TRUE | -0.376052101 | 0.016432856 | No Change | Significant Negative | TRUE | Cond1 DOWN + Cond2 Significant Negative | Core_DOWN | Cond1_DOWN |
| Cela2a | TRUE | -3.058893584 | 1 | No Change | Not Significant | FALSE | NA | NA | No Change | Not Detected | FALSE | Background = FALSE | Background = FALSE | Background = FALSE |
| Cela3b | TRUE | 0.176009995 | 1 | No Change | Not Significant | FALSE | NA | NA | No Change | Not Detected | FALSE | Background = FALSE | Background = FALSE | Background = FALSE |
| Celf1 | TRUE | 0.069565635 | 0.541253166 | No Change | Not Significant | TRUE | 0.02167544 | 0.763734926 | No Change | Not Significant | TRUE | Cond1 Not Significant + Cond2 Not Significant | None | None |
| Celf2 | TRUE | 0.164329055 | 0.756475084 | No Change | Not Significant | TRUE | 0.034306606 | 0.885371178 | No Change | Not Significant | TRUE | Cond1 Not Significant + Cond2 Not Significant | None | None |
| Celf3 | TRUE | 0.219679176 | 0.698946027 | No Change | Not Significant | TRUE | -0.069100689 | 0.865987203 | No Change | Not Significant | TRUE | Cond1 Not Significant + Cond2 Not Significant | None | None |
| Celf4 | TRUE | -0.262903676 | 0.805274537 | No Change | Not Significant | TRUE | -0.357442107 | 0.304713258 | No Change | Not Significant | TRUE | Cond1 Not Significant + Cond2 Not Significant | None | None |
| Celf5 | TRUE | -1.356624573 | 1.41E-05 | DOWN | DOWN | TRUE | -0.206681881 | 0.575843468 | No Change | Not Significant | TRUE | Cond1 DOWN + Cond2 Not Significant | Cond1_DOWN | Cond1_DOWN |
| Celf6 | TRUE | 0.66201581 | 0.129891785 | No Change | Not Significant | TRUE | -0.051212022 | 0.927643978 | No Change | Not Significant | TRUE | Cond1 Not Significant + Cond2 Not Significant | None | None |
| Celrr | TRUE | 2.45218749 | 1 | No Change | Not Significant | FALSE | NA | NA | No Change | Not Detected | FALSE | Background = FALSE | Background = FALSE | Background = FALSE |
| Celsr1 | TRUE | -0.04001813 | 0.818950015 | No Change | Not Significant | TRUE | 0.005956883 | 0.975106302 | No Change | Not Significant | TRUE | Cond1 Not Significant + Cond2 Not Significant | None | None |
| Celsr2 | TRUE | -0.117587558 | 0.772538316 | No Change | Not Significant | TRUE | -0.380660539 | 0.105513168 | No Change | Not Significant | TRUE | Cond1 Not Significant + Cond2 Not Significant | None | None |
| Celsr3 | TRUE | -0.154172415 | 0.888506311 | No Change | Not Significant | TRUE | 0.238126787 | 0.633814677 | No Change | Not Significant | TRUE | Cond1 Not Significant + Cond2 Not Significant | None | None |
| Cemip | TRUE | 0.552173621 | 0.747948226 | No Change | Not Significant | TRUE | 1.0348062 | 0.186591593 | No Change | Not Significant | TRUE | Cond1 Not Significant + Cond2 Not Significant | None | None |
| Cemip2 | TRUE | -0.163379362 | 0.197279169 | No Change | Not Significant | TRUE | -0.129599595 | 0.020709859 | No Change | Significant Negative | TRUE | Cond1 Not Significant + Cond2 Significant Negative | None | None |
| Cend1 | TRUE | 0.021061847 | 1 | No Change | Not Significant | FALSE | NA | NA | No Change | Not Detected | FALSE | Background = FALSE | Background = FALSE | Background = FALSE |
| Cenpa | TRUE | 0.797910241 | 1.27E-09 | UP | UP | TRUE | 0.063755451 | 0.68535506 | No Change | Not Significant | TRUE | Cond1 UP + Cond2 Not Significant | Cond1_UP | Cond1_UP |
| Cenpb | TRUE | -0.135814784 | 0.02823687 | No Change | Significant Negative | TRUE | -0.011980699 | 0.924226793 | No Change | Not Significant | TRUE | Cond1 Significant Negative + Cond2 Not Significant | None | None |
| Cenpc1 | TRUE | 0.278451186 | 0.032806559 | No Change | Significant Positive | TRUE | 0.104577181 | 0.361765545 | No Change | Not Significant | TRUE | Cond1 Significant Positive + Cond2 Not Significant | None | None |
| Cenpe | TRUE | 0.690159773 | 6.82E-05 | UP | UP | TRUE | -0.066552637 | 0.771707437 | No Change | Not Significant | TRUE | Cond1 UP + Cond2 Not Significant | Cond1_UP | Cond1_UP |
| Cenpf | TRUE | 0.673809703 | 0.000134753 | UP | UP | TRUE | -0.03806544 | 0.831904926 | No Change | Not Significant | TRUE | Cond1 UP + Cond2 Not Significant | Cond1_UP | Cond1_UP |
| Cenph | TRUE | 0.369132197 | 0.304738078 | No Change | Not Significant | TRUE | 0.1580372 | 0.58700794 | No Change | Not Significant | TRUE | Cond1 Not Significant + Cond2 Not Significant | None | None |
| Cenpi | TRUE | 0.785999121 | 0.000159134 | UP | UP | TRUE | 0.179207675 | 0.523831138 | No Change | Not Significant | TRUE | Cond1 UP + Cond2 Not Significant | Cond1_UP | Cond1_UP |
| Cenpj | TRUE | 0.211956891 | 0.265192684 | No Change | Not Significant | TRUE | 0.123922483 | 0.508185669 | No Change | Not Significant | TRUE | Cond1 Not Significant + Cond2 Not Significant | None | None |
| Cenpk | TRUE | 0.835550994 | 0.000153277 | UP | UP | TRUE | 0.271071643 | 0.21135596 | No Change | Not Significant | TRUE | Cond1 UP + Cond2 Not Significant | Cond1_UP | Cond1_UP |
| Cenpl | TRUE | 0.415274284 | 0.068863768 | No Change | Not Significant | TRUE | -0.081639684 | 0.708115319 | No Change | Not Significant | TRUE | Cond1 Not Significant + Cond2 Not Significant | None | None |
| Cenpm | TRUE | 0.746714064 | 0.000732064 | UP | UP | TRUE | -0.012408949 | 0.972704281 | No Change | Not Significant | TRUE | Cond1 UP + Cond2 Not Significant | Cond1_UP | Cond1_UP |
| Cenpn | TRUE | 0.574765665 | 0.045254367 | UP | UP | TRUE | 0.079955444 | 0.847823651 | No Change | Not Significant | TRUE | Cond1 UP + Cond2 Not Significant | Cond1_UP | Cond1_UP |
| Cenpo | TRUE | 0.196912148 | 0.25785531 | No Change | Not Significant | TRUE | -0.00420699 | 0.985688558 | No Change | Not Significant | TRUE | Cond1 Not Significant + Cond2 Not Significant | None | None |
| Cenpp | TRUE | 0.511673202 | 0.041791803 | UP | UP | TRUE | -0.280964033 | 0.267808218 | No Change | Not Significant | TRUE | Cond1 UP + Cond2 Not Significant | Cond1_UP | Cond1_UP |
| Cenpq | TRUE | 0.705969229 | 0.003689141 | UP | UP | TRUE | 0.144448179 | 0.529013894 | No Change | Not Significant | TRUE | Cond1 UP + Cond2 Not Significant | Cond1_UP | Cond1_UP |
| Cenps | TRUE | 0.384907365 | 0.30270362 | No Change | Not Significant | TRUE | 0.372522398 | 0.263862344 | No Change | Not Significant | TRUE | Cond1 Not Significant + Cond2 Not Significant | None | None |
| Cenpt | TRUE | 0.299419714 | 0.066267555 | No Change | Not Significant | TRUE | 0.151147543 | 0.485418672 | No Change | Not Significant | TRUE | Cond1 Not Significant + Cond2 Not Significant | None | None |
| Cenpu | TRUE | 0.528008918 | 0.050667362 | No Change | Not Significant | TRUE | 0.067833188 | 0.864247815 | No Change | Not Significant | TRUE | Cond1 Not Significant + Cond2 Not Significant | None | None |
| Cenpv | TRUE | -0.02670446 | 0.875081807 | No Change | Not Significant | TRUE | -0.032749546 | 0.800888833 | No Change | Not Significant | TRUE | Cond1 Not Significant + Cond2 Not Significant | None | None |
| Cenpw | TRUE | 1.828719621 | 6.18E-16 | UP | UP | TRUE | 1.097951315 | 2.73E-06 | UP | UP | TRUE | Cond1 UP + Cond2 UP | Core_UP | Core_UP |
| Cenpx | TRUE | 0.545023139 | 2.40E-07 | UP | UP | TRUE | 0.428780353 | 2.13E-07 | No Change | Significant Positive | TRUE | Cond1 UP + Cond2 Significant Positive | Core_UP | Cond1_UP |
| Cep104 | TRUE | 0.25672109 | 0.041117602 | No Change | Significant Positive | TRUE | 0.078197199 | 0.626292308 | No Change | Not Significant | TRUE | Cond1 Significant Positive + Cond2 Not Significant | None | None |
| Cep112 | TRUE | 1.040148282 | 0.005248535 | UP | UP | TRUE | 0.39776036 | 0.323030784 | No Change | Not Significant | TRUE | Cond1 UP + Cond2 Not Significant | Cond1_UP | Cond1_UP |
| Cep112it | TRUE | 2.239808779 | 1 | No Change | Not Significant | FALSE | NA | NA | No Change | Not Detected | FALSE | Background = FALSE | Background = FALSE | Background = FALSE |
| Cep120 | TRUE | -0.109663195 | 0.323295567 | No Change | Not Significant | TRUE | -0.123762408 | 0.264480943 | No Change | Not Significant | TRUE | Cond1 Not Significant + Cond2 Not Significant | None | None |
| Cep126 | TRUE | 0.292751161 | 0.288572541 | No Change | Not Significant | TRUE | 0.279273182 | 0.245400751 | No Change | Not Significant | TRUE | Cond1 Not Significant + Cond2 Not Significant | None | None |
| Cep128 | TRUE | 0.425902809 | 0.064265646 | No Change | Not Significant | TRUE | 0.055944943 | 0.808705755 | No Change | Not Significant | TRUE | Cond1 Not Significant + Cond2 Not Significant | None | None |
| Cep131 | TRUE | 1.218231392 | 8.16E-20 | UP | UP | TRUE | 0.534941229 | 9.66E-10 | UP | UP | TRUE | Cond1 UP + Cond2 UP | Core_UP | Core_UP |
| Cep135 | TRUE | 0.367392087 | 0.012618891 | No Change | Significant Positive | TRUE | -0.063275651 | 0.755288004 | No Change | Not Significant | TRUE | Cond1 Significant Positive + Cond2 Not Significant | None | None |
| Cep152 | TRUE | 0.128743451 | 0.465324602 | No Change | Not Significant | TRUE | -0.042113449 | 0.822000293 | No Change | Not Significant | TRUE | Cond1 Not Significant + Cond2 Not Significant | None | None |
| Cep162 | TRUE | 0.124579968 | 0.29613919 | No Change | Not Significant | TRUE | 0.059391766 | 0.672888818 | No Change | Not Significant | TRUE | Cond1 Not Significant + Cond2 Not Significant | None | None |
| Cep164 | TRUE | -0.139454471 | 0.211747172 | No Change | Not Significant | TRUE | 0.024488091 | 0.834200336 | No Change | Not Significant | TRUE | Cond1 Not Significant + Cond2 Not Significant | None | None |
| Cep170 | TRUE | 0.970867246 | 0.02891482 | UP | UP | TRUE | 0.486490662 | 0.174465906 | No Change | Not Significant | TRUE | Cond1 UP + Cond2 Not Significant | Cond1_UP | Cond1_UP |
| Cep170b | TRUE | -0.062341457 | 0.447960257 | No Change | Not Significant | TRUE | -0.033238658 | 0.765902723 | No Change | Not Significant | TRUE | Cond1 Not Significant + Cond2 Not Significant | None | None |
| Cep19 | TRUE | 0.034436265 | 0.825194181 | No Change | Not Significant | TRUE | 0.253274941 | 0.00054606 | No Change | Significant Positive | TRUE | Cond1 Not Significant + Cond2 Significant Positive | None | None |
| Cep192 | TRUE | 0.567637928 | 5.62E-05 | UP | UP | TRUE | 0.029005338 | 0.858410189 | No Change | Not Significant | TRUE | Cond1 UP + Cond2 Not Significant | Cond1_UP | Cond1_UP |
| Cep20 | TRUE | 0.167934176 | 0.109921135 | No Change | Not Significant | TRUE | 0.19378353 | 0.00596069 | No Change | Significant Positive | TRUE | Cond1 Not Significant + Cond2 Significant Positive | None | None |
| Cep250 | TRUE | 0.099576157 | 0.628183969 | No Change | Not Significant | TRUE | -0.079551031 | 0.550870027 | No Change | Not Significant | TRUE | Cond1 Not Significant + Cond2 Not Significant | None | None |
| Cep290 | TRUE | 0.554570361 | 8.98E-08 | UP | UP | TRUE | 0.330530885 | 0.017027773 | No Change | Significant Positive | TRUE | Cond1 UP + Cond2 Significant Positive | Core_UP | Cond1_UP |
| Cep295 | TRUE | 0.281560125 | 0.064474124 | No Change | Not Significant | TRUE | -0.06547475 | 0.751378251 | No Change | Not Significant | TRUE | Cond1 Not Significant + Cond2 Not Significant | None | None |
| Cep295nl | TRUE | -2.358989387 | 1 | No Change | Not Significant | FALSE | NA | NA | No Change | Not Detected | FALSE | Background = FALSE | Background = FALSE | Background = FALSE |
| Cep350 | TRUE | -0.007012866 | 0.968109605 | No Change | Not Significant | TRUE | -0.123206796 | 0.05836938 | No Change | Not Significant | TRUE | Cond1 Not Significant + Cond2 Not Significant | None | None |
| Cep41 | TRUE | 0.062348702 | 0.861143224 | No Change | Not Significant | TRUE | 0.08205331 | 0.767653434 | No Change | Not Significant | TRUE | Cond1 Not Significant + Cond2 Not Significant | None | None |
| Cep43 | TRUE | 0.26974494 | 0.058238694 | No Change | Not Significant | TRUE | -0.103844947 | 0.500746225 | No Change | Not Significant | TRUE | Cond1 Not Significant + Cond2 Not Significant | None | None |
| Cep44 | TRUE | 0.11749998 | 0.674169253 | No Change | Not Significant | TRUE | -0.055996471 | 0.820965922 | No Change | Not Significant | TRUE | Cond1 Not Significant + Cond2 Not Significant | None | None |
| Cep55 | TRUE | 0.71328156 | 8.11E-05 | UP | UP | TRUE | -0.020008204 | 0.93163806 | No Change | Not Significant | TRUE | Cond1 UP + Cond2 Not Significant | Cond1_UP | Cond1_UP |
| Cep57 | TRUE | 0.62877666 | 1.90E-07 | UP | UP | TRUE | 0.044256208 | 0.77637824 | No Change | Not Significant | TRUE | Cond1 UP + Cond2 Not Significant | Cond1_UP | Cond1_UP |
| Cep57l1 | TRUE | 0.328901725 | 0.216285499 | No Change | Not Significant | TRUE | 0.085327904 | 0.745625688 | No Change | Not Significant | TRUE | Cond1 Not Significant + Cond2 Not Significant | None | None |
| Cep63 | TRUE | -0.041135971 | 0.796376973 | No Change | Not Significant | TRUE | -0.055554043 | 0.606794107 | No Change | Not Significant | TRUE | Cond1 Not Significant + Cond2 Not Significant | None | None |
| Cep68 | TRUE | 0.12553171 | 0.430343308 | No Change | Not Significant | TRUE | -0.151352502 | 0.136174477 | No Change | Not Significant | TRUE | Cond1 Not Significant + Cond2 Not Significant | None | None |
| Cep70 | TRUE | -0.029385487 | 0.91020732 | No Change | Not Significant | TRUE | -0.12059684 | 0.503448119 | No Change | Not Significant | TRUE | Cond1 Not Significant + Cond2 Not Significant | None | None |
| Cep72 | TRUE | 0.141376907 | 0.512561987 | No Change | Not Significant | TRUE | -0.073783091 | 0.752522003 | No Change | Not Significant | TRUE | Cond1 Not Significant + Cond2 Not Significant | None | None |
| Cep76 | TRUE | 0.326024802 | 0.089783667 | No Change | Not Significant | TRUE | -0.143868353 | 0.415160239 | No Change | Not Significant | TRUE | Cond1 Not Significant + Cond2 Not Significant | None | None |
| Cep78 | TRUE | 0.30624331 | 0.229089299 | No Change | Not Significant | TRUE | 0.165248933 | 0.340981801 | No Change | Not Significant | TRUE | Cond1 Not Significant + Cond2 Not Significant | None | None |
| Cep83 | TRUE | 0.159512584 | 0.202844537 | No Change | Not Significant | TRUE | -0.008031807 | 0.970843006 | No Change | Not Significant | TRUE | Cond1 Not Significant + Cond2 Not Significant | None | None |
| Cep85 | TRUE | 0.38245962 | 8.30E-06 | No Change | Significant Positive | TRUE | 0.114655366 | 0.337846137 | No Change | Not Significant | TRUE | Cond1 Significant Positive + Cond2 Not Significant | None | None |
| Cep85l | TRUE | 1.782352283 | 1 | No Change | Not Significant | FALSE | NA | NA | No Change | Not Detected | FALSE | Background = FALSE | Background = FALSE | Background = FALSE |
| Cep89 | TRUE | 0.356809937 | 7.64E-05 | No Change | Significant Positive | TRUE | -0.004392567 | 0.980184745 | No Change | Not Significant | TRUE | Cond1 Significant Positive + Cond2 Not Significant | None | None |
| Cep95 | TRUE | -0.094126294 | 0.566016798 | No Change | Not Significant | TRUE | 0.006475667 | 0.983542327 | No Change | Not Significant | TRUE | Cond1 Not Significant + Cond2 Not Significant | None | None |
| Cep97 | TRUE | -0.351197761 | 0.084025676 | No Change | Not Significant | TRUE | -0.185407786 | 0.298245186 | No Change | Not Significant | TRUE | Cond1 Not Significant + Cond2 Not Significant | None | None |
| Cept1 | TRUE | 0.114645513 | 0.292413842 | No Change | Not Significant | TRUE | 0.001790117 | 0.990867082 | No Change | Not Significant | TRUE | Cond1 Not Significant + Cond2 Not Significant | None | None |
| Cercam | TRUE | 1.221558226 | 1 | No Change | Not Significant | FALSE | NA | NA | No Change | Not Detected | FALSE | Background = FALSE | Background = FALSE | Background = FALSE |
| Cerk | TRUE | 0.007645375 | 0.966840628 | No Change | Not Significant | TRUE | 0.098225312 | 0.298245186 | No Change | Not Significant | TRUE | Cond1 Not Significant + Cond2 Not Significant | None | None |
| Cerkl | TRUE | -0.792504828 | 1 | No Change | Not Significant | FALSE | NA | NA | No Change | Not Detected | FALSE | Background = FALSE | Background = FALSE | Background = FALSE |
| Cerox1 | TRUE | -0.353138341 | 1 | No Change | Not Significant | FALSE | NA | NA | No Change | Not Detected | FALSE | Background = FALSE | Background = FALSE | Background = FALSE |
| Cers1 | TRUE | 0.355499576 | 1 | No Change | Not Significant | FALSE | NA | NA | No Change | Not Detected | FALSE | Background = FALSE | Background = FALSE | Background = FALSE |
| Cers2 | TRUE | 0.024304618 | 0.865765398 | No Change | Not Significant | TRUE | -0.085698662 | 0.250247482 | No Change | Not Significant | TRUE | Cond1 Not Significant + Cond2 Not Significant | None | None |

Supplementary Table 3

| geneID | Cond1_DF_Detected | Cond1_DF_ValueCol | Cond1_DF_PadjCol | Cond1_DF_Cutoff | Cond1_DF_Cutoff_Specific.x | Cond2_DF_Detected | Cond2_DF_ValueCol | Cond2_DF_PadjCol | Cond2_DF_Cutoff | Cond2_DF_Cutoff_Specific | BG_Method | RG1_All | RG2_Significant | RG3_SignificantChange |
|---|---|---|---|---|---|---|---|---|---|---|---|---|---|---|
| Cers3 | TRUE | 0.958378092 | 0.102483165 | No Change | Not Significant | TRUE | 0.086422272 | 0.929613548 | No Change | Not Significant | TRUE | Cond1 Not Significant + Cond2 Not Significant | None | None |
| Cers4 | TRUE | -0.683809359 | 3.18E-06 | DOWN | DOWN | TRUE | -0.1991909 | 0.009610832 | No Change | Significant Negative | TRUE | Cond1 DOWN + Cond2 Significant Negative | Core_DOWN | Cond1_DOWN |
| Cers5 | TRUE | -0.696678289 | 7.84E-09 | DOWN | DOWN | TRUE | -0.22138991 | 0.042325937 | No Change | Significant Negative | TRUE | Cond1 DOWN + Cond2 Significant Negative | Core_DOWN | Cond1_DOWN |
| Cers6 | TRUE | -0.242953649 | 0.000875933 | No Change | Significant Negative | TRUE | -0.240125406 | 6.33E-13 | No Change | Significant Negative | TRUE | Cond1 Significant Negative + Cond2 Significant Negative | None | None |
| Cert1 | TRUE | 0.147919782 | 0.376049998 | No Change | Not Significant | TRUE | -0.020778248 | 0.817489503 | No Change | Not Significant | TRUE | Cond1 Not Significant + Cond2 Not Significant | None | None |
| Ces1b | TRUE | -2.440151183 | 1 | No Change | Not Significant | FALSE | NA | NA | No Change | Not Detected | FALSE | Background = FALSE | Background = FALSE | Background = FALSE |
| Ces1c | TRUE | -2.181689658 | 1 | No Change | Not Significant | TRUE | 0.429647106 | 0.674226502 | No Change | Not Significant | TRUE | Cond1 Not Significant + Cond2 Not Significant | None | None |
| Ces1d | TRUE | -1.116794538 | 0.000150204 | DOWN | DOWN | TRUE | -0.102260822 | 0.405874466 | No Change | Not Significant | TRUE | Cond1 DOWN + Cond2 Not Significant | Cond1_DOWN | Cond1_DOWN |
| Ces1e | TRUE | -0.14806101 | 0.404473864 | No Change | Not Significant | TRUE | -0.035022184 | 0.854251145 | No Change | Not Significant | TRUE | Cond1 Not Significant + Cond2 Not Significant | None | None |
| Ces1f | TRUE | -1.926624979 | 2.37E-31 | DOWN | DOWN | TRUE | -0.266688144 | 0.004378048 | No Change | Significant Negative | TRUE | Cond1 DOWN + Cond2 Significant Negative | Core_DOWN | Cond1_DOWN |
| Ces1g | TRUE | -3.33361884 | 8.33E-25 | DOWN | DOWN | TRUE | -0.977259409 | 1.41E-08 | DOWN | DOWN | TRUE | Cond1 DOWN + Cond2 DOWN | Core_DOWN | Core_DOWN |
| Ces1h | TRUE | 1.678194257 | 1 | No Change | Not Significant | FALSE | NA | NA | No Change | Not Detected | FALSE | Background = FALSE | Background = FALSE | Background = FALSE |
| Ces2a | TRUE | 0.082620146 | 0.811601896 | No Change | Not Significant | TRUE | 0.262268985 | 0.048102018 | No Change | Significant Positive | TRUE | Cond1 Not Significant + Cond2 Significant Positive | None | None |
| Ces2b | TRUE | -1.519518927 | 1.29E-06 | DOWN | DOWN | TRUE | -0.46554205 | 0.002410693 | No Change | Significant Negative | TRUE | Cond1 DOWN + Cond2 Significant Negative | Core_DOWN | Cond1_DOWN |
| [illegible] | | | | | | | | | | | | | | |

| geneID | Cond1_DF_ensembl_gene_id | Cond1_DF_entrezgene_id | Cond1_DF_gene_biotype | Cond1_DF_baseMean | Cond1_DF_log2FoldChange | Cond1_DF_lfcSE | Cond1_DF_stat | Cond1_DF_pvalue | Cond1_DF_padj | Cond1_DF_absLog2FC | Cond1_DF_Cutoff_Specific.y |
|---|---|---|---|---|---|---|---|---|---|---|---|
| Cers3 | ENSMUSG00000030510 | 545975 | protein_coding | 8.092370443 | 0.958378092 | 0.443303385 | -2.161901137 | 0.030625797 | 0.102483165 | 0.958378092 | Not Significant |
| Cers4 | ENSMUSG00000008206 | 67260 | protein_coding | 451.4739938 | -0.683809359 | 0.131594377 | 5.196341786 | 2.03E-07 | 3.18E-06 | 0.683809359 | DOWN |
| Cers5 | ENSMUSG00000023021 | 71949 | protein_coding | 339.1916331 | -0.696678289 | 0.110897792 | 6.28216555 | 3.34E-10 | 7.84E-09 | 0.696678289 | DOWN |
| Cers6 | ENSMUSG00000027035 | 241447 | protein_coding | 2359.595539 | -0.242953649 | 0.06252775 | 3.885533234 | 0.000102106 | 0.000875933 | 0.242953649 | Significant Negative |
| Cert1 | ENSMUSG00000021669 | 68018 | protein_coding | 820.7787678 | 0.147919782 | 0.110784093 | -1.335207767 | 0.181808356 | 0.376049998 | 0.147919782 | Not Significant |
| Ces1b | ENSMUSG00000078964 | 382044 | protein_coding | 2.73582639 | -2.440151183 | 0.984458944 | 2.478672368 | 0.013187238 | 1 | 2.440151183 | Not Significant |
| Ces1c | ENSMUSG00000057400 | 13884 | protein_coding | 1.789376869 | -2.181689658 | 1.001026044 | 2.179453443 | 0.029298 | 1 | 2.181689658 | Not Significant |
| Ces1d | ENSMUSG00000056973 | 104158 | protein_coding | 413.5835353 | -1.116794538 | 0.257078497 | 4.344177179 | 1.40E-05 | 0.000150204 | 1.116794538 | DOWN |
| Ces1e | ENSMUSG00000061959 | 13897 | protein_coding | 215.2919061 | -0.14806101 | 0.116170705 | 1.274512459 | 0.20248186 | 0.404473864 | 0.14806101 | Not Significant |
| Ces1f | ENSMUSG00000031725 | 234564 | protein_coding | 939.6115003 | -1.926624979 | 0.160280951 | 12.02029913 | 2.78E-33 | 2.37E-31 | 1.926624979 | DOWN |
| Ces1g | ENSMUSG00000057074 | 12623 | protein_coding | 342.3975464 | -3.33361884 | 0.311998218 | 10.6847368 | 1.20E-26 | 8.33E-25 | 3.33361884 | DOWN |
| Ces1h | ENSMUSG00000074156 | 75704 | protein_coding | 1.11941248 | 1.678194257 | 1.592499087 | -1.053811755 | 0.291969111 | 1 | 1.678194257 | Not Significant |
| Ces2a | ENSMUSG00000055730 | 102022 | protein_coding | 2071.791551 | 0.082620146 | 0.183594449 | -0.450014401 | 0.652700057 | 0.811601896 | 0.082620146 | Not Significant |
| Ces2b | ENSMUSG00000050097 | 234669 | protein_coding | 99.06141893 | -1.519518927 | 0.282762117 | 5.373841945 | 7.71E-08 | 1.29E-06 | 1.519518927 | DOWN |
| [illegible] | | | | | | | | | | | |

| geneID | Cond2_DF_ensembl_gene_id | Cond2_DF_entrezgene_id | Cond2_DF_gene_biotype | Cond2_DF_baseMean | Cond2_DF_log2FoldChange | Cond2_DF_lfcSE | Cond2_DF_stat | Cond2_DF_pvalue | Cond2_DF_padj | Cond2_DF_absLog2FC | RG1_Specific_Cond2 | RG1_Specific_Cond1 |
|---|---|---|---|---|---|---|---|---|---|---|---|---|
| Cers3 | ENSMUSG00000030510 | 545975 | protein_coding | 9.034044924 | 0.086422272 | 0.431361275 | 0.200347776 | 0.8412086 | 0.929613548 | 0.086422272 | Cond1 No Change + Cond2 Not Significant | Cond2 No Change + Cond1 Not Significant |
| Cers4 | ENSMUSG00000008206 | 67260 | protein_coding | 1061.6853 | -0.1991909 | 0.060254451 | -3.305828818 | 0.000946959 | 0.009610832 | 0.1991909 | Cond1 DOWN + Cond2 Significant Negative | Cond2 No Change + Cond1 DOWN |
| Cers5 | ENSMUSG00000023021 | 71949 | protein_coding | 735.7166809 | -0.22138991 | 0.080978319 | -2.733940539 | 0.006258134 | 0.042325937 | 0.22138991 | Cond1 DOWN + Cond2 Significant Negative | Cond2 No Change + Cond1 DOWN |
| Cers6 | ENSMUSG00000027035 | 241447 | protein_coding | 5197.434227 | -0.240125406 | 0.031041438 | -7.735640606 | 1.03E-14 | 6.33E-13 | 0.240125406 | Cond1 No Change + Cond2 Significant Negative | Cond2 No Change + Cond1 Significant Negative |
| Cert1 | ENSMUSG00000021669 | 68018 | protein_coding | 1555.974983 | -0.020778248 | 0.045411176 | -0.457558018 | 0.647270011 | 0.817489503 | 0.020778248 | Cond1 No Change + Cond2 Not Significant | Cond2 No Change + Cond1 Not Significant |
| Ces1b | NA | NA | NA | NA | NA | NA | NA | NA | NA | NA | Background = FALSE | Background = FALSE |
| Ces1c | ENSMUSG00000057400 | 13884 | protein_coding | 8.027832709 | 0.429647106 | 0.560275296 | 0.766849991 | 0.44317071 | 0.674226502 | 0.429647106 | Cond1 No Change + Cond2 Not Significant | Cond2 No Change + Cond1 Not Significant |
| Ces1d | ENSMUSG00000056973 | 104158 | protein_coding | 1236.788458 | -0.102260822 | 0.076047371 | -1.344698967 | 0.178722468 | 0.405874466 | 0.102260822 | Cond1 DOWN + Cond2 Not Significant | Cond2 No Change + Cond1 DOWN |
| Ces1e | ENSMUSG00000061959 | 13897 | protein_coding | 590.8899155 | -0.035022184 | 0.093064184 | -0.376322904 | 0.706676854 | 0.854251145 | 0.035022184 | Cond1 No Change + Cond2 Not Significant | Cond2 No Change + Cond1 Not Significant |
| Ces1f | ENSMUSG00000031725 | 234564 | protein_coding | 2096.41761 | -0.266688144 | 0.074851123 | -3.56291441 | 0.00036676 | 0.004378048 | 0.266688144 | Cond1 DOWN + Cond2 Significant Negative | Cond2 No Change + Cond1 DOWN |
| Ces1g | ENSMUSG00000057074 | 12623 | protein_coding | 1327.935083 | -0.977259409 | 0.155623473 | -6.279640161 | 3.39E-10 | 1.41E-08 | 0.977259409 | Cond1 DOWN + Cond2 DOWN | Cond2 DOWN + Cond1 DOWN |
| Ces1h | NA | NA | NA | NA | NA | NA | NA | NA | NA | NA | Background = FALSE | Background = FALSE |
| Ces2a | ENSMUSG00000055730 | 102022 | protein_coding | 1404.587618 | 0.262268985 | 0.098037504 | 2.67519035 | 0.007468683 | 0.048102018 | 0.262268985 | Cond1 No Change + Cond2 Significant Positive | Cond2 No Change + Cond1 Not Significant |
| Ces2b | ENSMUSG00000050097 | 234669 | protein_coding | 328.2826535 | -0.46554205 | 0.124339296 | -3.744126485 | 0.000181022 | 0.002410693 | 0.46554205 | Cond1 DOWN + Cond2 Significant Negative | Cond2 No Change + Cond1 DOWN |
| [illegible] | | | | | | | | | | | | |

| geneID | Cond1_DF_Detected | Cond1_DF_ValueCol | Cond1_DF_PadjCol | Cond1_DF_Cutoff | Cond1_DF_Cutoff_Specific.x | Cond2_DF_Detected | Cond2_DF_ValueCol | Cond2_DF_PadjCol | Cond2_DF_Cutoff | Cond2_DF_Cutoff_Specific | BG_Method | RG1_All |
|---|---|---|---|---|---|---|---|---|---|---|---|---|
| Ciao3 | TRUE | -0.070355689 | 0.475244062 | No Change | Not Significant | TRUE | 0.039109789 | 0.703210238 | No Change | Not Significant | TRUE | Cond1 Not Significant + Cond2 Not Significant |
| Ciapin1 | TRUE | 0.003662839 | 0.982416138 | No Change | Not Significant | TRUE | -0.00428067 | 0.983365559 | No Change | Not Significant | TRUE | Cond1 Not Significant + Cond2 Not Significant |
| Ciart | TRUE | -1.595751519 | 0.041509049 | DOWN | DOWN | TRUE | -2.250721792 | 3.10E-09 | DOWN | DOWN | TRUE | Cond1 DOWN + Cond2 DOWN |
| Cib1 | TRUE | 0.059526833 | 0.584928009 | No Change | Not Significant | TRUE | 0.190865091 | 0.029748403 | No Change | Significant Positive | TRUE | Cond1 Not Significant + Cond2 Significant Positive |
| Cib2 | TRUE | -0.001201816 | 0.998060837 | No Change | Not Significant | TRUE | 0.139322918 | 0.699625498 | No Change | Not Significant | TRUE | Cond1 Not Significant + Cond2 Not Significant |
| Cib3 | TRUE | -0.361309263 | 1 | No Change | Not Significant | FALSE | NA | NA | No Change | Not Detected | FALSE | Background = FALSE |
| Cibar1 | TRUE | 0.503752284 | 0.069848891 | No Change | Not Significant | TRUE | 0.274871277 | 0.188241916 | No Change | Not Significant | TRUE | Cond1 Not Significant + Cond2 Not Significant |
| Cibar2 | TRUE | 0.575623305 | 1 | No Change | Not Significant | FALSE | NA | NA | No Change | Not Detected | FALSE | Background = FALSE |
| Cic | TRUE | 0.510088394 | 0.006350049 | UP | UP | TRUE | -0.004847122 | 0.980585772 | No Change | Not Significant | TRUE | Cond1 UP + Cond2 Not Significant |
| Cidea | TRUE | -1.141977658 | 1 | No Change | Not Significant | FALSE | NA | NA | No Change | Not Detected | FALSE | Background = FALSE |
| Cideb | TRUE | -0.34416287 | 0.205991262 | No Change | Not Significant | TRUE | -0.019081668 | 0.94368516 | No Change | Not Significant | TRUE | Cond1 Not Significant + Cond2 Not Significant |
| Cidec | TRUE | 0.662755433 | 0.194226738 | No Change | Not Significant | TRUE | 0.120783318 | 0.698858175 | No Change | Not Significant | TRUE | Cond1 Not Significant + Cond2 Not Significant |
| Ciita | TRUE | 0.640447137 | 0.340247143 | No Change | Not Significant | FALSE | NA | NA | No Change | Not Detected | FALSE | Background = FALSE |
| Cilk1 | TRUE | 0.172358345 | 0.148948255 | No Change | Not Significant | TRUE | 0.113498529 | 0.157189965 | No Change | Not Significant | TRUE | Cond1 Not Significant + Cond2 Not Significant |
| Cilp | TRUE | -1.090286275 | 1 | No Change | Not Significant | FALSE | NA | NA | No Change | Not Detected | FALSE | Background = FALSE |
| Cilp2 | TRUE | 0.042557359 | 1 | No Change | Not Significant | FALSE | NA | NA | No Change | Not Detected | FALSE | Background = FALSE |
| Cinp | TRUE | 0.128636043 | 0.594533905 | No Change | Not Significant | TRUE | 0.043297287 | 0.789158883 | No Change | Not Significant | TRUE | Cond1 Not Significant + Cond2 Not Significant |
| Cip2a | TRUE | 0.824935044 | 0.000106469 | UP | UP | TRUE | 0.20706694 | 0.11303636 | No Change | Not Significant | TRUE | Cond1 UP + Cond2 Not Significant |
| Cipc | TRUE | -0.727524199 | 2.29E-06 | DOWN | DOWN | TRUE | -0.381074922 | 2.06E-07 | No Change | Significant Negative | TRUE | Cond1 DOWN + Cond2 Significant Negative |
| Cir1 | TRUE | -0.587283857 | 0.004768572 | DOWN | DOWN | TRUE | -0.165819163 | 0.34107542 | No Change | Not Significant | TRUE | Cond1 DOWN + Cond2 Not Significant |
| Cirbp | TRUE | 0.026144322 | 0.925637253 | No Change | Not Significant | TRUE | -0.37406941 | 0.00773893 | No Change | Significant Negative | TRUE | Cond1 Not Significant + Cond2 Significant Negative |
| Cirop | TRUE | 0.55124056 | 1 | No Change | Not Significant | FALSE | NA | NA | No Change | Not Detected | FALSE | Background = FALSE |
| Cisd1 | TRUE | -0.033098591 | 0.854853631 | No Change | Not Significant | TRUE | 0.126175726 | 0.193983147 | No Change | Not Significant | TRUE | Cond1 Not Significant + Cond2 Not Significant |
| Cisd2 | TRUE | 0.352007401 | 1.85E-07 | No Change | Significant Positive | TRUE | 0.048345957 | 0.491446343 | No Change | Not Significant | TRUE | Cond1 Significant Positive + Cond2 Not Significant |
| Cisd3 | TRUE | -0.615834858 | 8.30E-09 | DOWN | DOWN | TRUE | 0.00310569 | 0.979641635 | No Change | Not Significant | TRUE | Cond1 DOWN + Cond2 Not Significant |
| Cish | TRUE | -0.196682269 | 0.429214508 | No Change | Not Significant | TRUE | -0.080389371 | 0.724049267 | No Change | Not Significant | TRUE | Cond1 Not Significant + Cond2 Not Significant |
| Cit | TRUE | 1.495129073 | 1.68E-15 | UP | UP | TRUE | 0.181736139 | 0.171848571 | No Change | Not Significant | TRUE | Cond1 UP + Cond2 Not Significant |
| Cited1 | TRUE | -0.920942237 | 1 | No Change | Not Significant | FALSE | NA | NA | No Change | Not Detected | FALSE | Background = FALSE |
| Cited2 | TRUE | 0.031727834 | 0.933837244 | No Change | Not Significant | TRUE | -0.160351417 | 0.357173174 | No Change | Not Significant | TRUE | Cond1 Not Significant + Cond2 Not Significant |
| Cited4 | TRUE | 0.240655552 | 0.168040558 | No Change | Not Significant | TRUE | 0.200684671 | 0.295082324 | No Change | Not Significant | TRUE | Cond1 Not Significant + Cond2 Not Significant |
| Ciz1 | TRUE | -0.226227331 | 0.059173697 | No Change | Not Significant | TRUE | -0.118063526 | 0.121164388 | No Change | Not Significant | TRUE | Cond1 Not Significant + Cond2 Not Significant |
| Ckap2 | TRUE | 0.670434946 | 0.000464257 | UP | UP | TRUE | -0.106911282 | 0.569329485 | No Change | Not Significant | TRUE | Cond1 UP + Cond2 Not Significant |
| Ckap2l | TRUE | 0.808237041 | 2.05E-05 | UP | UP | TRUE | 0.115013527 | 0.573501586 | No Change | Not Significant | TRUE | Cond1 UP + Cond2 Not Significant |
| Ckap4 | TRUE | 0.941361207 | 2.82E-14 | UP | UP | TRUE | 0.280664692 | 0.054911988 | No Change | Not Significant | TRUE | Cond1 UP + Cond2 Not Significant |
| Ckap5 | TRUE | 0.326096937 | 0.011004052 | No Change | Significant Positive | TRUE | -0.011959338 | 0.935610703 | No Change | Not Significant | TRUE | Cond1 Significant Positive + Cond2 Not Significant |
| Ckb | TRUE | 1.553830894 | 1.63E-10 | UP | UP | TRUE | 0.524167194 | 0.008805649 | UP | UP | TRUE | Cond1 UP + Cond2 UP |
| Cklf | TRUE | 0.117375929 | 0.638714542 | No Change | Not Significant | TRUE | 0.013687914 | 0.959960814 | No Change | Not Significant | TRUE | Cond1 Not Significant + Cond2 Not Significant |
| Ckm | TRUE | 1.38112741 | 1 | No Change | Not Significant | FALSE | NA | NA | No Change | Not Detected | FALSE | Background = FALSE |
| Ckmt1 | TRUE | -0.206748307 | 0.408031364 | No Change | Not Significant | TRUE | 0.065674125 | 0.714113831 | No Change | Not Significant | TRUE | Cond1 Not Significant + Cond2 Not Significant |
| Ckmt2 | TRUE | 4.372820079 | 2.21E-182 | UP | UP | TRUE | 3.853417169 | 6.70E-133 | UP | UP | TRUE | Cond1 UP + Cond2 UP |
| Cks1b | TRUE | 1.292757517 | 2.45E-11 | UP | UP | TRUE | 0.415254129 | 0.041961584 | No Change | Significant Positive | TRUE | Cond1 UP + Cond2 Significant Positive |
| Cks2 | TRUE | -0.018177314 | 0.955211953 | No Change | Not Significant | TRUE | 0.133249272 | 0.205058028 | No Change | Not Significant | TRUE | Cond1 Not Significant + Cond2 Not Significant |
| Clasp1 | TRUE | 0.015218581 | 0.925637253 | No Change | Not Significant | TRUE | -0.000133487 | 0.999077457 | No Change | Not Significant | TRUE | Cond1 Not Significant + Cond2 Not Significant |
| Clasp2 | TRUE | 0.159335574 | 0.157297398 | No Change | Not Significant | TRUE | -0.101424943 | 0.278419092 | No Change | Not Significant | TRUE | Cond1 Not Significant + Cond2 Not Significant |
| Clasrp | TRUE | 0.253051485 | 0.255505462 | No Change | Not Significant | TRUE | -0.027778897 | 0.906028181 | No Change | Not Significant | TRUE | Cond1 Not Significant + Cond2 Not Significant |
| Clba1 | TRUE | 0.062800947 | 0.872440598 | No Change | Not Significant | TRUE | 0.409755911 | 0.104657804 | No Change | Not Significant | TRUE | Cond1 Not Significant + Cond2 Not Significant |
| Clca1 | TRUE | -1.195284189 | 0.008692083 | DOWN | DOWN | TRUE | 0.662444784 | 0.005267157 | UP | UP | TRUE | Cond1 DOWN + Cond2 UP |
| Clca2 | TRUE | -1.645417404 | 0.007425086 | DOWN | DOWN | TRUE | 0.633976099 | 0.260237414 | No Change | Not Significant | TRUE | Cond1 DOWN + Cond2 Not Significant |
| Clca3a1 | TRUE | 1.168566652 | 0.000114898 | UP | UP | TRUE | -0.542778239 | 0.128431439 | No Change | Not Significant | TRUE | Cond1 UP + Cond2 Not Significant |
| Clca3a2 | TRUE | 0.816244025 | 0.000331378 | UP | UP | TRUE | 0.144995522 | 0.566769091 | No Change | Not Significant | TRUE | Cond1 UP + Cond2 Not Significant |
| Clca3b | TRUE | 0.428701723 | 0.081360993 | No Change | Not Significant | TRUE | 0.13260466 | 0.388872616 | No Change | Not Significant | TRUE | Cond1 Not Significant + Cond2 Not Significant |
| Clca4a | TRUE | 0.337174935 | 0.559233675 | No Change | Not Significant | TRUE | -0.066249501 | 0.84768023 | No Change | Not Significant | TRUE | Cond1 Not Significant + Cond2 Not Significant |
| Clca4b | TRUE | 3.640561962 | 1.54E-13 | UP | UP | TRUE | 0.619967466 | 0.091425492 | No Change | Not Significant | TRUE | Cond1 UP + Cond2 Not Significant |
| Clcc1 | TRUE | 0.115804554 | 0.62627635 | No Change | Not Significant | TRUE | 0.027333246 | 0.837472006 | No Change | Not Significant | TRUE | Cond1 Not Significant + Cond2 Not Significant |
| Clcf1 | TRUE | 2.063995502 | 0.011264068 | UP | UP | FALSE | NA | NA | No Change | Not Detected | FALSE | Background = FALSE |
| Clcn1 | TRUE | 0.593807223 | 0.551855356 | No Change | Not Significant | TRUE | -0.003107555 | 0.996976705 | No Change | Not Significant | TRUE | Cond1 Not Significant + Cond2 Not Significant |
| Clcn2 | TRUE | -0.189943672 | 0.484112338 | No Change | Not Significant | TRUE | -0.055894302 | 0.708115319 | No Change | Not Significant | TRUE | Cond1 Not Significant + Cond2 Not Significant |
| Clcn3 | TRUE | 0.685873399 | 1.22E-33 | UP | UP | TRUE | 0.22168408 | 1.30E-06 | No Change | Significant Positive | TRUE | Cond1 UP + Cond2 Significant Positive |
| Clcn4 | TRUE | -0.038612373 | 0.806290266 | No Change | Not Significant | TRUE | 0.03692713 | 0.574280957 | No Change | Not Significant | TRUE | Cond1 Not Significant + Cond2 Not Significant |
| Clcn5 | TRUE | -0.263382488 | 0.044395495 | No Change | Significant Negative | TRUE | -0.316297645 | 5.65E-05 | No Change | Significant Negative | TRUE | Cond1 Significant Negative + Cond2 Significant Negative |
| Clcn6 | TRUE | -0.131222619 | 0.736796542 | No Change | Not Significant | TRUE | -0.284382613 | 0.176137379 | No Change | Not Significant | TRUE | Cond1 Not Significant + Cond2 Not Significant |
| Clcn7 | TRUE | -0.111843962 | 0.260339278 | No Change | Not Significant | TRUE | -0.073781572 | 0.488580666 | No Change | Not Significant | TRUE | Cond1 Not Significant + Cond2 Not Significant |
| Clcnka | TRUE | 1.039181163 | 1 | No Change | Not Significant | FALSE | NA | NA | No Change | Not Detected | FALSE | Background = FALSE |
| Clcnkb | TRUE | -1.413948679 | 1 | No Change | Not Significant | TRUE | 0.59083945 | 0.222950058 | No Change | Not Significant | TRUE | Cond1 Not Significant + Cond2 Not Significant |
| Cldn1 | TRUE | 1.088268146 | 1 | No Change | Not Significant | FALSE | NA | NA | No Change | Not Detected | FALSE | Background = FALSE |
| Cldn10 | TRUE | 0.47352596 | 1 | No Change | Not Significant | FALSE | NA | NA | No Change | Not Detected | FALSE | Background = FALSE |
| Cldn11 | TRUE | 1.361671607 | 1 | No Change | Not Significant | FALSE | NA | NA | No Change | Not Detected | FALSE | Background = FALSE |
| Cldn12 | TRUE | 0.099776418 | 0.671470886 | No Change | Not Significant | TRUE | 0.107134975 | 0.058770253 | No Change | Not Significant | TRUE | Cond1 Not Significant + Cond2 Not Significant |
| Cldn14 | TRUE | 0.232336736 | 0.512561987 | No Change | Not Significant | TRUE | -0.104176736 | 0.838995821 | No Change | Not Significant | TRUE | Cond1 Not Significant + Cond2 Not Significant |
| Cldn15 | TRUE | -0.107833568 | 0.548411142 | No Change | Not Significant | TRUE | 0.040972792 | 0.668045287 | No Change | Not Significant | TRUE | Cond1 Not Significant + Cond2 Not Significant |
| Cldn18 | TRUE | 1.023383311 | 1 | No Change | Not Significant | FALSE | NA | NA | No Change | Not Detected | FALSE | Background = FALSE |
| Cldn19 | TRUE | 1.246162585 | 1 | No Change | Not Significant | FALSE | NA | NA | No Change | Not Detected | FALSE | Background = FALSE |
| Cldn2 | TRUE | 0.311386101 | 0.322338142 | No Change | Not Significant | TRUE | -0.076611861 | 0.65186884 | No Change | Not Significant | TRUE | Cond1 Not Significant + Cond2 Not Significant |
| Cldn20 | TRUE | 0.152084204 | 1 | No Change | Not Significant | FALSE | NA | NA | No Change | Not Detected | FALSE | Background = FALSE |
| Cldn23 | TRUE | -0.272932919 | 0.415328565 | No Change | Not Significant | TRUE | -0.020289917 | 0.914534473 | No Change | Not Significant | TRUE | Cond1 Not Significant + Cond2 Not Significant |
| Cldn3 | TRUE | -0.047091336 | 0.803370142 | No Change | Not Significant | TRUE | 0.155588037 | 0.156629283 | No Change | Not Significant | TRUE | Cond1 Not Significant + Cond2 Not Significant |
| Cldn34b2 | TRUE | 0.60939749 | 1 | No Change | Not Significant | FALSE | NA | NA | No Change | Not Detected | FALSE | Background = FALSE |
| Cldn34c1 | TRUE | 1.871961866 | 0.09075569 | No Change | Not Significant | FALSE | NA | NA | No Change | Not Detected | FALSE | Background = FALSE |
| Cldn4 | TRUE | -0.534639453 | 0.133440901 | No Change | Not Significant | TRUE | 0.070098567 | 0.718207077 | No Change | Not Significant | TRUE | Cond1 Not Significant + Cond2 Not Significant |
| Cldn6 | TRUE | 0.961390733 | 1 | No Change | Not Significant | FALSE | NA | NA | No Change | Not Detected | FALSE | Background = FALSE |
| Cldn7 | TRUE | -0.039494916 | 0.780088572 | No Change | Not Significant | TRUE | 0.031978583 | 0.829882036 | No Change | Not Significant | TRUE | Cond1 Not Significant + Cond2 Not Significant |
| Cldn8 | TRUE | 0.254750674 | 0.363275924 | No Change | Not Significant | TRUE | 0.503611324 | 3.06E-05 | UP | UP | TRUE | Cond1 Not Significant + Cond2 UP |
| Cldn9 | TRUE | -0.101316685 | 1 | No Change | Not Significant | FALSE | NA | NA | No Change | Not Detected | FALSE | Background = FALSE |
| Cldnd1 | TRUE | -0.120434094 | 0.425467087 | No Change | Not Significant | TRUE | 0.08295654 | 0.493560011 | No Change | Not Significant | TRUE | Cond1 Not Significant + Cond2 Not Significant |
| Cldnd2 | TRUE | 2.373808506 | 1 | No Change | Not Significant | FALSE | NA | NA | No Change | Not Detected | FALSE | Background = FALSE |
| Clec10a | TRUE | 2.367967802 | 1 | No Change | Not Significant | FALSE | NA | NA | No Change | Not Detected | FALSE | Background = FALSE |
| Clec11a | TRUE | 0.964584379 | 0.004593536 | UP | UP | TRUE | 0.230628255 | 0.449133255 | No Change | Not Significant | TRUE | Cond1 UP + Cond2 Not Significant |
| Clec12a | TRUE | 1.321871466 | 1 | No Change | Not Significant | FALSE | NA | NA | No Change | Not Detected | FALSE | Background = FALSE |
| Clec14a | TRUE | -0.298587518 | 0.188104702 | No Change | Not Significant | TRUE | -0.094529378 | 0.481155234 | No Change | Not Significant | TRUE | Cond1 Not Significant + Cond2 Not Significant |
| Clec16a | TRUE | 0.110018766 | 0.268218365 | No Change | Not Significant | TRUE | 0.097074309 | 0.190759202 | No Change | Not Significant | TRUE | Cond1 Not Significant + Cond2 Not Significant |
| Clec18a | TRUE | 2.639638734 | 1 | No Change | Not Significant | FALSE | NA | NA | No Change | Not Detected | FALSE | Background = FALSE |
| Clec1a | TRUE | 3.455051354 | 1 | No Change | Not Significant | FALSE | NA | NA | No Change | Not Detected | FALSE | Background = FALSE |
| Clec2d | TRUE | -0.192533784 | 0.460660281 | No Change | Not Significant | TRUE | -0.244701093 | 0.013494603 | No Change | Significant Negative | TRUE | Cond1 Not Significant + Cond2 Significant Negative |
| Clec2e | TRUE | -0.295085333 | 0.532085855 | No Change | Not Significant | TRUE | -0.663917249 | 0.00104069 | DOWN | DOWN | TRUE | Cond1 Not Significant + Cond2 DOWN |
| Clec2f | TRUE | 2.059892812 | 1 | No Change | Not Significant | FALSE | NA | NA | No Change | Not Detected | FALSE | Background = FALSE |
| Clec2g | TRUE | 2.798263689 | 1 | No Change | Not Significant | FALSE | NA | NA | No Change | Not Detected | FALSE | Background = FALSE |
| Clec2h | TRUE | 0.047974793 | 0.849682329 | No Change | Not Significant | TRUE | 0.068974844 | 0.618232985 | No Change | Not Significant | TRUE | Cond1 Not Significant + Cond2 Not Significant |
| Clec2i | TRUE | 0.685734495 | 0.613316042 | No Change | Not Significant | FALSE | NA | NA | No Change | Not Detected | FALSE | Background = FALSE |
| Clec2l | TRUE | 1.545017955 | 1 | No Change | Not Significant | FALSE | NA | NA | No Change | Not Detected | FALSE | Background = FALSE |
| Clec2m | TRUE | 1.025959506 | 1 | No Change | Not Significant | FALSE | NA | NA | No Change | Not Detected | FALSE | Background = FALSE |
| Clec3b | TRUE | -0.015179713 | 1 | No Change | Not Significant | FALSE | NA | NA | No Change | Not Detected | FALSE | Background = FALSE |
| Clec4a1 | TRUE | -0.94474507 | 0.181090121 | No Change | Not Significant | TRUE | 0.277607633 | 0.730713836 | No Change | Not Significant | TRUE | Cond1 Not Significant + Cond2 Not Significant |
| Clec4a2 | TRUE | 0.52345303 | 1 | No Change | Not Significant | FALSE | NA | NA | No Change | Not Detected | FALSE | Background = FALSE |
| Clec4e | TRUE | 0.716084033 | 1 | No Change | Not Significant | FALSE | NA | NA | No Change | Not Detected | FALSE | Background = FALSE |
| Clec4g | TRUE | -0.76055164 | 1 | No Change | Not Significant | FALSE | NA | NA | No Change | Not Detected | FALSE | Background = FALSE |
| Clec7a | TRUE | -0.291376147 | 1 | No Change | Not Significant | FALSE | NA | NA | No Change | Not Detected | FALSE | Background = FALSE |
| Clgn | TRUE | 1.991358969 | 1 | No Change | Not Significant | FALSE | NA | NA | No Change | Not Detected | FALSE | Background = FALSE |
| Clhc1 | TRUE | 0.42111257 | 1 | No Change | Not Significant | FALSE | NA | NA | No Change | Not Detected | FALSE | Background = FALSE |
| Clic1 | TRUE | -0.08533874 | 0.377007156 | No Change | Not Significant | TRUE | 0.098675052 | 0.230421574 | No Change | Not Significant | TRUE | Cond1 Not Significant + Cond2 Not Significant |
| Clic3 | TRUE | 0.987438365 | 0.207264236 | No Change | Not Significant | TRUE | -0.113835536 | 0.890414429 | No Change | Not Significant | TRUE | Cond1 Not Significant + Cond2 Not Significant |
| Clic4 | TRUE | -0.161839735 | 0.31052626 | No Change | Not Significant | TRUE | 0.016757388 | 0.932292793 | No Change | Not Significant | TRUE | Cond1 Not Significant + Cond2 Not Significant |
| Clic5 | TRUE | -0.260753968 | 0.30074233 | No Change | Not Significant | TRUE | -0.265855113 | 0.10799004 | No Change | Not Significant | TRUE | Cond1 Not Significant + Cond2 Not Significant |
| Clic6 | TRUE | -0.08427539 | 0.809917796 | No Change | Not Significant | TRUE | 0.035337185 | 0.831606417 | No Change | Not Significant | TRUE | Cond1 Not Significant + Cond2 Not Significant |
| Clint1 | TRUE | -0.183368135 | 0.000580752 | No Change | Significant Negative | TRUE | -0.01235669 | 0.84457466 | No Change | Not Significant | TRUE | Cond1 Significant Negative + Cond2 Not Significant |
| Clip1 | TRUE | -0.2825768 | 0.001052445 | No Change | Significant Negative | TRUE | -0.001170575 | 0.994488063 | No Change | Not Significant | TRUE | Cond1 Significant Negative + Cond2 Not Significant |
| Clip2 | TRUE | 0.352747826 | 5.00E-05 | No Change | Significant Positive | TRUE | 0.295319489 | 3.01E-11 | No Change | Significant Positive | TRUE | Cond1 Significant Positive + Cond2 Significant Positive |
| Clip3 | TRUE | 0.190537368 | 1 | No Change | Not Significant | FALSE | NA | NA | No Change | Not Detected | FALSE | Background = FALSE |
| Clip4 | TRUE | 1.480995889 | 1 | No Change | Not Significant | FALSE | NA | NA | No Change | Not Detected | FALSE | Background = FALSE |
| Clk1 | TRUE | -0.200799445 | 0.292303079 | No Change | Not Significant | TRUE | -0.270784691 | 0.095778655 | No Change | Not Significant | TRUE | Cond1 Not Significant + Cond2 Not Significant |
| Clk2 | TRUE | -0.017700546 | 0.921642147 | No Change | Not Significant | TRUE | -0.202069253 | 0.082768143 | No Change | Not Significant | TRUE | Cond1 Not Significant + Cond2 Not Significant |
| Clk3 | TRUE | -0.076103174 | 0.441111962 | No Change | Not Significant | TRUE | -0.156216257 | 0.069808432 | No Change | Not Significant | TRUE | Cond1 Not Significant + Cond2 Not Significant |
| Clk4 | TRUE | -0.007045781 | 0.97587882 | No Change | Not Significant | TRUE | -0.315720096 | 0.018258611 | No Change | Significant Negative | TRUE | Cond1 Not Significant + Cond2 Significant Negative |
| Clmn | TRUE | -0.109223692 | 0.605068399 | No Change | Not Significant | TRUE | -0.097399713 | 0.332327332 | No Change | Not Significant | TRUE | Cond1 Not Significant + Cond2 Not Significant |
| Clmp | TRUE | 0.181351451 | 0.714354347 | No Change | Not Significant | TRUE | 0.061474225 | 0.893440495 | No Change | Not Significant | TRUE | Cond1 Not Significant + Cond2 Not Significant |
| Cln3 | TRUE | -0.354088317 | 0.000685437 | No Change | Significant Negative | TRUE | -0.300491529 | 0.000962527 | No Change | Significant Negative | TRUE | Cond1 Significant Negative + Cond2 Significant Negative |
| Cln5 | TRUE | -0.140903338 | 0.19938276 | No Change | Not Significant | TRUE | 0.045093422 | 0.72436245 | No Change | Not Significant | TRUE | Cond1 Not Significant + Cond2 Not Significant |
| Cln6 | TRUE | -0.158146758 | 0.419152975 | No Change | Not Significant | TRUE | -0.124975658 | 0.319001795 | No Change | Not Significant | TRUE | Cond1 Not Significant + Cond2 Not Significant |
| Cln8 | TRUE | 0.182325815 | 0.083878883 | No Change | Not Significant | TRUE | -0.073656686 | 0.5332246 | No Change | Not Significant | TRUE | Cond1 Not Significant + Cond2 Not Significant |
| Clnk | TRUE | 0.717331722 | 1 | No Change | Not Significant | FALSE | NA | NA | No Change | Not Detected | FALSE | Background = FALSE |
| Clns1a | TRUE | 0.036381845 | 0.866806221 | No Change | Not Significant | TRUE | -0.042963552 | 0.689357631 | No Change | Not Significant | TRUE | Cond1 Not Significant + Cond2 Not Significant |
| Clock | TRUE | -0.040685837 | 0.853197357 | No Change | Not Significant | TRUE | -0.149581555 | 0.082730576 | No Change | Not Significant | TRUE | Cond1 Not Significant + Cond2 Not Significant |
| Clp1 | TRUE | -0.01094104 | 0.957864571 | No Change | Not Significant | TRUE | -0.027849732 | 0.880070575 | No Change | Not Significant | TRUE | Cond1 Not Significant + Cond2 Not Significant |
| Clpb | TRUE | 0.053099824 | 0.669122175 | No Change | Not Significant | TRUE | 0.232962204 | 0.090105959 | No Change | Not Significant | TRUE | Cond1 Not Significant + Cond2 Not Significant |
| Clpp | TRUE | -0.11763216 | 0.338874866 | No Change | Not Significant | TRUE | 0.046985686 | 0.780976082 | No Change | Not Significant | TRUE | Cond1 Not Significant + Cond2 Not Significant |
| Clps | TRUE | -0.938701414 | 0.006667458 | DOWN | DOWN | TRUE | 0.325676095 | 0.052030743 | No Change | Not Significant | TRUE | Cond1 DOWN + Cond2 Not Significant |
| Clptm1 | TRUE | -0.182915722 | 0.020374669 | No Change | Significant Negative | TRUE | -0.066371263 | 0.616422248 | No Change | Not Significant | TRUE | Cond1 Significant Negative + Cond2 Not Significant |
| Clptm1l | TRUE | 0.736727029 | 1.31E-13 | UP | UP | TRUE | 0.407258442 | 2.46E-11 | No Change | Significant Positive | TRUE | Cond1 UP + Cond2 Significant Positive |
| Clpx | TRUE | -0.095831487 | 0.342262172 | No Change | Not Significant | TRUE | 0.014743219 | 0.896941755 | No Change | Not Significant | TRUE | Cond1 Not Significant + Cond2 Not Significant |
| Clrn2 | TRUE | 2.645497698 | 1 | No Change | Not Significant | FALSE | NA | NA | No Change | Not Detected | FALSE | Background = FALSE |
| Clrn3 | TRUE | -0.000761269 | 0.994433839 | No Change | Not Significant | TRUE | 0.022196248 | 0.833278687 | No Change | Not Significant | TRUE | Cond1 Not Significant + Cond2 Not Significant |
| Clspn | TRUE | 0.588582817 | 0.010922096 | UP | UP | TRUE | 0.187493967 | 0.609576053 | No Change | Not Significant | TRUE | Cond1 UP + Cond2 Not Significant |
| Clstn1 | TRUE | -0.132369401 | 0.078201839 | No Change | Not Significant | TRUE | -0.017932438 | 0.869255491 | No Change | Not Significant | TRUE | Cond1 Not Significant + Cond2 Not Significant |
| Clstn2 | TRUE | -0.63515752 | 1 | No Change | Not Significant | FALSE | NA | NA | No Change | Not Detected | FALSE | Background = FALSE |
| Clstn3 | TRUE | 0.453366472 | 0.871097624 | No Change | Not Significant | FALSE | NA | NA | No Change | Not Detected | FALSE | Background = FALSE |
| Clta | TRUE | -0.057686135 | 0.559233675 | No Change | Not Significant | TRUE | -0.020989682 | 0.837002876 | No Change | Not Significant | TRUE | Cond1 Not Significant + Cond2 Not Significant |
| Cltb | TRUE | 0.032747195 | 0.728048657 | No Change | Not Significant | TRUE | 0.082246788 | 0.280581712 | No Change | Not Significant | TRUE | Cond1 Not Significant + Cond2 Not Significant |
| Cltc | TRUE | 0.084985675 | 0.586288962 | No Change | Not Significant | TRUE | 0.05158223 | 0.500967327 | No Change | Not Significant | TRUE | Cond1 Not Significant + Cond2 Not Significant |
| Cltrn | TRUE | 1.186047621 | 1 | No Change | Not Significant | FALSE | NA | NA | No Change | Not Detected | FALSE | Background = FALSE |
| Clu | TRUE | -0.27685908 | 0.59260115 | No Change | Not Significant | TRUE | 0.072310952 | 0.850359085 | No Change | Not Significant | TRUE | Cond1 Not Significant + Cond2 Not Significant |
| Cluap1 | TRUE | 0.079669217 | 0.744656458 | No Change | Not Significant | TRUE | 0.109163401 | 0.624972092 | No Change | Not Significant | TRUE | Cond1 Not Significant + Cond2 Not Significant |
| Cluh | TRUE | -0.160190289 | 0.081727701 | No Change | Not Significant | TRUE | -0.000280286 | 0.998544326 | No Change | Not Significant | TRUE | Cond1 Not Significant + Cond2 Not Significant |
| Clvs1 | TRUE | 0.39347572 | 1 | No Change | Not Significant | FALSE | NA | NA | No Change | Not Detected | FALSE | Background = FALSE |
| Clvs2 | TRUE | 1.138948657 | 1 | No Change | Not Significant | FALSE | NA | NA | No Change | Not Detected | FALSE | Background = FALSE |
| Clybl | TRUE | -0.559641152 | 3.71E-10 | DOWN | DOWN | TRUE | -0.171188712 | 0.07724278 | No Change | Not Significant | TRUE | Cond1 DOWN + Cond2 Not Significant |
| Cma1 | TRUE | 1.620911491 | 0.065126762 | No Change | Not Significant | TRUE | 0.674433135 | 0.203987912 | No Change | Not Significant | TRUE | Cond1 Not Significant + Cond2 Not Significant |
| Cma2 | TRUE | -1.097806656 | 1 | No Change | Not Significant | FALSE | NA | NA | No Change | Not Detected | FALSE | Background = FALSE |
| Cmah | TRUE | -0.433267392 | 0.220951697 | No Change | Not Significant | TRUE | -0.07482571 | 0.836585484 | No Change | Not Significant | TRUE | Cond1 Not Significant + Cond2 Not Significant |
| Cmas | TRUE | 0.191449857 | 0.265233241 | No Change | Not Significant | TRUE | 0.173481331 | 0.010339898 | No Change | Significant Positive | TRUE | Cond1 Not Significant + Cond2 Significant Positive |
| Cmbl | TRUE | -0.197050982 | 0.135072906 | No Change | Not Significant | TRUE | 0.02171213 | 0.867367688 | No Change | Not Significant | TRUE | Cond1 Not Significant + Cond2 Not Significant |
| Cmc1 | TRUE | -0.319493567 | 0.019904115 | No Change | Significant Negative | TRUE | -0.119093119 | 0.263676126 | No Change | Not Significant | TRUE | Cond1 Significant Negative + Cond2 Not Significant |
| Cmc2 | TRUE | 0.121703439 | 0.479832481 | No Change | Not Significant | TRUE | 0.31411997 | 0.004683526 | No Change | Significant Positive | TRUE | Cond1 Not Significant + Cond2 Significant Positive |
| Cmc4 | TRUE | -0.041987745 | 0.929560416 | No Change | Not Significant | TRUE | -0.136496686 | 0.741606925 | No Change | Not Significant | TRUE | Cond1 Not Significant + Cond2 Not Significant |
| Cmip | TRUE | -0.09765765 | 0.696271072 | No Change | Not Significant | TRUE | -0.109357704 | 0.391964304 | No Change | Not Significant | TRUE | Cond1 Not Significant + Cond2 Not Significant |
| Cmklr1 | TRUE | -0.012170081 | 1 | No Change | Not Significant | FALSE | NA | NA | No Change | Not Detected | FALSE | Background = FALSE |
| Cmpk1 | TRUE | 0.011069918 | 0.954353007 | No Change | Not Significant | TRUE | 0.155359309 | 0.002142507 | No Change | Significant Positive | TRUE | Cond1 Not Significant + Cond2 Significant Positive |
| Cmpk2 | TRUE | 0.174040396 | 0.671509918 | No Change | Not Significant | TRUE | -0.104860333 | 0.672729777 | No Change | Not Significant | TRUE | Cond1 Not Significant + Cond2 Not Significant |
| Cmss1 | TRUE | 0.40157833 | 0.078035837 | No Change | Not Significant | TRUE | 0.075386174 | 0.755348304 | No Change | Not Significant | TRUE | Cond1 Not Significant + Cond2 Not Significant |
| Cmtm3 | TRUE | -0.184614238 | 0.832521206 | No Change | Not Significant | TRUE | 0.132644296 | 0.820984417 | No Change | Not Significant | TRUE | Cond1 Not Significant + Cond2 Not Significant |
| Cmtm4 | TRUE | -0.174697363 | 0.125813975 | No Change | Not Significant | TRUE | -0.031550727 | 0.690945086 | No Change | Not Significant | TRUE | Cond1 Not Significant + Cond2 Not Significant |
| Cmtm5 | TRUE | 1.188148867 | 1 | No Change | Not Significant | FALSE | NA | NA | No Change | Not Detected | FALSE | Background = FALSE |
| Cmtm6 | TRUE | -0.068092167 | 0.713191289 | No Change | Not Significant | TRUE | 0.061416749 | 0.360168963 | No Change | Not Significant | TRUE | Cond1 Not Significant + Cond2 Not Significant |
| Cmtm7 | TRUE | -0.05640198 | 0.914530359 | No Change | Not Significant | TRUE | 0.31781268 | 0.066299536 | No Change | Not Significant | TRUE | Cond1 Not Significant + Cond2 Not Significant |
| Cmtm8 | TRUE | -0.001139013 | 0.997384323 | No Change | Not Significant | TRUE | 0.474548258 | 0.01119127 | No Change | Significant Positive | TRUE | Cond1 Not Significant + Cond2 Significant Positive |
| Cmtr1 | TRUE | 0.281108642 | 0.305377683 | No Change | Not Significant | TRUE | -0.190360717 | 0.384596281 | No Change | Not Significant | TRUE | Cond1 Not Significant + Cond2 Not Significant |
| Cmtr2 | TRUE | 0.033451135 | 0.876856731 | No Change | Not Significant | TRUE | -0.062217957 | 0.682675952 | No Change | Not Significant | TRUE | Cond1 Not Significant + Cond2 Not Significant |
| Cmya5 | TRUE | 0.45430362 | 1 | No Change | Not Significant | FALSE | NA | NA | No Change | Not Detected | FALSE | Background = FALSE |
| CN725425 | TRUE | 0.684532733 | 1 | No Change | Not Significant | FALSE | NA | NA | No Change | Not Detected | FALSE | Background = FALSE |
| Cnbd2 | TRUE | 0.021736509 | 0.96968875 | No Change | Not Significant | TRUE | -0.166576383 | 0.617285476 | No Change | Not Significant | TRUE | Cond1 Not Significant + Cond2 Not Significant |
| Cnbp | TRUE | 0.019814758 | 0.924222269 | No Change | Not Significant | TRUE | 0.028726754 | 0.662000443 | No Change | Not Significant | TRUE | Cond1 Not Significant + Cond2 Not Significant |
| Cndp1 | TRUE | -0.891830348 | 1 | No Change | Not Significant | FALSE | NA | NA | No Change | Not Detected | FALSE | Background = FALSE |
| Cndp2 | TRUE | -0.113125767 | 0.378300667 | No Change | Not Significant | TRUE | -0.302091774 | 0.002187666 | No Change | Significant Negative | TRUE | Cond1 Not Significant + Cond2 Significant Negative |
| Cnep1r1 | TRUE | 0.263808343 | 0.043612621 | No Change | Significant Positive | TRUE | 0.050030183 | 0.722065604 | No Change | Not Significant | TRUE | Cond1 Significant Positive + Cond2 Not Significant |
| Cnga1 | TRUE | 1.315867317 | 1 | No Change | Not Significant | FALSE | NA | NA | No Change | Not Detected | FALSE | Background = FALSE |
| Cnga2 | TRUE | -0.709673445 | 1 | No Change | Not Significant | FALSE | NA | NA | No Change | Not Detected | FALSE | Background = FALSE |
| Cnga3 | TRUE | -0.358820897 | 1 | No Change | Not Significant | FALSE | NA | NA | No Change | Not Detected | FALSE | Background = FALSE |
| Cnga4 | TRUE | 1.044741107 | 1 | No Change | Not Significant | FALSE | NA | NA | No Change | Not Detected | FALSE | Background = FALSE |
| Cngb1 | TRUE | -0.553734208 | 0.876856731 | No Change | Not Significant | FALSE | NA | NA | No Change | Not Detected | FALSE | Background = FALSE |
| Cnih1 | TRUE | 0.251894288 | 0.015586518 | No Change | Significant Positive | TRUE | 0.16285009 | 0.023338386 | No Change | Significant Positive | TRUE | Cond1 Significant Positive + Cond2 Significant Positive |
| Cnih2 | TRUE | 0.191557042 | 0.740233019 | No Change | Not Significant | TRUE | -0.403983605 | 0.552491851 | No Change | Not Significant | TRUE | Cond1 Not Significant + Cond2 Not Significant |
| Cnih4 | TRUE | -0.37219464 | 0.00081486 | No Change | Significant Negative | TRUE | -0.116388232 | 0.14077647 | No Change | Not Significant | TRUE | Cond1 Significant Negative + Cond2 Not Significant |
| Cnksr1 | TRUE | 0.466984204 | 0.003327122 | No Change | Significant Positive | TRUE | 0.066285152 | 0.455719922 | No Change | Not Significant | TRUE | Cond1 Significant Positive + Cond2 Not Significant |
| Cnksr2 | TRUE | 0.102733846 | 1 | No Change | Not Significant | FALSE | NA | NA | No Change | Not Detected | FALSE | Background = FALSE |

Supplementary Table 3

| geneID | Cond1_DF_Detected | Cond1_DF_ValueCol | Cond1_DF_PadjCol | Cond1_DF_Cutoff | Cond1_DF_Cutoff_Specific.x | Cond2_D[illegible] | [illegible] |
|---|---|---|---|---|---|---|---|
| Cnksr3 | TRUE | -0.554244482 | 0.000754981 | DOWN | DOWN | TRUE | [illegible] |
| Cnmd | TRUE | -1.995936297 | 1 | No Change | Not Significant | FALSE | [illegible] |
| Cnn1 | TRUE | -0.273727896 | 1 | No Change | Not Significant | FALSE | [illegible] |
| Cnn2 | TRUE | 0.3829495 | 0.063634329 | No Change | Not Significant | TRUE | [illegible] |
| Cnn3 | TRUE | -0.417245677 | 0.023887859 | No Change | Significant Negative | TRUE | [illegible] |
| Cnnm1 | TRUE | -1.032811549 | 0.048307661 | DOWN | DOWN | TRUE | [illegible] |
| Cnnm2 | TRUE | 1.389694696 | 9.25E-25 | UP | UP | TRUE | [illegible] |
| Cnnm3 | TRUE | 0.164971308 | 0.131338332 | No Change | Not Significant | TRUE | [illegible] |
| Cnnm4 | TRUE | -0.103887441 | 0.45367543 | No Change | Not Significant | TRUE | [illegible] |
| Cnot1 | TRUE | 0.308936937 | 2.29E-05 | No Change | Significant Positive | TRUE | [illegible] |
| Cnot10 | TRUE | 0.101367657 | 0.169583213 | No Change | Not Significant | TRUE | [illegible] |
| Cnot11 | TRUE | 0.045796223 | 0.744703672 | No Change | Not Significant | TRUE | [illegible] |
| Cnot2 | TRUE | 0.020475391 | 0.892918574 | No Change | Not Significant | TRUE | [illegible] |
| Cnot3 | TRUE | 0.88729008 | 6.41E-09 | UP | UP | TRUE | [illegible] |
| Cnot4 | TRUE | -0.059151401 | 0.674995696 | No Change | Not Significant | TRUE | [illegible] |
| Cnot6 | TRUE | 0.330454269 | 0.003195929 | No Change | Significant Positive | TRUE | [illegible] |
| Cnot6l | TRUE | -0.223361095 | 0.220060838 | No Change | Not Significant | TRUE | [illegible] |
| Cnot7 | TRUE | 0.065161182 | 0.628584238 | No Change | Not Significant | TRUE | [illegible] |
| [illegible] | [illegible] | [illegible] | [illegible] | [illegible] | [illegible] | [illegible] | [illegible] |

| geneID | Cond1_DF_Detected | Cond1_DF_ValueCol | Cond1_DF_PadjCol | Cond1_DF_Cutoff | Cond1_DF_Cutoff_Specific.x | Cond2_DF_Detected |
|---|---|---|---|---|---|---|
| Cox7c | TRUE | -0.459314199 | 0.00035465 | No Change | Significant Negative | TRUE |
| Cox8a | TRUE | -0.247177347 | 0.008149473 | No Change | Significant Negative | TRUE |
| Cp | TRUE | 1.214256651 | 0.10685821 | No Change | Not Significant | FALSE |
| Cpa1 | TRUE | -2.616585831 | 1 | No Change | Not Significant | FALSE |
| Cpa2 | TRUE | 0.757549091 | 0.047864939 | UP | UP | TRUE |
| Cpa3 | TRUE | -0.14645552 | 1 | No Change | Not Significant | FALSE |
| Cpa4 | TRUE | -1.976174131 | 1 | No Change | Not Significant | FALSE |
| Cpa5 | TRUE | -0.023936136 | 1 | No Change | Not Significant | FALSE |
| Cpa6 | TRUE | 0.611545215 | 1 | No Change | Not Significant | FALSE |
| Cpb1 | TRUE | -3.666506859 | 1 | No Change | Not Significant | FALSE |
| Cpb2 | TRUE | -2.70406252 | 1 | No Change | Not Significant | TRUE |
| Cpd | TRUE | -0.008623041 | 0.977832194 | No Change | Not Significant | TRUE |
| Cpe | TRUE | -0.456128968 | 0.069446017 | No Change | Not Significant | TRUE |
| Cpeb1 | TRUE | -1.001113327 | 1 | No Change | Not Significant | FALSE |
| Cpeb2 | TRUE | 0.17720516 | 0.274255704 | No Change | Not Significant | TRUE |
| Cpeb3 | TRUE | -0.383744492 | 0.034924193 | No Change | Significant Negative | TRUE |
| Cpeb4 | TRUE | -0.120981286 | 0.302752759 | No Change | Not Significant | TRUE |
| Cped1 | TRUE | 0.262126885 | 0.722040793 | No Change | Not Significant | TRUE |
| Cplane1 | TRUE | 0.564715605 | 0.031766369 | UP | UP | TRUE |
| Cplane2 | TRUE | 0.098935843 | 0.904271659 | No Change | Not Significant | TRUE |
| Cplx1 | TRUE | -0.060137987 | 1 | No Change | Not Significant | FALSE |
| Cplx2 | TRUE | -0.132720122 | 0.674460666 | No Change | Not Significant | TRUE |
| Cplx3 | TRUE | 0.002520554 | 1 | No Change | Not Significant | FALSE |
| Cpm | TRUE | -0.190815389 | 0.265659566 | No Change | Not Significant | TRUE |
| Cpn1 | TRUE | 0.670747331 | 0.085735413 | No Change | Not Significant | TRUE |
| Cpn2 | TRUE | 1.111705476 | 0.031110715 | UP | UP | TRUE |
| Cpne1 | TRUE | -0.201573836 | 1 | No Change | Not Significant | FALSE |
| Cpne2 | TRUE | -0.013582262 | 0.953251304 | No Change | Not Significant | TRUE |
| Cpne3 | TRUE | -0.205168325 | 0.127638694 | No Change | Not Significant | TRUE |
| Cpne4 | TRUE | 3.092917108 | 1 | No Change | Not Significant | FALSE |
| Cpne5 | TRUE | -0.642308601 | 0.654912763 | No Change | Not Significant | FALSE |
| Cpne6 | TRUE | 1.113268318 | 1 | No Change | Not Significant | FALSE |
| Cpne7 | TRUE | -0.032641091 | 1 | No Change | Not Significant | FALSE |
| Cpne8 | TRUE | 0.332953162 | 0.000944559 | No Change | Significant Positive | TRUE |
| Cpne9 | TRUE | 0.862176842 | 0.69793377 | No Change | Not Significant | FALSE |
| Cpox | TRUE | 0.250062777 | 0.080540642 | No Change | Not Significant | TRUE |
| Cpped1 | TRUE | 0.580059138 | 1.23E-15 | UP | UP | TRUE |
| Cpq | TRUE | -0.626106737 | 0.003263111 | DOWN | DOWN | TRUE |
| Cps1 | TRUE | 0.437874025 | 0.093755613 | No Change | Not Significant | TRUE |
| Cpsf1 | TRUE | 0.204784486 | 0.01893658 | No Change | Significant Positive | TRUE |
| Cpsf2 | TRUE | 0.063674471 | 0.700292277 | No Change | Not Significant | TRUE |
| Cpsf3 | TRUE | 0.02073004 | 0.912523774 | No Change | Not Significant | TRUE |
| Cpsf4 | TRUE | 0.189225734 | 0.121313669 | No Change | Not Significant | TRUE |
| Cpsf4l | TRUE | 0.295406404 | 0.774787312 | No Change | Not Significant | FALSE |
| Cpsf6 | TRUE | 0.271602087 | 0.000260832 | No Change | Significant Positive | TRUE |
| Cpsf7 | TRUE | -0.003057727 | 0.980447529 | No Change | Not Significant | TRUE |
| Cpt1a | TRUE | -0.376751668 | 0.000104289 | No Change | Significant Negative | TRUE |
| Cpt1b | TRUE | 0.292127614 | 0.732754497 | No Change | Not Significant | FALSE |
| Cpt1c | TRUE | 4.30389438 | 1.34E-14 | UP | UP | TRUE |
| Cpt2 | TRUE | -0.092945958 | 0.377599236 | No Change | Not Significant | TRUE |
| Cptp | TRUE | 0.430032689 | 1.74E-07 | No Change | Significant Positive | TRUE |
| Cpvl | TRUE | -0.910128918 | 1 | No Change | Not Significant | TRUE |
| Cpxm1 | TRUE | 2.274936513 | 1 | No Change | Not Significant | FALSE |
| Cpxm2 | TRUE | -2.204652933 | 1 | No Change | Not Significant | FALSE |
| Cpz | TRUE | -0.382021156 | 1 | No Change | Not Significant | FALSE |
| Cr1l | TRUE | -0.41540833 | 0.00014838 | No Change | Significant Negative | TRUE |
| Cr2 | TRUE | -0.489086351 | 1 | No Change | Not Significant | FALSE |
| Cracd | TRUE | -0.180449318 | 0.266439513 | No Change | Not Significant | TRUE |
| Cracdl | TRUE | -0.349353152 | 0.001112705 | No Change | Significant Negative | TRUE |
| Cracr2a | TRUE | -0.496050193 | 0.026525797 | No Change | Significant Negative | TRUE |
| Cracr2b | TRUE | -0.11961301 | 0.720922137 | No Change | Not Significant | TRUE |
| Cradd | TRUE | -0.243428454 | 0.219498448 | No Change | Not Significant | TRUE |
| Cramp1 | TRUE | 0.031432357 | 0.915993052 | No Change | Not Significant | TRUE |
| Crat | TRUE | -0.419260572 | 1.27E-09 | No Change | Significant Negative | TRUE |
| Crb1 | TRUE | 0.637131204 | 1 | No Change | Not Significant | FALSE |
| Crb2 | TRUE | 0.057313272 | 1 | No Change | Not Significant | FALSE |
| Crb3 | TRUE | -0.228990498 | 0.084321363 | No Change | Not Significant | TRUE |
| Crbn | TRUE | -0.11815933 | 0.392885958 | No Change | Not Significant | TRUE |
| Crcp | TRUE | 0.35887156 | 6.93E-05 | No Change | Significant Positive | TRUE |
| Creb1 | TRUE | -0.070267291 | 0.704201299 | No Change | Not Significant | TRUE |
| Creb3 | TRUE | 0.177456754 | 0.045517455 | No Change | Significant Positive | TRUE |
| Creb3l1 | TRUE | 0.259004983 | 0.136567098 | No Change | Not Significant | TRUE |
| Creb3l2 | TRUE | 1.831449675 | 2.90E-48 | UP | UP | TRUE |
| Creb3l3 | TRUE | 1.380649509 | 0.02121487 | UP | UP | TRUE |
| Creb3l4 | TRUE | -0.020801755 | 0.966352899 | No Change | Not Significant | TRUE |
| Creb5 | TRUE | 0.132947593 | 0.977047797 | No Change | Not Significant | FALSE |
| Crebbp | TRUE | 0.084267748 | 0.696363286 | No Change | Not Significant | TRUE |
| Crebl2 | TRUE | -0.51579826 | 6.98E-07 | DOWN | DOWN | TRUE |
| Crebrf | TRUE | -0.664046989 | 0.000282016 | DOWN | DOWN | TRUE |
| Crebzf | TRUE | 0.206758492 | 0.056179837 | No Change | Not Significant | TRUE |
| Creg1 | TRUE | 0.558802346 | 6.18E-06 | UP | UP | TRUE |
| Creg2 | TRUE | 0.670662515 | 1 | No Change | Not Significant | FALSE |
| Creld1 | TRUE | 2.319294108 | 5.49E-96 | UP | UP | TRUE |
| Creld2 | TRUE | 4.29069154 | 1.28E-267 | UP | UP | TRUE |
| Crem | TRUE | 0.035102353 | 0.920873844 | No Change | Not Significant | TRUE |
| Crhr1 | TRUE | 0.753064844 | 1 | No Change | Not Significant | FALSE |
| Crhr2 | TRUE | 0.172974823 | 1 | No Change | Not Significant | FALSE |
| Crim1 | TRUE | -0.243320595 | 0.003829873 | No Change | Significant Negative | TRUE |
| Crip1 | TRUE | -0.9582824 | 1.47E-33 | DOWN | DOWN | TRUE |
| Crip2 | TRUE | -1.523506153 | 0.000153277 | DOWN | DOWN | TRUE |
| Crip3 | TRUE | 3.513912835 | 1 | No Change | Not Significant | FALSE |
| Cript | TRUE | -0.071379634 | 0.669812138 | No Change | Not Significant | TRUE |
| Crisp2 | TRUE | 2.375057318 | 1 | No Change | Not Significant | FALSE |
| Crisp3 | TRUE | -1.26819492 | 0.014751683 | DOWN | DOWN | TRUE |
| Crispld1 | TRUE | 0.176010528 | 1 | No Change | Not Significant | FALSE |
| Crispld2 | TRUE | 0.819694251 | 1 | No Change | Not Significant | FALSE |
| Crk | TRUE | 0.035483488 | 0.788502504 | No Change | Not Significant | TRUE |
| Crkl | TRUE | 0.008260695 | 0.942211125 | No Change | Not Significant | TRUE |
| Crlf1 | TRUE | -0.134841214 | 0.890541867 | No Change | Not Significant | TRUE |
| Crlf2 | TRUE | -0.102987395 | 0.793304622 | No Change | Not Significant | TRUE |
| Crlf3 | TRUE | -0.243676696 | 0.087113984 | No Change | Not Significant | TRUE |
| Crls1 | TRUE | 0.536962787 | 7.37E-07 | UP | UP | TRUE |
| Crmp1 | TRUE | 0.249984945 | 0.891996173 | No Change | Not Significant | FALSE |
| Crnkl1 | TRUE | 0.088969499 | 0.442885865 | No Change | Not Significant | TRUE |
| Crnn | TRUE | -0.707221292 | 1 | No Change | Not Significant | TRUE |
| Crocc | TRUE | -0.008597398 | 0.981095512 | No Change | Not Significant | TRUE |
| Crocc2 | TRUE | 0.410272984 | 0.682480636 | No Change | Not Significant | TRUE |
| Crot | TRUE | 0.250493667 | 0.002670749 | No Change | Significant Positive | TRUE |
| Crppa | TRUE | 0.067845313 | 0.883975099 | No Change | Not Significant | TRUE |
| Crtac1 | TRUE | 1.699497858 | 1 | No Change | Not Significant | FALSE |
| Crtam | TRUE | 0.510897655 | 1 | No Change | Not Significant | FALSE |
| Crtap | TRUE | -0.15368164 | 0.407353559 | No Change | Not Significant | TRUE |
| Crtc1 | TRUE | 0.133076042 | 0.571046919 | No Change | Not Significant | TRUE |
| Crtc2 | TRUE | -0.161967883 | 0.394941012 | No Change | Not Significant | TRUE |
| Crtc3 | TRUE | -0.075838714 | 0.687150319 | No Change | Not Significant | TRUE |
| Crxos | TRUE | 2.814719353 | 1 | No Change | Not Significant | FALSE |
| Cry1 | TRUE | 1.293845492 | 6.52E-06 | UP | UP | TRUE |
| Cry2 | TRUE | -0.011304959 | 0.979208448 | No Change | Not Significant | TRUE |
| Cryab | TRUE | -0.782106905 | 1 | No Change | Not Significant | FALSE |
| Cryba1 | TRUE | -0.225729281 | 1 | No Change | Not Significant | FALSE |
| Cryba2 | TRUE | 0.262361024 | 1 | No Change | Not Significant | TRUE |
| Cryba4 | TRUE | -0.989497186 | 1 | No Change | Not Significant | FALSE |
| Crybb1 | TRUE | 1.972646556 | 0.027492662 | UP | UP | FALSE |
| Crybb3 | TRUE | 0.147288446 | 0.898207529 | No Change | Not Significant | TRUE |
| Crybg1 | TRUE | 0.324050898 | 0.0043132 | No Change | Significant Positive | TRUE |
| Crybg2 | TRUE | -0.394969245 | 0.145821162 | No Change | Not Significant | TRUE |
| Crybg3 | TRUE | -0.151427048 | 0.128879463 | No Change | Not Significant | TRUE |
| Crygn | TRUE | -2.308656895 | 1 | No Change | Not Significant | FALSE |
| Cryl1 | TRUE | -0.386799961 | 8.96E-06 | No Change | Significant Negative | TRUE |
| Crym | TRUE | 0.052881732 | 0.884525396 | No Change | Not Significant | TRUE |
| Cryz | TRUE | -0.285283978 | 0.058004537 | No Change | Not Significant | TRUE |
| Cryzl1 | TRUE | -0.159841578 | 0.119334048 | No Change | Not Significant | TRUE |
| Cryzl2 | TRUE | -0.223260569 | 0.333845229 | No Change | Not Significant | TRUE |
| Cs | TRUE | -0.099126372 | 0.41130117 | No Change | Not Significant | TRUE |
| Csad | TRUE | -0.367923171 | 0.062458007 | No Change | Not Significant | TRUE |
| Csdc2 | TRUE | 1.11045444 | 1 | No Change | Not Significant | FALSE |
| Csde1 | TRUE | -0.0969835 | 0.488337435 | No Change | Not Significant | TRUE |
| Cse1l | TRUE | 0.349680453 | 0.057294003 | No Change | Not Significant | TRUE |
| Csf1 | TRUE | 0.318879481 | 0.767773712 | No Change | Not Significant | TRUE |
| Csf1r | TRUE | -0.141634131 | 0.908812719 | No Change | Not Significant | TRUE |
| Csf2ra | TRUE | 0.127493769 | 0.732676764 | No Change | Not Significant | TRUE |
| Csf2rb | TRUE | 0.173376172 | 0.932373279 | No Change | Not Significant | FALSE |
| Csf2rb2 | TRUE | 1.772790059 | 1 | No Change | Not Significant | FALSE |
| Csf3 | TRUE | 1.008190233 | 1 | No Change | Not Significant | FALSE |
| Csf3r | TRUE | 1.129838539 | 1 | No Change | Not Significant | FALSE |
| Csgalnact1 | TRUE | 1.243753377 | 1 | No Change | Not Significant | FALSE |
| Csgalnact2 | TRUE | 0.747535854 | 4.22E-08 | UP | UP | TRUE |
| Csk | TRUE | 0.027103422 | 0.827106157 | No Change | Not Significant | TRUE |
| Csl | TRUE | 1.240892662 | 1 | No Change | Not Significant | FALSE |
| Csmd1 | TRUE | 2.043782723 | 0.503546031 | No Change | Not Significant | FALSE |
| Csmd2 | TRUE | 0.188477542 | 0.969075596 | No Change | Not Significant | FALSE |
| Csmd3 | TRUE | 0.560991442 | 1 | No Change | Not Significant | FALSE |
| Csn1s2a | TRUE | 1.111548892 | 1 | No Change | Not Significant | FALSE |
| Csnk1a1 | TRUE | 0.027344422 | 0.771818402 | No Change | Not Significant | TRUE |
| Csnk1d | TRUE | 0.08701654 | 0.297863506 | No Change | Not Significant | TRUE |
| Csnk1e | TRUE | 0.103046206 | 0.665277305 | No Change | Not Significant | TRUE |
| Csnk1g1 | TRUE | 0.055802864 | 0.722040793 | No Change | Not Significant | TRUE |
| Csnk1g2 | TRUE | 0.032045671 | 0.767483504 | No Change | Not Significant | TRUE |
| Csnk1g3 | TRUE | -0.096742499 | 0.365932262 | No Change | Not Significant | TRUE |
| Csnk2a1 | TRUE | -0.366927986 | 9.17E-06 | No Change | Significant Negative | TRUE |
| Csnk2a2 | TRUE | -0.053029854 | 0.685206817 | No Change | Not Significant | TRUE |
| Csnk2b | TRUE | -0.047242881 | 0.622737045 | No Change | Not Significant | TRUE |
| Cspg4 | TRUE | 0.221914528 | 0.772317806 | No Change | Not Significant | TRUE |
| Cspg4b | TRUE | -0.065506437 | 1 | No Change | Not Significant | FALSE |
| Cspg5 | TRUE | -0.026972572 | 0.981497544 | No Change | Not Significant | TRUE |
| Cspp1 | TRUE | -0.11761941 | 0.49722072 | No Change | Not Significant | TRUE |
| Csrnp1 | TRUE | -0.684087076 | 0.045342919 | DOWN | DOWN | TRUE |
| Csrnp2 | TRUE | 0.062816033 | 0.878912418 | No Change | Not Significant | TRUE |
| Csrnp3 | TRUE | 0.693193961 | 1 | No Change | Not Significant | FALSE |
| Csrp1 | TRUE | 1.997140601 | 3.08E-52 | UP | UP | TRUE |
| Csrp2 | TRUE | -0.39862706 | 0.000339062 | No Change | Significant Negative | TRUE |
| Csrp3 | TRUE | 0.534451239 | 1 | No Change | Not Significant | FALSE |
| Cst3 | TRUE | -0.615528668 | 5.57E-07 | DOWN | DOWN | TRUE |
| Cst6 | TRUE | -0.663417721 | 0.19566778 | No Change | Not Significant | TRUE |
| Cst7 | TRUE | 0.324560635 | 1 | No Change | Not Significant | FALSE |
| Csta1 | TRUE | 0.473400164 | 0.313084659 | No Change | Not Significant | TRUE |
| Csta2 | TRUE | 0.214851854 | 0.892764273 | No Change | Not Significant | TRUE |
| Cstb | TRUE | -0.065092338 | 0.648455238 | No Change | Not Significant | TRUE |
| Cstf1 | TRUE | 0.155044453 | 0.362441146 | No Change | Not Significant | TRUE |
| Cstf2 | TRUE | 0.153305313 | 0.186963018 | No Change | Not Significant | TRUE |
| Cstf2t | TRUE | 0.010504451 | 0.950413359 | No Change | Not Significant | TRUE |
| Cstf3 | TRUE | 0.275812519 | 0.027885242 | No Change | Significant Positive | TRUE |
| Cstpp1 | TRUE | 0.153939345 | 0.585429376 | No Change | Not Significant | TRUE |
| Ct55 | TRUE | 0.261695311 | 1 | No Change | Not Significant | FALSE |
| Ctbp1 | TRUE | 0.028142367 | 0.833786242 | No Change | Not Significant | TRUE |
| Ctbp2 | TRUE | 0.072968638 | 0.37212295 | No Change | Not Significant | TRUE |
| Ctbs | TRUE | -0.11581193 | 0.661197851 | No Change | Not Significant | TRUE |
| Ctc1 | TRUE | 0.055431904 | 0.80301103 | No Change | Not Significant | TRUE |
| Ctcf | TRUE | 0.031658704 | 0.823884333 | No Change | Not Significant | TRUE |
| Ctcfl | TRUE | -0.440272828 | 0.429533455 | No Change | Not Significant | TRUE |
| Ctcflos | TRUE | -1.688395339 | 1 | No Change | Not Significant | FALSE |
| Ctdnep1 | TRUE | 0.251516192 | 0.117137463 | No Change | Not Significant | TRUE |

Supplementary Table 3

| geneID | Cond1_DF_Detected | Cond1_DF_ValueCol | Cond1_DF_PadjCol | Cond1_DF_Cutoff | Cond1_DF_Cutoff_Specific.x | Cond2_DF_Detected | Cond2_DF_ValueCol | Cond2_DF_PadjCol | Cond2_DF_Cutoff | Cond2_DF_Cutoff_Specific | BG_Method |
|---|---|---|---|---|---|---|---|---|---|---|---|
| Ctdp1 | TRUE | -0.082730852 | 0.535937072 | No Change | Not Significant | TRUE | -0.089370042 | 0.400571839 | No Change | Not Significant | TRUE |
| Ctdsp1 | TRUE | -0.089636795 | 0.411572816 | No Change | Not Significant | TRUE | -0.038133287 | 0.714747243 | No Change | Not Significant | TRUE |
| Ctdsp2 | TRUE | -0.115413094 | 0.249608668 | No Change | Not Significant | TRUE | -0.162596491 | 0.110958194 | No Change | Not Significant | TRUE |
| Ctdspl | TRUE | 0.246316386 | 0.015463527 | No Change | Significant Positive | TRUE | 0.087139275 | 0.209930404 | No Change | Not Significant | TRUE |
| Ctdspl2 | TRUE | -0.086807802 | 0.60160505 | No Change | Not Significant | TRUE | -0.212663667 | 0.040959773 | No Change | Significant Negative | TRUE |
| Ctf1 | TRUE | -0.304457004 | 0.496688848 | No Change | Not Significant | TRUE | 0.126896108 | 0.685788934 | No Change | Not Significant | TRUE |
| Ctf2 | TRUE | -1.665536768 | 1 | No Change | Not Significant | FALSE | NA | NA | No Change | Not Detected | FALSE |
| Cth | TRUE | -0.237203918 | 0.022621462 | No Change | Significant Negative | TRUE | -0.00460922 | 0.974178386 | No Change | Not Significant | TRUE |
| Cthrc1 | TRUE | 0.038091756 | 0.97108297 | No Change | Not Significant | FALSE | NA | NA | No Change | Not Detected | FALSE |
| Ctif | TRUE | 1.024441371 | 3.93E-11 | UP | UP | TRUE | 1.000065463 | 2.45E-24 | UP | UP | TRUE |
| Ctla2a | TRUE | -0.389518394 | 1 | No Change | Not Significant | FALSE | NA | NA | No Change | Not Detected | FALSE |
| Ctla4 | TRUE | 0.931539923 | 1 | No Change | Not Significant | FALSE | NA | NA | No Change | Not Detected | FALSE |
| Ctnna1 | TRUE | -0.125241774 | 0.295450802 | No Change | Not Significant | TRUE | 0.069616413 | 0.268368093 | No Change | Not Significant | TRUE |
| Ctnna2 | TRUE | -1.263074066 | 0.247378172 | No Change | Not Significant | TRUE | 0.131032797 | 0.893440495 | No Change | Not Significant | TRUE |
| Ctnna3 | TRUE | 0.831900577 | 1 | No Change | Not Significant | FALSE | NA | NA | No Change | Not Detected | FALSE |
| Ctnnal1 | TRUE | -0.205562702 | 0.374503142 | No Change | Not Significant | TRUE | -0.139371055 | 0.203588399 | No Change | Not Significant | TRUE |
| Ctnnb1 | TRUE | 0.107219554 | 0.04843157 | No Change | Significant Positive | TRUE | 0.144167153 | 0.001273135 | No Change | Significant Positive | TRUE |
| Ctnnbip1 | TRUE | -0.028056429 | 0.893150482 | No Change | Not Significant | TRUE | -0.011814996 | 0.937932225 | No Change | Not Significant | TRUE |
| Ctnnbl1 | TRUE | 0.098925557 | 0.37057407 | No Change | Not Significant | TRUE | 0.144495387 | 0.133152759 | No Change | Not Significant | TRUE |
| Ctnnd1 | TRUE | 0.043935492 | 0.77323163 | No Change | Not Significant | TRUE | 0.045231344 | 0.550111272 | No Change | Not Significant | TRUE |
| Ctnnd2 | TRUE | 0.316860209 | 0.824518194 | No Change | Not Significant | TRUE | 0.921894989 | 0.252369884 | No Change | Not Significant | TRUE |
| Ctns | TRUE | -0.268215736 | 0.114696833 | No Change | Not Significant | TRUE | -0.182801724 | 0.162264555 | No Change | Not Significant | TRUE |
| Ctps | TRUE | 0.227239417 | 0.279846882 | No Change | Not Significant | TRUE | -0.165314039 | 0.42427099 | No Change | Not Significant | TRUE |
| Ctps2 | TRUE | -0.312668638 | 0.038648666 | No Change | Significant Negative | TRUE | -0.154355554 | 0.090113624 | No Change | Not Significant | TRUE |
| Ctr9 | TRUE | -0.021169154 | 0.863385162 | No Change | Not Significant | TRUE | 0.05211946 | 0.533047399 | No Change | Not Significant | TRUE |
| Ctrb1 | TRUE | -2.328967293 | 1 | No Change | Not Significant | FALSE | NA | NA | No Change | Not Detected | FALSE |
| Ctrl | TRUE | 0.194619958 | 0.864656729 | No Change | Not Significant | TRUE | -0.51989747 | 0.468692446 | No Change | Not Significant | TRUE |
| Ctsa | TRUE | -0.139713124 | 0.028605439 | No Change | Significant Negative | TRUE | -0.01027971 | 0.929633324 | No Change | Not Significant | TRUE |
| Ctsb | TRUE | -0.394125291 | 2.22E-11 | No Change | Significant Negative | TRUE | -0.121462723 | 0.018994009 | No Change | Significant Negative | TRUE |
| Ctsc | TRUE | -0.22602808 | 0.057495825 | No Change | Not Significant | TRUE | -0.049557434 | 0.562506362 | No Change | Not Significant | TRUE |
| Ctsd | TRUE | -0.391853446 | 1.05E-05 | No Change | Significant Negative | TRUE | -0.033546272 | 0.716014372 | No Change | Not Significant | TRUE |
| Ctse | TRUE | -0.645210824 | 0.053943227 | No Change | Not Significant | TRUE | 0.039746411 | 0.903818244 | No Change | Not Significant | TRUE |
| Ctsf | TRUE | -1.115800778 | 2.51E-07 | DOWN | DOWN | TRUE | -0.268607349 | 0.061552292 | No Change | Not Significant | TRUE |
| Ctsh | TRUE | -0.35403182 | 0.000384378 | No Change | Significant Negative | TRUE | 0.149973632 | 0.041260849 | No Change | Significant Positive | TRUE |
| Ctsk | TRUE | 0.091327564 | 1 | No Change | Not Significant | FALSE | NA | NA | No Change | Not Detected | FALSE |
| Ctsl | TRUE | -0.504894193 | 6.99E-06 | DOWN | DOWN | TRUE | -0.273084124 | 0.000156868 | No Change | Significant Negative | TRUE |
| Ctso | TRUE | -0.179568632 | 0.193640076 | No Change | Not Significant | TRUE | -0.178100564 | 0.132451326 | No Change | Not Significant | TRUE |
| Ctss | TRUE | -0.64554174 | 2.16E-09 | DOWN | DOWN | TRUE | -0.409927723 | 4.57E-07 | No Change | Significant Negative | TRUE |
| Ctsw | TRUE | 0.183914416 | 0.78673795 | No Change | Not Significant | TRUE | 0.104072472 | 0.699528403 | No Change | Not Significant | TRUE |
| Ctsz | TRUE | -0.22922222 | 0.0002483 | No Change | Significant Negative | TRUE | -0.080501289 | 0.392580849 | No Change | Not Significant | TRUE |
| Cttn | TRUE | -0.064378775 | 0.622315002 | No Change | Not Significant | TRUE | 0.080486236 | 0.31618145 | No Change | Not Significant | TRUE |
| Cttnbp2 | TRUE | 0.371307988 | 0.221416423 | No Change | Not Significant | TRUE | -0.046359111 | 0.830822102 | No Change | Not Significant | TRUE |
| Cttnbp2nl | TRUE | -0.163278354 | 0.164593053 | No Change | Not Significant | TRUE | -0.052432759 | 0.657986673 | No Change | Not Significant | TRUE |
| Ctu1 | TRUE | -0.004391502 | 0.983565239 | No Change | Not Significant | TRUE | 0.085721425 | 0.577733884 | No Change | Not Significant | TRUE |
| Ctu2 | TRUE | 0.006235135 | 0.977515248 | No Change | Not Significant | TRUE | -0.007449946 | 0.965302141 | No Change | Not Significant | TRUE |
| Ctxn1 | TRUE | 0.584447873 | 0.53003414 | No Change | Not Significant | TRUE | -0.076086838 | 0.930889346 | No Change | Not Significant | TRUE |
| Ctxnd1 | TRUE | -0.707410548 | 0.000194686 | DOWN | DOWN | TRUE | -0.31253385 | 0.082452993 | No Change | Not Significant | TRUE |
| Cubn | TRUE | -0.798777366 | 0.07460396 | No Change | Not Significant | TRUE | -1.164119716 | 1.87E-05 | DOWN | DOWN | TRUE |
| Cuedc1 | TRUE | 0.015372014 | 0.964363534 | No Change | Not Significant | TRUE | -0.107713945 | 0.65828686 | No Change | Not Significant | TRUE |
| Cuedc2 | TRUE | -0.099377048 | 0.394964884 | No Change | Not Significant | TRUE | 0.016773001 | 0.915791177 | No Change | Not Significant | TRUE |
| Cul1 | TRUE | -0.133037951 | 0.088524117 | No Change | Not Significant | TRUE | 0.061233941 | 0.523452524 | No Change | Not Significant | TRUE |
| Cul2 | TRUE | 0.180377797 | 0.155922569 | No Change | Not Significant | TRUE | 0.062723972 | 0.517029701 | No Change | Not Significant | TRUE |
| Cul3 | TRUE | 0.006237504 | 0.971638507 | No Change | Not Significant | TRUE | -0.037231126 | 0.523508209 | No Change | Not Significant | TRUE |
| Cul4a | TRUE | 0.101256667 | 0.387604015 | No Change | Not Significant | TRUE | -0.026401693 | 0.707074879 | No Change | Not Significant | TRUE |
| Cul4b | TRUE | -0.013836281 | 0.931099354 | No Change | Not Significant | TRUE | -0.183210457 | 0.02835158 | No Change | Significant Negative | TRUE |
| Cul5 | TRUE | -0.00995262 | 0.965445424 | No Change | Not Significant | TRUE | -0.089764904 | 0.352354239 | No Change | Not Significant | TRUE |
| Cul7 | TRUE | 0.59277508 | 5.22E-06 | UP | UP | TRUE | 0.163829174 | 0.300613293 | No Change | Not Significant | TRUE |
| Cul9 | TRUE | -0.726153633 | 0.181269925 | No Change | Not Significant | TRUE | -0.46706758 | 0.082474622 | No Change | Not Significant | TRUE |
| Cuta | TRUE | 0.426591251 | 8.81E-05 | No Change | Significant Positive | TRUE | 0.17802425 | 0.013434123 | No Change | Significant Positive | TRUE |
| Cutal | TRUE | -0.196301507 | 0.294844608 | No Change | Not Significant | TRUE | 0.124362517 | 0.146403821 | No Change | Not Significant | TRUE |
| Cutc | TRUE | -0.469842636 | 0.029520727 | No Change | Significant Negative | TRUE | -0.225596951 | 0.077646289 | No Change | Not Significant | TRUE |
| Cux1 | TRUE | -0.335426486 | 5.67E-09 | No Change | Significant Negative | TRUE | -0.049358922 | 0.441566958 | No Change | Not Significant | TRUE |
| Cux2 | TRUE | 0.226316672 | 0.77804505 | No Change | Not Significant | TRUE | -0.577020041 | 0.133152759 | No Change | Not Significant | TRUE |
| Cwc15 | TRUE | 0.072731516 | 0.488030519 | No Change | Not Significant | TRUE | 0.072620532 | 0.34961375 | No Change | Not Significant | TRUE |
| Cwc22 | TRUE | 0.084673044 | 0.581717059 | No Change | Not Significant | TRUE | 0.056873939 | 0.657096531 | No Change | Not Significant | TRUE |
| Cwc25 | TRUE | -0.126868689 | 0.347646657 | No Change | Not Significant | TRUE | -0.076070272 | 0.487137296 | No Change | Not Significant | TRUE |
| Cwc27 | TRUE | 0.139412604 | 0.375643497 | No Change | Not Significant | TRUE | -0.029192653 | 0.861553924 | No Change | Not Significant | TRUE |
| Cwf19l1 | TRUE | 0.063973174 | 0.745058233 | No Change | Not Significant | TRUE | -0.122602088 | 0.392680814 | No Change | Not Significant | TRUE |
| Cwf19l2 | TRUE | -0.068214014 | 0.615862857 | No Change | Not Significant | TRUE | -0.088984105 | 0.510774618 | No Change | Not Significant | TRUE |
| Cwh43 | TRUE | -1.955044962 | 6.17E-24 | DOWN | DOWN | TRUE | -0.244089404 | 0.042017506 | No Change | Significant Negative | TRUE |
| Cx3cl1 | TRUE | -0.187521295 | 0.067704352 | No Change | Not Significant | TRUE | 0.122038606 | 0.345828328 | No Change | Not Significant | TRUE |
| Cx3cr1 | TRUE | 1.265325119 | 0.170406218 | No Change | Not Significant | FALSE | NA | NA | No Change | Not Detected | FALSE |
| Cxadr | TRUE | -0.274014119 | 0.002397361 | No Change | Significant Negative | TRUE | -0.09922747 | 0.210759777 | No Change | Not Significant | TRUE |
| Cxcl1 | TRUE | 0.595836128 | 0.400972961 | No Change | Not Significant | TRUE | -0.468917873 | 0.420477604 | No Change | Not Significant | TRUE |
| Cxcl10 | TRUE | 0.2336812 | 0.856549354 | No Change | Not Significant | TRUE | -1.43786284 | 0.012691889 | DOWN | DOWN | TRUE |
| Cxcl12 | TRUE | 0.199276268 | 1 | No Change | Not Significant | TRUE | -1.929794299 | 5.92E-06 | DOWN | DOWN | TRUE |
| Cxcl13 | TRUE | -0.097625161 | 1 | No Change | Not Significant | TRUE | 0.004392253 | 0.996471562 | No Change | Not Significant | TRUE |
| Cxcl14 | TRUE | 1.714569563 | 1 | No Change | Not Significant | FALSE | NA | NA | No Change | Not Detected | FALSE |
| Cxcl15 | TRUE | -0.94435704 | 1 | No Change | Not Significant | FALSE | NA | NA | No Change | Not Detected | FALSE |
| Cxcl16 | TRUE | -0.334500512 | 0.706691156 | No Change | Not Significant | TRUE | -0.808683356 | 0.003453319 | DOWN | DOWN | TRUE |
| Cxcl2 | TRUE | 3.307404296 | 1 | No Change | Not Significant | FALSE | NA | NA | No Change | Not Detected | FALSE |
| Cxcl3 | TRUE | 1.126282296 | 1 | No Change | Not Significant | FALSE | NA | NA | No Change | Not Detected | FALSE |
| Cxcl5 | TRUE | 1.118980437 | 0.125967345 | No Change | Not Significant | FALSE | NA | NA | No Change | Not Detected | FALSE |
| Cxcl9 | TRUE | 0.019806434 | 1 | No Change | Not Significant | FALSE | NA | NA | No Change | Not Detected | FALSE |
| Cxcr3 | TRUE | 0.555482437 | 0.564082441 | No Change | Not Significant | TRUE | 1.030513489 | 0.118807539 | No Change | Not Significant | TRUE |
| Cxcr4 | TRUE | -0.284864223 | 0.666174493 | No Change | Not Significant | TRUE | 0.419532491 | 0.365794955 | No Change | Not Significant | TRUE |
| Cxcr5 | TRUE | 0.420498878 | 0.716335759 | No Change | Not Significant | FALSE | NA | NA | No Change | Not Detected | FALSE |
| Cxcr6 | TRUE | -0.134994136 | 0.867615063 | No Change | Not Significant | TRUE | 0.443081161 | 0.302290895 | No Change | Not Significant | TRUE |
| Cxxc1 | TRUE | -0.016992625 | 0.884816185 | No Change | Not Significant | TRUE | 0.00533706 | 0.970843006 | No Change | Not Significant | TRUE |
| Cxxc4 | TRUE | 1.039667668 | 0.036961738 | UP | UP | TRUE | 0.366198704 | 0.24490649 | No Change | Not Significant | TRUE |
| Cxxc5 | TRUE | 0.133481403 | 0.616875898 | No Change | Not Significant | TRUE | -0.018377094 | 0.930889346 | No Change | Not Significant | TRUE |
| Cyb561 | TRUE | -0.026538283 | 0.910240646 | No Change | Not Significant | TRUE | 0.067669742 | 0.467843395 | No Change | Not Significant | TRUE |
| Cyb561a3 | TRUE | -0.367473755 | 6.49E-06 | No Change | Significant Negative | TRUE | -0.004969864 | 0.975483158 | No Change | Not Significant | TRUE |
| Cyb561d1 | TRUE | -0.486933506 | 4.86E-05 | No Change | Significant Negative | TRUE | -0.039611483 | 0.737950908 | No Change | Not Significant | TRUE |
| Cyb561d2 | TRUE | -0.177587151 | 0.259672367 | No Change | Not Significant | TRUE | 0.125739856 | 0.542414195 | No Change | Not Significant | TRUE |
| Cyb5a | TRUE | 0.108781453 | 0.273824144 | No Change | Not Significant | TRUE | 0.170483321 | 0.000116732 | No Change | Significant Positive | TRUE |
| Cyb5b | TRUE | 1.674686139 | 4.00E-104 | UP | UP | TRUE | 0.929782532 | 6.98E-43 | UP | UP | TRUE |
| Cyb5d1 | TRUE | -0.065959185 | 0.799444652 | No Change | Not Significant | TRUE | 0.100158663 | 0.585270749 | No Change | Not Significant | TRUE |
| Cyb5d2 | TRUE | -0.155109308 | 0.306361493 | No Change | Not Significant | TRUE | 0.111628886 | 0.456057983 | No Change | Not Significant | TRUE |
| Cyb5r1 | TRUE | -0.069445679 | 0.781981914 | No Change | Not Significant | TRUE | -0.093910889 | 0.668392242 | No Change | Not Significant | TRUE |
| Cyb5r2 | TRUE | 1.181372371 | 1 | No Change | Not Significant | FALSE | NA | NA | No Change | Not Detected | FALSE |
| Cyb5r3 | TRUE | 0.35195601 | 7.33E-13 | No Change | Significant Positive | TRUE | 0.181024377 | 0.00801981 | No Change | Significant Positive | TRUE |
| Cyb5r4 | TRUE | -0.191438113 | 0.032709605 | No Change | Significant Negative | TRUE | 0.037656244 | 0.647331425 | No Change | Not Significant | TRUE |
| Cyb5rl | TRUE | -0.100325673 | 0.725856654 | No Change | Not Significant | TRUE | 0.14470006 | 0.499179454 | No Change | Not Significant | TRUE |
| Cyba | TRUE | -2.5291 | 5.90E-64 | DOWN | DOWN | TRUE | -1.506440932 | 4.95E-102 | DOWN | DOWN | TRUE |
| Cybb | TRUE | -0.07484029 | 1 | No Change | Not Significant | FALSE | NA | NA | No Change | Not Detected | FALSE |
| Cybc1 | TRUE | -0.115552216 | 0.311717339 | No Change | Not Significant | TRUE | -0.070171815 | 0.589385665 | No Change | Not Significant | TRUE |
| Cybrd1 | TRUE | 1.486270984 | 1 | No Change | Not Significant | FALSE | NA | NA | No Change | Not Detected | FALSE |
| Cyc1 | TRUE | -0.098339805 | 0.345021291 | No Change | Not Significant | TRUE | 0.090614571 | 0.457022824 | No Change | Not Significant | TRUE |
| Cycs | TRUE | 0.056655075 | 0.779964063 | No Change | Not Significant | TRUE | 0.236135583 | 0.064133519 | No Change | Not Significant | TRUE |
| Cyct | TRUE | -1.197981398 | 1 | No Change | Not Significant | FALSE | NA | NA | No Change | Not Detected | FALSE |
| Cyfip1 | TRUE | -0.055966913 | 0.643957486 | No Change | Not Significant | TRUE | 0.022613271 | 0.767564688 | No Change | Not Significant | TRUE |
| Cyfip2 | TRUE | -0.021073963 | 0.973795719 | No Change | Not Significant | TRUE | 0.889600972 | 0.0020054 | UP | UP | TRUE |
| Cyhr1 | TRUE | -0.064764174 | 0.512903015 | No Change | Not Significant | TRUE | 0.010889099 | 0.910121497 | No Change | Not Significant | TRUE |
| Cylc2 | TRUE | -1.661403185 | 1 | No Change | Not Significant | FALSE | NA | NA | No Change | Not Detected | FALSE |
| Cyld | TRUE | 0.972709757 | 2.66E-43 | UP | UP | TRUE | 0.494145753 | 7.52E-20 | No Change | Significant Positive | TRUE |
| Cyp11a1 | TRUE | -1.608325729 | 1 | No Change | Not Significant | TRUE | -1.39301254 | 0.05258644 | No Change | Not Significant | TRUE |
| Cyp11b1 | TRUE | 3.505722 | 1 | No Change | Not Significant | FALSE | NA | NA | No Change | Not Detected | FALSE |
| Cyp17a1 | TRUE | -0.14384171 | 1 | No Change | Not Significant | FALSE | NA | NA | No Change | Not Detected | FALSE |
| Cyp19a1 | TRUE | -1.855903884 | 1 | No Change | Not Significant | FALSE | NA | NA | No Change | Not Detected | FALSE |
| Cyp1a1 | TRUE | 2.021657461 | 1 | No Change | Not Significant | FALSE | NA | NA | No Change | Not Detected | FALSE |
| Cyp20a1 | TRUE | 0.212746581 | 0.373991816 | No Change | Not Significant | TRUE | 0.109190613 | 0.321053395 | No Change | Not Significant | TRUE |
| Cyp21a1 | TRUE | 0.52340196 | 1 | No Change | Not Significant | FALSE | NA | NA | No Change | Not Detected | FALSE |
| Cyp26a1 | TRUE | 0.498589624 | 1 | No Change | Not Significant | FALSE | NA | NA | No Change | Not Detected | FALSE |
| Cyp26b1 | TRUE | 0.601011698 | 1 | No Change | Not Significant | FALSE | NA | NA | No Change | Not Detected | FALSE |
| Cyp26c1 | TRUE | -0.050403301 | 1 | No Change | Not Significant | FALSE | NA | NA | No Change | Not Detected | FALSE |
| Cyp27a1 | TRUE | -1.67242376 | 7.87E-15 | DOWN | DOWN | TRUE | -0.237258826 | 0.356748847 | No Change | Not Significant | TRUE |
| Cyp27b1 | TRUE | 0.975864231 | 1 | No Change | Not Significant | FALSE | NA | NA | No Change | Not Detected | FALSE |
| Cyp2a12 | TRUE | 0.061320056 | 1 | No Change | Not Significant | FALSE | NA | NA | No Change | Not Detected | FALSE |
| Cyp2a4 | TRUE | 0.004923469 | 1 | No Change | Not Significant | FALSE | NA | NA | No Change | Not Detected | FALSE |
| Cyp2a5 | TRUE | 0.606578239 | 1 | No Change | Not Significant | FALSE | NA | NA | No Change | Not Detected | FALSE |
| Cyp2ab1 | TRUE | -0.37122336 | 1 | No Change | Not Significant | FALSE | NA | NA | No Change | Not Detected | FALSE |
| Cyp2b10 | TRUE | 2.299619235 | 1 | No Change | Not Significant | TRUE | 0.977477124 | 0.415061056 | No Change | Not Significant | TRUE |
| Cyp2b19 | TRUE | -0.31541726 | 1 | No Change | Not Significant | FALSE | NA | NA | No Change | Not Detected | FALSE |
| Cyp2b23 | TRUE | 0.005693067 | 1 | No Change | Not Significant | FALSE | NA | NA | No Change | Not Detected | FALSE |
| Cyp2b9 | TRUE | 0.609860286 | 1 | No Change | Not Significant | FALSE | NA | NA | No Change | Not Detected | FALSE |
| Cyp2c23 | TRUE | -1.073332183 | 1 | No Change | Not Significant | TRUE | -0.737552944 | 0.374452893 | No Change | Not Significant | TRUE |
| Cyp2c40 | TRUE | 0.114111188 | 1 | No Change | Not Significant | FALSE | NA | NA | No Change | Not Detected | FALSE |
| Cyp2c55 | TRUE | -0.263253524 | 0.291600149 | No Change | Not Significant | TRUE | -0.163618234 | 0.386381076 | No Change | Not Significant | TRUE |
| Cyp2c65 | TRUE | -0.704725297 | 0.000933973 | DOWN | DOWN | TRUE | -0.427194064 | 1.01E-05 | No Change | Significant Negative | TRUE |
| Cyp2c66 | TRUE | -1.871767168 | 0.00120867 | DOWN | DOWN | TRUE | -0.257990837 | 0.220797621 | No Change | Not Significant | TRUE |
| Cyp2c67 | TRUE | 1.096178691 | 1 | No Change | Not Significant | FALSE | NA | NA | No Change | Not Detected | FALSE |
| Cyp2c68 | TRUE | -0.933341443 | 0.001056137 | DOWN | DOWN | TRUE | -0.671574065 | 1.69E-06 | DOWN | DOWN | TRUE |
| Cyp2c69 | TRUE | -1.165017536 | 0.000369764 | DOWN | DOWN | TRUE | -0.721027539 | 2.16E-05 | DOWN | DOWN | TRUE |
| Cyp2c70 | TRUE | 0.866854574 | 1 | No Change | Not Significant | FALSE | NA | NA | No Change | Not Detected | FALSE |
| Cyp2d10 | TRUE | -0.874775705 | 8.10E-05 | DOWN | DOWN | TRUE | -0.678832482 | 2.03E-06 | DOWN | DOWN | TRUE |
| Cyp2d11 | TRUE | 1.155641404 | 1 | No Change | Not Significant | FALSE | NA | NA | No Change | Not Detected | FALSE |
| Cyp2d12 | TRUE | -1.690796304 | 3.10E-05 | DOWN | DOWN | TRUE | -1.26910643 | 5.43E-07 | DOWN | DOWN | TRUE |
| Cyp2d13 | TRUE | -0.237349922 | 0.580790426 | No Change | Not Significant | TRUE | -0.814412191 | 9.82E-06 | DOWN | DOWN | TRUE |
| Cyp2d22 | TRUE | -0.562030869 | 0.012597287 | DOWN | DOWN | TRUE | -0.67173772 | 0.001567697 | DOWN | DOWN | TRUE |
| Cyp2d26 | TRUE | -0.71099169 | 2.10E-05 | DOWN | DOWN | TRUE | -0.338454727 | 0.086179968 | No Change | Not Significant | TRUE |
| Cyp2d34 | TRUE | -0.591985018 | 0.02346428 | DOWN | DOWN | TRUE | -0.738374145 | 2.63E-12 | DOWN | DOWN | TRUE |
| Cyp2d9 | TRUE | -0.831074683 | 0.078372637 | No Change | Not Significant | TRUE | -0.956213282 | 6.93E-06 | DOWN | DOWN | TRUE |
| Cyp2e1 | TRUE | -1.612844944 | 1 | No Change | Not Significant | FALSE | NA | NA | No Change | Not Detected | FALSE |
| Cyp2f2 | TRUE | -0.159703978 | 0.750868206 | No Change | Not Significant | TRUE | -0.202535538 | 0.298794445 | No Change | Not Significant | TRUE |
| Cyp2g1 | TRUE | 0.701620711 | 1 | No Change | Not Significant | FALSE | NA | NA | No Change | Not Detected | FALSE |
| Cyp2j11 | TRUE | 0.512838238 | 1 | No Change | Not Significant | FALSE | NA | NA | No Change | Not Detected | FALSE |
| Cyp2j13 | TRUE | 0.330788697 | 0.693930647 | No Change | Not Significant | TRUE | 0.287048199 | 0.712333599 | No Change | Not Significant | TRUE |
| Cyp2j5 | TRUE | 1.628366068 | 1 | No Change | Not Significant | FALSE | NA | NA | No Change | Not Detected | FALSE |
| Cyp2j6 | TRUE | 0.061920104 | 0.710135685 | No Change | Not Significant | TRUE | -0.024650198 | 0.805755951 | No Change | Not Significant | TRUE |
| Cyp2j9 | TRUE | -0.082975546 | 0.855204229 | No Change | Not Significant | TRUE | -0.143659404 | 0.726301064 | No Change | Not Significant | TRUE |
| Cyp2r1 | TRUE | 0.653684921 | 0.007657016 | UP | UP | TRUE | 0.583325194 | 0.001048252 | UP | UP | TRUE |
| Cyp2s1 | TRUE | -0.60903742 | 0.000290111 | DOWN | DOWN | TRUE | -0.232685544 | 0.007233294 | No Change | Significant Negative | TRUE |
| Cyp2t4 | TRUE | -0.368228148 | 1 | No Change | Not Significant | FALSE | NA | NA | No Change | Not Detected | FALSE |
| Cyp2u1 | TRUE | -0.108287819 | 0.882510888 | No Change | Not Significant | TRUE | -0.398865471 | 0.073403124 | No Change | Not Significant | TRUE |
| Cyp2w1 | TRUE | 0.414683309 | 0.329684129 | No Change | Not Significant | TRUE | -0.00617727 | 0.988590012 | No Change | Not Significant | TRUE |
| Cyp39a1 | TRUE | -0.33929377 | 0.055215758 | No Change | Not Significant | TRUE | -0.148259537 | 0.282652548 | No Change | Not Significant | TRUE |
| Cyp3a11 | TRUE | 0.041702782 | 0.982706396 | No Change | Not Significant | TRUE | 1.128729172 | 0.274499682 | No Change | Not Significant | TRUE |
| Cyp3a13 | TRUE | -0.687923692 | 6.88E-05 | DOWN | DOWN | TRUE | -0.305200553 | 0.007436649 | No Change | Significant Negative | TRUE |
| Cyp3a25 | TRUE | 0.333080698 | 0.630787089 | No Change | Not Significant | TRUE | 0.354939152 | 0.769561988 | No Change | Not Significant | TRUE |
| Cyp3a44 | TRUE | -0.903055216 | 0.228901189 | No Change | Not Significant | FALSE | NA | NA | No Change | Not Detected | FALSE |
| Cyp3a57 | TRUE | -0.959803648 | 1 | No Change | Not Significant | FALSE | NA | NA | No Change | Not Detected | FALSE |
| Cyp3a59 | TRUE | -0.58071659 | 1 | No Change | Not Significant | FALSE | NA | NA | No Change | Not Detected | FALSE |
| Cyp46a1 | TRUE | -0.394156911 | 1 | No Change | Not Significant | FALSE | NA | NA | No Change | Not Detected | FALSE |
| Cyp4a10 | TRUE | 0.497988766 | 1 | No Change | Not Significant | FALSE | NA | NA | No Change | Not Detected | FALSE |
| Cyp4a12a | TRUE | 0.541957066 | 1 | No Change | Not Significant | FALSE | NA | NA | No Change | Not Detected | FALSE |
| Cyp4a12b | TRUE | 0.125460556 | 1 | No Change | Not Significant | FALSE | NA | NA | No Change | Not Detected | FALSE |
| Cyp4a30b | TRUE | 0.170815812 | 1 | No Change | Not Significant | FALSE | NA | NA | No Change | Not Detected | FALSE |
| Cyp4a31 | TRUE | 0.048650852 | 1 | No Change | Not Significant | FALSE | NA | NA | No Change | Not Detected | FALSE |
| Cyp4a32 | TRUE | -0.167749584 | 1 | No Change | Not Significant | FALSE | NA | NA | No Change | Not Detected | FALSE |
| Cyp4b1 | TRUE | -3.768033417 | 3.91E-14 | DOWN | DOWN | TRUE | -1.654325263 | 4.72E-07 | DOWN | DOWN | TRUE |
| Cyp4f13 | TRUE | -0.367096703 | 0.0043368 | No Change | Significant Negative | TRUE | -0.200860886 | 0.036262298 | No Change | Significant Negative | TRUE |
| Cyp4f14 | TRUE | 0.160426508 | 0.607010336 | No Change | Not Significant | TRUE | -0.157338514 | 0.15130764 | No Change | Not Significant | TRUE |
| Cyp4f15 | TRUE | -0.224498508 | 1 | No Change | Not Significant | TRUE | -0.11551869 | 0.900132312 | No Change | Not Significant | TRUE |
| Cyp4f16 | TRUE | -0.28158371 | 0.013502881 | No Change | Significant Negative | TRUE | -0.161022708 | 0.194372111 | No Change | Not Significant | TRUE |
| Cyp4f17 | TRUE | -0.154294215 | 0.627713038 | No Change | Not Significant | TRUE | -0.070938262 | 0.828333496 | No Change | Not Significant | TRUE |
| Cyp4f18 | TRUE | 0.164771898 | 1 | No Change | Not Significant | FALSE | NA | NA | No Change | Not Detected | FALSE |
| Cyp4f37 | TRUE | -0.434832899 | 1 | No Change | Not Significant | FALSE | NA | NA | No Change | Not Detected | FALSE |
| Cyp4f39 | TRUE | 0.718004828 | 0.264732086 | No Change | Not Significant | TRUE | -0.550037895 | 0.503963985 | No Change | Not Significant | TRUE |
| Cyp4f40 | TRUE | 0.266322418 | 0.006461237 | No Change | Significant Positive | TRUE | 0.341362285 | 1.32E-05 | No Change | Significant Positive | TRUE |
| Cyp4v3 | TRUE | -0.699599762 | 1.42E-06 | DOWN | DOWN | TRUE | -0.344908533 | 0.080085937 | No Change | Not Significant | TRUE |
| Cyp4x1 | TRUE | -0.314601322 | 0.823849409 | No Change | Not Significant | TRUE | -0.244283631 | 0.737316445 | No Change | Not Significant | TRUE |

| geneID | Cond1_DF_Detected | Cond1_DF_ValueCol | Cond1_DF_PadjCol | Cond1_DF_Cutoff | Cond1_DF_Cutoff_Specific.x | Cond2_DF_Detected | Cond2_DF_ValueCol | Cond2_DF_PadjCol | Cond2_DF_Cutoff | Cond2_DF_Cutoff_Specific | BG_Method | RG1_All | RG2_Significant | RG3_SignificantChange | Cond1_DF_ensembl_gene_id | Cond1_DF_entrezgene_id | Cond1_DF_gene_biotype | Cond1_DF_baseMean |
|---|---|---|---|---|---|---|---|---|---|---|---|---|---|---|---|---|---|---|
| Cyp4x1os | TRUE | 0.206677864 | 1 | No Change | Not Significant | FALSE | NA | NA | No Change | Not Detected | FALSE | Background = FALSE | Background = FALSE | Background = FALSE | ENSMUSG00000086896 | 105244662 | lncRNA | 0.889061718 |
| Cyp51 | TRUE | 0.096155637 | 0.689777883 | No Change | Not Significant | TRUE | -0.045056294 | 0.847296219 | No Change | Not Significant | TRUE | Cond1 Not Significant + Cond2 Not Significant | None | None | ENSMUSG00000001467 | 13121 | protein_coding | 1493.539929 |
| Cyp7b1 | TRUE | 1.171675566 | 1 | No Change | Not Significant | FALSE | NA | NA | No Change | Not Detected | FALSE | Background = FALSE | Background = FALSE | Background = FALSE | ENSMUSG00000039519 | 13123 | protein_coding | 1.741779011 |
| Cyp8b1 | TRUE | 2.855657613 | 0.00425767 | UP | UP | FALSE | NA | NA | No Change | Not Detected | FALSE | Background = FALSE | Background = FALSE | Background = FALSE | ENSMUSG00000050445 | 13124 | protein_coding | 4.453082542 |
| Cypt1 | TRUE | -1.160285344 | 1 | No Change | Not Significant | FALSE | NA | NA | No Change | Not Detected | FALSE | Background = FALSE | Background = FALSE | Background = FALSE | ENSMUSG00000023257 | 66742 | protein_coding | 0.627707281 |
| Cyren | TRUE | 0.095670293 | 0.833066625 | No Change | Not Significant | TRUE | 0.423311773 | 0.228065548 | No Change | Not Significant | TRUE | Cond1 Not Significant + Cond2 Not Significant | None | None | ENSMUSG00000046806 | 78412 | protein_coding | 23.53843467 |
| Cyria | TRUE | -0.228145019 | 0.159457768 | No Change | Not Significant | TRUE | -0.162060904 | 0.493099444 | No Change | Not Significant | TRUE | Cond1 Not Significant + Cond2 Not Significant | None | None | ENSMUSG00000020589 | 76820 | protein_coding | 203.8355814 |
| Cyrib | TRUE | 0.226658909 | 0.010353962 | No Change | Significant Positive | TRUE | -0.025676005 | 0.851790685 | No Change | Not Significant | TRUE | Cond1 Significant Positive + Cond2 Not Significant | None | None | ENSMUSG00000022378 | 223601 | protein_coding | 398.2074156 |
| Cys1 | TRUE | 2.433837704 | 1 | No Change | Not Significant | FALSE | NA | NA | No Change | Not Detected | FALSE | Background = FALSE | Background = FALSE | Background = FALSE | ENSMUSG00000062563 | 12879 | protein_coding | 3.339574449 |
| Cysltr1 | TRUE | 0.442780331 | 1 | No Change | Not Significant | FALSE | NA | NA | No Change | Not Detected | FALSE | Background = FALSE | Background = FALSE | Background = FALSE | ENSMUSG00000052821 | 58861 | protein_coding | 4.094267078 |
| Cysltr2 | TRUE | 0.745826543 | 1 | No Change | Not Significant | FALSE | NA | NA | No Change | Not Detected | FALSE | Background = FALSE | Background = FALSE | Background = FALSE | ENSMUSG00000033470 | 70086 | protein_coding | 2.136066913 |
| Cysrt1 | TRUE | 0.739654525 | 1 | No Change | Not Significant | FALSE | NA | NA | No Change | Not Detected | FALSE | Background = FALSE | Background = FALSE | Background = FALSE | ENSMUSG00000036731 | 67859 | protein_coding | 1.570898927 |
| Cystm1 | TRUE | 0.015085864 | 0.94399288 | No Change | Not Significant | TRUE | 0.232109978 | 0.03683451 | No Change | Significant Positive | TRUE | Cond1 Not Significant + Cond2 Significant Positive | None | None | ENSMUSG00000046727 | 66060 | protein_coding | 3074.971537 |
| Cyth1 | TRUE | -0.048885926 | 0.736113301 | No Change | Not Significant | TRUE | 0.08803147 | 0.358198217 | No Change | Not Significant | TRUE | Cond1 Not Significant + Cond2 Not Significant | None | None | ENSMUSG00000017132 | 19157 | protein_coding | 668.8086033 |
| Cyth2 | TRUE | 0.33249873 | 0.000180437 | No Change | Significant Positive | TRUE | 0.026218965 | 0.790170193 | No Change | Not Significant | TRUE | Cond1 Significant Positive + Cond2 Not Significant | None | None | ENSMUSG00000003269 | 19158 | protein_coding | 703.7946589 |
| Cyth3 | TRUE | -0.272185123 | 0.011989004 | No Change | Significant Negative | TRUE | -0.113582193 | 0.352354239 | No Change | Not Significant | TRUE | Cond1 Significant Negative + Cond2 Not Significant | None | None | ENSMUSG00000018001 | 19159 | protein_coding | 195.2166562 |
| Cyth4 | TRUE | 0.285123282 | 0.768401324 | No Change | Not Significant | TRUE | 0.373222283 | 0.401621914 | No Change | Not Significant | TRUE | Cond1 Not Significant + Cond2 Not Significant | None | None | ENSMUSG00000018008 | 72318 | protein_coding | 15.43700199 |
| Cytip | TRUE | -0.927509643 | 5.45E-05 | DOWN | DOWN | TRUE | -0.006394898 | 0.98066383 | No Change | Not Significant | TRUE | Cond1 DOWN + Cond2 Not Significant | Cond1_DOWN | Cond1_DOWN | ENSMUSG00000026832 | 227929 | protein_coding | 57.45871774 |
| Cytl1 | TRUE | -1.27955412 | 1 | No Change | Not Significant | FALSE | NA | NA | No Change | Not Detected | FALSE | Background = FALSE | Background = FALSE | Background = FALSE | ENSMUSG00000062329 | 231162 | protein_coding | 1.140857146 |
| Cyyr1 | TRUE | 1.34891835 | 1 | No Change | Not Significant | FALSE | NA | NA | No Change | Not Detected | FALSE | Background = FALSE | Background = FALSE | Background = FALSE | ENSMUSG00000041134 | 224405 | protein_coding | 2.216381283 |
| Czib | TRUE | 0.573999054 | 1.50E-15 | UP | UP | TRUE | 0.447534571 | 1.86E-10 | No Change | Significant Positive | TRUE | Cond1 UP + Cond2 Significant Positive | Core_UP | Cond1_UP | ENSMUSG00000028608 | 74098 | protein_coding | 299.0997567 |
| D030025E07Rik | TRUE | -1.601539286 | 1 | No Change | Not Significant | FALSE | NA | NA | No Change | Not Detected | FALSE | Background = FALSE | Background = FALSE | Background = FALSE | ENSMUSG00000105816 | 402774 | lncRNA | 0.811085833 |
| D030025P21Rik | TRUE | -0.563556867 | 1 | No Change | Not Significant | FALSE | NA | NA | No Change | Not Detected | FALSE | Background = FALSE | Background = FALSE | Background = FALSE | ENSMUSG00000092130 | 100303738 | lncRNA | 1.583223529 |
| D030028A08Rik | TRUE | 1.031585129 | 7.78E-14 | UP | UP | TRUE | 0.335883694 | 0.013906591 | No Change | Significant Positive | TRUE | Cond1 UP + Cond2 Significant Positive | Core_UP | Cond1_UP | ENSMUSG00000078700 | 319371 | lncRNA | 150.8927474 |
| D030040B21Rik | TRUE | 0.574051576 | 1 | No Change | Not Significant | FALSE | NA | NA | No Change | Not Detected | FALSE | Background = FALSE | Background = FALSE | Background = FALSE | ENSMUSG00000097744 | 329101 | lncRNA | 0.858716145 |
| D030045P18Rik | TRUE | 0.176019727 | 1 | No Change | Not Significant | FALSE | NA | NA | No Change | Not Detected | FALSE | Background = FALSE | Background = FALSE | Background = FALSE | ENSMUSG00000110874 | 320363 | lncRNA | 1.395082835 |
| D030047H15Rik | TRUE | -0.57510283 | 1 | No Change | Not Significant | FALSE | NA | NA | No Change | Not Detected | FALSE | Background = FALSE | Background = FALSE | Background = FALSE | ENSMUSG00000085886 | 100037396 | lncRNA | 1.401581125 |
| D030056L22Rik | TRUE | 0.314871008 | 0.104544934 | No Change | Not Significant | TRUE | 0.021509111 | 0.914739662 | No Change | Not Significant | TRUE | Cond1 Not Significant + Cond2 Not Significant | None | None | ENSMUSG00000047044 | 225995 | protein_coding | 166.9608603 |
| D030068K23Rik | TRUE | 4.331839848 | 1 | No Change | Not Significant | FALSE | NA | NA | No Change | Not Detected | FALSE | Background = FALSE | Background = FALSE | Background = FALSE | ENSMUSG00000097393 | 628686 | lncRNA | 2.040943382 |
| D11Wsu47e | TRUE | -0.064995143 | 0.774805611 | No Change | Not Significant | TRUE | -0.052904802 | 0.794963793 | No Change | Not Significant | TRUE | Cond1 Not Significant + Cond2 Not Significant | None | None | ENSMUSG00000041623 | 276852 | protein_coding | 116.3223731 |
| D130009I18Rik | TRUE | 0.934309771 | 1 | No Change | Not Significant | FALSE | NA | NA | No Change | Not Detected | FALSE | Background = FALSE | Background = FALSE | Background = FALSE | ENSMUSG00000115432 | 320249 | lncRNA | 2.205453243 |
| D130020L05Rik | TRUE | -0.79781833 | 0.084706605 | No Change | Not Significant | TRUE | -0.025580858 | 0.962265867 | No Change | Not Significant | TRUE | Cond1 Not Significant + Cond2 Not Significant | None | None | ENSMUSG00000097121 | 319760 | lncRNA | 9.191067748 |
| D130040H23Rik | TRUE | -0.507739662 | 0.159769633 | No Change | Not Significant | TRUE | -0.149959722 | 0.768392944 | No Change | Not Significant | TRUE | Cond1 Not Significant + Cond2 Not Significant | None | None | ENSMUSG00000079038 | 211135 | protein_coding | 23.51296037 |
| D130043K22Rik | TRUE | -0.30456319 | 0.527501415 | No Change | Not Significant | TRUE | -0.174139301 | 0.658779673 | No Change | Not Significant | TRUE | Cond1 Not Significant + Cond2 Not Significant | None | None | ENSMUSG00000006711 | 210108 | protein_coding | 21.99730057 |
| D130058E05Rik | TRUE | -0.732796041 | 1 | No Change | Not Significant | FALSE | NA | NA | No Change | Not Detected | FALSE | Background = FALSE | Background = FALSE | Background = FALSE | ENSMUSG00000092627 | 102637806 | lncRNA | 0.984081372 |
| D130079A08Rik | TRUE | 3.217437259 | 1 | No Change | Not Significant | FALSE | NA | NA | No Change | Not Detected | FALSE | Background = FALSE | Background = FALSE | Background = FALSE | ENSMUSG00000115424 | 320671 | lncRNA | 1.897273497 |
| D16Ertd472e | TRUE | 2.825325856 | 4.80E-37 | UP | UP | TRUE | 1.029160445 | 0.001155432 | UP | UP | TRUE | Cond1 UP + Cond2 UP | Core_UP | Core_UP | ENSMUSG00000022864 | 67102 | protein_coding | 46.38016138 |
| D17H6S53E | TRUE | -0.207029614 | 0.00453877 | No Change | Significant Negative | TRUE | -0.042378651 | 0.692926016 | No Change | Not Significant | TRUE | Cond1 Significant Negative + Cond2 Not Significant | None | None | ENSMUSG00000043311 | 114585 | protein_coding | 401.9062859 |
| D2hgdh | TRUE | -0.128692712 | 0.280906233 | No Change | Not Significant | TRUE | -0.072362271 | 0.507292913 | No Change | Not Significant | TRUE | Cond1 Not Significant + Cond2 Not Significant | None | None | ENSMUSG00000073609 | 98314 | protein_coding | 211.2524378 |
| D330025C20Rik | TRUE | 1.138383346 | 1 | No Change | Not Significant | FALSE | NA | NA | No Change | Not Detected | FALSE | Background = FALSE | Background = FALSE | Background = FALSE | ENSMUSG00000102260 | 665054 | TEC | 1.524306224 |
| D330041H03Rik | TRUE | -0.22138433 | 0.704201299 | No Change | Not Significant | TRUE | -0.261700015 | 0.483669199 | No Change | Not Significant | TRUE | Cond1 Not Significant + Cond2 Not Significant | None | None | ENSMUSG00000073437 | 654822 | lncRNA | 19.76405829 |
| D330050G23Rik | TRUE | -0.757668725 | 1 | No Change | Not Significant | FALSE | NA | NA | No Change | Not Detected | FALSE | Background = FALSE | Background = FALSE | Background = FALSE | ENSMUSG00000085316 | 320975 | lncRNA | 0.90845457 |
| D3Ertd751e | TRUE | -0.066081347 | 0.834370416 | No Change | Not Significant | TRUE | 0.037843248 | 0.850189342 | No Change | Not Significant | TRUE | Cond1 Not Significant + Cond2 Not Significant | None | None | ENSMUSG00000025766 | 73852 | protein_coding | 109.685901 |
| D430019H16Rik | TRUE | 0.532209608 | 1 | No Change | Not Significant | FALSE | NA | NA | No Change | Not Detected | FALSE | Background = FALSE | Background = FALSE | Background = FALSE | ENSMUSG00000094910 | 268595 | protein_coding | 1.971240035 |
| D430020J02Rik | TRUE | -0.064162828 | 0.893161222 | No Change | Not Significant | TRUE | 0.092049359 | 0.82545938 | No Change | Not Significant | TRUE | Cond1 Not Significant + Cond2 Not Significant | None | None | ENSMUSG00000112980 | 319545 | lncRNA | 27.64794032 |
| D430041D05Rik | TRUE | 0.887869405 | 0.816921981 | No Change | Not Significant | FALSE | NA | NA | No Change | Not Detected | FALSE | Background = FALSE | Background = FALSE | Background = FALSE | ENSMUSG00000068373 | 241589 | protein_coding | 5.44791237 |
| D5Ertd579e | TRUE | -0.026040617 | 0.905779176 | No Change | Not Significant | TRUE | -0.286889589 | 0.000844425 | No Change | Significant Negative | TRUE | Cond1 Not Significant + Cond2 Significant Negative | None | None | ENSMUSG00000029190 | 320661 | protein_coding | 550.9171374 |
| D5Ertd615e | TRUE | 0.49787755 | 1 | No Change | Not Significant | FALSE | NA | NA | No Change | Not Detected | FALSE | Background = FALSE | Background = FALSE | Background = FALSE | ENSMUSG00000029092 | 52401 | lncRNA | 1.138702293 |
| D630003M21Rik | TRUE | 0.25515146 | 0.835769466 | No Change | Not Significant | TRUE | 0.39394917 | 0.574833929 | No Change | Not Significant | TRUE | Cond1 Not Significant + Cond2 Not Significant | None | None | ENSMUSG00000037813 | 228846 | protein_coding | 6.693502661 |
| D630023F18Rik | TRUE | 1.022015606 | 1 | No Change | Not Significant | FALSE | NA | NA | No Change | Not Detected | FALSE | Background = FALSE | Background = FALSE | Background = FALSE | ENSMUSG00000044816 | 98303 | protein_coding | 1.204243049 |
| D630024D03Rik | TRUE | -0.138505122 | 0.858172551 | No Change | Not Significant | TRUE | -0.026985581 | 0.972704281 | No Change | Not Significant | TRUE | Cond1 Not Significant + Cond2 Not Significant | None | None | ENSMUSG00000085772 | 414116 | lncRNA | 9.88362935 |
| D630033O11Rik | TRUE | -0.820620043 | 0.10409925 | No Change | Not Significant | TRUE | -0.735668674 | 0.000101583 | DOWN | DOWN | TRUE | Cond1 Not Significant + Cond2 DOWN | Cond2_DOWN | Cond2_DOWN | ENSMUSG00000095385 | 235302 | lncRNA | 20.20825997 |
| D630039A03Rik | TRUE | -0.311266161 | 0.294503539 | No Change | Not Significant | TRUE | -0.063540445 | 0.75798157 | No Change | Not Significant | TRUE | Cond1 Not Significant + Cond2 Not Significant | None | None | ENSMUSG00000052117 | 242484 | protein_coding | 493.4986947 |
| D630044L22Rik | TRUE | 0.084439837 | 1 | No Change | Not Significant | FALSE | NA | NA | No Change | Not Detected | FALSE | Background = FALSE | Background = FALSE | Background = FALSE | ENSMUSG00000097405 | 240054 | lncRNA | 1.607990219 |
| D630045J12Rik | TRUE | 0.487558809 | 0.409464624 | No Change | Not Significant | TRUE | -0.056364592 | 0.918422987 | No Change | Not Significant | TRUE | Cond1 Not Significant + Cond2 Not Significant | None | None | ENSMUSG00000063455 | 330286 | protein_coding | 20.76930276 |
| D6Ertd527e | TRUE | -0.577182737 | 0.647057325 | No Change | Not Significant | TRUE | -0.414904948 | 0.624972092 | No Change | Not Significant | TRUE | Cond1 Not Significant + Cond2 Not Significant | None | None | ENSMUSG00000090891 | 52372 | lncRNA | 4.802405877 |
| D6Wsu163e | TRUE | 0.04273884 | 0.855204229 | No Change | Not Significant | TRUE | 0.195406553 | 0.086330552 | No Change | Not Significant | TRUE | Cond1 Not Significant + Cond2 Not Significant | None | None | ENSMUSG00000030347 | 28040 | protein_coding | 164.7560529 |
| D730005E14Rik | TRUE | 0.029852925 | 0.966840628 | No Change | Not Significant | TRUE | -0.093345033 | 0.792648462 | No Change | Not Significant | TRUE | Cond1 Not Significant + Cond2 Not Significant | None | None | ENSMUSG00000115902 | 109361 | lncRNA | 28.75777974 |
| D7Ertd443e | TRUE | 2.70830429 | 1 | No Change | Not Significant | FALSE | NA | NA | No Change | Not Detected | FALSE | Background = FALSE | Background = FALSE | Background = FALSE | ENSMUSG00000030994 | 71007 | protein_coding | 3.147675411 |
| D830026I12Rik | TRUE | 0.609491162 | 1 | No Change | Not Significant | FALSE | NA | NA | No Change | Not Detected | FALSE | Background = FALSE | Background = FALSE | Background = FALSE | ENSMUSG00000085171 | 319682 | lncRNA | 1.208491621 |
| D8Ertd738e | TRUE | -0.050349732 | 0.522023104 | No Change | Not Significant | TRUE | 0.093357719 | 0.217970138 | No Change | Not Significant | TRUE | Cond1 Not Significant + Cond2 Not Significant | None | None | ENSMUSG00000019362 | 101966 | protein_coding | 741.5329195 |
| D930007P13Rik | TRUE | 2.764143018 | 1 | No Change | Not Significant | FALSE | NA | NA | No Change | Not Detected | FALSE | Background = FALSE | Background = FALSE | Background = FALSE | ENSMUSG00000097003 | 432989 | lncRNA | 1.622385447 |
| D930015M05Rik | TRUE | -1.105131125 | 0.071994506 | No Change | Not Significant | TRUE | -1.025889229 | 0.050937378 | No Change | Not Significant | TRUE | Cond1 Not Significant + Cond2 Not Significant | None | None | ENSMUSG00000084781 | 320507 | lncRNA | 8.377485496 |
| D930020B18Rik | TRUE | 0.296861095 | 0.629409554 | No Change | Not Significant | TRUE | 0.335619008 | 0.426540063 | No Change | Not Significant | TRUE | Cond1 Not Significant + Cond2 Not Significant | None | None | ENSMUSG00000047642 | 216393 | protein_coding | 21.42500458 |
| D930028M14Rik | TRUE | -0.674264019 | 1 | No Change | Not Significant | FALSE | NA | NA | No Change | Not Detected | FALSE | Background = FALSE | Background = FALSE | Background = FALSE | ENSMUSG00000074274 | 434147 | lncRNA | 1.725418167 |
| D930032P07Rik | TRUE | -0.99010684 | 1 | No Change | Not Significant | FALSE | NA | NA | No Change | Not Detected | FALSE | Background = FALSE | Background = FALSE | Background = FALSE | ENSMUSG00000097735 | 320176 | lncRNA | 0.751066119 |
| D930048N14Rik | TRUE | -1.198068452 | 0.001144911 | DOWN | DOWN | TRUE | -0.594880266 | 0.030313495 | DOWN | DOWN | TRUE | Cond1 DOWN + Cond2 DOWN | Core_DOWN | Core_DOWN | ENSMUSG00000052563 | 97775 | lncRNA | 19.75854223 |
| Daam1 | TRUE | -0.128737188 | 0.160822447 | No Change | Not Significant | TRUE | -0.156127547 | 0.011511368 | No Change | Significant Negative | TRUE | Cond1 Not Significant + Cond2 Significant Negative | None | None | ENSMUSG00000034574 | 208846 | protein_coding | 900.2476254 |
| Daam2 | TRUE | 0.236349465 | 0.802408631 | No Change | Not Significant | TRUE | -0.060862968 | 0.945241431 | No Change | Not Significant | TRUE | Cond1 Not Significant + Cond2 Not Significant | None | None | ENSMUSG00000040260 | 76441 | protein_coding | 10.98173043 |
| Dab1 | TRUE | -0.652376679 | 0.364292142 | No Change | Not Significant | TRUE | 0.474183666 | 0.314662099 | No Change | Not Significant | TRUE | Cond1 Not Significant + Cond2 Not Significant | None | None | ENSMUSG00000028519 | 13131 | protein_coding | 22.55843907 |
| Dab2 | TRUE | 0.015389577 | 0.99157486 | No Change | Not Significant | TRUE | -0.364327722 | 0.567390999 | No Change | Not Significant | TRUE | Cond1 Not Significant + Cond2 Not Significant | None | None | ENSMUSG00000022150 | 13132 | protein_coding | 7.416907053 |
| Dab2ip | TRUE | 0.005907888 | 0.957266079 | No Change | Not Significant | TRUE | -0.015406487 | 0.902954861 | No Change | Not Significant | TRUE | Cond1 Not Significant + Cond2 Not Significant | None | None | ENSMUSG00000026883 | 69601 | protein_coding | 1981.554412 |
| Dach1 | TRUE | -0.142707267 | 0.725185288 | No Change | Not Significant | TRUE | -0.184971009 | 0.437065467 | No Change | Not Significant | TRUE | Cond1 Not Significant + Cond2 Not Significant | None | None | ENSMUSG00000055639 | 13134 | protein_coding | 31.74633781 |
| Dach2 | TRUE | 0.061495585 | 0.94246215 | No Change | Not Significant | TRUE | -0.547425592 | 0.207110275 | No Change | Not Significant | TRUE | Cond1 Not Significant + Cond2 Not Significant | None | None | ENSMUSG00000025592 | 93837 | protein_coding | 7.424815713 |
| Dact1 | TRUE | -0.295941968 | 1 | No Change | Not Significant | FALSE | NA | NA | No Change | Not Detected | FALSE | Background = FALSE | Background = FALSE | Background = FALSE | ENSMUSG00000044548 | 59036 | protein_coding | 2.2809594 |
| Dact2 | TRUE | -0.165416307 | 0.450945654 | No Change | Not Significant | TRUE | 0.067006694 | 0.556656741 | No Change | Not Significant | TRUE | Cond1 Not Significant + Cond2 Not Significant | None | None | ENSMUSG00000048826 | 240025 | protein_coding | 320.0927784 |
| Dact3 | TRUE | 0.679847497 | 0.474068167 | No Change | Not Significant | FALSE | NA | NA | No Change | Not Detected | FALSE | Background = FALSE | Background = FALSE | Background = FALSE | ENSMUSG00000078794 | 629378 | protein_coding | 5.837009658 |
| Dad1 | TRUE | 0.117921429 | 0.559613498 | No Change | Not Significant | TRUE | 0.222948513 | 0.019489068 | No Change | Significant Positive | TRUE | Cond1 Not Significant + Cond2 Significant Positive | None | None | ENSMUSG00000022174 | 13135 | protein_coding | 828.5965287 |
| Dag1 | TRUE | -0.027715113 | 0.80314212 | No Change | Not Significant | TRUE | -0.089059446 | 0.336485813 | No Change | Not Significant | TRUE | Cond1 Not Significant + Cond2 Not Significant | None | None | ENSMUSG00000039952 | 13138 | protein_coding | 2232.536582 |
| Dagla | TRUE | -0.153711871 | 0.439074217 | No Change | Not Significant | TRUE | -0.03366864 | 0.865987203 | No Change | Not Significant | TRUE | Cond1 Not Significant + Cond2 Not Significant | None | None | ENSMUSG00000035735 | 269060 | protein_coding | 175.7919717 |
| Daglb | TRUE | 0.182872779 | 0.360481579 | No Change | Not Significant | TRUE | 0.09304166 | 0.485821117 | No Change | Not Significant | TRUE | Cond1 Not Significant + Cond2 Not Significant | None | None | ENSMUSG00000039206 | 231871 | protein_coding | 603.2413283 |
| Dalrd3 | TRUE | 1.249801693 | 4.17E-38 | UP | UP | TRUE | 0.685634837 | 1.74E-36 | UP | UP | TRUE | Cond1 UP + Cond2 UP | Core_UP | Core_UP | ENSMUSG00000019039 | 67789 | protein_coding | 638.3880739 |
| Dancr | TRUE | -0.131470466 | 0.83386713 | No Change | Not Significant | TRUE | -0.539218631 | 0.033016757 | DOWN | DOWN | TRUE | Cond1 Not Significant + Cond2 DOWN | Cond2_DOWN | Cond2_DOWN | ENSMUSG00000106943 | 70036 | lncRNA | 32.6243551 |
| Dand5 | TRUE | -0.072692751 | 0.879809661 | No Change | Not Significant | TRUE | 0.008880744 | 0.981205596 | No Change | Not Significant | TRUE | Cond1 Not Significant + Cond2 Not Significant | None | None | ENSMUSG00000053226 | 23863 | protein_coding | 27.28363634 |
| Dao | TRUE | 0.720995547 | 1 | No Change | Not Significant | FALSE | NA | NA | No Change | Not Detected | FALSE | Background = FALSE | Background = FALSE | Background = FALSE | ENSMUSG00000042096 | 13142 | protein_coding | 2.709141352 |
| Dap | TRUE | 0.412401096 | 3.15E-06 | No Change | Significant Positive | TRUE | 0.334994437 | 5.74E-05 | No Change | Significant Positive | TRUE | Cond1 Significant Positive + Cond2 Significant Positive | None | None | ENSMUSG00000039168 | 223453 | protein_coding | 490.0122428 |
| Dap3 | TRUE | -0.147483353 | 0.120977402 | No Change | Not Significant | TRUE | 0.006752024 | 0.954400543 | No Change | Not Significant | TRUE | Cond1 Not Significant + Cond2 Not Significant | None | None | ENSMUSG00000068921 | 65111 | protein_coding | 683.7040482 |
| Dapk1 | TRUE | -0.11999038 | 0.346538122 | No Change | Not Significant | TRUE | 0.019316191 | 0.91676947 | No Change | Not Significant | TRUE | Cond1 Not Significant + Cond2 Not Significant | None | None | ENSMUSG00000021559 | 69635 | protein_coding | 203.2765934 |
| Dapk2 | TRUE | -0.184321716 | 0.040902139 | No Change | Significant Negative | TRUE | -0.013246587 | 0.912178372 | No Change | Not Significant | TRUE | Cond1 Significant Negative + Cond2 Not Significant | None | None | ENSMUSG00000032380 | 13143 | protein_coding | 431.5321326 |
| Dapk3 | TRUE | 0.872676292 | 5.50E-23 | UP | UP | TRUE | 0.354968594 | 4.41E-09 | No Change | Significant Positive | TRUE | Cond1 UP + Cond2 Significant Positive | Core_UP | Cond1_UP | ENSMUSG00000034974 | 13144 | protein_coding | 517.7636994 |
| Dapl1 | TRUE | -0.524279594 | 1 | No Change | Not Significant | FALSE | NA | NA | No Change | Not Detected | FALSE | Background = FALSE | Background = FALSE | Background = FALSE | ENSMUSG00000026989 | 76747 | protein_coding | 0.858948479 |
| Dapp1 | TRUE | 0.347509934 | 0.185177119 | No Change | Not Significant | TRUE | -0.427764667 | 0.032713889 | No Change | Significant Negative | TRUE | Cond1 Not Significant + Cond2 Significant Negative | None | None | ENSMUSG00000028159 | 26377 | protein_coding | 116.7102726 |
| Dars | TRUE | -0.092606183 | 0.561689026 | No Change | Not Significant | TRUE | 0.003515696 | 0.981022852 | No Change | Not Significant | TRUE | Cond1 Not Significant + Cond2 Not Significant | None | None | ENSMUSG00000026356 | 226414 | protein_coding | 715.4848051 |
| Dars2 | TRUE | -0.09991662 | 0.640714813 | No Change | Not Significant | TRUE | 0.050776514 | 0.741375537 | No Change | Not Significant | TRUE | Cond1 Not Significant + Cond2 Not Significant | None | None | ENSMUSG00000026709 | 226539 | protein_coding | 310.5499348 |
| Daxx | TRUE | -0.113130192 | 0.286211349 | No Change | Not Significant | TRUE | -0.11608289 | 0.244581981 | No Change | Not Significant | TRUE | Cond1 Not Significant + Cond2 Not Significant | None | None | ENSMUSG00000002307 | 13163 | protein_coding | 317.9125459 |
| Dazap1 | TRUE | 0.194790937 | 0.070900795 | No Change | Not Significant | TRUE | 0.038054967 | 0.842728192 | No Change | Not Significant | TRUE | Cond1 Not Significant + Cond2 Not Significant | None | None | ENSMUSG00000069565 | 70248 | protein_coding | 901.1296809 |
| Dazap2 | TRUE | -0.077178556 | 0.356391801 | No Change | Not Significant | TRUE | 0.029746781 | 0.736749554 | No Change | Not Significant | TRUE | Cond1 Not Significant + Cond2 Not Significant | None | None | ENSMUSG00000000346 | 23994 | protein_coding | 2093.364858 |
| Dazl | TRUE | 1.266857763 | 1 | No Change | Not Significant | TRUE | -0.375814693 | 0.655860109 | No Change | Not Significant | TRUE | Cond1 Not Significant + Cond2 Not Significant | None | None | ENSMUSG00000010592 | 13164 | protein_coding | 3.708868644 |
| Dbf4 | TRUE | 0.551844212 | 0.000196012 | UP | UP | TRUE | 0.051333766 | 0.749012844 | No Change | Not Significant | TRUE | Cond1 UP + Cond2 Not Significant | Cond1_UP | Cond1_UP | ENSMUSG00000002297 | 27214 | protein_coding | 269.8232827 |
| Dbhos | TRUE | -1.688416624 | 1 | No Change | Not Significant | FALSE | NA | NA | No Change | Not Detected | FALSE | Background = FALSE | Background = FALSE | Background = FALSE | ENSMUSG00000085008 | 329371 | lncRNA | 1.442856055 |
| Dbi | TRUE | 0.226212963 | 0.0129987 | No Change | Significant Positive | TRUE | 0.160535521 | 0.004178688 | No Change | Significant Positive | TRUE | Cond1 Significant Positive + Cond2 Significant Positive | None | None | ENSMUSG00000026385 | 13167 | protein_coding | 2844.518478 |
| Dbil5 | TRUE | -0.197261975 | 1 | No Change | Not Significant | TRUE | 0.132208102 | 0.896972875 | No Change | Not Significant | TRUE | Cond1 Not Significant + Cond2 Not Significant | None | None | ENSMUSG00000038057 | 13168 | protein_coding | 2.730616878 |
| Dbn1 | TRUE | -0.114365012 | 0.904874646 | No Change | Not Significant | TRUE | 0.450270378 | 0.1947258 | No Change | Not Significant | TRUE | Cond1 Not Significant + Cond2 Not Significant | None | None | ENSMUSG00000034675 | 56320 | protein_coding | 12.64149407 |
| Dbndd1 | TRUE | 0.149487867 | 0.827422396 | No Change | Not Significant | TRUE | -0.298683897 | 0.487025714 | No Change | Not Significant | TRUE | Cond1 Not Significant + Cond2 Not Significant | None | None | ENSMUSG00000031970 | 72185 | protein_coding | 12.80893886 |
| Dbndd2 | TRUE | -0.095993311 | 0.801948109 | No Change | Not Significant | TRUE | -0.209377925 | 0.531418201 | No Change | Not Significant | TRUE | Cond1 Not Significant + Cond2 Not Significant | None | None | ENSMUSG00000017734 | 52840 | protein_coding | 36.28633515 |
| Dbnl | TRUE | -0.007079984 | 0.951341404 | No Change | Not Significant | TRUE | 0.106863436 | 0.247106483 | No Change | Not Significant | TRUE | Cond1 Not Significant + Cond2 Not Significant | None | None | ENSMUSG00000020476 | 13169 | protein_coding | 1323.534683 |
| Dbp | TRUE | -2.424805752 | 0.002774564 | DOWN | DOWN | TRUE | -1.961187426 | 2.10E-20 | DOWN | DOWN | TRUE | Cond1 DOWN + Cond2 DOWN | Core_DOWN | Core_DOWN | ENSMUSG00000059824 | 13170 | protein_coding | 103.2492282 |
| Dbpht2 | TRUE | -0.345383519 | 0.747120423 | No Change | Not Significant | TRUE | 0.325566824 | 0.609972551 | No Change | Not Significant | TRUE | Cond1 Not Significant + Cond2 Not Significant | None | None | ENSMUSG00000029878 | 386753 | lncRNA | 5.929207797 |
| Dbr1 | TRUE | 0.017767619 | 0.938415991 | No Change | Not Significant | TRUE | 0.050739526 | 0.684487275 | No Change | Not Significant | TRUE | Cond1 Not Significant + Cond2 Not Significant | None | None | ENSMUSG00000032469 | 83703 | protein_coding | 162.8266324 |
| Dbt | TRUE | -0.441389494 | 0.0011102 | No Change | Significant Negative | TRUE | -0.213767226 | 0.001620247 | No Change | Significant Negative | TRUE | Cond1 Significant Negative + Cond2 Significant Negative | None | None | ENSMUSG00000000340 | 13171 | protein_coding | 396.8491352 |
| Dbx1 | TRUE | -0.83683862 | 1 | No Change | Not Significant | FALSE | NA | NA | No Change | Not Detected | FALSE | Background = FALSE | Background = FALSE | Background = FALSE | ENSMUSG00000030507 | 13172 | protein_coding | 1.111564919 |
| Dbx2 | TRUE | 0.532854418 | 1 | No Change | Not Significant | FALSE | NA | NA | No Change | Not Detected | FALSE | Background = FALSE | Background = FALSE | Background = FALSE | ENSMUSG00000045608 | 223843 | protein_coding | 1.41702253 |
| Dcaf1 | TRUE | 0.121218801 | 0.225148876 | No Change | Not Significant | TRUE | 0.023463013 | 0.776292513 | No Change | Not Significant | TRUE | Cond1 Not Significant + Cond2 Not Significant | None | None | ENSMUSG00000040325 | 321006 | protein_coding | 700.3522082 |
| Dcaf10 | TRUE | 0.280249385 | 0.015447555 | No Change | Significant Positive | TRUE | 0.074115905 | 0.391491529 | No Change | Not Significant | TRUE | Cond1 Significant Positive + Cond2 Not Significant | None | None | ENSMUSG00000035572 | 242418 | protein_coding | 398.5828702 |
| Dcaf11 | TRUE | 0.046843559 | 0.628624818 | No Change | Not Significant | TRUE | -0.0170441 | 0.803190139 | No Change | Not Significant | TRUE | Cond1 Not Significant + Cond2 Not Significant | None | None | ENSMUSG00000022214 | 28199 | protein_coding | 1230.547606 |
| Dcaf12 | TRUE | -0.053059492 | 0.607042377 | No Change | Not Significant | TRUE | 0.036052418 | 0.750740503 | No Change | Not Significant | TRUE | Cond1 Not Significant + Cond2 Not Significant | None | None | ENSMUSG00000028436 | 68970 | protein_coding | 680.365107 |
| Dcaf12l1 | TRUE | -0.151706967 | 0.794593769 | No Change | Not Significant | TRUE | -0.067199252 | 0.850145091 | No Change | Not Significant | TRUE | Cond1 Not Significant + Cond2 Not Significant | None | None | ENSMUSG00000045284 | 245404 | protein_coding | 11.72648822 |
| Dcaf12l2 | TRUE | -0.431072745 | 1 | No Change | Not Significant | FALSE | NA | NA | No Change | Not Detected | FALSE | Background = FALSE | Background = FALSE | Background = FALSE | ENSMUSG00000050926 | 245403 | protein_coding | 1.173979019 |
| Dcaf13 | TRUE | 0.018367011 | 0.905779176 | No Change | Not Significant | TRUE | 0.044277559 | 0.708115319 | No Change | Not Significant | TRUE | Cond1 Not Significant + Cond2 Not Significant | None | None | ENSMUSG00000022300 | 223499 | protein_coding | 485.3927743 |
| Dcaf15 | TRUE | -0.057081304 | 0.669122175 | No Change | Not Significant | TRUE | 0.01610444 | 0.896941755 | No Change | Not Significant | TRUE | Cond1 Not Significant + Cond2 Not Significant | None | None | ENSMUSG00000037103 | 212123 | protein_coding | 506.2195272 |
| Dcaf17 | TRUE | -0.357287771 | 0.002533766 | No Change | Significant Negative | TRUE | -0.00903054 | 0.950733597 | No Change | Not Significant | TRUE | Cond1 Significant Negative + Cond2 Not Significant | None | None | ENSMUSG00000041966 | 75763 | protein_coding | 223.9542386 |
| Dcaf4 | TRUE | -0.114392342 | 0.570683233 | No Change | Not Significant | TRUE | -0.071352726 | 0.631515959 | No Change | Not Significant | TRUE | Cond1 Not Significant + Cond2 Not Significant | None | None | ENSMUSG00000021222 | 73828 | protein_coding | 150.676184 |
| Dcaf5 | TRUE | 0.289167671 | 1.39E-05 | No Change | Significant Positive | TRUE | 0.114680234 | 0.068138529 | No Change | Not Significant | TRUE | Cond1 Significant Positive + Cond2 Not Significant | None | None | ENSMUSG00000049106 | 320808 | protein_coding | 606.9068177 |
| Dcaf6 | TRUE | -0.241769659 | 0.012812723 | No Change | Significant Negative | TRUE | -0.359135424 | 5.45E-06 | No Change | Significant Negative | TRUE | Cond1 Significant Negative + Cond2 Significant Negative | None | None | ENSMUSG00000026571 | 74106 | protein_coding | 331.0316023 |
| Dcaf7 | TRUE | 0.112015514 | 0.477488842 | No Change | Not Significant | TRUE | -0.038515857 | 0.68117317 | No Change | Not Significant | TRUE | Cond1 Not Significant + Cond2 Not Significant | None | None | ENSMUSG00000049354 | 71833 | protein_coding | 1278.278334 |
| Dcaf8 | TRUE | -0.1425366 | 0.076884085 | No Change | Not Significant | TRUE | -0.021944884 | 0.74340107 | No Change | Not Significant | TRUE | Cond1 Not Significant + Cond2 Not Significant | None | None | ENSMUSG00000026554 | 98193 | protein_coding | 1267.788483 |
| Dcaf8l | TRUE | 0.649177342 | 1 | No Change | Not Significant | FALSE | NA | NA | No Change | Not Detected | FALSE | Background = FALSE | Background = FALSE | Background = FALSE | ENSMUSG00000035395 | 18630 | protein_coding | 1.290204042 |
| Dcakd | TRUE | 0.00457489 | 0.984756276 | No Change | Not Significant | TRUE | 0.098983014 | 0.549639002 | No Change | Not Significant | TRUE | Cond1 Not Significant + Cond2 Not Significant | None | None | ENSMUSG00000020935 | 68087 | protein_coding | 147.4528786 |
| Dcbld1 | TRUE | 0.350342809 | 0.000234331 | No Change | Significant Positive | TRUE | 0.155972137 | 0.081578678 | No Change | Not Significant | TRUE | Cond1 Significant Positive + Cond2 Not Significant | None | None | ENSMUSG00000019891 | 66686 | protein_coding | 415.0273793 |
| Dcbld2 | TRUE | 2.375864873 | 4.64E-115 | UP | UP | TRUE | 1.084738809 | 4.07E-12 | UP | UP | TRUE | Cond1 UP + Cond2 UP | Core_UP | Core_UP | ENSMUSG00000035107 | 73379 | protein_coding | 275.2317904 |
| Dcc | TRUE | 2.862261079 | 1 | No Change | Not Significant | FALSE | NA | NA | No Change | Not Detected | FALSE | Background = FALSE | Background = FALSE | Background = FALSE | ENSMUSG00000060534 | 13176 | protein_coding | 3.834657279 |
| Dcdc2a | TRUE | 0.917097499 | 0.042864306 | UP | UP | TRUE | 0.149391749 | 0.856166251 | No Change | Not Significant | TRUE | Cond1 UP + Cond2 Not Significant | Cond1_UP | Cond1_UP | ENSMUSG00000035910 | 195208 | protein_coding | 15.34025911 |
| Dcdc2b | TRUE | -0.266568397 | 0.711906345 | No Change | Not Significant | TRUE | -0.501808103 | 0.299347968 | No Change | Not Significant | TRUE | Cond1 Not Significant + Cond2 Not Significant | None | None | ENSMUSG00000078552 | 100504491 | protein_coding | 10.43441428 |
| Dchs1 | TRUE | 0.986682179 | 1 | No Change | Not Significant | FALSE | NA | NA | No Change | Not Detected | FALSE | Background = FALSE | Background = FALSE | Background = FALSE | ENSMUSG00000036862 | 233651 | protein_coding | 4.169660462 |
| Dchs2 | TRUE | 1.263166277 | 1 | No Change | Not Significant | FALSE | NA | NA | No Change | Not Detected | FALSE | Background = FALSE | Background = FALSE | Background = FALSE | ENSMUSG00000102692 | 100534287 | protein_coding | 3.704670798 |
| Dck | TRUE | 0.066582173 | 0.811137708 | No Change | Not Significant | TRUE | 0.079635969 | 0.667785655 | No Change | Not Significant | TRUE | Cond1 Not Significant + Cond2 Not Significant | None | None | ENSMUSG00000029366 | 13178 | protein_coding | 135.9592643 |
| Dclk1 | TRUE | -0.730229081 | 2.73E-05 | DOWN | DOWN | TRUE | 0.082326204 | 0.558485807 | No Change | Not Significant | TRUE | Cond1 DOWN + Cond2 Not Significant | Cond1_DOWN | Cond1_DOWN | ENSMUSG00000027797 | 13175 | protein_coding | 440.3316582 |
| Dclk2 | TRUE | 0.237078445 | 0.545970799 | No Change | Not Significant | TRUE | 0.443231287 | 0.05858417 | No Change | Not Significant | TRUE | Cond1 Not Significant + Cond2 Not Significant | None | None | ENSMUSG00000028078 | 70762 | protein_coding | 23.04028338 |
| Dclk3 | TRUE | -0.955859058 | 0.000157233 | DOWN | DOWN | TRUE | -0.157791909 | 0.601184005 | No Change | Not Significant | TRUE | Cond1 DOWN + Cond2 Not Significant | Cond1_DOWN | Cond1_DOWN | ENSMUSG00000032500 | 245038 | protein_coding | 40.14956701 |
| Dclre1a | TRUE | 0.049756818 | 0.736235501 | No Change | Not Significant | TRUE | -0.083454952 | 0.454403192 | No Change | Not Significant | TRUE | Cond1 Not Significant + Cond2 Not Significant | None | None | ENSMUSG00000025077 | 55947 | protein_coding | 210.0849668 |
| Dclre1b | TRUE | 0.070291785 | 0.69440476 | No Change | Not Significant | TRUE | 0.141971485 | 0.31570327 | No Change | Not Significant | TRUE | Cond1 Not Significant + Cond2 Not Significant | None | None | ENSMUSG00000027845 | 140917 | protein_coding | 119.6461029 |
| Dclre1c | TRUE | -0.042004155 | 0.860116914 | No Change | Not Significant | TRUE | -0.044346909 | 0.820125043 | No Change | Not Significant | TRUE | Cond1 Not Significant + Cond2 Not Significant | None | None | ENSMUSG00000026648 | 227525 | protein_coding | 95.8733748 |
| Dcn | TRUE | -0.180725958 | 0.901449255 | No Change | Not Significant | TRUE | 1.405947163 | 0.056167696 | No Change | Not Significant | TRUE | Cond1 Not Significant + Cond2 Not Significant | None | None | ENSMUSG00000019929 | 13179 | protein_coding | 4.849207902 |
| Dcp1a | TRUE | 0.022349482 | 0.870923726 | No Change | Not Significant | TRUE | -0.115718053 | 0.235215876 | No Change | Not Significant | TRUE | Cond1 Not Significant + Cond2 Not Significant | None | None | ENSMUSG00000021962 | 75901 | protein_coding | 330.7767239 |
| Dcp1b | TRUE | -0.41463763 | 0.047449241 | No Change | Significant Negative | TRUE | -0.166761202 | 0.133956365 | No Change | Not Significant | TRUE | Cond1 Significant Negative + Cond2 Not Significant | None | None | ENSMUSG00000041477 | 319618 | protein_coding | 148.6414247 |
| Dcp2 | TRUE | 0.011271539 | 0.957921881 | No Change | Not Significant | TRUE | -0.176273734 | 0.015184375 | No Change | Significant Negative | TRUE | Cond1 Not Significant + Cond2 Significant Negative | None | None | ENSMUSG00000024472 | 70640 | protein_coding | 752.9985476 |
| Dcps | TRUE | 0.599714219 | 1.95E-08 | UP | UP | TRUE | 0.541823726 | 1.73E-10 | UP | UP | TRUE | Cond1 UP + Cond2 UP | Core_UP | Core_UP | ENSMUSG00000032040 | 69305 | protein_coding | 257.9162321 |
| Dcst1 | TRUE | -0.050674431 | 0.938083652 | No Change | Not Significant | TRUE | 0.073044596 | 0.869327992 | No Change | Not Significant | TRUE | Cond1 Not Significant + Cond2 Not Significant | None | None | ENSMUSG00000042672 | 77772 | protein_coding | 24.96985472 |
| Dcst2 | TRUE | 0.367150158 | 0.25018373 | No Change | Not Significant | TRUE | -0.044956355 | 0.93820671 | No Change | Not Significant | TRUE | Cond1 Not Significant + Cond2 Not Significant | None | None | ENSMUSG00000109293 | 329702 | protein_coding | 25.33108856 |
| Dct | TRUE | 0.018116298 | 0.983632579 | No Change | Not Significant | TRUE | -0.13111564 | 0.800805946 | No Change | Not Significant | TRUE | Cond1 Not Significant + Cond2 Not Significant | None | None | ENSMUSG00000022129 | 13190 | protein_coding | 10.27179372 |
| Dctd | TRUE | 0.25050746 | 0.57042954 | No Change | Not Significant | TRUE | 0.238595487 | 0.35463345 | No Change | Not Significant | TRUE | Cond1 Not Significant + Cond2 Not Significant | None | None | ENSMUSG00000031562 | 320685 | protein_coding | 60.59424156 |
| Dctn1 | TRUE | 0.077893001 | 0.284857762 | No Change | Not Significant | TRUE | 0.123746938 | 0.199401327 | No Change | Not Significant | TRUE | Cond1 Not Significant + Cond2 Not Significant | None | None | ENSMUSG00000031865 | 13191 | protein_coding | 1687.350764 |
| Dctn2 | TRUE | -0.065901413 | 0.314808425 | No Change | Not Significant | TRUE | 0.057978942 | 0.45231664 | No Change | Not Significant | TRUE | Cond1 Not Significant + Cond2 Not Significant | None | None | ENSMUSG00000025410 | 69654 | protein_coding | 1478.18125 |
| Dctn3 | TRUE | -0.048694579 | 0.742355007 | No Change | Not Significant | TRUE | 0.15663653 | 0.082768143 | No Change | Not Significant | TRUE | Cond1 Not Significant + Cond2 Not Significant | None | None | ENSMUSG00000028447 | 53598 | protein_coding | 540.7918818 |
| Dctn4 | TRUE | -0.116671036 | 0.231605439 | No Change | Not Significant | TRUE | -0.017172059 | 0.82901007 | No Change | Not Significant | TRUE | Cond1 Not Significant + Cond2 Not Significant | None | None | ENSMUSG00000024603 | 67665 | protein_coding | 898.4175777 |
| Dctn5 | TRUE | 0.033870392 | 0.806325018 | No Change | Not Significant | TRUE | 0.123717723 | 0.088190332 | No Change | Not Significant | TRUE | Cond1 Not Significant + Cond2 Not Significant | None | None | ENSMUSG00000030868 | 59288 | protein_coding | 455.2974559 |
| Dctn6 | TRUE | -0.072203482 | 0.542231046 | No Change | Not Significant | TRUE | -0.039595038 | 0.705342483 | No Change | Not Significant | TRUE | Cond1 Not Significant + Cond2 Not Significant | None | None | ENSMUSG00000031516 | 22428 | protein_coding | 462.682918 |
| Dctpp1 | TRUE | -0.19629983 | 0.184932165 | No Change | Not Significant | TRUE | -0.044505787 | 0.723987816 | No Change | Not Significant | TRUE | Cond1 Not Significant + Cond2 Not Significant | None | None | ENSMUSG00000042462 | 66422 | protein_coding | 223.1751078 |
| Dcun1d1 | TRUE | -0.186902974 | 0.122282111 | No Change | Not Significant | TRUE | -0.096554595 | 0.220801611 | No Change | Not Significant | TRUE | Cond1 Not Significant + Cond2 Not Significant | None | None | ENSMUSG00000027708 | 114893 | protein_coding | 381.3928971 |
| Dcun1d2 | TRUE | -0.061492623 | 0.675987813 | No Change | Not Significant | TRUE | 0.053375227 | 0.640499152 | No Change | Not Significant | TRUE | Cond1 Not Significant + Cond2 Not Significant | None | None | ENSMUSG00000038506 | 102323 | protein_coding | 285.0660373 |
| Dcun1d3 | TRUE | 0.009436228 | 0.959501353 | No Change | Not Significant | TRUE | 0.002125296 | 0.993012764 | No Change | Not Significant | TRUE | Cond1 Not Significant + Cond2 Not Significant | None | None | ENSMUSG00000048787 | 233805 | protein_coding | 144.387298 |
| Dcun1d4 | TRUE | 1.447035708 | 2.49E-41 | UP | UP | TRUE | 0.638732258 | 1.58E-14 | UP | UP | TRUE | Cond1 UP + Cond2 UP | Core_UP | Core_UP | ENSMUSG00000051674 | 100737 | protein_coding | 228.4345946 |
| Dcun1d5 | TRUE | 0.096775869 | 0.37057407 | No Change | Not Significant | TRUE | 0.055569767 | 0.594351535 | No Change | Not Significant | TRUE | Cond1 Not Significant + Cond2 Not Significant | None | None | ENSMUSG00000032002 | 76863 | protein_coding | 401.9225579 |
| Dcx | TRUE | -0.253638579 | 1 | No Change | Not Significant | FALSE | NA | NA | No Change | Not Detected | FALSE | Background = FALSE | Background = FALSE | Background = FALSE | ENSMUSG00000031285 | 13193 | protein_coding | 4.36715505 |
| Dcxr | TRUE | -0.178453314 | 0.274998498 | No Change | Not Significant | TRUE | 0.053554456 | 0.715668065 | No Change | Not Significant | TRUE | Cond1 Not Significant + Cond2 Not Significant | None | None | ENSMUSG00000039450 | 67880 | protein_coding | 646.5563522 |
| Dda1 | TRUE | 0.512412498 | 3.14E-11 | UP | UP | TRUE | 0.17214613 | 0.09573909 | No Change | Not Significant | TRUE | Cond1 UP + Cond2 Not Significant | Cond1_UP | Cond1_UP | ENSMUSG00000074247 | 66498 | protein_coding | 534.250737 |
| Ddah1 | TRUE | 0.292301735 | 0.009870825 | No Change | Significant Positive | TRUE | 0.070063807 | 0.593278833 | No Change | Not Significant | TRUE | Cond1 Significant Positive + Cond2 Not Significant | None | None | ENSMUSG00000028194 | 69219 | protein_coding | 759.2474552 |
| Ddah2 | TRUE | -0.156019503 | 0.827931507 | No Change | Not Significant | TRUE | -0.313584245 | 0.426907135 | No Change | Not Significant | TRUE | Cond1 Not Significant + Cond2 Not Significant | None | None | ENSMUSG00000007039 | 51793 | protein_coding | 15.70547817 |
| Ddb1 | TRUE | 0.099989581 | 0.574143363 | No Change | Not Significant | TRUE | 0.094295268 | 0.172892408 | No Change | Not Significant | TRUE | Cond1 Not Significant + Cond2 Not Significant | None | None | ENSMUSG00000024740 | 13194 | protein_coding | 4802.556638 |
| Ddb2 | TRUE | -0.206984326 | 0.24534105 | No Change | Not Significant | TRUE | -0.453306693 | 0.010812233 | No Change | Significant Negative | TRUE | Cond1 Not Significant + Cond2 Significant Negative | None | None | ENSMUSG00000002109 | 107986 | protein_coding | 72.44921646 |
| Ddc | TRUE | -0.178487246 | 0.349520595 | No Change | Not Significant | TRUE | 0.217851216 | 0.045795479 | No Change | Significant Positive | TRUE | Cond1 Not Significant + Cond2 Significant Positive | None | None | ENSMUSG00000020182 | 13195 | protein_coding | 823.9024085 |
| Ddhd1 | TRUE | 0.158004082 | 0.120901073 | No Change | Not Significant | TRUE | -0.102363654 | 0.440293139 | No Change | Not Significant | TRUE | Cond1 Not Significant + Cond2 Not Significant | None | None | ENSMUSG00000037697 | 114874 | protein_coding | 202.3166707 |
| Ddhd2 | TRUE | -0.098815645 | 0.404039964 | No Change | Not Significant | TRUE | -0.149074523 | 0.213379724 | No Change | Not Significant | TRUE | Cond1 Not Significant + Cond2 Not Significant | None | None | ENSMUSG00000061313 | 72108 | protein_coding | 206.6101282 |
| Ddi1 | TRUE | -1.059268557 | 1 | No Change | Not Significant | TRUE | -0.973217822 | 0.224318083 | No Change | Not Significant | TRUE | Cond1 Not Significant + Cond2 Not Significant | None | None | ENSMUSG00000047619 | 71829 | protein_coding | 4.194305521 |
| Ddi2 | TRUE | 0.123297721 | 0.119675987 | No Change | Not Significant | TRUE | 0.030006891 | 0.766221312 | No Change | Not Significant | TRUE | Cond1 Not Significant + Cond2 Not Significant | None | None | ENSMUSG00000078515 | 68817 | protein_coding | 1399.41296 |
| Ddias | TRUE | 1.349093189 | 6.91E-09 | UP | UP | TRUE | 0.236282211 | 0.327353023 | No Change | Not Significant | TRUE | Cond1 UP + Cond2 Not Significant | Cond1_UP | Cond1_UP | ENSMUSG00000030641 | 74041 | protein_coding | 65.38817405 |
| Ddit3 | TRUE | 2.000643843 | 1.43E-28 | UP | UP | TRUE | 1.985240671 | 3.63E-112 | UP | UP | TRUE | Cond1 UP + Cond2 UP | Core_UP | Core_UP | ENSMUSG00000025408 | 13198 | protein_coding | 765.5283613 |
| Ddit4 | TRUE | 0.35097501 | 0.031711558 | No Change | Significant Positive | TRUE | 0.470247076 | 0.00107975 | No Change | Significant Positive | TRUE | Cond1 Significant Positive + Cond2 Significant Positive | None | None | ENSMUSG00000020108 | 74747 | protein_coding | 752.6388956 |
| Ddit4l | TRUE | 0.564700978 | 1 | No Change | Not Significant | TRUE | 0.090419093 | 0.948750818 | No Change | Not Significant | TRUE | Cond1 Not Significant + Cond2 Not Significant | None | None | ENSMUSG00000046818 | 73284 | protein_coding | 4.231561489 |
| Ddn | TRUE | 0.575495109 | 1 | No Change | Not Significant | FALSE | NA | NA | No Change | Not Detected | FALSE | Background = FALSE | Background = FALSE | Background = FALSE | ENSMUSG00000059213 | 13199 | protein_coding | 0.814631172 |
| Ddo | TRUE | 0.91452303 | 1 | No Change | Not Significant | FALSE | NA | NA | No Change | Not Detected | FALSE | Background = FALSE | Background = FALSE | Background = FALSE | ENSMUSG00000063428 | 70503 | protein_coding | 3.450322174 |
| Ddost | TRUE | 0.956099464 | 1.29E-16 | UP | UP | TRUE | 0.89030421 | 6.64E-30 | UP | UP | TRUE | Cond1 UP + Cond2 UP | Core_UP | Core_UP | ENSMUSG00000028757 | 13200 | protein_coding | 4763.674709 |
| Ddr1 | TRUE | -0.156590423 | 0.22515016 | No Change | Not Significant | TRUE | 0.063747477 | 0.685158105 | No Change | Not Significant | TRUE | Cond1 Not Significant + Cond2 Not Significant | None | None | ENSMUSG00000003534 | 12305 | protein_coding | 1570.915281 |
| Ddr2 | TRUE | 0.336472226 | 1 | No Change | Not Significant | FALSE | NA | NA | No Change | Not Detected | FALSE | Background = FALSE | Background = FALSE | Background = FALSE | ENSMUSG00000026674 | 18214 | protein_coding | 3.979259346 |
| Ddrgk1 | TRUE | -0.060889882 | 0.584169058 | No Change | Not Significant | TRUE | -0.030254201 | 0.680967534 | No Change | Not Significant | TRUE | Cond1 Not Significant + Cond2 Not Significant | None | None | ENSMUSG00000068290 | 77006 | protein_coding | 799.0359983 |
| Ddt | TRUE | 0.302655211 | 0.011213534 | No Change | Significant Positive | TRUE | 0.367984182 | 0.000611815 | No Change | Significant Positive | TRUE | Cond1 Significant Positive + Cond2 Significant Positive | None | None | ENSMUSG00000001666 | 13202 | protein_coding | 237.2367162 |
| Ddx1 | TRUE | 0.012190208 | 0.915295615 | No Change | Not Significant | TRUE | 0.049318333 | 0.604873354 | No Change | Not Significant | TRUE | Cond1 Not Significant + Cond2 Not Significant | None | None | ENSMUSG00000037149 | 104721 | protein_coding | 1343.195508 |
| Ddx10 | TRUE | 0.33702096 | 0.006694122 | No Change | Significant Positive | TRUE | 0.054924213 | 0.730184332 | No Change | Not Significant | TRUE | Cond1 Significant Positive + Cond2 Not Significant | None | None | ENSMUSG00000053289 | 77591 | protein_coding | 382.8725877 |
| Ddx11 | TRUE | 0.250841204 | 0.145350463 | No Change | Not Significant | TRUE | -0.024707316 | 0.921535618 | No Change | Not Significant | TRUE | Cond1 Not Significant + Cond2 Not Significant | None | None | ENSMUSG00000035842 | 320209 | protein_coding | 135.0800011 |
| Ddx17 | TRUE | -0.010994062 | 0.941973978 | No Change | Not Significant | TRUE | -0.051028951 | 0.536379174 | No Change | Not Significant | TRUE | Cond1 Not Significant + Cond2 Not Significant | None | None | ENSMUSG00000055065 | 67040 | protein_coding | 2704.206293 |
| Ddx18 | TRUE | 0.072702461 | 0.734239015 | No Change | Not Significant | TRUE | -0.016172962 | 0.942710088 | No Change | Not Significant | TRUE | Cond1 Not Significant + Cond2 Not Significant | None | None | ENSMUSG00000001674 | 66942 | protein_coding | 166.5279987 |
| Ddx19a | TRUE | 0.133698864 | 0.429893421 | No Change | Not Significant | TRUE | 0.062182506 | 0.609230875 | No Change | Not Significant | TRUE | Cond1 Not Significant + Cond2 Not Significant | None | None | ENSMUSG00000015023 | 13680 | protein_coding | 283.9410931 |

| geneID | Cond1_DF_Detected | Cond1_DF_ValueCol | Cond1_DF_PadjCol | Cond1_DF_Cutoff | Cond1_DF_Cutoff_Specific.x | Cond2_DF_Detected | Cond2_DF_ValueCol | Cond2_DF_PadjCol | Cond2_DF_Cutoff | Cond2_DF_Cutoff_Specific | BG_Method |
|---|---|---|---|---|---|---|---|---|---|---|---|
| Ddx19b | TRUE | -0.041089899 | 0.809917796 | No Change | Not Significant | TRUE | 0.035899729 | 0.732626853 | No Change | Not Significant | TRUE |
| Ddx20 | TRUE | -0.016096504 | 0.923798341 | No Change | Not Significant | TRUE | 0.023474827 | 0.872486041 | No Change | Not Significant | TRUE |
| Ddx21 | TRUE | -0.067901715 | 0.699692776 | No Change | Not Significant | TRUE | -0.099670647 | 0.350881819 | No Change | Not Significant | TRUE |
| Ddx23 | TRUE | 0.093433642 | 0.115594862 | No Change | Not Significant | TRUE | 0.067786327 | 0.27725095 | No Change | Not Significant | TRUE |
| Ddx24 | TRUE | -0.063789929 | 0.535990818 | No Change | Not Significant | TRUE | 0.009356728 | 0.930926537 | No Change | Not Significant | TRUE |
| Ddx25 | TRUE | -0.088300684 | 0.942424291 | No Change | Not Significant | FALSE | NA | NA | No Change | Not Detected | FALSE |
| Ddx27 | TRUE | -0.042119556 | 0.783067117 | No Change | Not Significant | TRUE | -0.023307944 | 0.851790951 | No Change | Not Significant | TRUE |
| Ddx28 | TRUE | -0.05132228 | 0.802698942 | No Change | Not Significant | TRUE | 0.019921523 | 0.916810722 | No Change | Not Significant | TRUE |
| Ddx31 | TRUE | 0.055175322 | 0.855204229 | No Change | Not Significant | TRUE | -0.073254301 | 0.706263486 | No Change | Not Significant | TRUE |
| Ddx39a | TRUE | 0.555343722 | 1.47E-07 | UP | UP | TRUE | 0.166811092 | 0.231492618 | No Change | Not Significant | TRUE |
| Ddx39b | TRUE | 0.134662622 | 0.155998286 | No Change | Not Significant | TRUE | 0.008868806 | 0.923349179 | No Change | Not Significant | TRUE |
| Ddx3x | TRUE | -0.012388067 | 0.942489774 | No Change | Not Significant | TRUE | -0.116769978 | 0.20675137 | No Change | Not Significant | TRUE |
| Ddx4 | TRUE | 1.135881427 | 1 | No Change | Not Significant | FALSE | NA | NA | No Change | Not Detected | FALSE |
| Ddx41 | TRUE | 0.02279054 | 0.878311377 | No Change | Not Significant | TRUE | 0.081126868 | 0.301772366 | No Change | Not Significant | TRUE |
| Ddx42 | TRUE | 0.009728521 | 0.946097348 | No Change | Not Significant | TRUE | -0.044761841 | 0.610334928 | No Change | Not Significant | TRUE |
| Ddx43 | TRUE | 1.849815186 | 3.21E-05 | UP | UP | TRUE | -0.152194114 | 0.76212257 | No Change | Not Significant | TRUE |
| Ddx46 | TRUE | 0.134544057 | 0.09071266 | No Change | Not Significant | TRUE | -0.085434804 | 0.337593648 | No Change | Not Significant | TRUE |
| Ddx47 | TRUE | -0.067958032 | 0.513092425 | No Change | Not Significant | TRUE | -0.060009706 | 0.537064913 | No Change | Not Significant | TRUE |
| Ddx49 | TRUE | 0.091265471 | 0.517476912 | No Change | Not Significant | TRUE | 0.039375673 | 0.772159116 | No Change | Not Significant | TRUE |
| Ddx5 | TRUE | -0.327018404 | 1.62E-07 | No Change | Significant Negative | TRUE | -0.148371295 | 0.024383381 | No Change | Significant Negative | TRUE |
| Ddx50 | TRUE | 0.848648336 | 6.55E-24 | UP | UP | TRUE | 0.545108748 | 2.22E-26 | UP | UP | TRUE |
| Ddx51 | TRUE | -0.08692926 | 0.706297653 | No Change | Not Significant | TRUE | -0.168817987 | 0.23074217 | No Change | Not Significant | TRUE |
| Ddx52 | TRUE | 0.162033713 | 0.170740454 | No Change | Not Significant | TRUE | -0.023430788 | 0.900451559 | No Change | Not Significant | TRUE |
| Ddx54 | TRUE | -0.022938897 | 0.811872956 | No Change | Not Significant | TRUE | 0.064543985 | 0.485822803 | No Change | Not Significant | TRUE |
| Ddx55 | TRUE | -0.019600975 | 0.927761944 | No Change | Not Significant | TRUE | -0.074335296 | 0.628080406 | No Change | Not Significant | TRUE |
| Ddx56 | TRUE | -0.033205966 | 0.87863969 | No Change | Not Significant | TRUE | -0.024979307 | 0.865841503 | No Change | Not Significant | TRUE |
| Ddx59 | TRUE | 0.061195165 | 0.874455706 | No Change | Not Significant | TRUE | 0.169935347 | 0.369310836 | No Change | Not Significant | TRUE |
| Ddx6 | TRUE | -0.033206768 | 0.852797822 | No Change | Not Significant | TRUE | -0.02078931 | 0.87479925 | No Change | Not Significant | TRUE |
| Ddx60 | TRUE | 0.371573418 | 0.639216792 | No Change | Not Significant | TRUE | -0.163932583 | 0.636981448 | No Change | Not Significant | TRUE |
| Deaf1 | TRUE | -0.187243465 | 0.223580206 | No Change | Not Significant | TRUE | -0.201985496 | 0.005413107 | No Change | Significant Negative | TRUE |
| Decr1 | TRUE | -0.318755893 | 0.011662382 | No Change | Significant Negative | TRUE | 0.031306602 | 0.79214131 | No Change | Not Significant | TRUE |
| Decr2 | TRUE | -0.164182188 | 0.053943227 | No Change | Not Significant | TRUE | 0.097558086 | 0.156316709 | No Change | Not Significant | TRUE |
| Dedd | TRUE | -0.10179726 | 0.634528665 | No Change | Not Significant | TRUE | -0.154996432 | 0.407223247 | No Change | Not Significant | TRUE |
| Dedd2 | TRUE | -0.189698026 | 0.422068056 | No Change | Not Significant | TRUE | -0.198698901 | 0.023776147 | No Change | Significant Negative | TRUE |
| Def6 | TRUE | 0.3652394 | 0.506958639 | No Change | Not Significant | TRUE | 0.065726463 | 0.918354815 | No Change | Not Significant | TRUE |
| Def8 | TRUE | 0.044150999 | 0.738518372 | No Change | Not Significant | TRUE | 0.096564578 | 0.328321636 | No Change | Not Significant | TRUE |
| Defa17 | FALSE | NA | NA | No Change | Not Detected | TRUE | 0.595857896 | 0.752000715 | No Change | Not Significant | FALSE |
| Defa21 | FALSE | NA | NA | No Change | Not Detected | TRUE | 1.505312643 | 0.435615504 | No Change | Not Significant | FALSE |
| Defa22 | FALSE | NA | NA | No Change | Not Detected | TRUE | 2.442262227 | 0.01394449 | UP | UP | FALSE |
| Defa24 | TRUE | 3.239752492 | 1 | No Change | Not Significant | TRUE | 0.815207808 | 0.691532821 | No Change | Not Significant | TRUE |
| Defa30 | TRUE | 2.738502405 | 1 | No Change | Not Significant | TRUE | 0.348786413 | 0.888612649 | No Change | Not Significant | TRUE |
| Defa38 | FALSE | NA | NA | No Change | Not Detected | TRUE | 0.716855891 | 0.715717596 | No Change | Not Significant | FALSE |
| Defa39 | FALSE | NA | NA | No Change | Not Detected | TRUE | 0.906848151 | 0.648598559 | No Change | Not Significant | FALSE |
| Defb37 | TRUE | 0.053138109 | 0.899929685 | No Change | Not Significant | TRUE | -0.33871156 | 0.29018928 | No Change | Not Significant | TRUE |
| Defb40 | TRUE | 0.853607398 | 1 | No Change | Not Significant | FALSE | NA | NA | No Change | Not Detected | FALSE |
| Defb42 | TRUE | 2.850701429 | 1 | No Change | Not Significant | FALSE | NA | NA | No Change | Not Detected | FALSE |
| Defb45 | TRUE | -0.481370283 | 0.504402765 | No Change | Not Significant | TRUE | -0.238867838 | 0.749854321 | No Change | Not Significant | TRUE |
| Degs1 | TRUE | 0.385305024 | 0.021273857 | No Change | Significant Positive | TRUE | 0.104370341 | 0.504092706 | No Change | Not Significant | TRUE |
| Degs2 | TRUE | 0.024954827 | 0.883975099 | No Change | Not Significant | TRUE | 0.088419681 | 0.357574711 | No Change | Not Significant | TRUE |
| Dek | TRUE | -0.015716336 | 0.958296749 | No Change | Not Significant | TRUE | -0.037986254 | 0.82145077 | No Change | Not Significant | TRUE |
| Dele1 | TRUE | -0.042224831 | 0.81729577 | No Change | Not Significant | TRUE | 0.041571542 | 0.673641376 | No Change | Not Significant | TRUE |
| Denn2b | TRUE | 0.164227303 | 0.1706035 | No Change | Not Significant | TRUE | 0.324738826 | 0.000320903 | No Change | Significant Positive | TRUE |
| Dennd10 | TRUE | 0.139564349 | 0.147459771 | No Change | Not Significant | TRUE | 0.128757652 | 0.381255129 | No Change | Not Significant | TRUE |
| Dennd11 | TRUE | -0.241064563 | 0.448589299 | No Change | Not Significant | TRUE | 0.202134546 | 0.000838976 | No Change | Significant Positive | TRUE |
| Dennd1a | TRUE | -0.039021584 | 0.839060823 | No Change | Not Significant | TRUE | 0.034463947 | 0.81580845 | No Change | Not Significant | TRUE |
| Dennd1b | TRUE | -0.172816984 | 0.233151171 | No Change | Not Significant | TRUE | -0.135603142 | 0.046962624 | No Change | Significant Negative | TRUE |
| Dennd1c | TRUE | -0.387759958 | 0.000787343 | No Change | Significant Negative | TRUE | -0.163784267 | 0.104068496 | No Change | Not Significant | TRUE |
| Dennd2a | TRUE | -0.00995431 | 0.995389267 | No Change | Not Significant | TRUE | 0.885838821 | 0.208682378 | No Change | Not Significant | TRUE |
| Dennd2c | TRUE | 0.144681679 | 0.728866518 | No Change | Not Significant | TRUE | -0.212414578 | 0.426951571 | No Change | Not Significant | TRUE |
| Dennd2d | TRUE | 0.215604586 | 0.075817398 | No Change | Not Significant | TRUE | 0.056769031 | 0.650098067 | No Change | Not Significant | TRUE |
| Dennd3 | TRUE | -0.132251671 | 0.337542702 | No Change | Not Significant | TRUE | 0.007721142 | 0.955322545 | No Change | Not Significant | TRUE |
| Dennd4a | TRUE | -0.072204126 | 0.735467298 | No Change | Not Significant | TRUE | -0.104422532 | 0.468724137 | No Change | Not Significant | TRUE |
| Dennd4b | TRUE | 0.130787054 | 0.8036518 | No Change | Not Significant | TRUE | -0.001587861 | 0.996791885 | No Change | Not Significant | TRUE |
| Dennd4c | TRUE | -0.236175374 | 0.002648228 | No Change | Significant Negative | TRUE | -0.016486977 | 0.827627866 | No Change | Not Significant | TRUE |
| Dennd5a | TRUE | 0.749302045 | 0.030010701 | UP | UP | TRUE | 0.277933459 | 0.3074195 | No Change | Not Significant | TRUE |
| Dennd5b | TRUE | 0.050652071 | 0.792160606 | No Change | Not Significant | TRUE | -0.05097585 | 0.718355583 | No Change | Not Significant | TRUE |
| Dennd6a | TRUE | -0.157111962 | 0.364391266 | No Change | Not Significant | TRUE | -0.026457915 | 0.800250831 | No Change | Not Significant | TRUE |
| Dennd6b | TRUE | -0.129936162 | 0.585225632 | No Change | Not Significant | TRUE | -0.207322788 | 0.167504953 | No Change | Not Significant | TRUE |
| Denr | TRUE | 0.269257583 | 0.003492164 | No Change | Significant Positive | TRUE | 0.135309279 | 0.134033812 | No Change | Not Significant | TRUE |
| Depdc1a | TRUE | 0.614533295 | 0.027682261 | UP | UP | TRUE | 0.111077161 | 0.708970987 | No Change | Not Significant | TRUE |
| Depdc1b | TRUE | 0.622139345 | 0.017134056 | UP | UP | TRUE | 0.055183693 | 0.901477241 | No Change | Not Significant | TRUE |
| Depdc5 | TRUE | -0.03071494 | 0.834370416 | No Change | Not Significant | TRUE | -0.108828763 | 0.167125066 | No Change | Not Significant | TRUE |
| Depdc7 | TRUE | -0.204311489 | 0.804106371 | No Change | Not Significant | TRUE | -0.225804932 | 0.65807782 | No Change | Not Significant | TRUE |
| Depp1 | TRUE | -1.163145511 | 0.035343465 | DOWN | DOWN | TRUE | -0.41372766 | 0.188241916 | No Change | Not Significant | TRUE |
| Deptor | TRUE | -0.181686584 | 0.383501006 | No Change | Not Significant | TRUE | -0.011124762 | 0.935951904 | No Change | Not Significant | TRUE |
| Dera | TRUE | 0.002121259 | 0.989055615 | No Change | Not Significant | TRUE | 0.018924551 | 0.883011233 | No Change | Not Significant | TRUE |
| Derl1 | TRUE | 0.665069758 | 5.76E-10 | UP | UP | TRUE | 0.470562721 | 6.54E-14 | No Change | Significant Positive | TRUE |
| Derl2 | TRUE | 1.199752212 | 1.76E-58 | UP | UP | TRUE | 0.952409784 | 8.82E-48 | UP | UP | TRUE |
| Derl3 | TRUE | 9.247893935 | 0 | UP | UP | TRUE | 7.442074117 | 1.72E-24 | UP | UP | TRUE |
| Des | TRUE | -0.996714308 | 1 | No Change | Not Significant | FALSE | NA | NA | No Change | Not Detected | FALSE |
| Desi1 | TRUE | -0.04100962 | 0.810888501 | No Change | Not Significant | TRUE | -0.185878643 | 0.049315416 | No Change | Significant Negative | TRUE |
| Desi2 | TRUE | 0.091201844 | 0.405906838 | No Change | Not Significant | TRUE | -0.056408816 | 0.585270749 | No Change | Not Significant | TRUE |
| Det1 | TRUE | 0.103246041 | 0.523380864 | No Change | Not Significant | TRUE | -0.018493738 | 0.928990077 | No Change | Not Significant | TRUE |
| Deup1 | TRUE | -0.823905113 | 1 | No Change | Not Significant | FALSE | NA | NA | No Change | Not Detected | FALSE |
| Dexi | TRUE | 0.428048998 | 0.004614708 | No Change | Significant Positive | TRUE | 0.241010508 | 0.144654187 | No Change | Not Significant | TRUE |
| Dffa | TRUE | -0.191814187 | 0.046167968 | No Change | Significant Negative | TRUE | -0.113073997 | 0.379596103 | No Change | Not Significant | TRUE |
| Dffb | TRUE | -0.164720326 | 0.381254943 | No Change | Not Significant | TRUE | -0.232512535 | 0.086102903 | No Change | Not Significant | TRUE |
| Dgat1 | TRUE | -0.139520578 | 0.410105269 | No Change | Not Significant | TRUE | -0.173100608 | 0.037056142 | No Change | Significant Negative | TRUE |
| Dgat2 | TRUE | 1.105499176 | 1.11E-08 | UP | UP | TRUE | 0.488778813 | 3.87E-07 | No Change | Significant Positive | TRUE |
| Dgat2l6 | TRUE | 1.138642844 | 1 | No Change | Not Significant | FALSE | NA | NA | No Change | Not Detected | FALSE |
| Dgcr2 | TRUE | -0.010207429 | 0.966286169 | No Change | Not Significant | TRUE | 0.011136397 | 0.880475742 | No Change | Not Significant | TRUE |
| Dgcr6 | TRUE | -0.220990277 | 0.028407451 | No Change | Significant Negative | TRUE | -0.084518937 | 0.485573395 | No Change | Not Significant | TRUE |
| Dgcr8 | TRUE | 0.162100157 | 0.084039656 | No Change | Not Significant | TRUE | -0.059007238 | 0.718418124 | No Change | Not Significant | TRUE |
| Dgka | TRUE | -0.304870513 | 0.042239714 | No Change | Significant Negative | TRUE | 0.015178228 | 0.911459024 | No Change | Not Significant | TRUE |
| Dgkb | TRUE | 0.410747998 | 1 | No Change | Not Significant | FALSE | NA | NA | No Change | Not Detected | FALSE |
| Dgkd | TRUE | -0.135831722 | 0.08602548 | No Change | Not Significant | TRUE | -0.059757102 | 0.352354239 | No Change | Not Significant | TRUE |
| Dgke | TRUE | 0.000402938 | 0.998249138 | No Change | Not Significant | TRUE | -0.099218223 | 0.438306743 | No Change | Not Significant | TRUE |
| Dgkg | TRUE | -0.164604129 | 0.668926246 | No Change | Not Significant | TRUE | -0.43659293 | 0.009146817 | No Change | Significant Negative | TRUE |
| Dgkh | TRUE | -0.609312179 | 0.000134753 | DOWN | DOWN | TRUE | -0.232566876 | 0.019391822 | No Change | Significant Negative | TRUE |
| Dgki | TRUE | -0.237398506 | 0.749146509 | No Change | Not Significant | TRUE | 0.1721224 | 0.706263486 | No Change | Not Significant | TRUE |
| Dgkk | TRUE | 2.094390115 | 1 | No Change | Not Significant | FALSE | NA | NA | No Change | Not Detected | FALSE |
| Dgkq | TRUE | -0.052187879 | 0.878895387 | No Change | Not Significant | TRUE | -0.229466278 | 0.067773928 | No Change | Not Significant | TRUE |
| Dgkz | TRUE | 0.035503631 | 0.726359126 | No Change | Not Significant | TRUE | 0.038070893 | 0.61248414 | No Change | Not Significant | TRUE |
| Dglucy | TRUE | 0.212248403 | 0.101632751 | No Change | Not Significant | TRUE | 0.265151638 | 4.62E-05 | No Change | Significant Positive | TRUE |
| Dguok | TRUE | 0.062403652 | 0.700835947 | No Change | Not Significant | TRUE | 0.089610769 | 0.45218843 | No Change | Not Significant | TRUE |
| Dhcr24 | TRUE | 0.027043337 | 0.86030736 | No Change | Not Significant | TRUE | 0.119597601 | 0.455009895 | No Change | Not Significant | TRUE |
| Dhcr7 | TRUE | -0.254984669 | 0.0190648 | No Change | Significant Negative | TRUE | -0.009941021 | 0.959153588 | No Change | Not Significant | TRUE |
| Dhdds | TRUE | 0.240868698 | 0.015327126 | No Change | Significant Positive | TRUE | 0.177801521 | 0.015196638 | No Change | Significant Positive | TRUE |
| Dhdh | TRUE | -0.263562419 | 0.657120605 | No Change | Not Significant | TRUE | -0.124461869 | 0.749854321 | No Change | Not Significant | TRUE |
| Dhfr | TRUE | 0.05894921 | 0.767282764 | No Change | Not Significant | TRUE | 0.109971232 | 0.604050305 | No Change | Not Significant | TRUE |
| Dhh | TRUE | 2.201512004 | 1 | No Change | Not Significant | FALSE | NA | NA | No Change | Not Detected | FALSE |
| Dhodh | TRUE | 0.116202641 | 0.625349723 | No Change | Not Significant | TRUE | 0.173320152 | 0.150979148 | No Change | Not Significant | TRUE |
| Dhps | TRUE | 0.107056447 | 0.454166041 | No Change | Not Significant | TRUE | -0.021631727 | 0.884485201 | No Change | Not Significant | TRUE |
| Dhrs1 | TRUE | -0.190312078 | 0.114659863 | No Change | Not Significant | TRUE | 0.042931473 | 0.762827428 | No Change | Not Significant | TRUE |
| Dhrs11 | TRUE | 0.888715601 | 4.76E-16 | UP | UP | TRUE | 0.763646259 | 2.42E-20 | UP | UP | TRUE |
| Dhrs13 | TRUE | -0.4464199 | 0.047310767 | No Change | Significant Negative | TRUE | -0.202582787 | 0.246155835 | No Change | Not Significant | TRUE |
| Dhrs13os | TRUE | -0.498380352 | 1 | No Change | Not Significant | FALSE | NA | NA | No Change | Not Detected | FALSE |
| Dhrs3 | TRUE | -0.734067566 | 7.56E-10 | DOWN | DOWN | TRUE | -0.391287625 | 8.62E-06 | No Change | Significant Negative | TRUE |
| Dhrs4 | TRUE | -0.141263478 | 0.324504542 | No Change | Not Significant | TRUE | 0.129914699 | 0.175422328 | No Change | Not Significant | TRUE |
| Dhrs7 | TRUE | -0.02656098 | 0.926816617 | No Change | Not Significant | TRUE | -0.031939164 | 0.832361742 | No Change | Not Significant | TRUE |
| Dhrs7b | TRUE | -0.188452186 | 0.022069064 | No Change | Significant Negative | TRUE | 0.123567591 | 0.257550033 | No Change | Not Significant | TRUE |
| Dhrs7c | TRUE | 1.126107492 | 1 | No Change | Not Significant | FALSE | NA | NA | No Change | Not Detected | FALSE |
| Dhrs9 | TRUE | 0.88284513 | 0.00357329 | UP | UP | TRUE | 1.076826809 | 5.87E-11 | UP | UP | TRUE |
| Dhtkd1 | TRUE | -0.190784915 | 1 | No Change | Not Significant | FALSE | NA | NA | No Change | Not Detected | FALSE |
| Dhx15 | TRUE | 0.115947462 | 0.386709513 | No Change | Not Significant | TRUE | -0.004303618 | 0.961085859 | No Change | Not Significant | TRUE |
| Dhx16 | TRUE | 0.04076287 | 0.693654751 | No Change | Not Significant | TRUE | 0.077152724 | 0.509217772 | No Change | Not Significant | TRUE |
| Dhx29 | TRUE | 0.413667927 | 0.003974816 | No Change | Significant Positive | TRUE | 0.282056096 | 0.015091528 | No Change | Significant Positive | TRUE |
| Dhx30 | TRUE | -0.006598053 | 0.963320302 | No Change | Not Significant | TRUE | 0.042530556 | 0.666166426 | No Change | Not Significant | TRUE |
| Dhx32 | TRUE | -0.023631275 | 0.882510888 | No Change | Not Significant | TRUE | 0.152882584 | 0.013434123 | No Change | Significant Positive | TRUE |
| Dhx33 | TRUE | 0.20141471 | 0.108625296 | No Change | Not Significant | TRUE | -0.078006162 | 0.51990697 | No Change | Not Significant | TRUE |
| Dhx34 | TRUE | -0.17808645 | 0.192028673 | No Change | Not Significant | TRUE | -0.00089372 | 0.996376861 | No Change | Not Significant | TRUE |
| Dhx35 | TRUE | -0.215632743 | 0.157015607 | No Change | Not Significant | TRUE | -0.08752314 | 0.336981111 | No Change | Not Significant | TRUE |
| Dhx36 | TRUE | -0.045010142 | 0.800507769 | No Change | Not Significant | TRUE | -0.023912705 | 0.832361742 | No Change | Not Significant | TRUE |
| Dhx37 | TRUE | 0.026067189 | 0.88791795 | No Change | Not Significant | TRUE | 0.011866022 | 0.945241431 | No Change | Not Significant | TRUE |
| Dhx38 | TRUE | 0.177393589 | 0.031812382 | No Change | Significant Positive | TRUE | 0.101498371 | 0.267808218 | No Change | Not Significant | TRUE |
| Dhx40 | TRUE | 0.181393691 | 0.164036653 | No Change | Not Significant | TRUE | 0.10566586 | 0.314662099 | No Change | Not Significant | TRUE |
| Dhx57 | TRUE | 0.211724449 | 0.008129295 | No Change | Significant Positive | TRUE | 0.074930181 | 0.486649916 | No Change | Not Significant | TRUE |
| Dhx58 | TRUE | -0.424373158 | 0.125929956 | No Change | Not Significant | TRUE | 0.075025012 | 0.803180854 | No Change | Not Significant | TRUE |
| Dhx8 | TRUE | 0.034514563 | 0.780311961 | No Change | Not Significant | TRUE | 0.026127793 | 0.780262005 | No Change | Not Significant | TRUE |
| Dhx9 | TRUE | 0.085650851 | 0.739162904 | No Change | Not Significant | TRUE | -0.166858881 | 0.17246929 | No Change | Not Significant | TRUE |
| Diablo | TRUE | -0.170000975 | 0.487629303 | No Change | Not Significant | TRUE | -0.2238795 | 0.258962499 | No Change | Not Significant | TRUE |
| Diaph1 | TRUE | -0.223706951 | 0.022343376 | No Change | Significant Negative | TRUE | -0.020633948 | 0.81829507 | No Change | Not Significant | TRUE |
| Diaph2 | TRUE | -0.357378827 | 0.000674132 | No Change | Significant Negative | TRUE | -0.327074852 | 0.000887414 | No Change | Significant Negative | TRUE |
| Diaph3 | TRUE | 0.935228457 | 1.54E-08 | UP | UP | TRUE | 0.126525477 | 0.604649053 | No Change | Not Significant | TRUE |
| Dicer1 | TRUE | 0.085588031 | 0.459757614 | No Change | Not Significant | TRUE | -0.018162876 | 0.863755116 | No Change | Not Significant | TRUE |
| Dido1 | TRUE | -0.015403445 | 0.91190529 | No Change | Not Significant | TRUE | -0.091083333 | 0.142359943 | No Change | Not Significant | TRUE |
| Dimt1 | TRUE | -0.251879036 | 0.301713259 | No Change | Not Significant | TRUE | -0.084189781 | 0.747901626 | No Change | Not Significant | TRUE |
| Dio1 | TRUE | -0.033457262 | 0.877003824 | No Change | Not Significant | TRUE | 0.119484545 | 0.235215876 | No Change | Not Significant | TRUE |
| Dio2 | TRUE | 1.119132346 | 1 | No Change | Not Significant | FALSE | NA | NA | No Change | Not Detected | FALSE |
| Dio3 | TRUE | -0.37657571 | 1 | No Change | Not Significant | FALSE | NA | NA | No Change | Not Detected | FALSE |
| Dio3os | TRUE | -0.987752801 | 0.171319292 | No Change | Not Significant | FALSE | NA | NA | No Change | Not Detected | FALSE |
| Dip2a | TRUE | -0.106610291 | 0.10695305 | No Change | Not Significant | TRUE | -0.080913689 | 0.410383236 | No Change | Not Significant | TRUE |
| Dip2b | TRUE | -0.178912128 | 0.141589694 | No Change | Not Significant | TRUE | -0.086227911 | 0.304905289 | No Change | Not Significant | TRUE |
| Dip2c | TRUE | -0.172997052 | 0.079188614 | No Change | Not Significant | TRUE | -0.235666546 | 0.001008172 | No Change | Significant Negative | TRUE |
| Dipk1a | TRUE | -0.114074408 | 0.801687789 | No Change | Not Significant | TRUE | 0.031438312 | 0.891466618 | No Change | Not Significant | TRUE |
| Dipk1b | TRUE | 0.042676433 | 0.913297021 | No Change | Not Significant | TRUE | -0.232968664 | 0.29018928 | No Change | Not Significant | TRUE |
| Dipk1c | TRUE | 0.836215055 | 0.17492772 | No Change | Not Significant | TRUE | 0.800392117 | 0.096931133 | No Change | Not Significant | TRUE |
| Dipk2a | TRUE | 0.253263853 | 0.110135672 | No Change | Not Significant | TRUE | 0.150173219 | 0.02180516 | No Change | Significant Positive | TRUE |
| Diras1 | TRUE | -1.882348289 | 1 | No Change | Not Significant | FALSE | NA | NA | No Change | Not Detected | FALSE |
| Diras2 | TRUE | 4.087707454 | 1 | No Change | Not Significant | FALSE | NA | NA | No Change | Not Detected | FALSE |
| Dis3 | TRUE | 0.2797226 | 0.161179189 | No Change | Not Significant | TRUE | -0.095145134 | 0.476450233 | No Change | Not Significant | TRUE |
| Dis3l | TRUE | -0.19924656 | 0.179403047 | No Change | Not Significant | TRUE | -0.008620749 | 0.945542552 | No Change | Not Significant | TRUE |
| Dis3l2 | TRUE | 0.078027354 | 0.574428953 | No Change | Not Significant | TRUE | -0.024129396 | 0.859442037 | No Change | Not Significant | TRUE |
| Disc1 | TRUE | -0.119284033 | 1 | No Change | Not Significant | FALSE | NA | NA | No Change | Not Detected | FALSE |
| Disp1 | TRUE | 0.155263497 | 0.332776929 | No Change | Not Significant | TRUE | -0.027120512 | 0.882337373 | No Change | Not Significant | TRUE |
| Disp2 | TRUE | 0.586623481 | 0.125648406 | No Change | Not Significant | TRUE | 0.233962928 | 0.29471814 | No Change | Not Significant | TRUE |
| Disp3 | TRUE | 0.937641839 | 1 | No Change | Not Significant | FALSE | NA | NA | No Change | Not Detected | FALSE |
| Dixdc1 | TRUE | -0.843020208 | 0.000869217 | DOWN | DOWN | TRUE | -0.027723978 | 0.930668864 | No Change | Not Significant | TRUE |
| Dkc1 | TRUE | 0.074898008 | 0.767077092 | No Change | Not Significant | TRUE | -0.102355044 | 0.500565425 | No Change | Not Significant | TRUE |
| Dkk2 | TRUE | 2.18049326 | 1 | No Change | Not Significant | FALSE | NA | NA | No Change | Not Detected | FALSE |
| Dkk3 | TRUE | 0.179285966 | 1 | No Change | Not Significant | FALSE | NA | NA | No Change | Not Detected | FALSE |
| Dkkl1 | TRUE | -1.888776311 | 0.001712088 | DOWN | DOWN | FALSE | NA | NA | No Change | Not Detected | FALSE |
| Dlat | TRUE | -0.225787931 | 0.07274578 | No Change | Not Significant | TRUE | 0.017233716 | 0.907439773 | No Change | Not Significant | TRUE |
| Dlc1 | TRUE | -1.87019999 | 0.009652762 | DOWN | DOWN | TRUE | -0.195186166 | 0.799032164 | No Change | Not Significant | TRUE |
| Dld | TRUE | -0.079869824 | 0.527501415 | No Change | Not Significant | TRUE | 0.031860566 | 0.74493282 | No Change | Not Significant | TRUE |
| Dlec1 | TRUE | 0.846796256 | 0.312959795 | No Change | Not Significant | TRUE | 0.604333272 | 0.352697729 | No Change | Not Significant | TRUE |
| Dlg1 | TRUE | -0.029389265 | 0.737366138 | No Change | Not Significant | TRUE | -0.016924756 | 0.836508911 | No Change | Not Significant | TRUE |
| Dlg2 | TRUE | -0.089922424 | 0.938919382 | No Change | Not Significant | TRUE | 0.444707649 | 0.395794258 | No Change | Not Significant | TRUE |
| Dlg3 | TRUE | -0.111650023 | 0.28762823 | No Change | Not Significant | TRUE | -0.005017136 | 0.962544624 | No Change | Not Significant | TRUE |
| Dlg4 | TRUE | 0.419656632 | 0.765756487 | No Change | Not Significant | TRUE | 0.235877681 | 0.737831675 | No Change | Not Significant | TRUE |
| Dlg5 | TRUE | 0.092386395 | 0.662536791 | No Change | Not Significant | TRUE | -0.043792544 | 0.763615826 | No Change | Not Significant | TRUE |
| Dlgap1 | TRUE | 0.090538825 | 0.958005484 | No Change | Not Significant | TRUE | 0.958835937 | 0.077285381 | No Change | Not Significant | TRUE |
| Dlgap2 | TRUE | 0.193720031 | 1 | No Change | Not Significant | FALSE | NA | NA | No Change | Not Detected | FALSE |
| Dlgap3 | TRUE | -0.814930409 | 0.448589299 | No Change | Not Significant | TRUE | -0.212484371 | 0.809423077 | No Change | Not Significant | TRUE |
| Dlgap4 | TRUE | 0.203990609 | 0.112998816 | No Change | Not Significant | TRUE | 0.068575337 | 0.65655478 | No Change | Not Significant | TRUE |
| Dlgap5 | TRUE | 0.673400227 | 6.42E-05 | UP | UP | TRUE | 0.148218648 | 0.244015888 | No Change | Not Significant | TRUE |
| Dlk1 | TRUE | -4.57398896 | 9.52E-14 | DOWN | DOWN | FALSE | NA | NA | No Change | Not Detected | FALSE |
| Dlk2 | TRUE | 0.297077112 | 1 | No Change | Not Significant | FALSE | NA | NA | No Change | Not Detected | FALSE |
| Dll1 | TRUE | -0.188128956 | 0.438743376 | No Change | Not Significant | TRUE | -0.122855935 | 0.523522715 | No Change | Not Significant | TRUE |
| Dll3 | TRUE | 0.258346906 | 0.822609935 | No Change | Not Significant | FALSE | NA | NA | No Change | Not Detected | FALSE |
| Dll4 | TRUE | 0.071298255 | 0.754850891 | No Change | Not Significant | TRUE | 0.098591005 | 0.668221346 | No Change | Not Significant | TRUE |
| Dlst | TRUE | -0.204672331 | 0.000130357 | No Change | Significant Negative | TRUE | 0.003440301 | 0.974004206 | No Change | Not Significant | TRUE |

| geneID | Cond1_DF_Detected | Cond1_DF_ValueCol | Cond1_DF_PadjCol | Cond1_DF_Cutoff | Cond1_DF_Cutoff_Specific.x | Cond2_DF_Detected | Cond2_DF_ValueCol |
|---|---|---|---|---|---|---|---|
| Dlx1 | TRUE | 1.126575062 | 1 | No Change | Not Significant | FALSE | NA |
| Dlx1as | TRUE | 0.172653526 | 1 | No Change | Not Significant | FALSE | NA |
| Dlx3 | TRUE | 0.887261424 | 1 | No Change | Not Significant | FALSE | NA |
| Dmac1 | TRUE | 0.086382785 | 0.69793377 | No Change | Not Significant | TRUE | 0.213802019 |
| Dmac2 | TRUE | -0.021792468 | 0.878750865 | No Change | Not Significant | TRUE | 0.023823418 |
| Dmac2l | TRUE | -0.024492227 | 0.923801556 | No Change | Not Significant | TRUE | -0.057873607 |
| Dmap1 | TRUE | -0.104754458 | 0.493649227 | No Change | Not Significant | TRUE | 0.094378048 |
| Dmbt1 | TRUE | -0.282472945 | 0.044864238 | No Change | Significant Negative | TRUE | -0.008646519 |
| Dmbx1 | TRUE | 0.698631533 | 1 | No Change | Not Significant | FALSE | NA |
| Dmd | TRUE | -0.065163213 | 0.716272317 | No Change | Not Significant | TRUE | 0.104964119 |
| Dmgdh | TRUE | 0.153378031 | 0.82411257 | No Change | Not Significant | TRUE | 0.144028574 |
| Dmkn | TRUE | -0.589143245 | 0.623441772 | No Change | Not Significant | FALSE | NA |
| Dmp1 | TRUE | -0.938138181 | 1 | No Change | Not Significant | FALSE | NA |
| Dmpk | TRUE | 0.296340018 | 0.298488621 | No Change | Not Significant | TRUE | -0.134302219 |
| Dmrt1 | TRUE | 1.125952056 | 1 | No Change | Not Significant | FALSE | NA |
| Dmrt1i | TRUE | 0.502545266 | 1 | No Change | Not Significant | FALSE | NA |
| Dmrta1 | TRUE | 0.490367274 | 0.50718864 | No Change | Not Significant | TRUE | 0.552407391 |
| Dmrta2 | TRUE | 0.1664749 | 0.894883064 | No Change | Not Significant | FALSE | NA |
| Dmrtb1 | TRUE | 0.607470915 | 1 | No Change | Not Significant | FALSE | NA |
| Dmrtc2 | TRUE | 0.00429568 | 1 | No Change | Not Significant | FALSE | NA |
| Dmtf1 | TRUE | -0.047225954 | 0.725511065 | No Change | Not Significant | TRUE | -0.122333206 |
| Dmtf1l | TRUE | 1.72643412 | 1 | No Change | Not Significant | FALSE | NA |
| Dmtn | TRUE | 0.85744874 | 0.318422618 | No Change | Not Significant | FALSE | NA |
| Dmwd | TRUE | 1.252016689 | 2.68E-05 | UP | UP | TRUE | 0.25081479 |
| Dmxl1 | TRUE | 1.15238916 | 9.41E-65 | UP | UP | TRUE | 0.144675175 |
| Dmxl2 | TRUE | -0.575338649 | 0.097491654 | No Change | Not Significant | TRUE | 0.012573969 |
| Dna2 | TRUE | 0.190515312 | 0.51911069 | No Change | Not Significant | TRUE | -0.057257611 |
| Dnaaf1 | TRUE | 0.195906286 | 0.720291626 | No Change | Not Significant | TRUE | 0.344891489 |
| Dnaaf10 | TRUE | 0.41601399 | 0.00621724 | No Change | Significant Positive | TRUE | 0.182481534 |
| Dnaaf2 | TRUE | -0.079568161 | 0.875244964 | No Change | Not Significant | TRUE | -0.099807439 |
| Dnaaf3 | TRUE | 1.749390357 | 0.007512139 | UP | UP | TRUE | 0.397601792 |
| Dnaaf5 | TRUE | 0.177050252 | 0.307109811 | No Change | Not Significant | TRUE | 0.00246135 |
| Dnaaf8 | TRUE | 0.209532359 | 0.70086955 | No Change | Not Significant | TRUE | 0.203023487 |
| Dnaaf9 | TRUE | -0.277009716 | 0.21050366 | No Change | Not Significant | TRUE | -0.047389369 |
| Dnah1 | TRUE | 0.959520353 | 0.556711723 | No Change | Not Significant | FALSE | NA |
| Dnah10 | TRUE | 2.390287275 | 0.000254828 | UP | UP | TRUE | 1.450480348 |
| Dnah11 | TRUE | 0.884201969 | 0.52766379 | No Change | Not Significant | TRUE | 0.802300269 |
| Dnah12 | TRUE | 0.589992711 | 1 | No Change | Not Significant | FALSE | NA |
| Dnah14 | TRUE | -0.201009636 | 0.890186044 | No Change | Not Significant | TRUE | -0.333484987 |
| Dnah17 | TRUE | 1.25007412 | 0.028549731 | UP | UP | TRUE | -0.403840489 |
| Dnah2 | TRUE | 0.299090581 | 0.794347303 | No Change | Not Significant | TRUE | 0.163968727 |
| Dnah3 | TRUE | -0.526749853 | 0.876790748 | No Change | Not Significant | FALSE | NA |
| Dnah5 | TRUE | -0.652361845 | 0.038404592 | DOWN | DOWN | TRUE | 0.026130532 |
| Dnah6 | TRUE | 0.893647282 | 0.756061621 | No Change | Not Significant | FALSE | NA |
| Dnah7a | TRUE | 0.541622678 | 1 | No Change | Not Significant | FALSE | NA |
| Dnah7b | TRUE | 0.65915375 | 0.300890309 | No Change | Not Significant | TRUE | 0.112852098 |
| Dnah7c | TRUE | 0.977453185 | 1 | No Change | Not Significant | FALSE | NA |
| Dnah8 | TRUE | -0.0471231 | 0.903013746 | No Change | Not Significant | TRUE | -0.05209125 |
| Dnah9 | TRUE | -0.069538975 | 0.937420397 | No Change | Not Significant | TRUE | -0.041462362 |
| Dnai1 | TRUE | -0.040247029 | 1 | No Change | Not Significant | FALSE | NA |
| Dnai2 | TRUE | 1.248816367 | 1 | No Change | Not Significant | FALSE | NA |
| Dnai3 | TRUE | -0.29010248 | 0.668687928 | No Change | Not Significant | TRUE | 0.153260531 |
| Dnai4 | TRUE | 0.648887577 | 0.049834789 | UP | UP | TRUE | 0.257372749 |
| Dnaja1 | TRUE | -0.217816695 | 0.267881142 | No Change | Not Significant | TRUE | -0.156301914 |
| Dnaja2 | TRUE | -0.048343849 | 0.732906636 | No Change | Not Significant | TRUE | -0.027630756 |
| Dnaja3 | TRUE | -0.0777377 | 0.546957754 | No Change | Not Significant | TRUE | 0.123070256 |
| Dnaja4 | TRUE | -0.090769039 | 0.832872319 | No Change | Not Significant | TRUE | 0.150263935 |
| Dnajb1 | TRUE | -0.370763676 | 0.072698514 | No Change | Not Significant | TRUE | 0.149265797 |
| Dnajb11 | TRUE | 2.304742478 | 6.02E-237 | UP | UP | TRUE | 2.111108898 |
| Dnajb12 | TRUE | 0.011419138 | 0.938415991 | No Change | Not Significant | TRUE | -0.005573064 |
| Dnajb13 | TRUE | -1.175555297 | 2.01E-05 | DOWN | DOWN | TRUE | -1.236355788 |
| Dnajb14 | TRUE | 0.115640608 | 0.415941821 | No Change | Not Significant | TRUE | 0.102119158 |
| Dnajb2 | TRUE | -0.01193962 | 0.956997404 | No Change | Not Significant | TRUE | 0.177528598 |
| Dnajb3 | TRUE | 0.244415257 | 0.683555242 | No Change | Not Significant | TRUE | -0.038107772 |
| Dnajb4 | TRUE | -0.248472362 | 0.149293713 | No Change | Not Significant | TRUE | -0.164257452 |
| Dnajb5 | TRUE | 1.481651878 | 8.14E-16 | UP | UP | TRUE | 0.626069493 |
| Dnajb6 | TRUE | -0.327659776 | 0.000716166 | No Change | Significant Negative | TRUE | -0.101641935 |
| Dnajb7 | TRUE | -0.085836778 | 0.932001143 | No Change | Not Significant | TRUE | -0.969715892 |
| Dnajb8 | TRUE | -0.407956891 | 1 | No Change | Not Significant | FALSE | NA |
| Dnajb9 | TRUE | 1.57992868 | 1.53E-71 | UP | UP | TRUE | 0.798429408 |
| Dnajc1 | TRUE | 0.134198479 | 0.3013667 | No Change | Not Significant | TRUE | 0.003211441 |
| Dnajc10 | TRUE | 0.646553531 | 0.001846439 | UP | UP | TRUE | 0.425769789 |
| Dnajc11 | TRUE | -0.106820343 | 0.327361658 | No Change | Not Significant | TRUE | 0.141498322 |
| Dnajc12 | TRUE | -0.187471245 | 0.510451645 | No Change | Not Significant | TRUE | 0.335708919 |
| Dnajc13 | TRUE | 0.085971647 | 0.38986575 | No Change | Not Significant | TRUE | 0.024397433 |
| Dnajc14 | TRUE | -0.144247371 | 0.00556186 | No Change | Significant Negative | TRUE | -0.067049994 |
| Dnajc15 | TRUE | 0.002082908 | 0.988088396 | No Change | Not Significant | TRUE | 0.12490322 |
| Dnajc16 | TRUE | -0.083749131 | 0.396311693 | No Change | Not Significant | TRUE | 0.051046723 |
| Dnajc17 | TRUE | -0.254347653 | 0.095640734 | No Change | Not Significant | TRUE | 0.015187535 |
| Dnajc18 | TRUE | 0.049033284 | 0.882319111 | No Change | Not Significant | TRUE | 0.136103753 |
| Dnajc19 | TRUE | -0.040643835 | 0.77680602 | No Change | Not Significant | TRUE | 0.03567288 |
| Dnajc2 | TRUE | -0.009864897 | 0.957266079 | No Change | Not Significant | TRUE | -0.122831885 |
| Dnajc21 | TRUE | 0.005455805 | 0.976016991 | No Change | Not Significant | TRUE | 0.039691214 |
| Dnajc22 | TRUE | 0.006994416 | 0.948996349 | No Change | Not Significant | TRUE | -0.00631637 |
| Dnajc24 | TRUE | 0.124214846 | 0.618855346 | No Change | Not Significant | TRUE | 0.204996488 |
| Dnajc25 | TRUE | 0.491775293 | 0.000316096 | No Change | Significant Positive | TRUE | 0.098351962 |
| Dnajc27 | TRUE | 0.286769818 | 0.522023104 | No Change | Not Significant | TRUE | -0.187327392 |
| Dnajc28 | TRUE | -0.645275281 | 0.006724263 | DOWN | DOWN | TRUE | -0.109685674 |
| Dnajc3 | TRUE | 2.507501229 | 7.90E-117 | UP | UP | TRUE | 1.981806734 |
| Dnajc30 | TRUE | -0.197632001 | 0.147967906 | No Change | Not Significant | TRUE | -0.07399976 |
| Dnajc4 | TRUE | -0.118255223 | 0.350538094 | No Change | Not Significant | TRUE | -0.019777686 |
| Dnajc5 | TRUE | 0.076733365 | 0.337876038 | No Change | Not Significant | TRUE | -0.007196878 |
| Dnajc6 | TRUE | 0.805730111 | 0.490623547 | No Change | Not Significant | TRUE | 0.318246664 |
| Dnajc7 | TRUE | -0.115263675 | 0.265200548 | No Change | Not Significant | TRUE | -0.094283937 |
| Dnajc8 | TRUE | -0.401302135 | 6.75E-07 | No Change | Significant Negative | TRUE | -0.203653752 |
| Dnajc9 | TRUE | 0.148414171 | 0.640805153 | No Change | Not Significant | TRUE | 0.169470033 |
| Dnal1 | TRUE | 0.339192921 | 0.4164509 | No Change | Not Significant | TRUE | 0.293044456 |
| Dnal4 | TRUE | 0.099833088 | 0.43912271 | No Change | Not Significant | TRUE | 0.10724542 |
| Dnali1 | TRUE | -0.791343041 | 1 | No Change | Not Significant | FALSE | NA |
| Dnase1 | TRUE | 0.275285559 | 0.647921517 | No Change | Not Significant | TRUE | -0.06998811 |
| Dnase1l1 | TRUE | -0.104649825 | 0.60223179 | No Change | Not Significant | TRUE | -0.003380789 |
| Dnase1l2 | TRUE | -0.617902131 | 0.161179189 | No Change | Not Significant | TRUE | -0.009007812 |
| Dnase1l3 | TRUE | 0.224572311 | 1 | No Change | Not Significant | FALSE | NA |
| Dnase2a | TRUE | -0.260077381 | 0.280913726 | No Change | Not Significant | TRUE | -0.001668937 |
| Dnase2b | TRUE | 2.746451838 | 1 | No Change | Not Significant | FALSE | NA |
| Dnd1 | TRUE | -0.00751441 | 0.994317649 | No Change | Not Significant | TRUE | 0.688225986 |
| Dner | TRUE | -0.195273738 | 0.904906279 | No Change | Not Significant | TRUE | -0.090834809 |
| Dnhd1 | TRUE | 0.259944861 | 0.797446101 | No Change | Not Significant | TRUE | -0.038934366 |
| Dnlz | TRUE | -0.274625287 | 0.012761429 | No Change | Significant Negative | TRUE | -0.033794993 |
| Dnm1 | TRUE | 0.440519662 | 0.437741278 | No Change | Not Significant | TRUE | -0.742164037 |
| Dnm1l | TRUE | 0.145106225 | 0.092148775 | No Change | Not Significant | TRUE | 0.079250085 |
| Dnm2 | TRUE | 0.182026993 | 0.166571051 | No Change | Not Significant | TRUE | -0.020429682 |
| Dnm3 | TRUE | 0.163119614 | 0.924166034 | No Change | Not Significant | FALSE | NA |
| Dnm3os | TRUE | 0.004898909 | 1 | No Change | Not Significant | FALSE | NA |
| Dnmbp | TRUE | -0.232977146 | 0.026393326 | No Change | Significant Negative | TRUE | -0.137982156 |
| Dnmt1 | TRUE | 0.087229584 | 0.725788994 | No Change | Not Significant | TRUE | 0.00801515 |
| Dnmt3a | TRUE | -0.214170615 | 0.05840978 | No Change | Not Significant | TRUE | -0.21265956 |
| Dnmt3aos | TRUE | -0.529259691 | 1 | No Change | Not Significant | FALSE | NA |
| Dnmt3b | TRUE | -0.107899985 | 0.799123664 | No Change | Not Significant | TRUE | 0.149889586 |
| Dnmt3l | TRUE | 0.511849205 | 1 | No Change | Not Significant | FALSE | NA |
| Dnpep | TRUE | 0.707757182 | 1.68E-24 | UP | UP | TRUE | 0.531655408 |
| Dnph1 | TRUE | -0.302354623 | 0.332817843 | No Change | Not Significant | TRUE | -0.57570903 |
| Dnttip1 | TRUE | 0.140996855 | 0.197148208 | No Change | Not Significant | TRUE | 0.09601431 |
| Dnttip2 | TRUE | -0.042430381 | 0.801630291 | No Change | Not Significant | TRUE | -0.09092489 |
| Doc2a | TRUE | 0.239025907 | 1 | No Change | Not Significant | FALSE | NA |
| Doc2b | TRUE | 0.031356704 | 1 | No Change | Not Significant | FALSE | NA |
| Doc2g | TRUE | 0.07272643 | 0.915204382 | No Change | Not Significant | TRUE | -0.298878665 |
| Dock1 | TRUE | -0.101691332 | 0.091363531 | No Change | Not Significant | TRUE | -0.067795732 |
| Dock10 | TRUE | -0.216642107 | 0.811870577 | No Change | Not Significant | TRUE | 0.285459572 |
| Dock11 | TRUE | -0.207000826 | 0.312060508 | No Change | Not Significant | TRUE | -0.188587615 |
| Dock2 | TRUE | 0.011563189 | 0.988088396 | No Change | Not Significant | TRUE | 0.517745751 |
| Dock3 | TRUE | 0.527468382 | 0.561130066 | No Change | Not Significant | TRUE | 0.360013803 |
| Dock4 | TRUE | -0.03444417 | 0.958199105 | No Change | Not Significant | TRUE | -0.273325057 |
| Dock5 | TRUE | -0.064996433 | 0.701521284 | No Change | Not Significant | TRUE | -0.04445215 |
| Dock6 | TRUE | 0.307697431 | 0.283960728 | No Change | Not Significant | TRUE | -0.118909239 |
| Dock7 | TRUE | 0.187003865 | 0.003959279 | No Change | Significant Positive | TRUE | -0.011535625 |
| Dock8 | TRUE | -0.431159217 | 1.77E-05 | No Change | Significant Negative | TRUE | -0.01237756 |
| Dock9 | TRUE | -0.132028323 | 0.26018046 | No Change | Not Significant | TRUE | -0.141207955 |
| Dohh | TRUE | 0.121196264 | 0.515232502 | No Change | Not Significant | TRUE | 0.084390343 |
| Dok1 | TRUE | -0.332319378 | 0.074952393 | No Change | Not Significant | TRUE | -0.20722985 |
| Dok2 | TRUE | -0.305369322 | 0.776544903 | No Change | Not Significant | TRUE | 0.725139862 |
| Dok3 | TRUE | 1.144354927 | 0.045481814 | UP | UP | TRUE | 0.623471014 |
| Dok4 | TRUE | -0.335872293 | 0.005445042 | No Change | Significant Negative | TRUE | -0.013273997 |
| Dok6 | TRUE | -1.16894898 | 1 | No Change | Not Significant | FALSE | NA |
| Dok7 | TRUE | -0.533036659 | 0.028755531 | DOWN | DOWN | TRUE | -0.15433492 |
| Dolk | TRUE | 0.102119786 | 0.384098334 | No Change | Not Significant | TRUE | 0.114283334 |
| Dolpp1 | TRUE | 0.858164341 | 5.65E-19 | UP | UP | TRUE | 0.178175927 |
| Donson | TRUE | 0.652821656 | 0.000415076 | UP | UP | TRUE | 0.025448633 |
| Dop1a | TRUE | -0.002335981 | 0.982347769 | No Change | Not Significant | TRUE | -0.106753892 |
| Dop1b | TRUE | -0.000727246 | 0.996939463 | No Change | Not Significant | TRUE | 0.058421479 |
| Dot1l | TRUE | 0.228836537 | 0.386709513 | No Change | Not Significant | TRUE | -0.195751908 |
| Dpagt1 | TRUE | 0.521307119 | 5.32E-11 | UP | UP | TRUE | 0.239961086 |
| Dpcd | TRUE | 0.1117364 | 0.406467904 | No Change | Not Significant | TRUE | 0.169873009 |
| Dpep1 | TRUE | -0.426998924 | 0.019150767 | No Change | Significant Negative | TRUE | -0.04935544 |
| Dpep2 | TRUE | 0.689446734 | 1 | No Change | Not Significant | FALSE | NA |
| Dpf1 | TRUE | -0.928771927 | 1 | No Change | Not Significant | FALSE | NA |
| Dpf2 | TRUE | -0.124304533 | 0.113038434 | No Change | Not Significant | TRUE | -0.055619614 |
| Dpf3 | TRUE | -0.085850713 | 0.884532517 | No Change | Not Significant | TRUE | -0.002471397 |
| Dph1 | TRUE | -0.32731663 | 0.027402344 | No Change | Significant Negative | TRUE | -0.126542121 |
| Dph2 | TRUE | 0.018604741 | 0.94451598 | No Change | Not Significant | TRUE | -0.033546047 |
| Dph3 | TRUE | -0.027610296 | 0.882690208 | No Change | Not Significant | TRUE | 0.041858882 |
| Dph5 | TRUE | -0.111358207 | 0.713786099 | No Change | Not Significant | TRUE | -0.021268377 |
| Dph6 | TRUE | 0.241763558 | 0.278322371 | No Change | Not Significant | TRUE | -0.092627134 |
| Dph7 | TRUE | 0.157685076 | 0.449849479 | No Change | Not Significant | TRUE | 0.055949955 |
| Dpm1 | TRUE | -0.184326674 | 0.474234146 | No Change | Not Significant | TRUE | -0.420740308 |
| Dpm2 | TRUE | 0.167492718 | 0.263984696 | No Change | Not Significant | TRUE | 0.121284502 |
| Dpm3 | TRUE | -0.313758526 | 0.007393398 | No Change | Significant Negative | TRUE | 0.096410354 |
| Dpp10 | TRUE | -0.124615777 | 1 | No Change | Not Significant | FALSE | NA |
| Dpp3 | TRUE | -0.047368559 | 0.698491194 | No Change | Not Significant | TRUE | 0.103348754 |
| Dpp4 | TRUE | 0.448495547 | 0.000109485 | No Change | Significant Positive | TRUE | 0.030637048 |
| Dpp6 | TRUE | 0.404882578 | 0.924222269 | No Change | Not Significant | FALSE | NA |
| Dpp7 | TRUE | -1.061779565 | 5.55E-27 | DOWN | DOWN | TRUE | -0.63213071 |
| Dpp8 | TRUE | -0.056454357 | 0.687965835 | No Change | Not Significant | TRUE | -0.006278571 |
| Dpp9 | TRUE | 0.373874438 | 0.000249836 | No Change | Significant Positive | TRUE | 0.29310449 |
| Dppa1 | TRUE | -0.428165315 | 1 | No Change | Not Significant | FALSE | NA |
| Dppa2 | TRUE | 0.604796954 | 1 | No Change | Not Significant | FALSE | NA |
| Dppa3 | TRUE | -2.020539754 | 1 | No Change | Not Significant | FALSE | NA |
| Dpt | TRUE | -1.526889254 | 1 | No Change | Not Significant | FALSE | NA |
| Dpy19l1 | TRUE | 2.094741287 | 3.07E-57 | UP | UP | TRUE | 1.022635063 |
| Dpy19l3 | TRUE | 0.003323706 | 0.989246416 | No Change | Not Significant | TRUE | -0.166224263 |
| Dpy19l4 | TRUE | 0.031145823 | 0.853579543 | No Change | Not Significant | TRUE | -0.072891452 |
| Dpy30 | TRUE | -0.075882907 | 0.447999484 | No Change | Not Significant | TRUE | -0.039457144 |
| Dpyd | TRUE | -0.586024374 | 4.40E-08 | DOWN | DOWN | TRUE | -0.243581055 |
| Dpys | TRUE | 0.094995913 | 1 | No Change | Not Significant | FALSE | NA |
| Dpysl2 | TRUE | 0.092049697 | 0.755650645 | No Change | Not Significant | TRUE | 0.236970805 |
| Dpysl3 | TRUE | -0.668799942 | 0.000888152 | DOWN | DOWN | TRUE | 0.062404523 |
| Dpysl4 | TRUE | 1.24919737 | 1 | No Change | Not Significant | FALSE | NA |
| Dpysl5 | TRUE | 0.221608535 | 0.927761944 | No Change | Not Significant | FALSE | NA |
| Dqx1 | TRUE | -0.145964937 | 0.336688663 | No Change | Not Significant | TRUE | -0.175552961 |
| Dr1 | TRUE | 0.084562741 | 0.568119977 | No Change | Not Significant | TRUE | -0.018707098 |
| Dram1 | TRUE | -0.573362416 | 1 | No Change | Not Significant | FALSE | NA |
| Dram2 | TRUE | 0.021806856 | 0.915295615 | No Change | Not Significant | TRUE | 0.003023299 |

Supplementary Table 3

| geneID | Cond1_DF_Detected | Cond1_DF_ValueCol | Cond1_DF_PadjCol | Cond1_DF_Cutoff |
|---|---|---|---|---|
| Drap1 | TRUE | 0.057944721 | 0.669812138 | No Change |
| Draxin | TRUE | -0.517626611 | 1 | No Change |
| Drc1 | TRUE | -1.241354517 | 0.006636053 | DOWN |
| Drc3 | TRUE | 0.407597803 | 0.182830803 | No Change |
| Drc7 | TRUE | 0.606312709 | 1 | No Change |
| Drd1 | TRUE | 1.138520494 | 1 | No Change |
| Drd2 | TRUE | -2.071360181 | 1 | No Change |
| Drd3 | TRUE | -0.326577474 | 0.697221779 | No Change |
| Drd4 | TRUE | -0.068691369 | 1 | No Change |
| Drg1 | TRUE | 0.129891196 | 0.356505326 | No Change |
| Drg2 | TRUE | 0.112639271 | 0.405734564 | No Change |
| Drosha | TRUE | 0.186696205 | 0.009870396 | No Change |
| Drp2 | TRUE | 1.578727141 | 1 | No Change |
| Dsc1 | TRUE | 0.097860256 | 1 | No Change |
| Dsc2 | TRUE | -0.098262823 | 0.551494032 | No Change |
| Dsc3 | TRUE | -0.364774476 | 1 | No Change |
| Dscam | TRUE | 1.523701076 | 0.270127107 | No Change |
| Dscaml1 | TRUE | -1.247281642 | 0.389463531 | No Change |
| Dscc1 | TRUE | 0.371988643 | 0.363647302 | No Change |
| Dse | TRUE | 0.694907274 | 1 | No Change |
| Dsel | TRUE | 0.553876144 | 0.000688653 | UP |
| Dsg1a | TRUE | 0.176025634 | 1 | No Change |
| Dsg1c | TRUE | 0.978897477 | 1 | No Change |
| Dsg2 | TRUE | -0.083266654 | 0.695624496 | No Change |
| Dsg3 | TRUE | -0.102571717 | 0.912240897 | No Change |
| Dsg4 | TRUE | -0.253049098 | 1 | No Change |
| Dsn1 | TRUE | 0.532999956 | 0.02379884 | UP |
| Dsp | TRUE | -0.301353385 | 0.014436949 | No Change |
| Dspp | TRUE | 0.176019553 | 1 | No Change |
| Dst | TRUE | -0.143124414 | 0.593868753 | No Change |
| Dstn | TRUE | 0.136957725 | 0.358767397 | No Change |
| Dstyk | TRUE | -0.3087111 | 0.004741185 | No Change |
| Dtd1 | TRUE | 0.018446517 | 0.936355518 | No Change |
| Dtd2 | TRUE | 0.073881297 | 0.699148013 | No Change |
| Dthd1 | TRUE | 0.095876308 | 1 | No Change |
| Dtl | TRUE | 0.35063343 | 0.147129207 | No Change |
| Dtna | TRUE | 0.302033553 | 0.908985828 | No Change |
| Dtnb | TRUE | 0.560275799 | 0.001413095 | UP |
| Dtnbp1 | TRUE | -0.085645151 | 0.657120605 | No Change |
| Dtwd1 | TRUE | 0.289313759 | 0.220499674 | No Change |
| Dtwd2 | TRUE | 0.200135044 | 0.615582315 | No Change |
| Dtx1 | TRUE | 0.519269587 | 0.429717907 | No Change |
| Dtx2 | TRUE | 0.057029675 | 0.789569374 | No Change |
| Dtx3 | TRUE | -0.869589677 | 0.000511803 | DOWN |
| Dtx3l | TRUE | -0.245512516 | 0.250425106 | No Change |
| Dtx4 | TRUE | 0.512270416 | 6.52E-05 | UP |
| Dtymk | TRUE | 0.510810865 | 0.001199022 | UP |
| Duox1 | TRUE | 0.564164703 | 0.249818224 | No Change |
| Duox2 | TRUE | 0.598252058 | 0.064232353 | No Change |
| Duoxa1 | TRUE | 0.356549314 | 0.618855346 | No Change |
| Duoxa2 | TRUE | 1.336060422 | 0.000397981 | UP |
| Dus1l | TRUE | -0.01088832 | 0.947202684 | No Change |
| Dus2 | TRUE | 0.029537336 | 0.912640622 | No Change |
| Dus3l | TRUE | 0.080428848 | 0.425536825 | No Change |
| Dus4l | TRUE | 0.113169026 | 0.732676764 | No Change |
| Dusp1 | TRUE | -0.479814061 | 0.037986091 | No Change |
| Dusp10 | TRUE | 0.839982418 | 0.005513702 | UP |
| Dusp11 | TRUE | 0.003089395 | 0.981701728 | No Change |
| Dusp12 | TRUE | 0.171532738 | 0.524360158 | No Change |
| Dusp13 | TRUE | -0.275016412 | 0.782086507 | No Change |
| Dusp14 | TRUE | 0.847410589 | 0.001109775 | UP |
| Dusp15 | TRUE | 1.21380942 | 1 | No Change |
| Dusp16 | TRUE | -0.215936183 | 0.189409212 | No Change |
| Dusp18 | TRUE | -0.22121872 | 0.115258628 | No Change |
| Dusp19 | TRUE | 0.088399733 | 0.683370901 | No Change |
| Dusp2 | TRUE | 1.378170869 | 0.053152102 | No Change |
| Dusp22 | TRUE | 0.573922108 | 0.027207712 | UP |
| Dusp23 | TRUE | 1.560770569 | 1.12E-41 | UP |
| Dusp26 | TRUE | 1.190007392 | 1 | No Change |
| Dusp27 | TRUE | -0.374105603 | 0.761222618 | No Change |
| Dusp28 | TRUE | -0.077617243 | 0.755320172 | No Change |
| Dusp3 | TRUE | -0.307201474 | 0.013089272 | No Change |
| Dusp4 | TRUE | -0.267940005 | 0.3013667 | No Change |
| Dusp5 | TRUE | -0.272105109 | 0.679736277 | No Change |
| Dusp6 | TRUE | 0.585678529 | 0.000545662 | UP |
| Dusp7 | TRUE | 0.255776407 | 0.262304772 | No Change |
| Dusp8 | TRUE | 0.269138306 | 0.37700811 | No Change |
| Dusp9 | TRUE | 0.564103837 | 0.292089426 | No Change |
| Dut | TRUE | 0.218862716 | 0.397518385 | No Change |
| Duxbl1 | TRUE | -1.259965774 | 1 | No Change |
| Duxf3 | TRUE | 0.523251474 | 1 | No Change |
| Dvl1 | TRUE | -0.025735671 | 0.897263004 | No Change |
| Dvl2 | TRUE | -0.088250779 | 0.69793377 | No Change |
| Dvl3 | TRUE | 0.237422529 | 0.347734431 | No Change |
| Dxo | TRUE | 0.082635177 | 0.488304745 | No Change |
| Dym | TRUE | 0.009389227 | 0.95811623 | No Change |
| Dync1h1 | TRUE | -0.000236844 | 0.998483602 | No Change |
| Dync1i1 | TRUE | 0.822266764 | 5.75E-05 | UP |
| Dync1i2 | TRUE | 0.092256529 | 0.329724247 | No Change |
| Dync1li1 | TRUE | 0.075731252 | 0.557076931 | No Change |
| Dync1li2 | TRUE | 0.017335951 | 0.916410806 | No Change |
| Dync2h1 | TRUE | 0.05514615 | 0.826265833 | No Change |
| Dync2i1 | TRUE | -0.086867136 | 0.699388858 | No Change |
| Dync2i2 | TRUE | 0.175033866 | 0.356436387 | No Change |
| Dync2li1 | TRUE | 0.522341618 | 0.173002023 | No Change |
| Dynll1 | TRUE | 0.345499564 | 9.54E-05 | No Change |
| Dynll2 | TRUE | 0.109407329 | 0.438122528 | No Change |
| Dynlrb1 | TRUE | -0.047396376 | 0.696686775 | No Change |
| Dynlt1a | TRUE | -0.065543253 | 0.872440598 | No Change |
| Dynlt1b | TRUE | -0.698202161 | 0.064246329 | No Change |
| Dynlt1c | TRUE | -0.430344196 | 0.358787591 | No Change |
| Dynlt1f | TRUE | 0.064519648 | 0.859382261 | No Change |
| Dynlt2b | TRUE | 0.146697733 | 0.735467298 | No Change |
| Dynlt3 | TRUE | 0.076247507 | 0.733377497 | No Change |
| Dynlt4 | TRUE | -0.00781399 | 0.989031323 | No Change |
| Dyrk1a | TRUE | 0.124065643 | 0.021615492 | No Change |
| Dyrk1b | TRUE | -0.340468081 | 0.230625736 | No Change |
| Dyrk2 | TRUE | -0.29240713 | 0.048327356 | No Change |
| Dyrk3 | TRUE | 0.294104743 | 0.537548029 | No Change |
| Dyrk4 | TRUE | -0.693741968 | 1 | No Change |
| Dysf | TRUE | 0.727573224 | 0.834370416 | No Change |
| Dytn | TRUE | 0.531585655 | 1 | No Change |
| Dzank1 | TRUE | -0.264109551 | 1 | No Change |
| Dzip1 | TRUE | 0.808872666 | 0.250788441 | No Change |
| Dzip1l | TRUE | -0.883189611 | 0.010564875 | DOWN |
| Dzip3 | TRUE | 0.539469407 | 0.000134753 | UP |
| E030013I19Rik | TRUE | 0.258010087 | 1 | No Change |
| E030018B13Rik | TRUE | -0.566976216 | 0.196696774 | No Change |
| E030030I06Rik | TRUE | 0.162017959 | 0.542047573 | No Change |
| E030044B06Rik | TRUE | 1.083040003 | 1 | No Change |
| E130006D01Rik | TRUE | 0.609618502 | 1 | No Change |
| E130008D07Rik | TRUE | -0.145818592 | 0.904254429 | No Change |
| E130018N17Rik | TRUE | -0.358477698 | 1 | No Change |
| E130018O15Rik | TRUE | -1.877823634 | 1 | No Change |
| E130102H24Rik | TRUE | -0.530033995 | 0.431111794 | No Change |
| E130201H02Rik | TRUE | 1.235167738 | 1 | No Change |
| E130215H24Rik | TRUE | 0.297899637 | 0.638701711 | No Change |
| E130307A14Rik | TRUE | 0.105969401 | 0.824348168 | No Change |
| E130308A19Rik | TRUE | 0.149300633 | 0.324750031 | No Change |
| E130310I04Rik | TRUE | 0.288730807 | 0.838607402 | No Change |
| E130311K13Rik | TRUE | 0.091811691 | 0.840340998 | No Change |
| E130317F20Rik | TRUE | 0.506302105 | 0.243576926 | No Change |
| E230013L22Rik | TRUE | 0.550082355 | 0.112356092 | No Change |
| E230016K23Rik | TRUE | -0.10204974 | 1 | No Change |
| E230016M11Rik | TRUE | -0.263807569 | 0.69281035 | No Change |
| E230025N22Rik | TRUE | 0.197785107 | 0.41642117 | No Change |
| E230029C05Rik | TRUE | -0.097033289 | 1 | No Change |
| E2f1 | TRUE | 0.822778533 | 0.00056319 | UP |
| E2f2 | TRUE | 1.203261947 | 5.76E-12 | UP |
| E2f3 | TRUE | 0.191383164 | 0.110941817 | No Change |
| E2f4 | TRUE | 0.135946588 | 0.09897928 | No Change |
| E2f5 | TRUE | 0.009657573 | 0.948731562 | No Change |
| E2f6 | TRUE | 0.127975903 | 0.490477055 | No Change |
| E2f7 | TRUE | 1.897761946 | 4.37E-17 | UP |
| E2f8 | TRUE | 0.502016328 | 0.066683451 | No Change |
| E330013P04Rik | TRUE | 0.825004905 | 1 | No Change |
| E330032C10Rik | TRUE | -0.536901519 | 1 | No Change |
| E430018J23Rik | TRUE | -0.293836313 | 0.016610875 | No Change |
| E430024P14Rik | TRUE | 0.096412519 | 0.878808271 | No Change |
| E4f1 | TRUE | -0.009944488 | 0.965802209 | No Change |
| Eaf1 | TRUE | 0.147253102 | 0.048232278 | No Change |
| Eaf2 | TRUE | 0.113232521 | 0.908481483 | No Change |
| Eapp | TRUE | -0.274406334 | 0.007009884 | No Change |
| Ears2 | TRUE | 0.23622927 | 0.196362629 | No Change |
| Ebag9 | TRUE | 0.365378522 | 0.001483889 | No Change |
| Ebf1 | TRUE | 0.36191929 | 0.929155653 | No Change |
| Ebf2 | TRUE | -1.181254602 | 1 | No Change |
| Ebf3 | TRUE | -0.169677051 | 0.877468906 | No Change |
| Ebf4 | TRUE | 0.93536173 | 1 | No Change |
| Ebi3 | TRUE | -0.072183023 | 1 | No Change |
| Ebna1bp2 | TRUE | -0.08068387 | 0.583549549 | No Change |
| Ebp | TRUE | 1.674542875 | 1.79E-63 | UP |
| Ebpl | TRUE | -0.0645994 | 0.79831751 | No Change |
| Ecd | TRUE | 0.104360966 | 0.367540047 | No Change |
| Ece1 | TRUE | -0.246221209 | 0.000632887 | No Change |
| Ece2 | TRUE | 1.896474723 | 1 | No Change |
| Ecel1 | TRUE | 1.937850432 | 1 | No Change |
| Ech1 | TRUE | -0.068751519 | 0.524504007 | No Change |
| Echdc1 | TRUE | -0.014075545 | 0.947202684 | No Change |
| Echdc2 | TRUE | -0.132203556 | 0.315341543 | No Change |
| Echdc3 | TRUE | -0.00267418 | 0.990774286 | No Change |
| Echs1 | TRUE | -0.562855336 | 3.50E-13 | DOWN |
| Eci1 | TRUE | -0.044353019 | 0.821176574 | No Change |
| Eci2 | TRUE | -0.082033637 | 0.471988664 | No Change |
| Eci3 | TRUE | -0.775927052 | 8.85E-10 | DOWN |
| Ecm1 | TRUE | 0.996754086 | 0.054382773 | No Change |
| Ecm2 | TRUE | 0.942085417 | 0.159434952 | No Change |
| Ecpas | TRUE | 0.003253207 | 0.985568894 | No Change |
| Ecscr | TRUE | 0.035553138 | 0.930749284 | No Change |
| Ecsit | TRUE | -0.015286207 | 0.938630585 | No Change |
| Ect2 | TRUE | 0.768012157 | 0.0001049 | UP |
| Ect2l | TRUE | 1.508894171 | 1 | No Change |
| Eda | TRUE | 0.62061009 | 0.715626628 | No Change |
| Eda2r | TRUE | 0.532216894 | 0.447999484 | No Change |
| Edar | TRUE | -1.310724405 | 1 | No Change |
| Edaradd | TRUE | 1.882179929 | 8.27E-06 | UP |
| Edc3 | TRUE | -0.006120192 | 0.975869522 | No Change |
| Edc4 | TRUE | 0.072372396 | 0.359584034 | No Change |
| Edem1 | TRUE | 1.957591901 | 1.45E-195 | UP |
| Edem2 | TRUE | 0.503062774 | 2.03E-08 | UP |
| Edem3 | TRUE | 0.404739527 | 0.00080104 | No Change |
| Edf1 | TRUE | -0.043169704 | 0.766711421 | No Change |

Supplementary Table 3

| geneID | Cond1_DF_Detected | Cond1_DF_ValueCol | Cond1_DF_PadjCol | Cond1_DF_Cutoff | Cond1_DF_Cutoff_Specific.x | Cond2_DF_Detected | Cond2_DF_ValueCol |
|---|---|---|---|---|---|---|---|
| Edil3 | TRUE | -2.240747123 | 1 | No Change | Not Significant | FALSE | NA |
| Edn1 | TRUE | -0.779291268 | 0.125633944 | No Change | Not Significant | TRUE | -0.626992195 |
| Edn2 | TRUE | -0.817004158 | 1.50E-05 | DOWN | DOWN | TRUE | -0.408891674 |
| Edn3 | TRUE | 0.155649413 | 0.767483504 | No Change | Not Significant | TRUE | 0.53878033 |
| Ednra | TRUE | 2.679482922 | 1 | No Change | Not Significant | FALSE | NA |
| Ednrb | TRUE | 0.073768977 | 1 | No Change | Not Significant | FALSE | NA |
| Edrf1 | TRUE | 0.30260664 | 0.003288476 | No Change | Significant Positive | TRUE | 0.096809213 |
| Eea1 | TRUE | 0.131389135 | 0.235970255 | No Change | Not Significant | TRUE | -0.019676013 |
| Eed | TRUE | 0.097725079 | 0.429954629 | No Change | Not Significant | TRUE | 0.027658654 |
| Eef1a1 | TRUE | -0.159397411 | 0.328982111 | No Change | Not Significant | TRUE | 0.007509912 |
| Eef1a2 | TRUE | 1.107311508 | 1 | No Change | Not Significant | FALSE | NA |
| Eef1akmt1 | TRUE | -0.017421873 | 0.935980063 | No Change | Not Significant | TRUE | 0.15297159 |
| Eef1akmt2 | TRUE | 0.314774741 | 0.00731937 | No Change | Significant Positive | TRUE | 0.137879525 |
| Eef1akmt3 | TRUE | 1.335518002 | 9.92E-05 | UP | UP | TRUE | 0.858207639 |
| Eef1akmt4 | TRUE | -0.126151102 | 0.712785998 | No Change | Not Significant | TRUE | -0.049919365 |
| Eef1aknmt | TRUE | -0.227507447 | 0.216495012 | No Change | Not Significant | TRUE | -0.306719689 |
| Eef1b2 | TRUE | -0.218693747 | 0.086284421 | No Change | Not Significant | TRUE | -0.094183988 |
| Eef1d | TRUE | 0.209293048 | 0.054259732 | No Change | Not Significant | TRUE | 0.122315878 |
| Eef1e1 | TRUE | 0.106536372 | 0.595844868 | No Change | Not Significant | TRUE | -0.115384526 |
| Eef1g | TRUE | -0.199441905 | 0.163952289 | No Change | Not Significant | TRUE | 0.061705019 |
| Eef2 | TRUE | -0.077284162 | 0.510848385 | No Change | Not Significant | TRUE | -0.048622895 |
| Eef2k | TRUE | -0.127532578 | 0.326885802 | No Change | Not Significant | TRUE | 0.099985236 |
| Eef2kmt | TRUE | -0.119614751 | 0.52065112 | No Change | Not Significant | TRUE | -0.021472175 |
| Eefsec | TRUE | 0.097199558 | 0.508005086 | No Change | Not Significant | TRUE | 0.25883264 |
| Eepd1 | TRUE | -0.169550877 | 0.590110273 | No Change | Not Significant | TRUE | -0.22142427 |
| Efcab1 | TRUE | -0.924080464 | 0.201525415 | No Change | Not Significant | TRUE | 0.351930032 |
| Efcab11 | TRUE | 1.230798197 | 1.49E-07 | UP | UP | TRUE | 0.308289852 |
| Efcab12 | TRUE | 0.544336616 | 0.446602599 | No Change | Not Significant | TRUE | 0.1515285 |
| Efcab14 | TRUE | -0.309869821 | 0.002331022 | No Change | Significant Negative | TRUE | -0.050767869 |
| Efcab15 | TRUE | 2.223069118 | 1 | No Change | Not Significant | FALSE | NA |
| Efcab2 | TRUE | -0.192699685 | 0.546769252 | No Change | Not Significant | TRUE | -0.145258856 |
| Efcab5 | TRUE | 0.094335114 | 0.751588072 | No Change | Not Significant | TRUE | 0.029803591 |
| Efcab6 | TRUE | 1.235209262 | 1 | No Change | Not Significant | FALSE | NA |
| Efcab7 | TRUE | 0.195567907 | 0.599541641 | No Change | Not Significant | TRUE | 0.077621902 |
| Efcab8 | TRUE | 1.574916656 | 0.480535393 | No Change | Not Significant | FALSE | NA |
| Efcab9 | TRUE | -1.595917759 | 1 | No Change | Not Significant | FALSE | NA |
| Efcc1 | TRUE | -0.393824995 | 1 | No Change | Not Significant | FALSE | NA |
| Efemp1 | TRUE | -0.243586078 | 1 | No Change | Not Significant | FALSE | NA |
| Efemp2 | TRUE | 0.563523196 | 0.141278128 | No Change | Not Significant | TRUE | 0.091972557 |
| Efhb | TRUE | -0.439327912 | 0.297019741 | No Change | Not Significant | TRUE | 0.431860398 |
| Efhc1 | TRUE | 1.705478032 | 0.000462789 | UP | UP | TRUE | 0.558707607 |
| Efhc2 | TRUE | 1.102592126 | 1 | No Change | Not Significant | FALSE | NA |
| Efhd1 | TRUE | 0.456594202 | 0.628507412 | No Change | Not Significant | TRUE | 0.0765026 |
| Efhd2 | TRUE | -0.21715897 | 0.036859691 | No Change | Significant Negative | TRUE | -0.010340418 |
| Efl1 | TRUE | -0.061598343 | 0.604102573 | No Change | Not Significant | TRUE | -0.017305975 |
| Efna1 | TRUE | -0.304931311 | 0.090795126 | No Change | Not Significant | TRUE | 0.120533584 |
| Efna2 | TRUE | 0.699955963 | 0.405848226 | No Change | Not Significant | FALSE | NA |
| Efna3 | TRUE | -0.537145893 | 0.435838332 | No Change | Not Significant | TRUE | -0.342697274 |
| Efna4 | TRUE | 0.262945115 | 0.180947387 | No Change | Not Significant | TRUE | 0.013939896 |
| Efna5 | TRUE | -1.837166519 | 1 | No Change | Not Significant | FALSE | NA |
| Efnb1 | TRUE | 0.017311088 | 0.930204755 | No Change | Not Significant | TRUE | 0.087763651 |
| Efnb2 | TRUE | 0.5481742 | 0.004596489 | UP | UP | TRUE | 0.115846689 |
| Efnb3 | TRUE | 0.763982366 | 0.42452042 | No Change | Not Significant | TRUE | 0.393819308 |
| Efr3a | TRUE | 0.056387225 | 0.71028091 | No Change | Not Significant | TRUE | -0.068313822 |
| Efr3b | TRUE | 0.842615219 | 0.108572726 | No Change | Not Significant | TRUE | 0.106362379 |
| Efs | TRUE | -0.152643317 | 0.908068261 | No Change | Not Significant | FALSE | NA |
| Eftud2 | TRUE | 0.240107607 | 0.049395172 | No Change | Significant Positive | TRUE | 0.065148518 |
| Egf | TRUE | -0.096909638 | 0.812167358 | No Change | Not Significant | TRUE | 0.215103244 |
| Egfl6 | TRUE | -0.142166392 | 1 | No Change | Not Significant | FALSE | NA |
| Egfl7 | TRUE | 0.357536595 | 0.602493175 | No Change | Not Significant | TRUE | -0.307023252 |
| Egfl8 | TRUE | -0.145140361 | 0.858172551 | No Change | Not Significant | TRUE | -0.681698652 |
| Egflam | TRUE | 0.425700475 | 0.83324102 | No Change | Not Significant | FALSE | NA |
| Egfr | TRUE | -0.210287316 | 0.01691664 | No Change | Significant Negative | TRUE | 0.001623348 |
| Egln1 | TRUE | -0.223366085 | 0.008692083 | No Change | Significant Negative | TRUE | -0.137030049 |
| Egln2 | TRUE | 0.069607827 | 0.367456023 | No Change | Not Significant | TRUE | 0.13676978 |
| Egln3 | TRUE | -0.660510356 | 0.016227616 | DOWN | DOWN | TRUE | 0.048927514 |
| Egr1 | TRUE | -0.137911946 | 0.695624496 | No Change | Not Significant | TRUE | 0.002376746 |
| Egr2 | TRUE | -0.11343386 | 0.86401456 | No Change | Not Significant | TRUE | -0.195934411 |
| Egr3 | TRUE | 0.492161909 | 0.649597228 | No Change | Not Significant | TRUE | -1.251913483 |
| Egr4 | TRUE | 1.703080131 | 1 | No Change | Not Significant | FALSE | NA |
| Ehbp1 | TRUE | 0.160686382 | 0.414545055 | No Change | Not Significant | TRUE | -0.12931987 |
| Ehbp1l1 | TRUE | 0.215798718 | 0.302752759 | No Change | Not Significant | TRUE | 0.154228284 |
| Ehd1 | TRUE | 0.265508081 | 0.007872301 | No Change | Significant Positive | TRUE | 0.167450743 |
| Ehd2 | TRUE | 0.048993582 | 0.975869522 | No Change | Not Significant | TRUE | 0.525625139 |
| Ehd3 | TRUE | 0.991488977 | 1 | No Change | Not Significant | FALSE | NA |
| Ehd4 | TRUE | 0.915940471 | 5.00E-19 | UP | UP | TRUE | 0.463958075 |
| Ehf | TRUE | -0.041211903 | 0.869465313 | No Change | Not Significant | TRUE | -0.074477483 |
| Ehhadh | TRUE | 0.024202596 | 0.878912418 | No Change | Not Significant | TRUE | 0.047026789 |
| Ehmt1 | TRUE | -0.040925912 | 0.748554316 | No Change | Not Significant | TRUE | -0.04372091 |
| Ehmt2 | TRUE | 0.055060149 | 0.245940261 | No Change | Not Significant | TRUE | -0.070348695 |
| Ei24 | TRUE | -0.16675286 | 0.173431717 | No Change | Not Significant | TRUE | -0.010319696 |
| Eid1 | TRUE | -0.360838848 | 0.208982974 | No Change | Not Significant | TRUE | -0.046638934 |
| Eid2 | TRUE | 0.563710593 | 0.260339278 | No Change | Not Significant | TRUE | 0.114436519 |
| Eid2b | TRUE | -0.127345766 | 0.736964526 | No Change | Not Significant | TRUE | 0.135705628 |
| Eid3 | TRUE | -0.507993576 | 0.092582579 | No Change | Not Significant | TRUE | -0.784459749 |
| Eif1 | TRUE | -0.251489454 | 0.061293105 | No Change | Not Significant | TRUE | -0.101892817 |
| Eif1a | TRUE | 0.574569815 | 3.58E-10 | UP | UP | TRUE | 0.333819344 |
| Eif1ad | TRUE | -0.004222943 | 0.983635967 | No Change | Not Significant | TRUE | 0.057541171 |
| Eif1ax | TRUE | -0.093296601 | 0.673341315 | No Change | Not Significant | TRUE | -0.029759124 |
| Eif1b | TRUE | 0.0861439 | 0.696186289 | No Change | Not Significant | TRUE | -0.094778663 |
| Eif2a | TRUE | 0.086003748 | 0.647320963 | No Change | Not Significant | TRUE | -0.114219833 |
| Eif2ak1 | TRUE | -0.046241246 | 0.720115363 | No Change | Not Significant | TRUE | 0.025939372 |
| Eif2ak2 | TRUE | 0.047081549 | 0.818950015 | No Change | Not Significant | TRUE | 0.03566993 |
| Eif2ak3 | TRUE | 0.915805282 | 1.80E-23 | UP | UP | TRUE | 0.409258965 |
| Eif2ak4 | TRUE | -0.02043919 | 0.915993052 | No Change | Not Significant | TRUE | 0.013945055 |
| Eif2b1 | TRUE | 0.202082711 | 0.086889644 | No Change | Not Significant | TRUE | 0.080353989 |
| Eif2b2 | TRUE | 0.169346511 | 0.080865411 | No Change | Not Significant | TRUE | 0.026122724 |
| Eif2b3 | TRUE | -0.10592928 | 0.698788598 | No Change | Not Significant | TRUE | -0.150690757 |
| Eif2b4 | TRUE | -0.104578941 | 0.466951438 | No Change | Not Significant | TRUE | 0.008185848 |
| Eif2b5 | TRUE | 0.065384126 | 0.667322041 | No Change | Not Significant | TRUE | 0.079058888 |
| Eif2d | TRUE | 0.078776554 | 0.404473864 | No Change | Not Significant | TRUE | 0.145727668 |
| Eif2s1 | TRUE | 0.120968618 | 0.318636429 | No Change | Not Significant | TRUE | 0.084557673 |
| Eif2s2 | TRUE | -0.213416593 | 0.236336338 | No Change | Not Significant | TRUE | -0.256341013 |
| Eif2s3x | TRUE | -0.34378638 | 0.026283536 | No Change | Significant Negative | TRUE | 0.037075708 |
| Eif3a | TRUE | 0.036588535 | 0.827372024 | No Change | Not Significant | TRUE | -0.044956911 |
| Eif3b | TRUE | 0.407328055 | 3.79E-07 | No Change | Significant Positive | TRUE | 0.118304063 |
| Eif3c | TRUE | -0.019489041 | 0.87503937 | No Change | Not Significant | TRUE | -0.076269668 |
| Eif3d | TRUE | 0.183095674 | 0.123758475 | No Change | Not Significant | TRUE | 0.064782901 |
| Eif3e | TRUE | -0.228775714 | 0.158163729 | No Change | Not Significant | TRUE | -0.144260595 |
| Eif3f | TRUE | -0.068725517 | 0.647921517 | No Change | Not Significant | TRUE | 0.007832665 |
| Eif3g | TRUE | 0.069040236 | 0.646027582 | No Change | Not Significant | TRUE | 0.084586283 |
| Eif3h | TRUE | -0.187589884 | 0.121629941 | No Change | Not Significant | TRUE | 0.000566799 |
| Eif3i | TRUE | -0.121479156 | 0.468911293 | No Change | Not Significant | TRUE | 0.039807169 |
| Eif3j1 | TRUE | -0.250683139 | 0.106468355 | No Change | Not Significant | TRUE | -0.062254701 |
| Eif3j2 | TRUE | 0.616203422 | 0.440150633 | No Change | Not Significant | TRUE | -0.428866145 |
| Eif3k | TRUE | 0.019010282 | 0.904795436 | No Change | Not Significant | TRUE | 0.016416956 |
| Eif3l | TRUE | -0.152230701 | 0.363163115 | No Change | Not Significant | TRUE | 0.014393068 |
| Eif3m | TRUE | -0.183104839 | 0.281607009 | No Change | Not Significant | TRUE | -0.039908411 |
| Eif4a1 | TRUE | -0.002639155 | 0.989584039 | No Change | Not Significant | TRUE | 0.073301574 |
| Eif4a2 | TRUE | -0.085814688 | 0.493582047 | No Change | Not Significant | TRUE | 0.090422901 |
| Eif4a3 | TRUE | 0.128105252 | 0.213086636 | No Change | Not Significant | TRUE | 0.060526604 |
| Eif4b | TRUE | -0.046322659 | 0.773864504 | No Change | Not Significant | TRUE | -0.10894416 |
| Eif4e | TRUE | 0.064827018 | 0.745748293 | No Change | Not Significant | TRUE | 0.033739781 |
| Eif4e1b | TRUE | -1.687093936 | 1 | No Change | Not Significant | FALSE | NA |
| Eif4e2 | TRUE | -0.096864931 | 0.404694914 | No Change | Not Significant | TRUE | -0.062410737 |
| Eif4e3 | TRUE | 0.113908239 | 0.809917796 | No Change | Not Significant | TRUE | 0.021972343 |
| Eif4ebp1 | TRUE | 0.313514057 | 0.027372809 | No Change | Significant Positive | TRUE | 0.109671625 |
| Eif4ebp2 | TRUE | -0.131321922 | 0.315287712 | No Change | Not Significant | TRUE | -0.122351161 |
| Eif4ebp3 | TRUE | -0.219422088 | 0.339181285 | No Change | Not Significant | TRUE | 0.158171359 |
| Eif4enif1 | TRUE | 0.103353276 | 0.20160769 | No Change | Not Significant | TRUE | -0.058508868 |
| Eif4g1 | TRUE | 0.242584003 | 0.000726214 | No Change | Significant Positive | TRUE | 0.158600525 |
| Eif4g2 | TRUE | 0.284799863 | 0.001000926 | No Change | Significant Positive | TRUE | 0.093606258 |
| Eif4g3 | TRUE | 0.269472181 | 0.000338705 | No Change | Significant Positive | TRUE | 0.143044502 |
| Eif4h | TRUE | -0.032504999 | 0.757643976 | No Change | Not Significant | TRUE | -0.059195588 |
| Eif5 | TRUE | 0.105506508 | 0.283788828 | No Change | Not Significant | TRUE | -0.113773438 |
| Eif5a | TRUE | 0.113114946 | 0.324941878 | No Change | Not Significant | TRUE | 0.120024772 |
| Eif5a2 | TRUE | 1.492319253 | 0.001219773 | UP | UP | TRUE | 0.799457449 |
| Eif5b | TRUE | 0.086399418 | 0.488518708 | No Change | Not Significant | TRUE | -0.019619441 |
| Eif6 | TRUE | -0.011768204 | 0.935658236 | No Change | Not Significant | TRUE | 0.077443991 |
| Eipr1 | TRUE | 0.164882076 | 0.130795377 | No Change | Not Significant | TRUE | 0.312094783 |
| Elac1 | TRUE | 0.146672168 | 0.462419363 | No Change | Not Significant | TRUE | -0.110370173 |
| Elac2 | TRUE | 0.075366903 | 0.732371878 | No Change | Not Significant | TRUE | 0.020345328 |
| Elapor1 | TRUE | -0.27679775 | 0.26676599 | No Change | Not Significant | TRUE | 0.092318053 |
| Elapor2 | TRUE | -0.975407897 | 0.163322332 | No Change | Not Significant | TRUE | -0.28115938 |
| Elavl1 | TRUE | 0.162226244 | 0.160666232 | No Change | Not Significant | TRUE | 0.190517682 |
| Elavl2 | TRUE | -0.544999166 | 1 | No Change | Not Significant | FALSE | NA |
| Elavl3 | TRUE | -0.242670161 | 0.909231489 | No Change | Not Significant | FALSE | NA |
| Elavl4 | TRUE | 0.582309075 | 1 | No Change | Not Significant | FALSE | NA |
| Eldr | TRUE | 0.253877432 | 0.493321208 | No Change | Not Significant | TRUE | -0.088676171 |
| Elf1 | TRUE | -0.127399497 | 0.168985435 | No Change | Not Significant | TRUE | -0.048107511 |
| Elf2 | TRUE | -0.19177291 | 0.028168424 | No Change | Significant Negative | TRUE | -0.160295776 |
| Elf3 | TRUE | -0.010470057 | 0.950848993 | No Change | Not Significant | TRUE | 0.170751903 |
| Elf4 | TRUE | -0.168549501 | 0.404694914 | No Change | Not Significant | TRUE | -0.078017594 |
| Elf5 | TRUE | -0.315896703 | 1 | No Change | Not Significant | FALSE | NA |
| Elfn2 | TRUE | 0.292239761 | 0.52454765 | No Change | Not Significant | TRUE | -0.093900865 |
| Elk1 | TRUE | 2.237651202 | 8.88E-35 | UP | UP | TRUE | 0.320439392 |
| Elk3 | TRUE | -0.323663063 | 0.30685209 | No Change | Not Significant | TRUE | 0.62518771 |
| Elk4 | TRUE | -0.0410376 | 0.742839333 | No Change | Not Significant | TRUE | -0.146634556 |
| Ell | TRUE | 0.161797163 | 0.530962336 | No Change | Not Significant | TRUE | 0.00787292 |
| Ell2 | TRUE | 0.329599082 | 0.000197035 | No Change | Significant Positive | TRUE | 0.062656726 |
| Ell3 | TRUE | -0.352696046 | 0.630552252 | No Change | Not Significant | TRUE | -0.541306413 |
| Elmo1 | TRUE | 0.111160457 | 0.704796705 | No Change | Not Significant | TRUE | -0.124048784 |
| Elmo2 | TRUE | 0.086501559 | 0.555809168 | No Change | Not Significant | TRUE | 0.003787102 |
| Elmo3 | TRUE | -0.051734674 | 0.642569972 | No Change | Not Significant | TRUE | -0.029106239 |
| Elmod1 | TRUE | 1.125916043 | 1 | No Change | Not Significant | FALSE | NA |
| Elmod2 | TRUE | 0.173510182 | 0.162665392 | No Change | Not Significant | TRUE | 0.00797479 |
| Elmod3 | TRUE | -0.23119745 | 0.092827346 | No Change | Not Significant | TRUE | 0.019552016 |
| Eln | TRUE | 0.66626444 | 0.807125838 | No Change | Not Significant | FALSE | NA |
| Eloa | TRUE | -0.042701079 | 0.696186289 | No Change | Not Significant | TRUE | -0.048512587 |
| Elob | TRUE | -0.130036941 | 0.222126524 | No Change | Not Significant | TRUE | 0.145741942 |
| Eloc | TRUE | 0.076307059 | 0.413054814 | No Change | Not Significant | TRUE | 0.079395489 |
| Elof1 | TRUE | 0.152101069 | 0.136567098 | No Change | Not Significant | TRUE | 0.262624177 |
| Elovl1 | TRUE | -0.029968959 | 0.825585716 | No Change | Not Significant | TRUE | 0.037600491 |
| Elovl2 | TRUE | 1.259802323 | 1 | No Change | Not Significant | FALSE | NA |
| Elovl3 | TRUE | -1.352096778 | 1 | No Change | Not Significant | FALSE | NA |
| Elovl4 | TRUE | 1.706124205 | 1 | No Change | Not Significant | FALSE | NA |
| Elovl5 | TRUE | -0.016305238 | 0.947906833 | No Change | Not Significant | TRUE | -0.124184993 |
| Elovl6 | TRUE | -0.095098631 | 0.55988186 | No Change | Not Significant | TRUE | 0.187374338 |
| Elovl7 | TRUE | -0.240859919 | 0.14799363 | No Change | Not Significant | TRUE | 0.17985223 |
| Elp1 | TRUE | 0.04390138 | 0.835998557 | No Change | Not Significant | TRUE | -0.117451506 |
| Elp2 | TRUE | 0.253220658 | 0.00535797 | No Change | Significant Positive | TRUE | 0.207378693 |
| Elp3 | TRUE | 0.297469239 | 0.026243183 | No Change | Significant Positive | TRUE | 0.127139327 |
| Elp4 | TRUE | 0.011493877 | 0.962530504 | No Change | Not Significant | TRUE | -0.032956461 |
| Elp5 | TRUE | 0.113549337 | 0.274864513 | No Change | Not Significant | TRUE | 0.053549372 |
| Elp6 | TRUE | 0.224568303 | 0.248992215 | No Change | Not Significant | TRUE | 0.156172653 |
| Emb | TRUE | 0.719954507 | 0.140291806 | No Change | Not Significant | TRUE | 0.297057797 |
| Emc1 | TRUE | 0.200644265 | 0.02939498 | No Change | Significant Positive | TRUE | 0.053998958 |
| Emc10 | TRUE | -0.160072055 | 0.054559819 | No Change | Not Significant | TRUE | 0.051476691 |
| Emc2 | TRUE | 0.053978908 | 0.72294031 | No Change | Not Significant | TRUE | 0.113341657 |
| Emc3 | TRUE | 0.245284029 | 0.008157095 | No Change | Significant Positive | TRUE | 0.135723554 |
| Emc4 | TRUE | -0.020404139 | 0.888506311 | No Change | Not Significant | TRUE | 0.082996114 |
| Emc6 | TRUE | 0.015804618 | 0.945659633 | No Change | Not Significant | TRUE | -0.006824462 |

Supplementary Table 3

| geneID | Cond1_DF_Detected | Cond1_DF_ValueCol | Cond1_DF_PadjCol | Cond1_DF_Cutoff | Cond1_DF_Cutoff_Specific.x | Cond2_DF_Detected | Cond2_DF_ValueCol | Cond2_DF_PadjCol | Cond2_DF_Cutoff | Cond2_DF_Cutoff_Specific |
|---|---|---|---|---|---|---|---|---|---|---|
| Emc7 | TRUE | 0.062229735 | 0.716210584 | No Change | Not Significant | TRUE | 0.103791456 | 0.189222442 | No Change | Not Significant |
| Emc8 | TRUE | 0.032667027 | 0.855204229 | No Change | Not Significant | TRUE | -0.029602231 | 0.829602333 | No Change | Not Significant |
| Emc9 | TRUE | -0.287824563 | 0.236536038 | No Change | Not Significant | TRUE | 0.116690268 | 0.669773628 | No Change | Not Significant |
| Emcn | TRUE | 1.126343937 | 1 | No Change | Not Significant | FALSE | NA | NA | No Change | Not Detected |
| Emd | TRUE | 0.186977924 | 0.148984107 | No Change | Not Significant | TRUE | 0.182436335 | 0.025378447 | No Change | Significant Positive |
| Eme1 | TRUE | 0.663876478 | 0.007373206 | UP | UP | TRUE | 0.411582593 | 0.084790715 | No Change | Not Significant |
| Eme2 | TRUE | 0.161170053 | 0.421098196 | No Change | Not Significant | TRUE | -0.078610407 | 0.749049145 | No Change | Not Significant |
| Emg1 | TRUE | 0.097154246 | 0.551855356 | No Change | Not Significant | TRUE | 0.198216986 | 0.003991974 | No Change | Significant Positive |
| Emid1 | TRUE | 0.056476664 | 0.970043107 | No Change | Not Significant | TRUE | 0.14706716 | 0.880021159 | No Change | Not Significant |
| Emilin1 | TRUE | 0.415574175 | 0.169937908 | No Change | Not Significant | TRUE | 0.668032696 | 0.006042973 | UP | UP |
| Emilin2 | TRUE | -0.148554024 | 0.933936484 | No Change | Not Significant | FALSE | NA | NA | No Change | Not Detected |
| Emilin3 | TRUE | 0.734449063 | 1 | No Change | Not Significant | FALSE | NA | NA | No Change | Not Detected |
| Eml1 | TRUE | 0.614906687 | 0.008537358 | UP | UP | TRUE | 0.386297897 | 0.059417614 | No Change | Not Significant |
| Eml2 | TRUE | -0.23333828 | 0.106287366 | No Change | Not Significant | TRUE | -0.056227539 | 0.720720873 | No Change | Not Significant |
| Eml3 | TRUE | 0.044332856 | 0.719334577 | No Change | Not Significant | TRUE | -0.148223611 | 0.141755849 | No Change | Not Significant |
| Eml4 | TRUE | -0.025412526 | 0.908409285 | No Change | Not Significant | TRUE | -0.069997285 | 0.301243853 | No Change | Not Significant |
| Eml5 | TRUE | 0.032921365 | 0.959505963 | No Change | Not Significant | TRUE | -0.415711653 | 0.212757025 | No Change | Not Significant |
| Eml6 | TRUE | 0.510011702 | 0.043909138 | UP | UP | TRUE | 0.31112326 | 0.043445662 | No Change | Significant Positive |
| Emp1 | TRUE | -0.259580256 | 0.419019832 | No Change | Not Significant | TRUE | -0.000855183 | 0.996976705 | No Change | Not Significant |
| Emp2 | TRUE | -0.6234834 | 3.42E-05 | DOWN | DOWN | TRUE | -0.638451778 | 1.63E-14 | DOWN | DOWN |
| Emp3 | TRUE | 1.652262873 | 0.051148256 | No Change | Not Significant | FALSE | NA | NA | No Change | Not Detected |
| Emsy | TRUE | 0.056358827 | 0.739498445 | No Change | Not Significant | TRUE | -0.100168006 | 0.365794955 | No Change | Not Significant |
| Emx2 | TRUE | 0.609569473 | 1 | No Change | Not Significant | FALSE | NA | NA | No Change | Not Detected |
| Emx2os | TRUE | -1.277066262 | 1 | No Change | Not Significant | FALSE | NA | NA | No Change | Not Detected |
| Enah | TRUE | 2.342642835 | 0.029218733 | UP | UP | FALSE | NA | NA | No Change | Not Detected |
| Enam | TRUE | 0.616165291 | 1 | No Change | Not Significant | FALSE | NA | NA | No Change | Not Detected |
| Enc1 | TRUE | -0.029723132 | 0.902394803 | No Change | Not Significant | TRUE | -0.204208597 | 0.141426796 | No Change | Not Significant |
| Endod1 | TRUE | -0.162888583 | 0.194381533 | No Change | Not Significant | TRUE | 0.297831253 | 0.009099034 | No Change | Significant Positive |
| Endog | TRUE | 0.236127773 | 0.211290803 | No Change | Not Significant | TRUE | -0.035140828 | 0.874754933 | No Change | Not Significant |
| Endou | TRUE | -1.677713795 | 1 | No Change | Not Significant | FALSE | NA | NA | No Change | Not Detected |
| Endov | TRUE | -0.207492062 | 0.117005071 | No Change | Not Significant | TRUE | -0.023975177 | 0.898056783 | No Change | Not Significant |
| Eng | TRUE | 0.766382352 | 0.715488638 | No Change | Not Significant | FALSE | NA | NA | No Change | Not Detected |
| Engase | TRUE | 1.147639622 | 1.49E-43 | UP | UP | TRUE | 0.189025298 | 0.004869416 | No Change | Significant Positive |
| Enho | TRUE | 1.65802145 | 0.099059146 | No Change | Not Significant | FALSE | NA | NA | No Change | Not Detected |
| Enkd1 | TRUE | 0.436191956 | 0.177458338 | No Change | Not Significant | TRUE | -0.030364043 | 0.95031765 | No Change | Not Significant |
| Enkur | TRUE | 0.082816995 | 0.932036168 | No Change | Not Significant | TRUE | 0.327830205 | 0.559864561 | No Change | Not Significant |
| Eno1 | TRUE | -0.036273618 | 0.878512436 | No Change | Not Significant | TRUE | 0.251876274 | 0.010029817 | No Change | Significant Positive |
| Eno1b | TRUE | 0.329946079 | 1 | No Change | Not Significant | FALSE | NA | NA | No Change | Not Detected |
| Eno2 | TRUE | 0.295268979 | 0.910297568 | No Change | Not Significant | FALSE | NA | NA | No Change | Not Detected |
| Eno3 | TRUE | -1.12447332 | 0.000200831 | DOWN | DOWN | TRUE | -0.494834384 | 0.079113565 | No Change | Not Significant |
| Eno4 | TRUE | 0.463692611 | 1 | No Change | Not Significant | FALSE | NA | NA | No Change | Not Detected |
| Enoph1 | TRUE | -0.183332984 | 0.125007242 | No Change | Not Significant | TRUE | 0.027843209 | 0.846888755 | No Change | Not Significant |
| Enox1 | TRUE | -0.453112849 | 0.736796542 | No Change | Not Significant | FALSE | NA | NA | No Change | Not Detected |
| Enox2 | TRUE | -0.128205667 | 0.34055723 | No Change | Not Significant | TRUE | -0.040377769 | 0.664948752 | No Change | Not Significant |
| Enpep | TRUE | -0.497725747 | 0.429396526 | No Change | Not Significant | TRUE | -0.404433172 | 0.294578028 | No Change | Not Significant |
| Enpp1 | TRUE | 0.762544857 | 4.98E-07 | UP | UP | TRUE | 0.712618678 | 5.68E-20 | UP | UP |
| Enpp2 | TRUE | -2.178395692 | 1 | No Change | Not Significant | FALSE | NA | NA | No Change | Not Detected |
| Enpp3 | TRUE | 0.801189222 | 0.038220989 | UP | UP | TRUE | 0.539966916 | 0.04380415 | UP | UP |
| Enpp4 | TRUE | -0.213765396 | 0.241953298 | No Change | Not Significant | TRUE | -0.198771838 | 0.062318117 | No Change | Not Significant |
| Enpp5 | TRUE | 0.138218184 | 0.577325459 | No Change | Not Significant | TRUE | 0.128525377 | 0.203825108 | No Change | Not Significant |
| Enpp6 | TRUE | 0.936693165 | 1 | No Change | Not Significant | FALSE | NA | NA | No Change | Not Detected |
| Enpp7 | TRUE | 0.1788072 | 0.940340033 | No Change | Not Significant | TRUE | 0.257948812 | 0.899522993 | No Change | Not Significant |
| Ensa | TRUE | 0.112298853 | 0.128987322 | No Change | Not Significant | TRUE | 0.082521455 | 0.058857178 | No Change | Not Significant |
| Entpd1 | TRUE | -0.173439466 | 0.908812719 | No Change | Not Significant | FALSE | NA | NA | No Change | Not Detected |
| Entpd2 | TRUE | -0.588167864 | 0.000106585 | DOWN | DOWN | TRUE | -0.326678131 | 0.002817596 | No Change | Significant Negative |
| Entpd3 | TRUE | -0.710390906 | 0.279846882 | No Change | Not Significant | TRUE | 0.029065099 | 0.973941151 | No Change | Not Significant |
| Entpd5 | TRUE | -0.385866626 | 0.000143449 | No Change | Significant Negative | TRUE | -0.247082221 | 0.001467526 | No Change | Significant Negative |
| Entpd6 | TRUE | 0.155121424 | 0.142788208 | No Change | Not Significant | TRUE | 0.196550825 | 0.00475227 | No Change | Significant Positive |
| Entpd7 | TRUE | 0.677968586 | 1.61E-07 | UP | UP | TRUE | 0.120351328 | 0.328198932 | No Change | Not Significant |
| Entpd8 | TRUE | -0.103135126 | 0.510066755 | No Change | Not Significant | TRUE | 0.084700627 | 0.43881778 | No Change | Not Significant |
| Entr1 | TRUE | 0.029986556 | 0.830576634 | No Change | Not Significant | TRUE | 0.026167652 | 0.801099657 | No Change | Not Significant |
| Entrep1 | TRUE | 0.319008233 | 0.026350675 | No Change | Significant Positive | TRUE | 0.037466244 | 0.761555988 | No Change | Not Significant |
| Entrep2 | TRUE | -0.564496836 | 0.080637284 | No Change | Not Significant | TRUE | -0.077381988 | 0.796788198 | No Change | Not Significant |
| Entrep3 | TRUE | 0.032215498 | 0.965151432 | No Change | Not Significant | TRUE | -0.0873982 | 0.824251158 | No Change | Not Significant |
| Eny2 | TRUE | 0.133002856 | 0.226282116 | No Change | Not Significant | TRUE | -0.010566382 | 0.933591927 | No Change | Not Significant |
| Eogt | TRUE | -0.045353385 | 0.795976331 | No Change | Not Significant | TRUE | 0.054915675 | 0.716067516 | No Change | Not Significant |
| Eola1 | TRUE | -0.073186747 | 0.644177805 | No Change | Not Significant | TRUE | 0.136107526 | 0.304322033 | No Change | Not Significant |
| Eomes | TRUE | -0.61196717 | 1 | No Change | Not Significant | FALSE | NA | NA | No Change | Not Detected |
| Ep300 | TRUE | -0.070133057 | 0.794593769 | No Change | Not Significant | TRUE | -0.248292531 | 0.026573931 | No Change | Significant Negative |
| Ep400 | TRUE | -0.006139831 | 0.976710073 | No Change | Not Significant | TRUE | -0.081558828 | 0.51227128 | No Change | Not Significant |
| Epas1 | TRUE | -0.372131699 | 0.036011679 | No Change | Significant Negative | TRUE | -0.095909412 | 0.314662099 | No Change | Not Significant |
| Epb41 | TRUE | 1.430082918 | 8.51E-131 | UP | UP | TRUE | 0.500736116 | 5.31E-24 | UP | UP |
| Epb41l1 | TRUE | -0.210715183 | 0.118476325 | No Change | Not Significant | TRUE | -0.094988442 | 0.429391175 | No Change | Not Significant |
| Epb41l2 | TRUE | 0.001912214 | 0.99157486 | No Change | Not Significant | TRUE | 0.029878522 | 0.795383546 | No Change | Not Significant |
| Epb41l3 | TRUE | 0.005703089 | 0.971417829 | No Change | Not Significant | TRUE | -0.055796132 | 0.357631975 | No Change | Not Significant |
| Epb41l4a | TRUE | -0.515180725 | 0.015962767 | DOWN | DOWN | TRUE | -0.208781258 | 0.067240721 | No Change | Not Significant |
| Epb41l4aos | TRUE | -0.514164761 | 0.000733845 | DOWN | DOWN | TRUE | -0.411568823 | 4.89E-05 | No Change | Significant Negative |
| Epb41l4b | TRUE | -0.216604453 | 0.073420411 | No Change | Not Significant | TRUE | 0.076504315 | 0.220072289 | No Change | Not Significant |
| Epb41l5 | TRUE | -0.102219515 | 0.535858528 | No Change | Not Significant | TRUE | -0.168621481 | 0.058437948 | No Change | Not Significant |
| Epb42 | TRUE | 0.481298027 | 1 | No Change | Not Significant | FALSE | NA | NA | No Change | Not Detected |
| Epc1 | TRUE | -0.047516328 | 0.791450739 | No Change | Not Significant | TRUE | -0.062848192 | 0.491745579 | No Change | Not Significant |
| Epc2 | TRUE | 0.135020704 | 0.384217774 | No Change | Not Significant | TRUE | 0.006888882 | 0.954463189 | No Change | Not Significant |
| Epcam | TRUE | -0.262666805 | 0.001253501 | No Change | Significant Negative | TRUE | -0.029549763 | 0.772616101 | No Change | Not Significant |
| Epdr1 | TRUE | 0.298334027 | 0.717449153 | No Change | Not Significant | TRUE | -0.346714979 | 0.44497561 | No Change | Not Significant |
| Epg5 | TRUE | 0.032887557 | 0.838326567 | No Change | Not Significant | TRUE | -0.082335421 | 0.309011907 | No Change | Not Significant |
| Epgn | TRUE | 0.64294922 | 1 | No Change | Not Significant | FALSE | NA | NA | No Change | Not Detected |
| Epha1 | TRUE | -0.345421699 | 0.048018124 | No Change | Significant Negative | TRUE | -0.262827171 | 0.007270796 | No Change | Significant Negative |
| Epha10 | TRUE | 0.19719824 | 1 | No Change | Not Significant | FALSE | NA | NA | No Change | Not Detected |
| Epha2 | TRUE | 0.434031298 | 0.288643819 | No Change | Not Significant | TRUE | -0.258237184 | 0.140386527 | No Change | Not Significant |
| Epha3 | TRUE | -0.211102461 | 1 | No Change | Not Significant | TRUE | -0.233183976 | 0.816434847 | No Change | Not Significant |
| Epha4 | TRUE | 0.108463443 | 0.675184655 | No Change | Not Significant | TRUE | 0.05907579 | 0.708241492 | No Change | Not Significant |
| Epha5 | TRUE | 1.480855931 | 0.426485613 | No Change | Not Significant | FALSE | NA | NA | No Change | Not Detected |
| Epha6 | TRUE | 1.69958101 | 1 | No Change | Not Significant | FALSE | NA | NA | No Change | Not Detected |
| Epha7 | TRUE | 0.488552136 | 0.356436387 | No Change | Not Significant | TRUE | -0.024693979 | 0.95873684 | No Change | Not Significant |
| Epha8 | TRUE | 1.795448077 | 1 | No Change | Not Significant | FALSE | NA | NA | No Change | Not Detected |
| Ephb1 | TRUE | -2.11929441 | 1 | No Change | Not Significant | FALSE | NA | NA | No Change | Not Detected |
| Ephb2 | TRUE | -0.030921328 | 0.876856731 | No Change | Not Significant | TRUE | -0.156558759 | 0.033802395 | No Change | Significant Negative |
| Ephb3 | TRUE | -0.020439559 | 0.940455052 | No Change | Not Significant | TRUE | -0.217893861 | 0.08873319 | No Change | Not Significant |
| Ephb4 | TRUE | -0.189944455 | 0.17492772 | No Change | Not Significant | TRUE | -0.133203883 | 0.165140974 | No Change | Not Significant |
| Ephb6 | TRUE | -0.618074239 | 0.132673445 | No Change | Not Significant | TRUE | 0.033099113 | 0.962265867 | No Change | Not Significant |
| Ephx1 | TRUE | 0.236594649 | 0.444677528 | No Change | Not Significant | TRUE | 0.094898139 | 0.483886312 | No Change | Not Significant |
| Ephx2 | TRUE | -0.272598269 | 0.03286063 | No Change | Significant Negative | TRUE | -0.306505019 | 0.006413818 | No Change | Significant Negative |
| Ephx3 | TRUE | 3.397864952 | 1 | No Change | Not Significant | FALSE | NA | NA | No Change | Not Detected |
| Ephx4 | TRUE | -0.455749266 | 0.033042863 | No Change | Significant Negative | TRUE | 0.045592091 | 0.898525697 | No Change | Not Significant |
| Epm2a | TRUE | -0.062097238 | 0.932166328 | No Change | Not Significant | TRUE | -0.095097673 | 0.818885486 | No Change | Not Significant |
| Epm2aip1 | TRUE | -0.006922837 | 0.973885877 | No Change | Not Significant | TRUE | -0.107924639 | 0.370189171 | No Change | Not Significant |
| Epn1 | TRUE | 0.055049933 | 0.548792208 | No Change | Not Significant | TRUE | 0.12246359 | 0.437382664 | No Change | Not Significant |
| Epn2 | TRUE | -0.132857634 | 0.416836029 | No Change | Not Significant | TRUE | -0.119319566 | 0.275847342 | No Change | Not Significant |
| Epn3 | TRUE | 0.134587517 | 0.258635202 | No Change | Not Significant | TRUE | 0.029951414 | 0.851559877 | No Change | Not Significant |
| Epop | TRUE | 0.45475179 | 0.507895701 | No Change | Not Significant | TRUE | -1.494104251 | 7.53E-08 | DOWN | DOWN |
| Epor | TRUE | -0.327321048 | 1 | No Change | Not Significant | FALSE | NA | NA | No Change | Not Detected |
| Epp13 | TRUE | 0.494518344 | 1 | No Change | Not Significant | FALSE | NA | NA | No Change | Not Detected |
| Eppin | TRUE | -1.295271625 | 1 | No Change | Not Significant | FALSE | NA | NA | No Change | Not Detected |
| Eprs | TRUE | 0.126237336 | 0.391427734 | No Change | Not Significant | TRUE | 0.025774122 | 0.859482312 | No Change | Not Significant |
| Eps15 | TRUE | -0.124482976 | 0.390165244 | No Change | Not Significant | TRUE | -0.093963604 | 0.34449158 | No Change | Not Significant |
| Eps15l1 | TRUE | -0.030542375 | 0.785464687 | No Change | Not Significant | TRUE | -0.027504225 | 0.808283189 | No Change | Not Significant |
| Eps8 | TRUE | 0.030376273 | 0.908812719 | No Change | Not Significant | TRUE | -0.243932014 | 0.014522485 | No Change | Significant Negative |
| Eps8l1 | TRUE | 0.15453759 | 0.702502904 | No Change | Not Significant | TRUE | 0.09149838 | 0.865987203 | No Change | Not Significant |
| Eps8l2 | TRUE | -0.220130961 | 0.438629566 | No Change | Not Significant | TRUE | -0.032511231 | 0.821648745 | No Change | Not Significant |
| Eps8l3 | TRUE | -0.156825862 | 0.408576247 | No Change | Not Significant | TRUE | -0.263217117 | 6.49E-05 | No Change | Significant Negative |
| Epsti1 | TRUE | -0.124540569 | 0.323295567 | No Change | Not Significant | TRUE | 0.114959519 | 0.277413181 | No Change | Not Significant |
| Epx | TRUE | -0.962323132 | 1 | No Change | Not Significant | FALSE | NA | NA | No Change | Not Detected |
| Eral1 | TRUE | -0.160608647 | 0.297380166 | No Change | Not Significant | TRUE | 0.069992884 | 0.588629676 | No Change | Not Significant |
| Erap1 | TRUE | -0.119865215 | 0.40541018 | No Change | Not Significant | TRUE | -0.119386654 | 0.140323897 | No Change | Not Significant |
| Erbb2 | TRUE | -0.227425714 | 0.148254918 | No Change | Not Significant | TRUE | -0.029216252 | 0.821762439 | No Change | Not Significant |
| Erbb3 | TRUE | -0.324708628 | 0.017334001 | No Change | Significant Negative | TRUE | -0.128965694 | 0.197514723 | No Change | Not Significant |
| Erbb4 | TRUE | 0.364785995 | 1 | No Change | Not Significant | FALSE | NA | NA | No Change | Not Detected |
| Erbin | TRUE | -0.225750849 | 0.073835702 | No Change | Not Significant | TRUE | -0.142892728 | 0.064598841 | No Change | Not Significant |
| Erc1 | TRUE | 0.102368436 | 0.370029503 | No Change | Not Significant | TRUE | 0.043258457 | 0.679535938 | No Change | Not Significant |
| Erc2 | TRUE | 1.459473638 | 0.2954831 | No Change | Not Significant | FALSE | NA | NA | No Change | Not Detected |
| Ercc1 | TRUE | 0.341558297 | 0.027440166 | No Change | Significant Positive | TRUE | 0.178574748 | 0.207110275 | No Change | Not Significant |
| Ercc2 | TRUE | 0.014880085 | 0.923801556 | No Change | Not Significant | TRUE | -0.049741102 | 0.697669633 | No Change | Not Significant |
| Ercc3 | TRUE | 0.035434447 | 0.806964824 | No Change | Not Significant | TRUE | 0.068138166 | 0.370570998 | No Change | Not Significant |
| Ercc4 | TRUE | -0.10628205 | 0.397106786 | No Change | Not Significant | TRUE | -0.098524652 | 0.402701636 | No Change | Not Significant |
| Ercc5 | TRUE | -0.058061519 | 0.720115363 | No Change | Not Significant | TRUE | -0.111341864 | 0.367724228 | No Change | Not Significant |
| Ercc6 | TRUE | -0.10076437 | 0.360359086 | No Change | Not Significant | TRUE | -0.182518252 | 0.149703476 | No Change | Not Significant |
| Ercc6l | TRUE | 0.61764252 | 0.011262661 | UP | UP | TRUE | -0.022651382 | 0.955523695 | No Change | Not Significant |
| Ercc6l2 | TRUE | 0.065567318 | 0.765654588 | No Change | Not Significant | TRUE | -0.113204903 | 0.302001032 | No Change | Not Significant |
| Ercc8 | TRUE | 0.063042579 | 0.787780401 | No Change | Not Significant | TRUE | -0.036615398 | 0.868306546 | No Change | Not Significant |
| Ereg | TRUE | 1.389298054 | 0.000561931 | UP | UP | TRUE | 0.156508239 | 0.713491312 | No Change | Not Significant |
| Erf | TRUE | -0.241689696 | 0.047685597 | No Change | Significant Negative | TRUE | -0.099589927 | 0.492727559 | No Change | Not Significant |
| Erfe | TRUE | 0.141465272 | 0.485271256 | No Change | Not Significant | TRUE | -0.092242458 | 0.594858267 | No Change | Not Significant |
| Erfl | TRUE | 1.277489581 | 1 | No Change | Not Significant | FALSE | NA | NA | No Change | Not Detected |
| Erg | TRUE | 1.789852734 | 1 | No Change | Not Significant | FALSE | NA | NA | No Change | Not Detected |
| Erg28 | TRUE | 0.797165457 | 2.32E-12 | UP | UP | TRUE | 0.276205322 | 0.000644642 | No Change | Significant Positive |
| Ergic1 | TRUE | 0.356610932 | 0.018819941 | No Change | Significant Positive | TRUE | 0.205277027 | 0.003688925 | No Change | Significant Positive |
| Ergic2 | TRUE | 0.367950831 | 4.67E-06 | No Change | Significant Positive | TRUE | 0.08178287 | 0.379001161 | No Change | Not Significant |
| Ergic3 | TRUE | -0.457682286 | 6.08E-09 | No Change | Significant Negative | TRUE | -0.101845848 | 0.20173823 | No Change | Not Significant |
| Erh | TRUE | -0.091196889 | 0.750868206 | No Change | Not Significant | TRUE | 0.12837483 | 0.406457009 | No Change | Not Significant |
| Eri1 | TRUE | 0.138298828 | 0.390726542 | No Change | Not Significant | TRUE | 0.065530259 | 0.644017631 | No Change | Not Significant |
| Eri2 | TRUE | 0.233097028 | 0.297863506 | No Change | Not Significant | TRUE | 0.015527718 | 0.960412417 | No Change | Not Significant |
| Eri3 | TRUE | 0.117799406 | 0.293933865 | No Change | Not Significant | TRUE | 0.135605914 | 0.178786609 | No Change | Not Significant |
| Erich1 | TRUE | -0.096919013 | 0.656764359 | No Change | Not Significant | TRUE | -0.022568929 | 0.930210801 | No Change | Not Significant |
| Erich2 | TRUE | 1.083888953 | 1 | No Change | Not Significant | FALSE | NA | NA | No Change | Not Detected |
| Erich3 | TRUE | 0.070079559 | 0.969802608 | No Change | Not Significant | FALSE | NA | NA | No Change | Not Detected |
| Erich4 | TRUE | 0.427615698 | 0.20160769 | No Change | Not Significant | TRUE | -0.395204498 | 0.095444865 | No Change | Not Significant |
| Erich5 | TRUE | 0.126308052 | 0.877286805 | No Change | Not Significant | TRUE | -0.397472291 | 0.463289311 | No Change | Not Significant |
| Erich6 | TRUE | 2.157317454 | 1 | No Change | Not Significant | FALSE | NA | NA | No Change | Not Detected |
| Erich6b | TRUE | 0.230063495 | 1 | No Change | Not Significant | FALSE | NA | NA | No Change | Not Detected |
| Erlec1 | TRUE | -0.217433186 | 0.277248109 | No Change | Not Significant | TRUE | -0.146215425 | 0.221867939 | No Change | Not Significant |
| Erlin1 | TRUE | 0.166389202 | 0.159701725 | No Change | Not Significant | TRUE | -0.223218497 | 0.001687476 | No Change | Significant Negative |
| Erlin2 | TRUE | -0.094255673 | 0.441179379 | No Change | Not Significant | TRUE | 0.06371924 | 0.15670896 | No Change | Not Significant |
| Ermap | TRUE | 1.114474874 | 1 | No Change | Not Significant | FALSE | NA | NA | No Change | Not Detected |
| Ermard | TRUE | -0.039283033 | 0.908714429 | No Change | Not Significant | TRUE | 0.073188708 | 0.737351049 | No Change | Not Significant |
| Ermn | TRUE | 0.493474007 | 0.40075936 | No Change | Not Significant | TRUE | 0.580668081 | 0.289522242 | No Change | Not Significant |
| Ermp1 | TRUE | -0.358088825 | 0.000104668 | No Change | Significant Negative | TRUE | -0.01697581 | 0.889849511 | No Change | Not Significant |
| Ern1 | TRUE | -0.002734335 | 0.984509162 | No Change | Not Significant | TRUE | -0.014897153 | 0.916810722 | No Change | Not Significant |
| Ern2 | TRUE | -0.075816135 | 0.79746811 | No Change | Not Significant | TRUE | 0.049944931 | 0.693679184 | No Change | Not Significant |
| Ero1a | TRUE | 0.329188136 | 0.000226707 | No Change | Significant Positive | TRUE | 0.098092995 | 0.339503983 | No Change | Not Significant |
| Ero1b | TRUE | 2.855756525 | 3.73E-132 | UP | UP | TRUE | 2.354847291 | 3.55E-161 | UP | UP |
| Erp27 | TRUE | 1.4516082 | 1 | No Change | Not Significant | FALSE | NA | NA | No Change | Not Detected |
| Erp29 | TRUE | 1.481045072 | 3.52E-137 | UP | UP | TRUE | 1.010528251 | 8.87E-82 | UP | UP |
| Erp44 | TRUE | 0.980774741 | 1.73E-29 | UP | UP | TRUE | 0.615158137 | 1.21E-26 | UP | UP |
| Errfi1 | TRUE | -0.01675059 | 0.968832408 | No Change | Not Significant | TRUE | -0.038868333 | 0.776097903 | No Change | Not Significant |
| Erv3 | TRUE | 2.597638802 | 1 | No Change | Not Significant | FALSE | NA | NA | No Change | Not Detected |
| Esam | TRUE | -1.060789067 | 0.045686082 | DOWN | DOWN | TRUE | -0.159582783 | 0.74719957 | No Change | Not Significant |
| Esco1 | TRUE | 0.044784593 | 0.793308598 | No Change | Not Significant | TRUE | 0.032241039 | 0.779660849 | No Change | Not Significant |
| Esco2 | TRUE | 0.664150522 | 0.014939401 | UP | UP | TRUE | 0.142706674 | 0.566769091 | No Change | Not Significant |
| Esd | TRUE | -0.165486692 | 0.106567719 | No Change | Not Significant | TRUE | 0.088046486 | 0.159045294 | No Change | Not Significant |
| Esf1 | TRUE | 0.141947683 | 0.513069369 | No Change | Not Significant | TRUE | -0.082187722 | 0.565705445 | No Change | Not Significant |
| Espl1 | TRUE | 1.041687552 | 1.13E-10 | UP | UP | TRUE | 0.028398243 | 0.900310808 | No Change | Not Significant |
| Espn | TRUE | -0.454883364 | 0.1525001 | No Change | Not Significant | TRUE | -0.483102022 | 4.35E-05 | No Change | Significant Negative |
| Espnl | TRUE | 2.80141191 | 0.002393317 | UP | UP | FALSE | NA | NA | No Change | Not Detected |
| Esr1 | TRUE | -0.399840883 | 0.403926995 | No Change | Not Significant | TRUE | 0.469794006 | 0.216846868 | No Change | Not Significant |
| Esr2 | TRUE | -0.173678646 | 0.683906563 | No Change | Not Significant | TRUE | 0.140447401 | 0.682675952 | No Change | Not Significant |
| Esrp1 | TRUE | 0.285134628 | 3.65E-06 | No Change | Significant Positive | TRUE | 0.08482125 | 0.094291987 | No Change | Not Significant |
| Esrp2 | TRUE | -0.19040145 | 0.018045571 | No Change | Significant Negative | TRUE | 0.103980914 | 0.213633392 | No Change | Not Significant |
| Esrra | TRUE | 0.086206116 | 0.564622747 | No Change | Not Significant | TRUE | 0.103594251 | 0.398306185 | No Change | Not Significant |
| Esrrb | TRUE | -2.31996059 | 1 | No Change | Not Significant | FALSE | NA | NA | No Change | Not Detected |
| Esrrg | TRUE | -0.987263703 | 0.000594601 | DOWN | DOWN | TRUE | -0.498332744 | 0.037542844 | No Change | Significant Negative |
| Ess2 | TRUE | 0.05037754 | 0.755621674 | No Change | Not Significant | TRUE | -0.057188279 | 0.692467352 | No Change | Not Significant |
| Esx1 | TRUE | 0.611520734 | 1 | No Change | Not Significant | FALSE | NA | NA | No Change | Not Detected |

| geneID | Cond1_DF_Detected | Cond1_DF_ValueCol | Cond1_DF_PadjCol | Cond1_DF_Cutoff | Cond1_DF_Cutoff_Specific.x | Cond2_DF_Detected | Cond2_DF_ValueCol | Cond2_DF_PadjCol | Cond2_DF_Cutoff | Cond2_DF_Cutoff_Specific | BG_Method | RG1_All |
|---|---|---|---|---|---|---|---|---|---|---|---|---|
| Esyt1 | TRUE | 0.309896286 | 0.015327126 | No Change | Significant Positive | TRUE | 0.000387152 | 0.997828443 | No Change | Not Significant | TRUE | Cond1 Significant Positive + Cond2 Not Significant |
| Esyt2 | TRUE | -0.086655095 | 0.393901406 | No Change | Not Significant | TRUE | -0.126506323 | 0.026787965 | No Change | Significant Negative | TRUE | Cond1 Not Significant + Cond2 Significant Negative |
| Esyt3 | TRUE | -0.406733526 | 0.788881899 | No Change | Not Significant | TRUE | -0.179060418 | 0.789158883 | No Change | Not Significant | TRUE | Cond1 Not Significant + Cond2 Not Significant |
| Etaa1 | TRUE | 0.06869181 | 0.822722373 | No Change | Not Significant | TRUE | -0.091755595 | 0.552028135 | No Change | Not Significant | TRUE | Cond1 Not Significant + Cond2 Not Significant |
| Etaa1os | TRUE | -0.437075845 | 0.549590181 | No Change | Not Significant | TRUE | 0.595397516 | 0.205867649 | No Change | Not Significant | TRUE | Cond1 Not Significant + Cond2 Not Significant |
| Etf1 | TRUE | 0.134092839 | 0.197442677 | No Change | Not Significant | TRUE | -0.010034306 | 0.919348861 | No Change | Not Significant | TRUE | Cond1 Not Significant + Cond2 Not Significant |
| Etfa | TRUE | -0.159964614 | 0.075235434 | No Change | Not Significant | TRUE | -0.083715794 | 0.180310883 | No Change | Not Significant | TRUE | Cond1 Not Significant + Cond2 Not Significant |
| Etfb | TRUE | 0.3001368 | 0.001180112 | No Change | Significant Positive | TRUE | 0.164806963 | 0.095081101 | No Change | Not Significant | TRUE | Cond1 Significant Positive + Cond2 Not Significant |
| Etfbkmt | TRUE | -0.399071296 | 0.011144808 | No Change | Significant Negative | TRUE | -0.075970381 | 0.628430063 | No Change | Not Significant | TRUE | Cond1 Significant Negative + Cond2 Not Significant |
| Etfdh | TRUE | -0.260719489 | 0.001412003 | No Change | Significant Negative | TRUE | 0.050111533 | 0.481643597 | No Change | Not Significant | TRUE | Cond1 Significant Negative + Cond2 Not Significant |
| Etfrf1 | TRUE | 0.093979141 | 0.561307404 | No Change | Not Significant | TRUE | 0.108632624 | 0.250536319 | No Change | Not Significant | TRUE | Cond1 Not Significant + Cond2 Not Significant |
| Ethe1 | TRUE | -0.665764782 | 1.03E-07 | DOWN | DOWN | TRUE | -0.100686877 | 0.461499638 | No Change | Not Significant | TRUE | Cond1 DOWN + Cond2 Not Significant |
| Etl4 | TRUE | 0.012355725 | 0.963320302 | No Change | Not Significant | TRUE | -0.083839798 | 0.481593365 | No Change | Not Significant | TRUE | Cond1 Not Significant + Cond2 Not Significant |
| Etnk1 | TRUE | 0.371642322 | 0.000632578 | No Change | Significant Positive | TRUE | 0.039166613 | 0.61867166 | No Change | Not Significant | TRUE | Cond1 Significant Positive + Cond2 Not Significant |
| Etnk2 | TRUE | -0.024939959 | 0.973795719 | No Change | Not Significant | TRUE | 0.354054461 | 0.434761962 | No Change | Not Significant | TRUE | Cond1 Not Significant + Cond2 Not Significant |
| Etnppl | TRUE | 1.05308989 | 1.93E-05 | UP | UP | TRUE | 0.471980094 | 0.001774894 | No Change | Significant Positive | TRUE | Cond1 UP + Cond2 Significant Positive |
| Ets1 | TRUE | 0.42819345 | 0.249900617 | No Change | Not Significant | TRUE | 0.559511655 | 0.01114091 | UP | UP | TRUE | Cond1 Not Significant + Cond2 UP |
| Ets2 | TRUE | 0.010307638 | 0.938630585 | No Change | Not Significant | TRUE | -0.068600402 | 0.221791752 | No Change | Not Significant | TRUE | Cond1 Not Significant + Cond2 Not Significant |
| Etv1 | TRUE | -0.306928895 | 0.683489308 | No Change | Not Significant | TRUE | 0.120138296 | 0.862768697 | No Change | Not Significant | TRUE | Cond1 Not Significant + Cond2 Not Significant |
| Etv3 | TRUE | 0.253097732 | 0.040777919 | No Change | Significant Positive | TRUE | 0.021745113 | 0.788682346 | No Change | Not Significant | TRUE | Cond1 Significant Positive + Cond2 Not Significant |
| Etv3l | TRUE | -1.389501579 | 1 | No Change | Not Significant | FALSE | NA | NA | No Change | Not Detected | FALSE | Background = FALSE |
| Etv4 | TRUE | 0.86964432 | 0.001846439 | UP | UP | TRUE | 0.420257887 | 0.057217045 | No Change | Not Significant | TRUE | Cond1 UP + Cond2 Not Significant |
| Etv5 | TRUE | -0.028626152 | 0.945659633 | No Change | Not Significant | TRUE | -0.154619404 | 0.435615504 | No Change | Not Significant | TRUE | Cond1 Not Significant + Cond2 Not Significant |
| Etv6 | TRUE | 0.188609481 | 0.145350463 | No Change | Not Significant | TRUE | -0.03812555 | 0.713271187 | No Change | Not Significant | TRUE | Cond1 Not Significant + Cond2 Not Significant |
| Eva1a | TRUE | -0.02076387 | 0.966128972 | No Change | Not Significant | TRUE | -0.336423033 | 0.301132615 | No Change | Not Significant | TRUE | Cond1 Not Significant + Cond2 Not Significant |
| Eva1b | TRUE | -0.260685031 | 0.570608171 | No Change | Not Significant | TRUE | 0.092770442 | 0.856166251 | No Change | Not Significant | TRUE | Cond1 Not Significant + Cond2 Not Significant |
| Eva1c | TRUE | -0.304980647 | 0.771232643 | No Change | Not Significant | TRUE | -0.209389981 | 0.789753 | No Change | Not Significant | TRUE | Cond1 Not Significant + Cond2 Not Significant |
| Evc | TRUE | -1.78517203 | 1 | No Change | Not Significant | FALSE | NA | NA | No Change | Not Detected | FALSE | Background = FALSE |
| Evc2 | TRUE | 0.578791019 | 1 | No Change | Not Significant | FALSE | NA | NA | No Change | Not Detected | FALSE | Background = FALSE |
| Evi2a | TRUE | -0.981349888 | 1 | No Change | Not Significant | TRUE | -0.320573959 | 0.734827388 | No Change | Not Significant | TRUE | Cond1 Not Significant + Cond2 Not Significant |
| Evi5 | TRUE | -0.050918729 | 0.708596728 | No Change | Not Significant | TRUE | -0.134893641 | 0.053478162 | No Change | Not Significant | TRUE | Cond1 Not Significant + Cond2 Not Significant |
| Evi5l | TRUE | 0.339960502 | 0.005488253 | No Change | Significant Positive | TRUE | 0.172167129 | 0.094722877 | No Change | Not Significant | TRUE | Cond1 Significant Positive + Cond2 Not Significant |
| Evl | TRUE | -0.294346368 | 0.340132673 | No Change | Not Significant | TRUE | -0.599700042 | 0.00791767 | DOWN | DOWN | TRUE | Cond1 Not Significant + Cond2 DOWN |
| Evpl | TRUE | -0.405824624 | 0.026309923 | No Change | Significant Negative | TRUE | 0.011369439 | 0.961246633 | No Change | Not Significant | TRUE | Cond1 Significant Negative + Cond2 Not Significant |
| Evx1 | TRUE | 0.323648812 | 0.367865096 | No Change | Not Significant | TRUE | 1.028668911 | 0.0153489 | UP | UP | TRUE | Cond1 Not Significant + Cond2 UP |
| Evx1os | TRUE | 0.566318329 | 0.276733031 | No Change | Not Significant | FALSE | NA | NA | No Change | Not Detected | FALSE | Background = FALSE |
| Evx2 | TRUE | 0.1486565 | 0.891264005 | No Change | Not Significant | TRUE | -0.031151105 | 0.975710693 | No Change | Not Significant | TRUE | Cond1 Not Significant + Cond2 Not Significant |
| Ewsr1 | TRUE | 0.031107711 | 0.80301103 | No Change | Not Significant | TRUE | -0.044476196 | 0.645753068 | No Change | Not Significant | TRUE | Cond1 Not Significant + Cond2 Not Significant |
| Exd1 | TRUE | 0.315472773 | 0.380697084 | No Change | Not Significant | TRUE | 0.658807984 | 0.026670568 | UP | UP | TRUE | Cond1 Not Significant + Cond2 UP |
| Exd2 | TRUE | 0.104264937 | 0.514332163 | No Change | Not Significant | TRUE | -0.015432286 | 0.909865152 | No Change | Not Significant | TRUE | Cond1 Not Significant + Cond2 Not Significant |
| Exo1 | TRUE | 0.559179316 | 0.009689999 | UP | UP | TRUE | 0.245473857 | 0.447440166 | No Change | Not Significant | TRUE | Cond1 UP + Cond2 Not Significant |
| Exo5 | TRUE | -0.032683113 | 0.853563561 | No Change | Not Significant | TRUE | -0.050646935 | 0.765233601 | No Change | Not Significant | TRUE | Cond1 Not Significant + Cond2 Not Significant |
| Exoc1 | TRUE | 0.093364045 | 0.405906838 | No Change | Not Significant | TRUE | -0.014122408 | 0.921722463 | No Change | Not Significant | TRUE | Cond1 Not Significant + Cond2 Not Significant |
| Exoc2 | TRUE | -0.166901082 | 0.302143144 | No Change | Not Significant | TRUE | -0.158543466 | 0.020439845 | No Change | Significant Negative | TRUE | Cond1 Not Significant + Cond2 Significant Negative |
| Exoc3 | TRUE | -0.031083932 | 0.822609935 | No Change | Not Significant | TRUE | 0.082677008 | 0.269652976 | No Change | Not Significant | TRUE | Cond1 Not Significant + Cond2 Not Significant |
| Exoc3l | TRUE | 0.367049993 | 0.601649715 | No Change | Not Significant | TRUE | 0.548250758 | 0.344951029 | No Change | Not Significant | TRUE | Cond1 Not Significant + Cond2 Not Significant |
| Exoc3l2 | TRUE | 0.179111933 | 1 | No Change | Not Significant | FALSE | NA | NA | No Change | Not Detected | FALSE | Background = FALSE |
| Exoc3l4 | TRUE | 0.898266752 | 0.006655171 | UP | UP | TRUE | 0.324714472 | 0.486955333 | No Change | Not Significant | TRUE | Cond1 UP + Cond2 Not Significant |
| Exoc4 | TRUE | 0.079218182 | 0.682033868 | No Change | Not Significant | TRUE | 0.139219012 | 0.13790318 | No Change | Not Significant | TRUE | Cond1 Not Significant + Cond2 Not Significant |
| Exoc5 | TRUE | 0.292799482 | 1.39E-05 | No Change | Significant Positive | TRUE | 0.062717664 | 0.390777655 | No Change | Not Significant | TRUE | Cond1 Significant Positive + Cond2 Not Significant |
| Exoc6 | TRUE | -0.082176162 | 0.696271072 | No Change | Not Significant | TRUE | -0.094916023 | 0.228015974 | No Change | Not Significant | TRUE | Cond1 Not Significant + Cond2 Not Significant |
| Exoc6b | TRUE | 0.072901256 | 0.656109195 | No Change | Not Significant | TRUE | -0.044110703 | 0.65828686 | No Change | Not Significant | TRUE | Cond1 Not Significant + Cond2 Not Significant |
| Exoc7 | TRUE | 0.247980608 | 0.000269538 | No Change | Significant Positive | TRUE | 0.16105953 | 0.001896124 | No Change | Significant Positive | TRUE | Cond1 Significant Positive + Cond2 Significant Positive |
| Exoc8 | TRUE | -0.081782213 | 0.439438221 | No Change | Not Significant | TRUE | -0.099984591 | 0.296257604 | No Change | Not Significant | TRUE | Cond1 Not Significant + Cond2 Not Significant |
| Exog | TRUE | 0.558622832 | 2.83E-05 | UP | UP | TRUE | 0.366374103 | 0.036667193 | No Change | Significant Positive | TRUE | Cond1 UP + Cond2 Significant Positive |
| Exosc1 | TRUE | 0.026545764 | 0.920280009 | No Change | Not Significant | TRUE | -0.02722551 | 0.917914545 | No Change | Not Significant | TRUE | Cond1 Not Significant + Cond2 Not Significant |
| Exosc10 | TRUE | -0.059051944 | 0.641714691 | No Change | Not Significant | TRUE | 0.024575567 | 0.864843268 | No Change | Not Significant | TRUE | Cond1 Not Significant + Cond2 Not Significant |
| Exosc2 | TRUE | 0.067566499 | 0.776435029 | No Change | Not Significant | TRUE | 0.016191839 | 0.925610506 | No Change | Not Significant | TRUE | Cond1 Not Significant + Cond2 Not Significant |
| Exosc3 | TRUE | 0.139393426 | 0.198651429 | No Change | Not Significant | TRUE | 0.07161386 | 0.530274756 | No Change | Not Significant | TRUE | Cond1 Not Significant + Cond2 Not Significant |
| Exosc4 | TRUE | 0.166658817 | 0.199902629 | No Change | Not Significant | TRUE | 0.15952505 | 0.228065548 | No Change | Not Significant | TRUE | Cond1 Not Significant + Cond2 Not Significant |
| Exosc5 | TRUE | -0.31094203 | 0.020269375 | No Change | Significant Negative | TRUE | 0.01492352 | 0.939561451 | No Change | Not Significant | TRUE | Cond1 Significant Negative + Cond2 Not Significant |
| Exosc7 | TRUE | 0.050447249 | 0.822722373 | No Change | Not Significant | TRUE | 0.043926573 | 0.759206347 | No Change | Not Significant | TRUE | Cond1 Not Significant + Cond2 Not Significant |
| Exosc8 | TRUE | 0.403362635 | 0.074589184 | No Change | Not Significant | TRUE | 0.066132144 | 0.73971984 | No Change | Not Significant | TRUE | Cond1 Not Significant + Cond2 Not Significant |
| Exosc9 | TRUE | -0.04939278 | 0.778956218 | No Change | Not Significant | TRUE | 0.015718061 | 0.917017949 | No Change | Not Significant | TRUE | Cond1 Not Significant + Cond2 Not Significant |
| Exph5 | TRUE | -0.35349995 | 0.004789552 | No Change | Significant Negative | TRUE | -0.304886059 | 0.005293007 | No Change | Significant Negative | TRUE | Cond1 Significant Negative + Cond2 Significant Negative |
| Ext1 | TRUE | 0.331589994 | 2.87E-05 | No Change | Significant Positive | TRUE | 0.177229731 | 0.042345703 | No Change | Significant Positive | TRUE | Cond1 Significant Positive + Cond2 Significant Positive |
| Ext2 | TRUE | 0.100006854 | 0.276057789 | No Change | Not Significant | TRUE | 0.096707689 | 0.244015888 | No Change | Not Significant | TRUE | Cond1 Not Significant + Cond2 Not Significant |
| Extl1 | TRUE | 0.211144467 | 0.728048657 | No Change | Not Significant | TRUE | -0.800518894 | 0.05831883 | No Change | Not Significant | TRUE | Cond1 Not Significant + Cond2 Not Significant |
| Extl2 | TRUE | 0.177165959 | 0.338919446 | No Change | Not Significant | TRUE | -0.019467602 | 0.929613548 | No Change | Not Significant | TRUE | Cond1 Not Significant + Cond2 Not Significant |
| Extl3 | TRUE | 0.006300884 | 0.972595566 | No Change | Not Significant | TRUE | -0.037776932 | 0.725570423 | No Change | Not Significant | TRUE | Cond1 Not Significant + Cond2 Not Significant |
| Eya1 | TRUE | -0.4284176 | 1 | No Change | Not Significant | FALSE | NA | NA | No Change | Not Detected | FALSE | Background = FALSE |
| Eya2 | TRUE | 0.581768439 | 0.152917797 | No Change | Not Significant | TRUE | 1.417391061 | 0.000381535 | UP | UP | TRUE | Cond1 Not Significant + Cond2 UP |
| Eya3 | TRUE | 0.06096203 | 0.595366473 | No Change | Not Significant | TRUE | 0.046809023 | 0.571645972 | No Change | Not Significant | TRUE | Cond1 Not Significant + Cond2 Not Significant |
| Eya4 | TRUE | 1.401733296 | 1 | No Change | Not Significant | FALSE | NA | NA | No Change | Not Detected | FALSE | Background = FALSE |
| Ezh1 | TRUE | -0.357921085 | 1.34E-06 | No Change | Significant Negative | TRUE | -0.109297676 | 0.086181037 | No Change | Not Significant | TRUE | Cond1 Significant Negative + Cond2 Not Significant |
| Ezh2 | TRUE | -0.034010117 | 0.856468824 | No Change | Not Significant | TRUE | -0.052039713 | 0.721608883 | No Change | Not Significant | TRUE | Cond1 Not Significant + Cond2 Not Significant |
| Ezr | TRUE | -0.259688605 | 0.049685066 | No Change | Significant Negative | TRUE | 0.040062963 | 0.791295959 | No Change | Not Significant | TRUE | Cond1 Significant Negative + Cond2 Not Significant |
| F10 | TRUE | 1.106337538 | 1 | No Change | Not Significant | FALSE | NA | NA | No Change | Not Detected | FALSE | Background = FALSE |
| F11 | TRUE | -0.236181783 | 1 | No Change | Not Significant | FALSE | NA | NA | No Change | Not Detected | FALSE | Background = FALSE |
| F11r | TRUE | -0.143913059 | 0.059503296 | No Change | Not Significant | TRUE | 0.091807021 | 0.141616335 | No Change | Not Significant | TRUE | Cond1 Not Significant + Cond2 Not Significant |
| F12 | TRUE | 0.180456164 | 1 | No Change | Not Significant | FALSE | NA | NA | No Change | Not Detected | FALSE | Background = FALSE |
| F13a1 | TRUE | 1.504490327 | 0.392188708 | No Change | Not Significant | FALSE | NA | NA | No Change | Not Detected | FALSE | Background = FALSE |
| F2 | TRUE | -0.577990849 | 0.477583041 | No Change | Not Significant | TRUE | -0.23567694 | 0.751266959 | No Change | Not Significant | TRUE | Cond1 Not Significant + Cond2 Not Significant |
| F2r | TRUE | 1.476879918 | 0.042442029 | UP | UP | TRUE | -0.079930466 | 0.942477935 | No Change | Not Significant | TRUE | Cond1 UP + Cond2 Not Significant |
| F2rl1 | TRUE | 0.114428297 | 0.289855686 | No Change | Not Significant | TRUE | 0.050858159 | 0.527617708 | No Change | Not Significant | TRUE | Cond1 Not Significant + Cond2 Not Significant |
| F2rl2 | TRUE | 0.037652298 | 0.947050968 | No Change | Not Significant | TRUE | -0.554105318 | 0.342694737 | No Change | Not Significant | TRUE | Cond1 Not Significant + Cond2 Not Significant |
| F2rl3 | TRUE | 1.616760303 | 1 | No Change | Not Significant | FALSE | NA | NA | No Change | Not Detected | FALSE | Background = FALSE |
| F3 | TRUE | -0.640420256 | 0.038576958 | DOWN | DOWN | TRUE | -0.29076401 | 0.014818642 | No Change | Significant Negative | TRUE | Cond1 DOWN + Cond2 Significant Negative |
| F420014N23Rik | TRUE | -0.248663811 | 0.749718386 | No Change | Not Significant | TRUE | -0.984159465 | 0.126769891 | No Change | Not Significant | TRUE | Cond1 Not Significant + Cond2 Not Significant |
| F5 | TRUE | -0.252624748 | 0.816921981 | No Change | Not Significant | TRUE | 0.663662259 | 0.219397086 | No Change | Not Significant | TRUE | Cond1 Not Significant + Cond2 Not Significant |
| F630040K05Rik | TRUE | -0.098940547 | 0.95803327 | No Change | Not Significant | TRUE | 0.385102792 | 0.719356203 | No Change | Not Significant | TRUE | Cond1 Not Significant + Cond2 Not Significant |
| F630042J09Rik | TRUE | -1.103014102 | 0.114635135 | No Change | Not Significant | TRUE | -0.189714682 | 0.769561988 | No Change | Not Significant | TRUE | Cond1 Not Significant + Cond2 Not Significant |
| F7 | TRUE | 1.137943524 | 1 | No Change | Not Significant | FALSE | NA | NA | No Change | Not Detected | FALSE | Background = FALSE |
| F730035M05Rik | TRUE | 1.720338855 | 1 | No Change | Not Significant | FALSE | NA | NA | No Change | Not Detected | FALSE | Background = FALSE |
| F730043M19Rik | TRUE | -0.60881501 | 1 | No Change | Not Significant | FALSE | NA | NA | No Change | Not Detected | FALSE | Background = FALSE |
| F730311O21Rik | TRUE | -1.71170909 | 1 | No Change | Not Significant | FALSE | NA | NA | No Change | Not Detected | FALSE | Background = FALSE |
| F8 | TRUE | 0.389275851 | 0.540015186 | No Change | Not Significant | TRUE | 0.160792689 | 0.755953522 | No Change | Not Significant | TRUE | Cond1 Not Significant + Cond2 Not Significant |
| F830016B08Rik | TRUE | 2.7462775 | 1 | No Change | Not Significant | FALSE | NA | NA | No Change | Not Detected | FALSE | Background = FALSE |
| F830045P16Rik | TRUE | 0.004283669 | 1 | No Change | Not Significant | FALSE | NA | NA | No Change | Not Detected | FALSE | Background = FALSE |
| F830208F22Rik | TRUE | -0.456991537 | 0.71414691 | No Change | Not Significant | TRUE | -1.021915466 | 0.194647414 | No Change | Not Significant | TRUE | Cond1 Not Significant + Cond2 Not Significant |
| F8a | TRUE | 0.186490528 | 0.214159123 | No Change | Not Significant | TRUE | 0.298233839 | 0.004976697 | No Change | Significant Positive | TRUE | Cond1 Not Significant + Cond2 Significant Positive |
| F930017D23Rik | TRUE | 0.210604882 | 1 | No Change | Not Significant | FALSE | NA | NA | No Change | Not Detected | FALSE | Background = FALSE |
| Fa2h | TRUE | 0.189889826 | 0.589852079 | No Change | Not Significant | TRUE | 0.141528859 | 0.357283876 | No Change | Not Significant | TRUE | Cond1 Not Significant + Cond2 Not Significant |
| Faah | TRUE | -0.223426399 | 0.048307661 | No Change | Significant Negative | TRUE | -0.220616972 | 0.000573713 | No Change | Significant Negative | TRUE | Cond1 Significant Negative + Cond2 Significant Negative |
| Faap100 | TRUE | -0.36743241 | 0.001406803 | No Change | Significant Negative | TRUE | -0.215023691 | 0.022022562 | No Change | Significant Negative | TRUE | Cond1 Significant Negative + Cond2 Significant Negative |
| Faap20 | TRUE | -0.340909081 | 0.003216222 | No Change | Significant Negative | TRUE | -0.091993956 | 0.510312204 | No Change | Not Significant | TRUE | Cond1 Significant Negative + Cond2 Not Significant |
| Faap24 | TRUE | 0.449223967 | 0.003393738 | No Change | Significant Positive | TRUE | 0.170945404 | 0.370743553 | No Change | Not Significant | TRUE | Cond1 Significant Positive + Cond2 Not Significant |
| Fabp1 | TRUE | -0.705644428 | 0.483654382 | No Change | Not Significant | TRUE | -0.052956757 | 0.979948241 | No Change | Not Significant | TRUE | Cond1 Not Significant + Cond2 Not Significant |
| Fabp2 | TRUE | 0.282729487 | 0.083216624 | No Change | Not Significant | TRUE | 0.221294012 | 0.032496922 | No Change | Significant Positive | TRUE | Cond1 Not Significant + Cond2 Significant Positive |
| Fabp3 | TRUE | -1.387040579 | 1 | No Change | Not Significant | FALSE | NA | NA | No Change | Not Detected | FALSE | Background = FALSE |
| Fabp5 | TRUE | -0.309765689 | 0.367316849 | No Change | Not Significant | TRUE | -0.221070496 | 0.328615354 | No Change | Not Significant | TRUE | Cond1 Not Significant + Cond2 Not Significant |
| Fabp6 | TRUE | 0.265855022 | 0.64392701 | No Change | Not Significant | TRUE | -1.93679198 | 1.72E-05 | DOWN | DOWN | TRUE | Cond1 Not Significant + Cond2 DOWN |
| Fadd | TRUE | -0.055569472 | 0.778759104 | No Change | Not Significant | TRUE | -0.006130736 | 0.971973978 | No Change | Not Significant | TRUE | Cond1 Not Significant + Cond2 Not Significant |
| Fads1 | TRUE | 0.927989838 | 6.42E-05 | UP | UP | TRUE | -0.01012782 | 0.96489645 | No Change | Not Significant | TRUE | Cond1 UP + Cond2 Not Significant |
| Fads2 | TRUE | -0.190202777 | 0.628507412 | No Change | Not Significant | TRUE | -0.133526471 | 0.381342548 | No Change | Not Significant | TRUE | Cond1 Not Significant + Cond2 Not Significant |
| Fads2b | TRUE | -0.619345334 | 1 | No Change | Not Significant | FALSE | NA | NA | No Change | Not Detected | FALSE | Background = FALSE |
| Fads3 | TRUE | 0.070109933 | 0.628822987 | No Change | Not Significant | TRUE | -0.194375834 | 0.195285263 | No Change | Not Significant | TRUE | Cond1 Not Significant + Cond2 Not Significant |
| Fads6 | TRUE | -0.460118197 | 0.38040956 | No Change | Not Significant | TRUE | -0.446357627 | 0.404955782 | No Change | Not Significant | TRUE | Cond1 Not Significant + Cond2 Not Significant |
| Faf1 | TRUE | 0.012285373 | 0.942424291 | No Change | Not Significant | TRUE | 0.039492311 | 0.73971984 | No Change | Not Significant | TRUE | Cond1 Not Significant + Cond2 Not Significant |
| Faf2 | TRUE | 0.187928364 | 0.077259526 | No Change | Not Significant | TRUE | 0.023931613 | 0.705835346 | No Change | Not Significant | TRUE | Cond1 Not Significant + Cond2 Not Significant |
| Fah | TRUE | -1.018137022 | 1.30E-07 | DOWN | DOWN | TRUE | -0.119865329 | 0.672975316 | No Change | Not Significant | TRUE | Cond1 DOWN + Cond2 Not Significant |
| Fahd1 | TRUE | -0.190106298 | 0.198286614 | No Change | Not Significant | TRUE | 0.102713305 | 0.337215278 | No Change | Not Significant | TRUE | Cond1 Not Significant + Cond2 Not Significant |
| Fahd2a | TRUE | -0.747796711 | 3.68E-10 | DOWN | DOWN | TRUE | -0.204420881 | 0.050786438 | No Change | Not Significant | TRUE | Cond1 DOWN + Cond2 Not Significant |
| Faim | TRUE | 0.076665374 | 0.717449153 | No Change | Not Significant | TRUE | -0.103381731 | 0.459085065 | No Change | Not Significant | TRUE | Cond1 Not Significant + Cond2 Not Significant |
| Faim2 | TRUE | 1.276768397 | 0.446602599 | No Change | Not Significant | FALSE | NA | NA | No Change | Not Detected | FALSE | Background = FALSE |
| Faiml | TRUE | -1.684239532 | 1 | No Change | Not Significant | FALSE | NA | NA | No Change | Not Detected | FALSE | Background = FALSE |
| Fam102a | TRUE | -0.208098 | 0.236336338 | No Change | Not Significant | TRUE | -0.03229166 | 0.853816014 | No Change | Not Significant | TRUE | Cond1 Not Significant + Cond2 Not Significant |
| Fam102b | TRUE | 0.488600574 | 8.28E-07 | No Change | Significant Positive | TRUE | 0.383146537 | 0.022022562 | No Change | Significant Positive | TRUE | Cond1 Significant Positive + Cond2 Significant Positive |
| Fam104a | TRUE | -0.060477996 | 0.761298159 | No Change | Not Significant | TRUE | -0.089743824 | 0.663421426 | No Change | Not Significant | TRUE | Cond1 Not Significant + Cond2 Not Significant |
| Fam107a | TRUE | 1.886446816 | 1 | No Change | Not Significant | FALSE | NA | NA | No Change | Not Detected | FALSE | Background = FALSE |
| Fam107b | TRUE | -0.237833673 | 0.195501402 | No Change | Not Significant | TRUE | -0.022333903 | 0.871991313 | No Change | Not Significant | TRUE | Cond1 Not Significant + Cond2 Not Significant |
| Fam110a | TRUE | 0.486156588 | 0.000113436 | No Change | Significant Positive | TRUE | 0.193590416 | 0.226895123 | No Change | Not Significant | TRUE | Cond1 Significant Positive + Cond2 Not Significant |
| Fam110b | TRUE | -0.209956646 | 0.709218761 | No Change | Not Significant | TRUE | -0.317108628 | 0.292453598 | No Change | Not Significant | TRUE | Cond1 Not Significant + Cond2 Not Significant |
| Fam110c | TRUE | -0.423701253 | 0.000155877 | No Change | Significant Negative | TRUE | -0.183926999 | 0.009794097 | No Change | Significant Negative | TRUE | Cond1 Significant Negative + Cond2 Significant Negative |
| Fam110d | TRUE | -0.490009715 | 1 | No Change | Not Significant | FALSE | NA | NA | No Change | Not Detected | FALSE | Background = FALSE |
| Fam111a | TRUE | 0.447590662 | 0.200815241 | No Change | Not Significant | TRUE | 0.254884888 | 0.322031193 | No Change | Not Significant | TRUE | Cond1 Not Significant + Cond2 Not Significant |
| Fam114a1 | TRUE | 0.150315233 | 0.253653789 | No Change | Not Significant | TRUE | 0.12086407 | 0.019455929 | No Change | Significant Positive | TRUE | Cond1 Not Significant + Cond2 Significant Positive |
| Fam114a2 | TRUE | -0.133804799 | 0.170333437 | No Change | Not Significant | TRUE | 0.030546342 | 0.750223246 | No Change | Not Significant | TRUE | Cond1 Not Significant + Cond2 Not Significant |
| Fam117a | TRUE | -0.321491147 | 0.184012672 | No Change | Not Significant | TRUE | 0.098217024 | 0.531357026 | No Change | Not Significant | TRUE | Cond1 Not Significant + Cond2 Not Significant |
| Fam117b | TRUE | 0.125385018 | 0.165344954 | No Change | Not Significant | TRUE | -0.069493858 | 0.388484989 | No Change | Not Significant | TRUE | Cond1 Not Significant + Cond2 Not Significant |
| Fam118a | TRUE | 0.688325738 | 5.03E-05 | UP | UP | TRUE | 0.107279729 | 0.484728722 | No Change | Not Significant | TRUE | Cond1 UP + Cond2 Not Significant |
| Fam118b | TRUE | -0.049495307 | 0.704868121 | No Change | Not Significant | TRUE | 0.012283833 | 0.934372006 | No Change | Not Significant | TRUE | Cond1 Not Significant + Cond2 Not Significant |
| Fam120a | TRUE | 0.128281167 | 0.035440956 | No Change | Significant Positive | TRUE | 0.044748805 | 0.596379092 | No Change | Not Significant | TRUE | Cond1 Significant Positive + Cond2 Not Significant |
| Fam120b | TRUE | 0.121739307 | 0.042377303 | No Change | Significant Positive | TRUE | 0.070329115 | 0.399424529 | No Change | Not Significant | TRUE | Cond1 Significant Positive + Cond2 Not Significant |
| Fam120c | TRUE | -0.097327135 | 0.699975951 | No Change | Not Significant | TRUE | 0.018104296 | 0.939855462 | No Change | Not Significant | TRUE | Cond1 Not Significant + Cond2 Not Significant |
| Fam122a | TRUE | 0.14860527 | 0.559860817 | No Change | Not Significant | TRUE | 0.193597694 | 0.236348478 | No Change | Not Significant | TRUE | Cond1 Not Significant + Cond2 Not Significant |
| Fam122b | TRUE | 0.383779122 | 0.134003711 | No Change | Not Significant | TRUE | -0.145927459 | 0.619086486 | No Change | Not Significant | TRUE | Cond1 Not Significant + Cond2 Not Significant |
| Fam124a | TRUE | -0.031657568 | 0.931679578 | No Change | Not Significant | TRUE | 0.382408075 | 0.002040458 | No Change | Significant Positive | TRUE | Cond1 Not Significant + Cond2 Significant Positive |
| Fam124b | TRUE | 1.574723239 | 1 | No Change | Not Significant | FALSE | NA | NA | No Change | Not Detected | FALSE | Background = FALSE |
| Fam131a | TRUE | -0.069591608 | 0.962004133 | No Change | Not Significant | TRUE | -0.988018934 | 0.041052542 | DOWN | DOWN | TRUE | Cond1 Not Significant + Cond2 DOWN |
| Fam131b | TRUE | -0.577063309 | 1 | No Change | Not Significant | FALSE | NA | NA | No Change | Not Detected | FALSE | Background = FALSE |
| Fam131c | TRUE | 0.596232356 | 0.559438733 | No Change | Not Significant | FALSE | NA | NA | No Change | Not Detected | FALSE | Background = FALSE |
| Fam133b | TRUE | 0.111528484 | 0.450443439 | No Change | Not Significant | TRUE | -0.016543945 | 0.929613548 | No Change | Not Significant | TRUE | Cond1 Not Significant + Cond2 Not Significant |
| Fam135a | TRUE | 0.06449214 | 0.588935008 | No Change | Not Significant | TRUE | -0.055368329 | 0.406831399 | No Change | Not Significant | TRUE | Cond1 Not Significant + Cond2 Not Significant |
| Fam135b | TRUE | 3.836297985 | 0.164631718 | No Change | Not Significant | FALSE | NA | NA | No Change | Not Detected | FALSE | Background = FALSE |
| Fam136a | TRUE | -0.146623179 | 0.439134108 | No Change | Not Significant | TRUE | 0.071075078 | 0.434133726 | No Change | Not Significant | TRUE | Cond1 Not Significant + Cond2 Not Significant |
| Fam13a | TRUE | 0.628312422 | 4.31E-07 | UP | UP | TRUE | 0.336572116 | 0.000908855 | No Change | Significant Positive | TRUE | Cond1 UP + Cond2 Significant Positive |
| Fam13b | TRUE | -0.129149796 | 0.309868458 | No Change | Not Significant | TRUE | -0.155616231 | 0.205361161 | No Change | Not Significant | TRUE | Cond1 Not Significant + Cond2 Not Significant |
| Fam13c | TRUE | -0.425416316 | 1 | No Change | Not Significant | FALSE | NA | NA | No Change | Not Detected | FALSE | Background = FALSE |
| Fam149a | TRUE | 0.477555724 | 0.003417675 | No Change | Significant Positive | TRUE | 0.595355991 | 0.000988977 | UP | UP | TRUE | Cond1 Significant Positive + Cond2 UP |
| Fam149b | TRUE | 0.008974215 | 0.941960269 | No Change | Not Significant | TRUE | 0.083362777 | 0.286054693 | No Change | Not Significant | TRUE | Cond1 Not Significant + Cond2 Not Significant |
| Fam151a | TRUE | -0.143795386 | 1 | No Change | Not Significant | FALSE | NA | NA | No Change | Not Detected | FALSE | Background = FALSE |
| Fam151b | TRUE | 0.201544158 | 0.506641059 | No Change | Not Significant | TRUE | 0.037532805 | 0.925325507 | No Change | Not Significant | TRUE | Cond1 Not Significant + Cond2 Not Significant |
| Fam161a | TRUE | -0.490754942 | 0.03273686 | No Change | Significant Negative | TRUE | 0.026661675 | 0.887703374 | No Change | Not Significant | TRUE | Cond1 Significant Negative + Cond2 Not Significant |
| Fam161b | TRUE | 0.12848565 | 0.720614094 | No Change | Not Significant | TRUE | 0.059793817 | 0.887756682 | No Change | Not Significant | TRUE | Cond1 Not Significant + Cond2 Not Significant |
| Fam162a | TRUE | -0.154857898 | 0.405453505 | No Change | Not Significant | TRUE | 0.13023726 | 0.302551116 | No Change | Not Significant | TRUE | Cond1 Not Significant + Cond2 Not Significant |
| Fam163a | TRUE | 0.944359006 | 1 | No Change | Not Significant | FALSE | NA | NA | No Change | Not Detected | FALSE | Background = FALSE |
| Fam163b | TRUE | -1.672344119 | 1 | No Change | Not Significant | FALSE | NA | NA | No Change | Not Detected | FALSE | Background = FALSE |
| Fam166a | TRUE | 0.852705991 | 0.199168782 | No Change | Not Significant | TRUE | -0.584941921 | 0.369310836 | No Change | Not Significant | TRUE | Cond1 Not Significant + Cond2 Not Significant |
| Fam166b | TRUE | -0.806588798 | 1 | No Change | Not Significant | FALSE | NA | NA | No Change | Not Detected | FALSE | Background = FALSE |
| Fam167a | TRUE | -0.522835259 | 0.483542311 | No Change | Not Significant | TRUE | -0.046724801 | 0.929981434 | No Change | Not Significant | TRUE | Cond1 Not Significant + Cond2 Not Significant |
| Fam167b | TRUE | -0.576253798 | 1 | No Change | Not Significant | FALSE | NA | NA | No Change | Not Detected | FALSE | Background = FALSE |
| Fam168a | TRUE | -0.223162447 | 0.13639879 | No Change | Not Significant | TRUE | -0.036749059 | 0.887577222 | No Change | Not Significant | TRUE | Cond1 Not Significant + Cond2 Not Significant |
| Fam168b | TRUE | -0.069818167 | 0.624327666 | No Change | Not Significant | TRUE | -0.039298373 | 0.495031103 | No Change | Not Significant | TRUE | Cond1 Not Significant + Cond2 Not Significant |
| Fam169a | TRUE | 0.30026026 | 0.709532566 | No Change | Not Significant | TRUE | 0.010390626 | 0.993012764 | No Change | Not Significant | TRUE | Cond1 Not Significant + Cond2 Not Significant |
| Fam169b | TRUE | 0.011369419 | 1 | No Change | Not Significant | FALSE | NA | NA | No Change | Not Detected | FALSE | Background = FALSE |
| Fam170b | TRUE | -0.958881852 | 1 | No Change | Not Significant | FALSE | NA | NA | No Change | Not Detected | FALSE | Background = FALSE |
| Fam171a1 | TRUE | -0.234081723 | 0.083972816 | No Change | Not Significant | TRUE | -0.030742607 | 0.841984797 | No Change | Not Significant | TRUE | Cond1 Not Significant + Cond2 Not Significant |
| Fam171a2 | TRUE | 0.323716638 | 0.71423395 | No Change | Not Significant | TRUE | -0.004029965 | 0.996791885 | No Change | Not Significant | TRUE | Cond1 Not Significant + Cond2 Not Significant |
| Fam171b | TRUE | 0.58962249 | 1 | No Change | Not Significant | FALSE | NA | NA | No Change | Not Detected | FALSE | Background = FALSE |
| Fam172a | TRUE | -0.103222116 | 0.449146195 | No Change | Not Significant | TRUE | -0.024103108 | 0.866650863 | No Change | Not Significant | TRUE | Cond1 Not Significant + Cond2 Not Significant |
| Fam174a | TRUE | -0.020940173 | 0.929465614 | No Change | Not Significant | TRUE | 0.089328323 | 0.440072685 | No Change | Not Significant | TRUE | Cond1 Not Significant + Cond2 Not Significant |
| Fam174b | TRUE | 1.239832475 | 6.63E-22 | UP | UP | TRUE | 0.979147659 | 7.89E-87 | UP | UP | TRUE | Cond1 UP + Cond2 UP |
| Fam174c | TRUE | -0.049157255 | 0.838533729 | No Change | Not Significant | TRUE | 0.231619888 | 0.146880196 | No Change | Not Significant | TRUE | Cond1 Not Significant + Cond2 Not Significant |
| Fam178b | TRUE | 0.8328237 | 1 | No Change | Not Significant | FALSE | NA | NA | No Change | Not Detected | FALSE | Background = FALSE |
| Fam183b | TRUE | 0.345803344 | 1 | No Change | Not Significant | TRUE | 0.34116675 | 0.558770915 | No Change | Not Significant | TRUE | Cond1 Not Significant + Cond2 Not Significant |
| Fam184a | TRUE | -0.174595324 | 1 | No Change | Not Significant | FALSE | NA | NA | No Change | Not Detected | FALSE | Background = FALSE |
| Fam184b | TRUE | 0.134198893 | 0.883589766 | No Change | Not Significant | TRUE | -0.374280042 | 0.399424529 | No Change | Not Significant | TRUE | Cond1 Not Significant + Cond2 Not Significant |
| Fam185a | TRUE | 0.365414807 | 0.006462165 | No Change | Significant Positive | TRUE | 0.12237055 | 0.407747138 | No Change | Not Significant | TRUE | Cond1 Significant Positive + Cond2 Not Significant |
| Fam186a | TRUE | -0.009888454 | 1 | No Change | Not Significant | FALSE | NA | NA | No Change | Not Detected | FALSE | Background = FALSE |
| Fam186b | TRUE | 1.156829676 | 1 | No Change | Not Significant | FALSE | NA | NA | No Change | Not Detected | FALSE | Background = FALSE |
| Fam187b | TRUE | -0.064274354 | 0.890125936 | No Change | Not Significant | TRUE | 0.25071099 | 0.378894236 | No Change | Not Significant | TRUE | Cond1 Not Significant + Cond2 Not Significant |

| geneID | Cond1_DF_Detected | Cond1_DF_ValueCol | Cond1_DF_PadjCol | Cond1_DF_Cutoff | Cond1_DF_Cutoff_Specific.x | Cond2_DF_Detected |
|---|---|---|---|---|---|---|
| Fam193a | TRUE | 0.044877696 | 0.79451259 | No Change | Not Significant | TRUE |
| Fam193b | TRUE | 0.071998913 | 0.822256556 | No Change | Not Significant | TRUE |
| Fam199x | TRUE | 0.227059536 | 0.197666177 | No Change | Not Significant | TRUE |
| Fam204a | TRUE | 0.194592319 | 0.154446119 | No Change | Not Significant | TRUE |
| Fam207a | TRUE | 0.156954947 | 0.085053259 | No Change | Not Significant | TRUE |
| Fam209 | TRUE | 1.701305958 | 1 | No Change | Not Significant | FALSE |
| Fam20a | TRUE | -0.21744222 | 1 | No Change | Not Significant | FALSE |
| Fam20b | TRUE | 0.074404107 | 0.493794621 | No Change | Not Significant | TRUE |
| Fam20c | TRUE | -1.55512212 | 0.16307245 | No Change | Not Significant | TRUE |
| Fam210a | TRUE | -0.163869172 | 0.28456734 | No Change | Not Significant | TRUE |
| Fam210b | TRUE | -0.660733136 | 1.01E-17 | DOWN | DOWN | TRUE |
| Fam216a | TRUE | 0.838462575 | 0.027169813 | UP | UP | TRUE |
| Fam217a | TRUE | 0.553925582 | 1 | No Change | Not Significant | FALSE |
| Fam217b | TRUE | 2.414020268 | 2.96E-27 | UP | UP | TRUE |
| Fam219a | TRUE | 0.846253661 | 0.006324576 | UP | UP | TRUE |
| Fam219b | TRUE | -0.167428877 | 0.292373241 | No Change | Not Significant | TRUE |
| Fam220a | TRUE | 0.438174238 | 0.61004314 | No Change | Not Significant | TRUE |
| Fam221a | TRUE | -0.473422634 | 0.014639914 | No Change | Significant Negative | TRUE |
| Fam221b | TRUE | 0.646121746 | 0.179322626 | No Change | Not Significant | TRUE |
| Fam222a | TRUE | 0.752740648 | 0.000487819 | UP | UP | TRUE |
| Fam222b | TRUE | 0.193948209 | 0.329600388 | No Change | Not Significant | TRUE |
| Fam227a | TRUE | -0.197538548 | 0.738667101 | No Change | Not Significant | TRUE |
| Fam227b | TRUE | 0.841863078 | 1 | No Change | Not Significant | FALSE |
| Fam228a | TRUE | 1.423202442 | 1 | No Change | Not Significant | FALSE |
| Fam228b | TRUE | -0.568205562 | 1 | No Change | Not Significant | FALSE |
| Fam229a | TRUE | 1.929741742 | 1 | No Change | Not Significant | FALSE |
| Fam229b | TRUE | 0.450307274 | 0.572963936 | No Change | Not Significant | FALSE |
| Fam234a | TRUE | -0.424675033 | 1.02E-10 | No Change | Significant Negative | TRUE |
| Fam234b | TRUE | 0.255391215 | 0.885199039 | No Change | Not Significant | TRUE |
| Fam241a | TRUE | -0.001839138 | 0.99357673 | No Change | Not Significant | TRUE |
| Fam241b | TRUE | -0.110812508 | 0.421254642 | No Change | Not Significant | TRUE |
| Fam243 | TRUE | -1.678951224 | 1 | No Change | Not Significant | FALSE |
| Fam24a | TRUE | 0.393942932 | 1 | No Change | Not Significant | FALSE |
| Fam32a | TRUE | -0.102547448 | 0.485210052 | No Change | Not Significant | TRUE |
| Fam3a | TRUE | 0.120371699 | 0.232815456 | No Change | Not Significant | TRUE |
| Fam3b | TRUE | -0.080998044 | 0.556711723 | No Change | Not Significant | TRUE |
| Fam3c | TRUE | 0.782354572 | 8.08E-06 | UP | UP | TRUE |
| Fam3d | TRUE | -0.113572816 | 0.728820184 | No Change | Not Significant | TRUE |
| Fam43a | TRUE | -0.984463149 | 0.038017517 | DOWN | DOWN | TRUE |
| Fam50a | TRUE | -0.038652073 | 0.85037506 | No Change | Not Significant | TRUE |
| Fam53a | TRUE | -0.058022654 | 0.733113936 | No Change | Not Significant | TRUE |
| Fam53b | TRUE | -0.208314742 | 0.090987984 | No Change | Not Significant | TRUE |
| Fam53c | TRUE | 0.251501755 | 0.024110197 | No Change | Significant Positive | TRUE |
| Fam72a | TRUE | 0.984316701 | 0.010831886 | UP | UP | TRUE |
| Fam76a | TRUE | -0.080060024 | 0.525167043 | No Change | Not Significant | TRUE |
| Fam76b | TRUE | 0.120955736 | 0.343912139 | No Change | Not Significant | TRUE |
| Fam78a | TRUE | -0.823949326 | 0.02103538 | DOWN | DOWN | TRUE |
| Fam78b | TRUE | -1.27901746 | 1 | No Change | Not Significant | FALSE |
| Fam81a | TRUE | 2.14292827 | 1 | No Change | Not Significant | FALSE |
| Fam81b | TRUE | -0.332893795 | 1 | No Change | Not Significant | FALSE |
| Fam83b | TRUE | -0.301958417 | 0.001601366 | No Change | Significant Negative | TRUE |
| Fam83c | TRUE | -0.272812866 | 0.088524117 | No Change | Not Significant | TRUE |
| Fam83d | TRUE | 0.452283151 | 0.022846718 | No Change | Significant Positive | TRUE |
| Fam83e | TRUE | -0.175057959 | 0.013704495 | No Change | Significant Negative | TRUE |
| Fam83f | TRUE | 0.183937295 | 0.039390404 | No Change | Significant Positive | TRUE |
| Fam83g | TRUE | -0.140242123 | 0.309858682 | No Change | Not Significant | TRUE |
| Fam83h | TRUE | -0.342203447 | 0.002381698 | No Change | Significant Negative | TRUE |
| Fam89a | TRUE | 0.500922036 | 0.532085855 | No Change | Not Significant | TRUE |
| Fam8a1 | TRUE | 0.165026026 | 0.166521981 | No Change | Not Significant | TRUE |
| Fam91a1 | TRUE | 0.02826489 | 0.898299072 | No Change | Not Significant | TRUE |
| Fam98a | TRUE | 0.232142081 | 0.133465102 | No Change | Not Significant | TRUE |
| Fam98b | TRUE | 0.049942596 | 0.81814452 | No Change | Not Significant | TRUE |
| Fam98c | TRUE | 0.90413415 | 1.09E-21 | UP | UP | TRUE |
| Fan1 | TRUE | -0.287151679 | 0.062762157 | No Change | Not Significant | TRUE |
| Fanca | TRUE | 0.619353788 | 0.018358264 | UP | UP | TRUE |
| Fancb | TRUE | 1.147873799 | 1.47E-06 | UP | UP | TRUE |
| Fancc | TRUE | -0.008435259 | 0.971883559 | No Change | Not Significant | TRUE |
| Fancd2 | TRUE | 0.781603972 | 0.000144113 | UP | UP | TRUE |
| Fance | TRUE | -0.068752066 | 0.708596728 | No Change | Not Significant | TRUE |
| Fancf | TRUE | 0.175255043 | 0.68833522 | No Change | Not Significant | TRUE |
| Fancg | TRUE | 0.094853144 | 0.647057325 | No Change | Not Significant | TRUE |
| Fanci | TRUE | 0.824528163 | 0.001853831 | UP | UP | TRUE |
| Fancl | TRUE | 0.134988823 | 0.519828028 | No Change | Not Significant | TRUE |
| Fancm | TRUE | 0.459563448 | 0.000952873 | No Change | Significant Positive | TRUE |
| Fank1 | TRUE | -0.252881425 | 0.762469572 | No Change | Not Significant | TRUE |
| Fap | TRUE | 0.97657379 | 5.88E-05 | UP | UP | TRUE |
| Far1 | TRUE | 2.696355798 | 9.26E-223 | UP | UP | TRUE |
| Far1os | TRUE | 2.37354346 | 1.81E-30 | UP | UP | TRUE |
| Far2 | TRUE | -0.016215803 | 0.957979661 | No Change | Not Significant | TRUE |
| Farp1 | TRUE | 0.265597705 | 0.004714752 | No Change | Significant Positive | TRUE |
| Farp2 | TRUE | -0.38895656 | 4.04E-05 | No Change | Significant Negative | TRUE |
| Fars2 | TRUE | -0.252955912 | 0.001176235 | No Change | Significant Negative | TRUE |
| Farsa | TRUE | 0.220791324 | 0.056074481 | No Change | Not Significant | TRUE |
| Farsb | TRUE | 0.194157525 | 0.176210159 | No Change | Not Significant | TRUE |
| Fas | TRUE | 0.672031196 | 0.000216378 | UP | UP | TRUE |
| Fasl | TRUE | -0.649880907 | 0.509468423 | No Change | Not Significant | TRUE |
| Fasn | TRUE | 0.884991564 | 7.19E-29 | UP | UP | TRUE |
| Fastk | TRUE | -0.026461362 | 0.853563561 | No Change | Not Significant | TRUE |
| Fastkd1 | TRUE | -0.329615412 | 0.005507949 | No Change | Significant Negative | TRUE |
| Fastkd2 | TRUE | 0.131250477 | 0.554128799 | No Change | Not Significant | TRUE |
| Fastkd3 | TRUE | 0.169560687 | 0.117439107 | No Change | Not Significant | TRUE |
| Fastkd5 | TRUE | -0.10071539 | 0.740505016 | No Change | Not Significant | TRUE |
| Fat1 | TRUE | -0.179665939 | 0.349981053 | No Change | Not Significant | TRUE |
| Fat2 | TRUE | 0.884051914 | 0.00179544 | UP | UP | TRUE |
| Fat3 | TRUE | 1.210281989 | 0.816884691 | No Change | Not Significant | FALSE |
| Fat4 | TRUE | 0.619711252 | 0.567081848 | No Change | Not Significant | TRUE |
| Fau | TRUE | -0.454229761 | 1.62E-05 | No Change | Significant Negative | TRUE |
| Faxc | TRUE | 0.733172212 | 0.347924739 | No Change | Not Significant | FALSE |
| Fbf1 | TRUE | 0.135027057 | 0.448702897 | No Change | Not Significant | TRUE |
| Fbh1 | TRUE | -0.134117749 | 0.093397187 | No Change | Not Significant | TRUE |
| Fbl | TRUE | -0.130829399 | 0.540682396 | No Change | Not Significant | TRUE |
| Fblim1 | TRUE | -0.324754597 | 0.000122946 | No Change | Significant Negative | TRUE |
| Fbln1 | TRUE | 1.484863979 | 9.56E-09 | UP | UP | TRUE |
| Fbln2 | TRUE | 1.013754555 | 1 | No Change | Not Significant | FALSE |
| Fbln5 | TRUE | 0.639394526 | 1 | No Change | Not Significant | FALSE |
| Fbn1 | TRUE | -1.710901044 | 1.01E-05 | DOWN | DOWN | TRUE |
| Fbn2 | TRUE | 0.503744998 | 0.822609935 | No Change | Not Significant | FALSE |
| Fbp1 | TRUE | -0.368086939 | 0.228942639 | No Change | Not Significant | TRUE |
| Fbp2 | TRUE | -0.106529392 | 0.347734431 | No Change | Not Significant | TRUE |
| Fbrs | TRUE | 0.263815366 | 0.372922396 | No Change | Not Significant | TRUE |
| Fbrsl1 | TRUE | -0.089875403 | 0.723107237 | No Change | Not Significant | TRUE |
| Fbxl12 | TRUE | 0.177416184 | 0.37341949 | No Change | Not Significant | TRUE |
| Fbxl12os | TRUE | -0.024478013 | 0.975869522 | No Change | Not Significant | TRUE |
| Fbxl13 | TRUE | -0.844673761 | 1 | No Change | Not Significant | FALSE |
| Fbxl14 | TRUE | 0.023705135 | 0.881056435 | No Change | Not Significant | TRUE |
| Fbxl15 | TRUE | -0.362447163 | 0.139280674 | No Change | Not Significant | TRUE |
| Fbxl16 | TRUE | -0.612309859 | 0.330166076 | No Change | Not Significant | TRUE |
| Fbxl17 | TRUE | -0.100688418 | 0.368933447 | No Change | Not Significant | TRUE |
| Fbxl18 | TRUE | -0.00221628 | 0.989008948 | No Change | Not Significant | TRUE |
| Fbxl19 | TRUE | 0.093584246 | 0.416065952 | No Change | Not Significant | TRUE |
| Fbxl2 | TRUE | -0.333708396 | 0.698788598 | No Change | Not Significant | TRUE |
| Fbxl20 | TRUE | -0.501437948 | 0.000958551 | DOWN | DOWN | TRUE |
| Fbxl21 | TRUE | -0.290007858 | 0.747996408 | No Change | Not Significant | TRUE |
| Fbxl22 | TRUE | 0.291363506 | 0.465324602 | No Change | Not Significant | TRUE |
| Fbxl3 | TRUE | -0.430407229 | 0.000113436 | No Change | Significant Negative | TRUE |
| Fbxl4 | TRUE | 0.012677035 | 0.957100023 | No Change | Not Significant | TRUE |
| Fbxl5 | TRUE | 0.334136804 | 7.13E-06 | No Change | Significant Positive | TRUE |
| Fbxl6 | TRUE | -0.247000629 | 0.03127124 | No Change | Significant Negative | TRUE |
| Fbxl7 | TRUE | -0.41852882 | 1 | No Change | Not Significant | FALSE |
| Fbxl8 | TRUE | -0.287241085 | 0.044513657 | No Change | Significant Negative | TRUE |
| Fbxl9 | TRUE | -0.241980938 | 0.284375933 | No Change | Not Significant | TRUE |
| Fbxo10 | TRUE | 0.236814524 | 0.405213026 | No Change | Not Significant | TRUE |
| Fbxo11 | TRUE | 0.083509159 | 0.603601732 | No Change | Not Significant | TRUE |
| Fbxo15 | TRUE | 2.447794607 | 1 | No Change | Not Significant | FALSE |
| Fbxo16 | TRUE | 2.669648191 | 3.82E-05 | UP | UP | TRUE |
| Fbxo17 | TRUE | 0.014603176 | 0.981387498 | No Change | Not Significant | TRUE |
| Fbxo2 | TRUE | 0.966532944 | 1 | No Change | Not Significant | FALSE |
| Fbxo21 | TRUE | 0.170309681 | 0.479629566 | No Change | Not Significant | TRUE |
| Fbxo22 | TRUE | -0.005363454 | 0.971883559 | No Change | Not Significant | TRUE |
| Fbxo24 | TRUE | -0.247817723 | 0.695055986 | No Change | Not Significant | TRUE |
| Fbxo25 | TRUE | -0.235829337 | 0.078282779 | No Change | Not Significant | TRUE |
| Fbxo27 | TRUE | -1.609386994 | 1 | No Change | Not Significant | FALSE |
| Fbxo28 | TRUE | 0.030190042 | 0.872440598 | No Change | Not Significant | TRUE |
| Fbxo3 | TRUE | -0.169492921 | 0.23816226 | No Change | Not Significant | TRUE |
| Fbxo30 | TRUE | -0.188832511 | 0.280659184 | No Change | Not Significant | TRUE |
| Fbxo31 | TRUE | -0.365511835 | 0.004347186 | No Change | Significant Negative | TRUE |
| Fbxo32 | TRUE | -1.033438972 | 1.14E-05 | DOWN | DOWN | TRUE |
| Fbxo33 | TRUE | 0.276874185 | 0.002788738 | No Change | Significant Positive | TRUE |
| Fbxo34 | TRUE | 0.118860047 | 0.468075728 | No Change | Not Significant | TRUE |
| Fbxo36 | TRUE | 0.385850604 | 0.031809007 | No Change | Significant Positive | TRUE |
| Fbxo38 | TRUE | 0.060597605 | 0.458272537 | No Change | Not Significant | TRUE |
| Fbxo4 | TRUE | 0.223496202 | 0.265192684 | No Change | Not Significant | TRUE |
| Fbxo40 | TRUE | -0.167710837 | 1 | No Change | Not Significant | FALSE |
| Fbxo41 | TRUE | 0.144108044 | 0.947328174 | No Change | Not Significant | FALSE |
| Fbxo42 | TRUE | 0.0207134 | 0.854899358 | No Change | Not Significant | TRUE |
| Fbxo43 | TRUE | 0.96571948 | 1 | No Change | Not Significant | FALSE |
| Fbxo44 | TRUE | -0.58187091 | 0.034924193 | DOWN | DOWN | TRUE |
| Fbxo45 | TRUE | 0.052315278 | 0.828386324 | No Change | Not Significant | TRUE |
| Fbxo46 | TRUE | -0.032511182 | 0.812675931 | No Change | Not Significant | TRUE |
| Fbxo47 | TRUE | 0.58738718 | 0.419497057 | No Change | Not Significant | TRUE |
| Fbxo48 | TRUE | 1.205929978 | 0.066917689 | No Change | Not Significant | TRUE |
| Fbxo5 | TRUE | 0.413797233 | 0.222484875 | No Change | Not Significant | TRUE |
| Fbxo6 | TRUE | -0.353158595 | 0.012311721 | No Change | Significant Negative | TRUE |
| Fbxo7 | TRUE | 0.032008134 | 0.888933284 | No Change | Not Significant | TRUE |
| Fbxo8 | TRUE | 0.079922805 | 0.494763373 | No Change | Not Significant | TRUE |
| Fbxo9 | TRUE | 0.136448193 | 0.16718219 | No Change | Not Significant | TRUE |
| Fbxw10 | TRUE | 1.252068254 | 0.080148748 | No Change | Not Significant | TRUE |
| Fbxw11 | TRUE | -0.121203179 | 0.225948198 | No Change | Not Significant | TRUE |
| Fbxw13 | TRUE | -0.434890619 | 1 | No Change | Not Significant | FALSE |
| Fbxw15 | TRUE | 1.442983445 | 1 | No Change | Not Significant | FALSE |
| Fbxw17 | TRUE | -0.378182632 | 0.062415953 | No Change | Not Significant | TRUE |
| Fbxw18 | TRUE | 0.35907496 | 1 | No Change | Not Significant | FALSE |
| Fbxw19 | TRUE | -1.209762248 | 1 | No Change | Not Significant | FALSE |
| Fbxw2 | TRUE | 0.129984703 | 0.107141645 | No Change | Not Significant | TRUE |
| Fbxw21 | TRUE | -0.731469516 | 1 | No Change | Not Significant | FALSE |
| Fbxw22 | TRUE | -0.744284082 | 0.266249604 | No Change | Not Significant | TRUE |
| Fbxw24 | TRUE | -0.715863867 | 1 | No Change | Not Significant | FALSE |
| Fbxw25 | TRUE | 1.726152933 | 1 | No Change | Not Significant | FALSE |
| Fbxw4 | TRUE | 0.164273921 | 0.554606509 | No Change | Not Significant | TRUE |
| Fbxw5 | TRUE | 0.05780426 | 0.618855346 | No Change | Not Significant | TRUE |
| Fbxw7 | TRUE | -0.095984172 | 0.389807821 | No Change | Not Significant | TRUE |
| Fbxw8 | TRUE | 0.265118971 | 0.000825955 | No Change | Significant Positive | TRUE |
| Fbxw9 | TRUE | 0.106874896 | 0.456768986 | No Change | Not Significant | TRUE |
| Fcamr | TRUE | 0.620654336 | 0.019227501 | UP | UP | TRUE |
| Fcer1g | TRUE | -0.028405548 | 0.972316965 | No Change | Not Significant | TRUE |
| Fcer2a | TRUE | -0.986683609 | 1 | No Change | Not Significant | FALSE |
| Fcf1 | TRUE | -0.140703634 | 0.467346556 | No Change | Not Significant | TRUE |
| Fcgbp | TRUE | -0.268258989 | 0.567700522 | No Change | Not Significant | TRUE |
| Fcgr1 | TRUE | 2.155812569 | 1 | No Change | Not Significant | FALSE |
| Fcgr2b | TRUE | 0.239950506 | 1 | No Change | Not Significant | FALSE |
| Fcgr3 | TRUE | 0.59395473 | 1 | No Change | Not Significant | FALSE |
| Fcgr4 | TRUE | 2.377806092 | 1 | No Change | Not Significant | FALSE |

Supplementary Table 3

| geneID | Cond1_DF_Detected | Cond1_DF_ValueCol | Cond1_DF_PadjCol | Cond1_DF_Cutoff | Cond1_DF_Cutoff_Specific.x | Cond2_DF_Detected |
|---|---|---|---|---|---|---|
| Fcgrt | TRUE | -0.588596695 | 1.13E-06 | DOWN | DOWN | TRUE |
| Fcho1 | TRUE | -0.187071256 | 0.617091033 | No Change | Not Significant | TRUE |
| Fcho2 | TRUE | -0.16756448 | 0.270067889 | No Change | Not Significant | TRUE |
| Fchsd1 | TRUE | -0.214466197 | 0.273253364 | No Change | Not Significant | TRUE |
| Fchsd2 | TRUE | -0.27488335 | 0.025095735 | No Change | Significant Negative | TRUE |
| Fcmr | TRUE | -1.848994113 | 1 | No Change | Not Significant | FALSE |
| Fcna | TRUE | -0.587894866 | 0.006961441 | DOWN | DOWN | TRUE |
| Fcnb | TRUE | 2.504712705 | 1 | No Change | Not Significant | FALSE |
| Fcor | TRUE | 0.497883308 | 1 | No Change | Not Significant | FALSE |
| Fcrl1 | TRUE | 1.057159491 | 1 | No Change | Not Significant | FALSE |
| Fcrl5 | TRUE | 0.916206348 | 1 | No Change | Not Significant | FALSE |
| Fcrla | TRUE | 1.395232939 | 1 | No Change | Not Significant | FALSE |
| Fcrlb | TRUE | 0.358622093 | 1 | No Change | Not Significant | FALSE |
| Fcrls | TRUE | 1.507009139 | 1 | No Change | Not Significant | FALSE |
| Fcsk | TRUE | 0.247056246 | 0.000786839 | No Change | Significant Positive | TRUE |
| Fdft1 | TRUE | 0.729386456 | 0.000440521 | UP | UP | TRUE |
| Fdps | TRUE | 1.650641319 | 9.59E-47 | UP | UP | TRUE |
| Fdx1 | TRUE | 0.522340887 | 3.64E-13 | UP | UP | TRUE |
| Fdx2 | TRUE | -0.319990589 | 1 | No Change | Not Significant | FALSE |
| Fdxacb1 | TRUE | -0.20993285 | 0.392885514 | No Change | Not Significant | TRUE |
| Fdxr | TRUE | -0.111454278 | 0.539389762 | No Change | Not Significant | TRUE |
| Fech | TRUE | -0.051868586 | 0.714893745 | No Change | Not Significant | TRUE |
| Fem1a | TRUE | -0.106408165 | 0.412396276 | No Change | Not Significant | TRUE |
| Fem1al | TRUE | -0.497317549 | 0.253780598 | No Change | Not Significant | TRUE |
| Fem1b | TRUE | -0.079279607 | 0.637333654 | No Change | Not Significant | TRUE |
| Fem1c | TRUE | -0.135564924 | 0.21130879 | No Change | Not Significant | TRUE |
| Fen1 | TRUE | 0.64444907 | 0.025782978 | UP | UP | TRUE |
| Fendrr | TRUE | -0.739928294 | 1 | No Change | Not Significant | FALSE |
| Fer | TRUE | 0.404584724 | 0.000157966 | No Change | Significant Positive | TRUE |
| Fer1l4 | TRUE | 0.891451016 | 0.004119773 | UP | UP | TRUE |
| Fer1l5 | TRUE | 0.314838993 | 0.404218578 | No Change | Not Significant | TRUE |
| Fer1l6 | TRUE | -0.689638183 | 0.009213125 | DOWN | DOWN | TRUE |
| Fermt1 | TRUE | 0.406625197 | 2.96E-06 | No Change | Significant Positive | TRUE |
| Fermt2 | TRUE | 0.53175057 | 1 | No Change | Not Significant | FALSE |
| Fermt3 | TRUE | -0.845895366 | 0.000298838 | DOWN | DOWN | TRUE |
| Fes | TRUE | -0.141604108 | 0.40732383 | No Change | Not Significant | TRUE |
| Fetub | TRUE | 0.841283981 | 1 | No Change | Not Significant | FALSE |
| Fev | TRUE | -0.44954904 | 1 | No Change | Not Significant | TRUE |
| Fez1 | TRUE | -0.764174255 | 1 | No Change | Not Significant | FALSE |
| Fez2 | TRUE | -0.283814893 | 0.029941356 | No Change | Significant Negative | TRUE |
| Fezf1 | TRUE | 5.005238748 | 3.57E-05 | UP | UP | FALSE |
| Fezf2 | TRUE | 2.406512542 | 1 | No Change | Not Significant | FALSE |
| Ffar1 | TRUE | -0.132709041 | 0.917066002 | No Change | Not Significant | TRUE |
| Ffar2 | TRUE | 0.271798773 | 0.340132673 | No Change | Not Significant | TRUE |
| Ffar3 | TRUE | -1.007173257 | 0.007727668 | DOWN | DOWN | TRUE |
| Ffar4 | TRUE | 0.151641968 | 0.722022907 | No Change | Not Significant | TRUE |
| Fga | TRUE | 2.196240183 | 1 | No Change | Not Significant | FALSE |
| Fgb | TRUE | 0.868392624 | 1 | No Change | Not Significant | FALSE |
| Fgd1 | TRUE | 0.757031003 | 0.003368135 | UP | UP | TRUE |
| Fgd2 | TRUE | -0.05004539 | 0.931236324 | No Change | Not Significant | TRUE |
| Fgd3 | TRUE | -0.128869497 | 0.509468423 | No Change | Not Significant | TRUE |
| Fgd4 | TRUE | -0.11066316 | 0.36280925 | No Change | Not Significant | TRUE |
| Fgd5 | TRUE | 0.817557059 | 1 | No Change | Not Significant | FALSE |
| Fgd6 | TRUE | -0.110361324 | 0.766128886 | No Change | Not Significant | TRUE |
| Fgf1 | TRUE | 0.409908226 | 0.053436476 | No Change | Not Significant | TRUE |
| Fgf10 | TRUE | 1.26652749 | 1 | No Change | Not Significant | FALSE |
| Fgf11 | TRUE | 0.313298631 | 0.510066755 | No Change | Not Significant | TRUE |
| Fgf12 | TRUE | -0.235057345 | 0.726753308 | No Change | Not Significant | TRUE |
| Fgf13 | TRUE | -0.666400007 | 1 | No Change | Not Significant | FALSE |
| Fgf14 | TRUE | 2.291718408 | 1 | No Change | Not Significant | FALSE |
| Fgf15 | TRUE | 0.47895882 | 1 | No Change | Not Significant | FALSE |
| Fgf17 | TRUE | -0.734715794 | 1 | No Change | Not Significant | FALSE |
| Fgf18 | TRUE | -0.898909891 | 1 | No Change | Not Significant | TRUE |
| Fgf2 | TRUE | 0.282893298 | 0.72618955 | No Change | Not Significant | TRUE |
| Fgf22 | TRUE | 0.718487953 | 1 | No Change | Not Significant | FALSE |
| Fgf6 | TRUE | -1.277286216 | 1 | No Change | Not Significant | FALSE |
| Fgf7 | TRUE | 0.563054826 | 1 | No Change | Not Significant | FALSE |
| Fgf9 | TRUE | -0.74336075 | 0.059503296 | No Change | Not Significant | TRUE |
| Fgfbp1 | TRUE | 0.207457459 | 0.1706035 | No Change | Not Significant | TRUE |
| Fgfbp3 | TRUE | -0.114985079 | 0.857956591 | No Change | Not Significant | TRUE |
| Fgfr1 | TRUE | -0.030644073 | 0.985766611 | No Change | Not Significant | FALSE |
| Fgfr1op2 | TRUE | 0.039845287 | 0.761475268 | No Change | Not Significant | TRUE |
| Fgfr2 | TRUE | -0.045193244 | 0.856426823 | No Change | Not Significant | TRUE |
| Fgfr3 | TRUE | -0.589893449 | 0.035343465 | DOWN | DOWN | TRUE |
| Fgfr4 | TRUE | 0.407225331 | 7.99E-06 | No Change | Significant Positive | TRUE |
| Fgfrl1 | TRUE | 0.182856205 | 0.37057407 | No Change | Not Significant | TRUE |
| Fggy | TRUE | -0.355782765 | 0.031346224 | No Change | Significant Negative | TRUE |
| Fgl1 | TRUE | -0.293897 | 0.513672903 | No Change | Not Significant | TRUE |
| Fgl2 | TRUE | -0.082852843 | 0.779177919 | No Change | Not Significant | TRUE |
| Fgr | TRUE | 0.309742593 | 0.534874425 | No Change | Not Significant | TRUE |
| Fh1 | TRUE | -0.073511809 | 0.563829035 | No Change | Not Significant | TRUE |
| Fhad1 | TRUE | 0.009495463 | 0.981474584 | No Change | Not Significant | TRUE |
| Fhad1os1 | TRUE | -0.904299492 | 1 | No Change | Not Significant | TRUE |
| Fhdc1 | TRUE | 0.157509124 | 0.707478587 | No Change | Not Significant | TRUE |
| Fhip1a | TRUE | 0.008981453 | 0.971381329 | No Change | Not Significant | TRUE |
| Fhip1b | TRUE | -0.105906149 | 0.395168001 | No Change | Not Significant | TRUE |
| Fhip2a | TRUE | 0.116869665 | 0.335676301 | No Change | Not Significant | TRUE |
| Fhip2b | TRUE | -0.130329577 | 0.443834541 | No Change | Not Significant | TRUE |
| Fhit | TRUE | 0.423700373 | 0.020879192 | No Change | Significant Positive | TRUE |
| Fhl1 | TRUE | -0.865110859 | 1.76E-05 | DOWN | DOWN | TRUE |
| Fhl2 | TRUE | -0.193584625 | 0.179528541 | No Change | Not Significant | TRUE |
| Fhl3 | TRUE | 0.009996746 | 0.990693936 | No Change | Not Significant | TRUE |
| Fhl4 | TRUE | 1.343979684 | 1 | No Change | Not Significant | FALSE |
| Fhod1 | TRUE | 0.373247786 | 0.029171357 | No Change | Significant Positive | TRUE |
| Fhod3 | TRUE | -0.877619708 | 1 | No Change | Not Significant | FALSE |
| Fibcd1 | TRUE | 1.789429852 | 1 | No Change | Not Significant | FALSE |
| Fibp | TRUE | -0.158520963 | 0.044259909 | No Change | Significant Negative | TRUE |
| Ficd | TRUE | 1.225347353 | 5.46E-27 | UP | UP | TRUE |
| Fig4 | TRUE | -0.024676458 | 0.881056435 | No Change | Not Significant | TRUE |
| Fign | TRUE | 0.392175428 | 0.788502504 | No Change | Not Significant | TRUE |
| Fignl1 | TRUE | 0.617537287 | 0.048978652 | UP | UP | TRUE |
| Fignl2 | TRUE | -0.312150931 | 0.473163139 | No Change | Not Significant | TRUE |
| Filip1 | TRUE | -2.390012578 | 1 | No Change | Not Significant | FALSE |
| Filip1l | TRUE | -0.259406357 | 0.316830918 | No Change | Not Significant | TRUE |
| Fip1l1 | TRUE | -0.001225636 | 0.990608762 | No Change | Not Significant | TRUE |
| Firre | TRUE | 0.297161967 | 0.130976957 | No Change | Not Significant | TRUE |
| Fis1 | TRUE | -0.046911992 | 0.720291626 | No Change | Not Significant | TRUE |
| Fitm2 | TRUE | -0.023412304 | 0.915629096 | No Change | Not Significant | TRUE |
| Fiz1 | TRUE | 0.204891219 | 0.020759714 | No Change | Significant Positive | TRUE |
| Fjx1 | TRUE | -0.53587768 | 1 | No Change | Not Significant | TRUE |
| Fkbp10 | TRUE | -0.307135473 | 1 | No Change | Not Significant | FALSE |
| Fkbp11 | TRUE | 2.304107505 | 7.91E-42 | UP | UP | TRUE |
| Fkbp14 | TRUE | 0.129199114 | 0.659260011 | No Change | Not Significant | TRUE |
| Fkbp15 | TRUE | -0.200036453 | 0.010183111 | No Change | Significant Negative | TRUE |
| Fkbp1a | TRUE | 0.038126608 | 0.826943172 | No Change | Not Significant | TRUE |
| Fkbp1b | TRUE | -0.646088512 | 0.169937908 | No Change | Not Significant | TRUE |
| Fkbp2 | TRUE | 0.748481867 | 5.98E-45 | UP | UP | TRUE |
| Fkbp3 | TRUE | -0.004672805 | 0.983100167 | No Change | Not Significant | TRUE |
| Fkbp4 | TRUE | 0.18793986 | 0.26324771 | No Change | Not Significant | TRUE |
| Fkbp5 | TRUE | -0.398096238 | 0.02983307 | No Change | Significant Negative | TRUE |
| Fkbp7 | TRUE | 0.830365917 | 0.001102492 | UP | UP | TRUE |
| Fkbp8 | TRUE | 0.100312008 | 0.216806333 | No Change | Not Significant | TRUE |
| Fkbp9 | TRUE | 0.117466263 | 0.67085817 | No Change | Not Significant | TRUE |
| Fkbpl | TRUE | 0.021658001 | 0.941067509 | No Change | Not Significant | TRUE |
| Fkrp | TRUE | 0.084340561 | 0.590040437 | No Change | Not Significant | TRUE |
| Fktn | TRUE | 0.081577299 | 0.762991261 | No Change | Not Significant | TRUE |
| Flacc1 | TRUE | -0.676813731 | 0.19999509 | No Change | Not Significant | TRUE |
| Flad1 | TRUE | -0.164321052 | 0.176502626 | No Change | Not Significant | TRUE |
| Flcn | TRUE | -0.599253067 | 5.53E-06 | DOWN | DOWN | TRUE |
| Flg | TRUE | 0.753081607 | 1 | No Change | Not Significant | FALSE |
| Flg2 | TRUE | 1.605992036 | 1 | No Change | Not Significant | FALSE |
| Fli1 | TRUE | -0.071818975 | 0.956641558 | No Change | Not Significant | TRUE |
| Flicr | TRUE | 0.143624926 | 1 | No Change | Not Significant | FALSE |
| Flii | TRUE | 0.003044211 | 0.981701728 | No Change | Not Significant | TRUE |
| Flna | TRUE | 0.107757987 | 0.582240877 | No Change | Not Significant | TRUE |
| Flnb | TRUE | -0.130571323 | 0.345760799 | No Change | Not Significant | TRUE |
| Flnc | TRUE | 0.507925078 | 0.846158956 | No Change | Not Significant | FALSE |
| Flot1 | TRUE | -0.05144612 | 0.784213744 | No Change | Not Significant | TRUE |
| Flot2 | TRUE | -0.036921611 | 0.877988085 | No Change | Not Significant | TRUE |
| Flrt1 | TRUE | -0.353208784 | 0.668047177 | No Change | Not Significant | TRUE |
| Flrt2 | TRUE | 0.145454923 | 0.736796542 | No Change | Not Significant | TRUE |
| Flrt3 | TRUE | 0.511167322 | 7.73E-05 | UP | UP | TRUE |
| Flt1 | TRUE | 0.469821249 | 0.885228942 | No Change | Not Significant | FALSE |
| Flt3 | TRUE | 1.774485732 | 1 | No Change | Not Significant | FALSE |
| Flt3l | TRUE | -0.923323817 | 0.052213508 | No Change | Not Significant | TRUE |
| Flt4 | TRUE | 3.465477832 | 1 | No Change | Not Significant | FALSE |
| Flvcr1 | TRUE | 0.871017881 | 6.45E-24 | UP | UP | TRUE |
| Flvcr2 | TRUE | 0.026574321 | 1 | No Change | Not Significant | FALSE |
| Flywch1 | TRUE | 0.362769133 | 5.97E-05 | No Change | Significant Positive | TRUE |
| Flywch2 | TRUE | 0.075695223 | 0.736796542 | No Change | Not Significant | TRUE |
| Fmc1 | TRUE | 0.014164972 | 0.952286247 | No Change | Not Significant | TRUE |
| Fmn1 | TRUE | 0.083409446 | 0.712111524 | No Change | Not Significant | TRUE |
| Fmn2 | TRUE | 3.082736865 | 5.24E-07 | UP | UP | TRUE |
| Fmnl1 | TRUE | 0.580371865 | 0.530096199 | No Change | Not Significant | TRUE |
| Fmnl2 | TRUE | 1.439968829 | 2.71E-09 | UP | UP | TRUE |
| Fmnl3 | TRUE | 0.334971772 | 0.472341459 | No Change | Not Significant | TRUE |
| Fmo1 | TRUE | -0.773329869 | 0.262814222 | No Change | Not Significant | TRUE |
| Fmo2 | TRUE | -0.138316445 | 0.882132403 | No Change | Not Significant | TRUE |
| Fmo4 | TRUE | -1.111644376 | 0.000102427 | DOWN | DOWN | TRUE |
| Fmo5 | TRUE | -0.739886122 | 0.044453504 | DOWN | DOWN | TRUE |
| Fmod | TRUE | 0.23518286 | 1 | No Change | Not Significant | FALSE |
| Fmr1 | TRUE | 0.122891754 | 0.30108314 | No Change | Not Significant | TRUE |
| Fmr1nb | TRUE | 1.565300093 | 1 | No Change | Not Significant | FALSE |
| Fn1 | TRUE | -0.081364919 | 0.909153704 | No Change | Not Significant | TRUE |
| Fn3k | TRUE | -1.12502843 | 0.136786828 | No Change | Not Significant | TRUE |
| Fn3krp | TRUE | -0.077251799 | 0.716335759 | No Change | Not Significant | TRUE |
| Fnbp1 | TRUE | -0.382803717 | 0.000810119 | No Change | Significant Negative | TRUE |
| Fnbp1l | TRUE | -0.115982013 | 0.242325103 | No Change | Not Significant | TRUE |
| Fnbp4 | TRUE | 0.11284666 | 0.53032725 | No Change | Not Significant | TRUE |
| Fndc1 | TRUE | 0.396148735 | 0.881056689 | No Change | Not Significant | FALSE |
| Fndc10 | TRUE | -0.240483407 | 0.464617944 | No Change | Not Significant | TRUE |
| Fndc11 | TRUE | 0.266048492 | 1 | No Change | Not Significant | FALSE |
| Fndc3a | TRUE | 0.236790242 | 0.001338605 | No Change | Significant Positive | TRUE |
| Fndc3b | TRUE | 0.903510838 | 3.69E-25 | UP | UP | TRUE |
| Fndc3c1 | TRUE | 0.072761562 | 1 | No Change | Not Significant | FALSE |
| Fndc4 | TRUE | 0.834881009 | 1 | No Change | Not Significant | FALSE |
| Fndc5 | TRUE | -1.354117352 | 2.35E-07 | DOWN | DOWN | TRUE |
| Fndc7 | TRUE | 3.23877555 | 1 | No Change | Not Significant | FALSE |
| Fndc8 | TRUE | 0.233242093 | 1 | No Change | Not Significant | TRUE |
| Fndc9 | TRUE | 2.185315 | 1 | No Change | Not Significant | FALSE |
| Fnip1 | TRUE | 0.260892133 | 0.079125787 | No Change | Not Significant | TRUE |
| Fnip2 | TRUE | -0.610884435 | 3.29E-05 | DOWN | DOWN | TRUE |
| Fnta | TRUE | 0.010370724 | 0.949998977 | No Change | Not Significant | TRUE |
| Fntb | TRUE | 0.099987792 | 0.422906972 | No Change | Not Significant | TRUE |
| Focad | TRUE | 0.24196797 | 0.042953833 | No Change | Significant Positive | TRUE |
| Folr1 | TRUE | 0.216040871 | 0.623763963 | No Change | Not Significant | TRUE |
| Fos | TRUE | -0.260515864 | 0.264383465 | No Change | Not Significant | TRUE |
| Fosb | TRUE | -0.448713739 | 0.001090541 | No Change | Significant Negative | TRUE |
| Fosl1 | TRUE | 0.006135872 | 0.993636457 | No Change | Not Significant | TRUE |
| Fosl2 | TRUE | 0.044721417 | 0.866982505 | No Change | Not Significant | TRUE |
| Foxa1 | TRUE | -0.001931471 | 0.99157486 | No Change | Not Significant | TRUE |
| Foxa2 | TRUE | 0.119831409 | 0.781981914 | No Change | Not Significant | TRUE |

Supplementary Table 3

| geneID | Cond1_DF_Detected | Cond1_DF_ValueCol | Cond1_DF_PadjCol | Cond1_DF_Cutoff | Cond1_DF_Cutoff_Specific.x | Cond2_DF_Detected | Cond2_DF_ValueCol | Cond2_DF_PadjCol | Cond2_DF_Cutoff | Cond2_DF_Cutoff_Specific | BG_Method | RG1_All |
|---|---|---|---|---|---|---|---|---|---|---|---|---|
| Foxa3 | TRUE | 0.161096529 | 0.556489079 | No Change | Not Significant | TRUE | 0.233648484 | 0.063259266 | No Change | Not Significant | TRUE | Cond1 Not Significant + Cond2 Not Significant |
| Foxb1 | TRUE | 0.522701262 | 1 | No Change | Not Significant | FALSE | NA | NA | No Change | Not Detected | FALSE | Background = FALSE |
| Foxc1 | TRUE | 0.287296977 | 0.626589558 | No Change | Not Significant | TRUE | -0.153108315 | 0.830161713 | No Change | Not Significant | TRUE | Cond1 Not Significant + Cond2 Not Significant |
| Foxd1 | TRUE | 1.251565036 | 1 | No Change | Not Significant | FALSE | NA | NA | No Change | Not Detected | FALSE | Background = FALSE |
| Foxd2 | TRUE | 0.214993114 | 0.108241146 | No Change | Not Significant | TRUE | -0.11877692 | 0.554719101 | No Change | Not Significant | TRUE | Cond1 Not Significant + Cond2 Not Significant |
| Foxd2os | TRUE | 0.006439132 | 0.975869522 | No Change | Not Significant | TRUE | -0.119470931 | 0.252438005 | No Change | Not Significant | TRUE | Cond1 Not Significant + Cond2 Not Significant |
| Foxd3 | TRUE | -0.864261797 | 1 | No Change | Not Significant | FALSE | NA | NA | No Change | Not Detected | FALSE | Background = FALSE |
| Foxe3 | TRUE | -0.003344915 | 1 | No Change | Not Significant | FALSE | NA | NA | No Change | Not Detected | FALSE | Background = FALSE |
| Foxf1 | TRUE | -1.61587138 | 1 | No Change | Not Significant | FALSE | NA | NA | No Change | Not Detected | FALSE | Background = FALSE |
| Foxg1 | TRUE | 0.005112629 | 1 | No Change | Not Significant | FALSE | NA | NA | No Change | Not Detected | FALSE | Background = FALSE |
| Foxi1 | TRUE | 2.480647131 | 1 | No Change | Not Significant | FALSE | NA | NA | No Change | Not Detected | FALSE | Background = FALSE |
| Foxj1 | TRUE | 0.044324184 | 1 | No Change | Not Significant | FALSE | NA | NA | No Change | Not Detected | FALSE | Background = FALSE |
| Foxj2 | TRUE | 0.007352731 | 0.978107631 | No Change | Not Significant | TRUE | -0.10016592 | 0.578448892 | No Change | Not Significant | TRUE | Cond1 Not Significant + Cond2 Not Significant |
| Foxj3 | TRUE | -0.107414385 | 0.47089684 | No Change | Not Significant | TRUE | -0.130071194 | 0.076460123 | No Change | Not Significant | TRUE | Cond1 Not Significant + Cond2 Not Significant |
| Foxk1 | TRUE | -0.080317304 | 0.607432671 | No Change | Not Significant | TRUE | -0.105723292 | 0.521098971 | No Change | Not Significant | TRUE | Cond1 Not Significant + Cond2 Not Significant |
| Foxk2 | TRUE | 1.047988767 | 1.24E-37 | UP | UP | TRUE | 0.530055888 | 1.71E-11 | UP | UP | TRUE | Cond1 UP + Cond2 UP |
| Foxl1 | TRUE | 0.49463165 | 1 | No Change | Not Significant | FALSE | NA | NA | No Change | Not Detected | FALSE | Background = FALSE |
| Foxl2 | TRUE | 0.611480109 | 1 | No Change | Not Significant | FALSE | NA | NA | No Change | Not Detected | FALSE | Background = FALSE |
| Foxm1 | TRUE | 0.876514164 | 1.10E-05 | UP | UP | TRUE | -0.033716795 | 0.898311362 | No Change | Not Significant | TRUE | Cond1 UP + Cond2 Not Significant |
| Foxn1 | TRUE | 0.426083349 | 0.458698364 | No Change | Not Significant | TRUE | 0.227883614 | 0.625831733 | No Change | Not Significant | TRUE | Cond1 Not Significant + Cond2 Not Significant |
| Foxn2 | TRUE | -0.204606586 | 0.254529872 | No Change | Not Significant | TRUE | -0.224245952 | 0.023394202 | No Change | Significant Negative | TRUE | Cond1 Not Significant + Cond2 Significant Negative |
| Foxn3 | TRUE | -0.421755796 | 5.21E-05 | No Change | Significant Negative | TRUE | -0.159871605 | 0.111759773 | No Change | Not Significant | TRUE | Cond1 Significant Negative + Cond2 Not Significant |
| Foxo1 | TRUE | -0.032026609 | 0.904098264 | No Change | Not Significant | TRUE | -0.171623128 | 0.08057857 | No Change | Not Significant | TRUE | Cond1 Not Significant + Cond2 Not Significant |
| Foxo3 | TRUE | -0.047124385 | 0.866982505 | No Change | Not Significant | TRUE | -0.07593379 | 0.406671386 | No Change | Not Significant | TRUE | Cond1 Not Significant + Cond2 Not Significant |
| Foxo4 | TRUE | -0.28081082 | 0.004853962 | No Change | Significant Negative | TRUE | -0.022090753 | 0.857670564 | No Change | Not Significant | TRUE | Cond1 Significant Negative + Cond2 Not Significant |
| Foxo6 | TRUE | -0.462381178 | 0.405213026 | No Change | Not Significant | TRUE | -0.321000156 | 0.299280839 | No Change | Not Significant | TRUE | Cond1 Not Significant + Cond2 Not Significant |
| Foxp1 | TRUE | -0.118234223 | 0.455456329 | No Change | Not Significant | TRUE | -0.087549243 | 0.359972154 | No Change | Not Significant | TRUE | Cond1 Not Significant + Cond2 Not Significant |

[illegible]

Supplementary Table 3

| geneID | Cond1_DF_Detected | Cond1_DF_ValueCol | Cond1_DF_PadjCol | Cond1_DF_Cutoff | Cond1_DF_Cutoff_Specific.x | Cond2_DF_Detected |
|---|---|---|---|---|---|---|
| Galm | TRUE | -0.175734473 | 0.015076179 | No Change | Significant Negative | TRUE |
| Galns | TRUE | -0.236173256 | 0.001167368 | No Change | Significant Negative | TRUE |
| Galnt1 | TRUE | 0.091246691 | 0.509468423 | No Change | Not Significant | TRUE |
| Galnt10 | TRUE | -0.109202698 | 0.630340996 | No Change | Not Significant | TRUE |
| Galnt11 | TRUE | -0.091291434 | 0.521287512 | No Change | Not Significant | TRUE |
| Galnt12 | TRUE | -0.027061741 | 0.900821606 | No Change | Not Significant | TRUE |
| Galnt13 | TRUE | 1.772255301 | 1 | No Change | Not Significant | FALSE |
| Galnt14 | TRUE | -0.137396105 | 1 | No Change | Not Significant | FALSE |
| Galnt15 | TRUE | 1.465996439 | 1 | No Change | Not Significant | FALSE |
| Galnt16 | TRUE | 1.638631185 | 1 | No Change | Not Significant | FALSE |
| Galnt17 | TRUE | -0.595562351 | 0.52648324 | No Change | Not Significant | TRUE |
| Galnt18 | TRUE | 0.896714807 | 0.041412525 | UP | UP | TRUE |
| Galnt2 | TRUE | 0.757551198 | 0.047764563 | UP | UP | TRUE |
| Galnt3 | TRUE | 0.504932721 | 0.000462865 | UP | UP | TRUE |
| Galnt4 | TRUE | -0.0522641 | 0.741922186 | No Change | Not Significant | TRUE |
| Galnt5 | TRUE | 0.162953911 | 0.698491194 | No Change | Not Significant | TRUE |
| Galnt6 | TRUE | -0.275804738 | 0.015207984 | No Change | Significant Negative | TRUE |
| Galnt7 | TRUE | 0.208262201 | 0.406008467 | No Change | Not Significant | TRUE |
| Galntl5 | TRUE | -0.427686945 | 1 | No Change | Not Significant | FALSE |
| Galntl6 | TRUE | 0.26417453 | 0.081931007 | No Change | Not Significant | TRUE |
| Galr1 | TRUE | -0.38935857 | 1 | No Change | Not Significant | TRUE |
| Galr2 | TRUE | 0.524761597 | 0.445131871 | No Change | Not Significant | TRUE |
| Galr3 | TRUE | 0.164656311 | 1 | No Change | Not Significant | FALSE |
| Galt | TRUE | -0.213932089 | 0.018266571 | No Change | Significant Negative | TRUE |
| Gamt | TRUE | -0.125547897 | 0.816056375 | No Change | Not Significant | TRUE |
| Gan | TRUE | -0.024397566 | 0.907217398 | No Change | Not Significant | TRUE |
| Ganab | TRUE | 0.257787275 | 0.028656498 | No Change | Significant Positive | TRUE |
| Ganc | TRUE | -0.131331587 | 0.334559671 | No Change | Not Significant | TRUE |
| Gap43 | TRUE | 2.767139052 | 1 | No Change | Not Significant | FALSE |
| Gapdh | TRUE | 0.066996259 | 0.801613541 | No Change | Not Significant | TRUE |
| Gapdhs | TRUE | -0.377192899 | 0.660786267 | No Change | Not Significant | FALSE |
| Gapvd1 | TRUE | -0.000151106 | 0.998483602 | No Change | Not Significant | TRUE |
| Gar1 | TRUE | 0.03110629 | 0.907147741 | No Change | Not Significant | TRUE |
| Garem1 | TRUE | -0.117049234 | 0.615439779 | No Change | Not Significant | TRUE |
| Garem2 | TRUE | 1.493682623 | 1 | No Change | Not Significant | FALSE |
| Garin1a | TRUE | 0.340578967 | 1 | No Change | Not Significant | FALSE |
| Garin1b | TRUE | 0.937657205 | 1 | No Change | Not Significant | FALSE |
| Garin3 | TRUE | 1.108383589 | 1 | No Change | Not Significant | FALSE |
| Garin4 | TRUE | 0.174859788 | 0.904107385 | No Change | Not Significant | TRUE |
| Garin5a | TRUE | 0.17090064 | 0.80301103 | No Change | Not Significant | TRUE |
| Garin5b | TRUE | 0.501900509 | 1 | No Change | Not Significant | FALSE |
| Garnl3 | TRUE | -1.424585141 | 0.073420411 | No Change | Not Significant | TRUE |
| Garre1 | TRUE | -0.277687105 | 0.019206682 | No Change | Significant Negative | TRUE |
| Gars | TRUE | 0.186627435 | 0.027890446 | No Change | Significant Positive | TRUE |
| Gart | TRUE | -0.109344088 | 0.623441772 | No Change | Not Significant | TRUE |
| Gas1 | TRUE | -0.062364763 | 1 | No Change | Not Significant | FALSE |
| Gas2 | TRUE | 0.245869253 | 0.368018849 | No Change | Not Significant | TRUE |
| Gas2l1 | TRUE | -0.392831179 | 0.001100095 | No Change | Significant Negative | TRUE |
| Gas2l2 | TRUE | 0.00621527 | 1 | No Change | Not Significant | FALSE |
| Gas2l3 | TRUE | 1.250982448 | 1.52E-18 | UP | UP | TRUE |
| Gas5 | TRUE | -0.123945791 | 0.283858253 | No Change | Not Significant | TRUE |
| Gas6 | TRUE | -0.441076375 | 0.0014822 | No Change | Significant Negative | TRUE |
| Gas7 | TRUE | -0.136260659 | 0.503462581 | No Change | Not Significant | TRUE |
| Gas8 | TRUE | 0.261236759 | 0.117137153 | No Change | Not Significant | TRUE |
| Gask1a | TRUE | -1.156578451 | 1 | No Change | Not Significant | FALSE |
| Gask1b | TRUE | 0.325040201 | 0.626929013 | No Change | Not Significant | TRUE |
| Gata1 | TRUE | 0.006340521 | 1 | No Change | Not Significant | FALSE |
| Gata2 | TRUE | -1.937533679 | 1 | No Change | Not Significant | FALSE |
| Gata3 | TRUE | 1.712323393 | 1 | No Change | Not Significant | FALSE |
| Gata4 | TRUE | -0.637158516 | 0.522874377 | No Change | Not Significant | TRUE |
| Gata5 | TRUE | 0.228823025 | 0.782288135 | No Change | Not Significant | TRUE |
| Gata5os | TRUE | -1.488677876 | 1 | No Change | Not Significant | FALSE |
| Gata6 | TRUE | -0.092158831 | 0.32396287 | No Change | Not Significant | TRUE |
| Gatad1 | TRUE | -0.03316914 | 0.800822518 | No Change | Not Significant | TRUE |
| Gatad2a | TRUE | 0.084615897 | 0.292373241 | No Change | Not Significant | TRUE |
| Gatad2b | TRUE | -0.069692455 | 0.730517043 | No Change | Not Significant | TRUE |
| Gatb | TRUE | -0.282902604 | 0.009444208 | No Change | Significant Negative | TRUE |
| Gatc | TRUE | -0.081948097 | 0.683706314 | No Change | Not Significant | TRUE |
| Gatd1 | TRUE | -0.031031429 | 0.859382261 | No Change | Not Significant | TRUE |
| Gatd3a | TRUE | -0.300643179 | 0.002244387 | No Change | Significant Negative | TRUE |
| Gatm | TRUE | 1.302213334 | 0.004096971 | UP | UP | TRUE |
| Gba | TRUE | 0.096696508 | 0.402137369 | No Change | Not Significant | TRUE |
| Gba2 | TRUE | -0.175653476 | 0.297415185 | No Change | Not Significant | TRUE |
| Gbe1 | TRUE | -0.1521273 | 0.338606949 | No Change | Not Significant | TRUE |
| Gbf1 | TRUE | 0.280540841 | 0.000429797 | No Change | Significant Positive | TRUE |
| Gbgt1 | TRUE | -1.120366474 | 0.126876666 | No Change | Not Significant | TRUE |
| Gbp10 | TRUE | 0.807870341 | 1 | No Change | Not Significant | FALSE |
| Gbp11 | TRUE | 1.947530208 | 1 | No Change | Not Significant | FALSE |
| Gbp2 | TRUE | -0.167739672 | 0.807553177 | No Change | Not Significant | FALSE |
| Gbp2b | TRUE | -0.003126751 | 1 | No Change | Not Significant | FALSE |
| Gbp3 | TRUE | 0.110497383 | 0.896761253 | No Change | Not Significant | FALSE |
| Gbp4 | TRUE | 0.172082969 | 0.896267126 | No Change | Not Significant | TRUE |
| Gbp5 | TRUE | -0.671134055 | 0.401690844 | No Change | Not Significant | TRUE |
| Gbp6 | TRUE | -0.62610467 | 0.401975042 | No Change | Not Significant | TRUE |
| Gbp7 | TRUE | -0.519233951 | 0.156808369 | No Change | Not Significant | TRUE |
| Gbp8 | TRUE | -0.416906976 | 0.647057325 | No Change | Not Significant | TRUE |
| Gbp9 | TRUE | 0.349006463 | 0.647232589 | No Change | Not Significant | TRUE |
| Gbx1 | TRUE | 0.336367624 | 1 | No Change | Not Significant | FALSE |
| Gca | TRUE | -0.072479971 | 0.690541222 | No Change | Not Significant | TRUE |
| Gcat | TRUE | 0.936088891 | 2.67E-09 | UP | UP | TRUE |
| Gcc1 | TRUE | -0.014806022 | 0.913297021 | No Change | Not Significant | TRUE |
| Gcc2 | TRUE | -0.054766271 | 0.761222618 | No Change | Not Significant | TRUE |
| Gcdh | TRUE | 0.559825317 | 7.39E-07 | UP | UP | TRUE |
| Gcfc2 | TRUE | 0.023415172 | 0.936825965 | No Change | Not Significant | TRUE |
| Gcg | TRUE | -0.448330828 | 0.004327722 | No Change | Significant Negative | TRUE |
| Gcgr | TRUE | 0.169203707 | 1 | No Change | Not Significant | FALSE |
| Gch1 | TRUE | -0.101646069 | 0.683156904 | No Change | Not Significant | TRUE |
| Gchfr | TRUE | 0.580029602 | 0.007756249 | UP | UP | TRUE |
| Gck | TRUE | -0.221623936 | 0.800507769 | No Change | Not Significant | TRUE |
| Gckr | TRUE | 1.313548046 | 1 | No Change | Not Significant | FALSE |
| Gclc | TRUE | 0.052777617 | 0.77652216 | No Change | Not Significant | TRUE |
| Gclm | TRUE | -0.28639703 | 0.003410008 | No Change | Significant Negative | TRUE |
| Gcm2 | TRUE | 1.110818634 | 1 | No Change | Not Significant | FALSE |
| Gcn1 | TRUE | 0.085754776 | 0.490619315 | No Change | Not Significant | TRUE |
| Gcnt1 | TRUE | -0.570754986 | 4.60E-07 | DOWN | DOWN | TRUE |
| Gcnt2 | TRUE | -0.314471705 | 0.512663892 | No Change | Not Significant | TRUE |
| Gcnt3 | TRUE | -0.229009389 | 0.265217693 | No Change | Not Significant | TRUE |
| Gcnt4 | TRUE | -0.820072591 | 9.96E-07 | DOWN | DOWN | TRUE |
| Gcnt7 | TRUE | -0.074234953 | 0.893375436 | No Change | Not Significant | TRUE |
| Gcsam | TRUE | 0.719682967 | 0.385300595 | No Change | Not Significant | TRUE |
| Gcsh | TRUE | 0.180474439 | 0.066769512 | No Change | Not Significant | TRUE |
| Gda | TRUE | -0.010865467 | 0.970517013 | No Change | Not Significant | TRUE |
| Gdap1 | TRUE | 0.259092501 | 1 | No Change | Not Significant | FALSE |
| Gdap1l1 | TRUE | -0.523842901 | 1 | No Change | Not Significant | FALSE |
| Gdap2 | TRUE | 0.683727153 | 5.13E-14 | UP | UP | TRUE |
| Gde1 | TRUE | -0.281578786 | 0.175022172 | No Change | Not Significant | TRUE |
| Gdf10 | TRUE | 0.609456085 | 1 | No Change | Not Significant | FALSE |
| Gdf11 | TRUE | 0.342422059 | 0.551684327 | No Change | Not Significant | TRUE |
| Gdf15 | TRUE | 0.770431739 | 0.001055133 | UP | UP | TRUE |
| Gdf5 | TRUE | -0.425057561 | 1 | No Change | Not Significant | FALSE |
| Gdf7 | TRUE | 0.520553101 | 1 | No Change | Not Significant | FALSE |
| Gdf9 | TRUE | -1.684854258 | 1 | No Change | Not Significant | FALSE |
| Gdi1 | TRUE | -0.149454096 | 0.059503296 | No Change | Not Significant | TRUE |
| Gdi2 | TRUE | 0.118373127 | 0.343330529 | No Change | Not Significant | TRUE |
| Gdnf | TRUE | 0.698601184 | 1 | No Change | Not Significant | FALSE |
| Gdpd1 | TRUE | -0.045173766 | 0.834766446 | No Change | Not Significant | TRUE |
| Gdpd2 | TRUE | -0.38295664 | 0.524764095 | No Change | Not Significant | TRUE |
| Gdpd3 | TRUE | 0.164217332 | 0.750140815 | No Change | Not Significant | TRUE |
| Gdpd5 | TRUE | -0.096488854 | 0.794916857 | No Change | Not Significant | TRUE |
| Gdpgp1 | TRUE | -0.314341631 | 0.036642767 | No Change | Significant Negative | TRUE |
| Gem | TRUE | -0.03185669 | 0.97587882 | No Change | Not Significant | TRUE |
| Gemin2 | TRUE | -0.022406353 | 0.936943344 | No Change | Not Significant | TRUE |
| Gemin4 | TRUE | 0.013874443 | 0.985442104 | No Change | Not Significant | TRUE |
| Gemin5 | TRUE | -0.056181658 | 0.812765484 | No Change | Not Significant | TRUE |
| Gemin6 | TRUE | 0.373210074 | 0.243645564 | No Change | Not Significant | TRUE |
| Gemin7 | TRUE | -0.141037557 | 0.198739711 | No Change | Not Significant | TRUE |
| Gemin8 | TRUE | -0.118486233 | 0.796376973 | No Change | Not Significant | TRUE |
| Gen1 | TRUE | 0.556944416 | 0.002788738 | UP | UP | TRUE |
| Get1 | TRUE | 0.162559758 | 0.475244062 | No Change | Not Significant | TRUE |
| Get3 | TRUE | 0.245551989 | 0.005454866 | No Change | Significant Positive | TRUE |
| Get4 | TRUE | 0.084564145 | 0.604102573 | No Change | Not Significant | TRUE |
| Gfap | TRUE | -1.909475485 | 1 | No Change | Not Significant | FALSE |
| Gfer | TRUE | 0.090594195 | 0.524504007 | No Change | Not Significant | TRUE |
| Gfi1 | TRUE | 0.016519843 | 0.966815843 | No Change | Not Significant | TRUE |
| Gfi1b | TRUE | -0.453853575 | 0.489845783 | No Change | Not Significant | TRUE |
| Gfm1 | TRUE | -0.179458292 | 0.13170242 | No Change | Not Significant | TRUE |
| Gfm2 | TRUE | -0.138100581 | 0.14626152 | No Change | Not Significant | TRUE |
| Gfod1 | TRUE | -0.640330516 | 0.022359199 | DOWN | DOWN | TRUE |
| Gfod2 | TRUE | -0.123412521 | 0.520301242 | No Change | Not Significant | TRUE |
| Gfpt1 | TRUE | -0.00489726 | 0.985876467 | No Change | Not Significant | TRUE |
| Gfpt2 | TRUE | 1.236551779 | 0.330514877 | No Change | Not Significant | TRUE |
| Gfra1 | TRUE | -0.399578926 | 1 | No Change | Not Significant | TRUE |
| Gfra2 | TRUE | 1.477486661 | 1 | No Change | Not Significant | FALSE |
| Gfra3 | TRUE | 0.132974263 | 0.797446101 | No Change | Not Significant | TRUE |
| Gfra4 | TRUE | 0.573749941 | 1 | No Change | Not Significant | FALSE |
| Gfus | TRUE | 0.007799894 | 0.965886893 | No Change | Not Significant | TRUE |
| Gfy | TRUE | -1.34954337 | 1 | No Change | Not Significant | FALSE |
| Gga1 | TRUE | 0.083584228 | 0.336792184 | No Change | Not Significant | TRUE |
| Gga2 | TRUE | 0.500352856 | 0.011044625 | UP | UP | TRUE |
| Gga3 | TRUE | 0.130308353 | 0.517311687 | No Change | Not Significant | TRUE |
| Ggact | TRUE | -0.298770069 | 9.48E-05 | No Change | Significant Negative | TRUE |
| Ggct | TRUE | 0.579543722 | 0.003689141 | UP | UP | TRUE |
| Ggcx | TRUE | 0.51314176 | 3.41E-08 | UP | UP | TRUE |
| Ggh | TRUE | -0.603473472 | 0.009680909 | DOWN | DOWN | TRUE |
| Ggn | TRUE | 0.30944396 | 0.812818663 | No Change | Not Significant | TRUE |
| Ggnbp1 | TRUE | -0.008486421 | 0.9902423 | No Change | Not Significant | TRUE |
| Ggnbp2 | TRUE | -0.066313716 | 0.387311023 | No Change | Not Significant | TRUE |
| Ggps1 | TRUE | -0.049291147 | 0.698491194 | No Change | Not Significant | TRUE |
| Ggt1 | TRUE | 0.932430644 | 0.129452831 | No Change | Not Significant | TRUE |
| Ggt5 | TRUE | 1.327704489 | 1 | No Change | Not Significant | FALSE |
| Ggt6 | TRUE | -0.213516817 | 0.102970942 | No Change | Not Significant | TRUE |
| Ggt7 | TRUE | 0.345580011 | 1 | No Change | Not Significant | FALSE |
| Ggta1 | TRUE | 0.211898521 | 0.891186777 | No Change | Not Significant | TRUE |
| Ghdc | TRUE | -0.140225178 | 0.255796319 | No Change | Not Significant | TRUE |
| Ghitm | TRUE | -0.206507975 | 0.000809147 | No Change | Significant Negative | TRUE |
| Ghr | TRUE | -0.317715381 | 0.095110269 | No Change | Not Significant | TRUE |
| Ghrh | TRUE | -0.73719167 | 1 | No Change | Not Significant | FALSE |
| Ghrhr | TRUE | 0.383483404 | 1 | No Change | Not Significant | FALSE |
| Ghrl | TRUE | 0.574555746 | 0.238311216 | No Change | Not Significant | TRUE |
| Gid4 | TRUE | -0.116318888 | 0.449590928 | No Change | Not Significant | TRUE |
| Gid8 | TRUE | -0.112760915 | 0.260514736 | No Change | Not Significant | TRUE |
| Gigyf1 | TRUE | -0.132923701 | 0.664871274 | No Change | Not Significant | TRUE |
| Gigyf2 | TRUE | -0.011178753 | 0.927761944 | No Change | Not Significant | TRUE |
| Gimap1 | TRUE | -0.08182311 | 0.877032235 | No Change | Not Significant | TRUE |
| Gimap3 | TRUE | 0.164951606 | 0.767282764 | No Change | Not Significant | TRUE |
| Gimap4 | TRUE | -0.061901439 | 0.93035707 | No Change | Not Significant | TRUE |
| Gimap5 | TRUE | 0.653095381 | 0.379085971 | No Change | Not Significant | TRUE |
| Gimap6 | TRUE | 0.059555951 | 0.931308041 | No Change | Not Significant | TRUE |
| Gimap7 | TRUE | 0.012199285 | 0.989913623 | No Change | Not Significant | TRUE |
| Gimap8 | TRUE | -0.125962717 | 0.751588072 | No Change | Not Significant | TRUE |
| Gimap9 | TRUE | 0.122201742 | 0.634755452 | No Change | Not Significant | TRUE |
| Gimd1 | TRUE | 0.607476643 | 1 | No Change | Not Significant | TRUE |

Supplementary Table 3

| geneID | Cond1_DF_Detected | Cond1_DF_ValueCol | Cond1_DF_PadjCol | Cond1_DF_Cutoff | Cond1_DF_Cutoff_Specific.x | Cond2_DF_Detected |
|---|---|---|---|---|---|---|
| Gin1 | TRUE | 0.061916074 | 0.801687789 | No Change | Not Significant | TRUE |
| Ginm1 | TRUE | -0.186622981 | 0.094598742 | No Change | Not Significant | TRUE |
| Gins1 | TRUE | 0.395348533 | 0.186051032 | No Change | Not Significant | TRUE |
| Gins2 | TRUE | 0.747439482 | 0.000125991 | UP | UP | TRUE |
| Gins3 | TRUE | 0.511262708 | 0.125499051 | No Change | Not Significant | TRUE |
| Gins4 | TRUE | 0.131208021 | 0.335284335 | No Change | Not Significant | TRUE |
| Gip | TRUE | -2.1170191 | 1 | No Change | Not Significant | TRUE |
| Gipc1 | TRUE | 1.173235741 | 2.17E-44 | UP | UP | TRUE |
| Gipc2 | TRUE | -0.451146109 | 5.08E-12 | No Change | Significant Negative | TRUE |
| Gipc3 | TRUE | 1.064726957 | 0.290347713 | No Change | Not Significant | FALSE |
| Gipr | TRUE | -0.50454266 | 0.626602117 | No Change | Not Significant | TRUE |
| Git1 | TRUE | -0.086779416 | 0.587211164 | No Change | Not Significant | TRUE |
| Git2 | TRUE | -0.177893981 | 0.304528653 | No Change | Not Significant | TRUE |
| Gja1 | TRUE | -0.507256316 | 0.131124269 | No Change | Not Significant | TRUE |
| Gja3 | TRUE | 0.891256148 | 1 | No Change | Not Significant | FALSE |
| Gja4 | TRUE | -0.711099432 | 1 | No Change | Not Significant | TRUE |
| Gja5 | TRUE | 0.482520714 | 1 | No Change | Not Significant | FALSE |
| Gja6 | TRUE | 1.273668807 | 1 | No Change | Not Significant | FALSE |
| Gja8 | TRUE | 0.409802034 | 1 | No Change | Not Significant | FALSE |
| Gjb1 | TRUE | 0.084031207 | 0.703090606 | No Change | Not Significant | TRUE |
| Gjb2 | TRUE | 0.247062026 | 0.183317181 | No Change | Not Significant | TRUE |
| Gjb3 | TRUE | -0.673217077 | 9.51E-06 | DOWN | DOWN | TRUE |
| Gjb4 | TRUE | -1.70857351 | 0.059481896 | No Change | Not Significant | TRUE |
| Gjb5 | TRUE | -2.840614381 | 3.32E-07 | DOWN | DOWN | TRUE |
| Gjc2 | TRUE | 0.269525444 | 1 | No Change | Not Significant | FALSE |
| Gjc3 | TRUE | -2.989140361 | 1 | No Change | Not Significant | FALSE |
| Gjd2 | TRUE | -1.874870124 | 1 | No Change | Not Significant | FALSE |
| Gjd3 | TRUE | -0.236784909 | 1 | No Change | Not Significant | FALSE |
| Gk | TRUE | -0.113919281 | 0.787858751 | No Change | Not Significant | TRUE |
| Gk5 | TRUE | -0.048567045 | 0.851944324 | No Change | Not Significant | TRUE |
| Gkap1 | TRUE | -0.014137623 | 0.976016991 | No Change | Not Significant | TRUE |
| Gla | TRUE | -0.077324265 | 0.630553153 | No Change | Not Significant | TRUE |
| Glb1 | TRUE | -0.27024638 | 0.036802352 | No Change | Significant Negative | TRUE |
| Glb1l | TRUE | -0.368204191 | 0.208654479 | No Change | Not Significant | TRUE |
| Glb1l2 | TRUE | -0.506915755 | 1 | No Change | Not Significant | FALSE |
| Glcci1 | TRUE | 0.617840667 | 0.117581482 | No Change | Not Significant | TRUE |
| Glce | TRUE | 0.213895075 | 0.183669899 | No Change | Not Significant | TRUE |
| Gldc | TRUE | 0.25964698 | 0.835534172 | No Change | Not Significant | TRUE |
| Gldn | TRUE | 3.114168152 | 1 | No Change | Not Significant | FALSE |
| Gle1 | TRUE | 0.244310141 | 0.146868346 | No Change | Not Significant | TRUE |
| Glg1 | TRUE | -0.126244425 | 0.274320379 | No Change | Not Significant | TRUE |
| Gli1 | TRUE | -1.19063743 | 1 | No Change | Not Significant | FALSE |
| Gli2 | TRUE | 1.224118591 | 1 | No Change | Not Significant | FALSE |
| Gli3 | TRUE | 1.200233499 | 0.371853021 | No Change | Not Significant | FALSE |
| Glipr1 | TRUE | 0.176218212 | 0.853563561 | No Change | Not Significant | TRUE |
| Glipr2 | TRUE | -0.555836272 | 0.079002436 | No Change | Not Significant | TRUE |
| Glis1 | TRUE | -1.144989295 | 1 | No Change | Not Significant | FALSE |
| Glis2 | TRUE | 0.195811174 | 0.768287864 | No Change | Not Significant | TRUE |
| Glis3 | TRUE | -0.174481752 | 0.370548786 | No Change | Not Significant | TRUE |
| Glmn | TRUE | 0.209542592 | 0.522795554 | No Change | Not Significant | TRUE |
| Glmp | TRUE | -0.325575765 | 3.66E-06 | No Change | Significant Negative | TRUE |
| Glo1 | TRUE | -0.090515557 | 0.48334534 | No Change | Not Significant | TRUE |
| Glod4 | TRUE | -0.098297037 | 0.59060209 | No Change | Not Significant | TRUE |
| Glod5 | TRUE | -0.672657069 | 6.98E-10 | DOWN | DOWN | TRUE |
| Glp1r | TRUE | 0.044339225 | 0.959274295 | No Change | Not Significant | TRUE |
| Glp2r | TRUE | 0.401155393 | 0.824897384 | No Change | Not Significant | TRUE |
| Glra1 | TRUE | 1.126067804 | 1 | No Change | Not Significant | FALSE |
| Glra3 | TRUE | -0.236173773 | 1 | No Change | Not Significant | FALSE |
| Glra4 | TRUE | 0.192332033 | 1 | No Change | Not Significant | FALSE |
| Glrb | TRUE | 2.37693728 | 1 | No Change | Not Significant | FALSE |
| Glrx | TRUE | -0.467243866 | 3.86E-07 | No Change | Significant Negative | TRUE |
| Glrx2 | TRUE | 0.105193699 | 0.492426209 | No Change | Not Significant | TRUE |
| Glrx3 | TRUE | -0.285526646 | 0.055893077 | No Change | Not Significant | TRUE |
| Glrx5 | TRUE | 0.242606887 | 0.031092878 | No Change | Significant Positive | TRUE |
| Gls | TRUE | -0.237292381 | 0.000292855 | No Change | Significant Negative | TRUE |
| Gls2 | TRUE | 0.622893965 | 0.173816344 | No Change | Not Significant | TRUE |
| Glt1d1 | TRUE | -0.191863439 | 0.338620816 | No Change | Not Significant | TRUE |
| Glt28d2 | TRUE | -0.063679242 | 0.851508089 | No Change | Not Significant | TRUE |
| Glt8d1 | TRUE | 0.313578353 | 0.070613276 | No Change | Not Significant | TRUE |
| Glt8d2 | TRUE | 0.198831088 | 0.844104331 | No Change | Not Significant | FALSE |
| Gltp | TRUE | 0.371393315 | 0.016227616 | No Change | Significant Positive | TRUE |
| Glud1 | TRUE | -0.113577449 | 0.339527301 | No Change | Not Significant | TRUE |
| Glul | TRUE | -0.427968029 | 0.054501706 | No Change | Not Significant | TRUE |
| Glyat | TRUE | -0.794286382 | 1 | No Change | Not Significant | FALSE |
| Glyatl3 | TRUE | -1.278598168 | 1 | No Change | Not Significant | FALSE |
| Glycam1 | TRUE | -0.719425423 | 1 | No Change | Not Significant | FALSE |
| Glyctk | TRUE | -0.308471618 | 0.483691716 | No Change | Not Significant | TRUE |
| Glyr1 | TRUE | 0.072748963 | 0.380783029 | No Change | Not Significant | TRUE |
| Gm10029 | TRUE | -0.830492162 | 1 | No Change | Not Significant | FALSE |
| Gm10030 | TRUE | -1.650015921 | 1 | No Change | Not Significant | FALSE |
| Gm10033 | TRUE | -0.186121636 | 0.408815356 | No Change | Not Significant | TRUE |
| Gm10037 | TRUE | -1.471512165 | 1 | No Change | Not Significant | FALSE |
| Gm10069 | TRUE | 0.804937656 | 0.214114094 | No Change | Not Significant | TRUE |
| Gm10110 | TRUE | 0.425796703 | 0.612475705 | No Change | Not Significant | TRUE |
| Gm10125 | TRUE | 0.55070705 | 1 | No Change | Not Significant | FALSE |
| Gm10130 | TRUE | -0.168708817 | 0.671470886 | No Change | Not Significant | TRUE |
| Gm10224 | TRUE | -0.303326585 | 1 | No Change | Not Significant | FALSE |
| Gm10244 | TRUE | -0.222342903 | 1 | No Change | Not Significant | FALSE |
| Gm10268 | TRUE | 2.016146467 | 1 | No Change | Not Significant | FALSE |
| Gm10277 | TRUE | -0.428995458 | 0.340684159 | No Change | Not Significant | TRUE |
| Gm10354 | TRUE | 1.125208798 | 1 | No Change | Not Significant | FALSE |
| Gm10364 | TRUE | 2.4776382 | 1 | No Change | Not Significant | FALSE |
| Gm10371 | TRUE | 0.946806439 | 1 | No Change | Not Significant | FALSE |
| Gm10373 | TRUE | 3.502981163 | 1 | No Change | Not Significant | FALSE |
| Gm10382 | TRUE | 0.276737404 | 0.447999484 | No Change | Not Significant | TRUE |
| Gm10384 | TRUE | -0.160051253 | 0.716728293 | No Change | Not Significant | TRUE |
| Gm10389 | TRUE | -0.149668163 | 0.902682351 | No Change | Not Significant | TRUE |
| Gm10390 | TRUE | -0.30761217 | 0.727166502 | No Change | Not Significant | TRUE |
| Gm10400 | TRUE | -0.202520675 | 0.733377497 | No Change | Not Significant | TRUE |
| Gm10415 | TRUE | 0.048513294 | 1 | No Change | Not Significant | FALSE |
| Gm10416 | TRUE | -0.506374562 | 1 | No Change | Not Significant | FALSE |
| Gm10420 | TRUE | 1.077718361 | 1 | No Change | Not Significant | FALSE |
| Gm10421 | TRUE | -1.041185052 | 1 | No Change | Not Significant | FALSE |
| Gm1043 | TRUE | 0.64106431 | 0.535752227 | No Change | Not Significant | TRUE |
| Gm10432 | TRUE | 0.264907021 | 0.851938182 | No Change | Not Significant | TRUE |
| Gm10433 | TRUE | 0.948988533 | 0.002138281 | UP | UP | TRUE |
| Gm10445 | TRUE | -1.932391573 | 1 | No Change | Not Significant | FALSE |
| Gm10451 | TRUE | -0.754872609 | 0.075366299 | No Change | Not Significant | TRUE |
| Gm10505 | TRUE | 1.251399223 | 1 | No Change | Not Significant | FALSE |
| Gm10509 | TRUE | -0.154620429 | 0.692354022 | No Change | Not Significant | TRUE |
| Gm10516 | TRUE | -0.25685421 | 0.151084803 | No Change | Not Significant | TRUE |
| Gm10532 | TRUE | 0.289559025 | 1 | No Change | Not Significant | FALSE |
| Gm10538 | TRUE | 4.380230648 | 1 | No Change | Not Significant | FALSE |
| Gm10548 | TRUE | 0.282034638 | 0.708596728 | No Change | Not Significant | TRUE |
| Gm10549 | TRUE | -0.432515945 | 1 | No Change | Not Significant | FALSE |
| Gm10575 | TRUE | -0.15656965 | 0.83386713 | No Change | Not Significant | TRUE |
| Gm10584 | TRUE | -0.609823365 | 1 | No Change | Not Significant | FALSE |
| Gm10612 | TRUE | -1.308492143 | 1 | No Change | Not Significant | FALSE |
| Gm10619 | TRUE | 0.886641803 | 0.169380754 | No Change | Not Significant | FALSE |
| Gm10636 | TRUE | -2.21994762 | 1 | No Change | Not Significant | FALSE |
| Gm10637 | TRUE | 2.604287209 | 1 | No Change | Not Significant | FALSE |
| Gm10644 | TRUE | 0.759749918 | 0.322298818 | No Change | Not Significant | TRUE |
| Gm10646 | TRUE | -1.032727423 | 1 | No Change | Not Significant | FALSE |
| Gm10653 | TRUE | -0.112845829 | 1 | No Change | Not Significant | FALSE |
| Gm10732 | TRUE | 0.208581517 | 0.721161093 | No Change | Not Significant | TRUE |
| Gm10767 | TRUE | -0.146669899 | 0.674995696 | No Change | Not Significant | TRUE |
| Gm10778 | TRUE | 0.346414964 | 1 | No Change | Not Significant | TRUE |
| Gm10785 | TRUE | -0.106586909 | 0.878912418 | No Change | Not Significant | TRUE |
| Gm10791 | TRUE | -0.23673454 | 0.714999514 | No Change | Not Significant | TRUE |
| Gm10804 | TRUE | -0.578146532 | 0.440150633 | No Change | Not Significant | TRUE |
| Gm10814 | TRUE | -0.103177514 | 0.930184975 | No Change | Not Significant | TRUE |
| Gm10825 | TRUE | 2.075940687 | 1 | No Change | Not Significant | FALSE |
| Gm10845 | TRUE | 1.18822375 | 1 | No Change | Not Significant | FALSE |
| Gm10865 | TRUE | 0.235641122 | 1 | No Change | Not Significant | FALSE |
| Gm10941 | TRUE | -0.405931714 | 1 | No Change | Not Significant | FALSE |
| Gm1110 | TRUE | 0.705828063 | 1 | No Change | Not Significant | FALSE |
| Gm11100 | TRUE | 1.694253631 | 1 | No Change | Not Significant | FALSE |
| Gm11110 | TRUE | -0.545862139 | 1 | No Change | Not Significant | TRUE |
| Gm11127 | TRUE | -1.252173552 | 1 | No Change | Not Significant | FALSE |
| Gm1113 | TRUE | 1.706285365 | 1 | No Change | Not Significant | FALSE |
| Gm11186 | TRUE | 2.695091457 | 1 | No Change | Not Significant | FALSE |
| Gm1123 | TRUE | -0.802903998 | 0.07689828 | No Change | Not Significant | TRUE |
| Gm11240 | TRUE | 1.712387185 | 1.88E-05 | UP | UP | TRUE |
| Gm11266 | TRUE | 0.264322206 | 1 | No Change | Not Significant | FALSE |
| Gm11423 | TRUE | -0.350985805 | 0.404039964 | No Change | Not Significant | TRUE |
| Gm11437 | TRUE | -0.5919031 | 0.020288204 | DOWN | DOWN | TRUE |
| Gm11513 | TRUE | -0.087819811 | 1 | No Change | Not Significant | FALSE |
| Gm11532 | TRUE | 1.181269444 | 1 | No Change | Not Significant | FALSE |
| Gm11696 | TRUE | 0.387050916 | 0.359584034 | No Change | Not Significant | TRUE |
| Gm11703 | TRUE | -0.102873761 | 1 | No Change | Not Significant | FALSE |
| Gm11714 | TRUE | -0.484460529 | 1 | No Change | Not Significant | FALSE |
| Gm11732 | TRUE | -0.428556401 | 1 | No Change | Not Significant | FALSE |
| Gm11747 | TRUE | 3.836795888 | 1.67E-05 | UP | UP | FALSE |
| Gm11767 | TRUE | -0.683087242 | 0.343481339 | No Change | Not Significant | TRUE |
| Gm11827 | TRUE | -0.692179427 | 1 | No Change | Not Significant | FALSE |
| Gm11832 | TRUE | 0.540584131 | 1 | No Change | Not Significant | FALSE |
| Gm11944 | TRUE | -0.404594709 | 1 | No Change | Not Significant | FALSE |
| Gm11964 | TRUE | 0.857269134 | 1 | No Change | Not Significant | FALSE |
| Gm11992 | TRUE | 0.825601471 | 0.037845599 | UP | UP | TRUE |
| Gm12108 | TRUE | -0.041119632 | 1 | No Change | Not Significant | FALSE |
| Gm12153 | TRUE | 0.402967328 | 1 | No Change | Not Significant | FALSE |
| Gm12185 | TRUE | 1.194531311 | 0.242240857 | No Change | Not Significant | FALSE |
| Gm12216 | TRUE | -0.057039889 | 0.943016089 | No Change | Not Significant | TRUE |
| Gm12248 | TRUE | -0.570325805 | 0.557777472 | No Change | Not Significant | TRUE |
| Gm12253 | TRUE | 1.447521761 | 1 | No Change | Not Significant | FALSE |
| Gm12258 | TRUE | 1.160413347 | 0.000209523 | UP | UP | TRUE |
| Gm12295 | TRUE | 0.500712722 | 1 | No Change | Not Significant | FALSE |
| Gm12296 | TRUE | 1.816185032 | 1 | No Change | Not Significant | FALSE |
| Gm12320 | TRUE | 0.02977079 | 0.951530343 | No Change | Not Significant | TRUE |
| Gm12359 | TRUE | -0.126608568 | 0.864094963 | No Change | Not Significant | TRUE |
| Gm12408 | TRUE | 0.62478894 | 0.06203487 | No Change | Not Significant | TRUE |
| Gm12409 | TRUE | -0.021335004 | 1 | No Change | Not Significant | FALSE |
| Gm12500 | TRUE | 0.794660878 | 0.14477215 | No Change | Not Significant | TRUE |
| Gm12522 | TRUE | 1.980889094 | 1 | No Change | Not Significant | FALSE |
| Gm12530 | TRUE | -1.626224128 | 1 | No Change | Not Significant | FALSE |
| Gm12592 | TRUE | 0.158648128 | 1 | No Change | Not Significant | FALSE |
| Gm12655 | TRUE | -0.015221604 | 1 | No Change | Not Significant | TRUE |
| Gm12669 | TRUE | 1.252864203 | 0.047144373 | UP | UP | FALSE |
| Gm12708 | TRUE | -0.739802039 | 1 | No Change | Not Significant | FALSE |
| Gm12712 | TRUE | 2.391500119 | 1 | No Change | Not Significant | FALSE |
| Gm12758 | TRUE | -0.091860709 | 0.941987155 | No Change | Not Significant | TRUE |
| Gm128 | TRUE | -0.101869059 | 1 | No Change | Not Significant | FALSE |
| Gm12816 | TRUE | 0.492109242 | 1 | No Change | Not Significant | FALSE |
| Gm12830 | TRUE | -0.273391961 | 0.691115487 | No Change | Not Significant | TRUE |
| Gm12866 | TRUE | 1.816136735 | 1 | No Change | Not Significant | FALSE |
| Gm12888 | TRUE | 0.297991747 | 1 | No Change | Not Significant | FALSE |
| Gm12992 | TRUE | -0.88962174 | 0.003846258 | DOWN | DOWN | TRUE |
| Gm13003 | TRUE | -0.567356641 | 1 | No Change | Not Significant | FALSE |
| Gm13032 | TRUE | -1.776593998 | 1 | No Change | Not Significant | FALSE |
| Gm13090 | TRUE | 1.507819475 | 1 | No Change | Not Significant | FALSE |
| Gm13199 | TRUE | 0.199122558 | 1 | No Change | Not Significant | FALSE |
| Gm13203 | TRUE | -0.42073824 | 0.660955754 | No Change | Not Significant | FALSE |

Supplementary Table 3

| geneID | Cond1_DF_Detected | Cond1_DF_ValueCol | Cond1_DF_PadjCol | Cond1_DF_Cutoff | Cond1_DF_Cutoff_Specific.x | Cond2_DF_Detected |
|---|---|---|---|---|---|---|
| Gm13238 | TRUE | -0.675491177 | 1 | No Change | Not Significant | FALSE |
| Gm13270 | TRUE | 0.297078903 | 0.753472451 | No Change | Not Significant | TRUE |
| Gm13415 | TRUE | -0.611806577 | 0.332892364 | No Change | Not Significant | TRUE |
| Gm13420 | TRUE | 0.231388363 | 0.828297243 | No Change | Not Significant | TRUE |
| Gm13421 | TRUE | 0.276239497 | 0.586981666 | No Change | Not Significant | TRUE |
| Gm13427 | TRUE | 1.098950187 | 1 | No Change | Not Significant | FALSE |
| Gm13470 | TRUE | 2.649169289 | 1 | No Change | Not Significant | FALSE |
| Gm13483 | TRUE | -0.283888703 | 1 | No Change | Not Significant | FALSE |
| Gm13544 | TRUE | -0.109568855 | 1 | No Change | Not Significant | FALSE |
| Gm13546 | TRUE | 1.131367261 | 1 | No Change | Not Significant | FALSE |
| Gm13572 | TRUE | 0.514543232 | 1 | No Change | Not Significant | TRUE |
| Gm13648 | TRUE | 0.134615101 | 0.758091673 | No Change | Not Significant | TRUE |
| Gm13652 | TRUE | 2.247026709 | 1 | No Change | Not Significant | FALSE |
| Gm13710 | TRUE | 0.128342114 | 1 | No Change | Not Significant | FALSE |
| Gm13807 | TRUE | 0.505754873 | 0.266628782 | No Change | Not Significant | TRUE |
| Gm13848 | TRUE | 3.075914829 | 1 | No Change | Not Significant | FALSE |
| Gm13889 | TRUE | -0.267261024 | 1 | No Change | Not Significant | FALSE |
| Gm13944 | TRUE | -0.777688981 | 1 | No Change | Not Significant | FALSE |
| Gm13981 | TRUE | 1.6646438 | 1 | No Change | Not Significant | FALSE |
| Gm13986 | TRUE | -1.168677992 | 0.039794231 | DOWN | DOWN | TRUE |
| Gm14023 | TRUE | -0.154339955 | 0.884482916 | No Change | Not Significant | TRUE |
| Gm14133 | TRUE | -0.286739242 | 1 | No Change | Not Significant | FALSE |
| Gm14137 | TRUE | 0.902850641 | 0.082552308 | No Change | Not Significant | TRUE |
| Gm14169 | TRUE | 0.176009277 | 1 | No Change | Not Significant | FALSE |
| Gm14204 | TRUE | 0.513276255 | 1 | No Change | Not Significant | FALSE |
| Gm14207 | TRUE | -0.487587963 | 0.075180916 | No Change | Not Significant | TRUE |
| Gm14267 | TRUE | 1.711609317 | 1 | No Change | Not Significant | FALSE |
| Gm14295 | TRUE | -0.342200973 | 0.469256898 | No Change | Not Significant | TRUE |
| Gm14296 | TRUE | 0.510921632 | 1 | No Change | Not Significant | FALSE |
| Gm14305 | TRUE | -0.697588185 | 0.01173204 | DOWN | DOWN | TRUE |
| Gm14317 | TRUE | 0.964046719 | 1 | No Change | Not Significant | FALSE |
| Gm14322 | TRUE | -0.693163296 | 1 | No Change | Not Significant | TRUE |
| Gm14325 | TRUE | -0.486725087 | 0.198945226 | No Change | Not Significant | TRUE |
| Gm14326 | TRUE | -0.202950656 | 0.596246587 | No Change | Not Significant | TRUE |
| Gm14379 | TRUE | 1.437009527 | 1 | No Change | Not Significant | FALSE |
| Gm14393 | TRUE | 0.03317613 | 1 | No Change | Not Significant | FALSE |
| Gm14399 | TRUE | -0.227065639 | 0.649798537 | No Change | Not Significant | TRUE |
| Gm14410 | TRUE | -0.713054723 | 0.047593622 | DOWN | DOWN | TRUE |
| Gm14419 | TRUE | -1.264025665 | 1 | No Change | Not Significant | FALSE |
| Gm14455 | TRUE | -1.132605876 | 1 | No Change | Not Significant | FALSE |
| Gm14496 | TRUE | -2.024152856 | 1 | No Change | Not Significant | FALSE |
| Gm14549 | TRUE | 0.684145136 | 1 | No Change | Not Significant | TRUE |
| Gm14569 | TRUE | 3.724276707 | 1 | No Change | Not Significant | FALSE |
| Gm14634 | TRUE | 1.208210341 | 1 | No Change | Not Significant | FALSE |
| Gm14635 | TRUE | 1.462847708 | 1 | No Change | Not Significant | FALSE |
| Gm14827 | TRUE | 1.86889453 | 1 | No Change | Not Significant | FALSE |
| Gm14858 | TRUE | -0.178627024 | 1 | No Change | Not Significant | FALSE |
| Gm14862 | TRUE | -1.031560513 | 1 | No Change | Not Significant | FALSE |
| Gm14963 | TRUE | 0.075908531 | 0.930374388 | No Change | Not Significant | TRUE |
| Gm14965 | TRUE | 1.186053269 | 1 | No Change | Not Significant | FALSE |
| Gm15050 | TRUE | 1.152847595 | 0.047864939 | UP | UP | TRUE |
| Gm15133 | TRUE | -0.321075954 | 0.629730716 | No Change | Not Significant | TRUE |
| Gm15232 | TRUE | -0.06545449 | 1 | No Change | Not Significant | FALSE |
| Gm15246 | TRUE | 1.684357323 | 1 | No Change | Not Significant | FALSE |
| Gm1527 | TRUE | 0.250471944 | 1 | No Change | Not Significant | FALSE |
| Gm15318 | TRUE | -0.509270857 | 1 | No Change | Not Significant | FALSE |
| Gm15326 | TRUE | -1.848257659 | 1 | No Change | Not Significant | FALSE |
| Gm15350 | TRUE | 0.390062999 | 1 | No Change | Not Significant | FALSE |
| Gm15413 | TRUE | 1.241807211 | 1 | No Change | Not Significant | FALSE |
| Gm15417 | TRUE | 0.209146019 | 0.472341459 | No Change | Not Significant | TRUE |
| Gm15441 | TRUE | -0.373956043 | 0.598604992 | No Change | Not Significant | TRUE |
| Gm15446 | TRUE | -0.328891661 | 0.166571051 | No Change | Not Significant | TRUE |
| Gm15455 | TRUE | 0.604386275 | 1 | No Change | Not Significant | FALSE |
| Gm15462 | TRUE | -0.544691053 | 0.12072884 | No Change | Not Significant | TRUE |
| Gm15501 | TRUE | 0.957525634 | 0.207566563 | No Change | Not Significant | TRUE |
| Gm15509 | TRUE | 2.679467287 | 1 | No Change | Not Significant | FALSE |
| Gm15510 | TRUE | -1.027656121 | 1 | No Change | Not Significant | FALSE |
| Gm15545 | TRUE | 0.042537816 | 0.939157889 | No Change | Not Significant | TRUE |
| Gm15559 | TRUE | -0.216214251 | 0.840342672 | No Change | Not Significant | TRUE |
| Gm15569 | TRUE | -0.213777194 | 1 | No Change | Not Significant | FALSE |
| Gm15612 | TRUE | -0.88259587 | 1 | No Change | Not Significant | FALSE |
| Gm15614 | TRUE | -0.004257881 | 1 | No Change | Not Significant | FALSE |
| Gm15638 | TRUE | -0.153081281 | 0.822609935 | No Change | Not Significant | TRUE |
| Gm15663 | TRUE | -0.172102161 | 0.676807614 | No Change | Not Significant | TRUE |
| Gm15682 | TRUE | -1.24901795 | 1 | No Change | Not Significant | FALSE |
| Gm15706 | TRUE | -0.20133178 | 0.468075728 | No Change | Not Significant | TRUE |
| Gm15708 | TRUE | -0.114433063 | 0.820143104 | No Change | Not Significant | TRUE |
| Gm15713 | TRUE | 0.276430578 | 1 | No Change | Not Significant | FALSE |
| Gm15737 | TRUE | -2.086205873 | 1 | No Change | Not Significant | FALSE |
| Gm15743 | TRUE | -0.978236137 | 0.065328114 | No Change | Not Significant | TRUE |
| Gm15756 | TRUE | 0.084141501 | 1 | No Change | Not Significant | FALSE |
| Gm15760 | TRUE | 0.017403314 | 0.978989985 | No Change | Not Significant | TRUE |
| Gm15787 | TRUE | -0.715375681 | 0.084361596 | No Change | Not Significant | TRUE |
| Gm15793 | TRUE | 0.196937996 | 0.840583034 | No Change | Not Significant | FALSE |
| Gm15816 | TRUE | 0.665426062 | 1 | No Change | Not Significant | FALSE |
| Gm15850 | TRUE | -1.382216818 | 1 | No Change | Not Significant | FALSE |
| Gm15880 | TRUE | -0.558399375 | 1 | No Change | Not Significant | FALSE |
| Gm15889 | TRUE | -0.26169447 | 0.795699228 | No Change | Not Significant | TRUE |
| Gm15910 | TRUE | -0.261151365 | 1 | No Change | Not Significant | TRUE |
| Gm15915 | TRUE | -1.409322879 | 1 | No Change | Not Significant | FALSE |
| Gm15941 | TRUE | -0.066689411 | 0.935658236 | No Change | Not Significant | TRUE |
| Gm15946 | TRUE | -0.137912922 | 1 | No Change | Not Significant | FALSE |
| Gm15972 | TRUE | -0.414267303 | 0.390140563 | No Change | Not Significant | TRUE |
| Gm1600 | TRUE | 1.318854328 | 1 | No Change | Not Significant | FALSE |
| Gm16015 | TRUE | -0.254339719 | 0.768390968 | No Change | Not Significant | TRUE |
| Gm16023 | TRUE | -0.183190557 | 0.687185565 | No Change | Not Significant | TRUE |
| Gm16090 | TRUE | 0.09030837 | 0.938415991 | No Change | Not Significant | FALSE |
| Gm16124 | TRUE | -0.185302015 | 0.836513744 | No Change | Not Significant | TRUE |
| Gm16136 | TRUE | -0.237023563 | 0.223580206 | No Change | Not Significant | TRUE |
| Gm16142 | TRUE | 0.176010272 | 1 | No Change | Not Significant | FALSE |
| Gm16143 | TRUE | -0.576060354 | 0.464129434 | No Change | Not Significant | TRUE |
| Gm16174 | TRUE | 0.456981812 | 0.279706106 | No Change | Not Significant | TRUE |
| Gm16244 | TRUE | 0.565195897 | 1 | No Change | Not Significant | FALSE |
| Gm16283 | TRUE | 0.083502453 | 1 | No Change | Not Significant | TRUE |
| Gm16287 | TRUE | -0.510568392 | 0.484112338 | No Change | Not Significant | TRUE |
| Gm16291 | TRUE | 1.237182731 | 1 | No Change | Not Significant | FALSE |
| Gm16299 | TRUE | -1.774042802 | 1 | No Change | Not Significant | FALSE |
| Gm16325 | TRUE | 0.320731476 | 1 | No Change | Not Significant | FALSE |
| Gm16364 | TRUE | 0.399576048 | 0.628274069 | No Change | Not Significant | TRUE |
| Gm16523 | TRUE | -0.09608863 | 1 | No Change | Not Significant | FALSE |
| Gm16537 | TRUE | 0.246567514 | 1 | No Change | Not Significant | FALSE |
| Gm16573 | TRUE | -0.117531255 | 0.934257579 | No Change | Not Significant | TRUE |
| Gm16576 | TRUE | -0.267773004 | 0.646068054 | No Change | Not Significant | TRUE |
| Gm16582 | TRUE | 1.128017972 | 1 | No Change | Not Significant | FALSE |
| Gm16617 | TRUE | 0.544087708 | 1 | No Change | Not Significant | FALSE |
| Gm16618 | TRUE | -0.330850672 | 1 | No Change | Not Significant | TRUE |
| Gm16638 | TRUE | 0.623434119 | 0.349409629 | No Change | Not Significant | TRUE |
| Gm16675 | TRUE | -0.191090598 | 0.702914252 | No Change | Not Significant | TRUE |
| Gm16685 | TRUE | 1.334652559 | 0.059061089 | No Change | Not Significant | TRUE |
| Gm1673 | TRUE | 2.592594912 | 6.62E-07 | UP | UP | FALSE |
| Gm16731 | TRUE | 0.243424062 | 0.726026078 | No Change | Not Significant | TRUE |
| Gm16740 | TRUE | 0.113140319 | 1 | No Change | Not Significant | TRUE |
| Gm16793 | TRUE | -0.095504279 | 1 | No Change | Not Significant | TRUE |
| Gm16833 | TRUE | 0.714626949 | 1 | No Change | Not Significant | FALSE |
| Gm16845 | TRUE | -0.516486036 | 0.321275859 | No Change | Not Significant | TRUE |
| Gm16853 | TRUE | -0.004852414 | 1 | No Change | Not Significant | FALSE |
| Gm16861 | TRUE | -0.204589542 | 0.750868206 | No Change | Not Significant | TRUE |
| Gm16863 | TRUE | -0.734238546 | 0.176609179 | No Change | Not Significant | TRUE |
| Gm16897 | TRUE | 0.375705714 | 0.628624818 | No Change | Not Significant | TRUE |
| Gm16973 | TRUE | -0.24879377 | 0.516613382 | No Change | Not Significant | TRUE |
| Gm16982 | TRUE | -0.311560839 | 0.630552252 | No Change | Not Significant | TRUE |
| Gm16998 | TRUE | 0.290797749 | 1 | No Change | Not Significant | FALSE |
| Gm17019 | TRUE | 1.189885147 | 1 | No Change | Not Significant | FALSE |
| Gm17039 | TRUE | -0.187484117 | 1 | No Change | Not Significant | FALSE |
| Gm17059 | TRUE | -0.482209316 | 1 | No Change | Not Significant | FALSE |
| Gm17067 | TRUE | -0.549169421 | 1 | No Change | Not Significant | FALSE |
| Gm17089 | TRUE | 0.113116873 | 0.935980063 | No Change | Not Significant | FALSE |
| Gm17227 | TRUE | -0.389228912 | 1 | No Change | Not Significant | FALSE |
| Gm17344 | TRUE | -0.757220093 | 1 | No Change | Not Significant | FALSE |
| Gm17359 | TRUE | -1.226871752 | 1 | No Change | Not Significant | FALSE |
| Gm17396 | TRUE | -1.601129965 | 1 | No Change | Not Significant | FALSE |
| Gm17399 | TRUE | -0.071480803 | 1 | No Change | Not Significant | FALSE |
| Gm17415 | TRUE | 1.046532765 | 0.000735343 | UP | UP | TRUE |
| Gm17484 | TRUE | -0.028712276 | 1 | No Change | Not Significant | FALSE |
| Gm17546 | TRUE | 2.312299902 | 1 | No Change | Not Significant | FALSE |
| Gm17634 | TRUE | -0.570239822 | 0.498301136 | No Change | Not Significant | TRUE |
| Gm17660 | TRUE | 1.941557954 | 1 | No Change | Not Significant | FALSE |
| Gm17733 | TRUE | 1.452666802 | 1 | No Change | Not Significant | FALSE |
| Gm17767 | TRUE | 2.923418399 | 3.84E-08 | UP | UP | TRUE |
| Gm17830 | TRUE | 2.862075234 | 1 | No Change | Not Significant | FALSE |
| Gm18409 | TRUE | -0.206662656 | 1 | No Change | Not Significant | FALSE |
| Gm19345 | TRUE | 1.806140241 | 1 | No Change | Not Significant | FALSE |
| Gm19410 | TRUE | 0.465267349 | 0.590110273 | No Change | Not Significant | TRUE |
| Gm19466 | TRUE | 0.629326914 | 1 | No Change | Not Significant | FALSE |
| Gm19500 | TRUE | -1.630155916 | 1 | No Change | Not Significant | FALSE |
| Gm19510 | TRUE | -0.812006855 | 1 | No Change | Not Significant | FALSE |
| Gm19519 | TRUE | -0.227903792 | 1 | No Change | Not Significant | FALSE |
| Gm19522 | TRUE | 0.110923618 | 0.885864136 | No Change | Not Significant | TRUE |
| Gm19557 | TRUE | -0.518939625 | 0.462014445 | No Change | Not Significant | TRUE |
| Gm19582 | TRUE | 0.141992706 | 1 | No Change | Not Significant | TRUE |
| Gm19589 | TRUE | 0.176019694 | 1 | No Change | Not Significant | FALSE |
| Gm19605 | TRUE | 0.748616688 | 9.12E-05 | UP | UP | TRUE |
| Gm1965 | TRUE | -1.914866615 | 7.61E-12 | DOWN | DOWN | TRUE |
| Gm19705 | TRUE | 0.413701185 | 1 | No Change | Not Significant | FALSE |
| Gm19710 | TRUE | -0.16164054 | 1 | No Change | Not Significant | FALSE |
| Gm1976 | TRUE | -0.127072234 | 0.735609498 | No Change | Not Significant | TRUE |
| Gm20052 | TRUE | 0.04533231 | 1 | No Change | Not Significant | FALSE |
| Gm20125 | TRUE | 1.694363094 | 0.33090651 | No Change | Not Significant | FALSE |
| Gm20275 | TRUE | -0.759355806 | 0.003290188 | DOWN | DOWN | TRUE |
| Gm20300 | TRUE | -0.148813043 | 0.432209358 | No Change | Not Significant | TRUE |
| Gm20324 | TRUE | 0.046161522 | 0.946708891 | No Change | Not Significant | TRUE |
| Gm20379 | TRUE | -0.276435203 | 1 | No Change | Not Significant | FALSE |
| Gm20498 | TRUE | 0.336817056 | 0.580678813 | No Change | Not Significant | TRUE |
| Gm20517 | TRUE | 0.969326668 | 0.040777919 | UP | UP | TRUE |
| Gm20556 | TRUE | -0.446138843 | 1 | No Change | Not Significant | FALSE |
| Gm20594 | TRUE | -0.03136418 | 1 | No Change | Not Significant | TRUE |
| Gm20597 | TRUE | -1.156997188 | 1 | No Change | Not Significant | FALSE |
| Gm20604 | TRUE | 0.184880188 | 0.448874722 | No Change | Not Significant | TRUE |
| Gm2061 | TRUE | 0.437466598 | 0.232355545 | No Change | Not Significant | TRUE |
| Gm20743 | TRUE | -0.602226083 | 1 | No Change | Not Significant | FALSE |
| Gm20744 | TRUE | 0.611484357 | 1 | No Change | Not Significant | FALSE |
| Gm20750 | TRUE | 0.906918411 | 1 | No Change | Not Significant | FALSE |
| Gm20753 | TRUE | -0.773583356 | 1 | No Change | Not Significant | FALSE |
| Gm20756 | TRUE | -0.643126742 | 1 | No Change | Not Significant | FALSE |
| Gm2093 | TRUE | -1.734872852 | 1 | No Change | Not Significant | FALSE |
| Gm20939 | TRUE | -0.347960265 | 0.297320449 | No Change | Not Significant | TRUE |
| Gm21057 | TRUE | 0.501399488 | 1 | No Change | Not Significant | TRUE |
| Gm21083 | TRUE | -0.887844969 | 1 | No Change | Not Significant | TRUE |
| Gm21190 | TRUE | -0.477079216 | 1 | No Change | Not Significant | TRUE |
| Gm21284 | TRUE | 0.623059425 | 1 | No Change | Not Significant | FALSE |
| Gm21388 | TRUE | 1.939937029 | 1 | No Change | Not Significant | FALSE |
| Gm21814 | TRUE | -0.434812763 | 0.535752227 | No Change | Not Significant | TRUE |

Supplementary Table 3

| geneID | Cond1_DF_Detected | Cond1_DF_ValueCol | Cond1_DF_PadjCol | Cond1_DF_Cutoff | Cond1_DF_Cutoff_Specific.x | Cond2_DF_Detected |
|---|---|---|---|---|---|---|
| Gm21847 | TRUE | -1.270105843 | 1 | No Change | Not Significant | FALSE |
| Gm21976 | TRUE | 1.20822665 | 1 | No Change | Not Significant | FALSE |
| Gm21992 | TRUE | 0.273166941 | 0.225445669 | No Change | Not Significant | TRUE |
| Gm22005 | TRUE | -1.49996275 | 1 | No Change | Not Significant | FALSE |
| Gm22009 | TRUE | -0.267648297 | 0.018648354 | No Change | Significant Negative | TRUE |
| Gm22039 | TRUE | -0.82491739 | 1 | No Change | Not Significant | FALSE |
| Gm22042 | TRUE | 3.002056631 | 1 | No Change | Not Significant | FALSE |
| Gm22043 | TRUE | 0.229463926 | 1 | No Change | Not Significant | FALSE |
| Gm22067 | TRUE | 2.157382834 | 1 | No Change | Not Significant | FALSE |
| Gm22079 | TRUE | 1.167142163 | 1 | No Change | Not Significant | FALSE |
| Gm22154 | TRUE | 0.150253357 | 0.685335967 | No Change | Not Significant | TRUE |
| Gm22200 | TRUE | 0.209359243 | 1 | No Change | Not Significant | FALSE |
| Gm22215 | TRUE | -0.139586877 | 1 | No Change | Not Significant | FALSE |
| Gm22270 | TRUE | 0.788358311 | 0.109549131 | No Change | Not Significant | TRUE |
| Gm22300 | TRUE | 0.502121167 | 1 | No Change | Not Significant | FALSE |
| Gm22303 | TRUE | 0.572079206 | 1 | No Change | Not Significant | FALSE |
| Gm22334 | TRUE | 0.316869955 | 1 | No Change | Not Significant | FALSE |
| Gm22353 | TRUE | -0.514968673 | 1 | No Change | Not Significant | FALSE |
| Gm22358 | TRUE | -0.254010285 | 1 | No Change | Not Significant | FALSE |
| Gm22362 | TRUE | -0.515082102 | 0.397814213 | No Change | Not Significant | TRUE |
| Gm22363 | TRUE | 1.666874867 | 1 | No Change | Not Significant | FALSE |
| Gm2237 | TRUE | -1.014885436 | 1 | No Change | Not Significant | TRUE |
| Gm22422 | TRUE | -0.012150737 | 0.982739526 | No Change | Not Significant | TRUE |
| Gm22442 | TRUE | -0.4839547 | 0.000673467 | No Change | Significant Negative | TRUE |
| Gm2245 | TRUE | 0.726659171 | 1 | No Change | Not Significant | FALSE |
| Gm22488 | TRUE | 1.660440712 | 1 | No Change | Not Significant | FALSE |
| Gm22499 | TRUE | 0.32125799 | 1 | No Change | Not Significant | FALSE |
| Gm22516 | TRUE | 0.653947575 | 1 | No Change | Not Significant | FALSE |
| Gm22571 | TRUE | -0.531382501 | 1 | No Change | Not Significant | FALSE |
| Gm22572 | TRUE | -0.255965213 | 1 | No Change | Not Significant | FALSE |
| Gm22573 | TRUE | -0.763009021 | 1 | No Change | Not Significant | FALSE |
| Gm22581 | TRUE | -0.328745788 | 1 | No Change | Not Significant | FALSE |
| Gm22595 | TRUE | -0.378459916 | 1 | No Change | Not Significant | FALSE |
| Gm22661 | TRUE | 1.266921841 | 1 | No Change | Not Significant | FALSE |
| Gm22728 | TRUE | -0.309796186 | 1 | No Change | Not Significant | FALSE |
| Gm22748 | TRUE | 0.032662317 | 0.938630585 | No Change | Not Significant | TRUE |
| Gm22753 | TRUE | -0.243669145 | 1 | No Change | Not Significant | FALSE |
| Gm22767 | TRUE | 0.32603211 | 0.641211258 | No Change | Not Significant | TRUE |
| Gm22772 | TRUE | 1.592772848 | 1 | No Change | Not Significant | FALSE |
| Gm22781 | TRUE | 0.885654256 | 1 | No Change | Not Significant | FALSE |
| Gm22786 | TRUE | 0.255374521 | 0.78673795 | No Change | Not Significant | TRUE |
| Gm22806 | TRUE | -0.287395315 | 1 | No Change | Not Significant | TRUE |
| Gm22840 | TRUE | 0.383646685 | 1 | No Change | Not Significant | FALSE |
| Gm22858 | TRUE | -0.565698695 | 1 | No Change | Not Significant | FALSE |
| Gm22888 | TRUE | -0.768504889 | 1 | No Change | Not Significant | FALSE |
| Gm22935 | TRUE | 1.1756056 | 1 | No Change | Not Significant | FALSE |
| Gm22960 | TRUE | 0.35151934 | 1 | No Change | Not Significant | FALSE |
| Gm22973 | TRUE | 1.246793204 | 1.55E-05 | UP | UP | TRUE |
| Gm22980 | TRUE | 0.174670336 | 1 | No Change | Not Significant | FALSE |
| Gm23105 | TRUE | 0.441162266 | 0.213539052 | No Change | Not Significant | TRUE |
| Gm23111 | TRUE | 2.224028278 | 1 | No Change | Not Significant | FALSE |
| Gm23119 | TRUE | -0.038079192 | 0.938630585 | No Change | Not Significant | TRUE |
| Gm23123 | TRUE | -0.712879617 | 1 | No Change | Not Significant | FALSE |
| Gm23169 | TRUE | 1.600986716 | 1 | No Change | Not Significant | FALSE |
| Gm23182 | TRUE | -0.33561454 | 1 | No Change | Not Significant | FALSE |
| Gm23199 | TRUE | 3.041684341 | 1 | No Change | Not Significant | FALSE |
| Gm23200 | TRUE | -1.032476832 | 1 | No Change | Not Significant | FALSE |
| Gm23201 | TRUE | -0.202936087 | 0.783873755 | No Change | Not Significant | TRUE |
| Gm23205 | TRUE | -0.604763763 | 1 | No Change | Not Significant | FALSE |
| Gm23206 | TRUE | -0.452575428 | 1 | No Change | Not Significant | FALSE |
| Gm23207 | TRUE | 0.094273203 | 1 | No Change | Not Significant | FALSE |
| Gm23238 | TRUE | -0.032102841 | 1 | No Change | Not Significant | FALSE |
| Gm23245 | TRUE | 0.208082124 | 0.800714821 | No Change | Not Significant | TRUE |
| Gm23297 | TRUE | -0.391507903 | 0.653742472 | No Change | Not Significant | TRUE |
| Gm23301 | TRUE | -0.029695604 | 0.975173098 | No Change | Not Significant | TRUE |
| Gm23318 | TRUE | 0.153466152 | 1 | No Change | Not Significant | FALSE |
| Gm23336 | TRUE | -2.147190622 | 1 | No Change | Not Significant | FALSE |
| Gm23346 | TRUE | 0.328608324 | 0.713884177 | No Change | Not Significant | FALSE |
| Gm23369 | TRUE | -0.67088004 | 1 | No Change | Not Significant | FALSE |
| Gm23442 | TRUE | 0.365572869 | 0.272838324 | No Change | Not Significant | TRUE |
| Gm23458 | TRUE | -0.622491999 | 1 | No Change | Not Significant | FALSE |
| Gm23547 | TRUE | -0.109174543 | 0.92126689 | No Change | Not Significant | FALSE |
| Gm23593 | TRUE | 0.005310503 | 1 | No Change | Not Significant | FALSE |
| Gm23608 | TRUE | -0.228812627 | 1 | No Change | Not Significant | FALSE |
| Gm23639 | TRUE | 0.439076428 | 0.314808425 | No Change | Not Significant | TRUE |
| Gm23656 | TRUE | 0.665754391 | 1 | No Change | Not Significant | FALSE |
| Gm23692 | TRUE | 0.366672309 | 1 | No Change | Not Significant | FALSE |
| Gm23722 | TRUE | -1.309404659 | 1 | No Change | Not Significant | FALSE |
| Gm23734 | TRUE | -0.394285306 | 0.429896633 | No Change | Not Significant | TRUE |
| Gm23747 | TRUE | -0.008876334 | 1 | No Change | Not Significant | FALSE |
| Gm23751 | TRUE | -0.323640972 | 1 | No Change | Not Significant | FALSE |
| Gm23787 | TRUE | -3.190571029 | 1 | No Change | Not Significant | FALSE |
| Gm23924 | TRUE | 0.971201145 | 1 | No Change | Not Significant | FALSE |
| Gm23940 | TRUE | -0.168485179 | 1 | No Change | Not Significant | FALSE |
| Gm23946 | TRUE | 0.814534465 | 0.127019194 | No Change | Not Significant | TRUE |
| Gm23951 | TRUE | -0.002661005 | 1 | No Change | Not Significant | FALSE |
| Gm23969 | TRUE | -0.034905137 | 0.971417829 | No Change | Not Significant | TRUE |
| Gm24031 | TRUE | -0.03849408 | 1 | No Change | Not Significant | FALSE |
| Gm24041 | TRUE | -0.261046832 | 1 | No Change | Not Significant | FALSE |
| Gm24119 | TRUE | -0.052895328 | 0.92158767 | No Change | Not Significant | TRUE |
| Gm24143 | TRUE | -1.949725517 | 1 | No Change | Not Significant | FALSE |
| Gm24145 | TRUE | 2.17764609 | 1 | No Change | Not Significant | FALSE |
| Gm24171 | TRUE | 2.888230242 | 1 | No Change | Not Significant | FALSE |
| Gm24207 | TRUE | 0.715602241 | 1 | No Change | Not Significant | FALSE |
| Gm24233 | TRUE | -0.090106975 | 0.930949982 | No Change | Not Significant | TRUE |
| Gm24252 | TRUE | -0.511873304 | 1 | No Change | Not Significant | FALSE |
| Gm24289 | TRUE | 0.180451529 | 1 | No Change | Not Significant | FALSE |
| Gm24313 | TRUE | 0.532987298 | 1 | No Change | Not Significant | FALSE |
| Gm24336 | TRUE | -0.334516564 | 0.00331551 | No Change | Significant Negative | TRUE |
| Gm24339 | TRUE | -0.089984546 | 0.736964526 | No Change | Not Significant | TRUE |
| Gm24400 | TRUE | -1.010788454 | 1 | No Change | Not Significant | FALSE |
| Gm24405 | TRUE | 0.214389654 | 1 | No Change | Not Significant | FALSE |
| Gm24440 | TRUE | 0.182675591 | 1 | No Change | Not Significant | FALSE |
| Gm24457 | TRUE | -0.38521708 | 1 | No Change | Not Significant | FALSE |
| Gm24461 | TRUE | 0.238332654 | 1 | No Change | Not Significant | FALSE |
| Gm24468 | TRUE | -0.016884576 | 1 | No Change | Not Significant | FALSE |
| Gm2447 | TRUE | 0.782942564 | 0.383501006 | No Change | Not Significant | FALSE |
| Gm24494 | TRUE | -0.192317492 | 0.776544903 | No Change | Not Significant | TRUE |
| Gm24507 | TRUE | 1.360243336 | 1 | No Change | Not Significant | FALSE |
| Gm24564 | TRUE | -3.521273579 | 1 | No Change | Not Significant | FALSE |
| Gm24582 | TRUE | -1.399136093 | 1 | No Change | Not Significant | FALSE |
| Gm24596 | TRUE | 0.251757321 | 1 | No Change | Not Significant | FALSE |
| Gm24607 | TRUE | 0.5788114 | 1 | No Change | Not Significant | FALSE |
| Gm24616 | TRUE | 0.809216592 | 0.181863858 | No Change | Not Significant | TRUE |
| Gm24620 | TRUE | 0.336675332 | 0.753386941 | No Change | Not Significant | TRUE |
| Gm24644 | TRUE | 0.377386716 | 1 | No Change | Not Significant | FALSE |
| Gm24671 | TRUE | 0.674119013 | 1 | No Change | Not Significant | FALSE |
| Gm24771 | TRUE | 0.344713418 | 1 | No Change | Not Significant | FALSE |
| Gm24837 | TRUE | 0.492951877 | 1 | No Change | Not Significant | FALSE |
| Gm24838 | TRUE | -2.602411195 | 1 | No Change | Not Significant | FALSE |
| Gm24867 | TRUE | 0.629322304 | 1 | No Change | Not Significant | FALSE |
| Gm24878 | TRUE | -0.503827488 | 1 | No Change | Not Significant | FALSE |
| Gm24879 | TRUE | 0.708201874 | 1 | No Change | Not Significant | FALSE |
| Gm24888 | TRUE | 0.235418516 | 1 | No Change | Not Significant | FALSE |
| Gm24924 | TRUE | 0.779017831 | 1 | No Change | Not Significant | FALSE |
| Gm24940 | TRUE | 0.104947673 | 1 | No Change | Not Significant | FALSE |
| Gm24978 | TRUE | -0.23121461 | 1 | No Change | Not Significant | FALSE |
| Gm24989 | TRUE | 0.793613855 | 1 | No Change | Not Significant | FALSE |
| Gm24992 | TRUE | -1.115995594 | 1 | No Change | Not Significant | FALSE |
| Gm25039 | TRUE | -1.444944969 | 1 | No Change | Not Significant | FALSE |
| Gm25117 | TRUE | -0.049743055 | 0.907147741 | No Change | Not Significant | TRUE |
| Gm25128 | TRUE | -0.166263377 | 0.861267215 | No Change | Not Significant | TRUE |
| Gm25133 | TRUE | 1.974405589 | 1 | No Change | Not Significant | FALSE |
| Gm25134 | TRUE | 2.577702347 | 1 | No Change | Not Significant | FALSE |
| Gm25147 | TRUE | -1.787372406 | 1 | No Change | Not Significant | FALSE |
| Gm25188 | TRUE | 0.088566162 | 1 | No Change | Not Significant | FALSE |
| Gm25189 | TRUE | 1.313997563 | 1 | No Change | Not Significant | FALSE |
| Gm25259 | TRUE | 0.875495501 | 1 | No Change | Not Significant | FALSE |
| Gm25290 | TRUE | -0.285658439 | 1 | No Change | Not Significant | FALSE |
| Gm25321 | TRUE | -0.179979246 | 1 | No Change | Not Significant | FALSE |
| Gm25323 | TRUE | -2.231512989 | 1 | No Change | Not Significant | FALSE |
| Gm25379 | TRUE | 0.408473279 | 1 | No Change | Not Significant | FALSE |
| Gm25394 | TRUE | 0.172212435 | 0.695624496 | No Change | Not Significant | TRUE |
| Gm25395 | TRUE | 0.019094175 | 0.971883559 | No Change | Not Significant | TRUE |
| Gm25410 | TRUE | -1.312277136 | 1 | No Change | Not Significant | FALSE |
| Gm25492 | TRUE | -0.048086371 | 0.924222269 | No Change | Not Significant | TRUE |
| Gm25517 | TRUE | 0.13368368 | 1 | No Change | Not Significant | FALSE |
| Gm25541 | TRUE | 0.218858431 | 0.703739933 | No Change | Not Significant | TRUE |
| Gm25579 | TRUE | 2.726379442 | 1 | No Change | Not Significant | FALSE |
| Gm25604 | TRUE | -1.823492891 | 1 | No Change | Not Significant | FALSE |
| Gm25630 | TRUE | -0.912751106 | 1 | No Change | Not Significant | FALSE |
| Gm25636 | TRUE | -0.429177642 | 0.519697021 | No Change | Not Significant | TRUE |
| Gm25683 | TRUE | -0.396427707 | 1 | No Change | Not Significant | FALSE |
| Gm25776 | TRUE | -0.103657309 | 0.854165577 | No Change | Not Significant | TRUE |
| Gm25777 | TRUE | 0.202643272 | 1 | No Change | Not Significant | FALSE |
| Gm25788 | TRUE | 0.253147144 | 0.612475705 | No Change | Not Significant | TRUE |
| Gm25813 | TRUE | 1.698522938 | 1 | No Change | Not Significant | FALSE |
| Gm25821 | TRUE | 1.888197479 | 1 | No Change | Not Significant | FALSE |
| Gm25827 | TRUE | 0.146979359 | 1 | No Change | Not Significant | FALSE |
| Gm25835 | TRUE | 0.042816834 | 0.90889719 | No Change | Not Significant | TRUE |
| Gm25848 | TRUE | 0.116877118 | 0.881386379 | No Change | Not Significant | TRUE |
| Gm25881 | TRUE | -0.323191781 | 1 | No Change | Not Significant | FALSE |
| Gm25897 | TRUE | -0.888372739 | 1 | No Change | Not Significant | FALSE |
| Gm25913 | TRUE | -0.154931699 | 1 | No Change | Not Significant | FALSE |
| Gm25926 | TRUE | -0.627638781 | 1 | No Change | Not Significant | FALSE |
| Gm25973 | TRUE | 0.740675743 | 1 | No Change | Not Significant | FALSE |
| Gm26020 | TRUE | 0.327672183 | 1 | No Change | Not Significant | FALSE |
| Gm26049 | TRUE | 0.036218233 | 1 | No Change | Not Significant | FALSE |
| Gm26064 | TRUE | 0.481223191 | 1 | No Change | Not Significant | FALSE |
| Gm26109 | TRUE | -0.382581941 | 1 | No Change | Not Significant | FALSE |
| Gm26110 | TRUE | 2.48896694 | 1 | No Change | Not Significant | FALSE |
| Gm26121 | TRUE | -1.379535719 | 1 | No Change | Not Significant | FALSE |
| Gm26165 | TRUE | 0.274930224 | 0.454613934 | No Change | Not Significant | TRUE |
| Gm26175 | TRUE | 0.430388218 | 0.590046157 | No Change | Not Significant | FALSE |
| Gm26202 | TRUE | 0.172226306 | 0.803678477 | No Change | Not Significant | TRUE |
| Gm26225 | TRUE | 0.381321403 | 0.627075992 | No Change | Not Significant | TRUE |
| Gm26244 | TRUE | 1.678864229 | 1 | No Change | Not Significant | FALSE |
| Gm26287 | TRUE | -1.051399812 | 1 | No Change | Not Significant | FALSE |
| Gm26343 | TRUE | -0.021756029 | 1 | No Change | Not Significant | FALSE |
| Gm26347 | TRUE | 0.651192431 | 1 | No Change | Not Significant | FALSE |
| Gm26387 | TRUE | 0.661947139 | 1 | No Change | Not Significant | FALSE |
| Gm26397 | TRUE | -0.020220335 | 0.970191419 | No Change | Not Significant | TRUE |
| Gm26403 | TRUE | -0.421417963 | 1 | No Change | Not Significant | FALSE |
| Gm26419 | TRUE | -1.695196852 | 1 | No Change | Not Significant | FALSE |
| Gm26424 | TRUE | -0.538221675 | 1 | No Change | Not Significant | FALSE |
| Gm26448 | TRUE | 0.380136693 | 0.169321114 | No Change | Not Significant | TRUE |
| Gm26457 | TRUE | -0.340483625 | 1 | No Change | Not Significant | FALSE |
| Gm26493 | TRUE | 0.703163892 | 0.410793982 | No Change | Not Significant | TRUE |
| Gm26520 | TRUE | -0.495588692 | 1 | No Change | Not Significant | FALSE |
| Gm26533 | TRUE | -0.575632906 | 1 | No Change | Not Significant | FALSE |
| Gm26544 | TRUE | 1.091902825 | 1 | No Change | Not Significant | FALSE |
| Gm26628 | TRUE | 0.364569232 | 0.710135685 | No Change | Not Significant | TRUE |

| geneID | Cond2_DF_ValueCol | Cond2_DF_PadjCol | Cond2_DF_Cutoff | Cond2_DF_Cutoff_Specific | BG_Method | RG1_All | RG2_Significant | RG3_SignificantChange | Cond1_DF_ensembl_gene_id | Cond1_DF_entrezgene_id | Cond1_DF_gene_biotype | Cond1_DF_baseMean | Cond1_DF_log2FoldChange | Cond1_DF_lfcSE | Cond1_DF_stat | Cond1_DF_pvalue | Cond1_DF_padj | Cond1_DF_absLog2FC | Cond1_DF_Cutoff_Specific.y | Cond2_DF_ensembl_gene_id | Cond2_DF_entrezgene_id | Cond2_DF_gene_biotype | Cond2_DF_baseMean | Cond2_DF_log2FoldChange | Cond2_DF_lfcSE | Cond2_DF_stat | Cond2_DF_pvalue | Cond2_DF_padj | Cond2_DF_absLog2FC | RG1_Specific_Cond2 | RG1_Specific_Cond1 |
|---|---|---|---|---|---|---|---|---|---|---|---|---|---|---|---|---|---|---|---|---|---|---|---|---|---|---|---|---|---|---|---|
| Gm21847 | NA | NA | No Change | Not Detected | FALSE | Background = FALSE | Background = FALSE | Background = FALSE | ENSMUSG00000094230 | 74062 | lncRNA | 1.402458198 | -1.270105843 | 2.002659978 | 0.63420943 | 0.525944154 | 1 | 1.270105843 | Not Significant | NA | NA | NA | NA | NA | NA | NA | NA | NA | NA | Background = FALSE | Background = FALSE |
| Gm21976 | NA | NA | No Change | Not Detected | FALSE | Background = FALSE | Background = FALSE | Background = FALSE | ENSMUSG00000096330 | 102637808 | transcribed_unprocessed_pseudogene | 1.711828582 | 1.20822665 | 1.552099988 | -0.778446401 | 0.436305893 | 1 | 1.20822665 | Not Significant | NA | NA | NA | NA | NA | NA | NA | NA | NA | NA | Background = FALSE | Background = FALSE |
| Gm21992 | 0.034626451 | 0.916289542 | No Change | Not Significant | TRUE | Cond1 Not Significant + Cond2 Not Significant | None | None | ENSMUSG00000096370 | 102902673 | protein_coding | 49.54391665 | 0.273166941 | 0.160233811 | -1.704802121 | 0.088231336 | 0.225445669 | 0.273166941 | Not Significant | ENSMUSG00000096370 | 102902673 | protein_coding | 84.4004912 | 0.034626451 | 0.147054491 | 0.235466804 | 0.813846441 | 0.916289542 | 0.034626451 | Cond1 No Change + Cond2 Not Significant | Cond2 No Change + Cond1 Not Significant |
| Gm22005 | NA | NA | No Change | Not Detected | FALSE | Background = FALSE | Background = FALSE | Background = FALSE | ENSMUSG00000075837 | 115489653 | snRNA | 0.926332705 | -1.49996275 | 1.2190344 | 1.230451536 | 0.218528065 | 1 | 1.49996275 | Not Significant | NA | NA | NA | NA | NA | NA | NA | NA | NA | NA | Background = FALSE | Background = FALSE |
| Gm22009 | -0.132223941 | 0.442487265 | No Change | Not Significant | TRUE | Cond1 Significant Negative + Cond2 Not Significant | None | None | ENSMUSG00000089417 | 115489854 | scaRNA | 1584.437012 | -0.267648297 | 0.092001192 | 2.909182933 | 0.003623748 | 0.018648354 | 0.267648297 | Significant Negative | ENSMUSG00000089417 | 115489854 | scaRNA | 2873.605295 | -0.132223941 | 0.104839766 | -1.261200265 | 0.207236703 | 0.442487265 | 0.132223941 | Cond1 No Change + Cond2 Not Significant | Cond2 No Change + Cond1 Significant Negative |
| Gm22039 | NA | NA | No Change | Not Detected | FALSE | Background = FALSE | Background = FALSE | Background = FALSE | ENSMUSG00000087903 | 115486273 | snoRNA | 0.567056251 | -0.82491739 | 1.466142426 | 0.562644785 | 0.573676789 | 1 | 0.82491739 | Not Significant | NA | NA | NA | NA | NA | NA | NA | NA | NA | NA | Background = FALSE | Background = FALSE |
| Gm22042 | NA | NA | No Change | Not Detected | FALSE | Background = FALSE | Background = FALSE | Background = FALSE | ENSMUSG00000064923 | 115490033 | snRNA | 1.356069965 | 3.002056631 | 1.732985456 | -1.732303419 | 0.083219554 | 1 | 3.002056631 | Not Significant | NA | NA | NA | NA | NA | NA | NA | NA | NA | NA | Background = FALSE | Background = FALSE |
| Gm22043 | NA | NA | No Change | Not Detected | FALSE | Background = FALSE | Background = FALSE | Background = FALSE | ENSMUSG00000064927 | 115489855 | snRNA | 2.181553871 | 0.229463926 | 0.783875279 | -0.292730148 | 0.769728424 | 1 | 0.229463926 | Not Significant | NA | NA | NA | NA | NA | NA | NA | NA | NA | NA | Background = FALSE | Background = FALSE |
| Gm22067 | NA | NA | No Change | Not Detected | FALSE | Background = FALSE | Background = FALSE | Background = FALSE | ENSMUSG00000084739 | 115488600 | snRNA | 1.344928801 | 2.157382834 | 1.338014743 | -1.612375981 | 0.106880148 | 1 | 2.157382834 | Not Significant | NA | NA | NA | NA | NA | NA | NA | NA | NA | NA | Background = FALSE | Background = FALSE |
| Gm22079 | NA | NA | No Change | Not Detected | FALSE | Background = FALSE | Background = FALSE | Background = FALSE | ENSMUSG00000088217 | 115488044 | snRNA | 2.013328377 | 1.167142163 | 0.837901353 | -1.39293505 | 0.163639432 | 1 | 1.167142163 | Not Significant | NA | NA | NA | NA | NA | NA | NA | NA | NA | NA | Background = FALSE | Background = FALSE |
| Gm22154 | 0.201113403 | 0.784827956 | No Change | Not Significant | TRUE | Cond1 Not Significant + Cond2 Not Significant | None | None | ENSMUSG00000065778 | 115490050 | snoRNA | 292.5540673 | 0.150253357 | 0.209449086 | -0.717374134 | 0.473143276 | 0.685335967 | 0.150253357 | Not Significant | ENSMUSG00000065778 | 115490050 | snoRNA | 551.9628916 | 0.201113403 | 0.382453207 | 0.525851005 | 0.598991733 | 0.784827956 | 0.201113403 | Cond1 No Change + Cond2 Not Significant | Cond2 No Change + Cond1 Not Significant |
| Gm22200 | NA | NA | No Change | Not Detected | FALSE | Background = FALSE | Background = FALSE | Background = FALSE | ENSMUSG00000089239 | 115489001 | snoRNA | 1.124106392 | 0.209359243 | 1.041889755 | -0.200941839 | 0.840744052 | 1 | 0.209359243 | Not Significant | NA | NA | NA | NA | NA | NA | NA | NA | NA | NA | Background = FALSE | Background = FALSE |
| Gm22215 | NA | NA | No Change | Not Detected | FALSE | Background = FALSE | Background = FALSE | Background = FALSE | ENSMUSG00000088183 | 115488890 | snRNA | 0.698573286 | -0.139586877 | 1.393595094 | 0.100163152 | 0.9202148 | 1 | 0.139586877 | Not Significant | NA | NA | NA | NA | NA | NA | NA | NA | NA | NA | Background = FALSE | Background = FALSE |

| geneID | Cond1_DF_Detected | Cond1_DF_ValueCol | Cond1_DF_PadjCol | Cond1_DF_Cutoff | Cond1_DF_Cutoff_Specific.x | Cond2_DF_Detected | Cond2_DF_ValueCol | Cond2_DF_PadjCol | Cond2_DF_Cutoff | Cond2_DF_Cutoff_Specific | BG_Method | RG1_All |
|---|---|---|---|---|---|---|---|---|---|---|---|---|
| Gm26633 | TRUE | 2.163836013 | 1 | No Change | Not Significant | FALSE | NA | NA | No Change | Not Detected | FALSE | Background = FALSE |
| Gm26644 | TRUE | 1.802502561 | 1 | No Change | Not Significant | FALSE | NA | NA | No Change | Not Detected | FALSE | Background = FALSE |
| Gm26685 | TRUE | -0.337792839 | 0.493664959 | No Change | Not Significant | TRUE | 0.28809143 | 0.4893628 | No Change | Not Significant | TRUE | Cond1 Not Significant + Cond2 Not Significant |
| Gm26782 | TRUE | -0.187607246 | 0.591883639 | No Change | Not Significant | TRUE | -0.082805667 | 0.787655348 | No Change | Not Significant | TRUE | Cond1 Not Significant + Cond2 Not Significant |
| Gm26787 | TRUE | 1.192116789 | 1 | No Change | Not Significant | FALSE | NA | NA | No Change | Not Detected | FALSE | Background = FALSE |
| Gm26788 | TRUE | -0.002166188 | 0.996966345 | No Change | Not Significant | TRUE | -0.423615033 | 0.421829944 | No Change | Not Significant | TRUE | Cond1 Not Significant + Cond2 Not Significant |
| Gm2694 | TRUE | -0.100267185 | 1 | No Change | Not Significant | FALSE | NA | NA | No Change | Not Detected | FALSE | Background = FALSE |
| Gm27011 | TRUE | 0.208577349 | 1 | No Change | Not Significant | TRUE | 0.491773857 | 0.469064473 | No Change | Not Significant | TRUE | Cond1 Not Significant + Cond2 Not Significant |
| Gm27217 | TRUE | -0.583617893 | 1 | No Change | Not Significant | FALSE | NA | NA | No Change | Not Detected | FALSE | Background = FALSE |
| Gm27343 | TRUE | 1.18378172 | 1 | No Change | Not Significant | FALSE | NA | NA | No Change | Not Detected | FALSE | Background = FALSE |
| Gm2762 | TRUE | -1.190240776 | 1 | No Change | Not Significant | FALSE | NA | NA | No Change | Not Detected | FALSE | Background = FALSE |
| Gm2788 | TRUE | 0.694743551 | 0.377599236 | No Change | Not Significant | TRUE | 0.344004595 | 0.551334926 | No Change | Not Significant | TRUE | Cond1 Not Significant + Cond2 Not Significant |
| Gm28040 | TRUE | -0.712226352 | 1 | No Change | Not Significant | FALSE | NA | NA | No Change | Not Detected | FALSE | Background = FALSE |
| Gm2808 | TRUE | -1.124255208 | 1 | No Change | Not Significant | FALSE | NA | NA | No Change | Not Detected | FALSE | Background = FALSE |
| Gm281 | TRUE | -1.526649531 | 1 | No Change | Not Significant | FALSE | NA | NA | No Change | Not Detected | FALSE | Background = FALSE |
| Gm28178 | TRUE | 0.609336272 | 1 | No Change | Not Significant | FALSE | NA | NA | No Change | Not Detected | FALSE | Background = FALSE |
| Gm28231 | TRUE | -0.989917373 | 1 | No Change | Not Significant | FALSE | NA | NA | No Change | Not Detected | FALSE | Background = FALSE |
| Gm28321 | TRUE | 1.294214421 | 1 | No Change | Not Significant | FALSE | NA | NA | No Change | Not Detected | FALSE | Background = FALSE |
| Gm28441 | TRUE | 1.188533176 | 1 | No Change | Not Significant | FALSE | NA | NA | No Change | Not Detected | FALSE | Background = FALSE |
| Gm28535 | TRUE | 0.547230313 | 1 | No Change | Not Significant | FALSE | NA | NA | No Change | Not Detected | FALSE | Background = FALSE |
| Gm28548 | TRUE | -0.790606981 | 1 | No Change | Not Significant | FALSE | NA | NA | No Change | Not Detected | FALSE | Background = FALSE |
| Gm28578 | TRUE | 1.396256738 | 1 | No Change | Not Significant | FALSE | NA | NA | No Change | Not Detected | FALSE | Background = FALSE |
| Gm28729 | TRUE | -0.4278909 | 1 | No Change | Not Significant | FALSE | NA | NA | No Change | Not Detected | FALSE | Background = FALSE |
| Gm28821 | TRUE | -0.551740533 | 1 | No Change | Not Significant | FALSE | NA | NA | No Change | Not Detected | FALSE | Background = FALSE |
| Gm29154 | TRUE | -0.873329638 | 1 | No Change | Not Significant | FALSE | NA | NA | No Change | Not Detected | FALSE | Background = FALSE |
| Gm29443 | TRUE | -0.421171803 | 1 | No Change | Not Significant | FALSE | NA | NA | No Change | Not Detected | FALSE | Background = FALSE |
| Gm29536 | TRUE | -0.245695568 | 1 | No Change | Not Significant | FALSE | NA | NA | No Change | Not Detected | FALSE | Background = FALSE |
| Gm29543 | TRUE | 0.253375255 | 1 | No Change | Not Significant | FALSE | NA | NA | No Change | Not Detected | FALSE | Background = FALSE |
| Gm29642 | TRUE | -0.5729566 | 1 | No Change | Not Significant | FALSE | NA | NA | No Change | Not Detected | FALSE | Background = FALSE |
| Gm29681 | TRUE | -0.020845445 | 1 | No Change | Not Significant | FALSE | NA | NA | No Change | Not Detected | FALSE | Background = FALSE |
| Gm29683 | TRUE | 1.126149062 | 1 | No Change | Not Significant | FALSE | NA | NA | No Change | Not Detected | FALSE | Background = FALSE |
| Gm29684 | TRUE | -0.271615637 | 1 | No Change | Not Significant | FALSE | NA | NA | No Change | Not Detected | FALSE | Background = FALSE |
| Gm29797 | TRUE | -1.534429571 | 1 | No Change | Not Significant | FALSE | NA | NA | No Change | Not Detected | FALSE | Background = FALSE |
| Gm29811 | TRUE | -0.605068704 | 1 | No Change | Not Significant | FALSE | NA | NA | No Change | Not Detected | FALSE | Background = FALSE |
| Gm29999 | TRUE | 0.16577657 | 1 | No Change | Not Significant | FALSE | NA | NA | No Change | Not Detected | FALSE | Background = FALSE |
| Gm2a | TRUE | -0.916925 | 8.92E-08 | DOWN | DOWN | TRUE | -0.389088993 | 0.003964431 | No Change | Significant Negative | TRUE | Cond1 DOWN + Cond2 Significant Negative |
| Gm30082 | TRUE | -0.659556872 | 1 | No Change | Not Significant | FALSE | NA | NA | No Change | Not Detected | FALSE | Background = FALSE |
| Gm30083 | TRUE | 0.004777584 | 1 | No Change | Not Significant | FALSE | NA | NA | No Change | Not Detected | FALSE | Background = FALSE |
| Gm30094 | TRUE | 2.140424283 | 1 | No Change | Not Significant | FALSE | NA | NA | No Change | Not Detected | FALSE | Background = FALSE |
| Gm30173 | TRUE | 0.843957941 | 1 | No Change | Not Significant | FALSE | NA | NA | No Change | Not Detected | FALSE | Background = FALSE |
| Gm30292 | TRUE | 1.735683327 | 1 | No Change | Not Significant | FALSE | NA | NA | No Change | Not Detected | FALSE | Background = FALSE |
| Gm30381 | TRUE | -1.987988258 | 1 | No Change | Not Significant | FALSE | NA | NA | No Change | Not Detected | FALSE | Background = FALSE |
| Gm30400 | TRUE | 0.574709928 | 1 | No Change | Not Significant | FALSE | NA | NA | No Change | Not Detected | FALSE | Background = FALSE |
| Gm30505 | TRUE | 2.97379987 | 0.000318174 | UP | UP | FALSE | NA | NA | No Change | Not Detected | FALSE | Background = FALSE |
| Gm30539 | TRUE | 4.876007816 | 0.00016086 | UP | UP | FALSE | NA | NA | No Change | Not Detected | FALSE | Background = FALSE |
| Gm3055 | TRUE | -0.324257985 | 0.702605511 | No Change | Not Significant | TRUE | 0.179661908 | 0.834153989 | No Change | Not Significant | TRUE | Cond1 Not Significant + Cond2 Not Significant |
| Gm30551 | TRUE | -0.753303576 | 1 | No Change | Not Significant | FALSE | NA | NA | No Change | Not Detected | FALSE | Background = FALSE |
| Gm30648 | TRUE | 1.291642925 | 1 | No Change | Not Significant | FALSE | NA | NA | No Change | Not Detected | FALSE | Background = FALSE |
| Gm30794 | TRUE | 0.552833786 | 1 | No Change | Not Significant | TRUE | -0.006614464 | 0.995768162 | No Change | Not Significant | TRUE | Cond1 Not Significant + Cond2 Not Significant |
| Gm3086 | TRUE | -1.429940603 | 1 | No Change | Not Significant | FALSE | NA | NA | No Change | Not Detected | FALSE | Background = FALSE |
| Gm30948 | TRUE | 0.138082061 | 1 | No Change | Not Significant | FALSE | NA | NA | No Change | Not Detected | FALSE | Background = FALSE |
| Gm30961 | TRUE | 0.054036263 | 1 | No Change | Not Significant | TRUE | -0.37452103 | 0.671427662 | No Change | Not Significant | TRUE | Cond1 Not Significant + Cond2 Not Significant |
| Gm31108 | TRUE | -1.210524524 | 1 | No Change | Not Significant | FALSE | NA | NA | No Change | Not Detected | FALSE | Background = FALSE |
| Gm31160 | TRUE | 0.141441557 | 0.812734264 | No Change | Not Significant | TRUE | -0.635929763 | 0.061835229 | No Change | Not Significant | TRUE | Cond1 Not Significant + Cond2 Not Significant |
| Gm31235 | TRUE | -0.477528797 | 0.36725457 | No Change | Not Significant | TRUE | 0.594071872 | 0.004783809 | UP | UP | TRUE | Cond1 Not Significant + Cond2 UP |
| Gm31333 | TRUE | -0.646713089 | 1 | No Change | Not Significant | FALSE | NA | NA | No Change | Not Detected | FALSE | Background = FALSE |
| Gm31508 | TRUE | 0.604125724 | 1 | No Change | Not Significant | FALSE | NA | NA | No Change | Not Detected | FALSE | Background = FALSE |
| Gm31510 | TRUE | -0.811424788 | 1 | No Change | Not Significant | FALSE | NA | NA | No Change | Not Detected | FALSE | Background = FALSE |
| Gm31517 | TRUE | 0.004278778 | 1 | No Change | Not Significant | FALSE | NA | NA | No Change | Not Detected | FALSE | Background = FALSE |
| Gm31562 | TRUE | 1.90180325 | 1 | No Change | Not Significant | FALSE | NA | NA | No Change | Not Detected | FALSE | Background = FALSE |
| Gm31592 | TRUE | 2.542699857 | 1 | No Change | Not Significant | FALSE | NA | NA | No Change | Not Detected | FALSE | Background = FALSE |
| Gm31728 | TRUE | 0.025335847 | 1 | No Change | Not Significant | FALSE | NA | NA | No Change | Not Detected | FALSE | Background = FALSE |
| Gm31763 | TRUE | -0.245868815 | 0.83310074 | No Change | Not Significant | TRUE | -0.425099179 | 0.682988492 | No Change | Not Significant | TRUE | Cond1 Not Significant + Cond2 Not Significant |
| Gm32014 | TRUE | -1.239686079 | 1 | No Change | Not Significant | FALSE | NA | NA | No Change | Not Detected | FALSE | Background = FALSE |
| Gm32090 | TRUE | 0.004301634 | 1 | No Change | Not Significant | FALSE | NA | NA | No Change | Not Detected | FALSE | Background = FALSE |
| Gm32123 | TRUE | -0.61327051 | 1 | No Change | Not Significant | FALSE | NA | NA | No Change | Not Detected | FALSE | Background = FALSE |
| Gm32200 | TRUE | 1.74690046 | 1 | No Change | Not Significant | FALSE | NA | NA | No Change | Not Detected | FALSE | Background = FALSE |
| Gm32282 | TRUE | -0.022795634 | 1 | No Change | Not Significant | TRUE | -0.078014331 | 0.943798397 | No Change | Not Significant | TRUE | Cond1 Not Significant + Cond2 Not Significant |
| Gm32391 | TRUE | 0.079212713 | 0.903803848 | No Change | Not Significant | TRUE | 0.123110074 | 0.82134757 | No Change | Not Significant | TRUE | Cond1 Not Significant + Cond2 Not Significant |
| Gm32592 | TRUE | 1.537619392 | 1 | No Change | Not Significant | FALSE | NA | NA | No Change | Not Detected | FALSE | Background = FALSE |
| Gm32618 | TRUE | -0.887351607 | 1 | No Change | Not Significant | FALSE | NA | NA | No Change | Not Detected | FALSE | Background = FALSE |
| Gm32624 | TRUE | 0.476172055 | 1 | No Change | Not Significant | FALSE | NA | NA | No Change | Not Detected | FALSE | Background = FALSE |
| Gm32687 | TRUE | -0.537825537 | 0.377733062 | No Change | Not Significant | TRUE | -0.286270223 | 0.569717873 | No Change | Not Significant | TRUE | Cond1 Not Significant + Cond2 Not Significant |
| Gm32742 | TRUE | -0.685222245 | 0.094313013 | No Change | Not Significant | TRUE | -0.14584228 | 0.786805165 | No Change | Not Significant | TRUE | Cond1 Not Significant + Cond2 Not Significant |
| Gm32828 | TRUE | -1.181428387 | 1 | No Change | Not Significant | FALSE | NA | NA | No Change | Not Detected | FALSE | Background = FALSE |
| Gm32856 | TRUE | 0.307195774 | 0.349988901 | No Change | Not Significant | TRUE | -0.022166274 | 0.962474368 | No Change | Not Significant | TRUE | Cond1 Not Significant + Cond2 Not Significant |
| Gm32926 | TRUE | -0.893874928 | 1 | No Change | Not Significant | FALSE | NA | NA | No Change | Not Detected | FALSE | Background = FALSE |
| Gm3294 | TRUE | -1.824151793 | 1 | No Change | Not Significant | FALSE | NA | NA | No Change | Not Detected | FALSE | Background = FALSE |
| Gm33035 | TRUE | 0.004913941 | 1 | No Change | Not Significant | FALSE | NA | NA | No Change | Not Detected | FALSE | Background = FALSE |
| Gm33054 | TRUE | -1.241731032 | 1 | No Change | Not Significant | FALSE | NA | NA | No Change | Not Detected | FALSE | Background = FALSE |
| Gm33104 | TRUE | 0.008449226 | 1 | No Change | Not Significant | FALSE | NA | NA | No Change | Not Detected | FALSE | Background = FALSE |
| Gm33148 | TRUE | 1.199610873 | 0.832085503 | No Change | Not Significant | FALSE | NA | NA | No Change | Not Detected | FALSE | Background = FALSE |
| Gm33175 | TRUE | 2.465236438 | 1 | No Change | Not Significant | FALSE | NA | NA | No Change | Not Detected | FALSE | Background = FALSE |
| Gm3325 | TRUE | -0.139190354 | 0.695624496 | No Change | Not Significant | TRUE | -0.217362098 | 0.457725814 | No Change | Not Significant | TRUE | Cond1 Not Significant + Cond2 Not Significant |
| Gm3336 | TRUE | -0.669694441 | 6.25E-13 | DOWN | DOWN | TRUE | 0.077938323 | 0.602381167 | No Change | Not Significant | TRUE | Cond1 DOWN + Cond2 Not Significant |
| Gm33543 | TRUE | 1.009284215 | 1 | No Change | Not Significant | FALSE | NA | NA | No Change | Not Detected | FALSE | Background = FALSE |
| Gm3363 | TRUE | 0.965089634 | 1 | No Change | Not Significant | FALSE | NA | NA | No Change | Not Detected | FALSE | Background = FALSE |
| Gm33677 | TRUE | -2.697642509 | 4.74E-08 | DOWN | DOWN | TRUE | -0.420258735 | 0.499743842 | No Change | Not Significant | TRUE | Cond1 DOWN + Cond2 Not Significant |
| Gm33989 | TRUE | -0.327623624 | 0.049685066 | No Change | Significant Negative | TRUE | -0.214577168 | 0.024315491 | No Change | Significant Negative | TRUE | Cond1 Significant Negative + Cond2 Significant Negative |
| Gm34005 | TRUE | 0.543076625 | 1 | No Change | Not Significant | FALSE | NA | NA | No Change | Not Detected | FALSE | Background = FALSE |
| Gm3411 | TRUE | 0.407412464 | 1 | No Change | Not Significant | TRUE | 0.66392614 | 0.395233324 | No Change | Not Significant | TRUE | Cond1 Not Significant + Cond2 Not Significant |
| Gm34121 | TRUE | -0.148611826 | 0.855204229 | No Change | Not Significant | TRUE | 0.109406728 | 0.862201627 | No Change | Not Significant | TRUE | Cond1 Not Significant + Cond2 Not Significant |
| Gm3435 | TRUE | -0.686471088 | 0.390530809 | No Change | Not Significant | TRUE | 0.278847771 | 0.645650963 | No Change | Not Significant | TRUE | Cond1 Not Significant + Cond2 Not Significant |
| Gm34408 | TRUE | -0.927042832 | 1 | No Change | Not Significant | FALSE | NA | NA | No Change | Not Detected | FALSE | Background = FALSE |
| Gm34574 | TRUE | -1.882236476 | 1 | No Change | Not Significant | FALSE | NA | NA | No Change | Not Detected | FALSE | Background = FALSE |
| Gm34589 | TRUE | 0.357325267 | 1 | No Change | Not Significant | FALSE | NA | NA | No Change | Not Detected | FALSE | Background = FALSE |
| Gm34643 | TRUE | 3.427245298 | 1 | No Change | Not Significant | FALSE | NA | NA | No Change | Not Detected | FALSE | Background = FALSE |
| Gm34785 | TRUE | -0.33984417 | 1 | No Change | Not Significant | FALSE | NA | NA | No Change | Not Detected | FALSE | Background = FALSE |
| Gm34821 | TRUE | 0.102600288 | 0.896636371 | No Change | Not Significant | TRUE | -0.336569712 | 0.50034619 | No Change | Not Significant | TRUE | Cond1 Not Significant + Cond2 Not Significant |
| Gm34882 | TRUE | -0.734658745 | 1 | No Change | Not Significant | TRUE | -0.465337324 | 0.573705524 | No Change | Not Significant | TRUE | Cond1 Not Significant + Cond2 Not Significant |
| Gm34885 | TRUE | 0.364340214 | 1 | No Change | Not Significant | FALSE | NA | NA | No Change | Not Detected | FALSE | Background = FALSE |
| Gm35021 | TRUE | 0.463900576 | 1 | No Change | Not Significant | FALSE | NA | NA | No Change | Not Detected | FALSE | Background = FALSE |
| Gm35166 | TRUE | -0.223702538 | 1 | No Change | Not Significant | FALSE | NA | NA | No Change | Not Detected | FALSE | Background = FALSE |
| Gm3519 | TRUE | 0.018195364 | 1 | No Change | Not Significant | FALSE | NA | NA | No Change | Not Detected | FALSE | Background = FALSE |
| Gm35191 | TRUE | 1.726125194 | 1 | No Change | Not Significant | FALSE | NA | NA | No Change | Not Detected | FALSE | Background = FALSE |
| Gm35279 | TRUE | 0.291226016 | 1 | No Change | Not Significant | FALSE | NA | NA | No Change | Not Detected | FALSE | Background = FALSE |
| Gm35315 | TRUE | 0.18709842 | 0.750140815 | No Change | Not Significant | TRUE | 0.320158257 | 0.299347968 | No Change | Not Significant | TRUE | Cond1 Not Significant + Cond2 Not Significant |
| Gm35339 | TRUE | 0.365572437 | 0.454124681 | No Change | Not Significant | TRUE | 0.215355216 | 0.62329844 | No Change | Not Significant | TRUE | Cond1 Not Significant + Cond2 Not Significant |
| Gm35392 | TRUE | 2.91851994 | 1 | No Change | Not Significant | FALSE | NA | NA | No Change | Not Detected | FALSE | Background = FALSE |
| Gm35507 | TRUE | -1.889119112 | 1 | No Change | Not Significant | FALSE | NA | NA | No Change | Not Detected | FALSE | Background = FALSE |
| Gm35549 | TRUE | 0.661463994 | 1 | No Change | Not Significant | FALSE | NA | NA | No Change | Not Detected | FALSE | Background = FALSE |
| Gm35554 | TRUE | 2.03695203 | 1 | No Change | Not Significant | FALSE | NA | NA | No Change | Not Detected | FALSE | Background = FALSE |
| Gm35715 | TRUE | -0.080035356 | 0.935174293 | No Change | Not Significant | TRUE | -0.481374301 | 0.584243096 | No Change | Not Significant | TRUE | Cond1 Not Significant + Cond2 Not Significant |
| Gm35725 | TRUE | 0.513577127 | 1 | No Change | Not Significant | FALSE | NA | NA | No Change | Not Detected | FALSE | Background = FALSE |
| Gm35853 | TRUE | -0.098877487 | 1 | No Change | Not Significant | FALSE | NA | NA | No Change | Not Detected | FALSE | Background = FALSE |
| Gm35909 | TRUE | 0.514569867 | 1 | No Change | Not Significant | FALSE | NA | NA | No Change | Not Detected | FALSE | Background = FALSE |
| Gm35974 | TRUE | -2.248304343 | 1 | No Change | Not Significant | FALSE | NA | NA | No Change | Not Detected | FALSE | Background = FALSE |
| Gm3604 | TRUE | -0.628958075 | 0.15977566 | No Change | Not Significant | TRUE | -0.202860937 | 0.536272425 | No Change | Not Significant | TRUE | Cond1 Not Significant + Cond2 Not Significant |
| Gm36266 | TRUE | -0.2611736 | 0.502676953 | No Change | Not Significant | TRUE | -0.142671668 | 0.639482053 | No Change | Not Significant | TRUE | Cond1 Not Significant + Cond2 Not Significant |
| Gm36278 | TRUE | 0.396939462 | 1 | No Change | Not Significant | FALSE | NA | NA | No Change | Not Detected | FALSE | Background = FALSE |
| Gm36283 | TRUE | -0.232289089 | 1 | No Change | Not Significant | FALSE | NA | NA | No Change | Not Detected | FALSE | Background = FALSE |
| Gm3636 | TRUE | -0.328495342 | 1 | No Change | Not Significant | TRUE | -0.100380017 | 0.930889346 | No Change | Not Significant | TRUE | Cond1 Not Significant + Cond2 Not Significant |
| Gm36582 | TRUE | 0.990838683 | 0.315055404 | No Change | Not Significant | TRUE | 0.281969448 | 0.725473655 | No Change | Not Significant | TRUE | Cond1 Not Significant + Cond2 Not Significant |
| Gm36595 | TRUE | 1.132689476 | 1 | No Change | Not Significant | FALSE | NA | NA | No Change | Not Detected | FALSE | Background = FALSE |
| Gm36608 | TRUE | 0.024717413 | 1 | No Change | Not Significant | FALSE | NA | NA | No Change | Not Detected | FALSE | Background = FALSE |
| Gm36756 | TRUE | 0.176289553 | 0.825096143 | No Change | Not Significant | TRUE | -0.599351234 | 0.574895496 | No Change | Not Significant | TRUE | Cond1 Not Significant + Cond2 Not Significant |
| Gm36864 | TRUE | 0.484884374 | 0.544366934 | No Change | Not Significant | TRUE | 0.490708787 | 0.538446559 | No Change | Not Significant | TRUE | Cond1 Not Significant + Cond2 Not Significant |
| Gm3693 | TRUE | 3.455364677 | 1 | No Change | Not Significant | FALSE | NA | NA | No Change | Not Detected | FALSE | Background = FALSE |
| Gm3716 | TRUE | -0.557510072 | 0.343200478 | No Change | Not Significant | TRUE | 0.212344524 | 0.771529581 | No Change | Not Significant | TRUE | Cond1 Not Significant + Cond2 Not Significant |
| Gm37240 | TRUE | 0.132223698 | 0.877032235 | No Change | Not Significant | TRUE | -0.103952404 | 0.892047592 | No Change | Not Significant | TRUE | Cond1 Not Significant + Cond2 Not Significant |
| Gm3764 | TRUE | -0.00856217 | 1 | No Change | Not Significant | FALSE | NA | NA | No Change | Not Detected | FALSE | Background = FALSE |
| Gm3776 | TRUE | 0.407966089 | 0.618256212 | No Change | Not Significant | TRUE | 0.214704672 | 0.765709584 | No Change | Not Significant | TRUE | Cond1 Not Significant + Cond2 Not Significant |
| Gm38404 | TRUE | 1.285355556 | 1 | No Change | Not Significant | FALSE | NA | NA | No Change | Not Detected | FALSE | Background = FALSE |
| Gm38409 | TRUE | 1.186512027 | 1 | No Change | Not Significant | TRUE | -0.026874237 | 0.979588542 | No Change | Not Significant | TRUE | Cond1 Not Significant + Cond2 Not Significant |
| Gm38410 | TRUE | 2.81927659 | 1 | No Change | Not Significant | FALSE | NA | NA | No Change | Not Detected | FALSE | Background = FALSE |
| Gm38412 | TRUE | -0.200233358 | 1 | No Change | Not Significant | FALSE | NA | NA | No Change | Not Detected | FALSE | Background = FALSE |
| Gm38413 | TRUE | 1.139010406 | 1 | No Change | Not Significant | FALSE | NA | NA | No Change | Not Detected | FALSE | Background = FALSE |
| Gm38416 | TRUE | 0.528455548 | 0.189251307 | No Change | Not Significant | TRUE | 0.28481671 | 0.410980449 | No Change | Not Significant | TRUE | Cond1 Not Significant + Cond2 Not Significant |
| Gm38509 | TRUE | -0.796645841 | 1 | No Change | Not Significant | FALSE | NA | NA | No Change | Not Detected | FALSE | Background = FALSE |
| Gm3854 | TRUE | -0.252267148 | 1 | No Change | Not Significant | FALSE | NA | NA | No Change | Not Detected | FALSE | Background = FALSE |
| Gm38574 | TRUE | -0.702678958 | 1 | No Change | Not Significant | FALSE | NA | NA | No Change | Not Detected | FALSE | Background = FALSE |
| Gm38576 | TRUE | -1.736379047 | 1 | No Change | Not Significant | FALSE | NA | NA | No Change | Not Detected | FALSE | Background = FALSE |
| Gm39043 | TRUE | 0.357821229 | 1 | No Change | Not Significant | FALSE | NA | NA | No Change | Not Detected | FALSE | Background = FALSE |
| Gm39079 | TRUE | 2.974169408 | 1 | No Change | Not Significant | FALSE | NA | NA | No Change | Not Detected | FALSE | Background = FALSE |
| Gm39307 | TRUE | 1.788397209 | 1 | No Change | Not Significant | FALSE | NA | NA | No Change | Not Detected | FALSE | Background = FALSE |
| Gm39822 | TRUE | 0.849780402 | 1 | No Change | Not Significant | FALSE | NA | NA | No Change | Not Detected | FALSE | Background = FALSE |
| Gm4013 | TRUE | -1.260162072 | 1 | No Change | Not Significant | FALSE | NA | NA | No Change | Not Detected | FALSE | Background = FALSE |
| Gm40155 | TRUE | 0.159777445 | 1 | No Change | Not Significant | FALSE | NA | NA | No Change | Not Detected | FALSE | Background = FALSE |
| Gm40614 | TRUE | -0.385425974 | 0.488906968 | No Change | Not Significant | TRUE | -0.026240674 | 0.967208345 | No Change | Not Significant | TRUE | Cond1 Not Significant + Cond2 Not Significant |
| Gm40617 | TRUE | -0.011331091 | 1 | No Change | Not Significant | FALSE | NA | NA | No Change | Not Detected | FALSE | Background = FALSE |
| Gm40909 | TRUE | 2.209802617 | 1 | No Change | Not Significant | FALSE | NA | NA | No Change | Not Detected | FALSE | Background = FALSE |
| Gm4131 | TRUE | -2.650755502 | 1 | No Change | Not Significant | FALSE | NA | NA | No Change | Not Detected | FALSE | Background = FALSE |
| Gm41584 | TRUE | -0.056681818 | 1 | No Change | Not Significant | TRUE | 0.104370031 | 0.926650089 | No Change | Not Significant | TRUE | Cond1 Not Significant + Cond2 Not Significant |
| Gm41609 | TRUE | -0.169264519 | 1 | No Change | Not Significant | FALSE | NA | NA | No Change | Not Detected | FALSE | Background = FALSE |
| Gm41611 | TRUE | 0.582362571 | 1 | No Change | Not Significant | FALSE | NA | NA | No Change | Not Detected | FALSE | Background = FALSE |
| Gm41790 | TRUE | -0.192081739 | 1 | No Change | Not Significant | FALSE | NA | NA | No Change | Not Detected | FALSE | Background = FALSE |
| Gm4211 | TRUE | 1.65750748 | 1 | No Change | Not Significant | FALSE | NA | NA | No Change | Not Detected | FALSE | Background = FALSE |
| Gm4221 | TRUE | 0.249219268 | 1 | No Change | Not Significant | FALSE | NA | NA | No Change | Not Detected | FALSE | Background = FALSE |
| Gm4265 | TRUE | 0.128326409 | 1 | No Change | Not Significant | FALSE | NA | NA | No Change | Not Detected | FALSE | Background = FALSE |
| Gm42788 | TRUE | -0.26072896 | 1 | No Change | Not Significant | FALSE | NA | NA | No Change | Not Detected | FALSE | Background = FALSE |
| Gm43050 | TRUE | -1.892045847 | 1 | No Change | Not Significant | FALSE | NA | NA | No Change | Not Detected | FALSE | Background = FALSE |
| Gm43247 | TRUE | 3.271859191 | 1 | No Change | Not Significant | FALSE | NA | NA | No Change | Not Detected | FALSE | Background = FALSE |
| Gm43391 | TRUE | 0.905333829 | 1 | No Change | Not Significant | FALSE | NA | NA | No Change | Not Detected | FALSE | Background = FALSE |
| Gm43403 | TRUE | 0.76046668 | 1 | No Change | Not Significant | FALSE | NA | NA | No Change | Not Detected | FALSE | Background = FALSE |
| Gm43868 | TRUE | -0.564437136 | 0.306022641 | No Change | Not Significant | TRUE | -0.725084435 | 0.160974426 | No Change | Not Significant | TRUE | Cond1 Not Significant + Cond2 Not Significant |
| Gm43915 | TRUE | 0.276235261 | 0.810260975 | No Change | Not Significant | TRUE | -0.222581945 | 0.813517602 | No Change | Not Significant | TRUE | Cond1 Not Significant + Cond2 Not Significant |
| Gm44126 | TRUE | 1.138814671 | 1 | No Change | Not Significant | FALSE | NA | NA | No Change | Not Detected | FALSE | Background = FALSE |
| Gm44148 | TRUE | 1.659992933 | 1 | No Change | Not Significant | FALSE | NA | NA | No Change | Not Detected | FALSE | Background = FALSE |
| Gm44851 | TRUE | -0.823506544 | 1 | No Change | Not Significant | FALSE | NA | NA | No Change | Not Detected | FALSE | Background = FALSE |
| Gm4489 | TRUE | -0.372010254 | 1 | No Change | Not Significant | FALSE | NA | NA | No Change | Not Detected | FALSE | Background = FALSE |
| Gm4544 | TRUE | 0.639765289 | 0.186383347 | No Change | Not Significant | TRUE | -0.20525956 | 0.720166783 | No Change | Not Significant | TRUE | Cond1 Not Significant + Cond2 Not Significant |
| Gm4566 | TRUE | 0.937127572 | 1 | No Change | Not Significant | FALSE | NA | NA | No Change | Not Detected | FALSE | Background = FALSE |
| Gm45799 | TRUE | 0.832422557 | 1 | No Change | Not Significant | FALSE | NA | NA | No Change | Not Detected | FALSE | Background = FALSE |
| Gm45861 | TRUE | 0.964101211 | 1 | No Change | Not Significant | FALSE | NA | NA | No Change | Not Detected | FALSE | Background = FALSE |
| Gm45871 | TRUE | -0.232119309 | 0.21833128 | No Change | Not Significant | TRUE | -0.081549327 | 0.65874317 | No Change | Not Significant | TRUE | Cond1 Not Significant + Cond2 Not Significant |
| Gm45902 | TRUE | 0.402198045 | 0.07969641 | No Change | Not Significant | TRUE | -0.156461108 | 0.672888818 | No Change | Not Significant | TRUE | Cond1 Not Significant + Cond2 Not Significant |
| Gm46123 | TRUE | 0.603896801 | 1 | No Change | Not Significant | FALSE | NA | NA | No Change | Not Detected | FALSE | Background = FALSE |
| Gm46234 | TRUE | -0.793691152 | 1 | No Change | Not Significant | FALSE | NA | NA | No Change | Not Detected | FALSE | Background = FALSE |
| Gm4632 | TRUE | 0.232779832 | 1 | No Change | Not Significant | FALSE | NA | NA | No Change | Not Detected | FALSE | Background = FALSE |
| Gm46355 | TRUE | 0.964211859 | 1 | No Change | Not Significant | FALSE | NA | NA | No Change | Not Detected | FALSE | Background = FALSE |
| Gm46392 | TRUE | 1.694049482 | 1 | No Change | Not Significant | FALSE | NA | NA | No Change | Not Detected | FALSE | Background = FALSE |
| Gm46430 | TRUE | -0.307653642 | 0.245826823 | No Change | Not Significant | TRUE | -0.006395 | 0.987251745 | No Change | Not Significant | TRUE | Cond1 Not Significant + Cond2 Not Significant |
| Gm4651 | TRUE | -0.285588798 | 0.557777472 | No Change | Not Significant | TRUE | -0.392164612 | 0.364719069 | No Change | Not Significant | TRUE | Cond1 Not Significant + Cond2 Not Significant |
| Gm46637 | TRUE | -0.046418157 | 1 | No Change | Not Significant | FALSE | NA | NA | No Change | Not Detected | FALSE | Background = FALSE |
| Gm4668 | TRUE | 0.230667468 | 1 | No Change | Not Significant | FALSE | NA | NA | No Change | Not Detected | FALSE | Background = FALSE |
| Gm4675 | TRUE | -0.15724788 | 0.816921981 | No Change | Not Significant | TRUE | -0.304114781 | 0.561627805 | No Change | Not Significant | TRUE | Cond1 Not Significant + Cond2 Not Significant |
| Gm47112 | TRUE | 0.532637789 | 0.606204104 | No Change | Not Significant | FALSE | NA | NA | No Change | Not Detected | FALSE | Background = FALSE |
| Gm4724 | TRUE | -2.123962033 | 1 | No Change | Not Significant | FALSE | NA | NA | No Change | Not Detected | FALSE | Background = FALSE |
| Gm47283 | TRUE | -0.072199419 | 0.96979553 | No Change | Not Significant | TRUE | -0.603944789 | 0.592953751 | No Change | Not Significant | TRUE | Cond1 Not Significant + Cond2 Not Significant |
| Gm47708 | TRUE | -1.210424796 | 0.006345534 | DOWN | DOWN | TRUE | -0.231946302 | 0.640499152 | No Change | Not Significant | TRUE | Cond1 DOWN + Cond2 Not Significant |
| Gm47792 | TRUE | 1.943642764 | 1 | No Change | Not Significant | FALSE | NA | NA | No Change | Not Detected | FALSE | Background = FALSE |
| Gm4787 | TRUE | -0.339154334 | 0.591017907 | No Change | Not Significant | TRUE | -0.440805045 | 0.430948755 | No Change | Not Significant | TRUE | Cond1 Not Significant + Cond2 Not Significant |

| geneID | Cond1_DF_Detected | Cond1_DF_ValueCol | Cond1_DF_PadjCol | Cond1_DF_Cutoff | Cond1_DF_Cutoff_Specific.x | Cond2_DF_Detected |
|---|---|---|---|---|---|---|
| Gm47985 | TRUE | -0.529372561 | 1 | No Change | Not Significant | FALSE |
| Gm4841 | TRUE | 1.281040714 | 1 | No Change | Not Significant | FALSE |
| Gm48485 | TRUE | -1.40578781 | 1 | No Change | Not Significant | FALSE |
| Gm48552 | TRUE | -0.68831078 | 1 | No Change | Not Significant | FALSE |
| Gm48623 | TRUE | -0.421612106 | 1 | No Change | Not Significant | FALSE |
| Gm48727 | TRUE | 1.310352523 | 0.047245724 | UP | UP | TRUE |
| Gm4876 | TRUE | 0.068494697 | 0.882510888 | No Change | Not Significant | TRUE |
| Gm4922 | TRUE | -1.89231043 | 1 | No Change | Not Significant | FALSE |
| Gm4924 | TRUE | -0.282994823 | 0.333254034 | No Change | Not Significant | TRUE |
| Gm4925 | TRUE | -0.643033905 | 0.159827185 | No Change | Not Significant | TRUE |
| Gm49336 | TRUE | 0.003976889 | 0.982229903 | No Change | Not Significant | TRUE |
| Gm4951 | TRUE | 1.25478849 | 1 | No Change | Not Significant | FALSE |
| Gm4952 | TRUE | -0.719381502 | 0.26676599 | No Change | Not Significant | TRUE |
| Gm4961 | TRUE | 0.446026964 | 0.618855346 | No Change | Not Significant | TRUE |
| Gm50340 | TRUE | -0.248159575 | 0.682320284 | No Change | Not Significant | TRUE |
| Gm5084 | TRUE | 0.00507117 | 1 | No Change | Not Significant | FALSE |
| Gm5086 | TRUE | -0.731865427 | 1 | No Change | Not Significant | FALSE |
| Gm5092 | TRUE | 1.083734455 | 1 | No Change | Not Significant | FALSE |
| Gm5095 | TRUE | 0.176017408 | 1 | No Change | Not Significant | FALSE |
| Gm5122 | TRUE | -1.152563293 | 1 | No Change | Not Significant | FALSE |
| Gm5124 | TRUE | 1.051752145 | 1 | No Change | Not Significant | FALSE |
| Gm5127 | TRUE | 0.17601115 | 1 | No Change | Not Significant | FALSE |
| Gm5129 | TRUE | 0.958048792 | 1 | No Change | Not Significant | FALSE |
| Gm5141 | TRUE | -0.3001414 | 0.541897747 | No Change | Not Significant | TRUE |
| Gm5142 | TRUE | 0.062008962 | 1 | No Change | Not Significant | FALSE |
| Gm5148 | TRUE | -0.124057468 | 0.84294636 | No Change | Not Significant | TRUE |
| Gm5165 | TRUE | -0.366557633 | 0.110042513 | No Change | Not Significant | TRUE |
| Gm5176 | TRUE | 1.122930086 | 1 | No Change | Not Significant | FALSE |
| Gm5177 | TRUE | 0.565429753 | 1 | No Change | Not Significant | FALSE |
| Gm527 | TRUE | -0.192365418 | 0.71205818 | No Change | Not Significant | TRUE |
| Gm52969 | TRUE | 0.322707696 | 1 | No Change | Not Significant | FALSE |
| Gm53058 | TRUE | -2.042177828 | 1 | No Change | Not Significant | FALSE |
| Gm5316 | TRUE | 0.604203923 | 1 | No Change | Not Significant | FALSE |
| Gm536 | TRUE | -0.953342058 | 1 | No Change | Not Significant | FALSE |
| Gm5414 | TRUE | -0.586630135 | 1 | No Change | Not Significant | FALSE |
| Gm5420 | TRUE | 0.990660471 | 0.254685181 | No Change | Not Significant | TRUE |
| Gm5431 | TRUE | -0.587054697 | 0.245852636 | No Change | Not Significant | TRUE |
| Gm5434 | TRUE | -0.69404751 | 1 | No Change | Not Significant | FALSE |
| Gm5441 | TRUE | 0.469637098 | 1 | No Change | Not Significant | FALSE |
| Gm5445 | TRUE | 1.359978428 | 1 | No Change | Not Significant | FALSE |
| Gm5460 | TRUE | -0.836693937 | 1 | No Change | Not Significant | FALSE |
| Gm5464 | TRUE | -0.410709652 | 1 | No Change | Not Significant | FALSE |
| Gm54719 | TRUE | -0.778992644 | 1 | No Change | Not Significant | FALSE |
| Gm5475 | TRUE | -0.553704743 | 0.607019295 | No Change | Not Significant | TRUE |
| Gm550 | TRUE | 1.285725628 | 1 | No Change | Not Significant | FALSE |
| Gm5523 | TRUE | 0.688863983 | 1 | No Change | Not Significant | FALSE |
| Gm5528 | TRUE | 1.341060391 | 1 | No Change | Not Significant | FALSE |
| Gm5535 | TRUE | 0.604257666 | 0.609455738 | No Change | Not Significant | FALSE |
| Gm5547 | TRUE | 1.099586072 | 1 | No Change | Not Significant | FALSE |
| Gm5553 | TRUE | -0.039732755 | 1 | No Change | Not Significant | FALSE |
| Gm55594 | TRUE | -0.443715301 | 0.785292483 | No Change | Not Significant | TRUE |
| Gm5577 | TRUE | 0.326917305 | 1 | No Change | Not Significant | FALSE |
| Gm5591 | TRUE | -1.705001228 | 1 | No Change | Not Significant | FALSE |
| Gm5592 | TRUE | 1.138682638 | 1 | No Change | Not Significant | FALSE |
| Gm5617 | TRUE | -0.543058475 | 0.004105122 | DOWN | DOWN | TRUE |
| Gm5637 | TRUE | 1.109095769 | 0.081330396 | No Change | Not Significant | FALSE |
| Gm572 | TRUE | -0.565232441 | 1 | No Change | Not Significant | FALSE |
| Gm5737 | TRUE | -1.268120971 | 1 | No Change | Not Significant | FALSE |
| Gm5766 | TRUE | -1.699022636 | 1 | No Change | Not Significant | FALSE |
| Gm5820 | TRUE | 2.326315868 | 1 | No Change | Not Significant | FALSE |
| Gm5833 | TRUE | 0.176009793 | 1 | No Change | Not Significant | FALSE |
| Gm5878 | TRUE | 0.878218646 | 1 | No Change | Not Significant | FALSE |
| Gm5893 | TRUE | 3.394628528 | 1 | No Change | Not Significant | FALSE |
| Gm5907 | TRUE | -0.736955739 | 1 | No Change | Not Significant | FALSE |
| Gm5914 | TRUE | -1.1112734 | 1 | No Change | Not Significant | FALSE |
| Gm6093 | TRUE | -0.355759482 | 1 | No Change | Not Significant | FALSE |
| Gm6109 | TRUE | 1.004038693 | 1 | No Change | Not Significant | FALSE |
| Gm6139 | TRUE | 3.484855576 | 1 | No Change | Not Significant | FALSE |
| Gm6145 | TRUE | 0.081107601 | 0.931471303 | No Change | Not Significant | TRUE |
| Gm6150 | TRUE | 1.070091284 | 1 | No Change | Not Significant | FALSE |
| Gm6195 | TRUE | 1.370201114 | 1 | No Change | Not Significant | FALSE |
| Gm6249 | TRUE | -0.694728932 | 1 | No Change | Not Significant | FALSE |
| Gm6329 | TRUE | 0.082172411 | 0.927743965 | No Change | Not Significant | TRUE |
| Gm6377 | TRUE | 1.17149212 | 1 | No Change | Not Significant | FALSE |
| Gm6559 | TRUE | 1.86910452 | 1 | No Change | Not Significant | FALSE |
| Gm6566 | TRUE | -0.434599514 | 0.579930477 | No Change | Not Significant | TRUE |
| Gm6567 | TRUE | -0.114761332 | 1 | No Change | Not Significant | FALSE |
| Gm6592 | TRUE | 2.272307913 | 1 | No Change | Not Significant | FALSE |
| Gm6639 | TRUE | 1.698703331 | 1 | No Change | Not Significant | FALSE |
| Gm6653 | TRUE | 0.887860094 | 1 | No Change | Not Significant | FALSE |
| Gm6712 | TRUE | -0.43160334 | 0.080385081 | No Change | Not Significant | TRUE |
| Gm6787 | TRUE | 0.528975272 | 1 | No Change | Not Significant | FALSE |
| Gm6878 | TRUE | 0.821627741 | 1 | No Change | Not Significant | FALSE |
| Gm6938 | TRUE | 0.224912598 | 1 | No Change | Not Significant | FALSE |
| Gm6961 | TRUE | 0.522911611 | 1 | No Change | Not Significant | FALSE |
| Gm6988 | TRUE | 0.975708232 | 1 | No Change | Not Significant | FALSE |
| Gm7008 | TRUE | 0.763345877 | 0.119049005 | No Change | Not Significant | TRUE |
| Gm7030 | TRUE | -1.300991763 | 1 | No Change | Not Significant | FALSE |
| Gm7072 | TRUE | -0.309444036 | 0.05718458 | No Change | Not Significant | TRUE |
| Gm7102 | TRUE | 0.341900338 | 1 | No Change | Not Significant | FALSE |
| Gm7135 | TRUE | -0.135661762 | 1 | No Change | Not Significant | FALSE |
| Gm7160 | TRUE | 0.003890691 | 0.992763569 | No Change | Not Significant | TRUE |
| Gm7457 | TRUE | 2.570592013 | 1 | No Change | Not Significant | FALSE |
| Gm7467 | TRUE | -0.027045576 | 0.963320302 | No Change | Not Significant | TRUE |
| Gm7579 | TRUE | 1.216541821 | 1 | No Change | Not Significant | FALSE |
| Gm7628 | TRUE | 0.316423756 | 1 | No Change | Not Significant | FALSE |
| Gm7694 | TRUE | -0.471555955 | 0.007756249 | No Change | Significant Negative | TRUE |
| Gm7854 | TRUE | -0.877493508 | 0.110042513 | No Change | Not Significant | TRUE |
| Gm7932 | TRUE | 0.043024715 | 0.941973978 | No Change | Not Significant | TRUE |
| Gm7972 | TRUE | 0.406331812 | 0.674823398 | No Change | Not Significant | FALSE |
| Gm8251 | TRUE | 0.641796044 | 0.438797199 | No Change | Not Significant | TRUE |
| Gm826 | TRUE | -0.293874231 | 1 | No Change | Not Significant | FALSE |
| Gm8369 | TRUE | 0.405436363 | 1 | No Change | Not Significant | FALSE |
| Gm8579 | TRUE | 0.903075423 | 1 | No Change | Not Significant | FALSE |
| Gm867 | TRUE | -0.483007423 | 1 | No Change | Not Significant | FALSE |
| Gm8817 | TRUE | 1.084478118 | 1 | No Change | Not Significant | FALSE |
| Gm884 | TRUE | 3.417462665 | 1 | No Change | Not Significant | FALSE |
| Gm8909 | TRUE | -0.292162539 | 0.656832884 | No Change | Not Significant | TRUE |
| Gm9182 | TRUE | 1.208387936 | 1 | No Change | Not Significant | FALSE |
| Gm9530 | TRUE | 1.027491299 | 0.284401928 | No Change | Not Significant | TRUE |
| Gm960 | TRUE | 0.200475922 | 0.898810919 | No Change | Not Significant | FALSE |
| Gm973 | TRUE | -1.173282438 | 1 | No Change | Not Significant | FALSE |
| Gm9758 | TRUE | 1.094934712 | 1 | No Change | Not Significant | FALSE |
| Gm9776 | TRUE | -0.364629863 | 0.232532135 | No Change | Not Significant | TRUE |
| Gm9828 | TRUE | 0.114351377 | 0.87943703 | No Change | Not Significant | TRUE |
| Gm9831 | TRUE | -0.383445196 | 0.772538316 | No Change | Not Significant | TRUE |
| Gm9866 | TRUE | 0.047997993 | 0.980506535 | No Change | Not Significant | FALSE |
| Gm9899 | TRUE | 0.049421649 | 1 | No Change | Not Significant | FALSE |
| Gm9917 | TRUE | -0.138419664 | 0.624001556 | No Change | Not Significant | TRUE |
| Gm9920 | TRUE | 0.234288352 | 0.459127576 | No Change | Not Significant | TRUE |
| Gm9926 | TRUE | -0.278392762 | 0.066267555 | No Change | Not Significant | TRUE |
| Gm9936 | TRUE | 2.32214194 | 1 | No Change | Not Significant | FALSE |
| Gm9949 | TRUE | -0.097691717 | 1 | No Change | Not Significant | TRUE |
| Gm9961 | TRUE | -0.134069603 | 1 | No Change | Not Significant | FALSE |
| Gm9968 | TRUE | 0.260895364 | 0.776893175 | No Change | Not Significant | TRUE |
| Gmcl1 | TRUE | -0.05646964 | 0.656764359 | No Change | Not Significant | TRUE |
| Gmds | TRUE | 0.523495274 | 2.55E-07 | UP | UP | TRUE |
| Gmeb1 | TRUE | -0.094200846 | 0.43912271 | No Change | Not Significant | TRUE |
| Gmeb2 | TRUE | -0.01573431 | 0.908812719 | No Change | Not Significant | TRUE |
| Gmfb | TRUE | -0.061433919 | 0.717835971 | No Change | Not Significant | TRUE |
| Gmfg | TRUE | 0.440410638 | 0.000703145 | No Change | Significant Positive | TRUE |
| Gmfg-ps | TRUE | 0.324116066 | 1 | No Change | Not Significant | FALSE |
| Gmip | TRUE | -0.389721487 | 0.003086183 | No Change | Significant Negative | TRUE |
| Gml2 | TRUE | 1.635441028 | 1 | No Change | Not Significant | FALSE |
| Gmnc | TRUE | 2.15297232 | 1 | No Change | Not Significant | FALSE |
| Gmnn | TRUE | 0.05596374 | 0.78770474 | No Change | Not Significant | TRUE |
| Gmppa | TRUE | 0.467421653 | 4.80E-09 | No Change | Significant Positive | TRUE |
| Gmppb | TRUE | 1.041276537 | 1.01E-26 | UP | UP | TRUE |
| Gmpr | TRUE | -0.707126804 | 0.001083461 | DOWN | DOWN | TRUE |
| Gmpr2 | TRUE | -0.179953792 | 0.553055905 | No Change | Not Significant | TRUE |
| Gmps | TRUE | 0.130975801 | 0.448874722 | No Change | Not Significant | TRUE |
| Gna11 | TRUE | -0.046009622 | 0.852575063 | No Change | Not Significant | TRUE |
| Gna12 | TRUE | 0.596628062 | 7.57E-12 | UP | UP | TRUE |
| Gna13 | TRUE | -0.132206396 | 0.090207431 | No Change | Not Significant | TRUE |
| Gna14 | TRUE | 0.188691319 | 0.54337393 | No Change | Not Significant | TRUE |
| Gna15 | TRUE | 0.802036857 | 1 | No Change | Not Significant | FALSE |
| Gnai1 | TRUE | -0.589613932 | 0.004518849 | DOWN | DOWN | TRUE |
| Gnai2 | TRUE | 0.078707682 | 0.445131871 | No Change | Not Significant | TRUE |
| Gnai3 | TRUE | 0.237352357 | 0.004448039 | No Change | Significant Positive | TRUE |
| Gnal | TRUE | -0.397322709 | 0.36280925 | No Change | Not Significant | TRUE |
| Gnao1 | TRUE | -0.639866091 | 0.194206788 | No Change | Not Significant | TRUE |
| Gnaq | TRUE | 0.057634367 | 0.604944412 | No Change | Not Significant | TRUE |
| Gnas | TRUE | 0.478133337 | 0.121338228 | No Change | Not Significant | TRUE |
| Gnasas1 | TRUE | -0.62440712 | 1 | No Change | Not Significant | FALSE |
| Gnat1 | TRUE | 1.008387162 | 1 | No Change | Not Significant | FALSE |
| Gnat2 | TRUE | 1.038008072 | 0.000602545 | UP | UP | TRUE |
| Gnat3 | TRUE | -0.840455148 | 0.002606475 | DOWN | DOWN | TRUE |
| Gnaz | TRUE | 0.143576367 | 0.838607402 | No Change | Not Significant | TRUE |
| Gnb1 | TRUE | 0.105957494 | 0.211992437 | No Change | Not Significant | TRUE |
| Gnb1l | TRUE | 0.338238164 | 0.157212375 | No Change | Not Significant | TRUE |
| Gnb2 | TRUE | 0.158826515 | 0.00832093 | No Change | Significant Positive | TRUE |
| Gnb3 | TRUE | 1.37850255 | 1 | No Change | Not Significant | FALSE |
| Gnb4 | TRUE | 0.164194263 | 0.80301103 | No Change | Not Significant | TRUE |
| Gnb5 | TRUE | -0.498836276 | 0.001087135 | No Change | Significant Negative | TRUE |
| Gne | TRUE | -0.143961874 | 0.515123206 | No Change | Not Significant | TRUE |
| Gng10 | TRUE | 0.116613596 | 0.874258149 | No Change | Not Significant | TRUE |
| Gng11 | TRUE | 0.052986986 | 0.75067522 | No Change | Not Significant | TRUE |
| Gng12 | TRUE | -0.384948004 | 2.83E-05 | No Change | Significant Negative | TRUE |
| Gng13 | TRUE | 0.893145496 | 1 | No Change | Not Significant | TRUE |
| Gng14 | TRUE | 0.130064266 | 1 | No Change | Not Significant | FALSE |
| Gng2 | TRUE | -0.130264722 | 0.846503532 | No Change | Not Significant | TRUE |
| Gng3 | TRUE | 0.435993152 | 1 | No Change | Not Significant | FALSE |
| Gng4 | TRUE | -0.418210736 | 0.552046615 | No Change | Not Significant | TRUE |
| Gng5 | TRUE | -0.504510291 | 0.000143449 | DOWN | DOWN | TRUE |
| Gng7 | TRUE | -0.64228231 | 0.375503154 | No Change | Not Significant | TRUE |
| Gng8 | TRUE | 3.673976391 | 1 | No Change | Not Significant | FALSE |
| Gngt1 | TRUE | 1.433507511 | 1 | No Change | Not Significant | FALSE |
| Gngt2 | TRUE | -0.806668866 | 0.015169415 | DOWN | DOWN | TRUE |
| Gnl1 | TRUE | -0.309436149 | 0.004469372 | No Change | Significant Negative | TRUE |
| Gnl2 | TRUE | 0.075791653 | 0.622662097 | No Change | Not Significant | TRUE |
| Gnl3 | TRUE | 0.047521426 | 0.858172551 | No Change | Not Significant | TRUE |
| Gnl3l | TRUE | 0.085797205 | 0.546769252 | No Change | Not Significant | TRUE |
| Gnmt | TRUE | 1.483097264 | 0.077202358 | No Change | Not Significant | FALSE |
| Gnpat | TRUE | 0.053525193 | 0.695624496 | No Change | Not Significant | TRUE |
| Gnpda1 | TRUE | -0.571897071 | 0.001381295 | DOWN | DOWN | TRUE |
| Gnpda2 | TRUE | -0.446021702 | 0.01154679 | No Change | Significant Negative | TRUE |
| Gnpnat1 | TRUE | 0.173812327 | 0.580790426 | No Change | Not Significant | TRUE |
| Gnptab | TRUE | -0.157036493 | 0.430343308 | No Change | Not Significant | TRUE |
| Gnptg | TRUE | -0.546261873 | 9.45E-07 | DOWN | DOWN | TRUE |
| Gnrh1 | TRUE | 0.925424042 | 0.146938331 | No Change | Not Significant | TRUE |
| Gns | TRUE | -0.165467197 | 0.00610432 | No Change | Significant Negative | TRUE |
| Golga1 | TRUE | 0.12675344 | 0.219013029 | No Change | Not Significant | TRUE |

| geneID | Cond1_DF_Detected | Cond1_DF_ValueCol | Cond1_DF_PadjCol | Cond1_DF_Cutoff | Cond1_DF_Cutoff_Specific.x | Cond2_DF_Detected | Cond2_DF_ValueCol | Cond2_DF_PadjCol | Cond2_DF_Cutoff | Cond2_DF_Cutoff_Specific | BG_Method | RG1_All |
|---|---|---|---|---|---|---|---|---|---|---|---|---|
| Golga2 | TRUE | 0.195443386 | 0.000238903 | No Change | Significant Positive | TRUE | 0.020237861 | 0.782426499 | No Change | Not Significant | TRUE | Cond1 Significant Positive + Cond2 Not Significant |
| Golga3 | TRUE | -0.15429926 | 0.283960728 | No Change | Not Significant | TRUE | -0.057003105 | 0.472500517 | No Change | Not Significant | TRUE | Cond1 Not Significant + Cond2 Not Significant |
| Golga4 | TRUE | 0.038165661 | 0.738779548 | No Change | Not Significant | TRUE | 0.010354162 | 0.876754335 | No Change | Not Significant | TRUE | Cond1 Not Significant + Cond2 Not Significant |
| Golga5 | TRUE | 0.166800903 | 0.108084245 | No Change | Not Significant | TRUE | 0.075267229 | 0.436583513 | No Change | Not Significant | TRUE | Cond1 Not Significant + Cond2 Not Significant |
| Golga7 | TRUE | -0.113116825 | 0.398802563 | No Change | Not Significant | TRUE | -0.097340489 | 0.29613801 | No Change | Not Significant | TRUE | Cond1 Not Significant + Cond2 Not Significant |
| Golga7b | TRUE | -0.033320622 | 1 | No Change | Not Significant | TRUE | -0.02063322 | 0.986105006 | No Change | Not Significant | TRUE | Cond1 Not Significant + Cond2 Not Significant |
| Golgb1 | TRUE | 0.247843936 | 0.000465269 | No Change | Significant Positive | TRUE | 0.120449049 | 0.013511107 | No Change | Significant Positive | TRUE | Cond1 Significant Positive + Cond2 Significant Positive |
| Golim4 | TRUE | 0.137688874 | 0.080642929 | No Change | Not Significant | TRUE | 0.048261796 | 0.593363209 | No Change | Not Significant | TRUE | Cond1 Not Significant + Cond2 Not Significant |
| Golm1 | TRUE | 0.330271326 | 0.009312644 | No Change | Significant Positive | TRUE | 0.252714814 | 0.000228758 | No Change | Significant Positive | TRUE | Cond1 Significant Positive + Cond2 Significant Positive |
| Golm2 | TRUE | -0.093363359 | 0.802698942 | No Change | Not Significant | TRUE | -0.05183845 | 0.745357832 | No Change | Not Significant | TRUE | Cond1 Not Significant + Cond2 Not Significant |
| Golph3 | TRUE | 0.012702154 | 0.932373279 | No Change | Not Significant | TRUE | 0.073693809 | 0.0501028 | No Change | Not Significant | TRUE | Cond1 Not Significant + Cond2 Not Significant |
| Golph3l | TRUE | -0.03483414 | 0.909608677 | No Change | Not Significant | TRUE | 0.053708236 | 0.487131402 | No Change | Not Significant | TRUE | Cond1 Not Significant + Cond2 Not Significant |
| [illegible] | [illegible] | [illegible] | [illegible] | [illegible] | [illegible] | [illegible] | [illegible] | [illegible] | [illegible] | [illegible] | [illegible] | [illegible] |

| geneID | RG2_Significant | RG3_SignificantChange | Cond1_DF_ensembl_gene_id | Cond1_DF_entrezgene_id | Cond1_DF_gene_biotype | Cond1_DF_baseMean | Cond1_DF_ log2FoldChange | Cond1_DF_lfcSE | Cond1_DF_stat | Cond1_DF_pvalue | Cond1_DF_ padj | Cond1_DF_absLog2FC | Cond1_DF_Cutoff_Specific.y |
|---|---|---|---|---|---|---|---|---|---|---|---|---|---|
| Golga2 | None | None | ENSMUSG00000002546 | 99412 | protein_coding | 1278.53553 | 0.195443386 | 0.046224763 | -4.228110077 | 2.36E-05 | 0.000238903 | 0.195443386 | Significant Positive |
| Golga3 | None | None | ENSMUSG00000029502 | 269682 | protein_coding | 1300.471166 | -0.15429926 | 0.099568277 | 1.549682938 | 0.121217635 | 0.283960728 | 0.15429926 | Not Significant |
| Golga4 | None | None | ENSMUSG00000038708 | 54214 | protein_coding | 3773.399457 | 0.038165661 | 0.063391172 | -0.60206587 | 0.547130292 | 0.738779548 | 0.038165661 | Not Significant |
| Golga5 | None | None | ENSMUSG00000021192 | 27277 | protein_coding | 734.3687518 | 0.166800903 | 0.078134721 | -2.134785919 | 0.032778501 | 0.108084245 | 0.166800903 | Not Significant |
| Golga7 | None | None | ENSMUSG00000015341 | 57437 | protein_coding | 418.4278076 | -0.113116825 | 0.087963419 | 1.285953025 | 0.198459456 | 0.398802563 | 0.113116825 | Not Significant |
| Golga7b | None | None | ENSMUSG00000042532 | 71146 | protein_coding | 3.172997258 | -0.033320622 | 0.613868866 | 0.054279707 | 0.956712317 | 1 | 0.033320622 | Not Significant |
| Golgb1 | None | None | ENSMUSG00000034243 | 224139 | protein_coding | 3597.660359 | 0.247843936 | 0.061099857 | -4.056375072 | 4.98E-05 | 0.000465269 | 0.247843936 | Significant Positive |
| Golim4 | None | None | ENSMUSG00000034109 | 73124 | protein_coding | 1055.517088 | 0.137688874 | 0.060355836 | -2.28128517 | 0.022531578 | 0.080642929 | 0.137688874 | Not Significant |
| Golm1 | None | None | ENSMUSG00000021556 | 105348 | protein_coding | 8577.456403 | 0.330271326 | 0.104551668 | -3.158929276 | 0.001583499 | 0.009312644 | 0.330271326 | Significant Positive |
| Golm2 | None | None | ENSMUSG00000060227 | 319996 | protein_coding | 233.903554 | -0.093363359 | 0.198385357 | 0.470616182 | 0.63791485 | 0.802698942 | 0.093363359 | Not Significant |
| Golph3 | None | None | ENSMUSG00000022200 | 66629 | protein_coding | 2056.407873 | 0.012702154 | 0.074065441 | -0.171499065 | 0.863831366 | 0.932373279 | 0.012702154 | Not Significant |
| Golph3l | None | None | ENSMUSG00000046519 | 229593 | protein_coding | 4145.26386 | -0.03483414 | 0.154086962 | 0.226068056 | 0.821148489 | 0.909608677 | 0.03483414 | Not Significant |
| [illegible] | [illegible] | [illegible] | [illegible] | [illegible] | [illegible] | [illegible] | [illegible] | [illegible] | [illegible] | [illegible] | [illegible] | [illegible] | [illegible] |

| geneID | Cond2_DF_ensembl_gene_id | Cond2_DF_entrezgene_id | Cond2_DF_gene_biotype | Cond2_DF_baseMean | Cond2_DF_ log2FoldChange | Cond2_DF_lfcSE | Cond2_DF_stat | Cond2_DF_pvalue | Cond2_DF_ padj | Cond2_DF_absLog2FC | RG1_Specific_Cond2 | RG1_Specific_Cond1 |
|---|---|---|---|---|---|---|---|---|---|---|---|---|
| Golga2 | ENSMUSG00000002546 | 99412 | protein_coding | 2136.176421 | 0.020237861 | 0.038110288 | 0.531034062 | 0.595395176 | 0.782426499 | 0.020237861 | Cond1 No Change + Cond2 Not Significant | Cond2 No Change + Cond1 Significant Positive |
| Golga3 | ENSMUSG00000029502 | 269682 | protein_coding | 2482.602904 | -0.057003105 | 0.047792979 | -1.192708767 | 0.232983451 | 0.472500517 | 0.057003105 | Cond1 No Change + Cond2 Not Significant | Cond2 No Change + Cond1 Not Significant |
| Golga4 | ENSMUSG00000038708 | 54214 | protein_coding | 6255.044664 | 0.010354162 | 0.031463074 | 0.329089323 | 0.742088174 | 0.876754335 | 0.010354162 | Cond1 No Change + Cond2 Not Significant | Cond2 No Change + Cond1 Not Significant |
| Golga5 | ENSMUSG00000021192 | 27277 | protein_coding | 1329.768012 | 0.075267229 | 0.059051477 | 1.274603667 | 0.202449559 | 0.436583513 | 0.075267229 | Cond1 No Change + Cond2 Not Significant | Cond2 No Change + Cond1 Not Significant |
| Golga7 | ENSMUSG00000015341 | 57437 | protein_coding | 785.4601452 | -0.097340489 | 0.059973433 | -1.623060152 | 0.104576547 | 0.29613801 | 0.097340489 | Cond1 No Change + Cond2 Not Significant | Cond2 No Change + Cond1 Not Significant |
| Golga7b | ENSMUSG00000042532 | 71146 | protein_coding | 6.534589551 | -0.02063322 | 0.475334103 | -0.043407826 | 0.965376439 | 0.986105006 | 0.02063322 | Cond1 No Change + Cond2 Not Significant | Cond2 No Change + Cond1 Not Significant |
| Golgb1 | ENSMUSG00000034243 | 224139 | protein_coding | 5209.74546 | 0.120449049 | 0.037800066 | 3.186477239 | 0.001440168 | 0.013511107 | 0.120449049 | Cond1 No Change + Cond2 Significant Positive | Cond2 No Change + Cond1 Significant Positive |
| Golim4 | ENSMUSG00000034109 | 73124 | protein_coding | 2047.713173 | 0.048261796 | 0.051746854 | 0.932651779 | 0.35099979 | 0.593363209 | 0.048261796 | Cond1 No Change + Cond2 Not Significant | Cond2 No Change + Cond1 Not Significant |
| Golm1 | ENSMUSG00000021556 | 105348 | protein_coding | 13970.74043 | 0.252714814 | 0.05770887 | 4.379132935 | 1.19E-05 | 0.000228758 | 0.252714814 | Cond1 No Change + Cond2 Significant Positive | Cond2 No Change + Cond1 Significant Positive |
| Golm2 | ENSMUSG00000060227 | 319996 | protein_coding | 406.0269443 | -0.05183845 | 0.084448802 | -0.613844712 | 0.539317948 | 0.745357832 | 0.05183845 | Cond1 No Change + Cond2 Not Significant | Cond2 No Change + Cond1 Not Significant |
| Golph3 | ENSMUSG00000022200 | 66629 | protein_coding | 3880.636828 | 0.073693809 | 0.02772924 | 2.6576209 | 0.007869436 | 0.0501028 | 0.073693809 | Cond1 No Change + Cond2 Not Significant | Cond2 No Change + Cond1 Not Significant |
| Golph3l | ENSMUSG00000046519 | 229593 | protein_coding | 8485.250843 | 0.053708236 | 0.046314303 | 1.159646853 | 0.246192617 | 0.487131402 | 0.053708236 | Cond1 No Change + Cond2 Not Significant | Cond2 No Change + Cond1 Not Significant |
| [illegible] | [illegible] | [illegible] | [illegible] | [illegible] | [illegible] | [illegible] | [illegible] | [illegible] | [illegible] | [illegible] | [illegible] | [illegible] |

| geneID | Cond1_DF_Detected | Cond1_DF_ValueCol | Cond1_DF_PadjCol | Cond1_DF_Cutoff | Cond1_DF_Cutoff_Specific.x | Cond2_DF_Detected | Cond2_DF_ValueCol |
|---|---|---|---|---|---|---|---|
| Grin2a | TRUE | 0.562239646 | 0.894265208 | No Change | Not Significant | FALSE | NA |
| Grin2b | TRUE | -0.419194825 | 0.770846581 | No Change | Not Significant | TRUE | -0.121668724 |
| Grin2c | TRUE | -0.094153306 | 1 | No Change | Not Significant | FALSE | NA |
| Grin2d | TRUE | -0.009844181 | 0.984770701 | No Change | Not Significant | TRUE | 0.015106749 |
| Grin3a | TRUE | -0.351916155 | 0.487146296 | No Change | Not Significant | TRUE | -1.113880863 |
| Grin3b | TRUE | 1.457565652 | 1 | No Change | Not Significant | FALSE | NA |
| Grina | TRUE | -0.39028532 | 0.015028634 | No Change | Significant Negative | TRUE | -0.218487483 |
| Grip1 | TRUE | -0.281466573 | 0.458287111 | No Change | Not Significant | TRUE | -0.15486918 |
| Grip1os1 | TRUE | -0.373557935 | 1 | No Change | Not Significant | FALSE | NA |
| Grip1os2 | TRUE | -1.202361699 | 1 | No Change | Not Significant | FALSE | NA |
| Grip2 | TRUE | -0.757109098 | 0.590046157 | No Change | Not Significant | TRUE | -0.997364513 |
| Gripap1 | TRUE | 0.019510153 | 0.931154836 | No Change | Not Significant | TRUE | 0.127453031 |
| Grk1 | TRUE | 1.012291144 | 1 | No Change | Not Significant | FALSE | NA |
| Grk2 | TRUE | 0.0475318 | 0.43599843 | No Change | Not Significant | TRUE | 0.056585386 |
| Grk3 | TRUE | -0.649403898 | 0.003576244 | DOWN | DOWN | TRUE | -0.322880946 |
| Grk4 | TRUE | -0.223985171 | 0.857084875 | No Change | Not Significant | TRUE | 0.474271609 |
| Grk5 | TRUE | 0.00711781 | 0.973367327 | No Change | Not Significant | TRUE | 0.132214338 |
| Grk6 | TRUE | -0.016868766 | 0.931236324 | No Change | Not Significant | TRUE | 0.036221758 |
| Grm1 | TRUE | 0.237379294 | 1 | No Change | Not Significant | FALSE | NA |
| Grm2 | TRUE | 0.609496082 | 1 | No Change | Not Significant | FALSE | NA |
| Grm4 | TRUE | -0.139110934 | 0.905233898 | No Change | Not Significant | TRUE | -0.5445822 |
| Grm5 | TRUE | 3.935968825 | 1 | No Change | Not Significant | FALSE | NA |
| Grm6 | TRUE | 0.966675221 | 1 | No Change | Not Significant | FALSE | NA |
| Grm7 | TRUE | 0.617017344 | 1 | No Change | Not Significant | FALSE | NA |
| Grm8 | TRUE | 1.899189798 | 1 | No Change | Not Significant | FALSE | NA |
| Grn | TRUE | -0.448419247 | 9.10E-12 | No Change | Significant Negative | TRUE | -0.169129898 |
| Grp | TRUE | 0.078788717 | 1 | No Change | Not Significant | FALSE | NA |

| geneID | Cond1_DF_Detected | Cond1_DF_ValueCol | Cond1_DF_PadjCol | Cond1_DF_Cutoff | Cond1_DF_Cutoff_Specific.x | Cond2_DF_Detected | Cond2_DF_ValueCol |
|---|---|---|---|---|---|---|---|
| H2ac7 | TRUE | 1.026333837 | 3.36E-05 | UP | UP | TRUE | 0.267688036 |
| H2ac8 | TRUE | 0.713094142 | 0.008065396 | UP | UP | TRUE | 0.258458906 |
| H2aj | TRUE | -0.072734076 | 0.616016681 | No Change | Not Significant | TRUE | 0.070478289 |
| H2al1m | TRUE | -0.110773252 | 1 | No Change | Not Significant | FALSE | NA |
| H2ax | TRUE | 1.007692222 | 9.64E-10 | UP | UP | TRUE | 0.335241227 |
| H2az1 | TRUE | 0.317482039 | 0.119688198 | No Change | Not Significant | TRUE | 0.142540942 |
| H2az2 | TRUE | -0.019302725 | 0.919173117 | No Change | Not Significant | TRUE | -0.053036828 |
| H2bc11 | TRUE | 0.82338366 | 0.003837987 | UP | UP | TRUE | 0.297442829 |
| H2bc12 | TRUE | 0.823893867 | 0.000369764 | UP | UP | TRUE | 0.28785902 |
| H2bc13 | TRUE | 0.430525053 | 0.114836207 | No Change | Not Significant | TRUE | 0.151518504 |
| H2bc14 | TRUE | 0.603353663 | 0.041285217 | UP | UP | TRUE | 0.313524612 |
| H2bc15 | TRUE | 0.349882429 | 0.199366082 | No Change | Not Significant | TRUE | 0.150628105 |
| H2bc21 | TRUE | 0.040267814 | 0.882363568 | No Change | Not Significant | TRUE | 0.229776082 |
| H2bc22 | TRUE | 0.505440086 | 0.134003711 | No Change | Not Significant | TRUE | 0.048008487 |
| H2bc23 | TRUE | -0.384562364 | 1 | No Change | Not Significant | FALSE | NA |
| H2bc24 | TRUE | -4.555701871 | 1 | No Change | Not Significant | FALSE | NA |
| H2bc26-ps | TRUE | 0.97613251 | 0.000398401 | UP | UP | TRUE | 0.158069512 |
| H2bc27 | TRUE | 1.013956342 | 0.003035203 | UP | UP | TRUE | 0.297366078 |
| H2bc3 | TRUE | 0.691172554 | 0.004789552 | UP | UP | TRUE | 0.123144154 |
| H2bc4 | TRUE | -0.014543961 | 0.919158483 | No Change | Not Significant | TRUE | 0.002913564 |
| H2bc6 | TRUE | 0.263444962 | 0.329724247 | No Change | Not Significant | TRUE | 0.152209397 |
| H2bc7 | TRUE | 0.811753814 | 0.000377208 | UP | UP | TRUE | 0.252748983 |
| H2bc8 | TRUE | 0.653836796 | 0.001254886 | UP | UP | TRUE | 0.092638205 |
| H2bc9 | TRUE | 0.711734107 | 0.002708917 | UP | UP | TRUE | 0.166584625 |
| H3c1 | TRUE | 0.921117764 | 1.99E-05 | UP | UP | TRUE | 0.329238618 |
| H3c10 | TRUE | 0.798153293 | 0.001385242 | UP | UP | TRUE | 0.210546284 |
| H3c11 | TRUE | 0.536716404 | 0.026785189 | UP | UP | TRUE | 0.157155746 |
| H3c15 | TRUE | 0.675248334 | 0.224444373 | No Change | Not Significant | TRUE | 0.608771304 |
| H3c2 | TRUE | 0.695142861 | 0.001949033 | UP | UP | TRUE | 0.185831023 |
| H3c3 | TRUE | 0.599528388 | 0.047749183 | UP | UP | TRUE | 0.227234845 |
| H3c4 | TRUE | 0.704687814 | 0.004647131 | UP | UP | TRUE | 0.264286206 |
| H3c6 | TRUE | 0.914352674 | 2.40E-07 | UP | UP | TRUE | 0.384405332 |
| H3c7 | TRUE | 0.419485542 | 0.107588657 | No Change | Not Significant | TRUE | 0.181515235 |
| H3c8 | TRUE | 0.609648058 | 0.022473218 | UP | UP | TRUE | 0.258797557 |
| H3f3a | TRUE | -0.566694806 | 0.000118988 | DOWN | DOWN | TRUE | -0.064380106 |
| H3f3b | TRUE | 0.010653336 | 0.959656087 | No Change | Not Significant | TRUE | 0.083404565 |
| H3f4 | TRUE | 1.935772068 | 1 | No Change | Not Significant | FALSE | NA |
| H4c1 | TRUE | 0.376576473 | 0.055018905 | No Change | Not Significant | TRUE | 0.123599908 |
| H4c11 | TRUE | 0.387032223 | 0.144852664 | No Change | Not Significant | TRUE | 0.322122178 |
| H4c12 | TRUE | 0.495966229 | 0.026283536 | No Change | Significant Positive | TRUE | 0.260621816 |
| H4c14 | TRUE | 0.605855018 | 0.002381578 | UP | UP | TRUE | 0.302609975 |
| H4c16 | TRUE | 0.252478734 | 0.047736459 | No Change | Significant Positive | TRUE | 0.07123465 |
| H4c17 | TRUE | -0.107329467 | 1 | No Change | Not Significant | FALSE | NA |
| H4c18 | TRUE | 0.28536644 | 0.196850041 | No Change | Not Significant | TRUE | 0.125808239 |
| H4c2 | TRUE | 0.788550487 | 0.000434422 | UP | UP | TRUE | 0.354235311 |
| H4c3 | TRUE | 0.18458529 | 0.253324166 | No Change | Not Significant | TRUE | 0.096591692 |
| H4c4 | TRUE | 0.404400776 | 0.010614744 | No Change | Significant Positive | TRUE | 0.222165539 |
| H4c6 | TRUE | 0.573575679 | 0.028137622 | UP | UP | TRUE | 0.223372487 |
| H4c8 | TRUE | 0.3372335 | 0.018508983 | No Change | Significant Positive | TRUE | 0.223017717 |
| H4c9 | TRUE | 0.15740548 | 0.563251236 | No Change | Not Significant | TRUE | 0.127220726 |
| H60b | TRUE | -0.74055595 | 1 | No Change | Not Significant | FALSE | NA |
| H60c | TRUE | 0.159247408 | 0.762395109 | No Change | Not Significant | TRUE | 1.114013727 |
| H6pd | TRUE | 0.29761894 | 0.061453081 | No Change | Not Significant | TRUE | 0.181223968 |
| Haao | TRUE | 0.612378877 | 0.68880035 | No Change | Not Significant | FALSE | NA |
| Habp2 | TRUE | -0.468630298 | 1 | No Change | Not Significant | TRUE | 0.025233479 |
| Habp4 | TRUE | 0.161058902 | 0.211618117 | No Change | Not Significant | TRUE | 0.082946817 |
| Hacd1 | TRUE | -0.280984393 | 0.10607158 | No Change | Not Significant | TRUE | -0.114123457 |
| Hacd2 | TRUE | -0.174645406 | 0.262189266 | No Change | Not Significant | TRUE | 0.084029586 |
| Hacd3 | TRUE | -0.368146147 | 0.015750587 | No Change | Significant Negative | TRUE | -0.17758342 |
| Hacd4 | TRUE | -0.484771825 | 0.05182755 | No Change | Not Significant | TRUE | -0.003880678 |
| Hace1 | TRUE | -0.015998138 | 0.92126689 | No Change | Not Significant | TRUE | 0.128598653 |
| Hacl1 | TRUE | -0.492670701 | 0.363688435 | No Change | Not Significant | TRUE | -0.17892832 |
| Hadh | TRUE | -0.440178053 | 2.11E-05 | No Change | Significant Negative | TRUE | -0.130525989 |
| Hadha | TRUE | -0.183964884 | 0.016920408 | No Change | Significant Negative | TRUE | 0.026362601 |
| Hadhb | TRUE | -0.659285547 | 3.59E-08 | DOWN | DOWN | TRUE | -0.009588851 |
| Hagh | TRUE | 0.370381466 | 5.65E-06 | No Change | Significant Positive | TRUE | 0.241102829 |
| Haghl | TRUE | -0.472633771 | 0.010565985 | No Change | Significant Negative | TRUE | -0.26076974 |
| Hal | TRUE | 0.327868004 | 1 | No Change | Not Significant | FALSE | NA |
| Hamp2 | TRUE | -1.497278567 | 1 | No Change | Not Significant | FALSE | NA |
| Hand1 | TRUE | -3.141029182 | 1 | No Change | Not Significant | FALSE | NA |
| Hand2os1 | TRUE | 1.57308725 | 0.632329199 | No Change | Not Significant | FALSE | NA |
| Hao1 | TRUE | 1.726431943 | 1 | No Change | Not Significant | FALSE | NA |
| Hao2 | TRUE | -0.779232826 | 8.50E-06 | DOWN | DOWN | TRUE | -0.062517881 |
| Hap1 | TRUE | -0.377551306 | 0.21167802 | No Change | Not Significant | TRUE | -0.056031112 |
| Hapln1 | TRUE | 1.428737819 | 0.000182029 | UP | UP | TRUE | 0.39177845 |
| Hapln2 | TRUE | 0.176019157 | 1 | No Change | Not Significant | FALSE | NA |
| Hapln3 | TRUE | -0.728246668 | 1 | No Change | Not Significant | FALSE | NA |
| Hapln4 | TRUE | 0.707464389 | 0.095316478 | No Change | Not Significant | TRUE | 1.092222657 |
| Harbi1 | TRUE | 0.182170754 | 0.270127107 | No Change | Not Significant | TRUE | -0.01132589 |
| Hars | TRUE | 0.174504142 | 0.12530374 | No Change | Not Significant | TRUE | 0.072059353 |
| Hars2 | TRUE | -0.192603783 | 0.092763808 | No Change | Not Significant | TRUE | -0.02531406 |
| Has1 | TRUE | 0.658199533 | 1 | No Change | Not Significant | FALSE | NA |
| Has2 | TRUE | 0.957222445 | 1 | No Change | Not Significant | FALSE | NA |
| Has2os | TRUE | -0.189594184 | 0.81231093 | No Change | Not Significant | TRUE | -0.35983492 |
| Has3 | TRUE | -0.43553238 | 0.463828671 | No Change | Not Significant | TRUE | -0.840144806 |
| Haspin | TRUE | 0.537757385 | 0.009897427 | UP | UP | TRUE | 0.035635709 |
| Hat1 | TRUE | 0.255247622 | 0.227592128 | No Change | Not Significant | TRUE | -0.002776559 |
| Haus1 | TRUE | 0.18108326 | 0.41642117 | No Change | Not Significant | TRUE | -0.000418266 |
| Haus2 | TRUE | -0.375386336 | 0.019710226 | No Change | Significant Negative | TRUE | -0.121605848 |
| Haus3 | TRUE | 0.385115226 | 0.023571307 | No Change | Significant Positive | TRUE | 0.049300457 |
| Haus4 | TRUE | 0.127267392 | 0.595133797 | No Change | Not Significant | TRUE | 0.086972221 |
| Haus5 | TRUE | -0.093270804 | 0.905485545 | No Change | Not Significant | TRUE | 0.004673898 |
| Haus6 | TRUE | 0.448176509 | 0.011401366 | No Change | Significant Positive | TRUE | -0.107762146 |
| Haus7 | TRUE | 0.738310518 | 5.88E-11 | UP | UP | TRUE | 0.194514641 |
| Haus8 | TRUE | -0.005395122 | 0.984509162 | No Change | Not Significant | TRUE | -0.001469142 |
| Havcr1 | TRUE | 2.880499286 | 1 | No Change | Not Significant | FALSE | NA |
| Havcr2 | TRUE | 1.533316849 | 1 | No Change | Not Significant | FALSE | NA |
| Hax1 | TRUE | -0.21860328 | 0.285320309 | No Change | Not Significant | TRUE | 0.221262109 |
| Hba-a1 | TRUE | -1.11603886 | 0.102914115 | No Change | Not Significant | TRUE | 0.420518492 |
| Hba-a2 | TRUE | -0.933618823 | 0.234124348 | No Change | Not Significant | TRUE | 0.324987898 |
| Hbb-bs | TRUE | -0.748975479 | 0.179449017 | No Change | Not Significant | TRUE | 0.443885017 |
| Hbb-bt | TRUE | -0.862253027 | 0.229589716 | No Change | Not Significant | TRUE | 0.523702693 |
| Hbegf | TRUE | -0.590601325 | 0.008491218 | DOWN | DOWN | TRUE | -0.162838396 |
| Hbp1 | TRUE | -0.210794757 | 0.305198626 | No Change | Not Significant | TRUE | -0.08847068 |
| Hbs1l | TRUE | 0.126487096 | 0.1731653 | No Change | Not Significant | TRUE | 0.083763501 |
| Hc | TRUE | -0.536689919 | 0.323295567 | No Change | Not Significant | TRUE | -0.565344067 |
| Hcar1 | TRUE | 1.178505623 | 1 | No Change | Not Significant | TRUE | -0.24744888 |
| Hcar2 | TRUE | -0.248815788 | 0.837421924 | No Change | Not Significant | TRUE | -0.175064492 |
| Hccs | TRUE | 0.159664382 | 0.053024211 | No Change | Not Significant | TRUE | -0.02511108 |
| Hcfc1 | TRUE | 0.297638565 | 0.000861774 | No Change | Significant Positive | TRUE | 0.041148605 |
| Hcfc1r1 | TRUE | -0.224546995 | 0.101276951 | No Change | Not Significant | TRUE | 0.078878535 |
| Hcfc2 | TRUE | 0.035689795 | 0.851829233 | No Change | Not Significant | TRUE | -0.014421328 |
| Hck | TRUE | -1.049626166 | 2.23E-07 | DOWN | DOWN | TRUE | 0.112469414 |
| Hcls1 | TRUE | -0.188991813 | 0.776544903 | No Change | Not Significant | TRUE | 0.164651397 |
| Hcn1 | TRUE | 1.10458982 | 0.655599676 | No Change | Not Significant | FALSE | NA |
| Hcn2 | TRUE | -0.600678683 | 0.003229253 | DOWN | DOWN | TRUE | -0.337243742 |
| Hcn3 | TRUE | 0.255559828 | 0.72280558 | No Change | Not Significant | TRUE | 0.375397997 |
| Hcn4 | TRUE | 0.069050852 | 0.965445424 | No Change | Not Significant | TRUE | -0.398514656 |
| Hcrtr1 | TRUE | 0.443131966 | 1 | No Change | Not Significant | FALSE | NA |
| Hcrtr2 | TRUE | -0.204989019 | 1 | No Change | Not Significant | FALSE | NA |
| Hcst | TRUE | -0.778403859 | 0.175901555 | No Change | Not Significant | TRUE | 0.10726657 |
| Hdac1 | TRUE | -0.061465602 | 0.720841738 | No Change | Not Significant | TRUE | -0.023149894 |
| Hdac10 | TRUE | -0.153659663 | 0.10685821 | No Change | Not Significant | TRUE | -0.013343565 |
| Hdac11 | TRUE | -0.556693217 | 1.30E-05 | DOWN | DOWN | TRUE | -0.203163184 |
| Hdac2 | TRUE | 0.121951241 | 0.302752759 | No Change | Not Significant | TRUE | 0.134145409 |
| Hdac3 | TRUE | 0.351510112 | 5.04E-07 | No Change | Significant Positive | TRUE | 0.149613839 |
| Hdac4 | TRUE | -0.139325935 | 0.456655966 | No Change | Not Significant | TRUE | -0.033140189 |
| Hdac5 | TRUE | -0.416716365 | 0.001685546 | No Change | Significant Negative | TRUE | -0.266227855 |
| Hdac6 | TRUE | 0.35627635 | 0.000916141 | No Change | Significant Positive | TRUE | 0.076207283 |
| Hdac7 | TRUE | 0.44317491 | 0.053833353 | No Change | Not Significant | TRUE | -0.204676748 |
| Hdac8 | TRUE | -0.303783708 | 0.058800778 | No Change | Not Significant | TRUE | -0.009148822 |
| Hdac9 | TRUE | -0.816184413 | 0.045361664 | DOWN | DOWN | TRUE | -0.589934282 |
| Hdc | TRUE | 0.703835459 | 1 | No Change | Not Significant | FALSE | NA |
| Hddc2 | TRUE | 0.261376926 | 0.116182363 | No Change | Not Significant | TRUE | 0.268359806 |
| Hddc3 | TRUE | -0.169440409 | 0.687965835 | No Change | Not Significant | TRUE | 0.104155896 |
| Hdgf | TRUE | -7.70E-05 | 0.999457543 | No Change | Not Significant | TRUE | -0.024788074 |
| Hdgfl2 | TRUE | 0.619734325 | 1.80E-08 | UP | UP | TRUE | 0.24792376 |
| Hdgfl3 | TRUE | -0.917410416 | 1 | No Change | Not Significant | FALSE | NA |
| Hdhd2 | TRUE | -0.242431836 | 0.169807667 | No Change | Not Significant | TRUE | -0.034603153 |
| Hdhd3 | TRUE | 0.23343732 | 0.199738463 | No Change | Not Significant | TRUE | 0.40020501 |
| Hdhd5 | TRUE | -0.075035311 | 0.64171609 | No Change | Not Significant | TRUE | 0.152530578 |
| Hdlbp | TRUE | 1.163137222 | 1.58E-64 | UP | UP | TRUE | 0.649408847 |
| Hdnr | TRUE | 3.194290096 | 1 | No Change | Not Significant | FALSE | NA |
| Hdx | TRUE | 0.696171344 | 1 | No Change | Not Significant | FALSE | NA |
| Heatr1 | TRUE | 0.019830254 | 0.947307136 | No Change | Not Significant | TRUE | 0.049077105 |
| Heatr3 | TRUE | -0.144328026 | 0.560532006 | No Change | Not Significant | TRUE | -0.100034534 |
| Heatr4 | TRUE | 0.615389379 | 1 | No Change | Not Significant | FALSE | NA |
| Heatr5a | TRUE | 0.702670673 | 1.69E-14 | UP | UP | TRUE | 0.322671309 |
| Heatr5b | TRUE | 0.087735563 | 0.451964647 | No Change | Not Significant | TRUE | 0.079548328 |
| Heatr6 | TRUE | 0.115139903 | 0.174007807 | No Change | Not Significant | TRUE | 0.152787161 |
| Heatr9 | TRUE | 0.452901766 | 1 | No Change | Not Significant | FALSE | NA |
| Hebp1 | TRUE | -0.271063955 | 0.797446101 | No Change | Not Significant | TRUE | -0.401987733 |
| Hebp2 | TRUE | 0.664026119 | 3.33E-08 | UP | UP | TRUE | 0.475961772 |
| Heca | TRUE | -0.300806345 | 0.005830986 | No Change | Significant Negative | TRUE | -0.110306094 |
| Hectd1 | TRUE | 0.251130333 | 0.000177661 | No Change | Significant Positive | TRUE | 0.21911421 |
| Hectd2 | TRUE | 0.885775373 | 0.214486414 | No Change | Not Significant | TRUE | 0.678959103 |
| Hectd2os | TRUE | 0.238188536 | 1 | No Change | Not Significant | FALSE | NA |
| Hectd3 | TRUE | -0.281874681 | 0.015417065 | No Change | Significant Negative | TRUE | -0.141640407 |
| Hectd4 | TRUE | 0.019860354 | 0.950621521 | No Change | Not Significant | TRUE | -0.038396959 |
| Hecw1 | TRUE | 0.01744059 | 1 | No Change | Not Significant | FALSE | NA |
| Hecw2 | TRUE | -0.208793391 | 0.906297546 | No Change | Not Significant | FALSE | NA |
| Heg1 | TRUE | 0.329208603 | 0.158736855 | No Change | Not Significant | TRUE | -0.060332753 |
| Helb | TRUE | 0.033353674 | 0.850705598 | No Change | Not Significant | TRUE | 0.14931088 |
| Hells | TRUE | 0.200789495 | 0.517488122 | No Change | Not Significant | TRUE | 0.058360566 |
| Helq | TRUE | -0.096977854 | 0.659125086 | No Change | Not Significant | TRUE | 0.154721698 |
| Helz | TRUE | -0.004095255 | 0.98618968 | No Change | Not Significant | TRUE | -0.129443226 |
| Helz2 | TRUE | 0.147618105 | 0.653309672 | No Change | Not Significant | TRUE | -0.075076356 |
| Hemgn | TRUE | -1.498533719 | 1 | No Change | Not Significant | FALSE | NA |
| Hemk1 | TRUE | -0.288632816 | 0.043322217 | No Change | Significant Negative | TRUE | -0.089229113 |
| Henmt1 | TRUE | 0.327789703 | 1 | No Change | Not Significant | FALSE | NA |
| Hepacam | TRUE | -0.95402362 | 0.342262172 | No Change | Not Significant | TRUE | -0.774775033 |
| Hepacam2 | TRUE | -0.061873139 | 0.9092783 | No Change | Not Significant | TRUE | 0.11085751 |
| Heph | TRUE | 0.74345289 | 6.08E-18 | UP | UP | TRUE | 0.249505241 |
| Hephl1 | TRUE | 0.333149048 | 1 | No Change | Not Significant | FALSE | NA |
| Herc1 | TRUE | -0.020000051 | 0.888933284 | No Change | Not Significant | TRUE | -0.123636363 |
| Herc2 | TRUE | 0.270520376 | 0.000589831 | No Change | Significant Positive | TRUE | 0.184328388 |
| Herc3 | TRUE | -0.096480381 | 0.546493327 | No Change | Not Significant | TRUE | -0.180999036 |
| Herc4 | TRUE | 0.034548751 | 0.822722373 | No Change | Not Significant | TRUE | -0.00246071 |
| Herc6 | TRUE | -0.371965456 | 0.470794255 | No Change | Not Significant | TRUE | -0.321697332 |
| Herpud1 | TRUE | 2.667464595 | 6.94E-251 | UP | UP | TRUE | 2.032888636 |
| Herpud2 | TRUE | 0.023764623 | 0.903431064 | No Change | Not Significant | TRUE | 0.076107447 |
| Hes1 | TRUE | 0.040505206 | 0.755664558 | No Change | Not Significant | TRUE | -0.237689577 |
| Hes2 | TRUE | 0.875382448 | 5.75E-09 | UP | UP | TRUE | 0.498337601 |
| Hes3 | TRUE | 0.6453065 | 1 | No Change | Not Significant | FALSE | NA |
| Hes5 | TRUE | -0.373139596 | 1 | No Change | Not Significant | FALSE | NA |
| Hes6 | TRUE | -0.251026462 | 0.164953548 | No Change | Not Significant | TRUE | -0.102191047 |
| Hes7 | TRUE | 0.318205469 | 1 | No Change | Not Significant | FALSE | NA |
| Hesx1 | TRUE | 1.095944839 | 0.368923116 | No Change | Not Significant | FALSE | NA |
| Hexa | TRUE | -0.597486339 | 5.94E-06 | DOWN | DOWN | TRUE | -0.060517539 |
| Hexb | TRUE | -0.712402786 | 0.0406597 | DOWN | DOWN | TRUE | -0.272951896 |
| Hexdc | TRUE | -0.253560193 | 0.102448537 | No Change | Not Significant | TRUE | -0.213609836 |
| Hexim1 | TRUE | 0.27747629 | 0.323442761 | No Change | Not Significant | TRUE | 0.198639847 |

| geneID | Cond1_DF_Detected | Cond1_DF_ValueCol | Cond1_DF_PadjCol | Cond1_DF_Cutoff | Cond1_DF_Cutoff_Specific.x | Cond2_DF_Detected |
|---|---|---|---|---|---|---|
| Hexim2 | TRUE | 0.362209931 | 0.204491718 | No Change | Not Significant | TRUE |
| Hey1 | TRUE | 2.895731197 | 0.001244042 | UP | UP | FALSE |
| Hey2 | TRUE | 6.282968506 | 6.67E-16 | UP | UP | TRUE |
| Heyl | TRUE | 0.252409205 | 1 | No Change | Not Significant | FALSE |
| Hfe | TRUE | -0.215655529 | 0.141278128 | No Change | Not Significant | TRUE |
| Hfm1 | TRUE | 1.068213355 | 1 | No Change | Not Significant | FALSE |
| Hgd | TRUE | 0.836459555 | 1 | No Change | Not Significant | FALSE |
| Hgf | TRUE | 1.269059236 | 0.139022943 | No Change | Not Significant | TRUE |
| Hgfac | TRUE | -0.150526403 | 0.606204104 | No Change | Not Significant | TRUE |
| Hgh1 | TRUE | 0.234111935 | 0.177826758 | No Change | Not Significant | TRUE |
| Hgs | TRUE | 0.053654723 | 0.703739933 | No Change | Not Significant | TRUE |
| Hgsnat | TRUE | -0.220409555 | 0.147109787 | No Change | Not Significant | TRUE |
| Hhat | TRUE | 0.056340337 | 0.879445484 | No Change | Not Significant | TRUE |
| Hhatl | TRUE | 1.186014542 | 1 | No Change | Not Significant | FALSE |
| Hhex | TRUE | -0.791146785 | 0.190837928 | No Change | Not Significant | TRUE |
| Hhip | TRUE | 2.328780537 | 1 | No Change | Not Significant | FALSE |
| Hhipl1 | TRUE | 0.486330355 | 1 | No Change | Not Significant | FALSE |
| Hhipl2 | TRUE | 2.4540206 | 1 | No Change | Not Significant | FALSE |
| Hibadh | TRUE | -0.471201004 | 8.68E-05 | No Change | Significant Negative | TRUE |
| Hibch | TRUE | -0.276357358 | 0.020119794 | No Change | Significant Negative | TRUE |
| Hic1 | TRUE | 0.20262558 | 0.823809965 | No Change | Not Significant | TRUE |
| Hic2 | TRUE | -0.132929216 | 0.384976843 | No Change | Not Significant | TRUE |
| Hid1 | TRUE | 0.125931446 | 0.722098569 | No Change | Not Significant | TRUE |
| Hif1a | TRUE | -0.194085906 | 0.467050404 | No Change | Not Significant | TRUE |
| Hif1an | TRUE | -0.032299062 | 0.791721985 | No Change | Not Significant | TRUE |
| Hif3a | TRUE | 0.173199069 | 0.555003602 | No Change | Not Significant | TRUE |
| Higd1a | TRUE | 0.140982549 | 0.480125333 | No Change | Not Significant | TRUE |
| Higd2a | TRUE | -0.546504134 | 1.42E-07 | DOWN | DOWN | TRUE |
| Hikeshi | TRUE | -0.111769038 | 0.586347377 | No Change | Not Significant | TRUE |
| Hilpda | TRUE | -0.095511281 | 0.607820554 | No Change | Not Significant | TRUE |
| Hinfp | TRUE | 0.19731082 | 0.108584543 | No Change | Not Significant | TRUE |
| Hint1 | TRUE | -0.091818325 | 0.560042619 | No Change | Not Significant | TRUE |
| Hint2 | TRUE | -0.099208085 | 0.274894315 | No Change | Not Significant | TRUE |
| Hint3 | TRUE | -0.138459229 | 0.155751645 | No Change | Not Significant | TRUE |
| Hip1 | TRUE | 1.022275715 | 0.483440174 | No Change | Not Significant | FALSE |
| Hip1r | TRUE | 0.134256759 | 0.036020338 | No Change | Significant Positive | TRUE |
| Hipk1 | TRUE | -0.116502989 | 0.06376765 | No Change | Not Significant | TRUE |
| Hipk2 | TRUE | -0.151172858 | 0.685675281 | No Change | Not Significant | TRUE |
| Hipk3 | TRUE | -0.018085311 | 0.897960915 | No Change | Not Significant | TRUE |
| Hipk4 | TRUE | 0.587968261 | 0.411419661 | No Change | Not Significant | TRUE |
| Hira | TRUE | 0.250933978 | 0.048232278 | No Change | Significant Positive | TRUE |
| Hirip3 | TRUE | 0.284166499 | 0.232532135 | No Change | Not Significant | TRUE |
| Hivep1 | TRUE | -0.155195792 | 0.313005269 | No Change | Not Significant | TRUE |
| Hivep2 | TRUE | 0.124051037 | 0.564822707 | No Change | Not Significant | TRUE |
| Hivep3 | TRUE | -0.071841607 | 0.816323585 | No Change | Not Significant | TRUE |
| Hjurp | TRUE | 0.216395705 | 0.017269937 | No Change | Significant Positive | TRUE |
| Hk1 | TRUE | -0.196722697 | 0.026363195 | No Change | Significant Negative | TRUE |
| Hk1os | TRUE | 0.361765089 | 0.662372535 | No Change | Not Significant | TRUE |
| Hk2 | TRUE | 0.835561713 | 8.39E-09 | UP | UP | TRUE |
| Hk3 | TRUE | -0.026798679 | 1 | No Change | Not Significant | FALSE |
| Hkdc1 | TRUE | -1.061656752 | 0.003240814 | DOWN | DOWN | TRUE |
| Hlcs | TRUE | -0.16042608 | 0.47491467 | No Change | Not Significant | TRUE |
| Hlf | TRUE | 0.398200479 | 1 | No Change | Not Significant | FALSE |
| Hltf | TRUE | 0.152642701 | 0.29840508 | No Change | Not Significant | TRUE |
| Hlx | TRUE | 0.886774714 | 1 | No Change | Not Significant | FALSE |
| Hmbox1 | TRUE | -0.049437431 | 0.818817302 | No Change | Not Significant | TRUE |
| Hmbs | TRUE | 1.43232264 | 6.67E-61 | UP | UP | TRUE |
| Hmces | TRUE | -0.438433935 | 0.002388097 | No Change | Significant Negative | TRUE |
| Hmcn1 | TRUE | 0.214412657 | 0.426993013 | No Change | Not Significant | TRUE |
| Hmcn2 | TRUE | 0.578022003 | 0.771555104 | No Change | Not Significant | FALSE |
| Hmg20a | TRUE | -0.083043465 | 0.561998621 | No Change | Not Significant | TRUE |
| Hmg20b | TRUE | 0.596362682 | 1.99E-06 | UP | UP | TRUE |
| Hmga1 | TRUE | -0.128801104 | 0.587772599 | No Change | Not Significant | TRUE |
| Hmga1b | TRUE | 0.915938278 | 0.127638694 | No Change | Not Significant | TRUE |
| Hmga2 | TRUE | -0.476212359 | 0.000932912 | No Change | Significant Negative | TRUE |
| Hmgb1 | TRUE | 0.145968861 | 0.53139398 | No Change | Not Significant | TRUE |
| Hmgb2 | TRUE | 0.182776853 | 0.438404222 | No Change | Not Significant | TRUE |
| Hmgb3 | TRUE | -0.048667171 | 0.847201146 | No Change | Not Significant | TRUE |
| Hmgcl | TRUE | -0.127759666 | 0.148611292 | No Change | Not Significant | TRUE |
| Hmgcll1 | TRUE | 0.176855391 | 0.855825916 | No Change | Not Significant | TRUE |
| Hmgcr | TRUE | 0.097416251 | 0.684381431 | No Change | Not Significant | TRUE |
| Hmgcs1 | TRUE | 0.977432719 | 3.14E-09 | UP | UP | TRUE |
| Hmgcs2 | TRUE | -0.41916398 | 0.005929802 | No Change | Significant Negative | TRUE |
| Hmgn1 | TRUE | 0.270745173 | 0.097670196 | No Change | Not Significant | TRUE |
| Hmgn2 | TRUE | 0.116261581 | 0.523793413 | No Change | Not Significant | TRUE |
| Hmgn3 | TRUE | -1.269921347 | 0.079007396 | No Change | Not Significant | TRUE |
| Hmgn5 | TRUE | -0.255563637 | 0.19654843 | No Change | Not Significant | TRUE |
| Hmgxb3 | TRUE | -0.167755075 | 0.027104593 | No Change | Significant Negative | TRUE |
| Hmgxb4 | TRUE | -0.020366307 | 0.911456552 | No Change | Not Significant | TRUE |
| Hmmr | TRUE | 0.681653547 | 0.001199022 | UP | UP | TRUE |
| Hmox1 | TRUE | 0.068022881 | 0.881056435 | No Change | Not Significant | TRUE |
| Hmox2 | TRUE | 0.077232424 | 0.628891312 | No Change | Not Significant | TRUE |
| Hmx2 | TRUE | -0.368930291 | 0.4551483 | No Change | Not Significant | TRUE |
| Hmx3 | TRUE | -0.368933831 | 0.429717907 | No Change | Not Significant | TRUE |
| Hnf1a | TRUE | 0.185506413 | 0.419407924 | No Change | Not Significant | TRUE |
| Hnf1b | TRUE | -0.029199673 | 0.908418149 | No Change | Not Significant | TRUE |
| Hnf4a | TRUE | -0.112770085 | 0.292047215 | No Change | Not Significant | TRUE |
| Hnf4aos | TRUE | -0.110642867 | 0.908120076 | No Change | Not Significant | TRUE |
| Hnf4g | TRUE | -0.224418672 | 0.083675806 | No Change | Not Significant | TRUE |
| Hnmt | TRUE | -0.500072962 | 0.412695527 | No Change | Not Significant | TRUE |
| Hnrnpa0 | TRUE | 0.395797622 | 5.17E-05 | No Change | Significant Positive | TRUE |
| Hnrnpa1 | TRUE | -0.48266035 | 3.55E-05 | No Change | Significant Negative | TRUE |
| Hnrnpa1l2-ps2 | TRUE | -0.984326775 | 1 | No Change | Not Significant | FALSE |
| Hnrnpa2b1 | TRUE | 0.036657159 | 0.801613541 | No Change | Not Significant | TRUE |
| Hnrnpa3 | TRUE | 0.521270281 | 3.23E-05 | UP | UP | TRUE |
| Hnrnpab | TRUE | 0.348964589 | 3.48E-06 | No Change | Significant Positive | TRUE |
| Hnrnpc | TRUE | -0.022546667 | 0.908714429 | No Change | Not Significant | TRUE |
| Hnrnpd | TRUE | 0.200349854 | 0.003951446 | No Change | Significant Positive | TRUE |
| Hnrnpdl | TRUE | 0.125774845 | 0.107787762 | No Change | Not Significant | TRUE |
| Hnrnpf | TRUE | -0.323087454 | 0.064232353 | No Change | Not Significant | TRUE |
| Hnrnph1 | TRUE | 0.215338231 | 0.000161703 | No Change | Significant Positive | TRUE |
| Hnrnph2 | TRUE | -0.077697522 | 0.728048657 | No Change | Not Significant | TRUE |
| Hnrnph3 | TRUE | -0.179254643 | 0.255796319 | No Change | Not Significant | TRUE |
| Hnrnpk | TRUE | 0.057246163 | 0.694394109 | No Change | Not Significant | TRUE |
| Hnrnpl | TRUE | 0.188758882 | 0.003917321 | No Change | Significant Positive | TRUE |
| Hnrnpll | TRUE | 0.101453637 | 0.238987471 | No Change | Not Significant | TRUE |
| Hnrnpm | TRUE | 0.215788923 | 0.012036117 | No Change | Significant Positive | TRUE |
| Hnrnpr | TRUE | 0.05846417 | 0.657836809 | No Change | Not Significant | TRUE |
| Hnrnpu | TRUE | 0.167562522 | 0.008791449 | No Change | Significant Positive | TRUE |
| Hnrnpul1 | TRUE | 0.141207473 | 0.009689537 | No Change | Significant Positive | TRUE |
| Hnrnpul2 | TRUE | 0.166475988 | 0.00731937 | No Change | Significant Positive | TRUE |
| Hoga1 | TRUE | 0.814151831 | 8.92E-05 | UP | UP | TRUE |
| Homer1 | TRUE | 0.019252799 | 0.913504709 | No Change | Not Significant | TRUE |
| Homer2 | TRUE | -0.052475194 | 0.781981914 | No Change | Not Significant | TRUE |
| Homer3 | TRUE | 0.545734606 | 0.19938276 | No Change | Not Significant | TRUE |
| Homez | TRUE | 0.601231002 | 2.23E-06 | UP | UP | TRUE |
| Hook1 | TRUE | 0.118286999 | 0.377733062 | No Change | Not Significant | TRUE |
| Hook2 | TRUE | -0.178026227 | 0.036437516 | No Change | Significant Negative | TRUE |
| Hook3 | TRUE | -0.110108377 | 0.361939775 | No Change | Not Significant | TRUE |
| Hopx | TRUE | 0.092887444 | 0.775716761 | No Change | Not Significant | TRUE |
| Hormad2 | TRUE | 0.107088192 | 1 | No Change | Not Significant | FALSE |
| Hotairm1 | TRUE | 2.764419192 | 8.50E-12 | UP | UP | TRUE |
| Hottip | TRUE | -0.02298329 | 0.957015366 | No Change | Not Significant | TRUE |
| Hoxa1 | TRUE | 5.076982057 | 3.31E-40 | UP | UP | TRUE |
| Hoxa10 | TRUE | -0.038809497 | 0.890125936 | No Change | Not Significant | TRUE |
| Hoxa11 | TRUE | 0.791592521 | 0.027440166 | UP | UP | TRUE |
| Hoxa11os | TRUE | -0.097953495 | 0.735467298 | No Change | Not Significant | TRUE |
| Hoxa13 | TRUE | -0.018610167 | 0.94399288 | No Change | Not Significant | TRUE |
| Hoxa2 | TRUE | 0.279495057 | 0.438111017 | No Change | Not Significant | TRUE |
| Hoxa3 | TRUE | -0.093797113 | 0.747162726 | No Change | Not Significant | TRUE |
| Hoxa4 | TRUE | -0.207424078 | 0.761180591 | No Change | Not Significant | TRUE |
| Hoxa5 | TRUE | 0.048837625 | 0.830017162 | No Change | Not Significant | TRUE |
| Hoxa6 | TRUE | 0.174316355 | 0.685206817 | No Change | Not Significant | TRUE |
| Hoxa7 | TRUE | 0.110932344 | 0.423608221 | No Change | Not Significant | TRUE |
| Hoxa9 | TRUE | -0.096970766 | 0.560708667 | No Change | Not Significant | TRUE |
| Hoxaas2 | TRUE | -0.179620915 | 0.761121093 | No Change | Not Significant | TRUE |
| Hoxaas3 | TRUE | 0.215336826 | 0.231983933 | No Change | Not Significant | TRUE |
| Hoxb13 | TRUE | -0.333371281 | 0.53655334 | No Change | Not Significant | TRUE |
| Hoxb2 | TRUE | 0.769742092 | 1 | No Change | Not Significant | TRUE |
| Hoxb3 | TRUE | -0.154816658 | 0.839976943 | No Change | Not Significant | TRUE |
| Hoxb4 | TRUE | -0.360280872 | 0.480866438 | No Change | Not Significant | TRUE |
| Hoxb5 | TRUE | -0.323697613 | 0.596644197 | No Change | Not Significant | TRUE |
| Hoxb5os | TRUE | -0.331280186 | 0.64833423 | No Change | Not Significant | TRUE |
| Hoxb6 | TRUE | -0.20637576 | 0.657892085 | No Change | Not Significant | TRUE |
| Hoxb7 | TRUE | 0.042522516 | 0.932166328 | No Change | Not Significant | TRUE |
| Hoxb8 | TRUE | -0.084901887 | 0.903013746 | No Change | Not Significant | TRUE |
| Hoxb9 | TRUE | 0.093278128 | 0.458272537 | No Change | Not Significant | TRUE |
| Hoxc13 | TRUE | 2.959253997 | 1 | No Change | Not Significant | FALSE |
| Hoxc4 | TRUE | 1.124943979 | 1 | No Change | Not Significant | FALSE |
| Hoxc5 | TRUE | 2.815461532 | 1 | No Change | Not Significant | FALSE |
| Hoxc6 | TRUE | -0.366378233 | 1 | No Change | Not Significant | FALSE |
| Hoxc8 | TRUE | 1.690130105 | 1 | No Change | Not Significant | FALSE |
| Hoxc9 | TRUE | 0.753080948 | 1 | No Change | Not Significant | FALSE |
| Hoxd1 | TRUE | 4.628251173 | 8.17E-07 | UP | UP | FALSE |
| Hoxd10 | TRUE | -0.341565452 | 0.648691985 | No Change | Not Significant | TRUE |
| Hoxd11 | TRUE | -0.040739057 | 0.961976016 | No Change | Not Significant | TRUE |
| Hoxd12 | TRUE | -0.191406363 | 0.835178936 | No Change | Not Significant | TRUE |
| Hoxd13 | TRUE | -0.323667915 | 0.665664739 | No Change | Not Significant | TRUE |
| Hoxd3 | TRUE | 0.830018519 | 1 | No Change | Not Significant | FALSE |
| Hoxd4 | TRUE | -0.144141378 | 0.829602026 | No Change | Not Significant | TRUE |
| Hoxd8 | TRUE | -0.120769856 | 0.855204229 | No Change | Not Significant | TRUE |
| Hoxd9 | TRUE | -0.707503725 | 1 | No Change | Not Significant | FALSE |
| Hp | TRUE | 1.571229055 | 1 | No Change | Not Significant | FALSE |
| Hp1bp3 | TRUE | 0.328268874 | 1.10E-07 | No Change | Significant Positive | TRUE |
| Hpca | TRUE | 0.680336069 | 0.115554504 | No Change | Not Significant | TRUE |
| Hpcal1 | TRUE | -0.290656035 | 0.000434691 | No Change | Significant Negative | TRUE |
| Hpcal4 | TRUE | -0.565128959 | 0.704201299 | No Change | Not Significant | TRUE |
| Hpd | TRUE | -0.264970043 | 0.550946551 | No Change | Not Significant | TRUE |
| Hpdl | TRUE | -0.584138069 | 0.020615447 | DOWN | DOWN | TRUE |
| Hpf1 | TRUE | 0.250466121 | 0.191813931 | No Change | Not Significant | TRUE |
| Hpgd | TRUE | -0.985941018 | 6.82E-17 | DOWN | DOWN | TRUE |
| Hpgds | TRUE | 0.109630084 | 0.674995696 | No Change | Not Significant | TRUE |
| Hpn | TRUE | -0.164014115 | 0.727184014 | No Change | Not Significant | TRUE |
| Hprt | TRUE | 0.136244781 | 0.389062992 | No Change | Not Significant | TRUE |
| Hps1 | TRUE | -0.210219103 | 0.132867477 | No Change | Not Significant | TRUE |
| Hps3 | TRUE | -0.152320069 | 0.431320242 | No Change | Not Significant | TRUE |
| Hps4 | TRUE | 0.169700712 | 0.025395551 | No Change | Significant Positive | TRUE |
| Hps5 | TRUE | 0.18727808 | 0.070214581 | No Change | Not Significant | TRUE |
| Hps6 | TRUE | -0.217042573 | 0.29613919 | No Change | Not Significant | TRUE |
| Hpse | TRUE | -0.004541391 | 0.983635967 | No Change | Not Significant | TRUE |
| Hpse2 | TRUE | 1.109713154 | 1 | No Change | Not Significant | FALSE |
| Hpx | TRUE | 0.638340145 | 1 | No Change | Not Significant | FALSE |
| Hr | TRUE | 0.221279592 | 0.081727701 | No Change | Not Significant | TRUE |
| Hras | TRUE | 0.354954551 | 4.21E-08 | No Change | Significant Positive | TRUE |
| Hrc | TRUE | -0.213504228 | 0.799444652 | No Change | Not Significant | TRUE |
| Hrct1 | TRUE | -1.081968297 | 1 | No Change | Not Significant | FALSE |
| Hrg | TRUE | -0.360585956 | 0.792979237 | No Change | Not Significant | TRUE |
| Hrh1 | TRUE | -0.344127542 | 0.157212621 | No Change | Not Significant | TRUE |
| Hrh2 | TRUE | 1.221696325 | 0.514264394 | No Change | Not Significant | FALSE |
| Hrh3 | TRUE | -1.305387711 | 1 | No Change | Not Significant | FALSE |
| Hrk | TRUE | -1.890499214 | 1 | No Change | Not Significant | FALSE |
| Hrob | TRUE | 0.566360625 | 0.048455198 | UP | UP | TRUE |

Supplementary Table 3

| geneID | Cond1_DF_Detected | Cond1_DF_ValueCol | Cond1_DF_PadjCol | Cond1_DF_Cutoff | Cond1_DF_Cutoff_Specific.x | Cond2_DF_Detected | Cond2_DF_ValueCol | [illegible] |
|---|---|---|---|---|---|---|---|---|
| Hs1bp3 | TRUE | -0.224892319 | 0.023359359 | No Change | Significant Negative | TRUE | 0.063198468 | [illegible] |
| Hs2st1 | TRUE | 0.176068944 | 0.11194538 | No Change | Not Significant | TRUE | -0.04444109 | [illegible] |
| Hs3st1 | TRUE | 0.087460763 | 0.626416691 | No Change | Not Significant | TRUE | -0.018656086 | [illegible] |
| Hs3st3a1 | TRUE | 0.270611999 | 0.619371535 | No Change | Not Significant | TRUE | 0.734455615 | [illegible] |
| Hs3st3b1 | TRUE | 3.460242948 | 2.45E-196 | UP | UP | TRUE | 2.835787146 | [illegible] |
| Hs3st4 | TRUE | -2.116946309 | 1 | No Change | Not Significant | FALSE | NA | NA |
| Hs3st5 | TRUE | -1.93326248 | 1 | No Change | Not Significant | FALSE | NA | NA |
| Hs3st6 | TRUE | -0.045708479 | 1 | No Change | Not Significant | FALSE | NA | NA |
| Hs6st1 | TRUE | 0.508625012 | 9.51E-08 | UP | UP | TRUE | 0.215025241 | [illegible] |
| Hs6st2 | TRUE | 0.974471121 | 1 | No Change | Not Significant | FALSE | NA | NA |
| Hsbp1 | TRUE | -0.066107773 | 0.594336299 | No Change | Not Significant | TRUE | 0.123243366 | [illegible] |
| Hsbp1l1 | TRUE | 0.310073751 | 0.31001526 | No Change | Not Significant | TRUE | 0.048456738 | [illegible] |
| Hscb | TRUE | -0.549263114 | 2.61E-05 | DOWN | DOWN | TRUE | -0.114890753 | [illegible] |
| Hsd11b1 | TRUE | 0.241461699 | 1 | No Change | Not Significant | FALSE | NA | NA |
| Hsd11b2 | TRUE | -0.122818186 | 0.73666903 | No Change | Not Significant | TRUE | 0.539172888 | [illegible] |
| Hsd17b1 | TRUE | 0.371606949 | 1 | No Change | Not Significant | FALSE | NA | NA |
| Hsd17b10 | TRUE | -0.347278546 | 0.00049841 | No Change | Significant Negative | TRUE | -0.076569135 | [illegible] |
| Hsd17b11 | TRUE | -0.341410768 | 0.000670367 | No Change | Significant Negative | TRUE | -0.081323198 | [illegible] |
| Hsd17b12 | TRUE | 0.353370118 | 0.004080362 | No Change | Significant Positive | TRUE | 0.095796001 | [illegible] |
| Hsd17b13 | TRUE | 0.193518789 | 0.43912271 | No Change | Not Significant | TRUE | 0.102535446 | [illegible] |
| Hsd17b14 | TRUE | -0.610693973 | 0.330029969 | No Change | Not Significant | TRUE | -0.742503335 | [illegible] |
| Hsd17b2 | TRUE | -0.541161357 | 0.003190048 | DOWN | DOWN | TRUE | -0.268190349 | [illegible] |
| Hsd17b4 | TRUE | -0.167678014 | 0.051560944 | No Change | Not Significant | TRUE | 0.088214679 | [illegible] |
| Hsd17b6 | TRUE | 0.484485058 | 1 | No Change | Not Significant | FALSE | NA | NA |
| [illegible] | [illegible] | [illegible] | [illegible] | [illegible] | [illegible] | [illegible] | [illegible] | [illegible] |

| geneID | Cond1_DF_Detected | Cond1_DF_ValueCol | Cond1_DF_PadjCol | Cond1_DF_Cutoff | Cond1_DF_Cutoff_Specific.x | Cond2_DF_Detected | Cond2_DF_ValueCol | Cond2_DF_PadjCol | Cond2_DF_Cutoff | Cond2_DF_Cutoff_Specific | BG_Method | RG1_All | RG2_Significant | RG3_SignificantChange | Cond1_DF_ensembl_gene_id | Cond1_DF_entrezgene_id | Cond1_DF_gene_biotype |
|---|---|---|---|---|---|---|---|---|---|---|---|---|---|---|---|---|---|
| Igdcc3 | TRUE | 1.29941813 | 1 | No Change | Not Significant | FALSE | NA | NA | No Change | Not Detected | FALSE | Background = FALSE | Background = FALSE | Background = FALSE | ENSMUSG00000032394 | 19289 | protein_coding |
| Igdcc4 | TRUE | 0.141467818 | 0.269506609 | No Change | Not Significant | TRUE | -0.138550683 | 0.122794781 | No Change | Not Significant | TRUE | Cond1 Not Significant + Cond2 Not Significant | None | None | ENSMUSG00000032816 | 56741 | protein_coding |
| Igf1 | TRUE | -0.236938311 | 1 | No Change | Not Significant | FALSE | NA | NA | No Change | Not Detected | FALSE | Background = FALSE | Background = FALSE | Background = FALSE | ENSMUSG00000020053 | 16000 | protein_coding |
| Igf1r | TRUE | -0.210690577 | 0.129309879 | No Change | Not Significant | TRUE | -0.150501391 | 0.242581587 | No Change | Not Significant | TRUE | Cond1 Not Significant + Cond2 Not Significant | None | None | ENSMUSG00000005533 | 16001 | protein_coding |
| Igf2 | TRUE | -0.427325 | 1 | No Change | Not Significant | FALSE | NA | NA | No Change | Not Detected | FALSE | Background = FALSE | Background = FALSE | Background = FALSE | ENSMUSG00000048583 | 16002 | protein_coding |
| Igf2bp1 | TRUE | -1.017506402 | 0.114834733 | No Change | Not Significant | TRUE | -0.812136165 | 0.032095218 | DOWN | DOWN | TRUE | Cond1 Not Significant + Cond2 DOWN | Cond2_DOWN | Cond2_DOWN | ENSMUSG00000013415 | 140486 | protein_coding |
| Igf2bp2 | TRUE | 0.085611314 | 0.761298159 | No Change | Not Significant | TRUE | -0.242542979 | 0.001327138 | No Change | Significant Negative | TRUE | Cond1 Not Significant + Cond2 Significant Negative | None | None | ENSMUSG00000033581 | 319765 | protein_coding |
| Igf2bp3 | TRUE | 4.3207522 | 1.34E-46 | UP | UP | TRUE | 0.44251195 | 0.40092091 | No Change | Not Significant | TRUE | Cond1 UP + Cond2 Not Significant | Cond1_UP | Cond1_UP | ENSMUSG00000029814 | 140488 | protein_coding |
| Igf2os | TRUE | 1.216269991 | 1 | No Change | Not Significant | FALSE | NA | NA | No Change | Not Detected | FALSE | Background = FALSE | Background = FALSE | Background = FALSE | ENSMUSG00000086266 | 111975 | lncRNA |
| Igf2r | TRUE | -0.186127996 | 0.289621372 | No Change | Not Significant | TRUE | -0.114862674 | 0.02017511 | No Change | Significant Negative | TRUE | Cond1 Not Significant + Cond2 Significant Negative | None | None | ENSMUSG00000023830 | 16004 | protein_coding |
| Igfals | TRUE | 0.015989234 | 0.976016284 | No Change | Not Significant | TRUE | -0.02095236 | 0.96169086 | No Change | Not Significant | TRUE | Cond1 Not Significant + Cond2 Not Significant | None | None | ENSMUSG00000046070 | 16005 | protein_coding |
| Igfbp2 | TRUE | 0.176019762 | 1 | No Change | Not Significant | FALSE | NA | NA | No Change | Not Detected | FALSE | Background = FALSE | Background = FALSE | Background = FALSE | ENSMUSG00000039323 | 16008 | protein_coding |
| Igfbp3 | TRUE | 1.078757978 | 0.186304828 | No Change | Not Significant | TRUE | 1.091532866 | 0.032705292 | UP | UP | TRUE | Cond1 Not Significant + Cond2 UP | Cond2_UP | Cond2_UP | ENSMUSG00000020427 | 16009 | protein_coding |
| Igfbp4 | TRUE | 1.504387411 | 1.15E-06 | UP | UP | TRUE | 0.157873083 | 0.756598137 | No Change | Not Significant | TRUE | Cond1 UP + Cond2 Not Significant | Cond1_UP | Cond1_UP | ENSMUSG00000017493 | 16010 | protein_coding |
| Igfbp5 | TRUE | -0.801514977 | 1 | No Change | Not Significant | TRUE | 2.534137136 | 0.001564246 | UP | UP | TRUE | Cond1 Not Significant + Cond2 UP | Cond2_UP | Cond2_UP | ENSMUSG00000026185 | 16011 | protein_coding |
| Igfbp6 | TRUE | 0.828263467 | 1 | No Change | Not Significant | FALSE | NA | NA | No Change | Not Detected | FALSE | Background = FALSE | Background = FALSE | Background = FALSE | ENSMUSG00000023046 | 16012 | protein_coding |
| Igfbp7 | TRUE | -0.925772055 | 1 | No Change | Not Significant | TRUE | 0.314401085 | 0.731850331 | No Change | Not Significant | TRUE | Cond1 Not Significant + Cond2 Not Significant | None | None | ENSMUSG00000036256 | 29817 | protein_coding |
| Igfbpl1 | TRUE | 0.270057249 | 1 | No Change | Not Significant | FALSE | NA | NA | No Change | Not Detected | FALSE | Background = FALSE | Background = FALSE | Background = FALSE | ENSMUSG00000035551 | 75426 | protein_coding |
| Igfn1 | TRUE | 0.357977556 | 0.820495762 | No Change | Not Significant | TRUE | 0.831465509 | 0.316071346 | No Change | Not Significant | TRUE | Cond1 Not Significant + Cond2 Not Significant | None | None | ENSMUSG00000051985 | 226438 | protein_coding |
| Ighmbp2 | TRUE | 0.232730988 | 0.07241317 | No Change | Not Significant | TRUE | -0.124590427 | 0.394052089 | No Change | Not Significant | TRUE | Cond1 Not Significant + Cond2 Not Significant | None | None | ENSMUSG00000024831 | 20589 | protein_coding |
| Igip | TRUE | 0.002638975 | 0.993692638 | No Change | Not Significant | TRUE | -0.028623631 | 0.886076667 | No Change | Not Significant | TRUE | Cond1 Not Significant + Cond2 Not Significant | None | None | ENSMUSG00000110185 | 109169 | protein_coding |
| Iglon5 | TRUE | -1.787632997 | 1 | No Change | Not Significant | FALSE | NA | NA | No Change | Not Detected | FALSE | Background = FALSE | Background = FALSE | Background = FALSE | ENSMUSG00000013367 | 210094 | protein_coding |
| Igsf1 | TRUE | -0.982004774 | 1 | No Change | Not Significant | FALSE | NA | NA | No Change | Not Detected | FALSE | Background = FALSE | Background = FALSE | Background = FALSE | ENSMUSG00000031111 | 209268 | protein_coding |
| Igsf10 | TRUE | -0.168458909 | 1 | No Change | Not Significant | FALSE | NA | NA | No Change | Not Detected | FALSE | Background = FALSE | Background = FALSE | Background = FALSE | ENSMUSG00000036334 | 242050 | protein_coding |
| Igsf11 | TRUE | 1.197544608 | 0.006161774 | UP | UP | TRUE | 0.360276914 | 0.381119039 | No Change | Not Significant | TRUE | Cond1 UP + Cond2 Not Significant | Cond1_UP | Cond1_UP | ENSMUSG00000022790 | 207683 | protein_coding |
| Igsf21 | TRUE | 0.655295163 | 1 | No Change | Not Significant | FALSE | NA | NA | No Change | Not Detected | FALSE | Background = FALSE | Background = FALSE | Background = FALSE | ENSMUSG00000040972 | 230868 | protein_coding |
| Igsf23 | TRUE | 0.163905892 | 0.464375257 | No Change | Not Significant | TRUE | 0.070214379 | 0.824909093 | No Change | Not Significant | TRUE | Cond1 Not Significant + Cond2 Not Significant | None | None | ENSMUSG00000040498 | 70080 | protein_coding |
| Igsf3 | TRUE | -0.328278663 | 0.050422608 | No Change | Not Significant | TRUE | -0.286350178 | 3.53E-05 | No Change | Significant Negative | TRUE | Cond1 Not Significant + Cond2 Significant Negative | None | None | ENSMUSG00000042035 | 78908 | protein_coding |
| Igsf5 | TRUE | 2.020057239 | 4.31E-181 | UP | UP | TRUE | 1.091935232 | 7.52E-34 | UP | UP | TRUE | Cond1 UP + Cond2 UP | Core_UP | Core_UP | ENSMUSG00000000159 | 72058 | protein_coding |
| Igsf6 | TRUE | -0.236072785 | 0.760617759 | No Change | Not Significant | TRUE | 0.097347746 | 0.916202738 | No Change | Not Significant | TRUE | Cond1 Not Significant + Cond2 Not Significant | None | None | ENSMUSG00000035004 | 80719 | protein_coding |
| Igsf8 | TRUE | -0.144008091 | 0.308134243 | No Change | Not Significant | TRUE | -0.052432356 | 0.718950524 | No Change | Not Significant | TRUE | Cond1 Not Significant + Cond2 Not Significant | None | None | ENSMUSG00000038034 | 140559 | protein_coding |
| Igsf9 | TRUE | 0.243829632 | 0.270265596 | No Change | Not Significant | TRUE | 0.212762589 | 0.115930543 | No Change | Not Significant | TRUE | Cond1 Not Significant + Cond2 Not Significant | None | None | ENSMUSG00000037995 | 93842 | protein_coding |
| Igsf9b | TRUE | 0.225058802 | 0.176210159 | No Change | Not Significant | TRUE | 0.42021769 | 0.017878316 | No Change | Significant Positive | TRUE | Cond1 Not Significant + Cond2 Significant Positive | None | None | ENSMUSG00000034275 | 235086 | protein_coding |
| Igtp | TRUE | 0.531183499 | 0.224469402 | No Change | Not Significant | FALSE | NA | NA | No Change | Not Detected | FALSE | Background = FALSE | Background = FALSE | Background = FALSE | ENSMUSG00000078853 | 16145 | protein_coding |
| Ihh | TRUE | -0.060693169 | 0.713797415 | No Change | Not Significant | TRUE | 0.049913921 | 0.762827428 | No Change | Not Significant | TRUE | Cond1 Not Significant + Cond2 Not Significant | None | None | ENSMUSG00000006538 | 16147 | protein_coding |
| Iho1 | TRUE | 1.602152381 | 1 | No Change | Not Significant | FALSE | NA | NA | No Change | Not Detected | FALSE | Background = FALSE | Background = FALSE | Background = FALSE | ENSMUSG00000047220 | 434438 | protein_coding |
| Iigp1 | TRUE | 1.213593757 | 0.253177382 | No Change | Not Significant | FALSE | NA | NA | No Change | Not Detected | FALSE | Background = FALSE | Background = FALSE | Background = FALSE | ENSMUSG00000054072 | 60440 | protein_coding |
| Ik | TRUE | 0.629751281 | 1.09E-39 | UP | UP | TRUE | 0.258365411 | 1.40E-05 | No Change | Significant Positive | TRUE | Cond1 UP + Cond2 Significant Positive | Core_UP | Cond1_UP | ENSMUSG00000024474 | 24010 | protein_coding |
| Ikbip | TRUE | 0.446698242 | 0.003857639 | No Change | Significant Positive | TRUE | 0.144738822 | 0.299347968 | No Change | Not Significant | TRUE | Cond1 Significant Positive + Cond2 Not Significant | None | None | ENSMUSG00000019975 | 67454 | protein_coding |
| Ikbkb | TRUE | 0.066949937 | 0.510000436 | No Change | Not Significant | TRUE | -0.065968838 | 0.443565039 | No Change | Not Significant | TRUE | Cond1 Not Significant + Cond2 Not Significant | None | None | ENSMUSG00000031537 | 16150 | protein_coding |
| Ikbke | TRUE | 1.613529755 | 8.78E-82 | UP | UP | TRUE | 1.198889768 | 5.61E-70 | UP | UP | TRUE | Cond1 UP + Cond2 UP | Core_UP | Core_UP | ENSMUSG00000042349 | 56489 | protein_coding |
| Ikbkg | TRUE | -0.055000016 | 0.669388475 | No Change | Not Significant | TRUE | 0.012607956 | 0.896853291 | No Change | Not Significant | TRUE | Cond1 Not Significant + Cond2 Not Significant | None | None | ENSMUSG00000004221 | 16151 | protein_coding |
| Ikzf1 | TRUE | 0.771194244 | 0.22115998 | No Change | Not Significant | TRUE | 0.354084275 | 0.414061984 | No Change | Not Significant | TRUE | Cond1 Not Significant + Cond2 Not Significant | None | None | ENSMUSG00000018654 | 22778 | protein_coding |
| Ikzf2 | TRUE | 0.146679679 | 0.539328021 | No Change | Not Significant | TRUE | 0.242589017 | 0.027819661 | No Change | Significant Positive | TRUE | Cond1 Not Significant + Cond2 Significant Positive | None | None | ENSMUSG00000025997 | 22779 | protein_coding |
| Ikzf3 | TRUE | -0.194372978 | 0.81021764 | No Change | Not Significant | TRUE | 0.071872248 | 0.88952542 | No Change | Not Significant | TRUE | Cond1 Not Significant + Cond2 Not Significant | None | None | ENSMUSG00000018168 | 22780 | protein_coding |
| Ikzf4 | TRUE | 0.078882238 | 0.966663747 | No Change | Not Significant | FALSE | NA | NA | No Change | Not Detected | FALSE | Background = FALSE | Background = FALSE | Background = FALSE | ENSMUSG00000002578 | 22781 | protein_coding |
| Ikzf5 | TRUE | -0.102550694 | 0.778864783 | No Change | Not Significant | TRUE | -0.083133322 | 0.813714227 | No Change | Not Significant | TRUE | Cond1 Not Significant + Cond2 Not Significant | None | None | ENSMUSG00000040167 | 67143 | protein_coding |
| Il10 | TRUE | 1.545476946 | 1 | No Change | Not Significant | FALSE | NA | NA | No Change | Not Detected | FALSE | Background = FALSE | Background = FALSE | Background = FALSE | ENSMUSG00000016529 | 16153 | protein_coding |
| Il10ra | TRUE | -0.383728671 | 1 | No Change | Not Significant | FALSE | NA | NA | No Change | Not Detected | FALSE | Background = FALSE | Background = FALSE | Background = FALSE | ENSMUSG00000032089 | 16154 | protein_coding |
| Il10rb | TRUE | -0.165515571 | 0.071194438 | No Change | Not Significant | TRUE | -0.026891985 | 0.816513895 | No Change | Not Significant | TRUE | Cond1 Not Significant + Cond2 Not Significant | None | None | ENSMUSG00000022969 | 16155 | protein_coding |
| Il11 | TRUE | 0.503922781 | 1 | No Change | Not Significant | FALSE | NA | NA | No Change | Not Detected | FALSE | Background = FALSE | Background = FALSE | Background = FALSE | ENSMUSG00000004371 | 16156 | protein_coding |
| Il11ra1 | TRUE | -0.162502395 | 0.447829813 | No Change | Not Significant | TRUE | -0.082608507 | 0.608127704 | No Change | Not Significant | TRUE | Cond1 Not Significant + Cond2 Not Significant | None | None | ENSMUSG00000073889 | 16157 | protein_coding |
| Il12a | TRUE | 1.170102447 | 1 | No Change | Not Significant | FALSE | NA | NA | No Change | Not Detected | FALSE | Background = FALSE | Background = FALSE | Background = FALSE | ENSMUSG00000027776 | 16159 | protein_coding |
| Il12b | TRUE | 0.176018703 | 1 | No Change | Not Significant | FALSE | NA | NA | No Change | Not Detected | FALSE | Background = FALSE | Background = FALSE | Background = FALSE | ENSMUSG00000004296 | 16160 | protein_coding |
| Il12rb1 | TRUE | 3.409782077 | 1.50E-09 | UP | UP | TRUE | 1.555579487 | 0.004508269 | UP | UP | TRUE | Cond1 UP + Cond2 UP | Core_UP | Core_UP | ENSMUSG00000000791 | 16161 | protein_coding |
| Il12rb2 | TRUE | -0.33455936 | 1 | No Change | Not Significant | FALSE | NA | NA | No Change | Not Detected | FALSE | Background = FALSE | Background = FALSE | Background = FALSE | ENSMUSG00000018341 | 16162 | protein_coding |
| Il13ra1 | TRUE | 0.248392818 | 0.342539221 | No Change | Not Significant | TRUE | 0.328585161 | 6.22E-08 | No Change | Significant Positive | TRUE | Cond1 Not Significant + Cond2 Significant Positive | None | None | ENSMUSG00000017057 | 16164 | protein_coding |
| Il15 | TRUE | -0.399597964 | 0.024770728 | No Change | Significant Negative | TRUE | -0.393034275 | 0.000509152 | No Change | Significant Negative | TRUE | Cond1 Significant Negative + Cond2 Significant Negative | None | None | ENSMUSG00000031712 | 16168 | protein_coding |
| Il15ra | TRUE | -0.342916562 | 0.008514609 | No Change | Significant Negative | TRUE | -0.229736413 | 0.273475636 | No Change | Not Significant | TRUE | Cond1 Significant Negative + Cond2 Not Significant | None | None | ENSMUSG00000023206 | 16169 | protein_coding |
| Il16 | TRUE | 0.14845343 | 0.856549354 | No Change | Not Significant | TRUE | 0.216244918 | 0.682675952 | No Change | Not Significant | TRUE | Cond1 Not Significant + Cond2 Not Significant | None | None | ENSMUSG00000001741 | 16170 | protein_coding |
| Il17c | TRUE | 0.003960856 | 0.995979059 | No Change | Not Significant | TRUE | -0.322567877 | 0.732548832 | No Change | Not Significant | TRUE | Cond1 Not Significant + Cond2 Not Significant | None | None | ENSMUSG00000046108 | 234836 | protein_coding |
| Il17d | TRUE | -0.265415575 | 0.544366934 | No Change | Not Significant | TRUE | -0.264732356 | 0.269652976 | No Change | Not Significant | TRUE | Cond1 Not Significant + Cond2 Not Significant | None | None | ENSMUSG00000050222 | 239114 | protein_coding |
| Il17f | TRUE | 0.267850176 | 1 | No Change | Not Significant | FALSE | NA | NA | No Change | Not Detected | FALSE | Background = FALSE | Background = FALSE | Background = FALSE | ENSMUSG00000041872 | 257630 | protein_coding |
| Il17ra | TRUE | -0.153343982 | 0.076491713 | No Change | Not Significant | TRUE | -0.195659896 | 0.027655107 | No Change | Significant Negative | TRUE | Cond1 Not Significant + Cond2 Significant Negative | None | None | ENSMUSG00000002897 | 16172 | protein_coding |
| Il17rb | TRUE | -0.378481184 | 0.296522686 | No Change | Not Significant | TRUE | -0.22036706 | 0.465617977 | No Change | Not Significant | TRUE | Cond1 Not Significant + Cond2 Not Significant | None | None | ENSMUSG00000015966 | 50905 | protein_coding |
| Il17rc | TRUE | -0.097676161 | 0.579526645 | No Change | Not Significant | TRUE | 0.038796117 | 0.816823161 | No Change | Not Significant | TRUE | Cond1 Not Significant + Cond2 Not Significant | None | None | ENSMUSG00000030281 | 171095 | protein_coding |
| Il17rd | TRUE | -0.062083096 | 0.86653427 | No Change | Not Significant | TRUE | -0.112562696 | 0.699935475 | No Change | Not Significant | TRUE | Cond1 Not Significant + Cond2 Not Significant | None | None | ENSMUSG00000040717 | 171463 | protein_coding |
| Il17re | TRUE | -0.058974973 | 0.769309474 | No Change | Not Significant | TRUE | 0.250312853 | 0.01118574 | No Change | Significant Positive | TRUE | Cond1 Not Significant + Cond2 Significant Positive | None | None | ENSMUSG00000043088 | 57890 | protein_coding |
| Il18 | TRUE | -0.614745025 | 0.027075673 | DOWN | DOWN | TRUE | -0.19215797 | 0.381977497 | No Change | Not Significant | TRUE | Cond1 DOWN + Cond2 Not Significant | Cond1_DOWN | Cond1_DOWN | ENSMUSG00000039217 | 16173 | protein_coding |
| Il18bp | TRUE | 0.057610149 | 0.898576349 | No Change | Not Significant | TRUE | -0.923177632 | 0.024532938 | DOWN | DOWN | TRUE | Cond1 Not Significant + Cond2 DOWN | Cond2_DOWN | Cond2_DOWN | ENSMUSG00000070427 | 16068 | protein_coding |
| Il18r1 | TRUE | 0.973780101 | 0.139328867 | No Change | Not Significant | TRUE | 0.571306942 | 0.256715961 | No Change | Not Significant | TRUE | Cond1 Not Significant + Cond2 Not Significant | None | None | ENSMUSG00000026070 | 16182 | protein_coding |
| Il18rap | TRUE | 0.467890195 | 0.749223673 | No Change | Not Significant | FALSE | NA | NA | No Change | Not Detected | FALSE | Background = FALSE | Background = FALSE | Background = FALSE | ENSMUSG00000026068 | 16174 | protein_coding |
| Il1a | TRUE | 0.495667151 | 1 | No Change | Not Significant | FALSE | NA | NA | No Change | Not Detected | FALSE | Background = FALSE | Background = FALSE | Background = FALSE | ENSMUSG00000027399 | 16175 | protein_coding |
| Il1b | TRUE | 1.952304339 | 0.096552424 | No Change | Not Significant | FALSE | NA | NA | No Change | Not Detected | FALSE | Background = FALSE | Background = FALSE | Background = FALSE | ENSMUSG00000027398 | 16176 | protein_coding |
| Il1r1 | TRUE | 0.143679373 | 0.511151353 | No Change | Not Significant | TRUE | 0.103507765 | 0.382604713 | No Change | Not Significant | TRUE | Cond1 Not Significant + Cond2 Not Significant | None | None | ENSMUSG00000026072 | 16177 | protein_coding |
| Il1r2 | TRUE | -0.065361826 | 1 | No Change | Not Significant | FALSE | NA | NA | No Change | Not Detected | FALSE | Background = FALSE | Background = FALSE | Background = FALSE | ENSMUSG00000026073 | 16178 | protein_coding |
| Il1rap | TRUE | 0.187461559 | 0.382529046 | No Change | Not Significant | TRUE | 0.0619793 | 0.7296319 | No Change | Not Significant | TRUE | Cond1 Not Significant + Cond2 Not Significant | None | None | ENSMUSG00000022514 | 16180 | protein_coding |
| Il1rapl1 | TRUE | 0.572487634 | 1 | No Change | Not Significant | FALSE | NA | NA | No Change | Not Detected | FALSE | Background = FALSE | Background = FALSE | Background = FALSE | ENSMUSG00000052372 | 331461 | protein_coding |
| Il1rapl2 | TRUE | 0.753064925 | 1 | No Change | Not Significant | FALSE | NA | NA | No Change | Not Detected | FALSE | Background = FALSE | Background = FALSE | Background = FALSE | ENSMUSG00000059203 | 60367 | protein_coding |
| Il1rl1 | TRUE | -0.15870246 | 1 | No Change | Not Significant | FALSE | NA | NA | No Change | Not Detected | FALSE | Background = FALSE | Background = FALSE | Background = FALSE | ENSMUSG00000026069 | 17082 | protein_coding |
| Il1rl2 | TRUE | -0.746598523 | 0.061535163 | No Change | Not Significant | TRUE | 0.31283987 | 0.495031103 | No Change | Not Significant | TRUE | Cond1 Not Significant + Cond2 Not Significant | None | None | ENSMUSG00000070942 | 107527 | protein_coding |
| Il1rn | TRUE | -0.933977923 | 0.000462865 | DOWN | DOWN | TRUE | -0.019151679 | 0.945960168 | No Change | Not Significant | TRUE | Cond1 DOWN + Cond2 Not Significant | Cond1_DOWN | Cond1_DOWN | ENSMUSG00000026981 | 16181 | protein_coding |
| Il20rb | TRUE | -0.280188043 | 0.630476965 | No Change | Not Significant | TRUE | -0.465869815 | 0.202823164 | No Change | Not Significant | TRUE | Cond1 Not Significant + Cond2 Not Significant | None | None | ENSMUSG00000044244 | 213208 | protein_coding |
| Il21 | TRUE | 3.454724777 | 1 | No Change | Not Significant | FALSE | NA | NA | No Change | Not Detected | FALSE | Background = FALSE | Background = FALSE | Background = FALSE | ENSMUSG00000027718 | 60505 | protein_coding |
| Il21r | TRUE | -0.110777557 | 0.912001512 | No Change | Not Significant | TRUE | 0.676368237 | 0.208682378 | No Change | Not Significant | TRUE | Cond1 Not Significant + Cond2 Not Significant | None | None | ENSMUSG00000030745 | 60504 | protein_coding |
| Il22ra1 | TRUE | 0.061454037 | 0.852797822 | No Change | Not Significant | TRUE | -0.035912029 | 0.75524772 | No Change | Not Significant | TRUE | Cond1 Not Significant + Cond2 Not Significant | None | None | ENSMUSG00000037157 | 230828 | protein_coding |
| Il22ra2 | TRUE | -1.417435554 | 1 | No Change | Not Significant | TRUE | 0.317658644 | 0.726170014 | No Change | Not Significant | TRUE | Cond1 Not Significant + Cond2 Not Significant | None | None | ENSMUSG00000039760 | 237310 | protein_coding |
| Il23a | TRUE | 0.932442036 | 1 | No Change | Not Significant | FALSE | NA | NA | No Change | Not Detected | FALSE | Background = FALSE | Background = FALSE | Background = FALSE | ENSMUSG00000025383 | 83430 | protein_coding |
| Il25 | TRUE | -1.018520451 | 1 | No Change | Not Significant | FALSE | NA | NA | No Change | Not Detected | FALSE | Background = FALSE | Background = FALSE | Background = FALSE | ENSMUSG00000040770 | 140806 | protein_coding |
| Il27ra | TRUE | 0.54342507 | 0.741566405 | No Change | Not Significant | TRUE | -0.03238044 | 0.970174281 | No Change | Not Significant | TRUE | Cond1 Not Significant + Cond2 Not Significant | None | None | ENSMUSG00000005465 | 50931 | protein_coding |
| Il2ra | TRUE | 1.130463219 | 1 | No Change | Not Significant | FALSE | NA | NA | No Change | Not Detected | FALSE | Background = FALSE | Background = FALSE | Background = FALSE | ENSMUSG00000026770 | 16184 | protein_coding |
| Il2rb | TRUE | 0.396071968 | 0.489835334 | No Change | Not Significant | TRUE | 0.425311402 | 0.017810477 | No Change | Significant Positive | TRUE | Cond1 Not Significant + Cond2 Significant Positive | None | None | ENSMUSG00000068227 | 16185 | protein_coding |
| Il2rg | TRUE | 0.821649756 | 0.350004803 | No Change | Not Significant | TRUE | 0.039241375 | 0.965011685 | No Change | Not Significant | TRUE | Cond1 Not Significant + Cond2 Not Significant | None | None | ENSMUSG00000031304 | 16186 | protein_coding |
| Il31ra | TRUE | -0.224764564 | 1 | No Change | Not Significant | FALSE | NA | NA | No Change | Not Detected | FALSE | Background = FALSE | Background = FALSE | Background = FALSE | ENSMUSG00000050377 | 218624 | protein_coding |
| Il33 | TRUE | 0.460083997 | 0.719844721 | No Change | Not Significant | TRUE | -0.40748639 | 0.511744838 | No Change | Not Significant | TRUE | Cond1 Not Significant + Cond2 Not Significant | None | None | ENSMUSG00000024810 | 77125 | protein_coding |
| Il34 | TRUE | -0.721594153 | 0.00707029 | DOWN | DOWN | TRUE | 0.065032865 | 0.880475742 | No Change | Not Significant | TRUE | Cond1 DOWN + Cond2 Not Significant | Cond1_DOWN | Cond1_DOWN | ENSMUSG00000031750 | 76527 | protein_coding |
| Il36rn | TRUE | -1.243701273 | 1 | No Change | Not Significant | FALSE | NA | NA | No Change | Not Detected | FALSE | Background = FALSE | Background = FALSE | Background = FALSE | ENSMUSG00000026983 | 54450 | protein_coding |
| Il3ra | TRUE | 1.359127637 | 0.00443976 | UP | UP | TRUE | 0.486807694 | 0.309576536 | No Change | Not Significant | TRUE | Cond1 UP + Cond2 Not Significant | Cond1_UP | Cond1_UP | ENSMUSG00000068758 | 16188 | protein_coding |
| Il4 | TRUE | 0.339143864 | 1 | No Change | Not Significant | TRUE | 0.561528911 | 0.601184005 | No Change | Not Significant | TRUE | Cond1 Not Significant + Cond2 Not Significant | None | None | ENSMUSG00000000869 | 16189 | protein_coding |
| Il4i1 | TRUE | -0.889929086 | 1 | No Change | Not Significant | FALSE | NA | NA | No Change | Not Detected | FALSE | Background = FALSE | Background = FALSE | Background = FALSE | ENSMUSG00000074141 | 14204 | protein_coding |
| Il4ra | TRUE | 0.341956837 | 0.020127052 | No Change | Significant Positive | TRUE | 0.092925509 | 0.320093959 | No Change | Not Significant | TRUE | Cond1 Significant Positive + Cond2 Not Significant | None | None | ENSMUSG00000030748 | 16190 | protein_coding |
| Il5 | TRUE | 0.607410622 | 1 | No Change | Not Significant | FALSE | NA | NA | No Change | Not Detected | FALSE | Background = FALSE | Background = FALSE | Background = FALSE | ENSMUSG00000036117 | 16191 | protein_coding |
| Il5ra | TRUE | 1.339631674 | 1 | No Change | Not Significant | FALSE | NA | NA | No Change | Not Detected | FALSE | Background = FALSE | Background = FALSE | Background = FALSE | ENSMUSG00000005364 | 16192 | protein_coding |
| Il6ra | TRUE | -0.227370779 | 0.718665364 | No Change | Not Significant | TRUE | -0.078616003 | 0.838364494 | No Change | Not Significant | TRUE | Cond1 Not Significant + Cond2 Not Significant | None | None | ENSMUSG00000027947 | 16194 | protein_coding |
| Il6st | TRUE | 0.655943921 | 1.64E-15 | UP | UP | TRUE | 0.346676251 | 2.46E-09 | No Change | Significant Positive | TRUE | Cond1 UP + Cond2 Significant Positive | Core_UP | Cond1_UP | ENSMUSG00000021756 | 16195 | protein_coding |
| Il7 | TRUE | 0.427540654 | 0.044806045 | No Change | Significant Positive | TRUE | 0.273755151 | 0.034576411 | No Change | Significant Positive | TRUE | Cond1 Significant Positive + Cond2 Significant Positive | None | None | ENSMUSG00000040329 | 16196 | protein_coding |
| Il7r | TRUE | -0.105611954 | 1 | No Change | Not Significant | FALSE | NA | NA | No Change | Not Detected | FALSE | Background = FALSE | Background = FALSE | Background = FALSE | ENSMUSG00000003882 | 16197 | protein_coding |
| Il9r | TRUE | -0.99642705 | 1 | No Change | Not Significant | FALSE | NA | NA | No Change | Not Detected | FALSE | Background = FALSE | Background = FALSE | Background = FALSE | ENSMUSG00000020279 | 16199 | protein_coding |
| Ildr1 | TRUE | 0.441977427 | 0.000192981 | No Change | Significant Positive | TRUE | 0.263759406 | 0.000149804 | No Change | Significant Positive | TRUE | Cond1 Significant Positive + Cond2 Significant Positive | None | None | ENSMUSG00000022900 | 106347 | protein_coding |
| Ildr2 | TRUE | 0.939251552 | 0.536652594 | No Change | Not Significant | TRUE | 1.180814152 | 0.118795321 | No Change | Not Significant | TRUE | Cond1 Not Significant + Cond2 Not Significant | None | None | ENSMUSG00000040612 | 100039795 | protein_coding |
| Ilf2 | TRUE | 0.283957736 | 0.025430219 | No Change | Significant Positive | TRUE | 0.012840184 | 0.923349179 | No Change | Not Significant | TRUE | Cond1 Significant Positive + Cond2 Not Significant | None | None | ENSMUSG00000001016 | 67781 | protein_coding |
| Ilf3 | TRUE | -0.046164298 | 0.816110122 | No Change | Not Significant | TRUE | -0.231291771 | 0.003319362 | No Change | Significant Negative | TRUE | Cond1 Not Significant + Cond2 Significant Negative | None | None | ENSMUSG00000032178 | 16201 | protein_coding |
| Ilk | TRUE | 0.391098554 | 0.392130341 | No Change | Not Significant | TRUE | -0.184820323 | 0.73971984 | No Change | Not Significant | TRUE | Cond1 Not Significant + Cond2 Not Significant | None | None | ENSMUSG00000030890 | 16202 | protein_coding |
| Ilkap | TRUE | -0.264480017 | 0.000647282 | No Change | Significant Negative | TRUE | -0.146848056 | 0.020731078 | No Change | Significant Negative | TRUE | Cond1 Significant Negative + Cond2 Significant Negative | None | None | ENSMUSG00000026309 | 67444 | protein_coding |
| Ilrun | TRUE | -0.055739553 | 0.398418975 | No Change | Not Significant | TRUE | 0.063373245 | 0.492478624 | No Change | Not Significant | TRUE | Cond1 Not Significant + Cond2 Not Significant | None | None | ENSMUSG00000056692 | 224647 | protein_coding |
| Ilvbl | TRUE | -0.28755931 | 0.000105168 | No Change | Significant Negative | TRUE | -0.01124185 | 0.889439786 | No Change | Not Significant | TRUE | Cond1 Significant Negative + Cond2 Not Significant | None | None | ENSMUSG00000032763 | 216136 | protein_coding |
| Immp1l | TRUE | -0.156679447 | 0.448045905 | No Change | Not Significant | TRUE | 0.065136794 | 0.57003615 | No Change | Not Significant | TRUE | Cond1 Not Significant + Cond2 Not Significant | None | None | ENSMUSG00000042670 | 66541 | protein_coding |
| Immp2l | TRUE | -0.523197476 | 0.003057453 | DOWN | DOWN | TRUE | -0.1370272 | 0.46714214 | No Change | Not Significant | TRUE | Cond1 DOWN + Cond2 Not Significant | Cond1_DOWN | Cond1_DOWN | ENSMUSG00000056899 | 93757 | protein_coding |
| Immt | TRUE | 0.313946739 | 1.62E-05 | No Change | Significant Positive | TRUE | 0.259625923 | 3.35E-06 | No Change | Significant Positive | TRUE | Cond1 Significant Positive + Cond2 Significant Positive | None | None | ENSMUSG00000052337 | 76614 | protein_coding |
| Imp3 | TRUE | -0.074144673 | 0.736456016 | No Change | Not Significant | TRUE | -0.111223192 | 0.318746628 | No Change | Not Significant | TRUE | Cond1 Not Significant + Cond2 Not Significant | None | None | ENSMUSG00000032288 | 102462 | protein_coding |
| Imp4 | TRUE | -0.087866425 | 0.527079777 | No Change | Not Significant | TRUE | 0.062654481 | 0.431211287 | No Change | Not Significant | TRUE | Cond1 Not Significant + Cond2 Not Significant | None | None | ENSMUSG00000026127 | 27993 | protein_coding |
| Impa1 | TRUE | 0.063160847 | 0.59579974 | No Change | Not Significant | TRUE | 0.091261492 | 0.21271653 | No Change | Not Significant | TRUE | Cond1 Not Significant + Cond2 Not Significant | None | None | ENSMUSG00000027531 | 55980 | protein_coding |
| Impa2 | TRUE | -0.301170544 | 0.041512301 | No Change | Significant Negative | TRUE | -0.033133777 | 0.820125043 | No Change | Not Significant | TRUE | Cond1 Significant Negative + Cond2 Not Significant | None | None | ENSMUSG00000024525 | 114663 | protein_coding |
| Impact | TRUE | -0.028785306 | 0.927574705 | No Change | Not Significant | TRUE | -0.356860492 | 0.054261734 | No Change | Not Significant | TRUE | Cond1 Not Significant + Cond2 Not Significant | None | None | ENSMUSG00000024423 | 16210 | protein_coding |
| Impdh1 | TRUE | -0.005981102 | 0.979508488 | No Change | Not Significant | TRUE | -0.083844343 | 0.44340091 | No Change | Not Significant | TRUE | Cond1 Not Significant + Cond2 Not Significant | None | None | ENSMUSG00000003500 | 23917 | protein_coding |
| Impdh2 | TRUE | -0.221384731 | 0.324596932 | No Change | Not Significant | TRUE | -0.090122 | 0.305187638 | No Change | Not Significant | TRUE | Cond1 Not Significant + Cond2 Not Significant | None | None | ENSMUSG00000062867 | 23918 | protein_coding |
| Impg1 | TRUE | 0.605248273 | 1 | No Change | Not Significant | FALSE | NA | NA | No Change | Not Detected | FALSE | Background = FALSE | Background = FALSE | Background = FALSE | ENSMUSG00000032343 | 63859 | protein_coding |
| Impg2 | TRUE | 1.149517962 | 0.186623428 | No Change | Not Significant | TRUE | -0.139030646 | 0.885839179 | No Change | Not Significant | TRUE | Cond1 Not Significant + Cond2 Not Significant | None | None | ENSMUSG00000035270 | 224224 | protein_coding |
| Ina | TRUE | -0.496494529 | 1 | No Change | Not Significant | FALSE | NA | NA | No Change | Not Detected | FALSE | Background = FALSE | Background = FALSE | Background = FALSE | ENSMUSG00000034336 | 226180 | protein_coding |
| Inafm1 | TRUE | -0.678508769 | 1.01E-07 | DOWN | DOWN | TRUE | -0.151965168 | 0.31618145 | No Change | Not Significant | TRUE | Cond1 DOWN + Cond2 Not Significant | Cond1_DOWN | Cond1_DOWN | ENSMUSG00000091811 | 66300 | protein_coding |
| Inafm2 | TRUE | -0.133805619 | 0.410708835 | No Change | Not Significant | TRUE | -0.106166494 | 0.350507776 | No Change | Not Significant | TRUE | Cond1 Not Significant + Cond2 Not Significant | None | None | ENSMUSG00000074918 | 100043272 | protein_coding |
| Inava | TRUE | -0.008664318 | 0.966128972 | No Change | Not Significant | TRUE | 0.045402256 | 0.699497125 | No Change | Not Significant | TRUE | Cond1 Not Significant + Cond2 Not Significant | None | None | ENSMUSG00000041605 | 67313 | protein_coding |
| Inca1 | TRUE | -0.081247231 | 0.906704483 | No Change | Not Significant | TRUE | -0.01083707 | 0.985900171 | No Change | Not Significant | TRUE | Cond1 Not Significant + Cond2 Not Significant | None | None | ENSMUSG00000057054 | 103844 | protein_coding |
| Incenp | TRUE | 0.526097888 | 0.001528454 | UP | UP | TRUE | 0.194632352 | 0.211105968 | No Change | Not Significant | TRUE | Cond1 UP + Cond2 Not Significant | Cond1_UP | Cond1_UP | ENSMUSG00000024660 | 16319 | protein_coding |
| Inf2 | TRUE | -0.314456481 | 0.172799808 | No Change | Not Significant | TRUE | -0.138601493 | 0.5061553 | No Change | Not Significant | TRUE | Cond1 Not Significant + Cond2 Not Significant | None | None | ENSMUSG00000037679 | 70435 | protein_coding |
| Ing1 | TRUE | 1.057661626 | 7.62E-19 | UP | UP | TRUE | 0.408988183 | 6.17E-15 | No Change | Significant Positive | TRUE | Cond1 UP + Cond2 Significant Positive | Core_UP | Cond1_UP | ENSMUSG00000045969 | 26356 | protein_coding |
| Ing2 | TRUE | -0.349014454 | 0.005227463 | No Change | Significant Negative | TRUE | -0.148221281 | 0.237154121 | No Change | Not Significant | TRUE | Cond1 Significant Negative + Cond2 Not Significant | None | None | ENSMUSG00000063049 | 69260 | protein_coding |
| Ing3 | TRUE | -0.027091206 | 0.917517221 | No Change | Not Significant | TRUE | -0.012309321 | 0.93968591 | No Change | Not Significant | TRUE | Cond1 Not Significant + Cond2 Not Significant | None | None | ENSMUSG00000029670 | 71777 | protein_coding |
| Ing4 | TRUE | 0.010230163 | 0.988088396 | No Change | Not Significant | TRUE | -0.186989974 | 0.730504162 | No Change | Not Significant | TRUE | Cond1 Not Significant + Cond2 Not Significant | None | None | ENSMUSG00000030330 | 28019 | protein_coding |
| Ing5 | TRUE | -0.149433585 | 0.316830918 | No Change | Not Significant | TRUE | -0.172650602 | 0.077108863 | No Change | Not Significant | TRUE | Cond1 Not Significant + Cond2 Not Significant | None | None | ENSMUSG00000026283 | 66262 | protein_coding |
| Inha | TRUE | 1.486002057 | 0.031886196 | UP | UP | FALSE | NA | NA | No Change | Not Detected | FALSE | Background = FALSE | Background = FALSE | Background = FALSE | ENSMUSG00000032968 | 16322 | protein_coding |
| Inhba | TRUE | 0.001378207 | 1 | No Change | Not Significant | FALSE | NA | NA | No Change | Not Detected | FALSE | Background = FALSE | Background = FALSE | Background = FALSE | ENSMUSG00000041324 | 16323 | protein_coding |
| Inhbb | TRUE | 0.914841623 | 1 | No Change | Not Significant | FALSE | NA | NA | No Change | Not Detected | FALSE | Background = FALSE | Background = FALSE | Background = FALSE | ENSMUSG00000037035 | 16324 | protein_coding |
| Inhbc | TRUE | -0.804998899 | 1 | No Change | Not Significant | FALSE | NA | NA | No Change | Not Detected | FALSE | Background = FALSE | Background = FALSE | Background = FALSE | ENSMUSG00000025405 | 16325 | protein_coding |
| Inhbe | TRUE | 0.531428888 | 1 | No Change | Not Significant | FALSE | NA | NA | No Change | Not Detected | FALSE | Background = FALSE | Background = FALSE | Background = FALSE | ENSMUSG00000047492 | 16326 | protein_coding |
| Inhca | TRUE | 0.058434591 | 1 | No Change | Not Significant | FALSE | NA | NA | No Change | Not Detected | FALSE | Background = FALSE | Background = FALSE | Background = FALSE | ENSMUSG00000033688 | 71775 | protein_coding |
| Inip | TRUE | 0.041449739 | 0.793440482 | No Change | Not Significant | TRUE | 0.064636858 | 0.571645972 | No Change | Not Significant | TRUE | Cond1 Not Significant + Cond2 Not Significant | None | None | ENSMUSG00000038544 | 66209 | protein_coding |
| Inka1 | TRUE | -0.254445992 | 1 | No Change | Not Significant | FALSE | NA | NA | No Change | Not Detected | FALSE | Background = FALSE | Background = FALSE | Background = FALSE | ENSMUSG00000042106 | 68176 | protein_coding |
| Inka2 | TRUE | 1.309648323 | 0.081458356 | No Change | Not Significant | TRUE | 1.028071465 | 0.117338412 | No Change | Not Significant | TRUE | Cond1 Not Significant + Cond2 Not Significant | None | None | ENSMUSG00000048458 | 109050 | protein_coding |
| Inmt | TRUE | -1.502168681 | 1 | No Change | Not Significant | FALSE | NA | NA | No Change | Not Detected | FALSE | Background = FALSE | Background = FALSE | Background = FALSE | ENSMUSG00000003477 | 21743 | protein_coding |
| Ino80 | TRUE | -0.204328061 | 0.039779225 | No Change | Significant Negative | TRUE | -0.269483385 | 0.000152227 | No Change | Significant Negative | TRUE | Cond1 Significant Negative + Cond2 Significant Negative | None | None | ENSMUSG00000034154 | 68142 | protein_coding |
| Ino80b | TRUE | -0.089888508 | 0.616333078 | No Change | Not Significant | TRUE | 0.10272661 | 0.504916665 | No Change | Not Significant | TRUE | Cond1 Not Significant + Cond2 Not Significant | None | None | ENSMUSG00000030034 | 70020 | protein_coding |
| Ino80c | TRUE | 0.209001466 | 0.350228168 | No Change | Not Significant | TRUE | 0.0498758 | 0.616422248 | No Change | Not Significant | TRUE | Cond1 Not Significant + Cond2 Not Significant | None | None | ENSMUSG00000047989 | 225280 | protein_coding |
| Ino80d | TRUE | 0.048466959 | 0.77103778 | No Change | Not Significant | TRUE | -0.15862338 | 0.067089927 | No Change | Not Significant | TRUE | Cond1 Not Significant + Cond2 Not Significant | None | None | ENSMUSG00000040865 | 227195 | protein_coding |
| Ino80dos | TRUE | 0.138259297 | 0.803282676 | No Change | Not Significant | TRUE | -0.583690836 | 0.101791489 | No Change | Not Significant | TRUE | Cond1 Not Significant + Cond2 Not Significant | None | None | ENSMUSG00000084799 | 100380944 | lncRNA |
| Ino80e | TRUE | 0.116375259 | 0.460059583 | No Change | Not Significant | TRUE | 0.133965526 | 0.214124788 | No Change | Not Significant | TRUE | Cond1 Not Significant + Cond2 Not Significant | None | None | ENSMUSG00000030689 | 233875 | protein_coding |
| Inpp1 | TRUE | 0.181690787 | 0.161615362 | No Change | Not Significant | TRUE | -0.026390792 | 0.864328625 | No Change | Not Significant | TRUE | Cond1 Not Significant + Cond2 Not Significant | None | None | ENSMUSG00000026102 | 16329 | protein_coding |
| Inpp4a | TRUE | -0.167433332 | 0.425861007 | No Change | Not Significant | TRUE | -0.123135472 | 0.444801234 | No Change | Not Significant | TRUE | Cond1 Not Significant + Cond2 Not Significant | None | None | ENSMUSG00000026113 | 269180 | protein_coding |
| Inpp4b | TRUE | 0.231197738 | 0.549974786 | No Change | Not Significant | TRUE | 0.149540582 | 0.160285476 | No Change | Not Significant | TRUE | Cond1 Not Significant + Cond2 Not Significant | None | None | ENSMUSG00000037940 | 234515 | protein_coding |
| Inpp5a | TRUE | -0.191026144 | 0.109773894 | No Change | Not Significant | TRUE | -0.143206427 | 0.044499775 | No Change | Significant Negative | TRUE | Cond1 Not Significant + Cond2 Significant Negative | None | None | ENSMUSG00000025477 | 212111 | protein_coding |
| Inpp5b | TRUE | 0.013336644 | 0.927421073 | No Change | Not Significant | TRUE | 0.140324324 | 0.064641 | No Change | Not Significant | TRUE | Cond1 Not Significant + Cond2 Not Significant | None | None | ENSMUSG00000028894 | 16330 | protein_coding |
| Inpp5d | TRUE | -0.358100587 | 0.227592128 | No Change | Not Significant | TRUE | 0.346442988 | 0.023082976 | No Change | Significant Positive | TRUE | Cond1 Not Significant + Cond2 Significant Positive | None | None | ENSMUSG00000026288 | 16331 | protein_coding |
| Inpp5e | TRUE | -0.272545729 | 0.053650817 | No Change | Not Significant | TRUE | -0.090077253 | 0.430673204 | No Change | Not Significant | TRUE | Cond1 Not Significant + Cond2 Not Significant | None | None | ENSMUSG00000026925 | 64436 | protein_coding |
| Inpp5f | TRUE | -0.345844839 | 0.031409224 | No Change | Significant Negative | TRUE | -0.056533862 | 0.730599032 | No Change | Not Significant | TRUE | Cond1 Significant Negative + Cond2 Not Significant | None | None | ENSMUSG00000042105 | 101490 | protein_coding |
| Inpp5j | TRUE | 0.060069566 | 0.893615254 | No Change | Not Significant | TRUE | -0.073810113 | 0.546718055 | No Change | Not Significant | TRUE | Cond1 Not Significant + Cond2 Not Significant | None | None | ENSMUSG00000034570 | 170835 | protein_coding |
| Inpp5k | TRUE | -0.203310899 | 0.06023867 | No Change | Not Significant | TRUE | -0.234069552 | 0.012740991 | No Change | Significant Negative | TRUE | Cond1 Not Significant + Cond2 Significant Negative | None | None | ENSMUSG00000006127 | 19062 | protein_coding |
| Inppl1 | TRUE | -0.063077663 | 0.564644176 | No Change | Not Significant | TRUE | -0.041965715 | 0.705835346 | No Change | Not Significant | TRUE | Cond1 Not Significant + Cond2 Not Significant | None | None | ENSMUSG00000032737 | 16332 | protein_coding |
| Ins2 | TRUE | 4.066791811 | 1 | No Change | Not Significant | FALSE | NA | NA | No Change | Not Detected | FALSE | Background = FALSE | Background = FALSE | Background = FALSE | ENSMUSG00000000215 | 16334 | protein_coding |
| Insc | TRUE | 0.992289325 | 0.000456675 | UP | UP | TRUE | 0.100893891 | 0.775222412 | No Change | Not Significant | TRUE | Cond1 UP + Cond2 Not Significant | Cond1_UP | Cond1_UP | ENSMUSG00000048782 | 233752 | protein_coding |
| Insig1 | TRUE | 0.011391345 | 0.975173098 | No Change | Not Significant | TRUE | 0.064269303 | 0.778738583 | No Change | Not Significant | TRUE | Cond1 Not Significant + Cond2 Not Significant | None | None | ENSMUSG00000045294 | 231070 | protein_coding |
| Insig2 | TRUE | 0.106635889 | 0.443879555 | No Change | Not Significant | TRUE | -0.002800177 | 0.984481423 | No Change | Not Significant | TRUE | Cond1 Not Significant + Cond2 Not Significant | None | None | ENSMUSG00000003721 | 72999 | protein_coding |
| Insl3 | TRUE | 0.660478101 | 1 | No Change | Not Significant | FALSE | NA | NA | No Change | Not Detected | FALSE | Background = FALSE | Background = FALSE | Background = FALSE | ENSMUSG00000079019 | 16336 | protein_coding |
| Insl5 | TRUE | -0.52854416 | 0.158286006 | No Change | Not Significant | TRUE | -0.087353316 | 0.753401915 | No Change | Not Significant | TRUE | Cond1 Not Significant + Cond2 Not Significant | None | None | ENSMUSG00000066090 | 23919 | protein_coding |
| Insl6 | TRUE | -0.271833052 | 0.408255959 | No Change | Not Significant | TRUE | 0.154588976 | 0.496173639 | No Change | Not Significant | TRUE | Cond1 Not Significant + Cond2 Not Significant | None | None | ENSMUSG00000050957 | 27356 | protein_coding |
| Insm1 | TRUE | 0.250862861 | 0.580790426 | No Change | Not Significant | TRUE | -0.04160265 | 0.9260086 | No Change | Not Significant | TRUE | Cond1 Not Significant + Cond2 Not Significant | None | None | ENSMUSG00000068154 | 53626 | protein_coding |
| Insm2 | TRUE | -1.987203215 | 1 | No Change | Not Significant | FALSE | NA | NA | No Change | Not Detected | FALSE | Background = FALSE | Background = FALSE | Background = FALSE | ENSMUSG00000045440 | 56856 | protein_coding |
| Insr | TRUE | -0.189775812 | 0.043863303 | No Change | Significant Negative | TRUE | -0.169563954 | 0.014803696 | No Change | Significant Negative | TRUE | Cond1 Significant Negative + Cond2 Significant Negative | None | None | ENSMUSG00000005534 | 16337 | protein_coding |
| Insrr | TRUE | -1.197216284 | 0.000206866 | DOWN | DOWN | TRUE | -0.054045773 | 0.825261698 | No Change | Not Significant | TRUE | Cond1 DOWN + Cond2 Not Significant | Cond1_DOWN | Cond1_DOWN | ENSMUSG00000005640 | 23920 | protein_coding |
| Insyn1 | TRUE | 2.516609316 | 1 | No Change | Not Significant | FALSE | NA | NA | No Change | Not Detected | FALSE | Background = FALSE | Background = FALSE | Background = FALSE | ENSMUSG00000066607 | 319477 | protein_coding |
| Insyn2a | TRUE | -0.360720444 | 0.322338142 | No Change | Not Significant | TRUE | -0.771289323 | 0.044203442 | DOWN | DOWN | TRUE | Cond1 Not Significant + Cond2 DOWN | Cond2_DOWN | Cond2_DOWN | ENSMUSG00000073805 | 627214 | protein_coding |
| Insyn2b | TRUE | -0.507463879 | 1 | No Change | Not Significant | FALSE | NA | NA | No Change | Not Detected | FALSE | Background = FALSE | Background = FALSE | Background = FALSE | ENSMUSG00000069911 | 574403 | protein_coding |
| Ints1 | TRUE | 0.313551424 | 0.000562777 | No Change | Significant Positive | TRUE | 0.129466685 | 0.46971763 | No Change | Not Significant | TRUE | Cond1 Significant Positive + Cond2 Not Significant | None | None | ENSMUSG00000029547 | 68510 | protein_coding |
| Ints10 | TRUE | 0.049234435 | 0.758091673 | No Change | Not Significant | TRUE | 0.194470829 | 0.164204113 | No Change | Not Significant | TRUE | Cond1 Not Significant + Cond2 Not Significant | None | None | ENSMUSG00000031864 | 70885 | protein_coding |
| Ints11 | TRUE | 0.521367499 | 2.41E-09 | UP | UP | TRUE | 0.339385247 | 2.57E-05 | No Change | Significant Positive | TRUE | Cond1 UP + Cond2 Significant Positive | Core_UP | Cond1_UP | ENSMUSG00000029034 | 71957 | protein_coding |
| Ints12 | TRUE | -0.00022356 | 0.998483602 | No Change | Not Significant | TRUE | -0.050984612 | 0.697134159 | No Change | Not Significant | TRUE | Cond1 Not Significant + Cond2 Not Significant | None | None | ENSMUSG00000028016 | 71793 | protein_coding |
| Ints13 | TRUE | 0.176621724 | 0.0542586 | No Change | Not Significant | TRUE | -0.031679953 | 0.781437157 | No Change | Not Significant | TRUE | Cond1 Not Significant + Cond2 Not Significant | None | None | ENSMUSG00000040250 | 71177 | protein_coding |
| Ints14 | TRUE | 0.168971093 | 0.17026467 | No Change | Not Significant | TRUE | 0.17462797 | 0.050384459 | No Change | Not Significant | TRUE | Cond1 Not Significant + Cond2 Not Significant | None | None | ENSMUSG00000034263 | 69882 | protein_coding |
| Ints15 | TRUE | 0.01951705 | 0.908146859 | No Change | Not Significant | TRUE | 0.005158947 | 0.983146794 | No Change | Not Significant | TRUE | Cond1 Not Significant + Cond2 Not Significant | None | None | ENSMUSG00000039244 | 231868 | protein_coding |
| Ints2 | TRUE | 0.449655412 | 0.001743841 | No Change | Significant Positive | TRUE | 0.378895052 | 0.005794729 | No Change | Significant Positive | TRUE | Cond1 Significant Positive + Cond2 Significant Positive | None | None | ENSMUSG00000018068 | 70422 | protein_coding |
| Ints3 | TRUE | 2.054884213 | 2.33E-133 | UP | UP | TRUE | 1.293673578 | 2.95E-54 | UP | UP | TRUE | Cond1 UP + Cond2 UP | Core_UP | Core_UP | ENSMUSG00000027933 | 229543 | protein_coding |
| Ints4 | TRUE | -0.018591758 | 0.919173117 | No Change | Not Significant | TRUE | 0.04328654 | 0.638789048 | No Change | Not Significant | TRUE | Cond1 Not Significant + Cond2 Not Significant | None | None | ENSMUSG00000025133 | 101861 | protein_coding |
| Ints5 | TRUE | 0.13078074 | 0.432354642 | No Change | Not Significant | TRUE | 0.057919582 | 0.762042164 | No Change | Not Significant | TRUE | Cond1 Not Significant + Cond2 Not Significant | None | None | ENSMUSG00000071652 | 109077 | protein_coding |

| geneID | Cond1_DF_Detected | Cond1_DF_ValueCol | Cond1_DF_PadjCol | Cond1_DF_Cutoff | Cond1_DF_Cutoff_Specific.x | Cond2_DF_Detected | Cond2_DF_ValueCol |
|---|---|---|---|---|---|---|---|
| Ints6 | TRUE | 0.364096697 | 0.002879422 | No Change | Significant Positive | TRUE | 0.109920317 |
| Ints6l | TRUE | -0.06757028 | 0.673998628 | No Change | Not Significant | TRUE | -0.138499313 |
| Ints7 | TRUE | 0.030598111 | 0.879513202 | No Change | Not Significant | TRUE | -0.016873425 |
| Ints8 | TRUE | 0.160158267 | 0.19557897 | No Change | Not Significant | TRUE | 0.005716057 |
| Ints9 | TRUE | -0.00715615 | 0.970043107 | No Change | Not Significant | TRUE | 0.01995759 |
| Intu | TRUE | 0.396808192 | 0.019107412 | No Change | Significant Positive | TRUE | -0.011134488 |
| Invs | TRUE | 0.655704547 | 7.71E-06 | UP | UP | TRUE | 0.415247656 |
| Ip6k1 | TRUE | 0.00502864 | 0.973795719 | No Change | Not Significant | TRUE | -0.056873558 |
| Ip6k2 | TRUE | -0.17071889 | 0.124671766 | No Change | Not Significant | TRUE | -0.252361722 |
| Ip6k3 | TRUE | -0.945389086 | 1 | No Change | Not Significant | FALSE | NA |
| Ipcef1 | TRUE | 0.447123242 | 0.407579707 | No Change | Not Significant | TRUE | 0.359702007 |
| Ipmk | TRUE | -0.185342078 | 0.196079582 | No Change | Not Significant | TRUE | 0.031227868 |
| Ipo11 | TRUE | 0.108071343 | 0.583719287 | No Change | Not Significant | TRUE | 0.019804962 |
| Ipo13 | TRUE | 0.066844711 | 0.67261003 | No Change | Not Significant | TRUE | 0.124150206 |
| Ipo4 | TRUE | 0.033673705 | 0.898463393 | No Change | Not Significant | TRUE | -0.156223129 |
| Ipo5 | TRUE | 0.947801492 | 2.41E-11 | UP | UP | TRUE | 0.509698445 |
| Ipo7 | TRUE | 0.057835365 | 0.786311623 | No Change | Not Significant | TRUE | -0.019951062 |
| Ipo8 | TRUE | -0.010924571 | 0.938630585 | No Change | Not Significant | TRUE | 0.061199586 |
| Ipo9 | TRUE | 0.293867801 | 0.094754639 | No Change | Not Significant | TRUE | 0.037352307 |
| Ipp | TRUE | -0.089662617 | 0.679601393 | No Change | Not Significant | TRUE | 0.011387126 |
| Ippk | TRUE | 0.128256942 | 0.546769252 | No Change | Not Significant | TRUE | 0.15497281 |
| [illegible] | [illegible] | [illegible] | [illegible] | [illegible] | [illegible] | [illegible] | [illegible] |

[illegible]

| geneID | Cond1_DF_Detected | Cond1_DF_ValueCol | Cond1_DF_PadjCol | Cond1_DF_Cutoff | Cond1_DF_Cutoff_Specific.x | Cond2_DF_Detected | Cond2_DF_ValueCol |
|---|---|---|---|---|---|---|---|
| Jpt1 | TRUE | -0.238362701 | 0.008896486 | No Change | Significant Negative | TRUE | -0.003117462 |
| Jpt2 | TRUE | 1.080039887 | 3.12E-21 | UP | UP | TRUE | 0.663832109 |
| Jpx | TRUE | -0.408847483 | 0.337678102 | No Change | Not Significant | TRUE | -0.287999148 |
| Jrk | TRUE | 0.008922725 | 0.977698054 | No Change | Not Significant | TRUE | 0.071462298 |
| Jrkl | TRUE | -0.315356696 | 0.017779903 | No Change | Significant Negative | TRUE | -0.080089178 |
| Jtb | TRUE | 0.019543598 | 0.930374388 | No Change | Not Significant | TRUE | 0.059848798 |
| Jun | TRUE | -0.101203975 | 0.60160505 | No Change | Not Significant | TRUE | 0.292267112 |
| Junb | TRUE | -0.355331694 | 0.100208451 | No Change | Not Significant | TRUE | -0.053110371 |
| Jund | TRUE | -0.062907297 | 0.825194181 | No Change | Not Significant | TRUE | 0.151066047 |
| Junos | TRUE | 0.036147875 | 0.9624373 | No Change | Not Significant | TRUE | -0.698578487 |
| Jup | TRUE | 0.04446186 | 0.821350533 | No Change | Not Significant | TRUE | 0.265620437 |
| Kalrn | TRUE | -0.339402219 | 0.084106216 | No Change | Not Significant | TRUE | -0.111839386 |
| Kank1 | TRUE | 0.272754803 | 0.112911971 | No Change | Not Significant | TRUE | 0.010813502 |
| Kank2 | TRUE | -0.54753588 | 0.009812883 | DOWN | DOWN | TRUE | -0.203833612 |
| Kank3 | TRUE | 0.163912064 | 0.624177234 | No Change | Not Significant | TRUE | -0.147344487 |
| Kank4 | TRUE | -0.691700084 | 0.402386658 | No Change | Not Significant | TRUE | -0.991689548 |
| Kank4os | TRUE | -1.075519595 | 1 | No Change | Not Significant | FALSE | NA |
| Kansl1 | TRUE | -0.113013012 | 0.305611042 | No Change | Not Significant | TRUE | -0.05949535 |
| Kansl1l | TRUE | -0.183283717 | 0.344285818 | No Change | Not Significant | TRUE | -0.259778473 |
| Kansl2 | TRUE | -0.033778157 | 0.885478012 | No Change | Not Significant | TRUE | -0.077391313 |
| Kansl3 | TRUE | -0.13706659 | 0.218595521 | No Change | Not Significant | TRUE | -0.130822449 |
| Kars | TRUE | -0.058813531 | 0.668257706 | No Change | Not Significant | TRUE | -0.075949448 |
| Kash5 | TRUE | -0.921523872 | 1 | No Change | Not Significant | FALSE | NA |
| Kat14 | TRUE | -0.047428779 | 0.669150322 | No Change | Not Significant | TRUE | -0.049776441 |
| Kat2a | TRUE | -0.214922585 | 0.139132959 | No Change | Not Significant | TRUE | -0.123568916 |
| Kat2b | TRUE | -0.090176727 | 0.53655334 | No Change | Not Significant | TRUE | -0.13994434 |
| Kat5 | TRUE | -0.069826651 | 0.607636589 | No Change | Not Significant | TRUE | -0.087631631 |
| Kat6a | TRUE | 0.049874132 | 0.768458962 | No Change | Not Significant | TRUE | -0.130026056 |
| Kat6b | TRUE | 0.41901964 | 7.11E-05 | No Change | Significant Positive | TRUE | 0.115898952 |
| Kat7 | TRUE | 0.143916953 | 0.070067307 | No Change | Not Significant | TRUE | -0.085793414 |
| Kat8 | TRUE | 0.102874521 | 0.438111017 | No Change | Not Significant | TRUE | 0.033839713 |
| Katna1 | TRUE | -0.056398034 | 0.64211714 | No Change | Not Significant | TRUE | 0.043104397 |
| Katnal1 | TRUE | 0.608407839 | 0.001255918 | UP | UP | TRUE | 0.4844129 |
| Katnal2 | TRUE | -0.140294461 | 0.915204382 | No Change | Not Significant | TRUE | -0.267605219 |
| Katnb1 | TRUE | 0.263184848 | 0.01219993 | No Change | Significant Positive | TRUE | 0.078012329 |
| Katnbl1 | TRUE | 0.474629343 | 0.000518229 | No Change | Significant Positive | TRUE | 0.060025729 |
| Katnip | TRUE | 0.008035023 | 0.937764615 | No Change | Not Significant | TRUE | -0.001481274 |
| Kazald1 | TRUE | -0.893457257 | 0.007657016 | DOWN | DOWN | TRUE | -0.352325998 |
| Kazn | TRUE | -0.496478459 | 0.003474964 | No Change | Significant Negative | TRUE | -0.379156089 |
| Kbtbd11 | TRUE | -0.081374952 | 0.719129457 | No Change | Not Significant | TRUE | 0.202599089 |
| Kbtbd12 | TRUE | 0.734030631 | 1 | No Change | Not Significant | FALSE | NA |
| Kbtbd13 | TRUE | 0.312956657 | 0.321034822 | No Change | Not Significant | TRUE | 0.076539808 |
| Kbtbd2 | TRUE | -0.160570809 | 0.174819917 | No Change | Not Significant | TRUE | -0.173198131 |
| Kbtbd3 | TRUE | 0.194964953 | 0.394941012 | No Change | Not Significant | TRUE | 0.012015116 |
| Kbtbd4 | TRUE | -0.166016781 | 0.440188363 | No Change | Not Significant | TRUE | 0.031267549 |
| Kbtbd6 | TRUE | 1.125895724 | 0.066324156 | No Change | Not Significant | TRUE | -0.240437886 |
| Kbtbd7 | TRUE | -0.087578661 | 0.612475705 | No Change | Not Significant | TRUE | -0.034191505 |
| Kbtbd8 | TRUE | 0.274713722 | 0.148948255 | No Change | Not Significant | TRUE | 0.230791347 |
| Kcmf1 | TRUE | 0.013754272 | 0.909752305 | No Change | Not Significant | TRUE | -0.052062153 |
| Kcna1 | TRUE | 0.176019553 | 1 | No Change | Not Significant | FALSE | NA |
| Kcna2 | TRUE | 2.991693874 | 1 | No Change | Not Significant | FALSE | NA |
| Kcna3 | TRUE | 1.358778294 | 1 | No Change | Not Significant | FALSE | NA |
| Kcna4 | TRUE | 1.064627075 | 1 | No Change | Not Significant | FALSE | NA |
| Kcna6 | TRUE | -0.751719705 | 1 | No Change | Not Significant | FALSE | NA |
| Kcna7 | TRUE | -0.329275573 | 1 | No Change | Not Significant | FALSE | NA |
| Kcnab1 | TRUE | 0.87976944 | 0.106381349 | No Change | Not Significant | TRUE | 0.586018203 |
| Kcnab2 | TRUE | 0.285639032 | 0.818817302 | No Change | Not Significant | TRUE | 0.40604036 |
| Kcnab3 | TRUE | -0.020054486 | 0.986602999 | No Change | Not Significant | TRUE | 0.157340694 |
| Kcnb1 | TRUE | -0.787476332 | 0.332667168 | No Change | Not Significant | TRUE | 0.054413892 |
| Kcnb2 | TRUE | -0.800238989 | 0.377007156 | No Change | Not Significant | TRUE | -0.206378319 |
| Kcnc1 | TRUE | -0.351774051 | 0.798287804 | No Change | Not Significant | TRUE | 0.521330091 |
| Kcnc2 | TRUE | 0.960275758 | 1 | No Change | Not Significant | FALSE | NA |
| Kcnc3 | TRUE | 3.261193207 | 1.18E-14 | UP | UP | TRUE | 1.078932153 |
| Kcnc4 | TRUE | 1.771908437 | 1 | No Change | Not Significant | FALSE | NA |
| Kcnd1 | TRUE | -0.56331929 | 0.548127746 | No Change | Not Significant | TRUE | -0.098257683 |
| Kcnd3 | TRUE | -0.298957849 | 0.796333696 | No Change | Not Significant | TRUE | 0.065195752 |
| Kcne1 | TRUE | 0.313603331 | 1 | No Change | Not Significant | FALSE | NA |
| Kcne1l | TRUE | -1.333296389 | 1 | No Change | Not Significant | FALSE | NA |
| Kcne2 | TRUE | -0.530592737 | 0.50295958 | No Change | Not Significant | TRUE | 0.325804032 |
| Kcne3 | TRUE | -0.326633126 | 0.258145624 | No Change | Not Significant | TRUE | -0.203696027 |
| Kcne4 | TRUE | -0.538873187 | 1 | No Change | Not Significant | FALSE | NA |
| Kcnf1 | TRUE | -0.674251787 | 0.012765795 | DOWN | DOWN | TRUE | -0.158476245 |
| Kcng2 | TRUE | -0.883927013 | 1 | No Change | Not Significant | FALSE | NA |
| Kcng3 | TRUE | -0.005321584 | 1 | No Change | Not Significant | TRUE | -1.198759677 |
| Kcng4 | TRUE | -0.421410452 | 1 | No Change | Not Significant | FALSE | NA |
| Kcnh1 | TRUE | -0.209118626 | 0.810851996 | No Change | Not Significant | TRUE | -0.303119788 |
| Kcnh2 | TRUE | -0.03043773 | 0.948996349 | No Change | Not Significant | TRUE | 0.056550166 |
| Kcnh3 | TRUE | -0.032926179 | 0.9092783 | No Change | Not Significant | TRUE | -0.509217299 |
| Kcnh4 | TRUE | 0.89565282 | 1 | No Change | Not Significant | FALSE | NA |
| Kcnh5 | TRUE | 1.189865565 | 1 | No Change | Not Significant | FALSE | NA |
| Kcnh6 | TRUE | 0.233619655 | 0.454872025 | No Change | Not Significant | TRUE | 0.021149409 |
| Kcnh7 | TRUE | -0.835632275 | 0.112936062 | No Change | Not Significant | TRUE | -0.270247284 |
| Kcnh8 | TRUE | -0.460373834 | 0.527698391 | No Change | Not Significant | TRUE | 0.100303006 |
| Kcnip1 | TRUE | 1.170568849 | 1 | No Change | Not Significant | FALSE | NA |
| Kcnip2 | TRUE | -0.279023329 | 0.858315628 | No Change | Not Significant | FALSE | NA |
| Kcnip3 | TRUE | -0.884593993 | 1 | No Change | Not Significant | FALSE | NA |
| Kcnip4 | TRUE | -0.284707939 | 0.766020418 | No Change | Not Significant | TRUE | -0.359922276 |
| Kcnj10 | TRUE | 1.8966475 | 1 | No Change | Not Significant | FALSE | NA |
| Kcnj11 | TRUE | -0.335156614 | 0.700180271 | No Change | Not Significant | TRUE | -0.222113906 |
| Kcnj13 | TRUE | -0.566974342 | 0.41130117 | No Change | Not Significant | TRUE | -0.46486282 |
| Kcnj14 | TRUE | 0.925178029 | 1 | No Change | Not Significant | FALSE | NA |
| Kcnj15 | TRUE | -1.187286212 | 1 | No Change | Not Significant | FALSE | NA |
| Kcnj16 | TRUE | -1.082939246 | 1 | No Change | Not Significant | FALSE | NA |
| Kcnj2 | TRUE | -0.77304442 | 0.06221406 | No Change | Not Significant | TRUE | -0.39671481 |
| Kcnj3 | TRUE | 0.006802836 | 1 | No Change | Not Significant | FALSE | NA |
| Kcnj5 | TRUE | 0.145207187 | 1 | No Change | Not Significant | FALSE | NA |
| Kcnj6 | TRUE | 1.651940298 | 1 | No Change | Not Significant | FALSE | NA |
| Kcnj8 | TRUE | -3.385334928 | 1 | No Change | Not Significant | TRUE | 0.625631054 |
| Kcnj9 | TRUE | 2.856955964 | 1 | No Change | Not Significant | FALSE | NA |
| Kcnk1 | TRUE | 0.242280872 | 0.049543507 | No Change | Significant Positive | TRUE | 0.276480184 |
| Kcnk10 | TRUE | -1.477889475 | 0.017169019 | DOWN | DOWN | TRUE | 0.739917 |
| Kcnk12 | TRUE | -0.688441877 | 1 | No Change | Not Significant | FALSE | NA |
| Kcnk13 | TRUE | 0.319385866 | 1 | No Change | Not Significant | FALSE | NA |
| Kcnk16 | TRUE | 0.643236407 | 0.587561035 | No Change | Not Significant | TRUE | 0.038944953 |
| Kcnk18 | TRUE | 2.851253078 | 1 | No Change | Not Significant | FALSE | NA |
| Kcnk2 | TRUE | 3.766619529 | 1 | No Change | Not Significant | FALSE | NA |
| Kcnk3 | TRUE | -0.586067569 | 0.465046296 | No Change | Not Significant | TRUE | 0.066187515 |
| Kcnk4 | TRUE | 0.120662689 | 0.907147741 | No Change | Not Significant | TRUE | 0.099607311 |
| Kcnk5 | TRUE | -0.496678342 | 0.000114755 | No Change | Significant Negative | TRUE | -0.033851234 |
| Kcnk6 | TRUE | 0.201257376 | 0.1706035 | No Change | Not Significant | TRUE | -0.040081954 |
| Kcnk7 | TRUE | -0.788808758 | 0.23910316 | No Change | Not Significant | TRUE | -0.122773412 |
| Kcnk9 | TRUE | 0.884268093 | 1 | No Change | Not Significant | FALSE | NA |
| Kcnma1 | TRUE | -0.421793642 | 0.151098 | No Change | Not Significant | TRUE | 0.169343947 |
| Kcnmb1 | TRUE | 2.500739607 | 1 | No Change | Not Significant | FALSE | NA |
| Kcnmb2 | TRUE | -0.895929959 | 1 | No Change | Not Significant | FALSE | NA |
| Kcnmb3 | TRUE | 0.124025873 | 1 | No Change | Not Significant | FALSE | NA |
| Kcnn1 | TRUE | -0.57921561 | 0.416114999 | No Change | Not Significant | TRUE | -0.3070875 |
| Kcnn2 | TRUE | -3.048209015 | 1 | No Change | Not Significant | FALSE | NA |
| Kcnn3 | TRUE | 0.008668059 | 1 | No Change | Not Significant | FALSE | NA |
| Kcnn4 | TRUE | -0.165637367 | 0.498301136 | No Change | Not Significant | TRUE | -0.017483453 |
| Kcnq1 | TRUE | -0.909804822 | 1.44E-19 | DOWN | DOWN | TRUE | -0.445671048 |
| Kcnq2 | TRUE | 1.47574415 | 0.561998621 | No Change | Not Significant | FALSE | NA |
| Kcnq3 | TRUE | 1.134495652 | 0.747162726 | No Change | Not Significant | FALSE | NA |
| Kcnq4 | TRUE | -1.113046098 | 0.259548816 | No Change | Not Significant | TRUE | -0.415418645 |
| Kcnq5 | TRUE | -1.500964582 | 1 | No Change | Not Significant | FALSE | NA |
| Kcnrg | TRUE | 0.011449021 | 1 | No Change | Not Significant | FALSE | NA |
| Kcns1 | TRUE | 1.823164588 | 1 | No Change | Not Significant | FALSE | NA |
| Kcns2 | TRUE | 0.702092941 | 1 | No Change | Not Significant | FALSE | NA |
| Kcns3 | TRUE | -1.838433926 | 1 | No Change | Not Significant | FALSE | NA |
| Kcnt1 | TRUE | 0.405521822 | 0.849668102 | No Change | Not Significant | FALSE | NA |
| Kcnt2 | TRUE | 0.835195026 | 1 | No Change | Not Significant | FALSE | NA |
| Kcnu1 | TRUE | -0.059571653 | 1 | No Change | Not Significant | TRUE | -0.687142161 |
| Kcnv1 | TRUE | 1.138682638 | 1 | No Change | Not Significant | FALSE | NA |
| Kcnv2 | TRUE | 0.552219762 | 0.499562412 | No Change | Not Significant | TRUE | 0.389193761 |
| Kcp | TRUE | 0.205546605 | 1 | No Change | Not Significant | FALSE | NA |
| Kctd1 | TRUE | -0.081790374 | 0.797713998 | No Change | Not Significant | TRUE | 0.215621352 |
| Kctd10 | TRUE | -0.074801162 | 0.592567182 | No Change | Not Significant | TRUE | 0.031072311 |
| Kctd11 | TRUE | -0.114094265 | 0.644012315 | No Change | Not Significant | TRUE | -0.091725622 |
| Kctd12 | TRUE | 0.252332132 | 0.40082018 | No Change | Not Significant | TRUE | 0.260736375 |
| Kctd12b | TRUE | -0.855116095 | 1 | No Change | Not Significant | TRUE | 0.398472642 |
| Kctd13 | TRUE | 0.140077824 | 0.60964446 | No Change | Not Significant | TRUE | -0.197635321 |
| Kctd14 | TRUE | 1.405995648 | 5.38E-18 | UP | UP | TRUE | 0.85650526 |
| Kctd15 | TRUE | -0.346411048 | 0.507895701 | No Change | Not Significant | TRUE | 0.087213379 |
| Kctd17 | TRUE | -0.298910656 | 0.158658236 | No Change | Not Significant | TRUE | -0.09171887 |
| Kctd18 | TRUE | -0.010049174 | 0.973795719 | No Change | Not Significant | TRUE | -0.138612501 |
| Kctd2 | TRUE | -0.048639268 | 0.756249943 | No Change | Not Significant | TRUE | -0.061334118 |
| Kctd20 | TRUE | 0.008159648 | 0.942360409 | No Change | Not Significant | TRUE | -0.101274469 |
| Kctd21 | TRUE | -0.50988532 | 0.002879422 | DOWN | DOWN | TRUE | -0.092569024 |
| Kctd3 | TRUE | 0.049878576 | 0.630552252 | No Change | Not Significant | TRUE | 0.001617417 |
| Kctd4 | TRUE | 0.118120261 | 0.874455706 | No Change | Not Significant | TRUE | 0.183428289 |
| Kctd5 | TRUE | 0.050565555 | 0.68930064 | No Change | Not Significant | TRUE | -0.009399514 |
| Kctd6 | TRUE | -0.212458934 | 0.183322799 | No Change | Not Significant | TRUE | -0.215165785 |
| Kctd7 | TRUE | 1.034920894 | 4.55E-10 | UP | UP | TRUE | 0.274885524 |
| Kctd9 | TRUE | 0.366007939 | 0.004051748 | No Change | Significant Positive | TRUE | 0.067163797 |
| Kdelr1 | TRUE | 0.137326653 | 0.270251212 | No Change | Not Significant | TRUE | 0.099356678 |
| Kdelr2 | TRUE | 0.60805151 | 1.05E-08 | UP | UP | TRUE | 0.349224024 |
| Kdelr3 | TRUE | 0.926542015 | 9.86E-09 | UP | UP | TRUE | 0.095123861 |
| Kdf1 | TRUE | -0.007706598 | 0.959486477 | No Change | Not Significant | TRUE | 0.061186235 |
| Kdm1a | TRUE | -0.000564145 | 0.995389267 | No Change | Not Significant | TRUE | -0.036233151 |
| Kdm1b | TRUE | -0.3057676 | 0.028864834 | No Change | Significant Negative | TRUE | -0.068791784 |
| Kdm2a | TRUE | -0.082828568 | 0.348151449 | No Change | Not Significant | TRUE | -0.089078511 |
| Kdm2b | TRUE | -0.184530422 | 0.139530465 | No Change | Not Significant | TRUE | 0.002259943 |
| Kdm3a | TRUE | -0.040945869 | 0.760164897 | No Change | Not Significant | TRUE | -0.118176142 |
| Kdm3b | TRUE | 0.650945843 | 6.21E-31 | UP | UP | TRUE | 0.402366757 |
| Kdm4a | TRUE | 0.218674428 | 0.009910776 | No Change | Significant Positive | TRUE | 0.069584601 |
| Kdm4b | TRUE | 0.646384095 | 4.45E-18 | UP | UP | TRUE | 0.15073448 |
| Kdm4c | TRUE | -0.008731003 | 0.948996349 | No Change | Not Significant | TRUE | -0.071109199 |
| Kdm4d | TRUE | 1.18001425 | 1 | No Change | Not Significant | FALSE | NA |
| Kdm5a | TRUE | -0.172566555 | 0.015940335 | No Change | Significant Negative | TRUE | -0.227111174 |
| Kdm5b | TRUE | -0.20861463 | 0.011209933 | No Change | Significant Negative | TRUE | -0.036929266 |
| Kdm5c | TRUE | 0.105344485 | 0.255796319 | No Change | Not Significant | TRUE | 0.123891695 |
| Kdm6a | TRUE | 0.564064706 | 2.07E-17 | UP | UP | TRUE | 0.167995903 |
| Kdm6b | TRUE | 0.214704055 | 0.654124055 | No Change | Not Significant | TRUE | -0.315152489 |
| Kdm7a | TRUE | 0.022447067 | 0.898891871 | No Change | Not Significant | TRUE | -0.056367844 |
| Kdm8 | TRUE | 0.087563344 | 0.800362194 | No Change | Not Significant | TRUE | -0.017532569 |
| Kdr | TRUE | -0.105236747 | 1 | No Change | Not Significant | FALSE | NA |
| Kdsr | TRUE | -0.4314308 | 0.003915665 | No Change | Significant Negative | TRUE | -0.10234948 |
| Keap1 | TRUE | 0.226855078 | 0.00627754 | No Change | Significant Positive | TRUE | 0.175568951 |
| Keg1 | TRUE | 0.604372832 | 1 | No Change | Not Significant | FALSE | NA |
| Kel | TRUE | 1.917557068 | 1 | No Change | Not Significant | FALSE | NA |
| Khdc1a | TRUE | -1.588670808 | 1 | No Change | Not Significant | FALSE | NA |
| Khdc3 | TRUE | -0.378473653 | 1 | No Change | Not Significant | FALSE | NA |
| Khdc4 | TRUE | 0.060384967 | 0.781739336 | No Change | Not Significant | TRUE | -0.219475561 |
| Khdrbs1 | TRUE | 0.158088623 | 0.098226787 | No Change | Not Significant | TRUE | -0.0472111 |
| Khdrbs2 | TRUE | 1.257611701 | 1 | No Change | Not Significant | FALSE | NA |
| Khdrbs3 | TRUE | 0.378545026 | 0.031096942 | No Change | Significant Positive | TRUE | 0.42658384 |
| Khk | TRUE | -0.412748484 | 0.007552486 | No Change | Significant Negative | TRUE | -0.190023593 |
| Khnyn | TRUE | -0.383377027 | 0.000440521 | No Change | Significant Negative | TRUE | -0.097886701 |
| Khsrp | TRUE | 0.983493967 | 1.55E-22 | UP | UP | TRUE | 0.332061094 |
| Kics2 | TRUE | -0.042626835 | 0.907147741 | No Change | Not Significant | TRUE | -0.044203723 |
| Kidins220 | TRUE | -0.255963493 | 0.000692995 | No Change | Significant Negative | TRUE | -0.154182626 |
| Kif11 | TRUE | 0.66976925 | 0.003254551 | UP | UP | TRUE | 0.054493777 |

Supplementary Table 3

| geneID | Cond1_DF_Detected | Cond1_DF_ValueCol | Cond1_DF_PadjCol | Cond1_DF_Cutoff | Cond1_DF_Cutoff_Specific.x | Cond2_DF_Detected | Cond2_DF_ValueCol | Cond2_DF_PadjCol | Cond2_DF_Cutoff | Cond2_DF_Cutoff_Specific | BG_Method |
|---|---|---|---|---|---|---|---|---|---|---|---|
| Kif12 | TRUE | 0.227919154 | 0.17162699 | No Change | Not Significant | TRUE | -0.406568742 | 0.001851734 | No Change | Significant Negative | TRUE |
| Kif13a | TRUE | -0.178578875 | 0.20691859 | No Change | Not Significant | TRUE | -0.029680686 | 0.76019675 | No Change | Not Significant | TRUE |
| Kif13b | TRUE | -0.272240509 | 0.01219993 | No Change | Significant Negative | TRUE | 0.022107778 | 0.853575129 | No Change | Not Significant | TRUE |
| Kif14 | TRUE | 0.845498246 | 2.24E-05 | UP | UP | TRUE | -0.036555199 | 0.895076992 | No Change | Not Significant | TRUE |
| Kif15 | TRUE | 0.711403928 | 0.000197903 | UP | UP | TRUE | -0.127440581 | 0.591564986 | No Change | Not Significant | TRUE |
| Kif16b | TRUE | -0.082234385 | 0.401975042 | No Change | Not Significant | TRUE | -0.003532403 | 0.967733155 | No Change | Not Significant | TRUE |
| Kif17 | TRUE | 0.951424786 | 0.462658024 | No Change | Not Significant | TRUE | 0.581979308 | 0.504916665 | No Change | Not Significant | TRUE |
| Kif18a | TRUE | 0.620899531 | 0.006350049 | UP | UP | TRUE | 0.076811704 | 0.789870733 | No Change | Not Significant | TRUE |
| Kif18b | TRUE | 0.739049054 | 0.000642047 | UP | UP | TRUE | 0.052499901 | 0.849391002 | No Change | Not Significant | TRUE |
| Kif19a | TRUE | -0.16672846 | 0.879422681 | No Change | Not Significant | TRUE | 0.117315802 | 0.836813383 | No Change | Not Significant | TRUE |
| Kif19b | TRUE | -1.048103914 | 1 | No Change | Not Significant | FALSE | NA | NA | No Change | Not Detected | FALSE |
| Kif1a | TRUE | -0.622793874 | 0.18810647 | No Change | Not Significant | TRUE | -0.041560283 | 0.930175172 | No Change | Not Significant | TRUE |
| Kif1b | TRUE | -0.100018059 | 0.270127107 | No Change | Not Significant | TRUE | -0.040112595 | 0.531747525 | No Change | Not Significant | TRUE |
| Kif1c | TRUE | -0.251947473 | 0.000334053 | No Change | Significant Negative | TRUE | -0.03726898 | 0.688774391 | No Change | Not Significant | TRUE |
| Kif20a | TRUE | 0.615902921 | 0.003200656 | UP | UP | TRUE | 0.018077877 | 0.93615636 | No Change | Not Significant | TRUE |
| Kif20b | TRUE | 0.730400234 | 0.000179112 | UP | UP | TRUE | 0.003796644 | 0.989174207 | No Change | Not Significant | TRUE |
| Kif21a | TRUE | 0.550300244 | 1.11E-06 | UP | UP | TRUE | 0.395257421 | 3.86E-05 | No Change | Significant Positive | TRUE |
| Kif21b | TRUE | 0.074173502 | 0.861253255 | No Change | Not Significant | TRUE | -0.248589799 | 0.040297853 | No Change | Significant Negative | TRUE |
| Kif22 | TRUE | 0.68239347 | 0.000563781 | UP | UP | TRUE | 0.139951825 | 0.378133941 | No Change | Not Significant | TRUE |
| Kif23 | TRUE | 0.651446664 | 8.87E-08 | UP | UP | TRUE | 0.073433885 | 0.653685981 | No Change | Not Significant | TRUE |
| Kif24 | TRUE | 0.79513207 | 2.64E-07 | UP | UP | TRUE | 0.199394599 | 0.31943494 | No Change | Not Significant | TRUE |
| Kif26a | TRUE | -1.175628621 | 0.400865192 | No Change | Not Significant | TRUE | -0.478300754 | 0.58240762 | No Change | Not Significant | TRUE |
| Kif26b | TRUE | -0.054861531 | 0.94900212 | No Change | Not Significant | TRUE | -0.335577312 | 0.455552165 | No Change | Not Significant | TRUE |
| Kif27 | TRUE | -0.210378264 | 0.68833522 | No Change | Not Significant | TRUE | -0.556596446 | 0.10129029 | No Change | Not Significant | TRUE |
| Kif28 | TRUE | -1.313023083 | 1 | No Change | Not Significant | FALSE | NA | NA | No Change | Not Detected | FALSE |
| Kif2a | TRUE | -0.089172253 | 0.56607413 | No Change | Not Significant | TRUE | -0.213652714 | 0.137687746 | No Change | Not Significant | TRUE |
| Kif2c | TRUE | 0.59260171 | 0.000216159 | UP | UP | TRUE | 0.031950852 | 0.902120505 | No Change | Not Significant | TRUE |
| Kif3a | TRUE | 0.182899557 | 0.35368873 | No Change | Not Significant | TRUE | 0.220627219 | 0.129304242 | No Change | Not Significant | TRUE |
| Kif3b | TRUE | 0.084925827 | 0.418758859 | No Change | Not Significant | TRUE | 0.08634493 | 0.405210667 | No Change | Not Significant | TRUE |
| Kif3c | TRUE | -0.346237137 | 0.406017465 | No Change | Not Significant | TRUE | -0.106500614 | 0.831653764 | No Change | Not Significant | TRUE |
| Kif4 | TRUE | 0.924811695 | 7.84E-09 | UP | UP | TRUE | 0.155581567 | 0.423163427 | No Change | Not Significant | TRUE |
| Kif5a | TRUE | 0.030517113 | 0.988831409 | No Change | Not Significant | FALSE | NA | NA | No Change | Not Detected | FALSE |
| Kif5b | TRUE | -0.001154204 | 0.993636457 | No Change | Not Significant | TRUE | 0.032697984 | 0.697472342 | No Change | Not Significant | TRUE |
| Kif5c | TRUE | 0.750374352 | 0.042030578 | UP | UP | TRUE | 0.816721499 | 0.003024955 | UP | UP | TRUE |
| Kif6 | TRUE | 1.691944921 | 1 | No Change | Not Significant | FALSE | NA | NA | No Change | Not Detected | FALSE |
| Kif7 | TRUE | 1.454037969 | 1 | No Change | Not Significant | FALSE | NA | NA | No Change | Not Detected | FALSE |
| Kif9 | TRUE | -0.107841425 | 0.819432282 | No Change | Not Significant | TRUE | 0.273120825 | 0.357616116 | No Change | Not Significant | TRUE |
| Kifap3 | TRUE | 0.032579899 | 0.873772754 | No Change | Not Significant | TRUE | 0.119625983 | 0.258860279 | No Change | Not Significant | TRUE |
| Kifbp | TRUE | 0.031606639 | 0.878912418 | No Change | Not Significant | TRUE | 0.08837406 | 0.487862729 | No Change | Not Significant | TRUE |
| Kifc1 | TRUE | 0.612047629 | 0.005165779 | UP | UP | TRUE | 0.162405343 | 0.484922952 | No Change | Not Significant | TRUE |
| Kifc2 | TRUE | -0.459052278 | 0.207507849 | No Change | Not Significant | TRUE | -0.345711216 | 0.117338412 | No Change | Not Significant | TRUE |
| Kifc3 | TRUE | 0.351166145 | 0.013622626 | No Change | Significant Positive | TRUE | 0.172855222 | 0.118917792 | No Change | Not Significant | TRUE |
| Kifc5b | TRUE | 0.738020856 | 0.002303836 | UP | UP | TRUE | 0.084846283 | 0.800888833 | No Change | Not Significant | TRUE |
| Kin | TRUE | 0.055493116 | 0.760811885 | No Change | Not Significant | TRUE | 0.083375041 | 0.545539031 | No Change | Not Significant | TRUE |
| Kirrel | TRUE | 0.04203288 | 0.976632343 | No Change | Not Significant | TRUE | 0.053438532 | 0.961718903 | No Change | Not Significant | TRUE |
| Kirrel2 | TRUE | 2.006208597 | 0.177264714 | No Change | Not Significant | FALSE | NA | NA | No Change | Not Detected | FALSE |
| Kirrel3 | TRUE | -0.514034302 | 0.309791228 | No Change | Not Significant | TRUE | 0.008623825 | 0.992312996 | No Change | Not Significant | TRUE |
| Kiss1r | TRUE | -0.275686987 | 0.569548832 | No Change | Not Significant | TRUE | -0.028448351 | 0.950656978 | No Change | Not Significant | TRUE |
| Kit | TRUE | -0.452503741 | 0.152129476 | No Change | Not Significant | TRUE | -0.026094604 | 0.883289673 | No Change | Not Significant | TRUE |
| Kitl | TRUE | -0.093557679 | 0.647539027 | No Change | Not Significant | TRUE | -0.038491684 | 0.688630936 | No Change | Not Significant | TRUE |
| Kiz | TRUE | -0.252256733 | 0.102738771 | No Change | Not Significant | TRUE | -0.164569212 | 0.167845772 | No Change | Not Significant | TRUE |
| Kl | TRUE | 0.086975651 | 1 | No Change | Not Significant | FALSE | NA | NA | No Change | Not Detected | FALSE |
| Klb | TRUE | -0.342947575 | 0.344813186 | No Change | Not Significant | TRUE | -0.202890798 | 0.420477604 | No Change | Not Significant | TRUE |
| Klc1 | TRUE | -0.045307723 | 0.816921981 | No Change | Not Significant | TRUE | -0.117443155 | 0.279854223 | No Change | Not Significant | TRUE |
| Klc2 | TRUE | 0.081525814 | 0.680564322 | No Change | Not Significant | TRUE | -0.075142561 | 0.561627805 | No Change | Not Significant | TRUE |
| Klc3 | TRUE | 0.178600657 | 0.724255209 | No Change | Not Significant | TRUE | 0.147699894 | 0.707879718 | No Change | Not Significant | TRUE |
| Klc4 | TRUE | -0.068373152 | 0.715750413 | No Change | Not Significant | TRUE | 0.126646867 | 0.140437951 | No Change | Not Significant | TRUE |
| Klf1 | TRUE | -0.110290175 | 0.894720656 | No Change | Not Significant | TRUE | -0.125552594 | 0.880070575 | No Change | Not Significant | TRUE |
| Klf10 | TRUE | 0.155675479 | 0.429285587 | No Change | Not Significant | TRUE | 0.023942226 | 0.936612161 | No Change | Not Significant | TRUE |
| Klf11 | TRUE | -0.775805632 | 0.001007326 | DOWN | DOWN | TRUE | -0.47718221 | 0.063172661 | No Change | Not Significant | TRUE |
| Klf12 | TRUE | 1.067585736 | 1.10E-15 | UP | UP | TRUE | 0.355033085 | 0.000105419 | No Change | Significant Positive | TRUE |
| Klf13 | TRUE | -0.369223238 | 2.22E-09 | No Change | Significant Negative | TRUE | -0.269254844 | 4.70E-06 | No Change | Significant Negative | TRUE |
| Klf14 | TRUE | -0.431021368 | 1 | No Change | Not Significant | FALSE | NA | NA | No Change | Not Detected | FALSE |
| Klf15 | TRUE | -2.428406813 | 0.00014909 | DOWN | DOWN | TRUE | -0.969765632 | 0.001697944 | DOWN | DOWN | TRUE |
| Klf16 | TRUE | -0.376046997 | 0.008896486 | No Change | Significant Negative | TRUE | -0.221726392 | 0.189222442 | No Change | Not Significant | TRUE |
| Klf17 | TRUE | -1.594409391 | 1 | No Change | Not Significant | FALSE | NA | NA | No Change | Not Detected | FALSE |
| Klf2 | TRUE | -0.788038073 | 0.000128314 | DOWN | DOWN | TRUE | 0.12119691 | 0.61633656 | No Change | Not Significant | TRUE |
| Klf3 | TRUE | -0.109008839 | 0.398279073 | No Change | Not Significant | TRUE | -0.065828869 | 0.481643597 | No Change | Not Significant | TRUE |
| Klf4 | TRUE | 0.077110614 | 0.762250234 | No Change | Not Significant | TRUE | 0.066453915 | 0.602198495 | No Change | Not Significant | TRUE |
| Klf5 | TRUE | -0.015324527 | 0.875049502 | No Change | Not Significant | TRUE | -0.017488871 | 0.90019746 | No Change | Not Significant | TRUE |
| Klf6 | TRUE | -0.04401657 | 0.900012841 | No Change | Not Significant | TRUE | 0.298857665 | 0.000182422 | No Change | Significant Positive | TRUE |
| Klf7 | TRUE | 0.237652152 | 0.101661214 | No Change | Not Significant | TRUE | 0.093103615 | 0.44497561 | No Change | Not Significant | TRUE |
| Klf8 | TRUE | -0.775871472 | 0.175901555 | No Change | Not Significant | TRUE | -0.363291613 | 0.315962272 | No Change | Not Significant | TRUE |
| Klf9 | TRUE | -0.430382996 | 0.007195824 | No Change | Significant Negative | TRUE | -0.245819607 | 0.0085163 | No Change | Significant Negative | TRUE |
| Klhdc1 | TRUE | -0.085276132 | 0.826943172 | No Change | Not Significant | TRUE | -0.005265537 | 0.987977748 | No Change | Not Significant | TRUE |
| Klhdc10 | TRUE | 0.150762931 | 0.093703879 | No Change | Not Significant | TRUE | 0.060702343 | 0.336624075 | No Change | Not Significant | TRUE |
| Klhdc2 | TRUE | -0.089788555 | 0.570683233 | No Change | Not Significant | TRUE | -0.092955996 | 0.310570612 | No Change | Not Significant | TRUE |
| Klhdc3 | TRUE | -0.378738331 | 0.000886497 | No Change | Significant Negative | TRUE | -0.184756288 | 0.091458234 | No Change | Not Significant | TRUE |
| Klhdc4 | TRUE | 0.281296752 | 0.02899794 | No Change | Significant Positive | TRUE | 0.235082606 | 0.106976995 | No Change | Not Significant | TRUE |
| Klhdc7a | TRUE | 0.016376416 | 0.973044134 | No Change | Not Significant | TRUE | 0.570171318 | 1.75E-06 | UP | UP | TRUE |
| Klhdc7b | TRUE | -1.011203112 | 1 | No Change | Not Significant | FALSE | NA | NA | No Change | Not Detected | FALSE |
| Klhdc8a | TRUE | -0.45916776 | 0.011271717 | No Change | Significant Negative | TRUE | -0.026957058 | 0.912727084 | No Change | Not Significant | TRUE |
| Klhdc8b | TRUE | -0.132740124 | 0.737121023 | No Change | Not Significant | TRUE | -0.053245894 | 0.880475742 | No Change | Not Significant | TRUE |
| Klhdc9 | TRUE | -0.23019347 | 0.760164897 | No Change | Not Significant | TRUE | -0.596048778 | 0.17246929 | No Change | Not Significant | TRUE |
| Klhl1 | TRUE | 1.310104526 | 1 | No Change | Not Significant | FALSE | NA | NA | No Change | Not Detected | FALSE |
| Klhl10 | TRUE | 0.087126778 | 1 | No Change | Not Significant | FALSE | NA | NA | No Change | Not Detected | FALSE |
| Klhl11 | TRUE | 0.185147526 | 0.172486883 | No Change | Not Significant | TRUE | -0.14405373 | 0.269714697 | No Change | Not Significant | TRUE |
| Klhl12 | TRUE | -0.099216367 | 0.409247045 | No Change | Not Significant | TRUE | -0.007601445 | 0.952805789 | No Change | Not Significant | TRUE |
| Klhl13 | TRUE | 0.449052102 | 0.270127107 | No Change | Not Significant | TRUE | 0.517811214 | 0.039352773 | UP | UP | TRUE |
| Klhl14 | TRUE | 1.049548395 | 1 | No Change | Not Significant | FALSE | NA | NA | No Change | Not Detected | FALSE |
| Klhl15 | TRUE | 0.022099092 | 0.931580268 | No Change | Not Significant | TRUE | -0.037759081 | 0.803018158 | No Change | Not Significant | TRUE |
| Klhl18 | TRUE | -0.137468516 | 0.196025521 | No Change | Not Significant | TRUE | 0.071012502 | 0.516979903 | No Change | Not Significant | TRUE |
| Klhl2 | TRUE | 0.012077255 | 0.908985828 | No Change | Not Significant | TRUE | 0.097644546 | 0.276314934 | No Change | Not Significant | TRUE |
| Klhl20 | TRUE | -0.047506375 | 0.77804505 | No Change | Not Significant | TRUE | -0.163372887 | 0.207235951 | No Change | Not Significant | TRUE |
| Klhl21 | TRUE | 0.756443518 | 1.68E-11 | UP | UP | TRUE | 0.22035765 | 0.035957058 | No Change | Significant Positive | TRUE |
| Klhl22 | TRUE | 0.06140126 | 0.653468095 | No Change | Not Significant | TRUE | 0.103392827 | 0.160521685 | No Change | Not Significant | TRUE |
| Klhl23 | TRUE | 0.44787949 | 0.207184962 | No Change | Not Significant | TRUE | 0.199990567 | 0.407223247 | No Change | Not Significant | TRUE |
| Klhl24 | TRUE | -0.491702114 | 0.08602548 | No Change | Not Significant | TRUE | -0.28106681 | 0.027013876 | No Change | Significant Negative | TRUE |
| Klhl25 | TRUE | 0.144607202 | 0.260075004 | No Change | Not Significant | TRUE | 0.107219315 | 0.395821216 | No Change | Not Significant | TRUE |
| Klhl26 | TRUE | -0.186132205 | 0.164355907 | No Change | Not Significant | TRUE | -0.214311169 | 0.042920997 | No Change | Significant Negative | TRUE |
| Klhl28 | TRUE | -0.1613404 | 0.406554718 | No Change | Not Significant | TRUE | -0.243486615 | 0.038300296 | No Change | Significant Negative | TRUE |
| Klhl29 | TRUE | 0.936787367 | 1 | No Change | Not Significant | FALSE | NA | NA | No Change | Not Detected | FALSE |
| Klhl3 | TRUE | -0.448341885 | 0.19999509 | No Change | Not Significant | TRUE | 0.133323979 | 0.708993627 | No Change | Not Significant | TRUE |
| Klhl30 | TRUE | 0.474833706 | 1 | No Change | Not Significant | FALSE | NA | NA | No Change | Not Detected | FALSE |
| Klhl31 | TRUE | 3.305540366 | 1.96E-05 | UP | UP | FALSE | NA | NA | No Change | Not Detected | FALSE |
| Klhl32 | TRUE | -0.23024918 | 1 | No Change | Not Significant | FALSE | NA | NA | No Change | Not Detected | FALSE |
| Klhl33 | TRUE | -0.379301387 | 1 | No Change | Not Significant | TRUE | 0.92528482 | 0.251280519 | No Change | Not Significant | TRUE |
| Klhl35 | TRUE | 0.333499315 | 0.750140815 | No Change | Not Significant | TRUE | -0.164183891 | 0.864654909 | No Change | Not Significant | TRUE |
| Klhl36 | TRUE | -0.41268764 | 0.00706591 | No Change | Significant Negative | TRUE | 0.005403564 | 0.984481423 | No Change | Not Significant | TRUE |
| Klhl4 | TRUE | -0.460517273 | 1 | No Change | Not Significant | FALSE | NA | NA | No Change | Not Detected | FALSE |
| Klhl40 | TRUE | 2.755708408 | 1 | No Change | Not Significant | FALSE | NA | NA | No Change | Not Detected | FALSE |
| Klhl41 | TRUE | -0.057988634 | 1 | No Change | Not Significant | FALSE | NA | NA | No Change | Not Detected | FALSE |
| Klhl42 | TRUE | -0.090454789 | 0.633873724 | No Change | Not Significant | TRUE | -0.078450845 | 0.556994886 | No Change | Not Significant | TRUE |
| Klhl5 | TRUE | 0.180503304 | 0.424858107 | No Change | Not Significant | TRUE | -0.028223626 | 0.892332662 | No Change | Not Significant | TRUE |
| Klhl6 | TRUE | 0.212325548 | 1 | No Change | Not Significant | TRUE | 0.952651089 | 0.117491746 | No Change | Not Significant | TRUE |
| Klhl7 | TRUE | 0.015357832 | 0.954353007 | No Change | Not Significant | TRUE | 0.100583068 | 0.395074475 | No Change | Not Significant | TRUE |
| Klhl8 | TRUE | 0.206305236 | 0.531259196 | No Change | Not Significant | TRUE | -0.029332831 | 0.910121497 | No Change | Not Significant | TRUE |
| Klhl9 | TRUE | 0.035973012 | 0.807443873 | No Change | Not Significant | TRUE | 0.05605799 | 0.431310511 | No Change | Not Significant | TRUE |
| Klk1 | TRUE | 0.867460145 | 3.26E-08 | UP | UP | TRUE | 0.932773909 | 6.22E-25 | UP | UP | TRUE |
| Klk10 | TRUE | 0.491312364 | 1 | No Change | Not Significant | FALSE | NA | NA | No Change | Not Detected | FALSE |
| Klk11 | TRUE | 2.226265413 | 1 | No Change | Not Significant | FALSE | NA | NA | No Change | Not Detected | FALSE |
| Klk12 | TRUE | 0.314797527 | 1 | No Change | Not Significant | FALSE | NA | NA | No Change | Not Detected | FALSE |
| Klk13 | TRUE | -0.344554053 | 1 | No Change | Not Significant | FALSE | NA | NA | No Change | Not Detected | FALSE |
| Klk15 | TRUE | -4.277574446 | 1 | No Change | Not Significant | TRUE | 0.370742148 | 0.671427662 | No Change | Not Significant | TRUE |
| Klk1b11 | TRUE | 2.760133999 | 1 | No Change | Not Significant | FALSE | NA | NA | No Change | Not Detected | FALSE |
| Klk1b16 | TRUE | 0.075355347 | 1 | No Change | Not Significant | FALSE | NA | NA | No Change | Not Detected | FALSE |
| Klk1b21 | TRUE | 0.383197788 | 1 | No Change | Not Significant | FALSE | NA | NA | No Change | Not Detected | FALSE |
| Klk1b3 | TRUE | -1.040056777 | 1 | No Change | Not Significant | FALSE | NA | NA | No Change | Not Detected | FALSE |
| Klk1b4 | TRUE | -0.851031913 | 0.165809177 | No Change | Not Significant | TRUE | -0.449350645 | 0.33118197 | No Change | Not Significant | TRUE |
| Klk1b5 | TRUE | 0.677859422 | 0.284460467 | No Change | Not Significant | TRUE | 0.43508624 | 0.335281809 | No Change | Not Significant | TRUE |
| Klk4 | TRUE | 1.233947612 | 1 | No Change | Not Significant | FALSE | NA | NA | No Change | Not Detected | FALSE |
| Klk6 | TRUE | 1.13903185 | 1 | No Change | Not Significant | FALSE | NA | NA | No Change | Not Detected | FALSE |
| Klk7 | TRUE | -0.899818639 | 1 | No Change | Not Significant | FALSE | NA | NA | No Change | Not Detected | FALSE |
| Klk8 | TRUE | 0.508051854 | 1 | No Change | Not Significant | FALSE | NA | NA | No Change | Not Detected | FALSE |
| Klk9 | TRUE | 0.007799139 | 1 | No Change | Not Significant | FALSE | NA | NA | No Change | Not Detected | FALSE |
| Klra1 | TRUE | 1.004058926 | 1 | No Change | Not Significant | FALSE | NA | NA | No Change | Not Detected | FALSE |
| Klra2 | TRUE | 1.934622735 | 1 | No Change | Not Significant | FALSE | NA | NA | No Change | Not Detected | FALSE |
| Klra5 | TRUE | -0.437938073 | 0.693738201 | No Change | Not Significant | FALSE | NA | NA | No Change | Not Detected | FALSE |
| Klra6 | TRUE | -0.467966747 | 1 | No Change | Not Significant | FALSE | NA | NA | No Change | Not Detected | FALSE |
| Klra7 | TRUE | 0.472829546 | 0.693375689 | No Change | Not Significant | TRUE | 0.808529445 | 0.044324634 | UP | UP | TRUE |
| Klra9 | TRUE | 1.181396493 | 1 | No Change | Not Significant | FALSE | NA | NA | No Change | Not Detected | FALSE |
| Klrb1 | TRUE | -0.408377706 | 0.686785939 | No Change | Not Significant | TRUE | 0.587437666 | 0.398908703 | No Change | Not Significant | TRUE |
| Klrb1a | TRUE | 0.055250317 | 0.966128972 | No Change | Not Significant | TRUE | 0.196495386 | 0.879811357 | No Change | Not Significant | TRUE |
| Klrb1b | TRUE | 0.546950286 | 1 | No Change | Not Significant | TRUE | 0.280987618 | 0.780262005 | No Change | Not Significant | TRUE |
| Klrb1c | TRUE | 0.234475128 | 0.83386713 | No Change | Not Significant | TRUE | 0.311999204 | 0.683797009 | No Change | Not Significant | TRUE |
| Klrb1f | TRUE | -0.171172317 | 1 | No Change | Not Significant | FALSE | NA | NA | No Change | Not Detected | FALSE |
| Klrc1 | TRUE | 0.611923538 | 1 | No Change | Not Significant | FALSE | NA | NA | No Change | Not Detected | FALSE |
| Klrc2 | TRUE | 0.918261555 | 1 | No Change | Not Significant | TRUE | 1.145870065 | 0.079378465 | No Change | Not Significant | TRUE |
| Klrd1 | TRUE | 0.601223253 | 0.227007559 | No Change | Not Significant | TRUE | 0.083986225 | 0.87485545 | No Change | Not Significant | TRUE |
| Klre1 | TRUE | 0.768127214 | 1 | No Change | Not Significant | TRUE | 1.189282631 | 0.137265172 | No Change | Not Significant | TRUE |
| Klrg1 | TRUE | 2.458022535 | 1 | No Change | Not Significant | FALSE | NA | NA | No Change | Not Detected | FALSE |
| Klrg2 | TRUE | 0.581999818 | 0.480125333 | No Change | Not Significant | TRUE | 0.258956817 | 0.769561988 | No Change | Not Significant | TRUE |
| Klrh1 | TRUE | -2.197977345 | 1 | No Change | Not Significant | FALSE | NA | NA | No Change | Not Detected | FALSE |
| Klri1 | TRUE | 0.188851216 | 1 | No Change | Not Significant | FALSE | NA | NA | No Change | Not Detected | FALSE |
| Klri2 | TRUE | 1.416073998 | 1 | No Change | Not Significant | FALSE | NA | NA | No Change | Not Detected | FALSE |
| Klrk1 | TRUE | -0.19180404 | 0.901300137 | No Change | Not Significant | TRUE | 0.277209046 | 0.62053716 | No Change | Not Significant | TRUE |
| Kmo | TRUE | -0.066013614 | 1 | No Change | Not Significant | FALSE | NA | NA | No Change | Not Detected | FALSE |
| Kmt2a | TRUE | 0.165118514 | 0.570180413 | No Change | Not Significant | TRUE | -0.221261451 | 0.084471059 | No Change | Not Significant | TRUE |
| Kmt2b | TRUE | 0.49232009 | 0.05352217 | No Change | Not Significant | TRUE | -0.029277554 | 0.843368224 | No Change | Not Significant | TRUE |
| Kmt2c | TRUE | 0.096185689 | 0.697463418 | No Change | Not Significant | TRUE | -0.122476838 | 0.162367908 | No Change | Not Significant | TRUE |
| Kmt2d | TRUE | 0.500422312 | 0.170942556 | No Change | Not Significant | TRUE | -0.108156138 | 0.530546055 | No Change | Not Significant | TRUE |
| Kmt2e | TRUE | -0.019967125 | 0.950352959 | No Change | Not Significant | TRUE | -0.198490714 | 0.104644605 | No Change | Not Significant | TRUE |
| Kmt5a | TRUE | 0.065986779 | 0.39045514 | No Change | Not Significant | TRUE | 0.109954475 | 0.178465808 | No Change | Not Significant | TRUE |
| Kmt5b | TRUE | -0.045884171 | 0.779660446 | No Change | Not Significant | TRUE | -0.085444159 | 0.254626651 | No Change | Not Significant | TRUE |
| Kmt5c | TRUE | 0.029063737 | 0.833786242 | No Change | Not Significant | TRUE | 0.043962067 | 0.682136666 | No Change | Not Significant | TRUE |
| Kncn | TRUE | 0.607060289 | 1 | No Change | Not Significant | FALSE | NA | NA | No Change | Not Detected | FALSE |
| Kndc1 | TRUE | 0.080216381 | 1 | No Change | Not Significant | FALSE | NA | NA | No Change | Not Detected | FALSE |
| Knl1 | TRUE | 0.739977036 | 0.000256367 | UP | UP | TRUE | -0.008793207 | 0.968383271 | No Change | Not Significant | TRUE |
| Knop1 | TRUE | -0.028996154 | 0.897242186 | No Change | Not Significant | TRUE | -0.054339129 | 0.645607651 | No Change | Not Significant | TRUE |
| Knstrn | TRUE | 0.719249197 | 7.60E-08 | UP | UP | TRUE | -0.047793213 | 0.81580845 | No Change | Not Significant | TRUE |
| Kntc1 | TRUE | 0.611412041 | 0.017730914 | UP | UP | TRUE | 0.179994088 | 0.434870954 | No Change | Not Significant | TRUE |
| Kpna1 | TRUE | 0.111182047 | 0.297320449 | No Change | Not Significant | TRUE | -0.000302912 | 0.997828443 | No Change | Not Significant | TRUE |
| Kpna2 | TRUE | -0.090030363 | 0.728820184 | No Change | Not Significant | TRUE | 0.033401994 | 0.758094025 | No Change | Not Significant | TRUE |
| Kpna3 | TRUE | 0.028148486 | 0.882690208 | No Change | Not Significant | TRUE | -0.045436281 | 0.633814677 | No Change | Not Significant | TRUE |
| Kpna4 | TRUE | -0.030340025 | 0.767930988 | No Change | Not Significant | TRUE | 0.044699931 | 0.500569859 | No Change | Not Significant | TRUE |
| Kpna6 | TRUE | -0.051805423 | 0.692354022 | No Change | Not Significant | TRUE | 0.020646375 | 0.81580845 | No Change | Not Significant | TRUE |
| Kpna7 | TRUE | -0.42545685 | 1 | No Change | Not Significant | FALSE | NA | NA | No Change | Not Detected | FALSE |
| Kpnb1 | TRUE | 0.774223651 | 5.15E-15 | UP | UP | TRUE | 0.208335077 | 0.00382426 | No Change | Significant Positive | TRUE |
| Kprp | TRUE | 0.611426782 | 1 | No Change | Not Significant | FALSE | NA | NA | No Change | Not Detected | FALSE |
| Kptn | TRUE | -0.244983635 | 0.045218819 | No Change | Significant Negative | TRUE | -0.050778632 | 0.741606925 | No Change | Not Significant | TRUE |
| Kras | TRUE | -0.092026371 | 0.270127107 | No Change | Not Significant | TRUE | -0.07591955 | 0.379596103 | No Change | Not Significant | TRUE |
| Krba1 | TRUE | 0.094772853 | 0.654986223 | No Change | Not Significant | TRUE | 0.399456863 | 1.32E-06 | No Change | Significant Positive | TRUE |
| Krcc1 | TRUE | 0.221526346 | 0.027917061 | No Change | Significant Positive | TRUE | 0.119048563 | 0.360790207 | No Change | Not Significant | TRUE |
| Kremen1 | TRUE | -0.333388009 | 0.003977173 | No Change | Significant Negative | TRUE | -0.126536441 | 0.242581587 | No Change | Not Significant | TRUE |
| Kremen2 | TRUE | -0.274967325 | 0.185023636 | No Change | Not Significant | TRUE | -0.219916689 | 0.228375593 | No Change | Not Significant | TRUE |
| Kri1 | TRUE | 0.032178704 | 0.891333845 | No Change | Not Significant | TRUE | -0.006466419 | 0.980775949 | No Change | Not Significant | TRUE |
| Krit1 | TRUE | -0.061152756 | 0.686291751 | No Change | Not Significant | TRUE | -0.083639042 | 0.595956629 | No Change | Not Significant | TRUE |
| Krr1 | TRUE | 0.187393365 | 0.19938276 | No Change | Not Significant | TRUE | 0.0043348 | 0.975710693 | No Change | Not Significant | TRUE |
| Krt1 | TRUE | -0.250252317 | 1 | No Change | Not Significant | FALSE | NA | NA | No Change | Not Detected | FALSE |
| Krt10 | TRUE | -0.530771881 | 0.098384506 | No Change | Not Significant | TRUE | -0.161299059 | 0.693047027 | No Change | Not Significant | TRUE |
| Krt12 | TRUE | -0.684400181 | 1 | No Change | Not Significant | FALSE | NA | NA | No Change | Not Detected | FALSE |
| Krt13 | TRUE | 1.223667455 | 1 | No Change | Not Significant | FALSE | NA | NA | No Change | Not Detected | FALSE |

Supplementary Table 3

| geneID | Cond1_DF_Detected | Cond1_DF_ValueCol | Cond1_DF_PadjCol | Cond1_DF_Cutoff | Cond1_DF_Cutoff_Specific.x | Cond2_DF_Detected | Cond2_DF_ValueCol | Cond2_DF_PadjCol | Cond2_DF_Cutoff | Cond2_DF_Cutoff_Specific | BG_Method | RG1_All |
|---|---|---|---|---|---|---|---|---|---|---|---|---|
| Krt14 | TRUE | -0.621421376 | 0.211290803 | No Change | Not Significant | TRUE | -0.002839186 | 0.996516146 | No Change | Not Significant | TRUE | Cond1 Not Significant + Cond2 Not Significant |
| Krt15 | TRUE | 0.62257051 | 0.442885865 | No Change | Not Significant | TRUE | -0.41171488 | 0.645614361 | No Change | Not Significant | TRUE | Cond1 Not Significant + Cond2 Not Significant |
| Krt16 | TRUE | 1.110058955 | 1 | No Change | Not Significant | FALSE | NA | NA | No Change | Not Detected | FALSE | Background = FALSE |
| Krt17 | TRUE | -0.842889762 | 1 | No Change | Not Significant | FALSE | NA | NA | No Change | Not Detected | FALSE | Background = FALSE |
| Krt18 | TRUE | -0.136909412 | 0.246352764 | No Change | Not Significant | TRUE | 0.242444932 | 1.33E-07 | No Change | Significant Positive | TRUE | Cond1 Not Significant + Cond2 Significant Positive |
| Krt19 | TRUE | -0.154497562 | 0.378300667 | No Change | Not Significant | TRUE | 0.355428024 | 0.002859216 | No Change | Significant Positive | TRUE | Cond1 Not Significant + Cond2 Significant Positive |
| Krt2 | TRUE | -1.186921824 | 1 | No Change | Not Significant | FALSE | NA | NA | No Change | Not Detected | FALSE | Background = FALSE |
| Krt20 | TRUE | 0.683740509 | 0.000397981 | UP | UP | TRUE | 0.402240332 | 0.001947167 | No Change | Significant Positive | TRUE | Cond1 UP + Cond2 Significant Positive |
| Krt222 | TRUE | -0.424577185 | 0.556110732 | No Change | Not Significant | TRUE | -0.533638784 | 0.10799004 | No Change | Not Significant | TRUE | Cond1 Not Significant + Cond2 Not Significant |
| Krt23 | TRUE | 0.320807655 | 0.130002609 | No Change | Not Significant | TRUE | 0.406745999 | 0.011636985 | No Change | Significant Positive | TRUE | Cond1 Not Significant + Cond2 Significant Positive |
| Krt24 | TRUE | 2.597766629 | 1 | No Change | Not Significant | FALSE | NA | NA | No Change | Not Detected | FALSE | Background = FALSE |
| Krt25 | TRUE | -1.351789474 | 1 | No Change | Not Significant | FALSE | NA | NA | No Change | Not Detected | FALSE | Background = FALSE |
| [illegible] | [illegible] | [illegible] | [illegible] | [illegible] | [illegible] | [illegible] | [illegible] | [illegible] | [illegible] | [illegible] | [illegible] | [illegible] |

| geneID | RG2_Significant | RG3_SignificantChange | Cond1_DF_ensembl_gene_id | Cond1_DF_entrezgene_id | Cond1_DF_gene_biotype | Cond1_DF_baseMean | Cond1_DF_log2FoldChange | Cond1_DF_lfcSE | Cond1_DF_stat | Cond1_DF_pvalue | Cond1_DF_padj | Cond1_DF_absLog2FC | Cond1_DF_Cutoff_Specific.y |
|---|---|---|---|---|---|---|---|---|---|---|---|---|---|
| Krt14 | None | None | ENSMUSG00000045545 | 16664 | protein_coding | 17.00490827 | -0.621421376 | 0.355812124 | 1.746487356 | 0.080726302 | 0.211290803 | 0.621421376 | Not Significant |
| Krt15 | None | None | ENSMUSG00000054146 | 16665 | protein_coding | 15.47004257 | 0.62257051 | 0.52252074 | -1.191475213 | 0.233467082 | 0.442885865 | 0.62257051 | Not Significant |
| Krt16 | Background = FALSE | Background = FALSE | ENSMUSG00000053797 | 16666 | protein_coding | 1.033283113 | 1.110058955 | 2.220281289 | -0.499963208 | 0.617100984 | 1 | 1.110058955 | Not Significant |
| Krt17 | Background = FALSE | Background = FALSE | ENSMUSG00000035557 | 16667 | protein_coding | 0.980486692 | -0.842889762 | 2.796621946 | 0.301395676 | 0.763112791 | 1 | 0.842889762 | Not Significant |
| Krt18 | None | None | ENSMUSG00000023043 | 16668 | protein_coding | 6615.610191 | -0.136909412 | 0.083078967 | 1.647943125 | 0.099364343 | 0.246352764 | 0.136909412 | Not Significant |
| Krt19 | None | None | ENSMUSG00000020911 | 16669 | protein_coding | 39271.83275 | -0.154497562 | 0.11614446 | 1.330218953 | 0.183446141 | 0.378300667 | 0.154497562 | Not Significant |
| Krt2 | Background = FALSE | Background = FALSE | ENSMUSG00000064201 | 16681 | protein_coding | 2.081342012 | -1.186921824 | 3.042661873 | 0.390093239 | 0.696467602 | 1 | 1.186921824 | Not Significant |
| Krt20 | Core_UP | Cond1_UP | ENSMUSG00000035775 | 66809 | protein_coding | 9465.065044 | 0.683740509 | 0.166824343 | -4.098565566 | 4.16E-05 | 0.000397981 | 0.683740509 | UP |
| Krt222 | None | None | ENSMUSG00000035849 | 268481 | protein_coding | 8.810325165 | -0.424577185 | 0.439287723 | 0.966512751 | 0.33378767 | 0.556110732 | 0.424577185 | Not Significant |
| Krt23 | None | None | ENSMUSG00000006777 | 94179 | protein_coding | 116.0193685 | 0.320807655 | 0.157552603 | -2.036193937 | 0.041730885 | 0.130002609 | 0.320807655 | Not Significant |
| Krt24 | Background = FALSE | Background = FALSE | ENSMUSG00000020913 | 75706 | protein_coding | 1.523960495 | 2.597766629 | 2.279346983 | -1.139697751 | 0.254412245 | 1 | 2.597766629 | Not Significant |
| Krt25 | Background = FALSE | Background = FALSE | ENSMUSG00000035831 | 70810 | protein_coding | 0.733329392 | -1.351789474 | 2.160411983 | 0.625709117 | 0.531505752 | 1 | 1.351789474 | Not Significant |
| [illegible] | [illegible] | [illegible] | [illegible] | [illegible] | [illegible] | [illegible] | [illegible] | [illegible] | [illegible] | [illegible] | [illegible] | [illegible] | [illegible] |

| geneID | Cond2_DF_ensembl_gene_id | Cond2_DF_entrezgene_id | Cond2_DF_gene_biotype | Cond2_DF_baseMean | Cond2_DF_log2FoldChange | Cond2_DF_lfcSE | Cond2_DF_stat | Cond2_DF_pvalue | Cond2_DF_padj | Cond2_DF_absLog2FC | RG1_Specific_Cond2 | RG1_Specific_Cond1 |
|---|---|---|---|---|---|---|---|---|---|---|---|---|
| Krt14 | ENSMUSG00000045545 | 16664 | protein_coding | 20.59932586 | -0.002839186 | 0.264923071 | -0.010717019 | 0.991449219 | 0.996516146 | 0.002839186 | Cond1 No Change + Cond2 Not Significant | Cond2 No Change + Cond1 Not Significant |
| Krt15 | ENSMUSG00000054146 | 16665 | protein_coding | 12.86842565 | -0.41171488 | 0.496378274 | -0.829437752 | 0.406856747 | 0.645614361 | 0.41171488 | Cond1 No Change + Cond2 Not Significant | Cond2 No Change + Cond1 Not Significant |
| Krt16 | NA | NA | NA | NA | NA | NA | NA | NA | NA | NA | Background = FALSE | Background = FALSE |
| Krt17 | NA | NA | NA | NA | NA | NA | NA | NA | NA | NA | Background = FALSE | Background = FALSE |
| Krt18 | ENSMUSG00000023043 | 16668 | protein_coding | 10734.52937 | 0.242444932 | 0.041066837 | 5.903667093 | 3.56E-09 | 1.33E-07 | 0.242444932 | Cond1 No Change + Cond2 Significant Positive | Cond2 No Change + Cond1 Not Significant |
| Krt19 | ENSMUSG00000020911 | 16669 | protein_coding | 68956.10272 | 0.355428024 | 0.096211645 | 3.694230812 | 0.000220553 | 0.002859216 | 0.355428024 | Cond1 No Change + Cond2 Significant Positive | Cond2 No Change + Cond1 Not Significant |
| Krt2 | NA | NA | NA | NA | NA | NA | NA | NA | NA | NA | Background = FALSE | Background = FALSE |
| Krt20 | ENSMUSG00000035775 | 66809 | protein_coding | 11721.72146 | 0.402240332 | 0.105557233 | 3.810637314 | 0.000138609 | 0.001947167 | 0.402240332 | Cond1 UP + Cond2 Significant Positive | Cond2 No Change + Cond1 UP |
| Krt222 | ENSMUSG00000035849 | 268481 | protein_coding | 39.58844253 | -0.533638784 | 0.234630104 | -2.274383274 | 0.022942952 | 0.10799004 | 0.533638784 | Cond1 No Change + Cond2 Not Significant | Cond2 No Change + Cond1 Not Significant |
| Krt23 | ENSMUSG00000006777 | 94179 | protein_coding | 195.5795993 | 0.406745999 | 0.125488845 | 3.241292078 | 0.001189892 | 0.011636985 | 0.406745999 | Cond1 No Change + Cond2 Significant Positive | Cond2 No Change + Cond1 Not Significant |
| Krt24 | NA | NA | NA | NA | NA | NA | NA | NA | NA | NA | Background = FALSE | Background = FALSE |
| Krt25 | NA | NA | NA | NA | NA | NA | NA | NA | NA | NA | Background = FALSE | Background = FALSE |
| [illegible] | [illegible] | [illegible] | [illegible] | [illegible] | [illegible] | [illegible] | [illegible] | [illegible] | [illegible] | [illegible] | [illegible] | [illegible] |

| geneID | Cond1_DF_Detected | Cond1_DF_ValueCol | Cond1_DF_PadjCol | Cond1_DF_Cutoff | Cond1_DF_Cutoff_Specific.x | Cond2_DF_Detected |
|---|---|---|---|---|---|---|
| Lhx3 | TRUE | 0.688571746 | 1 | No Change | Not Significant | TRUE |
| Lhx4 | TRUE | 2.423887142 | 1 | No Change | Not Significant | FALSE |
| Lhx5 | TRUE | 0.004293923 | 1 | No Change | Not Significant | FALSE |
| Lhx6 | TRUE | 0.176009083 | 1 | No Change | Not Significant | FALSE |
| Lhx9 | TRUE | 0.603343659 | 1 | No Change | Not Significant | FALSE |
| Lias | TRUE | -0.112064557 | 0.320998127 | No Change | Not Significant | TRUE |
| Lif | TRUE | 0.223402994 | 0.508294925 | No Change | Not Significant | TRUE |
| Lifr | TRUE | 0.032126577 | 0.952286247 | No Change | Not Significant | TRUE |
| Lig1 | TRUE | 0.072735172 | 0.793440482 | No Change | Not Significant | TRUE |
| Lig3 | TRUE | 0.085377459 | 0.595946987 | No Change | Not Significant | TRUE |
| Lig4 | TRUE | 0.127078846 | 0.299280888 | No Change | Not Significant | TRUE |
| Lilrb4a | TRUE | 2.413618952 | 1 | No Change | Not Significant | FALSE |
| Lilrb4b | TRUE | 0.399868314 | 1 | No Change | Not Significant | FALSE |
| Lima1 | TRUE | -0.202111738 | 0.106618553 | No Change | Not Significant | TRUE |
| Limch1 | TRUE | 0.498252014 | 0.058661192 | No Change | Not Significant | TRUE |
| Limd1 | TRUE | -0.110878435 | 0.264291499 | No Change | Not Significant | TRUE |
| Limd2 | TRUE | 0.078676228 | 0.80495599 | No Change | Not Significant | TRUE |
| Lime1 | TRUE | 0.705840509 | 1 | No Change | Not Significant | FALSE |
| Limk1 | TRUE | 0.394482481 | 0.088956393 | No Change | Not Significant | TRUE |
| Limk2 | TRUE | -0.17108866 | 0.188515765 | No Change | Not Significant | TRUE |
| Lims1 | TRUE | 0.028610992 | 0.932569867 | No Change | Not Significant | TRUE |
| Lims2 | TRUE | -0.032061663 | 0.981615003 | No Change | Not Significant | FALSE |
| Lin28a | TRUE | -0.481120496 | 1 | No Change | Not Significant | FALSE |
| Lin28b | TRUE | 1.602866662 | 1 | No Change | Not Significant | FALSE |
| Lin37 | TRUE | 0.01456435 | 0.947822351 | No Change | Not Significant | TRUE |
| Lin52 | TRUE | 0.744236466 | 2.41E-09 | UP | UP | TRUE |
| Lin7a | TRUE | -2.010177424 | 1 | No Change | Not Significant | TRUE |
| Lin7b | TRUE | 0.524773528 | 1 | No Change | Not Significant | FALSE |
| Lin7c | TRUE | -0.018996194 | 0.922491196 | No Change | Not Significant | TRUE |
| Lin9 | TRUE | 0.29187458 | 0.108626953 | No Change | Not Significant | TRUE |
| Lingo1 | TRUE | 1.380998427 | 1 | No Change | Not Significant | FALSE |
| Lingo2 | TRUE | 2.557594767 | 0.022335267 | UP | UP | FALSE |
| Lingo3 | TRUE | 1.205660165 | 1 | No Change | Not Significant | FALSE |
| Lingo4 | TRUE | 0.340146623 | 0.771232643 | No Change | Not Significant | TRUE |
| Lins1 | TRUE | 0.167724894 | 0.331035921 | No Change | Not Significant | TRUE |
| Lipa | TRUE | 2.010288107 | 2.35E-111 | UP | UP | TRUE |
| Lipc | TRUE | 0.191752716 | 0.852920113 | No Change | Not Significant | TRUE |
| Lipe | TRUE | -0.346352363 | 0.088285245 | No Change | Not Significant | TRUE |
| Lipf | TRUE | 0.753092189 | 1 | No Change | Not Significant | FALSE |
| Lipg | TRUE | -0.400148083 | 0.09052835 | No Change | Not Significant | TRUE |
| Liph | TRUE | -0.027920044 | 0.905233898 | No Change | Not Significant | TRUE |
| Lipk | TRUE | 0.13349536 | 0.907147741 | No Change | Not Significant | TRUE |
| Lipo2 | TRUE | 0.683327339 | 1 | No Change | Not Significant | FALSE |
| Lipo3 | TRUE | -0.27300293 | 0.265457875 | No Change | Not Significant | TRUE |
| Lipt1 | TRUE | -0.398561132 | 0.184012672 | No Change | Not Significant | TRUE |
| Lipt2 | TRUE | 0.232612012 | 0.363680341 | No Change | Not Significant | TRUE |
| Litaf | TRUE | -0.103205542 | 0.274894315 | No Change | Not Significant | TRUE |
| Litafd | TRUE | 1.910191893 | 1 | No Change | Not Significant | FALSE |
| Lix1 | TRUE | 0.14256569 | 0.881390987 | No Change | Not Significant | TRUE |
| Lix1l | TRUE | 0.058098962 | 0.959274295 | No Change | Not Significant | TRUE |
| Llcfc1 | TRUE | 2.17907174 | 1 | No Change | Not Significant | FALSE |
| Llgl1 | TRUE | -0.073768538 | 0.491020508 | No Change | Not Significant | TRUE |
| Llgl2 | TRUE | -0.075175955 | 0.524967146 | No Change | Not Significant | TRUE |
| Llph | TRUE | -0.084471881 | 0.724949186 | No Change | Not Significant | TRUE |
| Lman1 | TRUE | 0.574678953 | 0.001217281 | UP | UP | TRUE |
| Lman1l | TRUE | 0.646271909 | 0.640714813 | No Change | Not Significant | FALSE |
| Lman2 | TRUE | 0.740249406 | 5.83E-16 | UP | UP | TRUE |
| Lman2l | TRUE | 0.199794765 | 0.228911821 | No Change | Not Significant | TRUE |
| Lmbr1 | TRUE | -0.345495763 | 0.00613142 | No Change | Significant Negative | TRUE |
| Lmbr1l | TRUE | -0.271720106 | 0.156531285 | No Change | Not Significant | TRUE |
| Lmbrd1 | TRUE | -0.141151239 | 0.181983293 | No Change | Not Significant | TRUE |
| Lmbrd2 | TRUE | 0.82400734 | 2.85E-17 | UP | UP | TRUE |
| Lmcd1 | TRUE | 0.629680333 | 0.466691767 | No Change | Not Significant | TRUE |
| Lmf1 | TRUE | 1.503087321 | 5.35E-79 | UP | UP | TRUE |
| Lmf2 | TRUE | 0.190394311 | 0.174694151 | No Change | Not Significant | TRUE |
| Lmln | TRUE | -0.392090767 | 0.027189423 | No Change | Significant Negative | TRUE |
| Lmna | TRUE | -0.456558786 | 8.45E-07 | No Change | Significant Negative | TRUE |
| Lmnb1 | TRUE | 0.479803974 | 0.000626283 | No Change | Significant Positive | TRUE |
| Lmnb2 | TRUE | -0.15948353 | 0.117801163 | No Change | Not Significant | TRUE |
| Lmntd1 | TRUE | 1.356304939 | 1 | No Change | Not Significant | FALSE |
| Lmntd2 | TRUE | -0.159104902 | 0.680659303 | No Change | Not Significant | TRUE |
| Lmo1 | TRUE | 0.06646184 | 0.954311305 | No Change | Not Significant | TRUE |
| Lmo2 | TRUE | -0.185189524 | 0.816921981 | No Change | Not Significant | TRUE |
| Lmo3 | TRUE | -0.30632302 | 1 | No Change | Not Significant | FALSE |
| Lmo4 | TRUE | -0.164320079 | 0.133085576 | No Change | Not Significant | TRUE |
| Lmo7 | TRUE | 0.026562066 | 0.907710402 | No Change | Not Significant | TRUE |
| Lmod1 | TRUE | -1.203429859 | 1 | No Change | Not Significant | FALSE |
| Lmod2 | TRUE | 2.088936213 | 1 | No Change | Not Significant | FALSE |
| Lmod3 | TRUE | 0.75306431 | 1 | No Change | Not Significant | FALSE |
| Lmtk2 | TRUE | 0.109506812 | 0.294844608 | No Change | Not Significant | TRUE |
| Lmtk3 | TRUE | -0.663379117 | 0.546521427 | No Change | Not Significant | TRUE |
| Lmx1a | TRUE | -0.556879319 | 0.200044477 | No Change | Not Significant | TRUE |
| Lmx1b | TRUE | 0.852629933 | 1 | No Change | Not Significant | FALSE |
| Lncbate3 | TRUE | -1.263541308 | 1 | No Change | Not Significant | FALSE |
| Lncpint | TRUE | 0.543038536 | 0.059833793 | No Change | Not Significant | TRUE |
| Lnpep | TRUE | -0.050753731 | 0.773528957 | No Change | Not Significant | TRUE |
| Lnpk | TRUE | 0.390262478 | 0.009274475 | No Change | Significant Positive | TRUE |
| Lnx1 | TRUE | -0.408532995 | 0.043456274 | No Change | Significant Negative | TRUE |
| Lnx2 | TRUE | -0.031980142 | 0.797437818 | No Change | Not Significant | TRUE |
| Lonp1 | TRUE | -0.234269364 | 0.002907402 | No Change | Significant Negative | TRUE |
| Lonp2 | TRUE | -0.204405321 | 0.004126525 | No Change | Significant Negative | TRUE |
| Lonrf1 | TRUE | 0.169196489 | 0.560303 | No Change | Not Significant | TRUE |
| Lonrf2 | TRUE | -1.239691759 | 1 | No Change | Not Significant | FALSE |
| Lonrf3 | TRUE | 0.355292817 | 0.078139671 | No Change | Not Significant | TRUE |
| Lor | TRUE | 1.402764016 | 1 | No Change | Not Significant | TRUE |
| Loxhd1 | TRUE | 1.230875395 | 1 | No Change | Not Significant | FALSE |
| Loxl1 | TRUE | 0.676035745 | 1 | No Change | Not Significant | FALSE |
| Loxl2 | TRUE | -0.37109947 | 1 | No Change | Not Significant | FALSE |
| Loxl3 | TRUE | 1.507388832 | 0.001015447 | UP | UP | FALSE |
| Loxl4 | TRUE | -0.148609345 | 0.922115416 | No Change | Not Significant | TRUE |
| Lpar1 | TRUE | -0.147490881 | 0.1452832 | No Change | Not Significant | TRUE |
| Lpar2 | TRUE | -0.625675204 | 0.105793898 | No Change | Not Significant | TRUE |
| Lpar3 | TRUE | 1.802357487 | 0.011230894 | UP | UP | TRUE |
| Lpar5 | TRUE | 0.113568519 | 0.511613776 | No Change | Not Significant | TRUE |
| Lpar6 | TRUE | 0.089635477 | 0.587772599 | No Change | Not Significant | TRUE |
| Lpcat1 | TRUE | -0.163725881 | 0.525252463 | No Change | Not Significant | TRUE |
| Lpcat2 | TRUE | 0.116113613 | 0.730517043 | No Change | Not Significant | TRUE |
| Lpcat3 | TRUE | 0.191096024 | 0.035329044 | No Change | Significant Positive | TRUE |
| Lpcat4 | TRUE | 0.787475518 | 0.01218155 | UP | UP | TRUE |
| Lpgat1 | TRUE | 0.388119087 | 0.000137721 | No Change | Significant Positive | TRUE |
| Lpin1 | TRUE | 0.067164075 | 0.855640813 | No Change | Not Significant | TRUE |
| Lpin2 | TRUE | -0.093349617 | 0.833786242 | No Change | Not Significant | TRUE |
| Lpin3 | TRUE | 0.303753467 | 0.027610883 | No Change | Significant Positive | TRUE |
| Lpl | TRUE | -1.525640468 | 2.57E-15 | DOWN | DOWN | TRUE |
| Lpo | TRUE | 0.09284753 | 0.832872319 | No Change | Not Significant | TRUE |
| Lpp | TRUE | 0.00560402 | 0.980158095 | No Change | Not Significant | TRUE |
| Lpxn | TRUE | -0.074105982 | 0.89324602 | No Change | Not Significant | TRUE |
| Lrat | TRUE | -1.236684633 | 8.81E-14 | DOWN | DOWN | TRUE |
| Lratd1 | TRUE | 1.792746634 | 6.01E-32 | UP | UP | TRUE |
| Lratd2 | TRUE | 0.085441291 | 0.599520433 | No Change | Not Significant | TRUE |
| Lrba | TRUE | -0.098777694 | 0.515123206 | No Change | Not Significant | TRUE |
| Lrch1 | TRUE | -0.418659108 | 3.04E-05 | No Change | Significant Negative | TRUE |
| Lrch2 | TRUE | -0.658123968 | 1 | No Change | Not Significant | FALSE |
| Lrch3 | TRUE | -0.016309124 | 0.92805494 | No Change | Not Significant | TRUE |
| Lrch4 | TRUE | 0.286712022 | 0.829503357 | No Change | Not Significant | TRUE |
| Lrfn1 | TRUE | 2.086664679 | 0.00015813 | UP | UP | TRUE |
| Lrfn3 | TRUE | 0.146638729 | 0.696186289 | No Change | Not Significant | TRUE |
| Lrfn4 | TRUE | -0.318739351 | 0.015364319 | No Change | Significant Negative | TRUE |
| Lrfn5 | TRUE | 0.604778438 | 1 | No Change | Not Significant | FALSE |
| Lrg1 | TRUE | 2.82359387 | 0.000421772 | UP | UP | FALSE |
| Lrguk | TRUE | 0.43168399 | 0.8161622 | No Change | Not Significant | FALSE |
| Lrif1 | TRUE | 0.132546374 | 0.412028696 | No Change | Not Significant | TRUE |
| Lrig1 | TRUE | 0.159007014 | 0.487190988 | No Change | Not Significant | TRUE |
| Lrig2 | TRUE | -0.089539329 | 0.511503652 | No Change | Not Significant | TRUE |
| Lrig3 | TRUE | -0.244646108 | 0.074572727 | No Change | Not Significant | TRUE |
| Lrit1 | TRUE | -1.219807861 | 1 | No Change | Not Significant | FALSE |
| Lrit2 | TRUE | 0.874209496 | 1 | No Change | Not Significant | FALSE |
| Lrit3 | TRUE | -1.063750479 | 0.09528722 | No Change | Not Significant | TRUE |
| Lrmda | TRUE | -0.380737312 | 1 | No Change | Not Significant | FALSE |
| Lrp1 | TRUE | 0.199530824 | 0.582349713 | No Change | Not Significant | TRUE |
| Lrp10 | TRUE | 0.106347724 | 0.303828141 | No Change | Not Significant | TRUE |
| Lrp11 | TRUE | -0.517819641 | 0.134899501 | No Change | Not Significant | TRUE |
| Lrp12 | TRUE | -0.175887926 | 0.446089228 | No Change | Not Significant | TRUE |
| Lrp1b | TRUE | 0.940811512 | 0.8085485 | No Change | Not Significant | FALSE |
| Lrp2 | TRUE | 1.300846099 | 0.827904179 | No Change | Not Significant | FALSE |
| Lrp2bp | TRUE | -0.174683051 | 0.916826487 | No Change | Not Significant | FALSE |
| Lrp3 | TRUE | 0.578009842 | 0.504684233 | No Change | Not Significant | TRUE |
| Lrp4 | TRUE | -0.212780098 | 0.03390183 | No Change | Significant Negative | TRUE |
| Lrp5 | TRUE | -0.070180007 | 0.584758488 | No Change | Not Significant | TRUE |
| Lrp6 | TRUE | -0.097238143 | 0.269146142 | No Change | Not Significant | TRUE |
| Lrp8 | TRUE | 2.380478509 | 1.78E-37 | UP | UP | TRUE |
| Lrp8os2 | TRUE | 0.277540126 | 1 | No Change | Not Significant | FALSE |
| Lrp8os3 | TRUE | 1.112036257 | 1 | No Change | Not Significant | FALSE |
| Lrpap1 | TRUE | -0.411619941 | 7.99E-06 | No Change | Significant Negative | TRUE |
| Lrpprc | TRUE | -0.257255983 | 0.019034324 | No Change | Significant Negative | TRUE |
| Lrr1 | TRUE | 0.779290882 | 0.098851981 | No Change | Not Significant | TRUE |
| Lrrc1 | TRUE | 0.066839376 | 0.583801091 | No Change | Not Significant | TRUE |
| Lrrc10b | TRUE | 0.294798275 | 1 | No Change | Not Significant | FALSE |
| Lrrc14 | TRUE | 0.096638218 | 0.581581822 | No Change | Not Significant | TRUE |
| Lrrc15 | TRUE | -0.996420324 | 1 | No Change | Not Significant | FALSE |
| Lrrc17 | TRUE | 0.499449224 | 1 | No Change | Not Significant | FALSE |
| Lrrc18 | TRUE | -0.059556979 | 1 | No Change | Not Significant | FALSE |
| Lrrc19 | TRUE | -0.532925227 | 0.000391532 | DOWN | DOWN | TRUE |
| Lrrc20 | TRUE | 0.018065019 | 0.94399288 | No Change | Not Significant | TRUE |
| Lrrc23 | TRUE | -1.191492795 | 1 | No Change | Not Significant | FALSE |
| Lrrc24 | TRUE | -0.094536151 | 1 | No Change | Not Significant | FALSE |
| Lrrc25 | TRUE | 1.777352927 | 1 | No Change | Not Significant | FALSE |
| Lrrc26 | TRUE | -0.201206019 | 0.405213026 | No Change | Not Significant | TRUE |
| Lrrc27 | TRUE | -0.133385571 | 0.902595167 | No Change | Not Significant | TRUE |
| Lrrc28 | TRUE | 0.804404788 | 2.38E-07 | UP | UP | TRUE |
| Lrrc3 | TRUE | -0.900688253 | 0.24866598 | No Change | Not Significant | TRUE |
| Lrrc31 | TRUE | -0.345849376 | 0.219864981 | No Change | Not Significant | TRUE |
| Lrrc32 | TRUE | -2.021999258 | 1 | No Change | Not Significant | FALSE |
| Lrrc34 | TRUE | 1.07886645 | 1 | No Change | Not Significant | FALSE |
| Lrrc36 | TRUE | -1.434309532 | 1 | No Change | Not Significant | FALSE |
| Lrrc37a | TRUE | 1.829987391 | 1 | No Change | Not Significant | FALSE |
| Lrrc39 | TRUE | 0.515809416 | 0.554292124 | No Change | Not Significant | TRUE |
| Lrrc4 | TRUE | 0.582920693 | 0.25251889 | No Change | Not Significant | TRUE |
| Lrrc40 | TRUE | 0.210094154 | 0.1933428 | No Change | Not Significant | TRUE |
| Lrrc41 | TRUE | 0.260442749 | 0.019566455 | No Change | Significant Positive | TRUE |
| Lrrc42 | TRUE | -0.131517116 | 0.41642117 | No Change | Not Significant | TRUE |
| Lrrc43 | TRUE | 1.69759759 | 1 | No Change | Not Significant | FALSE |
| Lrrc45 | TRUE | 0.494027343 | 5.82E-07 | No Change | Significant Positive | TRUE |
| Lrrc46 | TRUE | -0.007998212 | 1 | No Change | Not Significant | FALSE |
| Lrrc47 | TRUE | 0.355201264 | 2.42E-06 | No Change | Significant Positive | TRUE |
| Lrrc49 | TRUE | 0.475097122 | 0.006582523 | No Change | Significant Positive | TRUE |
| Lrrc4b | TRUE | 1.12616829 | 1 | No Change | Not Significant | FALSE |
| Lrrc4c | TRUE | 0.523570038 | 1 | No Change | Not Significant | FALSE |
| Lrrc51 | TRUE | -0.45176234 | 0.068585239 | No Change | Not Significant | TRUE |
| Lrrc55 | TRUE | 1.249482307 | 1 | No Change | Not Significant | FALSE |
| Lrrc56 | TRUE | 0.102938429 | 0.620766699 | No Change | Not Significant | TRUE |
| Lrrc57 | TRUE | -0.421615767 | 0.00644975 | No Change | Significant Negative | TRUE |

| geneID | Cond1_DF_Detected | Cond1_DF_ValueCol | Cond1_DF_PadjCol | Cond1_DF_Cutoff | Cond1_DF_Cutoff_Specific.x |
|---|---|---|---|---|---|
| Lrrc58 | TRUE | 0.250850083 | 0.051198321 | No Change | Not Significant |
| Lrrc59 | TRUE | 0.588996746 | 1.94E-12 | UP | UP |
| Lrrc61 | TRUE | -0.290378518 | 0.198449829 | No Change | Not Significant |
| Lrrc66 | TRUE | 0.033463953 | 0.85879931 | No Change | Not Significant |
| Lrrc69 | TRUE | -0.659672398 | 1 | No Change | Not Significant |
| Lrrc7 | TRUE | 0.694548127 | 1 | No Change | Not Significant |
| Lrrc70 | TRUE | -0.138591338 | 0.900323208 | No Change | Not Significant |
| Lrrc71 | TRUE | 0.676374413 | 0.145821162 | No Change | Not Significant |
| Lrrc73 | TRUE | 0.7286128 | 0.589852079 | No Change | Not Significant |
| Lrrc74a | TRUE | -0.425304125 | 1 | No Change | Not Significant |
| Lrrc74b | TRUE | -0.758613426 | 1 | No Change | Not Significant |
| Lrrc75a | TRUE | 0.172385865 | 0.125530417 | No Change | Not Significant |
| Lrrc75b | TRUE | 1.464763071 | 0.028888694 | UP | UP |
| Lrrc8a | TRUE | -0.274350294 | 0.189251307 | No Change | Not Significant |
| Lrrc8b | TRUE | 0.324090969 | 0.0006509 | No Change | Significant Positive |
| Lrrc8c | TRUE | -0.144470543 | 0.877032235 | No Change | Not Significant |
| Lrrc8d | TRUE | 0.015697787 | 0.932932188 | No Change | Not Significant |
| Lrrc8e | TRUE | -0.997279514 | 8.30E-14 | DOWN | DOWN |
| Lrrc9 | TRUE | 3.101538723 | 1 | No Change | Not Significant |
| Lrrcc1 | TRUE | -0.144054457 | 0.559780524 | No Change | Not Significant |
| Lrrd1 | TRUE | 1.694227463 | 1 | No Change | Not Significant |
| Lrrfip1 | TRUE | -0.129414219 | 0.241894213 | No Change | Not Significant |
| Lrrfip2 | TRUE | -0.006634509 | 0.973904526 | No Change | Not Significant |
| Lrriq1 | TRUE | -0.257543301 | 1 | No Change | Not Significant |
| Lrriq3 | TRUE | -0.063433665 | 0.910171686 | No Change | Not Significant |
| Lrrk1 | TRUE | 0.187937915 | 0.025977317 | No Change | Significant Positive |
| Lrrk2 | TRUE | 0.30395797 | 0.470773677 | No Change | Not Significant |
| Lrrn2 | TRUE | 0.431950242 | 1 | No Change | Not Significant |
| Lrrn3 | TRUE | -0.716843402 | 0.105701154 | No Change | Not Significant |
| Lrrn4 | TRUE | 0.339620203 | 1 | No Change | Not Significant |
| Lrrn4cl | TRUE | 0.485921008 | 1 | No Change | Not Significant |
| Lrrtm1 | TRUE | 1.573849661 | 0.135678899 | No Change | Not Significant |
| Lrrtm2 | TRUE | 0.05072639 | 0.93371978 | No Change | Not Significant |
| Lrrtm3 | TRUE | -1.184325255 | 1 | No Change | Not Significant |
| Lrsam1 | TRUE | -0.037356609 | 0.833127005 | No Change | Not Significant |
| Lrtm1 | TRUE | 1.318958263 | 1 | No Change | Not Significant |
| Lrtm2 | TRUE | 0.065894449 | 1 | No Change | Not Significant |
| Lrwd1 | TRUE | 0.116938982 | 0.460735958 | No Change | Not Significant |
| Lsamp | TRUE | -0.231944788 | 0.975316352 | No Change | Not Significant |
| Lsg1 | TRUE | 0.098736864 | 0.362441146 | No Change | Not Significant |
| Lsm1 | TRUE | 0.026296129 | 0.855153117 | No Change | Not Significant |
| Lsm10 | TRUE | 0.074945543 | 0.735049182 | No Change | Not Significant |
| Lsm11 | TRUE | -0.082107958 | 0.736456016 | No Change | Not Significant |
| Lsm12 | TRUE | 0.112109648 | 0.410708835 | No Change | Not Significant |
| Lsm14a | TRUE | 0.160683743 | 0.118133785 | No Change | Not Significant |
| Lsm14b | TRUE | -0.232634373 | 0.018306921 | No Change | Significant Negative |
| Lsm2 | TRUE | 0.516023764 | 0.004567391 | UP | UP |
| Lsm3 | TRUE | 0.227690123 | 0.143668739 | No Change | Not Significant |
| Lsm4 | TRUE | 0.136666636 | 0.328779714 | No Change | Not Significant |
| Lsm5 | TRUE | 0.290448374 | 0.199909758 | No Change | Not Significant |
| Lsm6 | TRUE | 0.141894095 | 0.336996201 | No Change | Not Significant |
| Lsm7 | TRUE | -0.175438969 | 0.409951417 | No Change | Not Significant |
| Lsm8 | TRUE | 0.173554796 | 0.210915627 | No Change | Not Significant |
| Lsmem1 | TRUE | 2.89443152 | 1 | No Change | Not Significant |
| Lsmem2 | TRUE | -0.104708918 | 0.893302015 | No Change | Not Significant |
| Lsp1 | TRUE | 0.399657582 | 0.593065878 | No Change | Not Significant |
| Lsr | TRUE | 0.089748263 | 0.827951097 | No Change | Not Significant |
| Lss | TRUE | 0.354211154 | 0.004230202 | No Change | Significant Positive |
| Lst1 | TRUE | 0.664522581 | 1 | No Change | Not Significant |
| Lta | TRUE | 0.271370248 | 1 | No Change | Not Significant |
| Lta4h | TRUE | -0.127349993 | 0.195920934 | No Change | Not Significant |
| Ltb | TRUE | 0.09917413 | 0.930374388 | No Change | Not Significant |
| Ltb4r1 | TRUE | 1.332118186 | 1 | No Change | Not Significant |
| Ltb4r2 | TRUE | 0.318565136 | 0.710135685 | No Change | Not Significant |
| Ltbp1 | TRUE | 0.758745419 | 0.730517043 | No Change | Not Significant |
| Ltbp2 | TRUE | -0.494914792 | 0.736964526 | No Change | Not Significant |
| Ltbp3 | TRUE | 1.263996846 | 0.26018046 | No Change | Not Significant |
| Ltbp4 | TRUE | 0.152813908 | 0.868781842 | No Change | Not Significant |
| Ltbr | TRUE | 0.033481874 | 0.811870577 | No Change | Not Significant |
| Ltc4s | TRUE | -0.738415656 | 0.011948193 | DOWN | DOWN |
| Ltf | TRUE | 0.589356295 | 0.53139398 | No Change | Not Significant |
| Ltk | TRUE | -0.306331579 | 0.612497263 | No Change | Not Significant |
| Ltn1 | TRUE | 0.087634669 | 0.566362983 | No Change | Not Significant |
| LTO1 | TRUE | 0.030563314 | 0.856549354 | No Change | Not Significant |
| Ltv1 | TRUE | 0.099236674 | 0.571420483 | No Change | Not Significant |
| Luc7l | TRUE | 0.099822191 | 0.461422606 | No Change | Not Significant |
| Luc7l2 | TRUE | 0.291707299 | 0.00608931 | No Change | Significant Positive |
| Luc7l3 | TRUE | 0.053031482 | 0.691751611 | No Change | Not Significant |
| Lum | TRUE | -0.939013694 | 1 | No Change | Not Significant |
| Lurap1 | TRUE | 0.012076745 | 0.977047797 | No Change | Not Significant |
| Lurap1l | TRUE | -0.197558065 | 0.023543362 | No Change | Significant Negative |
| Luzp1 | TRUE | -0.230859779 | 0.001933622 | No Change | Significant Negative |
| Lvrn | TRUE | 0.044517408 | 1 | No Change | Not Significant |
| Lxn | TRUE | -0.06259451 | 0.771818402 | No Change | Not Significant |
| Ly6a | TRUE | -0.19919659 | 0.559633454 | No Change | Not Significant |
| Ly6c1 | TRUE | 0.48746065 | 0.537332461 | No Change | Not Significant |
| Ly6c2 | TRUE | 0.210619312 | 1 | No Change | Not Significant |
| Ly6d | TRUE | 1.847478298 | 1 | No Change | Not Significant |
| Ly6e | TRUE | -0.623876107 | 0.08026156 | No Change | Not Significant |
| Ly6g | TRUE | -0.845915893 | 0.029306374 | DOWN | DOWN |
| Ly6g2 | TRUE | 0.148406258 | 0.652884901 | No Change | Not Significant |
| Ly6g5b | TRUE | 2.40468178 | 1 | No Change | Not Significant |
| Ly6g6c | TRUE | -1.438722695 | 0.179926661 | No Change | Not Significant |
| Ly6g6d | TRUE | -0.506035273 | 0.027285164 | DOWN | DOWN |
| Ly6g6e | TRUE | -0.152819857 | 0.691465456 | No Change | Not Significant |
| Ly6g6f | TRUE | -0.917626737 | 0.004741185 | DOWN | DOWN |
| Ly6g6g | TRUE | 0.38554842 | 1 | No Change | Not Significant |
| Ly6h | TRUE | -1.697476823 | 1 | No Change | Not Significant |
| Ly6i | TRUE | 0.430878747 | 1 | No Change | Not Significant |
| Ly6k | TRUE | 0.01245404 | 1 | No Change | Not Significant |
| Ly6l | TRUE | -0.468603076 | 1 | No Change | Not Significant |
| Ly6m | TRUE | 0.058696855 | 0.952286247 | No Change | Not Significant |
| Ly75 | TRUE | -0.348131063 | 0.000352943 | No Change | Significant Negative |
| Ly86 | TRUE | -1.367667454 | 1 | No Change | Not Significant |
| Ly9 | TRUE | -0.224028974 | 0.867862447 | No Change | Not Significant |
| Ly96 | TRUE | -0.047572634 | 0.907746426 | No Change | Not Significant |
| Lyar | TRUE | 0.296220836 | 0.03900665 | No Change | Significant Positive |
| Lyg2 | TRUE | -0.430943803 | 1 | No Change | Not Significant |
| Lyl1 | TRUE | -0.134357609 | 0.878808271 | No Change | Not Significant |
| Lyn | TRUE | -0.028258807 | 0.864910546 | No Change | Not Significant |
| Lynx1 | TRUE | -0.192730023 | 0.415328565 | No Change | Not Significant |
| Lypd1 | TRUE | -0.242850518 | 1 | No Change | Not Significant |
| Lypd11 | TRUE | -0.593449832 | 1 | No Change | Not Significant |
| Lypd3 | TRUE | 1.533659827 | 1 | No Change | Not Significant |
| Lypd6 | TRUE | 0.088386276 | 1 | No Change | Not Significant |
| Lypd6b | TRUE | -0.10512821 | 0.844376272 | No Change | Not Significant |
| Lypd8 | TRUE | -0.424951336 | 0.000287573 | No Change | Significant Negative |
| Lypd8l | TRUE | 0.223458023 | 0.684298079 | No Change | Not Significant |
| Lypla1 | TRUE | 0.021696983 | 0.892925964 | No Change | Not Significant |
| Lypla2 | TRUE | -0.11649178 | 0.342262172 | No Change | Not Significant |
| Lyplal1 | TRUE | 0.286750292 | 0.113460169 | No Change | Not Significant |
| Lyrm1 | TRUE | 0.134490871 | 0.751901591 | No Change | Not Significant |
| Lyrm2 | TRUE | -0.095429241 | 0.690909666 | No Change | Not Significant |
| Lyrm4 | TRUE | -0.055597655 | 0.806964824 | No Change | Not Significant |
| Lyrm7 | TRUE | 0.258639057 | 0.288970255 | No Change | Not Significant |
| Lyrm9 | TRUE | -0.110337859 | 0.648857873 | No Change | Not Significant |
| Lysmd1 | TRUE | -0.43875895 | 0.201550674 | No Change | Not Significant |
| Lysmd2 | TRUE | 0.280890067 | 0.469401688 | No Change | Not Significant |
| Lysmd3 | TRUE | 0.376538697 | 2.87E-05 | No Change | Significant Positive |
| Lysmd4 | TRUE | 0.371468471 | 0.001253501 | No Change | Significant Positive |
| Lyst | TRUE | -0.041005409 | 0.813637356 | No Change | Not Significant |
| Lyve1 | TRUE | 0.389983214 | 0.232919564 | No Change | Not Significant |
| Lyz1 | TRUE | 1.211750338 | 1 | No Change | Not Significant |
| Lyz2 | TRUE | -0.198579455 | 0.852575063 | No Change | Not Significant |
| Lyzl4 | TRUE | 0.20800613 | 0.719935187 | No Change | Not Significant |
| Lzic | TRUE | -0.001291746 | 0.99304207 | No Change | Not Significant |
| Lztfl1 | TRUE | 0.262079705 | 0.022468565 | No Change | Significant Positive |
| Lztr1 | TRUE | -0.017469807 | 0.870063538 | No Change | Not Significant |
| Lzts1 | TRUE | 0.293670084 | 0.776186935 | No Change | Not Significant |
| Lzts2 | TRUE | -0.274238837 | 0.000493703 | No Change | Significant Negative |
| Lzts3 | TRUE | 0.094725775 | 0.331436869 | No Change | Not Significant |
| M1ap | TRUE | 0.385757696 | 0.597566653 | No Change | Not Significant |
| M5C1000I18Rik | TRUE | 0.752624423 | 0.319862266 | No Change | Not Significant |
| M6pr | TRUE | 0.32317392 | 0.006157833 | No Change | Significant Positive |
| Mab21l1 | TRUE | -1.304444185 | 1 | No Change | Not Significant |
| Mab21l2 | TRUE | -0.085455187 | 0.93121061 | No Change | Not Significant |
| Mab21l3 | TRUE | 1.980674387 | 0.048196656 | UP | UP |
| Mab21l4 | TRUE | -0.223357072 | 0.432685604 | No Change | Not Significant |
| Macc1 | TRUE | -0.144772081 | 0.690541222 | No Change | Not Significant |
| Macf1 | TRUE | 0.164499244 | 0.266215502 | No Change | Not Significant |
| Macir | TRUE | -0.362357196 | 0.001525421 | No Change | Significant Negative |
| Maco1 | TRUE | 0.194590276 | 0.252289573 | No Change | Not Significant |
| Macrod1 | TRUE | -1.020699659 | 1.17E-11 | DOWN | DOWN |
| Macrod2 | TRUE | 0.197100373 | 0.228436418 | No Change | Not Significant |
| Macroh2a1 | TRUE | 0.093015369 | 0.208999876 | No Change | Not Significant |
| Macroh2a2 | TRUE | 0.573274029 | 0.160445154 | No Change | Not Significant |
| Mad1l1 | TRUE | 0.480524586 | 4.36E-06 | No Change | Significant Positive |
| Mad2l1 | TRUE | 0.848555841 | 1.49E-06 | UP | UP |
| Mad2l1bp | TRUE | 0.163928727 | 0.327398864 | No Change | Not Significant |
| Mad2l2 | TRUE | -0.203044015 | 0.321507788 | No Change | Not Significant |
| Madd | TRUE | -0.076256626 | 0.470017696 | No Change | Not Significant |
| Maea | TRUE | 0.113439989 | 0.075334803 | No Change | Not Significant |
| Maf | TRUE | 1.137136474 | 0.005729834 | UP | UP |
| Maf1 | TRUE | -0.036764645 | 0.734843633 | No Change | Not Significant |
| Mafa | TRUE | 4.067167623 | 1 | No Change | Not Significant |
| Mafb | TRUE | 0.303673614 | 0.508545065 | No Change | Not Significant |
| Maff | TRUE | -0.32672142 | 0.368018849 | No Change | Not Significant |
| Mafg | TRUE | -0.257581208 | 0.067913025 | No Change | Not Significant |
| Mafk | TRUE | -0.218823186 | 0.37880229 | No Change | Not Significant |
| Mag | TRUE | -1.232184668 | 1 | No Change | Not Significant |
| Mageb3 | TRUE | 0.38075342 | 1 | No Change | Not Significant |
| Maged1 | TRUE | 1.247008094 | 1.24E-29 | UP | UP |
| Maged2 | TRUE | 1.826406771 | 1.94E-27 | UP | UP |
| Magee1 | TRUE | 0.459395617 | 0.322338142 | No Change | Not Significant |
| Magef1 | TRUE | -0.356357595 | 0.210915627 | No Change | Not Significant |
| Mageh1 | TRUE | 0.465195928 | 0.039582411 | No Change | Significant Positive |
| Magel2 | TRUE | 0.333895151 | 1 | No Change | Not Significant |
| Magi1 | TRUE | -0.013818769 | 0.92158767 | No Change | Not Significant |
| Magi2 | TRUE | -0.181876693 | 0.88791795 | No Change | Not Significant |
| Magi3 | TRUE | 0.139281 | 0.01440064 | No Change | Significant Positive |
| Magix | TRUE | -0.384947457 | 1 | No Change | Not Significant |
| Magoh | TRUE | 0.27492277 | 0.050254003 | No Change | Not Significant |
| Magohb | TRUE | 0.326968244 | 0.146863309 | No Change | Not Significant |
| Magt1 | TRUE | 0.835283714 | 2.25E-12 | UP | UP |
| Maip1 | TRUE | 0.082726463 | 0.589522317 | No Change | Not Significant |
| Majin | TRUE | 0.940030791 | 1 | No Change | Not Significant |
| Mak | TRUE | 0.284915268 | 1 | No Change | Not Significant |
| Mak16 | TRUE | 0.222305524 | 0.163589437 | No Change | Not Significant |
| Mal | TRUE | 0.065998262 | 0.882132403 | No Change | Not Significant |
| Mal2 | TRUE | -0.141463849 | 0.184167865 | No Change | Not Significant |
| Malat1 | TRUE | -0.106279632 | 0.679736277 | No Change | Not Significant |
| Mall | TRUE | -0.119540049 | 0.553861375 | No Change | Not Significant |

Supplementary Table 3

| geneID | Cond1_DF_Detected | Cond1_DF_ValueCol | Cond1_DF_PadjCol | Cond1_DF_Cutoff | Cond1_DF_Cutoff_Specific.x | Cond2_DF_Detected | Cond2_DF_ValueCol | Cond2_DF_PadjCol | Cond2_DF_Cutoff | Cond2_DF_Cutoff_Specific | BG_Method | RG1_All | RG2_Significant | RG3_SignificantChange | Cond1_DF_ensembl_gene_id | Cond1_DF_entrezgene_id | Cond1_DF_gene_biotype | Cond1_DF_baseMean | Cond1_DF_log2FoldChange | Cond1_DF_lfcSE | Cond1_DF_stat | Cond1_DF_pvalue | Cond1_DF_padj | Cond1_DF_absLog2FC | Cond1_DF_Cutoff_Specific.y | Cond2_DF_ensembl_gene_id | Cond2_DF_entrezgene_id | Cond2_DF_gene_biotype | Cond2_DF_baseMean | Cond2_DF_log2FoldChange | Cond2_DF_lfcSE | Cond2_DF_stat | Cond2_DF_pvalue | Cond2_DF_padj | Cond2_DF_absLog2FC | RG1_Specific_Cond2 | RG1_Specific_Cond1 |
|---|---|---|---|---|---|---|---|---|---|---|---|---|---|---|---|---|---|---|---|---|---|---|---|---|---|---|---|---|---|---|---|---|---|---|---|---|---|
| Malrd1 | TRUE | -0.465886769 | 0.111571269 | No Change | Not Significant | TRUE | -0.274198786 | 0.171444799 | No Change | Not Significant | TRUE | Cond1 Not Significant + Cond2 Not Significant | None | None | ENSMUSG00000075520 | 102635496 | protein_coding | 307.4158627 | -0.465886769 | 0.220056926 | 2.117119323 | 0.034249722 | 0.111571269 | 0.465886769 | Not Significant | ENSMUSG00000075520 | 102635496 | protein_coding | 1615.936714 | -0.274198786 | 0.136577784 | -2.007638275 | 0.044681744 | 0.171444799 | 0.274198786 | Cond1 No Change + Cond2 Not Significant | Cond2 No Change + Cond1 Not Significant |
| Malsu1 | TRUE | 0.037146804 | 0.826943172 | No Change | Not Significant | TRUE | 0.046366761 | 0.737831675 | No Change | Not Significant | TRUE | Cond1 Not Significant + Cond2 Not Significant | None | None | ENSMUSG00000029815 | 75593 | protein_coding | 164.0933818 | 0.037146804 | 0.089185976 | -0.416509469 | 0.67703724 | 0.826943172 | 0.037146804 | Not Significant | ENSMUSG00000029815 | 75593 | protein_coding | 308.1406713 | 0.046366761 | 0.073484585 | 0.630972611 | 0.528058433 | 0.737831675 | 0.046366761 | Cond1 No Change + Cond2 Not Significant | Cond2 No Change + Cond1 Not Significant |
| Malt1 | TRUE | 0.309882321 | 0.000919359 | No Change | Significant Positive | TRUE | -0.189230852 | 0.060891613 | No Change | Not Significant | TRUE | Cond1 Significant Positive + Cond2 Not Significant | None | None | ENSMUSG00000032688 | 240354 | protein_coding | 364.5250924 | 0.309882321 | 0.080016037 | -3.872752672 | 0.000107613 | 0.000919359 | 0.309882321 | Significant Positive | ENSMUSG00000032688 | 240354 | protein_coding | 744.0464504 | -0.189230852 | 0.073811734 | -2.563696062 | 0.010356419 | 0.060891613 | 0.189230852 | Cond1 No Change + Cond2 Not Significant | Cond2 No Change + Cond1 Significant Positive |
| Mamdc2 | TRUE | 3.115193563 | 1 | No Change | Not Significant | FALSE | NA | NA | No Change | Not Detected | FALSE | Background = FALSE | Background = FALSE | Background = FALSE | ENSMUSG00000033207 | 71738 | protein_coding | 1.297942174 | 3.115193563 | 3.510891757 | -0.887294106 | 0.374920578 | 1 | 3.115193563 | Not Significant | NA | NA | NA | NA | NA | NA | NA | NA | NA | NA | Background = FALSE | Background = FALSE |
| Mamdc4 | TRUE | 0.088094764 | 0.850751312 | No Change | Not Significant | TRUE | -0.4294887 | 0.104486659 | No Change | Not Significant | TRUE | Cond1 Not Significant + Cond2 Not Significant | None | None | ENSMUSG00000026941 | 381352 | protein_coding | 53.96107224 | 0.088094764 | 0.23944918 | -0.367905893 | 0.712943406 | 0.850751312 | 0.088094764 | Not Significant | ENSMUSG00000026941 | 381352 | protein_coding | 84.38987332 | -0.4294887 | 0.187163003 | -2.294730762 | 0.021748559 | 0.104486659 | 0.4294887 | Cond1 No Change + Cond2 Not Significant | Cond2 No Change + Cond1 Not Significant |
| Maml1 | TRUE | -0.251779992 | 0.120138597 | No Change | Not Significant | TRUE | -0.443430672 | 1.65E-05 | No Change | Significant Negative | TRUE | Cond1 Not Significant + Cond2 Significant Negative | None | None | ENSMUSG00000050567 | 103806 | protein_coding | 211.9123572 | -0.251779992 | 0.121130257 | 2.07858877 | 0.037655164 | 0.120138597 | 0.251779992 | Not Significant | ENSMUSG00000050567 | 103806 | protein_coding | 416.7854673 | -0.443430672 | 0.08908412 | -4.977662377 | 6.44E-07 | 1.65E-05 | 0.443430672 | Cond1 No Change + Cond2 Significant Negative | Cond2 No Change + Cond1 Not Significant |
| Maml2 | TRUE | -0.216729065 | 0.867757454 | No Change | Not Significant | TRUE | 0.266568354 | 0.741375537 | No Change | Not Significant | TRUE | Cond1 Not Significant + Cond2 Not Significant | None | None | ENSMUSG00000031925 | 270118 | protein_coding | 4.791131102 | -0.216729065 | 0.651353286 | 0.332736581 | 0.739333134 | 0.867757454 | 0.216729065 | Not Significant | ENSMUSG00000031925 | 270118 | protein_coding | 8.385737191 | 0.266568354 | 0.427996432 | 0.62282845 | 0.533397261 | 0.741375537 | 0.266568354 | Cond1 No Change + Cond2 Not Significant | Cond2 No Change + Cond1 Not Significant |
| Maml3 | TRUE | 0.111960158 | 0.408255959 | No Change | Not Significant | TRUE | -0.08277399 | 0.529611374 | No Change | Not Significant | TRUE | Cond1 Not Significant + Cond2 Not Significant | None | None | ENSMUSG00000061143 | 433586 | protein_coding | 314.9984307 | 0.111960158 | 0.088451362 | -1.265782187 | 0.205591074 | 0.408255959 | 0.111960158 | Not Significant | ENSMUSG00000061143 | 433586 | protein_coding | 573.3134804 | -0.08277399 | 0.077782256 | -1.064175751 | 0.28724909 | 0.529611374 | 0.08277399 | Cond1 No Change + Cond2 Not Significant | Cond2 No Change + Cond1 Not Significant |
| Mamld1 | TRUE | -0.283044729 | 0.566884219 | No Change | Not Significant | TRUE | 0.235360104 | 0.698577622 | No Change | Not Significant | TRUE | Cond1 Not Significant + Cond2 Not Significant | None | None | ENSMUSG00000059401 | 333639 | protein_coding | 13.40253352 | -0.283044729 | 0.299872341 | 0.943884079 | 0.345228888 | 0.566884219 | 0.283044729 | Not Significant | ENSMUSG00000059401 | 333639 | protein_coding | 15.43779359 | 0.235360104 | 0.329446958 | 0.714409707 | 0.474973871 | 0.698577622 | 0.235360104 | Cond1 No Change + Cond2 Not Significant | Cond2 No Change + Cond1 Not Significant |
| Mamstr | TRUE | 1.719682403 | 0.000207367 | UP | UP | TRUE | 0.094548793 | 0.880475742 | No Change | Not Significant | TRUE | Cond1 UP + Cond2 Not Significant | Cond1_UP | Cond1_UP | ENSMUSG00000042918 | 74490 | protein_coding | 21.79031266 | 1.719682403 | 0.403393592 | -4.263038471 | 2.02E-05 | 0.000207367 | 1.719682403 | UP | ENSMUSG00000042918 | 74490 | protein_coding | 16.77213961 | 0.094548793 | 0.296380066 | 0.319011986 | 0.749717424 | 0.880475742 | 0.094548793 | Cond1 UP + Cond2 Not Significant | Cond2 No Change + Cond1 UP |
| Man1a | TRUE | -0.247025764 | 0.001488577 | No Change | Significant Negative | TRUE | -0.078374306 | 0.302001032 | No Change | Not Significant | TRUE | Cond1 Significant Negative + Cond2 Not Significant | None | None | ENSMUSG00000003746 | 17155 | protein_coding | 1001.314946 | -0.247025764 | 0.066094094 | 3.737486192 | 0.000185869 | 0.001488577 | 0.247025764 | Significant Negative | ENSMUSG00000003746 | 17155 | protein_coding | 2465.111352 | -0.078374306 | 0.048865332 | -1.603883609 | 0.10873971 | 0.302001032 | 0.078374306 | Cond1 No Change + Cond2 Not Significant | Cond2 No Change + Cond1 Significant Negative |

Supplementary Table 3

| geneID | Cond1_DF_Detected | Cond1_DF_ValueCol | Cond1_DF_PadjCol | Cond1_DF_Cutoff |
|---|---|---|---|---|
| Mcmbp | TRUE | -0.07708786 | 0.398004294 | No Change |
| Mcmdc2 | TRUE | -0.190560289 | 0.899929685 | No Change |
| Mcoln1 | TRUE | -0.238786506 | 0.060935759 | No Change |
| Mcoln2 | TRUE | 0.112486818 | 0.656764359 | No Change |
| Mcoln3 | TRUE | 0.403349639 | 0.64425321 | No Change |
| Mcph1 | TRUE | 0.424533018 | 0.000844958 | No Change |
| Mcpt1 | TRUE | -0.400502756 | 0.677309474 | No Change |
| Mcpt2 | TRUE | -0.305526885 | 1 | No Change |
| Mcpt9 | TRUE | -2.511280455 | 1 | No Change |
| Mcrip1 | TRUE | -0.431594617 | 0.023285356 | No Change |
| Mcrip2 | TRUE | -0.635847541 | 9.57E-07 | DOWN |
| Mcrs1 | TRUE | -0.222960333 | 0.061495803 | No Change |
| Mctp1 | TRUE | -0.539664233 | 0.031903925 | DOWN |
| Mctp2 | TRUE | 0.647163144 | 5.46E-16 | UP |
| Mcts1 | TRUE | 0.175364306 | 0.176840786 | No Change |
| Mcts2 | TRUE | 0.009354106 | 0.976016991 | No Change |
| Mcu | TRUE | 0.053853093 | 0.614062569 | No Change |
| Mcub | TRUE | -0.59834082 | 0.023575839 | DOWN |
| Mcur1 | TRUE | -0.284784087 | 0.000647282 | No Change |
| Mdc1 | TRUE | 0.108639744 | 0.599511678 | No Change |
| Mdfi | TRUE | 0.455927113 | 0.585225632 | No Change |
| Mdfic | TRUE | 0.001907998 | 0.994724011 | No Change |
| Mdfic2 | TRUE | 0.416245099 | 1 | No Change |
| Mdga1 | TRUE | 0.060584822 | 0.959274295 | No Change |
| Mdga2 | TRUE | -0.559315516 | 1 | No Change |
| Mdh1 | TRUE | 1.247367001 | 2.73E-61 | UP |
| Mdh1b | TRUE | 1.512890776 | 1 | No Change |
| Mdh2 | TRUE | 0.113184491 | 0.321507788 | No Change |
| Mdk | TRUE | 0.079792592 | 0.943016089 | No Change |
| Mdm1 | TRUE | 1.793302828 | 2.30E-37 | UP |
| Mdm2 | TRUE | 0.054105887 | 0.736636318 | No Change |
| Mdm4 | TRUE | -0.031294017 | 0.839403501 | No Change |
| Mdn1 | TRUE | 0.102797169 | 0.635253445 | No Change |
| Mdp1 | TRUE | -0.038114162 | 0.841315261 | No Change |
| Me1 | TRUE | 0.332633958 | 0.225331394 | No Change |
| Me2 | TRUE | -0.106304017 | 0.50724071 | No Change |
| Me3 | TRUE | -0.15654281 | 0.859322531 | No Change |
| Mea1 | TRUE | -0.219540193 | 0.008308292 | No Change |
| Meaf6 | TRUE | -0.396618848 | 0.000193921 | No Change |
| Meak7 | TRUE | -0.022199311 | 0.927499565 | No Change |
| Mecom | TRUE | -0.313331381 | 0.0089816 | No Change |
| Mecp2 | TRUE | 0.002321396 | 0.98611318 | No Change |
| Mecr | TRUE | -0.055533145 | 0.827931507 | No Change |
| Med1 | TRUE | 0.144795428 | 0.009447848 | No Change |
| Med10 | TRUE | 0.058080722 | 0.65055862 | No Change |
| Med11 | TRUE | -0.032759723 | 0.886362513 | No Change |
| Med12 | TRUE | 0.007946554 | 0.966128972 | No Change |
| Med12l | TRUE | -0.216913132 | 0.723223689 | No Change |
| Med13 | TRUE | 0.090245199 | 0.380389352 | No Change |
| Med13l | TRUE | 0.12311756 | 0.570573694 | No Change |
| Med14 | TRUE | 0.412353565 | 0.000465269 | No Change |
| Med15 | TRUE | 0.01735429 | 0.930446023 | No Change |
| Med16 | TRUE | 0.011206505 | 0.956997404 | No Change |
| Med17 | TRUE | -0.140626553 | 0.246796371 | No Change |
| Med18 | TRUE | -0.066206484 | 0.761183901 | No Change |
| Med19 | TRUE | 0.196692533 | 0.063995287 | No Change |
| Med20 | TRUE | 0.113863998 | 0.397106786 | No Change |
| Med21 | TRUE | -0.001364518 | 0.994433839 | No Change |
| Med22 | TRUE | -0.230503783 | 0.167087933 | No Change |
| Med23 | TRUE | 0.026913402 | 0.855153117 | No Change |
| Med24 | TRUE | 0.139040917 | 0.137877355 | No Change |
| Med25 | TRUE | 0.095599878 | 0.394331932 | No Change |
| Med26 | TRUE | 0.081917136 | 0.643827591 | No Change |
| Med27 | TRUE | 0.250842986 | 0.063542317 | No Change |
| Med28 | TRUE | -0.159020027 | 0.190566644 | No Change |
| Med29 | TRUE | 0.019285778 | 0.942211125 | No Change |
| Med30 | TRUE | 0.116797053 | 0.582642414 | No Change |
| Med31 | TRUE | 0.094729703 | 0.720614094 | No Change |
| Med4 | TRUE | 0.038686139 | 0.835769466 | No Change |
| Med6 | TRUE | 0.184705161 | 0.233277344 | No Change |
| Med7 | TRUE | 0.042702409 | 0.84834929 | No Change |
| Med8 | TRUE | -0.140959726 | 0.140153435 | No Change |
| Med9 | TRUE | 0.315497312 | 0.021880184 | No Change |
| Med9os | TRUE | 1.03519968 | 1 | No Change |
| Medag | TRUE | -0.707063821 | 1 | No Change |
| Mef2a | TRUE | 0.123936123 | 0.237733773 | No Change |
| Mef2b | TRUE | -0.011509836 | 1 | No Change |
| Mef2c | TRUE | 1.256342981 | 0.649597228 | No Change |
| Mef2d | TRUE | 0.016068125 | 0.952874415 | No Change |
| Mefv | TRUE | -1.350050069 | 1 | No Change |
| Meg3 | TRUE | -4.175766366 | 1.86E-57 | DOWN |
| Megf10 | TRUE | 0.005749286 | 1 | No Change |
| Megf11 | TRUE | -0.250300596 | 0.498019388 | No Change |
| Megf6 | TRUE | -0.180511378 | 0.881056435 | No Change |
| Megf8 | TRUE | -0.053967189 | 0.761298159 | No Change |
| Megf9 | TRUE | 0.302775524 | 0.129592876 | No Change |
| Mei1 | TRUE | 1.016396951 | 1 | No Change |
| Mei4 | TRUE | 0.011525254 | 1 | No Change |
| Meig1 | TRUE | -0.304593136 | 1 | No Change |
| Meiob | TRUE | -0.685618257 | 1 | No Change |
| Meioc | TRUE | 0.322939255 | 1 | No Change |
| Meiosin | TRUE | 0.593304769 | 1 | No Change |
| Meis1 | TRUE | 1.093330825 | 1 | No Change |
| Meis2 | TRUE | 0.975309738 | 1 | No Change |
| Meis3 | TRUE | -0.47727088 | 0.02983307 | No Change |
| Melk | TRUE | 0.846576541 | 2.88E-05 | UP |
| Meltf | TRUE | -0.449908038 | 1 | No Change |
| Memo1 | TRUE | -0.161667423 | 0.138818348 | No Change |
| Men1 | TRUE | 0.109526443 | 0.541253166 | No Change |
| Meox1 | TRUE | 1.936826516 | 6.31E-07 | UP |
| Mep1a | TRUE | -0.57363312 | 1.06E-05 | DOWN |
| Mep1b | TRUE | -0.419936155 | 0.036593531 | No Change |
| Mepce | TRUE | 0.005114043 | 0.975869522 | No Change |
| Mertk | TRUE | -0.26490581 | 0.12190868 | No Change |
| Mesd | TRUE | 1.061246267 | 1.98E-24 | UP |
| Mesp2 | TRUE | 1.199060956 | 1 | No Change |
| Mest | TRUE | -0.497244424 | 0.449590928 | No Change |
| Met | TRUE | -0.29638196 | 0.004863751 | No Change |
| Metap1 | TRUE | 0.112915958 | 0.465324602 | No Change |
| Metap1d | TRUE | -0.116780087 | 0.450443439 | No Change |
| Metap2 | TRUE | 0.021326789 | 0.884145751 | No Change |
| Methig1 | TRUE | -2.962297115 | 1 | No Change |
| Metrn | TRUE | 0.435866018 | 0.23455206 | No Change |
| Metrnl | TRUE | 0.018171448 | 0.948731562 | No Change |
| Mettl1 | TRUE | 0.118805461 | 0.632451088 | No Change |
| Mettl14 | TRUE | -0.045072227 | 0.833904116 | No Change |
| Mettl15 | TRUE | -0.194344923 | 0.450382263 | No Change |
| Mettl16 | TRUE | 0.247894613 | 0.033210765 | No Change |
| Mettl17 | TRUE | -0.1849698 | 0.132446542 | No Change |
| Mettl18 | TRUE | 0.027248547 | 0.939901695 | No Change |
| Mettl2 | TRUE | -0.235331816 | 0.182841722 | No Change |
| Mettl21a | TRUE | 0.142637372 | 0.53655334 | No Change |
| Mettl21c | TRUE | 0.626539115 | 0.448589299 | No Change |
| Mettl22 | TRUE | -0.26050645 | 0.343804463 | No Change |
| Mettl23 | TRUE | -0.252551919 | 0.067342119 | No Change |
| Mettl24 | TRUE | 0.260257892 | 0.584685663 | No Change |
| Mettl25 | TRUE | 0.190499946 | 0.445943657 | No Change |
| Mettl26 | TRUE | -0.966139697 | 2.28E-39 | DOWN |
| Mettl27 | TRUE | -0.268131828 | 0.059578222 | No Change |
| Mettl3 | TRUE | -0.14022034 | 0.477601877 | No Change |
| Mettl4 | TRUE | -0.089420819 | 0.709931167 | No Change |
| Mettl5 | TRUE | -0.168940978 | 0.338979766 | No Change |
| Mettl6 | TRUE | 0.086938513 | 0.495777145 | No Change |
| Mettl7a1 | TRUE | -0.547496873 | 7.11E-05 | DOWN |
| Mettl7a2 | TRUE | -1.507073865 | 0.000418111 | DOWN |
| Mettl7a3 | TRUE | -1.091920857 | 0.036011679 | DOWN |
| Mettl7b | TRUE | -0.470090749 | 0.014644071 | No Change |
| Mettl8 | TRUE | 0.062569794 | 0.778258299 | No Change |
| Mettl9 | TRUE | 0.121541636 | 0.36973067 | No Change |
| Mex3a | TRUE | 0.149998997 | 0.677560974 | No Change |
| Mex3b | TRUE | 0.673855731 | 0.07201649 | No Change |
| Mex3c | TRUE | 0.18494024 | 0.032409794 | No Change |
| Mex3d | TRUE | -0.088824355 | 0.712111524 | No Change |
| Mexis | TRUE | -0.047478988 | 0.92788168 | No Change |
| Mfap1a | TRUE | -0.166251927 | 0.382601993 | No Change |
| Mfap1b | TRUE | -0.221626809 | 0.252989943 | No Change |
| Mfap2 | TRUE | 0.363339231 | 1 | No Change |
| Mfap3 | TRUE | 0.141790006 | 0.179403047 | No Change |
| Mfap3l | TRUE | -0.378614591 | 0.076601296 | No Change |
| Mfap4 | TRUE | -1.857721957 | 1 | No Change |
| Mfap5 | TRUE | -0.98878853 | 1 | No Change |
| Mff | TRUE | -0.053820558 | 0.671972423 | No Change |
| Mfge8 | TRUE | 1.505564026 | 9.73E-67 | UP |
| Mfhas1 | TRUE | 0.007835577 | 0.965818434 | No Change |
| Mfn1 | TRUE | 0.046851619 | 0.68930064 | No Change |
| Mfn2 | TRUE | 0.048967107 | 0.715988936 | No Change |
| Mfng | TRUE | 1.388906932 | 0.1532944 | No Change |
| Mfrp | TRUE | 1.589140364 | 0.020673719 | UP |
| Mfsd1 | TRUE | -0.197705104 | 0.098274011 | No Change |
| Mfsd10 | TRUE | 0.774678351 | 6.21E-12 | UP |
| Mfsd11 | TRUE | 0.461795869 | 5.07E-08 | No Change |
| Mfsd12 | TRUE | 1.849561526 | 1.41E-46 | UP |
| Mfsd13a | TRUE | -0.27037843 | 0.047677228 | No Change |
| Mfsd13b | TRUE | 0.693531729 | 0.293636687 | No Change |
| Mfsd14a | TRUE | 0.050849037 | 0.745972916 | No Change |
| Mfsd14b | TRUE | -0.067581404 | 0.431277012 | No Change |
| Mfsd2a | TRUE | 2.168458122 | 1.71E-29 | UP |
| Mfsd2b | TRUE | 0.613827405 | 0.328844638 | No Change |
| Mfsd3 | TRUE | -0.312790291 | 0.151424575 | No Change |
| Mfsd4a | TRUE | -0.141133389 | 0.652884901 | No Change |
| Mfsd4b1 | TRUE | 0.103830259 | 1 | No Change |
| Mfsd4b4 | TRUE | 0.492484066 | 0.112032105 | No Change |
| Mfsd4b5 | TRUE | 0.150292162 | 0.882510888 | No Change |
| Mfsd5 | TRUE | 0.183575234 | 0.083506033 | No Change |
| Mfsd6 | TRUE | -0.003228963 | 0.985840779 | No Change |
| Mfsd6l | TRUE | -0.038823255 | 0.830405926 | No Change |
| Mfsd7a | TRUE | 0.118345312 | 0.660786267 | No Change |
| Mfsd8 | TRUE | 0.031018764 | 0.891264005 | No Change |
| Mfsd9 | TRUE | -0.034002693 | 0.884510137 | No Change |
| Mga | TRUE | -0.162263617 | 0.163634492 | No Change |
| Mgam | TRUE | 0.122116725 | 0.407353559 | No Change |
| Mgarp | TRUE | 2.855639196 | 1 | No Change |
| Mgat1 | TRUE | -0.165244858 | 0.038640192 | No Change |
| Mgat2 | TRUE | 0.51069531 | 5.82E-08 | UP |
| Mgat3 | TRUE | 0.206885104 | 0.001419217 | No Change |
| Mgat4a | TRUE | -0.713087409 | 2.88E-05 | DOWN |
| Mgat4b | TRUE | 0.364188809 | 1.89E-08 | No Change |
| Mgat4c | TRUE | 0.509526724 | 0.000199648 | UP |
| Mgat5 | TRUE | -0.202537969 | 0.034412292 | No Change |
| Mgat5b | TRUE | -1.637941138 | 1 | No Change |
| Mgl2 | TRUE | 0.348574296 | 1 | No Change |
| Mgll | TRUE | 0.305269977 | 0.335154287 | No Change |

Supplementary Table 3

| geneID | Cond1_DF_Detected | Cond1_DF_ValueCol | Cond1_DF_PadjCol | Cond1_DF_Cutoff | Cond1_DF_Cutoff_Specific.x | Cond2_DF_Detected | Cond2_DF_ValueCol |
|---|---|---|---|---|---|---|---|
| Mgme1 | TRUE | -0.040941574 | 0.891196207 | No Change | Not Significant | TRUE | 0.063072997 |
| Mgmt | TRUE | -0.018997988 | 0.967478151 | No Change | Not Significant | TRUE | 0.091130699 |
| Mgp | TRUE | -0.284041886 | 1 | No Change | Not Significant | FALSE | NA |
| Mgrn1 | TRUE | -0.078181093 | 0.388417088 | No Change | Not Significant | TRUE | 0.038254211 |
| Mgst1 | TRUE | 0.076164051 | 0.646068054 | No Change | Not Significant | TRUE | -0.395795739 |
| Mgst2 | TRUE | -0.091247846 | 0.412914333 | No Change | Not Significant | TRUE | -0.059326714 |
| Mgst3 | TRUE | -0.469680091 | 8.36E-06 | No Change | Significant Negative | TRUE | 0.066445421 |
| Mia | TRUE | -0.08409649 | 0.921900136 | No Change | Not Significant | TRUE | 0.531707659 |
| Mia2 | TRUE | -0.046492238 | 0.643978573 | No Change | Not Significant | TRUE | -0.109597825 |
| Mia3 | TRUE | 0.281786686 | 0.164593053 | No Change | Not Significant | TRUE | 0.141740908 |
| Miat | TRUE | -0.535831858 | 0.67261003 | No Change | Not Significant | TRUE | -0.419836108 |
| Mib1 | TRUE | 0.202725377 | 0.036886891 | No Change | Significant Positive | TRUE | -0.00259509 |
| Mib2 | TRUE | -0.259604599 | 0.051357818 | No Change | Not Significant | TRUE | 0.029232572 |
| Mical1 | TRUE | 0.078862671 | 0.792979237 | No Change | Not Significant | TRUE | -0.148801496 |
| Mical2 | TRUE | -0.22071757 | 0.042179026 | No Change | Significant Negative | TRUE | -0.238965299 |
| Mical3 | TRUE | 0.773468916 | 0.000112689 | UP | UP | TRUE | 0.170645653 |
| Micall1 | TRUE | -0.215816589 | 0.104749157 | No Change | Not Significant | TRUE | -0.021612305 |
| Micall2 | TRUE | 0.38471319 | 0.025783888 | No Change | Significant Positive | TRUE | 0.056100646 |
| Micos10 | TRUE | -0.090474771 | 0.582491838 | No Change | Not Significant | TRUE | 0.03616618 |
| Micos13 | TRUE | -0.373741273 | 0.000602545 | No Change | Significant Negative | TRUE | 0.007600401 |
| Micu1 | TRUE | -0.380986705 | 2.80E-20 | No Change | Significant Negative | TRUE | -0.134095358 |
| Micu2 | TRUE | -0.123499749 | 0.19654843 | No Change | Not Significant | TRUE | -0.048719087 |
| Micu3 | TRUE | 0.306968967 | 0.535990818 | No Change | Not Significant | TRUE | -0.504220009 |
| Mid1 | TRUE | 0.058095514 | 0.907147741 | No Change | Not Significant | TRUE | -0.058351247 |
| Mid1ip1 | TRUE | 0.096857153 | 0.601768559 | No Change | Not Significant | TRUE | 0.251240301 |
| Mid2 | TRUE | 0.736201028 | 4.31E-19 | UP | UP | TRUE | 0.344249561 |
| Mideas | TRUE | 0.224464725 | 0.364427834 | No Change | Not Significant | TRUE | -0.020061701 |
| Midn | TRUE | 0.110570008 | 0.653940181 | No Change | Not Significant | TRUE | 0.059722041 |
| Mief1 | TRUE | 0.010952647 | 0.932569867 | No Change | Not Significant | TRUE | -0.013392306 |
| Mief2 | TRUE | 0.778155293 | 5.42E-14 | UP | UP | TRUE | 0.46917373 |
| Mien1 | TRUE | 0.151942232 | 0.224470391 | No Change | Not Significant | TRUE | 0.326475074 |
| Mier1 | TRUE | -0.017564331 | 0.93927028 | No Change | Not Significant | TRUE | -0.240472149 |
| Mier2 | TRUE | 0.657373216 | 3.48E-06 | UP | UP | TRUE | 0.139125176 |
| Mier3 | TRUE | -0.286453238 | 0.076382586 | No Change | Not Significant | TRUE | -0.406801966 |
| Mif | TRUE | 0.113891886 | 0.632329199 | No Change | Not Significant | TRUE | 0.394671369 |
| Mif4gd | TRUE | -0.332206987 | 0.000298838 | No Change | Significant Negative | TRUE | -0.035611644 |
| Miga1 | TRUE | -0.447827378 | 0.000421772 | No Change | Significant Negative | TRUE | -0.304940999 |
| Miga2 | TRUE | 0.123140639 | 0.295417576 | No Change | Not Significant | TRUE | 0.111235488 |
| Miip | TRUE | 0.156938057 | 0.226849015 | No Change | Not Significant | TRUE | -0.330601964 |
| Mill2 | TRUE | -0.120201425 | 0.736796542 | No Change | Not Significant | TRUE | -0.283243134 |
| Milr1 | TRUE | 0.268882492 | 0.784646895 | No Change | Not Significant | TRUE | 0.204810084 |
| Minar1 | TRUE | -0.808148857 | 1 | No Change | Not Significant | TRUE | 0.00562987 |
| Mindy1 | TRUE | -0.288582934 | 0.00151488 | No Change | Significant Negative | TRUE | -0.068585418 |
| Mindy2 | TRUE | -0.048405449 | 0.83310074 | No Change | Not Significant | TRUE | -0.063300739 |
| Mindy3 | TRUE | 0.066611907 | 0.5351691 | No Change | Not Significant | TRUE | 0.15713981 |
| Mindy4 | TRUE | 0.018886975 | 0.983482583 | No Change | Not Significant | TRUE | 0.667740053 |
| Mink1 | TRUE | -0.177741446 | 0.222084117 | No Change | Not Significant | TRUE | -0.00952658 |
| Minpp1 | TRUE | -0.027209297 | 0.908812719 | No Change | Not Significant | TRUE | -0.039429361 |
| Mios | TRUE | -0.112391783 | 0.401975042 | No Change | Not Significant | TRUE | -0.026413235 |
| Mip | TRUE | 0.894638867 | 1 | No Change | Not Significant | FALSE | NA |
| Mipep | TRUE | 0.030843846 | 0.873111243 | No Change | Not Significant | TRUE | 0.164824883 |
| Mipol1 | TRUE | -0.719164259 | 0.002788738 | DOWN | DOWN | TRUE | -0.267065367 |
| Mir100hg | TRUE | 0.262373999 | 0.929905153 | No Change | Not Significant | FALSE | NA |
| Mir10a | TRUE | -0.774890507 | 1 | No Change | Not Significant | FALSE | NA |
| Mir1188 | TRUE | -3.520060915 | 1 | No Change | Not Significant | FALSE | NA |
| Mir1190 | TRUE | 0.742350495 | 1 | No Change | Not Significant | FALSE | NA |
| Mir1192 | TRUE | -0.279976511 | 1 | No Change | Not Significant | TRUE | -1.233230265 |
| Mir1193 | TRUE | -2.7119888 | 1 | No Change | Not Significant | FALSE | NA |
| Mir1198 | TRUE | 0.82796429 | 1 | No Change | Not Significant | FALSE | NA |
| Mir124a-1hg | TRUE | -1.678386512 | 1 | No Change | Not Significant | FALSE | NA |
| Mir133a-1 | TRUE | -0.185079915 | 1 | No Change | Not Significant | FALSE | NA |
| Mir141 | TRUE | -0.787516947 | 1 | No Change | Not Significant | FALSE | NA |
| Mir148a | TRUE | -0.788245168 | 1 | No Change | Not Significant | FALSE | NA |
| Mir148b | TRUE | -0.480228289 | 1 | No Change | Not Significant | FALSE | NA |
| Mir150 | TRUE | -1.85534298 | 1 | No Change | Not Significant | FALSE | NA |
| Mir15a | TRUE | -1.353013989 | 1 | No Change | Not Significant | FALSE | NA |
| Mir181a-2 | TRUE | -1.275426418 | 1 | No Change | Not Significant | FALSE | NA |
| Mir182 | TRUE | 0.736889213 | 1 | No Change | Not Significant | FALSE | NA |
| Mir185 | TRUE | 0.606501033 | 1 | No Change | Not Significant | FALSE | NA |
| Mir186 | TRUE | -0.067290061 | 1 | No Change | Not Significant | FALSE | NA |
| Mir1898 | TRUE | -0.654632623 | 1 | No Change | Not Significant | FALSE | NA |
| Mir1904 | TRUE | 0.204456635 | 1 | No Change | Not Significant | FALSE | NA |
| Mir192 | TRUE | 0.390934888 | 0.604440039 | No Change | Not Significant | TRUE | 0.207055634 |
| Mir1927 | TRUE | -0.36877076 | 1 | No Change | Not Significant | FALSE | NA |
| Mir1931 | TRUE | -0.208989211 | 1 | No Change | Not Significant | FALSE | NA |
| Mir1932 | TRUE | 0.031321947 | 1 | No Change | Not Significant | FALSE | NA |
| Mir1933 | TRUE | 1.488759848 | 1 | No Change | Not Significant | FALSE | NA |
| Mir1936 | TRUE | 1.015485646 | 1 | No Change | Not Significant | FALSE | NA |
| Mir1938 | TRUE | -0.483955685 | 0.261778433 | No Change | Not Significant | TRUE | -0.483688288 |
| Mir193a | TRUE | -0.28424178 | 1 | No Change | Not Significant | FALSE | NA |
| Mir194-1 | TRUE | 0.141232963 | 1 | No Change | Not Significant | FALSE | NA |
| Mir194-2 | TRUE | 0.64503219 | 0.252397465 | No Change | Not Significant | TRUE | 0.124261827 |
| Mir1943 | TRUE | 0.031959333 | 1 | No Change | Not Significant | FALSE | NA |
| Mir1945 | TRUE | -1.698283058 | 1 | No Change | Not Significant | FALSE | NA |
| Mir1960 | TRUE | -1.014584629 | 1 | No Change | Not Significant | FALSE | NA |
| Mir1968 | TRUE | -1.313951572 | 0.03324812 | DOWN | DOWN | TRUE | 0.087361401 |
| Mir196a-1 | TRUE | -0.34404645 | 0.345012479 | No Change | Not Significant | TRUE | -0.064093227 |
| Mir1970 | TRUE | 0.799150847 | 1 | No Change | Not Significant | FALSE | NA |
| Mir1983 | TRUE | 0.695727226 | 1 | No Change | Not Significant | FALSE | NA |
| Mir199a-1 | TRUE | -0.219656101 | 1 | No Change | Not Significant | FALSE | NA |
| Mir200a | TRUE | -0.039785955 | 0.949794479 | No Change | Not Significant | TRUE | -0.226290599 |
| Mir200b | TRUE | 0.650993501 | 1 | No Change | Not Significant | FALSE | NA |
| Mir200c | TRUE | 1.573195802 | 1 | No Change | Not Significant | FALSE | NA |
| Mir203 | TRUE | -1.28526275 | 1 | No Change | Not Significant | FALSE | NA |
| Mir210 | TRUE | 0.16422262 | 1 | No Change | Not Significant | FALSE | NA |
| Mir212 | TRUE | -1.624004428 | 1 | No Change | Not Significant | FALSE | NA |
| Mir2137 | TRUE | 1.204326561 | 1 | No Change | Not Significant | FALSE | NA |
| Mir221 | TRUE | -0.579847218 | 1 | No Change | Not Significant | FALSE | NA |
| Mir22hg | TRUE | -0.524665042 | 0.000602988 | DOWN | DOWN | TRUE | -0.569913174 |
| Mir23a | TRUE | -0.676001431 | 0.058405732 | No Change | Not Significant | TRUE | -0.32025069 |
| Mir23b | TRUE | 0.848265384 | 1 | No Change | Not Significant | FALSE | NA |
| Mir24-2 | TRUE | 1.343537881 | 1 | No Change | Not Significant | FALSE | NA |
| Mir26a-1 | TRUE | 1.5634573 | 1 | No Change | Not Significant | FALSE | NA |
| Mir26a-2 | TRUE | 0.258802322 | 1 | No Change | Not Significant | FALSE | NA |
| Mir26b | TRUE | 0.605896151 | 1 | No Change | Not Significant | FALSE | NA |
| Mir27a | TRUE | -0.955578888 | 0.02077642 | DOWN | DOWN | TRUE | -0.252483488 |
| Mir28a | TRUE | -1.157564374 | 1 | No Change | Not Significant | FALSE | NA |
| Mir28c | TRUE | 0.557609492 | 1 | No Change | Not Significant | FALSE | NA |
| Mir29a | TRUE | 1.162624841 | 1 | No Change | Not Significant | TRUE | -0.084015978 |
| Mir301b | TRUE | 1.267997826 | 1 | No Change | Not Significant | FALSE | NA |
| Mir3058 | TRUE | -1.387245404 | 1 | No Change | Not Significant | FALSE | NA |
| Mir3060 | TRUE | -0.021662145 | 1 | No Change | Not Significant | FALSE | NA |
| Mir3075 | TRUE | -0.034383874 | 1 | No Change | Not Significant | FALSE | NA |
| Mir3081 | TRUE | 0.206824022 | 1 | No Change | Not Significant | FALSE | NA |
| Mir3091 | TRUE | -0.258984816 | 1 | No Change | Not Significant | FALSE | NA |
| Mir3092 | TRUE | 2.266345986 | 1 | No Change | Not Significant | FALSE | NA |
| Mir30a | TRUE | 0.536189771 | 1 | No Change | Not Significant | FALSE | NA |
| Mir30b | TRUE | -0.325306091 | 1 | No Change | Not Significant | FALSE | NA |
| Mir30c-1 | TRUE | 0.375042826 | 1 | No Change | Not Significant | FALSE | NA |
| Mir3105 | TRUE | -0.809404852 | 1 | No Change | Not Significant | FALSE | NA |
| Mir3109 | TRUE | -0.570251625 | 0.13974969 | No Change | Not Significant | TRUE | -0.645415552 |
| Mir320 | TRUE | 0.828807476 | 1 | No Change | Not Significant | FALSE | NA |
| Mir340 | TRUE | -1.58077567 | 1 | No Change | Not Significant | FALSE | NA |
| Mir341 | TRUE | -3.625453265 | 1 | No Change | Not Significant | FALSE | NA |
| Mir342 | TRUE | -0.375045051 | 1 | No Change | Not Significant | FALSE | NA |
| Mir343 | TRUE | 0.689070346 | 1 | No Change | Not Significant | FALSE | NA |
| Mir345 | TRUE | 0.932349776 | 1 | No Change | Not Significant | FALSE | NA |
| Mir3473a | TRUE | -0.6653201 | 1 | No Change | Not Significant | FALSE | NA |
| Mir3473e | TRUE | 0.329994827 | 1 | No Change | Not Significant | FALSE | NA |
| Mir3473g | TRUE | -0.787378117 | 1 | No Change | Not Significant | FALSE | NA |
| Mir3569 | TRUE | 1.646146288 | 1 | No Change | Not Significant | FALSE | NA |
| Mir375 | TRUE | -0.344142716 | 1 | No Change | Not Significant | FALSE | NA |
| Mir378b | TRUE | 0.338174944 | 1 | No Change | Not Significant | FALSE | NA |
| Mir423 | TRUE | -2.118775262 | 1 | No Change | Not Significant | FALSE | NA |
| Mir453 | TRUE | -3.726601777 | 1 | No Change | Not Significant | FALSE | NA |
| Mir466i | TRUE | -0.146570712 | 1 | No Change | Not Significant | FALSE | NA |
| Mir505 | TRUE | -0.441096104 | 1 | No Change | Not Significant | FALSE | NA |
| Mir5099 | TRUE | 2.582476486 | 1 | No Change | Not Significant | FALSE | NA |
| Mir5100 | TRUE | 0.176029121 | 1 | No Change | Not Significant | FALSE | NA |
| Mir5119 | TRUE | 0.43472486 | 1 | No Change | Not Significant | FALSE | NA |
| Mir5123 | TRUE | -0.396396717 | 1 | No Change | Not Significant | FALSE | NA |
| Mir5125 | TRUE | -0.023913029 | 0.973720842 | No Change | Not Significant | TRUE | -0.529100993 |
| Mir5128 | TRUE | -1.92398528 | 1 | No Change | Not Significant | FALSE | NA |
| Mir5135 | TRUE | -0.333547335 | 1 | No Change | Not Significant | FALSE | NA |
| Mir5136 | TRUE | 0.395151452 | 1 | No Change | Not Significant | FALSE | NA |
| Mir540 | TRUE | -3.415096596 | 1 | No Change | Not Significant | FALSE | NA |
| Mir5620 | TRUE | 1.164823929 | 1 | No Change | Not Significant | FALSE | NA |
| Mir5621 | TRUE | -1.386967519 | 1 | No Change | Not Significant | FALSE | NA |
| Mir5623 | TRUE | -1.936346 | 1 | No Change | Not Significant | FALSE | NA |
| Mir574 | TRUE | 0.141751176 | 1 | No Change | Not Significant | FALSE | NA |
| Mir599 | TRUE | 0.181223808 | 1 | No Change | Not Significant | FALSE | NA |
| Mir6236 | TRUE | 0.297110487 | 0.828402273 | No Change | Not Significant | TRUE | 0.431491452 |
| Mir6240 | TRUE | 4.708234746 | 0.00012735 | UP | UP | FALSE | NA |
| Mir6339 | TRUE | 0.324463581 | 1 | No Change | Not Significant | FALSE | NA |
| Mir6374 | TRUE | 0.048971346 | 1 | No Change | Not Significant | FALSE | NA |
| Mir6386 | TRUE | -0.101841474 | 0.901013699 | No Change | Not Significant | TRUE | -0.061639671 |
| Mir6395 | TRUE | 1.044246177 | 1 | No Change | Not Significant | FALSE | NA |
| Mir652 | TRUE | -0.247075957 | 1 | No Change | Not Significant | FALSE | NA |
| Mir6539 | TRUE | -0.32847835 | 1 | No Change | Not Significant | FALSE | NA |
| Mir654 | TRUE | -3.026472096 | 1 | No Change | Not Significant | FALSE | NA |
| Mir669n | TRUE | -1.768401455 | 1 | No Change | Not Significant | FALSE | NA |
| Mir674 | TRUE | 2.284359581 | 1 | No Change | Not Significant | FALSE | NA |
| Mir679 | TRUE | -3.825435563 | 1 | No Change | Not Significant | FALSE | NA |
| Mir687 | TRUE | 1.304343362 | 1 | No Change | Not Significant | FALSE | NA |
| Mir6897 | TRUE | 1.900073427 | 1 | No Change | Not Significant | FALSE | NA |
| Mir6909 | TRUE | -1.244729877 | 1 | No Change | Not Significant | FALSE | NA |
| Mir6919 | TRUE | -1.725202543 | 1 | No Change | Not Significant | FALSE | NA |
| Mir6939 | TRUE | -2.71723194 | 1 | No Change | Not Significant | FALSE | NA |
| Mir6942 | TRUE | 0.789824306 | 1 | No Change | Not Significant | FALSE | NA |
| Mir6946 | TRUE | 0.196329335 | 1 | No Change | Not Significant | FALSE | NA |
| Mir6951 | TRUE | -0.749998798 | 1 | No Change | Not Significant | FALSE | NA |
| Mir6955 | TRUE | -0.585396729 | 1 | No Change | Not Significant | FALSE | NA |
| Mir6959 | TRUE | 0.840944612 | 1 | No Change | Not Significant | FALSE | NA |
| Mir6962 | TRUE | -0.895239167 | 1 | No Change | Not Significant | FALSE | NA |
| Mir697 | TRUE | 0.089104948 | 1 | No Change | Not Significant | FALSE | NA |
| Mir6976 | TRUE | -0.351664531 | 1 | No Change | Not Significant | TRUE | -0.952465538 |
| Mir6997 | TRUE | -1.375935707 | 1 | No Change | Not Significant | FALSE | NA |
| Mir6998 | TRUE | 1.295541747 | 1 | No Change | Not Significant | FALSE | NA |
| Mir6999 | TRUE | 1.039126156 | 1 | No Change | Not Significant | FALSE | NA |
| Mir7-1 | TRUE | -0.569246782 | 0.462725114 | No Change | Not Significant | TRUE | -0.400689792 |
| Mir700 | TRUE | 0.240182063 | 1 | No Change | Not Significant | FALSE | NA |
| Mir7001 | TRUE | 1.064197432 | 1 | No Change | Not Significant | FALSE | NA |
| Mir7004 | TRUE | -0.446045312 | 1 | No Change | Not Significant | FALSE | NA |
| Mir7013 | TRUE | 0.733729113 | 1 | No Change | Not Significant | FALSE | NA |
| Mir7026 | TRUE | -1.646319427 | 1 | No Change | Not Significant | FALSE | NA |
| Mir7027 | TRUE | -1.462499108 | 1 | No Change | Not Significant | FALSE | NA |
| Mir703 | TRUE | -0.34109014 | 0.698491194 | No Change | Not Significant | TRUE | -0.019042575 |
| Mir7030 | TRUE | -0.245204787 | 1 | No Change | Not Significant | FALSE | NA |
| Mir7031 | TRUE | 1.368541916 | 1 | No Change | Not Significant | FALSE | NA |
| Mir704 | TRUE | 4.66228559 | 1 | No Change | Not Significant | FALSE | NA |
| Mir7041 | TRUE | -0.726398949 | 1 | No Change | Not Significant | FALSE | NA |
| Mir7042 | TRUE | -1.507267075 | 1 | No Change | Not Significant | FALSE | NA |

Supplementary Table 3

| geneID | Cond1_DF_Detected | Cond1_DF_ValueCol | Cond1_DF_PadjCol | Cond1_DF_Cutoff | Cond1_DF_Cutoff_Specific.x | Cond2_DF_Detected | Cond2_DF_ValueCol | Cond2_DF_PadjCol | Cond2_DF_Cutoff | Cond2_DF_Cutoff_Specific | BG_Method |
|---|---|---|---|---|---|---|---|---|---|---|---|
| Mir7048 | TRUE | -0.655373555 | 1 | No Change | Not Significant | FALSE | NA | NA | No Change | Not Detected | FALSE |
| Mir7051 | TRUE | 1.293319458 | 1 | No Change | Not Significant | FALSE | NA | NA | No Change | Not Detected | FALSE |
| Mir7052 | TRUE | 0.260285738 | 1 | No Change | Not Significant | FALSE | NA | NA | No Change | Not Detected | FALSE |
| Mir7054 | TRUE | -0.716588185 | 1 | No Change | Not Significant | FALSE | NA | NA | No Change | Not Detected | FALSE |
| Mir7089 | TRUE | -0.27945767 | 1 | No Change | Not Significant | FALSE | NA | NA | No Change | Not Detected | FALSE |
| Mir709 | TRUE | -1.316915071 | 1 | No Change | Not Significant | FALSE | NA | NA | No Change | Not Detected | FALSE |
| Mir7090 | TRUE | -0.415127297 | 1 | No Change | Not Significant | FALSE | NA | NA | No Change | Not Detected | FALSE |
| Mir7211 | TRUE | -0.908344037 | 0.165238877 | No Change | Not Significant | TRUE | -0.137857978 | 0.89645995 | No Change | Not Significant | TRUE |
| Mir7212 | TRUE | -0.691836813 | 1 | No Change | Not Significant | FALSE | NA | NA | No Change | Not Detected | FALSE |
| Mir7237 | TRUE | -1.020940057 | 1 | No Change | Not Significant | FALSE | NA | NA | No Change | Not Detected | FALSE |
| Mir744 | TRUE | -1.086848862 | 1 | No Change | Not Significant | FALSE | NA | NA | No Change | Not Detected | FALSE |
| Mir763 | TRUE | -0.240744467 | 1 | No Change | Not Significant | FALSE | NA | NA | No Change | Not Detected | FALSE |
| Mir7655 | TRUE | 0.790572641 | 1 | No Change | Not Significant | FALSE | NA | NA | No Change | Not Detected | FALSE |
| Mir7671 | TRUE | 0.986894554 | 1 | No Change | Not Significant | FALSE | NA | NA | No Change | Not Detected | FALSE |
| Mir7675 | TRUE | -1.066476856 | 1 | No Change | Not Significant | FALSE | NA | NA | No Change | Not Detected | FALSE |
| Mir7682 | TRUE | -1.891808405 | 1 | No Change | Not Significant | FALSE | NA | NA | No Change | Not Detected | FALSE |
| Mir7687 | TRUE | -0.018782232 | 1 | No Change | Not Significant | FALSE | NA | NA | No Change | Not Detected | FALSE |
| Mir8091 | TRUE | -0.184648679 | 0.857548131 | No Change | Not Significant | TRUE | -0.608395187 | 0.403959453 | No Change | Not Significant | TRUE |
| Mir8094 | TRUE | 0.31428815 | 1 | No Change | Not Significant | FALSE | NA | NA | No Change | Not Detected | FALSE |
| Mir8103 | TRUE | -0.611545011 | 1 | No Change | Not Significant | FALSE | NA | NA | No Change | Not Detected | FALSE |
| Mir8106 | TRUE | 0.490629303 | 1 | No Change | Not Significant | FALSE | NA | NA | No Change | Not Detected | FALSE |
| Mir8116 | TRUE | 0.061735789 | 1 | No Change | Not Significant | FALSE | NA | NA | No Change | Not Detected | FALSE |
| Mir8120 | TRUE | -1.678379569 | 1 | No Change | Not Significant | FALSE | NA | NA | No Change | Not Detected | FALSE |
| Mir96 | TRUE | -1.644944728 | 1 | No Change | Not Significant | FALSE | NA | NA | No Change | Not Detected | FALSE |
| Mir98 | TRUE | -0.373143504 | 1 | No Change | Not Significant | FALSE | NA | NA | No Change | Not Detected | FALSE |
| Mir99ahg | TRUE | 0.936906105 | 0.309726787 | No Change | Not Significant | TRUE | 0.987916201 | 0.066254458 | No Change | Not Significant | TRUE |
| Mirg | TRUE | -4.369007668 | 3.43E-25 | DOWN | DOWN | FALSE | NA | NA | No Change | Not Detected | FALSE |
| Mis12 | TRUE | 1.023423967 | 1.01E-16 | UP | UP | TRUE | 0.51301626 | 7.48E-08 | UP | UP | TRUE |
| Mis18a | TRUE | 0.684100018 | 0.000379723 | UP | UP | TRUE | 0.089925834 | 0.727325341 | No Change | Not Significant | TRUE |
| Mis18bp1 | TRUE | 0.514441893 | 0.023162411 | UP | UP | TRUE | -0.1233667 | 0.589851392 | No Change | Not Significant | TRUE |
| Misp | TRUE | 0.004452934 | 0.985568894 | No Change | Not Significant | TRUE | 0.248005693 | 0.011600354 | No Change | Significant Positive | TRUE |
| Misp3 | TRUE | -0.538655253 | 0.058854589 | No Change | Not Significant | TRUE | 0.087613199 | 0.737950908 | No Change | Not Significant | TRUE |
| Mitd1 | TRUE | 0.013199815 | 0.945840699 | No Change | Not Significant | TRUE | -0.10532328 | 0.469953022 | No Change | Not Significant | TRUE |
| Mitf | TRUE | 0.745375964 | 0.010174535 | UP | UP | TRUE | 0.677743371 | 0.002731844 | UP | UP | TRUE |
| Mixl1 | TRUE | -0.539502203 | 1 | No Change | Not Significant | FALSE | NA | NA | No Change | Not Detected | FALSE |
| Mki67 | TRUE | 0.635531071 | 0.000472967 | UP | UP | TRUE | 0.055757457 | 0.768361739 | No Change | Not Significant | TRUE |
| Mkks | TRUE | 0.028107364 | 0.932166328 | No Change | Not Significant | TRUE | 0.186945808 | 0.096539507 | No Change | Not Significant | TRUE |
| Mkln1 | TRUE | -0.176537717 | 0.001953624 | No Change | Significant Negative | TRUE | -0.029209741 | 0.783432973 | No Change | Not Significant | TRUE |
| Mknk1 | TRUE | 0.171824492 | 0.388417088 | No Change | Not Significant | TRUE | -0.120615244 | 0.428222446 | No Change | Not Significant | TRUE |
| Mknk2 | TRUE | -0.403256502 | 1.42E-06 | No Change | Significant Negative | TRUE | -0.111271935 | 0.233883785 | No Change | Not Significant | TRUE |
| Mkrn1 | TRUE | -0.42612796 | 3.02E-08 | No Change | Significant Negative | TRUE | -0.064346278 | 0.443185909 | No Change | Not Significant | TRUE |
| Mkrn2 | TRUE | -0.182567695 | 0.109497021 | No Change | Not Significant | TRUE | -0.094395165 | 0.357631975 | No Change | Not Significant | TRUE |
| Mkrn2os | TRUE | -0.222814579 | 0.085735413 | No Change | Not Significant | TRUE | 0.10657744 | 0.27921863 | No Change | Not Significant | TRUE |
| Mkrn3 | TRUE | -0.740938962 | 1 | No Change | Not Significant | FALSE | NA | NA | No Change | Not Detected | FALSE |
| Mks1 | TRUE | -0.058187892 | 0.783613584 | No Change | Not Significant | TRUE | -0.163231567 | 0.260228257 | No Change | Not Significant | TRUE |
| Mkx | TRUE | -0.037429829 | 0.888506311 | No Change | Not Significant | TRUE | 0.036139275 | 0.844292391 | No Change | Not Significant | TRUE |
| Mlana | TRUE | 1.061896477 | 1 | No Change | Not Significant | FALSE | NA | NA | No Change | Not Detected | FALSE |
| Mlc1 | TRUE | -0.427944429 | 1 | No Change | Not Significant | FALSE | NA | NA | No Change | Not Detected | FALSE |
| Mlec | TRUE | 0.572127174 | 5.38E-07 | UP | UP | TRUE | 0.609983246 | 1.07E-16 | UP | UP | TRUE |
| Mlf1 | TRUE | 0.820599083 | 1 | No Change | Not Significant | FALSE | NA | NA | No Change | Not Detected | FALSE |
| Mlf2 | TRUE | 0.200535082 | 0.01168152 | No Change | Significant Positive | TRUE | 0.106391309 | 0.391491529 | No Change | Not Significant | TRUE |
| Mlh1 | TRUE | 0.405876855 | 7.31E-06 | No Change | Significant Positive | TRUE | 0.04461973 | 0.761174286 | No Change | Not Significant | TRUE |
| Mlh3 | TRUE | 0.212711077 | 0.184147653 | No Change | Not Significant | TRUE | 0.052949271 | 0.709700079 | No Change | Not Significant | TRUE |
| Mlip | TRUE | -0.237604966 | 0.901189824 | No Change | Not Significant | TRUE | 0.52243434 | 0.515282046 | No Change | Not Significant | TRUE |
| Mlkl | TRUE | -0.279087006 | 0.014180725 | No Change | Significant Negative | TRUE | -0.222292389 | 0.101260467 | No Change | Not Significant | TRUE |
| Mllt1 | TRUE | 0.066053869 | 0.601975182 | No Change | Not Significant | TRUE | 0.044744178 | 0.700094114 | No Change | Not Significant | TRUE |
| Mllt10 | TRUE | -0.144725416 | 0.050180472 | No Change | Not Significant | TRUE | -0.097278676 | 0.226290026 | No Change | Not Significant | TRUE |
| Mllt11 | TRUE | -0.439631178 | 0.020879192 | No Change | Significant Negative | TRUE | -0.328821661 | 0.181275208 | No Change | Not Significant | TRUE |
| Mllt3 | TRUE | 0.017717716 | 0.932166328 | No Change | Not Significant | TRUE | -0.060923257 | 0.552220724 | No Change | Not Significant | TRUE |
| Mllt6 | TRUE | -0.348178642 | 0.00273381 | No Change | Significant Negative | TRUE | -0.318533172 | 6.24E-07 | No Change | Significant Negative | TRUE |
| Mlph | TRUE | 0.125531824 | 0.446602599 | No Change | Not Significant | TRUE | 0.166640702 | 0.086466472 | No Change | Not Significant | TRUE |
| Mlst8 | TRUE | 0.128639658 | 0.192986007 | No Change | Not Significant | TRUE | -0.011454022 | 0.930151569 | No Change | Not Significant | TRUE |
| Mlx | TRUE | -0.044321434 | 0.833638003 | No Change | Not Significant | TRUE | 0.112627273 | 0.179562869 | No Change | Not Significant | TRUE |
| Mlxip | TRUE | 0.053975657 | 0.736813469 | No Change | Not Significant | TRUE | 0.010440718 | 0.950733597 | No Change | Not Significant | TRUE |
| Mlxipl | TRUE | 0.067136597 | 0.866747322 | No Change | Not Significant | TRUE | -0.366706319 | 0.014456304 | No Change | Significant Negative | TRUE |
| Mlycd | TRUE | 0.060536959 | 0.56579982 | No Change | Not Significant | TRUE | 0.109647787 | 0.201440865 | No Change | Not Significant | TRUE |
| Mm2pr | TRUE | -0.299410194 | 1 | No Change | Not Significant | FALSE | NA | NA | No Change | Not Detected | FALSE |
| Mmaa | TRUE | -0.140513448 | 0.185483473 | No Change | Not Significant | TRUE | 0.094672232 | 0.444801234 | No Change | Not Significant | TRUE |
| Mmab | TRUE | -0.099655163 | 0.671162093 | No Change | Not Significant | TRUE | 0.002412084 | 0.993782244 | No Change | Not Significant | TRUE |
| Mmachc | TRUE | -0.092184279 | 0.664203441 | No Change | Not Significant | TRUE | 0.194502868 | 0.035604125 | No Change | Significant Positive | TRUE |
| Mmadhc | TRUE | 0.24356335 | 0.069204119 | No Change | Not Significant | TRUE | 0.201227773 | 0.001863973 | No Change | Significant Positive | TRUE |
| Mmd | TRUE | -0.152095721 | 0.566362983 | No Change | Not Significant | TRUE | -0.182598538 | 0.333871046 | No Change | Not Significant | TRUE |
| Mme | TRUE | 0.175899131 | 0.908409285 | No Change | Not Significant | TRUE | -0.417790627 | 0.352392829 | No Change | Not Significant | TRUE |
| Mmel1 | TRUE | 0.3393232 | 1 | No Change | Not Significant | FALSE | NA | NA | No Change | Not Detected | FALSE |
| Mmgt1 | TRUE | 0.461414286 | 0.001092699 | No Change | Significant Positive | TRUE | 0.167726968 | 0.040439555 | No Change | Significant Positive | TRUE |
| Mmgt2 | TRUE | 0.049019681 | 0.776081458 | No Change | Not Significant | TRUE | 0.171428586 | 0.051626789 | No Change | Not Significant | TRUE |
| Mmp11 | TRUE | 1.030768831 | 0.180532859 | No Change | Not Significant | TRUE | 0.135010235 | 0.872524487 | No Change | Not Significant | TRUE |
| Mmp12 | TRUE | -0.851981335 | 1 | No Change | Not Significant | FALSE | NA | NA | No Change | Not Detected | FALSE |
| Mmp14 | TRUE | 0.331557817 | 0.002004434 | No Change | Significant Positive | TRUE | 0.186639513 | 0.082978005 | No Change | Not Significant | TRUE |
| Mmp15 | TRUE | -0.562535022 | 9.57E-07 | DOWN | DOWN | TRUE | -0.044418687 | 0.797435496 | No Change | Not Significant | TRUE |
| Mmp16 | TRUE | 1.929269821 | 1 | No Change | Not Significant | FALSE | NA | NA | No Change | Not Detected | FALSE |
| Mmp17 | TRUE | 0.472811361 | 0.574847405 | No Change | Not Significant | FALSE | NA | NA | No Change | Not Detected | FALSE |
| Mmp19 | TRUE | -0.712763869 | 1 | No Change | Not Significant | FALSE | NA | NA | No Change | Not Detected | FALSE |
| Mmp2 | TRUE | -0.118846273 | 0.89873561 | No Change | Not Significant | TRUE | 0.605471647 | 0.340243435 | No Change | Not Significant | TRUE |
| Mmp20 | TRUE | 1.910997214 | 1 | No Change | Not Significant | FALSE | NA | NA | No Change | Not Detected | FALSE |
| Mmp21 | TRUE | 0.18535423 | 1 | No Change | Not Significant | FALSE | NA | NA | No Change | Not Detected | FALSE |
| Mmp23 | TRUE | 0.925558047 | 1 | No Change | Not Significant | FALSE | NA | NA | No Change | Not Detected | FALSE |
| Mmp24 | TRUE | -0.329554093 | 0.731167175 | No Change | Not Significant | TRUE | -0.179045554 | 0.81580845 | No Change | Not Significant | TRUE |
| Mmp25 | TRUE | -1.188458081 | 1 | No Change | Not Significant | FALSE | NA | NA | No Change | Not Detected | FALSE |
| Mmp28 | TRUE | -0.077782535 | 0.674875015 | No Change | Not Significant | TRUE | -0.010942685 | 0.955523695 | No Change | Not Significant | TRUE |
| Mmp7 | TRUE | 2.514588385 | 1.07E-14 | UP | UP | TRUE | 2.661532419 | 1.37E-38 | UP | UP | TRUE |
| Mmp8 | TRUE | 1.696703949 | 1 | No Change | Not Significant | FALSE | NA | NA | No Change | Not Detected | FALSE |
| Mmp9 | TRUE | 0.505237597 | 1 | No Change | Not Significant | FALSE | NA | NA | No Change | Not Detected | FALSE |
| Mmrn1 | TRUE | -1.668772642 | 1 | No Change | Not Significant | FALSE | NA | NA | No Change | Not Detected | FALSE |
| Mmrn2 | TRUE | 0.300269039 | 1 | No Change | Not Significant | FALSE | NA | NA | No Change | Not Detected | FALSE |
| Mms19 | TRUE | -0.151462642 | 0.148254918 | No Change | Not Significant | TRUE | -0.203772491 | 0.001113473 | No Change | Significant Negative | TRUE |
| Mms22l | TRUE | 0.54695205 | 0.00461924 | UP | UP | TRUE | 0.255423707 | 0.174452968 | No Change | Not Significant | TRUE |
| Mmut | TRUE | 0.236487191 | 0.047391714 | No Change | Significant Positive | TRUE | 0.128078751 | 0.048137725 | No Change | Significant Positive | TRUE |
| Mn1 | TRUE | -0.572598878 | 0.541897747 | No Change | Not Significant | TRUE | 0.239155636 | 0.67671134 | No Change | Not Significant | TRUE |
| Mnat1 | TRUE | -0.058726681 | 0.796810121 | No Change | Not Significant | TRUE | -0.057502296 | 0.715904735 | No Change | Not Significant | TRUE |
| Mnd1 | TRUE | -0.066603199 | 0.901569434 | No Change | Not Significant | TRUE | -0.025857832 | 0.955338492 | No Change | Not Significant | TRUE |
| Mnd1-ps | TRUE | 1.121896485 | 1 | No Change | Not Significant | FALSE | NA | NA | No Change | Not Detected | FALSE |
| Mndal | TRUE | -0.493502402 | 0.641211258 | No Change | Not Significant | TRUE | 0.721262831 | 0.231865576 | No Change | Not Significant | TRUE |
| Mns1 | TRUE | 0.779031703 | 0.006255193 | UP | UP | TRUE | 0.235236428 | 0.584118738 | No Change | Not Significant | TRUE |
| Mnt | TRUE | 0.096594361 | 0.662372535 | No Change | Not Significant | TRUE | -0.006208252 | 0.97886592 | No Change | Not Significant | TRUE |
| Mnx1 | TRUE | -1.025079076 | 0.005170946 | DOWN | DOWN | TRUE | -0.157592126 | 0.722065604 | No Change | Not Significant | TRUE |
| Moap1 | TRUE | 0.280776623 | 1 | No Change | Not Significant | TRUE | -0.164398821 | 0.856611363 | No Change | Not Significant | TRUE |
| Mob1a | TRUE | -0.117065781 | 0.501169049 | No Change | Not Significant | TRUE | -0.067408519 | 0.202990467 | No Change | Not Significant | TRUE |
| Mob1b | TRUE | 0.04308973 | 0.800362194 | No Change | Not Significant | TRUE | 0.04149861 | 0.66169488 | No Change | Not Significant | TRUE |
| Mob2 | TRUE | 0.137319925 | 0.033883971 | No Change | Significant Positive | TRUE | 0.083842755 | 0.132269137 | No Change | Not Significant | TRUE |
| Mob3a | TRUE | 0.149782392 | 0.046746214 | No Change | Significant Positive | TRUE | 0.226370724 | 0.016251873 | No Change | Significant Positive | TRUE |
| Mob3b | TRUE | 0.008552435 | 0.968634984 | No Change | Not Significant | TRUE | 0.002818683 | 0.989206294 | No Change | Not Significant | TRUE |
| Mob3c | TRUE | 0.048231634 | 0.795381772 | No Change | Not Significant | TRUE | 0.070943615 | 0.578888054 | No Change | Not Significant | TRUE |
| Mob4 | TRUE | 0.037215156 | 0.822609935 | No Change | Not Significant | TRUE | -0.011515134 | 0.913597373 | No Change | Not Significant | TRUE |
| Mobp | TRUE | 0.893784971 | 1 | No Change | Not Significant | FALSE | NA | NA | No Change | Not Detected | FALSE |
| Mocos | TRUE | -0.159224397 | 0.627409847 | No Change | Not Significant | TRUE | 0.043392099 | 0.881708496 | No Change | Not Significant | TRUE |
| Mocs1 | TRUE | -0.155465815 | 0.209433685 | No Change | Not Significant | TRUE | 0.022593475 | 0.892032325 | No Change | Not Significant | TRUE |
| Mocs2 | TRUE | -0.152211721 | 0.145669666 | No Change | Not Significant | TRUE | 0.007579278 | 0.949314078 | No Change | Not Significant | TRUE |
| Mocs3 | TRUE | 0.175842133 | 0.283633759 | No Change | Not Significant | TRUE | 0.045166936 | 0.843368224 | No Change | Not Significant | TRUE |
| Mog | TRUE | 3.244261967 | 1 | No Change | Not Significant | FALSE | NA | NA | No Change | Not Detected | FALSE |
| Mogat1 | TRUE | -0.59598025 | 1 | No Change | Not Significant | FALSE | NA | NA | No Change | Not Detected | FALSE |
| Mogat2 | TRUE | 0.47997555 | 0.330978129 | No Change | Not Significant | TRUE | 0.407387758 | 0.075318948 | No Change | Not Significant | TRUE |
| Mogs | TRUE | 0.661877126 | 1.21E-07 | UP | UP | TRUE | 0.407259571 | 7.72E-06 | No Change | Significant Positive | TRUE |
| Mok | TRUE | 0.004230274 | 0.995423402 | No Change | Not Significant | TRUE | -0.524877732 | 0.460551754 | No Change | Not Significant | TRUE |
| Mon1a | TRUE | 0.062579735 | 0.695624496 | No Change | Not Significant | TRUE | 0.161309185 | 0.121033168 | No Change | Not Significant | TRUE |
| Mon1b | TRUE | -0.072863931 | 0.647656102 | No Change | Not Significant | TRUE | 0.089126724 | 0.550870027 | No Change | Not Significant | TRUE |
| Mon2 | TRUE | 0.146179206 | 0.000913981 | No Change | Significant Positive | TRUE | 0.007005166 | 0.91448208 | No Change | Not Significant | TRUE |
| Morc1 | TRUE | 0.512262703 | 0.741309692 | No Change | Not Significant | FALSE | NA | NA | No Change | Not Detected | FALSE |
| Morc2a | TRUE | 0.041019985 | 0.75067522 | No Change | Not Significant | TRUE | -0.07677572 | 0.328198932 | No Change | Not Significant | TRUE |
| Morc2b | TRUE | 1.16228452 | 1 | No Change | Not Significant | FALSE | NA | NA | No Change | Not Detected | FALSE |
| Morc3 | TRUE | -0.028518169 | 0.823378739 | No Change | Not Significant | TRUE | -0.132073467 | 0.178786609 | No Change | Not Significant | TRUE |
| Morc4 | TRUE | 0.274486581 | 0.267599593 | No Change | Not Significant | TRUE | 0.253482926 | 0.154536441 | No Change | Not Significant | TRUE |
| Morf4l1 | TRUE | -0.344036446 | 0.020844259 | No Change | Significant Negative | TRUE | 0.076125396 | 0.210558459 | No Change | Not Significant | TRUE |
| Morf4l2 | TRUE | 0.826444388 | 5.20E-17 | UP | UP | TRUE | 0.533842318 | 4.70E-13 | UP | UP | TRUE |
| Morn1 | TRUE | -0.170458897 | 0.535859673 | No Change | Not Significant | TRUE | -0.229059865 | 0.293679015 | No Change | Not Significant | TRUE |
| Morn2 | TRUE | 0.256393602 | 0.36877688 | No Change | Not Significant | TRUE | 0.139109556 | 0.622373434 | No Change | Not Significant | TRUE |
| Morn3 | TRUE | 1.799007657 | 1 | No Change | Not Significant | FALSE | NA | NA | No Change | Not Detected | FALSE |
| Morn4 | TRUE | -0.434804829 | 1 | No Change | Not Significant | FALSE | NA | NA | No Change | Not Detected | FALSE |
| Morrbid | TRUE | -1.999139567 | 1 | No Change | Not Significant | FALSE | NA | NA | No Change | Not Detected | FALSE |
| Mosmo | TRUE | -0.129271852 | 0.568538439 | No Change | Not Significant | TRUE | -0.187816589 | 0.220005888 | No Change | Not Significant | TRUE |
| Mospd1 | TRUE | -0.157534991 | 0.389807821 | No Change | Not Significant | TRUE | -0.066438556 | 0.563409586 | No Change | Not Significant | TRUE |
| Mospd2 | TRUE | 0.281019103 | 0.014731479 | No Change | Significant Positive | TRUE | -0.047626401 | 0.749829352 | No Change | Not Significant | TRUE |
| Mospd3 | TRUE | 0.275952199 | 0.477220689 | No Change | Not Significant | TRUE | -0.043830235 | 0.911163158 | No Change | Not Significant | TRUE |
| Mov10 | TRUE | 0.700981112 | 0.016683177 | UP | UP | TRUE | -0.175184017 | 0.371487597 | No Change | Not Significant | TRUE |
| Mov10l1 | TRUE | 0.176018776 | 1 | No Change | Not Significant | FALSE | NA | NA | No Change | Not Detected | FALSE |
| Moxd1 | TRUE | 3.240806601 | 0.001763006 | UP | UP | FALSE | NA | NA | No Change | Not Detected | FALSE |
| Moxd2 | TRUE | 2.326914189 | 1 | No Change | Not Significant | FALSE | NA | NA | No Change | Not Detected | FALSE |
| Mpc1 | TRUE | -0.436137889 | 5.63E-05 | No Change | Significant Negative | TRUE | -0.024178774 | 0.878312994 | No Change | Not Significant | TRUE |
| Mpc2 | TRUE | -0.204788994 | 0.002381578 | No Change | Significant Negative | TRUE | 0.014694575 | 0.929706079 | No Change | Not Significant | TRUE |
| Mpdu1 | TRUE | -0.195410741 | 0.071455098 | No Change | Not Significant | TRUE | -0.090471322 | 0.165460865 | No Change | Not Significant | TRUE |
| Mpdz | TRUE | 0.671926585 | 0.015454578 | UP | UP | TRUE | 1.609526909 | 9.96E-06 | UP | UP | TRUE |
| Mpeg1 | TRUE | 0.903525304 | 1 | No Change | Not Significant | TRUE | -0.224418905 | 0.737165628 | No Change | Not Significant | TRUE |
| Mpg | TRUE | 0.083755843 | 0.713786099 | No Change | Not Significant | TRUE | -0.025317842 | 0.925602711 | No Change | Not Significant | TRUE |
| Mphosph10 | TRUE | 0.086168725 | 0.770846581 | No Change | Not Significant | TRUE | -0.089300355 | 0.521149121 | No Change | Not Significant | TRUE |
| Mphosph6 | TRUE | 0.376626745 | 0.002758114 | No Change | Significant Positive | TRUE | 0.143022215 | 0.292295199 | No Change | Not Significant | TRUE |
| Mphosph8 | TRUE | 0.372326902 | 0.00726647 | No Change | Significant Positive | TRUE | 0.233004722 | 0.086179968 | No Change | Not Significant | TRUE |
| Mphosph9 | TRUE | 0.470719327 | 0.008398372 | No Change | Significant Positive | TRUE | 0.033255569 | 0.853850569 | No Change | Not Significant | TRUE |
| Mpi | TRUE | 0.263807306 | 0.00164613 | No Change | Significant Positive | TRUE | 0.154695932 | 0.027572597 | No Change | Significant Positive | TRUE |
| Mpig6b | TRUE | -0.336646781 | 0.802306417 | No Change | Not Significant | TRUE | -0.683222464 | 0.25454282 | No Change | Not Significant | TRUE |
| Mpl | TRUE | -0.688658869 | 1 | No Change | Not Significant | FALSE | NA | NA | No Change | Not Detected | FALSE |
| Mplkip | TRUE | -0.226064428 | 0.082276233 | No Change | Not Significant | TRUE | -0.023893358 | 0.915791177 | No Change | Not Significant | TRUE |
| Mpnd | TRUE | 0.003682289 | 0.975869522 | No Change | Not Significant | TRUE | -0.009235861 | 0.942844904 | No Change | Not Significant | TRUE |
| Mpo | TRUE | -2.88398533 | 1 | No Change | Not Significant | FALSE | NA | NA | No Change | Not Detected | FALSE |
| Mpp1 | TRUE | -0.175122649 | 0.420143948 | No Change | Not Significant | TRUE | -0.042641623 | 0.797454855 | No Change | Not Significant | TRUE |
| Mpp2 | TRUE | 0.346097785 | 0.574134308 | No Change | Not Significant | TRUE | 0.355138446 | 0.413082963 | No Change | Not Significant | TRUE |
| Mpp3 | TRUE | 0.04093945 | 0.908812719 | No Change | Not Significant | TRUE | -0.165457293 | 0.416363961 | No Change | Not Significant | TRUE |
| Mpp4 | TRUE | 0.870450345 | 1 | No Change | Not Significant | FALSE | NA | NA | No Change | Not Detected | FALSE |
| Mpp7 | TRUE | -0.129696672 | 0.363688435 | No Change | Not Significant | TRUE | 0.08680784 | 0.536541439 | No Change | Not Significant | TRUE |
| Mppe1 | TRUE | -0.237848838 | 0.196529604 | No Change | Not Significant | TRUE | 0.215502334 | 0.012743984 | No Change | Significant Positive | TRUE |
| Mpped2 | TRUE | -0.428553829 | 1 | No Change | Not Significant | FALSE | NA | NA | No Change | Not Detected | FALSE |
| Mprip | TRUE | -0.001689492 | 0.992263312 | No Change | Not Significant | TRUE | -0.139788937 | 0.059031798 | No Change | Not Significant | TRUE |
| Mpst | TRUE | 0.422374913 | 0.068530286 | No Change | Not Significant | TRUE | 0.463899131 | 0.000297679 | No Change | Significant Positive | TRUE |
| Mptx1 | TRUE | -0.872531338 | 0.403987166 | No Change | Not Significant | TRUE | -0.00296231 | 0.993688498 | No Change | Not Significant | TRUE |
| Mptx2 | TRUE | -0.966091112 | 1 | No Change | Not Significant | FALSE | NA | NA | No Change | Not Detected | FALSE |
| Mpv17 | TRUE | -0.14953302 | 0.299596873 | No Change | Not Significant | TRUE | 0.048596811 | 0.761555988 | No Change | Not Significant | TRUE |
| Mpv17l | TRUE | 0.588269552 | 2.16E-05 | UP | UP | TRUE | 0.305270308 | 0.011052553 | No Change | Significant Positive | TRUE |
| Mpv17l2 | TRUE | 0.321776233 | 0.003236188 | No Change | Significant Positive | TRUE | 0.135098276 | 0.34107542 | No Change | Not Significant | TRUE |
| Mpz | TRUE | -1.189239056 | 1 | No Change | Not Significant | FALSE | NA | NA | No Change | Not Detected | FALSE |
| Mpzl1 | TRUE | 0.26158711 | 0.08629118 | No Change | Not Significant | TRUE | 0.146145181 | 0.333333335 | No Change | Not Significant | TRUE |
| Mpzl2 | TRUE | -0.590132845 | 0.000257174 | DOWN | DOWN | TRUE | -0.110016655 | 0.31154074 | No Change | Not Significant | TRUE |
| Mpzl3 | TRUE | -0.436384536 | 0.011092726 | No Change | Significant Negative | TRUE | -0.279811004 | 4.83E-05 | No Change | Significant Negative | TRUE |
| Mr1 | TRUE | 0.934627716 | 1 | No Change | Not Significant | FALSE | NA | NA | No Change | Not Detected | FALSE |
| Mrap2 | TRUE | 1.062779156 | 1 | No Change | Not Significant | FALSE | NA | NA | No Change | Not Detected | FALSE |
| Mras | TRUE | -0.591931532 | 0.181014958 | No Change | Not Significant | TRUE | -0.006780486 | 0.990489989 | No Change | Not Significant | TRUE |
| Mrc1 | TRUE | 0.455507691 | 0.481477497 | No Change | Not Significant | TRUE | 0.314093418 | 0.657004431 | No Change | Not Significant | TRUE |
| Mrc2 | TRUE | 3.800970518 | 1 | No Change | Not Significant | FALSE | NA | NA | No Change | Not Detected | FALSE |
| Mre11a | TRUE | 0.42552679 | 0.000162315 | No Change | Significant Positive | TRUE | 0.008777201 | 0.96026404 | No Change | Not Significant | TRUE |
| Mreg | TRUE | -0.236941855 | 0.761222618 | No Change | Not Significant | TRUE | -0.112471139 | 0.833307074 | No Change | Not Significant | TRUE |
| Mrfap1 | TRUE | -0.105999115 | 0.508065109 | No Change | Not Significant | TRUE | 0.130423601 | 0.194372111 | No Change | Not Significant | TRUE |
| Mrgbp | TRUE | -0.223547447 | 0.038648666 | No Change | Significant Negative | TRUE | -0.29799767 | 0.002230883 | No Change | Significant Negative | TRUE |
| Mrgpra6 | TRUE | 1.905376038 | 1 | No Change | Not Significant | FALSE | NA | NA | No Change | Not Detected | FALSE |
| Mrgprb1 | TRUE | -0.23619505 | 1 | No Change | Not Significant | FALSE | NA | NA | No Change | Not Detected | FALSE |

Supplementary Table 3

| geneID | Cond1_DF_Detected | Cond1_DF_ValueCol | Cond1_DF_PadjCol | Cond1_DF_Cutoff | Cond1_DF_Cutoff_Specific.x | Cond2_DF_Detected | Cond2_DF_ValueCol | Cond2_DF_PadjCol | Cond2_DF_Cutoff | Cond2_DF_Cutoff_Specific | BG_Method |
|---|---|---|---|---|---|---|---|---|---|---|---|
| Mrgpre | TRUE | -0.542932448 | 0.349330805 | No Change | Not Significant | TRUE | -0.25443315 | 0.658502858 | No Change | Not Significant | TRUE |
| Mrgprh | TRUE | 0.606360805 | 1 | No Change | Not Significant | FALSE | NA | NA | No Change | Not Detected | FALSE |
| Mri1 | TRUE | -0.044109369 | 0.816323585 | No Change | Not Significant | TRUE | -0.165629631 | 0.118917792 | No Change | Not Significant | TRUE |
| Mrln | TRUE | 0.88909289 | 1 | No Change | Not Significant | FALSE | NA | NA | No Change | Not Detected | FALSE |
| Mrm1 | TRUE | -0.117980817 | 0.582866455 | No Change | Not Significant | TRUE | -0.331147625 | 0.024042701 | No Change | Significant Negative | TRUE |
| Mrm2 | TRUE | -0.048217584 | 0.841035219 | No Change | Not Significant | TRUE | 0.085129971 | 0.678988684 | No Change | Not Significant | TRUE |
| Mrm3 | TRUE | -0.301197364 | 0.060682336 | No Change | Not Significant | TRUE | -0.019896112 | 0.930151569 | No Change | Not Significant | TRUE |
| Mrnip | TRUE | 0.202155634 | 0.691768261 | No Change | Not Significant | TRUE | -0.14671157 | 0.707465523 | No Change | Not Significant | TRUE |
| Mro | TRUE | -0.770684672 | 0.504312604 | No Change | Not Significant | TRUE | 0.158806312 | 0.836812138 | No Change | Not Significant | TRUE |
| Mroh1 | TRUE | 1.824102527 | 2.12E-117 | UP | UP | TRUE | 1.229443226 | 3.58E-142 | UP | UP | TRUE |
| Mroh2a | TRUE | 0.164861989 | 0.719334577 | No Change | Not Significant | TRUE | -0.096041348 | 0.67075219 | No Change | Not Significant | TRUE |
| Mroh2b | TRUE | 2.453442628 | 1 | No Change | Not Significant | FALSE | NA | NA | No Change | Not Detected | FALSE |
| Mroh3 | TRUE | -0.148411194 | 0.690104577 | No Change | Not Significant | TRUE | -0.288926687 | 0.391461735 | No Change | Not Significant | TRUE |
| Mroh4 | TRUE | 1.130555512 | 1 | No Change | Not Significant | FALSE | NA | NA | No Change | Not Detected | FALSE |
| Mroh5 | TRUE | 0.131186957 | 0.971883559 | No Change | Not Significant | FALSE | NA | NA | No Change | Not Detected | FALSE |
| Mroh6 | TRUE | 0.061150745 | 0.960078011 | No Change | Not Significant | TRUE | 0.121963672 | 0.790772957 | No Change | Not Significant | TRUE |
| Mroh7 | TRUE | 0.237349209 | 0.861267215 | No Change | Not Significant | TRUE | 0.067595378 | 0.943745489 | No Change | Not Significant | TRUE |
| Mroh8 | TRUE | 0.064963963 | 0.930446023 | No Change | Not Significant | TRUE | -0.514041632 | 0.245713848 | No Change | Not Significant | TRUE |
| Mrpl1 | TRUE | -0.057055291 | 0.751086942 | No Change | Not Significant | TRUE | 0.067778175 | 0.649442192 | No Change | Not Significant | TRUE |
| Mrpl10 | TRUE | -0.004424286 | 0.981434574 | No Change | Not Significant | TRUE | 0.041922016 | 0.679215578 | No Change | Not Significant | TRUE |
| Mrpl11 | TRUE | -0.015655312 | 0.910297568 | No Change | Not Significant | TRUE | 0.068180961 | 0.523522715 | No Change | Not Significant | TRUE |
| Mrpl12 | TRUE | -0.166458093 | 0.097034086 | No Change | Not Significant | TRUE | 0.103269775 | 0.243463775 | No Change | Not Significant | TRUE |
| Mrpl13 | TRUE | -0.043830918 | 0.789606655 | No Change | Not Significant | TRUE | 0.039891737 | 0.789753 | No Change | Not Significant | TRUE |
| Mrpl14 | TRUE | -0.108294409 | 0.391355356 | No Change | Not Significant | TRUE | 0.039558899 | 0.833463963 | No Change | Not Significant | TRUE |
| Mrpl15 | TRUE | -0.004784392 | 0.971638507 | No Change | Not Significant | TRUE | 0.174505602 | 0.005591547 | No Change | Significant Positive | TRUE |
| Mrpl16 | TRUE | 0.108351037 | 0.438019649 | No Change | Not Significant | TRUE | 0.10324499 | 0.393611906 | No Change | Not Significant | TRUE |
| Mrpl17 | TRUE | 0.10337485 | 0.510066755 | No Change | Not Significant | TRUE | 0.153194217 | 0.041146859 | No Change | Significant Positive | TRUE |
| Mrpl18 | TRUE | 0.178445449 | 0.084509808 | No Change | Not Significant | TRUE | 0.067522396 | 0.570343011 | No Change | Not Significant | TRUE |
| Mrpl19 | TRUE | -0.138653223 | 0.281393704 | No Change | Not Significant | TRUE | 0.024066371 | 0.886076667 | No Change | Not Significant | TRUE |
| Mrpl2 | TRUE | -0.068054784 | 0.610859762 | No Change | Not Significant | TRUE | -0.017473015 | 0.913844956 | No Change | Not Significant | TRUE |
| Mrpl20 | TRUE | 0.021817882 | 0.880144002 | No Change | Not Significant | TRUE | 0.085649631 | 0.444801234 | No Change | Not Significant | TRUE |
| Mrpl21 | TRUE | -0.078007358 | 0.577145813 | No Change | Not Significant | TRUE | 0.173532976 | 0.236142883 | No Change | Not Significant | TRUE |
| Mrpl22 | TRUE | -0.213305022 | 0.165344954 | No Change | Not Significant | TRUE | 0.166317098 | 0.176202916 | No Change | Not Significant | TRUE |
| Mrpl23 | TRUE | -0.483635253 | 0.022037337 | No Change | Significant Negative | TRUE | -0.052593239 | 0.826263572 | No Change | Not Significant | TRUE |
| Mrpl24 | TRUE | -0.162773681 | 0.009868582 | No Change | Significant Negative | TRUE | 0.005625076 | 0.956874924 | No Change | Not Significant | TRUE |
| Mrpl27 | TRUE | -0.116455745 | 0.377733062 | No Change | Not Significant | TRUE | 0.065412555 | 0.588629676 | No Change | Not Significant | TRUE |
| Mrpl28 | TRUE | 0.085064893 | 0.482492892 | No Change | Not Significant | TRUE | 0.117615945 | 0.20509246 | No Change | Not Significant | TRUE |
| Mrpl3 | TRUE | 0.031408059 | 0.830935108 | No Change | Not Significant | TRUE | 0.032899995 | 0.765987453 | No Change | Not Significant | TRUE |
| Mrpl30 | TRUE | -0.111164362 | 0.238987471 | No Change | Not Significant | TRUE | 0.026794904 | 0.772159116 | No Change | Not Significant | TRUE |
| Mrpl32 | TRUE | -0.055230924 | 0.721043477 | No Change | Not Significant | TRUE | -0.009550435 | 0.946674819 | No Change | Not Significant | TRUE |
| Mrpl33 | TRUE | 0.047222569 | 0.664930223 | No Change | Not Significant | TRUE | -0.013241138 | 0.915984616 | No Change | Not Significant | TRUE |
| Mrpl34 | TRUE | -0.089450532 | 0.60223179 | No Change | Not Significant | TRUE | 0.238814533 | 0.118795321 | No Change | Not Significant | TRUE |
| Mrpl35 | TRUE | -0.076840507 | 0.616875898 | No Change | Not Significant | TRUE | 0.103398039 | 0.287678519 | No Change | Not Significant | TRUE |
| Mrpl36 | TRUE | -0.001846181 | 0.990550804 | No Change | Not Significant | TRUE | 0.166242654 | 0.155879992 | No Change | Not Significant | TRUE |
| Mrpl37 | TRUE | -0.012083889 | 0.950671014 | No Change | Not Significant | TRUE | 0.065002025 | 0.625831733 | No Change | Not Significant | TRUE |
| Mrpl38 | TRUE | -0.13806635 | 0.435789505 | No Change | Not Significant | TRUE | 0.105083241 | 0.515530122 | No Change | Not Significant | TRUE |
| Mrpl39 | TRUE | -0.121257537 | 0.298680373 | No Change | Not Significant | TRUE | -0.044126334 | 0.687971202 | No Change | Not Significant | TRUE |
| Mrpl4 | TRUE | 0.058519788 | 0.676807614 | No Change | Not Significant | TRUE | 0.182459274 | 0.221007096 | No Change | Not Significant | TRUE |
| Mrpl40 | TRUE | -0.109858762 | 0.47814299 | No Change | Not Significant | TRUE | -0.011629426 | 0.930151569 | No Change | Not Significant | TRUE |
| Mrpl41 | TRUE | -0.170399382 | 0.194854619 | No Change | Not Significant | TRUE | 0.027309532 | 0.839028452 | No Change | Not Significant | TRUE |
| Mrpl42 | TRUE | -0.138406779 | 0.195652825 | No Change | Not Significant | TRUE | 0.106328449 | 0.247278021 | No Change | Not Significant | TRUE |

| geneID | Cond1_DF_Detected | Cond1_DF_ValueCol | Cond1_DF_PadjCol | Cond1_DF_Cutoff | Cond1_DF_Cutoff_Specific.x | Cond2_DF_Detected | Cond2_DF_ValueCol | Cond2_DF_PadjCol | Cond2_DF_Cutoff | Cond2_DF_Cutoff_Specific | BG_Method |
|---|---|---|---|---|---|---|---|---|---|---|---|
| Mthfs | TRUE | 0.078214456 | 0.884532517 | No Change | Not Significant | TRUE | 0.146791826 | 0.732526772 | No Change | Not Significant | TRUE |
| Mthfsd | TRUE | 0.186123325 | 0.106242441 | No Change | Not Significant | TRUE | 0.073703179 | 0.670824076 | No Change | Not Significant | TRUE |
| Mthfsl | TRUE | -0.449799404 | 0.011373612 | No Change | Significant Negative | TRUE | -0.026874759 | 0.901341033 | No Change | Not Significant | TRUE |
| Mtif2 | TRUE | -0.064514127 | 0.698491194 | No Change | Not Significant | TRUE | 0.063189636 | 0.547921576 | No Change | Not Significant | TRUE |
| Mtif3 | TRUE | 0.005610402 | 0.971840719 | No Change | Not Significant | TRUE | 0.100439354 | 0.448543447 | No Change | Not Significant | TRUE |
| Mtln | TRUE | -0.246458558 | 0.078315179 | No Change | Not Significant | TRUE | -0.024341342 | 0.884312567 | No Change | Not Significant | TRUE |
| Mtm1 | TRUE | 0.008187114 | 0.957266079 | No Change | Not Significant | TRUE | -0.079741196 | 0.349210912 | No Change | Not Significant | TRUE |
| Mtmr1 | TRUE | 0.038910263 | 0.840144679 | No Change | Not Significant | TRUE | 0.014172453 | 0.89315908 | No Change | Not Significant | TRUE |
| Mtmr10 | TRUE | -0.118101172 | 0.539479894 | No Change | Not Significant | TRUE | -0.030572998 | 0.803190139 | No Change | Not Significant | TRUE |
| Mtmr11 | TRUE | -0.04265778 | 0.896636371 | No Change | Not Significant | TRUE | 0.222614235 | 0.030908674 | No Change | Significant Positive | TRUE |
| Mtmr12 | TRUE | -0.094366858 | 0.270127107 | No Change | Not Significant | TRUE | -0.051439901 | 0.50318658 | No Change | Not Significant | TRUE |
| Mtmr14 | TRUE | 0.590609869 | 5.80E-16 | UP | UP | TRUE | 0.332925632 | 3.55E-09 | No Change | Significant Positive | TRUE |
| Mtmr2 | TRUE | 0.125622335 | 0.334228588 | No Change | Not Significant | TRUE | 0.037901784 | 0.719943844 | No Change | Not Significant | TRUE |
| Mtmr3 | TRUE | -0.195898851 | 0.016858279 | No Change | Significant Negative | TRUE | -0.057477966 | 0.406900734 | No Change | Not Significant | TRUE |
| Mtmr4 | TRUE | 0.600549347 | 1.25E-24 | UP | UP | TRUE | 0.726255528 | 3.25E-37 | UP | UP | TRUE |
| Mtmr6 | TRUE | -0.150946486 | 0.140312312 | No Change | Not Significant | TRUE | -0.070687962 | 0.285589823 | No Change | Not Significant | TRUE |
| Mtmr7 | TRUE | 0.418618779 | 0.000143449 | No Change | Significant Positive | TRUE | 0.09726939 | 0.394462093 | No Change | Not Significant | TRUE |
| Mtmr9 | TRUE | -0.113052217 | 0.474068167 | No Change | Not Significant | TRUE | 0.005122729 | 0.9774899 | No Change | Not Significant | TRUE |
| Mtnr1a | TRUE | -0.085537984 | 1 | No Change | Not Significant | FALSE | NA | NA | No Change | Not Detected | FALSE |
| Mto1 | TRUE | 0.13439791 | 0.308134243 | No Change | Not Significant | TRUE | 0.10565396 | 0.428764731 | No Change | Not Significant | TRUE |
| Mtor | TRUE | 0.145045602 | 0.031967742 | No Change | Significant Positive | TRUE | -0.002591451 | 0.981022852 | No Change | Not Significant | TRUE |
| Mtpap | TRUE | -0.025424452 | 0.894017228 | No Change | Not Significant | TRUE | -0.062637456 | 0.635629805 | No Change | Not Significant | TRUE |
| Mtpn | TRUE | 0.053372672 | 0.693558997 | No Change | Not Significant | TRUE | 0.101754226 | 0.07233951 | No Change | Not Significant | TRUE |
| Mtr | TRUE | 0.24301863 | 0.213630847 | No Change | Not Significant | TRUE | 0.044465113 | 0.823084402 | No Change | Not Significant | TRUE |
| Mtres1 | TRUE | -0.101565692 | 0.44640307 | No Change | Not Significant | TRUE | 0.03420632 | 0.820951722 | No Change | Not Significant | TRUE |
| Mtrex | TRUE | 0.162912161 | 0.173362261 | No Change | Not Significant | TRUE | 0.034845388 | 0.780262005 | No Change | Not Significant | TRUE |
| Mtrf1 | TRUE | -0.063254893 | 0.816921981 | No Change | Not Significant | TRUE | 0.07993111 | 0.718956777 | No Change | Not Significant | TRUE |
| Mtrf1l | TRUE | 0.016076243 | 0.942166544 | No Change | Not Significant | TRUE | -0.102196511 | 0.515514161 | No Change | Not Significant | TRUE |
| Mtrfr | TRUE | -0.274846038 | 0.164401504 | No Change | Not Significant | TRUE | -0.030360874 | 0.90358915 | No Change | Not Significant | TRUE |
| Mtrr | TRUE | 0.612812836 | 2.21E-06 | UP | UP | TRUE | 0.098333007 | 0.479144737 | No Change | Not Significant | TRUE |
| Mtss1 | TRUE | 0.01623178 | 0.905997055 | No Change | Not Significant | TRUE | 0.093445867 | 0.317445011 | No Change | Not Significant | TRUE |
| Mtss2 | TRUE | 0.141971894 | 0.935632034 | No Change | Not Significant | TRUE | -0.160523767 | 0.865580106 | No Change | Not Significant | TRUE |
| Mttp | TRUE | 0.626916371 | 0.228911821 | No Change | Not Significant | TRUE | 0.124771224 | 0.909275777 | No Change | Not Significant | TRUE |
| Mturn | TRUE | 0.677700404 | 0.107588657 | No Change | Not Significant | TRUE | 0.206949287 | 0.5332246 | No Change | Not Significant | TRUE |
| Mtus1 | TRUE | -0.371801121 | 9.12E-06 | No Change | Significant Negative | TRUE | -0.131661922 | 0.020319562 | No Change | Significant Negative | TRUE |
| Mtus2 | TRUE | 0.326498441 | 0.574354812 | No Change | Not Significant | TRUE | -0.561517954 | 0.000619329 | DOWN | DOWN | TRUE |
| Mtx1 | TRUE | 0.097479148 | 0.365696631 | No Change | Not Significant | TRUE | 0.03459677 | 0.793245503 | No Change | Not Significant | TRUE |
| Mtx2 | TRUE | -0.073532494 | 0.564917636 | No Change | Not Significant | TRUE | 0.093235624 | 0.219624067 | No Change | Not Significant | TRUE |
| Mtx3 | TRUE | -0.084503919 | 0.733377497 | No Change | Not Significant | TRUE | -0.170897911 | 0.177733885 | No Change | Not Significant | TRUE |
| Muc1 | TRUE | -0.459865791 | 0.123253915 | No Change | Not Significant | TRUE | -0.305299102 | 0.117491746 | No Change | Not Significant | TRUE |
| Muc13 | TRUE | -0.114360371 | 0.481401448 | No Change | Not Significant | TRUE | 0.060396153 | 0.3701874 | No Change | Not Significant | TRUE |
| Muc15 | TRUE | -0.428002642 | 1 | No Change | Not Significant | FALSE | NA | NA | No Change | Not Detected | FALSE |
| Muc16 | TRUE | 0.598254479 | 0.794172689 | No Change | Not Significant | TRUE | -0.258817247 | 0.78494916 | No Change | Not Significant | TRUE |
| Muc2 | TRUE | -1.282370647 | 1.63E-12 | DOWN | DOWN | TRUE | -0.586626949 | 2.51E-05 | DOWN | DOWN | TRUE |
| Muc20 | TRUE | 0.686103536 | 0.135006119 | No Change | Not Significant | TRUE | -0.200009078 | 0.785533185 | No Change | Not Significant | TRUE |
| Muc21 | TRUE | 0.890640229 | 0.793281643 | No Change | Not Significant | FALSE | NA | NA | No Change | Not Detected | FALSE |
| Muc3 | TRUE | -0.294300607 | 0.384388887 | No Change | Not Significant | TRUE | -0.222868563 | 0.061552292 | No Change | Not Significant | TRUE |
| Muc3a | TRUE | 0.223449498 | 0.699601077 | No Change | Not Significant | TRUE | -0.285557247 | 0.154385176 | No Change | Not Significant | TRUE |
| Muc5ac | TRUE | 0.792015668 | 0.848979902 | No Change | Not Significant | FALSE | NA | NA | No Change | Not Detected | FALSE |
| Muc5b | TRUE | 0.588996325 | 0.75345406 | No Change | Not Significant | FALSE | NA | NA | No Change | Not Detected | FALSE |
| Mucl2 | TRUE | 0.359735414 | 1 | No Change | Not Significant | FALSE | NA | NA | No Change | Not Detected | FALSE |
| Mucl3 | TRUE | -0.465534602 | 1 | No Change | Not Significant | FALSE | NA | NA | No Change | Not Detected | FALSE |
| Mug1 | TRUE | -1.174602832 | 1 | No Change | Not Significant | FALSE | NA | NA | No Change | Not Detected | FALSE |
| Mug2 | TRUE | 0.514766746 | 1 | No Change | Not Significant | FALSE | NA | NA | No Change | Not Detected | FALSE |
| Mul1 | TRUE | 0.066108239 | 0.486449404 | No Change | Not Significant | TRUE | 0.093681376 | 0.203825108 | No Change | Not Significant | TRUE |
| Mus81 | TRUE | 0.224330484 | 0.07241317 | No Change | Not Significant | TRUE | 0.096813102 | 0.420477604 | No Change | Not Significant | TRUE |
| Musk | TRUE | 1.652618261 | 1 | No Change | Not Significant | FALSE | NA | NA | No Change | Not Detected | FALSE |
| Mustn1 | TRUE | 0.485344954 | 1 | No Change | Not Significant | FALSE | NA | NA | No Change | Not Detected | FALSE |
| Mutyh | TRUE | 0.708388173 | 0.047391714 | UP | UP | TRUE | 0.254638361 | 0.480533622 | No Change | Not Significant | TRUE |
| Mvb12a | TRUE | -0.034008544 | 0.80663006 | No Change | Not Significant | TRUE | 0.125342573 | 0.286680207 | No Change | Not Significant | TRUE |
| Mvb12b | TRUE | 0.079687035 | 0.863385162 | No Change | Not Significant | TRUE | -0.402760868 | 0.114196664 | No Change | Not Significant | TRUE |
| Mvd | TRUE | 2.1157787 | 3.53E-52 | UP | UP | TRUE | 1.237639918 | 4.02E-20 | UP | UP | TRUE |
| Mvk | TRUE | 0.256875626 | 0.114758532 | No Change | Not Significant | TRUE | -0.001602989 | 0.995858831 | No Change | Not Significant | TRUE |
| Mvp | TRUE | -0.206183586 | 0.056473628 | No Change | Not Significant | TRUE | 0.037919828 | 0.737165628 | No Change | Not Significant | TRUE |
| Mx2 | TRUE | 0.112965071 | 0.904987729 | No Change | Not Significant | TRUE | -0.506531147 | 0.543683373 | No Change | Not Significant | TRUE |
| Mxd1 | TRUE | 0.367444627 | 0.174214948 | No Change | Not Significant | TRUE | -0.017939278 | 0.926650089 | No Change | Not Significant | TRUE |
| Mxd3 | TRUE | 0.958441155 | 0.000110204 | UP | UP | TRUE | -0.011186101 | 0.974690203 | No Change | Not Significant | TRUE |
| Mxd4 | TRUE | 0.526087807 | 3.29E-08 | UP | UP | TRUE | 0.255536784 | 0.000120676 | No Change | Significant Positive | TRUE |
| Mxi1 | TRUE | -0.089228796 | 0.720922137 | No Change | Not Significant | TRUE | 0.003795184 | 0.982192001 | No Change | Not Significant | TRUE |
| Mxra7 | TRUE | 0.249186016 | 0.444019633 | No Change | Not Significant | TRUE | -0.091613362 | 0.876632986 | No Change | Not Significant | TRUE |
| Mxra8 | TRUE | -0.618600201 | 1 | No Change | Not Significant | FALSE | NA | NA | No Change | Not Detected | FALSE |
| Myadm | TRUE | 0.130238842 | 0.129751344 | No Change | Not Significant | TRUE | 0.131130278 | 0.118482312 | No Change | Not Significant | TRUE |
| Myb | TRUE | 0.059873021 | 0.830405926 | No Change | Not Significant | TRUE | -0.000705347 | 0.995924128 | No Change | Not Significant | TRUE |
| Mybbp1a | TRUE | 0.026845184 | 0.913504709 | No Change | Not Significant | TRUE | -0.161252935 | 0.116216716 | No Change | Not Significant | TRUE |
| Mybl1 | TRUE | 0.638818225 | 0.054123578 | No Change | Not Significant | TRUE | 0.431178027 | 0.184123872 | No Change | Not Significant | TRUE |
| Mybl2 | TRUE | 0.081427824 | 0.816884691 | No Change | Not Significant | TRUE | -0.020153508 | 0.968789653 | No Change | Not Significant | TRUE |
| Mybpc1 | TRUE | -0.733071893 | 1 | No Change | Not Significant | FALSE | NA | NA | No Change | Not Detected | FALSE |
| Mybpc2 | TRUE | -0.456204822 | 1 | No Change | Not Significant | FALSE | NA | NA | No Change | Not Detected | FALSE |
| Mybpc3 | TRUE | 0.080487706 | 1 | No Change | Not Significant | FALSE | NA | NA | No Change | Not Detected | FALSE |
| Mybphl | TRUE | -0.861348063 | 1 | No Change | Not Significant | FALSE | NA | NA | No Change | Not Detected | FALSE |
| Myc | TRUE | 0.170352867 | 0.420617573 | No Change | Not Significant | TRUE | -0.093394222 | 0.523452524 | No Change | Not Significant | TRUE |
| Mycbp | TRUE | -0.09584154 | 0.555969435 | No Change | Not Significant | TRUE | -0.009989717 | 0.950733597 | No Change | Not Significant | TRUE |
| Mycbp2 | TRUE | 0.011048938 | 0.947202684 | No Change | Not Significant | TRUE | 0.064865024 | 0.546234922 | No Change | Not Significant | TRUE |
| Mycbpap | TRUE | -0.587676186 | 0.004591236 | DOWN | DOWN | TRUE | -0.384075301 | 0.063669058 | No Change | Not Significant | TRUE |
| Mycl | TRUE | -0.372279038 | 0.01556181 | No Change | Significant Negative | TRUE | -0.215109224 | 0.062200337 | No Change | Not Significant | TRUE |
| Mycn | TRUE | 0.221398125 | 0.653617891 | No Change | Not Significant | TRUE | 0.108293887 | 0.79296591 | No Change | Not Significant | TRUE |
| Myd88 | TRUE | -0.00117501 | 0.994042693 | No Change | Not Significant | TRUE | -0.025696049 | 0.803865895 | No Change | Not Significant | TRUE |
| Mydgf | TRUE | 0.891882137 | 2.16E-32 | UP | UP | TRUE | 0.675051347 | 1.26E-12 | UP | UP | TRUE |
| Myef2 | TRUE | 0.902434657 | 2.83E-09 | UP | UP | TRUE | 0.157154227 | 0.463780523 | No Change | Not Significant | TRUE |
| Myg1 | TRUE | 0.089087044 | 0.575163675 | No Change | Not Significant | TRUE | -0.059811773 | 0.649560515 | No Change | Not Significant | TRUE |
| Myh1 | TRUE | 0.532488595 | 1 | No Change | Not Significant | FALSE | NA | NA | No Change | Not Detected | FALSE |
| Myh10 | TRUE | 0.062001891 | 0.947050968 | No Change | Not Significant | TRUE | -0.537354059 | 0.233461352 | No Change | Not Significant | TRUE |
| Myh11 | TRUE | 0.776147317 | 0.560708667 | No Change | Not Significant | FALSE | NA | NA | No Change | Not Detected | FALSE |
| Myh13 | TRUE | -2.263295364 | 1 | No Change | Not Significant | FALSE | NA | NA | No Change | Not Detected | FALSE |
| Myh14 | TRUE | 0.276036414 | 0.236875615 | No Change | Not Significant | TRUE | 0.164139224 | 0.23362243 | No Change | Not Significant | TRUE |
| Myh15 | TRUE | 0.435301111 | 1 | No Change | Not Significant | FALSE | NA | NA | No Change | Not Detected | FALSE |
| Myh2 | TRUE | -1.205535809 | 1 | No Change | Not Significant | FALSE | NA | NA | No Change | Not Detected | FALSE |
| Myh3 | TRUE | 2.207408066 | 1 | No Change | Not Significant | FALSE | NA | NA | No Change | Not Detected | FALSE |
| Myh4 | TRUE | -1.244885241 | 1 | No Change | Not Significant | FALSE | NA | NA | No Change | Not Detected | FALSE |
| Myh6 | TRUE | 0.643021398 | 1 | No Change | Not Significant | FALSE | NA | NA | No Change | Not Detected | FALSE |
| Myh7 | TRUE | 0.523426532 | 0.810140405 | No Change | Not Significant | FALSE | NA | NA | No Change | Not Detected | FALSE |
| Myh7b | TRUE | 0.168046712 | 0.908812719 | No Change | Not Significant | FALSE | NA | NA | No Change | Not Detected | FALSE |
| Myh8 | TRUE | -0.791007773 | 1 | No Change | Not Significant | FALSE | NA | NA | No Change | Not Detected | FALSE |
| Myh9 | TRUE | 0.07150555 | 0.487195712 | No Change | Not Significant | TRUE | 0.060982838 | 0.568516308 | No Change | Not Significant | TRUE |
| Myl1 | TRUE | 1.189773279 | 1 | No Change | Not Significant | FALSE | NA | NA | No Change | Not Detected | FALSE |
| Myl12a | TRUE | -0.13660876 | 0.250669647 | No Change | Not Significant | TRUE | 0.010063324 | 0.933715176 | No Change | Not Significant | TRUE |
| Myl12b | TRUE | -0.020125991 | 0.930184975 | No Change | Not Significant | TRUE | 0.02920562 | 0.874869968 | No Change | Not Significant | TRUE |
| Myl3 | TRUE | 0.982393655 | 1 | No Change | Not Significant | FALSE | NA | NA | No Change | Not Detected | FALSE |
| Myl4 | TRUE | -0.068132319 | 1 | No Change | Not Significant | FALSE | NA | NA | No Change | Not Detected | FALSE |
| Myl6 | TRUE | -0.471235529 | 3.06E-07 | No Change | Significant Negative | TRUE | 0.021004684 | 0.774740391 | No Change | Not Significant | TRUE |
| Myl6b | TRUE | 0.378683869 | 0.408255959 | No Change | Not Significant | TRUE | -0.462297917 | 0.278626509 | No Change | Not Significant | TRUE |
| Myl7 | TRUE | -0.89851253 | 0.000310511 | DOWN | DOWN | TRUE | -0.382616376 | 0.399389438 | No Change | Not Significant | TRUE |
| Myl9 | TRUE | -0.331605523 | 1 | No Change | Not Significant | FALSE | NA | NA | No Change | Not Detected | FALSE |
| Mylip | TRUE | -0.146904962 | 0.252105692 | No Change | Not Significant | TRUE | -0.217933694 | 0.061938406 | No Change | Not Significant | TRUE |
| Mylk | TRUE | -0.160506778 | 0.436353234 | No Change | Not Significant | TRUE | -0.080171218 | 0.571636266 | No Change | Not Significant | TRUE |
| Mylk2 | TRUE | 2.253635255 | 1 | No Change | Not Significant | FALSE | NA | NA | No Change | Not Detected | FALSE |
| Mylk3 | TRUE | -1.832049159 | 1 | No Change | Not Significant | FALSE | NA | NA | No Change | Not Detected | FALSE |
| Mylk4 | TRUE | 0.882503618 | 1 | No Change | Not Significant | FALSE | NA | NA | No Change | Not Detected | FALSE |
| Mylpf | TRUE | 0.086801538 | 0.927761944 | No Change | Not Significant | TRUE | 0.125792485 | 0.793911553 | No Change | Not Significant | TRUE |
| Mynn | TRUE | -0.116451578 | 0.341473496 | No Change | Not Significant | TRUE | -0.097876477 | 0.371384959 | No Change | Not Significant | TRUE |
| Myo10 | TRUE | 0.04688266 | 0.665277305 | No Change | Not Significant | TRUE | -0.02172996 | 0.863755116 | No Change | Not Significant | TRUE |
| Myo15a | TRUE | 1.439644038 | 0.453944473 | No Change | Not Significant | FALSE | NA | NA | No Change | Not Detected | FALSE |
| Myo15b | TRUE | 0.193388031 | 0.618597629 | No Change | Not Significant | TRUE | -0.156323289 | 0.210995961 | No Change | Not Significant | TRUE |
| Myo16 | TRUE | -1.400991757 | 1 | No Change | Not Significant | FALSE | NA | NA | No Change | Not Detected | FALSE |
| Myo18a | TRUE | 0.050827413 | 0.85879931 | No Change | Not Significant | TRUE | -0.008487628 | 0.954467075 | No Change | Not Significant | TRUE |
| Myo18b | TRUE | 4.299491807 | 1.66E-26 | UP | UP | TRUE | 6.239168327 | 6.33E-33 | UP | UP | TRUE |
| Myo19 | TRUE | 0.008448878 | 0.98153379 | No Change | Not Significant | TRUE | -0.251638534 | 0.018182179 | No Change | Significant Negative | TRUE |
| Myo1a | TRUE | -0.549824885 | 0.000142578 | DOWN | DOWN | TRUE | -0.199005596 | 0.0051554 | No Change | Significant Negative | TRUE |
| Myo1b | TRUE | -0.066512338 | 0.720614094 | No Change | Not Significant | TRUE | -0.148260858 | 0.113433539 | No Change | Not Significant | TRUE |
| Myo1c | TRUE | -0.350337015 | 3.52E-10 | No Change | Significant Negative | TRUE | -0.104970532 | 0.137116178 | No Change | Not Significant | TRUE |
| Myo1d | TRUE | 0.187488349 | 0.017451579 | No Change | Significant Positive | TRUE | 0.196032922 | 0.012273523 | No Change | Significant Positive | TRUE |
| Myo1e | TRUE | -0.139149131 | 0.135006119 | No Change | Not Significant | TRUE | 0.01457361 | 0.915013749 | No Change | Not Significant | TRUE |
| Myo1f | TRUE | 0.148313366 | 0.846814623 | No Change | Not Significant | TRUE | 0.535258354 | 0.133735764 | No Change | Not Significant | TRUE |
| Myo1g | TRUE | 0.371154811 | 0.664082645 | No Change | Not Significant | TRUE | 0.124325713 | 0.853575129 | No Change | Not Significant | TRUE |
| Myo1h | TRUE | 0.148859218 | 0.846208704 | No Change | Not Significant | TRUE | -0.5939043 | 0.179789226 | No Change | Not Significant | TRUE |
| Myo3a | TRUE | -1.798091251 | 1.07E-21 | DOWN | DOWN | TRUE | -0.519398003 | 0.013126796 | DOWN | DOWN | TRUE |
| Myo3b | TRUE | -2.290872175 | 1 | No Change | Not Significant | FALSE | NA | NA | No Change | Not Detected | FALSE |
| Myo5a | TRUE | 0.409445697 | 0.322533618 | No Change | Not Significant | TRUE | 0.419440653 | 0.308074929 | No Change | Not Significant | TRUE |
| Myo5b | TRUE | -0.024001274 | 0.886212533 | No Change | Not Significant | TRUE | 0.098316266 | 0.102159742 | No Change | Not Significant | TRUE |
| Myo5c | TRUE | 0.085446586 | 0.600047814 | No Change | Not Significant | TRUE | 0.160576339 | 0.045316043 | No Change | Significant Positive | TRUE |
| Myo6 | TRUE | 0.016218638 | 0.908409285 | No Change | Not Significant | TRUE | -0.047473091 | 0.504916665 | No Change | Not Significant | TRUE |
| Myo7a | TRUE | 0.271646505 | 0.225018236 | No Change | Not Significant | TRUE | -0.002624858 | 0.989317668 | No Change | Not Significant | TRUE |
| Myo7b | TRUE | -0.18276335 | 0.472489189 | No Change | Not Significant | TRUE | -0.240563722 | 0.002933681 | No Change | Significant Negative | TRUE |
| Myo9a | TRUE | -0.303402546 | 0.064054228 | No Change | Not Significant | TRUE | -0.401627119 | 0.005446724 | No Change | Significant Negative | TRUE |
| Myo9b | TRUE | 0.120989033 | 0.472954995 | No Change | Not Significant | TRUE | -0.079521708 | 0.345432975 | No Change | Not Significant | TRUE |
| Myocd | TRUE | -1.562769459 | 1 | No Change | Not Significant | FALSE | NA | NA | No Change | Not Detected | FALSE |
| Myof | TRUE | 0.298748849 | 0.008692083 | No Change | Significant Positive | TRUE | 0.471138945 | 5.24E-09 | No Change | Significant Positive | TRUE |
| Myog | TRUE | -0.431800025 | 1 | No Change | Not Significant | FALSE | NA | NA | No Change | Not Detected | FALSE |
| Myom1 | TRUE | 0.718970636 | 0.07410998 | No Change | Not Significant | TRUE | -0.305647184 | 0.493560011 | No Change | Not Significant | TRUE |
| Myom2 | TRUE | 0.497745364 | 0.890653197 | No Change | Not Significant | FALSE | NA | NA | No Change | Not Detected | FALSE |
| Myom3 | TRUE | -0.110858157 | 0.876856731 | No Change | Not Significant | TRUE | -0.732527013 | 0.008422099 | DOWN | DOWN | TRUE |
| Myorg | TRUE | -0.196114635 | 0.388837505 | No Change | Not Significant | TRUE | 0.252372069 | 0.000588069 | No Change | Significant Positive | TRUE |
| Myot | TRUE | 1.904010835 | 1 | No Change | Not Significant | FALSE | NA | NA | No Change | Not Detected | FALSE |
| Myoz2 | TRUE | -0.525161794 | 1 | No Change | Not Significant | FALSE | NA | NA | No Change | Not Detected | FALSE |
| Myoz3 | TRUE | 0.954263278 | 1 | No Change | Not Significant | FALSE | NA | NA | No Change | Not Detected | FALSE |
| Mypn | TRUE | 1.408791457 | 1 | No Change | Not Significant | FALSE | NA | NA | No Change | Not Detected | FALSE |
| Mypop | TRUE | -0.417565205 | 0.363760214 | No Change | Not Significant | TRUE | -0.14652809 | 0.716156527 | No Change | Not Significant | TRUE |
| Myrf | TRUE | 0.406730402 | 0.810851996 | No Change | Not Significant | TRUE | 0.390204096 | 0.687894338 | No Change | Not Significant | TRUE |
| Myrfl | TRUE | -0.587516585 | 0.560514185 | No Change | Not Significant | TRUE | -0.764805379 | 0.076614782 | No Change | Not Significant | TRUE |
| Myrip | TRUE | -0.93296989 | 1.04E-06 | DOWN | DOWN | TRUE | -0.221272862 | 0.382237605 | No Change | Not Significant | TRUE |
| Mysm1 | TRUE | -0.066255862 | 0.566265254 | No Change | Not Significant | TRUE | -0.118631668 | 0.24490649 | No Change | Not Significant | TRUE |
| Myt1 | TRUE | -0.82811733 | 0.095392863 | No Change | Not Significant | TRUE | -0.172930858 | 0.668066015 | No Change | Not Significant | TRUE |
| Myt1l | TRUE | 1.286638922 | 0.729592345 | No Change | Not Significant | FALSE | NA | NA | No Change | Not Detected | FALSE |
| Myzap | TRUE | -0.113376109 | 0.808079957 | No Change | Not Significant | TRUE | 0.084358164 | 0.679126034 | No Change | Not Significant | TRUE |
| Mzf1 | TRUE | 0.010300221 | 0.9893404 | No Change | Not Significant | TRUE | 0.162616827 | 0.761252782 | No Change | Not Significant | TRUE |
| Mzt1 | TRUE | 0.338796292 | 0.00659513 | No Change | Significant Positive | TRUE | 0.069371682 | 0.516310954 | No Change | Not Significant | TRUE |
| Mzt2 | TRUE | -0.059803919 | 0.782626881 | No Change | Not Significant | TRUE | 0.010888529 | 0.952355017 | No Change | Not Significant | TRUE |
| n-R5s33 | TRUE | 1.718356607 | 1 | No Change | Not Significant | FALSE | NA | NA | No Change | Not Detected | FALSE |
| N4bp1 | TRUE | 0.054316843 | 0.690541222 | No Change | Not Significant | TRUE | -0.06899533 | 0.376690964 | No Change | Not Significant | TRUE |
| N4bp2 | TRUE | 1.159345913 | 1.07E-36 | UP | UP | TRUE | 0.51860713 | 3.83E-13 | UP | UP | TRUE |
| N4bp2l1 | TRUE | -1.163475781 | 0.000232062 | DOWN | DOWN | TRUE | -0.22942416 | 0.529236838 | No Change | Not Significant | TRUE |
| N4bp2l2 | TRUE | -0.048331813 | 0.684193826 | No Change | Not Significant | TRUE | -0.10200446 | 0.453522778 | No Change | Not Significant | TRUE |
| N4bp3 | TRUE | -0.869541783 | 9.74E-11 | DOWN | DOWN | TRUE | -0.334446642 | 0.016080083 | No Change | Significant Negative | TRUE |
| N6amt1 | TRUE | -0.317973672 | 0.027285164 | No Change | Significant Negative | TRUE | -0.222008507 | 0.149582964 | No Change | Not Significant | TRUE |
| Naa10 | TRUE | 0.116512663 | 0.487334112 | No Change | Not Significant | TRUE | 0.050669338 | 0.712754953 | No Change | Not Significant | TRUE |
| Naa12 | TRUE | 0.013962667 | 0.941973978 | No Change | Not Significant | TRUE | 0.01566572 | 0.898825528 | No Change | Not Significant | TRUE |
| Naa15 | TRUE | 0.130108747 | 0.420139706 | No Change | Not Significant | TRUE | -0.069872477 | 0.621060566 | No Change | Not Significant | TRUE |
| Naa16 | TRUE | 0.004683871 | 0.983635967 | No Change | Not Significant | TRUE | -0.102497903 | 0.500746225 | No Change | Not Significant | TRUE |
| Naa20 | TRUE | -0.189585912 | 0.081770074 | No Change | Not Significant | TRUE | 0.026145362 | 0.841671981 | No Change | Not Significant | TRUE |
| Naa25 | TRUE | 0.10666058 | 0.407409522 | No Change | Not Significant | TRUE | -0.09811257 | 0.410035436 | No Change | Not Significant | TRUE |
| Naa30 | TRUE | 0.040996796 | 0.717711842 | No Change | Not Significant | TRUE | -0.080216464 | 0.47644362 | No Change | Not Significant | TRUE |
| Naa35 | TRUE | 0.06984099 | 0.601173216 | No Change | Not Significant | TRUE | 0.139511684 | 0.056941378 | No Change | Not Significant | TRUE |
| Naa38 | TRUE | 0.07031446 | 0.714894653 | No Change | Not Significant | TRUE | 0.145791269 | 0.380615229 | No Change | Not Significant | TRUE |
| Naa40 | TRUE | -0.024874275 | 0.908812719 | No Change | Not Significant | TRUE | -0.0089611 | 0.952355017 | No Change | Not Significant | TRUE |
| Naa50 | TRUE | -0.035989854 | 0.860514985 | No Change | Not Significant | TRUE | 0.026432968 | 0.837472006 | No Change | Not Significant | TRUE |
| Naa60 | TRUE | -0.005831284 | 0.97354273 | No Change | Not Significant | TRUE | 0.03451585 | 0.762827428 | No Change | Not Significant | TRUE |
| Naa80 | TRUE | -0.203232771 | 0.198490252 | No Change | Not Significant | TRUE | 0.031059731 | 0.863217232 | No Change | Not Significant | TRUE |
| Naaa | TRUE | -0.407670322 | 0.033825714 | No Change | Significant Negative | TRUE | 0.028802217 | 0.863035664 | No Change | Not Significant | TRUE |
| Naalad2 | TRUE | -2.080708858 | 1 | No Change | Not Significant | FALSE | NA | NA | No Change | Not Detected | FALSE |
| Naaladl1 | TRUE | 3.800819753 | 2.10E-92 | UP | UP | TRUE | 0.647372364 | 9.93E-05 | UP | UP | TRUE |
| Naaladl2 | TRUE | -0.082305518 | 0.812494814 | No Change | Not Significant | TRUE | -0.063905599 | 0.733199598 | No Change | Not Significant | TRUE |
| Nab1 | TRUE | -0.181343356 | 0.199909758 | No Change | Not Significant | TRUE | -0.02383056 | 0.8024647 | No Change | Not Significant | TRUE |

Supplementary Table 3

| geneID | Cond1_DF_Detected | Cond1_DF_ValueCol | Cond1_DF_PadjCol | Cond1_DF_Cutoff | Cond1_DF_Cutoff_Specific.x | Cond2_DF_Detected | Cond2_DF_ValueCol | Cond2_DF_PadjCol | Cond2_DF_Cutoff | Cond2_DF_Cutoff_Specific | BG_Method | RG1_All | RG2_Significant | RG3_SignificantChange | Cond1_DF_ensembl_gene_id | Cond1_DF_entrezgene_id | Cond1_DF_gene_biotype |
|---|---|---|---|---|---|---|---|---|---|---|---|---|---|---|---|---|---|
| Nab2 | TRUE | 0.233872208 | 0.691562872 | No Change | Not Significant | TRUE | 0.168164448 | 0.78394657 | No Change | Not Significant | TRUE | Cond1 Not Significant + Cond2 Not Significant | None | None | ENSMUSG00000025402 | 17937 | protein_coding |
| Nabp1 | TRUE | 0.530220168 | 0.015927605 | UP | UP | TRUE | 0.171699513 | 0.13301537 | No Change | Not Significant | TRUE | Cond1 UP + Cond2 Not Significant | Cond1_UP | Cond1_UP | ENSMUSG00000026107 | 109019 | protein_coding |
| Nabp2 | TRUE | 0.151787312 | 0.22115998 | No Change | Not Significant | TRUE | 0.169701617 | 0.067221635 | No Change | Not Significant | TRUE | Cond1 Not Significant + Cond2 Not Significant | None | None | ENSMUSG00000025374 | 69917 | protein_coding |
| Naca | TRUE | -0.280372855 | 0.015492034 | No Change | Significant Negative | TRUE | -0.018136429 | 0.854394093 | No Change | Not Significant | TRUE | Cond1 Significant Negative + Cond2 Not Significant | None | None | ENSMUSG00000061315 | 17938 | protein_coding |
| Nacad | TRUE | -0.041679551 | 0.937028043 | No Change | Not Significant | TRUE | 0.746707406 | 0.001108091 | UP | UP | TRUE | Cond1 Not Significant + Cond2 UP | Cond2_UP | Cond2_UP | ENSMUSG00000041073 | 192950 | protein_coding |
| Nacc1 | TRUE | -0.033276826 | 0.785880315 | No Change | Not Significant | TRUE | 0.104780846 | 0.335814568 | No Change | Not Significant | TRUE | Cond1 Not Significant + Cond2 Not Significant | None | None | ENSMUSG00000001910 | 66830 | protein_coding |
| Nacc2 | TRUE | -0.126534121 | 0.306987369 | No Change | Not Significant | TRUE | 0.042660779 | 0.706609067 | No Change | Not Significant | TRUE | Cond1 Not Significant + Cond2 Not Significant | None | None | ENSMUSG00000026932 | 67991 | protein_coding |
| Nadk | TRUE | -0.261607421 | 6.90E-06 | No Change | Significant Negative | TRUE | 0.021293977 | 0.853157579 | No Change | Not Significant | TRUE | Cond1 Significant Negative + Cond2 Not Significant | None | None | ENSMUSG00000029063 | 192185 | protein_coding |
| Nadk2 | TRUE | 0.622028917 | 3.45E-09 | UP | UP | TRUE | 0.350774362 | 5.17E-07 | No Change | Significant Positive | TRUE | Cond1 UP + Cond2 Significant Positive | Core_UP | Cond1_UP | ENSMUSG00000022253 | 68646 | protein_coding |
| Nadsyn1 | TRUE | -0.628277989 | 5.19E-14 | DOWN | DOWN | TRUE | -0.153035181 | 0.018231711 | No Change | Significant Negative | TRUE | Cond1 DOWN + Cond2 Significant Negative | Core_DOWN | Cond1_DOWN | ENSMUSG00000031090 | 78914 | protein_coding |
| Nae1 | TRUE | -0.148489336 | 0.281753823 | No Change | Not Significant | TRUE | -0.132070467 | 0.139211338 | No Change | Not Significant | TRUE | Cond1 Not Significant + Cond2 Not Significant | None | None | ENSMUSG00000031878 | 234664 | protein_coding |
| Naf1 | TRUE | 0.146149947 | 0.514721638 | No Change | Not Significant | TRUE | 0.03481986 | 0.886815999 | No Change | Not Significant | TRUE | Cond1 Not Significant + Cond2 Not Significant | None | None | ENSMUSG00000014907 | 234344 | protein_coding |
| Naga | TRUE | 0.067500875 | 0.450443439 | No Change | Not Significant | TRUE | 0.109631541 | 0.213667404 | No Change | Not Significant | TRUE | Cond1 Not Significant + Cond2 Not Significant | None | None | ENSMUSG00000022453 | 17939 | protein_coding |
| Nagk | TRUE | -0.249019696 | 0.001453571 | No Change | Significant Negative | TRUE | -0.11591289 | 0.104705757 | No Change | Not Significant | TRUE | Cond1 Significant Negative + Cond2 Not Significant | None | None | ENSMUSG00000034744 | 56174 | protein_coding |
| Naglu | TRUE | 0.022328734 | 0.9092783 | No Change | Not Significant | TRUE | 0.275247677 | 0.030422534 | No Change | Significant Positive | TRUE | Cond1 Not Significant + Cond2 Significant Positive | None | None | ENSMUSG00000001751 | 27419 | protein_coding |
| Nagpa | TRUE | -0.154773928 | 0.31779121 | No Change | Not Significant | TRUE | 0.10969354 | 0.537823306 | No Change | Not Significant | TRUE | Cond1 Not Significant + Cond2 Not Significant | None | None | ENSMUSG00000023143 | 27426 | protein_coding |
| Nags | TRUE | -0.968948311 | 0.000714486 | DOWN | DOWN | TRUE | -0.509896245 | 0.00151806 | DOWN | DOWN | TRUE | Cond1 DOWN + Cond2 DOWN | Core_DOWN | Core_DOWN | ENSMUSG00000048217 | 217214 | protein_coding |

| geneID | Cond1_DF_baseMean | Cond1_DF_log2FoldChange | Cond1_DF_lfcSE | Cond1_DF_stat | Cond1_DF_pvalue | Cond1_DF_padj | Cond1_DF_absLog2FC | Cond1_DF_Cutoff_Specific.y | Cond2_DF_ensembl_gene_id | Cond2_DF_entrezgene_id | Cond2_DF_gene_biotype | Cond2_DF_baseMean | Cond2_DF_log2FoldChange | Cond2_DF_lfcSE | Cond2_DF_stat | Cond2_DF_pvalue | Cond2_DF_padj | Cond2_DF_absLog2FC | RG1_Specific_Cond2 | RG1_Specific_Cond1 |
|---|---|---|---|---|---|---|---|---|---|---|---|---|---|---|---|---|---|---|---|---|
| Nab2 | 14.34941495 | 0.233872208 | 0.331594795 | -0.705295172 | 0.480626564 | 0.691562872 | 0.233872208 | Not Significant | ENSMUSG00000025402 | 17937 | protein_coding | 14.49714278 | 0.168164448 | 0.318557057 | 0.527894279 | 0.597572716 | 0.78394657 | 0.168164448 | Cond1 No Change + Cond2 Not Significant | Cond2 No Change + Cond1 Not Significant |
| Nabp1 | 239.4180103 | 0.530220168 | 0.178747815 | -2.966302939 | 0.003014034 | 0.015927605 | 0.530220168 | UP | ENSMUSG00000026107 | 109019 | protein_coding | 341.6925097 | 0.171699513 | 0.079584749 | 2.1574424 | 0.030971211 | 0.13301537 | 0.171699513 | Cond1 UP + Cond2 Not Significant | Cond2 No Change + Cond1 UP |
| Nabp2 | 319.2694171 | 0.151787312 | 0.088353256 | -1.717959456 | 0.085804011 | 0.22115998 | 0.151787312 | Not Significant | ENSMUSG00000025374 | 69917 | protein_coding | 542.3347767 | 0.169701617 | 0.067397156 | 2.517934385 | 0.011804531 | 0.067221635 | 0.169701617 | Cond1 No Change + Cond2 Not Significant | Cond2 No Change + Cond1 Not Significant |
| Naca | 2724.967511 | -0.280372855 | 0.094192597 | 2.976591187 | 0.002914724 | 0.015492034 | 0.280372855 | Significant Negative | ENSMUSG00000061315 | 17938 | protein_coding | 5790.01091 | -0.018136429 | 0.048268901 | -0.375737348 | 0.707112169 | 0.854394093 | 0.018136429 | Cond1 No Change + Cond2 Not Significant | Cond2 No Change + Cond1 Significant Negative |
| Nacad | 73.23832666 | -0.041679551 | 0.259036406 | 0.16090229 | 0.872170359 | 0.937028043 | 0.041679551 | Not Significant | ENSMUSG00000041073 | 192950 | protein_coding | 63.18669415 | 0.746707406 | 0.187968022 | 3.972523612 | 7.11E-05 | 0.001108091 | 0.746707406 | Cond1 No Change + Cond2 UP | Cond2 UP + Cond1 Not Significant |
| Nacc1 | 1241.63056 | -0.033276826 | 0.065758911 | 0.506042833 | 0.612826581 | 0.785880315 | 0.033276826 | Not Significant | ENSMUSG00000001910 | 66830 | protein_coding | 2216.690742 | 0.104780846 | 0.06916042 | 1.515040628 | 0.129762116 | 0.335814568 | 0.104780846 | Cond1 No Change + Cond2 Not Significant | Cond2 No Change + Cond1 Not Significant |
| Nacc2 | 400.3580114 | -0.126534121 | 0.084797246 | 1.492196105 | 0.135647743 | 0.306987369 | 0.126534121 | Not Significant | ENSMUSG00000026932 | 67991 | protein_coding | 845.3330657 | 0.042660779 | 0.061141661 | 0.697736674 | 0.485341888 | 0.706609067 | 0.042660779 | Cond1 No Change + Cond2 Not Significant | Cond2 No Change + Cond1 Not Significant |
| Nadk | 2370.977621 | -0.261607421 | 0.051898485 | 5.040752602 | 4.64E-07 | 6.90E-06 | 0.261607421 | Significant Negative | ENSMUSG00000029063 | 192185 | protein_coding | 4438.145693 | 0.021293977 | 0.056159462 | 0.379169886 | 0.704561713 | 0.853157579 | 0.021293977 | Cond1 No Change + Cond2 Not Significant | Cond2 No Change + Cond1 Significant Negative |
| Nadk2 | 464.5988984 | 0.622028917 | 0.096979909 | -6.413997729 | 1.42E-10 | 3.45E-09 | 0.622028917 | UP | ENSMUSG00000022253 | 68646 | protein_coding | 814.2278451 | 0.350774362 | 0.061928763 | 5.66415904 | 1.48E-08 | 5.17E-07 | 0.350774362 | Cond1 UP + Cond2 Significant Positive | Cond2 No Change + Cond1 UP |
| Nadsyn1 | 493.4867996 | -0.628277989 | 0.078641172 | 7.989173754 | 1.36E-15 | 5.19E-14 | 0.628277989 | DOWN | ENSMUSG00000031090 | 78914 | protein_coding | 1280.109429 | -0.153035181 | 0.049754521 | -3.07580451 | 0.002099354 | 0.018231711 | 0.153035181 | Cond1 DOWN + Cond2 Significant Negative | Cond2 No Change + Cond1 DOWN |
| Nae1 | 277.1434762 | -0.148489336 | 0.095459801 | 1.555516934 | 0.119823005 | 0.281753823 | 0.148489336 | Not Significant | ENSMUSG00000031878 | 234664 | protein_coding | 617.0049755 | -0.132070467 | 0.061962841 | -2.131446275 | 0.033052394 | 0.139211338 | 0.132070467 | Cond1 No Change + Cond2 Not Significant | Cond2 No Change + Cond1 Not Significant |
| Naf1 | 92.9100874 | 0.146149947 | 0.139735408 | -1.045904899 | 0.295604942 | 0.514721638 | 0.146149947 | Not Significant | ENSMUSG00000014907 | 234344 | protein_coding | 157.0251814 | 0.03481986 | 0.114285923 | 0.304673219 | 0.760615049 | 0.886815999 | 0.03481986 | Cond1 No Change + Cond2 Not Significant | Cond2 No Change + Cond1 Not Significant |
| Naga | 1572.105303 | 0.067500875 | 0.05746042 | -1.174736895 | 0.240099994 | 0.450443439 | 0.067500875 | Not Significant | ENSMUSG00000022453 | 17939 | protein_coding | 2949.629593 | 0.109631541 | 0.058878079 | 1.862009481 | 0.062601754 | 0.213667404 | 0.109631541 | Cond1 No Change + Cond2 Not Significant | Cond2 No Change + Cond1 Not Significant |
| Nagk | 456.1158945 | -0.249019696 | 0.066510305 | 3.744076879 | 0.000181058 | 0.001453571 | 0.249019696 | Significant Negative | ENSMUSG00000034744 | 56174 | protein_coding | 964.4033226 | -0.11591289 | 0.050553057 | -2.292895751 | 0.021854008 | 0.104705757 | 0.11591289 | Cond1 No Change + Cond2 Not Significant | Cond2 No Change + Cond1 Significant Negative |
| Naglu | 376.0621458 | 0.022328734 | 0.098364693 | -0.226999477 | 0.820424149 | 0.9092783 | 0.022328734 | Not Significant | ENSMUSG00000001751 | 27419 | protein_coding | 699.7477329 | 0.275247677 | 0.09570022 | 2.87614465 | 0.004025652 | 0.030422534 | 0.275247677 | Cond1 No Change + Cond2 Significant Positive | Cond2 No Change + Cond1 Not Significant |
| Nagpa | 210.1540503 | -0.154773928 | 0.105485575 | 1.467252071 | 0.142307497 | 0.31779121 | 0.154773928 | Not Significant | ENSMUSG00000023143 | 27426 | protein_coding | 410.2612896 | 0.10969354 | 0.104696606 | 1.047727754 | 0.294764057 | 0.537823306 | 0.10969354 | Cond1 No Change + Cond2 Not Significant | Cond2 No Change + Cond1 Not Significant |
| Nags | 101.5841799 | -0.968948311 | 0.245779919 | 3.942341239 | 8.07E-05 | 0.000714486 | 0.968948311 | DOWN | ENSMUSG00000048217 | 217214 | protein_coding | 233.7784973 | -0.509896245 | 0.131246714 | -3.885021036 | 0.000102321 | 0.00151806 | 0.509896245 | Cond1 DOWN + Cond2 DOWN | Cond2 DOWN + Cond1 DOWN |

| geneID | Cond1_DF_Detected | Cond1_DF_ValueCol | Cond1_DF_PadjCol | Cond1_DF_Cutoff | Cond1_DF_Cutoff_Specific.x | Cond2_DF_Detected | Cond2_DF_ValueCol | Cond2_DF_PadjCol | Cond2_DF_Cutoff | Cond2_DF_Cutoff_Specific | BG_Method |
|---|---|---|---|---|---|---|---|---|---|---|---|
| Nefl | TRUE | 0.059484739 | 1 | No Change | Not Significant | TRUE | -0.084234628 | 0.93086312 | No Change | Not Significant | TRUE |
| Nefm | TRUE | 0.126939464 | 0.908714429 | No Change | Not Significant | TRUE | 0.070210356 | 0.940208703 | No Change | Not Significant | TRUE |
| Negr1 | TRUE | 0.713977144 | 0.722169631 | No Change | Not Significant | FALSE | NA | NA | No Change | Not Detected | FALSE |
| Neil1 | TRUE | -0.233821428 | 0.081077558 | No Change | Not Significant | TRUE | -0.020381617 | 0.917020505 | No Change | Not Significant | TRUE |
| Neil2 | TRUE | -0.640796509 | 1 | No Change | Not Significant | FALSE | NA | NA | No Change | Not Detected | FALSE |
| Neil3 | TRUE | 0.560784771 | 0.098234156 | No Change | Not Significant | TRUE | -0.407566315 | 0.122794781 | No Change | Not Significant | TRUE |
| Nek1 | TRUE | 0.086334605 | 0.566318105 | No Change | Not Significant | TRUE | -0.043166869 | 0.771101239 | No Change | Not Significant | TRUE |
| Nek10 | TRUE | 0.739633735 | 0.467050404 | No Change | Not Significant | TRUE | 1.259990946 | 0.068576338 | No Change | Not Significant | TRUE |
| Nek11 | TRUE | 0.605916899 | 0.465324602 | No Change | Not Significant | TRUE | 0.722524784 | 0.334104382 | No Change | Not Significant | TRUE |
| Nek2 | TRUE | 1.179542151 | 4.32E-14 | UP | UP | TRUE | 0.228350998 | 0.191597869 | No Change | Not Significant | TRUE |
| Nek3 | TRUE | -0.538188865 | 0.003784244 | DOWN | DOWN | TRUE | -0.0894494 | 0.573541103 | No Change | Not Significant | TRUE |
| Nek4 | TRUE | 0.213292478 | 0.123253915 | No Change | Not Significant | TRUE | 0.085373233 | 0.423163427 | No Change | Not Significant | TRUE |
| Nek5 | TRUE | -0.661665687 | 0.227794351 | No Change | Not Significant | TRUE | -0.020648177 | 0.971056929 | No Change | Not Significant | TRUE |
| Nek6 | TRUE | 0.043647733 | 0.780657323 | No Change | Not Significant | TRUE | 0.132596966 | 0.168649897 | No Change | Not Significant | TRUE |
| Nek7 | TRUE | 0.285002625 | 0.028422287 | No Change | Significant Positive | TRUE | 0.172046587 | 0.021014745 | No Change | Significant Positive | TRUE |
| Nek8 | TRUE | -0.520046707 | 1.84E-06 | DOWN | DOWN | TRUE | -0.116079828 | 0.393611906 | No Change | Not Significant | TRUE |
| Nek9 | TRUE | -0.102722942 | 0.144095106 | No Change | Not Significant | TRUE | -0.008510698 | 0.926479877 | No Change | Not Significant | TRUE |
| Nelfa | TRUE | 0.203579743 | 0.033677833 | No Change | Significant Positive | TRUE | 0.108563353 | 0.306818815 | No Change | Not Significant | TRUE |
| Nelfb | TRUE | -0.124437344 | 0.037065168 | No Change | Significant Negative | TRUE | 0.048231709 | 0.663709663 | No Change | Not Significant | TRUE |
| Nelfcd | TRUE | 0.017497389 | 0.897240511 | No Change | Not Significant | TRUE | 0.051561729 | 0.706043466 | No Change | Not Significant | TRUE |
| Nelfe | TRUE | 0.221210374 | 0.041864611 | No Change | Significant Positive | TRUE | 0.161757358 | 0.02531242 | No Change | Significant Positive | TRUE |
| Nell1 | TRUE | 0.968872108 | 1 | No Change | Not Significant | FALSE | NA | NA | No Change | Not Detected | FALSE |
| Nell2 | TRUE | -0.676671672 | 1 | No Change | Not Significant | FALSE | NA | NA | No Change | Not Detected | FALSE |
| Nemf | TRUE | 0.044485422 | 0.692354022 | No Change | Not Significant | TRUE | 0.019884161 | 0.892332662 | No Change | Not Significant | TRUE |
| Nemp1 | TRUE | 0.156241582 | 0.543156551 | No Change | Not Significant | TRUE | -0.070431606 | 0.644152928 | No Change | Not Significant | TRUE |
| Nemp2 | TRUE | -0.051151355 | 0.89369175 | No Change | Not Significant | TRUE | -0.229397766 | 0.444592741 | No Change | Not Significant | TRUE |
| Nenf | TRUE | -0.384776578 | 0.000694131 | No Change | Significant Negative | TRUE | -0.162837164 | 0.297255113 | No Change | Not Significant | TRUE |
| Neo1 | TRUE | -0.120660439 | 0.215774069 | No Change | Not Significant | TRUE | -0.171440494 | 0.009934207 | No Change | Significant Negative | TRUE |
| Nepro | TRUE | -0.027901499 | 0.888226437 | No Change | Not Significant | TRUE | -0.187981604 | 0.143220107 | No Change | Not Significant | TRUE |
| Nes | TRUE | 3.296137289 | 3.08E-13 | UP | UP | FALSE | NA | NA | No Change | Not Detected | FALSE |
| Net1 | TRUE | -0.270537662 | 0.001207955 | No Change | Significant Negative | TRUE | -0.022518526 | 0.893779608 | No Change | Not Significant | TRUE |
| Neto1 | TRUE | 1.242454041 | 1 | No Change | Not Significant | FALSE | NA | NA | No Change | Not Detected | FALSE |
| Neto2 | TRUE | 0.622118272 | 0.085571212 | No Change | Not Significant | TRUE | 0.05793062 | 0.880597382 | No Change | Not Significant | TRUE |
| Neu1 | TRUE | -0.03692206 | 0.920255229 | No Change | Not Significant | TRUE | -0.446332812 | 0.000645829 | No Change | Significant Negative | TRUE |
| Neu2 | TRUE | 0.370063176 | 0.314536999 | No Change | Not Significant | TRUE | -0.038799384 | 0.927227465 | No Change | Not Significant | TRUE |
| Neu3 | TRUE | -0.199631346 | 0.662922836 | No Change | Not Significant | TRUE | -0.629534203 | 0.019707576 | DOWN | DOWN | TRUE |
| Neu4 | TRUE | -0.879094113 | 1 | No Change | Not Significant | FALSE | NA | NA | No Change | Not Detected | FALSE |
| Neurl1a | TRUE | -0.674204907 | 0.009251404 | DOWN | DOWN | TRUE | -0.128955386 | 0.708292116 | No Change | Not Significant | TRUE |
| Neurl1b | TRUE | -0.521324639 | 1.46E-06 | DOWN | DOWN | TRUE | -0.336667691 | 0.000716702 | No Change | Significant Negative | TRUE |
| Neurl2 | TRUE | -0.473489351 | 1 | No Change | Not Significant | TRUE | -1.063514728 | 0.19482421 | No Change | Not Significant | TRUE |
| Neurl3 | TRUE | -0.050039541 | 0.816323585 | No Change | Not Significant | TRUE | -0.179642278 | 0.026841171 | No Change | Significant Negative | TRUE |
| Neurl4 | TRUE | -0.135330493 | 0.087793307 | No Change | Not Significant | TRUE | -0.024165533 | 0.873700342 | No Change | Not Significant | TRUE |
| Neurod1 | TRUE | -0.688459106 | 0.02823687 | DOWN | DOWN | TRUE | -0.139912521 | 0.696197974 | No Change | Not Significant | TRUE |
| Neurod2 | TRUE | 0.299073516 | 1 | No Change | Not Significant | FALSE | NA | NA | No Change | Not Detected | FALSE |
| Neurod4 | TRUE | 0.004903035 | 1 | No Change | Not Significant | FALSE | NA | NA | No Change | Not Detected | FALSE |
| Neurog3 | TRUE | 0.163773686 | 0.839403501 | No Change | Not Significant | TRUE | 0.098526626 | 0.853157579 | No Change | Not Significant | TRUE |
| Nexmif | TRUE | -0.179604336 | 0.750095737 | No Change | Not Significant | TRUE | 0.05380788 | 0.827608087 | No Change | Not Significant | TRUE |
| Nexn | TRUE | 0.189045893 | 0.734913884 | No Change | Not Significant | TRUE | -0.159368281 | 0.76862707 | No Change | Not Significant | TRUE |
| Nf1 | TRUE | 0.073385652 | 0.699975951 | No Change | Not Significant | TRUE | -0.10655586 | 0.463123925 | No Change | Not Significant | TRUE |
| Nf2 | TRUE | 0.016671686 | 0.867566194 | No Change | Not Significant | TRUE | 0.093632989 | 0.234551147 | No Change | Not Significant | TRUE |
| Nfam1 | TRUE | 1.372024828 | 0.228322063 | No Change | Not Significant | FALSE | NA | NA | No Change | Not Detected | FALSE |
| Nfasc | TRUE | 6.144157452 | 2.23E-71 | UP | UP | TRUE | 1.015261685 | 0.001203003 | UP | UP | TRUE |
| Nfat5 | TRUE | -0.328119322 | 0.193418659 | No Change | Not Significant | TRUE | -0.423322846 | 0.001163192 | No Change | Significant Negative | TRUE |
| Nfatc1 | TRUE | -0.146631426 | 0.285669361 | No Change | Not Significant | TRUE | 0.052801145 | 0.792924519 | No Change | Not Significant | TRUE |
| Nfatc2 | TRUE | -0.469996099 | 0.070745635 | No Change | Not Significant | TRUE | 0.106181578 | 0.767564688 | No Change | Not Significant | TRUE |
| Nfatc2ip | TRUE | 0.137715295 | 0.589852079 | No Change | Not Significant | TRUE | -0.025137909 | 0.936328291 | No Change | Not Significant | TRUE |
| Nfatc3 | TRUE | -0.037149118 | 0.675880563 | No Change | Not Significant | TRUE | -0.054207736 | 0.61867166 | No Change | Not Significant | TRUE |
| Nfatc4 | TRUE | 0.913160423 | 1 | No Change | Not Significant | FALSE | NA | NA | No Change | Not Detected | FALSE |
| Nfe2 | TRUE | 0.459223805 | 0.68833522 | No Change | Not Significant | TRUE | 0.295382234 | 0.691455871 | No Change | Not Significant | TRUE |
| Nfe2l1 | TRUE | 0.375839699 | 2.23E-09 | No Change | Significant Positive | TRUE | 0.229591655 | 5.10E-06 | No Change | Significant Positive | TRUE |
| Nfe2l2 | TRUE | -0.348219031 | 3.39E-05 | No Change | Significant Negative | TRUE | -0.101898412 | 0.229476957 | No Change | Not Significant | TRUE |
| Nfe2l3 | TRUE | -0.227731792 | 0.539052565 | No Change | Not Significant | TRUE | 0.122902652 | 0.693423019 | No Change | Not Significant | TRUE |
| Nfia | TRUE | 0.039206193 | 0.831662311 | No Change | Not Significant | TRUE | -0.096298891 | 0.447440166 | No Change | Not Significant | TRUE |
| Nfib | TRUE | 0.135911586 | 0.474208273 | No Change | Not Significant | TRUE | -0.089741899 | 0.414720716 | No Change | Not Significant | TRUE |
| Nfic | TRUE | -0.251842605 | 0.394112152 | No Change | Not Significant | TRUE | -0.346610455 | 0.001648627 | No Change | Significant Negative | TRUE |
| Nfil3 | TRUE | 0.95002452 | 0.005716696 | UP | UP | TRUE | 0.888827191 | 1.74E-06 | UP | UP | TRUE |
| Nfix | TRUE | -0.278852007 | 0.006384236 | No Change | Significant Negative | TRUE | -0.241497313 | 0.144401865 | No Change | Not Significant | TRUE |
| Nfkb1 | TRUE | -0.066425969 | 0.331061353 | No Change | Not Significant | TRUE | -0.014722639 | 0.839327919 | No Change | Not Significant | TRUE |
| Nfkb2 | TRUE | 0.203444516 | 0.050983705 | No Change | Not Significant | TRUE | -0.052377446 | 0.606684232 | No Change | Not Significant | TRUE |
| Nfkbia | TRUE | -0.022290147 | 0.964363534 | No Change | Not Significant | TRUE | -0.235641136 | 0.004729872 | No Change | Significant Negative | TRUE |
| Nfkbib | TRUE | 0.057722597 | 0.715798257 | No Change | Not Significant | TRUE | 0.059937923 | 0.768361739 | No Change | Not Significant | TRUE |
| Nfkbid | TRUE | 0.155487116 | 0.754128254 | No Change | Not Significant | TRUE | -0.252609834 | 0.379526132 | No Change | Not Significant | TRUE |
| Nfkbie | TRUE | -0.348082347 | 0.210845076 | No Change | Not Significant | TRUE | 0.057762297 | 0.849723168 | No Change | Not Significant | TRUE |
| Nfkbil1 | TRUE | 0.15434413 | 0.235970255 | No Change | Not Significant | TRUE | 0.034865424 | 0.892032325 | No Change | Not Significant | TRUE |
| Nfkbiz | TRUE | 0.50153355 | 0.004103449 | UP | UP | TRUE | -0.155172198 | 0.358583208 | No Change | Not Significant | TRUE |
| Nfrkb | TRUE | 0.086419484 | 0.439030384 | No Change | Not Significant | TRUE | -0.125642563 | 0.149664594 | No Change | Not Significant | TRUE |
| Nfs1 | TRUE | -0.324572356 | 0.000404346 | No Change | Significant Negative | TRUE | -0.088873092 | 0.273384904 | No Change | Not Significant | TRUE |
| Nfu1 | TRUE | -0.497880726 | 3.32E-06 | No Change | Significant Negative | TRUE | -0.336114265 | 0.000907852 | No Change | Significant Negative | TRUE |
| Nfx1 | TRUE | 0.056758308 | 0.728866518 | No Change | Not Significant | TRUE | -0.156021336 | 0.167263497 | No Change | Not Significant | TRUE |
| Nfxl1 | TRUE | 0.386626195 | 0.000169022 | No Change | Significant Positive | TRUE | 0.175682876 | 0.090113624 | No Change | Not Significant | TRUE |
| Nfya | TRUE | -0.047920122 | 0.750768863 | No Change | Not Significant | TRUE | 0.091650274 | 0.452109968 | No Change | Not Significant | TRUE |
| Nfyb | TRUE | 0.12591422 | 0.27724426 | No Change | Not Significant | TRUE | 0.087134665 | 0.405210667 | No Change | Not Significant | TRUE |
| Nfyc | TRUE | -0.009483748 | 0.959501353 | No Change | Not Significant | TRUE | 0.097638113 | 0.221624129 | No Change | Not Significant | TRUE |
| Ngb | TRUE | -0.743138469 | 1 | No Change | Not Significant | FALSE | NA | NA | No Change | Not Detected | FALSE |
| Ngdn | TRUE | 0.832229001 | 1.81E-29 | UP | UP | TRUE | 0.423124825 | 9.38E-14 | No Change | Significant Positive | TRUE |
| Ngef | TRUE | -0.176961836 | 0.12161285 | No Change | Not Significant | TRUE | -0.047329378 | 0.657152071 | No Change | Not Significant | TRUE |
| Ngf | TRUE | 0.533133096 | 1 | No Change | Not Significant | FALSE | NA | NA | No Change | Not Detected | FALSE |
| Ngfr | TRUE | 2.321363915 | 1 | No Change | Not Significant | FALSE | NA | NA | No Change | Not Detected | FALSE |
| Ngly1 | TRUE | -0.011651673 | 0.942430322 | No Change | Not Significant | TRUE | 0.039474684 | 0.630176941 | No Change | Not Significant | TRUE |
| Ngp | TRUE | 1.185907612 | 1 | No Change | Not Significant | FALSE | NA | NA | No Change | Not Detected | FALSE |
| Ngrn | TRUE | -0.055619525 | 0.767282764 | No Change | Not Significant | TRUE | 0.089015822 | 0.438338886 | No Change | Not Significant | TRUE |
| Nhej1 | TRUE | 0.048484036 | 0.875081807 | No Change | Not Significant | TRUE | 0.215616417 | 0.327154117 | No Change | Not Significant | TRUE |
| Nhlh1 | TRUE | -0.01147461 | 1 | No Change | Not Significant | FALSE | NA | NA | No Change | Not Detected | FALSE |
| Nhlrc1 | TRUE | -0.543798367 | 0.072556062 | No Change | Not Significant | TRUE | -0.172174278 | 0.502363316 | No Change | Not Significant | TRUE |
| Nhlrc2 | TRUE | -0.094979856 | 0.395804361 | No Change | Not Significant | TRUE | -0.211574517 | 0.000109362 | No Change | Significant Negative | TRUE |
| Nhlrc3 | TRUE | -0.015936325 | 0.924713357 | No Change | Not Significant | TRUE | 0.036130667 | 0.615647488 | No Change | Not Significant | TRUE |
| Nhp2 | TRUE | -0.134897981 | 0.390530809 | No Change | Not Significant | TRUE | -0.030365401 | 0.872420306 | No Change | Not Significant | TRUE |
| Nhs | TRUE | -0.188858939 | 0.760569151 | No Change | Not Significant | TRUE | -0.46719547 | 0.09546563 | No Change | Not Significant | TRUE |
| Nhsl1 | TRUE | 0.217431567 | 0.134498514 | No Change | Not Significant | TRUE | 0.175710923 | 0.019485867 | No Change | Significant Positive | TRUE |
| Nhsl2 | TRUE | 0.350535327 | 0.914187552 | No Change | Not Significant | FALSE | NA | NA | No Change | Not Detected | FALSE |
| Niban1 | TRUE | -0.08924032 | 0.523653558 | No Change | Not Significant | TRUE | -0.149922715 | 0.131915576 | No Change | Not Significant | TRUE |
| Niban2 | TRUE | -0.048526086 | 0.701603242 | No Change | Not Significant | TRUE | 0.149745263 | 0.109582031 | No Change | Not Significant | TRUE |
| Niban3 | TRUE | 0.084958069 | 0.912263113 | No Change | Not Significant | TRUE | 0.519378942 | 0.399060944 | No Change | Not Significant | TRUE |
| Nicn1 | TRUE | -0.228141558 | 0.189541363 | No Change | Not Significant | TRUE | 0.203061421 | 0.309125592 | No Change | Not Significant | TRUE |
| Nid1 | TRUE | 0.238227582 | 0.770846581 | No Change | Not Significant | TRUE | 1.369228283 | 0.019007112 | UP | UP | TRUE |
| Nid2 | TRUE | -0.098309335 | 0.690541222 | No Change | Not Significant | TRUE | -0.045224505 | 0.813747171 | No Change | Not Significant | TRUE |
| Nif3l1 | TRUE | 0.009479147 | 0.969552109 | No Change | Not Significant | TRUE | 0.120691861 | 0.20528446 | No Change | Not Significant | TRUE |
| Nifk | TRUE | -0.064900897 | 0.772689314 | No Change | Not Significant | TRUE | 0.008188754 | 0.961718903 | No Change | Not Significant | TRUE |
| Nim1k | TRUE | 0.508281003 | 0.53139398 | No Change | Not Significant | TRUE | 0.188341532 | 0.720023049 | No Change | Not Significant | TRUE |
| Nin | TRUE | -0.141710199 | 0.564520034 | No Change | Not Significant | TRUE | -0.055962907 | 0.737765595 | No Change | Not Significant | TRUE |
| Ninj1 | TRUE | -0.214267032 | 0.032851834 | No Change | Significant Negative | TRUE | 0.010080464 | 0.949475732 | No Change | Not Significant | TRUE |
| Ninj2 | TRUE | -1.076285516 | 0.139398299 | No Change | Not Significant | TRUE | 0.592220784 | 0.317074504 | No Change | Not Significant | TRUE |
| Ninl | TRUE | 0.090913909 | 0.833648609 | No Change | Not Significant | TRUE | 0.041216152 | 0.900662858 | No Change | Not Significant | TRUE |
| Nip7 | TRUE | -0.061919432 | 0.73666903 | No Change | Not Significant | TRUE | -0.093873251 | 0.554839644 | No Change | Not Significant | TRUE |
| Nipa1 | TRUE | -0.041442386 | 0.908409285 | No Change | Not Significant | TRUE | -0.066433457 | 0.832445548 | No Change | Not Significant | TRUE |
| Nipa2 | TRUE | 0.148346943 | 0.088668856 | No Change | Not Significant | TRUE | -0.030543438 | 0.755437307 | No Change | Not Significant | TRUE |
| Nipal1 | TRUE | -0.054953255 | 0.773824226 | No Change | Not Significant | TRUE | -0.071884416 | 0.525975748 | No Change | Not Significant | TRUE |
| Nipal2 | TRUE | 0.244148158 | 0.015594313 | No Change | Significant Positive | TRUE | 0.129641159 | 0.128328807 | No Change | Not Significant | TRUE |
| Nipal3 | TRUE | 0.314799236 | 1.88E-07 | No Change | Significant Positive | TRUE | 0.259213935 | 0.001424611 | No Change | Significant Positive | TRUE |
| Nipal4 | TRUE | 0.574209913 | 1 | No Change | Not Significant | FALSE | NA | NA | No Change | Not Detected | FALSE |
| Nipbl | TRUE | -0.095466018 | 0.48708267 | No Change | Not Significant | TRUE | -0.085364442 | 0.285546433 | No Change | Not Significant | TRUE |
| Nipsnap1 | TRUE | -0.297979209 | 0.007888281 | No Change | Significant Negative | TRUE | 0.012323111 | 0.915190139 | No Change | Not Significant | TRUE |
| Nipsnap2 | TRUE | -0.037873196 | 0.838743187 | No Change | Not Significant | TRUE | 0.023418947 | 0.827333871 | No Change | Not Significant | TRUE |
| Nipsnap3b | TRUE | 0.255923476 | 0.016585107 | No Change | Significant Positive | TRUE | 0.163243834 | 0.008657373 | No Change | Significant Positive | TRUE |
| Nisch | TRUE | -0.129050616 | 0.17504753 | No Change | Not Significant | TRUE | -0.087670627 | 0.304713258 | No Change | Not Significant | TRUE |
| Nit1 | TRUE | -0.203279563 | 0.028062449 | No Change | Significant Negative | TRUE | 0.019760545 | 0.8354677 | No Change | Not Significant | TRUE |
| Nit2 | TRUE | -0.499440768 | 0.000328678 | No Change | Significant Negative | TRUE | -0.134490495 | 0.267828024 | No Change | Not Significant | TRUE |
| Nkain1 | TRUE | 0.575226853 | 0.001502974 | UP | UP | TRUE | 0.106284886 | 0.616119646 | No Change | Not Significant | TRUE |
| Nkain3 | TRUE | -0.018392149 | 1 | No Change | Not Significant | FALSE | NA | NA | No Change | Not Detected | FALSE |
| Nkain4 | TRUE | 1.957364045 | 1 | No Change | Not Significant | FALSE | NA | NA | No Change | Not Detected | FALSE |
| Nkap | TRUE | -0.246231382 | 0.112051438 | No Change | Not Significant | TRUE | -0.0781592 | 0.495031103 | No Change | Not Significant | TRUE |
| Nkapd1 | TRUE | -0.176817636 | 0.423388606 | No Change | Not Significant | TRUE | -0.087876196 | 0.47621354 | No Change | Not Significant | TRUE |
| Nkapl | TRUE | -0.688594206 | 0.211378428 | No Change | Not Significant | TRUE | -0.432793103 | 0.553189591 | No Change | Not Significant | TRUE |
| Nkd1 | TRUE | -0.806444547 | 0.205647109 | No Change | Not Significant | TRUE | -0.909664294 | 0.009129074 | DOWN | DOWN | TRUE |
| Nkd2 | TRUE | 0.405792032 | 0.776544903 | No Change | Not Significant | FALSE | NA | NA | No Change | Not Detected | FALSE |
| Nkg7 | TRUE | 0.135763359 | 0.885199039 | No Change | Not Significant | TRUE | 0.602483136 | 0.143118696 | No Change | Not Significant | TRUE |
| Nkiras1 | TRUE | 0.021902842 | 0.930204755 | No Change | Not Significant | TRUE | -9.60E-05 | 0.999454275 | No Change | Not Significant | TRUE |
| Nkiras2 | TRUE | -0.081876693 | 0.566268598 | No Change | Not Significant | TRUE | 0.015163443 | 0.919509896 | No Change | Not Significant | TRUE |
| Nkpd1 | TRUE | -0.637824635 | 0.176657744 | No Change | Not Significant | TRUE | -0.087605073 | 0.923349179 | No Change | Not Significant | TRUE |
| Nkrf | TRUE | 0.052359824 | 0.877877519 | No Change | Not Significant | TRUE | 0.045319687 | 0.865314338 | No Change | Not Significant | TRUE |
| Nktr | TRUE | 1.143056506 | 1.89E-40 | UP | UP | TRUE | 0.661556486 | 1.19E-12 | UP | UP | TRUE |
| Nkx1-2 | TRUE | 1.131110074 | 1 | No Change | Not Significant | FALSE | NA | NA | No Change | Not Detected | FALSE |
| Nkx2-2 | TRUE | -0.812151927 | 0.144047837 | No Change | Not Significant | TRUE | 0.142969706 | 0.844374344 | No Change | Not Significant | TRUE |
| Nkx2-2os | TRUE | 1.127803748 | 1 | No Change | Not Significant | FALSE | NA | NA | No Change | Not Detected | FALSE |
| Nkx2-3 | TRUE | 1.135351955 | 1 | No Change | Not Significant | FALSE | NA | NA | No Change | Not Detected | FALSE |
| Nkx3-1 | TRUE | 1.126019775 | 1 | No Change | Not Significant | FALSE | NA | NA | No Change | Not Detected | FALSE |
| Nkx3-2 | TRUE | 0.753080655 | 1 | No Change | Not Significant | FALSE | NA | NA | No Change | Not Detected | FALSE |
| Nkx6-1 | TRUE | 1.063796885 | 1 | No Change | Not Significant | FALSE | NA | NA | No Change | Not Detected | FALSE |
| Nkx6-2 | TRUE | 1.19622767 | 1 | No Change | Not Significant | FALSE | NA | NA | No Change | Not Detected | FALSE |
| Nkx6-3 | TRUE | -1.05575962 | 0.145192693 | No Change | Not Significant | FALSE | NA | NA | No Change | Not Detected | FALSE |
| Nle1 | TRUE | 0.027841453 | 0.914497068 | No Change | Not Significant | TRUE | -0.134203303 | 0.413700119 | No Change | Not Significant | TRUE |
| Nlgn1 | TRUE | 0.557997456 | 1 | No Change | Not Significant | FALSE | NA | NA | No Change | Not Detected | FALSE |
| Nlgn2 | TRUE | 0.212713872 | 0.793088519 | No Change | Not Significant | TRUE | -0.64573411 | 0.113433539 | No Change | Not Significant | TRUE |
| Nlgn3 | TRUE | 1.098322756 | 1 | No Change | Not Significant | FALSE | NA | NA | No Change | Not Detected | FALSE |
| Nlk | TRUE | 0.313197746 | 0.015398144 | No Change | Significant Positive | TRUE | 0.165934388 | 0.210759777 | No Change | Not Significant | TRUE |
| Nln | TRUE | -0.281269231 | 0.000718512 | No Change | Significant Negative | TRUE | -0.069624712 | 0.303689326 | No Change | Not Significant | TRUE |
| Nlrc3 | TRUE | 0.393284643 | 0.684193826 | No Change | Not Significant | TRUE | -0.024009033 | 0.966069983 | No Change | Not Significant | TRUE |
| Nlrc4 | TRUE | -0.121349191 | 0.252139861 | No Change | Not Significant | TRUE | -0.078979881 | 0.318412428 | No Change | Not Significant | TRUE |
| Nlrc5 | TRUE | -0.097955839 | 0.897198344 | No Change | Not Significant | TRUE | -1.097131997 | 0.22777308 | No Change | Not Significant | TRUE |
| Nlrp10 | TRUE | -0.938338023 | 0.001014656 | DOWN | DOWN | TRUE | -0.358485177 | 0.134705162 | No Change | Not Significant | TRUE |
| Nlrp12 | TRUE | 3.547553688 | 1 | No Change | Not Significant | FALSE | NA | NA | No Change | Not Detected | FALSE |
| Nlrp1a | TRUE | 0.074748784 | 1 | No Change | Not Significant | FALSE | NA | NA | No Change | Not Detected | FALSE |
| Nlrp1b | TRUE | -0.88771264 | 5.17E-07 | DOWN | DOWN | TRUE | -0.221117343 | 0.098877846 | No Change | Not Significant | TRUE |
| Nlrp3 | TRUE | 1.276667741 | 1 | No Change | Not Significant | FALSE | NA | NA | No Change | Not Detected | FALSE |
| Nlrp4b | TRUE | 0.611520734 | 1 | No Change | Not Significant | FALSE | NA | NA | No Change | Not Detected | FALSE |
| Nlrp4f | TRUE | -0.427671646 | 1 | No Change | Not Significant | FALSE | NA | NA | No Change | Not Detected | FALSE |
| Nlrp5 | TRUE | 1.124480316 | 1 | No Change | Not Significant | FALSE | NA | NA | No Change | Not Detected | FALSE |
| Nlrp6 | TRUE | -0.241266475 | 0.108456517 | No Change | Not Significant | TRUE | -0.048527791 | 0.667195279 | No Change | Not Significant | TRUE |
| Nlrp9b | TRUE | 0.435134402 | 0.452315589 | No Change | Not Significant | TRUE | -0.270937818 | 0.373094934 | No Change | Not Significant | TRUE |
| Nlrx1 | TRUE | 0.043086199 | 0.878845311 | No Change | Not Significant | TRUE | 0.247634825 | 0.107008585 | No Change | Not Significant | TRUE |
| Nmb | TRUE | 0.851596353 | 0.254637565 | No Change | Not Significant | FALSE | NA | NA | No Change | Not Detected | FALSE |
| Nmbr | TRUE | 0.327877305 | 1 | No Change | Not Significant | FALSE | NA | NA | No Change | Not Detected | FALSE |
| Nmd3 | TRUE | 0.021569559 | 0.908700419 | No Change | Not Significant | TRUE | -0.109001087 | 0.449718678 | No Change | Not Significant | TRUE |
| Nme1 | TRUE | 0.290819314 | 0.112868558 | No Change | Not Significant | TRUE | 0.208740412 | 0.312196844 | No Change | Not Significant | TRUE |
| Nme2 | TRUE | 0.762905256 | 0.000174227 | UP | UP | TRUE | 0.168389687 | 0.513253429 | No Change | Not Significant | TRUE |
| Nme4 | TRUE | 0.118352001 | 0.67085817 | No Change | Not Significant | TRUE | -0.347056267 | 0.044292325 | No Change | Significant Negative | TRUE |
| Nme5 | TRUE | 0.741144604 | 1 | No Change | Not Significant | FALSE | NA | NA | No Change | Not Detected | FALSE |
| Nme6 | TRUE | 1.635022815 | 5.57E-27 | UP | UP | TRUE | 0.776224635 | 5.73E-12 | UP | UP | TRUE |
| Nme7 | TRUE | 0.478366609 | 0.001087135 | No Change | Significant Positive | TRUE | 0.138791304 | 0.407666398 | No Change | Not Significant | TRUE |
| Nme9 | TRUE | -0.407314113 | 0.732906636 | No Change | Not Significant | FALSE | NA | NA | No Change | Not Detected | FALSE |
| Nmi | TRUE | -0.163685435 | 0.242315905 | No Change | Not Significant | TRUE | 0.042801732 | 0.77864167 | No Change | Not Significant | TRUE |
| Nmnat1 | TRUE | -0.064153076 | 0.736796542 | No Change | Not Significant | TRUE | 0.009509649 | 0.960412417 | No Change | Not Significant | TRUE |
| Nmnat2 | TRUE | -0.982069418 | 0.340684159 | No Change | Not Significant | TRUE | 0.057590662 | 0.914534473 | No Change | Not Significant | TRUE |
| Nmnat3 | TRUE | -0.226351082 | 0.423784727 | No Change | Not Significant | TRUE | -0.01485157 | 0.945176551 | No Change | Not Significant | TRUE |
| Nmral1 | TRUE | 0.443454409 | 0.07241317 | No Change | Not Significant | TRUE | 0.451976854 | 0.087561384 | No Change | Not Significant | TRUE |
| Nmrk1 | TRUE | 0.214557823 | 0.40571187 | No Change | Not Significant | TRUE | 0.289057921 | 0.206112189 | No Change | Not Significant | TRUE |
| Nmrk2 | TRUE | 0.882488534 | 1 | No Change | Not Significant | FALSE | NA | NA | No Change | Not Detected | FALSE |
| Nmt1 | TRUE | 0.621851991 | 6.12E-18 | UP | UP | TRUE | 0.297106518 | 1.32E-06 | No Change | Significant Positive | TRUE |
| Nmt2 | TRUE | -0.134644982 | 0.291934232 | No Change | Not Significant | TRUE | -0.036426916 | 0.698573273 | No Change | Not Significant | TRUE |
| Nmur1 | TRUE | 2.594022889 | 1 | No Change | Not Significant | FALSE | NA | NA | No Change | Not Detected | FALSE |
| Nnat | TRUE | 0.200255877 | 0.804573016 | No Change | Not Significant | TRUE | -0.079701565 | 0.869327992 | No Change | Not Significant | TRUE |
| Nnt | TRUE | -0.15437655 | 0.833960201 | No Change | Not Significant | TRUE | 0.109658681 | 0.898825528 | No Change | Not Significant | TRUE |

| geneID | Cond1_DF_Detected | Cond1_DF_ValueCol | Cond1_DF_PadjCol | Cond1_DF_Cutoff | Cond1_DF_Cutoff_Specific.x | Cond2_DF_Detected |
|---|---|---|---|---|---|---|
| Noa1 | TRUE | -0.165840565 | 0.352762472 | No Change | Not Significant | TRUE |
| Nob1 | TRUE | -0.203709448 | 0.157556926 | No Change | Not Significant | TRUE |
| Nobox | TRUE | 0.413340268 | 1 | No Change | Not Significant | FALSE |
| Noc2l | TRUE | -0.004853522 | 0.980067227 | No Change | Not Significant | TRUE |
| Noc3l | TRUE | 0.03890479 | 0.882255147 | No Change | Not Significant | TRUE |
| Noc4l | TRUE | 0.290382613 | 0.030408957 | No Change | Significant Positive | TRUE |
| Noct | TRUE | -0.302504131 | 0.363275924 | No Change | Not Significant | TRUE |
| Nod1 | TRUE | 0.382983384 | 0.010863233 | No Change | Significant Positive | TRUE |
| Nod2 | TRUE | -0.819480454 | 1.49E-05 | DOWN | DOWN | TRUE |
| Nodal | TRUE | 1.726719411 | 1 | No Change | Not Significant | FALSE |
| Nol10 | TRUE | 0.106803025 | 0.680853016 | No Change | Not Significant | TRUE |
| Nol11 | TRUE | 0.163660072 | 0.446273789 | No Change | Not Significant | TRUE |
| Nol12 | TRUE | 0.132968882 | 0.398112257 | No Change | Not Significant | TRUE |
| Nol3 | TRUE | 0.261357411 | 0.34473544 | No Change | Not Significant | TRUE |
| Nol4 | TRUE | 0.231618473 | 1 | No Change | Not Significant | FALSE |
| Nol4l | TRUE | 0.216547925 | 0.283150481 | No Change | Not Significant | TRUE |
| Nol6 | TRUE | 0.137489578 | 0.398476941 | No Change | Not Significant | TRUE |
| Nol7 | TRUE | 0.244279695 | 0.026496643 | No Change | Significant Positive | TRUE |
| Nol8 | TRUE | 0.002628763 | 0.991590016 | No Change | Not Significant | TRUE |
| Nol9 | TRUE | 0.040559146 | 0.836513744 | No Change | Not Significant | TRUE |
| Nolc1 | TRUE | 0.125328134 | 0.552117058 | No Change | Not Significant | TRUE |
| Nom1 | TRUE | -0.008759657 | 0.965665343 | No Change | Not Significant | TRUE |
| Nomo1 | TRUE | 0.119157169 | 0.233671624 | No Change | Not Significant | TRUE |
| Nono | TRUE | -0.138575724 | 0.357709402 | No Change | Not Significant | TRUE |
| Nop10 | TRUE | -0.280081829 | 0.05145848 | No Change | Not Significant | TRUE |
| Nop14 | TRUE | 0.08365503 | 0.673388737 | No Change | Not Significant | TRUE |
| Nop16 | TRUE | 0.100878603 | 0.522421155 | No Change | Not Significant | TRUE |
| Nop2 | TRUE | 0.00838017 | 0.966352899 | No Change | Not Significant | TRUE |
| Nop53 | TRUE | -0.261471086 | 0.005835484 | No Change | Significant Negative | TRUE |
| Nop56 | TRUE | 0.026462379 | 0.907147741 | No Change | Not Significant | TRUE |
| Nop58 | TRUE | 0.111967589 | 0.412590307 | No Change | Not Significant | TRUE |
| Nop9 | TRUE | 0.106631506 | 0.603717953 | No Change | Not Significant | TRUE |
| Nopchap1 | TRUE | 0.773755075 | 8.11E-26 | UP | UP | TRUE |
| Norad | TRUE | 0.041790136 | 0.863168757 | No Change | Not Significant | TRUE |
| Nos1 | TRUE | 0.682122187 | 0.766833571 | No Change | Not Significant | FALSE |
| Nos1ap | TRUE | -0.198054579 | 0.628254175 | No Change | Not Significant | TRUE |
| Nos2 | TRUE | 1.966028867 | 0.015492034 | UP | UP | TRUE |
| Nos3 | TRUE | -0.618133974 | 1 | No Change | Not Significant | FALSE |
| Nosip | TRUE | -0.151538811 | 0.072388075 | No Change | Not Significant | TRUE |
| Nostrin | TRUE | 0.059216936 | 0.702568634 | No Change | Not Significant | TRUE |
| Notch1 | TRUE | 0.178926165 | 0.203670961 | No Change | Not Significant | TRUE |
| Notch2 | TRUE | 0.257209039 | 0.033167393 | No Change | Significant Positive | TRUE |
| Notch3 | TRUE | -0.469118693 | 0.781162199 | No Change | Not Significant | TRUE |
| Notch4 | TRUE | 0.254500311 | 0.268218365 | No Change | Not Significant | TRUE |
| Notum | TRUE | 1.526587289 | 1 | No Change | Not Significant | FALSE |
| Nova1 | TRUE | 2.269888519 | 1 | No Change | Not Significant | FALSE |
| Nova2 | TRUE | 0.418197266 | 0.799512805 | No Change | Not Significant | FALSE |
| Nox1 | TRUE | 0.594670989 | 0.051092662 | No Change | Not Significant | TRUE |
| Nox3 | TRUE | 0.753064977 | 1 | No Change | Not Significant | FALSE |
| Nox4 | TRUE | 1.890026489 | 1 | No Change | Not Significant | FALSE |
| Noxa1 | TRUE | 0.203528044 | 0.158138918 | No Change | Not Significant | TRUE |
| Noxo1 | TRUE | -0.056166049 | 0.882132403 | No Change | Not Significant | TRUE |
| Noxred1 | TRUE | 1.395870908 | 0.129368504 | No Change | Not Significant | FALSE |
| Npas1 | TRUE | 0.022516297 | 0.982587667 | No Change | Not Significant | TRUE |
| Npas2 | TRUE | 0.575570935 | 0.04294225 | UP | UP | TRUE |
| Npas3 | TRUE | -0.62293962 | 0.527928237 | No Change | Not Significant | TRUE |
| Npas4 | TRUE | 0.424507006 | 1 | No Change | Not Significant | FALSE |
| Npat | TRUE | 0.160458388 | 0.247269599 | No Change | Not Significant | TRUE |
| Npb | TRUE | -0.011374784 | 1 | No Change | Not Significant | TRUE |
| Npbwr1 | TRUE | 0.523295795 | 1 | No Change | Not Significant | FALSE |
| Npc1 | TRUE | -0.123126854 | 0.540502136 | No Change | Not Significant | TRUE |
| Npc1l1 | TRUE | -0.66562591 | 0.612813278 | No Change | Not Significant | TRUE |
| Npc2 | TRUE | -0.570144575 | 0.001371337 | DOWN | DOWN | TRUE |
| Npdc1 | TRUE | -0.38344183 | 7.68E-06 | No Change | Significant Negative | TRUE |
| Npepl1 | TRUE | -0.076293007 | 0.360508479 | No Change | Not Significant | TRUE |
| Npepps | TRUE | 0.019720117 | 0.832151567 | No Change | Not Significant | TRUE |
| Npff | TRUE | 0.385130425 | 1 | No Change | Not Significant | FALSE |
| Npffr1 | TRUE | -0.049839244 | 0.971404891 | No Change | Not Significant | FALSE |
| Nphp1 | TRUE | 0.590596561 | 0.00684621 | UP | UP | TRUE |
| Nphp3 | TRUE | 2.207919757 | 1.19E-65 | UP | UP | TRUE |
| Nphp4 | TRUE | 1.230992545 | 0.129891785 | No Change | Not Significant | FALSE |
| Nphs1 | TRUE | -1.47870466 | 1 | No Change | Not Significant | FALSE |
| Nphs2 | TRUE | -0.539196761 | 1 | No Change | Not Significant | FALSE |
| Npl | TRUE | -2.304508471 | 1 | No Change | Not Significant | FALSE |
| Nploc4 | TRUE | 0.166452471 | 0.051654946 | No Change | Not Significant | TRUE |
| Npm1 | TRUE | -0.168248369 | 0.389522022 | No Change | Not Significant | TRUE |
| Npm2 | TRUE | -1.105558784 | 1 | No Change | Not Significant | FALSE |
| Npm3 | TRUE | -0.333495105 | 0.086297303 | No Change | Not Significant | TRUE |
| Npnt | TRUE | -0.220121672 | 0.878750865 | No Change | Not Significant | TRUE |
| Nppc | TRUE | 1.643149699 | 1 | No Change | Not Significant | FALSE |
| Npr1 | TRUE | 1.696611654 | 0.000351256 | UP | UP | TRUE |
| Npr2 | TRUE | -0.269251144 | 1 | No Change | Not Significant | FALSE |
| Npr3 | TRUE | 0.69827775 | 1 | No Change | Not Significant | FALSE |
| Nprl2 | TRUE | -0.122435425 | 0.438111017 | No Change | Not Significant | TRUE |
| Nprl3 | TRUE | -0.101727108 | 0.616875898 | No Change | Not Significant | TRUE |
| Nps | TRUE | -0.773638031 | 1 | No Change | Not Significant | FALSE |
| Nptn | TRUE | -0.130805097 | 0.00775103 | No Change | Significant Negative | TRUE |
| Nptx1 | TRUE | -0.480659273 | 0.662290428 | No Change | Not Significant | TRUE |
| Nptx2 | TRUE | 0.443274809 | 0.527698391 | No Change | Not Significant | TRUE |
| Nptxr | TRUE | 0.387679682 | 0.72898503 | No Change | Not Significant | TRUE |
| Npy1r | TRUE | -0.008957873 | 0.988831409 | No Change | Not Significant | TRUE |
| Npy2r | TRUE | -0.307039448 | 0.740342018 | No Change | Not Significant | TRUE |
| Nqo1 | TRUE | -0.536760072 | 0.001107066 | DOWN | DOWN | TRUE |
| Nqo2 | TRUE | -0.131371773 | 0.406017465 | No Change | Not Significant | TRUE |
| Nr0b2 | TRUE | 0.363824673 | 1 | No Change | Not Significant | FALSE |
| Nr1d1 | TRUE | -2.054181308 | 0.001071245 | DOWN | DOWN | TRUE |
| Nr1d2 | TRUE | -1.852928612 | 1.81E-10 | DOWN | DOWN | TRUE |
| Nr1h2 | TRUE | -0.125849079 | 0.318637841 | No Change | Not Significant | TRUE |
| Nr1h3 | TRUE | -0.200752426 | 0.311384636 | No Change | Not Significant | TRUE |
| Nr1h4 | TRUE | 0.046544324 | 0.885228942 | No Change | Not Significant | TRUE |
| Nr1h5 | TRUE | -0.289913592 | 1 | No Change | Not Significant | FALSE |
| Nr1i2 | TRUE | -0.122893928 | 0.240948524 | No Change | Not Significant | TRUE |
| Nr1i3 | TRUE | 0.206984762 | 0.766020418 | No Change | Not Significant | TRUE |
| Nr2c1 | TRUE | 0.035216372 | 0.842528346 | No Change | Not Significant | TRUE |
| Nr2c2 | TRUE | -0.00511806 | 0.973990811 | No Change | Not Significant | TRUE |
| Nr2c2ap | TRUE | -0.093359538 | 0.816884691 | No Change | Not Significant | TRUE |
| Nr2e1 | TRUE | 3.455526097 | 1 | No Change | Not Significant | FALSE |
| Nr2e3 | TRUE | 0.403281118 | 0.170151917 | No Change | Not Significant | TRUE |
| Nr2f1 | TRUE | 1.725835269 | 1 | No Change | Not Significant | FALSE |
| Nr2f2 | TRUE | 0.012433848 | 0.962971397 | No Change | Not Significant | TRUE |
| Nr2f6 | TRUE | 0.177418388 | 0.210863618 | No Change | Not Significant | TRUE |
| Nr3c1 | TRUE | -0.105259843 | 0.186963018 | No Change | Not Significant | TRUE |
| Nr3c2 | TRUE | -0.455786415 | 2.10E-05 | No Change | Significant Negative | TRUE |
| Nr4a1 | TRUE | -0.659590838 | 0.026811304 | DOWN | DOWN | TRUE |
| Nr4a2 | TRUE | 0.684145793 | 0.000701882 | UP | UP | TRUE |
| Nr4a3 | TRUE | -0.266866655 | 0.813003196 | No Change | Not Significant | TRUE |
| Nr5a1 | TRUE | -1.655457959 | 1 | No Change | Not Significant | FALSE |
| Nr5a2 | TRUE | -0.205762785 | 0.011401366 | No Change | Significant Negative | TRUE |
| Nr6a1 | TRUE | -0.329288658 | 0.250862963 | No Change | Not Significant | TRUE |
| Nradd | TRUE | -0.589992978 | 0.128287829 | No Change | Not Significant | TRUE |
| Nrap | TRUE | -0.508316611 | 0.17331619 | No Change | Not Significant | TRUE |
| Nrarp | TRUE | 0.150862568 | 0.561491337 | No Change | Not Significant | TRUE |
| Nras | TRUE | -0.009369624 | 0.947050968 | No Change | Not Significant | TRUE |
| Nrbf2 | TRUE | -0.109867695 | 0.524360158 | No Change | Not Significant | TRUE |
| Nrbp1 | TRUE | -0.070174334 | 0.485621036 | No Change | Not Significant | TRUE |
| Nrbp2 | TRUE | -0.259830427 | 0.038193919 | No Change | Significant Negative | TRUE |
| Nrcam | TRUE | 2.148555058 | 0.270127107 | No Change | Not Significant | FALSE |
| Nrd1 | TRUE | 0.865141669 | 7.46E-31 | UP | UP | TRUE |
| Nrde2 | TRUE | 0.431224004 | 0.000254828 | No Change | Significant Positive | TRUE |
| Nrep | TRUE | -0.986072859 | 2.35E-09 | DOWN | DOWN | TRUE |
| Nrf1 | TRUE | 0.057683385 | 0.581669952 | No Change | Not Significant | TRUE |
| Nrg1 | TRUE | 0.286410199 | 0.907710402 | No Change | Not Significant | FALSE |
| Nrg2 | TRUE | -0.010523041 | 1 | No Change | Not Significant | FALSE |
| Nrg3 | TRUE | 0.81376651 | 1 | No Change | Not Significant | FALSE |
| Nrg4 | TRUE | -0.458523111 | 0.362486203 | No Change | Not Significant | TRUE |
| Nrgn | TRUE | -1.300353337 | 0.027492662 | DOWN | DOWN | TRUE |
| Nrip1 | TRUE | -0.206714916 | 0.009942925 | No Change | Significant Negative | TRUE |
| Nrip2 | TRUE | 0.424324004 | 0.528838189 | No Change | Not Significant | TRUE |
| Nrip3 | TRUE | -0.398691668 | 0.696497169 | No Change | Not Significant | TRUE |
| Nrk | TRUE | 2.156498531 | 0.102046874 | No Change | Not Significant | FALSE |
| Nrl | TRUE | 0.522351535 | 1 | No Change | Not Significant | FALSE |
| Nrm | TRUE | -0.134276027 | 0.703115827 | No Change | Not Significant | TRUE |
| Nrn1 | TRUE | -0.115594917 | 0.736796542 | No Change | Not Significant | TRUE |
| Nrp1 | TRUE | 0.218713719 | 0.437734172 | No Change | Not Significant | TRUE |
| Nrp2 | TRUE | -0.390953957 | 0.036708756 | No Change | Significant Negative | TRUE |
| Nrros | TRUE | 0.699136484 | 0.061791369 | No Change | Not Significant | TRUE |
| Nrsn1 | TRUE | -1.110320857 | 1 | No Change | Not Significant | FALSE |
| Nrsn2 | TRUE | 0.176012105 | 1 | No Change | Not Significant | FALSE |
| Nrtn | TRUE | 0.145798939 | 0.799512805 | No Change | Not Significant | TRUE |
| Nrxn1 | TRUE | -0.05221601 | 0.97136798 | No Change | Not Significant | TRUE |
| Nrxn2 | TRUE | 4.321630962 | 3.26E-15 | UP | UP | TRUE |
| Nrxn3 | TRUE | 0.830424244 | 1 | No Change | Not Significant | FALSE |
| Nsa2 | TRUE | -0.387886614 | 0.016359939 | No Change | Significant Negative | TRUE |
| Nscme3l | TRUE | 1.450356195 | 1 | No Change | Not Significant | FALSE |
| Nsd1 | TRUE | 0.049273017 | 0.632329199 | No Change | Not Significant | TRUE |
| Nsd2 | TRUE | 0.753945756 | 8.12E-17 | UP | UP | TRUE |
| Nsd3 | TRUE | 0.115487159 | 0.441179379 | No Change | Not Significant | TRUE |
| Nsdhl | TRUE | 0.239649309 | 0.051674706 | No Change | Not Significant | TRUE |
| Nsf | TRUE | 0.369241946 | 6.56E-05 | No Change | Significant Positive | TRUE |
| Nsfl1c | TRUE | 0.019250321 | 0.875081807 | No Change | Not Significant | TRUE |
| Nsg1 | TRUE | -0.636386418 | 1 | No Change | Not Significant | FALSE |
| Nsg2 | TRUE | 0.266338809 | 0.802698942 | No Change | Not Significant | TRUE |
| Nsl1 | TRUE | 0.921670427 | 0.000823203 | UP | UP | TRUE |
| Nsmaf | TRUE | -0.138076355 | 0.240996576 | No Change | Not Significant | TRUE |
| Nsmce1 | TRUE | 0.302643268 | 0.011638882 | No Change | Significant Positive | TRUE |
| Nsmce2 | TRUE | 0.21832304 | 0.075916739 | No Change | Not Significant | TRUE |
| Nsmce3 | TRUE | 0.163662119 | 0.302752759 | No Change | Not Significant | TRUE |
| Nsmce4a | TRUE | 0.018089231 | 0.918870578 | No Change | Not Significant | TRUE |
| Nsmf | TRUE | -0.384559589 | 0.000261416 | No Change | Significant Negative | TRUE |
| Nsrp1 | TRUE | 0.095488452 | 0.534985614 | No Change | Not Significant | TRUE |
| Nsun2 | TRUE | 0.037660883 | 0.839060823 | No Change | Not Significant | TRUE |
| Nsun3 | TRUE | -0.074851784 | 0.733662794 | No Change | Not Significant | TRUE |
| Nsun4 | TRUE | -0.210923407 | 0.142788208 | No Change | Not Significant | TRUE |
| Nsun5 | TRUE | 0.14808885 | 0.504312604 | No Change | Not Significant | TRUE |
| Nsun6 | TRUE | 0.222618592 | 0.131147943 | No Change | Not Significant | TRUE |
| Nsun7 | TRUE | 1.446011681 | 1 | No Change | Not Significant | FALSE |
| Nt5c | TRUE | -0.172349181 | 0.394567184 | No Change | Not Significant | TRUE |
| Nt5c1a | TRUE | -0.689556623 | 0.236536038 | No Change | Not Significant | TRUE |
| Nt5c1b | TRUE | 1.310147648 | 1 | No Change | Not Significant | FALSE |
| Nt5c2 | TRUE | 0.016571372 | 0.933981885 | No Change | Not Significant | TRUE |
| Nt5c3 | TRUE | -0.13339822 | 0.338448383 | No Change | Not Significant | TRUE |
| Nt5c3b | TRUE | -0.007201982 | 0.977515248 | No Change | Not Significant | TRUE |
| Nt5dc1 | TRUE | 0.011979355 | 0.947601039 | No Change | Not Significant | TRUE |
| Nt5dc2 | TRUE | 1.216262177 | 5.56E-06 | UP | UP | TRUE |
| Nt5dc3 | TRUE | -0.224800948 | 0.108084245 | No Change | Not Significant | TRUE |
| Nt5e | TRUE | -0.937884042 | 0.000105365 | DOWN | DOWN | TRUE |
| Nt5m | TRUE | -0.094950132 | 0.565545141 | No Change | Not Significant | TRUE |
| Ntan1 | TRUE | -0.232338195 | 0.004948633 | No Change | Significant Negative | TRUE |
| Ntf5 | TRUE | -1.150908368 | 1 | No Change | Not Significant | FALSE |
| Nthl1 | TRUE | -0.099678024 | 0.773317311 | No Change | Not Significant | TRUE |
| Ntm | TRUE | -0.046988221 | 1 | No Change | Not Significant | FALSE |
| Ntmt1 | TRUE | 0.102344118 | 0.345858144 | No Change | Not Significant | TRUE |

Supplementary Table 3

| geneID | Cond1_DF_Detected | Cond1_DF_ValueCol | Cond1_DF_PadjCol | Cond1_DF_Cutoff | Cond1_DF_Cutoff_Specific.x | Cond2_DF_Detected | Cond2_DF_ValueCol | Cond2_DF_PadjCol | Cond2_DF_Cutoff | Cond2_DF_Cutoff_Specific | BG_Method | RG1_All | RG2_Significant | RG3_SignificantChange | Cond1_DF_ensembl_gene_id | Cond1_DF_entrezgene_id | Cond1_DF_gene_biotype | Cond1_DF_baseMean | Cond1_DF_ log2FoldChange | Cond1_DF_lfcSE | Cond1_DF_stat | Cond1_DF_pvalue | Cond1_DF_ padj | Cond1_DF_absLog2FC | Cond1_DF_Cutoff_Specific.y | Cond2_DF_ensembl_gene_id | Cond2_DF_entrezgene_id | Cond2_DF_gene_biotype | Cond2_DF_baseMean | Cond2_DF_ log2FoldChange | Cond2_DF_lfcSE | Cond2_DF_stat | Cond2_DF_pvalue | Cond2_DF_ padj | Cond2_DF_absLog2FC | RG1_Specific_Cond2 | RG1_Specific_Cond1 |
|---|---|---|---|---|---|---|---|---|---|---|---|---|---|---|---|---|---|---|---|---|---|---|---|---|---|---|---|---|---|---|---|---|---|---|---|---|---|
| Ntn1 | TRUE | 0.23544405 | 0.88672609 | No Change | Not Significant | FALSE | NA | NA | No Change | Not Detected | FALSE | Background = FALSE | Background = FALSE | Background = FALSE | ENSMUSG00000020902 | 18208 | protein_coding | 5.303619582 | 0.23544405 | 0.81890679 | -0.287510194 | 0.773721698 | 0.88672609 | 0.23544405 | Not Significant | NA | NA | NA | NA | NA | NA | NA | NA | NA | NA | Background = FALSE | Background = FALSE |
| Ntn3 | TRUE | -0.314894761 | 1 | No Change | Not Significant | FALSE | NA | NA | No Change | Not Detected | FALSE | Background = FALSE | Background = FALSE | Background = FALSE | ENSMUSG00000117406 | 18209 | protein_coding | 3.962712511 | -0.314894761 | 0.861978746 | 0.365316155 | 0.714875421 | 1 | 0.314894761 | Not Significant | NA | NA | NA | NA | NA | NA | NA | NA | NA | NA | Background = FALSE | Background = FALSE |
| Ntn4 | TRUE | -1.758079437 | 1.35E-12 | DOWN | DOWN | TRUE | -0.708393565 | 1.94E-07 | DOWN | DOWN | TRUE | Cond1 DOWN + Cond2 DOWN | Core_DOWN | Core_DOWN | ENSMUSG00000020019 | 57764 | protein_coding | 72.20600955 | -1.758079437 | 0.232459418 | 7.562952083 | 3.94E-14 | 1.35E-12 | 1.758079437 | DOWN | ENSMUSG00000020019 | 57764 | protein_coding | 519.1718691 | -0.708393565 | 0.121311692 | -5.839450053 | 5.24E-09 | 1.94E-07 | 0.708393565 | Cond1 DOWN + Cond2 DOWN | Cond2 DOWN + Cond1 DOWN |
| Ntn5 | TRUE | 0.386193364 | 0.612426345 | No Change | Not Significant | FALSE | NA | NA | No Change | Not Detected | FALSE | Background = FALSE | Background = FALSE | Background = FALSE | ENSMUSG00000070564 | 243967 | protein_coding | 9.69480135 | 0.386193364 | 0.449216201 | -0.859704889 | 0.389951739 | 0.612426345 | 0.386193364 | Not Significant | NA | NA | NA | NA | NA | NA | NA | NA | NA | NA | Background = FALSE | Background = FALSE |
| Ntng1 | TRUE | -0.198904473 | 1 | No Change | Not Significant | FALSE | NA | NA | No Change | Not Detected | FALSE | Background = FALSE | Background = FALSE | Background = FALSE | ENSMUSG00000059857 | 80883 | protein_coding | 2.561819837 | -0.198904473 | 1.350431268 | 0.147289594 | 0.882903446 | 1 | 0.198904473 | Not Significant | NA | NA | NA | NA | NA | NA | NA | NA | NA | NA | Background = FALSE | Background = FALSE |
| Ntng2 | TRUE | 0.475135774 | 1 | No Change | Not Significant | FALSE | NA | NA | No Change | Not Detected | FALSE | Background = FALSE | Background = FALSE | Background = FALSE | ENSMUSG00000035513 | 171171 | protein_coding | 2.852251825 | 0.475135774 | 1.8115756 | -0.262277641 | 0.793107399 | 1 | 0.475135774 | Not Significant | NA | NA | NA | NA | NA | NA | NA | NA | NA | NA | Background = FALSE | Background = FALSE |
| Ntpcr | TRUE | -0.093915214 | 0.620277218 | No Change | Not Significant | TRUE | 0.040472179 | 0.768233148 | No Change | Not Significant | TRUE | Cond1 Not Significant + Cond2 Not Significant | None | None | ENSMUSG00000031851 | 66566 | protein_coding | 156.442917 | -0.093915214 | 0.111203024 | 0.844538306 | 0.398368668 | 0.620277218 | 0.093915214 | Not Significant | ENSMUSG00000031851 | 66566 | protein_coding | 339.779986 | 0.040472179 | 0.072130911 | 0.561093414 | 0.574733858 | 0.768233148 | 0.040472179 | Cond1 No Change + Cond2 Not Significant | Cond2 No Change + Cond1 Not Significant |
| Ntrk1 | TRUE | 1.326691038 | 1 | No Change | Not Significant | FALSE | NA | NA | No Change | Not Detected | FALSE | Background = FALSE | Background = FALSE | Background = FALSE | ENSMUSG00000028072 | 18211 | protein_coding | 1.408227333 | 1.326691038 | 1.608466067 | -0.824817549 | 0.409475182 | 1 | 1.326691038 | Not Significant | NA | NA | NA | NA | NA | NA | NA | NA | NA | NA | Background = FALSE | Background = FALSE |
| Ntrk2 | TRUE | 0.373409077 | 0.17492772 | No Change | Not Significant | TRUE | 0.065472394 | [illegible] | [illegible] | [illegible] | [illegible] | [illegible] | [illegible] | [illegible] | [illegible] | [illegible] | [illegible] | [illegible] | [illegible] | [illegible] | [illegible] | [illegible] | [illegible] | [illegible] | [illegible] | [illegible] | [illegible] | [illegible] | [illegible] | [illegible] | [illegible] | [illegible] | [illegible] | [illegible] | [illegible] | [illegible] | [illegible] |
| Ntrk3 | TRUE | 0.151337418 | 0.971381329 | No Change | Not Significant | FALSE | NA | [illegible] | [illegible] | [illegible] | [illegible] | [illegible] | [illegible] | [illegible] | [illegible] | [illegible] | [illegible] | [illegible] | [illegible] | [illegible] | [illegible] | [illegible] | [illegible] | [illegible] | [illegible] | [illegible] | [illegible] | [illegible] | [illegible] | [illegible] | [illegible] | [illegible] | [illegible] | [illegible] | [illegible] | [illegible] | [illegible] |
| Nts | TRUE | -0.350852711 | 0.512889463 | No Change | Not Significant | TRUE | -0.464102121 | [illegible] | [illegible] | [illegible] | [illegible] | [illegible] | [illegible] | [illegible] | [illegible] | [illegible] | [illegible] | [illegible] | [illegible] | [illegible] | [illegible] | [illegible] | [illegible] | [illegible] | [illegible] | [illegible] | [illegible] | [illegible] | [illegible] | [illegible] | [illegible] | [illegible] | [illegible] | [illegible] | [illegible] | [illegible] | [illegible] |
| Nuak1 | TRUE | 0.153832017 | 0.830450122 | No Change | Not Significant | TRUE | 0.309278033 | [illegible] | [illegible] | [illegible] | [illegible] | [illegible] | [illegible] | [illegible] | [illegible] | [illegible] | [illegible] | [illegible] | [illegible] | [illegible] | [illegible] | [illegible] | [illegible] | [illegible] | [illegible] | [illegible] | [illegible] | [illegible] | [illegible] | [illegible] | [illegible] | [illegible] | [illegible] | [illegible] | [illegible] | [illegible] | [illegible] |
| Nuak2 | TRUE | 0.289517925 | 0.055334304 | No Change | Not Significant | TRUE | -0.038241614 | [illegible] | [illegible] | [illegible] | [illegible] | [illegible] | [illegible] | [illegible] | [illegible] | [illegible] | [illegible] | [illegible] | [illegible] | [illegible] | [illegible] | [illegible] | [illegible] | [illegible] | [illegible] | [illegible] | [illegible] | [illegible] | [illegible] | [illegible] | [illegible] | [illegible] | [illegible] | [illegible] | [illegible] | [illegible] | [illegible] |
| Nub1 | TRUE | 0.01788478 | 0.880874346 | No Change | Not Significant | TRUE | -0.010907752 | [illegible] | [illegible] | [illegible] | [illegible] | [illegible] | [illegible] | [illegible] | [illegible] | [illegible] | [illegible] | [illegible] | [illegible] | [illegible] | [illegible] | [illegible] | [illegible] | [illegible] | [illegible] | [illegible] | [illegible] | [illegible] | [illegible] | [illegible] | [illegible] | [illegible] | [illegible] | [illegible] | [illegible] | [illegible] | [illegible] |
| Nubp1 | TRUE | 0.134110327 | 0.399325652 | No Change | Not Significant | TRUE | 0.059964037 | [illegible] | [illegible] | [illegible] | [illegible] | [illegible] | [illegible] | [illegible] | [illegible] | [illegible] | [illegible] | [illegible] | [illegible] | [illegible] | [illegible] | [illegible] | [illegible] | [illegible] | [illegible] | [illegible] | [illegible] | [illegible] | [illegible] | [illegible] | [illegible] | [illegible] | [illegible] | [illegible] | [illegible] | [illegible] | [illegible] |
| Nubp2 | TRUE | -0.032246658 | 0.837078181 | No Change | Not Significant | TRUE | 0.052459813 | [illegible] | [illegible] | [illegible] | [illegible] | [illegible] | [illegible] | [illegible] | [illegible] | [illegible] | [illegible] | [illegible] | [illegible] | [illegible] | [illegible] | [illegible] | [illegible] | [illegible] | [illegible] | [illegible] | [illegible] | [illegible] | [illegible] | [illegible] | [illegible] | [illegible] | [illegible] | [illegible] | [illegible] | [illegible] | [illegible] |
| Nubpl | TRUE | -0.04754577 | 0.863523565 | No Change | Not Significant | TRUE | -0.098877764 | [illegible] | [illegible] | [illegible] | [illegible] | [illegible] | [illegible] | [illegible] | [illegible] | [illegible] | [illegible] | [illegible] | [illegible] | [illegible] | [illegible] | [illegible] | [illegible] | [illegible] | [illegible] | [illegible] | [illegible] | [illegible] | [illegible] | [illegible] | [illegible] | [illegible] | [illegible] | [illegible] | [illegible] | [illegible] | [illegible] |
| Nucb1 | TRUE | 0.666561587 | 2.02E-54 | UP | UP | TRUE | 0.511672766 | [illegible] | [illegible] | [illegible] | [illegible] | [illegible] | [illegible] | [illegible] | [illegible] | [illegible] | [illegible] | [illegible] | [illegible] | [illegible] | [illegible] | [illegible] | [illegible] | [illegible] | [illegible] | [illegible] | [illegible] | [illegible] | [illegible] | [illegible] | [illegible] | [illegible] | [illegible] | [illegible] | [illegible] | [illegible] | [illegible] |

| geneID | Cond1_DF_Detected | Cond1_DF_ValueCol | Cond1_DF_PadjCol | Cond1_DF_Cutoff | Cond1_DF_Cutoff_Specific.x | Cond2_DF_Detected | Cond2_DF_ValueCol | Cond2_DF_PadjCol | Cond2_DF_Cutoff | Cond2_DF_Cutoff_Specific | BG_Method | RG1_All | RG2_Significant | RG3_SignificantChange | Cond1_DF_ensembl_gene_id | Cond1_DF_entrezgene_id | Cond1_DF_gene_biotype | Cond1_DF_baseMean | Cond1_DF_log2FoldChange | Cond1_DF_lfcSE | Cond1_DF_stat | Cond1_DF_pvalue | Cond1_DF_padj | Cond1_DF_absLog2FC | Cond1_DF_Cutoff_Specific.y | Cond2_DF_ensembl_gene_id | Cond2_DF_entrezgene_id | Cond2_DF_gene_biotype | Cond2_DF_baseMean | Cond2_DF_log2FoldChange | Cond2_DF_lfcSE | Cond2_DF_stat | Cond2_DF_pvalue | Cond2_DF_padj | Cond2_DF_absLog2FC | RG1_Specific_Cond2 | RG1_Specific_Cond1 |
|---|---|---|---|---|---|---|---|---|---|---|---|---|---|---|---|---|---|---|---|---|---|---|---|---|---|---|---|---|---|---|---|---|---|---|---|---|---|
| Or10a49 | TRUE | 1.38120202 | 1 | No Change | Not Significant | FALSE | NA | NA | No Change | Not Detected | FALSE | Background = FALSE | Background = FALSE | Background = FALSE | ENSMUSG00000066240 | 258136 | protein_coding | 1.110662649 | 1.38120202 | 2.686711337 | -0.514086497 | 0.607191518 | 1 | 1.38120202 | Not Significant | NA | NA | NA | NA | NA | NA | NA | NA | NA | NA | Background = FALSE | Background = FALSE |
| Or10a5 | TRUE | 3.642885794 | 1 | No Change | Not Significant | FALSE | NA | NA | No Change | Not Detected | FALSE | Background = FALSE | Background = FALSE | Background = FALSE | ENSMUSG00000073898 | 259036 | protein_coding | 1.331581821 | 3.642885794 | 2.030381896 | -1.794187488 | 0.072783249 | 1 | 3.642885794 | Not Significant | NA | NA | NA | NA | NA | NA | NA | NA | NA | NA | Background = FALSE | Background = FALSE |
| Or10aa1 | TRUE | 0.00571835 | 1 | No Change | Not Significant | FALSE | NA | NA | No Change | Not Detected | FALSE | Background = FALSE | Background = FALSE | Background = FALSE | ENSMUSG00000045381 | 258712 | protein_coding | 1.15172132 | 0.00571835 | 2.206371872 | -0.002591743 | 0.99793209 | 1 | 0.00571835 | Not Significant | NA | NA | NA | NA | NA | NA | NA | NA | NA | NA | Background = FALSE | Background = FALSE |
| Or10ab5 | TRUE | -0.754422907 | 1 | No Change | Not Significant | FALSE | NA | NA | No Change | Not Detected | FALSE | Background = FALSE | Background = FALSE | Background = FALSE | ENSMUSG00000049280 | 258369 | protein_coding | 0.847331937 | -0.754422907 | 2.231890207 | 0.338019722 | 0.735348326 | 1 | 0.754422907 | Not Significant | NA | NA | NA | NA | NA | NA | NA | NA | NA | NA | Background = FALSE | Background = FALSE |
| Or10ac1 | TRUE | 0.609882319 | 1 | No Change | Not Significant | FALSE | NA | NA | No Change | Not Detected | FALSE | Background = FALSE | Background = FALSE | Background = FALSE | ENSMUSG00000071494 | 546896 | protein_coding | 1.580721599 | 0.609882319 | 2.513103476 | -0.242680942 | 0.808252572 | 1 | 0.609882319 | Not Significant | NA | NA | NA | NA | NA | NA | NA | NA | NA | NA | Background = FALSE | Background = FALSE |
| Or10ad1b | TRUE | 0.53219246 | 1 | No Change | Not Significant | FALSE | NA | NA | No Change | Not Detected | FALSE | Background = FALSE | Background = FALSE | Background = FALSE | ENSMUSG00000059460 | 629524 | protein_coding | 1.923679423 | 0.53219246 | 3.02392225 | -0.175994095 | 0.860298582 | 1 | 0.53219246 | Not Significant | NA | NA | NA | NA | NA | NA | NA | NA | NA | NA | Background = FALSE | Background = FALSE |
| Or10ag2 | TRUE | -1.707211135 | 1 | No Change | Not Significant | FALSE | NA | NA | No Change | Not Detected | FALSE | Background = FALSE | Background = FALSE | Background = FALSE | ENSMUSG00000043274 | 258347 | protein_coding | 1.054714302 | -1.707211135 | 2.889551194 | 0.590822249 | 0.554639526 | 1 | 1.707211135 | Not Significant | NA | NA | NA | NA | NA | NA | NA | NA | NA | NA | Background = FALSE | Background = FALSE |
| Or10ag54 | TRUE | 0.005094936 | 1 | No Change | Not Significant | FALSE | NA | NA | No Change | Not Detected | FALSE | Background = FALSE | Background = FALSE | Background = FALSE | ENSMUSG00000070855 | 257875 | protein_coding | 2.244326701 | 0.005094936 | 2.464912278 | -0.002066985 | 0.998350786 | 1 | 0.005094936 | Not Significant | NA | NA | NA | NA | NA | NA | NA | NA | NA | NA | Background = FALSE | Background = FALSE |
| Or10ak16 | TRUE | 3.007526732 | 1 | No Change | Not Significant | FALSE | NA | NA | No Change | Not Detected | FALSE | Background = FALSE | Background = FALSE | Background = FALSE | ENSMUSG00000073768 | 258331 | protein_coding | 1.679047418 | 3.007526732 | 2.516899959 | -1.194932966 | 0.232113222 | 1 | 3.007526732 | Not Significant | NA | NA | NA | NA | NA | NA | NA | NA | NA | NA | Background = FALSE | Background = FALSE |
| Or10ak7 | TRUE | -0.78967306 | 1 | No Change | Not Significant | FALSE | NA | NA | No Change | Not Detected | FALSE | Background = FALSE | Background = FALSE | Background = FALSE | ENSMUSG00000111259 | 258394 | protein_coding | 1.593641439 | -0.78967306 | 2.363897956 | 0.334055477 | 0.738337696 | 1 | 0.78967306 | Not Significant | NA | NA | NA | NA | NA | NA | NA | NA | NA | NA | Background = FALSE | Background = FALSE |
| Or10al7 | TRUE | 2.851312973 | 1 | No Change | Not Significant | FALSE | NA | NA | No Change | Not Detected | FALSE | Background = FALSE | Background = FALSE | Background = FALSE | ENSMUSG00000081724 | 258324 | protein_coding | 1.462916313 | 2.851312973 | 3.02664268 | -0.942071224 | 0.346156176 | 1 | 2.851312973 | Not Significant | NA | NA | NA | NA | NA | NA | NA | NA | NA | NA | Background = FALSE | Background = FALSE |
| Or10am5 | TRUE | 3.186890288 | 1 | No Change | Not Significant | FALSE | NA | NA | No Change | Not Detected | FALSE | Background = FALSE | Background = FALSE | Background = FALSE | ENSMUSG00000048067 | 269862 | protein_coding | 1.963473853 | 3.186890288 | 2.351221812 | -1.355418817 | 0.175284158 | 1 | 3.186890288 | Not Significant | NA | NA | NA | NA | NA | NA | NA | NA | NA | NA | Background = FALSE | Background = FALSE |
| Or10c1 | TRUE | 0.753064876 | 1 | No Change | Not Significant | FALSE | NA | NA | No Change | Not Detected | FALSE | Background = FALSE | Background = FALSE | Background = FALSE | ENSMUSG00000049561 | 258506 | protein_coding | 1.506864841 | 0.753064876 | 3.175858456 | -0.237121675 | 0.812562398 | 1 | 0.753064876 | Not Significant | NA | NA | NA | NA | NA | NA | NA | NA | NA | NA | Background = FALSE | Background = FALSE |
| Or10d1 | TRUE | 0.523403065 | 1 | No Change | Not Significant | FALSE | NA | NA | No Change | Not Detected | FALSE | Background = FALSE | Background = FALSE | Background = FALSE | ENSMUSG00000059366 | 258501 | protein_coding | 1.241798554 | 0.523403065 | 2.652580406 | -0.197318454 | 0.843578339 | 1 | 0.523403065 | Not Significant | NA | NA | NA | NA | NA | NA | NA | NA | NA | NA | Background = FALSE | Background = FALSE |
| [illegible] | [illegible] | [illegible] | [illegible] | [illegible] | [illegible] | [illegible] | [illegible] | [illegible] | [illegible] | [illegible] | [illegible] | [illegible] | [illegible] | [illegible] | [illegible] | [illegible] | [illegible] | [illegible] | [illegible] | [illegible] | [illegible] | [illegible] | [illegible] | [illegible] | [illegible] | [illegible] | [illegible] | [illegible] | [illegible] | [illegible] | [illegible] | [illegible] | [illegible] | [illegible] | [illegible] | [illegible] | [illegible] |

| geneID | Cond1_DF_Detected | Cond1_DF_ValueCol | Cond1_DF_PadjCol | Cond1_DF_Cutoff | Cond1_DF_Cutoff_Specific.x | Cond2_DF_Detected | Cond2_DF_ValueCol | Cond2_DF_PadjCol | Cond2_DF_Cutoff | Cond2_DF_Cutoff_Specific | BG_Method | RG1_All | RG2_Significant | RG3_SignificantChange |
|---|---|---|---|---|---|---|---|---|---|---|---|---|---|---|
| Or52n20 | TRUE | -1.157533771 | 1 | No Change | Not Significant | FALSE | NA | NA | No Change | Not Detected | FALSE | Background = FALSE | Background = FALSE | Background = FALSE |
| Or52n4 | TRUE | -0.243790244 | 1 | No Change | Not Significant | FALSE | NA | NA | No Change | Not Detected | FALSE | Background = FALSE | Background = FALSE | Background = FALSE |
| Or52p1 | TRUE | 1.863801358 | 1 | No Change | Not Significant | FALSE | NA | NA | No Change | Not Detected | FALSE | Background = FALSE | Background = FALSE | Background = FALSE |
| Or52r1 | TRUE | 2.644748562 | 1 | No Change | Not Significant | FALSE | NA | NA | No Change | Not Detected | FALSE | Background = FALSE | Background = FALSE | Background = FALSE |
| Or52r1b | TRUE | 0.522284398 | 1 | No Change | Not Significant | FALSE | NA | NA | No Change | Not Detected | FALSE | Background = FALSE | Background = FALSE | Background = FALSE |
| Or52s1b | TRUE | 1.108501925 | 1 | No Change | Not Significant | FALSE | NA | NA | No Change | Not Detected | FALSE | Background = FALSE | Background = FALSE | Background = FALSE |
| Or52w1 | TRUE | 1.047075729 | 1 | No Change | Not Significant | FALSE | NA | NA | No Change | Not Detected | FALSE | Background = FALSE | Background = FALSE | Background = FALSE |
| Or55b10 | TRUE | 0.88104969 | 1 | No Change | Not Significant | FALSE | NA | NA | No Change | Not Detected | FALSE | Background = FALSE | Background = FALSE | Background = FALSE |
| Or55b3 | TRUE | 0.137231877 | 1 | No Change | Not Significant | FALSE | NA | NA | No Change | Not Detected | FALSE | Background = FALSE | Background = FALSE | Background = FALSE |
| Or55b4 | TRUE | -0.835362868 | 1 | No Change | Not Significant | FALSE | NA | NA | No Change | Not Detected | FALSE | Background = FALSE | Background = FALSE | Background = FALSE |
| Or56a3 | TRUE | -0.236181457 | 1 | No Change | Not Significant | FALSE | NA | NA | No Change | Not Detected | FALSE | Background = FALSE | Background = FALSE | Background = FALSE |
| Or56a4 | TRUE | 1.189960122 | 1 | No Change | Not Significant | FALSE | NA | NA | No Change | Not Detected | FALSE | Background = FALSE | Background = FALSE | Background = FALSE |

| geneID | Cond1_DF_ensembl_gene_id | Cond1_DF_entrezgene_id | Cond1_DF_gene_biotype | Cond1_DF_baseMean | Cond1_DF_log2FoldChange | Cond1_DF_lfcSE | Cond1_DF_stat | Cond1_DF_pvalue | Cond1_DF_padj | Cond1_DF_absLog2FC | Cond1_DF_Cutoff_Specific.y |
|---|---|---|---|---|---|---|---|---|---|---|---|
| Or52n20 | ENSMUSG00000073922 | 259052 | protein_coding | 2.005006497 | -1.157533771 | 2.294041467 | 0.504582758 | 0.613851922 | 1 | 1.157533771 | Not Significant |
| Or52n4 | ENSMUSG00000070421 | 259051 | protein_coding | 1.341653401 | -0.243790244 | 1.427930403 | 0.17072978 | 0.864436245 | 1 | 0.243790244 | Not Significant |
| Or52p1 | ENSMUSG00000073924 | 259078 | protein_coding | 1.648799992 | 1.863801358 | 2.684371975 | -0.694315607 | 0.487484305 | 1 | 1.863801358 | Not Significant |
| Or52r1 | ENSMUSG00000062142 | 259092 | protein_coding | 1.386828032 | 2.644748562 | 2.60556674 | -1.015037735 | 0.31008784 | 1 | 2.644748562 | Not Significant |
| Or52r1b | ENSMUSG00000073961 | 259055 | protein_coding | 1.298704097 | 0.522284398 | 2.613799216 | -0.199818102 | 0.841622844 | 1 | 0.522284398 | Not Significant |
| Or52s1b | ENSMUSG00000057461 | 258139 | protein_coding | 1.076326485 | 1.108501925 | 2.192897972 | -0.505496352 | 0.61321026 | 1 | 1.108501925 | Not Significant |
| Or52w1 | ENSMUSG00000073906 | 258352 | protein_coding | 0.830372288 | 1.047075729 | 1.523246305 | -0.687397518 | 0.491832262 | 1 | 1.047075729 | Not Significant |
| Or55b10 | ENSMUSG00000044824 | 258837 | protein_coding | 0.874700101 | 0.88104969 | 1.997307243 | -0.441118758 | 0.659127024 | 1 | 0.88104969 | Not Significant |
| Or55b3 | ENSMUSG00000044814 | 257947 | protein_coding | 1.131997778 | 0.137231877 | 0.967554186 | -0.141833789 | 0.887211294 | 1 | 0.137231877 | Not Significant |
| Or55b4 | ENSMUSG00000043925 | 257926 | protein_coding | 0.814797316 | -0.835362868 | 1.976574048 | 0.422631709 | 0.672563988 | 1 | 0.835362868 | Not Significant |
| Or56a3 | ENSMUSG00000096029 | 259046 | protein_coding | 0.897527938 | -0.236181457 | 2.67509266 | 0.088289075 | 0.929646922 | 1 | 0.236181457 | Not Significant |
| Or56a4 | ENSMUSG00000047225 | 244187 | protein_coding | 1.363459545 | 1.189960122 | 3.340475188 | -0.356224805 | 0.721672217 | 1 | 1.189960122 | Not Significant |

| geneID | Cond2_DF_ensembl_gene_id | Cond2_DF_entrezgene_id | Cond2_DF_gene_biotype | Cond2_DF_baseMean | Cond2_DF_log2FoldChange | Cond2_DF_lfcSE | Cond2_DF_stat | Cond2_DF_pvalue | Cond2_DF_padj | Cond2_DF_absLog2FC | RG1_Specific_Cond2 | RG1_Specific_Cond1 |
|---|---|---|---|---|---|---|---|---|---|---|---|---|
| Or52n20 | NA | NA | NA | NA | NA | NA | NA | NA | NA | NA | Background = FALSE | Background = FALSE |
| Or52n4 | NA | NA | NA | NA | NA | NA | NA | NA | NA | NA | Background = FALSE | Background = FALSE |
| Or52p1 | NA | NA | NA | NA | NA | NA | NA | NA | NA | NA | Background = FALSE | Background = FALSE |
| Or52r1 | NA | NA | NA | NA | NA | NA | NA | NA | NA | NA | Background = FALSE | Background = FALSE |
| Or52r1b | NA | NA | NA | NA | NA | NA | NA | NA | NA | NA | Background = FALSE | Background = FALSE |
| Or52s1b | NA | NA | NA | NA | NA | NA | NA | NA | NA | NA | Background = FALSE | Background = FALSE |
| Or52w1 | NA | NA | NA | NA | NA | NA | NA | NA | NA | NA | Background = FALSE | Background = FALSE |
| Or55b10 | NA | NA | NA | NA | NA | NA | NA | NA | NA | NA | Background = FALSE | Background = FALSE |
| Or55b3 | NA | NA | NA | NA | NA | NA | NA | NA | NA | NA | Background = FALSE | Background = FALSE |
| Or55b4 | NA | NA | NA | NA | NA | NA | NA | NA | NA | NA | Background = FALSE | Background = FALSE |
| Or56a3 | NA | NA | NA | NA | NA | NA | NA | NA | NA | NA | Background = FALSE | Background = FALSE |
| Or56a4 | NA | NA | NA | NA | NA | NA | NA | NA | NA | NA | Background = FALSE | Background = FALSE |

| geneID | Cond1_DF_Detected | Cond1_DF_ValueCol | Cond1_DF_PadjCol | Cond1_DF_Cutoff | Cond1_DF_Cutoff_Specific.x | Cond2_DF_Detected | Cond2_DF_ValueCol | Cond2_DF_PadjCol | Cond2_DF_Cutoff | Cond2_DF_Cutoff_Specific | BG_Method | RG1_All | RG2_Significant |
|---|---|---|---|---|---|---|---|---|---|---|---|---|---|
| P3h4 | TRUE | -0.248356801 | 0.388090095 | No Change | Not Significant | TRUE | 0.068794209 | 0.884953772 | No Change | Not Significant | TRUE | Cond1 Not Significant + Cond2 Not Significant | None |
| P3r3urf | TRUE | -0.622480635 | 1 | No Change | Not Significant | FALSE | NA | NA | No Change | Not Detected | FALSE | Background = FALSE | Background = FALSE |
| P4ha1 | TRUE | 0.106296831 | 0.498917274 | No Change | Not Significant | TRUE | 0.301108985 | 0.075680028 | No Change | Not Significant | TRUE | Cond1 Not Significant + Cond2 Not Significant | None |
| P4ha2 | TRUE | 1.696743554 | 1.29E-19 | UP | UP | TRUE | 1.040408424 | 2.10E-07 | UP | UP | TRUE | Cond1 UP + Cond2 UP | Core_UP |
| P4ha3 | TRUE | -0.811239627 | 1 | No Change | Not Significant | FALSE | NA | NA | No Change | Not Detected | FALSE | Background = FALSE | Background = FALSE |
| P4hb | TRUE | 2.113431818 | 1.12E-300 | UP | UP | TRUE | 1.210306569 | 1.68E-135 | UP | UP | TRUE | Cond1 UP + Cond2 UP | Core_UP |
| P4htm | TRUE | 1.129003225 | 0.21257502 | No Change | Not Significant | TRUE | 0.598043692 | 0.45055038 | No Change | Not Significant | TRUE | Cond1 Not Significant + Cond2 Not Significant | None |
| Pa2g4 | TRUE | 0.110723341 | 0.407364189 | No Change | Not Significant | TRUE | 0.010279857 | 0.94222547 | No Change | Not Significant | TRUE | Cond1 Not Significant + Cond2 Not Significant | None |
| Pabpc1 | TRUE | -0.109339784 | 0.48334534 | No Change | Not Significant | TRUE | -0.079036635 | 0.656821597 | No Change | Not Significant | TRUE | Cond1 Not Significant + Cond2 Not Significant | None |
| Pabpc1l | TRUE | 0.05255087 | 0.97354273 | No Change | Not Significant | TRUE | 0.392206334 | 0.640407651 | No Change | Not Significant | TRUE | Cond1 Not Significant + Cond2 Not Significant | None |
| Pabpc2 | TRUE | 0.939223255 | 1 | No Change | Not Significant | FALSE | NA | NA | No Change | Not Detected | FALSE | Background = FALSE | Background = FALSE |
| Pabpc4 | TRUE | -0.311887456 | 0.01173204 | No Change | Significant Negative | TRUE | -0.051998837 | 0.663421426 | No Change | Not Significant | TRUE | Cond1 Significant Negative + Cond2 Not Significant | None |
| Pabpc4l | TRUE | 0.522248131 | 1 | No Change | Not Significant | FALSE | NA | NA | No Change | Not Detected | FALSE | Background = FALSE | Background = FALSE |
| Pabpc5 | TRUE | 1.987224228 | 1 | No Change | Not Significant | FALSE | NA | NA | No Change | Not Detected | FALSE | Background = FALSE | Background = FALSE |
| Pabpn1 | TRUE | 0.466081556 | 0.049414074 | No Change | Significant Positive | TRUE | -0.085803339 | 0.61652794 | No Change | Not Significant | TRUE | Cond1 Significant Positive + Cond2 Not Significant | None |
| Pabpn1l | TRUE | -0.480472962 | 1 | No Change | Not Significant | FALSE | NA | NA | No Change | Not Detected | FALSE | Background = FALSE | Background = FALSE |
| Pacc1 | TRUE | -0.176243354 | 0.397106786 | No Change | Not Significant | TRUE | -0.030890641 | 0.907948215 | No Change | Not Significant | TRUE | Cond1 Not Significant + Cond2 Not Significant | None |
| Pacrg | TRUE | -1.107267657 | 1 | No Change | Not Significant | FALSE | NA | NA | No Change | Not Detected | FALSE | Background = FALSE | Background = FALSE |
| Pacs1 | TRUE | -0.386273991 | 0.008435285 | No Change | Significant Negative | TRUE | -0.127578134 | 0.409503276 | No Change | Not Significant | TRUE | Cond1 Significant Negative + Cond2 Not Significant | None |
| Pacs2 | TRUE | -0.132749499 | 0.051167007 | No Change | Not Significant | TRUE | -0.054410553 | 0.448799461 | No Change | Not Significant | TRUE | Cond1 Not Significant + Cond2 Not Significant | None |
| Pacsin1 | TRUE | -0.653652825 | 0.122282111 | No Change | Not Significant | TRUE | -0.488535194 | 0.258850238 | No Change | Not Significant | TRUE | Cond1 Not Significant + Cond2 Not Significant | None |
| Pacsin2 | TRUE | 0.735922133 | 1.22E-36 | UP | UP | TRUE | 0.400064428 | 4.35E-34 | No Change | Significant Positive | TRUE | Cond1 UP + Cond2 Significant Positive | Core_UP |
| Pacsin3 | TRUE | -0.295277783 | 0.097588853 | No Change | Not Significant | TRUE | 0.237396347 | 0.224854533 | No Change | Not Significant | TRUE | Cond1 Not Significant + Cond2 Not Significant | None |
| Padi1 | TRUE | 0.317630035 | 1 | No Change | Not Significant | FALSE | NA | NA | No Change | Not Detected | FALSE | Background = FALSE | Background = FALSE |
| Padi2 | TRUE | 0.055327217 | 0.884482916 | No Change | Not Significant | TRUE | -0.116254268 | 0.648598559 | No Change | Not Significant | TRUE | Cond1 Not Significant + Cond2 Not Significant | None |
| Padi4 | TRUE | 0.635110209 | 1 | No Change | Not Significant | FALSE | NA | NA | No Change | Not Detected | FALSE | Background = FALSE | Background = FALSE |
| Padi6 | TRUE | -0.430905498 | 1 | No Change | Not Significant | FALSE | NA | NA | No Change | Not Detected | FALSE | Background = FALSE | Background = FALSE |
| Paf1 | TRUE | 0.244992368 | 0.011044218 | No Change | Significant Positive | TRUE | 0.032870827 | 0.7673338 | No Change | Not Significant | TRUE | Cond1 Significant Positive + Cond2 Not Significant | None |
| Pafah1b1 | TRUE | 0.061429667 | 0.44767168 | No Change | Not Significant | TRUE | -0.043499469 | 0.4661143 | No Change | Not Significant | TRUE | Cond1 Not Significant + Cond2 Not Significant | None |
| Pafah1b2 | TRUE | 0.116760408 | 0.248835031 | No Change | Not Significant | TRUE | -0.017722249 | 0.84654428 | No Change | Not Significant | TRUE | Cond1 Not Significant + Cond2 Not Significant | None |
| Pafah1b3 | TRUE | 0.107040492 | 0.50522539 | No Change | Not Significant | TRUE | -0.053631625 | 0.732190524 | No Change | Not Significant | TRUE | Cond1 Not Significant + Cond2 Not Significant | None |
| Pafah2 | TRUE | -0.08896003 | 0.410345914 | No Change | Not Significant | TRUE | 0.024387803 | 0.880597382 | No Change | Not Significant | TRUE | Cond1 Not Significant + Cond2 Not Significant | None |
| Pag1 | TRUE | 0.008448826 | 0.973795719 | No Change | Not Significant | TRUE | -0.017014802 | 0.900006643 | No Change | Not Significant | TRUE | Cond1 Not Significant + Cond2 Not Significant | None |
| Paics | TRUE | -0.520580928 | 0.016937143 | DOWN | DOWN | TRUE | -0.225683598 | 0.019554576 | No Change | Significant Negative | TRUE | Cond1 DOWN + Cond2 Significant Negative | Core_DOWN |
| Paip1 | TRUE | 0.044856845 | 0.77323163 | No Change | Not Significant | TRUE | 0.031017359 | 0.698941369 | No Change | Not Significant | TRUE | Cond1 Not Significant + Cond2 Not Significant | None |
| Paip2 | TRUE | -0.221062455 | 0.146772084 | No Change | Not Significant | TRUE | 0.042925276 | 0.511015761 | No Change | Not Significant | TRUE | Cond1 Not Significant + Cond2 Not Significant | None |
| Paip2b | TRUE | 1.583098427 | 1.87E-72 | UP | UP | TRUE | 0.626717533 | 2.04E-17 | UP | UP | TRUE | Cond1 UP + Cond2 UP | Core_UP |
| Pak1 | TRUE | -0.014132872 | 0.9092783 | No Change | Not Significant | TRUE | 0.117797484 | 0.073503206 | No Change | Not Significant | TRUE | Cond1 Not Significant + Cond2 Not Significant | None |
| Pak1ip1 | TRUE | 0.090632618 | 0.483542311 | No Change | Not Significant | TRUE | 0.046870809 | 0.658779673 | No Change | Not Significant | TRUE | Cond1 Not Significant + Cond2 Not Significant | None |
| Pak2 | TRUE | -0.13412644 | 0.147459771 | No Change | Not Significant | TRUE | -0.131801057 | 0.050113226 | No Change | Not Significant | TRUE | Cond1 Not Significant + Cond2 Not Significant | None |
| Pak3 | TRUE | 0.509472072 | 0.837078181 | No Change | Not Significant | FALSE | NA | NA | No Change | Not Detected | FALSE | Background = FALSE | Background = FALSE |
| Pak4 | TRUE | 0.412332765 | 9.85E-10 | No Change | Significant Positive | TRUE | 0.180461789 | 0.055431266 | No Change | Not Significant | TRUE | Cond1 Significant Positive + Cond2 Not Significant | None |
| Pak5 | TRUE | 0.739587048 | 1 | No Change | Not Significant | FALSE | NA | NA | No Change | Not Detected | FALSE | Background = FALSE | Background = FALSE |
| Pak6 | TRUE | -0.306907482 | 0.360481579 | No Change | Not Significant | TRUE | -0.032516762 | 0.935604164 | No Change | Not Significant | TRUE | Cond1 Not Significant + Cond2 Not Significant | None |
| Pakap | TRUE | -0.279435872 | 0.585166009 | No Change | Not Significant | TRUE | -0.206496102 | 0.489027453 | No Change | Not Significant | TRUE | Cond1 Not Significant + Cond2 Not Significant | None |
| Palb2 | TRUE | 0.473754177 | 0.009275396 | No Change | Significant Positive | TRUE | -0.085488435 | 0.780262005 | No Change | Not Significant | TRUE | Cond1 Significant Positive + Cond2 Not Significant | None |
| Pald1 | TRUE | 0.26885565 | 0.580790426 | No Change | Not Significant | TRUE | 0.089592302 | 0.878396765 | No Change | Not Significant | TRUE | Cond1 Not Significant + Cond2 Not Significant | None |
| Palld | TRUE | 0.202510826 | 0.44350564 | No Change | Not Significant | TRUE | -0.15367435 | 0.439595026 | No Change | Not Significant | TRUE | Cond1 Not Significant + Cond2 Not Significant | None |
| Palm | TRUE | -0.081974743 | 0.900994567 | No Change | Not Significant | TRUE | -0.10876999 | 0.800250831 | No Change | Not Significant | TRUE | Cond1 Not Significant + Cond2 Not Significant | None |
| Palm3 | TRUE | 1.587685761 | 1 | No Change | Not Significant | FALSE | NA | NA | No Change | Not Detected | FALSE | Background = FALSE | Background = FALSE |
| Palmd | TRUE | 0.118444569 | 0.800822518 | No Change | Not Significant | TRUE | 0.04332589 | 0.930177636 | No Change | Not Significant | TRUE | Cond1 Not Significant + Cond2 Not Significant | None |
| Pals1 | TRUE | -0.057054123 | 0.642504934 | No Change | Not Significant | TRUE | -0.086591685 | 0.334230211 | No Change | Not Significant | TRUE | Cond1 Not Significant + Cond2 Not Significant | None |
| Pals2 | TRUE | 0.275434678 | 0.131303585 | No Change | Not Significant | TRUE | 0.165664456 | 0.314789716 | No Change | Not Significant | TRUE | Cond1 Not Significant + Cond2 Not Significant | None |
| Pam | TRUE | -0.275263022 | 0.236336338 | No Change | Not Significant | TRUE | -0.151268806 | 0.036869266 | No Change | Significant Negative | TRUE | Cond1 Not Significant + Cond2 Significant Negative | None |
| Pam16 | TRUE | -0.413026042 | 0.014104161 | No Change | Significant Negative | TRUE | 0.071316714 | 0.657014281 | No Change | Not Significant | TRUE | Cond1 Significant Negative + Cond2 Not Significant | None |
| Pamr1 | TRUE | -0.657741935 | 0.148382689 | No Change | Not Significant | TRUE | 0.286136735 | 0.484584527 | No Change | Not Significant | TRUE | Cond1 Not Significant + Cond2 Not Significant | None |
| Pan2 | TRUE | 0.60403047 | 7.13E-09 | UP | UP | TRUE | 0.345624222 | 1.98E-05 | No Change | Significant Positive | TRUE | Cond1 UP + Cond2 Significant Positive | Core_UP |
| Pan3 | TRUE | -0.004838029 | 0.962436702 | No Change | Not Significant | TRUE | -0.13597467 | 0.00500312 | No Change | Significant Negative | TRUE | Cond1 Not Significant + Cond2 Significant Negative | None |
| Panct2 | TRUE | 0.213961224 | 0.640973307 | No Change | Not Significant | TRUE | 0.465385687 | 0.018625542 | No Change | Significant Positive | TRUE | Cond1 Not Significant + Cond2 Significant Positive | None |
| Pank1 | TRUE | -0.357483776 | 0.000280811 | No Change | Significant Negative | TRUE | -0.076546218 | 0.583162735 | No Change | Not Significant | TRUE | Cond1 Significant Negative + Cond2 Not Significant | None |
| Pank2 | TRUE | 0.066969393 | 0.521072726 | No Change | Not Significant | TRUE | 0.090742612 | 0.370961375 | No Change | Not Significant | TRUE | Cond1 Not Significant + Cond2 Not Significant | None |
| Pank3 | TRUE | 0.061415995 | 0.714956306 | No Change | Not Significant | TRUE | -0.055694927 | 0.444801234 | No Change | Not Significant | TRUE | Cond1 Not Significant + Cond2 Not Significant | None |
| Pank4 | TRUE | -0.183495459 | 0.115758474 | No Change | Not Significant | TRUE | -0.136064688 | 0.258254223 | No Change | Not Significant | TRUE | Cond1 Not Significant + Cond2 Not Significant | None |
| Panx1 | TRUE | -0.292412637 | 0.015480494 | No Change | Significant Negative | TRUE | -0.172873908 | 0.10455009 | No Change | Not Significant | TRUE | Cond1 Significant Negative + Cond2 Not Significant | None |
| Panx2 | TRUE | 0.715493541 | 1 | No Change | Not Significant | FALSE | NA | NA | No Change | Not Detected | FALSE | Background = FALSE | Background = FALSE |
| Paox | TRUE | -0.828651482 | 6.25E-05 | DOWN | DOWN | TRUE | -0.329609877 | 0.00120322 | No Change | Significant Negative | TRUE | Cond1 DOWN + Cond2 Significant Negative | Core_DOWN |
| Papln | TRUE | 0.680509893 | 0.109621469 | No Change | Not Significant | TRUE | -0.237587778 | 0.513806497 | No Change | Not Significant | TRUE | Cond1 Not Significant + Cond2 Not Significant | None |
| Papola | TRUE | 0.046396586 | 0.743572273 | No Change | Not Significant | TRUE | 0.030979496 | 0.725207424 | No Change | Not Significant | TRUE | Cond1 Not Significant + Cond2 Not Significant | None |
| Papolg | TRUE | -0.303362071 | 0.000509035 | No Change | Significant Negative | TRUE | -0.099133824 | 0.369704242 | No Change | Not Significant | TRUE | Cond1 Significant Negative + Cond2 Not Significant | None |
| Pappa | TRUE | 1.135636128 | 0.214426852 | No Change | Not Significant | FALSE | NA | NA | No Change | Not Detected | FALSE | Background = FALSE | Background = FALSE |
| Pappa2 | TRUE | -0.081088709 | 0.849914408 | No Change | Not Significant | TRUE | 0.1124268 | 0.747864433 | No Change | Not Significant | TRUE | Cond1 Not Significant + Cond2 Not Significant | None |
| Papss1 | TRUE | 0.656151447 | 3.26E-24 | UP | UP | TRUE | 0.336159121 | 2.31E-12 | No Change | Significant Positive | TRUE | Cond1 UP + Cond2 Significant Positive | Core_UP |
| Papss2 | TRUE | -0.103269891 | 0.663778937 | No Change | Not Significant | TRUE | 0.037382941 | 0.724823423 | No Change | Not Significant | TRUE | Cond1 Not Significant + Cond2 Not Significant | None |
| Paqr3 | TRUE | 0.536135521 | 0.472341459 | No Change | Not Significant | TRUE | 0.016881214 | 0.984661462 | No Change | Not Significant | TRUE | Cond1 Not Significant + Cond2 Not Significant | None |
| Paqr4 | TRUE | 0.156730201 | 0.035388872 | No Change | Significant Positive | TRUE | 0.234921657 | 0.003904708 | No Change | Significant Positive | TRUE | Cond1 Significant Positive + Cond2 Significant Positive | None |
| Paqr5 | TRUE | -0.544481413 | 0.000402389 | DOWN | DOWN | TRUE | 0.025144295 | 0.826041967 | No Change | Not Significant | TRUE | Cond1 DOWN + Cond2 Not Significant | Cond1_DOWN |
| Paqr6 | TRUE | 2.486459092 | 1 | No Change | Not Significant | FALSE | NA | NA | No Change | Not Detected | FALSE | Background = FALSE | Background = FALSE |
| Paqr7 | TRUE | -0.556272197 | 0.01137907 | DOWN | DOWN | TRUE | -0.097510604 | 0.714326476 | No Change | Not Significant | TRUE | Cond1 DOWN + Cond2 Not Significant | Cond1_DOWN |
| Paqr8 | TRUE | 0.503905725 | 0.005268353 | UP | UP | TRUE | 0.359692174 | 0.001776055 | No Change | Significant Positive | TRUE | Cond1 UP + Cond2 Significant Positive | Core_UP |
| Paqr9 | TRUE | -1.130436016 | 1 | No Change | Not Significant | FALSE | NA | NA | No Change | Not Detected | FALSE | Background = FALSE | Background = FALSE |
| Pard3 | TRUE | -0.128903026 | 0.025397924 | No Change | Significant Negative | TRUE | -0.148843872 | 0.035735229 | No Change | Significant Negative | TRUE | Cond1 Significant Negative + Cond2 Significant Negative | None |
| Pard3b | TRUE | -0.229512675 | 0.013943178 | No Change | Significant Negative | TRUE | -0.098515182 | 0.326278724 | No Change | Not Significant | TRUE | Cond1 Significant Negative + Cond2 Not Significant | None |
| Pard6a | TRUE | -0.141056481 | 0.612497263 | No Change | Not Significant | TRUE | 0.034063765 | 0.916534992 | No Change | Not Significant | TRUE | Cond1 Not Significant + Cond2 Not Significant | None |
| Pard6b | TRUE | -0.287388739 | 0.283202471 | No Change | Not Significant | TRUE | -0.132549268 | 0.342321275 | No Change | Not Significant | TRUE | Cond1 Not Significant + Cond2 Not Significant | None |
| Pard6g | TRUE | -0.012585939 | 0.982706396 | No Change | Not Significant | TRUE | 0.225484376 | 0.716014372 | No Change | Not Significant | TRUE | Cond1 Not Significant + Cond2 Not Significant | None |
| Parg | TRUE | 0.051424391 | 0.684706793 | No Change | Not Significant | TRUE | 0.042544844 | 0.714326476 | No Change | Not Significant | TRUE | Cond1 Not Significant + Cond2 Not Significant | None |
| Park7 | TRUE | 0.100154769 | 0.44767168 | No Change | Not Significant | TRUE | 0.108531377 | 0.21029688 | No Change | Not Significant | TRUE | Cond1 Not Significant + Cond2 Not Significant | None |
| Parl | TRUE | -0.201471274 | 0.056736385 | No Change | Not Significant | TRUE | -0.123413242 | 0.091350728 | No Change | Not Significant | TRUE | Cond1 Not Significant + Cond2 Not Significant | None |
| Parm1 | TRUE | -0.285379801 | 0.003646336 | No Change | Significant Negative | TRUE | -0.085569379 | 0.487025714 | No Change | Not Significant | TRUE | Cond1 Significant Negative + Cond2 Not Significant | None |
| Parn | TRUE | 0.196175208 | 0.141461059 | No Change | Not Significant | TRUE | 0.048016999 | 0.701468541 | No Change | Not Significant | TRUE | Cond1 Not Significant + Cond2 Not Significant | None |
| Parp1 | TRUE | 1.036221179 | 2.60E-16 | UP | UP | TRUE | 0.537639882 | 1.82E-09 | UP | UP | TRUE | Cond1 UP + Cond2 UP | Core_UP |
| Parp10 | TRUE | -0.235210081 | 0.029064867 | No Change | Significant Negative | TRUE | -0.137065055 | 0.244881547 | No Change | Not Significant | TRUE | Cond1 Significant Negative + Cond2 Not Significant | None |
| Parp11 | TRUE | -0.06062911 | 0.80668897 | No Change | Not Significant | TRUE | -0.047129953 | 0.792823007 | No Change | Not Significant | TRUE | Cond1 Not Significant + Cond2 Not Significant | None |
| Parp12 | TRUE | -0.120017362 | 0.582711741 | No Change | Not Significant | TRUE | -0.076215675 | 0.715904735 | No Change | Not Significant | TRUE | Cond1 Not Significant + Cond2 Not Significant | None |
| Parp14 | TRUE | -0.192124859 | 0.524504007 | No Change | Not Significant | TRUE | -0.219731337 | 0.241032638 | No Change | Not Significant | TRUE | Cond1 Not Significant + Cond2 Not Significant | None |
| Parp16 | TRUE | 0.387835003 | 0.002788738 | No Change | Significant Positive | TRUE | 0.117445826 | 0.264813247 | No Change | Not Significant | TRUE | Cond1 Significant Positive + Cond2 Not Significant | None |
| Parp2 | TRUE | 0.006529404 | 0.969802608 | No Change | Not Significant | TRUE | -0.003656843 | 0.984637528 | No Change | Not Significant | TRUE | Cond1 Not Significant + Cond2 Not Significant | None |
| Parp3 | TRUE | -0.037913736 | 0.890125936 | No Change | Not Significant | TRUE | -0.104264731 | 0.783232176 | No Change | Not Significant | TRUE | Cond1 Not Significant + Cond2 Not Significant | None |
| Parp4 | TRUE | 0.053776561 | 0.929758221 | No Change | Not Significant | TRUE | -0.372550354 | 0.02917991 | No Change | Significant Negative | TRUE | Cond1 Not Significant + Cond2 Significant Negative | None |
| Parp6 | TRUE | -0.267210669 | 0.019227501 | No Change | Significant Negative | TRUE | -0.131753347 | 0.197440646 | No Change | Not Significant | TRUE | Cond1 Significant Negative + Cond2 Not Significant | None |
| Parp8 | TRUE | 1.511803912 | 0.110941817 | No Change | Not Significant | TRUE | 0.179892452 | 0.808283189 | No Change | Not Significant | TRUE | Cond1 Not Significant + Cond2 Not Significant | None |
| Parp9 | TRUE | -0.244278254 | 0.187361642 | No Change | Not Significant | TRUE | -0.122147681 | 0.610694985 | No Change | Not Significant | TRUE | Cond1 Not Significant + Cond2 Not Significant | None |
| Parpbp | TRUE | 0.821651173 | 9.77E-06 | UP | UP | TRUE | -0.008087062 | 0.980775949 | No Change | Not Significant | TRUE | Cond1 UP + Cond2 Not Significant | Cond1_UP |
| Pars2 | TRUE | -0.275026553 | 0.167325234 | No Change | Not Significant | TRUE | -0.150302963 | 0.379011242 | No Change | Not Significant | TRUE | Cond1 Not Significant + Cond2 Not Significant | None |
| Parva | TRUE | -0.136703295 | 0.441179379 | No Change | Not Significant | TRUE | 0.056430569 | 0.562649386 | No Change | Not Significant | TRUE | Cond1 Not Significant + Cond2 Not Significant | None |
| Parvaos | TRUE | 0.104097177 | 1 | No Change | Not Significant | FALSE | NA | NA | No Change | Not Detected | FALSE | Background = FALSE | Background = FALSE |
| Parvb | TRUE | -0.109804593 | 1 | No Change | Not Significant | FALSE | NA | NA | No Change | Not Detected | FALSE | Background = FALSE | Background = FALSE |
| Parvg | TRUE | 1.029974551 | 0.356592544 | No Change | Not Significant | TRUE | 1.035693967 | 0.048137725 | UP | UP | TRUE | Cond1 Not Significant + Cond2 UP | Cond2_UP |
| Pask | TRUE | 0.46487801 | 0.008514609 | No Change | Significant Positive | TRUE | 0.071962844 | 0.751487589 | No Change | Not Significant | TRUE | Cond1 Significant Positive + Cond2 Not Significant | None |
| Pate2 | TRUE | 0.338608699 | 1 | No Change | Not Significant | FALSE | NA | NA | No Change | Not Detected | FALSE | Background = FALSE | Background = FALSE |
| Pate3 | TRUE | 0.402746905 | 1 | No Change | Not Significant | FALSE | NA | NA | No Change | Not Detected | FALSE | Background = FALSE | Background = FALSE |
| Patj | TRUE | -0.260943477 | 0.000787953 | No Change | Significant Negative | TRUE | -0.036380672 | 0.578701569 | No Change | Not Significant | TRUE | Cond1 Significant Negative + Cond2 Not Significant | None |
| Patl1 | TRUE | 0.006077563 | 0.975869522 | No Change | Not Significant | TRUE | -0.125369244 | 0.408682037 | No Change | Not Significant | TRUE | Cond1 Not Significant + Cond2 Not Significant | None |

Supplementary Table 3

| geneID | Cond1_DF_Detected | Cond1_DF_ValueCol | Cond1_DF_PadjCol | Cond1_DF_Cutoff | Cond1_DF_Cutoff_Specific.x | Cond2_DF_Detected | Cond2_DF_ValueCol | Cond2_DF_PadjCol | Cond2_DF_Cutoff | Cond2_DF_Cutoff_Specific | BG_Method | RG1_All | RG2_Significant | RG3_SignificantChange |
|---|---|---|---|---|---|---|---|---|---|---|---|---|---|---|
| Pcdhga8 | TRUE | -0.487802845 | 1 | No Change | Not Significant | FALSE | NA | NA | No Change | Not Detected | FALSE | Background = FALSE | Background = FALSE | Background = FALSE |
| Pcdhga9 | TRUE | -0.105464497 | 0.859322531 | No Change | Not Significant | TRUE | 0.119570029 | 0.8642486 | No Change | Not Significant | TRUE | Cond1 Not Significant + Cond2 Not Significant | None | None |
| Pcdhgb1 | TRUE | 1.022930332 | 1 | No Change | Not Significant | FALSE | NA | NA | No Change | Not Detected | FALSE | Background = FALSE | Background = FALSE | Background = FALSE |
| Pcdhgb2 | TRUE | 0.098495431 | 0.904987729 | No Change | Not Significant | TRUE | 0.51744635 | 0.504916665 | No Change | Not Significant | TRUE | Cond1 Not Significant + Cond2 Not Significant | None | None |
| Pcdhgb4 | TRUE | -0.843404031 | 0.124359938 | No Change | Not Significant | TRUE | -0.568589803 | 0.323870851 | No Change | Not Significant | TRUE | Cond1 Not Significant + Cond2 Not Significant | None | None |
| Pcdhgb5 | TRUE | -0.273523263 | 0.619083741 | No Change | Not Significant | TRUE | 0.239627172 | 0.724201408 | No Change | Not Significant | TRUE | Cond1 Not Significant + Cond2 Not Significant | None | None |
| Pcdhgb6 | TRUE | 0.604734102 | 0.059944401 | No Change | Not Significant | TRUE | 0.143731982 | 0.679865123 | No Change | Not Significant | TRUE | Cond1 Not Significant + Cond2 Not Significant | None | None |
| Pcdhgb7 | TRUE | 0.757529337 | 0.101654269 | No Change | Not Significant | TRUE | 0.332479747 | 0.60618049 | No Change | Not Significant | TRUE | Cond1 Not Significant + Cond2 Not Significant | None | None |
| Pcdhgc3 | TRUE | 0.87752363 | 0.000184206 | UP | UP | TRUE | 0.346820662 | 0.080952136 | No Change | Not Significant | TRUE | Cond1 UP + Cond2 Not Significant | Cond1_UP | Cond1_UP |
| Pcdhgc4 | TRUE | 0.370757756 | 0.189607836 | No Change | Not Significant | TRUE | 0.150581183 | 0.432432462 | No Change | Not Significant | TRUE | Cond1 Not Significant + Cond2 Not Significant | None | None |
| Pcdhgc5 | TRUE | 0.266933356 | 0.624030976 | No Change | Not Significant | TRUE | 0.161399293 | 0.741098784 | No Change | Not Significant | TRUE | Cond1 Not Significant + Cond2 Not Significant | None | None |
| Pced1a | TRUE | -0.028835112 | 0.907710402 | No Change | Not Significant | TRUE | 0.041214715 | 0.843368224 | No Change | Not Significant | TRUE | Cond1 Not Significant + Cond2 Not Significant | None | None |

| geneID | Cond1_DF_ensembl_gene_id | Cond1_DF_entrezgene_id | Cond1_DF_gene_biotype | Cond1_DF_baseMean | Cond1_DF_log2FoldChange | Cond1_DF_lfcSE | Cond1_DF_stat | Cond1_DF_pvalue | Cond1_DF_padj | Cond1_DF_absLog2FC | Cond1_DF_Cutoff_Specific.y |
|---|---|---|---|---|---|---|---|---|---|---|---|
| Pcdhga8 | ENSMUSG00000103897 | 93716 | protein_coding | 4.337395447 | -0.487802845 | 0.552950631 | 0.882181551 | 0.377678637 | 1 | 0.487802845 | Not Significant |
| Pcdhga9 | ENSMUSG00000102440 | 93717 | protein_coding | 11.45871078 | -0.105464497 | 0.3029474 | 0.348128082 | 0.727743996 | 0.859322531 | 0.105464497 | Not Significant |
| Pcdhgb1 | ENSMUSG00000103037 | 93699 | protein_coding | 3.735897389 | 1.022930332 | 0.730380627 | -1.400544174 | 0.161350425 | 1 | 1.022930332 | Not Significant |
| Pcdhgb2 | ENSMUSG00000102748 | 93700 | protein_coding | 7.892418808 | 0.098495431 | 0.403786388 | -0.243929549 | 0.807285384 | 0.904987729 | 0.098495431 | Not Significant |
| Pcdhgb4 | ENSMUSG00000103585 | 93701 | protein_coding | 8.719215681 | -0.843404031 | 0.409226862 | 2.060969376 | 0.039305962 | 0.124359938 | 0.843404031 | Not Significant |
| Pcdhgb5 | ENSMUSG00000103749 | 93702 | protein_coding | 11.52110366 | -0.273523263 | 0.323047538 | 0.846696635 | 0.397164235 | 0.619083741 | 0.273523263 | Not Significant |
| Pcdhgb6 | ENSMUSG00000103088 | 93703 | protein_coding | 24.68370083 | 0.604734102 | 0.249831057 | -2.420572167 | 0.015496103 | 0.059944401 | 0.604734102 | Not Significant |
| Pcdhgb7 | ENSMUSG00000104063 | 93704 | protein_coding | 9.126358381 | 0.757529337 | 0.349630733 | -2.166655461 | 0.030261135 | 0.101654269 | 0.757529337 | Not Significant |
| Pcdhgc3 | ENSMUSG00000102918 | 93706 | protein_coding | 65.14849887 | 0.87752363 | 0.204399342 | -4.293182264 | 1.76E-05 | 0.000184206 | 0.87752363 | UP |
| Pcdhgc4 | ENSMUSG00000023036 | 93707 | protein_coding | 86.62277068 | 0.370757756 | 0.204137766 | -1.816213448 | 0.069337636 | 0.189607836 | 0.370757756 | Not Significant |
| Pcdhgc5 | ENSMUSG00000102543 | 93708 | protein_coding | 16.83475698 | 0.266933356 | 0.318924583 | -0.836979558 | 0.402604059 | 0.624030976 | 0.266933356 | Not Significant |
| Pced1a | ENSMUSG00000037773 | 319513 | protein_coding | 94.31679933 | -0.028835112 | 0.122571265 | 0.235251811 | 0.814013295 | 0.907710402 | 0.028835112 | Not Significant |

| geneID | Cond2_DF_ensembl_gene_id | Cond2_DF_entrezgene_id | Cond2_DF_gene_biotype | Cond2_DF_baseMean | Cond2_DF_log2FoldChange | Cond2_DF_lfcSE | Cond2_DF_stat | Cond2_DF_pvalue | Cond2_DF_padj | Cond2_DF_absLog2FC | RG1_Specific_Cond2 | RG1_Specific_Cond1 |
|---|---|---|---|---|---|---|---|---|---|---|---|---|
| Pcdhga8 | NA | NA | NA | NA | NA | NA | NA | NA | NA | NA | Background = FALSE | Background = FALSE |
| Pcdhga9 | ENSMUSG00000102440 | 93717 | protein_coding | 14.1312792 | 0.119570029 | 0.336503492 | 0.355330723 | 0.722341842 | 0.8642486 | 0.119570029 | Cond1 No Change + Cond2 Not Significant | Cond2 No Change + Cond1 Not Significant |
| Pcdhgb1 | NA | NA | NA | NA | NA | NA | NA | NA | NA | NA | Background = FALSE | Background = FALSE |
| Pcdhgb2 | ENSMUSG00000102748 | 93700 | protein_coding | 8.586033757 | 0.51744635 | 0.462707784 | 1.118300507 | 0.26343867 | 0.504916665 | 0.51744635 | Cond1 No Change + Cond2 Not Significant | Cond2 No Change + Cond1 Not Significant |
| Pcdhgb4 | ENSMUSG00000103585 | 93701 | protein_coding | 10.65943197 | -0.568589803 | 0.367787359 | -1.545974297 | 0.122110775 | 0.323870851 | 0.568589803 | Cond1 No Change + Cond2 Not Significant | Cond2 No Change + Cond1 Not Significant |
| Pcdhgb5 | ENSMUSG00000103749 | 93702 | protein_coding | 13.945775 | 0.239627172 | 0.362939422 | 0.660240133 | 0.509099742 | 0.724201408 | 0.239627172 | Cond1 No Change + Cond2 Not Significant | Cond2 No Change + Cond1 Not Significant |
| Pcdhgb6 | ENSMUSG00000103088 | 93703 | protein_coding | 38.08165121 | 0.143731982 | 0.190509921 | 0.7544593 | 0.450573475 | 0.679865123 | 0.143731982 | Cond1 No Change + Cond2 Not Significant | Cond2 No Change + Cond1 Not Significant |
| Pcdhgb7 | ENSMUSG00000104063 | 93704 | protein_coding | 10.95882673 | 0.332479747 | 0.366175132 | 0.907980137 | 0.363888716 | 0.60618049 | 0.332479747 | Cond1 No Change + Cond2 Not Significant | Cond2 No Change + Cond1 Not Significant |
| Pcdhgc3 | ENSMUSG00000102918 | 93706 | protein_coding | 98.3281417 | 0.346820662 | 0.142887428 | 2.427230061 | 0.015214604 | 0.080952136 | 0.346820662 | Cond1 UP + Cond2 Not Significant | Cond2 No Change + Cond1 UP |
| Pcdhgc4 | ENSMUSG00000023036 | 93707 | protein_coding | 178.8343881 | 0.150581183 | 0.117269903 | 1.284056508 | 0.199122184 | 0.432432462 | 0.150581183 | Cond1 No Change + Cond2 Not Significant | Cond2 No Change + Cond1 Not Significant |
| Pcdhgc5 | ENSMUSG00000102543 | 93708 | protein_coding | 23.97316375 | 0.161399293 | 0.258871195 | 0.623473359 | 0.532973504 | 0.741098784 | 0.161399293 | Cond1 No Change + Cond2 Not Significant | Cond2 No Change + Cond1 Not Significant |
| Pced1a | ENSMUSG00000037773 | 319513 | protein_coding | 176.9160356 | 0.041214715 | 0.102741867 | 0.4011482 | 0.688311015 | 0.843368224 | 0.041214715 | Cond1 No Change + Cond2 Not Significant | Cond2 No Change + Cond1 Not Significant |

| geneID | Cond1_DF_Detected | Cond1_DF_ValueCol | Cond1_DF_PadjCol | Cond1_DF_Cutoff | Cond1_DF_Cutoff_Specific.x | Cond2_DF_Detected | Cond2_DF_ValueCol | Cond2_DF_PadjCol | Cond2_DF_Cutoff | Cond2_DF_Cutoff_Specific |
|---|---|---|---|---|---|---|---|---|---|---|
| Pf4 | TRUE | 2.726948516 | 9.32E-10 | UP | UP | TRUE | 0.884968097 | 0.151794249 | No Change | Not Significant |
| Pfas | TRUE | -0.123984889 | 0.507895701 | No Change | Not Significant | TRUE | -0.03179887 | 0.829048013 | No Change | Not Significant |
| Pfdn1 | TRUE | 0.158037351 | 0.07322684 | No Change | Not Significant | TRUE | 0.118257811 | 0.17246929 | No Change | Not Significant |
| Pfdn2 | TRUE | -0.038542505 | 0.830017162 | No Change | Not Significant | TRUE | 0.168114713 | 0.221242374 | No Change | Not Significant |
| Pfdn4 | TRUE | 0.10610949 | 0.612056856 | No Change | Not Significant | TRUE | 0.13583288 | 0.479144737 | No Change | Not Significant |
| Pfdn5 | TRUE | -0.280222629 | 0.000552495 | No Change | Significant Negative | TRUE | -0.05277769 | 0.484078966 | No Change | Not Significant |
| Pfdn6 | TRUE | -0.039079299 | 0.770846581 | No Change | Not Significant | TRUE | 0.085393164 | 0.551334926 | No Change | Not Significant |
| Pfkfb1 | TRUE | 0.513141449 | 0.187297899 | No Change | Not Significant | TRUE | -0.251345611 | 0.525297065 | No Change | Not Significant |
| Pfkfb2 | TRUE | 0.489187412 | 1.79E-14 | No Change | Significant Positive | TRUE | 0.073648312 | 0.531357026 | No Change | Not Significant |
| Pfkfb3 | TRUE | 0.306242944 | 0.344460843 | No Change | Not Significant | TRUE | 0.455374646 | 0.10953031 | No Change | Not Significant |
| Pfkfb4 | TRUE | -0.308385675 | 0.09253023 | No Change | Not Significant | TRUE | -0.075185709 | 0.562649386 | No Change | Not Significant |
| Pfkl | TRUE | -0.243780343 | 0.052535985 | No Change | Not Significant | TRUE | 0.139644106 | 0.161750767 | No Change | Not Significant |
| Pfkm | TRUE | -0.471978446 | 0.001847862 | No Change | Significant Negative | TRUE | -0.169343207 | 0.167727999 | No Change | Not Significant |
| Pfkp | TRUE | -0.113283151 | 0.298087027 | No Change | Not Significant | TRUE | -0.093020613 | 0.484226665 | No Change | Not Significant |
| Pfn1 | TRUE | -0.084922422 | 0.565170522 | No Change | Not Significant | TRUE | 0.079339244 | 0.629397025 | No Change | Not Significant |
| Pfn2 | TRUE | 0.169057923 | 0.788881899 | No Change | Not Significant | TRUE | 0.019695808 | 0.97297734 | No Change | Not Significant |
| Pfn4 | TRUE | 0.150221459 | 1 | No Change | Not Significant | FALSE | NA | NA | No Change | Not Detected |
| Pga5 | TRUE | 0.753082542 | 1 | No Change | Not Significant | FALSE | NA | NA | No Change | Not Detected |
| Pgam1 | TRUE | -0.663293676 | 0.000245359 | DOWN | DOWN | TRUE | 0.171294486 | 0.180143349 | No Change | Not Significant |
| Pgam2 | TRUE | -0.224779337 | 0.669875536 | No Change | Not Significant | TRUE | -0.031891519 | 0.955322545 | No Change | Not Significant |
| Pgam5 | TRUE | 0.09032613 | 0.615578778 | No Change | Not Significant | TRUE | 0.08685203 | 0.474717773 | No Change | Not Significant |
| Pgap1 | TRUE | 0.148131484 | 0.169479714 | No Change | Not Significant | TRUE | 0.386157013 | 9.40E-10 | No Change | Significant Positive |
| Pgap2 | TRUE | -0.089074036 | 0.665664739 | No Change | Not Significant | TRUE | 0.072743277 | 0.425338507 | No Change | Not Significant |
| Pgap3 | TRUE | 0.374125464 | 0.000440521 | No Change | Significant Positive | TRUE | 0.230856425 | 0.006963823 | No Change | Significant Positive |
| Pgap4 | TRUE | -0.218721751 | 0.103308737 | No Change | Not Significant | TRUE | -0.008333452 | 0.950733597 | No Change | Not Significant |
| Pgap6 | TRUE | -0.166614852 | 0.023369313 | No Change | Significant Negative | TRUE | -0.043737078 | 0.656573069 | No Change | Not Significant |
| Pgbd1 | TRUE | 0.097408769 | 0.878912418 | No Change | Not Significant | TRUE | 0.062320845 | 0.915984616 | No Change | Not Significant |
| Pgbd5 | TRUE | 2.230887137 | 1 | No Change | Not Significant | FALSE | NA | NA | No Change | Not Detected |
| Pgc | TRUE | 0.022594638 | 1 | No Change | Not Significant | FALSE | NA | NA | No Change | Not Detected |
| Pgd | TRUE | 0.076550089 | 0.559438733 | No Change | Not Significant | TRUE | 0.147092235 | 0.158042883 | No Change | Not Significant |
| Pgf | TRUE | -0.187806878 | 1 | No Change | Not Significant | FALSE | NA | NA | No Change | Not Detected |
| Pgghg | TRUE | -0.05711895 | 0.716335759 | No Change | Not Significant | TRUE | -0.014821275 | 0.896141595 | No Change | Not Significant |
| Pggt1b | TRUE | -0.047554787 | 0.793440482 | No Change | Not Significant | TRUE | 0.058155367 | 0.593363209 | No Change | Not Significant |
| Pgk1 | TRUE | 0.65590269 | 8.36E-05 | UP | UP | TRUE | 0.61514463 | 2.31E-14 | UP | UP |
| Pgls | TRUE | 0.02827082 | 0.860514985 | No Change | Not Significant | TRUE | 0.095673649 | 0.407987666 | No Change | Not Significant |
| Pglyrp1 | TRUE | -0.181753777 | 0.094754639 | No Change | Not Significant | TRUE | 0.126695621 | 0.358198217 | No Change | Not Significant |
| Pglyrp2 | TRUE | 1.54295114 | 0.152928125 | No Change | Not Significant | TRUE | 0.09586742 | 0.916408517 | No Change | Not Significant |
| Pgm1 | TRUE | 0.459862032 | 0.000376608 | No Change | Significant Positive | TRUE | 0.321067501 | 7.14E-14 | No Change | Significant Positive |
| Pgm2 | TRUE | -0.40619376 | 9.01E-11 | No Change | Significant Negative | TRUE | -0.097650068 | 0.130971062 | No Change | Not Significant |
| Pgm2l1 | TRUE | -0.159203345 | 0.449052273 | No Change | Not Significant | TRUE | -0.032882959 | 0.822565162 | No Change | Not Significant |
| Pgm3 | TRUE | 0.193415723 | 0.216206632 | No Change | Not Significant | TRUE | 0.228396137 | 0.016796393 | No Change | Significant Positive |
| Pgm5 | TRUE | 1.830298862 | 0.057707786 | No Change | Not Significant | FALSE | NA | NA | No Change | Not Detected |
| Pgp | TRUE | 0.709649617 | 1.61E-15 | UP | UP | TRUE | 0.346778099 | 0.000682262 | No Change | Significant Positive |
| Pgpep1 | TRUE | -0.061835205 | 0.698491194 | No Change | Not Significant | TRUE | 0.083257765 | 0.264480943 | No Change | Not Significant |
| Pgpep1l | TRUE | -1.948582462 | 1 | No Change | Not Significant | FALSE | NA | NA | No Change | Not Detected |
| Pgr | TRUE | -1.514627453 | 1 | No Change | Not Significant | FALSE | NA | NA | No Change | Not Detected |
| Pgr15l | TRUE | 0.523792341 | 1 | No Change | Not Significant | FALSE | NA | NA | No Change | Not Detected |
| Pgrmc1 | TRUE | -0.333444916 | 1.17E-05 | No Change | Significant Negative | TRUE | -0.045199207 | 0.47743605 | No Change | Not Significant |
| Pgrmc2 | TRUE | 0.105258194 | 0.3984875 | No Change | Not Significant | TRUE | 0.085438517 | 0.523508209 | No Change | Not Significant |
| Pgs1 | TRUE | 0.320663934 | 0.004121541 | No Change | Significant Positive | TRUE | 0.079120533 | 0.423671625 | No Change | Not Significant |
| Phactr1 | TRUE | -0.055291844 | 0.988831409 | No Change | Not Significant | FALSE | NA | NA | No Change | Not Detected |
| Phactr2 | TRUE | -0.710429049 | 1.81E-08 | DOWN | DOWN | TRUE | -0.205304858 | 0.058099322 | No Change | Not Significant |
| Phactr3 | TRUE | -1.76958553 | 1 | No Change | Not Significant | FALSE | NA | NA | No Change | Not Detected |
| Phactr4 | TRUE | -0.137401817 | 0.499023496 | No Change | Not Significant | TRUE | -0.081757786 | 0.362726949 | No Change | Not Significant |
| Phaf1 | TRUE | -0.19396128 | 0.134811958 | No Change | Not Significant | TRUE | -0.147597681 | 0.164406979 | No Change | Not Significant |
| Phax | TRUE | -0.064596657 | 0.723107237 | No Change | Not Significant | TRUE | -0.083062282 | 0.389376941 | No Change | Not Significant |
| Phb | TRUE | -0.434729246 | 0.001648655 | No Change | Significant Negative | TRUE | 0.10804158 | 0.327433832 | No Change | Not Significant |
| Phb2 | TRUE | -0.078879829 | 0.436477048 | No Change | Not Significant | TRUE | 0.027499434 | 0.784827956 | No Change | Not Significant |
| Phc1 | TRUE | -0.129880725 | 0.548167261 | No Change | Not Significant | TRUE | -0.073842598 | 0.62920611 | No Change | Not Significant |
| Phc2 | TRUE | -0.079952463 | 0.302752759 | No Change | Not Significant | TRUE | -0.003732978 | 0.981205596 | No Change | Not Significant |
| Phc3 | TRUE | -0.063047429 | 0.714956306 | No Change | Not Significant | TRUE | -0.156497392 | 0.097944477 | No Change | Not Significant |
| Pheta1 | TRUE | 0.098348062 | 0.570284968 | No Change | Not Significant | TRUE | 0.108731556 | 0.293978751 | No Change | Not Significant |
| Pheta2 | TRUE | 0.238236169 | 0.733766095 | No Change | Not Significant | TRUE | 0.012385508 | 0.988537404 | No Change | Not Significant |
| Phex | TRUE | 0.617332913 | 1 | No Change | Not Significant | FALSE | NA | NA | No Change | Not Detected |
| Phf1 | TRUE | 0.072618682 | 0.816494106 | No Change | Not Significant | TRUE | -0.138534165 | 0.475463004 | No Change | Not Significant |
| Phf10 | TRUE | 0.251949314 | 0.012715494 | No Change | Significant Positive | TRUE | 0.067011707 | 0.536379174 | No Change | Not Significant |
| Phf11a | TRUE | 0.200356946 | 1 | No Change | Not Significant | FALSE | NA | NA | No Change | Not Detected |
| Phf11b | TRUE | -0.001382427 | 0.998067952 | No Change | Not Significant | TRUE | -0.335638839 | 0.383043628 | No Change | Not Significant |
| Phf11c | TRUE | 0.36014316 | 0.431271309 | No Change | Not Significant | TRUE | -0.096538371 | 0.872904276 | No Change | Not Significant |
| Phf11d | TRUE | -0.54383583 | 0.151449101 | No Change | Not Significant | TRUE | -0.151137568 | 0.623228806 | No Change | Not Significant |
| Phf12 | TRUE | 0.118560042 | 0.508545065 | No Change | Not Significant | TRUE | -0.128645709 | 0.176277123 | No Change | Not Significant |
| Phf13 | TRUE | 0.758029762 | 2.35E-09 | UP | UP | TRUE | 0.497837011 | 6.91E-06 | No Change | Significant Positive |
| Phf14 | TRUE | -0.15919458 | 0.230377922 | No Change | Not Significant | TRUE | -0.159668441 | 0.069915764 | No Change | Not Significant |
| Phf19 | TRUE | -0.233734691 | 0.430945001 | No Change | Not Significant | TRUE | -0.269218961 | 0.179979523 | No Change | Not Significant |
| Phf2 | TRUE | 0.192078361 | 0.051106386 | No Change | Not Significant | TRUE | -0.067574252 | 0.586511041 | No Change | Not Significant |
| Phf20 | TRUE | 0.506004464 | 0.000732396 | UP | UP | TRUE | 0.064405418 | 0.654207378 | No Change | Not Significant |
| Phf20l1 | TRUE | 0.075029504 | 0.447819239 | No Change | Not Significant | TRUE | -0.084927614 | 0.262563114 | No Change | Not Significant |
| Phf21a | TRUE | 0.009378697 | 0.971883559 | No Change | Not Significant | TRUE | 0.02812946 | 0.861553924 | No Change | Not Significant |
| Phf21b | TRUE | 0.067469343 | 0.809667817 | No Change | Not Significant | TRUE | -0.192167563 | 0.357032789 | No Change | Not Significant |
| Phf23 | TRUE | -0.030370139 | 0.872440598 | No Change | Not Significant | TRUE | -0.042820112 | 0.759659123 | No Change | Not Significant |
| Phf24 | TRUE | 0.555593914 | 1 | No Change | Not Significant | FALSE | NA | NA | No Change | Not Detected |
| Phf3 | TRUE | -0.01105393 | 0.931471303 | No Change | Not Significant | TRUE | -0.043596527 | 0.57990695 | No Change | Not Significant |
| Phf5a | TRUE | 0.087844644 | 0.637944042 | No Change | Not Significant | TRUE | 0.126764357 | 0.244589225 | No Change | Not Significant |
| Phf6 | TRUE | 0.36911082 | 0.030425148 | No Change | Significant Positive | TRUE | 0.145456067 | 0.121104897 | No Change | Not Significant |
| Phf7 | TRUE | -0.080504081 | 0.698215786 | No Change | Not Significant | TRUE | -0.211874019 | 0.119191767 | No Change | Not Significant |
| Phf8 | TRUE | 0.368073409 | 4.67E-07 | No Change | Significant Positive | TRUE | 0.244243746 | 0.000276807 | No Change | Significant Positive |
| Phgdh | TRUE | 2.231199673 | 2.84E-29 | UP | UP | TRUE | 1.119851407 | 4.39E-13 | UP | UP |
| Phgr1 | TRUE | -0.894531734 | 3.06E-13 | DOWN | DOWN | TRUE | -0.099696897 | 0.60759958 | No Change | Not Significant |
| Phip | TRUE | 0.117166738 | 0.349519674 | No Change | Not Significant | TRUE | -0.101663901 | 0.246739036 | No Change | Not Significant |
| Phka1 | TRUE | 0.128682725 | 0.319894352 | No Change | Not Significant | TRUE | 0.161162279 | 0.066299536 | No Change | Not Significant |
| Phka2 | TRUE | 0.191468665 | 0.204907671 | No Change | Not Significant | TRUE | -0.08978872 | 0.407666398 | No Change | Not Significant |
| Phkb | TRUE | -0.12909908 | 0.419775423 | No Change | Not Significant | TRUE | 0.078489936 | 0.309749988 | No Change | Not Significant |
| Phkg1 | TRUE | 0.799155089 | 0.667103975 | No Change | Not Significant | FALSE | NA | NA | No Change | Not Detected |
| Phkg2 | TRUE | 1.689228497 | 2.41E-67 | UP | UP | TRUE | 1.002930932 | 3.57E-33 | UP | UP |
| Phlda1 | TRUE | 0.347692886 | 0.183317181 | No Change | Not Significant | TRUE | -0.01167935 | 0.962443303 | No Change | Not Significant |
| Phlda2 | TRUE | -0.611364172 | 0.087305635 | No Change | Not Significant | TRUE | 0.050452633 | 0.936965584 | No Change | Not Significant |
| Phlda3 | TRUE | 0.989443335 | 0.00087081 | UP | UP | TRUE | 0.783098861 | 0.032617086 | UP | UP |
| Phldb1 | TRUE | -0.399220091 | 9.17E-06 | No Change | Significant Negative | TRUE | -0.115397769 | 0.290740541 | No Change | Not Significant |
| Phldb2 | TRUE | 0.040635919 | 0.870461483 | No Change | Not Significant | TRUE | -0.189899734 | 0.249990933 | No Change | Not Significant |
| Phldb3 | TRUE | -0.33203871 | 0.205917797 | No Change | Not Significant | TRUE | -0.143144196 | 0.415618234 | No Change | Not Significant |
| Phlpp1 | TRUE | -0.20448194 | 0.036230249 | No Change | Significant Negative | TRUE | -0.126799515 | 0.226923769 | No Change | Not Significant |
| Phlpp2 | TRUE | -0.355212281 | 0.130053252 | No Change | Not Significant | TRUE | -0.223790433 | 0.16134834 | No Change | Not Significant |
| Phospho1 | TRUE | 0.341914631 | 0.658275506 | No Change | Not Significant | TRUE | -0.395455969 | 0.345248914 | No Change | Not Significant |
| Phospho2 | TRUE | 0.004337741 | 0.984301824 | No Change | Not Significant | TRUE | 0.078904396 | 0.475376011 | No Change | Not Significant |
| Phox2a | TRUE | 0.616759946 | 1 | No Change | Not Significant | FALSE | NA | NA | No Change | Not Detected |
| Phox2b | TRUE | 1.649931959 | 1 | No Change | Not Significant | FALSE | NA | NA | No Change | Not Detected |
| Phpt1 | TRUE | 0.105522907 | 0.833389267 | No Change | Not Significant | TRUE | 0.226817993 | 0.528797792 | No Change | Not Significant |
| Phrf1 | TRUE | 0.153264582 | 0.243108963 | No Change | Not Significant | TRUE | 0.021965877 | 0.855350218 | No Change | Not Significant |
| Phtf1 | TRUE | 0.045875341 | 0.838607402 | No Change | Not Significant | TRUE | 0.078592047 | 0.519508601 | No Change | Not Significant |
| Phtf1os | TRUE | -0.066503815 | 1 | No Change | Not Significant | FALSE | NA | NA | No Change | Not Detected |
| Phtf2 | TRUE | 0.00182588 | 0.995401911 | No Change | Not Significant | TRUE | -0.134189408 | 0.589019803 | No Change | Not Significant |
| Phyh | TRUE | 0.566557303 | 1.69E-18 | UP | UP | TRUE | 0.445988645 | 2.58E-17 | No Change | Significant Positive |
| Phyhd1 | TRUE | 1.027627162 | 1 | No Change | Not Significant | FALSE | NA | NA | No Change | Not Detected |
| Phyhip | TRUE | 2.952805668 | 1 | No Change | Not Significant | FALSE | NA | NA | No Change | Not Detected |
| Phyhipl | TRUE | 0.62701474 | 1 | No Change | Not Significant | FALSE | NA | NA | No Change | Not Detected |
| Phykpl | TRUE | -0.00414275 | 0.984664559 | No Change | Not Significant | TRUE | -0.277502881 | 0.110140691 | No Change | Not Significant |
| Pi15 | TRUE | -0.428599756 | 1 | No Change | Not Significant | FALSE | NA | NA | No Change | Not Detected |
| Pi16 | TRUE | 1.007885888 | 1 | No Change | Not Significant | FALSE | NA | NA | No Change | Not Detected |
| Pi4k2a | TRUE | 0.58943222 | 8.28E-13 | UP | UP | TRUE | 0.255383592 | 0.003779053 | No Change | Significant Positive |
| Pi4k2b | TRUE | 0.263014818 | 0.000494524 | No Change | Significant Positive | TRUE | 0.08685914 | 0.067977533 | No Change | Not Significant |
| Pi4ka | TRUE | -0.16016862 | 0.068544401 | No Change | Not Significant | TRUE | -0.016985043 | 0.891725514 | No Change | Not Significant |
| Pi4kb | TRUE | -0.033993763 | 0.691324212 | No Change | Not Significant | TRUE | 0.034005003 | 0.726301064 | No Change | Not Significant |
| Pianp | TRUE | -0.490419903 | 1 | No Change | Not Significant | FALSE | NA | NA | No Change | Not Detected |
| Pias1 | TRUE | -0.109526903 | 0.294356962 | No Change | Not Significant | TRUE | -0.1516466 | 0.063224495 | No Change | Not Significant |
| Pias2 | TRUE | 0.043219029 | 0.782626881 | No Change | Not Significant | TRUE | -0.05733236 | 0.480533622 | No Change | Not Significant |
| Pias3 | TRUE | 0.004481135 | 0.986657316 | No Change | Not Significant | TRUE | -0.095279649 | 0.513977697 | No Change | Not Significant |
| Pias4 | TRUE | 0.026680563 | 0.859565127 | No Change | Not Significant | TRUE | -0.018232283 | 0.926650089 | No Change | Not Significant |
| Pibf1 | TRUE | 1.380945398 | 5.85E-69 | UP | UP | TRUE | 0.889712186 | 9.72E-20 | UP | UP |
| Picalm | TRUE | 0.053500216 | 0.43083733 | No Change | Not Significant | TRUE | -0.083926527 | 0.02955516 | No Change | Significant Negative |
| Pick1 | TRUE | 0.152727987 | 0.214305112 | No Change | Not Significant | TRUE | 0.147422677 | 0.275969898 | No Change | Not Significant |
| Pid1 | TRUE | 1.055212606 | 1 | No Change | Not Significant | FALSE | NA | NA | No Change | Not Detected |
| Pidd1 | TRUE | 0.276041931 | 0.267745952 | No Change | Not Significant | TRUE | 0.002386633 | 0.995550783 | No Change | Not Significant |
| Pierce1 | TRUE | 0.260141527 | 1 | No Change | Not Significant | FALSE | NA | NA | No Change | Not Detected |
| Pierce2 | TRUE | -0.134128737 | 0.612460672 | No Change | Not Significant | TRUE | 0.18640601 | 0.264317294 | No Change | Not Significant |
| Piezo1 | TRUE | -0.151574627 | 0.310179152 | No Change | Not Significant | TRUE | -0.161542683 | 0.32338554 | No Change | Not Significant |
| Piezo2 | TRUE | -0.583706057 | 0.332794609 | No Change | Not Significant | TRUE | -0.000295643 | 0.999195317 | No Change | Not Significant |
| Pif1 | TRUE | 0.852496082 | 2.97E-05 | UP | UP | TRUE | 0.062044851 | 0.836508911 | No Change | Not Significant |
| Piga | TRUE | 1.066284886 | 9.68E-35 | UP | UP | TRUE | 0.947589259 | 1.36E-32 | UP | UP |
| Pigb | TRUE | 0.149014322 | 0.462014445 | No Change | Not Significant | TRUE | 0.159332553 | 0.275588846 | No Change | Not Significant |
| Pigbos1 | TRUE | -0.105426074 | 0.632229811 | No Change | Not Significant | TRUE | 0.277572218 | 0.035899485 | No Change | Significant Positive |
| Pigc | TRUE | 0.178061439 | 0.224271688 | No Change | Not Significant | TRUE | 0.071185865 | 0.550544997 | No Change | Not Significant |
| Pigf | TRUE | -0.223394815 | 0.095391264 | No Change | Not Significant | TRUE | -0.10431209 | 0.38635024 | No Change | Not Significant |
| Pigg | TRUE | 0.026988559 | 0.877889692 | No Change | Not Significant | TRUE | -0.025741606 | 0.795599668 | No Change | Not Significant |
| Pigh | TRUE | -0.094360595 | 0.62019321 | No Change | Not Significant | TRUE | 0.148168817 | 0.336485813 | No Change | Not Significant |
| Pigk | TRUE | -0.058943045 | 0.720614094 | No Change | Not Significant | TRUE | 0.059770579 | 0.45050036 | No Change | Not Significant |
| Pigl | TRUE | -0.043643278 | 0.869560398 | No Change | Not Significant | TRUE | 0.104736302 | 0.592368513 | No Change | Not Significant |
| Pigm | TRUE | -0.039209929 | 0.794593745 | No Change | Not Significant | TRUE | -0.107139413 | 0.232615531 | No Change | Not Significant |
| Pign | TRUE | -0.054084415 | 0.676807614 | No Change | Not Significant | TRUE | -0.157381287 | 0.056752643 | No Change | Not Significant |
| Pigo | TRUE | 0.083103799 | 0.519676664 | No Change | Not Significant | TRUE | 0.115777294 | 0.366304868 | No Change | Not Significant |
| Pigp | TRUE | -0.333577769 | 0.00381346 | No Change | Significant Negative | TRUE | -0.197947801 | 0.118917792 | No Change | Not Significant |
| Pigq | TRUE | -0.085411872 | 0.356391801 | No Change | Not Significant | TRUE | 0.086035987 | 0.348184778 | No Change | Not Significant |
| Pigr | TRUE | -0.015534351 | 0.947202684 | No Change | Not Significant | TRUE | 0.108490865 | 0.197299838 | No Change | Not Significant |
| Pigs | TRUE | -0.176091603 | 0.103865546 | No Change | Not Significant | TRUE | -0.022662051 | 0.889251381 | No Change | Not Significant |
| Pigt | TRUE | 0.269246706 | 0.003030927 | No Change | Significant Positive | TRUE | 0.15895715 | 0.244881547 | No Change | Not Significant |
| Pigu | TRUE | 0.193375873 | 0.077538047 | No Change | Not Significant | TRUE | 0.047535282 | 0.659792516 | No Change | Not Significant |
| Pigv | TRUE | 0.057334736 | 0.676807614 | No Change | Not Significant | TRUE | 0.01390561 | 0.945572603 | No Change | Not Significant |
| Pigw | TRUE | 0.06839503 | 0.772538316 | No Change | Not Significant | TRUE | -0.032256007 | 0.910022729 | No Change | Not Significant |
| Pigx | TRUE | -0.149408463 | 0.136567098 | No Change | Not Significant | TRUE | -0.169031907 | 0.076012928 | No Change | Not Significant |
| Pigyl | TRUE | -0.083969951 | 0.727088376 | No Change | Not Significant | TRUE | 0.085685169 | 0.619192021 | No Change | Not Significant |
| Pigz | TRUE | -0.118132246 | 1 | No Change | Not Significant | FALSE | NA | NA | No Change | Not Detected |
| Pih1d1 | TRUE | 0.062607605 | 0.669122175 | No Change | Not Significant | TRUE | 0.129221819 | 0.31680248 | No Change | Not Significant |
| Pih1d2 | TRUE | 0.581261847 | 0.110624266 | No Change | Not Significant | TRUE | 0.615842221 | 0.057416794 | No Change | Not Significant |
| Pik3ap1 | TRUE | 3.609080189 | 2.12E-36 | UP | UP | TRUE | 0.760186359 | 0.006150272 | UP | UP |
| Pik3c2a | TRUE | 0.050436341 | 0.62645534 | No Change | Not Significant | TRUE | 0.049376505 | 0.527543906 | No Change | Not Significant |
| Pik3c2b | TRUE | -0.184539354 | 0.255480702 | No Change | Not Significant | TRUE | -0.290483197 | 3.88E-05 | No Change | Significant Negative |
| Pik3c2g | TRUE | -0.106086921 | 0.859322531 | No Change | Not Significant | TRUE | -0.230834815 | 0.665561246 | No Change | Not Significant |
| Pik3c3 | TRUE | -0.233160622 | 0.126157266 | No Change | Not Significant | TRUE | 0.00020322 | 0.998627072 | No Change | Not Significant |
| Pik3ca | TRUE | -0.072370836 | 0.671357918 | No Change | Not Significant | TRUE | -0.153157117 | 0.120633435 | No Change | Not Significant |
| Pik3cb | TRUE | 0.06054469 | 0.712245263 | No Change | Not Significant | TRUE | -0.002771492 | 0.988305042 | No Change | Not Significant |
| Pik3cd | TRUE | 0.042668937 | 0.951530343 | No Change | Not Significant | TRUE | 0.21139323 | 0.470848351 | No Change | Not Significant |
| Pik3cg | TRUE | -0.562926674 | 0.003837834 | DOWN | DOWN | TRUE | 0.313705962 | 0.0020054 | No Change | Significant Positive |
| Pik3ip1 | TRUE | -0.309987369 | 0.350225257 | No Change | Not Significant | TRUE | -0.579072268 | 0.003093734 | DOWN | DOWN |
| Pik3r1 | TRUE | -0.184474289 | 0.010614744 | No Change | Significant Negative | TRUE | -0.168529368 | 0.002731844 | No Change | Significant Negative |
| Pik3r2 | TRUE | 0.180341867 | 0.10878414 | No Change | Not Significant | TRUE | 0.04824657 | 0.708341774 | No Change | Not Significant |
| Pik3r3 | TRUE | 0.333096947 | 0.034782786 | No Change | Significant Positive | TRUE | 0.182158187 | 0.141755849 | No Change | Not Significant |
| Pik3r4 | TRUE | -0.07038589 | 0.624624492 | No Change | Not Significant | TRUE | 0.02051843 | 0.846659581 | No Change | Not Significant |
| Pik3r5 | TRUE | -0.856503145 | 0.000338705 | DOWN | DOWN | TRUE | -0.115731335 | 0.640407651 | No Change | Not Significant |
| Pik3r6 | TRUE | 1.038782641 | 1 | No Change | Not Significant | FALSE | NA | NA | No Change | Not Detected |
| Pikfyve | TRUE | -0.111374776 | 0.131176532 | No Change | Not Significant | TRUE | -0.082730682 | 0.286931191 | No Change | Not Significant |
| Pilra | TRUE | -0.557049385 | 1 | No Change | Not Significant | FALSE | NA | NA | No Change | Not Detected |
| Pilrb2 | TRUE | 1.638416066 | 1 | No Change | Not Significant | FALSE | NA | NA | No Change | Not Detected |
| Pim1 | TRUE | -0.051136138 | 0.874044129 | No Change | Not Significant | TRUE | -0.146336835 | 0.337846137 | No Change | Not Significant |
| Pim2 | TRUE | 3.051110609 | 5.95E-77 | UP | UP | TRUE | 1.911631424 | 1.62E-56 | UP | UP |
| Pim3 | TRUE | -0.445952152 | 0.004413704 | No Change | Significant Negative | TRUE | -0.521636408 | 0.002870449 | DOWN | DOWN |
| Pimreg | TRUE | 1.043536489 | 3.09E-05 | UP | UP | TRUE | 0.040044245 | 0.900451559 | No Change | Not Significant |
| Pin1 | TRUE | 1.655655389 | 8.27E-59 | UP | UP | TRUE | 0.810346773 | 1.01E-24 | UP | UP |
| Pin4 | TRUE | -0.607398007 | 0.006174237 | DOWN | DOWN | TRUE | 0.077782084 | 0.66096665 | No Change | Not Significant |
| Pink1 | TRUE | 0.369800761 | 0.00150081 | No Change | Significant Positive | TRUE | 0.207441369 | 0.012300224 | No Change | Significant Positive |
| Pinx1 | TRUE | 0.018871174 | 0.961855464 | No Change | Not Significant | TRUE | -0.031153389 | 0.916250984 | No Change | Not Significant |
| Pip4k2a | TRUE | 0.339574524 | 0.438341251 | No Change | Not Significant | TRUE | 0.394467015 | 0.240247585 | No Change | Not Significant |
| Pip4k2b | TRUE | 0.123300656 | 0.498558829 | No Change | Not Significant | TRUE | 0.040421614 | 0.81663734 | No Change | Not Significant |
| Pip4k2c | TRUE | -0.263571795 | 0.00025468 | No Change | Significant Negative | TRUE | 0.10872883 | 0.195649377 | No Change | Not Significant |

| geneID | Cond1_DF_Detected | Cond1_DF_ValueCol | Cond1_DF_PadjCol | Cond1_DF_Cutoff | Cond1_DF_Cutoff_Specific.x | Cond2_DF_Detected | Cond2_DF_ValueCol | Cond2_DF_PadjCol | Cond2_DF_Cutoff |
|---|---|---|---|---|---|---|---|---|---|
| Pip4p1 | TRUE | 0.004871196 | 0.97354273 | No Change | Not Significant | TRUE | -0.167583844 | 0.03065349 | No Change |
| Pip4p2 | TRUE | 0.006583543 | 0.971883559 | No Change | Not Significant | TRUE | 0.171227782 | 0.106296214 | No Change |
| Pip5k1a | TRUE | -0.137156237 | 0.226867192 | No Change | Not Significant | TRUE | -0.035300327 | 0.801920996 | No Change |
| Pip5k1b | TRUE | -0.034437815 | 0.784117555 | No Change | Not Significant | TRUE | -0.046175052 | 0.456057983 | No Change |
| Pip5k1c | TRUE | 0.220828368 | 0.00246612 | No Change | Significant Positive | TRUE | -0.068649223 | 0.402176854 | No Change |
| Pip5kl1 | TRUE | 0.332773483 | 1 | No Change | Not Significant | FALSE | NA | NA | No Change |
| Pipox | TRUE | 0.439060457 | 0.065598201 | No Change | Not Significant | TRUE | -0.814172605 | 2.23E-05 | DOWN |
| Pir | TRUE | 0.021854234 | 0.944306685 | No Change | Not Significant | TRUE | -0.249569715 | 0.212479973 | No Change |
| Pirb | TRUE | 2.104256686 | 1 | No Change | Not Significant | FALSE | NA | NA | No Change |
| Pirt | TRUE | -0.230798898 | 1 | No Change | Not Significant | FALSE | NA | NA | No Change |
| Pisd | TRUE | -0.091793861 | 0.512561987 | No Change | Not Significant | TRUE | 0.085310455 | 0.364802272 | No Change |
| Pithd1 | TRUE | 0.840196534 | 6.25E-16 | UP | UP | TRUE | 0.023500805 | 0.825416419 | No Change |
| Pitpna | TRUE | 0.028121098 | 0.780796859 | No Change | Not Significant | TRUE | 0.018161467 | 0.826041967 | No Change |
| Pitpnb | TRUE | 0.347152647 | 0.000475421 | No Change | Significant Positive | TRUE | 0.060978381 | 0.66862102 | No Change |
| Pitpnc1 | TRUE | -0.229870922 | 0.540266681 | No Change | Not Significant | TRUE | -0.070693995 | 0.841836579 | No Change |
| Pitpnm1 | TRUE | -0.13313691 | 0.672408259 | No Change | Not Significant | TRUE | -0.149473915 | 0.26456259 | No Change |
| Pitpnm2 | TRUE | -0.124438768 | 0.817662297 | No Change | Not Significant | TRUE | -0.452530303 | 0.019840672 | No Change |
| Pitpnm2os1 | TRUE | -0.466445507 | 1 | No Change | Not Significant | FALSE | NA | NA | No Change |
| Pitpnm2os2 | TRUE | 0.23893431 | 1 | No Change | Not Significant | FALSE | NA | NA | No Change |
| Pitpnm3 | TRUE | -0.025625957 | 0.931474719 | No Change | Not Significant | TRUE | -0.142998539 | 0.273529895 | No Change |
| Pitrm1 | TRUE | 0.332451084 | 0.001290307 | No Change | Significant Positive | TRUE | 0.14018935 | 0.116636001 | No Change |
| Pitx1 | TRUE | 0.233122813 | 1 | No Change | Not Significant | TRUE | 0.249620452 | 0.646554316 | No Change |
| Pitx2 | TRUE | 1.35931468 | 0.296850229 | No Change | Not Significant | TRUE | 0.345118499 | 0.232315871 | No Change |
| Piwil1 | TRUE | 2.589536514 | 1 | No Change | Not Significant | FALSE | NA | NA | No Change |
| Piwil2 | TRUE | 0.122973564 | 0.893150482 | No Change | Not Significant | TRUE | -0.243102743 | 0.75632376 | No Change |
| Piwil4 | TRUE | 0.780901194 | 0.000237545 | UP | UP | TRUE | 0.084897729 | 0.800805946 | No Change |
| Pja1 | TRUE | -0.070252011 | 0.583920353 | No Change | Not Significant | TRUE | 0.143065569 | 0.008056179 | No Change |
| Pja2 | TRUE | 1.094724871 | 1.69E-34 | UP | UP | TRUE | 0.598622151 | 1.31E-24 | UP |
| Pjvk | TRUE | -0.080825237 | 1 | No Change | Not Significant | FALSE | NA | NA | No Change |
| Pkd1 | TRUE | -0.031226144 | 0.877843331 | No Change | Not Significant | TRUE | -0.05714344 | 0.66295501 | No Change |
| Pkd1l1 | TRUE | 0.749413234 | 0.695624496 | No Change | Not Significant | FALSE | NA | NA | No Change |
| Pkd1l2 | TRUE | 0.735509992 | 1 | No Change | Not Significant | FALSE | NA | NA | No Change |
| Pkd1l3 | TRUE | -0.465668024 | 0.364445856 | No Change | Not Significant | TRUE | -0.220365401 | 0.673954486 | No Change |
| Pkd2 | TRUE | 0.053648959 | 0.823836911 | No Change | Not Significant | TRUE | -0.109350796 | 0.407666398 | No Change |
| Pkd2l1 | TRUE | 0.939420356 | 1 | No Change | Not Significant | FALSE | NA | NA | No Change |
| Pkd2l2 | TRUE | -0.181425422 | 0.366681641 | No Change | Not Significant | TRUE | -0.137758682 | 0.491535798 | No Change |
| Pkdcc | TRUE | 0.078213045 | 0.541493376 | No Change | Not Significant | TRUE | -0.006607031 | 0.96786874 | No Change |
| Pkdrej | TRUE | 0.629765167 | 0.005170946 | UP | UP | TRUE | 0.462692711 | 0.025297464 | No Change |
| Pkhd1 | TRUE | 0.400502636 | 0.002670749 | No Change | Significant Positive | TRUE | -0.029286546 | 0.89946133 | No Change |
| Pkhd1l1 | TRUE | 0.6010388 | 0.768458962 | No Change | Not Significant | FALSE | NA | NA | No Change |
| Pkia | TRUE | -0.076317538 | 0.840656682 | No Change | Not Significant | TRUE | 0.45324354 | 0.000258331 | No Change |
| Pkib | TRUE | 2.977799279 | 4.21E-34 | UP | UP | TRUE | 1.764184124 | 1.44E-12 | UP |
| Pkig | TRUE | 0.100553072 | 0.617417854 | No Change | Not Significant | TRUE | 0.130388577 | 0.205949343 | No Change |
| Pklr | TRUE | 0.080970502 | 0.493582047 | No Change | Not Significant | TRUE | 0.057686017 | 0.610106378 | No Change |
| Pkm | TRUE | -0.423037532 | 0.000445412 | No Change | Significant Negative | TRUE | -0.115110082 | 0.380084737 | No Change |
| Pkmyt1 | TRUE | 0.266111782 | 0.139280674 | No Change | Not Significant | TRUE | 0.004311928 | 0.988660819 | No Change |
| Pkn1 | TRUE | -0.219807619 | 0.001061844 | No Change | Significant Negative | TRUE | -0.142274012 | 0.108600285 | No Change |
| Pkn2 | TRUE | 0.022852622 | 0.832151567 | No Change | Not Significant | TRUE | -0.067698469 | 0.305372024 | No Change |
| Pkn3 | TRUE | 0.201393666 | 0.443634979 | No Change | Not Significant | TRUE | -0.21330239 | 0.218457165 | No Change |
| Pknox1 | TRUE | -0.041172028 | 0.84101582 | No Change | Not Significant | TRUE | 0.057823426 | 0.692786125 | No Change |
| Pknox2 | TRUE | -0.263383919 | 0.891560436 | No Change | Not Significant | FALSE | NA | NA | No Change |
| Pkp1 | TRUE | -0.22370453 | 0.291415309 | No Change | Not Significant | TRUE | 0.071758131 | 0.693047027 | No Change |
| Pkp2 | TRUE | -0.017693643 | 0.894394019 | No Change | Not Significant | TRUE | 0.061766744 | 0.510292007 | No Change |
| Pkp3 | TRUE | -0.080568224 | 0.684288244 | No Change | Not Significant | TRUE | 0.049683659 | 0.714476238 | No Change |
| Pkp4 | TRUE | 0.161337103 | 0.036683176 | No Change | Significant Positive | TRUE | 0.075188676 | 0.260026017 | No Change |
| Pla1a | TRUE | 1.115108744 | 1 | No Change | Not Significant | FALSE | NA | NA | No Change |
| Pla2g10 | TRUE | -0.304825594 | 0.129358637 | No Change | Not Significant | TRUE | 0.011442735 | 0.955322545 | No Change |
| Pla2g10os | TRUE | -0.549292782 | 0.011690393 | DOWN | DOWN | TRUE | -0.272303895 | 0.032623603 | No Change |
| Pla2g12a | TRUE | -0.286505298 | 0.182116478 | No Change | Not Significant | TRUE | -0.226243939 | 0.127299579 | No Change |
| Pla2g12b | TRUE | 0.673847463 | 0.136254601 | No Change | Not Significant | TRUE | 0.332560741 | 0.406691669 | No Change |
| Pla2g15 | TRUE | 0.351539495 | 0.010850695 | No Change | Significant Positive | TRUE | 0.244104554 | 0.130987701 | No Change |
| Pla2g1b | TRUE | 2.708893965 | 1 | No Change | Not Significant | FALSE | NA | NA | No Change |
| Pla2g2a | TRUE | 0.892005369 | 0.072289318 | No Change | Not Significant | TRUE | -0.00361564 | 0.996976705 | No Change |
| Pla2g2c | TRUE | 0.477786207 | 0.028888694 | No Change | Significant Positive | TRUE | 0.175339279 | 0.625632391 | No Change |
| Pla2g2d | TRUE | 1.60320806 | 1 | No Change | Not Significant | FALSE | NA | NA | No Change |
| Pla2g2e | TRUE | -0.096019963 | 1 | No Change | Not Significant | FALSE | NA | NA | No Change |
| Pla2g2f | TRUE | 0.128842355 | 0.872287153 | No Change | Not Significant | TRUE | 0.266904915 | 0.070827655 | No Change |
| Pla2g3 | TRUE | -0.020854421 | 0.948996349 | No Change | Not Significant | TRUE | -0.314939463 | 0.07559346 | No Change |
| Pla2g4a | TRUE | 0.056749219 | 0.864269275 | No Change | Not Significant | TRUE | 0.048509747 | 0.61867166 | No Change |
| Pla2g4b | TRUE | 0.582673604 | 0.483440174 | No Change | Not Significant | TRUE | -0.388270285 | 0.588620784 | No Change |
| Pla2g4c | FALSE | NA | NA | No Change | Not Detected | TRUE | 1.282856295 | 0.09418931 | No Change |
| Pla2g4d | TRUE | -1.101214934 | 1 | No Change | Not Significant | FALSE | NA | NA | No Change |
| Pla2g4e | TRUE | -1.804091334 | 1 | No Change | Not Significant | FALSE | NA | NA | No Change |
| Pla2g4f | TRUE | 0.215826651 | 0.497933561 | No Change | Not Significant | TRUE | 0.029365408 | 0.947331698 | No Change |
| Pla2g5 | TRUE | 1.719711858 | 1.50E-06 | UP | UP | TRUE | 0.464067845 | 0.101663099 | No Change |
| Pla2g6 | TRUE | 0.49619889 | 0.005843912 | No Change | Significant Positive | TRUE | 0.220879358 | 0.166920824 | No Change |
| Pla2g7 | TRUE | -1.58846117 | 0.049740685 | DOWN | DOWN | FALSE | NA | NA | No Change |
| Pla2r1 | TRUE | 1.609042273 | 1 | No Change | Not Significant | FALSE | NA | NA | No Change |
| Plaa | TRUE | 0.268179658 | 0.003202436 | No Change | Significant Positive | TRUE | 0.018873693 | 0.889849511 | No Change |
| Plaat3 | TRUE | -0.417444726 | 0.000870602 | No Change | Significant Negative | TRUE | -0.063956566 | 0.659618863 | No Change |
| Plaat5 | TRUE | 0.425595983 | 1 | No Change | Not Significant | FALSE | NA | NA | No Change |
| Plac8 | TRUE | -0.505043288 | 0.000317634 | DOWN | DOWN | TRUE | -0.017017528 | 0.936965584 | No Change |
| Plac8l1 | TRUE | 0.951392175 | 1 | No Change | Not Significant | FALSE | NA | NA | No Change |
| Plag1 | TRUE | -0.323655344 | 0.21047419 | No Change | Not Significant | TRUE | -0.278202361 | 0.281763913 | No Change |
| Plagl1 | TRUE | 0.55972028 | 0.094898539 | No Change | Not Significant | TRUE | 0.210206347 | 0.657586455 | No Change |
| Plagl2 | TRUE | 0.343181875 | 5.60E-06 | No Change | Significant Positive | TRUE | -0.06347308 | 0.669932492 | No Change |
| Plat | TRUE | 0.444688975 | 0.695624496 | No Change | Not Significant | FALSE | NA | NA | No Change |
| Platr10 | TRUE | 0.607206888 | 1 | No Change | Not Significant | FALSE | NA | NA | No Change |
| Platr21 | TRUE | 0.609787913 | 1 | No Change | Not Significant | FALSE | NA | NA | No Change |
| Platr26 | TRUE | -1.193284126 | 1 | No Change | Not Significant | FALSE | NA | NA | No Change |
| Platr27 | TRUE | -0.571127052 | 1 | No Change | Not Significant | FALSE | NA | NA | No Change |
| Platr4 | TRUE | -0.56801265 | 1 | No Change | Not Significant | FALSE | NA | NA | No Change |
| Platr8 | TRUE | -1.045495156 | 0.125530417 | No Change | Not Significant | TRUE | -0.900070647 | 0.261362607 | No Change |
| Platr9 | TRUE | -1.467125609 | 1 | No Change | Not Significant | FALSE | NA | NA | No Change |
| Plau | TRUE | 0.402841882 | 1 | No Change | Not Significant | FALSE | NA | NA | No Change |
| Plaur | TRUE | 0.706273731 | 0.091519937 | No Change | Not Significant | TRUE | 0.03784533 | 0.924270223 | No Change |
| Plb1 | TRUE | 0.703147441 | 0.635956548 | No Change | Not Significant | TRUE | 0.432828156 | 0.703236518 | No Change |
| Plbd1 | TRUE | -1.092954827 | 1.67E-06 | DOWN | DOWN | TRUE | -0.344458354 | 0.000138049 | No Change |
| Plbd2 | TRUE | -0.324324551 | 0.000180604 | No Change | Significant Negative | TRUE | -0.065255464 | 0.61867166 | No Change |
| Plcb1 | TRUE | -0.339167147 | 0.172130699 | No Change | Not Significant | TRUE | 0.073643825 | 0.709266447 | No Change |
| Plcb2 | TRUE | -0.617837852 | 0.004121541 | DOWN | DOWN | TRUE | -0.118771434 | 0.671631718 | No Change |
| Plcb3 | TRUE | -0.147009188 | 0.038236855 | No Change | Significant Negative | TRUE | -0.003069004 | 0.971391049 | No Change |
| Plcb4 | TRUE | 0.371680148 | 0.063995287 | No Change | Not Significant | TRUE | 0.203486289 | 0.154396187 | No Change |
| Plcd1 | TRUE | -0.040129547 | 0.885478012 | No Change | Not Significant | TRUE | -0.02140424 | 0.906052077 | No Change |
| Plcd3 | TRUE | -0.868692095 | 0.001752808 | DOWN | DOWN | TRUE | 0.162601218 | 0.773163315 | No Change |
| Plcd4 | TRUE | -0.475356803 | 0.631550268 | No Change | Not Significant | TRUE | -0.504584649 | 0.186021452 | No Change |
| Plce1 | TRUE | -0.152109621 | 0.183322799 | No Change | Not Significant | TRUE | -0.216174491 | 0.000224605 | No Change |
| Plcg1 | TRUE | -0.127388738 | 0.36983602 | No Change | Not Significant | TRUE | -0.153036226 | 0.060891613 | No Change |
| Plcg2 | TRUE | -0.756420364 | 9.17E-08 | DOWN | DOWN | TRUE | -0.105066525 | 0.473349371 | No Change |
| Plch1 | TRUE | 0.252789409 | 0.215060531 | No Change | Not Significant | TRUE | 0.236129393 | 0.027655107 | No Change |
| Plch2 | TRUE | 0.062961947 | 0.706397443 | No Change | Not Significant | TRUE | 0.044415576 | 0.706376429 | No Change |
| Plcl1 | TRUE | -0.677985861 | 0.301713259 | No Change | Not Significant | TRUE | -0.185912168 | 0.665280626 | No Change |
| Plcl2 | TRUE | -0.16980924 | 0.175865487 | No Change | Not Significant | TRUE | -0.006826999 | 0.958828915 | No Change |
| Plcxd1 | TRUE | -0.211312571 | 0.745058233 | No Change | Not Significant | TRUE | -0.316003514 | 0.548247546 | No Change |
| Plcxd2 | TRUE | 0.424377379 | 0.001159019 | No Change | Significant Positive | TRUE | 0.286444462 | 0.000298812 | No Change |
| Plcxd3 | TRUE | -0.170329931 | 1 | No Change | Not Significant | FALSE | NA | NA | No Change |
| Pld1 | TRUE | -0.278904581 | 0.00127605 | No Change | Significant Negative | TRUE | -0.046139602 | 0.584442043 | No Change |
| Pld2 | TRUE | 0.118481364 | 0.852650725 | No Change | Not Significant | TRUE | 0.388489146 | 0.165184357 | No Change |
| Pld3 | TRUE | -0.026411003 | 0.875092959 | No Change | Not Significant | TRUE | -0.010696368 | 0.950733597 | No Change |
| Pld4 | TRUE | 0.752684462 | 1 | No Change | Not Significant | FALSE | NA | NA | No Change |
| Pld5 | TRUE | 0.296267717 | 1 | No Change | Not Significant | TRUE | 0.539551817 | 0.327433832 | No Change |
| Pld6 | TRUE | -0.045173834 | 1 | No Change | Not Significant | TRUE | -0.855897979 | 0.260216003 | No Change |
| Plec | TRUE | -0.112577065 | 0.728820184 | No Change | Not Significant | TRUE | 0.053419854 | 0.803676612 | No Change |
| Plek | TRUE | 0.322841297 | 0.305048799 | No Change | Not Significant | TRUE | 0.125666816 | 0.720013604 | No Change |
| Plek2 | TRUE | -0.448957168 | 1.69E-05 | No Change | Significant Negative | TRUE | -0.008330771 | 0.967491161 | No Change |
| Plekha1 | TRUE | -0.286997142 | 0.020354233 | No Change | Significant Negative | TRUE | 0.016611594 | 0.917183108 | No Change |
| Plekha2 | TRUE | -0.165133198 | 0.228807636 | No Change | Not Significant | TRUE | 0.083715984 | 0.445219276 | No Change |
| Plekha3 | TRUE | -0.161996021 | 0.31327808 | No Change | Not Significant | TRUE | -0.100252437 | 0.291609013 | No Change |
| Plekha4 | TRUE | -0.646638811 | 0.467148794 | No Change | Not Significant | TRUE | -0.511523407 | 0.438439283 | No Change |
| Plekha5 | TRUE | 0.083499433 | 0.322004337 | No Change | Not Significant | TRUE | 0.022920392 | 0.824906092 | No Change |
| Plekha6 | TRUE | -0.040764323 | 0.70880782 | No Change | Not Significant | TRUE | 0.018752299 | 0.888754045 | No Change |
| Plekha7 | TRUE | 0.05258218 | 0.810140405 | No Change | Not Significant | TRUE | -0.043324005 | 0.724225297 | No Change |
| Plekha8 | TRUE | -0.209579068 | 0.000714089 | No Change | Significant Negative | TRUE | -0.296283703 | 1.18E-06 | No Change |
| Plekhb1 | TRUE | 0.203028532 | 0.762380457 | No Change | Not Significant | TRUE | -0.11456694 | 0.902745048 | No Change |
| Plekhb2 | TRUE | -0.061449204 | 0.438111017 | No Change | Not Significant | TRUE | 0.21616544 | 0.000301734 | No Change |
| Plekhd1 | TRUE | -0.334251802 | 0.312211636 | No Change | Not Significant | TRUE | -0.100225759 | 0.666799776 | No Change |
| Plekhd1os | TRUE | -0.074288736 | 0.902837235 | No Change | Not Significant | TRUE | 0.365511319 | 0.3590356 | No Change |
| Plekhf1 | TRUE | 0.740234207 | 0.050506026 | No Change | Not Significant | TRUE | 0.431505303 | 0.138573681 | No Change |
| Plekhf2 | TRUE | -0.034196606 | 0.724020576 | No Change | Not Significant | TRUE | -0.054199257 | 0.434385314 | No Change |
| Plekhg1 | TRUE | -0.301791238 | 0.01753991 | No Change | Significant Negative | TRUE | -0.050959743 | 0.678210988 | No Change |
| Plekhg2 | TRUE | 0.19364533 | 0.652884901 | No Change | Not Significant | TRUE | -0.0993448 | 0.60375884 | No Change |
| Plekhg3 | TRUE | -0.249850456 | 0.028864834 | No Change | Significant Negative | TRUE | -0.21101824 | 0.000746133 | No Change |
| Plekhg4 | TRUE | 0.960990363 | 0.260075004 | No Change | Not Significant | FALSE | NA | NA | No Change |
| Plekhg5 | TRUE | -0.286316801 | 0.03728716 | No Change | Significant Negative | TRUE | -0.194928563 | 0.108600285 | No Change |
| Plekhg6 | TRUE | -0.166865058 | 0.741922624 | No Change | Not Significant | TRUE | 0.07202724 | 0.750796946 | No Change |
| Plekhh1 | TRUE | -0.165005736 | 0.145896332 | No Change | Not Significant | TRUE | -0.092501693 | 0.340305926 | No Change |
| Plekhh2 | TRUE | 0.48027151 | 0.067726402 | No Change | Not Significant | TRUE | 0.588097022 | 0.051214509 | No Change |
| Plekhh3 | TRUE | -0.076501382 | 0.754784418 | No Change | Not Significant | TRUE | -0.143562268 | 0.368632313 | No Change |
| Plekhj1 | TRUE | -0.1451095 | 0.155400323 | No Change | Not Significant | TRUE | 0.139311928 | 0.242642163 | No Change |
| Plekhm1 | TRUE | -0.056285209 | 0.867716116 | No Change | Not Significant | TRUE | -0.079482774 | 0.488420788 | No Change |
| Plekhm2 | TRUE | -0.191657862 | 0.145821162 | No Change | Not Significant | TRUE | -0.056506605 | 0.64875856 | No Change |
| Plekhm3 | TRUE | -0.558260776 | 0.109351285 | No Change | Not Significant | TRUE | -0.161138221 | 0.488580666 | No Change |
| Plekhn1 | TRUE | -0.352485367 | 0.033071834 | No Change | Significant Negative | TRUE | -0.395017819 | 4.53E-06 | No Change |
| Plekho1 | TRUE | 0.187157055 | 0.712306522 | No Change | Not Significant | TRUE | 0.081572099 | 0.898825528 | No Change |
| Plekho2 | TRUE | -0.243238413 | 0.757617052 | No Change | Not Significant | TRUE | 0.173911005 | 0.699880814 | No Change |
| Plekhs1 | TRUE | 0.174394487 | 0.802947244 | No Change | Not Significant | TRUE | -0.259892896 | 0.666514881 | No Change |
| Plet1 | TRUE | -0.294412439 | 0.432027803 | No Change | Not Significant | TRUE | -0.153121638 | 0.521009858 | No Change |
| Plet1os | TRUE | -0.478255816 | 0.392717478 | No Change | Not Significant | TRUE | -0.092489789 | 0.853157579 | No Change |
| Plg | TRUE | 2.012945105 | 1 | No Change | Not Significant | FALSE | NA | NA | No Change |
| Plgrkt | TRUE | 0.058776888 | 0.716335759 | No Change | Not Significant | TRUE | -0.008023239 | 0.961718903 | No Change |
| Plin1 | TRUE | -0.0330824 | 1 | No Change | Not Significant | FALSE | NA | NA | No Change |
| Plin2 | TRUE | -0.586742052 | 1.17E-05 | DOWN | DOWN | TRUE | -0.013404193 | 0.929981434 | No Change |
| Plin3 | TRUE | 0.243331455 | 0.000396224 | No Change | Significant Positive | TRUE | 0.235296438 | 0.000176007 | No Change |
| Plin4 | TRUE | 0.162894302 | 0.894265208 | No Change | Not Significant | FALSE | NA | NA | No Change |
| Plin5 | TRUE | -0.796227926 | 0.367865096 | No Change | Not Significant | TRUE | -0.482107873 | 0.542138138 | No Change |
| Plk1 | TRUE | 0.309719941 | 0.09897928 | No Change | Not Significant | TRUE | 0.060347036 | 0.560212157 | No Change |
| Plk2 | TRUE | -0.327721486 | 0.157015607 | No Change | Not Significant | TRUE | -0.267288193 | 0.003321171 | No Change |
| Plk3 | TRUE | -0.401992116 | 0.141320893 | No Change | Not Significant | TRUE | -0.367480182 | 0.07953207 | No Change |
| Plk4 | TRUE | 0.839877049 | 2.75E-06 | UP | UP | TRUE | 0.12733784 | 0.493560011 | No Change |
| Plk5 | TRUE | -1.228137737 | 1 | No Change | Not Significant | TRUE | -0.457703732 | 0.637110813 | No Change |
| Pllp | TRUE | -0.239637798 | 0.159435243 | No Change | Not Significant | TRUE | 0.079273414 | 0.734525893 | No Change |
| Plod1 | TRUE | 0.35875691 | 0.059503296 | No Change | Not Significant | TRUE | 0.414196689 | 0.013723699 | No Change |
| Plod2 | TRUE | 2.044241093 | 3.06E-06 | UP | UP | TRUE | 1.538320601 | 4.64E-10 | UP |
| Plod3 | TRUE | 0.026757184 | 0.893788319 | No Change | Not Significant | TRUE | 0.08839554 | 0.445254566 | No Change |
| Plp1 | TRUE | -0.15304047 | 0.894265208 | No Change | Not Significant | TRUE | 0.046743947 | 0.956441676 | No Change |
| Plp2 | TRUE | -0.456933864 | 0.001154992 | No Change | Significant Negative | TRUE | -0.089028295 | 0.524516319 | No Change |
| Plpbp | TRUE | 0.86566634 | 2.52E-33 | UP | UP | TRUE | 0.37234129 | 1.67E-12 | No Change |
| Plpp1 | TRUE | 0.069268624 | 0.894265208 | No Change | Not Significant | TRUE | 0.092337994 | 0.523508209 | No Change |
| Plpp2 | TRUE | -0.047518007 | 0.806110415 | No Change | Not Significant | TRUE | 0.027494519 | 0.788784457 | No Change |
| Plpp3 | TRUE | -0.022926999 | 0.946592528 | No Change | Not Significant | TRUE | 0.089198541 | 0.671631718 | No Change |
| Plpp5 | TRUE | 0.168945861 | 0.06471615 | No Change | Not Significant | TRUE | 0.221510715 | 0.005414954 | No Change |
| Plpp6 | TRUE | 0.709911967 | 3.62E-21 | UP | UP | TRUE | 0.287145917 | 1.26E-08 | No Change |
| Plpp7 | TRUE | -1.442717587 | 1 | No Change | Not Significant | TRUE | -0.663815685 | 0.361643489 | No Change |
| Plppr1 | TRUE | 0.857958096 | 1 | No Change | Not Significant | FALSE | NA | NA | No Change |
| Plppr2 | TRUE | -0.039810783 | 0.947202684 | No Change | Not Significant | TRUE | -0.439989199 | 0.405224566 | No Change |
| Plppr3 | TRUE | 0.127722705 | 0.66852312 | No Change | Not Significant | TRUE | -0.338738676 | 0.298033498 | No Change |
| Plppr4 | TRUE | -0.42524228 | 1 | No Change | Not Significant | FALSE | NA | NA | No Change |
| Plppr5 | TRUE | -0.678334025 | 1 | No Change | Not Significant | FALSE | NA | NA | No Change |
| Plrg1 | TRUE | 0.142101763 | 0.17503883 | No Change | Not Significant | TRUE | 0.037412753 | 0.735943093 | No Change |
| Pls1 | TRUE | -0.019785213 | 0.9092783 | No Change | Not Significant | TRUE | 0.029876049 | 0.840526771 | No Change |
| Pls3 | TRUE | -0.202167731 | 0.109773894 | No Change | Not Significant | TRUE | 0.086263057 | 0.703879521 | No Change |
| Plscr1 | TRUE | -0.048432352 | 0.667939129 | No Change | Not Significant | TRUE | 0.000630915 | 0.994488063 | No Change |

Supplementary Table 3

| geneID | Cond1_DF_Detected | Cond1_DF_ValueCol | Cond1_DF_PadjCol | Cond1_DF_Cutoff | Cond1_DF_Cutoff_Specific.x | Cond2_DF_Detected |
|---|---|---|---|---|---|---|
| Plscr2 | TRUE | -1.109332403 | 0.012989531 | DOWN | DOWN | TRUE |
| Plscr3 | TRUE | -0.067403721 | 0.769577776 | No Change | Not Significant | TRUE |
| Plscr4 | TRUE | -0.526208512 | 0.250788441 | No Change | Not Significant | TRUE |
| Pltp | TRUE | 3.502336863 | 3.13E-12 | UP | UP | TRUE |
| Plut | TRUE | -0.590796393 | 0.540884146 | No Change | Not Significant | TRUE |
| Plvap | TRUE | 0.63603861 | 0.594037976 | No Change | Not Significant | FALSE |
| Plxdc1 | TRUE | 0.506927872 | 1 | No Change | Not Significant | FALSE |
| Plxdc2 | TRUE | 0.440264514 | 1 | No Change | Not Significant | FALSE |
| Plxna1 | TRUE | 0.270395221 | 0.012108371 | No Change | Significant Positive | TRUE |
| Plxna2 | TRUE | -0.269610799 | 0.043086943 | No Change | Significant Negative | TRUE |
| Plxna3 | TRUE | -0.462968882 | 0.023502823 | No Change | Significant Negative | TRUE |
| Plxna4 | TRUE | 0.487563046 | 0.876856731 | No Change | Not Significant | FALSE |
| Plxnb1 | TRUE | 0.638639882 | 0.000154732 | UP | UP | TRUE |
| Plxnb2 | TRUE | 0.082025033 | 0.476573341 | No Change | Not Significant | TRUE |
| Plxnb3 | TRUE | 0.451929709 | 1 | No Change | Not Significant | FALSE |
| Plxnc1 | TRUE | -0.26706617 | 0.795381772 | No Change | Not Significant | TRUE |
| Plxnd1 | TRUE | -0.331986626 | 0.41130117 | No Change | Not Significant | TRUE |
| Pm20d1 | TRUE | -0.566689389 | 0.000495149 | DOWN | DOWN | TRUE |
| Pm20d2 | TRUE | -0.164824177 | 0.659260011 | No Change | Not Significant | TRUE |
| Pmaip1 | TRUE | -0.790233511 | 0.000261952 | DOWN | DOWN | TRUE |
| Pmch | TRUE | 1.303899154 | 1 | No Change | Not Significant | FALSE |

[illegible]

| geneID | Cond1_DF_Detected | Cond1_DF_ValueCol | Cond1_DF_PadjCol | Cond1_DF_Cutoff | Cond1_DF_Cutoff_Specific.x | Cond2_DF_Detected | Cond2_DF_ValueCol |
|---|---|---|---|---|---|---|---|
| Ppib | TRUE | 1.535563757 | 4.86E-68 | UP | UP | TRUE | 1.002195685 |
| Ppic | TRUE | -1.395054202 | 0.000580034 | DOWN | DOWN | TRUE | -0.162113678 |
| Ppid | TRUE | -0.195615964 | 0.181090121 | No Change | Not Significant | TRUE | -0.19179069 |
| Ppie | TRUE | -0.161444766 | 0.475826228 | No Change | Not Significant | TRUE | 0.052634498 |
| Ppif | TRUE | -0.36377097 | 0.040712379 | No Change | Significant Negative | TRUE | -0.009243221 |
| Ppig | TRUE | 0.0126772 | 0.925637253 | No Change | Not Significant | TRUE | -0.064851124 |
| Ppih | TRUE | -0.010759792 | 0.973044134 | No Change | Not Significant | TRUE | 0.048164945 |
| Ppil1 | TRUE | 0.190254354 | 0.313984606 | No Change | Not Significant | TRUE | 0.161038923 |
| Ppil2 | TRUE | -0.06968577 | 0.261594876 | No Change | Not Significant | TRUE | 0.050926306 |
| Ppil3 | TRUE | 0.155530895 | 0.408887615 | No Change | Not Significant | TRUE | -0.062776471 |
| Ppil4 | TRUE | 0.030140672 | 0.839669485 | No Change | Not Significant | TRUE | -0.045412747 |
| Ppil6 | TRUE | -0.052541295 | 0.941010727 | No Change | Not Significant | TRUE | -0.372817534 |
| Ppip5k1 | TRUE | 1.221655722 | 1.18E-117 | UP | UP | TRUE | 0.668174954 |
| Ppip5k2 | TRUE | 0.177458431 | 0.10540525 | No Change | Not Significant | TRUE | 0.04332747 |
| Ppl | TRUE | -0.763349293 | 5.39E-10 | DOWN | DOWN | TRUE | -0.065981621 |
| Ppm1a | TRUE | -0.051908366 | 0.523653558 | No Change | Not Significant | TRUE | -0.028243871 |
| Ppm1b | TRUE | 0.168252461 | 0.028631939 | No Change | Significant Positive | TRUE | 0.122496733 |
| Ppm1d | TRUE | -0.205594074 | 0.11137126 | No Change | Not Significant | TRUE | -0.147432643 |
| Ppm1e | TRUE | 0.100369596 | 0.913504709 | No Change | Not Significant | TRUE | 0.614166735 |
| Ppm1f | TRUE | 0.151021222 | 0.532288131 | No Change | Not Significant | TRUE | 0.005158475 |
| Ppm1g | TRUE | 0.202479927 | 0.073420411 | No Change | Not Significant | TRUE | 0.063431298 |
| Ppm1h | TRUE | -0.20649176 | 0.000173843 | No Change | Significant Negative | TRUE | 0.019045096 |
| Ppm1j | TRUE | -0.562407619 | 0.271186338 | No Change | Not Significant | TRUE | -0.290668606 |
| Ppm1k | TRUE | -0.171200662 | 0.481173797 | No Change | Not Significant | TRUE | -0.05936382 |
| Ppm1l | TRUE | -0.178304405 | 0.284460467 | No Change | Not Significant | TRUE | 0.276206204 |
| Ppm1m | TRUE | -0.018310488 | 0.956044724 | No Change | Not Significant | TRUE | 0.040593067 |
| Ppm1n | TRUE | -0.075039042 | 1 | No Change | Not Significant | FALSE | NA |

Supplementary Table 3

| geneID | Cond1_DF_Detected | Cond1_DF_ValueCol | Cond1_DF_PadjCol | Cond1_DF_Cutoff | Cond1_DF_Cutoff_Specific.x | Cond2_DF_Detected | Cond2_DF_ValueCol | Cond2_DF_PadjCol | Cond2_DF_Cutoff | Cond2_DF_Cutoff_Specific | BG_Method |
|---|---|---|---|---|---|---|---|---|---|---|---|
| Prkg2 | TRUE | -0.405721418 | 0.0942345 | No Change | Not Significant | TRUE | -0.265961639 | 0.212494471 | No Change | Not Significant | TRUE |
| Prkn | TRUE | -0.646672762 | 0.184376923 | No Change | Not Significant | TRUE | -0.39675334 | 0.252239559 | No Change | Not Significant | TRUE |
| Prkra | TRUE | -0.249812214 | 0.028656498 | No Change | Significant Negative | TRUE | -0.080802614 | 0.461733499 | No Change | Not Significant | TRUE |
| Prkrip1 | TRUE | -0.212052646 | 0.095954019 | No Change | Not Significant | TRUE | 0.001478032 | 0.995445753 | No Change | Not Significant | TRUE |
| Prkx | TRUE | -0.249269734 | 0.087791464 | No Change | Not Significant | TRUE | -0.069928885 | 0.596118728 | No Change | Not Significant | TRUE |
| Prl2a1 | TRUE | 3.232987752 | 1 | No Change | Not Significant | FALSE | NA | NA | No Change | Not Detected | FALSE |
| Prl6a1 | TRUE | 0.113110437 | 1 | No Change | Not Significant | FALSE | NA | NA | No Change | Not Detected | FALSE |
| Prlh | TRUE | -0.14873706 | 1 | No Change | Not Significant | FALSE | NA | NA | No Change | Not Detected | FALSE |
| Prlr | TRUE | -1.043673282 | 8.76E-10 | DOWN | DOWN | TRUE | -0.265240753 | 0.003496865 | No Change | Significant Negative | TRUE |
| Prmt1 | TRUE | 0.484223374 | 1.88E-06 | No Change | Significant Positive | TRUE | 0.162897797 | 0.130372849 | No Change | Not Significant | TRUE |
| Prmt2 | TRUE | 0.008971286 | 0.975173098 | No Change | Not Significant | TRUE | 0.189342821 | 0.294235769 | No Change | Not Significant | TRUE |
| Prmt3 | TRUE | -0.133001107 | 0.572963936 | No Change | Not Significant | TRUE | -0.039313102 | 0.829602333 | No Change | Not Significant | TRUE |
| Prmt5 | TRUE | 0.312504328 | 0.049101931 | No Change | Significant Positive | TRUE | 0.14983586 | 0.18954598 | No Change | Not Significant | TRUE |
| Prmt6 | TRUE | -0.046300311 | 0.837885199 | No Change | Not Significant | TRUE | 0.072351803 | 0.65874317 | No Change | Not Significant | TRUE |
| Prmt7 | TRUE | -0.063076619 | 0.809117134 | No Change | Not Significant | TRUE | -0.071525957 | 0.606076959 | No Change | Not Significant | TRUE |
| Prmt8 | TRUE | -0.367817315 | 1 | No Change | Not Significant | FALSE | NA | NA | No Change | Not Detected | FALSE |
| Prmt9 | TRUE | 0.217392491 | 0.089747949 | No Change | Not Significant | TRUE | 0.098551517 | 0.4333093 | No Change | Not Significant | TRUE |
| Prnp | TRUE | -0.348750755 | 0.012282071 | No Change | Significant Negative | TRUE | 0.020401131 | 0.898695579 | No Change | Not Significant | TRUE |
| Prob1 | TRUE | 1.068446447 | 0.012714707 | UP | UP | TRUE | 0.244896132 | 0.666004719 | No Change | Not Significant | TRUE |
| Proc | TRUE | -0.630938839 | 0.010842012 | DOWN | DOWN | TRUE | -0.113393731 | 0.780331635 | No Change | Not Significant | TRUE |
| Proca1 | TRUE | -0.342376118 | 0.769209762 | No Change | Not Significant | TRUE | -0.260101764 | 0.813747171 | No Change | Not Significant | TRUE |
| Procr | TRUE | 0.903527141 | 1 | No Change | Not Significant | FALSE | NA | NA | No Change | Not Detected | FALSE |
| Prodh | TRUE | -0.59761599 | 4.70E-05 | DOWN | DOWN | TRUE | -0.424550477 | 0.000297303 | No Change | Significant Negative | TRUE |
| Prok1 | TRUE | 0.753081191 | 1 | No Change | Not Significant | FALSE | NA | NA | No Change | Not Detected | FALSE |
| Prok2 | TRUE | 1.158870617 | 1 | No Change | Not Significant | FALSE | NA | NA | No Change | Not Detected | FALSE |
| Prokr1 | TRUE | 2.287565719 | 1 | No Change | Not Significant | FALSE | NA | NA | No Change | Not Detected | FALSE |
| Prokr2 | TRUE | 1.143461477 | 1 | No Change | Not Significant | FALSE | NA | NA | No Change | Not Detected | FALSE |
| Prol1 | TRUE | 1.099242836 | 1 | No Change | Not Significant | FALSE | NA | NA | No Change | Not Detected | FALSE |
| Prom1 | TRUE | -0.323409648 | 0.020846249 | No Change | Significant Negative | TRUE | 0.06036985 | 0.713270579 | No Change | Not Significant | TRUE |
| Prom2 | TRUE | 0.213096806 | 1 | No Change | Not Significant | FALSE | NA | NA | No Change | Not Detected | FALSE |
| Prop1 | TRUE | 0.005722508 | 1 | No Change | Not Significant | FALSE | NA | NA | No Change | Not Detected | FALSE |
| Prorp | TRUE | -0.071354896 | 0.664688628 | No Change | Not Significant | TRUE | 0.013776189 | 0.935123077 | No Change | Not Significant | TRUE |
| Prorsd1 | TRUE | -0.0387402 | 0.882510888 | No Change | Not Significant | TRUE | 0.156391438 | 0.288516832 | No Change | Not Significant | TRUE |
| Pros1 | TRUE | 0.342300552 | 0.328390328 | No Change | Not Significant | TRUE | 0.461251586 | 0.001906168 | No Change | Significant Positive | TRUE |
| Proser1 | TRUE | 0.159094406 | 0.685318046 | No Change | Not Significant | TRUE | -0.108815065 | 0.678921982 | No Change | Not Significant | TRUE |
| Proser2 | TRUE | -0.507470304 | 0.000116906 | DOWN | DOWN | TRUE | -0.100192986 | 0.503509791 | No Change | Not Significant | TRUE |
| Proser3 | TRUE | 0.137302194 | 0.720291626 | No Change | Not Significant | TRUE | 0.180850358 | 0.491735724 | No Change | Not Significant | TRUE |
| Prox1 | TRUE | 0.570917658 | 0.074658557 | No Change | Not Significant | TRUE | 0.100394825 | 0.775454214 | No Change | Not Significant | TRUE |
| Prox2 | TRUE | -0.183759201 | 0.797440677 | No Change | Not Significant | TRUE | -0.073565774 | 0.918528635 | No Change | Not Significant | TRUE |
| Proz | TRUE | 0.140105451 | 0.776186935 | No Change | Not Significant | TRUE | -0.089883437 | 0.87419748 | No Change | Not Significant | TRUE |
| Prp2 | TRUE | 1.697698262 | 1 | No Change | Not Significant | FALSE | NA | NA | No Change | Not Detected | FALSE |
| Prpf18 | TRUE | 0.019513658 | 0.912143267 | No Change | Not Significant | TRUE | 0.025697499 | 0.814098293 | No Change | Not Significant | TRUE |
| Prpf19 | TRUE | 0.133235357 | 0.43083733 | No Change | Not Significant | TRUE | 0.134725601 | 0.120635733 | No Change | Not Significant | TRUE |
| Prpf3 | TRUE | 0.278151339 | 0.020855246 | No Change | Significant Positive | TRUE | 0.048169164 | 0.804068461 | No Change | Not Significant | TRUE |
| Prpf31 | TRUE | 0.180060287 | 0.113873419 | No Change | Not Significant | TRUE | 0.02657101 | 0.877650787 | No Change | Not Significant | TRUE |
| Prpf38a | TRUE | 0.050528735 | 0.720610174 | No Change | Not Significant | TRUE | 0.052155298 | 0.58688999 | No Change | Not Significant | TRUE |
| Prpf38b | TRUE | 0.464541548 | 8.69E-12 | No Change | Significant Positive | TRUE | 0.021208108 | 0.910363905 | No Change | Not Significant | TRUE |
| Prpf39 | TRUE | 0.164261422 | 0.298680373 | No Change | Not Significant | TRUE | -0.259470649 | 0.228944671 | No Change | Not Significant | TRUE |
| Prpf4 | TRUE | 0.139659853 | 0.396067528 | No Change | Not Significant | TRUE | 0.036165311 | 0.789908442 | No Change | Not Significant | TRUE |
| Prpf40a | TRUE | 0.278095914 | 0.000792278 | No Change | Significant Positive | TRUE | 0.051446553 | 0.594540465 | No Change | Not Significant | TRUE |
| Prpf40b | TRUE | -0.301362929 | 0.302183099 | No Change | Not Significant | TRUE | -0.059335675 | 0.784071601 | No Change | Not Significant | TRUE |
| Prpf4b | TRUE | 0.084101066 | 0.307249743 | No Change | Not Significant | TRUE | -0.053525745 | 0.622494568 | No Change | Not Significant | TRUE |
| Prpf6 | TRUE | 0.010088513 | 0.951530343 | No Change | Not Significant | TRUE | 0.041513865 | 0.682675952 | No Change | Not Significant | TRUE |
| Prpf8 | TRUE | 0.069759897 | 0.657874003 | No Change | Not Significant | TRUE | 0.002756997 | 0.986120526 | No Change | Not Significant | TRUE |
| Prph | TRUE | 3.019039839 | 1 | No Change | Not Significant | FALSE | NA | NA | No Change | Not Detected | FALSE |
| Prph2 | TRUE | -1.956777461 | 1 | No Change | Not Significant | FALSE | NA | NA | No Change | Not Detected | FALSE |
| Prps1 | TRUE | 0.307344335 | 0.168971808 | No Change | Not Significant | TRUE | 0.262873479 | 0.003473484 | No Change | Significant Positive | TRUE |
| Prps1l1 | TRUE | 0.015155022 | 1 | No Change | Not Significant | FALSE | NA | NA | No Change | Not Detected | FALSE |
| Prps1l3 | TRUE | 0.15212059 | 0.465710848 | No Change | Not Significant | TRUE | 0.075929136 | 0.489920593 | No Change | Not Significant | TRUE |
| Prps2 | TRUE | -0.212953285 | 0.334862894 | No Change | Not Significant | TRUE | -0.092873193 | 0.276898911 | No Change | Not Significant | TRUE |
| Prpsap1 | TRUE | -0.157454797 | 0.384369853 | No Change | Not Significant | TRUE | 0.043559988 | 0.775075241 | No Change | Not Significant | TRUE |
| Prpsap2 | TRUE | -0.193409315 | 0.21805685 | No Change | Not Significant | TRUE | 0.17830119 | 0.040239329 | No Change | Significant Positive | TRUE |
| Prr11 | TRUE | 0.649520891 | 0.009854484 | UP | UP | TRUE | -0.020384002 | 0.950733597 | No Change | Not Significant | TRUE |
| Prr12 | TRUE | 0.361542477 | 0.425016386 | No Change | Not Significant | TRUE | -0.20695107 | 0.4070526 | No Change | Not Significant | TRUE |
| Prr13 | TRUE | -0.096780977 | 0.581328932 | No Change | Not Significant | TRUE | 0.218871202 | 0.056965074 | No Change | Not Significant | TRUE |
| Prr14 | TRUE | 0.048007443 | 0.816884691 | No Change | Not Significant | TRUE | -0.039449632 | 0.709700079 | No Change | Not Significant | TRUE |
| Prr14l | TRUE | -0.153287546 | 0.07028111 | No Change | Not Significant | TRUE | -0.02235493 | 0.865987203 | No Change | Not Significant | TRUE |
| Prr15 | TRUE | 0.109540237 | 0.73156216 | No Change | Not Significant | TRUE | -0.041252005 | 0.844292391 | No Change | Not Significant | TRUE |
| Prr15l | TRUE | 0.074636867 | 0.674101639 | No Change | Not Significant | TRUE | 0.192608189 | 0.001644529 | No Change | Significant Positive | TRUE |
| Prr16 | TRUE | 0.179212321 | 0.853653903 | No Change | Not Significant | TRUE | 0.48831132 | 0.480740232 | No Change | Not Significant | TRUE |
| Prr18 | TRUE | -0.436286151 | 1 | No Change | Not Significant | TRUE | 0.231890601 | 0.813747171 | No Change | Not Significant | TRUE |
| Prr22 | TRUE | 1.332598275 | 0.044068147 | UP | UP | FALSE | NA | NA | No Change | Not Detected | FALSE |
| Prr29 | TRUE | 0.08870595 | 1 | No Change | Not Significant | FALSE | NA | NA | No Change | Not Detected | FALSE |
| Prr3 | TRUE | -0.186756041 | 0.56826406 | No Change | Not Significant | TRUE | -0.232953246 | 0.275342713 | No Change | Not Significant | TRUE |
| Prr30 | TRUE | 0.611278724 | 1 | No Change | Not Significant | FALSE | NA | NA | No Change | Not Detected | FALSE |
| Prr33 | TRUE | -1.015677768 | 1 | No Change | Not Significant | FALSE | NA | NA | No Change | Not Detected | FALSE |
| Prr36 | TRUE | -0.127260315 | 0.919582966 | No Change | Not Significant | TRUE | 0.083306605 | 0.929889314 | No Change | Not Significant | TRUE |
| Prr5 | TRUE | 1.601752306 | 3.54E-24 | UP | UP | TRUE | 0.245113868 | 0.02524933 | No Change | Significant Positive | TRUE |
| Prr5l | TRUE | -0.330690124 | 0.346538122 | No Change | Not Significant | TRUE | 0.096954262 | 0.706263486 | No Change | Not Significant | TRUE |
| Prr7 | TRUE | -0.314512755 | 0.697829595 | No Change | Not Significant | TRUE | -0.196440067 | 0.793255831 | No Change | Not Significant | TRUE |
| Prrc1 | TRUE | 0.399186394 | 1.63E-06 | No Change | Significant Positive | TRUE | 0.252500251 | 0.008266632 | No Change | Significant Positive | TRUE |
| Prrc2a | TRUE | 0.383742871 | 0.01632826 | No Change | Significant Positive | TRUE | 0.05370955 | 0.789765867 | No Change | Not Significant | TRUE |
| Prrc2b | TRUE | 0.220245947 | 0.012093437 | No Change | Significant Positive | TRUE | -0.123478904 | 0.254171925 | No Change | Not Significant | TRUE |
| Prrc2c | TRUE | 0.041389812 | 0.844600435 | No Change | Not Significant | TRUE | -0.121185754 | 0.196492651 | No Change | Not Significant | TRUE |
| Prrg1 | TRUE | 0.009354425 | 0.985415068 | No Change | Not Significant | TRUE | -0.467568427 | 0.044730048 | No Change | Significant Negative | TRUE |
| Prrg2 | TRUE | -0.028237607 | 0.888627112 | No Change | Not Significant | TRUE | 0.152739881 | 0.252562627 | No Change | Not Significant | TRUE |
| Prrg3 | TRUE | 1.213488638 | 1 | No Change | Not Significant | FALSE | NA | NA | No Change | Not Detected | FALSE |
| Prrg4 | TRUE | 0.127003099 | 0.869560398 | No Change | Not Significant | TRUE | 0.268514355 | 0.58688999 | No Change | Not Significant | TRUE |
| Prrt1 | TRUE | -0.642589663 | 1 | No Change | Not Significant | FALSE | NA | NA | No Change | Not Detected | FALSE |
| Prrt3 | TRUE | 0.602368027 | 0.300706943 | No Change | Not Significant | TRUE | 0.254209096 | 0.69880515 | No Change | Not Significant | TRUE |
| Prrt4 | TRUE | 2.863236197 | 1 | No Change | Not Significant | FALSE | NA | NA | No Change | Not Detected | FALSE |
| Prrx1 | TRUE | 0.772199795 | 1 | No Change | Not Significant | FALSE | NA | NA | No Change | Not Detected | FALSE |
| Prrx2 | TRUE | 1.730054115 | 1 | No Change | Not Significant | FALSE | NA | NA | No Change | Not Detected | FALSE |
| Prrxl1 | TRUE | 0.005685681 | 1 | No Change | Not Significant | FALSE | NA | NA | No Change | Not Detected | FALSE |
| Prss12 | TRUE | 0.794510139 | 0.358767397 | No Change | Not Significant | TRUE | -0.206008889 | 0.789053031 | No Change | Not Significant | TRUE |
| Prss16 | TRUE | 1.041026345 | 0.00457572 | UP | UP | TRUE | 0.832948166 | 0.038396884 | UP | UP | TRUE |
| Prss2 | TRUE | 0.60581203 | 1 | No Change | Not Significant | FALSE | NA | NA | No Change | Not Detected | FALSE |
| Prss22 | TRUE | 1.016690661 | 1 | No Change | Not Significant | FALSE | NA | NA | No Change | Not Detected | FALSE |
| Prss23 | TRUE | -0.8811276 | 0.001006726 | DOWN | DOWN | TRUE | -0.710520068 | 0.000200182 | DOWN | DOWN | TRUE |
| Prss27 | TRUE | 1.014806203 | 0.031640911 | UP | UP | TRUE | -0.253695737 | 0.761458074 | No Change | Not Significant | TRUE |
| Prss30 | TRUE | -0.299606637 | 0.344790726 | No Change | Not Significant | TRUE | -0.187306898 | 0.336096773 | No Change | Not Significant | TRUE |
| Prss32 | TRUE | -0.400520895 | 5.57E-05 | No Change | Significant Negative | TRUE | -0.005357824 | 0.972619969 | No Change | Not Significant | TRUE |
| Prss33 | TRUE | -0.009172701 | 1 | No Change | Not Significant | FALSE | NA | NA | No Change | Not Detected | FALSE |
| Prss35 | TRUE | 0.533297156 | 1 | No Change | Not Significant | FALSE | NA | NA | No Change | Not Detected | FALSE |
| Prss36 | TRUE | 0.142994366 | 0.695624496 | No Change | Not Significant | TRUE | 0.013483318 | 0.971253657 | No Change | Not Significant | TRUE |
| Prss37 | TRUE | 1.149806115 | 1 | No Change | Not Significant | FALSE | NA | NA | No Change | Not Detected | FALSE |
| Prss41 | TRUE | -0.519334748 | 0.38549753 | No Change | Not Significant | TRUE | -0.881032962 | 0.107506272 | No Change | Not Significant | TRUE |
| Prss43 | TRUE | 1.713591987 | 1 | No Change | Not Significant | FALSE | NA | NA | No Change | Not Detected | FALSE |
| Prss44 | TRUE | 0.546730673 | 1 | No Change | Not Significant | FALSE | NA | NA | No Change | Not Detected | FALSE |
| Prss46 | TRUE | 0.006548678 | 1 | No Change | Not Significant | FALSE | NA | NA | No Change | Not Detected | FALSE |
| Prss48 | TRUE | -0.462909157 | 0.513159947 | No Change | Not Significant | TRUE | -0.180667428 | 0.798512665 | No Change | Not Significant | TRUE |
| Prss50 | TRUE | 2.429177992 | 1 | No Change | Not Significant | FALSE | NA | NA | No Change | Not Detected | FALSE |
| Prss52 | TRUE | -0.946620721 | 1 | No Change | Not Significant | FALSE | NA | NA | No Change | Not Detected | FALSE |
| Prss53 | TRUE | 0.472823971 | 1 | No Change | Not Significant | TRUE | 0.151793082 | 0.859633975 | No Change | Not Significant | TRUE |
| Prss55 | TRUE | 0.004995012 | 1 | No Change | Not Significant | FALSE | NA | NA | No Change | Not Detected | FALSE |
| Prss56 | TRUE | -0.2361943 | 1 | No Change | Not Significant | FALSE | NA | NA | No Change | Not Detected | FALSE |
| Prss57 | TRUE | -1.815899003 | 1 | No Change | Not Significant | FALSE | NA | NA | No Change | Not Detected | FALSE |
| Prss8 | TRUE | 0.235835327 | 0.003837987 | No Change | Significant Positive | TRUE | 0.125262198 | 0.063591213 | No Change | Not Significant | TRUE |
| Prtg | TRUE | -0.562097334 | 1 | No Change | Not Significant | FALSE | NA | NA | No Change | Not Detected | FALSE |
| Prune1 | TRUE | -0.289216849 | 0.005227463 | No Change | Significant Negative | TRUE | -0.069739525 | 0.486120986 | No Change | Not Significant | TRUE |
| Prune2 | TRUE | 0.247037023 | 0.719474674 | No Change | Not Significant | TRUE | 0.150964956 | 0.76862707 | No Change | Not Significant | TRUE |
| Prx | TRUE | 0.579711663 | 0.002918524 | UP | UP | TRUE | 0.226808572 | 0.264527214 | No Change | Not Significant | TRUE |
| Prxl2a | TRUE | -0.00569601 | 0.985415068 | No Change | Not Significant | TRUE | 0.138607338 | 0.477141554 | No Change | Not Significant | TRUE |
| Prxl2b | TRUE | -0.53260639 | 0.00154796 | DOWN | DOWN | TRUE | -0.289410094 | 0.057550792 | No Change | Not Significant | TRUE |
| Prxl2c | TRUE | -0.414276826 | 0.008514609 | No Change | Significant Negative | TRUE | -0.128031059 | 0.197371991 | No Change | Not Significant | TRUE |
| Psap | TRUE | -0.542890657 | 2.37E-19 | DOWN | DOWN | TRUE | -0.161425785 | 0.000151003 | No Change | Significant Negative | TRUE |
| Psapl1 | TRUE | -0.309243197 | 1 | No Change | Not Significant | FALSE | NA | NA | No Change | Not Detected | FALSE |
| Psat1 | TRUE | 2.505342338 | 3.07E-47 | UP | UP | TRUE | 0.806477151 | 0.000175776 | UP | UP | TRUE |
| Psca | TRUE | -0.6626908 | 1 | No Change | Not Significant | FALSE | NA | NA | No Change | Not Detected | FALSE |
| Psd | TRUE | 0.275475078 | 0.234222517 | No Change | Not Significant | TRUE | 0.098853348 | 0.699497125 | No Change | Not Significant | TRUE |
| Psd2 | TRUE | 0.753055402 | 1 | No Change | Not Significant | FALSE | NA | NA | No Change | Not Detected | FALSE |
| Psd3 | TRUE | -0.114063772 | 0.283500399 | No Change | Not Significant | TRUE | -0.249598355 | 4.90E-05 | No Change | Significant Negative | TRUE |
| Psd4 | TRUE | -0.290240298 | 0.007094676 | No Change | Significant Negative | TRUE | -0.121317418 | 0.019496529 | No Change | Significant Negative | TRUE |
| Psen1 | TRUE | -0.022825967 | 0.819390913 | No Change | Not Significant | TRUE | 0.019370486 | 0.761818669 | No Change | Not Significant | TRUE |
| Psen2 | TRUE | -0.073974393 | 0.622382905 | No Change | Not Significant | TRUE | 0.041469878 | 0.794312578 | No Change | Not Significant | TRUE |
| Psenen | TRUE | -0.175641356 | 0.328033263 | No Change | Not Significant | TRUE | 0.136421939 | 0.179737215 | No Change | Not Significant | TRUE |
| Psg16 | TRUE | 0.893284555 | 1 | No Change | Not Significant | FALSE | NA | NA | No Change | Not Detected | FALSE |
| Psg17 | TRUE | 2.338145053 | 1 | No Change | Not Significant | FALSE | NA | NA | No Change | Not Detected | FALSE |
| Psg18 | TRUE | -2.295132616 | 1 | No Change | Not Significant | FALSE | NA | NA | No Change | Not Detected | FALSE |
| Psg19 | TRUE | 2.702518162 | 1 | No Change | Not Significant | FALSE | NA | NA | No Change | Not Detected | FALSE |
| Psg21 | TRUE | 0.543755776 | 1 | No Change | Not Significant | FALSE | NA | NA | No Change | Not Detected | FALSE |
| Psg22 | TRUE | 1.125963653 | 1 | No Change | Not Significant | FALSE | NA | NA | No Change | Not Detected | FALSE |
| Psg23 | TRUE | 0.52245806 | 1 | No Change | Not Significant | FALSE | NA | NA | No Change | Not Detected | FALSE |
| Psg25 | TRUE | -0.090938768 | 1 | No Change | Not Significant | FALSE | NA | NA | No Change | Not Detected | FALSE |
| Psg26 | TRUE | 1.23256742 | 1 | No Change | Not Significant | FALSE | NA | NA | No Change | Not Detected | FALSE |
| Psg28 | TRUE | 0.7285754 | 1 | No Change | Not Significant | TRUE | -1.426647173 | 0.245920034 | No Change | Not Significant | TRUE |
| Psg29 | TRUE | 1.186161563 | 1 | No Change | Not Significant | FALSE | NA | NA | No Change | Not Detected | FALSE |
| Psip1 | TRUE | 0.328638088 | 0.095392863 | No Change | Not Significant | TRUE | 0.068461392 | 0.625831733 | No Change | Not Significant | TRUE |
| Pskh1 | TRUE | -0.091181207 | 0.43633502 | No Change | Not Significant | TRUE | 0.034952185 | 0.771609416 | No Change | Not Significant | TRUE |
| Psma1 | TRUE | 0.223306131 | 0.071736644 | No Change | Not Significant | TRUE | 0.127417394 | 0.304713258 | No Change | Not Significant | TRUE |
| Psma2 | TRUE | 0.083459507 | 0.519260509 | No Change | Not Significant | TRUE | 0.119284181 | 0.12424678 | No Change | Not Significant | TRUE |
| Psma3 | TRUE | -0.156337342 | 0.344285818 | No Change | Not Significant | TRUE | 0.116031094 | 0.092745101 | No Change | Not Significant | TRUE |
| Psma4 | TRUE | 0.08552117 | 0.525632052 | No Change | Not Significant | TRUE | 0.146291432 | 0.10082379 | No Change | Not Significant | TRUE |
| Psma5 | TRUE | -0.150823881 | 0.529189296 | No Change | Not Significant | TRUE | 0.003589303 | 0.983542327 | No Change | Not Significant | TRUE |
| Psma6 | TRUE | 0.198852342 | 0.023571307 | No Change | Significant Positive | TRUE | 0.145669773 | 0.079524247 | No Change | Not Significant | TRUE |
| Psma7 | TRUE | 0.05492925 | 0.695624496 | No Change | Not Significant | TRUE | 0.036399636 | 0.681834975 | No Change | Not Significant | TRUE |
| Psma8 | TRUE | 0.367904381 | 1 | No Change | Not Significant | FALSE | NA | NA | No Change | Not Detected | FALSE |
| Psmb1 | TRUE | 0.059545846 | 0.717843966 | No Change | Not Significant | TRUE | 0.088926982 | 0.109582031 | No Change | Not Significant | TRUE |
| Psmb10 | TRUE | -0.108428406 | 0.240265168 | No Change | Not Significant | TRUE | -0.082094062 | 0.657067803 | No Change | Not Significant | TRUE |
| Psmb11 | TRUE | 1.539283735 | 1 | No Change | Not Significant | FALSE | NA | NA | No Change | Not Detected | FALSE |
| Psmb2 | TRUE | 0.181310364 | 0.042679767 | No Change | Significant Positive | TRUE | 0.118654475 | 0.149687228 | No Change | Not Significant | TRUE |
| Psmb3 | TRUE | 0.115730626 | 0.543328754 | No Change | Not Significant | TRUE | 0.236530472 | 0.001955285 | No Change | Significant Positive | TRUE |
| Psmb4 | TRUE | 0.128678614 | 0.230247597 | No Change | Not Significant | TRUE | 0.114131661 | 0.405715197 | No Change | Not Significant | TRUE |
| Psmb5 | TRUE | 0.047901874 | 0.818482437 | No Change | Not Significant | TRUE | 0.298932273 | 0.0020054 | No Change | Significant Positive | TRUE |
| Psmb6 | TRUE | 0.193478242 | 0.027610883 | No Change | Significant Positive | TRUE | 0.186814268 | 0.039036387 | No Change | Significant Positive | TRUE |
| Psmb7 | TRUE | 0.28808445 | 0.001731719 | No Change | Significant Positive | TRUE | 0.129775766 | 0.201440865 | No Change | Not Significant | TRUE |
| Psmb8 | TRUE | -0.558530861 | 0.011340379 | DOWN | DOWN | TRUE | -1.143660494 | 0.067692953 | No Change | Not Significant | TRUE |
| Psmb9 | TRUE | -0.51761319 | 0.006612235 | DOWN | DOWN | TRUE | -0.500652532 | 0.319914163 | No Change | Not Significant | TRUE |
| Psmc1 | TRUE | 0.288825077 | 0.003680621 | No Change | Significant Positive | TRUE | 0.099863642 | 0.240742704 | No Change | Not Significant | TRUE |
| Psmc2 | TRUE | 0.264309364 | 0.000602545 | No Change | Significant Positive | TRUE | 0.129697763 | 0.202153934 | No Change | Not Significant | TRUE |
| Psmc3 | TRUE | 0.237298826 | 0.011080142 | No Change | Significant Positive | TRUE | 0.16116143 | 0.077980613 | No Change | Not Significant | TRUE |
| Psmc3ip | TRUE | 0.437865057 | 0.208467978 | No Change | Not Significant | TRUE | -0.077084294 | 0.880597382 | No Change | Not Significant | TRUE |
| Psmc4 | TRUE | 0.192471022 | 0.014480556 | No Change | Significant Positive | TRUE | 0.190407983 | 0.041292666 | No Change | Significant Positive | TRUE |
| Psmc5 | TRUE | 0.14603488 | 0.258135524 | No Change | Not Significant | TRUE | 0.121970362 | 0.170760157 | No Change | Not Significant | TRUE |
| Psmc6 | TRUE | 0.038157726 | 0.807226927 | No Change | Not Significant | TRUE | 0.019030556 | 0.873729946 | No Change | Not Significant | TRUE |
| Psmd1 | TRUE | 0.375603187 | 8.60E-07 | No Change | Significant Positive | TRUE | 0.116916348 | 0.144210545 | No Change | Not Significant | TRUE |
| Psmd10 | TRUE | 0.020103837 | 0.907710402 | No Change | Not Significant | TRUE | 0.178378233 | 0.023286378 | No Change | Significant Positive | TRUE |
| Psmd11 | TRUE | 0.1902565 | 0.039961181 | No Change | Significant Positive | TRUE | 0.078138592 | 0.434666577 | No Change | Not Significant | TRUE |
| Psmd12 | TRUE | 0.160408311 | 0.179449017 | No Change | Not Significant | TRUE | 0.134399393 | 0.202129099 | No Change | Not Significant | TRUE |
| Psmd13 | TRUE | 0.213076246 | 0.005852887 | No Change | Significant Positive | TRUE | 0.136402357 | 0.201924436 | No Change | Not Significant | TRUE |
| Psmd14 | TRUE | 0.427909313 | 2.65E-07 | No Change | Significant Positive | TRUE | 0.191408897 | 0.143725313 | No Change | Not Significant | TRUE |
| Psmd2 | TRUE | 0.238840555 | 0.020312105 | No Change | Significant Positive | TRUE | 0.162561132 | 0.101348119 | No Change | Not Significant | TRUE |
| Psmd3 | TRUE | 0.338721871 | 4.37E-07 | No Change | Significant Positive | TRUE | 0.193355528 | 0.074800013 | No Change | Not Significant | TRUE |
| Psmd4 | TRUE | 0.099933448 | 0.164036653 | No Change | Not Significant | TRUE | 0.039900826 | 0.560212157 | No Change | Not Significant | TRUE |
| Psmd5 | TRUE | 0.17223714 | 0.010407861 | No Change | Significant Positive | TRUE | 0.14132311 | 0.04672709 | No Change | Significant Positive | TRUE |
| Psmd6 | TRUE | 0.186264494 | 0.093415051 | No Change | Not Significant | TRUE | 0.136546127 | 0.059943581 | No Change | Not Significant | TRUE |
| Psmd7 | TRUE | 0.303489312 | 5.89E-05 | No Change | Significant Positive | TRUE | 0.049514565 | 0.674900387 | No Change | Not Significant | TRUE |
| Psmd8 | TRUE | 0.039288085 | 0.722741011 | No Change | Not Significant | TRUE | 0.116231461 | 0.168799213 | No Change | Not Significant | TRUE |
| Psmd9 | TRUE | 0.08266454 | 0.595844868 | No Change | Not Significant | TRUE | 0.043412091 | 0.727325341 | No Change | Not Significant | TRUE |
| Psme1 | TRUE | -0.221013526 | 0.018490605 | No Change | Significant Negative | TRUE | -0.166086155 | 0.254568403 | No Change | Not Significant | TRUE |
| Psme2 | TRUE | -0.234254911 | 0.077290766 | No Change | Not Significant | TRUE | -0.053205952 | 0.785201968 | No Change | Not Significant | TRUE |
| Psme2b | TRUE | -0.195164037 | 0.548343875 | No Change | Not Significant | TRUE | 0.036538995 | 0.939855462 | No Change | Not Significant | TRUE |

Supplementary Table 3

| geneID | Cond1_DF_Detected | Cond1_DF_ValueCol | Cond1_DF_PadjCol | Cond1_DF_Cutoff | Cond1_DF_Cutoff_Specific.x | Cond2_DF_Detected | Cond2_DF_ValueCol | Cond2_DF_PadjCol | Cond2_DF_Cutoff | Cond2_DF_Cutoff_Specific | BG_Method |
|---|---|---|---|---|---|---|---|---|---|---|---|
| Psme3 | TRUE | 0.139915814 | 0.13918813 | No Change | Not Significant | TRUE | 0.031251142 | 0.79914261 | No Change | Not Significant | TRUE |
| Psme3ip1 | TRUE | 0.237611184 | 0.000636513 | No Change | Significant Positive | TRUE | 0.130785143 | 0.030809515 | No Change | Significant Positive | TRUE |
| Psme4 | TRUE | 0.147793898 | 0.204535912 | No Change | Not Significant | TRUE | 0.011372384 | 0.916408517 | No Change | Not Significant | TRUE |
| Psmf1 | TRUE | 0.022283897 | 0.866866795 | No Change | Not Significant | TRUE | 0.081164553 | 0.299280839 | No Change | Not Significant | TRUE |
| Psmg1 | TRUE | 0.001263429 | 0.994433839 | No Change | Not Significant | TRUE | 0.032121836 | 0.856166251 | No Change | Not Significant | TRUE |
| Psmg2 | TRUE | 0.029686351 | 0.923801556 | No Change | Not Significant | TRUE | 0.114053968 | 0.379061279 | No Change | Not Significant | TRUE |
| Psmg3 | TRUE | 0.104070987 | 0.599693731 | No Change | Not Significant | TRUE | 0.102271623 | 0.571134973 | No Change | Not Significant | TRUE |
| Psmg4 | TRUE | -0.125345998 | 0.610900332 | No Change | Not Significant | TRUE | -0.22256735 | 0.185048514 | No Change | Not Significant | TRUE |
| Psors1c2 | TRUE | 1.029340736 | 1 | No Change | Not Significant | FALSE | NA | NA | No Change | Not Detected | FALSE |
| Pspc1 | TRUE | 0.643123294 | 1.30E-11 | UP | UP | TRUE | 0.107589086 | 0.392724841 | No Change | Not Significant | TRUE |
| Psph | TRUE | 0.481605765 | 0.003676212 | No Change | Significant Positive | TRUE | 0.213214799 | 0.242489484 | No Change | Not Significant | TRUE |
| Pspn | TRUE | -0.696026783 | 0.280659184 | No Change | Not Significant | TRUE | 0.663215974 | 0.248514275 | No Change | Not Significant | TRUE |
| Psrc1 | TRUE | 0.423807377 | 0.123437405 | No Change | Not Significant | TRUE | -0.120164642 | 0.668045287 | No Change | Not Significant | TRUE |
| Pstk | TRUE | 0.337633179 | 0.005429865 | No Change | Significant Positive | TRUE | 0.20721354 | 0.044045718 | No Change | Significant Positive | TRUE |
| Pstpip1 | TRUE | -0.639639407 | 0.085349139 | No Change | Not Significant | TRUE | 0.048006436 | 0.923972613 | No Change | Not Significant | TRUE |
| Pstpip2 | TRUE | -0.318972698 | 0.022923163 | No Change | Significant Negative | TRUE | -0.168030221 | 0.353057236 | No Change | Not Significant | TRUE |
| Ptafr | TRUE | 2.086781472 | 1 | No Change | Not Significant | FALSE | NA | NA | No Change | Not Detected | FALSE |
| Ptar1 | TRUE | 0.161648619 | 0.364658188 | No Change | Not Significant | TRUE | 0.050267641 | 0.641658935 | No Change | Not Significant | TRUE |
| Ptbp1 | TRUE | 0.357607024 | 2.45E-07 | No Change | Significant Positive | TRUE | 0.124767292 | 0.062224986 | No Change | Not Significant | TRUE |
| Ptbp2 | TRUE | -0.092135697 | 0.746364189 | No Change | Not Significant | TRUE | -0.350664211 | 0.094931644 | No Change | Not Significant | TRUE |
| Ptbp3 | TRUE | -0.026840095 | 0.834763659 | No Change | Not Significant | TRUE | -0.113665584 | 0.198349916 | No Change | Not Significant | TRUE |
| Ptcd1 | TRUE | -0.056992393 | 0.683555242 | No Change | Not Significant | TRUE | 0.007556724 | 0.952330613 | No Change | Not Significant | TRUE |
| Ptcd2 | TRUE | 0.107060871 | 0.496167516 | No Change | Not Significant | TRUE | 0.128405612 | 0.031483955 | No Change | Significant Positive | TRUE |
| Ptcd3 | TRUE | -0.062151163 | 0.703739933 | No Change | Not Significant | TRUE | -0.11237452 | 0.308484011 | No Change | Not Significant | TRUE |
| Ptch1 | TRUE | -0.373668009 | 2.91E-05 | No Change | Significant Negative | TRUE | -0.301294805 | 0.00068978 | No Change | Significant Negative | TRUE |
| Ptch2 | TRUE | 0.70652013 | 0.606515707 | No Change | Not Significant | FALSE | NA | NA | No Change | Not Detected | FALSE |
| Ptchd4 | TRUE | 3.45400293 | 1 | No Change | Not Significant | FALSE | NA | NA | No Change | Not Detected | FALSE |
| Ptdss1 | TRUE | -0.111348059 | 0.22515016 | No Change | Not Significant | TRUE | 0.014570814 | 0.861383838 | No Change | Not Significant | TRUE |
| Ptdss2 | TRUE | -0.244170228 | 0.034050667 | No Change | Significant Negative | TRUE | -0.137934569 | 0.094069857 | No Change | Not Significant | TRUE |
| Pten | TRUE | -0.090188496 | 0.417180016 | No Change | Not Significant | TRUE | -0.148681897 | 0.00233347 | No Change | Significant Negative | TRUE |
| Pter | TRUE | -0.116738343 | 0.717206457 | No Change | Not Significant | TRUE | -0.019825437 | 0.925325507 | No Change | Not Significant | TRUE |
| Ptgdr2 | TRUE | 0.1925089 | 1 | No Change | Not Significant | FALSE | NA | NA | No Change | Not Detected | FALSE |
| Ptger1 | TRUE | -0.501025959 | 0.33362953 | No Change | Not Significant | TRUE | -0.502386054 | 0.299347968 | No Change | Not Significant | TRUE |
| Ptger2 | TRUE | 0.816401256 | 1 | No Change | Not Significant | FALSE | NA | NA | No Change | Not Detected | FALSE |
| Ptger3 | TRUE | -0.531663768 | 0.129774424 | No Change | Not Significant | TRUE | -0.201385059 | 0.279429875 | No Change | Not Significant | TRUE |
| Ptger4 | TRUE | -0.318260173 | 0.028104133 | No Change | Significant Negative | TRUE | -0.093699357 | 0.597225526 | No Change | Not Significant | TRUE |
| Ptges | TRUE | 1.718113539 | 7.79E-17 | UP | UP | TRUE | 1.634781032 | 1.83E-06 | UP | UP | TRUE |
| Ptges2 | TRUE | -0.372578599 | 0.000477499 | No Change | Significant Negative | TRUE | -0.124597103 | 0.203825108 | No Change | Not Significant | TRUE |
| Ptges3 | TRUE | -0.094095038 | 0.660473296 | No Change | Not Significant | TRUE | -0.004068695 | 0.97297734 | No Change | Not Significant | TRUE |
| Ptges3l | TRUE | -0.09395027 | 0.830902522 | No Change | Not Significant | TRUE | 0.690840626 | 0.010812233 | UP | UP | TRUE |
| Ptgfr | TRUE | -1.218943506 | 0.199909758 | No Change | Not Significant | TRUE | -0.011960119 | 0.989393511 | No Change | Not Significant | TRUE |
| Ptgfrn | TRUE | 0.087660114 | 0.276098635 | No Change | Not Significant | TRUE | 0.077750338 | 0.4712713 | No Change | Not Significant | TRUE |
| Ptgir | TRUE | 1.283809451 | 1 | No Change | Not Significant | FALSE | NA | NA | No Change | Not Detected | FALSE |
| Ptgis | TRUE | 1.520044328 | 1 | No Change | Not Significant | FALSE | NA | NA | No Change | Not Detected | FALSE |
| Ptgr1 | TRUE | -0.081500764 | 0.767483504 | No Change | Not Significant | TRUE | 0.222551426 | 0.051260837 | No Change | Not Significant | TRUE |
| Ptgr2 | TRUE | -0.209076631 | 0.03960484 | No Change | Significant Negative | TRUE | 0.021664151 | 0.830562128 | No Change | Not Significant | TRUE |
| Ptgs1 | TRUE | -0.686245094 | 0.011942524 | DOWN | DOWN | TRUE | 0.219005218 | 0.290200492 | No Change | Not Significant | TRUE |
| Ptgs2 | TRUE | 1.105462752 | 1 | No Change | Not Significant | FALSE | NA | NA | No Change | Not Detected | FALSE |
| Ptgs2os | TRUE | -1.003959845 | 1 | No Change | Not Significant | FALSE | NA | NA | No Change | Not Detected | FALSE |
| Pth1r | TRUE | 0.272001368 | 1 | No Change | Not Significant | FALSE | NA | NA | No Change | Not Detected | FALSE |
| Pth2r | TRUE | 1.109359464 | 1 | No Change | Not Significant | FALSE | NA | NA | No Change | Not Detected | FALSE |
| Pthlh | TRUE | 0.6182015 | 1 | No Change | Not Significant | FALSE | NA | NA | No Change | Not Detected | FALSE |
| Ptk2 | TRUE | 0.063683502 | 0.467839271 | No Change | Not Significant | TRUE | 0.100899101 | 0.133784989 | No Change | Not Significant | TRUE |
| Ptk2b | TRUE | -0.395746131 | 0.00811528 | No Change | Significant Negative | TRUE | -0.113333654 | 0.077775409 | No Change | Not Significant | TRUE |
| Ptk6 | TRUE | 0.315779698 | 0.352762472 | No Change | Not Significant | TRUE | 0.138837545 | 0.514223998 | No Change | Not Significant | TRUE |
| Ptk7 | TRUE | 0.06734848 | 0.733576122 | No Change | Not Significant | TRUE | -0.230112667 | 0.038253441 | No Change | Significant Negative | TRUE |
| Ptma | TRUE | 0.272740592 | 0.208481869 | No Change | Not Significant | TRUE | 0.45311205 | 9.60E-10 | No Change | Significant Positive | TRUE |
| Ptms | TRUE | 0.342824453 | 5.22E-07 | No Change | Significant Positive | TRUE | 0.147375554 | 0.34931326 | No Change | Not Significant | TRUE |
| Ptov1 | TRUE | 0.57598415 | 1.67E-13 | UP | UP | TRUE | 0.157266798 | 0.109264501 | No Change | Not Significant | TRUE |
| Ptp4a1 | TRUE | 0.012285901 | 0.959151705 | No Change | Not Significant | TRUE | -0.374603041 | 0.028308503 | No Change | Significant Negative | TRUE |
| Ptp4a2 | TRUE | 0.139943567 | 0.273747344 | No Change | Not Significant | TRUE | 0.04456096 | 0.480291336 | No Change | Not Significant | TRUE |
| Ptp4a3 | TRUE | -0.383030525 | 0.043086943 | No Change | Significant Negative | TRUE | -0.10758598 | 0.560212157 | No Change | Not Significant | TRUE |
| Ptpa | TRUE | 0.057704985 | 0.671255829 | No Change | Not Significant | TRUE | 0.076879718 | 0.433436829 | No Change | Not Significant | TRUE |
| Ptpdc1 | TRUE | 0.431246988 | 0.62300216 | No Change | Not Significant | TRUE | 0.063661532 | 0.91643725 | No Change | Not Significant | TRUE |
| Ptpmt1 | TRUE | -0.095626877 | 0.515205432 | No Change | Not Significant | TRUE | -0.135383309 | 0.156629283 | No Change | Not Significant | TRUE |
| Ptpn1 | TRUE | 0.126554552 | 0.309427758 | No Change | Not Significant | TRUE | -0.007174458 | 0.96489645 | No Change | Not Significant | TRUE |
| Ptpn11 | TRUE | 0.032754707 | 0.807324664 | No Change | Not Significant | TRUE | 0.068832257 | 0.375594388 | No Change | Not Significant | TRUE |
| Ptpn12 | TRUE | -0.061082831 | 0.669122175 | No Change | Not Significant | TRUE | -0.130769145 | 0.125530222 | No Change | Not Significant | TRUE |
| Ptpn13 | TRUE | -0.215384847 | 0.210900795 | No Change | Not Significant | TRUE | -0.032766183 | 0.820965922 | No Change | Not Significant | TRUE |
| Ptpn14 | TRUE | -0.110959643 | 0.665436557 | No Change | Not Significant | TRUE | -0.04625591 | 0.906911659 | No Change | Not Significant | TRUE |
| Ptpn18 | TRUE | -0.066524021 | 0.569827206 | No Change | Not Significant | TRUE | 0.103717146 | 0.162728957 | No Change | Not Significant | TRUE |
| Ptpn2 | TRUE | 0.275575619 | 0.001657697 | No Change | Significant Positive | TRUE | 0.104723667 | 0.214828885 | No Change | Not Significant | TRUE |
| Ptpn21 | TRUE | 0.048973918 | 0.817198338 | No Change | Not Significant | TRUE | -0.059483958 | 0.730504162 | No Change | Not Significant | TRUE |
| Ptpn22 | TRUE | 0.033538347 | 0.970043107 | No Change | Not Significant | TRUE | 0.242886868 | 0.580252873 | No Change | Not Significant | TRUE |
| Ptpn23 | TRUE | 0.327184347 | 0.206262665 | No Change | Not Significant | TRUE | -0.078681794 | 0.646860933 | No Change | Not Significant | TRUE |
| Ptpn3 | TRUE | -0.151782832 | 0.216936107 | No Change | Not Significant | TRUE | 0.041493495 | 0.720116015 | No Change | Not Significant | TRUE |
| Ptpn4 | TRUE | -0.135850484 | 0.393919217 | No Change | Not Significant | TRUE | -0.10863398 | 0.156571643 | No Change | Not Significant | TRUE |
| Ptpn5 | TRUE | -0.178256312 | 0.927198731 | No Change | Not Significant | TRUE | -0.219016785 | 0.839430978 | No Change | Not Significant | TRUE |
| Ptpn6 | TRUE | -0.242190736 | 0.032853604 | No Change | Significant Negative | TRUE | 0.113430341 | 0.227778986 | No Change | Not Significant | TRUE |
| Ptpn7 | TRUE | 0.490440165 | 0.722169631 | No Change | Not Significant | TRUE | 0.33304028 | 0.74493282 | No Change | Not Significant | TRUE |
| Ptpn9 | TRUE | 0.097919177 | 0.267419303 | No Change | Not Significant | TRUE | 0.058460432 | 0.487862729 | No Change | Not Significant | TRUE |
| Ptpra | TRUE | 0.018453983 | 0.882687823 | No Change | Not Significant | TRUE | -0.095955577 | 0.163100633 | No Change | Not Significant | TRUE |
| Ptprb | TRUE | -0.17940021 | 0.910240646 | No Change | Not Significant | TRUE | -0.338063324 | 0.649970343 | No Change | Not Significant | TRUE |
| Ptprc | TRUE | -0.184586818 | 0.518220794 | No Change | Not Significant | TRUE | 0.32779864 | 0.062860758 | No Change | Not Significant | TRUE |
| Ptprd | TRUE | -0.226648828 | 0.00846073 | No Change | Significant Negative | TRUE | -0.305530197 | 2.99E-06 | No Change | Significant Negative | TRUE |
| Ptpre | TRUE | -0.013017718 | 0.952450218 | No Change | Not Significant | TRUE | 0.055462839 | 0.742536306 | No Change | Not Significant | TRUE |
| Ptprf | TRUE | 0.046865271 | 0.581581822 | No Change | Not Significant | TRUE | -0.052636025 | 0.671885771 | No Change | Not Significant | TRUE |
| Ptprh | TRUE | -0.240053651 | 0.64620651 | No Change | Not Significant | TRUE | -0.326442627 | 0.002747673 | No Change | Significant Negative | TRUE |
| Ptprj | TRUE | -0.363008651 | 2.30E-05 | No Change | Significant Negative | TRUE | -0.316007137 | 0.000105429 | No Change | Significant Negative | TRUE |
| Ptprk | TRUE | -0.091447142 | 0.524250269 | No Change | Not Significant | TRUE | -0.065089827 | 0.317992618 | No Change | Not Significant | TRUE |
| Ptprm | TRUE | 0.554053145 | 0.032645166 | UP | UP | TRUE | 0.307916861 | 0.124668167 | No Change | Not Significant | TRUE |
| Ptprn | TRUE | -1.124548324 | 0.032709605 | DOWN | DOWN | TRUE | -0.240650552 | 0.501083584 | No Change | Not Significant | TRUE |
| Ptprn2 | TRUE | -0.511481213 | 0.040727249 | DOWN | DOWN | TRUE | 0.053948857 | 0.71518329 | No Change | Not Significant | TRUE |
| Ptpro | TRUE | 0.844764507 | 0.009881034 | UP | UP | TRUE | 0.452051171 | 0.034058175 | No Change | Significant Positive | TRUE |
| Ptprq | TRUE | -1.681115405 | 1 | No Change | Not Significant | FALSE | NA | NA | No Change | Not Detected | FALSE |
| Ptprr | TRUE | -0.329284976 | 0.417696153 | No Change | Not Significant | TRUE | -0.25474753 | 0.196571068 | No Change | Not Significant | TRUE |
| Ptprs | TRUE | -0.630476369 | 0.000232612 | DOWN | DOWN | TRUE | -0.465397552 | 3.29E-12 | No Change | Significant Negative | TRUE |
| Ptprt | TRUE | -0.749785017 | 0.015546587 | DOWN | DOWN | TRUE | -0.154349165 | 0.674900387 | No Change | Not Significant | TRUE |
| Ptpru | TRUE | -0.171761272 | 0.540704851 | No Change | Not Significant | TRUE | -0.268864462 | 0.305128872 | No Change | Not Significant | TRUE |
| Ptprv | TRUE | 1.963763043 | 1 | No Change | Not Significant | FALSE | NA | NA | No Change | Not Detected | FALSE |
| Ptprz1 | TRUE | 0.411786625 | 1 | No Change | Not Significant | FALSE | NA | NA | No Change | Not Detected | FALSE |
| Ptrh1 | TRUE | 0.99739694 | 7.57E-05 | UP | UP | TRUE | 0.439816577 | 0.003024955 | No Change | Significant Positive | TRUE |
| Ptrh2 | TRUE | 0.096932877 | 0.640714813 | No Change | Not Significant | TRUE | 0.02710693 | 0.869327992 | No Change | Not Significant | TRUE |
| Ptrhd1 | TRUE | 0.112947564 | 0.688162291 | No Change | Not Significant | TRUE | 0.00837101 | 0.974310135 | No Change | Not Significant | TRUE |
| Pts | TRUE | 0.13054717 | 0.309937719 | No Change | Not Significant | TRUE | 0.175498626 | 0.022015973 | No Change | Significant Positive | TRUE |
| Pttg1 | TRUE | 0.046791849 | 0.76356297 | No Change | Not Significant | TRUE | 0.145242455 | 0.149190319 | No Change | Not Significant | TRUE |
| Pttg1ip | TRUE | -0.193923967 | 0.088794097 | No Change | Not Significant | TRUE | -0.004533418 | 0.968095473 | No Change | Not Significant | TRUE |
| Puf60 | TRUE | 0.045878457 | 0.776186935 | No Change | Not Significant | TRUE | 0.116299153 | 0.252829537 | No Change | Not Significant | TRUE |
| Pum1 | TRUE | 0.060950128 | 0.460585845 | No Change | Not Significant | TRUE | -0.052465278 | 0.536379174 | No Change | Not Significant | TRUE |
| Pum2 | TRUE | 0.076308294 | 0.623781054 | No Change | Not Significant | TRUE | -0.075741244 | 0.329962116 | No Change | Not Significant | TRUE |
| Pum3 | TRUE | -0.108066435 | 0.509484153 | No Change | Not Significant | TRUE | -0.061053924 | 0.602834501 | No Change | Not Significant | TRUE |
| Pura | TRUE | -0.12643976 | 0.225215631 | No Change | Not Significant | TRUE | -0.032528413 | 0.78574508 | No Change | Not Significant | TRUE |
| Purb | TRUE | 0.316513189 | 0.021461314 | No Change | Significant Positive | TRUE | 0.174521968 | 0.009099034 | No Change | Significant Positive | TRUE |
| Purg | TRUE | -0.485040841 | 0.347033567 | No Change | Not Significant | TRUE | -0.074318614 | 0.883529645 | No Change | Not Significant | TRUE |
| Pus1 | TRUE | -0.184321559 | 0.30705241 | No Change | Not Significant | TRUE | 0.012808024 | 0.941878077 | No Change | Not Significant | TRUE |
| Pus10 | TRUE | 0.132059834 | 0.140963833 | No Change | Not Significant | TRUE | -0.032558069 | 0.833278687 | No Change | Not Significant | TRUE |
| Pus3 | TRUE | 1.916473964 | 1 | No Change | Not Significant | FALSE | NA | NA | No Change | Not Detected | FALSE |
| Pus7 | TRUE | 0.006486605 | 0.97686351 | No Change | Not Significant | TRUE | -0.005771113 | 0.979948241 | No Change | Not Significant | TRUE |
| Pus7l | TRUE | 0.22551766 | 0.510557756 | No Change | Not Significant | TRUE | 0.120449714 | 0.639523975 | No Change | Not Significant | TRUE |
| Pusl1 | TRUE | -0.340270623 | 0.15599306 | No Change | Not Significant | TRUE | -0.20350221 | 0.276680594 | No Change | Not Significant | TRUE |
| Pvalb | TRUE | 1.070357869 | 1 | No Change | Not Significant | FALSE | NA | NA | No Change | Not Detected | FALSE |
| Pvr | TRUE | -0.058599884 | 0.825971257 | No Change | Not Significant | TRUE | -0.318942259 | 0.041539585 | No Change | Significant Negative | TRUE |
| Pvrig | TRUE | 1.84312758 | 1 | No Change | Not Significant | FALSE | NA | NA | No Change | Not Detected | FALSE |
| Pvt1 | TRUE | 0.525929299 | 0.215982137 | No Change | Not Significant | TRUE | 0.625501011 | 0.110334952 | No Change | Not Significant | TRUE |
| Pwp1 | TRUE | 0.018433123 | 0.931471303 | No Change | Not Significant | TRUE | -0.025720456 | 0.848090631 | No Change | Not Significant | TRUE |
| Pwp2 | TRUE | -0.187656143 | 0.228807636 | No Change | Not Significant | TRUE | -0.228933365 | 0.017639786 | No Change | Significant Negative | TRUE |
| Pwwp2a | TRUE | 0.194429103 | 0.073869638 | No Change | Not Significant | TRUE | -0.102988218 | 0.437979805 | No Change | Not Significant | TRUE |
| Pwwp2b | TRUE | -0.045469352 | 0.794916857 | No Change | Not Significant | TRUE | 0.095490906 | 0.45050036 | No Change | Not Significant | TRUE |
| Pwwp3a | TRUE | -0.0906138 | 0.55548954 | No Change | Not Significant | TRUE | -0.07864046 | 0.355746355 | No Change | Not Significant | TRUE |
| Pwwp3b | TRUE | 0.51754662 | 0.456655966 | No Change | Not Significant | TRUE | 0.200231175 | 0.802084296 | No Change | Not Significant | TRUE |
| Pxdc1 | TRUE | 0.234608125 | 0.662854285 | No Change | Not Significant | TRUE | 0.440124438 | 0.096764517 | No Change | Not Significant | TRUE |
| Pxdn | TRUE | -0.299132342 | 0.810243834 | No Change | Not Significant | TRUE | 0.483484837 | 0.552526426 | No Change | Not Significant | TRUE |
| Pxk | TRUE | -0.191946335 | 0.028656498 | No Change | Significant Negative | TRUE | -0.06156009 | 0.399424529 | No Change | Not Significant | TRUE |
| Pxmp2 | TRUE | -0.157556578 | 0.5907413 | No Change | Not Significant | TRUE | -0.012058627 | 0.971967201 | No Change | Not Significant | TRUE |
| Pxmp4 | TRUE | 0.252365948 | 0.007373206 | No Change | Significant Positive | TRUE | 0.030283912 | 0.78841372 | No Change | Not Significant | TRUE |
| Pxn | TRUE | 0.0421598 | 0.742744607 | No Change | Not Significant | TRUE | 0.054684918 | 0.622645752 | No Change | Not Significant | TRUE |
| Pxt1 | TRUE | -1.006532126 | 1 | No Change | Not Significant | FALSE | NA | NA | No Change | Not Detected | FALSE |
| Pxylp1 | TRUE | -0.023380032 | 0.938630585 | No Change | Not Significant | TRUE | -0.023922539 | 0.836585484 | No Change | Not Significant | TRUE |
| Pycard | TRUE | 0.194780653 | 0.172631249 | No Change | Not Significant | TRUE | 0.395571152 | 9.16E-08 | No Change | Significant Positive | TRUE |
| Pycr1 | TRUE | 0.626967498 | 0.000232612 | UP | UP | TRUE | 0.104959768 | 0.71539272 | No Change | Not Significant | TRUE |
| Pycr2 | TRUE | 1.415453693 | 8.08E-59 | UP | UP | TRUE | 0.747454586 | 9.15E-17 | UP | UP | TRUE |
| Pycrl | TRUE | -0.108440908 | 0.412914333 | No Change | Not Significant | TRUE | 0.142881892 | 0.178864648 | No Change | Not Significant | TRUE |
| Pygb | TRUE | 0.245619338 | 0.092985996 | No Change | Not Significant | TRUE | 0.382500327 | 1.81E-07 | No Change | Significant Positive | TRUE |
| Pygl | TRUE | -0.688715445 | 2.24E-05 | DOWN | DOWN | TRUE | 0.169553697 | 0.303323933 | No Change | Not Significant | TRUE |
| Pygm | TRUE | 1.09596419 | 0.022923163 | UP | UP | TRUE | 0.148999856 | 0.864247815 | No Change | Not Significant | TRUE |
| Pygo1 | TRUE | -1.120501233 | 1 | No Change | Not Significant | FALSE | NA | NA | No Change | Not Detected | FALSE |
| Pygo2 | TRUE | 0.136764349 | 0.05583573 | No Change | Not Significant | TRUE | 0.006858589 | 0.966739549 | No Change | Not Significant | TRUE |
| Pym1 | TRUE | 0.013583248 | 0.933333191 | No Change | Not Significant | TRUE | 0.001026696 | 0.996376861 | No Change | Not Significant | TRUE |
| Pyroxd1 | TRUE | -0.069504034 | 0.761221133 | No Change | Not Significant | TRUE | -0.167318706 | 0.139211338 | No Change | Not Significant | TRUE |
| Pyroxd2 | TRUE | -0.48340776 | 0.000548043 | No Change | Significant Negative | TRUE | -0.181721298 | 0.377781372 | No Change | Not Significant | TRUE |
| Pyurf | TRUE | -0.17599588 | 0.171466333 | No Change | Not Significant | TRUE | 0.093671601 | 0.433436829 | No Change | Not Significant | TRUE |
| Pyy | TRUE | -0.396566613 | 0.154304393 | No Change | Not Significant | TRUE | -0.238328526 | 0.014762904 | No Change | Significant Negative | TRUE |
| Pzp | TRUE | 1.181376703 | 1 | No Change | Not Significant | FALSE | NA | NA | No Change | Not Detected | FALSE |
| Qars | TRUE | 0.425082825 | 3.40E-08 | No Change | Significant Positive | TRUE | 0.111721109 | 0.034738353 | No Change | Significant Positive | TRUE |
| Qdpr | TRUE | 0.587883296 | 4.24E-14 | UP | UP | TRUE | 0.342551808 | 1.16E-17 | No Change | Significant Positive | TRUE |
| Qki | TRUE | 0.461382129 | 0.508065109 | No Change | Not Significant | TRUE | 0.300886972 | 0.617215876 | No Change | Not Significant | TRUE |
| Qpct | TRUE | 0.089550597 | 0.923599766 | No Change | Not Significant | TRUE | 0.238549696 | 0.685788934 | No Change | Not Significant | TRUE |
| Qpctl | TRUE | 0.285163315 | 0.02767136 | No Change | Significant Positive | TRUE | 0.303508625 | 0.003141798 | No Change | Significant Positive | TRUE |
| Qprt | TRUE | 0.164470625 | 1 | No Change | Not Significant | FALSE | NA | NA | No Change | Not Detected | FALSE |
| Qrfp | TRUE | -0.379042677 | 0.76356297 | No Change | Not Significant | TRUE | -0.527801606 | 0.541946771 | No Change | Not Significant | TRUE |
| Qrich1 | TRUE | -0.013875186 | 0.882651278 | No Change | Not Significant | TRUE | -0.014182418 | 0.857064391 | No Change | Not Significant | TRUE |
| Qrich2 | TRUE | 1.776460859 | 2.25E-05 | UP | UP | TRUE | 0.521991759 | 0.449133255 | No Change | Not Significant | TRUE |
| Qrsl1 | TRUE | -0.233816149 | 0.073386004 | No Change | Not Significant | TRUE | 0.102207582 | 0.386825021 | No Change | Not Significant | TRUE |
| Qsox1 | TRUE | 0.012688605 | 0.976016991 | No Change | Not Significant | TRUE | 0.251104397 | 0.00107324 | No Change | Significant Positive | TRUE |
| Qsox2 | TRUE | 0.353977425 | 0.125701628 | No Change | Not Significant | TRUE | 0.092635482 | 0.375325502 | No Change | Not Significant | TRUE |
| Qtrt1 | TRUE | -0.14422675 | 0.456250391 | No Change | Not Significant | TRUE | -0.211259509 | 0.070685953 | No Change | Not Significant | TRUE |
| Qtrt2 | TRUE | -0.105962013 | 0.523269741 | No Change | Not Significant | TRUE | -0.279514439 | 0.066251304 | No Change | Not Significant | TRUE |
| R3hcc1 | TRUE | 0.054400137 | 0.751588072 | No Change | Not Significant | TRUE | -0.079066956 | 0.536541439 | No Change | Not Significant | TRUE |
| R3hcc1l | TRUE | -0.199963284 | 0.019336968 | No Change | Significant Negative | TRUE | -0.157383873 | 0.055950184 | No Change | Not Significant | TRUE |
| R3hdm1 | TRUE | 0.036589149 | 0.861359607 | No Change | Not Significant | TRUE | -0.059945728 | 0.702846794 | No Change | Not Significant | TRUE |
| R3hdm2 | TRUE | -0.019337212 | 0.945235545 | No Change | Not Significant | TRUE | -0.106364693 | 0.477271565 | No Change | Not Significant | TRUE |
| R3hdm4 | TRUE | -0.399492336 | 5.04E-07 | No Change | Significant Negative | TRUE | -0.062071583 | 0.65259636 | No Change | Not Significant | TRUE |
| R3hdml | TRUE | 2.257677207 | 1 | No Change | Not Significant | FALSE | NA | NA | No Change | Not Detected | FALSE |
| R74862 | TRUE | 0.408302088 | 0.713367573 | No Change | Not Significant | TRUE | -0.037325118 | 0.96964417 | No Change | Not Significant | TRUE |
| Rab10 | TRUE | 0.074947715 | 0.570683233 | No Change | Not Significant | TRUE | 0.096946861 | 0.143269254 | No Change | Not Significant | TRUE |
| Rab11a | TRUE | 0.00010618 | 0.999353427 | No Change | Not Significant | TRUE | 0.093278055 | 0.115148818 | No Change | Not Significant | TRUE |
| Rab11b | TRUE | -0.240007918 | 0.024640043 | No Change | Significant Negative | TRUE | 0.03237332 | 0.676462024 | No Change | Not Significant | TRUE |
| Rab11fip1 | TRUE | -0.306073352 | 0.000194686 | No Change | Significant Negative | TRUE | -0.079767768 | 0.192640198 | No Change | Not Significant | TRUE |
| Rab11fip2 | TRUE | -0.214303435 | 0.331061353 | No Change | Not Significant | TRUE | -0.137911079 | 0.468848355 | No Change | Not Significant | TRUE |
| Rab11fip3 | TRUE | 0.098381466 | 0.532495959 | No Change | Not Significant | TRUE | 0.061039901 | 0.59777452 | No Change | Not Significant | TRUE |
| Rab11fip4 | TRUE | -0.253449716 | 0.179258167 | No Change | Not Significant | TRUE | -0.289554551 | 1.20E-06 | No Change | Significant Negative | TRUE |
| Rab11fip4os1 | TRUE | 1.260024323 | 1 | No Change | Not Significant | FALSE | NA | NA | No Change | Not Detected | FALSE |
| Rab11fip4os2 | TRUE | -0.76954059 | 1 | No Change | Not Significant | FALSE | NA | NA | No Change | Not Detected | FALSE |
| Rab11fip5 | TRUE | -0.081868098 | 0.826766769 | No Change | Not Significant | TRUE | 0.002748451 | 0.994696524 | No Change | Not Significant | TRUE |
| Rab12 | TRUE | 0.131192375 | 0.56547568 | No Change | Not Significant | TRUE | -0.298496611 | 0.040151712 | No Change | Significant Negative | TRUE |
| Rab13 | TRUE | -0.182980407 | 0.191599363 | No Change | Not Significant | TRUE | 0.086992442 | 0.584131301 | No Change | Not Significant | TRUE |
| Rab14 | TRUE | -0.055241753 | 0.484149199 | No Change | Not Significant | TRUE | 0.021765536 | 0.740026398 | No Change | Not Significant | TRUE |
| Rab15 | TRUE | -0.011751116 | 0.97587882 | No Change | Not Significant | TRUE | 0.164615381 | 0.042790188 | No Change | Significant Positive | TRUE |
| Rab17 | TRUE | -0.690678672 | 0.000103273 | DOWN | DOWN | TRUE | -0.142054884 | 0.401707581 | No Change | Not Significant | TRUE |
| Rab18 | TRUE | 0.243661458 | 0.004152083 | No Change | Significant Positive | TRUE | 0.026918959 | 0.76190043 | No Change | Not Significant | TRUE |
| Rab19 | TRUE | -0.530564488 | 0.000476541 | DOWN | DOWN | TRUE | -0.14105066 | 0.280723736 | No Change | Not Significant | TRUE |
| Rab1a | TRUE | 0.456192869 | 7.94E-10 | No Change | Significant Positive | TRUE | 0.309672515 | 3.79E-10 | No Change | Significant Positive | TRUE |

[illegible]

Supplementary Table 3

| geneID | Cond1_DF_Detected | Cond1_DF_ValueCol | Cond1_DF_PadjCol | Cond1_DF_Cutoff | Cond1_DF_Cutoff_Specific.x | Cond2_DF_Detected |
|---|---|---|---|---|---|---|
| Rab1b | TRUE | 0.03111494 | 0.777615016 | No Change | Not Significant | TRUE |
| Rab20 | TRUE | 0.383266046 | 0.000592833 | No Change | Significant Positive | TRUE |
| Rab21 | TRUE | 0.125052294 | 0.235661079 | No Change | Not Significant | TRUE |
| Rab22a | TRUE | -0.014465097 | 0.922414938 | No Change | Not Significant | TRUE |
| Rab23 | TRUE | 0.04290368 | 0.895584805 | No Change | Not Significant | TRUE |
| Rab24 | TRUE | 0.071680861 | 0.722040793 | No Change | Not Significant | TRUE |
| Rab25 | TRUE | -0.292548452 | 0.001618275 | No Change | Significant Negative | TRUE |
| Rab26 | TRUE | 2.014125402 | 0.006956521 | UP | UP | FALSE |
| Rab26os | TRUE | 1.55530935 | 2.22E-05 | UP | UP | TRUE |
| Rab27a | TRUE | 0.281396592 | 0.20483749 | No Change | Not Significant | TRUE |
| Rab27b | TRUE | 0.160137711 | 0.601173216 | No Change | Not Significant | TRUE |
| Rab28 | TRUE | 0.117268377 | 0.282329367 | No Change | Not Significant | TRUE |
| Rab29 | TRUE | -0.297217402 | 0.103335112 | No Change | Not Significant | TRUE |
| Rab2a | TRUE | 0.152695905 | 0.314808425 | No Change | Not Significant | TRUE |
| Rab2b | TRUE | -0.009378499 | 0.989126875 | No Change | Not Significant | TRUE |
| Rab30 | TRUE | -0.795345059 | 0.19856618 | No Change | Not Significant | TRUE |
| Rab31 | TRUE | -0.169948066 | 0.450393623 | No Change | Not Significant | TRUE |
| Rab32 | TRUE | -0.156803125 | 0.69793377 | No Change | Not Significant | TRUE |
| Rab33a | TRUE | -0.476841013 | 1 | No Change | Not Significant | TRUE |
| Rab33b | TRUE | 0.048704502 | 0.813372119 | No Change | Not Significant | TRUE |
| Rab34 | TRUE | -0.089472058 | 1 | No Change | Not Significant | FALSE |
| Rab35 | TRUE | 0.0403834 | 0.771976939 | No Change | Not Significant | TRUE |
| Rab36 | TRUE | 2.892530458 | 2.64E-08 | UP | UP | TRUE |
| Rab37 | TRUE | -1.130496265 | 0.000903321 | DOWN | DOWN | TRUE |
| Rab38 | TRUE | 3.109626238 | 2.15E-77 | UP | UP | TRUE |
| Rab39b | TRUE | 1.739826812 | 1 | No Change | Not Significant | FALSE |
| Rab3a | TRUE | 0.582522966 | 5.63E-05 | UP | UP | TRUE |
| Rab3b | TRUE | 0.934226819 | 0.033659295 | UP | UP | TRUE |
| Rab3c | TRUE | -0.374487517 | 0.378653073 | No Change | Not Significant | TRUE |
| Rab3d | TRUE | 0.074250145 | 0.800627603 | No Change | Not Significant | TRUE |
| Rab3gap1 | TRUE | -0.111849565 | 0.405918699 | No Change | Not Significant | TRUE |
| Rab3gap2 | TRUE | 0.145483117 | 0.056578534 | No Change | Not Significant | TRUE |
| Rab3il1 | TRUE | 0.610058429 | 0.697627026 | No Change | Not Significant | FALSE |
| Rab3ip | TRUE | -0.1416793 | 0.26324771 | No Change | Not Significant | TRUE |
| Rab40b | TRUE | -0.79681305 | 0.368771559 | No Change | Not Significant | TRUE |
| Rab40c | TRUE | -0.055286397 | 0.653742472 | No Change | Not Significant | TRUE |
| Rab42 | TRUE | 0.517588778 | 0.6385535 | No Change | Not Significant | TRUE |
| Rab43 | TRUE | -0.174077476 | 0.601173216 | No Change | Not Significant | TRUE |
| Rab44 | TRUE | -0.423579577 | 1 | No Change | Not Significant | FALSE |
| Rab4a | TRUE | 0.127271171 | 0.484503153 | No Change | Not Significant | TRUE |
| Rab4b | TRUE | 0.171328702 | 0.617207172 | No Change | Not Significant | TRUE |
| Rab5a | TRUE | 0.140944967 | 0.116638837 | No Change | Not Significant | TRUE |
| Rab5b | TRUE | 0.056994421 | 0.743179634 | No Change | Not Significant | TRUE |
| Rab5c | TRUE | -0.023660149 | 0.86030736 | No Change | Not Significant | TRUE |
| Rab5if | TRUE | 0.38374015 | 0.00429793 | No Change | Significant Positive | TRUE |
| Rab6a | TRUE | 0.412851889 | 1.52E-09 | No Change | Significant Positive | TRUE |
| Rab6b | TRUE | 0.516862324 | 0.349700969 | No Change | Not Significant | TRUE |
| Rab7 | TRUE | -0.14464811 | 0.046796074 | No Change | Significant Negative | TRUE |
| Rab7b | TRUE | 0.80869267 | 1 | No Change | Not Significant | FALSE |
| Rab8a | TRUE | 0.257437932 | 4.01E-07 | No Change | Significant Positive | TRUE |
| Rab8b | TRUE | 0.172700427 | 0.259233994 | No Change | Not Significant | TRUE |
| Rab9 | TRUE | -0.154255467 | 0.14798755 | No Change | Not Significant | TRUE |
| Rabac1 | TRUE | 0.026589143 | 0.84294636 | No Change | Not Significant | TRUE |
| Rabep1 | TRUE | -0.179059982 | 0.201777892 | No Change | Not Significant | TRUE |
| Rabep2 | TRUE | -0.17581703 | 0.192028673 | No Change | Not Significant | TRUE |
| Rabepk | TRUE | 0.886602355 | 2.80E-36 | UP | UP | TRUE |
| Rabgap1 | TRUE | -0.074629653 | 0.566901476 | No Change | Not Significant | TRUE |
| Rabgap1l | TRUE | -0.27986703 | 0.002851946 | No Change | Significant Negative | TRUE |
| Rabgef1 | TRUE | -0.02487062 | 0.857652848 | No Change | Not Significant | TRUE |
| Rabggta | TRUE | -0.037058488 | 0.809917796 | No Change | Not Significant | TRUE |
| Rabggtb | TRUE | -0.103933569 | 0.455021685 | No Change | Not Significant | TRUE |
| Rabif | TRUE | 0.043777291 | 0.832872319 | No Change | Not Significant | TRUE |
| Rabl2 | TRUE | 0.482708549 | 0.00218877 | No Change | Significant Positive | TRUE |
| Rabl3 | TRUE | 0.075640297 | 0.687930892 | No Change | Not Significant | TRUE |
| Rabl6 | TRUE | 0.275901557 | 0.000359858 | No Change | Significant Positive | TRUE |
| Rac1 | TRUE | -0.046965088 | 0.670515281 | No Change | Not Significant | TRUE |
| Rac2 | TRUE | -0.73216623 | 0.003232956 | DOWN | DOWN | TRUE |
| Rac3 | TRUE | 0.06918545 | 0.785550184 | No Change | Not Significant | TRUE |
| Racgap1 | TRUE | 0.737301553 | 1.02E-05 | UP | UP | TRUE |
| Rack1 | TRUE | -0.158530305 | 0.285472293 | No Change | Not Significant | TRUE |
| Rad1 | TRUE | 0.136753108 | 0.506201691 | No Change | Not Significant | TRUE |
| Rad17 | TRUE | 0.124759208 | 0.28232699 | No Change | Not Significant | TRUE |
| Rad18 | TRUE | 0.661076796 | 0.000195468 | UP | UP | TRUE |
| Rad21 | TRUE | 0.303948725 | 0.012153013 | No Change | Significant Positive | TRUE |
| Rad23a | TRUE | -0.780733242 | 4.54E-20 | DOWN | DOWN | TRUE |
| Rad23b | TRUE | 0.285798149 | 0.00158386 | No Change | Significant Positive | TRUE |
| Rad50 | TRUE | 0.041636223 | 0.825194181 | No Change | Not Significant | TRUE |
| Rad51 | TRUE | 0.623652281 | 0.020744341 | UP | UP | TRUE |
| Rad51ap1 | TRUE | 0.577606627 | 0.06604638 | No Change | Not Significant | TRUE |
| Rad51ap2 | TRUE | 0.004954383 | 1 | No Change | Not Significant | FALSE |
| Rad51b | TRUE | 0.495773749 | 0.160039404 | No Change | Not Significant | TRUE |
| Rad51c | TRUE | 0.195677826 | 0.722040793 | No Change | Not Significant | TRUE |
| Rad51d | TRUE | -0.101662923 | 0.50827749 | No Change | Not Significant | TRUE |
| Rad52 | TRUE | -0.175027508 | 0.33564513 | No Change | Not Significant | TRUE |
| Rad54b | TRUE | 0.740981422 | 0.007288737 | UP | UP | TRUE |
| Rad54l | TRUE | 0.440134448 | 0.127940538 | No Change | Not Significant | TRUE |
| Rad54l2 | TRUE | -0.049953671 | 0.5825209 | No Change | Not Significant | TRUE |
| Rad9a | TRUE | 0.245919877 | 0.293636687 | No Change | Not Significant | TRUE |
| Rad9b | TRUE | -0.099349938 | 0.827422396 | No Change | Not Significant | TRUE |
| Radil | TRUE | 4.591896027 | 1 | No Change | Not Significant | FALSE |
| Radx | TRUE | -0.193004684 | 0.463479905 | No Change | Not Significant | TRUE |
| Rae1 | TRUE | 0.06628489 | 0.598887725 | No Change | Not Significant | TRUE |
| Raet1e | TRUE | -1.021362618 | 0.00034471 | DOWN | DOWN | TRUE |
| Raf1 | TRUE | 0.039522429 | 0.709123099 | No Change | Not Significant | TRUE |
| Rag1 | TRUE | 2.247946433 | 1 | No Change | Not Significant | FALSE |
| Rag2 | TRUE | 1.539687032 | 1 | No Change | Not Significant | FALSE |
| Rai1 | TRUE | 0.124276318 | 0.610103781 | No Change | Not Significant | TRUE |
| Rai14 | TRUE | 0.00391071 | 0.987234441 | No Change | Not Significant | TRUE |
| Rai2 | TRUE | -1.162377293 | 0.099855399 | No Change | Not Significant | TRUE |
| Rala | TRUE | 0.062539738 | 0.589852079 | No Change | Not Significant | TRUE |
| Ralb | TRUE | 0.967559894 | 4.17E-38 | UP | UP | TRUE |
| Ralbp1 | TRUE | -0.175364417 | 0.038706422 | No Change | Significant Negative | TRUE |
| Ralgapa1 | TRUE | 0.057858639 | 0.736573919 | No Change | Not Significant | TRUE |
| Ralgapa2 | TRUE | 0.00546321 | 0.973296906 | No Change | Not Significant | TRUE |
| Ralgapb | TRUE | -0.037604375 | 0.725715181 | No Change | Not Significant | TRUE |
| Ralgds | TRUE | 0.125135883 | 0.548343875 | No Change | Not Significant | TRUE |
| Ralgps1 | TRUE | 0.414733361 | 0.000754888 | No Change | Significant Positive | TRUE |
| Ralgps2 | TRUE | 0.190710583 | 0.002700918 | No Change | Significant Positive | TRUE |
| Raly | TRUE | 0.017881542 | 0.888775635 | No Change | Not Significant | TRUE |
| Ralyl | TRUE | 0.176010352 | 1 | No Change | Not Significant | FALSE |
| Ramac | TRUE | -0.143809191 | 0.433049501 | No Change | Not Significant | TRUE |
| Ramp1 | TRUE | -0.29497883 | 0.517476912 | No Change | Not Significant | TRUE |
| Ramp2 | TRUE | 0.239293354 | 1 | No Change | Not Significant | FALSE |
| Ramp3 | TRUE | -0.542625815 | 1 | No Change | Not Significant | TRUE |
| Ran | TRUE | 0.187539409 | 0.394975053 | No Change | Not Significant | TRUE |
| Ranbp1 | TRUE | 0.382948762 | 0.003214881 | No Change | Significant Positive | TRUE |
| Ranbp10 | TRUE | 0.003303517 | 0.984433119 | No Change | Not Significant | TRUE |
| Ranbp17 | TRUE | 0.588147826 | 0.257146025 | No Change | Not Significant | TRUE |
| Ranbp2 | TRUE | 0.020411815 | 0.905997055 | No Change | Not Significant | TRUE |
| Ranbp3 | TRUE | 0.086835264 | 0.252397465 | No Change | Not Significant | TRUE |
| Ranbp3l | TRUE | -2.28055445 | 1 | No Change | Not Significant | FALSE |
| Ranbp6 | TRUE | -0.037305766 | 0.819267308 | No Change | Not Significant | TRUE |
| Ranbp9 | TRUE | 0.001117371 | 0.99195007 | No Change | Not Significant | TRUE |
| Rangap1 | TRUE | 0.360585148 | 0.012608423 | No Change | Significant Positive | TRUE |
| Rangrf | TRUE | 0.369043629 | 1 | No Change | Not Significant | FALSE |
| Rap1a | TRUE | -0.131421529 | 0.425947586 | No Change | Not Significant | TRUE |
| Rap1b | TRUE | 0.159746657 | 0.109621469 | No Change | Not Significant | TRUE |
| Rap1gap | TRUE | -0.029453944 | 0.904600714 | No Change | Not Significant | TRUE |
| Rap1gap2 | TRUE | -0.346263225 | 2.70E-06 | No Change | Significant Negative | TRUE |
| Rap1gapos | TRUE | 0.761083792 | 0.008565982 | UP | UP | TRUE |
| Rap1gds1 | TRUE | -0.123585188 | 0.234124471 | No Change | Not Significant | TRUE |
| Rap2a | TRUE | 0.272261631 | 0.001439931 | No Change | Significant Positive | TRUE |
| Rap2b | TRUE | 0.598932904 | 0.045076681 | UP | UP | TRUE |
| Rap2c | TRUE | 0.031504161 | 0.82238042 | No Change | Not Significant | TRUE |
| Rapgef1 | TRUE | 0.112780096 | 0.18970918 | No Change | Not Significant | TRUE |
| Rapgef2 | TRUE | 0.824427974 | 5.01E-14 | UP | UP | TRUE |
| Rapgef3 | TRUE | 0.000520859 | 0.99837123 | No Change | Not Significant | TRUE |
| Rapgef4 | TRUE | -0.609527777 | 0.067539312 | No Change | Not Significant | TRUE |
| Rapgef5 | TRUE | 0.000212954 | 0.998492433 | No Change | Not Significant | TRUE |
| Rapgef6 | TRUE | 0.001574733 | 0.990883011 | No Change | Not Significant | TRUE |
| Rapgefl1 | TRUE | -0.406588354 | 0.006100279 | No Change | Significant Negative | TRUE |
| Raph1 | TRUE | 0.022789905 | 0.947786691 | No Change | Not Significant | TRUE |
| Rapsn | TRUE | 2.055925908 | 1 | No Change | Not Significant | FALSE |
| Rara | TRUE | -0.093472237 | 0.736522749 | No Change | Not Significant | TRUE |
| Rarb | TRUE | 1.437374105 | 1 | No Change | Not Significant | FALSE |
| Rarg | TRUE | -0.411080453 | 0.036642767 | No Change | Significant Negative | TRUE |
| Rarres1 | TRUE | 0.533830748 | 0.527150555 | No Change | Not Significant | FALSE |
| Rarres2 | TRUE | 0.342162104 | 1 | No Change | Not Significant | FALSE |
| Rars | TRUE | 0.165102278 | 0.053943227 | No Change | Not Significant | TRUE |
| Rars2 | TRUE | -0.166949868 | 0.224854857 | No Change | Not Significant | TRUE |
| Rasa1 | TRUE | 0.0253641 | 0.868161055 | No Change | Not Significant | TRUE |
| Rasa2 | TRUE | -0.300635324 | 1.32E-05 | No Change | Significant Negative | TRUE |
| Rasa3 | TRUE | 0.178837624 | 0.347924739 | No Change | Not Significant | TRUE |
| Rasa4 | TRUE | -0.378967558 | 0.033749795 | No Change | Significant Negative | TRUE |
| Rasal1 | TRUE | -0.493846566 | 0.695624496 | No Change | Not Significant | FALSE |
| Rasal2 | TRUE | -0.04053187 | 0.8379481 | No Change | Not Significant | TRUE |
| Rasal3 | TRUE | -0.047650937 | 0.956997404 | No Change | Not Significant | TRUE |
| Rasd1 | TRUE | 0.169918052 | 0.569001592 | No Change | Not Significant | TRUE |
| Rasd2 | TRUE | -0.353387711 | 0.417696153 | No Change | Not Significant | TRUE |
| Rasef | TRUE | 0.152181014 | 0.195041062 | No Change | Not Significant | TRUE |
| Rasgef1a | TRUE | 0.03571214 | 1 | No Change | Not Significant | TRUE |
| Rasgef1b | TRUE | -0.783195592 | 1.55E-05 | DOWN | DOWN | TRUE |
| Rasgef1c | TRUE | -2.119102208 | 1 | No Change | Not Significant | FALSE |
| Rasgrf1 | TRUE | 0.594201213 | 0.876856731 | No Change | Not Significant | FALSE |
| Rasgrf2 | TRUE | 0.313194295 | 0.089531187 | No Change | Not Significant | TRUE |
| Rasgrp1 | TRUE | -0.113506371 | 0.890186044 | No Change | Not Significant | TRUE |
| Rasgrp2 | TRUE | 0.751939502 | 0.606235185 | No Change | Not Significant | TRUE |
| Rasgrp3 | TRUE | -0.477833185 | 1 | No Change | Not Significant | FALSE |
| Rasgrp4 | TRUE | 1.133039009 | 0.272799628 | No Change | Not Significant | TRUE |
| Rasip1 | TRUE | 0.660565929 | 0.041791803 | UP | UP | TRUE |
| Rasl10a | TRUE | -0.415752372 | 1 | No Change | Not Significant | FALSE |
| Rasl10b | TRUE | 0.235705705 | 1 | No Change | Not Significant | FALSE |
| Rasl11a | TRUE | 2.728717738 | 1.44E-13 | UP | UP | TRUE |
| Rasl11b | TRUE | 1.649003205 | 1.06E-05 | UP | UP | TRUE |
| Rasl12 | TRUE | 0.888436555 | 0.124778744 | No Change | Not Significant | TRUE |
| Rasl2-9 | TRUE | -0.03504077 | 0.971762583 | No Change | Not Significant | TRUE |
| Rassf1 | TRUE | 0.095527919 | 0.707250275 | No Change | Not Significant | TRUE |
| Rassf10 | TRUE | -0.053846649 | 0.915629096 | No Change | Not Significant | TRUE |
| Rassf2 | TRUE | 0.482290007 | 0.5402916 | No Change | Not Significant | TRUE |
| Rassf3 | TRUE | -0.03059208 | 0.84101582 | No Change | Not Significant | TRUE |
| Rassf4 | TRUE | 0.203301273 | 0.466508229 | No Change | Not Significant | TRUE |
| Rassf5 | TRUE | 0.244073354 | 0.755154277 | No Change | Not Significant | TRUE |
| Rassf6 | TRUE | -0.179450175 | 0.310948291 | No Change | Not Significant | TRUE |
| Rassf7 | TRUE | -0.037325893 | 0.840298483 | No Change | Not Significant | TRUE |
| Rassf8 | TRUE | 0.398712208 | 0.350538094 | No Change | Not Significant | TRUE |
| Rassf9 | TRUE | -0.456358219 | 0.102451045 | No Change | Not Significant | TRUE |
| Raver2 | TRUE | 0.884457391 | 0.325550242 | No Change | Not Significant | TRUE |
| Rax | TRUE | 1.477811694 | 1 | No Change | Not Significant | FALSE |
| Rb1 | TRUE | -0.275463893 | 0.004469372 | No Change | Significant Negative | TRUE |
| Rb1cc1 | TRUE | -0.155329564 | 0.220621763 | No Change | Not Significant | TRUE |
| Rbak | TRUE | -0.08621808 | 0.690541222 | No Change | Not Significant | TRUE |
| Rbakdn | TRUE | -0.386701882 | 1 | No Change | Not Significant | FALSE |

| geneID | Cond1_DF_Detected | Cond1_DF_ValueCol | Cond1_DF_PadjCol | Cond1_DF_Cutoff | Cond1_DF_Cutoff_Specific.x | [illegible] |
|---|---|---|---|---|---|---|
| Rbbp4 | TRUE | 0.041942589 | 0.830769833 | No Change | Not Significant | [illegible] |
| Rbbp5 | TRUE | 0.05890086 | 0.648975432 | No Change | Not Significant | [illegible] |
| Rbbp6 | TRUE | 0.142265703 | 0.115831422 | No Change | Not Significant | [illegible] |
| Rbbp7 | TRUE | 0.11922535 | 0.429396526 | No Change | Not Significant | [illegible] |
| Rbbp8 | TRUE | 0.031161954 | 0.822609935 | No Change | Not Significant | [illegible] |
| Rbbp8nl | TRUE | 0.419971073 | 0.075129413 | No Change | Not Significant | [illegible] |
| Rbbp9 | TRUE | 0.004015954 | 0.986657316 | No Change | Not Significant | [illegible] |
| Rbck1 | TRUE | 0.261733392 | 1.13E-05 | No Change | Significant Positive | [illegible] |
| Rbfa | TRUE | -0.108698347 | 0.632451088 | No Change | Not Significant | [illegible] |
| Rbfaos | TRUE | 0.475835412 | 1 | No Change | Not Significant | [illegible] |
| Rbfox1 | TRUE | 0.176010904 | 1 | No Change | Not Significant | [illegible] |
| Rbfox2 | TRUE | 0.412781419 | 1.74E-07 | No Change | Significant Positive | [illegible] |
| Rbfox3 | TRUE | 2.862996577 | 0.13639879 | No Change | Not Significant | [illegible] |
| Rbis | TRUE | -0.170471728 | 0.284132491 | No Change | Not Significant | [illegible] |
| Rbks | TRUE | 0.225052316 | 0.19693237 | No Change | Not Significant | [illegible] |
| Rbl1 | TRUE | -0.075756947 | 0.64845799 | No Change | Not Significant | [illegible] |
| Rbl2 | TRUE | -0.265960896 | 0.004543659 | No Change | Significant Negative | [illegible] |
| Rbm10 | TRUE | 0.649415596 | 3.71E-17 | UP | UP | [illegible] |
| Rbm11 | TRUE | 1.131888598 | 0.228901189 | No Change | Not Significant | [illegible] |
| Rbm12 | TRUE | -0.012285136 | 0.955211953 | No Change | Not Significant | [illegible] |
| Rbm12b1 | TRUE | 0.333403821 | 0.463099872 | No Change | Not Significant | [illegible] |
| Rbm12b2 | TRUE | 0.037941789 | 0.90502442 | No Change | Not Significant | [illegible] |
| Rbm14 | TRUE | 0.444162972 | 6.36E-10 | No Change | Significant Positive | [illegible] |
| Rbm15 | TRUE | 0.21341483 | 0.00736753 | No Change | Significant Positive | [illegible] |
| Rbm15b | TRUE | 1.364934648 | 3.36E-53 | UP | UP | [illegible] |
| Rbm17 | TRUE | 0.081811758 | 0.552639068 | No Change | Not Significant | [illegible] |
| Rbm18 | TRUE | 0.136850286 | 0.299903938 | No Change | Not Significant | [illegible] |
| Rbm19 | TRUE | 0.002482288 | 0.990621616 | No Change | Not Significant | [illegible] |
| Rbm20 | TRUE | -0.210387907 | 0.356273302 | No Change | Not Significant | [illegible] |
| Rbm22 | TRUE | -0.112166691 | 0.208221097 | No Change | Not Significant | [illegible] |
| Rbm24 | TRUE | -0.04468744 | 1 | No Change | Not Significant | [illegible] |
| Rbm25 | TRUE | 0.185623132 | 0.026243183 | No Change | Significant Positive | [illegible] |
| Rbm26 | TRUE | 0.1281971 | 0.220951697 | No Change | Not Significant | [illegible] |
| Rbm27 | TRUE | 0.08681126 | 0.284063113 | No Change | Not Significant | [illegible] |
| Rbm28 | TRUE | 0.035330781 | 0.803282676 | No Change | Not Significant | [illegible] |
| Rbm3 | TRUE | 0.248781557 | 0.224907418 | No Change | Not Significant | [illegible] |
| Rbm33 | TRUE | 0.083615209 | 0.635394239 | No Change | Not Significant | [illegible] |
| Rbm34 | TRUE | -0.005686418 | 0.982587667 | No Change | Not Significant | [illegible] |
| Rbm38 | TRUE | -0.268231023 | 0.419407924 | No Change | Not Significant | [illegible] |
| Rbm39 | TRUE | 0.088299276 | 0.197983688 | No Change | Not Significant | [illegible] |
| Rbm3os | TRUE | -0.528980787 | 0.276478264 | No Change | Not Significant | [illegible] |
| Rbm4 | TRUE | 0.088365408 | 0.736098624 | No Change | Not Significant | [illegible] |
| Rbm41 | TRUE | -0.114287233 | 0.647625851 | No Change | Not Significant | [illegible] |
| Rbm42 | TRUE | -0.020004843 | 1 | No Change | Not Significant | [illegible] |
| Rbm43 | TRUE | -0.027532121 | 0.933837244 | No Change | Not Significant | [illegible] |
| Rbm44 | TRUE | -0.471634295 | 1 | No Change | Not Significant | [illegible] |
| Rbm45 | TRUE | -0.135070311 | 0.309858682 | No Change | Not Significant | [illegible] |
| Rbm46 | TRUE | 1.139196962 | 1 | No Change | Not Significant | [illegible] |
| Rbm46os | TRUE | -0.586158512 | 1 | No Change | Not Significant | [illegible] |
| Rbm47 | TRUE | 0.065825882 | 0.59609806 | No Change | Not Significant | [illegible] |
| Rbm48 | TRUE | -0.06900029 | 0.806666129 | No Change | Not Significant | [illegible] |
| Rbm4b | TRUE | 0.338826785 | 0.047662852 | No Change | Significant Positive | [illegible] |
| Rbm5 | TRUE | -0.030779711 | 0.783356356 | No Change | Not Significant | [illegible] |
| Rbm6 | TRUE | -0.117823004 | 0.102718051 | No Change | Not Significant | [illegible] |
| Rbm7 | TRUE | 0.046115593 | 0.741471864 | No Change | Not Significant | [illegible] |
| Rbm8a | TRUE | -0.009769357 | 0.947202684 | No Change | Not Significant | [illegible] |
| Rbms1 | TRUE | -0.165763687 | 0.264727397 | No Change | Not Significant | [illegible] |
| Rbms2 | TRUE | -0.064362498 | 0.674995696 | No Change | Not Significant | [illegible] |
| Rbms3 | TRUE | -0.773368007 | 0.012571198 | DOWN | DOWN | [illegible] |
| Rbmx | TRUE | 0.134833744 | 0.347734431 | No Change | Not Significant | [illegible] |
| Rbmx2 | TRUE | 0.314873842 | 0.275039552 | No Change | Not Significant | [illegible] |
| Rbmxl1 | TRUE | -0.016068035 | 0.959738982 | No Change | Not Significant | [illegible] |
| Rbp1 | TRUE | 1.589564337 | 0.038619645 | UP | UP | [illegible] |
| Rbp2 | TRUE | -1.13032031 | 0.028986639 | DOWN | DOWN | [illegible] |
| Rbp3 | TRUE | 3.214069076 | 1 | No Change | Not Significant | [illegible] |
| Rbp4 | TRUE | -0.54658956 | 0.065695873 | No Change | Not Significant | [illegible] |
| Rbp7 | TRUE | -0.879505238 | 0.052155522 | No Change | Not Significant | [illegible] |
| Rbpj | TRUE | -0.186624033 | 0.223449098 | No Change | Not Significant | [illegible] |
| Rbpjl | TRUE | 0.438554823 | 1 | No Change | Not Significant | [illegible] |
| Rbpms | TRUE | -0.708620316 | 0.065243941 | No Change | Not Significant | [illegible] |
| Rbpms2 | TRUE | 0.249823359 | 0.817346435 | No Change | Not Significant | [illegible] |
| Rbsn | TRUE | -0.006786905 | 0.96968875 | No Change | Not Significant | [illegible] |
| Rbx1 | TRUE | -0.07312623 | 0.68365582 | No Change | Not Significant | [illegible] |
| Rc3h1 | TRUE | 0.125420112 | 0.311742141 | No Change | Not Significant | [illegible] |
| Rc3h2 | TRUE | 0.186479356 | 0.028870044 | No Change | Significant Positive | [illegible] |
| Rcan1 | TRUE | 0.911665296 | 3.36E-38 | UP | UP | [illegible] |
| Rcan2 | TRUE | 0.930345112 | 1 | No Change | Not Significant | [illegible] |
| Rcan3 | TRUE | 0.30703113 | 0.02076044 | No Change | Significant Positive | [illegible] |
| Rcbtb1 | TRUE | 0.304323442 | 0.040790921 | No Change | Significant Positive | [illegible] |
| Rcbtb2 | TRUE | 0.252600335 | 0.06492078 | No Change | Not Significant | [illegible] |
| Rcc1 | TRUE | 0.395533607 | 7.43E-05 | No Change | Significant Positive | [illegible] |
| Rcc1l | TRUE | -0.12568036 | 0.597623629 | No Change | Not Significant | [illegible] |
| Rcc2 | TRUE | 0.057819851 | 0.859322531 | No Change | Not Significant | [illegible] |
| Rccd1 | TRUE | -0.276067487 | 0.282387155 | No Change | Not Significant | [illegible] |
| Rce1 | TRUE | -0.195168156 | 0.083756183 | No Change | Not Significant | [illegible] |
| Rchy1 | TRUE | 0.286672591 | 0.000751269 | No Change | Significant Positive | [illegible] |
| Rcl1 | TRUE | 0.786481933 | 7.37E-07 | UP | UP | [illegible] |
| Rcn1 | TRUE | 0.180977815 | 0.421530084 | No Change | Not Significant | [illegible] |
| Rcn2 | TRUE | 0.110764216 | 0.37057407 | No Change | Not Significant | [illegible] |
| Rcn3 | TRUE | -0.600874202 | 1 | No Change | Not Significant | [illegible] |
| Rcor1 | TRUE | 0.038002382 | 0.818950015 | No Change | Not Significant | [illegible] |
| Rcor2 | TRUE | 0.394508453 | 0.54358318 | No Change | Not Significant | [illegible] |
| Rcor3 | TRUE | -0.172223505 | 0.26676599 | No Change | Not Significant | [illegible] |
| Rcsd1 | TRUE | -0.135699935 | 0.91440345 | No Change | Not Significant | [illegible] |
| Rd3 | TRUE | -0.379881399 | 1 | No Change | Not Significant | [illegible] |
| Rdh1 | TRUE | 0.269895299 | 0.405947672 | No Change | Not Significant | [illegible] |
| Rdh10 | TRUE | -0.078015998 | 0.605788125 | No Change | Not Significant | [illegible] |
| Rdh11 | TRUE | 0.350066971 | 0.066596371 | No Change | Not Significant | [illegible] |
| Rdh12 | TRUE | -0.087743166 | 0.91440345 | No Change | Not Significant | [illegible] |
| Rdh13 | TRUE | -0.067474218 | 0.706297653 | No Change | Not Significant | [illegible] |
| Rdh14 | TRUE | 0.165881628 | 0.241136045 | No Change | Not Significant | [illegible] |
| Rdh16 | TRUE | -0.003654015 | 0.990621616 | No Change | Not Significant | [illegible] |
| Rdh16f2 | TRUE | -0.332348626 | 0.720257015 | No Change | Not Significant | [illegible] |
| Rdh19 | TRUE | -0.303788079 | 0.736895388 | No Change | Not Significant | [illegible] |
| Rdh5 | TRUE | -0.148934597 | 0.672242383 | No Change | Not Significant | [illegible] |
| Rdh7 | TRUE | -2.014574096 | 1 | No Change | Not Significant | [illegible] |
| Rdh9 | TRUE | 0.053644954 | 0.932166328 | No Change | Not Significant | [illegible] |
| Rdm1 | TRUE | -0.972595756 | 9.74E-19 | DOWN | DOWN | [illegible] |
| Rdx | TRUE | 0.236292504 | 0.719201828 | No Change | Not Significant | [illegible] |
| Rec114 | TRUE | 0.059680115 | 1 | No Change | Not Significant | [illegible] |
| Rec8 | TRUE | 0.414504439 | 0.418548305 | No Change | Not Significant | [illegible] |
| Reck | TRUE | -1.5185078 | 1 | No Change | Not Significant | [illegible] |
| Recql | TRUE | 0.16765257 | 0.198694494 | No Change | Not Significant | [illegible] |
| Recql4 | TRUE | 0.618975119 | 0.001004984 | UP | UP | [illegible] |
| Recql5 | TRUE | 0.013163109 | 0.964753099 | No Change | Not Significant | [illegible] |
| Reep1 | TRUE | 0.998569451 | 0.141758822 | No Change | Not Significant | [illegible] |
| Reep2 | TRUE | 0.018543014 | 1 | No Change | Not Significant | [illegible] |
| Reep3 | TRUE | -0.421705063 | 8.18E-08 | No Change | Significant Negative | [illegible] |
| Reep4 | TRUE | 0.240281594 | 0.010209199 | No Change | Significant Positive | [illegible] |
| Reep5 | TRUE | -0.080232445 | 0.714728777 | No Change | Not Significant | [illegible] |
| Reep6 | TRUE | -0.130285768 | 0.563829035 | No Change | Not Significant | [illegible] |
| Reg1 | TRUE | -0.616744231 | 0.388834077 | No Change | Not Significant | [illegible] |
| Reg3a | TRUE | 0.959509462 | 0.205062987 | No Change | Not Significant | [illegible] |
| Reg3b | TRUE | 1.129174827 | 0.009789581 | UP | UP | [illegible] |
| Reg3d | TRUE | 4.285612835 | 9.65E-07 | UP | UP | [illegible] |
| Reg3g | TRUE | 1.231647202 | 0.065583696 | No Change | Not Significant | [illegible] |
| Reg4 | TRUE | 0.27444871 | 0.772952576 | No Change | Not Significant | [illegible] |
| Rel | TRUE | -0.270230229 | 0.378441649 | No Change | Not Significant | [illegible] |
| Rela | TRUE | -0.110959475 | 0.215780426 | No Change | Not Significant | [illegible] |
| Relb | TRUE | -0.62891054 | 3.52E-10 | DOWN | DOWN | [illegible] |
| Relch | TRUE | 0.053793303 | 0.618912124 | No Change | Not Significant | [illegible] |
| Rell1 | TRUE | 0.422717325 | 1.54E-07 | No Change | Significant Positive | [illegible] |
| Rell2 | TRUE | -0.060931601 | 0.947833434 | No Change | Not Significant | [illegible] |
| Reln | TRUE | 0.225747796 | 0.966007248 | No Change | Not Significant | [illegible] |
| Relt | TRUE | 1.269546485 | 0.103308737 | No Change | Not Significant | [illegible] |
| Rem2 | TRUE | 0.724522061 | 0.271847534 | No Change | Not Significant | [illegible] |
| Ren1 | TRUE | 1.851075657 | 1 | No Change | Not Significant | [illegible] |
| Renbp | TRUE | -0.71923513 | 0.321916639 | No Change | Not Significant | [illegible] |
| Reno1 | TRUE | -0.526297273 | 0.065177848 | No Change | Not Significant | [illegible] |
| Rep15 | TRUE | 0.221836125 | 0.720922137 | No Change | Not Significant | [illegible] |
| Repin1 | TRUE | -0.319923343 | 0.20154342 | No Change | Not Significant | [illegible] |
| Reps1 | TRUE | 0.217424082 | 0.027880534 | No Change | Significant Positive | [illegible] |
| Reps2 | TRUE | 0.606960586 | 1.86E-08 | UP | UP | [illegible] |
| Rer1 | TRUE | -0.007849546 | 0.958483509 | No Change | Not Significant | [illegible] |
| Rere | TRUE | -0.12011419 | 0.666621321 | No Change | Not Significant | [illegible] |
| Rerg | TRUE | -0.999650084 | 0.127906199 | No Change | Not Significant | [illegible] |
| Resf1 | TRUE | 0.048958114 | 0.684625749 | No Change | Not Significant | [illegible] |
| Resp18 | TRUE | -1.478248205 | 8.26E-05 | DOWN | DOWN | [illegible] |
| Rest | TRUE | 0.041255038 | 0.807082912 | No Change | Not Significant | [illegible] |
| Ret | TRUE | 0.673916258 | 0.012721623 | UP | UP | [illegible] |
| Retn | TRUE | 3.249344393 | 1 | No Change | Not Significant | [illegible] |
| Retnla | TRUE | 2.572701581 | 1 | No Change | Not Significant | [illegible] |
| Retnlb | TRUE | 0.200509549 | 0.639652146 | No Change | Not Significant | [illegible] |
| Retreg1 | TRUE | -0.644461846 | 0.000602545 | DOWN | DOWN | [illegible] |
| Retreg2 | TRUE | -0.128003453 | 0.143502708 | No Change | Not Significant | [illegible] |
| Retreg3 | TRUE | -0.216533853 | 0.051707527 | No Change | Not Significant | [illegible] |
| Retsat | TRUE | 0.064408483 | 0.530813078 | No Change | Not Significant | [illegible] |
| Rev1 | TRUE | -0.04251491 | 0.873593484 | No Change | Not Significant | [illegible] |
| Rev3l | TRUE | 0.284323627 | 0.002350684 | No Change | Significant Positive | [illegible] |
| Rex1bd | TRUE | 0.128718247 | 0.459379022 | No Change | Not Significant | [illegible] |
| Rexo1 | TRUE | 0.070810711 | 0.633140737 | No Change | Not Significant | [illegible] |
| Rexo2 | TRUE | -0.16063478 | 0.357012648 | No Change | Not Significant | [illegible] |
| Rexo4 | TRUE | 0.158520777 | 0.224110872 | No Change | Not Significant | [illegible] |
| Rexo5 | TRUE | -0.097022951 | 0.714893745 | No Change | Not Significant | [illegible] |
| Rfc1 | TRUE | -0.040668887 | 0.843000612 | No Change | Not Significant | [illegible] |
| Rfc2 | TRUE | 0.014810812 | 0.945259194 | No Change | Not Significant | [illegible] |
| Rfc3 | TRUE | -0.057491289 | 0.827172038 | No Change | Not Significant | [illegible] |
| Rfc4 | TRUE | 0.260069865 | 0.27747708 | No Change | Not Significant | [illegible] |
| Rfc5 | TRUE | 0.324005288 | 0.121691433 | No Change | Not Significant | [illegible] |
| Rfesd | TRUE | -0.033297298 | 0.913225519 | No Change | Not Significant | [illegible] |
| Rffl | TRUE | -0.477713772 | 2.83E-05 | No Change | Significant Negative | [illegible] |
| Rfk | TRUE | -0.409658522 | 2.04E-05 | No Change | Significant Negative | [illegible] |
| Rflna | TRUE | -0.614968663 | 0.236336338 | No Change | Not Significant | [illegible] |
| Rflnb | TRUE | -0.401277487 | 0.003996721 | No Change | Significant Negative | [illegible] |
| Rfng | TRUE | 0.137269106 | 0.405696463 | No Change | Not Significant | [illegible] |
| Rft1 | TRUE | 0.331404405 | 0.008390955 | No Change | Significant Positive | [illegible] |
| Rftn1 | TRUE | 0.006475057 | 0.995423402 | No Change | Not Significant | [illegible] |
| Rftn2 | TRUE | -0.423819683 | 0.061425328 | No Change | Not Significant | [illegible] |
| Rfwd3 | TRUE | 0.120490053 | 0.461051436 | No Change | Not Significant | [illegible] |
| Rfx1 | TRUE | 0.275064467 | 0.094358402 | No Change | Not Significant | [illegible] |
| Rfx2 | TRUE | 0.001582597 | 0.99808463 | No Change | Not Significant | [illegible] |
| Rfx3 | TRUE | -0.055619553 | 0.816133095 | No Change | Not Significant | [illegible] |
| Rfx4 | TRUE | 0.531197576 | 1 | No Change | Not Significant | [illegible] |
| Rfx5 | TRUE | -0.010538784 | 0.958296749 | No Change | Not Significant | [illegible] |
| Rfx6 | TRUE | -0.165873451 | 0.825096143 | No Change | Not Significant | [illegible] |
| Rfx7 | TRUE | 0.469885054 | 3.29E-05 | No Change | Significant Positive | [illegible] |
| Rfxank | TRUE | -0.271474988 | 0.007703947 | No Change | Significant Negative | [illegible] |
| Rfxap | TRUE | 0.394555323 | 0.000830593 | No Change | Significant Positive | [illegible] |
| Rgcc | TRUE | 0.37511982 | 0.133608449 | No Change | Not Significant | [illegible] |
| Rgl1 | TRUE | 0.724411577 | 0.284375933 | No Change | Not Significant | [illegible] |
| Rgl2 | TRUE | -0.170539866 | 0.075644218 | No Change | Not Significant | [illegible] |
| Rgl3 | TRUE | 0.117066525 | 0.774449596 | No Change | Not Significant | [illegible] |

| geneID | Cond1_DF_Detected | Cond1_DF_ValueCol | Cond1_DF_PadjCol | Cond1_DF_Cutoff | Cond1_DF_Cutoff_Specific.x | Cond2_DF_Detected | Cond2_DF_ValueCol | Cond2_DF_PadjCol | Cond2_DF_Cutoff | Cond2_DF_Cutoff_Specific | BG_Method | RG1_All |
|---|---|---|---|---|---|---|---|---|---|---|---|---|
| Rgma | TRUE | 0.243266584 | 0.667221236 | No Change | Not Significant | TRUE | 0.189675872 | 0.673006845 | No Change | Not Significant | TRUE | Cond1 Not Significant + Cond2 Not Significant |
| Rgmb | TRUE | 0.101865072 | 0.559563944 | No Change | Not Significant | TRUE | -0.10179101 | 0.377840372 | No Change | Not Significant | TRUE | Cond1 Not Significant + Cond2 Not Significant |
| Rgn | TRUE | -0.220579854 | 0.619446591 | No Change | Not Significant | TRUE | 0.261013955 | 0.405350432 | No Change | Not Significant | TRUE | Cond1 Not Significant + Cond2 Not Significant |
| Rgp1 | TRUE | -0.020090829 | 0.908714429 | No Change | Not Significant | TRUE | -0.004813723 | 0.963947444 | No Change | Not Significant | TRUE | Cond1 Not Significant + Cond2 Not Significant |
| Rgr | TRUE | 1.459943037 | 1 | No Change | Not Significant | FALSE | NA | NA | No Change | Not Detected | FALSE | Background = FALSE |
| Rgs1 | TRUE | -0.092376724 | 0.859261888 | No Change | Not Significant | TRUE | 0.163541634 | 0.382898178 | No Change | Not Significant | TRUE | Cond1 Not Significant + Cond2 Not Significant |
| Rgs10 | TRUE | 0.505431459 | 0.472054853 | No Change | Not Significant | TRUE | 1.200601963 | 0.000491472 | UP | UP | TRUE | Cond1 Not Significant + Cond2 UP |
| Rgs11 | TRUE | -1.157942608 | 5.13E-05 | DOWN | DOWN | TRUE | -0.890913257 | 0.005058411 | DOWN | DOWN | TRUE | Cond1 DOWN + Cond2 DOWN |
| Rgs12 | TRUE | 0.209919485 | 0.408658758 | No Change | Not Significant | TRUE | -0.146638083 | 0.488954885 | No Change | Not Significant | TRUE | Cond1 Not Significant + Cond2 Not Significant |
| Rgs13 | TRUE | -0.873037886 | 8.98E-05 | DOWN | DOWN | TRUE | 0.198419163 | 0.275438171 | No Change | Not Significant | TRUE | Cond1 DOWN + Cond2 Not Significant |
| Rgs14 | TRUE | 0.982329999 | 9.77E-06 | UP | UP | TRUE | 0.105423131 | 0.729722934 | No Change | Not Significant | TRUE | Cond1 UP + Cond2 Not Significant |
| Rgs16 | TRUE | -0.284755502 | 0.588671613 | No Change | Not Significant | TRUE | 0.612641484 | 0.034453872 | UP | UP | TRUE | Cond1 Not Significant + Cond2 UP |
| Rgs17 | TRUE | -0.136684318 | 0.71379243 | No Change | Not Significant | TRUE | -0.134813023 | 0.452796783 | No Change | Not Significant | TRUE | Cond1 Not Significant + Cond2 Not Significant |
| Rgs19 | TRUE | 0.307855181 | 0.479304239 | No Change | Not Significant | TRUE | 0.233144603 | 0.455544189 | No Change | Not Significant | TRUE | Cond1 Not Significant + Cond2 Not Significant |
| Rgs2 | TRUE | 1.172183453 | 2.88E-16 | UP | UP | TRUE | 0.32388856 | 0.046747707 | No Change | Significant Positive | TRUE | Cond1 UP + Cond2 Significant Positive |
| Rgs20 | TRUE | 1.335433942 | 1 | No Change | Not Significant | FALSE | NA | NA | No Change | Not Detected | FALSE | Background = FALSE |
| Rgs22 | TRUE | -0.926915135 | 0.438111017 | No Change | Not Significant | TRUE | 0.116544366 | 0.903818244 | No Change | Not Significant | TRUE | Cond1 Not Significant + Cond2 Not Significant |
| Rgs3 | TRUE | 0.284539229 | 0.001976883 | No Change | Significant Positive | TRUE | 0.105962107 | 0.284213967 | No Change | Not Significant | TRUE | Cond1 Significant Positive + Cond2 Not Significant |
| Rgs4 | TRUE | 0.10600377 | 0.931099354 | No Change | Not Significant | TRUE | -0.226313913 | 0.827034368 | No Change | Not Significant | TRUE | Cond1 Not Significant + Cond2 Not Significant |
| Rgs5 | TRUE | -2.018579441 | 1 | No Change | Not Significant | FALSE | NA | NA | No Change | Not Detected | FALSE | Background = FALSE |
| Rgs6 | TRUE | 0.679586254 | 0.805274537 | No Change | Not Significant | FALSE | NA | NA | No Change | Not Detected | FALSE | Background = FALSE |
| Rgs7 | TRUE | -1.391578431 | 1 | No Change | Not Significant | TRUE | 0.004952926 | 0.995924128 | No Change | Not Significant | TRUE | Cond1 Not Significant + Cond2 Not Significant |
| Rgs7bp | TRUE | 1.264429286 | 0.114316197 | No Change | Not Significant | TRUE | 1.519193191 | 0.034603793 | UP | UP | TRUE | Cond1 Not Significant + Cond2 UP |
| Rgs8 | TRUE | -0.580010957 | 0.539333655 | No Change | Not Significant | TRUE | 0.158297318 | 0.800888833 | No Change | Not Significant | TRUE | Cond1 Not Significant + Cond2 Not Significant |
| Rgs9 | TRUE | 0.197148146 | 0.891754761 | No Change | Not Significant | FALSE | NA | NA | No Change | Not Detected | FALSE | Background = FALSE |
| Rgs9bp | TRUE | -1.571111221 | 0.021033325 | DOWN | DOWN | TRUE | 0.38387656 | 0.646488649 | No Change | Not Significant | TRUE | Cond1 DOWN + Cond2 Not Significant |
| Rgsl1 | TRUE | -0.784618321 | 0.533456622 | No Change | Not Significant | TRUE | -1.085257437 | 0.171421356 | No Change | Not Significant | TRUE | Cond1 Not Significant + Cond2 Not Significant |
| Rhag | TRUE | 0.448018642 | 1 | No Change | Not Significant | FALSE | NA | NA | No Change | Not Detected | FALSE | Background = FALSE |
| Rhbdd1 | TRUE | 0.025187809 | 0.849070124 | No Change | Not Significant | TRUE | 0.102868665 | 0.283793301 | No Change | Not Significant | TRUE | Cond1 Not Significant + Cond2 Not Significant |
| Rhbdd2 | TRUE | -0.125881057 | 0.456813522 | No Change | Not Significant | TRUE | -0.133034153 | 0.330057234 | No Change | Not Significant | TRUE | Cond1 Not Significant + Cond2 Not Significant |
| Rhbdd3 | TRUE | -0.37223745 | 0.007066978 | No Change | Significant Negative | TRUE | -0.191429088 | 0.147682543 | No Change | Not Significant | TRUE | Cond1 Significant Negative + Cond2 Not Significant |
| Rhbdf1 | TRUE | -0.171867201 | 0.509056399 | No Change | Not Significant | TRUE | -0.291641066 | 0.061911166 | No Change | Not Significant | TRUE | Cond1 Not Significant + Cond2 Not Significant |
| Rhbdf2 | TRUE | -0.353076302 | 0.003837834 | No Change | Significant Negative | TRUE | -0.213363004 | 0.035627054 | No Change | Significant Negative | TRUE | Cond1 Significant Negative + Cond2 Significant Negative |
| Rhbdl1 | TRUE | -0.059940983 | 0.931471303 | No Change | Not Significant | TRUE | -0.236139704 | 0.78841372 | No Change | Not Significant | TRUE | Cond1 Not Significant + Cond2 Not Significant |
| Rhbdl2 | TRUE | -0.043807791 | 0.826943172 | No Change | Not Significant | TRUE | 0.039208336 | 0.795599668 | No Change | Not Significant | TRUE | Cond1 Not Significant + Cond2 Not Significant |
| Rhbdl3 | TRUE | -1.127755409 | 3.28E-05 | DOWN | DOWN | TRUE | -0.307069664 | 0.198372469 | No Change | Not Significant | TRUE | Cond1 DOWN + Cond2 Not Significant |
| Rhbg | TRUE | -0.636272287 | 0.058119626 | No Change | Not Significant | TRUE | -0.150477556 | 0.688854258 | No Change | Not Significant | TRUE | Cond1 Not Significant + Cond2 Not Significant |
| Rhcg | TRUE | 0.123811418 | 1 | No Change | Not Significant | FALSE | NA | NA | No Change | Not Detected | FALSE | Background = FALSE |
| Rhd | TRUE | 0.209703173 | 1 | No Change | Not Significant | FALSE | NA | NA | No Change | Not Detected | FALSE | Background = FALSE |
| Rheb | TRUE | 0.276014289 | 6.62E-06 | No Change | Significant Positive | TRUE | -0.048148817 | 0.529611374 | No Change | Not Significant | TRUE | Cond1 Significant Positive + Cond2 Not Significant |
| Rhebl1 | TRUE | -0.306151179 | 0.612367665 | No Change | Not Significant | TRUE | -0.035214727 | 0.945176551 | No Change | Not Significant | TRUE | Cond1 Not Significant + Cond2 Not Significant |
| Rhno1 | TRUE | 0.160484055 | 0.465629861 | No Change | Not Significant | TRUE | -0.057358212 | 0.737129071 | No Change | Not Significant | TRUE | Cond1 Not Significant + Cond2 Not Significant |
| Rho | TRUE | 0.911831868 | 1 | No Change | Not Significant | FALSE | NA | NA | No Change | Not Detected | FALSE | Background = FALSE |
| Rhoa | TRUE | -0.122896041 | 0.361606551 | No Change | Not Significant | TRUE | 0.091398515 | 0.263635924 | No Change | Not Significant | TRUE | Cond1 Not Significant + Cond2 Not Significant |
| Rhob | TRUE | 0.034109265 | 0.903393958 | No Change | Not Significant | TRUE | 0.468296353 | 0.000976118 | No Change | Significant Positive | TRUE | Cond1 Not Significant + Cond2 Significant Positive |
| Rhobtb1 | TRUE | -0.685886305 | 0.008704007 | DOWN | DOWN | TRUE | -0.346630604 | 0.238816971 | No Change | Not Significant | TRUE | Cond1 DOWN + Cond2 Not Significant |
| Rhobtb2 | TRUE | -0.062271849 | 0.649802774 | No Change | Not Significant | TRUE | -0.027872929 | 0.846398669 | No Change | Not Significant | TRUE | Cond1 Not Significant + Cond2 Not Significant |
| Rhobtb3 | TRUE | 0.858384416 | 8.80E-11 | UP | UP | TRUE | 0.246083513 | 0.082474622 | No Change | Not Significant | TRUE | Cond1 UP + Cond2 Not Significant |
| Rhoc | TRUE | -0.401960489 | 0.105093391 | No Change | Not Significant | TRUE | -0.226667589 | 0.184123872 | No Change | Not Significant | TRUE | Cond1 Not Significant + Cond2 Not Significant |
| Rhod | TRUE | 0.331375315 | 0.361410598 | No Change | Not Significant | TRUE | 0.570147481 | 0.005660988 | UP | UP | TRUE | Cond1 Not Significant + Cond2 UP |
| Rhof | TRUE | 0.544968471 | 0.046796074 | UP | UP | TRUE | 0.327851327 | 0.004068924 | No Change | Significant Positive | TRUE | Cond1 UP + Cond2 Significant Positive |
| Rhog | TRUE | -0.078860687 | 0.638921152 | No Change | Not Significant | TRUE | 0.038193604 | 0.836813383 | No Change | Not Significant | TRUE | Cond1 Not Significant + Cond2 Not Significant |
| Rhoh | TRUE | 0.050042837 | 0.971883559 | No Change | Not Significant | TRUE | 0.403591372 | 0.424308967 | No Change | Not Significant | TRUE | Cond1 Not Significant + Cond2 Not Significant |
| Rhoj | TRUE | 1.66964858 | 1 | No Change | Not Significant | TRUE | 1.229964717 | 0.108400597 | No Change | Not Significant | TRUE | Cond1 Not Significant + Cond2 Not Significant |
| Rhoq | TRUE | 0.439630616 | 0.002484666 | No Change | Significant Positive | TRUE | -0.08491413 | 0.525210203 | No Change | Not Significant | TRUE | Cond1 Significant Positive + Cond2 Not Significant |
| Rhot1 | TRUE | 0.047519663 | 0.708596728 | No Change | Not Significant | TRUE | 0.029278086 | 0.80894922 | No Change | Not Significant | TRUE | Cond1 Not Significant + Cond2 Not Significant |
| Rhot2 | TRUE | -0.238129694 | 0.090795126 | No Change | Not Significant | TRUE | -0.030562419 | 0.756034291 | No Change | Not Significant | TRUE | Cond1 Not Significant + Cond2 Not Significant |
| Rhou | TRUE | -0.088935174 | 0.627248048 | No Change | Not Significant | TRUE | -0.017471374 | 0.923349179 | No Change | Not Significant | TRUE | Cond1 Not Significant + Cond2 Not Significant |
| Rhov | TRUE | 0.076293931 | 0.908812719 | No Change | Not Significant | TRUE | 0.020895376 | 0.975710693 | No Change | Not Significant | TRUE | Cond1 Not Significant + Cond2 Not Significant |
| Rhox2d | TRUE | 2.851937291 | 1 | No Change | Not Significant | FALSE | NA | NA | No Change | Not Detected | FALSE | Background = FALSE |
| Rhox9 | TRUE | 1.18831899 | 1 | No Change | Not Significant | FALSE | NA | NA | No Change | Not Detected | FALSE | Background = FALSE |
| Rhpn1 | TRUE | 0.247323032 | 1 | No Change | Not Significant | FALSE | NA | NA | No Change | Not Detected | FALSE | Background = FALSE |
| Rhpn2 | TRUE | -0.220836277 | 0.001688702 | No Change | Significant Negative | TRUE | -0.029353071 | 0.756983295 | No Change | Not Significant | TRUE | Cond1 Significant Negative + Cond2 Not Significant |
| Ribc1 | TRUE | -0.326099701 | 0.340684159 | No Change | Not Significant | TRUE | -0.227926375 | 0.42043505 | No Change | Not Significant | TRUE | Cond1 Not Significant + Cond2 Not Significant |
| Ribc2 | TRUE | 2.850701429 | 1 | No Change | Not Significant | FALSE | NA | NA | No Change | Not Detected | FALSE | Background = FALSE |
| Ric1 | TRUE | -0.098016529 | 0.503945908 | No Change | Not Significant | TRUE | -0.110240102 | 0.196492651 | No Change | Not Significant | TRUE | Cond1 Not Significant + Cond2 Not Significant |
| Ric3 | TRUE | -0.561225558 | 0.231529357 | No Change | Not Significant | TRUE | 0.233207405 | 0.598408466 | No Change | Not Significant | TRUE | Cond1 Not Significant + Cond2 Not Significant |
| Ric8a | TRUE | -0.100565581 | 0.26324771 | No Change | Not Significant | TRUE | -0.057841591 | 0.340947717 | No Change | Not Significant | TRUE | Cond1 Not Significant + Cond2 Not Significant |
| Ric8b | TRUE | 0.011660412 | 0.950394593 | No Change | Not Significant | TRUE | -0.029001349 | 0.817954527 | No Change | Not Significant | TRUE | Cond1 Not Significant + Cond2 Not Significant |
| Rictor | TRUE | -0.299151228 | 0.000124365 | No Change | Significant Negative | TRUE | -0.227379667 | 0.002243141 | No Change | Significant Negative | TRUE | Cond1 Significant Negative + Cond2 Significant Negative |
| Rida | TRUE | -0.102408367 | 0.581819665 | No Change | Not Significant | TRUE | 0.096109752 | 0.360218718 | No Change | Not Significant | TRUE | Cond1 Not Significant + Cond2 Not Significant |
| Rif1 | TRUE | -0.010298591 | 0.957864571 | No Change | Not Significant | TRUE | -0.131756877 | 0.221242374 | No Change | Not Significant | TRUE | Cond1 Not Significant + Cond2 Not Significant |
| Rigi | TRUE | -0.270275284 | 0.269245209 | No Change | Not Significant | TRUE | -0.157912105 | 0.297231022 | No Change | Not Significant | TRUE | Cond1 Not Significant + Cond2 Not Significant |
| Riiad1 | TRUE | -0.111519458 | 0.910171686 | No Change | Not Significant | FALSE | NA | NA | No Change | Not Detected | FALSE | Background = FALSE |
| Rilp | TRUE | -0.302840535 | 0.085735413 | No Change | Not Significant | TRUE | -0.09240423 | 0.529013894 | No Change | Not Significant | TRUE | Cond1 Not Significant + Cond2 Not Significant |
| Rilpl1 | TRUE | 0.61597314 | 0.109773894 | No Change | Not Significant | TRUE | -0.381178835 | 0.457721775 | No Change | Not Significant | TRUE | Cond1 Not Significant + Cond2 Not Significant |
| Rilpl2 | TRUE | 0.25398574 | 0.160507581 | No Change | Not Significant | TRUE | 0.0847055 | 0.599482386 | No Change | Not Significant | TRUE | Cond1 Not Significant + Cond2 Not Significant |
| Rimbp2 | TRUE | 0.266077247 | 0.578423233 | No Change | Not Significant | TRUE | -0.048216328 | 0.880475742 | No Change | Not Significant | TRUE | Cond1 Not Significant + Cond2 Not Significant |
| Rimbp3 | TRUE | -0.53602458 | 0.29413766 | No Change | Not Significant | TRUE | 0.267965942 | 0.526458884 | No Change | Not Significant | TRUE | Cond1 Not Significant + Cond2 Not Significant |
| Rimkla | TRUE | 0.095920559 | 0.890186044 | No Change | Not Significant | TRUE | 0.071599835 | 0.898056783 | No Change | Not Significant | TRUE | Cond1 Not Significant + Cond2 Not Significant |
| Rimklb | TRUE | 1.163856609 | 0.233671624 | No Change | Not Significant | FALSE | NA | NA | No Change | Not Detected | FALSE | Background = FALSE |
| Rimoc1 | TRUE | -0.376486691 | 0.002092344 | No Change | Significant Negative | TRUE | -0.346735137 | 0.00177444 | No Change | Significant Negative | TRUE | Cond1 Significant Negative + Cond2 Significant Negative |
| Rims1 | TRUE | 1.420724864 | 1 | No Change | Not Significant | TRUE | 0.407772191 | 0.614708431 | No Change | Not Significant | TRUE | Cond1 Not Significant + Cond2 Not Significant |
| Rims2 | TRUE | -0.944820178 | 0.155962044 | No Change | Not Significant | TRUE | -0.872494319 | 0.349163297 | No Change | Not Significant | TRUE | Cond1 Not Significant + Cond2 Not Significant |
| Rims3 | TRUE | -0.012122079 | 1 | No Change | Not Significant | FALSE | NA | NA | No Change | Not Detected | FALSE | Background = FALSE |
| Rims4 | TRUE | 0.084020043 | 0.893302015 | No Change | Not Significant | TRUE | -0.176590474 | 0.45050036 | No Change | Not Significant | TRUE | Cond1 Not Significant + Cond2 Not Significant |
| Rin1 | TRUE | -0.786167929 | 1.51E-05 | DOWN | DOWN | TRUE | -0.085001607 | 0.625831733 | No Change | Not Significant | TRUE | Cond1 DOWN + Cond2 Not Significant |
| Rin2 | TRUE | -0.344221389 | 1.49E-05 | No Change | Significant Negative | TRUE | -0.225855441 | 0.009476714 | No Change | Significant Negative | TRUE | Cond1 Significant Negative + Cond2 Significant Negative |
| Rin3 | TRUE | -0.167127467 | 0.716272317 | No Change | Not Significant | TRUE | 0.266871972 | 0.23731003 | No Change | Not Significant | TRUE | Cond1 Not Significant + Cond2 Not Significant |
| Ring1 | TRUE | -0.146263334 | 0.216212948 | No Change | Not Significant | TRUE | -0.062289888 | 0.629835803 | No Change | Not Significant | TRUE | Cond1 Not Significant + Cond2 Not Significant |
| Rinl | TRUE | 0.154276229 | 0.809117465 | No Change | Not Significant | TRUE | 0.198086184 | 0.663107066 | No Change | Not Significant | TRUE | Cond1 Not Significant + Cond2 Not Significant |
| Rint1 | TRUE | 0.312852097 | 0.005597504 | No Change | Significant Positive | TRUE | 0.172374221 | 0.020174917 | No Change | Significant Positive | TRUE | Cond1 Significant Positive + Cond2 Significant Positive |
| Riok1 | TRUE | 0.068686854 | 0.668047177 | No Change | Not Significant | TRUE | -0.101050907 | 0.521447386 | No Change | Not Significant | TRUE | Cond1 Not Significant + Cond2 Not Significant |
| Riok2 | TRUE | -0.026359277 | 0.908812719 | No Change | Not Significant | TRUE | -0.204134906 | 0.087104102 | No Change | Not Significant | TRUE | Cond1 Not Significant + Cond2 Not Significant |
| Riok3 | TRUE | 0.122375271 | 0.05194233 | No Change | Not Significant | TRUE | 0.095332886 | 0.033935297 | No Change | Significant Positive | TRUE | Cond1 Not Significant + Cond2 Significant Positive |
| Riox1 | TRUE | -0.10247479 | 0.526067129 | No Change | Not Significant | TRUE | -0.004198858 | 0.983923349 | No Change | Not Significant | TRUE | Cond1 Not Significant + Cond2 Not Significant |
| Riox2 | TRUE | -0.251950313 | 0.039805161 | No Change | Significant Negative | TRUE | 0.144981542 | 0.25893088 | No Change | Not Significant | TRUE | Cond1 Significant Negative + Cond2 Not Significant |
| Ripk1 | TRUE | -0.162564276 | 0.04769017 | No Change | Significant Negative | TRUE | -0.079266697 | 0.239447599 | No Change | Not Significant | TRUE | Cond1 Significant Negative + Cond2 Not Significant |
| Ripk2 | TRUE | 0.228061108 | 0.116216589 | No Change | Not Significant | TRUE | -0.165773317 | 0.29959893 | No Change | Not Significant | TRUE | Cond1 Not Significant + Cond2 Not Significant |
| Ripk3 | TRUE | 0.017678988 | 0.948680909 | No Change | Not Significant | TRUE | -0.227399809 | 0.183499374 | No Change | Not Significant | TRUE | Cond1 Not Significant + Cond2 Not Significant |
| Ripk4 | TRUE | -0.152418298 | 0.315242025 | No Change | Not Significant | TRUE | 0.118039022 | 0.125951884 | No Change | Not Significant | TRUE | Cond1 Not Significant + Cond2 Not Significant |
| Ripor1 | TRUE | -0.212969173 | 0.55048001 | No Change | Not Significant | TRUE | -0.098735291 | 0.695172778 | No Change | Not Significant | TRUE | Cond1 Not Significant + Cond2 Not Significant |
| Ripor2 | TRUE | 0.031135071 | 0.94517522 | No Change | Not Significant | TRUE | 0.225726923 | 0.420872061 | No Change | Not Significant | TRUE | Cond1 Not Significant + Cond2 Not Significant |
| Ripor3 | TRUE | -0.835678921 | 1 | No Change | Not Significant | FALSE | NA | NA | No Change | Not Detected | FALSE | Background = FALSE |
| Ripply1 | TRUE | -0.142283558 | 0.887804402 | No Change | Not Significant | TRUE | -0.137309818 | 0.880475742 | No Change | Not Significant | TRUE | Cond1 Not Significant + Cond2 Not Significant |
| Ripply3 | TRUE | -0.163713693 | 0.832307165 | No Change | Not Significant | TRUE | 0.226941819 | 0.792706869 | No Change | Not Significant | TRUE | Cond1 Not Significant + Cond2 Not Significant |
| Rit1 | TRUE | -0.191267786 | 0.019776625 | No Change | Significant Negative | TRUE | 0.047340151 | 0.562758791 | No Change | Not Significant | TRUE | Cond1 Significant Negative + Cond2 Not Significant |
| Rit2 | TRUE | 1.022349239 | 1 | No Change | Not Significant | FALSE | NA | NA | No Change | Not Detected | FALSE | Background = FALSE |
| Rita1 | TRUE | -0.048301087 | 0.821350533 | No Change | Not Significant | TRUE | 0.165762224 | 0.180362858 | No Change | Not Significant | TRUE | Cond1 Not Significant + Cond2 Not Significant |
| Rlbp1 | TRUE | -0.031716168 | 1 | No Change | Not Significant | FALSE | NA | NA | No Change | Not Detected | FALSE | Background = FALSE |
| Rlf | TRUE | 0.051741763 | 0.674995696 | No Change | Not Significant | TRUE | -0.109903419 | 0.34629961 | No Change | Not Significant | TRUE | Cond1 Not Significant + Cond2 Not Significant |
| Rlim | TRUE | 0.040870562 | 0.764064128 | No Change | Not Significant | TRUE | -0.019432014 | 0.798277593 | No Change | Not Significant | TRUE | Cond1 Not Significant + Cond2 Not Significant |
| Rln1 | TRUE | 2.659173826 | 1 | No Change | Not Significant | FALSE | NA | NA | No Change | Not Detected | FALSE | Background = FALSE |
| Rln3 | TRUE | 0.006624921 | 1 | No Change | Not Significant | FALSE | NA | NA | No Change | Not Detected | FALSE | Background = FALSE |
| Rmc1 | TRUE | -0.149489104 | 0.049834789 | No Change | Significant Negative | TRUE | 0.084487694 | 0.327122432 | No Change | Not Significant | TRUE | Cond1 Significant Negative + Cond2 Not Significant |
| Rmdn1 | TRUE | -0.151956063 | 0.343804463 | No Change | Not Significant | TRUE | -0.114013167 | 0.450343118 | No Change | Not Significant | TRUE | Cond1 Not Significant + Cond2 Not Significant |
| Rmdn2 | TRUE | 0.082052654 | 0.747162726 | No Change | Not Significant | TRUE | -0.02482815 | 0.934988261 | No Change | Not Significant | TRUE | Cond1 Not Significant + Cond2 Not Significant |
| Rmdn3 | TRUE | -0.383922202 | 0.000627847 | No Change | Significant Negative | TRUE | -0.014082821 | 0.924826781 | No Change | Not Significant | TRUE | Cond1 Significant Negative + Cond2 Not Significant |
| Rmi1 | TRUE | -0.111048941 | 0.528838189 | No Change | Not Significant | TRUE | -0.037779386 | 0.834309996 | No Change | Not Significant | TRUE | Cond1 Not Significant + Cond2 Not Significant |
| Rmi2 | TRUE | 0.078238102 | 0.891754761 | No Change | Not Significant | TRUE | 0.156764416 | 0.721828109 | No Change | Not Significant | TRUE | Cond1 Not Significant + Cond2 Not Significant |
| Rmnd1 | TRUE | -0.487500825 | 3.25E-06 | No Change | Significant Negative | TRUE | -0.070677869 | 0.561449273 | No Change | Not Significant | TRUE | Cond1 Significant Negative + Cond2 Not Significant |
| Rmnd5a | TRUE | 0.133649867 | 0.135669662 | No Change | Not Significant | TRUE | 0.086510105 | 0.088419803 | No Change | Not Significant | TRUE | Cond1 Not Significant + Cond2 Not Significant |
| Rmnd5b | TRUE | -0.302707093 | 0.02334356 | No Change | Significant Negative | TRUE | -0.040765576 | 0.760423283 | No Change | Not Significant | TRUE | Cond1 Significant Negative + Cond2 Not Significant |
| Rmst | TRUE | 1.137525318 | 1 | No Change | Not Significant | FALSE | NA | NA | No Change | Not Detected | FALSE | Background = FALSE |
| Rnase1 | TRUE | -0.962718244 | 1 | No Change | Not Significant | TRUE | -1.487136629 | 0.026790231 | DOWN | DOWN | TRUE | Cond1 Not Significant + Cond2 DOWN |
| Rnase10 | TRUE | 2.504628202 | 1 | No Change | Not Significant | FALSE | NA | NA | No Change | Not Detected | FALSE | Background = FALSE |
| Rnase12 | TRUE | 2.18143211 | 1 | No Change | Not Significant | FALSE | NA | NA | No Change | Not Detected | FALSE | Background = FALSE |
| Rnase13 | TRUE | 2.526564878 | 1 | No Change | Not Significant | FALSE | NA | NA | No Change | Not Detected | FALSE | Background = FALSE |
| Rnase4 | TRUE | -0.812448532 | 8.63E-12 | DOWN | DOWN | TRUE | -0.276215449 | 0.00064154 | No Change | Significant Negative | TRUE | Cond1 DOWN + Cond2 Significant Negative |
| Rnase6 | TRUE | 1.086885289 | 1 | No Change | Not Significant | FALSE | NA | NA | No Change | Not Detected | FALSE | Background = FALSE |
| Rnase9 | TRUE | 2.485086843 | 1 | No Change | Not Significant | FALSE | NA | NA | No Change | Not Detected | FALSE | Background = FALSE |
| Rnaseh1 | TRUE | -0.138105601 | 0.40124694 | No Change | Not Significant | TRUE | -0.043099951 | 0.803865895 | No Change | Not Significant | TRUE | Cond1 Not Significant + Cond2 Not Significant |
| Rnaseh2a | TRUE | -0.008887917 | 0.966007248 | No Change | Not Significant | TRUE | 0.017954117 | 0.882492633 | No Change | Not Significant | TRUE | Cond1 Not Significant + Cond2 Not Significant |
| Rnaseh2b | TRUE | 0.13734676 | 0.467346556 | No Change | Not Significant | TRUE | 0.016192445 | 0.937787345 | No Change | Not Significant | TRUE | Cond1 Not Significant + Cond2 Not Significant |
| Rnaseh2c | TRUE | 0.072099401 | 0.745551704 | No Change | Not Significant | TRUE | 0.002167329 | 0.994488063 | No Change | Not Significant | TRUE | Cond1 Not Significant + Cond2 Not Significant |
| Rnasek | TRUE | -0.286568966 | 0.746748674 | No Change | Not Significant | TRUE | -0.270609522 | 0.737001553 | No Change | Not Significant | TRUE | Cond1 Not Significant + Cond2 Not Significant |
| Rnasel | TRUE | -0.52715815 | 9.04E-05 | DOWN | DOWN | TRUE | -0.416233427 | 3.34E-06 | No Change | Significant Negative | TRUE | Cond1 DOWN + Cond2 Significant Negative |
| Rnaset2a | TRUE | -0.232874097 | 0.816530773 | No Change | Not Significant | TRUE | -0.095889101 | 0.931084542 | No Change | Not Significant | TRUE | Cond1 Not Significant + Cond2 Not Significant |
| Rnaset2b | TRUE | 0.116447158 | 0.89873561 | No Change | Not Significant | TRUE | 0.07767442 | 0.910363905 | No Change | Not Significant | TRUE | Cond1 Not Significant + Cond2 Not Significant |
| Rnd1 | TRUE | 0.920303649 | 0.380539475 | No Change | Not Significant | TRUE | -0.12601168 | 0.900576045 | No Change | Not Significant | TRUE | Cond1 Not Significant + Cond2 Not Significant |
| Rnd2 | TRUE | -0.515281634 | 0.331006534 | No Change | Not Significant | TRUE | -0.476229122 | 0.213181286 | No Change | Not Significant | TRUE | Cond1 Not Significant + Cond2 Not Significant |
| Rnd3 | TRUE | -0.208203945 | 0.416065952 | No Change | Not Significant | TRUE | -0.208216732 | 0.039783734 | No Change | Significant Negative | TRUE | Cond1 Not Significant + Cond2 Significant Negative |
| Rnf10 | TRUE | 0.050269601 | 0.744703672 | No Change | Not Significant | TRUE | 0.061917049 | 0.666779423 | No Change | Not Significant | TRUE | Cond1 Not Significant + Cond2 Not Significant |
| Rnf103 | TRUE | 0.118628188 | 0.293636687 | No Change | Not Significant | TRUE | 0.047536516 | 0.609389288 | No Change | Not Significant | TRUE | Cond1 Not Significant + Cond2 Not Significant |
| Rnf11 | TRUE | -0.080705496 | 0.654452039 | No Change | Not Significant | TRUE | 0.111665366 | 0.137199451 | No Change | Not Significant | TRUE | Cond1 Not Significant + Cond2 Not Significant |
| Rnf111 | TRUE | 0.035742675 | 0.763259485 | No Change | Not Significant | TRUE | -0.030002161 | 0.780786732 | No Change | Not Significant | TRUE | Cond1 Not Significant + Cond2 Not Significant |
| Rnf112 | TRUE | 0.303102066 | 1 | No Change | Not Significant | FALSE | NA | NA | No Change | Not Detected | FALSE | Background = FALSE |
| Rnf113a1 | TRUE | 0.037543065 | 0.930374388 | No Change | Not Significant | TRUE | 0.112375217 | 0.76841464 | No Change | Not Significant | TRUE | Cond1 Not Significant + Cond2 Not Significant |
| Rnf113a2 | TRUE | 0.43508864 | 0.001184764 | No Change | Significant Positive | TRUE | 0.113444056 | 0.469603637 | No Change | Not Significant | TRUE | Cond1 Significant Positive + Cond2 Not Significant |
| Rnf114 | TRUE | 0.263866543 | 9.45E-05 | No Change | Significant Positive | TRUE | 0.042147565 | 0.668067439 | No Change | Not Significant | TRUE | Cond1 Significant Positive + Cond2 Not Significant |
| Rnf115 | TRUE | 0.039455656 | 0.735049182 | No Change | Not Significant | TRUE | -0.006895923 | 0.959960814 | No Change | Not Significant | TRUE | Cond1 Not Significant + Cond2 Not Significant |
| Rnf121 | TRUE | 0.00210336 | 0.98618968 | No Change | Not Significant | TRUE | 0.00532126 | 0.966739549 | No Change | Not Significant | TRUE | Cond1 Not Significant + Cond2 Not Significant |
| Rnf122 | TRUE | -0.204811839 | 0.602075619 | No Change | Not Significant | TRUE | -0.090336738 | 0.779344838 | No Change | Not Significant | TRUE | Cond1 Not Significant + Cond2 Not Significant |
| Rnf123 | TRUE | -0.232823964 | 0.009122624 | No Change | Significant Negative | TRUE | -0.20154997 | 0.042345703 | No Change | Significant Negative | TRUE | Cond1 Significant Negative + Cond2 Significant Negative |
| Rnf125 | TRUE | -0.237530626 | 0.452145476 | No Change | Not Significant | TRUE | -0.035269747 | 0.912178372 | No Change | Not Significant | TRUE | Cond1 Not Significant + Cond2 Not Significant |
| Rnf126 | TRUE | 0.26474536 | 0.020123752 | No Change | Significant Positive | TRUE | 0.113978293 | 0.346341304 | No Change | Not Significant | TRUE | Cond1 Significant Positive + Cond2 Not Significant |
| Rnf128 | TRUE | -0.108143726 | 0.209820779 | No Change | Not Significant | TRUE | -0.101316709 | 0.181685836 | No Change | Not Significant | TRUE | Cond1 Not Significant + Cond2 Not Significant |
| Rnf13 | TRUE | -0.1828391 | 0.152668561 | No Change | Not Significant | TRUE | -0.102022082 | 0.253501793 | No Change | Not Significant | TRUE | Cond1 Not Significant + Cond2 Not Significant |
| Rnf130 | TRUE | -0.472623409 | 0.000437255 | No Change | Significant Negative | TRUE | -0.062871397 | 0.463123925 | No Change | Not Significant | TRUE | Cond1 Significant Negative + Cond2 Not Significant |
| Rnf133 | TRUE | 0.383896985 | 1 | No Change | Not Significant | FALSE | NA | NA | No Change | Not Detected | FALSE | Background = FALSE |
| Rnf135 | TRUE | -0.059266943 | 0.804573016 | No Change | Not Significant | TRUE | -0.000400349 | 0.998588735 | No Change | Not Significant | TRUE | Cond1 Not Significant + Cond2 Not Significant |
| Rnf138 | TRUE | -0.146196555 | 0.352113109 | No Change | Not Significant | TRUE | -0.077659672 | 0.496022219 | No Change | Not Significant | TRUE | Cond1 Not Significant + Cond2 Not Significant |
| Rnf139 | TRUE | -0.066386984 | 0.699749347 | No Change | Not Significant | TRUE | -0.088467997 | 0.335868401 | No Change | Not Significant | TRUE | Cond1 Not Significant + Cond2 Not Significant |
| Rnf14 | TRUE | -0.024073602 | 0.888933284 | No Change | Not Significant | TRUE | 0.05635692 | 0.354359599 | No Change | Not Significant | TRUE | Cond1 Not Significant + Cond2 Not Significant |
| Rnf141 | TRUE | 0.041114149 | 0.828381052 | No Change | Not Significant | TRUE | -0.07543804 | 0.336231563 | No Change | Not Significant | TRUE | Cond1 Not Significant + Cond2 Not Significant |
| Rnf144a | TRUE | 1.511757856 | 0.258312646 | No Change | Not Significant | FALSE | NA | NA | No Change | Not Detected | FALSE | Background = FALSE |
| Rnf144b | TRUE | -1.145133281 | 0.132607819 | No Change | Not Significant | TRUE | 0.762836462 | 0.295551199 | No Change | Not Significant | TRUE | Cond1 Not Significant + Cond2 Not Significant |
| Rnf145 | TRUE | 0.261634785 | 0.070067307 | No Change | Not Significant | TRUE | 0.061838575 | 0.615647488 | No Change | Not Significant | TRUE | Cond1 Not Significant + Cond2 Not Significant |
| Rnf146 | TRUE | 0.035068555 | 0.877032235 | No Change | Not Significant | TRUE | -0.028601506 | 0.856166251 | No Change | Not Significant | TRUE | Cond1 Not Significant + Cond2 Not Significant |
| Rnf148 | TRUE | 0.020507883 | 1 | No Change | Not Significant | FALSE | NA | NA | No Change | Not Detected | FALSE | Background = FALSE |
| Rnf149 | TRUE | -0.338211148 | 0.004789552 | No Change | Significant Negative | TRUE | -0.150535951 | 0.10680877 | No Change | Not Significant | TRUE | Cond1 Significant Negative + Cond2 Not Significant |
| Rnf150 | TRUE | 0.172581647 | 0.526136486 | No Change | Not Significant | TRUE | 0.033952305 | 0.889849511 | No Change | Not Significant | TRUE | Cond1 Not Significant + Cond2 Not Significant |
| Rnf151 | TRUE | 2.138904507 | 0.305377683 | No Change | Not Significant | FALSE | NA | NA | No Change | Not Detected | FALSE | Background = FALSE |
| Rnf152 | TRUE | -0.810218324 | 0.010862636 | DOWN | DOWN | TRUE | -0.1048647 | 0.576784962 | No Change | Not Significant | TRUE | Cond1 DOWN + Cond2 Not Significant |
| Rnf157 | TRUE | 1.53950685 | 6.16E-06 | UP | UP | TRUE | 0.675138398 | 0.013893501 | UP | UP | TRUE | Cond1 UP + Cond2 UP |
| Rnf165 | TRUE | -0.282264374 | 1 | No Change | Not Significant | FALSE | NA | NA | No Change | Not Detected | FALSE | Background = FALSE |
| Rnf166 | TRUE | 0.100151062 | 0.4551483 | No Change | Not Significant | TRUE | 0.092232569 | 0.528881227 | No Change | Not Significant | TRUE | Cond1 Not Significant + Cond2 Not Significant |
| Rnf167 | TRUE | -0.210534138 | 0.011930684 | No Change | Significant Negative | TRUE | -0.086093327 | 0.432432462 | No Change | Not Significant | TRUE | Cond1 Significant Negative + Cond2 Not Significant |
| Rnf168 | TRUE | -0.030738698 | 0.891061468 | No Change | Not Significant | TRUE | -0.026038099 | 0.821663954 | No Change | Not Significant | TRUE | Cond1 Not Significant + Cond2 Not Significant |
| Rnf169 | TRUE | -0.07792834 | 0.581328932 | No Change | Not Significant | TRUE | -0.022717433 | 0.849362552 | No Change | Not Significant | TRUE | Cond1 Not Significant + Cond2 Not Significant |
| Rnf17 | TRUE | 0.480237868 | 1 | No Change | Not Significant | FALSE | NA | NA | No Change | Not Detected | FALSE | Background = FALSE |
| Rnf170 | TRUE | 0.121181835 | 0.44769238 | No Change | Not Significant | TRUE | 0.070425328 | 0.5332246 | No Change | Not Significant | TRUE | Cond1 Not Significant + Cond2 Not Significant |
| Rnf180 | TRUE | -0.619559902 | 0.002815934 | DOWN | DOWN | TRUE | -0.219398938 | 0.203825108 | No Change | Not Significant | TRUE | Cond1 DOWN + Cond2 Not Significant |
| Rnf181 | TRUE | 0.487697626 | 1.33E-09 | No Change | Significant Positive | TRUE | 0.261472126 | 0.001219114 | No Change | Significant Positive | TRUE | Cond1 Significant Positive + Cond2 Significant Positive |
| Rnf183 | TRUE | 0.168398387 | 0.876856731 | No Change | Not Significant | TRUE | 1.230470807 | 0.038300296 | UP | UP | TRUE | Cond1 Not Significant + Cond2 UP |
| Rnf185 | TRUE | 0.067780642 | 0.503072723 | No Change | Not Significant | TRUE | 0.064416528 | 0.459085065 | No Change | Not Significant | TRUE | Cond1 Not Significant + Cond2 Not Significant |
| Rnf186 | TRUE | -0.061029613 | 0.781796866 | No Change | Not Significant | TRUE | -0.194866314 | 0.132728454 | No Change | Not Significant | TRUE | Cond1 Not Significant + Cond2 Not Significant |
| Rnf187 | TRUE | 0.23380682 | 0.017689676 | No Change | Significant Positive | TRUE | 0.05760207 | 0.64731555 | No Change | Not Significant | TRUE | Cond1 Significant Positive + Cond2 Not Significant |
| Rnf19a | TRUE | -0.172851916 | 0.290646979 | No Change | Not Significant | TRUE | -0.180491999 | 0.007846481 | No Change | Significant Negative | TRUE | Cond1 Not Significant + Cond2 Significant Negative |
| Rnf19b | TRUE | -0.007348579 | 0.978898029 | No Change | Not Significant | TRUE | -0.110317842 | 0.294196765 | No Change | Not Significant | TRUE | Cond1 Not Significant + Cond2 Not Significant |
| Rnf2 | TRUE | -0.03544895 | 0.833786242 | No Change | Not Significant | TRUE | -0.045007404 | 0.706867216 | No Change | Not Significant | TRUE | Cond1 Not Significant + Cond2 Not Significant |
| Rnf20 | TRUE | 0.005476486 | 0.971606814 | No Change | Not Significant | TRUE | 0.079448575 | 0.297926072 | No Change | Not Significant | TRUE | Cond1 Not Significant + Cond2 Not Significant |

| geneID | Cond1_DF_Detected | Cond1_DF_ValueCol | Cond1_DF_PadjCol | Cond1_DF_Cutoff | Cond1_DF_Cutoff_Specific.x | Cond2_DF_Detected | Cond2_DF_ValueCol |
|---|---|---|---|---|---|---|---|
| Rnf207 | TRUE | -0.090531864 | 1 | No Change | Not Significant | FALSE | NA |
| Rnf208 | TRUE | -0.443641531 | 0.611707822 | No Change | Not Significant | TRUE | -1.162431206 |
| Rnf212 | TRUE | -1.550448166 | 1.37E-05 | DOWN | DOWN | TRUE | -0.918880948 |
| Rnf212b | TRUE | 1.188112109 | 1 | No Change | Not Significant | FALSE | NA |
| Rnf213 | TRUE | 0.022838147 | 0.95200802 | No Change | Not Significant | TRUE | -0.104452486 |
| Rnf214 | TRUE | 0.146983789 | 0.221735064 | No Change | Not Significant | TRUE | -0.028408597 |
| Rnf215 | TRUE | -0.0760867 | 0.580632641 | No Change | Not Significant | TRUE | 0.001445959 |
| Rnf216 | TRUE | -0.141149551 | 0.151705224 | No Change | Not Significant | TRUE | -0.072497797 |
| Rnf217 | TRUE | 0.26525939 | 0.066679159 | No Change | Not Significant | TRUE | 0.239750282 |
| Rnf220 | TRUE | 0.102746836 | 0.395392153 | No Change | Not Significant | TRUE | -0.038194738 |
| Rnf222 | TRUE | 0.611169847 | 1 | No Change | Not Significant | FALSE | NA |
| Rnf223 | TRUE | 2.432406653 | 5.26E-09 | UP | UP | TRUE | 1.516161285 |
| Rnf224 | TRUE | 0.604092215 | 1 | No Change | Not Significant | FALSE | NA |
| Rnf225 | TRUE | 0.489095891 | 0.618855346 | No Change | Not Significant | TRUE | -0.382324761 |
| Rnf24 | TRUE | 0.486061153 | 0.186423523 | No Change | Not Significant | TRUE | 0.177240204 |
| Rnf25 | TRUE | -0.101923733 | 0.404864292 | No Change | Not Significant | TRUE | 0.035066012 |
| Rnf26 | TRUE | 1.140423057 | 1.09E-07 | UP | UP | TRUE | 0.511041837 |
| Rnf31 | TRUE | -0.11617086 | 0.652619695 | No Change | Not Significant | TRUE | -0.147293351 |
| Rnf32 | TRUE | -0.223588568 | 0.151405123 | No Change | Not Significant | TRUE | -0.266320861 |
| Rnf34 | TRUE | -0.034561919 | 0.8712288 | No Change | Not Significant | TRUE | -0.065798873 |
| Rnf38 | TRUE | 0.021573641 | 0.882690208 | No Change | Not Significant | TRUE | -0.118147277 |
| Rnf39 | TRUE | -0.603445817 | 3.36E-05 | DOWN | DOWN | TRUE | -0.308104615 |
| Rnf4 | TRUE | -0.008089478 | 0.937225143 | No Change | Not Significant | TRUE | -0.048310454 |
| Rnf40 | TRUE | 0.024135836 | 0.822256556 | No Change | Not Significant | TRUE | 0.035702232 |
| Rnf41 | TRUE | -0.037532683 | 0.789468115 | No Change | Not Significant | TRUE | -0.046379262 |
| Rnf43 | TRUE | -0.351375734 | 0.000434218 | No Change | Significant Negative | TRUE | -0.466906889 |
| Rnf44 | TRUE | -0.020354283 | 0.907147741 | No Change | Not Significant | TRUE | 0.060884118 |
| Rnf5 | TRUE | 0.754494379 | 1.31E-10 | UP | UP | TRUE | 0.200382421 |
| Rnf6 | TRUE | 0.082182997 | 0.342188672 | No Change | Not Significant | TRUE | 0.031428426 |
| Rnf7 | TRUE | -0.064408548 | 0.693738201 | No Change | Not Significant | TRUE | 0.108492009 |
| Rnf8 | TRUE | -0.238007866 | 0.222126524 | No Change | Not Significant | TRUE | -0.332584701 |
| Rnft1 | TRUE | 0.145684672 | 0.096986852 | No Change | Not Significant | TRUE | 0.070127899 |
| Rnft2 | TRUE | 1.095945607 | 0.026177574 | UP | UP | TRUE | 0.128959659 |
| Rngtt | TRUE | -0.074652007 | 0.575403714 | No Change | Not Significant | TRUE | 0.019708485 |
| Rnh1 | TRUE | 0.117007712 | 0.2820338 | No Change | Not Significant | TRUE | 0.24653169 |
| Rnls | TRUE | -0.416341212 | 0.1177122 | No Change | Not Significant | TRUE | 0.074220635 |
| Rnmt | TRUE | -0.038241026 | 0.864656729 | No Change | Not Significant | TRUE | -0.085529381 |
| Rnpc3 | TRUE | 0.004907738 | 0.979351528 | No Change | Not Significant | TRUE | -0.129927739 |
| Rnpep | TRUE | -0.036671933 | 0.829503357 | No Change | Not Significant | TRUE | 0.127311218 |
| Rnpepl1 | TRUE | -0.184672665 | 0.012806226 | No Change | Significant Negative | TRUE | 0.080235095 |
| Rnps1 | TRUE | 0.056437525 | 0.69829677 | No Change | Not Significant | TRUE | -0.061572381 |
| Rnu12 | TRUE | 0.540385968 | 0.426114561 | No Change | Not Significant | TRUE | 0.257008848 |
| Rny1 | TRUE | 1.070092384 | 0.001900364 | UP | UP | TRUE | 0.038181234 |
| Rny3 | TRUE | 0.647916849 | 0.105235002 | No Change | Not Significant | TRUE | 0.236680723 |
| Ro60 | TRUE | 0.107559139 | 0.544244621 | No Change | Not Significant | TRUE | 0.216728186 |
| Robo1 | TRUE | 0.068526001 | 0.919582966 | No Change | Not Significant | TRUE | 0.152838072 |
| Robo2 | TRUE | 0.332566963 | 0.379255294 | No Change | Not Significant | TRUE | 0.187515051 |
| Robo3 | TRUE | 0.270623254 | 1 | No Change | Not Significant | FALSE | NA |
| Robo4 | TRUE | -1.064969045 | 1 | No Change | Not Significant | FALSE | NA |
| Rock1 | TRUE | 0.048708981 | 0.690650055 | No Change | Not Significant | TRUE | -0.072893633 |
| Rock2 | TRUE | 0.22484787 | 0.05288871 | No Change | Not Significant | TRUE | -0.09445508 |
| Rogdi | TRUE | 0.282622183 | 0.157984593 | No Change | Not Significant | TRUE | 0.088114338 |
| Rom1 | TRUE | -0.089785096 | 0.875081807 | No Change | Not Significant | TRUE | 0.108863362 |
| Romo1 | TRUE | -0.367896222 | 0.01156526 | No Change | Significant Negative | TRUE | 0.018550332 |
| Ropn1l | TRUE | 0.286805303 | 1 | No Change | Not Significant | FALSE | NA |
| Ror1 | TRUE | -0.049783223 | 0.880273177 | No Change | Not Significant | TRUE | -0.000708577 |
| Ror2 | TRUE | 0.4905476 | 0.007373206 | No Change | Significant Positive | TRUE | 0.066474987 |
| Rora | TRUE | -0.823184713 | 0.216212948 | No Change | Not Significant | TRUE | -0.067191648 |
| Rorb | TRUE | 1.03142929 | 0.074253808 | No Change | Not Significant | TRUE | 0.225407099 |
| Rorc | TRUE | 0.814993003 | 0.000233941 | UP | UP | TRUE | 0.294330217 |
| Ros1 | TRUE | 1.031369965 | 0.125929956 | No Change | Not Significant | TRUE | 1.044843174 |
| Rp1 | TRUE | 1.856703098 | 1 | No Change | Not Significant | FALSE | NA |
| Rp1l1 | TRUE | -0.425289985 | 1 | No Change | Not Significant | FALSE | NA |
| Rp2 | TRUE | 0.81513182 | 5.37E-21 | UP | UP | TRUE | 0.444286358 |
| Rp9 | TRUE | -0.135811735 | 0.214493541 | No Change | Not Significant | TRUE | -0.055479034 |
| Rpa1 | TRUE | 0.324892458 | 0.088775474 | No Change | Not Significant | TRUE | 0.078936245 |
| Rpa2 | TRUE | 0.832862305 | 2.50E-05 | UP | UP | TRUE | 0.377875456 |
| Rpa3 | TRUE | 0.196037525 | 0.338979766 | No Change | Not Significant | TRUE | 0.105218935 |
| Rpain | TRUE | 0.694130771 | 8.58E-06 | UP | UP | TRUE | 0.281199311 |
| Rpap1 | TRUE | 0.133942796 | 0.319339495 | No Change | Not Significant | TRUE | 0.102798607 |
| Rpap2 | TRUE | -0.143280586 | 0.31779121 | No Change | Not Significant | TRUE | 0.03192769 |
| Rpap3 | TRUE | 0.03684929 | 0.882510888 | No Change | Not Significant | TRUE | -0.078979368 |
| Rpe | TRUE | 0.098472815 | 0.598740239 | No Change | Not Significant | TRUE | 0.218932409 |
| Rpe65 | TRUE | 0.522237567 | 1 | No Change | Not Significant | FALSE | NA |
| Rpf1 | TRUE | -0.102185402 | 0.508005086 | No Change | Not Significant | TRUE | -0.025911169 |
| Rpf2 | TRUE | -0.086935342 | 0.721840784 | No Change | Not Significant | TRUE | -0.053304906 |
| Rpgr | TRUE | 0.042483263 | 0.867333511 | No Change | Not Significant | TRUE | -0.151180025 |
| Rpgrip1 | TRUE | -0.342701509 | 0.708014489 | No Change | Not Significant | TRUE | 0.194661617 |
| Rpgrip1l | TRUE | 0.159861674 | 0.320998127 | No Change | Not Significant | TRUE | 0.191760379 |
| Rph3a | TRUE | 1.036920696 | 1 | No Change | Not Significant | FALSE | NA |
| Rph3al | TRUE | 0.470589922 | 0.023369313 | No Change | Significant Positive | TRUE | 0.207013725 |
| Rpia | TRUE | -0.255635863 | 0.05194233 | No Change | Not Significant | TRUE | -0.143871602 |
| Rpl10 | TRUE | -0.170755077 | 0.42858336 | No Change | Not Significant | TRUE | 0.215044065 |
| Rpl10a | TRUE | -0.411722483 | 0.007085436 | No Change | Significant Negative | TRUE | -0.067611243 |
| Rpl11 | TRUE | -0.351586576 | 0.037589453 | No Change | Significant Negative | TRUE | -0.002012401 |
| Rpl12 | TRUE | -0.341605305 | 0.04823163 | No Change | Significant Negative | TRUE | -0.066073871 |
| Rpl13 | TRUE | -0.355193453 | 0.017528923 | No Change | Significant Negative | TRUE | -0.011773789 |
| Rpl13-ps6 | TRUE | 1.483493716 | 1 | No Change | Not Significant | FALSE | NA |
| Rpl13a | TRUE | 0.223584741 | 0.031622347 | No Change | Significant Positive | TRUE | -0.060003242 |
| Rpl14 | TRUE | -0.199632438 | 0.241063164 | No Change | Not Significant | TRUE | -0.03223023 |
| Rpl15 | TRUE | -0.242956348 | 0.14663883 | No Change | Not Significant | TRUE | -0.007768434 |
| Rpl17 | TRUE | -0.28821999 | 0.041288574 | No Change | Significant Negative | TRUE | -0.009801983 |
| Rpl17-ps10 | TRUE | -0.767486617 | 1 | No Change | Not Significant | FALSE | NA |
| Rpl18 | TRUE | -0.26416624 | 0.042679767 | No Change | Significant Negative | TRUE | 0.011315896 |
| Rpl18a | TRUE | -0.442819122 | 0.003009871 | No Change | Significant Negative | TRUE | -0.057295544 |
| Rpl19 | TRUE | -0.443413852 | 0.003771026 | No Change | Significant Negative | TRUE | -0.003529862 |
| Rpl21 | TRUE | -0.291555507 | 0.050506026 | No Change | Not Significant | TRUE | -0.067269984 |
| Rpl22 | TRUE | -0.30771121 | 0.022923163 | No Change | Significant Negative | TRUE | -0.155568899 |
| Rpl22l1 | TRUE | 0.104483562 | 0.637415471 | No Change | Not Significant | TRUE | 0.037411119 |
| Rpl23 | TRUE | -0.324891988 | 0.03747705 | No Change | Significant Negative | TRUE | -0.065981742 |
| Rpl23a | TRUE | -0.294126466 | 0.04960305 | No Change | Significant Negative | TRUE | -0.078357852 |
| Rpl24 | TRUE | 0.022043859 | 0.942955032 | No Change | Not Significant | TRUE | 0.050239361 |
| Rpl26 | TRUE | -0.300292945 | 0.03389483 | No Change | Significant Negative | TRUE | -0.013930301 |
| Rpl27 | TRUE | -0.294252395 | 0.066917689 | No Change | Not Significant | TRUE | 0.03833573 |
| Rpl27a | TRUE | -0.268105641 | 0.071195349 | No Change | Not Significant | TRUE | -0.062054795 |
| Rpl28 | TRUE | -0.458245854 | 0.00156603 | No Change | Significant Negative | TRUE | -0.014260171 |
| Rpl29 | TRUE | -0.353580718 | 0.05194233 | No Change | Not Significant | TRUE | 0.022410562 |
| Rpl3 | TRUE | -0.318056981 | 0.053482492 | No Change | Not Significant | TRUE | -0.076403747 |
| Rpl30 | TRUE | -0.227138977 | 0.182183929 | No Change | Not Significant | TRUE | -0.033430158 |
| Rpl31 | TRUE | -0.608260287 | 0.000231089 | DOWN | DOWN | TRUE | -0.059917433 |
| Rpl31-ps12 | TRUE | 0.05962217 | 1 | No Change | Not Significant | FALSE | NA |
| Rpl32 | TRUE | -0.092603847 | 0.564974708 | No Change | Not Significant | TRUE | -0.053975317 |
| Rpl32l | TRUE | 0.146134235 | 0.819849053 | No Change | Not Significant | TRUE | -0.389244096 |
| Rpl34 | TRUE | -0.325253138 | 0.058674176 | No Change | Not Significant | TRUE | 0.015460329 |
| Rpl34-ps1 | TRUE | 0.69729104 | 1 | No Change | Not Significant | FALSE | NA |
| Rpl35 | TRUE | -0.423060183 | 0.011831403 | No Change | Significant Negative | TRUE | -0.043477861 |
| Rpl35a | TRUE | -0.260361813 | 0.045492347 | No Change | Significant Negative | TRUE | -0.016299107 |
| Rpl36 | TRUE | -0.358833261 | 0.00822343 | No Change | Significant Negative | TRUE | -0.061768588 |
| Rpl36a | TRUE | -0.38710609 | 0.006195065 | No Change | Significant Negative | TRUE | -0.091523223 |
| Rpl36al | TRUE | 0.442724948 | 0.036352985 | No Change | Significant Positive | TRUE | 0.207753029 |
| Rpl37 | TRUE | -0.342301202 | 0.010614744 | No Change | Significant Negative | TRUE | -0.029066293 |
| Rpl37a | TRUE | -0.280688302 | 0.038706422 | No Change | Significant Negative | TRUE | 0.020267931 |
| Rpl37rt | TRUE | 0.014792855 | 0.962478474 | No Change | Not Significant | TRUE | -0.097807174 |
| Rpl38 | TRUE | -0.37548697 | 0.015384668 | No Change | Significant Negative | TRUE | 0.02182384 |
| Rpl39 | TRUE | -0.562380217 | 0.009704716 | DOWN | DOWN | TRUE | -0.107137013 |
| Rpl3l | TRUE | -0.510303875 | 1 | No Change | Not Significant | FALSE | NA |
| Rpl4 | TRUE | -0.128908991 | 0.334133019 | No Change | Not Significant | TRUE | -0.08342418 |
| Rpl41 | TRUE | -0.223286823 | 0.103865546 | No Change | Not Significant | TRUE | 0.106106885 |
| Rpl5 | TRUE | -0.573032684 | 0.004258592 | DOWN | DOWN | TRUE | -0.069761572 |
| Rpl6 | TRUE | -0.261878671 | 0.105765439 | No Change | Not Significant | TRUE | -0.011499601 |
| Rpl6l | TRUE | 0.328164506 | 0.725771143 | No Change | Not Significant | TRUE | -0.457151567 |
| Rpl7 | TRUE | -0.295241172 | 0.053546803 | No Change | Not Significant | TRUE | -0.016557714 |
| Rpl7a | TRUE | -0.364298193 | 0.003410008 | No Change | Significant Negative | TRUE | -0.000394989 |
| Rpl7l1 | TRUE | 0.234200637 | 0.051642509 | No Change | Not Significant | TRUE | 0.170354087 |
| Rpl8 | TRUE | -0.105173886 | 0.442718029 | No Change | Not Significant | TRUE | 0.007463784 |
| Rpl9 | TRUE | -0.543976227 | 0.00123647 | DOWN | DOWN | TRUE | 0.006521279 |
| Rplp0 | TRUE | -0.070613456 | 0.73008549 | No Change | Not Significant | TRUE | 0.134776491 |
| Rplp1 | TRUE | -0.259050386 | 0.031962608 | No Change | Significant Negative | TRUE | -0.032635291 |
| Rplp2 | TRUE | -0.198513168 | 0.205793193 | No Change | Not Significant | TRUE | -0.007567894 |
| Rpn1 | TRUE | 1.065153387 | 6.66E-30 | UP | UP | TRUE | 0.909090167 |
| Rpn2 | TRUE | 1.15081477 | 1.08E-40 | UP | UP | TRUE | 0.843455592 |
| Rpp14 | TRUE | -0.040734364 | 0.883975099 | No Change | Not Significant | TRUE | -0.016775335 |
| Rpp21 | TRUE | -0.117961078 | 0.363480239 | No Change | Not Significant | TRUE | 0.050624377 |
| Rpp25 | TRUE | -0.835035541 | 0.065855474 | No Change | Not Significant | TRUE | -0.080873541 |
| Rpp25l | TRUE | -0.043548025 | 0.807082912 | No Change | Not Significant | TRUE | 0.066550515 |
| Rpp30 | TRUE | 0.309020721 | 0.109351285 | No Change | Not Significant | TRUE | 0.054919576 |
| Rpp38 | TRUE | 0.163293647 | 0.632329199 | No Change | Not Significant | TRUE | 0.036818089 |
| Rpp40 | TRUE | -0.280876633 | 0.483088189 | No Change | Not Significant | TRUE | -0.168013601 |
| Rpph1 | TRUE | 1.317071875 | 0.224929501 | No Change | Not Significant | TRUE | -0.040883423 |
| Rprd1a | TRUE | -0.148119612 | 0.416114999 | No Change | Not Significant | TRUE | -0.166173915 |
| Rprd1b | TRUE | -0.022315367 | 0.833786242 | No Change | Not Significant | TRUE | 0.007105867 |
| Rprd2 | TRUE | 0.046792797 | 0.789468115 | No Change | Not Significant | TRUE | -0.150637406 |
| Rprm | TRUE | 0.359977983 | 0.535446743 | No Change | Not Significant | TRUE | 0.382290023 |
| Rpml | TRUE | 0.455747102 | 1 | No Change | Not Significant | FALSE | NA |
| Rps10 | TRUE | 0.339197626 | 0.259543427 | No Change | Not Significant | TRUE | -0.178311553 |
| Rps11 | TRUE | -0.162334347 | 0.148948255 | No Change | Not Significant | TRUE | -0.001639166 |
| Rps12 | TRUE | -0.316378833 | 0.014342673 | No Change | Significant Negative | TRUE | -0.059332646 |
| Rps12-ps10 | TRUE | 1.958815428 | 1 | No Change | Not Significant | FALSE | NA |
| Rps12-ps24 | TRUE | 2.118912747 | 1 | No Change | Not Significant | FALSE | NA |
| Rps13 | TRUE | -0.321575563 | 0.057079639 | No Change | Not Significant | TRUE | 0.00721387 |
| Rps14 | TRUE | -0.278269296 | 0.012461318 | No Change | Significant Negative | TRUE | -0.002534027 |
| Rps15 | TRUE | -0.241257334 | 0.037046055 | No Change | Significant Negative | TRUE | -0.000801574 |
| Rps15a | TRUE | -0.229899931 | 0.23602289 | No Change | Not Significant | TRUE | -0.028019068 |
| Rps15a-ps5 | TRUE | -0.053485821 | 1 | No Change | Not Significant | FALSE | NA |
| Rps16 | TRUE | -0.377137828 | 0.079122383 | No Change | Not Significant | TRUE | 0.00380724 |
| Rps17 | TRUE | -0.376092653 | 0.007482994 | No Change | Significant Negative | TRUE | -0.027262186 |
| Rps18 | TRUE | -0.441755199 | 0.001589432 | No Change | Significant Negative | TRUE | -0.07580844 |
| Rps19 | TRUE | 0.119461886 | 0.509749949 | No Change | Not Significant | TRUE | 0.154359216 |
| Rps19-ps3 | TRUE | -0.278420033 | 0.695624496 | No Change | Not Significant | TRUE | -0.415112007 |
| Rps19bp1 | TRUE | 0.391424511 | 0.002810091 | No Change | Significant Positive | TRUE | 0.115067522 |
| Rps2 | TRUE | -0.063980306 | 0.729644779 | No Change | Not Significant | TRUE | 0.034990821 |
| Rps20 | TRUE | -0.205123907 | 0.139401698 | No Change | Not Significant | TRUE | -0.07145416 |
| Rps21 | TRUE | -0.225974219 | 0.19999509 | No Change | Not Significant | TRUE | -0.053580187 |
| Rps23 | TRUE | -0.347859962 | 0.020104181 | No Change | Significant Negative | TRUE | -0.050799038 |
| Rps23rg1 | TRUE | 0.829323784 | 0.021404668 | UP | UP | TRUE | 0.481252648 |
| Rps24 | TRUE | -0.377192592 | 0.007703947 | No Change | Significant Negative | TRUE | -0.063817472 |
| Rps25 | TRUE | -0.368139364 | 0.008312447 | No Change | Significant Negative | TRUE | -0.031076721 |
| Rps26 | TRUE | -0.252881204 | 0.041336308 | No Change | Significant Negative | TRUE | -0.048309994 |
| Rps27 | TRUE | -0.533207997 | 5.30E-06 | DOWN | DOWN | TRUE | -0.109807393 |
| Rps27a | TRUE | -0.346554033 | 0.065598201 | No Change | Not Significant | TRUE | 0.012450772 |
| Rps27l | TRUE | -0.00595958 | 0.977331011 | No Change | Not Significant | TRUE | 0.109800184 |
| Rps27rt | TRUE | 0.100493595 | 0.893606822 | No Change | Not Significant | TRUE | -0.133401062 |
| Rps28 | TRUE | -0.810081391 | 0.000341177 | DOWN | DOWN | TRUE | -0.043802027 |
| Rps29 | TRUE | -0.575773312 | 0.003154573 | DOWN | DOWN | TRUE | 0.028587536 |
| Rps3 | TRUE | -0.098769374 | 0.556313398 | No Change | Not Significant | TRUE | -0.01104179 |
| Rps3a1 | TRUE | -0.349629833 | 0.061038631 | No Change | Not Significant | TRUE | -0.027797102 |
| Rps4x | TRUE | -0.3202655 | 0.051375821 | No Change | Not Significant | TRUE | -0.026933377 |
| [illegible] | [illegible] | [illegible] | [illegible] | [illegible] | [illegible] | [illegible] | [illegible] |

Supplementary Table 3

| geneID | Cond1_DF_Detected | Cond1_DF_ValueCol | Cond1_DF_PadjCol | Cond1_DF_Cutoff | Cond1_DF_Cutoff_Specific.x | Cond2_DF_Detected | Cond2_DF_ValueCol | [illegible] |
|---|---|---|---|---|---|---|---|---|
| Rps6ka5 | TRUE | 0.412441705 | 0.119192842 | No Change | Not Significant | TRUE | 0.087004378 | [illegible] |
| Rps6ka6 | TRUE | 0.486045934 | 0.051921541 | No Change | Not Significant | TRUE | 0.248698347 | [illegible] |
| Rps6kb1 | TRUE | 0.175919614 | 0.069550933 | No Change | Not Significant | TRUE | 0.011999363 | [illegible] |
| Rps6kb2 | TRUE | 0.049541332 | 0.821333519 | No Change | Not Significant | TRUE | -0.080704942 | [illegible] |
| Rps6kc1 | TRUE | -0.002123631 | 0.986890148 | No Change | Not Significant | TRUE | -0.055260951 | [illegible] |
| Rps6kl1 | TRUE | 0.072649886 | 0.876856731 | No Change | Not Significant | TRUE | -0.536655239 | [illegible] |
| Rps7 | TRUE | -0.405628197 | 0.013074234 | No Change | Significant Negative | TRUE | -0.017174898 | [illegible] |
| Rps8 | TRUE | -0.256416246 | 0.090408616 | No Change | Not Significant | TRUE | -0.055399533 | [illegible] |
| Rps9 | TRUE | -0.164284977 | 0.191366503 | No Change | Not Significant | TRUE | -0.002960038 | [illegible] |
| Rpsa | TRUE | -0.393985027 | 0.022615701 | No Change | Significant Negative | TRUE | 0.013614023 | [illegible] |
| Rptor | TRUE | 0.192110089 | 0.049435028 | No Change | Significant Positive | TRUE | 0.134397631 | [illegible] |
| Rpusd1 | TRUE | 0.198895912 | 0.337710856 | No Change | Not Significant | TRUE | 0.018993444 | [illegible] |
| Rpusd2 | TRUE | 0.072330901 | 0.795381772 | No Change | Not Significant | TRUE | -0.173356117 | [illegible] |
| Rpusd3 | TRUE | 0.224566174 | 0.417785903 | No Change | Not Significant | TRUE | 0.171021325 | [illegible] |
| Rpusd4 | TRUE | -0.251072953 | 0.042442029 | No Change | Significant Negative | TRUE | 0.019678999 | [illegible] |
| Rrad | TRUE | -0.862494169 | 1 | No Change | Not Significant | FALSE | NA | NA |
| Rraga | TRUE | 0.01794139 | 0.926816617 | No Change | Not Significant | TRUE | 0.106834895 | [illegible] |
| Rragb | TRUE | 0.221143236 | 0.794593769 | No Change | Not Significant | TRUE | 0.177024762 | [illegible] |
| [illegible] | [illegible] | [illegible] | [illegible] | [illegible] | [illegible] | [illegible] | [illegible] | [illegible] |

Supplementary Table 3

| geneID | Cond1_DF_Detected | Cond1_DF_ValueCol | Cond1_DF_PadjCol | Cond1_DF_Cutoff | Cond1_DF_Cutoff_Specific.x | Cond2_DF_Detected | Cond2_DF_ValueCol | Cond2_DF_PadjCol | Cond2_DF_Cutoff | Cond2_DF_Cutoff_Specific | BG_Method | RG1_All | RG2_Significant | RG3_SignificantChange | Cond1_DF_ensembl_gene_id | Cond1_DF_entrezgene_id | Cond1_DF_gene_biotype | Cond1_DF_baseMean | Cond1_DF_log2FoldChange | Cond1_DF_lfcSE | Cond1_DF_stat | Cond1_DF_pvalue | Cond1_DF_padj | Cond1_DF_absLog2FC | Cond1_DF_Cutoff_Specific.y | Cond2_DF_ensembl_gene_id | Cond2_DF_entrezgene_id | Cond2_DF_gene_biotype | Cond2_DF_baseMean | Cond2_DF_log2FoldChange | Cond2_DF_lfcSE | Cond2_DF_stat | Cond2_DF_pvalue | Cond2_DF_padj | Cond2_DF_absLog2FC | RG1_Specific_Cond2 | RG1_Specific_Cond1 |
|---|---|---|---|---|---|---|---|---|---|---|---|---|---|---|---|---|---|---|---|---|---|---|---|---|---|---|---|---|---|---|---|---|---|---|---|---|---|
| Sapcd2 | TRUE | 1.241717499 | 2.24E-09 | UP | UP | TRUE | 0.222177187 | 0.179562869 | No Change | Not Significant | TRUE | Cond1 UP + Cond2 Not Significant | Cond1_UP | Cond1_UP | ENSMUSG00000026955 | 72080 | protein_coding | 131.2306225 | 1.241717499 | 0.191467215 | -6.485274766 | 8.86E-11 | 2.24E-09 | 1.241717499 | UP | ENSMUSG00000026955 | 72080 | protein_coding | 153.8877631 | 0.222177187 | 0.112268749 | 1.97897624 | 0.047818681 | 0.179562869 | 0.222177187 | Cond1 UP + Cond2 Not Significant | Cond2 No Change + Cond1 UP |
| Sar1a | TRUE | 0.388365974 | 0.003837834 | No Change | Significant Positive | TRUE | 0.23338365 | 8.08E-06 | No Change | Significant Positive | TRUE | Cond1 Significant Positive + Cond2 Significant Positive | None | None | ENSMUSG00000020088 | 20224 | protein_coding | 1404.553693 | 0.388365974 | 0.112391803 | -3.455465275 | 0.000549344 | 0.003837834 | 0.388365974 | Significant Positive | ENSMUSG00000020088 | 20224 | protein_coding | 2199.603288 | 0.23338365 | 0.045523047 | 5.126714114 | 2.95E-07 | 8.08E-06 | 0.23338365 | Cond1 No Change + Cond2 Significant Positive | Cond2 No Change + Cond1 Significant Positive |
| Sar1b | TRUE | 0.00933838 | 0.947202684 | No Change | Not Significant | TRUE | -0.001500016 | 0.987977748 | No Change | Not Significant | TRUE | Cond1 Not Significant + Cond2 Not Significant | None | None | ENSMUSG00000020386 | 66397 | protein_coding | 618.6740408 | 0.00933838 | 0.069215921 | -0.134916642 | 0.892677781 | 0.947202684 | 0.00933838 | Not Significant | ENSMUSG00000020386 | 66397 | protein_coding | 1188.098951 | -0.001500016 | 0.040168812 | -0.037342813 | 0.970211669 | 0.987977748 | 0.001500016 | Cond1 No Change + Cond2 Not Significant | Cond2 No Change + Cond1 Not Significant |
| Saraf | TRUE | 1.154226225 | 4.01E-69 | UP | UP | TRUE | 0.750635435 | 3.26E-38 | UP | UP | TRUE | Cond1 UP + Cond2 UP | Core_UP | Core_UP | ENSMUSG00000031532 | 67887 | protein_coding | 2457.699748 | 1.154226225 | 0.0645665 | -17.87654942 | 1.80E-71 | 4.01E-69 | 1.154226225 | UP | ENSMUSG00000031532 | 67887 | protein_coding | 3321.996148 | 0.750635435 | 0.056362547 | 13.31798295 | 1.82E-40 | 3.26E-38 | 0.750635435 | Cond1 UP + Cond2 UP | Cond2 UP + Cond1 UP |
| Sardh | TRUE | 0.95751099 | 3.01E-06 | UP | UP | TRUE | 0.644220929 | 3.53E-07 | UP | UP | TRUE | Cond1 UP + Cond2 UP | Core_UP | Core_UP | ENSMUSG00000009614 | 192166 | protein_coding | 113.1940832 | 0.95751099 | 0.183889971 | -5.206977751 | 1.92E-07 | 3.01E-06 | 0.95751099 | UP | ENSMUSG00000009614 | 192166 | protein_coding | 150.0573569 | 0.644220929 | 0.112378673 | 5.732590651 | 9.89E-09 | 3.53E-07 | 0.644220929 | Cond1 UP + Cond2 UP | Cond2 UP + Cond1 UP |
| Sarm1 | TRUE | -0.728251423 | 1 | No Change | Not Significant | FALSE | NA | NA | No Change | Not Detected | FALSE | Background = FALSE | Background = FALSE | Background = FALSE | ENSMUSG00000050132 | 237868 | protein_coding | 3.377446834 | -0.728251423 | 1.599798854 | 0.455214367 | 0.648955025 | 1 | 0.728251423 | Not Significant | NA | NA | NA | NA | NA | NA | NA | NA | NA | NA | Background = FALSE | Background = FALSE |
| Sarnp | TRUE | 0.701059762 | 7.80E-07 | UP | UP | TRUE | 0.841990225 | 2.50E-20 | UP | UP | TRUE | Cond1 UP + Cond2 UP | Core_UP | Core_UP | ENSMUSG00000078427 | 66118 | protein_coding | 368.4773167 | 0.701059762 | 0.1281801 | -5.469333877 | 4.52E-08 | 7.80E-07 | 0.701059762 | UP | ENSMUSG00000078427 | 66118 | protein_coding | 610.0104405 | 0.841990225 | 0.086663464 | 9.715630835 | 2.59E-22 | 2.50E-20 | 0.841990225 | Cond1 UP + Cond2 UP | Cond2 UP + Cond1 UP |
| Sars | TRUE | 0.334904228 | 1.33E-05 | No Change | Significant Positive | TRUE | 0.133201229 | 0.048835423 | No Change | Significant Positive | TRUE | Cond1 Significant Positive + Cond2 Significant Positive | None | None | ENSMUSG00000068739 | 20226 | protein_coding | 1417.324866 | 0.334904228 | 0.068358184 | -4.899255816 | 9.62E-07 | 1.33E-05 | 0.334904228 | Significant Positive | ENSMUSG00000068739 | 20226 | protein_coding | 2432.996049 | 0.133201229 | 0.049908498 | 2.668908814 | 0.007609811 | 0.048835423 | 0.133201229 | Cond1 No Change + Cond2 Significant Positive | Cond2 No Change + Cond1 Significant Positive |
| Sars2 | TRUE | 0.151932978 | 0.438295983 | No Change | Not Significant | TRUE | 0.082813981 | 0.655224931 | No Change | Not Significant | TRUE | Cond1 Not Significant + Cond2 Not Significant | None | None | ENSMUSG00000070699 | 71984 | protein_coding | 162.9186859 | 0.151932978 | 0.126494548 | -1.201102975 | 0.229711259 | 0.438295983 | 0.151932978 | Not Significant | ENSMUSG00000070699 | 71984 | protein_coding | 271.2656046 | 0.082813981 | 0.102125991 | 0.810900151 | 0.417423014 | 0.655224931 | 0.082813981 | Cond1 No Change + Cond2 Not Significant | Cond2 No Change + Cond1 Not Significant |
| Sart1 | TRUE | 0.023359414 | 0.81317775 | No Change | Not Significant | TRUE | 0.081764342 | 0.265315436 | No Change | Not Significant | TRUE | Cond1 Not Significant + Cond2 Not Significant | None | None | ENSMUSG00000039148 | 20227 | protein_coding | 699.9519779 | 0.023359414 | 0.052368501 | -0.446058476 | 0.655555007 | 0.81317775 | 0.023359414 | Not Significant | ENSMUSG00000039148 | 20227 | protein_coding | 1199.727599 | 0.081764342 | 0.047889012 | 1.707371661 | 0.087752994 | 0.265315436 | 0.081764342 | Cond1 No Change + Cond2 Not Significant | Cond2 No Change + Cond1 Not Significant |
| Sart3 | TRUE | 0.336166906 | 1.10E-05 | No Change | Significant Positive | TRUE | 0.138403157 | 0.114513814 | No Change | Not Significant | TRUE | Cond1 Significant Positive + Cond2 Not Significant | None | None | ENSMUSG00000018974 | 53890 | protein_coding | 533.4700765 | 0.336166906 | 0.068037858 | -4.9408802 | 7.78E-07 | 1.10E-05 | 0.336166906 | Significant Positive | ENSMUSG00000018974 | 53890 | protein_coding | 811.6565708 | 0.138403157 | 0.061715923 | 2.242584253 | 0.024923637 | 0.114513814 | 0.138403157 | Cond1 No Change + Cond2 Not Significant | Cond2 No Change + Cond1 Significant Positive |
| Sash1 | TRUE | -0.329252812 | 0.047749183 | No Change | Significant Negative | TRUE | -0.022606644 | 0.822565162 | No Change | Not Significant | TRUE | Cond1 Significant Negative + Cond2 Not Significant | None | None | ENSMUSG00000015305 | 70097 | protein_coding | 949.8679935 | -0.329252812 | 0.13037675 | 2.525395158 | 0.011556826 | 0.047749183 | 0.329252812 | Significant Negative | ENSMUSG00000015305 | 70097 | protein_coding | 2312.353634 | -0.022606644 | 0.050622268 | -0.446575102 | 0.655181876 | 0.822565162 | 0.022606644 | Cond1 No Change + Cond2 Not Significant | Cond2 No Change + Cond1 Significant Negative |
| [illegible] | [illegible] | [illegible] | [illegible] | [illegible] | [illegible] | [illegible] | [illegible] | [illegible] | [illegible] | [illegible] | [illegible] | [illegible] | [illegible] | [illegible] | [illegible] | [illegible] | [illegible] | [illegible] | [illegible] | [illegible] | [illegible] | [illegible] | [illegible] | [illegible] | [illegible] | [illegible] | [illegible] | [illegible] | [illegible] | [illegible] | [illegible] | [illegible] | [illegible] | [illegible] | [illegible] | [illegible] | [illegible] |

Supplementary Table 3

| geneID | Cond1_DF_Detected | Cond1_DF_ValueCol | Cond1_DF_PadjCol | Cond1_DF_Cutoff | Cond1_DF_Cutoff_Specific.x | Cond2_DF_Detected | Cond2_DF_ValueCol | Cond2_DF_PadjCol | Cond2_DF_Cutoff |
|---|---|---|---|---|---|---|---|---|---|
| Selenom | TRUE | 0.534515268 | 7.21E-07 | UP | UP | TRUE | 0.431283279 | 4.23E-06 | No Change |
| Selenon | TRUE | 0.175703725 | 0.664452584 | No Change | Not Significant | TRUE | 0.031282514 | 0.943679315 | No Change |
| Selenoo | TRUE | -0.336194924 | 2.88E-05 | No Change | Significant Negative | TRUE | -0.180437368 | 0.014823762 | No Change |
| Selenop | TRUE | -1.17778829 | 2.09E-09 | DOWN | DOWN | TRUE | -0.362122583 | 0.045838699 | No Change |
| Selenos | TRUE | 1.036688427 | 3.81E-23 | UP | UP | TRUE | 0.547759534 | 3.78E-41 | UP |
| Selenot | TRUE | 0.011107272 | 0.944306685 | No Change | Not Significant | TRUE | 0.177023133 | 0.000296401 | No Change |
| Selenov | TRUE | -1.243153468 | 1 | No Change | Not Significant | FALSE | NA | NA | No Change |
| Selenow | TRUE | -0.459776682 | 4.11E-11 | No Change | Significant Negative | TRUE | -0.245341169 | 1.71E-07 | No Change |
| Sell | TRUE | 0.220596102 | 0.908714429 | No Change | Not Significant | TRUE | 0.744055847 | 0.473794416 | No Change |
| Selp | TRUE | 0.71136946 | 1 | No Change | Not Significant | FALSE | NA | NA | No Change |
| Selplg | TRUE | 0.386026488 | 0.53622883 | No Change | Not Significant | TRUE | 0.643088574 | 0.030447057 | UP |
| Sem1 | TRUE | 0.039040554 | 0.751588072 | No Change | Not Significant | TRUE | 0.088078388 | 0.543683373 | No Change |
| Sema3a | TRUE | -1.588980932 | 1 | No Change | Not Significant | FALSE | NA | NA | No Change |
| Sema3b | TRUE | -0.033399405 | 0.797446101 | No Change | Not Significant | TRUE | -0.054290559 | 0.5285691 | No Change |
| Sema3c | TRUE | -0.169891974 | 0.379678597 | No Change | Not Significant | TRUE | -0.222957482 | 0.000108406 | No Change |
| Sema3d | TRUE | 1.135429577 | 0.137751634 | No Change | Not Significant | FALSE | NA | NA | No Change |
| Sema3e | TRUE | 2.340503899 | 0.001482597 | UP | UP | FALSE | NA | NA | No Change |
| Sema3f | TRUE | -0.591032177 | 0.112253043 | No Change | Not Significant | TRUE | -0.392417221 | 0.25741348 | No Change |
| [illegible] | [illegible] | [illegible] | [illegible] | [illegible] | [illegible] | [illegible] | [illegible] | [illegible] | [illegible] |

[illegible]

| geneID | Cond1_DF_Detected | Cond1_DF_ValueCol | Cond1_DF_PadjCol | Cond1_DF_Cutoff | Cond1_DF_Cutoff_Specific.x | Cond2_DF_Detected | Cond2_DF_ValueCol | Cond2_DF_PadjCol | Cond2_DF_Cutoff | Cond2_DF_Cutoff_Specific | BG_Method | RG1_All | RG2_Significant | RG3_SignificantChange | Cond1_DF_ensembl_gene_id | Cond1_DF_entrezgene_id | Cond1_DF_gene_biotype | Cond1_DF_baseMean | Cond1_DF_log2FoldChange | Cond1_DF_lfcSE | Cond1_DF_stat | Cond1_DF_pvalue | Cond1_DF_padj | Cond1_DF_absLog2FC | Cond1_DF_Cutoff_Specific.y | Cond2_DF_ensembl_gene_id | Cond2_DF_entrezgene_id | Cond2_DF_gene_biotype | Cond2_DF_baseMean | Cond2_DF_log2FoldChange | Cond2_DF_lfcSE | Cond2_DF_stat | Cond2_DF_pvalue | Cond2_DF_padj | Cond2_DF_absLog2FC | RG1_Specific_Cond2 | RG1_Specific_Cond1 |
|---|---|---|---|---|---|---|---|---|---|---|---|---|---|---|---|---|---|---|---|---|---|---|---|---|---|---|---|---|---|---|---|---|---|---|---|---|---|
| Sh3bp2 | TRUE | -0.07781803 | 0.769577776 | No Change | Not Significant | TRUE | -0.08629368 | 0.686760796 | No Change | Not Significant | TRUE | Cond1 Not Significant + Cond2 Not Significant | None | None | ENSMUSG00000054520 | 24055 | protein_coding | 83.24620012 | -0.07781803 | 0.143863359 | 0.540916263 | 0.588565301 | 0.769577776 | 0.07781803 | Not Significant | ENSMUSG00000054520 | 24055 | protein_coding | 138.0642733 | -0.08629368 | 0.116750276 | -0.739130416 | 0.459827809 | 0.686760796 | 0.08629368 | Cond1 No Change + Cond2 Not Significant | Cond2 No Change + Cond1 Not Significant |
| Sh3bp4 | TRUE | -0.219025198 | 0.195920934 | No Change | Not Significant | TRUE | -0.019053287 | 0.921867382 | No Change | Not Significant | TRUE | Cond1 Not Significant + Cond2 Not Significant | None | None | ENSMUSG00000036206 | 98402 | protein_coding | 225.6560132 | -0.219025198 | 0.121937802 | 1.796204242 | 0.07246204 | 0.195920934 | 0.219025198 | Not Significant | ENSMUSG00000036206 | 98402 | protein_coding | 352.1620808 | -0.019053287 | 0.085829158 | -0.221990841 | 0.824321013 | 0.921867382 | 0.019053287 | Cond1 No Change + Cond2 Not Significant | Cond2 No Change + Cond1 Not Significant |
| Sh3bp5 | TRUE | 0.000982382 | 0.996641735 | No Change | Not Significant | TRUE | 0.000180989 | 0.99936318 | No Change | Not Significant | TRUE | Cond1 Not Significant + Cond2 Not Significant | None | None | ENSMUSG00000021892 | 24056 | protein_coding | 69.2198607 | 0.000982382 | 0.13488677 | -0.007283011 | 0.99418905 | 0.996641735 | 0.000982382 | Not Significant | ENSMUSG00000021892 | 24056 | protein_coding | 160.9328122 | 0.000180989 | 0.13488544 | 0.001341797 | 0.998929401 | 0.99936318 | 0.000180989 | Cond1 No Change + Cond2 Not Significant | Cond2 No Change + Cond1 Not Significant |
| Sh3bp5l | TRUE | -0.03742073 | 0.744703672 | No Change | Not Significant | TRUE | 0.105347163 | 0.256181012 | No Change | Not Significant | TRUE | Cond1 Not Significant + Cond2 Not Significant | None | None | ENSMUSG00000013646 | 79566 | protein_coding | 428.9945128 | -0.03742073 | 0.063366107 | 0.590548046 | 0.554823284 | 0.744703672 | 0.03742073 | Not Significant | ENSMUSG00000013646 | 79566 | protein_coding | 863.519958 | 0.105347163 | 0.060760667 | 1.733805264 | 0.082952641 | 0.256181012 | 0.105347163 | Cond1 No Change + Cond2 Not Significant | Cond2 No Change + Cond1 Not Significant |
| Sh3d19 | TRUE | -0.132115188 | 0.05583573 | No Change | Not Significant | TRUE | -0.079903039 | 0.26624145 | No Change | Not Significant | TRUE | Cond1 Not Significant + Cond2 Not Significant | None | None | ENSMUSG00000028082 | 27059 | protein_coding | 1482.28052 | -0.132115188 | 0.053863892 | 2.452759757 | 0.014176498 | 0.05583573 | 0.132115188 | Not Significant | ENSMUSG00000028082 | 27059 | protein_coding | 2707.312705 | -0.079903039 | 0.046856199 | -1.705282133 | 0.088141818 | 0.26624145 | 0.079903039 | Cond1 No Change + Cond2 Not Significant | Cond2 No Change + Cond1 Not Significant |
| Sh3d21 | TRUE | -0.408800062 | 0.13890357 | No Change | Not Significant | TRUE | 0.198908508 | 0.189536742 | No Change | Not Significant | TRUE | Cond1 Not Significant + Cond2 Not Significant | None | None | ENSMUSG00000073758 | 66938 | protein_coding | 538.27075 | -0.408800062 | 0.204380557 | 2.000190565 | 0.04547969 | 0.13890357 | 0.408800062 | Not Significant | ENSMUSG00000073758 | 66938 | protein_coding | 967.5019705 | 0.198908508 | 0.102295604 | 1.944448258 | 0.051841419 | 0.189536742 | 0.198908508 | Cond1 No Change + Cond2 Not Significant | Cond2 No Change + Cond1 Not Significant |
| Sh3gl1 | TRUE | 0.054452555 | 0.662983366 | No Change | Not Significant | TRUE | 0.089214493 | 0.336982889 | No Change | Not Significant | TRUE | Cond1 Not Significant + Cond2 Not Significant | None | None | ENSMUSG00000003200 | 20405 | protein_coding | 438.6154944 | 0.054452555 | 0.071419804 | -0.762429351 | 0.445803794 | 0.662983366 | 0.054452555 | Not Significant | ENSMUSG00000003200 | 20405 | protein_coding | 769.9976403 | 0.089214493 | 0.05909094 | 1.509782944 | 0.131098819 | 0.336982889 | 0.089214493 | Cond1 No Change + Cond2 Not Significant | Cond2 No Change + Cond1 Not Significant |
| Sh3gl2 | TRUE | -0.194611615 | 0.3894353 | No Change | Not Significant | TRUE | -0.09386989 | 0.553781446 | No Change | Not Significant | TRUE | Cond1 Not Significant + Cond2 Not Significant | None | None | ENSMUSG00000028488 | 20404 | protein_coding | 159.573154 | -0.194611615 | 0.149046845 | 1.305707716 | 0.191651973 | 0.3894353 | 0.194611615 | Not Significant | ENSMUSG00000028488 | 20404 | protein_coding | 310.3601731 | -0.09386989 | 0.092490679 | -1.014911895 | 0.310147828 | 0.553781446 | 0.09386989 | Cond1 No Change + Cond2 Not Significant | Cond2 No Change + Cond1 Not Significant |

| geneID | Cond1_DF_Detected | Cond1_DF_ValueCol | Cond1_DF_PadjCol | Cond1_DF_Cutoff | Cond1_DF_Cutoff_Specific.x | Cond2_DF_Detected | Cond2_DF_ValueCol | Cond2_DF_PadjCol | Cond2_DF_Cutoff | Cond2_DF_Cutoff_Specific | BG_Method | RG1_All | RG2_Significant | RG3_SignificantChange | Cond1_DF_ensembl_gene_id | Cond1_DF_entrezgene_id | Cond1_DF_gene_biotype | Cond1_DF_baseMean | Cond1_DF_log2FoldChange | Cond1_DF_lfcSE | Cond1_DF_stat | Cond1_DF_pvalue | Cond1_DF_padj | Cond1_DF_absLog2FC | Cond1_DF_Cutoff_Specific.y | Cond2_DF_ensembl_gene_id | Cond2_DF_entrezgene_id | Cond2_DF_gene_biotype | Cond2_DF_baseMean | Cond2_DF_log2FoldChange | Cond2_DF_lfcSE | Cond2_DF_stat | Cond2_DF_pvalue | Cond2_DF_padj | Cond2_DF_absLog2FC | RG1_Specific_Cond2 | RG1_Specific_Cond1 |
|---|---|---|---|---|---|---|---|---|---|---|---|---|---|---|---|---|---|---|---|---|---|---|---|---|---|---|---|---|---|---|---|---|---|---|---|---|---|
| Slc18b1 | TRUE | -0.077798016 | 0.630052795 | No Change | Not Significant | TRUE | 0.284814124 | 0.002161701 | No Change | Significant Positive | TRUE | Cond1 Not Significant + Cond2 Significant Positive | None | None | ENSMUSG00000037455 | 76306 | protein_coding | 321.0670197 | -0.077798016 | 0.094318868 | 0.824840436 | 0.409462186 | 0.630052795 | 0.077798016 | Not Significant | ENSMUSG00000037455 | 76306 | protein_coding | 538.5881787 | 0.284814124 | 0.075398382 | 3.777456712 | 0.000158438 | 0.002161701 | 0.284814124 | Cond1 No Change + Cond2 Significant Positive | Cond2 No Change + Cond1 Not Significant |
| Slc19a1 | TRUE | 0.111449673 | 0.633723045 | No Change | Not Significant | TRUE | -0.134016809 | 0.286052251 | No Change | Not Significant | TRUE | Cond1 Not Significant + Cond2 Not Significant | None | None | ENSMUSG00000001436 | 20509 | protein_coding | 111.996956 | 0.111449673 | 0.13631875 | -0.81756672 | 0.413604637 | 0.633723045 | 0.111449673 | Not Significant | ENSMUSG00000001436 | 20509 | protein_coding | 232.326863 | -0.134016809 | 0.081211293 | -1.650223798 | 0.098897171 | 0.286052251 | 0.134016809 | Cond1 No Change + Cond2 Not Significant | Cond2 No Change + Cond1 Not Significant |
| Slc19a2 | TRUE | -0.033532372 | 0.893615254 | No Change | Not Significant | TRUE | -0.114070375 | 0.505545025 | No Change | Not Significant | TRUE | Cond1 Not Significant + Cond2 Not Significant | None | None | ENSMUSG00000040918 | 116914 | protein_coding | 111.0741671 | -0.033532372 | 0.124407416 | 0.269536763 | 0.787516656 | 0.893615254 | 0.033532372 | Not Significant | ENSMUSG00000040918 | 116914 | protein_coding | 229.4757077 | -0.114070375 | 0.102152395 | -1.116668634 | 0.264136034 | 0.505545025 | 0.114070375 | Cond1 No Change + Cond2 Not Significant | Cond2 No Change + Cond1 Not Significant |
| Slc19a3 | TRUE | -0.048924273 | 1 | No Change | Not Significant | FALSE | NA | NA | No Change | Not Detected | FALSE | Background = FALSE | Background = FALSE | Background = FALSE | ENSMUSG00000038496 | 80721 | protein_coding | 2.434640418 | -0.048924273 | 0.683765647 | 0.07155123 | 0.942959054 | 1 | 0.048924273 | Not Significant | NA | NA | NA | NA | NA | NA | NA | NA | NA | NA | Background = FALSE | Background = FALSE |
| Slc1a1 | TRUE | -0.707788677 | 0.002014251 | DOWN | DOWN | TRUE | -0.117899445 | 0.516713029 | No Change | Not Significant | TRUE | Cond1 DOWN + Cond2 Not Significant | Cond1_DOWN | Cond1_DOWN | ENSMUSG00000024935 | 20510 | protein_coding | 352.5273427 | -0.707788677 | 0.193895665 | 3.650358443 | 0.000261875 | 0.002014251 | 0.707788677 | DOWN | ENSMUSG00000024935 | 20510 | protein_coding | 793.7857887 | -0.117899445 | 0.107817085 | -1.093513567 | 0.274168375 | 0.516713029 | 0.117899445 | Cond1 DOWN + Cond2 Not Significant | Cond2 No Change + Cond1 DOWN |
| Slc1a2 | TRUE | 0.279765404 | 0.788881899 | No Change | Not Significant | TRUE | -0.340766334 | 0.50123327 | No Change | Not Significant | TRUE | Cond1 Not Significant + Cond2 Not Significant | None | None | ENSMUSG00000005089 | 20511 | protein_coding | 12.82068931 | 0.279765404 | 0.559349707 | -0.500161886 | 0.616961093 | 0.788881899 | 0.279765404 | Not Significant | ENSMUSG00000005089 | 20511 | protein_coding | 17.17061031 | -0.340766334 | 0.302379606 | -1.126948801 | 0.259764129 | 0.50123327 | 0.340766334 | Cond1 No Change + Cond2 Not Significant | Cond2 No Change + Cond1 Not Significant |
| Slc1a3 | TRUE | -1.703176661 | 0.008564059 | DOWN | DOWN | TRUE | -0.276160557 | 0.50123327 | No Change | Not Significant | TRUE | Cond1 DOWN + Cond2 Not Significant | Cond1_DOWN | Cond1_DOWN | ENSMUSG00000005360 | 20512 | protein_coding | 13.01389393 | -1.703176661 | 0.534144604 | 3.188605947 | 0.001429606 | 0.008564059 | 1.703176661 | DOWN | ENSMUSG00000005360 | 20512 | protein_coding | 28.30894583 | -0.276160557 | 0.24500006 | -1.127185673 | 0.259663987 | 0.50123327 | 0.276160557 | Cond1 DOWN + Cond2 Not Significant | Cond2 No Change + Cond1 DOWN |
| Slc1a4 | TRUE | 0.148875054 | 0.387358014 | No Change | Not Significant | TRUE | -0.15326256 | 0.225933166 | No Change | Not Significant | TRUE | Cond1 Not Significant + Cond2 Not Significant | None | None | ENSMUSG00000020142 | 55963 | protein_coding | 545.5931425 | 0.148875054 | 0.113609599 | -1.310409122 | 0.190057469 | 0.387358014 | 0.148875054 | Not Significant | ENSMUSG00000020142 | 55963 | protein_coding | 771.5973436 | -0.15326256 | 0.0839939 | -1.824686786 | 0.068048309 | 0.225933166 | 0.15326256 | Cond1 No Change + Cond2 Not Significant | Cond2 No Change + Cond1 Not Significant |
| Slc1a5 | TRUE | -0.13055567 | 0.298275885 | No Change | Not Significant | TRUE | -0.334897828 | 3.46E-08 | No Change | Significant Negative | TRUE | Cond1 Not Significant + Cond2 Significant Negative | None | None | ENSMUSG00000001918 | 20514 | protein_coding | 1456.892046 | -0.13055567 | 0.086297732 | 1.512851682 | 0.130317338 | 0.298275885 | 0.13055567 | Not Significant | ENSMUSG00000001918 | 20514 | protein_coding | 3309.369402 | -0.334897828 | 0.054613303 | -6.13216574 | 8.67E-10 | 3.46E-08 | 0.334897828 | Cond1 No Change + Cond2 Significant Negative | Cond2 No Change + Cond1 Not Significant |
| Slc1a6 | TRUE | -0.668906048 | 1 | No Change | Not Significant | FALSE | NA | NA | No Change | Not Detected | FALSE | Background = FALSE | Background = FALSE | Background = FALSE | ENSMUSG00000005357 | 20513 | protein_coding | 1.245792686 | -0.668906048 | 2.00840731 | 0.333052984 | 0.739094288 | 1 | 0.668906048 | Not Significant | NA | NA | NA | NA | NA | NA | NA | NA | NA | NA | Background = FALSE | Background = FALSE |
| Slc20a1 | TRUE | 1.119804598 | 5.11E-07 | UP | UP | TRUE | 0.00641526 | 0.976480713 | No Change | Not Significant | TRUE | Cond1 UP + Cond2 Not Significant | Cond1_UP | Cond1_UP | ENSMUSG00000027397 | 20515 | protein_coding | 5439.840287 | 1.119804598 | 0.201770312 | -5.549897727 | 2.86E-08 | 5.11E-07 | 1.119804598 | UP | ENSMUSG00000027397 | 20515 | protein_coding | 8494.564857 | 0.00641526 | 0.093499332 | 0.068612894 | 0.945297755 | 0.976480713 | 0.00641526 | Cond1 UP + Cond2 Not Significant | Cond2 No Change + Cond1 UP |
| Slc20a2 | TRUE | -0.198884607 | 0.202452247 | No Change | Not Significant | TRUE | -0.092658075 | 0.432738515 | No Change | Not Significant | TRUE | Cond1 Not Significant + Cond2 Not Significant | None | None | ENSMUSG00000037656 | 20516 | protein_coding | 232.252308 | -0.198884607 | 0.112099304 | 1.774182353 | 0.076032992 | 0.202452247 | 0.198884607 | Not Significant | ENSMUSG00000037656 | 20516 | protein_coding | 506.5359681 | -0.092658075 | 0.072191693 | -1.283500517 | 0.199316779 | 0.432738515 | 0.092658075 | Cond1 No Change + Cond2 Not Significant | Cond2 No Change + Cond1 Not Significant |
| Slc22a1 | TRUE | -0.619845144 | 1.47E-12 | DOWN | DOWN | TRUE | -0.105253847 | 0.259255851 | No Change | Not Significant | TRUE | Cond1 DOWN + Cond2 Not Significant | Cond1_DOWN | Cond1_DOWN | ENSMUSG00000023829 | 20517 | protein_coding | 333.641529 | -0.619845144 | 0.082093192 | 7.550506079 | 4.34E-14 | 1.47E-12 | 0.619845144 | DOWN | ENSMUSG00000023829 | 20517 | protein_coding | 894.6073677 | -0.105253847 | 0.06101979 | -1.724913309 | 0.084543097 | 0.259255851 | 0.105253847 | Cond1 DOWN + Cond2 Not Significant | Cond2 No Change + Cond1 DOWN |
| [illegible] | [illegible] | [illegible] | [illegible] | [illegible] | [illegible] | [illegible] | [illegible] | [illegible] | [illegible] | [illegible] | [illegible] | [illegible] | [illegible] | [illegible] | [illegible] | [illegible] | [illegible] | [illegible] | [illegible] | [illegible] | [illegible] | [illegible] | [illegible] | [illegible] | [illegible] | [illegible] | [illegible] | [illegible] | [illegible] | [illegible] | [illegible] | [illegible] | [illegible] | [illegible] | [illegible] | [illegible] | [illegible] |

| geneID | Cond1_DF_Detected | Cond1_DF_ValueCol | Cond1_DF_PadjCol | Cond1_DF_Cutoff | Cond1_DF_Cutoff_Specific.x | Cond2_DF_Detected | Cond2_DF_ValueCol | Cond2_DF_PadjCol | Cond2_DF_Cutoff | Cond2_DF_Cutoff_Specific | BG_Method | RG1_All | RG2_Significant | RG3_SignificantChange | Cond1_DF_ensembl_gene_id | Cond1_DF_entrezgene_id | Cond1_DF_gene_biotype | Cond1_DF_baseMean | Cond1_DF_log2FoldChange | Cond1_DF_lfcSE |
|---|---|---|---|---|---|---|---|---|---|---|---|---|---|---|---|---|---|---|---|---|
| Slc39a11 | TRUE | 0.274952073 | 0.028942592 | No Change | Significant Positive | TRUE | 0.341801313 | 0.00078781 | No Change | Significant Positive | TRUE | Cond1 Significant Positive + Cond2 Significant Positive | None | None | ENSMUSG00000041654 | 69806 | protein_coding | 1622.606275 | 0.274952073 | 0.100512099 |
| Slc39a12 | TRUE | -0.579581062 | 1 | No Change | Not Significant | FALSE | NA | NA | No Change | Not Detected | FALSE | Background = FALSE | Background = FALSE | Background = FALSE | ENSMUSG00000036949 | 277468 | protein_coding | 1.509730784 | -0.579581062 | 2.081280083 |
| Slc39a13 | TRUE | 0.113584677 | 0.622315002 | No Change | Not Significant | TRUE | -0.081214601 | 0.721674201 | No Change | Not Significant | TRUE | Cond1 Not Significant + Cond2 Not Significant | None | None | ENSMUSG00000002105 | 68427 | protein_coding | 75.74068633 | 0.113584677 | 0.135044328 |
| Slc39a14 | TRUE | 1.387311401 | 2.80E-26 | UP | UP | TRUE | 0.370594038 | 9.57E-13 | No Change | Significant Positive | TRUE | Cond1 UP + Cond2 Significant Positive | Core_UP | Cond1_UP | ENSMUSG00000022094 | 213053 | protein_coding | 983.5419524 | 1.387311401 | 0.126131894 |
| Slc39a2 | TRUE | 0.121299615 | 0.921900136 | No Change | Not Significant | TRUE | 0.230026802 | 0.80962603 | No Change | Not Significant | TRUE | Cond1 Not Significant + Cond2 Not Significant | None | None | ENSMUSG00000072572 | 214922 | protein_coding | 5.398220294 | 0.121299615 | 0.609390043 |
| Slc39a3 | TRUE | 0.040736839 | 0.812494814 | No Change | Not Significant | TRUE | 0.123306052 | 0.335820819 | No Change | Not Significant | TRUE | Cond1 Not Significant + Cond2 Not Significant | None | None | ENSMUSG00000046822 | 106947 | protein_coding | 396.0290589 | 0.040736839 | 0.091006069 |
| Slc39a4 | TRUE | 0.341123246 | 0.057657592 | No Change | Not Significant | TRUE | -0.021456631 | 0.930175172 | No Change | Not Significant | TRUE | Cond1 Not Significant + Cond2 Not Significant | None | None | ENSMUSG00000063354 | 72027 | protein_coding | 2083.852109 | 0.341123246 | 0.139934015 |
| Slc39a5 | TRUE | -0.140179494 | 0.213539052 | No Change | Not Significant | TRUE | -0.058694761 | 0.679535938 | No Change | Not Significant | TRUE | Cond1 Not Significant + Cond2 Not Significant | None | None | ENSMUSG00000039878 | 72002 | protein_coding | 1179.049675 | -0.140179494 | 0.080578096 |
| Slc39a6 | TRUE | 0.577963723 | 0.009312046 | UP | UP | TRUE | 0.357098455 | 0.091124885 | No Change | Not Significant | TRUE | Cond1 UP + Cond2 Not Significant | Cond1_UP | Cond1_UP | ENSMUSG00000024270 | 106957 | protein_coding | 44.524232 | 0.577963723 | 0.182951109 |
| Slc39a7 | TRUE | 0.553030599 | 0.000180273 | UP | UP | TRUE | 0.225423134 | 0.170791501 | No Change | Not Significant | TRUE | Cond1 UP + Cond2 Not Significant | Cond1_UP | Cond1_UP | ENSMUSG00000024327 | 14977 | protein_coding | 229.5242044 | 0.553030599 | 0.128643051 |
| Slc39a8 | TRUE | 0.100159145 | 0.681791678 | No Change | Not Significant | TRUE | -0.038454649 | 0.73174917 | No Change | Not Significant | TRUE | Cond1 Not Significant + Cond2 Not Significant | None | None | ENSMUSG00000053897 | 67547 | protein_coding | 528.3273163 | 0.100159145 | 0.138066891 |
| Slc39a9 | TRUE | 0.068443509 | 0.559860817 | No Change | Not Significant | TRUE | 0.058930601 | 0.420477604 | No Change | Not Significant | TRUE | Cond1 Not Significant + Cond2 Not Significant | None | None | ENSMUSG00000048833 | 328133 | protein_coding | 1829.84863 | 0.068443509 | 0.071461275 |
| Slc3a1 | TRUE | 0.542286752 | 0.056508103 | No Change | Not Significant | TRUE | -0.232645901 | 0.331093889 | No Change | Not Significant | TRUE | Cond1 Not Significant + Cond2 Not Significant | None | None | ENSMUSG00000024131 | 20532 | protein_coding | 428.6073139 | 0.542286752 | 0.221653509 |
| Slc3a2 | TRUE | 0.204563725 | 0.019203423 | No Change | Significant Positive | TRUE | -0.157658774 | 0.007881807 | No Change | Significant Negative | TRUE | Cond1 Significant Positive + Cond2 Significant Negative | None | None | ENSMUSG00000010095 | 17254 | protein_coding | 546.909011 | 0.204563725 | 0.070591325 |
| Slc40a1 | TRUE | -0.962772625 | 4.31E-13 | DOWN | DOWN | TRUE | -0.287275047 | 0.097082061 | No Change | Not Significant | TRUE | Cond1 DOWN + Cond2 Not Significant | Cond1_DOWN | Cond1_DOWN | ENSMUSG00000025993 | 53945 | protein_coding | 2723.548767 | -0.962772625 | 0.12480765 |
| Slc41a1 | TRUE | -0.113774259 | 0.621033677 | No Change | Not Significant | TRUE | -0.013555249 | 0.927812059 | No Change | Not Significant | TRUE | Cond1 Not Significant + Cond2 Not Significant | None | None | ENSMUSG00000013275 | 98396 | protein_coding | 180.3469236 | -0.113774259 | 0.134910327 |
| Slc41a2 | TRUE | 0.30876327 | 0.175901555 | No Change | Not Significant | TRUE | 0.178719261 | 0.028790463 | No Change | Significant Positive | TRUE | Cond1 Not Significant + Cond2 Significant Positive | None | None | ENSMUSG00000034591 | 338365 | protein_coding | 283.8871807 | 0.30876327 | 0.165821388 |
| Slc41a3 | TRUE | 0.211791925 | 0.567902926 | No Change | Not Significant | TRUE | -0.138443895 | 0.649970343 | No Change | Not Significant | TRUE | Cond1 Not Significant + Cond2 Not Significant | None | None | ENSMUSG00000030089 | 71699 | protein_coding | 25.89017749 | 0.211791925 | 0.224817672 |
| Slc43a1 | TRUE | 0.775252327 | 0.000311984 | UP | UP | TRUE | 0.149611326 | 0.472718587 | No Change | Not Significant | TRUE | Cond1 UP + Cond2 Not Significant | Cond1_UP | Cond1_UP | ENSMUSG00000027075 | 72401 | protein_coding | 75.47221184 | 0.775252327 | 0.186306675 |
| Slc43a2 | TRUE | 0.072687287 | 0.842918326 | No Change | Not Significant | TRUE | 0.026683498 | 0.927834878 | No Change | Not Significant | TRUE | Cond1 Not Significant + Cond2 Not Significant | None | None | ENSMUSG00000038178 | 215113 | protein_coding | 154.6380287 | 0.072687287 | 0.190101289 |
| Slc43a3 | TRUE | -0.393513263 | 0.59846773 | No Change | Not Significant | TRUE | 0.21277957 | 0.786125192 | No Change | Not Significant | TRUE | Cond1 Not Significant + Cond2 Not Significant | None | None | ENSMUSG00000027074 | 58207 | protein_coding | 14.50789535 | -0.393513263 | 0.444729071 |
| Slc44a1 | TRUE | -0.300187531 | 0.011857691 | No Change | Significant Negative | TRUE | -0.147060075 | 0.044211463 | No Change | Significant Negative | TRUE | Cond1 Significant Negative + Cond2 Significant Negative | None | None | ENSMUSG00000028412 | 100434 | protein_coding | 4697.03714 | -0.300187531 | 0.097648495 |
| Slc44a2 | TRUE | -0.145434454 | 0.37057407 | No Change | Not Significant | TRUE | 0.008645031 | 0.937561241 | No Change | Not Significant | TRUE | Cond1 Not Significant + Cond2 Not Significant | None | None | ENSMUSG00000057193 | 68682 | protein_coding | 345.6675785 | -0.145434454 | 0.108034029 |
| Slc44a3 | TRUE | -0.150355741 | 0.460363236 | No Change | Not Significant | TRUE | -0.078061315 | 0.330006473 | No Change | Not Significant | TRUE | Cond1 Not Significant + Cond2 Not Significant | None | None | ENSMUSG00000039865 | 213603 | protein_coding | 951.0497586 | -0.150355741 | 0.130174916 |
| Slc44a4 | TRUE | -0.32878964 | 5.32E-06 | No Change | Significant Negative | TRUE | -0.068115908 | 0.523508209 | No Change | Not Significant | TRUE | Cond1 Significant Negative + Cond2 Not Significant | None | None | ENSMUSG00000007034 | 70129 | protein_coding | 4978.067283 | -0.32878964 | 0.06454245 |
| Slc44a5 | TRUE | 0.123137285 | 1 | No Change | Not Significant | TRUE | -1.027579096 | 0.156498963 | No Change | Not Significant | TRUE | Cond1 Not Significant + Cond2 Not Significant | None | None | ENSMUSG00000028360 | 242259 | protein_coding | 3.988389921 | 0.123137285 | 1.028455698 |
| Slc45a1 | TRUE | 1.792506424 | 1 | No Change | Not Significant | FALSE | NA | NA | No Change | Not Detected | FALSE | Background = FALSE | Background = FALSE | Background = FALSE | ENSMUSG00000039838 | 242773 | protein_coding | 2.947341486 | 1.792506424 | 1.574727739 |
| Slc45a3 | TRUE | -0.125626928 | 0.630552252 | No Change | Not Significant | TRUE | 0.308561135 | 0.04029418 | No Change | Significant Positive | TRUE | Cond1 Not Significant + Cond2 Significant Positive | None | None | ENSMUSG00000026435 | 212980 | protein_coding | 428.4746985 | -0.125626928 | 0.15254239 |
| Slc45a4 | TRUE | 0.251495018 | 0.025463175 | No Change | Significant Positive | TRUE | 0.062661787 | 0.563769199 | No Change | Not Significant | TRUE | Cond1 Significant Positive + Cond2 Not Significant | None | None | ENSMUSG00000079020 | 106068 | protein_coding | 350.0184222 | 0.251495018 | 0.090143571 |
| Slc46a1 | TRUE | -0.582589573 | 0.012189964 | DOWN | DOWN | TRUE | 0.068019212 | 0.840526771 | No Change | Not Significant | TRUE | Cond1 DOWN + Cond2 Not Significant | Cond1_DOWN | Cond1_DOWN | ENSMUSG00000020829 | 52466 | protein_coding | 164.98649 | -0.582589573 | 0.190204775 |
| Slc46a2 | TRUE | 1.706333729 | 1 | No Change | Not Significant | FALSE | NA | NA | No Change | Not Detected | FALSE | Background = FALSE | Background = FALSE | Background = FALSE | ENSMUSG00000028386 | 30936 | protein_coding | 1.545880134 | 1.706333729 | 1.393206313 |
| Slc46a3 | TRUE | 0.06415446 | 0.905779176 | No Change | Not Significant | TRUE | -0.032674397 | 0.938113902 | No Change | Not Significant | TRUE | Cond1 Not Significant + Cond2 Not Significant | None | None | ENSMUSG00000029650 | 71706 | protein_coding | 80.90939854 | 0.06415446 | 0.265741767 |
| Slc47a1 | TRUE | 0.087561559 | 0.906704483 | No Change | Not Significant | TRUE | -0.029797053 | 0.951128072 | No Change | Not Significant | TRUE | Cond1 Not Significant + Cond2 Not Significant | None | None | ENSMUSG00000010122 | 67473 | protein_coding | 22.98406562 | 0.087561559 | 0.367254457 |
| Slc47a2 | TRUE | 2.074607811 | 1 | No Change | Not Significant | FALSE | NA | NA | No Change | Not Detected | FALSE | Background = FALSE | Background = FALSE | Background = FALSE | ENSMUSG00000069855 | 380701 | protein_coding | 1.252686932 | 2.074607811 | 2.177547235 |
| Slc48a1 | TRUE | -0.071655155 | 0.698491194 | No Change | Not Significant | TRUE | -0.006522163 | 0.962265867 | No Change | Not Significant | TRUE | Cond1 Not Significant + Cond2 Not Significant | None | None | ENSMUSG00000081534 | 67739 | protein_coding | 277.6326829 | -0.071655155 | 0.104190545 |
| Slc49a4 | TRUE | -0.074214473 | 0.616875898 | No Change | Not Significant | TRUE | -0.04680286 | 0.703236518 | No Change | Not Significant | TRUE | Cond1 Not Significant + Cond2 Not Significant | None | None | ENSMUSG00000022848 | 224132 | protein_coding | 481.2973534 | -0.074214473 | 0.087141073 |
| Slc4a1 | TRUE | -0.011594929 | 0.994279305 | No Change | Not Significant | FALSE | NA | NA | No Change | Not Detected | FALSE | Background = FALSE | Background = FALSE | Background = FALSE | ENSMUSG00000006574 | 20533 | protein_coding | 7.440871256 | -0.011594929 | 0.919993543 |
| Slc4a10 | TRUE | 1.193339323 | 1 | No Change | Not Significant | FALSE | NA | NA | No Change | Not Detected | FALSE | Background = FALSE | Background = FALSE | Background = FALSE | ENSMUSG00000026904 | 94229 | protein_coding | 2.567275284 | 1.193339323 | 1.133089196 |
| Slc4a11 | TRUE | 0.374024715 | 0.789053391 | No Change | Not Significant | TRUE | 0.477329753 | 0.485573395 | No Change | Not Significant | TRUE | Cond1 Not Significant + Cond2 Not Significant | None | None | ENSMUSG00000074796 | 269356 | protein_coding | 5.207805302 | 0.374024715 | 0.748385872 |
| Slc4a1ap | TRUE | 0.073842236 | 0.539389762 | No Change | Not Significant | TRUE | -0.007491604 | 0.960412417 | No Change | Not Significant | TRUE | Cond1 Not Significant + Cond2 Not Significant | None | None | ENSMUSG00000029141 | 20534 | protein_coding | 233.4924963 | 0.073842236 | 0.073952941 |
| Slc4a2 | TRUE | -0.120491158 | 0.583801091 | No Change | Not Significant | TRUE | -0.171124513 | 0.103384952 | No Change | Not Significant | TRUE | Cond1 Not Significant + Cond2 Not Significant | None | None | ENSMUSG00000028962 | 20535 | protein_coding | 2677.040284 | -0.120491158 | 0.132016723 |
| Slc4a3 | TRUE | -0.21498554 | 0.827306109 | No Change | Not Significant | TRUE | 0.202312739 | 0.799662139 | No Change | Not Significant | TRUE | Cond1 Not Significant + Cond2 Not Significant | None | None | ENSMUSG00000006576 | 20536 | protein_coding | 13.22576273 | -0.21498554 | 0.517232477 |
| Slc4a4 | TRUE | -0.666046466 | 3.02E-14 | DOWN | DOWN | TRUE | -0.171696782 | 0.106731485 | No Change | Not Significant | TRUE | Cond1 DOWN + Cond2 Not Significant | Cond1_DOWN | Cond1_DOWN | ENSMUSG00000060961 | 54403 | protein_coding | 1349.922103 | -0.666046466 | 0.082639428 |
| Slc4a5 | TRUE | 1.173971074 | 0.350538094 | No Change | Not Significant | TRUE | -0.153373365 | 0.853011339 | No Change | Not Significant | TRUE | Cond1 Not Significant + Cond2 Not Significant | None | None | ENSMUSG00000068323 | 232156 | protein_coding | 8.157313142 | 1.173971074 | 0.84401368 |
| Slc4a7 | TRUE | 0.394831665 | 0.051167007 | No Change | Not Significant | TRUE | 0.416207243 | 2.55E-12 | No Change | Significant Positive | TRUE | Cond1 Not Significant + Cond2 Significant Positive | None | None | ENSMUSG00000021733 | 218756 | protein_coding | 2889.599175 | 0.394831665 | 0.158388243 |
| Slc4a8 | TRUE | 0.487591769 | 0.324596932 | No Change | Not Significant | TRUE | 0.006417865 | 0.991109854 | No Change | Not Significant | TRUE | Cond1 Not Significant + Cond2 Not Significant | None | None | ENSMUSG00000023032 | 59033 | protein_coding | 31.15778632 | 0.487591769 | 0.336037729 |
| Slc4a9 | TRUE | -0.125462538 | 0.914162556 | No Change | Not Significant | TRUE | -0.285211554 | 0.775689464 | No Change | Not Significant | TRUE | Cond1 Not Significant + Cond2 Not Significant | None | None | ENSMUSG00000024485 | 240215 | protein_coding | 5.911253614 | -0.125462538 | 0.583849004 |
| Slc50a1 | TRUE | -0.301172524 | 0.17492772 | No Change | Not Significant | TRUE | -0.060911595 | 0.644516088 | No Change | Not Significant | TRUE | Cond1 Not Significant + Cond2 Not Significant | None | None | ENSMUSG00000027953 | 19729 | protein_coding | 417.2787955 | -0.301172524 | 0.161442068 |
| Slc5a1 | TRUE | -0.458183881 | 0.243645564 | No Change | Not Significant | TRUE | -0.359369355 | 0.039304177 | No Change | Significant Negative | TRUE | Cond1 Not Significant + Cond2 Significant Negative | None | None | ENSMUSG00000035699 | 106407 | protein_coding | 562.9237994 | -0.458183881 | 0.276894457 |
| Slc51b | TRUE | 0.542537598 | 0.017604641 | UP | UP | TRUE | -0.235544001 | 0.028426826 | No Change | Significant Negative | TRUE | Cond1 UP + Cond2 Significant Negative | Opposite | Cond1_UP | ENSMUSG00000053862 | 330962 | protein_coding | 500.6281086 | 0.542537598 | 0.185144008 |
| Slc52a2 | TRUE | -0.13760615 | 0.552639068 | No Change | Not Significant | TRUE | -0.141989788 | 0.101895312 | No Change | Not Significant | TRUE | Cond1 Not Significant + Cond2 Not Significant | None | None | ENSMUSG00000022560 | 52710 | protein_coding | 239.673431 | -0.13760615 | 0.141389217 |
| Slc52a3 | TRUE | -0.024453735 | 0.896829747 | No Change | Not Significant | TRUE | 0.059629446 | 0.682136666 | No Change | Not Significant | TRUE | Cond1 Not Significant + Cond2 Not Significant | None | None | ENSMUSG00000027463 | 69698 | protein_coding | 493.0956375 | -0.024453735 | 0.09308978 |
| Slc5a1 | TRUE | 0.176083509 | 0.034445098 | No Change | Significant Positive | TRUE | -0.096053679 | 0.106976995 | No Change | Not Significant | TRUE | Cond1 Significant Positive + Cond2 Not Significant | None | None | ENSMUSG00000011034 | 20537 | protein_coding | 2802.657482 | 0.176083509 | 0.066054419 |
| Slc5a10 | TRUE | 0.423119598 | 0.773864504 | No Change | Not Significant | TRUE | -0.346132362 | 0.689357631 | No Change | Not Significant | TRUE | Cond1 Not Significant + Cond2 Not Significant | None | None | ENSMUSG00000042371 | 109342 | protein_coding | 5.357167305 | 0.423119598 | 0.797136874 |
| Slc5a11 | TRUE | 2.309067048 | 1 | No Change | Not Significant | TRUE | -0.12383809 | 0.937787345 | No Change | Not Significant | TRUE | Cond1 Not Significant + Cond2 Not Significant | None | None | ENSMUSG00000030769 | 233836 | protein_coding | 4.045484028 | 2.309067048 | 1.29718221 |
| Slc5a12 | TRUE | 0.167899258 | 0.962213622 | No Change | Not Significant | TRUE | 0.483144436 | 0.755247648 | No Change | Not Significant | TRUE | Cond1 Not Significant + Cond2 Not Significant | None | None | ENSMUSG00000041644 | 241612 | protein_coding | 4.957144405 | 0.167899258 | 1.706013127 |
| Slc5a2 | TRUE | 0.373301133 | 1 | No Change | Not Significant | FALSE | NA | NA | No Change | Not Detected | FALSE | Background = FALSE | Background = FALSE | Background = FALSE | ENSMUSG00000030781 | 246787 | protein_coding | 2.995941002 | 0.373301133 | 0.931483376 |
| Slc5a3 | TRUE | -0.488684188 | 0.002018623 | No Change | Significant Negative | TRUE | -0.085730863 | 0.598521667 | No Change | Not Significant | TRUE | Cond1 Significant Negative + Cond2 Not Significant | None | None | ENSMUSG00000089774 | 53881 | protein_coding | 1060.367175 | -0.488684188 | 0.133902133 |
| Slc5a4a | TRUE | 1.640813392 | 0.09822375 | No Change | Not Significant | TRUE | -0.090062796 | 0.939855462 | No Change | Not Significant | TRUE | Cond1 Not Significant + Cond2 Not Significant | None | None | ENSMUSG00000020229 | 64452 | protein_coding | 4.635481713 | 1.640813392 | 0.75127936 |
| Slc5a4b | TRUE | 0.097555095 | 0.936356422 | No Change | Not Significant | TRUE | 0.440976494 | 0.616661588 | No Change | Not Significant | TRUE | Cond1 Not Significant + Cond2 Not Significant | None | None | ENSMUSG00000020226 | 64454 | protein_coding | 9.386331724 | 0.097555095 | 0.600427605 |
| Slc5a5 | TRUE | 0.478489045 | 0.588390676 | No Change | Not Significant | FALSE | NA | NA | No Change | Not Detected | FALSE | Background = FALSE | Background = FALSE | Background = FALSE | ENSMUSG00000000792 | 114479 | protein_coding | 6.212422916 | 0.478489045 | 0.529059428 |
| Slc5a6 | TRUE | -0.142106661 | 0.656764359 | No Change | Not Significant | TRUE | -0.10090552 | 0.445602229 | No Change | Not Significant | TRUE | Cond1 Not Significant + Cond2 Not Significant | None | None | ENSMUSG00000006641 | 330064 | protein_coding | 358.654102 | -0.142106661 | 0.18346632 |
| Slc5a7 | TRUE | -1.187814625 | 1 | No Change | Not Significant | FALSE | NA | NA | No Change | Not Detected | FALSE | Background = FALSE | Background = FALSE | Background = FALSE | ENSMUSG00000023945 | 63993 | protein_coding | 3.132386009 | -1.187814625 | 1.158049288 |
| Slc5a8 | TRUE | -0.367856439 | 0.038748264 | No Change | Significant Negative | TRUE | -0.116446218 | 0.488015969 | No Change | Not Significant | TRUE | Cond1 Significant Negative + Cond2 Not Significant | None | None | ENSMUSG00000020062 | 216225 | protein_coding | 4559.823663 | -0.367856439 | 0.140637397 |
| Slc5a9 | TRUE | 0.72465246 | 2.34E-06 | UP | UP | TRUE | -0.023993281 | 0.930926537 | No Change | Not Significant | TRUE | Cond1 UP + Cond2 Not Significant | Cond1_UP | Cond1_UP | ENSMUSG00000028544 | 230612 | protein_coding | 312.6270619 | 0.72465246 | 0.137822407 |
| Slc66a1 | TRUE | 0.053477986 | 0.837254214 | No Change | Not Significant | TRUE | -0.207718625 | 0.221421603 | No Change | Not Significant | TRUE | Cond1 Not Significant + Cond2 Not Significant | None | None | ENSMUSG00000028744 | 212555 | protein_coding | 80.02297489 | 0.053477986 | 0.135673942 |
| Slc66a2 | TRUE | -0.008358132 | 0.948896349 | No Change | Not Significant | TRUE | 0.032104755 | 0.73971984 | No Change | Not Significant | TRUE | Cond1 Not Significant + Cond2 Not Significant | None | None | ENSMUSG00000034006 | 66943 | protein_coding | 632.3857677 | -0.008358132 | 0.064046479 |
| Slc6a1 | TRUE | 1.727679261 | 0.170249866 | No Change | Not Significant | FALSE | NA | NA | No Change | Not Detected | FALSE | Background = FALSE | Background = FALSE | Background = FALSE | ENSMUSG00000030310 | 232333 | protein_coding | 6.445095464 | 1.727679261 | 0.917593807 |
| Slc6a11 | TRUE | 0.176008836 | 1 | No Change | Not Significant | FALSE | NA | NA | No Change | Not Detected | FALSE | Background = FALSE | Background = FALSE | Background = FALSE | ENSMUSG00000030307 | 243616 | protein_coding | 2.04742851 | 0.176008836 | 2.687610432 |
| Slc6a12 | TRUE | -0.855559377 | 1 | No Change | Not Significant | TRUE | -1.116643893 | 0.002687827 | DOWN | DOWN | TRUE | Cond1 Not Significant + Cond2 DOWN | Cond2_DOWN | Cond2_DOWN | ENSMUSG00000030109 | 14411 | protein_coding | 4.11555343 | -0.855559377 | 1.228589073 |

| geneID | Cond1_DF_Detected | Cond1_DF_ValueCol |
|---|---|---|
| Smcr8 | TRUE | 0.165390786 |
| Smdt1 | TRUE | 0.22233244 |
| Smg1 | TRUE | 0.252378721 |
| Smg5 | TRUE | 0.203957351 |
| Smg6 | TRUE | -0.13808157 |
| Smg7 | TRUE | 0.141418787 |
| Smg8 | TRUE | 0.313318037 |
| Smg9 | TRUE | 0.026580454 |
| Smgc | TRUE | 2.568849921 |
| Smim1 | TRUE | 0.303887818 |
| Smim10l1 | TRUE | 0.084108821 |
| Smim10l2a | TRUE | 0.301878233 |
| Smim11 | TRUE | -0.225466172 |
| Smim12 | TRUE | -0.026254346 |
| Smim13 | TRUE | -0.204991241 |
| Smim14 | TRUE | 0.090836138 |
| Smim15 | TRUE | -0.019834591 |
| Smim17 | TRUE | -1.007487242 |
| Smim18 | TRUE | -0.320127289 |
| Smim19 | TRUE | -1.24E-05 |
| Smim20 | TRUE | 0.05812298 |
| Smim22 | TRUE | -0.499669921 |
| Smim24 | TRUE | -1.295617344 |
| Smim26 | TRUE | -0.224378272 |
| Smim27 | TRUE | -0.17930853 |
| Smim29 | TRUE | -0.60947472 |
| Smim3 | TRUE | 0.30705683 |
| Smim30 | TRUE | -0.003464967 |
| Smim31 | TRUE | -0.036046764 |
| Smim32 | TRUE | -0.042612387 |
| Smim33 | TRUE | 0.526494829 |
| Smim35 | TRUE | 0.325534315 |
| Smim36 | TRUE | -0.202471634 |
| Smim38 | TRUE | -1.01107479 |
| Smim4 | TRUE | 0.002623413 |
| Smim41 | TRUE | 0.637463961 |
| Smim43 | TRUE | 0.349816756 |
| Smim45 | TRUE | -1.479193358 |
| Smim5 | TRUE | -0.544754562 |
| Smim6 | TRUE | -0.13539381 |
| Smim7 | TRUE | 0.077214431 |
| Smim8 | TRUE | 0.095670924 |
| Smn1 | TRUE | 0.297911938 |
| Smndc1 | TRUE | -0.154920355 |
| Smo | TRUE | 0.209003491 |
| Smoc1 | TRUE | 0.739957025 |
| Smoc2 | TRUE | 0.684906305 |
| Smok2a | TRUE | 2.739184389 |
| Smok2b | TRUE | 0.383184039 |
| Smox | TRUE | -0.277338309 |
| Smpd1 | TRUE | -0.021641978 |
| Smpd2 | TRUE | -0.175555228 |
| Smpd3 | TRUE | 0.17136173 |
| Smpd4 | TRUE | 0.356660698 |
| Smpd5 | TRUE | -0.332413846 |
| Smpdl3a | TRUE | -1.049860095 |
| Smpdl3b | TRUE | 0.24596875 |
| Smpx | TRUE | 0.470258404 |
| Sms | TRUE | -0.247142637 |
| Smtn | TRUE | 1.537415836 |
| Smtnl2 | TRUE | -0.162189666 |
| Smu1 | TRUE | 0.144013782 |
| Smug1 | TRUE | -0.040295407 |
| Smurf1 | TRUE | 0.034321688 |
| Smurf2 | TRUE | -0.195260785 |
| Smyd1 | TRUE | -0.866298295 |
| Smyd2 | TRUE | -0.093523672 |
| Smyd3 | TRUE | -0.415305598 |
| Smyd4 | TRUE | -0.118057828 |
| Smyd5 | TRUE | 0.298996428 |
| Snai1 | TRUE | 4.616014404 |
| Snai3 | TRUE | 0.236709171 |
| Snap23 | TRUE | 0.646703301 |
| Snap25 | TRUE | 0.530573616 |
| Snap29 | TRUE | 0.029869816 |
| Snap47 | TRUE | 0.006495774 |
| Snap91 | TRUE | -0.108598233 |
| Snapc1 | TRUE | 0.073279998 |
| Snapc2 | TRUE | -0.062131293 |
| Snapc3 | TRUE | -0.086838266 |
| Snapc4 | TRUE | 0.102614407 |
| Snapc5 | TRUE | -0.364277342 |
| Snapin | TRUE | -0.16065561 |
| Snca | TRUE | 1.381159212 |
| Sncaip | TRUE | -0.729945512 |
| Sncb | TRUE | -1.324200569 |
| Snd1 | TRUE | 0.620253078 |
| Sned1 | TRUE | -0.326181896 |
| Snf8 | TRUE | -0.130230157 |
| Snhg10 | TRUE | -0.228644344 |
| Snhg11 | TRUE | 0.777843895 |
| Snhg12 | TRUE | -0.250630292 |
| Snhg14 | TRUE | -0.416559237 |
| Snhg16 | TRUE | -0.097119863 |
| Snhg17 | TRUE | -0.044531457 |
| Snhg18 | TRUE | -0.04243755 |
| Snhg20 | TRUE | -0.157261829 |
| Snhg3 | TRUE | 0.088154168 |
| Snhg4 | TRUE | -0.18483802 |
| Snhg8 | TRUE | -0.023738594 |
| Snip1 | TRUE | -0.058379754 |
| Snn | TRUE | 1.093110381 |
| Snora17 | TRUE | 0.715808121 |
| Snora21 | TRUE | 0.235401744 |
| Snora23 | TRUE | 1.069993549 |
| Snora47 | TRUE | 0.079247055 |
| Snora57 | TRUE | 0.480731066 |
| Snora69 | TRUE | 1.108873608 |
| Snorc | TRUE | 0.555035832 |
| Snord118 | TRUE | 0.730752057 |
| Snord13 | TRUE | 0.295717737 |
| Snord138 | TRUE | -0.631051825 |
| Snord15a | TRUE | -0.247107515 |
| Snord17 | TRUE | -0.001302503 |
| Snord23 | TRUE | 0.591675737 |
| Snord32a | TRUE | 0.695830019 |
| Snord34 | TRUE | 0.682707826 |
| Snord35a | TRUE | 0.709690897 |
| Snord55 | TRUE | 0.200286192 |
| Snord59a | TRUE | 0.358975826 |
| Snord7 | TRUE | 0.153174558 |
| Snord73a | TRUE | 0.603899616 |
| Snord73b | TRUE | 0.136606542 |
| Snord8 | TRUE | -0.040926452 |
| Snord82 | TRUE | 0.314654047 |
| Snord92 | TRUE | 0.403406466 |
| Snord94 | TRUE | 0.113909207 |
| Snph | TRUE | -0.246688606 |
| Snrk | TRUE | -0.046117388 |
| Snrnp200 | TRUE | 0.079173056 |
| Snrnp25 | TRUE | -0.425458636 |
| Snrnp27 | TRUE | 0.152219365 |
| Snrnp35 | TRUE | 0.074103994 |
| Snrnp40 | TRUE | 0.467791684 |
| Snrnp48 | TRUE | 0.392071961 |
| Snrnp70 | TRUE | 0.693055886 |
| Snrpa | TRUE | 0.698754122 |
| Snrpa1 | TRUE | 0.48901604 |
| Snrpb | TRUE | 0.217990473 |
| Snrpb2 | TRUE | 0.167545761 |
| Snrpc | TRUE | -0.030330513 |
| Snrpd1 | TRUE | 0.226856357 |
| Snrpd2 | TRUE | -0.240000183 |
| Snrpd3 | TRUE | 0.229273077 |
| Snrpe | TRUE | -0.112602412 |
| Snrpf | TRUE | 0.133690852 |
| Snrpg | TRUE | -0.044297035 |
| Snrpn | TRUE | -0.363446474 |
| Snta1 | TRUE | 0.257212276 |
| Sntb1 | TRUE | 1.234797897 |
| Sntb2 | TRUE | 0.10412869 |
| Sntg1 | TRUE | 0.871409962 |
| Sntg2 | TRUE | 1.881174622 |
| Snu13 | TRUE | -0.335847375 |
| Snupn | TRUE | -0.050512862 |
| Snw1 | TRUE | -0.048153245 |
| Snx1 | TRUE | -0.206881826 |
| Snx10 | TRUE | -0.307094572 |
| Snx11 | TRUE | 0.221908695 |
| Snx12 | TRUE | -0.01998145 |
| Snx13 | TRUE | 0.02985685 |
| Snx14 | TRUE | -0.119972533 |
| Snx15 | TRUE | -0.221301584 |
| Snx16 | TRUE | -0.136944335 |
| Snx17 | TRUE | 0.072538072 |
| Snx18 | TRUE | -0.021315632 |
| Snx19 | TRUE | 0.083014614 |
| Snx2 | TRUE | -0.17401915 |
| Snx20 | TRUE | 0.471883486 |
| Snx21 | TRUE | 0.136517893 |
| Snx22 | TRUE | 1.28669393 |
| Snx24 | TRUE | 0.060425642 |
| Snx25 | TRUE | -0.088155482 |
| Snx27 | TRUE | -0.075605554 |
| Snx29 | TRUE | -0.099860303 |
| Snx3 | TRUE | 0.066530456 |
| Snx30 | TRUE | -0.047276657 |
| Snx31 | TRUE | -1.014241654 |
| Snx32 | TRUE | -0.102368693 |
| Snx33 | TRUE | -0.1675095 |
| Snx4 | TRUE | -0.06266725 |
| Snx5 | TRUE | -0.261062845 |
| Snx6 | TRUE | -0.045959911 |
| Snx7 | TRUE | -0.13772874 |
| Snx8 | TRUE | 0.365497185 |
| Snx9 | TRUE | -0.210181756 |
| Soat1 | TRUE | -0.209833715 |
| Soat2 | TRUE | 1.530451805 |
| Sobp | TRUE | -1.103276618 |
| Socs1 | TRUE | 0.493836298 |
| Socs2 | TRUE | -0.387527001 |
| Socs3 | TRUE | -0.172990601 |

| geneID | Cond1_DF_PadjCol | Cond1_DF_Cutoff | Cond1_DF_Cutoff_Specific.x | Cond2_DF_Detected | Cond2_DF_ValueCol | Cond2_DF_PadjCol | Cond2_DF_Cutoff | Cond2_DF_Cutoff_Specific | BG_Method | RG1_All |
|---|---|---|---|---|---|---|---|---|---|---|
| Smcr8 | 0.123236664 | No Change | Not Significant | TRUE | -0.072557944 | 0.326892409 | No Change | Not Significant | TRUE | Cond1 Not Significant + Cond2 Not Significant |
| Smdt1 | 0.040715581 | No Change | Significant Positive | TRUE | 0.166871259 | 0.17573461 | No Change | Not Significant | TRUE | Cond1 Significant Positive + Cond2 Not Significant |
| Smg1 | 0.149272747 | No Change | Not Significant | TRUE | -0.091473808 | 0.26456259 | No Change | Not Significant | TRUE | Cond1 Not Significant + Cond2 Not Significant |
| Smg5 | 0.001633842 | No Change | Significant Positive | TRUE | 0.032624411 | 0.784048864 | No Change | Not Significant | TRUE | Cond1 Significant Positive + Cond2 Not Significant |
| Smg6 | 0.274864513 | No Change | Not Significant | TRUE | -0.155605946 | 0.022791902 | No Change | Significant Negative | TRUE | Cond1 Not Significant + Cond2 Significant Negative |
| Smg7 | 0.097089608 | No Change | Not Significant | TRUE | 0.011954445 | 0.898301545 | No Change | Not Significant | TRUE | Cond1 Not Significant + Cond2 Not Significant |
| Smg8 | 0.007596149 | No Change | Significant Positive | TRUE | 0.089014734 | 0.476068282 | No Change | Not Significant | TRUE | Cond1 Significant Positive + Cond2 Not Significant |
| Smg9 | 0.888933284 | No Change | Not Significant | TRUE | 0.043817544 | 0.794327773 | No Change | Not Significant | TRUE | Cond1 Not Significant + Cond2 Not Significant |
| Smgc | 1 | No Change | Not Significant | FALSE | NA | NA | No Change | Not Detected | FALSE | Background = FALSE |
| Smim1 | 0.628624818 | No Change | Not Significant | TRUE | -0.024159526 | 0.970920986 | No Change | Not Significant | TRUE | Cond1 Not Significant + Cond2 Not Significant |
| Smim10l1 | 0.579378227 | No Change | Not Significant | TRUE | 0.160678355 | 3.44E-05 | No Change | Significant Positive | TRUE | Cond1 Not Significant + Cond2 Significant Positive |
| Smim10l2a | 0.496825672 | No Change | Not Significant | TRUE | 0.352693137 | 0.20889399 | No Change | Not Significant | TRUE | Cond1 Not Significant + Cond2 Not Significant |
| Smim11 | 0.097546045 | No Change | Not Significant | TRUE | 0.152878015 | 0.114712014 | No Change | Not Significant | TRUE | Cond1 Not Significant + Cond2 Not Significant |
| Smim12 | 0.880245376 | No Change | Not Significant | TRUE | 0.087520556 | 0.606833732 | No Change | Not Significant | TRUE | Cond1 Not Significant + Cond2 Not Significant |
| Smim13 | 0.161560315 | No Change | Not Significant | TRUE | -0.213364306 | 0.105292377 | No Change | Not Significant | TRUE | Cond1 Not Significant + Cond2 Not Significant |
| Smim14 | 0.685335967 | No Change | Not Significant | TRUE | 0.054938841 | 0.377781372 | No Change | Not Significant | TRUE | Cond1 Not Significant + Cond2 Not Significant |
| Smim15 | 0.884245488 | No Change | Not Significant | TRUE | 0.112681967 | 0.112975905 | No Change | Not Significant | TRUE | Cond1 Not Significant + Cond2 Not Significant |
| Smim17 | 1 | No Change | Not Significant | FALSE | NA | NA | No Change | Not Detected | FALSE | Background = FALSE |

Supplementary Table 3

| geneID | Cond1_DF_Detected | Cond1_DF_ValueCol | Cond1_DF_PadjCol | Cond1_DF_Cutoff | Cond1_DF_Cutoff_Specific.x | Cond2_DF_Detected | Cond2_DF_ValueCol | Cond2_DF_PadjCol | Cond2_DF_Cutoff | Cond2_DF_Cutoff_Specific | BG_Method | RG1_All |
|---|---|---|---|---|---|---|---|---|---|---|---|---|
| Socs4 | TRUE | -0.008933254 | 0.947705869 | No Change | Not Significant | TRUE | -0.045168797 | 0.54999264 | No Change | Not Significant | TRUE | Cond1 Not Significant + Cond2 Not Significant |
| Socs5 | TRUE | 0.010920228 | 0.971381329 | No Change | Not Significant | TRUE | 0.010709545 | 0.955895949 | No Change | Not Significant | TRUE | Cond1 Not Significant + Cond2 Not Significant |
| Socs6 | TRUE | -0.023707059 | 0.869043726 | No Change | Not Significant | TRUE | 0.096421165 | 0.064914238 | No Change | Not Significant | TRUE | Cond1 Not Significant + Cond2 Not Significant |
| Socs7 | TRUE | -0.019278244 | 0.926816617 | No Change | Not Significant | TRUE | -0.083335364 | 0.375480655 | No Change | Not Significant | TRUE | Cond1 Not Significant + Cond2 Not Significant |
| Sod1 | TRUE | -0.065742618 | 0.536227262 | No Change | Not Significant | TRUE | -0.069643084 | 0.416513477 | No Change | Not Significant | TRUE | Cond1 Not Significant + Cond2 Not Significant |
| Sod2 | TRUE | -0.209705163 | 0.01632826 | No Change | Significant Negative | TRUE | 0.092258713 | 0.276898911 | No Change | Not Significant | TRUE | Cond1 Significant Negative + Cond2 Not Significant |
| Sod3 | TRUE | 0.686956963 | 0.372872861 | No Change | Not Significant | TRUE | 0.38485523 | 0.603998706 | No Change | Not Significant | TRUE | Cond1 Not Significant + Cond2 Not Significant |
| Soga1 | TRUE | 0.40920219 | 0.339025906 | No Change | Not Significant | TRUE | -0.030039548 | 0.925920981 | No Change | Not Significant | TRUE | Cond1 Not Significant + Cond2 Not Significant |
| Soga3 | TRUE | 0.725883553 | 1 | No Change | Not Significant | FALSE | NA | NA | No Change | Not Detected | FALSE | Background = FALSE |
| Sohlh2 | TRUE | -0.845309728 | 1 | No Change | Not Significant | FALSE | NA | NA | No Change | Not Detected | FALSE | Background = FALSE |
| Son | TRUE | 0.017121281 | 0.88093126 | No Change | Not Significant | TRUE | -0.11115393 | 0.002365286 | No Change | Significant Negative | TRUE | Cond1 Not Significant + Cond2 Significant Negative |
| Sorbs1 | TRUE | 0.193045869 | 0.680659303 | No Change | Not Significant | TRUE | -0.00482459 | 0.984744632 | No Change | Not Significant | TRUE | Cond1 Not Significant + Cond2 Not Significant |
| Sorbs2 | TRUE | 0.004701927 | 0.987234441 | No Change | Not Significant | TRUE | -0.150355807 | 0.140900901 | No Change | Not Significant | TRUE | Cond1 Not Significant + Cond2 Not Significant |
| Sorbs2os | TRUE | 0.432574454 | 0.599649878 | No Change | Not Significant | TRUE | -0.782074639 | 0.256750769 | No Change | Not Significant | TRUE | Cond1 Not Significant + Cond2 Not Significant |
| Sorbs3 | TRUE | -0.813059834 | 0.011044625 | DOWN | DOWN | TRUE | 0.211926722 | 0.529168268 | No Change | Not Significant | TRUE | Cond1 DOWN + Cond2 Not Significant |
| Sorcs1 | TRUE | -0.327877411 | 1 | No Change | Not Significant | FALSE | NA | NA | No Change | Not Detected | FALSE | Background = FALSE |
| Sorcs2 | TRUE | 0.204757325 | 0.440496352 | No Change | Not Significant | TRUE | -0.178487181 | 0.380799766 | No Change | Not Significant | TRUE | Cond1 Not Significant + Cond2 Not Significant |
| Sorcs3 | TRUE | 0.887401022 | 0.420598049 | No Change | Not Significant | TRUE | -0.442148657 | 0.257723064 | No Change | Not Significant | TRUE | Cond1 Not Significant + Cond2 Not Significant |
| Sord | TRUE | -0.83089097 | 3.99E-28 | DOWN | DOWN | TRUE | -0.522307696 | 1.34E-22 | DOWN | DOWN | TRUE | Cond1 DOWN + Cond2 DOWN |
| Sorl1 | TRUE | -0.479479594 | 6.20E-06 | No Change | Significant Negative | TRUE | -0.076893832 | 0.529168268 | No Change | Not Significant | TRUE | Cond1 Significant Negative + Cond2 Not Significant |
| Sort1 | TRUE | -0.248021799 | 0.017451579 | No Change | Significant Negative | TRUE | -0.005026595 | 0.958041702 | No Change | Not Significant | TRUE | Cond1 Significant Negative + Cond2 Not Significant |
| Sos1 | TRUE | 0.068817842 | 0.652196761 | No Change | Not Significant | TRUE | -0.102800081 | 0.342598955 | No Change | Not Significant | TRUE | Cond1 Not Significant + Cond2 Not Significant |
| Sos2 | TRUE | -0.181338418 | 0.092985996 | No Change | Not Significant | TRUE | -0.187014611 | 0.00158722 | No Change | Significant Negative | TRUE | Cond1 Not Significant + Cond2 Significant Negative |
| Sowaha | TRUE | -0.74609658 | 0.000487336 | DOWN | DOWN | TRUE | 0.006486606 | 0.978324393 | No Change | Not Significant | TRUE | Cond1 DOWN + Cond2 Not Significant |
| Sowahb | TRUE | -0.087280416 | 0.684542733 | No Change | Not Significant | TRUE | -0.176099388 | 0.015264121 | No Change | Significant Negative | TRUE | Cond1 Not Significant + Cond2 Significant Negative |
| Sowahc | TRUE | -0.327467688 | 0.003307101 | No Change | Significant Negative | TRUE | -0.232360506 | 0.009823815 | No Change | Significant Negative | TRUE | Cond1 Significant Negative + Cond2 Significant Negative |
| Sox1 | TRUE | 2.120602075 | 1 | No Change | Not Significant | FALSE | NA | NA | No Change | Not Detected | FALSE | Background = FALSE |
| Sox10 | TRUE | 1.97282262 | 1 | No Change | Not Significant | FALSE | NA | NA | No Change | Not Detected | FALSE | Background = FALSE |
| Sox11 | TRUE | 2.906239593 | 1 | No Change | Not Significant | FALSE | NA | NA | No Change | Not Detected | FALSE | Background = FALSE |
| Sox12 | TRUE | 2.823591774 | 1.03E-20 | UP | UP | TRUE | 1.309800718 | 1.23E-05 | UP | UP | TRUE | Cond1 UP + Cond2 UP |
| Sox13 | TRUE | 0.063016424 | 0.786724259 | No Change | Not Significant | TRUE | -0.131965615 | 0.234551147 | No Change | Not Significant | TRUE | Cond1 Not Significant + Cond2 Not Significant |
| Sox15 | TRUE | -0.22599297 | 1 | No Change | Not Significant | FALSE | NA | NA | No Change | Not Detected | FALSE | Background = FALSE |
| Sox17 | TRUE | 1.714807483 | 1 | No Change | Not Significant | FALSE | NA | NA | No Change | Not Detected | FALSE | Background = FALSE |
| Sox1ot | TRUE | 1.683586674 | 0.000968688 | UP | UP | TRUE | 0.639833772 | 0.267564435 | No Change | Not Significant | TRUE | Cond1 UP + Cond2 Not Significant |
| Sox21 | TRUE | 0.240271925 | 0.376052415 | No Change | Not Significant | TRUE | 0.025755337 | 0.945542552 | No Change | Not Significant | TRUE | Cond1 Not Significant + Cond2 Not Significant |
| Sox2ot | TRUE | 3.548386848 | 1 | No Change | Not Significant | FALSE | NA | NA | No Change | Not Detected | FALSE | Background = FALSE |
| Sox30 | TRUE | 2.106010415 | 1 | No Change | Not Significant | FALSE | NA | NA | No Change | Not Detected | FALSE | Background = FALSE |
| Sox4 | TRUE | 0.138116618 | 0.2494703 | No Change | Not Significant | TRUE | -0.033256984 | 0.800888833 | No Change | Not Significant | TRUE | Cond1 Not Significant + Cond2 Not Significant |
| Sox5 | TRUE | 0.298798081 | 0.766436139 | No Change | Not Significant | TRUE | 0.128350811 | 0.861383838 | No Change | Not Significant | TRUE | Cond1 Not Significant + Cond2 Not Significant |
| Sox6 | TRUE | 0.074469665 | 0.850752177 | No Change | Not Significant | TRUE | -0.381022342 | 0.079418761 | No Change | Not Significant | TRUE | Cond1 Not Significant + Cond2 Not Significant |
| Sox6os | TRUE | 0.292023816 | 1 | No Change | Not Significant | FALSE | NA | NA | No Change | Not Detected | FALSE | Background = FALSE |
| Sox7 | TRUE | -2.060849111 | 1 | No Change | Not Significant | FALSE | NA | NA | No Change | Not Detected | FALSE | Background = FALSE |
| Sox8 | TRUE | -0.17431704 | 1 | No Change | Not Significant | FALSE | NA | NA | No Change | Not Detected | FALSE | Background = FALSE |
| Sox9 | TRUE | 0.060118921 | 0.760117839 | No Change | Not Significant | TRUE | -0.029944657 | 0.893742272 | No Change | Not Significant | TRUE | Cond1 Not Significant + Cond2 Not Significant |
| Sp1 | TRUE | 0.004346101 | 0.978107631 | No Change | Not Significant | TRUE | -0.065079014 | 0.44497561 | No Change | Not Significant | TRUE | Cond1 Not Significant + Cond2 Not Significant |
| Sp100 | TRUE | -0.748641141 | 0.107730217 | No Change | Not Significant | TRUE | -0.218908231 | 0.619709567 | No Change | Not Significant | TRUE | Cond1 Not Significant + Cond2 Not Significant |
| Sp110 | TRUE | -0.318664831 | 0.817346435 | No Change | Not Significant | TRUE | -0.142803512 | 0.805987676 | No Change | Not Significant | TRUE | Cond1 Not Significant + Cond2 Not Significant |
| Sp140 | TRUE | -1.55503074 | 0.001489031 | DOWN | DOWN | TRUE | -0.532140381 | 0.306958481 | No Change | Not Significant | TRUE | Cond1 DOWN + Cond2 Not Significant |
| Sp2 | TRUE | 0.261572026 | 0.443634979 | No Change | Not Significant | TRUE | 0.137577236 | 0.529021743 | No Change | Not Significant | TRUE | Cond1 Not Significant + Cond2 Not Significant |
| Sp3 | TRUE | -0.064858456 | 0.540878801 | No Change | Not Significant | TRUE | 0.007326108 | 0.935610703 | No Change | Not Significant | TRUE | Cond1 Not Significant + Cond2 Not Significant |
| Sp3os | TRUE | 0.151040136 | 0.685174282 | No Change | Not Significant | TRUE | -0.139794521 | 0.661841071 | No Change | Not Significant | TRUE | Cond1 Not Significant + Cond2 Not Significant |
| Sp4 | TRUE | -0.366818087 | 0.008323805 | No Change | Significant Negative | TRUE | -0.158296535 | 0.323423905 | No Change | Not Significant | TRUE | Cond1 Significant Negative + Cond2 Not Significant |
| Sp5 | TRUE | 2.102335219 | 1.45E-05 | UP | UP | TRUE | 1.013249493 | 0.036330185 | UP | UP | TRUE | Cond1 UP + Cond2 UP |
| Sp7 | TRUE | 1.90180325 | 1 | No Change | Not Significant | FALSE | NA | NA | No Change | Not Detected | FALSE | Background = FALSE |
| Sp9 | TRUE | 1.221029954 | 0.113250188 | No Change | Not Significant | FALSE | NA | NA | No Change | Not Detected | FALSE | Background = FALSE |
| Spa17 | TRUE | 0.12415305 | 0.837885199 | No Change | Not Significant | TRUE | 0.551559462 | 0.187205038 | No Change | Not Significant | TRUE | Cond1 Not Significant + Cond2 Not Significant |
| Spaar | TRUE | 0.846003123 | 1 | No Change | Not Significant | FALSE | NA | NA | No Change | Not Detected | FALSE | Background = FALSE |
| Spaca1 | TRUE | 1.979825483 | 1 | No Change | Not Significant | FALSE | NA | NA | No Change | Not Detected | FALSE | Background = FALSE |
| Spaca3 | TRUE | 3.277256133 | 1 | No Change | Not Significant | FALSE | NA | NA | No Change | Not Detected | FALSE | Background = FALSE |
| Spaca4 | TRUE | -0.291677068 | 0.310491768 | No Change | Not Significant | TRUE | -0.563976344 | 0.001930175 | DOWN | DOWN | TRUE | Cond1 Not Significant + Cond2 DOWN |
| Spaca6 | TRUE | -0.130750905 | 0.887843124 | No Change | Not Significant | TRUE | 0.199981833 | 0.828333496 | No Change | Not Significant | TRUE | Cond1 Not Significant + Cond2 Not Significant |
| Spaca7 | TRUE | 0.69474401 | 1 | No Change | Not Significant | FALSE | NA | NA | No Change | Not Detected | FALSE | Background = FALSE |
| Spaca7b | TRUE | -1.885829728 | 1 | No Change | Not Significant | FALSE | NA | NA | No Change | Not Detected | FALSE | Background = FALSE |
| Spaca9 | TRUE | 0.023038858 | 1 | No Change | Not Significant | FALSE | NA | NA | No Change | Not Detected | FALSE | Background = FALSE |
| Spacdr | TRUE | -0.195376404 | 0.180584617 | No Change | Not Significant | TRUE | -0.003566595 | 0.989317668 | No Change | Not Significant | TRUE | Cond1 Not Significant + Cond2 Not Significant |
| Spag1 | TRUE | 0.475652166 | 0.170406218 | No Change | Not Significant | TRUE | -0.104148081 | 0.835024575 | No Change | Not Significant | TRUE | Cond1 Not Significant + Cond2 Not Significant |
| Spag16 | TRUE | 2.323477309 | 1 | No Change | Not Significant | FALSE | NA | NA | No Change | Not Detected | FALSE | Background = FALSE |
| Spag17 | TRUE | 1.828327058 | 2.71E-07 | UP | UP | TRUE | 0.486842434 | 0.334110714 | No Change | Not Significant | TRUE | Cond1 UP + Cond2 Not Significant |
| Spag4 | TRUE | 0.740488015 | 0.494519255 | No Change | Not Significant | FALSE | NA | NA | No Change | Not Detected | FALSE | Background = FALSE |
| Spag5 | TRUE | 0.69667876 | 5.06E-06 | UP | UP | TRUE | 0.038544153 | 0.856292985 | No Change | Not Significant | TRUE | Cond1 UP + Cond2 Not Significant |
| Spag6 | TRUE | -0.072655037 | 1 | No Change | Not Significant | FALSE | NA | NA | No Change | Not Detected | FALSE | Background = FALSE |
| Spag6l | TRUE | -0.442895326 | 1 | No Change | Not Significant | FALSE | NA | NA | No Change | Not Detected | FALSE | Background = FALSE |
| Spag7 | TRUE | 0.02044203 | 0.91001573 | No Change | Not Significant | TRUE | 0.101261159 | 0.20232926 | No Change | Not Significant | TRUE | Cond1 Not Significant + Cond2 Not Significant |
| Spag8 | TRUE | -0.545366518 | 1 | No Change | Not Significant | FALSE | NA | NA | No Change | Not Detected | FALSE | Background = FALSE |
| Spag9 | TRUE | -0.08732851 | 0.56607413 | No Change | Not Significant | TRUE | -0.125136874 | 0.03448615 | No Change | Significant Negative | TRUE | Cond1 Not Significant + Cond2 Significant Negative |
| Sparc | TRUE | 0.425387635 | 0.727166502 | No Change | Not Significant | TRUE | 2.149235893 | 8.18E-05 | UP | UP | TRUE | Cond1 Not Significant + Cond2 UP |
| Sparcl1 | TRUE | 0.732545491 | 1 | No Change | Not Significant | FALSE | NA | NA | No Change | Not Detected | FALSE | Background = FALSE |
| Spast | TRUE | -0.003591132 | 0.982740683 | No Change | Not Significant | TRUE | -0.111217239 | 0.082474622 | No Change | Not Significant | TRUE | Cond1 Not Significant + Cond2 Not Significant |
| Spata1 | TRUE | -0.403372461 | 0.376665775 | No Change | Not Significant | TRUE | -0.207725307 | 0.584131301 | No Change | Not Significant | TRUE | Cond1 Not Significant + Cond2 Not Significant |
| Spata13 | TRUE | -0.35418316 | 0.005188088 | No Change | Significant Negative | TRUE | -0.281981482 | 1.06E-07 | No Change | Significant Negative | TRUE | Cond1 Significant Negative + Cond2 Significant Negative |
| Spata17 | TRUE | -0.303048782 | 0.616464692 | No Change | Not Significant | TRUE | -0.085861224 | 0.886610448 | No Change | Not Significant | TRUE | Cond1 Not Significant + Cond2 Not Significant |
| Spata18 | TRUE | -1.704813955 | 1 | No Change | Not Significant | FALSE | NA | NA | No Change | Not Detected | FALSE | Background = FALSE |
| Spata2 | TRUE | -0.258643938 | 0.000261409 | No Change | Significant Negative | TRUE | -0.118040225 | 0.225910228 | No Change | Not Significant | TRUE | Cond1 Significant Negative + Cond2 Not Significant |
| Spata20 | TRUE | 0.125827252 | 1 | No Change | Not Significant | FALSE | NA | NA | No Change | Not Detected | FALSE | Background = FALSE |
| Spata21 | TRUE | 0.737819138 | 0.519448318 | No Change | Not Significant | TRUE | -0.373321596 | 0.693047027 | No Change | Not Significant | TRUE | Cond1 Not Significant + Cond2 Not Significant |
| Spata22 | TRUE | -1.284935711 | 1 | No Change | Not Significant | FALSE | NA | NA | No Change | Not Detected | FALSE | Background = FALSE |
| Spata24 | TRUE | 1.454309395 | 0.000475973 | UP | UP | TRUE | 0.51186825 | 0.248174588 | No Change | Not Significant | TRUE | Cond1 UP + Cond2 Not Significant |
| Spata2l | TRUE | -0.557777911 | 0.000267672 | DOWN | DOWN | TRUE | -0.261711355 | 0.030298572 | No Change | Significant Negative | TRUE | Cond1 DOWN + Cond2 Significant Negative |
| Spata3 | TRUE | -0.694495665 | 1 | No Change | Not Significant | FALSE | NA | NA | No Change | Not Detected | FALSE | Background = FALSE |
| Spata31d1a | TRUE | 2.150362598 | 1 | No Change | Not Significant | FALSE | NA | NA | No Change | Not Detected | FALSE | Background = FALSE |
| Spata31d1b | TRUE | -0.363032201 | 1 | No Change | Not Significant | FALSE | NA | NA | No Change | Not Detected | FALSE | Background = FALSE |
| Spata31d1d | TRUE | 1.566407838 | 1 | No Change | Not Significant | FALSE | NA | NA | No Change | Not Detected | FALSE | Background = FALSE |
| Spata33 | TRUE | -0.36587796 | 0.720791305 | No Change | Not Significant | TRUE | 0.18571928 | 0.805989706 | No Change | Not Significant | TRUE | Cond1 Not Significant + Cond2 Not Significant |
| Spata46 | TRUE | -0.875483492 | 0.322533618 | No Change | Not Significant | FALSE | NA | NA | No Change | Not Detected | FALSE | Background = FALSE |
| Spata5 | TRUE | -0.207697153 | 0.265751013 | No Change | Not Significant | TRUE | -0.103799143 | 0.482991121 | No Change | Not Significant | TRUE | Cond1 Not Significant + Cond2 Not Significant |
| Spata6 | TRUE | -0.233148978 | 0.441303539 | No Change | Not Significant | TRUE | 0.118813303 | 0.676953924 | No Change | Not Significant | TRUE | Cond1 Not Significant + Cond2 Not Significant |
| Spata6l | TRUE | 1.129651021 | 0.001642193 | UP | UP | TRUE | 0.266317468 | 0.415618234 | No Change | Not Significant | TRUE | Cond1 UP + Cond2 Not Significant |
| Spata7 | TRUE | -0.232383858 | 0.255121545 | No Change | Not Significant | TRUE | -0.086546231 | 0.744805917 | No Change | Not Significant | TRUE | Cond1 Not Significant + Cond2 Not Significant |
| Spata9 | TRUE | 0.022180583 | 1 | No Change | Not Significant | TRUE | 0.186567614 | 0.847142841 | No Change | Not Significant | TRUE | Cond1 Not Significant + Cond2 Not Significant |
| Spatc1 | TRUE | 1.178829447 | 1 | No Change | Not Significant | FALSE | NA | NA | No Change | Not Detected | FALSE | Background = FALSE |
| Spatc1l | TRUE | 0.961753691 | 1 | No Change | Not Significant | FALSE | NA | NA | No Change | Not Detected | FALSE | Background = FALSE |
| Spats1 | TRUE | 0.44279073 | 0.482454733 | No Change | Not Significant | FALSE | NA | NA | No Change | Not Detected | FALSE | Background = FALSE |
| Spats2 | TRUE | 0.104433152 | 0.559203284 | No Change | Not Significant | TRUE | 0.005708931 | 0.975397839 | No Change | Not Significant | TRUE | Cond1 Not Significant + Cond2 Not Significant |
| Spats2l | TRUE | -0.156145043 | 0.695300529 | No Change | Not Significant | TRUE | 0.05348531 | 0.888663035 | No Change | Not Significant | TRUE | Cond1 Not Significant + Cond2 Not Significant |
| Spc24 | TRUE | 0.715621535 | 8.17E-05 | UP | UP | TRUE | 0.112174329 | 0.572083958 | No Change | Not Significant | TRUE | Cond1 UP + Cond2 Not Significant |
| Spc25 | TRUE | 0.825928893 | 0.000108339 | UP | UP | TRUE | 0.277769858 | 0.079396931 | No Change | Not Significant | TRUE | Cond1 UP + Cond2 Not Significant |
| Spcs1 | TRUE | 0.338678296 | 0.00461816 | No Change | Significant Positive | TRUE | 0.206725292 | 0.006954202 | No Change | Significant Positive | TRUE | Cond1 Significant Positive + Cond2 Significant Positive |
| Spcs2 | TRUE | 0.887327469 | 1.07E-14 | UP | UP | TRUE | 0.678327726 | 7.04E-33 | UP | UP | TRUE | Cond1 UP + Cond2 UP |
| Spcs3 | TRUE | 1.38179639 | 4.50E-28 | UP | UP | TRUE | 1.03870666 | 2.41E-80 | UP | UP | TRUE | Cond1 UP + Cond2 UP |
| Spdef | TRUE | 0.290029529 | 0.23299599 | No Change | Not Significant | TRUE | 0.111757748 | 0.478917898 | No Change | Not Significant | TRUE | Cond1 Not Significant + Cond2 Not Significant |
| Spdl1 | TRUE | 0.768058449 | 0.000434207 | UP | UP | TRUE | 0.287514313 | 0.229201544 | No Change | Not Significant | TRUE | Cond1 UP + Cond2 Not Significant |
| Spdya | TRUE | 0.00586697 | 1 | No Change | Not Significant | FALSE | NA | NA | No Change | Not Detected | FALSE | Background = FALSE |
| Spdye4b | TRUE | 0.607430715 | 1 | No Change | Not Significant | FALSE | NA | NA | No Change | Not Detected | FALSE | Background = FALSE |
| Specc1 | TRUE | -0.286519446 | 0.017356745 | No Change | Significant Negative | TRUE | -0.0592308 | 0.527473335 | No Change | Not Significant | TRUE | Cond1 Significant Negative + Cond2 Not Significant |
| Specc1l | TRUE | -0.027944287 | 0.918959391 | No Change | Not Significant | TRUE | -0.110022932 | 0.288630668 | No Change | Not Significant | TRUE | Cond1 Not Significant + Cond2 Not Significant |
| Speer3 | TRUE | -1.249233464 | 1 | No Change | Not Significant | FALSE | NA | NA | No Change | Not Detected | FALSE | Background = FALSE |
| Speer4a | TRUE | 1.411869924 | 1 | No Change | Not Significant | FALSE | NA | NA | No Change | Not Detected | FALSE | Background = FALSE |
| Speer4cos | TRUE | 1.110965784 | 1 | No Change | Not Significant | FALSE | NA | NA | No Change | Not Detected | FALSE | Background = FALSE |
| Speer4f2 | TRUE | 0.043036239 | 1 | No Change | Not Significant | FALSE | NA | NA | No Change | Not Detected | FALSE | Background = FALSE |
| Speer5-ps1 | TRUE | 2.477627628 | 1 | No Change | Not Significant | FALSE | NA | NA | No Change | Not Detected | FALSE | Background = FALSE |
| Spef1 | TRUE | -0.135802938 | 0.633934205 | No Change | Not Significant | TRUE | 0.111316265 | 0.692404394 | No Change | Not Significant | TRUE | Cond1 Not Significant + Cond2 Not Significant |
| Spef2 | TRUE | -0.707580343 | 1 | No Change | Not Significant | FALSE | NA | NA | No Change | Not Detected | FALSE | Background = FALSE |
| Speg | TRUE | -0.251661373 | 0.933837244 | No Change | Not Significant | FALSE | NA | NA | No Change | Not Detected | FALSE | Background = FALSE |
| Spem2 | TRUE | 0.968345084 | 1 | No Change | Not Significant | FALSE | NA | NA | No Change | Not Detected | FALSE | Background = FALSE |
| Spem3 | TRUE | -0.176233663 | 1 | No Change | Not Significant | FALSE | NA | NA | No Change | Not Detected | FALSE | Background = FALSE |
| Spen | TRUE | 0.146443952 | 0.642978943 | No Change | Not Significant | TRUE | -0.115105035 | 0.4893628 | No Change | Not Significant | TRUE | Cond1 Not Significant + Cond2 Not Significant |
| Spg11 | TRUE | -0.09733751 | 0.167999957 | No Change | Not Significant | TRUE | -0.034689225 | 0.606076959 | No Change | Not Significant | TRUE | Cond1 Not Significant + Cond2 Not Significant |
| Spg20 | TRUE | 0.358771191 | 0.015401876 | No Change | Significant Positive | TRUE | 0.104581662 | 0.408949491 | No Change | Not Significant | TRUE | Cond1 Significant Positive + Cond2 Not Significant |
| Spg21 | TRUE | -0.051953249 | 0.751050176 | No Change | Not Significant | TRUE | 0.06055846 | 0.425472739 | No Change | Not Significant | TRUE | Cond1 Not Significant + Cond2 Not Significant |
| Spg7 | TRUE | -0.367742173 | 8.95E-06 | No Change | Significant Negative | TRUE | -0.166627963 | 0.00035792 | No Change | Significant Negative | TRUE | Cond1 Significant Negative + Cond2 Significant Negative |
| Sphk1 | TRUE | -0.410966841 | 1 | No Change | Not Significant | FALSE | NA | NA | No Change | Not Detected | FALSE | Background = FALSE |
| Sphk2 | TRUE | -0.093965992 | 0.560935494 | No Change | Not Significant | TRUE | 0.044782755 | 0.755247648 | No Change | Not Significant | TRUE | Cond1 Not Significant + Cond2 Not Significant |
| Sphkap | TRUE | -0.348241529 | 1 | No Change | Not Significant | FALSE | NA | NA | No Change | Not Detected | FALSE | Background = FALSE |
| Spi1 | TRUE | 1.037104012 | 1 | No Change | Not Significant | FALSE | NA | NA | No Change | Not Detected | FALSE | Background = FALSE |
| Spib | TRUE | -0.711271608 | 0.000377711 | DOWN | DOWN | TRUE | 0.172798042 | 0.35758991 | No Change | Not Significant | TRUE | Cond1 DOWN + Cond2 Not Significant |
| Spice1 | TRUE | -0.181772024 | 0.15592012 | No Change | Not Significant | TRUE | 0.076976745 | 0.58240762 | No Change | Not Significant | TRUE | Cond1 Not Significant + Cond2 Not Significant |
| Spidr | TRUE | 0.079129305 | 0.667366241 | No Change | Not Significant | TRUE | 0.000211538 | 0.999077457 | No Change | Not Significant | TRUE | Cond1 Not Significant + Cond2 Not Significant |
| Spin1 | TRUE | -0.087139987 | 0.461770971 | No Change | Not Significant | TRUE | -0.039214282 | 0.666012107 | No Change | Not Significant | TRUE | Cond1 Not Significant + Cond2 Not Significant |
| Spin2c | TRUE | 1.931255725 | 3.95E-09 | UP | UP | TRUE | 1.220965099 | 0.002281792 | UP | UP | TRUE | Cond1 UP + Cond2 UP |
| Spin4 | TRUE | 0.07723982 | 0.890125936 | No Change | Not Significant | TRUE | 0.20486304 | 0.654811627 | No Change | Not Significant | TRUE | Cond1 Not Significant + Cond2 Not Significant |
| Spindoc | TRUE | 0.988863189 | 1.36E-16 | UP | UP | TRUE | 0.387356958 | 2.61E-06 | No Change | Significant Positive | TRUE | Cond1 UP + Cond2 Significant Positive |
| Spink1 | TRUE | -0.196978643 | 0.773528957 | No Change | Not Significant | TRUE | 0.22221153 | 0.185896496 | No Change | Not Significant | TRUE | Cond1 Not Significant + Cond2 Not Significant |
| Spink10 | TRUE | 0.995247095 | 1 | No Change | Not Significant | FALSE | NA | NA | No Change | Not Detected | FALSE | Background = FALSE |
| Spink2 | TRUE | -0.825759144 | 1 | No Change | Not Significant | FALSE | NA | NA | No Change | Not Detected | FALSE | Background = FALSE |
| Spink4 | TRUE | 0.538533378 | 0.136998766 | No Change | Not Significant | TRUE | 0.885962386 | 1.50E-23 | UP | UP | TRUE | Cond1 Not Significant + Cond2 UP |
| Spink5 | TRUE | 0.901246433 | 1 | No Change | Not Significant | TRUE | 0.342869836 | 0.749029274 | No Change | Not Significant | TRUE | Cond1 Not Significant + Cond2 Not Significant |
| Spint1 | TRUE | -0.366739034 | 0.000281155 | No Change | Significant Negative | TRUE | -0.139203787 | 0.073314226 | No Change | Not Significant | TRUE | Cond1 Significant Negative + Cond2 Not Significant |
| Spint2 | TRUE | -0.300778371 | 0.001074484 | No Change | Significant Negative | TRUE | -0.080189531 | 0.523508209 | No Change | Not Significant | TRUE | Cond1 Significant Negative + Cond2 Not Significant |
| Spire1 | TRUE | -0.032360539 | 0.941973978 | No Change | Not Significant | TRUE | 0.12197994 | 0.593968278 | No Change | Not Significant | TRUE | Cond1 Not Significant + Cond2 Not Significant |
| Spire2 | TRUE | 9.66E-05 | 0.999457543 | No Change | Not Significant | TRUE | 0.106928943 | 0.230847912 | No Change | Not Significant | TRUE | Cond1 Not Significant + Cond2 Not Significant |
| Spn | TRUE | -0.220292071 | 0.764339772 | No Change | Not Significant | TRUE | 0.224365608 | 0.679535938 | No Change | Not Significant | TRUE | Cond1 Not Significant + Cond2 Not Significant |
| Spn-ps | TRUE | 0.176010528 | 1 | No Change | Not Significant | FALSE | NA | NA | No Change | Not Detected | FALSE | Background = FALSE |
| Spns1 | TRUE | 0.059595994 | 0.695624496 | No Change | Not Significant | TRUE | 0.050431023 | 0.760578285 | No Change | Not Significant | TRUE | Cond1 Not Significant + Cond2 Not Significant |
| Spns2 | TRUE | -0.610026426 | 0.048151079 | DOWN | DOWN | TRUE | -0.198186801 | 0.563525822 | No Change | Not Significant | TRUE | Cond1 DOWN + Cond2 Not Significant |
| Spns3 | TRUE | -0.566211581 | 1 | No Change | Not Significant | FALSE | NA | NA | No Change | Not Detected | FALSE | Background = FALSE |
| Spock1 | TRUE | -0.438238364 | 0.776186935 | No Change | Not Significant | FALSE | NA | NA | No Change | Not Detected | FALSE | Background = FALSE |
| Spock2 | TRUE | -0.10548779 | 0.908700419 | No Change | Not Significant | TRUE | -0.071890262 | 0.914739662 | No Change | Not Significant | TRUE | Cond1 Not Significant + Cond2 Not Significant |
| Spon1 | TRUE | 0.27578355 | 0.708014489 | No Change | Not Significant | TRUE | -0.131695439 | 0.84565495 | No Change | Not Significant | TRUE | Cond1 Not Significant + Cond2 Not Significant |
| Spon2 | TRUE | -0.053651654 | 0.915629096 | No Change | Not Significant | TRUE | 0.357983897 | 0.178786609 | No Change | Not Significant | TRUE | Cond1 Not Significant + Cond2 Not Significant |
| Spop | TRUE | -0.283830657 | 0.025001604 | No Change | Significant Negative | TRUE | 0.053759532 | 0.532196551 | No Change | Not Significant | TRUE | Cond1 Significant Negative + Cond2 Not Significant |
| Spopl | TRUE | -0.264894251 | 0.01801348 | No Change | Significant Negative | TRUE | -0.251568738 | 0.002308089 | No Change | Significant Negative | TRUE | Cond1 Significant Negative + Cond2 Significant Negative |
| Spout1 | TRUE | 0.173887909 | 0.230377922 | No Change | Not Significant | TRUE | 0.038267041 | 0.768186995 | No Change | Not Significant | TRUE | Cond1 Not Significant + Cond2 Not Significant |
| Spp1 | TRUE | 0.965265726 | 1 | No Change | Not Significant | FALSE | NA | NA | No Change | Not Detected | FALSE | Background = FALSE |
| Sppl2a | TRUE | -0.067880437 | 0.736796542 | No Change | Not Significant | TRUE | 0.05399622 | 0.627358857 | No Change | Not Significant | TRUE | Cond1 Not Significant + Cond2 Not Significant |
| Sppl2b | TRUE | 0.030604855 | 0.802408631 | No Change | Not Significant | TRUE | -0.028342467 | 0.849281085 | No Change | Not Significant | TRUE | Cond1 Not Significant + Cond2 Not Significant |
| Sppl3 | TRUE | -0.024353007 | 0.824971081 | No Change | Not Significant | TRUE | -0.049938767 | 0.624972092 | No Change | Not Significant | TRUE | Cond1 Not Significant + Cond2 Not Significant |
| Spr | TRUE | -0.051781055 | 0.717711842 | No Change | Not Significant | TRUE | 0.129662849 | 0.194579678 | No Change | Not Significant | TRUE | Cond1 Not Significant + Cond2 Not Significant |
| Spred1 | TRUE | 0.182070272 | 0.332817843 | No Change | Not Significant | TRUE | -0.002187157 | 0.987804745 | No Change | Not Significant | TRUE | Cond1 Not Significant + Cond2 Not Significant |
| Spred2 | TRUE | -0.005354374 | 0.981434574 | No Change | Not Significant | TRUE | 0.054797079 | 0.774890517 | No Change | Not Significant | TRUE | Cond1 Not Significant + Cond2 Not Significant |
| Spred3 | TRUE | 0.379994896 | 0.712606767 | No Change | Not Significant | TRUE | -0.008526121 | 0.995445753 | No Change | Not Significant | TRUE | Cond1 Not Significant + Cond2 Not Significant |
| Spring1 | TRUE | 0.383093724 | 0.004563174 | No Change | Significant Positive | TRUE | 0.18574915 | 0.07724278 | No Change | Not Significant | TRUE | Cond1 Significant Positive + Cond2 Not Significant |
| Sprn | TRUE | -0.830262047 | 1 | No Change | Not Significant | FALSE | NA | NA | No Change | Not Detected | FALSE | Background = FALSE |
| Sprr1a | TRUE | 1.547902425 | 0.011942524 | UP | UP | TRUE | 1.101001936 | 6.43E-05 | UP | UP | TRUE | Cond1 UP + Cond2 UP |
| Sprr2a1 | TRUE | 0.37693391 | 0.803370142 | No Change | Not Significant | TRUE | -0.277004941 | 0.852303638 | No Change | Not Significant | TRUE | Cond1 Not Significant + Cond2 Not Significant |
| Sprr2a2 | TRUE | -0.345943638 | 0.620277218 | No Change | Not Significant | FALSE | NA | NA | No Change | Not Detected | FALSE | Background = FALSE |
| Sprr2a3 | TRUE | -0.071866267 | 0.893615254 | No Change | Not Significant | TRUE | 0.679163961 | 0.010812233 | UP | UP | TRUE | Cond1 Not Significant + Cond2 UP |
| Sprr2b | TRUE | 0.592196747 | 1 | No Change | Not Significant | FALSE | NA | NA | No Change | Not Detected | FALSE | Background = FALSE |
| Sprr2h | TRUE | 2.93974439 | 1 | No Change | Not Significant | FALSE | NA | NA | No Change | Not Detected | FALSE | Background = FALSE |
| Sprtn | TRUE | -0.190867477 | 0.114480981 | No Change | Not Significant | TRUE | -0.14571561 | 0.142368697 | No Change | Not Significant | TRUE | Cond1 Not Significant + Cond2 Not Significant |
| Spry1 | TRUE | -0.128338896 | 0.509010748 | No Change | Not Significant | TRUE | -0.199648229 | 0.200957912 | No Change | Not Significant | TRUE | Cond1 Not Significant + Cond2 Not Significant |
| Spry2 | TRUE | -0.227944023 | 0.208221097 | No Change | Not Significant | TRUE | -0.07801814 | 0.4893628 | No Change | Not Significant | TRUE | Cond1 Not Significant + Cond2 Not Significant |
| Spry4 | TRUE | -0.062857895 | 0.905678041 | No Change | Not Significant | TRUE | -0.283714753 | 0.423163427 | No Change | Not Significant | TRUE | Cond1 Not Significant + Cond2 Not Significant |
| Spryd3 | TRUE | 0.520896648 | 2.53E-14 | UP | UP | TRUE | 0.404991128 | 1.54E-08 | No Change | Significant Positive | TRUE | Cond1 UP + Cond2 Significant Positive |
| Spryd4 | TRUE | -0.121172183 | 0.647057325 | No Change | Not Significant | TRUE | 0.024167038 | 0.898825528 | No Change | Not Significant | TRUE | Cond1 Not Significant + Cond2 Not Significant |
| Spryd7 | TRUE | -0.165036259 | 0.226286043 | No Change | Not Significant | TRUE | -0.06214845 | 0.695023769 | No Change | Not Significant | TRUE | Cond1 Not Significant + Cond2 Not Significant |
| Spsb1 | TRUE | -1.343072901 | 1.23E-10 | DOWN | DOWN | TRUE | 0.041606737 | 0.925277452 | No Change | Not Significant | TRUE | Cond1 DOWN + Cond2 Not Significant |
| Spsb2 | TRUE | -0.172239348 | 0.32409141 | No Change | Not Significant | TRUE | -0.032856701 | 0.834153989 | No Change | Not Significant | TRUE | Cond1 Not Significant + Cond2 Not Significant |
| Spsb3 | TRUE | 0.112294181 | 0.557631454 | No Change | Not Significant | TRUE | -0.047206389 | 0.858612846 | No Change | Not Significant | TRUE | Cond1 Not Significant + Cond2 Not Significant |
| Spsb4 | TRUE | -0.744951603 | 0.022441881 | DOWN | DOWN | TRUE | -0.208684416 | 0.484894029 | No Change | Not Significant | TRUE | Cond1 DOWN + Cond2 Not Significant |
| Spta1 | TRUE | -1.357739883 | 0.138995748 | No Change | Not Significant | TRUE | 0.707733167 | 0.225459333 | No Change | Not Significant | TRUE | Cond1 Not Significant + Cond2 Not Significant |
| Sptan1 | TRUE | -0.092311379 | 0.190422043 | No Change | Not Significant | TRUE | 0.000600789 | 0.995768162 | No Change | Not Significant | TRUE | Cond1 Not Significant + Cond2 Not Significant |
| Sptb | TRUE | -0.208103374 | 0.702910017 | No Change | Not Significant | TRUE | -0.252163949 | 0.254690744 | No Change | Not Significant | TRUE | Cond1 Not Significant + Cond2 Not Significant |

Supplementary Table 3

| geneID | Cond1_DF_Detected | Cond1_DF_ValueCol | Cond1_DF_PadjCol | Cond1_DF_Cutoff | Cond1_DF_Cutoff_Specific.x |
|---|---|---|---|---|---|
| Sptbn1 | TRUE | -0.174854227 | 0.020064348 | No Change | Significant Negative |
| Sptbn2 | TRUE | -0.42676438 | 0.039545746 | No Change | Significant Negative |
| Sptbn4 | TRUE | 2.443241539 | 6.50E-19 | UP | UP |
| Sptbn5 | TRUE | 1.644642133 | 0.00156603 | UP | UP |
| Sptlc1 | TRUE | 0.126697493 | 0.289400593 | No Change | Not Significant |
| Sptlc2 | TRUE | 0.423421325 | 1.05E-13 | No Change | Significant Positive |
| Sptlc3 | TRUE | 0.609300338 | 1 | No Change | Not Significant |
| Sptssa | TRUE | 0.722456142 | 4.73E-17 | UP | UP |
| Sptssb | TRUE | -0.522907799 | 0.047353826 | DOWN | DOWN |
| Spty2d1 | TRUE | 0.179886373 | 0.075334803 | No Change | Not Significant |
| Spx | TRUE | 0.166723789 | 1 | No Change | Not Significant |
| Sqle | TRUE | 0.172412637 | 0.462725114 | No Change | Not Significant |
| Sqor | TRUE | -0.490602079 | 5.14E-08 | No Change | Significant Negative |
| Sqstm1 | TRUE | 0.039642848 | 0.871840133 | No Change | Not Significant |
| Sra1 | TRUE | 0.000208183 | 0.998483602 | No Change | Not Significant |
| Srbd1 | TRUE | 0.260098263 | 0.06023867 | No Change | Not Significant |
| Src | TRUE | 0.001730339 | 0.99157486 | No Change | Not Significant |
| Srcap | TRUE | 0.292921331 | 0.300075862 | No Change | Not Significant |
| Srcin1 | TRUE | 1.783261259 | 0.084025676 | No Change | Not Significant |
| Srd5a1 | TRUE | 0.080097728 | 0.776035105 | No Change | Not Significant |
| Srd5a2 | TRUE | -0.423570324 | 0.419775423 | No Change | Not Significant |
| Srd5a3 | TRUE | -0.018178256 | 0.931722347 | No Change | Not Significant |
| Srebf1 | TRUE | 0.345332054 | 0.005694853 | No Change | Significant Positive |
| Srebf2 | TRUE | -0.1106203 | 0.162548412 | No Change | Not Significant |
| Srek1 | TRUE | 0.0689099 | 0.647057325 | No Change | Not Significant |
| Srek1ip1 | TRUE | -0.154500631 | 0.25251889 | No Change | Not Significant |
| Srf | TRUE | 0.343718029 | 0.013458042 | No Change | Significant Positive |
| Srfbp1 | TRUE | -0.070781633 | 0.69793377 | No Change | Not Significant |
| Srgap1 | TRUE | 0.038893656 | 0.890030875 | No Change | Not Significant |
| Srgap2 | TRUE | 0.633715224 | 6.28E-08 | UP | UP |
| Srgap3 | TRUE | -0.507682509 | 0.027715628 | DOWN | DOWN |
| Srgn | TRUE | 0.069264202 | 0.949716026 | No Change | Not Significant |
| Sri | TRUE | -0.217829092 | 0.053374514 | No Change | Not Significant |
| Srl | TRUE | 0.470860619 | 0.00949063 | No Change | Significant Positive |
| Srm | TRUE | 0.998393584 | 1.46E-09 | UP | UP |
| Srms | TRUE | -0.111063911 | 0.911456552 | No Change | Not Significant |
| Srp14 | TRUE | 0.078226981 | 0.643957486 | No Change | Not Significant |
| Srp19 | TRUE | 0.290424732 | 0.008943393 | No Change | Significant Positive |
| Srp54a | TRUE | -0.013947643 | 0.973795719 | No Change | Not Significant |
| Srp54b | TRUE | -0.099995834 | 1 | No Change | Not Significant |
| Srp54c | TRUE | 0.134867704 | 0.60719955 | No Change | Not Significant |
| Srp68 | TRUE | 0.142314847 | 0.114836207 | No Change | Not Significant |
| Srp72 | TRUE | 0.025863753 | 0.901569434 | No Change | Not Significant |
| Srp9 | TRUE | 0.491421245 | 3.74E-08 | No Change | Significant Positive |
| Srpk1 | TRUE | 0.000922545 | 0.994183621 | No Change | Not Significant |
| Srpk2 | TRUE | 0.005893837 | 0.976016991 | No Change | Not Significant |
| Srpk3 | TRUE | 0.220679545 | 0.794172689 | No Change | Not Significant |
| Srpr | TRUE | 0.139427072 | 0.29575977 | No Change | Not Significant |
| Srprb | TRUE | 0.33524281 | 0.00890944 | No Change | Significant Positive |
| Srpx2 | TRUE | -1.589164766 | 0.00221215 | DOWN | DOWN |
| Srr | TRUE | -0.053934066 | 0.740730056 | No Change | Not Significant |
| Srrd | TRUE | 0.78413154 | 5.73E-08 | UP | UP |
| Srrm1 | TRUE | 0.081720786 | 0.55512192 | No Change | Not Significant |
| Srrm2 | TRUE | 0.071088608 | 0.819511083 | No Change | Not Significant |
| Srrm3 | TRUE | 0.860489351 | 0.632329199 | No Change | Not Significant |
| Srrm4 | TRUE | -0.76527639 | 1 | No Change | Not Significant |
| Srrm4os | TRUE | 3.422263582 | 1 | No Change | Not Significant |
| Srrt | TRUE | 0.469244548 | 3.00E-11 | No Change | Significant Positive |
| Srsf1 | TRUE | 0.368957906 | 0.004647131 | No Change | Significant Positive |
| Srsf10 | TRUE | 0.133427899 | 0.515610058 | No Change | Not Significant |
| Srsf11 | TRUE | 0.182698537 | 0.020927689 | No Change | Significant Positive |
| Srsf12 | TRUE | 0.49745976 | 1 | No Change | Not Significant |
| Srsf2 | TRUE | 0.365132936 | 5.09E-08 | No Change | Significant Positive |
| Srsf3 | TRUE | -0.001799904 | 0.989337077 | No Change | Not Significant |
| Srsf4 | TRUE | -0.097487416 | 0.52811293 | No Change | Not Significant |
| Srsf5 | TRUE | -0.119557357 | 0.319067066 | No Change | Not Significant |
| Srsf6 | TRUE | 0.060420672 | 0.595257447 | No Change | Not Significant |
| Srsf7 | TRUE | 0.395706161 | 1.39E-05 | No Change | Significant Positive |
| Srsf9 | TRUE | 0.145772993 | 0.255086991 | No Change | Not Significant |
| Srxn1 | TRUE | -0.139009811 | 0.475244062 | No Change | Not Significant |
| Ss18 | TRUE | 0.226801209 | 0.037634187 | No Change | Significant Positive |
| Ss18l1 | TRUE | -0.057199303 | 0.77680602 | No Change | Not Significant |
| Ss18l2 | TRUE | 0.388395202 | 0.003550608 | No Change | Significant Positive |
| Ssb | TRUE | 0.248247857 | 0.040294527 | No Change | Significant Positive |
| Ssbp1 | TRUE | 0.193253449 | 0.22143558 | No Change | Not Significant |
| Ssbp2 | TRUE | -0.750447022 | 0.037851103 | DOWN | DOWN |
| Ssbp3 | TRUE | 0.182139263 | 0.178321807 | No Change | Not Significant |
| Ssbp4 | TRUE | 0.152833773 | 0.47603234 | No Change | Not Significant |
| Ssc4d | TRUE | 0.289184993 | 0.468471767 | No Change | Not Significant |
| Ssc5d | TRUE | 0.221682627 | 0.908714429 | No Change | Not Significant |
| Ssh1 | TRUE | 0.125996664 | 0.467106114 | No Change | Not Significant |
| Ssh2 | TRUE | 0.366342818 | 1.52E-05 | No Change | Significant Positive |
| Ssh3 | TRUE | -0.148401398 | 0.050745295 | No Change | Not Significant |
| Ssmem1 | TRUE | 0.183333759 | 1 | No Change | Not Significant |
| Ssna1 | TRUE | -0.088775865 | 0.659125086 | No Change | Not Significant |
| Sspn | TRUE | -0.846591811 | 1 | No Change | Not Significant |
| Sspo | TRUE | -0.141532377 | 0.750060917 | No Change | Not Significant |
| Ssr1 | TRUE | 0.376163407 | 0.020064348 | No Change | Significant Positive |
| Ssr2 | TRUE | 0.16001605 | 0.268614824 | No Change | Not Significant |
| Ssr3 | TRUE | 1.438327976 | 7.72E-48 | UP | UP |
| Ssr4 | TRUE | 0.552283634 | 9.98E-10 | UP | UP |
| Ssrp1 | TRUE | 0.13935266 | 0.359501207 | No Change | Not Significant |
| Sst | TRUE | -0.521098357 | 0.018374318 | DOWN | DOWN |
| Sstr1 | TRUE | -2.058046773 | 4.94E-17 | DOWN | DOWN |
| Sstr2 | TRUE | -0.913204287 | 0.010565985 | DOWN | DOWN |
| Sstr3 | TRUE | 0.669615663 | 0.187078687 | No Change | Not Significant |
| Sstr5 | TRUE | -0.301664632 | 0.716728293 | No Change | Not Significant |
| Ssu2 | TRUE | -0.431220179 | 1 | No Change | Not Significant |
| Ssu72 | TRUE | -0.017556003 | 0.889135053 | No Change | Not Significant |
| Ssx2ip | TRUE | 0.339158903 | 8.86E-06 | No Change | Significant Positive |
| St13 | TRUE | -0.075104917 | 0.647057325 | No Change | Not Significant |
| St14 | TRUE | 0.048136601 | 0.702914252 | No Change | Not Significant |
| St18 | TRUE | -1.037112138 | 0.000143007 | DOWN | DOWN |
| St3gal1 | TRUE | 0.549809583 | 0.320178792 | No Change | Not Significant |
| St3gal2 | TRUE | -0.227901892 | 0.788172589 | No Change | Not Significant |
| St3gal3 | TRUE | -0.582591085 | 3.57E-05 | DOWN | DOWN |
| St3gal4 | TRUE | -0.86388726 | 0.006446644 | DOWN | DOWN |
| St3gal5 | TRUE | 0.701585407 | 0.559233675 | No Change | Not Significant |
| St3gal6 | TRUE | -0.490825459 | 0.170971405 | No Change | Not Significant |
| St6gal1 | TRUE | -0.25290226 | 0.704201299 | No Change | Not Significant |
| St6gal2 | TRUE | -0.569471219 | 1 | No Change | Not Significant |
| St6galnac1 | TRUE | 2.593412692 | 1 | No Change | Not Significant |
| St6galnac2 | TRUE | -0.470468499 | 0.000623516 | No Change | Significant Negative |
| St6galnac3 | TRUE | 0.093176513 | 1 | No Change | Not Significant |
| St6galnac4 | TRUE | 1.034643947 | 7.32E-12 | UP | UP |
| St6galnac5 | TRUE | -0.548788999 | 1 | No Change | Not Significant |
| St6galnac6 | TRUE | 0.157564674 | 0.210845076 | No Change | Not Significant |
| St7 | TRUE | 0.162210006 | 0.42426293 | No Change | Not Significant |
| St7l | TRUE | -0.126927442 | 0.130773645 | No Change | Not Significant |
| St8sia1 | TRUE | 1.088038813 | 0.461664802 | No Change | Not Significant |
| St8sia2 | TRUE | -0.600112377 | 1 | No Change | Not Significant |
| St8sia3 | TRUE | 0.572585409 | 1 | No Change | Not Significant |
| St8sia4 | TRUE | 0.442553511 | 0.674995696 | No Change | Not Significant |
| St8sia5 | TRUE | -1.09961008 | 0.074195482 | No Change | Not Significant |
| St8sia6 | TRUE | -0.393803154 | 1 | No Change | Not Significant |
| Stab1 | TRUE | 1.982919276 | 0.231252996 | No Change | Not Significant |
| Stab2 | TRUE | 0.549458023 | 0.79831751 | No Change | Not Significant |
| Stac | TRUE | -0.836827619 | 1 | No Change | Not Significant |
| Stac2 | TRUE | 1.228324303 | 1 | No Change | Not Significant |
| Stac3 | TRUE | 0.17551007 | 0.874258149 | No Change | Not Significant |
| Stag1 | TRUE | 0.374090899 | 0.000227524 | No Change | Significant Positive |
| Stag2 | TRUE | -0.207737079 | 0.0473047 | No Change | Significant Negative |
| Stag3 | TRUE | 0.201640609 | 1 | No Change | Not Significant |
| Stam | TRUE | -0.023782587 | 0.879445484 | No Change | Not Significant |
| Stam2 | TRUE | 0.012586483 | 0.925637253 | No Change | Not Significant |
| Stambp | TRUE | 0.049829363 | 0.82238042 | No Change | Not Significant |
| Stambpl1 | TRUE | 0.060676547 | 0.71414691 | No Change | Not Significant |
| Stap1 | TRUE | 0.955510688 | 1 | No Change | Not Significant |
| Stap2 | TRUE | -0.198559053 | 0.131848679 | No Change | Not Significant |
| Star | TRUE | -0.115178679 | 0.775818184 | No Change | Not Significant |
| Stard10 | TRUE | -0.027705043 | 0.812734264 | No Change | Not Significant |
| Stard13 | TRUE | 0.191110107 | 0.032851834 | No Change | Significant Positive |
| Stard3 | TRUE | 0.057624068 | 0.648871201 | No Change | Not Significant |
| Stard3nl | TRUE | 0.07690232 | 0.472341459 | No Change | Not Significant |
| Stard4 | TRUE | 0.936429016 | 2.63E-19 | UP | UP |
| Stard5 | TRUE | -0.133863187 | 0.48334534 | No Change | Not Significant |
| Stard6 | TRUE | 1.068018184 | 1 | No Change | Not Significant |
| Stard7 | TRUE | 0.014413072 | 0.926810134 | No Change | Not Significant |
| Stard8 | TRUE | -0.708807823 | 0.000759025 | DOWN | DOWN |
| Stard9 | TRUE | 0.452526152 | 0.722022907 | No Change | Not Significant |
| Stat1 | TRUE | 0.079010309 | 0.770846581 | No Change | Not Significant |
| Stat2 | TRUE | 0.739313993 | 0.000156354 | UP | UP |
| Stat3 | TRUE | 0.253605191 | 0.021650799 | No Change | Significant Positive |
| Stat4 | TRUE | 1.297457492 | 0.122583491 | No Change | Not Significant |
| Stat5a | TRUE | -0.151716931 | 0.282675077 | No Change | Not Significant |
| Stat5b | TRUE | -0.426376201 | 6.51E-08 | No Change | Significant Negative |
| Stat6 | TRUE | -0.337793239 | 1.77E-09 | No Change | Significant Negative |
| Stau1 | TRUE | -0.023195197 | 0.872440598 | No Change | Not Significant |
| Stau2 | TRUE | 0.060375179 | 0.714689206 | No Change | Not Significant |
| Stbd1 | TRUE | -0.607747782 | 0.50683356 | No Change | Not Significant |
| Stc1 | TRUE | 2.354935918 | 1 | No Change | Not Significant |
| Stc2 | TRUE | 0.522853157 | 0.000602545 | UP | UP |
| Steap1 | TRUE | 0.369644266 | 0.163322332 | No Change | Not Significant |
| Steap2 | TRUE | 0.048347362 | 0.801687789 | No Change | Not Significant |
| Steap3 | TRUE | -0.214164316 | 0.037106194 | No Change | Significant Negative |
| Steap4 | TRUE | 0.465989625 | 0.207507849 | No Change | Not Significant |
| Steep1 | TRUE | 0.037450363 | 0.857956591 | No Change | Not Significant |
| Stfa3 | TRUE | -0.372038226 | 1 | No Change | Not Significant |
| Stil | TRUE | 0.724403745 | 7.04E-09 | UP | UP |
| Stim1 | TRUE | -0.273772599 | 0.081578077 | No Change | Not Significant |
| Stim2 | TRUE | -0.143645678 | 0.467148794 | No Change | Not Significant |
| Stimate | TRUE | 0.060355174 | 0.754583735 | No Change | Not Significant |
| Sting1 | TRUE | 0.48014115 | 0.084474747 | No Change | Not Significant |
| Stip1 | TRUE | 0.380276057 | 0.011575143 | No Change | Significant Positive |
| Stk10 | TRUE | -0.167846688 | 0.71356425 | No Change | Not Significant |
| Stk11 | TRUE | 0.144135979 | 0.114637957 | No Change | Not Significant |
| Stk11ip | TRUE | -0.020220892 | 0.894720656 | No Change | Not Significant |
| Stk16 | TRUE | 0.073332435 | 0.519697021 | No Change | Not Significant |
| Stk17b | TRUE | -0.306580081 | 0.168010721 | No Change | Not Significant |
| Stk19 | TRUE | -0.098192667 | 0.580947662 | No Change | Not Significant |
| Stk24 | TRUE | 0.408569297 | 2.72E-09 | No Change | Significant Positive |
| Stk25 | TRUE | -0.209617363 | 0.097993747 | No Change | Not Significant |
| Stk26 | TRUE | 0.054066297 | 0.938406345 | No Change | Not Significant |
| Stk3 | TRUE | -0.045874478 | 0.723506743 | No Change | Not Significant |
| Stk32a | TRUE | -1.103318249 | 0.04269855 | DOWN | DOWN |
| Stk32b | TRUE | -2.11233539 | 1 | No Change | Not Significant |
| Stk32c | TRUE | 1.570988283 | 1 | No Change | Not Significant |
| Stk35 | TRUE | 0.048134155 | 0.715485319 | No Change | Not Significant |
| Stk36 | TRUE | 0.426763775 | 0.499023496 | No Change | Not Significant |
| Stk38 | TRUE | -0.14490986 | 0.036559585 | No Change | Significant Negative |
| Stk38l | TRUE | -0.183941685 | 0.375235922 | No Change | Not Significant |
| Stk39 | TRUE | -0.0418334 | 0.873310067 | No Change | Not Significant |

| geneID | Cond2_DF_Detected | Cond2_DF_ValueCol | Cond2_DF_PadjCol | Cond2_DF_Cutoff | Cond2_DF_Cutoff_Specific | BG_Method | RG1_All | RG2_Significant | RG3_SignificantChange |
|---|---|---|---|---|---|---|---|---|---|
| Sptbn1 | TRUE | -0.096828237 | 0.040237686 | No Change | Significant Negative | TRUE | Cond1 Significant Negative + Cond2 Significant Negative | None | None |
| Sptbn2 | TRUE | -0.270204595 | 0.012985416 | No Change | Significant Negative | TRUE | Cond1 Significant Negative + Cond2 Significant Negative | None | None |
| Sptbn4 | TRUE | 0.476632381 | 0.119015238 | No Change | Not Significant | TRUE | Cond1 UP + Cond2 Not Significant | Cond1_UP | Cond1_UP |
| Sptbn5 | TRUE | 0.501046686 | 0.15749077 | No Change | Not Significant | TRUE | Cond1 UP + Cond2 Not Significant | Cond1_UP | Cond1_UP |
| Sptlc1 | TRUE | 0.101499111 | 0.116930286 | No Change | Not Significant | TRUE | Cond1 Not Significant + Cond2 Not Significant | None | None |
| Sptlc2 | TRUE | 0.106744545 | 0.061486489 | No Change | Not Significant | TRUE | Cond1 Significant Positive + Cond2 Not Significant | None | None |
| Sptlc3 | FALSE | NA | NA | No Change | Not Detected | FALSE | Background = FALSE | Background = FALSE | Background = FALSE |
| Sptssa | TRUE | 0.322451171 | 1.13E-05 | No Change | Significant Positive | TRUE | Cond1 UP + Cond2 Significant Positive | Core_UP | Cond1_UP |
| Sptssb | TRUE | 0.264921463 | 0.084538762 | No Change | Not Significant | TRUE | Cond1 DOWN + Cond2 Not Significant | Cond1_DOWN | Cond1_DOWN |
| Spty2d1 | TRUE | 0.075534749 | 0.290503186 | No Change | Not Significant | TRUE | Cond1 Not Significant + Cond2 Not Significant | None | None |
| Spx | FALSE | NA | NA | No Change | Not Detected | FALSE | Background = FALSE | Background = FALSE | Background = FALSE |
| Sqle | TRUE | -0.074209507 | 0.75210185 | No Change | Not Significant | TRUE | Cond1 Not Significant + Cond2 Not Significant | None | None |
| [illegible] | [illegible] | [illegible] | [illegible] | [illegible] | [illegible] | [illegible] | [illegible] | [illegible] | [illegible] |

| geneID | Cond1_DF_ensembl_gene_id | Cond1_DF_entrezgene_id | Cond1_DF_gene_biotype | Cond1_DF_baseMean | Cond1_DF_log2FoldChange | Cond1_DF_lfcSE | Cond1_DF_stat | Cond1_DF_pvalue | Cond1_DF_padj | Cond1_DF_absLog2FC | Cond1_DF_Cutoff_Specific.y |
|---|---|---|---|---|---|---|---|---|---|---|---|
| Sptbn1 | ENSMUSG00000020315 | 20742 | protein_coding | 9189.480466 | -0.174854227 | 0.060696866 | 2.880778519 | 0.003966943 | 0.020064348 | 0.174854227 | Significant Negative |
| Sptbn2 | ENSMUSG00000067889 | 20743 | protein_coding | 223.3785657 | -0.42676438 | 0.16364717 | 2.607832323 | 0.009111758 | 0.039545746 | 0.42676438 | Significant Negative |
| Sptbn4 | ENSMUSG00000011751 | 80297 | protein_coding | 100.9281533 | 2.443241539 | 0.262331932 | -9.313549904 | 1.24E-20 | 6.50E-19 | 2.443241539 | UP |
| Sptbn5 | ENSMUSG00000074899 | 640524 | protein_coding | 44.67714294 | 1.644642133 | 0.441909957 | -3.721667971 | 0.000197911 | 0.00156603 | 1.644642133 | UP |
| Sptlc1 | ENSMUSG00000021468 | 268656 | protein_coding | 761.1247772 | 0.126697493 | 0.082506431 | -1.535607495 | 0.124634675 | 0.289400593 | 0.126697493 | Not Significant |
| Sptlc2 | ENSMUSG00000021036 | 20773 | protein_coding | 3490.325014 | 0.423421325 | 0.053607084 | -7.898607685 | 2.82E-15 | 1.05E-13 | 0.423421325 | Significant Positive |
| Sptlc3 | ENSMUSG00000039092 | 228677 | protein_coding | 1.160807392 | 0.609300338 | 2.413558415 | -0.252448971 | 0.800694052 | 1 | 0.609300338 | Not Significant |
| Sptssa | ENSMUSG00000044408 | 104725 | protein_coding | 692.0618823 | 0.722456142 | 0.081766978 | -8.835548977 | 9.96E-19 | 4.73E-17 | 0.722456142 | UP |
| Sptssb | ENSMUSG00000043461 | 66183 | protein_coding | 1079.122323 | -0.522907799 | 0.206713361 | 2.529627488 | 0.011418368 | 0.047353826 | 0.522907799 | DOWN |
| Spty2d1 | ENSMUSG00000049516 | 101685 | protein_coding | 605.8779915 | 0.179886373 | 0.077761923 | -2.313296363 | 0.020706351 | 0.075334803 | 0.179886373 | Not Significant |
| Spx | ENSMUSG00000071112 | 319552 | protein_coding | 2.455548208 | 0.166723789 | 0.808225378 | -0.206283783 | 0.836569245 | 1 | 0.166723789 | Not Significant |
| Sqle | ENSMUSG00000022351 | 20775 | protein_coding | 1321.294961 | 0.172412637 | 0.150028017 | -1.149202927 | 0.250472314 | 0.462725114 | 0.172412637 | Not Significant |
| [illegible] | [illegible] | [illegible] | [illegible] | [illegible] | [illegible] | [illegible] | [illegible] | [illegible] | [illegible] | [illegible] | [illegible] |

| geneID | Cond2_DF_ensembl_gene_id | Cond2_DF_entrezgene_id | Cond2_DF_gene_biotype | Cond2_DF_baseMean | Cond2_DF_log2FoldChange | Cond2_DF_lfcSE | Cond2_DF_stat | Cond2_DF_pvalue | Cond2_DF_padj | Cond2_DF_absLog2FC | RG1_Specific_Cond2 | RG1_Specific_Cond1 |
|---|---|---|---|---|---|---|---|---|---|---|---|---|
| Sptbn1 | ENSMUSG00000020315 | 20742 | protein_coding | 17802.06153 | -0.096828237 | 0.035130209 | -2.756267046 | 0.005846524 | 0.040237686 | 0.096828237 | Cond1 No Change + Cond2 Significant Negative | Cond2 No Change + Cond1 Significant Negative |
| Sptbn2 | ENSMUSG00000067889 | 20743 | protein_coding | 510.7586274 | -0.270204595 | 0.084422182 | -3.200635052 | 0.001371251 | 0.012985416 | 0.270204595 | Cond1 No Change + Cond2 Significant Negative | Cond2 No Change + Cond1 Significant Negative |
| Sptbn4 | ENSMUSG00000011751 | 80297 | protein_coding | 30.09540084 | 0.476632381 | 0.214853728 | 2.218404055 | 0.026527295 | 0.119015238 | 0.476632381 | Cond1 UP + Cond2 Not Significant | Cond2 No Change + Cond1 UP |
| Sptbn5 | ENSMUSG00000074899 | 640524 | protein_coding | 26.53575933 | 0.501046686 | 0.243534914 | 2.057391602 | 0.039648572 | 0.15749077 | 0.501046686 | Cond1 UP + Cond2 Not Significant | Cond2 No Change + Cond1 UP |
| Sptlc1 | ENSMUSG00000021468 | 268656 | protein_coding | 1517.191384 | 0.101499111 | 0.045497871 | 2.23085409 | 0.025690796 | 0.116930286 | 0.101499111 | Cond1 No Change + Cond2 Not Significant | Cond2 No Change + Cond1 Not Significant |
| Sptlc2 | ENSMUSG00000021036 | 20773 | protein_coding | 5561.487687 | 0.106744545 | 0.041708634 | 2.559291337 | 0.01048858 | 0.061486489 | 0.106744545 | Cond1 No Change + Cond2 Not Significant | Cond2 No Change + Cond1 Significant Positive |
| Sptlc3 | NA | NA | NA | NA | NA | NA | NA | NA | NA | NA | Background = FALSE | Background = FALSE |
| Sptssa | ENSMUSG00000044408 | 104725 | protein_coding | 1202.95962 | 0.322451171 | 0.063766315 | 5.056763467 | 4.26E-07 | 1.13E-05 | 0.322451171 | Cond1 UP + Cond2 Significant Positive | Cond2 No Change + Cond1 UP |
| Sptssb | ENSMUSG00000043461 | 66183 | protein_coding | 710.9696801 | 0.264921463 | 0.110088551 | 2.406439729 | 0.016108862 | 0.084538762 | 0.264921463 | Cond1 DOWN + Cond2 Not Significant | Cond2 No Change + Cond1 DOWN |
| Spty2d1 | ENSMUSG00000049516 | 101685 | protein_coding | 1110.451053 | 0.075534749 | 0.046130233 | 1.637423963 | 0.101541915 | 0.290503186 | 0.075534749 | Cond1 No Change + Cond2 Not Significant | Cond2 No Change + Cond1 Not Significant |
| Spx | NA | NA | NA | NA | NA | NA | NA | NA | NA | NA | Background = FALSE | Background = FALSE |
| Sqle | ENSMUSG00000022351 | 20775 | protein_coding | 3135.131492 | -0.074209507 | 0.124489032 | -0.59611281 | 0.551099867 | 0.75210185 | 0.074209507 | Cond1 No Change + Cond2 Not Significant | Cond2 No Change + Cond1 Not Significant |
| [illegible] | [illegible] | [illegible] | [illegible] | [illegible] | [illegible] | [illegible] | [illegible] | [illegible] | [illegible] | [illegible] | [illegible] | [illegible] |

| geneID | Cond1_DF_Detected | Cond1_DF_ValueCol | Cond1_DF_PadjCol | Cond1_DF_Cutoff | Cond1_DF_Cutoff_Specific.x | Cond2_DF_Detected | Cond2_DF_ValueCol | Cond2_DF_PadjCol | Cond2_DF_Cutoff | Cond2_DF_Cutoff_Specific | BG_Method | RG1_All | RG2_Significant | RG3_SignificantChange | Cond1_DF_ensembl_gene_id | Cond1_DF_entrezgene_id | Cond1_DF_gene_biotype | Cond1_DF_baseMean |
|---|---|---|---|---|---|---|---|---|---|---|---|---|---|---|---|---|---|---|
| Stk4 | TRUE | -0.238222266 | 0.019529117 | No Change | Significant Negative | TRUE | -0.065472087 | 0.345547355 | No Change | Not Significant | TRUE | Cond1 Significant Negative + Cond2 Not Significant | None | None | ENSMUSG00000018209 | 58231 | protein_coding | 702.0451212 |
| Stk40 | TRUE | 0.064197434 | 0.695380179 | No Change | Not Significant | TRUE | 0.026065799 | 0.861383838 | No Change | Not Significant | TRUE | Cond1 Not Significant + Cond2 Not Significant | None | None | ENSMUSG00000042608 | 74178 | protein_coding | 544.4584298 |
| Stkld1 | TRUE | -0.357102687 | 0.723645701 | No Change | Not Significant | TRUE | -0.284689182 | 0.777011446 | No Change | Not Significant | TRUE | Cond1 Not Significant + Cond2 Not Significant | None | None | ENSMUSG00000049897 | 279029 | protein_coding | 5.319311173 |
| Stmn1 | TRUE | -0.064729844 | 0.861253255 | No Change | Not Significant | TRUE | -0.069190441 | 0.721608883 | No Change | Not Significant | TRUE | Cond1 Not Significant + Cond2 Not Significant | None | None | ENSMUSG00000028832 | 16765 | protein_coding | 432.9105832 |
| Stmn2 | TRUE | -1.18856102 | 1 | No Change | Not Significant | FALSE | NA | NA | No Change | Not Detected | FALSE | Background = FALSE | Background = FALSE | Background = FALSE | ENSMUSG00000027500 | 20257 | protein_coding | 1.10551655 |
| Stmn3 | TRUE | -1.632379905 | 1 | No Change | Not Significant | FALSE | NA | NA | No Change | Not Detected | FALSE | Background = FALSE | Background = FALSE | Background = FALSE | ENSMUSG00000027581 | 20262 | protein_coding | 1.184027352 |
| Stmn4 | TRUE | -2.589492777 | 1 | No Change | Not Significant | FALSE | NA | NA | No Change | Not Detected | FALSE | Background = FALSE | Background = FALSE | Background = FALSE | ENSMUSG00000022044 | 56471 | protein_coding | 1.939248618 |
| Stmnd1 | TRUE | -1.177684599 | 1 | No Change | Not Significant | FALSE | NA | NA | No Change | Not Detected | FALSE | Background = FALSE | Background = FALSE | Background = FALSE | ENSMUSG00000063529 | 380842 | protein_coding | 1.194880588 |
| Stn1 | TRUE | 0.130119279 | 0.532085855 | No Change | Not Significant | TRUE | 0.027447513 | 0.898963943 | No Change | Not Significant | TRUE | Cond1 Not Significant + Cond2 Not Significant | None | None | ENSMUSG00000042694 | 108689 | protein_coding | 106.601017 |
| Stom | TRUE | -0.482105248 | 0.070692483 | No Change | Not Significant | TRUE | -0.589124985 | 5.40E-05 | DOWN | DOWN | TRUE | Cond1 Not Significant + Cond2 DOWN | Cond2_DOWN | Cond2_DOWN | ENSMUSG00000026880 | 13830 | protein_coding | 409.2167108 |
| Stoml1 | TRUE | -0.113669354 | 0.559233675 | No Change | Not Significant | TRUE | 0.063577506 | 0.687468885 | No Change | Not Significant | TRUE | Cond1 Not Significant + Cond2 Not Significant | None | None | ENSMUSG00000032333 | 69106 | protein_coding | 128.4566268 |
| Stoml2 | TRUE | -0.067688858 | 0.646395058 | No Change | Not Significant | TRUE | 0.070177885 | 0.429486734 | No Change | Not Significant | TRUE | Cond1 Not Significant + Cond2 Not Significant | None | None | ENSMUSG00000028455 | 66592 | protein_coding | 506.2256191 |
| Stoml3 | TRUE | 0.210626998 | 1 | No Change | Not Significant | FALSE | NA | NA | No Change | Not Detected | FALSE | Background = FALSE | Background = FALSE | Background = FALSE | ENSMUSG00000027744 | 229277 | protein_coding | 1.461318199 |
| Ston1 | TRUE | -0.15384153 | 0.669122175 | No Change | Not Significant | TRUE | -0.296254681 | 0.207368721 | No Change | Not Significant | TRUE | Cond1 Not Significant + Cond2 Not Significant | None | None | ENSMUSG00000033855 | 77057 | protein_coding | 57.27315809 |
| Ston2 | TRUE | -0.15908998 | 0.527754626 | No Change | Not Significant | TRUE | 0.039068712 | 0.809382719 | No Change | Not Significant | TRUE | Cond1 Not Significant + Cond2 Not Significant | None | None | ENSMUSG00000020961 | 108800 | protein_coding | 401.0276201 |
| Stox1 | TRUE | 0.286718592 | 0.721840784 | No Change | Not Significant | TRUE | 0.571181662 | 0.491645551 | No Change | Not Significant | TRUE | Cond1 Not Significant + Cond2 Not Significant | None | None | ENSMUSG00000036923 | 216021 | protein_coding | 5.85741463 |
| Stox2 | TRUE | 2.930234456 | 4.79E-239 | UP | UP | TRUE | 1.425593182 | 6.35E-42 | UP | UP | TRUE | Cond1 UP + Cond2 UP | Core_UP | Core_UP | ENSMUSG00000038143 | 71069 | protein_coding | 479.3872002 |
| Stpg1 | TRUE | -0.551389485 | 0.324596932 | No Change | Not Significant | TRUE | 0.242550165 | 0.692926016 | No Change | Not Significant | TRUE | Cond1 Not Significant + Cond2 Not Significant | None | None | ENSMUSG00000028801 | 78806 | protein_coding | 11.46335694 |
| [illegible] | [illegible] | [illegible] | [illegible] | [illegible] | [illegible] | [illegible] | [illegible] | [illegible] | [illegible] | [illegible] | [illegible] | [illegible] | [illegible] | [illegible] | [illegible] | [illegible] | [illegible] | [illegible] |

| geneID | Cond1_DF_log2FoldChange | Cond1_DF_lfcSE | Cond1_DF_stat | Cond1_DF_pvalue | Cond1_DF_padj | Cond1_DF_absLog2FC | Cond1_DF_Cutoff_Specific.y | Cond2_DF_ensembl_gene_id | Cond2_DF_entrezgene_id | Cond2_DF_gene_biotype | Cond2_DF_baseMean | Cond2_DF_log2FoldChange | Cond2_DF_lfcSE | Cond2_DF_stat | Cond2_DF_pvalue | Cond2_DF_padj | Cond2_DF_absLog2FC | RG1_Specific_Cond2 | RG1_Specific_Cond1 |
|---|---|---|---|---|---|---|---|---|---|---|---|---|---|---|---|---|---|---|---|
| Stk4 | -0.238222266 | 0.08238673 | 2.891512572 | 0.003833923 | 0.019529117 | 0.238222266 | Significant Negative | ENSMUSG00000018209 | 58231 | protein_coding | 1444.8088 | -0.065472087 | 0.043972945 | -1.488917493 | 0.136509097 | 0.345547355 | 0.065472087 | Cond1 No Change + Cond2 Not Significant | Cond2 No Change + Cond1 Significant Negative |
| Stk40 | 0.064197434 | 0.092029766 | -0.697572503 | 0.485444582 | 0.695380179 | 0.064197434 | Not Significant | ENSMUSG00000042608 | 74178 | protein_coding | 908.2833889 | 0.026065799 | 0.071936626 | 0.362343918 | 0.717095043 | 0.861383838 | 0.026065799 | Cond1 No Change + Cond2 Not Significant | Cond2 No Change + Cond1 Not Significant |
| Stkld1 | -0.357102687 | 0.562094579 | 0.635307118 | 0.525228132 | 0.723645701 | 0.357102687 | Not Significant | ENSMUSG00000049897 | 279029 | protein_coding | 7.360199135 | -0.284689182 | 0.523869681 | -0.543435118 | 0.586830252 | 0.777011446 | 0.284689182 | Cond1 No Change + Cond2 Not Significant | Cond2 No Change + Cond1 Not Significant |
| Stmn1 | -0.064729844 | 0.18799083 | 0.344324478 | 0.730602278 | 0.861253255 | 0.064729844 | Not Significant | ENSMUSG00000028832 | 16765 | protein_coding | 982.3978169 | -0.069190441 | 0.103940002 | -0.665676732 | 0.505617749 | 0.721608883 | 0.069190441 | Cond1 No Change + Cond2 Not Significant | Cond2 No Change + Cond1 Not Significant |
| Stmn2 | -1.18856102 | 1.776148778 | 0.66917875 | 0.50338146 | 1 | 1.18856102 | Not Significant | NA | NA | NA | NA | NA | NA | NA | NA | NA | NA | Background = FALSE | Background = FALSE |
| Stmn3 | -1.632379905 | 1.185641956 | 1.376789929 | 0.168577205 | 1 | 1.632379905 | Not Significant | NA | NA | NA | NA | NA | NA | NA | NA | NA | NA | Background = FALSE | Background = FALSE |
| Stmn4 | -2.589492777 | 2.201967021 | 1.175990718 | 0.239598585 | 1 | 2.589492777 | Not Significant | NA | NA | NA | NA | NA | NA | NA | NA | NA | NA | Background = FALSE | Background = FALSE |
| Stmnd1 | -1.177684599 | 1.192616037 | 0.987480096 | 0.323407336 | 1 | 1.177684599 | Not Significant | NA | NA | NA | NA | NA | NA | NA | NA | NA | NA | Background = FALSE | Background = FALSE |
| Stn1 | 0.130119279 | 0.128482197 | -1.012741704 | 0.311183552 | 0.532085855 | 0.130119279 | Not Significant | ENSMUSG00000042694 | 108689 | protein_coding | 208.3077358 | 0.027447513 | 0.099853047 | 0.274879076 | 0.783409143 | 0.898963943 | 0.027447513 | Cond1 No Change + Cond2 Not Significant | Cond2 No Change + Cond1 Not Significant |
| Stom | -0.482105248 | 0.205640535 | 2.344407671 | 0.019057328 | 0.070692483 | 0.482105248 | Not Significant | ENSMUSG00000026880 | 13830 | protein_coding | 635.0133498 | -0.589124985 | 0.124900731 | -4.716745692 | 2.40E-06 | 5.40E-05 | 0.589124985 | Cond1 No Change + Cond2 DOWN | Cond2 DOWN + Cond1 Not Significant |
| Stoml1 | -0.113669354 | 0.118321317 | 0.960683641 | 0.336711259 | 0.559233675 | 0.113669354 | Not Significant | ENSMUSG00000032333 | 69106 | protein_coding | 281.3910126 | 0.063577506 | 0.086205849 | 0.737508035 | 0.460813459 | 0.687468885 | 0.063577506 | Cond1 No Change + Cond2 Not Significant | Cond2 No Change + Cond1 Not Significant |
| Stoml2 | -0.067688858 | 0.085201507 | 0.794456111 | 0.426929949 | 0.646395058 | 0.067688858 | Not Significant | ENSMUSG00000028455 | 66592 | protein_coding | 927.4103576 | 0.070177885 | 0.054397748 | 1.290088069 | 0.197020082 | 0.429486734 | 0.070177885 | Cond1 No Change + Cond2 Not Significant | Cond2 No Change + Cond1 Not Significant |
| Stoml3 | 0.210626998 | 2.071065045 | -0.101699847 | 0.918994924 | 1 | 0.210626998 | Not Significant | NA | NA | NA | NA | NA | NA | NA | NA | NA | NA | Background = FALSE | Background = FALSE |
| Ston1 | -0.15384153 | 0.205089367 | 0.750119486 | 0.453182745 | 0.669122175 | 0.15384153 | Not Significant | ENSMUSG00000033855 | 77057 | protein_coding | 110.4921082 | -0.296254681 | 0.157300462 | -1.883368156 | 0.059650493 | 0.207368721 | 0.296254681 | Cond1 No Change + Cond2 Not Significant | Cond2 No Change + Cond1 Not Significant |
| Ston2 | -0.15908998 | 0.155835463 | 1.020884313 | 0.307309252 | 0.527754626 | 0.15908998 | Not Significant | ENSMUSG00000020961 | 108800 | protein_coding | 701.9688585 | 0.039068712 | 0.082315347 | 0.474622447 | 0.635056104 | 0.809382719 | 0.039068712 | Cond1 No Change + Cond2 Not Significant | Cond2 No Change + Cond1 Not Significant |
| Stox1 | 0.286718592 | 0.448465678 | -0.639332296 | 0.522606783 | 0.721840784 | 0.286718592 | Not Significant | ENSMUSG00000036923 | 216021 | protein_coding | 5.977161413 | 0.571181662 | 0.497186002 | 1.148828928 | 0.250626528 | 0.491645551 | 0.571181662 | Cond1 No Change + Cond2 Not Significant | Cond2 No Change + Cond1 Not Significant |
| Stox2 | 2.930234456 | 0.088206578 | -33.22013528 | 5.51E-242 | 4.79E-239 | 2.930234456 | UP | ENSMUSG00000038143 | 71069 | protein_coding | 305.0061851 | 1.425593182 | 0.102180768 | 13.95167804 | 3.07E-44 | 6.35E-42 | 1.425593182 | Cond1 UP + Cond2 UP | Cond2 UP + Cond1 UP |
| Stpg1 | -0.551389485 | 0.380004833 | 1.451006508 | 0.146778048 | 0.324596932 | 0.551389485 | Not Significant | ENSMUSG00000028801 | 78806 | protein_coding | 16.46261041 | 0.242550165 | 0.333939463 | 0.726329744 | 0.467636649 | 0.692926016 | 0.242550165 | Cond1 No Change + Cond2 Not Significant | Cond2 No Change + Cond1 Not Significant |
| [illegible] | [illegible] | [illegible] | [illegible] | [illegible] | [illegible] | [illegible] | [illegible] | [illegible] | [illegible] | [illegible] | [illegible] | [illegible] | [illegible] | [illegible] | [illegible] | [illegible] | [illegible] | [illegible] | [illegible] |

| geneID | Cond1_DF_Detected | Cond1_DF_ValueCol | Cond1_DF_PadjCol | Cond1_DF_Cutoff | Cond1_DF_Cutoff_Specific.x | Cond2_DF_Detected | Cond2_DF_ValueCol | Cond2_DF_PadjCol | Cond2_DF_Cutoff | Cond2_DF_Cutoff_Specific | BG_Method |
|---|---|---|---|---|---|---|---|---|---|---|---|
| Syt9 | TRUE | 1.259744827 | 1 | No Change | Not Significant | FALSE | NA | NA | No Change | Not Detected | FALSE |
| Sytl1 | TRUE | -0.399213284 | 0.05132314 | No Change | Not Significant | TRUE | -0.011611926 | 0.951039003 | No Change | Not Significant | TRUE |
| Sytl2 | TRUE | -0.17417268 | 0.579378227 | No Change | Not Significant | TRUE | 0.112784757 | 0.263070646 | No Change | Not Significant | TRUE |
| Sytl3 | TRUE | -0.214174434 | 0.734651013 | No Change | Not Significant | TRUE | -0.305433823 | 0.331056319 | No Change | Not Significant | TRUE |
| Sytl4 | TRUE | -0.249616128 | 0.131923303 | No Change | Not Significant | TRUE | 0.096309692 | 0.405350432 | No Change | Not Significant | TRUE |
| Sytl5 | TRUE | 0.979379169 | 0.007064806 | UP | UP | TRUE | 0.239300229 | 0.488954885 | No Change | Not Significant | TRUE |
| Syvn1 | TRUE | 2.305526002 | 6.87E-55 | UP | UP | TRUE | 1.480926223 | 2.26E-100 | UP | UP | TRUE |
| Szrd1 | TRUE | 0.023044213 | 0.855153117 | No Change | Not Significant | TRUE | 0.133232631 | 0.12038719 | No Change | Not Significant | TRUE |
| Szt2 | TRUE | 0.171821339 | 0.396311693 | No Change | Not Significant | TRUE | 0.024949745 | 0.889541232 | No Change | Not Significant | TRUE |
| T2 | TRUE | -0.612971421 | 1 | No Change | Not Significant | FALSE | NA | NA | No Change | Not Detected | FALSE |
| Tab1 | TRUE | -0.130031592 | 0.309078852 | No Change | Not Significant | TRUE | -0.046440832 | 0.628768653 | No Change | Not Significant | TRUE |
| Tab2 | TRUE | 0.139853493 | 0.069749255 | No Change | Not Significant | TRUE | -0.03029978 | 0.718418124 | No Change | Not Significant | TRUE |
| Tab3 | TRUE | 0.15048345 | 0.273724003 | No Change | Not Significant | TRUE | -0.046162947 | 0.775075241 | No Change | Not Significant | TRUE |
| Tac1 | TRUE | -0.045491951 | 0.913446239 | No Change | Not Significant | TRUE | 0.038642983 | 0.914739662 | No Change | Not Significant | TRUE |
| Tac2 | TRUE | 1.77978055 | 1 | No Change | Not Significant | FALSE | NA | NA | No Change | Not Detected | FALSE |
| Tac4 | TRUE | -0.194850801 | 1 | No Change | Not Significant | FALSE | NA | NA | No Change | Not Detected | FALSE |
| Tacc1 | TRUE | 0.576863487 | 0.002234472 | UP | UP | TRUE | 0.078002675 | 0.680801421 | No Change | Not Significant | TRUE |
| Tacc2 | TRUE | -0.423441562 | 0.045227455 | No Change | Significant Negative | TRUE | -0.049584554 | 0.708115319 | No Change | Not Significant | TRUE |
| Tacc3 | TRUE | 0.829187605 | 6.64E-07 | UP | UP | TRUE | 0.134730051 | 0.410775296 | No Change | Not Significant | TRUE |
| Taco1 | TRUE | -0.224732268 | 0.10695305 | No Change | Not Significant | TRUE | 0.053865899 | 0.755288004 | No Change | Not Significant | TRUE |
| Tacr1 | TRUE | 0.93761448 | 1 | No Change | Not Significant | FALSE | NA | NA | No Change | Not Detected | FALSE |
| Tacr2 | TRUE | 4.323780638 | 3.25E-05 | UP | UP | FALSE | NA | NA | No Change | Not Detected | FALSE |
| Tada1 | TRUE | -0.026230709 | 0.906158059 | No Change | Not Significant | TRUE | 0.014083193 | 0.940565051 | No Change | Not Significant | TRUE |
| Tada2a | TRUE | 0.044670688 | 0.844104331 | No Change | Not Significant | TRUE | -0.070505092 | 0.700254387 | No Change | Not Significant | TRUE |
| Tada2b | TRUE | 0.030902571 | 0.880874346 | No Change | Not Significant | TRUE | -0.024681566 | 0.84654428 | No Change | Not Significant | TRUE |
| Tada3 | TRUE | 0.020424733 | 0.899452571 | No Change | Not Significant | TRUE | 0.087726565 | 0.421109924 | No Change | Not Significant | TRUE |
| Taf1 | TRUE | 0.114833907 | 0.432209358 | No Change | Not Significant | TRUE | -0.034431082 | 0.713490595 | No Change | Not Significant | TRUE |
| Taf10 | TRUE | 0.099874568 | 0.499023496 | No Change | Not Significant | TRUE | 0.062819379 | 0.677665469 | No Change | Not Significant | TRUE |
| Taf11 | TRUE | -0.024964718 | 0.882510888 | No Change | Not Significant | TRUE | 0.018128783 | 0.917020505 | No Change | Not Significant | TRUE |
| Taf12 | TRUE | 0.118007568 | 0.31192207 | No Change | Not Significant | TRUE | 0.243284231 | 0.002907732 | No Change | Significant Positive | TRUE |
| Taf13 | TRUE | 0.651204042 | 3.61E-12 | UP | UP | TRUE | 0.150672321 | 0.284518548 | No Change | Not Significant | TRUE |
| Taf15 | TRUE | 0.185038753 | 0.070049242 | No Change | Not Significant | TRUE | 0.088730997 | 0.302685656 | No Change | Not Significant | TRUE |
| Taf1a | TRUE | -0.160384564 | 0.350247335 | No Change | Not Significant | TRUE | -0.014082114 | 0.961596444 | No Change | Not Significant | TRUE |
| Taf1b | TRUE | 0.128609813 | 0.668047177 | No Change | Not Significant | TRUE | 0.059811714 | 0.82591596 | No Change | Not Significant | TRUE |
| Taf1c | TRUE | 0.184940097 | 0.055659328 | No Change | Not Significant | TRUE | 0.081430787 | 0.625286182 | No Change | Not Significant | TRUE |
| Taf1d | TRUE | -0.277595716 | 0.040914089 | No Change | Significant Negative | TRUE | -0.421388032 | 0.003096681 | No Change | Significant Negative | TRUE |
| Taf2 | TRUE | 0.210957105 | 0.006239 | No Change | Significant Positive | TRUE | -0.032453381 | 0.789053031 | No Change | Not Significant | TRUE |
| Taf3 | TRUE | -0.079141408 | 0.739192192 | No Change | Not Significant | TRUE | -0.081067825 | 0.691634539 | No Change | Not Significant | TRUE |
| Taf4 | TRUE | 0.006641053 | 0.973795719 | No Change | Not Significant | TRUE | -0.111435743 | 0.177906613 | No Change | Not Significant | TRUE |
| Taf4b | TRUE | 0.161264525 | 0.524146449 | No Change | Not Significant | TRUE | -0.121926446 | 0.488633619 | No Change | Not Significant | TRUE |
| Taf5 | TRUE | 0.204040164 | 0.331061353 | No Change | Not Significant | TRUE | -0.21757983 | 0.086330552 | No Change | Not Significant | TRUE |
| Taf5l | TRUE | -0.107961682 | 0.168040558 | No Change | Not Significant | TRUE | -0.044970759 | 0.584243096 | No Change | Not Significant | TRUE |
| Taf6 | TRUE | 0.005886869 | 0.97587882 | No Change | Not Significant | TRUE | 0.021332921 | 0.912178372 | No Change | Not Significant | TRUE |
| Taf6l | TRUE | 0.474185589 | 0.010174535 | No Change | Significant Positive | TRUE | 0.116614441 | 0.487131402 | No Change | Not Significant | TRUE |
| Taf7 | TRUE | -0.23582262 | 0.096042838 | No Change | Not Significant | TRUE | -0.23175173 | 0.149108639 | No Change | Not Significant | TRUE |
| Taf8 | TRUE | 0.179860769 | 0.13918813 | No Change | Not Significant | TRUE | 0.053698178 | 0.624972092 | No Change | Not Significant | TRUE |
| Taf9 | TRUE | -0.211122395 | 1 | No Change | Not Significant | FALSE | NA | NA | No Change | Not Detected | FALSE |
| Taf9b | TRUE | 0.197398864 | 0.462725114 | No Change | Not Significant | TRUE | 0.175280283 | 0.352075628 | No Change | Not Significant | TRUE |
| Tafa1 | TRUE | -0.827809935 | 0.326976379 | No Change | Not Significant | TRUE | -0.013589642 | 0.991618025 | No Change | Not Significant | TRUE |
| Tafa2 | TRUE | 0.947459468 | 1 | No Change | Not Significant | FALSE | NA | NA | No Change | Not Detected | FALSE |
| Tafa3 | TRUE | -0.373855918 | 0.732538277 | No Change | Not Significant | TRUE | -0.168803919 | 0.840526771 | No Change | Not Significant | TRUE |
| Tafa4 | TRUE | 3.03441491 | 1 | No Change | Not Significant | FALSE | NA | NA | No Change | Not Detected | FALSE |
| Tafazzin | TRUE | -0.140733863 | 0.078410101 | No Change | Not Significant | TRUE | 0.015745521 | 0.875013582 | No Change | Not Significant | TRUE |
| Tagap | TRUE | -0.536442181 | 0.090822909 | No Change | Not Significant | TRUE | 0.021379662 | 0.961295463 | No Change | Not Significant | TRUE |
| Tagap1 | TRUE | -0.033428158 | 0.913504709 | No Change | Not Significant | TRUE | 0.010449543 | 0.967142108 | No Change | Not Significant | TRUE |
| Tagln | TRUE | 0.324298969 | 0.483440174 | No Change | Not Significant | TRUE | -0.07431353 | 0.879750811 | No Change | Not Significant | TRUE |
| Tagln2 | TRUE | -0.265071697 | 0.000602545 | No Change | Significant Negative | TRUE | 0.278547302 | 0.030422534 | No Change | Significant Positive | TRUE |
| Tagln3 | TRUE | 1.346798899 | 1 | No Change | Not Significant | FALSE | NA | NA | No Change | Not Detected | FALSE |
| Tal1 | TRUE | 1.195108172 | 1 | No Change | Not Significant | FALSE | NA | NA | No Change | Not Detected | FALSE |
| Tal2 | TRUE | -1.439919656 | 1 | No Change | Not Significant | FALSE | NA | NA | No Change | Not Detected | FALSE |
| Taldo1 | TRUE | -0.207428904 | 0.003617932 | No Change | Significant Negative | TRUE | 0.070805752 | 0.53329119 | No Change | Not Significant | TRUE |
| Tamalin | TRUE | 0.062735671 | 0.899834189 | No Change | Not Significant | TRUE | -0.269087503 | 0.247934382 | No Change | Not Significant | TRUE |
| Tamm41 | TRUE | 0.231052467 | 0.116153464 | No Change | Not Significant | TRUE | -0.151655397 | 0.205867649 | No Change | Not Significant | TRUE |
| Tanc1 | TRUE | -0.053414515 | 0.625216588 | No Change | Not Significant | TRUE | -0.144929167 | 0.016534966 | No Change | Significant Negative | TRUE |
| Tanc2 | TRUE | 0.027509125 | 0.898808084 | No Change | Not Significant | TRUE | -0.201555676 | 0.225933166 | No Change | Not Significant | TRUE |
| Tango2 | TRUE | 0.210400062 | 0.067342119 | No Change | Not Significant | TRUE | 0.155333486 | 0.125530222 | No Change | Not Significant | TRUE |
| Tango6 | TRUE | -0.109740156 | 0.660786267 | No Change | Not Significant | TRUE | -0.148668586 | 0.406744622 | No Change | Not Significant | TRUE |
| Tank | TRUE | 0.010641928 | 0.954311305 | No Change | Not Significant | TRUE | -0.017710367 | 0.92587823 | No Change | Not Significant | TRUE |
| Taok1 | TRUE | 0.109935622 | 0.141690444 | No Change | Not Significant | TRUE | -0.081116227 | 0.088190332 | No Change | Not Significant | TRUE |
| Taok2 | TRUE | 0.072873564 | 0.571706158 | No Change | Not Significant | TRUE | -0.046980671 | 0.727964612 | No Change | Not Significant | TRUE |
| Taok3 | TRUE | 0.047738536 | 0.817346435 | No Change | Not Significant | TRUE | 0.022148477 | 0.81580845 | No Change | Not Significant | TRUE |
| Tap1 | TRUE | -0.327598158 | 0.281287196 | No Change | Not Significant | TRUE | -0.685103245 | 0.158163814 | No Change | Not Significant | TRUE |
| Tap2 | TRUE | -0.414388914 | 9.24E-05 | No Change | Significant Negative | TRUE | -0.262556243 | 0.206435622 | No Change | Not Significant | TRUE |
| Tapbp | TRUE | -0.486059838 | 3.93E-14 | No Change | Significant Negative | TRUE | -0.214550882 | 0.239967558 | No Change | Not Significant | TRUE |
| Tapbpl | TRUE | 0.289407673 | 0.011158438 | No Change | Significant Positive | TRUE | -0.064412735 | 0.718582863 | No Change | Not Significant | TRUE |
| Tapt1 | TRUE | 0.069603202 | 0.589522317 | No Change | Not Significant | TRUE | -0.132310433 | 0.087561384 | No Change | Not Significant | TRUE |
| Tarbp1 | TRUE | -0.254197095 | 0.173816344 | No Change | Not Significant | TRUE | -0.410639869 | 0.002351219 | No Change | Significant Negative | TRUE |
| Tarbp2 | TRUE | 0.028624042 | 0.907710402 | No Change | Not Significant | TRUE | 0.070758706 | 0.664948752 | No Change | Not Significant | TRUE |
| Tardbp | TRUE | 0.107621515 | 0.431773126 | No Change | Not Significant | TRUE | 0.203840254 | 0.022244465 | No Change | Significant Positive | TRUE |
| Tars | TRUE | 0.255698025 | 0.043279016 | No Change | Significant Positive | TRUE | 0.177140007 | 0.012163642 | No Change | Significant Positive | TRUE |
| Tars2 | TRUE | -0.129056064 | 0.500444242 | No Change | Not Significant | TRUE | 0.051082065 | 0.719943844 | No Change | Not Significant | TRUE |
| Tarsl2 | TRUE | -0.000637517 | 0.998249138 | No Change | Not Significant | TRUE | -0.138685392 | 0.31807442 | No Change | Not Significant | TRUE |
| Tas1r1 | TRUE | 0.517132158 | 0.555630454 | No Change | Not Significant | FALSE | NA | NA | No Change | Not Detected | FALSE |
| Tas1r3 | TRUE | 0.391801048 | 0.27902128 | No Change | Not Significant | TRUE | -0.218601378 | 0.552028135 | No Change | Not Significant | TRUE |
| Tasl | TRUE | 0.242938041 | 1 | No Change | Not Significant | FALSE | NA | NA | No Change | Not Detected | FALSE |
| Tasor | TRUE | -0.066610822 | 0.636091497 | No Change | Not Significant | TRUE | -0.164048553 | 0.079926167 | No Change | Not Significant | TRUE |
| Tasor2 | TRUE | -0.07127605 | 0.652026162 | No Change | Not Significant | TRUE | -0.047023493 | 0.737190295 | No Change | Not Significant | TRUE |
| Tasp1 | TRUE | 0.287493419 | 0.013987474 | No Change | Significant Positive | TRUE | 0.234350613 | 0.125723897 | No Change | Not Significant | TRUE |
| Tat | TRUE | 0.845930388 | 0.018707218 | UP | UP | TRUE | 0.054397568 | 0.838364494 | No Change | Not Significant | TRUE |
| Tatdn1 | TRUE | 0.172936226 | 0.235970255 | No Change | Not Significant | TRUE | -0.156851581 | 0.376993804 | No Change | Not Significant | TRUE |
| Tatdn2 | TRUE | 0.189079595 | 0.015214774 | No Change | Significant Positive | TRUE | -0.030917973 | 0.706263486 | No Change | Not Significant | TRUE |
| Tatdn3 | TRUE | 0.123685226 | 0.630279329 | No Change | Not Significant | TRUE | -0.033283615 | 0.916470201 | No Change | Not Significant | TRUE |
| Tax1bp1 | TRUE | 0.116229797 | 0.47841828 | No Change | Not Significant | TRUE | -0.130188279 | 0.201924436 | No Change | Not Significant | TRUE |
| Tax1bp3 | TRUE | -0.521853896 | 0.000154732 | DOWN | DOWN | TRUE | 0.071041876 | 0.621784722 | No Change | Not Significant | TRUE |
| Tbata | TRUE | 0.523248649 | 1 | No Change | Not Significant | FALSE | NA | NA | No Change | Not Detected | FALSE |
| Tbc1d1 | TRUE | 0.588132746 | 1.91E-09 | UP | UP | TRUE | 0.229516999 | 0.015154147 | No Change | Significant Positive | TRUE |
| Tbc1d10a | TRUE | 1.007618313 | 9.77E-26 | UP | UP | TRUE | 0.289242644 | 0.001092885 | No Change | Significant Positive | TRUE |
| Tbc1d10b | TRUE | 0.010327065 | 0.947365784 | No Change | Not Significant | TRUE | 0.011681286 | 0.950733597 | No Change | Not Significant | TRUE |
| Tbc1d10c | TRUE | -0.220036688 | 0.822845898 | No Change | Not Significant | TRUE | -0.255331676 | 0.708970987 | No Change | Not Significant | TRUE |
| Tbc1d12 | TRUE | 0.197749349 | 0.31052626 | No Change | Not Significant | TRUE | -0.088486779 | 0.550870027 | No Change | Not Significant | TRUE |
| Tbc1d13 | TRUE | -0.068770352 | 0.604102573 | No Change | Not Significant | TRUE | 0.079492394 | 0.437979805 | No Change | Not Significant | TRUE |
| Tbc1d14 | TRUE | -0.010378651 | 0.947705869 | No Change | Not Significant | TRUE | 0.0379774 | 0.678988684 | No Change | Not Significant | TRUE |
| Tbc1d15 | TRUE | -0.02540319 | 0.932166328 | No Change | Not Significant | TRUE | -0.12736551 | 0.328925409 | No Change | Not Significant | TRUE |
| Tbc1d16 | TRUE | -0.645180207 | 0.002935352 | DOWN | DOWN | TRUE | -0.267894169 | 0.0358309 | No Change | Significant Negative | TRUE |
| Tbc1d17 | TRUE | -0.237386142 | 0.00466172 | No Change | Significant Negative | TRUE | -0.067874251 | 0.495031103 | No Change | Not Significant | TRUE |
| Tbc1d19 | TRUE | 0.230634623 | 0.429954629 | No Change | Not Significant | TRUE | 0.154668886 | 0.428764731 | No Change | Not Significant | TRUE |
| Tbc1d2 | TRUE | 0.12237575 | 0.467157647 | No Change | Not Significant | TRUE | -0.01385606 | 0.93708084 | No Change | Not Significant | TRUE |
| Tbc1d20 | TRUE | 0.026183089 | 0.839976943 | No Change | Not Significant | TRUE | -0.051491203 | 0.484584527 | No Change | Not Significant | TRUE |
| Tbc1d22a | TRUE | -0.131111521 | 0.342763083 | No Change | Not Significant | TRUE | -0.015456956 | 0.914534473 | No Change | Not Significant | TRUE |
| Tbc1d22b | TRUE | 0.105614135 | 0.414545055 | No Change | Not Significant | TRUE | 0.119010938 | 0.256348065 | No Change | Not Significant | TRUE |
| Tbc1d22bos | TRUE | 0.093466233 | 1 | No Change | Not Significant | TRUE | -0.507115413 | 0.554568301 | No Change | Not Significant | TRUE |
| Tbc1d23 | TRUE | 0.213775139 | 0.069955433 | No Change | Not Significant | TRUE | -0.033879551 | 0.822476285 | No Change | Not Significant | TRUE |
| Tbc1d24 | TRUE | -0.610226133 | 0.031251999 | DOWN | DOWN | TRUE | -0.217802478 | 0.052462144 | No Change | Not Significant | TRUE |
| Tbc1d25 | TRUE | -0.106900275 | 0.588694 | No Change | Not Significant | TRUE | -0.008255495 | 0.968095473 | No Change | Not Significant | TRUE |
| Tbc1d2b | TRUE | -0.055550977 | 0.826798642 | No Change | Not Significant | TRUE | 0.04396021 | 0.789158883 | No Change | Not Significant | TRUE |
| Tbc1d30 | TRUE | -0.150668286 | 0.188104702 | No Change | Not Significant | TRUE | -0.037911384 | 0.746586685 | No Change | Not Significant | TRUE |
| Tbc1d31 | TRUE | 0.538566674 | 0.006667458 | UP | UP | TRUE | 0.173114636 | 0.317992618 | No Change | Not Significant | TRUE |
| Tbc1d32 | TRUE | -0.228263217 | 0.002672795 | No Change | Significant Negative | TRUE | -0.009939585 | 0.935592329 | No Change | Not Significant | TRUE |
| Tbc1d4 | TRUE | 0.015131838 | 0.939378465 | No Change | Not Significant | TRUE | -0.109197363 | 0.328615354 | No Change | Not Significant | TRUE |
| Tbc1d5 | TRUE | -0.179325013 | 0.083023198 | No Change | Not Significant | TRUE | -0.025022498 | 0.821648745 | No Change | Not Significant | TRUE |
| Tbc1d7 | TRUE | 0.234372434 | 0.141882067 | No Change | Not Significant | TRUE | 0.298168469 | 0.113115816 | No Change | Not Significant | TRUE |
| Tbc1d8 | TRUE | -0.010793067 | 0.968172466 | No Change | Not Significant | TRUE | -0.046554288 | 0.740026398 | No Change | Not Significant | TRUE |
| Tbc1d8b | TRUE | 0.204548559 | 0.171354072 | No Change | Not Significant | TRUE | 0.196900145 | 0.001539685 | No Change | Significant Positive | TRUE |
| Tbc1d9 | TRUE | 1.368201876 | 6.48E-07 | UP | UP | TRUE | 0.392349095 | 0.01586756 | No Change | Significant Positive | TRUE |
| Tbc1d9b | TRUE | -0.031253524 | 0.815856848 | No Change | Not Significant | TRUE | 0.034589051 | 0.694009261 | No Change | Not Significant | TRUE |
| Tbca | TRUE | -0.165274441 | 0.231775376 | No Change | Not Significant | TRUE | -0.008809951 | 0.939033668 | No Change | Not Significant | TRUE |
| Tbcb | TRUE | 1.673441515 | 3.43E-96 | UP | UP | TRUE | 1.498413291 | 3.78E-85 | UP | UP | TRUE |
| Tbcc | TRUE | 0.164448104 | 0.236076969 | No Change | Not Significant | TRUE | 0.134969917 | 0.245557111 | No Change | Not Significant | TRUE |
| Tbccd1 | TRUE | 1.109836739 | 9.33E-23 | UP | UP | TRUE | 0.364936097 | 7.54E-06 | No Change | Significant Positive | TRUE |
| Tbcd | TRUE | 0.020052876 | 0.908714429 | No Change | Not Significant | TRUE | 0.048286527 | 0.588629676 | No Change | Not Significant | TRUE |
| Tbce | TRUE | -0.0416114 | 0.841825268 | No Change | Not Significant | TRUE | -0.194440684 | 0.09661718 | No Change | Not Significant | TRUE |
| Tbcel | TRUE | -0.94252204 | 1.94E-22 | DOWN | DOWN | TRUE | -0.509151917 | 1.11E-05 | DOWN | DOWN | TRUE |
| Tbck | TRUE | -0.043889696 | 0.734843633 | No Change | Not Significant | TRUE | -0.010532376 | 0.921867382 | No Change | Not Significant | TRUE |
| Tbk1 | TRUE | -0.046944077 | 0.690909666 | No Change | Not Significant | TRUE | -0.155562727 | 0.081578678 | No Change | Not Significant | TRUE |
| Tbkbp1 | TRUE | -1.355016289 | 0.239895137 | No Change | Not Significant | FALSE | NA | NA | No Change | Not Detected | FALSE |
| Tbl1x | TRUE | -0.126901547 | 0.215537515 | No Change | Not Significant | TRUE | -0.064807642 | 0.445849236 | No Change | Not Significant | TRUE |
| Tbl1xr1 | TRUE | 0.118455264 | 0.295635135 | No Change | Not Significant | TRUE | 0.053294731 | 0.483875678 | No Change | Not Significant | TRUE |
| Tbl2 | TRUE | 0.413921531 | 0.001766577 | No Change | Significant Positive | TRUE | 0.200913713 | 0.03683451 | No Change | Significant Positive | TRUE |
| Tbl3 | TRUE | -0.039909113 | 0.813424841 | No Change | Not Significant | TRUE | -0.066879979 | 0.605769124 | No Change | Not Significant | TRUE |
| Tbp | TRUE | 0.037987586 | 0.845574994 | No Change | Not Significant | TRUE | -0.109240167 | 0.435188726 | No Change | Not Significant | TRUE |
| Tbpl1 | TRUE | 0.199172937 | 0.219864981 | No Change | Not Significant | TRUE | 0.068451501 | 0.571937821 | No Change | Not Significant | TRUE |
| Tbr1 | TRUE | 0.371169709 | 1 | No Change | Not Significant | FALSE | NA | NA | No Change | Not Detected | FALSE |
| Tbrg1 | TRUE | 0.220848082 | 0.029831627 | No Change | Significant Positive | TRUE | 0.17065668 | 0.001073278 | No Change | Significant Positive | TRUE |
| Tbrg4 | TRUE | -0.006804982 | 0.948996349 | No Change | Not Significant | TRUE | 0.084416864 | 0.380724711 | No Change | Not Significant | TRUE |
| Tbx1 | TRUE | -2.059808424 | 1 | No Change | Not Significant | FALSE | NA | NA | No Change | Not Detected | FALSE |
| Tbx10 | TRUE | 2.126269847 | 1 | No Change | Not Significant | FALSE | NA | NA | No Change | Not Detected | FALSE |
| Tbx15 | TRUE | -0.003963924 | 1 | No Change | Not Significant | FALSE | NA | NA | No Change | Not Detected | FALSE |
| Tbx19 | TRUE | 0.969179695 | 1 | No Change | Not Significant | FALSE | NA | NA | No Change | Not Detected | FALSE |
| Tbx2 | TRUE | 0.322110222 | 1 | No Change | Not Significant | FALSE | NA | NA | No Change | Not Detected | FALSE |
| Tbx20 | TRUE | 1.328530785 | 1 | No Change | Not Significant | FALSE | NA | NA | No Change | Not Detected | FALSE |
| Tbx21 | TRUE | 1.072024528 | 0.332693773 | No Change | Not Significant | TRUE | 0.866597955 | 0.143636725 | No Change | Not Significant | TRUE |
| Tbx22 | TRUE | 0.005729717 | 1 | No Change | Not Significant | FALSE | NA | NA | No Change | Not Detected | FALSE |
| Tbx3 | TRUE | 0.143844022 | 0.776435029 | No Change | Not Significant | TRUE | -0.519617742 | 0.000293943 | DOWN | DOWN | TRUE |
| Tbx3os1 | TRUE | 0.203096321 | 1 | No Change | Not Significant | FALSE | NA | NA | No Change | Not Detected | FALSE |
| Tbx3os2 | TRUE | -0.48465604 | 1 | No Change | Not Significant | FALSE | NA | NA | No Change | Not Detected | FALSE |
| Tbx4 | TRUE | 1.189983526 | 1 | No Change | Not Significant | FALSE | NA | NA | No Change | Not Detected | FALSE |
| Tbx5 | TRUE | -1.18738031 | 1 | No Change | Not Significant | FALSE | NA | NA | No Change | Not Detected | FALSE |
| Tbx6 | TRUE | -0.172933351 | 0.857652848 | No Change | Not Significant | TRUE | -0.424292766 | 0.522078519 | No Change | Not Significant | TRUE |
| Tbxas1 | TRUE | -0.578994238 | 0.452145476 | No Change | Not Significant | TRUE | -0.057399373 | 0.949475732 | No Change | Not Significant | TRUE |
| Tc2n | TRUE | 0.592089927 | 1.24E-07 | UP | UP | TRUE | 0.281800934 | 1.08E-05 | No Change | Significant Positive | TRUE |
| Tcaf1 | TRUE | 0.02519433 | 0.889481359 | No Change | Not Significant | TRUE | 0.10635629 | 0.401740784 | No Change | Not Significant | TRUE |
| Tcaf2 | TRUE | -0.427202432 | 0.03924437 | No Change | Significant Negative | TRUE | 0.035579419 | 0.749029274 | No Change | Not Significant | TRUE |
| Tcaim | TRUE | 0.071400061 | 0.738779548 | No Change | Not Significant | TRUE | 0.159557579 | 0.259710803 | No Change | Not Significant | TRUE |
| Tcam1 | TRUE | 0.896858759 | 1 | No Change | Not Significant | FALSE | NA | NA | No Change | Not Detected | FALSE |
| Tcap | TRUE | 3.090445101 | 1 | No Change | Not Significant | FALSE | NA | NA | No Change | Not Detected | FALSE |
| Tcea1 | TRUE | 0.150453218 | 0.264518622 | No Change | Not Significant | TRUE | 0.150839211 | 0.105379648 | No Change | Not Significant | TRUE |
| Tcea2 | TRUE | -0.066462149 | 0.895242955 | No Change | Not Significant | TRUE | -0.688055879 | 0.026557202 | DOWN | DOWN | TRUE |
| Tcea3 | TRUE | -0.520275318 | 1.20E-11 | DOWN | DOWN | TRUE | -0.219936089 | 0.047758664 | No Change | Significant Negative | TRUE |
| Tceal1 | TRUE | -0.600144985 | 0.158286006 | No Change | Not Significant | TRUE | 0.077150013 | 0.886076667 | No Change | Not Significant | TRUE |
| Tceal3 | TRUE | -0.61780601 | 1 | No Change | Not Significant | FALSE | NA | NA | No Change | Not Detected | FALSE |
| Tceal5 | TRUE | -2.115599068 | 1 | No Change | Not Significant | FALSE | NA | NA | No Change | Not Detected | FALSE |
| Tceal8 | TRUE | 0.304484462 | 0.119750527 | No Change | Not Significant | TRUE | 0.108897316 | 0.60723078 | No Change | Not Significant | TRUE |
| Tceal9 | TRUE | 0.183540381 | 0.351137093 | No Change | Not Significant | TRUE | 0.055495501 | 0.666254386 | No Change | Not Significant | TRUE |
| Tceanc | TRUE | -0.098448193 | 0.752577549 | No Change | Not Significant | TRUE | -0.063653016 | 0.768361739 | No Change | Not Significant | TRUE |
| Tceanc2 | TRUE | -0.206523212 | 0.076654886 | No Change | Not Significant | TRUE | -0.034944652 | 0.788968011 | No Change | Not Significant | TRUE |
| Tcerg1 | TRUE | 0.423452835 | 1.71E-05 | No Change | Significant Positive | TRUE | -0.056319211 | 0.638789048 | No Change | Not Significant | TRUE |
| Tcf12 | TRUE | 0.082617352 | 0.261665898 | No Change | Not Significant | TRUE | -0.091068378 | 0.27082801 | No Change | Not Significant | TRUE |
| Tcf19 | TRUE | 0.424011127 | 0.130987801 | No Change | Not Significant | TRUE | 0.288814124 | 0.256750769 | No Change | Not Significant | TRUE |
| Tcf20 | TRUE | -0.186628557 | 2.65E-05 | No Change | Significant Negative | TRUE | -0.084254903 | 0.099626908 | No Change | Not Significant | TRUE |
| Tcf23 | TRUE | 0.207715901 | 0.484040442 | No Change | Not Significant | TRUE | 0.054567551 | 0.845418668 | No Change | Not Significant | TRUE |
| Tcf24 | TRUE | 0.890246541 | 0.062098357 | No Change | Not Significant | TRUE | 0.307117744 | 0.447821073 | No Change | Not Significant | TRUE |
| Tcf25 | TRUE | 0.067110102 | 0.134681337 | No Change | Not Significant | TRUE | -0.035937236 | 0.546174109 | No Change | Not Significant | TRUE |
| Tcf3 | TRUE | 0.53868721 | 3.96E-10 | UP | UP | TRUE | 0.073795555 | 0.548046454 | No Change | Not Significant | TRUE |
| Tcf4 | TRUE | 0.135397243 | 0.331843045 | No Change | Not Significant | TRUE | -0.021517507 | 0.889388602 | No Change | Not Significant | TRUE |
| Tcf7 | TRUE | -0.546414731 | 0.060155346 | No Change | Not Significant | TRUE | -0.194403195 | 0.463022048 | No Change | Not Significant | TRUE |
| Tcf7l1 | TRUE | -0.054929635 | 0.890125936 | No Change | Not Significant | TRUE | 0.180924821 | 0.500565425 | No Change | Not Significant | TRUE |
| Tcf7l2 | TRUE | 0.005450128 | 0.988983332 | No Change | Not Significant | TRUE | -0.145489161 | 0.245659762 | No Change | Not Significant | TRUE |
| Tchh | TRUE | 0.804877404 | 0.028073652 | UP | UP | TRUE | 0.572346629 | 0.061495716 | No Change | Not Significant | TRUE |
| Tchp | TRUE | 0.241984867 | 0.292089426 | No Change | Not Significant | TRUE | -0.159480492 | 0.512797437 | No Change | Not Significant | TRUE |
| Tcim | TRUE | -0.272622025 | 0.10016904 | No Change | Not Significant | TRUE | 0.136161642 | 0.082768143 | No Change | Not Significant | TRUE |
| Tcirg1 | TRUE | -0.556270107 | 0.002539175 | DOWN | DOWN | TRUE | -0.525663581 | 7.79E-13 | DOWN | DOWN | TRUE |
| Tcn2 | TRUE | -0.486957032 | 1.52E-07 | No Change | Significant Negative | TRUE | 0.016998026 | 0.880695457 | No Change | Not Significant | TRUE |

Supplementary Table 3

| geneID | Cond1_DF_Detected | Cond1_DF_ValueCol | Cond1_DF_PadjCol | Cond1_DF_Cutoff | Cond1_DF_Cutoff_Specific.x | Cond2_DF_Detected |
|---|---|---|---|---|---|---|
| Tcof1 | TRUE | 0.34861853 | 0.007701301 | No Change | Significant Positive | TRUE |
| Tcp1 | TRUE | 0.029301821 | 0.891264005 | No Change | Not Significant | TRUE |
| Tcp10a | TRUE | -0.428252751 | 1 | No Change | Not Significant | FALSE |
| Tcp10b | TRUE | -0.649266125 | 1 | No Change | Not Significant | FALSE |
| Tcp10c | TRUE | 1.183851457 | 1 | No Change | Not Significant | FALSE |
| Tcp11 | TRUE | -0.567563051 | 0.578423233 | No Change | Not Significant | TRUE |
| Tcp11l1 | TRUE | 0.224839263 | 0.852489762 | No Change | Not Significant | FALSE |
| Tcp11l2 | TRUE | -0.359002722 | 0.207979566 | No Change | Not Significant | TRUE |
| Tcp11x2 | TRUE | -0.843032531 | 1 | No Change | Not Significant | FALSE |
| Tcta | TRUE | 0.033813907 | 0.838124245 | No Change | Not Significant | TRUE |
| Tcte1 | TRUE | -1.884324805 | 1 | No Change | Not Significant | FALSE |
| Tcte2 | TRUE | -0.008933736 | 0.98994327 | No Change | Not Significant | TRUE |
| Tctn1 | TRUE | -0.116310211 | 0.725980089 | No Change | Not Significant | TRUE |
| Tctn3 | TRUE | 0.0441477 | 0.886362513 | No Change | Not Significant | TRUE |
| Tdg | TRUE | -0.0863714 | 0.622737045 | No Change | Not Significant | TRUE |
| Tdh | TRUE | -0.777251667 | 1 | No Change | Not Significant | FALSE |
| Tdo2 | TRUE | -1.583198028 | 0.008633007 | DOWN | DOWN | TRUE |
| Tdp1 | TRUE | 0.192686022 | 0.451758428 | No Change | Not Significant | TRUE |
| Tdp2 | TRUE | -0.061161461 | 0.672740084 | No Change | Not Significant | TRUE |
| Tdpoz9 | TRUE | 0.073729475 | 1 | No Change | Not Significant | FALSE |
| Tdrd1 | TRUE | -0.722718153 | 0.021981059 | DOWN | DOWN | TRUE |
| Tdrd12 | TRUE | 0.057948541 | 0.931908663 | No Change | Not Significant | TRUE |
| Tdrd3 | TRUE | 0.010356888 | 0.93820896 | No Change | Not Significant | TRUE |
| Tdrd5 | TRUE | -0.725452455 | 1 | No Change | Not Significant | FALSE |
| Tdrd6 | TRUE | 0.604606687 | 1 | No Change | Not Significant | FALSE |
| Tdrd7 | TRUE | -0.000857793 | 0.994433839 | No Change | Not Significant | TRUE |
| Tdrd9 | TRUE | 1.719050937 | 1 | No Change | Not Significant | FALSE |
| Tdrkh | TRUE | 1.126280298 | 0.086352522 | No Change | Not Significant | FALSE |
| Tdrp | TRUE | -0.9455264 | 0.351529573 | No Change | Not Significant | FALSE |
| Tead1 | TRUE | -0.138488099 | 0.646395058 | No Change | Not Significant | TRUE |
| Tead2 | TRUE | 0.141407164 | 0.65700846 | No Change | Not Significant | TRUE |
| Tead3 | TRUE | 0.069282088 | 0.7765798 | No Change | Not Significant | TRUE |
| Tead4 | TRUE | 0.196920407 | 0.432209358 | No Change | Not Significant | TRUE |
| Tec | TRUE | -0.448841359 | 9.80E-05 | No Change | Significant Negative | TRUE |
| Tecpr1 | TRUE | 0.437856356 | 1.29E-12 | No Change | Significant Positive | TRUE |
| Tecpr2 | TRUE | -0.148921648 | 0.151293715 | No Change | Not Significant | TRUE |
| Tecr | TRUE | 0.114090558 | 0.185177119 | No Change | Not Significant | TRUE |
| Tecta | TRUE | -0.325928564 | 1 | No Change | Not Significant | FALSE |
| Tectb | TRUE | 0.589064346 | 1 | No Change | Not Significant | FALSE |
| Tedc1 | TRUE | 0.585293518 | 0.001633842 | UP | UP | TRUE |
| Tedc2 | TRUE | -0.146147613 | 0.577906578 | No Change | Not Significant | TRUE |
| Teddm1a | TRUE | 0.462645232 | 1 | No Change | Not Significant | FALSE |
| Teddm2 | TRUE | 0.504208949 | 0.354149765 | No Change | Not Significant | TRUE |
| Tef | TRUE | -1.075105086 | 0.095014843 | No Change | Not Significant | TRUE |
| Tefm | TRUE | -0.13297915 | 0.60223179 | No Change | Not Significant | TRUE |
| Tek | TRUE | -1.02794468 | 0.141461059 | No Change | Not Significant | TRUE |
| Tekt1 | TRUE | -1.650667186 | 1 | No Change | Not Significant | FALSE |
| Tekt2 | TRUE | 1.255356072 | 1 | No Change | Not Significant | FALSE |
| Tekt3 | TRUE | -0.046119424 | 1 | No Change | Not Significant | FALSE |
| Tekt4 | TRUE | 1.463829371 | 1 | No Change | Not Significant | FALSE |
| Tekt5 | TRUE | -0.873051686 | 1 | No Change | Not Significant | FALSE |
| Tektip1 | TRUE | 0.028781523 | 0.970587024 | No Change | Not Significant | TRUE |
| Telo2 | TRUE | 0.196286987 | 0.30108314 | No Change | Not Significant | TRUE |
| Ten1 | TRUE | 0.0466692 | 0.824189967 | No Change | Not Significant | TRUE |
| Tenm1 | TRUE | -0.22214601 | 1 | No Change | Not Significant | FALSE |
| Tenm2 | TRUE | -0.921013558 | 1 | No Change | Not Significant | FALSE |
| Tenm3 | TRUE | -0.188518534 | 0.927743965 | No Change | Not Significant | FALSE |
| Tenm4 | TRUE | -0.033034674 | 0.915935963 | No Change | Not Significant | TRUE |
| Tent2 | TRUE | 0.061380938 | 0.710428845 | No Change | Not Significant | TRUE |
| Tent4a | TRUE | 0.290473902 | 0.000506309 | No Change | Significant Positive | TRUE |
| Tent4b | TRUE | -0.039332791 | 0.736796542 | No Change | Not Significant | TRUE |
| Tent5a | TRUE | -0.325252827 | 0.003584189 | No Change | Significant Negative | TRUE |
| Tent5b | TRUE | 1.744553054 | 1.34E-11 | UP | UP | TRUE |
| Tent5c | TRUE | -0.134603939 | 0.791152348 | No Change | Not Significant | TRUE |
| Tent5d | TRUE | 1.149881521 | 1 | No Change | Not Significant | FALSE |
| Tep1 | TRUE | -0.215507074 | 0.011900366 | No Change | Significant Negative | TRUE |
| Tepp | TRUE | 1.713294283 | 1 | No Change | Not Significant | FALSE |
| Tepsin | TRUE | 0.306980166 | 0.003403673 | No Change | Significant Positive | TRUE |
| Terb1 | TRUE | 0.005879139 | 1 | No Change | Not Significant | FALSE |
| Terf1 | TRUE | 0.348717967 | 0.08100219 | No Change | Not Significant | TRUE |
| Terf2 | TRUE | 0.033437942 | 0.837421924 | No Change | Not Significant | TRUE |
| Terf2ip | TRUE | 0.03838255 | 0.865628713 | No Change | Not Significant | TRUE |
| Tert | TRUE | 0.255003196 | 0.360508479 | No Change | Not Significant | TRUE |
| Tes | TRUE | 0.929949386 | 5.86E-71 | UP | UP | TRUE |
| Tesc | TRUE | 0.640951883 | 0.063163803 | No Change | Not Significant | TRUE |
| Tesk1 | TRUE | -0.019287735 | 0.923798341 | No Change | Not Significant | TRUE |
| Tesk2 | TRUE | 0.06073987 | 0.804696828 | No Change | Not Significant | TRUE |
| Tesmin | TRUE | 0.782094265 | 0.359143126 | No Change | Not Significant | TRUE |
| Tespa1 | TRUE | 0.16204845 | 1 | No Change | Not Significant | TRUE |
| Tet1 | TRUE | -0.512342663 | 0.131848679 | No Change | Not Significant | TRUE |
| Tet2 | TRUE | -0.220848987 | 0.201270862 | No Change | Not Significant | TRUE |
| Tet3 | TRUE | -0.102941168 | 0.580790426 | No Change | Not Significant | TRUE |
| Tex10 | TRUE | -0.155329035 | 0.352113109 | No Change | Not Significant | TRUE |
| Tex11 | TRUE | 0.20506807 | 1 | No Change | Not Significant | FALSE |
| Tex12 | TRUE | -0.192424504 | 0.753386941 | No Change | Not Significant | TRUE |
| Tex13a | TRUE | -1.691445467 | 1 | No Change | Not Significant | FALSE |
| Tex13b | TRUE | 1.682932149 | 1 | No Change | Not Significant | FALSE |
| Tex14 | TRUE | 1.730554086 | 0.059503296 | No Change | Not Significant | TRUE |
| Tex15 | TRUE | 1.271517038 | 0.184012672 | No Change | Not Significant | TRUE |
| Tex19.1 | TRUE | 0.048201861 | 1 | No Change | Not Significant | FALSE |
| Tex2 | TRUE | -0.019691637 | 0.881383982 | No Change | Not Significant | TRUE |
| Tex22 | TRUE | 0.165917784 | 1 | No Change | Not Significant | FALSE |
| Tex26 | TRUE | -2.00420288 | 1 | No Change | Not Significant | FALSE |
| Tex261 | TRUE | -0.101597236 | 0.289400593 | No Change | Not Significant | TRUE |
| Tex264 | TRUE | -0.443043409 | 2.09E-10 | No Change | Significant Negative | TRUE |
| Tex29 | TRUE | 0.327326124 | 1 | No Change | Not Significant | FALSE |
| Tex30 | TRUE | 0.356877088 | 0.116818048 | No Change | Not Significant | TRUE |
| Tex33 | TRUE | -0.194459857 | 1 | No Change | Not Significant | FALSE |
| Tex38 | TRUE | 1.977302665 | 1 | No Change | Not Significant | FALSE |
| Tex45 | TRUE | 2.517929105 | 1 | No Change | Not Significant | FALSE |
| Tex48 | TRUE | 0.793401284 | 1 | No Change | Not Significant | FALSE |
| Tex49 | TRUE | -0.239267745 | 0.593868753 | No Change | Not Significant | TRUE |
| Tex51 | TRUE | 0.604891481 | 1 | No Change | Not Significant | FALSE |
| Tex53 | TRUE | 2.389306901 | 1 | No Change | Not Significant | FALSE |
| Tex54 | TRUE | 0.490433924 | 1 | No Change | Not Significant | FALSE |
| Tex56 | TRUE | -0.960358877 | 1 | No Change | Not Significant | FALSE |
| Tex9 | TRUE | 0.128461956 | 0.698215786 | No Change | Not Significant | TRUE |
| Tfam | TRUE | -0.102042716 | 0.456505337 | No Change | Not Significant | TRUE |
| Tfap2a | TRUE | -1.16027404 | 1 | No Change | Not Significant | FALSE |
| Tfap2b | TRUE | 0.376800034 | 1 | No Change | Not Significant | FALSE |
| Tfap2c | TRUE | -0.633207558 | 1 | No Change | Not Significant | FALSE |
| Tfap2e | TRUE | 2.322938793 | 1 | No Change | Not Significant | FALSE |
| Tfap4 | TRUE | -0.090814079 | 0.669812138 | No Change | Not Significant | TRUE |
| Tfb1m | TRUE | 0.543874835 | 0.001786002 | UP | UP | TRUE |
| Tfb2m | TRUE | -0.334314597 | 0.015750587 | No Change | Significant Negative | TRUE |
| Tfcp2 | TRUE | 0.664405729 | 2.62E-11 | UP | UP | TRUE |
| Tfcp2l1 | TRUE | -0.312557686 | 0.01240278 | No Change | Significant Negative | TRUE |
| Tfdp1 | TRUE | 0.214222661 | 0.207790664 | No Change | Not Significant | TRUE |
| Tfdp2 | TRUE | 0.205773809 | 0.128287829 | No Change | Not Significant | TRUE |
| Tfe3 | TRUE | -0.000688296 | 0.995778769 | No Change | Not Significant | TRUE |
| Tfeb | TRUE | -0.258696578 | 0.098824478 | No Change | Not Significant | TRUE |
| Tfec | TRUE | 0.456878508 | 1 | No Change | Not Significant | FALSE |
| Tff2 | TRUE | 0.016538551 | 0.988831409 | No Change | Not Significant | TRUE |
| Tff3 | TRUE | -0.133194113 | 0.825194181 | No Change | Not Significant | TRUE |
| Tfg | TRUE | 1.395751208 | 2.12E-91 | UP | UP | TRUE |
| Tfip11 | TRUE | 0.590470796 | 8.09E-20 | UP | UP | TRUE |
| Tfpi | TRUE | 0.05021247 | 0.910297568 | No Change | Not Significant | TRUE |
| Tfpi2 | TRUE | 0.845948693 | 2.57E-11 | UP | UP | TRUE |
| Tfpt | TRUE | -0.066398877 | 0.704656555 | No Change | Not Significant | TRUE |
| Tfr2 | TRUE | -0.367345285 | 1 | No Change | Not Significant | FALSE |
| Tfrc | TRUE | 0.363321044 | 0.181269925 | No Change | Not Significant | TRUE |
| Tg | TRUE | -2.830101356 | 1 | No Change | Not Significant | FALSE |
| Tgds | TRUE | 0.272027242 | 0.043873796 | No Change | Significant Positive | TRUE |
| Tgfa | TRUE | -0.176390707 | 0.088763622 | No Change | Not Significant | TRUE |
| Tgfb1 | TRUE | -1.249659055 | 0.024649528 | DOWN | DOWN | TRUE |
| Tgfb1i1 | TRUE | -0.150128869 | 0.940676072 | No Change | Not Significant | FALSE |
| Tgfb2 | TRUE | 0.07988608 | 1 | No Change | Not Significant | FALSE |
| Tgfb3 | TRUE | 0.622333711 | 0.501395265 | No Change | Not Significant | FALSE |
| Tgfbi | TRUE | -0.925286928 | 0.060682336 | No Change | Not Significant | TRUE |
| Tgfbr1 | TRUE | 0.299333795 | 0.021806863 | No Change | Significant Positive | TRUE |
| Tgfbr2 | TRUE | -0.090217474 | 0.606155692 | No Change | Not Significant | TRUE |
| Tgfbr3 | TRUE | 1.443911404 | 1 | No Change | Not Significant | FALSE |
| Tgfbr3l | TRUE | -0.515704215 | 0.502997128 | No Change | Not Significant | TRUE |
| Tgfbrap1 | TRUE | -0.041259513 | 0.749146509 | No Change | Not Significant | TRUE |
| Tgif1 | TRUE | -0.263247305 | 0.038706422 | No Change | Significant Negative | TRUE |
| Tgif2 | TRUE | -0.268092956 | 0.776544903 | No Change | Not Significant | TRUE |
| Tgm1 | TRUE | -0.611037511 | 1 | No Change | Not Significant | FALSE |
| Tgm2 | TRUE | 1.145795117 | 1.22E-08 | UP | UP | TRUE |
| Tgm3 | TRUE | -1.161717891 | 0.003708381 | DOWN | DOWN | TRUE |
| Tgm4 | TRUE | -0.035782645 | 0.966352899 | No Change | Not Significant | TRUE |
| Tgm5 | TRUE | 0.585748457 | 0.652783039 | No Change | Not Significant | TRUE |
| Tgm6 | TRUE | 0.501207938 | 1 | No Change | Not Significant | FALSE |
| Tgm7 | TRUE | 0.32771691 | 0.825194181 | No Change | Not Significant | TRUE |
| Tgoln1 | TRUE | -0.460470368 | 8.37E-05 | No Change | Significant Negative | TRUE |
| Tgs1 | TRUE | 0.001434609 | 0.994190961 | No Change | Not Significant | TRUE |
| Tgtp1 | TRUE | 0.505678027 | 1 | No Change | Not Significant | FALSE |
| Tgtp2 | TRUE | 0.307740669 | 0.784402584 | No Change | Not Significant | TRUE |
| Th | TRUE | -0.340409452 | 1 | No Change | Not Significant | FALSE |
| Tha1 | TRUE | -0.593429597 | 9.04E-06 | DOWN | DOWN | TRUE |
| Thada | TRUE | 0.087949953 | 0.683786987 | No Change | Not Significant | TRUE |
| Thap1 | TRUE | 0.225046941 | 0.176210159 | No Change | Not Significant | TRUE |
| Thap11 | TRUE | -0.093899262 | 0.547073822 | No Change | Not Significant | TRUE |
| Thap12 | TRUE | -0.00373743 | 0.984868386 | No Change | Not Significant | TRUE |
| Thap2 | TRUE | 0.104995771 | 0.630279329 | No Change | Not Significant | TRUE |
| Thap3 | TRUE | 2.738998392 | 1.44E-87 | UP | UP | TRUE |
| Thap4 | TRUE | -0.25381308 | 0.001515379 | No Change | Significant Negative | TRUE |
| Thap7 | TRUE | 0.020734925 | 0.91001573 | No Change | Not Significant | TRUE |
| Thbd | TRUE | -1.016596215 | 1 | No Change | Not Significant | TRUE |
| Thbs1 | TRUE | -0.755254636 | 0.008870554 | DOWN | DOWN | TRUE |
| Thbs2 | TRUE | 0.530563254 | 1 | No Change | Not Significant | FALSE |
| Thbs3 | TRUE | -0.487785561 | 0.352762472 | No Change | Not Significant | TRUE |
| Thbs4 | TRUE | -0.328465371 | 0.350538094 | No Change | Not Significant | TRUE |
| Theg | TRUE | 0.006367339 | 1 | No Change | Not Significant | FALSE |
| Thegl | TRUE | 3.112721026 | 1 | No Change | Not Significant | FALSE |
| Them4 | TRUE | -0.211301966 | 0.261594876 | No Change | Not Significant | TRUE |
| Them5 | TRUE | 1.656179979 | 1 | No Change | Not Significant | FALSE |
| Them6 | TRUE | 0.136504276 | 0.78655252 | No Change | Not Significant | TRUE |
| Them7 | TRUE | 2.572121689 | 1 | No Change | Not Significant | FALSE |
| Themis | TRUE | 0.481545056 | 1 | No Change | Not Significant | TRUE |
| Themis2 | TRUE | 0.539902022 | 1 | No Change | Not Significant | FALSE |
| Themis3 | TRUE | -0.058034106 | 0.79831751 | No Change | Not Significant | TRUE |
| Thg1l | TRUE | 0.079029534 | 0.779964063 | No Change | Not Significant | TRUE |
| Thnsl1 | TRUE | -0.316326043 | 0.027548077 | No Change | Significant Negative | TRUE |
| Thnsl2 | TRUE | 0.101218754 | 0.664930223 | No Change | Not Significant | TRUE |
| Thoc1 | TRUE | 0.134555609 | 0.337697097 | No Change | Not Significant | TRUE |
| Thoc2 | TRUE | 0.091772245 | 0.343609993 | No Change | Not Significant | TRUE |
| Thoc2l | TRUE | -0.13299648 | 0.601975182 | No Change | Not Significant | TRUE |
| Thoc3 | TRUE | 0.002567986 | 0.990427908 | No Change | Not Significant | TRUE |
| Thoc5 | TRUE | -0.087321949 | 0.611719756 | No Change | Not Significant | TRUE |
| Thoc6 | TRUE | 0.158395807 | 0.50453867 | No Change | Not Significant | TRUE |
| Thoc7 | TRUE | -0.196636839 | 0.024436932 | No Change | Significant Negative | TRUE |
| Thop1 | TRUE | 1.884272595 | 3.48E-89 | UP | UP | TRUE |

[illegible]

Supplementary Table 3

| geneID | Cond1_DF_Detected | Cond1_DF_ValueCol | Cond1_DF_PadjCol | Cond1_DF_Cutoff | Cond1_DF_Cutoff_Specific.x | Cond2_DF_Detected | Cond2_DF_ValueCol | Cond2_DF_PadjCol | Cond2_DF_Cutoff | Cond2_DF_Cutoff_Specific | BG_Method | RG1_All | RG2_Significant | RG3_SignificantChange | Cond1_DF_ensembl_gene_id | Cond1_DF_entrezgene_id | Cond1_DF_gene_biotype | Cond1_DF_baseMean |
|---|---|---|---|---|---|---|---|---|---|---|---|---|---|---|---|---|---|---|
| Thpo | TRUE | 0.194673451 | 0.735271124 | No Change | Not Significant | TRUE | 0.114336476 | 0.885066476 | No Change | Not Significant | TRUE | Cond1 Not Significant + Cond2 Not Significant | None | None | ENSMUSG00000022847 | 21832 | protein_coding | 13.50957582 |
| Thra | TRUE | -0.407737595 | 0.001828517 | No Change | Significant Negative | TRUE | -0.165356252 | 0.059625892 | No Change | Not Significant | TRUE | Cond1 Significant Negative + Cond2 Not Significant | None | None | ENSMUSG00000058756 | 21833 | protein_coding | 528.8281767 |
| Thrap3 | TRUE | 0.004776233 | 0.977538039 | No Change | Not Significant | TRUE | 0.067602351 | 0.271210133 | No Change | Not Significant | TRUE | Cond1 Not Significant + Cond2 Not Significant | None | None | ENSMUSG00000043962 | 230753 | protein_coding | 1391.323634 |
| Thrb | TRUE | -0.116073158 | 0.22586956 | No Change | Not Significant | TRUE | -0.214225834 | 0.007859568 | No Change | Significant Negative | TRUE | Cond1 Not Significant + Cond2 Significant Negative | None | None | ENSMUSG00000021779 | 21834 | protein_coding | 421.219062 |
| Thrsp | TRUE | -0.087365452 | 1 | No Change | Not Significant | FALSE | NA | NA | No Change | Not Detected | FALSE | Background = FALSE | Background = FALSE | Background = FALSE | ENSMUSG00000035686 | 21835 | protein_coding | 1.204472807 |
| Thsd1 | TRUE | 0.286961766 | 1 | No Change | Not Significant | FALSE | NA | NA | No Change | Not Detected | FALSE | Background = FALSE | Background = FALSE | Background = FALSE | ENSMUSG00000031480 | 56229 | protein_coding | 3.374912589 |
| Thsd4 | TRUE | -0.420332611 | 0.025839819 | No Change | Significant Negative | TRUE | -0.473974632 | 2.43E-07 | No Change | Significant Negative | TRUE | Cond1 Significant Negative + Cond2 Significant Negative | None | None | ENSMUSG00000032289 | 207596 | protein_coding | 414.198872 |
| Thsd7a | TRUE | -1.318107703 | 0.476857637 | No Change | Not Significant | TRUE | -0.883563819 | 0.192705082 | No Change | Not Significant | TRUE | Cond1 Not Significant + Cond2 Not Significant | None | None | ENSMUSG00000032625 | 330267 | protein_coding | 5.362482199 |
| Thsd7b | TRUE | 0.506436241 | 1 | No Change | Not Significant | FALSE | NA | NA | No Change | Not Detected | FALSE | Background = FALSE | Background = FALSE | Background = FALSE | ENSMUSG00000042581 | 210417 | protein_coding | 2.741922176 |
| Thtpa | TRUE | 0.016628122 | 0.947544143 | No Change | Not Significant | TRUE | 0.093398292 | 0.609949562 | No Change | Not Significant | TRUE | Cond1 Not Significant + Cond2 Not Significant | None | None | ENSMUSG00000045691 | 105663 | protein_coding | 92.86672598 |
| Thumpd1 | TRUE | 0.04717634 | 0.828120806 | No Change | Not Significant | TRUE | -0.060891914 | 0.5061553 | No Change | Not Significant | TRUE | Cond1 Not Significant + Cond2 Not Significant | None | None | ENSMUSG00000030942 | 233802 | protein_coding | 347.9742012 |
| Thumpd2 | TRUE | -0.179831394 | 0.622191478 | No Change | Not Significant | TRUE | -0.249189769 | 0.303863905 | No Change | Not Significant | TRUE | Cond1 Not Significant + Cond2 Not Significant | None | None | ENSMUSG00000024246 | 72167 | protein_coding | 33.75852895 |
| Thumpd3 | TRUE | -0.114112816 | 0.301146282 | No Change | Not Significant | TRUE | -0.087434703 | 0.477271565 | No Change | Not Significant | TRUE | Cond1 Not Significant + Cond2 Not Significant | None | None | ENSMUSG00000030264 | 14911 | protein_coding | 611.7116579 |
| [illegible] | [illegible] | [illegible] | [illegible] | [illegible] | [illegible] | [illegible] | [illegible] | [illegible] | [illegible] | [illegible] | [illegible] | [illegible] | [illegible] | [illegible] | [illegible] | [illegible] | [illegible] | [illegible] |

| geneID | Cond1_DF_ log2FoldChange | Cond1_DF_lfcSE | Cond1_DF_stat | Cond1_DF_pvalue | Cond1_DF_ padj | Cond1_DF_absLog2FC | Cond1_DF_Cutoff_Specific.y | Cond2_DF_ensembl_gene_id | Cond2_DF_entrezgene_id | Cond2_DF_gene_biotype | Cond2_DF_baseMean | Cond2_DF_ log2FoldChange | Cond2_DF_lfcSE | Cond2_DF_stat | Cond2_DF_pvalue | Cond2_DF_ padj | Cond2_DF_absLog2FC | RG1_Specific_Cond2 | RG1_Specific_Cond1 |
|---|---|---|---|---|---|---|---|---|---|---|---|---|---|---|---|---|---|---|---|
| Thpo | 0.194673451 | 0.318677514 | -0.610879157 | 0.541279584 | 0.735271124 | 0.194673451 | Not Significant | ENSMUSG00000022847 | 21832 | protein_coding | 11.13436587 | 0.114336476 | 0.370078861 | 0.308951653 | 0.757358304 | 0.885066476 | 0.114336476 | Cond1 No Change + Cond2 Not Significant | Cond2 No Change + Cond1 Not Significant |
| Thra | -0.407737595 | 0.110869363 | 3.677639923 | 0.000235402 | 0.001828517 | 0.407737595 | Significant Negative | ENSMUSG00000058756 | 21833 | protein_coding | 1237.285819 | -0.165356252 | 0.064270355 | -2.57282306 | 0.010087276 | 0.059625892 | 0.165356252 | Cond1 No Change + Cond2 Not Significant | Cond2 No Change + Cond1 Significant Negative |
| Thrap3 | 0.004776233 | 0.084724529 | -0.056373675 | 0.955044128 | 0.977538039 | 0.004776233 | Not Significant | ENSMUSG00000043962 | 230753 | protein_coding | 2608.140387 | 0.067602351 | 0.03997233 | 1.691228701 | 0.090793131 | 0.271210133 | 0.067602351 | Cond1 No Change + Cond2 Not Significant | Cond2 No Change + Cond1 Not Significant |
| Thrb | -0.116073158 | 0.068133653 | 1.703609794 | 0.088454011 | 0.22586956 | 0.116073158 | Not Significant | ENSMUSG00000021779 | 21834 | protein_coding | 884.0975818 | -0.214225834 | 0.063489273 | -3.374205167 | 0.000740292 | 0.007859568 | 0.214225834 | Cond1 No Change + Cond2 Significant Negative | Cond2 No Change + Cond1 Not Significant |
| Thrsp | -0.087365452 | 2.016817188 | 0.043318479 | 0.965447661 | 1 | 0.087365452 | Not Significant | NA | NA | NA | NA | NA | NA | NA | NA | NA | NA | Background = FALSE | Background = FALSE |
| Thsd1 | 0.286961766 | 0.853984513 | -0.336026897 | 0.736850584 | 1 | 0.286961766 | Not Significant | NA | NA | NA | NA | NA | NA | NA | NA | NA | NA | Background = FALSE | Background = FALSE |
| Thsd4 | -0.420332611 | 0.150984236 | 2.783950317 | 0.005370126 | 0.025839819 | 0.420332611 | Significant Negative | ENSMUSG00000032289 | 207596 | protein_coding | 614.1251644 | -0.473974632 | 0.08174575 | -5.798156225 | 6.70E-09 | 2.43E-07 | 0.473974632 | Cond1 No Change + Cond2 Significant Negative | Cond2 No Change + Cond1 Significant Negative |
| Thsd7a | -1.318107703 | 1.176952443 | 1.11993285 | 0.262742378 | 0.476857637 | 1.318107703 | Not Significant | ENSMUSG00000032625 | 330267 | protein_coding | 10.86352259 | -0.883563819 | 0.456936989 | -1.933666653 | 0.053154126 | 0.192705082 | 0.883563819 | Cond1 No Change + Cond2 Not Significant | Cond2 No Change + Cond1 Not Significant |
| Thsd7b | 0.506436241 | 2.376770433 | -0.213077474 | 0.831266534 | 1 | 0.506436241 | Not Significant | NA | NA | NA | NA | NA | NA | NA | NA | NA | NA | Background = FALSE | Background = FALSE |
| Thtpa | 0.016628122 | 0.124197443 | -0.133884576 | 0.893493847 | 0.947544143 | 0.016628122 | Not Significant | ENSMUSG00000045691 | 105663 | protein_coding | 186.0464166 | 0.093398292 | 0.103644375 | 0.901141932 | 0.36751286 | 0.609949562 | 0.093398292 | Cond1 No Change + Cond2 Not Significant | Cond2 No Change + Cond1 Not Significant |
| Thumpd1 | 0.04717634 | 0.114165543 | -0.41322748 | 0.679439956 | 0.828120806 | 0.04717634 | Not Significant | ENSMUSG00000030942 | 233802 | protein_coding | 658.742652 | -0.060891914 | 0.054580733 | -1.115630185 | 0.264580466 | 0.5061553 | 0.060891914 | Cond1 No Change + Cond2 Not Significant | Cond2 No Change + Cond1 Not Significant |
| Thumpd2 | -0.179831394 | 0.213678278 | 0.841598855 | 0.40001253 | 0.622191478 | 0.179831394 | Not Significant | ENSMUSG00000024246 | 72167 | protein_coding | 66.30931518 | -0.249189769 | 0.155828444 | -1.59912891 | 0.109791962 | 0.303863905 | 0.249189769 | Cond1 No Change + Cond2 Not Significant | Cond2 No Change + Cond1 Not Significant |
| Thumpd3 | -0.114112816 | 0.075757161 | 1.506297407 | 0.131990843 | 0.301146282 | 0.114112816 | Not Significant | ENSMUSG00000030264 | 14911 | protein_coding | 1093.257982 | -0.087434703 | 0.07399182 | -1.181680663 | 0.237332434 | 0.477271565 | 0.087434703 | Cond1 No Change + Cond2 Not Significant | Cond2 No Change + Cond1 Not Significant |
| [illegible] | [illegible] | [illegible] | [illegible] | [illegible] | [illegible] | [illegible] | [illegible] | [illegible] | [illegible] | [illegible] | [illegible] | [illegible] | [illegible] | [illegible] | [illegible] | [illegible] | [illegible] | [illegible] | [illegible] |

Supplementary Table 3

| geneID | Cond1_DF_Detected | Cond1_DF_ValueCol | Cond1_DF_PadjCol | Cond1_DF_Cutoff | Cond1_DF_Cutoff_Specific.x | Cond2_DF_Detected | Cond2_DF_ValueCol |
|---|---|---|---|---|---|---|---|
| Tmem139 | TRUE | -0.28112956 | 0.156906987 | No Change | Not Significant | TRUE | 0.450880647 |
| Tmem140 | TRUE | -0.319803166 | 0.648871201 | No Change | Not Significant | TRUE | -0.178980469 |
| Tmem141 | TRUE | -0.269155431 | 0.054614131 | No Change | Not Significant | TRUE | 0.080268541 |
| Tmem143 | TRUE | -0.197885238 | 0.193614755 | No Change | Not Significant | TRUE | 0.070819056 |
| Tmem144 | TRUE | -0.084641179 | 0.778302948 | No Change | Not Significant | TRUE | -0.269354211 |
| Tmem145 | TRUE | 1.981393479 | 1 | No Change | Not Significant | FALSE | NA |
| Tmem147 | TRUE | -0.219638218 | 0.143734593 | No Change | Not Significant | TRUE | -0.082832164 |
| Tmem14a | TRUE | -0.417110704 | 0.22572726 | No Change | Not Significant | TRUE | 0.060489897 |
| Tmem14c | TRUE | -0.121589799 | 0.370076028 | No Change | Not Significant | TRUE | 0.156749483 |
| Tmem150a | TRUE | -0.42212035 | 0.612813278 | No Change | Not Significant | TRUE | -0.326953854 |
| Tmem150b | TRUE | -0.103410617 | 0.624336681 | No Change | Not Significant | TRUE | -0.028212076 |
| Tmem150c | TRUE | 0.597471716 | 1 | No Change | Not Significant | FALSE | NA |
| Tmem151a | TRUE | 0.417005729 | 0.394331932 | No Change | Not Significant | TRUE | 0.211695972 |
| Tmem151b | TRUE | 0.974568529 | 1 | No Change | Not Significant | FALSE | NA |
| Tmem154 | TRUE | 0.80414709 | 0.154444113 | No Change | Not Significant | TRUE | -0.164802483 |
| Tmem156 | TRUE | -1.335395988 | 1 | No Change | Not Significant | FALSE | NA |
| Tmem158 | TRUE | 1.403701197 | 0.004936679 | UP | UP | TRUE | 0.728002725 |
| Tmem159 | TRUE | -0.082121767 | 0.765104392 | No Change | Not Significant | TRUE | 0.025761491 |
| Tmem160 | TRUE | -0.207931635 | 0.260075004 | No Change | Not Significant | TRUE | -0.121840693 |
| Tmem161a | TRUE | -0.101295018 | 0.385300595 | No Change | Not Significant | TRUE | 0.049012021 |
| Tmem161b | TRUE | 0.306867181 | 0.003214427 | No Change | Significant Positive | TRUE | 0.151858162 |
| Tmem163 | TRUE | -1.228395303 | 0.131303585 | No Change | Not Significant | TRUE | -0.105740633 |
| Tmem164 | TRUE | -0.175497181 | 0.108093077 | No Change | Not Significant | TRUE | -0.077103314 |
| Tmem165 | TRUE | 0.745192944 | 7.57E-12 | UP | UP | TRUE | 0.193131788 |
| [illegible] | [illegible] | [illegible] | [illegible] | [illegible] | [illegible] | [illegible] | [illegible] |

[illegible]

| geneID | Cond1_DF_Detected | Cond1_DF_ValueCol | Cond1_DF_PadjCol | Cond1_DF_Cutoff | Cond1_DF_Cutoff_Specific.x | Cond2_DF_Detected | [illegible] |
|---|---|---|---|---|---|---|---|
| Tmod4 | TRUE | -0.369326391 | 0.695624496 | No Change | Not Significant | TRUE | [illegible] |
| Tmpo | TRUE | 0.453574005 | 0.007482994 | No Change | Significant Positive | TRUE | [illegible] |
| Tmppe | TRUE | -0.266479309 | 0.067952726 | No Change | Not Significant | TRUE | [illegible] |
| Tmprss11a | TRUE | 2.161332341 | 1 | No Change | Not Significant | FALSE | [illegible] |
| Tmprss11b | TRUE | 0.176024601 | 1 | No Change | Not Significant | FALSE | [illegible] |
| Tmprss11d | TRUE | 0.845470933 | 1 | No Change | Not Significant | FALSE | [illegible] |
| Tmprss11e | TRUE | 1.652429091 | 1 | No Change | Not Significant | FALSE | [illegible] |
| Tmprss11g | TRUE | 2.457270557 | 1 | No Change | Not Significant | FALSE | [illegible] |
| Tmprss12 | TRUE | 0.094644394 | 1 | No Change | Not Significant | FALSE | [illegible] |
| Tmprss13 | TRUE | -0.655587529 | 0.255970313 | No Change | Not Significant | TRUE | [illegible] |
| Tmprss15 | TRUE | 2.584731255 | 1 | No Change | Not Significant | FALSE | [illegible] |
| Tmprss2 | TRUE | -0.279479413 | 0.184932165 | No Change | Not Significant | TRUE | [illegible] |
| Tmprss3 | TRUE | -0.75327634 | 1 | No Change | Not Significant | FALSE | [illegible] |
| Tmprss4 | TRUE | 0.259533107 | 2.03E-06 | No Change | Significant Positive | TRUE | [illegible] |
| Tmprss5 | TRUE | 1.293268614 | 1 | No Change | Not Significant | FALSE | [illegible] |
| Tmprss6 | TRUE | 0.115431761 | 0.937028043 | No Change | Not Significant | FALSE | [illegible] |
| Tmprss7 | TRUE | 0.40487886 | 0.373498571 | No Change | Not Significant | TRUE | [illegible] |
| Tmprss9 | TRUE | -0.041978167 | 1 | No Change | Not Significant | FALSE | [illegible] |
| Tmsb10 | TRUE | 0.026449728 | 0.890072723 | No Change | Not Significant | TRUE | [illegible] |
| Tmsb15b1 | TRUE | 1.993279538 | 0.004397156 | UP | UP | TRUE | [illegible] |
| Tmsb15b2 | TRUE | 1.539293299 | 1 | No Change | Not Significant | FALSE | [illegible] |
| Tmsb15l | TRUE | 1.625101251 | 1 | No Change | Not Significant | FALSE | [illegible] |
| Tmsb4x | TRUE | -0.119621311 | 0.258193749 | No Change | Not Significant | TRUE | [illegible] |
| Tmtc1 | TRUE | 0.783941547 | 0.453835428 | No Change | Not Significant | TRUE | [illegible] |
| Tmtc2 | TRUE | -0.11629577 | 0.361449656 | No Change | Not Significant | TRUE | [illegible] |
| Tmtc3 | TRUE | 1.137726409 | 8.99E-20 | UP | UP | TRUE | [illegible] |
| Tmtc4 | TRUE | 0.091282593 | 0.562302999 | No Change | Not Significant | TRUE | [illegible] |
| Tmub1 | TRUE | 0.004962895 | 0.97354273 | No Change | Not Significant | TRUE | [illegible] |
| Tmub2 | TRUE | 0.075294381 | 0.33115426 | No Change | Not Significant | TRUE | [illegible] |
| Tmx1 | TRUE | 0.208391728 | 0.214493541 | No Change | Not Significant | TRUE | [illegible] |
| Tmx2 | TRUE | 0.02255415 | 0.913317606 | No Change | Not Significant | TRUE | [illegible] |
| Tmx3 | TRUE | -0.014736298 | 0.937394946 | No Change | Not Significant | TRUE | [illegible] |
| Tmx4 | TRUE | 0.266527235 | 0.322533618 | No Change | Not Significant | TRUE | [illegible] |
| Tnc | TRUE | 0.566471238 | 0.67278137 | No Change | Not Significant | FALSE | [illegible] |
| Tnf | TRUE | 0.36948065 | 0.601768559 | No Change | Not Significant | TRUE | [illegible] |
| Tnfaip1 | TRUE | 0.069120306 | 0.580211292 | No Change | Not Significant | TRUE | [illegible] |
| Tnfaip2 | TRUE | 1.634960263 | 0.028888694 | UP | UP | TRUE | [illegible] |
| Tnfaip3 | TRUE | -0.11044017 | 0.795976331 | No Change | Not Significant | TRUE | [illegible] |
| Tnfaip8 | TRUE | -0.171705298 | 0.580790426 | No Change | Not Significant | TRUE | [illegible] |
| Tnfaip8l1 | TRUE | 0.114711712 | 0.564082441 | No Change | Not Significant | TRUE | [illegible] |
| Tnfaip8l2 | TRUE | 0.823804893 | 1 | No Change | Not Significant | FALSE | [illegible] |
| Tnfaip8l3 | TRUE | 0.573664545 | 1 | No Change | Not Significant | FALSE | [illegible] |
| Tnfrsf10b | TRUE | -0.391308826 | 0.097419765 | No Change | Not Significant | TRUE | [illegible] |
| Tnfrsf11a | TRUE | 0.030256876 | 0.884245212 | No Change | Not Significant | TRUE | [illegible] |
| Tnfrsf11b | TRUE | 0.668129811 | 0.190353587 | No Change | Not Significant | TRUE | [illegible] |
| Tnfrsf12a | TRUE | 0.081905157 | 0.826426838 | No Change | Not Significant | TRUE | [illegible] |
| Tnfrsf13b | TRUE | 0.418415659 | 1 | No Change | Not Significant | FALSE | [illegible] |
| Tnfrsf13c | TRUE | 2.285709147 | 1 | No Change | Not Significant | FALSE | [illegible] |
| Tnfrsf14 | TRUE | -0.347184124 | 0.037639301 | No Change | Significant Negative | TRUE | [illegible] |
| Tnfrsf18 | TRUE | 0.334393219 | 0.296685693 | No Change | Not Significant | TRUE | [illegible] |
| Tnfrsf19 | TRUE | 0.154040064 | 0.734530441 | No Change | Not Significant | TRUE | [illegible] |
| Tnfrsf1a | TRUE | 0.281813301 | 0.011855051 | No Change | Significant Positive | TRUE | [illegible] |
| Tnfrsf1b | TRUE | -0.143042196 | 0.525029915 | No Change | Not Significant | TRUE | [illegible] |
| Tnfrsf21 | TRUE | -0.199497964 | 0.073733764 | No Change | Not Significant | TRUE | [illegible] |
| Tnfrsf22 | TRUE | -0.119498811 | 0.93035707 | No Change | Not Significant | TRUE | [illegible] |
| Tnfrsf23 | TRUE | -0.598609813 | 0.175858569 | No Change | Not Significant | TRUE | [illegible] |
| Tnfrsf25 | TRUE | -0.66810726 | 0.341264466 | No Change | Not Significant | TRUE | [illegible] |
| Tnfrsf26 | TRUE | -0.149297217 | 1 | No Change | Not Significant | FALSE | [illegible] |
| Tnfrsf4 | TRUE | 0.041252848 | 0.970559847 | No Change | Not Significant | TRUE | [illegible] |
| Tnfrsf8 | TRUE | 1.220317078 | 0.01100288 | UP | UP | TRUE | [illegible] |
| Tnfrsf9 | TRUE | 0.638700319 | 0.532495959 | No Change | Not Significant | FALSE | [illegible] |
| Tnfsf10 | TRUE | -0.218770409 | 0.36023903 | No Change | Not Significant | TRUE | [illegible] |
| Tnfsf12 | TRUE | -0.373626051 | 0.724255209 | No Change | Not Significant | TRUE | [illegible] |
| Tnfsf13 | TRUE | 0.056916211 | 0.794593769 | No Change | Not Significant | TRUE | [illegible] |
| Tnfsf13b | TRUE | 0.023278894 | 0.954867376 | No Change | Not Significant | TRUE | [illegible] |
| Tnfsf14 | TRUE | 0.426423076 | 1 | No Change | Not Significant | FALSE | [illegible] |
| Tnfsf15 | TRUE | -1.009490378 | 0.000845894 | DOWN | DOWN | TRUE | [illegible] |
| Tnfsf8 | TRUE | -0.798742015 | 1 | No Change | Not Significant | FALSE | [illegible] |
| Tnfsf9 | TRUE | -0.200619626 | 1 | No Change | Not Significant | TRUE | [illegible] |
| Tnfsfm13 | TRUE | 0.051049192 | 0.937028043 | No Change | Not Significant | TRUE | [illegible] |
| Tnik | TRUE | -0.387879843 | 0.000926263 | No Change | Significant Negative | TRUE | [illegible] |
| Tnip1 | TRUE | -0.347907761 | 0.045254367 | No Change | Significant Negative | TRUE | [illegible] |
| Tnip2 | TRUE | 0.298459575 | 0.015925715 | No Change | Significant Positive | TRUE | [illegible] |
| Tnip3 | TRUE | 0.49133867 | 0.130180251 | No Change | Not Significant | TRUE | [illegible] |
| Tnk1 | TRUE | -0.10767586 | 0.470794255 | No Change | Not Significant | TRUE | [illegible] |
| Tnk2 | TRUE | -0.004916283 | 0.972316965 | No Change | Not Significant | TRUE | [illegible] |
| Tnk2os | TRUE | -0.621755725 | 1 | No Change | Not Significant | FALSE | [illegible] |
| Tnks | TRUE | 0.053615906 | 0.709931167 | No Change | Not Significant | TRUE | [illegible] |
| Tnks1bp1 | TRUE | -0.252188643 | 0.001134417 | No Change | Significant Negative | TRUE | [illegible] |
| Tnks2 | TRUE | 0.125065092 | 0.239040298 | No Change | Not Significant | TRUE | [illegible] |
| Tnn | TRUE | 1.887874966 | 1 | No Change | Not Significant | FALSE | [illegible] |
| Tnnc1 | TRUE | -1.304404379 | 1 | No Change | Not Significant | FALSE | [illegible] |
| Tnni1 | TRUE | -0.704847077 | 0.012707239 | DOWN | DOWN | TRUE | [illegible] |
| Tnni2 | TRUE | 1.62776654 | 1 | No Change | Not Significant | FALSE | [illegible] |
| Tnni3 | TRUE | 0.678355474 | 1 | No Change | Not Significant | FALSE | [illegible] |
| Tnni3k | TRUE | 0.822403492 | 1 | No Change | Not Significant | FALSE | [illegible] |
| Tnnt1 | TRUE | -0.161913455 | 0.806814922 | No Change | Not Significant | TRUE | [illegible] |
| Tnnt2 | TRUE | -0.341055711 | 0.565422739 | No Change | Not Significant | TRUE | [illegible] |
| Tnnt3 | TRUE | 1.186097182 | 1 | No Change | Not Significant | FALSE | [illegible] |
| Tnpo1 | TRUE | 0.03866323 | 0.800822518 | No Change | Not Significant | TRUE | [illegible] |
| Tnpo2 | TRUE | 0.068527928 | 0.369037513 | No Change | Not Significant | TRUE | [illegible] |
| Tnpo3 | TRUE | 0.211917761 | 0.036011679 | No Change | Significant Positive | TRUE | [illegible] |
| Tnr | TRUE | -1.182667262 | 0.393348786 | No Change | Not Significant | TRUE | [illegible] |
| Tnrc18 | TRUE | 0.081135179 | 0.79763624 | No Change | Not Significant | TRUE | [illegible] |
| Tnrc6a | TRUE | -0.13370557 | 0.383265272 | No Change | Not Significant | TRUE | [illegible] |
| Tnrc6b | TRUE | 0.250983823 | 0.016594879 | No Change | Significant Positive | TRUE | [illegible] |
| Tnrc6c | TRUE | 0.033991502 | 0.799065123 | No Change | Not Significant | TRUE | [illegible] |
| Tns1 | TRUE | 0.675183528 | 0.727211667 | No Change | Not Significant | TRUE | [illegible] |
| Tns2 | TRUE | 1.773785721 | 4.85E-52 | UP | UP | TRUE | [illegible] |
| Tns3 | TRUE | -0.412545758 | 7.89E-08 | No Change | Significant Negative | TRUE | [illegible] |
| Tns4 | TRUE | -0.406723963 | 0.12652989 | No Change | Not Significant | TRUE | [illegible] |
| Tnxb | TRUE | 0.03872054 | 0.980697519 | No Change | Not Significant | TRUE | [illegible] |
| Tob1 | TRUE | -0.148443941 | 0.398335908 | No Change | Not Significant | TRUE | [illegible] |
| Tob2 | TRUE | -0.227833951 | 0.066534014 | No Change | Not Significant | TRUE | [illegible] |
| Toe1 | TRUE | 0.445860824 | 0.000244236 | No Change | Significant Positive | TRUE | [illegible] |
| Togaram1 | TRUE | -0.020675644 | 0.90234117 | No Change | Not Significant | TRUE | [illegible] |
| Togaram2 | TRUE | 3.787723339 | 1 | No Change | Not Significant | FALSE | [illegible] |
| Tollip | TRUE | -0.104375461 | 0.141461059 | No Change | Not Significant | TRUE | [illegible] |
| Tom1 | TRUE | -1.09325751 | 1.04E-13 | DOWN | DOWN | TRUE | [illegible] |
| Tom1l1 | TRUE | -0.198516232 | 0.009870396 | No Change | Significant Negative | TRUE | [illegible] |
| Tom1l2 | TRUE | 0.195770464 | 0.318770942 | No Change | Not Significant | TRUE | [illegible] |
| Tomm20 | TRUE | -0.367567977 | 0.014241781 | No Change | Significant Negative | TRUE | [illegible] |
| Tomm22 | TRUE | -0.161576683 | 0.194652614 | No Change | Not Significant | TRUE | [illegible] |
| Tomm34 | TRUE | -0.342639135 | 0.000412515 | No Change | Significant Negative | TRUE | [illegible] |
| Tomm40 | TRUE | 0.437179665 | 2.17E-05 | No Change | Significant Positive | TRUE | [illegible] |
| Tomm40l | TRUE | -0.009198506 | 0.970043107 | No Change | Not Significant | TRUE | [illegible] |
| Tomm5 | TRUE | -0.301605186 | 0.028162478 | No Change | Significant Negative | TRUE | [illegible] |
| Tomm6 | TRUE | 2.105559045 | 1 | No Change | Not Significant | FALSE | [illegible] |
| Tomm6os | TRUE | -0.108824458 | 0.919169532 | No Change | Not Significant | TRUE | [illegible] |
| Tomm7 | TRUE | -0.298167841 | 0.000236946 | No Change | Significant Negative | TRUE | [illegible] |
| Tomm70a | TRUE | 0.143057634 | 0.208654479 | No Change | Not Significant | TRUE | [illegible] |
| Tomt | TRUE | -0.269841857 | 0.781418386 | No Change | Not Significant | TRUE | [illegible] |
| Tonsl | TRUE | 0.292495762 | 0.05273746 | No Change | Not Significant | TRUE | [illegible] |
| Top1 | TRUE | 0.443452829 | 2.01E-12 | No Change | Significant Positive | TRUE | [illegible] |
| Top1mt | TRUE | -0.423332295 | 0.028080198 | No Change | Significant Negative | TRUE | [illegible] |
| Top2a | TRUE | 1.36788918 | 5.00E-11 | UP | UP | TRUE | [illegible] |
| Top2b | TRUE | -0.18782445 | 0.185483473 | No Change | Not Significant | TRUE | [illegible] |
| Top3a | TRUE | 0.070157103 | 0.694394109 | No Change | Not Significant | TRUE | [illegible] |
| Top3b | TRUE | 0.023567597 | 0.810197705 | No Change | Not Significant | TRUE | [illegible] |
| Topaz1 | TRUE | -1.923698605 | 1 | No Change | Not Significant | FALSE | [illegible] |
| Topbp1 | TRUE | 0.414152072 | 0.02338927 | No Change | Significant Positive | TRUE | [illegible] |
| Topors | TRUE | -0.195504927 | 0.007575843 | No Change | Significant Negative | TRUE | [illegible] |
| Tor1a | TRUE | 0.125522773 | 0.211442486 | No Change | Not Significant | TRUE | [illegible] |
| Tor1aip1 | TRUE | -0.112962648 | 0.448589299 | No Change | Not Significant | TRUE | [illegible] |
| Tor1aip2 | TRUE | 0.218441622 | 0.011213534 | No Change | Significant Positive | TRUE | [illegible] |
| Tor1b | TRUE | -0.10201792 | 0.301713259 | No Change | Not Significant | TRUE | [illegible] |
| Tor2a | TRUE | 0.017519695 | 0.918959391 | No Change | Not Significant | TRUE | [illegible] |
| Tor3a | TRUE | -0.38152458 | 0.005884119 | No Change | Significant Negative | TRUE | [illegible] |
| Tor4a | TRUE | 0.076863667 | 0.605068399 | No Change | Not Significant | TRUE | [illegible] |
| Tox | TRUE | -0.003330158 | 0.987811867 | No Change | Not Significant | TRUE | [illegible] |
| Tox2 | TRUE | 0.805782095 | 1 | No Change | Not Significant | FALSE | [illegible] |
| Tox3 | TRUE | -0.190359934 | 0.105418633 | No Change | Not Significant | TRUE | [illegible] |
| Tox4 | TRUE | 0.04097033 | 0.687930892 | No Change | Not Significant | TRUE | [illegible] |
| Tpbg | TRUE | -0.095879838 | 0.772328749 | No Change | Not Significant | TRUE | [illegible] |
| Tpbgl | TRUE | -0.434164969 | 1 | No Change | Not Significant | FALSE | [illegible] |
| Tpcn1 | TRUE | 0.130255164 | 0.104013109 | No Change | Not Significant | TRUE | [illegible] |
| Tpcn2 | TRUE | 0.336971253 | 0.083204714 | No Change | Not Significant | TRUE | [illegible] |
| Tpd52 | TRUE | 0.172925983 | 0.212151185 | No Change | Not Significant | TRUE | [illegible] |
| Tpd52l1 | TRUE | 0.413547683 | 0.101319853 | No Change | Not Significant | TRUE | [illegible] |
| Tpd52l2 | TRUE | -0.008082954 | 0.95132126 | No Change | Not Significant | TRUE | [illegible] |
| Tpgs1 | TRUE | -0.070851502 | 0.785464687 | No Change | Not Significant | TRUE | [illegible] |
| Tpgs2 | TRUE | 0.058820115 | 0.861504353 | No Change | Not Significant | TRUE | [illegible] |
| Tph1 | TRUE | -1.372544829 | 3.34E-05 | DOWN | DOWN | TRUE | [illegible] |
| Tph2 | TRUE | 3.508829846 | 1 | No Change | Not Significant | FALSE | [illegible] |
| Tpi1 | TRUE | -0.144261699 | 0.050511589 | No Change | Not Significant | TRUE | [illegible] |
| Tpk1 | TRUE | -0.143236148 | 0.15967971 | No Change | Not Significant | TRUE | [illegible] |
| Tpm1 | TRUE | -0.31878597 | 8.30E-06 | No Change | Significant Negative | TRUE | [illegible] |
| Tpm2 | TRUE | -0.216059168 | 0.730731431 | No Change | Not Significant | TRUE | [illegible] |
| Tpm3 | TRUE | 0.173027899 | 0.181579513 | No Change | Not Significant | TRUE | [illegible] |
| Tpm4 | TRUE | -0.239012743 | 0.125611281 | No Change | Not Significant | TRUE | [illegible] |
| Tpmt | TRUE | -0.451275875 | 3.10E-05 | No Change | Significant Negative | TRUE | [illegible] |
| Tpo | TRUE | 1.278355093 | 1 | No Change | Not Significant | FALSE | [illegible] |
| Tpp1 | TRUE | -0.437643545 | 1.34E-09 | No Change | Significant Negative | TRUE | [illegible] |
| Tpp2 | TRUE | 0.868185121 | 3.14E-48 | UP | UP | TRUE | [illegible] |
| Tppp | TRUE | -0.987291325 | 8.69E-16 | DOWN | DOWN | TRUE | [illegible] |
| Tppp3 | TRUE | -1.23506993 | 8.94E-11 | DOWN | DOWN | TRUE | [illegible] |
| Tpr | TRUE | -0.051009976 | 0.762991261 | No Change | Not Significant | TRUE | [illegible] |
| Tpra1 | TRUE | -0.409121075 | 2.62E-11 | No Change | Significant Negative | TRUE | [illegible] |
| Tprg | TRUE | 0.720976388 | 1 | No Change | Not Significant | FALSE | [illegible] |
| Tprgl | TRUE | -0.325689518 | 3.47E-05 | No Change | Significant Negative | TRUE | [illegible] |
| Tprkb | TRUE | -0.260202647 | 0.043322217 | No Change | Significant Negative | TRUE | [illegible] |
| Tprn | TRUE | 0.246818892 | 0.107730217 | No Change | Not Significant | TRUE | [illegible] |
| Tpsb2 | TRUE | 0.369668109 | 1 | No Change | Not Significant | FALSE | [illegible] |
| Tpsg1 | TRUE | -0.64927211 | 0.032592144 | DOWN | DOWN | TRUE | [illegible] |
| Tpst1 | TRUE | 2.511594419 | 4.50E-53 | UP | UP | TRUE | [illegible] |
| Tpst2 | TRUE | 0.351725731 | 0.017328863 | No Change | Significant Positive | TRUE | [illegible] |
| Tpt1 | TRUE | -0.425796484 | 0.000129591 | No Change | Significant Negative | TRUE | [illegible] |
| Tpte | TRUE | -1.196166348 | 1 | No Change | Not Significant | FALSE | [illegible] |
| Tpx2 | TRUE | 0.80776252 | 4.48E-06 | UP | UP | TRUE | [illegible] |
| Tra2a | TRUE | 0.224106756 | 0.002893353 | No Change | Significant Positive | TRUE | [illegible] |
| Tra2b | TRUE | 0.012803723 | 0.940676072 | No Change | Not Significant | TRUE | [illegible] |
| Trabd | TRUE | 0.821830893 | 9.45E-29 | UP | UP | TRUE | [illegible] |
| Trabd2b | TRUE | -0.052010479 | 0.936825965 | No Change | Not Significant | TRUE | [illegible] |
| Tradd | TRUE | -0.144168096 | 0.228322063 | No Change | Not Significant | TRUE | [illegible] |
| Traf1 | TRUE | -1.50315082 | 7.83E-15 | DOWN | DOWN | TRUE | [illegible] |
| Traf2 | TRUE | 0.352063563 | 0.000850344 | No Change | Significant Positive | TRUE | [illegible] |
| Traf3 | TRUE | 0.024443397 | 0.910384401 | No Change | Not Significant | TRUE | [illegible] |
| Traf3ip1 | TRUE | 0.070594894 | 0.703146268 | No Change | Not Significant | TRUE | [illegible] |
| Traf3ip2 | TRUE | 0.0414376 | 0.784979105 | No Change | Not Significant | TRUE | [illegible] |
| Traf3ip3 | TRUE | 0.068284394 | 0.951341404 | No Change | Not Significant | TRUE | [illegible] |
| Traf4 | TRUE | -0.221346285 | 0.02696225 | No Change | Significant Negative | TRUE | [illegible] |
| Traf5 | TRUE | -0.197412391 | 0.532085855 | No Change | Not Significant | TRUE | [illegible] |

Supplementary Table 3

| geneID | Cond1_DF_Detected | Cond1_DF_ValueCol | Cond1_DF_PadjCol | Cond1_DF_Cutoff | Cond1_DF_Cutoff_Specific.x | Cond2_DF_Detected | Cond2_DF_ValueCol | Cond2_DF_PadjCol | Cond2_DF_Cutoff | Cond2_DF_Cutoff_Specific | BG_Method |
|---|---|---|---|---|---|---|---|---|---|---|---|
| Traf6 | TRUE | 0.142434844 | 0.196079582 | No Change | Not Significant | TRUE | -0.003366071 | 0.974755742 | No Change | Not Significant | TRUE |
| Traf7 | TRUE | 0.723122986 | 2.23E-28 | UP | UP | TRUE | 0.149317469 | 0.013485044 | No Change | Significant Positive | TRUE |
| Trafd1 | TRUE | -0.014039949 | 0.955778515 | No Change | Not Significant | TRUE | -0.305870102 | 0.015483388 | No Change | Significant Negative | TRUE |
| Traip | TRUE | 0.839908926 | 0.005762227 | UP | UP | TRUE | 0.279218923 | 0.302001032 | No Change | Not Significant | TRUE |
| Trak1 | TRUE | -0.3927503 | 0.00166012 | No Change | Significant Negative | TRUE | -0.078051279 | 0.453693043 | No Change | Not Significant | TRUE |
| Trak2 | TRUE | -0.134983299 | 0.365337006 | No Change | Not Significant | TRUE | -0.171324383 | 0.015372629 | No Change | Significant Negative | TRUE |
| Tram1 | TRUE | 0.761235499 | 6.66E-18 | UP | UP | TRUE | 0.480227942 | 2.81E-40 | No Change | Significant Positive | TRUE |
| Tram2 | TRUE | 0.307991391 | 0.551537722 | No Change | Not Significant | TRUE | -0.319395099 | 0.263676126 | No Change | Not Significant | TRUE |
| Trank1 | TRUE | 0.754303953 | 0.553734049 | No Change | Not Significant | TRUE | 0.571317181 | 0.389376941 | No Change | Not Significant | TRUE |
| Trap1 | TRUE | -0.283101573 | 0.008569101 | No Change | Significant Negative | TRUE | -0.032558206 | 0.749029274 | No Change | Not Significant | TRUE |
| Trappc1 | TRUE | 0.101828932 | 0.446443342 | No Change | Not Significant | TRUE | 0.095895319 | 0.331045348 | No Change | Not Significant | TRUE |
| Trappc10 | TRUE | 0.158524524 | 0.064265646 | No Change | Not Significant | TRUE | 0.057500295 | 0.326278724 | No Change | Not Significant | TRUE |
| Trappc11 | TRUE | 0.291088313 | 0.000195572 | No Change | Significant Positive | TRUE | 0.139622133 | 0.037252106 | No Change | Significant Positive | TRUE |
| Trappc12 | TRUE | -0.100933818 | 0.309427758 | No Change | Not Significant | TRUE | 0.053838987 | 0.531418201 | No Change | Not Significant | TRUE |
| Trappc13 | TRUE | 0.015675958 | 0.913026208 | No Change | Not Significant | TRUE | -0.058592987 | 0.576322837 | No Change | Not Significant | TRUE |
| Trappc14 | TRUE | -0.084047818 | 0.541897747 | No Change | Not Significant | TRUE | -0.102449211 | 0.428914823 | No Change | Not Significant | TRUE |
| Trappc2 | TRUE | -0.255306066 | 0.322533618 | No Change | Not Significant | TRUE | -0.111710274 | 0.740026398 | No Change | Not Significant | TRUE |
| Trappc2l | TRUE | -0.267710744 | 0.014029681 | No Change | Significant Negative | TRUE | -0.042270872 | 0.744499039 | No Change | Not Significant | TRUE |
| Trappc3 | TRUE | -0.081713428 | 0.381372755 | No Change | Not Significant | TRUE | 0.036351274 | 0.736439527 | No Change | Not Significant | TRUE |
| Trappc4 | TRUE | 0.010131836 | 0.947692153 | No Change | Not Significant | TRUE | 0.148932338 | 0.10995238 | No Change | Not Significant | TRUE |
| Trappc5 | TRUE | 0.041922706 | 0.760117839 | No Change | Not Significant | TRUE | 0.138068012 | 0.071196124 | No Change | Not Significant | TRUE |
| Trappc6a | TRUE | 0.151315055 | 0.224312683 | No Change | Not Significant | TRUE | 0.267211774 | 0.021439236 | No Change | Significant Positive | TRUE |
| Trappc6b | TRUE | 0.028892651 | 0.843587573 | No Change | Not Significant | TRUE | 0.063743324 | 0.502716586 | No Change | Not Significant | TRUE |
| Trappc8 | TRUE | 0.025556398 | 0.839060823 | No Change | Not Significant | TRUE | 0.016220829 | 0.856611363 | No Change | Not Significant | TRUE |
| Trappc9 | TRUE | -0.001342385 | 0.991234621 | No Change | Not Significant | TRUE | 0.096302333 | 0.356724723 | No Change | Not Significant | TRUE |
| Trarg1 | TRUE | -1.450210411 | 1 | No Change | Not Significant | FALSE | NA | NA | No Change | Not Detected | FALSE |
| Trat1 | TRUE | -0.444044077 | 1 | No Change | Not Significant | TRUE | 1.139422986 | 0.114590169 | No Change | Not Significant | TRUE |
| Trcg1 | TRUE | -0.377466739 | 1 | No Change | Not Significant | FALSE | NA | NA | No Change | Not Detected | FALSE |
| Trdmt1 | TRUE | -0.713854819 | 0.007010715 | DOWN | DOWN | TRUE | -0.446992154 | 0.00876128 | No Change | Significant Negative | TRUE |
| Trdn | TRUE | -0.236177031 | 1 | No Change | Not Significant | FALSE | NA | NA | No Change | Not Detected | FALSE |
| Treh | TRUE | 4.05579021 | 1.32E-43 | UP | UP | TRUE | 1.943586016 | 3.46E-13 | UP | UP | TRUE |
| Trem1 | TRUE | 0.555368883 | 1 | No Change | Not Significant | FALSE | NA | NA | No Change | Not Detected | FALSE |
| Trem2 | TRUE | 3.142978889 | 1 | No Change | Not Significant | FALSE | NA | NA | No Change | Not Detected | FALSE |
| Treml1 | TRUE | -0.861609348 | 1 | No Change | Not Significant | FALSE | NA | NA | No Change | Not Detected | FALSE |
| Treml2 | TRUE | 0.315226323 | 0.884582288 | No Change | Not Significant | FALSE | NA | NA | No Change | Not Detected | FALSE |
| Treml4 | TRUE | 0.57251034 | 1 | No Change | Not Significant | FALSE | NA | NA | No Change | Not Detected | FALSE |
| Trerf1 | TRUE | 0.529344127 | 0.735609498 | No Change | Not Significant | TRUE | 0.36421211 | 0.685928116 | No Change | Not Significant | TRUE |
| Trex2 | TRUE | -1.37509257 | 1 | No Change | Not Significant | FALSE | NA | NA | No Change | Not Detected | FALSE |
| Trf | TRUE | 1.266750057 | 0.042192071 | UP | UP | TRUE | 0.696389469 | 0.009325517 | UP | UP | TRUE |
| Trh | TRUE | 1.188326988 | 1 | No Change | Not Significant | FALSE | NA | NA | No Change | Not Detected | FALSE |
| Trhde | TRUE | 3.882054496 | 1 | No Change | Not Significant | FALSE | NA | NA | No Change | Not Detected | FALSE |
| Trhr | TRUE | -0.428634943 | 1 | No Change | Not Significant | FALSE | NA | NA | No Change | Not Detected | FALSE |
| Triap1 | TRUE | -0.233787832 | 0.064634295 | No Change | Not Significant | TRUE | 0.042561491 | 0.780129105 | No Change | Not Significant | TRUE |
| Trib1 | TRUE | 0.047563414 | 0.889721317 | No Change | Not Significant | TRUE | 0.227773127 | 0.000210953 | No Change | Significant Positive | TRUE |
| Trib2 | TRUE | 0.229835835 | 0.423711673 | No Change | Not Significant | TRUE | 0.226348963 | 0.380751845 | No Change | Not Significant | TRUE |
| Trib3 | TRUE | 5.837363353 | 3.01E-39 | UP | UP | TRUE | 3.339878378 | 8.07E-27 | UP | UP | TRUE |
| Tril | TRUE | 1.024883515 | 1 | No Change | Not Significant | FALSE | NA | NA | No Change | Not Detected | FALSE |
| Trim10 | TRUE | 1.714977473 | 1 | No Change | Not Significant | FALSE | NA | NA | No Change | Not Detected | FALSE |
| Trim11 | TRUE | -0.03636857 | 0.838339763 | No Change | Not Significant | TRUE | -0.077493812 | 0.426539942 | No Change | Not Significant | TRUE |
| Trim12a | TRUE | -0.382146648 | 0.068336322 | No Change | Not Significant | TRUE | -0.208127334 | 0.215277659 | No Change | Not Significant | TRUE |
| Trim12c | TRUE | -0.493748641 | 0.000848668 | No Change | Significant Negative | TRUE | -0.02106087 | 0.904736983 | No Change | Not Significant | TRUE |
| Trim13 | TRUE | -0.202054945 | 0.454822267 | No Change | Not Significant | TRUE | 0.146593744 | 0.492145102 | No Change | Not Significant | TRUE |
| Trim14 | TRUE | -0.2099016 | 0.058209168 | No Change | Not Significant | TRUE | 0.069682355 | 0.592096117 | No Change | Not Significant | TRUE |
| Trim15 | TRUE | 0.677927972 | 0.040476816 | UP | UP | TRUE | -0.056250141 | 0.781437157 | No Change | Not Significant | TRUE |
| Trim16 | TRUE | -0.150585904 | 0.271607716 | No Change | Not Significant | TRUE | -0.071092463 | 0.470931774 | No Change | Not Significant | TRUE |
| Trim17 | TRUE | 0.628256228 | 0.482609908 | No Change | Not Significant | TRUE | -0.240789649 | 0.790773109 | No Change | Not Significant | TRUE |
| Trim2 | TRUE | -0.215274243 | 0.025873521 | No Change | Significant Negative | TRUE | -0.017780001 | 0.771609416 | No Change | Not Significant | TRUE |
| Trim21 | TRUE | -0.19751841 | 0.178888658 | No Change | Not Significant | TRUE | -0.151461429 | 0.360385206 | No Change | Not Significant | TRUE |
| Trim23 | TRUE | 0.035948594 | 0.859565127 | No Change | Not Significant | TRUE | -0.150934981 | 0.366529497 | No Change | Not Significant | TRUE |
| Trim24 | TRUE | 0.329125603 | 1.51E-08 | No Change | Significant Positive | TRUE | 0.015641113 | 0.909183504 | No Change | Not Significant | TRUE |
| Trim25 | TRUE | -0.077291639 | 0.728048657 | No Change | Not Significant | TRUE | 0.04114853 | 0.525210203 | No Change | Not Significant | TRUE |
| Trim26 | TRUE | 0.079543119 | 0.445367775 | No Change | Not Significant | TRUE | -0.001008389 | 0.994488063 | No Change | Not Significant | TRUE |
| Trim27 | TRUE | 0.200130512 | 0.045254367 | No Change | Significant Positive | TRUE | 0.029252778 | 0.776873083 | No Change | Not Significant | TRUE |
| Trim28 | TRUE | 0.219274124 | 0.081579713 | No Change | Not Significant | TRUE | -0.017835447 | 0.855350218 | No Change | Not Significant | TRUE |
| Trim29 | TRUE | -0.488965175 | 0.466951438 | No Change | Not Significant | TRUE | -0.380132726 | 0.573356749 | No Change | Not Significant | TRUE |
| Trim3 | TRUE | -0.157559259 | 0.591274514 | No Change | Not Significant | TRUE | -0.06263372 | 0.752092411 | No Change | Not Significant | TRUE |
| Trim30a | TRUE | -0.139957002 | 0.856549354 | No Change | Not Significant | TRUE | -0.062722623 | 0.890725386 | No Change | Not Significant | TRUE |
| Trim30b | TRUE | 0.322614971 | 1 | No Change | Not Significant | FALSE | NA | NA | No Change | Not Detected | FALSE |
| Trim30d | TRUE | -0.494571577 | 0.019107412 | No Change | Significant Negative | TRUE | -0.296697509 | 0.120556652 | No Change | Not Significant | TRUE |
| Trim31 | TRUE | 0.493320339 | 1.76E-09 | No Change | Significant Positive | TRUE | 0.245201174 | 0.012784915 | No Change | Significant Positive | TRUE |
| Trim32 | TRUE | 0.585274623 | 1.16E-08 | UP | UP | TRUE | 0.414787574 | 4.05E-07 | No Change | Significant Positive | TRUE |
| Trim33 | TRUE | 0.011272042 | 0.914187552 | No Change | Not Significant | TRUE | -0.08456679 | 0.334558164 | No Change | Not Significant | TRUE |
| Trim34a | TRUE | -0.507121452 | 0.030092813 | DOWN | DOWN | TRUE | 0.030923916 | 0.892331737 | No Change | Not Significant | TRUE |
| Trim34b | TRUE | -0.552372834 | 0.44122879 | No Change | Not Significant | TRUE | 0.125173222 | 0.862409121 | No Change | Not Significant | TRUE |
| Trim35 | TRUE | 0.551869803 | 0.000348044 | UP | UP | TRUE | 0.349703978 | 0.003318073 | No Change | Significant Positive | TRUE |
| Trim36 | TRUE | -0.679962767 | 5.70E-09 | DOWN | DOWN | TRUE | -0.368700507 | 1.59E-07 | No Change | Significant Negative | TRUE |
| Trim37 | TRUE | 0.55058312 | 0.000536429 | UP | UP | TRUE | 0.191703496 | 0.170737422 | No Change | Not Significant | TRUE |
| Trim38 | TRUE | -1.458358196 | 4.63E-05 | DOWN | DOWN | TRUE | -0.010592717 | 0.984093538 | No Change | Not Significant | TRUE |
| Trim39 | TRUE | 0.116114806 | 0.495289271 | No Change | Not Significant | TRUE | -0.161066593 | 0.301043575 | No Change | Not Significant | TRUE |
| Trim40 | TRUE | 0.826453357 | 0.006936888 | UP | UP | TRUE | -0.169789862 | 0.474717773 | No Change | Not Significant | TRUE |
| Trim41 | TRUE | 0.047344629 | 0.644974953 | No Change | Not Significant | TRUE | 0.009827476 | 0.93615636 | No Change | Not Significant | TRUE |
| Trim43a | TRUE | 0.350361804 | 1 | No Change | Not Significant | FALSE | NA | NA | No Change | Not Detected | FALSE |
| Trim43c | TRUE | -0.708585903 | 1 | No Change | Not Significant | FALSE | NA | NA | No Change | Not Detected | FALSE |
| Trim44 | TRUE | -0.215903311 | 0.248746733 | No Change | Not Significant | TRUE | -0.090615189 | 0.353156981 | No Change | Not Significant | TRUE |
| Trim45 | TRUE | 0.33043343 | 0.154934252 | No Change | Not Significant | TRUE | 0.14642419 | 0.589027996 | No Change | Not Significant | TRUE |
| Trim46 | TRUE | -0.136527749 | 0.683156904 | No Change | Not Significant | TRUE | 0.218862487 | 0.247744036 | No Change | Not Significant | TRUE |
| Trim47 | TRUE | 0.53410827 | 0.106687692 | No Change | Not Significant | TRUE | 0.757602756 | 0.010257788 | UP | UP | TRUE |
| Trim5 | TRUE | -1.051174323 | 0.028250624 | DOWN | DOWN | TRUE | -0.624249455 | 0.087225321 | No Change | Not Significant | TRUE |
| Trim56 | TRUE | -0.063087466 | 0.594533905 | No Change | Not Significant | TRUE | 0.051033881 | 0.699728324 | No Change | Not Significant | TRUE |
| Trim58 | TRUE | 0.609434461 | 1 | No Change | Not Significant | FALSE | NA | NA | No Change | Not Detected | FALSE |
| Trim59 | TRUE | 0.707844549 | 1.22E-06 | UP | UP | TRUE | 0.087572995 | 0.649970343 | No Change | Not Significant | TRUE |
| Trim6 | TRUE | -0.298646566 | 0.101663614 | No Change | Not Significant | TRUE | 0.233125862 | 0.417073819 | No Change | Not Significant | TRUE |
| Trim62 | TRUE | -0.261775564 | 0.398975857 | No Change | Not Significant | TRUE | 0.0152605 | 0.959446759 | No Change | Not Significant | TRUE |
| Trim65 | TRUE | -0.033056967 | 0.88672609 | No Change | Not Significant | TRUE | 0.051901708 | 0.780422622 | No Change | Not Significant | TRUE |
| Trim66 | TRUE | -0.412327019 | 0.890758364 | No Change | Not Significant | FALSE | NA | NA | No Change | Not Detected | FALSE |
| Trim67 | TRUE | -0.088143735 | 1 | No Change | Not Significant | FALSE | NA | NA | No Change | Not Detected | FALSE |
| Trim68 | TRUE | 1.145814899 | 1.94E-22 | UP | UP | TRUE | 0.761147438 | 2.74E-16 | UP | UP | TRUE |
| Trim69 | TRUE | 1.482180746 | 1 | No Change | Not Significant | FALSE | NA | NA | No Change | Not Detected | FALSE |
| Trim7 | TRUE | -0.264021182 | 0.542303804 | No Change | Not Significant | TRUE | -0.302711129 | 0.252369884 | No Change | Not Significant | TRUE |
| Trim71 | TRUE | 0.919146257 | 0.359891361 | No Change | Not Significant | FALSE | NA | NA | No Change | Not Detected | FALSE |
| Trim72 | TRUE | 0.068452154 | 0.856795141 | No Change | Not Significant | TRUE | 0.537502518 | 0.000896291 | UP | UP | TRUE |
| Trim8 | TRUE | -0.056436427 | 0.746166522 | No Change | Not Significant | TRUE | 0.022753312 | 0.910278712 | No Change | Not Significant | TRUE |
| Trim80 | TRUE | 0.047217835 | 1 | No Change | Not Significant | FALSE | NA | NA | No Change | Not Detected | FALSE |
| Trim9 | TRUE | -0.603726686 | 0.529343875 | No Change | Not Significant | TRUE | -0.224953916 | 0.768546546 | No Change | Not Significant | TRUE |
| Triml1 | TRUE | -0.734087227 | 1 | No Change | Not Significant | FALSE | NA | NA | No Change | Not Detected | FALSE |
| Trio | TRUE | 0.255867475 | 0.286211349 | No Change | Not Significant | TRUE | -0.075438624 | 0.37913729 | No Change | Not Significant | TRUE |
| Triobp | TRUE | 0.10994406 | 0.08105111 | No Change | Not Significant | TRUE | 0.078620071 | 0.515483555 | No Change | Not Significant | TRUE |
| Trip10 | TRUE | -0.404301997 | 0.001385242 | No Change | Significant Negative | TRUE | -0.154594699 | 0.088092517 | No Change | Not Significant | TRUE |
| Trip11 | TRUE | 0.129547622 | 0.188061933 | No Change | Not Significant | TRUE | 0.05949174 | 0.433436829 | No Change | Not Significant | TRUE |
| Trip12 | TRUE | -0.041606352 | 0.637283772 | No Change | Not Significant | TRUE | -0.056884653 | 0.272007465 | No Change | Not Significant | TRUE |
| Trip13 | TRUE | 0.796745829 | 8.50E-05 | UP | UP | TRUE | 0.204165841 | 0.375078854 | No Change | Not Significant | TRUE |
| Trip4 | TRUE | 0.223567771 | 0.081931007 | No Change | Not Significant | TRUE | 0.168513004 | 0.048515603 | No Change | Significant Positive | TRUE |
| Trip6 | TRUE | 0.148747681 | 0.49283969 | No Change | Not Significant | TRUE | -0.609077054 | 3.97E-06 | DOWN | DOWN | TRUE |
| Triqk | TRUE | 0.147957296 | 0.617453792 | No Change | Not Significant | TRUE | -0.004748623 | 0.98474512 | No Change | Not Significant | TRUE |
| Trir | TRUE | 0.036561438 | 0.80301103 | No Change | Not Significant | TRUE | 0.05144631 | 0.727325341 | No Change | Not Significant | TRUE |
| Trit1 | TRUE | 0.010574981 | 0.967739018 | No Change | Not Significant | TRUE | -0.066214958 | 0.657926757 | No Change | Not Significant | TRUE |
| Trmo | TRUE | -0.136730783 | 0.640714813 | No Change | Not Significant | TRUE | 0.002705286 | 0.992886774 | No Change | Not Significant | TRUE |
| Trmt1 | TRUE | -0.289310082 | 0.009099247 | No Change | Significant Negative | TRUE | -0.159345155 | 0.016779727 | No Change | Significant Negative | TRUE |
| Trmt10a | TRUE | 0.568862593 | 0.010614744 | UP | UP | TRUE | 0.05775902 | 0.84654428 | No Change | Not Significant | TRUE |
| Trmt10b | TRUE | 0.044939354 | 0.825096143 | No Change | Not Significant | TRUE | -0.001073061 | 0.996369812 | No Change | Not Significant | TRUE |
| Trmt10c | TRUE | -0.202917557 | 0.199909758 | No Change | Not Significant | TRUE | -0.109165936 | 0.350881819 | No Change | Not Significant | TRUE |
| Trmt11 | TRUE | 0.079152254 | 0.661197851 | No Change | Not Significant | TRUE | 0.107234283 | 0.438066569 | No Change | Not Significant | TRUE |
| Trmt112 | TRUE | -0.078981785 | 0.728500002 | No Change | Not Significant | TRUE | -0.032178092 | 0.862386247 | No Change | Not Significant | TRUE |
| Trmt12 | TRUE | -0.029994987 | 0.924722861 | No Change | Not Significant | TRUE | 0.120548429 | 0.533617367 | No Change | Not Significant | TRUE |
| Trmt13 | TRUE | 0.052608674 | 0.856549354 | No Change | Not Significant | TRUE | -0.221479859 | 0.420211186 | No Change | Not Significant | TRUE |
| Trmt1l | TRUE | -0.017523958 | 0.881386379 | No Change | Not Significant | TRUE | -0.032115563 | 0.838143427 | No Change | Not Significant | TRUE |
| Trmt2a | TRUE | 0.176715969 | 0.106039571 | No Change | Not Significant | TRUE | 0.005989835 | 0.967733155 | No Change | Not Significant | TRUE |
| Trmt2b | TRUE | -0.112843963 | 0.295101944 | No Change | Not Significant | TRUE | 0.011967114 | 0.928042853 | No Change | Not Significant | TRUE |
| Trmt44 | TRUE | 0.015258087 | 0.956997404 | No Change | Not Significant | TRUE | -0.028065558 | 0.916752358 | No Change | Not Significant | TRUE |
| Trmt5 | TRUE | 0.028382547 | 0.917959327 | No Change | Not Significant | TRUE | -0.033884836 | 0.87299831 | No Change | Not Significant | TRUE |
| Trmt6 | TRUE | 0.557458166 | 2.93E-07 | UP | UP | TRUE | 0.04225769 | 0.811381567 | No Change | Not Significant | TRUE |
| Trmt61a | TRUE | -0.022143405 | 0.95630475 | No Change | Not Significant | TRUE | -0.081253918 | 0.742878353 | No Change | Not Significant | TRUE |
| Trmt61b | TRUE | 0.094988042 | 0.686659782 | No Change | Not Significant | TRUE | 0.028531102 | 0.914754428 | No Change | Not Significant | TRUE |
| Trmt9b | TRUE | -0.86052575 | 0.062193652 | No Change | Not Significant | TRUE | -0.415300402 | 0.20352728 | No Change | Not Significant | TRUE |
| Trmu | TRUE | -0.183610443 | 0.214101896 | No Change | Not Significant | TRUE | 0.004587323 | 0.983146794 | No Change | Not Significant | TRUE |
| Trnau1ap | TRUE | 0.148677489 | 0.26786833 | No Change | Not Significant | TRUE | -0.007540297 | 0.955338492 | No Change | Not Significant | TRUE |
| Trnp1 | TRUE | 1.025049646 | 1 | No Change | Not Significant | FALSE | NA | NA | No Change | Not Detected | FALSE |
| Trnt1 | TRUE | 0.026489297 | 0.897079532 | No Change | Not Significant | TRUE | -0.181523428 | 0.137199451 | No Change | Not Significant | TRUE |
| Tro | TRUE | -0.408967407 | 1 | No Change | Not Significant | FALSE | NA | NA | No Change | Not Detected | FALSE |
| Troap | TRUE | 0.798095327 | 0.00077231 | UP | UP | TRUE | 0.072029071 | 0.831366151 | No Change | Not Significant | TRUE |
| Trp53 | TRUE | -0.074556251 | 0.62645577 | No Change | Not Significant | TRUE | -0.02155678 | 0.904781532 | No Change | Not Significant | TRUE |
| Trp53bp1 | TRUE | 0.089979206 | 0.50901339 | No Change | Not Significant | TRUE | -0.019735828 | 0.87479925 | No Change | Not Significant | TRUE |
| Trp53bp2 | TRUE | -0.127366449 | 0.068347858 | No Change | Not Significant | TRUE | 0.065203456 | 0.504916665 | No Change | Not Significant | TRUE |
| Trp53cor1 | TRUE | 0.934282916 | 1 | No Change | Not Significant | FALSE | NA | NA | No Change | Not Detected | FALSE |
| Trp53i11 | TRUE | 0.28242601 | 0.071374882 | No Change | Not Significant | TRUE | 0.180856585 | 0.147231287 | No Change | Not Significant | TRUE |
| Trp53i13 | TRUE | -0.16445423 | 0.625349723 | No Change | Not Significant | TRUE | -0.10972103 | 0.669842781 | No Change | Not Significant | TRUE |
| Trp53inp1 | TRUE | -0.441026455 | 0.013886585 | No Change | Significant Negative | TRUE | -0.091238493 | 0.586900547 | No Change | Not Significant | TRUE |
| Trp53inp2 | TRUE | 0.011569362 | 0.968697092 | No Change | Not Significant | TRUE | 0.153434009 | 0.107552177 | No Change | Not Significant | TRUE |
| Trp53rka | TRUE | 0.010928475 | 0.959151705 | No Change | Not Significant | TRUE | -0.056996691 | 0.784048864 | No Change | Not Significant | TRUE |
| Trp53rkb | TRUE | 0.018993881 | 0.961855464 | No Change | Not Significant | TRUE | -0.088840622 | 0.758094025 | No Change | Not Significant | TRUE |
| Trp53tg5 | TRUE | 0.91613651 | 1 | No Change | Not Significant | FALSE | NA | NA | No Change | Not Detected | FALSE |
| Trp63 | TRUE | 1.573767554 | 1 | No Change | Not Significant | FALSE | NA | NA | No Change | Not Detected | FALSE |
| Trp73 | TRUE | -0.913422601 | 1 | No Change | Not Significant | FALSE | NA | NA | No Change | Not Detected | FALSE |
| Trpc1 | TRUE | 0.011053501 | 0.987905762 | No Change | Not Significant | TRUE | -0.069792734 | 0.90924234 | No Change | Not Significant | TRUE |
| Trpc3 | TRUE | 0.824005884 | 1 | No Change | Not Significant | FALSE | NA | NA | No Change | Not Detected | FALSE |
| Trpc4 | TRUE | 0.056526052 | 1 | No Change | Not Significant | FALSE | NA | NA | No Change | Not Detected | FALSE |
| Trpc4ap | TRUE | 0.103950863 | 0.184376923 | No Change | Not Significant | TRUE | 0.04757323 | 0.511930778 | No Change | Not Significant | TRUE |
| Trpc5 | TRUE | -1.052590348 | 1 | No Change | Not Significant | FALSE | NA | NA | No Change | Not Detected | FALSE |
| Trpc6 | TRUE | -1.899902658 | 1 | No Change | Not Significant | FALSE | NA | NA | No Change | Not Detected | FALSE |
| Trpc7 | TRUE | 1.061401138 | 1 | No Change | Not Significant | FALSE | NA | NA | No Change | Not Detected | FALSE |
| Trpm1 | TRUE | 0.364507488 | 0.913297021 | No Change | Not Significant | FALSE | NA | NA | No Change | Not Detected | FALSE |
| Trpm2 | TRUE | 0.792160215 | 0.661290522 | No Change | Not Significant | TRUE | -0.076723433 | 0.927834878 | No Change | Not Significant | TRUE |
| Trpm3 | TRUE | 0.011209403 | 0.998060837 | No Change | Not Significant | FALSE | NA | NA | No Change | Not Detected | FALSE |
| Trpm4 | TRUE | -0.118248885 | 0.557080699 | No Change | Not Significant | TRUE | -0.091790749 | 0.553733449 | No Change | Not Significant | TRUE |
| Trpm5 | TRUE | -0.773240639 | 3.78E-05 | DOWN | DOWN | TRUE | 0.011326344 | 0.954463189 | No Change | Not Significant | TRUE |
| Trpm6 | TRUE | 0.563489158 | 0.036254161 | UP | UP | TRUE | -0.007436124 | 0.970931898 | No Change | Not Significant | TRUE |
| Trpm7 | TRUE | -0.015165258 | 0.931474719 | No Change | Not Significant | TRUE | -0.130283993 | 0.107171866 | No Change | Not Significant | TRUE |
| Trpm8 | TRUE | -0.921916708 | 1 | No Change | Not Significant | FALSE | NA | NA | No Change | Not Detected | FALSE |
| Trps1 | TRUE | 0.362109116 | 1 | No Change | Not Significant | FALSE | NA | NA | No Change | Not Detected | FALSE |
| Trpt1 | TRUE | -0.192234852 | 0.3224445 | No Change | Not Significant | TRUE | -0.096635145 | 0.644152928 | No Change | Not Significant | TRUE |
| Trpv1 | TRUE | -0.459110636 | 0.559233675 | No Change | Not Significant | TRUE | -0.155536873 | 0.84094949 | No Change | Not Significant | TRUE |
| Trpv2 | TRUE | 0.124246983 | 0.946057182 | No Change | Not Significant | FALSE | NA | NA | No Change | Not Detected | FALSE |
| Trpv3 | TRUE | -0.924358444 | 0.003550608 | DOWN | DOWN | TRUE | -0.328359139 | 0.094722877 | No Change | Not Significant | TRUE |
| Trpv4 | TRUE | 2.573554024 | 0.045227455 | UP | UP | FALSE | NA | NA | No Change | Not Detected | FALSE |
| Trpv5 | TRUE | 1.198801452 | 1 | No Change | Not Significant | FALSE | NA | NA | No Change | Not Detected | FALSE |
| Trpv6 | TRUE | -0.767245018 | 0.101750358 | No Change | Not Significant | TRUE | 0.130390552 | 0.876765146 | No Change | Not Significant | TRUE |
| Trrap | TRUE | 0.219275372 | 0.027040753 | No Change | Significant Positive | TRUE | 0.016900355 | 0.910796813 | No Change | Not Significant | TRUE |
| Trub1 | TRUE | -0.153545974 | 0.465961592 | No Change | Not Significant | TRUE | -0.060949642 | 0.70267522 | No Change | Not Significant | TRUE |
| Trub2 | TRUE | -0.268647349 | 0.039959466 | No Change | Significant Negative | TRUE | -0.156163949 | 0.248328382 | No Change | Not Significant | TRUE |
| Try4 | TRUE | -2.021452192 | 1 | No Change | Not Significant | FALSE | NA | NA | No Change | Not Detected | FALSE |
| Try5 | TRUE | 0.144117814 | 1 | No Change | Not Significant | FALSE | NA | NA | No Change | Not Detected | FALSE |
| Tsacc | TRUE | -0.431236851 | 0.299128777 | No Change | Not Significant | TRUE | -0.101515388 | 0.818405307 | No Change | Not Significant | TRUE |
| Tsc1 | TRUE | 0.012660113 | 0.92805494 | No Change | Not Significant | TRUE | -0.076360686 | 0.389715871 | No Change | Not Significant | TRUE |
| Tsc2 | TRUE | -0.084403674 | 0.277378268 | No Change | Not Significant | TRUE | -0.023712553 | 0.768227276 | No Change | Not Significant | TRUE |
| Tsc22d1 | TRUE | -0.160414272 | 0.062245517 | No Change | Not Significant | TRUE | 0.173254877 | 0.001671239 | No Change | Significant Positive | TRUE |
| Tsc22d2 | TRUE | 0.153979681 | 0.514332163 | No Change | Not Significant | TRUE | -0.154943899 | 0.314327481 | No Change | Not Significant | TRUE |
| Tsc22d3 | TRUE | -0.04064785 | 0.859322531 | No Change | Not Significant | TRUE | 0.571145826 | 1.25E-09 | UP | UP | TRUE |
| Tsen15 | TRUE | 0.104392778 | 0.746218772 | No Change | Not Significant | TRUE | 0.047798854 | 0.845589334 | No Change | Not Significant | TRUE |
| Tsen2 | TRUE | 0.02337026 | 0.954081249 | No Change | Not Significant | TRUE | 0.167952452 | 0.45050036 | No Change | Not Significant | TRUE |
| Tsen34 | TRUE | 0.056544466 | 0.714893745 | No Change | Not Significant | TRUE | 0.006665259 | 0.959446759 | No Change | Not Significant | TRUE |
| Tsen54 | TRUE | -0.058700559 | 0.703739933 | No Change | Not Significant | TRUE | -0.087771978 | 0.491447666 | No Change | Not Significant | TRUE |
| Tsfm | TRUE | -0.074273414 | 0.580790426 | No Change | Not Significant | TRUE | 0.073418353 | 0.694009261 | No Change | Not Significant | TRUE |

Supplementary Table 3

| geneID | Cond1_DF_Detected | Cond1_DF_ValueCol | Cond1_DF_PadjCol | Cond1_DF_Cutoff | Cond1_DF_Cutoff_Specific.x | Cond2_DF_Detected |
|---|---|---|---|---|---|---|
| Tsg101 | TRUE | -0.149878845 | 0.123393648 | No Change | Not Significant | TRUE |
| Tsga10 | TRUE | -0.243057407 | 0.295987343 | No Change | Not Significant | TRUE |
| Tsga10ip | TRUE | -0.155817219 | 1 | No Change | Not Significant | FALSE |
| Tsga13 | TRUE | 0.004940645 | 1 | No Change | Not Significant | FALSE |
| Tshb | TRUE | -0.879648951 | 1 | No Change | Not Significant | FALSE |
| Tshr | TRUE | 1.175977866 | 1 | No Change | Not Significant | FALSE |
| Tshz1 | TRUE | -0.13714312 | 0.109497021 | No Change | Not Significant | TRUE |
| Tshz2 | TRUE | -0.118189928 | 0.429893421 | No Change | Not Significant | TRUE |
| Tshz3 | TRUE | -0.279093086 | 1 | No Change | Not Significant | FALSE |
| Tsix | TRUE | -1.067387222 | 1 | No Change | Not Significant | FALSE |
| Tsks | TRUE | 4.064381306 | 1 | No Change | Not Significant | FALSE |
| Tsku | TRUE | -0.825561487 | 0.003220245 | DOWN | DOWN | TRUE |
| Tslp | TRUE | -0.795754367 | 1 | No Change | Not Significant | FALSE |
| Tsn | TRUE | 0.133207702 | 0.363760214 | No Change | Not Significant | TRUE |
| Tsnax | TRUE | -0.105471187 | 0.48334534 | No Change | Not Significant | TRUE |
| Tsnaxip1 | TRUE | 1.021507129 | 1 | No Change | Not Significant | FALSE |
| Tspan1 | TRUE | -0.211514293 | 0.05461877 | No Change | Not Significant | TRUE |
| Tspan10 | TRUE | 1.068710871 | 1 | No Change | Not Significant | FALSE |
| Tspan11 | TRUE | 0.827598697 | 1 | No Change | Not Significant | FALSE |
| Tspan12 | TRUE | 0.028749978 | 0.895862069 | No Change | Not Significant | TRUE |
| Tspan13 | TRUE | 0.204359956 | 0.457189099 | No Change | Not Significant | TRUE |
| Tspan14 | TRUE | 0.055367825 | 0.834370416 | No Change | Not Significant | TRUE |
| Tspan15 | TRUE | -0.261794868 | 0.014482058 | No Change | Significant Negative | TRUE |
| Tspan17 | TRUE | -1.264995287 | 4.68E-18 | DOWN | DOWN | TRUE |
| Tspan18 | TRUE | -0.298806777 | 0.812734264 | No Change | Not Significant | TRUE |
| Tspan2 | TRUE | -1.174150591 | 1 | No Change | Not Significant | FALSE |
| Tspan3 | TRUE | 0.110024268 | 0.5825209 | No Change | Not Significant | TRUE |
| Tspan31 | TRUE | 0.639019193 | 1.13E-08 | UP | UP | TRUE |
| Tspan32 | TRUE | 2.46870981 | 0.185243988 | No Change | Not Significant | FALSE |
| Tspan33 | TRUE | 0.115114444 | 0.728048657 | No Change | Not Significant | TRUE |
| Tspan4 | TRUE | 0.373866554 | 0.427441699 | No Change | Not Significant | TRUE |
| Tspan5 | TRUE | 0.743071238 | 0.000702799 | UP | UP | TRUE |
| Tspan6 | TRUE | -0.578804103 | 0.006385957 | DOWN | DOWN | TRUE |
| Tspan7 | TRUE | -0.298894797 | 0.00024793 | No Change | Significant Negative | TRUE |
| Tspan8 | TRUE | -0.533258525 | 2.99E-08 | DOWN | DOWN | TRUE |
| Tspan9 | TRUE | 0.487955868 | 0.019469435 | No Change | Significant Positive | TRUE |
| Tspear | TRUE | -0.521827552 | 0.465690304 | No Change | Not Significant | TRUE |
| Tspo | TRUE | -0.935116904 | 1.87E-17 | DOWN | DOWN | TRUE |
| Tspo2 | TRUE | -0.325849092 | 1 | No Change | Not Significant | FALSE |
| Tspoap1 | TRUE | 0.798509669 | 0.522421155 | No Change | Not Significant | TRUE |
| Tspyl1 | TRUE | 0.090697981 | 0.41124734 | No Change | Not Significant | TRUE |
| Tspyl2 | TRUE | 2.032960203 | 5.47E-38 | UP | UP | TRUE |
| Tspyl3 | TRUE | 0.495405745 | 0.507809662 | No Change | Not Significant | TRUE |
| Tspyl4 | TRUE | -0.087685452 | 0.896636371 | No Change | Not Significant | TRUE |
| Tspyl5 | TRUE | -0.113474161 | 0.894017228 | No Change | Not Significant | TRUE |
| Tsr1 | TRUE | -0.073911507 | 0.659125086 | No Change | Not Significant | TRUE |
| Tsr2 | TRUE | 0.102020548 | 0.714390744 | No Change | Not Significant | TRUE |
| Tsr3 | TRUE | 0.300167798 | 0.060109302 | No Change | Not Significant | TRUE |
| Tssc4 | TRUE | 0.806488763 | 2.26E-12 | UP | UP | TRUE |
| Tssk1 | TRUE | -0.176367481 | 0.907710402 | No Change | Not Significant | FALSE |
| Tssk2 | TRUE | -0.529883082 | 1 | No Change | Not Significant | FALSE |
| Tssk3 | TRUE | -0.343101692 | 1 | No Change | Not Significant | FALSE |
| Tssk4 | TRUE | 0.441470647 | 0.204058016 | No Change | Not Significant | TRUE |
| Tssk5 | TRUE | 0.068603908 | 1 | No Change | Not Significant | FALSE |
| Tssk6 | TRUE | -0.278521165 | 0.522023104 | No Change | Not Significant | TRUE |
| Tst | TRUE | -0.185352884 | 0.152129476 | No Change | Not Significant | TRUE |
| Tstd1 | TRUE | -0.059182376 | 0.783067117 | No Change | Not Significant | TRUE |
| Tstd2 | TRUE | 0.054603527 | 0.695624496 | No Change | Not Significant | TRUE |
| Tstd3 | TRUE | -0.295888761 | 0.195389167 | No Change | Not Significant | TRUE |
| Tsx | TRUE | 2.416128035 | 1 | No Change | Not Significant | FALSE |
| Ttbk1 | TRUE | 0.410928984 | 0.878808271 | No Change | Not Significant | FALSE |
| Ttbk2 | TRUE | -0.057683385 | 0.855725733 | No Change | Not Significant | TRUE |
| Ttc1 | TRUE | -0.016761612 | 0.908409285 | No Change | Not Significant | TRUE |
| Ttc12 | TRUE | 0.817411464 | 1 | No Change | Not Significant | FALSE |
| Ttc13 | TRUE | 0.308060764 | 0.000661879 | No Change | Significant Positive | TRUE |
| Ttc14 | TRUE | 0.051436245 | 0.758644032 | No Change | Not Significant | TRUE |
| Ttc16 | TRUE | -0.002559799 | 0.998249138 | No Change | Not Significant | TRUE |
| Ttc17 | TRUE | 0.583396273 | 2.13E-24 | UP | UP | TRUE |
| Ttc19 | TRUE | -0.086867349 | 0.524360158 | No Change | Not Significant | TRUE |
| Ttc21a | TRUE | 0.544729276 | 0.566901476 | No Change | Not Significant | TRUE |
| Ttc21b | TRUE | -0.010534006 | 0.966400364 | No Change | Not Significant | TRUE |
| Ttc22 | TRUE | 0.358113362 | 0.055871658 | No Change | Not Significant | TRUE |
| Ttc23 | TRUE | 0.211430048 | 0.048509931 | No Change | Significant Positive | TRUE |
| Ttc24 | TRUE | 1.147935066 | 1 | No Change | Not Significant | FALSE |
| Ttc26 | TRUE | 1.028678067 | 6.80E-05 | UP | UP | TRUE |
| Ttc27 | TRUE | -0.073578972 | 0.802764652 | No Change | Not Significant | TRUE |
| Ttc28 | TRUE | 0.593665151 | 0.703138729 | No Change | Not Significant | TRUE |
| Ttc29 | TRUE | 0.611375929 | 1 | No Change | Not Significant | FALSE |
| Ttc3 | TRUE | -0.078603092 | 0.62645534 | No Change | Not Significant | TRUE |
| Ttc30a1 | TRUE | -0.109730398 | 0.755800484 | No Change | Not Significant | TRUE |
| Ttc30a2 | TRUE | 0.008898676 | 1 | No Change | Not Significant | FALSE |
| Ttc30b | TRUE | -0.372067206 | 0.025985072 | No Change | Significant Negative | TRUE |
| Ttc32 | TRUE | 0.093771418 | 0.687930892 | No Change | Not Significant | TRUE |
| Ttc33 | TRUE | -0.263103331 | 0.058719535 | No Change | Not Significant | TRUE |
| Ttc34 | TRUE | 0.219302825 | 1 | No Change | Not Significant | FALSE |
| Ttc36 | TRUE | -0.780288156 | 0.32611438 | No Change | Not Significant | TRUE |
| Ttc37 | TRUE | 0.173333367 | 0.246388083 | No Change | Not Significant | TRUE |
| Ttc38 | TRUE | 0.147685263 | 0.286482691 | No Change | Not Significant | TRUE |
| Ttc39a | TRUE | 0.232845173 | 0.295198531 | No Change | Not Significant | TRUE |
| Ttc39b | TRUE | -0.238384944 | 0.040464115 | No Change | Significant Negative | TRUE |
| Ttc39c | TRUE | 0.004744183 | 0.993283417 | No Change | Not Significant | TRUE |
| Ttc39d | TRUE | 1.927578355 | 1 | No Change | Not Significant | FALSE |
| Ttc4 | TRUE | 0.156981598 | 0.044782795 | No Change | Significant Positive | TRUE |
| Ttc41 | TRUE | 1.957895381 | 4.41E-17 | UP | UP | TRUE |
| Ttc5 | TRUE | -0.037987029 | 0.823378739 | No Change | Not Significant | TRUE |
| Ttc6 | TRUE | 0.087931677 | 0.774560557 | No Change | Not Significant | TRUE |
| Ttc7 | TRUE | 0.015457046 | 0.897130018 | No Change | Not Significant | TRUE |
| Ttc7b | TRUE | 0.823325488 | 5.56E-05 | UP | UP | TRUE |
| Ttc8 | TRUE | 0.672749058 | 0.066630388 | No Change | Not Significant | TRUE |
| Ttc9 | TRUE | -0.076894186 | 1 | No Change | Not Significant | TRUE |
| Ttc9b | TRUE | -0.932056829 | 1 | No Change | Not Significant | FALSE |
| Ttc9c | TRUE | 0.26802445 | 0.007349514 | No Change | Significant Positive | TRUE |
| Ttf1 | TRUE | 0.009440753 | 0.94348537 | No Change | Not Significant | TRUE |
| Ttf2 | TRUE | 0.249560693 | 0.184592245 | No Change | Not Significant | TRUE |
| Tti1 | TRUE | -0.032707053 | 0.89588517 | No Change | Not Significant | TRUE |
| Tti2 | TRUE | 0.004141045 | 0.98178904 | No Change | Not Significant | TRUE |
| Ttk | TRUE | 0.949546046 | 1.14E-05 | UP | UP | TRUE |
| Ttl | TRUE | -0.033327432 | 0.892918574 | No Change | Not Significant | TRUE |
| Ttll1 | TRUE | -0.523509333 | 0.015303887 | DOWN | DOWN | TRUE |
| Ttll10 | TRUE | -0.425335724 | 0.130972952 | No Change | Not Significant | TRUE |
| Ttll11 | TRUE | 0.100379667 | 0.884053089 | No Change | Not Significant | TRUE |
| Ttll12 | TRUE | 0.075245009 | 0.68565173 | No Change | Not Significant | TRUE |
| Ttll13 | TRUE | 0.288635389 | 0.744666516 | No Change | Not Significant | TRUE |
| Ttll3 | TRUE | 0.51296032 | 0.236098206 | No Change | Not Significant | TRUE |
| Ttll4 | TRUE | 0.045266841 | 0.78655252 | No Change | Not Significant | TRUE |
| Ttll5 | TRUE | 0.449024949 | 1.74E-06 | No Change | Significant Positive | TRUE |
| Ttll6 | TRUE | -0.069980892 | 0.757928885 | No Change | Not Significant | TRUE |
| Ttll7 | TRUE | 0.526987429 | 0.001280348 | UP | UP | TRUE |
| Ttll8 | TRUE | -1.83029565 | 1 | No Change | Not Significant | FALSE |
| Ttll9 | TRUE | -0.457094242 | 1 | No Change | Not Significant | FALSE |
| Ttn | TRUE | 0.07865088 | 0.965445424 | No Change | Not Significant | TRUE |
| Ttpa | TRUE | 1.888397246 | 3.92E-16 | UP | UP | TRUE |
| Ttpal | TRUE | 0.197481588 | 0.01536058 | No Change | Significant Positive | TRUE |
| Ttr | TRUE | -0.133172329 | 0.867980313 | No Change | Not Significant | TRUE |
| Ttyh1 | TRUE | -0.234370351 | 0.490866888 | No Change | Not Significant | TRUE |
| Ttyh2 | TRUE | -0.090448272 | 0.733766095 | No Change | Not Significant | TRUE |
| Ttyh3 | TRUE | 0.163405228 | 0.417979787 | No Change | Not Significant | TRUE |
| Tub | TRUE | 0.668985978 | 1 | No Change | Not Significant | FALSE |
| Tuba1a | TRUE | -0.513972688 | 0.025735772 | DOWN | DOWN | TRUE |
| Tuba1b | TRUE | 0.026131731 | 0.957921881 | No Change | Not Significant | TRUE |
| Tuba1c | TRUE | 0.11080861 | 0.561116348 | No Change | Not Significant | TRUE |
| Tuba3a | TRUE | 1.464830046 | 1 | No Change | Not Significant | FALSE |
| Tuba3b | TRUE | 1.188192682 | 1 | No Change | Not Significant | FALSE |
| Tuba4a | TRUE | 0.072095135 | 0.698213329 | No Change | Not Significant | TRUE |
| Tuba8 | TRUE | 2.959832988 | 1 | No Change | Not Significant | FALSE |
| Tubal3 | TRUE | -0.710688962 | 6.87E-10 | DOWN | DOWN | TRUE |
| Tubb1 | TRUE | 1.965823582 | 1 | No Change | Not Significant | FALSE |
| Tubb2a | TRUE | -0.818704134 | 0.009870396 | DOWN | DOWN | TRUE |
| Tubb2b | TRUE | -1.109314908 | 0.071982309 | No Change | Not Significant | TRUE |
| Tubb3 | TRUE | 3.689688608 | 0.000333855 | UP | UP | FALSE |
| Tubb4a | TRUE | 0.493257067 | 1 | No Change | Not Significant | FALSE |
| Tubb4b | TRUE | 0.214218861 | 0.09253023 | No Change | Not Significant | TRUE |
| Tubb5 | TRUE | 0.877286824 | 1.82E-07 | UP | UP | TRUE |
| Tubb6 | TRUE | -0.386666957 | 0.03690111 | No Change | Significant Negative | TRUE |
| Tubd1 | TRUE | 0.031248598 | 0.927761944 | No Change | Not Significant | TRUE |
| Tube1 | TRUE | 0.316607861 | 0.251954286 | No Change | Not Significant | TRUE |
| Tubg1 | TRUE | 0.33189879 | 0.07190116 | No Change | Not Significant | TRUE |
| Tubg2 | TRUE | 0.083453624 | 0.910493405 | No Change | Not Significant | TRUE |
| Tubgcp2 | TRUE | 0.151210126 | 0.330166076 | No Change | Not Significant | TRUE |
| Tubgcp3 | TRUE | -0.111976306 | 0.267099369 | No Change | Not Significant | TRUE |
| Tubgcp4 | TRUE | -0.187165731 | 0.080560978 | No Change | Not Significant | TRUE |
| Tubgcp5 | TRUE | 0.095804183 | 0.597546855 | No Change | Not Significant | TRUE |
| Tubgcp6 | TRUE | 0.06001989 | 0.73470834 | No Change | Not Significant | TRUE |
| Tufm | TRUE | -0.594394325 | 7.45E-05 | DOWN | DOWN | TRUE |
| Tuft1 | TRUE | -0.51642958 | 0.000464149 | DOWN | DOWN | TRUE |
| Tulp1 | TRUE | 1.025749071 | 1 | No Change | Not Significant | FALSE |
| Tulp2 | TRUE | 0.795796329 | 0.204938354 | No Change | Not Significant | TRUE |
| Tulp3 | TRUE | 0.486286036 | 0.186607563 | No Change | Not Significant | TRUE |
| Tulp4 | TRUE | -0.062024402 | 0.56547568 | No Change | Not Significant | TRUE |
| Tusc1 | TRUE | 0.047938046 | 0.911460908 | No Change | Not Significant | TRUE |
| Tusc2 | TRUE | 0.189947267 | 0.07607843 | No Change | Not Significant | TRUE |
| Tut1 | TRUE | 0.100872213 | 0.425500655 | No Change | Not Significant | TRUE |
| Tut4 | TRUE | -0.137246971 | 0.343942789 | No Change | Not Significant | TRUE |
| Tut7 | TRUE | 0.064363657 | 0.422162995 | No Change | Not Significant | TRUE |
| Tvp23a | TRUE | 0.008104718 | 0.99222255 | No Change | Not Significant | TRUE |
| Tvp23b | TRUE | 0.114997891 | 0.535752227 | No Change | Not Significant | TRUE |
| Twf1 | TRUE | 0.448198056 | 1.23E-08 | No Change | Significant Positive | TRUE |
| Twf2 | TRUE | 0.22744504 | 0.066614215 | No Change | Not Significant | TRUE |
| Twnk | TRUE | -0.28074595 | 0.004704636 | No Change | Significant Negative | TRUE |
| Twsg1 | TRUE | -0.069225681 | 0.762564348 | No Change | Not Significant | TRUE |
| Txk | TRUE | 0.118604285 | 0.927286869 | No Change | Not Significant | TRUE |
| Txlna | TRUE | 0.5161024 | 3.00E-13 | UP | UP | TRUE |
| Txlnb | TRUE | 0.385879457 | 1 | No Change | Not Significant | FALSE |
| Txlng | TRUE | -0.181808272 | 0.065976358 | No Change | Not Significant | TRUE |
| Txn1 | TRUE | -0.296362763 | 0.004614708 | No Change | Significant Negative | TRUE |
| Txn2 | TRUE | -0.073004039 | 0.458233276 | No Change | Not Significant | TRUE |
| Txndc11 | TRUE | 0.889811171 | 9.95E-27 | UP | UP | TRUE |
| Txndc12 | TRUE | -0.102127446 | 0.561689026 | No Change | Not Significant | TRUE |
| Txndc15 | TRUE | -0.166673964 | 0.082503558 | No Change | Not Significant | TRUE |
| Txndc16 | TRUE | -0.00773743 | 0.984509162 | No Change | Not Significant | TRUE |
| Txndc17 | TRUE | -0.165386368 | 0.056890248 | No Change | Not Significant | TRUE |
| Txndc2 | TRUE | -0.424310931 | 1 | No Change | Not Significant | FALSE |
| Txndc5 | TRUE | 0.228551931 | 0.239468239 | No Change | Not Significant | TRUE |
| Txndc9 | TRUE | -0.063028424 | 0.618855346 | No Change | Not Significant | TRUE |
| Txnip | TRUE | -0.43362476 | 0.000809062 | No Change | Significant Negative | TRUE |
| Txnl1 | TRUE | 0.354098756 | 0.013647086 | No Change | Significant Positive | TRUE |
| Txnl4a | TRUE | 0.062490468 | 0.750770075 | No Change | Not Significant | TRUE |
| Txnl4b | TRUE | 0.140557058 | 0.490619315 | No Change | Not Significant | TRUE |
| Txnrd1 | TRUE | -0.108097001 | 0.262880883 | No Change | Not Significant | TRUE |
| Txnrd2 | TRUE | -0.165251588 | 0.249608668 | No Change | Not Significant | TRUE |
| Txnrd3 | TRUE | -0.590978359 | 0.000716166 | DOWN | DOWN | TRUE |
| Tyk2 | TRUE | 0.758586458 | 3.90E-70 | UP | UP | TRUE |

Supplementary Table 3

| geneID | Cond1_DF_Detected | Cond1_DF_ValueCol | Cond1_DF_PadjCol | Cond1_DF_Cutoff | Cond1_DF_Cutoff_Specific.x | Cond2_DF_Detected | Cond2_DF_ValueCol | Cond2_DF_PadjCol | Cond2_DF_Cutoff | Cond2_DF_Cutoff_Specific | BG_Method |
|---|---|---|---|---|---|---|---|---|---|---|---|
| Tymp | TRUE | -0.527562445 | 0.043340911 | DOWN | DOWN | TRUE | -0.085678956 | 0.741395712 | No Change | Not Significant | TRUE |
| Tyms | TRUE | 0.230809853 | 0.324759105 | No Change | Not Significant | TRUE | 0.221206095 | 0.137270396 | No Change | Not Significant | TRUE |
| Tyro3 | TRUE | -0.367174751 | 0.045273886 | No Change | Significant Negative | TRUE | 0.094987818 | 0.567946164 | No Change | Not Significant | TRUE |
| Tyrobp | TRUE | 0.69656227 | 0.368253661 | No Change | Not Significant | TRUE | 0.409113816 | 0.500729794 | No Change | Not Significant | TRUE |
| Tysnd1 | TRUE | 0.303997092 | 0.007027417 | No Change | Significant Positive | TRUE | 0.244709112 | 0.039815194 | No Change | Significant Positive | TRUE |
| Tyw1 | TRUE | 0.02476003 | 0.885199039 | No Change | Not Significant | TRUE | -0.045108219 | 0.699994557 | No Change | Not Significant | TRUE |
| Tyw3 | TRUE | -0.030563584 | 0.951530343 | No Change | Not Significant | TRUE | 0.093179995 | 0.765238104 | No Change | Not Significant | TRUE |
| Tyw5 | TRUE | 0.2933697 | 0.139478188 | No Change | Not Significant | TRUE | 0.053379518 | 0.83633117 | No Change | Not Significant | TRUE |
| U2af1 | TRUE | -0.147076656 | 0.481307968 | No Change | Not Significant | TRUE | 0.035196856 | 0.780262005 | No Change | Not Significant | TRUE |
| U2af1l4 | TRUE | -0.168661187 | 0.872440598 | No Change | Not Significant | TRUE | 0.033980415 | 0.974493399 | No Change | Not Significant | TRUE |
| U2af2 | TRUE | 0.197333245 | 0.034686086 | No Change | Significant Positive | TRUE | 0.128716984 | 0.110184505 | No Change | Not Significant | TRUE |
| U2surp | TRUE | -0.137873643 | 0.128477424 | No Change | Not Significant | TRUE | -0.08665238 | 0.40304403 | No Change | Not Significant | TRUE |
| U90926 | TRUE | 0.215179619 | 1 | No Change | Not Significant | FALSE | NA | NA | No Change | Not Detected | FALSE |
| Uaca | TRUE | 0.257203664 | 0.001204311 | No Change | Significant Positive | TRUE | 0.04118369 | 0.657926757 | No Change | Not Significant | TRUE |
| Uap1 | TRUE | 0.514886443 | 3.47E-14 | UP | UP | TRUE | 0.158353033 | 0.002370426 | No Change | Significant Positive | TRUE |
| Uap1l1 | TRUE | -0.383141486 | 0.007322266 | No Change | Significant Negative | TRUE | -0.260858134 | 0.006056005 | No Change | Significant Negative | TRUE |
| Uba1 | TRUE | 0.194351526 | 0.027443223 | No Change | Significant Positive | TRUE | 0.182085708 | 0.032477324 | No Change | Significant Positive | TRUE |
| Uba2 | TRUE | 0.07273834 | 0.579347577 | No Change | Not Significant | TRUE | 0.045754966 | 0.629623876 | No Change | Not Significant | TRUE |
| Uba3 | TRUE | -0.095504751 | 0.294670784 | No Change | Not Significant | TRUE | -0.093358319 | 0.171818006 | No Change | Not Significant | TRUE |
| Uba5 | TRUE | 0.520026901 | 3.29E-07 | UP | UP | TRUE | 0.455742859 | 8.08E-06 | No Change | Significant Positive | TRUE |
| Uba52 | TRUE | 0.623509656 | 0.000161703 | UP | UP | TRUE | -0.028666324 | 0.93513139 | No Change | Not Significant | TRUE |
| Uba6 | TRUE | 0.2697567 | 0.031386624 | No Change | Significant Positive | TRUE | 0.102296374 | 0.292221615 | No Change | Not Significant | TRUE |
| Uba7 | TRUE | -0.111817972 | 0.556711723 | No Change | Not Significant | TRUE | -0.196666552 | 0.142262102 | No Change | Not Significant | TRUE |
| Ubac1 | TRUE | -0.142113898 | 0.236336338 | No Change | Not Significant | TRUE | 0.008754317 | 0.952805789 | No Change | Not Significant | TRUE |
| Ubac2 | TRUE | -0.012440761 | 0.908812719 | No Change | Not Significant | TRUE | 7.12E-05 | 0.999454275 | No Change | Not Significant | TRUE |
| Ubald1 | TRUE | -0.017857415 | 0.947202684 | No Change | Not Significant | TRUE | -0.334550106 | 0.042967024 | No Change | Significant Negative | TRUE |
| Ubald2 | TRUE | 1.112699959 | 3.61E-09 | UP | UP | TRUE | 0.725125543 | 1.82E-10 | UP | UP | TRUE |
| Ubap1 | TRUE | -0.091158433 | 0.294806545 | No Change | Not Significant | TRUE | 0.021225803 | 0.786805165 | No Change | Not Significant | TRUE |
| Ubap1l | TRUE | 0.5703913 | 1 | No Change | Not Significant | FALSE | NA | NA | No Change | Not Detected | FALSE |
| Ubap2 | TRUE | 0.038228781 | 0.824518194 | No Change | Not Significant | TRUE | -0.038361316 | 0.775689464 | No Change | Not Significant | TRUE |
| Ubap2l | TRUE | 0.006992239 | 0.955501531 | No Change | Not Significant | TRUE | 0.00042534 | 0.996814475 | No Change | Not Significant | TRUE |
| Ubash3a | TRUE | 0.392543432 | 0.87755486 | No Change | Not Significant | FALSE | NA | NA | No Change | Not Detected | FALSE |
| Ubash3b | TRUE | -0.001811379 | 0.99621657 | No Change | Not Significant | TRUE | 0.445070692 | 0.084879315 | No Change | Not Significant | TRUE |
| Ubb | TRUE | 0.172890592 | 0.149272747 | No Change | Not Significant | TRUE | 0.205139998 | 0.122653508 | No Change | Not Significant | TRUE |
| Ubb-ps | FALSE | NA | NA | No Change | Not Detected | TRUE | 1.031599414 | 0.33296048 | No Change | Not Significant | FALSE |
| Ubc | TRUE | -0.119945677 | 0.373495661 | No Change | Not Significant | TRUE | 0.141642991 | 0.414426406 | No Change | Not Significant | TRUE |
| Ubd | TRUE | -1.589533426 | 1 | No Change | Not Significant | TRUE | -1.117286552 | 0.304759215 | No Change | Not Significant | TRUE |
| Ube2a | TRUE | 0.013509571 | 0.93927028 | No Change | Not Significant | TRUE | 0.067980419 | 0.611593275 | No Change | Not Significant | TRUE |
| Ube2b | TRUE | -0.193907755 | 0.038324307 | No Change | Significant Negative | TRUE | -0.122165073 | 0.079254935 | No Change | Not Significant | TRUE |
| Ube2c | TRUE | 1.462138743 | 3.09E-18 | UP | UP | TRUE | 0.391673944 | 0.000274493 | No Change | Significant Positive | TRUE |
| Ube2cbp | TRUE | -0.19975783 | 0.595844868 | No Change | Not Significant | TRUE | -0.190317611 | 0.417346835 | No Change | Not Significant | TRUE |
| Ube2d1 | TRUE | -0.059281539 | 0.801613541 | No Change | Not Significant | TRUE | -0.078357696 | 0.638672592 | No Change | Not Significant | TRUE |
| Ube2d2a | TRUE | -0.013285755 | 0.930184975 | No Change | Not Significant | TRUE | 0.087168824 | 0.273701805 | No Change | Not Significant | TRUE |
| Ube2d2b | TRUE | -0.674261056 | 1 | No Change | Not Significant | FALSE | NA | NA | No Change | Not Detected | FALSE |
| Ube2d3 | TRUE | 0.031144826 | 0.723223689 | No Change | Not Significant | TRUE | -0.02063439 | 0.799982033 | No Change | Not Significant | TRUE |
| Ube2e1 | TRUE | 0.335850655 | 0.002930061 | No Change | Significant Positive | TRUE | 0.046254914 | 0.740616938 | No Change | Not Significant | TRUE |
| Ube2e2 | TRUE | 0.409561494 | 0.006462165 | No Change | Significant Positive | TRUE | 0.008253982 | 0.97153938 | No Change | Not Significant | TRUE |
| Ube2e3 | TRUE | 0.303438357 | 0.0089185 | No Change | Significant Positive | TRUE | 0.12241416 | 0.154660479 | No Change | Not Significant | TRUE |
| Ube2f | TRUE | 0.003909755 | 0.976016991 | No Change | Not Significant | TRUE | 0.109598514 | 0.32795219 | No Change | Not Significant | TRUE |
| Ube2g1 | TRUE | 0.01788768 | 0.877843331 | No Change | Not Significant | TRUE | -0.101520683 | 0.10100479 | No Change | Not Significant | TRUE |
| Ube2g2 | TRUE | 0.30391006 | 0.008622498 | No Change | Significant Positive | TRUE | 0.216915806 | 0.011447596 | No Change | Significant Positive | TRUE |
| Ube2h | TRUE | 0.035295936 | 0.821398208 | No Change | Not Significant | TRUE | 0.046044293 | 0.542488541 | No Change | Not Significant | TRUE |
| Ube2i | TRUE | 0.035613905 | 0.841086565 | No Change | Not Significant | TRUE | -0.112302596 | 0.297761537 | No Change | Not Significant | TRUE |
| Ube2j1 | TRUE | 0.384676223 | 6.55E-08 | No Change | Significant Positive | TRUE | 0.1684846 | 0.001773678 | No Change | Significant Positive | TRUE |
| Ube2j2 | TRUE | -0.119096763 | 0.26091246 | No Change | Not Significant | TRUE | 0.063937894 | 0.537837327 | No Change | Not Significant | TRUE |
| Ube2k | TRUE | 0.081733647 | 0.367893 | No Change | Not Significant | TRUE | 0.05459042 | 0.567389191 | No Change | Not Significant | TRUE |
| Ube2l3 | TRUE | -0.039313043 | 0.770506411 | No Change | Not Significant | TRUE | 0.230710508 | 0.001454132 | No Change | Significant Positive | TRUE |
| Ube2l6 | TRUE | -0.358318851 | 0.283161202 | No Change | Not Significant | TRUE | -0.247235313 | 0.304322033 | No Change | Not Significant | TRUE |
| Ube2m | TRUE | 0.234340819 | 0.086566519 | No Change | Not Significant | TRUE | 0.157362489 | 0.319001795 | No Change | Not Significant | TRUE |

[illegible]

| geneID | Cond1_DF_Detected | Cond1_DF_ValueCol | Cond1_DF_PadjCol | Cond1_DF_Cutoff | Cond1_DF_Cutoff_Specific.x | Cond2_DF_Detected | Cond2_DF_ValueCol | Cond2_DF_PadjCol | Cond2_DF_Cutoff | Cond2_DF_Cutoff_Specific | BG_Method | RG1_All | RG2_Significant | RG3_SignificantChange |
|---|---|---|---|---|---|---|---|---|---|---|---|---|---|---|
| Upk3a | TRUE | 2.13019944 | 1 | No Change | Not Significant | FALSE | NA | NA | No Change | Not Detected | FALSE | Background = FALSE | Background = FALSE | Background = FALSE |
| Upk3b | TRUE | -1.774337348 | 1 | No Change | Not Significant | FALSE | NA | NA | No Change | Not Detected | FALSE | Background = FALSE | Background = FALSE | Background = FALSE |
| Upk3bl | TRUE | 2.356717939 | 1 | No Change | Not Significant | FALSE | NA | NA | No Change | Not Detected | FALSE | Background = FALSE | Background = FALSE | Background = FALSE |
| Upp1 | TRUE | -0.753128231 | 0.004145606 | DOWN | DOWN | TRUE | -0.805738977 | 0.171777386 | No Change | Not Significant | TRUE | Cond1 DOWN + Cond2 Not Significant | Cond1_DOWN | Cond1_DOWN |
| Upp2 | TRUE | -0.216686317 | 0.584928009 | No Change | Not Significant | TRUE | 0.416566907 | 0.085315693 | No Change | Not Significant | TRUE | Cond1 Not Significant + Cond2 Not Significant | None | None |
| Uprt | TRUE | -0.073501791 | 0.691115487 | No Change | Not Significant | TRUE | -0.043976962 | 0.789401467 | No Change | Not Significant | TRUE | Cond1 Not Significant + Cond2 Not Significant | None | None |
| Uqcc1 | TRUE | -0.061336181 | 0.698494442 | No Change | Not Significant | TRUE | 0.044331967 | 0.584025646 | No Change | Not Significant | TRUE | Cond1 Not Significant + Cond2 Not Significant | None | None |
| Uqcc2 | TRUE | -0.193097982 | 0.174107671 | No Change | Not Significant | TRUE | 0.047737309 | 0.751487589 | No Change | Not Significant | TRUE | Cond1 Not Significant + Cond2 Not Significant | None | None |
| Uqcc3 | TRUE | -0.287059784 | 0.039999889 | No Change | Significant Negative | TRUE | -0.071645023 | 0.625831733 | No Change | Not Significant | TRUE | Cond1 Significant Negative + Cond2 Not Significant | None | None |
| Uqcr10 | TRUE | -0.188600188 | 0.114725166 | No Change | Not Significant | TRUE | 0.120645336 | 0.317252001 | No Change | Not Significant | TRUE | Cond1 Not Significant + Cond2 Not Significant | None | None |
| Uqcr11 | TRUE | -0.198526172 | 0.103991793 | No Change | Not Significant | TRUE | 0.178645542 | 0.122594973 | No Change | Not Significant | TRUE | Cond1 Not Significant + Cond2 Not Significant | None | None |
| Uqcrb | TRUE | -0.322614932 | 0.003520597 | No Change | Significant Negative | TRUE | 0.054710579 | 0.428222446 | No Change | Not Significant | TRUE | Cond1 Significant Negative + Cond2 Not Significant | None | None |
| [illegible] | [illegible] | [illegible] | [illegible] | [illegible] | [illegible] | [illegible] | [illegible] | [illegible] | [illegible] | [illegible] | [illegible] | [illegible] | [illegible] | [illegible] |

| geneID | Cond1_DF_ensembl_gene_id | Cond1_DF_entrezgene_id | Cond1_DF_gene_biotype | Cond1_DF_baseMean | Cond1_DF_ log2FoldChange | Cond1_DF_lfcSE | Cond1_DF_stat | Cond1_DF_pvalue | Cond1_DF_ padj | Cond1_DF_absLog2FC | Cond1_DF_Cutoff_Specific.y | Cond2_DF_ensembl_gene_id | Cond2_DF_entrezgene_id | Cond2_DF_gene_biotype |
|---|---|---|---|---|---|---|---|---|---|---|---|---|---|---|
| Upk3a | ENSMUSG00000022435 | 22270 | protein_coding | 1.862724539 | 2.13019944 | 1.768020749 | -1.204849796 | 0.228261292 | 1 | 2.13019944 | Not Significant | NA | NA | NA |
| Upk3b | ENSMUSG00000042985 | 100647 | protein_coding | 1.982125444 | -1.774337348 | 1.418233607 | 1.251089623 | 0.210901781 | 1 | 1.774337348 | Not Significant | NA | NA | NA |
| Upk3bl | ENSMUSG00000006143 | 69665 | protein_coding | 1.159790034 | 2.356717939 | 1.680382397 | -1.402489066 | 0.160769253 | 1 | 2.356717939 | Not Significant | NA | NA | NA |
| Upp1 | ENSMUSG00000020407 | 22271 | protein_coding | 210.1279515 | -0.753128231 | 0.219553836 | 3.430266785 | 0.000602988 | 0.004145606 | 0.753128231 | DOWN | ENSMUSG00000020407 | 22271 | protein_coding |
| Upp2 | ENSMUSG00000026839 | 76654 | protein_coding | 41.83729662 | -0.216686317 | 0.238000018 | 0.910446642 | 0.36258701 | 0.584928009 | 0.216686317 | Not Significant | ENSMUSG00000026839 | 76654 | protein_coding |
| Uprt | ENSMUSG00000073016 | 331487 | protein_coding | 229.5660483 | -0.073501791 | 0.104068154 | 0.706285137 | 0.480010834 | 0.691115487 | 0.073501791 | Not Significant | ENSMUSG00000073016 | 331487 | protein_coding |
| Uqcc1 | ENSMUSG00000005882 | 56046 | protein_coding | 642.8517209 | -0.061336181 | 0.089203118 | 0.687601313 | 0.491703881 | 0.698494442 | 0.061336181 | Not Significant | ENSMUSG00000005882 | 56046 | protein_coding |
| Uqcc2 | ENSMUSG00000024208 | 67267 | protein_coding | 348.4102897 | -0.193097982 | 0.103341817 | 1.868536736 | 0.061687295 | 0.174107671 | 0.193097982 | Not Significant | ENSMUSG00000024208 | 67267 | protein_coding |
| Uqcc3 | ENSMUSG00000071654 | 107197 | protein_coding | 209.430851 | -0.287059784 | 0.110298485 | 2.602572328 | 0.00925273 | 0.039999889 | 0.287059784 | Significant Negative | ENSMUSG00000071654 | 107197 | protein_coding |
| Uqcr10 | ENSMUSG00000059534 | 66152 | protein_coding | 2293.749251 | -0.188600188 | 0.089702442 | 2.10250896 | 0.035508716 | 0.114725166 | 0.188600188 | Not Significant | ENSMUSG00000059534 | 66152 | protein_coding |
| Uqcr11 | ENSMUSG00000020163 | 66594 | protein_coding | 2359.194875 | -0.198526172 | 0.092135448 | 2.154720863 | 0.031183684 | 0.103991793 | 0.198526172 | Not Significant | ENSMUSG00000020163 | 66594 | protein_coding |
| Uqcrb | ENSMUSG00000021520 | 67530 | protein_coding | 2463.672785 | -0.322614932 | 0.092620161 | 3.48320417 | 0.00049545 | 0.003520597 | 0.322614932 | Significant Negative | ENSMUSG00000021520 | 67530 | protein_coding |
| [illegible] | [illegible] | [illegible] | [illegible] | [illegible] | [illegible] | [illegible] | [illegible] | [illegible] | [illegible] | [illegible] | [illegible] | [illegible] | [illegible] | [illegible] |

| geneID | Cond2_DF_baseMean | Cond2_DF_ log2FoldChange | Cond2_DF_lfcSE | Cond2_DF_stat | Cond2_DF_pvalue | Cond2_DF_ padj | Cond2_DF_absLog2FC | RG1_Specific_Cond2 | RG1_Specific_Cond1 |
|---|---|---|---|---|---|---|---|---|---|
| Upk3a | NA | NA | NA | NA | NA | NA | NA | Background = FALSE | Background = FALSE |
| Upk3b | NA | NA | NA | NA | NA | NA | NA | Background = FALSE | Background = FALSE |
| Upk3bl | NA | NA | NA | NA | NA | NA | NA | Background = FALSE | Background = FALSE |
| Upp1 | 875.718231 | -0.805738977 | 0.401559613 | -2.006523939 | 0.044800377 | 0.171777386 | 0.805738977 | Cond1 DOWN + Cond2 Not Significant | Cond2 No Change + Cond1 DOWN |
| Upp2 | 78.40533309 | 0.416566907 | 0.173440231 | 2.401789389 | 0.016315099 | 0.085315693 | 0.416566907 | Cond1 No Change + Cond2 Not Significant | Cond2 No Change + Cond1 Not Significant |
| Uprt | 462.0945202 | -0.043976962 | 0.085174174 | -0.516318035 | 0.605632302 | 0.789401467 | 0.043976962 | Cond1 No Change + Cond2 Not Significant | Cond2 No Change + Cond1 Not Significant |
| Uqcc1 | 1217.321358 | 0.044331967 | 0.04656759 | 0.951991865 | 0.341101108 | 0.584025646 | 0.044331967 | Cond1 No Change + Cond2 Not Significant | Cond2 No Change + Cond1 Not Significant |
| Uqcc2 | 702.7759192 | 0.047737309 | 0.079800184 | 0.598210515 | 0.549699476 | 0.751487589 | 0.047737309 | Cond1 No Change + Cond2 Not Significant | Cond2 No Change + Cond1 Not Significant |
| Uqcc3 | 517.4169361 | -0.071645023 | 0.082575635 | -0.867629095 | 0.385597413 | 0.625831733 | 0.071645023 | Cond1 No Change + Cond2 Not Significant | Cond2 No Change + Cond1 Significant Negative |
| Uqcr10 | 5164.142001 | 0.120645336 | 0.077138679 | 1.564005728 | 0.117816223 | 0.317252001 | 0.120645336 | Cond1 No Change + Cond2 Not Significant | Cond2 No Change + Cond1 Not Significant |
| Uqcr11 | 5338.463357 | 0.178645542 | 0.081095884 | 2.202892826 | 0.027602303 | 0.122594973 | 0.178645542 | Cond1 No Change + Cond2 Not Significant | Cond2 No Change + Cond1 Not Significant |
| Uqcrb | 5442.090595 | 0.054710579 | 0.042285663 | 1.293832815 | 0.195723175 | 0.428222446 | 0.054710579 | Cond1 No Change + Cond2 Not Significant | Cond2 No Change + Cond1 Significant Negative |
| [illegible] | [illegible] | [illegible] | [illegible] | [illegible] | [illegible] | [illegible] | [illegible] | [illegible] | [illegible] |

| geneID | Cond1_DF_Detected | Cond1_DF_ValueCol | Cond1_DF_PadjCol | Cond1_DF_Cutoff |
|---|---|---|---|---|
| Vmn1r226 | TRUE | 0.493953082 | 1 | No Change |
| Vmn1r227 | TRUE | -0.839498973 | 1 | No Change |
| Vmn1r229 | TRUE | -0.862351878 | 1 | No Change |
| Vmn1r230 | TRUE | 1.318322364 | 1 | No Change |
| Vmn1r234 | TRUE | 1.860728975 | 1 | No Change |
| Vmn1r28 | TRUE | -1.242695851 | 1 | No Change |
| Vmn1r29 | TRUE | 0.532759063 | 1 | No Change |
| Vmn1r32 | TRUE | 2.494522379 | 1 | No Change |
| Vmn1r39 | TRUE | -2.073017988 | 1 | No Change |
| Vmn1r4 | TRUE | -0.262808786 | 1 | No Change |
| Vmn1r43 | TRUE | -1.313915695 | 1 | No Change |
| Vmn1r44 | TRUE | -2.482423969 | 1 | No Change |
| Vmn1r45 | TRUE | 0.611500265 | 1 | No Change |
| Vmn1r46 | TRUE | 0.905140064 | 1 | No Change |
| Vmn1r49 | TRUE | 0.004960812 | 1 | No Change |
| Vmn1r50 | TRUE | 3.249360954 | 1 | No Change |
| Vmn1r51 | TRUE | 0.224826849 | 1 | No Change |
| Vmn1r52 | TRUE | 1.931241477 | 1 | No Change |
| Vmn1r55 | TRUE | 0.99794293 | 1 | No Change |
| Vmn1r58 | TRUE | -1.351214375 | 1 | No Change |
| Vmn1r59 | TRUE | 2.852361715 | 1 | No Change |
| Vmn1r6 | TRUE | 1.126419763 | 1 | No Change |
| Vmn1r60 | TRUE | 1.999557035 | 1 | No Change |
| Vmn1r61 | TRUE | 0.523413292 | 1 | No Change |
| Vmn1r68 | TRUE | 2.120520709 | 1 | No Change |
| Vmn1r71 | TRUE | 1.126460431 | 1 | No Change |
| Vmn1r74 | TRUE | -0.272243953 | 1 | No Change |
| Vmn1r75 | TRUE | 2.217322124 | 1 | No Change |
| Vmn1r76 | TRUE | 2.337000462 | 1 | No Change |
| Vmn1r77 | TRUE | -1.787413755 | 1 | No Change |
| Vmn1r78 | TRUE | -0.842962086 | 1 | No Change |
| Vmn1r79 | TRUE | 1.862605678 | 1 | No Change |
| Vmn1r80 | TRUE | -1.792124438 | 1 | No Change |
| Vmn1r81 | TRUE | 2.367254853 | 1 | No Change |
| Vmn1r82 | TRUE | 2.598371764 | 1 | No Change |
| Vmn1r85 | TRUE | 0.224163717 | 1 | No Change |
| Vmn1r86 | TRUE | 2.657012505 | 1 | No Change |
| Vmn1r87 | TRUE | -0.334180396 | 1 | No Change |
| Vmn1r88 | TRUE | -1.660833345 | 1 | No Change |
| Vmn1r9 | TRUE | 1.189927792 | 1 | No Change |
| Vmn1r90 | TRUE | -2.575331535 | 1 | No Change |
| Vmn1r91 | TRUE | 3.624601194 | 1 | No Change |
| Vmn2r1 | TRUE | -0.456743966 | 1 | No Change |
| Vmn2r10 | TRUE | 1.813472748 | 1 | No Change |
| Vmn2r100 | TRUE | 1.139495317 | 1 | No Change |
| Vmn2r102 | TRUE | 0.327555553 | 1 | No Change |
| Vmn2r104 | TRUE | 0.005834303 | 1 | No Change |
| Vmn2r105 | TRUE | 0.463887774 | 1 | No Change |
| Vmn2r107 | TRUE | 1.138520494 | 1 | No Change |
| Vmn2r108 | TRUE | 3.18949291 | 1 | No Change |
| Vmn2r11 | TRUE | 0.609523642 | 1 | No Change |
| Vmn2r111 | TRUE | -1.690361931 | 1 | No Change |
| Vmn2r112 | TRUE | -1.078810251 | 1 | No Change |
| Vmn2r113 | TRUE | -1.204086316 | 1 | No Change |
| Vmn2r114 | TRUE | 0.07460139 | 1 | No Change |
| Vmn2r115 | TRUE | 0.706310531 | 1 | No Change |
| Vmn2r116 | TRUE | -0.498982911 | 1 | No Change |
| Vmn2r117 | TRUE | 2.502079956 | 1 | No Change |
| Vmn2r118 | TRUE | 0.00507117 | 1 | No Change |
| Vmn2r120 | TRUE | 0.535088639 | 0.923234456 | No Change |
| Vmn2r121 | TRUE | 0.16630737 | 1 | No Change |
| Vmn2r124 | TRUE | -0.086488948 | 1 | No Change |
| Vmn2r13 | TRUE | 0.5331027 | 1 | No Change |
| Vmn2r14 | TRUE | -1.277819923 | 1 | No Change |
| Vmn2r15 | TRUE | -2.116757596 | 1 | No Change |
| Vmn2r16 | TRUE | -0.304010144 | 1 | No Change |
| Vmn2r17 | TRUE | 2.739951092 | 1 | No Change |
| Vmn2r18 | TRUE | 0.573337695 | 1 | No Change |
| Vmn2r2 | TRUE | -0.828300908 | 1 | No Change |
| Vmn2r20 | TRUE | 0.876866601 | 0.826766769 | No Change |
| Vmn2r23 | TRUE | 2.951668584 | 1 | No Change |
| Vmn2r25 | TRUE | 3.813378746 | 1 | No Change |
| Vmn2r26 | TRUE | -0.716570326 | 1 | No Change |
| Vmn2r27 | TRUE | 1.188228368 | 1 | No Change |
| Vmn2r28 | TRUE | 2.502049708 | 1 | No Change |
| Vmn2r29 | TRUE | -1.662582196 | 1 | No Change |
| Vmn2r3 | TRUE | 2.974902894 | 1 | No Change |
| Vmn2r31 | TRUE | -0.840316958 | 1 | No Change |
| Vmn2r4 | TRUE | 0.803971406 | 1 | No Change |
| Vmn2r5 | TRUE | 2.457219744 | 1 | No Change |
| Vmn2r53 | TRUE | 1.348644659 | 0.722098569 | No Change |
| Vmn2r56 | TRUE | 2.45271743 | 1 | No Change |
| Vmn2r57 | TRUE | -1.257027768 | 1 | No Change |
| Vmn2r59 | TRUE | -0.325572464 | 1 | No Change |
| Vmn2r6 | TRUE | 3.535143065 | 1 | No Change |
| Vmn2r60 | TRUE | 2.695336073 | 1 | No Change |
| Vmn2r63 | TRUE | -0.027577402 | 1 | No Change |
| Vmn2r65 | TRUE | -2.004490308 | 1 | No Change |
| Vmn2r66 | TRUE | 0.176009164 | 1 | No Change |
| Vmn2r67 | TRUE | -0.905728555 | 1 | No Change |
| Vmn2r69 | TRUE | 1.014302054 | 1 | No Change |
| Vmn2r7 | TRUE | 1.090796954 | 1 | No Change |
| Vmn2r74 | TRUE | -1.618722037 | 1 | No Change |
| Vmn2r75 | TRUE | -0.839228664 | 1 | No Change |
| Vmn2r76 | TRUE | 1.099472178 | 0.735735661 | No Change |
| Vmn2r77 | TRUE | -0.842855656 | 1 | No Change |
| Vmn2r78 | TRUE | 2.850605432 | 1 | No Change |
| Vmn2r79 | TRUE | -1.811200482 | 1 | No Change |
| Vmn2r8 | TRUE | 3.793663065 | 1 | No Change |
| Vmn2r82 | TRUE | 0.004983316 | 1 | No Change |
| Vmn2r83 | TRUE | -0.072485024 | 1 | No Change |
| Vmn2r84 | TRUE | -1.239944091 | 1 | No Change |
| Vmn2r85 | TRUE | 2.507241405 | 1 | No Change |
| Vmn2r86 | TRUE | 0.004304003 | 1 | No Change |
| Vmn2r88 | TRUE | 0.532875078 | 1 | No Change |
| Vmn2r89 | TRUE | -0.714519352 | 1 | No Change |
| Vmn2r9 | TRUE | 0.522883946 | 1 | No Change |
| Vmn2r90 | TRUE | -0.84047043 | 1 | No Change |
| Vmn2r92 | TRUE | 0.511093042 | 1 | No Change |
| Vmn2r93 | TRUE | 1.110242233 | 1 | No Change |
| Vmn2r94 | TRUE | -1.488972246 | 1 | No Change |
| Vmn2r95 | TRUE | -2.453315439 | 1 | No Change |
| Vmn2r96 | TRUE | 1.715561361 | 1 | No Change |
| Vmn2r97 | TRUE | 0.051360728 | 1 | No Change |
| Vmn2r99 | TRUE | 0.004984744 | 1 | No Change |
| Vmp1 | TRUE | 0.084705934 | 0.606155692 | No Change |
| Vnn1 | TRUE | -0.703707486 | 0.030225443 | DOWN |
| Vopp1 | TRUE | 0.004352234 | 0.984116541 | No Change |
| Vpreb1 | TRUE | 0.045328285 | 1 | No Change |
| Vpreb3 | TRUE | 2.7830779 | 1 | No Change |
| Vps11 | TRUE | 0.251036541 | 0.003679413 | No Change |
| Vps13a | TRUE | 0.117614369 | 0.26767214 | No Change |
| Vps13b | TRUE | -0.0309433 | 0.827845828 | No Change |
| Vps13c | TRUE | -0.088034521 | 0.736456016 | No Change |
| Vps13d | TRUE | 0.020404811 | 0.926816617 | No Change |
| Vps16 | TRUE | -0.095694264 | 0.281411586 | No Change |
| Vps18 | TRUE | -0.114780349 | 0.349482473 | No Change |
| Vps25 | TRUE | -0.28321466 | 0.649494967 | No Change |
| Vps26a | TRUE | -0.259915418 | 6.52E-05 | No Change |
| Vps26b | TRUE | 0.047328472 | 0.728066081 | No Change |
| Vps26c | TRUE | -0.02543624 | 0.80597323 | No Change |
| Vps28 | TRUE | -0.176871092 | 0.016858279 | No Change |
| Vps29 | TRUE | 0.003438003 | 0.979202348 | No Change |
| Vps33a | TRUE | 0.102972337 | 0.345496868 | No Change |
| Vps33b | TRUE | -0.016195859 | 0.919794636 | No Change |
| Vps35 | TRUE | 0.107551381 | 0.395770858 | No Change |
| Vps35l | TRUE | -0.116845196 | 0.108013659 | No Change |
| Vps36 | TRUE | -0.065750776 | 0.591834816 | No Change |
| Vps37a | TRUE | -0.077910813 | 0.419497057 | No Change |
| Vps37b | TRUE | 0.200015287 | 0.383856501 | No Change |
| Vps37c | TRUE | 0.260129223 | 0.050530713 | No Change |
| Vps37d | TRUE | -0.247907228 | 0.700667401 | No Change |
| Vps39 | TRUE | -0.094756948 | 0.418308017 | No Change |
| Vps41 | TRUE | -0.087832254 | 0.439030384 | No Change |
| Vps45 | TRUE | 0.098420604 | 0.443108941 | No Change |
| Vps4a | TRUE | 0.002373671 | 0.988088396 | No Change |
| Vps4b | TRUE | 0.026052428 | 0.792833975 | No Change |
| Vps50 | TRUE | 0.148030648 | 0.254671525 | No Change |
| Vps51 | TRUE | -0.11433075 | 0.208527092 | No Change |
| Vps52 | TRUE | 0.089617978 | 0.454166041 | No Change |
| Vps53 | TRUE | 0.052866334 | 0.591883639 | No Change |
| Vps54 | TRUE | -0.035569578 | 0.811818066 | No Change |
| Vps72 | TRUE | -0.066879209 | 0.541378279 | No Change |
| Vps8 | TRUE | 0.046756465 | 0.728048657 | No Change |
| Vps9d1 | TRUE | -0.164787235 | 0.15977566 | No Change |
| Vrk1 | TRUE | 0.493533219 | 0.004819991 | No Change |
| Vrk2 | TRUE | 0.060360134 | 0.783550764 | No Change |
| Vrk3 | TRUE | -0.082358149 | 0.475244062 | No Change |
| Vsig1 | TRUE | 0.211923331 | 0.700749306 | No Change |
| Vsig10 | TRUE | -0.388271857 | 1.03E-07 | No Change |
| Vsig10l | TRUE | 0.28318488 | 0.612475705 | No Change |
| Vsig2 | TRUE | -1.443213287 | 5.86E-21 | DOWN |
| Vsig8 | TRUE | 0.492267962 | 0.64425321 | No Change |
| Vsir | TRUE | 0.414086236 | 0.56579982 | No Change |
| Vstm2a | TRUE | -1.514616261 | 1 | No Change |
| Vstm2l | TRUE | -0.972144085 | 0.034419117 | DOWN |
| Vstm4 | TRUE | -0.236172321 | 1 | No Change |
| Vstm5 | TRUE | -1.153349091 | 2.69E-12 | DOWN |
| Vsx1 | TRUE | 2.245633944 | 1 | No Change |
| Vta1 | TRUE | 0.209502614 | 0.057630735 | No Change |
| Vtcn1 | TRUE | 0.377622921 | 0.567534275 | No Change |
| Vti1a | TRUE | 0.048452385 | 0.712495114 | No Change |
| Vti1b | TRUE | -0.225748829 | 0.0065599 | No Change |
| Vwa1 | TRUE | -0.37065897 | 0.660955754 | No Change |
| Vwa2 | TRUE | 0.223264401 | 0.776544903 | No Change |
| Vwa3a | TRUE | 0.506369295 | 1 | No Change |
| Vwa3b | TRUE | -0.191426123 | 0.733113936 | No Change |
| Vwa5a | TRUE | -0.343184019 | 0.010467953 | No Change |
| Vwa5b1 | TRUE | 0.392115453 | 0.899007849 | No Change |
| Vwa5b2 | TRUE | -0.020222688 | 0.975399666 | No Change |
| Vwa7 | TRUE | -0.260451487 | 0.80301103 | No Change |
| Vwa8 | TRUE | -0.105933378 | 0.415010021 | No Change |
| Vwc2 | TRUE | 2.696470522 | 1 | No Change |
| Vwc2l | TRUE | -1.24353889 | 1 | No Change |
| Vwce | TRUE | -0.836916392 | 0.592205336 | No Change |
| Vwde | TRUE | 1.108894451 | 1 | No Change |
| Vwf | TRUE | 0.570943865 | 0.111570191 | No Change |
| Wac | TRUE | 0.010141492 | 0.932166328 | No Change |
| Wapl | TRUE | 0.103252645 | 0.250425106 | No Change |
| Wars1 | TRUE | 1.330417026 | 2.00E-40 | UP |
| Wars2 | TRUE | -0.204580529 | 0.147853975 | No Change |
| Was | TRUE | 0.664997785 | 0.526861229 | No Change |

| geneID | Cond1_DF_Detected | Cond1_DF_ValueCol | Cond1_DF_PadjCol | Cond1_DF_Cutoff | Cond1_DF_Cutoff_Specific.x | Cond2_DF_Detected | Cond2_DF_ValueCol | Cond2_DF_PadjCol | Cond2_DF_Cutoff | Cond2_DF_Cutoff_Specific | BG_Method | RG1_All |
|---|---|---|---|---|---|---|---|---|---|---|---|---|
| Wasf1 | TRUE | 0.728032456 | 1 | No Change | Not Significant | FALSE | NA | NA | No Change | Not Detected | FALSE | Background = FALSE |
| Wasf2 | TRUE | 0.066778902 | 0.54788666 | No Change | Not Significant | TRUE | 0.020264575 | 0.891415233 | No Change | Not Significant | TRUE | Cond1 Not Significant + Cond2 Not Significant |
| Wasf3 | TRUE | 1.110273449 | 0.626138614 | No Change | Not Significant | FALSE | NA | NA | No Change | Not Detected | FALSE | Background = FALSE |
| Washc1 | TRUE | -0.127546581 | 0.157814254 | No Change | Not Significant | TRUE | 0.064202442 | 0.535833676 | No Change | Not Significant | TRUE | Cond1 Not Significant + Cond2 Not Significant |
| Washc2 | TRUE | -0.107255342 | 0.388133006 | No Change | Not Significant | TRUE | -0.061544858 | 0.33815124 | No Change | Not Significant | TRUE | Cond1 Not Significant + Cond2 Not Significant |
| Washc3 | TRUE | 0.003543076 | 0.982706396 | No Change | Not Significant | TRUE | 0.11712269 | 0.181262767 | No Change | Not Significant | TRUE | Cond1 Not Significant + Cond2 Not Significant |
| Washc4 | TRUE | 0.252771906 | 0.009886119 | No Change | Significant Positive | TRUE | 0.105778783 | 0.249525136 | No Change | Not Significant | TRUE | Cond1 Significant Positive + Cond2 Not Significant |
| Washc5 | TRUE | 0.023284061 | 0.869666246 | No Change | Not Significant | TRUE | -0.054223704 | 0.604050305 | No Change | Not Significant | TRUE | Cond1 Not Significant + Cond2 Not Significant |
| Wasl | TRUE | 0.013017536 | 0.931649419 | No Change | Not Significant | TRUE | 0.006781836 | 0.952773623 | No Change | Not Significant | TRUE | Cond1 Not Significant + Cond2 Not Significant |
| Wbp1 | TRUE | 1.073003377 | 3.42E-63 | UP | UP | TRUE | 0.679571199 | 1.54E-26 | UP | UP | TRUE | Cond1 UP + Cond2 UP |
| Wbp11 | TRUE | 0.170303702 | 0.112749861 | No Change | Not Significant | TRUE | 0.07409043 | 0.443013562 | No Change | Not Significant | TRUE | Cond1 Not Significant + Cond2 Not Significant |
| Wbp1l | TRUE | 0.017657055 | 0.878912418 | No Change | Not Significant | TRUE | 0.041367036 | 0.634775849 | No Change | Not Significant | TRUE | Cond1 Not Significant + Cond2 Not Significant |
| [illegible] | [illegible] | [illegible] | [illegible] | [illegible] | [illegible] | [illegible] | [illegible] | [illegible] | [illegible] | [illegible] | [illegible] | [illegible] |

| geneID | RG2_Significant | RG3_SignificantChange | Cond1_DF_ensembl_gene_id | Cond1_DF_entrezgene_id | Cond1_DF_gene_biotype | Cond1_DF_baseMean | Cond1_DF_log2FoldChange | Cond1_DF_lfcSE | Cond1_DF_stat | Cond1_DF_pvalue | Cond1_DF_padj | Cond1_DF_absLog2FC | Cond1_DF_Cutoff_Specific.y |
|---|---|---|---|---|---|---|---|---|---|---|---|---|---|
| Wasf1 | Background = FALSE | Background = FALSE | ENSMUSG00000019831 | 83767 | protein_coding | 2.002579354 | 0.728032456 | 1.650939757 | -0.440980631 | 0.659227019 | 1 | 0.728032456 | Not Significant |
| Wasf2 | None | None | ENSMUSG00000028868 | 242687 | protein_coding | 1230.209667 | 0.066778902 | 0.067974683 | -0.982408437 | 0.325898671 | 0.54788666 | 0.066778902 | Not Significant |
| Wasf3 | Background = FALSE | Background = FALSE | ENSMUSG00000029636 | 245880 | protein_coding | 6.001062622 | 1.110273449 | 1.333020795 | -0.832900322 | 0.404900948 | 0.626138614 | 1.110273449 | Not Significant |
| Washc1 | None | None | ENSMUSG00000024101 | 68767 | protein_coding | 279.9942724 | -0.127546581 | 0.06616407 | 1.927731809 | 0.053888493 | 0.157814254 | 0.127546581 | Not Significant |
| Washc2 | None | None | ENSMUSG00000024104 | 28006 | protein_coding | 569.1278381 | -0.107255342 | 0.081965993 | 1.308534653 | 0.190692029 | 0.388133006 | 0.107255342 | Not Significant |
| Washc3 | None | None | ENSMUSG00000020056 | 67282 | protein_coding | 272.9817775 | 0.003543076 | 0.081970585 | -0.043223744 | 0.965523178 | 0.982706396 | 0.003543076 | Not Significant |
| Washc4 | None | None | ENSMUSG00000034560 | 319277 | protein_coding | 441.4067678 | 0.252771906 | 0.080542719 | -3.138358235 | 0.001698971 | 0.009886119 | 0.252771906 | Significant Positive |
| Washc5 | None | None | ENSMUSG00000022350 | 223593 | protein_coding | 560.9633071 | 0.023284061 | 0.070733802 | -0.329178692 | 0.742020628 | 0.869666246 | 0.023284061 | Not Significant |
| Wasl | None | None | ENSMUSG00000029684 | 73178 | protein_coding | 821.3664904 | 0.013017536 | 0.074492075 | -0.174750615 | 0.861275595 | 0.931649419 | 0.013017536 | Not Significant |
| Wbp1 | Core_UP | Core_UP | ENSMUSG00000030035 | 22377 | protein_coding | 1160.275275 | 1.073003377 | 0.062792611 | -17.08805166 | 1.82E-65 | 3.42E-63 | 1.073003377 | UP |
| Wbp11 | None | None | ENSMUSG00000030216 | 60321 | protein_coding | 560.6834918 | 0.170303702 | 0.080641149 | -2.111870983 | 0.034697517 | 0.112749861 | 0.170303702 | Not Significant |
| Wbp1l | None | None | ENSMUSG00000047731 | 226178 | protein_coding | 912.2921443 | 0.017657055 | 0.05747504 | -0.307212576 | 0.758681574 | 0.878912418 | 0.017657055 | Not Significant |
| [illegible] | [illegible] | [illegible] | [illegible] | [illegible] | [illegible] | [illegible] | [illegible] | [illegible] | [illegible] | [illegible] | [illegible] | [illegible] | [illegible] |

| geneID | Cond2_DF_ensembl_gene_id | Cond2_DF_entrezgene_id | Cond2_DF_gene_biotype | Cond2_DF_baseMean | Cond2_DF_log2FoldChange | Cond2_DF_lfcSE | Cond2_DF_stat | Cond2_DF_pvalue | Cond2_DF_padj | Cond2_DF_absLog2FC | RG1_Specific_Cond2 | RG1_Specific_Cond1 |
|---|---|---|---|---|---|---|---|---|---|---|---|---|
| Wasf1 | NA | NA | NA | NA | NA | NA | NA | NA | NA | NA | Background = FALSE | Background = FALSE |
| Wasf2 | ENSMUSG00000028868 | 242687 | protein_coding | 2324.991726 | 0.020264575 | 0.068792549 | 0.294575139 | 0.768318456 | 0.891415233 | 0.020264575 | Cond1 No Change + Cond2 Not Significant | Cond2 No Change + Cond1 Not Significant |
| Wasf3 | NA | NA | NA | NA | NA | NA | NA | NA | NA | NA | Background = FALSE | Background = FALSE |
| Washc1 | ENSMUSG00000024101 | 68767 | protein_coding | 598.1209256 | 0.064202442 | 0.061048284 | 1.051666612 | 0.292952534 | 0.535833676 | 0.064202442 | Cond1 No Change + Cond2 Not Significant | Cond2 No Change + Cond1 Not Significant |
| Washc2 | ENSMUSG00000024104 | 28006 | protein_coding | 1221.980652 | -0.061544858 | 0.040840937 | -1.5069404 | 0.131825935 | 0.33815124 | 0.061544858 | Cond1 No Change + Cond2 Not Significant | Cond2 No Change + Cond1 Not Significant |
| Washc3 | ENSMUSG00000020056 | 67282 | protein_coding | 532.0488683 | 0.11712269 | 0.059356198 | 1.973217516 | 0.048470786 | 0.181262767 | 0.11712269 | Cond1 No Change + Cond2 Not Significant | Cond2 No Change + Cond1 Not Significant |
| Washc4 | ENSMUSG00000034560 | 319277 | protein_coding | 761.3809536 | 0.105778783 | 0.060339285 | 1.75306657 | 0.079590581 | 0.249525136 | 0.105778783 | Cond1 No Change + Cond2 Not Significant | Cond2 No Change + Cond1 Significant Positive |
| Washc5 | ENSMUSG00000022350 | 223593 | protein_coding | 1167.963105 | -0.054223704 | 0.059338437 | -0.91380404 | 0.360819823 | 0.604050305 | 0.054223704 | Cond1 No Change + Cond2 Not Significant | Cond2 No Change + Cond1 Not Significant |
| Wasl | ENSMUSG00000029684 | 73178 | protein_coding | 1416.503896 | 0.006781836 | 0.0510299 | 0.132899252 | 0.894273058 | 0.952773623 | 0.006781836 | Cond1 No Change + Cond2 Not Significant | Cond2 No Change + Cond1 Not Significant |
| Wbp1 | ENSMUSG00000030035 | 22377 | protein_coding | 1893.766168 | 0.679571199 | 0.061217614 | 11.10090962 | 1.24E-28 | 1.54E-26 | 0.679571199 | Cond1 UP + Cond2 UP | Cond2 UP + Cond1 UP |
| Wbp11 | ENSMUSG00000030216 | 60321 | protein_coding | 883.0821427 | 0.07409043 | 0.058813382 | 1.25975462 | 0.207757895 | 0.443013562 | 0.07409043 | Cond1 No Change + Cond2 Not Significant | Cond2 No Change + Cond1 Not Significant |
| Wbp1l | ENSMUSG00000047731 | 226178 | protein_coding | 1665.521099 | 0.041367036 | 0.04868917 | 0.849614741 | 0.395539314 | 0.634775849 | 0.041367036 | Cond1 No Change + Cond2 Not Significant | Cond2 No Change + Cond1 Not Significant |
| [illegible] | [illegible] | [illegible] | [illegible] | [illegible] | [illegible] | [illegible] | [illegible] | [illegible] | [illegible] | [illegible] | [illegible] | [illegible] |

| geneID | Cond1_DF_Detected | Cond1_DF_ValueCol | Cond1_DF_PadjCol | Cond1_DF_Cutoff | Cond1_DF_Cutoff_Specific.x | Cond2_DF_Detected | Cond2_DF_ValueCol | Cond2_DF_PadjCol | Cond2_DF_Cutoff | Cond2_DF_Cutoff_Specific | BG_Method | RG1_All | RG2_Significant | RG3_SignificantChange |
|---|---|---|---|---|---|---|---|---|---|---|---|---|---|---|
| Xrcc4 | TRUE | 0.00876372 | 0.971787076 | No Change | Not Significant | TRUE | 0.008292962 | 0.967733155 | No Change | Not Significant | TRUE | Cond1 Not Significant + Cond2 Not Significant | None | None |
| Xrcc5 | TRUE | -0.150302657 | 0.398762803 | No Change | Not Significant | TRUE | -0.040400755 | 0.753561 | No Change | Not Significant | TRUE | Cond1 Not Significant + Cond2 Not Significant | None | None |
| Xrcc6 | TRUE | -0.03396037 | 0.884277416 | No Change | Not Significant | TRUE | 0.013532448 | 0.938452729 | No Change | Not Significant | TRUE | Cond1 Not Significant + Cond2 Not Significant | None | None |
| Xrn1 | TRUE | -0.11691987 | 0.419000785 | No Change | Not Significant | TRUE | -0.250580164 | 0.003277783 | No Change | Significant Negative | TRUE | Cond1 Not Significant + Cond2 Significant Negative | None | None |
| Xrn2 | TRUE | -0.10967475 | 0.123253915 | No Change | Not Significant | TRUE | -0.208285948 | 0.014251137 | No Change | Significant Negative | TRUE | Cond1 Not Significant + Cond2 Significant Negative | None | None |
| Xrra1 | TRUE | 0.786231082 | 1 | No Change | Not Significant | FALSE | NA | NA | No Change | Not Detected | FALSE | Background = FALSE | Background = FALSE | Background = FALSE |
| Xxylt1 | TRUE | 0.07171652 | 0.742718368 | No Change | Not Significant | TRUE | 0.009218478 | 0.957280308 | No Change | Not Significant | TRUE | Cond1 Not Significant + Cond2 Not Significant | None | None |
| Xylb | TRUE | -0.098827827 | 0.695380179 | No Change | Not Significant | TRUE | -0.115694421 | 0.650456717 | No Change | Not Significant | TRUE | Cond1 Not Significant + Cond2 Not Significant | None | None |
| Xylt1 | TRUE | -0.023978244 | 0.961084696 | No Change | Not Significant | TRUE | 0.214963806 | 0.162728786 | No Change | Not Significant | TRUE | Cond1 Not Significant + Cond2 Not Significant | None | None |
| Xylt2 | TRUE | -0.066956935 | 0.752299084 | No Change | Not Significant | TRUE | -0.020636008 | 0.910121497 | No Change | Not Significant | TRUE | Cond1 Not Significant + Cond2 Not Significant | None | None |
| Yae1d1 | TRUE | -0.202396138 | 0.099059146 | No Change | Not Significant | TRUE | -0.085890167 | 0.514726356 | No Change | Not Significant | TRUE | Cond1 Not Significant + Cond2 Not Significant | None | None |
| Yaf2 | TRUE | 0.125359905 | 0.472212364 | No Change | Not Significant | TRUE | 0.048392957 | 0.800888833 | No Change | Not Significant | TRUE | Cond1 Not Significant + Cond2 Not Significant | None | None |

| geneID | Cond1_DF_ensembl_gene_id | Cond1_DF_entrezgene_id | Cond1_DF_gene_biotype | Cond1_DF_baseMean | Cond1_DF_log2FoldChange | Cond1_DF_lfcSE | Cond1_DF_stat | Cond1_DF_pvalue | Cond1_DF_padj | Cond1_DF_absLog2FC | Cond1_DF_Cutoff_Specific.y |
|---|---|---|---|---|---|---|---|---|---|---|---|
| Xrcc4 | ENSMUSG00000021615 | 108138 | protein_coding | 99.6666059 | 0.00876372 | 0.119143263 | -0.073556153 | 0.941363562 | 0.971787076 | 0.00876372 | Not Significant |
| Xrcc5 | ENSMUSG00000026187 | 22596 | protein_coding | 289.7646833 | -0.150302657 | 0.116868923 | 1.286078911 | 0.198415523 | 0.398762803 | 0.150302657 | Not Significant |
| Xrcc6 | ENSMUSG00000022471 | 14375 | protein_coding | 297.1039015 | -0.03396037 | 0.11568634 | 0.293555577 | 0.769097525 | 0.884277416 | 0.03396037 | Not Significant |
| Xrn1 | ENSMUSG00000032410 | 24127 | protein_coding | 587.0815179 | -0.11691987 | 0.094088007 | 1.24266497 | 0.213991317 | 0.419000785 | 0.11691987 | Not Significant |
| Xrn2 | ENSMUSG00000027433 | 24128 | protein_coding | 907.6336118 | -0.10967475 | 0.053097663 | 2.065528783 | 0.038872995 | 0.123253915 | 0.10967475 | Not Significant |
| Xrra1 | ENSMUSG00000035211 | 446101 | protein_coding | 4.196301237 | 0.786231082 | 0.834337989 | -0.942341224 | 0.346017969 | 1 | 0.786231082 | Not Significant |
| Xxylt1 | ENSMUSG00000047434 | 268880 | protein_coding | 161.8929539 | 0.07171652 | 0.120586714 | -0.594729865 | 0.552024055 | 0.742718368 | 0.07171652 | Not Significant |
| Xylb | ENSMUSG00000035769 | 102448 | protein_coding | 60.77354562 | -0.098827827 | 0.141652954 | 0.697675721 | 0.485380015 | 0.695380179 | 0.098827827 | Not Significant |
| Xylt1 | ENSMUSG00000030657 | 233781 | protein_coding | 142.1495528 | -0.023978244 | 0.237869193 | 0.100804326 | 0.919705792 | 0.961084696 | 0.023978244 | Not Significant |
| Xylt2 | ENSMUSG00000020868 | 217119 | protein_coding | 213.1859992 | -0.066956935 | 0.116557127 | 0.574455947 | 0.565659303 | 0.752299084 | 0.066956935 | Not Significant |
| Yae1d1 | ENSMUSG00000075054 | 67008 | protein_coding | 144.0469453 | -0.202396138 | 0.09288543 | 2.178986915 | 0.02933264 | 0.099059146 | 0.202396138 | Not Significant |
| Yaf2 | ENSMUSG00000022634 | 67057 | protein_coding | 105.2669957 | 0.125359905 | 0.111009171 | -1.129275211 | 0.258781755 | 0.472212364 | 0.125359905 | Not Significant |

| geneID | Cond2_DF_ensembl_gene_id | Cond2_DF_entrezgene_id | Cond2_DF_gene_biotype | Cond2_DF_baseMean | Cond2_DF_log2FoldChange | Cond2_DF_lfcSE | Cond2_DF_stat | Cond2_DF_pvalue | Cond2_DF_padj | Cond2_DF_absLog2FC | RG1_Specific_Cond2 | RG1_Specific_Cond1 |
|---|---|---|---|---|---|---|---|---|---|---|---|---|
| Xrcc4 | ENSMUSG00000021615 | 108138 | protein_coding | 182.6197827 | 0.008292962 | 0.089311335 | 0.092854524 | 0.926019134 | 0.967733155 | 0.008292962 | Cond1 No Change + Cond2 Not Significant | Cond2 No Change + Cond1 Not Significant |
| Xrcc5 | ENSMUSG00000026187 | 22596 | protein_coding | 647.2918501 | -0.040400755 | 0.068134619 | -0.592954885 | 0.553211345 | 0.753561 | 0.040400755 | Cond1 No Change + Cond2 Not Significant | Cond2 No Change + Cond1 Not Significant |
| Xrcc6 | ENSMUSG00000022471 | 14375 | protein_coding | 538.2495155 | 0.013532448 | 0.079260615 | 0.170733569 | 0.864433266 | 0.938452729 | 0.013532448 | Cond1 No Change + Cond2 Not Significant | Cond2 No Change + Cond1 Not Significant |
| Xrn1 | ENSMUSG00000032410 | 24127 | protein_coding | 1334.432863 | -0.250580164 | 0.068609839 | -3.652248231 | 0.000259954 | 0.003277783 | 0.250580164 | Cond1 No Change + Cond2 Significant Negative | Cond2 No Change + Cond1 Not Significant |
| Xrn2 | ENSMUSG00000027433 | 24128 | protein_coding | 1823.156281 | -0.208285948 | 0.065746985 | -3.167992387 | 0.001534955 | 0.014251137 | 0.208285948 | Cond1 No Change + Cond2 Significant Negative | Cond2 No Change + Cond1 Not Significant |
| Xrra1 | NA | NA | NA | NA | NA | NA | NA | NA | NA | NA | Background = FALSE | Background = FALSE |
| Xxylt1 | ENSMUSG00000047434 | 268880 | protein_coding | 281.6160468 | 0.009218478 | 0.075614458 | 0.121914222 | 0.902966952 | 0.957280308 | 0.009218478 | Cond1 No Change + Cond2 Not Significant | Cond2 No Change + Cond1 Not Significant |
| Xylb | ENSMUSG00000035769 | 102448 | protein_coding | 113.7003527 | -0.115694421 | 0.141258672 | -0.819025263 | 0.412772 | 0.650456717 | 0.115694421 | Cond1 No Change + Cond2 Not Significant | Cond2 No Change + Cond1 Not Significant |
| Xylt1 | ENSMUSG00000030657 | 233781 | protein_coding | 232.7638287 | 0.214963806 | 0.105453036 | 2.038479066 | 0.041502046 | 0.162728786 | 0.214963806 | Cond1 No Change + Cond2 Not Significant | Cond2 No Change + Cond1 Not Significant |
| Xylt2 | ENSMUSG00000020868 | 217119 | protein_coding | 393.1031172 | -0.020636008 | 0.082447116 | -0.250293878 | 0.80236009 | 0.910121497 | 0.020636008 | Cond1 No Change + Cond2 Not Significant | Cond2 No Change + Cond1 Not Significant |
| Yae1d1 | ENSMUSG00000075054 | 67008 | protein_coding | 275.47079 | -0.085890167 | 0.078223 | -1.098016778 | 0.272197163 | 0.514726356 | 0.085890167 | Cond1 No Change + Cond2 Not Significant | Cond2 No Change + Cond1 Not Significant |
| Yaf2 | ENSMUSG00000022634 | 67057 | protein_coding | 202.9601066 | 0.048392957 | 0.098388233 | 0.49185716 | 0.622820325 | 0.800888833 | 0.048392957 | Cond1 No Change + Cond2 Not Significant | Cond2 No Change + Cond1 Not Significant |

Supplementary Table 3

| geneID | Cond1_DF_Detected | Cond1_DF_ValueCol | Cond1_DF_PadjCol | Cond1_DF_Cutoff | Cond1_DF_Cutoff_Specific.x | Cond2_DF_Detected | Cond2_DF_ValueCol | Cond2_DF_PadjCol | Cond2_DF_Cutoff | Cond2_DF_Cutoff_Specific | BG_Method | RG1_All | RG2_Significant | RG3_SignificantChange | Cond1_DF_ensembl_gene_id | Cond1_DF_entrezgene_id | Cond1_DF_gene_biotype | Cond1_DF_baseMean | Cond1_DF_log2FoldChange | Cond1_DF_lfcSE | Cond1_DF_stat | Cond1_DF_pvalue | Cond1_DF_padj | Cond1_DF_absLog2FC | Cond1_DF_Cutoff_Specific.y | Cond2_DF_ensembl_gene_id | Cond2_DF_entrezgene_id | Cond2_DF_gene_biotype | Cond2_DF_baseMean | Cond2_DF_log2FoldChange | Cond2_DF_lfcSE | Cond2_DF_stat | Cond2_DF_pvalue | Cond2_DF_padj | Cond2_DF_absLog2FC | RG1_Specific_Cond2 | RG1_Specific_Cond1 |
|---|---|---|---|---|---|---|---|---|---|---|---|---|---|---|---|---|---|---|---|---|---|---|---|---|---|---|---|---|---|---|---|---|---|---|---|---|---|
| Zfp111 | TRUE | -0.136846968 | 0.467151421 | No Change | Not Significant | TRUE | -0.137870393 | 0.275714286 | No Change | Not Significant | TRUE | Cond1 Not Significant + Cond2 Not Significant | None | None | ENSMUSG00000087598 | 56707 | protein_coding | 138.9887755 | -0.136846968 | 0.12011895 | 1.13926211 | 0.254593847 | 0.467151421 | 0.136846968 | Not Significant | ENSMUSG00000087598 | 56707 | protein_coding | 258.0933699 | -0.137870393 | 0.082124788 | -1.678791458 | 0.093192694 | 0.275714286 | 0.137870393 | Cond1 No Change + Cond2 Not Significant | Cond2 No Change + Cond1 Not Significant |
| Zfp112 | TRUE | -0.110661266 | 0.817399333 | No Change | Not Significant | TRUE | -0.133388834 | 0.737165747 | No Change | Not Significant | TRUE | Cond1 Not Significant + Cond2 Not Significant | None | None | ENSMUSG00000052675 | 57745 | protein_coding | 26.77296208 | -0.110661266 | 0.253558033 | 0.43643368 | 0.662522107 | 0.817399333 | 0.110661266 | Not Significant | ENSMUSG00000052675 | 57745 | protein_coding | 42.37743302 | -0.133388834 | 0.210926112 | -0.632396023 | 0.527128132 | 0.737165747 | 0.133388834 | Cond1 No Change + Cond2 Not Significant | Cond2 No Change + Cond1 Not Significant |
| Zfp113 | TRUE | -0.195834674 | 0.109753236 | No Change | Not Significant | TRUE | -0.142711633 | 0.31618145 | No Change | Not Significant | TRUE | Cond1 Not Significant + Cond2 Not Significant | None | None | ENSMUSG00000037007 | 56314 | protein_coding | 168.0004753 | -0.195834674 | 0.092117391 | 2.125925109 | 0.033509496 | 0.109753236 | 0.195834674 | Not Significant | ENSMUSG00000037007 | 56314 | protein_coding | 362.16028 | -0.142711633 | 0.091047408 | -1.567443117 | 0.117011152 | 0.31618145 | 0.142711633 | Cond1 No Change + Cond2 Not Significant | Cond2 No Change + Cond1 Not Significant |
| Zfp114 | TRUE | -0.265394098 | 1 | No Change | Not Significant | FALSE | NA | NA | No Change | Not Detected | FALSE | Background = FALSE | Background = FALSE | Background = FALSE | ENSMUSG00000068962 | 232966 | protein_coding | 2.436509382 | -0.265394098 | 0.774630555 | 0.34260732 | 0.731893894 | 1 | 0.265394098 | Not Significant | NA | NA | NA | NA | NA | NA | NA | NA | NA | NA | Background = FALSE | Background = FALSE |
| Zfp119a | TRUE | -0.224433701 | 0.269245209 | No Change | Not Significant | TRUE | -0.06101545 | 0.797655914 | No Change | Not Significant | TRUE | Cond1 Not Significant + Cond2 Not Significant | None | None | ENSMUSG00000057835 | 104349 | protein_coding | 49.92334411 | -0.224433701 | 0.141466441 | 1.586480151 | 0.112630431 | 0.269245209 | 0.224433701 | Not Significant | ENSMUSG00000057835 | 104349 | protein_coding | 119.1471963 | -0.06101545 | 0.122477754 | -0.498175774 | 0.618360156 | 0.797655914 | 0.06101545 | Cond1 No Change + Cond2 Not Significant | Cond2 No Change + Cond1 Not Significant |
| Zfp119b | TRUE | -0.331328164 | 0.210665378 | No Change | Not Significant | TRUE | -0.062410023 | 0.831002845 | No Change | Not Significant | TRUE | Cond1 Not Significant + Cond2 Not Significant | None | None | ENSMUSG00000062101 | 240120 | protein_coding | 35.81965475 | -0.331328164 | 0.189470677 | 1.748704179 | 0.080342167 | 0.210665378 | 0.331328164 | Not Significant | ENSMUSG00000062101 | 240120 | protein_coding | 82.34435645 | -0.062410023 | 0.145857519 | -0.427883479 | 0.668735956 | 0.831002845 | 0.062410023 | Cond1 No Change + Cond2 Not Significant | Cond2 No Change + Cond1 Not Significant |
| Zfp12 | TRUE | -0.079471768 | 0.678429586 | No Change | Not Significant | TRUE | -0.053245186 | 0.660009298 | No Change | Not Significant | TRUE | Cond1 Not Significant + Cond2 Not Significant | None | None | ENSMUSG00000029587 | 231866 | protein_coding | 191.7825272 | -0.079471768 | 0.108732255 | 0.73089414 | 0.464843815 | 0.678429586 | 0.079471768 | Not Significant | ENSMUSG00000029587 | 231866 | protein_coding | 437.562556 | -0.053245186 | 0.066620208 | -0.799234764 | 0.424154298 | 0.660009298 | 0.053245186 | Cond1 No Change + Cond2 Not Significant | Cond2 No Change + Cond1 Not Significant |
| Zfp120 | TRUE | -0.081601253 | 0.644361593 | No Change | Not Significant | TRUE | -0.169427496 | 0.118807539 | No Change | Not Significant | TRUE | Cond1 Not Significant + Cond2 Not Significant | None | None | ENSMUSG00000068134 | 104348 | protein_coding | 193.560182 | -0.081601253 | 0.10225911 | 0.79798517 | 0.424879096 | 0.644361593 | 0.081601253 | Not Significant | ENSMUSG00000068134 | 104348 | protein_coding | 434.1675761 | -0.169427496 | 0.076303399 | -2.220444952 | 0.026388579 | 0.118807539 | 0.169427496 | Cond1 No Change + Cond2 Not Significant | Cond2 No Change + Cond1 Not Significant |
| Zfp128 | TRUE | 0.012332229 | 0.978805568 | No Change | Not Significant | TRUE | 0.037763347 | 0.933520082 | No Change | Not Significant | TRUE | Cond1 Not Significant + Cond2 Not Significant | None | None | ENSMUSG00000060397 | 243833 | protein_coding | 21.12053501 | 0.012332229 | 0.228685995 | -0.053926474 | 0.956993744 | 0.978805568 | 0.012332229 | Not Significant | ENSMUSG00000060397 | 243833 | protein_coding | 31.90224583 | 0.037763347 | 0.201999781 | 0.186947466 | 0.851701823 | 0.933520082 | 0.037763347 | Cond1 No Change + Cond2 Not Significant | Cond2 No Change + Cond1 Not Significant |
| Zfp13 | TRUE | 0.132588839 | 0.665277305 | No Change | Not Significant | TRUE | -0.154000241 | 0.508053902 | No Change | Not Significant | TRUE | Cond1 Not Significant + Cond2 Not Significant | None | None | ENSMUSG00000062012 | 22654 | protein_coding | 40.79942229 | 0.132588839 | 0.174883347 | -0.758155887 | 0.448357666 | 0.665277305 | 0.132588839 | Not Significant | ENSMUSG00000062012 | 22654 | protein_coding | 76.31216762 | -0.154000241 | 0.13846258 | -1.112215595 | 0.266045464 | 0.508053902 | 0.154000241 | Cond1 No Change + Cond2 Not Significant | Cond2 No Change + Cond1 Not Significant |
| Zfp131 | TRUE | 0.025659682 | 0.875081807 | No Change | Not Significant | TRUE | -0.182025591 | 0.110275809 | No Change | Not Significant | TRUE | Cond1 Not Significant + Cond2 Not Significant | None | None | ENSMUSG00000094870 | 72465 | protein_coding | 322.0304232 | 0.025659682 | 0.080981008 | -0.316860497 | 0.751349458 | 0.875081807 | 0.025659682 | Not Significant | ENSMUSG00000094870 | 72465 | protein_coding | 535.2761777 | -0.182025591 | 0.080445221 | -2.262727223 | 0.02365251 | 0.110275809 | 0.182025591 | Cond1 No Change + Cond2 Not Significant | Cond2 No Change + Cond1 Not Significant |
| Zfp133-ps | TRUE | -0.209009054 | 0.811601896 | No Change | Not Significant | TRUE | 0.366340986 | 0.593903542 | No Change | Not Significant | TRUE | Cond1 Not Significant + Cond2 Not Significant | None | None | ENSMUSG00000083674 | 668917 | transcribed_unitary_pseudogene | 6.779029773 | -0.209009054 | 0.464483465 | 0.449981688 | 0.652723644 | 0.811601896 | 0.209009054 | Not Significant | ENSMUSG00000083674 | 668917 | transcribed_unitary_pseudogene | 11.6463539 | 0.366340986 | 0.393239681 | 0.931597202 | 0.35154473 | 0.593903542 | 0.366340986 | Cond1 No Change + Cond2 Not Significant | Cond2 No Change + Cond1 Not Significant |

| geneID | Cond1_DF_Detected | Cond1_DF_ValueCol | Cond1_DF_PadjCol | Cond1_DF_Cutoff | Cond1_DF_Cutoff_Specific.x | Cond2_DF_Detected | Cond2_DF_ValueCol |
|---|---|---|---|---|---|---|---|
| Zfp618 | TRUE | -0.098065834 | 0.649489062 | No Change | Not Significant | TRUE | -0.055182037 |
| Zfp619 | TRUE | -0.166352933 | 0.64845799 | No Change | Not Significant | TRUE | -0.168034082 |
| Zfp62 | TRUE | -0.234280574 | 0.030425148 | No Change | Significant Negative | TRUE | -0.166740566 |
| Zfp622 | TRUE | -0.12921977 | 0.224854857 | No Change | Not Significant | TRUE | -0.164360016 |
| Zfp623 | TRUE | -0.18700087 | 0.296522686 | No Change | Not Significant | TRUE | -0.153700811 |
| Zfp626 | TRUE | -0.235846754 | 0.086166827 | No Change | Not Significant | TRUE | -0.167745746 |
| Zfp628 | TRUE | 0.147244921 | 0.510066755 | No Change | Not Significant | TRUE | 0.011370127 |
| Zfp629 | TRUE | -0.279701255 | 0.042953833 | No Change | Significant Negative | TRUE | -0.264312659 |
| Zfp637 | TRUE | -0.020399088 | 0.914175515 | No Change | Not Significant | TRUE | 0.114218264 |
| Zfp638 | TRUE | 0.056507038 | 0.731167175 | No Change | Not Significant | TRUE | 0.014141789 |
| Zfp639 | TRUE | -0.073156674 | 0.648260298 | No Change | Not Significant | TRUE | -0.173639443 |
| Zfp64 | TRUE | -0.052080748 | 0.773008398 | No Change | Not Significant | TRUE | 0.043344377 |
| Zfp641 | TRUE | 0.50595657 | 0.622275328 | No Change | Not Significant | TRUE | 0.629388782 |
| Zfp644 | TRUE | -0.198712684 | 0.055555275 | No Change | Not Significant | TRUE | -0.095325873 |
| Zfp646 | TRUE | 0.037953364 | 0.551855356 | No Change | Not Significant | TRUE | -0.014147511 |
| Zfp647 | TRUE | -0.574765446 | 0.390165244 | No Change | Not Significant | TRUE | -0.548099443 |
| Zfp648 | TRUE | 0.697276486 | 1 | No Change | Not Significant | FALSE | NA |
| Zfp65 | TRUE | -0.268593678 | 0.208654479 | No Change | Not Significant | TRUE | -0.132417317 |
| Zfp651 | TRUE | 2.084936086 | 0.004614708 | UP | UP | TRUE | 1.278335492 |
| Zfp652 | TRUE | 0.140478469 | 0.260126811 | No Change | Not Significant | TRUE | -0.037217088 |
| Zfp653 | TRUE | 0.099307603 | 0.747162726 | No Change | Not Significant | TRUE | -0.177863235 |
| Zfp654 | TRUE | 0.296238497 | 0.040685989 | No Change | Significant Positive | TRUE | 0.00557657 |
| Zfp655 | TRUE | -0.269107521 | 0.018535691 | No Change | Significant Negative | TRUE | -0.203810422 |
| Zfp658 | TRUE | -0.187578712 | 0.585225632 | No Change | Not Significant | TRUE | -0.062305607 |
| Zfp660 | TRUE | -0.017858784 | 1 | No Change | Not Significant | FALSE | NA |
| Zfp661 | TRUE | -0.193876613 | 0.364391266 | No Change | Not Significant | TRUE | 0.038411472 |
| Zfp663 | TRUE | -0.071450831 | 1 | No Change | Not Significant | FALSE | NA |

[illegible]

Supplementary Table 3

| geneID | Cond1_DF_Detected | Cond1_DF_ValueCol | Cond1_DF_PadjCol | Cond1_DF_Cutoff | Cond1_DF_Cutoff_Specific.x | Cond2_DF_Detected | Cond2_DF_ValueCol | Cond2_DF_PadjCol | Cond2_DF_Cutoff | Cond2_DF_Cutoff_Specific | BG_Method |
|---|---|---|---|---|---|---|---|---|---|---|---|
| Zfyve26 | TRUE | 0.132450996 | 0.45758404 | No Change | Not Significant | TRUE | -0.078011842 | 0.441678211 | No Change | Not Significant | TRUE |
| Zfyve27 | TRUE | -0.193054713 | 0.008606748 | No Change | Significant Negative | TRUE | -0.088482829 | 0.383544922 | No Change | Not Significant | TRUE |
| Zfyve28 | TRUE | 0.079076046 | 1 | No Change | Not Significant | FALSE | NA | NA | No Change | Not Detected | FALSE |
| Zfyve9 | TRUE | -0.308385659 | 0.000149154 | No Change | Significant Negative | TRUE | -0.18422642 | 0.007100804 | No Change | Significant Negative | TRUE |
| Zg16 | TRUE | 0.279760853 | 0.638103523 | No Change | Not Significant | TRUE | 0.556720745 | 1.40E-09 | UP | UP | TRUE |
| Zgpat | TRUE | -0.139426343 | 0.128762902 | No Change | Not Significant | TRUE | -0.047957688 | 0.655860109 | No Change | Not Significant | TRUE |
| Zgrf1 | TRUE | 0.068065409 | 0.88791795 | No Change | Not Significant | TRUE | 0.064530906 | 0.87317221 | No Change | Not Significant | TRUE |
| Zhx1 | TRUE | -0.097273246 | 0.472369953 | No Change | Not Significant | TRUE | -0.100676968 | 0.254568403 | No Change | Not Significant | TRUE |
| Zhx2 | TRUE | -0.083739055 | 0.532495959 | No Change | Not Significant | TRUE | 0.114339437 | 0.29784969 | No Change | Not Significant | TRUE |
| Zhx3 | TRUE | 0.377713306 | 0.009308568 | No Change | Significant Positive | TRUE | -0.005897987 | 0.96881133 | No Change | Not Significant | TRUE |
| Zic1 | TRUE | -1.504027329 | 1 | No Change | Not Significant | FALSE | NA | NA | No Change | Not Detected | FALSE |
| Zic3 | TRUE | 0.46280127 | 1 | No Change | Not Significant | FALSE | NA | NA | No Change | Not Detected | FALSE |
| Zic4 | TRUE | 1.138682638 | 1 | No Change | Not Significant | FALSE | NA | NA | No Change | Not Detected | FALSE |
| Zic5 | TRUE | 0.196198178 | 1 | No Change | Not Significant | FALSE | NA | NA | No Change | Not Detected | FALSE |
| Zik1 | TRUE | 0.141982838 | 1 | No Change | Not Significant | FALSE | NA | NA | No Change | Not Detected | FALSE |
| Zim1 | TRUE | 2.537534922 | 1 | No Change | Not Significant | FALSE | NA | NA | No Change | Not Detected | FALSE |
| Zkscan1 | TRUE | -0.255064076 | 0.047391714 | No Change | Significant Negative | TRUE | -0.100383908 | 0.234551147 | No Change | Not Significant | TRUE |
| Zkscan14 | TRUE | -0.358433428 | 0.047830343 | No Change | Significant Negative | TRUE | -0.13684261 | 0.499365134 | No Change | Not Significant | TRUE |
| Zkscan16 | TRUE | 0.452167249 | 1 | No Change | Not Significant | FALSE | NA | NA | No Change | Not Detected | FALSE |
| Zkscan17 | TRUE | 0.223837065 | 0.280974277 | No Change | Not Significant | TRUE | -0.317283429 | 0.002345091 | No Change | Significant Negative | TRUE |
| Zkscan2 | TRUE | 0.034076653 | 1 | No Change | Not Significant | FALSE | NA | NA | No Change | Not Detected | FALSE |
| Zkscan3 | TRUE | 0.079405655 | 0.379181439 | No Change | Not Significant | TRUE | 0.14114423 | 0.132040471 | No Change | Not Significant | TRUE |
| Zkscan4 | TRUE | -0.076189848 | 0.935658236 | No Change | Not Significant | TRUE | 0.145085352 | 0.827097835 | No Change | Not Significant | TRUE |
| Zkscan5 | TRUE | -0.134609502 | 0.340965701 | No Change | Not Significant | TRUE | 0.018051662 | 0.914739662 | No Change | Not Significant | TRUE |
| Zkscan6 | TRUE | -0.141489984 | 0.240948524 | No Change | Not Significant | TRUE | 0.042056284 | 0.742852833 | No Change | Not Significant | TRUE |
| Zkscan7 | TRUE | -0.207092189 | 0.070735054 | No Change | Not Significant | TRUE | -0.121055685 | 0.41301159 | No Change | Not Significant | TRUE |
| Zkscan8 | TRUE | -0.455062337 | 0.015825683 | No Change | Significant Negative | TRUE | -0.278100488 | 0.011052553 | No Change | Significant Negative | TRUE |
| Zmat1 | TRUE | 0.067100169 | 0.91001573 | No Change | Not Significant | TRUE | 0.324822099 | 0.482850861 | No Change | Not Significant | TRUE |
| Zmat2 | TRUE | 0.014834619 | 0.883975099 | No Change | Not Significant | TRUE | 0.070362816 | 0.374523994 | No Change | Not Significant | TRUE |
| Zmat3 | TRUE | 0.025465434 | 0.909231489 | No Change | Not Significant | TRUE | 0.015699372 | 0.957171506 | No Change | Not Significant | TRUE |
| Zmat4 | TRUE | -0.471481987 | 0.573211901 | No Change | Not Significant | TRUE | -0.222724213 | 0.659950229 | No Change | Not Significant | TRUE |
| Zmat5 | TRUE | -0.172424996 | 0.30685209 | No Change | Not Significant | TRUE | 0.041397726 | 0.808283189 | No Change | Not Significant | TRUE |
| Zmiz1 | TRUE | -0.047269506 | 0.938083652 | No Change | Not Significant | TRUE | -0.252984105 | 0.007978132 | No Change | Significant Negative | TRUE |
| Zmiz2 | TRUE | -0.097286875 | 0.645752753 | No Change | Not Significant | TRUE | -0.054584338 | 0.750752059 | No Change | Not Significant | TRUE |
| Zmpste24 | TRUE | 0.124663252 | 0.334272057 | No Change | Not Significant | TRUE | 0.086428171 | 0.272341441 | No Change | Not Significant | TRUE |
| Zmym1 | TRUE | 0.380697276 | 0.14581122 | No Change | Not Significant | TRUE | -0.036610866 | 0.890725386 | No Change | Not Significant | TRUE |
| Zmym2 | TRUE | -0.1159501 | 0.303828141 | No Change | Not Significant | TRUE | -0.142052977 | 0.078555143 | No Change | Not Significant | TRUE |
| Zmym3 | TRUE | -0.005163266 | 0.980506535 | No Change | Not Significant | TRUE | -0.137900301 | 0.105936335 | No Change | Not Significant | TRUE |
| Zmym4 | TRUE | -0.092505817 | 0.456768986 | No Change | Not Significant | TRUE | -0.266356297 | 0.031797108 | No Change | Significant Negative | TRUE |
| Zmym5 | TRUE | -0.083484068 | 0.524764095 | No Change | Not Significant | TRUE | -0.138249484 | 0.358198217 | No Change | Not Significant | TRUE |
| Zmym6 | TRUE | -0.512217309 | 0.000391737 | DOWN | DOWN | TRUE | -0.394143501 | 0.004775176 | No Change | Significant Negative | TRUE |
| Zmynd10 | TRUE | 0.44218702 | 1 | No Change | Not Significant | FALSE | NA | NA | No Change | Not Detected | FALSE |
| Zmynd11 | TRUE | 0.023636953 | 0.787654245 | No Change | Not Significant | TRUE | -0.078770593 | 0.26640858 | No Change | Not Significant | TRUE |
| Zmynd12 | TRUE | 0.943086626 | 0.22515016 | No Change | Not Significant | TRUE | 0.172012978 | 0.809845517 | No Change | Not Significant | TRUE |
| Zmynd15 | TRUE | 1.163703358 | 0.491701663 | No Change | Not Significant | FALSE | NA | NA | No Change | Not Detected | FALSE |
| Zmynd19 | TRUE | 0.570366321 | 4.90E-05 | UP | UP | TRUE | 0.269807448 | 0.006617243 | No Change | Significant Positive | TRUE |
| Zmynd8 | TRUE | -0.049357596 | 0.696047947 | No Change | Not Significant | TRUE | 0.008980416 | 0.930210801 | No Change | Not Significant | TRUE |
| Znfx1 | TRUE | -0.18629565 | 0.244320693 | No Change | Not Significant | TRUE | -0.106357965 | 0.227937517 | No Change | Not Significant | TRUE |
| Znhit1 | TRUE | 0.006826791 | 0.97354273 | No Change | Not Significant | TRUE | 0.133572277 | 0.393219762 | No Change | Not Significant | TRUE |
| Znhit2 | TRUE | 0.259735632 | 0.146868346 | No Change | Not Significant | TRUE | 0.193800488 | 0.211953026 | No Change | Not Significant | TRUE |
| Znhit3 | TRUE | 0.130014742 | 0.614686552 | No Change | Not Significant | TRUE | 0.047103173 | 0.861407542 | No Change | Not Significant | TRUE |
| Znhit6 | TRUE | -0.056159059 | 0.869560398 | No Change | Not Significant | TRUE | 0.095299113 | 0.658144381 | No Change | Not Significant | TRUE |
| Znrd2 | TRUE | 1.061652254 | 1 | No Change | Not Significant | FALSE | NA | NA | No Change | Not Detected | FALSE |
| Znrf1 | TRUE | 0.130642706 | 0.298368931 | No Change | Not Significant | TRUE | 0.042567935 | 0.756975167 | No Change | Not Significant | TRUE |
| Znrf2 | TRUE | -0.189580124 | 0.032806559 | No Change | Significant Negative | TRUE | -0.164307588 | 0.005996667 | No Change | Significant Negative | TRUE |
| Znrf3 | TRUE | -0.214977172 | 0.236872902 | No Change | Not Significant | TRUE | -0.287815276 | 0.03084881 | No Change | Significant Negative | TRUE |
| Zp1 | TRUE | -0.50950679 | 1 | No Change | Not Significant | TRUE | 0.353287648 | 0.632267311 | No Change | Not Significant | TRUE |
| Zp2 | TRUE | -0.103404622 | 0.938415991 | No Change | Not Significant | FALSE | NA | NA | No Change | Not Detected | FALSE |
| Zpbp2 | TRUE | -1.23186727 | 1 | No Change | Not Significant | FALSE | NA | NA | No Change | Not Detected | FALSE |
| Zpld2 | TRUE | -0.752194537 | 1 | No Change | Not Significant | FALSE | NA | NA | No Change | Not Detected | FALSE |
| Zpr1 | TRUE | -0.151262874 | 0.353403803 | No Change | Not Significant | TRUE | -0.11578679 | 0.20964847 | No Change | Not Significant | TRUE |
| Zranb1 | TRUE | -0.123402115 | 0.352762472 | No Change | Not Significant | TRUE | -0.097740711 | 0.27725095 | No Change | Not Significant | TRUE |
| Zranb2 | TRUE | -0.008287956 | 0.966049475 | No Change | Not Significant | TRUE | 0.13689921 | 0.218015518 | No Change | Not Significant | TRUE |
| Zranb3 | TRUE | 0.418656907 | 0.069313561 | No Change | Not Significant | TRUE | 0.070398327 | 0.751266959 | No Change | Not Significant | TRUE |
| Zrsr2 | TRUE | 0.057151045 | 0.650053783 | No Change | Not Significant | TRUE | 0.015485158 | 0.913844956 | No Change | Not Significant | TRUE |
| Zscan10 | TRUE | 0.960847928 | 1 | No Change | Not Significant | FALSE | NA | NA | No Change | Not Detected | FALSE |
| Zscan12 | TRUE | -0.035544558 | 0.927281893 | No Change | Not Significant | TRUE | -0.043850654 | 0.856166251 | No Change | Not Significant | TRUE |
| Zscan18 | TRUE | 1.040685346 | 0.109658699 | No Change | Not Significant | TRUE | 0.32371056 | 0.706263486 | No Change | Not Significant | TRUE |
| Zscan2 | TRUE | -0.12115792 | 0.751588072 | No Change | Not Significant | TRUE | -0.043960322 | 0.890414429 | No Change | Not Significant | TRUE |
| Zscan20 | TRUE | -0.190382776 | 0.270127107 | No Change | Not Significant | TRUE | 0.015267903 | 0.943767347 | No Change | Not Significant | TRUE |
| Zscan21 | TRUE | -0.005181551 | 0.980506535 | No Change | Not Significant | TRUE | -0.072214029 | 0.579426898 | No Change | Not Significant | TRUE |
| Zscan22 | TRUE | 0.40564216 | 0.025963765 | No Change | Significant Positive | TRUE | 0.125022599 | 0.576562634 | No Change | Not Significant | TRUE |
| Zscan25 | TRUE | -0.005173196 | 0.981111951 | No Change | Not Significant | TRUE | -0.112771364 | 0.34868951 | No Change | Not Significant | TRUE |
| Zscan26 | TRUE | -0.193072506 | 0.079002436 | No Change | Not Significant | TRUE | -0.201011959 | 0.134058533 | No Change | Not Significant | TRUE |
| Zscan29 | TRUE | -0.230043323 | 0.022291098 | No Change | Significant Negative | TRUE | -0.144314767 | 0.037436109 | No Change | Significant Negative | TRUE |
| Zswim1 | TRUE | 0.469334349 | 0.003190048 | No Change | Significant Positive | TRUE | 0.240695172 | 0.131915576 | No Change | Not Significant | TRUE |
| Zswim3 | TRUE | -0.02951894 | 0.913717368 | No Change | Not Significant | TRUE | 0.03583183 | 0.886968072 | No Change | Not Significant | TRUE |
| Zswim4 | TRUE | 0.084786377 | 0.778258299 | No Change | Not Significant | TRUE | -0.277972733 | 0.001614979 | No Change | Significant Negative | TRUE |
| Zswim5 | TRUE | -0.243145437 | 0.043464619 | No Change | Significant Negative | TRUE | -0.204648335 | 0.008441125 | No Change | Significant Negative | TRUE |
| Zswim6 | TRUE | -0.107552582 | 0.622394781 | No Change | Not Significant | TRUE | -0.138390133 | 0.188695979 | No Change | Not Significant | TRUE |
| Zswim7 | TRUE | -0.199054215 | 0.358402373 | No Change | Not Significant | TRUE | -0.103554197 | 0.560212157 | No Change | Not Significant | TRUE |
| Zswim8 | TRUE | 0.10743396 | 0.460883095 | No Change | Not Significant | TRUE | -0.157983007 | 0.16048917 | No Change | Not Significant | TRUE |
| Zswim9 | TRUE | -0.049512787 | 0.878912418 | No Change | Not Significant | TRUE | 0.080046226 | 0.769561988 | No Change | Not Significant | TRUE |
| Zup1 | TRUE | -0.373556329 | 0.000226486 | No Change | Significant Negative | TRUE | -0.259386991 | 0.040905563 | No Change | Significant Negative | TRUE |
| Zw10 | TRUE | 0.056599846 | 0.690104577 | No Change | Not Significant | TRUE | 0.180726297 | 0.008303236 | No Change | Significant Positive | TRUE |
| Zwilch | TRUE | 0.92085516 | 7.70E-08 | UP | UP | TRUE | 0.119877471 | 0.583344264 | No Change | Not Significant | TRUE |
| Zwint | TRUE | 0.53111505 | 1.09E-10 | UP | UP | TRUE | 0.038937189 | 0.818686277 | No Change | Not Significant | TRUE |
| Zxdb | TRUE | -0.141200252 | 0.667666133 | No Change | Not Significant | TRUE | -0.027206937 | 0.923060252 | No Change | Not Significant | TRUE |
| Zxdc | TRUE | -0.146363094 | 0.26704356 | No Change | Not Significant | TRUE | -0.133708218 | 0.102159742 | No Change | Not Significant | TRUE |
| Zyg11a | TRUE | -0.086502189 | 0.951530343 | No Change | Not Significant | TRUE | -1.09199946 | 0.052253619 | No Change | Not Significant | TRUE |
| Zyg11b | TRUE | -0.031317745 | 0.805935268 | No Change | Not Significant | TRUE | 0.028161196 | 0.73971984 | No Change | Not Significant | TRUE |
| Zyx | TRUE | -0.266370601 | 0.099059146 | No Change | Not Significant | TRUE | -0.302439055 | 0.034530138 | No Change | Significant Negative | TRUE |
| Zzef1 | TRUE | 0.167539874 | 0.181269925 | No Change | Not Significant | TRUE | -0.009275707 | 0.91832199 | No Change | Not Significant | TRUE |
| Zzz3 | TRUE | -0.012832101 | 0.938406345 | No Change | Not Significant | TRUE | -0.061679852 | 0.452365863 | No Change | Not Significant | TRUE |

| geneID | RG1_All | RG2_Significant | RG3_SignificantChange | Cond1_DF_ensembl_gene_id | Cond1_DF_entrezgene_id | Cond1_DF_gene_biotype |
|---|---|---|---|---|---|---|
| Zfyve26 | Cond1 Not Significant + Cond2 Not Significant | None | None | ENSMUSG00000066440 | 211978 | protein_coding |
| Zfyve27 | Cond1 Significant Negative + Cond2 Not Significant | None | None | ENSMUSG00000018820 | 319740 | protein_coding |
| Zfyve28 | Background = FALSE | Background = FALSE | Background = FALSE | ENSMUSG00000037224 | 231125 | protein_coding |
| Zfyve9 | Cond1 Significant Negative + Cond2 Significant Negative | None | None | ENSMUSG00000034557 | 230597 | protein_coding |
| Zg16 | Cond1 Not Significant + Cond2 UP | Cond2_UP | Cond2_UP | ENSMUSG00000049350 | 69036 | protein_coding |
| Zgpat | Cond1 Not Significant + Cond2 Not Significant | None | None | ENSMUSG00000027582 | 229007 | protein_coding |
| Zgrf1 | Cond1 Not Significant + Cond2 Not Significant | None | None | ENSMUSG00000051278 | 71643 | protein_coding |
| Zhx1 | Cond1 Not Significant + Cond2 Not Significant | None | None | ENSMUSG00000022361 | 22770 | protein_coding |
| Zhx2 | Cond1 Not Significant + Cond2 Not Significant | None | None | ENSMUSG00000071757 | 387609 | protein_coding |
| Zhx3 | Cond1 Significant Positive + Cond2 Not Significant | None | None | ENSMUSG00000035877 | 320799 | protein_coding |
| Zic1 | Background = FALSE | Background = FALSE | Background = FALSE | ENSMUSG00000032368 | 22771 | protein_coding |
| Zic3 | Background = FALSE | Background = FALSE | Background = FALSE | ENSMUSG00000067860 | 22773 | protein_coding |
| Zic4 | Background = FALSE | Background = FALSE | Background = FALSE | ENSMUSG00000036972 | 22774 | protein_coding |
| Zic5 | Background = FALSE | Background = FALSE | Background = FALSE | ENSMUSG00000041703 | 65100 | protein_coding |
| Zik1 | Background = FALSE | Background = FALSE | Background = FALSE | ENSMUSG00000030393 | 22775 | protein_coding |
| Zim1 | Background = FALSE | Background = FALSE | Background = FALSE | ENSMUSG00000002266 | 22776 | protein_coding |
| Zkscan1 | Cond1 Significant Negative + Cond2 Not Significant | None | None | ENSMUSG00000029729 | 74570 | protein_coding |
| Zkscan14 | Cond1 Significant Negative + Cond2 Not Significant | None | None | ENSMUSG00000029627 | 67235 | protein_coding |
| Zkscan16 | Background = FALSE | Background = FALSE | Background = FALSE | ENSMUSG00000038630 | 100041581 | protein_coding |
| Zkscan17 | Cond1 Not Significant + Cond2 Significant Negative | None | None | ENSMUSG00000020472 | 268417 | protein_coding |
| Zkscan2 | Background = FALSE | Background = FALSE | Background = FALSE | ENSMUSG00000030757 | 210162 | protein_coding |
| Zkscan3 | Cond1 Not Significant + Cond2 Not Significant | None | None | ENSMUSG00000021327 | 72739 | protein_coding |
| Zkscan4 | Cond1 Not Significant + Cond2 Not Significant | None | None | ENSMUSG00000054931 | 544922 | protein_coding |
| Zkscan5 | Cond1 Not Significant + Cond2 Not Significant | None | None | ENSMUSG00000055991 | 22757 | protein_coding |
| Zkscan6 | Cond1 Not Significant + Cond2 Not Significant | None | None | ENSMUSG00000018347 | 52712 | protein_coding |
| Zkscan7 | Cond1 Not Significant + Cond2 Not Significant | None | None | ENSMUSG00000063488 | 382118 | protein_coding |
| Zkscan8 | Cond1 Significant Negative + Cond2 Significant Negative | None | None | ENSMUSG00000063894 | 93681 | protein_coding |
| Zmat1 | Cond1 Not Significant + Cond2 Not Significant | None | None | ENSMUSG00000052676 | 215693 | protein_coding |
| Zmat2 | Cond1 Not Significant + Cond2 Not Significant | None | None | ENSMUSG00000001383 | 66492 | protein_coding |
| Zmat3 | Cond1 Not Significant + Cond2 Not Significant | None | None | ENSMUSG00000027663 | 22401 | protein_coding |
| Zmat4 | Cond1 Not Significant + Cond2 Not Significant | None | None | ENSMUSG00000037492 | 320158 | protein_coding |
| Zmat5 | Cond1 Not Significant + Cond2 Not Significant | None | None | ENSMUSG00000009076 | 67178 | protein_coding |
| Zmiz1 | Cond1 Not Significant + Cond2 Significant Negative | None | None | ENSMUSG00000007817 | 328365 | protein_coding |
| Zmiz2 | Cond1 Not Significant + Cond2 Not Significant | None | None | ENSMUSG00000041164 | 52915 | protein_coding |
| Zmpste24 | Cond1 Not Significant + Cond2 Not Significant | None | None | ENSMUSG00000043207 | 230709 | protein_coding |
| Zmym1 | Cond1 Not Significant + Cond2 Not Significant | None | None | ENSMUSG00000043872 | 68310 | protein_coding |
| Zmym2 | Cond1 Not Significant + Cond2 Not Significant | None | None | ENSMUSG00000021945 | 76007 | protein_coding |
| Zmym3 | Cond1 Not Significant + Cond2 Not Significant | None | None | ENSMUSG00000031310 | 56364 | protein_coding |
| Zmym4 | Cond1 Not Significant + Cond2 Significant Negative | None | None | ENSMUSG00000042446 | 67785 | protein_coding |
| Zmym5 | Cond1 Not Significant + Cond2 Not Significant | None | None | ENSMUSG00000040123 | 219105 | protein_coding |
| Zmym6 | Cond1 DOWN + Cond2 Significant Negative | Core_DOWN | Cond1_DOWN | ENSMUSG00000042408 | 100177 | protein_coding |
| Zmynd10 | Background = FALSE | Background = FALSE | Background = FALSE | ENSMUSG00000010044 | 114602 | protein_coding |
| Zmynd11 | Cond1 Not Significant + Cond2 Not Significant | None | None | ENSMUSG00000021156 | 66505 | protein_coding |
| Zmynd12 | Cond1 Not Significant + Cond2 Not Significant | None | None | ENSMUSG00000070806 | 332934 | protein_coding |
| Zmynd15 | Background = FALSE | Background = FALSE | Background = FALSE | ENSMUSG00000040829 | 574428 | protein_coding |
| Zmynd19 | Cond1 UP + Cond2 Significant Positive | Core_UP | Cond1_UP | ENSMUSG00000026974 | 67187 | protein_coding |
| Zmynd8 | Cond1 Not Significant + Cond2 Not Significant | None | None | ENSMUSG00000039671 | 228880 | protein_coding |
| Znfx1 | Cond1 Not Significant + Cond2 Not Significant | None | None | ENSMUSG00000039501 | 98999 | protein_coding |
| Znhit1 | Cond1 Not Significant + Cond2 Not Significant | None | None | ENSMUSG00000059518 | 70103 | protein_coding |
| Znhit2 | Cond1 Not Significant + Cond2 Not Significant | None | None | ENSMUSG00000075227 | 29805 | protein_coding |
| Znhit3 | Cond1 Not Significant + Cond2 Not Significant | None | None | ENSMUSG00000020526 | 448850 | protein_coding |
| Znhit6 | Cond1 Not Significant + Cond2 Not Significant | None | None | ENSMUSG00000074182 | 229937 | protein_coding |
| Znrd2 | Background = FALSE | Background = FALSE | Background = FALSE | ENSMUSG00000079478 | 56390 | protein_coding |
| Znrf1 | Cond1 Not Significant + Cond2 Not Significant | None | None | ENSMUSG00000033545 | 170737 | protein_coding |
| Znrf2 | Cond1 Significant Negative + Cond2 Significant Negative | None | None | ENSMUSG00000058446 | 387524 | protein_coding |
| Znrf3 | Cond1 Not Significant + Cond2 Significant Negative | None | None | ENSMUSG00000041961 | 407821 | protein_coding |
| Zp1 | Cond1 Not Significant + Cond2 Not Significant | None | None | ENSMUSG00000024734 | 22786 | protein_coding |
| Zp2 | Background = FALSE | Background = FALSE | Background = FALSE | ENSMUSG00000030911 | 22787 | protein_coding |
| Zpbp2 | Background = FALSE | Background = FALSE | Background = FALSE | ENSMUSG00000017195 | 69376 | protein_coding |
| Zpld2 | Background = FALSE | Background = FALSE | Background = FALSE | ENSMUSG00000073747 | 665186 | protein_coding |
| Zpr1 | Cond1 Not Significant + Cond2 Not Significant | None | None | ENSMUSG00000032078 | 22687 | protein_coding |
| Zranb1 | Cond1 Not Significant + Cond2 Not Significant | None | None | ENSMUSG00000030967 | 360216 | protein_coding |
| Zranb2 | Cond1 Not Significant + Cond2 Not Significant | None | None | ENSMUSG00000028180 | 53861 | protein_coding |
| Zranb3 | Cond1 Not Significant + Cond2 Not Significant | None | None | ENSMUSG00000036086 | 226409 | protein_coding |
| Zrsr2 | Cond1 Not Significant + Cond2 Not Significant | None | None | ENSMUSG00000031370 | 22184 | protein_coding |
| Zscan10 | Background = FALSE | Background = FALSE | Background = FALSE | ENSMUSG00000023902 | 332221 | protein_coding |
| Zscan12 | Cond1 Not Significant + Cond2 Not Significant | None | None | ENSMUSG00000036721 | 22758 | protein_coding |
| Zscan18 | Cond1 Not Significant + Cond2 Not Significant | None | None | ENSMUSG00000070822 | 232875 | protein_coding |
| Zscan2 | Cond1 Not Significant + Cond2 Not Significant | None | None | ENSMUSG00000038797 | 22691 | protein_coding |
| Zscan20 | Cond1 Not Significant + Cond2 Not Significant | None | None | ENSMUSG00000061894 | 269585 | protein_coding |
| Zscan21 | Cond1 Not Significant + Cond2 Not Significant | None | None | ENSMUSG00000037017 | 22697 | protein_coding |
| Zscan22 | Cond1 Significant Positive + Cond2 Not Significant | None | None | ENSMUSG00000054715 | 232878 | protein_coding |
| Zscan25 | Cond1 Not Significant + Cond2 Not Significant | None | None | ENSMUSG00000070420 | 666311 | protein_coding |
| Zscan26 | Cond1 Not Significant + Cond2 Not Significant | None | None | ENSMUSG00000022228 | 432731 | protein_coding |
| Zscan29 | Cond1 Significant Negative + Cond2 Significant Negative | None | None | ENSMUSG00000050619 | 99334 | protein_coding |
| Zswim1 | Cond1 Significant Positive + Cond2 Not Significant | None | None | ENSMUSG00000017764 | 71971 | protein_coding |
| Zswim3 | Cond1 Not Significant + Cond2 Not Significant | None | None | ENSMUSG00000045822 | 67538 | protein_coding |
| Zswim4 | Cond1 Not Significant + Cond2 Significant Negative | None | None | ENSMUSG00000035671 | 212168 | protein_coding |
| Zswim5 | Cond1 Significant Negative + Cond2 Significant Negative | None | None | ENSMUSG00000033948 | 74464 | protein_coding |
| Zswim6 | Cond1 Not Significant + Cond2 Not Significant | None | None | ENSMUSG00000032846 | 67263 | protein_coding |
| Zswim7 | Cond1 Not Significant + Cond2 Not Significant | None | None | ENSMUSG00000014243 | 69747 | protein_coding |
| Zswim8 | Cond1 Not Significant + Cond2 Not Significant | None | None | ENSMUSG00000021819 | 268721 | protein_coding |
| Zswim9 | Cond1 Not Significant + Cond2 Not Significant | None | None | ENSMUSG00000070814 | 321008 | protein_coding |
| Zup1 | Cond1 Significant Negative + Cond2 Significant Negative | None | None | ENSMUSG00000039531 | 72580 | protein_coding |
| Zw10 | Cond1 Not Significant + Cond2 Significant Positive | None | None | ENSMUSG00000032264 | 26951 | protein_coding |
| Zwilch | Cond1 UP + Cond2 Not Significant | Cond1_UP | Cond1_UP | ENSMUSG00000032400 | 68014 | protein_coding |
| Zwint | Cond1 UP + Cond2 Not Significant | Cond1_UP | Cond1_UP | ENSMUSG00000019923 | 52696 | protein_coding |
| Zxdb | Cond1 Not Significant + Cond2 Not Significant | None | None | ENSMUSG00000073062 | 668166 | protein_coding |
| Zxdc | Cond1 Not Significant + Cond2 Not Significant | None | None | ENSMUSG00000034430 | 80292 | protein_coding |
| Zyg11a | Cond1 Not Significant + Cond2 Not Significant | None | None | ENSMUSG00000034645 | 230590 | protein_coding |
| Zyg11b | Cond1 Not Significant + Cond2 Not Significant | None | None | ENSMUSG00000034636 | 414872 | protein_coding |
| Zyx | Cond1 Not Significant + Cond2 Significant Negative | None | None | ENSMUSG00000029860 | 22793 | protein_coding |
| Zzef1 | Cond1 Not Significant + Cond2 Not Significant | None | None | ENSMUSG00000055670 | 195018 | protein_coding |
| Zzz3 | Cond1 Not Significant + Cond2 Not Significant | None | None | ENSMUSG00000039068 | 108946 | protein_coding |

| geneID | Cond1_DF_baseMean | Cond1_DF_log2FoldChange | Cond1_DF_lfcSE | Cond1_DF_stat | Cond1_DF_pvalue | Cond1_DF_padj | Cond1_DF_absLog2FC | Cond1_DF_Cutoff_Specific.y | Cond2_DF_ensembl_gene_id | Cond2_DF_entrezgene_id | Cond2_DF_gene_biotype |
|---|---|---|---|---|---|---|---|---|---|---|---|
| Zfyve26 | 333.4367724 | 0.132450996 | 0.114100482 | -1.160827658 | 0.245711993 | 0.45758404 | 0.132450996 | Not Significant | ENSMUSG00000066440 | 211978 | protein_coding |
| Zfyve27 | 366.1668607 | -0.193054713 | 0.060598969 | 3.185775544 | 0.001443666 | 0.008606748 | 0.193054713 | Significant Negative | ENSMUSG00000018820 | 319740 | protein_coding |
| Zfyve28 | 3.076241437 | 0.079076046 | 1.821232139 | -0.043418983 | 0.965367546 | 1 | 0.079076046 | Not Significant | NA | NA | NA |
| Zfyve9 | 415.1646789 | -0.308385659 | 0.070958448 | 4.346003435 | 1.39E-05 | 0.000149154 | 0.308385659 | Significant Negative | ENSMUSG00000034557 | 230597 | protein_coding |
| Zg16 | 28919.21281 | 0.279760853 | 0.345492335 | -0.809745469 | 0.418086479 | 0.638103523 | 0.279760853 | Not Significant | ENSMUSG00000049350 | 69036 | protein_coding |
| Zgpat | 388.260569 | -0.139426343 | 0.068280005 | 2.041979118 | 0.041153603 | 0.128762902 | 0.139426343 | Not Significant | ENSMUSG00000027582 | 229007 | protein_coding |
| Zgrf1 | 55.05004615 | 0.068065409 | 0.238787815 | -0.285045572 | 0.775609228 | 0.88791795 | 0.068065409 | Not Significant | ENSMUSG00000051278 | 71643 | protein_coding |
| Zhx1 | 403.1193151 | -0.097273246 | 0.086189616 | 1.128595888 | 0.259068346 | 0.472369953 | 0.097273246 | Not Significant | ENSMUSG00000022361 | 22770 | protein_coding |
| Zhx2 | 188.4805382 | -0.083739055 | 0.082776127 | 1.011632913 | 0.311713603 | 0.532495959 | 0.083739055 | Not Significant | ENSMUSG00000071757 | 387609 | protein_coding |
| Zhx3 | 215.7609004 | 0.377713306 | 0.119553424 | -3.15936837 | 0.001581115 | 0.009308568 | 0.377713306 | Significant Positive | ENSMUSG00000035877 | 320799 | protein_coding |
| Zic1 | 2.223568625 | -1.504027329 | 2.344681671 | 0.641463337 | 0.521221694 | 1 | 1.504027329 | Not Significant | NA | NA | NA |
| Zic3 | 1.206198641 | 0.46280127 | 2.277154727 | -0.203236637 | 0.838950078 | 1 | 0.46280127 | Not Significant | NA | NA | NA |
| Zic4 | 1.43331246 | 1.138682638 | 2.971621624 | -0.383185608 | 0.701582148 | 1 | 1.138682638 | Not Significant | NA | NA | NA |
| Zic5 | 1.093696463 | 0.196198178 | 1.843358759 | -0.106435156 | 0.915237101 | 1 | 0.196198178 | Not Significant | NA | NA | NA |
| Zik1 | 4.115317907 | 0.141982838 | 0.754099517 | -0.188281301 | 0.850656143 | 1 | 0.141982838 | Not Significant | NA | NA | NA |
| Zim1 | 3.629382414 | 2.537534922 | 3.387829635 | -0.749014914 | 0.453848217 | 1 | 2.537534922 | Not Significant | NA | NA | NA |
| Zkscan1 | 1425.653857 | -0.255064076 | 0.100852267 | 2.529086204 | 0.011435994 | 0.047391714 | 0.255064076 | Significant Negative | ENSMUSG00000029729 | 74570 | protein_coding |
| Zkscan14 | 95.98420759 | -0.358433428 | 0.141985361 | 2.524439318 | 0.011588301 | 0.047830343 | 0.358433428 | Significant Negative | ENSMUSG00000029627 | 67235 | protein_coding |
| Zkscan16 | 1.072622222 | 0.452167249 | 1.784841166 | -0.253337528 | 0.800007399 | 1 | 0.452167249 | Not Significant | NA | NA | NA |
| Zkscan17 | 185.4149093 | 0.223837065 | 0.143723609 | -1.557413332 | 0.119372383 | 0.280974277 | 0.223837065 | Not Significant | ENSMUSG00000020472 | 268417 | protein_coding |
| Zkscan2 | 2.228054327 | 0.034076653 | 2.429485077 | -0.014026286 | 0.98880901 | 1 | 0.034076653 | Not Significant | NA | NA | NA |
| Zkscan3 | 635.4172376 | 0.079405655 | 0.059788539 | -1.328108299 | 0.184142332 | 0.379181439 | 0.079405655 | Not Significant | ENSMUSG00000021327 | 72739 | protein_coding |
| Zkscan4 | 7.231578324 | -0.076189848 | 0.464695302 | 0.163956571 | 0.869765332 | 0.935658236 | 0.076189848 | Not Significant | ENSMUSG00000054931 | 544922 | protein_coding |
| Zkscan5 | 178.1272042 | -0.134609502 | 0.095247317 | 1.41326292 | 0.157578432 | 0.340965701 | 0.134609502 | Not Significant | ENSMUSG00000055991 | 22757 | protein_coding |
| Zkscan6 | 224.7153379 | -0.141489984 | 0.085149977 | 1.66165616 | 0.096581736 | 0.240948524 | 0.141489984 | Not Significant | ENSMUSG00000018347 | 52712 | protein_coding |
| Zkscan7 | 170.4805137 | -0.207092189 | 0.088349776 | 2.344003541 | 0.01907799 | 0.070735054 | 0.207092189 | Not Significant | ENSMUSG00000063488 | 382118 | protein_coding |
| Zkscan8 | 187.2335421 | -0.455062337 | 0.153283676 | 2.968759297 | 0.002990047 | 0.015825683 | 0.455062337 | Significant Negative | ENSMUSG00000063894 | 93681 | protein_coding |
| Zmat1 | 14.26073181 | 0.067100169 | 0.297847318 | -0.225283778 | 0.821758519 | 0.91001573 | 0.067100169 | Not Significant | ENSMUSG00000052676 | 215693 | protein_coding |
| Zmat2 | 516.7085864 | 0.014834619 | 0.050359418 | -0.29457488 | 0.768318654 | 0.883975099 | 0.014834619 | Not Significant | ENSMUSG00000001383 | 66492 | protein_coding |
| Zmat3 | 207.815639 | 0.025465434 | 0.111960651 | -0.227449858 | 0.820073955 | 0.909231489 | 0.025465434 | Not Significant | ENSMUSG00000027663 | 22401 | protein_coding |
| Zmat4 | 13.51560238 | -0.471481987 | 0.505673073 | 0.932384998 | 0.351137595 | 0.573211901 | 0.471481987 | Not Significant | ENSMUSG00000037492 | 320158 | protein_coding |
| Zmat5 | 111.8835088 | -0.172424996 | 0.115509578 | 1.49273332 | 0.135507008 | 0.30685209 | 0.172424996 | Not Significant | ENSMUSG00000009076 | 67178 | protein_coding |
| Zmiz1 | 825.7428542 | -0.047269506 | 0.298233177 | 0.158498483 | 0.874064016 | 0.938083652 | 0.047269506 | Not Significant | ENSMUSG00000007817 | 328365 | protein_coding |
| Zmiz2 | 482.1686083 | -0.097286875 | 0.122229754 | 0.795934474 | 0.426070122 | 0.645752753 | 0.097286875 | Not Significant | ENSMUSG00000041164 | 52915 | protein_coding |
| Zmpste24 | 588.4330586 | 0.124663252 | 0.087288949 | -1.42816763 | 0.153243615 | 0.334272057 | 0.124663252 | Not Significant | ENSMUSG00000043207 | 230709 | protein_coding |
| Zmym1 | 57.05730672 | 0.380697276 | 0.192995832 | -1.972567347 | 0.048544876 | 0.14581122 | 0.380697276 | Not Significant | ENSMUSG00000043872 | 68310 | protein_coding |
| Zmym2 | 777.057143 | -0.1159501 | 0.077328042 | 1.499457328 | 0.133755031 | 0.303828141 | 0.1159501 | Not Significant | ENSMUSG00000021945 | 76007 | protein_coding |
| Zmym3 | 260.5321973 | -0.005163266 | 0.102853979 | 0.050199963 | 0.959963041 | 0.980506535 | 0.005163266 | Not Significant | ENSMUSG00000031310 | 56364 | protein_coding |
| Zmym4 | 247.137144 | -0.092505817 | 0.079586569 | 1.16232951 | 0.245101644 | 0.456768986 | 0.092505817 | Not Significant | ENSMUSG00000042446 | 67785 | protein_coding |
| Zmym5 | 332.9204503 | -0.083484068 | 0.081316382 | 1.026657432 | 0.3045818 | 0.524764095 | 0.083484068 | Not Significant | ENSMUSG00000040123 | 219105 | protein_coding |
| Zmym6 | 134.3919918 | -0.512217309 | 0.124842858 | 4.102896364 | 4.08E-05 | 0.000391737 | 0.512217309 | DOWN | ENSMUSG00000042408 | 100177 | protein_coding |
| Zmynd10 | 3.300740004 | 0.44218702 | 0.954294101 | -0.46336556 | 0.643102361 | 1 | 0.44218702 | Not Significant | NA | NA | NA |
| Zmynd11 | 572.5057062 | 0.023636953 | 0.04701495 | -0.502753962 | 0.615137265 | 0.787654245 | 0.023636953 | Not Significant | ENSMUSG00000021156 | 66505 | protein_coding |
| Zmynd12 | 7.591215054 | 0.943086626 | 0.552850986 | -1.705860439 | 0.088034067 | 0.22515016 | 0.943086626 | Not Significant | ENSMUSG00000070806 | 332934 | protein_coding |
| Zmynd15 | 4.395792338 | 1.163703358 | 1.066298432 | -1.091348654 | 0.275119496 | 0.491701663 | 1.163703358 | Not Significant | NA | NA | NA |
| Zmynd19 | 235.5425989 | 0.570366321 | 0.123696051 | -4.611030971 | 4.01E-06 | 4.90E-05 | 0.570366321 | UP | ENSMUSG00000026974 | 67187 | protein_coding |
| Zmynd8 | 1037.212499 | -0.049357596 | 0.071101014 | 0.694189759 | 0.487563214 | 0.696047947 | 0.049357596 | Not Significant | ENSMUSG00000039671 | 228880 | protein_coding |
| Znfx1 | 778.126088 | -0.18629565 | 0.112694952 | 1.65309667 | 0.098311191 | 0.244320693 | 0.18629565 | Not Significant | ENSMUSG00000039501 | 98999 | protein_coding |
| Znhit1 | 294.015806 | 0.006826791 | 0.099291014 | -0.068755372 | 0.945184342 | 0.97354273 | 0.006826791 | Not Significant | ENSMUSG00000059518 | 70103 | protein_coding |
| Znhit2 | 194.5982238 | 0.259735632 | 0.131947076 | -1.968483426 | 0.049012441 | 0.146868346 | 0.259735632 | Not Significant | ENSMUSG00000075227 | 29805 | protein_coding |
| Znhit3 | 63.69480418 | 0.130014742 | 0.151918568 | -0.855818633 | 0.392098107 | 0.614686552 | 0.130014742 | Not Significant | ENSMUSG00000020526 | 448850 | protein_coding |
| Znhit6 | 64.53388172 | -0.056159059 | 0.170500143 | 0.329378371 | 0.741869714 | 0.869560398 | 0.056159059 | Not Significant | ENSMUSG00000074182 | 229937 | protein_coding |
| Znrd2 | 1.554802687 | 1.061652254 | 0.942175918 | -1.126808946 | 0.259823267 | 1 | 1.061652254 | Not Significant | NA | NA | NA |
| Znrf1 | 190.064011 | 0.130642706 | 0.086368456 | -1.512620606 | 0.130376057 | 0.298368931 | 0.130642706 | Not Significant | ENSMUSG00000033545 | 170737 | protein_coding |
| Znrf2 | 962.0838925 | -0.189580124 | 0.070595156 | 2.685455139 | 0.00724311 | 0.032806559 | 0.189580124 | Significant Negative | ENSMUSG00000058446 | 387524 | protein_coding |
| Znrf3 | 161.9108852 | -0.214977172 | 0.128558501 | 1.672212803 | 0.094482367 | 0.236872902 | 0.214977172 | Not Significant | ENSMUSG00000041961 | 407821 | protein_coding |
| Zp1 | 3.889098907 | -0.50950679 | 0.885199012 | 0.575584454 | 0.564896091 | 1 | 0.50950679 | Not Significant | ENSMUSG00000024734 | 22786 | protein_coding |
| Zp2 | 5.418931895 | -0.103404622 | 0.658441335 | 0.157044548 | 0.875209739 | 0.938415991 | 0.103404622 | Not Significant | NA | NA | NA |
| Zpbp2 | 1.450859149 | -1.23186727 | 1.152904387 | 1.068490401 | 0.285299358 | 1 | 1.23186727 | Not Significant | NA | NA | NA |
| Zpld2 | 1.729529425 | -0.752194537 | 3.05833442 | 0.24594908 | 0.80572165 | 1 | 0.752194537 | Not Significant | NA | NA | NA |
| Zpr1 | 323.6301268 | -0.151262874 | 0.109213036 | 1.385025813 | 0.166044575 | 0.353403803 | 0.151262874 | Not Significant | ENSMUSG00000032078 | 22687 | protein_coding |
| Zranb1 | 628.0893306 | -0.123402115 | 0.08902304 | 1.386181765 | 0.165691408 | 0.352762472 | 0.123402115 | Not Significant | ENSMUSG00000030967 | 360216 | protein_coding |
| Zranb2 | 606.6898468 | -0.008287956 | 0.092870269 | 0.089242298 | 0.928889351 | 0.966049475 | 0.008287956 | Not Significant | ENSMUSG00000028180 | 53861 | protein_coding |
| Zranb3 | 67.09382733 | 0.418656907 | 0.17788128 | -2.353574845 | 0.018593864 | 0.069313561 | 0.418656907 | Not Significant | ENSMUSG00000036086 | 226409 | protein_coding |
| Zrsr2 | 242.5704049 | 0.057151045 | 0.072685253 | -0.786281159 | 0.431702788 | 0.650053783 | 0.057151045 | Not Significant | ENSMUSG00000031370 | 22184 | protein_coding |
| Zscan10 | 1.848399832 | 0.960847928 | 1.575285764 | -0.609951508 | 0.54189393 | 1 | 0.960847928 | Not Significant | NA | NA | NA |
| Zscan12 | 74.02585578 | -0.035544558 | 0.189226891 | 0.187840944 | 0.851001338 | 0.927281893 | 0.035544558 | Not Significant | ENSMUSG00000036721 | 22758 | protein_coding |
| Zscan18 | 7.950824682 | 1.040685346 | 0.489404607 | -2.126431447 | 0.033467352 | 0.109658699 | 1.040685346 | Not Significant | ENSMUSG00000070822 | 232875 | protein_coding |
| Zscan2 | 33.10194188 | -0.12115792 | 0.210436665 | 0.575745295 | 0.564787355 | 0.751588072 | 0.12115792 | Not Significant | ENSMUSG00000038797 | 22691 | protein_coding |
| Zscan20 | 93.18035012 | -0.190382776 | 0.120212052 | 1.583724534 | 0.113256419 | 0.270127107 | 0.190382776 | Not Significant | ENSMUSG00000061894 | 269585 | protein_coding |
| Zscan21 | 212.0206503 | -0.005181551 | 0.103095303 | 0.050259817 | 0.959915345 | 0.980506535 | 0.005181551 | Not Significant | ENSMUSG00000037017 | 22697 | protein_coding |
| Zscan22 | 84.03098568 | 0.40564216 | 0.145803556 | -2.78211431 | 0.005400601 | 0.025963765 | 0.40564216 | Significant Positive | ENSMUSG00000054715 | 232878 | protein_coding |
| Zscan25 | 139.3103244 | -0.005173196 | 0.10685669 | 0.048412465 | 0.961387525 | 0.981111951 | 0.005173196 | Not Significant | ENSMUSG00000070420 | 666311 | protein_coding |
| Zscan26 | 414.0563171 | -0.193072506 | 0.084277391 | 2.290916975 | 0.021968216 | 0.079002436 | 0.193072506 | Not Significant | ENSMUSG00000022228 | 432731 | protein_coding |
| Zscan29 | 334.144663 | -0.230043323 | 0.080968017 | 2.841162869 | 0.004494935 | 0.022291098 | 0.230043323 | Significant Negative | ENSMUSG00000050619 | 99334 | protein_coding |
| Zswim1 | 117.9526382 | 0.469334349 | 0.133521737 | -3.51504077 | 0.000439686 | 0.003190048 | 0.469334349 | Significant Positive | ENSMUSG00000017764 | 71971 | protein_coding |
| Zswim3 | 77.55914867 | -0.02951894 | 0.136947711 | 0.215548984 | 0.829339326 | 0.913717368 | 0.02951894 | Not Significant | ENSMUSG00000045822 | 67538 | protein_coding |
| Zswim4 | 228.9599354 | 0.084786377 | 0.162616833 | -0.521387454 | 0.602096889 | 0.778258299 | 0.084786377 | Not Significant | ENSMUSG00000035671 | 212168 | protein_coding |
| Zswim5 | 165.9298045 | -0.243145437 | 0.094730573 | 2.566705014 | 0.010266991 | 0.043464619 | 0.243145437 | Significant Negative | ENSMUSG00000033948 | 74464 | protein_coding |
| Zswim6 | 290.8133656 | -0.107552582 | 0.127934594 | 0.840684123 | 0.400524917 | 0.622394781 | 0.107552582 | Not Significant | ENSMUSG00000032846 | 67263 | protein_coding |
| Zswim7 | 57.47589174 | -0.199054215 | 0.144788669 | 1.374791382 | 0.169196131 | 0.358402373 | 0.199054215 | Not Significant | ENSMUSG00000014243 | 69747 | protein_coding |
| Zswim8 | 558.7977277 | 0.10743396 | 0.09313172 | -1.153570021 | 0.248676495 | 0.460883095 | 0.10743396 | Not Significant | ENSMUSG00000021819 | 268721 | protein_coding |
| Zswim9 | 59.71020104 | -0.049512787 | 0.161145928 | 0.307254349 | 0.758649781 | 0.878912418 | 0.049512787 | Not Significant | ENSMUSG00000070814 | 321008 | protein_coding |
| Zup1 | 167.9050686 | -0.373556329 | 0.088066233 | 4.241765726 | 2.22E-05 | 0.000226486 | 0.373556329 | Significant Negative | ENSMUSG00000039531 | 72580 | protein_coding |
| Zw10 | 345.8537793 | 0.056599846 | 0.079867209 | -0.708674402 | 0.478526554 | 0.690104577 | 0.056599846 | Not Significant | ENSMUSG00000032264 | 26951 | protein_coding |
| Zwilch | 107.7737516 | 0.92085516 | 0.156210727 | -5.894954703 | 3.75E-09 | 7.70E-08 | 0.92085516 | UP | ENSMUSG00000032400 | 68014 | protein_coding |
| Zwint | 386.914665 | 0.53111505 | 0.076462237 | -6.946109265 | 3.75E-12 | 1.09E-10 | 0.53111505 | UP | ENSMUSG00000019923 | 52696 | protein_coding |
| Zxdb | 49.24936466 | -0.141200252 | 0.187390343 | 0.753508688 | 0.451144292 | 0.667666133 | 0.141200252 | Not Significant | ENSMUSG00000073062 | 668166 | protein_coding |
| Zxdc | 279.5139735 | -0.146363094 | 0.091915664 | 1.592362904 | 0.111303186 | 0.26704356 | 0.146363094 | Not Significant | ENSMUSG00000034430 | 80292 | protein_coding |
| Zyg11a | 5.359491047 | -0.086502189 | 0.703925806 | 0.122885378 | 0.902197865 | 0.951530343 | 0.086502189 | Not Significant | ENSMUSG00000034645 | 230590 | protein_coding |
| Zyg11b | 965.9856194 | -0.031317745 | 0.067732098 | 0.462376716 | 0.643811194 | 0.805935268 | 0.031317745 | Not Significant | ENSMUSG00000034636 | 414872 | protein_coding |
| Zyx | 523.582034 | -0.266370601 | 0.122243214 | 2.179021577 | 0.029330065 | 0.099059146 | 0.266370601 | Not Significant | ENSMUSG00000029860 | 22793 | protein_coding |
| Zzef1 | 2593.609642 | 0.167539874 | 0.090851951 | -1.8440977 | 0.065168903 | 0.181269925 | 0.167539874 | Not Significant | ENSMUSG00000055670 | 195018 | protein_coding |
| Zzz3 | 709.3471847 | -0.012832101 | 0.081304141 | 0.157828387 | 0.87459203 | 0.938406345 | 0.012832101 | Not Significant | ENSMUSG00000039068 | 108946 | protein_coding |

| geneID | Cond2_DF_baseMean | Cond2_DF_log2FoldChange | Cond2_DF_lfcSE | Cond2_DF_stat | Cond2_DF_pvalue | Cond2_DF_padj | Cond2_DF_absLog2FC | RG1_Specific_Cond2 | RG1_Specific_Cond1 |
|---|---|---|---|---|---|---|---|---|---|
| Zfyve26 | 583.4170965 | -0.078011842 | 0.06179247 | -1.262481374 | 0.206775624 | 0.441678211 | 0.078011842 | Cond1 No Change + Cond2 Not Significant | Cond2 No Change + Cond1 Not Significant |
| Zfyve27 | 779.2007931 | -0.088482829 | 0.063355529 | -1.396607841 | 0.162531531 | 0.383544922 | 0.088482829 | Cond1 No Change + Cond2 Not Significant | Cond2 No Change + Cond1 Significant Negative |
| Zfyve28 | NA | NA | NA | NA | NA | NA | NA | Background = FALSE | Background = FALSE |
| Zfyve9 | 1034.698187 | -0.18422642 | 0.054024156 | -3.410074913 | 0.00064945 | 0.007100804 | 0.18422642 | Cond1 No Change + Cond2 Significant Negative | Cond2 No Change + Cond1 Significant Negative |
| Zg16 | 90928.39472 | 0.556720745 | 0.083796487 | 6.643724167 | 3.06E-11 | 1.40E-09 | 0.556720745 | Cond1 No Change + Cond2 UP | Cond2 UP + Cond1 Not Significant |
| Zgpat | 646.173453 | -0.047957688 | 0.059244451 | -0.809488265 | 0.41823435 | 0.655860109 | 0.047957688 | Cond1 No Change + Cond2 Not Significant | Cond2 No Change + Cond1 Not Significant |
| Zgrf1 | 86.82887421 | 0.064530906 | 0.191500023 | 0.336975972 | 0.736135014 | 0.87317221 | 0.064530906 | Cond1 No Change + Cond2 Not Significant | Cond2 No Change + Cond1 Not Significant |
| Zhx1 | 964.4948513 | -0.100676968 | 0.057926656 | -1.738007583 | 0.082209478 | 0.254568403 | 0.100676968 | Cond1 No Change + Cond2 Not Significant | Cond2 No Change + Cond1 Not Significant |
| Zhx2 | 317.2680636 | 0.114339437 | 0.070676812 | 1.617778628 | 0.105710314 | 0.29784969 | 0.114339437 | Cond1 No Change + Cond2 Not Significant | Cond2 No Change + Cond1 Not Significant |
| Zhx3 | 424.8978733 | -0.005897987 | 0.066619674 | -0.088532211 | 0.929453684 | 0.96881133 | 0.005897987 | Cond1 No Change + Cond2 Not Significant | Cond2 No Change + Cond1 Significant Positive |
| Zic1 | NA | NA | NA | NA | NA | NA | NA | Background = FALSE | Background = FALSE |
| Zic3 | NA | NA | NA | NA | NA | NA | NA | Background = FALSE | Background = FALSE |
| Zic4 | NA | NA | NA | NA | NA | NA | NA | Background = FALSE | Background = FALSE |
| Zic5 | NA | NA | NA | NA | NA | NA | NA | Background = FALSE | Background = FALSE |
| Zik1 | NA | NA | NA | NA | NA | NA | NA | Background = FALSE | Background = FALSE |
| Zim1 | NA | NA | NA | NA | NA | NA | NA | Background = FALSE | Background = FALSE |
| Zkscan1 | 2902.083401 | -0.100383908 | 0.055831829 | -1.797969191 | 0.07218189 | 0.234551147 | 0.100383908 | Cond1 No Change + Cond2 Not Significant | Cond2 No Change + Cond1 Significant Negative |
| Zkscan14 | 259.3398744 | -0.13684261 | 0.120851837 | -1.132317171 | 0.257501114 | 0.499365134 | 0.13684261 | Cond1 No Change + Cond2 Not Significant | Cond2 No Change + Cond1 Significant Negative |
| Zkscan16 | NA | NA | NA | NA | NA | NA | NA | Background = FALSE | Background = FALSE |
| Zkscan17 | 362.7926074 | -0.317283429 | 0.084543114 | -3.752918667 | 0.000174788 | 0.002345091 | 0.317283429 | Cond1 No Change + Cond2 Significant Negative | Cond2 No Change + Cond1 Not Significant |
| Zkscan2 | NA | NA | NA | NA | NA | NA | NA | Background = FALSE | Background = FALSE |
| Zkscan3 | 1004.047403 | 0.14114423 | 0.065278951 | 2.162170616 | 0.030605028 | 0.132040471 | 0.14114423 | Cond1 No Change + Cond2 Not Significant | Cond2 No Change + Cond1 Not Significant |
| Zkscan4 | 16.82274518 | 0.145085352 | 0.331625014 | 0.437498217 | 0.661750069 | 0.827097835 | 0.145085352 | Cond1 No Change + Cond2 Not Significant | Cond2 No Change + Cond1 Not Significant |
| Zkscan5 | 343.1848433 | 0.018051662 | 0.075193723 | 0.24006874 | 0.810276968 | 0.914739662 | 0.018051662 | Cond1 No Change + Cond2 Not Significant | Cond2 No Change + Cond1 Not Significant |
| Zkscan6 | 429.9570381 | 0.042056284 | 0.067961473 | 0.618825378 | 0.536031402 | 0.742852833 | 0.042056284 | Cond1 No Change + Cond2 Not Significant | Cond2 No Change + Cond1 Not Significant |
| Zkscan7 | 391.2935747 | -0.121055685 | 0.091184788 | -1.327586408 | 0.184314777 | 0.41301159 | 0.121055685 | Cond1 No Change + Cond2 Not Significant | Cond2 No Change + Cond1 Not Significant |
| Zkscan8 | 395.431613 | -0.278100488 | 0.08533095 | -3.259081132 | 0.001117737 | 0.011052553 | 0.278100488 | Cond1 No Change + Cond2 Significant Negative | Cond2 No Change + Cond1 Significant Negative |
| Zmat1 | 21.19169905 | 0.324822099 | 0.277577661 | 1.170202594 | 0.241919449 | 0.482850861 | 0.324822099 | Cond1 No Change + Cond2 Not Significant | Cond2 No Change + Cond1 Not Significant |
| Zmat2 | 961.3995351 | 0.070362816 | 0.049602871 | 1.418523039 | 0.156038117 | 0.374523994 | 0.070362816 | Cond1 No Change + Cond2 Not Significant | Cond2 No Change + Cond1 Not Significant |
| Zmat3 | 359.8715644 | 0.015699372 | 0.128479392 | 0.122193695 | 0.90274562 | 0.957171506 | 0.015699372 | Cond1 No Change + Cond2 Not Significant | Cond2 No Change + Cond1 Not Significant |
| Zmat4 | 49.88283932 | -0.222724213 | 0.278624392 | -0.799370836 | 0.424075416 | 0.659950229 | 0.222724213 | Cond1 No Change + Cond2 Not Significant | Cond2 No Change + Cond1 Not Significant |
| Zmat5 | 240.1403864 | 0.041397726 | 0.086860346 | 0.476600977 | 0.633646288 | 0.808283189 | 0.041397726 | Cond1 No Change + Cond2 Not Significant | Cond2 No Change + Cond1 Not Significant |
| Zmiz1 | 1530.742433 | -0.252984105 | 0.075092586 | -3.368962466 | 0.000754517 | 0.007978132 | 0.252984105 | Cond1 No Change + Cond2 Significant Negative | Cond2 No Change + Cond1 Not Significant |
| Zmiz2 | 777.4122677 | -0.054584338 | 0.090799082 | -0.601155177 | 0.547736636 | 0.750752059 | 0.054584338 | Cond1 No Change + Cond2 Not Significant | Cond2 No Change + Cond1 Not Significant |
| Zmpste24 | 1044.735578 | 0.086428171 | 0.051185465 | 1.688529581 | 0.091309616 | 0.272341441 | 0.086428171 | Cond1 No Change + Cond2 Not Significant | Cond2 No Change + Cond1 Not Significant |
| Zmym1 | 92.86367388 | -0.036610866 | 0.123735261 | -0.295880623 | 0.76732125 | 0.890725386 | 0.036610866 | Cond1 No Change + Cond2 Not Significant | Cond2 No Change + Cond1 Not Significant |
| Zmym2 | 1511.200437 | -0.142052977 | 0.058149017 | -2.442912763 | 0.014569259 | 0.078555143 | 0.142052977 | Cond1 No Change + Cond2 Not Significant | Cond2 No Change + Cond1 Not Significant |
| Zmym3 | 518.3210291 | -0.137900301 | 0.060309521 | -2.28654281 | 0.022222523 | 0.105936335 | 0.137900301 | Cond1 No Change + Cond2 Not Significant | Cond2 No Change + Cond1 Not Significant |
| Zmym4 | 495.766588 | -0.266356297 | 0.093224328 | -2.857154385 | 0.004274579 | 0.031797108 | 0.266356297 | Cond1 No Change + Cond2 Significant Negative | Cond2 No Change + Cond1 Not Significant |
| Zmym5 | 682.3068614 | -0.138249484 | 0.094951299 | -1.456004133 | 0.145391477 | 0.358198217 | 0.138249484 | Cond1 No Change + Cond2 Not Significant | Cond2 No Change + Cond1 Not Significant |
| Zmym6 | 297.4706813 | -0.394143501 | 0.111544785 | -3.533500045 | 0.000410096 | 0.004775176 | 0.394143501 | Cond1 DOWN + Cond2 Significant Negative | Cond2 No Change + Cond1 DOWN |
| Zmynd10 | NA | NA | NA | NA | NA | NA | NA | Background = FALSE | Background = FALSE |
| Zmynd11 | 1158.18267 | -0.078770593 | 0.046204957 | -1.704808279 | 0.088230187 | 0.26640858 | 0.078770593 | Cond1 No Change + Cond2 Not Significant | Cond2 No Change + Cond1 Not Significant |
| Zmynd12 | 14.27203808 | 0.172012978 | 0.363699406 | 0.472953697 | 0.636246214 | 0.809845517 | 0.172012978 | Cond1 No Change + Cond2 Not Significant | Cond2 No Change + Cond1 Not Significant |
| Zmynd15 | NA | NA | NA | NA | NA | NA | NA | Background = FALSE | Background = FALSE |
| Zmynd19 | 340.397109 | 0.269807448 | 0.078593099 | 3.432966131 | 0.000597017 | 0.006617243 | 0.269807448 | Cond1 UP + Cond2 Significant Positive | Cond2 No Change + Cond1 UP |
| Zmynd8 | 1578.427575 | 0.008980416 | 0.045788228 | 0.196129352 | 0.844508923 | 0.930210801 | 0.008980416 | Cond1 No Change + Cond2 Not Significant | Cond2 No Change + Cond1 Not Significant |
| Znfx1 | 1477.633342 | -0.106357965 | 0.058486614 | -1.818500963 | 0.0689876 | 0.227937517 | 0.106357965 | Cond1 No Change + Cond2 Not Significant | Cond2 No Change + Cond1 Not Significant |
| Znhit1 | 492.7889635 | 0.133572277 | 0.097118274 | 1.375356788 | 0.169020859 | 0.393219762 | 0.133572277 | Cond1 No Change + Cond2 Not Significant | Cond2 No Change + Cond1 Not Significant |
| Znhit2 | 359.8308531 | 0.193800488 | 0.103724913 | 1.868408298 | 0.061705181 | 0.211953026 | 0.193800488 | Cond1 No Change + Cond2 Not Significant | Cond2 No Change + Cond1 Not Significant |
| Znhit3 | 100.2568881 | 0.047103173 | 0.130092593 | 0.362074209 | 0.717296576 | 0.861407542 | 0.047103173 | Cond1 No Change + Cond2 Not Significant | Cond2 No Change + Cond1 Not Significant |
| Znhit6 | 104.864322 | 0.095299113 | 0.11855413 | 0.803844739 | 0.421486648 | 0.658144381 | 0.095299113 | Cond1 No Change + Cond2 Not Significant | Cond2 No Change + Cond1 Not Significant |
| Znrd2 | NA | NA | NA | NA | NA | NA | NA | Background = FALSE | Background = FALSE |
| Znrf1 | 401.77805 | 0.042567935 | 0.072743161 | 0.585181263 | 0.558425842 | 0.756975167 | 0.042567935 | Cond1 No Change + Cond2 Not Significant | Cond2 No Change + Cond1 Not Significant |
| Znrf2 | 2381.226926 | -0.164307588 | 0.047451703 | -3.462627819 | 0.000534928 | 0.005996667 | 0.164307588 | Cond1 No Change + Cond2 Significant Negative | Cond2 No Change + Cond1 Significant Negative |
| Znrf3 | 386.2756673 | -0.287815276 | 0.100315725 | -2.869094318 | 0.00411649 | 0.03084881 | 0.287815276 | Cond1 No Change + Cond2 Significant Negative | Cond2 No Change + Cond1 Not Significant |
| Zp1 | 10.06743462 | 0.353287648 | 0.4132162 | 0.854970467 | 0.392567498 | 0.632267311 | 0.353287648 | Cond1 No Change + Cond2 Not Significant | Cond2 No Change + Cond1 Not Significant |
| Zp2 | NA | NA | NA | NA | NA | NA | NA | Background = FALSE | Background = FALSE |
| Zpbp2 | NA | NA | NA | NA | NA | NA | NA | Background = FALSE | Background = FALSE |
| Zpld2 | NA | NA | NA | NA | NA | NA | NA | Background = FALSE | Background = FALSE |
| Zpr1 | 627.336632 | -0.11578679 | 0.061737658 | -1.87546455 | 0.060728842 | 0.20964847 | 0.11578679 | Cond1 No Change + Cond2 Not Significant | Cond2 No Change + Cond1 Not Significant |
| Zranb1 | 1348.252433 | -0.097740711 | 0.058368717 | -1.674539314 | 0.094024655 | 0.27725095 | 0.097740711 | Cond1 No Change + Cond2 Not Significant | Cond2 No Change + Cond1 Not Significant |
| Zranb2 | 1082.267197 | 0.13689921 | 0.074036737 | 1.84907136 | 0.064447509 | 0.218015518 | 0.13689921 | Cond1 No Change + Cond2 Not Significant | Cond2 No Change + Cond1 Not Significant |
| Zranb3 | 103.6845357 | 0.070398327 | 0.117461115 | 0.599333042 | 0.548950818 | 0.751266959 | 0.070398327 | Cond1 No Change + Cond2 Not Significant | Cond2 No Change + Cond1 Not Significant |
| Zrsr2 | 437.5160725 | 0.015485158 | 0.063959843 | 0.242107512 | 0.808696858 | 0.913844956 | 0.015485158 | Cond1 No Change + Cond2 Not Significant | Cond2 No Change + Cond1 Not Significant |
| Zscan10 | NA | NA | NA | NA | NA | NA | NA | Background = FALSE | Background = FALSE |
| Zscan12 | 128.6066407 | -0.043850654 | 0.117616728 | -0.372826681 | 0.709277444 | 0.856166251 | 0.043850654 | Cond1 No Change + Cond2 Not Significant | Cond2 No Change + Cond1 Not Significant |
| Zscan18 | 7.304793562 | 0.32371056 | 0.463297836 | 0.698709416 | 0.484733648 | 0.706263486 | 0.32371056 | Cond1 No Change + Cond2 Not Significant | Cond2 No Change + Cond1 Not Significant |
| Zscan2 | 71.51568947 | -0.043960322 | 0.148052526 | -0.296923821 | 0.76652467 | 0.890414429 | 0.043960322 | Cond1 No Change + Cond2 Not Significant | Cond2 No Change + Cond1 Not Significant |
| Zscan20 | 182.5730863 | 0.015267903 | 0.097151549 | 0.157155531 | 0.875122274 | 0.943767347 | 0.015267903 | Cond1 No Change + Cond2 Not Significant | Cond2 No Change + Cond1 Not Significant |
| Zscan21 | 414.9582595 | -0.072214029 | 0.075043349 | -0.962297525 | 0.335900172 | 0.579426898 | 0.072214029 | Cond1 No Change + Cond2 Not Significant | Cond2 No Change + Cond1 Not Significant |
| Zscan22 | 118.7483404 | 0.125022599 | 0.129202571 | 0.967647923 | 0.333220235 | 0.576562634 | 0.125022599 | Cond1 No Change + Cond2 Not Significant | Cond2 No Change + Cond1 Significant Positive |
| Zscan25 | 262.7870236 | -0.112771364 | 0.076142924 | -1.481048507 | 0.138593648 | 0.34868951 | 0.112771364 | Cond1 No Change + Cond2 Not Significant | Cond2 No Change + Cond1 Not Significant |
| Zscan26 | 815.9763847 | -0.201011959 | 0.093396151 | -2.152250992 | 0.031377591 | 0.134058533 | 0.201011959 | Cond1 No Change + Cond2 Not Significant | Cond2 No Change + Cond1 Not Significant |
| Zscan29 | 723.6371644 | -0.144314767 | 0.051797878 | -2.78611349 | 0.005334419 | 0.037436109 | 0.144314767 | Cond1 No Change + Cond2 Significant Negative | Cond2 No Change + Cond1 Significant Negative |
| Zswim1 | 193.4772098 | 0.240695172 | 0.111287667 | 2.162819816 | 0.030555041 | 0.131915576 | 0.240695172 | Cond1 No Change + Cond2 Not Significant | Cond2 No Change + Cond1 Significant Positive |
| Zswim3 | 118.9946016 | 0.03583183 | 0.117729381 | 0.304357584 | 0.760855479 | 0.886968072 | 0.03583183 | Cond1 No Change + Cond2 Not Significant | Cond2 No Change + Cond1 Not Significant |
| Zswim4 | 417.9395745 | -0.277972733 | 0.071882193 | -3.867059712 | 0.000110155 | 0.001614979 | 0.277972733 | Cond1 No Change + Cond2 Significant Negative | Cond2 No Change + Cond1 Not Significant |
| Zswim5 | 388.4984927 | -0.204648335 | 0.061095834 | -3.34962832 | 0.000809201 | 0.008441125 | 0.204648335 | Cond1 No Change + Cond2 Significant Negative | Cond2 No Change + Cond1 Significant Negative |
| Zswim6 | 556.9697921 | -0.138390133 | 0.071041426 | -1.9480202 | 0.051412546 | 0.188695979 | 0.138390133 | Cond1 No Change + Cond2 Not Significant | Cond2 No Change + Cond1 Not Significant |
| Zswim7 | 137.4601393 | -0.103554197 | 0.103310985 | -1.002354169 | 0.316172569 | 0.560212157 | 0.103554197 | Cond1 No Change + Cond2 Not Significant | Cond2 No Change + Cond1 Not Significant |
| Zswim8 | 1026.490705 | -0.157983007 | 0.077187033 | -2.046755785 | 0.040682069 | 0.16048917 | 0.157983007 | Cond1 No Change + Cond2 Not Significant | Cond2 No Change + Cond1 Not Significant |
| Zswim9 | 87.28893798 | 0.080046226 | 0.143462885 | 0.557957731 | 0.576873248 | 0.769561988 | 0.080046226 | Cond1 No Change + Cond2 Not Significant | Cond2 No Change + Cond1 Not Significant |
| Zup1 | 340.875373 | -0.259386991 | 0.094353412 | -2.749100261 | 0.00597591 | 0.040905563 | 0.259386991 | Cond1 No Change + Cond2 Significant Negative | Cond2 No Change + Cond1 Significant Negative |
| Zw10 | 576.7191269 | 0.180726297 | 0.053863511 | 3.355263962 | 0.000792893 | 0.008303236 | 0.180726297 | Cond1 No Change + Cond2 Significant Positive | Cond2 No Change + Cond1 Not Significant |
| Zwilch | 127.6281628 | 0.119877471 | 0.125781346 | 0.953062396 | 0.340558457 | 0.583344264 | 0.119877471 | Cond1 UP + Cond2 Not Significant | Cond2 No Change + Cond1 UP |
| Zwint | 673.3679052 | 0.038937189 | 0.085515704 | 0.455322093 | 0.648877534 | 0.818686277 | 0.038937189 | Cond1 UP + Cond2 Not Significant | Cond2 No Change + Cond1 UP |
| Zxdb | 100.2353003 | -0.027206937 | 0.124437924 | -0.218638629 | 0.826931559 | 0.923060252 | 0.027206937 | Cond1 No Change + Cond2 Not Significant | Cond2 No Change + Cond1 Not Significant |
| Zxdc | 584.0646787 | -0.133708218 | 0.058015544 | -2.304696468 | 0.021183579 | 0.102159742 | 0.133708218 | Cond1 No Change + Cond2 Not Significant | Cond2 No Change + Cond1 Not Significant |
| Zyg11a | 12.02743039 | -1.09199946 | 0.414085606 | -2.637134553 | 0.008360966 | 0.052253619 | 1.09199946 | Cond1 No Change + Cond2 Not Significant | Cond2 No Change + Cond1 Not Significant |
| Zyg11b | 1883.760765 | 0.028161196 | 0.044927663 | 0.626811948 | 0.53078251 | 0.73971984 | 0.028161196 | Cond1 No Change + Cond2 Not Significant | Cond2 No Change + Cond1 Not Significant |
| Zyx | 802.6401604 | -0.302439055 | 0.107237536 | -2.820272323 | 0.004798291 | 0.034530138 | 0.302439055 | Cond1 No Change + Cond2 Significant Negative | Cond2 No Change + Cond1 Not Significant |
| Zzef1 | 4640.831356 | -0.009275707 | 0.04034434 | -0.229913469 | 0.81815901 | 0.91832199 | 0.009275707 | Cond1 No Change + Cond2 Not Significant | Cond2 No Change + Cond1 Not Significant |
| Zzz3 | 1490.920055 | -0.061679852 | 0.049903333 | -1.235986636 | 0.216463531 | 0.452365863 | 0.061679852 | Cond1 No Change + Cond2 Not Significant | Cond2 No Change + Cond1 Not Significant |

Supplementary Table 3

| Cond1 | Cond1_Detailed | Cond2 | Cond2_Detailed | RG1_All | RG2_Significant | RG3_SignificantChange |
|---|---|---|---|---|---|---|
| DOWN | DOWN | DOWN | DOWN | Cond1 DOWN + Cond2 DOWN | Core_DOWN | Core_DOWN |
| DOWN | DOWN | Not Detected | Not Detected | Cond1 DOWN + Cond2 Not Detected | Cond1_DOWN | Cond1_DOWN |
| DOWN | DOWN | No Change | Not Significant | Cond1 DOWN + Cond2 Not Significant | Cond1_DOWN | Cond1_DOWN |
| DOWN | DOWN | No Change | Significant Negative | Cond1 DOWN + Cond2 Significant Negative | Core_DOWN | Cond1_DOWN |
| DOWN | DOWN | No Change | Significant Positive | Cond1 DOWN + Cond2 Significant Positive | Opposite | Cond1_DOWN |
| DOWN | DOWN | UP | UP | Cond1 DOWN + Cond2 UP | Opposite | Opposite |
| UP | UP | DOWN | DOWN | Cond1 UP + Cond2 DOWN | Opposite | Opposite |
| UP | UP | Not Detected | Not Detected | Cond1 UP + Cond2 Not Detected | Cond1_UP | Cond1_UP |
| UP | UP | No Change | Not Significant | Cond1 UP + Cond2 Not Significant | Cond1_UP | Cond1_UP |
| UP | UP | No Change | Significant Negative | Cond1 UP + Cond2 Significant Negative | Opposite | Cond1_UP |
| UP | UP | No Change | Significant Positive | Cond1 UP + Cond2 Significant Positive | Core_UP | Cond1_UP |
| UP | UP | UP | UP | Cond1 UP + Cond2 UP | Core_UP | Core_UP |
| Not Detected | Not Detected | DOWN | DOWN | Cond1 Not Detected + Cond2 DOWN | Cond2_DOWN | Cond2_DOWN |
| Not Detected | Not Detected | Not Detected | Not Detected | Cond1 Not Detected + Cond2 Not Detected | None | None |
| Not Detected | Not Detected | No Change | Not Significant | Cond1 Not Detected + Cond2 Not Significant | None | None |
| Not Detected | Not Detected | No Change | Significant Negative | Cond1 Not Detected + Cond2 Significant Negative | None | None |
| Not Detected | Not Detected | No Change | Significant Positive | Cond1 Not Detected + Cond2 Significant Positive | None | None |
| Not Detected | Not Detected | UP | UP | Cond1 Not Detected + Cond2 UP | Cond2_UP | Cond2_UP |
| No Change | Significant Negative | DOWN | DOWN | Cond1 Significant Negative + Cond2 DOWN | Core_DOWN | Cond2_DOWN |
| No Change | Significant Negative | Not Detected | Not Detected | Cond1 Significant Negative + Cond2 Not Detected | None | None |
| No Change | Significant Negative | No Change | Not Significant | Cond1 Significant Negative + Cond2 Not Significant | None | None |
| No Change | Significant Negative | No Change | Significant Negative | Cond1 Significant Negative + Cond2 Significant Negative | None | None |
| No Change | Significant Negative | No Change | Significant Positive | Cond1 Significant Negative + Cond2 Significant Positive | None | None |
| No Change | Significant Negative | UP | UP | Cond1 Significant Negative + Cond2 UP | Opposite | Cond2_UP |
| No Change | Significant Positive | DOWN | DOWN | Cond1 Significant Positive + Cond2 DOWN | Opposite | Cond2_DOWN |
| No Change | Significant Positive | Not Detected | Not Detected | Cond1 Significant Positive + Cond2 Not Detected | None | None |
| No Change | Significant Positive | No Change | Not Significant | Cond1 Significant Positive + Cond2 Not Significant | None | None |
| No Change | Significant Positive | No Change | Significant Negative | Cond1 Significant Positive + Cond2 Significant Negative | None | None |
| No Change | Significant Positive | No Change | Significant Positive | Cond1 Significant Positive + Cond2 Significant Positive | None | None |
| No Change | Significant Positive | UP | UP | Cond1 Significant Positive + Cond2 UP | Core_UP | Cond2_UP |
| No Change | Not Significant | DOWN | DOWN | Cond1 Not Significant + Cond2 DOWN | Cond2_DOWN | Cond2_DOWN |
| No Change | Not Significant | Not Detected | Not Detected | Cond1 Not Significant + Cond2 Not Detected | None | None |

Supplementary Table 3

| Cond1 | Cond1_Detailed | Cond2 | Cond2_Detailed | RG1_All | RG2_Significant | RG3_SignificantChange |
|---|---|---|---|---|---|---|
| No Change | Not Significant | No Change | Not Significant | Cond1 Not Significant + Cond2 Not Significant | None | None |
| No Change | Not Significant | No Change | Significant Negative | Cond1 Not Significant + Cond2 Significant Negative | None | None |
| No Change | Not Significant | No Change | Significant Positive | Cond1 Not Significant + Cond2 Significant Positive | None | None |
| No Change | Not Significant | UP | UP | Cond1 Not Significant + Cond2 UP | Cond1_UP | Cond1_UP |
