## Supplementary Table 4 for "ATF6 activation alters colonic lipid metabolism causing tumor-associated microbial adaptation"

| ID | Description | GeneRatio | BgRatio | pvalue | p.adjust | qvalue | geneID | Count | Genes_in_Pa | Percentage_of_Pathway_detected |
|---|---|---|---|---|---|---|---|---|---|---|
| KEGG_DRUG_METABOLISM_CYTOCHROME_P450 | DRUG_METABOLISM_CYTOCHROME_P450 | May-33 | 46/3594 | 5.00E-05 | 0.00204812 | 0.00184042 | Aldh3b1/Cyp2d10/Cyp2d12/Cyp2d22/Cyp2d34 | 5 | 84 | 5.95 |
| KEGG_CIRCADIAN_RHYTHM_MAMMAL | CIRCADIAN_RHYTHM_MAMMAL | Feb-33 | Dec-94 | 0.00509532 | 0.10445415 | 0.09386124 | Nr1d1/Per3 | 2 | 13 | 15.38 |
| KEGG_PROXIMAL_TUBULE_BICARBONATE_RECLAMATION | PROXIMAL_TUBULE_BICARBONATE_RECLAMATION | Feb-33 | 15/3594 | 0.00796788 | 0.10889434 | 0.09785114 | Aqp1/Car4 | 2 | 23 | 8.7 |
| KEGG_HISTIDINE_METABOLISM | HISTIDINE_METABOLISM | Feb-33 | 22/3594 | 0.01684116 | 0.1380975 | 0.12409274 | Aldh3b1/Aspa | 2 | 33 | 6.06 |
| KEGG_O_GLYCAN_BIOSYNTHESIS | O_GLYCAN_BIOSYNTHESIS | Feb-33 | 22/3594 | 0.01684116 | 0.1380975 | 0.12409274 | B3gnt6/Gcnt4 | 2 | 29 | 6.9 |
| KEGG_ABC_TRANSPORTERS | ABC_TRANSPORTERS | Feb-33 | 30/3594 | 0.03030072 | 0.20705491 | 0.18605704 | Abcb1a/Abcc8 | 2 | 43 | 4.65 |
| KEGG_AXON_GUIDANCE | AXON_GUIDANCE | Mar-33 | 101/3594 | 0.06396417 | 0.33750074 | 0.30327409 | Ablim2/Ntn4/Sema7a | 3 | 129 | 2.33 |
| KEGG_LEISHMANIA_INFECTION | LEISHMANIA_INFECTION | Feb-33 | 46/3594 | 0.0658538 | 0.33750074 | 0.30327409 | Cyba/Tlr2 | 2 | 70 | 2.86 |
| KEGG_ECM_RECEPTOR_INTERACTION | ECM_RECEPTOR_INTERACTION | Feb-33 | 58/3594 | 0.09832503 | 0.44792512 | 0.40250011 | Lamb1/Thbs1 | 2 | 83 | 2.41 |
| KEGG_PHENYLALANINE_METABOLISM | PHENYLALANINE_METABOLISM | Jan-33 | 14/3594 | 0.121357 | 0.49756371 | 0.44710475 | Aldh3b1 | 1 | 21 | 4.76 |
| KEGG_GLYCOSPHINGOLIPID_BIOSYNTHESIS_LACTO_AND_NEOLACTO_SERIES | GLYCOSPHINGOLIPID_BIOSYNTHESIS_LACTO_AND_NEOLACTO_SERIES | Jan-33 | 18/3594 | 0.15332191 | 0.50819567 | 0.45665849 | St3gal4 | 1 | 23 | 4.35 |
| KEGG_NITROGEN_METABOLISM | NITROGEN_METABOLISM | Jan-33 | 19/3594 | 0.16113521 | 0.50819567 | 0.45665849 | Car4 | 1 | 23 | 4.35 |
| KEGG_SMALL_CELL_LUNG_CANCER | SMALL_CELL_LUNG_CANCER | Feb-33 | 75/3594 | 0.15017155 | 0.50819567 | 0.45665849 | Lamb1/Traf1 | 2 | 82 | 2.44 |
| KEGG_ALANINE_ASPARTATE_AND_GLUTAMATE_METABOLISM | ALANINE_ASPARTATE_AND_GLUTAMATE_METABOLISM | Jan-33 | 25/3594 | 0.20656741 | 0.52659715 | 0.47319385 | Aspa | 1 | 33 | 3.03 |
| KEGG_ARGININE_AND_PROLINE_METABOLISM | ARGININE_AND_PROLINE_METABOLISM | Jan-33 | 46/3594 | 0.34754509 | 0.52659715 | 0.47319385 | Nags | 1 | 59 | 1.69 |
| KEGG_ARRHYTHMOGENIC_RIGHT_VENTRICULAR_CARDIOMYOPATHY_ARVC | ARRHYTHMOGENIC_RIGHT_VENTRICULAR_CARDIOMYOPATHY_ARVC | Jan-33 | 48/3594 | 0.35962732 | 0.52659715 | 0.47319385 | Cacnb3 | 1 | 74 | 1.35 |
| KEGG_BLADDER_CANCER | BLADDER_CANCER | Jan-33 | 37/3594 | 0.29038396 | 0.52659715 | 0.47319385 | Thbs1 | 1 | 42 | 2.38 |
| KEGG_CARDIAC_MUSCLE_CONTRACTION | CARDIAC_MUSCLE_CONTRACTION | Jan-33 | 45/3594 | 0.34142136 | 0.52659715 | 0.47319385 | Cacnb3 | 1 | 79 | 1.27 |
| KEGG_DRUG_METABOLISM_OTHER_ENZYMES | DRUG_METABOLISM_OTHER_ENZYMES | Jan-33 | 37/3594 | 0.29038396 | 0.52659715 | 0.47319385 | Ces1g | 1 | 73 | 1.37 |
| KEGG_FRUCTOSE_AND_MANNOSE_METABOLISM | FRUCTOSE_AND_MANNOSE_METABOLISM | Jan-33 | 35/3594 | 0.27703725 | 0.52659715 | 0.47319385 | Sord | 1 | 38 | 2.63 |
| KEGG_GLYCOLYSIS_GLUCONEOGENESIS | GLYCOLYSIS_GLUCONEOGENESIS | Jan-33 | 48/3594 | 0.35962732 | 0.52659715 | 0.47319385 | Aldh3b1 | 1 | 67 | 1.49 |
| KEGG_HYPERTROPHIC_CARDIOMYOPATHY_HCM | HYPERTROPHIC_CARDIOMYOPATHY_HCM | Jan-33 | 48/3594 | 0.35962732 | 0.52659715 | 0.47319385 | Cacnb3 | 1 | 84 | 1.19 |
| KEGG_METABOLISM_OF_XENOBIOTICS_BY_CYTOCHROME_P450 | METABOLISM_OF_XENOBIOTICS_BY_CYTOCHROME_P450 | Jan-33 | 37/3594 | 0.29038396 | 0.52659715 | 0.47319385 | Aldh3b1 | 1 | 67 | 1.49 |
| KEGG_OXIDATIVE_PHOSPHORYLATION | OXIDATIVE_PHOSPHORYLATION | Feb-33 | 111/3594 | 0.2713629 | 0.52659715 | 0.47319385 | Atp12a/Tcirg1 | 2 | 132 | 1.52 |
| KEGG_STARCH_AND_SUCROSE_METABOLISM | STARCH_AND_SUCROSE_METABOLISM | Jan-33 | 32/3594 | 0.25655967 | 0.52659715 | 0.47319385 | Sis | 1 | 52 | 1.92 |
| KEGG_TYPE_II_DIABETES_MELLITUS | TYPE_II_DIABETES_MELLITUS | Jan-33 | 40/3594 | 0.3099571 | 0.52659715 | 0.47319385 | Abcc8 | 1 | 48 | 2.08 |
| KEGG_TYROSINE_METABOLISM | TYROSINE_METABOLISM | Jan-33 | 27/3594 | 0.22117423 | 0.52659715 | 0.47319385 | Aldh3b1 | 1 | 44 | 2.27 |
| KEGG_VIBRIO_CHOLERAE_INFECTION | VIBRIO_CHOLERAE_INFECTION | Jan-33 | 46/3594 | 0.34754509 | 0.52659715 | 0.47319385 | Tcirg1 | 1 | 52 | 1.92 |
| KEGG_DILATED_CARDIOMYOPATHY | DILATED_CARDIOMYOPATHY | Jan-33 | 53/3594 | 0.38889144 | 0.53148496 | 0.47758597 | Cacnb3 | 1 | 90 | 1.11 |
| KEGG_EPITHELIAL_CELL_SIGNALING_IN_HELICOBACTER_PYLORI_INFECTION | EPITHELIAL_CELL_SIGNALING_IN_HELICOBACTER_PYLORI_INFECTION | Jan-33 | 52/3594 | 0.38314433 | 0.53148496 | 0.47758597 | Tcirg1 | 1 | 64 | 1.56 |
| KEGG_FOCAL_ADHESION | FOCAL_ADHESION | Feb-33 | 152/3594 | 0.41018717 | 0.53801475 | 0.48345356 | Lamb1/Thbs1 | 2 | 195 | 1.03 |
| KEGG_P53_SIGNALING_PATHWAY | P53_SIGNALING_PATHWAY | Jan-33 | 61/3594 | 0.43303626 | 0.53801475 | 0.48345356 | Thbs1 | 1 | 66 | 1.52 |
| KEGG_TGF_BETA_SIGNALING_PATHWAY | TGF_BETA_SIGNALING_PATHWAY | Jan-33 | 61/3594 | 0.43303626 | 0.53801475 | 0.48345356 | Thbs1 | 1 | 83 | 1.2 |
| KEGG_TOLL_LIKE_RECEPTOR_SIGNALING_PATHWAY | TOLL_LIKE_RECEPTOR_SIGNALING_PATHWAY | Jan-33 | 68/3594 | 0.4691134 | 0.56569557 | 0.50832722 | Tlr2 | 1 | 267 | 0.37 |
| KEGG_NEUROACTIVE_LIGAND_RECEPTOR_INTERACTION | NEUROACTIVE_LIGAND_RECEPTOR_INTERACTION | Jan-33 | 84/3594 | 0.54341446 | 0.63657122 | 0.57201522 | Sstr1 | 1 | 313 | 0.32 |
| KEGG_LEUKOCYTE_TRANSENDOTHELIAL_MIGRATION | LEUKOCYTE_TRANSENDOTHELIAL_MIGRATION | Jan-33 | 88/3594 | 0.5603517 | 0.63735349 | 0.57271815 | Cyba | 1 | 112 | 0.89 |
| KEGG_MAPK_SIGNALING_PATHWAY | MAPK_SIGNALING_PATHWAY | Feb-33 | 207/3594 | 0.57517266 | 0.63735349 | 0.57271815 | Cacna1h/Cacnb3 | 2 | 266 | 0.75 |
| KEGG_CALCIUM_SIGNALING_PATHWAY | CALCIUM_SIGNALING_PATHWAY | Jan-33 | 103/3594 | 0.61860539 | 0.66528212 | 0.59781448 | Cacna1h | 1 | 173 | 0.58 |
| KEGG_LYSOSOME | LYSOSOME | Jan-33 | 107/3594 | 0.63282933 | 0.66528212 | 0.59781448 | Tcirg1 | 1 | 118 | 0.85 |
| KEGG_CYTOKINE_CYTOKINE_RECEPTOR_INTERACTION | CYTOKINE_CYTOKINE_RECEPTOR_INTERACTION | Jan-33 | 112/3594 | 0.64988719 | 0.66613437 | 0.59858031 | Tnfsf15 | 1 | 460 | 0.22 |
| KEGG_PATHWAYS_IN_CANCER | PATHWAYS_IN_CANCER | Feb-33 | 262/3594 | 0.70566063 | 0.70566063 | 0.63409812 | Lamb1/Traf1 | 2 | 340 | 0.59 |
| KEGG_ACUTE_MYELOID_LEUKEMIA | ACUTE_MYELOID_LEUKEMIA | NA | NA | NA | NA | NA | NA | 0 | 59 | 0 |
| KEGG_ADHERENS_JUNCTION | ADHERENS_JUNCTION | NA | NA | NA | NA | NA | NA | 0 | 73 | 0 |
| KEGG_ADIPOCYTOKINE_SIGNALING_PATHWAY | ADIPOCYTOKINE_SIGNALING_PATHWAY | NA | NA | NA | NA | NA | NA | 0 | 67 | 0 |
| KEGG_ALDOSTERONE_REGULATED_SODIUM_REABSORPTION | ALDOSTERONE_REGULATED_SODIUM_REABSORPTION | NA | NA | NA | NA | NA | NA | 0 | 43 | 0 |
| KEGG_ALLOGRAFT_REJECTION | ALLOGRAFT_REJECTION | NA | NA | NA | NA | NA | NA | 0 | 37 | 0 |
| KEGG_ALPHA_LINOLENIC_ACID_METABOLISM | ALPHA_LINOLENIC_ACID_METABOLISM | NA | NA | NA | NA | NA | NA | 0 | 18 | 0 |
| KEGG_ALZHEIMERS_DISEASE | ALZHEIMERS_DISEASE | NA | NA | NA | NA | NA | NA | 0 | 165 | 0 |
| KEGG_AMINO_SUGAR_AND_NUCLEOTIDE_SUGAR_METABOLISM | AMINO_SUGAR_AND_NUCLEOTIDE_SUGAR_METABOLISM | NA | NA | NA | NA | NA | NA | 0 | 48 | 0 |
| KEGG_AMINOACYL_TRNA_BIOSYNTHESIS | AMINOACYL_TRNA_BIOSYNTHESIS | NA | NA | NA | NA | NA | NA | 0 | 41 | 0 |
| KEGG_AMYOTROPHIC_LATERAL_SCLEROSIS_ALS | AMYOTROPHIC_LATERAL_SCLEROSIS_ALS | NA | NA | NA | NA | NA | NA | 0 | 51 | 0 |
| KEGG_ANTIGEN_PROCESSING_AND_PRESENTATION | ANTIGEN_PROCESSING_AND_PRESENTATION | NA | NA | NA | NA | NA | NA | 0 | 279 | 0 |
| KEGG_APOPTOSIS | APOPTOSIS | NA | NA | NA | NA | NA | NA | 0 | 89 | 0 |
| KEGG_ARACHIDONIC_ACID_METABOLISM | ARACHIDONIC_ACID_METABOLISM | NA | NA | NA | NA | NA | NA | 0 | 60 | 0 |
| KEGG_ASCORBATE_AND_ALDARATE_METABOLISM | ASCORBATE_AND_ALDARATE_METABOLISM | NA | NA | NA | NA | NA | NA | 0 | 31 | 0 |
| KEGG_ASTHMA | ASTHMA | NA | NA | NA | NA | NA | NA | 0 | 32 | 0 |
| KEGG_AUTOIMMUNE_THYROID_DISEASE | AUTOIMMUNE_THYROID_DISEASE | NA | NA | NA | NA | NA | NA | 0 | 221 | 0 |
| KEGG_B_CELL_RECEPTOR_SIGNALING_PATHWAY | B_CELL_RECEPTOR_SIGNALING_PATHWAY | NA | NA | NA | NA | NA | NA | 0 | 90 | 0 |
| KEGG_BASAL_CELL_CARCINOMA | BASAL_CELL_CARCINOMA | NA | NA | NA | NA | NA | NA | 0 | 55 | 0 |
| KEGG_BASAL_TRANSCRIPTION_FACTORS | BASAL_TRANSCRIPTION_FACTORS | NA | NA | NA | NA | NA | NA | 0 | 38 | 0 |
| KEGG_BASE_EXCISION_REPAIR | BASE_EXCISION_REPAIR | NA | NA | NA | NA | NA | NA | 0 | 32 | 0 |
| KEGG_BETA_ALANINE_METABOLISM | BETA_ALANINE_METABOLISM | NA | NA | NA | NA | NA | NA | 0 | 22 | 0 |
| KEGG_BIOSYNTHESIS_OF_UNSATURATED_FATTY_ACIDS | BIOSYNTHESIS_OF_UNSATURATED_FATTY_ACIDS | NA | NA | NA | NA | NA | NA | 0 | 29 | 0 |
| KEGG_BUTANOATE_METABOLISM | BUTANOATE_METABOLISM | NA | NA | NA | NA | NA | NA | 0 | 38 | 0 |
| KEGG_CELL_ADHESION_MOLECULES_CAMS | CELL_ADHESION_MOLECULES_CAMS | NA | NA | NA | NA | NA | NA | 0 | 129 | 0 |
| KEGG_CELL_CYCLE | CELL_CYCLE | NA | NA | NA | NA | NA | NA | 0 | 124 | 0 |
| KEGG_CHEMOKINE_SIGNALING_PATHWAY | CHEMOKINE_SIGNALING_PATHWAY | NA | NA | NA | NA | NA | NA | 0 | 178 | 0 |
| KEGG_CHRONIC_MYELOID_LEUKEMIA | CHRONIC_MYELOID_LEUKEMIA | NA | NA | NA | NA | NA | NA | 0 | 72 | 0 |
| KEGG_CITRATE_CYCLE_TCA_CYCLE | CITRATE_CYCLE_TCA_CYCLE | NA | NA | NA | NA | NA | NA | 0 | 31 | 0 |
| KEGG_COLORECTAL_CANCER | COLORECTAL_CANCER | NA | NA | NA | NA | NA | NA | 0 | 64 | 0 |
| KEGG_COMPLEMENT_AND_COAGULATION_CASCADES | COMPLEMENT_AND_COAGULATION_CASCADES | NA | NA | NA | NA | NA | NA | 0 | 74 | 0 |
| KEGG_CYSTEINE_AND_METHIONINE_METABOLISM | CYSTEINE_AND_METHIONINE_METABOLISM | NA | NA | NA | NA | NA | NA | 0 | 38 | 0 |
| KEGG_CYTOSOLIC_DNA_SENSING_PATHWAY | CYTOSOLIC_DNA_SENSING_PATHWAY | NA | NA | NA | NA | NA | NA | 0 | 221 | 0 |
| KEGG_DNA_REPLICATION | DNA_REPLICATION | NA | NA | NA | NA | NA | NA | 0 | 34 | 0 |
| KEGG_DORSO_VENTRAL_AXIS_FORMATION | DORSO_VENTRAL_AXIS_FORMATION | NA | NA | NA | NA | NA | NA | 0 | 21 | 0 |
| KEGG_ENDOCYTOSIS | ENDOCYTOSIS | NA | NA | NA | NA | NA | NA | 0 | 184 | 0 |
| KEGG_ENDOMETRIAL_CANCER | ENDOMETRIAL_CANCER | NA | NA | NA | NA | NA | NA | 0 | 52 | 0 |
| KEGG_ERBB_SIGNALING_PATHWAY | ERBB_SIGNALING_PATHWAY | NA | NA | NA | NA | NA | NA | 0 | 86 | 0 |
| KEGG_ETHER_LIPID_METABOLISM | ETHER_LIPID_METABOLISM | NA | NA | NA | NA | NA | NA | 0 | 32 | 0 |
| KEGG_FATTY_ACID_METABOLISM | FATTY_ACID_METABOLISM | NA | NA | NA | NA | NA | NA | 0 | 51 | 0 |
| KEGG_FC_EPSILON_RI_SIGNALING_PATHWAY | FC_EPSILON_RI_SIGNALING_PATHWAY | NA | NA | NA | NA | NA | NA | 0 | 78 | 0 |
| KEGG_FC_GAMMA_R_MEDIATED_PHAGOCYTOSIS | FC_GAMMA_R_MEDIATED_PHAGOCYTOSIS | NA | NA | NA | NA | NA | NA | 0 | 99 | 0 |
| KEGG_FOLATE_BIOSYNTHESIS | FOLATE_BIOSYNTHESIS | NA | NA | NA | NA | NA | NA | 0 | 17 | 0 |
| KEGG_GALACTOSE_METABOLISM | GALACTOSE_METABOLISM | NA | NA | NA | NA | NA | NA | 0 | 26 | 0 |
| KEGG_GAP_JUNCTION | GAP_JUNCTION | NA | NA | NA | NA | NA | NA | 0 | 84 | 0 |
| KEGG_GLIOMA | GLIOMA | NA | NA | NA | NA | NA | NA | 0 | 61 | 0 |
| KEGG_GLUTATHIONE_METABOLISM | GLUTATHIONE_METABOLISM | NA | NA | NA | NA | NA | NA | 0 | 48 | 0 |
| KEGG_GLYCEROLIPID_METABOLISM | GLYCEROLIPID_METABOLISM | NA | NA | NA | NA | NA | NA | 0 | 49 | 0 |
| KEGG_GLYCEROPHOSPHOLIPID_METABOLISM | GLYCEROPHOSPHOLIPID_METABOLISM | NA | NA | NA | NA | NA | NA | 0 | 76 | 0 |
| KEGG_GLYCINE_SERINE_AND_THREONINE_METABOLISM | GLYCINE_SERINE_AND_THREONINE_METABOLISM | NA | NA | NA | NA | NA | NA | 0 | 32 | 0 |
| KEGG_GLYCOSAMINOGLYCAN_BIOSYNTHESIS_CHONDROITIN_SULFATE | GLYCOSAMINOGLYCAN_BIOSYNTHESIS_CHONDROITIN_SULFATE | NA | NA | NA | NA | NA | NA | 0 | 22 | 0 |
| KEGG_GLYCOSAMINOGLYCAN_BIOSYNTHESIS_HEPARAN_SULFATE | GLYCOSAMINOGLYCAN_BIOSYNTHESIS_HEPARAN_SULFATE | NA | NA | NA | NA | NA | NA | 0 | 26 | 0 |
| KEGG_GLYCOSAMINOGLYCAN_BIOSYNTHESIS_KERATAN_SULFATE | GLYCOSAMINOGLYCAN_BIOSYNTHESIS_KERATAN_SULFATE | NA | NA | NA | NA | NA | NA | 0 | 14 | 0 |
| KEGG_GLYCOSAMINOGLYCAN_DEGRADATION | GLYCOSAMINOGLYCAN_DEGRADATION | NA | NA | NA | NA | NA | NA | 0 | 22 | 0 |
| KEGG_GLYCOSPHINGOLIPID_BIOSYNTHESIS_GANGLIO_SERIES | GLYCOSPHINGOLIPID_BIOSYNTHESIS_GANGLIO_SERIES | NA | NA | NA | NA | NA | NA | 0 | 15 | 0 |
| KEGG_GLYCOSPHINGOLIPID_BIOSYNTHESIS_GLOBO_SERIES | GLYCOSPHINGOLIPID_BIOSYNTHESIS_GLOBO_SERIES | NA | NA | NA | NA | NA | NA | 0 | 14 | 0 |
| KEGG_GLYCOSYLPHOSPHATIDYLINOSITOL_GPI_ANCHOR_BIOSYNTHESIS | GLYCOSYLPHOSPHATIDYLINOSITOL_GPI_ANCHOR_BIOSYNTHESIS | NA | NA | NA | NA | NA | NA | 0 | 24 | 0 |
| KEGG_GLYOXYLATE_AND_DICARBOXYLATE_METABOLISM | GLYOXYLATE_AND_DICARBOXYLATE_METABOLISM | NA | NA | NA | NA | NA | NA | 0 | 16 | 0 |
| KEGG_GNRH_SIGNALING_PATHWAY | GNRH_SIGNALING_PATHWAY | NA | NA | NA | NA | NA | NA | 0 | 95 | 0 |
| KEGG_GRAFT_VERSUS_HOST_DISEASE | GRAFT_VERSUS_HOST_DISEASE | NA | NA | NA | NA | NA | NA | 0 | 50 | 0 |
| KEGG_HEDGEHOG_SIGNALING_PATHWAY | HEDGEHOG_SIGNALING_PATHWAY | NA | NA | NA | NA | NA | NA | 0 | 52 | 0 |
| KEGG_HEMATOPOIETIC_CELL_LINEAGE | HEMATOPOIETIC_CELL_LINEAGE | NA | NA | NA | NA | NA | NA | 0 | 88 | 0 |
| KEGG_HOMOLOGOUS_RECOMBINATION | HOMOLOGOUS_RECOMBINATION | NA | NA | NA | NA | NA | NA | 0 | 26 | 0 |
| KEGG_HUNTINGTONS_DISEASE | HUNTINGTONS_DISEASE | NA | NA | NA | NA | NA | NA | 0 | 175 | 0 |
| KEGG_INOSITOL_PHOSPHATE_METABOLISM | INOSITOL_PHOSPHATE_METABOLISM | NA | NA | NA | NA | NA | NA | 0 | 53 | 0 |
| KEGG_INSULIN_SIGNALING_PATHWAY | INSULIN_SIGNALING_PATHWAY | NA | NA | NA | NA | NA | NA | 0 | 137 | 0 |
| KEGG_INTESTINAL_IMMUNE_NETWORK_FOR_IGA_PRODUCTION | INTESTINAL_IMMUNE_NETWORK_FOR_IGA_PRODUCTION | NA | NA | NA | NA | NA | NA | 0 | 46 | 0 |
| KEGG_JAK_STAT_SIGNALING_PATHWAY | JAK_STAT_SIGNALING_PATHWAY | NA | NA | NA | NA | NA | NA | 0 | 353 | 0 |
| KEGG_LIMONENE_AND_PINENE_DEGRADATION | LIMONENE_AND_PINENE_DEGRADATION | NA | NA | NA | NA | NA | NA | 0 | 10 | 0 |
| KEGG_LINOLEIC_ACID_METABOLISM | LINOLEIC_ACID_METABOLISM | NA | NA | NA | NA | NA | NA | 0 | 28 | 0 |
| KEGG_LONG_TERM_DEPRESSION | LONG_TERM_DEPRESSION | NA | NA | NA | NA | NA | NA | 0 | 67 | 0 |
| KEGG_LONG_TERM_POTENTIATION | LONG_TERM_POTENTIATION | NA | NA | NA | NA | NA | NA | 0 | 68 | 0 |
| KEGG_LYSINE_DEGRADATION | LYSINE_DEGRADATION | NA | NA | NA | NA | NA | NA | 0 | 42 | 0 |
| KEGG_MATURITY_ONSET_DIABETES_OF_THE_YOUNG | MATURITY_ONSET_DIABETES_OF_THE_YOUNG | NA | NA | NA | NA | NA | NA | 0 | 25 | 0 |
| KEGG_MELANOGENESIS | MELANOGENESIS | NA | NA | NA | NA | NA | NA | 0 | 97 | 0 |
| KEGG_MELANOMA | MELANOMA | NA | NA | NA | NA | NA | NA | 0 | 69 | 0 |
| KEGG_MISMATCH_REPAIR | MISMATCH_REPAIR | NA | NA | NA | NA | NA | NA | 0 | 21 | 0 |
| KEGG_MTOR_SIGNALING_PATHWAY | MTOR_SIGNALING_PATHWAY | NA | NA | NA | NA | NA | NA | 0 | 52 | 0 |
| KEGG_N_GLYCAN_BIOSYNTHESIS | N_GLYCAN_BIOSYNTHESIS | NA | NA | NA | NA | NA | NA | 0 | 46 | 0 |
| KEGG_NATURAL_KILLER_CELL_MEDIATED_CYTOTOXICITY | NATURAL_KILLER_CELL_MEDIATED_CYTOTOXICITY | NA | NA | NA | NA | NA | NA | 0 | 316 | 0 |
| KEGG_NEUROTROPHIN_SIGNALING_PATHWAY | NEUROTROPHIN_SIGNALING_PATHWAY | NA | NA | NA | NA | NA | NA | 0 | 126 | 0 |
| KEGG_NICOTINATE_AND_NICOTINAMIDE_METABOLISM | NICOTINATE_AND_NICOTINAMIDE_METABOLISM | NA | NA | NA | NA | NA | NA | 0 | 26 | 0 |
| KEGG_NOD_LIKE_RECEPTOR_SIGNALING_PATHWAY | NOD_LIKE_RECEPTOR_SIGNALING_PATHWAY | NA | NA | NA | NA | NA | NA | 0 | 52 | 0 |
| KEGG_NON_HOMOLOGOUS_END_JOINING | NON_HOMOLOGOUS_END_JOINING | NA | NA | NA | NA | NA | NA | 0 | 13 | 0 |
| KEGG_NON_SMALL_CELL_LUNG_CANCER | NON_SMALL_CELL_LUNG_CANCER | NA | NA | NA | NA | NA | NA | 0 | 53 | 0 |
| KEGG_NOTCH_SIGNALING_PATHWAY | NOTCH_SIGNALING_PATHWAY | NA | NA | NA | NA | NA | NA | 0 | 46 | 0 |
| KEGG_NUCLEOTIDE_EXCISION_REPAIR | NUCLEOTIDE_EXCISION_REPAIR | NA | NA | NA | NA | NA | NA | 0 | 42 | 0 |
| KEGG_OLFACTORY_TRANSDUCTION | OLFACTORY_TRANSDUCTION | NA | NA | NA | NA | NA | NA | 0 | 650 | 0 |
| KEGG_ONE_CARBON_POOL_BY_FOLATE | ONE_CARBON_POOL_BY_FOLATE | NA | NA | NA | NA | NA | NA | 0 | 18 | 0 |
| KEGG_OOCYTE_MEIOSIS | OOCYTE_MEIOSIS | NA | NA | NA | NA | NA | NA | 0 | 111 | 0 |
| KEGG_OTHER_GLYCAN_DEGRADATION | OTHER_GLYCAN_DEGRADATION | NA | NA | NA | NA | NA | NA | 0 | 16 | 0 |
| KEGG_PANCREATIC_CANCER | PANCREATIC_CANCER | NA | NA | NA | NA | NA | NA | 0 | 71 | 0 |
| KEGG_PANTOTHENATE_AND_COA_BIOSYNTHESIS | PANTOTHENATE_AND_COA_BIOSYNTHESIS | NA | NA | NA | NA | NA | NA | 0 | 15 | 0 |
| KEGG_PARKINSONS_DISEASE | PARKINSONS_DISEASE | NA | NA | NA | NA | NA | NA | 0 | 126 | 0 |
| KEGG_PATHOGENIC_ESCHERICHIA_COLI_INFECTION | PATHOGENIC_ESCHERICHIA_COLI_INFECTION | NA | NA | NA | NA | NA | NA | 0 | 52 | 0 |
| KEGG_PENTOSE_AND_GLUCURONATE_INTERCONVERSIONS | PENTOSE_AND_GLUCURONATE_INTERCONVERSIONS | NA | NA | NA | NA | NA | NA | 0 | 34 | 0 |
| KEGG_PENTOSE_PHOSPHATE_PATHWAY | PENTOSE_PHOSPHATE_PATHWAY | NA | NA | NA | NA | NA | NA | 0 | 31 | 0 |
| KEGG_PEROXISOME | PEROXISOME | NA | NA | NA | NA | NA | NA | 0 | 80 | 0 |
| KEGG_PHOSPHATIDYLINOSITOL_SIGNALING_SYSTEM | PHOSPHATIDYLINOSITOL_SIGNALING_SYSTEM | NA | NA | NA | NA | NA | NA | 0 | 75 | 0 |
| KEGG_PORPHYRIN_AND_CHLOROPHYLL_METABOLISM | PORPHYRIN_AND_CHLOROPHYLL_METABOLISM | NA | NA | NA | NA | NA | NA | 0 | 47 | 0 |
| KEGG_PPAR_SIGNALING_PATHWAY | PPAR_SIGNALING_PATHWAY | NA | NA | NA | NA | NA | NA | 0 | 78 | 0 |
| KEGG_PRIMARY_BILE_ACID_BIOSYNTHESIS | PRIMARY_BILE_ACID_BIOSYNTHESIS | NA | NA | NA | NA | NA | NA | 0 | 15 | 0 |
| KEGG_PRIMARY_IMMUNODEFICIENCY | PRIMARY_IMMUNODEFICIENCY | NA | NA | NA | NA | NA | NA | 0 | 38 | 0 |
| KEGG_PRION_DISEASES | PRION_DISEASES | NA | NA | NA | NA | NA | NA | 0 | 34 | 0 |
| KEGG_PROGESTERONE_MEDIATED_OOCYTE_MATURATION | PROGESTERONE_MEDIATED_OOCYTE_MATURATION | NA | NA | NA | NA | NA | NA | 0 | 84 | 0 |
| KEGG_PROPANOATE_METABOLISM | PROPANOATE_METABOLISM | NA | NA | NA | NA | NA | NA | 0 | 32 | 0 |
| KEGG_PROSTATE_CANCER | PROSTATE_CANCER | NA | NA | NA | NA | NA | NA | 0 | 103 | 0 |
| KEGG_PROTEASOME | PROTEASOME | NA | NA | NA | NA | NA | NA | 0 | 45 | 0 |
| KEGG_PROTEIN_EXPORT | PROTEIN_EXPORT | NA | NA | NA | NA | NA | NA | 0 | 26 | 0 |
| KEGG_PURINE_METABOLISM | PURINE_METABOLISM | NA | NA | NA | NA | NA | NA | 0 | 158 | 0 |
| KEGG_PYRIMIDINE_METABOLISM | PYRIMIDINE_METABOLISM | NA | NA | NA | NA | NA | NA | 0 | 96 | 0 |
| KEGG_PYRUVATE_METABOLISM | PYRUVATE_METABOLISM | NA | NA | NA | NA | NA | NA | 0 | 39 | 0 |
| KEGG_REGULATION_OF_ACTIN_CYTOSKELETON | REGULATION_OF_ACTIN_CYTOSKELETON | NA | NA | NA | NA | NA | NA | 0 | 211 | 0 |
| KEGG_REGULATION_OF_AUTOPHAGY | REGULATION_OF_AUTOPHAGY | NA | NA | NA | NA | NA | NA | 0 | 206 | 0 |
| KEGG_RENAL_CELL_CARCINOMA | RENAL_CELL_CARCINOMA | NA | NA | NA | NA | NA | NA | 0 | 70 | 0 |
| KEGG_RENIN_ANGIOTENSIN_SYSTEM | RENIN_ANGIOTENSIN_SYSTEM | NA | NA | NA | NA | NA | NA | 0 | 19 | 0 |
| KEGG_RETINOL_METABOLISM | RETINOL_METABOLISM | NA | NA | NA | NA | NA | NA | 0 | 81 | 0 |
| KEGG_RIBOFLAVIN_METABOLISM | RIBOFLAVIN_METABOLISM | NA | NA | NA | NA | NA | NA | 0 | 16 | 0 |
| KEGG_RIBOSOME | RIBOSOME | NA | NA | NA | NA | NA | NA | 0 | 74 | 0 |
| KEGG_RIG_I_LIKE_RECEPTOR_SIGNALING_PATHWAY | RIG_I_LIKE_RECEPTOR_SIGNALING_PATHWAY | NA | NA | NA | NA | NA | NA | 0 | 238 | 0 |
| KEGG_RNA_DEGRADATION | RNA_DEGRADATION | NA | NA | NA | NA | NA | NA | 0 | 58 | 0 |
| KEGG_RNA_POLYMERASE | RNA_POLYMERASE | NA | NA | NA | NA | NA | NA | 0 | 27 | 0 |
| KEGG_SELENOAMINO_ACID_METABOLISM | SELENOAMINO_ACID_METABOLISM | NA | NA | NA | NA | NA | NA | 0 | 27 | 0 |
| KEGG_SNARE_INTERACTIONS_IN_VESICULAR_TRANSPORT | SNARE_INTERACTIONS_IN_VESICULAR_TRANSPORT | NA | NA | NA | NA | NA | NA | 0 | 34 | 0 |
| KEGG_SPHINGOLIPID_METABOLISM | SPHINGOLIPID_METABOLISM | NA | NA | NA | NA | NA | NA | 0 | 40 | 0 |
| KEGG_SPLICEOSOME | SPLICEOSOME | NA | NA | NA | NA | NA | NA | 0 | 154 | 0 |
| KEGG_STEROID_BIOSYNTHESIS | STEROID_BIOSYNTHESIS | NA | NA | NA | NA | NA | NA | 0 | 17 | 0 |
| KEGG_STEROID_HORMONE_BIOSYNTHESIS | STEROID_HORMONE_BIOSYNTHESIS | NA | NA | NA | NA | NA | NA | 0 | 69 | 0 |
| KEGG_SULFUR_METABOLISM | SULFUR_METABOLISM | NA | NA | NA | NA | NA | NA | 0 | 14 | 0 |
| KEGG_SYSTEMIC_LUPUS_ERYTHEMATOSUS | SYSTEMIC_LUPUS_ERYTHEMATOSUS | NA | NA | NA | NA | NA | NA | 0 | 105 | 0 |
| KEGG_T_CELL_RECEPTOR_SIGNALING_PATHWAY | T_CELL_RECEPTOR_SIGNALING_PATHWAY | NA | NA | NA | NA | NA | NA | 0 | 108 | 0 |
| KEGG_TASTE_TRANSDUCTION | TASTE_TRANSDUCTION | NA | NA | NA | NA | NA | NA | 0 | 69 | 0 |
| KEGG_TAURINE_AND_HYPOTAURINE_METABOLISM | TAURINE_AND_HYPOTAURINE_METABOLISM | NA | NA | NA | NA | NA | NA | 0 | 10 | 0 |
| KEGG_TERPENOID_BACKBONE_BIOSYNTHESIS | TERPENOID_BACKBONE_BIOSYNTHESIS | NA | NA | NA | NA | NA | NA | 0 | 17 | 0 |
| KEGG_THYROID_CANCER | THYROID_CANCER | NA | NA | NA | NA | NA | NA | 0 | 31 | 0 |
| KEGG_TIGHT_JUNCTION | TIGHT_JUNCTION | NA | NA | NA | NA | NA | NA | 0 | 125 | 0 |
| KEGG_TRYPTOPHAN_METABOLISM | TRYPTOPHAN_METABOLISM | NA | NA | NA | NA | NA | NA | 0 | 45 | 0 |
| KEGG_TYPE_I_DIABETES_MELLITUS | TYPE_I_DIABETES_MELLITUS | NA | NA | NA | NA | NA | NA | 0 | 44 | 0 |
| KEGG_UBIQUITIN_MEDIATED_PROTEOLYSIS | UBIQUITIN_MEDIATED_PROTEOLYSIS | NA | NA | NA | NA | NA | NA | 0 | 136 | 0 |
| KEGG_VALINE_LEUCINE_AND_ISOLEUCINE_BIOSYNTHESIS | VALINE_LEUCINE_AND_ISOLEUCINE_BIOSYNTHESIS | NA | NA | NA | NA | NA | NA | 0 | 11 | 0 |
| KEGG_VALINE_LEUCINE_AND_ISOLEUCINE_DEGRADATION | VALINE_LEUCINE_AND_ISOLEUCINE_DEGRADATION | NA | NA | NA | NA | NA | NA | 0 | 48 | 0 |
| KEGG_VASCULAR_SMOOTH_MUSCLE_CONTRACTION | VASCULAR_SMOOTH_MUSCLE_CONTRACTION | NA | NA | NA | NA | NA | NA | 0 | 119 | 0 |
| KEGG_VASOPRESSIN_REGULATED_WATER_REABSORPTION | VASOPRESSIN_REGULATED_WATER_REABSORPTION | NA | NA | NA | NA | NA | NA | 0 | 44 | 0 |
| KEGG_VEGF_SIGNALING_PATHWAY | VEGF_SIGNALING_PATHWAY | NA | NA | NA | NA | NA | NA | 0 | 74 | 0 |
| KEGG_VIRAL_MYOCARDITIS | VIRAL_MYOCARDITIS | NA | NA | NA | NA | NA | NA | 0 | 70 | 0 |
| KEGG_WNT_SIGNALING_PATHWAY | WNT_SIGNALING_PATHWAY | NA | NA | NA | NA | NA | NA | 0 | 151 | 0 |

| ID | Description | GeneRatio | BgRatio | pvalue | p.adjust | qvalue | geneID | Count | Genes_in_Pa | Percentage_of_Pathway_detected |
|---|---|---|---|---|---|---|---|---|---|---|
| KEGG_DRUG_METABOLISM_OTHER_ENZYMES | DRUG_METABOLISM_OTHER_ENZYMES | 11/182 | 37/3594 | 1.12E-06 | 0.00016255 | 0.00014632 | Ces1d/Ces2b/Ces2c/Ces2g/Cyp3a13/Dpyd/Tymp/Ugt1a6a/Ugt1a7c/Upb1/Upp1 | 11 | 73 | 15.07 |
| KEGG_BETA_ALANINE_METABOLISM | BETA_ALANINE_METABOLISM | 5/182 | 18/3594 | 0.00157419 | 0.06683015 | 0.06015926 | Acadm/Aldh2/Dpyd/Echs1/Upb1 | 5 | 22 | 22.73 |
| KEGG_DRUG_METABOLISM_CYTOCHROME_P450 | DRUG_METABOLISM_CYTOCHROME_P450 | 8/182 | 46/3594 | 0.00184359 | 0.06683015 | 0.06015926 | Cyp2d26/Cyp3a13/Fmo4/Fmo5/Gstp2/Maoa/Ugt1a6a/Ugt1a7c | 8 | 84 | 9.52 |
| KEGG_TRYPTOPHAN_METABOLISM | TRYPTOPHAN_METABOLISM | 6/182 | 25/3594 | 0.00122671 | 0.06683015 | 0.06015926 | Aldh2/Aoc1/Echs1/Maoa/Tdo2/Tph1 | 6 | 45 | 13.33 |
| KEGG_ASCORBATE_AND_ALDARATE_METABOLISM | ASCORBATE_AND_ALDARATE_METABOLISM | 4/182 | 13/3594 | 0.00317238 | 0.07981694 | 0.07184973 | Aldh2/Ugdh/Ugt1a6a/Ugt1a7c | 4 | 31 | 12.9 |
| KEGG_STEROID_HORMONE_BIOSYNTHESIS | STEROID_HORMONE_BIOSYNTHESIS | 5/182 | 21/3594 | 0.00330277 | 0.07981694 | 0.07184973 | Cyp3a13/Hsd17b2/Hsd3b3/Ugt1a6a/Ugt1a7c | 5 | 69 | 7.25 |
| KEGG_LEUKOCYTE_TRANSENDOTHELIAL_MIGRATION | LEUKOCYTE_TRANSENDOTHELIAL_MIGRATION | 11/182 | 88/3594 | 0.0043556 | 0.08466231 | 0.07621144 | Actg1/Esam/Gnai1/Mapk12/Myl7/Pecam1/Pik3cg/Pik3r5/Plcg2/Rac2/Vav1 | 11 | 112 | 9.82 |
| KEGG_RETINOL_METABOLISM | RETINOL_METABOLISM | 6/182 | 32/3594 | 0.00467102 | 0.08466231 | 0.07621144 | Aldh1a1/Cyp3a13/Dhrs3/Lrat/Ugt1a6a/Ugt1a7c | 6 | 81 | 7.41 |
| KEGG_BUTANOATE_METABOLISM | BUTANOATE_METABOLISM | 5/182 | 28/3594 | 0.01196585 | 0.19278316 | 0.17353983 | Akr1b8/Aldh2/Bdh1/Echs1/Oxct1 | 5 | 38 | 13.16 |
| KEGG_TASTE_TRANSDUCTION | TASTE_TRANSDUCTION | 4/182 | 20/3594 | 0.0162826 | 0.19674808 | 0.17710898 | Gnat3/Plcb2/Scnn1a/Trpm5 | 4 | 69 | 5.8 |
| KEGG_TYPE_I_DIABETES_MELLITUS | TYPE_I_DIABETES_MELLITUS | 3/182 | Nov-94 | 0.01559184 | 0.19674808 | 0.17710898 | H2-DMb1/Ptprn/Ptprn2 | 3 | 44 | 6.82 |
| KEGG_VEGF_SIGNALING_PATHWAY | VEGF_SIGNALING_PATHWAY | 8/182 | 64/3594 | 0.01437746 | 0.19674808 | 0.17710898 | Chp2/Hspb1/Mapk12/Pik3cg/Pik3r5/Plcg2/Ppp3cc/Rac2 | 8 | 74 | 10.81 |
| KEGG_ARGININE_AND_PROLINE_METABOLISM | ARGININE_AND_PROLINE_METABOLISM | 6/182 | 46/3594 | 0.0268985 | 0.27859156 | 0.25078297 | Aldh2/Aoc1/Ass1/Maoa/Otc/Prodh | 6 | 59 | 10.17 |
| KEGG_PANTOTHENATE_AND_COA_BIOSYNTHESIS | PANTOTHENATE_AND_COA_BIOSYNTHESIS | 3/182 | 13/3594 | 0.02508362 | 0.27859156 | 0.25078297 | Dpyd/Upb1/Vnn1 | 3 | 15 | 20 |
| KEGG_FATTY_ACID_METABOLISM | FATTY_ACID_METABOLISM | 5/182 | 36/3594 | 0.03315424 | 0.31603871 | 0.2844922 | Acadm/Aldh2/Echs1/Eci3/Hadhb | 5 | 51 | 9.8 |
| KEGG_METABOLISM_OF_XENOBIOTICS_BY_CYTOCHROME_P450 | METABOLISM_OF_XENOBIOTICS_BY_CYTOCHROME_P450 | 5/182 | 37/3594 | 0.03681669 | 0.31603871 | 0.2844922 | Cyp2s1/Cyp3a13/Gstp2/Ugt1a6a/Ugt1a7c | 5 | 67 | 7.46 |
| KEGG_OTHER_GLYCAN_DEGRADATION | OTHER_GLYCAN_DEGRADATION | 3/182 | 15/3594 | 0.03705281 | 0.31603871 | 0.2844922 | Hexa/Hexb/Man2b2 | 3 | 16 | 18.75 |
| KEGG_GAP_JUNCTION | GAP_JUNCTION | 7/182 | 64/3594 | 0.04118912 | 0.31863626 | 0.28683046 | Gnai1/Pdgfd/Pdgfrb/Plcb2/Tuba1a/Tubal3/Tubb2a | 7 | 84 | 8.33 |
| KEGG_PENTOSE_AND_GLUCURONATE_INTERCONVERSIONS | PENTOSE_AND_GLUCURONATE_INTERCONVERSIONS | 3/182 | 16/3594 | 0.04394983 | 0.31863626 | 0.28683046 | Ugdh/Ugt1a6a/Ugt1a7c | 3 | 34 | 8.82 |
| KEGG_REGULATION_OF_ACTIN_CYTOSKELETON | REGULATION_OF_ACTIN_CYTOSKELETON | 14/182 | 168/3594 | 0.04310648 | 0.31863626 | 0.28683046 | Actg1/Chrm3/Fgfr3/Insrr/Itgb8/Myl7/Pdgfd/Pdgfrb/Pik3cg/Pik3r5/Rac2/Rras/Scin/Vav1 | 14 | 211 | 6.64 |
| KEGG_AMINO_SUGAR_AND_NUCLEOTIDE_SUGAR_METABOLISM | AMINO_SUGAR_AND_NUCLEOTIDE_SUGAR_METABOLISM | 5/182 | 41/3594 | 0.05388297 | 0.32554292 | 0.29304771 | Amdhd2/Gnpda1/Hexa/Hexb/Ugdh | 5 | 48 | 10.42 |
| KEGG_B_CELL_RECEPTOR_SIGNALING_PATHWAY | B_CELL_RECEPTOR_SIGNALING_PATHWAY | 7/182 | 66/3594 | 0.0474848 | 0.32554292 | 0.29304771 | Chp2/Pik3cg/Pik3r5/Plcg2/Ppp3cc/Rac2/Vav1 | 7 | 90 | 7.78 |
| KEGG_NATURAL_KILLER_CELL_MEDIATED_CYTOTOXICITY | NATURAL_KILLER_CELL_MEDIATED_CYTOTOXICITY | 8/182 | 82/3594 | 0.05372916 | 0.32554292 | 0.29304771 | Chp2/Ifngr2/Pik3cg/Pik3r5/Plcg2/Ppp3cc/Rac2/Vav1 | 8 | 316 | 2.53 |
| KEGG_VALINE_LEUCINE_AND_ISOLEUCINE_DEGRADATION | VALINE_LEUCINE_AND_ISOLEUCINE_DEGRADATION | 5/182 | 41/3594 | 0.05388297 | 0.32554292 | 0.29304771 | Acadm/Aldh2/Echs1/Hadhb/Oxct1 | 5 | 48 | 10.42 |
| KEGG_COMPLEMENT_AND_COAGULATION_CASCADES | COMPLEMENT_AND_COAGULATION_CASCADES | 4/182 | 29/3594 | 0.05614019 | 0.32561308 | 0.29311087 | C1ra/Cd55/Cd59a/F3 | 4 | 74 | 5.41 |
| KEGG_ALDOSTERONE_REGULATED_SODIUM_REABSORPTION | ALDOSTERONE_REGULATED_SODIUM_REABSORPTION | 4/182 | 31/3594 | 0.06883766 | 0.36968372 | 0.33278244 | Fxyd4/Pik3cg/Pik3r5/Scnn1a | 4 | 43 | 9.3 |
| KEGG_VIRAL_MYOCARDITIS | VIRAL_MYOCARDITIS | 4/182 | 31/3594 | 0.06883766 | 0.36968372 | 0.33278244 | Actg1/Cd55/H2-DMb1/Rac2 | 4 | 70 | 5.71 |
| KEGG_CALCIUM_SIGNALING_PATHWAY | CALCIUM_SIGNALING_PATHWAY | 9/182 | 103/3594 | 0.07481467 | 0.37343194 | 0.33615652 | Chp2/Chrm3/Pdgfrb/Plcb2/Plcd3/Plcg2/Ppp3cc/Slc8a1/Vdac3 | 9 | 173 | 5.2 |
| KEGG_MATURITY_ONSET_DIABETES_OF_THE_YOUNG | MATURITY_ONSET_DIABETES_OF_THE_YOUNG | 3/182 | 20/3594 | 0.07726178 | 0.37343194 | 0.33615652 | Iapp/Mnx1/Neurod1 | 3 | 25 | 12 |
| KEGG_STARCH_AND_SUCROSE_METABOLISM | STARCH_AND_SUCROSE_METABOLISM | 4/182 | 32/3594 | 0.07569614 | 0.37343194 | 0.33615652 | Pygl/Ugdh/Ugt1a6a/Ugt1a7c | 4 | 52 | 7.69 |
| KEGG_LIMONENE_AND_PINENE_DEGRADATION | LIMONENE_AND_PINENE_DEGRADATION | 2/182 | Oct-94 | 0.08782746 | 0.39796819 | 0.3582436 | Aldh2/Echs1 | 2 | 10 | 20 |
| KEGG_NICOTINATE_AND_NICOTINAMIDE_METABOLISM | NICOTINATE_AND_NICOTINAMIDE_METABOLISM | 3/182 | 21/3594 | 0.08691597 | 0.39796819 | 0.3582436 | Nadsyn1/Nt5e/Pnp2 | 3 | 26 | 11.54 |
| KEGG_HYPERTROPHIC_CARDIOMYOPATHY_HCM | HYPERTROPHIC_CARDIOMYOPATHY_HCM | 5/182 | 48/3594 | 0.09311259 | 0.40913106 | 0.36829221 | Actg1/Itgb8/Prkab2/Slc8a1/Tgfb1 | 5 | 84 | 5.95 |
| KEGG_HISTIDINE_METABOLISM | HISTIDINE_METABOLISM | 3/182 | 22/3594 | 0.09705181 | 0.41389744 | 0.37258281 | Aldh2/Aoc1/Maoa | 3 | 33 | 9.09 |
| KEGG_ANTIGEN_PROCESSING_AND_PRESENTATION | ANTIGEN_PROCESSING_AND_PRESENTATION | 4/182 | 36/3594 | 0.10636212 | 0.42871733 | 0.3859234 | B2m/Ctss/H2-DMb1/Ifi30 | 4 | 279 | 1.43 |
| KEGG_PPAR_SIGNALING_PATHWAY | PPAR_SIGNALING_PATHWAY | 5/182 | 50/3594 | 0.10644016 | 0.42871733 | 0.3859234 | Acadm/Cyp27a1/Lpl/Pck1/Ppara | 5 | 78 | 6.41 |
| KEGG_FC_EPSILON_RI_SIGNALING_PATHWAY | FC_EPSILON_RI_SIGNALING_PATHWAY | 6/182 | 67/3594 | 0.12127493 | 0.45089395 | 0.40588639 | Mapk12/Pik3cg/Pik3r5/Plcg2/Rac2/Vav1 | 6 | 78 | 7.69 |
| KEGG_GLYCOSPHINGOLIPID_BIOSYNTHESIS_GANGLIO_SERIES | GLYCOSPHINGOLIPID_BIOSYNTHESIS_GANGLIO_SERIES | 2/182 | Dec-94 | 0.12065035 | 0.45089395 | 0.40588639 | Hexa/Hexb | 2 | 15 | 13.33 |
| KEGG_GLYCOSPHINGOLIPID_BIOSYNTHESIS_GLOBO_SERIES | GLYCOSPHINGOLIPID_BIOSYNTHESIS_GLOBO_SERIES | 2/182 | Dec-94 | 0.12065035 | 0.45089395 | 0.40588639 | Hexa/Hexb | 2 | 14 | 14.29 |
| KEGG_CHEMOKINE_SIGNALING_PATHWAY | CHEMOKINE_SIGNALING_PATHWAY | 10/182 | 132/3594 | 0.12936475 | 0.4689472 | 0.42213759 | Gnai1/Gngt2/Grk3/Hck/Pik3cg/Pik3r5/Plcb2/Prex1/Rac2/Vav1 | 10 | 178 | 5.62 |
| KEGG_FC_GAMMA_R_MEDIATED_PHAGOCYTOSIS | FC_GAMMA_R_MEDIATED_PHAGOCYTOSIS | 7/182 | 85/3594 | 0.13690859 | 0.48418891 | 0.43585789 | Hck/Pik3cg/Pik3r5/Plcg2/Rac2/Scin/Vav1 | 7 | 99 | 7.07 |
| KEGG_PHOSPHATIDYLINOSITOL_SIGNALING_SYSTEM | PHOSPHATIDYLINOSITOL_SIGNALING_SYSTEM | 6/182 | 70/3594 | 0.14119165 | 0.48744735 | 0.43879108 | Dgkh/Pik3cg/Pik3r5/Plcb2/Plcd3/Plcg2 | 6 | 75 | 8 |
| KEGG_PHENYLALANINE_METABOLISM | PHENYLALANINE_METABOLISM | 2/182 | 14/3594 | 0.15590369 | 0.52572175 | 0.47324499 | Maoa/Prdx6 | 2 | 21 | 9.52 |
| KEGG_GLYCEROLIPID_METABOLISM | GLYCEROLIPID_METABOLISM | 4/182 | 43/3594 | 0.17093516 | 0.52619425 | 0.47367032 | Agk/Aldh2/Dgkh/Lpl | 4 | 49 | 8.16 |
| KEGG_LYSOSOME | LYSOSOME | 8/182 | 107/3594 | 0.17251483 | 0.52619425 | 0.47367032 | Ctsf/Ctss/Gm2a/Gnptg/Hexa/Hexb/Npc2/Psap | 8 | 118 | 6.78 |
| KEGG_NOD_LIKE_RECEPTOR_SIGNALING_PATHWAY | NOD_LIKE_RECEPTOR_SIGNALING_PATHWAY | 4/182 | 42/3594 | 0.16097681 | 0.52619425 | 0.47367032 | Il18/Mapk12/Nlrp1b/Nod2 | 4 | 52 | 7.69 |
| KEGG_PORPHYRIN_AND_CHLOROPHYLL_METABOLISM | PORPHYRIN_AND_CHLOROPHYLL_METABOLISM | 3/182 | 28/3594 | 0.16643541 | 0.52619425 | 0.47367032 | Fth1/Ugt1a6a/Ugt1a7c | 3 | 47 | 6.38 |
| KEGG_PROXIMAL_TUBULE_BICARBONATE_RECLAMATION | PROXIMAL_TUBULE_BICARBONATE_RECLAMATION | 2/182 | 15/3594 | 0.17418844 | 0.52619425 | 0.47367032 | Pck1/Slc4a4 | 2 | 23 | 8.7 |
| KEGG_PROPANOATE_METABOLISM | PROPANOATE_METABOLISM | 3/182 | 29/3594 | 0.17916178 | 0.5301726 | 0.47725156 | Acadm/Aldh2/Echs1 | 3 | 32 | 9.38 |
| KEGG_HEMATOPOIETIC_CELL_LINEAGE | HEMATOPOIETIC_CELL_LINEAGE | 4/182 | 45/3594 | 0.19146927 | 0.55269113 | 0.49752232 | Cd55/Cd59a/Cd9/Siglecf | 4 | 88 | 4.55 |
| KEGG_LEISHMANIA_INFECTION | LEISHMANIA_INFECTION | 4/182 | 46/3594 | 0.20201814 | 0.55269113 | 0.49752232 | H2-DMb1/Ifngr2/Mapk12/Tgfb1 | 4 | 70 | 5.71 |
| KEGG_PATHOGENIC_ESCHERICHIA_COLI_INFECTION | PATHOGENIC_ESCHERICHIA_COLI_INFECTION | 4/182 | 46/3594 | 0.20201814 | 0.55269113 | 0.49752232 | Actg1/Tuba1a/Tubal3/Tubb2a | 4 | 52 | 7.69 |
| KEGG_VIBRIO_CHOLERAE_INFECTION | VIBRIO_CHOLERAE_INFECTION | 4/182 | 46/3594 | 0.20201814 | 0.55269113 | 0.49752232 | Actg1/Atp6v0e2/Kcnq1/Plcg2 | 4 | 52 | 7.69 |
| KEGG_CELL_ADHESION_MOLECULES_CAMS | CELL_ADHESION_MOLECULES_CAMS | 5/182 | 65/3594 | 0.23114954 | 0.57787386 | 0.52019135 | Cntnap1/Esam/H2-DMb1/Itgb8/Pecam1 | 5 | 129 | 3.88 |
| KEGG_GLYCOLYSIS_GLUCONEOGENESIS | GLYCOLYSIS_GLUCONEOGENESIS | 4/182 | 48/3594 | 0.22361175 | 0.57787386 | 0.52019135 | Aldh2/Eno3/Pck1/Pgam1 | 4 | 67 | 5.97 |
| KEGG_GLYCOSAMINOGLYCAN_DEGRADATION | GLYCOSAMINOGLYCAN_DEGRADATION | 2/182 | 18/3594 | 0.23065919 | 0.57787386 | 0.52019135 | Hexa/Hexb | 2 | 22 | 9.09 |
| KEGG_GLYCOSPHINGOLIPID_BIOSYNTHESIS_LACTO_AND_NEOLACTO_SERIES | GLYCOSPHINGOLIPID_BIOSYNTHESIS_LACTO_AND_NEOLACTO_SERIES | 2/182 | 18/3594 | 0.23065919 | 0.57787386 | 0.52019135 | B3galt2/St3gal3 | 2 | 23 | 8.7 |
| KEGG_RENAL_CELL_CARCINOMA | RENAL_CELL_CARCINOMA | 5/182 | 64/3594 | 0.22174062 | 0.57787386 | 0.52019135 | Egln3/Flcn/Pik3cg/Pik3r5/Tgfb1 | 5 | 70 | 7.14 |
| KEGG_LINOLEIC_ACID_METABOLISM | LINOLEIC_ACID_METABOLISM | 2/182 | 19/3594 | 0.24978205 | 0.59616696 | 0.53665846 | Akr1b8/Cyp3a13 | 2 | 28 | 7.14 |
| KEGG_MELANOMA | MELANOMA | 4/182 | 50/3594 | 0.24577628 | 0.59616696 | 0.53665846 | Pdgfd/Pdgfrb/Pik3cg/Pik3r5 | 4 | 69 | 5.8 |
| KEGG_PROSTATE_CANCER | PROSTATE_CANCER | 6/182 | 84/3594 | 0.25080127 | 0.59616696 | 0.53665846 | Gstp2/Insrr/Pdgfd/Pdgfrb/Pik3cg/Pik3r5 | 6 | 103 | 5.83 |
| KEGG_INOSITOL_PHOSPHATE_METABOLISM | INOSITOL_PHOSPHATE_METABOLISM | 4/182 | 51/3594 | 0.25703908 | 0.60113978 | 0.5411349 | Pik3cg/Plcb2/Plcd3/Plcg2 | 4 | 53 | 7.55 |
| KEGG_ARACHIDONIC_ACID_METABOLISM | ARACHIDONIC_ACID_METABOLISM | 3/182 | 36/3594 | 0.27392103 | 0.60566678 | 0.54521002 | Alox5/Cbr3/Ptgs1 | 3 | 60 | 5 |
| KEGG_DILATED_CARDIOMYOPATHY | DILATED_CARDIOMYOPATHY | 4/182 | 53/3594 | 0.27985982 | 0.60566678 | 0.54521002 | Actg1/Itgb8/Slc8a1/Tgfb1 | 4 | 90 | 4.44 |
| KEGG_EPITHELIAL_CELL_SIGNALING_IN_HELICOBACTER_PYLORI_INFECTION | EPITHELIAL_CELL_SIGNALING_IN_HELICOBACTER_PYLORI_INFECTION | 4/182 | 52/3594 | 0.26840459 | 0.60566678 | 0.54521002 | Atp6v0e2/Hbegf/Mapk12/Plcg2 | 4 | 64 | 6.25 |
| KEGG_INTESTINAL_IMMUNE_NETWORK_FOR_IGA_PRODUCTION | INTESTINAL_IMMUNE_NETWORK_FOR_IGA_PRODUCTION | 2/182 | 20/3594 | 0.26895097 | 0.60566678 | 0.54521002 | H2-DMb1/Tgfb1 | 2 | 46 | 4.35 |
| KEGG_PYRIMIDINE_METABOLISM | PYRIMIDINE_METABOLISM | 6/182 | 87/3594 | 0.27698467 | 0.60566678 | 0.54521002 | Dpyd/Nt5e/Pnp2/Tymp/Upb1/Upp1 | 6 | 96 | 6.25 |
| KEGG_ADIPOCYTOKINE_SIGNALING_PATHWAY | ADIPOCYTOKINE_SIGNALING_PATHWAY | 4/182 | 55/3594 | 0.30298852 | 0.63671501 | 0.57315907 | Pck1/Ppara/Ppargc1a/Prkab2 | 4 | 67 | 5.97 |
| KEGG_GLIOMA | GLIOMA | 4/182 | 55/3594 | 0.30298852 | 0.63671501 | 0.57315907 | Pdgfrb/Pik3cg/Pik3r5/Plcg2 | 4 | 61 | 6.56 |
| KEGG_OLFACTORY_TRANSDUCTION | OLFACTORY_TRANSDUCTION | 2/182 | 23/3594 | 0.32622587 | 0.66623593 | 0.59973325 | Clca2/Grk3 | 2 | 650 | 0.31 |
| KEGG_T_CELL_RECEPTOR_SIGNALING_PATHWAY | T_CELL_RECEPTOR_SIGNALING_PATHWAY | 6/182 | 92/3594 | 0.32190771 | 0.66623593 | 0.59973325 | Chp2/Mapk12/Pik3cg/Pik3r5/Ppp3cc/Vav1 | 6 | 108 | 5.56 |
| KEGG_COLORECTAL_CANCER | COLORECTAL_CANCER | 4/182 | 58/3594 | 0.33804322 | 0.68078148 | 0.61282689 | Pik3cg/Pik3r5/Rac2/Tgfb1 | 4 | 64 | 6.25 |
| KEGG_GLYCINE_SERINE_AND_THREONINE_METABOLISM | GLYCINE_SERINE_AND_THREONINE_METABOLISM | 2/182 | 24/3594 | 0.34509545 | 0.68546357 | 0.61704161 | Chdh/Maoa | 2 | 32 | 6.25 |
| KEGG_ALANINE_ASPARTATE_AND_GLUTAMATE_METABOLISM | ALANINE_ASPARTATE_AND_GLUTAMATE_METABOLISM | 2/182 | 25/3594 | 0.36379268 | 0.70333251 | 0.63312691 | Ass1/Gpt2 | 2 | 33 | 6.06 |
| KEGG_FOCAL_ADHESION | FOCAL_ADHESION | 9/182 | 152/3594 | 0.36302161 | 0.70333251 | 0.63312691 | Actg1/Itgb8/Myl7/Pdgfd/Pdgfrb/Pik3cg/Pik3r5/Rac2/Vav1 | 9 | 195 | 4.62 |
| KEGG_AMYOTROPHIC_LATERAL_SCLEROSIS_ALS | AMYOTROPHIC_LATERAL_SCLEROSIS_ALS | 3/182 | 43/3594 | 0.37266004 | 0.71099612 | 0.64002555 | Chp2/Mapk12/Ppp3cc | 3 | 51 | 5.88 |
| KEGG_SYSTEMIC_LUPUS_ERYTHEMATOSUS | SYSTEMIC_LUPUS_ERYTHEMATOSUS | 3/182 | 44/3594 | 0.38667635 | 0.72815676 | 0.65547324 | C1ra/H2-DMb1/H3f3a | 3 | 105 | 2.86 |
| KEGG_CYSTEINE_AND_METHIONINE_METABOLISM | CYSTEINE_AND_METHIONINE_METABOLISM | 2/182 | 27/3594 | 0.40055014 | 0.73096724 | 0.65800317 | Ahcyl2/Trdmt1 | 2 | 38 | 5.26 |
| KEGG_FOLATE_BIOSYNTHESIS | FOLATE_BIOSYNTHESIS | 1/182 | Oct-94 | 0.40568196 | 0.73096724 | 0.65800317 | Ggh | 1 | 17 | 5.88 |
| KEGG_PANCREATIC_CANCER | PANCREATIC_CANCER | 4/182 | 64/3594 | 0.40833342 | 0.73096724 | 0.65800317 | Pik3cg/Pik3r5/Rac2/Tgfb1 | 4 | 71 | 5.63 |
| KEGG_TYROSINE_METABOLISM | TYROSINE_METABOLISM | 2/182 | 27/3594 | 0.40055014 | 0.73096724 | 0.65800317 | Fah/Maoa | 2 | 44 | 4.55 |
| KEGG_GLYCOSAMINOGLYCAN_BIOSYNTHESIS_KERATAN_SULFATE | GLYCOSAMINOGLYCAN_BIOSYNTHESIS_KERATAN_SULFATE | 1/182 | Nov-94 | 0.43586217 | 0.77073188 | 0.69379858 | St3gal3 | 1 | 14 | 7.14 |
| KEGG_ARRHYTHMOGENIC_RIGHT_VENTRICULAR_CARDIOMYOPATHY_ARVC | ARRHYTHMOGENIC_RIGHT_VENTRICULAR_CARDIOMYOPATHY_ARVC | 3/182 | 48/3594 | 0.44187181 | 0.77194472 | 0.69489035 | Actg1/Itgb8/Slc8a1 | 3 | 74 | 4.05 |
| KEGG_RENIN_ANGIOTENSIN_SYSTEM | RENIN_ANGIOTENSIN_SYSTEM | 1/182 | Dec-94 | 0.46451779 | 0.80184618 | 0.72180709 | Ace2 | 1 | 19 | 5.26 |
| KEGG_MAPK_SIGNALING_PATHWAY | MAPK_SIGNALING_PATHWAY | 11/182 | 207/3594 | 0.48025682 | 0.81267453 | 0.73155456 | Chp2/Fgfr3/Hspb1/Mapk12/Nr4a1/Pdgfrb/Ppp3cc/Rac2/Relb/Rras/Tgfb1 | 11 | 266 | 4.14 |
| KEGG_NON_SMALL_CELL_LUNG_CANCER | NON_SMALL_CELL_LUNG_CANCER | 3/182 | 51/3594 | 0.48200006 | 0.81267453 | 0.73155456 | Pik3cg/Pik3r5/Plcg2 | 3 | 53 | 5.66 |
| KEGG_APOPTOSIS | APOPTOSIS | 4/182 | 74/3594 | 0.52124724 | 0.8126973 | 0.73157507 | Chp2/Pik3cg/Pik3r5/Ppp3cc | 4 | 89 | 4.49 |
| KEGG_GLYCOSAMINOGLYCAN_BIOSYNTHESIS_CHONDROITIN_SULFATE | GLYCOSAMINOGLYCAN_BIOSYNTHESIS_CHONDROITIN_SULFATE | 1/182 | 14/3594 | 0.51755787 | 0.8126973 | 0.73157507 | Chst15 | 1 | 22 | 4.55 |
| KEGG_GLYOXYLATE_AND_DICARBOXYLATE_METABOLISM | GLYOXYLATE_AND_DICARBOXYLATE_METABOLISM | 1/182 | 14/3594 | 0.51755787 | 0.8126973 | 0.73157507 | Hao2 | 1 | 16 | 6.25 |
| KEGG_PEROXISOME | PEROXISOME | 4/182 | 74/3594 | 0.52124724 | 0.8126973 | 0.73157507 | Eci3/Hao2/Paox/Pex16 | 4 | 80 | 5 |
| KEGG_PROGESTERONE_MEDIATED_OOCYTE_MATURATION | PROGESTERONE_MEDIATED_OOCYTE_MATURATION | 4/182 | 74/3594 | 0.52124724 | 0.8126973 | 0.73157507 | Gnai1/Mapk12/Pik3cg/Pik3r5 | 4 | 84 | 4.76 |
| KEGG_PURINE_METABOLISM | PURINE_METABOLISM | 7/182 | 131/3594 | 0.49872994 | 0.8126973 | 0.73157507 | Entpd2/Gmpr/Nt5e/Paics/Pde7a/Pde8b/Pnp2 | 7 | 158 | 4.43 |
| KEGG_PYRUVATE_METABOLISM | PYRUVATE_METABOLISM | 2/182 | 34/3594 | 0.52026588 | 0.8126973 | 0.73157507 | Aldh2/Pck1 | 2 | 39 | 5.13 |
| KEGG_ALZHEIMERS_DISEASE | ALZHEIMERS_DISEASE | 7/182 | 140/3594 | 0.57085153 | 0.83514076 | 0.75177825 | Atp5g2/Bace2/Chp2/Cox6b2/Lpl/Plcb2/Ppp3cc | 7 | 165 | 4.24 |
| KEGG_AXON_GUIDANCE | AXON_GUIDANCE | 5/182 | 101/3594 | 0.58703069 | 0.83514076 | 0.75177825 | Chp2/Gnai1/Ppp3cc/Rac2/Srgap3 | 5 | 129 | 3.88 |
| KEGG_BLADDER_CANCER | BLADDER_CANCER | 2/182 | 37/3594 | 0.56651811 | 0.83514076 | 0.75177825 | Fgfr3/Tymp | 2 | 42 | 4.76 |
| KEGG_ERBB_SIGNALING_PATHWAY | ERBB_SIGNALING_PATHWAY | 4/182 | 78/3594 | 0.56358178 | 0.83514076 | 0.75177825 | Hbegf/Pik3cg/Pik3r5/Plcg2 | 4 | 86 | 4.65 |
| KEGG_INSULIN_SIGNALING_PATHWAY | INSULIN_SIGNALING_PATHWAY | 6/182 | 119/3594 | 0.56549017 | 0.83514076 | 0.75177825 | Pck1/Pik3cg/Pik3r5/Ppargc1a/Prkab2/Pygl | 6 | 137 | 4.38 |
| KEGG_LONG_TERM_POTENTIATION | LONG_TERM_POTENTIATION | 3/182 | 57/3594 | 0.55785614 | 0.83514076 | 0.75177825 | Chp2/Plcb2/Ppp3cc | 3 | 68 | 4.41 |
| KEGG_LYSINE_DEGRADATION | LYSINE_DEGRADATION | 2/182 | 38/3594 | 0.58121033 | 0.83514076 | 0.75177825 | Aldh2/Echs1 | 2 | 42 | 4.76 |
| KEGG_ONE_CARBON_POOL_BY_FOLATE | ONE_CARBON_POOL_BY_FOLATE | 1/182 | 17/3594 | 0.58747833 | 0.83514076 | 0.75177825 | Aldh1l1 | 1 | 18 | 5.56 |
| KEGG_WNT_SIGNALING_PATHWAY | WNT_SIGNALING_PATHWAY | 6/182 | 121/3594 | 0.58216013 | 0.83514076 | 0.75177825 | Chp2/Frat2/Plcb2/Ppp3cc/Rac2/Wif1 | 6 | 151 | 3.97 |
| KEGG_P53_SIGNALING_PATHWAY | P53_SIGNALING_PATHWAY | 3/182 | 61/3594 | 0.60459801 | 0.84168208 | 0.75766662 | Adgrb1/Pmaip1/Serpinb5 | 3 | 66 | 4.55 |
| KEGG_TGF_BETA_SIGNALING_PATHWAY | TGF_BETA_SIGNALING_PATHWAY | 3/182 | 61/3594 | 0.60459801 | 0.84168208 | 0.75766662 | Acvr1c/Lefty1/Tgfb1 | 3 | 83 | 3.61 |
| KEGG_TYPE_II_DIABETES_MELLITUS | TYPE_II_DIABETES_MELLITUS | 2/182 | 40/3594 | 0.60949392 | 0.84168208 | 0.75766662 | Pik3cg/Pik3r5 | 2 | 48 | 4.17 |
| KEGG_NEUROACTIVE_LIGAND_RECEPTOR_INTERACTION | NEUROACTIVE_LIGAND_RECEPTOR_INTERACTION | 4/182 | 84/3594 | 0.62308375 | 0.8464671 | 0.76197402 | Chrm3/Prlr/Sstr2/Tspo | 4 | 313 | 1.28 |
| KEGG_NOTCH_SIGNALING_PATHWAY | NOTCH_SIGNALING_PATHWAY | 2/182 | 42/3594 | 0.63630975 | 0.8464671 | 0.76197402 | Cir1/Dtx3 | 2 | 46 | 4.35 |
| KEGG_PROTEASOME | PROTEASOME | 2/182 | 42/3594 | 0.63630975 | 0.8464671 | 0.76197402 | Psmb8/Psmb9 | 2 | 45 | 4.44 |
| KEGG_RIBOSOME | RIBOSOME | 3/182 | 63/3594 | 0.62671997 | 0.8464671 | 0.76197402 | Rpl39/Rps28/Rps29 | 3 | 74 | 4.05 |
| KEGG_ADHERENS_JUNCTION | ADHERENS_JUNCTION | 3/182 | 65/3594 | 0.64798489 | 0.85022595 | 0.76535766 | Actg1/Nectin4/Rac2 | 3 | 73 | 4.11 |
| KEGG_JAK_STAT_SIGNALING_PATHWAY | JAK_STAT_SIGNALING_PATHWAY | 4/182 | 87/3594 | 0.65086262 | 0.85022595 | 0.76535766 | Ifngr2/Pik3cg/Pik3r5/Prlr | 4 | 353 | 1.13 |
| KEGG_CARDIAC_MUSCLE_CONTRACTION | CARDIAC_MUSCLE_CONTRACTION | 2/182 | 45/3594 | 0.67381234 | 0.85254168 | 0.76744224 | Cox6b2/Slc8a1 | 2 | 79 | 2.53 |
| KEGG_CHRONIC_MYELOID_LEUKEMIA | CHRONIC_MYELOID_LEUKEMIA | 3/182 | 69/3594 | 0.68791294 | 0.85254168 | 0.76744224 | Pik3cg/Pik3r5/Tgfb1 | 3 | 72 | 4.17 |
| KEGG_CYTOKINE_CYTOKINE_RECEPTOR_INTERACTION | CYTOKINE_CYTOKINE_RECEPTOR_INTERACTION | 5/182 | 112/3594 | 0.67894808 | 0.85254168 | 0.76744224 | Ifngr2/Il18/Pdgfrb/Prlr/Tgfb1 | 5 | 460 | 1.09 |
| KEGG_MTOR_SIGNALING_PATHWAY | MTOR_SIGNALING_PATHWAY | 2/182 | 46/3594 | 0.68560037 | 0.85254168 | 0.76744224 | Pik3cg/Pik3r5 | 2 | 52 | 3.85 |
| KEGG_O_GLYCAN_BIOSYNTHESIS | O_GLYCAN_BIOSYNTHESIS | 1/182 | 22/3594 | 0.68232297 | 0.85254168 | 0.76744224 | Gcnt1 | 1 | 29 | 3.45 |
| KEGG_TOLL_LIKE_RECEPTOR_SIGNALING_PATHWAY | TOLL_LIKE_RECEPTOR_SIGNALING_PATHWAY | 3/182 | 68/3594 | 0.67825685 | 0.85254168 | 0.76744224 | Mapk12/Pik3cg/Pik3r5 | 3 | 267 | 1.12 |
| KEGG_LONG_TERM_DEPRESSION | LONG_TERM_DEPRESSION | 2/182 | 48/3594 | 0.70813326 | 0.85547006 | 0.77007831 | Gnai1/Plcb2 | 2 | 67 | 2.99 |
| KEGG_PROTEIN_EXPORT | PROTEIN_EXPORT | 1/182 | 24/3594 | 0.71387501 | 0.85547006 | 0.77007831 | Immp2l | 1 | 26 | 3.85 |
| KEGG_SELENOAMINO_ACID_METABOLISM | SELENOAMINO_ACID_METABOLISM | 1/182 | 24/3594 | 0.71387501 | 0.85547006 | 0.77007831 | Ahcyl2 | 1 | 27 | 3.7 |
| KEGG_TIGHT_JUNCTION | TIGHT_JUNCTION | 4/182 | 94/3594 | 0.71027329 | 0.85547006 | 0.77007831 | Actg1/Gnai1/Myl7/Rras | 4 | 125 | 3.2 |
| KEGG_ENDOMETRIAL_CANCER | ENDOMETRIAL_CANCER | 2/182 | 51/3594 | 0.7394044 | 0.87880032 | 0.79107978 | Pik3cg/Pik3r5 | 2 | 52 | 3.85 |
| KEGG_ACUTE_MYELOID_LEUKEMIA | ACUTE_MYELOID_LEUKEMIA | 2/182 | 56/3594 | 0.78518898 | 0.91324819 | 0.82208911 | Pik3cg/Pik3r5 | 2 | 59 | 3.39 |
| KEGG_CITRATE_CYCLE_TCA_CYCLE | CITRATE_CYCLE_TCA_CYCLE | 1/182 | 29/3594 | 0.77977516 | 0.91324819 | 0.82208911 | Pck1 | 1 | 31 | 3.23 |
| KEGG_GNRH_SIGNALING_PATHWAY | GNRH_SIGNALING_PATHWAY | 3/182 | 81/3594 | 0.78728292 | 0.91324819 | 0.82208911 | Hbegf/Mapk12/Plcb2 | 3 | 95 | 3.16 |
| KEGG_CYTOSOLIC_DNA_SENSING_PATHWAY | CYTOSOLIC_DNA_SENSING_PATHWAY | 1/182 | 33/3594 | 0.82143481 | 0.93255599 | 0.83946964 | Il18 | 1 | 221 | 0.45 |
| KEGG_NEUROTROPHIN_SIGNALING_PATHWAY | NEUROTROPHIN_SIGNALING_PATHWAY | 4/182 | 109/3594 | 0.81212619 | 0.93255599 | 0.83946964 | Mapk12/Pik3cg/Pik3r5/Plcg2 | 4 | 126 | 3.17 |
| KEGG_OXIDATIVE_PHOSPHORYLATION | OXIDATIVE_PHOSPHORYLATION | 4/182 | 111/3594 | 0.82322184 | 0.93255599 | 0.83946964 | Atp5g2/Atp6v0e2/Cox6b2/Lhpp | 4 | 132 | 3.03 |
| KEGG_FRUCTOSE_AND_MANNOSE_METABOLISM | FRUCTOSE_AND_MANNOSE_METABOLISM | 1/182 | 35/3594 | 0.83922346 | 0.94331319 | 0.84915307 | Akr1b8 | 1 | 38 | 2.63 |
| KEGG_OOCYTE_MEIOSIS | OOCYTE_MEIOSIS | 3/182 | 92/3594 | 0.85401118 | 0.95255093 | 0.85746872 | Chp2/Mapk12/Ppp3cc | 3 | 111 | 2.7 |
| KEGG_GLUTATHIONE_METABOLISM | GLUTATHIONE_METABOLISM | 1/182 | 44/3594 | 0.89981258 | 0.95356553 | 0.85838203 | Gstp2 | 1 | 48 | 2.08 |
| KEGG_GLYCEROPHOSPHOLIPID_METABOLISM | GLYCEROPHOSPHOLIPID_METABOLISM | 2/182 | 70/3594 | 0.87792641 | 0.95356553 | 0.85838203 | Dgkh/Gpd1 | 2 | 76 | 2.63 |
| KEGG_HUNTINGTONS_DISEASE | HUNTINGTONS_DISEASE | 5/182 | 150/3594 | 0.88638992 | 0.95356553 | 0.85838203 | Atp5g2/Cox6b2/Plcb2/Ppargc1a/Vdac3 | 5 | 175 | 2.86 |
| KEGG_MELANOGENESIS | MELANOGENESIS | 2/182 | 74/3594 | 0.89669849 | 0.95356553 | 0.85838203 | Gnai1/Plcb2 | 2 | 97 | 2.06 |
| KEGG_N_GLYCAN_BIOSYNTHESIS | N_GLYCAN_BIOSYNTHESIS | 1/182 | 43/3594 | 0.89440026 | 0.95356553 | 0.85838203 | Mgat4a | 1 | 46 | 2.17 |
| KEGG_NUCLEOTIDE_EXCISION_REPAIR | NUCLEOTIDE_EXCISION_REPAIR | 1/182 | 42/3594 | 0.88869725 | 0.95356553 | 0.85838203 | Rad23a | 1 | 42 | 2.38 |
| KEGG_SMALL_CELL_LUNG_CANCER | SMALL_CELL_LUNG_CANCER | 2/182 | 75/3594 | 0.90095501 | 0.95356553 | 0.85838203 | Pik3cg/Pik3r5 | 2 | 82 | 2.44 |
| KEGG_PARKINSONS_DISEASE | PARKINSONS_DISEASE | 3/182 | 107/3594 | 0.91535366 | 0.95486532 | 0.85955209 | Atp5g2/Cox6b2/Vdac3 | 3 | 126 | 2.38 |
| KEGG_RIG_I_LIKE_RECEPTOR_SIGNALING_PATHWAY | RIG_I_LIKE_RECEPTOR_SIGNALING_PATHWAY | 1/182 | 47/3594 | 0.91444911 | 0.95486532 | 0.85955209 | Mapk12 | 1 | 238 | 0.42 |
| KEGG_PATHWAYS_IN_CANCER | PATHWAYS_IN_CANCER | 9/182 | 262/3594 | 0.9251802 | 0.95822235 | 0.86257402 | Egln3/Fgfr3/Gstp2/Pdgfrb/Pik3cg/Pik3r5/Plcg2/Rac2/Tgfb1 | 9 | 340 | 2.65 |
| KEGG_RNA_DEGRADATION | RNA_DEGRADATION | 1/182 | 54/3594 | 0.94085031 | 0.9675411 | 0.87096259 | Eno3 | 1 | 58 | 1.72 |
| KEGG_ECM_RECEPTOR_INTERACTION | ECM_RECEPTOR_INTERACTION | 1/182 | 58/3594 | 0.95211249 | 0.97222754 | 0.87518124 | Itgb8 | 1 | 83 | 1.2 |
| KEGG_VASCULAR_SMOOTH_MUSCLE_CONTRACTION | VASCULAR_SMOOTH_MUSCLE_CONTRACTION | 1/182 | 77/3594 | 0.98249977 | 0.99624103 | 0.89679773 | Plcb2 | 1 | 119 | 0.84 |
| KEGG_CELL_CYCLE | CELL_CYCLE | 1/182 | 118/3594 | 0.99804415 | 0.99804415 | 0.89842087 | Tgfb1 | 1 | 124 | 0.81 |
| KEGG_ENDOCYTOSIS | ENDOCYTOSIS | 2/182 | 152/3594 | 0.99704684 | 0.99804415 | 0.89842087 | Fgfr3/Grk3 | 2 | 184 | 1.09 |
| KEGG_ABC_TRANSPORTERS | ABC_TRANSPORTERS | NA | NA | NA | NA | NA | NA | 0 | 43 | 0 |
| KEGG_ALLOGRAFT_REJECTION | ALLOGRAFT_REJECTION | NA | NA | NA | NA | NA | NA | 0 | 37 | 0 |
| KEGG_ALPHA_LINOLENIC_ACID_METABOLISM | ALPHA_LINOLENIC_ACID_METABOLISM | NA | NA | NA | NA | NA | NA | 0 | 18 | 0 |
| KEGG_AMINOACYL_TRNA_BIOSYNTHESIS | AMINOACYL_TRNA_BIOSYNTHESIS | NA | NA | NA | NA | NA | NA | 0 | 41 | 0 |
| KEGG_ASTHMA | ASTHMA | NA | NA | NA | NA | NA | NA | 0 | 32 | 0 |
| KEGG_AUTOIMMUNE_THYROID_DISEASE | AUTOIMMUNE_THYROID_DISEASE | NA | NA | NA | NA | NA | NA | 0 | 221 | 0 |
| KEGG_BASAL_CELL_CARCINOMA | BASAL_CELL_CARCINOMA | NA | NA | NA | NA | NA | NA | 0 | 55 | 0 |
| KEGG_BASAL_TRANSCRIPTION_FACTORS | BASAL_TRANSCRIPTION_FACTORS | NA | NA | NA | NA | NA | NA | 0 | 38 | 0 |
| KEGG_BASE_EXCISION_REPAIR | BASE_EXCISION_REPAIR | NA | NA | NA | NA | NA | NA | 0 | 32 | 0 |
| KEGG_BIOSYNTHESIS_OF_UNSATURATED_FATTY_ACIDS | BIOSYNTHESIS_OF_UNSATURATED_FATTY_ACIDS | NA | NA | NA | NA | NA | NA | 0 | 29 | 0 |
| KEGG_CIRCADIAN_RHYTHM_MAMMAL | CIRCADIAN_RHYTHM_MAMMAL | NA | NA | NA | NA | NA | NA | 0 | 13 | 0 |
| KEGG_DNA_REPLICATION | DNA_REPLICATION | NA | NA | NA | NA | NA | NA | 0 | 34 | 0 |
| KEGG_DORSO_VENTRAL_AXIS_FORMATION | DORSO_VENTRAL_AXIS_FORMATION | NA | NA | NA | NA | NA | NA | 0 | 21 | 0 |
| KEGG_ETHER_LIPID_METABOLISM | ETHER_LIPID_METABOLISM | NA | NA | NA | NA | NA | NA | 0 | 32 | 0 |
| KEGG_GALACTOSE_METABOLISM | GALACTOSE_METABOLISM | NA | NA | NA | NA | NA | NA | 0 | 26 | 0 |
| KEGG_GLYCOSAMINOGLYCAN_BIOSYNTHESIS_HEPARAN_SULFATE | GLYCOSAMINOGLYCAN_BIOSYNTHESIS_HEPARAN_SULFATE | NA | NA | NA | NA | NA | NA | 0 | 26 | 0 |
| KEGG_GLYCOSYLPHOSPHATIDYLINOSITOL_GPI_ANCHOR_BIOSYNTHESIS | GLYCOSYLPHOSPHATIDYLINOSITOL_GPI_ANCHOR_BIOSYNTHESIS | NA | NA | NA | NA | NA | NA | 0 | 24 | 0 |
| KEGG_GRAFT_VERSUS_HOST_DISEASE | GRAFT_VERSUS_HOST_DISEASE | NA | NA | NA | NA | NA | NA | 0 | 50 | 0 |
| KEGG_HEDGEHOG_SIGNALING_PATHWAY | HEDGEHOG_SIGNALING_PATHWAY | NA | NA | NA | NA | NA | NA | 0 | 52 | 0 |
| KEGG_HOMOLOGOUS_RECOMBINATION | HOMOLOGOUS_RECOMBINATION | NA | NA | NA | NA | NA | NA | 0 | 26 | 0 |
| KEGG_MISMATCH_REPAIR | MISMATCH_REPAIR | NA | NA | NA | NA | NA | NA | 0 | 21 | 0 |
| KEGG_NITROGEN_METABOLISM | NITROGEN_METABOLISM | NA | NA | NA | NA | NA | NA | 0 | 23 | 0 |
| KEGG_NON_HOMOLOGOUS_END_JOINING | NON_HOMOLOGOUS_END_JOINING | NA | NA | NA | NA | NA | NA | 0 | 13 | 0 |
| KEGG_PENTOSE_PHOSPHATE_PATHWAY | PENTOSE_PHOSPHATE_PATHWAY | NA | NA | NA | NA | NA | NA | 0 | 31 | 0 |
| KEGG_PRIMARY_BILE_ACID_BIOSYNTHESIS | PRIMARY_BILE_ACID_BIOSYNTHESIS | NA | NA | NA | NA | NA | NA | 0 | 15 | 0 |
| KEGG_PRIMARY_IMMUNODEFICIENCY | PRIMARY_IMMUNODEFICIENCY | NA | NA | NA | NA | NA | NA | 0 | 38 | 0 |
| KEGG_PRION_DISEASES | PRION_DISEASES | NA | NA | NA | NA | NA | NA | 0 | 34 | 0 |
| KEGG_REGULATION_OF_AUTOPHAGY | REGULATION_OF_AUTOPHAGY | NA | NA | NA | NA | NA | NA | 0 | 206 | 0 |
| KEGG_RIBOFLAVIN_METABOLISM | RIBOFLAVIN_METABOLISM | NA | NA | NA | NA | NA | NA | 0 | 16 | 0 |
| KEGG_RNA_POLYMERASE | RNA_POLYMERASE | NA | NA | NA | NA | NA | NA | 0 | 27 | 0 |
| KEGG_SNARE_INTERACTIONS_IN_VESICULAR_TRANSPORT | SNARE_INTERACTIONS_IN_VESICULAR_TRANSPORT | NA | NA | NA | NA | NA | NA | 0 | 34 | 0 |
| KEGG_SPHINGOLIPID_METABOLISM | SPHINGOLIPID_METABOLISM | NA | NA | NA | NA | NA | NA | 0 | 40 | 0 |
| KEGG_SPLICEOSOME | SPLICEOSOME | NA | NA | NA | NA | NA | NA | 0 | 154 | 0 |
| KEGG_STEROID_BIOSYNTHESIS | STEROID_BIOSYNTHESIS | NA | NA | NA | NA | NA | NA | 0 | 17 | 0 |
| KEGG_SULFUR_METABOLISM | SULFUR_METABOLISM | NA | NA | NA | NA | NA | NA | 0 | 14 | 0 |
| KEGG_TAURINE_AND_HYPOTAURINE_METABOLISM | TAURINE_AND_HYPOTAURINE_METABOLISM | NA | NA | NA | NA | NA | NA | 0 | 10 | 0 |
| KEGG_TERPENOID_BACKBONE_BIOSYNTHESIS | TERPENOID_BACKBONE_BIOSYNTHESIS | NA | NA | NA | NA | NA | NA | 0 | 17 | 0 |
| KEGG_THYROID_CANCER | THYROID_CANCER | NA | NA | NA | NA | NA | NA | 0 | 31 | 0 |
| KEGG_UBIQUITIN_MEDIATED_PROTEOLYSIS | UBIQUITIN_MEDIATED_PROTEOLYSIS | NA | NA | NA | NA | NA | NA | 0 | 136 | 0 |
| KEGG_VALINE_LEUCINE_AND_ISOLEUCINE_BIOSYNTHESIS | VALINE_LEUCINE_AND_ISOLEUCINE_BIOSYNTHESIS | NA | NA | NA | NA | NA | NA | 0 | 11 | 0 |
| KEGG_VASOPRESSIN_REGULATED_WATER_REABSORPTION | VASOPRESSIN_REGULATED_WATER_REABSORPTION | NA | NA | NA | NA | NA | NA | 0 | 44 | 0 |

| ID | Description | GeneRatio | BgRatio | pvalue | p.adjust | qvalue | geneID | Count | Genes_in_Pathway | Percentage_of_Pathway_detected |
|---|---|---|---|---|---|---|---|---|---|---|
| KEGG_SYSTEMIC_LUPUS_ERYTHEMATOSUS | SYSTEMIC_LUPUS_ERYTHEMATOSUS | 12/223 | 44/3594 | 8.68E-06 | 0.0013017 | 0.00126973 | H2ac15/H2ac20/H2ac25/H2ax/H2bc14/H2bc26-ps/H2bc27/H2bc8/H3c1/H3c10/H3c8/H4c14 | 12 | 105 | 11.43 |
| KEGG_CELL_CYCLE | CELL_CYCLE | 20/223 | 118/3594 | 2.67E-05 | 0.00200279 | 0.0019536 | Bub1/Bub1b/Ccna2/Ccnb1/Ccnb2/Ccnd1/Cdc20/Cdc25c/Cdc45/Cdc7/Cdk1/Cdkn1c/Dbf4/E2f1/Espl1/Gadd45b/Gadd45g/Mad2l1/Skp2/Ttk | 20 | 124 | 16.13 |
| KEGG_HOMOLOGOUS_RECOMBINATION | HOMOLOGOUS_RECOMBINATION | 6/223 | 26/3594 | 0.00429464 | 0.21473211 | 0.20945799 | Blm/Eme1/Rad51/Rad54b/Rpa2/Xrcc3 | 6 | 26 | 23.08 |
| KEGG_ARGININE_AND_PROLINE_METABOLISM | ARGININE_AND_PROLINE_METABOLISM | 8/223 | 46/3594 | 0.00648906 | 0.22224411 | 0.21678548 | Agmat/Gatm/Got1/Got2/Nos2/Oat/Pycr1/Srm | 8 | 59 | 13.56 |
| KEGG_GALACTOSE_METABOLISM | GALACTOSE_METABOLISM | 5/223 | 23/3594 | 0.01177756 | 0.22224411 | 0.21678548 | Akr1b3/B4galt2/Galk1/Hk2/Ugp2 | 5 | 26 | 19.23 |
| KEGG_GLYCOSAMINOGLYCAN_BIOSYNTHESIS_CHONDROITIN_SULFATE | GLYCOSAMINOGLYCAN_BIOSYNTHESIS_CHONDROITIN_SULFATE | 4/223 | 14/3594 | 0.0088107 | 0.22224411 | 0.21678548 | Chpf/Chst11/Chsy1/Csgalnact2 | 4 | 22 | 18.18 |
| KEGG_OOCYTE_MEIOSIS | OOCYTE_MEIOSIS | 12/223 | 92/3594 | 0.01049002 | 0.22224411 | 0.21678548 | Adcy3/Aurka/Bub1/Ccnb1/Ccnb2/Cdc20/Cdc25c/Cdk1/Espl1/Mad2l1/Prkaca/Sgo1 | 12 | 111 | 10.81 |
| KEGG_P53_SIGNALING_PATHWAY | P53_SIGNALING_PATHWAY | 9/223 | 61/3594 | 0.01185302 | 0.22224411 | 0.21678548 | Ccnb1/Ccnb2/Ccnd1/Ccng1/Cdk1/Fas/Gadd45b/Gadd45g/Gtse1 | 9 | 66 | 13.64 |
| KEGG_PROGESTERONE_MEDIATED_OOCYTE_MATURATION | PROGESTERONE_MEDIATED_OOCYTE_MATURATION | 10/223 | 74/3594 | 0.01488405 | 0.24806752 | 0.24197464 | Adcy3/Bub1/Ccna2/Ccnb1/Ccnb2/Cdc25c/Cdk1/Mad2l1/Mapk9/Prkaca | 10 | 84 | 11.9 |
| KEGG_N_GLYCAN_BIOSYNTHESIS | N_GLYCAN_BIOSYNTHESIS | 6/223 | 43/3594 | 0.04733499 | 0.64498338 | 0.62914169 | Alg8/B4galt2/Dolpp1/Dpagt1/Mgat2/Mogs | 6 | 46 | 13.04 |
| KEGG_O_GLYCAN_BIOSYNTHESIS | O_GLYCAN_BIOSYNTHESIS | 4/223 | 22/3594 | 0.0436469 | 0.64498338 | 0.62914169 | C1galt1c1/Galnt18/Galnt2/Galnt3 | 4 | 29 | 13.79 |
| KEGG_PHENYLALANINE_METABOLISM | PHENYLALANINE_METABOLISM | 3/223 | 14/3594 | 0.05159867 | 0.64498338 | 0.62914169 | Got1/Got2/Tat | 3 | 21 | 14.29 |
| KEGG_CYSTEINE_AND_METHIONINE_METABOLISM | CYSTEINE_AND_METHIONINE_METABOLISM | 4/223 | 27/3594 | 0.08244847 | 0.85447105 | 0.83348404 | Got1/Got2/Srm/Tat | 4 | 38 | 10.53 |
| KEGG_LYSINE_DEGRADATION | LYSINE_DEGRADATION | 5/223 | 38/3594 | 0.08309579 | 0.85447105 | 0.83348404 | Gcdh/Nsd2/Setd1b/Suv39h1/Suv39h2 | 5 | 42 | 11.9 |
| KEGG_PEROXISOME | PEROXISOME | 8/223 | 74/3594 | 0.08544711 | 0.85447105 | 0.83348404 | Acaa1b/Acox2/Cat/Mpv17l/Nos2/Pex2/Phyh/Xdh | 8 | 80 | 10 |
| KEGG_GLYCOSPHINGOLIPID_BIOSYNTHESIS_LACTO_AND_NEOLACTO_SERIES | GLYCOSPHINGOLIPID_BIOSYNTHESIS_LACTO_AND_NEOLACTO_SERIES | 3/223 | 18/3594 | 0.09657499 | 0.90539056 | 0.8831529 | B4galt2/Fut1/Fut2 | 3 | 23 | 13.04 |
| KEGG_ABC_TRANSPORTERS | ABC_TRANSPORTERS | 1/223 | 30/3594 | 0.85481448 | 0.99938151 | 0.97483529 | Abca7 | 1 | 43 | 2.33 |
| KEGG_ACUTE_MYELOID_LEUKEMIA | ACUTE_MYELOID_LEUKEMIA | 1/223 | 56/3594 | 0.97310478 | 0.99938151 | 0.97483529 | Ccnd1 | 1 | 59 | 1.69 |
| KEGG_ADHERENS_JUNCTION | ADHERENS_JUNCTION | 1/223 | 65/3594 | 0.98504079 | 0.99938151 | 0.97483529 | Ptprm | 1 | 73 | 1.37 |
| KEGG_ADIPOCYTOKINE_SIGNALING_PATHWAY | ADIPOCYTOKINE_SIGNALING_PATHWAY | 1/223 | 55/3594 | 0.97129608 | 0.99938151 | 0.97483529 | Mapk9 | 1 | 67 | 1.49 |
| KEGG_ALANINE_ASPARTATE_AND_GLUTAMATE_METABOLISM | ALANINE_ASPARTATE_AND_GLUTAMATE_METABOLISM | 3/223 | 25/3594 | 0.19980844 | 0.99938151 | 0.97483529 | Asns/Got1/Got2 | 3 | 33 | 9.09 |
| KEGG_ALPHA_LINOLENIC_ACID_METABOLISM | ALPHA_LINOLENIC_ACID_METABOLISM | 1/223 | 14/3594 | 0.5928071 | 0.99938151 | 0.97483529 | Pla2g5 | 1 | 18 | 5.56 |
| KEGG_ALZHEIMERS_DISEASE | ALZHEIMERS_DISEASE | 2/223 | 140/3594 | 0.99886991 | 0.99938151 | 0.97483529 | Eif2ak3/Fas | 2 | 165 | 1.21 |
| KEGG_AMINO_SUGAR_AND_NUCLEOTIDE_SUGAR_METABOLISM | AMINO_SUGAR_AND_NUCLEOTIDE_SUGAR_METABOLISM | 4/223 | 41/3594 | 0.247916 | 0.99938151 | 0.97483529 | Galk1/Hk2/Uap1/Ugp2 | 4 | 48 | 8.33 |
| KEGG_AMINOACYL_TRNA_BIOSYNTHESIS | AMINOACYL_TRNA_BIOSYNTHESIS | 1/223 | 41/3594 | 0.92874725 | 0.99938151 | 0.97483529 | Aars | 1 | 41 | 2.44 |
| KEGG_AMYOTROPHIC_LATERAL_SCLEROSIS_ALS | AMYOTROPHIC_LATERAL_SCLEROSIS_ALS | 2/223 | 43/3594 | 0.75713049 | 0.99938151 | 0.97483529 | Cat/Derl1 | 2 | 51 | 3.92 |
| KEGG_APOPTOSIS | APOPTOSIS | 3/223 | 74/3594 | 0.84802626 | 0.99938151 | 0.97483529 | Exog/Fas/Prkaca | 3 | 89 | 3.37 |
| KEGG_ARACHIDONIC_ACID_METABOLISM | ARACHIDONIC_ACID_METABOLISM | 1/223 | 36/3594 | 0.90150054 | 0.99938151 | 0.97483529 | Pla2g5 | 1 | 60 | 1.67 |
| KEGG_AXON_GUIDANCE | AXON_GUIDANCE | 4/223 | 101/3594 | 0.88259148 | 0.99938151 | 0.97483529 | Efnb2/Plxnb1/Sema4c/Srgap2 | 4 | 129 | 3.1 |
| KEGG_B_CELL_RECEPTOR_SIGNALING_PATHWAY | B_CELL_RECEPTOR_SIGNALING_PATHWAY | 1/223 | 66/3594 | 0.98598607 | 0.99938151 | 0.97483529 | Ifitm3 | 1 | 90 | 1.11 |
| KEGG_BASAL_CELL_CARCINOMA | BASAL_CELL_CARCINOMA | 3/223 | 35/3594 | 0.37124591 | 0.99938151 | 0.97483529 | Fzd2/Fzd3/Fzd7 | 3 | 55 | 5.45 |
| KEGG_BASAL_TRANSCRIPTION_FACTORS | BASAL_TRANSCRIPTION_FACTORS | 1/223 | 31/3594 | 0.86389876 | 0.99938151 | 0.97483529 | Taf13 | 1 | 38 | 2.63 |
| KEGG_BASE_EXCISION_REPAIR | BASE_EXCISION_REPAIR | 4/223 | 31/3594 | 0.12236714 | 0.99938151 | 0.97483529 | Apex1/Fen1/Mbd4/Mutyh | 4 | 32 | 12.5 |
| KEGG_BETA_ALANINE_METABOLISM | BETA_ALANINE_METABOLISM | 1/223 | 18/3594 | 0.685207 | 0.99938151 | 0.97483529 | Srm | 1 | 22 | 4.55 |
| KEGG_BIOSYNTHESIS_OF_UNSATURATED_FATTY_ACIDS | BIOSYNTHESIS_OF_UNSATURATED_FATTY_ACIDS | 3/223 | 22/3594 | 0.15258213 | 0.99938151 | 0.97483529 | Acaa1b/Acot1/Fads1 | 3 | 29 | 10.34 |
| KEGG_BLADDER_CANCER | BLADDER_CANCER | 3/223 | 37/3594 | 0.40564261 | 0.99938151 | 0.97483529 | Ccnd1/Dapk3/E2f1 | 3 | 42 | 7.14 |
| KEGG_BUTANOATE_METABOLISM | BUTANOATE_METABOLISM | 1/223 | 28/3594 | 0.83479569 | 0.99938151 | 0.97483529 | Hmgcs1 | 1 | 38 | 2.63 |
| KEGG_CALCIUM_SIGNALING_PATHWAY | CALCIUM_SIGNALING_PATHWAY | 5/223 | 103/3594 | 0.77676967 | 0.99938151 | 0.97483529 | Adcy3/Atp2b1/F2r/Nos2/Prkaca | 5 | 173 | 2.89 |
| KEGG_CELL_ADHESION_MOLECULES_CAMS | CELL_ADHESION_MOLECULES_CAMS | 3/223 | 65/3594 | 0.77866375 | 0.99938151 | 0.97483529 | Itgam/Ptprm/Vcan | 3 | 129 | 2.33 |
| KEGG_CHEMOKINE_SIGNALING_PATHWAY | CHEMOKINE_SIGNALING_PATHWAY | 7/223 | 132/3594 | 0.72221815 | 0.99938151 | 0.97483529 | Adcy3/Ccl9/Ccr1/Prkaca/Prkcz/Stat2/Tiam2 | 7 | 178 | 3.93 |
| KEGG_CHRONIC_MYELOID_LEUKEMIA | CHRONIC_MYELOID_LEUKEMIA | 2/223 | 69/3594 | 0.9348125 | 0.99938151 | 0.97483529 | Ccnd1/E2f1 | 2 | 72 | 2.78 |
| KEGG_CIRCADIAN_RHYTHM_MAMMAL | CIRCADIAN_RHYTHM_MAMMAL | 1/223 | Dec-94 | 0.53693772 | 0.99938151 | 0.97483529 | Per1 | 1 | 13 | 7.69 |
| KEGG_COLORECTAL_CANCER | COLORECTAL_CANCER | 4/223 | 58/3594 | 0.48955441 | 0.99938151 | 0.97483529 | Appl1/Birc5/Ccnd1/Mapk9 | 4 | 64 | 6.25 |
| KEGG_COMPLEMENT_AND_COAGULATION_CASCADES | COMPLEMENT_AND_COAGULATION_CASCADES | 2/223 | 29/3594 | 0.54552601 | 0.99938151 | 0.97483529 | F2r/Serpina1b | 2 | 74 | 2.7 |
| KEGG_CYTOKINE_CYTOKINE_RECEPTOR_INTERACTION | CYTOKINE_CYTOKINE_RECEPTOR_INTERACTION | 7/223 | 112/3594 | 0.54947703 | 0.99938151 | 0.97483529 | Ccl9/Ccr1/Fas/Ifnar1/Il6st/Pdgfc/Tnfrsf8 | 7 | 460 | 1.52 |
| KEGG_CYTOSOLIC_DNA_SENSING_PATHWAY | CYTOSOLIC_DNA_SENSING_PATHWAY | 1/223 | 33/3594 | 0.88040439 | 0.99938151 | 0.97483529 | Polr3k | 1 | 221 | 0.45 |
| KEGG_DILATED_CARDIOMYOPATHY | DILATED_CARDIOMYOPATHY | 2/223 | 53/3594 | 0.85088898 | 0.99938151 | 0.97483529 | Adcy3/Prkaca | 2 | 90 | 2.22 |
| KEGG_DNA_REPLICATION | DNA_REPLICATION | 2/223 | 34/3594 | 0.63325356 | 0.99938151 | 0.97483529 | Fen1/Rpa2 | 2 | 34 | 5.88 |
| KEGG_DORSO_VENTRAL_AXIS_FORMATION | DORSO_VENTRAL_AXIS_FORMATION | 1/223 | 19/3594 | 0.70483755 | 0.99938151 | 0.97483529 | Piwil4 | 1 | 21 | 4.76 |
| KEGG_DRUG_METABOLISM_CYTOCHROME_P450 | DRUG_METABOLISM_CYTOCHROME_P450 | 1/223 | 46/3594 | 0.94848131 | 0.99938151 | 0.97483529 | Gstm6 | 1 | 84 | 1.19 |
| KEGG_DRUG_METABOLISM_OTHER_ENZYMES | DRUG_METABOLISM_OTHER_ENZYMES | 2/223 | 37/3594 | 0.67925386 | 0.99938151 | 0.97483529 | Tk1/Xdh | 2 | 73 | 2.74 |
| KEGG_ECM_RECEPTOR_INTERACTION | ECM_RECEPTOR_INTERACTION | 1/223 | 58/3594 | 0.97638879 | 0.99938151 | 0.97483529 | Hmmr | 1 | 83 | 1.2 |
| KEGG_ENDOCYTOSIS | ENDOCYTOSIS | 4/223 | 152/3594 | 0.98791934 | 0.99938151 | 0.97483529 | Ehd4/F2r/Prkcz/Ret | 4 | 184 | 2.17 |
| KEGG_ENDOMETRIAL_CANCER | ENDOMETRIAL_CANCER | 2/223 | 51/3594 | 0.83526191 | 0.99938151 | 0.97483529 | Ccnd1/Elk1 | 2 | 52 | 3.85 |
| KEGG_EPITHELIAL_CELL_SIGNALING_IN_HELICOBACTER_PYLORI_INFECTION | EPITHELIAL_CELL_SIGNALING_IN_HELICOBACTER_PYLORI_INFECTION | 1/223 | 52/3594 | 0.9651109 | 0.99938151 | 0.97483529 | Mapk9 | 1 | 64 | 1.56 |
| KEGG_ERBB_SIGNALING_PATHWAY | ERBB_SIGNALING_PATHWAY | 3/223 | 78/3594 | 0.87227602 | 0.99938151 | 0.97483529 | Elk1/Ereg/Mapk9 | 3 | 86 | 3.49 |
| KEGG_ETHER_LIPID_METABOLISM | ETHER_LIPID_METABOLISM | 2/223 | 25/3594 | 0.46554143 | 0.99938151 | 0.97483529 | Lpcat4/Pla2g5 | 2 | 32 | 6.25 |
| KEGG_FATTY_ACID_METABOLISM | FATTY_ACID_METABOLISM | 2/223 | 36/3594 | 0.66446406 | 0.99938151 | 0.97483529 | Acaa1b/Gcdh | 2 | 51 | 3.92 |
| KEGG_FC_EPSILON_RI_SIGNALING_PATHWAY | FC_EPSILON_RI_SIGNALING_PATHWAY | 2/223 | 67/3594 | 0.9275232 | 0.99938151 | 0.97483529 | Mapk9/Pla2g5 | 2 | 78 | 2.56 |
| KEGG_FC_GAMMA_R_MEDIATED_PHAGOCYTOSIS | FC_GAMMA_R_MEDIATED_PHAGOCYTOSIS | 1/223 | 85/3594 | 0.9959603 | 0.99938151 | 0.97483529 | Marcks | 1 | 99 | 1.01 |
| KEGG_FOCAL_ADHESION | FOCAL_ADHESION | 4/223 | 152/3594 | 0.98791934 | 0.99938151 | 0.97483529 | Ccnd1/Elk1/Mapk9/Pdgfc | 4 | 195 | 2.05 |
| KEGG_FOLATE_BIOSYNTHESIS | FOLATE_BIOSYNTHESIS | 1/223 | Oct-94 | 0.47344175 | 0.99938151 | 0.97483529 | Qdpr | 1 | 17 | 5.88 |
| KEGG_FRUCTOSE_AND_MANNOSE_METABOLISM | FRUCTOSE_AND_MANNOSE_METABOLISM | 2/223 | 35/3594 | 0.64913281 | 0.99938151 | 0.97483529 | Akr1b3/Hk2 | 2 | 38 | 5.26 |
| KEGG_GAP_JUNCTION | GAP_JUNCTION | 5/223 | 64/3594 | 0.36508829 | 0.99938151 | 0.97483529 | Adcy3/Cdk1/Pdgfc/Prkaca/Tubb5 | 5 | 84 | 5.95 |
| KEGG_GLIOMA | GLIOMA | 2/223 | 55/3594 | 0.86515995 | 0.99938151 | 0.97483529 | Ccnd1/E2f1 | 2 | 61 | 3.28 |
| KEGG_GLUTATHIONE_METABOLISM | GLUTATHIONE_METABOLISM | 3/223 | 44/3594 | 0.52044446 | 0.99938151 | 0.97483529 | Ggct/Gstm6/Srm | 3 | 48 | 6.25 |
| KEGG_GLYCEROLIPID_METABOLISM | GLYCEROLIPID_METABOLISM | 3/223 | 43/3594 | 0.50473647 | 0.99938151 | 0.97483529 | Agpat3/Akr1b3/Dgat2 | 3 | 49 | 6.12 |
| KEGG_GLYCEROPHOSPHOLIPID_METABOLISM | GLYCEROPHOSPHOLIPID_METABOLISM | 5/223 | 70/3594 | 0.44056676 | 0.99938151 | 0.97483529 | Agpat3/Chka/Crls1/Lpcat4/Pla2g5 | 5 | 76 | 6.58 |
| KEGG_GLYCINE_SERINE_AND_THREONINE_METABOLISM | GLYCINE_SERINE_AND_THREONINE_METABOLISM | 2/223 | 24/3594 | 0.44428279 | 0.99938151 | 0.97483529 | Gatm/Gcat | 2 | 32 | 6.25 |
| KEGG_GLYCOLYSIS_GLUCONEOGENESIS | GLYCOLYSIS_GLUCONEOGENESIS | 1/223 | 48/3594 | 0.95475478 | 0.99938151 | 0.97483529 | Hk2 | 1 | 67 | 1.49 |
| KEGG_GLYCOSAMINOGLYCAN_BIOSYNTHESIS_HEPARAN_SULFATE | GLYCOSAMINOGLYCAN_BIOSYNTHESIS_HEPARAN_SULFATE | 1/223 | 19/3594 | 0.70483755 | 0.99938151 | 0.97483529 | Hs6st1 | 1 | 26 | 3.85 |
| KEGG_GLYCOSAMINOGLYCAN_BIOSYNTHESIS_KERATAN_SULFATE | GLYCOSAMINOGLYCAN_BIOSYNTHESIS_KERATAN_SULFATE | 1/223 | Nov-94 | 0.50620472 | 0.99938151 | 0.97483529 | B4galt2 | 1 | 14 | 7.14 |
| KEGG_GLYCOSPHINGOLIPID_BIOSYNTHESIS_GANGLIO_SERIES | GLYCOSPHINGOLIPID_BIOSYNTHESIS_GANGLIO_SERIES | 1/223 | Dec-94 | 0.53693772 | 0.99938151 | 0.97483529 | St6galnac4 | 1 | 15 | 6.67 |
| KEGG_GLYCOSPHINGOLIPID_BIOSYNTHESIS_GLOBO_SERIES | GLYCOSPHINGOLIPID_BIOSYNTHESIS_GLOBO_SERIES | 2/223 | Dec-94 | 0.1681417 | 0.99938151 | 0.97483529 | Fut1/Fut2 | 2 | 14 | 14.29 |
| KEGG_GLYOXYLATE_AND_DICARBOXYLATE_METABOLISM | GLYOXYLATE_AND_DICARBOXYLATE_METABOLISM | 2/223 | 14/3594 | 0.21423169 | 0.99938151 | 0.97483529 | Mthfd2/Pgp | 2 | 16 | 12.5 |
| KEGG_GNRH_SIGNALING_PATHWAY | GNRH_SIGNALING_PATHWAY | 7/223 | 81/3594 | 0.23511536 | 0.99938151 | 0.97483529 | Adcy3/Atf4/Elk1/Map3k3/Mapk9/Pla2g5/Prkaca | 7 | 95 | 7.37 |
| KEGG_HEDGEHOG_SIGNALING_PATHWAY | HEDGEHOG_SIGNALING_PATHWAY | 1/223 | 27/3594 | 0.82377878 | 0.99938151 | 0.97483529 | Prkaca | 1 | 52 | 1.92 |
| KEGG_HEMATOPOIETIC_CELL_LINEAGE | HEMATOPOIETIC_CELL_LINEAGE | 1/223 | 45/3594 | 0.94502711 | 0.99938151 | 0.97483529 | Itgam | 1 | 88 | 1.14 |
| KEGG_HUNTINGTONS_DISEASE | HUNTINGTONS_DISEASE | 2/223 | 150/3594 | 0.99938151 | 0.99938151 | 0.97483529 | Polr2i/Tgm2 | 2 | 175 | 1.14 |
| KEGG_INOSITOL_PHOSPHATE_METABOLISM | INOSITOL_PHOSPHATE_METABOLISM | 1/223 | 51/3594 | 0.96276744 | 0.99938151 | 0.97483529 | Aldh6a1 | 1 | 53 | 1.89 |
| KEGG_INSULIN_SIGNALING_PATHWAY | INSULIN_SIGNALING_PATHWAY | 8/223 | 119/3594 | 0.46087672 | 0.99938151 | 0.97483529 | Elk1/Fasn/Gys1/Hk2/Mapk9/Prkaca/Prkcz/Pygm | 8 | 137 | 5.84 |
| KEGG_JAK_STAT_SIGNALING_PATHWAY | JAK_STAT_SIGNALING_PATHWAY | 5/223 | 87/3594 | 0.63628792 | 0.99938151 | 0.97483529 | Ccnd1/Ifnar1/Il6st/Stat2/Tyk2 | 5 | 353 | 1.42 |
| KEGG_LEISHMANIA_INFECTION | LEISHMANIA_INFECTION | 3/223 | 46/3594 | 0.5510334 | 0.99938151 | 0.97483529 | Elk1/Itgam/Nos2 | 3 | 70 | 4.29 |
| KEGG_LEUKOCYTE_TRANSENDOTHELIAL_MIGRATION | LEUKOCYTE_TRANSENDOTHELIAL_MIGRATION | 1/223 | 88/3594 | 0.99668276 | 0.99938151 | 0.97483529 | Itgam | 1 | 112 | 0.89 |
| KEGG_LINOLEIC_ACID_METABOLISM | LINOLEIC_ACID_METABOLISM | 1/223 | 19/3594 | 0.70483755 | 0.99938151 | 0.97483529 | Pla2g5 | 1 | 28 | 3.57 |
| KEGG_LONG_TERM_DEPRESSION | LONG_TERM_DEPRESSION | 2/223 | 48/3594 | 0.80905537 | 0.99938151 | 0.97483529 | Gna12/Pla2g5 | 2 | 67 | 2.99 |
| KEGG_LONG_TERM_POTENTIATION | LONG_TERM_POTENTIATION | 3/223 | 57/3594 | 0.69704201 | 0.99938151 | 0.97483529 | Atf4/Ppp1r1a/Prkaca | 3 | 68 | 4.41 |
| KEGG_LYSOSOME | LYSOSOME | 2/223 | 107/3594 | 0.99211947 | 0.99938151 | 0.97483529 | Ap3d1/Gga2 | 2 | 118 | 1.69 |
| KEGG_MAPK_SIGNALING_PATHWAY | MAPK_SIGNALING_PATHWAY | 15/223 | 207/3594 | 0.30139543 | 0.99938151 | 0.97483529 | Atf2/Atf4/Dusp14/Dusp6/Elk1/Fas/Gadd45b/Gadd45g/Gna12/Map3k3/Mapk8ip3/Mapk9/Pla2g5/Prkaca/Rapgef2 | 15 | 266 | 5.64 |
| KEGG_MELANOGENESIS | MELANOGENESIS | 5/223 | 74/3594 | 0.48979965 | 0.99938151 | 0.97483529 | Adcy3/Fzd2/Fzd3/Fzd7/Prkaca | 5 | 97 | 5.15 |
| KEGG_MELANOMA | MELANOMA | 3/223 | 50/3594 | 0.60862929 | 0.99938151 | 0.97483529 | Ccnd1/E2f1/Pdgfc | 3 | 69 | 4.35 |
| KEGG_METABOLISM_OF_XENOBIOTICS_BY_CYTOCHROME_P450 | METABOLISM_OF_XENOBIOTICS_BY_CYTOCHROME_P450 | 1/223 | 37/3594 | 0.90767406 | 0.99938151 | 0.97483529 | Gstm6 | 1 | 67 | 1.49 |
| KEGG_MISMATCH_REPAIR | MISMATCH_REPAIR | 2/223 | 21/3594 | 0.37789118 | 0.99938151 | 0.97483529 | Exo1/Rpa2 | 2 | 21 | 9.52 |
| KEGG_NATURAL_KILLER_CELL_MEDIATED_CYTOTOXICITY | NATURAL_KILLER_CELL_MEDIATED_CYTOTOXICITY | 2/223 | 82/3594 | 0.96774319 | 0.99938151 | 0.97483529 | Fas/Ifnar1 | 2 | 316 | 0.63 |
| KEGG_NEUROACTIVE_LIGAND_RECEPTOR_INTERACTION | NEUROACTIVE_LIGAND_RECEPTOR_INTERACTION | 3/223 | 84/3594 | 0.9022695 | 0.99938151 | 0.97483529 | F2r/Lpar3/P2ry4 | 3 | 313 | 0.96 |
| KEGG_NEUROTROPHIN_SIGNALING_PATHWAY | NEUROTROPHIN_SIGNALING_PATHWAY | 4/223 | 109/3594 | 0.91534621 | 0.99938151 | 0.97483529 | Atf4/Bex3/Map3k3/Mapk9 | 4 | 126 | 3.17 |
| KEGG_NITROGEN_METABOLISM | NITROGEN_METABOLISM | 1/223 | 19/3594 | 0.70483755 | 0.99938151 | 0.97483529 | Asns | 1 | 23 | 4.35 |
| KEGG_NOD_LIKE_RECEPTOR_SIGNALING_PATHWAY | NOD_LIKE_RECEPTOR_SIGNALING_PATHWAY | 1/223 | 42/3594 | 0.93321935 | 0.99938151 | 0.97483529 | Mapk9 | 1 | 52 | 1.92 |
| KEGG_NON_HOMOLOGOUS_END_JOINING | NON_HOMOLOGOUS_END_JOINING | 1/223 | Dec-94 | 0.53693772 | 0.99938151 | 0.97483529 | Fen1 | 1 | 13 | 7.69 |
| KEGG_NON_SMALL_CELL_LUNG_CANCER | NON_SMALL_CELL_LUNG_CANCER | 2/223 | 51/3594 | 0.83526191 | 0.99938151 | 0.97483529 | Ccnd1/E2f1 | 2 | 53 | 3.77 |
| KEGG_NOTCH_SIGNALING_PATHWAY | NOTCH_SIGNALING_PATHWAY | 1/223 | 42/3594 | 0.93321935 | 0.99938151 | 0.97483529 | Dtx4 | 1 | 46 | 2.17 |
| KEGG_NUCLEOTIDE_EXCISION_REPAIR | NUCLEOTIDE_EXCISION_REPAIR | 1/223 | 42/3594 | 0.93321935 | 0.99938151 | 0.97483529 | Rpa2 | 1 | 42 | 2.38 |
| KEGG_OLFACTORY_TRANSDUCTION | OLFACTORY_TRANSDUCTION | 3/223 | 23/3594 | 0.16791601 | 0.99938151 | 0.97483529 | Adcy3/Clca4b/Prkaca | 3 | 650 | 0.46 |
| KEGG_ONE_CARBON_POOL_BY_FOLATE | ONE_CARBON_POOL_BY_FOLATE | 1/223 | 17/3594 | 0.66427711 | 0.99938151 | 0.97483529 | Mthfd2 | 1 | 18 | 5.56 |
| KEGG_OTHER_GLYCAN_DEGRADATION | OTHER_GLYCAN_DEGRADATION | 1/223 | 15/3594 | 0.61817135 | 0.99938151 | 0.97483529 | Engase | 1 | 16 | 6.25 |
| KEGG_PANCREATIC_CANCER | PANCREATIC_CANCER | 4/223 | 64/3594 | 0.56865341 | 0.99938151 | 0.97483529 | Ccnd1/E2f1/Mapk9/Rad51 | 4 | 71 | 5.63 |
| KEGG_PATHOGENIC_ESCHERICHIA_COLI_INFECTION | PATHOGENIC_ESCHERICHIA_COLI_INFECTION | 2/223 | 46/3594 | 0.78958811 | 0.99938151 | 0.97483529 | Ncl/Tubb5 | 2 | 52 | 3.85 |
| KEGG_PATHWAYS_IN_CANCER | PATHWAYS_IN_CANCER | 16/223 | 262/3594 | 0.56701945 | 0.99938151 | 0.97483529 | Appl1/Arnt2/Birc5/Ccnd1/Cks1b/Dapk3/E2f1/Fas/Fzd2/Fzd3/Fzd7/Mapk9/Nos2/Rad51/Ret/Skp2 | 16 | 340 | 4.71 |
| KEGG_PENTOSE_AND_GLUCURONATE_INTERCONVERSIONS | PENTOSE_AND_GLUCURONATE_INTERCONVERSIONS | 2/223 | 16/3594 | 0.26130719 | 0.99938151 | 0.97483529 | Akr1b3/Ugp2 | 2 | 34 | 5.88 |
| KEGG_PORPHYRIN_AND_CHLOROPHYLL_METABOLISM | PORPHYRIN_AND_CHLOROPHYLL_METABOLISM | 2/223 | 28/3594 | 0.5263221 | 0.99938151 | 0.97483529 | Blvra/Ppox | 2 | 47 | 4.26 |
| KEGG_PPAR_SIGNALING_PATHWAY | PPAR_SIGNALING_PATHWAY | 3/223 | 50/3594 | 0.60862929 | 0.99938151 | 0.97483529 | Acaa1b/Acox2/Pltp | 3 | 78 | 3.85 |
| KEGG_PRION_DISEASES | PRION_DISEASES | 2/223 | 23/3594 | 0.42256553 | 0.99938151 | 0.97483529 | Elk1/Prkaca | 2 | 34 | 5.88 |
| KEGG_PROPANOATE_METABOLISM | PROPANOATE_METABOLISM | 1/223 | 29/3594 | 0.84512675 | 0.99938151 | 0.97483529 | Aldh6a1 | 1 | 32 | 3.12 |
| KEGG_PROSTATE_CANCER | PROSTATE_CANCER | 4/223 | 84/3594 | 0.77587296 | 0.99938151 | 0.97483529 | Atf4/Ccnd1/E2f1/Pdgfc | 4 | 103 | 3.88 |
| KEGG_PROTEIN_EXPORT | PROTEIN_EXPORT | 2/223 | 24/3594 | 0.44428279 | 0.99938151 | 0.97483529 | Arxes2/Sec63 | 2 | 26 | 7.69 |
| KEGG_PURINE_METABOLISM | PURINE_METABOLISM | 9/223 | 131/3594 | 0.42605087 | 0.99938151 | 0.97483529 | Adcy3/Ampd2/Nme2/Nudt2/Papss1/Pde7b/Polr2i/Polr3k/Xdh | 9 | 158 | 5.7 |
| KEGG_PYRIMIDINE_METABOLISM | PYRIMIDINE_METABOLISM | 6/223 | 87/3594 | 0.45646683 | 0.99938151 | 0.97483529 | Dtymk/Nme2/Nudt2/Polr2i/Polr3k/Tk1 | 6 | 96 | 6.25 |
| KEGG_PYRUVATE_METABOLISM | PYRUVATE_METABOLISM | 1/223 | 34/3594 | 0.88789381 | 0.99938151 | 0.97483529 | Akr1b3 | 1 | 39 | 2.56 |
| KEGG_REGULATION_OF_ACTIN_CYTOSKELETON | REGULATION_OF_ACTIN_CYTOSKELETON | 9/223 | 168/3594 | 0.72678873 | 0.99938151 | 0.97483529 | Abi2/Diaph3/F2r/Fgd1/Gna12/Iqgap3/Itgam/Pdgfc/Tiam2 | 9 | 211 | 4.27 |
| KEGG_RENAL_CELL_CARCINOMA | RENAL_CELL_CARCINOMA | 1/223 | 64/3594 | 0.98403205 | 0.99938151 | 0.97483529 | Arnt2 | 1 | 70 | 1.43 |
| KEGG_RETINOL_METABOLISM | RETINOL_METABOLISM | 1/223 | 32/3594 | 0.87241702 | 0.99938151 | 0.97483529 | Dgat2 | 1 | 81 | 1.23 |
| KEGG_RIBOSOME | RIBOSOME | 1/223 | 63/3594 | 0.98295561 | 0.99938151 | 0.97483529 | Uba52 | 1 | 74 | 1.35 |
| KEGG_RIG_I_LIKE_RECEPTOR_SIGNALING_PATHWAY | RIG_I_LIKE_RECEPTOR_SIGNALING_PATHWAY | 2/223 | 47/3594 | 0.79953013 | 0.99938151 | 0.97483529 | Cyld/Mapk9 | 2 | 238 | 0.84 |
| KEGG_RNA_DEGRADATION | RNA_DEGRADATION | 2/223 | 54/3594 | 0.85818774 | 0.99938151 | 0.97483529 | Cnot3/Lsm2 | 2 | 58 | 3.45 |
| KEGG_RNA_POLYMERASE | RNA_POLYMERASE | 2/223 | 25/3594 | 0.46554143 | 0.99938151 | 0.97483529 | Polr2i/Polr3k | 2 | 27 | 7.41 |
| KEGG_SELENOAMINO_ACID_METABOLISM | SELENOAMINO_ACID_METABOLISM | 1/223 | 24/3594 | 0.78614469 | 0.99938151 | 0.97483529 | Papss1 | 1 | 27 | 3.7 |
| KEGG_SMALL_CELL_LUNG_CANCER | SMALL_CELL_LUNG_CANCER | 5/223 | 75/3594 | 0.5018774 | 0.99938151 | 0.97483529 | Ccnd1/Cks1b/E2f1/Nos2/Skp2 | 5 | 82 | 6.1 |
| KEGG_SNARE_INTERACTIONS_IN_VESICULAR_TRANSPORT | SNARE_INTERACTIONS_IN_VESICULAR_TRANSPORT | 2/223 | 31/3594 | 0.5822861 | 0.99938151 | 0.97483529 | Snap23/Stx2 | 2 | 34 | 5.88 |
| KEGG_SPLICEOSOME | SPLICEOSOME | 3/223 | 120/3594 | 0.98289574 | 0.99938151 | 0.97483529 | Lsm2/Snrnp70/Snrpa | 3 | 154 | 1.95 |
| KEGG_STARCH_AND_SUCROSE_METABOLISM | STARCH_AND_SUCROSE_METABOLISM | 4/223 | 32/3594 | 0.13343923 | 0.99938151 | 0.97483529 | Gys1/Hk2/Pygm/Ugp2 | 4 | 52 | 7.69 |
| KEGG_STEROID_BIOSYNTHESIS | STEROID_BIOSYNTHESIS | 2/223 | 15/3594 | 0.2377076 | 0.99938151 | 0.97483529 | Fdft1/Soat2 | 2 | 17 | 11.76 |
| KEGG_T_CELL_RECEPTOR_SIGNALING_PATHWAY | T_CELL_RECEPTOR_SIGNALING_PATHWAY | 1/223 | 92/3594 | 0.99744987 | 0.99938151 | 0.97483529 | Mapk9 | 1 | 108 | 0.93 |
| KEGG_TASTE_TRANSDUCTION | TASTE_TRANSDUCTION | 1/223 | 20/3594 | 0.72324908 | 0.99938151 | 0.97483529 | Prkaca | 1 | 69 | 1.45 |
| KEGG_TERPENOID_BACKBONE_BIOSYNTHESIS | TERPENOID_BACKBONE_BIOSYNTHESIS | 1/223 | 14/3594 | 0.5928071 | 0.99938151 | 0.97483529 | Hmgcs1 | 1 | 17 | 5.88 |
| KEGG_TGF_BETA_SIGNALING_PATHWAY | TGF_BETA_SIGNALING_PATHWAY | 2/223 | 61/3594 | 0.90080081 | 0.99938151 | 0.97483529 | Id3/Id4 | 2 | 83 | 2.41 |
| KEGG_THYROID_CANCER | THYROID_CANCER | 2/223 | 28/3594 | 0.5263221 | 0.99938151 | 0.97483529 | Ccnd1/Ret | 2 | 31 | 6.45 |
| KEGG_TIGHT_JUNCTION | TIGHT_JUNCTION | 3/223 | 94/3594 | 0.93845963 | 0.99938151 | 0.97483529 | Amotl1/Prkcz/Rab3b | 3 | 125 | 2.4 |
| KEGG_TOLL_LIKE_RECEPTOR_SIGNALING_PATHWAY | TOLL_LIKE_RECEPTOR_SIGNALING_PATHWAY | 4/223 | 68/3594 | 0.6174607 | 0.99938151 | 0.97483529 | Ifnar1/Lbp/Mapk9/Ticam1 | 4 | 267 | 1.5 |
| KEGG_TRYPTOPHAN_METABOLISM | TRYPTOPHAN_METABOLISM | 2/223 | 25/3594 | 0.46554143 | 0.99938151 | 0.97483529 | Cat/Gcdh | 2 | 45 | 4.44 |
| KEGG_TYPE_I_DIABETES_MELLITUS | TYPE_I_DIABETES_MELLITUS | 2/223 | Nov-94 | 0.14581204 | 0.99938151 | 0.97483529 | Fas/Ica1 | 2 | 44 | 4.55 |
| KEGG_TYPE_II_DIABETES_MELLITUS | TYPE_II_DIABETES_MELLITUS | 3/223 | 40/3594 | 0.45612225 | 0.99938151 | 0.97483529 | Hk2/Mapk9/Prkcz | 3 | 48 | 6.25 |
| KEGG_TYROSINE_METABOLISM | TYROSINE_METABOLISM | 3/223 | 27/3594 | 0.23299003 | 0.99938151 | 0.97483529 | Got1/Got2/Tat | 3 | 44 | 6.82 |
| KEGG_UBIQUITIN_MEDIATED_PROTEOLYSIS | UBIQUITIN_MEDIATED_PROTEOLYSIS | 8/223 | 129/3594 | 0.55461427 | 0.99938151 | 0.97483529 | Brca1/Cdc20/Cul7/Skp2/Trim32/Trim37/Ube2c/Ube2s | 8 | 136 | 5.88 |
| KEGG_VALINE_LEUCINE_AND_ISOLEUCINE_DEGRADATION | VALINE_LEUCINE_AND_ISOLEUCINE_DEGRADATION | 3/223 | 41/3594 | 0.47255959 | 0.99938151 | 0.97483529 | Acaa1b/Aldh6a1/Hmgcs1 | 3 | 48 | 6.25 |
| KEGG_VASCULAR_SMOOTH_MUSCLE_CONTRACTION | VASCULAR_SMOOTH_MUSCLE_CONTRACTION | 4/223 | 77/3594 | 0.7139822 | 0.99938151 | 0.97483529 | Adcy3/Gna12/Pla2g5/Prkaca | 4 | 119 | 3.36 |
| KEGG_VASOPRESSIN_REGULATED_WATER_REABSORPTION | VASOPRESSIN_REGULATED_WATER_REABSORPTION | 3/223 | 38/3594 | 0.42264235 | 0.99938151 | 0.97483529 | Adcy3/Dync1i1/Prkaca | 3 | 44 | 6.82 |
| KEGG_VEGF_SIGNALING_PATHWAY | VEGF_SIGNALING_PATHWAY | 1/223 | 64/3594 | 0.98403205 | 0.99938151 | 0.97483529 | Pla2g5 | 1 | 74 | 1.35 |
| KEGG_VIBRIO_CHOLERAE_INFECTION | VIBRIO_CHOLERAE_INFECTION | 4/223 | 46/3594 | 0.31869106 | 0.99938151 | 0.97483529 | Adcy3/Kdelr2/Kdelr3/Prkaca | 4 | 52 | 7.69 |
| KEGG_VIRAL_MYOCARDITIS | VIRAL_MYOCARDITIS | 1/223 | 31/3594 | 0.86389876 | 0.99938151 | 0.97483529 | Ccnd1 | 1 | 70 | 1.43 |
| KEGG_WNT_SIGNALING_PATHWAY | WNT_SIGNALING_PATHWAY | 7/223 | 121/3594 | 0.63301772 | 0.99938151 | 0.97483529 | Ccnd1/Cxxc4/Fzd2/Fzd3/Fzd7/Mapk9/Prkaca | 7 | 151 | 4.64 |
| KEGG_ALDOSTERONE_REGULATED_SODIUM_REABSORPTION | ALDOSTERONE_REGULATED_SODIUM_REABSORPTION | NA | NA | NA | NA | NA | NA | 0 | 43 | 0 |
| KEGG_ALLOGRAFT_REJECTION | ALLOGRAFT_REJECTION | NA | NA | NA | NA | NA | NA | 0 | 37 | 0 |
| KEGG_ANTIGEN_PROCESSING_AND_PRESENTATION | ANTIGEN_PROCESSING_AND_PRESENTATION | NA | NA | NA | NA | NA | NA | 0 | 279 | 0 |
| KEGG_ARRHYTHMOGENIC_RIGHT_VENTRICULAR_CARDIOMYOPATHY_ARVC | ARRHYTHMOGENIC_RIGHT_VENTRICULAR_CARDIOMYOPATHY_ARVC | NA | NA | NA | NA | NA | NA | 0 | 74 | 0 |
| KEGG_ASCORBATE_AND_ALDARATE_METABOLISM | ASCORBATE_AND_ALDARATE_METABOLISM | NA | NA | NA | NA | NA | NA | 0 | 31 | 0 |
| KEGG_ASTHMA | ASTHMA | NA | NA | NA | NA | NA | NA | 0 | 32 | 0 |
| KEGG_AUTOIMMUNE_THYROID_DISEASE | AUTOIMMUNE_THYROID_DISEASE | NA | NA | NA | NA | NA | NA | 0 | 221 | 0 |
| KEGG_CARDIAC_MUSCLE_CONTRACTION | CARDIAC_MUSCLE_CONTRACTION | NA | NA | NA | NA | NA | NA | 0 | 79 | 0 |
| KEGG_CITRATE_CYCLE_TCA_CYCLE | CITRATE_CYCLE_TCA_CYCLE | NA | NA | NA | NA | NA | NA | 0 | 31 | 0 |
| KEGG_GLYCOSAMINOGLYCAN_DEGRADATION | GLYCOSAMINOGLYCAN_DEGRADATION | NA | NA | NA | NA | NA | NA | 0 | 22 | 0 |
| KEGG_GLYCOSYLPHOSPHATIDYLINOSITOL_GPI_ANCHOR_BIOSYNTHESIS | GLYCOSYLPHOSPHATIDYLINOSITOL_GPI_ANCHOR_BIOSYNTHESIS | NA | NA | NA | NA | NA | NA | 0 | 24 | 0 |
| KEGG_GRAFT_VERSUS_HOST_DISEASE | GRAFT_VERSUS_HOST_DISEASE | NA | NA | NA | NA | NA | NA | 0 | 50 | 0 |
| KEGG_HISTIDINE_METABOLISM | HISTIDINE_METABOLISM | NA | NA | NA | NA | NA | NA | 0 | 33 | 0 |
| KEGG_HYPERTROPHIC_CARDIOMYOPATHY_HCM | HYPERTROPHIC_CARDIOMYOPATHY_HCM | NA | NA | NA | NA | NA | NA | 0 | 84 | 0 |
| KEGG_INTESTINAL_IMMUNE_NETWORK_FOR_IGA_PRODUCTION | INTESTINAL_IMMUNE_NETWORK_FOR_IGA_PRODUCTION | NA | NA | NA | NA | NA | NA | 0 | 46 | 0 |
| KEGG_LIMONENE_AND_PINENE_DEGRADATION | LIMONENE_AND_PINENE_DEGRADATION | NA | NA | NA | NA | NA | NA | 0 | 10 | 0 |
| KEGG_MATURITY_ONSET_DIABETES_OF_THE_YOUNG | MATURITY_ONSET_DIABETES_OF_THE_YOUNG | NA | NA | NA | NA | NA | NA | 0 | 25 | 0 |
| KEGG_MTOR_SIGNALING_PATHWAY | MTOR_SIGNALING_PATHWAY | NA | NA | NA | NA | NA | NA | 0 | 52 | 0 |
| KEGG_NICOTINATE_AND_NICOTINAMIDE_METABOLISM | NICOTINATE_AND_NICOTINAMIDE_METABOLISM | NA | NA | NA | NA | NA | NA | 0 | 26 | 0 |
| KEGG_OXIDATIVE_PHOSPHORYLATION | OXIDATIVE_PHOSPHORYLATION | NA | NA | NA | NA | NA | NA | 0 | 132 | 0 |
| KEGG_PANTOTHENATE_AND_COA_BIOSYNTHESIS | PANTOTHENATE_AND_COA_BIOSYNTHESIS | NA | NA | NA | NA | NA | NA | 0 | 15 | 0 |
| KEGG_PARKINSONS_DISEASE | PARKINSONS_DISEASE | NA | NA | NA | NA | NA | NA | 0 | 126 | 0 |
| KEGG_PENTOSE_PHOSPHATE_PATHWAY | PENTOSE_PHOSPHATE_PATHWAY | NA | NA | NA | NA | NA | NA | 0 | 31 | 0 |
| KEGG_PHOSPHATIDYLINOSITOL_SIGNALING_SYSTEM | PHOSPHATIDYLINOSITOL_SIGNALING_SYSTEM | NA | NA | NA | NA | NA | NA | 0 | 75 | 0 |
| KEGG_PRIMARY_BILE_ACID_BIOSYNTHESIS | PRIMARY_BILE_ACID_BIOSYNTHESIS | NA | NA | NA | NA | NA | NA | 0 | 15 | 0 |
| KEGG_PRIMARY_IMMUNODEFICIENCY | PRIMARY_IMMUNODEFICIENCY | NA | NA | NA | NA | NA | NA | 0 | 38 | 0 |
| KEGG_PROTEASOME | PROTEASOME | NA | NA | NA | NA | NA | NA | 0 | 45 | 0 |
| KEGG_PROXIMAL_TUBULE_BICARBONATE_RECLAMATION | PROXIMAL_TUBULE_BICARBONATE_RECLAMATION | NA | NA | NA | NA | NA | NA | 0 | 23 | 0 |
| KEGG_REGULATION_OF_AUTOPHAGY | REGULATION_OF_AUTOPHAGY | NA | NA | NA | NA | NA | NA | 0 | 206 | 0 |
| KEGG_RENIN_ANGIOTENSIN_SYSTEM | RENIN_ANGIOTENSIN_SYSTEM | NA | NA | NA | NA | NA | NA | 0 | 19 | 0 |
| KEGG_RIBOFLAVIN_METABOLISM | RIBOFLAVIN_METABOLISM | NA | NA | NA | NA | NA | NA | 0 | 16 | 0 |
| KEGG_SPHINGOLIPID_METABOLISM | SPHINGOLIPID_METABOLISM | NA | NA | NA | NA | NA | NA | 0 | 40 | 0 |
| KEGG_STEROID_HORMONE_BIOSYNTHESIS | STEROID_HORMONE_BIOSYNTHESIS | NA | NA | NA | NA | NA | NA | 0 | 69 | 0 |
| KEGG_SULFUR_METABOLISM | SULFUR_METABOLISM | NA | NA | NA | NA | NA | NA | 0 | 14 | 0 |
| KEGG_TAURINE_AND_HYPOTAURINE_METABOLISM | TAURINE_AND_HYPOTAURINE_METABOLISM | NA | NA | NA | NA | NA | NA | 0 | 10 | 0 |
| KEGG_VALINE_LEUCINE_AND_ISOLEUCINE_BIOSYNTHESIS | VALINE_LEUCINE_AND_ISOLEUCINE_BIOSYNTHESIS | NA | NA | NA | NA | NA | NA | 0 | 11 | 0 |
