## Supplementary Table 5 for "ATF6 activation alters colonic lipid metabolism causing tumor-associated microbial adaptation"

| TF | gene | GeneRatio | BgRatio | pvalue | p.adjust | qvalue | DetectedTargets | Count | TF_targets | Percentage_of_TFtargets_detected |
|---|---|---|---|---|---|---|---|---|---|---|
| Atf6 | Atp2a2 | 8/200 | Oct-37 | 2.00E-11 | 3.42E-09 | 3.39E-09 | Atp2a2/Calr/Ddit3/Dnajc3/Hsp90b1/Hspa5/Sel1l/Xbp1 | 8 | 10 | 80 |
| Atf4 | Ccdc9 | 9/200 | 88/6837 | 0.00101415 | 0.08670941 | 0.08593546 | Abcg1/Chac1/Ddit3/Hdlbp/Herpud1/P4hb/Phgdh/Trib3/Wars1 | 9 | 110 | 8.18 |
| Bcl6 | Ldlr | 5/200 | 42/6837 | 0.00719231 | 0.30839756 | 0.30564485 | Dusp22/Gmds/Osbpl10/Prdm1/Ubald2 | 5 | 46 | 10.87 |
| Irf3 | Hgs | 6/200 | 59/6837 | 0.00721398 | 0.30839756 | 0.30564485 | Abcb9/Cep131/Cnpy3/Foxk2/Mvd/Wbp1 | 6 | 66 | 9.09 |
| Srebf2 | Pcyt2 | 7/200 | 82/6837 | 0.0098365 | 0.33640847 | 0.33340574 | Abca1/Ebp/Fdps/Foxk2/Gipc1/Mvd/Sardh | 7 | 90 | 7.78 |
| Nfyb | Txnip | 6/200 | 73/6837 | 0.01949566 | 0.55562627 | 0.55066685 | Epb41/Lipa/Mlec/Nucb2/Pdia4/Vcp | 6 | 76 | 7.89 |
| Cebpg | Edem1 | 4/200 | 41/6837 | 0.03094894 | 0.63554315 | 0.62987041 | Atp2a2/Edem1/Nfil3/Trib3 | 4 | 46 | 8.7 |
| Lef1 | Cdx1 | 4/200 | 42/6837 | 0.03344964 | 0.63554315 | 0.62987041 | Ctif/Mitf/Mmp7/Myo18b | 4 | 54 | 7.41 |
| Nr1h3 | Rnf145 | 2/200 | Oct-37 | 0.03282782 | 0.63554315 | 0.62987041 | Abca1/Abcg1 | 2 | 12 | 16.67 |
| Spib | Bank1 | 4/200 | 44/6837 | 0.03879544 | 0.66340203 | 0.65748062 | Gmds/Pik3ap1/Prdm1/Xbp1 | 4 | 58 | 6.9 |
| Nr1h2 | Fasn | 2/200 | 13/6837 | 0.05373649 | 0.83535814 | 0.82790188 | Abca1/Abcg1 | 2 | 13 | 15.38 |
| Ar | Klk1b11 | 8/200 | 187/6837 | 0.18137095 | 0.96740996 | 0.95877503 | Atf6/Cdc6/Fam174b/Grb10/Klk1/Lipa/Map2/Pin1 | 8 | 222 | 3.6 |
| Arnt | Eno1 | 1/200 | 54/6837 | 0.8000216 | 0.96740996 | 0.95877503 | Nfasc | 1 | 63 | 1.59 |
| Arntl | Cry2 | 1/200 | 17/6837 | 0.39668338 | 0.96740996 | 0.95877503 | Cry1 | 1 | 20 | 5 |
| Atf1 | Il10 | 2/200 | 80/6837 | 0.68454933 | 0.96740996 | 0.95877503 | Creb3l2/Pdxk | 2 | 91 | 2.2 |
| Atf2 | Ncoa7 | 2/200 | 52/6837 | 0.45242631 | 0.96740996 | 0.95877503 | Cdk14/Dusp10 | 2 | 62 | 3.23 |
| Atf3 | Vgll4 | 2/200 | 48/6837 | 0.41211622 | 0.96740996 | 0.95877503 | Ddit3/Gdf15 | 2 | 50 | 4 |
| Bach1 | Piezo1 | 2/200 | 79/6837 | 0.6778458 | 0.96740996 | 0.95877503 | Bcat2/Slc7a11 | 2 | 84 | 2.38 |
| Bach2 | Swap70 | 2/200 | 44/6837 | 0.3704092 | 0.96740996 | 0.95877503 | Osbpl10/Siah2 | 2 | 49 | 4.08 |
| Batf | Apbb1ip | 1/200 | 44/6837 | 0.73031559 | 0.96740996 | 0.95877503 | Pik3ap1 | 1 | 49 | 2.04 |
| Bcl11a | Cacna1e | 1/200 | 29/6837 | 0.57801059 | 0.96740996 | 0.95877503 | Pik3ap1 | 1 | 34 | 2.94 |
| Bhlhe40 | Creb5 | 1/200 | 53/6837 | 0.79394692 | 0.96740996 | 0.95877503 | Gdf15 | 1 | 57 | 1.75 |
| Cdx2 | Sipa1l1 | 1/200 | 49/6837 | 0.76775641 | 0.96740996 | 0.95877503 | Agr2 | 1 | 52 | 1.92 |
| Cebpa | Fstl5 | 21/200 | 788/6837 | 0.71044304 | 0.96740996 | 0.95877503 | Arntl/Atp2a2/Ccne1/Csrp1/Dusp10/Dusp22/E2f2/Enpp3/Gmds/Grb10/Ipo5/Myo18b/Nbeal2/Nfil3/Npas2/Nucb2/Pik3ap1/Ppib/Septin11/Sil1/Xbp1 | 21 | 959 | 2.19 |
| Cebpb | Smarca2 | 3/200 | 62/6837 | 0.27200131 | 0.96740996 | 0.95877503 | Atp2a2/Csrp1/Xbp1 | 3 | 85 | 3.53 |
| Cebpd | Lep | 3/200 | 61/6837 | 0.26402392 | 0.96740996 | 0.95877503 | Atp2a2/Nfil3/Ubald2 | 3 | 71 | 4.23 |
| Clock | Bhlhe40 | 1/200 | 41/6837 | 0.70502866 | 0.96740996 | 0.95877503 | Cry1 | 1 | 51 | 1.96 |
| Creb1 | Pold2 | 2/200 | 74/6837 | 0.64258931 | 0.96740996 | 0.95877503 | Creb3l2/Mitf | 2 | 117 | 1.71 |
| Creb3l1 | Nrsn2 | 1/200 | 17/6837 | 0.39668338 | 0.96740996 | 0.95877503 | Serpinh1 | 1 | 20 | 5 |
| Crem | Pde4b | 1/200 | 44/6837 | 0.73031559 | 0.96740996 | 0.95877503 | Creb3l2 | 1 | 50 | 2 |
| Ctcf | Snx18 | 18/200 | 729/6837 | 0.81164611 | 0.96740996 | 0.95877503 | Arhgef17/Arntl/Ccdc149/Cdk14/Ctif/Gmds/Grb10/Hspa5/Mroh1/Nfasc/Nrxn2/Parp1/Pdia6/Rpn1/Rps6ka2/Sardh/Ubfd1/Wfs1 | 18 | 936 | 1.92 |
| Ctcfl | Mast1 | 1/200 | 39/6837 | 0.68687377 | 0.96740996 | 0.95877503 | Nrxn2 | 1 | 53 | 1.89 |
| Cux1 | Fto | 1/200 | 37/6837 | 0.6676074 | 0.96740996 | 0.95877503 | Gmds | 1 | 40 | 2.5 |
| E2f1 | Rbpj | 18/200 | 576/6837 | 0.42041584 | 0.96740996 | 0.95877503 | Abca1/Arhgef17/Canx/Ccne1/Cdc6/Hdlbp/Herpud1/Hspa5/Il12rb1/Isyna1/N4bp2/Pdxk/Prdm1/Serp1/Sox12/Tspyl2/Ung/Vcp | 18 | 675 | 2.67 |
| E2f2 | Dock9 | 1/200 | 22/6837 | 0.48013076 | 0.96740996 | 0.95877503 | Cdc6 | 1 | 22 | 4.55 |
| E2f3 | E2f1 | 1/200 | Nov-37 | 0.27878753 | 0.96740996 | 0.95877503 | Cdc6 | 1 | 14 | 7.14 |
| E2f4 | Mfsd12 | 22/200 | 695/6837 | 0.38018341 | 0.96740996 | 0.95877503 | Abcg1/Atp8b2/Calr/Canx/Ccdc134/Cdc6/Derl2/E2f2/Hmbs/Mcfd2/Mfsd12/Mis12/Mlec/P4ha2/Pclaf/Pdia3/Pdxk/Pim2/Thop1/Trib3/Ung/Vcp | 22 | 743 | 2.96 |
| E2f6 | E2f1 | 2/200 | 59/6837 | 0.51901026 | 0.96740996 | 0.95877503 | Mroh1/Trib3 | 2 | 67 | 2.99 |
| Ebf1 | Il16 | 1/200 | 67/6837 | 0.86452546 | 0.96740996 | 0.95877503 | Arhgef17 | 1 | 74 | 1.35 |
| Egr1 | Map3k11 | 25/200 | 819/6837 | 0.44203066 | 0.96740996 | 0.95877503 | Arhgef17/Arid3a/Atp2a2/Cacna2d1/Creb3l2/Derl3/Dusp22/Dusp23/Fkbp2/Gdf15/Kcnc3/Lmf1/Mroh1/Olfm1/Pdxk/Ppa1/Ptges/Rpn1/Rps6ka2/Septin11/Slc30a3/Sox12/Tfpi2/Thop1/Trib3 | 25 | 976 | 2.56 |
| Elf3 | Grhl2 | 2/200 | 49/6837 | 0.42233416 | 0.96740996 | 0.95877503 | Atp2a2/Gmds | 2 | 52 | 3.85 |
| Elk1 | Ly96 | 2/200 | 106/6837 | 0.82199533 | 0.96740996 | 0.95877503 | Fkbp2/Tmem41b | 2 | 113 | 1.77 |
| Erg | Dab2ip | 4/200 | 78/6837 | 0.1938765 | 0.96740996 | 0.95877503 | Abcg1/Fam174b/Grb10/Pik3ap1 | 4 | 89 | 4.49 |
| Esr1 | Sipa1l2 | 14/200 | 579/6837 | 0.8104345 | 0.96740996 | 0.95877503 | Abcb9/Cacna2d1/Cad/Dhrs9/E2f2/Fam174b/Gmds/Isg20/Nfasc/P4ha2/Pkib/Scnn1b/Sil1/Syt8 | 14 | 722 | 1.94 |
| Esr2 | Dgke | 4/200 | 59/6837 | 0.0933376 | 0.96740996 | 0.95877503 | Anxa6/Isg20/Mtmr4/P4ha2 | 4 | 74 | 5.41 |
| Esrra | Slc22a23 | 2/200 | 64/6837 | 0.56317705 | 0.96740996 | 0.95877503 | Capn8/Trib3 | 2 | 69 | 2.9 |
| Ets1 | 2210016F16f | 21/200 | 837/6837 | 0.80690417 | 0.96740996 | 0.95877503 | Abcb9/Abcg1/Arhgef17/Arid3a/Canx/Creb3l2/Csrp1/Ctif/Dalrd3/Ddost/Dnajb9/Maged1/Mlec/Mtmr4/Nucb1/Parp1/Rps6ka2/Septin11/Tfg/Tmem41b/Wars1 | 21 | 934 | 2.25 |
| Etv4 | Sub1 | 3/200 | 55/6837 | 0.21695729 | 0.96740996 | 0.95877503 | Mmp7/Tns2/Trib3 | 3 | 59 | 5.08 |
| Fos | Clip4 | 17/200 | 686/6837 | 0.80073822 | 0.96740996 | 0.95877503 | Arhgef10/Ddit3/Dusp10/Fkbp2/Gmds/Ints3/Isg20/Klk1/Lipa/Map2/Mfsd12/Mmp7/Mtmr4/Mydgf/Nucb2/Osbpl10/Rpn1 | 17 | 847 | 2.01 |
| Foxa1 | Eya2 | 22/200 | 674/6837 | 0.32491884 | 0.96740996 | 0.95877503 | Abca1/Agr2/Arhgef10/Cacna2d1/Cd101/Cdk14/Dnpep/Epb41/Gdf15/Gmds/Grb10/Ibtk/Igsf5/Klk1/Map2/Mitf/Nucb2/Osbpl10/Paip2b/Pkib/Scly/Sil1 | 22 | 851 | 2.59 |
| Foxa2 | Adgrl2 | 2/200 | 42/6837 | 0.34911857 | 0.96740996 | 0.95877503 | Mitf/Plod2 | 2 | 49 | 4.08 |
| Foxm1 | Cenpa | 1/200 | 83/6837 | 0.91619881 | 0.96740996 | 0.95877503 | Cdc6 | 1 | 92 | 1.09 |
| Gata2 | Dock10 | 17/200 | 542/6837 | 0.41862854 | 0.96740996 | 0.95877503 | Atp8b2/Cdk14/Dusp10/Fdps/Gmds/Grb10/Map2/Mcfd2/Mitf/Mpdz/Nfasc/Nucb2/Osbpl10/Pkib/Rps6ka2/Serpine2/Sil1 | 17 | 676 | 2.51 |
| Gata3 | Csgalnact1 | 15/200 | 596/6837 | 0.76788612 | 0.96740996 | 0.95877503 | Arntl/Cacna2d1/Dusp10/Dusp22/Epb41/Fbxo16/Fmn2/Gmds/Lipa/Map2/Mitf/Mpdz/Nucb2/Osbpl10/Sil1 | 15 | 760 | 1.97 |
| Gata6 | Stard13 | 1/200 | 49/6837 | 0.76775641 | 0.96740996 | 0.95877503 | Pkib | 1 | 52 | 1.92 |
| Hbp1 | Klhdc2 | 1/200 | 38/6837 | 0.67738365 | 0.96740996 | 0.95877503 | Pja2 | 1 | 38 | 2.63 |
| Hif1a | Ldhb | 7/200 | 208/6837 | 0.40759316 | 0.96740996 | 0.95877503 | Abca1/Grb10/Hsp90b1/Klk1/P4ha2/Pdk3/Pgk1 | 7 | 269 | 2.6 |
| Hnf4g | Atp11a | 1/200 | 49/6837 | 0.76775641 | 0.96740996 | 0.95877503 | Creb3l3 | 1 | 58 | 1.72 |
| Hsf1 | Gna12 | 4/200 | 76/6837 | 0.18200501 | 0.96740996 | 0.95877503 | Bag3/Mroh1/Pdxk/Serpinh1 | 4 | 83 | 4.82 |
| Ikzf1 | Plb1 | 2/200 | 33/6837 | 0.25121438 | 0.96740996 | 0.95877503 | Pik3ap1/Trib3 | 2 | 40 | 5 |
| Irf4 | Fcer2 | 2/200 | 56/6837 | 0.49112871 | 0.96740996 | 0.95877503 | Pik3ap1/Txndc11 | 2 | 65 | 3.08 |
| Irf9 | Runx1 | 1/200 | 44/6837 | 0.73031559 | 0.96740996 | 0.95877503 | Wars1 | 1 | 46 | 2.17 |
| Jun | Plaur | 5/200 | 119/6837 | 0.26858102 | 0.96740996 | 0.95877503 | Ddit3/Dusp10/Mmp7/Sprr1a/Tfpi2 | 5 | 185 | 2.7 |
| Junb | Rtn4 | 1/200 | 48/6837 | 0.76070697 | 0.96740996 | 0.95877503 | Gmds | 1 | 50 | 2 |
| Jund | Itgav | 1/200 | 66/6837 | 0.86040206 | 0.96740996 | 0.95877503 | Dusp10 | 1 | 75 | 1.33 |
| Klf6 | Ehd1 | 2/200 | 51/6837 | 0.44249247 | 0.96740996 | 0.95877503 | Dusp10/Wfs1 | 2 | 52 | 3.85 |
| Klf9 | Tns1 | 2/200 | 63/6837 | 0.55457872 | 0.96740996 | 0.95877503 | Pcbp3/Tns2 | 2 | 68 | 2.94 |
| Lyl1 | Inpp5d | 1/200 | 72/6837 | 0.8833938 | 0.96740996 | 0.95877503 | Chst12 | 1 | 84 | 1.19 |
| Mafb | Snx29 | 1/200 | 44/6837 | 0.73031559 | 0.96740996 | 0.95877503 | Prdm1 | 1 | 53 | 1.89 |
| Maff | Jmjd1c | 1/200 | 39/6837 | 0.68687377 | 0.96740996 | 0.95877503 | Cdk14 | 1 | 43 | 2.33 |
| Mafg | Stau2 | 1/200 | 32/6837 | 0.61412449 | 0.96740996 | 0.95877503 | Ctif | 1 | 37 | 2.7 |
| Mafk | Foxk1 | 2/200 | 40/6837 | 0.32759388 | 0.96740996 | 0.95877503 | Cdk14/Ctif | 2 | 50 | 4 |
| Max | Serf2 | 2/200 | 50/6837 | 0.43246071 | 0.96740996 | 0.95877503 | Nrxn2/Rps6ka2 | 2 | 53 | 3.77 |
| Maz | Kdm2b | 4/200 | 67/6837 | 0.13202429 | 0.96740996 | 0.95877503 | Mfsd12/Pdxk/Serpinh1/Sox12 | 4 | 70 | 5.71 |
| Mbd1 | Exoc7 | 2/200 | 40/6837 | 0.32759388 | 0.96740996 | 0.95877503 | Foxk2/Thop1 | 2 | 43 | 4.65 |
| Mef2b | Gnas | 1/200 | 27/6837 | 0.55208585 | 0.96740996 | 0.95877503 | Pik3ap1 | 1 | 36 | 2.78 |
| Mef2c | Ank3 | 2/200 | 62/6837 | 0.54586209 | 0.96740996 | 0.95877503 | Cdk14/Nucb2 | 2 | 70 | 2.86 |
| Meis2 | Fam184a | 1/200 | 51/6837 | 0.78124131 | 0.96740996 | 0.95877503 | Mitf | 1 | 60 | 1.67 |
| Mnt | Rara | 2/200 | 54/6837 | 0.47198898 | 0.96740996 | 0.95877503 | Foxk2/Pdxk | 2 | 56 | 3.57 |
| Mxi1 | Rusc1 | 4/200 | 74/6837 | 0.17038853 | 0.96740996 | 0.95877503 | Arid3a/Ctif/Epb41/Pdxk | 4 | 77 | 5.19 |
| Myc | Gnl3 | 10/200 | 403/6837 | 0.75026255 | 0.96740996 | 0.95877503 | Cad/Ccne1/Gmds/Herpud1/Hmbs/Isg20/Nfil3/Parp1/Pdxk/Rps6ka2 | 10 | 447 | 2.24 |
| Nfe2 | Sema4d | 1/200 | 44/6837 | 0.73031559 | 0.96740996 | 0.95877503 | Pik3ap1 | 1 | 47 | 2.13 |
| Nfe2l1 | Parn | 1/200 | 29/6837 | 0.57801059 | 0.96740996 | 0.95877503 | Arntl | 1 | 30 | 3.33 |
| Nfe2l2 | Abcb9 | 1/200 | 70/6837 | 0.87618253 | 0.96740996 | 0.95877503 | Abcb9 | 1 | 72 | 1.39 |
| Nfic | Svil | 2/200 | 94/6837 | 0.7668834 | 0.96740996 | 0.95877503 | Ctif/Tfpi2 | 2 | 108 | 1.85 |
| Nfkb1 | Icam1 | 3/200 | 175/6837 | 0.89173273 | 0.96740996 | 0.95877503 | Klk1/Ralb/Tfpi2 | 3 | 251 | 1.2 |
| Nfya | Dnajb1 | 4/200 | 61/6837 | 0.10243745 | 0.96740996 | 0.95877503 | Ap3m2/Copz1/Ddit3/Fdps | 4 | 70 | 5.71 |
| Nr2f1 | Abca2 | 3/200 | 51/6837 | 0.18668259 | 0.96740996 | 0.95877503 | Ctif/Nrxn2/Pdxk | 3 | 66 | 4.55 |
| Nr2f2 | Fbln1 | 2/200 | 68/6837 | 0.59637794 | 0.96740996 | 0.95877503 | Mitf/Osbpl10 | 2 | 84 | 2.38 |
| Nr2f6 | Ptprf | 1/200 | 27/6837 | 0.55208585 | 0.96740996 | 0.95877503 | Osbpl10 | 1 | 32 | 3.12 |
| Nr3c1 | Mbnl2 | 1/200 | 70/6837 | 0.87618253 | 0.96740996 | 0.95877503 | Mitf | 1 | 76 | 1.32 |
| Nr4a1 | Hsd3b1 | 2/200 | 36/6837 | 0.28405297 | 0.96740996 | 0.95877503 | Creb3l2/Ctif | 2 | 41 | 4.88 |
| Nr5a2 | Txnrd1 | 1/200 | 28/6837 | 0.56524045 | 0.96740996 | 0.95877503 | Scnn1b | 1 | 31 | 3.23 |
| Nrf1 | Cic | 3/200 | 73/6837 | 0.36061023 | 0.96740996 | 0.95877503 | Pkib/Serp1/Tbcb | 3 | 77 | 3.9 |
| Onecut1 | Dyrk1a | 2/200 | 40/6837 | 0.32759388 | 0.96740996 | 0.95877503 | Cacna2d1/Mitf | 2 | 52 | 3.85 |
| Pax5 | Dusp10 | 3/200 | 42/6837 | 0.12366029 | 0.96740996 | 0.95877503 | Dusp10/Prdm1/Xbp1 | 3 | 53 | 5.66 |
| Pax6 | Pdlim5 | 1/200 | 34/6837 | 0.63647468 | 0.96740996 | 0.95877503 | Mitf | 1 | 43 | 2.33 |
| Pbx2 | Bcor | 2/200 | 38/6837 | 0.30588604 | 0.96740996 | 0.95877503 | Mitf/Nfasc | 2 | 44 | 4.55 |
| Pbx3 | Pde4d | 2/200 | 46/6837 | 0.39142129 | 0.96740996 | 0.95877503 | Cacna2d1/Rps6ka2 | 2 | 51 | 3.92 |
| Pgr | Sipa1l1 | 3/200 | 67/6837 | 0.31219912 | 0.96740996 | 0.95877503 | Creb3l2/Klk1/Nfasc | 3 | 89 | 3.37 |
| Pknox1 | Lims1 | 1/200 | 28/6837 | 0.56524045 | 0.96740996 | 0.95877503 | Sil1 | 1 | 35 | 2.86 |
| Pou2f2 | Kdm4b | 1/200 | 69/6837 | 0.87241221 | 0.96740996 | 0.95877503 | Cdk14 | 1 | 86 | 1.16 |
| Pou5f1 | Cobl | 1/200 | 59/6837 | 0.82782352 | 0.96740996 | 0.95877503 | Osbpl10 | 1 | 72 | 1.39 |
| Ppara | Pdha1 | 1/200 | 26/6837 | 0.53853528 | 0.96740996 | 0.95877503 | Lipa | 1 | 37 | 2.7 |
| Pparg | Palmd | 2/200 | 84/6837 | 0.71023201 | 0.96740996 | 0.95877503 | Atp2a2/Lipa | 2 | 104 | 1.92 |
| Prdm14 | Rtl4 | 9/200 | 227/6837 | 0.21993993 | 0.96740996 | 0.95877503 | Arhgef17/Arntl/Bcat2/Fam174b/Gmds/Grb10/Npr1/Rps6ka2/Ttpa | 9 | 306 | 2.94 |
| Prox1 | Add3 | 1/200 | 23/6837 | 0.49538738 | 0.96740996 | 0.95877503 | Osbpl10 | 1 | 26 | 3.85 |
| Rbpj | Snx25 | 2/200 | 52/6837 | 0.45242631 | 0.96740996 | 0.95877503 | Hey2/Pik3ap1 | 2 | 59 | 3.39 |
| Rel | Cd40lg | 1/200 | Oct-37 | 0.25702165 | 0.96740996 | 0.95877503 | Csrp1 | 1 | 23 | 4.35 |
| Rela | Cdk6 | 4/200 | 180/6837 | 0.77826802 | 0.96740996 | 0.95877503 | Csrp1/Dusp22/Klk1/Trib3 | 4 | 245 | 1.63 |
| Relb | Ccl12 | 1/200 | 58/6837 | 0.8225894 | 0.96740996 | 0.95877503 | Dusp22 | 1 | 74 | 1.35 |
| Rest | Acap3 | 1/200 | 40/6837 | 0.69608607 | 0.96740996 | 0.95877503 | Nfasc | 1 | 67 | 1.49 |
| Rfx5 | Arvcf | 4/200 | 64/6837 | 0.11681712 | 0.96740996 | 0.95877503 | Bcat2/Cdk2ap2/Sox12/Tmem231 | 4 | 80 | 5 |
| Runx3 | Clnk | 1/200 | 80/6837 | 0.90829305 | 0.96740996 | 0.95877503 | Arntl | 1 | 97 | 1.03 |
| Rxra | Sult2b1 | 1/200 | 67/6837 | 0.86452546 | 0.96740996 | 0.95877503 | Cpt1c | 1 | 78 | 1.28 |
| Six5 | Rfx1 | 1/200 | 21/6837 | 0.46441525 | 0.96740996 | 0.95877503 | Bcat2 | 1 | 23 | 4.35 |
| Smad1 | Spp1 | 1/200 | 13/6837 | 0.32043406 | 0.96740996 | 0.95877503 | Gdf15 | 1 | 17 | 5.88 |
| Smad3 | Afap1 | 1/200 | 78/6837 | 0.9026146 | 0.96740996 | 0.95877503 | Il12rb1 | 1 | 96 | 1.04 |
| Smad5 | Nr3c1 | 1/200 | Oct-37 | 0.25702165 | 0.96740996 | 0.95877503 | Hnrnpa3 | 1 | 10 | 10 |
| Snai2 | Agpat3 | 1/200 | 62/6837 | 0.84262169 | 0.96740996 | 0.95877503 | C1qtnf1 | 1 | 66 | 1.52 |
| Sox10 | Sh3kbp1 | 2/200 | 35/6837 | 0.27310966 | 0.96740996 | 0.95877503 | Col11a2/Mitf | 2 | 45 | 4.44 |
| Sox2 | Ccdc91 | 7/200 | 291/6837 | 0.75489158 | 0.96740996 | 0.95877503 | Cacna2d1/Ccdc88a/Map2/Mpdz/Pcbp3/Pdia4/Prdm5 | 7 | 398 | 1.76 |
| Sox6 | Sox5 | 2/200 | 29/6837 | 0.20767642 | 0.96740996 | 0.95877503 | Dusp10/Nfasc | 2 | 32 | 6.25 |
| Sox9 | Chd2 | 3/200 | 67/6837 | 0.31219912 | 0.96740996 | 0.95877503 | Dcun1d4/Epb41/Serp1 | 3 | 74 | 4.05 |
| Sp1 | Pnoc | 6/200 | 296/6837 | 0.87135632 | 0.96740996 | 0.95877503 | Arid3a/Atp2a2/Cenpw/Klk1/Tfpi2/Ung | 6 | 394 | 1.52 |
| Sp3 | Igfbp3 | 2/200 | 54/6837 | 0.47198898 | 0.96740996 | 0.95877503 | Abca1/Klk1 | 2 | 91 | 2.2 |
| Sp4 | Nfyc | 3/200 | 52/6837 | 0.19414654 | 0.96740996 | 0.95877503 | Hnrnpa3/Hyou1/Kcnc3 | 3 | 59 | 5.08 |
| Spi1 | Vcl | 12/200 | 506/6837 | 0.81582057 | 0.96740996 | 0.95877503 | Abcg1/Arntl/Ctif/Dhrs9/Dusp10/Dusp22/Herpud1/Nrxn2/Pik3ap1/Ppib/Rps6ka2/Sil1 | 12 | 585 | 2.05 |
| Srebf1 | Rab26 | 2/200 | 67/6837 | 0.5882572 | 0.96740996 | 0.95877503 | Fdps/Mvd | 2 | 71 | 2.82 |
| Srf | Mthfr | 3/200 | 100/6837 | 0.56455278 | 0.96740996 | 0.95877503 | Arhgef17/Csrp1/Ctif | 3 | 109 | 2.75 |
| Stat1 | Pitpnb | 19/200 | 845/6837 | 0.91649365 | 0.96740996 | 0.95877503 | Agtrap/C1qtnf1/Cacna2d1/Ddx50/Dusp10/Dusp22/Gdf15/Gmppb/Hoxa1/Il12rb1/Isg20/Lipa/Mitf/Pgk1/Phgdh/Pik3ap1/Rps6ka2/Tns2/Wars1 | 19 | 1014 | 1.87 |
| Stat2 | Atp13a1 | 1/200 | 74/6837 | 0.89018705 | 0.96740996 | 0.95877503 | Lipa | 1 | 80 | 1.25 |
| Stat3 | Cdc42ep4 | 3/200 | 140/6837 | 0.78286658 | 0.96740996 | 0.95877503 | Csrp1/Dusp22/Foxk2 | 3 | 171 | 1.75 |
| Stat5a | Bcl2l1 | 2/200 | 56/6837 | 0.49112871 | 0.96740996 | 0.95877503 | Ctif/Dusp22 | 2 | 66 | 3.03 |
| Stat5b | Crtc3 | 1/200 | 66/6837 | 0.86040206 | 0.96740996 | 0.95877503 | Dusp22 | 1 | 78 | 1.28 |
| Tal1 | Plcb1 | 8/200 | 364/6837 | 0.84372738 | 0.96740996 | 0.95877503 | Creb3l2/Ctif/Dusp10/Gmds/Grb10/Pik3ap1/Rps6ka2/Sil1 | 8 | 416 | 1.92 |
| Tbp | Serpine1 | 1/200 | 37/6837 | 0.6676074 | 0.96740996 | 0.95877503 | Smtn | 1 | 46 | 2.17 |
| Tcf7l2 | Stxbp5 | 3/200 | 66/6837 | 0.30413159 | 0.96740996 | 0.95877503 | Mmp7/Sil1/Sp5 | 3 | 74 | 4.05 |
| Tead1 | Osbpl9 | 3/200 | 116/6837 | 0.66525287 | 0.96740996 | 0.95877503 | Cacna2d1/Fmn2/Gmds | 3 | 129 | 2.33 |
| Tead2 | Fam89a | 2/200 | 26/6837 | 0.17561686 | 0.96740996 | 0.95877503 | Atp8b2/Dnajb5 | 2 | 29 | 6.9 |
| Tead4 | Tgm2 | 1/200 | 67/6837 | 0.86452546 | 0.96740996 | 0.95877503 | Gmds | 1 | 73 | 1.37 |
| Tfap2a | Pold2 | 2/200 | 97/6837 | 0.78190321 | 0.96740996 | 0.95877503 | Nfasc/Scnn1b | 2 | 125 | 1.6 |
| Tfap2c | Dpf3 | 18/200 | 737/6837 | 0.82556799 | 0.96740996 | 0.95877503 | Abcg1/Capn8/Ctif/E2f2/Fam174b/Foxk2/Gmds/Grb10/Isg20/Nbeal2/Nfasc/Npas2/Osbpl10/Pcbp3/Pdia6/Scnn1b/Sil1/Tns2 | 18 | 890 | 2.02 |
| Tfdp1 | Gapdh | 4/200 | 66/6837 | 0.12686683 | 0.96740996 | 0.95877503 | Ccne1/Pclaf/Pdia4/Trib3 | 4 | 69 | 5.8 |
| Thap1 | Gak | 1/200 | 51/6837 | 0.78124131 | 0.96740996 | 0.95877503 | Trabd | 1 | 55 | 1.82 |
| Thap11 | Atl3 | 4/200 | 112/6837 | 0.41577431 | 0.96740996 | 0.95877503 | Erp44/Hsp90b1/Mitf/Sel1l | 4 | 114 | 3.51 |
| Tp53 | Tent4a | 10/200 | 483/6837 | 0.9081901 | 0.96740996 | 0.95877503 | Abcb9/Arid3a/Atp2a2/E2f7/Gdf15/Klk1/Pcbp3/Phlda3/Prdm1/Syt8 | 10 | 582 | 1.72 |
| Tp63 | Rin3 | 3/200 | 76/6837 | 0.38466779 | 0.96740996 | 0.95877503 | Arntl/Grb10/Syt8 | 3 | 90 | 3.33 |
| Tp73 | Megf6 | 1/200 | 36/6837 | 0.65753642 | 0.96740996 | 0.95877503 | Foxk2 | 1 | 41 | 2.44 |
| Twist1 | Map4k4 | 1/200 | 55/6837 | 0.80591806 | 0.96740996 | 0.95877503 | Creb3l2 | 1 | 67 | 1.49 |
| Usf1 | Stard13 | 3/200 | 89/6837 | 0.48564901 | 0.96740996 | 0.95877503 | Abca1/Isg20/Serpine2 | 3 | 103 | 2.91 |
| Usf2 | Serpine2 | 3/200 | 103/6837 | 0.58476837 | 0.96740996 | 0.95877503 | Abca1/Agtrap/Serpine2 | 3 | 113 | 2.65 |
| Vdr | Stap2 | 5/200 | 210/6837 | 0.74192578 | 0.96740996 | 0.95877503 | Agtrap/Arntl/Bag3/Calr/Ubald2 | 5 | 244 | 2.05 |
| Yy1 | Mrps10 | 5/200 | 91/6837 | 0.12722073 | 0.96740996 | 0.95877503 | Fdps/Hdlbp/Hspa5/Manf/Rnf26 | 5 | 99 | 5.05 |
| Zbtb33 | Ehd1 | 1/200 | 83/6837 | 0.91619881 | 0.96740996 | 0.95877503 | Chst12 | 1 | 88 | 1.14 |
| Zbtb7a | Cdc42ep4 | 1/200 | 82/6837 | 0.91364195 | 0.96740996 | 0.95877503 | Obsl1 | 1 | 90 | 1.11 |
| Zeb2 | Pip4k2a | 1/200 | 46/6837 | 0.74596309 | 0.96740996 | 0.95877503 | Creb3l2 | 1 | 53 | 1.89 |
| Zfp384 | Kansl1 | 2/200 | 61/6837 | 0.53702787 | 0.96740996 | 0.95877503 | Herpud1/Nde1 | 2 | 65 | 3.08 |
| Zfx | Zbtb7a | 1/200 | 53/6837 | 0.79394692 | 0.96740996 | 0.95877503 | Derl2 | 1 | 54 | 1.85 |
| Zkscan1 | Ripor1 | 1/200 | 40/6837 | 0.69608607 | 0.96740996 | 0.95877503 | Nfasc | 1 | 46 | 2.17 |
| Znf143 | Dnajb6 | 2/200 | 47/6837 | 0.4018106 | 0.96740996 | 0.95877503 | Ctif/Nucb1 | 2 | 51 | 3.92 |
| Znf263 | Tcf3 | 12/200 | 351/6837 | 0.33033045 | 0.96740996 | 0.95877503 | Arhgef17/Bag3/Nfasc/Npas2/Nrxn2/Pcbp3/Pdxk/Rps6ka2/Sardh/Shisa7/Stt3b/Susd4 | 12 | 432 | 2.78 |
| Hnf4a | Kcnab2 | 17/200 | 788/6837 | 0.93454569 | 0.97011147 | 0.96145243 | Bcat2/Dnah10/Ebp/Gdf15/Gmds/Grb10/Lrp8/Mcfd2/N4bp2/Nfasc/Npas2/Osbpl10/Pik3ap1/Sardh/Sil1/Surf4/Unc5b | 17 | 907 | 1.87 |
| Mef2a | Pam | 1/200 | 92/6837 | 0.93607247 | 0.97011147 | 0.96145243 | Pik3ap1 | 1 | 102 | 0.98 |
| Zeb1 | Pip4k2a | 1/200 | 92/6837 | 0.93607247 | 0.97011147 | 0.96145243 | Csrp1 | 1 | 96 | 1.04 |
| Fli1 | Sh2b3 | 1/200 | 97/6837 | 0.94500709 | 0.97189469 | 0.96321973 | Foxk2 | 1 | 107 | 0.93 |
| Foxo1 | Aff1 | 1/200 | 105/6837 | 0.956789 | 0.97189469 | 0.96321973 | Ctif | 1 | 122 | 0.82 |
| Rara | Rarb | 1/200 | 108/6837 | 0.9605275 | 0.97189469 | 0.96321973 | Hoxa1 | 1 | 122 | 0.82 |
| Runx1 | Efna5 | 5/200 | 306/6837 | 0.94962833 | 0.97189469 | 0.96321973 | Dusp10/Gmds/Klk1/Pik3ap1/Sil1 | 5 | 380 | 1.32 |
| Tcf3 | Mad1l1 | 1/200 | 114/6837 | 0.96706621 | 0.97275483 | 0.96407219 | Trabd | 1 | 125 | 0.8 |
| Mitf | Csmd1 | 10/200 | 574/6837 | 0.97720202 | 0.97720202 | 0.96847969 | Agtrap/Calr/Ccdc149/Gmds/Grb10/Phgdh/Rps6ka2/Sil1/Slc33a1/Tmem116 | 10 | 660 | 1.52 |
| Adnp | Atf7ip | NA | NA | NA | NA | NA | NA | 0 | 4 | 0 |
| Ahr | Arhgap15 | NA | NA | NA | NA | NA | NA | 0 | 43 | 0 |
| Arid2 | Cdk17 | NA | NA | NA | NA | NA | NA | 0 | 30 | 0 |
| Arid3a | Ptk2b | NA | NA | NA | NA | NA | NA | 0 | 23 | 0 |
| Arntl2 | Serpine1 | NA | NA | NA | NA | NA | NA | 0 | 1 | 0 |
| Ascl1 | Cplx2 | NA | NA | NA | NA | NA | NA | 0 | 54 | 0 |
| Atf5 | Disc1 | NA | NA | NA | NA | NA | NA | 0 | 1 | 0 |
| Atoh1 | Hes6 | NA | NA | NA | NA | NA | NA | 0 | 1 | 0 |
| Bhlhe22 | Elf4 | NA | NA | NA | NA | NA | NA | 0 | 21 | 0 |
| Cdx1 | Ada | NA | NA | NA | NA | NA | NA | 0 | 3 | 0 |
| Cebpe | Myc | NA | NA | NA | NA | NA | NA | 0 | 6 | 0 |
| Cebpz | Col11a1 | NA | NA | NA | NA | NA | NA | 0 | 3 | 0 |
| Creb3 | Utp11 | NA | NA | NA | NA | NA | NA | 0 | 25 | 0 |
| Creb3l2 | Cpt1c | NA | NA | NA | NA | NA | NA | 0 | 1 | 0 |
| Creb5 | Pou1f1 | NA | NA | NA | NA | NA | NA | 0 | 1 | 0 |
| Ddit3 | Trib3 | NA | NA | NA | NA | NA | NA | 0 | 2 | 0 |
| Deaf1 | Rac3 | NA | NA | NA | NA | NA | NA | 0 | 8 | 0 |
| Dnmt1 | Sfrp5 | NA | NA | NA | NA | NA | NA | 0 | 4 | 0 |
| E2f5 | Jade1 | NA | NA | NA | NA | NA | NA | 0 | 12 | 0 |
| E2f7 | Hnrnpd | NA | NA | NA | NA | NA | NA | 0 | 29 | 0 |
| E2f8 | E2f1 | NA | NA | NA | NA | NA | NA | 0 | 1 | 0 |
| E4f1 | Ccna2 | NA | NA | NA | NA | NA | NA | 0 | 1 | 0 |
| Egr2 | Spint1 | NA | NA | NA | NA | NA | NA | 0 | 4 | 0 |
| Egr3 | Nab2 | NA | NA | NA | NA | NA | NA | 0 | 1 | 0 |
| Ehf | Myh14 | NA | NA | NA | NA | NA | NA | 0 | 19 | 0 |
| Elf1 | Nedd4l | NA | NA | NA | NA | NA | NA | 0 | 50 | 0 |
| Elf2 | Vcp | NA | NA | NA | NA | NA | NA | 0 | 1 | 0 |
| Elf5 | Rbm34 | NA | NA | NA | NA | NA | NA | 0 | 19 | 0 |
| Elk3 | Fos | NA | NA | NA | NA | NA | NA | 0 | 1 | 0 |
| Elk4 | Scaf11 | NA | NA | NA | NA | NA | NA | 0 | 14 | 0 |
| En1 | Utrn | NA | NA | NA | NA | NA | NA | 0 | 1 | 0 |
| Eomes | Astn2 | NA | NA | NA | NA | NA | NA | 0 | 21 | 0 |
| Epas1 | Asap1 | NA | NA | NA | NA | NA | NA | 0 | 80 | 0 |
| Erf | Ets2 | NA | NA | NA | NA | NA | NA | 0 | 1 | 0 |
| Ets2 | Msr1 | NA | NA | NA | NA | NA | NA | 0 | 31 | 0 |
| Etv1 | Mzf1 | NA | NA | NA | NA | NA | NA | 0 | 29 | 0 |
| Etv3 | Nfkb1 | NA | NA | NA | NA | NA | NA | 0 | 1 | 0 |

| | | | | | | | | | | |
|---|---|---|---|---|---|---|---|---|---|---|
| Etv6 | Cap1 | NA | NA | NA | NA | NA | NA | 0 | 6 | 0 |
| Fosb | Runx2 | NA | NA | NA | NA | NA | NA | 0 | 1 | 0 |
| Fosl1 | Skt | NA | NA | NA | NA | NA | NA | 0 | 53 | 0 |
| Fosl2 | Zbtb20 | NA | NA | NA | NA | NA | NA | 0 | 79 | 0 |
| Foxa3 | Cyp2c38 | NA | NA | NA | NA | NA | NA | 0 | 3 | 0 |
| Foxj2 | Crb1 | NA | NA | NA | NA | NA | NA | 0 | 36 | 0 |
| Foxk1 | Cald1 | NA | NA | NA | NA | NA | NA | 0 | 8 | 0 |
| Foxk2 | Foxk1 | NA | NA | NA | NA | NA | NA | 0 | 50 | 0 |
| Foxl2 | Cxcl2 | NA | NA | NA | NA | NA | NA | 0 | 20 | 0 |
| Foxo3 | Pink1 | NA | NA | NA | NA | NA | NA | 0 | 40 | 0 |
| Foxo4 | Cdkn1a | NA | NA | NA | NA | NA | NA | 0 | 15 | 0 |
| Foxo6 | Fbxo25 | NA | NA | NA | NA | NA | NA | 0 | 1 | 0 |
| Foxp2 | Gnas | NA | NA | NA | NA | NA | NA | 0 | 35 | 0 |
| Foxp3 | Il2ra | NA | NA | NA | NA | NA | NA | 0 | 3 | 0 |
| Gabpa | Efr3a | NA | NA | NA | NA | NA | NA | 0 | 55 | 0 |
| Gata1 | Xpo7 | NA | NA | NA | NA | NA | NA | 0 | 64 | 0 |
| Gata4 | Cyp17a1 | NA | NA | NA | NA | NA | NA | 0 | 10 | 0 |
| Gcm2 | Pth | NA | NA | NA | NA | NA | NA | 0 | 1 | 0 |
| Gfi1 | Bax | NA | NA | NA | NA | NA | NA | 0 | 2 | 0 |
| Gfi1b | Ahcyl2 | NA | NA | NA | NA | NA | NA | 0 | 28 | 0 |
| Gli1 | Egr2 | NA | NA | NA | NA | NA | NA | 0 | 7 | 0 |
| Gli2 | Efemp1 | NA | NA | NA | NA | NA | NA | 0 | 16 | 0 |
| Gli3 | Wnt7b | NA | NA | NA | NA | NA | NA | 0 | 3 | 0 |
| Grhl2 | Tiam2 | NA | NA | NA | NA | NA | NA | 0 | 79 | 0 |
| Gtf2i | Sirt1 | NA | NA | NA | NA | NA | NA | 0 | 1 | 0 |
| Hes1 | Snx6 | NA | NA | NA | NA | NA | NA | 0 | 6 | 0 |
| Hey1 | Kdr | NA | NA | NA | NA | NA | NA | 0 | 1 | 0 |
| Hey2 | Gata6 | NA | NA | NA | NA | NA | NA | 0 | 2 | 0 |
| Hhex | Wscd1 | NA | NA | NA | NA | NA | NA | 0 | 11 | 0 |
| Hic1 | Fgfbp1 | NA | NA | NA | NA | NA | NA | 0 | 7 | 0 |
| Hinfp | Gpn1 | NA | NA | NA | NA | NA | NA | 0 | 16 | 0 |
| Hlf | Runx1 | NA | NA | NA | NA | NA | NA | 0 | 2 | 0 |
| Hmbox1 | Cyth1 | NA | NA | NA | NA | NA | NA | 0 | 26 | 0 |
| Hmga1 | Cdc25a | NA | NA | NA | NA | NA | NA | 0 | 1 | 0 |
| Hnf1a | Fgl1 | NA | NA | NA | NA | NA | NA | 0 | 23 | 0 |
| Hnf1b | Tmco5a | NA | NA | NA | NA | NA | NA | 0 | 37 | 0 |
| Hoxa10 | Emx2 | NA | NA | NA | NA | NA | NA | 0 | 4 | 0 |
| Hoxa11 | Hoxa10 | NA | NA | NA | NA | NA | NA | 0 | 1 | 0 |
| Hoxa4 | Th | NA | NA | NA | NA | NA | NA | 0 | 1 | 0 |
| Hoxa5 | Th | NA | NA | NA | NA | NA | NA | 0 | 4 | 0 |
| Hoxa9 | Tle4 | NA | NA | NA | NA | NA | NA | 0 | 22 | 0 |
| Hoxb13 | Ctnnal1 | NA | NA | NA | NA | NA | NA | 0 | 46 | 0 |
| Hoxb3 | Otx2 | NA | NA | NA | NA | NA | NA | 0 | 1 | 0 |
| Hoxb7 | Ppp5c | NA | NA | NA | NA | NA | NA | 0 | 5 | 0 |
| Hoxc11 | Lct | NA | NA | NA | NA | NA | NA | 0 | 2 | 0 |
| Hoxc8 | Tnfrsf11b | NA | NA | NA | NA | NA | NA | 0 | 1 | 0 |
| Hoxd3 | Itgb1 | NA | NA | NA | NA | NA | NA | 0 | 3 | 0 |
| Hsf2 | Mapk8 | NA | NA | NA | NA | NA | NA | 0 | 3 | 0 |
| Insm1 | Neurod1 | NA | NA | NA | NA | NA | NA | 0 | 3 | 0 |
| Irf1 | Tap1 | NA | NA | NA | NA | NA | NA | 0 | 77 | 0 |
| Irf2 | Tasor2 | NA | NA | NA | NA | NA | NA | 0 | 67 | 0 |
| Irf5 | Tnf | NA | NA | NA | NA | NA | NA | 0 | 1 | 0 |
| Irf7 | Ifnb1 | NA | NA | NA | NA | NA | NA | 0 | 2 | 0 |
| Irf8 | Tlr4 | NA | NA | NA | NA | NA | NA | 0 | 8 | 0 |
| Kdm5b | Prrc2b | NA | NA | NA | NA | NA | NA | 0 | 17 | 0 |
| Klf1 | Snap47 | NA | NA | NA | NA | NA | NA | 0 | 50 | 0 |
| Klf10 | Cdkn1a | NA | NA | NA | NA | NA | NA | 0 | 1 | 0 |
| Klf11 | Pdx1 | NA | NA | NA | NA | NA | NA | 0 | 2 | 0 |
| Klf12 | Tfap2a | NA | NA | NA | NA | NA | NA | 0 | 1 | 0 |
| Klf13 | Dnajb1 | NA | NA | NA | NA | NA | NA | 0 | 31 | 0 |
| Klf15 | Lrp5 | NA | NA | NA | NA | NA | NA | 0 | 1 | 0 |
| Klf2 | Thbd | NA | NA | NA | NA | NA | NA | 0 | 2 | 0 |
| Klf3 | Cast | NA | NA | NA | NA | NA | NA | 0 | 13 | 0 |
| Klf4 | Gpa33 | NA | NA | NA | NA | NA | NA | 0 | 49 | 0 |
| Klf5 | Actn1 | NA | NA | NA | NA | NA | NA | 0 | 80 | 0 |
| Kmt2b | Supt5h | NA | NA | NA | NA | NA | NA | 0 | 8 | 0 |
| Lhx2 | Sox6 | NA | NA | NA | NA | NA | NA | 0 | 10 | 0 |
| Lhx3 | Fshb | NA | NA | NA | NA | NA | NA | 0 | 3 | 0 |
| Lhx4 | Pou1f1 | NA | NA | NA | NA | NA | NA | 0 | 4 | 0 |
| Maf | Galm | NA | NA | NA | NA | NA | NA | 0 | 38 | 0 |
| Mafa | Ins2 | NA | NA | NA | NA | NA | NA | 0 | 1 | 0 |
| Mbd2 | Tyk2 | NA | NA | NA | NA | NA | NA | 0 | 32 | 0 |
| Mbd3 | Esr1 | NA | NA | NA | NA | NA | NA | 0 | 1 | 0 |
| Mecp2 | Esr1 | NA | NA | NA | NA | NA | NA | 0 | 1 | 0 |
| Mef2d | Sost | NA | NA | NA | NA | NA | NA | 0 | 4 | 0 |
| Meis1 | Rere | NA | NA | NA | NA | NA | NA | 0 | 54 | 0 |
| Msc | Mmp2 | NA | NA | NA | NA | NA | NA | 0 | 2 | 0 |
| Mxd1 | Odc1 | NA | NA | NA | NA | NA | NA | 0 | 1 | 0 |
| Mxd4 | Odc1 | NA | NA | NA | NA | NA | NA | 0 | 1 | 0 |
| Myb | Vgll4 | NA | NA | NA | NA | NA | NA | 0 | 65 | 0 |
| Mybl1 | Ncl | NA | NA | NA | NA | NA | NA | 0 | 2 | 0 |
| Mybl2 | Cenpf | NA | NA | NA | NA | NA | NA | 0 | 25 | 0 |
| Mycn | Rxra | NA | NA | NA | NA | NA | NA | 0 | 64 | 0 |
| Myf6 | Acacb | NA | NA | NA | NA | NA | NA | 0 | 1 | 0 |
| Mynn | Tra2b | NA | NA | NA | NA | NA | NA | 0 | 4 | 0 |
| Myod1 | Cuedc1 | NA | NA | NA | NA | NA | NA | 0 | 46 | 0 |
| Myog | Acacb | NA | NA | NA | NA | NA | NA | 0 | 1 | 0 |
| Mzf1 | Myb | NA | NA | NA | NA | NA | NA | 0 | 3 | 0 |
| Nanog | Rabgap1l | NA | NA | NA | NA | NA | NA | 0 | 48 | 0 |
| Ncoa1 | Hipk1 | NA | NA | NA | NA | NA | NA | 0 | 20 | 0 |
| Ncoa2 | Daam1 | NA | NA | NA | NA | NA | NA | 0 | 18 | 0 |
| Ncoa3 | Arhgap10 | NA | NA | NA | NA | NA | NA | 0 | 23 | 0 |
| Neurod1 | Ctsb | NA | NA | NA | NA | NA | NA | 0 | 35 | 0 |
| Neurog3 | Neurod1 | NA | NA | NA | NA | NA | NA | 0 | 1 | 0 |
| Nfat5 | Tnf | NA | NA | NA | NA | NA | NA | 0 | 2 | 0 |
| Nfatc1 | Tmem131l | NA | NA | NA | NA | NA | NA | 0 | 34 | 0 |
| Nfatc2 | Cdk4 | NA | NA | NA | NA | NA | NA | 0 | 10 | 0 |
| Nfatc3 | Rcan1 | NA | NA | NA | NA | NA | NA | 0 | 2 | 0 |
| Nfatc4 | Trpc6 | NA | NA | NA | NA | NA | NA | 0 | 1 | 0 |
| Nfia | Tp53 | NA | NA | NA | NA | NA | NA | 0 | 3 | 0 |
| Nfil3 | Il1b | NA | NA | NA | NA | NA | NA | 0 | 1 | 0 |
| Nfkb2 | Wnt10a | NA | NA | NA | NA | NA | NA | 0 | 62 | 0 |
| Nfyc | P04224 | NA | NA | NA | NA | NA | NA | 0 | 3 | 0 |
| Nkx2-1 | Sftpb | NA | NA | NA | NA | NA | NA | 0 | 1 | 0 |
| Nkx2-5 | Gata4 | NA | NA | NA | NA | NA | NA | 0 | 2 | 0 |
| Nme2 | Myc | NA | NA | NA | NA | NA | NA | 0 | 1 | 0 |
| Npas2 | Cry2 | NA | NA | NA | NA | NA | NA | 0 | 4 | 0 |
| Nr0b1 | Nr5a1 | NA | NA | NA | NA | NA | NA | 0 | 2 | 0 |
| Nr1d1 | Arntl | NA | NA | NA | NA | NA | NA | 0 | 1 | 0 |
| Nr1h4 | Nr1i2 | NA | NA | NA | NA | NA | NA | 0 | 7 | 0 |
| Nr1i2 | Vamp7 | NA | NA | NA | NA | NA | NA | 0 | 4 | 0 |
| Nr1i3 | Ugt1a1 | NA | NA | NA | NA | NA | NA | 0 | 1 | 0 |
| Nr2c2 | Ankib1 | NA | NA | NA | NA | NA | NA | 0 | 84 | 0 |
| Nr2e1 | Pax2 | NA | NA | NA | NA | NA | NA | 0 | 3 | 0 |
| Nr3c2 | Atp1b1 | NA | NA | NA | NA | NA | NA | 0 | 2 | 0 |
| Nr5a1 | Cyp19a1 | NA | NA | NA | NA | NA | NA | 0 | 12 | 0 |
| Nrf1 | Znf12 | NA | NA | NA | NA | NA | NA | 0 | 9 | 0 |
| Nrl | Rho | NA | NA | NA | NA | NA | NA | 0 | 1 | 0 |
| Onecut2 | Sytl4 | NA | NA | NA | NA | NA | NA | 0 | 1 | 0 |
| Otx2 | Creb5 | NA | NA | NA | NA | NA | NA | 0 | 23 | 0 |
| Pax1 | Pdgfra | NA | NA | NA | NA | NA | NA | 0 | 1 | 0 |
| Pax2 | Tp53 | NA | NA | NA | NA | NA | NA | 0 | 2 | 0 |
| Pax3 | Mitf | NA | NA | NA | NA | NA | NA | 0 | 3 | 0 |
| Pax4 | Ins2 | NA | NA | NA | NA | NA | NA | 0 | 1 | 0 |
| Pax8 | Tp53 | NA | NA | NA | NA | NA | NA | 0 | 11 | 0 |
| Pbx1 | Vcp | NA | NA | NA | NA | NA | NA | 0 | 4 | 0 |
| Pcgf2 | Hoxa7 | NA | NA | NA | NA | NA | NA | 0 | 1 | 0 |
| Pdx1 | Fto | NA | NA | NA | NA | NA | NA | 0 | 50 | 0 |
| Pitx1 | Fshb | NA | NA | NA | NA | NA | NA | 0 | 1 | 0 |
| Pitx2 | Fshb | NA | NA | NA | NA | NA | NA | 0 | 1 | 0 |
| Plag1 | Abcc6 | NA | NA | NA | NA | NA | NA | 0 | 1 | 0 |
| Pou1f1 | Ghr | NA | NA | NA | NA | NA | NA | 0 | 1 | 0 |
| Pou2f1 | Tnrc6c | NA | NA | NA | NA | NA | NA | 0 | 61 | 0 |
| Pou2f3 | Sprr2d | NA | NA | NA | NA | NA | NA | 0 | 1 | 0 |
| Pou3f1 | Gnrh1 | NA | NA | NA | NA | NA | NA | 0 | 1 | 0 |
| Pou3f2 | Kcnn3 | NA | NA | NA | NA | NA | NA | 0 | 2 | 0 |
| Pou4f2 | Kcnip4 | NA | NA | NA | NA | NA | NA | 0 | 28 | 0 |
| Ppard | Sat1 | NA | NA | NA | NA | NA | NA | 0 | 4 | 0 |
| Prdm1 | Mecom | NA | NA | NA | NA | NA | NA | 0 | 40 | 0 |
| Pura | Ar | NA | NA | NA | NA | NA | NA | 0 | 1 | 0 |
| Rarb | Adh5 | NA | NA | NA | NA | NA | NA | 0 | 5 | 0 |
| Rarg | Greb1 | NA | NA | NA | NA | NA | NA | 0 | 8 | 0 |
| Rfx1 | Fgf1 | NA | NA | NA | NA | NA | NA | 0 | 40 | 0 |
| Rfx2 | Ttc30b | NA | NA | NA | NA | NA | NA | 0 | 20 | 0 |
| Rora | Tuba1b | NA | NA | NA | NA | NA | NA | 0 | 3 | 0 |
| Rorc | Il22b | NA | NA | NA | NA | NA | NA | 0 | 3 | 0 |
| Rreb1 | Klk1b5 | NA | NA | NA | NA | NA | NA | 0 | 14 | 0 |
| Runx2 | Pbx1 | NA | NA | NA | NA | NA | NA | 0 | 65 | 0 |
| Rxrb | Angptl4 | NA | NA | NA | NA | NA | NA | 0 | 6 | 0 |
| Rxrg | Plin5 | NA | NA | NA | NA | NA | NA | 0 | 6 | 0 |
| Sall4 | Zeb1 | NA | NA | NA | NA | NA | NA | 0 | 3 | 0 |
| Satb1 | Tmem245 | NA | NA | NA | NA | NA | NA | 0 | 9 | 0 |
| Six2 | Col13a1 | NA | NA | NA | NA | NA | NA | 0 | 39 | 0 |
| Ski | Smad7 | NA | NA | NA | NA | NA | NA | 0 | 1 | 0 |
| Skil | Smad7 | NA | NA | NA | NA | NA | NA | 0 | 1 | 0 |
| Smad4 | Met | NA | NA | NA | NA | NA | NA | 0 | 75 | 0 |
| Snai1 | Cdh1 | NA | NA | NA | NA | NA | NA | 0 | 5 | 0 |
| Snapc4 | Znf335 | NA | NA | NA | NA | NA | NA | 0 | 29 | 0 |
| Sox11 | Col28a1 | NA | NA | NA | NA | NA | NA | 0 | 29 | 0 |
| Sox13 | Jag1 | NA | NA | NA | NA | NA | NA | 0 | 45 | 0 |
| Sox4 | Ncam1 | NA | NA | NA | NA | NA | NA | 0 | 3 | 0 |
| Sox8 | Cldn23 | NA | NA | NA | NA | NA | NA | 0 | 1 | 0 |
| Sp2 | Usp16 | NA | NA | NA | NA | NA | NA | 0 | 22 | 0 |
| Spdef | Agr2 | NA | NA | NA | NA | NA | NA | 0 | 1 | 0 |
| Stat4 | Prf1 | NA | NA | NA | NA | NA | NA | 0 | 13 | 0 |
| Stat6 | Sidt1 | NA | NA | NA | NA | NA | NA | 0 | 13 | 0 |
| Tbx2 | Cdkn2a | NA | NA | NA | NA | NA | NA | 0 | 1 | 0 |
| Tbx21 | Gas7 | NA | NA | NA | NA | NA | NA | 0 | 76 | 0 |
| Tbx5 | Cdkn2a | NA | NA | NA | NA | NA | NA | 0 | 2 | 0 |
| Tcf12 | Lpp | NA | NA | NA | NA | NA | NA | 0 | 78 | 0 |
| Tcf4 | Vim | NA | NA | NA | NA | NA | NA | 0 | 60 | 0 |
| Tcf7 | Lef1 | NA | NA | NA | NA | NA | NA | 0 | 58 | 0 |
| Tcf7l1 | Nkd1 | NA | NA | NA | NA | NA | NA | 0 | 5 | 0 |
| Tcfl5 | Cd2 | NA | NA | NA | NA | NA | NA | 0 | 1 | 0 |
| Tfap2b | Drd1 | NA | NA | NA | NA | NA | NA | 0 | 2 | 0 |
| Tfap4 | Pemt | NA | NA | NA | NA | NA | NA | 0 | 40 | 0 |
| Tfcp2 | Tf | NA | NA | NA | NA | NA | NA | 0 | 1 | 0 |
| Tgif2 | Septin9 | NA | NA | NA | NA | NA | NA | 0 | 19 | 0 |
| Thra | Cga | NA | NA | NA | NA | NA | NA | 0 | 1 | 0 |
| Thrb | Myh6 | NA | NA | NA | NA | NA | NA | 0 | 4 | 0 |
| Tlx1 | Aldh1a1 | NA | NA | NA | NA | NA | NA | 0 | 1 | 0 |
| Trps1 | Runx2 | NA | NA | NA | NA | NA | NA | 0 | 1 | 0 |
| Twist2 | Srpx | NA | NA | NA | NA | NA | NA | 0 | 3 | 0 |
| Wt1 | Cdh1 | NA | NA | NA | NA | NA | NA | 0 | 30 | 0 |
| Xbp1 | Rapgef4 | NA | NA | NA | NA | NA | NA | 0 | 16 | 0 |
| Ybx1 | Egfr | NA | NA | NA | NA | NA | NA | 0 | 6 | 0 |
| Zbtb11 | Uspl1 | NA | NA | NA | NA | NA | NA | 0 | 22 | 0 |
| Zbtb16 | Ccna2 | NA | NA | NA | NA | NA | NA | 0 | 1 | 0 |
| Zbtb17 | Cdkn2b | NA | NA | NA | NA | NA | NA | 0 | 1 | 0 |
| Zc3h8 | Has3 | NA | NA | NA | NA | NA | NA | 0 | 6 | 0 |
| Zfp202 | Abca1 | NA | NA | NA | NA | NA | NA | 0 | 2 | 0 |
| Zfp217 | Arid5b | NA | NA | NA | NA | NA | NA | 0 | 10 | 0 |
| Zfp644 | Epc1 | NA | NA | NA | NA | NA | NA | 0 | 7 | 0 |
| Zhx1 | Atf1 | NA | NA | NA | NA | NA | NA | 0 | 9 | 0 |
| Zhx2 | Arid1b | NA | NA | NA | NA | NA | NA | 0 | 8 | 0 |
| Zic1 | Apoe | NA | NA | NA | NA | NA | NA | 0 | 1 | 0 |
| Zic2 | Apoe | NA | NA | NA | NA | NA | NA | 0 | 2 | 0 |
| Znf148 | Irs2 | NA | NA | NA | NA | NA | NA | 0 | 2 | 0 |
| Znf24 | Csmd2 | NA | NA | NA | NA | NA | NA | 0 | 18 | 0 |
| Znf382 | Hmga2 | NA | NA | NA | NA | NA | NA | 0 | 1 | 0 |
| Znf592 | Fbxl18 | NA | NA | NA | NA | NA | NA | 0 | 14 | 0 |
| Znf639 | Ahctf1 | NA | NA | NA | NA | NA | NA | 0 | 28 | 0 |
| Znf740 | Pdgfa | NA | NA | NA | NA | NA | NA | 0 | 36 | 0 |

| TF | gene | GeneRatio | BgRatio | pvalue | p.adjust | qvalue | DetectedTargets | Count | TF_targets | Percentage_of_TFtargets_detected |
|---|---|---|---|---|---|---|---|---|---|---|
| Arntl | Sema4d | Apr-48 | 17/6837 | 4.76E-06 | 0.00041893 | 0.00038586 | Dbp/Nr1d1/Nr1d2/Per3 | 4 | 20 | 20 |
| Clock | Fam91a1 | Feb-48 | 41/6837 | 0.03326541 | 0.3512416 | 0.323512 | Nr1d2/Per3 | 2 | 51 | 3.92 |
| Foxo1 | Timp3 | Mar-48 | 105/6837 | 0.03697136 | 0.3512416 | 0.323512 | Abcb1a/Hdac9/Timp3 | 3 | 122 | 2.46 |
| Hif1a | Cxcr4 | May-48 | 208/6837 | 0.01470668 | 0.3512416 | 0.323512 | Abcb1a/Aqp1/Bnip3/Ciart/Tlr2 | 5 | 269 | 1.86 |
| Hnf4a | Sec14l1 | Oct-48 | 788/6837 | 0.0439052 | 0.3512416 | 0.323512 | Abcb1a/Aldh3b1/Cacna1h/Cnnm1/Fnip2/Lhfpl2/Nags/Nr1d1/Sun2/Tcirg1 | 10 | 907 | 1.1 |
| Meis1 | Lsamp | Feb-48 | 46/6837 | 0.04107172 | 0.3512416 | 0.323512 | Fbn1/Hdac9 | 2 | 54 | 3.7 |
| Nanog | Tcf4 | Feb-48 | 37/6837 | 0.02749991 | 0.3512416 | 0.323512 | Hdac9/Lamb1 | 2 | 48 | 4.17 |
| Rela | Maml2 | Apr-48 | 180/6837 | 0.03679153 | 0.3512416 | 0.323512 | Abcb1a/Bnip3/Saa1/Traf1 | 4 | 245 | 1.63 |
| Usf2 | Zbtb40 | Mar-48 | 103/6837 | 0.03521763 | 0.3512416 | 0.323512 | Dbp/Klf15/Thbs1 | 3 | 113 | 2.65 |
| Wt1 | Ccn2 | Feb-48 | 21/6837 | 0.00930976 | 0.3512416 | 0.323512 | Abcb1a/Thbs1 | 2 | 30 | 6.67 |
| Zkscan1 | Aldh3b1 | Feb-48 | 40/6837 | 0.03178277 | 0.3512416 | 0.323512 | Ablim2/Aldh3b1 | 2 | 46 | 4.35 |
| Gata3 | Malrd1 | Aug-48 | 596/6837 | 0.05358672 | 0.38343434 | 0.3531632 | Ablim2/Asb11/Atp12a/Fnip2/Gcnt4/Hdac9/Lhfpl2/Nbl1 | 8 | 760 | 1.05 |
| Twist1 | Hpcal1 | Feb-48 | 55/6837 | 0.05664371 | 0.38343434 | 0.3531632 | Hdac9/Lamb1 | 2 | 67 | 2.99 |
| Cebpb | Magi1 | Feb-48 | 62/6837 | 0.06995369 | 0.41039499 | 0.37799538 | Abcb1a/Saa1 | 2 | 85 | 2.35 |
| Mef2c | Tbl1xr1 | Feb-48 | 62/6837 | 0.06995369 | 0.41039499 | 0.37799538 | Hdac9/Nrep | 2 | 70 | 2.86 |
| Tead4 | Tgm2 | Feb-48 | 67/6837 | 0.0800263 | 0.44014464 | 0.40539638 | Ablim2/Hdac9 | 2 | 73 | 2.74 |
| Nr3c1 | Robo1 | Feb-48 | 70/6837 | 0.08627557 | 0.44660296 | 0.41134483 | Bglap3/Hdac9 | 2 | 76 | 2.63 |
| Esr1 | Kctd3 | Jul-48 | 579/6837 | 0.10744752 | 0.47276908 | 0.43544521 | Ablim2/Adamts9/Emp2/Lhfpl2/Macrod1/Prss23/Tcirg1 | 7 | 722 | 0.97 |
| Hinfp | Atf7ip2 | Jan-48 | 16/6837 | 0.10671573 | 0.47276908 | 0.43544521 | Sun2 | 1 | 16 | 6.25 |
| Rfx2 | Mapre2 | Jan-48 | 15/6837 | 0.10038599 | 0.47276908 | 0.43544521 | Tbcel | 1 | 20 | 5 |
| Kdm5b | Rreb1 | Jan-48 | 17/6837 | 0.11300186 | 0.47353159 | 0.43614751 | Lhfpl2 | 1 | 17 | 5.88 |
| Nfkb1 | Cdk4 | Mar-48 | 175/6837 | 0.12391432 | 0.4799613 | 0.44206962 | Abcb1a/Tlr2/Traf1 | 3 | 251 | 1.2 |
| Tgif2 | Tcirg1 | Jan-48 | 19/6837 | 0.12544443 | 0.4799613 | 0.44206962 | Tcirg1 | 1 | 19 | 5.26 |
| Mef2a | Bcas3 | Feb-48 | 92/6837 | 0.13596903 | 0.49855309 | 0.45919364 | Hdac9/Lhfpl2 | 2 | 102 | 1.96 |
| Prox1 | Arhgap42 | Jan-48 | 23/6837 | 0.14981902 | 0.50707975 | 0.46704714 | Abcb1a | 1 | 26 | 3.85 |
| Tfap2a | Nr3c1 | Feb-48 | 97/6837 | 0.14802286 | 0.50707975 | 0.46704714 | Abcb1a/Lamb1 | 2 | 125 | 1.6 |
| Mef2b | Tox2 | Jan-48 | 27/6837 | 0.17352803 | 0.56557284 | 0.52092235 | Hdac9 | 1 | 36 | 2.78 |
| Ahr | Vgll4 | Jan-48 | 35/6837 | 0.21902006 | 0.58633672 | 0.54004698 | Lhfpl2 | 1 | 43 | 2.33 |
| Bhlhe40 | Trappc9 | Jan-48 | 53/6837 | 0.31259917 | 0.58633672 | 0.54004698 | Nr1d1 | 1 | 57 | 1.75 |
| Cdx2 | Capza2 | Jan-48 | 49/6837 | 0.2928028 | 0.58633672 | 0.54004698 | Abcb1a | 1 | 52 | 1.92 |
| Cebpg | Trib3 | Jan-48 | 41/6837 | 0.25152096 | 0.58633672 | 0.54004698 | Hdac9 | 1 | 46 | 2.17 |
| Ctcfl | Nfe2l2 | Jan-48 | 39/6837 | 0.24083728 | 0.58633672 | 0.54004698 | Cacna1h | 1 | 53 | 1.89 |
| Egr1 | Numbl | Aug-48 | 819/6837 | 0.21075707 | 0.58633672 | 0.54004698 | Abcb1a/Cacna1h/Pdlim4/Slc37a2/Sun2/Syt2/Thbs1/Timp3 | 8 | 976 | 0.82 |
| Foxa2 | Tln2 | Jan-48 | 42/6837 | 0.25680745 | 0.58633672 | 0.54004698 | Hdac9 | 1 | 49 | 2.04 |
| Foxo3 | Jak1 | Jan-48 | 35/6837 | 0.21902006 | 0.58633672 | 0.54004698 | Abcb1a | 1 | 40 | 2.5 |
| Hnf4g | Hsd17b2 | Jan-48 | 49/6837 | 0.2928028 | 0.58633672 | 0.54004698 | Aldh3b1 | 1 | 58 | 1.72 |
| Klf6 | Lhfpl2 | Jan-48 | 51/6837 | 0.30276978 | 0.58633672 | 0.54004698 | Lhfpl2 | 1 | 52 | 1.92 |
| Max | Ppp2r2c | Jan-48 | 50/6837 | 0.2978036 | 0.58633672 | 0.54004698 | Ablim2 | 1 | 53 | 1.89 |
| Myc | Tbcd | Apr-48 | 403/6837 | 0.31315711 | 0.58633672 | 0.54004698 | Dbp/Nr1d1/Tcirg1/Thbs1 | 4 | 447 | 0.89 |
| Nfkb2 | Nek6 | Jan-48 | 48/6837 | 0.28776712 | 0.58633672 | 0.54004698 | Traf1 | 1 | 62 | 1.61 |
| Pax6 | Gck | Jan-48 | 34/6837 | 0.21347053 | 0.58633672 | 0.54004698 | Hdac9 | 1 | 43 | 2.33 |
| Pdx1 | Lhfpl2 | Jan-48 | 45/6837 | 0.27244852 | 0.58633672 | 0.54004698 | Lhfpl2 | 1 | 50 | 2 |
| Rbpj | Slc12a8 | Jan-48 | 52/6837 | 0.30770155 | 0.58633672 | 0.54004698 | Pex5l | 1 | 59 | 1.69 |
| Rest | Nxph1 | Jan-48 | 40/6837 | 0.24619766 | 0.58633672 | 0.54004698 | Cacna1h | 1 | 67 | 1.49 |
| Sox10 | Mbnl2 | Jan-48 | 35/6837 | 0.21902006 | 0.58633672 | 0.54004698 | Hdac9 | 1 | 45 | 2.22 |
| Tcf4 | Phactr2 | Jan-48 | 52/6837 | 0.30770155 | 0.58633672 | 0.54004698 | Abcb1a | 1 | 60 | 1.67 |
| Zeb2 | Adgrb3 | Jan-48 | 46/6837 | 0.27759023 | 0.58633672 | 0.54004698 | Lhfpl2 | 1 | 53 | 1.89 |
| Ar | Banf1 | Feb-48 | 187/6837 | 0.37960965 | 0.58993628 | 0.54336236 | Hdac9/Sord | 2 | 222 | 0.9 |
| Fos | Rras2 | Jun-48 | 686/6837 | 0.34971613 | 0.58993628 | 0.54336236 | Hdac9/Hkdc1/Lhfpl2/Nbl1/Nr1d1/Tmem44 | 6 | 847 | 0.71 |
| Gata2 | Il1rl1 | May-48 | 542/6837 | 0.33125318 | 0.58993628 | 0.54336236 | Fnip2/Hdac9/Hkdc1/Lhfpl2/Tlr2 | 5 | 676 | 0.74 |
| Jund | Ndrg1 | Jan-48 | 66/6837 | 0.37326271 | 0.58993628 | 0.54336236 | Bglap3 | 1 | 75 | 1.33 |
| Nfya | Pnrc1 | Jan-48 | 61/6837 | 0.35057772 | 0.58993628 | 0.54336236 | Abcb1a | 1 | 70 | 1.43 |
| Nr2f2 | Synj2 | Jan-48 | 68/6837 | 0.38211782 | 0.58993628 | 0.54336236 | Bglap3 | 1 | 84 | 1.19 |
| Relb | Tmtc2 | Jan-48 | 58/6837 | 0.33658287 | 0.58993628 | 0.54336236 | Abcb1a | 1 | 74 | 1.35 |
| Rxra | Cyp7a1 | Jan-48 | 67/6837 | 0.37770569 | 0.58993628 | 0.54336236 | Nrep | 1 | 78 | 1.28 |
| Smad4 | Gadd45b | Jan-48 | 59/6837 | 0.34128032 | 0.58993628 | 0.54336236 | Bglap3 | 1 | 75 | 1.33 |
| Tcf7l2 | Plcb4 | Jan-48 | 66/6837 | 0.37326271 | 0.58993628 | 0.54336236 | Abcb1a | 1 | 74 | 1.35 |
| Bach1 | Dst | Jan-48 | 79/6837 | 0.42867446 | 0.60744892 | 0.55949243 | Lhfpl2 | 1 | 84 | 1.19 |
| Foxm1 | Lama2 | Jan-48 | 83/6837 | 0.44473767 | 0.60744892 | 0.55949243 | Hdac9 | 1 | 92 | 1.09 |
| Pparg | Serpine1 | Jan-48 | 84/6837 | 0.44868386 | 0.60744892 | 0.55949243 | Nr1d1 | 1 | 104 | 0.96 |
| Runx3 | Arhgap10 | Jan-48 | 80/6837 | 0.43273241 | 0.60744892 | 0.55949243 | Hdac9 | 1 | 97 | 1.03 |
| Smad3 | Mylk | Jan-48 | 78/6837 | 0.42458809 | 0.60744892 | 0.55949243 | Bglap3 | 1 | 96 | 1.04 |
| Tfap2c | Mideas | Jun-48 | 737/6837 | 0.41622286 | 0.60744892 | 0.55949243 | Ablim2/Cbx7/Lhfpl2/Prss23/Rtn4r/Tcirg1 | 6 | 890 | 0.67 |
| Tp63 | Ston2 | Jan-48 | 76/6837 | 0.41632931 | 0.60744892 | 0.55949243 | Ablim2 | 1 | 90 | 1.11 |
| Vdr | Bag3 | Feb-48 | 210/6837 | 0.436577 | 0.60744892 | 0.55949243 | Bglap3/Lhfpl2 | 2 | 244 | 0.82 |
| Tal1 | Plekho1 | Mar-48 | 364/6837 | 0.474653 | 0.62342483 | 0.57420708 | Fnip2/Hdac9/Lhfpl2 | 3 | 416 | 0.72 |
| Usf1 | Ceacam2 | Jan-48 | 89/6837 | 0.46800658 | 0.62342483 | 0.57420708 | Ablim2 | 1 | 103 | 0.97 |
| Nfic | Pold2 | Jan-48 | 94/6837 | 0.48666569 | 0.62980266 | 0.5800814 | Nrep | 1 | 108 | 0.93 |
| Srf | Acta1 | Jan-48 | 100/6837 | 0.50821204 | 0.64815449 | 0.5969844 | Sun2 | 1 | 109 | 0.92 |
| Rara | Apoa1 | Jan-48 | 108/6837 | 0.53556802 | 0.67328551 | 0.62013139 | Macrod1 | 1 | 122 | 0.82 |
| Stat1 | Hk1 | Jun-48 | 845/6837 | 0.55395947 | 0.686195 | 0.63202171 | Fnip2/Hdac9/Lhfpl2/Sstr1/Thbs1/Traf1 | 6 | 1014 | 0.59 |
| Tead1 | Schip1 | Jan-48 | 116/6837 | 0.56143227 | 0.686195 | 0.63202171 | Hdac9 | 1 | 129 | 0.78 |
| Jun | Ccl12 | Jan-48 | 119/6837 | 0.57076318 | 0.68804329 | 0.63372408 | Abcb1a | 1 | 185 | 0.54 |
| Sox2 | Col14a1 | Feb-48 | 291/6837 | 0.61246811 | 0.72834046 | 0.6708399 | Hdac9/Nrep | 2 | 398 | 0.5 |
| Sp1 | Nfkb2 | Feb-48 | 296/6837 | 0.62189411 | 0.72968909 | 0.67208206 | Abcb1a/Macrod1 | 2 | 394 | 0.51 |
| Stat3 | Plec | Jan-48 | 140/6837 | 0.63085109 | 0.73045916 | 0.67279133 | Saa1 | 1 | 171 | 0.58 |
| Runx1 | Tlk1 | Feb-48 | 306/6837 | 0.64023184 | 0.73169353 | 0.67392825 | Fnip2/Hdac9 | 2 | 380 | 0.53 |
| Spi1 | Nrxn2 | Mar-48 | 506/6837 | 0.7004876 | 0.79029371 | 0.7279021 | Hdac9/Lhfpl2/Macrod1 | 3 | 585 | 0.51 |
| Znf263 | Plcg2 | Feb-48 | 351/6837 | 0.714409 | 0.79579737 | 0.73297126 | Lhfpl2/Rtn4r | 2 | 432 | 0.46 |
| E2f1 | Ap4m1 | Mar-48 | 576/6837 | 0.78204812 | 0.84963253 | 0.78255627 | Bnip3/Sstr1/Thbs1 | 3 | 675 | 0.44 |
| Mitf | Cops7a | Mar-48 | 574/6837 | 0.77998802 | 0.84963253 | 0.78255627 | Ablim2/Fnip2/Lhfpl2 | 3 | 660 | 0.45 |
| Prdm14 | Arntl | Jan-48 | 227/6837 | 0.80336994 | 0.86215311 | 0.79408839 | Lamb1 | 1 | 306 | 0.33 |
| Cebpa | Pcnx1 | Apr-48 | 788/6837 | 0.81995333 | 0.86934811 | 0.80071536 | Abcb1a/Fnip2/Lamb1/Saa1 | 4 | 959 | 0.42 |
| Ets1 | Rpl3 | Apr-48 | 837/6837 | 0.85572838 | 0.88557159 | 0.81565804 | Abcb1a/Adamts9/Lhfpl2/Thbs1 | 4 | 934 | 0.43 |
| Foxa1 | Focad | Mar-48 | 674/6837 | 0.86544496 | 0.88557159 | 0.81565804 | Hdac9/Lhfpl2/Macrod1 | 3 | 851 | 0.35 |
| Tp53 | Adrb2 | Feb-48 | 483/6837 | 0.86287647 | 0.88557159 | 0.81565804 | Abcb1a/Thbs1 | 2 | 582 | 0.34 |
| Ctcf | Elovl2 | Feb-48 | 729/6837 | 0.97036451 | 0.98151812 | 0.90402985 | Hdac9/Lhfpl2 | 2 | 936 | 0.21 |
| E2f4 | Lmo4 | Jan-48 | 695/6837 | 0.9942829 | 0.9942829 | 0.91578689 | St3gal4 | 1 | 743 | 0.13 |
| Adnp | Atf7ip | NA | NA | NA | NA | NA | NA | 0 | 4 | 0 |
| Arid2 | Cdk17 | NA | NA | NA | NA | NA | NA | 0 | 30 | 0 |
| Arid3a | Ptk2b | NA | NA | NA | NA | NA | NA | 0 | 23 | 0 |
| Arnt | Uvrag | NA | NA | NA | NA | NA | NA | 0 | 63 | 0 |
| Arntl2 | Serpine1 | NA | NA | NA | NA | NA | NA | 0 | 1 | 0 |
| Ascl1 | Cplx2 | NA | NA | NA | NA | NA | NA | 0 | 54 | 0 |
| Atf1 | Ntrk3 | NA | NA | NA | NA | NA | NA | 0 | 91 | 0 |
| Atf2 | Msi2 | NA | NA | NA | NA | NA | NA | 0 | 62 | 0 |
| Atf3 | Creb5 | NA | NA | NA | NA | NA | NA | 0 | 50 | 0 |
| Atf4 | Hrk | NA | NA | NA | NA | NA | NA | 0 | 110 | 0 |
| Atf5 | Disc1 | NA | NA | NA | NA | NA | NA | 0 | 1 | 0 |
| Atf6 | Ddit3 | NA | NA | NA | NA | NA | NA | 0 | 10 | 0 |
| Atoh1 | Hes6 | NA | NA | NA | NA | NA | NA | 0 | 1 | 0 |
| Bach2 | Tuba4a | NA | NA | NA | NA | NA | NA | 0 | 49 | 0 |
| Batf | Dennd1a | NA | NA | NA | NA | NA | NA | 0 | 49 | 0 |
| Bcl11a | Elmo1 | NA | NA | NA | NA | NA | NA | 0 | 34 | 0 |
| Bcl6 | St3gal1 | NA | NA | NA | NA | NA | NA | 0 | 46 | 0 |
| Bhlhe22 | Elf4 | NA | NA | NA | NA | NA | NA | 0 | 21 | 0 |
| Cdx1 | Ada | NA | NA | NA | NA | NA | NA | 0 | 3 | 0 |
| Cebpd | Itgb5 | NA | NA | NA | NA | NA | NA | 0 | 71 | 0 |
| Cebpe | Myc | NA | NA | NA | NA | NA | NA | 0 | 6 | 0 |
| Cebpz | Col11a1 | NA | NA | NA | NA | NA | NA | 0 | 3 | 0 |
| Creb1 | Slc19a1 | NA | NA | NA | NA | NA | NA | 0 | 117 | 0 |
| Creb3 | Utp11 | NA | NA | NA | NA | NA | NA | 0 | 25 | 0 |
| Creb3l1 | Plec | NA | NA | NA | NA | NA | NA | 0 | 20 | 0 |
| Creb3l2 | Cpt1c | NA | NA | NA | NA | NA | NA | 0 | 1 | 0 |
| Creb5 | Pou1f1 | NA | NA | NA | NA | NA | NA | 0 | 1 | 0 |
| Crem | Esyt2 | NA | NA | NA | NA | NA | NA | 0 | 50 | 0 |
| Cux1 | Srcin1 | NA | NA | NA | NA | NA | NA | 0 | 40 | 0 |
| Ddit3 | Trib3 | NA | NA | NA | NA | NA | NA | 0 | 2 | 0 |
| Deaf1 | Rac3 | NA | NA | NA | NA | NA | NA | 0 | 8 | 0 |
| Dnmt1 | Sfrp5 | NA | NA | NA | NA | NA | NA | 0 | 4 | 0 |
| E2f2 | E2f1 | NA | NA | NA | NA | NA | NA | 0 | 22 | 0 |
| E2f3 | Aurka | NA | NA | NA | NA | NA | NA | 0 | 14 | 0 |
| E2f5 | Jade1 | NA | NA | NA | NA | NA | NA | 0 | 12 | 0 |
| E2f6 | Slc9a3 | NA | NA | NA | NA | NA | NA | 0 | 67 | 0 |
| E2f7 | Hnrnpd | NA | NA | NA | NA | NA | NA | 0 | 29 | 0 |
| E2f8 | E2f1 | NA | NA | NA | NA | NA | NA | 0 | 1 | 0 |
| E4f1 | Ccna2 | NA | NA | NA | NA | NA | NA | 0 | 1 | 0 |
| Ebf1 | Rbm47 | NA | NA | NA | NA | NA | NA | 0 | 74 | 0 |
| Egr2 | Spint1 | NA | NA | NA | NA | NA | NA | 0 | 4 | 0 |
| Egr3 | Nab2 | NA | NA | NA | NA | NA | NA | 0 | 1 | 0 |
| Ehf | Tmc5 | NA | NA | NA | NA | NA | NA | 0 | 19 | 0 |
| Elf1 | Nedd4l | NA | NA | NA | NA | NA | NA | 0 | 50 | 0 |
| Elf2 | Vcp | NA | NA | NA | NA | NA | NA | 0 | 1 | 0 |
| Elf3 | Sprr2d | NA | NA | NA | NA | NA | NA | 0 | 52 | 0 |
| Elf5 | Rbm34 | NA | NA | NA | NA | NA | NA | 0 | 19 | 0 |
| Elk1 | Tagln | NA | NA | NA | NA | NA | NA | 0 | 113 | 0 |
| Elk3 | Fos | NA | NA | NA | NA | NA | NA | 0 | 1 | 0 |
| Elk4 | Krt8 | NA | NA | NA | NA | NA | NA | 0 | 14 | 0 |
| En1 | Utrn | NA | NA | NA | NA | NA | NA | 0 | 1 | 0 |
| Eomes | Ksr1 | NA | NA | NA | NA | NA | NA | 0 | 21 | 0 |
| Epas1 | Serpine1 | NA | NA | NA | NA | NA | NA | 0 | 80 | 0 |
| Erf | Ets2 | NA | NA | NA | NA | NA | NA | 0 | 1 | 0 |
| Erg | Pik3ap1 | NA | NA | NA | NA | NA | NA | 0 | 89 | 0 |
| Esr2 | Adamtsl5 | NA | NA | NA | NA | NA | NA | 0 | 74 | 0 |
| Esrra | Slc9a1 | NA | NA | NA | NA | NA | NA | 0 | 69 | 0 |
| Ets2 | Il5 | NA | NA | NA | NA | NA | NA | 0 | 31 | 0 |
| Etv1 | Limch1 | NA | NA | NA | NA | NA | NA | 0 | 29 | 0 |
| Etv3 | Nfkb1 | NA | NA | NA | NA | NA | NA | 0 | 1 | 0 |
| Etv4 | H6pd | NA | NA | NA | NA | NA | NA | 0 | 59 | 0 |
| Etv6 | Cap1 | NA | NA | NA | NA | NA | NA | 0 | 6 | 0 |
| Fli1 | Hdac7 | NA | NA | NA | NA | NA | NA | 0 | 107 | 0 |
| Fosb | Runx2 | NA | NA | NA | NA | NA | NA | 0 | 1 | 0 |
| Fosl1 | Il1r1 | NA | NA | NA | NA | NA | NA | 0 | 53 | 0 |
| Fosl2 | Cast | NA | NA | NA | NA | NA | NA | 0 | 79 | 0 |
| Foxa3 | Cyp2c38 | NA | NA | NA | NA | NA | NA | 0 | 3 | 0 |
| Foxj2 | Crb1 | NA | NA | NA | NA | NA | NA | 0 | 36 | 0 |
| Foxk1 | Cald1 | NA | NA | NA | NA | NA | NA | 0 | 8 | 0 |
| Foxk2 | Foxk1 | NA | NA | NA | NA | NA | NA | 0 | 50 | 0 |
| Foxl2 | Bcl2a1 | NA | NA | NA | NA | NA | NA | 0 | 20 | 0 |
| Foxo4 | Cdkn1a | NA | NA | NA | NA | NA | NA | 0 | 15 | 0 |
| Foxo6 | Fbxo25 | NA | NA | NA | NA | NA | NA | 0 | 1 | 0 |
| Foxp2 | Gnas | NA | NA | NA | NA | NA | NA | 0 | 35 | 0 |
| Foxp3 | Il2ra | NA | NA | NA | NA | NA | NA | 0 | 3 | 0 |
| Gabpa | Efr3a | NA | NA | NA | NA | NA | NA | 0 | 55 | 0 |
| Gata1 | Dst | NA | NA | NA | NA | NA | NA | 0 | 64 | 0 |
| Gata4 | Cyp17a1 | NA | NA | NA | NA | NA | NA | 0 | 10 | 0 |
| Gata6 | Zfp36l1 | NA | NA | NA | NA | NA | NA | 0 | 52 | 0 |
| Gcm2 | Pth | NA | NA | NA | NA | NA | NA | 0 | 1 | 0 |
| Gfi1 | Bax | NA | NA | NA | NA | NA | NA | 0 | 2 | 0 |
| Gfi1b | Plb1 | NA | NA | NA | NA | NA | NA | 0 | 28 | 0 |
| Gli1 | Egr2 | NA | NA | NA | NA | NA | NA | 0 | 7 | 0 |
| Gli2 | Efemp1 | NA | NA | NA | NA | NA | NA | 0 | 16 | 0 |
| Gli3 | Ptch2 | NA | NA | NA | NA | NA | NA | 0 | 3 | 0 |
| Grhl2 | Slco3a1 | NA | NA | NA | NA | NA | NA | 0 | 79 | 0 |
| Gtf2i | Sirt1 | NA | NA | NA | NA | NA | NA | 0 | 1 | 0 |
| Hbp1 | P3h4 | NA | NA | NA | NA | NA | NA | 0 | 38 | 0 |
| Hes1 | Snx6 | NA | NA | NA | NA | NA | NA | 0 | 6 | 0 |
| Hey1 | Kdr | NA | NA | NA | NA | NA | NA | 0 | 1 | 0 |
| Hey2 | Gata6 | NA | NA | NA | NA | NA | NA | 0 | 2 | 0 |
| Hhex | Wscd1 | NA | NA | NA | NA | NA | NA | 0 | 11 | 0 |
| Hic1 | Fgfbp1 | NA | NA | NA | NA | NA | NA | 0 | 7 | 0 |
| Hlf | Dbp | NA | NA | NA | NA | NA | NA | 0 | 2 | 0 |
| Hmbox1 | Cyth1 | NA | NA | NA | NA | NA | NA | 0 | 26 | 0 |
| Hmga1 | Cdc25a | NA | NA | NA | NA | NA | NA | 0 | 1 | 0 |
| Hnf1a | Ugt1a1 | NA | NA | NA | NA | NA | NA | 0 | 23 | 0 |
| Hnf1b | Wdr72 | NA | NA | NA | NA | NA | NA | 0 | 37 | 0 |
| Hoxa10 | Emx2 | NA | NA | NA | NA | NA | NA | 0 | 4 | 0 |
| Hoxa11 | Hoxa10 | NA | NA | NA | NA | NA | NA | 0 | 1 | 0 |
| Hoxa4 | Th | NA | NA | NA | NA | NA | NA | 0 | 1 | 0 |
| Hoxa5 | Th | NA | NA | NA | NA | NA | NA | 0 | 4 | 0 |
| Hoxa9 | Tle4 | NA | NA | NA | NA | NA | NA | 0 | 22 | 0 |
| Hoxb13 | Ctnnal1 | NA | NA | NA | NA | NA | NA | 0 | 46 | 0 |
| Hoxb3 | Otx2 | NA | NA | NA | NA | NA | NA | 0 | 1 | 0 |
| Hoxb7 | Fgf2 | NA | NA | NA | NA | NA | NA | 0 | 5 | 0 |
| Hoxc11 | Eef1a1 | NA | NA | NA | NA | NA | NA | 0 | 2 | 0 |
| Hoxc8 | Tnfrsf11b | NA | NA | NA | NA | NA | NA | 0 | 1 | 0 |
| Hoxd3 | Itgb1 | NA | NA | NA | NA | NA | NA | 0 | 3 | 0 |
| Hsf1 | Hsp90aa1 | NA | NA | NA | NA | NA | NA | 0 | 83 | 0 |
| Hsf2 | Mapk8 | NA | NA | NA | NA | NA | NA | 0 | 3 | 0 |
| Ikzf1 | Nabp1 | NA | NA | NA | NA | NA | NA | 0 | 40 | 0 |
| Insm1 | Neurod1 | NA | NA | NA | NA | NA | NA | 0 | 3 | 0 |
| Irf1 | Tap1 | NA | NA | NA | NA | NA | NA | 0 | 77 | 0 |
| Irf2 | Il15ra | NA | NA | NA | NA | NA | NA | 0 | 67 | 0 |
| Irf3 | Hgs | NA | NA | NA | NA | NA | NA | 0 | 66 | 0 |
| Irf4 | Samd4a | NA | NA | NA | NA | NA | NA | 0 | 65 | 0 |
| Irf5 | Tnf | NA | NA | NA | NA | NA | NA | 0 | 1 | 0 |
| Irf7 | Ifnb1 | NA | NA | NA | NA | NA | NA | 0 | 2 | 0 |
| Irf8 | Cd68 | NA | NA | NA | NA | NA | NA | 0 | 8 | 0 |

| | | | | | | | | | | |
|---|---|---|---|---|---|---|---|---|---|---|
| Irf9 | Runx1 | NA | NA | NA | NA | NA | NA | 0 | 46 | 0 |
| Junb | Ampd3 | NA | NA | NA | NA | NA | NA | 0 | 50 | 0 |
| Klf1 | Pkm | NA | NA | NA | NA | NA | NA | 0 | 50 | 0 |
| Klf10 | Cdkn1a | NA | NA | NA | NA | NA | NA | 0 | 1 | 0 |
| Klf11 | Pdx1 | NA | NA | NA | NA | NA | NA | 0 | 2 | 0 |
| Klf12 | Tfap2a | NA | NA | NA | NA | NA | NA | 0 | 1 | 0 |
| Klf13 | Dnajb1 | NA | NA | NA | NA | NA | NA | 0 | 31 | 0 |
| Klf15 | Lrp5 | NA | NA | NA | NA | NA | NA | 0 | 1 | 0 |
| Klf2 | Thbd | NA | NA | NA | NA | NA | NA | 0 | 2 | 0 |
| Klf3 | Cast | NA | NA | NA | NA | NA | NA | 0 | 13 | 0 |
| Klf4 | Gse1 | NA | NA | NA | NA | NA | NA | 0 | 49 | 0 |
| Klf5 | Myo1c | NA | NA | NA | NA | NA | NA | 0 | 80 | 0 |
| Klf9 | Nfic | NA | NA | NA | NA | NA | NA | 0 | 68 | 0 |
| Kmt2b | Supt5h | NA | NA | NA | NA | NA | NA | 0 | 8 | 0 |
| Lef1 | Rnf43 | NA | NA | NA | NA | NA | NA | 0 | 54 | 0 |
| Lhx2 | Sox6 | NA | NA | NA | NA | NA | NA | 0 | 10 | 0 |
| Lhx3 | Fshb | NA | NA | NA | NA | NA | NA | 0 | 3 | 0 |
| Lhx4 | Pou1f1 | NA | NA | NA | NA | NA | NA | 0 | 4 | 0 |
| Lyl1 | Dysf | NA | NA | NA | NA | NA | NA | 0 | 84 | 0 |
| Maf | Fgd3 | NA | NA | NA | NA | NA | NA | 0 | 38 | 0 |
| Mafa | Ins2 | NA | NA | NA | NA | NA | NA | 0 | 1 | 0 |
| Mafb | Adamts2 | NA | NA | NA | NA | NA | NA | 0 | 53 | 0 |
| Maff | Tiam2 | NA | NA | NA | NA | NA | NA | 0 | 43 | 0 |
| Mafg | Hdac4 | NA | NA | NA | NA | NA | NA | 0 | 37 | 0 |
| Mafk | Scn8a | NA | NA | NA | NA | NA | NA | 0 | 50 | 0 |
| Maz | Hnrnpf | NA | NA | NA | NA | NA | NA | 0 | 70 | 0 |
| Mbd1 | Dazap1 | NA | NA | NA | NA | NA | NA | 0 | 43 | 0 |
| Mbd2 | Tyk2 | NA | NA | NA | NA | NA | NA | 0 | 32 | 0 |
| Mbd3 | Esr1 | NA | NA | NA | NA | NA | NA | 0 | 1 | 0 |
| Mecp2 | Esr1 | NA | NA | NA | NA | NA | NA | 0 | 1 | 0 |
| Mef2d | Atm | NA | NA | NA | NA | NA | NA | 0 | 4 | 0 |
| Meis2 | Hk1 | NA | NA | NA | NA | NA | NA | 0 | 60 | 0 |
| Mnt | Ap3d1 | NA | NA | NA | NA | NA | NA | 0 | 56 | 0 |
| Msc | Mmp2 | NA | NA | NA | NA | NA | NA | 0 | 2 | 0 |
| Mxd1 | Odc1 | NA | NA | NA | NA | NA | NA | 0 | 1 | 0 |
| Mxd4 | Odc1 | NA | NA | NA | NA | NA | NA | 0 | 1 | 0 |
| Mxi1 | Klf10 | NA | NA | NA | NA | NA | NA | 0 | 77 | 0 |
| Myb | Vgll4 | NA | NA | NA | NA | NA | NA | 0 | 65 | 0 |
| Mybl1 | Ncl | NA | NA | NA | NA | NA | NA | 0 | 2 | 0 |
| Mybl2 | Top2a | NA | NA | NA | NA | NA | NA | 0 | 25 | 0 |
| Mycn | Clip2 | NA | NA | NA | NA | NA | NA | 0 | 64 | 0 |
| Myf6 | Acacb | NA | NA | NA | NA | NA | NA | 0 | 1 | 0 |
| Mynn | Thumpd3 | NA | NA | NA | NA | NA | NA | 0 | 4 | 0 |
| Myod1 | Fyn | NA | NA | NA | NA | NA | NA | 0 | 46 | 0 |
| Myog | Acacb | NA | NA | NA | NA | NA | NA | 0 | 1 | 0 |
| Mzf1 | Myb | NA | NA | NA | NA | NA | NA | 0 | 3 | 0 |
| Ncoa1 | Mapkap1 | NA | NA | NA | NA | NA | NA | 0 | 20 | 0 |
| Ncoa2 | Vps13b | NA | NA | NA | NA | NA | NA | 0 | 18 | 0 |
| Ncoa3 | Asap1 | NA | NA | NA | NA | NA | NA | 0 | 23 | 0 |
| Neurod1 | Scrt1 | NA | NA | NA | NA | NA | NA | 0 | 35 | 0 |
| Neurog3 | Neurod1 | NA | NA | NA | NA | NA | NA | 0 | 1 | 0 |
| Nfat5 | Tnf | NA | NA | NA | NA | NA | NA | 0 | 2 | 0 |
| Nfatc1 | Chd2 | NA | NA | NA | NA | NA | NA | 0 | 34 | 0 |
| Nfatc2 | Il4 | NA | NA | NA | NA | NA | NA | 0 | 10 | 0 |
| Nfatc3 | Rcan1 | NA | NA | NA | NA | NA | NA | 0 | 2 | 0 |
| Nfatc4 | Trpc6 | NA | NA | NA | NA | NA | NA | 0 | 1 | 0 |
| Nfe2 | Plb1 | NA | NA | NA | NA | NA | NA | 0 | 47 | 0 |
| Nfe2l1 | Sqstm1 | NA | NA | NA | NA | NA | NA | 0 | 30 | 0 |
| Nfe2l2 | Fhad1 | NA | NA | NA | NA | NA | NA | 0 | 72 | 0 |
| Nfia | Tp53 | NA | NA | NA | NA | NA | NA | 0 | 3 | 0 |
| Nfil3 | Il1b | NA | NA | NA | NA | NA | NA | 0 | 1 | 0 |
| Nfyb | Rhof | NA | NA | NA | NA | NA | NA | 0 | 76 | 0 |
| Nfyc | P04224 | NA | NA | NA | NA | NA | NA | 0 | 3 | 0 |
| Nkx2-1 | Sftpb | NA | NA | NA | NA | NA | NA | 0 | 1 | 0 |
| Nkx2-5 | Gata4 | NA | NA | NA | NA | NA | NA | 0 | 2 | 0 |
| Nme2 | Myc | NA | NA | NA | NA | NA | NA | 0 | 1 | 0 |
| Npas2 | Per2 | NA | NA | NA | NA | NA | NA | 0 | 4 | 0 |
| Nr0b1 | Nr5a1 | NA | NA | NA | NA | NA | NA | 0 | 2 | 0 |
| Nr1d1 | Arntl | NA | NA | NA | NA | NA | NA | 0 | 1 | 0 |
| Nr1h2 | Cdc42ep4 | NA | NA | NA | NA | NA | NA | 0 | 13 | 0 |
| Nr1h3 | Abcd1 | NA | NA | NA | NA | NA | NA | 0 | 12 | 0 |
| Nr1h4 | Nr1i2 | NA | NA | NA | NA | NA | NA | 0 | 7 | 0 |
| Nr1i2 | Ttc21a | NA | NA | NA | NA | NA | NA | 0 | 4 | 0 |
| Nr1i3 | Ugt1a1 | NA | NA | NA | NA | NA | NA | 0 | 1 | 0 |
| Nr2c2 | Ankib1 | NA | NA | NA | NA | NA | NA | 0 | 84 | 0 |
| Nr2e1 | Pax2 | NA | NA | NA | NA | NA | NA | 0 | 3 | 0 |
| Nr2f1 | Tnk2 | NA | NA | NA | NA | NA | NA | 0 | 66 | 0 |
| Nr2f6 | Plcd3 | NA | NA | NA | NA | NA | NA | 0 | 32 | 0 |
| Nr3c2 | Scnn1a | NA | NA | NA | NA | NA | NA | 0 | 2 | 0 |
| Nr4a1 | Rnf220 | NA | NA | NA | NA | NA | NA | 0 | 41 | 0 |
| Nr5a1 | Cyp19a1 | NA | NA | NA | NA | NA | NA | 0 | 12 | 0 |
| Nr5a2 | Ppm1d | NA | NA | NA | NA | NA | NA | 0 | 31 | 0 |
| Nrf1 | Pkib | NA | NA | NA | NA | NA | NA | 0 | 77 | 0 |
| Nrif1 | Zfp397 | NA | NA | NA | NA | NA | NA | 0 | 9 | 0 |
| Nrl | Rho | NA | NA | NA | NA | NA | NA | 0 | 1 | 0 |
| Onecut1 | Rapgef4 | NA | NA | NA | NA | NA | NA | 0 | 52 | 0 |
| Onecut2 | Sytl4 | NA | NA | NA | NA | NA | NA | 0 | 1 | 0 |
| Otx2 | Pde1a | NA | NA | NA | NA | NA | NA | 0 | 23 | 0 |
| Pax1 | Pdgfra | NA | NA | NA | NA | NA | NA | 0 | 1 | 0 |
| Pax2 | Tp53 | NA | NA | NA | NA | NA | NA | 0 | 2 | 0 |
| Pax3 | Mitf | NA | NA | NA | NA | NA | NA | 0 | 3 | 0 |
| Pax4 | Ins2 | NA | NA | NA | NA | NA | NA | 0 | 1 | 0 |
| Pax5 | Cd22 | NA | NA | NA | NA | NA | NA | 0 | 53 | 0 |
| Pax8 | Wt1 | NA | NA | NA | NA | NA | NA | 0 | 11 | 0 |
| Pbx1 | Cyp17a1 | NA | NA | NA | NA | NA | NA | 0 | 4 | 0 |
| Pbx2 | Sox6 | NA | NA | NA | NA | NA | NA | 0 | 44 | 0 |
| Pbx3 | Galnt2 | NA | NA | NA | NA | NA | NA | 0 | 51 | 0 |
| Pcgf2 | Hoxa7 | NA | NA | NA | NA | NA | NA | 0 | 1 | 0 |
| Pgr | Klk1b24 | NA | NA | NA | NA | NA | NA | 0 | 89 | 0 |
| Pitx1 | Fshb | NA | NA | NA | NA | NA | NA | 0 | 1 | 0 |
| Pitx2 | Fshb | NA | NA | NA | NA | NA | NA | 0 | 1 | 0 |
| Pknox1 | Astn2 | NA | NA | NA | NA | NA | NA | 0 | 35 | 0 |
| Plag1 | Abcc6 | NA | NA | NA | NA | NA | NA | 0 | 1 | 0 |
| Pou1f1 | Ghr | NA | NA | NA | NA | NA | NA | 0 | 1 | 0 |
| Pou2f1 | Tnrc6c | NA | NA | NA | NA | NA | NA | 0 | 61 | 0 |
| Pou2f2 | Myh9 | NA | NA | NA | NA | NA | NA | 0 | 86 | 0 |
| Pou2f3 | Sprr2d | NA | NA | NA | NA | NA | NA | 0 | 1 | 0 |
| Pou3f1 | Gnrh1 | NA | NA | NA | NA | NA | NA | 0 | 1 | 0 |
| Pou3f2 | Kcnn3 | NA | NA | NA | NA | NA | NA | 0 | 2 | 0 |
| Pou4f2 | Kcnip4 | NA | NA | NA | NA | NA | NA | 0 | 28 | 0 |
| Pou5f1 | Tnrc6b | NA | NA | NA | NA | NA | NA | 0 | 72 | 0 |
| Ppara | Apoa1 | NA | NA | NA | NA | NA | NA | 0 | 37 | 0 |
| Ppard | Cpt1b | NA | NA | NA | NA | NA | NA | 0 | 4 | 0 |
| Prdm1 | Ifit2 | NA | NA | NA | NA | NA | NA | 0 | 40 | 0 |
| Pura | Ar | NA | NA | NA | NA | NA | NA | 0 | 1 | 0 |
| Rarb | Rarg | NA | NA | NA | NA | NA | NA | 0 | 5 | 0 |
| Rarg | Rarb | NA | NA | NA | NA | NA | NA | 0 | 8 | 0 |
| Rel | Icam1 | NA | NA | NA | NA | NA | NA | 0 | 23 | 0 |
| Rfx1 | Fgf1 | NA | NA | NA | NA | NA | NA | 0 | 40 | 0 |
| Rfx5 | Iqcg | NA | NA | NA | NA | NA | NA | 0 | 80 | 0 |
| Rora | Tuba1b | NA | NA | NA | NA | NA | NA | 0 | 3 | 0 |
| Rorc | Il22b | NA | NA | NA | NA | NA | NA | 0 | 3 | 0 |
| Rreb1 | Klk1b16 | NA | NA | NA | NA | NA | NA | 0 | 14 | 0 |
| Runx2 | Dock10 | NA | NA | NA | NA | NA | NA | 0 | 65 | 0 |
| Rxrb | Angptl4 | NA | NA | NA | NA | NA | NA | 0 | 6 | 0 |
| Rxrg | Plin5 | NA | NA | NA | NA | NA | NA | 0 | 6 | 0 |
| Sall4 | Zeb1 | NA | NA | NA | NA | NA | NA | 0 | 3 | 0 |
| Satb1 | Tmem245 | NA | NA | NA | NA | NA | NA | 0 | 9 | 0 |
| Six2 | Itpr1 | NA | NA | NA | NA | NA | NA | 0 | 39 | 0 |
| Six5 | Tcea2 | NA | NA | NA | NA | NA | NA | 0 | 23 | 0 |
| Ski | Smad7 | NA | NA | NA | NA | NA | NA | 0 | 1 | 0 |
| Skil | Smad7 | NA | NA | NA | NA | NA | NA | 0 | 1 | 0 |
| Smad1 | Tnfrsf11b | NA | NA | NA | NA | NA | NA | 0 | 17 | 0 |
| Smad5 | Sp3 | NA | NA | NA | NA | NA | NA | 0 | 10 | 0 |
| Snai1 | Hbegf | NA | NA | NA | NA | NA | NA | 0 | 5 | 0 |
| Snai2 | Ank1 | NA | NA | NA | NA | NA | NA | 0 | 66 | 0 |
| Snapc4 | Pask | NA | NA | NA | NA | NA | NA | 0 | 29 | 0 |
| Sox11 | Col28a1 | NA | NA | NA | NA | NA | NA | 0 | 29 | 0 |
| Sox13 | Cbx2 | NA | NA | NA | NA | NA | NA | 0 | 45 | 0 |
| Sox4 | Ncam1 | NA | NA | NA | NA | NA | NA | 0 | 3 | 0 |
| Sox6 | Zeb2 | NA | NA | NA | NA | NA | NA | 0 | 32 | 0 |
| Sox8 | Cldn23 | NA | NA | NA | NA | NA | NA | 0 | 1 | 0 |
| Sox9 | Ceacam2 | NA | NA | NA | NA | NA | NA | 0 | 74 | 0 |
| Sp2 | Usp16 | NA | NA | NA | NA | NA | NA | 0 | 22 | 0 |
| Sp3 | Klk1b9 | NA | NA | NA | NA | NA | NA | 0 | 91 | 0 |
| Sp4 | Thra | NA | NA | NA | NA | NA | NA | 0 | 59 | 0 |
| Spdef | Agr2 | NA | NA | NA | NA | NA | NA | 0 | 1 | 0 |
| Spib | Gmds | NA | NA | NA | NA | NA | NA | 0 | 58 | 0 |
| Srebf1 | Lrp1 | NA | NA | NA | NA | NA | NA | 0 | 71 | 0 |
| Srebf2 | Pcsk9 | NA | NA | NA | NA | NA | NA | 0 | 90 | 0 |
| Stat2 | Zfp37 | NA | NA | NA | NA | NA | NA | 0 | 80 | 0 |
| Stat4 | Socs3 | NA | NA | NA | NA | NA | NA | 0 | 13 | 0 |
| Stat5a | Dennd3 | NA | NA | NA | NA | NA | NA | 0 | 66 | 0 |
| Stat5b | Svil | NA | NA | NA | NA | NA | NA | 0 | 78 | 0 |
| Stat6 | Itga2b | NA | NA | NA | NA | NA | NA | 0 | 13 | 0 |
| Tbp | Prrc2b | NA | NA | NA | NA | NA | NA | 0 | 46 | 0 |
| Tbx2 | Cdkn2a | NA | NA | NA | NA | NA | NA | 0 | 1 | 0 |
| Tbx21 | Inpp5d | NA | NA | NA | NA | NA | NA | 0 | 76 | 0 |
| Tbx5 | Nppa | NA | NA | NA | NA | NA | NA | 0 | 2 | 0 |
| Tcf12 | Sertad2 | NA | NA | NA | NA | NA | NA | 0 | 78 | 0 |
| Tcf3 | Med15 | NA | NA | NA | NA | NA | NA | 0 | 125 | 0 |
| Tcf7 | Pde7a | NA | NA | NA | NA | NA | NA | 0 | 58 | 0 |
| Tcf7l1 | Nkd1 | NA | NA | NA | NA | NA | NA | 0 | 5 | 0 |
| Tcfl5 | Cd2 | NA | NA | NA | NA | NA | NA | 0 | 1 | 0 |
| Tead2 | Plekha4 | NA | NA | NA | NA | NA | NA | 0 | 29 | 0 |
| Tfap2b | Ptgds | NA | NA | NA | NA | NA | NA | 0 | 2 | 0 |
| Tfap4 | Pemt | NA | NA | NA | NA | NA | NA | 0 | 40 | 0 |
| Tfcp2 | Tf | NA | NA | NA | NA | NA | NA | 0 | 1 | 0 |
| Tfdp1 | Tlcd1 | NA | NA | NA | NA | NA | NA | 0 | 69 | 0 |
| Thap1 | Znf276 | NA | NA | NA | NA | NA | NA | 0 | 55 | 0 |
| Thap11 | Wbp2 | NA | NA | NA | NA | NA | NA | 0 | 114 | 0 |
| Thra | Cga | NA | NA | NA | NA | NA | NA | 0 | 1 | 0 |
| Thrb | Myh6 | NA | NA | NA | NA | NA | NA | 0 | 4 | 0 |
| Tlx1 | Aldh1a1 | NA | NA | NA | NA | NA | NA | 0 | 1 | 0 |
| Tp73 | Syne1 | NA | NA | NA | NA | NA | NA | 0 | 41 | 0 |
| Trps1 | Runx2 | NA | NA | NA | NA | NA | NA | 0 | 1 | 0 |
| Twist2 | Mmp2 | NA | NA | NA | NA | NA | NA | 0 | 3 | 0 |
| Xbp1 | Cacnb2 | NA | NA | NA | NA | NA | NA | 0 | 16 | 0 |
| Ybx1 | Rbbp8 | NA | NA | NA | NA | NA | NA | 0 | 6 | 0 |
| Yy1 | Hsf1 | NA | NA | NA | NA | NA | NA | 0 | 99 | 0 |
| Zbtb11 | Uspl1 | NA | NA | NA | NA | NA | NA | 0 | 22 | 0 |
| Zbtb16 | Ccna2 | NA | NA | NA | NA | NA | NA | 0 | 1 | 0 |
| Zbtb17 | Cdkn2b | NA | NA | NA | NA | NA | NA | 0 | 1 | 0 |
| Zbtb33 | Lrsam1 | NA | NA | NA | NA | NA | NA | 0 | 88 | 0 |
| Zbtb7a | Comt | NA | NA | NA | NA | NA | NA | 0 | 90 | 0 |
| Zc3h8 | Mdm4 | NA | NA | NA | NA | NA | NA | 0 | 6 | 0 |
| Zeb1 | Col4a3 | NA | NA | NA | NA | NA | NA | 0 | 96 | 0 |
| Zfp202 | Abca1 | NA | NA | NA | NA | NA | NA | 0 | 2 | 0 |
| Zfp217 | Arid5b | NA | NA | NA | NA | NA | NA | 0 | 10 | 0 |
| Zfp384 | Ankrd11 | NA | NA | NA | NA | NA | NA | 0 | 65 | 0 |
| Zfp644 | Epc1 | NA | NA | NA | NA | NA | NA | 0 | 7 | 0 |
| Zfx | Zfp398 | NA | NA | NA | NA | NA | NA | 0 | 54 | 0 |
| Zhx1 | Chd9 | NA | NA | NA | NA | NA | NA | 0 | 9 | 0 |
| Zhx2 | Ndrg1 | NA | NA | NA | NA | NA | NA | 0 | 8 | 0 |
| Zic1 | Apoe | NA | NA | NA | NA | NA | NA | 0 | 1 | 0 |
| Zic2 | Apoe | NA | NA | NA | NA | NA | NA | 0 | 2 | 0 |
| Znf143 | Sptbn1 | NA | NA | NA | NA | NA | NA | 0 | 51 | 0 |
| Znf148 | Irs2 | NA | NA | NA | NA | NA | NA | 0 | 2 | 0 |
| Znf24 | Tbxas1 | NA | NA | NA | NA | NA | NA | 0 | 18 | 0 |
| Znf382 | Hmga2 | NA | NA | NA | NA | NA | NA | 0 | 1 | 0 |
| Znf592 | Pcif1 | NA | NA | NA | NA | NA | NA | 0 | 14 | 0 |
| Znf639 | Ahctf1 | NA | NA | NA | NA | NA | NA | 0 | 28 | 0 |
| Znf740 | Pdgfa | NA | NA | NA | NA | NA | NA | 0 | 36 | 0 |

| TF | gene | GeneRatio | BgRatio | pvalue | p.adjust | qvalue | DetectedTargets | Count | TF_targets | Percentage_of_TFtargets_detected |
|---|---|---|---|---|---|---|---|---|---|---|
| E2f4 | Irf2bp2 | 99/428 | 695/6837 | 4.13E-16 | 9.09E-14 | 9.09E-14 | Abca7/Agpat3/Anln/Asf1b/Aspm/Atad2/Aurkb/Birc5/Blm/Brca1/Brip1/Ccna2/Ccnb1/Ccnd1/Ccnf/Cdc25c/Cdc45/Cdc7/Cdca2/Cdk1/Cenpa/Cenpe/Cenpm/Cep57/Cic/Cit/Clspn/Creg1/Dbf4/Ddx39a/Depdc1b/Dlgap5/E2f1/Elk1/Exo1/Fanca/Fasn/Fdft1/Fen1/Foxm1/Gen1/H2ax/H2bc14/H3c11/Hmmr/Id3/Ier5/Ing1/Iqgap3/Katnal1/Kd | 99 | 743 | 13.32 |
| Foxm1 | Brca2 | 29/428 | 83/6837 | 4.18E-15 | 4.60E-13 | 4.60E-13 | Anln/Aspm/Aurkb/Birc5/Brip1/Ccnb1/Ccnd1/Cdc25c/Cdk1/Cenpa/Cenpf/Cit/Cks1b/Dtnb/Ect2/H2ac11/H2ac13/H2ac4/H2ac7/H2ac8/Hmmr/Kif11/Kif14/Kif23/Ncapd2/Ncaph/Prr11/Skp2/Ube2s | 29 | 92 | 31.52 |
| Nfyb | Bfsp1 | 12/428 | 73/6837 | 0.0017352 | 0.12724798 | 0.12724798 | Brsk2/Ccnb1/Ccnb2/Cdk1/Eif2ak3/Fads1/Fdft1/Fzd2/Gna12/Ier5/Rhof/Tbc1d31 | 12 | 76 | 15.79 |
| Cebpd | Srcin1 | 10/428 | 61/6837 | 0.00420275 | 0.23115102 | 0.23115102 | Atf7ip2/Bcl3/Ccm2/Ccnd1/Exoc3l4/Fads1/Gadd45g/Gramd1a/Rrbp1/Tgm2 | 10 | 71 | 14.08 |
| E2f1 | Top2a | 50/428 | 576/6837 | 0.00999926 | 0.37317281 | 0.37317281 | Asf1b/Atad2/Aurka/B9d1/Birc5/Blm/Blvra/Brca1/Ccna2/Ccnd1/Cdc45/Cdk1/Cdkn1c/Chka/Dbf4/Ect2/Etv4/Fanca/Fancd2/Fndc3b/Foxm1/Id3/Ift80/Jmjd1c/Kif18a/Kif23/Kif24/Kif2c/Kpnb1/Mcm10/Mcm8/Mical3/Ncapg2/Ndc80/Nek2/Nr4a2/Pdk4/Per1/Phf13/Plk4/Ptpro/Racgap1/Rad18/Rad51/Sertad3/Skp2/Smc4/Snn/Tk1/Top2a | 50 | 675 | 7.41 |
| E2f2 | Plk4 | 5/428 | 22/6837 | 0.01017744 | 0.37317281 | 0.37317281 | Asf1b/E2f1/Foxm1/Mcm10/Plk4 | 5 | 22 | 22.73 |
| Myc | Notch4 | 36/428 | 403/6837 | 0.01821452 | 0.57245645 | 0.57245645 | Agpat3/Ampd2/Apex1/Atf4/Bcl3/Brca1/Ccna2/Ccnb1/Ccnd1/Cdc25c/E2f1/Eif5a2/Elk1/Entpd7/Fam13a/Fasn/Foxm1/Fut1/Id3/Ier5/Kdm4b/Khsrp/Mst1/Ncl/Nme2/Per1/Prkcz/Qdpr/Rad51/Rgs2/Slc25a28/Slc39a7/Srm/Tnfrsf8/Uba52/Ube2c | 36 | 447 | 8.05 |
| Smad5 | Dmtf1 | 3/428 | Oct-37 | 0.02100291 | 0.57758015 | 0.57758015 | Id3/Rhobtb3/Smc4 | 3 | 10 | 30 |
| E2f3 | Tp73 | 3/428 | Nov-37 | 0.02756273 | 0.64870707 | 0.64870707 | Aurka/Ccnb1/E2f1 | 3 | 14 | 21.43 |
| Irf3 | Tmem160 | 8/428 | 59/6837 | 0.02948669 | 0.64870707 | 0.64870707 | Abca7/Cdca3/Cnot3/Ddx39a/Slc25a28/Tcf3/Tyk2/Xrcc3 | 8 | 66 | 12.12 |
| Ahr | Dock4 | 1/428 | 35/6837 | 0.89652596 | 0.99638415 | 0.99638415 | Trim37 | 1 | 43 | 2.33 |
| Ar | Rbpms | 6/428 | 187/6837 | 0.97995328 | 0.99638415 | 0.99638415 | Cdkl5/Fam13a/Hk2/Kdm4b/Ptprm/Tacc1 | 6 | 222 | 2.7 |
| Arid2 | Bbx | 1/428 | 29/6837 | 0.84721214 | 0.99638415 | 0.99638415 | Jmjd1c | 1 | 30 | 3.33 |
| Arid3a | Stk10 | 1/428 | 20/6837 | 0.72604056 | 0.99638415 | 0.99638415 | Cic | 1 | 23 | 4.35 |
| Arnt | Mid1 | 1/428 | 54/6837 | 0.96995099 | 0.99638415 | 0.99638415 | Fndc3b | 1 | 63 | 1.59 |
| Arntl | Cerk | 1/428 | 17/6837 | 0.66723175 | 0.99638415 | 0.99638415 | Per1 | 1 | 20 | 5 |
| Ascl1 | Arhgef10l | 1/428 | 46/6837 | 0.94940296 | 0.99638415 | 0.99638415 | Fam13a | 1 | 54 | 1.85 |
| Atf1 | Stat3 | 8/428 | 80/6837 | 0.12612177 | 0.99638415 | 0.99638415 | Apex1/Brca1/Nsd2/Per1/Slc20a1/Snd1/Top2a/Tspan31 | 8 | 91 | 8.79 |
| Atf2 | Msi2 | 5/428 | 52/6837 | 0.22423515 | 0.99638415 | 0.99638415 | Ccnd1/Fndc3b/Nr4a2/Slc39a10/Snd1 | 5 | 62 | 8.06 |
| Atf3 | Klf6 | 5/428 | 48/6837 | 0.17935312 | 0.99638415 | 0.99638415 | Asns/Gna12/Per1/Rcan1/Snd1 | 5 | 50 | 10 |
| Atf4 | Vldlr | 8/428 | 88/6837 | 0.18426167 | 0.99638415 | 0.99638415 | Asns/Cbx4/Fut1/Got1/Hspb6/Nupr1/Pycr1/Slc6a9 | 8 | 110 | 7.27 |
| Bach1 | Coro1c | 7/428 | 79/6837 | 0.22386712 | 0.99638415 | 0.99638415 | Bicdl1/Chsy1/Fndc3b/Klhl21/Msantd3/Prr5/Tnfaip2 | 7 | 84 | 8.33 |
| Bach2 | Daam1 | 3/428 | 44/6837 | 0.52606432 | 0.99638415 | 0.99638415 | Jmjd1c/Klhl21/Slc16a6 | 3 | 49 | 6.12 |
| Batf | Pik3ap1 | 4/428 | 44/6837 | 0.29583974 | 0.99638415 | 0.99638415 | Apbb1ip/Ier5/Nabp1/Tnfrsf8 | 4 | 49 | 8.16 |
| Bcl11a | Klhl6 | 1/428 | 29/6837 | 0.84721214 | 0.99638415 | 0.99638415 | Dtnb | 1 | 34 | 2.94 |
| Bcl6 | Eipr1 | 1/428 | 42/6837 | 0.93435989 | 0.99638415 | 0.99638415 | Tcf3 | 1 | 46 | 2.17 |
| Bhlhe22 | Cdyl | 2/428 | 17/6837 | 0.28850161 | 0.99638415 | 0.99638415 | Ap3d1/Pcdhgc3 | 2 | 21 | 9.52 |
| Bhlhe40 | Xxylt1 | 5/428 | 53/6837 | 0.23595042 | 0.99638415 | 0.99638415 | Bcl3/Gna12/Jmjd1c/Mical3/Slc7a5 | 5 | 57 | 8.77 |
| Cdx2 | Vav3 | 1/428 | 49/6837 | 0.95838051 | 0.99638415 | 0.99638415 | Svil | 1 | 52 | 1.92 |
| Cebpa | Ak4 | 51/428 | 788/6837 | 0.42076142 | 0.99638415 | 0.99638415 | Acox2/Agpat3/Apbb1ip/Atf7ip2/Ccnb1/Ccnd1/Cdkl5/Dapk3/Ddx39a/Efhc1/Eml6/Exoc3l4/Fads1/Fam13a/Fanci/Fas/Fasn/Fndc3b/Gadd45b/Galnt2/Gna12/Gpc6/Gramd1a/Heatr5a/Jmjd1c/Kdm4b/Lbp/Mad2l1/Nabp1/Pacsin2/Paqr8/Pla2g5/Pom121/Rapgef2/Rcan1/Rhobtb3/Scd1/Sntb1/Svil/Tacc1/Tbc1d1/Tgm2/Tiam2/Tk1/Tle1/ | 51 | 959 | 5.32 |
| Cebpb | Bcr | 2/428 | 62/6837 | 0.90761384 | 0.99638415 | 0.99638415 | Asns/Gadd45g | 2 | 85 | 2.35 |
| Cebpg | Trib3 | 1/428 | 41/6837 | 0.92994814 | 0.99638415 | 0.99638415 | Tbc1d31 | 1 | 46 | 2.17 |
| Clock | Nr1d2 | 3/428 | 41/6837 | 0.47818143 | 0.99638415 | 0.99638415 | Per1/Smoc2/Sptbn4 | 3 | 51 | 5.88 |
| Creb1 | Msi2 | 7/428 | 74/6837 | 0.17920111 | 0.99638415 | 0.99638415 | Brca1/Ccnd1/Clcn3/Nr4a2/Per1/Slc20a1/Snd1 | 7 | 117 | 5.98 |
| Creb3 | Dtnbp1 | 1/428 | 24/6837 | 0.78864184 | 0.99638415 | 0.99638415 | Galnt2 | 1 | 25 | 4 |
| Creb3l1 | Asph | 2/428 | 17/6837 | 0.28850161 | 0.99638415 | 0.99638415 | Asph/Fndc3b | 2 | 20 | 10 |
| Crem | Ern1 | 4/428 | 44/6837 | 0.29583974 | 0.99638415 | 0.99638415 | Ccna2/Nr4a2/Prkcz/Snd1 | 4 | 50 | 8 |
| Ctcf | Arl8b | 36/428 | 729/6837 | 0.95306482 | 0.99638415 | 0.99638415 | Ackr3/Alkbh2/Arhgap39/Bmper/Cbx4/Ccnd1/Dync1i1/Ect2/Efhc1/Eml1/Fam13a/Fam3c/Fam98c/Fndc3b/Gadd45g/Haus7/Hebp2/Kdm4b/Lrrc59/Mapk8ip3/Mcm10/Mical3/Nkain1/Nos2/Nsd2/Pacsin2/Prkcz/Prss27/Ptprm/Rrbp1/Sptbn4/Svil/Tacc1/Tbc1d1/Tiam2/Zc3h7a | 36 | 936 | 3.85 |
| Ctcfl | Gad1 | 3/428 | 39/6837 | 0.44504548 | 0.99638415 | 0.99638415 | Cbx4/Mapk8ip3/Svil | 3 | 53 | 5.66 |
| Cux1 | Fto | 5/428 | 37/6837 | 0.07860863 | 0.99638415 | 0.99638415 | Depdc1b/Ect2/Etv4/Racgap1/Tbc1d1 | 5 | 40 | 12.5 |
| E2f6 | Piezo1 | 3/428 | 59/6837 | 0.72394802 | 0.99638415 | 0.99638415 | Brca1/E2f1/Tcf3 | 3 | 67 | 4.48 |
| E2f7 | Cd44 | 1/428 | 27/6837 | 0.82603025 | 0.99638415 | 0.99638415 | E2f1 | 1 | 29 | 3.45 |
| Ebf1 | Ankrd44 | 5/428 | 67/6837 | 0.41035765 | 0.99638415 | 0.99638415 | Ackr3/Pacsin2/Tacc1/Tcf3/Tnfrsf8 | 5 | 74 | 6.76 |
| Egr1 | Capzb | 53/428 | 819/6837 | 0.41858212 | 0.99638415 | 0.99638415 | Agpat3/Anxa5/Ap3d1/Apex1/Atad2/Bicdl1/Cat/Cbx4/Ccm2/Ccnd1/Chtf18/Cic/Cpped1/Cyld/Dmwd/Elk1/Etv4/Fam118a/Fam13a/Fas/Fasn/Fzd2/Gadd45b/Hes2/Igf2bp3/Igfbp4/Incenp/Kdelr3/Kdm4b/Klhl21/Kpnb1/Mical3/Mov10/Msantd3/Nr4a2/P3h1/Pacsin2/Pdgfc/Per1/Prkaca/Prkcz/Ptov1/Ret/Rhof/Setx/Sptbn4/Tbc1d10a/Tcf | 53 | 976 | 5.43 |
| Elf1 | Lsm4 | 2/428 | 47/6837 | 0.80271551 | 0.99638415 | 0.99638415 | Fndc3b/Tacc1 | 2 | 50 | 4 |
| Elf3 | Septin9 | 1/428 | 49/6837 | 0.95838051 | 0.99638415 | 0.99638415 | Hk2 | 1 | 52 | 1.92 |
| Elk1 | Ier2 | 3/428 | 106/6837 | 0.96620317 | 0.99638415 | 0.99638415 | Arid4a/Cdc45/Mapk8ip3 | 3 | 113 | 2.65 |
| Epas1 | Pdgfb | 6/428 | 66/6837 | 0.23036794 | 0.99638415 | 0.99638415 | Chsy1/Dnah17/Fam13a/Slc7a5/Tgm2/Tle1 | 6 | 80 | 7.5 |
| Erg | Ptk2 | 4/428 | 78/6837 | 0.72864266 | 0.99638415 | 0.99638415 | Amotl1/Chsy1/Pacsin2/Ptprm | 4 | 89 | 4.49 |
| Esr1 | Bmper | 34/428 | 579/6837 | 0.68282623 | 0.99638415 | 0.99638415 | Amotl1/Atp2b1/Bcl3/Blm/Bmper/Brip1/Camta1/Ccnd1/Cdk18/Cep112/Cul7/E2f1/Fbln1/Fndc3b/Frmd5/Gadd45g/Hk2/Hs6st1/Igfbp4/Kdm4b/Mctp2/Nkain1/Otud6b/Pacsin2/Ret/Rhobtb3/Rrbp1/Sec24d/Snd1/Svil/Tacc1/Tbc1d1/Tle1/Trim37 | 34 | 722 | 4.71 |
| Esrra | Synpo | 2/428 | 64/6837 | 0.91678674 | 0.99638415 | 0.99638415 | Bcl3/Tgm2 | 2 | 69 | 2.9 |
| Ets1 | Gpaa1 | 54/428 | 837/6837 | 0.42688423 | 0.99638415 | 0.99638415 | Abi2/Ampd2/Arid4a/Atf4/Bcl3/Ccnd1/Cdc45/Cic/Cip2a/Csgalnact2/Dapk3/Depdc1b/Dusp6/Efcab11/Efhc1/Eme1/Etv4/Fas/Fasn/Fndc3b/Gadd45b/Gna12/Gramd1a/Id3/Ier5/Igf2bp3/Ing1/Jmjd1c/Kif4/Kifc1/Klhl21/Kpnb1/Mical3/Mtmr14/Nr4a2/Per1/Pex2/Pf4/Prkaca/Rad51/Rhof/Sertad3/Svil/Tacc1/Taf13/Tfcp2/Tnfrsf8/Trim35/T | 54 | 934 | 5.78 |
| Ets2 | Anpep | 2/428 | 17/6837 | 0.28850161 | 0.99638415 | 0.99638415 | Brca1/Cdk1 | 2 | 31 | 6.45 |
| Etv1 | Birc6 | 2/428 | 27/6837 | 0.5110695 | 0.99638415 | 0.99638415 | Cdk18/Dusp6 | 2 | 29 | 6.9 |
| Etv4 | Trib3 | 2/428 | 55/6837 | 0.86756376 | 0.99638415 | 0.99638415 | Ap3d1/Pacsin2 | 2 | 59 | 3.39 |
| Fos | Tnip1 | 38/428 | 686/6837 | 0.81630408 | 0.99638415 | 0.99638415 | Ccnd1/Cdca2/Cenpm/Cep112/Cfp/Col7a1/Dtnb/Efhc1/Engase/F2r/Fam13a/Fas/Fasn/Fndc3b/Fut2/Galnt2/Gna12/Heph/Hk2/Ints11/Map7d1/Msantd3/Nabp1/Nos2/Pacsin2/Pi4k2a/Prr5/Ptprm/Rnft2/Snd1/Sptbn5/Svil/Tbc1d1/Tbc1d31/Tent5b/Tiam2/Ticrr/Zc3h7a | 38 | 847 | 4.49 |
| Fosl2 | Pfkfb3 | 6/428 | 72/6837 | 0.29495484 | 0.99638415 | 0.99638415 | Brca1/Etv4/Nos2/Slc7a5/Tgm2/Tiam2 | 6 | 79 | 7.59 |
| Foxa1 | Luzp2 | 40/428 | 674/6837 | 0.66825878 | 0.99638415 | 0.99638415 | Amotl1/Bmper/Brca1/Brip1/Cdkl5/Cenpi/Cep112/Dpagt1/Fam13a/Fdft1/Fndc3b/H2ac11/H2ac13/H2ac4/H2ac7/H2ac8/Ica1/Il6st/Jmjd1c/Kdm6a/Kif21a/Mgat4c/Nbas/Nup210l/Papss1/Pbx3/Pcdh17/Pspc1/Ptprm/Ret/Snd1/Spag17/Svil/Tbc1d1/Tiam2/Tle1/Tmem98/Ttc7b/Uap1/Zc3h12c | 40 | 851 | 4.7 |
| Foxa2 | Mitf | 4/428 | 42/6837 | 0.2673215 | 0.99638415 | 0.99638415 | Fam13a/Fndc3b/Svil/Tbc1d1 | 4 | 49 | 8.16 |
| Foxj2 | Fam81b | 3/428 | 30/6837 | 0.28905432 | 0.99638415 | 0.99638415 | Arid4a/Fam13a/Per1 | 3 | 36 | 8.33 |
| Foxk2 | Ankrd11 | 3/428 | 49/6837 | 0.60012969 | 0.99638415 | 0.99638415 | Atf7ip2/Fdft1/Snap23 | 3 | 50 | 6 |
| Foxo1 | Negr1 | 8/428 | 105/6837 | 0.33526 | 0.99638415 | 0.99638415 | Ccnb1/Ccnb2/Ccnd1/Cdkn1c/Fam13a/Pdk4/Ptprm/Tacc1 | 8 | 122 | 6.56 |
| Foxo3 | Ar | 3/428 | 35/6837 | 0.37656605 | 0.99638415 | 0.99638415 | Cat/Ccnd1/Dusp6 | 3 | 40 | 7.5 |
| Gabpa | Ccdc171 | 1/428 | 52/6837 | 0.9657682 | 0.99638415 | 0.99638415 | Brca1 | 1 | 55 | 1.82 |
| Gata1 | Mxi1 | 6/428 | 54/6837 | 0.1195116 | 0.99638415 | 0.99638415 | Amotl1/Galnt2/Gna12/Pacsin2/Pf4/Ppox | 6 | 64 | 9.38 |
| Gata2 | Aff3 | 26/428 | 542/6837 | 0.94469324 | 0.99638415 | 0.99638415 | Adamts18/Amotl1/Apex1/Ccm2/Chsy1/Cntn3/Dtnb/Efhc1/Eml1/Fam13a/Fdft1/Fignl1/Fndc3b/Frmd5/Galnt2/Gna12/Jmjd1c/Kdm6a/Nbas/Pacsin2/Ptprm/Rapgef2/Sec24d/Snd1/Svil/Tiam2 | 26 | 676 | 3.85 |
| Gata3 | Ap1m1 | 34/428 | 596/6837 | 0.74618425 | 0.99638415 | 0.99638415 | Amotl1/Arnt2/Atp2b1/Bmper/Brip1/Cenpp/Cep112/Dtnb/Efhc1/Eml1/Etv4/Fam13a/Fdft1/Fndc3b/Frmd5/Galnt2/Gramd1a/Ier5/Kif21a/Mical3/Nadk2/Pacsin2/Pbx3/Pde7b/Rhobtb3/Sec24d/Snd1/Svil/Tacc1/Tbc1d1/Tle1/Tnfrsf8/Trim37/Unc13b | 34 | 760 | 4.47 |
| Gata6 | Krt18 | 4/428 | 49/6837 | 0.36829365 | 0.99638415 | 0.99638415 | Atp2b1/Galnt2/Svil/Tbc1d1 | 4 | 52 | 7.69 |
| Gfi1b | Ahcyl2 | 1/428 | 24/6837 | 0.78864184 | 0.99638415 | 0.99638415 | Snd1 | 1 | 28 | 3.57 |
| Grhl2 | Spidr | 4/428 | 67/6837 | 0.61177336 | 0.99638415 | 0.99638415 | Fat2/Pacsin2/Slc35f2/Tiam2 | 4 | 79 | 5.06 |
| Hbp1 | Ercc1 | 2/428 | 38/6837 | 0.69753944 | 0.99638415 | 0.99638415 | Atad2/Jmjd1c | 2 | 38 | 5.26 |
| Hif1a | Bach1 | 13/428 | 208/6837 | 0.54451496 | 0.99638415 | 0.99638415 | Aurka/Birc5/Brsk2/Ccnd1/Chsy1/Fndc3b/Foxm1/Gys1/Hk2/Kdm4b/Nme2/Nos2/Stc2 | 13 | 269 | 4.83 |
| Hinfp | C2cd2l | 2/428 | 16/6837 | 0.26469727 | 0.99638415 | 0.99638415 | Atf7ip2/Gpn1 | 2 | 16 | 12.5 |
| Hmbox1 | Adcy10 | 1/428 | 25/6837 | 0.80191959 | 0.99638415 | 0.99638415 | Asph | 1 | 26 | 3.85 |
| Hnf4a | St3gal1 | 47/428 | 788/6837 | 0.66567788 | 0.99638415 | 0.99638415 | Acox2/Adamts18/Agpat3/Aldh6a1/Ampd2/Atp2b1/Bcl3/Camk2n1/Card14/Donson/Efhc1/Eml6/Etv4/Fam13a/Fam222a/Fdft1/Fndc3b/Galnt2/Gas2l3/Gna12/Igf2bp3/Kif21a/Knl1/Lgals2/Mical3/Nadk2/Nek2/Nfkbiz/Nsl1/Pacsin2/Pde7b/Prr5/Ptprm/Rapgef2/Rcan1/Rhobtb3/Rrbp1/Scd1/Serpina1b/Slc7a5/Sntb1/Soat2/Svil/Tbc1d1/ | 47 | 907 | 5.18 |
| Hnf4g | Ncoa4 | 1/428 | 49/6837 | 0.95838051 | 0.99638415 | 0.99638415 | Serpina1b | 1 | 58 | 1.72 |
| Hoxa9 | Ghr | 1/428 | 19/6837 | 0.70769086 | 0.99638415 | 0.99638415 | Spag17 | 1 | 22 | 4.55 |
| Hoxb13 | Psd3 | 2/428 | 37/6837 | 0.68336198 | 0.99638415 | 0.99638415 | Snd1/Svil | 2 | 46 | 4.35 |
| Hsf1 | Mcf2l | 4/428 | 76/6837 | 0.70953989 | 0.99638415 | 0.99638415 | Dnah17/Engase/Gna12/Ier5 | 4 | 83 | 4.82 |
| Ikzf1 | Chst11 | 2/428 | 33/6837 | 0.62118575 | 0.99638415 | 0.99638415 | Chst11/Nabp1 | 2 | 40 | 5 |
| Irf1 | Etv6 | 3/428 | 63/6837 | 0.76430397 | 0.99638415 | 0.99638415 | Bcl3/H2-Q7/Nfkbiz | 3 | 77 | 3.9 |
| Irf2 | Ube2l6 | 5/428 | 61/6837 | 0.33432855 | 0.99638415 | 0.99638415 | Fam13a/Fanca/Ing1/Per1/Sema4c | 5 | 67 | 7.46 |
| Irf4 | Nlrc5 | 2/428 | 56/6837 | 0.87412952 | 0.99638415 | 0.99638415 | Dtnb/Nfkbiz | 2 | 65 | 3.08 |
| Irf9 | Rtn4 | 1/428 | 44/6837 | 0.94236905 | 0.99638415 | 0.99638415 | Dnah17 | 1 | 46 | 2.17 |
| Jun | Vcam1 | 9/428 | 119/6837 | 0.32781533 | 0.99638415 | 0.99638415 | Brca1/Ccnd1/Fas/Gna12/Ier5/Igf2bp3/Lbp/Nabp1/Nos2 | 9 | 185 | 4.86 |
| Junb | Egfr | 3/428 | 48/6837 | 0.58593565 | 0.99638415 | 0.99638415 | Gna12/Ier5/Tiam2 | 3 | 50 | 6 |
| Jund | Klf6 | 4/428 | 66/6837 | 0.59974854 | 0.99638415 | 0.99638415 | Gna12/Ier5/Igf2bp3/Nos2 | 4 | 75 | 5.33 |
| Klf1 | Chid1 | 1/428 | 46/6837 | 0.94940296 | 0.99638415 | 0.99638415 | Klhl21 | 1 | 50 | 2 |
| Klf13 | Eml2 | 4/428 | 29/6837 | 0.1044227 | 0.99638415 | 0.99638415 | Ap3d1/Fasn/Klhl21/Ptov1 | 4 | 31 | 12.9 |
| Klf4 | Mylk | 4/428 | 47/6837 | 0.33922527 | 0.99638415 | 0.99638415 | Birc5/Ccnd1/Mical3/Rcan1 | 4 | 49 | 8.16 |
| Klf5 | Rxra | 4/428 | 78/6837 | 0.72864266 | 0.99638415 | 0.99638415 | Birc5/Fasn/Prkcz/Tcf3 | 4 | 80 | 5 |
| Klf6 | Macf1 | 3/428 | 51/6837 | 0.62754591 | 0.99638415 | 0.99638415 | Ier5/Nos2/Tacc1 | 3 | 52 | 5.77 |
| Klf9 | Gprc5b | 3/428 | 63/6837 | 0.76430397 | 0.99638415 | 0.99638415 | Ampd2/Per1/Tacc1 | 3 | 68 | 4.41 |
| Lef1 | Lgi1 | 4/428 | 42/6837 | 0.2673215 | 0.99638415 | 0.99638415 | Birc5/Ccnd1/Slc25a28/Tcf3 | 4 | 54 | 7.41 |
| Lyl1 | Ccm2 | 2/428 | 72/6837 | 0.94558611 | 0.99638415 | 0.99638415 | Ccm2/Prr5 | 2 | 84 | 2.38 |
| Mafb | Zfp407 | 3/428 | 44/6837 | 0.52606432 | 0.99638415 | 0.99638415 | Jmjd1c/Nbas/Tent5b | 3 | 53 | 5.66 |
| Maff | Tbc1d14 | 2/428 | 39/6837 | 0.71118776 | 0.99638415 | 0.99638415 | Jmjd1c/Tiam2 | 2 | 43 | 4.65 |
| Mafg | Scin | 3/428 | 32/6837 | 0.32408962 | 0.99638415 | 0.99638415 | Cep112/Nabp1/Tiam2 | 3 | 37 | 8.11 |
| Mafk | Tiam2 | 3/428 | 40/6837 | 0.46172342 | 0.99638415 | 0.99638415 | Cep112/Gna12/Tiam2 | 3 | 50 | 6 |
| Max | Rap1gap2 | 5/428 | 50/6837 | 0.20136692 | 0.99638415 | 0.99638415 | Agpat3/Ap3d1/Ccnd1/Kdm4b/Prkcz | 5 | 53 | 9.43 |
| Maz | Pdxk | 3/428 | 67/6837 | 0.79974161 | 0.99638415 | 0.99638415 | Agpat3/Ap3d1/Tcf3 | 3 | 70 | 4.29 |
| Mbd1 | Mc1r | 1/428 | 40/6837 | 0.92524062 | 0.99638415 | 0.99638415 | Kdm4b | 1 | 43 | 2.33 |
| Mbd2 | Espn | 2/428 | 17/6837 | 0.28850161 | 0.99638415 | 0.99638415 | Prkcz/Tyk2 | 2 | 32 | 6.25 |
| Mef2a | Ncoa3 | 2/428 | 92/6837 | 0.98186871 | 0.99638415 | 0.99638415 | Jmjd1c/Svil | 2 | 102 | 1.96 |
| Mef2b | Klf6 | 1/428 | 27/6837 | 0.82603025 | 0.99638415 | 0.99638415 | Tcf3 | 1 | 36 | 2.78 |
| Mef2c | Erc1 | 1/428 | 62/6837 | 0.98216557 | 0.99638415 | 0.99638415 | Cdkl5 | 1 | 70 | 1.43 |
| Meis1 | Slc8a1 | 2/428 | 46/6837 | 0.79287334 | 0.99638415 | 0.99638415 | Dtnb/Gna12 | 2 | 54 | 3.7 |
| Meis2 | Cnot10 | 2/428 | 51/6837 | 0.83804922 | 0.99638415 | 0.99638415 | Fam13a/Tiam2 | 2 | 60 | 3.33 |
| Mitf | Commd4 | 24/428 | 574/6837 | 0.99046743 | 0.99638415 | 0.99638415 | Anln/Anxa5/Apex1/Cdkl3/Depdc1a/Dtnb/Etv4/Fam13a/Fndc3b/Foxm1/Ica1/Kdm4b/Kdm6a/Lin52/Mical3/Pacsin2/Pltp/Ptprm/Rcan1/Sgo1/Snd1/Svil/Tacc1/Tbc1d1 | 24 | 660 | 3.64 |
| Mnt | Greb1 | 2/428 | 54/6837 | 0.86068532 | 0.99638415 | 0.99638415 | Ap3d1/Kdm4b | 2 | 56 | 3.57 |
| Mxi1 | Stk32c | 6/428 | 74/6837 | 0.3172718 | 0.99638415 | 0.99638415 | Ampd2/Ap3d1/Atad2/Fam13a/Ing1/Kdm4b | 6 | 77 | 7.79 |
| Myb | Gstm2 | 3/428 | 56/6837 | 0.69028831 | 0.99638415 | 0.99638415 | Mctp2/Slc39a14/Tk1 | 3 | 65 | 4.62 |
| Mybl2 | Fam169a | 4/428 | 23/6837 | 0.05202091 | 0.99638415 | 0.99638415 | Cdkn3/Cenpf/Depdc1b/Top2a | 4 | 25 | 16 |
| Mycn | Diaph1 | 3/428 | 56/6837 | 0.69028831 | 0.99638415 | 0.99638415 | Ampd2/Mcm8/Prr5 | 3 | 64 | 4.69 |
| Myod1 | Actc1 | 3/428 | 41/6837 | 0.47818143 | 0.99638415 | 0.99638415 | Prr5/Svil/Tbc1d1 | 3 | 46 | 6.52 |
| Nanog | Cdk6 | 5/428 | 37/6837 | 0.07860863 | 0.99638415 | 0.99638415 | Ccnd1/Gpc6/Igf2bp3/Jmjd1c/Tiam2 | 5 | 48 | 10.42 |
| Nfe2 | Mical2 | 3/428 | 44/6837 | 0.52606432 | 0.99638415 | 0.99638415 | Gna12/Nabp1/Prr5 | 3 | 47 | 6.38 |
| Nfic | Gsn | 3/428 | 94/6837 | 0.93976926 | 0.99638415 | 0.99638415 | Dtnb/Got1/Svil | 3 | 108 | 2.78 |
| Nfkb1 | Edn1 | 11/428 | 175/6837 | 0.53989105 | 0.99638415 | 0.99638415 | Bcl3/Birc5/Brca1/Ccnd1/Fas/Fndc3b/Gadd45g/Gna12/Mical3/Nos2/Slc35f2 | 11 | 251 | 4.38 |
| Nfkb2 | Rara | 3/428 | 48/6837 | 0.58593565 | 0.99638415 | 0.99638415 | Bcl3/Fndc3b/Snd1 | 3 | 62 | 4.84 |
| Nfya | H2-Oa | 5/428 | 61/6837 | 0.33432855 | 0.99638415 | 0.99638415 | Brsk2/Eif2ak3/Gna12/Rhof/Tbc1d31 | 5 | 70 | 7.14 |
| Nr1h2 | Srebf1 | 2/428 | 13/6837 | 0.19372938 | 0.99638415 | 0.99638415 | Fasn/Tbc1d1 | 2 | 13 | 15.38 |
| Nr1h3 | Srebf1 | 1/428 | Oct-37 | 0.47633223 | 0.99638415 | 0.99638415 | Fasn | 1 | 12 | 8.33 |
| Nr2c2 | Usp37 | 2/428 | 78/6837 | 0.96067114 | 0.99638415 | 0.99638415 | Parpbp/Trip13 | 2 | 84 | 2.38 |
| Nr2f1 | Pepd | 3/428 | 51/6837 | 0.62754591 | 0.99638415 | 0.99638415 | Agpat3/Brsk2/Fbln1 | 3 | 66 | 4.55 |
| Nr2f2 | Hnf4a | 5/428 | 68/6837 | 0.42297217 | 0.99638415 | 0.99638415 | Atp2b1/Fbln1/Gna12/Pacsin2/Tbc1d1 | 5 | 84 | 5.95 |
| Nr2f6 | Aff1 | 1/428 | 27/6837 | 0.82603025 | 0.99638415 | 0.99638415 | Agpat3 | 1 | 32 | 3.12 |
| Nr3c1 | Bglap3 | 6/428 | 70/6837 | 0.27297315 | 0.99638415 | 0.99638415 | Atf2/Pacsin2/Tacc1/Tat/Tbc1d1/Tle1 | 6 | 76 | 7.89 |
| Nr4a1 | Parvb | 1/428 | 36/6837 | 0.90303682 | 0.99638415 | 0.99638415 | E2f1 | 1 | 41 | 2.44 |
| Nr5a2 | Map3k14 | 1/428 | 28/6837 | 0.83696403 | 0.99638415 | 0.99638415 | Bmper | 1 | 31 | 3.23 |
| Nrf1 | Rsrp1 | 5/428 | 73/6837 | 0.48508976 | 0.99638415 | 0.99638415 | B9d1/Cic/Fads1/Ing1/Per1 | 5 | 77 | 6.49 |
| Onecut1 | Kiaa1109 | 1/428 | 40/6837 | 0.92524062 | 0.99638415 | 0.99638415 | Diaph3 | 1 | 52 | 1.92 |
| Otx2 | Xirp2 | 2/428 | 15/6837 | 0.24089409 | 0.99638415 | 0.99638415 | Adamts18/Diaph3 | 2 | 23 | 8.7 |
| Pax5 | Dusp10 | 2/428 | 42/6837 | 0.74906675 | 0.99638415 | 0.99638415 | Mical3/Per1 | 2 | 53 | 3.77 |
| Pax6 | Plcb1 | 1/428 | 34/6837 | 0.889579 | 0.99638415 | 0.99638415 | Frmd5 | 1 | 43 | 2.33 |
| Pbx2 | Arhgef1 | 5/428 | 38/6837 | 0.08610689 | 0.99638415 | 0.99638415 | Galnt2/Nbas/Pacsin2/Tacc1/Tiam2 | 5 | 44 | 11.36 |
| Pbx3 | Csnk1d | 4/428 | 46/6837 | 0.32471291 | 0.99638415 | 0.99638415 | Fndc3b/Galnt2/Pacsin2/Tiam2 | 4 | 51 | 7.84 |
| Pdx1 | Daam1 | 2/428 | 45/6837 | 0.78260074 | 0.99638415 | 0.99638415 | Dync1i1/Nbas | 2 | 50 | 4 |
| Pgr | Runx1 | 3/428 | 67/6837 | 0.79974161 | 0.99638415 | 0.99638415 | Ccnd1/Rcl1/Tacc1 | 3 | 89 | 3.37 |
| Pknox1 | Wnt3 | 3/428 | 28/6837 | 0.25433719 | 0.99638415 | 0.99638415 | Fam13a/Galnt2/Mical3 | 3 | 35 | 8.57 |
| Pou2f1 | Rictor | 2/428 | 42/6837 | 0.74906675 | 0.99638415 | 0.99638415 | Ccnd1/Uggt2 | 2 | 61 | 3.28 |
| Pou2f2 | Cd226 | 1/428 | 69/6837 | 0.98870769 | 0.99638415 | 0.99638415 | Kdm4b | 1 | 86 | 1.16 |
| Pou4f2 | Ctnnd2 | 2/428 | 24/6837 | 0.44866829 | 0.99638415 | 0.99638415 | Ccnd1/Fam13a | 2 | 28 | 7.14 |
| Pou5f1 | Galnt10 | 4/428 | 59/6837 | 0.50975316 | 0.99638415 | 0.99638415 | Gpc6/Igf2bp3/Jmjd1c/Tiam2 | 4 | 72 | 5.56 |
| Ppara | Mmp12 | 3/428 | 26/6837 | 0.22028719 | 0.99638415 | 0.99638415 | Acox2/Ccnd1/Hmgcs1 | 3 | 37 | 8.11 |
| Pparg | Mmp9 | 4/428 | 84/6837 | 0.78029345 | 0.99638415 | 0.99638415 | Ccnd1/Dtx4/Ptprm/Tacc1 | 4 | 104 | 3.85 |
| Rara | Crabp2 | 10/428 | 108/6837 | 0.13748142 | 0.99638415 | 0.99638415 | Bcl3/Cdr2/Chsy1/Cic/Gramd1a/Khsrp/Kpnb1/Prkaca/Scd1/Tnfaip2 | 10 | 122 | 8.2 |
| Rbpj | Cep85l | 3/428 | 52/6837 | 0.64076053 | 0.99638415 | 0.99638415 | Fmnl2/Fndc3b/Gna12 | 3 | 59 | 5.08 |
| Rela | Il17c | 11/428 | 180/6837 | 0.57842056 | 0.99638415 | 0.99638415 | Bcl3/Birc5/Ccnd1/Col7a1/Fndc3b/Gadd45b/Gadd45g/Gramd1a/Nfkbiz/Nos2/Pacsin2 | 11 | 245 | 4.49 |
| Relb | Rasgrp1 | 3/428 | 58/6837 | 0.71305676 | 0.99638415 | 0.99638415 | Bcl3/Fndc3b/Slc39a14 | 3 | 74 | 4.05 |
| Rest | Gsn | 2/428 | 40/6837 | 0.72431747 | 0.99638415 | 0.99638415 | Prss27/Sptbn4 | 2 | 67 | 2.99 |
| Rfx1 | Stk32c | 2/428 | 35/6837 | 0.65338226 | 0.99638415 | 0.99638415 | Cep112/Swsap1 | 2 | 40 | 5 |
| Rfx5 | Sult2b1 | 3/428 | 64/6837 | 0.77361186 | 0.99638415 | 0.99638415 | Eml1/Per1/Rhof | 3 | 80 | 3.75 |
| Runx1 | Arhgap10 | 16/428 | 306/6837 | 0.80955115 | 0.99638415 | 0.99638415 | Agpat3/Atg13/Ccm2/Fndc3b/Galnt2/Jmjd1c/Kdm4b/Mad2l1/Mctp2/Nbas/Rorc/Snd1/Svil/Tbc[illegible] | 16 | 380 | 4.21 |
| Runx2 | Mid1 | 6/428 | 51/6837 | 0.09708383 | 0.99638415 | 0.99638415 | Nabp1/Pacsin2/Ptprm/Svil/Tiam2/Tspan5 | 6 | 65 | 9.23 |
| Runx3 | Cd226 | 2/428 | 80/6837 | 0.96474153 | 0.99638415 | 0.99638415 | Pacsin2/Tiam2 | 2 | 97 | 2.06 |
| Rxra | Plcb1 | 2/428 | 67/6837 | 0.92895706 | 0.99638415 | 0.99638415 | Galnt2/Pdgfc | 2 | 78 | 2.56 |
| Six2 | Tbc1d1 | 2/428 | 28/6837 | 0.53081814 | 0.99638415 | 0.99638415 | Fbln1/Tbc1d1 | 2 | 39 | 5.13 |
| Six5 | Fuz | 1/428 | 21/6837 | 0.74324089 | 0.99638415 | 0.99638415 | Mamstr | 1 | 23 | 4.35 |
| Smad1 | Stk32b | 1/428 | 13/6837 | 0.56878913 | 0.99638415 | 0.99638415 | Id3 | 1 | 17 | 5.88 |
| Smad3 | Pxn | 5/428 | 78/6837 | 0.54472186 | 0.99638415 | 0.99638415 | Col7a1/Fndc3b/Nabp1/Tbc1d1/Tiam2 | 5 | 96 | 5.21 |
| Smad4 | Brwd1 | 5/428 | 59/6837 | 0.30917724 | 0.99638415 | 0.99638415 | Col7a1/Gadd45b/Id3/Map3k3/Mctp2 | 5 | 75 | 6.67 |
| Snai2 | Adap1 | 3/428 | 62/6837 | 0.75468828 | 0.99638415 | 0.99638415 | Agpat3/Kdm4b/Nsd2 | 3 | 66 | 4.55 |
| Sox10 | Mbp | 2/428 | 35/6837 | 0.65338226 | 0.99638415 | 0.99638415 | Arnt2/Ret | 2 | 45 | 4.44 |
| Sox13 | Rcan1 | 3/428 | 39/6837 | 0.44504548 | 0.99638415 | 0.99638415 | Dusp6/Rcan1/Serpina1b | 3 | 45 | 6.67 |
| Sox2 | Naaladl2 | 14/428 | 291/6837 | 0.88138448 | 0.99638415 | 0.99638415 | Ackr3/Ccnd1/Cep112/Diaph3/Fam13a/Fndc3b/Jmjd1c/Nek2/Pbx3/Plxnb1/Ptprm/Rapgef2/Rtl6/Tnfrsf8 | 14 | 398 | 3.52 |
| Sox9 | Hapln1 | 3/428 | 67/6837 | 0.79974161 | 0.99638415 | 0.99638415 | Clcn3/Hapln1/Nmt1 | 3 | 74 | 4.05 |
| Sp1 | Cd163 | 20/428 | 296/6837 | 0.39406803 | 0.99638415 | 0.99638415 | Apex1/Atf2/Birc5/Brca1/Cat/Ccnd1/Cdc25c/Cdk1/Cic/Col7a1/E2f1/F2r/Fas/Igfbp4/Kif2c/Pdgfc/Per1/Skp2/Slc7a5/Tnfrsf8 | 20 | 394 | 5.08 |
| Sp3 | Plat | 3/428 | 54/6837 | 0.66619174 | 0.99638415 | 0.99638415 | Ccnd1/F2r/Ret | 3 | 91 | 3.3 |
| Sp4 | Kcnip2 | 2/428 | 52/6837 | 0.84594076 | 0.99638415 | 0.99638415 | Ccnd1/Sptbn4 | 2 | 59 | 3.39 |
| Spi1 | Wipf1 | 27/428 | 506/6837 | 0.83823764 | 0.99638415 | 0.99638415 | Ampd2/Apbb1ip/Cdc7/Cdkl3/Chst11/Dtnb/Fam13a/Fndc3b/Galnt2/H2bc12/Hk2/Ier5/Ifnar1/Jmjd1c/Kif18b/Mctp2/Nabp1/Nbas/Pacsin2/Rcan1/Rrbp1/Scamp4/Snd1/Svil/Tacc1/Tbc1d1/Tiam2 | 27 | 585 | 4.62 |
| Spib | Cd40 | 1/428 | 44/6837 | 0.94236905 | 0.99638415 | 0.99638415 | Dtnb | 1 | 58 | 1.72 |
| Srebf1 | Scd1 | 7/428 | 67/6837 | 0.12408583 | 0.99638415 | 0.99638415 | Cic/Fasn/Fdft1/Hk2/Hmgcs1/Mical3/Scd1 | 7 | 71 | 9.86 |
| Srebf2 | Dnah1 | 6/428 | 82/6837 | 0.40818867 | 0.99638415 | 0.99638415 | Ampd2/Fads1/Fasn/Fdft1/Sec16a/Stard4 | 6 | 90 | 6.67 |
| Srf | Fermt2 | 5/428 | 100/6837 | 0.75917382 | 0.99638415 | 0.99638415 | Dapk3/Gadd45g/Svil/Tacc1/Tbc1d1 | 5 | 109 | 4.59 |
| Stat1 | Hivep3 | 49/428 | 845/6837 | 0.74463855 | 0.99638415 | 0.99638415 | Ap3d1/Bcl3/Blvra/Cables2/Ccnd1/Ccr1/Cdc7/Cdkl3/Cenpf/Dtnb/Edem2/Eif2ak3/Exo1/Fas/Fndc3b/Galnt3/Gen1/Gna12/Gramd1a/Hk2/Ier5/Ifitm3/Il6st/Jmjd1c/Kif21a/Nabp1/Nfkbiz/Nos2/Nr4a2/Pacsin2/Rapgef2/Rasip1/Rhof/Sertad3/Smc4/Snd1/Stat2/Tacc1/Tbc1d1/Tcf3/Tgm2/Tiam2/Ticam1/Tnfrsf8/Trim40/Ttc17/Ttc26/Ttc7b/ | 49 | 1014 | 4.83 |
| Stat2 | Pim1 | 1/428 | 74/6837 | 0.99185513 | 0.99638415 | 0.99638415 | Mov10 | 1 | 80 | 1.25 |
| Stat3 | Cd55b | 10/428 | 140/6837 | 0.38005019 | 0.99638415 | 0.99638415 | Bcl3/Birc5/Ccnd1/Fas/Fndc3b/Galnt2/Ifnar1/Il6st/Lbp/Tgm2 | 10 | 171 | 5.85 |
| Stat5a | Ccnd1 | 6/428 | 56/6837 | 0.13576868 | 0.99638415 | 0.99638415 | Ccnd1/Chst11/Il6st/Tcf3/Tiam2/Tnfrsf8 | 6 | 66 | 9.09 |
| Stat5b | Rapgef1 | 4/428 | 66/6837 | 0.59974854 | 0.99638415 | 0.99638415 | Ccnd1/Il6st/Pacsin2/Svil | 4 | 78 | 5.13 |
| Tal1 | Cdk5rap2 | 18/428 | 364/6837 | 0.88286909 | 0.99638415 | 0.99638415 | Chst11/Chsy1/Fam13a/Galnt2/Gna12/Ier5/Kdm4b/Nbas/Pacsin2/Prr5/Ptprm/Rapgef2/Snd1/Svil/Tacc1/Tbc1d1/Tiam2/Ttc7b | 18 | 416 | 4.33 |
| Tbp | St6gal1 | 1/428 | 37/6837 | 0.9091389 | 0.99638415 | 0.99638415 | Ier5 | 1 | 46 | 2.17 |
| Tbx21 | Arhgap25 | 1/428 | 66/6837 | 0.98626369 | 0.99638415 | 0.99638415 | Tnfrsf8 | 1 | 76 | 1.32 |
| Tcf12 | Prickle2 | 4/428 | 72/6837 | 0.66845766 | 0.99638415 | 0.99638415 | Fndc3b/Svil/Tbc1d1/Tiam2 | 4 | 78 | 5.13 |
| Tcf3 | Brf1 | 5/428 | 114/6837 | 0.84993932 | 0.99638415 | 0.99638415 | Ccm2/Prkcz/Prr5/Scamp4/Tbc1d1 | 5 | 125 | 4 |
| Tcf4 | Arid5b | 3/428 | 52/6837 | 0.64076053 | 0.99638415 | 0.99638415 | Ccnd1/Cep112/Slc25a28 | 3 | 60 | 5 |
| Tcf7 | Dmd | 5/428 | 47/6837 | 0.16870502 | 0.99638415 | 0.99638415 | Birc5/Ccnd1/Cpped1/Dtnb/Tcf3 | 5 | 58 | 8.62 |
| Tcf7l2 | Asap1 | 4/428 | 66/6837 | 0.59974854 | 0.99638415 | 0.99638415 | Birc5/Ccnd1/Rhobtb3/Tiam2 | 4 | 74 | 5.41 |
| Tead1 | Lamc2 | 8/428 | 116/6837 | 0.44169616 | 0.99638415 | 0.99638415 | Arnt2/Eml1/Pdgfc/Ptprm/Rapgef2/Svil/Tacc1/Tiam2 | 8 | 129 | 6.2 |
| Tead2 | Vps37b | 2/428 | 26/6837 | 0.49078329 | 0.99638415 | 0.99638415 | Galnt2/Jmjd1c | 2 | 29 | 6.9 |
| Tead4 | Rai14 | 5/428 | 67/6837 | 0.41035765 | 0.99638415 | 0.99638415 | Fndc3b/Galnt2/Pdgfc/Tgm2/Tiam2 | 5 | 73 | 6.85 |
| Tfap2a | Th | 9/428 | 97/6837 | 0.15211674 | 0.99638415 | 0.99638415 | Atf2/Bicdl1/Ccnb1/F2r/Fas/Galnt3/Hk2/Oat/Pacsin2 | 9 | 125 | 7.2 |
| Tfap2c | Pacsin2 | 43/428 | 737/6837 | 0.71672779 | 0.99638415 | 0.99638415 | Agpat3/Ampd2/Asns/Bmper/Brip1/Ccnf/Cep112/Cyld/Efhc1/Eml1/Fads1/Fas/Fbln1/Fdft1/Fgd1/Fndc3b/Galnt2/Gna12/Hk2/Hs6st1/Ier5/Kdm4b/Lpcat4/Mical3/Nkain1/Nr4a2/Pacsin2/Pbk/Pi4k2a/Piwil4/Ret/Rrbp1/Sec16a/Sertad3/Slc7a5/Smc2/Snd1/Stc2/Svil/Tbc1d1/Top2a/Tspan31/Ttc7b | 43 | 890 | 4.83 |
| Tfap4 | Dennd1a | 5/428 | 37/6837 | 0.07860863 | 0.99638415 | 0.99638415 | Agpat3/Gna12/Prkcz/Svil/Tiam2 | 5 | 40 | 12.5 |
| Tfdp1 | Tchp | 5/428 | 66/6837 | 0.39770573 | 0.99638415 | 0.99638415 | Anp32e/Cdk1/Ckap2l/Kpnb1/Ncaph | 5 | 69 | 7.25 |
| Tgif2 | Asxl1 | 1/428 | 19/6837 | 0.70769086 | 0.99638415 | 0.99638415 | Dpagt1 | 1 | 19 | 5.26 |
| Thap1 | Bcor | 1/428 | 51/6837 | 0.96346383 | 0.99638415 | 0.99638415 | Gramd1a | 1 | 55 | 1.82 |
| Thap11 | Cnih4 | 6/428 | 112/6837 | 0.71111186 | 0.99638415 | 0.99638415 | Bub1b/Invs/Lmbrd2/Nmt1/Skp2/Tbc1d31 | 6 | 114 | 5.26 |
| Tp53 | Fam183b | 34/428 | 483/6837 | 0.25769753 | 0.99638415 | 0.99638415 | Amotl1/Bcl3/Birc5/Bmper/Brca1/Ccna2/Ccnb1/Ccnd1/Ccng1/Cdc25c/Cdk1/Cic/E2f1/Fas/Fbln1/Frmd5/Gadd45g/Gna12/Gpc6/Ier5/Ing1/Mki67/Nos2/Pdgfc/Pi4k2a/Prr5/Rad51/Slc39a14/Svil/Tbc1d31/Tle1/Tmem182/Trim32/Vcan | 34 | 582 | 5.84 |
| Tp63 | Itgb6 | 2/428 | 76/6837 | 0.95615287 | 0.99638415 | 0.99638415 | Fat2/Svil | 2 | 90 | 2.22 |
| Tp73 | Pou2af1 | 1/428 | 36/6837 | 0.90303682 | 0.99638415 | 0.99638415 | Cdk1 | 1 | 41 | 2.44 |
| Twist1 | Plxnd1 | 4/428 | 55/6837 | 0.45446527 | 0.99638415 | 0.99638415 | Fbln1/Igfbp4/Kdm4b/Mgat4c | 4 | 67 | 5.97 |
| Usf1 | Cyp19a1 | 9/428 | 89/6837 | 0.10347256 | 0.99638415 | 0.99638415 | Apex1/Ccnb1/Ccnd1/Dnah17/Fasn/Id4/Pf4/Sptbn4/Tbc1d1 | 9 | 103 | 8.74 |
| Usf2 | Fis1 | 5/428 | 103/6837 | 0.78152249 | 0.99638415 | 0.99638415 | Atf4/Borcs6/Brca1/Pf4/Sptbn4 | 5 | 113 | 4.42 |
| Vdr | Cox5a | 13/428 | 210/6837 | 0.55881335 | 0.99638415 | 0.99638415 | Alkbh2/Apex1/Atf4/Bmper/Cbx4/Chst11/Gna12/Id3/Kpnb1/Prkaca/Ptprm/Rnf5/Tpx2 | 13 | 244 | 5.33 |
| Xbp1 | Frmd4b | 2/428 | Nov-37 | 0.14803312 | 0.99638415 | 0.99638415 | Amotl1/Snd1 | 2 | 16 | 12.5 |
| Yy1 | Smad7 | 4/428 | 91/6837 | 0.83036975 | 0.99638415 | 0.99638415 | Ap3d1/Fas/Nos2/Ttc17 | 4 | 99 | 4.04 |
| Zbtb33 | Smg7 | 5/428 | 83/6837 | 0.60090084 | 0.99638415 | 0.99638415 | Dmxl1/Mms22l/Smc4/Spdl1/Suv39h1 | 5 | 88 | 5.68 |
| Zbtb7a | Ncor2 | 5/428 | 82/6837 | 0.58997744 | 0.99638415 | 0.99638415 | Ap3d1/Cic/Hs6st1/Khsrp/Zfpm1 | 5 | 90 | 5.56 |
| Zeb1 | Mark2 | 3/428 | 92/6837 | 0.93382993 | 0.99638415 | 0.99638415 | Abca7/Nsd2/Tbc1d1 | 3 | 96 | 3.12 |
| Zeb2 | Susd1 | 2/428 | 46/6837 | 0.79287334 | 0.99638415 | 0.99638415 | Creg1/Fndc3b | 2 | 53 | 3.77 |
| Zfp217 | Arid5b | 1/428 | Oct-37 | 0.47633223 | 0.99638415 | 0.99638415 | Fndc3b | 1 | 10 | 10 |
| Zfp384 | Cacna1a | 6/428 | 61/6837 | 0.18055704 | 0.99638415 | 0.99638415 | Ap3d1/Dennd5a/Fdft1/Igf2bp3/Jmjd1c/Racgap1 | 6 | 65 | 9.23 |
| Zfx | Setd5 | 2/428 | 53/6837 | 0.85348186 | 0.99638415 | 0.99638415 | Klhl21/Nsd2 | 2 | 54 | 3.7 |
| Zkscan1 | Spag9 | 3/428 | 40/6837 | 0.46172342 | 0.99638415 | 0.99638415 | Prr5/Ticam1/Ttc7b | 3 | 46 | 6.52 |
| Znf143 | Nucb1 | 2/428 | 47/6837 | 0.80271551 | 0.99638415 | 0.99638415 | Esco2/Fndc3b | 2 | 51 | 3.92 |
| Znf263 | Cfap46 | 19/428 | 351/6837 | 0.7806512 | 0.99638415 | 0.99638415 | Abca7/Adcy3/Agpat3/Cdk18/Cic/Dnah17/Fam13a/Igf2bp3/Mark1/Mical3/Prkcz/Prr5/Ret/Setd1b/Smoc2/Snd1/Tbc1d1/Tcf3/Vcan | 19 | 432 | 4.4 |
| Znf639 | Sin3a | 3/428 | 26/6837 | 0.22028719 | 0.99638415 | 0.99638415 | Mapk9/Slc30a5/Zc3h7a | 3 | 28 | 10.71 |
| Znf740 | Map1a | 2/428 | 34/6837 | 0.63756419 | 0.99638415 | 0.99638415 | Rhof/Setd1b | 2 | 36 | 5.56 |
| Prdm14 | Nr4a1 | 6/428 | 227/6837 | 0.99650711 | 0.99650711 | 0.99650711 | Agpat3/Hes2/Kdm4b/Map7d1/Tbc1d9/Tent5b | 6 | 306 | 1.96 |
| Adnp | Atf7ip | NA | NA | NA | NA | NA | NA | 0 | 4 | 0 |
| Arntl2 | Serpine1 | NA | NA | NA | NA | NA | NA | 0 | 1 | 0 |
| Atf5 | Disc1 | NA | NA | NA | NA | NA | NA | 0 | 1 | 0 |
| Atf6 | Ddit3 | NA | NA | NA | NA | NA | NA | 0 | 10 | 0 |
| Atoh1 | Hes6 | NA | NA | NA | NA | NA | NA | 0 | 1 | 0 |
| Cdx1 | Ada | NA | NA | NA | NA | NA | NA | 0 | 3 | 0 |
| Cebpe | Myc | NA | NA | NA | NA | NA | NA | 0 | 6 | 0 |
| Cebpz | Col11a1 | NA | NA | NA | NA | NA | NA | 0 | 3 | 0 |
| Creb3l2 | Cpt1c | NA | NA | NA | NA | NA | NA | 0 | 1 | 0 |
| Creb5 | Pou1f1 | NA | NA | NA | NA | NA | NA | 0 | 1 | 0 |
| Ddit3 | Trib3 | NA | NA | NA | NA | NA | NA | 0 | 2 | 0 |
| Deaf1 | Rac3 | NA | NA | NA | NA | NA | NA | 0 | 8 | 0 |
| Dnmt1 | Sfrp5 | NA | NA | NA | NA | NA | NA | 0 | 4 | 0 |
| E2f5 | Jade1 | NA | NA | NA | NA | NA | NA | 0 | 12 | 0 |
| E2f8 | E2f1 | NA | NA | NA | NA | NA | NA | 0 | 1 | 0 |
| E4f1 | Ccna2 | NA | NA | NA | NA | NA | NA | 0 | 1 | 0 |
| Egr2 | Spint1 | NA | NA | NA | NA | NA | NA | 0 | 4 | 0 |
| Egr3 | Nab2 | NA | NA | NA | NA | NA | NA | 0 | 1 | 0 |
| Ehf | Limd1 | NA | NA | NA | NA | NA | NA | 0 | 19 | 0 |
| Elf2 | Vcp | NA | NA | NA | NA | NA | NA | 0 | 1 | 0 |
| Elf5 | Ffar4 | NA | NA | NA | NA | NA | NA | 0 | 19 | 0 |
| Elk3 | Fos | NA | NA | NA | NA | NA | NA | 0 | 1 | 0 |
| Elk4 | Krt8 | NA | NA | NA | NA | NA | NA | 0 | 14 | 0 |
| En1 | Utrn | NA | NA | NA | NA | NA | NA | 0 | 1 | 0 |
| Eomes | Csmd3 | NA | NA | NA | NA | NA | NA | 0 | 21 | 0 |
| Erf | Ets2 | NA | NA | NA | NA | NA | NA | 0 | 1 | 0 |
| Esr2 | Adamtsl5 | NA | NA | NA | NA | NA | NA | 0 | 74 | 0 |
| Etv3 | Nfkb1 | NA | NA | NA | NA | NA | NA | 0 | 1 | 0 |
| Etv6 | Hivep2 | NA | NA | NA | NA | NA | NA | 0 | 6 | 0 |
| Fli1 | Ets1 | NA | NA | NA | NA | NA | NA | 0 | 107 | 0 |
| Fosb | Runx2 | NA | NA | NA | NA | NA | NA | 0 | 1 | 0 |
| Fosl1 | Egfr | NA | NA | NA | NA | NA | NA | 0 | 53 | 0 |
| Foxa3 | Cyp2c38 | NA | NA | NA | NA | NA | NA | 0 | 3 | 0 |
| Foxk1 | Cald1 | NA | NA | NA | NA | NA | NA | 0 | 8 | 0 |
| Foxl2 | Cxcl1 | NA | NA | NA | NA | NA | NA | 0 | 20 | 0 |
| Foxo4 | Tnfsf10 | NA | NA | NA | NA | NA | NA | 0 | 15 | 0 |
| Foxo6 | Fbxo25 | NA | NA | NA | NA | NA | NA | 0 | 1 | 0 |
| Foxp2 | Gnas | NA | NA | NA | NA | NA | NA | 0 | 35 | 0 |
| Foxp3 | Il2ra | NA | NA | NA | NA | NA | NA | 0 | 3 | 0 |
| Gata4 | Cldn14 | NA | NA | NA | NA | NA | NA | 0 | 10 | 0 |
| Gcm2 | Pth | NA | NA | NA | NA | NA | NA | 0 | 1 | 0 |
| Gfi1 | Bax | NA | NA | NA | NA | NA | NA | 0 | 2 | 0 |
| Gli1 | Egr2 | NA | NA | NA | NA | NA | NA | 0 | 7 | 0 |
| Gli2 | Efemp1 | NA | NA | NA | NA | NA | NA | 0 | 16 | 0 |
| Gli3 | Ptch2 | NA | NA | NA | NA | NA | NA | 0 | 3 | 0 |
| Gtf2i | Sirt1 | NA | NA | NA | NA | NA | NA | 0 | 1 | 0 |
| Hes1 | Snx6 | NA | NA | NA | NA | NA | NA | 0 | 6 | 0 |
| Hey1 | Kdr | NA | NA | NA | NA | NA | NA | 0 | 1 | 0 |
| Hey2 | Acta2 | NA | NA | NA | NA | NA | NA | 0 | 2 | 0 |
| Hhex | Elmo1 | NA | NA | NA | NA | NA | NA | 0 | 11 | 0 |
| Hic1 | Sirt1 | NA | NA | NA | NA | NA | NA | 0 | 7 | 0 |
| Hlf | Runx1 | NA | NA | NA | NA | NA | NA | 0 | 2 | 0 |
| Hmga1 | Cdc25a | NA | NA | NA | NA | NA | NA | 0 | 1 | 0 |
| Hnf1a | Ugt1a9 | NA | NA | NA | NA | NA | NA | 0 | 23 | 0 |
| Hnf1b | Wdr72 | NA | NA | NA | NA | NA | NA | 0 | 37 | 0 |
| Hoxa10 | Emx2 | NA | NA | NA | NA | NA | NA | 0 | 4 | 0 |
| Hoxa11 | Hoxa10 | NA | NA | NA | NA | NA | NA | 0 | 1 | 0 |
| Hoxa4 | Th | NA | NA | NA | NA | NA | NA | 0 | 1 | 0 |
| Hoxa5 | Th | NA | NA | NA | NA | NA | NA | 0 | 4 | 0 |
| Hoxb3 | Otx2 | NA | NA | NA | NA | NA | NA | 0 | 1 | 0 |
| Hoxb7 | Ppfibp2 | NA | NA | NA | NA | NA | NA | 0 | 5 | 0 |
| Hoxc11 | Lct | NA | NA | NA | NA | NA | NA | 0 | 2 | 0 |
| Hoxc8 | Tnfrsf11b | NA | NA | NA | NA | NA | NA | 0 | 1 | 0 |
| Hoxd3 | Itgb1 | NA | NA | NA | NA | NA | NA | 0 | 3 | 0 |
| Hsf2 | Mapk8 | NA | NA | NA | NA | NA | NA | 0 | 3 | 0 |
| Insm1 | Neurod1 | NA | NA | NA | NA | NA | NA | 0 | 3 | 0 |
| Irf5 | Tnf | NA | NA | NA | NA | NA | NA | 0 | 1 | 0 |
| Irf7 | Ifnb1 | NA | NA | NA | NA | NA | NA | 0 | 2 | 0 |
| Irf8 | Nf1 | NA | NA | NA | NA | NA | NA | 0 | 8 | 0 |
| Kdm5b | Prrc2b | NA | NA | NA | NA | NA | NA | 0 | 17 | 0 |
| Klf10 | Cdkn1a | NA | NA | NA | NA | NA | NA | 0 | 1 | 0 |
| Klf11 | Ins2 | NA | NA | NA | NA | NA | NA | 0 | 2 | 0 |
| Klf12 | Tfap2a | NA | NA | NA | NA | NA | NA | 0 | 1 | 0 |
| Klf15 | Lrp5 | NA | NA | NA | NA | NA | NA | 0 | 1 | 0 |
| Klf2 | Thbd | NA | NA | NA | NA | NA | NA | 0 | 2 | 0 |
| Klf3 | Cast | NA | NA | NA | NA | NA | NA | 0 | 13 | 0 |
| Kmt2b | Supt5h | NA | NA | NA | NA | NA | NA | 0 | 8 | 0 |
| Lhx2 | Sox6 | NA | NA | NA | NA | NA | NA | 0 | 10 | 0 |
| Lhx3 | Tshb | NA | NA | NA | NA | NA | NA | 0 | 3 | 0 |
| Lhx4 | Fshb | NA | NA | NA | NA | NA | NA | 0 | 4 | 0 |
| Maf | Osbpl3 | NA | NA | NA | NA | NA | NA | 0 | 38 | 0 |
| Mafa | Ins2 | NA | NA | NA | NA | NA | NA | 0 | 1 | 0 |
| Mbd3 | Esr1 | NA | NA | NA | NA | NA | NA | 0 | 1 | 0 |
| Mecp2 | Esr1 | NA | NA | NA | NA | NA | NA | 0 | 1 | 0 |
| Mef2d | Atf3 | NA | NA | NA | NA | NA | NA | 0 | 4 | 0 |
| Msc | Mmp2 | NA | NA | NA | NA | NA | NA | 0 | 2 | 0 |
| Mxd1 | Odc1 | NA | NA | NA | NA | NA | NA | 0 | 1 | 0 |
| Mxd4 | Odc1 | NA | NA | NA | NA | NA | NA | 0 | 1 | 0 |
| Mybl1 | Ncl | NA | NA | NA | NA | NA | NA | 0 | 2 | 0 |
| Myf6 | Acacb | NA | NA | NA | NA | NA | NA | 0 | 1 | 0 |
| Mynn | Thumpd3 | NA | NA | NA | NA | NA | NA | 0 | 4 | 0 |
| Myog | Acacb | NA | NA | NA | NA | NA | NA | 0 | 1 | 0 |
| Mzf1 | Myb | NA | NA | NA | NA | NA | NA | 0 | 3 | 0 |
| Ncoa1 | Asap1 | NA | NA | NA | NA | NA | NA | 0 | 20 | 0 |
| Ncoa2 | Daam1 | NA | NA | NA | NA | NA | NA | 0 | 18 | 0 |
| Ncoa3 | Arhgap10 | NA | NA | NA | NA | NA | NA | 0 | 23 | 0 |
| Neurod1 | Ctsb | NA | NA | NA | NA | NA | NA | 0 | 35 | 0 |
| Neurog3 | Neurod1 | NA | NA | NA | NA | NA | NA | 0 | 1 | 0 |
| Nfat5 | Tnf | NA | NA | NA | NA | NA | NA | 0 | 2 | 0 |
| Nfatc1 | Tmem131l | NA | NA | NA | NA | NA | NA | 0 | 34 | 0 |
| Nfatc2 | Il4 | NA | NA | NA | NA | NA | NA | 0 | 10 | 0 |
| Nfatc3 | Rcan1 | NA | NA | NA | NA | NA | NA | 0 | 2 | 0 |
| Nfatc4 | Trpc6 | NA | NA | NA | NA | NA | NA | 0 | 1 | 0 |
| Nfe2l1 | Sqstm1 | NA | NA | NA | NA | NA | NA | 0 | 30 | 0 |
| Nfe2l2 | Gatad2a | NA | NA | NA | NA | NA | NA | 0 | 72 | 0 |
| Nfia | Tp53 | NA | NA | NA | NA | NA | NA | 0 | 3 | 0 |
| Nfil3 | Il1b | NA | NA | NA | NA | NA | NA | 0 | 1 | 0 |
| Nfyc | Fdps | NA | NA | NA | NA | NA | NA | 0 | 3 | 0 |
| Nkx2-1 | Sftpb | NA | NA | NA | NA | NA | NA | 0 | 1 | 0 |
| Nkx2-5 | Gata4 | NA | NA | NA | NA | NA | NA | 0 | 2 | 0 |
| Nme2 | Myc | NA | NA | NA | NA | NA | NA | 0 | 1 | 0 |
| Npas2 | Cry2 | NA | NA | NA | NA | NA | NA | 0 | 4 | 0 |
| Nr0b1 | Nr5a1 | NA | NA | NA | NA | NA | NA | 0 | 2 | 0 |
| Nr1d1 | Arntl | NA | NA | NA | NA | NA | NA | 0 | 1 | 0 |
| Nr1h4 | Nr1i2 | NA | NA | NA | NA | NA | NA | 0 | 7 | 0 |
| Nr1i2 | Ttc21a | NA | NA | NA | NA | NA | NA | 0 | 4 | 0 |
| Nr1i3 | Ugt1a1 | NA | NA | NA | NA | NA | NA | 0 | 1 | 0 |
| Nr2e1 | Pax2 | NA | NA | NA | NA | NA | NA | 0 | 3 | 0 |
| Nr3c2 | Atp1b1 | NA | NA | NA | NA | NA | NA | 0 | 2 | 0 |
| Nr5a1 | Mc2r | NA | NA | NA | NA | NA | NA | 0 | 12 | 0 |
| Nrif1 | Tnfaip8 | NA | NA | NA | NA | NA | NA | 0 | 9 | 0 |
| Nrl | Rho | NA | NA | NA | NA | NA | NA | 0 | 1 | 0 |
| Onecut2 | Sytl4 | NA | NA | NA | NA | NA | NA | 0 | 1 | 0 |
| Pax1 | Pdgfra | NA | NA | NA | NA | NA | NA | 0 | 1 | 0 |
| Pax2 | Tp53 | NA | NA | NA | NA | NA | NA | 0 | 2 | 0 |
| Pax3 | Mitf | NA | NA | NA | NA | NA | NA | 0 | 3 | 0 |
| Pax4 | Ins2 | NA | NA | NA | NA | NA | NA | 0 | 1 | 0 |
| Pax8 | Tpo | NA | NA | NA | NA | NA | NA | 0 | 11 | 0 |
| Pbx1 | Il10 | NA | NA | NA | NA | NA | NA | 0 | 4 | 0 |
| Pcgf2 | Hoxa7 | NA | NA | NA | NA | NA | NA | 0 | 1 | 0 |
| Pitx1 | Fshb | NA | NA | NA | NA | NA | NA | 0 | 1 | 0 |
| Pitx2 | Fshb | NA | NA | NA | NA | NA | NA | 0 | 1 | 0 |
| Plag1 | Abcc6 | NA | NA | NA | NA | NA | NA | 0 | 1 | 0 |
| Pou1f1 | Ghr | NA | NA | NA | NA | NA | NA | 0 | 1 | 0 |
| Pou2f3 | Sprr2d | NA | NA | NA | NA | NA | NA | 0 | 1 | 0 |
| Pou3f1 | Gnrh1 | NA | NA | NA | NA | NA | NA | 0 | 1 | 0 |
| Pou3f2 | Kcnn3 | NA | NA | NA | NA | NA | NA | 0 | 2 | 0 |
| Ppard | Sat1 | NA | NA | NA | NA | NA | NA | 0 | 4 | 0 |
| Prdm1 | Nfix | NA | NA | NA | NA | NA | NA | 0 | 40 | 0 |
| Prox1 | Arhgap42 | NA | NA | NA | NA | NA | NA | 0 | 26 | 0 |
| Pura | Ar | NA | NA | NA | NA | NA | NA | 0 | 1 | 0 |
| Rarb | Adh5 | NA | NA | NA | NA | NA | NA | 0 | 5 | 0 |
| Rarg | Greb1 | NA | NA | NA | NA | NA | NA | 0 | 8 | 0 |
| Rel | Csf2 | NA | NA | NA | NA | NA | NA | 0 | 23 | 0 |
| Rfx2 | Ttc30b | NA | NA | NA | NA | NA | NA | 0 | 20 | 0 |
| Rora | Tuba1b | NA | NA | NA | NA | NA | NA | 0 | 3 | 0 |
| Rorc | Il22b | NA | NA | NA | NA | NA | NA | 0 | 3 | 0 |
| Rreb1 | Klk1b5 | NA | NA | NA | NA | NA | NA | 0 | 14 | 0 |
| Rxrb | Angptl4 | NA | NA | NA | NA | NA | NA | 0 | 6 | 0 |
| Rxrg | Plin5 | NA | NA | NA | NA | NA | NA | 0 | 6 | 0 |
| Sall4 | Zeb1 | NA | NA | NA | NA | NA | NA | 0 | 3 | 0 |
| Satb1 | Tmem245 | NA | NA | NA | NA | NA | NA | 0 | 9 | 0 |
| Ski | Smad7 | NA | NA | NA | NA | NA | NA | 0 | 1 | 0 |
| Skil | Smad7 | NA | NA | NA | NA | NA | NA | 0 | 1 | 0 |
| Snai1 | Thbs1 | NA | NA | NA | NA | NA | NA | 0 | 5 | 0 |
| Snapc4 | Dlk2 | NA | NA | NA | NA | NA | NA | 0 | 29 | 0 |
| Sox11 | Gria2 | NA | NA | NA | NA | NA | NA | 0 | 29 | 0 |
| Sox4 | Dicer1 | NA | NA | NA | NA | NA | NA | 0 | 3 | 0 |
| Sox6 | Aplf | NA | NA | NA | NA | NA | NA | 0 | 32 | 0 |
| Sox8 | Cldn23 | NA | NA | NA | NA | NA | NA | 0 | 1 | 0 |
| Sp2 | Mon1b | NA | NA | NA | NA | NA | NA | 0 | 22 | 0 |
| Spdef | Agr2 | NA | NA | NA | NA | NA | NA | 0 | 1 | 0 |
| Stat4 | Prf1 | NA | NA | NA | NA | NA | NA | 0 | 13 | 0 |
| Stat6 | Sidt1 | NA | NA | NA | NA | NA | NA | 0 | 13 | 0 |
| Tbx2 | Cdkn2a | NA | NA | NA | NA | NA | NA | 0 | 1 | 0 |
| Tbx5 | Nppa | NA | NA | NA | NA | NA | NA | 0 | 2 | 0 |
| Tcf7l1 | Nkd1 | NA | NA | NA | NA | NA | NA | 0 | 5 | 0 |
| Tcfl5 | Cd2 | NA | NA | NA | NA | NA | NA | 0 | 1 | 0 |
| Tfap2b | Drd1 | NA | NA | NA | NA | NA | NA | 0 | 2 | 0 |
| Tfcp2 | Tf | NA | NA | NA | NA | NA | NA | 0 | 1 | 0 |
| Thra | Cga | NA | NA | NA | NA | NA | NA | 0 | 1 | 0 |
| Thrb | Atp2a2 | NA | NA | NA | NA | NA | NA | 0 | 4 | 0 |
| Tlx1 | Aldh1a1 | NA | NA | NA | NA | NA | NA | 0 | 1 | 0 |
| Trps1 | Runx2 | NA | NA | NA | NA | NA | NA | 0 | 1 | 0 |
| Twist2 | Mmp2 | NA | NA | NA | NA | NA | NA | 0 | 3 | 0 |
| Wt1 | Cdh1 | NA | NA | NA | NA | NA | NA | 0 | 30 | 0 |
| Ybx1 | Kpna2 | NA | NA | NA | NA | NA | NA | 0 | 6 | 0 |
| Zbtb11 | Uspl1 | NA | NA | NA | NA | NA | NA | 0 | 22 | 0 |
| Zbtb16 | Ccna2 | NA | NA | NA | NA | NA | NA | 0 | 1 | 0 |
| Zbtb17 | Cdkn2b | NA | NA | NA | NA | NA | NA | 0 | 1 | 0 |
| Zc3h8 | Pdcd11 | NA | NA | NA | NA | NA | NA | 0 | 6 | 0 |
| Zfp202 | Abca1 | NA | NA | NA | NA | NA | NA | 0 | 2 | 0 |
| Zfp644 | Epc1 | NA | NA | NA | NA | NA | NA | 0 | 7 | 0 |
