## Supplementary Figures 1 and 2 for "ATF6 activation alters colonic lipid metabolism causing tumor-associated microbial adaptation"

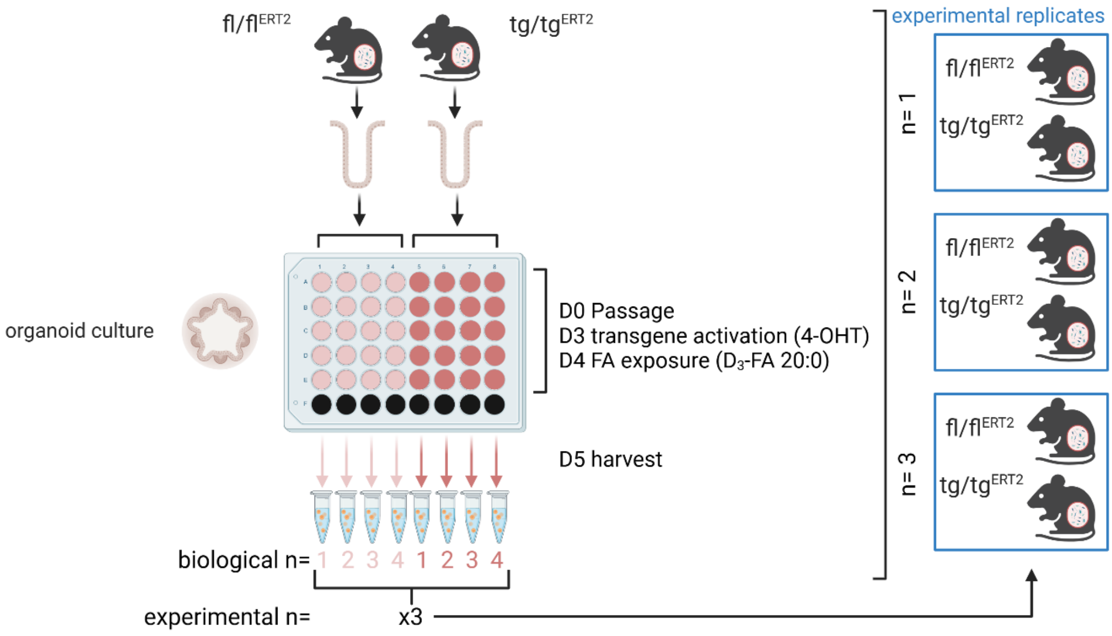


**Supplementary Figure 1. Scheme depicting organoid experiment.** Crypts from donor mice were isolated to generate 4 x 5 organoid cultures. The biological induction of the nATF6 transgene was performed after organoid formation through the application of tamoxifen to the respective 20 wells. After tamoxifen-induced nATF6 expression, all 20 wells were exposed to the isotope labelled fatty acid. Five wells were pooled to allow for enough material to determine fatty acid elongation in 4 different (biological) replicates per mouse. The setup was performed in three replicates with different ATF6fl/fl;Vil-CreERT2 and ATF6tg/tg;Vil-CreERT2 donor mice for organoid cultivation.

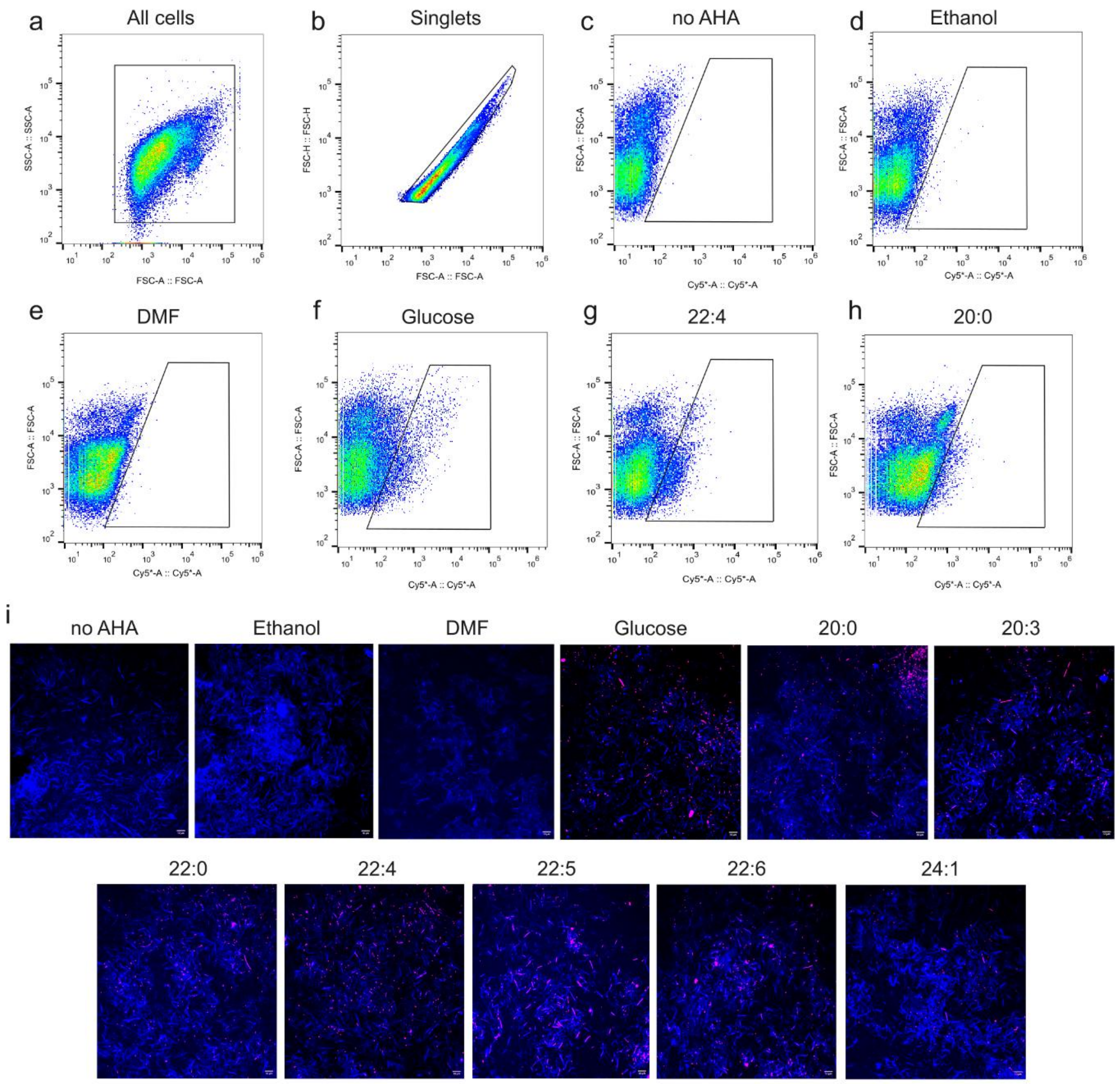


**Supplementary Figure 2. Gating strategy and representative microscopic images of the cecal microbiota of fl/fl mice**. Bacteria were sorted with FACS Melody cell sorter. The sorting gate was utilized to collect the active fraction after incubation with different long chain fatty acids (LCFAs). **a,** Bacteria were visualized using the forward scatter (FSC) and side scatter (SSC). **b,** Doublet discrimination was conducted to exclude doublets from analysis, preventing false positives in the sorted fraction. **c,** The negative control no AHA, which did not contain AHA, was used to set the gate position for BONCAT-positive cells. **d, e,** Ethanol and DMF were included as solvent controls for 20:3, 22:4, 22:5, 22:6, 24:1 and 20:0, 22:0, respectively. **f,** Positive control with glucose was included in every experiment to verify the accuracy of the click reaction and active cell labeling. All four controls were included in every sorting experiment. Example of Cy5-labeled cells in the presence of **g,** 22:4 and h, 20:0. Translationally active (BONCAT-positive) cells are represented as density plots. i, The images display BONCAT-positive (Cy5-labeled bacterial cells) following 6 hours of exposure to long-chain fatty acids (LCFAs) in fl/fl mice. Glucose was used as positive control to confirm the success of the BONCAT and click reaction, no AHA condition was used as a negative control to ensure that no background noise was present in the Cy5 channel. Ethanol and DMF were used as solvent controls for LCFAs 20:3, 22:4, 22:5, 22:6, 24:1, and 20:0, 22:0, respectively. Translationally active bacterial cells are shown in pink (BONCAT positive signal), while all cells are shown in blue (DAPI). The images are displayed as merged channels, with a scale bar of 10 µm.
