## Supplementary Table 6 for "ATF6 activation alters colonic lipid metabolism causing tumor-associated microbial adaptation"

| Id | Feature | Network |
|---|---|---|
| 1 | Michler's ketone | Mucosal |
| 2 | myo−Inositol−1−phosphate | Mucosal |
| 3 | FAHFA 22:6/3:0 | Mucosal |
| 4 | SM 12:2;2O/2:0 | Mucosal |
| 5 | DG 8:0_16:3 | Mucosal |
| 6 | SL 10:0;O/13:0;O | Mucosal |
| 7 | SL 10:0;O/22:2;O | Mucosal |
| 8 | SL 10:1;O/19:2 | Mucosal |
| 9 | TG 8:0_8:0_16:0 | Mucosal |
| 10 | DG O−8:0_24:5 | Mucosal |
| 11 | IV/VI/LV/VL | Mucosal |
| 12 | IV/VI/LV/VL | Mucosal |
| 13 | IT/TI/LT/TL | Mucosal |
| 14 | DG 40:4 | Mucosal |
| 15 | IL/LI | Mucosal |
| 16 | Zotu61; MuribaculaceaeRIAY | Mucosal |
| 17 | Zotu61; MuribaculaceaeRIAY | Mucosal |
| 18 | Zotu154; Lachnoclostridium | Mucosal |
| 19 | Zotu488; Lachnoclostridium | Mucosal |
| 20 | Zotu153; Oscillibacter | Mucosal |
| 21 | Zotu328; Acutalibacter | Mucosal |
| 22 | Zotu497; LachnospiraceaeNK4A136grc | Mucosal |
| 23 | Zotu343; Alistipes | Mucosal |
| 24 | Zotu274; Oscillibacter | Mucosal |
| 1 | Michler's ketone | Luminal |
| 2 | myo−Inositol−1−phosphate | Luminal |
| 3 | FAHFA 22:6/3:0 | Luminal |
| 4 | SM 12:2;2O/2:0 | Luminal |
| 5 | DG 8:0_16:3 | Luminal |
| 6 | SL 10:0;O/13:0;O | Luminal |
| 7 | SL 10:0;O/22:2;O | Luminal |
| 8 | SL 10:1;O/19:2 | Luminal |
| 9 | TG 8:0_8:0_16:0 | Luminal |
| 10 | DG O−8:0_24:5 | Luminal |
| 11 | IV/VI/LV/VL | Luminal |
| 12 | IV/VI/LV/VL | Luminal |
| 13 | IT/TI/LT/TL | Luminal |
| 14 | DG 40:4 | Luminal |
| 15 | IL/LI | Luminal |
| 16 | PA 6:0_22:5 | Luminal |
| 17 | FAHFA 3:0/20:4 | Luminal |
| 18 | PA 3:0_18:4 | Luminal |
| 19 | IT/TI/LT/TL | Luminal |
| 20 | Zotu343; Alistipes | Luminal |
| 21 | Zotu271; MuribaculaceaeRIAY | Luminal |
| 22 | Zotu61; MuribaculaceaeRIAY | Luminal |
