## Supplementary Table 7 for "ATF6 activation alters colonic lipid metabolism causing tumor-associated microbial adaptation"

| | | | | |
|---|---|---|---|---|
| C3:0 | Propionic acid | 73.02898 | C3:0_Propionic acid_73.02897943252 | |
| C4:0 | Butyric acid | 87.04463 | C4:0_Butyric acid_87.04462949698 | |
| C3:0 | Hydroxy-Propionic acid | 89.02392 | C3:0_Hydroxy-Propionic acid_89.02391908432 | |
| C5:0 | Valeric acid | 101.0603 | C5:0_Valeric acid_101.06027956144 | |
| C4:0 | Hydroxy-Butyric acid | 103.0396 | C4:0_Hydroxy-Butyric acid_103.03956914878 | |
| C6:0 | Caproic acid | 115.0759 | C6:0_Caproic acid_115.075929625899 | |
| C5:0 | Hydroxy-Valeric acid | 117.0552 | C5:0_Hydroxy-Valeric acid_117.05521921324 | |
| C7:0 | Enanthic acid | 129.0916 | C7:0_Enanthic acid_129.09157969036 | |
| C6:0 | Hydroxy-Caproic acid | 131.0709 | C6:0_Hydroxy-Caproic acid_131.0708692777 | |
| C8:0 | Caprylic acid | 143.1072 | C8:0_Caprylic acid_143.10722975482 | |
| C7:0 | Hydroxy-Enanthic acid | 145.0865 | C7:0_Hydroxy-Enanthic acid_145.08651934216 | |
| C9:0 | Pelargonic acid | 157.1229 | C9:0_Pelargonic acid_157.12287981928 | |
| C8:0 | Hydroxy-Caprylic acid | 159.1022 | C8:0_Hydroxy-Caprylic acid_159.10216940662 | |
| C10:0 | Capric acid | 171.1385 | C10:0_Capric acid_171.13852988374 | |
| C9:0 | Hydroxy-Pelargonic acid | 173.1178 | C9:0_Hydroxy-Pelargonic acid_173.11781947108 | |
| C11:0 | Undecylic acid | 185.1542 | C11:0_Undecylic acid_185.154179948199 | |
| C10:0 | Hydroxy-Capric acid | 187.1335 | C10:0_Hydroxy-Capric acid_187.13346953554 | |
| C12:0 | Lauric acid | 199.1698 | C12:0_Lauric acid_199.16983001266 | |
| C11:0 | Hydroxy-Undecylic acid | 201.1491 | C11:0_Hydroxy-Undecylic acid_201.1491196 | |
| C13:0 | Tridecylic acid | 213.1855 | C13:0_Tridecylic acid_213.18548007712 | |
| C12:0 | Hydroxy-Lauric acid | 215.1648 | C12:0_Hydroxy-Lauric acid_215.16476966446 | |
| C14:1 | Myristoleic acid | 225.1855 | C14:1_Myristoleic acid_225.1854 | |
| C14:0 | Myristic acid | 227.2011 | C14:0_Myristic acid_227.20113014158 | |
| C13:0 | Hydroxy-Tridecylic acid | 229.1804 | C13:0_Hydroxy-Tridecylic acid_229.18041972892 | |
| C14:1 | Hydroxy-Myristoleic acid | 241.1804 | C14:1_Hydroxy-Myristoleic acid_241.18041972892 | |
| C15:0 | Pentadecylic acid | 241.2168 | C15:0_Pentadecylic acid_241.21678020604 | |
| C14:0 | Hydroxy-Myristic acid | 243.1961 | C14:0_Hydroxy-Myristic acid_243.19606979338 | |
| C16:1 | Sapienic acid | 253.2168 | C16:1_Sapienic acid_Palmitoleic acid_253.21678020604 | |
| C16:0 | Palmitic acid | 255.2324 | C16:0_Palmitic acid_255.2324302705 | |
| C15:0 | Hydroxy-Pentadecylic acid | 257.2117 | C15:0_Hydroxy-Pentadecylic acid_257.21171985784 | |
| C16:1 | Hydroxy-Sapienic aci | 269.2117 | C16:1_Hydroxy-Sapienic aci_Hydroxy-Palmitoleic acid_269.21171985784 | |
| C17:0 | Margaric acid | 269.2481 | C17:0_Margaric acid_269.24808033496 | |
| C16:0 | Hydroxy-Palmitic acid | 271.2274 | C16:0_Hydroxy-Palmitic acid_271.2273699223 | |
| C18:4 | Octadecatetraenoic acid | 275.2011 | C18:4_Octadecatetraenoic acid_Stearidonic acid_Eicosatetraenoic acid_275.20113014158 | 275.2011 |
| C18:3 | SeveralPossibleIsomers | 277.2168 | C18:3_SeveralPossibleIsomers_277.21678020604 | |
| C18:2 | Linoleic acid | 279.2324 | C18:2_Linoleic acid_Rumenic acid or Bovinic acid_279.2324302705 | |
| C18:1 | Oleic acid | 281.2481 | C18:1_Oleic acid_Vaccenic acid_Elaidic acid_281.24808033496 | 281.2481 |
| C18:0 | Stearic acid | 283.2637 | C18:0_Stearic acid_283.26373039942 | |
| C17:0 | Hydroxy-Margaric acid | 285.243 | C17:0_Hydroxy-Margaric acid_285.24301998676 | |
| C18:4 | Hydroxy-Stearidonic acid | 291.1961 | C18:4_Hydroxy-Stearidonic acid_Hydroxy-Octadecatetraenoic acid_Eicosatetraenoic acid_291.19606979338 | 291.1961 |
| C18:3 | SeveralPossibleIsomers | 293.2117 | C18:3_SeveralPossibleIsomers_293.21171985784 | |
| C18:2 | Hydroxy-Rumenic acid | 295.2274 | C18:2_Hydroxy-Rumenic acid_Hydroxy-Linoleic acid_Bovinic acid_295.2273699223 | 295.2274 |
| C18:1 | Hydroxy-Oleic acid | 297.243 | C18:1_Hydroxy-Oleic acid_Hydroxy-Elaidic acid_Hydroxy-Vaccenic acid_297.24301998676 | 297.243 |
| C19:0 | Nonadecylic acid | 297.2794 | C19:0_Nonadecylic acid_297.279380463879 | |
| C18:0 | Hydroxy-Stearic acid | 299.2587 | C18:0_Hydroxy-Stearic acid_299.25867005122 | |
| C20:5 | Bosseopentaenoic acid | 301.2168 | C20:5_Bosseopentaenoic acid_Eicosapentanoic acid_301.21678020604 | |
| C20:4 | Arachidonic acid or Eicosatetra | 303.2324 | C20:4_Arachidonic acid or Eicosatetraenoic acid_303.2324302705 | |
| C20:3 | Mead acid | 305.2481 | C20:3_Mead acid_Dihomo-linolenic acid_305.24808033496 | |
| C20:2 | Eicosadienoid acid | 307.2637 | C20:2_Eicosadienoid acid _307.26373039942 | |
| C20:1 | Eicosenoic acid | 309.2794 | C20:1_Eicosenoic acid _309.279380463879 | |
| C20:0 | Arachidic acid | 311.295 | C20:0_Arachidic acid_311.29503052834 | |
| C19:0 | Hydroxy-Nonadecylic acid | 313.2743 | C19:0_Hydroxy-Nonadecylic acid_313.27432011568 | |
| C20:5 | Hydroxy-Bosseopentaenoic aci | 317.2117 | C20:5_Hydroxy-Bosseopentaenoic acid_Hydroxy-Eicosapentanoic acid_317.21171985784 | |
| C20:4 | Hydroxy-Arachidonic acid or Ei | 319.2274 | C20:4_Hydroxy-Arachidonic acid or Eicosatetraenoic acid_319.2273699223 | |
| C20:3 | Hydroxy-Mead acid | 321.243 | C20:3_Hydroxy-Mead acid_Hydroxy-Dihomo-linolenic acid_321.24301998676 | |
| C20:2 | Hydroxy-Eicosadienoid acid | 323.2587 | C20:2_Hydroxy-Eicosadienoid acid _323.25867005122 | |
| C20:1 | Hydroxy-Eicosenoic acid | 325.2743 | C20:1_Hydroxy-Eicosenoic acid_325.27432011568 | |
| C21:0 | Heneicosylic acid | 325.3107 | C21:0_Heneicosylic acid_325.3106805928 | |
| C22:6 | Cervonic acid | 327.2324 | C22:6_Cervonic acid_Docosahexaenoic acid_327.2324302705 | |
| C20:0 | Hydroxy-Arachidic acid | 327.29 | C20:0_Hydroxy-Arachidic acid_327.289970180139 | |
| C22:5 | Ozubondo acid | 329.2481 | C22:5_Ozubondo acid_Sardine acid_Docosapentanoic acid_329.24808033496 | 329.2481 |
| C22:4 | Docosatetraenoic acid | 331.2637 | C22:4_Docosatetraenoic acid_Adrenic acid_331.26373039942 | |
| C22:2 | Docosadienoic acid | 335.295 | C22:2_Docosadienoic acid_335.29503052834 | |
| C22:1 | Erucic acid | 337.3107 | C22:1_Erucic acid_337.3106805928 | |
| C22:0 | Behenic acid | 339.3263 | C22:0_Behenic acid_339.32633065726 | |
| C21:0 | Hydroxy-Heneicosylic acid | 341.3056 | C21:0_Hydroxy-Heneicosylic acid_341.3056202446 | |
| C22:6 | Hydroxy-docosahexaenoic acid | 343.2274 | C22:6_Hydroxy-docosahexaenoic acid__Hydroxy-Cervonic acid_343.2273699223 | 343.2274 |
| C22:5 | Hydroxy-Docosapentanoic acid | 345.243 | C22:5_Hydroxy-Docosapentanoic acid_Hydroxy-Ozubondo acid_Hydroxy-Sardine acid_345.24301998676 | 345.243 |
| C22:4 | Hydroxy-Docosatetraenoic acid | 347.2587 | C22:4_Hydroxy-Docosatetraenoic acid_Hydroxy-Adrenic acid_347.25867005122 | |
| C22:2 | Hydroxy-Docosadienoic acid | 351.29 | C22:2_Hydroxy-Docosadienoic acid_351.289970180139 | |
| C22:1 | Hydroxy-Erucic acid | 353.3056 | C22:1_Hydroxy-Erucic acid_353.3056202446 | |
| C23:0 | Tricosylic acid | 353.342 | C23:0_Tricosylic acid_353.34198072172 | |
| C24:6 | Herring acid | 355.2637 | C24:6_Herring acid_355.26373039942 | |
| C22:0 | Hydroxy-Behenic acid | 355.3213 | C22:0_Hydroxy-Behenic acid_355.32127030906 | |
| C24:5 | Tetracosapentaenoic acid | 357.2794 | C24:5_Tetracosapentaenoic acid_357.279380463879 | |
| C24:1 | Nervonic acid | 365.342 | C24:1_Nervonic acid_365.34198072172 | |
| C24:0 | Lignoceric acid | 367.3576 | C24:0_Lignoceric acid_367.35763078618 | |
| C23:0 | Hydroxy-Tricosylic acid | 369.3369 | C23:0_Hydroxy-Tricosylic acid_369.33692037352 | |
| C24:6 | Hydroxy-Herring acid | 371.2587 | C24:6_Hydroxy-Herring acid_371.25867005122 | |
| C24:5 | Hydroxy-Tetracosapentaenoic a | 373.2743 | C24:5_Hydroxy-Tetracosapentaenoic acid_373.27432011568 | |
| C24:1 | Hydroxy-Nervonic acid | 381.3369 | C24:1_Hydroxy-Nervonic acid_381.33692037352 | |
| C25:0 | Pentacosylic acid | 381.3733 | C25:0_Pentacosylic acid_381.37328085064 | |
| C24:0 | Hydroxy-Lignoceric acid | 383.3526 | C24:0_Hydroxy-Lignoceric acid_383.35257043798 | |
| C26:0 | Cerotic acid | 395.3889 | C26:0_Cerotic acid_395.388930915099 | |
| C25:0 | Hydroxy-Pentacosylic acid | 397.3682 | C25:0_Hydroxy-Pentacosylic acid_397.36822050244 | |
| C27:0 | Carboceric acid | 409.4046 | C27:0_Carboceric acid_409.40458097956 | |
| C26:0 | Hydroxy-Cerotic acid | 411.3839 | C26:0_Hydroxy-Cerotic acid_411.3838705669 | |
| C28:0 | Montanic acid | 423.4202 | C28:0_Montanic acid_423.42023104402 | |
| C27:0 | Hydroxy-Carboceric acid | 425.3995 | C27:0_Hydroxy-Carboceric acid_425.399520631359 | |
| C29:0 | Nonacosylic acid | 437.4359 | C29:0_Nonacosylic acid_437.43588110848 | |
| C28:0 | Hydroxy-Montanic acid | 439.4152 | C28:0_Hydroxy-Montanic acid_439.41517069582 | |
| C30:0 | Melissic acid | 451.4515 | C30:0_Melissic acid_451.45153117294 | |
| C29:0 | Hydroxy-Nonacosylic acid | 453.4308 | C29:0_Hydroxy-Nonacosylic acid_453.43082076028 | |
| C31:0 | Hentriacontylic acid | 465.4672 | C31:0_Hentriacontylic acid_465.4671812374 | |
| C30:0 | Hydroxy-Melissic acid | 467.4465 | C30:0_Hydroxy-Melissic acid_467.44647082474 | |
| C32:0 | Lacceroic acid | 479.4828 | C32:0_Lacceroic acid_479.48283130186 | |
| C31:0 | Hydroxy-Hentriacontylic acid | 481.4621 | C31:0_Hydroxy-Hentriacontylic acid_481.4621208892 | |
| C33:0 | Psyllic acid | 493.4985 | C33:0_Psyllic acid_493.498481366319 | |
| C32:0 | Hydroxy-Lacceroic acid | 495.4778 | C32:0_Hydroxy-Lacceroic acid_495.47777095366 | |
| C34:0 | Geddic acid | 507.5141 | C34:0_Geddic acid_507.51413143078 | |
| C33:0 | Hydroxy-Psyllic acid | 509.4934 | C33:0_Hydroxy-Psyllic acid_509.49342101812 | |
| C35:0 | Ceroplastic acid | 521.5298 | C35:0_Ceroplastic acid_521.52978149524 | |
| C34:0 | Hydroxy-Geddic acid | 523.5091 | C34:0_Hydroxy-Geddic acid_523.50907108258 | |
| C36:0 | Hexatriacontylic acid | 535.5454 | C36:0_Hexatriacontylic acid_535.5454315597 | |
| C35:0 | Hydroxy-Ceroplastic acid | 537.5247 | C35:0_Hydroxy-Ceroplastic acid_537.52472114704 | |
| C37:0 | Heptatriacontylic acid | 549.5611 | C37:0_Heptatriacontylic acid_549.56108162416 | |
| C36:0 | Hydroxy-Hexatriacontylic acid | 551.5404 | C36:0_Hydroxy-Hexatriacontylic acid_551.5403712115 | |
| C38:0 | Octatriacontylic acid | 563.5767 | C38:0_Octatriacontylic acid_563.57673168862 | |
| C37:0 | Hydroxy-Heptatriacontylic acid | 565.556 | C37:0_Hydroxy-Heptatriacontylic acid_565.55602127596 | |
| C39:0 | Nonatriacontylic acid | 577.5924 | C39:0_Nonatriacontylic acid_577.59238175308 | |
| C38:0 | Hydroxy-Octatriacontylic acid | 579.5717 | C38:0_Hydroxy-Octatriacontylic acid_579.57167134042 | |
| C40:0 | Tetracontylic acid | 591.608 | C40:0_Tetracontylic acid_591.60803181754 | |
| C39:0 | Hydroxy-Nonatriacontylic acid | 593.5873 | C39:0_Hydroxy-Nonatriacontylic acid_593.58732140488 | |
| C40:0 | Hydroxy-Tetracontylic acid | 607.603 | C40:0_Hydroxy-Tetracontylic acid_607.602971469339 | |
